# Integrating spatial and single-cell multi-omics analysis of induced pluripotent stem cell-derived cervical adenocarcinoma model

**DOI:** 10.64898/2026.05.01.722143

**Authors:** Saki Kamata, Ayumi Taguchi, Hitoshi Iuchi, Yuji Ikeda, Rie Maruyama, Yoko Nakanishi, Toshihiro Sugi, Yuki Okuma, Osamu Kobayashi, Naoko Tomita, Daisuke Yoshimoto, Lifang Wang, Nozomi Moritsugu, Chitose Takahashi, Michihira Tagami, Hiroko Matsunaga, Toshitsugu Okayama, Ri-ichiroh Manabe, Kazuma Kiyotani, Kazuho Ikeo, Yasushi Okazaki, Tohru Kiyono, Shinobu Masuda, Michiaki Hamada, Haruko Takeyama, Kei Kawana

## Abstract

Human papillomavirus 18 (HPV18) preferentially infects cervical stem cell-like cells and is strongly associated with adenocarcinoma. However, the mechanisms underlying differentiation into cervical adenocarcinoma remain unclear due to the lack of appropriate experimental models. We aimed to establish a model of HPV18-associated cervical adenocarcinoma and elucidate its molecular and cellular differentiation mechanisms. HPV18 E6/E7 were introduced into induced pluripotent stem cell-derived reserve cell-like cells (iRCs) to generate tumor models. Spatial transcriptomics and single-cell multi-omics analyses were performed to integrate histological and molecular data. A distinct component (Gland_A) exhibited morphological and immunohistochemical features of cervical adenocarcinoma and was efficiently induced in iRC-18 tumors. Gland_A showed increased chromatin accessibility and elevated expression of FOXA1, FOXA2, and ALDH1A1. Analysis of clinical samples confirmed enrichment of ALDH1A1 in HPV-associated adenocarcinomas. This model recapitulates key features of HPV18-associated cervical adenocarcinoma and provides insights into its differentiation mechanisms.

## Introduction

In developed countries, the incidence of cervical cancer among young individuals has decreased owing to the widespread implementation of human papillomavirus (HPV) vaccination^1^. However, incomplete implementation in low- and middle-income countries hinders eradication efforts. The molecular mechanisms of HPV-associated carcinogenesis are well established and shared across HPV-associated cancers.

Most cervical cancers (> 95%) are caused by HPV infection, particularly types 16 and 18, which account for approximately 70% of detectable HPV genotypes in cervical cancer^2^. Cells in the squamocolumnar junction (SCJ) of the uterine cervix are considered the origin of HPV-associated cervical cancer^3^. After infecting SCJ cells, HPV replicates in parallel with the differentiation of stratified squamous epithelium, establishing persistent infection through cycles of latency and replication^4^. The HPV genome, approximately 8,000 base pairs in circular double-stranded DNA, encodes six early region genes (E1, E2, E4, E5, E6, and E7) involved in viral replication. The E6 protein inactivates p53, whereas E7 inactivates Rb^5^. During persistent HPV infection, the HPV genome integrates into the human genome, leading to continuous expression of E6 and E7 and to genetic changes such as oncogene amplification and chromosomal rearrangements^5,6^. Through cycles of progression and regression, HPV-infected cells eventually develop into cervical cancer^7^.

HPV16 is implicated in approximately half of all cervical cancers, whereas HPV18 accounts for 10–20%. Squamous cell carcinoma represents about 70% of HPV-related cancers, although adenocarcinomas also occur and comprise over 20% of cases. Adenocarcinomas have a worse prognosis than squamous cell carcinoma^8,9^. In cervical adenocarcinoma, the prevalence of HPV16 and HPV18 is nearly equivalent (49.8% and 45.3%, respectively) and largely independent of histological subtype or geographic region. Notably, HPV18 prevalence in adenocarcinoma is approximately three times higher than in squamous cell carcinoma^10^.

The molecular mechanisms underlying squamous cell and glandular differentiation remain unclear^11^. A detailed investigation of how HPV18 drives adenocarcinoma development is therefore warranted.

HPV18 is frequently detected in adenocarcinoma and exhibits distinctive characteristics among high-risk HPVs; its associated lesions are less commonly identified at precancerous stages^12^. Recent studies have examined the cellular origin of HPV18 and its carcinogenic properties. HPV18-infected cells uniquely retain stem cell pluripotency, targeting undifferentiated stem cell-like cells^11,13,14^. The HPV18 genome integrates into the host genome at the early stages of infection, and precancerous lesions often lack expression of the capsid protein L1^15,16^. HPV18-positive cells remain pluripotent and can differentiate into multiple tissue types even after viral integration^17,18^. Because HPV18 maintains stem cell–like characteristics, establishing differentiation models from stem cells is crucial to elucidate adenocarcinoma development. However, due to the lack of suitable models, the intracellular regulatory mechanisms governing differentiation into cervical adenocarcinoma remain poorly understood.

In a previous study, we established induced reserve cell-like cells (iRCs) from induced pluripotent stem (iPS) cells^19^. These iRCs expressed markers of stem cells, SCJ, and Müllerian duct-derived cells markers, and were capable of differentiating into glandular and stratified squamous epithelium in 3D culture. In the present study, we generated a mouse model using HPV18 E6/E7-transduced iRCs and investigated the molecular mechanisms of HPV18-induced adenocarcinomas using spatial and single-cell multi-omics analyses.

## Results

### HPV16/18 oncogenes increase tumorigenic potential in iRCs

HPV16 and HPV18 E6 and E7 were introduced into iRCs to generate iRCs expressing HPV16 or HPV18 E6/E7 (iRC-16 and iRC-18) (Fig. 1a and Supplementary Fig. 1). A colony formation assay showed that iRC-16 and iRC-18 formed significantly more colonies than control iRCs treated with retrovirus alone (iRC-cont) (Fig. 1b). Proliferation curves indicated that iRC-16 and iRC-18 proliferated significantly faster than iRC-cont (Fig. 1c). In mouse tumor models, the engraftment rates of iRC-16 and iRC-18 were 100%, whereas that of iRC-cont was 60%, suggesting that HPV16/18 E6/E7 expression enhances tumor engraftment in vivo (Fig. 1e).

**Fig. 1:**
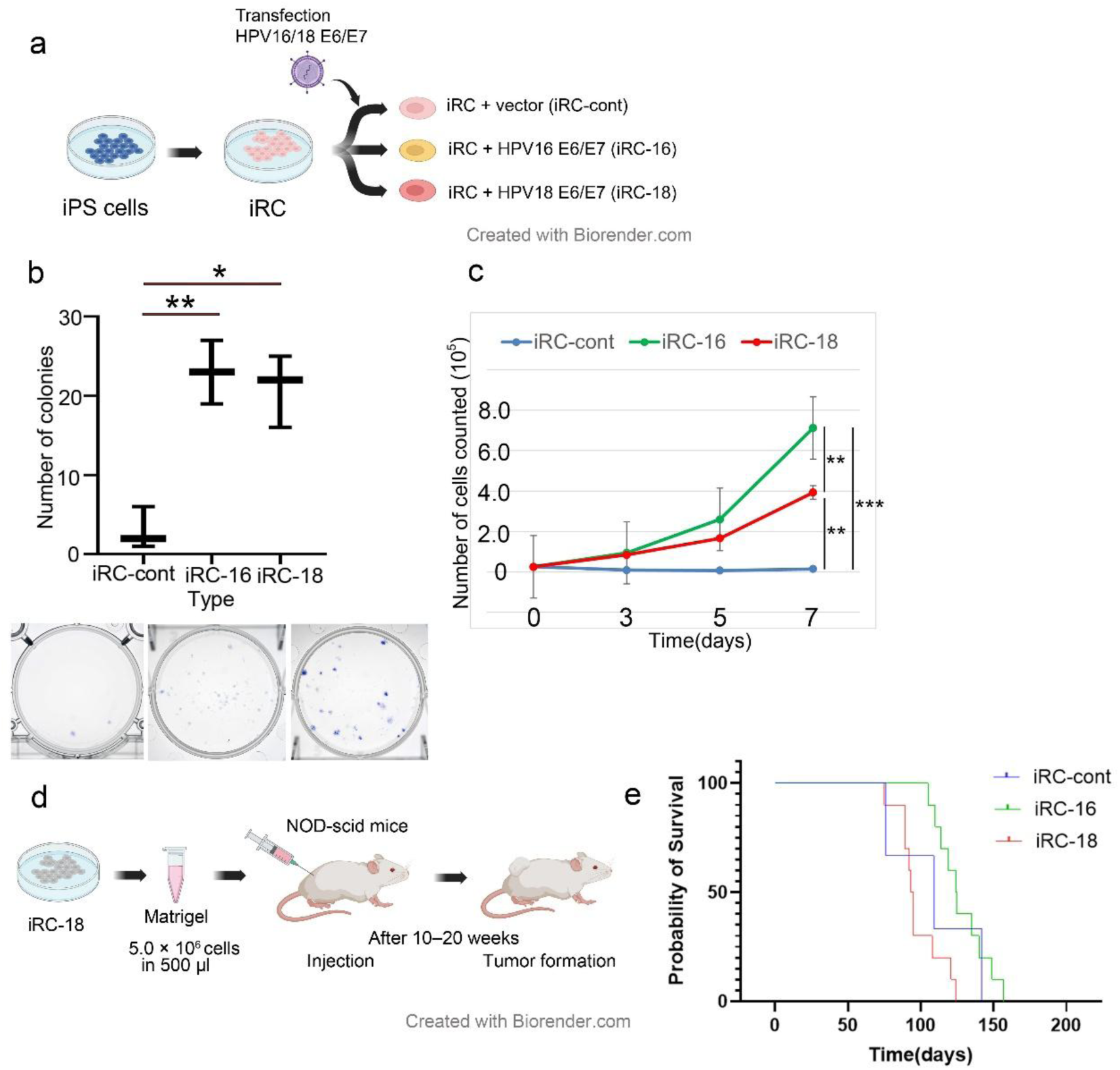
Establishment of HPV18 E6/E7-transduced iRC (iRC-18). (**a**) Flowchart illustrating the establishment of iRC-18 from iPS cells. iRCs were generated from iPS cells under specific culture conditions. On day 6, primary iRCs were inoculated with retroviral fluid of LXSN (iRC-cont) or LXSN-18E6E7 (iRC-18s). (**b**) Colony formation assay of iRC-cont, iRC-16, and iRC-18. Cells were seeded at 100 cells/dish and incubated for 2 weeks. Colony numbers were counted. One-way ANOVA followed by Tukey’s test. *P < 0.05, **P < 0.01. (**C**) Cell proliferation assay of iRC-cont, iRC-16, and iRC-18. Cells (passage 7–9) were seeded at 2.5 × 10^4^ cells/dish, and cell numbers were counted on days 3, 5, and 7. Statistical comparisons were performed for day 7. One-way ANOVA followed by Tukey’s test. **P < 0.01, ***P < 0.001. (**d**) Flowchart of the establishment of iRC mouse tumor models. iRC-cont, iRC-16, and iRC-18 (5.0 × 10^5^ cells/mouse) were subcutaneously injected with 500 μL Matrigel into NOD SCID mice (n = 3, 10, and 10 respectively). Tumors developed 10–20 weeks after injection. (**e**) Kaplan–Meier curves for iRC-cont (n = 3), iRC-16 (n = 10), and iRC-18 (n = 10) tumors, where the event was defined as the time from transplantation to when the tumor reached a maximum diameter of 2.0 cm (sacrifice time). iRC; induced reserve cell-like, iPS; induced pluripotent stem; NOD SCID, non-obese diabetic severe combined immunodeficiency.

### iRC-derived tumors comprise undifferentiated and glandular components

Tumors derived from iRC-18 were pathologically classified into undifferentiated (Undiff) and glandular components (Gland_A and Gland_B) (Fig. 2a). Undifferentiated tumors displayed partial neural differentiation, with CD56 expressed in approximately 40% of cells, and synaptophysin and chromogranin A expressed in 5% and 1% of cells, respectively, while being negative for p63 (data not shown). Gland_A tumors showed well-differentiated glandular lesions with secretions and were periodic acid–Schiff (PAS)-positive. In contrast, Gland_B tumors exhibited a papillary growth pattern, were moderately differentiated, and were PAS-negative.

**Fig. 2:**
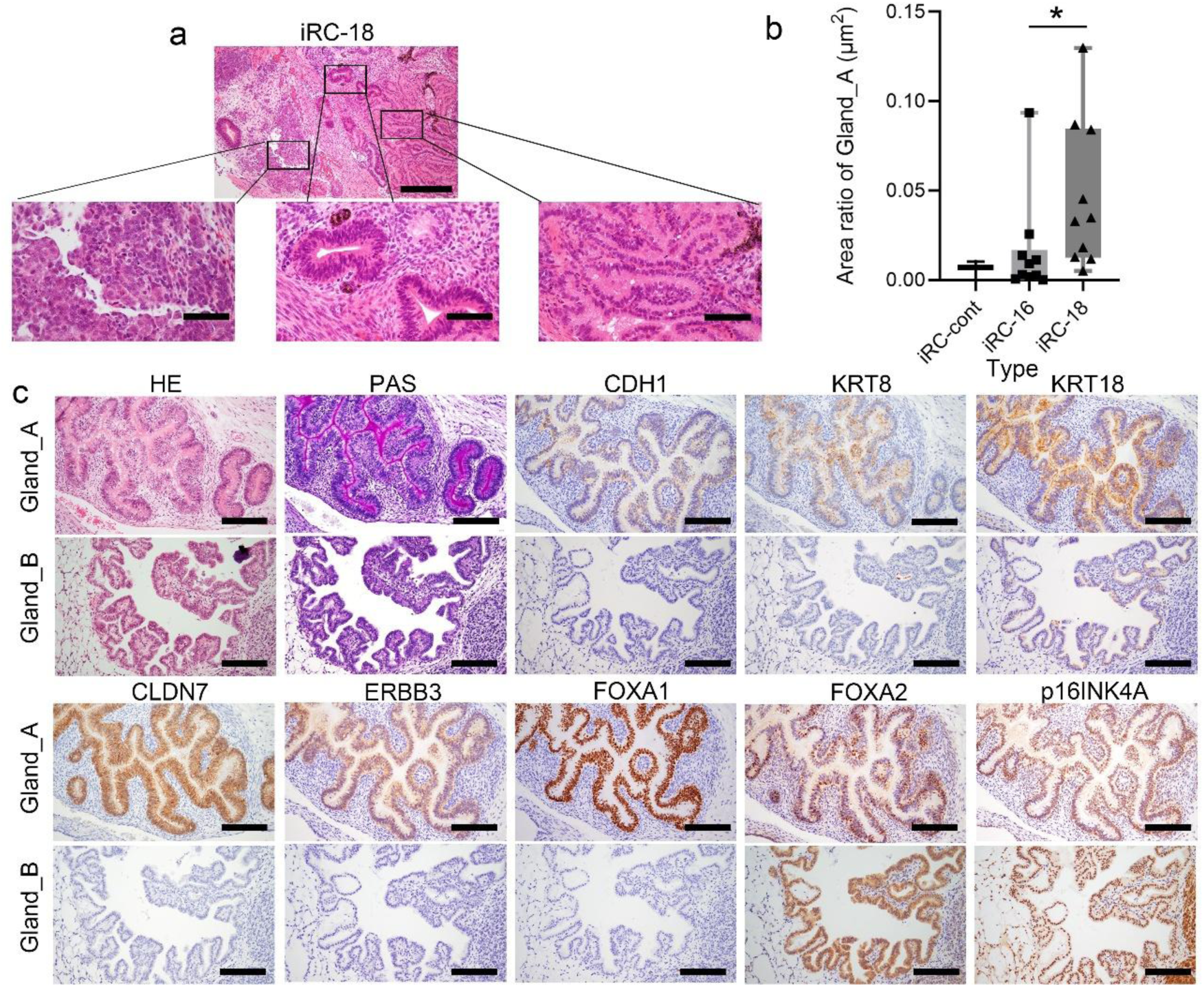
Pathological examination of iRC-18 tumors. (**a**) H&E staining of iRC-18 tumors. The upper image shows H&E staining (magnification, 100×). The lower left, middle, and right panels represent Undiff, Gland_A, and Gland_B, respectively (magnification, 400×). Scale bar indicates 200 μm (upper panel) and 50 μm (lower panels). (**b**) Area ratio of Gland_A in iRC-cont (n = 3), iRC-16 (n = 10), and iRC-18 (n = 10). Kruskal–Wallis test followed by Steel–Dwass test. *P < 0.05. (**c**) Representative staining images of Gland_A and Gland_B. The upper panel shows H&E, PAS, and immunostaining for Gland_A, whereas the lower panel shows staining for Gland_B (magnification, 200×). From upper left to lower right, H&E, PAS, CDH1, KRT8, KRT18, CLDN7, ERBB3, FOXA1, FOXA2, and p16INK4A staining. FOXA1 and FOXA2 were considered positive when nuclear staining was observed, and p16INK4A was considered positive when nuclear or cytoplasmic staining was present. Membrane staining for CDH1, KRT8, KRT18, CLDN7, and ERBB3 was considered positive.

The proportion of Gland_A tumors was significantly higher in iRC-18 than in iRC-16 (p = 0.04). However, no significant difference was observed compared with iRC-cont, likely due to the small sample size (Fig. 2b and Supplementary Fig. 2a).

Immunohistological analysis revealed that Gland_A was positive for cadherin 1 (CDH1), whereas Gland_B was not. Gland_A also expressed keratin 8 (KRT8), KRT18, claudin 7 (CLDN7), Erb-b2 receptor tyrosine kinase 3 (ERBB3), FOXA1, and FOXA2, which are typical markers of cervical adenocarcinoma^20^ (Fig. 2c, and Supplementary Table 1).

In normal cervix immunostaining, KRT8 and KRT18 were strongly positive; FOXA1 and FOXA2 were positive; and CDH1, CLDN7, ERBB3, and p16INK4A were weakly positive (Supplementary Fig. 2b, and Supplementary Table 1). In contrast, Gland_B lacked these markers (FOXA2 in Gland_B was considered negative because staining was only cytoplasmic) (Fig. 2c and Supplementary Table 1). We subsequently assessed p16INK4A expression in each tumor component. Cyclin-dependent kinase inhibitor 2A (CDKN2A; p16INK4A) was positive in Gland_A, Gland_B, and Undiff tissues (Fig. 2c and Supplementary Table 1). Quantitative analysis showed that CDH1, KRT8, KRT18, CLDN7, ERBB3, FOXA1, and FOXA2 were more highly expressed in Gland_A than in Gland_B and Undiff, whereas p16INK4A expression did not differ significantly among the groups (Supplementary Fig. 3). Given that Gland_A exhibited immunohistochemical features characteristic of HPV-associated adenocarcinoma, and that iRC-18 showed high induction efficiency of Gland_A, these findings suggest that introduction of HPV18 E6/E7 into cervical stem cells can drive the development of HPV-associated adenocarcinoma.

### Distinct gene expression profiles for tumor components

To characterize the molecular features of the two glandular structures and the undifferentiated component in iRC-18 tumors, we performed microdissection-based spatial transcriptome analysis (Fig. 3a). Principal component analysis and hierarchical clustering confirmed distinct gene expression patterns across tissue types (Fig. 3b, c, and Supplementary Fig. 4a). Differentially expressed gene (DEG) analysis revealed 2,460, 3,799, and 3,047 significantly different genes between Undiff vs. Gland_A, Undiff vs. Gland_B, and Gland_A vs. B, respectively (Fig. 3d, Supplementary Fig. 4b, and Supplementary Tables 2–4). Gland_A showed elevated expression of cervical adenocarcinoma markers, including FOXA2, KRT8, KRT18, mucin 1, CLDN4, and CLDN7. Both Undiff and Gland_A exhibited higher CDKN2A (p16INK4A) expression than Gland_B (Supplementary Fig. 4c). Kyoto Encyclopedia of Genes and Genomes (KEGG) pathway analysis showed that Gland_A was enriched in “glycerolipid metabolism” and “fat digestion and absorption,” whereas Gland_B was enriched in genes related to the “Wnt” and “TGF-β” pathways. The Undiff component showed elevated expression of genes involved in “DNA replication,” “cell cycle,” and “synaptic vesicle cycle pathways,” suggesting high proliferative and neural-like characteristics (Supplementary Tables 5–10). Gene Ontology (GO) analysis further revealed that Gland_A was associated with lipid metabolism, Gland_B with cilium-related processes, and the Undiff group with nerve-related functions (Supplementary Tables 11–16).

**Fig. 3:**
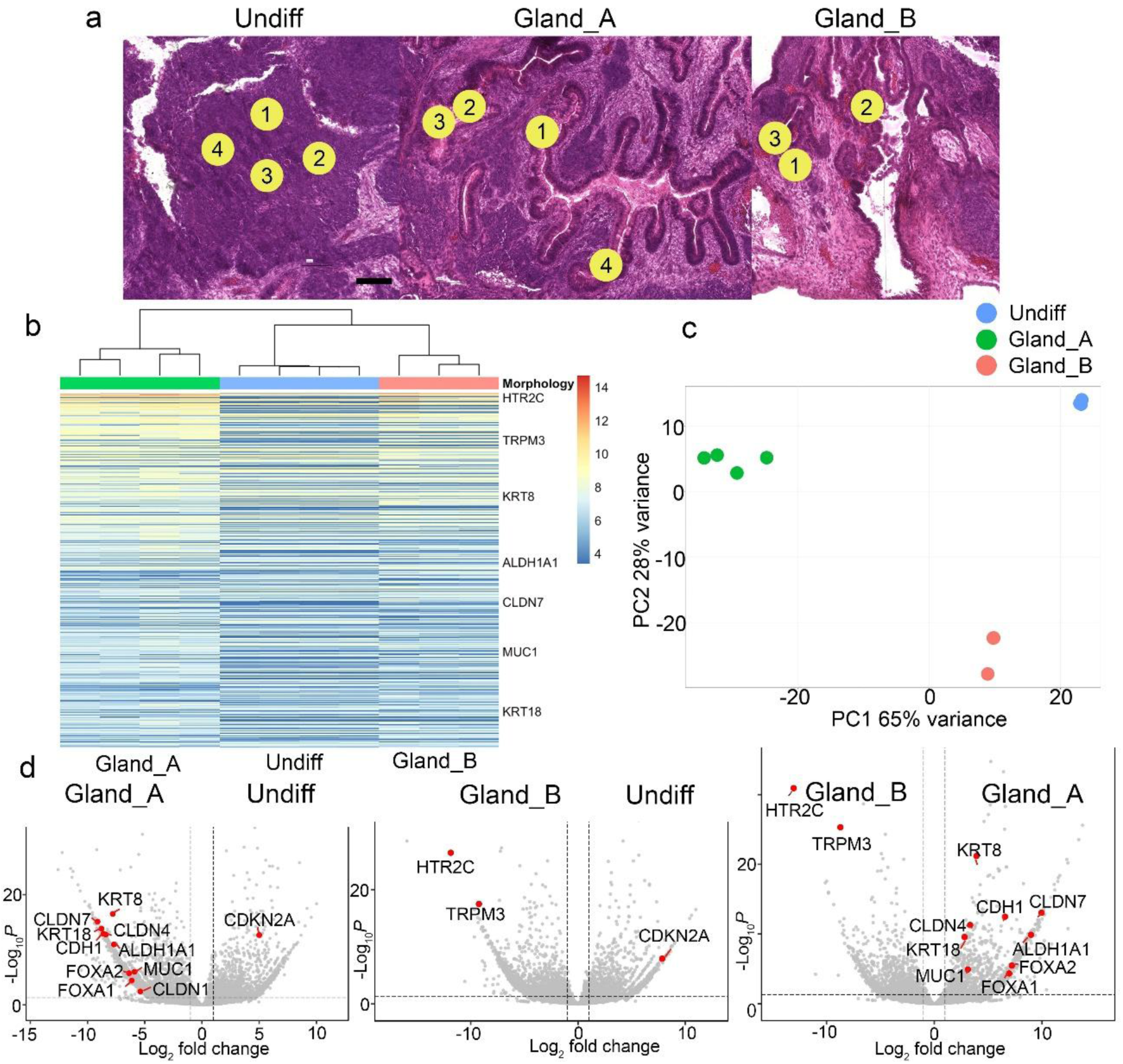
Spatial transcriptome analysis of iRC-18 tumors. (**a**) Microdissection was performed using 100-μm circular regions at three representative sites for Undiff, Gland_A, and Gland_B. (**b**) Heatmap of microdissection-based spatial transcriptomics. The 500 most variable genes based on RNA sequencing of Undiff, Gland_A, and Gland_B. (**c**) Principal component analysis (PCA) of spatial transcriptomics data. PCA plot of Undiff, Gland_A, and Gland_B samples. (**d**) Volcano plots of DEGs among the three tissue types (Undiff, Gland_A, and Gland_B). The left, middle, and right panels show comparisons between Undiff vs. Gland_A, Undiff vs. Gland_B, and Gland_A vs. Gland_B. Fold change > 1, P < 0.05. The horizontal dashed line indicates an adjusted P value of 0.05 (Wald test in DESeq2 with Benjamini–Hochberg correction), and the vertical dashed lines indicate log_2_FC of −1 and 1. Each dot represents a gene, with selected candidate genes highlighted in red. KRT, keratin; CLDN, claudin; ALDH, aldehyde dehydrogenase; MUC, mucin; HTR, 5-hydroxytryptamine receptor; TRPM; transient receptor potential cation channel subfamily M member; CDH, cadherin.

### Single-cell RNA sequencing (scRNA-seq) reveals HPV18-positive cell-enrichment in Gland_A

To elucidate intracellular mechanisms of cell differentiation at single-cell resolution, we conducted single-cell multi-omics analysis of iRC-18 tumors. Clustering of scRNA-seq data identified six distinct clusters, which were named based on their gene expression profiles: Cluster 0, extracellular matrix (ECM)/invasion; Cluster 1, oncogenic/ECM-remodeling; Cluster 2, neuronal/synaptic; Cluster 3, mesenchymal/epithelial–mesenchymal transition (EMT)-like; Cluster 4, neuro-ciliated/stem-like; and Cluster 5, neuroendocrine/secretory (Fig. 4a and Supplementary Table 17).

**Fig. 4:**
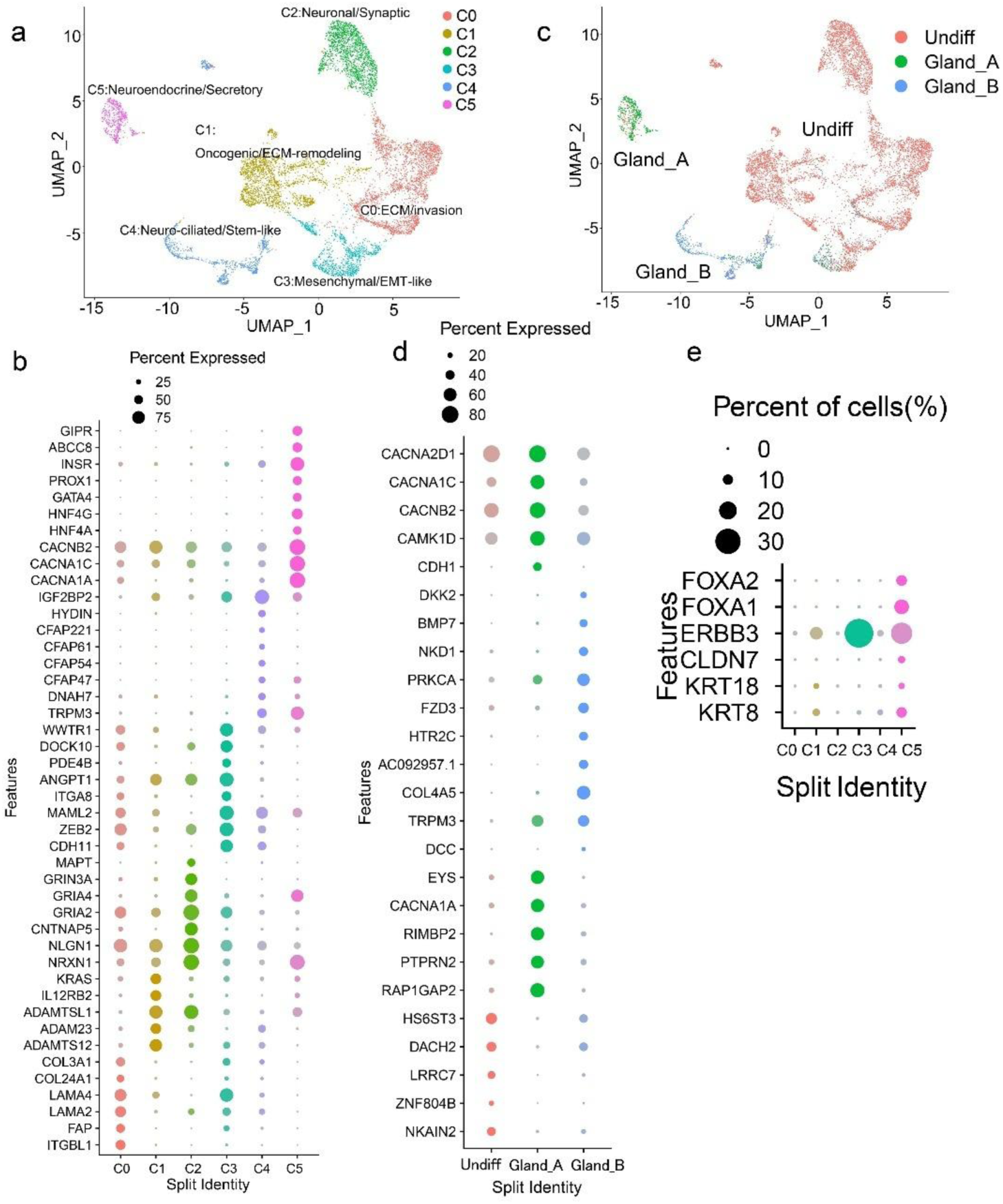
Single-cell RNA-seq analysis of iRC-18 tumors. (**a**) UMAP of single-cell RNA-seq data. Single-cell ATAC-seq and gene expression analyses were performed on iRC-18 tumors from two mice. After batch correction, dimensionality reduction was performed based on the RNA-seq data. (**b**) Heatmap of single-cell RNA-seq data based on gene expression-defined clusters. (**c**) Spatial integration of single-cell and microdissection-based RNA-seq data. Single-cell classification was performed using SingleR, with microdissection-based RNA-seq data as a reference to categorize each cell into Undiff (red), Gland_A (green), or Gland_B (blue). (**d**) Heatmap of single-cell RNA-seq data based on clusters derived from spatial integration of single-cell and microdissection-based RNA-seq data. (**e**) Heatmap of genes associated with glandular structures and cervical adenocarcinoma. Genes related to cervical adenocarcinoma (FOXA2, FOXA1, and ERBB3) and glandular structures (CLDN7, KRT18, and KRT8) were used to construct the heatmap based on gene expression-based clusters. C, cluster; ECM, extracellular matrix; EMT, epithelial–mesenchymal transition; ITG, integrin; FAP, fibroblast activation protein; LAMA, laminin subunit alpha; COL, collagen; ADAMTS, a disintegrin-like and metalloproteinase with thrombospondin; IL, interleukin; KRAS, Kirsten rat sarcoma viral oncogene homolog; NRXN, neurexin; NLGN, neuroligin; CNTNAP, contactin-associated protein; GRIA, glutamate ionotropic receptor AMPA; GRIN, glutamate ionotropic receptor NMDA; MAPT, microtubule associated protein tau; CDH, cadherin; ZEB, zinc finger E-box binding homeobox; MAML, mastermind like; ANGPT, angiopoietin; PDE, phosphodiesterase; DOCK, dedicator of cytokinesis; WWTR, WW domain containing transcription regulator, TRPM, transient receptor potential cation channel subfamily M member; DNAH, dynein axonemal heavy chain; CFAP, cilia- and flagella-associated protein; HYDIN, HYDIN axonemal central pair apparatus protein; IG, insulin like growth factor; CACN, calcium voltage-gated channel; HNF, hepatocyte nuclear factor; PROX, prospero homeobox; INSR, insulin receptor; ABCC, ATP binding cassette subfamily C; GIPR, gastric inhibitory polypeptide receptor; NKAIN, sodium/potassium transporting ATPase interacting; ZNF, Zinc finger protein; LRRC, leucine rich repeat containing; DACH, dachshund family transcription; HS6SST3, heparan sulfate 6-O-sulfotransferase 3; RAP1GAP, RAP1 GTPase activating protein; PTPR, protein tyrosine phosphatase receptor; RIMBP, RIMS binding protein; EYS, EGF-like photoreceptor maintenance factor; DCC, DCC netrin 1 receptor; HTR, 5-hydroxytryptamine receptor; FZD, frizzled class receptor; PRKCA, protein kinase C alpha; NKD, NKD inhibitor of Wnt signaling pathway; BMP, bone morphogenetic protein; DKK, dickkopf Wnt signaling pathway inhibitor; CAMK, calcium/calmodulin dependent protein kinase; KRT, keratin; CLDN, claudin; ERBB, Erb-B2 receptor tyrosine kinase; FOX, Forkhead box.

Cluster 2 showed upregulation of neuronal genes, including core synaptic adhesion genes (NRXN1, NLGN1, CNTNAP5). Cluster 4 showed increased expression of cilia-related genes, such as TRPM3, DNAH7, and CFAP family genes (CFAP47, CFAP54, CFAP61, CFAP221), as well as HYDIN. Cluster 5 showed increased expression of calcium channel genes (CACNA1A, CACNA1C, CACNB2) and endocrine/metabolic genes (HNF4A, HNF4G, GATA4, PROX1) (Fig. 4b and Supplementary Table 17).

SingleR integration annotated 7,524 cells as Undiff, 659 as Gland_A, and 863 as Gland_B (Fig. 4c and Supplementary Fig. 5a). Analysis of the correspondence between tissue types and clusters revealed that Undiff corresponded to the neuronal/synaptic cluster (Cluster 2), Gland_B to the neuro-ciliated/stem-like cluster (Cluster 4), and Gland_A to the neuroendocrine/secretory cluster (Cluster 5). Based on the spatially annotated cells, we examined the characteristic gene expression profiles of the Undiff, Gland_A, and Gland_B components. Undiff showed increased expression of neuronal-related genes. Gland_B was characterized by upregulation of Wnt signaling, TGF-β/BMP-related molecules, and cancer-associated genes, including MAPK10 and ERBB4. Gland_A exhibited high expression of calcium channel genes, including CACNA1A, CACNA1B, CACNA1C, CACNA2D1, CAMK1D, and CAMK2D, along with elevated CDH1 expression (Fig. 4d and Supplementary Table 17). In this study, iRC-18 tumors from two mice were analyzed; however, a cell population corresponding to Gland_A was detected in only one tumor (Supplementary Fig. 5b).

We next examined HPV-derived gene expression and found that HPV18 E6 and E7 were sporadically expressed in Clusters 0, 1, and 2, with marked enrichment in Cluster 5 (Supplementary Fig. 5c). These four HPV-positive clusters showed strong CDKN2A (p16INK4A) expression (Supplementary Fig. 5d). In situ hybridization further demonstrated that Gland A and Undiff were particularly positive for HPV18 E7 (Supplementary Fig. 5e–g). These findings suggest that Gland_A represents HPV-positive adenocarcinoma characterized by enhanced calcium channel activity and endocrine metabolism, Gland_B represents glandular structures with enhanced ciliated-related gene expression and Wnt and TGF-β/BMP signaling, and Undiff represents HPV-positive components with neural-like features.

We subsequently evaluated the expression of HPV-positive cervical adenocarcinoma genes (FOXA1, FOXA2, and ERBB3)^20^ and glandular structure-related genes (KRT8, KRT18, and CLDN7)^21–23^. KRT8 and KRT18 were highly expressed in Clusters 1 and 5, whereas FOXA1, FOXA2, and CLDN7 were specifically enriched in Cluster 5 (Gland_A, Fig. 4e and Supplementary Fig. 5h).

### Enhanced chromatin accessibility in Gland_A for HPV-associated genes

Integrated analysis of microdissection-based spatial transcriptomics and single-cell RNA-seq enabled identification of cells corresponding to each histological type at the single-cell resolution. Dimensionality reduction was performed using single-cell assay for transposase-accessible chromatin with high-throughput sequencing (ATAC-seq) data. Gene expression-based clusters (Clusters 0, 1, 2, 3, 4, 5, and 6 in Fig. 4a) were recapitulated in the ATAC-based Uniform Manifold Approximation and Projection (UMAP) (Fig. 5a and Supplementary Fig. 6a for iRC-18-1 and iRC-18-2, respectively). Cells from each cluster grouped closely in ATAC-based UMAP space (Fig. 5a, b, and Supplementary Fig. 6a, b).

**Fig. 5:**
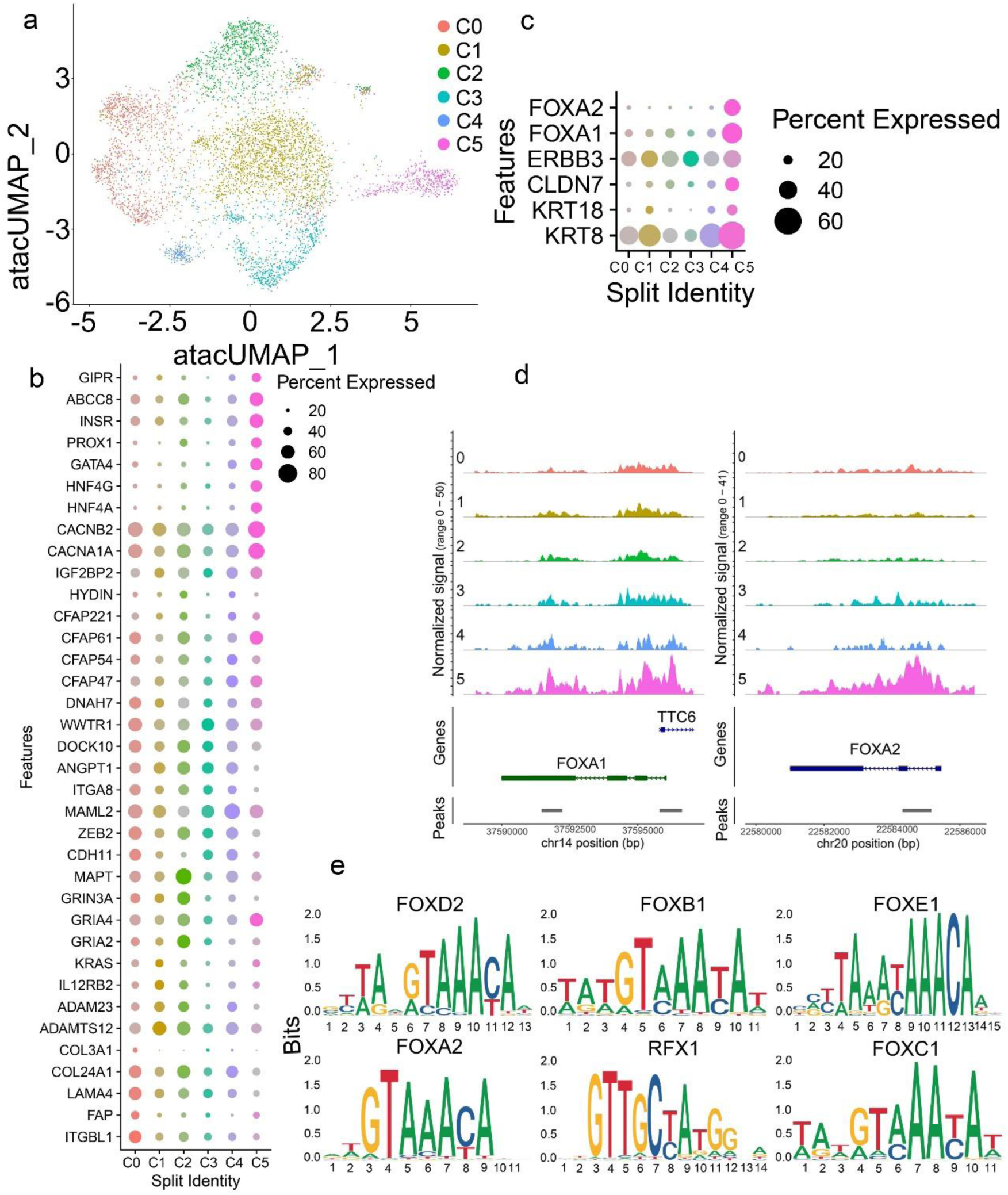
Single-cell ATAC-seq analysis of the iRC-18-1 tumor. (**a**) UMAP of single-cell ATAC-seq data. Single-cell ATAC-seq and gene expression analyses were performed on iRC-18-1 tumors from two mice. Dimensionality reduction was performed based on ATAC-seq data from the iRC-18-1 tumor. Cluster identities were assigned according to gene expression-based analyses. (**b**) Heatmap of single-cell ATAC-seq data based on gene expression-defined clusters. Candidate genes were selected for clusters 0–5. Some genes (CACNA1A, TRPM3, PDE4B, CNTNAP5, NLGN1, NRXN1, ADAMTSL1, and LAMA2) were not detected in the ATAC-seq dataset. (**c**) ATAC-seq heatmap of genes associated with glandular structures and cervical adenocarcinoma. Genes related to cervical adenocarcinoma (FOXA2, FOXA1, and ERBB3) and glandular structures (CLDN7, KRT18, and KRT8) were used to construct the heatmap based on gene expression-based clusters. Chromatin accessibility profiles at the FOXA1 and FOXA2 loci, visualized using the Seurat CoveragePlot function. (**e**) Top six genes identified from motif analysis of HPV-positive genes highly expressed in Gland_A. C, cluster; ITG, integrin; FAP, fibroblast activation protein; LAMA, laminin subunit alpha; COL, collagen; ADAMTS, a disintegrin-like and metalloproteinase with thrombospondin; IL, interleukin; KRAS, Kirsten rat sarcoma viral oncogene homolog; GRIA, glutamate ionotropic receptor AMPA; GRIN, glutamate ionotropic receptor NMDA; MAPT, microtubule associated protein tau; CDH, cadherin; ZEB, zinc finger E-box binding homeobox; MAML, mastermind like; ANGPT, angiopoietin; DOCK, dedicator of cytokinesis; WWTR, WW domain containing transcription regulator; DNAH, dynein axonemal heavy chain; CFAP, cilia- and flagella-associated protein; HYDIN, HYDIN axonemal central pair apparatus protein; IG, insulin like growth factor; CACN, calcium voltage-gated channel; HNF, hepatocyte nuclear factor; PROX, prospero homeobox; INSR, insulin receptor; ABCC, ATP binding cassette subfamily C; GIPR, gastric inhibitory polypeptide receptor; KRT, keratin; CLDN, claudin; ERBB, Erb-B2 receptor tyrosine kinase; FOX, Forkhead box; RFX, Regulatory factor X.

Changes in chromatin accessibility were less pronounced than changes in gene expression (Fig. 5b and Supplementary Fig. 6b). However, accessibility of HPV-positive cervical adenocarcinoma genes (FOXA2 and FOXA1) was specifically increased in Cluster 5 (Fig. 5c, d). Motif enrichment analysis of HPV-positive cells in the Gland_A cluster showed that the top enriched motifs predominantly belonged to the FOX and RFX families. Notably, the FOXA1 (p = 3.24 × 10^-8^) and FOXA2 (p = 3.27 × 10^-10^) motifs were significantly enriched, suggesting that forkhead transcription factors play key roles in regulating chromatin accessibility in these cells (Fig. 5e and Supplementary Table 18).

### Gene expression profiles of Gland_A and Gland_B

To clarify the clinical characteristics of each histological component, we performed gene set enrichment analysis using The Cancer Genome Atlas (TCGA) cervical cancer dataset (Supplementary Table 19), based on the characteristic gene sets of Undiff, Gland_A, and Gland_B clusters (Clusters 2, 5, and 4, respectively). Gene sets representing HPV18-positive adenocarcinoma (HPV18-ADC), other-HPV-positive adenocarcinoma (other-HPV-ADC), HPV-negative adenocarcinoma (HPV-negative ADC), HPV18-positive squamous cell carcinoma (HPV18-SCC), other-HPV-positive squamous cell carcinoma (other-HPV-SCC), and HPV-negative squamous cell carcinoma (HPV-negative SCC) were compared. Clusters 2 and 4 showed minimal enrichment across all groups but were relatively higher in HPV-negative SCC (Fig. 6a, c and Supplementary Fig. 7a, b). In contrast, Cluster 5 gene sets were strongly expressed in HPV-positive cervical adenocarcinoma (Fig. 6b, Supplementary Fig. 7a, b). These enrichment scores were not associated with cervical cancer patient prognosis (Supplementary Fig. 7c–e).

**Fig. 6:**
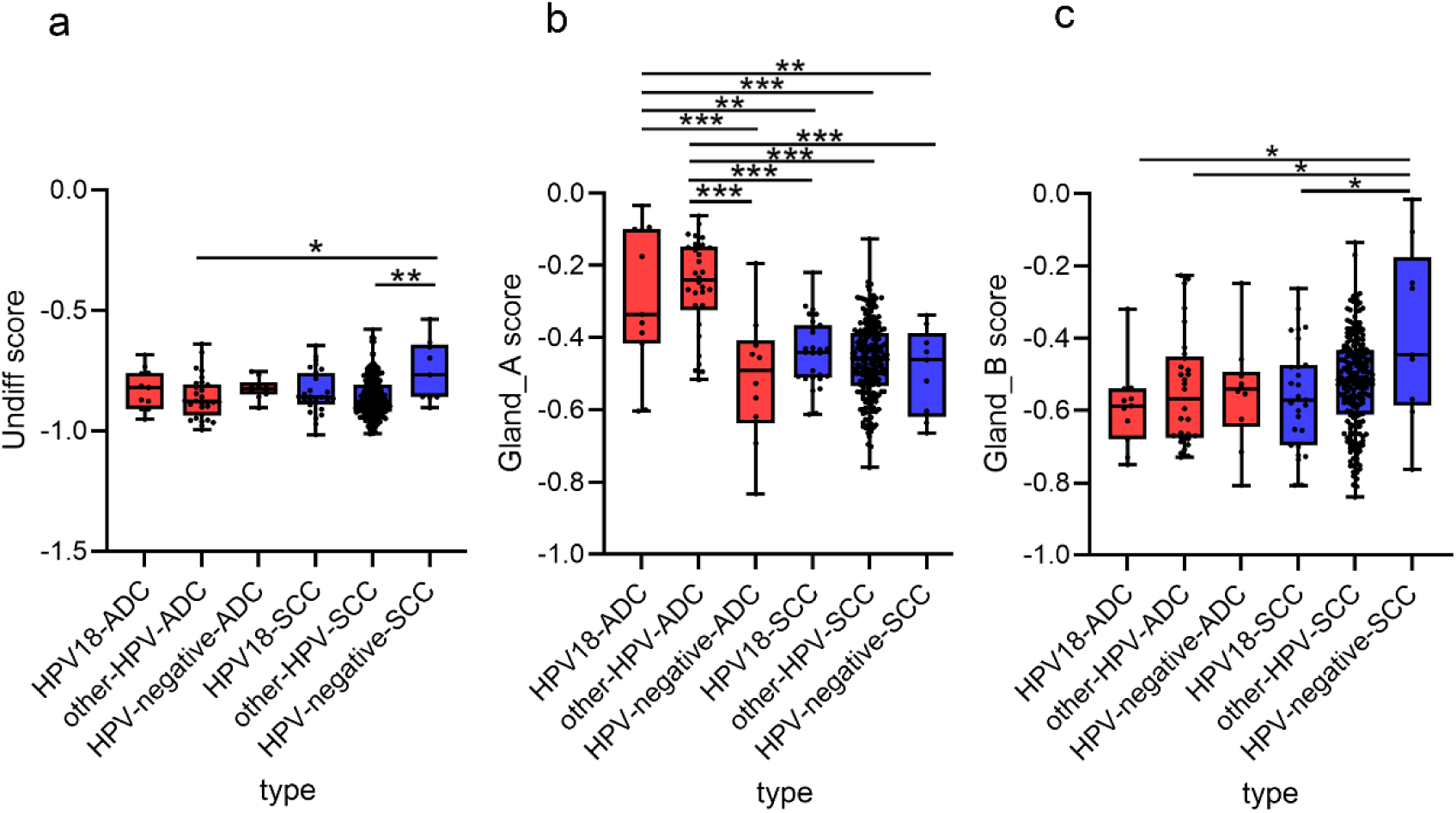
Characterization of Undiff, Gland_A, and Gland_B based on TCGA database. Gene set enrichment analysis was performed using the TCGA cervical cancer dataset (n = 299), based on characteristic gene sets (top 10 genes) from the Undiff, Gland_A, and Gland_B clusters (Clusters 2, 5, and 4, respectively). Gene set scores were compared according to histological type and HPV status. (**a**) Cluster 2 (Undiff) score; (**b**) Cluster 5 (Gland_A) score; and (**c**) Cluster 4 (Gland_B) score. “HPV18” indicates patients infected exclusively with HPV18 infection only; “other-HPV” indicates patients infected with HPV types other than HPV type 18; and “HPV-negative” indicates patients without HPV infection. The "normal" category was excluded from the analysis. Each dot represents gene expression for an individual patient. ADC, cervical adenocarcinoma; SCC, cervical squamous cell carcinoma. *P < 0.05, **P < 0.01, ***P < 0.001.

### Histological variation in stem cell markers expression

Spatial transcriptome analysis showed that ALDH1A1, a stem cell marker, was upregulated in Gland_A cells (Supplementary Tables 2 and 4). Stem cell markers such as ALDH1A1, CD44, PROM1, and SRY-box transcription factor 2 (SOX2) are linked to drug and radiotherapy resistance in cervical cancer^24–26^; however, their expression varies by histological type. Using iPS-derived reserve cells, we examined the regulation of stem cell marker expression during histological differentiation.

Expression patterns differed by histology (Fig. 7a): leucine-rich repeat-containing G protein-coupled receptor 5 (LGR5) was enriched in Cluster 2 (Undiff), ALDH1A1 was specific to Cluster 5 (Gland_A), and integrin subunit alpha 6 (ITGA6) and PROM1 were prominent in Cluster 4 (Gland_B) (Fig. 7a). Additionally, PROM1 was highly expressed in glandular components (Gland_A and Gland_B), whereas POU class 5 homeobox 1 (POU5F1) expression was downregulated in Gland_A (Fig. 7a). Changes in chromatin accessibility were more pronounced for ALDH1A in Gland_A and PROM1 in Gland_B, but remained less distinct than changes in gene expression (Fig. 7a, b).

**Fig. 7:**
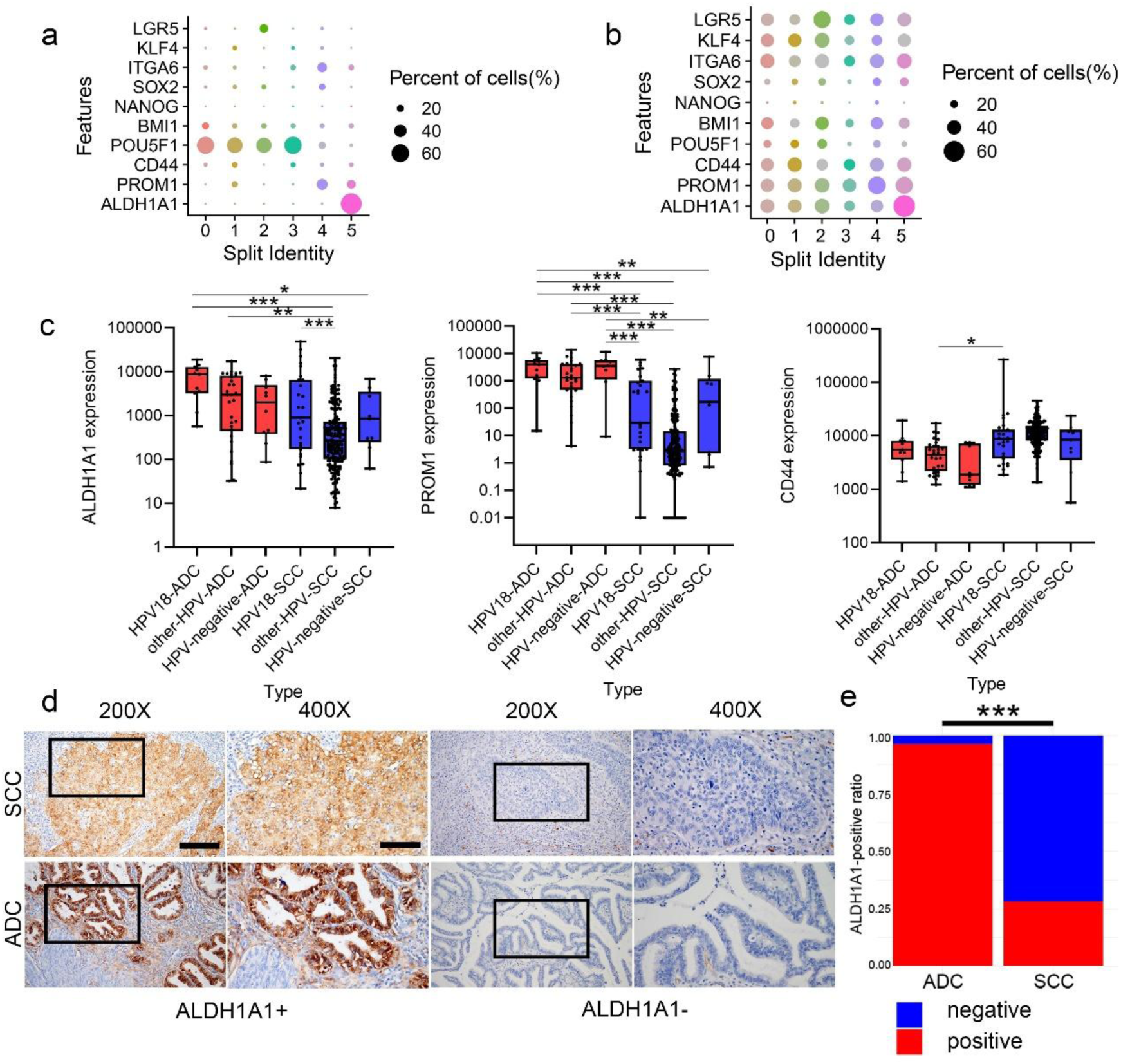
Heterogeneous expression of cancer stem cell markers. (**a**) Heatmap of expression of stem cell markers by gene expression-based clusters. (**b**) Heatmap of chromatin accessibility of stem cell markers using gene expression-based clusters. (**c**) Expression of cancer stem cell markers based on TCGA dataset. Expression of each cancer stem cell marker was compared by histology and HPV status based on TCGA dataset. The left, middle, and right panels represent ALDH1A1, PROM1, and CD44 expression, respectively. “HPV18” are patients infected with HPV18 infection only; “other-HPV” are patients infected with HPV other than HPV type 18; “HPV-negative” are patients without HPV infection. Dots in the graph represent gene expression for each patient. The vertical axis was transformed on a log10 scale after adding 0.01 to all values. *P < 0.05, **P < 0.01, ***P < 0.001. (**d**) Immunostaining for ALDH1A1 in clinical samples of cervical cancer. ALDH1A1 IHC was performed on 78 cervical cancer samples. Representative images are shown. The area squared at 200× magnification was enlarged and imaged at 400×. ALDH1A1 was determined to be positive for cytoplasmic staining. Scale bar, 100 μm. (d) Differences in ALDH1A1 positivity by cervical cancer histology. ALDH1A1 positivity was compared between squamous cell carcinomas and adenocarcinomas. Pearson’s Chi-square test with Yates continuity correction was used. ADC, n = 28; SCC, n = 50. *P < 0.05, **P < 0.01, ***P < 0.001. ADC, cervical adenocarcinoma; ALDH, aldehyde dehydrogenase; PROM1 (CD133), prominin 1; POU5F, POU class 5 homeobox; BMI1, BMI1 proto-oncogene, polycomb ring finger; NANOG, Nanog homeobox; SOX, SRY-box transcription factor; ITGA6 (CD49f), integrin subunit alpha 6; KLF, KLF transcription factor; LGR, leucine-rich repeat-containing G protein-coupled receptor; SCC, cervical squamous cell carcinoma.

We next evaluated the relationship between stem cell marker expression, histological type, and HPV genotypes using the TCGA database. Consistent with the findings in iRC-18 tumors, ALDH1A1 was highly expressed in HPV18-positive adenocarcinoma, whereas PROM1 expression was elevated in adenocarcinoma regardless of HPV status. In contrast, CD44 was predominantly expressed in SCC (Fig. 7c). These results suggest that stem cell marker expression profiles vary according to HPV status, genotype, and histological type.

Immunohistochemical analysis of clinical cervical cancer samples (50 SCCs and 28 adenocarcinomas) showed that 77 specimens (99%) were p16INK4A-positive. The proportion of ALDH1A1-positive cases was significantly higher in adenocarcinomas than in SCCs (96% vs. 28%, P = 2.6 × 10^8^) (Fig. 7d, e, and Supplementary Table 20), supporting ALDH1A1 as a marker of HPV-associated adenocarcinoma. Finally, shRNA-mediated knockdown of *ALDH1A1* in iRC-18 cells reduced *FOXA1* expression and showed a decreasing trend in *FOXA2* expression; however, this did not reach statistical significance (p = 0.1), suggesting a potential trend (Supplementary Fig. 8).

## Discussion

In this study, we generated a model of HPV-related cervical adenocarcinoma by transducing HPV18 oncogenes E6 and E7 into iRCs. Using spatial transcriptomics and single-cell multi-omics analyses, we successfully integrated tissue-type information at single-cell resolution and showed that HPV18-positive adenocarcinomas are associated with increased chromatin accessibility and elevated FOXA1 and FOXA2 expression. Furthermore, we investigated the relationship between stem cell marker expression, histological type, and HPV infection in cervical cancer.

Previous models of HPV-associated cervical cancer, such as HeLa cells, are limited in their ability to evaluate HPV-induced adenocarcinoma due to their differentiated state^27,28^. More recent models using SCJ organoids or SCJ-derived cell lines often require additional oncogenes, such as *KRAS*, alongside HPV genes, making it challenging to isolate HPV-driven histological differentiation^29,30^. To overcome these limitations, we developed a model using iRCs. The pluripotent nature of iRCs provides a suitable system for assessing HPV-induced histological differentiation. Although iRC-18 tumors are heterogeneous, they reproducibly generate adenocarcinoma-like lesions, including Gland_A structures, which exhibit features of HPV-associated adenocarcinomas. This enables investigation of HPV-associated cervical adenocarcinoma development. Additionally, this model shows greater reproducibility compared with patient-derived models.

iRC-18 tumors were heterogeneous, comprising undifferentiated structures (Undiff) and two glandular types (Gland_A and Gland_B). By integrating spatial transcriptomic and single-cell multi-omics analyses, we simultaneously evaluated gene expression and chromatin accessibility for each histological type at single-cell resolution. This analysis showed that the gene expression profile of Gland_A closely resembled that of HPV-associated adenocarcinomas, with elevated expression of HPV-associated genes such as FOXA1 and FOXA2 accompanied by chromatin remodeling. In addition to FOXA1 and FOXA2, the Wnt-β–catenin pathway is crucial for glandular differentiation in the cervix during development^22^. Wnt-related genes, including MAPK10, Dickkopf Wnt signaling pathway inhibitor 2, and NKD inhibitor of WNT signaling pathway 1, were highly expressed in the Gland_B cluster, suggesting that Gland_B cells differentiate into glandular structures via activation of the Wnt pathway. Increased expression of the Wnt receptor frizzled class receptor 3 in Gland_B was also associated with changes in chromatin accessibility (Fig. 5b), indicating that histological changes in the uterine cervix are accompanied by chromatin remodeling. In addition, single-cell analysis revealed upregulation of ion channel-related genes, such as CACNA1A, in Gland_A. Calcium channels are known to be upregulated in various cancers, including leukemia, ovarian cancer, and sarcoma^31^, and are involved in proliferation and drug resistance^32,33^. These findings suggest that increased calcium channel expression may contribute to HPV-associated cervical adenocarcinoma development.

Stem cell markers such as ALDH1, CD44, PROM1, and p63 are recognized in cervical cancer; however, their histological distribution and relationship with HPV infection remain unclear. Analysis of TCGA cervical cancer data showed that stem cell marker expression varies by histological type and HPV status. Specifically, ALDH1A1 expression was higher in HPV18-positive adenocarcinomas, PROM1 expression was elevated in adenocarcinomas regardless of HPV status, and CD44 expression was higher in SCCs. The iRCs used in this study exhibit stem cell–like properties and provide a suitable model for investigating regulation of stem cell marker expression during histological differentiation. In iRC-18 tumors, the expression patterns of stem cell markers across histological subtypes were consistent with TCGA data: ALDH1A1 was highly expressed in HPV-positive glandular structure (Gland_A), and PROM1 was highly expressed in glandular structures (Gland_A and Gland_B) regardless of HPV status. These findings indicate that stem cell marker expression varies accordingly to histological type and HPV status, even within cells of the same origin. ALDH1A1, which is characteristic of HPV-associated adenocarcinoma, has been reported to regulate pathways such as ubiquitin specific peptidase 28–MYC, hypoxia-inducible factor 1 subunit alpha–vascular endothelial growth factor, and Wnt–β-catenin signaling, and to contribute to cell differentiation, proliferation, and drug resistance through metabolic regulation^34^.

Our results suggest that ALDH1A1 is a marker preferentially associated with adenocarcinoma lesions, which may reflect its role in adenocarcinoma differentiation rather than being specific to HPV18-driven carcinogenesis^35^. Future studies should explore ALDH1A1’s role in cervical adenocarcinoma development and its potential as a therapeutic target to enhance understanding and inform treatment strategies.

This study has several limitations. First, only HPV18 E6/E7 were introduced into iRCs, and therefore the model does not recapitulate early stages of HPV infection, such as viral entry, episomal maintenance, or early gene expression. Instead, it primarily reflects cellular and histological changes associated with constitutive E6/E7 expression following viral integration, which is characteristic of HPV18-related carcinogenesis. Second, despite the use of a forced-expression system, heterogeneity in HPV expression was observed in iRC-18-derived tumors, likely due to reduced long terminal repeat promoter activity in a subset of cells. This heterogeneity may partially reflect features of HPV-associated cancers. Third, tumor formation in control iRC-cont cells indicates that HPV is not strictly required for tumorigenesis in this model; rather, HPV18 appears to specifically contribute to the induction of adenocarcinoma-like structures. Fourth, squamous-like differentiation was not observed in vivo, suggesting that lineage specification of iRC-like cells depends on microenvironmental cues that may not be fully recapitulated in the mouse model. Finally, as this study focused on HPV18, further studies are required to determine whether these findings are generalizable to other HPV types.

In conclusion, HPV-associated cervical adenocarcinoma was successfully induced by transducing HPV18 E6 and E7 into iRCs. Spatially integrated multi-omics analysis revealed the expression profiles and chromatin remodeling associated with HPV-associated cervical adenocarcinoma. This model has the potential for elucidate the carcinogenic mechanisms of HPV-associated cervical adenocarcinoma and to support the development of future therapeutic strategies.

## Methods

### Construction and production of retroviral vectors

The destination vectors pCLXSN-DEST and pCMSCVpuro-DEST were constructed by inserting a modified RfA cassette containing chloramphenicol-resistant and ccdB genes, enclosed by attR1 and attR2 sequences (Thermo Fisher Scientific, Waltham, MA, USA), between the EcoRI and BglII sites of pCLXSN (IMGENEX, Novus Biologicals LLC., Centennial, CO, USA). HPV16 E6E7 (16E6E7) and HPV18 E6E7 (18E6E7) segments were recombined into pDONR201 (Thermo Fisher Scientific) using the BP reaction and subsequently into retroviral vectors using the LR reaction (Thermo Fisher Scientific) to generate pCLXSN-16E6E7 and pCLXSN-18E6E7. Recombinant retroviruses were produced as described previously^36^. Briefly, retroviruses were generated by polyethylenimine Max-mediated triple transfection of the retroviral vector and packaging constructs, pCL-GagPol and pEF-10A1, into HEK293T cells (American Type Culture Collection, Manassas, VA, USA), and the culture medium was harvested at 60–72 h post-transfection^37^.

### Cell culture and generation of cell lines

We purchased iPS cells (201B7) from RIKEN BRC (Tokyo, Japan)^38,39^. As reported previously, iRCs were generated from iPS cells under specific culture conditions. iRCs, confirmed to be positive for both stem cell and reserve cell markers, were cultured in keratinocyte serum-free medium (KcSFM; Thermo Fisher Scientific) containing 10 μM Y-27632 (Wako, Tokyo, Japan) on dishes coated with collagen IV (Astena Holdings Co., Ltd., Tokyo, Japan).

On day 6, primary iRCs were harvested and seeded on 6-well plates coated with collagen type IV at a density of 5.0 × 10^4^ cells/well and cultured in KcSFM supplemented with 10 μM Y-27632. On the same day, cells were inoculated with retroviral supernatants of LXSN (iRC-cont), LXSN-16E6E7(iRC-16), or LXSN-18E6E7 (iRC-18) in the presence of polybrene (4 μg/mL) at a multiplicity of infection of approximately 1. Infected cells were selected with 100 μg/mL G418 until all mock-infected cells died. G418-resistant cells were propagated and used for subsequent experiments (Fig. 1a).

### Colony formation assay

iRCs (iRC-cont, iRC-16, and iRC-18) were seeded in 6-well plates (100 cells per well) and incubated at 37 °C with 5% CO_2_ concentration for 2 weeks. Colonies were stained with Giemsa stain, and colony numbers were counted to estimate viability. Each experiment was performed in triplicate.

### Cell proliferation assay

iRC-cont, iRC-16, and iRC-18 (passages 7–9) were seeded in triplicate at 2.5 x 10^4^ cells per dish, and cell numbers were counted on days 3, 5, and 7. The average cell count was calculated to generate proliferation curves.

### In vivo mouse tumor formation assay

Immunocompromised mice (NOD.CB17-Prkdc^scid^/J, females, 5 weeks old; The Jackson Laboratory Japan, Inc., Kanagawa, Japan) were used. iRC-cont, iRC-16, and iRC-18 (passages 8–14) were subcutaneously injected with 500 μL Matrigel (Corning). Tumor growth was monitored weekly, and tumor length (L) and width (W) were measured twice weekly using calipers. Tumor volume (TV) was calculated using the formula: TV = (L × W^2^)/2. Mice were euthanized when tumors reached 20 mm in diameter. Mice without tumor engraftment after 6 months were considered negative for tumor formation (Fig. 1d).

### Immunohistochemistry of mouse tumors and clinical cervical cancer specimens

Mouse tumor tissues were fixed in 4% paraformaldehyde (Wako) and embedded in paraffin. Hematoxylin and eosin (H&E) and PAS staining (Muto Pure Chemicals Co., Ltd., Tokyo, Japan) were used for pathological evaluation.

For immunohistochemistry (IHC), 4-μm sections were cut and mounted on silane-coated glass slides. Antigen retrieval was performed via dewaxing and washing. Sections were then treated with primary antibodies (Supplementary Table 21). Next, membranes were incubated with secondary antibodies (Simple Stain MAX-PO Rabbit, Nichirei Biosciences Inc., Tokyo, Japan) at room temperature (20–25 °C) and counter-stained with 3,3’-diaminobenzidine (DAB, Nichirei Biosciences Inc.). FOXA1 and FOXA2 were considered positive when nuclear staining was observed, whereas p16INK4A positively was defined as nuclear or cytoplasmic staining. Membrane staining for CDH1, KRT8, KRT18, CLDN7, and ERBB3 was considered positive.

Clinical cervical cancer specimens were processed similarly. Pathological diagnosis was based on H&E staining. IHC for ALDH1A1 and p16INK4A was performed as described above.

### Spatial gene expression analysis

Three to four spots (100-µm diameter per punched region) were prepared for iRC-18-1 and iRC-18-2 tumors (Fig. 3a).

Formalin-fixed, paraffin-embedded tissue samples stained with H&E were covered with Cryofilm (#CF-FS103, Cryofilm type 2C(10) 2.0 cm, SECTION-LAB Co., Hiroshima, Japan) and sectioned into 10-μm-thick slices. Subsequently, they were attached to glass slides using super cryo embedding medium (SECTION-LAB Co.) as the encapsulant. Regions corresponding to each histological type were punched using a microtissue collection device with a 100 µm-inner diameter needle^40^. Microtissue sections were dispensed into 99.5% ethanol at the time of sampling and stored intact at –80 °C. A vacuum evaporator was used to completely remove ethanol during cDNA library preparation prior to subsequent reactions.

RNA-seq library construction was performed as previously described^41^. Amplified cDNA products were purified with 0.8 × volume of AMPure XP beads (Beckman Coulter, Brea, CA, USA). Purified cDNA was used for sequencing library preparation using a Nextera XT DNA Library Prep Kit (Illumina). Libraries were sequenced with 50-bp paired-end reads on an Illumina NextSeq 2000.

We trimmed adapter sequences from all reads using Flexbar (ver. 3.5.0). Trimmed reads were aligned to the Ensembl human reference genome (GRCh38 ver.98), using Hisat2 (ver. 2.1.0) with default parameters. Gene expression levels (expressed as TPM) were calculated using Stringtie (ver. 2.1.7) with a transcriptome reference obtained from Ensembl. To compare expression levels, we normalized the counts of mapped reads using the median-of-ratios method in DESeq2.

The data included log_2_ fold changes, Wald test P-values, and Benjamini–Hochberg adjusted P-values for each DEG. KEGG pathway enrichment analysis was performed using DAVID 2021 (https://david.ncifcrf.gov/).

### Nuclei extraction and library preparation for single-cell multi-omics

To simultaneously measure gene expression and chromatin accessibility in the same cell, single-cell ATAC-seq and gene expression analyses were conducted using the 10x Genomics Single Cell Multiome ATAC + Gene Expression platform, according to the manufacturer’s instructions. Briefly, nuclei were extracted from flash-frozen xenografts by Dounce homogenization (ten strokes with pestle A followed by ten strokes with pestle B) in cold NP40 lysis buffer, stained with 7-AAD (Nacalai Tesque, Kyoto, Japan), and sorted using a FACSAria III (BD Biosciences, New Jersey, NJ, USA), following the 10x Genomics protocol (CG000375_DemonstratedProtocol_NucleiIsolationComplexSample_ATAC_GEX_ Sequencing_RevA). Nuclei were sorted at the FACS Core Facility of RIKEN IMS. The sorted nuclei were processed to generate single-cell gene expression and single-cell ATAC libraries using a Chromium Next GEM Single-Cell Multiome ATAC + Gene Expression Reagent Bundle and a Chromium controller (10x Genomics), according to the manufacturer’s instructions. The resulting libraries were sequenced using paired-end reads on a NextSeq 2000 system using NextSeq P3 reagents (Illumina) at the RIKEN IMS sequencing platform. Read lengths were as follows: 28 bp for Read1, 10 bp for i7 Index, 10 bp for i5 Index, and 90 bp for Read2 for the single-cell gene expression library, and 50 bp for Read1N, 8 bp for i7 Index, 24 bp for i5 Index, and 49 bp for Read2N for the single-cell ATAC library.

### scRNA and scATAC processing

Raw reads from the single-cell gene expression and single-cell ATAC libraries for each tissue were processed and aligned using Cell Ranger ARC software (v2.0.0, 10x Genomics) against a combined reference including the 10x Genomics human genome (refdata-cellranger-arc-GRCh38-2020-A-2.0.0), the HPV18 reference genome (GenBank accession number X05015), and the 10x Genomics mouse genome (refdata-cellranger-arc-mm10-2020-A-2.0.0), using default parameters. Reads were mapped to both mouse and human genomes, and cells with more 50% of reads mapped to the human genome were retained for analysis.

R software (version 4.1.2; https://www.R-project.org/) was used as the analytical platform for both scRNA-seq and scATAC-seq data. Data processing was performed using Seurat (v.4.3.0)^42^ for scRNA-seq and Signac (v.1.10.0)^43^ for scATAC-seq. Cells with UMI counts per cell ≤ 2,000 or ≥ 15,000, ≥ 2 nucleosome signals, or a transcription start site enrichment score ≤ 1 were excluded. RNA expression data from the two mice were independently normalized and scaled using the SCTransform (vst.flavour = "v2") function. Subsequently, dimensionality reduction was performed using the RunPCA (npcs = 30) and RunUMAP (dims = 1:30) functions with variable genes. A set of 3000 genes was selected for integration using the SelectIntegrationFeatures function, and integration anchors were determined using the FindIntegrationAnchors function. These anchors were subsequently used to merge the datasets using the IntegrateData function.

For scATAC data, nucleosome signal and transcription start site enrichment scores were calculated using the NucleosomeSignal and TSSEnrichment functions, respectively. Data were then normalized using the RunTFIDF function. Dimensionality reduction was performed using the RunSVD function with features identified by the FindTopFeatures function, followed by projection into two dimensions using the RunUMAP (reduction = "lsi,” dims = 2:30) function. Clustering was performed similarly to scRNA-seq using the FindNeighbours (reduction = ‘lsi,’ dims = 2:30) and FindClusters (resolution = 0.4) functions.

Motif position frequency matrices were obtained from the JASPAR2020 database (CORE, vertebrates) and added to the ATAC assay using the AddMotifs function with the hg38 reference genome. Differentially accessible peaks were identified using the FindAllMarkers function (test.use = "LR"), and enrichment of transcription factor binding motifs in cluster-specific peaks was assessed using the FindMotifs function. Representative enriched motifs were visualized with MotifPlot.

Single-cell classification was achieved using SingleR (v. 1.6.1)^44^ with default parameters, using bulk RNA-seq data as a reference to categorize each cell into Undiff, Gland_A, or Gland_B. Differential gene expression analysis for each cluster was performed using the FindAllMarkers (test.use = "MAST") function.

### TCGA data analysis

Transcriptome profiling and prognostic data for cervical cancer from TCGA (TCGA-CESC) were downloaded using the R package (TCGAbiolinks). A total of 299 cervical carcinoma samples, including 247 squamous cell carcinomas and 52 adenocarcinomas, were analyzed. HPV status was determined using RNA-seq BAM files from TCGA-CESC with VirTect. Gene set enrichment analysis was performed using the R package IOBR to calculate gene set expression scores for each cluster in TCGA-CESC samples, based on the top 10 marker genes specific to Clusters 2, 5, and 4 identified in our single-cell sequencing data. Student’s *t*-test was used to compare gene set expression scores across samples with different HPV types within each cluster.

### Statistical analysis

Colony formation and cell proliferation assays were performed with three technical and biological replicates. One-way ANOVA followed by Tukey test was used for statistical analysis. For tumor area comparisons, the Kruskal–Wallis test followed by the Steel–Dwass test was used.

For TCGA analysis, Student’s *t*-test was used to compare gene set expression scores across samples with different HPV types within each cluster. Survival analysis was conducted using R packages (survival and survminer) based on prognostic data. Patients were classified into two groups within each cluster according to the median gene set expression score. Survival rates were compared using Kaplan–Meier curves and the log-rank test.

For comparisons among multiple groups (e.g., CDKN2A gene expression and TCGA analysis), we used one-way ANOVA tests for continuous outcome variables, followed by Bonferroni correction for pairwise comparisons. Pearson’s Chi-square test with Yates’ continuity correction was used to compare ALDH1A1 positivity between SCCs and ADCs by immunostaining. Statistical analyses were performed using GraphPad Prism (version 10.2.2) and EZR (version 1.55)^45^. In all analyses, P-values < 0.05 were considered statistically significant.

## Study approval

All in vivo experiments were approved by the Institutional Review Board of Nihon University.

This study included patients (n = 78) with cervical cancer treated at Nihon University Itabashi Hospital between 2013 and 2021. All samples were collected with written informed consent in accordance with the Declaration of Helsinki, and the study was approved by the Ethics Committee of Nihon University (approval number: RK-170711-6).

## Data availability

The sequencing datasets produced in this study are available in the following databases:

- DDBJ Sequence Read Archive (DRA https://www.ddbj.nig.ac.jp/dra/index.html) Bioinformation and DDBJ Center accession number DRR555131-DRR555141 and DRR587192-DRR587195
- Genomic Expression Archive (GEA https://www.ddbj.nig.ac.jp/gea/index.html), Bioinformation and DDBJ Center accession number E-GEAD-805.

## Author contributions

S.K. performed all in vitro and in vivo experiments, interpreted the bioinformatics data, and contributed to the writing of the paper. A.T. contributed to designing the research studies, interpreting the data, and writing the manuscript. H.I., T.O., K.I., and M.H. performed bioinformatics (scRNA and ATAC) analyses. Y.I. designed the study. R. Maruyama and T.S. supported the in vitro and in vivo experiments. Y.N. and S.M. evaluated the pathological specimens. Y. Okuma sampled the ALDH1A patient cohort. O.K. guided research. N.T. performed paperwork and laboratory assistance. D.Y. contributed to the stem cell marker analysis. L.W. and K. Kiyotani analyzed TCGA data. N.M. performed single-cell experiments. C.T. and M.T. processed and aligned raw data using cellranger-arc. H.M. and H.T. analyzed the microdissection-based spatial gene expression. R. Manabe designed and performed single-cell experiments. Y. Okazaki administered single-cell experiments. T.K. established the iRC-cont, iRC-16, and iRC-18. K. Kawana conceptualized the study, designed the experiments, and wrote the manuscript.

## Acknowledgments

We thank Toyoharu Jige for preparing pathological specimens, Rina Kawatake for assistance with mouse experiments, Chiho Kohno for performing gene transfer experiments, Hiroko Kinoshita for assistance with sequence library preparation, and Yoshiyuki Ishii for valuable assistance with the experiments. Moreover, we are grateful to Dr. Shinya Yamanaka at the University of Kyoto for kindly providing the human iPSC line (201B7) used in this study. This study was financially supported by a grant to Ayumi Taguchi from AMED (grant numbers: 22wm0325014h0003 and 23wm0325057h0001) and to Kei Kawana from JSPS KAKENHI (grant number: 21K09553). This study was partially supported by the Research Support Project for Life Science and Drug Discovery (BINDS) from AMED (grant number: JP23ama121055).

