## Supplementary Figures and Methods for "Integrating spatial and single-cell multi-omics analysis of induced pluripotent stem cell-derived cervical adenocarcinoma model"

### Supplementary Methods

#### *Western blot analysis*

Whole-cell protein extracts were used for immunoblotting. Antibodies against CDH1, GAPDH, HPV18 E6, and p16INK4A (presented in Supplementary Table 22) were used as probes. Horseradish peroxidase-conjugated anti-mouse and anti-rabbit (#7074S; Cell Signaling Technology, Danvers, MA, USA) immunoglobulins were used as secondary antibodies. An anti-HPV16 E6 monoclonal antibody (clone 46A4), generated in-house and raised against the N-terminal 16 amino acids, was used as a probe. The LAS3000 charge-coupled device imaging system (Fujifilm Co. Ltd., Tokyo, Japan) was employed for detecting proteins, which were visualized using Lumi-light plus western blotting substrate (Roche Diagnostics, Basel, Switzerland).

#### *RNA in situ hybridization*

In situ hybridization for HPV18 E7 RNA was performed using the RNAscope® 2.5 HD

Duplex Reagent Kit (Advanced Cell Diagnostics, a brand of Bio-Techne Corporation, Newark, CA, USA) according to the manufacturer's instructions. Probes were selected for HPV18 E7 (#463471-C2) and labeled with Fast Red. A positive diagnosis was established by the detection of one or more red dots in the cytoplasm.

*TCGA data analysis*

Transcriptome and clinical data for cervical cancer (TCGA-CESC) were obtained using TCGAbiolinks. A total of 299 samples (247 squamous cell carcinomas and 52 adenocarcinomas) were analyzed. HPV status was determined from RNA-seq BAM files using VirTect. Gene Set Enrichment Analysis (GSEA) was performed using IOBR to calculate gene set expression scores based on the top 10 marker genes for Clusters 2, 5, and 4 identified in single-cell data. Differences in gene set expression scores across HPV types were assessed using Student's t-test.

*Short hairpin RNA (shRNA)-mediated targeting of ALDH1A1*

Knockdown of *ALDH1A1* was achieved using pLKO.1-puro shRNA lentiviral particles obtained from the MISSION$^{TM}$ shRNA library (Merck KGaA, Darmstadt, Germany). The shRNA construct (TRCN0000276459) was tested together with a non-targeting (scrambled) shRNA (SHC202V-1EA) control. In brief, cells were plated in 6-well plates ($2 \times 10^4$ cells/well) and incubated overnight with the appropriate lentiviral particles (MOI = 2) in the presence of hexadimethrine bromide (final concentration, 8 μg/mL). Cells were then selected with 0.2 μg/mL puromycin for 2 weeks, and resistant colonies were picked, expanded, and assayed for knockdown efficiency.

*Quantitative real-time PCR (qPCR)*

Total RNA was extracted from cells using the miRNeasy Mini Kit (QIAGEN, Hilden, Germany) and complementary DNA (cDNA) was synthesized using the High-Capacity cDNA Reverse Transcription Kit (Thermo Fisher Scientific, Waltham, MA, USA) according to the manufacturers' instructions. Quantitative real-time PCR was performed using SYBR Green chemistry (Cat# 1725121; Bio-Rad, Hercules, CA, USA) on a QuantStudio 3 system (Thermo Fisher Scientific, Waltham, MA, USA).

The gene expression levels of *ALDH1A1*, *FOXA1*, and *FOXA2* were analyzed using the following primers:

ALDH1A1-F: 5′-GCACGCCAGACTTACCTGTC-3′,

ALDH1A1-R: 5′-CCTCCTCAGTTGCAGGATTAAAG-3′;

FOXA1-F: 5′-GAAGGGCATGAAACCAGCGA-3′,

FOXA1-R: 5′-TCATGTTGCCGCTCGTAGTC-3′;

FOXA2-F: 5′-GCCCCAACAAGATGCTGAC-3′,

FOXA2-R: 5′-CACCTTCAGGAAACAGTCGTTG-3′.

ACTB-F：5'- TCACCCACACTGTGCCCATCTACGA -3'

ACTB-R：5'- CAGCGGAACCGCTCATTGCCAATGG -3'

PCR reactions were performed in triplicate. Relative gene expression levels were calculated using the 2^(-ΔCt) method and normalized to an internal control gene (β-actin, ACTB). Statistical analyses were performed using ΔCt values. ΔCt = Ct(target) – Ct(ACTB). Undetermined Ct values were assigned a value of 40.

**Supplementary Tables**

**Supplementary Table 1.** Immunostaining results of mouse specimens (Excel file).

The transcription factors FOXA1 and FOXA2 were considered positive if the nucleus was stained, and p16INK4A was evaluated as positive in samples with nuclear and/or cytoplasmic staining. CDH1, KRT8, KRT18, CLDN7, and ERBB3 were considered positive when the membrane was stained. H&E, hematoxylin & eosin; PAS, periodic acid–Schiff; CDH, cadherin; KRT, keratin; CLDN, claudin; ERBB, Erb-B2 receptor tyrosine kinase; FOX, forkhead box.

**Supplementary Tables 2–4.** Comparative list of gene expression in undifferentiated tumor (Undiff), Gland_A, and Gland_B tissue components assessed via spatial gene expression analysis. (Excel file).

**2**. In the Undiff vs. Gland_A comparison, the gene group highly expressed in Undiff had a positive log value, and that in Gland_A had a negative log value.

**3**. In the Undiff vs. Gland_B comparison, the gene group highly expressed in Undiff had a positive log value, and that in Gland_B had a negative log value.

**4**. In the Gland_A vs. Gland_B comparison, the gene group highly expressed in Gland_A had a positive log value, whereas that in Gland_B had a negative log value.

Lists of gene groups with an absolute $\log_2$ value greater than 1.

KEGG, Kyoto Encyclopedia of Genes and Genomes.

**Supplementary Tables 5–10.** Comparative list of KEGG pathways in undifferentiated tumors (Undiff), Gland_A, and Gland_B tissue components assessed via microdissection (Excel file).

**5**. Upregulated pathways in Undiff compared to those in Gland_A.

**6**. Upregulated pathways in Gland_A compared to those in Undiff.

**7**. Upregulated pathways in Undiff compared to those in Gland_B.

**8**. Upregulated pathways in Gland_B compared to those in Undiff.

**9**. Upregulated pathways in Gland_A compared to those in Gland_B.

**10**. Upregulated pathways in Gland_B compared to those in Gland_A.

The analysis was performed using DAVID (version 2021) (https://davidbioinformatics.nih.gov/home.jsp). Statistical significance was set at $P < 0.05$.

KEGG, Kyoto Encyclopedia of Genes and Genomes.

**Supplementary Tables 11–16.** Comparative list of GO pathways in undifferentiated tumors (Undiff), Gland_A, and Gland_B tissue components via microdissection (Excel file).

11. Upregulated pathways in Undiff compared to those in Gland_A.

12. Upregulated pathways in Gland_A compared to those in Undiff.

13. Upregulated pathways in Undiff compared to those in Gland_B.

14. Upregulated pathways in Gland_B compared to those in Undiff.

15. Upregulated pathways in Gland_A compared to those in Gland_B.

16. Upregulated pathways in Gland_B compared to those in Gland_A.

The analysis was performed using DAVID (version 2021) (https://davidbioinformatics.nih.gov/home.jsp). Statistical significance was set at $P < 0.05$.

GO, Gene Ontology.

**Supplementary Table 17**. Marker genes for each cluster in single-cell analysis (Excel file). For each cluster (0–5) versus all clusters, genes with an absolute value > 0.25 in average log2 fold change were defined as marker genes.

**Supplementary Table 18.** List of genes identified through motif analysis of HPV-positive genes highly expressed in Gland_A (Excel file).

List of genes with adjusted P-values < 0.05.

**Supplementary Table 19.** Patient information for TCGA (Excel file).

“Adenocarcinoma” contains “Endocervical type of adenocarcinoma,” “Mucinous adenocarcinoma,” “Endocervical Adenocarcinoma of the usual type,” and “Endometrioid adenocarcinoma of the endocervix.”

“other-HPV” includes HPV types 16, 26, 30, 31, 33, 35, 39, 45, 51, 52, 56, 58, 59, 68, 69, 70, and 73, and their mixed infections. Mixed infections with HPV18 and other HPV types were also included in the “other HPV” group.

TCGA, The Cancer Genome Atlas.

**Supplementary Table 20.** Patient information for ALDH1A1 immunostaining (Excel file).

Squamous cell carcinoma (SCC) includes keratinizing, non-keratinizing, and microinvasive squamous cell carcinomas. Adenocarcinoma (ADC) includes endocervical, mucinous, and gastric-type adenocarcinomas.

**Supplementary Table 21.** Information on antibodies used for immunostaining (Excel file).

CDH, cadherin; KRT, keratin; CLDN, claudin; ERBB, Erb-B2 receptor tyrosine kinase; FOX, forkhead box; ALDH, aldehyde dehydrogenase; SYP, synaptophysin; CHGA, chromogranin A.

**Supplementary Table 22.** Information on antibodies used for immunoblotting (Excel file).

CDH, cadherin; HPV, human papillomavirus; GAPDH, glyceraldehyde-3-phosphate dehydrogenase.

**Supplementary Figures**

**Supplementary Figure 1.** Western blot analysis of iRC-cont, iRC-16, and iRC-18.

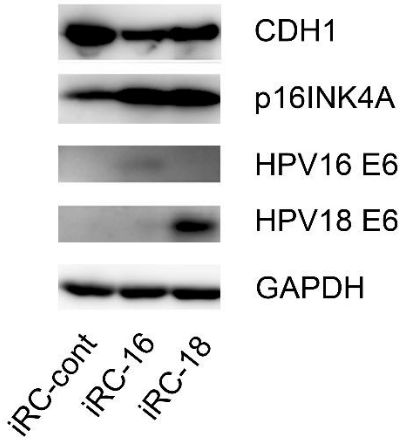


CDH, cadherin; HPV, human papillomavirus; GAPDH, glyceraldehyde-3-phosphate dehydrogenase.

**Supplementary Figure 2.**

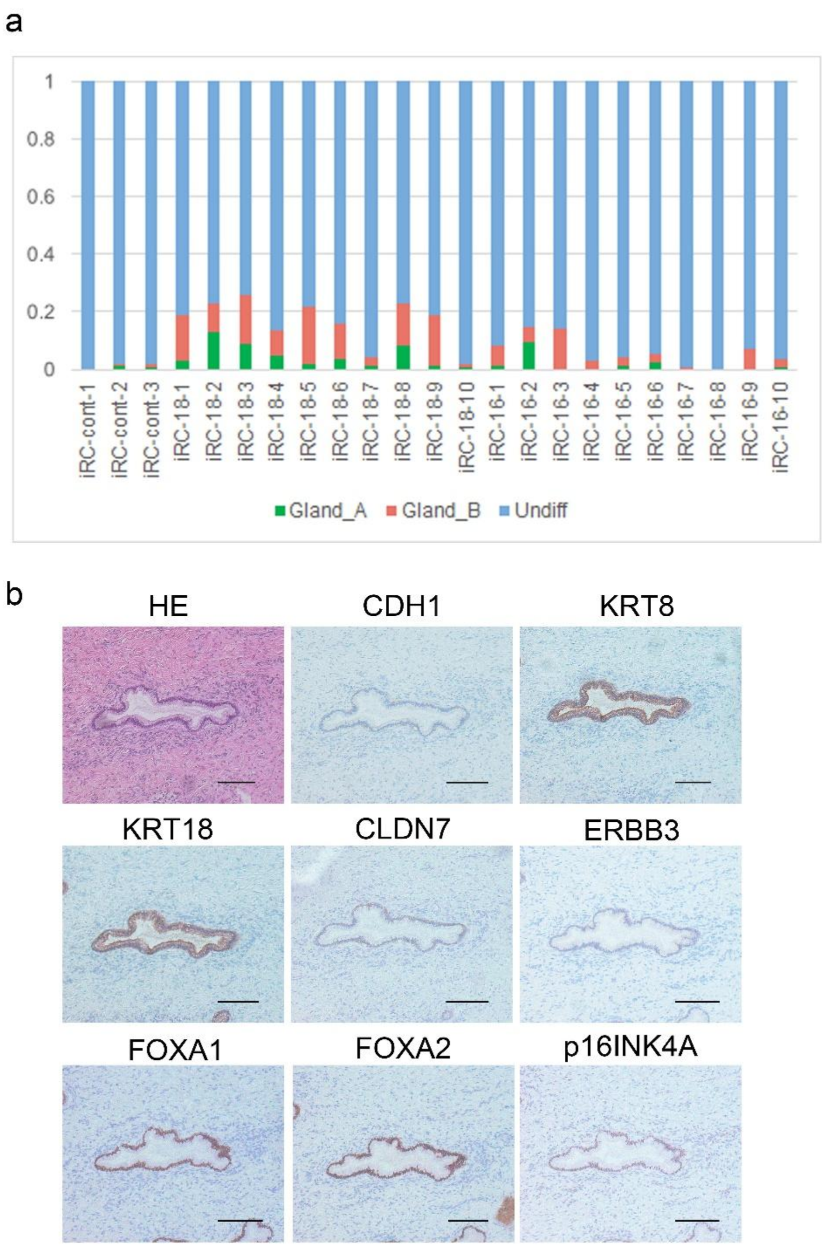


**a.** Proportions of Gland_A, Gland_B, and Undiff tissue in iRC-cont, iRC-16, and iRC-18 tumors. **b.** Representative panel of normal cervical glandular tissue. Supplementary Table

21 contains antibody information. CDH, cadherin; KRT, keratin; CLDN, claudin; ERBB, Erb-B2 receptor tyrosine kinase; FOX, forkhead box. Scale bar = 100 μm.

**Supplementary Figure 3.** Intensity of antibody immunostaining in Figure 2c.

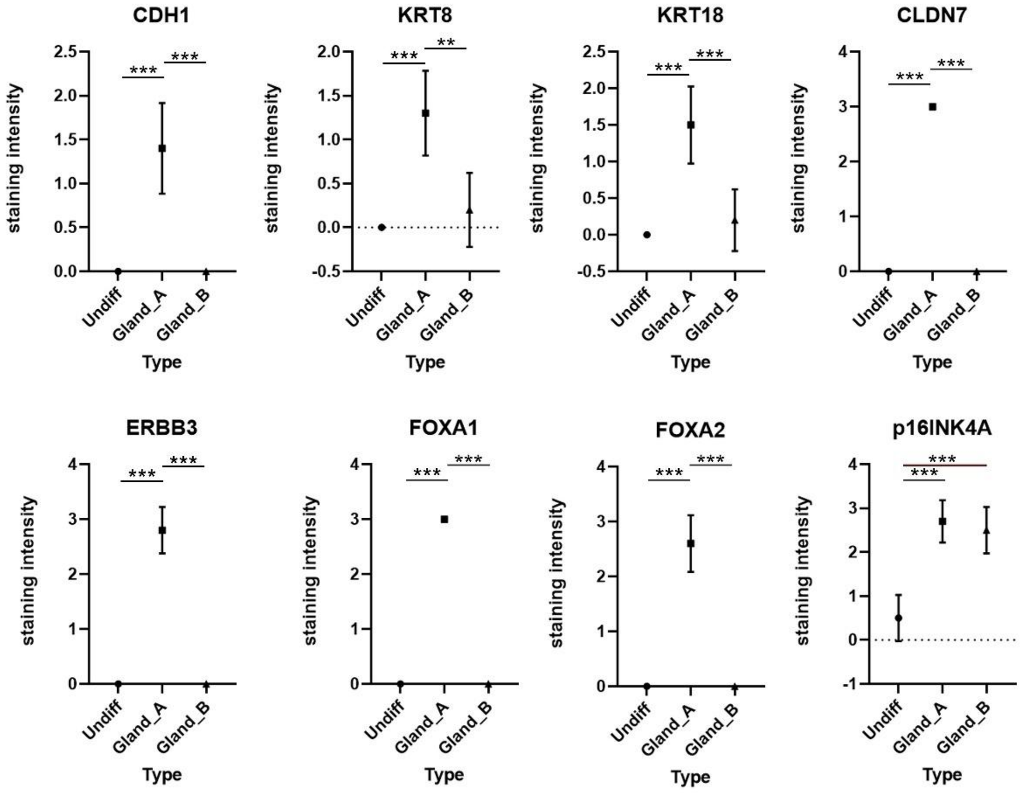


Column plot representing immunofluorescence staining intensity. In iRC-18 tumors (n = 10), three structures (Gland_A, Gland_B, and Undiff) were randomly selected from each case and evaluated for staining intensity on a scale of 0 to 3. Kruskal–Wallis test followed by Steel–Dwass tests. **P < 0.01, ***P < 0.001. CDH, cadherin; KRT, keratin; CLDN, claudin; ERBB, Erb-B2 receptor tyrosine kinase; FOX, forkhead box.

**Supplementary Figure 4.** Spatial transcriptomic characterization of Undiff, Gland_A, and Gland_B clusters

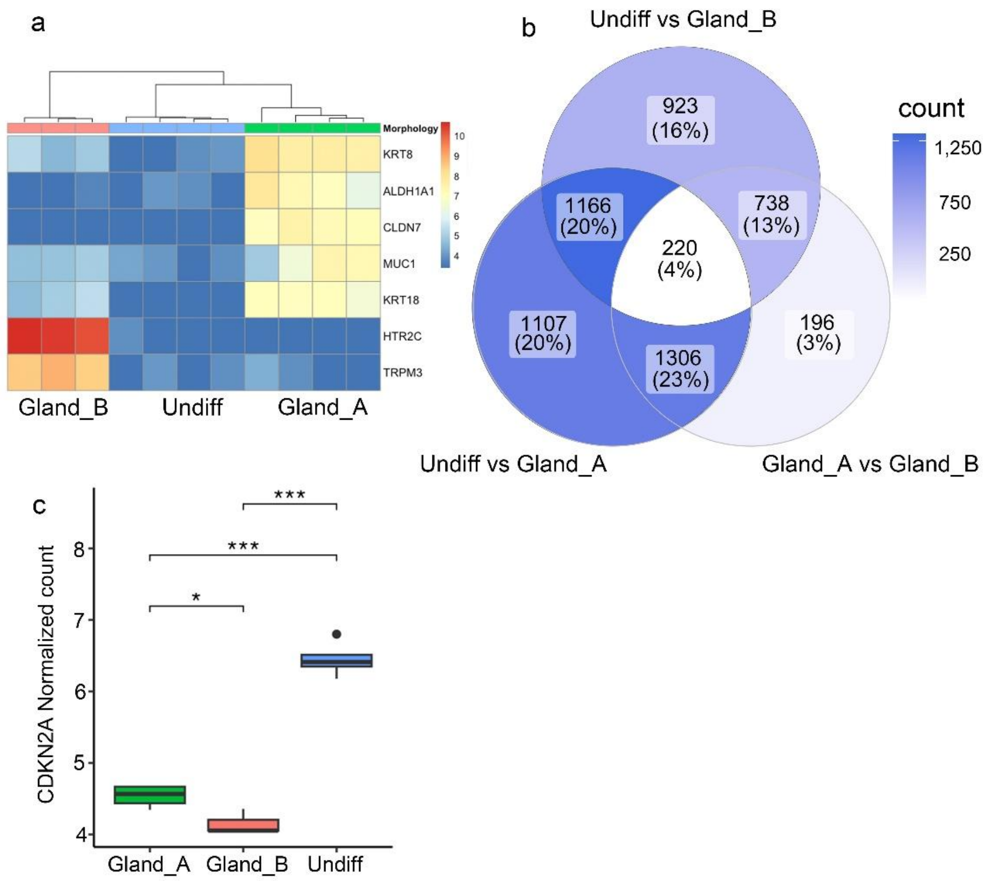


**a**. Heatmap of spatial transcriptomics analysis based on microdissection. Heatmap of candidate genes based on RNA sequencing of Undiff, Gland_A, and Gland_B. KRT, keratin; ALDH, aldehyde dehydrogenase; CLDN, claudin; MUC, mucin; HTR, 5-hydroxytryptamine receptor; TRPM, transient receptor potential cation channel subfamily M member. **b**. Venn diagram showing the number of significant genes in spatial gene expression analysis among Undiff, Gland_A, and Gland_B tissue components. Dark blue indicates a high number of significant genes, and white indicates a low number. **c**. Boxplot focused on the *CDKN2A* gene in spatial gene expression analysis. n = 3–4; one-way ANOVA followed by Bonferroni tests. *P < 0.05, **P < 0.01, ***P < 0.001. CDKN2A, cyclin-dependent kinase inhibitor 2A**.**

**Supplementary Figure 5**. Supplementary data for single-cell RNA-seq.

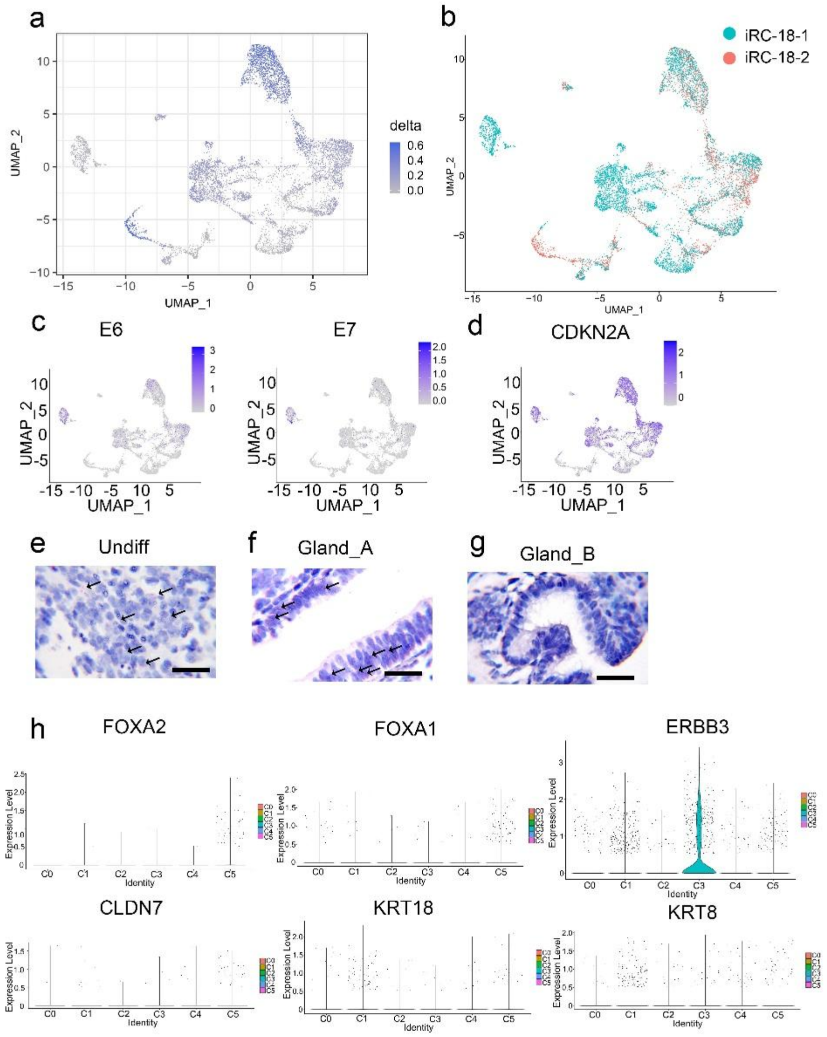


**a**. Single-cell analysis. This analysis was performed to identify the clusters that are more representative of Undiff tissue based on their relative functions. **b.** Distribution of cells in UMAP plots of iRC-18-1 and iRC-18-2. **c**. Expression of HPV18 E6 and E7. HPV18 E6 and

E7 expression is reflected in gene expression-based UMAP. **d.** Expression of *CDKN2A* (p16INK4A). *CDKN2A* expression is reflected in gene expression-based UMAP. CDKN2A, cyclin-dependent kinase inhibitor 2A. **e**. HPV18 E7 RNA ISH of Undiff. **f.** HPV18 E7 RNA ISH of Gland_A. **g**. HPV18 E7 RNA ISH of Gland_B. A positive diagnosis was established when one or more red dots were detected in the cytoplasm. Scale bar = 20 μm. PAS, periodic acid–Schiff; CDH, cadherin; KRT, keratin; CLDN, claudin; ERBB, Erb-B2 receptor tyrosine kinase. **h**. Violin plots for candidate genes from clusters 0 to 5. FOX, forkhead box; ERBB, Erb-B2 receptor tyrosine kinase; KRT, keratin; CLDN, claudin.

**Supplementary Figure 6.** Single-cell ATAC-seq of the iRC-18-2 tumor.

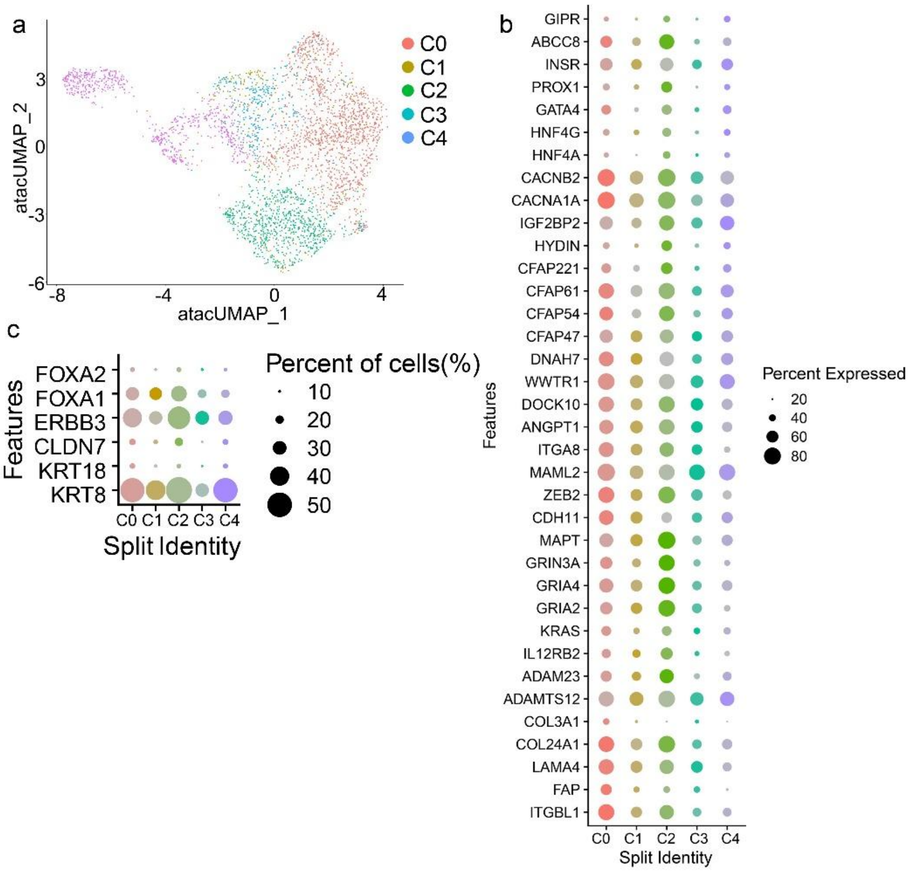


**a**. UMAP of ATAC-seq for single-cell analysis. Single-cell ATAC-seq and gene expression analyses were performed on iRC-18 tumors from two mice. Dimensionality reduction was performed based on ATAC-seq data of the iRC-18-2 tumor. Cluster numbers were assigned as determined using gene expression-based analyses. **b**. Heatmap of single-cell ATAC-seq data using gene expression-based clusters. Candidate genes were extracted and constructed for clusters 0 through 5. Some genes (*CACNA1A, TRPM3, PDE4B*, *CNTNAP5, NLGN1, NRXN1, ADAMTSL1*, and *LAMA2)* were not detected in the ATAC-seq dataset. **c**. ATAC-seq heatmap of genes related to glandular structures and cervical adenocarcinoma. Genes related to the development of cervical adenocarcinoma (*FOXA2, FOXA*1, and *ERBB3*), and those related to glandular structures (*CLDN7, KRT18, and KRT8*) were selected to construct a heatmap. The heatmap was constructed using gene expression-based clusters. C, cluster; ITG, integrin; FAP, fibroblast activation protein; LAMA, laminin subunit alpha; COL, collagen; ADAMTS, a disintegrin-like and metalloproteinase with thrombospondin; IL, interleukin; KRAS, Kirsten rat sarcoma viral oncogene homolog; GRIA, glutamate ionotropic receptor AMPA; GRIN, glutamate ionotropic receptor NMDA; MAPT, microtubule associated protein tau; CDH, cadherin; ZEB, zinc finger E-box binding homeobox; MAML, mastermind like; ANGPT, angiopoietin; DOCK, dedicator of cytokinesis; WWTR, WW domain containing transcription regulator; DNAH, dynein axonemal heavy chain; CFAP, cilia- and flagella-associated protein; HYDIN, HYDIN axonemal central pair apparatus protein; IG, insulin like growth factor; CACN, calcium voltage-gated channel; HNF, hepatocyte nuclear factor; PROX, prospero homeobox; INSR, insulin receptor; ABCC, ATP binding cassette subfamily C; GIPR, gastric inhibitory polypeptide receptor; KRT, keratin; CLDN, claudin; ERBB, Erb-B2 receptor tyrosine kinase; FOX, forkhead box.

**Supplementary Figure 7.** Data projected to TCGA for the highly expressed genes in Clusters 2, 5, and 4 of single-cell analysis.

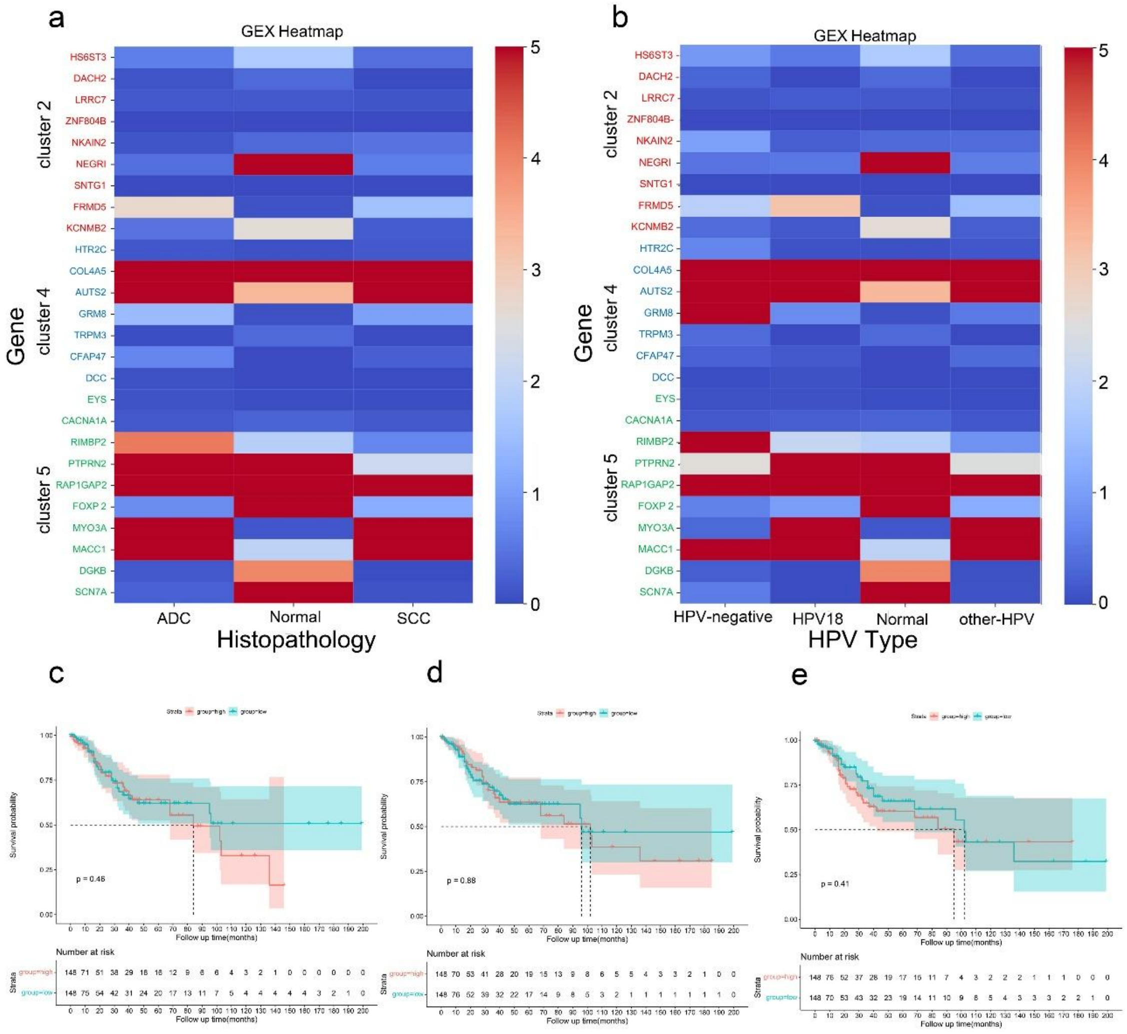


**a**. Distribution of the highly expressed genes of Clusters 2, 5, and 4 by histological type.

The top ten genes in Clusters 2, 5, and 4 were scored and compared with gene expression levels in TCGA patients. “Adenocarcinoma” includes “Endocervical type of adenocarcinoma,” “Mucinous adenocarcinoma,” “Endocervical Adenocarcinoma of the usual type,” and “Endometrioid adenocarcinoma of the endocervix.” **b.** Distribution of the highly expressed genes in Clusters 2, 5, and 4, stratified into neg, HPV18, normal, and other HPV groups. Neg, patients without HPV infection; HPV18, patients infected with HPV18 only; normal, normal tissue of the cervix; other HPV, patients infected with HPV other than the HPV type 18. Moreover, “other HPV” includes HPV types 16, 26, 30, 31, 33, 35, 39, 45, 51, 52, 56, 58, 59, 68, 69, 70, and 73, and their mixed infections. Mixed infections with HPV18 and other HPV types were also included in the “other HPV” group. **c.** Overall survival analysis of high- and low-gene expression groups in squamous cell carcinoma and adenocarcinoma in Clusters 2, 5, and 4. The left, middle, and right panels represent Clusters 2, 5, and 4, respectively. Survival curves were created using Kaplan–Meier analysis, and $P < 0.05$ (log-rank test). Survival probabilities were compared based on the median expression value.

**Supplementary Figure 8.** Comparison of relative RNA expression levels by RT-PCR for sh-*ALDH1A1*.

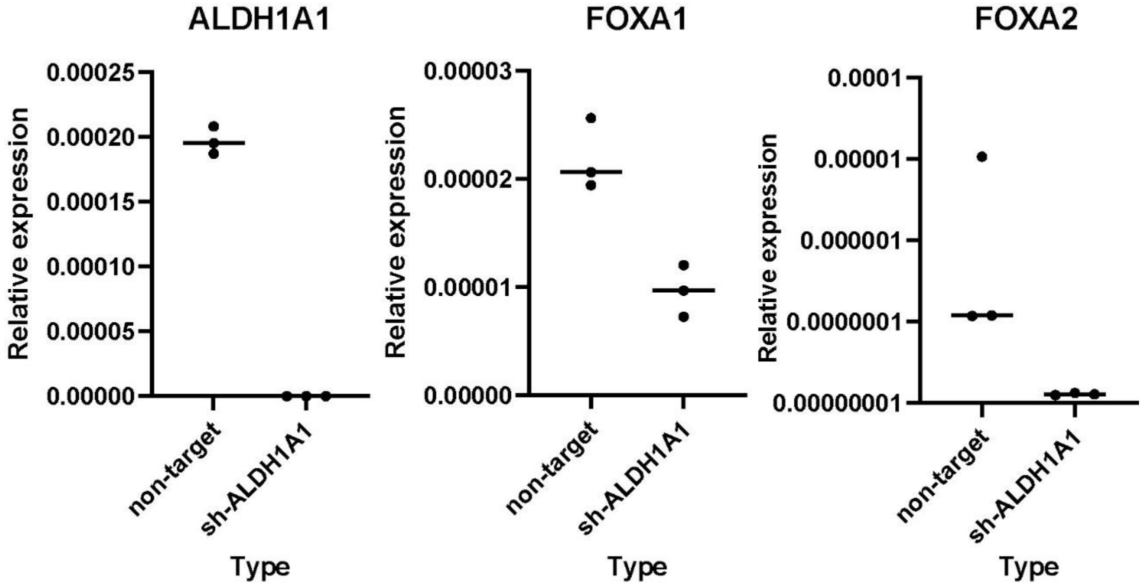


Expression levels are presented as 2^(-ΔCt) values normalized to ACTB. Each dot represents an independent experiment (n = 3). Data are shown as the median.

Due to low expression levels, FOXA2 expression is presented on a logarithmic scale.

Statistical analyses were performed using the Mann–Whitney U test.

ALDH, aldehyde dehydrogenase; FOX, forkhead box; ACTB, β-actin.
