## Supplementary Tables for "Integrating spatial and single-cell multi-omics analysis of induced pluripotent stem cell-derived cervical adenocarcinoma model"

TableS1

| Immunostaining results | | | | | | | | | |
|---|---|---|---|---|---|---|---|---|---|
| | PAS | CDH1 | KRT8 | KRT18 | CLDN7 | ERBB3 | FOXA1 | FOXA2 | p16INK4A |
| Gland_A | + | ++ | + | + | ++ | + | ++ | + | + |
| Gland_B | - | - | - | - | - | - | - | - | + |
| Cervical glandular epithelium | | ±〜+ | ++ | ++ | ±〜+ | ±〜+ | + | + | ±〜+ |

TableS2

| Genes upregulated in Undiff compared to Gland_A | | | | | | | |
|---|---|---|---|---|---|---|---|
| **gene ID** | **gene name** | **baseMean** | **log2FoldChange** | **lfcSE** | **stat** | **p-value** | **adjusted p-value** |
| 1253 | NEFL | 78.8660084 | 10.02369764 | 1.0539 | 9.51117 | 1.89E-21 | 5.26E-19 |
| 675 | NCAN | 57.3136667 | 9.554619696 | 1.0795 | 8.8513 | 8.65E-19 | 1.56E-16 |
| 47 | CYP4Z1 | 116.345268 | 9.185843848 | 0.9895 | 9.28355 | 1.64E-20 | 3.82E-18 |
| 1101 | CPLX2 | 39.2625567 | 9.018869547 | 1.0725 | 8.40923 | 4.13E-17 | 5.10E-15 |
| 1243 | MIR124-1HG | 30.740167 | 8.665354948 | 1.0989 | 7.88583 | 3.12E-15 | 2.57E-13 |
| 848 | MAFB | 143.803567 | 8.528055857 | 0.7594 | 11.2298 | 2.91E-29 | 2.18E-26 |
| 819 | KIF1A | 71.3528859 | 8.458513305 | 0.9828 | 8.60629 | 7.55E-18 | 1.08E-15 |
| 835 | SNAP25 | 24.7092831 | 8.367101143 | 1.1207 | 7.46629 | 8.25E-14 | 5.26E-12 |
| 129 | SYT14 | 22.9951872 | 8.256592379 | 1.1189 | 7.37937 | 1.59E-13 | 9.67E-12 |
| 411 | NKX2-1 | 22.8405124 | 8.235402351 | 1.1221 | 7.33949 | 2.14E-13 | 1.27E-11 |
| 548 | P2RX1 | 19.8821362 | 8.040675002 | 1.0951 | 7.3421 | 2.10E-13 | 1.25E-11 |
| 1274 | EYA1 | 19.61116 | 8.030938568 | 1.1098 | 7.23612 | 4.62E-13 | 2.55E-11 |
| 484 | MINAR1 | 18.6435802 | 7.960491583 | 1.1168 | 7.12813 | 1.02E-12 | 5.22E-11 |
| 1208 | DLX6 | 16.6002891 | 7.786193953 | 1.112 | 7.00181 | 2.53E-12 | 1.22E-10 |
| 1055 | CARTPT | 15.4675527 | 7.682519339 | 1.1217 | 6.84916 | 7.43E-12 | 3.27E-10 |
| 181 | LBX1 | 15.5479551 | 7.673445178 | 1.1268 | 6.81008 | 9.75E-12 | 4.20E-10 |
| 510 | HS3ST4 | 14.1882545 | 7.55991405 | 1.1269 | 6.70854 | 1.97E-11 | 7.75E-10 |
| 972 | PCOLCE2 | 23.1410879 | 7.541174677 | 1.0986 | 6.86419 | 6.69E-12 | 2.99E-10 |
| 291 | LINC02470 | 13.653017 | 7.498577522 | 1.172 | 6.39829 | 1.57E-10 | 4.97E-09 |
| 273 | IGSF9B | 12.4198953 | 7.346913202 | 1.1387 | 6.45228 | 1.10E-10 | 3.68E-09 |
| 1223 | CTTNBP2 | 20.05893 | 7.317960751 | 1.106 | 6.61686 | 3.67E-11 | 1.37E-09 |
| 1099 | FBLL1 | 11.5757702 | 7.251904862 | 1.1604 | 6.2493 | 4.12E-10 | 1.19E-08 |
| 218 | LMO1 | 11.6731787 | 7.248095511 | 1.1663 | 6.21437 | 5.15E-10 | 1.44E-08 |
| 986 | PEX5L | 29.991459 | 7.232953249 | 1.0225 | 7.07347 | 1.51E-12 | 7.62E-11 |
| 435 | TUNAR | 110.405534 | 7.210431045 | 1.3927 | 5.17725 | 2.25E-07 | 3.47E-06 |
| 509 | PRKCB | 18.6127189 | 7.206823689 | 1.1139 | 6.46973 | 9.82E-11 | 3.32E-09 |
| 1400 | DACH2 | 28.3989103 | 7.153841596 | 1.0138 | 7.05617 | 1.71E-12 | 8.57E-11 |
| 698 | ATP1A3 | 17.9524869 | 7.152460756 | 1.1088 | 6.45057 | 1.11E-10 | 3.69E-09 |
| 1007 | ENSG00000286318 | 17.7452675 | 7.151243776 | 1.1017 | 6.49093 | 8.53E-11 | 2.95E-09 |
| 1360 | CACNA1B | 10.8206921 | 7.149388219 | 1.182 | 6.04858 | 1.46E-09 | 3.69E-08 |
| 188 | CFAP58 | 10.6252725 | 7.143881686 | 1.1607 | 6.15499 | 7.51E-10 | 2.01E-08 |
| 1105 | AACSP1 | 10.2779233 | 7.109493818 | 1.2272 | 5.79332 | 6.90E-09 | 1.51E-07 |
| 903 | SULT4A1 | 17.1276214 | 7.106132648 | 1.1058 | 6.4262 | 1.31E-10 | 4.24E-09 |
| 277 | GLB1L3 | 16.9835071 | 7.089982208 | 1.1248 | 6.30356 | 2.91E-10 | 8.62E-09 |
| 1305 | SCRT1 | 10.2817701 | 7.082690669 | 1.1595 | 6.10847 | 1.01E-09 | 2.62E-08 |

| | | | | | | | |
|---|---|---|---|---|---|---|---|
| 80 | KCNA2 | 16.697188 | 7.071971922 | 1.1138 | 6.34928 | 2.16E-10 | 6.60E-09 |
| 1209 | DLX5 | 25.951494 | 7.01180171 | 1.0138 | 6.91666 | 4.62E-12 | 2.14E-10 |
| 770 | ENSG00000236885 | 16.0161166 | 7.002512125 | 1.1241 | 6.22967 | 4.67E-10 | 1.32E-08 |
| 850 | GDAP1L1 | 9.20607152 | 6.913184593 | 1.1771 | 5.87285 | 4.28E-09 | 9.75E-08 |
| 699 | TMEM145 | 9.07802852 | 6.906233617 | 1.1721 | 5.89205 | 3.81E-09 | 8.78E-08 |
| 1219 | VGF | 14.0177947 | 6.79758969 | 1.1247 | 6.044 | 1.50E-09 | 3.79E-08 |
| 31 | HPCA | 8.27518512 | 6.780760635 | 1.2077 | 5.61448 | 1.97E-08 | 3.94E-07 |
| 369 | GPR12 | 8.03633028 | 6.750569312 | 1.2092 | 5.58289 | 2.37E-08 | 4.67E-07 |
| 1211 | TMEM130 | 8.03700796 | 6.741598886 | 1.196 | 5.63668 | 1.73E-08 | 3.51E-07 |
| 124 | CNTN2 | 13.3199059 | 6.732411144 | 1.1452 | 5.87864 | 4.14E-09 | 9.46E-08 |
| 1312 | SLC24A2 | 30.5394081 | 6.710273551 | 0.8997 | 7.45871 | 8.74E-14 | 5.55E-12 |
| 1293 | TMEM74 | 12.8779776 | 6.674253369 | 1.1293 | 5.9102 | 3.42E-09 | 7.93E-08 |
| 341 | SLC6A15 | 12.8349796 | 6.671717422 | 1.1464 | 5.81986 | 5.89E-09 | 1.31E-07 |
| 1395 | NALF2 | 7.46694294 | 6.650582088 | 1.2379 | 5.37234 | 7.77E-08 | 1.34E-06 |
| 1040 | ACSL1 | 68.3327872 | 6.641744029 | 0.6037 | 11.0026 | 3.71E-28 | 2.30E-25 |
| 1217 | ACTL6B | 7.58813027 | 6.63883909 | 1.2042 | 5.51297 | 3.53E-08 | 6.67E-07 |
| 264 | POU2F3 | 7.53634069 | 6.635363547 | 1.2467 | 5.32255 | 1.02E-07 | 1.73E-06 |
| 201 | EBF3 | 20.3529808 | 6.634581403 | 1.0253 | 6.47117 | 9.72E-11 | 3.29E-09 |
| 190 | SORCS1 | 7.41756302 | 6.623900362 | 1.2003 | 5.51846 | 3.42E-08 | 6.49E-07 |
| 1406 | TCEAL5 | 57.4469178 | 6.622336617 | 0.6481 | 10.2182 | 1.64E-24 | 6.32E-22 |
| 1009 | KCNIP4-IT1 | 7.51587522 | 6.619790878 | 1.218 | 5.43512 | 5.48E-08 | 9.84E-07 |
| 1207 | DLX6-AS1 | 28.3822337 | 6.610280751 | 0.9108 | 7.25793 | 3.93E-13 | 2.18E-11 |
| 1087 | PPP2R2B | 7.27064327 | 6.607869429 | 1.2316 | 5.36517 | 8.09E-08 | 1.39E-06 |
| 748 | LRRTM4 | 7.42018009 | 6.600272739 | 1.2285 | 5.37268 | 7.76E-08 | 1.34E-06 |
| 722 | KLHL29 | 27.9312102 | 6.596099917 | 0.9185 | 7.18101 | 6.92E-13 | 3.66E-11 |
| 442 | TMEM179 | 7.23715825 | 6.583255998 | 1.2036 | 5.46957 | 4.51E-08 | 8.27E-07 |
| 398 | MYO16 | 7.11400557 | 6.579259569 | 1.2347 | 5.32852 | 9.90E-08 | 1.68E-06 |
| 308 | CCDC184 | 7.3315657 | 6.578244801 | 1.2498 | 5.26363 | 1.41E-07 | 2.29E-06 |
| 99 | SYT11 | 19.0618407 | 6.577887192 | 1.0332 | 6.3663 | 1.94E-10 | 6.00E-09 |
| 971 | CLSTN2 | 7.22504249 | 6.577497051 | 1.2114 | 5.4296 | 5.65E-08 | 1.01E-06 |
| 1249 | EGR3 | 11.555819 | 6.53682004 | 1.1428 | 5.72003 | 1.07E-08 | 2.25E-07 |
| 862 | SLCO4A1 | 7.00630016 | 6.518231737 | 1.2694 | 5.13496 | 2.82E-07 | 4.28E-06 |
| 1082 | HBEGF | 34.473302 | 6.487455255 | 0.8149 | 7.9607 | 1.71E-15 | 1.48E-13 |
| 48 | CYP4A22 | 6.84809284 | 6.473110939 | 1.2438 | 5.20419 | 1.95E-07 | 3.05E-06 |
| 1078 | SPOCK1 | 18.2813012 | 6.460775047 | 1.0335 | 6.2516 | 4.06E-10 | 1.17E-08 |
| 1187 | DPY19L2P1 | 6.5122444 | 6.446914443 | 1.2691 | 5.07995 | 3.78E-07 | 5.54E-06 |
| 208 | CEND1 | 6.64077676 | 6.442203957 | 1.2281 | 5.24552 | 1.56E-07 | 2.5E-06 |
| 956 | TAGLN3 | 17.3499772 | 6.439705219 | 1.0453 | 6.16065 | 7.24E-10 | 1.94E-08 |
| 543 | ENSG00000256982 | 6.53961414 | 6.43827868 | 1.272 | 5.06162 | 4.16E-07 | 6.03E-06 |

| | | | | | | | |
|---|---|---|---|---|---|---|---|
| 64 | ERICH3 | 10.5024357 | 6.39983935 | 1.2164 | 5.2612 | 1.43E-07 | 2.32E-06 |
| 187 | INA | 63.334515 | 6.372860815 | 1.1451 | 5.56553 | 2.61E-08 | 5.11E-07 |
| 1252 | NEFM | 63.0137527 | 6.369477718 | 1.1659 | 5.46315 | 4.68E-08 | 8.54E-07 |
| 455 | RASGRP1 | 16.7589168 | 6.366261997 | 1.056 | 6.02874 | 1.65E-09 | 4.13E-08 |
| 489 | SH3GL3 | 10.2363944 | 6.358347969 | 1.1523 | 5.51815 | 3.43E-08 | 6.50E-07 |
| 1329 | GRIN3A | 16.380181 | 6.351008302 | 1.0422 | 6.0941 | 1.10E-09 | 2.83E-08 |
| 839 | SYNDIG1 | 10.238804 | 6.339662469 | 1.1784 | 5.37976 | 7.46E-08 | 1.3E-06 |
| 350 | FOXN4 | 6.20970244 | 6.336414249 | 1.3169 | 4.81174 | 1.5E-06 | 1.88E-05 |
| 547 | P2RX5 | 5.98949928 | 6.328250023 | 1.2672 | 4.99399 | 5.91E-07 | 8.31E-06 |
| 1412 | FRMPD3 | 6.07310957 | 6.32348624 | 1.2421 | 5.09081 | 3.57E-07 | 5.27E-06 |
| 1012 | KCTD8 | 5.93978053 | 6.298250127 | 1.2418 | 5.07176 | 3.94E-07 | 5.76E-06 |
| 1315 | LINC01241 | 5.94362838 | 6.288867795 | 1.2412 | 5.06668 | 4.05E-07 | 5.9E-06 |
| 344 | ANO4 | 44.0224401 | 6.276976233 | 0.7208 | 8.70895 | 3.07E-18 | 4.79E-16 |
| 498 | FBXL16 | 5.80712916 | 6.257565014 | 1.2464 | 5.02049 | 5.15E-07 | 7.35E-06 |
| 556 | ATP1B2 | 5.6783516 | 6.24663548 | 1.2716 | 4.91233 | 9.00E-07 | 1.21E-05 |
| 329 | B4GALNT1 | 5.77269634 | 6.240939186 | 1.3237 | 4.7146 | 2.4E-06 | 2.92E-05 |
| 278 | B3GAT1 | 9.39051139 | 6.229832084 | 1.1613 | 5.36459 | 8.11E-08 | 1.39E-06 |
| 665 | ELAVL3 | 43.0206813 | 6.170343843 | 0.6618 | 9.32396 | 1.12E-20 | 2.80E-18 |
| 395 | HS6ST3 | 65.3847173 | 6.167956915 | 0.5567 | 11.0789 | 1.59E-28 | 1.13E-25 |
| 881 | CECR7 | 5.33872299 | 6.162868368 | 1.2805 | 4.81272 | 1.5E-06 | 1.88E-05 |
| 1235 | TMEM178B | 5.35117759 | 6.162775406 | 1.2803 | 4.8137 | 1.5E-06 | 1.87E-05 |
| 803 | PTPRN | 8.90028543 | 6.16216971 | 1.1896 | 5.18 | 2.22E-07 | 3.43E-06 |
| 39 | POU3F1 | 8.95477284 | 6.158366484 | 1.2146 | 5.07049 | 3.97E-07 | 5.79E-06 |
| 709 | HAS1 | 5.29096958 | 6.151529144 | 1.2825 | 4.79633 | 1.6E-06 | 2.02E-05 |
| 1136 | PTCHD4 | 5.27992722 | 6.135610942 | 1.3043 | 4.70429 | 2.5E-06 | 3.05E-05 |
| 527 | GNAO1 | 46.4046799 | 6.097276856 | 0.6325 | 9.64008 | 5.41E-22 | 1.64E-19 |
| 949 | CADM2 | 5.11662217 | 6.067343312 | 1.2886 | 4.70862 | 2.5E-06 | 2.99E-05 |
| 397 | FGF14 | 13.6226787 | 6.066633739 | 1.0879 | 5.57667 | 2.45E-08 | 4.82E-07 |
| 367 | ENSG00000243008 | 5.04036875 | 6.061920175 | 1.3081 | 4.63414 | 3.6E-06 | 4.12E-05 |
| 532 | SLC9A5 | 8.3431075 | 6.061087794 | 1.1938 | 5.07728 | 3.83E-07 | 5.61E-06 |
| 204 | LINC01166 | 5.0998044 | 6.054897368 | 1.2906 | 4.6915 | 2.7E-06 | 3.21E-05 |
| 1290 | RIMS2 | 13.2732235 | 6.048508522 | 1.0582 | 5.7158 | 1.09E-08 | 2.30E-07 |
| 1072 | LINC01170 | 4.99223871 | 6.040444846 | 1.3666 | 4.42005 | 9.9E-06 | 0.000101 |
| 433 | UNC79 | 13.5616806 | 6.037988076 | 1.0671 | 5.65832 | 1.53E-08 | 3.15E-07 |
| 300 | CNTN1 | 18.8216345 | 6.033858172 | 0.9502 | 6.34987 | 2.15E-10 | 6.59E-09 |
| 655 | ATCAY | 12.982011 | 6.030317505 | 1.0884 | 5.54058 | 3.01E-08 | 5.80E-07 |
| 925 | MYRIP | 13.1756489 | 6.010571055 | 1.0878 | 5.52532 | 3.29E-08 | 6.27E-07 |
| 767 | THSD7B | 4.91401994 | 6.008424181 | 1.3219 | 4.54519 | 5.5E-06 | 5.97E-05 |
| 296 | RASSF8-AS1 | 8.06899405 | 6.004292406 | 1.2043 | 4.98578 | 6.17E-07 | 8.62E-06 |

| | | | | | | | |
|---|---|---|---|---|---|---|---|
| 271 | FEZ1 | 49.6677534 | 5.978945054 | 0.5826 | 10.2627 | 1.04E-24 | 4.22E-22 |
| 868 | MYT1 | 7.94430133 | 5.962593066 | 1.1925 | 4.99989 | 5.74E-07 | 8.08E-06 |
| 49 | TAL1 | 4.81537971 | 5.958960224 | 1.56 | 3.81976 | 0.00013 | 0.001012 |
| 752 | ITPRIPL1 | 4.68041335 | 5.955434135 | 1.2928 | 4.60668 | 4.1E-06 | 4.63E-05 |
| 1291 | ENSG00000254615 | 7.78584985 | 5.953898716 | 1.2016 | 4.95504 | 7.23E-07 | 9.94E-06 |
| 1418 | DOCK11 | 29.5757008 | 5.947466961 | 0.7728 | 7.69588 | 1.41E-14 | 1.06E-12 |
| 1348 | OLFM1 | 12.354919 | 5.938842475 | 1.0774 | 5.51232 | 3.54E-08 | 6.69E-07 |
| 10 | DISP3 | 4.55701548 | 5.927739664 | 1.3395 | 4.42524 | 9.6E-06 | 9.94E-05 |
| 935 | CAMKV | 63.7191891 | 5.911169363 | 0.5353 | 11.0436 | 2.35E-28 | 1.60E-25 |
| 345 | SNX5P2 | 4.43245554 | 5.863087339 | 1.3587 | 4.31517 | 1.6E-05 | 0.000156 |
| 287 | ENO2 | 59.455733 | 5.852584903 | 0.9525 | 6.14443 | 8.03E-10 | 2.12E-08 |
| 467 | TMOD2 | 7.37370498 | 5.848273122 | 1.2261 | 4.76992 | 1.8E-06 | 2.27E-05 |
| 654 | ZFR2 | 7.31859471 | 5.846678045 | 1.2053 | 4.85085 | 1.2E-06 | 1.59E-05 |
| 423 | RTN1 | 22.0584564 | 5.844481508 | 0.8291 | 7.04931 | 1.80E-12 | 8.94E-11 |
| 864 | EEF1A2 | 7.28093901 | 5.843288649 | 1.2226 | 4.7794 | 1.8E-06 | 2.18E-05 |
| 4 | FNDC10 | 4.35822331 | 5.827582377 | 1.3246 | 4.39938 | 1.1E-05 | 0.000111 |
| 156 | MKX | 17.2758992 | 5.798269537 | 0.9321 | 6.22062 | 4.95E-10 | 1.40E-08 |
| 1358 | GRIN1 | 7.02495848 | 5.784201738 | 1.2163 | 4.75562 | 2E-06 | 2.42E-05 |
| 407 | JPH4 | 16.0401793 | 5.782005277 | 0.9438 | 6.12641 | 8.99E-10 | 2.36E-08 |
| 1283 | CALB1 | 21.5112706 | 5.779954498 | 0.8461 | 6.8309 | 8.44E-12 | 3.66E-10 |
| 179 | CRTAC1 | 4.07511224 | 5.778388521 | 1.4273 | 4.04849 | 5.2E-05 | 0.000441 |
| 314 | POU6F1 | 7.04915595 | 5.770606212 | 1.2305 | 4.68974 | 2.7E-06 | 3.23E-05 |
| 229 | FJX1 | 4.09943741 | 5.768246217 | 1.333 | 4.32739 | 1.5E-05 | 0.000148 |
| 576 | SEZ6 | 6.85713508 | 5.751174851 | 1.2135 | 4.73944 | 2.1E-06 | 2.6E-05 |
| 908 | PANX2 | 4.05076112 | 5.746566962 | 1.3771 | 4.17283 | 3E-05 | 0.000273 |
| 1421 | RAB33A | 3.98922869 | 5.737267033 | 1.3504 | 4.24864 | 2.2E-05 | 0.000203 |
| 691 | MAP4K1 | 6.72489509 | 5.711650214 | 1.2844 | 4.44704 | 8.7E-06 | 9.05E-05 |
| 393 | POU4F1 | 11.0110703 | 5.711386697 | 1.0809 | 5.28401 | 1.26E-07 | 2.08E-06 |
| 138 | SUSD4 | 24.7663652 | 5.694238108 | 0.7876 | 7.22949 | 4.85E-13 | 2.66E-11 |
| 629 | TMEM200C | 3.83225216 | 5.691890961 | 1.4048 | 4.0518 | 5.1E-05 | 0.000435 |
| 102 | MIR9-1HG | 10.5488183 | 5.688798729 | 1.0921 | 5.20884 | 1.90E-07 | 2.99E-06 |
| 301 | ENSG00000274943 | 3.99350902 | 5.680901421 | 1.4073 | 4.03682 | 5.4E-05 | 0.000461 |
| 1206 | DYNC1I1 | 24.5029692 | 5.678408365 | 0.7604 | 7.46726 | 8.19E-14 | 5.25E-12 |
| 504 | SEC14L5 | 3.93124203 | 5.666472493 | 1.3749 | 4.12149 | 3.8E-05 | 0.000334 |
| 694 | DLL3 | 14.5771438 | 5.654367648 | 0.9525 | 5.93659 | 2.91E-09 | 6.81E-08 |
| 749 | CTNNA2 | 3.7745107 | 5.650271581 | 1.3841 | 4.08237 | 4.5E-05 | 0.000389 |
| 930 | CSPG5 | 3.80319903 | 5.648863992 | 1.3595 | 4.15496 | 3.3E-05 | 0.000293 |
| 480 | PSTPIP1 | 3.75609238 | 5.646871644 | 1.44 | 3.92137 | 8.8E-05 | 0.000704 |
| 117 | COLGALT2 | 6.22483677 | 5.642492456 | 1.2463 | 4.52735 | 6E-06 | 6.45E-05 |

| | | | | | | | |
|---|---|---|---|---|---|---|---|
| 177 | SLIT1 | 14.3418797 | 5.640557555 | 0.9708 | 5.81014 | 6.24E-09 | 1.38E-07 |
| 1191 | GCK | 6.26014741 | 5.631733128 | 1.2623 | 4.46163 | 8.1E-06 | 8.51E-05 |
| 486 | AP3B2 | 18.8744104 | 5.626127691 | 0.8587 | 6.55205 | 5.68E-11 | 2.02E-09 |
| 561 | HES7 | 3.67198586 | 5.621259208 | 1.4283 | 3.93568 | 8.3E-05 | 0.000669 |
| 615 | TCAM1P | 3.740348 | 5.620692953 | 1.3597 | 4.13378 | 3.6E-05 | 0.000318 |
| 584 | LHX1 | 3.71670052 | 5.617964071 | 1.4371 | 3.90928 | 9.3E-05 | 0.000733 |
| 1048 | CDH10 | 6.03756141 | 5.600527125 | 1.2429 | 4.50605 | 6.6E-06 | 7.09E-05 |
| 115 | NMNAT2 | 6.08305151 | 5.598170875 | 1.2437 | 4.50137 | 6.8E-06 | 7.23E-05 |
| 182 | LBX1-AS1 | 23.1470285 | 5.597193648 | 0.8247 | 6.78686 | 1.15E-11 | 4.87E-10 |
| 1279 | CRISPLD1 | 64.101703 | 5.578955868 | 0.4949 | 11.2736 | 1.77E-29 | 1.40E-26 |
| 997 | RNF212 | 22.9641365 | 5.573402503 | 0.7604 | 7.32923 | 2.31E-13 | 1.35E-11 |
| 1125 | HLA-DPB1 | 6.12425438 | 5.568407554 | 1.2735 | 4.37244 | 1.2E-05 | 0.000123 |
| 325 | NXPH4 | 3.56470212 | 5.567174602 | 1.4404 | 3.86506 | 0.00011 | 0.000863 |
| 1270 | BHLHE22 | 18.1171477 | 5.562408141 | 0.8578 | 6.48442 | 8.91E-11 | 3.05E-09 |
| 893 | ENSG00000166897 | 3.53986164 | 5.555066566 | 1.4014 | 3.96402 | 7.4E-05 | 0.000601 |
| 1281 | STMN2 | 46.22985 | 5.552179794 | 1.3351 | 4.1585 | 3.2E-05 | 0.000289 |
| 151 | SFMBT2 | 3.57060896 | 5.546515454 | 1.3804 | 4.01806 | 5.9E-05 | 0.000495 |
| 916 | GRIP2 | 5.88194023 | 5.543854518 | 1.2309 | 4.50386 | 6.7E-06 | 7.15E-05 |
| 800 | CDK5R2 | 3.54822319 | 5.53915111 | 1.4014 | 3.9525 | 7.7E-05 | 0.000628 |
| 577 | ANKRD13B | 18.5952827 | 5.532502672 | 0.8348 | 6.6275 | 3.41E-11 | 1.28E-09 |
| 811 | DNER | 31.6267446 | 5.522421463 | 0.6611 | 8.35301 | 6.65E-17 | 7.70E-15 |
| 941 | CACNA2D3 | 3.47980416 | 5.518813513 | 1.4166 | 3.89578 | 9.8E-05 | 0.00077 |
| 479 | ISL2 | 5.89207464 | 5.517020016 | 1.2649 | 4.36177 | 1.3E-05 | 0.000129 |
| 517 | ASPHD1 | 14.1342369 | 5.50738117 | 0.9499 | 5.79795 | 6.71E-09 | 1.48E-07 |
| 1338 | PTGS1 | 9.51622589 | 5.49841758 | 1.1263 | 4.88173 | 1.1E-06 | 1.38E-05 |
| 1314 | ELAVL2 | 55.6956857 | 5.49013386 | 1.1325 | 4.84795 | 1.2E-06 | 1.61E-05 |
| 744 | DYSF | 3.42569131 | 5.489337323 | 1.4681 | 3.73899 | 0.00018 | 0.001336 |
| 596 | RUNDC3A | 21.6204394 | 5.48815583 | 0.781 | 7.02733 | 2.11E-12 | 1.04E-10 |
| 761 | INHBB | 18.6839438 | 5.48399173 | 0.8577 | 6.39356 | 1.62E-10 | 5.11E-09 |
| 751 | DUSP2 | 3.33494339 | 5.48083938 | 1.4266 | 3.84184 | 0.00012 | 0.000938 |
| 863 | KCNQ2 | 21.2886142 | 5.473656519 | 0.7843 | 6.97891 | 2.97E-12 | 1.42E-10 |
| 74 | COL11A1 | 33.5775547 | 5.463997273 | 0.6391 | 8.55 | 1.23E-17 | 1.72E-15 |
| 1083 | PCDHB16 | 3.31812562 | 5.463400057 | 1.3894 | 3.93232 | 8.4E-05 | 0.000677 |
| 1262 | ENSG00000279099 | 3.26406965 | 5.455283675 | 1.4124 | 3.86241 | 0.00011 | 0.000871 |
| 683 | FXYD7 | 3.31846446 | 5.450141961 | 1.3919 | 3.91549 | 9E-05 | 0.000719 |
| 762 | CNTNAP5 | 3.32495533 | 5.448329349 | 1.4166 | 3.84599 | 0.00012 | 0.000925 |
| 822 | SCRT2 | 3.22732082 | 5.432153678 | 1.4344 | 3.78701 | 0.00015 | 0.00113 |
| 1095 | EBF1 | 8.63197282 | 5.42987983 | 1.1204 | 4.84647 | 1.3E-06 | 1.62E-05 |
| 42 | CFAP57 | 5.50954925 | 5.426422055 | 1.2498 | 4.34185 | 1.4E-05 | 0.00014 |

| | | | | | | | |
|---|---|---|---|---|---|---|---|
| 1015 | NMU | 29.4708463 | 5.425586241 | 0.6634 | 8.17839 | 2.88E-16 | 2.90E-14 |
| 610 | PPM1E | 3.26508616 | 5.41603111 | 1.3956 | 3.88073 | 0.0001 | 0.000814 |
| 643 | SLC14A1 | 3.21348491 | 5.41456064 | 1.4163 | 3.8229 | 0.00013 | 0.001003 |
| 1123 | ATP6V1G2 | 3.28781078 | 5.412858094 | 1.4182 | 3.81682 | 0.00014 | 0.001023 |
| 30 | FNDC5 | 12.4885717 | 5.403823483 | 0.9821 | 5.50248 | 3.74E-08 | 7.03E-07 |
| 780 | CHN1 | 78.1674255 | 5.399793735 | 0.428 | 12.6175 | 1.69E-36 | 2.67E-33 |
| 360 | ENSG00000274191 | 3.20819886 | 5.385719646 | 1.4169 | 3.80107 | 0.00014 | 0.001077 |
| 85 | NHLH2 | 3.0829872 | 5.383536986 | 1.4991 | 3.59126 | 0.00033 | 0.00222 |
| 408 | CARMIL3 | 55.370656 | 5.379276599 | 0.4994 | 10.7725 | 4.64E-27 | 2.36E-24 |
| 795 | UNC80 | 5.36022345 | 5.362848578 | 1.3211 | 4.05941 | 4.9E-05 | 0.000422 |
| 674 | TMEM59L | 3.14328884 | 5.359316789 | 1.4148 | 3.78797 | 0.00015 | 0.001127 |
| 745 | FBXO41 | 8.86677262 | 5.358796799 | 1.1229 | 4.77246 | 1.8E-06 | 2.25E-05 |
| 454 | GJD2 | 3.0330492 | 5.351412825 | 1.6827 | 3.18029 | 0.00147 | 0.008 |
| 833 | LAMP5 | 11.9447652 | 5.346523448 | 1.0096 | 5.29556 | 1.19E-07 | 1.97E-06 |
| 26 | NKAIN1 | 3.16967299 | 5.338352901 | 1.5035 | 3.55061 | 0.00038 | 0.002541 |
| 377 | DCLK1 | 35.9294217 | 5.335339629 | 0.5829 | 9.1534 | 5.52E-20 | 1.23E-17 |
| 381 | ENOX1 | 2.98132835 | 5.333940951 | 1.5099 | 3.53265 | 0.00041 | 0.002691 |
| 1248 | LZTS1 | 5.09221001 | 5.329645273 | 1.2736 | 4.18456 | 2.9E-05 | 0.000262 |
| 1210 | NPTX2 | 19.0999612 | 5.318386863 | 0.7812 | 6.80812 | 9.89E-12 | 4.25E-10 |
| 1328 | PLPPR1 | 2.9524517 | 5.318291304 | 1.6332 | 3.25632 | 0.00113 | 0.006356 |
| 1320 | IGFBPL1 | 30.4594748 | 5.309826328 | 1.2004 | 4.42343 | 9.7E-06 | 0.0001 |
| 768 | KIF5C | 28.5554903 | 5.309459867 | 0.6631 | 8.00682 | 1.18E-15 | 1.05E-13 |
| 801 | FEV | 23.061082 | 5.295064098 | 0.7049 | 7.51211 | 5.82E-14 | 3.81E-12 |
| 1004 | HS3ST1 | 27.0811035 | 5.287431952 | 0.663 | 7.97468 | 1.53E-15 | 1.35E-13 |
| 1254 | EBF2 | 2.98698999 | 5.287018696 | 1.6573 | 3.19018 | 0.00142 | 0.007776 |
| 1075 | ACSL6 | 2.9702291 | 5.283342872 | 1.43 | 3.69464 | 0.00022 | 0.001558 |
| 392 | ENSG00000271776 | 2.89941225 | 5.280848248 | 1.4794 | 3.56954 | 0.00036 | 0.002389 |
| 1181 | ENSG00000253308 | 3.01097631 | 5.270108965 | 1.4778 | 3.56623 | 0.00036 | 0.002412 |
| 747 | SEMA4F | 2.82297051 | 5.256173736 | 1.5516 | 3.38758 | 0.00071 | 0.004274 |
| 630 | L3MBTL4 | 2.82801136 | 5.254581439 | 1.4902 | 3.52619 | 0.00042 | 0.00275 |
| 906 | PRR34-AS1 | 2.87047872 | 5.246403864 | 1.4661 | 3.57846 | 0.00035 | 0.002317 |
| 923 | ARPP21 | 15.9466832 | 5.233780358 | 0.8822 | 5.93267 | 2.98E-09 | 6.96E-08 |
| 189 | SORCS3 | 2.81676159 | 5.222614256 | 1.4867 | 3.5129 | 0.00044 | 0.002876 |
| 1403 | TCEAL2 | 4.74172735 | 5.220408307 | 1.2743 | 4.09654 | 4.2E-05 | 0.000369 |
| 513 | ENSG00000246465 | 2.84671163 | 5.2165803 | 1.5087 | 3.45758 | 0.00055 | 0.003441 |
| 946 | GPR27 | 2.80258684 | 5.216108841 | 1.5162 | 3.44022 | 0.00058 | 0.003638 |
| 1164 | PPP1R14C | 35.6075425 | 5.210804874 | 0.5909 | 8.818 | 1.17E-18 | 1.96E-16 |
| 1415 | CHRDL1 | 2.75045843 | 5.208530504 | 1.6685 | 3.12178 | 0.0018 | 0.009492 |
| 126 | KLHDC8A | 4.64839703 | 5.208189807 | 1.2962 | 4.01798 | 5.9E-05 | 0.000495 |

| | | | | | | | |
|---|---|---|---|---|---|---|---|
| 1057 | FOXD1 | 4.72253458 | 5.198508811 | 1.277 | 4.07083 | 4.7E-05 | 0.000405 |
| 616 | CD79B | 2.77703089 | 5.195294455 | 1.4991 | 3.46568 | 0.00053 | 0.003356 |
| 865 | STMN3 | 26.1725516 | 5.172986555 | 0.6624 | 7.80909 | 5.76E-15 | 4.50E-13 |
| 1127 | PACSIN1 | 4.6208369 | 5.170568314 | 1.2846 | 4.02509 | 5.7E-05 | 0.000481 |
| 1310 | SH3GL2 | 2.81288778 | 5.165874611 | 1.5379 | 3.35902 | 0.00078 | 0.004681 |
| 160 | FRMPD2 | 2.6717125 | 5.165761095 | 1.5198 | 3.39889 | 0.00068 | 0.004124 |
| 687 | ZFP82 | 4.60021011 | 5.163159467 | 1.3199 | 3.91174 | 9.2E-05 | 0.000727 |
| 1224 | KCND2 | 2.75286808 | 5.160870998 | 1.7012 | 3.03362 | 0.00242 | 0.012228 |
| 78 | ATXN7L2 | 4.46857839 | 5.157056027 | 1.2954 | 3.98099 | 6.9E-05 | 0.000569 |
| 106 | NHLH1 | 4.62472358 | 5.153161643 | 1.3171 | 3.91243 | 9.1E-05 | 0.000726 |
| 910 | MAPK8IP2 | 14.1667488 | 5.144010743 | 0.8654 | 5.944 | 2.78E-09 | 6.56E-08 |
| 203 | DPYSL4 | 16.7298579 | 5.142588588 | 0.8334 | 6.17091 | 6.79E-10 | 1.84E-08 |
| 114 | RGS16 | 10.4179427 | 5.140582875 | 1.0005 | 5.13779 | 2.78E-07 | 4.22E-06 |
| 496 | ST8SIA2 | 2.64071998 | 5.136592495 | 1.5403 | 3.33485 | 0.00085 | 0.005048 |
| 1268 | ST18 | 39.7148245 | 5.136469726 | 0.604 | 8.50446 | 1.82E-17 | 2.40E-15 |
| 1250 | STC1 | 10.7591978 | 5.132695677 | 1.1339 | 4.52672 | 6E-06 | 6.47E-05 |
| 198 | PLPP4 | 2.67888105 | 5.132469307 | 1.4618 | 3.51099 | 0.00045 | 0.002891 |
| 786 | ITGA4 | 34.3267152 | 5.130050753 | 0.5596 | 9.16699 | 4.86E-20 | 1.10E-17 |
| 1061 | LINC00461 | 7.1563252 | 5.112433882 | 1.1427 | 4.47413 | 7.7E-06 | 8.08E-05 |
| 778 | SP9 | 4.49597905 | 5.110221286 | 1.3675 | 3.7369 | 0.00019 | 0.001344 |
| 961 | KLF15 | 2.59945745 | 5.109320419 | 1.6834 | 3.03513 | 0.0024 | 0.012183 |
| 1184 | NEUROD6 | 2.57005365 | 5.107126793 | 1.5441 | 3.30755 | 0.00094 | 0.00546 |
| 583 | LHX1-DT | 4.32943614 | 5.103204585 | 1.3331 | 3.8281 | 0.00013 | 0.000986 |
| 491 | MIR9-3HG | 2.56793778 | 5.085961622 | 1.4912 | 3.41062 | 0.00065 | 0.003984 |
| 1194 | SRRM3 | 16.8349998 | 5.077853608 | 0.813 | 6.24606 | 4.21E-10 | 1.20E-08 |
| 285 | PIANP | 2.48466633 | 5.0723872 | 1.6722 | 3.03339 | 0.00242 | 0.012233 |
| 963 | KBTBD12 | 2.62353741 | 5.069458592 | 1.7646 | 2.87287 | 0.00407 | 0.018952 |
| 1255 | PNMA2 | 26.3918394 | 5.063642637 | 0.6508 | 7.78053 | 7.22E-15 | 5.52E-13 |
| 16 | IGSF21 | 2.57980834 | 5.060644525 | 1.5391 | 3.28804 | 0.00101 | 0.005784 |
| 339 | SYT1 | 16.1979608 | 5.058442831 | 0.8018 | 6.3092 | 2.80E-10 | 8.36E-09 |
| 673 | KCNN1 | 2.4977986 | 5.050501312 | 1.5309 | 3.29904 | 0.00097 | 0.005598 |
| 75 | WDR47 | 9.87317449 | 5.047705918 | 0.9873 | 5.11279 | 3.17E-07 | 4.73E-06 |
| 340 | PPFIA2 | 2.41452715 | 5.035116398 | 1.689 | 2.98107 | 0.00287 | 0.014134 |
| 666 | MAST1 | 9.50083526 | 5.033192733 | 1.0443 | 4.81949 | 1.4E-06 | 1.83E-05 |
| 37 | MANEAL | 9.96497879 | 5.03275988 | 1.0091 | 4.9875 | 6.12E-07 | 8.56E-06 |
| 905 | LINC00899 | 2.48396269 | 5.027757611 | 1.5708 | 3.20084 | 0.00137 | 0.007546 |
| 1182 | EVX1 | 2.43442042 | 5.013901742 | 1.531 | 3.27494 | 0.00106 | 0.006021 |
| 1385 | SYN1 | 2.38002561 | 5.01048717 | 1.6729 | 2.99505 | 0.00274 | 0.013605 |
| 915 | WNT7A | 2.4069938 | 5.009116288 | 1.5623 | 3.20622 | 0.00134 | 0.007424 |

| | | | | | | | |
|---|---|---|---|---|---|---|---|
| 1313 | CDKN2A | 26.7396811 | 5.007389754 | 0.6341 | 7.89743 | 2.85E-15 | 2.38E-13 |
| 904 | PHF21B | 18.4588775 | 4.997251884 | 0.7481 | 6.67966 | 2.39E-11 | 9.28E-10 |
| 730 | GALNT14 | 6.55962257 | 4.995932936 | 1.1847 | 4.21718 | 2.5E-05 | 0.000229 |
| 96 | CHRNB2 | 4.04285882 | 4.994760613 | 1.3598 | 3.67309 | 0.00024 | 0.001676 |
| 482 | RASGRF1 | 2.40733264 | 4.993024843 | 1.5905 | 3.13932 | 0.00169 | 0.009025 |
| 1056 | MAP1B | 560.413476 | 4.990240856 | 1.0295 | 4.84743 | 1.3E-06 | 1.61E-05 |
| 579 | RHBDL3 | 2.46099288 | 4.988290766 | 1.5531 | 3.21187 | 0.00132 | 0.007297 |
| 334 | KCNMB4 | 4.03670678 | 4.985522282 | 1.3144 | 3.79296 | 0.00015 | 0.001108 |
| 1289 | ENSG00000271830 | 3.94224244 | 4.982509339 | 1.3394 | 3.7199 | 0.0002 | 0.00143 |
| 470 | MYO5A | 12.1175756 | 4.968561809 | 0.9102 | 5.45855 | 4.80E-08 | 8.75E-07 |
| 266 | TECTA | 2.31022527 | 4.964304947 | 1.5683 | 3.16533 | 0.00155 | 0.008356 |
| 1430 | ARHGEF6 | 19.1329951 | 4.96126705 | 0.7666 | 6.47187 | 9.68E-11 | 3.29E-09 |
| 268 | SCN3B | 2.45047767 | 4.957790565 | 1.6393 | 3.02441 | 0.00249 | 0.012522 |
| 841 | XKR7 | 2.36163822 | 4.957620776 | 1.539 | 3.22133 | 0.00128 | 0.007084 |
| 1103 | F12 | 3.9201795 | 4.945274639 | 1.3347 | 3.70503 | 0.00021 | 0.001502 |
| 1246 | LPL | 14.5041246 | 4.939465194 | 0.8882 | 5.56126 | 2.68E-08 | 5.21E-07 |
| 63 | NEGR1 | 22.9621049 | 4.938954049 | 0.7086 | 6.97009 | 3.17E-12 | 1.51E-10 |
| 912 | LHFPL4 | 8.73615979 | 4.938282771 | 1.0659 | 4.63309 | 3.6E-06 | 4.14E-05 |
| 834 | PAK5 | 20.2864978 | 4.932228279 | 0.7332 | 6.72659 | 1.74E-11 | 6.94E-10 |
| 1157 | SLC35D3 | 2.26751272 | 4.930317669 | 1.5755 | 3.12937 | 0.00175 | 0.009278 |
| 250 | ENSG00000278989 | 2.27400359 | 4.926873887 | 1.5911 | 3.09654 | 0.00196 | 0.010198 |
| 624 | RBFOX3 | 6.68639971 | 4.921817195 | 1.2011 | 4.09789 | 4.2E-05 | 0.000368 |
| 224 | KCNC1 | 2.26486969 | 4.912000083 | 1.5306 | 3.20922 | 0.00133 | 0.007356 |
| 227 | KIAA1549L | 11.6846088 | 4.908408253 | 0.9117 | 5.38378 | 7.29E-08 | 1.27E-06 |
| 754 | LONRF2 | 3.79579604 | 4.908036764 | 1.3613 | 3.60537 | 0.00031 | 0.002113 |
| 1090 | MYOZ3 | 6.08555064 | 4.899311035 | 1.184 | 4.13805 | 3.5E-05 | 0.000314 |
| 692 | CCER2 | 20.5001646 | 4.896046255 | 0.7112 | 6.88385 | 5.83E-12 | 2.62E-10 |
| 466 | SCG3 | 18.3359415 | 4.890956669 | 0.76 | 6.43538 | 1.23E-10 | 4.03E-09 |
| 451 | OTUD7A | 3.80789375 | 4.887855056 | 1.3653 | 3.58019 | 0.00034 | 0.002303 |
| 821 | TCF15 | 2.25103378 | 4.884461773 | 1.5293 | 3.194 | 0.0014 | 0.007683 |
| 1297 | SYBU-AS1 | 2.26905638 | 4.872670206 | 1.6139 | 3.01914 | 0.00253 | 0.012706 |
| 331 | C12orf56 | 2.18227592 | 4.871799316 | 1.5812 | 3.08113 | 0.00206 | 0.010679 |
| 1060 | EDIL3-DT | 5.78101755 | 4.868477637 | 1.279 | 3.80645 | 0.00014 | 0.001059 |
| 95 | SHE | 5.86036249 | 4.862724549 | 1.1988 | 4.05644 | 5E-05 | 0.000427 |
| 1434 | AFF2 | 17.4237058 | 4.862713251 | 0.7623 | 6.37876 | 1.79E-10 | 5.58E-09 |
| 979 | SHOX2 | 69.695196 | 4.858124011 | 0.415 | 11.7072 | 1.17E-31 | 1.39E-28 |
| 72 | PLPPR5 | 3.61937472 | 4.85363699 | 1.3529 | 3.58753 | 0.00033 | 0.002247 |
| 173 | NRG3 | 3.73432355 | 4.84741985 | 1.4809 | 3.27337 | 0.00106 | 0.006048 |
| 279 | IQSEC3 | 2.21739824 | 4.844980339 | 1.8482 | 2.62143 | 0.00876 | 0.035498 |

| | | | | | | | |
|---|---|---|---|---|---|---|---|
| 109 | CD247 | 3.60646169 | 4.84163216 | 1.4055 | 3.44482 | 0.00057 | 0.003584 |
| 1002 | STK32B | 13.5441484 | 4.84043352 | 0.8357 | 5.79182 | 6.96E-09 | 1.52E-07 |
| 46 | CYP4X1 | 30.2173806 | 4.829267198 | 0.6193 | 7.79839 | 6.27E-15 | 4.87E-13 |
| 349 | ALDH1L2 | 13.5277532 | 4.820880639 | 0.8682 | 5.55302 | 2.81E-08 | 5.43E-07 |
| 323 | HSD17B6 | 2.12591576 | 4.820525546 | 1.5836 | 3.04393 | 0.00234 | 0.011876 |
| 1034 | GRIA2 | 102.917615 | 4.818190927 | 0.9603 | 5.01721 | 5.24E-07 | 7.45E-06 |
| 671 | ANKLE1 | 8.23525917 | 4.808624598 | 1.0521 | 4.57053 | 4.9E-06 | 5.39E-05 |
| 831 | LRRN4 | 2.13221832 | 4.802938124 | 1.5585 | 3.08176 | 0.00206 | 0.01066 |
| 1045 | UBE2QL1 | 2.09531896 | 4.802056242 | 1.5685 | 3.06153 | 0.0022 | 0.011313 |
| 737 | NRXN1 | 27.7646842 | 4.796908088 | 1.1685 | 4.10527 | 4E-05 | 0.000357 |
| 409 | COCH | 35.2935492 | 4.788691541 | 0.529 | 9.05198 | 1.40E-19 | 2.81E-17 |
| 693 | LRFN1 | 3.57820583 | 4.786016119 | 1.3538 | 3.53521 | 0.00041 | 0.002677 |
| 1146 | RIPPLY2 | 2.04928571 | 4.781523267 | 1.599 | 2.99029 | 0.00279 | 0.013777 |
| 889 | NEFH | 20.3478347 | 4.775179966 | 0.7717 | 6.18781 | 6.10E-10 | 1.67E-08 |
| 1402 | PCDH19 | 2.06902846 | 4.774355622 | 1.8043 | 2.64615 | 0.00814 | 0.033477 |
| 144 | GNG4 | 50.8372396 | 4.771703091 | 0.4636 | 10.292 | 7.66E-25 | 3.20E-22 |
| 858 | DOK5 | 13.0036366 | 4.77089421 | 0.8674 | 5.50049 | 3.79E-08 | 7.09E-07 |
| 353 | DTX1 | 2.132896 | 4.769962639 | 1.6688 | 2.85838 | 0.00426 | 0.019659 |
| 512 | SBK1 | 17.7529489 | 4.762614931 | 0.8078 | 5.89602 | 3.72E-09 | 8.59E-08 |
| 860 | CDH4 | 5.44480329 | 4.754149397 | 1.2223 | 3.88945 | 0.0001 | 0.00079 |
| 1264 | ANK1 | 7.8704673 | 4.751533121 | 1.0506 | 4.52276 | 6.1E-06 | 6.58E-05 |
| 1204 | DPY19L2P4 | 5.49766626 | 4.750373907 | 1.1994 | 3.96054 | 7.5E-05 | 0.00061 |
| 60 | DNAJC6 | 21.4510272 | 4.745110958 | 0.6449 | 7.35813 | 1.87E-13 | 1.12E-11 |
| 20 | RUNX3 | 2.04996338 | 4.740660157 | 1.6091 | 2.9461 | 0.00322 | 0.01556 |
| 708 | SHANK1 | 2.04698152 | 4.740153298 | 1.6392 | 2.89183 | 0.00383 | 0.018062 |
| 1167 | LPAL2 | 1.9923984 | 4.731934474 | 1.7 | 2.7835 | 0.00538 | 0.023965 |
| 178 | ARHGAP19-SLIT1 | 1.97299449 | 4.718865669 | 1.7107 | 2.75842 | 0.00581 | 0.025477 |
| 1011 | SHISA3 | 20.0830612 | 4.718668226 | 0.7109 | 6.63789 | 3.18E-11 | 1.21E-09 |
| 338 | NAV3 | 30.4893832 | 4.716031194 | 0.5643 | 8.35769 | 6.40E-17 | 7.46E-15 |
| 607 | TMEM100 | 10.1919728 | 4.713364602 | 0.9583 | 4.91832 | 8.73E-07 | 1.18E-05 |
| 1257 | STMN4 | 2.02229156 | 4.697744556 | 1.8269 | 2.57149 | 0.01013 | 0.039803 |
| 1142 | RIMS1 | 7.57583943 | 4.697509326 | 1.0331 | 4.54713 | 5.4E-06 | 5.93E-05 |
| 270 | NRGN | 14.9313929 | 4.693112864 | 0.7578 | 6.19346 | 5.89E-10 | 1.63E-08 |
| 25 | PTPRU | 23.2000202 | 4.689419757 | 0.6165 | 7.60675 | 2.81E-14 | 1.94E-12 |
| 842 | ENSG00000278012 | 1.99360323 | 4.687107868 | 1.6799 | 2.79016 | 0.00527 | 0.023551 |
| 962 | PODXL2 | 8.03868059 | 4.686341986 | 1.0329 | 4.5372 | 5.7E-06 | 6.19E-05 |
| 431 | ZDHHC22 | 31.3964151 | 4.682550075 | 1.0709 | 4.37234 | 1.2E-05 | 0.000123 |
| 1302 | FAM135B | 3.34058209 | 4.680636985 | 1.4016 | 3.33939 | 0.00084 | 0.004975 |
| 647 | CDH7 | 7.97924671 | 4.672824119 | 1.0458 | 4.46824 | 7.9E-06 | 8.29E-05 |

| | | | | | | | |
|---|---|---|---|---|---|---|---|
| 269 | TMEM225 | 1.95704271 | 4.661211897 | 1.6612 | 2.80591 | 0.00502 | 0.022642 |
| 362 | DNAH10 | 5.22587264 | 4.661026041 | 1.2206 | 3.81859 | 0.00013 | 0.001016 |
| 1301 | KCNQ3 | 1.90544145 | 4.655509068 | 1.6053 | 2.90007 | 0.00373 | 0.017667 |
| 1441 | ATP2B3 | 1.9715563 | 4.649023984 | 1.6613 | 2.79835 | 0.00514 | 0.023035 |
| 573 | LINC02693 | 3.19873688 | 4.648583263 | 1.4729 | 3.15614 | 0.0016 | 0.00859 |
| 535 | CARMIL2 | 1.89424857 | 4.648009231 | 1.6234 | 2.8631 | 0.0042 | 0.019451 |
| 760 | EN1 | 7.3088792 | 4.647509733 | 1.0956 | 4.24203 | 2.2E-05 | 0.000208 |
| 856 | ENSG00000227705 | 1.85138548 | 4.641583508 | 1.6571 | 2.80107 | 0.00509 | 0.022885 |
| 735 | KCNK12 | 3.26985887 | 4.630590124 | 1.4867 | 3.11468 | 0.00184 | 0.009677 |
| 1278 | GDAP1 | 76.1648406 | 4.628627086 | 0.3725 | 12.4242 | 1.93E-35 | 2.75E-32 |
| 1137 | TFAP2B | 22.4127604 | 4.617818209 | 0.6512 | 7.09112 | 1.33E-12 | 6.73E-11 |
| 558 | DNAH2 | 8.80221526 | 4.605294376 | 1.0518 | 4.3783 | 1.2E-05 | 0.00012 |
| 289 | LINC00937 | 1.82933855 | 4.604380569 | 1.6443 | 2.80027 | 0.00511 | 0.022913 |
| 587 | EPOP | 3.14920644 | 4.596360289 | 1.4511 | 3.16743 | 0.00154 | 0.008309 |
| 924 | SCN5A | 3.12257709 | 4.587840156 | 1.4027 | 3.27082 | 0.00107 | 0.006094 |
| 1003 | CRMP1 | 28.3041101 | 4.573749239 | 0.5803 | 7.88187 | 3.23E-15 | 2.64E-13 |
| 632 | APCDD1 | 18.2420933 | 4.568793397 | 0.6887 | 6.63382 | 3.27E-11 | 1.24E-09 |
| 434 | PPP4R4 | 1.80430975 | 4.5641131 | 1.7047 | 2.67734 | 0.00742 | 0.031039 |
| 1316 | ANKRD18B | 1.86393372 | 4.564059732 | 1.7248 | 2.6461 | 0.00814 | 0.033477 |
| 17 | ENSG00000280222 | 1.78398297 | 4.532786594 | 1.6567 | 2.73605 | 0.00622 | 0.026926 |
| 603 | NXPH3 | 1.78398297 | 4.532786594 | 1.6567 | 2.73605 | 0.00622 | 0.026926 |
| 1286 | SLC26A7 | 1.71018425 | 4.527961343 | 1.7051 | 2.65561 | 0.00792 | 0.032698 |
| 781 | HOXD3 | 1.69899137 | 4.51856293 | 1.733 | 2.60743 | 0.00912 | 0.036699 |
| 1124 | HLA-DPA1 | 1.72171598 | 4.513151216 | 1.6641 | 2.71208 | 0.00669 | 0.028597 |
| 965 | GATA2 | 1.73026584 | 4.495208533 | 1.7018 | 2.6414 | 0.00826 | 0.033867 |
| 108 | FAM78B | 11.2150671 | 4.493534557 | 0.8285 | 5.4238 | 5.83E-08 | 1.03E-06 |
| 884 | RTN4R | 4.52472588 | 4.48922939 | 1.2738 | 3.52432 | 0.00042 | 0.002768 |
| 446 | GABRB3 | 8.61410207 | 4.487489148 | 0.9997 | 4.48875 | 7.2E-06 | 7.61E-05 |
| 36 | GRIK3 | 6.29968039 | 4.480983587 | 1.1911 | 3.762 | 0.00017 | 0.001233 |
| 1032 | DCLK2 | 42.7151253 | 4.479326486 | 0.4594 | 9.75036 | 1.84E-22 | 6.08E-20 |
| 987 | ENSG00000287645 | 1.66150797 | 4.478191229 | 1.6956 | 2.64099 | 0.00827 | 0.033897 |
| 448 | ENTREP2 | 22.9271499 | 4.475251202 | 0.6693 | 6.68663 | 2.28E-11 | 8.92E-10 |
| 802 | CRYBA2 | 19.8475617 | 4.472861007 | 0.6635 | 6.74123 | 1.57E-11 | 6.37E-10 |
| 867 | RGS19 | 2.76145466 | 4.468414407 | 1.5855 | 2.81833 | 0.00483 | 0.021898 |
| 1070 | CDO1 | 4.87634051 | 4.463747147 | 1.3114 | 3.40383 | 0.00066 | 0.00406 |
| 1352 | DIPK1B | 1.62177728 | 4.430691741 | 1.7828 | 2.48525 | 0.01295 | 0.04868 |
| 999 | NAT8L | 2.65979581 | 4.414338051 | 1.5633 | 2.82382 | 0.00475 | 0.021574 |
| 180 | CNNM1 | 4.79949618 | 4.409678955 | 1.2535 | 3.5179 | 0.00043 | 0.002827 |
| 976 | MBNL1-AS1 | 2.68606827 | 4.401749874 | 1.4606 | 3.01357 | 0.00258 | 0.012919 |

| | | | | | | | |
|---|---|---|---|---|---|---|---|
| 595 | LINC01976 | 1.62562513 | 4.396536095 | 1.6936 | 2.59591 | 0.00943 | 0.037623 |
| 620 | MGAT5B | 4.21590148 | 4.394592346 | 1.3842 | 3.17493 | 0.0015 | 0.008127 |
| 166 | FAM241B | 6.24011022 | 4.381772995 | 1.0965 | 3.99607 | 6.4E-05 | 0.000538 |
| 909 | MAPK12 | 8.41005396 | 4.376037106 | 0.976 | 4.4837 | 7.3E-06 | 7.76E-05 |
| 743 | ADD2 | 16.3884413 | 4.375261312 | 0.7327 | 5.97154 | 2.35E-09 | 5.67E-08 |
| 304 | NELL2 | 47.2270736 | 4.372974176 | 0.4865 | 8.98844 | 2.51E-19 | 4.74E-17 |
| 1439 | PNMA6A | 8.83021594 | 4.367133927 | 0.9657 | 4.52204 | 6.1E-06 | 6.59E-05 |
| 711 | UBE2S | 2.69231395 | 4.351882884 | 1.4604 | 2.97986 | 0.00288 | 0.01418 |
| 1008 | ENSG00000248343 | 1.51732488 | 4.348782422 | 1.7367 | 2.50401 | 0.01228 | 0.046688 |
| 1106 | FLT4 | 19.0908574 | 4.346713081 | 1.3206 | 3.29144 | 0.001 | 0.005736 |
| 483 | ANKRD34C | 1.50911385 | 4.341293708 | 1.7479 | 2.4837 | 0.013 | 0.04888 |
| 375 | NBEA | 7.48812197 | 4.327619453 | 1.064 | 4.06714 | 4.8E-05 | 0.000411 |
| 1225 | PTPRZ1 | 40.8654117 | 4.313011822 | 0.5131 | 8.40623 | 4.23E-17 | 5.19E-15 |
| 738 | ENSG00000286412 | 2.65344924 | 4.303522392 | 1.5525 | 2.772 | 0.00557 | 0.024628 |
| 659 | PTPRS | 4.11810237 | 4.299318981 | 1.3069 | 3.28968 | 0.001 | 0.00576 |
| 872 | TIAM1 | 2.44730334 | 4.280026296 | 1.4846 | 2.88296 | 0.00394 | 0.018465 |
| 807 | COL4A4 | 2.54879288 | 4.273162126 | 1.5275 | 2.79741 | 0.00515 | 0.023088 |
| 983 | LINC02082 | 2.48758401 | 4.262116835 | 1.5282 | 2.78906 | 0.00529 | 0.023617 |
| 514 | PRRT2 | 21.8482372 | 4.258601954 | 0.5916 | 7.1986 | 6.08E-13 | 3.28E-11 |
| 193 | ATRNL1 | 7.25810198 | 4.250934381 | 0.9859 | 4.31184 | 1.6E-05 | 0.000158 |
| 1240 | PRKAG2-AS1 | 13.6268794 | 4.242236929 | 0.7348 | 5.77299 | 7.79E-09 | 1.69E-07 |
| 1145 | SNAP91 | 4.36627468 | 4.240134891 | 1.3626 | 3.11188 | 0.00186 | 0.009754 |
| 1353 | LINC02908 | 5.5539234 | 4.237291576 | 1.1625 | 3.64499 | 0.00027 | 0.001847 |
| 112 | BRINP2 | 2.53476866 | 4.235130068 | 1.48 | 2.86149 | 0.00422 | 0.019512 |
| 1340 | STXBP1 | 39.1065286 | 4.235057228 | 0.4618 | 9.17024 | 4.72E-20 | 1.08E-17 |
| 404 | CDH24 | 19.8623001 | 4.231458734 | 0.6559 | 6.45124 | 1.11E-10 | 3.69E-09 |
| 1155 | HEY2 | 2.53463378 | 4.228775725 | 1.5643 | 2.70326 | 0.00687 | 0.029174 |
| 932 | CELSR3 | 32.1334266 | 4.227581344 | 0.5672 | 7.45351 | 9.09E-14 | 5.67E-12 |
| 832 | ENSG00000278192 | 5.2874578 | 4.22169317 | 1.175 | 3.59288 | 0.00033 | 0.002209 |
| 524 | ENSG00000275191 | 2.51500273 | 4.219543737 | 1.56 | 2.7048 | 0.00683 | 0.02907 |
| 1444 | RAB39B | 4.17254386 | 4.21401318 | 1.2952 | 3.25354 | 0.00114 | 0.006406 |
| 525 | IRX3 | 12.2461879 | 4.213234435 | 0.8399 | 5.01612 | 5.27E-07 | 7.49E-06 |
| 447 | APBA2 | 20.7236682 | 4.205058875 | 0.6364 | 6.60743 | 3.91E-11 | 1.45E-09 |
| 1041 | PLEKHG4B | 2.3629734 | 4.203472543 | 1.4871 | 2.8266 | 0.0047 | 0.0214 |
| 779 | CHRNA1 | 10.9215171 | 4.202648111 | 0.8671 | 4.8469 | 1.3E-06 | 1.61E-05 |
| 1188 | AMPH | 12.2817017 | 4.200712543 | 0.8751 | 4.80006 | 1.6E-06 | 1.99E-05 |
| 1180 | SNX10 | 5.69909212 | 4.20011529 | 1.1217 | 3.74435 | 0.00018 | 0.001309 |
| 1391 | NUDT11 | 17.749447 | 4.190239849 | 0.6405 | 6.54205 | 6.07E-11 | 2.15E-09 |
| 1022 | NKX6-1 | 5.52436703 | 4.181527628 | 1.0701 | 3.90771 | 9.3E-05 | 0.000738 |

| | | | | | | | |
|---|---|---|---|---|---|---|---|
| 970 | RBP1 | 135.267126 | 4.180888136 | 0.2682 | 15.5899 | 8.52E-55 | 1.21E-50 |
| 902 | SEPTIN3 | 6.9368188 | 4.171192326 | 1.0315 | 4.04378 | 5.3E-05 | 0.000449 |
| 426 | TTC9 | 5.38807213 | 4.169281083 | 1.1503 | 3.62444 | 0.00029 | 0.001983 |
| 968 | SLC35G2 | 6.78916449 | 4.162298256 | 1.0631 | 3.91533 | 9E-05 | 0.000719 |
| 1143 | KCNQ5 | 4.17667367 | 4.159193135 | 1.3611 | 3.05571 | 0.00225 | 0.011488 |
| 1130 | GLP1R | 4.02339791 | 4.158503229 | 1.2677 | 3.28035 | 0.00104 | 0.005923 |
| 246 | SPTBN2 | 9.14427084 | 4.154599491 | 0.8778 | 4.73278 | 2.2E-06 | 2.68E-05 |
| 568 | FOXO3B | 2.34970348 | 4.153759614 | 1.4715 | 2.82275 | 0.00476 | 0.021626 |
| 376 | ENSG00000287650 | 2.25444396 | 4.150930932 | 1.5303 | 2.71248 | 0.00668 | 0.028584 |
| 917 | FGD5 | 19.7851248 | 4.146089477 | 0.6872 | 6.03366 | 1.60E-09 | 4.01E-08 |
| 1047 | BASP1 | 75.3606755 | 4.135745024 | 0.3973 | 10.4091 | 2.25E-25 | 1.00E-22 |
| 511 | ENSG00000283662 | 4.05148656 | 4.13243314 | 1.365 | 3.02734 | 0.00247 | 0.012436 |
| 1370 | RAI2 | 3.70069949 | 4.130124238 | 1.2881 | 3.20641 | 0.00134 | 0.007422 |
| 158 | ZNF25 | 3.55296855 | 4.129986074 | 1.3163 | 3.1375 | 0.0017 | 0.009071 |
| 1405 | BEX1 | 31.5411038 | 4.124217813 | 0.6155 | 6.70028 | 2.08E-11 | 8.17E-10 |
| 98 | MTX1 | 2.28777532 | 4.119477207 | 1.5027 | 2.74134 | 0.00612 | 0.026602 |
| 1228 | CCDC136 | 34.0117839 | 4.109640144 | 0.5061 | 8.11988 | 4.67E-16 | 4.52E-14 |
| 1043 | IRX2 | 24.3005333 | 4.103242414 | 1.5488 | 2.64928 | 0.00807 | 0.033231 |
| 750 | ANKRD36BP2 | 2.19583651 | 4.096507599 | 1.5025 | 2.72644 | 0.0064 | 0.027564 |
| 1347 | SLC2A6 | 3.42280968 | 4.096074464 | 1.3489 | 3.03651 | 0.00239 | 0.012141 |
| 936 | CACNA2D2 | 6.57774546 | 4.093510914 | 1.0557 | 3.87765 | 0.00011 | 0.000824 |
| 969 | MRAS | 16.6286826 | 4.092353977 | 0.7284 | 5.61802 | 1.93E-08 | 3.87E-07 |
| 794 | MAP2 | 20.4827467 | 4.077492628 | 0.6002 | 6.7937 | 1.09E-11 | 4.65E-10 |
| 1027 | ENSG00000251259 | 2.14216026 | 4.071021581 | 1.5705 | 2.5922 | 0.00954 | 0.037903 |
| 303 | ADAMTS20 | 2.22188182 | 4.069496676 | 1.5726 | 2.58776 | 0.00966 | 0.038278 |
| 1242 | GS1-24F4.2 | 3.85967603 | 4.069437338 | 1.3418 | 3.03274 | 0.00242 | 0.012255 |
| 18 | FAM43B | 3.67072349 | 4.066910235 | 1.3583 | 2.99411 | 0.00275 | 0.013632 |
| 1152 | SLC35F1 | 2.21969513 | 4.064520616 | 1.496 | 2.71684 | 0.00659 | 0.02824 |
| 957 | GAP43 | 6.32538166 | 4.062822878 | 1.0518 | 3.86258 | 0.00011 | 0.000871 |
| 1390 | NUDT10 | 14.1335626 | 4.059079332 | 0.7342 | 5.52832 | 3.23E-08 | 6.18E-07 |
| 1327 | CAVIN4 | 6.91831595 | 4.057590303 | 1.0069 | 4.02973 | 5.6E-05 | 0.000474 |
| 492 | MESP1 | 3.76894922 | 4.041631235 | 1.2729 | 3.17506 | 0.0015 | 0.008127 |
| 1189 | BLVRA | 7.80568433 | 4.033866439 | 1.0632 | 3.79412 | 0.00015 | 0.001103 |
| 81 | ENSG00000259834 | 2.10818587 | 4.029816344 | 1.5446 | 2.60896 | 0.00908 | 0.036597 |
| 146 | AKT3 | 7.70250777 | 4.026534296 | 0.9768 | 4.12231 | 3.8E-05 | 0.000334 |
| 960 | STXBP5L | 3.91514969 | 4.022267366 | 1.4217 | 2.82914 | 0.00467 | 0.021259 |
| 286 | GNB3 | 11.3443917 | 4.021817375 | 0.772 | 5.20938 | 1.89E-07 | 2.98E-06 |
| 937 | GRM2 | 2.2635022 | 4.010648935 | 1.559 | 2.57252 | 0.0101 | 0.039695 |
| 523 | NETO2 | 24.6326918 | 4.006088104 | 0.5783 | 6.92757 | 4.28E-12 | 2.00E-10 |

| | | | | | | | |
|---|---|---|---|---|---|---|---|
| 836 | RPS19P1 | 2.15489681 | 4.003119575 | 1.5573 | 2.57061 | 0.01015 | 0.039867 |
| 169 | CDH23 | 7.65529032 | 4.001975426 | 0.9709 | 4.12202 | 3.8E-05 | 0.000334 |
| 1006 | SLIT2 | 39.2049996 | 3.992352537 | 0.5488 | 7.2749 | 3.47E-13 | 1.97E-11 |
| 685 | APLP1 | 10.5338351 | 3.986063944 | 0.8364 | 4.76548 | 1.9E-06 | 2.32E-05 |
| 672 | UNC13A | 16.2958415 | 3.985924241 | 0.6549 | 6.0866 | 1.15E-09 | 2.95E-08 |
| 720 | TRIB2 | 53.5155663 | 3.985042146 | 0.4125 | 9.66135 | 4.40E-22 | 1.39E-19 |
| 878 | PCBP3 | 10.505326 | 3.984946516 | 0.8366 | 4.76342 | 1.9E-06 | 2.34E-05 |
| 438 | BEGAIN | 8.01292389 | 3.981917343 | 0.9096 | 4.37751 | 1.2E-05 | 0.000121 |
| 695 | EID2B | 2.16116567 | 3.976780342 | 1.528 | 2.60262 | 0.00925 | 0.037113 |
| 1366 | CLCN4 | 4.60370186 | 3.976138386 | 1.1845 | 3.35689 | 0.00079 | 0.004713 |
| 1183 | MTURN | 41.690273 | 3.96987019 | 0.4859 | 8.16964 | 3.09E-16 | 3.08E-14 |
| 110 | DNM3OS | 54.5556661 | 3.968728194 | 0.4777 | 8.30829 | 9.71E-17 | 1.11E-14 |
| 449 | NSMCE3 | 71.9591602 | 3.954533442 | 0.3692 | 10.7123 | 8.91E-27 | 4.37E-24 |
| 274 | LINC02731 | 1.9666458 | 3.950440867 | 1.553 | 2.54369 | 0.01097 | 0.042423 |
| 697 | TTC9B | 8.83596937 | 3.948738671 | 0.9437 | 4.18415 | 2.9E-05 | 0.000262 |
| 1226 | GPR37 | 6.89621966 | 3.94586606 | 1.0526 | 3.74866 | 0.00018 | 0.00129 |
| 1113 | TMEM170B | 6.83151671 | 3.944671808 | 0.965 | 4.08762 | 4.4E-05 | 0.000382 |
| 808 | COL4A3 | 3.46387695 | 3.935692756 | 1.3286 | 2.96222 | 0.00305 | 0.014884 |
| 753 | ENSG00000277701 | 1.96641661 | 3.925018657 | 1.5336 | 2.55941 | 0.01049 | 0.040964 |
| 1169 | HPAT5 | 21.2289994 | 3.920673972 | 0.6565 | 5.97251 | 2.34E-09 | 5.64E-08 |
| 1401 | PABPC5 | 6.70638131 | 3.915988834 | 0.9973 | 3.92674 | 8.6E-05 | 0.00069 |
| 1147 | SOBP | 31.3315931 | 3.913446773 | 0.4878 | 8.02302 | 1.03E-15 | 9.41E-14 |
| 154 | CCDC3 | 3.24008267 | 3.883907187 | 1.436 | 2.70474 | 0.00684 | 0.02907 |
| 783 | PDE11A | 7.38685528 | 3.879281518 | 0.9604 | 4.03929 | 5.4E-05 | 0.000456 |
| 348 | CHST11 | 2.94758872 | 3.873450284 | 1.5217 | 2.54545 | 0.01091 | 0.042278 |
| 1110 | TUBB2B | 62.0351988 | 3.87034651 | 0.9581 | 4.03951 | 5.4E-05 | 0.000456 |
| 897 | SYNGR1 | 12.5317144 | 3.865860982 | 0.8084 | 4.78238 | 1.7E-06 | 2.16E-05 |
| 1162 | EPM2A | 4.96273534 | 3.865248004 | 1.2095 | 3.19582 | 0.00139 | 0.007646 |
| 1233 | CHRM2 | 4.90047006 | 3.85222351 | 1.1692 | 3.29479 | 0.00098 | 0.005675 |
| 317 | SCN8A | 17.9487057 | 3.845318167 | 0.6827 | 5.63289 | 1.77E-08 | 3.58E-07 |
| 424 | SIX4 | 3.28364729 | 3.844805182 | 1.4073 | 2.73196 | 0.0063 | 0.027172 |
| 676 | VSTM2B | 5.88367833 | 3.837409334 | 1.0161 | 3.77676 | 0.00016 | 0.00117 |
| 61 | IL12RB2 | 3.11328538 | 3.833250963 | 1.3394 | 2.86181 | 0.00421 | 0.019499 |
| 125 | TMCC2 | 2.86976573 | 3.822633969 | 1.5247 | 2.50718 | 0.01217 | 0.046387 |
| 1438 | PNMA3 | 5.14129227 | 3.822234561 | 1.2555 | 3.04437 | 0.00233 | 0.011867 |
| 1098 | TENM2 | 8.58484547 | 3.816491963 | 0.8789 | 4.34211 | 1.4E-05 | 0.00014 |
| 920 | KCNH8 | 5.39201042 | 3.795600372 | 1.1317 | 3.35398 | 0.0008 | 0.004752 |
| 820 | AGXT | 2.79366282 | 3.785553434 | 1.4446 | 2.62053 | 0.00878 | 0.03557 |
| 104 | KIRREL1 | 6.75949933 | 3.781852243 | 0.9569 | 3.95201 | 7.7E-05 | 0.000629 |

| | | | | | | | |
|---|---|---|---|---|---|---|---|
| 1251 | ENSG00000272163 | 3.15628197 | 3.780547914 | 1.4599 | 2.58963 | 0.00961 | 0.038155 |
| 661 | ZNF561-AS1 | 3.16464352 | 3.777894261 | 1.4131 | 2.67341 | 0.00751 | 0.031331 |
| 235 | VWCE | 4.7390752 | 3.774140609 | 1.1322 | 3.33348 | 0.00086 | 0.005065 |
| 152 | CELF2 | 54.7117737 | 3.770808926 | 0.3715 | 10.1514 | 3.27E-24 | 1.22E-21 |
| 211 | BRSK2 | 14.1307465 | 3.764276998 | 0.6972 | 5.39951 | 6.68E-08 | 1.17E-06 |
| 1321 | APBA1 | 6.36352868 | 3.759294329 | 1.0464 | 3.59255 | 0.00033 | 0.002211 |
| 1044 | IRX2-DT | 4.02574448 | 3.745214075 | 1.2933 | 2.89593 | 0.00378 | 0.017872 |
| 1323 | ENSG00000224945 | 2.76672557 | 3.742146918 | 1.4732 | 2.54022 | 0.01108 | 0.042787 |
| 283 | TSPAN9 | 11.3943881 | 3.738369129 | 0.7311 | 5.11335 | 3.16E-07 | 4.72E-06 |
| 1389 | PPP1R3F | 7.05131452 | 3.737162301 | 0.9443 | 3.95744 | 7.6E-05 | 0.000617 |
| 996 | CPLX1 | 5.48904738 | 3.736050121 | 1.0162 | 3.67651 | 0.00024 | 0.001658 |
| 855 | TSHZ2 | 65.3767285 | 3.730869125 | 0.4366 | 8.5444 | 1.29E-17 | 1.77E-15 |
| 153 | SEC61A2 | 5.07879192 | 3.724170252 | 1.1284 | 3.30048 | 0.00097 | 0.005577 |
| 726 | DPYSL5 | 14.5199289 | 3.717968361 | 0.8774 | 4.23727 | 2.3E-05 | 0.000211 |
| 1133 | TTBK1 | 7.5391654 | 3.71085491 | 0.9856 | 3.76501 | 0.00017 | 0.001221 |
| 379 | RGCC | 3.76667662 | 3.702974247 | 1.2811 | 2.89056 | 0.00385 | 0.018119 |
| 93 | MLLT11 | 28.3047523 | 3.69737026 | 0.5473 | 6.75562 | 1.42E-11 | 5.84E-10 |
| 678 | ZNF536 | 23.918275 | 3.688865207 | 0.558 | 6.61086 | 3.82E-11 | 1.42E-09 |
| 309 | TUBA1A | 293.577161 | 3.686915205 | 0.3192 | 11.5509 | 7.30E-31 | 6.93E-28 |
| 1234 | KIAA1549 | 13.1326853 | 3.683875596 | 0.6946 | 5.30377 | 1.13E-07 | 1.89E-06 |
| 633 | GNAL | 2.82482488 | 3.679604931 | 1.3643 | 2.69702 | 0.007 | 0.029637 |
| 228 | SLC1A2 | 9.53789073 | 3.679218988 | 0.8369 | 4.39601 | 1.1E-05 | 0.000112 |
| 1046 | MIR4458HG | 3.87956719 | 3.673160313 | 1.1988 | 3.06408 | 0.00218 | 0.011229 |
| 520 | DOC2A | 2.65313927 | 3.670245804 | 1.4087 | 2.60537 | 0.00918 | 0.036869 |
| 416 | TRIM9 | 4.00442436 | 3.665967671 | 1.2248 | 2.9932 | 0.00276 | 0.013664 |
| 216 | TUB | 4.03345153 | 3.653130797 | 1.2188 | 2.99728 | 0.00272 | 0.01352 |
| 1261 | UNC5D | 21.5844779 | 3.649438806 | 0.5343 | 6.83084 | 8.44E-12 | 3.66E-10 |
| 1093 | GRIA1 | 13.4316887 | 3.645073011 | 0.7992 | 4.56097 | 5.1E-06 | 5.61E-05 |
| 581 | ASIC2 | 2.817125 | 3.634631583 | 1.3592 | 2.6741 | 0.00749 | 0.031285 |
| 252 | FAT3 | 10.381245 | 3.631092427 | 0.7847 | 4.62738 | 3.7E-06 | 4.23E-05 |
| 450 | ENSG00000259690 | 11.832732 | 3.625891052 | 0.7558 | 4.79722 | 1.6E-06 | 2.01E-05 |
| 704 | CPT1C | 9.75277311 | 3.615671743 | 0.751 | 4.81434 | 1.5E-06 | 1.87E-05 |
| 660 | PRR36 | 3.56073196 | 3.612750558 | 1.3015 | 2.77591 | 0.0055 | 0.024417 |
| 1108 | FOXF2 | 27.7141531 | 3.599359496 | 0.5073 | 7.09547 | 1.29E-12 | 6.55E-11 |
| 1033 | DCHS2 | 5.22663832 | 3.598863235 | 1.0865 | 3.31244 | 0.00092 | 0.005379 |
| 955 | DPPA4 | 71.9613633 | 3.594950165 | 0.4701 | 7.64774 | 2.05E-14 | 1.47E-12 |
| 1317 | DNAJB5 | 2.49259679 | 3.594251235 | 1.4459 | 2.48579 | 0.01293 | 0.048619 |
| 335 | LGR5 | 4.29773086 | 3.591405209 | 1.154 | 3.11213 | 0.00186 | 0.00975 |
| 162 | OGDHL | 22.1476455 | 3.586784857 | 0.5815 | 6.16811 | 6.91E-10 | 1.87E-08 |

| | | | | | | | |
|---|---|---|---|---|---|---|---|
| 1377 | ENSG00000237994 | 2.52071709 | 3.58416324 | 1.3721 | 2.61225 | 0.00899 | 0.036319 |
| 1100 | STC2 | 15.8028718 | 3.58323505 | 0.6985 | 5.12959 | 2.90E-07 | 4.38E-06 |
| 657 | FSD1 | 14.9230355 | 3.580550425 | 0.6626 | 5.40354 | 6.53E-08 | 1.15E-06 |
| 1244 | MTMR7 | 8.9215312 | 3.576710231 | 0.8315 | 4.30153 | 1.7E-05 | 0.000164 |
| 130 | SERTAD4-AS1 | 5.45501334 | 3.567225089 | 1.0469 | 3.40744 | 0.00066 | 0.004023 |
| 533 | PLEKHG4 | 5.68438689 | 3.563563027 | 1.005 | 3.5457 | 0.00039 | 0.002584 |
| 689 | ZNF382 | 4.5826104 | 3.561443219 | 1.0911 | 3.26418 | 0.0011 | 0.006219 |
| 462 | FRMD5 | 5.88054885 | 3.551928778 | 0.986 | 3.60241 | 0.00032 | 0.002136 |
| 84 | LINC01357 | 2.6071659 | 3.54616714 | 1.3642 | 2.59949 | 0.00934 | 0.037379 |
| 157 | ZNF248 | 4.60572258 | 3.543254753 | 1.1497 | 3.08196 | 0.00206 | 0.010657 |
| 417 | GPR137C | 9.94553569 | 3.528396335 | 0.7777 | 4.53676 | 5.7E-06 | 6.2E-05 |
| 1374 | CNKSR2 | 16.9586444 | 3.528108821 | 0.7388 | 4.77569 | 1.8E-06 | 2.22E-05 |
| 729 | CLIP4 | 10.3315313 | 3.522166152 | 0.7771 | 4.53233 | 5.8E-06 | 6.32E-05 |
| 536 | ENSG00000262141 | 3.69409566 | 3.519212314 | 1.2985 | 2.71031 | 0.00672 | 0.028715 |
| 444 | JAG2 | 14.1715219 | 3.51914456 | 0.7199 | 4.88868 | 1E-06 | 1.34E-05 |
| 759 | DPP10-AS1 | 16.2057072 | 3.500029034 | 0.6615 | 5.29114 | 1.22E-07 | 2.01E-06 |
| 1192 | CAMK2B | 6.8753648 | 3.494705604 | 0.8969 | 3.89657 | 9.8E-05 | 0.000768 |
| 493 | ZNF710-AS1 | 12.4564912 | 3.493144334 | 0.6804 | 5.13365 | 2.84E-07 | 4.3E-06 |
| 265 | TLCD5 | 4.92819186 | 3.491376517 | 1.0712 | 3.25921 | 0.00112 | 0.006307 |
| 793 | ZDBF2 | 4.33790451 | 3.486415611 | 1.1542 | 3.02053 | 0.00252 | 0.012657 |
| 875 | RIPPLY3 | 5.2413384 | 3.47310962 | 1.161 | 2.99146 | 0.00278 | 0.013733 |
| 297 | RASSF8 | 16.8740031 | 3.472941882 | 0.6014 | 5.77514 | 7.69E-09 | 1.68E-07 |
| 879 | ENSG00000275139 | 6.12055067 | 3.470638666 | 1.1428 | 3.03702 | 0.00239 | 0.012126 |
| 1238 | ATP6V0E2 | 13.890319 | 3.459499932 | 0.6289 | 5.50101 | 3.78E-08 | 7.08E-07 |
| 1175 | SP4 | 3.51649474 | 3.458218669 | 1.3875 | 2.49234 | 0.01269 | 0.04795 |
| 351 | FAM216A | 12.5661076 | 3.452138346 | 0.729 | 4.73557 | 2.2E-06 | 2.65E-05 |
| 716 | GRASLND | 3.50318598 | 3.452110518 | 1.2658 | 2.72711 | 0.00639 | 0.027524 |
| 560 | KCNAB3 | 18.2262817 | 3.451278326 | 0.607 | 5.68569 | 1.30E-08 | 2.71E-07 |
| 601 | TBKBP1 | 3.51772307 | 3.440413601 | 1.2365 | 2.78241 | 0.0054 | 0.02403 |
| 76 | CELSR2 | 10.820197 | 3.439989146 | 0.8161 | 4.21526 | 2.5E-05 | 0.000231 |
| 701 | CADM4 | 5.55130781 | 3.429350006 | 1.0522 | 3.25924 | 0.00112 | 0.006307 |
| 764 | ARHGEF4 | 8.16946661 | 3.424044263 | 0.869 | 3.94028 | 8.1E-05 | 0.000657 |
| 230 | LDLRAD3 | 3.89946387 | 3.421822238 | 1.2681 | 2.69833 | 0.00697 | 0.029539 |
| 1017 | ARL9 | 3.14775889 | 3.407328767 | 1.3194 | 2.58246 | 0.00981 | 0.038795 |
| 1271 | CYP7B1 | 6.06814411 | 3.404572527 | 1.1217 | 3.03507 | 0.0024 | 0.012183 |
| 943 | C3orf14 | 15.24919 | 3.404143287 | 0.626 | 5.43816 | 5.38E-08 | 9.69E-07 |
| 1149 | WASF1 | 23.2111588 | 3.402055283 | 0.5492 | 6.19416 | 5.86E-10 | 1.62E-08 |
| 1372 | MAP7D2 | 15.5841765 | 3.40139197 | 0.6976 | 4.87596 | 1.1E-06 | 1.42E-05 |
| 715 | SOX11 | 54.7171371 | 3.397531261 | 1.0235 | 3.31957 | 0.0009 | 0.005271 |

| | | | | | | | |
|---|---|---|---|---|---|---|---|
| 727 | CGREF1 | 5.92714808 | 3.389147858 | 0.9921 | 3.41606 | 0.00064 | 0.003924 |
| 934 | BSN | 9.25855001 | 3.384526344 | 0.824 | 4.10745 | 4E-05 | 0.000354 |
| 958 | LSAMP | 16.9485539 | 3.381501618 | 0.6477 | 5.22048 | 1.78E-07 | 2.83E-06 |
| 1300 | PVT1 | 10.9866671 | 3.380936074 | 0.7701 | 4.39003 | 1.1E-05 | 0.000115 |
| 1013 | SLC10A4 | 4.62929308 | 3.376584718 | 1.1079 | 3.04771 | 0.00231 | 0.011748 |
| 464 | MYEF2 | 17.8977842 | 3.373667749 | 0.5537 | 6.09301 | 1.11E-09 | 2.85E-08 |
| 1154 | CLVS2 | 6.95094882 | 3.36919822 | 1.0598 | 3.17918 | 0.00148 | 0.008025 |
| 315 | ENSG00000278126 | 7.56251747 | 3.35652885 | 0.8659 | 3.87648 | 0.00011 | 0.000827 |
| 1378 | SYTL5 | 17.052559 | 3.352971621 | 0.6622 | 5.06318 | 4.12E-07 | 6E-06 |
| 1351 | GPSM1 | 9.13793329 | 3.34994348 | 0.8529 | 3.92759 | 8.6E-05 | 0.000689 |
| 896 | NPTXR | 7.80866148 | 3.349417163 | 0.843 | 3.97331 | 7.1E-05 | 0.000583 |
| 123 | NFASC | 11.3859981 | 3.347470332 | 0.7184 | 4.65939 | 3.2E-06 | 3.7E-05 |
| 742 | CNRIP1 | 10.1038899 | 3.346473921 | 0.8214 | 4.0739 | 4.6E-05 | 0.000401 |
| 1079 | REEP2 | 10.0460092 | 3.344991297 | 0.8027 | 4.16696 | 3.1E-05 | 0.00028 |
| 372 | KATNAL1 | 3.52795147 | 3.34485222 | 1.2255 | 2.72936 | 0.00635 | 0.027379 |
| 1216 | TFR2 | 12.1530914 | 3.344344621 | 0.7728 | 4.32761 | 1.5E-05 | 0.000148 |
| 894 | MFNG | 19.252804 | 3.343316785 | 0.5271 | 6.34277 | 2.26E-10 | 6.83E-09 |
| 256 | GUCY1A2 | 7.69173823 | 3.338620532 | 0.9028 | 3.69795 | 0.00022 | 0.001541 |
| 387 | CNMD | 3.82208378 | 3.336876841 | 1.2383 | 2.69483 | 0.00704 | 0.029771 |
| 570 | USP32P3 | 3.97526522 | 3.334783308 | 1.1599 | 2.87516 | 0.00404 | 0.01884 |
| 567 | ZNF286B | 3.21895336 | 3.328223749 | 1.3421 | 2.47992 | 0.01314 | 0.049295 |
| 217 | RIC3 | 21.7620073 | 3.325877268 | 0.5233 | 6.35597 | 2.07E-10 | 6.36E-09 |
| 921 | ZNF385D | 14.5073868 | 3.322904668 | 0.7055 | 4.70976 | 2.5E-06 | 2.98E-05 |
| 948 | CNTN3 | 20.2519924 | 3.322067465 | 0.5971 | 5.56405 | 2.64E-08 | 5.14E-07 |
| 86 | SLC22A15 | 3.00189653 | 3.318521793 | 1.2615 | 2.6306 | 0.00852 | 0.034733 |
| 79 | GSTM3 | 5.67870399 | 3.318250952 | 0.999 | 3.32149 | 0.0009 | 0.005242 |
| 259 | NPAT | 7.50851202 | 3.316837569 | 0.8589 | 3.86184 | 0.00011 | 0.000873 |
| 1116 | RIPOR2 | 12.037759 | 3.314563935 | 0.7811 | 4.24368 | 2.2E-05 | 0.000206 |
| 502 | ZSCAN32 | 3.54784475 | 3.313377831 | 1.2084 | 2.74197 | 0.00611 | 0.026559 |
| 790 | TMEM237 | 9.51300911 | 3.312072503 | 0.8378 | 3.95336 | 7.7E-05 | 0.000627 |
| 1010 | UCHL1 | 45.9233995 | 3.310988779 | 1.147 | 2.88664 | 0.00389 | 0.018293 |
| 1038 | SAP30 | 12.343734 | 3.303066244 | 0.6708 | 4.92423 | 8.47E-07 | 1.15E-05 |
| 378 | SMAD9 | 20.0327019 | 3.302771853 | 0.5724 | 5.77001 | 7.93E-09 | 1.72E-07 |
| 66 | TTLL7 | 14.5123768 | 3.301919937 | 0.6115 | 5.39985 | 6.67E-08 | 1.17E-06 |
| 205 | B4GALNT4 | 16.0997145 | 3.2954496 | 0.5585 | 5.90042 | 3.63E-09 | 8.39E-08 |
| 1280 | PKIA | 13.2334102 | 3.287315373 | 0.7579 | 4.33766 | 1.4E-05 | 0.000142 |
| 806 | DOCK10 | 8.98336428 | 3.284660667 | 0.9808 | 3.34883 | 0.00081 | 0.00483 |
| 1001 | NSG1 | 12.253673 | 3.283176221 | 0.7053 | 4.65469 | 3.2E-06 | 3.77E-05 |
| 213 | TRIM22 | 15.7939769 | 3.274312173 | 0.7099 | 4.61222 | 4E-06 | 4.51E-05 |

| | | | | | | | |
|---|---|---|---|---|---|---|---|
| 196 | INPP5F | 54.8156823 | 3.273391877 | 0.3762 | 8.70141 | 3.28E-18 | 5.07E-16 |
| 1230 | STRIP2 | 7.51077971 | 3.266542471 | 0.9539 | 3.42437 | 0.00062 | 0.003821 |
| 945 | FRMD4B | 13.8420402 | 3.258323592 | 0.6002 | 5.42839 | 5.69E-08 | 1.01E-06 |
| 766 | CXCR4 | 55.7799091 | 3.257385216 | 0.4847 | 6.72061 | 1.81E-11 | 7.19E-10 |
| 796 | TMEM169 | 2.82253422 | 3.254083671 | 1.2744 | 2.55346 | 0.01067 | 0.041549 |
| 530 | NDRG4 | 12.1261506 | 3.249666183 | 0.7257 | 4.47789 | 7.5E-06 | 7.94E-05 |
| 1173 | VWDE | 7.27687271 | 3.247927433 | 0.8843 | 3.67305 | 0.00024 | 0.001676 |
| 461 | WDR76 | 14.074874 | 3.245248093 | 0.6549 | 4.95516 | 7.23E-07 | 9.94E-06 |
| 145 | CEP170 | 12.0559317 | 3.239700118 | 0.6366 | 5.0894 | 3.59E-07 | 5.3E-06 |
| 1428 | MAP7D3 | 17.9610683 | 3.239150398 | 0.5791 | 5.59314 | 2.23E-08 | 4.42E-07 |
| 740 | EFEMP1 | 11.8391576 | 3.231900055 | 0.7642 | 4.229 | 2.3E-05 | 0.000218 |
| 707 | C19orf81 | 9.57024033 | 3.210738286 | 0.716 | 4.48421 | 7.3E-06 | 7.75E-05 |
| 330 | DPY19L2 | 3.6151593 | 3.210315145 | 1.1748 | 2.73261 | 0.00628 | 0.027135 |
| 1354 | NPDC1 | 39.1410363 | 3.209450451 | 0.3881 | 8.27027 | 1.34E-16 | 1.46E-14 |
| 1053 | NLN | 13.1553624 | 3.207737852 | 0.6505 | 4.93119 | 8.17E-07 | 1.11E-05 |
| 1282 | MMP16 | 10.3335724 | 3.206515728 | 0.7139 | 4.49157 | 7.1E-06 | 7.53E-05 |
| 723 | WDCP | 5.96182596 | 3.205625981 | 0.9347 | 3.42941 | 0.0006 | 0.003764 |
| 1202 | RUNDC3B | 5.18825642 | 3.19884333 | 1.0469 | 3.05562 | 0.00225 | 0.011488 |
| 575 | PROCA1 | 3.08085155 | 3.195140193 | 1.2699 | 2.51615 | 0.01186 | 0.045369 |
| 474 | MEGF11 | 4.49915727 | 3.189951612 | 1.1053 | 2.886 | 0.0039 | 0.018324 |
| 755 | CHST10 | 6.30615231 | 3.189602663 | 0.9974 | 3.19783 | 0.00138 | 0.007605 |
| 1407 | BEX2 | 52.8014078 | 3.187889143 | 0.3608 | 8.83644 | 9.88E-19 | 1.74E-16 |
| 712 | ENSG00000176593 | 4.80553586 | 3.185452758 | 1.0665 | 2.98675 | 0.00282 | 0.013907 |
| 534 | RIPOR1 | 6.55753268 | 3.182366496 | 0.8652 | 3.67802 | 0.00024 | 0.00165 |
| 1427 | FHL1 | 84.7983191 | 3.181763963 | 0.4697 | 6.77404 | 1.25E-11 | 5.26E-10 |
| 929 | TMEM158 | 12.8144414 | 3.175183119 | 0.6501 | 4.88432 | 1E-06 | 1.37E-05 |
| 784 | OSBPL6 | 5.61193492 | 3.16443799 | 1.0004 | 3.16306 | 0.00156 | 0.008409 |
| 519 | ENSG00000279789 | 7.81292776 | 3.160942447 | 0.862 | 3.66704 | 0.00025 | 0.00171 |
| 321 | PDE1B | 10.4791451 | 3.16065303 | 0.8143 | 3.8813 | 0.0001 | 0.000814 |
| 787 | NEUROD1 | 98.1741774 | 3.158350661 | 0.8386 | 3.76636 | 0.00017 | 0.001216 |
| 823 | FAM110A | 5.28993654 | 3.15128032 | 1.0031 | 3.1414 | 0.00168 | 0.008965 |
| 1172 | THSD7A | 7.03117944 | 3.151089321 | 0.9167 | 3.43729 | 0.00059 | 0.003666 |
| 887 | EMID1 | 8.5482869 | 3.144731072 | 0.7501 | 4.19236 | 2.8E-05 | 0.000254 |
| 1205 | PEG10 | 29.1159711 | 3.137091716 | 0.4477 | 7.00726 | 2.43E-12 | 1.18E-10 |
| 782 | HAGLR | 5.62217249 | 3.132215258 | 0.9206 | 3.40237 | 0.00067 | 0.004077 |
| 1119 | NOP56P1 | 4.31066101 | 3.128603903 | 1.0516 | 2.97506 | 0.00293 | 0.014368 |
| 292 | FAM234B | 7.50434612 | 3.113673489 | 0.9886 | 3.14953 | 0.00164 | 0.008752 |
| 73 | PLPPR4 | 4.97678658 | 3.110681953 | 1.1088 | 2.80537 | 0.00503 | 0.022668 |
| 926 | EIF1B-AS1 | 4.13117893 | 3.109582417 | 1.2348 | 2.51821 | 0.0118 | 0.045153 |

| | | | | | | | |
|---|---|---|---|---|---|---|---|
| 366 | SACS | 17.5703854 | 3.106954374 | 0.6614 | 4.69729 | 2.6E-06 | 3.13E-05 |
| 15 | CROCC | 34.8113395 | 3.105304806 | 0.4193 | 7.40524 | 1.31E-13 | 8.06E-12 |
| 1148 | SESN1 | 23.4194131 | 3.092238692 | 0.485 | 6.37624 | 1.81E-10 | 5.65E-09 |
| 1028 | PAPSS1 | 11.1751907 | 3.088608647 | 0.6568 | 4.70231 | 2.6E-06 | 3.07E-05 |
| 215 | DCHS1 | 7.081233 | 3.080052339 | 0.8705 | 3.53841 | 0.0004 | 0.00265 |
| 938 | PCBP4 | 25.139873 | 3.078721035 | 0.4988 | 6.17253 | 6.72E-10 | 1.82E-08 |
| 913 | ATP2B2 | 6.4647051 | 3.075137039 | 0.9576 | 3.21118 | 0.00132 | 0.007311 |
| 390 | TBC1D4 | 11.858 | 3.070903715 | 0.7743 | 3.96594 | 7.3E-05 | 0.000598 |
| 1134 | ENSG00000231881 | 7.16822264 | 3.068353577 | 1.0261 | 2.99027 | 0.00279 | 0.013777 |
| 888 | RFPL1S | 4.30530415 | 3.066774151 | 1.0126 | 3.02869 | 0.00246 | 0.012394 |
| 746 | MTHFD2 | 37.8195959 | 3.066635161 | 0.4533 | 6.76458 | 1.34E-11 | 5.58E-10 |
| 1068 | MCC | 15.7530475 | 3.066368239 | 0.6273 | 4.88802 | 1E-06 | 1.35E-05 |
| 1064 | MCTP1 | 10.5869047 | 3.062107319 | 0.7722 | 3.96529 | 7.3E-05 | 0.000599 |
| 1097 | GABRB2 | 10.2160338 | 3.061557096 | 0.7194 | 4.25559 | 2.1E-05 | 0.000198 |
| 977 | P2RY1 | 50.5867797 | 3.05649475 | 0.4284 | 7.13385 | 9.76E-13 | 5.05E-11 |
| 644 | RNF165 | 16.3934363 | 3.051794065 | 0.67 | 4.55471 | 5.2E-06 | 5.75E-05 |
| 667 | NFIX | 62.670907 | 3.05001635 | 0.3976 | 7.67038 | 1.71E-14 | 1.25E-12 |
| 953 | TMEM45A | 9.88547741 | 3.050015581 | 0.8345 | 3.65509 | 0.00026 | 0.001781 |
| 413 | LRFN5 | 4.89394778 | 3.042696926 | 1.0695 | 2.84499 | 0.00444 | 0.020373 |
| 243 | NRXN2 | 6.67285369 | 3.040771544 | 0.866 | 3.51111 | 0.00045 | 0.002891 |
| 1417 | LRCH2 | 10.990709 | 3.038447126 | 0.7452 | 4.07757 | 4.6E-05 | 0.000396 |
| 82 | LRIF1 | 10.1550527 | 3.033643877 | 0.7601 | 3.99116 | 6.6E-05 | 0.000547 |
| 485 | ARNT2 | 6.84020469 | 3.033158803 | 1.0528 | 2.88112 | 0.00396 | 0.01853 |
| 131 | SERTAD4 | 12.171967 | 3.030076558 | 0.6786 | 4.46508 | 8E-06 | 8.41E-05 |
| 27 | ADGRB2 | 15.5089866 | 3.01408359 | 0.5879 | 5.12729 | 2.94E-07 | 4.42E-06 |
| 1030 | RNF150 | 3.8615961 | 3.008735212 | 1.1468 | 2.62352 | 0.0087 | 0.03533 |
| 1029 | HSPA4L | 7.85485327 | 3.007877919 | 0.8654 | 3.47584 | 0.00051 | 0.003243 |
| 585 | ARHGAP23 | 7.73503599 | 2.999068068 | 0.8111 | 3.69743 | 0.00022 | 0.001544 |
| 89 | LIX1L | 9.16271465 | 2.997207629 | 0.7113 | 4.21377 | 2.5E-05 | 0.000232 |
| 1025 | UNC5C | 5.06983175 | 2.997073979 | 0.9944 | 3.01399 | 0.00258 | 0.012906 |
| 499 | NPW | 10.272651 | 2.990996408 | 0.7743 | 3.86274 | 0.00011 | 0.000871 |
| 773 | GALNT13 | 3.21760882 | 2.98041903 | 1.1784 | 2.52914 | 0.01143 | 0.043948 |
| 586 | SRCIN1 | 4.34208977 | 2.97757518 | 1.0571 | 2.81672 | 0.00485 | 0.022 |
| 1200 | PCLO | 32.8907808 | 2.974804447 | 0.4838 | 6.14894 | 7.80E-10 | 2.07E-08 |
| 1260 | DUSP26 | 5.55220902 | 2.973255788 | 1.1496 | 2.58629 | 0.0097 | 0.03841 |
| 147 | KIF26B | 3.73280972 | 2.972389328 | 1.1362 | 2.61615 | 0.00889 | 0.035988 |
| 1199 | CACNA2D1 | 12.6564832 | 2.962475529 | 0.6594 | 4.49263 | 7E-06 | 7.5E-05 |
| 724 | KIF3C | 9.57208246 | 2.961526243 | 0.8479 | 3.49298 | 0.00048 | 0.003066 |
| 241 | RCOR2 | 8.13685212 | 2.952127159 | 0.8041 | 3.67126 | 0.00024 | 0.001685 |

| | | | | | | | |
|---|---|---|---|---|---|---|---|
| 225 | SERGEF | 10.4192448 | 2.95112284 | 0.6968 | 4.23524 | 2.3E-05 | 0.000213 |
| 1179 | KLHL7 | 29.6072337 | 2.950135539 | 0.425 | 6.9418 | 3.87E-12 | 1.83E-10 |
| 336 | PHLDA1 | 19.8300629 | 2.948580982 | 0.5492 | 5.36886 | 7.92E-08 | 1.37E-06 |
| 541 | CDT1 | 30.5426637 | 2.946261413 | 0.464 | 6.35035 | 2.15E-10 | 6.59E-09 |
| 405 | SLC22A17 | 20.3192074 | 2.944798299 | 0.5106 | 5.76782 | 8.03E-09 | 1.74E-07 |
| 500 | ABCA3 | 12.1644326 | 2.934393497 | 0.7236 | 4.05526 | 5E-05 | 0.000429 |
| 1122 | TCF19 | 18.095379 | 2.933381817 | 0.5296 | 5.53887 | 3.04E-08 | 5.84E-07 |
| 55 | LRP8 | 10.1120871 | 2.930612056 | 0.9017 | 3.25013 | 0.00115 | 0.00647 |
| 374 | FRY | 6.24799505 | 2.917507127 | 0.9371 | 3.11347 | 0.00185 | 0.009713 |
| 1135 | SUPT3H | 3.80659915 | 2.917401303 | 1.1224 | 2.59934 | 0.00934 | 0.037385 |
| 765 | MCM6 | 70.0257746 | 2.912756952 | 0.4237 | 6.8741 | 6.24E-12 | 2.80E-10 |
| 1220 | SH2B2 | 3.84559837 | 2.910478254 | 1.1024 | 2.64007 | 0.00829 | 0.03396 |
| 373 | B3GLCT | 13.0121741 | 2.906416979 | 0.6674 | 4.35457 | 1.3E-05 | 0.000133 |
| 690 | DPF1 | 7.36013913 | 2.903917153 | 0.9395 | 3.09089 | 0.002 | 0.010375 |
| 640 | CDH2 | 19.9621929 | 2.899104755 | 0.6993 | 4.14567 | 3.4E-05 | 0.000304 |
| 1411 | RADX | 31.3883737 | 2.897916108 | 0.4891 | 5.92533 | 3.12E-09 | 7.26E-08 |
| 436 | C14orf132 | 7.29561872 | 2.896653056 | 0.856 | 3.38389 | 0.00071 | 0.00433 |
| 1398 | PHKA1 | 15.6753425 | 2.889352871 | 0.6488 | 4.45351 | 8.4E-06 | 8.82E-05 |
| 1435 | MAMLD1 | 14.2503194 | 2.876357159 | 0.7064 | 4.07183 | 4.7E-05 | 0.000404 |
| 1285 | NECAB1 | 7.91128543 | 2.874339799 | 0.9045 | 3.17784 | 0.00148 | 0.008056 |
| 1363 | NLGN4X | 19.8868542 | 2.873219404 | 0.5227 | 5.49739 | 3.85E-08 | 7.20E-07 |
| 33 | NCDN | 18.6641852 | 2.872765323 | 0.5801 | 4.95257 | 7.32E-07 | 1.01E-05 |
| 1141 | OGFRL1 | 16.8993325 | 2.87005755 | 0.6198 | 4.63068 | 3.6E-06 | 4.17E-05 |
| 171 | VCL | 169.216354 | 2.864350466 | 0.3198 | 8.95635 | 3.36E-19 | 6.20E-17 |
| 257 | ELMOD1 | 17.2409193 | 2.86129858 | 0.7601 | 3.76425 | 0.00017 | 0.001224 |
| 1423 | HS6ST2 | 10.5987431 | 2.861290826 | 0.6643 | 4.30732 | 1.7E-05 | 0.000161 |
| 282 | NRIP2 | 10.3921026 | 2.854912678 | 0.8195 | 3.48383 | 0.00049 | 0.003163 |
| 127 | SLC41A1 | 4.18486007 | 2.852715575 | 1.0972 | 2.60003 | 0.00932 | 0.037341 |
| 1371 | SCML2 | 6.34990404 | 2.848872108 | 0.9274 | 3.07176 | 0.00213 | 0.010976 |
| 1380 | TSPAN7 | 72.7342744 | 2.845620466 | 0.3481 | 8.17358 | 2.99E-16 | 3.00E-14 |
| 725 | GAREM2 | 4.95692917 | 2.839498429 | 0.9637 | 2.94649 | 0.00321 | 0.015545 |
| 918 | RFTN1 | 6.74718315 | 2.837944345 | 0.8834 | 3.21239 | 0.00132 | 0.007292 |
| 1177 | SNHG26 | 13.1602456 | 2.836349559 | 0.7601 | 3.73155 | 0.00019 | 0.001371 |
| 34 | C1orf216 | 5.58193685 | 2.819266216 | 0.9443 | 2.9857 | 0.00283 | 0.013936 |
| 608 | NOG | 5.35021525 | 2.817524692 | 0.9407 | 2.99524 | 0.00274 | 0.013602 |
| 472 | UBAP1L | 5.64152583 | 2.816987207 | 1.0304 | 2.73394 | 0.00626 | 0.02705 |
| 837 | INSM1 | 31.6997219 | 2.81600569 | 0.4443 | 6.3384 | 2.32E-10 | 7.01E-09 |
| 542 | CBFA2T3 | 9.19755746 | 2.809401595 | 1.0047 | 2.79615 | 0.00517 | 0.02317 |
| 636 | ZNF519 | 21.7996205 | 2.807943118 | 0.5713 | 4.91493 | 8.88E-07 | 1.19E-05 |

| | | | | | | | |
|---|---|---|---|---|---|---|---|
| 1322 | DAPK1 | 13.9897898 | 2.801456557 | 0.7037 | 3.98102 | 6.9E-05 | 0.000569 |
| 580 | CDK5R1 | 8.80307793 | 2.800690557 | 1.0643 | 2.63136 | 0.0085 | 0.034673 |
| 1287 | RUNX1T1 | 36.7938631 | 2.7978582 | 0.3641 | 7.68461 | 1.53E-14 | 1.15E-12 |
| 83 | INKA2 | 6.59248485 | 2.793170399 | 0.9509 | 2.93753 | 0.00331 | 0.015915 |
| 816 | RAMP1 | 11.0102741 | 2.782094791 | 0.7567 | 3.6764 | 0.00024 | 0.001658 |
| 1231 | LINC-PINT | 5.48617091 | 2.781656379 | 1.0511 | 2.64641 | 0.00814 | 0.033475 |
| 982 | EGFEM1P | 12.3252907 | 2.781510424 | 0.728 | 3.82065 | 0.00013 | 0.00101 |
| 1161 | HYMAI | 5.13771666 | 2.780485662 | 1.0273 | 2.7066 | 0.0068 | 0.02896 |
| 142 | PGBD5 | 29.25917 | 2.780255302 | 0.5118 | 5.43207 | 5.57E-08 | 9.98E-07 |
| 22 | STMN1 | 398.428511 | 2.775618572 | 0.2653 | 10.4623 | 1.29E-25 | 5.90E-23 |
| 1063 | NR2F1 | 16.9274489 | 2.774623219 | 0.6641 | 4.17795 | 2.9E-05 | 0.000268 |
| 219 | DENND5A | 13.1520346 | 2.77247061 | 0.6518 | 4.2533 | 2.1E-05 | 0.000199 |
| 1440 | ZNF275 | 9.70222782 | 2.768129473 | 0.785 | 3.52634 | 0.00042 | 0.002749 |
| 526 | CRNDE | 15.6976472 | 2.767720068 | 0.7116 | 3.88917 | 0.0001 | 0.00079 |
| 298 | TSPAN11 | 6.51676108 | 2.764408875 | 0.8155 | 3.38977 | 0.0007 | 0.004245 |
| 1196 | MAGI2 | 10.7400938 | 2.764196617 | 0.6466 | 4.27479 | 1.9E-05 | 0.000183 |
| 355 | BICDL1 | 9.14904732 | 2.759356997 | 0.8108 | 3.40328 | 0.00067 | 0.004065 |
| 1059 | EDIL3 | 23.032697 | 2.753937155 | 0.5591 | 4.92581 | 8.40E-07 | 1.14E-05 |
| 1388 | SYP | 34.6620622 | 2.750802117 | 0.4281 | 6.42607 | 1.31E-10 | 4.24E-09 |
| 1144 | CD109 | 4.20803897 | 2.747870017 | 1.0667 | 2.57593 | 0.01 | 0.039414 |
| 1160 | PLAGL1 | 89.9371092 | 2.739373928 | 0.3099 | 8.83818 | 9.73E-19 | 1.73E-16 |
| 594 | ARL4D | 6.25102682 | 2.736899389 | 0.8592 | 3.18551 | 0.00145 | 0.007882 |
| 328 | DTX3 | 18.1476987 | 2.732803557 | 0.6634 | 4.11924 | 3.8E-05 | 0.000337 |
| 1114 | GFOD1 | 5.73815288 | 2.72897088 | 0.9708 | 2.81093 | 0.00494 | 0.022357 |
| 984 | CLDN11 | 7.59968601 | 2.725762363 | 0.9008 | 3.02587 | 0.00248 | 0.012475 |
| 1365 | SHROOM2 | 23.6274556 | 2.718534454 | 0.4762 | 5.70837 | 1.14E-08 | 2.39E-07 |
| 1369 | AP1S2 | 49.2821515 | 2.715541343 | 0.3278 | 8.28538 | 1.18E-16 | 1.33E-14 |
| 876 | PRDM15 | 9.09640947 | 2.709977932 | 0.7065 | 3.83561 | 0.00013 | 0.000959 |
| 134 | VASH2 | 8.69014199 | 2.705714832 | 0.8552 | 3.16366 | 0.00156 | 0.008395 |
| 54 | ZYG11A | 8.32924319 | 2.704227022 | 0.8673 | 3.11802 | 0.00182 | 0.0096 |
| 944 | ADAMTS9 | 19.0459568 | 2.694484838 | 0.5257 | 5.12586 | 2.96E-07 | 4.45E-06 |
| 139 | MIXL1 | 4.31732408 | 2.691133389 | 1.0177 | 2.6444 | 0.00818 | 0.033615 |
| 1000 | GRK4 | 8.36750949 | 2.690810241 | 0.782 | 3.4408 | 0.00058 | 0.003631 |
| 775 | SP5 | 7.31260062 | 2.688657136 | 0.8931 | 3.01051 | 0.00261 | 0.013041 |
| 1168 | QKI | 45.2976337 | 2.687843796 | 0.4049 | 6.63824 | 3.17E-11 | 1.21E-09 |
| 545 | MC1R | 6.52312072 | 2.683748737 | 0.9148 | 2.93382 | 0.00335 | 0.016074 |
| 1024 | NAP1L5 | 10.0598276 | 2.683715131 | 0.6592 | 4.07094 | 4.7E-05 | 0.000405 |
| 12 | PDPN | 23.0609586 | 2.681350059 | 0.5343 | 5.01867 | 5.20E-07 | 7.41E-06 |
| 358 | BCL7A | 19.6365906 | 2.679151932 | 0.6058 | 4.42262 | 9.8E-06 | 0.0001 |

| | | | | | | | |
|---|---|---|---|---|---|---|---|
| 851 | TOMM34 | 24.5712503 | 2.677498945 | 0.441 | 6.07184 | 1.26E-09 | 3.21E-08 |
| 1088 | DPYSL3 | 41.0616655 | 2.67590302 | 0.4565 | 5.86203 | 4.57E-09 | 1.04E-07 |
| 967 | PLXND1 | 15.8881961 | 2.675316073 | 0.6276 | 4.26258 | 2E-05 | 0.000193 |
| 469 | GNB5 | 8.08236555 | 2.67518038 | 0.7891 | 3.39014 | 0.0007 | 0.004241 |
| 167 | LRRC20 | 5.17592129 | 2.663883295 | 0.9102 | 2.92684 | 0.00342 | 0.016404 |
| 900 | C22orf46P | 4.74723187 | 2.662349219 | 1.0509 | 2.53344 | 0.01129 | 0.043507 |
| 1318 | UNC13B | 118.71832 | 2.661327606 | 0.2817 | 9.44682 | 3.49E-21 | 9.37E-19 |
| 825 | EBF4 | 18.3180435 | 2.657253993 | 0.6805 | 3.90491 | 9.4E-05 | 0.000745 |
| 234 | TMEM132A | 5.69216221 | 2.656317942 | 0.8888 | 2.98876 | 0.0028 | 0.013836 |
| 582 | RASL10B | 6.71805385 | 2.656307756 | 0.9515 | 2.79164 | 0.00524 | 0.023459 |
| 642 | CELF4 | 40.8894833 | 2.656243708 | 0.4278 | 6.20961 | 5.31E-10 | 1.48E-08 |
| 137 | MARK1 | 9.70924735 | 2.654771819 | 0.766 | 3.46593 | 0.00053 | 0.003355 |
| 113 | CACNA1E | 8.19965601 | 2.653557427 | 0.9836 | 2.69793 | 0.00698 | 0.029565 |
| 1158 | ARFGEF3 | 24.3315614 | 2.648922351 | 0.4928 | 5.37562 | 7.63E-08 | 1.32E-06 |
| 1084 | FCHSD1 | 4.83554418 | 2.647644878 | 1.0046 | 2.63558 | 0.0084 | 0.034334 |
| 311 | NCKAP5L | 6.66322465 | 2.643880214 | 0.8148 | 3.24464 | 0.00118 | 0.006581 |
| 826 | CPXM1 | 21.7791943 | 2.64374389 | 0.5713 | 4.62738 | 3.7E-06 | 4.23E-05 |
| 430 | VASH1 | 10.2480767 | 2.639856984 | 0.6845 | 3.85673 | 0.00011 | 0.000889 |
| 769 | LYPD6 | 6.83494546 | 2.633670129 | 0.9413 | 2.79801 | 0.00514 | 0.023051 |
| 815 | SH3BP4 | 12.6897221 | 2.62986389 | 0.6158 | 4.27086 | 1.9E-05 | 0.000186 |
| 670 | PKN1 | 6.19353492 | 2.628450578 | 0.8858 | 2.9674 | 0.003 | 0.014683 |
| 65 | ST6GALNAC5 | 4.93508136 | 2.626399386 | 0.9505 | 2.76322 | 0.00572 | 0.025214 |
| 388 | DACH1 | 33.828917 | 2.620458562 | 0.3832 | 6.83845 | 8.01E-12 | 3.50E-10 |
| 886 | KREMEN1 | 6.04996006 | 2.619815565 | 0.9824 | 2.66676 | 0.00766 | 0.031883 |
| 419 | RPL13AP3 | 4.44554714 | 2.61325397 | 0.9762 | 2.67694 | 0.00743 | 0.031067 |
| 1170 | FSCN1 | 34.908647 | 2.611229047 | 0.5137 | 5.08303 | 3.71E-07 | 5.46E-06 |
| 1292 | ANGPT1 | 11.6841406 | 2.610108671 | 0.6926 | 3.76878 | 0.00016 | 0.001206 |
| 200 | CHST15 | 4.57418819 | 2.605857573 | 1.0066 | 2.58865 | 0.00964 | 0.038211 |
| 319 | LINC02381 | 9.05422086 | 2.602658495 | 0.6791 | 3.83261 | 0.00013 | 0.00097 |
| 175 | EXOC6 | 10.4223714 | 2.598278577 | 0.7083 | 3.66848 | 0.00024 | 0.001702 |
| 1379 | SRPX | 18.3281616 | 2.596707207 | 0.5553 | 4.67643 | 2.9E-06 | 3.43E-05 |
| 1404 | TMSB15A | 20.8361643 | 2.596648021 | 0.523 | 4.96478 | 6.88E-07 | 9.51E-06 |
| 598 | GJC1 | 30.1944951 | 2.593509622 | 0.4677 | 5.54563 | 2.93E-08 | 5.65E-07 |
| 62 | PTGER3 | 10.7551682 | 2.589545958 | 0.7161 | 3.61633 | 0.0003 | 0.002036 |
| 159 | RPS19P7 | 7.61062282 | 2.586251129 | 0.7597 | 3.40449 | 0.00066 | 0.004056 |
| 1215 | PCOLCE | 28.3455247 | 2.582664249 | 0.5627 | 4.5897 | 4.4E-06 | 4.98E-05 |
| 627 | RAC3 | 7.27391994 | 2.575268884 | 0.7872 | 3.27138 | 0.00107 | 0.006085 |
| 1332 | SVEP1 | 5.48881177 | 2.567507261 | 0.9203 | 2.78988 | 0.00527 | 0.023564 |
| 458 | CHST14 | 8.4964856 | 2.562568998 | 0.7443 | 3.44315 | 0.00057 | 0.003605 |

| | | | | | | | |
|---|---|---|---|---|---|---|---|
| 476 | LARP6 | 6.78993995 | 2.56031148 | 0.9733 | 2.63058 | 0.00852 | 0.034733 |
| 324 | NAB2 | 9.23074023 | 2.55870343 | 0.7419 | 3.44886 | 0.00056 | 0.00354 |
| 756 | MAP4K4 | 79.1716879 | 2.55533318 | 0.2965 | 8.61693 | 6.88E-18 | 1.02E-15 |
| 1381 | JADE3 | 7.73743976 | 2.553518558 | 0.7875 | 3.24259 | 0.00118 | 0.00662 |
| 847 | SOGA1 | 36.3322435 | 2.544986581 | 0.5117 | 4.9738 | 6.57E-07 | 9.13E-06 |
| 361 | CDK2AP1 | 63.1762486 | 2.539976318 | 0.3349 | 7.5842 | 3.35E-14 | 2.27E-12 |
| 995 | PDE6B | 17.9179105 | 2.536776396 | 0.5001 | 5.07217 | 3.93E-07 | 5.76E-06 |
| 1256 | DPYSL2 | 73.8812475 | 2.536745716 | 0.3621 | 7.00645 | 2.44E-12 | 1.19E-10 |
| 1102 | ARL10 | 18.5314962 | 2.531923761 | 0.5501 | 4.60285 | 4.2E-06 | 4.71E-05 |
| 552 | PIMREG | 13.6811212 | 2.527602877 | 0.7005 | 3.6082 | 0.00031 | 0.002093 |
| 686 | CLIP3 | 14.5864604 | 2.520242824 | 0.6348 | 3.97041 | 7.2E-05 | 0.000589 |
| 1343 | CERCAM | 8.90053681 | 2.519327276 | 0.6823 | 3.6923 | 0.00022 | 0.001572 |
| 302 | PRICKLE1 | 14.9532345 | 2.518709869 | 0.7108 | 3.54355 | 0.00039 | 0.002603 |
| 128 | SRGAP2 | 7.78803258 | 2.518094051 | 0.7997 | 3.14864 | 0.00164 | 0.008768 |
| 1074 | ADAMTS19 | 4.93208049 | 2.512242206 | 0.9477 | 2.6508 | 0.00803 | 0.033109 |
| 710 | ZNF331 | 10.1169556 | 2.508800278 | 0.7585 | 3.30756 | 0.00094 | 0.00546 |
| 133 | DTL | 17.6916108 | 2.508483808 | 0.5049 | 4.96829 | 6.75E-07 | 9.36E-06 |
| 634 | PRELID3A | 6.06294042 | 2.507811677 | 0.8439 | 2.97173 | 0.00296 | 0.014505 |
| 1442 | ARHGAP4 | 28.0387359 | 2.499737338 | 0.5719 | 4.37068 | 1.2E-05 | 0.000124 |
| 1073 | LMNB1 | 31.056331 | 2.496860263 | 0.3817 | 6.54202 | 6.07E-11 | 2.15E-09 |
| 244 | SAC3D1 | 6.4285272 | 2.49355823 | 0.8133 | 3.06605 | 0.00217 | 0.011163 |
| 1111 | PXDC1 | 22.329186 | 2.489831588 | 0.4791 | 5.19667 | 2.03E-07 | 3.16E-06 |
| 1085 | ARAP3 | 12.1299015 | 2.483764281 | 0.6249 | 3.97475 | 7E-05 | 0.00058 |
| 293 | LMO3 | 7.00202249 | 2.483190944 | 0.8389 | 2.96 | 0.00308 | 0.014963 |
| 901 | TNFRSF13C | 5.24282852 | 2.482134867 | 0.9531 | 2.60431 | 0.00921 | 0.036972 |
| 882 | HIRA | 5.57391971 | 2.481351539 | 0.9458 | 2.62356 | 0.0087 | 0.03533 |
| 316 | SLC4A8 | 7.66129639 | 2.475012471 | 0.7007 | 3.53203 | 0.00041 | 0.002695 |
| 506 | BMERB1 | 8.01660986 | 2.474961197 | 0.749 | 3.30433 | 0.00095 | 0.005512 |
| 501 | TEDC2 | 7.47434625 | 2.473082538 | 0.7421 | 3.33247 | 0.00086 | 0.005079 |
| 1232 | AKR1B1 | 18.9261596 | 2.472490152 | 0.5695 | 4.34177 | 1.4E-05 | 0.00014 |
| 412 | ENSG00000259121 | 4.35515712 | 2.469839339 | 0.9269 | 2.66466 | 0.00771 | 0.032064 |
| 394 | OBI1 | 22.771348 | 2.469658783 | 0.4991 | 4.94865 | 7.47E-07 | 1.02E-05 |
| 505 | EMP2 | 11.7801068 | 2.469289612 | 0.6308 | 3.91445 | 9.1E-05 | 0.000721 |
| 306 | VDR | 41.4953842 | 2.468084542 | 0.4233 | 5.83002 | 5.54E-09 | 1.24E-07 |
| 843 | E2F1 | 27.3728584 | 2.466886918 | 0.4925 | 5.00937 | 5.46E-07 | 7.74E-06 |
| 1026 | TSPAN5 | 8.63392562 | 2.465335993 | 0.723 | 3.40997 | 0.00065 | 0.003992 |
| 1150 | FYN | 13.0229523 | 2.462726709 | 0.6757 | 3.64465 | 0.00027 | 0.001849 |
| 1236 | TCAF1 | 13.7916539 | 2.461015252 | 0.581 | 4.23567 | 2.3E-05 | 0.000213 |
| 8 | KIF1B | 35.1211221 | 2.46028852 | 0.4384 | 5.61151 | 2.01E-08 | 4.01E-07 |

| | | | | | | | |
|---|---|---|---|---|---|---|---|
| 1077 | CXCL14 | 45.6267943 | 2.459602486 | 0.3225 | 7.62708 | 2.40E-14 | 1.69E-12 |
| 487 | HOMER2 | 9.78985475 | 2.457095871 | 0.7377 | 3.33062 | 0.00087 | 0.005102 |
| 717 | RNF144A | 12.7294938 | 2.443890061 | 0.6249 | 3.91069 | 9.2E-05 | 0.000729 |
| 942 | PTPRG | 14.5187942 | 2.427525444 | 0.5451 | 4.45361 | 8.4E-06 | 8.82E-05 |
| 593 | RND2 | 9.20758927 | 2.421227384 | 0.895 | 2.70534 | 0.00682 | 0.029044 |
| 155 | PIP4K2A | 13.2400542 | 2.420791259 | 0.6303 | 3.84052 | 0.00012 | 0.000942 |
| 870 | JAM2 | 4.97401825 | 2.414938037 | 0.9643 | 2.50439 | 0.01227 | 0.046656 |
| 1349 | CAMSAP1 | 14.7775736 | 2.411278963 | 0.6351 | 3.79665 | 0.00015 | 0.001092 |
| 389 | MZT1 | 17.2771481 | 2.408286068 | 0.5191 | 4.63949 | 3.5E-06 | 4.03E-05 |
| 706 | POLD1 | 10.7465239 | 2.405023239 | 0.6011 | 4.00102 | 6.3E-05 | 0.000528 |
| 1341 | BBLN | 28.452253 | 2.402026225 | 0.413 | 5.81582 | 6.03E-09 | 1.34E-07 |
| 391 | SLAIN1 | 5.86961624 | 2.401103348 | 0.9364 | 2.56431 | 0.01034 | 0.0405 |
| 299 | BICD1 | 14.1610956 | 2.397540563 | 0.5552 | 4.31813 | 1.6E-05 | 0.000154 |
| 481 | LINGO1 | 5.69105938 | 2.396459616 | 0.8629 | 2.77724 | 0.00548 | 0.02434 |
| 1052 | RAB3C | 4.99847697 | 2.390546838 | 0.9392 | 2.5452 | 0.01092 | 0.042286 |
| 645 | TCF4 | 96.7327805 | 2.389238924 | 0.2706 | 8.83026 | 1.04E-18 | 1.79E-16 |
| 195 | HSPA12A | 11.7376577 | 2.387649954 | 0.8087 | 2.95242 | 0.00315 | 0.015292 |
| 1368 | GPM6B | 19.0715679 | 2.384162698 | 0.6422 | 3.71259 | 0.00021 | 0.001463 |
| 950 | ST3GAL6 | 8.12883672 | 2.381839356 | 0.8004 | 2.97571 | 0.00292 | 0.014348 |
| 702 | KPTN | 7.9081496 | 2.379938025 | 0.7092 | 3.35577 | 0.00079 | 0.00473 |
| 668 | BRME1 | 7.71430355 | 2.377876181 | 0.8694 | 2.73494 | 0.00624 | 0.027001 |
| 529 | CFAP20 | 46.4619442 | 2.37567353 | 0.3987 | 5.95878 | 2.54E-09 | 6.07E-08 |
| 1049 | RAI14 | 17.5034427 | 2.375016622 | 0.6301 | 3.76906 | 0.00016 | 0.001205 |
| 1422 | MBNL3 | 46.7967433 | 2.373717031 | 0.36 | 6.59405 | 4.28E-11 | 1.57E-09 |
| 830 | ENSG00000275632 | 12.0887458 | 2.372839931 | 0.5894 | 4.02558 | 5.7E-05 | 0.000481 |
| 1376 | EEF1B2P3 | 4.37295771 | 2.369553304 | 0.9522 | 2.4885 | 0.01283 | 0.048389 |
| 327 | KIF5A | 6.51967564 | 2.363594905 | 0.9288 | 2.54467 | 0.01094 | 0.042327 |
| 1386 | PIM2 | 32.0032938 | 2.358732418 | 0.3843 | 6.1376 | 8.38E-10 | 2.21E-08 |
| 460 | MAP1A | 6.036328 | 2.358533571 | 0.9487 | 2.48616 | 0.01291 | 0.048582 |
| 1335 | WHRN | 5.65568804 | 2.353598059 | 0.8978 | 2.6216 | 0.00875 | 0.035498 |
| 1042 | LPCAT1 | 19.1378349 | 2.351115564 | 0.6083 | 3.86503 | 0.00011 | 0.000863 |
| 194 | GFRA1 | 6.49976919 | 2.350580672 | 0.8325 | 2.82343 | 0.00475 | 0.021593 |
| 24 | RPA2 | 29.0332954 | 2.345068563 | 0.4309 | 5.44263 | 5.25E-08 | 9.48E-07 |
| 385 | FAM124A | 5.86593855 | 2.342434396 | 0.9417 | 2.48747 | 0.01287 | 0.048441 |
| 539 | TERF2IP | 43.4066817 | 2.341950862 | 0.3464 | 6.76039 | 1.38E-11 | 5.71E-10 |
| 1178 | HYCC1 | 19.4296651 | 2.341603987 | 0.5348 | 4.37816 | 1.2E-05 | 0.00012 |
| 1375 | POLA1 | 16.5810334 | 2.334473437 | 0.5306 | 4.39953 | 1.1E-05 | 0.000111 |
| 1413 | PRPS1 | 41.1315988 | 2.331713898 | 0.3259 | 7.15556 | 8.33E-13 | 4.33E-11 |
| 736 | ENSG00000230979 | 9.63802157 | 2.331462806 | 0.71 | 3.28383 | 0.00102 | 0.005862 |

| | | | | | | | |
|---|---|---|---|---|---|---|---|
| 354 | SRRM4 | 7.87007191 | 2.329267031 | 0.7919 | 2.94147 | 0.00327 | 0.015746 |
| 618 | KCNJ2 | 10.7976401 | 2.324752962 | 0.6325 | 3.6756 | 0.00024 | 0.001661 |
| 518 | KCTD13 | 32.2539479 | 2.324085175 | 0.3768 | 6.16768 | 6.93E-10 | 1.87E-08 |
| 733 | RHOQ | 20.2093369 | 2.323984956 | 0.621 | 3.74211 | 0.00018 | 0.00132 |
| 1344 | PKN3 | 6.60914057 | 2.320507399 | 0.8415 | 2.75759 | 0.00582 | 0.025518 |
| 254 | CEP126 | 9.78700488 | 2.319533223 | 0.7656 | 3.02966 | 0.00245 | 0.012363 |
| 1023 | PKD2 | 11.9247596 | 2.319079429 | 0.6564 | 3.53323 | 0.00041 | 0.002689 |
| 516 | SEZ6L2 | 34.1019703 | 2.316565714 | 0.3853 | 6.01293 | 1.82E-09 | 4.50E-08 |
| 1 | KLHL17 | 12.2916922 | 2.315216293 | 0.5743 | 4.03128 | 5.5E-05 | 0.000471 |
| 871 | GABPA | 12.3194141 | 2.285134225 | 0.6415 | 3.56226 | 0.00037 | 0.002446 |
| 68 | LRRC8B | 11.2056954 | 2.283238602 | 0.7503 | 3.04329 | 0.00234 | 0.011897 |
| 92 | CTSK | 8.40926319 | 2.281337747 | 0.6768 | 3.37074 | 0.00075 | 0.004517 |
| 333 | IRAK3 | 6.45373929 | 2.281331054 | 0.8105 | 2.81458 | 0.00488 | 0.022141 |
| 1066 | APC | 36.2990071 | 2.279344936 | 0.5866 | 3.88577 | 0.0001 | 0.000801 |
| 1129 | FKBP5 | 10.7018778 | 2.269868372 | 0.7094 | 3.1997 | 0.00138 | 0.00757 |
| 280 | CCDC77 | 5.7209088 | 2.269856985 | 0.8497 | 2.67143 | 0.00755 | 0.03148 |
| 368 | CDK8 | 9.04860854 | 2.269770187 | 0.6554 | 3.46338 | 0.00053 | 0.003378 |
| 1107 | EXOC2 | 24.1635982 | 2.269661344 | 0.5056 | 4.48864 | 7.2E-06 | 7.61E-05 |
| 168 | PALD1 | 14.7024226 | 2.265074863 | 0.5924 | 3.8236 | 0.00013 | 0.001 |
| 420 | PELI2 | 8.86160082 | 2.261221205 | 0.734 | 3.08065 | 0.00207 | 0.010692 |
| 980 | RSRC1 | 19.9396358 | 2.258102232 | 0.4955 | 4.55738 | 5.2E-06 | 5.69E-05 |
| 791 | SUMO1 | 11.7569885 | 2.250126542 | 0.6078 | 3.70212 | 0.00021 | 0.001518 |
| 818 | GPC1 | 13.6085261 | 2.245364352 | 0.5859 | 3.8321 | 0.00013 | 0.000972 |
| 69 | CDC7 | 25.6077059 | 2.244650345 | 0.5536 | 4.05451 | 5E-05 | 0.00043 |
| 57 | DAB1 | 15.4941927 | 2.239186309 | 0.7512 | 2.98072 | 0.00288 | 0.014145 |
| 105 | IGSF8 | 12.9060252 | 2.236540648 | 0.665 | 3.36329 | 0.00077 | 0.004619 |
| 1350 | QSOX2 | 21.9250646 | 2.235671127 | 0.5252 | 4.2567 | 2.1E-05 | 0.000197 |
| 14 | NBPF1 | 20.4025005 | 2.23471798 | 0.5256 | 4.2517 | 2.1E-05 | 0.0002 |
| 363 | STX2 | 7.31812561 | 2.233795485 | 0.7461 | 2.99403 | 0.00275 | 0.013632 |
| 1416 | PAK3 | 16.8113804 | 2.227375019 | 0.728 | 3.05965 | 0.00222 | 0.011369 |
| 1306 | PTPRD | 33.9239702 | 2.226100366 | 0.375 | 5.93675 | 2.91E-09 | 6.81E-08 |
| 951 | CMSS1 | 22.4739648 | 2.225946338 | 0.5227 | 4.25861 | 2.1E-05 | 0.000196 |
| 1432 | FGF13 | 13.439324 | 2.224752997 | 0.7535 | 2.95267 | 0.00315 | 0.015285 |
| 1298 | ENPP2 | 6.99515885 | 2.224315834 | 0.8212 | 2.70847 | 0.00676 | 0.02884 |
| 732 | TMEM178A | 47.3915517 | 2.220382015 | 0.5598 | 3.9667 | 7.3E-05 | 0.000597 |
| 384 | RCBTB1 | 10.8381669 | 2.217687861 | 0.7602 | 2.9172 | 0.00353 | 0.016859 |
| 428 | RPS6KL1 | 14.1500867 | 2.213898591 | 0.601 | 3.68389 | 0.00023 | 0.001615 |
| 473 | IGDCC3 | 10.732353 | 2.204206082 | 0.7989 | 2.75908 | 0.0058 | 0.025449 |
| 700 | ZNF428 | 19.7511075 | 2.202907485 | 0.4637 | 4.75048 | 2E-06 | 2.47E-05 |

| | | | | | | | |
|---|---|---|---|---|---|---|---|
| 805 | EPHA4 | 20.5653622 | 2.202488922 | 0.5288 | 4.16501 | 3.1E-05 | 0.000282 |
| 97 | TRIM46 | 6.79957839 | 2.199991692 | 0.7899 | 2.78505 | 0.00535 | 0.023866 |
| 1419 | SEPTIN6 | 10.1620046 | 2.198863242 | 0.6676 | 3.29366 | 0.00099 | 0.005693 |
| 1345 | MED27 | 7.13030424 | 2.192236591 | 0.7714 | 2.8418 | 0.00449 | 0.020532 |
| 679 | TSHZ3 | 22.5649241 | 2.191674302 | 0.5441 | 4.02776 | 5.6E-05 | 0.000477 |
| 1112 | ENSG00000270504 | 6.54684876 | 2.189905596 | 0.8724 | 2.51025 | 0.01206 | 0.046048 |
| 90 | SV2A | 11.9960978 | 2.189690357 | 0.6324 | 3.46254 | 0.00054 | 0.003386 |
| 611 | TRIM37 | 12.7383175 | 2.186919997 | 0.6236 | 3.50685 | 0.00045 | 0.002928 |
| 639 | RBBP8 | 22.3644219 | 2.180622923 | 0.4672 | 4.66722 | 3.1E-06 | 3.57E-05 |
| 650 | SHC2 | 20.4070989 | 2.179862657 | 0.4718 | 4.61999 | 3.8E-06 | 4.36E-05 |
| 141 | H2AC25 | 9.89659223 | 2.178979329 | 0.6724 | 3.24067 | 0.00119 | 0.006655 |
| 43 | KIF2C | 26.6114739 | 2.176305427 | 0.4356 | 4.99564 | 5.86E-07 | 8.24E-06 |
| 71 | PTBP2 | 31.1716275 | 2.158186251 | 0.374 | 5.77022 | 7.92E-09 | 1.72E-07 |
| 1307 | NFIB | 55.9632195 | 2.155792767 | 0.3819 | 5.6451 | 1.65E-08 | 3.36E-07 |
| 406 | EFS | 8.08737596 | 2.154672886 | 0.859 | 2.5083 | 0.01213 | 0.046265 |
| 1193 | CASTOR2 | 11.6466263 | 2.15335774 | 0.6887 | 3.12667 | 0.00177 | 0.009349 |
| 696 | MAP3K10 | 5.62468071 | 2.151327533 | 0.806 | 2.66898 | 0.00761 | 0.031701 |
| 253 | SLC36A4 | 6.98215539 | 2.146630811 | 0.7581 | 2.83144 | 0.00463 | 0.021148 |
| 1410 | TCEAL3 | 17.9608971 | 2.142650466 | 0.5489 | 3.90333 | 9.5E-05 | 0.00075 |
| 1121 | TUBB | 359.904396 | 2.142415201 | 0.1878 | 11.4086 | 3.79E-30 | 3.17E-27 |
| 284 | IFFO1 | 12.816177 | 2.141792464 | 0.5719 | 3.74522 | 0.00018 | 0.001306 |
| 121 | UBE2T | 16.6376636 | 2.137761279 | 0.6773 | 3.1561 | 0.0016 | 0.00859 |
| 989 | ABCC5 | 15.6656114 | 2.137072503 | 0.5476 | 3.90268 | 9.5E-05 | 0.000751 |
| 1018 | ADGRL3 | 12.8933214 | 2.136326049 | 0.5879 | 3.63366 | 0.00028 | 0.00192 |
| 569 | SPECC1 | 11.595276 | 2.125516419 | 0.6077 | 3.49785 | 0.00047 | 0.003019 |
| 288 | C12orf57 | 57.5143796 | 2.114483176 | 0.3066 | 6.89568 | 5.36E-12 | 2.44E-10 |
| 488 | HDGFL3 | 23.7560969 | 2.11259526 | 0.5897 | 3.58265 | 0.00034 | 0.002284 |
| 1019 | DCK | 8.45806308 | 2.111001194 | 0.6729 | 3.13737 | 0.0017 | 0.009072 |
| 70 | CNN3 | 41.1384886 | 2.110679024 | 0.453 | 4.65943 | 3.2E-06 | 3.7E-05 |
| 1304 | BOP1 | 24.082472 | 2.107086215 | 0.5161 | 4.08271 | 4.5E-05 | 0.000389 |
| 1284 | TMEM64 | 7.57157575 | 2.101891089 | 0.7231 | 2.90682 | 0.00365 | 0.017335 |
| 1408 | BEX3 | 110.151794 | 2.100928221 | 0.2379 | 8.83246 | 1.02E-18 | 1.78E-16 |
| 1397 | KIF4A | 14.683458 | 2.09925502 | 0.5734 | 3.66098 | 0.00025 | 0.001746 |
| 1222 | PRKAR2B | 13.9919291 | 2.098969075 | 0.7101 | 2.95604 | 0.00312 | 0.015144 |
| 777 | PDK1 | 26.2626571 | 2.097473878 | 0.5147 | 4.07527 | 4.6E-05 | 0.000399 |
| 410 | STRN3 | 30.7355906 | 2.09685829 | 0.4582 | 4.57661 | 4.7E-06 | 5.25E-05 |
| 648 | DOK6 | 8.24005883 | 2.094774466 | 0.8071 | 2.59537 | 0.00945 | 0.037648 |
| 1394 | FAM120C | 5.2008108 | 2.092590718 | 0.8249 | 2.53692 | 0.01118 | 0.043112 |
| 550 | ENO3 | 6.89755647 | 2.089115272 | 0.7718 | 2.70687 | 0.00679 | 0.028953 |

| | | | | | | | |
|---|---|---|---|---|---|---|---|
| 600 | DCAKD | 14.3342954 | 2.08621624 | 0.5203 | 4.00993 | 6.1E-05 | 0.000509 |
| 658 | ENSG00000280239 | 8.35142407 | 2.085482254 | 0.7741 | 2.69422 | 0.00706 | 0.029808 |
| 919 | SATB1 | 64.7320964 | 2.0851014 | 0.3016 | 6.91458 | 4.69E-12 | 2.17E-10 |
| 1203 | ADAM22 | 9.4250744 | 2.083894913 | 0.7232 | 2.88166 | 0.00396 | 0.018511 |
| 1361 | PLCXD1 | 28.9302915 | 2.07992185 | 0.3959 | 5.25344 | 1.49E-07 | 2.41E-06 |
| 267 | MIR100HG | 16.4334362 | 2.077460224 | 0.6042 | 3.4384 | 0.00059 | 0.003654 |
| 788 | SLC39A10 | 17.3020831 | 2.068741524 | 0.4866 | 4.25131 | 2.1E-05 | 0.000201 |
| 1104 | DBN1 | 25.5576929 | 2.06818766 | 0.5545 | 3.7299 | 0.00019 | 0.001379 |
| 120 | NAV1 | 16.4807421 | 2.065916458 | 0.5474 | 3.77434 | 0.00016 | 0.001181 |
| 1393 | SMC1A | 69.8067517 | 2.065844839 | 0.2814 | 7.34205 | 2.10E-13 | 1.25E-11 |
| 100 | MEX3A | 36.4592219 | 2.063371211 | 0.3814 | 5.40971 | 6.31E-08 | 1.11E-06 |
| 1069 | CCDC112 | 8.55268259 | 2.062507285 | 0.765 | 2.69597 | 0.00702 | 0.029705 |
| 140 | PARP1 | 56.8613457 | 2.061130559 | 0.3329 | 6.19189 | 5.94E-10 | 1.64E-08 |
| 538 | MTSS2 | 15.1499319 | 2.059985352 | 0.6542 | 3.14904 | 0.00164 | 0.00876 |
| 1356 | UAP1L1 | 12.6445796 | 2.054517794 | 0.7208 | 2.85042 | 0.00437 | 0.020087 |
| 184 | PSD | 14.9016575 | 2.050493728 | 0.7295 | 2.81075 | 0.00494 | 0.022363 |
| 1020 | PAQR3 | 13.8685964 | 2.048687284 | 0.5142 | 3.98446 | 6.8E-05 | 0.000562 |
| 731 | MAP4K3 | 19.8806804 | 2.042680713 | 0.5078 | 4.02227 | 5.8E-05 | 0.000486 |
| 1153 | HSF2 | 16.5503889 | 2.028286969 | 0.5317 | 3.81442 | 0.00014 | 0.00103 |
| 814 | ARL4C | 8.54794327 | 2.025521545 | 0.7605 | 2.66328 | 0.00774 | 0.032148 |
| 828 | RNF24 | 8.6353887 | 2.024199479 | 0.703 | 2.87937 | 0.00398 | 0.018621 |
| 214 | APBB1 | 18.0583361 | 2.021882802 | 0.4837 | 4.18005 | 2.9E-05 | 0.000266 |
| 150 | PFKFB3 | 27.1421161 | 2.018914085 | 0.5981 | 3.37572 | 0.00074 | 0.004447 |
| 1197 | MAGI2-AS3 | 14.2785214 | 2.015677805 | 0.6085 | 3.31247 | 0.00092 | 0.005379 |
| 892 | FOXRED2 | 19.5397733 | 2.01401067 | 0.4959 | 4.06131 | 4.9E-05 | 0.00042 |
| 88 | PHGDH | 43.5297334 | 2.013934911 | 0.3262 | 6.17414 | 6.65E-10 | 1.81E-08 |
| 631 | RAB31 | 15.9723355 | 2.011586739 | 0.7769 | 2.58939 | 0.00961 | 0.038161 |
| 612 | SKA2 | 50.5628508 | 2.011065771 | 0.3068 | 6.55554 | 5.54E-11 | 1.98E-09 |
| 249 | PPME1 | 18.2573696 | 2.010879957 | 0.4866 | 4.1323 | 3.6E-05 | 0.00032 |
| 1201 | ABCB1 | 9.81510541 | 2.010727063 | 0.7703 | 2.61016 | 0.00905 | 0.036489 |
| 1067 | DCP2 | 34.9642049 | 2.010441747 | 0.431 | 4.66406 | 3.1E-06 | 3.62E-05 |
| 810 | PID1 | 18.1545786 | 2.009040349 | 0.4729 | 4.24857 | 2.2E-05 | 0.000203 |
| 1277 | JPH1 | 17.2064046 | 2.008855889 | 0.6921 | 2.90242 | 0.0037 | 0.017552 |
| 797 | LINC01963 | 8.51173813 | 1.997692873 | 0.7418 | 2.69316 | 0.00708 | 0.029893 |
| 465 | DUT | 51.6194271 | 1.996064605 | 0.3359 | 5.9423 | 2.81E-09 | 6.61E-08 |
| 255 | DCUN1D5 | 22.704907 | 1.992902786 | 0.4622 | 4.31146 | 1.6E-05 | 0.000158 |
| 684 | FXYD5 | 13.0419382 | 1.990982581 | 0.768 | 2.59235 | 0.00953 | 0.037898 |
| 56 | PRKAA2 | 6.30837407 | 1.988682086 | 0.7939 | 2.50499 | 0.01225 | 0.046625 |
| 422 | DACT1 | 11.0086831 | 1.984020341 | 0.7388 | 2.68537 | 0.00724 | 0.030464 |

| | | | | | | | |
|---|---|---|---|---|---|---|---|
| 1309 | CNTLN | 9.37100893 | 1.978928925 | 0.7283 | 2.71735 | 0.00658 | 0.028213 |
| 94 | SEMA6C | 18.7080538 | 1.978333693 | 0.4825 | 4.09986 | 4.1E-05 | 0.000365 |
| 1021 | ENSG00000273156 | 7.3447809 | 1.977898335 | 0.7951 | 2.48764 | 0.01286 | 0.048437 |
| 1429 | HTATSF1 | 42.8511266 | 1.976989533 | 0.4107 | 4.81382 | 1.5E-06 | 1.87E-05 |
| 1039 | CENPU | 32.5212665 | 1.964682874 | 0.3911 | 5.02406 | 5.06E-07 | 7.25E-06 |
| 192 | TRUB1 | 21.2828249 | 1.961452656 | 0.5325 | 3.68379 | 0.00023 | 0.001615 |
| 1294 | NUDCD1 | 14.2977209 | 1.96042362 | 0.6416 | 3.05562 | 0.00225 | 0.011488 |
| 591 | NT5C3B | 21.7971498 | 1.960168831 | 0.439 | 4.46463 | 8E-06 | 8.42E-05 |
| 1330 | FSD1L | 9.42302698 | 1.959578579 | 0.745 | 2.63021 | 0.00853 | 0.034761 |
| 540 | PIEZO1 | 11.3076752 | 1.959292892 | 0.5738 | 3.41477 | 0.00064 | 0.00394 |
| 741 | CCT4 | 8.49109315 | 1.959116497 | 0.7354 | 2.66404 | 0.00772 | 0.0321 |
| 122 | PPP1R12B | 38.1904025 | 1.954295944 | 0.54 | 3.61898 | 0.0003 | 0.002019 |
| 688 | ZNF566 | 9.55860717 | 1.947443552 | 0.6616 | 2.94343 | 0.00325 | 0.015673 |
| 170 | FUT11 | 20.1864581 | 1.946070646 | 0.5881 | 3.30904 | 0.00094 | 0.005436 |
| 907 | CERK | 42.3520596 | 1.945485795 | 0.4162 | 4.67456 | 2.9E-06 | 3.46E-05 |
| 1031 | HHIP | 14.780828 | 1.942518621 | 0.5625 | 3.45311 | 0.00055 | 0.00349 |
| 220 | WEE1 | 48.9007851 | 1.941871368 | 0.3697 | 5.25286 | 1.50E-07 | 2.41E-06 |
| 522 | VKORC1 | 24.0754395 | 1.938143052 | 0.4479 | 4.32722 | 1.5E-05 | 0.000148 |
| 232 | SLC43A3 | 13.468351 | 1.934023507 | 0.6119 | 3.16049 | 0.00158 | 0.008477 |
| 1414 | ACSL4 | 25.5453923 | 1.9322101 | 0.3831 | 5.04327 | 4.58E-07 | 6.6E-06 |
| 508 | CCP110 | 8.90465482 | 1.932010574 | 0.7269 | 2.65774 | 0.00787 | 0.032568 |
| 757 | TTL | 31.46543 | 1.931598818 | 0.4184 | 4.61704 | 3.9E-06 | 4.42E-05 |
| 1131 | BYSL | 10.4309261 | 1.931168934 | 0.7096 | 2.72137 | 0.0065 | 0.027914 |
| 981 | ENSG00000269888 | 10.3454299 | 1.928281069 | 0.6325 | 3.04864 | 0.0023 | 0.011716 |
| 233 | FAM111B | 15.234503 | 1.924345453 | 0.5249 | 3.6658 | 0.00025 | 0.001718 |
| 247 | AIP | 15.2486442 | 1.921211078 | 0.5082 | 3.78026 | 0.00016 | 0.001156 |
| 1245 | SLC7A2 | 7.85671711 | 1.918711613 | 0.7428 | 2.58293 | 0.0098 | 0.038764 |
| 939 | DUSP7 | 10.0082464 | 1.916452525 | 0.7728 | 2.47975 | 0.01315 | 0.049295 |
| 1118 | ZNF184 | 8.07989952 | 1.914583411 | 0.7623 | 2.5117 | 0.01201 | 0.045883 |
| 1295 | ENY2 | 41.8382524 | 1.914566495 | 0.4357 | 4.39442 | 1.1E-05 | 0.000113 |
| 1426 | SLC9A6 | 11.0924854 | 1.905143387 | 0.6656 | 2.86229 | 0.00421 | 0.019482 |
| 1276 | ELOC | 36.3220417 | 1.903764515 | 0.3765 | 5.05614 | 4.28E-07 | 6.19E-06 |
| 477 | LOXL1 | 15.020379 | 1.903108976 | 0.5133 | 3.70781 | 0.00021 | 0.001487 |
| 911 | SRGAP3 | 9.44803773 | 1.90243502 | 0.6128 | 3.10425 | 0.00191 | 0.00998 |
| 1299 | ATAD2 | 21.3206233 | 1.891420644 | 0.4914 | 3.84905 | 0.00012 | 0.000917 |
| 176 | HELLS | 71.3572647 | 1.891055351 | 0.4147 | 4.55993 | 5.1E-06 | 5.63E-05 |
| 1221 | CCDC71L | 17.2343302 | 1.889684633 | 0.4868 | 3.88169 | 0.0001 | 0.000813 |
| 275 | JAM3 | 9.21016691 | 1.887096274 | 0.7202 | 2.62031 | 0.00879 | 0.035583 |
| 310 | FMNL3 | 8.65957135 | 1.886813535 | 0.7097 | 2.65866 | 0.00785 | 0.032517 |

| | | | | | | | |
|---|---|---|---|---|---|---|---|
| 494 | IDH2 | 81.9425624 | 1.885279983 | 0.3412 | 5.52472 | 3.30E-08 | 6.28E-07 |
| 662 | PIN1 | 34.5268838 | 1.883523635 | 0.4047 | 4.65376 | 3.3E-06 | 3.78E-05 |
| 571 | CCDC144NL-AS1 | 21.4576957 | 1.882222304 | 0.5449 | 3.45418 | 0.00055 | 0.00348 |
| 272 | ETS1 | 9.92409213 | 1.880222945 | 0.7301 | 2.57534 | 0.01001 | 0.039453 |
| 1159 | HIVEP2 | 12.1620715 | 1.876211326 | 0.5654 | 3.31834 | 0.00091 | 0.00529 |
| 606 | UTP18 | 11.993975 | 1.872123079 | 0.6804 | 2.75133 | 0.00594 | 0.025947 |
| 1359 | ENSG00000284976 | 19.9651971 | 1.870383196 | 0.529 | 3.53564 | 0.00041 | 0.002674 |
| 604 | SPAG9 | 81.4644877 | 1.868863496 | 0.257 | 7.2718 | 3.55E-13 | 2.01E-11 |
| 356 | MSI1 | 7.87033555 | 1.865925491 | 0.7342 | 2.54146 | 0.01104 | 0.042671 |
| 1241 | LMBR1 | 11.3884291 | 1.853163352 | 0.6179 | 2.99907 | 0.00271 | 0.013464 |
| 13 | PRDM2 | 36.2682415 | 1.849255471 | 0.3591 | 5.14978 | 2.61E-07 | 3.98E-06 |
| 772 | FMNL2 | 71.5997017 | 1.844287175 | 0.2978 | 6.19207 | 5.94E-10 | 1.64E-08 |
| 993 | FBXO45 | 11.520906 | 1.842762083 | 0.5945 | 3.09948 | 0.00194 | 0.010116 |
| 445 | MTA1 | 12.9117139 | 1.841220876 | 0.5385 | 3.41913 | 0.00063 | 0.003885 |
| 927 | KIF15 | 13.4279922 | 1.840050777 | 0.5525 | 3.33021 | 0.00087 | 0.005107 |
| 429 | NEK9 | 17.2424662 | 1.835912765 | 0.5339 | 3.43877 | 0.00058 | 0.003652 |
| 23 | CEP85 | 8.46991075 | 1.834312632 | 0.6961 | 2.63524 | 0.00841 | 0.034358 |
| 371 | SLC7A1 | 12.6323874 | 1.834184759 | 0.5666 | 3.23693 | 0.00121 | 0.00674 |
| 812 | PDE6D | 10.2762291 | 1.832751559 | 0.6716 | 2.72904 | 0.00635 | 0.027397 |
| 551 | RABEP1 | 9.35875022 | 1.83166816 | 0.6194 | 2.95737 | 0.0031 | 0.015084 |
| 1275 | STAU2 | 11.5488535 | 1.828893735 | 0.6626 | 2.76 | 0.00578 | 0.025409 |
| 1342 | DNM1 | 22.7328525 | 1.824925317 | 0.5487 | 3.32619 | 0.00088 | 0.005173 |
| 988 | YEATS2 | 32.9554763 | 1.823987123 | 0.4196 | 4.34672 | 1.4E-05 | 0.000137 |
| 5 | ACOT7 | 21.6776342 | 1.82255382 | 0.418 | 4.36049 | 1.3E-05 | 0.00013 |
| 954 | CEP97 | 9.55170139 | 1.820100969 | 0.6687 | 2.72168 | 0.0065 | 0.027897 |
| 437 | VRK1 | 17.8515621 | 1.819083295 | 0.4843 | 3.75582 | 0.00017 | 0.001258 |
| 619 | UNK | 21.2546176 | 1.818822538 | 0.4457 | 4.08094 | 4.5E-05 | 0.000391 |
| 432 | FOXN3 | 50.8474157 | 1.810718141 | 0.3761 | 4.81482 | 1.5E-06 | 1.86E-05 |
| 1239 | AGAP3 | 43.7389004 | 1.810321999 | 0.4073 | 4.44428 | 8.8E-06 | 9.14E-05 |
| 1214 | GPC2 | 12.2019804 | 1.808446661 | 0.6135 | 2.94794 | 0.0032 | 0.015483 |
| 838 | NAPB | 16.4722059 | 1.808403094 | 0.5499 | 3.28851 | 0.00101 | 0.00578 |
| 789 | PGAP1 | 15.0882083 | 1.80680386 | 0.5858 | 3.08457 | 0.00204 | 0.010579 |
| 11 | MAD2L2 | 18.8033273 | 1.803953517 | 0.5449 | 3.31072 | 0.00093 | 0.005407 |
| 52 | COA7 | 9.25993761 | 1.802301531 | 0.6951 | 2.59279 | 0.00952 | 0.03787 |
| 118 | TSEN15 | 8.1854238 | 1.799692489 | 0.7235 | 2.4876 | 0.01286 | 0.048437 |
| 399 | IRS2 | 57.6046887 | 1.798930748 | 0.3704 | 4.857 | 1.2E-06 | 1.54E-05 |
| 1334 | ZNF618 | 12.867003 | 1.797527175 | 0.5386 | 3.33738 | 0.00085 | 0.005006 |
| 1092 | ANXA6 | 31.565094 | 1.797521646 | 0.4281 | 4.19862 | 2.7E-05 | 0.000247 |
| 400 | CHAMP1 | 8.66669559 | 1.796175325 | 0.629 | 2.85565 | 0.00429 | 0.01981 |

| | | | | | | | |
|---|---|---|---|---|---|---|---|
| 623 | TIMP2 | 59.6587104 | 1.796097623 | 0.432 | 4.15722 | 3.2E-05 | 0.000291 |
| 263 | MCAM | 16.060433 | 1.79521719 | 0.6801 | 2.63978 | 0.0083 | 0.033979 |
| 2 | B3GALT6 | 8.73820743 | 1.794189726 | 0.6944 | 2.58387 | 0.00977 | 0.038669 |
| 1247 | ATP6V1B2 | 32.8540803 | 1.789216477 | 0.3898 | 4.59052 | 4.4E-06 | 4.96E-05 |
| 992 | RNF168 | 23.670336 | 1.788898298 | 0.4095 | 4.36809 | 1.3E-05 | 0.000126 |
| 877 | RRP1B | 13.1025542 | 1.787696093 | 0.6677 | 2.67748 | 0.00742 | 0.031035 |
| 9 | SRM | 51.6595108 | 1.784364805 | 0.3187 | 5.59967 | 2.15E-08 | 4.27E-07 |
| 1190 | AEBP1 | 20.9121526 | 1.783666192 | 0.4381 | 4.0716 | 4.7E-05 | 0.000404 |
| 1094 | CYFIP2 | 10.8376332 | 1.78217065 | 0.5758 | 3.095 | 0.00197 | 0.010244 |
| 1273 | SNHG6 | 49.5452658 | 1.781388935 | 0.3156 | 5.64448 | 1.66E-08 | 3.36E-07 |
| 58 | NFIA | 92.7848549 | 1.78008101 | 0.2635 | 6.75627 | 1.42E-11 | 5.84E-10 |
| 682 | GRAMD1A | 24.4237365 | 1.777732578 | 0.4864 | 3.65522 | 0.00026 | 0.001781 |
| 452 | SCG5 | 12.5610217 | 1.776418454 | 0.5726 | 3.10252 | 0.00192 | 0.010026 |
| 1259 | FZD3 | 17.6377946 | 1.767902441 | 0.5698 | 3.10246 | 0.00192 | 0.010026 |
| 652 | TCF3 | 49.7157156 | 1.765723159 | 0.3713 | 4.75588 | 2E-06 | 2.41E-05 |
| 210 | TSPAN4 | 16.5890882 | 1.76383374 | 0.5953 | 2.96298 | 0.00305 | 0.014853 |
| 53 | RPS13P2 | 32.2499605 | 1.757162479 | 0.3646 | 4.81944 | 1.4E-06 | 1.83E-05 |
| 549 | ARRB2 | 22.6606763 | 1.755459877 | 0.4206 | 4.17393 | 3E-05 | 0.000272 |
| 1424 | HPRT1 | 36.2115376 | 1.754496633 | 0.3916 | 4.47985 | 7.5E-06 | 7.88E-05 |
| 885 | MIAT | 21.7230059 | 1.753960622 | 0.5762 | 3.04404 | 0.00233 | 0.011876 |
| 891 | RBFOX2 | 57.1967688 | 1.75253377 | 0.3816 | 4.59204 | 4.4E-06 | 4.94E-05 |
| 792 | ABI2 | 33.8142763 | 1.752288742 | 0.3675 | 4.76778 | 1.9E-06 | 2.29E-05 |
| 136 | RPS15AP12 | 8.90362301 | 1.751451558 | 0.707 | 2.47725 | 0.01324 | 0.049575 |
| 221 | RPL23AP65 | 16.3053514 | 1.749392785 | 0.5138 | 3.40503 | 0.00066 | 0.004051 |
| 91 | PLEKHO1 | 14.3199727 | 1.748486282 | 0.5533 | 3.16034 | 0.00158 | 0.008478 |
| 574 | TMEM97 | 21.7758612 | 1.747844806 | 0.5221 | 3.34783 | 0.00081 | 0.004846 |
| 312 | RACGAP1 | 11.558842 | 1.747710832 | 0.5593 | 3.12461 | 0.00178 | 0.009408 |
| 590 | FKBP10 | 7.88888054 | 1.743365162 | 0.6868 | 2.53835 | 0.01114 | 0.04296 |
| 1139 | TMEM14A | 18.4213275 | 1.739713041 | 0.54 | 3.22185 | 0.00127 | 0.007074 |
| 638 | MIB1 | 33.4922684 | 1.731536481 | 0.4038 | 4.28859 | 1.8E-05 | 0.000173 |
| 824 | NOP56 | 78.0657669 | 1.730376108 | 0.3591 | 4.81877 | 1.4E-06 | 1.83E-05 |
| 646 | ZNF532 | 15.0478782 | 1.72759783 | 0.5089 | 3.39464 | 0.00069 | 0.004181 |
| 163 | ZWINT | 34.2439009 | 1.725453005 | 0.4785 | 3.60596 | 0.00031 | 0.00211 |
| 295 | BCAT1 | 19.8964331 | 1.723174677 | 0.4525 | 3.80784 | 0.00014 | 0.001056 |
| 132 | TRAF5 | 32.3319033 | 1.720924252 | 0.5342 | 3.22124 | 0.00128 | 0.007084 |
| 1076 | KIF3A | 13.423675 | 1.719602982 | 0.5747 | 2.99228 | 0.00277 | 0.013701 |
| 29 | MARCKSL1 | 113.97851 | 1.717854039 | 0.2593 | 6.62508 | 3.47E-11 | 1.30E-09 |
| 931 | CDC25A | 12.1670505 | 1.712039086 | 0.5965 | 2.86996 | 0.00411 | 0.019097 |
| 87 | WDR3 | 25.1376416 | 1.708784795 | 0.4545 | 3.7599 | 0.00017 | 0.001241 |

| | | | | | | | |
|---|---|---|---|---|---|---|---|
| 260 | ATM | 24.196821 | 1.707119758 | 0.4026 | 4.23998 | 2.2E-05 | 0.000209 |
| 1346 | RALGDS | 18.2685439 | 1.705375355 | 0.6226 | 2.73919 | 0.00616 | 0.026727 |
| 1080 | SNHG4 | 19.646609 | 1.70422739 | 0.6414 | 2.65692 | 0.00789 | 0.032629 |
| 457 | KNSTRN | 13.5258767 | 1.703535906 | 0.6823 | 2.49678 | 0.01253 | 0.047439 |
| 427 | PNMA1 | 20.7386778 | 1.703287645 | 0.5982 | 2.84725 | 0.00441 | 0.020249 |
| 651 | MEX3D | 14.8171832 | 1.70312342 | 0.5077 | 3.35427 | 0.0008 | 0.00475 |
| 663 | DNMT1 | 56.981313 | 1.70178436 | 0.4227 | 4.02585 | 5.7E-05 | 0.000481 |
| 1014 | DANCR | 45.4755048 | 1.69902996 | 0.3181 | 5.34066 | 9.26E-08 | 1.58E-06 |
| 609 | DGKE | 9.71576417 | 1.698615088 | 0.6541 | 2.59675 | 0.00941 | 0.037562 |
| 621 | TNRC6C | 17.8489027 | 1.690440451 | 0.5865 | 2.88202 | 0.00395 | 0.018501 |
| 1186 | DPY19L1 | 15.2494191 | 1.690434098 | 0.4895 | 3.45363 | 0.00055 | 0.003484 |
| 281 | ERC1 | 9.33564689 | 1.674914043 | 0.6769 | 2.47449 | 0.01334 | 0.049908 |
| 990 | RFC4 | 20.0898417 | 1.673885868 | 0.4458 | 3.75498 | 0.00017 | 0.001261 |
| 985 | USP13 | 12.6325974 | 1.67182925 | 0.6629 | 2.52186 | 0.01167 | 0.044736 |
| 564 | MYH10 | 39.4569702 | 1.668525931 | 0.3555 | 4.69405 | 2.7E-06 | 3.18E-05 |
| 172 | ZMIZ1 | 37.9225439 | 1.663414875 | 0.3654 | 4.55247 | 5.3E-06 | 5.8E-05 |
| 503 | PPL | 17.1969629 | 1.663195489 | 0.5968 | 2.78687 | 0.00532 | 0.02374 |
| 852 | NCOA3 | 11.2154097 | 1.6621111 | 0.6555 | 2.53559 | 0.01123 | 0.043263 |
| 463 | CTDSPL2 | 20.8515699 | 1.661569112 | 0.4466 | 3.72023 | 0.0002 | 0.001429 |
| 914 | NUP210 | 29.2958611 | 1.655657562 | 0.4915 | 3.36844 | 0.00076 | 0.004551 |
| 67 | RBMXL1 | 9.15418252 | 1.644160176 | 0.6368 | 2.58211 | 0.00982 | 0.038824 |
| 1198 | GNAI1 | 18.3651055 | 1.643318628 | 0.5688 | 2.88933 | 0.00386 | 0.018154 |
| 1062 | ARRDC3 | 12.0744367 | 1.642988142 | 0.6618 | 2.48259 | 0.01304 | 0.049006 |
| 3 | ACAP3 | 33.293561 | 1.642913123 | 0.4494 | 3.65613 | 0.00026 | 0.001777 |
| 572 | USP22 | 45.3638132 | 1.642228911 | 0.3207 | 5.12034 | 3.05E-07 | 4.57E-06 |
| 1174 | BZW2 | 36.6880837 | 1.636628847 | 0.4181 | 3.91431 | 9.1E-05 | 0.000721 |
| 1212 | PTCD1 | 9.77027187 | 1.633066645 | 0.6296 | 2.59391 | 0.00949 | 0.037768 |
| 521 | YPEL3 | 24.3793226 | 1.630952303 | 0.446 | 3.65675 | 0.00026 | 0.001773 |
| 1387 | PRAF2 | 16.170915 | 1.629376431 | 0.5811 | 2.80396 | 0.00505 | 0.022738 |
| 1156 | ENSG00000260418 | 12.6126631 | 1.622004853 | 0.5863 | 2.76658 | 0.00566 | 0.024979 |
| 418 | SOCS4 | 12.1833475 | 1.620686961 | 0.5408 | 2.99688 | 0.00273 | 0.013533 |
| 1058 | BTF3 | 102.124105 | 1.617453742 | 0.2377 | 6.80387 | 1.02E-11 | 4.36E-10 |
| 869 | HSPA13 | 12.2717059 | 1.613882455 | 0.61 | 2.64561 | 0.00815 | 0.033516 |
| 845 | CEP250 | 16.3153364 | 1.613615149 | 0.4924 | 3.27726 | 0.00105 | 0.005981 |
| 261 | NCAM1 | 17.8589252 | 1.613039539 | 0.6429 | 2.50899 | 0.01211 | 0.0462 |
| 1436 | MTMR1 | 35.8424733 | 1.60924757 | 0.3907 | 4.11844 | 3.8E-05 | 0.000338 |
| 456 | BUB1B | 21.5589979 | 1.602115719 | 0.5924 | 2.7046 | 0.00684 | 0.029073 |
| 185 | CUEDC2 | 22.2802548 | 1.600328087 | 0.4785 | 3.34446 | 0.00082 | 0.004891 |
| 1319 | TPM2 | 51.0542683 | 1.598112612 | 0.5116 | 3.12401 | 0.00178 | 0.009424 |

| | | | | | | | |
|---|---|---|---|---|---|---|---|
| 207 | DEAF1 | 22.2879489 | 1.597942402 | 0.4416 | 3.61883 | 0.0003 | 0.002019 |
| 1096 | PTTG1 | 46.9586298 | 1.597186609 | 0.3755 | 4.25356 | 2.1E-05 | 0.000199 |
| 415 | NIN | 16.5267599 | 1.59296013 | 0.5269 | 3.02338 | 0.0025 | 0.012551 |
| 41 | EBNA1BP2 | 35.0343623 | 1.591416614 | 0.3383 | 4.70393 | 2.6E-06 | 3.05E-05 |
| 1109 | TUBB2A | 54.8449703 | 1.587948257 | 0.2895 | 5.48492 | 4.14E-08 | 7.64E-07 |
| 237 | FADS1 | 24.1115252 | 1.584485973 | 0.4534 | 3.49484 | 0.00047 | 0.003046 |
| 1383 | RBM10 | 29.3661917 | 1.583080461 | 0.3913 | 4.04543 | 5.2E-05 | 0.000446 |
| 622 | CYTH1 | 16.7916601 | 1.582651807 | 0.481 | 3.29044 | 0.001 | 0.005749 |
| 563 | PFAS | 28.0378543 | 1.581803187 | 0.4818 | 3.28345 | 0.00103 | 0.005868 |
| 669 | SAMD1 | 18.1400912 | 1.577786156 | 0.4951 | 3.1867 | 0.00144 | 0.007855 |
| 1382 | NDUFB11 | 40.6383854 | 1.565221498 | 0.3839 | 4.07762 | 4.5E-05 | 0.000396 |
| 1176 | RAPGEF5 | 20.621157 | 1.558175022 | 0.5374 | 2.89972 | 0.00373 | 0.01768 |
| 1213 | MCM7 | 82.2616714 | 1.555793501 | 0.3433 | 4.53249 | 5.8E-06 | 6.32E-05 |
| 305 | HDAC7 | 21.4035589 | 1.553524435 | 0.5299 | 2.9316 | 0.00337 | 0.016179 |
| 1051 | MOCS2 | 19.2667377 | 1.552985401 | 0.497 | 3.12463 | 0.00178 | 0.009408 |
| 402 | LINC00641 | 15.1974074 | 1.546619864 | 0.593 | 2.60796 | 0.00911 | 0.036674 |
| 880 | PCNT | 28.9378367 | 1.54563536 | 0.3873 | 3.99074 | 6.6E-05 | 0.000548 |
| 116 | ARPC5 | 59.5055373 | 1.545302154 | 0.2966 | 5.21046 | 1.88E-07 | 2.97E-06 |
| 1443 | GDI1 | 53.0725235 | 1.542597282 | 0.3528 | 4.37264 | 1.2E-05 | 0.000123 |
| 322 | NABP2 | 29.84334 | 1.542489562 | 0.4078 | 3.78244 | 0.00016 | 0.001148 |
| 1054 | CENPH | 11.6607335 | 1.539219504 | 0.5737 | 2.6829 | 0.0073 | 0.030609 |
| 1005 | DCAF16 | 40.8979174 | 1.535323839 | 0.3753 | 4.09038 | 4.3E-05 | 0.000378 |
| 186 | ARL3 | 13.3059903 | 1.530511465 | 0.6124 | 2.49938 | 0.01244 | 0.047193 |
| 248 | CCND1 | 20.5203655 | 1.530149825 | 0.4553 | 3.36057 | 0.00078 | 0.004657 |
| 119 | CAMSAP2 | 14.7118594 | 1.52844657 | 0.5559 | 2.74935 | 0.00597 | 0.026088 |
| 370 | GTF3A | 18.0068192 | 1.52660777 | 0.549 | 2.78052 | 0.00543 | 0.024141 |
| 206 | HRAS | 16.9987222 | 1.523706278 | 0.4522 | 3.36921 | 0.00075 | 0.00454 |
| 1081 | PFDN1 | 11.9005316 | 1.520423441 | 0.5947 | 2.55654 | 0.01057 | 0.04124 |
| 318 | EIF4B | 21.65386 | 1.51910491 | 0.4336 | 3.50387 | 0.00046 | 0.002958 |
| 1336 | PHF19 | 22.5858423 | 1.517106201 | 0.4164 | 3.64343 | 0.00027 | 0.001857 |
| 857 | PFDN4 | 17.0324396 | 1.516827512 | 0.5163 | 2.93782 | 0.00331 | 0.015906 |
| 515 | CDIPT | 35.7524391 | 1.51426093 | 0.3654 | 4.14407 | 3.4E-05 | 0.000306 |
| 599 | KIF18B | 24.1237001 | 1.51350098 | 0.4732 | 3.19851 | 0.00138 | 0.007593 |
| 346 | TXNRD1 | 18.2561906 | 1.512139351 | 0.4745 | 3.18708 | 0.00144 | 0.007851 |
| 238 | CSKMT | 18.1386251 | 1.510184254 | 0.5817 | 2.59629 | 0.00942 | 0.037602 |
| 1357 | DPP7 | 12.1428888 | 1.50836421 | 0.5684 | 2.65378 | 0.00796 | 0.032848 |
| 758 | DPP10 | 11.1036201 | 1.505785684 | 0.5962 | 2.52545 | 0.01156 | 0.044317 |
| 290 | RIMKLB | 13.9532346 | 1.505605472 | 0.512 | 2.9409 | 0.00327 | 0.01577 |
| 242 | VEGFB | 38.9684723 | 1.505279038 | 0.3933 | 3.82709 | 0.00013 | 0.000989 |

| | | | | | | | |
|---|---|---|---|---|---|---|---|
| 1333 | ZNF483 | 15.8557258 | 1.500555228 | 0.5128 | 2.92622 | 0.00343 | 0.016428 |
| 443 | CDCA4 | 14.7501956 | 1.499714654 | 0.4962 | 3.02232 | 0.00251 | 0.012586 |
| 628 | EPB41L3 | 16.2858201 | 1.495972616 | 0.4867 | 3.07379 | 0.00211 | 0.010913 |
| 553 | NLGN2 | 18.1765977 | 1.489479747 | 0.4777 | 3.11785 | 0.00182 | 0.009601 |
| 382 | MED4 | 23.8923515 | 1.486544594 | 0.4948 | 3.00463 | 0.00266 | 0.013258 |
| 161 | MAPK8 | 29.3853304 | 1.486118475 | 0.4126 | 3.60173 | 0.00032 | 0.00214 |
| 459 | MGA | 19.8375578 | 1.483029011 | 0.4362 | 3.39971 | 0.00067 | 0.004113 |
| 1016 | PPAT | 19.7299299 | 1.480946485 | 0.5012 | 2.95459 | 0.00313 | 0.015204 |
| 1425 | MMGT1 | 17.597414 | 1.477838867 | 0.469 | 3.15097 | 0.00163 | 0.008716 |
| 734 | MSH2 | 23.7181352 | 1.474796795 | 0.5082 | 2.90193 | 0.00371 | 0.017574 |
| 1392 | GPR173 | 28.4959809 | 1.471852143 | 0.5402 | 2.72458 | 0.00644 | 0.027686 |
| 165 | ADO | 17.0039043 | 1.470546287 | 0.4784 | 3.07381 | 0.00211 | 0.010913 |
| 149 | PITRM1 | 25.3202427 | 1.469159037 | 0.4207 | 3.49208 | 0.00048 | 0.003074 |
| 164 | CDK1 | 59.5145467 | 1.466651235 | 0.3557 | 4.12283 | 3.7E-05 | 0.000333 |
| 1035 | C4orf46 | 13.5562444 | 1.463599162 | 0.5186 | 2.82202 | 0.00477 | 0.021668 |
| 1265 | PRKDC | 52.6931125 | 1.463263491 | 0.294 | 4.97756 | 6.44E-07 | 8.97E-06 |
| 236 | FEN1 | 19.0462962 | 1.462399506 | 0.5013 | 2.9171 | 0.00353 | 0.016859 |
| 1325 | WNK2 | 25.7906892 | 1.457502157 | 0.5042 | 2.89054 | 0.00385 | 0.018119 |
| 947 | PDZRN3 | 19.3277388 | 1.455168613 | 0.4579 | 3.17786 | 0.00148 | 0.008056 |
| 107 | PFDN2 | 21.479964 | 1.452868753 | 0.464 | 3.13142 | 0.00174 | 0.009227 |
| 1355 | SAPCD2 | 24.5833901 | 1.445774097 | 0.4068 | 3.55438 | 0.00038 | 0.002514 |
| 174 | RPP30 | 16.7726979 | 1.442656599 | 0.527 | 2.73759 | 0.00619 | 0.026841 |
| 846 | DSN1 | 23.3340429 | 1.441856833 | 0.4913 | 2.93508 | 0.00333 | 0.016015 |
| 337 | NAP1L1 | 122.049751 | 1.437685491 | 0.3256 | 4.4158 | 1E-05 | 0.000103 |
| 978 | GMPS | 16.3732263 | 1.43038751 | 0.5489 | 2.60579 | 0.00917 | 0.036834 |
| 1165 | ARMT1 | 16.0194304 | 1.418380869 | 0.5542 | 2.55928 | 0.01049 | 0.040968 |
| 1420 | ZBTB33 | 19.4904089 | 1.417212635 | 0.4782 | 2.96343 | 0.00304 | 0.014841 |
| 840 | NINL | 15.115784 | 1.416719756 | 0.5502 | 2.57495 | 0.01003 | 0.039461 |
| 1120 | MDC1 | 14.3327099 | 1.415039112 | 0.5125 | 2.76114 | 0.00576 | 0.025342 |
| 565 | RPL13P12 | 26.523964 | 1.413928683 | 0.3986 | 3.54754 | 0.00039 | 0.002569 |
| 714 | RNASEH1 | 15.4762672 | 1.413723888 | 0.5048 | 2.80037 | 0.0051 | 0.022913 |
| 763 | POLR2D | 21.9804494 | 1.411747525 | 0.4541 | 3.10868 | 0.00188 | 0.009853 |
| 895 | CSNK1E | 34.0617302 | 1.411647976 | 0.346 | 4.08018 | 4.5E-05 | 0.000392 |
| 653 | AMH | 19.9065234 | 1.411637393 | 0.4218 | 3.34656 | 0.00082 | 0.004858 |
| 1384 | USP11 | 81.5852327 | 1.409853718 | 0.3754 | 3.7561 | 0.00017 | 0.001258 |
| 50 | BTF3L4 | 15.2259107 | 1.408124733 | 0.5018 | 2.80591 | 0.00502 | 0.022642 |
| 103 | NES | 31.2572613 | 1.407192515 | 0.5209 | 2.70139 | 0.00691 | 0.029312 |
| 251 | ME3 | 15.5389299 | 1.406322535 | 0.5362 | 2.62268 | 0.00872 | 0.035407 |
| 35 | CLSPN | 29.6150859 | 1.40608118 | 0.4686 | 3.00065 | 0.00269 | 0.013404 |

| | | | | | | | |
|---|---|---|---|---|---|---|---|
| 991 | OPA1 | 34.7268366 | 1.40426025 | 0.3936 | 3.56811 | 0.00036 | 0.002399 |
| 677 | URI1 | 23.3864522 | 1.403444991 | 0.4136 | 3.39302 | 0.00069 | 0.004202 |
| 605 | NME1 | 65.0770658 | 1.403179883 | 0.2977 | 4.71289 | 2.4E-06 | 2.94E-05 |
| 1171 | GLCCI1 | 17.3491194 | 1.402691276 | 0.5574 | 2.51629 | 0.01186 | 0.045363 |
| 222 | TEAD1 | 40.8436084 | 1.402327899 | 0.3901 | 3.59495 | 0.00032 | 0.002194 |
| 425 | FUT8 | 18.9724647 | 1.401290049 | 0.5344 | 2.62221 | 0.00874 | 0.035445 |
| 719 | ODC1 | 48.537736 | 1.400014186 | 0.3668 | 3.81655 | 0.00014 | 0.001023 |
| 148 | ZNF692 | 30.9756495 | 1.398450443 | 0.3689 | 3.79135 | 0.00015 | 0.001114 |
| 705 | NUP62 | 39.6244322 | 1.393022521 | 0.398 | 3.49994 | 0.00047 | 0.002999 |
| 739 | CCDC88A | 37.6915564 | 1.392312363 | 0.3665 | 3.79892 | 0.00015 | 0.001085 |
| 347 | RPL18AP3 | 19.7036207 | 1.390452382 | 0.419 | 3.31889 | 0.0009 | 0.005282 |
| 478 | PTPN9 | 13.990985 | 1.384653485 | 0.5306 | 2.60968 | 0.00906 | 0.036531 |
| 1037 | HMGB2 | 92.6818282 | 1.383662102 | 0.2572 | 5.38031 | 7.44E-08 | 1.29E-06 |
| 849 | MYBL2 | 18.2678809 | 1.383287997 | 0.485 | 2.85186 | 0.00435 | 0.020016 |
| 1050 | NIPBL-DT | 30.9474032 | 1.382524573 | 0.4587 | 3.01401 | 0.00258 | 0.012906 |
| 32 | ZNF362 | 17.7380052 | 1.38249939 | 0.4593 | 3.00985 | 0.00261 | 0.01306 |
| 544 | FANCA | 13.5859973 | 1.38189465 | 0.5157 | 2.67989 | 0.00736 | 0.03084 |
| 1272 | PDE7A | 16.0591593 | 1.380627374 | 0.4704 | 2.93522 | 0.00333 | 0.016013 |
| 975 | MBNL1 | 62.4614065 | 1.379269572 | 0.2898 | 4.7595 | 1.9E-06 | 2.38E-05 |
| 320 | CBX5 | 71.8757252 | 1.374913548 | 0.3092 | 4.44608 | 8.7E-06 | 9.08E-05 |
| 421 | EXOC5 | 25.4372944 | 1.366131922 | 0.3938 | 3.46945 | 0.00052 | 0.003315 |
| 1138 | MCM3 | 55.2938597 | 1.365361124 | 0.3086 | 4.42423 | 9.7E-06 | 9.98E-05 |
| 664 | ILF3 | 158.170432 | 1.363756027 | 0.2377 | 5.73694 | 9.64E-09 | 2.06E-07 |
| 866 | TCEA2 | 15.5824974 | 1.36082878 | 0.5211 | 2.61141 | 0.00902 | 0.036382 |
| 554 | POLR2A | 19.3615996 | 1.358201495 | 0.4708 | 2.88461 | 0.00392 | 0.01838 |
| 829 | PCNA | 59.5036207 | 1.354799392 | 0.3427 | 3.95382 | 7.7E-05 | 0.000626 |
| 1269 | CHD7 | 55.6140214 | 1.353197004 | 0.3021 | 4.47993 | 7.5E-06 | 7.88E-05 |
| 626 | ALYREF | 25.4195109 | 1.349849078 | 0.4012 | 3.36452 | 0.00077 | 0.004601 |
| 1071 | ENSG00000185641 | 55.9806104 | 1.349665076 | 0.3488 | 3.86918 | 0.00011 | 0.00085 |
| 933 | NCKIPSD | 19.0249908 | 1.346986665 | 0.4878 | 2.76161 | 0.00575 | 0.025315 |
| 1151 | HDAC2 | 58.9084937 | 1.345275558 | 0.288 | 4.67074 | 3E-06 | 3.52E-05 |
| 555 | FXR2 | 19.4791082 | 1.342959313 | 0.4557 | 2.94676 | 0.00321 | 0.015538 |
| 785 | UBE2E3 | 43.107621 | 1.341597491 | 0.3224 | 4.16147 | 3.2E-05 | 0.000286 |
| 641 | SLC39A6 | 24.104715 | 1.340916816 | 0.4833 | 2.77442 | 0.00553 | 0.024505 |
| 294 | LDHB | 218.558763 | 1.339008827 | 0.2455 | 5.4549 | 4.90E-08 | 8.91E-07 |
| 597 | GPATCH8 | 19.0427849 | 1.336781669 | 0.4488 | 2.97874 | 0.00289 | 0.014227 |
| 922 | CRTAP | 18.1531914 | 1.332679299 | 0.4654 | 2.86349 | 0.00419 | 0.01944 |
| 1036 | CPE | 39.1533774 | 1.328198584 | 0.3707 | 3.58308 | 0.00034 | 0.002282 |
| 396 | IPO5 | 44.8357675 | 1.327710949 | 0.3306 | 4.01602 | 5.9E-05 | 0.000498 |

| | | | | | | | |
|---|---|---|---|---|---|---|---|
| 1331 | PALM2AKAP2 | 19.5400529 | 1.326558404 | 0.4637 | 2.86106 | 0.00422 | 0.019533 |
| 898 | PHF5A | 15.5153847 | 1.323282076 | 0.514 | 2.57431 | 0.01004 | 0.039513 |
| 307 | PFKM | 38.0955277 | 1.322255656 | 0.4226 | 3.12886 | 0.00175 | 0.009287 |
| 352 | RPL6 | 55.2956588 | 1.319114301 | 0.3159 | 4.17609 | 3E-05 | 0.00027 |
| 77 | PSRC1 | 21.8286895 | 1.316099156 | 0.4549 | 2.8929 | 0.00382 | 0.018021 |
| 1303 | AGO2 | 29.2477285 | 1.312347499 | 0.3713 | 3.53492 | 0.00041 | 0.002679 |
| 1339 | STRBP | 29.966675 | 1.306413555 | 0.3955 | 3.30343 | 0.00096 | 0.005523 |
| 602 | CBX1 | 34.0898589 | 1.302793592 | 0.339 | 3.84331 | 0.00012 | 0.000934 |
| 899 | SNU13 | 72.8244882 | 1.299455835 | 0.2743 | 4.73798 | 2.2E-06 | 2.62E-05 |
| 613 | DHX40 | 22.0722023 | 1.297412903 | 0.4536 | 2.8604 | 0.00423 | 0.01956 |
| 681 | UBA2 | 42.767065 | 1.297228721 | 0.3247 | 3.9947 | 6.5E-05 | 0.00054 |
| 1367 | PRPS2 | 35.5983453 | 1.291785336 | 0.3554 | 3.63518 | 0.00028 | 0.001912 |
| 1089 | GRPEL2 | 14.3001328 | 1.291461401 | 0.5206 | 2.48053 | 0.01312 | 0.049239 |
| 589 | TOP2A | 107.551362 | 1.287747452 | 0.2886 | 4.46232 | 8.1E-06 | 8.5E-05 |
| 359 | KNTC1 | 22.8171316 | 1.287343519 | 0.4025 | 3.19876 | 0.00138 | 0.007589 |
| 890 | MCM5 | 30.3554116 | 1.287311794 | 0.495 | 2.60087 | 0.0093 | 0.037265 |
| 38 | UTP11 | 22.554166 | 1.285687832 | 0.4209 | 3.05439 | 0.00226 | 0.011516 |
| 453 | GOLGA8B | 50.3975386 | 1.283422206 | 0.468 | 2.74245 | 0.0061 | 0.026529 |
| 332 | XPOT | 18.0906173 | 1.281732074 | 0.513 | 2.49846 | 0.01247 | 0.047303 |
| 6 | CAMTA1 | 25.969344 | 1.280684249 | 0.4085 | 3.13489 | 0.00172 | 0.009142 |
| 326 | SHMT2 | 35.6853904 | 1.276872864 | 0.3992 | 3.1982 | 0.00138 | 0.007598 |
| 44 | NASP | 66.5483072 | 1.273580689 | 0.3391 | 3.75556 | 0.00017 | 0.001259 |
| 798 | IGFBP2 | 123.994035 | 1.273298795 | 0.2668 | 4.77308 | 1.8E-06 | 2.24E-05 |
| 1373 | EIF1AX | 50.4464953 | 1.271642859 | 0.304 | 4.18347 | 2.9E-05 | 0.000263 |
| 1399 | ZNF711 | 38.5224917 | 1.269812905 | 0.4485 | 2.83126 | 0.00464 | 0.021153 |
| 844 | RPL31P2 | 44.1231297 | 1.268237196 | 0.311 | 4.07731 | 4.6E-05 | 0.000396 |
| 973 | COMMD2 | 21.2120372 | 1.267779558 | 0.4256 | 2.9786 | 0.0029 | 0.014228 |
| 625 | GAA | 15.8532216 | 1.265165611 | 0.4953 | 2.5542 | 0.01064 | 0.041479 |
| 1308 | PSIP1 | 47.2087017 | 1.264030079 | 0.3492 | 3.61993 | 0.00029 | 0.002012 |
| 1431 | RBMX | 145.029228 | 1.263820492 | 0.2269 | 5.56913 | 2.56E-08 | 5.02E-07 |
| 1326 | ANP32B | 80.0806101 | 1.259692338 | 0.2762 | 4.56112 | 5.1E-06 | 5.61E-05 |
| 817 | COPS9 | 45.0844239 | 1.258816398 | 0.2874 | 4.37956 | 1.2E-05 | 0.00012 |
| 468 | ENSG00000242327 | 23.9152224 | 1.254531564 | 0.4925 | 2.54703 | 0.01086 | 0.042156 |
| 364 | POLE | 23.4948346 | 1.253277808 | 0.4234 | 2.96005 | 0.00308 | 0.014963 |
| 804 | OBSL1 | 24.8544964 | 1.252475447 | 0.4459 | 2.80879 | 0.00497 | 0.022492 |
| 1258 | PBK | 26.0190293 | 1.244597369 | 0.4118 | 3.02244 | 0.00251 | 0.012586 |
| 1364 | TBL1X | 38.6223829 | 1.242472786 | 0.3229 | 3.84777 | 0.00012 | 0.00092 |
| 562 | VAMP2 | 21.4541649 | 1.236754541 | 0.4231 | 2.92328 | 0.00346 | 0.016573 |
| 490 | FANCI | 31.663332 | 1.236155248 | 0.4079 | 3.03066 | 0.00244 | 0.012327 |

| | | | | | | | |
|---|---|---|---|---|---|---|---|
| 1337 | STOM | 31.4435661 | 1.235781748 | 0.4163 | 2.96833 | 0.00299 | 0.014647 |
| 874 | DONSON | 19.6109796 | 1.234663845 | 0.4767 | 2.5901 | 0.00959 | 0.038114 |
| 883 | RANBP1 | 49.7060717 | 1.232190505 | 0.3242 | 3.80114 | 0.00014 | 0.001077 |
| 342 | VEZT | 42.5446079 | 1.231340274 | 0.3562 | 3.45696 | 0.00055 | 0.003448 |
| 1218 | POP7 | 19.8192602 | 1.231273332 | 0.4264 | 2.8876 | 0.00388 | 0.018243 |
| 441 | KLC1 | 35.1888113 | 1.231158451 | 0.3906 | 3.1519 | 0.00162 | 0.008694 |
| 365 | PSPC1 | 20.5039578 | 1.229456763 | 0.4714 | 2.60806 | 0.00911 | 0.036673 |
| 853 | CSE1L | 38.2228349 | 1.228114084 | 0.3563 | 3.44678 | 0.00057 | 0.003561 |
| 495 | PRC1 | 33.2174027 | 1.226299461 | 0.4417 | 2.77636 | 0.0055 | 0.024398 |
| 528 | NUP93 | 21.2890704 | 1.224396479 | 0.4475 | 2.73615 | 0.00622 | 0.026926 |
| 1163 | SAMD5 | 34.1719056 | 1.2237387 | 0.3457 | 3.53985 | 0.0004 | 0.002638 |
| 1437 | HMGB3 | 50.1255528 | 1.222645467 | 0.3584 | 3.41175 | 0.00065 | 0.003973 |
| 1132 | KLHDC3 | 16.900828 | 1.218555735 | 0.4786 | 2.54614 | 0.01089 | 0.042218 |
| 617 | MAP2K6 | 35.0801833 | 1.214179899 | 0.4334 | 2.80178 | 0.00508 | 0.022848 |
| 928 | CDCP1 | 50.6854033 | 1.214115621 | 0.3072 | 3.95254 | 7.7E-05 | 0.000628 |
| 873 | GART | 23.7859632 | 1.213195589 | 0.4361 | 2.78203 | 0.0054 | 0.024051 |
| 439 | PPP2R5C | 49.9338941 | 1.209498282 | 0.3048 | 3.96756 | 7.3E-05 | 0.000595 |
| 728 | PPM1G | 49.4664182 | 1.209458707 | 0.3443 | 3.5132 | 0.00044 | 0.002874 |
| 1396 | PDZD11 | 22.6478709 | 1.204048256 | 0.4732 | 2.54468 | 0.01094 | 0.042327 |
| 1267 | UBE2V2 | 32.8862136 | 1.203926299 | 0.3505 | 3.43527 | 0.00059 | 0.003692 |
| 199 | BUB3 | 71.0091184 | 1.203890932 | 0.2545 | 4.73069 | 2.2E-06 | 2.71E-05 |
| 223 | RPS13 | 364.558761 | 1.202386739 | 0.1863 | 6.45294 | 1.10E-10 | 3.68E-09 |
| 1266 | MCM4 | 54.0851402 | 1.202352053 | 0.3197 | 3.76087 | 0.00017 | 0.001237 |
| 1126 | HMGA1 | 137.621521 | 1.202165619 | 0.3007 | 3.99771 | 6.4E-05 | 0.000534 |
| 197 | MCMBP | 25.4642773 | 1.201012163 | 0.4297 | 2.79499 | 0.00519 | 0.023246 |
| 1065 | NREP | 30.8035526 | 1.195899044 | 0.3596 | 3.32525 | 0.00088 | 0.005188 |
| 383 | FNDC3A | 26.0831799 | 1.195399003 | 0.4137 | 2.88944 | 0.00386 | 0.018154 |
| 1324 | IARS1 | 25.9569861 | 1.195050828 | 0.4153 | 2.87727 | 0.00401 | 0.018727 |
| 380 | DGKH | 30.2081031 | 1.193910688 | 0.3652 | 3.26948 | 0.00108 | 0.006118 |
| 357 | KDM2B | 21.7351063 | 1.192675443 | 0.4407 | 2.70606 | 0.00681 | 0.028994 |
| 245 | DRAP1 | 15.0674111 | 1.192060665 | 0.4786 | 2.49073 | 0.01275 | 0.048135 |
| 313 | CERS5 | 17.3942346 | 1.191446445 | 0.4771 | 2.49719 | 0.01252 | 0.047411 |
| 258 | CUL5 | 17.7461498 | 1.189692044 | 0.4516 | 2.63412 | 0.00844 | 0.034452 |
| 1128 | ENSG00000219023 | 27.9204722 | 1.188064412 | 0.3717 | 3.19663 | 0.00139 | 0.007631 |
| 1229 | ATP6V1F | 39.3444 | 1.186648571 | 0.3629 | 3.27014 | 0.00107 | 0.006107 |
| 859 | FAM210B | 22.2947851 | 1.185457802 | 0.4032 | 2.93976 | 0.00328 | 0.015812 |
| 854 | ADNP | 52.0049467 | 1.185382004 | 0.3307 | 3.5847 | 0.00034 | 0.002269 |
| 471 | PCLAF | 32.9291464 | 1.185009323 | 0.4597 | 2.5777 | 0.00995 | 0.039246 |
| 226 | QSER1 | 41.8150292 | 1.18131143 | 0.4094 | 2.88531 | 0.00391 | 0.018352 |

| | | | | | | | |
|---|---|---|---|---|---|---|---|
| 1296 | SYBU | 20.1154377 | 1.181252725 | 0.4517 | 2.61538 | 0.00891 | 0.036045 |
| 191 | SMC3 | 29.302261 | 1.177050156 | 0.3537 | 3.32812 | 0.00087 | 0.005142 |
| 143 | TOMM20 | 107.093381 | 1.174035089 | 0.2386 | 4.91965 | 8.67E-07 | 1.17E-05 |
| 1117 | H4C3 | 33.0156202 | 1.172023528 | 0.3719 | 3.15172 | 0.00162 | 0.008696 |
| 656 | EEF2 | 623.587465 | 1.171353146 | 0.1796 | 6.52302 | 6.89E-11 | 2.41E-09 |
| 1227 | IMPDH1 | 26.2567163 | 1.170943503 | 0.394 | 2.97196 | 0.00296 | 0.014499 |
| 1409 | TCEAL4 | 36.7853031 | 1.16965471 | 0.3414 | 3.42628 | 0.00061 | 0.003801 |
| 718 | YWHAQ | 69.6000237 | 1.164266339 | 0.273 | 4.26527 | 2E-05 | 0.00019 |
| 827 | CDC25B | 24.3317555 | 1.160711528 | 0.407 | 2.85157 | 0.00435 | 0.020021 |
| 403 | SUPT16H | 39.3131765 | 1.158684797 | 0.3186 | 3.63728 | 0.00028 | 0.001898 |
| 809 | MFF | 31.5613038 | 1.158141727 | 0.3453 | 3.35391 | 0.0008 | 0.004752 |
| 507 | RPS15A | 166.410865 | 1.157792914 | 0.1963 | 5.89664 | 3.71E-09 | 8.57E-08 |
| 1433 | FMR1 | 33.844557 | 1.1529887 | 0.4634 | 2.48785 | 0.01285 | 0.048429 |
| 966 | H1-10 | 68.9149577 | 1.151890191 | 0.3153 | 3.65284 | 0.00026 | 0.001796 |
| 592 | ACLY | 110.768937 | 1.150921433 | 0.2083 | 5.52425 | 3.31E-08 | 6.29E-07 |
| 1166 | TCP1 | 16.0945226 | 1.149543885 | 0.454 | 2.53193 | 0.01134 | 0.043612 |
| 537 | EXOSC6 | 33.1168929 | 1.147160139 | 0.3494 | 3.28305 | 0.00103 | 0.005874 |
| 721 | DDX1 | 55.9570486 | 1.144673468 | 0.3215 | 3.56051 | 0.00037 | 0.002462 |
| 1263 | EIF4EBP1 | 21.9046482 | 1.143825306 | 0.4614 | 2.47902 | 0.01317 | 0.049377 |
| 1115 | NUP153 | 22.9827057 | 1.142317657 | 0.407 | 2.80652 | 0.00501 | 0.022615 |
| 566 | TOP3A | 20.25633 | 1.141678207 | 0.4581 | 2.49232 | 0.01269 | 0.04795 |
| 1288 | ZNF706 | 28.702616 | 1.138568719 | 0.4003 | 2.84437 | 0.00445 | 0.0204 |
| 703 | RPS11 | 701.287348 | 1.135123425 | 0.1705 | 6.65567 | 2.82E-11 | 1.09E-09 |
| 964 | RUVBL1 | 20.5777865 | 1.134936026 | 0.409 | 2.77492 | 0.00552 | 0.024476 |
| 994 | PAK2 | 17.3573329 | 1.132725057 | 0.4389 | 2.58108 | 0.00985 | 0.038929 |
| 637 | SNRPD1 | 43.191934 | 1.131743996 | 0.3776 | 2.99754 | 0.00272 | 0.013518 |
| 343 | METAP2 | 52.1219275 | 1.128908742 | 0.2813 | 4.01343 | 6E-05 | 0.000503 |
| 998 | SLBP | 28.0489652 | 1.127439082 | 0.3809 | 2.95996 | 0.00308 | 0.014963 |
| 1091 | DCTN4 | 23.1782194 | 1.126845438 | 0.4199 | 2.68388 | 0.00728 | 0.030536 |
| 635 | SEH1L | 38.2374779 | 1.124122563 | 0.3821 | 2.94163 | 0.00326 | 0.015743 |
| 614 | DCAF7 | 74.5180975 | 1.123905714 | 0.2567 | 4.37874 | 1.2E-05 | 0.00012 |
| 1140 | ELOVL5 | 26.3006028 | 1.120882292 | 0.4222 | 2.65469 | 0.00794 | 0.032778 |
| 51 | TUT4 | 25.6303086 | 1.12082065 | 0.4309 | 2.60083 | 0.0093 | 0.037265 |
| 578 | SUZ12 | 21.8973814 | 1.117197909 | 0.4029 | 2.77257 | 0.00556 | 0.024592 |
| 440 | DYNC1H1 | 83.0869439 | 1.10977487 | 0.2337 | 4.74918 | 2E-06 | 2.49E-05 |
| 276 | VPS26B | 19.4033339 | 1.109473568 | 0.433 | 2.56238 | 0.0104 | 0.040671 |
| 262 | KMT2A | 40.6038238 | 1.105447152 | 0.3207 | 3.44667 | 0.00057 | 0.003561 |
| 28 | KHDRBS1 | 79.8477415 | 1.101024965 | 0.2521 | 4.36784 | 1.3E-05 | 0.000126 |
| 1311 | RRAGA | 31.959662 | 1.100470955 | 0.3516 | 3.12951 | 0.00175 | 0.009277 |

| | | | | | | | |
|---|---|---|---|---|---|---|---|
| 7 | PARK7 | 66.1740146 | 1.099480785 | 0.2691 | 4.08536 | 4.4E-05 | 0.000386 |
| 386 | CKAP2 | 25.0701508 | 1.099478444 | 0.4134 | 2.6599 | 0.00782 | 0.032426 |
| 19 | HNRNPR | 50.5943056 | 1.089425718 | 0.318 | 3.42586 | 0.00061 | 0.003805 |
| 231 | MDK | 61.9888988 | 1.083803682 | 0.276 | 3.92742 | 8.6E-05 | 0.000689 |
| 959 | FSTL1 | 53.943823 | 1.083776846 | 0.2851 | 3.80164 | 0.00014 | 0.001076 |
| 531 | GOT2 | 70.3132332 | 1.083147678 | 0.3096 | 3.4981 | 0.00047 | 0.003018 |
| 202 | GLRX3 | 24.2590223 | 1.075558312 | 0.3958 | 2.71751 | 0.00658 | 0.028208 |
| 940 | PBRM1 | 34.1579581 | 1.068711892 | 0.3853 | 2.77387 | 0.00554 | 0.024517 |
| 239 | SNHG1 | 83.2369129 | 1.06805225 | 0.2752 | 3.88099 | 0.0001 | 0.000814 |
| 557 | TP53 | 29.6097558 | 1.066008935 | 0.3863 | 2.75985 | 0.00578 | 0.025412 |
| 212 | RRM1 | 54.8391952 | 1.062871833 | 0.3008 | 3.53369 | 0.00041 | 0.002686 |
| 713 | UBE2M | 30.768193 | 1.062471059 | 0.4185 | 2.53868 | 0.01113 | 0.042942 |
| 401 | APEX1 | 33.4611186 | 1.055068148 | 0.4137 | 2.55038 | 0.01076 | 0.041845 |
| 183 | NOLC1 | 46.5740988 | 1.05362993 | 0.4063 | 2.59293 | 0.00952 | 0.037865 |
| 588 | CASC3 | 32.8268857 | 1.051813466 | 0.3906 | 2.69274 | 0.00709 | 0.029923 |
| 1185 | KBTBD2 | 24.9097863 | 1.051531499 | 0.4196 | 2.50591 | 0.01221 | 0.046529 |
| 799 | CNOT9 | 43.4640465 | 1.050927083 | 0.2887 | 3.64 | 0.00027 | 0.001879 |
| 414 | MIS18BP1 | 42.5919988 | 1.050814774 | 0.3337 | 3.14937 | 0.00164 | 0.008754 |
| 771 | RIF1 | 35.4555735 | 1.049612833 | 0.3761 | 2.79106 | 0.00525 | 0.023494 |
| 952 | TBC1D23 | 31.3101984 | 1.049466646 | 0.3781 | 2.77531 | 0.00551 | 0.024454 |
| 209 | RPLP2 | 525.340941 | 1.04649329 | 0.1673 | 6.25351 | 4.01E-10 | 1.16E-08 |
| 475 | ANP32A | 31.877838 | 1.043103306 | 0.3806 | 2.74064 | 0.00613 | 0.02665 |
| 1086 | DELE1 | 40.6105393 | 1.041863941 | 0.3214 | 3.24197 | 0.00119 | 0.006632 |
| 59 | USP1 | 36.2872564 | 1.036236011 | 0.3921 | 2.64246 | 0.00823 | 0.033779 |
| 45 | UQCRH | 195.212098 | 1.034515671 | 0.2437 | 4.24468 | 2.2E-05 | 0.000206 |
| 40 | CTPS1 | 27.1487036 | 1.033935043 | 0.4156 | 2.48795 | 0.01285 | 0.048428 |
| 649 | TXNL4A | 58.7034616 | 1.033481132 | 0.2632 | 3.927 | 8.6E-05 | 0.00069 |
| 1195 | YWHAG | 69.5810952 | 1.03311518 | 0.299 | 3.45515 | 0.00055 | 0.003469 |
| 774 | BAZ2B | 53.8264036 | 1.030922741 | 0.3217 | 3.20495 | 0.00135 | 0.007451 |
| 559 | NAA38 | 35.6279385 | 1.028276066 | 0.3646 | 2.8202 | 0.0048 | 0.021777 |
| 240 | RTN3 | 57.1443703 | 1.025521826 | 0.2785 | 3.68195 | 0.00023 | 0.001626 |
| 546 | RPA1 | 29.0056181 | 1.022655799 | 0.4102 | 2.49327 | 0.01266 | 0.04786 |
| 813 | DGKD | 37.4723605 | 1.021423709 | 0.3439 | 2.9699 | 0.00298 | 0.014587 |
| 776 | RPS15P4 | 53.5661606 | 1.016667601 | 0.3665 | 2.77407 | 0.00554 | 0.02451 |
| 680 | GPI | 82.056992 | 1.016657551 | 0.3129 | 3.24943 | 0.00116 | 0.006483 |
| 861 | RPS21 | 470.278549 | 1.014281747 | 0.1624 | 6.24438 | 4.25E-10 | 1.21E-08 |
| 1362 | SLC25A6 | 304.604729 | 1.012237896 | 0.1894 | 5.34335 | 9.12E-08 | 1.56E-06 |
| 21 | SELENON | 36.2413136 | 1.011367224 | 0.3729 | 2.71211 | 0.00669 | 0.028597 |
| 135 | CENPF | 48.7134764 | 1.01075946 | 0.3902 | 2.5903 | 0.00959 | 0.038102 |

| | | | | | | | |
|---|---|---|---|---|---|---|---|
| 111 | CACYBP | 38.709405 | 1.009517779 | 0.3503 | 2.88166 | 0.00396 | 0.018511 |
| 1237 | EZH2 | 30.7205484 | 1.009140458 | 0.3402 | 2.96625 | 0.00301 | 0.014731 |
| 974 | PFN2 | 67.6736275 | 1.007749885 | 0.3728 | 2.70317 | 0.00687 | 0.029174 |
| 101 | CCT3 | 75.4657518 | 1.002927902 | 0.2859 | 3.50752 | 0.00045 | 0.002922 |
| 497 | IGF1R | 28.7482627 | 1.001921166 | 0.358 | 2.79847 | 0.00513 | 0.023033 |
| 2087 | MT-TY | 34.0927339 | -1.012735726 | 0.405 | -2.5006 | 0.0124 | 0.047054 |
| 1059 | CAPNS1 | 69.4082731 | -1.013167172 | 0.3002 | -3.3746 | 0.00074 | 0.004464 |
| 2093 | MT-CYB | 988.932202 | -1.017306986 | 0.1961 | -5.1884 | 2.1E-07 | 3.29E-06 |
| 635 | CNIH1 | 53.8459477 | -1.021174863 | 0.3973 | -2.5705 | 0.01015 | 0.039867 |
| 1091 | SELENOW | 72.9733478 | -1.023889486 | 0.2975 | -3.4411 | 0.00058 | 0.003629 |
| 187 | MPZL1 | 22.5745314 | -1.024136036 | 0.4116 | -2.4884 | 0.01283 | 0.048389 |
| 309 | GSTO1 | 44.552379 | -1.024191018 | 0.3302 | -3.1018 | 0.00192 | 0.010042 |
| 1484 | PCGF3 | 38.7004854 | -1.025244358 | 0.3386 | -3.0281 | 0.00246 | 0.012408 |
| 1163 | MOB1A | 32.8233304 | -1.025452512 | 0.3661 | -2.8007 | 0.0051 | 0.022899 |
| 1672 | PANK3 | 26.9568582 | -1.043936876 | 0.409 | -2.5525 | 0.0107 | 0.041616 |
| 1997 | VPS28 | 50.1785034 | -1.049754432 | 0.2979 | -3.5238 | 0.00043 | 0.002771 |
| 860 | NCOR1 | 57.2618969 | -1.053497749 | 0.2858 | -3.6867 | 0.00023 | 0.0016 |
| 1235 | PNKD | 41.7749101 | -1.054544055 | 0.3078 | -3.4263 | 0.00061 | 0.003801 |
| 700 | SERF2 | 201.275143 | -1.06510157 | 0.1887 | -5.6451 | 1.7E-08 | 3.36E-07 |
| 1371 | MRPS25 | 42.1548418 | -1.066822348 | 0.3712 | -2.8737 | 0.00406 | 0.018915 |
| 2168 | ERCC00074 | 19736.7572 | -1.068715379 | 0.307 | -3.4809 | 0.0005 | 0.00319 |
| 592 | TSC22D1 | 91.6391829 | -1.071868904 | 0.4282 | -2.5034 | 0.0123 | 0.046732 |
| 1943 | SARAF | 27.1269464 | -1.075063051 | 0.4324 | -2.4862 | 0.01291 | 0.048582 |
| 1917 | REPIN1 | 46.2105389 | -1.075095889 | 0.287 | -3.7455 | 0.00018 | 0.001305 |
| 1124 | ROCK2 | 26.0461876 | -1.079665265 | 0.403 | -2.6792 | 0.00738 | 0.030889 |
| 173 | TAGLN2 | 36.4960637 | -1.082341022 | 0.3823 | -2.831 | 0.00464 | 0.021164 |
| 1889 | ZNHIT1 | 27.219747 | -1.083324445 | 0.4054 | -2.672 | 0.00754 | 0.03144 |
| 332 | MTG1 | 26.3961293 | -1.086829291 | 0.3745 | -2.9024 | 0.0037 | 0.017552 |
| 1009 | ZBTB7A | 51.7748321 | -1.089712649 | 0.3562 | -3.0596 | 0.00222 | 0.011369 |
| 2084 | MT-RNR2 | 2563.65137 | -1.092280124 | 0.1757 | -6.2161 | 5.1E-10 | 1.43E-08 |
| 1131 | PTRHD1 | 21.2216393 | -1.09433495 | 0.437 | -2.5039 | 0.01228 | 0.046688 |
| 1287 | EIF6 | 45.2406275 | -1.098042926 | 0.3375 | -3.2535 | 0.00114 | 0.006406 |
| 2103 | PDK3 | 34.2962476 | -1.099175598 | 0.4177 | -2.6314 | 0.0085 | 0.034673 |
| 2096 | CD99 | 30.1905445 | -1.104268681 | 0.3838 | -2.8775 | 0.00401 | 0.01872 |
| 345 | CD151 | 59.4129191 | -1.10832497 | 0.352 | -3.1486 | 0.00164 | 0.008768 |
| 1913 | MKRN1 | 25.7913696 | -1.111350909 | 0.4316 | -2.575 | 0.01002 | 0.039461 |
| 133 | RAP1A | 21.6718069 | -1.11376521 | 0.4349 | -2.5609 | 0.01044 | 0.040819 |
| 2085 | MT-ND1 | 213.628528 | -1.114453359 | 0.223 | -4.9982 | 5.8E-07 | 8.14E-06 |
| 619 | DCAF11 | 24.0406415 | -1.114672626 | 0.4101 | -2.7182 | 0.00656 | 0.028168 |

| | | | | | | | |
|---|---|---|---|---|---|---|---|
| 1211 | ATP5MC3 | 151.576993 | -1.114964219 | 0.2874 | -3.8789 | 0.0001 | 0.00082 |
| 2075 | TPRN | 24.2439528 | -1.116434054 | 0.3903 | -2.8601 | 0.00423 | 0.019564 |
| 1706 | HLA-A | 239.938997 | -1.118011954 | 0.2437 | -4.5882 | 4.5E-06 | 5.01E-05 |
| 2147 | TMEM164 | 28.279909 | -1.11868472 | 0.3878 | -2.8849 | 0.00391 | 0.018367 |
| 38 | CDC42 | 86.8721638 | -1.121475019 | 0.3946 | -2.842 | 0.00448 | 0.020526 |
| 1775 | CD24 | 123.417522 | -1.131893313 | 0.2945 | -3.8437 | 0.00012 | 0.000933 |
| 660 | CALM1 | 140.504307 | -1.133760268 | 0.2485 | -4.5631 | 5E-06 | 5.57E-05 |
| 1294 | SERINC3 | 24.1190284 | -1.134672895 | 0.3965 | -2.8619 | 0.00421 | 0.019499 |
| 1877 | LAMTOR4 | 66.9035301 | -1.13512907 | 0.3232 | -3.5117 | 0.00045 | 0.002885 |
| 1530 | SEC31A | 67.4562041 | -1.137005228 | 0.3785 | -3.0038 | 0.00267 | 0.01329 |
| 1285 | CHMP4B | 49.9773544 | -1.138593542 | 0.3136 | -3.6308 | 0.00028 | 0.00194 |
| 1062 | ECH1 | 51.9952966 | -1.139040041 | 0.3439 | -3.3125 | 0.00092 | 0.005379 |
| 722 | DPP8 | 24.6831416 | -1.141294048 | 0.4495 | -2.5387 | 0.01112 | 0.042942 |
| 1830 | TSPAN13 | 36.4309516 | -1.141891175 | 0.4344 | -2.6285 | 0.00858 | 0.034904 |
| 322 | NSMCE4A | 20.2147654 | -1.143197974 | 0.4442 | -2.5739 | 0.01006 | 0.039549 |
| 1329 | GUCD1 | 28.0452614 | -1.144480711 | 0.4374 | -2.6163 | 0.00889 | 0.035985 |
| 968 | ENOSF1 | 35.0720358 | -1.148085844 | 0.3985 | -2.8809 | 0.00397 | 0.018538 |
| 2090 | MT-ND3 | 584.174408 | -1.152439771 | 0.2007 | -5.7432 | 9.3E-09 | 2E-07 |
| 129 | AMPD2 | 22.8114447 | -1.154042568 | 0.437 | -2.6408 | 0.00827 | 0.033911 |
| 1210 | CIR1 | 22.3070555 | -1.155410799 | 0.4052 | -2.8516 | 0.00435 | 0.020021 |
| 426 | NDUFC2 | 53.7568147 | -1.160124716 | 0.3047 | -3.8069 | 0.00014 | 0.001059 |
| 2056 | HSPA5 | 59.7485183 | -1.160801449 | 0.3205 | -3.6214 | 0.00029 | 0.002004 |
| 1320 | PDXK | 26.2147117 | -1.161186879 | 0.4327 | -2.6833 | 0.00729 | 0.030583 |
| 161 | TPM3 | 21.9167544 | -1.162357043 | 0.459 | -2.5325 | 0.01133 | 0.043586 |
| 1364 | PLXNB2 | 34.949844 | -1.164863095 | 0.4007 | -2.9067 | 0.00365 | 0.017335 |
| 1271 | MAVS | 35.9024263 | -1.170447142 | 0.3574 | -3.2752 | 0.00106 | 0.006019 |
| 852 | TXNDC17 | 60.657341 | -1.175725053 | 0.3879 | -3.0313 | 0.00243 | 0.012303 |
| 1388 | SHISA5 | 78.585744 | -1.177335771 | 0.2504 | -4.7026 | 2.6E-06 | 3.07E-05 |
| 1891 | CUX1 | 31.4914634 | -1.180938429 | 0.414 | -2.8528 | 0.00433 | 0.019966 |
| 1374 | STT3B | 41.7458669 | -1.182581826 | 0.3572 | -3.3103 | 0.00093 | 0.005413 |
| 1258 | LRRFIP1 | 30.6301873 | -1.183630413 | 0.3858 | -3.0679 | 0.00216 | 0.011104 |
| 241 | IDI1 | 45.5803392 | -1.184339275 | 0.3814 | -3.1054 | 0.0019 | 0.009944 |
| 2026 | VPS13A | 44.4129273 | -1.186091664 | 0.4571 | -2.595 | 0.00946 | 0.03766 |
| 1125 | LAPTM4A | 61.4379946 | -1.188877013 | 0.2874 | -4.1366 | 3.5E-05 | 0.000315 |
| 2091 | MT-ND5 | 147.039576 | -1.190081311 | 0.3075 | -3.8708 | 0.00011 | 0.000845 |
| 1118 | SH3YL1 | 26.2149005 | -1.190972644 | 0.4395 | -2.7098 | 0.00673 | 0.028747 |
| 647 | SYNJ2BP | 31.9113096 | -1.194503771 | 0.4484 | -2.6637 | 0.00773 | 0.032123 |
| 2118 | MAGED2 | 47.5393446 | -1.197343922 | 0.3552 | -3.3709 | 0.00075 | 0.004516 |
| 22 | PEX14 | 14.5986766 | -1.198215132 | 0.4765 | -2.5146 | 0.01192 | 0.045529 |

| | | | | | | | |
|---|---|---|---|---|---|---|---|
| 1152 | RTN4 | 108.928334 | -1.19861741 | 0.2519 | -4.7583 | 2E-06 | 2.39E-05 |
| 1492 | ADD1 | 43.1898125 | -1.199949956 | 0.341 | -3.5192 | 0.00043 | 0.002814 |
| 310 | XPNPEP1 | 31.8329537 | -1.203836762 | 0.4421 | -2.7228 | 0.00647 | 0.027821 |
| 879 | GGNBP2 | 34.4083925 | -1.210364656 | 0.4485 | -2.6986 | 0.00696 | 0.029531 |
| 1190 | DBI | 97.2187436 | -1.211478048 | 0.2199 | -5.5093 | 3.6E-08 | 6.79E-07 |
| 730 | CLK3 | 26.1824322 | -1.213826731 | 0.4581 | -2.65 | 0.00805 | 0.033182 |
| 2126 | MED12 | 27.169937 | -1.222007075 | 0.4432 | -2.7573 | 0.00583 | 0.025533 |
| 1169 | TGOLN2 | 31.0174747 | -1.225199502 | 0.4155 | -2.9487 | 0.00319 | 0.015451 |
| 750 | PGAP6 | 24.7730451 | -1.229017821 | 0.4018 | -3.0584 | 0.00222 | 0.011405 |
| 356 | TMEM41B | 25.2179061 | -1.230054088 | 0.3881 | -3.1694 | 0.00153 | 0.008256 |
| 1671 | CCNG1 | 60.7501572 | -1.230615745 | 0.4585 | -2.684 | 0.00727 | 0.030532 |
| 832 | SPIRE2 | 36.9799184 | -1.23141067 | 0.4305 | -2.8602 | 0.00423 | 0.019564 |
| 1175 | TMEM131 | 17.2594215 | -1.233840304 | 0.4941 | -2.4972 | 0.01252 | 0.047411 |
| 1093 | KDELR1 | 30.1628387 | -1.242073865 | 0.3518 | -3.5308 | 0.00041 | 0.002706 |
| 608 | LAMP1 | 54.0140032 | -1.242511023 | 0.3128 | -3.9722 | 7.1E-05 | 0.000586 |
| 540 | DYRK2 | 19.8598973 | -1.243050632 | 0.4308 | -2.8856 | 0.00391 | 0.018343 |
| 1219 | HIBCH | 30.7454046 | -1.243206847 | 0.4645 | -2.6765 | 0.00744 | 0.031099 |
| 896 | COA3 | 27.6146059 | -1.243254847 | 0.3901 | -3.1869 | 0.00144 | 0.007854 |
| 845 | ZZEF1 | 22.767503 | -1.24707821 | 0.4225 | -2.9514 | 0.00316 | 0.015331 |
| 44 | PNRC2 | 38.431712 | -1.250214633 | 0.4816 | -2.5962 | 0.00943 | 0.037604 |
| 1478 | TFRC | 50.2634546 | -1.250766933 | 0.3849 | -3.2494 | 0.00116 | 0.006483 |
| 1352 | TRIOBP | 23.7464197 | -1.252734315 | 0.447 | -2.8023 | 0.00507 | 0.022817 |
| 2166 | ERCC00046 | 2790.14175 | -1.256098852 | 0.4829 | -2.6011 | 0.00929 | 0.037256 |
| 929 | AKAP1 | 51.4885623 | -1.258907179 | 0.3678 | -3.423 | 0.00062 | 0.003838 |
| 575 | MLXIP | 54.2686322 | -1.26221976 | 0.3626 | -3.4809 | 0.0005 | 0.00319 |
| 1116 | EPN1 | 30.0308405 | -1.262602132 | 0.3999 | -3.1571 | 0.00159 | 0.008571 |
| 577 | TMED2 | 38.8920381 | -1.264362536 | 0.3575 | -3.5368 | 0.0004 | 0.002663 |
| 2107 | CASK | 55.9816528 | -1.270855469 | 0.3247 | -3.9142 | 9.1E-05 | 0.000721 |
| 302 | SCD | 76.458802 | -1.271075231 | 0.4925 | -2.5808 | 0.00986 | 0.03895 |
| 1908 | AHCYL2 | 29.1596434 | -1.272338041 | 0.3922 | -3.244 | 0.00118 | 0.006593 |
| 863 | SREBF1 | 18.6145 | -1.273209133 | 0.4992 | -2.5503 | 0.01076 | 0.041847 |
| 1270 | DDRGK1 | 19.898087 | -1.276541176 | 0.4202 | -3.0382 | 0.00238 | 0.012083 |
| 233 | CNIH4 | 25.4269712 | -1.279052135 | 0.4162 | -3.0734 | 0.00212 | 0.010923 |
| 372 | HIPK3 | 20.7478165 | -1.286424212 | 0.519 | -2.4785 | 0.01319 | 0.049428 |
| 2048 | UGCG | 18.716709 | -1.287317675 | 0.4784 | -2.6907 | 0.00713 | 0.030056 |
| 1035 | ILVBL | 37.0373108 | -1.288398683 | 0.3751 | -3.4346 | 0.00059 | 0.003698 |
| 1646 | SAR1B | 33.7475861 | -1.290520174 | 0.4598 | -2.8069 | 0.005 | 0.022598 |
| 1967 | RMDN1 | 17.6682699 | -1.290591605 | 0.5177 | -2.493 | 0.01267 | 0.047881 |
| 49 | SYF2 | 24.0966487 | -1.29142049 | 0.434 | -2.9756 | 0.00292 | 0.014348 |

| | | | | | | | |
|---|---|---|---|---|---|---|---|
| 128 | PSMA5 | 43.2975974 | -1.292768355 | 0.415 | -3.1151 | 0.00184 | 0.009673 |
| 2031 | FAM120A | 54.9089246 | -1.294037085 | 0.3262 | -3.9674 | 7.3E-05 | 0.000595 |
| 633 | ERO1A | 31.6892764 | -1.29607986 | 0.3926 | -3.3016 | 0.00096 | 0.005556 |
| 1370 | SEC13 | 38.8066677 | -1.299297517 | 0.3964 | -3.2778 | 0.00105 | 0.005972 |
| 1652 | PURA | 26.5770238 | -1.302355081 | 0.4766 | -2.7323 | 0.00629 | 0.02715 |
| 464 | SC5D | 20.2816003 | -1.305432922 | 0.4683 | -2.7874 | 0.00531 | 0.023723 |
| 242 | KLF6 | 31.3971701 | -1.306784112 | 0.4596 | -2.8436 | 0.00446 | 0.020445 |
| 1994 | GRINA | 20.692297 | -1.306995347 | 0.4477 | -2.9193 | 0.00351 | 0.016763 |
| 623 | GMPR2 | 19.9153892 | -1.309406469 | 0.4779 | -2.7401 | 0.00614 | 0.026675 |
| 1496 | LAP3 | 21.3413632 | -1.310505439 | 0.5081 | -2.5791 | 0.0099 | 0.039104 |
| 767 | LITAF | 16.3683861 | -1.311872038 | 0.4971 | -2.6393 | 0.00831 | 0.034018 |
| 265 | JMJD1C | 18.2832806 | -1.312440067 | 0.4674 | -2.8082 | 0.00498 | 0.022519 |
| 471 | STT3A | 25.3508897 | -1.31453752 | 0.4213 | -3.1203 | 0.00181 | 0.009531 |
| 1607 | DAP | 17.6392094 | -1.316038664 | 0.49 | -2.6861 | 0.00723 | 0.030417 |
| 1282 | ACSS1 | 17.2361038 | -1.32039387 | 0.4871 | -2.7107 | 0.00671 | 0.028692 |
| 1196 | PTPN18 | 35.4510637 | -1.320627614 | 0.4147 | -3.1848 | 0.00145 | 0.007896 |
| 1988 | LY6E | 29.4694886 | -1.322785717 | 0.399 | -3.3154 | 0.00092 | 0.005337 |
| 1014 | MYDGF | 20.102837 | -1.326648356 | 0.492 | -2.6963 | 0.00701 | 0.029696 |
| 857 | SAT2 | 48.0369947 | -1.333203254 | 0.4382 | -3.0424 | 0.00235 | 0.011929 |
| 2083 | MT-RNR1 | 162.913246 | -1.333496074 | 0.2102 | -6.3449 | 2.2E-10 | 6.75E-09 |
| 2167 | ERCC00062 | 29.3142706 | -1.335066389 | 0.5067 | -2.6351 | 0.00841 | 0.034364 |
| 1876 | ZKSCAN1 | 52.6635339 | -1.336495541 | 0.4034 | -3.3128 | 0.00092 | 0.005377 |
| 1090 | SLC1A5 | 25.5802859 | -1.337546276 | 0.3975 | -3.3645 | 0.00077 | 0.004601 |
| 1593 | MSMO1 | 26.6946656 | -1.337658592 | 0.468 | -2.858 | 0.00426 | 0.01968 |
| 516 | DAZAP2 | 55.3389182 | -1.344366356 | 0.2833 | -4.7449 | 2.1E-06 | 2.54E-05 |
| 1572 | PGRMC2 | 49.0050131 | -1.344616533 | 0.2926 | -4.5954 | 4.3E-06 | 4.86E-05 |
| 1555 | OSTC | 41.4716375 | -1.344888416 | 0.429 | -3.1346 | 0.00172 | 0.009145 |
| 398 | CAPN1 | 45.0968285 | -1.346451846 | 0.3321 | -4.0542 | 5E-05 | 0.000431 |
| 894 | STAT3 | 27.6927215 | -1.348650993 | 0.4375 | -3.0829 | 0.00205 | 0.010628 |
| 1580 | ZNF827 | 21.0983069 | -1.349098036 | 0.4696 | -2.8726 | 0.00407 | 0.018961 |
| 184 | TMCO1 | 36.9386299 | -1.349659388 | 0.3529 | -3.8247 | 0.00013 | 0.000997 |
| 650 | NUMB | 25.9737464 | -1.352916603 | 0.4356 | -3.1059 | 0.0019 | 0.009933 |
| 557 | ARL1 | 27.1409934 | -1.353049386 | 0.4005 | -3.3782 | 0.00073 | 0.004412 |
| 725 | UACA | 20.1341691 | -1.353599302 | 0.4644 | -2.9146 | 0.00356 | 0.016972 |
| 964 | CSNK1D | 60.3520165 | -1.355306125 | 0.3277 | -4.136 | 3.5E-05 | 0.000316 |
| 2063 | SURF4 | 31.766387 | -1.358254726 | 0.3988 | -3.4057 | 0.00066 | 0.004044 |
| 843 | NCBP3 | 26.190608 | -1.3611612 | 0.4851 | -2.8058 | 0.00502 | 0.022642 |
| 1494 | WDR1 | 56.0537415 | -1.361971425 | 0.3739 | -3.6429 | 0.00027 | 0.001859 |
| 1156 | RAB1A | 78.7516969 | -1.363276342 | 0.3713 | -3.6717 | 0.00024 | 0.001684 |

| | | | | | | | |
|---|---|---|---|---|---|---|---|
| 1198 | RAB3GAP1 | 21.3338601 | -1.364542795 | 0.4248 | -3.212 | 0.00132 | 0.007297 |
| 1266 | RBCK1 | 33.6525588 | -1.365004248 | 0.3629 | -3.7618 | 0.00017 | 0.001233 |
| 996 | BSG | 78.2623094 | -1.366617444 | 0.2784 | -4.9094 | 9.1E-07 | 1.22E-05 |
| 1765 | PGM3 | 18.9507271 | -1.367915186 | 0.5368 | -2.5485 | 0.01082 | 0.041993 |
| 1751 | FBXO9 | 27.9943058 | -1.368375259 | 0.46 | -2.9744 | 0.00294 | 0.014389 |
| 115 | PKN2 | 29.104655 | -1.372709372 | 0.4941 | -2.7784 | 0.00546 | 0.024284 |
| 902 | TMUB2 | 20.148687 | -1.372898481 | 0.5086 | -2.6993 | 0.00695 | 0.029478 |
| 1123 | HPCAL1 | 19.3177434 | -1.374701229 | 0.4833 | -2.8444 | 0.00445 | 0.0204 |
| 1865 | GATAD1 | 26.1735388 | -1.377893547 | 0.5155 | -2.6732 | 0.00751 | 0.031344 |
| 2110 | ARAF | 22.1677208 | -1.379160828 | 0.4952 | -2.7849 | 0.00535 | 0.023871 |
| 2113 | HDAC6 | 33.7242169 | -1.380138063 | 0.4123 | -3.3472 | 0.00082 | 0.00485 |
| 1345 | MPST | 28.1078759 | -1.381379906 | 0.4478 | -3.0846 | 0.00204 | 0.010579 |
| 1962 | IMPA1 | 20.6512703 | -1.383214516 | 0.495 | -2.7943 | 0.0052 | 0.023286 |
| 2144 | RBM41 | 15.2740018 | -1.384296483 | 0.5211 | -2.6563 | 0.0079 | 0.032661 |
| 1482 | DLG1 | 36.5498762 | -1.387919053 | 0.3436 | -4.0396 | 5.4E-05 | 0.000456 |
| 1873 | BAIAP2L1 | 21.7360115 | -1.387934685 | 0.546 | -2.5421 | 0.01102 | 0.0426 |
| 413 | KMT5B | 16.5184032 | -1.389806023 | 0.5219 | -2.6628 | 0.00775 | 0.032162 |
| 2114 | SLC35A2 | 17.6035836 | -1.390608251 | 0.5306 | -2.6207 | 0.00877 | 0.035562 |
| 179 | TSTD1 | 21.7718392 | -1.391070632 | 0.5593 | -2.4871 | 0.01288 | 0.048479 |
| 1485 | GAK | 43.9900747 | -1.391200206 | 0.3948 | -3.5242 | 0.00042 | 0.002768 |
| 1582 | LRBA | 19.0621154 | -1.392368289 | 0.5497 | -2.5331 | 0.01131 | 0.043539 |
| 1633 | HSD17B4 | 30.3106073 | -1.394631759 | 0.4282 | -3.2566 | 0.00113 | 0.006352 |
| 1276 | DSTN | 78.8093511 | -1.400212736 | 0.2476 | -5.6553 | 1.6E-08 | 3.2E-07 |
| 2088 | MT-CO1 | 250.860546 | -1.402657212 | 0.2133 | -6.576 | 4.8E-11 | 1.75E-09 |
| 407 | CDK2AP2 | 17.4908012 | -1.403955289 | 0.5134 | -2.7347 | 0.00624 | 0.027009 |
| 1650 | CTNNA1 | 46.3676469 | -1.404419336 | 0.3489 | -4.0256 | 5.7E-05 | 0.000481 |
| 323 | PLEKHA1 | 35.5144157 | -1.405536047 | 0.359 | -3.9155 | 9E-05 | 0.000719 |
| 1824 | KDELR2 | 63.1546519 | -1.406359008 | 0.3514 | -4.0016 | 6.3E-05 | 0.000527 |
| 737 | ABHD17C | 31.2965869 | -1.411606026 | 0.3754 | -3.7598 | 0.00017 | 0.001241 |
| 204 | RNPEP | 19.3606678 | -1.412043075 | 0.4941 | -2.8576 | 0.00427 | 0.019698 |
| 2112 | EBP | 33.7827291 | -1.412520122 | 0.3518 | -4.0149 | 5.9E-05 | 0.0005 |
| 1803 | AIG1 | 22.8053312 | -1.418867665 | 0.5338 | -2.6581 | 0.00786 | 0.032542 |
| 5 | SDF4 | 40.8404394 | -1.419168413 | 0.3728 | -3.8065 | 0.00014 | 0.001059 |
| 1721 | PSMB8 | 31.3174493 | -1.42337323 | 0.4248 | -3.3505 | 0.00081 | 0.004805 |
| 495 | ATF7IP | 19.0778482 | -1.424523032 | 0.4619 | -3.0842 | 0.00204 | 0.010588 |
| 114 | LMO4 | 62.5596047 | -1.425410527 | 0.3361 | -4.2417 | 2.2E-05 | 0.000208 |
| 1512 | EXOC1 | 13.9137063 | -1.428260262 | 0.5701 | -2.5055 | 0.01223 | 0.046577 |
| 81 | AKR1A1 | 43.172392 | -1.428685113 | 0.3677 | -3.8853 | 0.0001 | 0.000802 |
| 731 | CSK | 17.0683042 | -1.434293588 | 0.5672 | -2.5289 | 0.01144 | 0.043959 |

| | | | | | | | |
|---|---|---|---|---|---|---|---|
| 1399 | PRKCD | 17.9341449 | -1.434685078 | 0.4725 | -3.0363 | 0.00239 | 0.012141 |
| 1921 | FASTK | 14.8398121 | -1.435353665 | 0.5336 | -2.6901 | 0.00714 | 0.030099 |
| 1012 | SH3GL1 | 25.3824743 | -1.435437567 | 0.4306 | -3.3338 | 0.00086 | 0.005062 |
| 369 | LGR4 | 27.9282943 | -1.437417483 | 0.4753 | -3.0244 | 0.00249 | 0.012522 |
| 2060 | ASS1 | 32.4045205 | -1.440558376 | 0.459 | -3.1388 | 0.0017 | 0.009038 |
| 2128 | OGT | 82.4122111 | -1.44218743 | 0.29 | -4.9728 | 6.6E-07 | 9.17E-06 |
| 1423 | FAM162A | 40.756035 | -1.445392239 | 0.4237 | -3.4116 | 0.00065 | 0.003974 |
| 1442 | XRN1 | 17.3304055 | -1.447159038 | 0.5007 | -2.8904 | 0.00385 | 0.018119 |
| 2165 | ERCC00043 | 308.17754 | -1.447480985 | 0.3825 | -3.7841 | 0.00015 | 0.001141 |
| 962 | SIRT7 | 26.2080275 | -1.44951358 | 0.397 | -3.6508 | 0.00026 | 0.001808 |
| 1154 | UGP2 | 28.4639417 | -1.449830729 | 0.426 | -3.4035 | 0.00067 | 0.004063 |
| 368 | CCDC34 | 28.7634973 | -1.455205825 | 0.3948 | -3.6857 | 0.00023 | 0.001606 |
| 1213 | DNAJC10 | 53.9033684 | -1.462909056 | 0.5841 | -2.5048 | 0.01225 | 0.046642 |
| 1713 | HLA-B | 119.204836 | -1.464465979 | 0.2993 | -4.893 | 9.9E-07 | 1.32E-05 |
| 1578 | GAB1 | 13.4066376 | -1.466767547 | 0.5435 | -2.6985 | 0.00696 | 0.029531 |
| 1140 | EIF2AK2 | 26.4280969 | -1.468608177 | 0.4011 | -3.6613 | 0.00025 | 0.001745 |
| 1842 | GARS1-DT | 18.1457338 | -1.468881078 | 0.5086 | -2.8882 | 0.00387 | 0.018213 |
| 246 | NET1 | 58.5780194 | -1.469805754 | 0.3857 | -3.8105 | 0.00014 | 0.001045 |
| 1934 | FHIP2B | 19.5230379 | -1.472634715 | 0.4646 | -3.1696 | 0.00153 | 0.008253 |
| 1768 | RARS2 | 22.1886707 | -1.474659729 | 0.434 | -3.3975 | 0.00068 | 0.004143 |
| 1812 | CEP43 | 16.3829345 | -1.477307237 | 0.5878 | -2.5131 | 0.01197 | 0.04571 |
| 2070 | PHPT1 | 20.249545 | -1.483933629 | 0.5515 | -2.6909 | 0.00713 | 0.030055 |
| 120 | TMED5 | 25.3344364 | -1.484107388 | 0.411 | -3.6112 | 0.0003 | 0.002075 |
| 1554 | HADH | 49.8059026 | -1.484568267 | 0.3864 | -3.8422 | 0.00012 | 0.000938 |
| 319 | CCDC186 | 23.6763574 | -1.48544396 | 0.4597 | -3.2312 | 0.00123 | 0.006861 |
| 947 | NT5C | 15.3668521 | -1.488460177 | 0.5665 | -2.6275 | 0.0086 | 0.034956 |
| 1060 | ACTN4 | 35.2166774 | -1.490925256 | 0.4002 | -3.7251 | 0.0002 | 0.001404 |
| 2164 | BCAP31 | 14.4662862 | -1.492719777 | 0.5722 | -2.6088 | 0.00909 | 0.036603 |
| 2136 | DIAPH2 | 15.1374781 | -1.495166688 | 0.5392 | -2.7727 | 0.00556 | 0.024589 |
| 404 | SSH3 | 15.5232637 | -1.496412098 | 0.603 | -2.4818 | 0.01307 | 0.04909 |
| 1223 | SPATS2L | 29.820946 | -1.499689628 | 0.3784 | -3.9636 | 7.4E-05 | 0.000602 |
| 558 | HSP90B1 | 116.660998 | -1.506918191 | 0.2482 | -6.0719 | 1.3E-09 | 3.21E-08 |
| 1599 | SDHA | 18.9013792 | -1.509354407 | 0.4657 | -3.241 | 0.00119 | 0.00665 |
| 1097 | NUCB1 | 28.4313067 | -1.509796165 | 0.4154 | -3.6342 | 0.00028 | 0.001917 |
| 1384 | HIGD1A | 91.1135144 | -1.510592818 | 0.3564 | -4.2385 | 2.2E-05 | 0.00021 |
| 550 | ATP2B1 | 19.9855645 | -1.513168714 | 0.4986 | -3.0346 | 0.00241 | 0.012194 |
| 1176 | UNC50 | 14.2894245 | -1.514873529 | 0.5286 | -2.8658 | 0.00416 | 0.019324 |
| 253 | COMMD3 | 22.0097047 | -1.515703359 | 0.5565 | -2.7236 | 0.00646 | 0.027763 |
| 1691 | SSR1 | 34.6018056 | -1.516269583 | 0.3487 | -4.3482 | 1.4E-05 | 0.000136 |

| | | | | | | | |
|---|---|---|---|---|---|---|---|
| 1385 | ZDHHC3 | 18.5040585 | -1.516407035 | 0.4859 | -3.1207 | 0.0018 | 0.009522 |
| 2016 | TMEM8B | 19.1173388 | -1.517653323 | 0.5653 | -2.6848 | 0.00726 | 0.030499 |
| 61 | ATP5IF1 | 139.800886 | -1.518505944 | 0.2159 | -7.0339 | 2E-12 | 9.95E-11 |
| 86 | CMPK1 | 30.4471763 | -1.520207445 | 0.4235 | -3.59 | 0.00033 | 0.002229 |
| 534 | LRP1 | 34.2902693 | -1.520504676 | 0.3461 | -4.3932 | 1.1E-05 | 0.000113 |
| 829 | ACSF3 | 13.3182984 | -1.52195642 | 0.5854 | -2.5999 | 0.00933 | 0.037348 |
| 960 | RNF213 | 28.0302567 | -1.52410726 | 0.4248 | -3.588 | 0.00033 | 0.002244 |
| 817 | MAF | 53.3951678 | -1.526364111 | 0.5555 | -2.7476 | 0.006 | 0.026218 |
| 2072 | FBXW5 | 39.415822 | -1.527148842 | 0.3565 | -4.284 | 1.8E-05 | 0.000176 |
| 699 | PDIA3 | 44.0305755 | -1.528748181 | 0.3329 | -4.5916 | 4.4E-06 | 4.94E-05 |
| 1601 | SLC9A3-AS1 | 101.78023 | -1.530495886 | 0.2792 | -5.4818 | 4.2E-08 | 7.76E-07 |
| 1509 | FRYL | 48.3569565 | -1.531912249 | 0.3622 | -4.2293 | 2.3E-05 | 0.000218 |
| 2154 | ELF4 | 12.8587019 | -1.531931577 | 0.6127 | -2.5002 | 0.01241 | 0.047091 |
| 400 | MALAT1 | 2302.54401 | -1.534994746 | 0.2252 | -6.8173 | 9.3E-12 | 4.01E-10 |
| 708 | TMOD3 | 24.1717191 | -1.537149415 | 0.4076 | -3.771 | 0.00016 | 0.001196 |
| 1912 | SLC37A3 | 17.6793104 | -1.53981232 | 0.5887 | -2.6158 | 0.0089 | 0.036011 |
| 3 | C1orf159 | 10.7337437 | -1.540714146 | 0.59 | -2.6114 | 0.00902 | 0.036382 |
| 1167 | TMSB10 | 270.163405 | -1.540840057 | 0.3377 | -4.5624 | 5.1E-06 | 5.58E-05 |
| 1440 | RNF7 | 32.8206529 | -1.541181054 | 0.4337 | -3.5534 | 0.00038 | 0.00252 |
| 1436 | DNAJC13 | 12.4742536 | -1.542047612 | 0.5787 | -2.6645 | 0.00771 | 0.03207 |
| 24 | EFHD2 | 17.9136664 | -1.544367089 | 0.5228 | -2.954 | 0.00314 | 0.015222 |
| 943 | NHERF1 | 54.9382812 | -1.547458029 | 0.4164 | -3.716 | 0.0002 | 0.001448 |
| 1752 | GCLC | 18.0140276 | -1.547775016 | 0.5595 | -2.7663 | 0.00567 | 0.024985 |
| 10 | RER1 | 30.8982395 | -1.549438792 | 0.3644 | -4.2517 | 2.1E-05 | 0.0002 |
| 189 | DCAF6 | 14.4361351 | -1.550249336 | 0.5889 | -2.6325 | 0.00847 | 0.034593 |
| 751 | MCRIP2 | 22.1061647 | -1.556050814 | 0.4717 | -3.2986 | 0.00097 | 0.005602 |
| 591 | DNAJC15 | 32.7936109 | -1.55926628 | 0.3728 | -4.1829 | 2.9E-05 | 0.000263 |
| 194 | QSOX1 | 34.1522263 | -1.560830515 | 0.4979 | -3.1348 | 0.00172 | 0.009142 |
| 1230 | IDH1 | 38.1050956 | -1.566337753 | 0.4117 | -3.8049 | 0.00014 | 0.001064 |
| 1857 | TMEM120A | 25.0695257 | -1.567723341 | 0.5521 | -2.8395 | 0.00452 | 0.02066 |
| 665 | TRIP11 | 14.747271 | -1.572990841 | 0.612 | -2.5702 | 0.01016 | 0.039897 |
| 1954 | GGH | 17.0339497 | -1.574409214 | 0.5935 | -2.6525 | 0.00799 | 0.032959 |
| 1947 | ERLIN2 | 14.9920443 | -1.577959175 | 0.549 | -2.8742 | 0.00405 | 0.018894 |
| 503 | CMAS | 33.8039441 | -1.579405066 | 0.4177 | -3.7814 | 0.00016 | 0.001152 |
| 1409 | CHMP2B | 20.4788451 | -1.582344176 | 0.5787 | -2.7345 | 0.00625 | 0.02702 |
| 888 | ORMDL3 | 22.9858161 | -1.587200034 | 0.4635 | -3.4246 | 0.00062 | 0.00382 |
| 701 | B2M | 378.68829 | -1.589145306 | 0.1961 | -8.1055 | 5.3E-16 | 5.02E-14 |
| 1864 | AKAP9 | 86.5300098 | -1.589667175 | 0.3573 | -4.4488 | 8.6E-06 | 9E-05 |
| 1155 | AFTPH | 17.3153227 | -1.589678665 | 0.5294 | -3.0029 | 0.00267 | 0.013318 |

| | | | | | | | |
|---|---|---|---|---|---|---|---|
| 1025 | DNM2 | 52.6642707 | -1.591429072 | 0.3471 | -4.5845 | 4.6E-06 | 5.08E-05 |
| 186 | CREG1 | 14.2060714 | -1.591659499 | 0.5071 | -3.1385 | 0.0017 | 0.009045 |
| 1740 | DNPH1 | 39.8688229 | -1.592125214 | 0.3668 | -4.3405 | 1.4E-05 | 0.00014 |
| 1911 | PARP12 | 16.5236338 | -1.59610412 | 0.5788 | -2.7576 | 0.00582 | 0.025518 |
| 786 | FBRS | 18.0541458 | -1.598260707 | 0.4928 | -3.2433 | 0.00118 | 0.006606 |
| 687 | LPCAT4 | 18.6281459 | -1.607841704 | 0.4823 | -3.3337 | 0.00086 | 0.005062 |
| 990 | RELCH | 15.5894264 | -1.610030288 | 0.5165 | -3.1174 | 0.00182 | 0.009612 |
| 785 | MAPK3 | 53.0135126 | -1.611216587 | 0.3329 | -4.8403 | 1.3E-06 | 1.66E-05 |
| 831 | VPS9D1 | 12.2108794 | -1.612591919 | 0.6252 | -2.5795 | 0.00989 | 0.039074 |
| 816 | MON1B | 16.6585966 | -1.613764533 | 0.6129 | -2.6329 | 0.00847 | 0.034571 |
| 555 | GOLGA2P5 | 27.6753711 | -1.614405322 | 0.6193 | -2.6068 | 0.00914 | 0.036761 |
| 1680 | TBC1D9B | 20.5973673 | -1.616949124 | 0.473 | -3.4186 | 0.00063 | 0.003889 |
| 1004 | SPPL2B | 33.7360984 | -1.617590847 | 0.3818 | -4.2365 | 2.3E-05 | 0.000212 |
| 684 | CYFIP1 | 29.071835 | -1.621698756 | 0.396 | -4.0956 | 4.2E-05 | 0.00037 |
| 1600 | EXOC3 | 17.8881243 | -1.622215245 | 0.5307 | -3.0565 | 0.00224 | 0.011468 |
| 1153 | AHSA2P | 33.7680655 | -1.623616122 | 0.3919 | -4.1428 | 3.4E-05 | 0.000308 |
| 546 | PAWR | 32.1000621 | -1.624175506 | 0.4375 | -3.7126 | 0.00021 | 0.001463 |
| 1574 | NDUFC1 | 43.5469028 | -1.627735367 | 0.4297 | -3.7879 | 0.00015 | 0.001127 |
| 1696 | MCUR1 | 18.6511736 | -1.627926873 | 0.4479 | -3.6348 | 0.00028 | 0.001914 |
| 989 | MALT1 | 10.9914208 | -1.628259936 | 0.6379 | -2.5527 | 0.01069 | 0.041607 |
| 1189 | SLC35F5 | 26.4262942 | -1.628651082 | 0.4831 | -3.3712 | 0.00075 | 0.004513 |
| 96 | HOOK1 | 18.6208003 | -1.628976425 | 0.4772 | -3.4136 | 0.00064 | 0.003955 |
| 1491 | TNIP2 | 9.6656066 | -1.631963884 | 0.6145 | -2.6559 | 0.00791 | 0.03269 |
| 1242 | GMPPA | 13.5838683 | -1.632244067 | 0.6032 | -2.706 | 0.00681 | 0.028994 |
| 1809 | EZR | 56.9706933 | -1.632641921 | 0.4144 | -3.9393 | 8.2E-05 | 0.000659 |
| 232 | CAPN2 | 33.1782877 | -1.632688064 | 0.3697 | -4.4159 | 1E-05 | 0.000103 |
| 658 | GALC | 21.5440806 | -1.63489397 | 0.5905 | -2.7686 | 0.00563 | 0.024853 |
| 1942 | CCDC25 | 11.8595461 | -1.638157591 | 0.6096 | -2.6874 | 0.0072 | 0.030322 |
| 278 | TSPAN14 | 20.1589544 | -1.640663002 | 0.5266 | -3.1159 | 0.00183 | 0.009656 |
| 434 | SRSF8 | 18.3304065 | -1.641082727 | 0.5541 | -2.9617 | 0.00306 | 0.014903 |
| 865 | ALDH3A2 | 27.3583775 | -1.641274675 | 0.4918 | -3.3371 | 0.00085 | 0.005009 |
| 752 | WDR90 | 25.8028084 | -1.643438442 | 0.4385 | -3.7483 | 0.00018 | 0.001292 |
| 1778 | MFSD4B | 17.1327948 | -1.647534168 | 0.5443 | -3.0266 | 0.00247 | 0.012457 |
| 347 | TOLLIP | 12.0373014 | -1.647663585 | 0.5754 | -2.8635 | 0.00419 | 0.01944 |
| 688 | SPRED1 | 12.964355 | -1.648062309 | 0.66 | -2.497 | 0.01253 | 0.047426 |
| 1932 | ASAH1 | 41.6472837 | -1.650059464 | 0.3551 | -4.6466 | 3.4E-06 | 3.9E-05 |
| 541 | RAP1B | 32.9958544 | -1.654582989 | 0.4276 | -3.8694 | 0.00011 | 0.00085 |
| 77 | PTPRF | 101.584071 | -1.655129937 | 0.2961 | -5.5902 | 2.3E-08 | 4.49E-07 |
| 952 | ACOX1 | 18.2367221 | -1.655833403 | 0.584 | -2.8351 | 0.00458 | 0.020925 |

| | | | | | | | |
|---|---|---|---|---|---|---|---|
| 280 | CDHR1 | 22.4947479 | -1.656413404 | 0.5649 | -2.932 | 0.00337 | 0.016162 |
| 1571 | ANXA5 | 21.4712615 | -1.658033686 | 0.553 | -2.9983 | 0.00271 | 0.013493 |
| 1323 | PTTG1IP | 44.7662829 | -1.658207023 | 0.3198 | -5.1854 | 2.2E-07 | 3.34E-06 |
| 247 | OPTN | 24.5826323 | -1.661052015 | 0.4499 | -3.6922 | 0.00022 | 0.001572 |
| 183 | MGST3 | 48.1176906 | -1.661261222 | 0.3318 | -5.0062 | 5.6E-07 | 7.87E-06 |
| 301 | ERLIN1 | 11.6762039 | -1.662303696 | 0.566 | -2.9371 | 0.00331 | 0.015929 |
| 1677 | LMAN2 | 24.9812639 | -1.664617018 | 0.4285 | -3.8847 | 0.0001 | 0.000803 |
| 570 | MLEC | 72.5392045 | -1.670535383 | 0.2798 | -5.9704 | 2.4E-09 | 5.7E-08 |
| 808 | CMTR2 | 14.8804354 | -1.670541987 | 0.5217 | -3.202 | 0.00136 | 0.00752 |
| 295 | PDLIM1 | 95.8773148 | -1.670564841 | 0.2728 | -6.1236 | 9.1E-10 | 2.4E-08 |
| 112 | DDAH1 | 63.254923 | -1.67541622 | 0.3104 | -5.3973 | 6.8E-08 | 1.19E-06 |
| 614 | NDRG2 | 24.2238331 | -1.675583685 | 0.5262 | -3.1842 | 0.00145 | 0.007907 |
| 150 | GOLPH3L | 15.3085716 | -1.67622645 | 0.58 | -2.8902 | 0.00385 | 0.018124 |
| 1212 | ITPRID2 | 20.1309217 | -1.676977984 | 0.4738 | -3.5397 | 0.0004 | 0.002638 |
| 102 | JAK1 | 19.6131086 | -1.677506924 | 0.4455 | -3.7653 | 0.00017 | 0.00122 |
| 1407 | FOXP1 | 47.2844762 | -1.678284873 | 0.3697 | -4.5398 | 5.6E-06 | 6.12E-05 |
| 1502 | UGDH | 31.380531 | -1.682967995 | 0.3781 | -4.4511 | 8.5E-06 | 8.92E-05 |
| 539 | HMGA2-AS1 | 10.4297802 | -1.686436421 | 0.6728 | -2.5065 | 0.01219 | 0.046459 |
| 970 | MYL12B | 54.7975529 | -1.687132327 | 0.329 | -5.1273 | 2.9E-07 | 4.42E-06 |
| 134 | MOV10 | 15.203709 | -1.688820581 | 0.6123 | -2.7583 | 0.00581 | 0.025479 |
| 1895 | SYPL1 | 30.5151058 | -1.692447571 | 0.525 | -3.224 | 0.00126 | 0.007027 |
| 1788 | HINT3 | 12.6972427 | -1.692458067 | 0.6014 | -2.8144 | 0.00489 | 0.022144 |
| 846 | ANKFY1 | 10.1912688 | -1.693155338 | 0.6827 | -2.48 | 0.01314 | 0.049295 |
| 830 | CDK10 | 36.0750327 | -1.69345075 | 0.4451 | -3.8051 | 0.00014 | 0.001064 |
| 640 | PLEKHG3 | 21.6218916 | -1.696055508 | 0.5044 | -3.3628 | 0.00077 | 0.004624 |
| 2095 | AKAP17A | 23.7015835 | -1.696703158 | 0.4756 | -3.5678 | 0.00036 | 0.0024 |
| 1938 | DOCK5 | 18.5233787 | -1.697321245 | 0.616 | -2.7552 | 0.00587 | 0.025681 |
| 1297 | ZMYND8 | 21.9299664 | -1.698297518 | 0.4686 | -3.6239 | 0.00029 | 0.001987 |
| 56 | RPS6KA1 | 18.9430313 | -1.701857841 | 0.588 | -2.8942 | 0.0038 | 0.017951 |
| 1923 | DYNC2I1 | 18.0565911 | -1.702893935 | 0.4592 | -3.7084 | 0.00021 | 0.001484 |
| 754 | FAHD1 | 16.1793732 | -1.704563902 | 0.5199 | -3.2786 | 0.00104 | 0.005958 |
| 2092 | MT-ND6 | 406.21143 | -1.707844757 | 0.2352 | -7.2623 | 3.8E-13 | 2.13E-11 |
| 384 | CTNND1 | 63.0336106 | -1.708588646 | 0.3488 | -4.8992 | 9.6E-07 | 1.28E-05 |
| 1613 | PRKAA1 | 22.6155853 | -1.710896836 | 0.4383 | -3.9032 | 9.5E-05 | 0.00075 |
| 100 | ALG6 | 9.40515139 | -1.71194368 | 0.6899 | -2.4814 | 0.01309 | 0.049133 |
| 1032 | JUNB | 8.41195546 | -1.711969199 | 0.6795 | -2.5194 | 0.01176 | 0.045016 |
| 1305 | SLC17A9 | 29.1696395 | -1.716723327 | 0.4058 | -4.2301 | 2.3E-05 | 0.000218 |
| 1441 | TFDP2 | 18.9757754 | -1.718293389 | 0.4806 | -3.575 | 0.00035 | 0.002345 |
| 1110 | MBOAT7 | 24.0430244 | -1.722203205 | 0.424 | -4.0619 | 4.9E-05 | 0.000419 |

| | | | | | | | |
|---|---|---|---|---|---|---|---|
| 631 | ARF6 | 41.5109218 | -1.726842673 | 0.3906 | -4.4215 | 9.8E-06 | 0.000101 |
| 1958 | RDH10 | 14.9450564 | -1.726898693 | 0.5942 | -2.906 | 0.00366 | 0.017369 |
| 1415 | CBLB | 8.53955412 | -1.72857843 | 0.6975 | -2.4782 | 0.01321 | 0.049458 |
| 416 | DHCR7 | 20.6575228 | -1.730041389 | 0.6219 | -2.7819 | 0.0054 | 0.024052 |
| 1309 | TMEM50B | 14.0132646 | -1.730753235 | 0.6242 | -2.7729 | 0.00556 | 0.024586 |
| 1488 | C4orf48 | 17.1995817 | -1.731366417 | 0.5947 | -2.9112 | 0.0036 | 0.017124 |
| 1681 | ZFP62 | 11.8128532 | -1.737007042 | 0.6716 | -2.5865 | 0.0097 | 0.038406 |
| 1688 | ECI2 | 21.5235552 | -1.737027975 | 0.4252 | -4.0857 | 4.4E-05 | 0.000385 |
| 1387 | PLXNB1 | 39.3507902 | -1.739088057 | 0.3563 | -4.8809 | 1.1E-06 | 1.39E-05 |
| 1181 | GCC2 | 30.5020563 | -1.739856167 | 0.4659 | -3.7343 | 0.00019 | 0.001357 |
| 91 | TMEM59 | 51.7034205 | -1.741894708 | 0.3583 | -4.861 | 1.2E-06 | 1.51E-05 |
| 1849 | AUTS2 | 19.1181349 | -1.742051337 | 0.5229 | -3.3315 | 0.00086 | 0.005091 |
| 1955 | SGK3 | 10.5068841 | -1.745561484 | 0.7033 | -2.4821 | 0.01306 | 0.049063 |
| 1611 | NADK2 | 17.8033985 | -1.745761905 | 0.5441 | -3.2083 | 0.00134 | 0.007376 |
| 473 | APLP2 | 155.555196 | -1.749900413 | 0.2601 | -6.7271 | 1.7E-11 | 6.93E-10 |
| 718 | RPS27L | 67.010154 | -1.753848803 | 0.278 | -6.3088 | 2.8E-10 | 8.37E-09 |
| 957 | LGALS3BP | 121.432692 | -1.75413964 | 0.2772 | -6.3292 | 2.5E-10 | 7.4E-09 |
| 1104 | EMC10 | 12.3863692 | -1.75628644 | 0.6467 | -2.7159 | 0.00661 | 0.028311 |
| 593 | TPT1-AS1 | 21.1582856 | -1.75771229 | 0.4945 | -3.5548 | 0.00038 | 0.002511 |
| 1586 | TRIM2 | 41.2243565 | -1.75798345 | 0.3578 | -4.9136 | 8.9E-07 | 1.2E-05 |
| 1321 | CSTB | 38.9559715 | -1.758420841 | 0.3693 | -4.7613 | 1.9E-06 | 2.36E-05 |
| 653 | TMED10 | 47.2891144 | -1.758454406 | 0.412 | -4.2677 | 2E-05 | 0.000189 |
| 366 | HTATIP2 | 16.107878 | -1.75920315 | 0.5293 | -3.3233 | 0.00089 | 0.005218 |
| 928 | TRIM25 | 25.9718066 | -1.76076233 | 0.4152 | -4.2404 | 2.2E-05 | 0.000209 |
| 1548 | DNAJB14 | 15.8869491 | -1.766743682 | 0.5445 | -3.2447 | 0.00118 | 0.006581 |
| 1130 | NCOA1 | 15.6101629 | -1.767548337 | 0.5043 | -3.5046 | 0.00046 | 0.002951 |
| 1566 | UGT8 | 30.3967016 | -1.769769125 | 0.5114 | -3.4609 | 0.00054 | 0.003402 |
| 1464 | ALG3 | 17.1146375 | -1.770750018 | 0.5601 | -3.1613 | 0.00157 | 0.008456 |
| 1619 | SMIM15 | 8.21655976 | -1.771660298 | 0.7062 | -2.5088 | 0.01211 | 0.046211 |
| 2089 | MT-CO2 | 719.77367 | -1.776011035 | 0.2118 | -8.3866 | 5E-17 | 6.03E-15 |
| 1953 | CHCHD7 | 14.5884617 | -1.776430902 | 0.5261 | -3.3769 | 0.00073 | 0.004432 |
| 276 | USP54 | 14.8936619 | -1.78249066 | 0.5614 | -3.1749 | 0.0015 | 0.008127 |
| 856 | TMEM256 | 17.520173 | -1.785684578 | 0.5521 | -3.2344 | 0.00122 | 0.006794 |
| 1462 | TBL1XR1 | 47.0219203 | -1.7896678 | 0.3687 | -4.8538 | 1.2E-06 | 1.57E-05 |
| 2064 | RXRA | 17.9054017 | -1.790184861 | 0.5115 | -3.4998 | 0.00047 | 0.003 |
| 480 | CD9 | 53.0476921 | -1.791429969 | 0.3828 | -4.6798 | 2.9E-06 | 3.38E-05 |
| 1229 | FZD5 | 12.8419604 | -1.794096516 | 0.597 | -3.0049 | 0.00266 | 0.013249 |
| 1781 | NT5DC1 | 12.6577994 | -1.794829217 | 0.7002 | -2.5633 | 0.01037 | 0.040572 |
| 937 | ۸P27P1-BPTFP1-KP | 23.4817675 | -1.796389309 | 0.5515 | -3.2573 | 0.00112 | 0.006344 |

| | | | | | | | |
|---|---|---|---|---|---|---|---|
| 719 | CA12 | 37.7803473 | -1.802238808 | 0.4206 | -4.2849 | 1.8E-05 | 0.000176 |
| 584 | UBL3 | 15.453744 | -1.804228413 | 0.5297 | -3.406 | 0.00066 | 0.004042 |
| 755 | MSRB1 | 10.5086222 | -1.805111704 | 0.6807 | -2.652 | 0.008 | 0.033006 |
| 1898 | LAMB1 | 47.3296581 | -1.808210343 | 0.316 | -5.7229 | 1E-08 | 2.22E-07 |
| 1817 | COX19 | 16.0904673 | -1.811703039 | 0.5299 | -3.4187 | 0.00063 | 0.003889 |
| 1300 | PARD6B | 10.233396 | -1.814860011 | 0.6413 | -2.8301 | 0.00465 | 0.021207 |
| 1789 | ARHGAP18 | 15.3541062 | -1.815896696 | 0.6099 | -2.9773 | 0.00291 | 0.014285 |
| 697 | TMEM87A | 25.3842676 | -1.818031957 | 0.4386 | -4.145 | 3.4E-05 | 0.000305 |
| 621 | PSME2 | 63.0692404 | -1.819695913 | 0.289 | -6.2975 | 3E-10 | 8.93E-09 |
| 106 | WLS | 14.3604614 | -1.825780918 | 0.551 | -3.3133 | 0.00092 | 0.005372 |
| 869 | FAM222B | 10.5121076 | -1.826367852 | 0.671 | -2.7218 | 0.00649 | 0.027891 |
| 405 | CORO1B | 13.2396434 | -1.828318562 | 0.572 | -3.1962 | 0.00139 | 0.007639 |
| 904 | HEXIM1 | 7.96916366 | -1.828465756 | 0.7141 | -2.5606 | 0.01045 | 0.040841 |
| 2105 | DMD | 12.1986424 | -1.832765366 | 0.6227 | -2.9431 | 0.00325 | 0.015686 |
| 1269 | LZTS3 | 9.95031001 | -1.836222728 | 0.6735 | -2.7263 | 0.0064 | 0.027564 |
| 941 | SLC39A11 | 9.8743184 | -1.838499514 | 0.6565 | -2.8007 | 0.0051 | 0.022899 |
| 1620 | PIK3R1 | 11.9172991 | -1.845518197 | 0.6219 | -2.9674 | 0.003 | 0.014683 |
| 1381 | GORASP1 | 10.9555124 | -1.845816164 | 0.6539 | -2.823 | 0.00476 | 0.021617 |
| 477 | B4GALNT3 | 12.917548 | -1.847977702 | 0.7114 | -2.5977 | 0.00938 | 0.037491 |
| 83 | GPBP1L1 | 25.946618 | -1.848817196 | 0.3991 | -4.6329 | 3.6E-06 | 4.14E-05 |
| 1002 | MKNK2 | 23.6517621 | -1.851064078 | 0.4447 | -4.1622 | 3.2E-05 | 0.000285 |
| 963 | SLC16A3 | 23.9618224 | -1.859828313 | 0.7354 | -2.529 | 0.01144 | 0.043948 |
| 2059 | TBC1D13 | 8.70793504 | -1.860371021 | 0.693 | -2.6847 | 0.00726 | 0.030499 |
| 1439 | ZBTB38 | 16.5266771 | -1.86046665 | 0.4978 | -3.7374 | 0.00019 | 0.001342 |
| 2055 | GSN | 20.9666357 | -1.860899427 | 0.5537 | -3.3606 | 0.00078 | 0.004657 |
| 240 | SCCPDH | 12.4448352 | -1.86162709 | 0.643 | -2.8953 | 0.00379 | 0.017895 |
| 1132 | TMEM214 | 22.4695225 | -1.866890775 | 0.5587 | -3.3413 | 0.00083 | 0.004942 |
| 1447 | SELENOT | 15.6032077 | -1.867639668 | 0.6592 | -2.833 | 0.00461 | 0.021052 |
| 130 | GSTM4 | 13.2681683 | -1.868284529 | 0.6131 | -3.0473 | 0.00231 | 0.011759 |
| 678 | ANKRD9 | 17.5942703 | -1.870498036 | 0.5095 | -3.6711 | 0.00024 | 0.001686 |
| 1126 | SDC1 | 26.6879179 | -1.871737135 | 0.4302 | -4.3504 | 1.4E-05 | 0.000135 |
| 18 | VAMP3 | 23.9199861 | -1.872141207 | 0.4752 | -3.9398 | 8.2E-05 | 0.000658 |
| 1376 | CMTM6 | 11.1258439 | -1.875582612 | 0.6419 | -2.922 | 0.00348 | 0.016634 |
| 2045 | PTPN3 | 17.4198481 | -1.878396929 | 0.6551 | -2.8675 | 0.00414 | 0.019231 |
| 745 | IQGAP1 | 27.3764812 | -1.87870166 | 0.4011 | -4.6844 | 2.8E-06 | 3.32E-05 |
| 197 | EDEM3 | 25.257095 | -1.882331558 | 0.4965 | -3.7914 | 0.00015 | 0.001114 |
| 589 | ELF1 | 12.6725854 | -1.882854048 | 0.5861 | -3.2127 | 0.00131 | 0.007286 |
| 1145 | EML4 | 38.6038645 | -1.887138249 | 0.4277 | -4.4123 | 1E-05 | 0.000105 |
| 1326 | CHCHD10 | 71.5527876 | -1.894864702 | 0.3183 | -5.9531 | 2.6E-09 | 6.23E-08 |

| | | | | | | | |
|---|---|---|---|---|---|---|---|
| 1049 | IPO5P1 | 15.0512652 | -1.895682628 | 0.5037 | -3.7638 | 0.00017 | 0.001226 |
| 668 | CHGA | 95.9296658 | -1.899373031 | 0.6814 | -2.7874 | 0.00531 | 0.023723 |
| 842 | TAX1BP3 | 14.7020072 | -1.905806574 | 0.5699 | -3.3443 | 0.00083 | 0.004892 |
| 682 | INF2 | 24.1645033 | -1.906783888 | 0.4862 | -3.9219 | 8.8E-05 | 0.000703 |
| 381 | ACP2 | 13.2651161 | -1.910689716 | 0.7312 | -2.613 | 0.00897 | 0.036267 |
| 2071 | MAMDC4 | 21.6276381 | -1.912804101 | 0.5979 | -3.1994 | 0.00138 | 0.007575 |
| 1355 | APOBEC3C | 7.25909649 | -1.91511862 | 0.7477 | -2.5613 | 0.01043 | 0.040791 |
| 1731 | STK38 | 18.8447045 | -1.924759534 | 0.5065 | -3.7999 | 0.00014 | 0.001082 |
| 983 | ACAA2 | 49.4257484 | -1.925800993 | 0.4008 | -4.8052 | 1.5E-06 | 1.94E-05 |
| 39 | LUZP1 | 15.9859175 | -1.926610948 | 0.5623 | -3.4265 | 0.00061 | 0.003801 |
| 1165 | SUCLG1 | 62.7857006 | -1.927417358 | 0.3064 | -6.2912 | 3.2E-10 | 9.26E-09 |
| 521 | CSAD | 17.1734591 | -1.93286517 | 0.5274 | -3.6648 | 0.00025 | 0.001724 |
| 974 | ABHD3 | 13.3404853 | -1.933466367 | 0.5937 | -3.2569 | 0.00113 | 0.006349 |
| 1334 | ASCC2 | 10.4400159 | -1.938816216 | 0.6843 | -2.8332 | 0.00461 | 0.021048 |
| 811 | ZFHX3 | 13.4347234 | -1.939659002 | 0.6655 | -2.9145 | 0.00356 | 0.016972 |
| 50 | TMEM50A | 19.7747682 | -1.940414095 | 0.5065 | -3.831 | 0.00013 | 0.000976 |
| 2015 | GALT | 10.834217 | -1.944366073 | 0.6188 | -3.1421 | 0.00168 | 0.008946 |
| 729 | NPTN | 44.6718989 | -1.944979783 | 0.4682 | -4.154 | 3.3E-05 | 0.000294 |
| 2050 | HSDL2 | 12.2616906 | -1.946604713 | 0.6286 | -3.0966 | 0.00196 | 0.010198 |
| 1003 | PLEKHJ1 | 52.0925195 | -1.94865145 | 0.3507 | -5.556 | 2.8E-08 | 5.36E-07 |
| 1750 | CILK1 | 11.8891163 | -1.949314459 | 0.7096 | -2.7471 | 0.00601 | 0.026226 |
| 1637 | CDC42SE2 | 15.4123061 | -1.952840662 | 0.5254 | -3.7167 | 0.0002 | 0.001445 |
| 2116 | CLCN5 | 16.4482283 | -1.954626524 | 0.6009 | -3.2527 | 0.00114 | 0.006419 |
| 822 | ZDHHC7 | 19.8486338 | -1.960292148 | 0.5094 | -3.848 | 0.00012 | 0.00092 |
| 1322 | PFKL | 61.4564884 | -1.960315516 | 0.3709 | -5.286 | 1.3E-07 | 2.06E-06 |
| 624 | DHRS1 | 8.67912663 | -1.960450452 | 0.6778 | -2.8924 | 0.00382 | 0.018042 |
| 1031 | HOOK2 | 19.2081078 | -1.966591215 | 0.5044 | -3.8985 | 9.7E-05 | 0.000763 |
| 537 | GNS | 24.9640932 | -1.967364731 | 0.5567 | -3.534 | 0.00041 | 0.002685 |
| 195 | LAMC1 | 11.3131494 | -1.968516196 | 0.7328 | -2.6864 | 0.00722 | 0.03041 |
| 7 | CPTP | 7.02948219 | -1.973884343 | 0.7665 | -2.5754 | 0.01001 | 0.039453 |
| 296 | ALDH18A1 | 48.0005915 | -1.974956434 | 0.3448 | -5.7278 | 1E-08 | 2.16E-07 |
| 406 | TMEM134 | 9.54418404 | -1.978035817 | 0.735 | -2.6911 | 0.00712 | 0.030041 |
| 1377 | GLB1 | 18.3528238 | -1.978098882 | 0.5532 | -3.5757 | 0.00035 | 0.00234 |
| 560 | GLTP | 8.34692068 | -1.97862249 | 0.7901 | -2.5044 | 0.01227 | 0.046656 |
| 1814 | ERMARD | 10.4222048 | -1.981740734 | 0.6808 | -2.9111 | 0.0036 | 0.017124 |
| 709 | RAB27A | 9.8886163 | -1.98219822 | 0.7494 | -2.6452 | 0.00816 | 0.033549 |
| 606 | TM9SF2 | 53.7642793 | -1.986426634 | 0.316 | -6.2868 | 3.2E-10 | 9.49E-09 |
| 2018 | EBLN3P | 26.8932414 | -1.990292136 | 0.4874 | -4.0835 | 4.4E-05 | 0.000388 |
| 669 | ITPK1 | 18.4119542 | -1.992043894 | 0.5052 | -3.9428 | 8.1E-05 | 0.000652 |

| | | | | | | | |
|---|---|---|---|---|---|---|---|
| 188 | MPC2 | 46.4390202 | -1.995496845 | 0.3638 | -5.4854 | 4.1E-08 | 7.63E-07 |
| 875 | TMEM98 | 28.6241948 | -1.998322756 | 0.583 | -3.4279 | 0.00061 | 0.003783 |
| 594 | ITM2B | 80.2059632 | -2.000898805 | 0.301 | -6.6481 | 3E-11 | 1.14E-09 |
| 2099 | MID1 | 15.8378557 | -2.001311969 | 0.6033 | -3.3172 | 0.00091 | 0.00531 |
| 484 | LTBR | 21.2009236 | -2.001328589 | 0.503 | -3.9791 | 6.9E-05 | 0.000572 |
| 1079 | PVR | 14.9417811 | -2.005380935 | 0.5677 | -3.5328 | 0.00041 | 0.002691 |
| 1159 | PCYOX1 | 16.1968713 | -2.008037347 | 0.535 | -3.7535 | 0.00017 | 0.001268 |
| 1879 | TSC22D4 | 11.8932416 | -2.01046814 | 0.6194 | -3.246 | 0.00117 | 0.006555 |
| 2100 | TMSB4X | 143.335206 | -2.012104175 | 0.3039 | -6.6212 | 3.6E-11 | 1.33E-09 |
| 1596 | GALNT7 | 22.2514087 | -2.016222355 | 0.4327 | -4.6592 | 3.2E-06 | 3.7E-05 |
| 190 | FMO1 | 8.50252133 | -2.016250749 | 0.799 | -2.5236 | 0.01162 | 0.044525 |
| 446 | FDX1 | 15.5985366 | -2.022854324 | 0.5426 | -3.7279 | 0.00019 | 0.001389 |
| 850 | KIF1C | 32.0342984 | -2.023536188 | 0.429 | -4.7169 | 2.4E-06 | 2.89E-05 |
| 1259 | RNPEPL1 | 16.1341212 | -2.025715856 | 0.5592 | -3.6224 | 0.00029 | 0.001997 |
| 1890 | CLDN15 | 20.4475413 | -2.027773506 | 0.5056 | -4.0105 | 6.1E-05 | 0.000509 |
| 226 | LPGAT1 | 17.1581525 | -2.028500883 | 0.6434 | -3.1529 | 0.00162 | 0.008672 |
| 1760 | SLC17A5 | 8.6730859 | -2.029216043 | 0.6846 | -2.9641 | 0.00304 | 0.014817 |
| 993 | CNDP2 | 40.5785669 | -2.031505471 | 0.4802 | -4.2303 | 2.3E-05 | 0.000218 |
| 749 | FAM234A | 19.0212311 | -2.032904655 | 0.5159 | -3.9405 | 8.1E-05 | 0.000657 |
| 969 | MYL12A | 47.616477 | -2.036055546 | 0.3328 | -6.1175 | 9.5E-10 | 2.48E-08 |
| 1351 | LGALS1 | 16.5797515 | -2.039575038 | 0.6463 | -3.1557 | 0.0016 | 0.008597 |
| 1069 | ITPKC | 7.5280901 | -2.042864151 | 0.7353 | -2.7781 | 0.00547 | 0.024284 |
| 1936 | LOXL2 | 6.41104035 | -2.043156854 | 0.7763 | -2.6318 | 0.00849 | 0.034645 |
| 1350 | SH3BP1 | 12.9950468 | -2.047783731 | 0.6008 | -3.4085 | 0.00065 | 0.004009 |
| 1772 | PREP | 26.6298548 | -2.050276301 | 0.4252 | -4.822 | 1.4E-06 | 1.81E-05 |
| 1144 | PKDCC | 12.2959303 | -2.052329577 | 0.575 | -3.5695 | 0.00036 | 0.002389 |
| 2160 | MOSPD1 | 9.43878766 | -2.055849144 | 0.7113 | -2.8904 | 0.00385 | 0.018119 |
| 141 | IGSF3 | 18.9638733 | -2.066504365 | 0.5411 | -3.8193 | 0.00013 | 0.001014 |
| 1402 | FLNB | 33.8200105 | -2.074704933 | 0.4424 | -4.6892 | 2.7E-06 | 3.24E-05 |
| 601 | SPRY2 | 9.97481882 | -2.077187706 | 0.7207 | -2.882 | 0.00395 | 0.018501 |
| 1856 | POR | 15.8040055 | -2.079333593 | 0.5954 | -3.4922 | 0.00048 | 0.003074 |
| 1179 | NCK2 | 18.8001763 | -2.081607073 | 0.5955 | -3.4955 | 0.00047 | 0.003041 |
| 1343 | MYH9 | 57.805342 | -2.084195961 | 0.3289 | -6.3373 | 2.3E-10 | 7.05E-09 |
| 182 | UHMK1 | 20.9408225 | -2.085749058 | 0.5381 | -3.8764 | 0.00011 | 0.000827 |
| 898 | NBR1 | 30.5518645 | -2.087165929 | 0.48 | -4.3487 | 1.4E-05 | 0.000136 |
| 1366 | ARSA | 18.0042358 | -2.089908836 | 0.5339 | -3.9148 | 9E-05 | 0.00072 |
| 262 | IPMK | 6.14660304 | -2.093705319 | 0.8459 | -2.475 | 0.01332 | 0.049843 |
| 274 | MCU | 8.52512844 | -2.096485974 | 0.6908 | -3.0349 | 0.00241 | 0.012184 |
| 191 | RABGAP1L | 18.4694641 | -2.102148218 | 0.5596 | -3.7568 | 0.00017 | 0.001255 |

| | | | | | | | |
|---|---|---|---|---|---|---|---|
| 498 | H2AJ | 8.01324391 | -2.103220195 | 0.7781 | -2.7032 | 0.00687 | 0.029174 |
| 674 | CLMN | 15.8058804 | -2.103525396 | 0.6336 | -3.3198 | 0.0009 | 0.005268 |
| 1039 | OCEL1 | 10.1818004 | -2.103673756 | 0.7625 | -2.7588 | 0.0058 | 0.025455 |
| 220 | CD46 | 56.4186577 | -2.105475101 | 0.3376 | -6.2367 | 4.5E-10 | 1.27E-08 |
| 1739 | KLC4 | 16.6828811 | -2.107447149 | 0.5575 | -3.7803 | 0.00016 | 0.001156 |
| 1367 | SUMF1 | 8.47413145 | -2.112705408 | 0.6806 | -3.1041 | 0.00191 | 0.00998 |
| 170 | LMNA | 20.1339525 | -2.112826455 | 0.4873 | -4.3355 | 1.5E-05 | 0.000143 |
| 738 | MESD | 20.1757849 | -2.11510865 | 0.49 | -4.3167 | 1.6E-05 | 0.000155 |
| 936 | TEX2 | 12.8501666 | -2.116775223 | 0.6734 | -3.1436 | 0.00167 | 0.008905 |
| 1973 | RIDA | 9.51894427 | -2.117467456 | 0.7059 | -2.9999 | 0.0027 | 0.013434 |
| 861 | PIGL | 6.34092604 | -2.121037956 | 0.7934 | -2.6734 | 0.00751 | 0.031331 |
| 1119 | ADI1 | 29.351855 | -2.122094401 | 0.386 | -5.4971 | 3.9E-08 | 7.2E-07 |
| 1743 | TNFRSF21 | 7.97763039 | -2.123594847 | 0.7825 | -2.7139 | 0.00665 | 0.028473 |
| 68 | SH3D21 | 10.4207291 | -2.123782095 | 0.7845 | -2.7072 | 0.00678 | 0.028931 |
| 2133 | HMGN5 | 11.099438 | -2.128451887 | 0.7662 | -2.7781 | 0.00547 | 0.024284 |
| 1107 | ZNF600 | 10.7361362 | -2.128702118 | 0.6877 | -3.0955 | 0.00196 | 0.010231 |
| 1397 | ABHD14B | 18.8238192 | -2.130922794 | 0.5766 | -3.696 | 0.00022 | 0.001552 |
| 377 | ACCS | 14.3346108 | -2.13106867 | 0.6543 | -3.2572 | 0.00113 | 0.006344 |
| 1861 | CROT | 11.5114929 | -2.131421241 | 0.6338 | -3.3628 | 0.00077 | 0.004624 |
| 1283 | PYGB | 38.9130092 | -2.135143282 | 0.4534 | -4.7088 | 2.5E-06 | 2.99E-05 |
| 1807 | SNX9 | 7.29094845 | -2.136926998 | 0.7985 | -2.6762 | 0.00745 | 0.031108 |
| 37 | ECE1 | 14.0433195 | -2.13746053 | 0.5656 | -3.7789 | 0.00016 | 0.001161 |
| 698 | SNAP23 | 11.6586659 | -2.13868482 | 0.8483 | -2.5212 | 0.0117 | 0.044809 |
| 1298 | SULF2 | 24.2189938 | -2.143086086 | 0.4533 | -4.7275 | 2.3E-06 | 2.75E-05 |
| 1486 | DGKQ | 17.8870214 | -2.144116043 | 0.6132 | -3.4964 | 0.00047 | 0.003033 |
| 927 | TOB1 | 23.1807182 | -2.144476308 | 0.5648 | -3.7968 | 0.00015 | 0.001092 |
| 2079 | EXD3 | 7.51497171 | -2.147664478 | 0.8522 | -2.5202 | 0.01173 | 0.044928 |
| 1949 | PLPP5 | 21.5211509 | -2.150173011 | 0.458 | -4.6951 | 2.7E-06 | 3.16E-05 |
| 1302 | ZNF217 | 9.88319984 | -2.15398066 | 0.6685 | -3.2222 | 0.00127 | 0.007068 |
| 524 | CD63 | 136.652912 | -2.155144296 | 0.2398 | -8.9875 | 2.5E-19 | 4.74E-17 |
| 611 | TEP1 | 13.6887994 | -2.155777698 | 0.5903 | -3.6522 | 0.00026 | 0.0018 |
| 707 | SPPL2A | 19.2505971 | -2.156289669 | 0.4761 | -4.5289 | 5.9E-06 | 6.41E-05 |
| 1359 | PACSIN2 | 14.5239245 | -2.159117134 | 0.5706 | -3.7841 | 0.00015 | 0.001141 |
| 41 | GALE | 34.0798619 | -2.159357124 | 0.4177 | -5.17 | 2.3E-07 | 3.6E-06 |
| 1799 | ABRACL | 29.9583286 | -2.162208594 | 0.4234 | -5.1067 | 3.3E-07 | 4.87E-06 |
| 538 | HMGA2 | 23.7568304 | -2.162824663 | 0.6097 | -3.5474 | 0.00039 | 0.00257 |
| 1770 | TSTD3 | 7.60546127 | -2.168166569 | 0.8584 | -2.5257 | 0.01155 | 0.0443 |
| 1392 | GMPPB | 8.39257577 | -2.170254851 | 0.8391 | -2.5864 | 0.0097 | 0.038409 |
| 1281 | CST3 | 234.049792 | -2.170982734 | 0.2511 | -8.6459 | 5.3E-18 | 8.08E-16 |

| | | | | | | | |
|---|---|---|---|---|---|---|---|
| 1538 | NUDT9 | 9.32544584 | -2.172221106 | 0.7422 | -2.9268 | 0.00342 | 0.016404 |
| 166 | SLC50A1 | 18.8476683 | -2.172734938 | 0.532 | -4.0842 | 4.4E-05 | 0.000387 |
| 553 | NUDT4 | 12.6297351 | -2.175871392 | 0.6226 | -3.4949 | 0.00047 | 0.003046 |
| 2148 | AMOT | 15.5154912 | -2.176560284 | 0.5687 | -3.8271 | 0.00013 | 0.000989 |
| 1712 | HLA-C | 72.7341181 | -2.178263764 | 0.3225 | -6.7553 | 1.4E-11 | 5.84E-10 |
| 764 | NAGPA | 6.81559252 | -2.179262586 | 0.8524 | -2.5567 | 0.01057 | 0.04124 |
| 387 | STX3 | 15.7771743 | -2.179763724 | 0.588 | -3.7069 | 0.00021 | 0.001492 |
| 348 | CTSD | 42.7870754 | -2.190244854 | 0.3509 | -6.2422 | 4.3E-10 | 1.23E-08 |
| 642 | PALS1 | 14.9590309 | -2.193081281 | 0.6243 | -3.5127 | 0.00044 | 0.002877 |
| 1862 | STEAP1 | 9.53425703 | -2.19647584 | 0.7434 | -2.9545 | 0.00313 | 0.015204 |
| 1673 | ATP6V0E1 | 20.376683 | -2.198599767 | 0.4699 | -4.6794 | 2.9E-06 | 3.39E-05 |
| 32 | NBL1 | 30.2375583 | -2.204147752 | 0.4731 | -4.6591 | 3.2E-06 | 3.7E-05 |
| 784 | TMEM219 | 19.0478622 | -2.20438251 | 0.4993 | -4.4149 | 1E-05 | 0.000104 |
| 1022 | STXBP2 | 11.8942142 | -2.20444266 | 0.6536 | -3.3729 | 0.00074 | 0.004488 |
| 126 | STXBP3 | 10.786123 | -2.204717379 | 0.6719 | -3.2815 | 0.00103 | 0.0059 |
| 720 | DAPK2 | 5.61748167 | -2.208928586 | 0.8485 | -2.6032 | 0.00923 | 0.037077 |
| 838 | MYO1C | 24.8632951 | -2.215233321 | 0.4917 | -4.5055 | 6.6E-06 | 7.1E-05 |
| 98 | PATJ | 12.100208 | -2.216005575 | 0.601 | -3.6874 | 0.00023 | 0.001597 |
| 1984 | MTSS1 | 15.6405396 | -2.219926414 | 0.6434 | -3.4504 | 0.00056 | 0.003522 |
| 1825 | C1GALT1 | 10.3236783 | -2.220477508 | 0.6817 | -3.2573 | 0.00112 | 0.006344 |
| 1971 | TP53INP1 | 9.04195698 | -2.22414315 | 0.8102 | -2.7452 | 0.00605 | 0.026361 |
| 994 | HSBP1L1 | 27.2370494 | -2.227392073 | 0.4562 | -4.8821 | 1E-06 | 1.38E-05 |
| 333 | BET1L | 12.1999334 | -2.227953766 | 0.6656 | -3.3475 | 0.00082 | 0.004849 |
| 2078 | NRARP | 20.7479335 | -2.23739344 | 0.5213 | -4.2919 | 1.8E-05 | 0.000171 |
| 2082 | ARRDC1 | 12.136748 | -2.237876737 | 0.6885 | -3.2502 | 0.00115 | 0.00647 |
| 511 | TMBIM6 | 164.586265 | -2.239426538 | 0.2184 | -10.253 | 1.1E-24 | 4.52E-22 |
| 1182 | SOWAHC | 7.36667491 | -2.244356964 | 0.8225 | -2.7288 | 0.00636 | 0.027403 |
| 1914 | GSTK1 | 38.0373728 | -2.245329533 | 0.3611 | -6.2176 | 5E-10 | 1.42E-08 |
| 1874 | ZNF789 | 12.5253217 | -2.252276245 | 0.5977 | -3.7684 | 0.00016 | 0.001207 |
| 1098 | FTL | 255.001839 | -2.252903214 | 0.2767 | -8.1407 | 3.9E-16 | 3.83E-14 |
| 1275 | KIF16B | 7.27754776 | -2.253194274 | 0.8183 | -2.7536 | 0.00589 | 0.025794 |
| 1272 | PLCB4 | 30.1653718 | -2.255987488 | 0.4221 | -5.3451 | 9E-08 | 1.54E-06 |
| 603 | DNAJC3 | 10.921596 | -2.263616017 | 0.6721 | -3.368 | 0.00076 | 0.004552 |
| 1006 | GNA11 | 43.204542 | -2.270460012 | 0.3721 | -6.1025 | 1E-09 | 2.71E-08 |
| 428 | PICALM | 28.1458897 | -2.271758528 | 0.4551 | -4.9917 | 6E-07 | 8.39E-06 |
| 683 | CEP170B | 20.538971 | -2.272980302 | 0.5979 | -3.8017 | 0.00014 | 0.001076 |
| 326 | LHPP | 6.59981286 | -2.278624336 | 0.9059 | -2.5154 | 0.01189 | 0.045452 |
| 1390 | LAMB2 | 10.136525 | -2.279141391 | 0.6532 | -3.4892 | 0.00048 | 0.003102 |
| 1935 | SLC39A14 | 32.2899987 | -2.281617408 | 0.5242 | -4.3525 | 1.3E-05 | 0.000134 |

| | | | | | | | |
|---|---|---|---|---|---|---|---|
| 306 | NEURL1 | 12.5659892 | -2.290661126 | 0.6942 | -3.2996 | 0.00097 | 0.005592 |
| 238 | NID1 | 15.3189589 | -2.292489355 | 0.7615 | -3.0104 | 0.00261 | 0.013041 |
| 104 | SLC35D1 | 11.3201796 | -2.294573513 | 0.6818 | -3.3657 | 0.00076 | 0.004585 |
| 325 | OAT | 38.4185575 | -2.29691412 | 0.6264 | -3.6671 | 0.00025 | 0.00171 |
| 160 | S100A13 | 7.41956167 | -2.299934332 | 0.8404 | -2.7366 | 0.00621 | 0.026904 |
| 2029 | GOLM1 | 143.736817 | -2.3039765 | 0.2321 | -9.925 | 3.2E-23 | 1.12E-20 |
| 1737 | TFEB | 7.0727427 | -2.305812721 | 0.9077 | -2.5402 | 0.01108 | 0.042787 |
| 768 | CPPED1 | 7.04093445 | -2.306078194 | 0.8649 | -2.6662 | 0.00767 | 0.031929 |
| 1157 | GFPT1 | 43.1218239 | -2.307338238 | 0.3556 | -6.4891 | 8.6E-11 | 2.97E-09 |
| 585 | SPART | 6.88847017 | -2.308262628 | 0.7939 | -2.9075 | 0.00364 | 0.017306 |
| 1459 | FNDC3B | 12.037674 | -2.309982803 | 0.6478 | -3.5661 | 0.00036 | 0.002412 |
| 1427 | PDIA5 | 8.94793339 | -2.310147066 | 0.8569 | -2.6958 | 0.00702 | 0.02971 |
| 799 | CMTM4 | 15.2761742 | -2.313328765 | 0.6233 | -3.7116 | 0.00021 | 0.001467 |
| 70 | MYCBP | 12.5077567 | -2.314965765 | 0.6631 | -3.4911 | 0.00048 | 0.003083 |
| 1678 | PHYKPL | 33.7457198 | -2.318821908 | 0.5026 | -4.6133 | 4E-06 | 4.49E-05 |
| 522 | PRR13 | 49.7966366 | -2.320285263 | 0.3383 | -6.8587 | 6.9E-12 | 3.1E-10 |
| 1289 | BLCAP | 28.09091 | -2.322664877 | 0.438 | -5.3031 | 1.1E-07 | 1.9E-06 |
| 73 | PPCS | 25.9345482 | -2.32340805 | 0.4435 | -5.2387 | 1.6E-07 | 2.58E-06 |
| 1795 | IFNGR1 | 13.750551 | -2.328879052 | 0.6237 | -3.734 | 0.00019 | 0.001358 |
| 1241 | DNPEP | 16.2243457 | -2.329098087 | 0.5301 | -4.3934 | 1.1E-05 | 0.000113 |
| 2131 | TAF9B | 5.9090488 | -2.329147075 | 0.8968 | -2.5971 | 0.0094 | 0.037546 |
| 1687 | RIPK1 | 7.13655285 | -2.334381922 | 0.8458 | -2.7601 | 0.00578 | 0.025409 |
| 933 | CYB561 | 8.08902282 | -2.337464751 | 0.8516 | -2.7449 | 0.00605 | 0.026372 |
| 1263 | FARP2 | 10.8012813 | -2.339312038 | 0.6574 | -3.5582 | 0.00037 | 0.00248 |
| 1278 | RALGAPA2 | 20.1979176 | -2.341200533 | 0.5187 | -4.5139 | 6.4E-06 | 6.84E-05 |
| 549 | POC1B | 15.5488907 | -2.346185208 | 0.7812 | -3.0034 | 0.00267 | 0.013302 |
| 389 | TMEM109 | 6.2468955 | -2.352706748 | 0.8266 | -2.8463 | 0.00442 | 0.020294 |
| 429 | TMEM135 | 9.60556456 | -2.354893638 | 0.752 | -3.1315 | 0.00174 | 0.009227 |
| 1769 | UFL1 | 13.3777218 | -2.355670714 | 0.7524 | -3.1309 | 0.00174 | 0.009239 |
| 298 | CUTC | 6.64856994 | -2.356468291 | 0.9189 | -2.5645 | 0.01033 | 0.040492 |
| 2057 | NIBAN2 | 30.3480187 | -2.357226844 | 0.4516 | -5.2193 | 1.8E-07 | 2.84E-06 |
| 491 | DUSP16 | 16.466868 | -2.361994652 | 0.6126 | -3.8556 | 0.00012 | 0.000893 |
| 361 | RRAS2 | 15.1923881 | -2.3625381 | 0.8604 | -2.7459 | 0.00603 | 0.026315 |
| 1992 | PLEC | 19.7924427 | -2.366851339 | 0.6222 | -3.8043 | 0.00014 | 0.001066 |
| 123 | TLCD4 | 10.0801612 | -2.36812386 | 0.8435 | -2.8073 | 0.005 | 0.022572 |
| 1822 | GNA12 | 6.21387128 | -2.368716417 | 0.8655 | -2.7369 | 0.0062 | 0.026892 |
| 944 | FDXR | 5.00355463 | -2.369157482 | 0.9413 | -2.517 | 0.01184 | 0.045289 |
| 2065 | COL5A1 | 21.276865 | -2.369934111 | 0.6755 | -3.5083 | 0.00045 | 0.002917 |
| 1957 | LACTB2 | 10.1855571 | -2.373649861 | 0.7132 | -3.3284 | 0.00087 | 0.005139 |

| | | | | | | | |
|---|---|---|---|---|---|---|---|
| 1567 | SEC24D | 13.1279896 | -2.37679663 | 0.6177 | -3.8481 | 0.00012 | 0.00092 |
| 1727 | TEAD3 | 13.2876696 | -2.377240165 | 0.7676 | -3.0971 | 0.00195 | 0.010187 |
| 620 | PSME1 | 78.3574836 | -2.37777999 | 0.3105 | -7.658 | 1.9E-14 | 1.37E-12 |
| 227 | PPP2R5A | 27.4539276 | -2.38071284 | 0.6171 | -3.8579 | 0.00011 | 0.000886 |
| 723 | SMAD3 | 10.8412849 | -2.389766852 | 0.695 | -3.4384 | 0.00059 | 0.003654 |
| 284 | LIPA | 12.7402954 | -2.390471613 | 0.6932 | -3.4485 | 0.00056 | 0.003543 |
| 735 | TSPAN3 | 223.094579 | -2.395696911 | 0.3497 | -6.8502 | 7.4E-12 | 3.26E-10 |
| 1510 | PDGFRA | 8.50722124 | -2.395986772 | 0.7292 | -3.2856 | 0.00102 | 0.005828 |
| 1793 | STX7 | 29.1319829 | -2.397853397 | 0.5062 | -4.7373 | 2.2E-06 | 2.63E-05 |
| 1829 | ANKMY2 | 9.31140152 | -2.398033533 | 0.7425 | -3.2298 | 0.00124 | 0.006889 |
| 390 | TKFC | 21.9542126 | -2.400475545 | 0.5187 | -4.628 | 3.7E-06 | 4.23E-05 |
| 1306 | HELZ2 | 5.87283169 | -2.402129908 | 0.8568 | -2.8035 | 0.00506 | 0.022765 |
| 903 | GRN | 57.5424288 | -2.402829125 | 0.3765 | -6.3818 | 1.8E-10 | 5.48E-09 |
| 1114 | ZNF524 | 5.84122605 | -2.4181902 | 0.9309 | -2.5978 | 0.00938 | 0.037491 |
| 1192 | RALB | 10.9934982 | -2.423635113 | 0.708 | -3.4234 | 0.00062 | 0.003833 |
| 493 | HEBP1 | 12.2692439 | -2.42642151 | 0.8021 | -3.0252 | 0.00248 | 0.012499 |
| 1667 | SPARC | 22.9035516 | -2.434693549 | 0.6813 | -3.5736 | 0.00035 | 0.002356 |
| 1639 | SLC22A5 | 9.58179871 | -2.435472906 | 0.6994 | -3.4824 | 0.0005 | 0.003177 |
| 732 | SCAMP2 | 20.5438459 | -2.439312999 | 0.5279 | -4.6209 | 3.8E-06 | 4.35E-05 |
| 728 | NEO1 | 19.8776164 | -2.439984749 | 0.6078 | -4.0146 | 6E-05 | 0.000501 |
| 2094 | MT-TP | 21.5962118 | -2.446028741 | 0.462 | -5.2944 | 1.2E-07 | 1.98E-06 |
| 1444 | PLOD2 | 21.45824 | -2.449783708 | 0.7164 | -3.4194 | 0.00063 | 0.003884 |
| 810 | PHLPP2 | 7.34896414 | -2.450875899 | 0.8545 | -2.868 | 0.00413 | 0.019206 |
| 1080 | CEACAM19 | 9.63759285 | -2.451118275 | 0.7733 | -3.1696 | 0.00153 | 0.008253 |
| 136 | LRIG2 | 18.4990457 | -2.451499383 | 0.7098 | -3.4539 | 0.00055 | 0.003482 |
| 1382 | TRAK1 | 21.8660488 | -2.452188885 | 0.4718 | -5.1975 | 2E-07 | 3.15E-06 |
| 140 | CD58 | 6.67002468 | -2.459525397 | 0.8774 | -2.8031 | 0.00506 | 0.022779 |
| 1086 | VASP | 8.02283419 | -2.469930885 | 0.7474 | -3.3046 | 0.00095 | 0.005508 |
| 313 | SHOC2 | 8.8240975 | -2.471969899 | 0.8207 | -3.012 | 0.0026 | 0.012982 |
| 790 | TENT4B | 20.6742881 | -2.473937607 | 0.6606 | -3.745 | 0.00018 | 0.001307 |
| 1928 | CTSB | 40.212414 | -2.474539069 | 0.3572 | -6.9271 | 4.3E-12 | 2E-10 |
| 1668 | CLINT1 | 29.9374271 | -2.479197607 | 0.4402 | -5.6316 | 1.8E-08 | 3.6E-07 |
| 13 | PRXL2B | 14.0416583 | -2.481501596 | 0.566 | -4.3845 | 1.2E-05 | 0.000117 |
| 1679 | MGAT4B | 91.4184624 | -2.482854286 | 0.3992 | -6.2189 | 5E-10 | 1.41E-08 |
| 1419 | ZDHHC23 | 6.1793414 | -2.483408684 | 0.9838 | -2.5243 | 0.01159 | 0.044448 |
| 1880 | AGFG2 | 5.94522626 | -2.4900269 | 0.9077 | -2.7433 | 0.00608 | 0.026477 |
| 2169 | ERCC00165 | 45.3612221 | -2.496720482 | 0.9575 | -2.6076 | 0.00912 | 0.036686 |
| 847 | MED11 | 5.32765718 | -2.527921327 | 0.9627 | -2.6258 | 0.00865 | 0.035117 |
| 1645 | JADE2 | 6.15298623 | -2.528627749 | 0.9512 | -2.6584 | 0.00785 | 0.032525 |

| | | | | | | | |
|---|---|---|---|---|---|---|---|
| 439 | BIRC2 | 20.6034924 | -2.529302374 | 0.6049 | -4.1816 | 2.9E-05 | 0.000264 |
| 681 | PPP1R13B | 6.7239001 | -2.530308168 | 0.887 | -2.8526 | 0.00434 | 0.019978 |
| 52 | PAFAH2 | 7.25980002 | -2.532463246 | 0.8053 | -3.1447 | 0.00166 | 0.008875 |
| 1228 | KLF7 | 7.66991518 | -2.534116495 | 0.8135 | -3.1149 | 0.00184 | 0.009673 |
| 794 | PLLP | 8.71026024 | -2.540625661 | 0.7586 | -3.3492 | 0.00081 | 0.004826 |
| 1170 | RETSAT | 22.383033 | -2.546594259 | 0.5069 | -5.0242 | 5.1E-07 | 7.25E-06 |
| 304 | SFXN3 | 16.3525856 | -2.549076426 | 0.5995 | -4.2523 | 2.1E-05 | 0.0002 |
| 353 | TPP1 | 28.2204234 | -2.550084799 | 0.5314 | -4.7989 | 1.6E-06 | 1.99E-05 |
| 828 | CYBA | 30.488315 | -2.551605567 | 0.5686 | -4.4877 | 7.2E-06 | 7.64E-05 |
| 1826 | ICA1 | 12.7208004 | -2.55435476 | 0.6283 | -4.0654 | 4.8E-05 | 0.000414 |
| 612 | ANG | 8.00171667 | -2.556862294 | 0.8118 | -3.1496 | 0.00163 | 0.008752 |
| 1420 | B4GALT4 | 7.7477914 | -2.559206587 | 0.9201 | -2.7815 | 0.00541 | 0.024074 |
| 1858 | DTX2 | 9.57197096 | -2.561111373 | 0.7604 | -3.3681 | 0.00076 | 0.004552 |
| 340 | RASSF7 | 23.3280906 | -2.563234108 | 0.489 | -5.2418 | 1.6E-07 | 2.54E-06 |
| 272 | TSPAN15 | 6.90332076 | -2.563999904 | 0.8348 | -3.0713 | 0.00213 | 0.010984 |
| 1888 | PLOD3 | 21.4704007 | -2.567990813 | 0.486 | -5.2844 | 1.3E-07 | 2.07E-06 |
| 1133 | EMILIN1 | 11.9502725 | -2.569098457 | 0.8728 | -2.9434 | 0.00325 | 0.015673 |
| 748 | LRRC28 | 12.2704698 | -2.574380449 | 0.7031 | -3.6614 | 0.00025 | 0.001745 |
| 872 | CPD | 17.8766836 | -2.575397478 | 0.6524 | -3.9475 | 7.9E-05 | 0.00064 |
| 858 | TMEM220 | 10.580229 | -2.575588526 | 0.7598 | -3.3897 | 0.0007 | 0.004245 |
| 266 | REEP3 | 15.397158 | -2.58266453 | 0.5937 | -4.35 | 1.4E-05 | 0.000135 |
| 836 | C17orf97 | 5.58026834 | -2.582718937 | 0.9636 | -2.6802 | 0.00736 | 0.030816 |
| 2049 | PTBP3 | 26.8738466 | -2.583117832 | 0.5227 | -4.9422 | 7.7E-07 | 1.05E-05 |
| 1896 | BCAP29 | 10.3683542 | -2.592860934 | 0.7185 | -3.6088 | 0.00031 | 0.002089 |
| 1379 | CTDSPL | 13.250163 | -2.594498339 | 0.7055 | -3.6773 | 0.00024 | 0.001653 |
| 694 | CHP1 | 45.6804079 | -2.59953218 | 0.3677 | -7.0702 | 1.5E-12 | 7.77E-11 |
| 1026 | DOCK6 | 12.1592059 | -2.601292217 | 0.6851 | -3.7969 | 0.00015 | 0.001092 |
| 1255 | COL6A3 | 11.7591268 | -2.606584947 | 0.7982 | -3.2657 | 0.00109 | 0.006189 |
| 1986 | EFR3A | 11.5207412 | -2.610405504 | 0.6276 | -4.1592 | 3.2E-05 | 0.000289 |
| 2101 | RPS6KA3 | 39.1017359 | -2.610983777 | 0.3906 | -6.6852 | 2.3E-11 | 8.99E-10 |
| 62 | ENSG00000270605 | 13.4231353 | -2.614317968 | 0.7843 | -3.3332 | 0.00086 | 0.005068 |
| 1378 | GOLGA4 | 65.9947282 | -2.622370083 | 0.4181 | -6.2715 | 3.6E-10 | 1.04E-08 |
| 1264 | D2HGDH | 28.2489487 | -2.629892956 | 0.5112 | -5.1445 | 2.7E-07 | 4.09E-06 |
| 1 | MTCO1P12 | 10.073712 | -2.642339553 | 1.0495 | -2.5176 | 0.01181 | 0.045213 |
| 1549 | SLC39A8 | 12.0395821 | -2.646048744 | 0.6607 | -4.0052 | 6.2E-05 | 0.000519 |
| 2023 | PCSK5 | 17.2723441 | -2.652487555 | 0.6147 | -4.3154 | 1.6E-05 | 0.000156 |
| 1338 | TIMP3 | 7.69264564 | -2.656305292 | 0.8784 | -3.024 | 0.00249 | 0.012534 |
| 59 | SLC9A1 | 8.95506411 | -2.657432245 | 0.8628 | -3.0802 | 0.00207 | 0.010706 |
| 559 | MVK | 6.81906322 | -2.663727545 | 0.9172 | -2.9043 | 0.00368 | 0.017457 |

| | | | | | | | |
|---|---|---|---|---|---|---|---|
| 1542 | FAM13A | 23.8627958 | -2.664084148 | 0.505 | -5.2757 | 1.3E-07 | 2.15E-06 |
| 1835 | ITGB8 | 19.6685335 | -2.669118447 | 0.7396 | -3.6087 | 0.00031 | 0.002089 |
| 1800 | HECA | 11.8209068 | -2.672076862 | 0.7687 | -3.4763 | 0.00051 | 0.00324 |
| 1356 | CBX7 | 5.3494683 | -2.679502128 | 0.98 | -2.7342 | 0.00625 | 0.02704 |
| 1665 | SYNPO | 7.9803315 | -2.679812439 | 0.8949 | -2.9944 | 0.00275 | 0.013626 |
| 35 | CAMK2N1 | 77.0874541 | -2.683733535 | 0.3891 | -6.8974 | 5.3E-12 | 2.41E-10 |
| 1187 | TMEM87B | 9.31921966 | -2.701009751 | 0.7605 | -3.5516 | 0.00038 | 0.002535 |
| 527 | SUOX | 12.2906767 | -2.702230276 | 0.6655 | -4.0605 | 4.9E-05 | 0.000421 |
| 1463 | B3GNT5 | 6.09556232 | -2.705766476 | 0.9849 | -2.7471 | 0.00601 | 0.026226 |
| 1221 | NABP1 | 21.4413047 | -2.710169858 | 0.6636 | -4.0842 | 4.4E-05 | 0.000387 |
| 622 | IRF9 | 11.978202 | -2.711266509 | 0.7346 | -3.6907 | 0.00022 | 0.00158 |
| 2137 | TSPAN6 | 45.6021842 | -2.716362698 | 0.3644 | -7.4552 | 9E-14 | 5.67E-12 |
| 873 | RNF135 | 9.1571543 | -2.717515924 | 0.7747 | -3.508 | 0.00045 | 0.00292 |
| 1589 | CTSO | 5.03492725 | -2.71824202 | 1.0624 | -2.5586 | 0.01051 | 0.041026 |
| 797 | MMP15 | 29.4505871 | -2.72055389 | 0.474 | -5.739 | 9.5E-09 | 2.04E-07 |
| 1284 | NOL4L | 33.7540624 | -2.726414405 | 0.4573 | -5.9624 | 2.5E-09 | 5.95E-08 |
| 101 | RAVER2 | 17.2336549 | -2.72964868 | 0.5718 | -4.7738 | 1.8E-06 | 2.24E-05 |
| 789 | PYCARD | 5.88451585 | -2.730843856 | 0.9929 | -2.7503 | 0.00595 | 0.026017 |
| 487 | C1RL | 10.0234435 | -2.73602283 | 0.7231 | -3.7836 | 0.00015 | 0.001143 |
| 1434 | MGLL | 36.0928357 | -2.736919804 | 0.502 | -5.4521 | 5E-08 | 9.04E-07 |
| 1627 | GLRX | 21.4040627 | -2.740407565 | 0.5637 | -4.8619 | 1.2E-06 | 1.51E-05 |
| 1736 | FOXP4 | 14.9543084 | -2.741673028 | 0.6594 | -4.158 | 3.2E-05 | 0.00029 |
| 506 | RESF1 | 19.0122908 | -2.744152973 | 0.601 | -4.5658 | 5E-06 | 5.5E-05 |
| 1332 | TPST2 | 7.96213338 | -2.744160149 | 0.9342 | -2.9374 | 0.00331 | 0.015915 |
| 329 | PWWP2B | 11.3996378 | -2.75038688 | 0.6994 | -3.9324 | 8.4E-05 | 0.000677 |
| 507 | FGD4 | 6.36312082 | -2.750747956 | 0.9518 | -2.8901 | 0.00385 | 0.018124 |
| 648 | PSEN1 | 25.279512 | -2.752537372 | 0.4926 | -5.588 | 2.3E-08 | 4.54E-07 |
| 1220 | MFSD6 | 5.48651109 | -2.756104529 | 1.0411 | -2.6472 | 0.00812 | 0.033405 |
| 1878 | TRAPPC14 | 13.7504481 | -2.756961815 | 0.6535 | -4.219 | 2.5E-05 | 0.000228 |
| 1717 | NEU1 | 23.6912312 | -2.759193653 | 0.5588 | -4.938 | 7.9E-07 | 1.07E-05 |
| 1569 | USP53 | 18.2177641 | -2.760385029 | 0.5713 | -4.8314 | 1.4E-06 | 1.73E-05 |
| 119 | EVI5 | 15.1383741 | -2.760907658 | 0.5868 | -4.7053 | 2.5E-06 | 3.03E-05 |
| 854 | ACADVL | 75.2836841 | -2.761017312 | 0.328 | -8.4174 | 3.8E-17 | 4.8E-15 |
| 562 | SH2B3 | 9.55277178 | -2.762714952 | 0.7971 | -3.4658 | 0.00053 | 0.003355 |
| 198 | DENND1B | 7.0668218 | -2.764046078 | 0.9034 | -3.0596 | 0.00222 | 0.011369 |
| 574 | ORAI1 | 9.85086451 | -2.765073356 | 0.7222 | -3.8287 | 0.00013 | 0.000984 |
| 1506 | ATP8A1 | 10.0518556 | -2.767448627 | 0.8377 | -3.3035 | 0.00095 | 0.005523 |
| 1564 | CAMK2D | 30.8617599 | -2.768538358 | 0.5799 | -4.7738 | 1.8E-06 | 2.24E-05 |
| 1960 | TPD52 | 41.9373625 | -2.770137116 | 0.3944 | -7.0231 | 2.2E-12 | 1.06E-10 |

| | | | | | | | |
|---|---|---|---|---|---|---|---|
| 2036 | TRIM14 | 13.9209105 | -2.770596621 | 0.6228 | -4.4488 | 8.6E-06 | 9E-05 |
| 775 | RRN3P3 | 4.01689752 | -2.770828783 | 1.1102 | -2.4958 | 0.01257 | 0.047556 |
| 766 | ATF7IP2 | 18.9916821 | -2.771873789 | 0.5646 | -4.9092 | 9.1E-07 | 1.22E-05 |
| 1634 | SNX24 | 8.26942997 | -2.772630242 | 0.9298 | -2.9819 | 0.00286 | 0.014099 |
| 181 | NIT1 | 5.79161722 | -2.773377204 | 1.029 | -2.6953 | 0.00703 | 0.029748 |
| 807 | FCSK | 10.4050045 | -2.774087957 | 0.7501 | -3.6985 | 0.00022 | 0.001539 |
| 583 | SLC46A3 | 15.2856243 | -2.774151811 | 0.7737 | -3.5857 | 0.00034 | 0.002261 |
| 566 | TPCN1 | 19.2113416 | -2.775698874 | 0.5342 | -5.196 | 2E-07 | 3.17E-06 |
| 576 | TMED2-DT | 5.0299895 | -2.776282648 | 1.0241 | -2.711 | 0.00671 | 0.028669 |
| 992 | ZCCHC2 | 10.1507567 | -2.777746716 | 0.8217 | -3.3804 | 0.00072 | 0.004379 |
| 351 | TRIM5 | 5.35004451 | -2.777830424 | 1.0162 | -2.7335 | 0.00627 | 0.02707 |
| 646 | SUSD6 | 10.0600306 | -2.784046456 | 0.8235 | -3.3807 | 0.00072 | 0.004377 |
| 373 | CAT | 10.0931648 | -2.784497326 | 0.841 | -3.311 | 0.00093 | 0.005404 |
| 139 | ATP1A1 | 102.832146 | -2.784684271 | 0.2952 | -9.4323 | 4E-21 | 1.06E-18 |
| 1237 | CTDSP1 | 22.3113522 | -2.787545218 | 0.5009 | -5.5649 | 2.6E-08 | 5.13E-07 |
| 231 | CAPN8 | 6.94639414 | -2.789646779 | 0.8962 | -3.1127 | 0.00185 | 0.009736 |
| 1716 | CLIC1 | 40.3491121 | -2.791500575 | 0.4082 | -6.8385 | 8E-12 | 3.5E-10 |
| 1400 | CHDH | 14.839209 | -2.792286629 | 0.6223 | -4.487 | 7.2E-06 | 7.66E-05 |
| 291 | PLCE1 | 12.8862236 | -2.807472467 | 0.636 | -4.4144 | 1E-05 | 0.000104 |
| 264 | ARID5B | 12.2951274 | -2.80917472 | 0.6602 | -4.2548 | 2.1E-05 | 0.000198 |
| 1744 | CD2AP | 25.2955105 | -2.811416238 | 0.544 | -5.1685 | 2.4E-07 | 3.63E-06 |
| 1640 | IRF1-AS1 | 4.68512129 | -2.814596605 | 1.0703 | -2.6298 | 0.00854 | 0.034788 |
| 1757 | LMBRD1 | 18.530591 | -2.817093286 | 0.6557 | -4.2964 | 1.7E-05 | 0.000168 |
| 1424 | PARP9 | 15.1138186 | -2.822372398 | 0.58 | -4.8661 | 1.1E-06 | 1.48E-05 |
| 1544 | PDLIM5 | 19.352456 | -2.82453965 | 0.534 | -5.2896 | 1.2E-07 | 2.02E-06 |
| 1160 | SPR | 12.4236299 | -2.833623319 | 0.7217 | -3.9263 | 8.6E-05 | 0.000691 |
| 1910 | CALD1 | 17.3599438 | -2.835820612 | 0.8184 | -3.4652 | 0.00053 | 0.003358 |
| 2129 | PBDC1 | 27.5302986 | -2.836965793 | 0.4827 | -5.8773 | 4.2E-09 | 9.52E-08 |
| 8 | MXRA8 | 5.29955357 | -2.841636875 | 1.0421 | -2.7267 | 0.0064 | 0.027548 |
| 1927 | XKR6 | 4.39251654 | -2.847226591 | 1.0834 | -2.6281 | 0.00859 | 0.034941 |
| 981 | GALNT1 | 27.9594107 | -2.854813274 | 0.4832 | -5.9083 | 3.5E-09 | 8.01E-08 |
| 1232 | FN1 | 27.5078381 | -2.861707614 | 0.5909 | -4.8433 | 1.3E-06 | 1.64E-05 |
| 168 | THBS3 | 9.54602256 | -2.862491307 | 0.8677 | -3.2989 | 0.00097 | 0.005598 |
| 1666 | SLC36A1 | 19.755092 | -2.862543135 | 0.5848 | -4.8947 | 9.8E-07 | 1.31E-05 |
| 2130 | MAGT1 | 13.4285582 | -2.86333494 | 0.6837 | -4.1882 | 2.8E-05 | 0.000258 |
| 753 | LMF1 | 4.60508451 | -2.865532575 | 1.1043 | -2.595 | 0.00946 | 0.03766 |
| 1389 | SLC26A6 | 16.4217774 | -2.866825305 | 0.5798 | -4.9449 | 7.6E-07 | 1.04E-05 |
| 1699 | TDP2 | 44.6259737 | -2.867517052 | 0.4451 | -6.4425 | 1.2E-10 | 3.88E-09 |
| 561 | PHETA1 | 4.23106981 | -2.871986359 | 1.119 | -2.5666 | 0.01027 | 0.040265 |

| | | | | | | | |
|---|---|---|---|---|---|---|---|
| 1443 | PLS1 | 22.6744864 | -2.874387986 | 0.4829 | -5.9529 | 2.6E-09 | 6.23E-08 |
| 277 | ANXA11 | 28.461028 | -2.877916496 | 0.4533 | -6.3482 | 2.2E-10 | 6.64E-09 |
| 1561 | PITX2 | 22.425845 | -2.881309214 | 0.5224 | -5.5151 | 3.5E-08 | 6.6E-07 |
| 1733 | KCNK5 | 6.60398099 | -2.884596224 | 0.9057 | -3.185 | 0.00145 | 0.007893 |
| 1802 | ADGRG6 | 5.37591007 | -2.888135241 | 1.0427 | -2.77 | 0.00561 | 0.024765 |
| 741 | AKAP13 | 8.8836859 | -2.892863729 | 0.7857 | -3.6817 | 0.00023 | 0.001627 |
| 1886 | TRIM56 | 20.2646084 | -2.896897283 | 0.5485 | -5.2818 | 1.3E-07 | 2.09E-06 |
| 740 | PDE8A | 19.8406643 | -2.898317743 | 0.5441 | -5.3272 | 1E-07 | 1.69E-06 |
| 2039 | GALNT12 | 23.3155262 | -2.902771123 | 0.5529 | -5.2501 | 1.5E-07 | 2.45E-06 |
| 211 | CDK18 | 12.4355669 | -2.905454909 | 0.7563 | -3.8415 | 0.00012 | 0.000939 |
| 124 | SLC35A3 | 20.4473621 | -2.908821379 | 0.5481 | -5.3072 | 1.1E-07 | 1.87E-06 |
| 1446 | PLSCR1 | 24.4148427 | -2.908921298 | 0.5586 | -5.2078 | 1.9E-07 | 3E-06 |
| 1956 | TRAM1 | 52.1530024 | -2.911997553 | 0.4855 | -5.9976 | 2E-09 | 4.89E-08 |
| 423 | SERPINH1 | 12.8542964 | -2.916440684 | 0.701 | -4.1603 | 3.2E-05 | 0.000287 |
| 661 | DGLUCY | 11.2143346 | -2.918437762 | 0.7158 | -4.0771 | 4.6E-05 | 0.000396 |
| 733 | NEIL1 | 11.0041922 | -2.918488387 | 0.8201 | -3.5588 | 0.00037 | 0.002475 |
| 1122 | ID2 | 19.9126666 | -2.920603395 | 0.724 | -4.0339 | 5.5E-05 | 0.000466 |
| 451 | C11orf71 | 6.44572697 | -2.921257834 | 0.8978 | -3.254 | 0.00114 | 0.006401 |
| 193 | RALGPS2 | 9.0762145 | -2.926285278 | 0.7776 | -3.7631 | 0.00017 | 0.001229 |
| 1425 | DTX3L | 15.4763444 | -2.927036719 | 0.5883 | -4.9756 | 6.5E-07 | 9.06E-06 |
| 795 | DOK4 | 11.3786789 | -2.928341537 | 0.7154 | -4.0931 | 4.3E-05 | 0.000374 |
| 940 | LINC00511 | 4.46981684 | -2.928445254 | 1.1831 | -2.4753 | 0.01331 | 0.049828 |
| 1040 | NR2F6 | 56.109385 | -2.929111216 | 0.3489 | -8.3955 | 4.6E-17 | 5.64E-15 |
| 1811 | RNASET2 | 12.3554495 | -2.932603081 | 0.6671 | -4.396 | 1.1E-05 | 0.000112 |
| 649 | PAPLN | 5.6445378 | -2.935745461 | 1.0434 | -2.8136 | 0.0049 | 0.022188 |
| 1056 | LGI4 | 5.12408245 | -2.936396558 | 1.0051 | -2.9216 | 0.00348 | 0.016653 |
| 456 | SIDT2 | 6.68349625 | -2.936554454 | 0.8699 | -3.3758 | 0.00074 | 0.004447 |
| 2111 | TIMP1 | 15.3772806 | -2.93935147 | 0.7515 | -3.9113 | 9.2E-05 | 0.000728 |
| 401 | EHBP1L1 | 8.38340897 | -2.941502631 | 0.902 | -3.2612 | 0.00111 | 0.006271 |
| 1622 | HEXB | 15.399656 | -2.943190456 | 0.5998 | -4.9069 | 9.3E-07 | 1.24E-05 |
| 2033 | SLC35D2 | 21.6837513 | -2.944459544 | 0.6136 | -4.799 | 1.6E-06 | 1.99E-05 |
| 1990 | FAM83H | 12.2024693 | -2.945408657 | 0.7258 | -4.0581 | 4.9E-05 | 0.000424 |
| 1759 | FAM135A | 20.4357868 | -2.954265927 | 0.6121 | -4.8264 | 1.4E-06 | 1.77E-05 |
| 430 | NAALAD2 | 6.03580338 | -2.956939503 | 0.9643 | -3.0666 | 0.00217 | 0.011148 |
| 1470 | ST6GAL1 | 17.8773667 | -2.962675653 | 0.7726 | -3.8346 | 0.00013 | 0.000963 |
| 971 | TGIF1 | 6.66329031 | -2.96505104 | 0.9093 | -3.2609 | 0.00111 | 0.006274 |
| 1776 | CD164 | 142.058742 | -2.972676607 | 0.3916 | -7.5911 | 3.2E-14 | 2.17E-12 |
| 1623 | IQGAP2 | 14.4042446 | -2.973239172 | 0.6183 | -4.809 | 1.5E-06 | 1.91E-05 |
| 2047 | PTGR1 | 36.5943656 | -2.975347256 | 0.5542 | -5.369 | 7.9E-08 | 1.37E-06 |

| | | | | | | | |
|---|---|---|---|---|---|---|---|
| 288 | FGFBP3 | 7.72817595 | -2.977958521 | 1.1179 | -2.664 | 0.00772 | 0.0321 |
| 762 | DNASE1 | 43.1727553 | -2.981324975 | 0.4762 | -6.2606 | 3.8E-10 | 1.11E-08 |
| 1999 | SLC1A1 | 10.3300964 | -2.984677774 | 0.7613 | -3.9207 | 8.8E-05 | 0.000706 |
| 2054 | CUTALP | 4.55550098 | -2.985978399 | 1.1615 | -2.5708 | 0.01015 | 0.039858 |
| 1536 | AFF1 | 11.3734988 | -2.99144273 | 0.7235 | -4.1345 | 3.6E-05 | 0.000318 |
| 1085 | PPP1R13L | 6.16852656 | -2.997606254 | 0.9068 | -3.3056 | 0.00095 | 0.005492 |
| 200 | INAVA | 4.94251433 | -2.999023727 | 1.0408 | -2.8815 | 0.00396 | 0.018512 |
| 417 | NADSYN1 | 26.4235883 | -3.010307632 | 0.4851 | -6.2056 | 5.4E-10 | 1.52E-08 |
| 1233 | IGFBP5 | 4.67213963 | -3.019634458 | 1.207 | -2.5019 | 0.01235 | 0.046914 |
| 1483 | BDH1 | 25.0848548 | -3.02174079 | 0.5021 | -6.0178 | 1.8E-09 | 4.39E-08 |
| 2005 | CDKN2B | 8.68073587 | -3.022153227 | 0.9266 | -3.2615 | 0.00111 | 0.006266 |
| 1968 | DECR1 | 19.554089 | -3.026444207 | 0.5697 | -5.3127 | 1.1E-07 | 1.82E-06 |
| 219 | CD55 | 6.07947732 | -3.034263775 | 1.0825 | -2.8031 | 0.00506 | 0.022779 |
| 2034 | CTSV | 3.78042114 | -3.039682384 | 1.1879 | -2.5589 | 0.0105 | 0.041007 |
| 1357 | FAM83F | 6.9990732 | -3.04776913 | 0.9311 | -3.2733 | 0.00106 | 0.006048 |
| 1108 | ZNF415 | 12.2284389 | -3.05370068 | 0.9978 | -3.0604 | 0.00221 | 0.01135 |
| 827 | MTHFSD | 4.8506735 | -3.056952951 | 1.0687 | -2.8605 | 0.00423 | 0.01956 |
| 667 | LGMN | 13.7082917 | -3.060878902 | 0.7119 | -4.2998 | 1.7E-05 | 0.000165 |
| 210 | NUAK2 | 3.6592385 | -3.067312535 | 1.2229 | -2.5082 | 0.01214 | 0.04627 |
| 627 | NYNRIN | 6.53322669 | -3.068014016 | 0.9673 | -3.1718 | 0.00152 | 0.008205 |
| 1941 | CLU | 39.8379579 | -3.071730132 | 0.4996 | -6.1479 | 7.9E-10 | 2.08E-08 |
| 710 | CCPG1 | 8.08077215 | -3.079036007 | 0.9905 | -3.1086 | 0.00188 | 0.009853 |
| 370 | PRRG4 | 7.93763164 | -3.081096099 | 0.9082 | -3.3925 | 0.00069 | 0.004207 |
| 675 | GSKIP | 10.9504709 | -3.081522402 | 0.7336 | -4.2007 | 2.7E-05 | 0.000245 |
| 1240 | STK16 | 3.8554179 | -3.082842677 | 1.2361 | -2.4941 | 0.01263 | 0.047767 |
| 1715 | DDAH2 | 15.7649115 | -3.088094645 | 0.6259 | -4.9342 | 8E-07 | 1.09E-05 |
| 1728 | MAPK13 | 32.7746736 | -3.089409263 | 0.4211 | -7.3365 | 2.2E-13 | 1.29E-11 |
| 2124 | FOXO4 | 4.16955974 | -3.096217469 | 1.1418 | -2.7118 | 0.00669 | 0.028616 |
| 1333 | XBP1 | 42.9130166 | -3.096424468 | 0.486 | -6.3713 | 1.9E-10 | 5.82E-09 |
| 705 | FBN1 | 6.16022382 | -3.097402357 | 1.1694 | -2.6488 | 0.00808 | 0.033267 |
| 1128 | TP53I3 | 14.8483054 | -3.098263629 | 0.7294 | -4.2477 | 2.2E-05 | 0.000203 |
| 2097 | ARSL | 12.7433716 | -3.098522766 | 0.6831 | -4.5357 | 5.7E-06 | 6.23E-05 |
| 1428 | KALRN | 12.5373976 | -3.099257833 | 0.7555 | -4.1021 | 4.1E-05 | 0.000361 |
| 54 | CNKSR1 | 9.80862055 | -3.101528081 | 0.8205 | -3.7799 | 0.00016 | 0.001157 |
| 2041 | TMEM38B | 12.2851052 | -3.103196419 | 0.7013 | -4.4249 | 9.6E-06 | 9.95E-05 |
| 2008 | ACO1 | 14.2475649 | -3.10857293 | 0.6625 | -4.6925 | 2.7E-06 | 3.2E-05 |
| 1638 | MIR3936HG | 4.90756328 | -3.113951354 | 1.0766 | -2.8923 | 0.00382 | 0.018042 |
| 931 | VMP1 | 59.4684794 | -3.114497357 | 0.4178 | -7.4543 | 9E-14 | 5.67E-12 |
| 422 | ARRB1 | 8.48576623 | -3.115189219 | 0.9946 | -3.132 | 0.00174 | 0.009215 |

| | | | | | | | |
|---|---|---|---|---|---|---|---|
| 1998 | PPP1R16A | 35.0185558 | -3.12478505 | 0.5115 | -6.1088 | 1E-09 | 2.62E-08 |
| 734 | SNX33 | 7.44105797 | -3.126412562 | 0.9739 | -3.2103 | 0.00133 | 0.007332 |
| 961 | PCYT2 | 11.4536452 | -3.13399215 | 0.727 | -4.3109 | 1.6E-05 | 0.000158 |
| 1933 | DMTN | 10.4448043 | -3.135724087 | 0.919 | -3.4121 | 0.00064 | 0.00397 |
| 1541 | HERC6 | 5.00926211 | -3.14622519 | 1.207 | -2.6066 | 0.00914 | 0.036768 |
| 2155 | ENOX2 | 6.28370248 | -3.150930662 | 1.1057 | -2.8497 | 0.00438 | 0.020122 |
| 1860 | TP53TG1 | 4.20603274 | -3.154055763 | 1.2135 | -2.5992 | 0.00934 | 0.037391 |
| 1653 | CYSTM1 | 13.0563673 | -3.157886672 | 0.67 | -4.7133 | 2.4E-06 | 2.93E-05 |
| 66 | KIAA1522 | 17.2814327 | -3.159877402 | 0.5681 | -5.5625 | 2.7E-08 | 5.18E-07 |
| 290 | SLC35G1 | 26.4103405 | -3.1695241 | 0.8306 | -3.8162 | 0.00014 | 0.001024 |
| 1106 | ETFB | 14.1650654 | -3.187484417 | 0.6997 | -4.5552 | 5.2E-06 | 5.74E-05 |
| 1625 | LHFPL2 | 11.3608094 | -3.188245044 | 0.7944 | -4.0132 | 6E-05 | 0.000503 |
| 1777 | SMPD2 | 5.28827141 | -3.194816206 | 1.0459 | -3.0545 | 0.00225 | 0.011516 |
| 973 | IMPA2 | 23.0624319 | -3.196444217 | 0.5685 | -5.6226 | 1.9E-08 | 3.77E-07 |
| 568 | PXN | 10.5273099 | -3.198207302 | 0.7911 | -4.043 | 5.3E-05 | 0.00045 |
| 958 | ENGASE | 11.5540787 | -3.198395754 | 0.7658 | -4.1763 | 3E-05 | 0.00027 |
| 1362 | PPARA | 13.4670261 | -3.21268443 | 0.714 | -4.4994 | 6.8E-06 | 7.28E-05 |
| 482 | TNFRSF1A | 22.047292 | -3.221582643 | 0.6218 | -5.1807 | 2.2E-07 | 3.42E-06 |
| 178 | F11R | 20.0653667 | -3.228293814 | 0.5714 | -5.6498 | 1.6E-08 | 3.29E-07 |
| 924 | SPATA20 | 27.0986229 | -3.235476918 | 0.4799 | -6.7416 | 1.6E-11 | 6.37E-10 |
| 338 | SIGIRR | 10.2294268 | -3.237660008 | 0.7519 | -4.3062 | 1.7E-05 | 0.000161 |
| 352 | FHIP1B | 27.7706792 | -3.237796437 | 0.5434 | -5.9582 | 2.6E-09 | 6.08E-08 |
| 866 | DHRS7B | 7.32669055 | -3.239723911 | 0.955 | -3.3925 | 0.00069 | 0.004207 |
| 465 | SORL1 | 18.2040009 | -3.239725494 | 0.629 | -5.1505 | 2.6E-07 | 3.97E-06 |
| 2044 | EPB41L4B | 17.763552 | -3.241344624 | 0.6499 | -4.9877 | 6.1E-07 | 8.55E-06 |
| 2000 | JAK2 | 4.26650643 | -3.241709215 | 1.2339 | -2.6272 | 0.00861 | 0.034981 |
| 870 | ABHD15 | 4.22353589 | -3.245783016 | 1.1395 | -2.8484 | 0.00439 | 0.020191 |
| 1518 | SLC4A4 | 6.05103922 | -3.246910675 | 1.0323 | -3.1453 | 0.00166 | 0.00886 |
| 1308 | HUNK | 5.82542998 | -3.25452085 | 1.0664 | -3.0517 | 0.00228 | 0.011604 |
| 2007 | MOB3B | 8.12798 | -3.259610139 | 0.9414 | -3.4625 | 0.00054 | 0.003386 |
| 1556 | CASP6 | 9.07920652 | -3.262735968 | 0.8425 | -3.8727 | 0.00011 | 0.000839 |
| 610 | GAS6 | 19.5576425 | -3.265326743 | 0.6857 | -4.762 | 1.9E-06 | 2.35E-05 |
| 1174 | CNNM4 | 10.8586554 | -3.271119648 | 0.7577 | -4.317 | 1.6E-05 | 0.000155 |
| 2098 | PUDP | 13.399636 | -3.271632338 | 0.704 | -4.6471 | 3.4E-06 | 3.9E-05 |
| 394 | PLCB3 | 17.717408 | -3.273613272 | 0.5816 | -5.6291 | 1.8E-08 | 3.65E-07 |
| 108 | NEXN | 6.96429018 | -3.278533748 | 1.0254 | -3.1975 | 0.00139 | 0.007612 |
| 162 | PBXIP1 | 6.67045338 | -3.280430246 | 0.974 | -3.3682 | 0.00076 | 0.004552 |
| 918 | CALCOCO2 | 26.3279826 | -3.284136066 | 0.5421 | -6.0585 | 1.4E-09 | 3.48E-08 |
| 1644 | SHROOM1 | 10.7771979 | -3.297404352 | 0.7818 | -4.2176 | 2.5E-05 | 0.000229 |

| | | | | | | | |
|---|---|---|---|---|---|---|---|
| 1224 | CFLAR | 20.8027686 | -3.309950691 | 0.6632 | -4.9909 | 6E-07 | 8.42E-06 |
| 1722 | HSD17B8 | 4.49666414 | -3.310309011 | 1.1216 | -2.9514 | 0.00316 | 0.015331 |
| 1250 | C2orf72 | 25.661863 | -3.310501622 | 0.5509 | -6.0096 | 1.9E-09 | 4.58E-08 |
| 344 | CRACR2B | 18.7839495 | -3.312183206 | 0.6677 | -4.9605 | 7E-07 | 9.69E-06 |
| 1534 | MAPK10 | 5.55747222 | -3.324693733 | 1.0697 | -3.108 | 0.00188 | 0.00987 |
| 371 | DEPDC7 | 9.50706967 | -3.327608262 | 0.843 | -3.9471 | 7.9E-05 | 0.000641 |
| 2149 | PLS3 | 37.6373929 | -3.334216907 | 0.445 | -7.4923 | 6.8E-14 | 4.4E-12 |
| 1253 | SNORC | 7.49115135 | -3.33515822 | 0.8961 | -3.722 | 0.0002 | 0.001419 |
| 2134 | RPS6KA6 | 11.5478491 | -3.335272699 | 0.7307 | -4.5647 | 5E-06 | 5.53E-05 |
| 1100 | FCGRT | 113.897474 | -3.341766592 | 0.3485 | -9.5903 | 8.8E-22 | 2.5E-19 |
| 2035 | TDRD7 | 5.94159601 | -3.345660273 | 1.0471 | -3.1951 | 0.0014 | 0.007657 |
| 630 | MIA2 | 22.4577316 | -3.357581993 | 0.647 | -5.1891 | 2.1E-07 | 3.29E-06 |
| 1435 | ASTE1 | 4.90557747 | -3.362080225 | 1.1917 | -2.8212 | 0.00478 | 0.021713 |
| 229 | C1orf115 | 13.2354374 | -3.362632246 | 0.772 | -4.356 | 1.3E-05 | 0.000132 |
| 1961 | PAG1 | 21.3632379 | -3.369637095 | 0.6024 | -5.5938 | 2.2E-08 | 4.41E-07 |
| 1177 | MGAT4A | 10.6891326 | -3.373981002 | 0.8264 | -4.0828 | 4.4E-05 | 0.000389 |
| 163 | ZBTB7B | 3.34230856 | -3.379304371 | 1.3271 | -2.5464 | 0.01088 | 0.042208 |
| 1761 | MYO6 | 57.3856126 | -3.379619005 | 0.3734 | -9.0507 | 1.4E-19 | 2.81E-17 |
| 69 | ZC3H12A | 4.45851523 | -3.379849172 | 1.1241 | -3.0066 | 0.00264 | 0.01319 |
| 1993 | PARP10 | 3.30545634 | -3.38322185 | 1.3113 | -2.58 | 0.00988 | 0.039028 |
| 2069 | ENSG00000273066 | 21.3257115 | -3.383394697 | 0.6004 | -5.6351 | 1.7E-08 | 3.54E-07 |
| 880 | DHRS11 | 53.1631017 | -3.385552435 | 0.4137 | -8.1834 | 2.8E-16 | 2.8E-14 |
| 267 | LINC01515 | 3.10798312 | -3.386548527 | 1.3641 | -2.4827 | 0.01304 | 0.049006 |
| 1473 | LPP | 52.0807646 | -3.386657925 | 0.4226 | -8.0131 | 1.1E-15 | 1.01E-13 |
| 250 | ST8SIA6-AS1 | 3.33969493 | -3.410544143 | 1.3487 | -2.5287 | 0.01145 | 0.043962 |
| 563 | ALDH2 | 36.6662426 | -3.413542755 | 0.4306 | -7.9265 | 2.3E-15 | 1.92E-13 |
| 29 | MST1P2 | 5.06692616 | -3.421799736 | 1.2455 | -2.7474 | 0.00601 | 0.026226 |
| 1974 | NIPAL2 | 8.66098938 | -3.42257483 | 0.9951 | -3.4394 | 0.00058 | 0.003646 |
| 547 | KITLG | 4.62217891 | -3.428852412 | 1.177 | -2.9133 | 0.00358 | 0.017019 |
| 1964 | CA13 | 7.50617095 | -3.430361569 | 1.0584 | -3.2412 | 0.00119 | 0.006648 |
| 652 | NPC2 | 18.8077551 | -3.433502709 | 0.6021 | -5.7024 | 1.2E-08 | 2.46E-07 |
| 1820 | LFNG | 8.96505406 | -3.437983591 | 0.947 | -3.6304 | 0.00028 | 0.001942 |
| 1801 | CITED2 | 6.39058423 | -3.439492509 | 1.0078 | -3.4128 | 0.00064 | 0.003963 |
| 359 | MICAL2 | 11.3349842 | -3.442279917 | 0.974 | -3.5343 | 0.00041 | 0.002682 |
| 536 | SRGAP1 | 6.39975355 | -3.450968729 | 0.9768 | -3.533 | 0.00041 | 0.002689 |
| 1021 | CAMSAP3 | 23.6035751 | -3.458959751 | 0.5889 | -5.8738 | 4.3E-09 | 9.71E-08 |
| 1393 | UBA7 | 7.85978736 | -3.463332925 | 0.9536 | -3.6318 | 0.00028 | 0.001933 |
| 281 | PAPSS2 | 56.3070782 | -3.472769385 | 0.3607 | -9.6274 | 6.1E-22 | 1.81E-19 |
| 1503 | SMIM14 | 26.2802367 | -3.473378092 | 0.615 | -5.6476 | 1.6E-08 | 3.33E-07 |

| | | | | | | | |
|---|---|---|---|---|---|---|---|
| 743 | SEMA4B | 11.416991 | -3.483610568 | 0.7595 | -4.5869 | 4.5E-06 | 5.04E-05 |
| 1626 | SERINC5 | 10.8536773 | -3.488052185 | 0.8069 | -4.3228 | 1.5E-05 | 0.000151 |
| 862 | LRRC75A | 3.56316833 | -3.488772614 | 1.2732 | -2.7402 | 0.00614 | 0.026675 |
| 1762 | SH3BGRL2 | 23.6860851 | -3.490759519 | 0.5703 | -6.1207 | 9.3E-10 | 2.44E-08 |
| 1084 | NECTIN2 | 22.2327521 | -3.491468796 | 0.564 | -6.1902 | 6E-10 | 1.65E-08 |
| 1527 | ANTXR2 | 4.73623224 | -3.494698239 | 1.2174 | -2.8706 | 0.0041 | 0.019068 |
| 1766 | ME1 | 6.43834756 | -3.505842499 | 1.191 | -2.9437 | 0.00324 | 0.01567 |
| 317 | CASP7 | 8.52715073 | -3.506468596 | 0.8821 | -3.9751 | 7E-05 | 0.00058 |
| 887 | GSDMB | 32.7266022 | -3.507090223 | 0.5476 | -6.4043 | 1.5E-10 | 4.81E-09 |
| 526 | PMEL | 3.5887705 | -3.513205451 | 1.3798 | -2.5463 | 0.01089 | 0.042216 |
| 1466 | TMEM41A | 8.15774427 | -3.515331094 | 0.9018 | -3.8982 | 9.7E-05 | 0.000764 |
| 1558 | EGF | 3.57759216 | -3.531405441 | 1.344 | -2.6275 | 0.0086 | 0.034956 |
| 1234 | TNS1 | 5.31650823 | -3.532127653 | 1.2255 | -2.8823 | 0.00395 | 0.018499 |
| 616 | LRP10 | 33.7197871 | -3.534483773 | 0.5408 | -6.5356 | 6.3E-11 | 2.23E-09 |
| 467 | GRAMD1B | 4.91808744 | -3.538044296 | 1.1929 | -2.966 | 0.00302 | 0.014739 |
| 920 | ITGA3 | 14.064842 | -3.539423549 | 0.7957 | -4.4481 | 8.7E-06 | 9.02E-05 |
| 1103 | SPIB | 11.6627387 | -3.540140385 | 0.9883 | -3.5821 | 0.00034 | 0.002288 |
| 1068 | LTBP4 | 25.8804222 | -3.544403104 | 0.6053 | -5.8555 | 4.8E-09 | 1.07E-07 |
| 632 | PTGDR | 5.05950108 | -3.554241843 | 1.1903 | -2.9861 | 0.00283 | 0.013932 |
| 1723 | TAPBP | 46.3615155 | -3.55822429 | 0.4944 | -7.1967 | 6.2E-13 | 3.31E-11 |
| 1121 | RSAD2 | 3.53351441 | -3.570183215 | 1.2925 | -2.7623 | 0.00574 | 0.025273 |
| 74 | ERMAP | 3.50642157 | -3.573718816 | 1.3256 | -2.696 | 0.00702 | 0.029705 |
| 1693 | GCNT2 | 5.27127072 | -3.574567551 | 1.1506 | -3.1067 | 0.00189 | 0.009908 |
| 206 | BTG2 | 19.4186954 | -3.57634302 | 0.6312 | -5.6657 | 1.5E-08 | 3.03E-07 |
| 946 | SLC16A5 | 7.21693043 | -3.581962731 | 1.1006 | -3.2546 | 0.00114 | 0.00639 |
| 1248 | SP100 | 12.4708591 | -3.582705291 | 0.806 | -4.4452 | 8.8E-06 | 9.11E-05 |
| 1102 | RRAS | 6.51844956 | -3.583480276 | 1.1268 | -3.1802 | 0.00147 | 0.008 |
| 746 | FAM174B | 3.50868595 | -3.588928772 | 1.3083 | -2.7433 | 0.00608 | 0.026477 |
| 923 | ACSF2 | 5.27933911 | -3.601554349 | 1.1041 | -3.262 | 0.00111 | 0.006256 |
| 2156 | IGSF1 | 28.8753632 | -3.605830114 | 0.6197 | -5.8189 | 5.9E-09 | 1.32E-07 |
| 1431 | SLC12A8 | 3.53912868 | -3.607025561 | 1.3548 | -2.6625 | 0.00776 | 0.032186 |
| 820 | WFDC1 | 3.65649599 | -3.614499846 | 1.3168 | -2.7449 | 0.00605 | 0.026372 |
| 1398 | GLYCTK | 28.8044068 | -3.626602371 | 0.5511 | -6.5803 | 4.7E-11 | 1.71E-09 |
| 259 | RASSF4 | 11.1997746 | -3.626679551 | 0.8065 | -4.497 | 6.9E-06 | 7.36E-05 |
| 2161 | INTS6L | 13.7359551 | -3.634306957 | 0.7466 | -4.868 | 1.1E-06 | 1.47E-05 |
| 787 | ORAI3 | 3.8275766 | -3.635431794 | 1.2977 | -2.8013 | 0.00509 | 0.022872 |
| 716 | C2CD4B | 3.9046861 | -3.636268959 | 1.317 | -2.7611 | 0.00576 | 0.025342 |
| 826 | FOXF1 | 3.89919038 | -3.639333408 | 1.2561 | -2.8973 | 0.00376 | 0.017802 |
| 939 | SOX9 | 6.92572829 | -3.64216956 | 1.0251 | -3.5528 | 0.00038 | 0.002525 |

| | | | | | | | |
|---|---|---|---|---|---|---|---|
| 447 | COLCA1 | 5.66375717 | -3.643151368 | 1.2015 | -3.0321 | 0.00243 | 0.012278 |
| 1929 | DLC1 | 7.83274876 | -3.649144178 | 1.1777 | -3.0986 | 0.00194 | 0.010143 |
| 1632 | SEMA6A | 20.1174732 | -3.655426056 | 0.6053 | -6.0393 | 1.5E-09 | 3.88E-08 |
| 425 | CAPN5 | 32.4039764 | -3.657844623 | 0.5019 | -7.2879 | 3.1E-13 | 1.81E-11 |
| 542 | CPM | 3.82310096 | -3.659873415 | 1.3101 | -2.7936 | 0.00521 | 0.023326 |
| 60 | IFI6 | 78.9989786 | -3.662400813 | 0.4195 | -8.7314 | 2.5E-18 | 4.02E-16 |
| 2122 | EDA | 7.23277201 | -3.669033647 | 0.9554 | -3.8404 | 0.00012 | 0.000942 |
| 1610 | SPEF2 | 3.65617963 | -3.670595947 | 1.3901 | -2.6405 | 0.00828 | 0.033923 |
| 479 | DYRK4 | 5.69258922 | -3.673043319 | 1.1054 | -3.323 | 0.00089 | 0.005222 |
| 427 | SYTL2 | 33.4792922 | -3.68059018 | 0.5578 | -6.5979 | 4.2E-11 | 1.54E-09 |
| 225 | SLC30A1 | 17.4030055 | -3.681095503 | 0.6541 | -5.6281 | 1.8E-08 | 3.66E-07 |
| 209 | PIK3C2B | 11.661334 | -3.690224547 | 0.8308 | -4.4419 | 8.9E-06 | 9.24E-05 |
| 893 | DHX58 | 3.64577705 | -3.708822661 | 1.4907 | -2.488 | 0.01285 | 0.048428 |
| 1710 | HLA-E | 38.9174723 | -3.709251342 | 0.4499 | -8.244 | 1.7E-16 | 1.77E-14 |
| 1631 | MAN2A1 | 16.9800922 | -3.714580598 | 0.6568 | -5.6558 | 1.6E-08 | 3.19E-07 |
| 42 | HMGCL | 11.4983989 | -3.716743254 | 0.7885 | -4.7137 | 2.4E-06 | 2.93E-05 |
| 305 | NFKB2 | 2.34875692 | -3.719547067 | 1.4717 | -2.5275 | 0.01149 | 0.044087 |
| 656 | ADCK1 | 5.63233889 | -3.725993522 | 1.1219 | -3.3212 | 0.0009 | 0.005245 |
| 28 | FBLIM1 | 22.6771753 | -3.726424601 | 0.5799 | -6.4265 | 1.3E-10 | 4.24E-09 |
| 1636 | GRAMD2B | 7.63331746 | -3.726784304 | 1.01 | -3.69 | 0.00022 | 0.001584 |
| 1044 | PGPEP1 | 24.4758154 | -3.726826502 | 0.6029 | -6.1814 | 6.4E-10 | 1.73E-08 |
| 1711 | DDR1 | 32.4943508 | -3.727885433 | 0.4786 | -7.7892 | 6.7E-15 | 5.18E-13 |
| 1565 | ARSJ | 2.42727214 | -3.734135538 | 1.4664 | -2.5465 | 0.01088 | 0.042208 |
| 385 | FAM111A | 22.0857492 | -3.734818981 | 0.6089 | -6.1334 | 8.6E-10 | 2.27E-08 |
| 998 | ARID3A | 58.2712133 | -3.736026888 | 0.4089 | -9.1369 | 6.4E-20 | 1.41E-17 |
| 350 | TRIM21 | 7.72051171 | -3.738598583 | 0.9533 | -3.9215 | 8.8E-05 | 0.000704 |
| 1531 | PLAC8 | 14.2759948 | -3.738975016 | 0.7451 | -5.018 | 5.2E-07 | 7.43E-06 |
| 834 | RPH3AL | 2.40787543 | -3.743984189 | 1.4784 | -2.5324 | 0.01133 | 0.043586 |
| 837 | ABR | 7.97548985 | -3.746765159 | 0.9843 | -3.8065 | 0.00014 | 0.001059 |
| 2046 | LPAR1 | 2.44377617 | -3.747165147 | 1.4718 | -2.546 | 0.0109 | 0.042221 |
| 1587 | TLR2 | 2.44123915 | -3.747659619 | 1.48 | -2.5322 | 0.01133 | 0.043587 |
| 1368 | BHLHE40 | 12.0935807 | -3.756466793 | 0.9055 | -4.1487 | 3.3E-05 | 0.000301 |
| 604 | MBNL2 | 14.6889767 | -3.762829397 | 0.7683 | -4.8975 | 9.7E-07 | 1.29E-05 |
| 1360 | PRR5 | 5.55755867 | -3.763113339 | 1.1536 | -3.2621 | 0.00111 | 0.006256 |
| 1977 | EXT1 | 8.0679967 | -3.763482522 | 0.9185 | -4.0973 | 4.2E-05 | 0.000368 |
| 213 | RAB29 | 11.7457215 | -3.769381041 | 0.8728 | -4.3185 | 1.6E-05 | 0.000154 |
| 307 | ENSG00000273108 | 3.95295879 | -3.769607903 | 1.2635 | -2.9835 | 0.00285 | 0.01403 |
| 1900 | FOXP2 | 6.8324335 | -3.77765691 | 1.1634 | -3.2471 | 0.00117 | 0.006531 |
| 1794 | SGK1 | 2.52468464 | -3.785227976 | 1.4749 | -2.5665 | 0.01027 | 0.040268 |

| | | | | | | | |
|---|---|---|---|---|---|---|---|
| 1537 | HSD17B11 | 42.5820299 | -3.787718696 | 0.4956 | -7.6421 | 2.1E-14 | 1.52E-12 |
| 1391 | MST1 | 23.6547567 | -3.788552873 | 0.7434 | -5.0965 | 3.5E-07 | 5.13E-06 |
| 1787 | SMPDL3A | 4.4537633 | -3.799495318 | 1.3179 | -2.883 | 0.00394 | 0.018465 |
| 1129 | ITSN2 | 17.2235314 | -3.801781027 | 0.7515 | -5.0591 | 4.2E-07 | 6.11E-06 |
| 2086 | MT-TI | 2.58329063 | -3.808135727 | 1.4937 | -2.5494 | 0.01079 | 0.041913 |
| 1606 | CMBL | 35.2819109 | -3.811092462 | 0.6068 | -6.2808 | 3.4E-10 | 9.84E-09 |
| 1412 | ABI3BP | 4.32707464 | -3.816291582 | 1.3117 | -2.9093 | 0.00362 | 0.017214 |
| 2138 | NOX1 | 13.8015822 | -3.824900381 | 0.8108 | -4.7176 | 2.4E-06 | 2.88E-05 |
| 1194 | MAP3K2-DT | 2.52636557 | -3.830262929 | 1.5 | -2.5534 | 0.01067 | 0.041549 |
| 1989 | GSDMD | 30.0399256 | -3.833262575 | 0.5671 | -6.7598 | 1.4E-11 | 5.71E-10 |
| 2077 | TOR4A | 4.34926835 | -3.835325247 | 1.2621 | -3.0388 | 0.00238 | 0.012064 |
| 803 | ELMO3 | 11.7492155 | -3.840023421 | 0.8225 | -4.669 | 3E-06 | 3.54E-05 |
| 956 | SYNGR2 | 25.5884718 | -3.840899751 | 0.6294 | -6.1023 | 1E-09 | 2.71E-08 |
| 1432 | LINC02614 | 2.53394367 | -3.841032495 | 1.4618 | -2.6276 | 0.0086 | 0.034956 |
| 2017 | GNE | 3.98302488 | -3.847451031 | 1.4553 | -2.6438 | 0.0082 | 0.033667 |
| 530 | RBMS2 | 4.16572444 | -3.848405468 | 1.3401 | -2.8718 | 0.00408 | 0.019002 |
| 1698 | ALDH5A1 | 4.40321374 | -3.850825695 | 1.2607 | -3.0546 | 0.00225 | 0.011516 |
| 1331 | ENSG00000279110 | 4.44881927 | -3.855705642 | 1.2752 | -3.0236 | 0.0025 | 0.012545 |
| 258 | CXCL12 | 7.90722678 | -3.856273453 | 1.2153 | -3.173 | 0.00151 | 0.008176 |
| 174 | IGSF9 | 17.8007125 | -3.858472244 | 0.7153 | -5.3942 | 6.9E-08 | 1.21E-06 |
| 676 | DEGS2 | 2.53887394 | -3.860333781 | 1.4633 | -2.638 | 0.00834 | 0.034117 |
| 1694 | NEDD9 | 2.61348413 | -3.865636096 | 1.4631 | -2.642 | 0.00824 | 0.033812 |
| 2158 | GPC4 | 8.15525314 | -3.868920574 | 0.9846 | -3.9294 | 8.5E-05 | 0.000684 |
| 92 | ACOT11 | 15.1564546 | -3.871114795 | 0.7277 | -5.3198 | 1E-07 | 1.76E-06 |
| 393 | PLAAT3 | 4.5615531 | -3.876670266 | 1.3659 | -2.8381 | 0.00454 | 0.020744 |
| 1579 | MMAA | 4.30411593 | -3.880120747 | 1.3536 | -2.8666 | 0.00415 | 0.019284 |
| 921 | COL1A1 | 55.1123822 | -3.887069242 | 1.0266 | -3.7863 | 0.00015 | 0.001132 |
| 157 | S100A6 | 241.225641 | -3.8906311 | 0.2643 | -14.722 | 4.6E-49 | 3.29E-45 |
| 142 | TENT5C | 4.2844712 | -3.891032693 | 1.4646 | -2.6567 | 0.00789 | 0.032634 |
| 1685 | GMDS-DT | 8.33361765 | -3.891922083 | 1.1231 | -3.4652 | 0.00053 | 0.003358 |
| 1214 | ENSG00000287129 | 2.71462224 | -3.90004706 | 1.5278 | -2.5528 | 0.01069 | 0.041604 |
| 1828 | ETV1 | 4.45026013 | -3.908272313 | 1.2801 | -3.053 | 0.00227 | 0.011559 |
| 21 | NMNAT1 | 4.45328485 | -3.908660722 | 1.276 | -3.0632 | 0.00219 | 0.011254 |
| 1296 | SLC2A10 | 2.62485439 | -3.909676942 | 1.5064 | -2.5953 | 0.00945 | 0.037648 |
| 89 | ELAVL4 | 2.74532728 | -3.910989024 | 1.5061 | -2.5968 | 0.00941 | 0.037562 |
| 770 | ABCC6 | 13.9861411 | -3.911798114 | 0.8592 | -4.5531 | 5.3E-06 | 5.79E-05 |
| 1726 | DEF6 | 2.66469879 | -3.912557475 | 1.5075 | -2.5953 | 0.00945 | 0.037648 |
| 286 | IFIT1 | 8.73148662 | -3.91492491 | 0.9126 | -4.29 | 1.8E-05 | 0.000172 |
| 312 | PDCD4 | 38.2756744 | -3.915413148 | 0.5116 | -7.6528 | 2E-14 | 1.42E-12 |

| | | | | | | | |
|---|---|---|---|---|---|---|---|
| 105 | GNG12 | 32.181932 | -3.927063557 | 0.4996 | -7.8599 | 3.8E-15 | 3.09E-13 |
| 1500 | KLF3 | 26.2887122 | -3.931641704 | 0.5939 | -6.6202 | 3.6E-11 | 1.34E-09 |
| 1171 | CAPG | 8.24552232 | -3.943652864 | 0.9918 | -3.9761 | 7E-05 | 0.000578 |
| 1892 | LRRC17 | 2.6988357 | -3.945271566 | 1.5475 | -2.5495 | 0.01079 | 0.041913 |
| 801 | CES2 | 56.5813154 | -3.945949033 | 0.4499 | -8.7709 | 1.8E-18 | 2.86E-16 |
| 1871 | PON2 | 26.1961338 | -3.94706252 | 0.5508 | -7.1665 | 7.7E-13 | 4.01E-11 |
| 617 | ENSG00000215271 | 2.74322544 | -3.953982567 | 1.4758 | -2.6792 | 0.00738 | 0.030889 |
| 1753 | LRRC1 | 8.72627974 | -3.956638726 | 0.9666 | -4.0933 | 4.3E-05 | 0.000374 |
| 1262 | ANO7 | 2.92127908 | -3.959919263 | 1.4313 | -2.7666 | 0.00566 | 0.024979 |
| 1948 | RAB11FIP1 | 4.53704777 | -3.963588022 | 1.3236 | -2.9946 | 0.00275 | 0.013622 |
| 1660 | CARMN | 16.3234913 | -3.969385646 | 0.9226 | -4.3023 | 1.7E-05 | 0.000164 |
| 2024 | GCNT1 | 34.5711878 | -3.971962673 | 0.5472 | -7.259 | 3.9E-13 | 2.17E-11 |
| 1418 | PHLDB2 | 6.83730904 | -3.975611852 | 1.0979 | -3.621 | 0.00029 | 0.002005 |
| 263 | ANK3 | 14.389122 | -3.992678184 | 0.8186 | -4.8775 | 1.1E-06 | 1.41E-05 |
| 1475 | LINC02924 | 2.85695535 | -3.995351839 | 1.402 | -2.8498 | 0.00437 | 0.02012 |
| 1577 | INPP4B | 2.80266966 | -3.997486654 | 1.4893 | -2.6842 | 0.00727 | 0.030526 |
| 686 | PWRN1 | 2.96691717 | -4.000258861 | 1.4496 | -2.7595 | 0.00579 | 0.025428 |
| 16 | TPRG1L | 15.8108805 | -4.001947178 | 0.6971 | -5.7405 | 9.4E-09 | 2.03E-07 |
| 531 | TAC3 | 4.71401528 | -4.002572729 | 1.2849 | -3.1151 | 0.00184 | 0.009673 |
| 1137 | FOSL2 | 11.6237938 | -4.004821918 | 0.8629 | -4.6413 | 3.5E-06 | 4E-05 |
| 1819 | TMEM184A | 10.6394639 | -4.006716639 | 0.9866 | -4.061 | 4.9E-05 | 0.00042 |
| 1395 | SEMA3B | 18.2912855 | -4.009449514 | 0.6966 | -5.7554 | 8.6E-09 | 1.86E-07 |
| 1096 | PPP1R15A | 4.75388449 | -4.011408991 | 1.2248 | -3.2751 | 0.00106 | 0.00602 |
| 840 | TLCD2 | 13.9736868 | -4.017317762 | 0.8152 | -4.928 | 8.3E-07 | 1.13E-05 |
| 1295 | SDC4 | 13.0743376 | -4.017745802 | 0.8425 | -4.7688 | 1.9E-06 | 2.28E-05 |
| 1480 | PIGZ | 16.3434411 | -4.019287151 | 0.732 | -5.4908 | 4E-08 | 7.43E-07 |
| 51 | LDLRAP1 | 2.88474879 | -4.020961674 | 1.4937 | -2.692 | 0.0071 | 0.029983 |
| 781 | QPRT | 36.2157968 | -4.021870376 | 0.5482 | -7.3364 | 2.2E-13 | 1.29E-11 |
| 1036 | CYP4F3 | 2.94812839 | -4.022343402 | 1.4611 | -2.7529 | 0.00591 | 0.025831 |
| 1149 | EPAS1 | 15.6518121 | -4.022621025 | 0.7769 | -5.1778 | 2.2E-07 | 3.47E-06 |
| 1628 | CAST | 28.2630338 | -4.025191159 | 0.6444 | -6.2464 | 4.2E-10 | 1.2E-08 |
| 311 | ADD3 | 40.0191933 | -4.030972879 | 0.5286 | -7.6256 | 2.4E-14 | 1.7E-12 |
| 2010 | B4GALT1 | 20.6211453 | -4.034018127 | 0.6665 | -6.0526 | 1.4E-09 | 3.61E-08 |
| 1394 | MST1R | 10.9668 | -4.034238317 | 0.9073 | -4.4464 | 8.7E-06 | 9.07E-05 |
| 1747 | PAQR8 | 6.83212215 | -4.034598213 | 1.0561 | -3.8204 | 0.00013 | 0.00101 |
| 651 | ENTPD5 | 4.88421957 | -4.036276072 | 1.2174 | -3.3156 | 0.00091 | 0.005336 |
| 1158 | ANXA4 | 57.0803112 | -4.038345605 | 0.4301 | -9.3896 | 6E-21 | 1.53E-18 |
| 1584 | SH3D19 | 27.0519595 | -4.044430188 | 0.6279 | -6.4408 | 1.2E-10 | 3.91E-09 |
| 935 | ERN1 | 5.17390074 | -4.046427886 | 1.2298 | -3.2904 | 0.001 | 0.005749 |

| | | | | | | | |
|---|---|---|---|---|---|---|---|
| 382 | NR1H3 | 11.5543316 | -4.047315473 | 0.8684 | -4.6608 | 3.2E-06 | 3.68E-05 |
| 637 | CCDC198 | 6.78026975 | -4.05163837 | 1.1681 | -3.4687 | 0.00052 | 0.003323 |
| 977 | LAMA3 | 2.93419538 | -4.053667447 | 1.409 | -2.8771 | 0.00401 | 0.018733 |
| 2163 | BGN | 4.73612638 | -4.055513966 | 1.4167 | -2.8626 | 0.0042 | 0.019466 |
| 1950 | PLAT | 5.18007161 | -4.057851752 | 1.4356 | -2.8265 | 0.00471 | 0.0214 |
| 1852 | ABHD11 | 9.11493618 | -4.059879503 | 0.9524 | -4.263 | 2E-05 | 0.000192 |
| 341 | IRF7 | 12.1035071 | -4.062012823 | 0.8152 | -4.983 | 6.3E-07 | 8.74E-06 |
| 2117 | IQSEC2 | 5.15820648 | -4.063709383 | 1.1973 | -3.3941 | 0.00069 | 0.004187 |
| 386 | DTX4 | 19.7451814 | -4.06445772 | 0.7488 | -5.4281 | 5.7E-08 | 1.01E-06 |
| 1705 | H2AC6 | 7.05001694 | -4.067245662 | 1.0575 | -3.8463 | 0.00012 | 0.000924 |
| 1301 | NFATC2 | 3.0056171 | -4.068092094 | 1.3904 | -2.9259 | 0.00344 | 0.016441 |
| 492 | GPRC5A | 13.0473755 | -4.07375695 | 0.8233 | -4.9479 | 7.5E-07 | 1.03E-05 |
| 1848 | ASL | 37.6664227 | -4.074630773 | 0.5493 | -7.4183 | 1.2E-13 | 7.34E-12 |
| 1146 | OXER1 | 6.92711051 | -4.075074882 | 1.1018 | -3.6984 | 0.00022 | 0.001539 |
| 711 | MYZAP | 4.77206224 | -4.081503014 | 1.2391 | -3.2939 | 0.00099 | 0.005691 |
| 269 | TET1 | 3.05267725 | -4.083565767 | 1.3888 | -2.9403 | 0.00328 | 0.01579 |
| 1030 | ZNF443 | 3.09875284 | -4.085066792 | 1.5738 | -2.5956 | 0.00944 | 0.037641 |
| 1078 | ETHE1 | 38.2979069 | -4.085384081 | 0.5072 | -8.0551 | 7.9E-16 | 7.39E-14 |
| 886 | GRB7 | 7.76204297 | -4.094366764 | 1.0249 | -3.9949 | 6.5E-05 | 0.00054 |
| 2115 | ENSG00000270012 | 3.13620822 | -4.100830612 | 1.5051 | -2.7247 | 0.00644 | 0.027686 |
| 1585 | FHDC1 | 5.32363768 | -4.102074432 | 1.2837 | -3.1955 | 0.0014 | 0.007648 |
| 2 | ISG15 | 22.0788941 | -4.102492907 | 0.6134 | -6.6877 | 2.3E-11 | 8.88E-10 |
| 1277 | RRBP1 | 49.4303186 | -4.104524926 | 0.4505 | -9.1102 | 8.2E-20 | 1.75E-17 |
| 618 | PCK2 | 45.7737766 | -4.111151113 | 0.5226 | -7.8669 | 3.6E-15 | 2.94E-13 |
| 1299 | CEBPB | 3.20042332 | -4.115783646 | 1.3774 | -2.988 | 0.00281 | 0.013858 |
| 1592 | SMIM31 | 3.0774537 | -4.119833715 | 1.4205 | -2.9002 | 0.00373 | 0.017666 |
| 528 | ERBB3 | 57.4428391 | -4.120478274 | 0.4151 | -9.9275 | 3.2E-23 | 1.12E-20 |
| 303 | SEMA4G | 4.99763515 | -4.123830227 | 1.2376 | -3.3321 | 0.00086 | 0.005081 |
| 1048 | ZNF493 | 1.79945106 | -4.127733108 | 1.6415 | -2.5146 | 0.01192 | 0.045529 |
| 1855 | ELN | 3.16270563 | -4.127879696 | 1.5527 | -2.6585 | 0.00785 | 0.032524 |
| 1499 | ARAP2 | 3.15886365 | -4.128054684 | 1.6197 | -2.5486 | 0.01082 | 0.041993 |
| 1456 | LRRC31 | 1.82341774 | -4.130181239 | 1.6191 | -2.5509 | 0.01074 | 0.041791 |
| 578 | RFLNA | 14.6863884 | -4.130296293 | 0.7666 | -5.3878 | 7.1E-08 | 1.25E-06 |
| 1786 | GJA1 | 5.4050076 | -4.132028008 | 1.2979 | -3.1836 | 0.00145 | 0.007918 |
| 1498 | PCDH7 | 5.15568198 | -4.132808686 | 1.3647 | -3.0283 | 0.00246 | 0.012405 |
| 1735 | FOXP4-AS1 | 1.86737489 | -4.135287213 | 1.66 | -2.4912 | 0.01273 | 0.048091 |
| 132 | DENND2D | 10.0209611 | -4.142121119 | 0.9003 | -4.601 | 4.2E-06 | 4.74E-05 |
| 1674 | CPEB4 | 8.2493981 | -4.145977774 | 1.0539 | -3.9338 | 8.4E-05 | 0.000674 |
| 1363 | TTC38 | 18.0208593 | -4.150839487 | 0.7232 | -5.7393 | 9.5E-09 | 2.04E-07 |

| | | | | | | | |
|---|---|---|---|---|---|---|---|
| 871 | SLC6A4 | 1.83657756 | -4.153965317 | 1.6289 | -2.5501 | 0.01077 | 0.041856 |
| 243 | AKR1C1 | 1.85940818 | -4.155662917 | 1.6191 | -2.5666 | 0.01027 | 0.040265 |
| 1972 | MATN2 | 3.19446549 | -4.157446011 | 1.5972 | -2.603 | 0.00924 | 0.037086 |
| 1054 | CEBPA-DT | 1.85248782 | -4.161141842 | 1.6328 | -2.5485 | 0.01082 | 0.041993 |
| 1511 | KDR | 1.87475041 | -4.163483906 | 1.6471 | -2.5278 | 0.01148 | 0.04407 |
| 1532 | GPAT3 | 1.83282622 | -4.169068958 | 1.6668 | -2.5013 | 0.01238 | 0.046981 |
| 859 | MYOCD | 5.3440703 | -4.170350965 | 1.5286 | -2.7282 | 0.00637 | 0.027443 |
| 1495 | PROM1 | 15.0778239 | -4.173889649 | 0.8313 | -5.0208 | 5.1E-07 | 7.35E-06 |
| 137 | BCL2L15 | 15.7236859 | -4.176805425 | 0.7518 | -5.5557 | 2.8E-08 | 5.37E-07 |
| 31 | ARHGEF10L | 35.7131893 | -4.177000035 | 0.5482 | -7.6199 | 2.5E-14 | 1.76E-12 |
| 844 | ATP2A3 | 7.39203809 | -4.177275529 | 1.163 | -3.5918 | 0.00033 | 0.002216 |
| 1870 | PPP1R9A | 5.48483718 | -4.179798783 | 1.2075 | -3.4614 | 0.00054 | 0.003397 |
| 107 | ST6GALNAC3 | 1.8742721 | -4.180041388 | 1.6091 | -2.5977 | 0.00939 | 0.037491 |
| 85 | FAAH | 12.8936616 | -4.180130577 | 0.8037 | -5.201 | 2E-07 | 3.1E-06 |
| 43 | FUCA1 | 10.5142305 | -4.180713882 | 0.9141 | -4.5736 | 4.8E-06 | 5.32E-05 |
| 1260 | GPR35 | 10.4528111 | -4.180772658 | 0.9077 | -4.606 | 4.1E-06 | 4.64E-05 |
| 1307 | NRIP1 | 7.71365835 | -4.186235014 | 1.022 | -4.096 | 4.2E-05 | 0.00037 |
| 1120 | CMPK2 | 5.25045691 | -4.189404865 | 1.3639 | -3.0715 | 0.00213 | 0.01098 |
| 1358 | PHETA2 | 1.8908401 | -4.205559697 | 1.5827 | -2.6573 | 0.00788 | 0.032603 |
| 1905 | CADPS2 | 3.27207097 | -4.205886286 | 1.4514 | -2.8979 | 0.00376 | 0.017778 |
| 1700 | CARMIL1 | 16.1796179 | -4.206584366 | 0.7339 | -5.7318 | 9.9E-09 | 2.12E-07 |
| 1034 | ADGRE5 | 9.93966821 | -4.207734504 | 0.9425 | -4.4642 | 8E-06 | 8.43E-05 |
| 885 | ERBB2 | 12.7729358 | -4.220841006 | 0.8497 | -4.9677 | 6.8E-07 | 9.38E-06 |
| 40 | TCEA3 | 10.2326407 | -4.22265907 | 1.0683 | -3.9528 | 7.7E-05 | 0.000628 |
| 1945 | ENSG00000253414 | 1.92990985 | -4.223382918 | 1.6911 | -2.4974 | 0.01251 | 0.047401 |
| 80 | BTBD19 | 1.95046835 | -4.223474359 | 1.6179 | -2.6105 | 0.00904 | 0.036467 |
| 1602 | SLC12A7 | 25.8380915 | -4.223546429 | 0.6542 | -6.4563 | 1.1E-10 | 3.62E-09 |
| 360 | SPON1 | 10.792597 | -4.225252711 | 0.926 | -4.5628 | 5E-06 | 5.57E-05 |
| 573 | P2RX4 | 17.7808218 | -4.227448767 | 0.8423 | -5.0187 | 5.2E-07 | 7.41E-06 |
| 282 | RNLS | 3.32230014 | -4.227470987 | 1.4159 | -2.9857 | 0.00283 | 0.013936 |
| 1851 | ENSG00000274080 | 1.92569448 | -4.227709295 | 1.6897 | -2.502 | 0.01235 | 0.046911 |
| 1325 | LINC01637 | 1.94966116 | -4.230813489 | 1.5912 | -2.6589 | 0.00784 | 0.032507 |
| 1369 | IL17RC | 1.94966116 | -4.230813489 | 1.5912 | -2.6589 | 0.00784 | 0.032507 |
| 127 | ELAPOR1 | 5.53456911 | -4.232347481 | 1.2413 | -3.4095 | 0.00065 | 0.003995 |
| 644 | TMEM229B | 1.97941215 | -4.233484786 | 1.6997 | -2.4907 | 0.01275 | 0.048135 |
| 1033 | MISP3 | 1.90773697 | -4.237379428 | 1.6217 | -2.6129 | 0.00898 | 0.036268 |
| 462 | USP2 | 8.01370767 | -4.241588733 | 1.1126 | -3.8123 | 0.00014 | 0.001039 |
| 1821 | IQCE | 5.65988626 | -4.242061983 | 1.2156 | -3.4897 | 0.00048 | 0.003097 |
| 1115 | ZNF784 | 3.29141938 | -4.248519461 | 1.3763 | -3.0869 | 0.00202 | 0.010503 |

| | | | | | | | |
|---|---|---|---|---|---|---|---|
| 399 | NEAT1 | 816.514286 | -4.249059405 | 0.5282 | -8.0451 | 8.6E-16 | 7.91E-14 |
| 2028 | NTRK2 | 7.65607039 | -4.251752744 | 1.0721 | -3.9656 | 7.3E-05 | 0.000599 |
| 443 | CASP5 | 1.97598967 | -4.252494792 | 1.607 | -2.6462 | 0.00814 | 0.033477 |
| 239 | CHRM3-AS2 | 3.38401852 | -4.254170659 | 1.5287 | -2.7829 | 0.00539 | 0.023999 |
| 1833 | LINC02889 | 2.01767471 | -4.255328451 | 1.6253 | -2.6182 | 0.00884 | 0.035793 |
| 1719 | C2 | 5.71423792 | -4.260314322 | 1.3366 | -3.1875 | 0.00143 | 0.007844 |
| 882 | LASP1 | 3.32797785 | -4.262779022 | 1.3741 | -3.1023 | 0.00192 | 0.010029 |
| 835 | RPH3AL-AS2 | 1.96769409 | -4.264916792 | 1.575 | -2.7079 | 0.00677 | 0.028878 |
| 1354 | KDELR3 | 5.93563866 | -4.266531337 | 1.2078 | -3.5324 | 0.00041 | 0.002692 |
| 1926 | PPP1R3B | 8.613921 | -4.266598253 | 1.0199 | -4.1832 | 2.9E-05 | 0.000263 |
| 476 | B3GAT1-DT | 5.63458775 | -4.267331739 | 1.2504 | -3.4129 | 0.00064 | 0.003963 |
| 1319 | SLC37A1 | 23.2859956 | -4.268544837 | 0.6637 | -6.4312 | 1.3E-10 | 4.13E-09 |
| 1982 | SNTB1 | 8.10170337 | -4.275093791 | 1.0074 | -4.2437 | 2.2E-05 | 0.000206 |
| 135 | RHOC | 57.0726106 | -4.276123802 | 0.4428 | -9.6569 | 4.6E-22 | 1.42E-19 |
| 1312 | RUNX1 | 3.48509839 | -4.276337215 | 1.3665 | -3.1295 | 0.00175 | 0.009277 |
| 402 | RIN1 | 3.35597017 | -4.279662937 | 1.4431 | -2.9657 | 0.00302 | 0.014749 |
| 458 | SCN4B | 2.02561826 | -4.280689922 | 1.5887 | -2.6944 | 0.00705 | 0.029798 |
| 1216 | COL3A1 | 121.571915 | -4.29595947 | 0.7875 | -5.4549 | 4.9E-08 | 8.91E-07 |
| 1184 | MALL | 2.0057175 | -4.297637756 | 1.5687 | -2.7396 | 0.00615 | 0.026705 |
| 1200 | KYNU | 2.06160871 | -4.306285788 | 1.6641 | -2.5878 | 0.00966 | 0.038278 |
| 1975 | STK3 | 3.54508861 | -4.308366478 | 1.4553 | -2.9605 | 0.00307 | 0.014954 |
| 815 | LDHD | 2.01968451 | -4.309796986 | 1.5991 | -2.6951 | 0.00704 | 0.029755 |
| 215 | DYRK3 | 3.61204899 | -4.313917897 | 1.7232 | -2.5034 | 0.0123 | 0.046732 |
| 1629 | ERAP2 | 8.57687648 | -4.315883091 | 1.1544 | -3.7385 | 0.00019 | 0.001337 |
| 287 | PCGF5 | 8.85000498 | -4.321023495 | 1.0177 | -4.2459 | 2.2E-05 | 0.000205 |
| 533 | STAT6 | 37.1588313 | -4.323739254 | 0.5135 | -8.4194 | 3.8E-17 | 4.77E-15 |
| 523 | METTL7B | 2.12986142 | -4.326301175 | 1.6803 | -2.5748 | 0.01003 | 0.03947 |
| 1690 | RREB1 | 25.7074718 | -4.326355423 | 0.6029 | -7.1758 | 7.2E-13 | 3.79E-11 |
| 1755 | FAM83B | 2.13132634 | -4.327314447 | 1.5902 | -2.7212 | 0.0065 | 0.027921 |
| 1135 | KRTCAP3 | 13.9731214 | -4.332248785 | 0.8091 | -5.3543 | 8.6E-08 | 1.47E-06 |
| 900 | DUSP3 | 8.62162336 | -4.333636982 | 1.0758 | -4.0283 | 5.6E-05 | 0.000476 |
| 494 | EMP1 | 2.0446378 | -4.33389525 | 1.6467 | -2.6319 | 0.00849 | 0.034645 |
| 47 | IFNLR1 | 2.13222324 | -4.334553992 | 1.5726 | -2.7562 | 0.00585 | 0.025608 |
| 1708 | TRIM10 | 2.12995113 | -4.33563517 | 1.6113 | -2.6908 | 0.00713 | 0.030056 |
| 1023 | SHFL | 15.0472507 | -4.335955458 | 0.7993 | -5.4248 | 5.8E-08 | 1.03E-06 |
| 17 | ESPN | 3.65860859 | -4.336918611 | 1.4329 | -3.0267 | 0.00247 | 0.012457 |
| 1714 | MICA | 2.08575481 | -4.338200446 | 1.6289 | -2.6632 | 0.00774 | 0.032148 |
| 316 | RPS15AP30 | 6.06468764 | -4.340141013 | 1.214 | -3.5752 | 0.00035 | 0.002344 |
| 432 | FUT4 | 11.1335213 | -4.345150852 | 1.0097 | -4.3035 | 1.7E-05 | 0.000163 |

| | | | | | | | |
|---|---|---|---|---|---|---|---|
| 1836 | HOTAIRM1 | 2.09417668 | -4.348250536 | 1.6482 | -2.6382 | 0.00833 | 0.034107 |
| 821 | ATP2C2 | 25.2044682 | -4.350808475 | 0.6652 | -6.5406 | 6.1E-11 | 2.16E-09 |
| 489 | STYK1 | 2.08257149 | -4.352451234 | 1.5843 | -2.7472 | 0.00601 | 0.026226 |
| 643 | PLEK2 | 3.50714115 | -4.35331219 | 1.3811 | -3.1522 | 0.00162 | 0.008689 |
| 773 | GP2 | 2.11506406 | -4.357161229 | 1.5905 | -2.7396 | 0.00615 | 0.026705 |
| 1328 | ADORA2A-AS1 | 2.12903107 | -4.369695757 | 1.7178 | -2.5438 | 0.01096 | 0.042415 |
| 2051 | BSPRY | 3.61643259 | -4.371415371 | 1.3373 | -3.269 | 0.00108 | 0.006127 |
| 2123 | GDPD2 | 8.55826769 | -4.371464275 | 1.0817 | -4.0413 | 5.3E-05 | 0.000453 |
| 55 | CRYBG2 | 2.13243924 | -4.371895089 | 1.7176 | -2.5453 | 0.01092 | 0.04228 |
| 744 | CIB1 | 43.7942614 | -4.377001854 | 0.4711 | -9.2909 | 1.5E-20 | 3.63E-18 |
| 1535 | PTPN13 | 3.64898912 | -4.378690563 | 1.3208 | -3.3153 | 0.00092 | 0.005338 |
| 339 | ANO9 | 80.7322905 | -4.379042065 | 0.481 | -9.104 | 8.7E-20 | 1.82E-17 |
| 1790 | TMEM200A | 3.63905548 | -4.379442962 | 1.3308 | -3.2909 | 0.001 | 0.005745 |
| 1810 | RSPH3 | 3.77203158 | -4.38094348 | 1.376 | -3.1839 | 0.00145 | 0.007915 |
| 1162 | ACTG2 | 17.0036566 | -4.383265854 | 1.5044 | -2.9137 | 0.00357 | 0.01701 |
| 212 | SLC45A3 | 2.14058537 | -4.384074738 | 1.5366 | -2.8531 | 0.00433 | 0.01996 |
| 1016 | CRB3 | 3.71001834 | -4.386236831 | 1.3339 | -3.2884 | 0.00101 | 0.00578 |
| 383 | AGBL2 | 2.19728378 | -4.388416286 | 1.6606 | -2.6427 | 0.00823 | 0.033769 |
| 543 | ENSG00000274979 | 2.25872899 | -4.391995446 | 1.7133 | -2.5635 | 0.01036 | 0.040561 |
| 409 | LINC02754 | 6.39070949 | -4.392517575 | 1.2311 | -3.5679 | 0.00036 | 0.0024 |
| 440 | MMP1 | 2.2201144 | -4.395062841 | 1.698 | -2.5883 | 0.00964 | 0.038238 |
| 965 | RAB40B | 8.63629316 | -4.395536077 | 1.1078 | -3.9678 | 7.3E-05 | 0.000595 |
| 798 | CDH11 | 3.61714545 | -4.39729706 | 1.38 | -3.1865 | 0.00144 | 0.007859 |
| 876 | SLFN5 | 2.20770202 | -4.397999797 | 1.6035 | -2.7427 | 0.00609 | 0.026514 |
| 1348 | CDC42EP1 | 8.70870037 | -4.400784424 | 1.0821 | -4.0668 | 4.8E-05 | 0.000411 |
| 1265 | NEU4 | 2.13432275 | -4.401524505 | 1.5865 | -2.7743 | 0.00553 | 0.024505 |
| 666 | RIN3 | 2.15544929 | -4.404855086 | 1.5372 | -2.8654 | 0.00416 | 0.019339 |
| 1907 | TSPAN33 | 2.14715371 | -4.406290356 | 1.7158 | -2.5681 | 0.01022 | 0.040113 |
| 1816 | ADAP1 | 9.13218877 | -4.411448929 | 0.9943 | -4.4365 | 9.1E-06 | 9.46E-05 |
| 890 | ENSG00000234477 | 2.22370199 | -4.417016422 | 1.5516 | -2.8468 | 0.00442 | 0.02027 |
| 1684 | GMDS | 23.8999635 | -4.418774547 | 0.6543 | -6.7535 | 1.4E-11 | 5.9E-10 |
| 355 | CYB5R2 | 2.28821765 | -4.427683361 | 1.6297 | -2.7168 | 0.00659 | 0.02824 |
| 736 | CTSH | 18.3702788 | -4.435099138 | 0.801 | -5.5372 | 3.1E-08 | 5.89E-07 |
| 328 | PTPRE | 2.23970196 | -4.437694902 | 1.5346 | -2.8918 | 0.00383 | 0.018062 |
| 1642 | CCNI2 | 2.22994146 | -4.440260279 | 1.5323 | -2.8977 | 0.00376 | 0.017781 |
| 934 | ACE | 12.5235063 | -4.441093164 | 1.1041 | -4.0223 | 5.8E-05 | 0.000486 |
| 164 | EFNA4 | 3.71373646 | -4.442121019 | 1.4183 | -3.132 | 0.00174 | 0.009215 |
| 1204 | LINC01806 | 2.27276255 | -4.44317833 | 1.5707 | -2.8289 | 0.00467 | 0.021271 |
| 1560 | PANCR | 2.23254244 | -4.448782571 | 1.5501 | -2.8701 | 0.0041 | 0.019096 |

| | | | | | | | |
|---|---|---|---|---|---|---|---|
| 82 | CCDC17 | 2.27479551 | -4.449480262 | 1.5054 | -2.9557 | 0.00312 | 0.015154 |
| 1551 | CXXC4 | 2.3184238 | -4.451527896 | 1.6092 | -2.7663 | 0.00567 | 0.024985 |
| 1604 | SEMA5A | 3.88935 | -4.453378269 | 1.3408 | -3.3215 | 0.0009 | 0.005242 |
| 1513 | REST | 12.2000682 | -4.457579594 | 0.8873 | -5.0237 | 5.1E-07 | 7.25E-06 |
| 1207 | SCN9A | 2.2551339 | -4.458040717 | 1.5276 | -2.9184 | 0.00352 | 0.016808 |
| 214 | IKBKE | 3.7775775 | -4.461723904 | 1.5408 | -2.8957 | 0.00378 | 0.017879 |
| 1796 | TNFAIP3 | 2.31704858 | -4.465002411 | 1.7633 | -2.5322 | 0.01133 | 0.043587 |
| 466 | CLMP | 3.86243197 | -4.466306684 | 1.6024 | -2.7872 | 0.00532 | 0.023724 |
| 502 | PYROXD1 | 3.91579502 | -4.468085642 | 1.3549 | -3.2977 | 0.00097 | 0.005619 |
| 256 | SVIL | 10.1904185 | -4.471876918 | 0.9954 | -4.4925 | 7E-06 | 7.5E-05 |
| 375 | CD44 | 6.42238671 | -4.473008794 | 1.205 | -3.7121 | 0.00021 | 0.001465 |
| 450 | BCO2 | 3.95587857 | -4.482266522 | 1.6696 | -2.6846 | 0.00726 | 0.030502 |
| 1336 | TCN2 | 9.47589689 | -4.483532547 | 1.0214 | -4.3897 | 1.1E-05 | 0.000115 |
| 199 | NR5A2 | 13.4041439 | -4.497284178 | 0.981 | -4.5843 | 4.6E-06 | 5.08E-05 |
| 544 | PTPRB | 2.32076247 | -4.501308322 | 1.5915 | -2.8284 | 0.00468 | 0.021293 |
| 1845 | TNS3 | 15.8240838 | -4.504061188 | 0.8406 | -5.3581 | 8.4E-08 | 1.44E-06 |
| 470 | VSIG2 | 12.0610491 | -4.504987713 | 1.0276 | -4.3842 | 1.2E-05 | 0.000118 |
| 639 | PPP1R36 | 2.31687597 | -4.509435583 | 1.6345 | -2.759 | 0.0058 | 0.025451 |
| 1015 | FUT3 | 2.33207761 | -4.514049016 | 1.7434 | -2.5893 | 0.00962 | 0.038161 |
| 1823 | CARD11 | 2.3026555 | -4.51465954 | 1.7375 | -2.5984 | 0.00937 | 0.037461 |
| 1925 | CLDN23 | 2.36788864 | -4.516622318 | 1.7619 | -2.5635 | 0.01036 | 0.040561 |
| 249 | ST8SIA6 | 2.37684196 | -4.518757051 | 1.5784 | -2.8628 | 0.0042 | 0.019462 |
| 63 | SESN2 | 10.0523364 | -4.520620259 | 1.0742 | -4.2085 | 2.6E-05 | 0.000237 |
| 1805 | ZC3H12D | 2.43126824 | -4.522465471 | 1.5983 | -2.8295 | 0.00466 | 0.021244 |
| 567 | SLC8B1 | 3.99221356 | -4.526485352 | 1.3167 | -3.4378 | 0.00059 | 0.003661 |
| 1150 | EPCAM-DT | 2.3874008 | -4.526796694 | 1.4843 | -3.0498 | 0.00229 | 0.011676 |
| 180 | KLHDC9 | 2.41885587 | -4.534078243 | 1.7184 | -2.6385 | 0.00833 | 0.03409 |
| 586 | POSTN | 16.3506716 | -4.53436053 | 0.8912 | -5.0881 | 3.6E-07 | 5.33E-06 |
| 2104 | ARX | 2.39933485 | -4.534389477 | 1.7594 | -2.5772 | 0.00996 | 0.039294 |
| 712 | ALDH1A2 | 2.42419843 | -4.53986861 | 1.5044 | -3.0178 | 0.00255 | 0.012759 |
| 587 | FREM2 | 2.36090854 | -4.548486603 | 1.5589 | -2.9178 | 0.00353 | 0.016834 |
| 912 | HOXB7 | 6.49075851 | -4.548982036 | 1.1921 | -3.816 | 0.00014 | 0.001024 |
| 349 | SLC22A18 | 22.8259718 | -4.550806124 | 0.6862 | -6.6317 | 3.3E-11 | 1.25E-09 |
| 883 | LASP1NB | 2.36350952 | -4.551718197 | 1.6207 | -2.8086 | 0.00498 | 0.022501 |
| 431 | SMCO4 | 6.70229785 | -4.552197425 | 1.2112 | -3.7585 | 0.00017 | 0.001247 |
| 436 | MAML2 | 2.38146704 | -4.552262422 | 1.5078 | -3.0192 | 0.00253 | 0.012706 |
| 1225 | CASP10 | 10.0933165 | -4.55264747 | 1.0137 | -4.491 | 7.1E-06 | 7.55E-05 |
| 1209 | ITGA6 | 15.8497961 | -4.557573452 | 0.8737 | -5.2162 | 1.8E-07 | 2.89E-06 |
| 727 | GRAMD2A | 2.43238114 | -4.557791211 | 1.7116 | -2.6629 | 0.00775 | 0.032162 |

| | | | | | | | |
|---|---|---|---|---|---|---|---|
| 411 | ALDH3B1 | 4.11490194 | -4.558000654 | 1.3277 | -3.4331 | 0.0006 | 0.003715 |
| 945 | FADS6 | 4.2059213 | -4.560695292 | 1.3455 | -3.3895 | 0.0007 | 0.004246 |
| 1408 | ZNF717 | 2.48349782 | -4.562474116 | 1.722 | -2.6495 | 0.00806 | 0.03322 |
| 833 | GAS8 | 2.44052727 | -4.566238555 | 1.4768 | -3.0919 | 0.00199 | 0.010344 |
| 1481 | MELTF | 9.95598221 | -4.566380306 | 1.1114 | -4.1087 | 4E-05 | 0.000352 |
| 986 | RAB27B | 2.49424493 | -4.56778511 | 1.5299 | -2.9858 | 0.00283 | 0.013936 |
| 1615 | CCDC152 | 2.50731505 | -4.570525667 | 1.5105 | -3.0259 | 0.00248 | 0.012475 |
| 955 | TMC6 | 6.7129178 | -4.571451457 | 1.2403 | -3.6856 | 0.00023 | 0.001606 |
| 1738 | C6orf132 | 6.8544655 | -4.573209139 | 1.2008 | -3.8086 | 0.00014 | 0.001053 |
| 1089 | HIF3A | 6.67939235 | -4.575516713 | 1.2935 | -3.5373 | 0.0004 | 0.00266 |
| 1522 | LINC02562 | 2.53877012 | -4.5758846 | 1.5706 | -2.9134 | 0.00358 | 0.017019 |
| 1840 | HOXA10 | 4.10702494 | -4.576105388 | 1.3375 | -3.4214 | 0.00062 | 0.003857 |
| 1342 | APOL1 | 2.40981965 | -4.579076477 | 1.7048 | -2.686 | 0.00723 | 0.030417 |
| 1946 | ZNF703 | 16.8321125 | -4.587845961 | 0.7846 | -5.8476 | 5E-09 | 1.12E-07 |
| 629 | SSTR1 | 2.49910918 | -4.592929771 | 1.4898 | -3.0829 | 0.00205 | 0.010628 |
| 877 | MMP28 | 2.43354717 | -4.595156799 | 1.5969 | -2.8776 | 0.00401 | 0.01872 |
| 283 | FAS | 4.26258827 | -4.597767772 | 1.3516 | -3.4016 | 0.00067 | 0.004086 |
| 2002 | FREM1 | 4.20680813 | -4.601155372 | 1.3841 | -3.3244 | 0.00089 | 0.0052 |
| 671 | SERPINA10 | 2.53728203 | -4.601291551 | 1.5311 | -3.0052 | 0.00265 | 0.013243 |
| 1906 | SLC13A1 | 2.48446127 | -4.601599489 | 1.7139 | -2.6848 | 0.00726 | 0.030499 |
| 1138 | CAPN13 | 2.49194966 | -4.60302203 | 1.506 | -3.0565 | 0.00224 | 0.011468 |
| 1449 | MME | 7.09525065 | -4.605206298 | 1.2464 | -3.6948 | 0.00022 | 0.001558 |
| 706 | SLC27A2 | 7.03269021 | -4.607427487 | 1.1943 | -3.8579 | 0.00011 | 0.000886 |
| 2068 | TMEM141 | 59.0176029 | -4.607556357 | 0.4668 | -9.8706 | 5.6E-23 | 1.89E-20 |
| 1553 | SGMS2 | 6.88420014 | -4.608795067 | 1.2706 | -3.6274 | 0.00029 | 0.001963 |
| 1944 | RBPMS | 7.13604249 | -4.610047666 | 1.1985 | -3.8464 | 0.00012 | 0.000924 |
| 597 | NEK3 | 6.98225813 | -4.610292834 | 1.2611 | -3.6558 | 0.00026 | 0.001778 |
| 453 | NXPE4 | 6.64423415 | -4.612644623 | 1.4078 | -3.2766 | 0.00105 | 0.005993 |
| 1471 | RTP4 | 2.61952491 | -4.614487965 | 1.7272 | -2.6716 | 0.00755 | 0.031473 |
| 1401 | WNT5A | 2.5287473 | -4.615714443 | 1.5505 | -2.977 | 0.00291 | 0.014293 |
| 2011 | SPINK4 | 2.58091032 | -4.62004936 | 1.8563 | -2.4889 | 0.01282 | 0.048369 |
| 1490 | RN7SL589P | 2.52737208 | -4.620072628 | 1.6162 | -2.8587 | 0.00425 | 0.019648 |
| 1543 | ATOH1 | 2.56166729 | -4.620557039 | 1.4595 | -3.1658 | 0.00155 | 0.00835 |
| 289 | MYOF | 2.48918492 | -4.622780622 | 1.6664 | -2.7741 | 0.00554 | 0.02451 |
| 146 | PDZK1 | 2.6426844 | -4.62296938 | 1.7091 | -2.7049 | 0.00683 | 0.02907 |
| 1991 | EPPK1 | 4.35398616 | -4.623360205 | 1.332 | -3.471 | 0.00052 | 0.003298 |
| 985 | MRO | 2.61122933 | -4.624414148 | 1.8063 | -2.5602 | 0.01046 | 0.040881 |
| 1756 | BMP5 | 2.61099017 | -4.624821515 | 1.7548 | -2.6356 | 0.0084 | 0.034334 |
| 769 | MYH11 | 14.8079956 | -4.625768931 | 1.6801 | -2.7533 | 0.0059 | 0.025805 |

| | | | | | | | |
|---|---|---|---|---|---|---|---|
| 499 | ERP27 | 2.56256419 | -4.628283311 | 1.4761 | -3.1355 | 0.00172 | 0.009128 |
| 1405 | FEZF2 | 2.49211478 | -4.628503589 | 1.7422 | -2.6567 | 0.00789 | 0.032634 |
| 1254 | NGEF | 2.53110912 | -4.629844166 | 1.4804 | -3.1275 | 0.00176 | 0.009326 |
| 602 | SLITRK6 | 2.54417924 | -4.632128336 | 1.5753 | -2.9404 | 0.00328 | 0.015788 |
| 917 | HOXB13 | 2.5118661 | -4.634733706 | 1.7786 | -2.6059 | 0.00916 | 0.036833 |
| 1113 | TMEM238 | 17.0264663 | -4.636184806 | 0.7953 | -5.8296 | 5.6E-09 | 1.24E-07 |
| 1655 | ARHGAP26 | 13.6989756 | -4.636476004 | 0.8637 | -5.3679 | 8E-08 | 1.37E-06 |
| 1396 | HYAL1 | 4.29338123 | -4.637786571 | 1.3026 | -3.5603 | 0.00037 | 0.002462 |
| 1249 | ITM2C | 139.932192 | -4.640756769 | 0.3336 | -13.911 | 5.4E-44 | 1.93E-40 |
| 505 | REP15 | 2.6193523 | -4.64134568 | 1.498 | -3.0984 | 0.00195 | 0.010146 |
| 1562 | FAM241A | 2.52891242 | -4.646387637 | 1.6866 | -2.7549 | 0.00587 | 0.0257 |
| 1792 | AKAP7 | 7.18013533 | -4.647104421 | 1.1922 | -3.8979 | 9.7E-05 | 0.000764 |
| 950 | SMIM5 | 2.62810305 | -4.658883577 | 1.5791 | -2.9504 | 0.00317 | 0.015377 |
| 9 | VWA1 | 10.2783471 | -4.660612425 | 1.0662 | -4.3713 | 1.2E-05 | 0.000124 |
| 1186 | MERTK | 2.58309954 | -4.66367288 | 1.5246 | -3.0589 | 0.00222 | 0.011394 |
| 504 | SSPN | 2.65680768 | -4.66641787 | 1.4975 | -3.116 | 0.00183 | 0.009653 |
| 1451 | ENSG00000272087 | 2.64207006 | -4.670217363 | 1.6399 | -2.848 | 0.0044 | 0.020211 |
| 2022 | NMRK1 | 10.8862641 | -4.67547196 | 1.0362 | -4.5122 | 6.4E-06 | 6.89E-05 |
| 2157 | FIRRE | 11.4378471 | -4.676170019 | 1.0182 | -4.5924 | 4.4E-06 | 4.93E-05 |
| 1341 | APOL2 | 7.26019658 | -4.679638313 | 1.1785 | -3.9708 | 7.2E-05 | 0.000589 |
| 414 | CPT1A | 49.4757955 | -4.681592075 | 0.5739 | -8.1578 | 3.4E-16 | 3.35E-14 |
| 598 | PCDH17 | 2.74902706 | -4.686704425 | 1.5436 | -3.0363 | 0.00239 | 0.012141 |
| 78 | TMEM53 | 4.4922916 | -4.687627194 | 1.3262 | -3.5348 | 0.00041 | 0.002679 |
| 365 | LDHA | 182.292977 | -4.68785926 | 0.6605 | -7.0974 | 1.3E-12 | 6.48E-11 |
| 1702 | SLC17A1 | 2.68985397 | -4.690754004 | 1.5311 | -3.0636 | 0.00219 | 0.011243 |
| 1545 | ADH6 | 4.49927594 | -4.691195634 | 1.2718 | -3.6885 | 0.00023 | 0.001592 |
| 1029 | ZNF799 | 2.7399243 | -4.696337261 | 1.5106 | -3.1089 | 0.00188 | 0.00985 |
| 1037 | CYP4F12 | 2.67800962 | -4.697963975 | 1.5082 | -3.115 | 0.00184 | 0.009673 |
| 248 | ITGA8 | 2.68539945 | -4.699480003 | 1.5729 | -2.9878 | 0.00281 | 0.013864 |
| 1909 | AKR1B10 | 4.66479865 | -4.700511607 | 1.6762 | -2.8043 | 0.00504 | 0.022728 |
| 1208 | NOSTRIN | 7.47351941 | -4.703473712 | 1.1831 | -3.9755 | 7E-05 | 0.000579 |
| 1863 | STEAP2 | 14.2119917 | -4.703816882 | 0.8637 | -5.4459 | 5.2E-08 | 9.34E-07 |
| 1020 | INSR | 18.0867612 | -4.704350284 | 0.8215 | -5.7268 | 1E-08 | 2.17E-07 |
| 1782 | RFX6 | 2.73097098 | -4.704463102 | 1.7201 | -2.7351 | 0.00624 | 0.027 |
| 1010 | CREB3L3 | 4.77987327 | -4.704655479 | 1.4858 | -3.1663 | 0.00154 | 0.008337 |
| 103 | LINC01359 | 4.65896658 | -4.712916619 | 1.3399 | -3.5174 | 0.00044 | 0.002831 |
| 579 | GJB2 | 2.71115992 | -4.713705433 | 1.4675 | -3.2121 | 0.00132 | 0.007297 |
| 1041 | BST2 | 7.38374674 | -4.715646991 | 1.2105 | -3.8957 | 9.8E-05 | 0.00077 |
| 2043 | ABITRAM | 4.63777864 | -4.717418608 | 1.3583 | -3.4731 | 0.00051 | 0.003274 |

| | | | | | | | |
|---|---|---|---|---|---|---|---|
| 1915 | EPHA1 | 2.70707971 | -4.718407624 | 1.4786 | -3.1912 | 0.00142 | 0.007752 |
| 1453 | ARL14 | 2.78021982 | -4.719829875 | 1.4829 | -3.1827 | 0.00146 | 0.007936 |
| 1976 | GRHL2 | 2.72014982 | -4.722084739 | 1.5337 | -3.0788 | 0.00208 | 0.010746 |
| 659 | EFCAB11 | 4.77234392 | -4.728883352 | 1.3543 | -3.4918 | 0.00048 | 0.003075 |
| 1745 | C6orf141 | 4.74993704 | -4.730954473 | 1.3015 | -3.6351 | 0.00028 | 0.001912 |
| 1469 | IGF2BP2-AS1 | 2.69432961 | -4.732365279 | 1.5391 | -3.0747 | 0.00211 | 0.010888 |
| 569 | ENSG00000286067 | 2.7311361 | -4.732373923 | 1.4779 | -3.2021 | 0.00136 | 0.007519 |
| 1774 | CRYBG1 | 22.4789719 | -4.733017985 | 0.7266 | -6.5136 | 7.3E-11 | 2.55E-09 |
| 273 | VSIR | 2.74079804 | -4.734358682 | 1.4511 | -3.2627 | 0.0011 | 0.006249 |
| 1183 | LINC01123 | 2.84667873 | -4.736763018 | 1.6221 | -2.9201 | 0.0035 | 0.016727 |
| 1148 | ABCG8 | 2.82978186 | -4.736932066 | 1.8473 | -2.5642 | 0.01034 | 0.0405 |
| 1001 | ADAMTSL5 | 4.7362643 | -4.740137761 | 1.3384 | -3.5416 | 0.0004 | 0.002621 |
| 6 | C1QTNF12 | 7.84157376 | -4.742249473 | 1.2158 | -3.9004 | 9.6E-05 | 0.000758 |
| 330 | PRAP1 | 2.76953926 | -4.74854821 | 1.6025 | -2.9633 | 0.00304 | 0.014843 |
| 693 | SPINT1 | 25.4347123 | -4.750408322 | 0.7362 | -6.4523 | 1.1E-10 | 3.68E-09 |
| 1027 | ZNF823 | 4.75298433 | -4.761058738 | 1.4209 | -3.3508 | 0.00081 | 0.004801 |
| 967 | COLEC12 | 2.75575166 | -4.761330985 | 1.4721 | -3.2343 | 0.00122 | 0.006794 |
| 257 | MAP3K8 | 2.85196155 | -4.763529045 | 1.5416 | -3.0901 | 0.002 | 0.0104 |
| 1791 | SMLR1 | 4.88332801 | -4.76504497 | 1.5509 | -3.0725 | 0.00212 | 0.010951 |
| 1267 | TMEM74B | 7.78327012 | -4.770062115 | 1.1873 | -4.0174 | 5.9E-05 | 0.000496 |
| 71 | RHBDL2 | 4.67647911 | -4.772501278 | 1.5429 | -3.0933 | 0.00198 | 0.0103 |
| 2073 | C8G | 4.73735404 | -4.773820125 | 1.5131 | -3.1551 | 0.0016 | 0.008613 |
| 1922 | SHH | 2.84716386 | -4.780040146 | 1.4769 | -3.2365 | 0.00121 | 0.006748 |
| 437 | ARHGAP42 | 5.01685414 | -4.781625948 | 1.2976 | -3.685 | 0.00023 | 0.001609 |
| 1939 | PTK2B | 7.89900929 | -4.782579434 | 1.2025 | -3.9771 | 7E-05 | 0.000577 |
| 1771 | GRIK2 | 2.86364215 | -4.783885428 | 1.4262 | -3.3544 | 0.0008 | 0.00475 |
| 514 | ENSG00000257298 | 4.95337597 | -4.784893619 | 1.341 | -3.5682 | 0.00036 | 0.002399 |
| 1951 | CEBPD | 11.1909283 | -4.786526402 | 1.001 | -4.7817 | 1.7E-06 | 2.16E-05 |
| 2081 | ENTPD8 | 2.82218741 | -4.786532992 | 1.6774 | -2.8535 | 0.00432 | 0.019939 |
| 234 | ENSG00000242861 | 7.92346082 | -4.791284856 | 1.158 | -4.1377 | 3.5E-05 | 0.000314 |
| 673 | SERPINA4 | 2.82308431 | -4.791317315 | 1.6438 | -2.9148 | 0.00356 | 0.016972 |
| 171 | BCAN-AS1 | 2.90906423 | -4.792932226 | 1.4456 | -3.3156 | 0.00091 | 0.005336 |
| 726 | THSD4 | 2.85397135 | -4.798149939 | 1.4316 | -3.3516 | 0.0008 | 0.004789 |
| 275 | PLA2G12B | 2.84932311 | -4.799738449 | 1.5394 | -3.118 | 0.00182 | 0.0096 |
| 358 | IRAG1 | 4.79655969 | -4.799748056 | 1.5135 | -3.1713 | 0.00152 | 0.008214 |
| 1867 | SAMD9L | 2.85657234 | -4.805451399 | 1.4243 | -3.3739 | 0.00074 | 0.004472 |
| 1445 | PLSCR4 | 2.89825738 | -4.805628873 | 1.4364 | -3.3457 | 0.00082 | 0.004871 |
| 2139 | SERPINA7 | 2.96642037 | -4.807651591 | 1.4782 | -3.2523 | 0.00114 | 0.006426 |
| 461 | TREH | 2.88792884 | -4.807758076 | 1.7002 | -2.8277 | 0.00469 | 0.021331 |

| | | | | | | | |
|---|---|---|---|---|---|---|---|
| 1624 | F2RL1 | 12.5408334 | -4.808591621 | 0.9553 | -5.0336 | 4.8E-07 | 6.94E-06 |
| 1523 | FAM47E | 2.97244384 | -4.808750326 | 1.5731 | -3.0569 | 0.00224 | 0.01146 |
| 67 | TMEM54 | 22.2973771 | -4.814023554 | 0.886 | -5.4334 | 5.5E-08 | 9.92E-07 |
| 1724 | MLN | 2.92573624 | -4.814374529 | 1.7923 | -2.6862 | 0.00723 | 0.030414 |
| 984 | MYO5B | 15.291907 | -4.814401288 | 0.8685 | -5.5434 | 3E-08 | 5.71E-07 |
| 1450 | LXN | 2.87665252 | -4.814611579 | 1.4896 | -3.2322 | 0.00123 | 0.00684 |
| 953 | EVPL | 7.93182348 | -4.81997426 | 1.2959 | -3.7195 | 0.0002 | 0.001431 |
| 721 | SLC51B | 2.98071626 | -4.822165455 | 1.4754 | -3.2684 | 0.00108 | 0.006134 |
| 420 | P2RY6 | 2.94607787 | -4.83390259 | 1.7395 | -2.7789 | 0.00545 | 0.024256 |
| 1869 | COL1A2 | 35.0392091 | -4.837046621 | 1.2166 | -3.976 | 7E-05 | 0.000578 |
| 689 | THBS1 | 8.58178523 | -4.84041775 | 1.2071 | -4.01 | 6.1E-05 | 0.000509 |
| 270 | STOX1 | 5.08694404 | -4.841456372 | 1.2665 | -3.8227 | 0.00013 | 0.001003 |
| 1573 | SLC7A11 | 2.9493366 | -4.846699923 | 1.5353 | -3.1569 | 0.00159 | 0.008572 |
| 293 | CYP2C19 | 3.05532129 | -4.847029312 | 1.622 | -2.9884 | 0.0028 | 0.013847 |
| 138 | OLFML3 | 4.95107487 | -4.848288937 | 1.4439 | -3.3577 | 0.00079 | 0.004701 |
| 1489 | ZFYVE28 | 5.1625388 | -4.850379599 | 1.3455 | -3.6048 | 0.00031 | 0.002117 |
| 1525 | FRAS1 | 2.95658583 | -4.851260482 | 1.4349 | -3.3809 | 0.00072 | 0.004375 |
| 1540 | ABCG2 | 3.07806221 | -4.858396207 | 1.7824 | -2.7258 | 0.00642 | 0.027603 |
| 14 | PRDM16 | 3.11698251 | -4.864119168 | 1.5675 | -3.1031 | 0.00192 | 0.010012 |
| 2004 | BNC2 | 2.95998515 | -4.864574075 | 1.4787 | -3.2897 | 0.001 | 0.00576 |
| 268 | PBLD | 27.874359 | -4.864818911 | 0.8447 | -5.7593 | 8.4E-09 | 1.82E-07 |
| 1161 | NAT8 | 2.96542628 | -4.870164029 | 1.5064 | -3.2329 | 0.00123 | 0.006824 |
| 2074 | LCN12 | 5.25615802 | -4.874063782 | 1.35 | -3.6105 | 0.00031 | 0.002078 |
| 378 | CD82 | 3.00540723 | -4.875632272 | 1.4093 | -3.4596 | 0.00054 | 0.003417 |
| 2006 | LRRC19 | 21.2675294 | -4.878900937 | 0.7671 | -6.3599 | 2E-10 | 6.23E-09 |
| 23 | TNFRSF1B | 2.99824771 | -4.881072635 | 1.4758 | -3.3074 | 0.00094 | 0.005461 |
| 79 | BEST4 | 5.09800052 | -4.885137464 | 1.3927 | -3.5076 | 0.00045 | 0.002922 |
| 1764 | TPBG | 3.01344935 | -4.885186426 | 1.5238 | -3.2059 | 0.00135 | 0.007429 |
| 1730 | ETV7 | 5.26013072 | -4.885393089 | 1.3391 | -3.6483 | 0.00026 | 0.001824 |
| 995 | PLPP2 | 12.5043024 | -4.886819439 | 1.0193 | -4.7943 | 1.6E-06 | 2.03E-05 |
| 654 | FOS | 5.10891222 | -4.886877504 | 1.3702 | -3.5665 | 0.00036 | 0.002411 |
| 25 | ENSG00000272510 | 3.0575471 | -4.889842524 | 1.496 | -3.2687 | 0.00108 | 0.006131 |
| 818 | CDYL2 | 3.14810871 | -4.890406564 | 1.8426 | -2.654 | 0.00795 | 0.032832 |
| 595 | PHF11 | 16.5684121 | -4.890460304 | 0.8345 | -5.8604 | 4.6E-09 | 1.04E-07 |
| 1520 | RASSF6 | 8.54233962 | -4.895334865 | 1.1633 | -4.2082 | 2.6E-05 | 0.000238 |
| 824 | LINC01082 | 3.00907568 | -4.896043544 | 1.7404 | -2.8131 | 0.00491 | 0.022211 |
| 36 | CDA | 3.00330023 | -4.896664497 | 1.5958 | -3.0685 | 0.00215 | 0.011085 |
| 518 | KRT7 | 3.16849461 | -4.904488571 | 1.6063 | -3.0533 | 0.00226 | 0.011554 |
| 588 | FOXO1 | 3.10557017 | -4.905385146 | 1.4922 | -3.2874 | 0.00101 | 0.005795 |

| | | | | | | | |
|---|---|---|---|---|---|---|---|
| 1246 | SP110 | 3.04806458 | -4.907039444 | 1.4571 | -3.3676 | 0.00076 | 0.004556 |
| 2052 | HDHD3 | 44.1041157 | -4.913789171 | 0.5992 | -8.201 | 2.4E-16 | 2.46E-14 |
| 554 | TMCC3 | 5.31380723 | -4.918528799 | 1.338 | -3.676 | 0.00024 | 0.001659 |
| 897 | IFI35 | 8.59227875 | -4.922170456 | 1.1145 | -4.4165 | 1E-05 | 0.000103 |
| 1539 | SPARCL1 | 8.5349569 | -4.923487809 | 1.2474 | -3.947 | 7.9E-05 | 0.000641 |
| 1827 | SCIN | 5.53716552 | -4.923538782 | 1.3697 | -3.5946 | 0.00032 | 0.002196 |
| 1617 | IL6ST | 12.7382964 | -4.925722043 | 0.995 | -4.9504 | 7.4E-07 | 1.02E-05 |
| 551 | LUM | 5.22364615 | -4.936918089 | 1.5109 | -3.2675 | 0.00109 | 0.006152 |
| 758 | ENSG00000205890 | 3.1887765 | -4.937515712 | 1.4103 | -3.5011 | 0.00046 | 0.002988 |
| 391 | MYRF | 9.03473643 | -4.946107186 | 1.1774 | -4.2007 | 2.7E-05 | 0.000245 |
| 1758 | COL9A1 | 3.22387004 | -4.946389584 | 1.4055 | -3.5194 | 0.00043 | 0.002814 |
| 914 | HOXB-AS4 | 3.15829918 | -4.952581122 | 1.7432 | -2.8411 | 0.0045 | 0.020569 |
| 1052 | LINC01224 | 3.21727855 | -4.955577059 | 1.4407 | -3.4398 | 0.00058 | 0.003642 |
| 1437 | TF | 5.55946635 | -4.956551811 | 1.4919 | -3.3223 | 0.00089 | 0.005232 |
| 1779 | MROCKI | 3.29098669 | -4.958033737 | 1.436 | -3.4527 | 0.00056 | 0.003493 |
| 1047 | ZNF682 | 3.31977422 | -4.959161016 | 1.7444 | -2.8429 | 0.00447 | 0.020477 |
| 1588 | FGA | 3.26570453 | -4.961454997 | 1.7926 | -2.7678 | 0.00564 | 0.024912 |
| 1843 | AOAH | 3.209551 | -4.962222736 | 1.4862 | -3.3388 | 0.00084 | 0.004982 |
| 525 | DGKA | 12.7268269 | -4.962238575 | 1.0131 | -4.8983 | 9.7E-07 | 1.29E-05 |
| 252 | MALRD1 | 3.31178436 | -4.964371662 | 1.5248 | -3.2558 | 0.00113 | 0.006365 |
| 1780 | FRK | 5.5567996 | -4.966020977 | 1.3188 | -3.7656 | 0.00017 | 0.00122 |
| 1881 | EPHB4 | 5.45389561 | -4.967722657 | 1.2567 | -3.9529 | 7.7E-05 | 0.000628 |
| 906 | WNT3 | 3.26450874 | -4.968812463 | 1.6529 | -3.0061 | 0.00265 | 0.013206 |
| 1215 | TFPI | 9.33522612 | -4.971961994 | 1.2541 | -3.9647 | 7.3E-05 | 0.0006 |
| 19 | SLC2A5 | 5.58525971 | -4.972381521 | 1.3806 | -3.6016 | 0.00032 | 0.00214 |
| 1327 | SUSD2 | 3.28733936 | -4.973911699 | 1.6532 | -3.0086 | 0.00262 | 0.013107 |
| 1901 | MDFIC | 3.20689028 | -4.979494629 | 1.4374 | -3.4642 | 0.00053 | 0.003369 |
| 1818 | MICALL2 | 17.3658424 | -4.985958154 | 0.9148 | -5.4502 | 5E-08 | 9.13E-07 |
| 118 | GBP2 | 5.4662014 | -4.986605265 | 1.248 | -3.9958 | 6.4E-05 | 0.000538 |
| 1568 | SYNPO2 | 3.2218439 | -4.995942972 | 1.5039 | -3.322 | 0.00089 | 0.005236 |
| 463 | OAF | 17.892371 | -4.99915298 | 0.891 | -5.611 | 2E-08 | 4.01E-07 |
| 1784 | ENSG00000253194 | 3.36044201 | -5.003392739 | 1.4007 | -3.572 | 0.00035 | 0.00237 |
| 1559 | ENPEP | 5.69757807 | -5.009238222 | 1.3458 | -3.7222 | 0.0002 | 0.001419 |
| 1649 | SMIM32 | 3.26067449 | -5.013058662 | 1.4764 | -3.3954 | 0.00069 | 0.004171 |
| 772 | TMC5 | 13.7785917 | -5.014810583 | 0.9677 | -5.1824 | 2.2E-07 | 3.4E-06 |
| 1583 | MAB21L2 | 3.33438263 | -5.018666586 | 1.5189 | -3.3042 | 0.00095 | 0.005513 |
| 1641 | IRF1 | 9.30726086 | -5.019474761 | 1.1407 | -4.4005 | 1.1E-05 | 0.00011 |
| 255 | MPP7 | 5.68807239 | -5.026992624 | 1.2496 | -4.0228 | 5.8E-05 | 0.000486 |
| 1940 | EPHX2 | 47.7809531 | -5.031374693 | 0.5853 | -8.5968 | 8.2E-18 | 1.17E-15 |

| | | | | | | | |
|---|---|---|---|---|---|---|---|
| 1894 | LHFPL3-AS2 | 3.43143629 | -5.031404149 | 1.5868 | -3.1708 | 0.00152 | 0.008225 |
| 1279 | LINC00261 | 5.69270633 | -5.03434187 | 1.2771 | -3.942 | 8.1E-05 | 0.000653 |
| 853 | ALOX12P2 | 3.40034892 | -5.042778085 | 1.5938 | -3.164 | 0.00156 | 0.008388 |
| 938 | PRKCA | 13.9109375 | -5.042857817 | 1.0272 | -4.9095 | 9.1E-07 | 1.22E-05 |
| 1461 | NAALADL2 | 3.41753583 | -5.046488157 | 1.4495 | -3.4815 | 0.0005 | 0.003186 |
| 459 | MPZL3 | 3.38399692 | -5.05245781 | 1.3762 | -3.6713 | 0.00024 | 0.001685 |
| 1872 | PDK4 | 3.43908095 | -5.060450381 | 1.4179 | -3.5689 | 0.00036 | 0.002394 |
| 1346 | C1QTNF6 | 5.82757421 | -5.066445077 | 1.2568 | -4.0312 | 5.6E-05 | 0.000471 |
| 655 | FLVCR2 | 3.50643675 | -5.067891452 | 1.4549 | -3.4833 | 0.0005 | 0.003168 |
| 2150 | DANT2 | 3.57072211 | -5.069569377 | 1.4725 | -3.4429 | 0.00058 | 0.003606 |
| 2037 | CORO2A | 13.9547862 | -5.076150068 | 0.9847 | -5.1552 | 2.5E-07 | 3.88E-06 |
| 1689 | PPP1R3G | 3.52641293 | -5.07973729 | 1.4074 | -3.6094 | 0.00031 | 0.002086 |
| 1245 | CCL20 | 3.49358264 | -5.083358229 | 1.4077 | -3.611 | 0.0003 | 0.002075 |
| 1217 | COL5A2 | 9.50480706 | -5.083605087 | 1.1729 | -4.3341 | 1.5E-05 | 0.000144 |
| 488 | APOBEC1 | 3.6006891 | -5.085282092 | 1.4323 | -3.5505 | 0.00038 | 0.002541 |
| 1222 | CAVIN2 | 3.46117093 | -5.088844881 | 1.6041 | -3.1725 | 0.00151 | 0.008188 |
| 761 | CASP16P | 3.53175549 | -5.091112443 | 1.4957 | -3.4039 | 0.00066 | 0.00406 |
| 813 | MLKL | 3.45402027 | -5.092278719 | 1.4435 | -3.5278 | 0.00042 | 0.002735 |
| 1630 | SLCO4C1 | 3.53208436 | -5.094025136 | 1.4474 | -3.5195 | 0.00043 | 0.002814 |
| 926 | ANKRD40CL | 6.1582281 | -5.100335108 | 1.3037 | -3.9123 | 9.1E-05 | 0.000726 |
| 930 | RNF43 | 3.47490764 | -5.100700533 | 1.3971 | -3.6509 | 0.00026 | 0.001808 |
| 1576 | MAML3 | 3.5229816 | -5.101768463 | 1.4937 | -3.4156 | 0.00064 | 0.003929 |
| 20 | SPSB1 | 3.48603451 | -5.103596476 | 1.4491 | -3.5219 | 0.00043 | 0.002789 |
| 792 | MT1F | 9.84529537 | -5.108606141 | 1.3425 | -3.8054 | 0.00014 | 0.001063 |
| 2142 | TBC1D8B | 10.1400168 | -5.108871265 | 1.145 | -4.462 | 8.1E-06 | 8.5E-05 |
| 1058 | DMKN | 3.6182437 | -5.110613167 | 1.438 | -3.554 | 0.00038 | 0.002516 |
| 442 | CASP4 | 3.63221072 | -5.114300544 | 1.3771 | -3.7138 | 0.0002 | 0.001458 |
| 1046 | TM6SF2 | 6.12489634 | -5.118696999 | 1.2369 | -4.1382 | 3.5E-05 | 0.000314 |
| 27 | TMEM82 | 3.56962945 | -5.120097897 | 1.379 | -3.7128 | 0.0002 | 0.001463 |
| 1838 | HOXA5 | 3.57278961 | -5.126986719 | 1.6646 | -3.0799 | 0.00207 | 0.01071 |
| 1533 | CDS1 | 29.7669991 | -5.129632549 | 0.7548 | -6.7962 | 1.1E-11 | 4.59E-10 |
| 1083 | ENSG00000285128 | 3.59837067 | -5.129677686 | 1.5064 | -3.4052 | 0.00066 | 0.00405 |
| 228 | MIR215 | 3.71296551 | -5.129853645 | 1.4147 | -3.6261 | 0.00029 | 0.001972 |
| 613 | RNASE1 | 14.2413922 | -5.13343902 | 0.9673 | -5.3067 | 1.1E-07 | 1.87E-06 |
| 1479 | SLC51A | 10.6776478 | -5.136786502 | 1.1962 | -4.2942 | 1.8E-05 | 0.000169 |
| 509 | CPNE8 | 3.56800077 | -5.137133366 | 1.4466 | -3.5511 | 0.00038 | 0.002538 |
| 122 | SLC44A3 | 14.5687799 | -5.137845999 | 0.9631 | -5.3346 | 9.6E-08 | 1.63E-06 |
| 552 | DCN | 19.4311378 | -5.141068496 | 0.9899 | -5.1935 | 2.1E-07 | 3.21E-06 |
| 1057 | LSR | 24.2020816 | -5.142326939 | 0.7524 | -6.8349 | 8.2E-12 | 3.58E-10 |

| | | | | | | | |
|---|---|---|---|---|---|---|---|
| 907 | SP6 | 3.64114088 | -5.142827434 | 1.3583 | -3.7863 | 0.00015 | 0.001132 |
| 1893 | LHFPL3 | 3.74026496 | -5.14886985 | 1.7666 | -2.9145 | 0.00356 | 0.016972 |
| 2021 | TRPM6 | 3.70035056 | -5.150995082 | 1.6132 | -3.193 | 0.00141 | 0.007707 |
| 1344 | TST | 39.4954016 | -5.151647261 | 0.6065 | -8.4946 | 2E-17 | 2.57E-15 |
| 1837 | HOXA3 | 3.61795823 | -5.152200681 | 1.5453 | -3.3342 | 0.00086 | 0.005058 |
| 2120 | ZC4H2 | 3.7085864 | -5.157683162 | 1.4596 | -3.5336 | 0.00041 | 0.002686 |
| 1581 | NR3C2 | 3.72449666 | -5.158887625 | 1.3761 | -3.7489 | 0.00018 | 0.00129 |
| 125 | VAV3 | 14.443259 | -5.161008052 | 0.9768 | -5.2836 | 1.3E-07 | 2.08E-06 |
| 315 | TCF7L2 | 35.6775745 | -5.161895862 | 0.6491 | -7.9527 | 1.8E-15 | 1.57E-13 |
| 1813 | FRMD1 | 3.67077014 | -5.164278383 | 1.5428 | -3.3473 | 0.00082 | 0.00485 |
| 99 | L1TD1 | 3.64599083 | -5.166141197 | 1.4862 | -3.4762 | 0.00051 | 0.00324 |
| 1931 | MTUS1 | 3.68450685 | -5.169733958 | 1.3617 | -3.7966 | 0.00015 | 0.001092 |
| 1608 | AGXT2 | 3.73860427 | -5.172552905 | 1.653 | -3.1292 | 0.00175 | 0.009278 |
| 1806 | PLEKHG1 | 3.8138759 | -5.174598765 | 1.437 | -3.6011 | 0.00032 | 0.002144 |
| 1426 | PARP14 | 24.8009758 | -5.174652097 | 0.7849 | -6.5925 | 4.3E-11 | 1.59E-09 |
| 64 | SERINC2 | 30.3458141 | -5.175885459 | 0.6833 | -7.5748 | 3.6E-14 | 2.42E-12 |
| 1798 | NHSL1 | 10.3170946 | -5.176307279 | 1.1333 | -4.5675 | 4.9E-06 | 5.46E-05 |
| 1720 | HLA-DRB1 | 3.73096643 | -5.179018309 | 1.5542 | -3.3323 | 0.00086 | 0.005081 |
| 2132 | CHMP1B2P | 3.80696984 | -5.179073048 | 1.6206 | -3.1957 | 0.00139 | 0.007646 |
| 1081 | BCL3 | 10.1403056 | -5.18393474 | 1.144 | -4.5316 | 5.9E-06 | 6.33E-05 |
| 1373 | TGFBR2 | 6.24198174 | -5.19251897 | 1.2473 | -4.1631 | 3.1E-05 | 0.000284 |
| 412 | TCIRG1 | 3.69499165 | -5.192544427 | 1.5514 | -3.347 | 0.00082 | 0.004853 |
| 1563 | ALPK1 | 10.4275629 | -5.192563599 | 1.1184 | -4.6427 | 3.4E-06 | 3.97E-05 |
| 1066 | SERTAD1 | 3.81418161 | -5.19611486 | 1.3579 | -3.8267 | 0.00013 | 0.000989 |
| 626 | NFATC4 | 3.73569008 | -5.197051132 | 1.3813 | -3.7624 | 0.00017 | 0.001231 |
| 715 | ANXA2 | 89.1865768 | -5.199195531 | 0.4262 | -12.199 | 3.2E-34 | 4.08E-31 |
| 724 | PAQR5 | 6.26417574 | -5.204237444 | 1.3074 | -3.9805 | 6.9E-05 | 0.00057 |
| 1134 | KHK | 10.4769811 | -5.206689054 | 1.1706 | -4.4478 | 8.7E-06 | 9.03E-05 |
| 441 | DYNC2H1 | 3.86935534 | -5.212565955 | 1.358 | -3.8383 | 0.00012 | 0.00095 |
| 260 | ALOX5 | 3.91958398 | -5.213770952 | 1.5489 | -3.366 | 0.00076 | 0.004581 |
| 53 | TRIM63 | 3.84518609 | -5.21957955 | 1.5009 | -3.4775 | 0.00051 | 0.003227 |
| 2003 | TTC39B | 25.5903895 | -5.21991109 | 0.8092 | -6.4511 | 1.1E-10 | 3.69E-09 |
| 1746 | ENSG00000235122 | 3.95945206 | -5.223647138 | 1.4729 | -3.5466 | 0.00039 | 0.002576 |
| 1365 | TYMP | 3.84892313 | -5.223763173 | 1.4558 | -3.5881 | 0.00033 | 0.002244 |
| 1335 | GAL3ST1 | 3.81557794 | -5.227682322 | 1.4437 | -3.6212 | 0.00029 | 0.002005 |
| 1883 | ENSG00000274993 | 3.81433787 | -5.235312252 | 1.3761 | -3.8044 | 0.00014 | 0.001066 |
| 93 | MROH7 | 3.88022875 | -5.23578419 | 1.3252 | -3.9508 | 7.8E-05 | 0.000632 |
| 802 | CES3 | 6.58225034 | -5.238933201 | 1.234 | -4.2455 | 2.2E-05 | 0.000205 |
| 2119 | ENSG00000227486 | 3.98041485 | -5.242951623 | 1.4454 | -3.6274 | 0.00029 | 0.001963 |

| | | | | | | | |
|---|---|---|---|---|---|---|---|
| 1095 | FUT2 | 20.4554479 | -5.246380756 | 0.8369 | -6.269 | 3.6E-10 | 1.06E-08 |
| 1261 | ENSG00000219159 | 3.85203241 | -5.248036445 | 1.3714 | -3.8269 | 0.00013 | 0.000989 |
| 1375 | CMTM8 | 3.97164095 | -5.248154805 | 1.3435 | -3.9064 | 9.4E-05 | 0.000741 |
| 221 | MIR205HG | 4.03379478 | -5.253850812 | 1.4522 | -3.6179 | 0.0003 | 0.002025 |
| 657 | FLRT2 | 3.94070847 | -5.256173702 | 1.7296 | -3.0389 | 0.00237 | 0.012063 |
| 362 | SOX6 | 11.6874921 | -5.260352703 | 1.2899 | -4.0781 | 4.5E-05 | 0.000395 |
| 1433 | CHST13 | 4.08035293 | -5.261648619 | 1.438 | -3.6591 | 0.00025 | 0.001758 |
| 196 | LAMC2 | 6.66645305 | -5.263510095 | 1.2593 | -4.1797 | 2.9E-05 | 0.000266 |
| 331 | FUOM | 6.78250709 | -5.264765537 | 1.2475 | -4.2203 | 2.4E-05 | 0.000227 |
| 1310 | LINC00649 | 4.00851944 | -5.277928834 | 1.593 | -3.3132 | 0.00092 | 0.005372 |
| 48 | RCAN3 | 6.8538534 | -5.278069254 | 1.3629 | -3.8728 | 0.00011 | 0.000839 |
| 959 | ENPP7 | 4.0479778 | -5.278903672 | 1.5308 | -3.4484 | 0.00056 | 0.003544 |
| 449 | IL18 | 3.95398029 | -5.280216672 | 1.521 | -3.4714 | 0.00052 | 0.003293 |
| 245 | TUBAL3 | 4.07194448 | -5.28298673 | 1.5259 | -3.4622 | 0.00054 | 0.003389 |
| 1773 | PRDM1 | 4.13950286 | -5.284568214 | 1.4761 | -3.5802 | 0.00034 | 0.002303 |
| 444 | CASP1 | 6.91815073 | -5.287451475 | 1.2367 | -4.2755 | 1.9E-05 | 0.000183 |
| 717 | TLN2 | 6.68266553 | -5.289448745 | 1.2322 | -4.2927 | 1.8E-05 | 0.00017 |
| 1018 | TRIP10 | 6.80660687 | -5.300814092 | 1.232 | -4.3026 | 1.7E-05 | 0.000164 |
| 2058 | LCN2 | 4.03077454 | -5.317435739 | 1.5526 | -3.4248 | 0.00062 | 0.003819 |
| 915 | LINC03057 | 4.05034644 | -5.320289001 | 1.4229 | -3.7391 | 0.00018 | 0.001335 |
| 1340 | APOL6 | 17.950539 | -5.321989681 | 0.9809 | -5.4257 | 5.8E-08 | 1.02E-06 |
| 380 | CREB3L1 | 11.1788065 | -5.32331509 | 1.1432 | -4.6567 | 3.2E-06 | 3.74E-05 |
| 418 | ENSG00000285933 | 4.19846589 | -5.329249602 | 1.3363 | -3.9881 | 6.7E-05 | 0.000553 |
| 236 | AGT | 7.01986771 | -5.33240082 | 1.2918 | -4.1279 | 3.7E-05 | 0.000326 |
| 472 | TMEM45B | 49.9666878 | -5.333253543 | 0.584 | -9.1328 | 6.7E-20 | 1.44E-17 |
| 2140 | PWWP3B | 4.12972601 | -5.335783709 | 1.4356 | -3.7169 | 0.0002 | 0.001445 |
| 1515 | UGT2B7 | 4.2026724 | -5.337262518 | 1.3435 | -3.9726 | 7.1E-05 | 0.000585 |
| 396 | BATF2 | 4.27434758 | -5.34349459 | 1.4336 | -3.7274 | 0.00019 | 0.001392 |
| 1686 | SERPINB1 | 51.0143752 | -5.343820721 | 0.5747 | -9.2986 | 1.4E-20 | 3.43E-18 |
| 1227 | PARD3B | 4.15572565 | -5.345429875 | 1.3389 | -3.9925 | 6.5E-05 | 0.000545 |
| 1142 | RMDN2 | 4.30545063 | -5.357567 | 1.4041 | -3.8156 | 0.00014 | 0.001026 |
| 1411 | ADGRG7 | 7.15351921 | -5.367029526 | 1.278 | -4.1996 | 2.7E-05 | 0.000246 |
| 147 | TXNIP | 67.2323032 | -5.375701089 | 0.5909 | -9.0982 | 9.2E-20 | 1.89E-17 |
| 535 | GLI1 | 4.20106484 | -5.378321151 | 1.4577 | -3.6895 | 0.00022 | 0.001586 |
| 97 | CYP2J2 | 4.32116055 | -5.380442594 | 1.5461 | -3.48 | 0.0005 | 0.0032 |
| 1474 | CLDN1 | 4.42720499 | -5.380495463 | 1.5773 | -3.4111 | 0.00065 | 0.003979 |
| 1729 | BNIP5 | 4.30751355 | -5.382102811 | 1.4401 | -3.7374 | 0.00019 | 0.001342 |
| 1178 | SLC9A2 | 7.16402459 | -5.385799236 | 1.2358 | -4.3582 | 1.3E-05 | 0.000131 |
| 1916 | ARHGEF5 | 4.33217456 | -5.393349054 | 1.3112 | -4.1132 | 3.9E-05 | 0.000346 |

| | | | | | | | |
|---|---|---|---|---|---|---|---|
| 415 | LINC02747 | 4.41829507 | -5.3959065 | 1.4025 | -3.8473 | 0.00012 | 0.000921 |
| 2009 | RIGI | 7.45983606 | -5.39926745 | 1.272 | -4.2446 | 2.2E-05 | 0.000206 |
| 1136 | RBKS | 4.4714447 | -5.40105626 | 1.413 | -3.8224 | 0.00013 | 0.001003 |
| 1050 | ZNF91 | 7.50616623 | -5.401954879 | 1.2684 | -4.2588 | 2.1E-05 | 0.000196 |
| 756 | TNFRSF12A | 4.36020715 | -5.403261033 | 1.3583 | -3.9781 | 6.9E-05 | 0.000575 |
| 1505 | NSUN7 | 7.45115188 | -5.403709859 | 1.2325 | -4.3845 | 1.2E-05 | 0.000117 |
| 320 | ABLIM1 | 17.9479532 | -5.406936431 | 0.9821 | -5.5056 | 3.7E-08 | 6.92E-07 |
| 1202 | PLA2R1 | 12.1187266 | -5.410691995 | 1.1113 | -4.869 | 1.1E-06 | 1.46E-05 |
| 1839 | HOXA9 | 12.2238718 | -5.410839218 | 1.0902 | -4.963 | 6.9E-07 | 9.58E-06 |
| 600 | EDNRB | 4.48094289 | -5.413802094 | 1.3326 | -4.0625 | 4.9E-05 | 0.000418 |
| 788 | PRSS8 | 4.36879959 | -5.421876276 | 1.4987 | -3.6178 | 0.0003 | 0.002025 |
| 1709 | TRIM15 | 4.49694287 | -5.423859639 | 1.3299 | -4.0783 | 4.5E-05 | 0.000395 |
| 851 | SLC52A1 | 4.45539842 | -5.42513037 | 1.4453 | -3.7535 | 0.00017 | 0.001268 |
| 664 | FBLN5 | 4.43937529 | -5.430194679 | 1.6508 | -3.2893 | 0.001 | 0.005765 |
| 1985 | LRATD2 | 4.57414889 | -5.437739694 | 1.3663 | -3.98 | 6.9E-05 | 0.000571 |
| 1866 | SAMD9 | 4.49351153 | -5.443272835 | 1.3719 | -3.9676 | 7.3E-05 | 0.000595 |
| 2109 | MAOB | 19.9110725 | -5.445600211 | 1.1127 | -4.8939 | 9.9E-07 | 1.31E-05 |
| 1256 | MLPH | 4.5949234 | -5.45168103 | 1.3388 | -4.0719 | 4.7E-05 | 0.000404 |
| 1995 | DGAT1 | 90.2046228 | -5.455087504 | 0.8653 | -6.3046 | 2.9E-10 | 8.58E-09 |
| 695 | ITPKA | 12.4971929 | -5.455404601 | 1.118 | -4.8798 | 1.1E-06 | 1.39E-05 |
| 475 | ADAMTS15 | 7.56891282 | -5.455640078 | 1.1982 | -4.5532 | 5.3E-06 | 5.78E-05 |
| 814 | FA2H | 4.50576016 | -5.456076224 | 1.4686 | -3.7153 | 0.0002 | 0.001452 |
| 1924 | PRAG1 | 4.57251135 | -5.458073603 | 1.2915 | -4.2263 | 2.4E-05 | 0.000221 |
| 1930 | MSR1 | 7.86516155 | -5.459230162 | 1.3651 | -3.9992 | 6.4E-05 | 0.000531 |
| 1521 | CXCL5 | 4.47970964 | -5.460608659 | 1.3309 | -4.103 | 4.1E-05 | 0.00036 |
| 376 | PRR5L | 7.5040894 | -5.463379815 | 1.2836 | -4.2563 | 2.1E-05 | 0.000197 |
| 823 | IRF8 | 4.63076439 | -5.469568296 | 1.3035 | -4.196 | 2.7E-05 | 0.00025 |
| 1205 | DPP4 | 24.3182122 | -5.470769439 | 0.8233 | -6.6452 | 3E-11 | 1.15E-09 |
| 172 | IFI16 | 12.5670316 | -5.470950147 | 1.101 | -4.969 | 6.7E-07 | 9.33E-06 |
| 1274 | FLRT3 | 4.73518015 | -5.476362304 | 1.431 | -3.827 | 0.00013 | 0.000989 |
| 1147 | HAAO | 4.61474127 | -5.480207009 | 1.3113 | -4.1793 | 2.9E-05 | 0.000266 |
| 565 | OAS2 | 4.60086396 | -5.481339726 | 1.2995 | -4.2182 | 2.5E-05 | 0.000229 |
| 1188 | PSD4 | 12.5540609 | -5.490561001 | 1.1158 | -4.921 | 8.6E-07 | 1.16E-05 |
| 1042 | BISPR | 4.72979127 | -5.490595843 | 1.3219 | -4.1537 | 3.3E-05 | 0.000295 |
| 1247 | SP140L | 4.60823948 | -5.492426646 | 1.3091 | -4.1956 | 2.7E-05 | 0.00025 |
| 949 | MYO15B | 32.8858164 | -5.505649882 | 0.8121 | -6.7797 | 1.2E-11 | 5.1E-10 |
| 713 | MYO1E | 7.91548379 | -5.505709885 | 1.2008 | -4.5851 | 4.5E-06 | 5.07E-05 |
| 1197 | GPR39 | 4.78941953 | -5.506343183 | 1.3846 | -3.9768 | 7E-05 | 0.000577 |
| 702 | GATM | 12.924934 | -5.507781263 | 1.0874 | -5.065 | 4.1E-07 | 5.95E-06 |

| | | | | | | | |
|---|---|---|---|---|---|---|---|
| 2076 | STPG3-AS1 | 4.75521402 | -5.509975022 | 1.4047 | -3.9225 | 8.8E-05 | 0.000702 |
| 357 | ADM | 4.63008487 | -5.514151085 | 1.4497 | -3.8036 | 0.00014 | 0.001068 |
| 864 | MFAP4 | 7.8671769 | -5.519692365 | 1.3594 | -4.0603 | 4.9E-05 | 0.000421 |
| 203 | LMOD1 | 4.66365919 | -5.521587394 | 1.5732 | -3.5097 | 0.00045 | 0.002903 |
| 336 | IFITM3 | 82.3864083 | -5.524718343 | 0.4812 | -11.482 | 1.6E-30 | 1.45E-27 |
| 1785 | MAN1A1 | 13.3368177 | -5.529238355 | 1.0933 | -5.0573 | 4.3E-07 | 6.16E-06 |
| 2001 | ERMP1 | 8.02133291 | -5.535998296 | 1.1977 | -4.6223 | 3.8E-06 | 4.32E-05 |
| 1853 | CLDN3 | 13.1357095 | -5.539111263 | 1.0738 | -5.1582 | 2.5E-07 | 3.82E-06 |
| 176 | CD244 | 4.71580792 | -5.541620387 | 1.3421 | -4.1292 | 3.6E-05 | 0.000325 |
| 1072 | ERICH4 | 4.90876576 | -5.543659467 | 1.5033 | -3.6877 | 0.00023 | 0.001596 |
| 1013 | LRG1 | 4.88050967 | -5.546822107 | 1.2933 | -4.289 | 1.8E-05 | 0.000173 |
| 410 | UNC93B1 | 4.78316372 | -5.549037328 | 1.2854 | -4.3171 | 1.6E-05 | 0.000155 |
| 121 | SLC44A3-AS1 | 4.91551351 | -5.549215221 | 1.3245 | -4.1898 | 2.8E-05 | 0.000256 |
| 1959 | HNF4G | 8.24419006 | -5.553176505 | 1.2153 | -4.5695 | 4.9E-06 | 5.42E-05 |
| 1005 | TLE2 | 8.08839074 | -5.554901969 | 1.2148 | -4.5727 | 4.8E-06 | 5.34E-05 |
| 1088 | FOXA3 | 4.84449607 | -5.556659953 | 1.2728 | -4.3655 | 1.3E-05 | 0.000127 |
| 760 | ENSG00000262370 | 4.86220013 | -5.566347271 | 1.2915 | -4.3101 | 1.6E-05 | 0.000159 |
| 1605 | SNHG18 | 4.88512046 | -5.574222278 | 1.2776 | -4.3631 | 1.3E-05 | 0.000128 |
| 1614 | C6 | 5.09567036 | -5.586315781 | 1.5726 | -3.5522 | 0.00038 | 0.00253 |
| 1654 | SPRY4 | 5.00267288 | -5.588059951 | 1.2905 | -4.3301 | 1.5E-05 | 0.000147 |
| 849 | GLTPD2 | 5.05733833 | -5.596103214 | 1.4288 | -3.9167 | 9E-05 | 0.000716 |
| 848 | TM4SF5 | 5.08708931 | -5.598537338 | 1.4287 | -3.9188 | 8.9E-05 | 0.000711 |
| 445 | CARD16 | 5.04993965 | -5.599388092 | 1.4001 | -3.9992 | 6.4E-05 | 0.000531 |
| 1380 | VILL | 13.6941805 | -5.613724945 | 1.0726 | -5.2339 | 1.7E-07 | 2.64E-06 |
| 662 | ENSG00000260711 | 5.04030768 | -5.613763221 | 1.2754 | -4.4017 | 1.1E-05 | 0.00011 |
| 1314 | FAM3B | 5.21648152 | -5.623622818 | 1.6493 | -3.4098 | 0.00065 | 0.003993 |
| 374 | EHF | 5.06809226 | -5.632766553 | 1.2973 | -4.3419 | 1.4E-05 | 0.00014 |
| 1979 | TNFRSF11B | 5.14264417 | -5.6342827 | 1.3038 | -4.3213 | 1.6E-05 | 0.000152 |
| 696 | PLA2G4F | 5.06992264 | -5.636904861 | 1.3711 | -4.1112 | 3.9E-05 | 0.000349 |
| 76 | C1orf210 | 5.16418928 | -5.643609233 | 1.2831 | -4.3983 | 1.1E-05 | 0.000111 |
| 2066 | AGPAT2 | 56.3476472 | -5.651242305 | 0.5813 | -9.7216 | 2.4E-22 | 7.88E-20 |
| 1875 | CYP3A5 | 45.7163101 | -5.65130452 | 1.0318 | -5.4771 | 4.3E-08 | 7.95E-07 |
| 2027 | RASEF | 5.12892313 | -5.660199014 | 1.3282 | -4.2617 | 2E-05 | 0.000193 |
| 625 | RIPK3 | 5.19944795 | -5.671687661 | 1.2663 | -4.4789 | 7.5E-06 | 7.91E-05 |
| 408 | ACY3 | 5.32690148 | -5.685300599 | 1.3793 | -4.122 | 3.8E-05 | 0.000334 |
| 1286 | ACSS2 | 28.4360933 | -5.686038127 | 0.7931 | -7.1698 | 7.5E-13 | 3.93E-11 |
| 2151 | KLHL13 | 9.07305514 | -5.689164523 | 1.2001 | -4.7404 | 2.1E-06 | 2.59E-05 |
| 334 | PGGHG | 27.9071989 | -5.689175943 | 0.8876 | -6.4097 | 1.5E-10 | 4.67E-09 |
| 1664 | CD74 | 5.38525308 | -5.689378241 | 1.2643 | -4.5001 | 6.8E-06 | 7.27E-05 |

| | | | | | | | |
|---|---|---|---|---|---|---|---|
| 1635 | PPIC | 14.7466239 | -5.693335041 | 1.0852 | -5.2463 | 1.6E-07 | 2.49E-06 |
| 1203 | ITGB6 | 5.47203134 | -5.699859931 | 1.3317 | -4.28 | 1.9E-05 | 0.000179 |
| 469 | SIAE | 9.07587728 | -5.716065213 | 1.1725 | -4.8751 | 1.1E-06 | 1.42E-05 |
| 1661 | SLC26A2 | 150.306348 | -5.718912347 | 0.421 | -13.585 | 4.9E-42 | 9.97E-39 |
| 878 | CCL15 | 5.35014319 | -5.720925763 | 1.4515 | -3.9414 | 8.1E-05 | 0.000655 |
| 966 | METRNL | 9.16643082 | -5.724605188 | 1.1723 | -4.8834 | 1E-06 | 1.37E-05 |
| 1981 | DEPTOR | 5.5256518 | -5.733785678 | 1.2668 | -4.5262 | 6E-06 | 6.48E-05 |
| 192 | CRYZL2P | 15.0835772 | -5.739263158 | 1.1874 | -4.8334 | 1.3E-06 | 1.72E-05 |
| 403 | C11orf86 | 5.62257031 | -5.739738225 | 1.3329 | -4.3063 | 1.7E-05 | 0.000161 |
| 2152 | IL13RA1 | 14.9671739 | -5.744685154 | 1.0866 | -5.2867 | 1.2E-07 | 2.05E-06 |
| 580 | CRYL1 | 29.5588724 | -5.747802122 | 0.8618 | -6.6693 | 2.6E-11 | 9.93E-10 |
| 1508 | NIPAL1 | 5.52003806 | -5.754838334 | 1.2626 | -4.558 | 5.2E-06 | 5.67E-05 |
| 913 | HOXB9 | 5.47868189 | -5.755108667 | 1.2828 | -4.4864 | 7.2E-06 | 7.68E-05 |
| 1704 | HFE | 5.58710158 | -5.757699181 | 1.3523 | -4.2577 | 2.1E-05 | 0.000196 |
| 433 | PIWIL4 | 5.73976615 | -5.761285403 | 1.4434 | -3.9916 | 6.6E-05 | 0.000546 |
| 1963 | CHMP4C | 5.58591464 | -5.762928493 | 1.2537 | -4.5968 | 4.3E-06 | 4.84E-05 |
| 1767 | NT5E | 9.30874501 | -5.774889324 | 1.1745 | -4.9171 | 8.8E-07 | 1.18E-05 |
| 747 | MCTP2 | 5.66367984 | -5.777335923 | 1.2437 | -4.6453 | 3.4E-06 | 3.92E-05 |
| 1732 | CDKN1A | 29.7033335 | -5.778247435 | 0.8196 | -7.0504 | 1.8E-12 | 8.9E-11 |
| 2106 | DYNLT3 | 5.6600182 | -5.788770148 | 1.2451 | -4.6492 | 3.3E-06 | 3.86E-05 |
| 155 | S100A11 | 45.2714567 | -5.790039117 | 0.7447 | -7.7748 | 7.6E-15 | 5.75E-13 |
| 1643 | SOWAHA | 15.7822219 | -5.792510794 | 1.1522 | -5.0276 | 5E-07 | 7.15E-06 |
| 901 | NAGS | 5.82420281 | -5.797327756 | 1.4003 | -4.1401 | 3.5E-05 | 0.000311 |
| 1477 | MUC4 | 5.65090113 | -5.798818177 | 1.3058 | -4.4407 | 9E-06 | 9.28E-05 |
| 765 | ABAT | 9.73293688 | -5.799412339 | 1.1871 | -4.8854 | 1E-06 | 1.36E-05 |
| 1676 | FGFR4 | 9.51345681 | -5.802386189 | 1.1933 | -4.8623 | 1.2E-06 | 1.5E-05 |
| 1808 | SYNJ2 | 9.74602446 | -5.805378569 | 1.2034 | -4.8243 | 1.4E-06 | 1.79E-05 |
| 2102 | SAT1 | 184.271988 | -5.8055356 | 0.4013 | -14.466 | 2E-47 | 9.42E-44 |
| 1028 | ZNF44 | 5.8090834 | -5.825939774 | 1.3225 | -4.4051 | 1.1E-05 | 0.000108 |
| 1101 | RCN3 | 9.81124953 | -5.826900492 | 1.2121 | -4.8072 | 1.5E-06 | 1.92E-05 |
| 1651 | SLC23A1 | 5.8688611 | -5.828542074 | 1.3655 | -4.2686 | 2E-05 | 0.000188 |
| 94 | TTC22 | 9.69132521 | -5.830945716 | 1.1603 | -5.0256 | 5E-07 | 7.21E-06 |
| 1429 | ITGB5 | 5.8665381 | -5.832876155 | 1.2385 | -4.7097 | 2.5E-06 | 2.98E-05 |
| 1019 | SH2D3A | 9.87005114 | -5.835451812 | 1.1597 | -5.0319 | 4.9E-07 | 6.99E-06 |
| 1529 | PRKG2 | 10.023567 | -5.835864716 | 1.3631 | -4.2815 | 1.9E-05 | 0.000178 |
| 1076 | CEACAM1 | 23.3265186 | -5.836708275 | 0.9403 | -6.2072 | 5.4E-10 | 1.5E-08 |
| 343 | EPS8L2 | 47.5330922 | -5.839511099 | 0.7036 | -8.2996 | 1E-16 | 1.19E-14 |
| 367 | SLC5A12 | 16.1965511 | -5.848647232 | 1.1834 | -4.9424 | 7.7E-07 | 1.05E-05 |
| 117 | GBP1 | 6.09266191 | -5.851115547 | 1.2859 | -4.5503 | 5.4E-06 | 5.85E-05 |

| | | | | | | | |
|---|---|---|---|---|---|---|---|
| 972 | LAMA1 | 10.0973968 | -5.863674732 | 1.9246 | -3.0466 | 0.00231 | 0.011782 |
| 1657 | SH3RF2 | 16.2839953 | -5.867592455 | 1.0925 | -5.3706 | 7.8E-08 | 1.36E-06 |
| 867 | LGALS9 | 16.3661692 | -5.867700885 | 1.0692 | -5.4877 | 4.1E-08 | 7.55E-07 |
| 251 | TMEM236 | 10.279258 | -5.870316237 | 1.2725 | -4.6133 | 4E-06 | 4.49E-05 |
| 90 | TTC39A | 5.99571139 | -5.870747406 | 1.2342 | -4.7569 | 2E-06 | 2.4E-05 |
| 1552 | NPNT | 6.04956964 | -5.870903857 | 1.3093 | -4.4842 | 7.3E-06 | 7.75E-05 |
| 812 | NPIPB15 | 6.04429363 | -5.883592478 | 1.2685 | -4.6382 | 3.5E-06 | 4.05E-05 |
| 167 | MUC1 | 42.1101375 | -5.884600477 | 1.0861 | -5.4182 | 6E-08 | 1.06E-06 |
| 1064 | LGALS14 | 6.00438809 | -5.889566718 | 1.3143 | -4.4812 | 7.4E-06 | 7.84E-05 |
| 908 | PRR15L | 49.7200383 | -5.899770716 | 0.6888 | -8.5648 | 1.1E-17 | 1.52E-15 |
| 1045 | GDF15 | 6.06818627 | -5.901926254 | 1.2881 | -4.582 | 4.6E-06 | 5.13E-05 |
| 285 | IFIT3 | 6.35652365 | -5.9125467 | 1.3075 | -4.5221 | 6.1E-06 | 6.59E-05 |
| 663 | TC2N | 10.3518862 | -5.920284589 | 1.2229 | -4.8412 | 1.3E-06 | 1.66E-05 |
| 515 | METTL7A | 17.0847564 | -5.920918499 | 1.0518 | -5.6291 | 1.8E-08 | 3.65E-07 |
| 1517 | SULT1E1 | 10.610424 | -5.923573347 | 1.2814 | -4.6226 | 3.8E-06 | 4.32E-05 |
| 909 | HOXB3 | 6.15331269 | -5.926626755 | 1.3143 | -4.5092 | 6.5E-06 | 6.99E-05 |
| 481 | PLEKHG6 | 17.5436666 | -5.932932099 | 1.1117 | -5.337 | 9.4E-08 | 1.61E-06 |
| 1557 | CFI | 6.44322787 | -5.934725066 | 1.3057 | -4.5453 | 5.5E-06 | 5.97E-05 |
| 1252 | ALPI | 6.31170414 | -5.93821354 | 1.4089 | -4.2147 | 2.5E-05 | 0.000231 |
| 379 | SYT13 | 6.27509478 | -5.938224245 | 1.2298 | -4.8288 | 1.4E-06 | 1.75E-05 |
| 1868 | HEPACAM2 | 6.34198657 | -5.938616172 | 1.3275 | -4.4736 | 7.7E-06 | 8.09E-05 |
| 1087 | GIPR | 6.29047584 | -5.944035854 | 1.3407 | -4.4335 | 9.3E-06 | 9.58E-05 |
| 485 | LPAR5 | 6.4010325 | -5.95003414 | 1.2223 | -4.8679 | 1.1E-06 | 1.47E-05 |
| 1193 | TFCP2L1 | 10.5205112 | -5.951451386 | 1.1675 | -5.0975 | 3.4E-07 | 5.11E-06 |
| 75 | TMEM125 | 6.42806963 | -5.957015523 | 1.267 | -4.7016 | 2.6E-06 | 3.08E-05 |
| 1168 | TCF7L1 | 6.48462743 | -5.957103931 | 1.2347 | -4.8248 | 1.4E-06 | 1.79E-05 |
| 839 | RILP | 6.37557773 | -5.959003015 | 1.2877 | -4.6275 | 3.7E-06 | 4.23E-05 |
| 208 | GOLT1A | 6.42318222 | -5.960609536 | 1.2649 | -4.7122 | 2.5E-06 | 2.94E-05 |
| 1185 | MTLN | 6.49542543 | -5.961809458 | 1.2825 | -4.6487 | 3.3E-06 | 3.87E-05 |
| 804 | ESRP2 | 6.45690055 | -5.96772302 | 1.3024 | -4.5823 | 4.6E-06 | 5.12E-05 |
| 1476 | HES1 | 17.5287712 | -5.971159419 | 1.0464 | -5.7063 | 1.2E-08 | 2.41E-07 |
| 1598 | SORBS2 | 6.64917974 | -5.975145661 | 1.289 | -4.6356 | 3.6E-06 | 4.1E-05 |
| 916 | LINC02086 | 10.8551264 | -5.978611309 | 1.2519 | -4.7756 | 1.8E-06 | 2.22E-05 |
| 438 | YAP1 | 10.7154116 | -5.980569013 | 1.1671 | -5.1241 | 3E-07 | 4.48E-06 |
| 1595 | DDX60L | 6.73229636 | -5.991134463 | 1.3078 | -4.5809 | 4.6E-06 | 5.15E-05 |
| 905 | ARHGAP27 | 35.1718538 | -5.99243759 | 0.8021 | -7.4705 | 8E-14 | 5.14E-12 |
| 899 | ETV4 | 6.61292835 | -6.005156528 | 1.3689 | -4.3868 | 1.2E-05 | 0.000116 |
| 1656 | PRELID2 | 6.56660051 | -6.007096556 | 1.2264 | -4.898 | 9.7E-07 | 1.29E-05 |
| 874 | MYO1D | 26.4172438 | -6.010396678 | 0.9212 | -6.5249 | 6.8E-11 | 2.39E-09 |

| | | | | | | | |
|---|---|---|---|---|---|---|---|
| 26 | AGMAT | 11.1646548 | -6.018290053 | 1.135 | -5.3027 | 1.1E-07 | 1.9E-06 |
| 1038 | HSH2D | 6.73095024 | -6.01963615 | 1.235 | -4.8743 | 1.1E-06 | 1.43E-05 |
| 207 | FMOD | 6.86539359 | -6.019753919 | 1.4046 | -4.2859 | 1.8E-05 | 0.000175 |
| 1094 | FAM83E | 6.65866159 | -6.025437595 | 1.2318 | -4.8917 | 1E-06 | 1.32E-05 |
| 421 | KCNE3 | 6.64070407 | -6.026784434 | 1.2445 | -4.8426 | 1.3E-06 | 1.64E-05 |
| 468 | VWA5A | 11.36353 | -6.030129481 | 1.1598 | -5.1991 | 2E-07 | 3.13E-06 |
| 1847 | COBL | 6.84923444 | -6.030691664 | 1.3517 | -4.4617 | 8.1E-06 | 8.51E-05 |
| 87 | FOXD2 | 6.6336254 | -6.032053934 | 1.3151 | -4.5868 | 4.5E-06 | 5.04E-05 |
| 1304 | PCK1 | 6.70784844 | -6.035587996 | 1.3167 | -4.5837 | 4.6E-06 | 5.09E-05 |
| 691 | PPP1R14D | 6.72385727 | -6.039004696 | 1.2641 | -4.7773 | 1.8E-06 | 2.2E-05 |
| 1291 | SGK2 | 6.75768847 | -6.039165201 | 1.2071 | -5.0032 | 5.6E-07 | 7.98E-06 |
| 224 | IRF6 | 6.87978513 | -6.050119999 | 1.224 | -4.9429 | 7.7E-07 | 1.05E-05 |
| 1330 | KIAA1671 | 6.7923934 | -6.051362674 | 1.2417 | -4.8734 | 1.1E-06 | 1.43E-05 |
| 911 | HOXB6 | 6.75981795 | -6.054961893 | 1.4151 | -4.2789 | 1.9E-05 | 0.00018 |
| 1937 | ADAM28 | 7.02256746 | -6.0583106 | 1.3018 | -4.6538 | 3.3E-06 | 3.78E-05 |
| 1406 | SUCLG2 | 11.4043201 | -6.059621153 | 1.1398 | -5.3166 | 1.1E-07 | 1.78E-06 |
| 1465 | EHHADH | 7.01189575 | -6.060482964 | 1.2894 | -4.7003 | 2.6E-06 | 3.09E-05 |
| 448 | POU2AF3 | 6.96465672 | -6.064158536 | 1.3326 | -4.5506 | 5.4E-06 | 5.85E-05 |
| 1206 | GALNT3 | 6.87473942 | -6.06655531 | 1.2342 | -4.9154 | 8.9E-07 | 1.19E-05 |
| 645 | ZFP36L1 | 19.1058924 | -6.069206433 | 1.0588 | -5.7319 | 9.9E-09 | 2.12E-07 |
| 2038 | TBC1D2 | 6.93433109 | -6.070538809 | 1.2409 | -4.892 | 1E-06 | 1.32E-05 |
| 1117 | FAM110C | 7.02408464 | -6.073900677 | 1.2272 | -4.9496 | 7.4E-07 | 1.02E-05 |
| 609 | GRTP1 | 11.6549052 | -6.082653282 | 1.2253 | -4.964 | 6.9E-07 | 9.54E-06 |
| 910 | HOXB5 | 6.90223395 | -6.083263619 | 1.2796 | -4.754 | 2E-06 | 2.43E-05 |
| 1590 | GASK1B | 11.9014669 | -6.086089786 | 1.1464 | -5.309 | 1.1E-07 | 1.85E-06 |
| 987 | CCDC68 | 7.02449436 | -6.086179986 | 1.2172 | -5.0003 | 5.7E-07 | 8.07E-06 |
| 1659 | SPINK5 | 6.91593321 | -6.092750476 | 1.3433 | -4.5357 | 5.7E-06 | 6.23E-05 |
| 1846 | DDC | 28.3016767 | -6.094784233 | 0.9827 | -6.2021 | 5.6E-10 | 1.55E-08 |
| 895 | WNK4 | 11.6517612 | -6.098106575 | 1.1862 | -5.1411 | 2.7E-07 | 4.15E-06 |
| 778 | IL4R | 7.00483957 | -6.108871144 | 1.2838 | -4.7583 | 2E-06 | 2.39E-05 |
| 841 | MIR22HG | 7.3339074 | -6.11392183 | 1.3409 | -4.5597 | 5.1E-06 | 5.63E-05 |
| 15 | ARHGEF16 | 19.2001413 | -6.115002279 | 1.0694 | -5.7182 | 1.1E-08 | 2.27E-07 |
| 978 | DSC2 | 28.865349 | -6.115692719 | 0.8923 | -6.8536 | 7.2E-12 | 3.19E-10 |
| 1996 | SLC39A4 | 19.5804881 | -6.124646299 | 1.194 | -5.1295 | 2.9E-07 | 4.38E-06 |
| 2162 | GABRE | 7.32484803 | -6.127351429 | 1.2446 | -4.9232 | 8.5E-07 | 1.15E-05 |
| 680 | EXOC3L4 | 7.35516704 | -6.128505539 | 1.2502 | -4.9021 | 9.5E-07 | 1.27E-05 |
| 757 | BICDL2 | 7.19505291 | -6.128551658 | 2.1583 | -2.8396 | 0.00452 | 0.02066 |
| 628 | FOXA1 | 7.30576875 | -6.129251763 | 1.3235 | -4.6309 | 3.6E-06 | 4.17E-05 |
| 1455 | MECOM | 19.7525529 | -6.131618214 | 1.0369 | -5.9135 | 3.3E-09 | 7.79E-08 |

| | | | | | | | |
|---|---|---|---|---|---|---|---|
| 513 | LIMA1 | 38.1282133 | -6.1326275 | 0.7868 | -7.7948 | 6.5E-15 | 4.99E-13 |
| 1071 | CYP2S1 | 7.33611004 | -6.136187683 | 1.3107 | -4.6816 | 2.8E-06 | 3.36E-05 |
| 1372 | THRB | 7.41374213 | -6.138373713 | 1.4158 | -4.3356 | 1.5E-05 | 0.000143 |
| 1011 | STAP2 | 12.1511847 | -6.139887172 | 1.1276 | -5.445 | 5.2E-08 | 9.37E-07 |
| 774 | ANKS4B | 12.1631187 | -6.140694162 | 1.1318 | -5.4258 | 5.8E-08 | 1.02E-06 |
| 997 | MISP | 29.272951 | -6.145560406 | 0.9246 | -6.6469 | 3E-11 | 1.14E-09 |
| 1725 | SPDEF | 7.22333897 | -6.145798118 | 1.2614 | -4.8721 | 1.1E-06 | 1.44E-05 |
| 1063 | ZFP36 | 12.2346796 | -6.153092469 | 1.1393 | -5.4009 | 6.6E-08 | 1.17E-06 |
| 1324 | COMT | 29.4360787 | -6.154102414 | 0.9144 | -6.7302 | 1.7E-11 | 6.81E-10 |
| 1859 | SEMA3C | 12.108445 | -6.154553238 | 1.1742 | -5.2413 | 1.6E-07 | 2.55E-06 |
| 45 | MYOM3 | 7.41669223 | -6.156700033 | 1.2847 | -4.7922 | 1.6E-06 | 2.05E-05 |
| 590 | EPSTI1 | 12.4909744 | -6.160134748 | 1.1638 | -5.293 | 1.2E-07 | 1.99E-06 |
| 237 | KCNK1 | 7.30540924 | -6.162374331 | 1.2272 | -5.0213 | 5.1E-07 | 7.34E-06 |
| 1141 | QPCT | 7.35358176 | -6.163465059 | 1.2086 | -5.0997 | 3.4E-07 | 5.05E-06 |
| 1422 | HGD | 7.56926708 | -6.163738425 | 1.392 | -4.4279 | 9.5E-06 | 9.83E-05 |
| 892 | JUP | 40.5256308 | -6.163862482 | 0.7872 | -7.8297 | 4.9E-15 | 3.87E-13 |
| 1251 | B3GNT7 | 7.52845374 | -6.176582532 | 1.2179 | -5.0714 | 3.9E-07 | 5.77E-06 |
| 793 | MT1G | 22.2329213 | -6.180409989 | 2.1104 | -2.9285 | 0.00341 | 0.01633 |
| 1410 | PROS1 | 12.6172813 | -6.185359865 | 1.129 | -5.4789 | 4.3E-08 | 7.88E-07 |
| 1386 | SLC6A20 | 7.62389303 | -6.187586925 | 1.2849 | -4.8158 | 1.5E-06 | 1.86E-05 |
| 318 | PLEKHS1 | 7.66104269 | -6.188980616 | 1.3169 | -4.6996 | 2.6E-06 | 3.1E-05 |
| 607 | F7 | 7.60778019 | -6.191366438 | 1.3393 | -4.623 | 3.8E-06 | 4.32E-05 |
| 460 | MPZL2 | 7.71206217 | -6.20064307 | 1.2117 | -5.1173 | 3.1E-07 | 4.64E-06 |
| 1783 | GOPC | 12.6831275 | -6.202946521 | 1.1431 | -5.4265 | 5.7E-08 | 1.02E-06 |
| 483 | SCNN1A | 7.67022768 | -6.205537022 | 1.207 | -5.1412 | 2.7E-07 | 4.15E-06 |
| 714 | GCNT3 | 12.6042822 | -6.206065403 | 1.1404 | -5.442 | 5.3E-08 | 9.5E-07 |
| 490 | BCL2L14 | 7.72090262 | -6.208634852 | 1.2012 | -5.1687 | 2.4E-07 | 3.62E-06 |
| 948 | LLGL2 | 82.2281501 | -6.21203752 | 0.5632 | -11.03 | 2.7E-28 | 1.77E-25 |
| 1051 | ENSG00000267934 | 7.75390347 | -6.221461798 | 1.2166 | -5.114 | 3.2E-07 | 4.71E-06 |
| 95 | PCSK9 | 7.61812576 | -6.221616155 | 1.2874 | -4.8328 | 1.3E-06 | 1.72E-05 |
| 548 | DUSP6 | 12.9218182 | -6.222374477 | 1.1898 | -5.2299 | 1.7E-07 | 2.7E-06 |
| 335 | IFITM2 | 31.0711666 | -6.230197414 | 0.8894 | -7.0047 | 2.5E-12 | 1.2E-10 |
| 65 | TINAGL1 | 7.70251018 | -6.240035446 | 1.2327 | -5.0621 | 4.1E-07 | 6.03E-06 |
| 596 | ATP7B | 8.12114348 | -6.267258811 | 1.2643 | -4.9572 | 7.2E-07 | 9.85E-06 |
| 868 | SLC13A2 | 8.03385489 | -6.273939784 | 1.1952 | -5.2494 | 1.5E-07 | 2.45E-06 |
| 254 | KIAA1217 | 13.3814444 | -6.276099726 | 1.1487 | -5.4639 | 4.7E-08 | 8.52E-07 |
| 2080 | NOXA1 | 8.02828659 | -6.28460795 | 1.2262 | -5.1252 | 3E-07 | 4.46E-06 |
| 148 | FMO5 | 13.5711592 | -6.28584675 | 1.1943 | -5.2631 | 1.4E-07 | 2.3E-06 |
| 1804 | RAB32 | 8.21590868 | -6.294001482 | 1.2234 | -5.1445 | 2.7E-07 | 4.09E-06 |

| | | | | | | | |
|---|---|---|---|---|---|---|---|
| 976 | GATA6 | 22.2123401 | -6.295080439 | 1.0587 | -5.9459 | 2.7E-09 | 6.5E-08 |
| 1292 | R3HDML-AS1 | 8.13485499 | -6.297113975 | 1.1975 | -5.2586 | 1.5E-07 | 2.34E-06 |
| 1575 | MGST2 | 21.9166366 | -6.297498894 | 1.0337 | -6.092 | 1.1E-09 | 2.86E-08 |
| 435 | ENDOD1 | 8.19931752 | -6.299064219 | 1.2315 | -5.115 | 3.1E-07 | 4.69E-06 |
| 1952 | PLAG1 | 8.01067906 | -6.299628939 | 1.2651 | -4.9795 | 6.4E-07 | 8.89E-06 |
| 496 | PLBD1 | 8.09453631 | -6.299895482 | 1.1933 | -5.2794 | 1.3E-07 | 2.11E-06 |
| 1524 | SHROOM3 | 13.8676893 | -6.302970427 | 1.152 | -5.4716 | 4.5E-08 | 8.19E-07 |
| 218 | PFKFB2 | 22.1868139 | -6.303324103 | 1.055 | -5.9746 | 2.3E-09 | 5.58E-08 |
| 1416 | HHLA2 | 8.25856603 | -6.304784792 | 1.2102 | -5.2097 | 1.9E-07 | 2.98E-06 |
| 615 | SLC7A7 | 13.6729968 | -6.306415179 | 1.2261 | -5.1434 | 2.7E-07 | 4.11E-06 |
| 2061 | CELP | 8.1846376 | -6.313210603 | 1.4659 | -4.3068 | 1.7E-05 | 0.000161 |
| 881 | HNF1B | 8.16545656 | -6.321020829 | 1.1907 | -5.3087 | 1.1E-07 | 1.85E-06 |
| 217 | C1orf116 | 8.42878977 | -6.323081178 | 1.2334 | -5.1264 | 3E-07 | 4.44E-06 |
| 46 | IL22RA1 | 8.35402643 | -6.32717916 | 1.2327 | -5.1328 | 2.9E-07 | 4.32E-06 |
| 999 | EFNA2 | 8.22654974 | -6.328070251 | 1.1816 | -5.3556 | 8.5E-08 | 1.46E-06 |
| 1507 | GABRA2 | 8.26084496 | -6.3280727 | 1.1821 | -5.3534 | 8.6E-08 | 1.48E-06 |
| 1231 | CPS1 | 56.0492692 | -6.332762359 | 0.8067 | -7.8499 | 4.2E-15 | 3.31E-13 |
| 919 | IGF2BP1 | 22.5927281 | -6.333224207 | 1.0274 | -6.1641 | 7.1E-10 | 1.91E-08 |
| 57 | SFN | 13.6775806 | -6.336321703 | 1.208 | -5.2455 | 1.6E-07 | 2.5E-06 |
| 2042 | KLF4 | 14.016672 | -6.341821403 | 1.1538 | -5.4967 | 3.9E-08 | 7.21E-07 |
| 582 | URAD | 8.35350179 | -6.346724943 | 1.269 | -5.0013 | 5.7E-07 | 8.04E-06 |
| 486 | MIR200CHG | 8.30584642 | -6.350802135 | 1.2198 | -5.2064 | 1.9E-07 | 3.02E-06 |
| 564 | OAS1 | 45.4178548 | -6.353048174 | 0.8062 | -7.8801 | 3.3E-15 | 2.66E-13 |
| 1707 | TRIM31 | 8.41464266 | -6.358464662 | 1.202 | -5.2897 | 1.2E-07 | 2.02E-06 |
| 819 | HSD17B2 | 8.67983486 | -6.364181839 | 1.2724 | -5.0016 | 5.7E-07 | 8.04E-06 |
| 1594 | DDX60 | 8.40343834 | -6.36794757 | 1.2514 | -5.0888 | 3.6E-07 | 5.31E-06 |
| 1280 | FOXA2 | 8.51456644 | -6.371148174 | 1.2017 | -5.3017 | 1.1E-07 | 1.91E-06 |
| 2012 | AQP7 | 8.70073566 | -6.381827261 | 1.3578 | -4.7001 | 2.6E-06 | 3.09E-05 |
| 1695 | ADTRP | 8.69327724 | -6.382330523 | 1.2491 | -5.1096 | 3.2E-07 | 4.8E-06 |
| 363 | PLEKHA7 | 8.61596059 | -6.384878793 | 1.1762 | -5.4285 | 5.7E-08 | 1.01E-06 |
| 109 | GIPC2 | 14.4990006 | -6.385766724 | 1.163 | -5.491 | 4E-08 | 7.43E-07 |
| 1902 | TFEC | 8.80511483 | -6.389790324 | 1.3936 | -4.585 | 4.5E-06 | 5.07E-05 |
| 806 | NQO1 | 8.57363348 | -6.394644003 | 1.2258 | -5.2169 | 1.8E-07 | 2.88E-06 |
| 1844 | NPC1L1 | 8.82335628 | -6.402147953 | 1.2121 | -5.282 | 1.3E-07 | 2.09E-06 |
| 1413 | NFKBIZ | 70.5450991 | -6.411046641 | 0.7777 | -8.2436 | 1.7E-16 | 1.77E-14 |
| 419 | STARD10 | 34.8779115 | -6.414780419 | 0.8924 | -7.1886 | 6.5E-13 | 3.48E-11 |
| 1092 | SULT2A1 | 8.88578673 | -6.425423818 | 2.4785 | -2.5924 | 0.00953 | 0.037898 |
| 497 | GUCY2C | 8.87299168 | -6.428531777 | 1.1715 | -5.4873 | 4.1E-08 | 7.56E-07 |
| 517 | ACVRL1 | 8.74990092 | -6.430061954 | 1.2672 | -5.0743 | 3.9E-07 | 5.7E-06 |

| | | | | | | | |
|---|---|---|---|---|---|---|---|
| 153 | CGN | 24.2902468 | -6.430147766 | 1.0451 | -6.1524 | 7.6E-10 | 2.04E-08 |
| 1616 | SELENOP | 8.97753373 | -6.430765237 | 1.2281 | -5.2364 | 1.6E-07 | 2.61E-06 |
| 692 | SPINT1-AS1 | 8.8959486 | -6.433549868 | 1.2221 | -5.2641 | 1.4E-07 | 2.29E-06 |
| 1082 | CBLC | 8.85046677 | -6.437314577 | 1.2118 | -5.3121 | 1.1E-07 | 1.82E-06 |
| 1164 | DQX1 | 9.1628837 | -6.446577422 | 1.2658 | -5.0928 | 3.5E-07 | 5.22E-06 |
| 1111 | CDC42EP5 | 14.9726127 | -6.449293961 | 1.1406 | -5.6545 | 1.6E-08 | 3.21E-07 |
| 1528 | BMP3 | 8.96000161 | -6.456250424 | 1.1777 | -5.4822 | 4.2E-08 | 7.75E-07 |
| 779 | APOBR | 8.96216086 | -6.456779693 | 1.2471 | -5.1775 | 2.2E-07 | 3.47E-06 |
| 1682 | BTNL8 | 15.5413261 | -6.480454865 | 1.2413 | -5.2205 | 1.8E-07 | 2.83E-06 |
| 2030 | SYK | 9.36804406 | -6.481627234 | 1.1926 | -5.435 | 5.5E-08 | 9.84E-07 |
| 1239 | IHH | 9.14741907 | -6.486218416 | 1.2226 | -5.3054 | 1.1E-07 | 1.88E-06 |
| 2025 | PRUNE2 | 9.35737916 | -6.504750901 | 1.1597 | -5.6091 | 2E-08 | 4.05E-07 |
| 1191 | TMEM37 | 9.46916159 | -6.510062041 | 1.1991 | -5.4291 | 5.7E-08 | 1.01E-06 |
| 244 | AKR1C3 | 9.55831053 | -6.510741357 | 1.1844 | -5.497 | 3.9E-08 | 7.2E-07 |
| 1099 | TRPM4 | 15.5050854 | -6.515987256 | 1.1424 | -5.7037 | 1.2E-08 | 2.45E-07 |
| 512 | GPD1 | 9.4921185 | -6.516563078 | 1.2279 | -5.3069 | 1.1E-07 | 1.87E-06 |
| 1603 | TERLR1 | 9.72498326 | -6.538495388 | 1.3319 | -4.9093 | 9.1E-07 | 1.22E-05 |
| 1347 | CARD10 | 16.1856309 | -6.54797526 | 1.1274 | -5.8078 | 6.3E-09 | 1.4E-07 |
| 1978 | SAMD12 | 9.75595933 | -6.564297311 | 1.1564 | -5.6767 | 1.4E-08 | 2.85E-07 |
| 1647 | PITX1 | 16.2710453 | -6.570186499 | 1.1924 | -5.5101 | 3.6E-08 | 6.76E-07 |
| 1339 | ISX | 9.92475473 | -6.576036958 | 1.1841 | -5.5538 | 2.8E-08 | 5.42E-07 |
| 30 | PADI2 | 52.1257207 | -6.588804856 | 0.7795 | -8.453 | 2.8E-17 | 3.6E-15 |
| 501 | PDE3A | 10.0640405 | -6.589314043 | 1.1674 | -5.6445 | 1.7E-08 | 3.36E-07 |
| 1273 | SPTLC3 | 10.1557891 | -6.590858263 | 1.2578 | -5.2399 | 1.6E-07 | 2.57E-06 |
| 1609 | PRLR | 10.0949785 | -6.591914117 | 1.2106 | -5.4453 | 5.2E-08 | 9.36E-07 |
| 634 | BMP4 | 9.91469328 | -6.599273321 | 1.2786 | -5.1613 | 2.5E-07 | 3.76E-06 |
| 1648 | TGFBI | 9.90317917 | -6.605365896 | 1.3567 | -4.8688 | 1.1E-06 | 1.46E-05 |
| 1797 | PERP | 16.7749973 | -6.605664955 | 1.1026 | -5.9908 | 2.1E-09 | 5.08E-08 |
| 1663 | SLC6A7 | 10.1409522 | -6.617030647 | 1.1722 | -5.6449 | 1.7E-08 | 3.36E-07 |
| 1268 | SDCBP2 | 16.8120543 | -6.623652994 | 1.1325 | -5.8488 | 5E-09 | 1.11E-07 |
| 951 | ITGB4 | 10.0458044 | -6.626539589 | 1.1878 | -5.5789 | 2.4E-08 | 4.77E-07 |
| 1467 | LIPH | 10.3609554 | -6.648253923 | 1.1498 | -5.7823 | 7.4E-09 | 1.61E-07 |
| 1311 | CLIC6 | 10.2948254 | -6.650745762 | 1.1519 | -5.7737 | 7.8E-09 | 1.69E-07 |
| 165 | EFNA1 | 41.0447437 | -6.651660259 | 0.9079 | -7.3261 | 2.4E-13 | 1.38E-11 |
| 1448 | AADAC | 10.4303823 | -6.652699866 | 1.9586 | -3.3966 | 0.00068 | 0.004155 |
| 1077 | CD177 | 10.407968 | -6.660837756 | 2.2638 | -2.9424 | 0.00326 | 0.015717 |
| 297 | BLNK | 10.5522071 | -6.664023309 | 1.202 | -5.5443 | 3E-08 | 5.69E-07 |
| 500 | MGST1 | 55.3617678 | -6.670553041 | 0.8403 | -7.9385 | 2E-15 | 1.75E-13 |
| 222 | LAMB3 | 10.5135551 | -6.674693232 | 1.221 | -5.4666 | 4.6E-08 | 8.4E-07 |

| | | | | | | | |
|---|---|---|---|---|---|---|---|
| 2135 | POF1B | 29.0110901 | -6.693203027 | 1.0207 | -6.5576 | 5.5E-11 | 1.95E-09 |
| 1383 | VIPR1 | 17.8598121 | -6.693883792 | 1.1416 | -5.8638 | 4.5E-09 | 1.03E-07 |
| 982 | LIPG | 10.61413 | -6.696831393 | 1.1647 | -5.7499 | 8.9E-09 | 1.92E-07 |
| 1361 | UPK3A | 10.7529561 | -6.700749055 | 1.3187 | -5.0813 | 3.7E-07 | 5.51E-06 |
| 1226 | CASP8 | 10.9537566 | -6.701259752 | 1.2453 | -5.3812 | 7.4E-08 | 1.29E-06 |
| 2145 | COL4A6 | 10.6560433 | -6.70440954 | 1.3408 | -5.0003 | 5.7E-07 | 8.07E-06 |
| 975 | GATA6-AS1 | 18.1375038 | -6.706002424 | 1.2743 | -5.2625 | 1.4E-07 | 2.3E-06 |
| 988 | ATP8B1 | 17.824261 | -6.706604924 | 1.1104 | -6.0399 | 1.5E-09 | 3.88E-08 |
| 638 | TMEM30B | 17.9681628 | -6.706754085 | 1.114 | -6.0206 | 1.7E-09 | 4.32E-08 |
| 1303 | BCAS1 | 18.0311361 | -6.714908059 | 1.1112 | -6.0429 | 1.5E-09 | 3.81E-08 |
| 1754 | TINAG | 10.9223606 | -6.715232019 | 1.1461 | -5.8594 | 4.6E-09 | 1.05E-07 |
| 771 | ABCC6P1 | 11.0766007 | -6.716110845 | 1.2461 | -5.3896 | 7.1E-08 | 1.24E-06 |
| 1457 | GPR160 | 18.030216 | -6.718748996 | 1.1169 | -6.0155 | 1.8E-09 | 4.44E-08 |
| 1017 | C3 | 11.1123103 | -6.744404185 | 1.3678 | -4.9307 | 8.2E-07 | 1.11E-05 |
| 1884 | MUC12 | 11.0551676 | -6.749700071 | 2.0534 | -3.287 | 0.00101 | 0.0058 |
| 152 | SELENBP1 | 30.747914 | -6.764446018 | 1.017 | -6.6516 | 2.9E-11 | 1.11E-09 |
| 800 | CA7 | 11.253493 | -6.764825407 | 1.2803 | -5.2838 | 1.3E-07 | 2.08E-06 |
| 763 | SMIM22 | 18.4392672 | -6.765112131 | 1.126 | -6.0079 | 1.9E-09 | 4.62E-08 |
| 942 | GPRC5C | 18.4781272 | -6.767965962 | 1.1105 | -6.0943 | 1.1E-09 | 2.83E-08 |
| 739 | ALPK3 | 11.4301034 | -6.772563355 | 1.2307 | -5.5031 | 3.7E-08 | 7.01E-07 |
| 1885 | MUC17 | 11.3153636 | -6.778734873 | 1.1842 | -5.7243 | 1E-08 | 2.2E-07 |
| 1236 | TMBIM1 | 18.6685252 | -6.781280484 | 1.1378 | -5.9599 | 2.5E-09 | 6.03E-08 |
| 1550 | BDH2 | 11.4508824 | -6.78413693 | 1.1946 | -5.6789 | 1.4E-08 | 2.82E-07 |
| 1618 | ELOVL7 | 11.6427139 | -6.794301763 | 1.3523 | -5.0241 | 5.1E-07 | 7.25E-06 |
| 2053 | KIF12 | 11.5442658 | -6.812120338 | 1.1922 | -5.714 | 1.1E-08 | 2.32E-07 |
| 1621 | MARVELD2 | 11.7490694 | -6.819810852 | 1.1486 | -5.9374 | 2.9E-09 | 6.8E-08 |
| 321 | PNLIPRP2 | 11.6675285 | -6.828546548 | 1.1765 | -5.804 | 6.5E-09 | 1.43E-07 |
| 776 | ERN2 | 11.5449915 | -6.829089208 | 1.2338 | -5.5351 | 3.1E-08 | 5.95E-07 |
| 1404 | FAM3D | 11.6105548 | -6.835498412 | 1.2941 | -5.2822 | 1.3E-07 | 2.09E-06 |
| 1965 | CA1 | 11.8229814 | -6.838215683 | 1.2843 | -5.3243 | 1E-07 | 1.72E-06 |
| 1421 | NR1I2 | 12.0144423 | -6.839839488 | 1.17 | -5.8458 | 5E-09 | 1.13E-07 |
| 1591 | TMEM144 | 11.8118646 | -6.844255922 | 1.1623 | -5.8887 | 3.9E-09 | 8.93E-08 |
| 230 | MTARC2 | 12.015607 | -6.8482436 | 1.302 | -5.2597 | 1.4E-07 | 2.33E-06 |
| 58 | KDF1 | 12.0093592 | -6.856406881 | 1.1462 | -5.982 | 2.2E-09 | 5.35E-08 |
| 143 | HSD3B2 | 12.1521124 | -6.861175416 | 1.3174 | -5.2083 | 1.9E-07 | 3E-06 |
| 1337 | SLC5A1 | 31.3517333 | -6.865614276 | 1.4862 | -4.6197 | 3.8E-06 | 4.37E-05 |
| 1675 | CDHR2 | 12.089866 | -6.86620752 | 1.1728 | -5.8544 | 4.8E-09 | 1.08E-07 |
| 1024 | AP1M2 | 12.134326 | -6.872118421 | 1.1413 | -6.0214 | 1.7E-09 | 4.31E-08 |
| 1043 | B3GNT3 | 12.0519248 | -6.881038243 | 1.1511 | -5.9776 | 2.3E-09 | 5.49E-08 |

| | | | | | | | |
|---|---|---|---|---|---|---|---|
| 1053 | SLC7A9 | 12.2553197 | -6.888931801 | 1.3025 | -5.2892 | 1.2E-07 | 2.03E-06 |
| 1501 | FAM114A1 | 12.4380335 | -6.901398168 | 1.1506 | -5.9982 | 2E-09 | 4.88E-08 |
| 1741 | CLIC5 | 12.5786212 | -6.902075892 | 1.2122 | -5.6936 | 1.2E-08 | 2.59E-07 |
| 685 | NDN | 12.2460689 | -6.902524361 | 1.1811 | -5.8442 | 5.1E-09 | 1.14E-07 |
| 1417 | C3orf85 | 12.4946422 | -6.90478434 | 1.1897 | -5.8039 | 6.5E-09 | 1.43E-07 |
| 1414 | ALCAM | 12.6964861 | -6.920521916 | 1.1644 | -5.9436 | 2.8E-09 | 6.57E-08 |
| 1763 | TENT5A | 12.6315587 | -6.926473091 | 1.1351 | -6.1019 | 1E-09 | 2.71E-08 |
| 1257 | RAB17 | 12.506725 | -6.929811455 | 1.1527 | -6.012 | 1.8E-09 | 4.52E-08 |
| 677 | DIO3OS | 20.8842874 | -6.935832632 | 1.139 | -6.0895 | 1.1E-09 | 2.9E-08 |
| 1318 | TFF1 | 12.6215781 | -6.936320111 | 1.3142 | -5.2779 | 1.3E-07 | 2.13E-06 |
| 1841 | PRR15 | 12.9002576 | -6.938471216 | 1.2156 | -5.7077 | 1.1E-08 | 2.4E-07 |
| 1403 | ACOX2 | 13.0199327 | -6.951703303 | 1.2272 | -5.6648 | 1.5E-08 | 3.04E-07 |
| 354 | PPFIBP2 | 12.7511732 | -6.954195465 | 1.1409 | -6.0952 | 1.1E-09 | 2.82E-08 |
| 796 | ADGRG1 | 12.7267193 | -6.95765626 | 1.1567 | -6.015 | 1.8E-09 | 4.45E-08 |
| 1244 | SLC19A3 | 13.0918741 | -6.97024026 | 1.162 | -5.9985 | 2E-09 | 4.88E-08 |
| 1317 | TFF3 | 170.348261 | -6.976656562 | 0.509 | -13.708 | 9.1E-43 | 2.55E-39 |
| 1055 | HPN | 21.7724457 | -6.988075189 | 1.192 | -5.8627 | 4.6E-09 | 1.03E-07 |
| 510 | DNAJC22 | 13.2802205 | -6.98819573 | 1.1709 | -5.9683 | 2.4E-09 | 5.76E-08 |
| 1815 | PDGFA | 13.2244236 | -6.996133183 | 1.1749 | -5.9546 | 2.6E-09 | 6.19E-08 |
| 223 | G0S2 | 13.2843112 | -7.013872405 | 1.2506 | -5.6086 | 2E-08 | 4.06E-07 |
| 149 | MTMR11 | 55.6011656 | -7.020168476 | 0.8895 | -7.8922 | 3E-15 | 2.47E-13 |
| 1526 | ANXA3 | 13.4027097 | -7.025970283 | 1.1778 | -5.9651 | 2.4E-09 | 5.86E-08 |
| 2032 | FBP1 | 13.57806 | -7.035020654 | 1.1811 | -5.9562 | 2.6E-09 | 6.14E-08 |
| 2014 | MYORG | 13.6572846 | -7.040546823 | 1.1355 | -6.2001 | 5.6E-10 | 1.56E-08 |
| 424 | MOGAT2 | 13.6785817 | -7.048409252 | 1.1468 | -6.1464 | 7.9E-10 | 2.1E-08 |
| 922 | TMEM92 | 13.7379565 | -7.055311051 | 1.1299 | -6.2442 | 4.3E-10 | 1.21E-08 |
| 395 | MIR194-2HG | 37.5910872 | -7.057273828 | 1.0417 | -6.7749 | 1.2E-11 | 5.24E-10 |
| 397 | NAALADL1 | 13.9056668 | -7.069117418 | 1.2673 | -5.5779 | 2.4E-08 | 4.79E-07 |
| 1139 | XDH | 14.0313279 | -7.083498002 | 1.2483 | -5.6745 | 1.4E-08 | 2.88E-07 |
| 1201 | GALNT5 | 14.1435413 | -7.098464193 | 1.1323 | -6.269 | 3.6E-10 | 1.06E-08 |
| 1612 | DAB2 | 24.2076199 | -7.12959545 | 1.1317 | -6.2998 | 3E-10 | 8.81E-09 |
| 780 | SULT1A1 | 39.2858808 | -7.133564119 | 1.0292 | -6.9312 | 4.2E-12 | 1.95E-10 |
| 1966 | CA2 | 14.2999162 | -7.134886113 | 1.231 | -5.7959 | 6.8E-09 | 1.49E-07 |
| 1987 | NDRG1 | 112.761484 | -7.141118921 | 0.6554 | -10.896 | 1.2E-27 | 6.61E-25 |
| 1899 | TMEM168 | 14.567634 | -7.144924389 | 1.1161 | -6.4018 | 1.5E-10 | 4.87E-09 |
| 1514 | UGT2A3 | 14.5142497 | -7.147055597 | 1.2147 | -5.8837 | 4E-09 | 9.19E-08 |
| 1497 | GBA3 | 14.9325019 | -7.161230622 | 1.189 | -6.0229 | 1.7E-09 | 4.28E-08 |
| 1067 | BLVRB | 14.8871852 | -7.161731876 | 1.1286 | -6.3457 | 2.2E-10 | 6.73E-09 |
| 1112 | PTPRH | 15.0539334 | -7.174235645 | 1.1489 | -6.2445 | 4.3E-10 | 1.21E-08 |

| | | | | | | | |
|---|---|---|---|---|---|---|---|
| 1748 | GSTA2 | 15.0041703 | -7.179722503 | 1.7788 | -4.0363 | 5.4E-05 | 0.000462 |
| 1887 | MOGAT3 | 15.0031524 | -7.185107779 | 1.1193 | -6.4192 | 1.4E-10 | 4.42E-09 |
| 759 | IL32 | 15.0798412 | -7.186716341 | 1.1417 | -6.2949 | 3.1E-10 | 9.06E-09 |
| 308 | COL17A1 | 15.168751 | -7.187331581 | 1.1217 | -6.4075 | 1.5E-10 | 4.73E-09 |
| 1703 | TRIM38 | 25.3396797 | -7.192592209 | 1.1438 | -6.2882 | 3.2E-10 | 9.42E-09 |
| 88 | SLC5A9 | 15.3353774 | -7.209154667 | 1.2007 | -6.004 | 1.9E-09 | 4.72E-08 |
| 116 | GBP3 | 15.474043 | -7.21428752 | 1.2033 | -5.9953 | 2E-09 | 4.95E-08 |
| 299 | ABCC2 | 15.3513078 | -7.215466292 | 2.6914 | -2.6809 | 0.00734 | 0.030763 |
| 111 | IFI44 | 15.6129797 | -7.217286807 | 1.2254 | -5.8899 | 3.9E-09 | 8.88E-08 |
| 202 | CSRP1 | 15.2932454 | -7.222927876 | 1.1327 | -6.3768 | 1.8E-10 | 5.64E-09 |
| 571 | ACADS | 25.4716726 | -7.228489553 | 1.1143 | -6.4871 | 8.7E-11 | 3E-09 |
| 2125 | IL2RG | 15.5636932 | -7.242259438 | 1.1902 | -6.0851 | 1.2E-09 | 2.97E-08 |
| 1570 | FABP2 | 26.3210028 | -7.255356038 | 1.2236 | -5.9295 | 3E-09 | 7.09E-08 |
| 1670 | ATP10B | 15.6204427 | -7.259130046 | 1.1322 | -6.4116 | 1.4E-10 | 4.63E-09 |
| 1199 | HNMT | 15.8298014 | -7.265455134 | 1.1178 | -6.4997 | 8E-11 | 2.79E-09 |
| 2146 | COL4A5 | 15.7829355 | -7.267530635 | 1.1631 | -6.2485 | 4.1E-10 | 1.19E-08 |
| 33 | RNF186 | 16.0666028 | -7.289779949 | 1.1208 | -6.5042 | 7.8E-11 | 2.71E-09 |
| 825 | FENDRR | 16.202299 | -7.30657957 | 1.1405 | -6.4065 | 1.5E-10 | 4.75E-09 |
| 1697 | MBOAT1 | 27.4577477 | -7.319457094 | 1.0823 | -6.7626 | 1.4E-11 | 5.64E-10 |
| 452 | NXPE1 | 16.3552012 | -7.328449765 | 1.1528 | -6.3572 | 2.1E-10 | 6.33E-09 |
| 235 | TMEM63A | 16.7013335 | -7.353121669 | 1.1484 | -6.4028 | 1.5E-10 | 4.84E-09 |
| 145 | REG4 | 46.5956968 | -7.363132821 | 1.093 | -6.7368 | 1.6E-11 | 6.54E-10 |
| 1832 | AGR3 | 28.582431 | -7.368863021 | 1.1486 | -6.4155 | 1.4E-10 | 4.52E-09 |
| 110 | IFI44L | 28.7706162 | -7.379405983 | 1.1595 | -6.3645 | 2E-10 | 6.06E-09 |
| 300 | DNMBP | 17.1938072 | -7.386479088 | 1.1404 | -6.4772 | 9.3E-11 | 3.18E-09 |
| 1070 | CYP2B6 | 17.2988264 | -7.390607562 | 1.1339 | -6.5176 | 7.1E-11 | 2.49E-09 |
| 2143 | CLDN2 | 17.5004975 | -7.399679647 | 1.1687 | -6.3314 | 2.4E-10 | 7.31E-09 |
| 11 | TNFRSF14-AS1 | 17.5412259 | -7.402319711 | 1.1525 | -6.4231 | 1.3E-10 | 4.32E-09 |
| 1597 | HPGD | 17.9109435 | -7.416254346 | 1.1999 | -6.1809 | 6.4E-10 | 1.74E-08 |
| 704 | SQOR | 17.4145946 | -7.4184646 | 1.1493 | -6.455 | 1.1E-10 | 3.64E-09 |
| 279 | GPR15LG | 17.4790959 | -7.422973586 | 1.1452 | -6.4817 | 9.1E-11 | 3.09E-09 |
| 1903 | TES | 18.0474454 | -7.441149168 | 1.1084 | -6.7132 | 1.9E-11 | 7.53E-10 |
| 201 | LAD1 | 17.749878 | -7.443968042 | 1.1338 | -6.5652 | 5.2E-11 | 1.88E-09 |
| 572 | HNF1A-AS1 | 30.0116926 | -7.449362343 | 1.0784 | -6.9075 | 4.9E-12 | 2.27E-10 |
| 185 | GPA33 | 17.897218 | -7.450334483 | 1.1352 | -6.5633 | 5.3E-11 | 1.89E-09 |
| 327 | CLRN3 | 18.076056 | -7.451522611 | 1.132 | -6.5824 | 4.6E-11 | 1.69E-09 |
| 388 | MS4A8 | 18.3671102 | -7.454616029 | 1.2117 | -6.1522 | 7.6E-10 | 2.04E-08 |
| 1315 | MX1 | 49.4959401 | -7.46247705 | 1.0089 | -7.397 | 1.4E-13 | 8.51E-12 |
| 1008 | TJP3 | 18.7142934 | -7.510953186 | 1.1095 | -6.7696 | 1.3E-11 | 5.4E-10 |

| | | | | | | | |
|---|---|---|---|---|---|---|---|
| 1288 | MYL9 | 18.7835641 | -7.513532962 | 1.6218 | -4.6328 | 3.6E-06 | 4.14E-05 |
| 34 | PLA2G2A | 51.0187507 | -7.521065291 | 1.0842 | -6.9368 | 4E-12 | 1.89E-10 |
| 175 | LINC01133 | 19.2298071 | -7.547451333 | 1.0952 | -6.8911 | 5.5E-12 | 2.51E-10 |
| 337 | PKP3 | 19.6688323 | -7.577621073 | 1.0978 | -6.9025 | 5.1E-12 | 2.34E-10 |
| 1718 | SLC44A4 | 32.4508019 | -7.582603443 | 1.0792 | -7.0265 | 2.1E-12 | 1.04E-10 |
| 791 | MT1E | 19.9099832 | -7.596188072 | 1.771 | -4.2893 | 1.8E-05 | 0.000173 |
| 1166 | TRABD2A | 19.7830267 | -7.596490198 | 1.1295 | -6.7258 | 1.7E-11 | 6.96E-10 |
| 884 | PPP1R1B | 19.7342964 | -7.597274311 | 1.1548 | -6.5789 | 4.7E-11 | 1.72E-09 |
| 1546 | ADH1C | 20.2283603 | -7.608602354 | 1.16 | -6.5591 | 5.4E-11 | 1.94E-09 |
| 1683 | BTNL3 | 20.3194957 | -7.609390628 | 1.1591 | -6.5649 | 5.2E-11 | 1.88E-09 |
| 1313 | B3GALT5 | 20.3056443 | -7.616220587 | 1.1235 | -6.7793 | 1.2E-11 | 5.1E-10 |
| 1458 | PLD1 | 20.5056065 | -7.623571975 | 1.1842 | -6.4378 | 1.2E-10 | 3.97E-09 |
| 556 | NR1H4 | 20.5258959 | -7.62630801 | 1.2391 | -6.1547 | 7.5E-10 | 2.01E-08 |
| 294 | CYP2C9 | 20.396213 | -7.631498091 | 1.1365 | -6.7152 | 1.9E-11 | 7.44E-10 |
| 1834 | MACC1 | 20.9235469 | -7.643976315 | 1.1728 | -6.5175 | 7.1E-11 | 2.49E-09 |
| 1243 | TM4SF20 | 20.8096286 | -7.645394108 | 1.1968 | -6.3884 | 1.7E-10 | 5.26E-09 |
| 1904 | CFTR | 20.4184174 | -7.64599716 | 1.1629 | -6.5749 | 4.9E-11 | 1.76E-09 |
| 156 | S100A9 | 20.7299479 | -7.654960126 | 1.5692 | -4.8784 | 1.1E-06 | 1.4E-05 |
| 261 | A1CF | 21.2329706 | -7.667469862 | 1.1187 | -6.854 | 7.2E-12 | 3.19E-10 |
| 508 | PKP2 | 21.232582 | -7.668183153 | 1.1106 | -6.9046 | 5E-12 | 2.31E-10 |
| 1195 | MYO7B | 20.9087948 | -7.671226185 | 1.1389 | -6.7357 | 1.6E-11 | 6.57E-10 |
| 1460 | TNFSF10 | 21.4138332 | -7.685961197 | 1.1394 | -6.7455 | 1.5E-11 | 6.22E-10 |
| 2020 | ALDH1A1 | 58.1171693 | -7.687336866 | 1.0481 | -7.3348 | 2.2E-13 | 1.3E-11 |
| 1970 | ESRP1 | 21.3522387 | -7.694440133 | 1.1096 | -6.9347 | 4.1E-12 | 1.91E-10 |
| 392 | AHNAK | 36.1645432 | -7.714076128 | 1.1041 | -6.9867 | 2.8E-12 | 1.35E-10 |
| 1983 | ANXA13 | 36.8202116 | -7.751117008 | 1.7572 | -4.411 | 1E-05 | 0.000105 |
| 809 | MARVELD3 | 22.4705293 | -7.754532851 | 1.103 | -7.0303 | 2.1E-12 | 1.02E-10 |
| 519 | KRT8 | 90.2770876 | -7.795496608 | 0.8653 | -9.0086 | 2.1E-19 | 4.01E-17 |
| 158 | S100A16 | 23.2780397 | -7.82368489 | 1.0863 | -7.2024 | 5.9E-13 | 3.21E-11 |
| 605 | SLC15A1 | 23.6465334 | -7.827086155 | 1.1866 | -6.596 | 4.2E-11 | 1.56E-09 |
| 454 | APOC3 | 23.4451252 | -7.828149188 | 1.799 | -4.3514 | 1.4E-05 | 0.000135 |
| 4 | ENSG00000272141 | 23.9891448 | -7.868034261 | 1.0933 | -7.1964 | 6.2E-13 | 3.31E-11 |
| 1143 | GALM | 24.4514729 | -7.87949757 | 1.0929 | -7.21 | 5.6E-13 | 3.05E-11 |
| 177 | ITLN1 | 24.4176717 | -7.883122688 | 1.0843 | -7.2703 | 3.6E-13 | 2.03E-11 |
| 2013 | PRSS3 | 40.3707653 | -7.88685024 | 1.0693 | -7.3761 | 1.6E-13 | 9.87E-12 |
| 1487 | FGFR3 | 24.7476285 | -7.912254377 | 1.1489 | -6.8866 | 5.7E-12 | 2.58E-10 |
| 159 | S100A14 | 25.0162316 | -7.927796372 | 1.1158 | -7.1051 | 1.2E-12 | 6.15E-11 |
| 581 | CDX2 | 25.3783948 | -7.944683152 | 1.0929 | -7.2695 | 3.6E-13 | 2.03E-11 |
| 271 | HKDC1 | 25.6559701 | -7.961310873 | 1.0966 | -7.2603 | 3.9E-13 | 2.15E-11 |

| | | | | | | | |
|---|---|---|---|---|---|---|---|
| 991 | TNFRSF11A | 26.0236452 | -7.975328645 | 1.093 | -7.2966 | 3E-13 | 1.7E-11 |
| 2127 | GJB1 | 26.3556214 | -7.994254565 | 1.1067 | -7.2236 | 5.1E-13 | 2.77E-11 |
| 1734 | UNC5CL | 26.8701837 | -8.018682962 | 1.1532 | -6.9534 | 3.6E-12 | 1.69E-10 |
| 2141 | RNF128 | 45.0762085 | -8.03287653 | 1.1198 | -7.1734 | 7.3E-13 | 3.84E-11 |
| 151 | CTSS | 27.169846 | -8.043384624 | 1.0791 | -7.4536 | 9.1E-14 | 5.67E-12 |
| 979 | DSG2 | 27.2126478 | -8.056730193 | 1.1302 | -7.1285 | 1E-12 | 5.22E-11 |
| 478 | CCND2 | 76.1845432 | -8.080716612 | 1.0023 | -8.0621 | 7.5E-16 | 7.02E-14 |
| 1316 | TMPRSS2 | 28.1142614 | -8.10059157 | 1.0832 | -7.4787 | 7.5E-14 | 4.85E-12 |
| 1897 | SLC26A3 | 28.5850561 | -8.124038796 | 1.0795 | -7.5256 | 5.2E-14 | 3.49E-12 |
| 1516 | SULT1B1 | 28.7066499 | -8.124322678 | 1.0831 | -7.5007 | 6.3E-14 | 4.14E-12 |
| 1180 | SULT1C2 | 29.4100164 | -8.15148847 | 1.6938 | -4.8124 | 1.5E-06 | 1.88E-05 |
| 1109 | TMC4 | 29.6999426 | -8.164033679 | 1.1028 | -7.4028 | 1.3E-13 | 8.18E-12 |
| 1172 | VAMP8 | 30.0531034 | -8.191998093 | 1.0749 | -7.621 | 2.5E-14 | 1.75E-12 |
| 169 | RAB25 | 30.053818 | -8.195765846 | 1.1228 | -7.2991 | 2.9E-13 | 1.68E-11 |
| 1290 | TGM2 | 30.4309206 | -8.198861934 | 1.1142 | -7.3585 | 1.9E-13 | 1.12E-11 |
| 1000 | REEP6 | 30.5033362 | -8.19908674 | 1.1761 | -6.9717 | 3.1E-12 | 1.49E-10 |
| 292 | CYP2C18 | 30.5121914 | -8.20319419 | 1.4523 | -5.6482 | 1.6E-08 | 3.32E-07 |
| 1980 | MAL2 | 30.5866732 | -8.213660579 | 1.0765 | -7.6298 | 2.4E-14 | 1.66E-12 |
| 1105 | KLK1 | 30.6170342 | -8.213894097 | 1.0701 | -7.6758 | 1.6E-14 | 1.21E-12 |
| 932 | CA4 | 31.0858748 | -8.24326887 | 1.0782 | -7.6452 | 2.1E-14 | 1.49E-12 |
| 783 | MVP | 31.2691964 | -8.248116332 | 1.0977 | -7.5138 | 5.7E-14 | 3.8E-12 |
| 1504 | RBM47 | 85.1115979 | -8.254684123 | 1.0061 | -8.2044 | 2.3E-16 | 2.41E-14 |
| 314 | ACSL5 | 52.4250023 | -8.264784586 | 1.0873 | -7.6013 | 2.9E-14 | 2.01E-12 |
| 1468 | IGF2BP2 | 32.0200264 | -8.27460741 | 1.0773 | -7.6812 | 1.6E-14 | 1.17E-12 |
| 2019 | GDA | 32.1782248 | -8.27644602 | 1.1506 | -7.1934 | 6.3E-13 | 3.37E-11 |
| 72 | GUCA2A | 32.2144029 | -8.286863189 | 1.6267 | -5.0944 | 3.5E-07 | 5.18E-06 |
| 144 | HMGCS2 | 86.4167 | -8.289577253 | 1.0279 | -8.0643 | 7.4E-16 | 6.94E-14 |
| 1218 | SLC40A1 | 33.5653355 | -8.330276986 | 1.1088 | -7.5128 | 5.8E-14 | 3.81E-12 |
| 703 | C15orf48 | 33.5733543 | -8.359212712 | 1.161 | -7.2 | 6E-13 | 3.26E-11 |
| 1701 | SLC17A4 | 34.625294 | -8.380428697 | 1.0915 | -7.6777 | 1.6E-14 | 1.2E-12 |
| 1493 | S100P | 34.3947768 | -8.394745045 | 1.1113 | -7.5538 | 4.2E-14 | 2.82E-12 |
| 1854 | CLDN4 | 57.2252687 | -8.400218714 | 1.0611 | -7.9168 | 2.4E-15 | 2.05E-13 |
| 1547 | MTTP | 36.2663252 | -8.455988989 | 1.8836 | -4.4892 | 7.2E-06 | 7.61E-05 |
| 1692 | DSP | 37.3870221 | -8.489509147 | 1.1189 | -7.5874 | 3.3E-14 | 2.22E-12 |
| 529 | SLC39A5 | 38.1195797 | -8.536009359 | 1.1108 | -7.6844 | 1.5E-14 | 1.15E-12 |
| 1293 | HNF4A | 38.5083623 | -8.541400745 | 1.066 | -8.0124 | 1.1E-15 | 1.01E-13 |
| 2153 | XPNPEP2 | 39.1219361 | -8.566947478 | 1.3884 | -6.1704 | 6.8E-10 | 1.84E-08 |
| 925 | ABCC3 | 39.1320951 | -8.578402987 | 1.0972 | -7.8187 | 5.3E-15 | 4.19E-13 |
| 954 | ST6GALNAC1 | 66.0935505 | -8.590888662 | 1.0784 | -7.9666 | 1.6E-15 | 1.42E-13 |

| | | | | | | | |
|---|---|---|---|---|---|---|---|
| 805 | CDH1 | 40.4353295 | -8.603198539 | 1.0792 | -7.9718 | 1.6E-15 | 1.37E-13 |
| 364 | USH1C | 40.1292489 | -8.608250722 | 1.122 | -7.6723 | 1.7E-14 | 1.24E-12 |
| 1353 | BAIAP2L2 | 42.0866248 | -8.670911493 | 1.1041 | -7.8533 | 4.1E-15 | 3.24E-13 |
| 457 | TMPRSS4 | 42.7316926 | -8.687439523 | 1.0647 | -8.1599 | 3.4E-16 | 3.31E-14 |
| 12 | TNFRSF14 | 44.0910291 | -8.737502673 | 1.078 | -8.1054 | 5.3E-16 | 5.02E-14 |
| 532 | MYO1A | 44.5312417 | -8.755681137 | 1.0876 | -8.0505 | 8.2E-16 | 7.62E-14 |
| 455 | APOA1 | 119.026251 | -8.756406614 | 1.8708 | -4.6806 | 2.9E-06 | 3.37E-05 |
| 1127 | APOB | 73.8377459 | -8.758886812 | 1.5847 | -5.5271 | 3.3E-08 | 6.22E-07 |
| 599 | KLF5 | 75.1261207 | -8.779085678 | 1.0488 | -8.3706 | 5.7E-17 | 6.74E-15 |
| 1882 | MUC3A | 45.5644216 | -8.779112518 | 1.1087 | -7.9185 | 2.4E-15 | 2.03E-13 |
| 1662 | CDX1 | 45.2913674 | -8.78191959 | 1.0618 | -8.2709 | 1.3E-16 | 1.46E-14 |
| 520 | KRT18 | 45.5990452 | -8.783952593 | 1.0651 | -8.2471 | 1.6E-16 | 1.75E-14 |
| 131 | EPS8L3 | 45.7962596 | -8.803108088 | 1.0713 | -8.2172 | 2.1E-16 | 2.18E-14 |
| 1452 | RARRES1 | 76.4757775 | -8.81449121 | 1.0753 | -8.1969 | 2.5E-16 | 2.52E-14 |
| 1349 | LGALS2 | 47.4507638 | -8.845018184 | 1.0567 | -8.3707 | 5.7E-17 | 6.74E-15 |
| 1007 | SMIM24 | 47.8569991 | -8.854449994 | 1.109 | -7.984 | 1.4E-15 | 1.26E-13 |
| 1920 | AOC1 | 47.8111114 | -8.857931026 | 1.0575 | -8.3763 | 5.5E-17 | 6.53E-15 |
| 2062 | ABO | 47.8419635 | -8.864798906 | 1.0735 | -8.258 | 1.5E-16 | 1.61E-14 |
| 889 | KRT20 | 49.2398147 | -8.886811295 | 1.1261 | -7.8914 | 3E-15 | 2.47E-13 |
| 891 | KRT19 | 80.8062383 | -8.8964111 | 1.0753 | -8.2734 | 1.3E-16 | 1.45E-14 |
| 1073 | CEACAM7 | 84.081087 | -8.948466521 | 1.4611 | -6.1247 | 9.1E-10 | 2.39E-08 |
| 679 | AMN | 51.5292015 | -8.966092552 | 1.1077 | -8.0941 | 5.8E-16 | 5.47E-14 |
| 1454 | SI | 52.1827868 | -8.981582951 | 1.5557 | -5.7735 | 7.8E-09 | 1.69E-07 |
| 1238 | VIL1 | 55.7420016 | -9.073234757 | 1.0678 | -8.497 | 1.9E-17 | 2.54E-15 |
| 782 | ZG16 | 56.7383875 | -9.101456833 | 1.4025 | -6.4893 | 8.6E-11 | 2.97E-09 |
| 855 | CLDN7 | 57.3805016 | -9.122204063 | 1.0574 | -8.6273 | 6.3E-18 | 9.41E-16 |
| 1850 | MLXIPL | 59.4491559 | -9.172857987 | 1.1128 | -8.2428 | 1.7E-16 | 1.77E-14 |
| 1919 | TMEM176A | 98.523709 | -9.175168236 | 1.0652 | -8.6137 | 7.1E-18 | 1.03E-15 |
| 742 | ANPEP | 319.217686 | -9.189073248 | 0.7855 | -11.698 | 1.3E-31 | 1.43E-28 |
| 777 | CHP2 | 60.4979813 | -9.191377938 | 1.1102 | -8.2788 | 1.2E-16 | 1.39E-14 |
| 2108 | MAOA | 62.7450197 | -9.240748938 | 1.0906 | -8.4731 | 2.4E-17 | 3.06E-15 |
| 1438 | RBP2 | 104.319874 | -9.259035322 | 1.9828 | -4.6696 | 3E-06 | 3.54E-05 |
| 672 | SERPINA1 | 104.537007 | -9.262456253 | 1.4662 | -6.3172 | 2.7E-10 | 7.98E-09 |
| 1969 | CDH17 | 105.726806 | -9.271117656 | 1.0663 | -8.6947 | 3.5E-18 | 5.32E-16 |
| 342 | CDHR5 | 64.1852363 | -9.282339287 | 1.2755 | -7.2774 | 3.4E-13 | 1.94E-11 |
| 1742 | MEP1A | 64.8858787 | -9.29090091 | 1.0883 | -8.537 | 1.4E-17 | 1.87E-15 |
| 1918 | TMEM176B | 65.6353755 | -9.310081445 | 1.0577 | -8.8026 | 1.3E-18 | 2.21E-16 |
| 2159 | GPC3 | 66.0177438 | -9.318174725 | 1.0815 | -8.6162 | 6.9E-18 | 1.02E-15 |
| 474 | ST14 | 66.0517112 | -9.330817426 | 1.0698 | -8.7224 | 2.7E-18 | 4.31E-16 |

| | | | | | | | |
|---|---|---|---|---|---|---|---|
| 641 | GPX2 | 67.0264608 | -9.352809767 | 1.0989 | -8.5112 | 1.7E-17 | 2.29E-15 |
| 2121 | HEPH | 68.5866869 | -9.370452729 | 1.0627 | -8.8174 | 1.2E-18 | 1.96E-16 |
| 1074 | CEACAM5 | 68.3448117 | -9.379121582 | 1.1003 | -8.5244 | 1.5E-17 | 2.06E-15 |
| 346 | MUC2 | 70.056487 | -9.410121129 | 1.1013 | -8.5448 | 1.3E-17 | 1.77E-15 |
| 1519 | AFP | 72.3964671 | -9.454544685 | 1.9622 | -4.8185 | 1.4E-06 | 1.83E-05 |
| 1749 | GSTA1 | 72.4709424 | -9.45503702 | 1.467 | -6.4451 | 1.2E-10 | 3.82E-09 |
| 1075 | CEACAM6 | 76.1180362 | -9.527390033 | 1.0847 | -8.7833 | 1.6E-18 | 2.6E-16 |
| 1065 | FCGBP | 77.3794328 | -9.552700073 | 1.0681 | -8.944 | 3.8E-19 | 6.85E-17 |
| 1831 | AGR2 | 128.899191 | -9.567297943 | 1.0543 | -9.075 | 1.1E-19 | 2.31E-17 |
| 980 | TTR | 78.283118 | -9.567743801 | 1.4972 | -6.3906 | 1.7E-10 | 5.2E-09 |
| 1472 | SST | 79.4793597 | -9.590471232 | 3.0609 | -3.1332 | 0.00173 | 0.009184 |
| 636 | LGALS3 | 540.240283 | -9.619164429 | 0.7024 | -13.696 | 1.1E-42 | 2.55E-39 |
| 1430 | MUC13 | 136.528445 | -9.646953048 | 1.0701 | -9.0149 | 2E-19 | 3.84E-17 |
| 1061 | LGALS4 | 462.921355 | -9.746014423 | 0.7533 | -12.938 | 2.8E-38 | 4.91E-35 |
| 205 | ELF3 | 147.351284 | -9.761527154 | 1.0485 | -9.3103 | 1.3E-20 | 3.13E-18 |
| 670 | IFI27 | 154.229813 | -9.824322166 | 1.0382 | -9.4632 | 3E-21 | 8.17E-19 |
| 84 | TSPAN1 | 97.5770463 | -9.892835706 | 1.0532 | -9.3933 | 5.8E-21 | 1.5E-18 |
| 2067 | LCN15 | 112.890338 | -10.09695765 | 1.5114 | -6.6805 | 2.4E-11 | 9.25E-10 |
| 154 | S100A10 | 204.116517 | -10.22420642 | 1.0646 | -9.6036 | 7.7E-22 | 2.24E-19 |
| 1669 | FABP6 | 152.439875 | -10.52981571 | 1.8961 | -5.5534 | 2.8E-08 | 5.42E-07 |
| 1658 | SPINK1 | 164.168071 | -10.64118452 | 1.0567 | -10.071 | 7.5E-24 | 2.72E-21 |
| 113 | CLCA1 | 187.198299 | -10.8256027 | 1.0473 | -10.336 | 4.8E-25 | 2.08E-22 |
| 216 | PIGR | 249.975796 | -11.24286242 | 1.5089 | -7.4512 | 9.3E-14 | 5.75E-12 |
| 545 | TSPAN8 | 254.695561 | -11.26893954 | 1.0356 | -10.881 | 1.4E-27 | 7.47E-25 |
| 1151 | EPCAM | 270.228877 | -11.3521493 | 1.0677 | -10.633 | 2.1E-26 | 9.97E-24 |
| 690 | PHGR1 | 293.485536 | -11.47567339 | 1.0436 | -10.996 | 4E-28 | 2.36E-25 |
| 2040 | ALDOB | 320.89558 | -11.60353043 | 1.536 | -7.5543 | 4.2E-14 | 2.82E-12 |
| 1173 | FABP1 | 805.356852 | -12.20946444 | 1.0516 | -11.611 | 3.6E-31 | 3.69E-28 |
| 324 | DMBT1 | 630.02651 | -12.57741701 | 1.1475 | -10.961 | 5.9E-28 | 3.36E-25 |

TableS3

| Genes upregulated in Undiff compared to Gland_B | | | | | | | |
|---|---|---|---|---|---|---|---|
| gene ID | gene name | baseMean | log2FoldChange | lfcSE | stat | p-value | adjusted p-value |
| 45 | CYP4Z1 | 146.63877 | 10.9805468 | 1.21099 | 9.06745 | 1.22E-19 | 3.23E-17 |
| 610 | NEUROD1 | 111.42639 | 10.57953157 | 1.20167 | 8.804 | 1.32E-18 | 3.31E-16 |
| 1011 | NEFL | 99.430059 | 10.41061791 | 1.20467 | 8.64192 | 5.53E-18 | 1.15E-15 |
| 860 | CPLX2 | 49.570996 | 9.408014046 | 1.2211 | 7.70453 | 1.31E-14 | 1.72E-12 |
| 311 | ZDHHC22 | 38.225312 | 9.029601242 | 1.22228 | 7.38751 | 1.50E-13 | 1.66E-11 |
| 44 | CYP4X1 | 36.975197 | 8.981129524 | 1.2289 | 7.30828 | 2.71E-13 | 2.80E-11 |
| 974 | DLX5 | 32.566631 | 8.803600812 | 1.22946 | 7.16053 | 8.04E-13 | 7.43E-11 |
| 398 | GNAO1 | 57.919909 | 8.682310186 | 1.21395 | 7.1521 | 8.55E-13 | 7.85E-11 |
| 755 | P2RY1 | 57.366944 | 8.66253623 | 1.22621 | 7.06449 | 1.61E-12 | 1.36E-10 |
| 294 | NKX2-1 | 28.816368 | 8.623437255 | 1.26389 | 6.82292 | 8.92E-12 | 6.47E-10 |
| 936 | HPAT5 | 25.265812 | 8.448196345 | 1.27501 | 6.62599 | 3.45E-11 | 2.13E-09 |
| 423 | P2RX1 | 25.086833 | 8.429871782 | 1.23966 | 6.80017 | 1.04E-11 | 7.46E-10 |
| 619 | CRYBA2 | 24.044291 | 8.371151525 | 1.25687 | 6.66032 | 2.73E-11 | 1.75E-09 |
| 711 | FGD5 | 23.856271 | 8.360678068 | 1.25423 | 6.666 | 2.63E-11 | 1.70E-09 |
| 756 | SHOX2 | 85.537271 | 8.33840219 | 1.05353 | 7.91475 | 2.48E-15 | 3.63E-13 |
| 782 | ENSG00000286318 | 22.303638 | 8.262397869 | 1.24715 | 6.62502 | 3.47E-11 | 2.14E-09 |
| 538 | ATP1A3 | 22.494107 | 8.260561957 | 1.25338 | 6.59061 | 4.38E-11 | 2.64E-09 |
| 690 | MFNG | 22.206299 | 8.242786731 | 1.25213 | 6.58304 | 4.61E-11 | 2.75E-09 |
| 204 | GLB1L3 | 21.321322 | 8.200357437 | 1.26799 | 6.4672 | 9.98E-11 | 5.62E-09 |
| 119 | MKX | 21.250667 | 8.180269617 | 1.2568 | 6.50879 | 7.58E-11 | 4.41E-09 |
| 973 | DLX6 | 20.949978 | 8.176544144 | 1.25412 | 6.51972 | 7.04E-11 | 4.11E-09 |
| 1073 | GRIN3A | 20.486124 | 8.141926448 | 1.25188 | 6.50373 | 7.84E-11 | 4.54E-09 |
| 517 | NCAN | 72.361789 | 8.088689306 | 1.07796 | 7.50371 | 6.20E-14 | 7.25E-12 |
| 643 | FERMT1 | 19.117827 | 8.046194683 | 1.27961 | 6.28802 | 3.22E-10 | 1.61E-08 |
| 137 | SLIT1 | 17.835874 | 7.951761054 | 1.27116 | 6.25549 | 3.96E-10 | 1.95E-08 |
| 209 | LINC02470 | 17.256453 | 7.889584617 | 1.30758 | 6.03375 | 1.60E-09 | 6.86E-08 |
| 776 | STK32B | 16.635796 | 7.856939454 | 1.28259 | 6.12585 | 9.02E-10 | 4.06E-08 |
| 1060 | CDKN2A | 32.721031 | 7.843886072 | 1.23756 | 6.33817 | 2.33E-10 | 1.23E-08 |
| 250 | ALDH1L2 | 16.592572 | 7.817925041 | 1.27726 | 6.12088 | 9.31E-10 | 4.18E-08 |
| 999 | PRKAG2-AS1 | 16.281372 | 7.812482079 | 1.27406 | 6.13195 | 8.68E-10 | 3.95E-08 |
| 618 | FEV | 28.553123 | 7.660642288 | 1.23622 | 6.19681 | 5.76E-10 | 2.71E-08 |
| 658 | MAFB | 181.64677 | 7.623516891 | 0.59102 | 12.8989 | 4.57E-38 | 1.57E-34 |
| 278 | POU4F1 | 13.604275 | 7.541359797 | 1.28689 | 5.86014 | 4.62E-09 | 1.75E-07 |
| 1021 | ST18 | 48.474609 | 7.528581356 | 1.06572 | 7.06435 | 1.61E-12 | 1.36E-10 |
| 314 | CHGA | 25.574558 | 7.503464031 | 1.24985 | 6.00349 | 1.93E-09 | 8.11E-08 |
| 866 | AACSP1 | 12.972698 | 7.502199418 | 1.35528 | 5.53554 | 3.10E-08 | 9.48E-07 |

| | | | | | | | |
|---|---|---|---|---|---|---|---|
| 819 | ADAMTS16-DT | 12.594808 | 7.438107491 | 1.29767 | 5.73188 | 9.93E-09 | 3.47E-07 |
| 1061 | ELAVL2 | 68.839628 | 7.433367963 | 0.87486 | 8.49664 | 1.95E-17 | 3.90E-15 |
| 786 | SHISA3 | 24.658887 | 7.421763887 | 1.24638 | 5.95465 | 2.61E-09 | 1.04E-07 |
| 379 | PRKCB | 23.487919 | 7.353244727 | 1.25654 | 5.85199 | 4.86E-09 | 1.82E-07 |
| 742 | DPPA4 | 84.659275 | 7.32185969 | 0.77586 | 9.43707 | 3.83E-21 | 1.54E-18 |
| 486 | RNF125 | 11.323755 | 7.303256329 | 1.34154 | 5.44394 | 5.21E-08 | 1.5E-06 |
| 779 | ENSG00000287778 | 11.482266 | 7.287316986 | 1.31417 | 5.54518 | 2.94E-08 | 9.07E-07 |
| 620 | PTPRN | 11.101332 | 7.276258407 | 1.32786 | 5.47968 | 4.26E-08 | 1.2E-06 |
| 736 | ROBO2 | 11.173936 | 7.259269024 | 1.32581 | 5.47536 | 4.37E-08 | 1.3E-06 |
| 706 | LHFPL4 | 10.758007 | 7.232582989 | 1.33625 | 5.41261 | 6.21E-08 | 1.7E-06 |
| 29 | HPCA | 10.453142 | 7.171783715 | 1.33664 | 5.36555 | 8.07E-08 | 2.2E-06 |
| 593 | ENSG00000236885 | 20.171693 | 7.1523835 | 1.26543 | 5.65215 | 1.58E-08 | 5.31E-07 |
| 266 | GPR12 | 10.158764 | 7.144492556 | 1.33736 | 5.34225 | 9.18E-08 | 2.4E-06 |
| 830 | CARTPT | 19.619715 | 7.110913972 | 1.26048 | 5.64145 | 1.69E-08 | 5.60E-07 |
| 718 | ARPP21 | 19.579678 | 7.076252988 | 1.28477 | 5.50781 | 3.63E-08 | 1.1E-06 |
| 673 | MYT1 | 9.9215411 | 7.069969338 | 1.3302 | 5.31496 | 1.07E-07 | 2.8E-06 |
| 1101 | SYTL5 | 19.489252 | 7.068655853 | 1.27508 | 5.54368 | 2.96E-08 | 9.13E-07 |
| 649 | INSM1 | 35.513418 | 7.051980822 | 1.08535 | 6.4974 | 8.17E-11 | 4.67E-09 |
| 980 | ACTL6B | 9.5743855 | 7.025939936 | 1.33281 | 5.27151 | 1.35E-07 | 3.4E-06 |
| 548 | NLRP2 | 9.4652556 | 7.014085804 | 1.33695 | 5.24635 | 1.55E-07 | 3.9E-06 |
| 582 | EN1 | 8.9300428 | 6.927791371 | 1.35859 | 5.09924 | 3.41E-07 | 7.6E-06 |
| 1091 | GRIN1 | 8.7545157 | 6.89070764 | 1.35241 | 5.09513 | 3.49E-07 | 7.8E-06 |
| 315 | UNC79 | 16.904255 | 6.880736471 | 1.27654 | 5.39013 | 7.04E-08 | 2E-06 |
| 587 | TMEM163 | 8.8101666 | 6.870902438 | 1.41412 | 4.85877 | 1.2E-06 | 2.3E-05 |
| 971 | DYNC1I1 | 30.640973 | 6.867565934 | 1.08553 | 6.32647 | 2.51E-10 | 1.31E-08 |
| 107 | SUSD4 | 30.983187 | 6.866555936 | 1.08962 | 6.30179 | 2.94E-10 | 1.50E-08 |
| 46 | CYP4A22 | 8.622661 | 6.854687891 | 1.36699 | 5.01444 | 5.32E-07 | 1.1E-05 |
| 417 | ENSG00000256982 | 8.2578275 | 6.828666397 | 1.39313 | 4.90168 | 9.50E-07 | 1.9E-05 |
| 988 | GPR37 | 8.1270481 | 6.808985982 | 1.3652 | 4.98754 | 6.12E-07 | 1.3E-05 |
| 708 | PPARG | 7.9732434 | 6.784086288 | 1.35329 | 5.01304 | 5.36E-07 | 1.1E-05 |
| 143 | LBX1-AS1 | 28.878695 | 6.754481258 | 1.10936 | 6.08861 | 1.14E-09 | 5.03E-08 |
| 32 | GRIK3 | 7.6712044 | 6.752406755 | 1.43201 | 4.71533 | 2.4E-06 | 4.3E-05 |
| 918 | CLVS2 | 8.0801397 | 6.749488511 | 1.39699 | 4.83146 | 1.4E-06 | 2.6E-05 |
| 609 | ITGA4 | 42.546237 | 6.74259282 | 0.88886 | 7.58566 | 3.31E-14 | 4.00E-12 |
| 823 | CDH10 | 7.529524 | 6.713697646 | 1.37823 | 4.87126 | 1.1E-06 | 2.2E-05 |
| 136 | SORBS1 | 15.083248 | 6.705356474 | 1.2795 | 5.24062 | 1.60E-07 | 4E-06 |
| 645 | LAMP5 | 14.872279 | 6.677738727 | 1.29387 | 5.16107 | 2.46E-07 | 5.7E-06 |
| 857 | FBLL1 | 14.694823 | 6.676780356 | 1.29183 | 5.16848 | 2.36E-07 | 5.5E-06 |
| 148 | INA | 79.628631 | 6.655067219 | 0.65224 | 10.2035 | 1.91E-24 | 1.26E-21 |

| | | | | | | | |
|---|---|---|---|---|---|---|---|
| 710 | GRIP2 | 7.3311219 | 6.654949308 | 1.36702 | 4.86822 | 1.1E-06 | 2.2E-05 |
| 1010 | NEFM | 79.282015 | 6.651171181 | 0.65585 | 10.1414 | 3.62E-24 | 2.14E-21 |
| 835 | EDIL3-DT | 7.1229064 | 6.645925354 | 1.4523 | 4.57613 | 4.7E-06 | 7.6E-05 |
| 77 | SHE | 7.2189556 | 6.640976101 | 1.40308 | 4.73315 | 2.2E-06 | 4E-05 |
| 351 | ISL2 | 7.3319202 | 6.623413202 | 1.39955 | 4.73253 | 2.2E-06 | 4E-05 |
| 1003 | MIR124-1HG | 39.071894 | 6.617122267 | 0.92279 | 7.17081 | 7.46E-13 | 6.99E-11 |
| 400 | NLRC5 | 13.663665 | 6.592138022 | 1.31484 | 5.01363 | 5.34E-07 | 1.1E-05 |
| 149 | CFAP58 | 13.520111 | 6.573431738 | 1.29206 | 5.08756 | 3.63E-07 | 8.1E-06 |
| 533 | CCER2 | 25.288745 | 6.564436596 | 1.09002 | 6.02231 | 1.72E-09 | 7.34E-08 |
| 681 | CECR7 | 6.74905 | 6.557208233 | 1.39812 | 4.69003 | 2.7E-06 | 4.7E-05 |
| 72 | TBX15 | 24.691914 | 6.515298951 | 1.11482 | 5.84428 | 5.09E-09 | 1.89E-07 |
| 794 | NKX6-1 | 6.6382012 | 6.510371719 | 1.38552 | 4.69888 | 2.6E-06 | 4.5E-05 |
| 605 | CHRNA1 | 13.230903 | 6.509611806 | 1.30795 | 4.97694 | 6.46E-07 | 1.3E-05 |
| 608 | HAGLR | 6.3511727 | 6.467013837 | 1.42771 | 4.52965 | 5.9E-06 | 9.2E-05 |
| 583 | INHBB | 23.161824 | 6.460236441 | 1.11208 | 5.80914 | 6.28E-09 | 2.31E-07 |
| 678 | PCBP3 | 12.43044 | 6.459651354 | 1.30126 | 4.96417 | 6.90E-07 | 1.4E-05 |
| 316 | TUNAR | 139.45484 | 6.44836313 | 0.467 | 13.8079 | 2.28E-43 | 1.57E-39 |
| 37 | MFSD2A | 6.1795442 | 6.442157528 | 1.48507 | 4.33796 | 1.4E-05 | 0.0002 |
| 159 | LINC01166 | 6.4302694 | 6.439371059 | 1.40761 | 4.57467 | 4.8E-06 | 7.7E-05 |
| 34 | MANEAL | 12.295274 | 6.438636707 | 1.31298 | 4.90382 | 9.40E-07 | 1.9E-05 |
| 714 | KCNH8 | 6.4097097 | 6.432935426 | 1.41628 | 4.54215 | 5.6E-06 | 8.8E-05 |
| 841 | LINC01170 | 6.2965264 | 6.427789485 | 1.47535 | 4.35679 | 1.3E-05 | 0.00019 |
| 810 | DCHS2 | 6.0826115 | 6.415019359 | 1.4334 | 4.47539 | 7.6E-06 | 0.00012 |
| 559 | NRXN1 | 33.916632 | 6.396638177 | 0.90661 | 7.05554 | 1.72E-12 | 1.43E-10 |
| 590 | THSD7B | 6.1860465 | 6.392282476 | 1.43525 | 4.45377 | 8.4E-06 | 0.00013 |
| 58 | PTGER3 | 11.778415 | 6.381581099 | 1.34408 | 4.74791 | 2.1E-06 | 3.7E-05 |
| 870 | FOXF2 | 32.862613 | 6.364422623 | 0.92213 | 6.9019 | 5.13E-12 | 3.89E-10 |
| 47 | TAL1 | 6.0547669 | 6.335829939 | 1.64243 | 3.85758 | 0.00011 | 0.00121 |
| 68 | KCNA2 | 21.191973 | 6.327319801 | 1.11051 | 5.69765 | 1.21E-08 | 4.18E-07 |
| 766 | VWA5B2 | 5.6938176 | 6.323099203 | 1.47353 | 4.29112 | 1.8E-05 | 0.00024 |
| 647 | SNAP25 | 31.478322 | 6.32091019 | 0.94687 | 6.67558 | 2.46E-11 | 1.63E-09 |
| 63 | PLPPR4 | 5.6199247 | 6.270686096 | 1.47556 | 4.24971 | 2.1E-05 | 0.00029 |
| 604 | SP9 | 5.568242 | 6.215047771 | 1.50116 | 4.14017 | 3.5E-05 | 0.00044 |
| 290 | CARMIL3 | 69.213231 | 6.212839241 | 0.6 | 10.3547 | 3.98E-25 | 2.89E-22 |
| 622 | DOCK10 | 10.357983 | 6.209516025 | 1.36579 | 4.54647 | 5.5E-06 | 8.7E-05 |
| 142 | LBX1 | 19.812763 | 6.202026805 | 1.1203 | 5.53606 | 3.09E-08 | 9.48E-07 |
| 516 | UNC13A | 19.549732 | 6.186375061 | 1.11898 | 5.5286 | 3.23E-08 | 9.76E-07 |
| 59 | NEGR1 | 28.549327 | 6.179998409 | 0.92761 | 6.66229 | 2.70E-11 | 1.74E-09 |
| 416 | CBFA2T3 | 10.382716 | 6.16901107 | 1.32819 | 4.64467 | 3.4E-06 | 5.6E-05 |

| | | | | | | | |
|---|---|---|---|---|---|---|---|
| 1124 | RAB33A | 5.0366901 | 6.129341069 | 1.45931 | 4.20018 | 2.7E-05 | 0.00035 |
| 624 | PID1 | 18.491503 | 6.120796044 | 1.11706 | 5.47937 | 4.27E-08 | 1.2E-06 |
| 1036 | ENSG00000271830 | 4.8859947 | 6.096303442 | 1.4734 | 4.13758 | 3.5E-05 | 0.00044 |
| 199 | POU2F3 | 9.6109183 | 6.062510996 | 1.36332 | 4.44688 | 8.7E-06 | 0.00013 |
| 123 | ZNF488 | 4.7901518 | 6.052964891 | 1.46938 | 4.11939 | 3.8E-05 | 0.00047 |
| 388 | ASPHD1 | 17.504058 | 6.041740909 | 1.12433 | 5.37365 | 7.72E-08 | 2.1E-06 |
| 382 | ENSG00000283662 | 4.6867213 | 6.039247296 | 1.51698 | 3.9811 | 6.9E-05 | 0.00078 |
| 1028 | JPH1 | 17.317366 | 6.020635125 | 1.11852 | 5.38267 | 7.34E-08 | 2E-06 |
| 959 | VSTM2A | 4.7086713 | 6.019069804 | 1.45846 | 4.12699 | 3.7E-05 | 0.00046 |
| 432 | HES7 | 4.6383665 | 6.014369465 | 1.52929 | 3.93279 | 8.4E-05 | 0.00092 |
| 62 | PLPPR5 | 4.4791584 | 5.967357522 | 1.48738 | 4.01199 | 6E-05 | 0.0007 |
| 719 | MYRIP | 16.557006 | 5.953064214 | 1.15282 | 5.16389 | 2.42E-07 | 5.6E-06 |
| 88 | CD247 | 4.4518824 | 5.952005395 | 1.54078 | 3.86298 | 0.00011 | 0.00119 |
| 363 | SYNM | 4.4459598 | 5.944663561 | 1.48745 | 3.99654 | 6.4E-05 | 0.00074 |
| 689 | ENSG00000166897 | 4.4613391 | 5.943673363 | 1.50503 | 3.9492 | 7.8E-05 | 0.00087 |
| 214 | CNTN1 | 23.788029 | 5.911981077 | 0.94012 | 6.28855 | 3.20E-10 | 1.61E-08 |
| 438 | SEZ6 | 8.543475 | 5.901701135 | 1.343 | 4.39441 | 1.1E-05 | 0.00016 |
| 868 | FLT4 | 23.32671 | 5.846651438 | 0.93542 | 6.25032 | 4.10E-10 | 1.99E-08 |
| 1025 | BHLHE22 | 22.786732 | 5.836238558 | 0.92547 | 6.30625 | 2.86E-10 | 1.48E-08 |
| 808 | HHIP | 15.11926 | 5.834000523 | 1.13818 | 5.12574 | 2.96E-07 | 6.7E-06 |
| 585 | CNTNAP5 | 4.1872288 | 5.833311048 | 1.51815 | 3.84237 | 0.00012 | 0.00127 |
| 633 | SCRT2 | 4.0779487 | 5.825120367 | 1.53358 | 3.79838 | 0.00015 | 0.00148 |
| 348 | HCN4 | 4.2666215 | 5.814963217 | 1.62465 | 3.57922 | 0.00034 | 0.00303 |
| 310 | RPS6KL1 | 15.051136 | 5.809998706 | 1.15068 | 5.04917 | 4.44E-07 | 9.7E-06 |
| 220 | VDR | 44.752405 | 5.807904341 | 0.66783 | 8.69673 | 3.42E-18 | 7.72E-16 |
| 489 | SLC14A1 | 4.057093 | 5.804769704 | 1.51752 | 3.82518 | 0.00013 | 0.00135 |
| 955 | GCK | 7.8964692 | 5.779053882 | 1.38561 | 4.17077 | 3E-05 | 0.00039 |
| 422 | P2RX5 | 7.6750227 | 5.762150792 | 1.37615 | 4.18715 | 2.8E-05 | 0.00036 |
| 979 | TFR2 | 14.175688 | 5.75330345 | 1.14899 | 5.00729 | 5.52E-07 | 1.2E-05 |
| 972 | DLX6-AS1 | 36.069751 | 5.744237377 | 0.7383 | 7.78031 | 7.23E-15 | 1.02E-12 |
| 558 | KCNK12 | 3.9155584 | 5.743031379 | 1.62692 | 3.53001 | 0.00042 | 0.00352 |
| 632 | ENSG00000280119 | 3.8526393 | 5.725974799 | 1.56546 | 3.6577 | 0.00025 | 0.00235 |
| 679 | ENSG00000275139 | 7.2737355 | 5.691569051 | 1.43645 | 3.96225 | 7.4E-05 | 0.00083 |
| 892 | GRM4 | 3.8891535 | 5.686621629 | 1.60426 | 3.5447 | 0.00039 | 0.00337 |
| 367 | FBXL16 | 7.420213 | 5.68345412 | 1.35959 | 4.18027 | 2.9E-05 | 0.00037 |
| 87 | POU2F1-DT | 3.7319603 | 5.682563948 | 1.52213 | 3.73329 | 0.00019 | 0.00184 |
| 277 | ENSG00000271776 | 3.6590654 | 5.673061535 | 1.57425 | 3.60366 | 0.00031 | 0.0028 |
| 596 | GALNT13 | 3.6408728 | 5.672317545 | 1.5792 | 3.59189 | 0.00033 | 0.00291 |
| 1012 | EBF2 | 3.7688434 | 5.672130575 | 1.72391 | 3.29028 | 0.001 | 0.00722 |

| | | | | | | | |
|---|---|---|---|---|---|---|---|
| 729 | CAMKV | 80.402407 | 5.664539581 | 0.50064 | 11.3146 | 1.11E-29 | 1.39E-26 |
| 526 | FXYD5 | 13.51551 | 5.655100146 | 1.17612 | 4.80828 | 1.5E-06 | 2.8E-05 |
| 55 | IL12RB2 | 3.6903219 | 5.652604254 | 1.56421 | 3.61371 | 0.0003 | 0.00271 |
| 480 | L3MBTL4 | 3.5746214 | 5.650221928 | 1.5814 | 3.57292 | 0.00035 | 0.00308 |
| 949 | ENSG00000253308 | 3.7869631 | 5.64725215 | 1.5735 | 3.58898 | 0.00033 | 0.00293 |
| 1029 | GDAP1 | 93.882656 | 5.641446663 | 0.4494 | 12.5532 | 3.82E-36 | 1.05E-32 |
| 518 | VSTM2B | 7.0542031 | 5.635705205 | 1.37435 | 4.10063 | 4.1E-05 | 0.0005 |
| 418 | CPNE7 | 3.5055567 | 5.6262456 | 1.59367 | 3.53036 | 0.00041 | 0.00351 |
| 1040 | ANGPT1 | 12.819881 | 5.602898245 | 1.16168 | 4.82311 | 1.4E-06 | 2.7E-05 |
| 460 | KCNH6 | 3.5177539 | 5.598976167 | 1.55062 | 3.6108 | 0.00031 | 0.00274 |
| 883 | HLA-F | 6.964433 | 5.597886067 | 1.41178 | 3.96514 | 7.3E-05 | 0.00083 |
| 763 | PEX5L | 38.240844 | 5.58858102 | 0.70059 | 7.97691 | 1.50E-15 | 2.32E-13 |
| 812 | GRIA2 | 127.50668 | 5.575750593 | 0.88459 | 6.30321 | 2.92E-10 | 1.49E-08 |
| 807 | HHIP-AS1 | 7.0947751 | 5.574830268 | 1.43158 | 3.89419 | 9.9E-05 | 0.00106 |
| 901 | PTCHD4 | 6.77034 | 5.565910219 | 1.40694 | 3.95605 | 7.6E-05 | 0.00085 |
| 744 | SIDT1 | 3.481446 | 5.559474066 | 1.63705 | 3.39604 | 0.00068 | 0.00528 |
| 404 | HSF4 | 3.4152675 | 5.550998247 | 1.55456 | 3.57079 | 0.00036 | 0.0031 |
| 631 | AGXT | 3.3460526 | 5.548823355 | 1.61509 | 3.43561 | 0.00059 | 0.00468 |
| 986 | KCND2 | 3.4647166 | 5.541160427 | 1.76422 | 3.14086 | 0.00168 | 0.0112 |
| 642 | ENSG00000278192 | 6.4941678 | 5.533318605 | 1.43984 | 3.84301 | 0.00012 | 0.00127 |
| 917 | HS3ST5 | 3.3033243 | 5.515657021 | 1.61455 | 3.41622 | 0.00063 | 0.00497 |
| 844 | CDC25C | 3.3130771 | 5.514019441 | 1.59319 | 3.46099 | 0.00054 | 0.00434 |
| 380 | HS3ST4 | 18.203481 | 5.505893741 | 0.95759 | 5.74976 | 8.94E-09 | 3.17E-07 |
| 704 | CNTN6 | 3.204093 | 5.502798727 | 1.73186 | 3.17739 | 0.00149 | 0.0101 |
| 952 | NEUROD6 | 3.252006 | 5.502195352 | 1.63129 | 3.3729 | 0.00074 | 0.00565 |
| 761 | SLC7A14 | 3.2245797 | 5.501140848 | 1.73956 | 3.16238 | 0.00156 | 0.01057 |
| 222 | KCNH3 | 3.2796554 | 5.489043101 | 1.56682 | 3.50331 | 0.00046 | 0.00382 |
| 264 | ENSG00000243008 | 6.4600674 | 5.488929754 | 1.40874 | 3.89634 | 9.8E-05 | 0.00105 |
| 331 | ARHGAP11B | 6.5242524 | 5.48207309 | 1.57592 | 3.47865 | 0.0005 | 0.00411 |
| 981 | VGF | 17.760032 | 5.476271659 | 0.95288 | 5.74708 | 9.08E-09 | 3.21E-07 |
| 984 | ENSG00000287829 | 3.1677851 | 5.471841737 | 1.61061 | 3.39737 | 0.00068 | 0.00526 |
| 881 | SCGN | 3.2285602 | 5.470166018 | 1.56891 | 3.48661 | 0.00049 | 0.00401 |
| 856 | GABRB2 | 11.679457 | 5.451958886 | 1.17785 | 4.62874 | 3.7E-06 | 6.1E-05 |
| 777 | CRMP1 | 34.575497 | 5.443316632 | 0.70977 | 7.66918 | 1.73E-14 | 2.23E-12 |
| 339 | SCG3 | 22.697784 | 5.434719697 | 0.8668 | 6.26985 | 3.61E-10 | 1.79E-08 |
| 238 | PPFIA2 | 3.0533247 | 5.434070035 | 1.74757 | 3.10951 | 0.00187 | 0.01222 |
| 391 | DOC2A | 3.1591995 | 5.430321758 | 1.5824 | 3.43169 | 0.0006 | 0.00474 |
| 950 | EVX1 | 3.0775688 | 5.405508589 | 1.62016 | 3.3364 | 0.00085 | 0.00631 |
| 778 | HS3ST1 | 33.966421 | 5.400683866 | 0.68077 | 7.93324 | 2.14E-15 | 3.16E-13 |

| | | | | | | | |
|---|---|---|---|---|---|---|---|
| 947 | SKAP2 | 3.0898388 | 5.392385238 | 1.67266 | 3.22383 | 0.00126 | 0.00882 |
| 355 | CTXND1 | 5.9357342 | 5.376845549 | 1.44868 | 3.71155 | 0.00021 | 0.00197 |
| 695 | SULT4A1 | 21.924783 | 5.3642909 | 0.83428 | 6.42983 | 1.28E-10 | 7.13E-09 |
| 1058 | SLC24A2 | 38.962203 | 5.348818747 | 0.6719 | 7.96071 | 1.71E-15 | 2.59E-13 |
| 832 | FOXD1 | 5.9069762 | 5.345234099 | 1.39963 | 3.81904 | 0.00013 | 0.00138 |
| 652 | XKR7 | 2.9781188 | 5.344451284 | 1.62687 | 3.28512 | 0.00102 | 0.00733 |
| 377 | ENSG00000280000 | 3.0814764 | 5.342587415 | 1.65915 | 3.22008 | 0.00128 | 0.00891 |
| 303 | GCH1 | 5.9289991 | 5.337031206 | 1.42859 | 3.73588 | 0.00019 | 0.00183 |
| 407 | HSD11B2 | 3.0181111 | 5.336052797 | 1.66543 | 3.20401 | 0.00136 | 0.00938 |
| 53 | DAB1 | 16.321623 | 5.334418814 | 0.96883 | 5.50603 | 3.67E-08 | 1.1E-06 |
| 928 | HYMAI | 5.791983 | 5.312146588 | 1.41332 | 3.75862 | 0.00017 | 0.00169 |
| 141 | CNNM1 | 5.8611979 | 5.311858859 | 1.42205 | 3.73534 | 0.00019 | 0.00183 |
| 817 | TERT | 2.8907886 | 5.293885125 | 1.64024 | 3.22751 | 0.00125 | 0.00874 |
| 1044 | SYBU-AS1 | 2.8521866 | 5.251007851 | 1.69538 | 3.09724 | 0.00195 | 0.01263 |
| 246 | ANO4 | 56.086631 | 5.24346112 | 0.55621 | 9.42706 | 4.22E-21 | 1.61E-18 |
| 994 | FAM131B | 10.126574 | 5.236687926 | 1.1936 | 4.38732 | 1.1E-05 | 0.00017 |
| 1009 | EGR3 | 14.772693 | 5.225691362 | 0.98525 | 5.3039 | 1.13E-07 | 3E-06 |
| 818 | IRX2 | 29.636831 | 5.195548243 | 0.7126 | 7.29093 | 3.08E-13 | 3.10E-11 |
| 196 | ELMOD1 | 19.668032 | 5.190488836 | 0.84221 | 6.1629 | 7.14E-10 | 3.30E-08 |
| 753 | PCOLCE2 | 29.751741 | 5.189365589 | 0.73796 | 7.03204 | 2.04E-12 | 1.67E-10 |
| 641 | LRRN4 | 2.6889251 | 5.188674307 | 1.64425 | 3.15565 | 0.0016 | 0.01076 |
| 801 | ENSG00000251259 | 2.6129938 | 5.18425315 | 1.71646 | 3.02031 | 0.00253 | 0.01553 |
| 909 | RIPPLY2 | 2.592211 | 5.176440685 | 1.67935 | 3.08241 | 0.00205 | 0.01319 |
| 354 | MINAR1 | 24.072278 | 5.166185115 | 0.77966 | 6.62618 | 3.44E-11 | 2.13E-09 |
| 281 | HS6ST3 | 82.929691 | 5.157758359 | 0.46543 | 11.0816 | 1.54E-28 | 1.77E-25 |
| 69 | ENSG00000259834 | 2.5685421 | 5.142050653 | 1.69016 | 3.04235 | 0.00235 | 0.01469 |
| 79 | CHRNB2 | 5.0983413 | 5.141782068 | 1.47292 | 3.49087 | 0.00048 | 0.00396 |
| 284 | MYO16 | 9.1728798 | 5.137164998 | 1.22505 | 4.19344 | 2.7E-05 | 0.00036 |
| 784 | KCNIP4-IT1 | 9.6798068 | 5.133060643 | 1.20139 | 4.27259 | 1.9E-05 | 0.00026 |
| 15 | RUNX3 | 2.5864387 | 5.125891089 | 1.6911 | 3.03111 | 0.00244 | 0.01514 |
| 86 | FAM78B | 13.894995 | 5.12033321 | 0.97397 | 5.25717 | 1.46E-07 | 3.7E-06 |
| 340 | TMOD2 | 9.3765631 | 5.11655876 | 1.22814 | 4.16611 | 3.1E-05 | 0.00039 |
| 1093 | CACNA1B | 13.909097 | 5.115512562 | 1.02568 | 4.98741 | 6.12E-07 | 1.3E-05 |
| 138 | ARHGAP19-SLIT1 | 2.4953512 | 5.112231667 | 1.7849 | 2.86416 | 0.00418 | 0.0233 |
| 308 | ENSG00000258490 | 2.4899477 | 5.104174206 | 1.66838 | 3.05935 | 0.00222 | 0.01403 |
| 420 | CCDC92B | 5.0079018 | 5.101997104 | 1.43108 | 3.56513 | 0.00036 | 0.00316 |
| 865 | F12 | 4.9565835 | 5.094643897 | 1.44998 | 3.51361 | 0.00044 | 0.0037 |
| 221 | CCDC184 | 9.4434677 | 5.093643865 | 1.22057 | 4.17316 | 3E-05 | 0.00038 |
| 376 | SEC14L5 | 5.0478213 | 5.08205633 | 1.46105 | 3.47837 | 0.0005 | 0.00411 |

| | | | | | | | |
|---|---|---|---|---|---|---|---|
| 97 | SYT2 | 2.4111303 | 5.07209038 | 1.72995 | 2.93193 | 0.00337 | 0.01951 |
| 905 | ADGRB3 | 2.4537128 | 5.069399221 | 1.70532 | 2.9727 | 0.00295 | 0.01755 |
| 653 | ENSG00000278012 | 2.5116746 | 5.068438795 | 1.75644 | 2.88564 | 0.00391 | 0.02195 |
| 447 | KCNH4 | 2.3875342 | 5.061675482 | 1.71745 | 2.9472 | 0.00321 | 0.0188 |
| 76 | C2CD4D | 2.4965184 | 5.057403334 | 1.71659 | 2.94619 | 0.00322 | 0.01883 |
| 669 | SLCO4A1 | 9.0253997 | 5.041260466 | 1.25237 | 4.02537 | 5.7E-05 | 0.00067 |
| 349 | RPP25 | 8.8122376 | 5.040332062 | 1.22077 | 4.12882 | 3.6E-05 | 0.00045 |
| 40 | HPDL | 8.8738331 | 5.039775203 | 1.23712 | 4.07381 | 4.6E-05 | 0.00056 |
| 1052 | SCRT1 | 13.249471 | 5.039247094 | 0.98472 | 5.11744 | 3.10E-07 | 7E-06 |
| 179 | SYT7 | 2.3874614 | 5.035935165 | 1.68031 | 2.99703 | 0.00273 | 0.0165 |
| 906 | B3GAT2 | 2.4148876 | 5.025178299 | 1.74114 | 2.88614 | 0.0039 | 0.02194 |
| 750 | ESYT3 | 2.3183967 | 5.000156302 | 1.74695 | 2.86223 | 0.00421 | 0.02341 |
| 232 | NXPH4 | 4.5918456 | 4.996459753 | 1.51417 | 3.2998 | 0.00097 | 0.00702 |
| 1123 | APLN | 2.3542628 | 4.992305443 | 1.74276 | 2.86461 | 0.00418 | 0.02328 |
| 863 | GPRIN1 | 2.3278535 | 4.983876704 | 1.91598 | 2.60121 | 0.00929 | 0.04386 |
| 239 | EPYC | 2.2368388 | 4.981964618 | 1.89442 | 2.62981 | 0.00854 | 0.04111 |
| 504 | ATCAY | 16.580755 | 4.979386592 | 0.89073 | 5.59024 | 2.27E-08 | 7.26E-07 |
| 962 | SRRM3 | 21.117604 | 4.977940986 | 0.79842 | 6.23473 | 4.53E-10 | 2.17E-08 |
| 529 | PPP1R14A | 8.1568975 | 4.967064388 | 1.29254 | 3.84288 | 0.00012 | 0.00127 |
| 617 | CDK5R2 | 4.5735664 | 4.961708109 | 1.4814 | 3.34933 | 0.00081 | 0.00607 |
| 471 | ENSG00000267568 | 2.3277806 | 4.953924842 | 1.77717 | 2.78753 | 0.00531 | 0.02841 |
| 190 | KLHL35 | 2.2092668 | 4.950063739 | 1.76022 | 2.81219 | 0.00492 | 0.02674 |
| 226 | HOXC8 | 4.4457975 | 4.945735375 | 1.45991 | 3.38769 | 0.0007 | 0.00541 |
| 654 | ENSG00000275576 | 2.3126244 | 4.944027253 | 1.76173 | 2.80634 | 0.00501 | 0.02711 |
| 497 | NETO1 | 2.2520724 | 4.939752646 | 1.89597 | 2.60539 | 0.00918 | 0.0435 |
| 797 | GPRIN3 | 12.368065 | 4.934951046 | 1.02506 | 4.81429 | 1.5E-06 | 2.8E-05 |
| 1137 | PNCK | 33.545602 | 4.934454701 | 0.66476 | 7.42286 | 1.15E-13 | 1.28E-11 |
| 113 | SIPA1L2 | 28.443786 | 4.913949274 | 0.66843 | 7.35151 | 1.96E-13 | 2.11E-11 |
| 607 | HOXD3 | 2.1430154 | 4.911626659 | 1.8049 | 2.72128 | 0.0065 | 0.03333 |
| 200 | ENSG00000255219 | 2.2766125 | 4.910663129 | 1.91898 | 2.55899 | 0.0105 | 0.04837 |
| 888 | HLA-DPA1 | 2.1704416 | 4.901769359 | 1.73997 | 2.81716 | 0.00485 | 0.02638 |
| 464 | ENSG00000279917 | 2.1250459 | 4.900498777 | 1.82806 | 2.68071 | 0.00735 | 0.03658 |
| 206 | NRIP2 | 11.969573 | 4.89485205 | 1.01621 | 4.81679 | 1.5E-06 | 2.8E-05 |
| 572 | FAM178B | 2.1370928 | 4.893022598 | 1.75571 | 2.78693 | 0.00532 | 0.02846 |
| 746 | GATA2 | 2.1798984 | 4.878918905 | 1.77484 | 2.74893 | 0.00598 | 0.03119 |
| 764 | ENSG00000287645 | 2.1011538 | 4.872805838 | 1.76785 | 2.75634 | 0.00585 | 0.03068 |
| 1031 | STMN2 | 58.609778 | 4.867007493 | 0.48046 | 10.1298 | 4.07E-24 | 2.25E-21 |
| 320 | MEG8 | 4.2715834 | 4.865306806 | 1.50717 | 3.2281 | 0.00125 | 0.00873 |
| 1089 | SAPCD2 | 23.490493 | 4.865199129 | 0.7166 | 6.78931 | 1.13E-11 | 7.96E-10 |

| | | | | | | | |
|---|---|---|---|---|---|---|---|
| 510 | ELAVL3 | 54.913884 | 4.862310138 | 0.49969 | 9.73059 | 2.23E-22 | 1.06E-19 |
| 127 | TMEM26 | 7.4587965 | 4.833774181 | 1.35076 | 3.57856 | 0.00035 | 0.00303 |
| 1049 | FAM135B | 4.1060279 | 4.833176372 | 1.51288 | 3.19469 | 0.0014 | 0.00962 |
| 36 | POU3F1 | 11.503876 | 4.822262372 | 1.07041 | 4.50504 | 6.6E-06 | 0.0001 |
| 54 | DNAJC6 | 26.857541 | 4.821469722 | 0.65639 | 7.34543 | 2.05E-13 | 2.19E-11 |
| 33 | EPHA10 | 4.1736104 | 4.81856206 | 1.50759 | 3.1962 | 0.00139 | 0.00958 |
| 395 | IRX3 | 15.186822 | 4.809700559 | 0.88797 | 5.41651 | 6.08E-08 | 1.7E-06 |
| 820 | ADAMTS16 | 4.1054357 | 4.809467287 | 1.47594 | 3.25858 | 0.00112 | 0.00797 |
| 1068 | ENSG00000228430 | 2.1438137 | 4.805841136 | 1.86312 | 2.57946 | 0.0099 | 0.0462 |
| 144 | KCNIP2 | 7.3757662 | 4.791741867 | 1.2283 | 3.90113 | 9.6E-05 | 0.00104 |
| 964 | CACNA2D1 | 14.674863 | 4.776069065 | 0.8929 | 5.34892 | 8.85E-08 | 2.4E-06 |
| 931 | PPP1R14C | 45.188112 | 4.774374391 | 0.5484 | 8.70602 | 3.15E-18 | 7.23E-16 |
| 122 | ENSG00000259869 | 1.9988177 | 4.750508432 | 1.81979 | 2.61047 | 0.00904 | 0.043 |
| 783 | ENSG00000248343 | 1.9172598 | 4.74280556 | 1.80541 | 2.627 | 0.00861 | 0.04141 |
| 1065 | PAX5 | 1.9144465 | 4.739416774 | 1.86195 | 2.54541 | 0.01091 | 0.04984 |
| 353 | ANKRD34C | 1.9049169 | 4.73413689 | 1.81637 | 2.60637 | 0.00915 | 0.04341 |
| 726 | CELSR3 | 39.674442 | 4.730199893 | 0.54013 | 8.75752 | 2.00E-18 | 4.83E-16 |
| 330 | CHRFAM7A | 1.8992903 | 4.730088744 | 1.85085 | 2.55563 | 0.0106 | 0.04872 |
| 546 | HAS1 | 6.8695665 | 4.712685159 | 1.26729 | 3.71873 | 0.0002 | 0.00194 |
| 522 | TSHZ3 | 24.275789 | 4.700470082 | 0.70506 | 6.66673 | 2.62E-11 | 1.70E-09 |
| 723 | CDCP1 | 46.119172 | 4.700464434 | 0.51112 | 9.1964 | 3.70E-20 | 1.21E-17 |
| 133 | CDHR1 | 7.1112602 | 4.693475486 | 1.27113 | 3.69237 | 0.00022 | 0.00211 |
| 902 | TFAP2B | 28.003353 | 4.691012135 | 0.6994 | 6.70715 | 1.98E-11 | 1.33E-09 |
| 451 | PNPO | 3.6438198 | 4.679755477 | 1.54621 | 3.02659 | 0.00247 | 0.01529 |
| 995 | FAM131B-AS1 | 1.8507124 | 4.678074258 | 1.83533 | 2.5489 | 0.01081 | 0.04941 |
| 435 | KCNJ12 | 1.8660917 | 4.676633303 | 1.8317 | 2.55317 | 0.01067 | 0.04897 |
| 78 | TDRD10 | 1.9143008 | 4.674119472 | 1.83346 | 2.54934 | 0.01079 | 0.04937 |
| 170 | ENSG00000240881 | 1.9143008 | 4.674119472 | 1.83346 | 2.54934 | 0.01079 | 0.04937 |
| 788 | NMU | 37.447714 | 4.659797711 | 0.55052 | 8.46441 | 2.57E-17 | 5.07E-15 |
| 463 | CD79B | 3.6092713 | 4.622726169 | 1.5576 | 2.96786 | 0.003 | 0.01779 |
| 1050 | ARC | 3.4843156 | 4.600368292 | 1.58805 | 2.89686 | 0.00377 | 0.02136 |
| 853 | GRIA1 | 16.253157 | 4.594489537 | 0.81524 | 5.63573 | 1.74E-08 | 5.72E-07 |
| 1019 | ANK1 | 9.9542741 | 4.583768245 | 1.01664 | 4.50872 | 6.5E-06 | 0.0001 |
| 1057 | SH3GL2 | 3.6367749 | 4.573858195 | 1.59172 | 2.87352 | 0.00406 | 0.02272 |
| 262 | FGF9 | 3.4977014 | 4.560390607 | 1.6351 | 2.78906 | 0.00529 | 0.02829 |
| 397 | IRX5 | 3.2891171 | 4.530704555 | 1.59597 | 2.83884 | 0.00453 | 0.02495 |
| 185 | RCOR2 | 9.2953081 | 4.511318954 | 1.02179 | 4.41512 | 1E-05 | 0.00015 |
| 427 | PIMREG | 15.425474 | 4.500280704 | 0.8518 | 5.28327 | 1.27E-07 | 3.2E-06 |
| 101 | KLHDC8A | 5.919324 | 4.476839742 | 1.27665 | 3.5067 | 0.00045 | 0.00378 |

| | | | | | | | |
|---|---|---|---|---|---|---|---|
| 697 | LINC00899 | 3.2265501 | 4.449206541 | 1.61263 | 2.75897 | 0.0058 | 0.0305 |
| 894 | PACSIN1 | 5.9118465 | 4.444646621 | 1.28681 | 3.45401 | 0.00055 | 0.00444 |
| 164 | LMO1 | 15.214665 | 4.443935653 | 0.85222 | 5.21452 | 1.84E-07 | 4.5E-06 |
| 560 | ENSG00000286412 | 3.288283 | 4.441995462 | 1.6528 | 2.68756 | 0.0072 | 0.03606 |
| 1120 | DOCK11 | 38.062863 | 4.436770975 | 0.5471 | 8.10958 | 5.08E-16 | 8.64E-14 |
| 173 | SLC1A2 | 11.533851 | 4.428532749 | 0.96268 | 4.6002 | 4.2E-06 | 6.9E-05 |
| 282 | FGF14 | 17.570031 | 4.417686635 | 0.7837 | 5.63699 | 1.73E-08 | 5.69E-07 |
| 178 | VWCE | 5.6341245 | 4.405623258 | 1.2944 | 3.40361 | 0.00067 | 0.00517 |
| 241 | FGD6 | 8.8430049 | 4.401544698 | 1.07234 | 4.10461 | 4E-05 | 0.0005 |
| 685 | NEFH | 25.840922 | 4.396028841 | 0.64264 | 6.84053 | 7.89E-12 | 5.78E-10 |
| 625 | C2orf72 | 3.1114977 | 4.395309173 | 1.5892 | 2.76574 | 0.00568 | 0.02998 |
| 646 | PAK5 | 25.803928 | 4.391245029 | 0.62509 | 7.02503 | 2.14E-12 | 1.73E-10 |
| 421 | RAP1GAP2 | 5.9377093 | 4.389551447 | 1.33384 | 3.29091 | 0.001 | 0.00721 |
| 275 | LMO7 | 8.6384766 | 4.388365572 | 1.04226 | 4.21044 | 2.5E-05 | 0.00033 |
| 91 | BRINP2 | 3.078372 | 4.381751432 | 1.58622 | 2.76239 | 0.00574 | 0.03025 |
| 715 | ZNF385D | 17.106362 | 4.370182243 | 0.77242 | 5.65775 | 1.53E-08 | 5.19E-07 |
| 1069 | SHC3 | 30.693321 | 4.367384235 | 0.68038 | 6.41906 | 1.37E-10 | 7.50E-09 |
| 924 | SLC35D3 | 2.9665017 | 4.365648024 | 1.61321 | 2.70618 | 0.00681 | 0.03452 |
| 443 | LHX1-DT | 5.5777007 | 4.363662839 | 1.30831 | 3.33535 | 0.00085 | 0.00633 |
| 3 | FNDC10 | 5.6848371 | 4.361657621 | 1.28129 | 3.40411 | 0.00066 | 0.00516 |
| 168 | KCNC1 | 2.9608022 | 4.341829831 | 1.57769 | 2.75201 | 0.00592 | 0.03099 |
| 537 | TTC9B | 10.985105 | 4.290931189 | 0.9754 | 4.39917 | 1.1E-05 | 0.00016 |
| 966 | ABCB1 | 10.29686 | 4.248088128 | 0.97943 | 4.33731 | 1.4E-05 | 0.0002 |
| 244 | NTN4 | 5.1828598 | 4.246221301 | 1.35756 | 3.12783 | 0.00176 | 0.0116 |
| 923 | MAP3K5 | 7.6373714 | 4.240939314 | 1.15704 | 3.66533 | 0.00025 | 0.0023 |
| 942 | RAPGEF5 | 20.456247 | 4.240247637 | 0.66274 | 6.39803 | 1.57E-10 | 8.47E-09 |
| 693 | ENSG00000273096 | 2.8858924 | 4.239404223 | 1.65578 | 2.56037 | 0.01046 | 0.04824 |
| 1090 | UAP1L1 | 13.181474 | 4.229800539 | 0.84444 | 5.00901 | 5.47E-07 | 1.2E-05 |
| 352 | PSTPIP1 | 4.9325409 | 4.217582619 | 1.41377 | 2.98323 | 0.00285 | 0.01713 |
| 922 | ENSG00000260418 | 12.54714 | 4.204073892 | 0.8885 | 4.73165 | 2.2E-06 | 4E-05 |
| 1030 | CRISPLD1 | 82.774712 | 4.201828935 | 0.35497 | 11.8372 | 2.51E-32 | 4.93E-29 |
| 684 | RFPL1S | 5.1208095 | 4.200466135 | 1.30477 | 3.21932 | 0.00128 | 0.00893 |
| 215 | ENSG00000274943 | 5.226596 | 4.194506503 | 1.3756 | 3.04922 | 0.00229 | 0.01442 |
| 172 | KIAA1549L | 14.932497 | 4.173639981 | 0.7924 | 5.26706 | 1.39E-07 | 3.5E-06 |
| 248 | RNY1P16 | 2.6536391 | 4.173197255 | 1.61952 | 2.57681 | 0.00997 | 0.04646 |
| 775 | NSG1 | 14.534556 | 4.153251929 | 0.78679 | 5.27874 | 1.30E-07 | 3.3E-06 |
| 665 | SALL4 | 9.5815294 | 4.136282106 | 0.96949 | 4.26643 | 2E-05 | 0.00027 |
| 880 | RIPOR2 | 14.38263 | 4.122842545 | 0.82534 | 4.99533 | 5.87E-07 | 1.2E-05 |
| 1118 | RADX | 36.075511 | 4.119221985 | 1.12657 | 3.65643 | 0.00026 | 0.00236 |

| | | | | | | | |
|---|---|---|---|---|---|---|---|
| 948 | HOXA13 | 9.4950919 | 4.115586032 | 0.99668 | 4.12932 | 3.6E-05 | 0.00045 |
| 969 | DPY19L2P4 | 7.0413742 | 4.07832322 | 1.11098 | 3.67093 | 0.00024 | 0.00226 |
| 738 | ST3GAL6 | 8.9251443 | 4.018097502 | 1.02683 | 3.9131 | 9.1E-05 | 0.001 |
| 563 | DYSF | 4.5241411 | 4.012224306 | 1.42937 | 2.80698 | 0.005 | 0.02708 |
| 987 | PTPRZ1 | 51.684286 | 4.008590475 | 0.51328 | 7.80972 | 5.73E-15 | 8.17E-13 |
| 733 | CACNA1D | 4.526212 | 4.000123514 | 1.44837 | 2.76181 | 0.00575 | 0.03028 |
| 846 | HBEGF | 45.119078 | 3.996066357 | 0.5226 | 7.64644 | 2.07E-14 | 2.57E-12 |
| 325 | GABRB3 | 10.959451 | 3.992329284 | 0.86789 | 4.60003 | 4.2E-06 | 6.9E-05 |
| 158 | EBF3 | 26.548394 | 3.991525 | 0.5797 | 6.88549 | 5.76E-12 | 4.34E-10 |
| 938 | THSD7A | 8.4191762 | 3.978682653 | 1.06852 | 3.72354 | 0.0002 | 0.0019 |
| 259 | DNAH10 | 6.7027874 | 3.97686623 | 1.10146 | 3.61054 | 0.00031 | 0.00274 |
| 847 | PCDHB16 | 4.4001665 | 3.973298255 | 1.35043 | 2.94225 | 0.00326 | 0.019 |
| 525 | FXYD7 | 4.3972803 | 3.956156358 | 1.35312 | 2.92374 | 0.00346 | 0.0199 |
| 112 | PGBD5 | 34.051762 | 3.950501965 | 0.51242 | 7.70951 | 1.26E-14 | 1.67E-12 |
| 1105 | SLC38A5 | 8.4185523 | 3.946873298 | 0.97782 | 4.0364 | 5.4E-05 | 0.00064 |
| 1072 | CAVIN4 | 8.6100572 | 3.946158725 | 0.99312 | 3.97351 | 7.1E-05 | 0.0008 |
| 324 | JAG2 | 17.344568 | 3.932073642 | 0.70396 | 5.58564 | 2.33E-08 | 7.39E-07 |
| 535 | DLL3 | 19.024314 | 3.927656927 | 0.76986 | 5.1018 | 3.36E-07 | 7.6E-06 |
| 515 | ANKLE1 | 10.58062 | 3.924773176 | 0.87633 | 4.47866 | 7.5E-06 | 0.00011 |
| 1132 | ARHGEF6 | 24.481615 | 3.922480171 | 0.67643 | 5.79883 | 6.68E-09 | 2.44E-07 |
| 887 | ATP6V1G2 | 4.3478716 | 3.91865829 | 1.34874 | 2.90542 | 0.00367 | 0.02088 |
| 1077 | SVEP1 | 6.3207313 | 3.916697966 | 1.1378 | 3.44236 | 0.00058 | 0.00459 |
| 615 | SATB2-AS1 | 10.566369 | 3.909228592 | 0.89323 | 4.3765 | 1.2E-05 | 0.00017 |
| 935 | SLC22A3 | 10.252405 | 3.901998355 | 0.88439 | 4.4121 | 1E-05 | 0.00015 |
| 369 | NPW | 12.23863 | 3.899110959 | 0.82923 | 4.70207 | 2.6E-06 | 4.5E-05 |
| 24 | NKAIN1 | 4.1698951 | 3.890097392 | 1.46761 | 2.65064 | 0.00803 | 0.03916 |
| 660 | GDAP1L1 | 12.172154 | 3.87643304 | 0.81704 | 4.74451 | 2.1E-06 | 3.8E-05 |
| 318 | BEGAIN | 10.017991 | 3.862135257 | 0.88709 | 4.35372 | 1.3E-05 | 0.00019 |
| 360 | TICRR | 5.7524338 | 3.861396283 | 1.23668 | 3.12238 | 0.00179 | 0.01177 |
| 1062 | LINC01241 | 7.8622925 | 3.854987188 | 1.02295 | 3.76848 | 0.00016 | 0.00163 |
| 798 | UNC5C | 6.0341277 | 3.846359405 | 1.15875 | 3.3194 | 0.0009 | 0.00662 |
| 444 | EPOP | 4.0653429 | 3.835553147 | 1.44886 | 2.64729 | 0.00811 | 0.03943 |
| 933 | ZDHHC14 | 5.838639 | 3.829980738 | 1.12387 | 3.40786 | 0.00065 | 0.00511 |
| 1064 | UNC13B | 136.47398 | 3.822269225 | 0.29497 | 12.9581 | 2.11E-38 | 9.71E-35 |
| 247 | SNX5P2 | 5.8693248 | 3.818344817 | 1.17248 | 3.25664 | 0.00113 | 0.00801 |
| 1037 | RIMS2 | 17.447498 | 3.818298602 | 0.6858 | 5.56764 | 2.58E-08 | 8.07E-07 |
| 1125 | LINC01201 | 3.8439641 | 3.817250286 | 1.41196 | 2.70351 | 0.00686 | 0.0347 |
| 335 | WDR76 | 17.050956 | 3.80594236 | 0.72196 | 5.27169 | 1.35E-07 | 3.4E-06 |
| 269 | DCLK1 | 46.670454 | 3.805584528 | 0.41564 | 9.15591 | 5.39E-20 | 1.65E-17 |

| | | | | | | | |
|---|---|---|---|---|---|---|---|
| 672 | EEF1A2 | 9.5625207 | 3.802573935 | 0.93624 | 4.06156 | 4.9E-05 | 0.00058 |
| 125 | PRKG1 | 6.0352376 | 3.796527398 | 1.16738 | 3.25218 | 0.00115 | 0.00813 |
| 234 | B4GALNT1 | 7.6385427 | 3.794497246 | 1.07672 | 3.52412 | 0.00042 | 0.00358 |
| 291 | COCH | 45.415801 | 3.786328755 | 0.43402 | 8.72394 | 2.69E-18 | 6.38E-16 |
| 650 | SYNDIG1 | 13.516765 | 3.772137928 | 0.83213 | 4.53312 | 5.8E-06 | 9.1E-05 |
| 48 | ORC1 | 5.651028 | 3.765055989 | 1.20101 | 3.13491 | 0.00172 | 0.01138 |
| 762 | NLGN1 | 9.2481257 | 3.764815921 | 0.99889 | 3.76901 | 0.00016 | 0.00163 |
| 670 | SLC17A9 | 9.2324445 | 3.762467796 | 0.95788 | 3.92793 | 8.6E-05 | 0.00094 |
| 1138 | ARHGAP4 | 31.656303 | 3.752843385 | 0.55184 | 6.80057 | 1.04E-11 | 7.46E-10 |
| 470 | MGAT5B | 5.4285646 | 3.750020585 | 1.26104 | 2.97375 | 0.00294 | 0.01751 |
| 89 | DNM3OS | 68.576019 | 3.740913268 | 0.52382 | 7.14166 | 9.22E-13 | 8.42E-11 |
| 384 | ENSG00000246465 | 3.7867328 | 3.721447199 | 1.41924 | 2.62214 | 0.00874 | 0.04185 |
| 197 | NCAM1 | 17.737589 | 3.706958359 | 0.75628 | 4.90157 | 9.51E-07 | 1.9E-05 |
| 346 | MEGF11 | 5.4819942 | 3.706003828 | 1.17968 | 3.14154 | 0.00168 | 0.01119 |
| 289 | JPH4 | 21.05707 | 3.698567701 | 0.69969 | 5.28597 | 1.25E-07 | 3.2E-06 |
| 205 | B3GAT1 | 12.427781 | 3.694759567 | 0.77141 | 4.78959 | 1.7E-06 | 3.1E-05 |
| 975 | NPTX2 | 25.019987 | 3.686686845 | 0.62741 | 5.87604 | 4.20E-09 | 1.60E-07 |
| 100 | CNTN2 | 17.572474 | 3.668076286 | 0.66668 | 5.502 | 3.76E-08 | 1.1E-06 |
| 225 | HOXC6 | 5.2612171 | 3.667949746 | 1.15041 | 3.18839 | 0.00143 | 0.00979 |
| 448 | RUNDC3A | 28.360215 | 3.667830065 | 0.54907 | 6.68013 | 2.39E-11 | 1.59E-09 |
| 725 | CDC25A | 12.488014 | 3.652386536 | 0.83185 | 4.39067 | 1.1E-05 | 0.00016 |
| 743 | TAGLN3 | 23.027448 | 3.647521707 | 1.43074 | 2.54939 | 0.01079 | 0.04937 |
| 990 | CCDC136 | 43.045016 | 3.645082017 | 0.43748 | 8.33194 | 7.95E-17 | 1.48E-14 |
| 1098 | MAP7D2 | 18.758389 | 3.633445689 | 0.66963 | 5.42605 | 5.76E-08 | 1.6E-06 |
| 216 | PRICKLE1 | 17.248075 | 3.63342953 | 0.65692 | 5.531 | 3.18E-08 | 9.67E-07 |
| 228 | LINC02381 | 10.554499 | 3.632754064 | 0.85334 | 4.25712 | 2.1E-05 | 0.00028 |
| 507 | PTPRS | 5.284942 | 3.626391503 | 1.18521 | 3.05971 | 0.00222 | 0.01402 |
| 1134 | MAMLD1 | 16.964945 | 3.625687887 | 0.6976 | 5.1974 | 2.02E-07 | 4.9E-06 |
| 161 | DUSP8 | 8.5514303 | 3.625073357 | 0.9997 | 3.62618 | 0.00029 | 0.00261 |
| 671 | KCNQ2 | 27.95494 | 3.614644582 | 0.67122 | 5.38518 | 7.24E-08 | 2E-06 |
| 580 | DPP10-AS1 | 19.847704 | 3.613643758 | 0.67749 | 5.33391 | 9.61E-08 | 2.6E-06 |
| 683 | EMID1 | 10.302579 | 3.604474519 | 0.93834 | 3.84132 | 0.00012 | 0.00128 |
| 908 | KCNQ5 | 5.2729407 | 3.598508106 | 1.29433 | 2.78022 | 0.00543 | 0.02889 |
| 495 | CDH7 | 10.293251 | 3.594105246 | 0.94853 | 3.78913 | 0.00015 | 0.00152 |
| 1102 | SRPX | 21.354697 | 3.583216149 | 0.5903 | 6.0702 | 1.28E-09 | 5.54E-08 |
| 332 | OTUD7A | 4.9471144 | 3.578404542 | 1.20583 | 2.96759 | 0.003 | 0.0178 |
| 394 | TOX3 | 18.296551 | 3.57182857 | 0.66494 | 5.37167 | 7.80E-08 | 2.1E-06 |
| 115 | GNG4 | 65.687917 | 3.570418296 | 0.42202 | 8.46033 | 2.67E-17 | 5.18E-15 |
| 1085 | OLFM1 | 16.373453 | 3.553533681 | 0.69872 | 5.08579 | 3.66E-07 | 8.2E-06 |

| | | | | | | | |
|---|---|---|---|---|---|---|---|
| 889 | HLA-DPB1 | 7.9745335 | 3.549360629 | 1.05426 | 3.36669 | 0.00076 | 0.00577 |
| 337 | SEMA6D | 27.515058 | 3.53407841 | 0.53625 | 6.59033 | 4.39E-11 | 2.64E-09 |
| 556 | TMEM178A | 51.897254 | 3.526249384 | 0.37185 | 9.48299 | 2.47E-21 | 1.03E-18 |
| 51 | ZYG11A | 9.462237 | 3.520662184 | 0.90202 | 3.90307 | 9.5E-05 | 0.00103 |
| 406 | SLC9A5 | 11.078107 | 3.515880764 | 0.83027 | 4.23461 | 2.3E-05 | 0.0003 |
| 236 | NAV3 | 39.28638 | 3.512328302 | 0.47763 | 7.35359 | 1.93E-13 | 2.09E-11 |
| 334 | DISP2 | 4.65244 | 3.511418528 | 1.23062 | 2.85336 | 0.00433 | 0.02399 |
| 319 | MEG3 | 42.819797 | 3.504852538 | 0.49565 | 7.07128 | 1.54E-12 | 1.32E-10 |
| 543 | PPFIA3 | 6.1812808 | 3.487915066 | 1.07011 | 3.2594 | 0.00112 | 0.00796 |
| 811 | GUCY1B1 | 4.9214512 | 3.480030434 | 1.2747 | 2.73009 | 0.00633 | 0.03263 |
| 390 | ENSG00000279789 | 9.6103949 | 3.478347979 | 0.86644 | 4.01452 | 6E-05 | 0.00069 |
| 793 | ENSG00000273156 | 7.8394912 | 3.477580288 | 0.94361 | 3.6854 | 0.00023 | 0.00215 |
| 366 | RHBDL1 | 6.4290475 | 3.476150889 | 1.01154 | 3.4365 | 0.00059 | 0.00468 |
| 203 | ETS1 | 10.579371 | 3.473018747 | 0.90416 | 3.84117 | 0.00012 | 0.00128 |
| 118 | PIP4K2A | 15.20207 | 3.443251405 | 0.75683 | 4.54956 | 5.4E-06 | 8.6E-05 |
| 553 | GAREM2 | 5.9618942 | 3.432550172 | 1.05362 | 3.25786 | 0.00112 | 0.00799 |
| 253 | TMEM116 | 6.1125915 | 3.422630659 | 1.08371 | 3.15824 | 0.00159 | 0.01069 |
| 821 | OTULINL | 11.556087 | 3.410276095 | 0.83372 | 4.09041 | 4.3E-05 | 0.00052 |
| 177 | FAM111B | 16.283903 | 3.396696257 | 0.708 | 4.79758 | 1.6E-06 | 3E-05 |
| 1043 | SYBU | 18.878125 | 3.38898947 | 0.60778 | 5.57602 | 2.46E-08 | 7.74E-07 |
| 336 | FRMD5 | 7.3794451 | 3.385795324 | 0.96855 | 3.49573 | 0.00047 | 0.0039 |
| 1115 | DACH2 | 38.288518 | 3.384544494 | 0.44797 | 7.55528 | 4.18E-14 | 4.98E-12 |
| 968 | ADAM22 | 10.209718 | 3.371966864 | 0.90776 | 3.71461 | 0.0002 | 0.00196 |
| 698 | MIRLET7BHG | 4.3562202 | 3.356207301 | 1.21286 | 2.76718 | 0.00565 | 0.02989 |
| 552 | LRATD1 | 42.914681 | 3.354416062 | 0.44906 | 7.46988 | 8.03E-14 | 9.25E-12 |
| 270 | DGKH | 28.693443 | 3.347534924 | 0.58591 | 5.71343 | 1.11E-08 | 3.83E-07 |
| 589 | CXCR4 | 68.607401 | 3.34731789 | 0.80528 | 4.15673 | 3.2E-05 | 0.00041 |
| 201 | NRGN | 19.603033 | 3.333558272 | 0.61802 | 5.39394 | 6.89E-08 | 1.9E-06 |
| 17 | TRNP1 | 8.2157461 | 3.333451073 | 0.97867 | 3.4061 | 0.00066 | 0.00514 |
| 542 | BLOC1S3 | 4.3903861 | 3.332914751 | 1.2098 | 2.75492 | 0.00587 | 0.03076 |
| 907 | RIMS1 | 9.9784942 | 3.322914952 | 0.784 | 4.23841 | 2.3E-05 | 0.0003 |
| 1005 | MTMR7 | 11.162903 | 3.315684973 | 0.77875 | 4.25773 | 2.1E-05 | 0.00028 |
| 592 | KIF5C | 37.600263 | 3.30904729 | 0.44862 | 7.37605 | 1.63E-13 | 1.80E-11 |
| 140 | CRTAC1 | 5.5619116 | 3.307129204 | 1.23638 | 2.67486 | 0.00748 | 0.03709 |
| 473 | RBFOX3 | 8.7516425 | 3.303205317 | 0.93126 | 3.54703 | 0.00039 | 0.00334 |
| 701 | PANX2 | 5.5035757 | 3.298395706 | 1.10959 | 2.97263 | 0.00295 | 0.01755 |
| 699 | GTSE1 | 17.745208 | 3.297288718 | 0.70657 | 4.66662 | 3.1E-06 | 5.2E-05 |
| 574 | ANKRD36B | 9.5396417 | 3.290938822 | 0.83364 | 3.94767 | 7.9E-05 | 0.00088 |
| 1023 | TOX | 6.8831897 | 3.289044254 | 1.07447 | 3.0611 | 0.00221 | 0.01397 |

| | | | | | | | |
|---|---|---|---|---|---|---|---|
| 323 | TMEM179 | 9.8473869 | 3.28898442 | 0.81901 | 4.01579 | 5.9E-05 | 0.00069 |
| 623 | COL4A3 | 4.3486748 | 3.286431508 | 1.24612 | 2.63733 | 0.00836 | 0.04035 |
| 210 | LMO3 | 8.100425 | 3.279001625 | 0.93656 | 3.50111 | 0.00046 | 0.00384 |
| 288 | CDH24 | 25.791829 | 3.26826631 | 0.53401 | 6.12025 | 9.34E-10 | 4.18E-08 |
| 540 | TMEM145 | 12.348173 | 3.261743282 | 0.7554 | 4.31792 | 1.6E-05 | 0.00022 |
| 1033 | RUNX1T1 | 43.801157 | 3.250386271 | 0.9554 | 3.4021 | 0.00067 | 0.00519 |
| 850 | DPYSL3 | 48.762499 | 3.24654426 | 0.48645 | 6.67401 | 2.49E-11 | 1.64E-09 |
| 1078 | SNX30 | 6.6903006 | 3.239560831 | 1.00639 | 3.21899 | 0.00129 | 0.00893 |
| 613 | MYO1B | 26.950904 | 3.234867166 | 0.57366 | 5.63899 | 1.71E-08 | 5.64E-07 |
| 1016 | DUSP26 | 6.699874 | 3.229029551 | 1.1963 | 2.69918 | 0.00695 | 0.03506 |
| 545 | C19orf81 | 11.822326 | 3.228069664 | 0.72768 | 4.43613 | 9.2E-06 | 0.00014 |
| 965 | PCLO | 40.011691 | 3.225204859 | 0.44186 | 7.29908 | 2.90E-13 | 2.96E-11 |
| 834 | EDIL3 | 27.462667 | 3.221471736 | 0.52115 | 6.18145 | 6.35E-10 | 2.97E-08 |
| 531 | MAP4K1 | 8.9814717 | 3.219703581 | 1.04749 | 3.07372 | 0.00211 | 0.01348 |
| 493 | ONECUT2 | 3.9698842 | 3.219271765 | 1.2247 | 2.62862 | 0.00857 | 0.04124 |
| 662 | NCOA3 | 11.739267 | 3.216367988 | 0.82374 | 3.90457 | 9.4E-05 | 0.00103 |
| 1096 | STS | 14.589671 | 3.21580393 | 0.65136 | 4.93703 | 7.93E-07 | 1.6E-05 |
| 227 | HOXC4 | 10.401205 | 3.210502388 | 0.89587 | 3.58366 | 0.00034 | 0.00299 |
| 925 | ARFGEF3 | 28.81894 | 3.209567727 | 0.50385 | 6.37005 | 1.89E-10 | 1.01E-08 |
| 1038 | ENSG00000254615 | 10.515665 | 3.20399511 | 0.86191 | 3.71731 | 0.0002 | 0.00194 |
| 272 | DACH1 | 39.727417 | 3.201373823 | 0.45999 | 6.95961 | 3.41E-12 | 2.63E-10 |
| 255 | SRRM4 | 8.9235509 | 3.191289427 | 0.8744 | 3.64968 | 0.00026 | 0.00242 |
| 488 | CELF4 | 48.224323 | 3.183593654 | 0.53437 | 5.95767 | 2.56E-09 | 1.03E-07 |
| 963 | GSAP | 5.090319 | 3.1785816 | 1.0838 | 2.93282 | 0.00336 | 0.01947 |
| 120 | RPS12P16 | 6.3811155 | 3.173483586 | 1.14557 | 2.77022 | 0.0056 | 0.02969 |
| 109 | MIXL1 | 5.1097569 | 3.164984803 | 1.16704 | 2.71199 | 0.00669 | 0.03408 |
| 914 | MICAL1 | 29.627604 | 3.150919238 | 0.47703 | 6.60531 | 3.97E-11 | 2.42E-09 |
| 213 | ENSG00000276115 | 4.8921724 | 3.143831909 | 1.11974 | 2.80765 | 0.00499 | 0.02704 |
| 462 | TCAM1P | 5.1191408 | 3.142642091 | 1.15942 | 2.71053 | 0.00672 | 0.03418 |
| 9 | NBPF1 | 22.792696 | 3.13988101 | 0.5796 | 5.41737 | 6.05E-08 | 1.7E-06 |
| 1117 | TCEAL5 | 78.03801 | 3.13071752 | 0.3088 | 10.1385 | 3.73E-24 | 2.14E-21 |
| 815 | NEIL3 | 11.446358 | 3.130365336 | 0.81432 | 3.84415 | 0.00012 | 0.00127 |
| 836 | RHOBTB3 | 28.315846 | 3.124262785 | 0.53622 | 5.82645 | 5.66E-09 | 2.09E-07 |
| 720 | EIF1B-AS1 | 5.0673916 | 3.121325701 | 1.13544 | 2.74899 | 0.00598 | 0.03119 |
| 668 | CDH4 | 7.2686508 | 3.109691298 | 0.93717 | 3.31816 | 0.00091 | 0.00664 |
| 328 | NSMCE3 | 92.881155 | 3.106358471 | 0.292 | 10.6381 | 1.98E-26 | 1.61E-23 |
| 1013 | PNMA2 | 35.19424 | 3.102765837 | 0.46054 | 6.7372 | 1.61E-11 | 1.13E-09 |
| 5 | PER3 | 11.164242 | 3.101548618 | 0.82537 | 3.75777 | 0.00017 | 0.00169 |
| 842 | LMNB1 | 36.492933 | 3.088257771 | 0.5921 | 5.2158 | 1.83E-07 | 4.5E-06 |

| | | | | | | | |
|---|---|---|---|---|---|---|---|
| 494 | NEDD4L | 32.117508 | 3.086211562 | 0.48969 | 6.30235 | 2.93E-10 | 1.50E-08 |
| 534 | LRFN1 | 4.7816145 | 3.085221206 | 1.18633 | 2.60063 | 0.00931 | 0.04392 |
| 383 | SBK1 | 23.660172 | 3.075297544 | 0.58871 | 5.2238 | 1.75E-07 | 4.3E-06 |
| 326 | APBA2 | 27.343277 | 3.0591609 | 0.49062 | 6.23523 | 4.51E-10 | 2.17E-08 |
| 577 | MAP4K4 | 93.539466 | 3.048419233 | 0.29802 | 10.2287 | 1.47E-24 | 1.02E-21 |
| 1053 | RECQL4 | 12.021619 | 3.043437752 | 0.74645 | 4.0772 | 4.6E-05 | 0.00055 |
| 787 | SLC10A4 | 5.7976396 | 3.033748127 | 1.02991 | 2.94564 | 0.00322 | 0.01886 |
| 898 | TTBK1 | 9.7555417 | 3.033126246 | 0.82732 | 3.66619 | 0.00025 | 0.00229 |
| 675 | DOP1B | 17.200737 | 3.028896532 | 0.63335 | 4.78236 | 1.7E-06 | 3.2E-05 |
| 478 | NDC80 | 17.28818 | 3.024538401 | 0.65741 | 4.60067 | 4.2E-06 | 6.9E-05 |
| 666 | TSHZ2 | 84.374452 | 3.020464509 | 0.31006 | 9.7414 | 2.01E-22 | 9.88E-20 |
| 378 | PLK1 | 18.32341 | 3.019256631 | 0.64522 | 4.67942 | 2.9E-06 | 4.9E-05 |
| 1015 | PBK | 25.253114 | 3.01865267 | 0.63666 | 4.7414 | 2.1E-06 | 3.8E-05 |
| 358 | SH3GL3 | 14.043367 | 3.018559511 | 0.64602 | 4.67258 | 3E-06 | 5E-05 |
| 745 | PODXL2 | 10.597056 | 3.017849546 | 0.79975 | 3.77351 | 0.00016 | 0.00161 |
| 804 | UGT8 | 9.3293573 | 3.016355863 | 0.82103 | 3.67388 | 0.00024 | 0.00224 |
| 921 | ENPP1 | 15.239391 | 3.01090586 | 0.67871 | 4.43624 | 9.2E-06 | 0.00014 |
| 568 | LRRTM4 | 10.189521 | 2.997832414 | 0.84413 | 3.55138 | 0.00038 | 0.0033 |
| 265 | CDK8 | 10.481266 | 2.993818355 | 0.76223 | 3.92773 | 8.6E-05 | 0.00094 |
| 16 | STMN1 | 479.91604 | 2.978495554 | 0.67993 | 4.38062 | 1.2E-05 | 0.00017 |
| 771 | RNF212 | 31.228729 | 2.977869347 | 0.48034 | 6.19955 | 5.66E-10 | 2.67E-08 |
| 419 | FANCA | 13.513782 | 2.974351495 | 0.72148 | 4.12256 | 3.7E-05 | 0.00046 |
| 831 | MAP1B | 751.68115 | 2.961987558 | 0.18445 | 16.0584 | 4.99E-58 | 6.88E-54 |
| 229 | PDE1B | 13.186565 | 2.947679995 | 0.81253 | 3.62776 | 0.00029 | 0.0026 |
| 927 | PLAGL1 | 107.99561 | 2.94564654 | 0.32268 | 9.12866 | 6.93E-20 | 2.03E-17 |
| 237 | SYT1 | 21.868949 | 2.940536596 | 0.62751 | 4.68605 | 2.8E-06 | 4.8E-05 |
| 983 | CCDC71L | 18.742893 | 2.935135386 | 0.56781 | 5.16919 | 2.35E-07 | 5.5E-06 |
| 735 | ADAMTS9 | 22.917965 | 2.931711465 | 0.534 | 5.49011 | 4.02E-08 | 1.2E-06 |
| 389 | KCTD13 | 37.343619 | 2.931634746 | 0.40938 | 7.16118 | 8.00E-13 | 7.43E-11 |
| 634 | FAM110A | 6.666895 | 2.931632109 | 0.97539 | 3.0056 | 0.00265 | 0.01613 |
| 94 | ASPM | 27.127651 | 2.93026342 | 0.59326 | 4.93922 | 7.84E-07 | 1.6E-05 |
| 1048 | KHDRBS3 | 8.7072615 | 2.924527186 | 0.83381 | 3.50745 | 0.00045 | 0.00377 |
| 570 | ITPRIPL1 | 6.4819067 | 2.921677316 | 0.93942 | 3.11008 | 0.00187 | 0.0122 |
| 104 | SERTAD4 | 15.083664 | 2.919024825 | 0.72506 | 4.02593 | 5.7E-05 | 0.00067 |
| 791 | NAAA | 17.962407 | 2.918837756 | 0.70859 | 4.11925 | 3.8E-05 | 0.00047 |
| 434 | SPECC1 | 13.249164 | 2.917848395 | 0.67037 | 4.35259 | 1.3E-05 | 0.00019 |
| 1127 | HS6ST2 | 13.062095 | 2.911606139 | 0.66288 | 4.39232 | 1.1E-05 | 0.00016 |
| 872 | TUBB2B | 80.76252 | 2.905581016 | 0.45496 | 6.3865 | 1.70E-10 | 9.10E-09 |
| 843 | SPOCK1 | 24.984281 | 2.903715083 | 0.59743 | 4.86038 | 1.2E-06 | 2.3E-05 |

| | | | | | | | |
|---|---|---|---|---|---|---|---|
| 439 | ANKRD13B | 25.191453 | 2.903704626 | 0.61079 | 4.75401 | 2E-06 | 3.6E-05 |
| 1112 | KIF4A | 16.630551 | 2.902598775 | 0.69538 | 4.17413 | 3E-05 | 0.00038 |
| 640 | ENSG00000275632 | 14.052964 | 2.901240131 | 0.68358 | 4.2442 | 2.2E-05 | 0.00029 |
| 741 | CIP2A | 12.794753 | 2.888310374 | 0.79467 | 3.63463 | 0.00028 | 0.00254 |
| 530 | KCNK6 | 6.4210793 | 2.886591328 | 1.12742 | 2.56036 | 0.01046 | 0.04824 |
| 300 | TRIM9 | 5.1383098 | 2.870794524 | 1.09604 | 2.61925 | 0.00881 | 0.04217 |
| 897 | GLP1R | 5.2018982 | 2.870257545 | 1.05556 | 2.71919 | 0.00654 | 0.03349 |
| 157 | MKI67 | 65.080537 | 2.862184979 | 0.43038 | 6.65034 | 2.92E-11 | 1.86E-09 |
| 770 | CPLX1 | 7.0893583 | 2.861835558 | 0.96078 | 2.97865 | 0.0029 | 0.01729 |
| 614 | SATB2 | 18.858155 | 2.860765118 | 0.57528 | 4.97286 | 6.60E-07 | 1.4E-05 |
| 202 | FEZ1 | 68.109162 | 2.857582444 | 0.33034 | 8.6504 | 5.13E-18 | 1.11E-15 |
| 372 | ENSG00000260293 | 7.2289373 | 2.856370281 | 0.95387 | 2.99449 | 0.00275 | 0.01659 |
| 333 | RYR3 | 9.2750048 | 2.846390734 | 0.83686 | 3.40128 | 0.00067 | 0.00521 |
| 606 | CHN1 | 106.78165 | 2.827046493 | 0.25596 | 11.0447 | 2.33E-28 | 2.29E-25 |
| 414 | GSE1 | 33.05215 | 2.825427639 | 0.49894 | 5.66285 | 1.49E-08 | 5.05E-07 |
| 212 | ENSG00000177359 | 9.1570372 | 2.81271541 | 0.83566 | 3.36584 | 0.00076 | 0.00579 |
| 298 | L2HGDH | 5.1208477 | 2.804069139 | 1.08106 | 2.59382 | 0.00949 | 0.04455 |
| 490 | RNF165 | 20.688746 | 2.803266773 | 0.63951 | 4.38346 | 1.2E-05 | 0.00017 |
| 1046 | MYC | 7.8715016 | 2.796636801 | 1.02055 | 2.74032 | 0.00614 | 0.03181 |
| 1126 | MBNL3 | 55.12407 | 2.79382225 | 0.39461 | 7.08003 | 1.44E-12 | 1.25E-10 |
| 717 | TRIM71 | 7.0386092 | 2.793658145 | 0.92764 | 3.01158 | 0.0026 | 0.01588 |
| 429 | DNAH2 | 11.587063 | 2.787376797 | 0.81566 | 3.41731 | 0.00063 | 0.00496 |
| 1055 | GLDC | 4.8682665 | 2.783988203 | 1.07253 | 2.59572 | 0.00944 | 0.04438 |
| 895 | FANCE | 8.2153823 | 2.773467787 | 0.81966 | 3.38368 | 0.00072 | 0.00548 |
| 102 | PLXNA2 | 14.055651 | 2.772805802 | 0.74568 | 3.71848 | 0.0002 | 0.00194 |
| 327 | ENTREP2 | 30.993206 | 2.765548999 | 0.4268 | 6.47976 | 9.19E-11 | 5.21E-09 |
| 1110 | NALF2 | 10.488674 | 2.762735186 | 0.81748 | 3.37957 | 0.00073 | 0.00554 |
| 484 | ZNF519 | 26.582093 | 2.75147921 | 0.53141 | 5.17767 | 2.25E-07 | 5.3E-06 |
| 674 | GABPA | 14.533312 | 2.750416311 | 0.61568 | 4.46726 | 7.9E-06 | 0.00012 |
| 235 | IRAK3 | 7.5818563 | 2.727886033 | 0.9015 | 3.02594 | 0.00248 | 0.01531 |
| 1087 | GPSM1 | 11.763774 | 2.719520097 | 0.73096 | 3.72048 | 0.0002 | 0.00192 |
| 39 | KIF2C | 30.772421 | 2.708474275 | 0.57718 | 4.69259 | 2.7E-06 | 4.7E-05 |
| 702 | MAPK12 | 11.282036 | 2.707310159 | 0.76711 | 3.52922 | 0.00042 | 0.00352 |
| 758 | TRIM59 | 8.735334 | 2.69718159 | 0.82143 | 3.28353 | 0.00103 | 0.00737 |
| 677 | WDR4 | 5.915199 | 2.696780069 | 1.01067 | 2.6683 | 0.00762 | 0.03759 |
| 345 | IGDCC3 | 12.541845 | 2.692455387 | 0.8451 | 3.18597 | 0.00144 | 0.00985 |
| 472 | CYTH1 | 17.713612 | 2.691858535 | 0.55933 | 4.81263 | 1.5E-06 | 2.8E-05 |
| 106 | VASH2 | 10.482671 | 2.689490661 | 0.71326 | 3.7707 | 0.00016 | 0.00162 |
| 1082 | PTGS1 | 13.10065 | 2.677823372 | 0.73506 | 3.64301 | 0.00027 | 0.00247 |

| | | | | | | | |
|---|---|---|---|---|---|---|---|
| 967 | RUNDC3B | 6.5519195 | 2.673713423 | 0.92635 | 2.88628 | 0.0039 | 0.02194 |
| 900 | ENSG00000231881 | 8.9124808 | 2.660253118 | 1.00423 | 2.64906 | 0.00807 | 0.03933 |
| 128 | LRRC20 | 6.4060362 | 2.6601158 | 0.89388 | 2.97592 | 0.00292 | 0.01742 |
| 211 | ITPR2 | 9.0735558 | 2.659717709 | 0.85151 | 3.12353 | 0.00179 | 0.01173 |
| 655 | E2F1 | 32.945864 | 2.649809837 | 0.60956 | 4.34709 | 1.4E-05 | 0.00019 |
| 415 | CDT1 | 38.377687 | 2.642700649 | 0.43454 | 6.08158 | 1.19E-09 | 5.22E-08 |
| 521 | ZNF536 | 31.463625 | 2.63752514 | 0.47237 | 5.5836 | 2.36E-08 | 7.45E-07 |
| 591 | ZEB2 | 15.261235 | 2.632325147 | 0.69427 | 3.79148 | 0.00015 | 0.00151 |
| 939 | VWDE | 9.2764237 | 2.628878392 | 0.88001 | 2.98734 | 0.00281 | 0.01692 |
| 588 | MCM6 | 86.80905 | 2.626468408 | 0.85364 | 3.07677 | 0.00209 | 0.01339 |
| 92 | CACNA1E | 10.112524 | 2.626088373 | 0.84672 | 3.10148 | 0.00193 | 0.01248 |
| 996 | TCAF1 | 16.528179 | 2.618811334 | 0.60993 | 4.29366 | 1.8E-05 | 0.00024 |
| 554 | DPYSL5 | 19.240784 | 2.618203434 | 0.68053 | 3.84732 | 0.00012 | 0.00125 |
| 147 | MFSD13A | 8.4108572 | 2.615359147 | 0.93513 | 2.79678 | 0.00516 | 0.02773 |
| 387 | SEZ6L2 | 40.176588 | 2.615161861 | 0.47936 | 5.45551 | 4.88E-08 | 1.4E-06 |
| 30 | CLSPN | 30.469384 | 2.612987643 | 0.637 | 4.10202 | 4.1E-05 | 0.0005 |
| 481 | MTCL1 | 10.535999 | 2.610623902 | 0.84007 | 3.10762 | 0.00189 | 0.01228 |
| 707 | OGG1 | 6.1791726 | 2.610348001 | 0.93045 | 2.80545 | 0.00502 | 0.02715 |
| 991 | CHRM2 | 6.4679855 | 2.604726947 | 0.93714 | 2.77944 | 0.00545 | 0.02893 |
| 302 | GPR137C | 13.13918 | 2.600298845 | 0.6684 | 3.89033 | 0.0001 | 0.00107 |
| 74 | MLLT11 | 37.387584 | 2.595951466 | 0.40731 | 6.37339 | 1.85E-10 | 9.88E-09 |
| 329 | ENSG00000259690 | 15.682098 | 2.5956716 | 0.72308 | 3.58975 | 0.00033 | 0.00293 |
| 1002 | MCPH1 | 7.9407695 | 2.589971881 | 0.79528 | 3.25669 | 0.00113 | 0.00801 |
| 455 | TMEM100 | 13.99827 | 2.587239836 | 0.67461 | 3.83516 | 0.00013 | 0.00131 |
| 116 | PFKFB3 | 30.920217 | 2.586762469 | 0.61797 | 4.18587 | 2.8E-05 | 0.00037 |
| 188 | SPTBN2 | 12.309941 | 2.577298222 | 0.7064 | 3.64851 | 0.00026 | 0.00242 |
| 859 | STC2 | 20.670291 | 2.575357613 | 0.59352 | 4.33916 | 1.4E-05 | 0.0002 |
| 891 | KIFC1 | 12.932628 | 2.568574154 | 0.76979 | 3.33671 | 0.00085 | 0.00631 |
| 67 | KCNC4 | 10.906708 | 2.567672172 | 0.80715 | 3.18115 | 0.00147 | 0.00998 |
| 941 | TSPAN13 | 16.047277 | 2.567548129 | 0.62741 | 4.09227 | 4.3E-05 | 0.00052 |
| 482 | APCDD1 | 25.071748 | 2.567183755 | 0.54691 | 4.69396 | 2.7E-06 | 4.6E-05 |
| 267 | FRY | 7.9269127 | 2.56666359 | 0.84584 | 3.03444 | 0.00241 | 0.01501 |
| 426 | ENO3 | 7.8728128 | 2.564873307 | 0.79665 | 3.21955 | 0.00128 | 0.00892 |
| 70 | LRIF1 | 12.932831 | 2.561565129 | 0.7205 | 3.55526 | 0.00038 | 0.00325 |
| 1074 | SMC2 | 37.103466 | 2.561469973 | 0.57453 | 4.4584 | 8.3E-06 | 0.00012 |
| 826 | MOCS2 | 20.146917 | 2.556943092 | 0.5829 | 4.38661 | 1.2E-05 | 0.00017 |
| 760 | CLDN11 | 9.537602 | 2.552084817 | 0.96143 | 2.65447 | 0.00794 | 0.03885 |
| 527 | WDR62 | 7.8330977 | 2.551984056 | 0.83848 | 3.04357 | 0.00234 | 0.01464 |
| 852 | MYOZ3 | 8.4222683 | 2.544003274 | 0.89225 | 2.85121 | 0.00436 | 0.02411 |

| | | | | | | | |
|---|---|---|---|---|---|---|---|
| 544 | POLD1 | 12.946322 | 2.543665152 | 0.63482 | 4.00688 | 6.2E-05 | 0.00071 |
| 993 | TMEM178B | 7.6323507 | 2.54014478 | 0.88794 | 2.86073 | 0.00423 | 0.0235 |
| 1080 | PHF19 | 23.890459 | 2.537726317 | 0.53156 | 4.77414 | 1.8E-06 | 3.3E-05 |
| 528 | DPF1 | 9.3643619 | 2.533443356 | 0.75621 | 3.35017 | 0.00081 | 0.00606 |
| 1084 | EXOSC2 | 11.491279 | 2.531177914 | 0.73391 | 3.44889 | 0.00056 | 0.0045 |
| 1136 | PNMA6A | 11.942635 | 2.525039314 | 0.72855 | 3.46585 | 0.00053 | 0.00427 |
| 93 | ARPC5 | 63.391232 | 2.523341742 | 0.42731 | 5.90517 | 3.52E-09 | 1.36E-07 |
| 1045 | ATAD2 | 23.993192 | 2.501842049 | 0.58955 | 4.24364 | 2.2E-05 | 0.00029 |
| 186 | MACROD1 | 7.7486444 | 2.501251962 | 0.90384 | 2.76736 | 0.00565 | 0.02989 |
| 95 | UBE2T | 19.640657 | 2.500574768 | 0.64002 | 3.90703 | 9.3E-05 | 0.00102 |
| 43 | PIK3R3 | 11.065615 | 2.499961533 | 0.70211 | 3.56066 | 0.00037 | 0.0032 |
| 644 | BMP2 | 12.488124 | 2.489445547 | 0.6894 | 3.61103 | 0.0003 | 0.00273 |
| 343 | PIF1 | 20.147138 | 2.481389578 | 0.75271 | 3.29659 | 0.00098 | 0.00708 |
| 309 | DCAF4 | 10.409217 | 2.479843472 | 0.76296 | 3.2503 | 0.00115 | 0.00817 |
| 595 | FMNL2 | 80.525643 | 2.474465218 | 0.36802 | 6.72367 | 1.77E-11 | 1.20E-09 |
| 454 | ENSG00000264895 | 7.5895384 | 2.474146415 | 0.84522 | 2.92721 | 0.00342 | 0.01975 |
| 296 | RN7SL1 | 6.5284735 | 2.472741411 | 0.87322 | 2.83175 | 0.00463 | 0.0254 |
| 532 | NFKBIB | 5.7631723 | 2.470457629 | 0.89993 | 2.74516 | 0.00605 | 0.03144 |
| 134 | PAPSS2 | 6.7408205 | 2.470406912 | 0.88467 | 2.79246 | 0.00523 | 0.02808 |
| 436 | TMEM97 | 24.301272 | 2.467897245 | 0.47965 | 5.14521 | 2.67E-07 | 6.1E-06 |
| 13 | E2F2 | 10.851294 | 2.463336155 | 0.79934 | 3.08169 | 0.00206 | 0.01322 |
| 871 | TUBB2A | 58.642127 | 2.459766597 | 0.43836 | 5.61135 | 2.01E-08 | 6.48E-07 |
| 1007 | LPL | 20.246919 | 2.456960873 | 0.68167 | 3.60431 | 0.00031 | 0.0028 |
| 217 | SLC38A1 | 48.727491 | 2.455032075 | 0.41731 | 5.88304 | 4.03E-09 | 1.55E-07 |
| 951 | MTURN | 55.996385 | 2.444495297 | 0.36369 | 6.72135 | 1.80E-11 | 1.22E-09 |
| 56 | DEPDC1 | 9.5829898 | 2.442318918 | 0.81672 | 2.99041 | 0.00279 | 0.01677 |
| 876 | TMEM170B | 9.0989078 | 2.438296185 | 0.74376 | 3.27835 | 0.00104 | 0.00749 |
| 83 | NHLH1 | 6.4367914 | 2.434550038 | 0.95134 | 2.55908 | 0.0105 | 0.04837 |
| 555 | CGREF1 | 7.9099349 | 2.433502562 | 0.83169 | 2.92599 | 0.00343 | 0.0198 |
| 774 | GRK4 | 10.588614 | 2.432588484 | 0.72425 | 3.35879 | 0.00078 | 0.0059 |
| 913 | SESN1 | 30.056505 | 2.429641404 | 0.55552 | 4.37365 | 1.2E-05 | 0.00017 |
| 628 | HJURP | 18.104289 | 2.427202588 | 0.66999 | 3.62277 | 0.00029 | 0.00264 |
| 22 | EPB41 | 8.0708829 | 2.422173595 | 0.89351 | 2.71084 | 0.00671 | 0.03418 |
| 2 | ACAP3 | 36.601953 | 2.421471035 | 0.40881 | 5.92325 | 3.16E-09 | 1.25E-07 |
| 105 | DTL | 21.451213 | 2.421397044 | 0.65214 | 3.713 | 0.0002 | 0.00196 |
| 449 | GJC1 | 37.467445 | 2.421112627 | 0.46947 | 5.15713 | 2.51E-07 | 5.8E-06 |
| 840 | MCC | 20.319915 | 2.417278537 | 0.52538 | 4.60098 | 4.2E-06 | 6.9E-05 |
| 19 | RCC1 | 8.6426135 | 2.409957484 | 0.77411 | 3.1132 | 0.00185 | 0.0121 |
| 257 | BCL7A | 24.518755 | 2.408801265 | 0.57593 | 4.18246 | 2.9E-05 | 0.00037 |

| | | | | | | | |
|---|---|---|---|---|---|---|---|
| 1083 | STXBP1 | 53.521077 | 2.408469828 | 0.40309 | 5.97508 | 2.30E-09 | 9.46E-08 |
| 740 | TMEM45A | 12.824266 | 2.399512429 | 0.73577 | 3.26125 | 0.00111 | 0.00791 |
| 171 | LINC00294 | 7.8736133 | 2.393438909 | 0.79296 | 3.01834 | 0.00254 | 0.0156 |
| 150 | SORCS1 | 10.68681 | 2.390816413 | 0.76637 | 3.11966 | 0.00181 | 0.01187 |
| 1001 | NCAPG2 | 19.827443 | 2.389029333 | 0.68231 | 3.50139 | 0.00046 | 0.00384 |
| 42 | MAST2 | 19.680162 | 2.379449243 | 0.5474 | 4.34684 | 1.4E-05 | 0.00019 |
| 280 | MIR17HG | 12.697837 | 2.375518188 | 0.80796 | 2.94013 | 0.00328 | 0.01911 |
| 14 | GALE | 9.0772552 | 2.372413543 | 0.82819 | 2.86457 | 0.00418 | 0.02328 |
| 769 | PDE6B | 21.975942 | 2.364972336 | 0.52923 | 4.46873 | 7.9E-06 | 0.00012 |
| 410 | RFWD3 | 27.227257 | 2.363414786 | 0.58765 | 4.02184 | 5.8E-05 | 0.00067 |
| 576 | AFF3 | 9.5355956 | 2.36333769 | 0.83882 | 2.81745 | 0.00484 | 0.02637 |
| 160 | BRSK2 | 19.066972 | 2.363143582 | 0.63338 | 3.73102 | 0.00019 | 0.00186 |
| 737 | ROBO1 | 10.041861 | 2.358521549 | 0.85901 | 2.74563 | 0.00604 | 0.03141 |
| 550 | RRM2 | 45.985214 | 2.358199183 | 0.54382 | 4.33638 | 1.4E-05 | 0.0002 |
| 512 | BRME1 | 9.438414 | 2.357922318 | 0.89409 | 2.63722 | 0.00836 | 0.04035 |
| 809 | DCLK2 | 59.165751 | 2.357141688 | 0.33798 | 6.97427 | 3.07E-12 | 2.41E-10 |
| 785 | C4orf19 | 19.806406 | 2.354478072 | 0.53561 | 4.39585 | 1.1E-05 | 0.00016 |
| 1059 | MLLT3 | 12.491331 | 2.351946288 | 0.65419 | 3.59518 | 0.00032 | 0.00288 |
| 813 | HMGB2 | 97.288096 | 2.351239339 | 0.41892 | 5.61263 | 1.99E-08 | 6.45E-07 |
| 691 | ENSG00000279738 | 7.7208953 | 2.34019347 | 0.89826 | 2.60524 | 0.00918 | 0.0435 |
| 245 | GAS2L3 | 21.332994 | 2.332931848 | 0.58564 | 3.98354 | 6.8E-05 | 0.00077 |
| 734 | C3orf14 | 20.249243 | 2.331732979 | 0.58457 | 3.98877 | 6.6E-05 | 0.00076 |
| 505 | FSD1 | 20.186871 | 2.325542972 | 0.63082 | 3.68652 | 0.00023 | 0.00215 |
| 396 | CRNDE | 20.064599 | 2.325046256 | 0.63879 | 3.63975 | 0.00027 | 0.00249 |
| 219 | AMIGO2 | 7.4982841 | 2.324526171 | 0.8576 | 2.71051 | 0.00672 | 0.03418 |
| 297 | LRR1 | 14.961379 | 2.324514821 | 0.57175 | 4.06563 | 4.8E-05 | 0.00057 |
| 748 | TOPBP1 | 34.73934 | 2.318647503 | 0.52022 | 4.45706 | 8.3E-06 | 0.00012 |
| 739 | TBC1D23 | 30.457135 | 2.315512891 | 0.4329 | 5.34889 | 8.85E-08 | 2.4E-06 |
| 299 | NIN | 18.254255 | 2.311591821 | 0.5754 | 4.01734 | 5.9E-05 | 0.00068 |
| 441 | CDK5R1 | 10.963732 | 2.311104864 | 0.77252 | 2.99166 | 0.00277 | 0.01672 |
| 985 | CTTNBP2 | 29.00566 | 2.307463754 | 0.50654 | 4.55535 | 5.2E-06 | 8.4E-05 |
| 998 | PRKAG2 | 5.9545267 | 2.306339596 | 0.90259 | 2.55525 | 0.01061 | 0.04875 |
| 131 | COMTD1 | 12.137462 | 2.301533072 | 0.65351 | 3.52183 | 0.00043 | 0.0036 |
| 187 | CDCA5 | 12.736877 | 2.298770961 | 0.80619 | 2.85139 | 0.00435 | 0.02411 |
| 491 | SKA1 | 14.136526 | 2.297963111 | 0.75824 | 3.03065 | 0.00244 | 0.01516 |
| 496 | DOK6 | 9.8296003 | 2.297071871 | 0.79703 | 2.88205 | 0.00395 | 0.02218 |
| 223 | POU6F1 | 10.046818 | 2.293802269 | 0.74161 | 3.09301 | 0.00198 | 0.01279 |
| 757 | RSRC1 | 24.102844 | 2.288840296 | 0.49733 | 4.60227 | 4.2E-06 | 6.8E-05 |
| 503 | ZFR2 | 10.55055 | 2.27477143 | 0.85292 | 2.66703 | 0.00765 | 0.0377 |

| | | | | | | | |
|---|---|---|---|---|---|---|---|
| 189 | FCHSD2 | 10.775913 | 2.274180283 | 0.72463 | 3.13838 | 0.0017 | 0.01128 |
| 896 | FKBP5 | 13.084335 | 2.274084007 | 0.77263 | 2.94329 | 0.00325 | 0.01898 |
| 912 | SOBP | 42.834318 | 2.272849343 | 0.38158 | 5.95646 | 2.58E-09 | 1.04E-07 |
| 747 | PLXND1 | 20.071643 | 2.270619259 | 0.60476 | 3.75458 | 0.00017 | 0.00171 |
| 600 | KLHL23 | 23.675141 | 2.269603 | 0.55045 | 4.12319 | 3.7E-05 | 0.00046 |
| 224 | SCN8A | 24.447509 | 2.268599628 | 0.51226 | 4.42857 | 9.5E-06 | 0.00014 |
| 1099 | CNKSR2 | 23.055042 | 2.258999968 | 0.4677 | 4.83001 | 1.4E-06 | 2.6E-05 |
| 356 | ABHD17C | 12.66684 | 2.255539215 | 0.77622 | 2.9058 | 0.00366 | 0.02086 |
| 867 | ZNF354B | 7.9127019 | 2.241265629 | 0.87042 | 2.57491 | 0.01003 | 0.04668 |
| 139 | AVPI1 | 8.0388744 | 2.240023965 | 0.8185 | 2.73675 | 0.0062 | 0.0321 |
| 705 | RAD18 | 12.571869 | 2.238901045 | 0.73736 | 3.03637 | 0.00239 | 0.01493 |
| 712 | EAF1 | 10.166086 | 2.232995259 | 0.75096 | 2.97353 | 0.00294 | 0.01752 |
| 1047 | PVT1 | 14.711503 | 2.232983816 | 0.60151 | 3.7123 | 0.00021 | 0.00197 |
| 551 | TRIB2 | 74.138561 | 2.227367588 | 0.36013 | 6.18498 | 6.21E-10 | 2.91E-08 |
| 875 | ENSG00000270504 | 7.6370079 | 2.222595619 | 0.83627 | 2.65776 | 0.00787 | 0.03856 |
| 49 | COA7 | 10.658079 | 2.221734216 | 0.74931 | 2.96504 | 0.00303 | 0.01792 |
| 956 | CAMK2B | 9.3146484 | 2.204451439 | 0.83469 | 2.64106 | 0.00826 | 0.04006 |
| 362 | PRC1 | 34.496185 | 2.20204614 | 0.54142 | 4.06715 | 4.8E-05 | 0.00057 |
| 52 | ECHDC2 | 71.279299 | 2.197589767 | 0.31131 | 7.05924 | 1.67E-12 | 1.40E-10 |
| 364 | NPRL3 | 18.184049 | 2.194411402 | 0.56107 | 3.91115 | 9.2E-05 | 0.001 |
| 26 | TMEM39B | 9.6341558 | 2.191600553 | 0.7072 | 3.09899 | 0.00194 | 0.01257 |
| 99 | NFASC | 15.162716 | 2.190508134 | 0.56746 | 3.86017 | 0.00011 | 0.0012 |
| 1056 | PTPRD | 40.945657 | 2.189492025 | 0.4429 | 4.94356 | 7.67E-07 | 1.6E-05 |
| 954 | DPY19L2P1 | 9.5907632 | 2.188227625 | 0.74182 | 2.94982 | 0.00318 | 0.01865 |
| 295 | MIS18BP1 | 41.82956 | 2.183299105 | 0.56316 | 3.87685 | 0.00011 | 0.00113 |
| 845 | SNHG4 | 22.264388 | 2.182268889 | 0.58202 | 3.74945 | 0.00018 | 0.00174 |
| 450 | KIF18B | 26.575561 | 2.175547093 | 0.62521 | 3.47972 | 0.0005 | 0.0041 |
| 1116 | TMSB15A | 26.614969 | 2.173548306 | 0.59524 | 3.65153 | 0.00026 | 0.0024 |
| 566 | MTHFD2 | 50.191243 | 2.163858464 | 0.32799 | 6.59729 | 4.19E-11 | 2.54E-09 |
| 375 | PPL | 18.944171 | 2.163404214 | 0.64827 | 3.33718 | 0.00085 | 0.0063 |
| 499 | SHC2 | 24.434144 | 2.15813266 | 0.4646 | 4.64511 | 3.4E-06 | 5.6E-05 |
| 301 | TXNDC16 | 8.6762281 | 2.156581876 | 0.81189 | 2.65626 | 0.0079 | 0.03865 |
| 344 | ANKDD1A | 8.4300027 | 2.15631139 | 0.81666 | 2.64042 | 0.00828 | 0.04007 |
| 459 | PRR11 | 13.869378 | 2.155868998 | 0.80681 | 2.67209 | 0.00754 | 0.0373 |
| 357 | AP3B2 | 27.379498 | 2.151110061 | 0.49088 | 4.38218 | 1.2E-05 | 0.00017 |
| 146 | PSD | 17.905892 | 2.133736568 | 0.56829 | 3.75464 | 0.00017 | 0.00171 |
| 412 | ENSG00000261061 | 8.1446039 | 2.129196825 | 0.7516 | 2.83288 | 0.00461 | 0.02535 |
| 598 | BAZ2B | 53.631202 | 2.117060076 | 0.37574 | 5.63436 | 1.76E-08 | 5.75E-07 |
| 402 | GINS3 | 6.6277654 | 2.108778469 | 0.82203 | 2.56533 | 0.01031 | 0.04775 |

| | | | | | | | |
|---|---|---|---|---|---|---|---|
| 754 | MBNL1 | 66.857085 | 2.107913838 | 0.35959 | 5.86194 | 4.58E-09 | 1.73E-07 |
| 60 | ERICH3 | 15.383897 | 2.104163187 | 0.82014 | 2.56563 | 0.0103 | 0.04772 |
| 932 | ARMT1 | 17.475936 | 2.103933789 | 0.5848 | 3.59771 | 0.00032 | 0.00286 |
| 1086 | QSOX2 | 26.633322 | 2.103743111 | 0.50255 | 4.18614 | 2.8E-05 | 0.00037 |
| 82 | IGSF8 | 15.822992 | 2.103328896 | 0.57634 | 3.64947 | 0.00026 | 0.00242 |
| 629 | HES6 | 31.890353 | 2.099074431 | 0.43412 | 4.83519 | 1.3E-06 | 2.5E-05 |
| 601 | PDK1 | 31.685671 | 2.097519584 | 0.49707 | 4.21975 | 2.4E-05 | 0.00032 |
| 571 | NCAPH | 14.217793 | 2.097217321 | 0.72958 | 2.87454 | 0.00405 | 0.02267 |
| 192 | ME3 | 16.691969 | 2.093979608 | 0.70716 | 2.96113 | 0.00307 | 0.01808 |
| 1095 | NLGN4X | 25.689523 | 2.087009144 | 0.59731 | 3.494 | 0.00048 | 0.00392 |
| 198 | THY1 | 11.546735 | 2.085741631 | 0.72965 | 2.85855 | 0.00426 | 0.02363 |
| 1017 | UNC5D | 29.635661 | 2.079559923 | 0.41875 | 4.96606 | 6.83E-07 | 1.4E-05 |
| 240 | BTG1 | 106.45487 | 2.072963832 | 0.26372 | 7.86041 | 3.83E-15 | 5.55E-13 |
| 838 | APC | 44.260584 | 2.071665324 | 0.34942 | 5.9288 | 3.05E-09 | 1.21E-07 |
| 242 | VEZT | 44.090094 | 2.068740722 | 0.37538 | 5.51101 | 3.57E-08 | 1.1E-06 |
| 317 | VRK1 | 20.696659 | 2.067854534 | 0.6362 | 3.25033 | 0.00115 | 0.00817 |
| 292 | STRN3 | 36.739463 | 2.064324907 | 0.40133 | 5.14367 | 2.69E-07 | 6.2E-06 |
| 1114 | ZNF711 | 39.987627 | 2.060707661 | 0.41463 | 4.96999 | 6.70E-07 | 1.4E-05 |
| 130 | VCL | 222.40731 | 2.055730895 | 0.22373 | 9.18862 | 3.98E-20 | 1.28E-17 |
| 191 | PAK1 | 36.279834 | 2.052210024 | 0.38723 | 5.29974 | 1.16E-07 | 3E-06 |
| 1076 | PALM2AKAP2 | 20.635351 | 2.051229133 | 0.51279 | 4.00014 | 6.3E-05 | 0.00073 |
| 520 | URI1 | 25.454134 | 2.045094413 | 0.4908 | 4.16686 | 3.1E-05 | 0.00039 |
| 1131 | HTATSF1 | 50.698139 | 2.0402952 | 0.34391 | 5.93259 | 2.98E-09 | 1.19E-07 |
| 393 | NETO2 | 34.811677 | 2.039529951 | 0.43384 | 4.70115 | 2.6E-06 | 4.5E-05 |
| 307 | FUT8 | 20.312745 | 2.034504282 | 0.58473 | 3.47937 | 0.0005 | 0.0041 |
| 977 | GPC2 | 14.343399 | 2.033354017 | 0.69874 | 2.91005 | 0.00361 | 0.02066 |
| 828 | NLN | 17.887632 | 2.032797341 | 0.50081 | 4.05905 | 4.9E-05 | 0.00059 |
| 731 | DUSP7 | 11.94107 | 2.032453262 | 0.74918 | 2.71289 | 0.00667 | 0.03402 |
| 465 | MAP2K6 | 36.570432 | 2.030662462 | 0.43366 | 4.6826 | 2.8E-06 | 4.8E-05 |
| 1119 | ACSL4 | 30.112849 | 2.029932022 | 0.39928 | 5.08401 | 3.70E-07 | 8.2E-06 |
| 676 | PRDM15 | 11.880404 | 2.029282356 | 0.66928 | 3.03205 | 0.00243 | 0.01511 |
| 312 | FOXN3 | 58.851382 | 2.029096103 | 0.38308 | 5.29674 | 1.18E-07 | 3.1E-06 |
| 110 | PARP1 | 68.221334 | 2.024726775 | 0.33838 | 5.98357 | 2.18E-09 | 9.06E-08 |
| 573 | ANKRD36 | 17.716768 | 2.023550836 | 0.72061 | 2.80809 | 0.00498 | 0.02702 |
| 722 | KIF15 | 15.785791 | 2.020527942 | 0.68756 | 2.93868 | 0.0033 | 0.01917 |
| 193 | FAT3 | 14.401557 | 2.014997265 | 0.74433 | 2.70711 | 0.00679 | 0.03445 |
| 829 | CENPH | 13.090894 | 2.007220893 | 0.60342 | 3.3264 | 0.00088 | 0.0065 |
| 474 | GAA | 16.716128 | 2.00504134 | 0.53797 | 3.72707 | 0.00019 | 0.00188 |
| 483 | SEH1L | 39.230504 | 2.003922166 | 0.38512 | 5.20338 | 1.96E-07 | 4.7E-06 |

| | | | | | | | |
|---|---|---|---|---|---|---|---|
| 1103 | TSPAN7 | 96.000403 | 2.002950591 | 0.76589 | 2.61519 | 0.00892 | 0.04254 |
| 11 | NBL1 | 7.9970175 | 2.002414014 | 0.77101 | 2.59714 | 0.0094 | 0.04424 |
| 827 | FST | 17.384355 | 1.999681807 | 0.60663 | 3.29639 | 0.00098 | 0.00708 |
| 1094 | PLCXD1 | 35.144407 | 1.997881802 | 0.50973 | 3.91946 | 8.9E-05 | 0.00097 |
| 602 | CDCA7 | 35.025232 | 1.996699285 | 0.42124 | 4.74004 | 2.1E-06 | 3.8E-05 |
| 1066 | IGFBPL1 | 44.721723 | 1.995633469 | 0.52862 | 3.77518 | 0.00016 | 0.0016 |
| 61 | PTBP2 | 37.943508 | 1.990311535 | 0.48543 | 4.10007 | 4.1E-05 | 0.0005 |
| 468 | UNK | 25.040276 | 1.985560355 | 0.52293 | 3.79697 | 0.00015 | 0.00148 |
| 893 | HMGA1 | 144.35294 | 1.985487736 | 0.38559 | 5.14919 | 2.62E-07 | 6E-06 |
| 154 | INPP5F | 75.143484 | 1.979087844 | 0.32579 | 6.07477 | 1.24E-09 | 5.43E-08 |
| 960 | LANCL2 | 11.798828 | 1.974443196 | 0.62823 | 3.14286 | 0.00167 | 0.01115 |
| 1071 | ANP32B | 85.512329 | 1.968204273 | 0.25425 | 7.74124 | 9.85E-15 | 1.33E-12 |
| 911 | MMS22L | 14.724477 | 1.963462927 | 0.60613 | 3.23935 | 0.0012 | 0.00844 |
| 1034 | PDP1 | 20.698564 | 1.96041953 | 0.48703 | 4.02525 | 5.7E-05 | 0.00067 |
| 165 | WEE1 | 57.801104 | 1.957454938 | 0.34675 | 5.64514 | 1.65E-08 | 5.49E-07 |
| 882 | H4C3 | 34.565344 | 1.954606832 | 0.48933 | 3.99444 | 6.5E-05 | 0.00074 |
| 565 | FBXO41 | 12.825301 | 1.945473473 | 0.58546 | 3.32296 | 0.00089 | 0.00656 |
| 233 | SHMT2 | 38.379461 | 1.944054674 | 0.37242 | 5.22003 | 1.79E-07 | 4.4E-06 |
| 283 | TEX30 | 13.938172 | 1.943527472 | 0.59817 | 3.24912 | 0.00116 | 0.00819 |
| 359 | FANCI | 33.632721 | 1.938067929 | 0.56752 | 3.41495 | 0.00064 | 0.00499 |
| 371 | CCNF | 16.741776 | 1.937847196 | 0.69674 | 2.78132 | 0.00541 | 0.02882 |
| 66 | SORT1 | 13.136225 | 1.937119534 | 0.61652 | 3.142 | 0.00168 | 0.01118 |
| 437 | SPAG5 | 23.740624 | 1.935374234 | 0.65762 | 2.94301 | 0.00325 | 0.01899 |
| 752 | CLSTN2 | 10.914709 | 1.934479836 | 0.64042 | 3.02064 | 0.00252 | 0.01552 |
| 803 | ELOVL6 | 10.014696 | 1.924406509 | 0.69974 | 2.75017 | 0.00596 | 0.03112 |
| 562 | ADD2 | 23.730199 | 1.916903194 | 0.50984 | 3.75978 | 0.00017 | 0.00168 |
| 373 | CORO7 | 13.517157 | 1.916578069 | 0.63573 | 3.01476 | 0.00257 | 0.01575 |
| 23 | PTPRU | 33.874533 | 1.915395074 | 0.41007 | 4.67089 | 3E-06 | 5.1E-05 |
| 919 | NCOA7 | 16.6929 | 1.915302119 | 0.74523 | 2.57008 | 0.01017 | 0.04723 |
| 920 | PTPRK | 26.540171 | 1.910582953 | 0.48098 | 3.97229 | 7.1E-05 | 0.0008 |
| 1054 | C8orf82 | 11.856997 | 1.905685693 | 0.7094 | 2.68635 | 0.00722 | 0.03613 |
| 800 | CENPE | 20.745728 | 1.901643604 | 0.6357 | 2.99143 | 0.00278 | 0.01672 |
| 873 | PSMG4 | 15.236642 | 1.900928094 | 0.59868 | 3.17518 | 0.0015 | 0.01015 |
| 899 | ZNF318 | 10.857983 | 1.895137921 | 0.68512 | 2.76616 | 0.00567 | 0.02995 |
| 822 | ANKH | 12.535145 | 1.894770669 | 0.60223 | 3.14627 | 0.00165 | 0.01104 |
| 700 | CERK | 50.716034 | 1.894378667 | 0.33143 | 5.71578 | 1.09E-08 | 3.80E-07 |
| 659 | MYBL2 | 20.234854 | 1.893811746 | 0.65873 | 2.87496 | 0.00404 | 0.02265 |
| 630 | KIF1A | 107.77736 | 1.892969913 | 0.30104 | 6.28817 | 3.21E-10 | 1.61E-08 |
| 802 | TET2 | 21.655159 | 1.891800113 | 0.46757 | 4.04598 | 5.2E-05 | 0.00062 |

| | | | | | | | |
|---|---|---|---|---|---|---|---|
| 982 | NAMPT | 18.673408 | 1.890829264 | 0.49697 | 3.8047 | 0.00014 | 0.00144 |
| 878 | E2F3 | 11.180145 | 1.889668535 | 0.65267 | 2.8953 | 0.00379 | 0.02143 |
| 12 | CAMK2N1 | 15.825918 | 1.879016613 | 0.70689 | 2.65813 | 0.00786 | 0.03854 |
| 466 | KCNJ2 | 13.734906 | 1.878531731 | 0.59591 | 3.15239 | 0.00162 | 0.01085 |
| 930 | SAMD5 | 36.455458 | 1.874615955 | 0.63032 | 2.97408 | 0.00294 | 0.0175 |
| 796 | NAP1L5 | 13.094057 | 1.873783613 | 0.59404 | 3.15432 | 0.00161 | 0.01079 |
| 1032 | MMP16 | 14.125269 | 1.871804238 | 0.62027 | 3.01772 | 0.00255 | 0.01562 |
| 694 | SREBF2 | 65.128212 | 1.865888669 | 0.30441 | 6.12957 | 8.81E-10 | 3.99E-08 |
| 749 | MRAS | 24.034145 | 1.861309342 | 0.4705 | 3.95599 | 7.6E-05 | 0.00085 |
| 445 | CDC6 | 15.970636 | 1.85954303 | 0.65634 | 2.83321 | 0.00461 | 0.02533 |
| 904 | ENSG00000216775 | 13.598167 | 1.852344754 | 0.62371 | 2.96986 | 0.00298 | 0.01769 |
| 549 | SOX11 | 76.439024 | 1.848161057 | 0.39471 | 4.6823 | 2.8E-06 | 4.8E-05 |
| 64 | SASS6 | 17.503869 | 1.847238167 | 0.54001 | 3.42075 | 0.00062 | 0.00491 |
| 440 | ATAD5 | 15.891214 | 1.847197965 | 0.58913 | 3.13546 | 0.00172 | 0.01137 |
| 651 | TPX2 | 26.428761 | 1.845998639 | 0.64254 | 2.87298 | 0.00407 | 0.02275 |
| 635 | NOP56 | 91.986145 | 1.844514219 | 0.26343 | 7.00184 | 2.53E-12 | 1.99E-10 |
| 713 | SATB1 | 79.713721 | 1.840919323 | 0.25477 | 7.22579 | 4.98E-13 | 4.84E-11 |
| 479 | EPB41L3 | 18.146024 | 1.837291493 | 0.51964 | 3.53572 | 0.00041 | 0.00346 |
| 976 | MCM7 | 94.377974 | 1.830336425 | 0.42361 | 4.32081 | 1.6E-05 | 0.00021 |
| 806 | MAD2L1 | 29.583027 | 1.829311123 | 0.62556 | 2.92429 | 0.00345 | 0.01987 |
| 368 | MRPS34 | 26.570848 | 1.829127681 | 0.44216 | 4.13683 | 3.5E-05 | 0.00044 |
| 851 | GRPEL2 | 15.408523 | 1.820194362 | 0.60332 | 3.01696 | 0.00255 | 0.01564 |
| 926 | HIVEP2 | 14.725701 | 1.81825105 | 0.60263 | 3.01719 | 0.00255 | 0.01563 |
| 639 | MCM8 | 21.752144 | 1.816684224 | 0.48749 | 3.72661 | 0.00019 | 0.00189 |
| 519 | CCNE1 | 11.771727 | 1.814316678 | 0.68537 | 2.6472 | 0.00812 | 0.03943 |
| 252 | FAM216A | 17.82307 | 1.808956172 | 0.53214 | 3.39939 | 0.00068 | 0.00524 |
| 476 | RAC3 | 9.601199 | 1.806038768 | 0.69986 | 2.58059 | 0.00986 | 0.04608 |
| 839 | DCP2 | 43.195348 | 1.803995008 | 0.35477 | 5.08492 | 3.68E-07 | 8.2E-06 |
| 934 | TMEM181 | 26.345812 | 1.803048434 | 0.49292 | 3.65791 | 0.00025 | 0.00235 |
| 1024 | CHD7 | 61.744182 | 1.800095457 | 0.3399 | 5.29596 | 1.18E-07 | 3.1E-06 |
| 855 | PTTG1 | 54.309412 | 1.799275745 | 0.46223 | 3.89261 | 9.9E-05 | 0.00107 |
| 230 | PA2G4 | 21.896739 | 1.795734197 | 0.49865 | 3.6012 | 0.00032 | 0.00283 |
| 260 | BRI3BP | 19.054674 | 1.792195228 | 0.58942 | 3.04061 | 0.00236 | 0.01476 |
| 431 | KCNAB3 | 25.604422 | 1.79202533 | 0.52608 | 3.40638 | 0.00066 | 0.00514 |
| 703 | MAPK8IP2 | 21.091875 | 1.789153538 | 0.54219 | 3.29984 | 0.00097 | 0.00702 |
| 425 | ARRB2 | 26.767361 | 1.787750107 | 0.42862 | 4.17097 | 3E-05 | 0.00039 |
| 271 | CKAP2 | 26.429238 | 1.787282259 | 0.49953 | 3.5779 | 0.00035 | 0.00303 |
| 861 | ARL10 | 24.055388 | 1.786937787 | 0.45 | 3.97098 | 7.2E-05 | 0.00081 |
| 575 | MRPL30 | 18.084815 | 1.7799363 | 0.53571 | 3.3226 | 0.00089 | 0.00657 |

| | | | | | | | |
|---|---|---|---|---|---|---|---|
| 854 | EBF1 | 13.076207 | 1.777916087 | 0.62912 | 2.82602 | 0.00471 | 0.02578 |
| 446 | ACLY | 117.68538 | 1.776988718 | 0.24985 | 7.11222 | 1.14E-12 | 1.03E-10 |
| 579 | DPP10 | 12.452701 | 1.775981743 | 0.61866 | 2.87068 | 0.0041 | 0.02286 |
| 273 | MZT1 | 22.560153 | 1.775236211 | 0.50414 | 3.5213 | 0.00043 | 0.0036 |
| 792 | CNOT6L | 21.587869 | 1.773112952 | 0.50311 | 3.52434 | 0.00042 | 0.00358 |
| 886 | TCF19 | 24.461347 | 1.768675828 | 0.67003 | 2.6397 | 0.0083 | 0.0401 |
| 293 | HECTD1 | 51.834958 | 1.758716943 | 0.33289 | 5.28323 | 1.27E-07 | 3.2E-06 |
| 958 | CCM2 | 18.931891 | 1.756093435 | 0.49629 | 3.53844 | 0.0004 | 0.00343 |
| 687 | RBFOX2 | 68.110262 | 1.753722312 | 0.33632 | 5.21443 | 1.84E-07 | 4.5E-06 |
| 274 | TBC1D4 | 16.616833 | 1.752387998 | 0.52102 | 3.36341 | 0.00077 | 0.00583 |
| 759 | ENSG00000269888 | 12.815088 | 1.747135099 | 0.59952 | 2.91422 | 0.00357 | 0.02043 |
| 405 | MATCAP1 | 13.222964 | 1.746553732 | 0.68399 | 2.55349 | 0.01066 | 0.04894 |
| 167 | ENSG00000260196 | 15.025833 | 1.744690682 | 0.56547 | 3.08536 | 0.00203 | 0.01307 |
| 386 | PAGR1 | 10.38261 | 1.744631415 | 0.6598 | 2.6442 | 0.00819 | 0.03973 |
| 597 | GPD2 | 14.420144 | 1.743688658 | 0.6691 | 2.60603 | 0.00916 | 0.04344 |
| 953 | NT5C3A | 19.692699 | 1.743253343 | 0.48986 | 3.55869 | 0.00037 | 0.00321 |
| 442 | MYO19 | 27.777259 | 1.739171116 | 0.46578 | 3.73392 | 0.00019 | 0.00184 |
| 181 | FADS2 | 30.942804 | 1.738742969 | 0.43746 | 3.97464 | 7E-05 | 0.0008 |
| 279 | OBI1 | 30.097632 | 1.732842323 | 0.40397 | 4.28956 | 1.8E-05 | 0.00024 |
| 174 | MDK | 65.339309 | 1.72917115 | 0.46851 | 3.69078 | 0.00022 | 0.00211 |
| 124 | OGDHL | 31.868223 | 1.726435001 | 0.52739 | 3.27355 | 0.00106 | 0.00761 |
| 727 | NCKIPSD | 21.17767 | 1.723068072 | 0.56902 | 3.02816 | 0.00246 | 0.01523 |
| 599 | CERS6 | 16.194551 | 1.717825776 | 0.62537 | 2.74689 | 0.00602 | 0.03133 |
| 1051 | BOP1 | 30.476487 | 1.715527459 | 0.46407 | 3.6967 | 0.00022 | 0.00208 |
| 637 | CDC25B | 26.322539 | 1.713881982 | 0.49135 | 3.4881 | 0.00049 | 0.00399 |
| 1122 | UPF3B | 35.578785 | 1.709619634 | 0.48786 | 3.50432 | 0.00046 | 0.00381 |
| 126 | ZWINT | 41.037925 | 1.706244901 | 0.60201 | 2.83425 | 0.00459 | 0.02526 |
| 195 | DCUN1D5 | 28.078996 | 1.699290583 | 0.40535 | 4.19216 | 2.8E-05 | 0.00036 |
| 869 | EXOC2 | 31.087109 | 1.697500942 | 0.44485 | 3.81588 | 0.00014 | 0.00139 |
| 721 | ANO10 | 21.252828 | 1.696169001 | 0.52434 | 3.23487 | 0.00122 | 0.00855 |
| 915 | TRAF3IP2 | 14.204969 | 1.69015318 | 0.6294 | 2.68533 | 0.00725 | 0.03621 |
| 1129 | FHL1 | 119.39043 | 1.677575329 | 0.26687 | 6.28603 | 3.26E-10 | 1.63E-08 |
| 65 | PSRC1 | 24.539653 | 1.670198857 | 0.5159 | 3.23744 | 0.00121 | 0.00849 |
| 96 | PPP1R12B | 47.639262 | 1.667367765 | 0.40584 | 4.10839 | 4E-05 | 0.00049 |
| 103 | SYT14 | 35.923362 | 1.667237458 | 0.47055 | 3.54318 | 0.0004 | 0.00338 |
| 8 | PRDM2 | 44.083978 | 1.666029409 | 0.39393 | 4.22925 | 2.3E-05 | 0.00031 |
| 1108 | NUDT11 | 26.447738 | 1.664214 | 0.47453 | 3.5071 | 0.00045 | 0.00377 |
| 276 | KCTD12 | 86.38496 | 1.660800091 | 0.31822 | 5.21897 | 1.80E-07 | 4.4E-06 |
| 57 | LRRC40 | 11.700556 | 1.660404019 | 0.64997 | 2.5546 | 0.01063 | 0.04882 |

| | | | | | | | |
|---|---|---|---|---|---|---|---|
| 1139 | UBL4A | 10.940641 | 1.659736118 | 0.65029 | 2.55232 | 0.0107 | 0.04907 |
| 163 | RRM1 | 58.088669 | 1.651941972 | 0.46357 | 3.5635 | 0.00037 | 0.00317 |
| 506 | CHAF1A | 23.413979 | 1.650685324 | 0.52802 | 3.12621 | 0.00177 | 0.01164 |
| 467 | JPT1 | 70.953392 | 1.649260925 | 0.36518 | 4.51626 | 6.3E-06 | 9.8E-05 |
| 989 | IMPDH1 | 28.649662 | 1.64723313 | 0.42329 | 3.89147 | 1E-04 | 0.00107 |
| 1133 | FMR1 | 36.123103 | 1.646129989 | 0.45431 | 3.62337 | 0.00029 | 0.00263 |
| 1113 | PHKA1 | 21.283086 | 1.641325664 | 0.53416 | 3.07273 | 0.00212 | 0.01351 |
| 508 | DNMT1 | 68.584026 | 1.641311918 | 0.28587 | 5.74148 | 9.39E-09 | 3.31E-07 |
| 500 | TCF3 | 60.505814 | 1.637610581 | 0.36712 | 4.46073 | 8.2E-06 | 0.00012 |
| 688 | FOXRED2 | 24.662039 | 1.632578302 | 0.49055 | 3.32802 | 0.00087 | 0.00647 |
| 696 | PHF21B | 28.144499 | 1.631277512 | 0.43037 | 3.7904 | 0.00015 | 0.00152 |
| 814 | SAP30 | 17.595913 | 1.631062203 | 0.62134 | 2.62506 | 0.00866 | 0.0416 |
| 1075 | SLC44A1 | 37.805414 | 1.630726305 | 0.39741 | 4.10337 | 4.1E-05 | 0.0005 |
| 392 | SEPHS2 | 16.919735 | 1.63017616 | 0.54119 | 3.0122 | 0.00259 | 0.01586 |
| 1081 | STOM | 34.4885 | 1.621692195 | 0.47202 | 3.43565 | 0.00059 | 0.00468 |
| 231 | NABP2 | 34.716915 | 1.615912431 | 0.38621 | 4.18405 | 2.9E-05 | 0.00037 |
| 341 | CCNB2 | 15.968224 | 1.613938003 | 0.6212 | 2.59808 | 0.00937 | 0.04414 |
| 453 | SPAG9 | 100.85491 | 1.6109845 | 0.24221 | 6.65108 | 2.91E-11 | 1.86E-09 |
| 943 | SNHG26 | 18.078253 | 1.607859791 | 0.55898 | 2.87642 | 0.00402 | 0.02257 |
| 937 | PHF14 | 46.580154 | 1.599860671 | 0.38034 | 4.20641 | 2.6E-05 | 0.00034 |
| 1121 | SEPTIN6 | 13.350006 | 1.597640589 | 0.58433 | 2.73413 | 0.00625 | 0.03231 |
| 1039 | OXR1 | 32.381215 | 1.596689273 | 0.36762 | 4.34332 | 1.4E-05 | 0.0002 |
| 117 | HSPA14 | 13.66355 | 1.594558587 | 0.59051 | 2.7003 | 0.00693 | 0.03498 |
| 1000 | PAXIP1 | 15.489635 | 1.591047845 | 0.60004 | 2.65156 | 0.00801 | 0.03913 |
| 84 | PFDN2 | 25.111027 | 1.586414382 | 0.5158 | 3.07566 | 0.0021 | 0.01344 |
| 1027 | TMEM70 | 17.833275 | 1.583348181 | 0.52055 | 3.04167 | 0.00235 | 0.01472 |
| 768 | RNF168 | 29.173729 | 1.582313609 | 0.4585 | 3.45103 | 0.00056 | 0.00448 |
| 31 | MRPS15 | 18.9745 | 1.57894397 | 0.48771 | 3.2375 | 0.00121 | 0.00849 |
| 251 | CORO1C | 21.619866 | 1.572263692 | 0.51127 | 3.07524 | 0.0021 | 0.01344 |
| 129 | FUT11 | 25.444952 | 1.563416966 | 0.59967 | 2.60715 | 0.00913 | 0.04337 |
| 890 | DAXX | 20.129526 | 1.560204642 | 0.51499 | 3.02961 | 0.00245 | 0.01518 |
| 385 | KIF22 | 27.538168 | 1.560028872 | 0.52303 | 2.98266 | 0.00286 | 0.01714 |
| 1097 | SHROOM2 | 32.464005 | 1.551841638 | 0.45038 | 3.44566 | 0.00057 | 0.00455 |
| 27 | HDAC1 | 25.835088 | 1.550024521 | 0.42082 | 3.68335 | 0.00023 | 0.00217 |
| 401 | CFAP20 | 62.096613 | 1.546873053 | 0.3149 | 4.91233 | 9.00E-07 | 1.8E-05 |
| 781 | DCAF16 | 48.563353 | 1.544386024 | 0.32236 | 4.79086 | 1.7E-06 | 3.1E-05 |
| 169 | SERGEF | 14.799706 | 1.543579373 | 0.57846 | 2.66844 | 0.00762 | 0.03759 |
| 433 | PFAS | 33.208072 | 1.543195077 | 0.43535 | 3.54472 | 0.00039 | 0.00337 |
| 992 | KIAA1549 | 19.387562 | 1.542155889 | 0.56769 | 2.71657 | 0.0066 | 0.03374 |

| | | | | | | | |
|---|---|---|---|---|---|---|---|
| 1100 | POLA1 | 21.807557 | 1.538912658 | 0.55011 | 2.79748 | 0.00515 | 0.02768 |
| 910 | CASP8AP2 | 28.023754 | 1.538667572 | 0.43392 | 3.54594 | 0.00039 | 0.00335 |
| 799 | RAP1GDS1 | 24.79965 | 1.538626963 | 0.46267 | 3.32551 | 0.00088 | 0.00651 |
| 1014 | DPYSL2 | 100.02035 | 1.537113447 | 0.43442 | 3.53829 | 0.0004 | 0.00343 |
| 1020 | MCM4 | 59.629224 | 1.534402754 | 0.47403 | 3.23693 | 0.00121 | 0.0085 |
| 477 | FASN | 52.6218 | 1.533563276 | 0.32598 | 4.70451 | 2.5E-06 | 4.5E-05 |
| 627 | PDE6D | 12.976533 | 1.530535859 | 0.56389 | 2.71425 | 0.00664 | 0.03393 |
| 1026 | EYA1 | 31.277681 | 1.525502575 | 0.44666 | 3.41535 | 0.00064 | 0.00499 |
| 287 | SUPT16H | 43.335967 | 1.522749169 | 0.41157 | 3.69984 | 0.00022 | 0.00206 |
| 547 | ZNF160 | 17.98676 | 1.522500441 | 0.50633 | 3.00695 | 0.00264 | 0.01607 |
| 511 | NFIX | 89.710618 | 1.521668244 | 0.28165 | 5.4027 | 6.56E-08 | 1.8E-06 |
| 156 | BUB3 | 78.7682 | 1.517289205 | 0.31837 | 4.76585 | 1.9E-06 | 3.5E-05 |
| 638 | PCNA | 68.632808 | 1.509033492 | 0.33858 | 4.45695 | 8.3E-06 | 0.00012 |
| 514 | DDX39A | 43.894339 | 1.50667511 | 0.44399 | 3.3935 | 0.00069 | 0.00532 |
| 469 | RNF157 | 19.367378 | 1.505245697 | 0.57405 | 2.62217 | 0.00874 | 0.04185 |
| 584 | TSN | 25.893258 | 1.503950942 | 0.44762 | 3.35989 | 0.00078 | 0.00588 |
| 541 | ZNF428 | 26.002679 | 1.503597112 | 0.42243 | 3.55942 | 0.00037 | 0.00321 |
| 513 | SAMD1 | 21.941212 | 1.49098505 | 0.45129 | 3.3038 | 0.00095 | 0.00693 |
| 772 | SLBP | 30.60873 | 1.489992548 | 0.48165 | 3.09354 | 0.00198 | 0.01277 |
| 1130 | MAP7D3 | 25.836139 | 1.489557767 | 0.44524 | 3.34555 | 0.00082 | 0.00614 |
| 1107 | SYP | 48.068196 | 1.488651688 | 0.3519 | 4.23029 | 2.3E-05 | 0.00031 |
| 492 | TCF4 | 129.70143 | 1.483575808 | 0.30525 | 4.86025 | 1.2E-06 | 2.3E-05 |
| 509 | LDLR | 15.384111 | 1.482973688 | 0.58274 | 2.54483 | 0.01093 | 0.04989 |
| 864 | MXD3 | 23.071717 | 1.482938513 | 0.55545 | 2.66978 | 0.00759 | 0.03748 |
| 18 | RPA2 | 39.098466 | 1.47949178 | 0.3615 | 4.09263 | 4.3E-05 | 0.00052 |
| 1092 | ENSG00000284976 | 25.27235 | 1.478802176 | 0.45197 | 3.27188 | 0.00107 | 0.00765 |
| 208 | C12orf57 | 75.081091 | 1.477318347 | 0.27543 | 5.36363 | 8.16E-08 | 2.2E-06 |
| 458 | SKA2 | 64.625613 | 1.476362526 | 0.28955 | 5.09884 | 3.42E-07 | 7.6E-06 |
| 656 | SOGA1 | 49.95094 | 1.47322097 | 0.34147 | 4.31433 | 1.6E-05 | 0.00022 |
| 1070 | CKS2 | 39.993748 | 1.470289459 | 0.5193 | 2.8313 | 0.00464 | 0.02542 |
| 28 | MARCKSL1 | 140.44405 | 1.470264778 | 0.38291 | 3.83969 | 0.00012 | 0.00129 |
| 1128 | HPRT1 | 44.686835 | 1.470240447 | 0.38475 | 3.82133 | 0.00013 | 0.00137 |
| 1106 | PIM2 | 42.734199 | 1.468566543 | 0.32626 | 4.50124 | 6.8E-06 | 0.0001 |
| 1041 | NUDCD1 | 18.390135 | 1.465278021 | 0.5006 | 2.92704 | 0.00342 | 0.01976 |
| 498 | TSHZ1 | 21.302323 | 1.463102293 | 0.48381 | 3.02416 | 0.00249 | 0.01538 |
| 1111 | PDZD11 | 25.425548 | 1.459959734 | 0.48034 | 3.03945 | 0.00237 | 0.01479 |
| 1 | KLHL17 | 16.484431 | 1.45841697 | 0.5646 | 2.58308 | 0.00979 | 0.04581 |
| 180 | FEN1 | 22.702533 | 1.456828797 | 0.50061 | 2.9101 | 0.00361 | 0.02066 |
| 877 | DEK | 133.38873 | 1.456490183 | 0.25563 | 5.69758 | 1.22E-08 | 4.18E-07 |

| | | | | | | | |
|---|---|---|---|---|---|---|---|
| 564 | RAB11FIP5 | 17.269091 | 1.454379464 | 0.51921 | 2.80114 | 0.00509 | 0.02746 |
| 1042 | ENY2 | 53.690469 | 1.437164883 | 0.34019 | 4.22462 | 2.4E-05 | 0.00032 |
| 268 | PDS5B | 22.934071 | 1.433672832 | 0.45241 | 3.16894 | 0.00153 | 0.01035 |
| 970 | PEG10 | 42.230965 | 1.430929702 | 0.4022 | 3.55775 | 0.00037 | 0.00322 |
| 501 | SF3A2 | 22.625063 | 1.430502088 | 0.49054 | 2.91619 | 0.00354 | 0.02033 |
| 1004 | FDFT1 | 36.271393 | 1.428204321 | 0.42217 | 3.38299 | 0.00072 | 0.00548 |
| 263 | PARP4 | 15.863535 | 1.427904674 | 0.55951 | 2.55207 | 0.01071 | 0.04909 |
| 286 | PARP2 | 26.323742 | 1.427519619 | 0.48761 | 2.92759 | 0.00342 | 0.01974 |
| 249 | TDG | 22.519262 | 1.424781491 | 0.47956 | 2.971 | 0.00297 | 0.01763 |
| 916 | HDAC2 | 68.361534 | 1.422985595 | 0.29839 | 4.76895 | 1.9E-06 | 3.4E-05 |
| 557 | MSH2 | 28.438482 | 1.4213081 | 0.41365 | 3.43605 | 0.00059 | 0.00468 |
| 350 | SCAMP5 | 33.014238 | 1.41687144 | 0.40748 | 3.47717 | 0.00051 | 0.00412 |
| 1109 | SMC1A | 90.898994 | 1.415051609 | 0.23527 | 6.01454 | 1.80E-09 | 7.67E-08 |
| 207 | ENO2 | 94.845042 | 1.40559056 | 0.28387 | 4.95157 | 7.36E-07 | 1.5E-05 |
| 636 | VPS16 | 20.7449 | 1.405435472 | 0.45311 | 3.10174 | 0.00192 | 0.01248 |
| 256 | CAMKK2 | 21.091982 | 1.404461206 | 0.46306 | 3.03299 | 0.00242 | 0.01507 |
| 108 | LBR | 23.36998 | 1.400760213 | 0.52158 | 2.68558 | 0.00724 | 0.0362 |
| 218 | SLC38A2 | 56.380703 | 1.394854235 | 0.47988 | 2.9067 | 0.00365 | 0.02082 |
| 816 | ACSL1 | 109.88868 | 1.393510191 | 0.32174 | 4.33115 | 1.5E-05 | 0.00021 |
| 342 | USP3 | 27.151187 | 1.392117704 | 0.41906 | 3.32198 | 0.00089 | 0.00658 |
| 80 | RUSC1 | 30.21446 | 1.391914772 | 0.40348 | 3.44976 | 0.00056 | 0.00449 |
| 166 | RPL23AP65 | 20.696347 | 1.390916283 | 0.46023 | 3.02221 | 0.00251 | 0.01546 |
| 321 | DYNC1H1 | 92.241146 | 1.381174581 | 0.25206 | 5.47954 | 4.26E-08 | 1.2E-06 |
| 837 | EPB41L4A-AS1 | 28.932224 | 1.374522071 | 0.46138 | 2.97915 | 0.00289 | 0.01727 |
| 789 | PPAT | 23.319467 | 1.374188979 | 0.4493 | 3.05854 | 0.00222 | 0.01404 |
| 612 | STAT1 | 47.60327 | 1.373117931 | 0.35232 | 3.89735 | 9.7E-05 | 0.00105 |
| 81 | MEX3A | 48.153103 | 1.372197802 | 0.42056 | 3.26278 | 0.0011 | 0.00787 |
| 667 | PRELID3B | 24.732607 | 1.36926068 | 0.46686 | 2.93292 | 0.00336 | 0.01947 |
| 1008 | ATP6V1B2 | 40.993499 | 1.36829699 | 0.39136 | 3.49629 | 0.00047 | 0.00389 |
| 456 | AKAP1 | 24.993343 | 1.367701977 | 0.45097 | 3.03279 | 0.00242 | 0.01508 |
| 730 | PCBP4 | 36.852663 | 1.365304734 | 0.37969 | 3.59586 | 0.00032 | 0.00287 |
| 1079 | FBXW2 | 51.283761 | 1.358674842 | 0.30793 | 4.41235 | 1E-05 | 0.00015 |
| 98 | PLEKHA6 | 45.976141 | 1.358670658 | 0.37313 | 3.64132 | 0.00027 | 0.00248 |
| 399 | NUDT21 | 29.719295 | 1.356179473 | 0.4251 | 3.19024 | 0.00142 | 0.00975 |
| 716 | SLC4A7 | 26.369558 | 1.351455932 | 0.49536 | 2.72825 | 0.00637 | 0.03275 |
| 85 | PBX1 | 50.263794 | 1.34885473 | 0.36593 | 3.68608 | 0.00023 | 0.00215 |
| 428 | POLR2A | 22.987453 | 1.345868199 | 0.46998 | 2.86368 | 0.00419 | 0.02333 |
| 903 | MCM3 | 65.133521 | 1.344878072 | 0.38123 | 3.52772 | 0.00042 | 0.00354 |
| 833 | HMGCR | 22.023222 | 1.344633562 | 0.45514 | 2.95432 | 0.00313 | 0.01842 |

| | | | | | | | |
|---|---|---|---|---|---|---|---|
| 7 | EXOSC10 | 25.083347 | 1.339238921 | 0.42635 | 3.14119 | 0.00168 | 0.01119 |
| 194 | TAF1D | 28.837122 | 1.330523097 | 0.42084 | 3.16161 | 0.00157 | 0.01059 |
| 874 | PXDC1 | 30.686899 | 1.328703857 | 0.45882 | 2.89591 | 0.00378 | 0.02141 |
| 261 | PSPC1 | 23.786534 | 1.320267186 | 0.46685 | 2.828 | 0.00468 | 0.02565 |
| 879 | GMNN | 32.998378 | 1.318591685 | 0.43514 | 3.03026 | 0.00244 | 0.01516 |
| 411 | TERF2IP | 59.57683 | 1.3163336 | 0.32131 | 4.09676 | 4.2E-05 | 0.00051 |
| 978 | PCOLCE | 40.255829 | 1.315191721 | 0.48521 | 2.71054 | 0.00672 | 0.03418 |
| 75 | SEMA6C | 24.689732 | 1.313284739 | 0.46111 | 2.84809 | 0.0044 | 0.02429 |
| 686 | YWHAH | 43.916346 | 1.304576514 | 0.40509 | 3.22049 | 0.00128 | 0.0089 |
| 581 | DDX18 | 35.322177 | 1.304460473 | 0.36326 | 3.59094 | 0.00033 | 0.00292 |
| 682 | YDJC | 26.179074 | 1.301173342 | 0.42148 | 3.08715 | 0.00202 | 0.01301 |
| 1063 | C9orf72 | 23.691707 | 1.298419457 | 0.49173 | 2.64051 | 0.00828 | 0.04007 |
| 825 | NIPBL-DT | 36.325264 | 1.297961077 | 0.41639 | 3.11714 | 0.00183 | 0.01196 |
| 381 | XPO6 | 34.658484 | 1.297751157 | 0.383 | 3.38843 | 0.0007 | 0.0054 |
| 724 | TMEM158 | 19.196375 | 1.296957827 | 0.50384 | 2.57414 | 0.01005 | 0.04677 |
| 1006 | PSD3 | 18.082477 | 1.290201 | 0.50578 | 2.5509 | 0.01074 | 0.04924 |
| 586 | POLR2D | 26.041196 | 1.285140571 | 0.4297 | 2.99077 | 0.00278 | 0.01675 |
| 692 | GTPBP1 | 23.092435 | 1.281786445 | 0.44209 | 2.89939 | 0.00374 | 0.0212 |
| 71 | TTF2 | 20.29797 | 1.281652427 | 0.45151 | 2.83859 | 0.00453 | 0.02496 |
| 626 | PTMA | 162.12023 | 1.280712477 | 0.27448 | 4.6659 | 3.1E-06 | 5.2E-05 |
| 183 | CSKMT | 22.507754 | 1.279014258 | 0.48604 | 2.63152 | 0.0085 | 0.04092 |
| 135 | HELLS | 92.938473 | 1.27783285 | 0.41149 | 3.1054 | 0.0019 | 0.01236 |
| 21 | SNHG12 | 26.234485 | 1.271703754 | 0.46823 | 2.71596 | 0.00661 | 0.03379 |
| 155 | MCMBP | 29.050359 | 1.269540775 | 0.47274 | 2.6855 | 0.00724 | 0.0362 |
| 175 | CKAP5 | 40.206676 | 1.269472761 | 0.35498 | 3.57614 | 0.00035 | 0.00305 |
| 176 | ZFP91 | 22.617969 | 1.267959668 | 0.45883 | 2.76345 | 0.00572 | 0.03016 |
| 487 | RNF138 | 22.056188 | 1.261142023 | 0.46341 | 2.72146 | 0.0065 | 0.03332 |
| 408 | EXOSC6 | 37.807531 | 1.259018093 | 0.35492 | 3.54738 | 0.00039 | 0.00334 |
| 885 | TUBB | 486.95818 | 1.255600288 | 0.24951 | 5.03221 | 4.85E-07 | 1E-05 |
| 374 | UBN1 | 27.137542 | 1.253678328 | 0.40651 | 3.08397 | 0.00204 | 0.01313 |
| 780 | QDPR | 19.951392 | 1.251278338 | 0.47811 | 2.61712 | 0.00887 | 0.04233 |
| 765 | YEATS2 | 43.006641 | 1.250593795 | 0.3854 | 3.24494 | 0.00117 | 0.00829 |
| 709 | NUP210 | 37.049096 | 1.244300834 | 0.39908 | 3.11792 | 0.00182 | 0.01194 |
| 664 | ZFAS1 | 111.69947 | 1.242285987 | 0.26437 | 4.699 | 2.6E-06 | 4.5E-05 |
| 1135 | HMGB3 | 58.329692 | 1.237355926 | 0.36839 | 3.35878 | 0.00078 | 0.0059 |
| 313 | PPP4R3A | 25.10478 | 1.235073804 | 0.42751 | 2.88901 | 0.00386 | 0.02176 |
| 285 | IRS2 | 74.768244 | 1.233584955 | 0.31647 | 3.89789 | 9.7E-05 | 0.00105 |
| 41 | NASP | 78.463619 | 1.230819367 | 0.38135 | 3.22751 | 0.00125 | 0.00874 |
| 162 | ZNF195 | 24.931309 | 1.229308628 | 0.47146 | 2.60748 | 0.00912 | 0.04336 |

| | | | | | | | |
|---|---|---|---|---|---|---|---|
| 536 | EID2 | 20.794842 | 1.2195777 | 0.46042 | 2.64881 | 0.00808 | 0.03933 |
| 1035 | ZNF706 | 32.628321 | 1.214742419 | 0.39552 | 3.07129 | 0.00213 | 0.01356 |
| 805 | SNHG8 | 104.32798 | 1.212823081 | 0.23367 | 5.19036 | 2.10E-07 | 5E-06 |
| 751 | RBP1 | 214.93347 | 1.207723547 | 0.27356 | 4.41491 | 1E-05 | 0.00015 |
| 38 | CTPS1 | 30.575965 | 1.207260319 | 0.3887 | 3.10589 | 0.0019 | 0.01234 |
| 594 | RIF1 | 39.683591 | 1.206080366 | 0.35768 | 3.37191 | 0.00075 | 0.00567 |
| 661 | TOMM34 | 35.359666 | 1.206054384 | 0.38287 | 3.15007 | 0.00163 | 0.01093 |
| 305 | MTHFD1 | 37.509574 | 1.204423085 | 0.40005 | 3.0107 | 0.00261 | 0.01591 |
| 153 | TRUB1 | 28.627537 | 1.20083033 | 0.39457 | 3.04338 | 0.00234 | 0.01465 |
| 111 | CDC42BPA | 44.486044 | 1.199861131 | 0.34278 | 3.50042 | 0.00046 | 0.00385 |
| 616 | LANCL1 | 41.525299 | 1.198966159 | 0.34714 | 3.45388 | 0.00055 | 0.00444 |
| 145 | NOLC1 | 52.789998 | 1.198861228 | 0.34522 | 3.47279 | 0.00052 | 0.00418 |
| 524 | UBA2 | 51.020885 | 1.197493815 | 0.33614 | 3.56254 | 0.00037 | 0.00318 |
| 539 | PAFAH1B3 | 39.813858 | 1.19683354 | 0.37754 | 3.17005 | 0.00152 | 0.01031 |
| 306 | MAX | 34.907448 | 1.196406296 | 0.39863 | 3.00131 | 0.00269 | 0.01631 |
| 1022 | BPNT2 | 36.919359 | 1.196367611 | 0.38972 | 3.06979 | 0.00214 | 0.01361 |
| 151 | MXI1 | 25.359755 | 1.19512985 | 0.4295 | 2.78264 | 0.00539 | 0.02872 |
| 621 | EPHA4 | 28.345324 | 1.19486802 | 0.43595 | 2.74087 | 0.00613 | 0.03179 |
| 795 | WDFY3 | 22.701918 | 1.193083171 | 0.44382 | 2.68823 | 0.00718 | 0.03601 |
| 611 | GLS | 72.72332 | 1.192875197 | 0.28786 | 4.14394 | 3.4E-05 | 0.00043 |
| 485 | SNRPD1 | 49.914112 | 1.192755528 | 0.38278 | 3.11606 | 0.00183 | 0.012 |
| 569 | IMMT | 28.944168 | 1.191153086 | 0.40603 | 2.93366 | 0.00335 | 0.01943 |
| 732 | PBRM1 | 39.295523 | 1.189488116 | 0.38029 | 3.12783 | 0.00176 | 0.0116 |
| 961 | GUSB | 27.081208 | 1.187738178 | 0.41653 | 2.85149 | 0.00435 | 0.02411 |
| 35 | UTP11 | 26.799836 | 1.186390595 | 0.46006 | 2.57879 | 0.00991 | 0.04625 |
| 361 | IDH2 | 108.67107 | 1.184683486 | 0.22688 | 5.22165 | 1.77E-07 | 4.4E-06 |
| 561 | PPP3R1 | 27.802346 | 1.179498758 | 0.457 | 2.58098 | 0.00985 | 0.04606 |
| 1104 | RBM10 | 37.363996 | 1.175422277 | 0.3508 | 3.35066 | 0.00081 | 0.00605 |
| 152 | SMC3 | 33.975272 | 1.174522975 | 0.38517 | 3.04935 | 0.00229 | 0.01442 |
| 475 | ALYREF | 30.807972 | 1.173589624 | 0.39293 | 2.9868 | 0.00282 | 0.01694 |
| 944 | KLHL7 | 43.956814 | 1.173004207 | 0.34315 | 3.4183 | 0.00063 | 0.00495 |
| 424 | PELP1 | 33.543267 | 1.168356398 | 0.3727 | 3.13486 | 0.00172 | 0.01138 |
| 790 | PAICS | 99.282139 | 1.163605831 | 0.27199 | 4.2782 | 1.9E-05 | 0.00025 |
| 457 | MTMR4 | 33.326196 | 1.161915575 | 0.38504 | 3.01761 | 0.00255 | 0.01562 |
| 997 | AGAP3 | 57.730839 | 1.160956826 | 0.33638 | 3.45131 | 0.00056 | 0.00447 |
| 957 | ZMIZ2 | 44.55431 | 1.155843328 | 0.36156 | 3.19679 | 0.00139 | 0.00956 |
| 1018 | GPAT4 | 23.744683 | 1.151517048 | 0.42013 | 2.74086 | 0.00613 | 0.03179 |
| 663 | CSE1L | 45.50753 | 1.147324444 | 0.34787 | 3.29814 | 0.00097 | 0.00705 |
| 338 | DUT | 70.108856 | 1.145765456 | 0.31147 | 3.67862 | 0.00023 | 0.0022 |

| | | | | | | | |
|---|---|---|---|---|---|---|---|
| 25 | KHDRBS1 | 92.390257 | 1.144850739 | 0.24639 | 4.64652 | 3.4E-06 | 5.6E-05 |
| 680 | PCNT | 36.236145 | 1.136577029 | 0.35504 | 3.20127 | 0.00137 | 0.00946 |
| 773 | NSD2 | 50.595746 | 1.135731927 | 0.41697 | 2.72375 | 0.00645 | 0.03312 |
| 502 | LMNB2 | 38.909024 | 1.135717955 | 0.41978 | 2.70552 | 0.00682 | 0.03457 |
| 728 | IMPDH2 | 55.049779 | 1.134368644 | 0.32614 | 3.47811 | 0.0005 | 0.00411 |
| 452 | SPOP | 22.437387 | 1.134145855 | 0.43949 | 2.58058 | 0.00986 | 0.04608 |
| 258 | CDK2AP1 | 91.364095 | 1.133449841 | 0.2781 | 4.07572 | 4.6E-05 | 0.00055 |
| 884 | HLA-A | 127.76558 | 1.131767773 | 0.34851 | 3.24749 | 0.00116 | 0.00823 |
| 767 | OPA1 | 42.383628 | 1.128362415 | 0.35703 | 3.16046 | 0.00158 | 0.01063 |
| 182 | FADS1 | 30.671105 | 1.128015695 | 0.39592 | 2.8491 | 0.00438 | 0.02425 |
| 365 | CAPN15 | 27.766689 | 1.123740715 | 0.43337 | 2.59302 | 0.00951 | 0.04464 |
| 849 | TCERG1 | 77.270714 | 1.123275462 | 0.31332 | 3.58504 | 0.00034 | 0.00298 |
| 824 | TARS1 | 37.164305 | 1.120520928 | 0.35482 | 3.15797 | 0.00159 | 0.01069 |
| 10 | CROCCP2 | 81.379712 | 1.117224861 | 0.27684 | 4.03569 | 5.4E-05 | 0.00064 |
| 114 | TOMM20 | 125.72954 | 1.11295453 | 0.2357 | 4.72188 | 2.3E-06 | 4.1E-05 |
| 945 | CYCS | 87.319282 | 1.112823266 | 0.24883 | 4.47229 | 7.7E-06 | 0.00012 |
| 940 | BZW2 | 47.44429 | 1.112424036 | 0.38291 | 2.90515 | 0.00367 | 0.02089 |
| 50 | RPS13P2 | 42.508481 | 1.111131353 | 0.33064 | 3.3605 | 0.00078 | 0.00588 |
| 304 | RTN1 | 37.090148 | 1.110322522 | 0.40139 | 2.76622 | 0.00567 | 0.02995 |
| 370 | SNHG19 | 28.581899 | 1.10778833 | 0.41029 | 2.7 | 0.00693 | 0.035 |
| 848 | DELE1 | 46.510958 | 1.104305545 | 0.32138 | 3.43609 | 0.00059 | 0.00468 |
| 184 | SNHG1 | 96.228898 | 1.098465637 | 0.28398 | 3.86806 | 0.00011 | 0.00116 |
| 657 | SNHG17 | 35.80092 | 1.098076589 | 0.37482 | 2.92962 | 0.00339 | 0.01963 |
| 90 | CACYBP | 43.645869 | 1.09734036 | 0.34299 | 3.19931 | 0.00138 | 0.0095 |
| 461 | DCAF7 | 86.832796 | 1.095166375 | 0.26278 | 4.16761 | 3.1E-05 | 0.00039 |
| 121 | ZNF33B | 22.960528 | 1.082429378 | 0.41529 | 2.60641 | 0.00915 | 0.04341 |
| 243 | METAP2 | 61.141066 | 1.078741233 | 0.28651 | 3.7651 | 0.00017 | 0.00165 |
| 929 | SF3B5 | 43.43971 | 1.073981037 | 0.34614 | 3.1027 | 0.00192 | 0.01245 |
| 322 | CKB | 136.84182 | 1.071015227 | 0.2366 | 4.52665 | 6E-06 | 9.3E-05 |
| 4 | ACOT7 | 28.892041 | 1.070433978 | 0.41185 | 2.59908 | 0.00935 | 0.04404 |
| 413 | USP10 | 41.247335 | 1.068015642 | 0.33408 | 3.19689 | 0.00139 | 0.00956 |
| 1067 | PTAR1 | 37.653507 | 1.066899934 | 0.39048 | 2.73227 | 0.00629 | 0.03246 |
| 1088 | NPDC1 | 60.914507 | 1.059827647 | 0.3624 | 2.9245 | 0.00345 | 0.01987 |
| 409 | SF3B3 | 52.594872 | 1.057523859 | 0.31747 | 3.33109 | 0.00087 | 0.0064 |
| 567 | RTKN | 36.728852 | 1.05679556 | 0.39458 | 2.67831 | 0.0074 | 0.0368 |
| 648 | SNX5 | 50.285192 | 1.054225863 | 0.39667 | 2.65772 | 0.00787 | 0.03856 |
| 403 | GOT2 | 82.528112 | 1.051259624 | 0.28645 | 3.6699 | 0.00024 | 0.00227 |
| 430 | CHD3 | 42.844827 | 1.048462718 | 0.3967 | 2.64297 | 0.00822 | 0.03986 |
| 523 | GPI | 93.991648 | 1.047395609 | 0.28958 | 3.61689 | 0.0003 | 0.00268 |

| | | | | | | | |
|---|---|---|---|---|---|---|---|
| 347 | ANP32A | 37.185229 | 1.042172064 | 0.39798 | 2.61864 | 0.00883 | 0.04221 |
| 858 | ERGIC1 | 33.494172 | 1.032117374 | 0.38616 | 2.67279 | 0.00752 | 0.03724 |
| 73 | PHGDH | 59.972396 | 1.029597294 | 0.37963 | 2.71212 | 0.00669 | 0.03408 |
| 603 | OLA1 | 53.766115 | 1.026835412 | 0.30922 | 3.32076 | 0.0009 | 0.0066 |
| 6 | KIF1B | 51.67521 | 1.024018973 | 0.38282 | 2.67493 | 0.00747 | 0.03709 |
| 20 | SNHG3 | 86.913794 | 1.023824416 | 0.30795 | 3.32467 | 0.00089 | 0.00653 |
| 946 | CBX3 | 153.8824 | 1.020319566 | 0.23038 | 4.42882 | 9.5E-06 | 0.00014 |
| 862 | RNF44 | 50.609736 | 1.017253413 | 0.3592 | 2.83196 | 0.00463 | 0.02539 |
| 254 | PEBP1 | 155.63102 | 1.015438143 | 0.20934 | 4.8506 | 1.2E-06 | 2.4E-05 |
| 578 | TTL | 43.57387 | 1.010762845 | 0.3661 | 2.76089 | 0.00576 | 0.03035 |
| 132 | ZMIZ1 | 50.454553 | 1.00373294 | 0.316 | 3.1764 | 0.00149 | 0.01012 |
| 351 | CD9 | 37.815103 | -1.000581291 | 0.33837 | -2.957 | 0.00311 | 0.01829 |
| 2 | SDF4 | 35.117418 | -1.002534254 | 0.37103 | -2.7021 | 0.00689 | 0.03484 |
| 119 | TAGLN2 | 37.364666 | -1.006975595 | 0.37941 | -2.6541 | 0.00795 | 0.03886 |
| 1139 | DLG1 | 31.958645 | -1.020712662 | 0.36977 | -2.7604 | 0.00577 | 0.03038 |
| 1008 | MAPK1 | 31.182963 | -1.020739129 | 0.3791 | -2.6926 | 0.00709 | 0.03564 |
| 438 | SLC7A1 | 31.269831 | -1.021616578 | 0.36671 | -2.7859 | 0.00534 | 0.02851 |
| 1400 | TMED4 | 38.279282 | -1.03232218 | 0.36725 | -2.811 | 0.00494 | 0.0268 |
| 1608 | MT-ND6 | 304.84119 | -1.036262436 | 0.27581 | -3.7571 | 0.00017 | 0.0017 |
| 278 | HSD17B12 | 42.636886 | -1.043339099 | 0.3522 | -2.9623 | 0.00305 | 0.01804 |
| 478 | CNIH1 | 56.651502 | -1.044441238 | 0.3317 | -3.1487 | 0.00164 | 0.01096 |
| 1289 | RNF145 | 29.137353 | -1.045449093 | 0.38352 | -2.7259 | 0.00641 | 0.03291 |
| 945 | NAA20 | 35.034897 | -1.054204266 | 0.37331 | -2.8239 | 0.00474 | 0.02591 |
| 94 | ATP1A1 | 42.007945 | -1.055806395 | 0.387 | -2.7282 | 0.00637 | 0.03275 |
| 1123 | NDUFB5 | 72.023144 | -1.056494071 | 0.34401 | -3.0711 | 0.00213 | 0.01356 |
| 1477 | PPP2CB | 30.971143 | -1.059226348 | 0.4064 | -2.6064 | 0.00915 | 0.04341 |
| 1535 | GLIS3 | 29.132305 | -1.062936363 | 0.41074 | -2.5878 | 0.00966 | 0.04519 |
| 1632 | ATP6AP2 | 80.249917 | -1.063282689 | 0.27614 | -3.8505 | 0.00012 | 0.00124 |
| 193 | CCDC6 | 28.253299 | -1.069630525 | 0.41038 | -2.6064 | 0.00915 | 0.04341 |
| 347 | WNK1 | 53.432379 | -1.070073129 | 0.40008 | -2.6746 | 0.00748 | 0.0371 |
| 743 | GPX4 | 49.809711 | -1.071999992 | 0.34926 | -3.0694 | 0.00215 | 0.01362 |
| 1642 | PRAF2 | 40.168665 | -1.072126301 | 0.35122 | -3.0525 | 0.00227 | 0.01429 |
| 1457 | REPIN1 | 48.607596 | -1.076526006 | 0.31 | -3.4726 | 0.00052 | 0.00418 |
| 486 | SYNE2 | 50.210407 | -1.079274078 | 0.36207 | -2.9808 | 0.00287 | 0.01721 |
| 733 | TPGS2 | 78.242851 | -1.083085012 | 0.3086 | -3.5097 | 0.00045 | 0.00375 |
| 14 | CLSTN1 | 43.451879 | -1.098423067 | 0.3452 | -3.182 | 0.00146 | 0.00996 |
| 973 | APP | 110.83191 | -1.099946758 | 0.24369 | -4.5137 | 6.4E-06 | 9.9E-05 |
| 784 | TMEM147 | 39.883762 | -1.10541372 | 0.36826 | -3.0017 | 0.00268 | 0.0163 |
| 721 | MYL12B | 42.830755 | -1.105957664 | 0.40513 | -2.7299 | 0.00634 | 0.03264 |

| | | | | | | | |
|---|---|---|---|---|---|---|---|
| 1107 | COPB2 | 46.892787 | -1.107575368 | 0.31626 | -3.5021 | 0.00046 | 0.00383 |
| 992 | CSTB | 29.515605 | -1.108274492 | 0.40757 | -2.7192 | 0.00654 | 0.03349 |
| 74 | DDAH1 | 50.28556 | -1.122703062 | 0.43863 | -2.5596 | 0.01048 | 0.04833 |
| 786 | CAPNS1 | 76.808953 | -1.123811467 | 0.28229 | -3.981 | 6.9E-05 | 0.00078 |
| 1288 | CYFIP2 | 28.186275 | -1.124058147 | 0.42034 | -2.6742 | 0.00749 | 0.03711 |
| 636 | CDK10 | 28.401599 | -1.124360413 | 0.43606 | -2.5785 | 0.00992 | 0.04627 |
| 1044 | BRK1 | 54.65802 | -1.124427288 | 0.41003 | -2.7423 | 0.0061 | 0.0317 |
| 493 | PNMA1 | 53.268942 | -1.125632833 | 0.30119 | -3.7372 | 0.00019 | 0.00182 |
| 921 | RETREG2 | 29.765929 | -1.128397352 | 0.42632 | -2.6468 | 0.00813 | 0.03946 |
| 55 | TMEM59 | 39.445085 | -1.128996869 | 0.35282 | -3.1999 | 0.00137 | 0.00949 |
| 432 | ULK1 | 26.321597 | -1.135252098 | 0.42459 | -2.6738 | 0.0075 | 0.03714 |
| 1607 | MT-CO3 | 1305.4959 | -1.137024572 | 0.20342 | -5.5895 | 2.3E-08 | 7.3E-07 |
| 327 | RDX | 32.371971 | -1.137206003 | 0.37394 | -3.0412 | 0.00236 | 0.01474 |
| 1115 | PFN2 | 150.418 | -1.138349344 | 0.28874 | -3.9425 | 8.1E-05 | 0.00089 |
| 335 | ARCN1 | 28.186519 | -1.144021365 | 0.39575 | -2.8908 | 0.00384 | 0.02168 |
| 1623 | PRDX4 | 57.588709 | -1.14666593 | 0.31704 | -3.6167 | 0.0003 | 0.00268 |
| 249 | TSPAN4 | 42.95314 | -1.147024684 | 0.4108 | -2.7922 | 0.00524 | 0.02809 |
| 1702 | ATP11C | 68.901708 | -1.149092416 | 0.34182 | -3.3616 | 0.00077 | 0.00586 |
| 389 | LRP1 | 30.060171 | -1.164131805 | 0.37453 | -3.1082 | 0.00188 | 0.01226 |
| 1654 | MSN | 32.747635 | -1.164685473 | 0.4541 | -2.5648 | 0.01032 | 0.0478 |
| 1560 | TLE4 | 27.155462 | -1.167996279 | 0.42709 | -2.7348 | 0.00624 | 0.03226 |
| 1418 | AKAP9 | 72.595595 | -1.168794579 | 0.294 | -3.9755 | 7E-05 | 0.0008 |
| 1147 | WDR1 | 53.295752 | -1.170426729 | 0.39058 | -2.9966 | 0.00273 | 0.01651 |
| 1403 | TMEM248 | 34.959292 | -1.179779268 | 0.42815 | -2.7555 | 0.00586 | 0.03073 |
| 991 | NDUFV3 | 31.11138 | -1.186801937 | 0.44199 | -2.6852 | 0.00725 | 0.03621 |
| 852 | RTN4 | 112.25797 | -1.187107608 | 0.22488 | -5.2789 | 1.3E-07 | 3.3E-06 |
| 294 | TRMT112 | 48.090826 | -1.188170219 | 0.34014 | -3.4932 | 0.00048 | 0.00393 |
| 1428 | ZNHIT1 | 29.675919 | -1.192669407 | 0.45867 | -2.6003 | 0.00932 | 0.04393 |
| 528 | TJP1 | 26.497217 | -1.196418035 | 0.45821 | -2.6111 | 0.00903 | 0.04295 |
| 878 | ARHGEF4 | 25.213096 | -1.197014774 | 0.43734 | -2.737 | 0.0062 | 0.03209 |
| 893 | GORASP2 | 35.144335 | -1.202774592 | 0.37155 | -3.2372 | 0.00121 | 0.00849 |
| 1611 | AKAP17A | 19.020923 | -1.207210174 | 0.46949 | -2.5713 | 0.01013 | 0.04709 |
| 830 | LAPTM4A | 64.521028 | -1.208506701 | 0.29589 | -4.0844 | 4.4E-05 | 0.00053 |
| 1298 | TMED9 | 54.224244 | -1.213589798 | 0.34288 | -3.5394 | 0.0004 | 0.00342 |
| 43 | AKIRIN1 | 36.74151 | -1.216222165 | 0.37159 | -3.273 | 0.00106 | 0.00762 |
| 1644 | MAGED1 | 131.41981 | -1.220129407 | 0.24226 | -5.0364 | 4.7E-07 | 1E-05 |
| 423 | MSI1 | 21.494461 | -1.223132454 | 0.46836 | -2.6115 | 0.00901 | 0.04292 |
| 690 | TSPOAP1 | 27.198281 | -1.229104699 | 0.44996 | -2.7316 | 0.0063 | 0.03251 |
| 1606 | MT-CO2 | 566.32025 | -1.229538003 | 0.2814 | -4.3694 | 1.2E-05 | 0.00018 |

| | | | | | | | |
|---|---|---|---|---|---|---|---|
| 1688 | OCRL | 23.513 | -1.22987974 | 0.45973 | -2.6752 | 0.00747 | 0.03707 |
| 1499 | NECAB1 | 24.68622 | -1.231521467 | 0.41729 | -2.9512 | 0.00317 | 0.01858 |
| 923 | OBSL1 | 61.173693 | -1.232259695 | 0.30601 | -4.0269 | 5.7E-05 | 0.00066 |
| 698 | PRKAR1A | 84.03436 | -1.233174594 | 0.30179 | -4.0862 | 4.4E-05 | 0.00053 |
| 1045 | SEC13 | 39.05694 | -1.239046662 | 0.37459 | -3.3077 | 0.00094 | 0.00684 |
| 856 | RAB1A | 76.541941 | -1.240007271 | 0.27352 | -4.5336 | 5.8E-06 | 9.1E-05 |
| 261 | CTR9 | 20.887359 | -1.240023407 | 0.45672 | -2.715 | 0.00663 | 0.03387 |
| 795 | RABAC1 | 28.655486 | -1.240273592 | 0.44783 | -2.7695 | 0.00561 | 0.02973 |
| 1328 | DNPH1 | 34.660071 | -1.242693212 | 0.42435 | -2.9284 | 0.00341 | 0.01969 |
| 825 | CHMP2A | 27.005249 | -1.245842388 | 0.45194 | -2.7566 | 0.00584 | 0.03067 |
| 977 | SOD1 | 113.19395 | -1.247124437 | 0.27922 | -4.4665 | 8E-06 | 0.00012 |
| 305 | GSTP1 | 170.81134 | -1.248131129 | 0.26645 | -4.6843 | 2.8E-06 | 4.8E-05 |
| 1011 | CABIN1 | 20.926694 | -1.260390677 | 0.48245 | -2.6125 | 0.00899 | 0.04285 |
| 18 | PLOD1 | 21.4962 | -1.26523211 | 0.45162 | -2.8016 | 0.00509 | 0.02745 |
| 1071 | DAG1 | 34.428265 | -1.271225439 | 0.36942 | -3.4411 | 0.00058 | 0.00461 |
| 1587 | COQ4 | 33.058316 | -1.271923144 | 0.39878 | -3.1895 | 0.00142 | 0.00976 |
| 834 | SNX17 | 25.733609 | -1.273791009 | 0.46478 | -2.7406 | 0.00613 | 0.03181 |
| 1355 | SERINC1 | 55.664624 | -1.275026034 | 0.31061 | -4.1049 | 4E-05 | 0.0005 |
| 1348 | GTF3C6 | 26.806016 | -1.278616505 | 0.47853 | -2.672 | 0.00754 | 0.0373 |
| 664 | STARD3 | 22.971035 | -1.280747218 | 0.46484 | -2.7553 | 0.00586 | 0.03074 |
| 1687 | SMARCA1 | 48.697093 | -1.281539197 | 0.38686 | -3.3127 | 0.00092 | 0.00674 |
| 465 | ARHGEF40 | 25.704373 | -1.281971944 | 0.46061 | -2.7832 | 0.00538 | 0.02868 |
| 1209 | SLC25A4 | 30.941152 | -1.285049885 | 0.48281 | -2.6616 | 0.00778 | 0.03818 |
| 1452 | HIPK2 | 29.634849 | -1.286995092 | 0.42069 | -3.0593 | 0.00222 | 0.01403 |
| 747 | SPPL2B | 29.790464 | -1.287244962 | 0.41923 | -3.0705 | 0.00214 | 0.01359 |
| 673 | GRN | 32.83181 | -1.295532145 | 0.44554 | -2.9078 | 0.00364 | 0.02078 |
| 802 | ERCC1 | 23.236502 | -1.302197499 | 0.44703 | -2.913 | 0.00358 | 0.0205 |
| 1305 | SSR1 | 32.018459 | -1.302405563 | 0.38968 | -3.3422 | 0.00083 | 0.0062 |
| 535 | GOLM2 | 29.75372 | -1.314956296 | 0.42429 | -3.0992 | 0.00194 | 0.01257 |
| 293 | OTUB1 | 33.978008 | -1.315286847 | 0.3892 | -3.3795 | 0.00073 | 0.00554 |
| 466 | DAD1 | 68.827929 | -1.315603458 | 0.27967 | -4.7041 | 2.6E-06 | 4.5E-05 |
| 1466 | ASAH1 | 36.2888 | -1.318092256 | 0.37116 | -3.5513 | 0.00038 | 0.0033 |
| 415 | TMEM263 | 21.470523 | -1.32042068 | 0.47953 | -2.7536 | 0.00589 | 0.03088 |
| 968 | STX16 | 39.340044 | -1.325601412 | 0.43841 | -3.0237 | 0.0025 | 0.01539 |
| 146 | DSTYK | 22.857736 | -1.32740859 | 0.47325 | -2.8049 | 0.00503 | 0.02719 |
| 230 | RAB11FIP2 | 21.931985 | -1.328514448 | 0.49361 | -2.6914 | 0.00712 | 0.03575 |
| 209 | TNKS2 | 29.509786 | -1.332979154 | 0.41314 | -3.2265 | 0.00125 | 0.00876 |
| 688 | MSI2 | 60.74637 | -1.333601258 | 0.32752 | -4.0718 | 4.7E-05 | 0.00056 |
| 53 | RNF11 | 19.953979 | -1.3397001 | 0.50693 | -2.6428 | 0.00822 | 0.03988 |

| | | | | | | | |
|---|---|---|---|---|---|---|---|
| 679 | NFE2L1 | 22.426556 | -1.339757935 | 0.45354 | -2.954 | 0.00314 | 0.01843 |
| 386 | STAT2 | 23.953474 | -1.342149926 | 0.48595 | -2.7619 | 0.00575 | 0.03028 |
| 1508 | EMC2 | 28.009344 | -1.346827483 | 0.48675 | -2.767 | 0.00566 | 0.0299 |
| 390 | MBD6 | 21.657154 | -1.347345281 | 0.49775 | -2.7069 | 0.00679 | 0.03446 |
| 1624 | ACOT9 | 25.157433 | -1.34996875 | 0.44324 | -3.0457 | 0.00232 | 0.01456 |
| 1659 | SLC16A2 | 20.358729 | -1.352930927 | 0.51086 | -2.6484 | 0.00809 | 0.03936 |
| 941 | DSTN | 79.936482 | -1.355732758 | 0.29484 | -4.5982 | 4.3E-06 | 6.9E-05 |
| 227 | PDZD8 | 18.452626 | -1.358737144 | 0.51772 | -2.6245 | 0.00868 | 0.04166 |
| 833 | ATRAID | 26.054238 | -1.359012883 | 0.46027 | -2.9526 | 0.00315 | 0.01851 |
| 1600 | TUBB4B | 36.505948 | -1.360098107 | 0.42986 | -3.1641 | 0.00156 | 0.01051 |
| 706 | WBP2 | 25.75835 | -1.360104143 | 0.46866 | -2.9021 | 0.00371 | 0.02106 |
| 414 | HSP90B1 | 111.75078 | -1.361150874 | 0.23672 | -5.7501 | 8.9E-09 | 3.2E-07 |
| 1054 | GOLGA4 | 33.764333 | -1.361936846 | 0.40617 | -3.3531 | 0.0008 | 0.006 |
| 659 | COPRS | 25.906389 | -1.363906783 | 0.45591 | -2.9916 | 0.00278 | 0.01672 |
| 1300 | TBC1D9B | 18.894005 | -1.372103376 | 0.52871 | -2.5952 | 0.00945 | 0.04442 |
| 137 | GLUL | 80.870198 | -1.373927742 | 0.28926 | -4.7498 | 2E-06 | 3.7E-05 |
| 1198 | SCOC | 36.124679 | -1.380364979 | 0.37892 | -3.6429 | 0.00027 | 0.00247 |
| 1669 | FAM199X | 19.813343 | -1.380556423 | 0.46604 | -2.9623 | 0.00305 | 0.01804 |
| 851 | ERLEC1 | 26.714823 | -1.383092101 | 0.4662 | -2.9668 | 0.00301 | 0.01784 |
| 938 | BTBD3 | 27.426725 | -1.387133165 | 0.41442 | -3.3471 | 0.00082 | 0.00611 |
| 666 | ORMDL3 | 21.372815 | -1.390375567 | 0.46405 | -2.9962 | 0.00273 | 0.01653 |
| 811 | NUCB1 | 28.076423 | -1.398582064 | 0.39827 | -3.5116 | 0.00045 | 0.00372 |
| 1303 | ECI2 | 18.652608 | -1.399991175 | 0.48239 | -2.9022 | 0.00371 | 0.02106 |
| 490 | ACTN1 | 28.753988 | -1.401132213 | 0.49151 | -2.8506 | 0.00436 | 0.02415 |
| 215 | ENTPD7 | 14.779681 | -1.401521014 | 0.53895 | -2.6005 | 0.00931 | 0.04392 |
| 949 | NOL4L | 16.362945 | -1.409585612 | 0.53758 | -2.6221 | 0.00874 | 0.04185 |
| 1480 | FGFR1 | 29.650798 | -1.411118083 | 0.4473 | -3.1547 | 0.00161 | 0.01079 |
| 946 | APMAP | 35.755992 | -1.412726711 | 0.37801 | -3.7373 | 0.00019 | 0.00182 |
| 1639 | HDAC6 | 35.475945 | -1.41462061 | 0.38067 | -3.7162 | 0.0002 | 0.00195 |
| 1274 | CTNNA1 | 48.4674 | -1.415252203 | 0.31777 | -4.4536 | 8.4E-06 | 0.00013 |
| 1641 | SLC35A2 | 18.458107 | -1.41733427 | 0.54728 | -2.5898 | 0.0096 | 0.04502 |
| 1594 | SURF4 | 33.636051 | -1.418437805 | 0.42899 | -3.3064 | 0.00094 | 0.00687 |
| 551 | UACA | 21.380792 | -1.418441624 | 0.51761 | -2.7404 | 0.00614 | 0.03181 |
| 1022 | IFT27 | 17.881536 | -1.431870302 | 0.54963 | -2.6052 | 0.00918 | 0.0435 |
| 1380 | CEP43 | 16.294145 | -1.432396486 | 0.53416 | -2.6816 | 0.00733 | 0.03652 |
| 634 | COTL1 | 51.121991 | -1.434243119 | 0.34726 | -4.1301 | 3.6E-05 | 0.00045 |
| 385 | CD63 | 96.152917 | -1.436547386 | 0.25969 | -5.5317 | 3.2E-08 | 9.6E-07 |
| 623 | BCAR1 | 28.837443 | -1.440445006 | 0.42459 | -3.3925 | 0.00069 | 0.00533 |
| 375 | ACVR1B | 19.548165 | -1.44251579 | 0.5087 | -2.8357 | 0.00457 | 0.02517 |

| | | | | | | | |
|---|---|---|---|---|---|---|---|
| 398 | OSBPL8 | 35.753036 | -1.443496413 | 0.37135 | -3.8871 | 0.0001 | 0.00108 |
| 1295 | LMAN2 | 23.132172 | -1.444330846 | 0.45915 | -3.1456 | 0.00166 | 0.01106 |
| 1550 | EBLN3P | 20.770603 | -1.451183476 | 0.49642 | -2.9233 | 0.00346 | 0.01992 |
| 9 | VAMP3 | 19.99767 | -1.451927452 | 0.52596 | -2.7605 | 0.00577 | 0.03038 |
| 167 | GALNT2 | 28.629754 | -1.453551812 | 0.48053 | -3.0249 | 0.00249 | 0.01535 |
| 291 | SLC3A2 | 30.760082 | -1.454383325 | 0.39619 | -3.671 | 0.00024 | 0.00226 |
| 336 | CENATAC | 26.145034 | -1.456275834 | 0.45847 | -3.1764 | 0.00149 | 0.01012 |
| 1230 | NADK2 | 15.58858 | -1.459348167 | 0.54281 | -2.6885 | 0.00718 | 0.03601 |
| 1345 | CD164 | 61.850452 | -1.460035376 | 0.31347 | -4.6577 | 3.2E-06 | 5.3E-05 |
| 1121 | PHC3 | 17.512196 | -1.466060342 | 0.50109 | -2.9257 | 0.00344 | 0.0198 |
| 534 | SERF2 | 252.3659 | -1.467023943 | 0.21802 | -6.7288 | 1.7E-11 | 1.2E-09 |
| 1712 | ATP6AP1 | 16.48668 | -1.467502219 | 0.52087 | -2.8174 | 0.00484 | 0.02637 |
| 1002 | PRMT2 | 40.645811 | -1.467710163 | 0.34702 | -4.2295 | 2.3E-05 | 0.00031 |
| 734 | ME2 | 57.201385 | -1.468255065 | 0.50522 | -2.9062 | 0.00366 | 0.02084 |
| 716 | CSNK1D | 65.812406 | -1.468401323 | 0.31328 | -4.6871 | 2.8E-06 | 4.8E-05 |
| 220 | TRIM8 | 29.093635 | -1.471243876 | 0.48257 | -3.0488 | 0.0023 | 0.01444 |
| 860 | WDR54 | 20.70982 | -1.47654366 | 0.50529 | -2.9222 | 0.00348 | 0.01998 |
| 850 | ASB3 | 18.616712 | -1.482131401 | 0.55779 | -2.6571 | 0.00788 | 0.03859 |
| 533 | PDIA3 | 44.149215 | -1.486525552 | 0.34018 | -4.3699 | 1.2E-05 | 0.00018 |
| 1612 | CD99 | 37.644841 | -1.491072644 | 0.40506 | -3.6811 | 0.00023 | 0.00218 |
| 592 | TMEM219 | 13.291993 | -1.491393666 | 0.55878 | -2.669 | 0.00761 | 0.03755 |
| 1650 | NBDY | 19.263131 | -1.493417506 | 0.53728 | -2.7796 | 0.00544 | 0.02893 |
| 1696 | RTL8C | 96.174355 | -1.49467906 | 0.26499 | -5.6405 | 1.7E-08 | 5.6E-07 |
| 1268 | C5orf15 | 26.077656 | -1.497287389 | 0.4517 | -3.3148 | 0.00092 | 0.0067 |
| 416 | CRY1 | 19.252226 | -1.498449707 | 0.53381 | -2.8071 | 0.005 | 0.02708 |
| 162 | CAPN2 | 31.921494 | -1.499430057 | 0.42018 | -3.5686 | 0.00036 | 0.00312 |
| 1638 | WDR13 | 29.587182 | -1.501037232 | 0.4134 | -3.631 | 0.00028 | 0.00257 |
| 319 | SRSF8 | 17.487574 | -1.501968503 | 0.53911 | -2.786 | 0.00534 | 0.02851 |
| 1605 | MT-CO1 | 272.00805 | -1.504823706 | 0.29039 | -5.1821 | 2.2E-07 | 5.2E-06 |
| 511 | CCDC88C | 17.021022 | -1.514242199 | 0.52911 | -2.8619 | 0.00421 | 0.02342 |
| 1340 | HMGN3 | 80.589206 | -1.514367726 | 0.28864 | -5.2466 | 1.5E-07 | 3.9E-06 |
| 1572 | PTGR1 | 16.237859 | -1.518708957 | 0.58951 | -2.5762 | 0.00999 | 0.04652 |
| 1142 | LRPAP1 | 39.463115 | -1.521419385 | 0.37763 | -4.0288 | 5.6E-05 | 0.00066 |
| 153 | CD46 | 42.062842 | -1.524883083 | 0.37939 | -4.0193 | 5.8E-05 | 0.00068 |
| 446 | TSC22D1 | 118.00326 | -1.530446947 | 0.27529 | -5.5594 | 2.7E-08 | 8.4E-07 |
| 554 | NPTN | 36.425164 | -1.532287575 | 0.45243 | -3.3868 | 0.00071 | 0.00542 |
| 1099 | HEG1 | 17.478954 | -1.534829566 | 0.56817 | -2.7013 | 0.00691 | 0.03489 |
| 437 | ENSG00000182957 | 17.162376 | -1.537909912 | 0.50927 | -3.0198 | 0.00253 | 0.01554 |
| 1374 | CNKSR3 | 15.127627 | -1.539597253 | 0.59479 | -2.5885 | 0.00964 | 0.04516 |

| | | | | | | | |
|---|---|---|---|---|---|---|---|
| 134 | RABGAP1L | 13.807496 | -1.542076128 | 0.5731 | -2.6908 | 0.00713 | 0.0358 |
| 1359 | SLC18B1 | 17.061979 | -1.543311007 | 0.53995 | -2.8582 | 0.00426 | 0.02364 |
| 631 | HSBP1 | 112.69471 | -1.547838387 | 0.25253 | -6.1292 | 8.8E-10 | 4E-08 |
| 1468 | SORBS3 | 14.439883 | -1.551033336 | 0.60529 | -2.5625 | 0.01039 | 0.04803 |
| 993 | PTTG1IP | 43.504403 | -1.554433129 | 0.34337 | -4.527 | 6E-06 | 9.3E-05 |
| 391 | CTDSP2 | 22.50416 | -1.559687337 | 0.49457 | -3.1536 | 0.00161 | 0.01081 |
| 1101 | RPN1 | 30.192366 | -1.56170109 | 0.42297 | -3.6923 | 0.00022 | 0.00211 |
| 268 | NUCB2 | 16.202531 | -1.562411539 | 0.59171 | -2.6405 | 0.00828 | 0.04007 |
| 746 | CSNK1G2 | 22.408858 | -1.57508953 | 0.47083 | -3.3454 | 0.00082 | 0.00614 |
| 598 | AMFR | 13.372504 | -1.576212805 | 0.56561 | -2.7868 | 0.00532 | 0.02846 |
| 1622 | RPS6KA3 | 22.38549 | -1.582762605 | 0.44429 | -3.5625 | 0.00037 | 0.00318 |
| 125 | UHMK1 | 16.130099 | -1.586193676 | 0.52345 | -3.0302 | 0.00244 | 0.01516 |
| 904 | SPATS2L | 31.88314 | -1.58671199 | 0.37794 | -4.1983 | 2.7E-05 | 0.00035 |
| 1591 | SH3GLB2 | 33.067947 | -1.588983735 | 0.37755 | -4.2087 | 2.6E-05 | 0.00034 |
| 1196 | NDUFC1 | 43.496606 | -1.589602444 | 0.37461 | -4.2433 | 2.2E-05 | 0.00029 |
| 808 | CYTH2 | 41.330121 | -1.589602872 | 0.37752 | -4.2107 | 2.5E-05 | 0.00033 |
| 1409 | GTF2IRD1 | 14.284392 | -1.59593074 | 0.58465 | -2.7297 | 0.00634 | 0.03264 |
| 735 | POLI | 16.870688 | -1.597018128 | 0.53662 | -2.976 | 0.00292 | 0.01742 |
| 626 | MAF | 56.71595 | -1.600735203 | 0.43203 | -3.7051 | 0.00021 | 0.00202 |
| 1032 | TSPO | 22.346017 | -1.601774217 | 0.5021 | -3.1901 | 0.00142 | 0.00975 |
| 807 | KDELR1 | 37.356857 | -1.608329521 | 0.35822 | -4.4898 | 7.1E-06 | 0.00011 |
| 1443 | CALU | 69.905162 | -1.614900161 | 0.33232 | -4.8595 | 1.2E-06 | 2.3E-05 |
| 829 | MBOAT2 | 26.308091 | -1.615315984 | 0.48916 | -3.3022 | 0.00096 | 0.00696 |
| 296 | EHD1 | 13.875678 | -1.617481403 | 0.61019 | -2.6508 | 0.00803 | 0.03916 |
| 1575 | WHRN | 19.4076 | -1.626754918 | 0.52946 | -3.0725 | 0.00212 | 0.01351 |
| 135 | RASAL2 | 15.997443 | -1.626810843 | 0.55371 | -2.938 | 0.0033 | 0.01921 |
| 309 | ARHGEF17 | 22.880146 | -1.631997272 | 0.4861 | -3.3574 | 0.00079 | 0.00592 |
| 556 | LOXL1 | 49.397177 | -1.635227625 | 0.34408 | -4.7524 | 2E-06 | 3.7E-05 |
| 559 | TSPAN3 | 149.44449 | -1.635667318 | 0.20945 | -7.8093 | 5.7E-15 | 8.2E-13 |
| 835 | IFT172 | 15.457294 | -1.637604056 | 0.58104 | -2.8184 | 0.00483 | 0.02632 |
| 1678 | TMEM164 | 37.437834 | -1.637920436 | 0.45552 | -3.5957 | 0.00032 | 0.00287 |
| 1264 | FBN2 | 29.294165 | -1.640307676 | 0.48406 | -3.3887 | 0.0007 | 0.00539 |
| 63 | LEPROT | 26.584537 | -1.644763436 | 0.54566 | -3.0143 | 0.00258 | 0.01576 |
| 248 | POLR2L | 48.179943 | -1.647179102 | 0.39697 | -4.1494 | 3.3E-05 | 0.00042 |
| 942 | RRBP1 | 11.648645 | -1.652591332 | 0.61886 | -2.6704 | 0.00758 | 0.03747 |
| 978 | CFAP298 | 20.756373 | -1.65363107 | 0.55231 | -2.994 | 0.00275 | 0.01661 |
| 715 | CCDC57 | 16.487939 | -1.654011079 | 0.55091 | -3.0023 | 0.00268 | 0.01627 |
| 420 | ALDH2 | 13.464056 | -1.659218374 | 0.61157 | -2.7131 | 0.00667 | 0.03402 |
| 382 | CALCOCO1 | 18.849579 | -1.660635561 | 0.57383 | -2.894 | 0.0038 | 0.0215 |

| | | | | | | | |
|---|---|---|---|---|---|---|---|
| 395 | TMBIM4 | 16.780077 | -1.663375295 | 0.51987 | -3.1996 | 0.00138 | 0.0095 |
| 1671 | MORC4 | 15.801795 | -1.67276605 | 0.56212 | -2.9758 | 0.00292 | 0.01742 |
| 1618 | HCCS | 10.913254 | -1.674888954 | 0.65529 | -2.556 | 0.01059 | 0.04871 |
| 251 | CD81 | 111.11121 | -1.675362682 | 0.28561 | -5.8659 | 4.5E-09 | 1.7E-07 |
| 1571 | TMEM245 | 33.599408 | -1.681990576 | 0.38845 | -4.33 | 1.5E-05 | 0.00021 |
| 537 | SPPL2A | 15.060687 | -1.682791236 | 0.61285 | -2.7458 | 0.00604 | 0.03141 |
| 561 | MESD | 16.209687 | -1.683811099 | 0.57455 | -2.9307 | 0.00338 | 0.01957 |
| 1395 | GARS1-DT | 20.426706 | -1.685353015 | 0.52949 | -3.183 | 0.00146 | 0.00993 |
| 1494 | NCOA2 | 12.935051 | -1.68910683 | 0.62969 | -2.6824 | 0.00731 | 0.03646 |
| 312 | UVRAG | 21.019173 | -1.68973375 | 0.5977 | -2.8271 | 0.0047 | 0.02571 |
| 93 | HIPK1 | 25.937697 | -1.693114228 | 0.4291 | -3.9458 | 8E-05 | 0.00088 |
| 1458 | SMARCD3 | 16.146728 | -1.693816176 | 0.54693 | -3.097 | 0.00196 | 0.01264 |
| 298 | SCYL1 | 17.765359 | -1.694597427 | 0.52594 | -3.222 | 0.00127 | 0.00886 |
| 1113 | WWTR1 | 13.071074 | -1.695367571 | 0.64003 | -2.6489 | 0.00808 | 0.03933 |
| 163 | FBXO28 | 11.013131 | -1.695579124 | 0.65963 | -2.5705 | 0.01016 | 0.04718 |
| 671 | NBR1 | 25.165259 | -1.700040016 | 0.49514 | -3.4335 | 0.0006 | 0.00471 |
| 325 | TMEM123 | 33.118863 | -1.700283727 | 0.39473 | -4.3075 | 1.7E-05 | 0.00023 |
| 575 | JMJD8 | 15.874898 | -1.702184638 | 0.60102 | -2.8322 | 0.00462 | 0.02539 |
| 657 | ALDH3A2 | 28.487048 | -1.704050211 | 0.51937 | -3.281 | 0.00103 | 0.00743 |
| 159 | MARK1 | 35.983931 | -1.710043415 | 0.4447 | -3.8454 | 0.00012 | 0.00126 |
| 963 | PARD6B | 9.7637971 | -1.711225692 | 0.66947 | -2.5561 | 0.01059 | 0.04871 |
| 843 | THADA | 13.435196 | -1.711317819 | 0.61311 | -2.7912 | 0.00525 | 0.02814 |
| 1674 | PSMD10 | 27.930367 | -1.713093894 | 0.42554 | -4.0257 | 5.7E-05 | 0.00067 |
| 1080 | FLNB | 28.538174 | -1.722669684 | 0.45473 | -3.7883 | 0.00015 | 0.00153 |
| 820 | NAT14 | 12.32437 | -1.723098905 | 0.66792 | -2.5798 | 0.00989 | 0.04617 |
| 1096 | FSTL1 | 162.15288 | -1.72378841 | 0.21699 | -7.9439 | 2E-15 | 2.9E-13 |
| 459 | TM9SF2 | 47.345033 | -1.728308087 | 0.32711 | -5.2836 | 1.3E-07 | 3.2E-06 |
| 1050 | NGLY1 | 31.574058 | -1.728606447 | 0.38732 | -4.463 | 8.1E-06 | 0.00012 |
| 848 | MCFD2 | 26.851466 | -1.729092787 | 0.46183 | -3.744 | 0.00018 | 0.00178 |
| 745 | SCAMP4 | 12.723383 | -1.742420851 | 0.63188 | -2.7575 | 0.00582 | 0.03062 |
| 1604 | MT-ND1 | 298.8848 | -1.743894135 | 0.33134 | -5.2632 | 1.4E-07 | 3.5E-06 |
| 285 | TNKS1BP1 | 19.214974 | -1.744321342 | 0.54642 | -3.1923 | 0.00141 | 0.00969 |
| 1585 | CDK9 | 12.436255 | -1.745062999 | 0.65821 | -2.6512 | 0.00802 | 0.03915 |
| 1444 | UBE2H | 29.163068 | -1.746416177 | 0.4711 | -3.7071 | 0.00021 | 0.002 |
| 616 | PRMT7 | 25.18835 | -1.74832727 | 0.46883 | -3.7291 | 0.00019 | 0.00187 |
| 691 | VMP1 | 26.912687 | -1.749411431 | 0.60526 | -2.8903 | 0.00385 | 0.02169 |
| 273 | RCN1 | 22.839029 | -1.75680242 | 0.466 | -3.7699 | 0.00016 | 0.00163 |
| 1052 | CMTM6 | 10.8026 | -1.757980786 | 0.6564 | -2.6782 | 0.0074 | 0.0368 |
| 498 | TMED10 | 47.482825 | -1.758775865 | 0.33788 | -5.2053 | 1.9E-07 | 4.7E-06 |

| | | | | | | | |
|---|---|---|---|---|---|---|---|
| 1389 | HDAC9 | 13.090947 | -1.759857849 | 0.64894 | -2.7119 | 0.00669 | 0.03408 |
| 513 | IFI27L1 | 23.501569 | -1.776595152 | 0.49599 | -3.582 | 0.00034 | 0.003 |
| 306 | KMT5B | 20.494661 | -1.780792012 | 0.49766 | -3.5783 | 0.00035 | 0.00303 |
| 838 | YPEL5 | 27.180723 | -1.782328185 | 0.43481 | -4.0991 | 4.1E-05 | 0.00051 |
| 854 | AHSA2P | 36.994958 | -1.782862463 | 0.36765 | -4.8493 | 1.2E-06 | 2.4E-05 |
| 1474 | FZD3 | 62.644718 | -1.783519653 | 0.32347 | -5.5137 | 3.5E-08 | 1.1E-06 |
| 491 | PSEN1 | 14.727621 | -1.785865672 | 0.59635 | -2.9947 | 0.00275 | 0.01659 |
| 204 | ANXA11 | 15.490996 | -1.786231227 | 0.55635 | -3.2106 | 0.00132 | 0.00917 |
| 558 | SCAMP2 | 14.82399 | -1.787383578 | 0.63453 | -2.8169 | 0.00485 | 0.02639 |
| 186 | ZEB1 | 12.479292 | -1.790485607 | 0.67937 | -2.6355 | 0.0084 | 0.0405 |
| 1574 | RGS3 | 10.727583 | -1.792438795 | 0.64042 | -2.7989 | 0.00513 | 0.02761 |
| 412 | ARL1 | 34.723512 | -1.794096278 | 0.40178 | -4.4654 | 8E-06 | 0.00012 |
| 1046 | ATP2B2 | 26.465209 | -1.79455334 | 0.46847 | -3.8307 | 0.00013 | 0.00133 |
| 1199 | TBC1D9 | 13.33006 | -1.799162399 | 0.59112 | -3.0436 | 0.00234 | 0.01464 |
| 54 | GPX7 | 21.228593 | -1.799197579 | 0.46471 | -3.8717 | 0.00011 | 0.00115 |
| 809 | PLEKHA4 | 14.70763 | -1.79920469 | 0.59042 | -3.0473 | 0.00231 | 0.0145 |
| 1602 | NSMF | 40.192239 | -1.79970222 | 0.35588 | -5.057 | 4.3E-07 | 9.4E-06 |
| 1647 | FAM104B | 12.229129 | -1.801435795 | 0.61928 | -2.9089 | 0.00363 | 0.02071 |
| 1689 | IGSF1 | 10.019302 | -1.810007395 | 0.65811 | -2.7503 | 0.00595 | 0.03111 |
| 77 | SELENOF | 36.623284 | -1.813352012 | 0.39856 | -4.5497 | 5.4E-06 | 8.6E-05 |
| 76 | ODF2L | 14.47342 | -1.815923956 | 0.64437 | -2.8182 | 0.00483 | 0.02633 |
| 543 | ZNF280D | 10.160282 | -1.81844223 | 0.7004 | -2.5963 | 0.00942 | 0.04434 |
| 519 | EML1 | 13.940778 | -1.82294045 | 0.64427 | -2.8295 | 0.00466 | 0.02555 |
| 1013 | ZNRF3 | 11.910587 | -1.826518702 | 0.62209 | -2.9361 | 0.00332 | 0.0193 |
| 1299 | MGAT4B | 63.652316 | -1.82779444 | 0.36533 | -5.0031 | 5.6E-07 | 1.2E-05 |
| 1658 | ZMYM3 | 27.174029 | -1.828347804 | 0.46156 | -3.9612 | 7.5E-05 | 0.00084 |
| 81 | TMED5 | 30.732747 | -1.833958031 | 0.52794 | -3.4738 | 0.00051 | 0.00417 |
| 264 | RRAS2 | 11.396917 | -1.840750165 | 0.70807 | -2.5997 | 0.00933 | 0.044 |
| 1588 | ODF2 | 26.812351 | -1.84129471 | 0.46062 | -3.9974 | 6.4E-05 | 0.00073 |
| 1213 | PLEKHG4B | 10.709518 | -1.842299104 | 0.64784 | -2.8437 | 0.00446 | 0.0246 |
| 371 | TMBIM6 | 133.50789 | -1.842482785 | 0.21463 | -8.5844 | 9.1E-18 | 1.9E-15 |
| 1273 | EGR1 | 17.340674 | -1.842833907 | 0.69786 | -2.6407 | 0.00827 | 0.04007 |
| 913 | FN1 | 15.47202 | -1.846104972 | 0.63393 | -2.9121 | 0.00359 | 0.02055 |
| 38 | ADPRS | 23.367479 | -1.850383508 | 0.50782 | -3.6438 | 0.00027 | 0.00246 |
| 304 | PITPNM1 | 11.097722 | -1.851049646 | 0.67375 | -2.7474 | 0.00601 | 0.0313 |
| 732 | GALNT1 | 15.662056 | -1.853202589 | 0.56552 | -3.277 | 0.00105 | 0.00752 |
| 214 | TCTN3 | 14.498719 | -1.857730227 | 0.5917 | -3.1397 | 0.00169 | 0.01124 |
| 234 | ATE1 | 12.160032 | -1.860332274 | 0.60796 | -3.06 | 0.00221 | 0.01402 |
| 1438 | AASS | 25.708322 | -1.861905315 | 0.49162 | -3.7873 | 0.00015 | 0.00153 |

| | | | | | | | |
|---|---|---|---|---|---|---|---|
| 1491 | SGK3 | 11.344812 | -1.868061148 | 0.69077 | -2.7043 | 0.00684 | 0.03466 |
| 217 | SFXN3 | 11.116194 | -1.869150611 | 0.68008 | -2.7484 | 0.00599 | 0.03122 |
| 88 | AMPD2 | 33.727631 | -1.871079845 | 0.40621 | -4.6062 | 4.1E-06 | 6.7E-05 |
| 1404 | AUTS2 | 20.728104 | -1.874185214 | 0.56429 | -3.3213 | 0.0009 | 0.00659 |
| 876 | SLC35F5 | 30.241007 | -1.876689617 | 0.47888 | -3.9189 | 8.9E-05 | 0.00097 |
| 439 | UBL3 | 16.109723 | -1.879151262 | 0.56319 | -3.3366 | 0.00085 | 0.00631 |
| 1067 | SHISA5 | 114.81711 | -1.882584309 | 0.43372 | -4.3405 | 1.4E-05 | 0.0002 |
| 1370 | MTHFD1L | 12.718293 | -1.887072858 | 0.62534 | -3.0177 | 0.00255 | 0.01562 |
| 1525 | GSDMD | 9.6336956 | -1.888835605 | 0.7087 | -2.6652 | 0.00769 | 0.03785 |
| 1573 | HSDL2 | 11.902516 | -1.891110455 | 0.65129 | -2.9037 | 0.00369 | 0.02097 |
| 150 | SLC41A1 | 18.386602 | -1.892817254 | 0.59375 | -3.1879 | 0.00143 | 0.0098 |
| 903 | PGAP1 | 57.126456 | -1.904935579 | 0.39105 | -4.8713 | 1.1E-06 | 2.2E-05 |
| 1307 | TMEM14C | 34.918554 | -1.906260008 | 0.52692 | -3.6178 | 0.0003 | 0.00268 |
| 695 | TEX2 | 11.422171 | -1.906336425 | 0.71278 | -2.6745 | 0.00748 | 0.0371 |
| 364 | STK38L | 14.453657 | -1.907649318 | 0.61571 | -3.0983 | 0.00195 | 0.01259 |
| 403 | ATP2B1 | 24.716455 | -1.908526938 | 0.50476 | -3.781 | 0.00016 | 0.00157 |
| 429 | SLC15A4 | 26.22522 | -1.909174619 | 0.49896 | -3.8263 | 0.00013 | 0.00135 |
| 253 | STIM1 | 9.6237905 | -1.912122278 | 0.68041 | -2.8102 | 0.00495 | 0.02685 |
| 532 | ZNF106 | 32.128164 | -1.913023417 | 0.50725 | -3.7713 | 0.00016 | 0.00162 |
| 658 | CPD | 12.276938 | -1.919269523 | 0.66839 | -2.8715 | 0.00409 | 0.02283 |
| 106 | S100A6 | 74.321794 | -1.920233644 | 0.38388 | -5.0022 | 5.7E-07 | 1.2E-05 |
| 566 | MFGE8 | 19.317226 | -1.924548201 | 0.56534 | -3.4042 | 0.00066 | 0.00516 |
| 814 | MED25 | 28.594967 | -1.924650164 | 0.40756 | -4.7223 | 2.3E-06 | 4.1E-05 |
| 1000 | COL6A2 | 43.293191 | -1.925939595 | 0.5356 | -3.5959 | 0.00032 | 0.00287 |
| 1252 | ADGRV1 | 24.698619 | -1.930225369 | 0.48536 | -3.9769 | 7E-05 | 0.00079 |
| 884 | TANC1 | 23.962548 | -1.930444256 | 0.54683 | -3.5303 | 0.00042 | 0.00351 |
| 1130 | LPP | 22.146841 | -1.934355039 | 0.56253 | -3.4387 | 0.00058 | 0.00464 |
| 463 | TMCO3 | 10.311562 | -1.934473654 | 0.74329 | -2.6026 | 0.00925 | 0.04371 |
| 1673 | RBM41 | 20.396461 | -1.935938102 | 0.60214 | -3.2151 | 0.0013 | 0.00904 |
| 1648 | MAGEH1 | 18.252207 | -1.937065896 | 0.54851 | -3.5315 | 0.00041 | 0.00351 |
| 317 | PICALM | 23.301225 | -1.937409103 | 0.46028 | -4.2092 | 2.6E-05 | 0.00034 |
| 1652 | ARHGEF9 | 13.107258 | -1.941556341 | 0.61775 | -3.1429 | 0.00167 | 0.01115 |
| 1162 | COMMD8 | 14.734398 | -1.942615511 | 0.59841 | -3.2463 | 0.00117 | 0.00825 |
| 720 | MYL12A | 45.610968 | -1.945818787 | 0.38151 | -5.1003 | 3.4E-07 | 7.6E-06 |
| 1104 | NPHP3 | 10.59508 | -1.948919231 | 0.6703 | -2.9075 | 0.00364 | 0.02079 |
| 1301 | MGAT1 | 14.18702 | -1.953329527 | 0.5629 | -3.4701 | 0.00052 | 0.00422 |
| 1363 | AHI1 | 21.166971 | -1.964548601 | 0.62686 | -3.134 | 0.00172 | 0.01141 |
| 944 | RIN2 | 18.593587 | -1.964967062 | 0.54298 | -3.6189 | 0.0003 | 0.00267 |
| 922 | DNAJB2 | 15.717843 | -1.9662569 | 0.56954 | -3.4523 | 0.00056 | 0.00447 |

| | | | | | | | |
|---|---|---|---|---|---|---|---|
| 1460 | DYNC2I1 | 20.947375 | -1.966949307 | 0.48036 | -4.0947 | 4.2E-05 | 0.00051 |
| 924 | SERPINE2 | 11.611454 | -1.96717023 | 0.72636 | -2.7083 | 0.00676 | 0.03434 |
| 1636 | ARAF | 30.581374 | -1.973449967 | 0.44156 | -4.4692 | 7.8E-06 | 0.00012 |
| 243 | RIC8A | 13.648924 | -1.974049365 | 0.69963 | -2.8216 | 0.00478 | 0.02608 |
| 1191 | CAMK2D | 19.523161 | -1.974744916 | 0.58992 | -3.3475 | 0.00082 | 0.0061 |
| 1435 | SMIM30 | 10.849371 | -1.975378552 | 0.7383 | -2.6756 | 0.00746 | 0.03706 |
| 726 | IMPACT | 11.554692 | -1.976166537 | 0.69528 | -2.8423 | 0.00448 | 0.0247 |
| 200 | PSAP | 112.1479 | -1.977005387 | 0.27372 | -7.2229 | 5.1E-13 | 4.9E-11 |
| 34 | IFI6 | 29.287413 | -1.977225689 | 0.56364 | -3.508 | 0.00045 | 0.00377 |
| 30 | TMEM50A | 20.315323 | -1.97747943 | 0.53792 | -3.6761 | 0.00024 | 0.00222 |
| 969 | NPEPL1 | 16.885059 | -1.980339699 | 0.55971 | -3.5381 | 0.0004 | 0.00343 |
| 470 | SLC22A17 | 90.706286 | -1.980836162 | 0.3056 | -6.4819 | 9.1E-11 | 5.2E-09 |
| 1371 | AKAP12 | 35.935168 | -1.982922342 | 0.55982 | -3.5421 | 0.0004 | 0.00339 |
| 283 | CSTPP1 | 13.287853 | -1.982933832 | 0.61474 | -3.2256 | 0.00126 | 0.00878 |
| 503 | POMT2 | 9.0814485 | -1.98395275 | 0.74301 | -2.6702 | 0.00758 | 0.03747 |
| 955 | BLCAP | 23.400988 | -1.989926212 | 0.50273 | -3.9583 | 7.5E-05 | 0.00084 |
| 1475 | SARAF | 44.164058 | -1.990693421 | 0.38554 | -5.1634 | 2.4E-07 | 5.6E-06 |
| 1547 | GALT | 11.316912 | -1.990919611 | 0.64664 | -3.0789 | 0.00208 | 0.01331 |
| 275 | CD59 | 50.74945 | -2.003423835 | 0.42025 | -4.7672 | 1.9E-06 | 3.4E-05 |
| 1312 | ATXN1 | 9.9384777 | -2.00943048 | 0.68659 | -2.9267 | 0.00343 | 0.01977 |
| 1534 | RFX3 | 21.244199 | -2.015825528 | 0.52694 | -3.8255 | 0.00013 | 0.00135 |
| 260 | RNF141 | 14.614673 | -2.016127202 | 0.58426 | -3.4507 | 0.00056 | 0.00448 |
| 284 | SLC39A13 | 26.219503 | -2.018355924 | 0.55866 | -3.6129 | 0.0003 | 0.00272 |
| 827 | PXDN | 36.500145 | -2.021847759 | 0.41943 | -4.8205 | 1.4E-06 | 2.7E-05 |
| 906 | ALS2 | 14.263156 | -2.02387207 | 0.65809 | -3.0754 | 0.0021 | 0.01344 |
| 1478 | ERLIN2 | 19.101856 | -2.024919422 | 0.53265 | -3.8016 | 0.00014 | 0.00146 |
| 254 | FHIP1B | 13.652257 | -2.028212907 | 0.59687 | -3.3981 | 0.00068 | 0.00525 |
| 917 | CTDSP1 | 14.677003 | -2.029885428 | 0.56945 | -3.5646 | 0.00036 | 0.00316 |
| 815 | VRK3 | 14.20706 | -2.033924151 | 0.7626 | -2.6671 | 0.00765 | 0.0377 |
| 1490 | GGH | 22.164635 | -2.034174429 | 0.55035 | -3.6962 | 0.00022 | 0.00208 |
| 637 | VPS9D1 | 15.454724 | -2.036248944 | 0.62798 | -3.2425 | 0.00118 | 0.00836 |
| 1114 | RNF13 | 11.368444 | -2.036906919 | 0.68527 | -2.9724 | 0.00295 | 0.01756 |
| 1615 | PNPLA4 | 11.438251 | -2.039836651 | 0.64902 | -3.1429 | 0.00167 | 0.01115 |
| 1565 | CTSL | 13.358795 | -2.04093639 | 0.62513 | -3.2648 | 0.0011 | 0.00782 |
| 595 | TENT4B | 16.052798 | -2.041010799 | 0.6389 | -3.1946 | 0.0014 | 0.00962 |
| 789 | PLD3 | 63.598223 | -2.042565 | 0.30331 | -6.7342 | 1.6E-11 | 1.1E-09 |
| 59 | PATJ | 11.11268 | -2.044354832 | 0.66901 | -3.0558 | 0.00224 | 0.01416 |
| 344 | APLP2 | 183.24334 | -2.046372304 | 0.22207 | -9.2149 | 3.1E-20 | 1E-17 |
| 964 | BCAS4 | 9.6976854 | -2.051006494 | 0.72584 | -2.8257 | 0.00472 | 0.02579 |

| | | | | | | | |
|---|---|---|---|---|---|---|---|
| 1247 | SCAMP1-AS1 | 9.3417096 | -2.052283787 | 0.7528 | -2.7262 | 0.00641 | 0.0329 |
| 516 | CLMN | 15.756779 | -2.054136566 | 0.66827 | -3.0738 | 0.00211 | 0.01348 |
| 457 | ABCC4 | 10.984384 | -2.054485809 | 0.68532 | -2.9978 | 0.00272 | 0.01646 |
| 1194 | FGF2 | 8.2952695 | -2.055592646 | 0.8035 | -2.5583 | 0.01052 | 0.04845 |
| 1664 | ARMCX2 | 37.000783 | -2.055601193 | 0.46568 | -4.4142 | 1E-05 | 0.00015 |
| 593 | FBRS | 23.319713 | -2.05633972 | 0.52767 | -3.897 | 9.7E-05 | 0.00105 |
| 1270 | CXCL14 | 200.90964 | -2.056633187 | 0.31905 | -6.4462 | 1.1E-10 | 6.4E-09 |
| 1291 | ATP6V0E1 | 18.961811 | -2.057194269 | 0.61296 | -3.3562 | 0.00079 | 0.00594 |
| 67 | GADD45A | 9.9082264 | -2.060086316 | 0.8094 | -2.5452 | 0.01092 | 0.04985 |
| 599 | BBS2 | 11.89288 | -2.064236852 | 0.71364 | -2.8926 | 0.00382 | 0.02158 |
| 1068 | P4HTM | 18.673307 | -2.064811363 | 0.54134 | -3.8143 | 0.00014 | 0.0014 |
| 1024 | KCTD17 | 14.303737 | -2.065054279 | 0.57638 | -3.5828 | 0.00034 | 0.00299 |
| 35 | COL16A1 | 9.5374717 | -2.066576638 | 0.80748 | -2.5593 | 0.01049 | 0.04836 |
| 1459 | GALNT11 | 37.28295 | -2.066703216 | 0.62731 | -3.2945 | 0.00099 | 0.00712 |
| 574 | WDR90 | 32.685353 | -2.066740639 | 0.38974 | -5.3029 | 1.1E-07 | 3E-06 |
| 689 | CUEDC1 | 14.01451 | -2.067985036 | 0.58992 | -3.5055 | 0.00046 | 0.00379 |
| 1504 | RIDA | 9.1893305 | -2.068467109 | 0.79687 | -2.5957 | 0.00944 | 0.04438 |
| 867 | UNC50 | 19.695892 | -2.072451551 | 0.51062 | -4.0587 | 4.9E-05 | 0.00059 |
| 864 | ST3GAL5 | 11.115013 | -2.075659986 | 0.77869 | -2.6656 | 0.00769 | 0.03783 |
| 1225 | CDH6 | 19.903944 | -2.077436631 | 0.56015 | -3.7087 | 0.00021 | 0.00199 |
| 1245 | F2R | 9.2586721 | -2.077542825 | 0.79339 | -2.6186 | 0.00883 | 0.04221 |
| 853 | CFAP36 | 14.372342 | -2.081132201 | 0.64685 | -3.2173 | 0.00129 | 0.00898 |
| 1381 | ERMARD | 10.914788 | -2.082686557 | 0.68205 | -3.0536 | 0.00226 | 0.01426 |
| 936 | TMEM230 | 22.842458 | -2.084142483 | 0.52577 | -3.964 | 7.4E-05 | 0.00083 |
| 1195 | SPRY1 | 12.087209 | -2.084826875 | 0.66365 | -3.1415 | 0.00168 | 0.01119 |
| 742 | BSG | 116.01424 | -2.085313717 | 0.27602 | -7.555 | 4.2E-14 | 5E-12 |
| 642 | PITPNA | 13.277843 | -2.088395387 | 0.64816 | -3.222 | 0.00127 | 0.00886 |
| 418 | IFT81 | 10.572942 | -2.090684956 | 0.75856 | -2.7561 | 0.00585 | 0.03068 |
| 719 | LPIN2 | 18.256873 | -2.094295715 | 0.55573 | -3.7686 | 0.00016 | 0.00163 |
| 1259 | LOX | 18.364408 | -2.094524227 | 0.64491 | -3.2478 | 0.00116 | 0.00822 |
| 785 | PSENEN | 36.926307 | -2.097096979 | 0.47581 | -4.4074 | 1E-05 | 0.00015 |
| 1567 | NINJ1 | 10.834045 | -2.099428396 | 0.71381 | -2.9412 | 0.00327 | 0.01905 |
| 315 | PRCP | 19.140849 | -2.100984802 | 0.52939 | -3.9687 | 7.2E-05 | 0.00081 |
| 381 | CSAD | 19.134637 | -2.107212759 | 0.55001 | -3.8312 | 0.00013 | 0.00133 |
| 648 | ATP1B2 | 30.960078 | -2.107268737 | 0.46407 | -4.5408 | 5.6E-06 | 8.8E-05 |
| 760 | TMEM205 | 33.378036 | -2.107313743 | 0.4206 | -5.0103 | 5.4E-07 | 1.2E-05 |
| 751 | MLLT1 | 52.964451 | -2.111736771 | 0.33786 | -6.2504 | 4.1E-10 | 2E-08 |
| 1667 | TCEAL7 | 12.48408 | -2.113675976 | 0.76331 | -2.7691 | 0.00562 | 0.02976 |
| 1531 | PLEC | 17.335662 | -2.114671819 | 0.6333 | -3.3391 | 0.00084 | 0.00626 |

| | | | | | | | |
|---|---|---|---|---|---|---|---|
| 434 | IFT88 | 8.4877649 | -2.116497304 | 0.77613 | -2.727 | 0.00639 | 0.03283 |
| 590 | QPRT | 11.532465 | -2.119520242 | 0.76875 | -2.7571 | 0.00583 | 0.03064 |
| 858 | PCYOX1 | 17.17018 | -2.122886192 | 0.5835 | -3.6382 | 0.00027 | 0.0025 |
| 122 | PEA15 | 29.087249 | -2.123772689 | 0.47914 | -4.4325 | 9.3E-06 | 0.00014 |
| 300 | EFEMP2 | 10.096743 | -2.124122145 | 0.7121 | -2.9829 | 0.00286 | 0.01714 |
| 1529 | NRBP2 | 43.672268 | -2.132915835 | 0.34304 | -6.2176 | 5E-10 | 2.4E-08 |
| 223 | PDCD4 | 12.504122 | -2.133904398 | 0.67738 | -3.1502 | 0.00163 | 0.01092 |
| 1411 | POR | 16.669426 | -2.13772127 | 0.53928 | -3.9641 | 7.4E-05 | 0.00083 |
| 12 | H6PD | 8.065406 | -2.137930108 | 0.80474 | -2.6567 | 0.00789 | 0.03862 |
| 905 | CFLAR | 10.393332 | -2.137936414 | 0.7617 | -2.8068 | 0.005 | 0.02708 |
| 33 | AHDC1 | 12.87768 | -2.143482514 | 0.61427 | -3.4895 | 0.00048 | 0.00397 |
| 488 | ATP6V1D | 13.776981 | -2.147106685 | 0.60056 | -3.5752 | 0.00035 | 0.00305 |
| 1105 | AMOTL2 | 19.531056 | -2.15090498 | 0.5706 | -3.7695 | 0.00016 | 0.00163 |
| 1481 | ADAM9 | 15.367821 | -2.151817921 | 0.5947 | -3.6183 | 0.0003 | 0.00267 |
| 1532 | PPP1R16A | 19.875127 | -2.159532252 | 0.53459 | -4.0396 | 5.4E-05 | 0.00063 |
| 572 | ARRDC4 | 13.289325 | -2.161571704 | 0.6995 | -3.0902 | 0.002 | 0.01289 |
| 919 | TTLL4 | 29.84796 | -2.162490685 | 0.58497 | -3.6967 | 0.00022 | 0.00208 |
| 1495 | TRAM1 | 33.539093 | -2.162961529 | 0.43378 | -4.9863 | 6.2E-07 | 1.3E-05 |
| 515 | IFI27L2 | 11.06842 | -2.164682825 | 0.80552 | -2.6873 | 0.0072 | 0.03606 |
| 1337 | FAM135A | 13.130168 | -2.165966285 | 0.78836 | -2.7474 | 0.00601 | 0.0313 |
| 1141 | ENSG00000288380 | 8.2886559 | -2.168384285 | 0.75564 | -2.8696 | 0.00411 | 0.02293 |
| 1464 | TRMT9B | 25.92541 | -2.169879553 | 0.77171 | -2.8118 | 0.00493 | 0.02676 |
| 1619 | TMSB4X | 157.77991 | -2.171482901 | 0.34605 | -6.2751 | 3.5E-10 | 1.7E-08 |
| 1066 | PLXNB1 | 50.554915 | -2.173359093 | 0.34351 | -6.327 | 2.5E-10 | 1.3E-08 |
| 1589 | ZER1 | 11.081969 | -2.175436595 | 0.73448 | -2.9619 | 0.00306 | 0.01805 |
| 280 | ACCS | 14.769071 | -2.176187245 | 0.67389 | -3.2293 | 0.00124 | 0.0087 |
| 553 | NEO1 | 17.180016 | -2.176617516 | 0.61016 | -3.5673 | 0.00036 | 0.00314 |
| 1661 | HMGN5 | 11.26458 | -2.176737539 | 0.73399 | -2.9656 | 0.00302 | 0.0179 |
| 495 | ALDH6A1 | 11.311088 | -2.178449444 | 0.69997 | -3.1122 | 0.00186 | 0.01213 |
| 384 | ITGA7 | 22.73695 | -2.182487485 | 0.52111 | -4.1881 | 2.8E-05 | 0.00036 |
| 502 | IRF2BPL | 28.565537 | -2.187838398 | 0.47378 | -4.6178 | 3.9E-06 | 6.3E-05 |
| 1697 | ZNF449 | 12.673973 | -2.192589689 | 0.70189 | -3.1239 | 0.00179 | 0.01173 |
| 1711 | FLNA | 58.687529 | -2.193942063 | 0.36658 | -5.9848 | 2.2E-09 | 9E-08 |
| 1432 | LAMB1 | 58.962058 | -2.200001517 | 0.31316 | -7.0252 | 2.1E-12 | 1.7E-10 |
| 930 | IQCA1 | 16.88247 | -2.204210849 | 0.56669 | -3.8896 | 0.0001 | 0.00108 |
| 597 | RPGRIP1L | 7.8885155 | -2.204684511 | 0.80424 | -2.7413 | 0.00612 | 0.03179 |
| 1584 | NIBAN2 | 28.267263 | -2.20569318 | 0.52541 | -4.198 | 2.7E-05 | 0.00035 |
| 1426 | TRIM56 | 13.340479 | -2.212907591 | 0.64069 | -3.454 | 0.00055 | 0.00444 |
| 1021 | MYH9 | 62.957802 | -2.215424635 | 0.39912 | -5.5508 | 2.8E-08 | 8.8E-07 |

| | | | | | | | |
|---|---|---|---|---|---|---|---|
| 797 | CEACAM19 | 8.4300203 | -2.219748742 | 0.7783 | -2.8521 | 0.00434 | 0.02408 |
| 552 | BBS4 | 17.973826 | -2.21976269 | 0.62168 | -3.5706 | 0.00036 | 0.0031 |
| 358 | HEBP1 | 10.8609 | -2.236537059 | 0.72897 | -3.0681 | 0.00215 | 0.01367 |
| 931 | COL6A3 | 9.6427665 | -2.237739347 | 0.83636 | -2.6756 | 0.00746 | 0.03706 |
| 966 | ZNF217 | 10.341122 | -2.240860958 | 0.70374 | -3.1842 | 0.00145 | 0.00989 |
| 1152 | PACRGL | 7.5985147 | -2.244740226 | 0.85036 | -2.6397 | 0.0083 | 0.0401 |
| 1484 | SLC20A2 | 14.542002 | -2.24924806 | 0.71848 | -3.1306 | 0.00174 | 0.01153 |
| 676 | HEXIM1 | 10.099863 | -2.251597015 | 0.69776 | -3.2269 | 0.00125 | 0.00875 |
| 1609 | MT-TP | 19.290553 | -2.260013232 | 0.52327 | -4.319 | 1.6E-05 | 0.00022 |
| 1586 | AK1 | 7.6981685 | -2.266550304 | 0.82523 | -2.7466 | 0.00602 | 0.03135 |
| 115 | LMNA | 22.306586 | -2.267247012 | 0.50847 | -4.4589 | 8.2E-06 | 0.00012 |
| 274 | HIPK3 | 34.723151 | -2.269043101 | 0.4382 | -5.1781 | 2.2E-07 | 5.3E-06 |
| 1122 | FNDC3B | 11.87089 | -2.269619789 | 0.77333 | -2.9349 | 0.00334 | 0.01937 |
| 158 | RAB3GAP2 | 56.898314 | -2.271473903 | 0.43825 | -5.183 | 2.2E-07 | 5.2E-06 |
| 1492 | CSPP1 | 27.400835 | -2.273872371 | 0.54621 | -4.163 | 3.1E-05 | 0.0004 |
| 1023 | TST | 6.4779162 | -2.275862222 | 0.84972 | -2.6784 | 0.0074 | 0.0368 |
| 730 | B4GALT6 | 12.844536 | -2.279563285 | 0.73245 | -3.1123 | 0.00186 | 0.01213 |
| 573 | FAM234A | 22.175073 | -2.291904489 | 0.49906 | -4.5925 | 4.4E-06 | 7.1E-05 |
| 510 | DGLUCY | 7.7676276 | -2.295847305 | 0.86022 | -2.6689 | 0.00761 | 0.03755 |
| 1470 | TNFRSF10B | 24.573147 | -2.298127725 | 0.76352 | -3.0099 | 0.00261 | 0.01594 |
| 1506 | FZD6 | 24.803166 | -2.298303995 | 0.4908 | -4.6828 | 2.8E-06 | 4.8E-05 |
| 271 | METTL15 | 7.5407472 | -2.303749789 | 0.81583 | -2.8238 | 0.00475 | 0.02591 |
| 320 | SESN3 | 14.497301 | -2.309113728 | 0.60546 | -3.8138 | 0.00014 | 0.0014 |
| 1304 | RREB1 | 7.1901734 | -2.30925066 | 0.8795 | -2.6256 | 0.00865 | 0.04154 |
| 82 | GCLM | 11.686693 | -2.312365128 | 0.65498 | -3.5305 | 0.00041 | 0.00351 |
| 456 | GPC6 | 7.6643179 | -2.312874643 | 0.83532 | -2.7689 | 0.00563 | 0.02977 |
| 1410 | RHBDD2 | 31.742845 | -2.313770094 | 0.44297 | -5.2233 | 1.8E-07 | 4.3E-06 |
| 303 | TMEM134 | 11.694239 | -2.32155128 | 0.84428 | -2.7497 | 0.00596 | 0.03115 |
| 8 | NPHP4 | 12.708847 | -2.325555006 | 0.67509 | -3.4448 | 0.00057 | 0.00456 |
| 1061 | TMEM42 | 9.642108 | -2.326165119 | 0.78452 | -2.9651 | 0.00303 | 0.01792 |
| 57 | PLPP3 | 10.871746 | -2.329062947 | 0.71309 | -3.2661 | 0.00109 | 0.00779 |
| 935 | SLC23A2 | 25.264308 | -2.330090627 | 0.52875 | -4.4068 | 1E-05 | 0.00015 |
| 207 | PANK1 | 8.472755 | -2.3318165 | 0.86369 | -2.6998 | 0.00694 | 0.03501 |
| 80 | EVI5 | 11.859865 | -2.332810057 | 0.69113 | -3.3754 | 0.00074 | 0.00562 |
| 1037 | PLXNB2 | 65.578707 | -2.346880934 | 0.35769 | -6.5613 | 5.3E-11 | 3.2E-09 |
| 1580 | DENND1A | 9.3152414 | -2.356733477 | 0.74843 | -3.1489 | 0.00164 | 0.01096 |
| 171 | SCCPDH | 16.399087 | -2.370006361 | 0.6631 | -3.5741 | 0.00035 | 0.00306 |
| 61 | CACHD1 | 35.988687 | -2.370587193 | 0.39979 | -5.9296 | 3E-09 | 1.2E-07 |
| 1222 | DAP | 30.547502 | -2.371490186 | 0.42547 | -5.5738 | 2.5E-08 | 7.8E-07 |

| | | | | | | | |
|---|---|---|---|---|---|---|---|
| 1202 | MAP9 | 26.718765 | -2.37475306 | 0.50376 | -4.714 | 2.4E-06 | 4.3E-05 |
| 78 | HS2ST1 | 15.517888 | -2.380891101 | 0.59431 | -4.0062 | 6.2E-05 | 0.00071 |
| 151 | ENSG00000286619 | 18.07286 | -2.382907433 | 0.54162 | -4.3996 | 1.1E-05 | 0.00016 |
| 1463 | CTSB | 38.017363 | -2.386085801 | 0.37113 | -6.4292 | 1.3E-10 | 7.1E-09 |
| 975 | ADAMTS1 | 21.674108 | -2.386551221 | 0.49284 | -4.8425 | 1.3E-06 | 2.5E-05 |
| 419 | TCTN1 | 16.669473 | -2.388829817 | 0.71641 | -3.3345 | 0.00085 | 0.00635 |
| 1347 | AK9 | 8.8169472 | -2.390378614 | 0.78425 | -3.048 | 0.0023 | 0.01447 |
| 170 | EFCAB2 | 8.9255038 | -2.397605929 | 0.75002 | -3.1967 | 0.00139 | 0.00956 |
| 621 | ZFHX3 | 17.482965 | -2.401794962 | 0.60616 | -3.9623 | 7.4E-05 | 0.00083 |
| 957 | PKIG | 8.6496002 | -2.411286562 | 0.88262 | -2.732 | 0.0063 | 0.03248 |
| 999 | COL6A1 | 156.9155 | -2.41325687 | 0.2908 | -8.2987 | 1.1E-16 | 1.9E-14 |
| 1479 | PLPP5 | 24.920234 | -2.415379115 | 0.50455 | -4.7872 | 1.7E-06 | 3.1E-05 |
| 985 | GET1 | 26.112353 | -2.416148864 | 0.61889 | -3.904 | 9.5E-05 | 0.00103 |
| 1041 | RABL2B | 18.02256 | -2.417965926 | 0.62091 | -3.8942 | 9.9E-05 | 0.00106 |
| 39 | COL8A2 | 7.6663336 | -2.420357176 | 0.9142 | -2.6475 | 0.00811 | 0.03943 |
| 447 | ITM2B | 101.9714 | -2.431547027 | 0.26083 | -9.3223 | 1.1E-20 | 4E-18 |
| 980 | TMEM50B | 20.35001 | -2.431885279 | 0.51671 | -4.7065 | 2.5E-06 | 4.4E-05 |
| 500 | IFT43 | 16.019034 | -2.434992037 | 0.60488 | -4.0256 | 5.7E-05 | 0.00067 |
| 1631 | RPGR | 42.928557 | -2.439045317 | 0.46792 | -5.2125 | 1.9E-07 | 4.5E-06 |
| 1578 | GSN | 29.49467 | -2.441181835 | 0.48022 | -5.0834 | 3.7E-07 | 8.2E-06 |
| 1079 | IL17RD | 29.367891 | -2.444258779 | 0.52372 | -4.6671 | 3.1E-06 | 5.2E-05 |
| 357 | C1RL | 8.2706287 | -2.44629281 | 0.82639 | -2.9602 | 0.00307 | 0.01812 |
| 388 | STAT6 | 11.623114 | -2.448989006 | 0.80719 | -3.034 | 0.00241 | 0.01503 |
| 1412 | HSPB1 | 46.064579 | -2.451747514 | 0.3468 | -7.0697 | 1.6E-12 | 1.3E-10 |
| 1192 | SEC24D | 13.715937 | -2.454990766 | 0.61385 | -3.9993 | 6.4E-05 | 0.00073 |
| 996 | LINC00205 | 11.130493 | -2.459780803 | 0.81666 | -3.012 | 0.0026 | 0.01586 |
| 407 | CEP83 | 19.322835 | -2.467422104 | 0.54639 | -4.5159 | 6.3E-06 | 9.8E-05 |
| 896 | TTN | 10.516401 | -2.469099274 | 0.72318 | -3.4142 | 0.00064 | 0.005 |
| 62 | JAK1 | 30.414803 | -2.469927581 | 0.49312 | -5.0088 | 5.5E-07 | 1.2E-05 |
| 1665 | ZMAT1 | 8.8145986 | -2.473879757 | 0.84149 | -2.9399 | 0.00328 | 0.01912 |
| 890 | BBS5 | 10.581937 | -2.477032968 | 0.72992 | -3.3935 | 0.00069 | 0.00532 |
| 1242 | MAST4 | 9.9438129 | -2.477806161 | 0.90777 | -2.7296 | 0.00634 | 0.03265 |
| 1385 | SLC29A4 | 8.7647443 | -2.480799281 | 0.75201 | -3.2989 | 0.00097 | 0.00704 |
| 203 | TMEM254 | 9.873731 | -2.483115575 | 0.8289 | -2.9957 | 0.00274 | 0.01655 |
| 780 | ENSG00000268362 | 11.205754 | -2.484651228 | 0.73556 | -3.3779 | 0.00073 | 0.00557 |
| 196 | REEP3 | 14.523946 | -2.491697273 | 0.64312 | -3.8744 | 0.00011 | 0.00114 |
| 1375 | SNX9 | 9.0874754 | -2.496457415 | 0.79081 | -3.1568 | 0.00159 | 0.01072 |
| 1036 | PPARA | 8.4819658 | -2.499736361 | 0.84951 | -2.9426 | 0.00325 | 0.019 |
| 692 | MRC2 | 15.559259 | -2.501971871 | 0.59129 | -4.2314 | 2.3E-05 | 0.00031 |

| | | | | | | | |
|---|---|---|---|---|---|---|---|
| 1322 | TAPBP | 23.780818 | -2.502075725 | 0.65021 | -3.8481 | 0.00012 | 0.00125 |
| 1431 | BCAP29 | 9.5956871 | -2.507666946 | 0.73633 | -3.4056 | 0.00066 | 0.00514 |
| 1076 | DOCK3 | 14.95999 | -2.510302347 | 0.64718 | -3.8789 | 0.0001 | 0.00112 |
| 680 | CALCOCO2 | 16.369192 | -2.513319865 | 0.59264 | -4.2409 | 2.2E-05 | 0.0003 |
| 26 | HMGCL | 5.5278257 | -2.518712995 | 0.98748 | -2.5506 | 0.01075 | 0.04926 |
| 796 | GRIK5 | 8.6403658 | -2.527009408 | 0.8637 | -2.9258 | 0.00344 | 0.0198 |
| 1357 | LAMA2 | 7.5967935 | -2.531553283 | 0.82032 | -3.0861 | 0.00203 | 0.01305 |
| 571 | SLCO3A1 | 12.630557 | -2.532999116 | 0.74852 | -3.384 | 0.00071 | 0.00548 |
| 1244 | IQGAP2 | 10.983427 | -2.533249203 | 0.73094 | -3.4658 | 0.00053 | 0.00427 |
| 1250 | ZFYVE16 | 16.258722 | -2.533351654 | 0.61515 | -4.1183 | 3.8E-05 | 0.00047 |
| 1488 | FAM110B | 5.6118878 | -2.534599215 | 0.98713 | -2.5676 | 0.01024 | 0.0475 |
| 768 | SLC27A1 | 10.737263 | -2.547082187 | 0.72023 | -3.5365 | 0.00041 | 0.00345 |
| 136 | RALGPS2 | 7.2543492 | -2.548069125 | 0.84165 | -3.0275 | 0.00247 | 0.01525 |
| 98 | NOTCH2 | 24.970404 | -2.548913217 | 0.5902 | -4.3187 | 1.6E-05 | 0.00022 |
| 393 | GNS | 34.359211 | -2.548944948 | 0.46685 | -5.4599 | 4.8E-08 | 1.4E-06 |
| 396 | MDM1 | 13.330024 | -2.551677145 | 0.75567 | -3.3767 | 0.00073 | 0.0056 |
| 174 | CAMK1D | 16.037274 | -2.552636527 | 0.71393 | -3.5755 | 0.00035 | 0.00305 |
| 1149 | CC2D2A | 14.19926 | -2.552702527 | 0.8047 | -3.1722 | 0.00151 | 0.01024 |
| 1208 | TENM3 | 10.382517 | -2.553551263 | 0.74284 | -3.4376 | 0.00059 | 0.00466 |
| 346 | GLB1L2 | 15.53533 | -2.561245591 | 0.67495 | -3.7947 | 0.00015 | 0.0015 |
| 765 | NOTCH3 | 14.487891 | -2.562302908 | 0.69773 | -3.6723 | 0.00024 | 0.00225 |
| 1700 | TM9SF5P | 14.140486 | -2.562826052 | 0.69552 | -3.6848 | 0.00023 | 0.00215 |
| 1473 | SCARA3 | 10.962695 | -2.566892913 | 0.76099 | -3.3731 | 0.00074 | 0.00565 |
| 687 | STXBP4 | 7.9478406 | -2.568317894 | 0.84474 | -3.0404 | 0.00236 | 0.01476 |
| 560 | CTSH | 5.5755881 | -2.56922681 | 0.94812 | -2.7098 | 0.00673 | 0.03423 |
| 399 | CSRP2 | 9.699478 | -2.572558671 | 0.77997 | -3.2983 | 0.00097 | 0.00705 |
| 1254 | CAST | 11.32767 | -2.579044675 | 0.88724 | -2.9068 | 0.00365 | 0.02082 |
| 800 | NECTIN2 | 12.285642 | -2.582118643 | 0.7287 | -3.5434 | 0.00039 | 0.00338 |
| 889 | STK39 | 35.794327 | -2.585186003 | 0.41971 | -6.1595 | 7.3E-10 | 3.4E-08 |
| 492 | DNAL1 | 9.6090489 | -2.58823327 | 0.76024 | -3.4045 | 0.00066 | 0.00516 |
| 1540 | IFT74 | 8.4441336 | -2.593087096 | 0.92909 | -2.791 | 0.00525 | 0.02814 |
| 981 | MRPS6 | 94.704335 | -2.608974145 | 0.27049 | -9.6453 | 5.1E-22 | 2.4E-19 |
| 505 | DIO2 | 11.226723 | -2.611195886 | 0.84751 | -3.081 | 0.00206 | 0.01323 |
| 109 | S100A13 | 8.9193771 | -2.615024908 | 0.83548 | -3.13 | 0.00175 | 0.01154 |
| 138 | LAMC1 | 16.253365 | -2.615287928 | 0.68481 | -3.819 | 0.00013 | 0.00138 |
| 1502 | TP53INP1 | 11.216386 | -2.627889344 | 0.90839 | -2.8929 | 0.00382 | 0.02157 |
| 1693 | ENSG00000283638 | 5.555212 | -2.631291758 | 0.97247 | -2.7058 | 0.00681 | 0.03455 |
| 1599 | AGPAT2 | 8.3367936 | -2.632247589 | 0.88979 | -2.9583 | 0.00309 | 0.01823 |
| 1453 | DENND2A | 7.0550731 | -2.634573883 | 0.9138 | -2.8831 | 0.00394 | 0.02212 |

| | | | | | | | |
|---|---|---|---|---|---|---|---|
| 222 | ADD3 | 16.768138 | -2.636524386 | 0.63839 | -4.13 | 3.6E-05 | 0.00045 |
| 23 | IFFO2 | 6.7594679 | -2.637204739 | 0.87807 | -3.0034 | 0.00267 | 0.01623 |
| 1026 | LGALS1 | 23.435925 | -2.637919278 | 0.50038 | -5.2718 | 1.4E-07 | 3.4E-06 |
| 307 | CPT1A | 13.03466 | -2.642028736 | 0.6806 | -3.8819 | 0.0001 | 0.00111 |
| 926 | ITM2C | 38.309367 | -2.646902042 | 0.45636 | -5.8 | 6.6E-09 | 2.4E-07 |
| 538 | MYO5C | 13.181004 | -2.650664422 | 0.68058 | -3.8947 | 9.8E-05 | 0.00106 |
| 897 | FAM171B | 6.9916758 | -2.653159807 | 0.88209 | -3.0078 | 0.00263 | 0.01604 |
| 1505 | SPAG1 | 10.428958 | -2.657537002 | 0.80365 | -3.3068 | 0.00094 | 0.00686 |
| 406 | PLXNC1 | 7.9731245 | -2.664956664 | 0.89768 | -2.9687 | 0.00299 | 0.01775 |
| 1386 | C1GALT1 | 13.246059 | -2.669282136 | 0.79116 | -3.3739 | 0.00074 | 0.00564 |
| 256 | TPP1 | 29.646903 | -2.670641963 | 0.46814 | -5.7048 | 1.2E-08 | 4E-07 |
| 1695 | MOSPD1 | 13.096868 | -2.672493338 | 0.75418 | -3.5436 | 0.00039 | 0.00338 |
| 793 | LTBP4 | 15.126165 | -2.682702033 | 0.6971 | -3.8484 | 0.00012 | 0.00125 |
| 1521 | PTP4A3 | 19.933744 | -2.684499396 | 0.59465 | -4.5144 | 6.3E-06 | 9.8E-05 |
| 464 | GAS6 | 14.202699 | -2.686436405 | 0.85814 | -3.1305 | 0.00174 | 0.01153 |
| 1257 | SEMA6A | 10.645233 | -2.686777644 | 0.98695 | -2.7223 | 0.00648 | 0.03325 |
| 202 | ZNF503 | 13.714012 | -2.689247225 | 0.9368 | -2.8707 | 0.0041 | 0.02286 |
| 250 | CTSD | 57.334918 | -2.699204741 | 0.45265 | -5.9631 | 2.5E-09 | 1E-07 |
| 588 | KATNIP | 14.42453 | -2.702924885 | 0.75288 | -3.5901 | 0.00033 | 0.00293 |
| 579 | CLUAP1 | 11.037242 | -2.706063249 | 0.69885 | -3.8721 | 0.00011 | 0.00115 |
| 995 | POFUT2 | 21.503766 | -2.708232134 | 0.58546 | -4.6258 | 3.7E-06 | 6.1E-05 |
| 582 | LITAF | 35.648587 | -2.713556775 | 0.48761 | -5.565 | 2.6E-08 | 8.2E-07 |
| 542 | MNS1 | 5.040119 | -2.716219822 | 1.05745 | -2.5687 | 0.01021 | 0.04739 |
| 757 | ADAMTS10 | 10.344154 | -2.73723226 | 0.81429 | -3.3615 | 0.00078 | 0.00586 |
| 1319 | DDAH2 | 12.492592 | -2.737716978 | 0.68443 | -4 | 6.3E-05 | 0.00073 |
| 1029 | CBY1 | 21.697539 | -2.740749431 | 0.68553 | -3.998 | 6.4E-05 | 0.00073 |
| 1269 | TCF7 | 13.581576 | -2.743504483 | 0.70679 | -3.8816 | 0.0001 | 0.00111 |
| 1512 | ENPP2 | 42.95824 | -2.748312308 | 0.59447 | -4.6231 | 3.8E-06 | 6.2E-05 |
| 602 | CPNE2 | 19.643957 | -2.74870445 | 0.54202 | -5.0712 | 4E-07 | 8.7E-06 |
| 1030 | DNAL4 | 7.9327452 | -2.767244624 | 0.90481 | -3.0584 | 0.00223 | 0.01404 |
| 224 | TCF7L2 | 7.6125888 | -2.767638532 | 0.84919 | -3.2592 | 0.00112 | 0.00796 |
| 740 | C2CD4C | 7.0089385 | -2.768839774 | 0.93699 | -2.955 | 0.00313 | 0.01839 |
| 1518 | EFR3A | 12.455863 | -2.769005 | 0.65657 | -4.2174 | 2.5E-05 | 0.00032 |
| 113 | MUC1 | 5.4300583 | -2.772428798 | 1.02802 | -2.6969 | 0.007 | 0.03524 |
| 316 | ANKRD42 | 11.830732 | -2.772779394 | 0.70073 | -3.957 | 7.6E-05 | 0.00085 |
| 1323 | SYNGAP1 | 19.256197 | -2.773804782 | 0.59956 | -4.6264 | 3.7E-06 | 6.1E-05 |
| 1276 | CXXC5 | 19.619558 | -2.786602373 | 0.61889 | -4.5026 | 6.7E-06 | 0.0001 |
| 1117 | MLF1 | 20.698354 | -2.791076607 | 0.56902 | -4.9051 | 9.3E-07 | 1.9E-05 |
| 1660 | MAGT1 | 12.664182 | -2.792321113 | 0.68308 | -4.0878 | 4.4E-05 | 0.00053 |

| | | | | | | | |
|---|---|---|---|---|---|---|---|
| 1498 | DECR1 | 16.807785 | -2.801224299 | 0.64549 | -4.3397 | 1.4E-05 | 0.0002 |
| 943 | DZANK1 | 11.904722 | -2.806517326 | 0.72654 | -3.8629 | 0.00011 | 0.00119 |
| 1074 | SEMA3B | 8.3399819 | -2.811705028 | 0.83322 | -3.3745 | 0.00074 | 0.00563 |
| 1566 | CDK20 | 9.0030557 | -2.814412028 | 0.84581 | -3.3275 | 0.00088 | 0.00648 |
| 1350 | FAM229B | 16.778412 | -2.815975171 | 0.59642 | -4.7215 | 2.3E-06 | 4.1E-05 |
| 441 | SPART | 9.1588225 | -2.816145898 | 0.79929 | -3.5233 | 0.00043 | 0.00358 |
| 710 | MXRA7 | 9.5042085 | -2.819768152 | 0.86842 | -3.247 | 0.00117 | 0.00824 |
| 1335 | LMBRD1 | 18.102401 | -2.822326643 | 0.66779 | -4.2264 | 2.4E-05 | 0.00031 |
| 366 | FGD4 | 6.4652137 | -2.825567387 | 1.00926 | -2.7996 | 0.00512 | 0.02756 |
| 1646 | MAGED2 | 117.26936 | -2.838284303 | 0.25645 | -11.068 | 1.8E-28 | 1.9E-25 |
| 1542 | ACO1 | 11.696746 | -2.84136276 | 0.79427 | -3.5773 | 0.00035 | 0.00304 |
| 369 | TMEM117 | 9.8627995 | -2.841379676 | 1.01563 | -2.7977 | 0.00515 | 0.02768 |
| 693 | TANC2 | 14.599945 | -2.841863983 | 0.67429 | -4.2146 | 2.5E-05 | 0.00033 |
| 1527 | FAM83H | 11.61629 | -2.84669985 | 0.85863 | -3.3154 | 0.00092 | 0.00669 |
| 685 | SPATA20 | 20.754094 | -2.847046913 | 0.59308 | -4.8004 | 1.6E-06 | 3E-05 |
| 392 | GIHCG | 13.675731 | -2.853245038 | 0.67005 | -4.2583 | 2.1E-05 | 0.00028 |
| 287 | FAM111A | 12.609446 | -2.853326367 | 0.74935 | -3.8078 | 0.00014 | 0.00143 |
| 1596 | COL5A1 | 28.330489 | -2.858123775 | 0.58642 | -4.8739 | 1.1E-06 | 2.2E-05 |
| 286 | SERPING1 | 21.895929 | -2.86035545 | 0.58599 | -4.8813 | 1.1E-06 | 2.1E-05 |
| 206 | LIPA | 16.405341 | -2.861578842 | 0.57863 | -4.9455 | 7.6E-07 | 1.6E-05 |
| 1465 | TUSC3 | 70.371481 | -2.863505991 | 0.37383 | -7.66 | 1.9E-14 | 2.4E-12 |
| 1205 | PALLD | 36.831567 | -2.863639169 | 0.54635 | -5.2414 | 1.6E-07 | 3.9E-06 |
| 1469 | PDLIM2 | 11.187753 | -2.865006707 | 0.83177 | -3.4445 | 0.00057 | 0.00456 |
| 487 | PALS1 | 21.661592 | -2.86788943 | 0.54101 | -5.301 | 1.2E-07 | 3E-06 |
| 225 | CASP7 | 5.7358735 | -2.869347635 | 1.01229 | -2.8345 | 0.00459 | 0.02525 |
| 781 | WTIP | 6.3153829 | -2.872610937 | 0.99995 | -2.8727 | 0.00407 | 0.02276 |
| 247 | IRF7 | 5.7184539 | -2.877903509 | 1.02559 | -2.8061 | 0.00501 | 0.02712 |
| 711 | SYNGR2 | 13.956117 | -2.884617538 | 0.74387 | -3.8779 | 0.00011 | 0.00112 |
| 1266 | SLC22A5 | 12.21374 | -2.886104019 | 0.77638 | -3.7174 | 0.0002 | 0.00194 |
| 970 | LAMA5 | 61.038798 | -2.894556168 | 0.33493 | -8.6422 | 5.5E-18 | 1.2E-15 |
| 1155 | STIM2 | 38.595561 | -2.895429521 | 0.40834 | -7.0907 | 1.3E-12 | 1.2E-10 |
| 476 | MIA2 | 16.460711 | -2.898930317 | 0.6103 | -4.75 | 2E-06 | 3.7E-05 |
| 1190 | TIFA | 5.5124344 | -2.900635573 | 1.13781 | -2.5493 | 0.01079 | 0.04937 |
| 277 | IFTAP | 8.6280672 | -2.902506987 | 0.83001 | -3.497 | 0.00047 | 0.00389 |
| 411 | SLC17A8 | 17.9346 | -2.903529526 | 0.71501 | -4.0608 | 4.9E-05 | 0.00058 |
| 1256 | CDO1 | 40.109255 | -2.90671848 | 0.52021 | -5.5875 | 2.3E-08 | 7.3E-07 |
| 849 | STON1 | 6.136594 | -2.911457668 | 1.10408 | -2.637 | 0.00836 | 0.04036 |
| 1366 | CITED2 | 4.4057932 | -2.918268825 | 1.1421 | -2.5552 | 0.01061 | 0.04875 |
| 1699 | INTS6L | 8.667136 | -2.924519468 | 0.85322 | -3.4276 | 0.00061 | 0.0048 |

| | | | | | | | |
|---|---|---|---|---|---|---|---|
| 479 | SAMD4A | 5.8291033 | -2.938887846 | 1.12374 | -2.6153 | 0.00892 | 0.04254 |
| 124 | ADAMTS4 | 4.4764776 | -2.93904064 | 1.10369 | -2.6629 | 0.00775 | 0.03807 |
| 1265 | PDLIM4 | 22.29427 | -2.940924004 | 0.55462 | -5.3026 | 1.1E-07 | 3E-06 |
| 142 | CYB5R1 | 10.841926 | -2.942739499 | 0.83395 | -3.5287 | 0.00042 | 0.00353 |
| 994 | ADARB1 | 18.914068 | -2.949515399 | 0.70705 | -4.1716 | 3E-05 | 0.00039 |
| 842 | MORN2 | 22.793095 | -2.953058921 | 0.55558 | -5.3153 | 1.1E-07 | 2.8E-06 |
| 655 | B9D1 | 22.567041 | -2.963408142 | 0.58692 | -5.0491 | 4.4E-07 | 9.7E-06 |
| 238 | MGMT | 7.6050981 | -2.969002183 | 0.96586 | -3.0739 | 0.00211 | 0.01348 |
| 232 | BAG3 | 5.6106799 | -2.969660455 | 1.01707 | -2.9198 | 0.0035 | 0.02013 |
| 894 | CYBRD1 | 12.729493 | -2.971926109 | 0.73086 | -4.0663 | 4.8E-05 | 0.00057 |
| 772 | ISYNA1 | 33.121584 | -2.978743919 | 0.45592 | -6.5334 | 6.4E-11 | 3.8E-09 |
| 85 | F3 | 7.9080971 | -2.979301348 | 1.01296 | -2.9412 | 0.00327 | 0.01905 |
| 213 | PLCE1 | 14.197988 | -2.983653806 | 0.66056 | -4.5169 | 6.3E-06 | 9.8E-05 |
| 187 | FZD8 | 41.363804 | -2.987632297 | 0.51111 | -5.8454 | 5.1E-09 | 1.9E-07 |
| 508 | TTC8 | 8.5769931 | -2.993362911 | 0.99145 | -3.0192 | 0.00253 | 0.01557 |
| 548 | SMAD3 | 15.118856 | -2.997000668 | 0.64455 | -4.6498 | 3.3E-06 | 5.5E-05 |
| 694 | CYB561 | 12.187395 | -3.006556708 | 0.78927 | -3.8093 | 0.00014 | 0.00142 |
| 1106 | DZIP1L | 8.0326434 | -3.01290532 | 0.93773 | -3.213 | 0.00131 | 0.00911 |
| 1590 | PHYHD1 | 5.2808287 | -3.025915258 | 1.13723 | -2.6608 | 0.0078 | 0.03825 |
| 647 | ACADVL | 86.7956 | -3.026935297 | 0.30165 | -10.035 | 1.1E-23 | 5.5E-21 |
| 640 | NXN | 14.480468 | -3.028340448 | 0.66266 | -4.57 | 4.9E-06 | 7.8E-05 |
| 1413 | CCDC146 | 11.736862 | -3.031608123 | 0.86743 | -3.4949 | 0.00047 | 0.00391 |
| 857 | ANTXR1 | 26.029514 | -3.033930594 | 0.51256 | -5.9192 | 3.2E-09 | 1.3E-07 |
| 467 | LRP10 | 24.660976 | -3.035929144 | 0.6402 | -4.7421 | 2.1E-06 | 3.8E-05 |
| 201 | SPOCK2 | 6.9111259 | -3.036793022 | 1.15995 | -2.618 | 0.00884 | 0.04225 |
| 1035 | RIBC2 | 5.5523277 | -3.042484723 | 1.08957 | -2.7924 | 0.00523 | 0.02808 |
| 1193 | ANXA5 | 47.245256 | -3.044257205 | 0.46065 | -6.6086 | 3.9E-11 | 2.4E-09 |
| 883 | ACVR1 | 5.3348183 | -3.050952138 | 1.10694 | -2.7562 | 0.00585 | 0.03068 |
| 1159 | WDR19 | 15.49634 | -3.054739863 | 0.63267 | -4.8283 | 1.4E-06 | 2.6E-05 |
| 523 | CLBA1 | 9.3766281 | -3.055369298 | 0.9222 | -3.3131 | 0.00092 | 0.00673 |
| 1617 | MID1 | 29.498172 | -3.055791553 | 0.56419 | -5.4163 | 6.1E-08 | 1.7E-06 |
| 1176 | SCD5 | 9.9437671 | -3.056549187 | 0.76916 | -3.9739 | 7.1E-05 | 0.0008 |
| 72 | NEXN | 5.9123963 | -3.057738926 | 1.05573 | -2.8963 | 0.00378 | 0.02139 |
| 92 | RHOC | 25.440367 | -3.058216549 | 0.53103 | -5.7591 | 8.5E-09 | 3E-07 |
| 1399 | MATCAP2 | 7.4031023 | -3.058611464 | 0.92354 | -3.3118 | 0.00093 | 0.00675 |
| 1562 | KIF27 | 6.3767759 | -3.068397 | 0.94922 | -3.2326 | 0.00123 | 0.00861 |
| 1577 | CUTALP | 4.6301262 | -3.088166203 | 1.17088 | -2.6375 | 0.00835 | 0.04035 |
| 373 | LIMA1 | 4.7605202 | -3.088563676 | 1.14235 | -2.7037 | 0.00686 | 0.0347 |
| 817 | MYADM | 15.907486 | -3.090421458 | 0.68067 | -4.5403 | 5.6E-06 | 8.9E-05 |

| | | | | | | | |
|---|---|---|---|---|---|---|---|
| 759 | SHFL | 6.9250746 | -3.091648039 | 1.05193 | -2.939 | 0.00329 | 0.01916 |
| 1063 | LZTFL1 | 15.453102 | -3.094387646 | 0.66497 | -4.6534 | 3.3E-06 | 5.4E-05 |
| 911 | CPS1 | 6.5147529 | -3.095648616 | 1.02009 | -3.0347 | 0.00241 | 0.01501 |
| 823 | ZNF776 | 8.8487146 | -3.107647753 | 0.84722 | -3.6681 | 0.00024 | 0.00228 |
| 145 | LRRN2 | 13.725512 | -3.110346905 | 0.66342 | -4.6884 | 2.8E-06 | 4.7E-05 |
| 1595 | BRD3OS | 9.9224582 | -3.119679504 | 0.77605 | -4.02 | 5.8E-05 | 0.00068 |
| 299 | LTBP3 | 15.886469 | -3.124416 | 0.6567 | -4.7577 | 2E-06 | 3.6E-05 |
| 1451 | PTN | 9.2753483 | -3.127372655 | 0.87735 | -3.5646 | 0.00036 | 0.00316 |
| 918 | STK36 | 41.581723 | -3.127943273 | 0.55768 | -5.6088 | 2E-08 | 6.6E-07 |
| 660 | TMEM98 | 54.715177 | -3.131643601 | 0.48744 | -6.4247 | 1.3E-10 | 7.3E-09 |
| 1625 | SAT1 | 30.932495 | -3.14093485 | 0.52646 | -5.9661 | 2.4E-09 | 1E-07 |
| 45 | CCDC30 | 10.213593 | -3.141032133 | 0.82254 | -3.8187 | 0.00013 | 0.00138 |
| 1321 | ENSG00000272217 | 4.1417023 | -3.142260511 | 1.19545 | -2.6285 | 0.00858 | 0.04124 |
| 725 | OSBPL1A | 14.444766 | -3.144837639 | 0.6656 | -4.7248 | 2.3E-06 | 4.1E-05 |
| 555 | LOXL1-AS1 | 5.4518232 | -3.147844733 | 1.14419 | -2.7512 | 0.00594 | 0.03106 |
| 804 | INAFM1 | 16.489543 | -3.152173334 | 0.6436 | -4.8977 | 9.7E-07 | 1.9E-05 |
| 565 | AKAP13 | 10.316921 | -3.155939709 | 0.82643 | -3.8188 | 0.00013 | 0.00138 |
| 1048 | SH3BP5-AS1 | 5.3388409 | -3.171928816 | 1.09671 | -2.8922 | 0.00383 | 0.0216 |
| 1709 | ZNF185 | 6.8700058 | -3.173265028 | 0.98231 | -3.2304 | 0.00124 | 0.00867 |
| 1246 | F2RL1 | 4.6359609 | -3.175400778 | 1.21904 | -2.6048 | 0.00919 | 0.04351 |
| 352 | TNFRSF1A | 21.214345 | -3.199119584 | 0.67858 | -4.7144 | 2.4E-06 | 4.3E-05 |
| 1163 | ATP10D | 6.6557671 | -3.199999113 | 1.02826 | -3.1121 | 0.00186 | 0.01213 |
| 639 | C17orf97 | 7.8925794 | -3.200851261 | 0.93664 | -3.4174 | 0.00063 | 0.00496 |
| 1031 | NAGA | 3.973031 | -3.201416259 | 1.24856 | -2.5641 | 0.01034 | 0.04789 |
| 1582 | MVB12B | 8.7471581 | -3.207998587 | 0.88046 | -3.6436 | 0.00027 | 0.00246 |
| 1072 | UBA7 | 6.5612689 | -3.214856501 | 1.09678 | -2.9312 | 0.00338 | 0.01955 |
| 1217 | SLC12A7 | 13.044636 | -3.217360858 | 0.74081 | -4.343 | 1.4E-05 | 0.0002 |
| 331 | SIDT2 | 7.7826105 | -3.219179472 | 0.94527 | -3.4056 | 0.00066 | 0.00514 |
| 40 | OSCP1 | 10.120803 | -3.223754107 | 0.91206 | -3.5346 | 0.00041 | 0.00347 |
| 1018 | SELENOM | 9.2673181 | -3.22578606 | 0.86392 | -3.7339 | 0.00019 | 0.00184 |
| 1201 | SH3D19 | 15.342271 | -3.227154146 | 0.74946 | -4.3059 | 1.7E-05 | 0.00023 |
| 1538 | CDKN2B | 9.5163836 | -3.231271511 | 0.97304 | -3.3208 | 0.0009 | 0.0066 |
| 1251 | SSBP2 | 20.466588 | -3.244482611 | 0.6041 | -5.3708 | 7.8E-08 | 2.1E-06 |
| 1351 | NT5DC1 | 28.590426 | -3.249805028 | 0.782 | -4.1558 | 3.2E-05 | 0.00041 |
| 1614 | PUDP | 12.785224 | -3.251800597 | 0.71919 | -4.5215 | 6.1E-06 | 9.6E-05 |
| 99 | TXNIP | 16.243509 | -3.259813347 | 0.69875 | -4.6652 | 3.1E-06 | 5.2E-05 |
| 60 | EFCAB7 | 7.9780552 | -3.260029879 | 1.00338 | -3.2491 | 0.00116 | 0.00819 |
| 191 | SLC16A9 | 35.397617 | -3.260079234 | 0.4528 | -7.1999 | 6E-13 | 5.7E-11 |
| 361 | MGP | 7.2448194 | -3.262362674 | 1.02329 | -3.1881 | 0.00143 | 0.0098 |

| | | | | | | | |
|---|---|---|---|---|---|---|---|
| 1042 | ITPR1 | 6.6212015 | -3.262582964 | 0.97456 | -3.3478 | 0.00081 | 0.0061 |
| 1248 | LHFPL2 | 11.719662 | -3.26459389 | 0.78892 | -4.1381 | 3.5E-05 | 0.00044 |
| 1178 | CDS1 | 8.5881405 | -3.271551156 | 0.97164 | -3.367 | 0.00076 | 0.00577 |
| 166 | OBSCN-AS1 | 8.7945134 | -3.278643932 | 0.88788 | -3.6927 | 0.00022 | 0.0021 |
| 741 | TPGS1 | 5.5163263 | -3.281930653 | 1.13279 | -2.8972 | 0.00376 | 0.02134 |
| 1302 | SERPINB1 | 12.822164 | -3.283684526 | 0.70207 | -4.6771 | 2.9E-06 | 4.9E-05 |
| 1286 | GALNT10 | 16.299961 | -3.284028954 | 0.71937 | -4.5651 | 5E-06 | 8E-05 |
| 775 | HOMER3 | 5.3254978 | -3.287403613 | 1.14979 | -2.8591 | 0.00425 | 0.0236 |
| 1515 | TRMT12 | 4.074176 | -3.288141823 | 1.26297 | -2.6035 | 0.00923 | 0.04362 |
| 683 | ITGA3 | 12.20445 | -3.290560754 | 0.92001 | -3.5766 | 0.00035 | 0.00304 |
| 832 | SLC5A6 | 45.861582 | -3.291585868 | 0.50647 | -6.499 | 8.1E-11 | 4.6E-09 |
| 929 | ARL4C | 73.177792 | -3.296528707 | 0.32559 | -10.125 | 4.3E-24 | 2.3E-21 |
| 1379 | PRR18 | 4.48483 | -3.298055265 | 1.17932 | -2.7966 | 0.00516 | 0.02774 |
| 1558 | NMRK1 | 4.2356313 | -3.305575113 | 1.23603 | -2.6744 | 0.00749 | 0.0371 |
| 354 | LRRC23 | 4.1272516 | -3.309525162 | 1.23336 | -2.6833 | 0.00729 | 0.03638 |
| 68 | GNG12 | 20.70935 | -3.310726408 | 0.62171 | -5.3252 | 1E-07 | 2.7E-06 |
| 568 | PEX11A | 4.129658 | -3.312171089 | 1.26657 | -2.6151 | 0.00892 | 0.04254 |
| 982 | SLC5A3 | 61.350299 | -3.317369818 | 0.39832 | -8.3285 | 8.2E-17 | 1.5E-14 |
| 1390 | ITGB8 | 27.518031 | -3.317693789 | 0.64048 | -5.18 | 2.2E-07 | 5.3E-06 |
| 828 | ID2 | 24.839521 | -3.324668227 | 0.74739 | -4.4484 | 8.7E-06 | 0.00013 |
| 1373 | SYNE1 | 16.904552 | -3.325804788 | 0.71361 | -4.6605 | 3.2E-06 | 5.3E-05 |
| 1514 | TMEM65 | 16.997858 | -3.328890248 | 0.64853 | -5.133 | 2.9E-07 | 6.5E-06 |
| 1541 | MOB3B | 8.3210712 | -3.337273707 | 0.98176 | -3.3993 | 0.00068 | 0.00524 |
| 314 | FAM181B | 9.6784475 | -3.343880976 | 0.86656 | -3.8588 | 0.00011 | 0.0012 |
| 343 | ROBO3 | 7.0915493 | -3.351350841 | 0.99745 | -3.3599 | 0.00078 | 0.00588 |
| 707 | FBF1 | 5.0253807 | -3.351892639 | 1.18445 | -2.8299 | 0.00466 | 0.02552 |
| 1124 | SOX2-OT | 22.237384 | -3.352444929 | 0.76731 | -4.3691 | 1.2E-05 | 0.00018 |
| 844 | DYNC2LI1 | 13.848713 | -3.358627979 | 0.85353 | -3.935 | 8.3E-05 | 0.00092 |
| 1277 | CYSTM1 | 14.420898 | -3.365599262 | 0.69345 | -4.8534 | 1.2E-06 | 2.3E-05 |
| 615 | SLC7A6 | 70.677358 | -3.367116508 | 0.36468 | -9.2332 | 2.6E-20 | 9.1E-18 |
| 1448 | COPG2IT1 | 11.708226 | -3.368834541 | 0.85136 | -3.957 | 7.6E-05 | 0.00085 |
| 1456 | RARRES2 | 39.783856 | -3.370895694 | 0.47458 | -7.1029 | 1.2E-12 | 1.1E-10 |
| 714 | SLC26A11 | 17.173441 | -3.377685605 | 0.64151 | -5.2652 | 1.4E-07 | 3.5E-06 |
| 84 | ARHGAP29 | 12.098393 | -3.390616498 | 0.77441 | -4.3783 | 1.2E-05 | 0.00017 |
| 701 | DNAI2 | 5.7695761 | -3.391905542 | 1.27279 | -2.6649 | 0.0077 | 0.03787 |
| 184 | BAMBI | 13.638967 | -3.400268358 | 0.69995 | -4.8579 | 1.2E-06 | 2.3E-05 |
| 1296 | PRR7 | 5.965359 | -3.405704674 | 1.0726 | -3.1752 | 0.0015 | 0.01015 |
| 1015 | SEC14L2 | 10.519448 | -3.4061087 | 0.93439 | -3.6453 | 0.00027 | 0.00245 |
| 120 | IGSF9 | 12.64395 | -3.406606576 | 0.83174 | -4.0958 | 4.2E-05 | 0.00051 |

| | | | | | | | |
|---|---|---|---|---|---|---|---|
| 877 | CFAP221 | 5.9701864 | -3.408420776 | 1.33299 | -2.557 | 0.01056 | 0.04862 |
| 1239 | MAP3K1 | 8.009461 | -3.418810741 | 0.95963 | -3.5626 | 0.00037 | 0.00318 |
| 48 | CCDC24 | 7.2667066 | -3.420570438 | 1.00802 | -3.3934 | 0.00069 | 0.00532 |
| 188 | RASGEF1A | 11.425242 | -3.429309205 | 0.87733 | -3.9088 | 9.3E-05 | 0.00101 |
| 297 | NEAT1 | 467.69196 | -3.431366268 | 0.63713 | -5.3857 | 7.2E-08 | 2E-06 |
| 1206 | MFAP3L | 8.4119137 | -3.433141518 | 0.97864 | -3.5081 | 0.00045 | 0.00377 |
| 349 | DYRK4 | 4.8937117 | -3.441868381 | 1.17633 | -2.9259 | 0.00343 | 0.0198 |
| 1616 | GPR143 | 4.6114095 | -3.442313851 | 1.19749 | -2.8746 | 0.00405 | 0.02267 |
| 1543 | B4GALT1 | 13.840654 | -3.466546183 | 0.70444 | -4.921 | 8.6E-07 | 1.7E-05 |
| 1663 | TSPAN6 | 70.96065 | -3.469222351 | 0.45472 | -7.6293 | 2.4E-14 | 2.9E-12 |
| 473 | NYNRIN | 8.2959288 | -3.491939763 | 0.91337 | -3.8232 | 0.00013 | 0.00136 |
| 1496 | ZFHX4 | 6.5578348 | -3.499445761 | 1.03416 | -3.3838 | 0.00071 | 0.00548 |
| 801 | APOE | 13.827482 | -3.505811855 | 0.72066 | -4.8647 | 1.1E-06 | 2.2E-05 |
| 1423 | PON2 | 18.995487 | -3.507125714 | 0.6203 | -5.6539 | 1.6E-08 | 5.3E-07 |
| 1346 | SMPD2 | 6.3312764 | -3.513500737 | 1.22203 | -2.8751 | 0.00404 | 0.02265 |
| 632 | NECAB2 | 6.4235639 | -3.515233106 | 1.10367 | -3.185 | 0.00145 | 0.00987 |
| 178 | PLXDC2 | 9.3500979 | -3.521319527 | 0.88964 | -3.9582 | 7.6E-05 | 0.00084 |
| 239 | LRRC27 | 4.8513959 | -3.521646247 | 1.18352 | -2.9756 | 0.00292 | 0.01743 |
| 506 | STON2 | 15.141963 | -3.523129257 | 0.71305 | -4.9409 | 7.8E-07 | 1.6E-05 |
| 1181 | AFF1 | 15.489224 | -3.523179833 | 0.70854 | -4.9725 | 6.6E-07 | 1.4E-05 |
| 1434 | GPR85 | 4.9061532 | -3.524530815 | 1.35807 | -2.5953 | 0.00945 | 0.04442 |
| 782 | LSR | 8.1312772 | -3.536732524 | 0.92775 | -3.8122 | 0.00014 | 0.00141 |
| 567 | ABHD2 | 45.071855 | -3.541338379 | 0.53755 | -6.5879 | 4.5E-11 | 2.7E-09 |
| 1150 | PROM1 | 9.8751999 | -3.549866213 | 0.88241 | -4.0229 | 5.7E-05 | 0.00067 |
| 1704 | LDOC1 | 6.2796269 | -3.557424611 | 1.06408 | -3.3432 | 0.00083 | 0.00618 |
| 436 | MIPEP | 6.343717 | -3.558153833 | 1.06472 | -3.3419 | 0.00083 | 0.00621 |
| 1211 | PDLIM3 | 10.551244 | -3.565347716 | 1.08782 | -3.2775 | 0.00105 | 0.00751 |
| 1315 | CARMIL1 | 10.555652 | -3.567200831 | 0.9554 | -3.7337 | 0.00019 | 0.00184 |
| 1645 | IQSEC2 | 3.6824485 | -3.569275486 | 1.38389 | -2.5792 | 0.0099 | 0.04622 |
| 4 | VWA1 | 4.9316373 | -3.571642788 | 1.17237 | -3.0465 | 0.00232 | 0.01453 |
| 147 | NUAK2 | 5.0025615 | -3.575064519 | 1.29898 | -2.7522 | 0.00592 | 0.03098 |
| 233 | FGFR2 | 49.206564 | -3.576562591 | 0.44747 | -7.9928 | 1.3E-15 | 2.1E-13 |
| 103 | CGN | 3.4343131 | -3.578685007 | 1.39581 | -2.5639 | 0.01035 | 0.04789 |
| 1613 | MXRA5 | 7.512002 | -3.582159968 | 1.05492 | -3.3957 | 0.00068 | 0.00529 |
| 755 | CTXN1 | 13.80464 | -3.582373657 | 0.76595 | -4.677 | 2.9E-06 | 4.9E-05 |
| 15 | NMNAT1 | 3.4390543 | -3.585385087 | 1.36707 | -2.6227 | 0.00872 | 0.04182 |
| 885 | RBMS1 | 20.345976 | -3.585756622 | 0.5996 | -5.9802 | 2.2E-09 | 9.2E-08 |
| 576 | METRN | 6.7459024 | -3.587766617 | 1.03372 | -3.4707 | 0.00052 | 0.00421 |
| 665 | ERBB2 | 8.169025 | -3.590987196 | 1.02607 | -3.4997 | 0.00047 | 0.00385 |

| | | | | | | | |
|---|---|---|---|---|---|---|---|
| 788 | DYRK1B | 6.4685625 | -3.592112053 | 1.08252 | -3.3183 | 0.00091 | 0.00664 |
| 1320 | HSD17B8 | 5.3128433 | -3.598123023 | 1.16781 | -3.0811 | 0.00206 | 0.01323 |
| 310 | SERPINH1 | 19.41976 | -3.607352836 | 0.64095 | -5.6281 | 1.8E-08 | 5.9E-07 |
| 612 | ZDHHC1 | 3.4888812 | -3.607573755 | 1.40897 | -2.5604 | 0.01045 | 0.04824 |
| 1564 | GAS1 | 3.4888812 | -3.607573755 | 1.40897 | -2.5604 | 0.01045 | 0.04824 |
| 1267 | IRF1 | 3.5624757 | -3.608207805 | 1.4029 | -2.572 | 0.01011 | 0.04702 |
| 563 | NMB | 5.0182235 | -3.609901589 | 1.31474 | -2.7457 | 0.00604 | 0.03141 |
| 1017 | OSBP2 | 6.6725256 | -3.611917415 | 1.12922 | -3.1986 | 0.00138 | 0.00952 |
| 1485 | CEBPD | 4.8508017 | -3.612111867 | 1.23746 | -2.919 | 0.00351 | 0.02017 |
| 682 | FAM117A | 5.0225753 | -3.620269575 | 1.21082 | -2.9899 | 0.00279 | 0.01679 |
| 1461 | TDRP | 14.642737 | -3.625384694 | 0.77262 | -4.6923 | 2.7E-06 | 4.7E-05 |
| 798 | BCL3 | 3.4025369 | -3.627489866 | 1.42427 | -2.5469 | 0.01087 | 0.04968 |
| 270 | NAV2 | 11.230974 | -3.628020163 | 0.83882 | -4.3251 | 1.5E-05 | 0.00021 |
| 25 | ID3 | 41.047449 | -3.630983702 | 0.4977 | -7.2955 | 3E-13 | 3E-11 |
| 451 | NEK3 | 3.5496543 | -3.6365227 | 1.40389 | -2.5903 | 0.00959 | 0.04496 |
| 1500 | TMEM67 | 15.670195 | -3.645685441 | 0.74624 | -4.8854 | 1E-06 | 2E-05 |
| 1583 | ANGPTL2 | 5.3317297 | -3.653277453 | 1.15905 | -3.152 | 0.00162 | 0.01087 |
| 641 | ABR | 7.4665805 | -3.664452104 | 0.98373 | -3.7251 | 0.0002 | 0.00189 |
| 1668 | RAB40A | 5.2393519 | -3.672162671 | 1.23388 | -2.9761 | 0.00292 | 0.01742 |
| 27 | FUCA1 | 7.4876556 | -3.677759792 | 1.00465 | -3.6607 | 0.00025 | 0.00233 |
| 1203 | CTSO | 8.749348 | -3.685063462 | 1.02924 | -3.5804 | 0.00034 | 0.00302 |
| 1554 | CFAP95 | 6.0816843 | -3.687264284 | 1.18429 | -3.1135 | 0.00185 | 0.01209 |
| 1235 | CDC20B | 7.3787403 | -3.69593104 | 1.2786 | -2.8906 | 0.00384 | 0.02168 |
| 154 | SLC30A1 | 17.140365 | -3.697916661 | 0.71522 | -5.1703 | 2.3E-07 | 5.5E-06 |
| 609 | TK2 | 3.6362436 | -3.698804403 | 1.34551 | -2.749 | 0.00598 | 0.03119 |
| 652 | GAS7 | 9.3875882 | -3.715151193 | 1.0019 | -3.7081 | 0.00021 | 0.002 |
| 750 | RFX2 | 12.865919 | -3.720006746 | 0.8045 | -4.624 | 3.8E-06 | 6.2E-05 |
| 1313 | RBM24 | 5.210842 | -3.724483964 | 1.22967 | -3.0289 | 0.00245 | 0.0152 |
| 614 | PSKH1 | 4.0317636 | -3.727171865 | 1.35178 | -2.7572 | 0.00583 | 0.03064 |
| 127 | RGS5 | 15.233323 | -3.732985338 | 0.76648 | -4.8703 | 1.1E-06 | 2.2E-05 |
| 675 | PLCD3 | 3.9066606 | -3.734538349 | 1.35771 | -2.7506 | 0.00595 | 0.0311 |
| 1549 | GNE | 3.6556834 | -3.734858569 | 1.39608 | -2.6752 | 0.00747 | 0.03707 |
| 1214 | TPPP | 3.7886554 | -3.73530863 | 1.37331 | -2.7199 | 0.00653 | 0.03344 |
| 1153 | PPARGC1A | 9.1296001 | -3.739537714 | 1.03228 | -3.6226 | 0.00029 | 0.00264 |
| 504 | ADCK1 | 5.4378335 | -3.739589032 | 1.15949 | -3.2252 | 0.00126 | 0.00878 |
| 971 | COL9A3 | 7.1106395 | -3.753023009 | 1.15107 | -3.2605 | 0.00111 | 0.00793 |
| 1097 | PARP14 | 9.1848137 | -3.754524492 | 0.96154 | -3.9047 | 9.4E-05 | 0.00103 |
| 895 | ITGA6 | 9.1490211 | -3.75914888 | 0.92556 | -4.0615 | 4.9E-05 | 0.00058 |
| 36 | KIAA1522 | 24.4267 | -3.760281206 | 0.5871 | -6.4049 | 1.5E-10 | 8.2E-09 |

| | | | | | | | |
|---|---|---|---|---|---|---|---|
| 17 | AGTRAP | 14.683984 | -3.762084536 | 0.74568 | -5.0452 | 4.5E-07 | 9.8E-06 |
| 1140 | IQCG | 15.001336 | -3.772850816 | 0.75055 | -5.0268 | 5E-07 | 1.1E-05 |
| 197 | LINC01515 | 3.8829389 | -3.773974015 | 1.34512 | -2.8057 | 0.00502 | 0.02715 |
| 570 | CIB1 | 28.474822 | -3.774977235 | 0.55367 | -6.8181 | 9.2E-12 | 6.7E-10 |
| 1392 | OSBPL3 | 9.6871226 | -3.78072977 | 0.95741 | -3.9489 | 7.9E-05 | 0.00087 |
| 920 | CFAP65 | 10.481889 | -3.783087304 | 0.94508 | -4.0029 | 6.3E-05 | 0.00072 |
| 244 | PGGHG | 7.5025846 | -3.786311167 | 1.00624 | -3.7628 | 0.00017 | 0.00167 |
| 1089 | RIOX2 | 4.1538133 | -3.791052945 | 1.46336 | -2.5906 | 0.00958 | 0.04493 |
| 1129 | ETV5 | 5.6257583 | -3.793694012 | 1.15454 | -3.2859 | 0.00102 | 0.00732 |
| 699 | SOX9 | 7.6061086 | -3.802865579 | 1.00158 | -3.7969 | 0.00015 | 0.00148 |
| 868 | TSGA10 | 3.8232292 | -3.805143667 | 1.429 | -2.6628 | 0.00775 | 0.03807 |
| 722 | TGIF1 | 11.117966 | -3.814791561 | 0.9955 | -3.832 | 0.00013 | 0.00132 |
| 1633 | MAOB | 6.586342 | -3.818544777 | 1.17852 | -3.2401 | 0.00119 | 0.00842 |
| 972 | MIR99AHG | 17.560866 | -3.828003884 | 0.77916 | -4.913 | 9E-07 | 1.8E-05 |
| 1419 | CDK6 | 7.7776709 | -3.828238206 | 1.01743 | -3.7626 | 0.00017 | 0.00167 |
| 194 | ANK3 | 12.292946 | -3.830701405 | 0.81975 | -4.673 | 3E-06 | 5E-05 |
| 425 | CFAP251 | 12.09335 | -3.836181356 | 0.90232 | -4.2515 | 2.1E-05 | 0.00028 |
| 311 | MAP6 | 19.366751 | -3.838306833 | 0.71105 | -5.3981 | 6.7E-08 | 1.9E-06 |
| 1034 | SHISAL1 | 6.2576332 | -3.85041251 | 1.11108 | -3.4655 | 0.00053 | 0.00427 |
| 279 | ENSG00000246250 | 7.8291078 | -3.853710459 | 1.07527 | -3.584 | 0.00034 | 0.00299 |
| 269 | LDHA | 99.305534 | -3.85513591 | 0.41972 | -9.1849 | 4.1E-20 | 1.3E-17 |
| 338 | OAF | 8.2288639 | -3.859813898 | 1.05267 | -3.6667 | 0.00025 | 0.00229 |
| 1708 | CETN2 | 35.936012 | -3.866257344 | 0.54549 | -7.0877 | 1.4E-12 | 1.2E-10 |
| 557 | STRA6 | 7.7305206 | -3.872803727 | 1.01447 | -3.8176 | 0.00013 | 0.00139 |
| 1116 | PLCH1 | 4.0814777 | -3.88023489 | 1.33294 | -2.911 | 0.0036 | 0.02061 |
| 1057 | CSRNP1 | 4.4521699 | -3.88640699 | 1.37105 | -2.8346 | 0.00459 | 0.02525 |
| 1324 | ZBTB9 | 61.458821 | -3.893313408 | 0.48525 | -8.0234 | 1E-15 | 1.6E-13 |
| 449 | PHF11 | 8.3727422 | -3.904819007 | 0.963 | -4.0548 | 5E-05 | 0.0006 |
| 668 | JUP | 9.3614991 | -3.905655703 | 0.97414 | -4.0093 | 6.1E-05 | 0.00071 |
| 97 | HMGCS2 | 4.2294627 | -3.907480607 | 1.53471 | -2.5461 | 0.01089 | 0.04976 |
| 176 | VIM | 169.88921 | -3.909319082 | 0.37062 | -10.548 | 5.2E-26 | 4E-23 |
| 818 | CACNG8 | 6.1543551 | -3.911292664 | 1.10792 | -3.5303 | 0.00042 | 0.00351 |
| 377 | KRT8 | 6.010801 | -3.915876186 | 1.16345 | -3.3657 | 0.00076 | 0.00579 |
| 617 | SMPD3 | 4.2637442 | -3.916933997 | 1.34868 | -2.9043 | 0.00368 | 0.02094 |
| 1649 | USP51 | 4.325021 | -3.917615327 | 1.30367 | -3.0051 | 0.00266 | 0.01615 |
| 909 | NRP2 | 12.130274 | -3.918488849 | 0.93334 | -4.1984 | 2.7E-05 | 0.00035 |
| 813 | RCN3 | 2.6231058 | -3.932130236 | 1.53126 | -2.5679 | 0.01023 | 0.04748 |
| 496 | NPC2 | 24.989614 | -3.933052488 | 0.56643 | -6.9436 | 3.8E-12 | 2.9E-10 |
| 1056 | TTC21A | 4.7808788 | -3.9397973 | 1.35864 | -2.8998 | 0.00373 | 0.02119 |

| | | | | | | | |
|---|---|---|---|---|---|---|---|
| 549 | PAQR5 | 2.6250799 | -3.942651438 | 1.52641 | -2.583 | 0.0098 | 0.04581 |
| 1292 | CPEB4 | 7.1477101 | -3.945441003 | 1.18813 | -3.3207 | 0.0009 | 0.0066 |
| 1333 | ENSG00000227885 | 2.7694734 | -3.948796975 | 1.53131 | -2.5787 | 0.00992 | 0.04625 |
| 1581 | NEK6 | 6.5186265 | -3.949337743 | 1.10856 | -3.5626 | 0.00037 | 0.00318 |
| 1603 | ENSG00000278817 | 2.6397323 | -3.950200177 | 1.52631 | -2.5881 | 0.00965 | 0.04518 |
| 168 | FAM89A | 18.310653 | -3.953311045 | 0.8397 | -4.708 | 2.5E-06 | 4.4E-05 |
| 950 | BPIFB4 | 4.5763365 | -3.964103485 | 1.52193 | -2.6046 | 0.0092 | 0.04351 |
| 1053 | FBXL2 | 7.160945 | -3.965944766 | 1.09928 | -3.6078 | 0.00031 | 0.00276 |
| 1640 | PCSK1N | 56.821158 | -3.966067433 | 0.43428 | -9.1325 | 6.7E-20 | 2E-17 |
| 101 | MINDY1 | 4.6306585 | -3.968950316 | 1.41164 | -2.8116 | 0.00493 | 0.02677 |
| 1279 | NR3C1 | 6.3012106 | -3.969426167 | 1.13891 | -3.4853 | 0.00049 | 0.00403 |
| 531 | BMF | 6.6881685 | -3.97186518 | 1.09531 | -3.6262 | 0.00029 | 0.00261 |
| 387 | RBMS2 | 4.4102895 | -3.972563831 | 1.35378 | -2.9344 | 0.00334 | 0.01939 |
| 379 | TNS2 | 4.6810387 | -3.973805563 | 1.27849 | -3.1082 | 0.00188 | 0.01226 |
| 1472 | CLU | 68.968539 | -3.993023239 | 0.45387 | -8.7977 | 1.4E-18 | 3.4E-16 |
| 342 | SPA17 | 15.998905 | -3.993153115 | 0.839 | -4.7594 | 1.9E-06 | 3.6E-05 |
| 1344 | ARMC2 | 4.6634296 | -3.996935019 | 1.46539 | -2.7276 | 0.00638 | 0.03279 |
| 754 | INSR | 10.930648 | -3.99809383 | 0.90232 | -4.4309 | 9.4E-06 | 0.00014 |
| 1157 | KLF3 | 26.333465 | -3.998656271 | 0.60297 | -6.6316 | 3.3E-11 | 2.1E-09 |
| 953 | FER1L4 | 9.4181618 | -4.004213442 | 0.94692 | -4.2287 | 2.4E-05 | 0.00031 |
| 1362 | SGK1 | 2.73609 | -4.012885642 | 1.5696 | -2.5566 | 0.01057 | 0.04865 |
| 1690 | FIRRE | 7.2952904 | -4.017707358 | 1.09016 | -3.6854 | 0.00023 | 0.00215 |
| 1331 | NFKBIE | 4.3782208 | -4.018709009 | 1.48308 | -2.7097 | 0.00673 | 0.03423 |
| 959 | SDC4 | 12.693495 | -4.020741423 | 0.93404 | -4.3047 | 1.7E-05 | 0.00023 |
| 907 | FZD7 | 4.539797 | -4.021476606 | 1.3716 | -2.932 | 0.00337 | 0.01951 |
| 370 | COL2A1 | 17.823241 | -4.023588347 | 0.78402 | -5.132 | 2.9E-07 | 6.5E-06 |
| 1326 | MDGA1 | 2.7543981 | -4.027428665 | 1.53565 | -2.6226 | 0.00873 | 0.04182 |
| 812 | TRPM4 | 2.7828574 | -4.028701943 | 1.54216 | -2.6124 | 0.00899 | 0.04285 |
| 1476 | RBPMS | 4.7926273 | -4.034307238 | 1.2888 | -3.1303 | 0.00175 | 0.01153 |
| 1365 | NHSL1 | 4.5843329 | -4.036135286 | 1.33138 | -3.0316 | 0.00243 | 0.01512 |
| 334 | IFT46 | 11.684865 | -4.043011534 | 0.86236 | -4.6883 | 2.8E-06 | 4.7E-05 |
| 458 | MBNL2 | 17.068155 | -4.044860593 | 0.77841 | -5.1963 | 2E-07 | 4.9E-06 |
| 1394 | LINC01176 | 2.8563089 | -4.061008045 | 1.53089 | -2.6527 | 0.00798 | 0.03901 |
| 218 | DPCD | 16.346637 | -4.063489329 | 0.73401 | -5.536 | 3.1E-08 | 9.5E-07 |
| 1249 | SERINC5 | 15.289484 | -4.064895607 | 0.78407 | -5.1843 | 2.2E-07 | 5.2E-06 |
| 107 | S100A4 | 5.1325001 | -4.065087089 | 1.32727 | -3.0627 | 0.00219 | 0.01391 |
| 932 | SPEF1 | 8.9409188 | -4.065475985 | 1.0996 | -3.6972 | 0.00022 | 0.00208 |
| 69 | WLS | 54.583199 | -4.068444786 | 0.43431 | -9.3675 | 7.4E-21 | 2.8E-18 |
| 112 | EFNA4 | 2.8435336 | -4.071169717 | 1.55142 | -2.6242 | 0.00869 | 0.04167 |

| | | | | | | | |
|---|---|---|---|---|---|---|---|
| 372 | AQP5 | 4.7766316 | -4.074586905 | 1.26902 | -3.2108 | 0.00132 | 0.00917 |
| 822 | ZNF667-AS1 | 2.8729353 | -4.077740708 | 1.53248 | -2.6609 | 0.00779 | 0.03825 |
| 1510 | EXT1 | 9.5062037 | -4.080667308 | 0.94545 | -4.3161 | 1.6E-05 | 0.00022 |
| 1146 | ENSG00000287117 | 3.0173289 | -4.083231113 | 1.51122 | -2.7019 | 0.00689 | 0.03484 |
| 365 | PTHLH | 4.7437457 | -4.084489433 | 1.36006 | -3.0032 | 0.00267 | 0.01623 |
| 1316 | TRIM38 | 2.908949 | -4.086574942 | 1.55726 | -2.6242 | 0.00869 | 0.04167 |
| 499 | FOS | 2.8908365 | -4.091309079 | 1.52464 | -2.6835 | 0.00729 | 0.03638 |
| 32 | UBXN11 | 23.730759 | -4.092001888 | 0.69259 | -5.9082 | 3.5E-09 | 1.3E-07 |
| 663 | GAS2L2 | 4.7730506 | -4.095332741 | 1.47582 | -2.775 | 0.00552 | 0.02932 |
| 1091 | IFT57 | 15.005624 | -4.115151672 | 0.78011 | -5.2751 | 1.3E-07 | 3.4E-06 |
| 837 | FOSL2 | 12.098209 | -4.118412267 | 0.93514 | -4.404 | 1.1E-05 | 0.00015 |
| 208 | PCGF5 | 7.6039616 | -4.120404306 | 1.09896 | -3.7494 | 0.00018 | 0.00174 |
| 600 | MT2A | 7.186335 | -4.123378807 | 1.11756 | -3.6896 | 0.00022 | 0.00212 |
| 1519 | DNAAF11 | 13.990544 | -4.126502702 | 0.86815 | -4.7532 | 2E-06 | 3.6E-05 |
| 337 | USP2 | 7.161604 | -4.131058329 | 1.15282 | -3.5834 | 0.00034 | 0.00299 |
| 1489 | ENSG00000274956 | 3.0094661 | -4.143426392 | 1.49517 | -2.7712 | 0.00558 | 0.02961 |
| 1621 | PIR | 3.0208698 | -4.143858767 | 1.53208 | -2.7047 | 0.00684 | 0.03463 |
| 1628 | TMEM47 | 7.3890301 | -4.144294379 | 1.11159 | -3.7282 | 0.00019 | 0.00188 |
| 1094 | ZBTB20 | 10.239271 | -4.154693831 | 0.95586 | -4.3466 | 1.4E-05 | 0.00019 |
| 139 | LAMC2 | 3.1501357 | -4.155051438 | 1.48327 | -2.8013 | 0.00509 | 0.02746 |
| 1656 | EDA2R | 3.0121649 | -4.155759542 | 1.60574 | -2.5881 | 0.00965 | 0.04518 |
| 362 | MGST1 | 9.4806068 | -4.156789415 | 1.0762 | -3.8625 | 0.00011 | 0.00119 |
| 219 | NFKB2 | 3.0241933 | -4.158175279 | 1.46977 | -2.8291 | 0.00467 | 0.02557 |
| 173 | ECHDC3 | 3.0223401 | -4.160840981 | 1.63523 | -2.5445 | 0.01094 | 0.0499 |
| 161 | DISP1 | 3.0188054 | -4.162542183 | 1.47994 | -2.8126 | 0.00491 | 0.02671 |
| 984 | RUNX1 | 3.1650521 | -4.166545817 | 1.52061 | -2.7401 | 0.00614 | 0.03183 |
| 1651 | LINC01278 | 3.0850346 | -4.177455988 | 1.48357 | -2.8158 | 0.00487 | 0.02647 |
| 1334 | LRRC1 | 9.6455109 | -4.179439988 | 1.03009 | -4.0574 | 5E-05 | 0.00059 |
| 1670 | TBC1D8B | 5.2265541 | -4.181138552 | 1.36216 | -3.0695 | 0.00214 | 0.01362 |
| 915 | TNS1 | 7.7939605 | -4.183801167 | 1.05964 | -3.9483 | 7.9E-05 | 0.00087 |
| 494 | BBOF1 | 10.029983 | -4.184321341 | 0.97543 | -4.2897 | 1.8E-05 | 0.00024 |
| 265 | SOX6 | 5.5287717 | -4.189394789 | 1.28829 | -3.2519 | 0.00115 | 0.00813 |
| 455 | SLITRK5 | 5.5287717 | -4.189394789 | 1.28829 | -3.2519 | 0.00115 | 0.00813 |
| 756 | MARCHF2 | 5.275706 | -4.190060444 | 1.27534 | -3.2855 | 0.00102 | 0.00733 |
| 672 | DUSP3 | 7.4722529 | -4.196979439 | 1.0754 | -3.9027 | 9.5E-05 | 0.00103 |
| 1483 | PLAT | 5.4830642 | -4.227163679 | 1.24932 | -3.3836 | 0.00072 | 0.00548 |
| 1597 | PPP1R26-AS1 | 3.1822347 | -4.237290893 | 1.52242 | -2.7833 | 0.00538 | 0.02868 |
| 653 | MYH3 | 3.1779323 | -4.241826828 | 1.49767 | -2.8323 | 0.00462 | 0.02538 |
| 100 | CA14 | 12.601348 | -4.244302631 | 0.91666 | -4.6302 | 3.7E-06 | 6E-05 |

| | | | | | | | |
|---|---|---|---|---|---|---|---|
| 1553 | ENTREP1 | 5.4518882 | -4.246212375 | 1.35651 | -3.1303 | 0.00175 | 0.01153 |
| 712 | SOCS3 | 5.4155138 | -4.247757577 | 1.27413 | -3.3339 | 0.00086 | 0.00636 |
| 322 | ARHGAP42 | 3.3604104 | -4.256584751 | 1.46797 | -2.8996 | 0.00374 | 0.0212 |
| 1135 | PLAAT1 | 3.3671939 | -4.259440846 | 1.43823 | -2.9616 | 0.00306 | 0.01806 |
| 5 | CFAP74 | 3.2768582 | -4.269872421 | 1.5012 | -2.8443 | 0.00445 | 0.02456 |
| 355 | C1S | 24.062422 | -4.279370869 | 0.62553 | -6.8412 | 7.9E-12 | 5.8E-10 |
| 148 | BLACAT1 | 3.4181987 | -4.285517317 | 1.43798 | -2.9802 | 0.00288 | 0.01723 |
| 242 | FUOM | 3.3147776 | -4.297418694 | 1.45845 | -2.9466 | 0.00321 | 0.01882 |
| 697 | PRKCA | 7.9776815 | -4.298709379 | 1.07726 | -3.9904 | 6.6E-05 | 0.00075 |
| 902 | C2orf66 | 5.3813773 | -4.301387206 | 1.27932 | -3.3622 | 0.00077 | 0.00585 |
| 1382 | MICALL2 | 10.66005 | -4.312570297 | 0.97144 | -4.4393 | 9E-06 | 0.00013 |
| 1339 | COL12A1 | 3.3610412 | -4.316324172 | 1.44724 | -2.9824 | 0.00286 | 0.01715 |
| 155 | PTPN14 | 7.7569234 | -4.317910936 | 1.11701 | -3.8656 | 0.00011 | 0.00117 |
| 1425 | EPHB4 | 3.345742 | -4.320735081 | 1.4489 | -2.9821 | 0.00286 | 0.01715 |
| 1627 | DMD | 54.198627 | -4.322117077 | 0.51481 | -8.3956 | 4.6E-17 | 8.8E-15 |
| 1183 | PDLIM5 | 48.600819 | -4.327015916 | 0.49637 | -8.7173 | 2.8E-18 | 6.7E-16 |
| 1156 | PCDH7 | 5.6534117 | -4.328973539 | 1.35768 | -3.1885 | 0.00143 | 0.00979 |
| 1349 | SLC16A10 | 32.80628 | -4.335200675 | 0.56733 | -7.6414 | 2.1E-14 | 2.6E-12 |
| 989 | MX1 | 5.6934418 | -4.336672064 | 1.38602 | -3.1289 | 0.00175 | 0.01157 |
| 1666 | BEX5 | 3.4109605 | -4.340969764 | 1.57008 | -2.7648 | 0.0057 | 0.03005 |
| 1197 | MGST2 | 5.572937 | -4.351190205 | 1.26491 | -3.4399 | 0.00058 | 0.00462 |
| 649 | EFNB3 | 8.1470121 | -4.358065306 | 1.08985 | -3.9988 | 6.4E-05 | 0.00073 |
| 1372 | CCDC170 | 13.542579 | -4.360244618 | 0.97356 | -4.4786 | 7.5E-06 | 0.00011 |
| 428 | TCTN2 | 5.8156844 | -4.360333322 | 1.26722 | -3.4409 | 0.00058 | 0.00461 |
| 1446 | MEST | 71.92368 | -4.363224501 | 0.42963 | -10.156 | 3.1E-24 | 2E-21 |
| 1401 | TNS3 | 13.761568 | -4.364102454 | 0.86492 | -5.0457 | 4.5E-07 | 9.8E-06 |
| 703 | GPRC5C | 3.4758039 | -4.370670748 | 1.43294 | -3.0501 | 0.00229 | 0.01439 |
| 1563 | NTRK2 | 7.9056254 | -4.372874817 | 1.28692 | -3.3979 | 0.00068 | 0.00525 |
| 613 | TSNAXIP1 | 5.676175 | -4.382010587 | 1.31516 | -3.3319 | 0.00086 | 0.00639 |
| 313 | TENM4 | 3.5190827 | -4.387604972 | 1.44843 | -3.0292 | 0.00245 | 0.01519 |
| 1691 | GPC4 | 11.257763 | -4.398247809 | 0.98724 | -4.4551 | 8.4E-06 | 0.00013 |
| 861 | DNAH6 | 6.0122918 | -4.407876247 | 1.24429 | -3.5425 | 0.0004 | 0.00338 |
| 771 | SSBP4 | 12.294811 | -4.412744873 | 0.9314 | -4.7378 | 2.2E-06 | 3.9E-05 |
| 810 | PPP1R15A | 5.891809 | -4.413613193 | 1.2797 | -3.4489 | 0.00056 | 0.0045 |
| 767 | CPAMD8 | 2.0815747 | -4.414439019 | 1.68551 | -2.619 | 0.00882 | 0.04218 |
| 769 | JAK3 | 6.1172117 | -4.422454306 | 1.22845 | -3.6 | 0.00032 | 0.00284 |
| 1429 | IFT22 | 14.587665 | -4.425845067 | 0.90379 | -4.897 | 9.7E-07 | 1.9E-05 |
| 866 | FER1L5 | 2.1112865 | -4.433174432 | 1.70372 | -2.6021 | 0.00927 | 0.04376 |
| 916 | TMBIM1 | 3.6466667 | -4.433191883 | 1.4135 | -3.1363 | 0.00171 | 0.01135 |

| | | | | | | | |
|---|---|---|---|---|---|---|---|
| 442 | CCNA1 | 2.1161707 | -4.436201521 | 1.70329 | -2.6045 | 0.0092 | 0.04351 |
| 462 | ZIC2 | 58.655298 | -4.437464732 | 0.48652 | -9.1209 | 7.5E-20 | 2.1E-17 |
| 1338 | LINC00472 | 31.003087 | -4.438482606 | 0.61547 | -7.2115 | 5.5E-13 | 5.3E-11 |
| 350 | NTF3 | 2.112636 | -4.439680164 | 1.72825 | -2.5689 | 0.0102 | 0.04737 |
| 91 | WNT2B | 11.755737 | -4.440947067 | 1.14087 | -3.8926 | 9.9E-05 | 0.00107 |
| 601 | MT1F | 6.085621 | -4.44402594 | 1.3208 | -3.3646 | 0.00077 | 0.00581 |
| 580 | VASN | 6.0090494 | -4.445782131 | 1.25514 | -3.542 | 0.0004 | 0.00339 |
| 195 | ARID5B | 33.521103 | -4.453445647 | 0.57495 | -7.7459 | 9.5E-15 | 1.3E-12 |
| 1038 | DENND6B | 11.682572 | -4.456310507 | 0.91488 | -4.8709 | 1.1E-06 | 2.2E-05 |
| 1215 | ZDHHC11 | 5.9033115 | -4.457897871 | 1.27537 | -3.4954 | 0.00047 | 0.0039 |
| 899 | COL3A1 | 129.72872 | -4.462945375 | 0.78525 | -5.6834 | 1.3E-08 | 4.5E-07 |
| 1537 | BNC2 | 2.1521161 | -4.464220294 | 1.67451 | -2.666 | 0.00768 | 0.0378 |
| 1131 | P3H2 | 6.5748539 | -4.467839024 | 1.26603 | -3.529 | 0.00042 | 0.00352 |
| 974 | CYYR1 | 2.1588996 | -4.468411127 | 1.68802 | -2.6471 | 0.00812 | 0.03943 |
| 1551 | CNTNAP3 | 2.160392 | -4.469302031 | 1.65591 | -2.699 | 0.00695 | 0.03507 |
| 51 | SPATA6 | 3.7191821 | -4.470648777 | 1.39995 | -3.1934 | 0.00141 | 0.00965 |
| 497 | LTBP2 | 6.4922442 | -4.47224547 | 1.39762 | -3.1999 | 0.00137 | 0.00949 |
| 1069 | LAMB2 | 39.07135 | -4.472971608 | 0.51469 | -8.6907 | 3.6E-18 | 8E-16 |
| 622 | CLEC18B | 3.7482693 | -4.478440054 | 1.43249 | -3.1263 | 0.00177 | 0.01164 |
| 1151 | LDB2 | 6.151596 | -4.479535801 | 1.2401 | -3.6122 | 0.0003 | 0.00272 |
| 933 | PRNP | 72.113198 | -4.481745813 | 0.5538 | -8.0927 | 5.8E-16 | 9.7E-14 |
| 484 | ARMH4 | 3.7518787 | -4.482498391 | 1.41893 | -3.1591 | 0.00158 | 0.01066 |
| 413 | DRAM1 | 5.9867489 | -4.484140779 | 1.33881 | -3.3494 | 0.00081 | 0.00607 |
| 859 | SLC4A5 | 6.1745243 | -4.484910009 | 1.2524 | -3.581 | 0.00034 | 0.00301 |
| 1686 | GRIA3 | 2.187119 | -4.485673491 | 1.75046 | -2.5626 | 0.01039 | 0.04803 |
| 1317 | DDR1 | 51.686699 | -4.48649956 | 0.47554 | -9.4346 | 3.9E-21 | 1.5E-18 |
| 1601 | NOXA1 | 2.1901038 | -4.487245965 | 1.66564 | -2.694 | 0.00706 | 0.03549 |
| 1228 | ENSG00000279737 | 2.1998721 | -4.493049858 | 1.66491 | -2.6987 | 0.00696 | 0.03508 |
| 696 | CEP112 | 6.2063994 | -4.494391779 | 1.25307 | -3.5867 | 0.00033 | 0.00296 |
| 589 | SULT1A1 | 6.214804 | -4.498455583 | 1.27951 | -3.5158 | 0.00044 | 0.00367 |
| 1522 | LYNX1 | 3.8140695 | -4.502225032 | 1.41257 | -3.1873 | 0.00144 | 0.00981 |
| 986 | B3GALT5 | 2.2143816 | -4.507185266 | 1.7003 | -2.6508 | 0.00803 | 0.03916 |
| 21 | MFAP2 | 2.2246998 | -4.508085571 | 1.70222 | -2.6484 | 0.00809 | 0.03936 |
| 737 | LINC00305 | 2.2295839 | -4.510974199 | 1.70183 | -2.6507 | 0.00803 | 0.03916 |
| 539 | DNAAF4 | 3.8213346 | -4.511358649 | 1.4108 | -3.1977 | 0.00139 | 0.00954 |
| 536 | FBN1 | 13.457092 | -4.511416835 | 0.90689 | -4.9746 | 6.5E-07 | 1.4E-05 |
| 1455 | ZNF425 | 2.2245568 | -4.513234891 | 1.72435 | -2.6174 | 0.00886 | 0.04231 |
| 450 | NEK5 | 15.953751 | -4.516516402 | 0.88174 | -5.1223 | 3E-07 | 6.9E-06 |
| 302 | RHOD | 6.3110375 | -4.51957785 | 1.32016 | -3.4235 | 0.00062 | 0.00487 |

| | | | | | | | |
|---|---|---|---|---|---|---|---|
| 1297 | PDLIM7 | 49.95546 | -4.521658874 | 0.5086 | -8.8904 | 6.1E-19 | 1.6E-16 |
| 1610 | CSF2RA | 2.2540047 | -4.525661142 | 1.74448 | -2.5943 | 0.00948 | 0.04451 |
| 979 | C21orf62 | 2.2474851 | -4.526947925 | 1.65531 | -2.7348 | 0.00624 | 0.03226 |
| 774 | CERS1 | 2.2489775 | -4.527741754 | 1.6599 | -2.7277 | 0.00638 | 0.03278 |
| 564 | ALPK3 | 2.2504699 | -4.528419832 | 1.72427 | -2.6263 | 0.00863 | 0.04148 |
| 198 | ENSG00000260400 | 2.2625445 | -4.535760394 | 1.69889 | -2.6698 | 0.00759 | 0.03748 |
| 522 | MOK | 20.796929 | -4.537193456 | 0.76166 | -5.957 | 2.6E-09 | 1E-07 |
| 777 | ZNF429 | 2.2659362 | -4.537744047 | 1.74303 | -2.6034 | 0.00923 | 0.04363 |
| 19 | FHAD1 | 3.9060468 | -4.538415765 | 1.40715 | -3.2253 | 0.00126 | 0.00878 |
| 787 | ZFP36 | 3.9325099 | -4.54441052 | 1.45329 | -3.127 | 0.00177 | 0.01162 |
| 1449 | CALD1 | 50.445523 | -4.547142284 | 0.95869 | -4.7431 | 2.1E-06 | 3.8E-05 |
| 773 | COMP | 2.2937487 | -4.553824556 | 1.66513 | -2.7348 | 0.00624 | 0.03226 |
| 871 | SULT1C4 | 9.4540033 | -4.555114263 | 1.12605 | -4.0452 | 5.2E-05 | 0.00062 |
| 1177 | GPAT3 | 2.2967335 | -4.555370476 | 1.64619 | -2.7672 | 0.00565 | 0.02989 |
| 591 | MVP | 2.2982259 | -4.556279093 | 1.69006 | -2.6959 | 0.00702 | 0.03532 |
| 330 | CADM1 | 19.217745 | -4.560654946 | 0.77104 | -5.9149 | 3.3E-09 | 1.3E-07 |
| 1520 | ENSG00000279518 | 2.3162701 | -4.566795359 | 1.72977 | -2.6401 | 0.00829 | 0.04009 |
| 749 | CAPS | 16.271428 | -4.567425125 | 0.97629 | -4.6783 | 2.9E-06 | 4.9E-05 |
| 152 | DYRK3 | 3.9668946 | -4.569222906 | 1.4057 | -3.2505 | 0.00115 | 0.00817 |
| 289 | TMEM109 | 25.026104 | -4.572936496 | 0.65582 | -6.9728 | 3.1E-12 | 2.4E-10 |
| 863 | VAMP8 | 2.3313295 | -4.575245293 | 1.67069 | -2.7385 | 0.00617 | 0.03196 |
| 1569 | TBC1D2 | 2.324403 | -4.576335253 | 1.68665 | -2.7133 | 0.00666 | 0.03402 |
| 1511 | MAL2 | 2.3377061 | -4.579054181 | 1.71496 | -2.6701 | 0.00758 | 0.03747 |
| 836 | RBKS | 2.3311865 | -4.580424925 | 1.64473 | -2.7849 | 0.00535 | 0.02856 |
| 237 | PTPRE | 2.3410978 | -4.580885107 | 1.69103 | -2.7089 | 0.00675 | 0.0343 |
| 262 | ZBED5-AS1 | 4.1457632 | -4.581026123 | 1.64762 | -2.7804 | 0.00543 | 0.02889 |
| 318 | SMCO4 | 6.5789574 | -4.583444897 | 1.23056 | -3.7247 | 0.0002 | 0.0019 |
| 13 | PIK3CD | 2.3394624 | -4.585114971 | 1.63849 | -2.7984 | 0.00514 | 0.02764 |
| 530 | FSIP1 | 2.3493737 | -4.585678706 | 1.70168 | -2.6948 | 0.00704 | 0.03542 |
| 123 | KLHDC9 | 2.3413617 | -4.586066442 | 1.70916 | -2.6832 | 0.00729 | 0.03638 |
| 1165 | PDGFRA | 32.368836 | -4.590486965 | 0.63429 | -7.2373 | 4.6E-13 | 4.5E-11 |
| 353 | GPR162 | 9.6309346 | -4.593680167 | 1.04983 | -4.3757 | 1.2E-05 | 0.00017 |
| 965 | NFATC2 | 4.0527813 | -4.59591648 | 1.48882 | -3.0869 | 0.00202 | 0.01302 |
| 1014 | CCDC157 | 7.1398778 | -4.601746496 | 1.24497 | -3.6963 | 0.00022 | 0.00208 |
| 892 | ENSG00000239467 | 6.8240856 | -4.602140843 | 1.22428 | -3.7591 | 0.00017 | 0.00169 |
| 914 | IGFBP5 | 12.9485 | -4.608654335 | 0.95817 | -4.8098 | 1.5E-06 | 2.8E-05 |
| 976 | ADAMTS5 | 4.1090024 | -4.611621172 | 1.45169 | -3.1767 | 0.00149 | 0.01012 |
| 321 | MAML2 | 2.3872184 | -4.61166403 | 1.6191 | -2.8483 | 0.0044 | 0.02429 |
| 1281 | PRELID2 | 2.3872184 | -4.61166403 | 1.6191 | -2.8483 | 0.0044 | 0.02429 |

| | | | | | | | |
|---|---|---|---|---|---|---|---|
| 1417 | FZD1 | 9.9120367 | -4.613444283 | 1.02306 | -4.5094 | 6.5E-06 | 0.0001 |
| 594 | ORAI3 | 6.90738 | -4.621791929 | 1.2128 | -3.8108 | 0.00014 | 0.00141 |
| 1657 | GDPD2 | 9.7535871 | -4.622382304 | 1.12059 | -4.1249 | 3.7E-05 | 0.00046 |
| 1065 | KIF9 | 20.568405 | -4.625351324 | 0.75039 | -6.1639 | 7.1E-10 | 3.3E-08 |
| 1261 | PPIC | 6.7667107 | -4.626195194 | 1.22342 | -3.7814 | 0.00016 | 0.00157 |
| 514 | IFI27 | 4.1608287 | -4.629035908 | 1.47127 | -3.1463 | 0.00165 | 0.01104 |
| 947 | ID1 | 47.154685 | -4.636639412 | 1.43335 | -3.2348 | 0.00122 | 0.00855 |
| 1182 | SPARCL1 | 6.8223356 | -4.637511587 | 1.23025 | -3.7696 | 0.00016 | 0.00163 |
| 1593 | CFAP77 | 6.7393589 | -4.639744802 | 1.31754 | -3.5215 | 0.00043 | 0.0036 |
| 448 | CAB39L | 10.432426 | -4.640252609 | 1.02329 | -4.5347 | 5.8E-06 | 9.1E-05 |
| 90 | KCND3 | 4.1903976 | -4.641259812 | 1.40119 | -3.3124 | 0.00093 | 0.00674 |
| 1016 | RPL13AP26 | 4.1625629 | -4.642999652 | 1.39105 | -3.3378 | 0.00084 | 0.00629 |
| 562 | CFAP161 | 2.4393087 | -4.644813973 | 1.75004 | -2.6541 | 0.00795 | 0.03886 |
| 958 | KCNK15 | 4.1816926 | -4.649781746 | 1.4828 | -3.1358 | 0.00171 | 0.01136 |
| 778 | ZNF91 | 4.3333734 | -4.650473918 | 1.39703 | -3.3288 | 0.00087 | 0.00645 |
| 1561 | ENSG00000231616 | 2.4608876 | -4.65260025 | 1.75101 | -2.6571 | 0.00788 | 0.03859 |
| 1368 | ADGB | 2.4694274 | -4.661452604 | 1.60985 | -2.8956 | 0.00378 | 0.02142 |
| 267 | OR7E14P | 2.488964 | -4.671717731 | 1.60164 | -2.9168 | 0.00354 | 0.0203 |
| 380 | IGFBP6 | 2.5040234 | -4.679806902 | 1.62 | -2.8888 | 0.00387 | 0.02177 |
| 744 | REEP6 | 2.5055158 | -4.680486336 | 1.60539 | -2.9155 | 0.00355 | 0.02037 |
| 1204 | PDGFC | 10.461926 | -4.682992781 | 1.03881 | -4.508 | 6.5E-06 | 0.0001 |
| 1544 | MYORG | 2.5122993 | -4.684247846 | 1.63668 | -2.862 | 0.00421 | 0.02342 |
| 409 | CFAP54 | 2.5137917 | -4.684889112 | 1.61438 | -2.902 | 0.00371 | 0.02106 |
| 1682 | DANT2 | 2.5171834 | -4.686784154 | 1.63189 | -2.872 | 0.00408 | 0.0228 |
| 628 | CDYL2 | 2.5190828 | -4.688177344 | 1.71234 | -2.7379 | 0.00618 | 0.03201 |
| 1559 | PRUNE2 | 2.5121563 | -4.688453344 | 1.73412 | -2.7036 | 0.00686 | 0.0347 |
| 1 | LINC02593 | 2.5205752 | -4.688726442 | 1.66011 | -2.8243 | 0.00474 | 0.02589 |
| 1020 | APOL2 | 7.071226 | -4.693670481 | 1.19004 | -3.9441 | 8E-05 | 0.00089 |
| 1509 | TRPS1 | 4.2982116 | -4.695832046 | 1.40901 | -3.3327 | 0.00086 | 0.00638 |
| 1447 | ENSG00000270953 | 2.5264018 | -4.696007688 | 1.68252 | -2.791 | 0.00525 | 0.02814 |
| 175 | RPP38-DT | 2.5371269 | -4.697914685 | 1.75207 | -2.6814 | 0.00733 | 0.03653 |
| 1070 | KLHDC8B | 28.063667 | -4.699502403 | 0.62917 | -7.4694 | 8.1E-14 | 9.3E-12 |
| 363 | DNAI7 | 4.3284733 | -4.702325634 | 1.36286 | -3.4503 | 0.00056 | 0.00448 |
| 1406 | CLDN3 | 7.2650857 | -4.702975046 | 1.20982 | -3.8873 | 0.0001 | 0.00108 |
| 205 | DYDC2 | 2.5513725 | -4.704797687 | 1.66485 | -2.826 | 0.00471 | 0.02578 |
| 586 | ENSG00000261596 | 2.544303 | -4.708744476 | 1.78864 | -2.6326 | 0.00847 | 0.04082 |
| 1224 | BASP1-AS1 | 2.5679242 | -4.71374153 | 1.69971 | -2.7733 | 0.00555 | 0.02945 |
| 1033 | EFCAB6 | 2.5628971 | -4.715261426 | 1.5822 | -2.9802 | 0.00288 | 0.01723 |
| 241 | KNDC1 | 2.5692737 | -4.718445802 | 1.61617 | -2.9195 | 0.00351 | 0.02014 |

| | | | | | | | |
|---|---|---|---|---|---|---|---|
| 1051 | TGFBR2 | 4.3958628 | -4.718734465 | 1.40269 | -3.3641 | 0.00077 | 0.00582 |
| 37 | TEKT2 | 4.3962697 | -4.718841456 | 1.38978 | -3.3954 | 0.00069 | 0.00529 |
| 763 | ZNF491 | 2.5810843 | -4.721039296 | 1.75621 | -2.6882 | 0.00718 | 0.03601 |
| 547 | TLN2 | 4.3976191 | -4.722284699 | 1.35065 | -3.4963 | 0.00047 | 0.00389 |
| 1643 | CENPVL3 | 4.4019534 | -4.726189612 | 1.43521 | -3.293 | 0.00099 | 0.00716 |
| 1552 | BANCR | 7.0607647 | -4.728576144 | 1.2728 | -3.7151 | 0.0002 | 0.00195 |
| 1216 | NKD2 | 2.5856825 | -4.731389008 | 1.6368 | -2.8906 | 0.00384 | 0.02168 |
| 131 | CCDC181 | 4.4052483 | -4.732420498 | 1.37106 | -3.4516 | 0.00056 | 0.00447 |
| 339 | JHY | 2.605769 | -4.737105615 | 1.58882 | -2.9815 | 0.00287 | 0.01718 |
| 16 | FBXO2 | 2.605626 | -4.741630545 | 1.63127 | -2.9067 | 0.00365 | 0.02082 |
| 577 | BAIAP3 | 2.6219138 | -4.745353849 | 1.63114 | -2.9092 | 0.00362 | 0.0207 |
| 803 | HIF3A | 7.2096625 | -4.746376212 | 1.29593 | -3.6625 | 0.00025 | 0.00232 |
| 29 | RCAN3 | 4.4900034 | -4.753620762 | 1.36605 | -3.4798 | 0.0005 | 0.0041 |
| 410 | RMST | 18.127306 | -4.753998987 | 0.81234 | -5.8522 | 4.9E-09 | 1.8E-07 |
| 452 | PCDH8 | 4.5132639 | -4.755249093 | 1.42722 | -3.3318 | 0.00086 | 0.00639 |
| 383 | ITGA5 | 2.6368302 | -4.75750709 | 1.5786 | -3.0137 | 0.00258 | 0.01578 |
| 172 | PRKCQ | 2.6451061 | -4.761640552 | 1.57597 | -3.0214 | 0.00252 | 0.01549 |
| 258 | CYB5R2 | 2.6484979 | -4.763332819 | 1.58769 | -3.0002 | 0.0027 | 0.01636 |
| 546 | ANXA2 | 65.574686 | -4.772738821 | 0.5254 | -9.084 | 1E-19 | 2.9E-17 |
| 762 | ODAD3 | 2.6714262 | -4.774634517 | 1.56405 | -3.0527 | 0.00227 | 0.01429 |
| 46 | CLDN19 | 2.6729186 | -4.775313446 | 1.56766 | -3.0461 | 0.00232 | 0.01454 |
| 678 | COPZ2 | 2.687571 | -4.782511605 | 1.60363 | -2.9823 | 0.00286 | 0.01715 |
| 1077 | TNNC1 | 2.6913697 | -4.784522457 | 1.58931 | -3.0104 | 0.00261 | 0.01592 |
| 1219 | SEMA5A | 4.7309944 | -4.785823234 | 1.36833 | -3.4976 | 0.00047 | 0.00388 |
| 190 | TMEM72 | 2.7008741 | -4.786267977 | 1.78125 | -2.687 | 0.00721 | 0.03608 |
| 1226 | PRLR | 2.7057582 | -4.788867349 | 1.80218 | -2.6573 | 0.00788 | 0.03859 |
| 761 | EPOR | 4.5958654 | -4.789898786 | 1.40937 | -3.3986 | 0.00068 | 0.00525 |
| 667 | KRT17 | 2.7022235 | -4.789981793 | 1.71994 | -2.785 | 0.00535 | 0.02856 |
| 408 | ELK3 | 4.6058515 | -4.793837015 | 1.37652 | -3.4826 | 0.0005 | 0.00406 |
| 1120 | WDR49 | 2.7123987 | -4.794641472 | 1.57847 | -3.0375 | 0.00239 | 0.01488 |
| 1127 | EPHB3 | 7.571721 | -4.795047927 | 1.1988 | -3.9999 | 6.3E-05 | 0.00073 |
| 483 | ENSG00000259969 | 2.7138911 | -4.795462592 | 1.61235 | -2.9742 | 0.00294 | 0.0175 |
| 471 | MYH7 | 2.7172829 | -4.797047531 | 1.58847 | -3.0199 | 0.00253 | 0.01554 |
| 1025 | C1QTNF6 | 4.6262019 | -4.799908972 | 1.3802 | -3.4777 | 0.00051 | 0.00411 |
| 527 | OCA2 | 2.7152406 | -4.799978202 | 1.71353 | -2.8012 | 0.00509 | 0.02746 |
| 424 | MORN3 | 4.6722699 | -4.803268319 | 1.44844 | -3.3162 | 0.00091 | 0.00667 |
| 1169 | BTC | 2.7304429 | -4.803502711 | 1.5984 | -3.0052 | 0.00265 | 0.01615 |
| 295 | PYGM | 2.735734 | -4.806331364 | 1.62332 | -2.9608 | 0.00307 | 0.01809 |
| 705 | ITGB4 | 2.7406181 | -4.808688531 | 1.62231 | -2.9641 | 0.00304 | 0.01796 |

| | | | | | | | |
|---|---|---|---|---|---|---|---|
| 1287 | FAXDC2 | 2.7366765 | -4.810676467 | 1.65061 | -2.9145 | 0.00356 | 0.02042 |
| 1396 | MINDY4 | 2.7552706 | -4.815806738 | 1.62928 | -2.9558 | 0.00312 | 0.01836 |
| 1555 | MAMDC2 | 2.756763 | -4.816486078 | 1.62955 | -2.9557 | 0.00312 | 0.01836 |
| 1420 | GNG11 | 2.7682876 | -4.826092735 | 1.54675 | -3.1202 | 0.00181 | 0.01186 |
| 1148 | LINC01085 | 2.7750711 | -4.829381832 | 1.56049 | -3.0948 | 0.00197 | 0.01272 |
| 281 | CD82 | 2.7764205 | -4.833848027 | 1.64617 | -2.9364 | 0.00332 | 0.01929 |
| 1393 | TRIL | 8.078318 | -4.843557271 | 1.32149 | -3.6652 | 0.00025 | 0.0023 |
| 541 | TEX9 | 7.6660825 | -4.844279338 | 1.20237 | -4.029 | 5.6E-05 | 0.00066 |
| 738 | DSEL | 4.7746181 | -4.845524962 | 1.42933 | -3.3901 | 0.0007 | 0.00538 |
| 951 | SNTA1 | 7.945668 | -4.850512715 | 1.18945 | -4.078 | 4.5E-05 | 0.00055 |
| 1467 | PHYHIP | 2.8209278 | -4.850931544 | 1.56382 | -3.102 | 0.00192 | 0.01247 |
| 144 | PRELP | 2.8224202 | -4.851652401 | 1.57753 | -3.0755 | 0.0021 | 0.01344 |
| 1004 | TXNRD2 | 4.7898522 | -4.854036575 | 1.33044 | -3.6485 | 0.00026 | 0.00242 |
| 1356 | TPD52L1 | 29.701745 | -4.854091518 | 1.32285 | -3.6694 | 0.00024 | 0.00227 |
| 545 | MYO1E | 4.9593133 | -4.857568882 | 1.3773 | -3.5269 | 0.00042 | 0.00355 |
| 1579 | MORN5 | 2.8355802 | -4.857885816 | 1.588 | -3.0591 | 0.00222 | 0.01403 |
| 1086 | ZNF717 | 2.8358442 | -4.862284908 | 1.57947 | -3.0784 | 0.00208 | 0.01333 |
| 1306 | DSP | 2.8453485 | -4.862640642 | 1.62043 | -3.0008 | 0.00269 | 0.01633 |
| 753 | TRIP10 | 4.8277938 | -4.863321541 | 1.3428 | -3.6218 | 0.00029 | 0.00264 |
| 1200 | NR3C2 | 2.8553808 | -4.871337316 | 1.53466 | -3.1742 | 0.0015 | 0.01018 |
| 669 | ODAD4 | 8.1479991 | -4.872860699 | 1.18678 | -4.106 | 4E-05 | 0.0005 |
| 1310 | CD83 | 15.571522 | -4.880263322 | 0.90632 | -5.3847 | 7.3E-08 | 2E-06 |
| 1186 | TACR3 | 2.8818438 | -4.880436313 | 1.70465 | -2.863 | 0.0042 | 0.02337 |
| 44 | COL9A2 | 132.89201 | -4.881193841 | 0.45249 | -10.787 | 3.9E-27 | 3.4E-24 |
| 126 | DDR2 | 8.4844056 | -4.881764498 | 1.18266 | -4.1278 | 3.7E-05 | 0.00046 |
| 1398 | BBS9 | 2.886585 | -4.885574676 | 1.53255 | -3.1879 | 0.00143 | 0.0098 |
| 64 | PDE4B | 2.8869919 | -4.885978173 | 1.57992 | -3.0925 | 0.00198 | 0.0128 |
| 1010 | C22orf15 | 2.8884843 | -4.886558054 | 1.54909 | -3.1545 | 0.00161 | 0.01079 |
| 474 | LINC01551 | 2.881151 | -4.886683827 | 1.64892 | -2.9636 | 0.00304 | 0.01798 |
| 1568 | CTSV | 11.878549 | -4.889210412 | 1.07652 | -4.5417 | 5.6E-06 | 8.8E-05 |
| 140 | PTPRC | 2.8963533 | -4.890013099 | 1.54331 | -3.1685 | 0.00153 | 0.01036 |
| 1445 | CPA4 | 2.8928186 | -4.892507207 | 1.5766 | -3.1032 | 0.00191 | 0.01244 |
| 11 | MIR34AHG | 8.221215 | -4.892589651 | 1.43373 | -3.4125 | 0.00064 | 0.00503 |
| 1662 | PCDH11X | 2.9031368 | -4.8931457 | 1.53884 | -3.1798 | 0.00147 | 0.01003 |
| 10 | SLC2A5 | 5.066064 | -4.895378018 | 1.33398 | -3.6698 | 0.00024 | 0.00227 |
| 824 | C19orf18 | 2.9059786 | -4.898544893 | 1.56692 | -3.1262 | 0.00177 | 0.01164 |
| 1367 | RAB32 | 2.9225304 | -4.906167646 | 1.54014 | -3.1855 | 0.00144 | 0.00986 |
| 1059 | CCDC13-AS1 | 2.9508928 | -4.91512843 | 1.59865 | -3.0746 | 0.00211 | 0.01346 |
| 670 | CAVIN1 | 16.880923 | -4.917580192 | 0.88183 | -5.5766 | 2.5E-08 | 7.7E-07 |

| | | | | | | | |
|---|---|---|---|---|---|---|---|
| 618 | CDH3 | 2.9655452 | -4.921788085 | 1.60903 | -3.0589 | 0.00222 | 0.01403 |
| 1318 | MICA | 2.9655452 | -4.921788085 | 1.60903 | -3.0589 | 0.00222 | 0.01403 |
| 952 | GDF5 | 2.9639099 | -4.924733543 | 1.52211 | -3.2355 | 0.00121 | 0.00853 |
| 1283 | PDGFRB | 24.280867 | -4.929578994 | 0.72541 | -6.7956 | 1.1E-11 | 7.7E-10 |
| 799 | BCAM | 2.9838534 | -4.933645207 | 1.5481 | -3.1869 | 0.00144 | 0.00982 |
| 1634 | EFHC2 | 15.604177 | -4.946045399 | 0.97834 | -5.0556 | 4.3E-07 | 9.4E-06 |
| 1278 | SPRY4 | 3.0214342 | -4.950067334 | 1.53565 | -3.2234 | 0.00127 | 0.00883 |
| 1437 | TES | 3.0210272 | -4.950082852 | 1.60147 | -3.091 | 0.002 | 0.01287 |
| 1327 | TREM1 | 3.0259114 | -4.952384834 | 1.63645 | -3.0263 | 0.00248 | 0.0153 |
| 997 | ENSG00000273796 | 3.0291602 | -4.957323869 | 1.55971 | -3.1784 | 0.00148 | 0.01007 |
| 1698 | SMIM10L2A | 5.1632923 | -4.959364101 | 1.32373 | -3.7465 | 0.00018 | 0.00176 |
| 1517 | PCAT1 | 3.0458549 | -4.960886922 | 1.56991 | -3.16 | 0.00158 | 0.01064 |
| 75 | CCN1 | 12.899729 | -4.963339543 | 1.1484 | -4.3219 | 1.5E-05 | 0.00021 |
| 1258 | TNFAIP8 | 5.182422 | -4.964995383 | 1.43595 | -3.4576 | 0.00054 | 0.00439 |
| 183 | ENKUR | 5.3238307 | -4.966701274 | 1.33718 | -3.7143 | 0.0002 | 0.00196 |
| 402 | DUSP6 | 5.1911048 | -4.966745814 | 1.34216 | -3.7006 | 0.00022 | 0.00205 |
| 962 | CEBPB | 5.3236878 | -4.969240081 | 1.31132 | -3.7895 | 0.00015 | 0.00152 |
| 1043 | LMCD1 | 3.0519455 | -4.970344616 | 1.64294 | -3.0253 | 0.00248 | 0.01534 |
| 42 | RSPO1 | 3.0557442 | -4.971967993 | 1.66872 | -2.9795 | 0.00289 | 0.01726 |
| 1570 | ABITRAM | 5.3699514 | -4.981137481 | 1.30606 | -3.8139 | 0.00014 | 0.0014 |
| 356 | C1R | 26.325734 | -4.982494102 | 0.7015 | -7.1026 | 1.2E-12 | 1.1E-10 |
| 1530 | EPPK1 | 5.3844608 | -4.987617343 | 1.30717 | -3.8156 | 0.00014 | 0.00139 |
| 1161 | NSUN7 | 5.2677511 | -4.989654455 | 1.40133 | -3.5607 | 0.00037 | 0.0032 |
| 1166 | REST | 16.573888 | -4.989660775 | 0.88692 | -5.6258 | 1.8E-08 | 6E-07 |
| 1332 | EFHC1 | 72.622515 | -4.990430188 | 0.5202 | -9.5933 | 8.5E-22 | 3.8E-19 |
| 1232 | OSMR | 3.1219513 | -5.000901815 | 1.66986 | -2.9948 | 0.00275 | 0.01659 |
| 426 | LRRC43 | 3.1393246 | -5.004991071 | 1.60022 | -3.1277 | 0.00176 | 0.0116 |
| 1027 | KDELR3 | 9.0847245 | -5.01300064 | 1.17106 | -4.2807 | 1.9E-05 | 0.00025 |
| 1329 | LRRC73 | 3.1508493 | -5.013293072 | 1.51446 | -3.3103 | 0.00093 | 0.00679 |
| 1145 | SORCS2 | 8.7572506 | -5.01430034 | 1.16835 | -4.2918 | 1.8E-05 | 0.00024 |
| 374 | METTL7A | 9.0204914 | -5.019610845 | 1.15543 | -4.3444 | 1.4E-05 | 0.0002 |
| 873 | MTLN | 3.1653587 | -5.022627676 | 1.58995 | -3.159 | 0.00158 | 0.01066 |
| 417 | ABTB3 | 3.18898 | -5.026381313 | 1.62647 | -3.0904 | 0.002 | 0.01289 |
| 709 | RHBDF2 | 3.201997 | -5.034520918 | 1.49921 | -3.3581 | 0.00078 | 0.00591 |
| 1238 | IL6ST | 13.190277 | -5.036155741 | 0.98544 | -5.1106 | 3.2E-07 | 7.3E-06 |
| 1134 | OSTN | 3.2332012 | -5.047441507 | 1.52169 | -3.317 | 0.00091 | 0.00666 |
| 405 | DCN | 17.619421 | -5.048725037 | 0.98959 | -5.1018 | 3.4E-07 | 7.6E-06 |
| 1462 | FAM167A | 3.2403917 | -5.051060302 | 1.61439 | -3.1288 | 0.00176 | 0.01157 |
| 1353 | GOPC | 5.4967948 | -5.051848531 | 1.35178 | -3.7372 | 0.00019 | 0.00182 |

| | | | | | | | |
|---|---|---|---|---|---|---|---|
| 1407 | CLDN4 | 5.5141681 | -5.053831255 | 1.35664 | -3.7253 | 0.0002 | 0.00189 |
| 662 | SLFN5 | 3.2447259 | -5.055928613 | 1.48255 | -3.4103 | 0.00065 | 0.00507 |
| 1290 | KCNIP1 | 5.5154636 | -5.05874098 | 1.3279 | -3.8096 | 0.00014 | 0.00142 |
| 509 | EFCAB11 | 5.5317514 | -5.060704509 | 1.36276 | -3.7136 | 0.0002 | 0.00196 |
| 95 | TENT5C | 8.8616443 | -5.060906279 | 1.1767 | -4.3009 | 1.7E-05 | 0.00023 |
| 475 | NPAS3 | 3.2627701 | -5.063181756 | 1.47911 | -3.4231 | 0.00062 | 0.00487 |
| 52 | DMRTA2 | 3.2626271 | -5.066572661 | 1.54202 | -3.2857 | 0.00102 | 0.00733 |
| 117 | ACKR1 | 3.2902966 | -5.079698107 | 1.67989 | -3.0238 | 0.0025 | 0.01539 |
| 333 | MPZL2 | 3.310933 | -5.082782731 | 1.53402 | -3.3134 | 0.00092 | 0.00673 |
| 192 | MRLN | 3.3283063 | -5.087985389 | 1.72904 | -2.9427 | 0.00325 | 0.019 |
| 1007 | LRRC74B | 3.3255855 | -5.088551786 | 1.51962 | -3.3486 | 0.00081 | 0.00608 |
| 610 | RRAD | 5.6392027 | -5.089144548 | 1.56597 | -3.2498 | 0.00115 | 0.00818 |
| 1391 | GPNMB | 14.139234 | -5.089717082 | 1.03788 | -4.9039 | 9.4E-07 | 1.9E-05 |
| 443 | FREM2 | 3.3481069 | -5.097509391 | 1.53619 | -3.3183 | 0.00091 | 0.00664 |
| 961 | SLC2A10 | 5.6553716 | -5.098538793 | 1.34416 | -3.7931 | 0.00015 | 0.0015 |
| 1513 | SNTB1 | 13.500764 | -5.099842416 | 0.98148 | -5.1961 | 2E-07 | 4.9E-06 |
| 427 | RILPL2 | 3.353255 | -5.103014932 | 1.47135 | -3.4682 | 0.00052 | 0.00424 |
| 779 | ENSG00000267934 | 3.353255 | -5.103014932 | 1.47135 | -3.4682 | 0.00052 | 0.00424 |
| 440 | KL | 3.3661511 | -5.104885471 | 1.56334 | -3.2654 | 0.00109 | 0.00781 |
| 185 | MAP3K8 | 3.3611239 | -5.10599232 | 1.49434 | -3.4169 | 0.00063 | 0.00497 |
| 1397 | AQP1 | 27.836384 | -5.106038551 | 0.89011 | -5.7364 | 9.7E-09 | 3.4E-07 |
| 1263 | C5orf63 | 3.3615309 | -5.106261957 | 1.47319 | -3.4661 | 0.00053 | 0.00427 |
| 1154 | SOD3 | 3.3729345 | -5.107555424 | 1.55275 | -3.2894 | 0.001 | 0.00724 |
| 102 | SELENBP1 | 9.8134672 | -5.110393388 | 1.15829 | -4.412 | 1E-05 | 0.00015 |
| 988 | BACE2 | 3.3795751 | -5.113270283 | 1.469 | -3.4808 | 0.0005 | 0.00408 |
| 937 | ANKEF1 | 3.3882579 | -5.116922732 | 1.54306 | -3.3161 | 0.00091 | 0.00667 |
| 1369 | PLEKHG1 | 3.3893433 | -5.117032016 | 1.46905 | -3.4832 | 0.0005 | 0.00405 |
| 468 | AJUBA | 9.5650963 | -5.118330708 | 1.15943 | -4.4145 | 1E-05 | 0.00015 |
| 401 | LRRIQ1 | 46.451443 | -5.119682596 | 0.56441 | -9.0709 | 1.2E-19 | 3.2E-17 |
| 1705 | GABRE | 3.3995186 | -5.121142841 | 1.49343 | -3.4291 | 0.00061 | 0.00478 |
| 1383 | IQCE | 9.7413187 | -5.123047857 | 1.14078 | -4.4908 | 7.1E-06 | 0.00011 |
| 525 | MAGEL2 | 3.4155204 | -5.130892016 | 1.47164 | -3.4865 | 0.00049 | 0.00401 |
| 605 | DRC7 | 3.4254317 | -5.131065135 | 1.4899 | -3.4439 | 0.00057 | 0.00457 |
| 263 | DKK3 | 28.089932 | -5.133759669 | 0.73096 | -7.0233 | 2.2E-12 | 1.7E-10 |
| 739 | NFATC1 | 3.4322152 | -5.133771834 | 1.4952 | -3.4335 | 0.0006 | 0.00471 |
| 1084 | SUCLG2 | 5.7919772 | -5.135072091 | 1.28767 | -3.9879 | 6.7E-05 | 0.00076 |
| 130 | MAEL | 3.4375063 | -5.136272713 | 1.56056 | -3.2913 | 0.001 | 0.0072 |
| 656 | MFAP4 | 5.8386698 | -5.138263288 | 1.28654 | -3.9939 | 6.5E-05 | 0.00074 |
| 826 | FAM110C | 3.4452323 | -5.142302201 | 1.46126 | -3.5191 | 0.00043 | 0.00363 |

| | | | | | | | |
|---|---|---|---|---|---|---|---|
| 368 | PDZRN4 | 3.4662613 | -5.150205541 | 1.48597 | -3.4659 | 0.00053 | 0.00427 |
| 1416 | ENSG00000234459 | 3.4627266 | -5.151954177 | 1.53675 | -3.3525 | 0.0008 | 0.00601 |
| 684 | COL1A1 | 123.50561 | -5.154290891 | 0.85067 | -6.0591 | 1.4E-09 | 5.9E-08 |
| 1179 | MAPK10 | 18.547026 | -5.154590544 | 0.86211 | -5.979 | 2.2E-09 | 9.3E-08 |
| 521 | DIO3OS | 6.0300726 | -5.15845134 | 1.4491 | -3.5598 | 0.00037 | 0.00321 |
| 332 | FXYD6 | 14.413331 | -5.163620609 | 1.0239 | -5.0431 | 4.6E-07 | 9.9E-06 |
| 1655 | HEPH | 3.5072338 | -5.165952513 | 1.54982 | -3.3333 | 0.00086 | 0.00637 |
| 1060 | ZNF662 | 3.5060053 | -5.16883879 | 1.47784 | -3.4976 | 0.00047 | 0.00388 |
| 569 | WDR93 | 3.5222932 | -5.171434166 | 1.48308 | -3.487 | 0.00049 | 0.00401 |
| 517 | AK7 | 6.0893609 | -5.171551667 | 1.29193 | -4.003 | 6.3E-05 | 0.00072 |
| 1653 | ZC4H2 | 3.517266 | -5.17308205 | 1.45351 | -3.559 | 0.00037 | 0.00321 |
| 79 | GBP1 | 3.5416868 | -5.182012985 | 1.51092 | -3.4297 | 0.0006 | 0.00477 |
| 156 | SPATA17 | 6.1668287 | -5.189314303 | 1.31899 | -3.9343 | 8.3E-05 | 0.00092 |
| 49 | BEST4 | 6.0367268 | -5.193100926 | 1.45143 | -3.5779 | 0.00035 | 0.00303 |
| 1523 | LY6H | 3.5716626 | -5.196467336 | 1.50613 | -3.4502 | 0.00056 | 0.00448 |
| 454 | EDNRB | 3.5880934 | -5.196630724 | 1.54845 | -3.356 | 0.00079 | 0.00594 |
| 108 | S100A16 | 3.6156199 | -5.212636867 | 1.48887 | -3.5011 | 0.00046 | 0.00384 |
| 791 | SERTAD1 | 3.6370558 | -5.220503945 | 1.471 | -3.5489 | 0.00039 | 0.00332 |
| 776 | ZNF493 | 3.6408545 | -5.222037304 | 1.45126 | -3.5983 | 0.00032 | 0.00285 |
| 252 | ART5 | 3.6522582 | -5.222920995 | 1.48745 | -3.5113 | 0.00045 | 0.00373 |
| 105 | S100A11 | 29.655397 | -5.223292847 | 0.77036 | -6.7803 | 1.2E-11 | 8.4E-10 |
| 790 | PRX | 3.647638 | -5.224506257 | 1.48137 | -3.5268 | 0.00042 | 0.00355 |
| 748 | GNG7 | 3.6451887 | -5.226519921 | 1.49636 | -3.4928 | 0.00048 | 0.00393 |
| 875 | RABL2A | 14.838478 | -5.230336912 | 1.05326 | -4.9658 | 6.8E-07 | 1.4E-05 |
| 1361 | SLC2A12 | 3.6781713 | -5.232975045 | 1.57805 | -3.3161 | 0.00091 | 0.00667 |
| 1342 | CFAP206 | 3.6720587 | -5.233234509 | 1.43989 | -3.6345 | 0.00028 | 0.00254 |
| 397 | CAPS2 | 3.6882036 | -5.23893966 | 1.44673 | -3.6212 | 0.00029 | 0.00265 |
| 1143 | MSX1 | 42.120336 | -5.24244325 | 0.63771 | -8.2207 | 2E-16 | 3.6E-14 |
| 948 | TTLL9 | 3.7081471 | -5.246089448 | 1.44484 | -3.6309 | 0.00028 | 0.00257 |
| 540 | PRTG | 15.270615 | -5.254510822 | 1.08045 | -4.8633 | 1.2E-06 | 2.2E-05 |
| 518 | CYP46A1 | 3.7455849 | -5.262625669 | 1.4426 | -3.648 | 0.00026 | 0.00243 |
| 1471 | PTK2B | 10.365944 | -5.263087356 | 1.25976 | -4.1778 | 2.9E-05 | 0.00038 |
| 1294 | HRH2 | 3.7671638 | -5.267338537 | 1.49469 | -3.524 | 0.00043 | 0.00358 |
| 96 | SPAG17 | 10.540609 | -5.268167763 | 1.15573 | -4.5583 | 5.2E-06 | 8.2E-05 |
| 821 | ZNF667 | 3.7621367 | -5.268444194 | 1.43069 | -3.6824 | 0.00023 | 0.00217 |
| 400 | ACSS3 | 3.7625436 | -5.268696709 | 1.44378 | -3.6492 | 0.00026 | 0.00242 |
| 1384 | SDK1 | 3.7704126 | -5.27133754 | 1.42787 | -3.6917 | 0.00022 | 0.00211 |
| 645 | WSCD1 | 6.3578148 | -5.273399713 | 1.28645 | -4.0992 | 4.1E-05 | 0.00051 |
| 133 | ANKRD45 | 3.7786885 | -5.274226322 | 1.42703 | -3.6959 | 0.00022 | 0.00208 |

| | | | | | | | |
|---|---|---|---|---|---|---|---|
| 855 | OTX1 | 3.7804448 | -5.277939584 | 1.48941 | -3.5436 | 0.00039 | 0.00338 |
| 1126 | SOX2 | 21.47975 | -5.289202249 | 0.83325 | -6.3477 | 2.2E-10 | 1.2E-08 |
| 1189 | LEF1 | 3.8234597 | -5.290029898 | 1.46778 | -3.6041 | 0.00031 | 0.0028 |
| 723 | PTPRM | 6.6132912 | -5.294726039 | 1.35587 | -3.905 | 9.4E-05 | 0.00103 |
| 879 | CCDC74A | 10.511649 | -5.297027675 | 1.12587 | -4.7048 | 2.5E-06 | 4.4E-05 |
| 783 | DMKN | 3.8364767 | -5.297580454 | 1.44868 | -3.6568 | 0.00026 | 0.00236 |
| 1111 | TM4SF18 | 3.8423034 | -5.302330489 | 1.47615 | -3.592 | 0.00033 | 0.00291 |
| 1293 | MSX2 | 10.693551 | -5.30898675 | 1.17195 | -4.53 | 5.9E-06 | 9.2E-05 |
| 1125 | ENSG00000241231 | 3.867538 | -5.31093094 | 1.5082 | -3.5214 | 0.00043 | 0.0036 |
| 199 | STOX1 | 6.6957642 | -5.314763392 | 1.26638 | -4.1968 | 2.7E-05 | 0.00035 |
| 1055 | DLEC1 | 3.8971288 | -5.315980729 | 1.53207 | -3.4698 | 0.00052 | 0.00422 |
| 841 | GALM | 3.8953504 | -5.32076977 | 1.43756 | -3.7013 | 0.00021 | 0.00205 |
| 1039 | ODF3B | 3.9215496 | -5.324699664 | 1.56812 | -3.3956 | 0.00068 | 0.00529 |
| 1364 | PERP | 6.6282138 | -5.326453148 | 1.281 | -4.1581 | 3.2E-05 | 0.00041 |
| 1536 | ERMP1 | 6.6485421 | -5.337376804 | 1.27372 | -4.1904 | 2.8E-05 | 0.00036 |
| 110 | KCNN3 | 3.9545101 | -5.337693611 | 1.45856 | -3.6596 | 0.00025 | 0.00234 |
| 367 | CPNE8 | 3.9612936 | -5.340134327 | 1.47291 | -3.6256 | 0.00029 | 0.00261 |
| 1255 | TRIM36 | 3.9645424 | -5.34398461 | 1.41582 | -3.7745 | 0.00016 | 0.00161 |
| 529 | THBS1 | 11.487638 | -5.346075862 | 1.18979 | -4.4933 | 7E-06 | 0.00011 |
| 1422 | PPP1R9A | 11.217093 | -5.351979844 | 1.13449 | -4.7175 | 2.4E-06 | 4.2E-05 |
| 1188 | SGMS2 | 10.995954 | -5.352024871 | 1.32322 | -4.0447 | 5.2E-05 | 0.00062 |
| 1144 | WFS1 | 4.0053719 | -5.360591762 | 1.41522 | -3.7878 | 0.00015 | 0.00153 |
| 485 | RHOJ | 4.0333274 | -5.366685916 | 1.43392 | -3.7427 | 0.00018 | 0.00179 |
| 1680 | SNORA35 | 4.0249085 | -5.36693936 | 1.41129 | -3.8028 | 0.00014 | 0.00145 |
| 31 | LDLRAP1 | 6.7924887 | -5.367249362 | 1.35182 | -3.9704 | 7.2E-05 | 0.00081 |
| 1343 | ANKRD6 | 4.0444451 | -5.373235135 | 1.44305 | -3.7235 | 0.0002 | 0.0019 |
| 453 | PCDH17 | 4.0512286 | -5.375497501 | 1.40819 | -3.8173 | 0.00013 | 0.00139 |
| 189 | ZNF487 | 6.8389701 | -5.37603534 | 1.25657 | -4.2783 | 1.9E-05 | 0.00025 |
| 1260 | SNCAIP | 6.8627806 | -5.376196629 | 1.33716 | -4.0206 | 5.8E-05 | 0.00068 |
| 607 | CCDC113 | 4.0546203 | -5.376618286 | 1.40139 | -3.8366 | 0.00012 | 0.0013 |
| 149 | SLC45A3 | 4.0550273 | -5.376849294 | 1.42288 | -3.7789 | 0.00016 | 0.00158 |
| 1358 | CCN2 | 57.624552 | -5.380241199 | 1.15417 | -4.6616 | 3.1E-06 | 5.3E-05 |
| 1545 | C9orf24 | 27.535438 | -5.384626986 | 1.37145 | -3.9262 | 8.6E-05 | 0.00095 |
| 421 | CFAP73 | 4.0908517 | -5.386027194 | 1.48951 | -3.616 | 0.0003 | 0.00269 |
| 323 | CFAP300 | 6.9178557 | -5.388085028 | 1.27205 | -4.2357 | 2.3E-05 | 0.0003 |
| 272 | DCDC1 | 4.1255686 | -5.402568748 | 1.41657 | -3.8138 | 0.00014 | 0.0014 |
| 650 | TMEM88 | 4.1467406 | -5.406408083 | 1.43325 | -3.7721 | 0.00016 | 0.00162 |
| 606 | KIFC3 | 4.1398142 | -5.406986827 | 1.41533 | -3.8203 | 0.00013 | 0.00138 |
| 1164 | SPATA18 | 7.0065159 | -5.410658763 | 1.27691 | -4.2373 | 2.3E-05 | 0.0003 |

| | | | | | | | |
|---|---|---|---|---|---|---|---|
| 1637 | TIMP1 | 73.676651 | -5.414701363 | 0.4967 | -10.901 | 1.1E-27 | 1E-24 |
| 1093 | CCDC191 | 7.1530728 | -5.415137148 | 1.25957 | -4.2992 | 1.7E-05 | 0.00023 |
| 1170 | CDKL2 | 4.1676266 | -5.415842562 | 1.45338 | -3.7264 | 0.00019 | 0.00189 |
| 422 | HSPB8 | 4.195989 | -5.422619013 | 1.48555 | -3.6502 | 0.00026 | 0.00242 |
| 111 | PBXIP1 | 27.465967 | -5.422993432 | 0.77842 | -6.9667 | 3.2E-12 | 2.5E-10 |
| 512 | FBLN5 | 4.2042649 | -5.425402192 | 1.49883 | -3.6197 | 0.00029 | 0.00266 |
| 480 | LGALS3 | 29.853627 | -5.425763487 | 0.88224 | -6.15 | 7.7E-10 | 3.5E-08 |
| 1019 | TIMP3 | 45.118439 | -5.425958464 | 0.6545 | -8.2902 | 1.1E-16 | 2E-14 |
| 1626 | ARX | 4.2008512 | -5.430748207 | 1.57793 | -3.4417 | 0.00058 | 0.0046 |
| 603 | CCDC102A | 4.2220232 | -5.435882956 | 1.39781 | -3.8889 | 0.0001 | 0.00108 |
| 624 | TMEM231 | 23.208984 | -5.438277072 | 0.83633 | -6.5026 | 7.9E-11 | 4.6E-09 |
| 635 | CIBAR2 | 4.2436021 | -5.439822359 | 1.42505 | -3.8173 | 0.00013 | 0.00139 |
| 116 | CRABP2 | 28.549721 | -5.440901048 | 0.76782 | -7.0862 | 1.4E-12 | 1.2E-10 |
| 1487 | PLAG1 | 4.2472578 | -5.443832142 | 1.40927 | -3.8629 | 0.00011 | 0.00119 |
| 1090 | ALCAM | 4.2823893 | -5.456991356 | 1.49136 | -3.6591 | 0.00025 | 0.00234 |
| 847 | EPAS1 | 36.86934 | -5.459810156 | 0.68549 | -7.9648 | 1.7E-15 | 2.5E-13 |
| 179 | NEBL | 4.3057246 | -5.464593462 | 1.40034 | -3.9023 | 9.5E-05 | 0.00103 |
| 430 | GLT1D1 | 4.3156358 | -5.464903184 | 1.43093 | -3.8191 | 0.00013 | 0.00138 |
| 1681 | PLS3 | 145.12107 | -5.46558707 | 0.76616 | -7.1338 | 9.8E-13 | 8.9E-11 |
| 1103 | NEK11 | 11.889806 | -5.466206353 | 1.15689 | -4.7249 | 2.3E-06 | 4.1E-05 |
| 340 | GRAMD1B | 17.105816 | -5.469957771 | 1.0028 | -5.4547 | 4.9E-08 | 1.4E-06 |
| 56 | MROH7 | 4.3484755 | -5.474081565 | 1.59538 | -3.4312 | 0.0006 | 0.00475 |
| 231 | LINC02944 | 4.360407 | -5.478703755 | 1.42774 | -3.8373 | 0.00012 | 0.0013 |
| 1012 | KIAA1671 | 4.3929607 | -5.491035339 | 1.39457 | -3.9374 | 8.2E-05 | 0.00091 |
| 1040 | SHANK3 | 7.4083743 | -5.497232781 | 1.25512 | -4.3799 | 1.2E-05 | 0.00017 |
| 1064 | PTH1R | 4.4184592 | -5.50142948 | 1.49869 | -3.6708 | 0.00024 | 0.00226 |
| 839 | LTBP1 | 12.551129 | -5.504434995 | 1.13177 | -4.8635 | 1.2E-06 | 2.2E-05 |
| 888 | CSRNP3 | 4.4392243 | -5.504988186 | 1.40311 | -3.9234 | 8.7E-05 | 0.00096 |
| 550 | KIF23-AS1 | 4.4797899 | -5.517904585 | 1.5194 | -3.6316 | 0.00028 | 0.00256 |
| 792 | BLVRB | 4.4751697 | -5.518259636 | 1.37629 | -4.0095 | 6.1E-05 | 0.00071 |
| 1138 | HES1 | 12.122447 | -5.518512097 | 1.26195 | -4.373 | 1.2E-05 | 0.00017 |
| 1102 | EFCAB12 | 4.4888796 | -5.52056445 | 1.48639 | -3.7141 | 0.0002 | 0.00196 |
| 1387 | ENSG00000226690 | 4.505967 | -5.527663002 | 1.48779 | -3.7153 | 0.0002 | 0.00195 |
| 1005 | COMT | 18.298438 | -5.532150518 | 0.95642 | -5.7842 | 7.3E-09 | 2.7E-07 |
| 435 | LATS2 | 7.7142578 | -5.552634203 | 1.28546 | -4.3196 | 1.6E-05 | 0.00022 |
| 865 | TEKT4 | 4.615046 | -5.559994964 | 1.47418 | -3.7716 | 0.00016 | 0.00162 |
| 764 | PODNL1 | 4.606891 | -5.562026499 | 1.37951 | -4.0319 | 5.5E-05 | 0.00065 |
| 164 | ITPKB | 7.9309712 | -5.567447943 | 1.38423 | -4.0221 | 5.8E-05 | 0.00067 |
| 1630 | DYNLT3 | 4.6478636 | -5.573657207 | 1.35797 | -4.1044 | 4.1E-05 | 0.0005 |

| | | | | | | | |
|---|---|---|---|---|---|---|---|
| 1376 | RSPH3 | 7.8410204 | -5.574324673 | 1.22868 | -4.5369 | 5.7E-06 | 9E-05 |
| 255 | CAVIN3 | 4.6722843 | -5.580493481 | 1.36485 | -4.0887 | 4.3E-05 | 0.00052 |
| 1414 | SEMA3C | 7.8210327 | -5.580677212 | 1.31385 | -4.2476 | 2.2E-05 | 0.00029 |
| 444 | LHFPL6 | 12.64848 | -5.583116065 | 1.14728 | -4.8664 | 1.1E-06 | 2.2E-05 |
| 900 | COL5A2 | 12.864746 | -5.583869896 | 1.10024 | -5.0751 | 3.9E-07 | 8.6E-06 |
| 627 | DYNLRB2 | 4.7002398 | -5.586533344 | 1.44851 | -3.8567 | 0.00011 | 0.00121 |
| 681 | IGF2BP1 | 12.996301 | -5.589266989 | 1.09493 | -5.1047 | 3.3E-07 | 7.5E-06 |
| 1503 | MATN2 | 7.8949878 | -5.590825947 | 1.22467 | -4.5652 | 5E-06 | 8E-05 |
| 235 | CPXM2 | 18.75653 | -5.592016681 | 1.01054 | -5.5337 | 3.1E-08 | 9.6E-07 |
| 1284 | CD74 | 4.718141 | -5.593531201 | 1.40138 | -3.9914 | 6.6E-05 | 0.00075 |
| 1439 | FEZF1 | 4.7173195 | -5.595832425 | 1.41842 | -3.9451 | 8E-05 | 0.00088 |
| 700 | SDK2 | 4.7500161 | -5.604996664 | 1.37129 | -4.0874 | 4.4E-05 | 0.00053 |
| 216 | DNMBP | 4.7642616 | -5.608815369 | 1.35882 | -4.1277 | 3.7E-05 | 0.00046 |
| 1073 | SEMA3F | 4.7740299 | -5.611520709 | 1.37054 | -4.0944 | 4.2E-05 | 0.00051 |
| 1234 | CCDC152 | 4.7839412 | -5.612302827 | 1.43433 | -3.9128 | 9.1E-05 | 0.001 |
| 345 | NTM | 4.8285694 | -5.626580031 | 1.36977 | -4.1077 | 4E-05 | 0.00049 |
| 704 | SMIM5 | 4.834682 | -5.627399422 | 1.53693 | -3.6615 | 0.00025 | 0.00233 |
| 816 | LRRC4B | 4.8417294 | -5.63023901 | 1.37695 | -4.0889 | 4.3E-05 | 0.00052 |
| 939 | SPTLC3 | 4.8367023 | -5.631341114 | 1.34532 | -4.1859 | 2.8E-05 | 0.00037 |
| 1092 | CCDC80 | 8.2138137 | -5.643203229 | 1.22111 | -4.6214 | 3.8E-06 | 6.2E-05 |
| 71 | ST6GALNAC3 | 4.8977613 | -5.646094684 | 1.43101 | -3.9455 | 8E-05 | 0.00088 |
| 1243 | MARVELD2 | 4.9141701 | -5.652308309 | 1.35945 | -4.1578 | 3.2E-05 | 0.00041 |
| 20 | EPHA2 | 4.9248809 | -5.657527242 | 1.39973 | -4.0419 | 5.3E-05 | 0.00063 |
| 22 | KLHDC7A | 4.9400833 | -5.659343077 | 1.38023 | -4.1003 | 4.1E-05 | 0.0005 |
| 282 | PRDM11 | 8.3916777 | -5.672578114 | 1.26485 | -4.4848 | 7.3E-06 | 0.00011 |
| 3 | MXRA8 | 33.254512 | -5.67349849 | 0.77127 | -7.356 | 1.9E-13 | 2.1E-11 |
| 1311 | GMPR | 8.4422755 | -5.681882476 | 1.24741 | -4.555 | 5.2E-06 | 8.4E-05 |
| 501 | TGFB3 | 13.723996 | -5.682234273 | 1.13066 | -5.0256 | 5E-07 | 1.1E-05 |
| 1262 | GRAMD2B | 27.319205 | -5.69896957 | 0.91323 | -6.2404 | 4.4E-10 | 2.1E-08 |
| 630 | HSD17B2 | 5.1074861 | -5.708180268 | 1.37238 | -4.1593 | 3.2E-05 | 0.0004 |
| 908 | PARD3B | 5.1180683 | -5.710979928 | 1.35926 | -4.2015 | 2.7E-05 | 0.00035 |
| 717 | METRNL | 8.607779 | -5.711074052 | 1.26202 | -4.5253 | 6E-06 | 9.4E-05 |
| 1168 | IGFBP7-AS1 | 5.1203745 | -5.712122142 | 1.45731 | -3.9196 | 8.9E-05 | 0.00097 |
| 1325 | CDKN1A | 25.382257 | -5.71264033 | 1.44172 | -3.9624 | 7.4E-05 | 0.00083 |
| 431 | RIMBP2 | 5.1603902 | -5.724088496 | 1.32834 | -4.3092 | 1.6E-05 | 0.00022 |
| 1576 | TNC | 5.1727144 | -5.732098006 | 1.49934 | -3.8231 | 0.00013 | 0.00136 |
| 1314 | ID4 | 14.24065 | -5.73406758 | 1.10326 | -5.1974 | 2E-07 | 4.9E-06 |
| 240 | CFAP46 | 8.7970247 | -5.743013527 | 1.22973 | -4.6701 | 3E-06 | 5.1E-05 |
| 165 | WNT9A | 5.2605224 | -5.748805979 | 1.47711 | -3.8919 | 9.9E-05 | 0.00107 |

| | | | | | | | |
|---|---|---|---|---|---|---|---|
| 1280 | KCTD16 | 5.2804659 | -5.754669221 | 1.55878 | -3.6918 | 0.00022 | 0.00211 |
| 881 | LRP1B | 5.2774591 | -5.758275736 | 1.38827 | -4.1478 | 3.4E-05 | 0.00042 |
| 819 | DNAAF3 | 5.3031083 | -5.762444313 | 1.34431 | -4.2865 | 1.8E-05 | 0.00025 |
| 1175 | LINC01088 | 5.4728174 | -5.808608675 | 1.32105 | -4.397 | 1.1E-05 | 0.00016 |
| 874 | MERTK | 5.4893692 | -5.812639132 | 1.33013 | -4.37 | 1.2E-05 | 0.00018 |
| 1210 | C4orf47 | 14.869125 | -5.821227821 | 1.15975 | -5.0194 | 5.2E-07 | 1.1E-05 |
| 940 | FLRT3 | 5.5638522 | -5.831283657 | 1.43278 | -4.0699 | 4.7E-05 | 0.00056 |
| 898 | GULP1 | 15.216819 | -5.832387945 | 1.08319 | -5.3845 | 7.3E-08 | 2E-06 |
| 1629 | CFAP47 | 5.5763414 | -5.833529563 | 1.53737 | -3.7945 | 0.00015 | 0.0015 |
| 28 | STPG1 | 5.5692719 | -5.833806834 | 1.31792 | -4.4265 | 9.6E-06 | 0.00014 |
| 1180 | PTPN13 | 9.3115495 | -5.833926217 | 1.19852 | -4.8676 | 1.1E-06 | 2.2E-05 |
| 1119 | SERPINI2 | 5.6093019 | -5.842016793 | 1.40285 | -4.1644 | 3.1E-05 | 0.0004 |
| 460 | ENSG00000286757 | 5.6518878 | -5.8555084 | 1.3113 | -4.4654 | 8E-06 | 0.00012 |
| 643 | MIR22HG | 5.7248641 | -5.876388648 | 1.34858 | -4.3575 | 1.3E-05 | 0.00019 |
| 288 | MS4A8 | 5.7535125 | -5.878207151 | 1.42476 | -4.1258 | 3.7E-05 | 0.00046 |
| 1227 | SPEF2 | 15.552174 | -5.881411436 | 1.11154 | -5.2912 | 1.2E-07 | 3.1E-06 |
| 292 | PLAAT3 | 16.000438 | -5.885710819 | 1.09574 | -5.3715 | 7.8E-08 | 2.1E-06 |
| 583 | XYLT1 | 5.8130571 | -5.894825794 | 1.32259 | -4.457 | 8.3E-06 | 0.00012 |
| 728 | DSG2 | 5.8110148 | -5.896412181 | 1.29819 | -4.542 | 5.6E-06 | 8.8E-05 |
| 1049 | EFHB | 5.842098 | -5.900307276 | 1.40498 | -4.1996 | 2.7E-05 | 0.00035 |
| 1285 | SPARC | 209.87971 | -5.909036747 | 0.93513 | -6.319 | 2.6E-10 | 1.4E-08 |
| 1635 | CHST7 | 5.8542792 | -5.909142572 | 1.42304 | -4.1525 | 3.3E-05 | 0.00041 |
| 1229 | CAPSL | 5.9028711 | -5.915557614 | 1.36584 | -4.3311 | 1.5E-05 | 0.00021 |
| 157 | SLC30A10 | 5.9046274 | -5.917407773 | 1.3052 | -4.5337 | 5.8E-06 | 9.1E-05 |
| 629 | PKD1L2 | 5.9010927 | -5.918741623 | 1.29722 | -4.5626 | 5.1E-06 | 8.1E-05 |
| 1058 | CCK | 5.9217071 | -5.924488736 | 1.50781 | -3.9292 | 8.5E-05 | 0.00094 |
| 1528 | IQANK1 | 10.162839 | -5.939003775 | 1.19306 | -4.9779 | 6.4E-07 | 1.3E-05 |
| 1592 | PTGES | 6.0195331 | -5.944952473 | 1.30963 | -4.5394 | 5.6E-06 | 8.9E-05 |
| 378 | KRT18 | 6.0045947 | -5.945630835 | 1.31881 | -4.5083 | 6.5E-06 | 0.0001 |
| 752 | C3 | 6.0445037 | -5.949591878 | 1.4011 | -4.2464 | 2.2E-05 | 0.00029 |
| 702 | BTBD17 | 6.0790777 | -5.962016835 | 1.31271 | -4.5418 | 5.6E-06 | 8.8E-05 |
| 1441 | CADPS2 | 10.199973 | -5.962293856 | 1.25445 | -4.7529 | 2E-06 | 3.7E-05 |
| 1221 | ROPN1L | 6.1265697 | -5.970272847 | 1.30366 | -4.5796 | 4.7E-06 | 7.5E-05 |
| 1433 | TMEM168 | 6.1280621 | -5.970585785 | 1.30541 | -4.5737 | 4.8E-06 | 7.7E-05 |
| 1454 | CNTNAP2 | 24.58599 | -5.970992 | 0.94694 | -6.3056 | 2.9E-10 | 1.5E-08 |
| 1710 | BGN | 16.86747 | -5.98121777 | 1.11551 | -5.3619 | 8.2E-08 | 2.2E-06 |
| 404 | LUM | 10.318287 | -5.983935072 | 1.26003 | -4.749 | 2E-06 | 3.7E-05 |
| 674 | FZD2 | 6.1937194 | -5.986578456 | 1.29185 | -4.6341 | 3.6E-06 | 5.9E-05 |
| 1377 | PACRG | 6.2246596 | -5.992191866 | 1.38502 | -4.3264 | 1.5E-05 | 0.00021 |

| | | | | | | | |
|---|---|---|---|---|---|---|---|
| 89 | PIFO | 49.523037 | -5.998865654 | 1.71331 | -3.5013 | 0.00046 | 0.00384 |
| 1598 | PIERCE1 | 10.579882 | -6.013398089 | 1.21508 | -4.949 | 7.5E-07 | 1.5E-05 |
| 713 | CCDC40 | 17.033865 | -6.015560248 | 1.11803 | -5.3805 | 7.4E-08 | 2E-06 |
| 869 | POU3F3 | 6.3242123 | -6.019410178 | 1.41838 | -4.2439 | 2.2E-05 | 0.00029 |
| 341 | VWA5A | 10.880309 | -6.056885491 | 1.22488 | -4.9449 | 7.6E-07 | 1.6E-05 |
| 677 | LRRC46 | 6.5521462 | -6.065993235 | 1.37529 | -4.4107 | 1E-05 | 0.00015 |
| 1421 | COL1A2 | 77.415452 | -6.070025491 | 1.23305 | -4.9228 | 8.5E-07 | 1.7E-05 |
| 212 | ENSG00000280660 | 6.573967 | -6.074750582 | 1.33552 | -4.5486 | 5.4E-06 | 8.6E-05 |
| 1112 | TM4SF1 | 6.5876627 | -6.079284392 | 1.2942 | -4.6973 | 2.6E-06 | 4.6E-05 |
| 301 | CTSF | 6.6472293 | -6.089396813 | 1.26576 | -4.8109 | 1.5E-06 | 2.8E-05 |
| 1424 | PDK4 | 6.6883448 | -6.096090446 | 1.31384 | -4.6399 | 3.5E-06 | 5.8E-05 |
| 73 | DNAI3 | 6.7165642 | -6.102021332 | 1.33804 | -4.5604 | 5.1E-06 | 8.2E-05 |
| 1081 | ACOX2 | 6.7161353 | -6.106659036 | 1.28412 | -4.7555 | 2E-06 | 3.6E-05 |
| 766 | NWD1 | 11.247737 | -6.107239002 | 1.37609 | -4.4381 | 9.1E-06 | 0.00013 |
| 638 | GAS8 | 6.7450476 | -6.109214444 | 1.33103 | -4.5898 | 4.4E-06 | 7.2E-05 |
| 1087 | EPHA3 | 6.732951 | -6.110228645 | 1.48002 | -4.1285 | 3.7E-05 | 0.00045 |
| 880 | MAP3K19 | 6.7918467 | -6.119607256 | 1.26876 | -4.8233 | 1.4E-06 | 2.7E-05 |
| 1088 | PROS1 | 11.495564 | -6.1222665 | 1.1919 | -5.1366 | 2.8E-07 | 6.4E-06 |
| 276 | CD44 | 18.843414 | -6.123268245 | 1.15993 | -5.279 | 1.3E-07 | 3.3E-06 |
| 736 | RAX | 6.8139535 | -6.125829346 | 1.31438 | -4.6606 | 3.2E-06 | 5.3E-05 |
| 1620 | EGFL6 | 6.8427085 | -6.133124531 | 1.27091 | -4.8258 | 1.4E-06 | 2.6E-05 |
| 805 | C5AR1 | 11.502165 | -6.139824693 | 1.2268 | -5.0048 | 5.6E-07 | 1.2E-05 |
| 132 | F5 | 6.913008 | -6.143319466 | 1.42721 | -4.3044 | 1.7E-05 | 0.00023 |
| 182 | KIAA1217 | 11.625656 | -6.156785075 | 1.18501 | -5.1955 | 2E-07 | 4.9E-06 |
| 1009 | RAB36 | 18.854689 | -6.160366007 | 1.07799 | -5.7147 | 1.1E-08 | 3.8E-07 |
| 1133 | GMNC | 7.0018575 | -6.16219723 | 1.32092 | -4.6651 | 3.1E-06 | 5.2E-05 |
| 1001 | S100B | 7.0314263 | -6.169059197 | 1.28503 | -4.8007 | 1.6E-06 | 3E-05 |
| 359 | EMP1 | 7.0231284 | -6.17208739 | 1.3135 | -4.6989 | 2.6E-06 | 4.5E-05 |
| 1172 | SHROOM3 | 11.883681 | -6.1725843 | 1.16345 | -5.3054 | 1.1E-07 | 3E-06 |
| 160 | MTARC2 | 7.0726628 | -6.178373191 | 1.25795 | -4.9114 | 9E-07 | 1.8E-05 |
| 901 | DNAH7 | 7.1003323 | -6.185283707 | 1.25758 | -4.9184 | 8.7E-07 | 1.8E-05 |
| 1360 | EYA4 | 7.1499876 | -6.194465256 | 1.25076 | -4.9526 | 7.3E-07 | 1.5E-05 |
| 1692 | GPC3 | 7.1501086 | -6.197290545 | 1.33801 | -4.6317 | 3.6E-06 | 6E-05 |
| 1539 | LINC01239 | 7.1691173 | -6.198048314 | 1.2887 | -4.8095 | 1.5E-06 | 2.8E-05 |
| 1683 | IL13RA1 | 19.479279 | -6.209372218 | 1.09242 | -5.684 | 1.3E-08 | 4.5E-07 |
| 1354 | GJA1 | 20.197605 | -6.223498271 | 1.05794 | -5.8827 | 4E-09 | 1.5E-07 |
| 86 | PALMD | 7.3111569 | -6.225617488 | 1.25872 | -4.946 | 7.6E-07 | 1.5E-05 |
| 445 | FOXO1 | 7.4040987 | -6.242576856 | 1.40582 | -4.4405 | 9E-06 | 0.00013 |
| 1330 | RSPH9 | 7.4388156 | -6.250277698 | 1.27844 | -4.889 | 1E-06 | 2E-05 |

| | | | | | | | |
|---|---|---|---|---|---|---|---|
| 1236 | GPX8 | 12.488266 | -6.261918612 | 1.16597 | -5.3706 | 7.8E-08 | 2.1E-06 |
| 928 | KCNJ13 | 12.668891 | -6.266005314 | 1.16095 | -5.3973 | 6.8E-08 | 1.9E-06 |
| 1136 | ATP13A5 | 7.5286296 | -6.26754756 | 1.42803 | -4.389 | 1.1E-05 | 0.00016 |
| 236 | FANK1 | 7.5720294 | -6.276782812 | 1.24652 | -5.0354 | 4.8E-07 | 1E-05 |
| 724 | LDLRAD4 | 7.6286112 | -6.285942707 | 1.31946 | -4.764 | 1.9E-06 | 3.5E-05 |
| 1440 | FEZF1-AS1 | 7.618007 | -6.288070404 | 1.24694 | -5.0428 | 4.6E-07 | 9.9E-06 |
| 1685 | AKAP14 | 7.6492332 | -6.289543325 | 1.32097 | -4.7613 | 1.9E-06 | 3.5E-05 |
| 348 | CCND2 | 21.423316 | -6.294539992 | 1.0789 | -5.8342 | 5.4E-09 | 2E-07 |
| 58 | FYB2 | 7.7574764 | -6.311922173 | 1.24873 | -5.0547 | 4.3E-07 | 9.4E-06 |
| 1075 | ZMYND10 | 7.7791983 | -6.313729344 | 1.33341 | -4.735 | 2.2E-06 | 3.9E-05 |
| 585 | ENSG00000259871 | 7.8633912 | -6.33307938 | 2.35668 | -2.6873 | 0.0072 | 0.03606 |
| 1158 | FAM114A1 | 7.9155178 | -6.341683919 | 1.2289 | -5.1604 | 2.5E-07 | 5.7E-06 |
| 925 | DAW1 | 13.290239 | -6.348102162 | 1.17107 | -5.4208 | 5.9E-08 | 1.7E-06 |
| 50 | BTBD19 | 7.9659947 | -6.348948032 | 1.29049 | -4.9198 | 8.7E-07 | 1.7E-05 |
| 731 | DTNA | 13.407789 | -6.363405794 | 1.15213 | -5.5232 | 3.3E-08 | 1E-06 |
| 998 | COL18A1 | 8.0892972 | -6.37336192 | 1.23317 | -5.1683 | 2.4E-07 | 5.5E-06 |
| 1282 | SPINK5 | 8.2185836 | -6.39590476 | 1.25156 | -5.1103 | 3.2E-07 | 7.3E-06 |
| 1220 | SNHG18 | 8.2926674 | -6.407023045 | 1.32974 | -4.8183 | 1.4E-06 | 2.7E-05 |
| 794 | AXL | 13.791821 | -6.407364937 | 1.16419 | -5.5037 | 3.7E-08 | 1.1E-06 |
| 862 | TCF7L1 | 8.3088046 | -6.410343156 | 1.24402 | -5.1529 | 2.6E-07 | 5.9E-06 |
| 604 | ADGRG1 | 8.2995498 | -6.413006066 | 1.34638 | -4.7631 | 1.9E-06 | 3.5E-05 |
| 1684 | ZCCHC12 | 8.310275 | -6.413830066 | 1.35559 | -4.7314 | 2.2E-06 | 4E-05 |
| 1240 | PLK2 | 8.3370317 | -6.414471503 | 1.35184 | -4.745 | 2.1E-06 | 3.8E-05 |
| 1501 | GEM | 8.3695853 | -6.420926969 | 1.39909 | -4.5894 | 4.4E-06 | 7.2E-05 |
| 257 | PPFIBP2 | 8.3801675 | -6.421883264 | 1.27028 | -5.0555 | 4.3E-07 | 9.4E-06 |
| 1083 | FEZF2 | 8.3555817 | -6.422557471 | 1.30194 | -4.9331 | 8.1E-07 | 1.6E-05 |
| 1173 | FRAS1 | 8.4869325 | -6.440618891 | 1.31668 | -4.8916 | 1E-06 | 2E-05 |
| 912 | ERBB4 | 8.5156875 | -6.445693015 | 1.23577 | -5.2159 | 1.8E-07 | 4.5E-06 |
| 70 | RPE65 | 8.5315464 | -6.450976675 | 1.22015 | -5.287 | 1.2E-07 | 3.2E-06 |
| 1118 | ZBBX | 8.5528613 | -6.45150309 | 1.24579 | -5.1786 | 2.2E-07 | 5.3E-06 |
| 360 | PLBD1 | 8.558974 | -6.452095431 | 1.32935 | -4.8536 | 1.2E-06 | 2.3E-05 |
| 24 | UBXN10 | 8.6493159 | -6.467971716 | 1.27646 | -5.0671 | 4E-07 | 8.9E-06 |
| 1352 | RSPH4A | 8.6512152 | -6.468106474 | 1.2541 | -5.1576 | 2.5E-07 | 5.8E-06 |
| 1694 | CCDC160 | 8.6615334 | -6.468947706 | 1.28878 | -5.0194 | 5.2E-07 | 1.1E-05 |
| 1402 | C7orf57 | 8.6711587 | -6.471159453 | 1.24944 | -5.1792 | 2.2E-07 | 5.3E-06 |
| 524 | CRIP2 | 34.337277 | -6.481257263 | 0.92285 | -7.0231 | 2.2E-12 | 1.7E-10 |
| 1703 | LINC00632 | 35.221723 | -6.486507085 | 0.89388 | -7.2566 | 4E-13 | 3.9E-11 |
| 1507 | RSPO2 | 24.112597 | -6.487477331 | 1.12651 | -5.7589 | 8.5E-09 | 3E-07 |
| 1275 | STING1 | 14.634599 | -6.489656943 | 1.25294 | -5.1795 | 2.2E-07 | 5.3E-06 |

| | | | | | | | |
|---|---|---|---|---|---|---|---|
| 1533 | DMRT3 | 8.7715329 | -6.490361027 | 1.21144 | -5.3575 | 8.4E-08 | 2.3E-06 |
| 180 | SPAG6 | 8.9465627 | -6.515966685 | 1.39142 | -4.683 | 2.8E-06 | 4.8E-05 |
| 141 | CSRP1 | 9.0370256 | -6.532394333 | 1.23446 | -5.2917 | 1.2E-07 | 3.1E-06 |
| 66 | DNAI4 | 9.0618533 | -6.536015647 | 1.22539 | -5.3338 | 9.6E-08 | 2.6E-06 |
| 469 | SLC7A8 | 15.093642 | -6.536221075 | 1.14117 | -5.7277 | 1E-08 | 3.6E-07 |
| 1212 | SORBS2 | 9.0621172 | -6.537432494 | 1.20579 | -5.4217 | 5.9E-08 | 1.7E-06 |
| 1436 | MDFIC | 9.1375647 | -6.546414323 | 1.27394 | -5.1387 | 2.8E-07 | 6.3E-06 |
| 1672 | CLDN2 | 9.1674195 | -6.550765957 | 1.34789 | -4.86 | 1.2E-06 | 2.3E-05 |
| 1415 | CFAP69 | 15.343443 | -6.560689034 | 1.16805 | -5.6168 | 1.9E-08 | 6.3E-07 |
| 259 | STK33 | 15.570142 | -6.568218576 | 1.16179 | -5.6535 | 1.6E-08 | 5.3E-07 |
| 872 | NPHP1 | 15.574477 | -6.569046445 | 1.14061 | -5.7592 | 8.4E-09 | 3E-07 |
| 1078 | WNT5A | 9.3255599 | -6.580033858 | 1.26877 | -5.1861 | 2.1E-07 | 5.1E-06 |
| 1430 | COL26A1 | 9.3605628 | -6.584926886 | 1.30475 | -5.0469 | 4.5E-07 | 9.8E-06 |
| 1223 | DNAH5 | 9.4753475 | -6.600300998 | 1.21657 | -5.4253 | 5.8E-08 | 1.7E-06 |
| 806 | ODAD1 | 25.786441 | -6.602835377 | 1.08352 | -6.0939 | 1.1E-09 | 4.9E-08 |
| 433 | ENSG00000274373 | 9.7344636 | -6.638449374 | 1.21208 | -5.4769 | 4.3E-08 | 1.3E-06 |
| 118 | CFAP45 | 9.7769286 | -6.644366023 | 1.22596 | -5.4197 | 6E-08 | 1.7E-06 |
| 934 | PRND | 9.7964652 | -6.647152532 | 1.23751 | -5.3714 | 7.8E-08 | 2.1E-06 |
| 1442 | GRM8 | 9.8560098 | -6.657228975 | 1.19566 | -5.5678 | 2.6E-08 | 8.1E-07 |
| 520 | DLK1 | 40.286174 | -6.660313292 | 0.88647 | -7.5133 | 5.8E-14 | 6.8E-12 |
| 581 | ABAT | 16.422607 | -6.660901084 | 1.13413 | -5.8731 | 4.3E-09 | 1.6E-07 |
| 266 | PLEKHA7 | 9.9412036 | -6.669805511 | 1.19325 | -5.5896 | 2.3E-08 | 7.3E-07 |
| 47 | CFAP144 | 9.9741862 | -6.672669454 | 1.25976 | -5.2968 | 1.2E-07 | 3.1E-06 |
| 596 | CHD9NB | 9.990474 | -6.67489514 | 1.33239 | -5.0097 | 5.5E-07 | 1.2E-05 |
| 1309 | RNF182 | 10.028561 | -6.683884405 | 1.1969 | -5.5843 | 2.3E-08 | 7.4E-07 |
| 1006 | AIFM3 | 10.17118 | -6.702856163 | 2.12849 | -3.1491 | 0.00164 | 0.01096 |
| 1706 | CSAG1 | 10.175492 | -6.703127733 | 1.22496 | -5.4721 | 4.4E-08 | 1.3E-06 |
| 1047 | FBLN2 | 10.240042 | -6.715614404 | 1.23643 | -5.4315 | 5.6E-08 | 1.6E-06 |
| 1701 | ZIC3 | 10.278579 | -6.719819977 | 1.21971 | -5.5093 | 3.6E-08 | 1.1E-06 |
| 1408 | ELN | 17.328051 | -6.724867216 | 1.21172 | -5.5499 | 2.9E-08 | 8.9E-07 |
| 654 | DRC3 | 10.39907 | -6.733485937 | 1.24503 | -5.4083 | 6.4E-08 | 1.8E-06 |
| 482 | OTX2-AS1 | 10.399334 | -6.734165779 | 1.20238 | -5.6007 | 2.1E-08 | 6.9E-07 |
| 1218 | CFAP90 | 10.468383 | -6.744018147 | 1.19374 | -5.6495 | 1.6E-08 | 5.4E-07 |
| 226 | ABLIM1 | 42.731629 | -6.750244276 | 0.96317 | -7.0084 | 2.4E-12 | 1.9E-10 |
| 1171 | FAM47E | 10.522251 | -6.75038598 | 1.25335 | -5.3859 | 7.2E-08 | 2E-06 |
| 1388 | SOSTDC1 | 10.620334 | -6.763628918 | 1.22838 | -5.5062 | 3.7E-08 | 1.1E-06 |
| 228 | EMX2OS | 10.618698 | -6.764142977 | 1.19947 | -5.6393 | 1.7E-08 | 5.6E-07 |
| 578 | TNFRSF12A | 10.628452 | -6.767355338 | 1.23495 | -5.4798 | 4.3E-08 | 1.2E-06 |
| 1308 | NEDD9 | 17.747771 | -6.770306381 | 1.12894 | -5.997 | 2E-09 | 8.4E-08 |

| | | | | | | | |
|---|---|---|---|---|---|---|---|
| 611 | TPPP3 | 29.295537 | -6.77305742 | 1.07657 | -6.2913 | 3.1E-10 | 1.6E-08 |
| 1132 | CLDN1 | 10.914724 | -6.803062983 | 1.29774 | -5.2423 | 1.6E-07 | 3.9E-06 |
| 121 | ATP1A2 | 10.892302 | -6.804254219 | 1.19969 | -5.6717 | 1.4E-08 | 4.8E-07 |
| 646 | TEKT1 | 10.956775 | -6.808411734 | 1.23122 | -5.5298 | 3.2E-08 | 9.7E-07 |
| 840 | CYP1B1 | 10.98063 | -6.814033408 | 2.27273 | -2.9982 | 0.00272 | 0.01646 |
| 394 | WIF1 | 10.995568 | -6.815750511 | 2.15848 | -3.1577 | 0.00159 | 0.0107 |
| 608 | CDH11 | 18.286409 | -6.819654152 | 1.17978 | -5.7804 | 7.5E-09 | 2.7E-07 |
| 1187 | DKK2 | 18.40083 | -6.82387095 | 1.12885 | -6.045 | 1.5E-09 | 6.4E-08 |
| 845 | LINC01833 | 11.062297 | -6.824859755 | 1.18701 | -5.7496 | 8.9E-09 | 3.2E-07 |
| 1524 | RHPN1 | 18.432442 | -6.826148728 | 1.12184 | -6.0848 | 1.2E-09 | 5.1E-08 |
| 1546 | DNAI1 | 11.153218 | -6.833980886 | 1.28633 | -5.3128 | 1.1E-07 | 2.8E-06 |
| 507 | FLRT2 | 11.150882 | -6.836473167 | 1.18197 | -5.784 | 7.3E-09 | 2.7E-07 |
| 1098 | ITGB5 | 11.184807 | -6.839332754 | 1.18421 | -5.7754 | 7.7E-09 | 2.8E-07 |
| 1707 | MAGEA12 | 11.32534 | -6.858981736 | 1.19489 | -5.7403 | 9.5E-09 | 3.3E-07 |
| 169 | KCNK1 | 11.414212 | -6.868552326 | 1.21436 | -5.6561 | 1.5E-08 | 5.2E-07 |
| 891 | CFAP210 | 11.497785 | -6.878388446 | 1.20858 | -5.6913 | 1.3E-08 | 4.3E-07 |
| 489 | ZFP36L1 | 31.775584 | -6.897505658 | 1.03392 | -6.6712 | 2.5E-11 | 1.7E-09 |
| 620 | HYDIN | 11.659368 | -6.897991197 | 1.2704 | -5.4298 | 5.6E-08 | 1.6E-06 |
| 619 | NQO1 | 11.669379 | -6.901712375 | 1.16801 | -5.9089 | 3.4E-09 | 1.3E-07 |
| 1184 | DDIT4L | 11.80257 | -6.920300463 | 1.28473 | -5.3866 | 7.2E-08 | 2E-06 |
| 210 | MYOF | 11.95263 | -6.937259672 | 1.2004 | -5.7791 | 7.5E-09 | 2.7E-07 |
| 129 | ILDR2 | 11.979371 | -6.938551892 | 1.18524 | -5.8542 | 4.8E-09 | 1.8E-07 |
| 584 | SYT17 | 12.074597 | -6.95095527 | 1.18139 | -5.8837 | 4E-09 | 1.5E-07 |
| 1427 | SERPINE1 | 12.161518 | -6.961941279 | 2.00863 | -3.466 | 0.00053 | 0.00427 |
| 1253 | FAM81B | 12.23416 | -6.967626704 | 1.25837 | -5.537 | 3.1E-08 | 9.5E-07 |
| 1095 | CFAP91 | 12.263172 | -6.972363012 | 1.16932 | -5.9628 | 2.5E-09 | 1E-07 |
| 1241 | ADAMTS6 | 12.295725 | -6.976819745 | 1.16128 | -6.0079 | 1.9E-09 | 7.9E-08 |
| 927 | ECEL1 | 33.982251 | -7.008907406 | 1.18488 | -5.9153 | 3.3E-09 | 1.3E-07 |
| 910 | MDH1B | 12.602191 | -7.011426566 | 1.18582 | -5.9127 | 3.4E-09 | 1.3E-07 |
| 651 | CFAP52 | 12.624055 | -7.012683688 | 1.27376 | -5.5055 | 3.7E-08 | 1.1E-06 |
| 324 | YAP1 | 21.18622 | -7.028362346 | 1.11165 | -6.3225 | 2.6E-10 | 1.3E-08 |
| 870 | ECRG4 | 12.777213 | -7.030134547 | 1.24934 | -5.6271 | 1.8E-08 | 6E-07 |
| 114 | THBS3-AS1 | 12.75629 | -7.030173824 | 1.17953 | -5.9602 | 2.5E-09 | 1E-07 |
| 1272 | NME5 | 12.832673 | -7.038063714 | 1.15938 | -6.0705 | 1.3E-09 | 5.5E-08 |
| 328 | HOATZ | 12.878958 | -7.041564992 | 1.24688 | -5.6473 | 1.6E-08 | 5.4E-07 |
| 727 | ZNF521 | 13.153497 | -7.07537936 | 1.16493 | -6.0736 | 1.3E-09 | 5.5E-08 |
| 1028 | FAM227A | 13.25118 | -7.085095064 | 1.1579 | -6.1189 | 9.4E-10 | 4.2E-08 |
| 1082 | FAM107A | 13.267196 | -7.085878322 | 1.16345 | -6.0904 | 1.1E-09 | 5E-08 |
| 1110 | CP | 22.130389 | -7.091981444 | 1.17943 | -6.0131 | 1.8E-09 | 7.7E-08 |

| | | | | | | | |
|---|---|---|---|---|---|---|---|
| 1174 | ANXA3 | 13.469587 | -7.109404023 | 1.15591 | -6.1505 | 7.7E-10 | 3.5E-08 |
| 1062 | SLC6A20 | 13.601067 | -7.120956814 | 1.19586 | -5.9547 | 2.6E-09 | 1E-07 |
| 1493 | SULF1 | 127.38805 | -7.124430466 | 0.61433 | -11.597 | 4.3E-31 | 7.3E-28 |
| 1233 | DAB2 | 22.993371 | -7.139067825 | 1.15057 | -6.2048 | 5.5E-10 | 2.6E-08 |
| 376 | KRT7 | 13.774095 | -7.141444286 | 2.37393 | -3.0083 | 0.00263 | 0.01602 |
| 472 | NFATC4 | 13.837082 | -7.149038873 | 1.18959 | -6.0097 | 1.9E-09 | 7.9E-08 |
| 887 | SLC38A11 | 38.461764 | -7.185244004 | 1.02418 | -7.0156 | 2.3E-12 | 1.8E-10 |
| 329 | CRYAB | 14.215688 | -7.189181877 | 1.23139 | -5.8383 | 5.3E-09 | 2E-07 |
| 1100 | CFAP100 | 14.420873 | -7.205070524 | 1.21682 | -5.9212 | 3.2E-09 | 1.3E-07 |
| 481 | OTX2 | 118.66147 | -7.229907572 | 0.62987 | -11.478 | 1.7E-30 | 2.6E-27 |
| 644 | SERPINF1 | 74.117934 | -7.232623724 | 1.07402 | -6.7341 | 1.6E-11 | 1.1E-09 |
| 1378 | C6orf118 | 14.996621 | -7.261339687 | 1.21839 | -5.9598 | 2.5E-09 | 1E-07 |
| 1405 | GALNT17 | 15.377962 | -7.298170873 | 1.26689 | -5.7607 | 8.4E-09 | 3E-07 |
| 960 | PLTP | 25.557984 | -7.301152301 | 1.14062 | -6.401 | 1.5E-10 | 8.3E-09 |
| 246 | IFITM3 | 263.25505 | -7.313759205 | 0.76861 | -9.5156 | 1.8E-21 | 7.8E-19 |
| 1548 | SPAG8 | 25.87792 | -7.319999971 | 1.15932 | -6.314 | 2.7E-10 | 1.4E-08 |
| 1185 | BDH2 | 15.953667 | -7.35338737 | 1.13557 | -6.4755 | 9.5E-11 | 5.3E-09 |
| 1085 | MITF | 26.767083 | -7.369954394 | 1.09681 | -6.7195 | 1.8E-11 | 1.2E-09 |
| 221 | CFAP43 | 16.219053 | -7.374577151 | 1.18117 | -6.2435 | 4.3E-10 | 2.1E-08 |
| 1450 | SLC13A4 | 83.989502 | -7.378558071 | 1.48205 | -4.9786 | 6.4E-07 | 1.3E-05 |
| 290 | AHNAK | 26.999307 | -7.383842616 | 1.10655 | -6.6729 | 2.5E-11 | 1.6E-09 |
| 128 | LMX1A | 16.32239 | -7.386594054 | 1.13203 | -6.5251 | 6.8E-11 | 4E-09 |
| 229 | EMX2 | 16.408519 | -7.394916022 | 1.20407 | -6.1416 | 8.2E-10 | 3.7E-08 |
| 886 | SLC4A10 | 16.523454 | -7.402803941 | 1.1526 | -6.4227 | 1.3E-10 | 7.3E-09 |
| 177 | SLC39A12 | 16.680575 | -7.415003786 | 1.19302 | -6.2153 | 5.1E-10 | 2.4E-08 |
| 104 | S100A10 | 27.586434 | -7.416494011 | 1.18149 | -6.2772 | 3.4E-10 | 1.7E-08 |
| 1516 | LRATD2 | 16.721918 | -7.420649794 | 1.15471 | -6.4264 | 1.3E-10 | 7.2E-09 |
| 1231 | SLC1A3 | 16.913107 | -7.435042362 | 1.24109 | -5.9907 | 2.1E-09 | 8.7E-08 |
| 990 | RSPH1 | 17.255224 | -7.464181342 | 1.16358 | -6.4148 | 1.4E-10 | 7.7E-09 |
| 83 | ABCA4 | 17.301202 | -7.468613648 | 1.13911 | -6.5565 | 5.5E-11 | 3.3E-09 |
| 143 | CHI3L1 | 17.393806 | -7.478043799 | 1.78936 | -4.1792 | 2.9E-05 | 0.00037 |
| 87 | CFAP276 | 17.529543 | -7.486679368 | 1.20134 | -6.232 | 4.6E-10 | 2.2E-08 |
| 967 | BMP7 | 17.739381 | -7.505104896 | 1.13172 | -6.6316 | 3.3E-11 | 2.1E-09 |
| 1160 | RBM47 | 48.067901 | -7.513040131 | 1.01009 | -7.438 | 1E-13 | 1.2E-11 |
| 326 | DYNC2H1 | 18.088968 | -7.534128632 | 1.13633 | -6.6303 | 3.4E-11 | 2.1E-09 |
| 7 | PRDM16 | 18.13441 | -7.538175445 | 1.11933 | -6.7345 | 1.6E-11 | 1.1E-09 |
| 831 | DRC1 | 18.292231 | -7.548117587 | 1.20717 | -6.2527 | 4E-10 | 2E-08 |
| 1128 | IGF2BP2 | 18.343742 | -7.553998961 | 1.12248 | -6.7297 | 1.7E-11 | 1.2E-09 |
| 1137 | ATP13A4 | 19.922294 | -7.67252362 | 1.11604 | -6.8748 | 6.2E-12 | 4.7E-10 |

| | | | | | | | |
|---|---|---|---|---|---|---|---|
| 1237 | CCNO | 20.330739 | -7.703685538 | 1.16256 | -6.6265 | 3.4E-11 | 2.1E-09 |
| 544 | CGNL1 | 20.532361 | -7.716035632 | 1.12475 | -6.8602 | 6.9E-12 | 5.1E-10 |
| 633 | DNAAF1 | 34.846807 | -7.750949806 | 1.12155 | -6.9109 | 4.8E-12 | 3.7E-10 |
| 211 | LGI1 | 21.921231 | -7.812083868 | 1.10597 | -7.0636 | 1.6E-12 | 1.4E-10 |
| 587 | VWA3A | 21.963597 | -7.812497637 | 1.14057 | -6.8496 | 7.4E-12 | 5.5E-10 |
| 1336 | COL9A1 | 22.174059 | -7.827659655 | 2.35939 | -3.3177 | 0.00091 | 0.00665 |
| 65 | DYNLT5 | 22.613166 | -7.855785748 | 1.66723 | -4.7119 | 2.5E-06 | 4.3E-05 |
| 770 | GDF15 | 22.707886 | -7.862015001 | 2.55226 | -3.0804 | 0.00207 | 0.01326 |
| 245 | IFITM2 | 91.755827 | -7.885279169 | 0.86895 | -9.0745 | 1.1E-19 | 3.1E-17 |
| 181 | ARMC3 | 23.570919 | -7.914154233 | 1.15919 | -6.8273 | 8.7E-12 | 6.3E-10 |
| 308 | FOLR1 | 23.804965 | -7.92995669 | 1.91736 | -4.1359 | 3.5E-05 | 0.00044 |
| 1108 | ZIC4 | 25.253743 | -8.015349658 | 1.09582 | -7.3145 | 2.6E-13 | 2.7E-11 |
| 758 | COL5A3 | 25.28849 | -8.016430152 | 1.11456 | -7.1925 | 6.4E-13 | 6E-11 |
| 41 | DNALI1 | 25.518051 | -8.028768095 | 1.14014 | -7.0419 | 1.9E-12 | 1.6E-10 |
| 6 | PRDM16-DT | 25.677948 | -8.039729333 | 1.09926 | -7.3137 | 2.6E-13 | 2.7E-11 |
| 846 | SIX3 | 25.6934 | -8.041393505 | 1.09896 | -7.3173 | 2.5E-13 | 2.7E-11 |
| 882 | GALNT5 | 28.157166 | -8.171503753 | 1.09743 | -7.4461 | 9.6E-14 | 1.1E-11 |
| 461 | ZIC5 | 28.897289 | -8.21076205 | 1.12435 | -7.3027 | 2.8E-13 | 2.9E-11 |
| 1486 | CLXN | 31.744753 | -8.343915631 | 1.14564 | -7.2832 | 3.3E-13 | 3.3E-11 |
| 956 | NNAT | 31.762197 | -8.346107804 | 1.14984 | -7.2585 | 3.9E-13 | 3.9E-11 |
| 526 | NDN | 32.567478 | -8.381963147 | 1.08113 | -7.753 | 9E-15 | 1.2E-12 |
| 1675 | COL4A6 | 33.232318 | -8.412445874 | 1.09876 | -7.6563 | 1.9E-14 | 2.4E-12 |
| 718 | COLEC12 | 33.70418 | -8.431408977 | 1.08586 | -7.7647 | 8.2E-15 | 1.1E-12 |
| 1167 | IGFBP7 | 337.54859 | -8.491403822 | 1.82059 | -4.6641 | 3.1E-06 | 5.2E-05 |
| 708 | FOXJ1 | 36.35827 | -8.540099247 | 1.10388 | -7.7365 | 1E-14 | 1.4E-12 |
| 1557 | ANXA1 | 37.677285 | -8.592049423 | 1.12058 | -7.6675 | 1.8E-14 | 2.2E-12 |
| 1109 | ZIC1 | 63.767893 | -8.627777084 | 1.07252 | -8.0444 | 8.7E-16 | 1.4E-13 |
| 661 | CCL2 | 39.067821 | -8.644875828 | 1.48923 | -5.8049 | 6.4E-09 | 2.4E-07 |
| 1482 | SFRP1 | 65.506471 | -8.665502592 | 2.15012 | -4.0302 | 5.6E-05 | 0.00066 |
| 1271 | TGFBI | 41.178147 | -8.721334355 | 1.44472 | -6.0367 | 1.6E-09 | 6.8E-08 |
| 477 | BMP4 | 41.656968 | -8.73732871 | 1.59032 | -5.4941 | 3.9E-08 | 1.2E-06 |
| 954 | MYL9 | 43.040859 | -8.784838352 | 1.09412 | -8.0292 | 9.8E-16 | 1.6E-13 |
| 1497 | CA2 | 44.797048 | -8.841351568 | 1.10724 | -7.985 | 1.4E-15 | 2.2E-13 |
| 987 | PCP4 | 45.294291 | -8.859166401 | 1.07859 | -8.2137 | 2.1E-16 | 3.7E-14 |
| 1003 | CLDN5 | 46.975528 | -8.910673214 | 1.44502 | -6.1665 | 7E-10 | 3.3E-08 |
| 1341 | TPBG | 48.489889 | -8.957695378 | 1.09104 | -8.2102 | 2.2E-16 | 3.8E-14 |
| 625 | VAT1L | 50.539035 | -9.015322252 | 1.1119 | -8.108 | 5.1E-16 | 8.6E-14 |
| 1676 | COL4A5 | 54.838588 | -9.133946893 | 1.06222 | -8.5989 | 8E-18 | 1.7E-15 |
| 1556 | TRPM3 | 158.47946 | -9.23555342 | 0.98931 | -9.3354 | 1E-20 | 3.6E-18 |

| | | | | | | | |
|---|---|---|---|---|---|---|---|
| 1526 | MAPK15 | 61.790729 | -9.305471197 | 1.10314 | -8.4354 | 3.3E-17 | 6.3E-15 |
| 1677 | ENSG00000261409 | 63.869985 | -9.354929354 | 1.07801 | -8.678 | 4E-18 | 8.8E-16 |
| 1207 | GPM6A | 77.899422 | -9.639991605 | 1.06431 | -9.0575 | 1.3E-19 | 3.5E-17 |
| 983 | CLIC6 | 79.265039 | -9.665527256 | 1.25692 | -7.6899 | 1.5E-14 | 1.9E-12 |
| 686 | WFIKKN2 | 146.26427 | -10.54949098 | 2.36646 | -4.4579 | 8.3E-06 | 0.00012 |
| 1679 | HTR2C | 595.47993 | -11.85311959 | 1.03708 | -11.429 | 3E-30 | 4.1E-27 |
| 729 | TTR | 6137.5126 | -15.94058083 | 1.3414 | -11.884 | 1.4E-32 | 3.3E-29 |

TableS4

| Genes upregulated in Gland_A compared to Gland_B | | | | | | | |
|---|---|---|---|---|---|---|---|
| gene ID | gene name | baseMean | log2FoldChange | lfcSE | stat | p-value | adjusted p-value |
| 694 | FABP1 | 1210.37191 | 13.75188221 | 1.22169 | 11.2564 | 2.15E-29 | 2.69E-26 |
| 195 | DMBT1 | 947.996678 | 13.40020237 | 1.24072 | 10.8003 | 3.43E-27 | 3.15E-24 |
| 1218 | ALDOB | 475.982654 | 12.40523698 | 1.67575 | 7.4028 | 1.33E-13 | 1.70E-11 |
| 429 | ANPEP | 474.540759 | 12.40069558 | 1.23532 | 10.0384 | 1.03E-23 | 6.77E-21 |
| 402 | PHGR1 | 438.856783 | 12.28897158 | 1.2101 | 10.1553 | 3.14E-24 | 2.29E-21 |
| 72 | CLCA1 | 280.86512 | 11.64393717 | 1.21536 | 9.58068 | 9.64E-22 | 4.28E-19 |
| 800 | TFF3 | 253.580685 | 11.49949302 | 1.21124 | 9.49397 | 2.22E-21 | 9.28E-19 |
| 990 | SPINK1 | 245.917429 | 11.45631532 | 1.22435 | 9.35704 | 8.20E-21 | 3.05E-18 |
| 862 | MUC13 | 203.102412 | 11.17689496 | 1.23634 | 9.04028 | 1.56E-19 | 5.00E-17 |
| 315 | TSPAN8 | 381.796397 | 11.12421106 | 1.20278 | 9.24875 | 2.27E-20 | 7.82E-18 |
| 1237 | LCN15 | 167.741147 | 10.90127924 | 1.65245 | 6.59706 | 4.19E-11 | 2.71E-09 |
| 1182 | CDH17 | 158.303107 | 10.81063615 | 1.23418 | 8.75937 | 1.96E-18 | 5.63E-16 |
| 392 | SERPINA1 | 154.513745 | 10.78201161 | 1.69574 | 6.3583 | 2.04E-10 | 1.14E-08 |
| 891 | SST | 119.647977 | 10.41354234 | 3.2209 | 3.23312 | 0.00122 | 0.00998 |
| 664 | APOB | 109.38836 | 10.28297736 | 1.85236 | 5.55129 | 2.84E-08 | 8.89E-07 |
| 1055 | GSTA1 | 108.233367 | 10.26646462 | 1.61371 | 6.36201 | 1.99E-10 | 1.12E-08 |
| 209 | MUC2 | 105.272935 | 10.23024616 | 1.26886 | 8.06255 | 7.47E-16 | 1.47E-13 |
| 628 | CEACAM5 | 103.216428 | 10.20582831 | 1.2694 | 8.0399 | 8.99E-16 | 1.74E-13 |
| 133 | PIGR | 373.484455 | 10.19069748 | 1.50539 | 6.76949 | 1.29E-11 | 9.47E-10 |
| 380 | GPX2 | 101.009724 | 10.17636078 | 1.26744 | 8.02905 | 9.82E-16 | 1.88E-13 |
| 1103 | AGR2 | 193.700273 | 10.14814765 | 1.22322 | 8.29624 | 1.07E-16 | 2.39E-14 |
| 274 | ST14 | 99.0106787 | 10.14684 | 1.23682 | 8.20398 | 2.33E-16 | 4.93E-14 |
| 264 | APOA1 | 175.983273 | 10.00674324 | 2.00541 | 4.98988 | 6.04E-07 | 1.3E-05 |
| 453 | CHP2 | 90.0435468 | 9.998566925 | 1.2743 | 7.84631 | 4.28E-15 | 7.56E-13 |
| 504 | CLDN7 | 85.8641982 | 9.936657231 | 1.22346 | 8.12178 | 4.59E-16 | 9.30E-14 |
| 459 | ZG16 | 85.1459232 | 9.919796726 | 1.55532 | 6.37799 | 1.79E-10 | 1.03E-08 |
| 879 | SI | 77.477019 | 9.784586462 | 1.69581 | 5.76986 | 7.93E-09 | 2.96E-07 |
| 19 | PLA2G2A | 76.6954628 | 9.779792354 | 1.30827 | 7.47534 | 7.70E-14 | 1.03E-11 |
| 396 | AMN | 76.6316406 | 9.772063677 | 1.27105 | 7.68821 | 1.49E-14 | 2.39E-12 |
| 517 | KRT20 | 73.9031176 | 9.706135883 | 1.29192 | 7.51296 | 5.78E-14 | 7.88E-12 |
| 525 | PRR15L | 73.4315405 | 9.702333707 | 1.24873 | 7.76973 | 7.87E-15 | 1.29E-12 |
| 1235 | ABO | 71.5426782 | 9.677882722 | 1.23896 | 7.81132 | 5.66E-15 | 9.62E-13 |
| 587 | SMIM24 | 71.1968661 | 9.661076826 | 1.27215 | 7.59431 | 3.09E-14 | 4.59E-12 |
| 91 | REG4 | 69.6955777 | 9.623958521 | 1.31586 | 7.31381 | 2.60E-13 | 2.93E-11 |
| 82 | EPS8L3 | 68.5823364 | 9.617929529 | 1.23754 | 7.77183 | 7.74E-15 | 1.28E-12 |
| 993 | CDX1 | 67.8005283 | 9.596817197 | 1.22792 | 7.81551 | 5.47E-15 | 9.42E-13 |

| | | | | | | | |
|---|---|---|---|---|---|---|---|
| 1268 | RNF128 | 67.3599689 | 9.571375721 | 1.28666 | 7.43892 | 1.02E-13 | 1.32E-11 |
| 312 | MYO1A | 66.4276052 | 9.566072918 | 1.2517 | 7.64248 | 2.13E-14 | 3.26E-12 |
| 620 | LGALS4 | 692.440816 | 9.506923507 | 0.65015 | 14.6227 | 2.01E-48 | 2.77E-44 |
| 519 | KRT19 | 121.698442 | 9.479454872 | 1.24423 | 7.61874 | 2.56E-14 | 3.88E-12 |
| 387 | CHGA | 113.153579 | 9.366780562 | 1.51231 | 6.19367 | 5.88E-10 | 2.90E-08 |
| 1271 | XPNPEP2 | 57.9258155 | 9.365370693 | 1.32973 | 7.04307 | 1.88E-12 | 1.66E-10 |
| 783 | HNF4A | 57.6946423 | 9.358322305 | 1.23234 | 7.59394 | 3.10E-14 | 4.59E-12 |
| 310 | SLC39A5 | 56.8099208 | 9.344939713 | 1.27403 | 7.33495 | 2.22E-13 | 2.61E-11 |
| 923 | AFP | 108.011735 | 9.300971977 | 1.94126 | 4.79121 | 1.7E-06 | 3.2E-05 |
| 1189 | ANXA13 | 55.0061967 | 9.289311326 | 2.08317 | 4.45923 | 8.2E-06 | 0.00013 |
| 906 | S100P | 51.7660216 | 9.216413263 | 1.27773 | 7.21313 | 5.47E-13 | 5.66E-11 |
| 1019 | SLC17A4 | 51.781047 | 9.195012916 | 1.25605 | 7.32055 | 2.47E-13 | 2.83E-11 |
| 551 | ST6GALNAC1 | 99.3400769 | 9.173037259 | 1.24615 | 7.3611 | 1.82E-13 | 2.24E-11 |
| 1051 | MEP1A | 96.9085133 | 9.139761749 | 1.25212 | 7.29941 | 2.89E-13 | 3.13E-11 |
| 1035 | SLC44A4 | 48.4777602 | 9.119635242 | 1.24635 | 7.3171 | 2.53E-13 | 2.88E-11 |
| 45 | GUCA2A | 47.5143842 | 9.080588137 | 1.76099 | 5.15652 | 2.52E-07 | 6.1E-06 |
| 811 | SLC5A1 | 46.4626711 | 9.044297582 | 1.71815 | 5.26398 | 1.41E-07 | 3.7E-06 |
| 647 | KLK1 | 45.8851822 | 9.03073739 | 1.23653 | 7.3033 | 2.81E-13 | 3.09E-11 |
| 109 | RAB25 | 44.9190732 | 9.008693135 | 1.28635 | 7.00329 | 2.50E-12 | 2.15E-10 |
| 172 | CYP2C18 | 45.6783058 | 9.008088741 | 1.35856 | 6.63062 | 3.34E-11 | 2.23E-09 |
| 332 | HNF1A-AS1 | 44.7103105 | 8.986348823 | 1.24475 | 7.21941 | 5.22E-13 | 5.45E-11 |
| 650 | TMC4 | 44.3050048 | 8.974735946 | 1.26588 | 7.0897 | 1.34E-12 | 1.23E-10 |
| 1206 | ALDH1A1 | 86.550685 | 8.972807593 | 1.26849 | 7.0736 | 1.51E-12 | 1.37E-10 |
| 1145 | SLC26A3 | 42.8958948 | 8.941595976 | 1.24625 | 7.17477 | 7.24E-13 | 7.22E-11 |
| 920 | SULT1B1 | 42.9210248 | 8.937841477 | 1.24795 | 7.16203 | 7.95E-13 | 7.82E-11 |
| 740 | VIL1 | 83.4675645 | 8.925535376 | 1.23251 | 7.24178 | 4.43E-13 | 4.65E-11 |
| 799 | TMPRSS2 | 42.1267894 | 8.916257941 | 1.24898 | 7.13882 | 9.41E-13 | 9.00E-11 |
| 1104 | AGR3 | 42.7684287 | 8.91122382 | 1.31535 | 6.77477 | 1.25E-11 | 9.22E-10 |
| 867 | RBP2 | 155.590124 | 8.875167354 | 2.12082 | 4.18479 | 2.9E-05 | 0.00039 |
| 95 | CTSS | 40.8324614 | 8.863618043 | 1.24648 | 7.11091 | 1.15E-12 | 1.07E-10 |
| 1263 | GJB1 | 39.3253358 | 8.805356212 | 1.26959 | 6.93558 | 4.05E-12 | 3.26E-10 |
| 951 | FABP2 | 39.0592657 | 8.78519069 | 1.38832 | 6.32792 | 2.48E-10 | 1.36E-08 |
| 340 | CDX2 | 37.9123321 | 8.757200481 | 1.25686 | 6.96754 | 3.23E-12 | 2.69E-10 |
| 102 | S100A14 | 37.3639067 | 8.740006856 | 1.27887 | 6.83415 | 8.25E-12 | 6.35E-10 |
| 1157 | AOC1 | 71.8122307 | 8.713411673 | 1.22391 | 7.11935 | 1.08E-12 | 1.02E-10 |
| 113 | ITLN1 | 36.5813417 | 8.699799905 | 1.24994 | 6.96019 | 3.40E-12 | 2.82E-10 |
| 818 | LGALS2 | 71.1983254 | 8.699692638 | 1.22267 | 7.11533 | 1.12E-12 | 1.05E-10 |
| 2 | ENSG00000272141 | 35.8695671 | 8.681497714 | 1.25778 | 6.90225 | 5.12E-12 | 4.03E-10 |
| 627 | CEACAM7 | 127.515062 | 8.638782617 | 1.55658 | 5.54985 | 2.86E-08 | 8.93E-07 |

| | | | | | | | |
|---|---|---|---|---|---|---|---|
| 263 | APOC3 | 34.5249437 | 8.619550273 | 1.93483 | 4.45494 | 8.4E-06 | 0.00013 |
| 265 | TMPRSS4 | 64.2635937 | 8.545753363 | 1.23126 | 6.94066 | 3.90E-12 | 3.16E-10 |
| 39 | TMEM54 | 32.3639897 | 8.538819017 | 1.38739 | 6.15461 | 7.53E-10 | 3.61E-08 |
| 820 | BAIAP2L2 | 62.7752598 | 8.51689136 | 1.26565 | 6.72927 | 1.71E-11 | 1.22E-09 |
| 615 | HPN | 32.131227 | 8.511685414 | 1.35492 | 6.28207 | 3.34E-10 | 1.76E-08 |
| 1183 | ESRP1 | 31.9783332 | 8.510132557 | 1.27369 | 6.68147 | 2.37E-11 | 1.64E-09 |
| 99 | S100A9 | 30.9705372 | 8.474494514 | 1.38823 | 6.10451 | 1.03E-09 | 4.85E-08 |
| 1148 | CFTR | 30.7504554 | 8.468955083 | 1.32614 | 6.38619 | 1.70E-10 | 9.88E-09 |
| 1202 | PRSS3 | 60.3306649 | 8.460798742 | 1.23456 | 6.85327 | 7.22E-12 | 5.61E-10 |
| 212 | USH1C | 59.8157958 | 8.452966143 | 1.28259 | 6.59052 | 4.38E-11 | 2.82E-09 |
| 744 | TM4SF20 | 30.9400911 | 8.450712369 | 1.35501 | 6.23664 | 4.47E-10 | 2.32E-08 |
| 939 | ADH1C | 30.2715735 | 8.4231339 | 1.32137 | 6.37454 | 1.84E-10 | 1.04E-08 |
| 1004 | BTNL3 | 30.2680997 | 8.417676242 | 1.31908 | 6.38146 | 1.75E-10 | 1.01E-08 |
| 201 | PKP3 | 29.4843081 | 8.394724259 | 1.26331 | 6.64502 | 3.03E-11 | 2.06E-09 |
| 1194 | SLC39A4 | 28.6020053 | 8.344261802 | 1.39764 | 5.97025 | 2.37E-09 | 1.05E-07 |
| 54 | TSPAN1 | 146.780112 | 8.267171665 | 0.89162 | 9.27212 | 1.82E-20 | 6.44E-18 |
| 125 | LAD1 | 26.6965688 | 8.264899308 | 1.29849 | 6.365 | 1.95E-10 | 1.10E-08 |
| 197 | CLRN3 | 26.94066 | 8.261003039 | 1.29333 | 6.3874 | 1.69E-10 | 9.85E-09 |
| 378 | TMEM30B | 26.8161811 | 8.248800672 | 1.28193 | 6.43466 | 1.24E-10 | 7.34E-09 |
| 168 | GPR15LG | 26.2973833 | 8.244436583 | 1.30924 | 6.29711 | 3.03E-10 | 1.63E-08 |
| 831 | VIPR1 | 26.4348934 | 8.225566875 | 1.30611 | 6.29777 | 3.02E-10 | 1.63E-08 |
| 414 | C15orf48 | 50.5650812 | 8.217644709 | 1.32175 | 6.21723 | 5.06E-10 | 2.53E-08 |
| 4 | TNFRSF14-AS1 | 26.1104391 | 8.2098985 | 1.31239 | 6.25569 | 3.96E-10 | 2.07E-08 |
| 261 | NXPE1 | 24.5818399 | 8.14862393 | 1.31598 | 6.19204 | 5.94E-10 | 2.92E-08 |
| 1121 | MLXIPL | 88.5897487 | 8.119755972 | 1.13173 | 7.17467 | 7.25E-13 | 7.22E-11 |
| 18 | RNF186 | 24.0123475 | 8.102932088 | 1.28401 | 6.31067 | 2.78E-10 | 1.51E-08 |
| 998 | ATP10B | 23.4939891 | 8.07994921 | 1.29711 | 6.22917 | 4.69E-10 | 2.39E-08 |
| 981 | SOWAHA | 23.2090314 | 8.026144463 | 1.36026 | 5.90046 | 3.62E-09 | 1.49E-07 |
| 898 | PIGZ | 23.1477237 | 8.023693319 | 1.29833 | 6.18 | 6.41E-10 | 3.13E-08 |
| 1140 | MOGAT3 | 22.4321902 | 7.998954399 | 1.28287 | 6.23522 | 4.51E-10 | 2.33E-08 |
| 1054 | GSTA2 | 22.3617561 | 7.987878879 | 1.92078 | 4.15867 | 3.2E-05 | 0.00043 |
| 653 | PTPRH | 22.4807803 | 7.986354471 | 1.31006 | 6.09617 | 1.09E-09 | 5.09E-08 |
| 1265 | POF1B | 43.254472 | 7.982428458 | 1.24242 | 6.42488 | 1.32E-10 | 7.80E-09 |
| 578 | MISP | 43.1929295 | 7.977139642 | 1.2604 | 6.32904 | 2.47E-10 | 1.35E-08 |
| 908 | GBA3 | 22.2800135 | 7.971468641 | 1.3478 | 5.91444 | 3.33E-09 | 1.40E-07 |
| 918 | UGT2A3 | 21.6058428 | 7.955198411 | 1.37163 | 5.79981 | 6.64E-09 | 2.55E-07 |
| 15 | PADI2 | 77.524262 | 7.937335079 | 1.07938 | 7.35361 | 1.93E-13 | 2.35E-11 |
| 189 | ACSL5 | 78.1741883 | 7.937070336 | 1.10231 | 7.20038 | 6.00E-13 | 6.08E-11 |
| 629 | CEACAM6 | 115.190015 | 7.90910342 | 0.93498 | 8.45913 | 2.69E-17 | 6.29E-15 |

| | | | | | | | |
|---|---|---|---|---|---|---|---|
| 358 | RNASE1 | 20.9349531 | 7.895643218 | 1.29313 | 6.10584 | 1.02E-09 | 4.82E-08 |
| 336 | RFLNA | 20.8664277 | 7.890386943 | 1.3048 | 6.04721 | 1.47E-09 | 6.83E-08 |
| 669 | XDH | 20.7665241 | 7.882923207 | 1.40267 | 5.61994 | 1.91E-08 | 6.34E-07 |
| 248 | MOGAT2 | 20.4189158 | 7.859993072 | 1.30809 | 6.00875 | 1.87E-09 | 8.44E-08 |
| 1212 | FBP1 | 20.2011052 | 7.8418889 | 1.33952 | 5.85424 | 4.79E-09 | 1.90E-07 |
| 93 | FMO5 | 20.0990621 | 7.818948979 | 1.35931 | 5.75213 | 8.81E-09 | 3.24E-07 |
| 165 | HKDC1 | 38.4297184 | 7.811812073 | 1.25862 | 6.20667 | 5.41E-10 | 2.69E-08 |
| 801 | TFF1 | 19.1644362 | 7.771093495 | 1.46948 | 5.28831 | 1.23E-07 | 3.3E-06 |
| 1112 | PRR15 | 19.3596104 | 7.756374143 | 1.37442 | 5.64337 | 1.67E-08 | 5.66E-07 |
| 756 | RAB17 | 18.674981 | 7.741838201 | 1.31404 | 5.89161 | 3.82E-09 | 1.57E-07 |
| 1050 | CLIC5 | 18.9023347 | 7.721971496 | 1.3717 | 5.62951 | 1.81E-08 | 6.04E-07 |
| 851 | C3orf85 | 18.6377428 | 7.71481524 | 1.34835 | 5.72168 | 1.05E-08 | 3.73E-07 |
| 407 | ITPKA | 18.1177762 | 7.702037331 | 1.33976 | 5.74883 | 8.99E-09 | 3.26E-07 |
| 345 | EPSTI1 | 18.5421983 | 7.699188453 | 1.33092 | 5.78485 | 7.26E-09 | 2.74E-07 |
| 609 | B3GNT3 | 18.0231059 | 7.694999924 | 1.31335 | 5.85907 | 4.65E-09 | 1.87E-07 |
| 451 | ANKS4B | 18.0225757 | 7.677318948 | 1.29951 | 5.90786 | 3.47E-09 | 1.44E-07 |
| 89 | HSD3B2 | 18.2062944 | 7.67616173 | 1.47022 | 5.22109 | 1.78E-07 | 4.5E-06 |
| 31 | KDF1 | 17.9694258 | 7.671422694 | 1.30881 | 5.86138 | 4.59E-09 | 1.86E-07 |
| 844 | FAM3D | 17.5523642 | 7.664626124 | 1.44958 | 5.28747 | 1.24E-07 | 3.3E-06 |
| 751 | C2orf72 | 34.9441527 | 7.662759978 | 1.27658 | 6.00257 | 1.94E-09 | 8.68E-08 |
| 855 | NR1I2 | 18.0247051 | 7.658097608 | 1.33178 | 5.75027 | 8.91E-09 | 3.25E-07 |
| 541 | ACE | 17.598419 | 7.647053737 | 1.50385 | 5.085 | 3.68E-07 | 8.5E-06 |
| 194 | PNLIPRP2 | 17.4690347 | 7.643864941 | 1.33727 | 5.71601 | 1.09E-08 | 3.84E-07 |
| 206 | CDHR5 | 95.7134316 | 7.634723657 | 0.9766 | 7.81766 | 5.38E-15 | 9.38E-13 |
| 521 | WNK4 | 17.2085913 | 7.629056968 | 1.35197 | 5.64292 | 1.67E-08 | 5.66E-07 |
| 564 | GATA6 | 33.0097712 | 7.572605542 | 1.27292 | 5.949 | 2.70E-09 | 1.16E-07 |
| 471 | CA7 | 16.6468395 | 7.563605898 | 1.433 | 5.27817 | 1.30E-07 | 3.4E-06 |
| 766 | FERMT1 | 16.7853336 | 7.559241139 | 1.32518 | 5.70431 | 1.17E-08 | 4.10E-07 |
| 11 | AGMAT | 16.5680405 | 7.557834947 | 1.30418 | 5.79509 | 6.83E-09 | 2.59E-07 |
| 1056 | TINAG | 16.3750144 | 7.532799258 | 1.30978 | 5.75119 | 8.86E-09 | 3.25E-07 |
| 708 | MYO7B | 31.2731137 | 7.518587418 | 1.2969 | 5.79737 | 6.74E-09 | 2.57E-07 |
| 677 | EPCAM | 406.82939 | 7.500641241 | 0.54622 | 13.732 | 6.53E-43 | 4.50E-39 |
| 1106 | MACC1 | 31.4851231 | 7.498959093 | 1.33013 | 5.63776 | 1.72E-08 | 5.77E-07 |
| 707 | TFCP2L1 | 15.562731 | 7.488321218 | 1.33612 | 5.60453 | 2.09E-08 | 6.78E-07 |
| 631 | CD177 | 15.6519138 | 7.482855481 | 2.36539 | 3.16348 | 0.00156 | 0.01228 |
| 174 | CYP2C9 | 30.5110291 | 7.478682372 | 1.29451 | 5.77723 | 7.59E-09 | 2.86E-07 |
| 320 | NR1H4 | 30.5770497 | 7.466880988 | 1.38928 | 5.37463 | 7.67E-08 | 2.2E-06 |
| 873 | AADAC | 15.579415 | 7.463815494 | 2.10445 | 3.54669 | 0.00039 | 0.00378 |
| 921 | SULT1E1 | 15.6374208 | 7.45393 | 1.44718 | 5.15067 | 2.60E-07 | 6.3E-06 |

| | | | | | | | |
|---|---|---|---|---|---|---|---|
| 231 | MIR194-2HG | 55.9112411 | 7.445337275 | 1.11387 | 6.68423 | 2.32E-11 | 1.61E-09 |
| 940 | MTTP | 54.3063774 | 7.427056006 | 1.68677 | 4.40312 | 1.1E-05 | 0.00017 |
| 1098 | TMEM184A | 15.0084364 | 7.42011437 | 1.37568 | 5.39376 | 6.90E-08 | 2E-06 |
| 158 | TMEM236 | 15.1648284 | 7.404662371 | 1.43904 | 5.14556 | 2.67E-07 | 6.4E-06 |
| 812 | ISX | 14.8079403 | 7.386935433 | 1.34373 | 5.49735 | 3.86E-08 | 1.2E-06 |
| 723 | NEUROD1 | 14.714607 | 7.385761399 | 1.51691 | 4.86895 | 1.1E-06 | 2.3E-05 |
| 116 | GPA33 | 26.9437232 | 7.305573445 | 1.29522 | 5.64043 | 1.70E-08 | 5.71E-07 |
| 1211 | SYK | 14.0535137 | 7.299505835 | 1.35355 | 5.39285 | 6.93E-08 | 2E-06 |
| 563 | GATA6-AS1 | 27.0562525 | 7.278862705 | 1.43117 | 5.08597 | 3.66E-07 | 8.4E-06 |
| 929 | BMP3 | 13.4371757 | 7.273862728 | 1.34016 | 5.42759 | 5.71E-08 | 1.7E-06 |
| 456 | APOBR | 13.3975226 | 7.270990413 | 1.40342 | 5.18089 | 2.21E-07 | 5.5E-06 |
| 684 | DQX1 | 13.68357 | 7.257297239 | 1.42088 | 5.10759 | 3.26E-07 | 7.7E-06 |
| 287 | GUCY2C | 13.2713336 | 7.242899851 | 1.33359 | 5.43112 | 5.60E-08 | 1.6E-06 |
| 639 | SULT2A1 | 13.2627336 | 7.235642509 | 2.58823 | 2.79559 | 0.00518 | 0.03251 |
| 1115 | NPC1L1 | 13.2118735 | 7.217664643 | 1.37135 | 5.26317 | 1.42E-07 | 3.7E-06 |
| 1095 | ADAP1 | 12.9769113 | 7.214236778 | 1.34326 | 5.37071 | 7.84E-08 | 2.2E-06 |
| 827 | PPARG | 12.8779626 | 7.209897756 | 1.39498 | 5.16846 | 2.36E-07 | 5.8E-06 |
| 1146 | TFEC | 13.154047 | 7.200038957 | 1.54182 | 4.66983 | 3E-06 | 5.5E-05 |
| 774 | FOXA2 | 12.8407206 | 7.196095233 | 1.36412 | 5.27528 | 1.33E-07 | 3.5E-06 |
| 1027 | TRIM31 | 12.6886697 | 7.183125534 | 1.36447 | 5.26441 | 1.41E-07 | 3.7E-06 |
| 283 | MIR200CHG | 12.4371983 | 7.166701223 | 1.37885 | 5.19761 | 2.02E-07 | 5.1E-06 |
| 1136 | MUC3A | 68.2616349 | 7.163701802 | 0.96873 | 7.39495 | 1.41E-13 | 1.79E-11 |
| 341 | URAD | 12.4044848 | 7.151705373 | 1.42277 | 5.0266 | 4.99E-07 | 1.1E-05 |
| 914 | GABRA2 | 12.4242923 | 7.149345646 | 1.34554 | 5.31338 | 1.08E-07 | 2.9E-06 |
| 134 | C1orf116 | 12.6442578 | 7.14046196 | 1.3918 | 5.13037 | 2.89E-07 | 6.9E-06 |
| 514 | HNF1B | 12.2357564 | 7.13761347 | 1.35236 | 5.27789 | 1.31E-07 | 3.4E-06 |
| 26 | IL22RA1 | 12.4607444 | 7.137179421 | 1.38971 | 5.13574 | 2.81E-07 | 6.8E-06 |
| 924 | RASSF6 | 12.3296236 | 7.137102734 | 1.38919 | 5.13761 | 2.78E-07 | 6.7E-06 |
| 850 | HHLA2 | 12.3834359 | 7.122200699 | 1.3704 | 5.19717 | 2.02E-07 | 5.1E-06 |
| 1234 | CELP | 12.051879 | 7.106034043 | 1.60913 | 4.41606 | 1E-05 | 0.00016 |
| 510 | SLC13A2 | 12.0214451 | 7.088610381 | 1.35599 | 5.22764 | 1.72E-07 | 4.4E-06 |
| 690 | CAPG | 11.7410609 | 7.086777851 | 1.38774 | 5.10671 | 3.28E-07 | 7.7E-06 |
| 1176 | HNF4G | 11.933524 | 7.082687928 | 1.38446 | 5.11586 | 3.12E-07 | 7.4E-06 |
| 44 | GUCA2B | 11.6964839 | 7.05608185 | 1.59937 | 4.41178 | 1E-05 | 0.00016 |
| 1003 | BTNL8 | 23.1776851 | 7.052051449 | 1.3986 | 5.04221 | 4.60E-07 | 1E-05 |
| 192 | PLEKHS1 | 11.4391984 | 6.999099429 | 1.46922 | 4.76383 | 1.9E-06 | 3.7E-05 |
| 1164 | MSR1 | 11.4126657 | 6.993064311 | 1.53625 | 4.55205 | 5.3E-06 | 9E-05 |
| 753 | B3GNT7 | 11.2748541 | 6.992114952 | 1.37762 | 5.0755 | 3.86E-07 | 8.9E-06 |
| 856 | HGD | 11.3819718 | 6.982926737 | 1.54151 | 4.52993 | 5.9E-06 | 9.9E-05 |

| | | | | | | | |
|---|---|---|---|---|---|---|---|
| 79 | VAV3 | 21.29962 | 6.971643172 | 1.30755 | 5.33184 | 9.72E-08 | 2.7E-06 |
| 25 | MYOM3 | 11.0192103 | 6.961488796 | 1.43832 | 4.84 | 1.3E-06 | 2.6E-05 |
| 1042 | SPDEF | 10.7733267 | 6.957061271 | 1.4171 | 4.90938 | 9.14E-07 | 1.9E-05 |
| 436 | BICDL2 | 10.7234855 | 6.949329767 | 1.54524 | 4.49726 | 6.9E-06 | 0.00011 |
| 397 | EXOC3L4 | 11.0125216 | 6.943070195 | 1.40745 | 4.93308 | 8.09E-07 | 1.7E-05 |
| 497 | ATP2A3 | 10.5879917 | 6.941895446 | 1.51047 | 4.59585 | 4.3E-06 | 7.6E-05 |
| 367 | FOXA1 | 10.9076522 | 6.939686132 | 1.47536 | 4.70372 | 2.6E-06 | 4.7E-05 |
| 697 | SLC9A2 | 10.5340409 | 6.920358017 | 1.40635 | 4.92078 | 8.62E-07 | 1.8E-05 |
| 847 | ADGRG7 | 10.4496903 | 6.891572947 | 1.44639 | 4.76468 | 1.9E-06 | 3.6E-05 |
| 527 | HOXB6 | 10.2028008 | 6.882218587 | 1.56247 | 4.4047 | 1.1E-05 | 0.00016 |
| 149 | AGT | 10.2382615 | 6.857207812 | 1.46069 | 4.69449 | 2.7E-06 | 4.9E-05 |
| 403 | PPP1R14D | 10.0546988 | 6.853628041 | 1.42077 | 4.82388 | 1.4E-06 | 2.8E-05 |
| 640 | FAM83E | 9.99867749 | 6.844680604 | 1.39204 | 4.91701 | 8.79E-07 | 1.8E-05 |
| 298 | DNAJC22 | 20.0276476 | 6.842154399 | 1.32683 | 5.15675 | 2.51E-07 | 6.1E-06 |
| 606 | HSH2D | 10.0959835 | 6.837613062 | 1.39467 | 4.90267 | 9.45E-07 | 2E-05 |
| 718 | DPP4 | 35.9545844 | 6.815072645 | 1.1054 | 6.16523 | 7.04E-10 | 3.41E-08 |
| 262 | NXPE4 | 9.69995128 | 6.813741756 | 1.60227 | 4.25256 | 2.1E-05 | 0.0003 |
| 123 | NR5A2 | 19.3119641 | 6.781132994 | 1.38913 | 4.88158 | 1.1E-06 | 2.2E-05 |
| 129 | GOLT1A | 9.55738249 | 6.768293743 | 1.42071 | 4.76403 | 1.9E-06 | 3.7E-05 |
| 282 | LPAR5 | 9.57832578 | 6.764818168 | 1.3827 | 4.89246 | 9.96E-07 | 2E-05 |
| 528 | HOXB7 | 9.41976139 | 6.764004523 | 1.40112 | 4.82757 | 1.4E-06 | 2.8E-05 |
| 376 | CCDC198 | 9.60593599 | 6.758992997 | 1.45123 | 4.65741 | 3.2E-06 | 5.8E-05 |
| 1131 | HEPACAM2 | 9.47813716 | 6.751172111 | 1.47996 | 4.56171 | 5.1E-06 | 8.7E-05 |
| 754 | ALPI | 9.32733891 | 6.736348412 | 1.5556 | 4.3304 | 1.5E-05 | 0.00022 |
| 621 | LGALS14 | 9.03891794 | 6.713199363 | 1.46904 | 4.56979 | 4.9E-06 | 8.4E-05 |
| 1000 | CDHR2 | 18.1327364 | 6.713095385 | 1.32686 | 5.05938 | 4.21E-07 | 9.5E-06 |
| 1181 | CA1 | 18.0111878 | 6.706023741 | 1.42956 | 4.69098 | 2.7E-06 | 5E-05 |
| 1155 | TMEM176B | 98.8610303 | 6.689623109 | 0.67599 | 9.89599 | 4.33E-23 | 2.71E-20 |
| 60 | TTC39A | 8.98484824 | 6.687500584 | 1.39464 | 4.79513 | 1.6E-06 | 3.2E-05 |
| 877 | RARRES1 | 114.7786 | 6.686097401 | 0.67682 | 9.87862 | 5.15E-23 | 3.09E-20 |
| 145 | CAPN8 | 8.89660707 | 6.68367998 | 1.41684 | 4.71731 | 2.4E-06 | 4.5E-05 |
| 1249 | ARSL | 17.3793493 | 6.64581088 | 1.30194 | 5.10456 | 3.32E-07 | 7.8E-06 |
| 536 | ANKRD40CL | 8.90185683 | 6.64137216 | 1.47968 | 4.48838 | 7.2E-06 | 0.00012 |
| 986 | SLC23A1 | 8.69117225 | 6.629360476 | 1.5149 | 4.37612 | 1.2E-05 | 0.00018 |
| 997 | FABP6 | 226.396946 | 6.59050094 | 1.7167 | 3.83904 | 0.00012 | 0.00141 |
| 1191 | NDRG1 | 169.049906 | 6.587691902 | 0.55912 | 11.7823 | 4.82E-32 | 8.29E-29 |
| 773 | LINC00261 | 8.38542618 | 6.577737415 | 1.45268 | 4.52802 | 6E-06 | 9.9E-05 |
| 529 | HOXB9 | 8.23345756 | 6.575897366 | 1.44066 | 4.56452 | 5E-06 | 8.6E-05 |
| 915 | NIPAL1 | 8.27056998 | 6.571431721 | 1.42139 | 4.62325 | 3.8E-06 | 6.7E-05 |

| | | | | | | | |
|---|---|---|---|---|---|---|---|
| 477 | CDH1 | 61.0614225 | 6.559406684 | 0.82547 | 7.94626 | 1.92E-15 | 3.48E-13 |
| 641 | FUT2 | 30.2919129 | 6.557601897 | 1.10593 | 5.9295 | 3.04E-09 | 1.29E-07 |
| 1188 | DEPTOR | 8.32312913 | 6.557001444 | 1.42653 | 4.59648 | 4.3E-06 | 7.5E-05 |
| 237 | C11orf86 | 8.39169288 | 6.549387581 | 1.48539 | 4.4092 | 1E-05 | 0.00016 |
| 512 | CCL15 | 8.02065682 | 6.54006032 | 1.59167 | 4.10892 | 4E-05 | 0.00053 |
| 825 | UPK3A | 16.0011547 | 6.534793909 | 1.45742 | 4.48382 | 7.3E-06 | 0.00012 |
| 137 | LAMB3 | 15.7475407 | 6.519211337 | 1.36946 | 4.76044 | 1.9E-06 | 3.7E-05 |
| 1156 | TMEM176A | 148.060526 | 6.515039347 | 0.5866 | 11.1065 | 1.17E-28 | 1.24E-25 |
| 716 | ITGB6 | 8.17417685 | 6.510700122 | 1.48471 | 4.38515 | 1.2E-05 | 0.00018 |
| 276 | B3GAT1-DT | 8.10926581 | 6.508984909 | 1.46239 | 4.45092 | 8.6E-06 | 0.00014 |
| 364 | RIPK3 | 7.80207821 | 6.490356928 | 1.42586 | 4.5519 | 5.3E-06 | 9E-05 |
| 239 | ACY3 | 7.89911358 | 6.487636254 | 1.52847 | 4.24454 | 2.2E-05 | 0.00031 |
| 994 | SLC6A7 | 15.2640873 | 6.467262386 | 1.32692 | 4.87387 | 1.1E-06 | 2.2E-05 |
| 506 | MYOCD | 7.6108115 | 6.463599752 | 1.738 | 3.71898 | 0.0002 | 0.00214 |
| 219 | EHF | 7.60711316 | 6.451653545 | 1.45454 | 4.43552 | 9.2E-06 | 0.00014 |
| 409 | PLA2G4F | 7.55655128 | 6.447783586 | 1.52079 | 4.23977 | 2.2E-05 | 0.00031 |
| 797 | FAM3B | 7.84634475 | 6.441917969 | 1.78656 | 3.60577 | 0.00031 | 0.00313 |
| 384 | ENSG00000260711 | 7.57802692 | 6.434550407 | 1.4348 | 4.48464 | 7.3E-06 | 0.00012 |
| 127 | ELF3 | 222.299246 | 6.416593654 | 0.50368 | 12.7394 | 3.57E-37 | 1.23E-33 |
| 969 | C6 | 7.66114146 | 6.403889291 | 1.71333 | 3.73768 | 0.00019 | 0.00201 |
| 498 | TM4SF5 | 7.56637567 | 6.403061331 | 1.57566 | 4.06373 | 4.8E-05 | 0.00063 |
| 259 | CARD16 | 7.48186947 | 6.399927658 | 1.54866 | 4.13257 | 3.6E-05 | 0.00048 |
| 43 | MFSD2A | 7.32662497 | 6.397007357 | 1.47678 | 4.33174 | 1.5E-05 | 0.00022 |
| 499 | GLTPD2 | 7.49855819 | 6.396980358 | 1.57599 | 4.05904 | 4.9E-05 | 0.00064 |
| 439 | ENSG00000262370 | 7.28854499 | 6.383888136 | 1.44951 | 4.40416 | 1.1E-05 | 0.00016 |
| 965 | TERLR1 | 14.5951624 | 6.380884519 | 1.46965 | 4.34177 | 1.4E-05 | 0.00021 |
| 637 | FOXA3 | 7.25471958 | 6.372686206 | 1.43222 | 4.44953 | 8.6E-06 | 0.00014 |
| 592 | LRG1 | 7.33215816 | 6.366200416 | 1.45138 | 4.3863 | 1.2E-05 | 0.00018 |
| 112 | CD244 | 7.06757742 | 6.359466854 | 1.49596 | 4.25108 | 2.1E-05 | 0.0003 |
| 626 | ERICH4 | 7.31977339 | 6.351046565 | 1.64638 | 3.85758 | 0.00011 | 0.00132 |
| 372 | PTGDR | 6.9938393 | 6.343544085 | 1.52578 | 4.15757 | 3.2E-05 | 0.00044 |
| 169 | CDHR1 | 25.9379597 | 6.339382258 | 1.19574 | 5.30164 | 1.15E-07 | 3.1E-06 |
| 742 | IHH | 13.7667548 | 6.336366801 | 1.3712 | 4.62104 | 3.8E-06 | 6.8E-05 |
| 1161 | MNX1 | 6.93300717 | 6.323700088 | 1.49273 | 4.23633 | 2.3E-05 | 0.00032 |
| 711 | GPR39 | 7.15254646 | 6.316634708 | 1.53515 | 4.11466 | 3.9E-05 | 0.00052 |
| 1246 | STPG3-AS1 | 7.07474218 | 6.315778516 | 1.55339 | 4.06581 | 4.8E-05 | 0.00062 |
| 1076 | SMLR1 | 7.1473832 | 6.304647869 | 1.73271 | 3.6386 | 0.00027 | 0.0028 |
| 608 | BISPR | 7.07807482 | 6.304456791 | 1.47726 | 4.26768 | 2E-05 | 0.00028 |
| 475 | HSD11B2 | 6.75943338 | 6.29795665 | 1.55266 | 4.05623 | 5E-05 | 0.00065 |

| | | | | | | | |
|---|---|---|---|---|---|---|---|
| 1242 | C8G | 6.83673083 | 6.29229746 | 1.69605 | 3.70998 | 0.00021 | 0.00221 |
| 675 | HAAO | 6.88173776 | 6.290554691 | 1.46715 | 4.28759 | 1.8E-05 | 0.00026 |
| 485 | IRF8 | 6.94046648 | 6.285791905 | 1.46091 | 4.30266 | 1.7E-05 | 0.00025 |
| 481 | FA2H | 6.68770528 | 6.260908325 | 1.61362 | 3.88004 | 0.0001 | 0.00122 |
| 1107 | SKAP2 | 6.86889227 | 6.248335435 | 1.57624 | 3.96408 | 7.4E-05 | 0.00091 |
| 590 | CREB3L3 | 6.79453953 | 6.237595528 | 1.67117 | 3.73248 | 0.00019 | 0.00204 |
| 1029 | TRIM15 | 6.72404967 | 6.236958244 | 1.48514 | 4.19957 | 2.7E-05 | 0.00037 |
| 464 | PRSS8 | 6.50493659 | 6.231773197 | 1.63618 | 3.80872 | 0.00014 | 0.00156 |
| 938 | ADH6 | 6.57682023 | 6.228514413 | 1.45218 | 4.28909 | 1.8E-05 | 0.00026 |
| 1013 | ADTRP | 13.0448618 | 6.226174243 | 1.39446 | 4.46494 | 8E-06 | 0.00013 |
| 1044 | BNIP5 | 6.49087299 | 6.205609508 | 1.58781 | 3.90829 | 9.3E-05 | 0.00111 |
| 214 | SLC5A12 | 23.8966704 | 6.204280292 | 1.2401 | 5.00305 | 5.64E-07 | 1.2E-05 |
| 540 | CA4 | 47.0208974 | 6.203839542 | 0.82472 | 7.52234 | 5.38E-14 | 7.41E-12 |
| 630 | CEACAM1 | 34.9526759 | 6.190264592 | 0.96583 | 6.40926 | 1.46E-10 | 8.60E-09 |
| 567 | RNF125 | 6.31590332 | 6.187947951 | 1.56136 | 3.96318 | 7.4E-05 | 0.00091 |
| 59 | SLC5A9 | 22.9864084 | 6.175329586 | 1.2261 | 5.03655 | 4.74E-07 | 1.1E-05 |
| 251 | SYTL2 | 46.5780437 | 6.172423316 | 0.85043 | 7.25804 | 3.93E-13 | 4.19E-11 |
| 782 | R3HDML-AS1 | 12.3509883 | 6.155221509 | 1.35113 | 4.55559 | 5.2E-06 | 8.9E-05 |
| 244 | ENSG00000285933 | 6.31406158 | 6.150120736 | 1.49278 | 4.11991 | 3.8E-05 | 0.00051 |
| 919 | UGT2B7 | 6.2666678 | 6.147289976 | 1.49815 | 4.10324 | 4.1E-05 | 0.00054 |
| 1233 | LCN2 | 6.05747038 | 6.140178616 | 1.68813 | 3.63727 | 0.00028 | 0.00281 |
| 531 | LINC03057 | 6.04698786 | 6.133554376 | 1.57159 | 3.90278 | 9.5E-05 | 0.00113 |
| 63 | PCSK9 | 11.6538253 | 6.092160443 | 1.43207 | 4.25409 | 2.1E-05 | 0.0003 |
| 546 | FADS6 | 6.06402052 | 6.09136716 | 1.52772 | 3.98722 | 6.7E-05 | 0.00084 |
| 260 | IL18 | 5.89048125 | 6.091020049 | 1.65795 | 3.67383 | 0.00024 | 0.00249 |
| 155 | TUBAL3 | 6.03129432 | 6.082265074 | 1.66951 | 3.64315 | 0.00027 | 0.00276 |
| 752 | ENSG00000261829 | 6.04048922 | 6.079703628 | 1.57175 | 3.86812 | 0.00011 | 0.00127 |
| 553 | ENPP7 | 5.9878478 | 6.076713774 | 1.67361 | 3.6309 | 0.00028 | 0.00287 |
| 864 | CHST13 | 6.11621397 | 6.075783976 | 1.58687 | 3.82877 | 0.00013 | 0.00146 |
| 760 | ENSG00000219159 | 5.78167572 | 6.067811078 | 1.52571 | 3.97704 | 7E-05 | 0.00087 |
| 136 | MIR205HG | 6.02901363 | 6.064407776 | 1.59981 | 3.79071 | 0.00015 | 0.00166 |
| 68 | GIPC2 | 21.7346992 | 6.061753004 | 1.18846 | 5.10053 | 3.39E-07 | 7.9E-06 |
| 1137 | ENSG00000274993 | 5.72914045 | 6.055993894 | 1.53026 | 3.95749 | 7.6E-05 | 0.00093 |
| 1117 | DDC | 42.359508 | 6.055203674 | 0.948 | 6.38737 | 1.69E-10 | 9.85E-09 |
| 622 | FCGBP | 117.114234 | 6.042188044 | 0.62038 | 9.73948 | 2.05E-22 | 1.08E-19 |
| 162 | A1CF | 32.1934294 | 6.029027059 | 0.97766 | 6.16682 | 6.97E-10 | 3.39E-08 |
| 683 | LBX2-AS1 | 5.83484049 | 6.025027587 | 1.83423 | 3.28478 | 0.00102 | 0.00853 |
| 28 | TRIM63 | 5.67704413 | 6.016657208 | 1.64612 | 3.65505 | 0.00026 | 0.00266 |
| 427 | SLC28A1 | 5.7634892 | 6.015663158 | 1.88177 | 3.19681 | 0.00139 | 0.01112 |

| | | | | | | | |
|---|---|---|---|---|---|---|---|
| 907 | ENSG00000287778 | 5.76633725 | 6.003890757 | 1.61749 | 3.71186 | 0.00021 | 0.00219 |
| 309 | ERBB3 | 82.1445391 | 5.997278228 | 0.61671 | 9.72457 | 2.37E-22 | 1.16E-19 |
| 1216 | CORO2A | 20.6407771 | 5.996039828 | 1.17088 | 5.12099 | 3.04E-07 | 7.3E-06 |
| 720 | NOSTRIN | 11.0212608 | 5.988665432 | 1.40164 | 4.27261 | 1.9E-05 | 0.00028 |
| 1225 | C9orf152 | 5.53440044 | 5.982012449 | 1.52141 | 3.93189 | 8.4E-05 | 0.00101 |
| 1037 | HLA-DRB1 | 5.51237069 | 5.977033609 | 1.69653 | 3.52309 | 0.00043 | 0.00409 |
| 667 | KRTCAP3 | 20.2293405 | 5.975077713 | 1.15923 | 5.15435 | 2.55E-07 | 6.2E-06 |
| 968 | AGXT2 | 5.52168449 | 5.968645382 | 1.78447 | 3.34477 | 0.00082 | 0.00716 |
| 524 | SP6 | 5.48509166 | 5.966211225 | 1.51505 | 3.93796 | 8.2E-05 | 0.00099 |
| 1142 | LHFPL3 | 5.58501171 | 5.956654271 | 1.89267 | 3.14723 | 0.00165 | 0.01283 |
| 526 | HOXB5 | 10.5255142 | 5.946413325 | 1.42308 | 4.17854 | 2.9E-05 | 0.0004 |
| 143 | MIR215 | 5.56789678 | 5.945368047 | 1.56626 | 3.79589 | 0.00015 | 0.00163 |
| 652 | NLRP2 | 5.31361016 | 5.934652383 | 1.57596 | 3.76574 | 0.00017 | 0.00183 |
| 12 | TMEM82 | 5.3217186 | 5.93072225 | 1.53323 | 3.86813 | 0.00011 | 0.00127 |
| 357 | ANG | 10.429344 | 5.930534001 | 1.37259 | 4.3207 | 1.6E-05 | 0.00023 |
| 636 | ENSG00000285128 | 5.32130515 | 5.929596694 | 1.65156 | 3.59029 | 0.00033 | 0.0033 |
| 1109 | HOXA5 | 5.28554086 | 5.927740214 | 1.79556 | 3.30134 | 0.00096 | 0.00813 |
| 170 | PAPSS2 | 78.4142602 | 5.920818992 | 0.60302 | 9.81865 | 9.36E-23 | 5.15E-20 |
| 594 | CRB3 | 5.33840732 | 5.919821858 | 1.52113 | 3.89172 | 1E-04 | 0.00117 |
| 419 | C2CD4B | 5.37671235 | 5.916509555 | 1.57832 | 3.74861 | 0.00018 | 0.00194 |
| 1230 | BSPRY | 5.24351199 | 5.914173839 | 1.5269 | 3.87333 | 0.00011 | 0.00125 |
| 480 | MLKL | 5.18660325 | 5.913158764 | 1.59323 | 3.71142 | 0.00021 | 0.00219 |
| 1239 | ENSG00000273066 | 29.1740371 | 5.911996182 | 0.94893 | 6.23019 | 4.66E-10 | 2.38E-08 |
| 284 | APOBEC1 | 5.40644984 | 5.902440378 | 1.58306 | 3.7285 | 0.00019 | 0.00207 |
| 1134 | CYP3A5 | 67.4724121 | 5.901900477 | 1.2012 | 4.91333 | 8.95E-07 | 1.9E-05 |
| 978 | SLCO4C1 | 5.24352474 | 5.898576303 | 1.59686 | 3.69386 | 0.00022 | 0.00233 |
| 579 | CFD | 5.21462691 | 5.897604108 | 1.52069 | 3.87823 | 0.00011 | 0.00123 |
| 381 | PLEK2 | 5.1146851 | 5.894538134 | 1.57062 | 3.75301 | 0.00017 | 0.00191 |
| 440 | CASP16P | 5.23261601 | 5.89262072 | 1.6425 | 3.5876 | 0.00033 | 0.00332 |
| 176 | SORBS1 | 10.2425343 | 5.890974834 | 1.46821 | 4.01236 | 6E-05 | 0.00076 |
| 789 | PCK1 | 10.0871614 | 5.880872821 | 1.45217 | 4.04973 | 5.1E-05 | 0.00066 |
| 442 | SMIM22 | 27.7628402 | 5.851228626 | 0.98021 | 5.96938 | 2.38E-09 | 1.05E-07 |
| 1143 | LHFPL3-AS2 | 5.08687747 | 5.831288886 | 1.72879 | 3.37305 | 0.00074 | 0.00657 |
| 985 | SMIM32 | 4.88474767 | 5.830920312 | 1.62464 | 3.58905 | 0.00033 | 0.00331 |
| 883 | GPR160 | 27.0809882 | 5.827993785 | 0.98319 | 5.92766 | 3.07E-09 | 1.30E-07 |
| 452 | ERN2 | 17.5810524 | 5.81654895 | 1.26273 | 4.60632 | 4.1E-06 | 7.2E-05 |
| 228 | SYT7 | 4.88912114 | 5.816381656 | 1.62468 | 3.58001 | 0.00034 | 0.00339 |
| 415 | SQOR | 26.5510991 | 5.790483486 | 1.00596 | 5.75616 | 8.61E-09 | 3.17E-07 |
| 105 | EFNA1 | 62.1369631 | 5.790079303 | 0.69042 | 8.38629 | 5.02E-17 | 1.13E-14 |

| | | | | | | | |
|---|---|---|---|---|---|---|---|
| 572 | ATP8B1 | 26.7528299 | 5.787634328 | 0.97192 | 5.95487 | 2.60E-09 | 1.13E-07 |
| 618 | PPP1R14A | 17.5038665 | 5.785006448 | 1.19082 | 4.858 | 1.2E-06 | 2.4E-05 |
| 1069 | MROCKI | 4.92937779 | 5.772166896 | 1.58762 | 3.63573 | 0.00028 | 0.00282 |
| 1114 | AOAH | 4.77180648 | 5.76920751 | 1.63456 | 3.52952 | 0.00042 | 0.004 |
| 956 | FGA | 4.84229634 | 5.760625679 | 1.91878 | 3.00223 | 0.00268 | 0.01903 |
| 467 | NLRC5 | 9.17863761 | 5.757770726 | 1.49108 | 3.86148 | 0.00011 | 0.0013 |
| 437 | ENSG00000205890 | 4.79244246 | 5.757635888 | 1.56475 | 3.6796 | 0.00023 | 0.00245 |
| 624 | CYP2B6 | 26.1674259 | 5.756764565 | 0.9974 | 5.77176 | 7.84E-09 | 2.94E-07 |
| 21 | CDA | 4.54849771 | 5.729137014 | 1.73707 | 3.29815 | 0.00097 | 0.0082 |
| 486 | LINC01082 | 4.50811315 | 5.71484263 | 1.86897 | 3.05775 | 0.00223 | 0.01642 |
| 408 | LTK | 4.49487597 | 5.711992688 | 1.6466 | 3.46895 | 0.00052 | 0.00486 |
| 10 | ENSG00000272510 | 4.5363141 | 5.693938365 | 1.64441 | 3.4626 | 0.00053 | 0.00496 |
| 9 | TNFRSF1B | 4.4727512 | 5.693333813 | 1.62507 | 3.50343 | 0.00046 | 0.00436 |
| 681 | NAT8 | 4.45939512 | 5.69300432 | 1.65424 | 3.44147 | 0.00058 | 0.00531 |
| 699 | LINC01594 | 4.57680691 | 5.682158318 | 1.90808 | 2.97794 | 0.0029 | 0.02038 |
| 269 | SORL1 | 24.7507641 | 5.675112072 | 0.97631 | 5.8128 | 6.14E-09 | 2.37E-07 |
| 1046 | KCNK5 | 8.9357132 | 5.669310772 | 1.39048 | 4.07723 | 4.6E-05 | 0.0006 |
| 173 | CYP2C19 | 4.59318833 | 5.6643933 | 1.76288 | 3.21314 | 0.00131 | 0.01058 |
| 1065 | CRYBG1 | 33.0697018 | 5.659153266 | 0.86069 | 6.57512 | 4.86E-11 | 3.08E-09 |
| 959 | SMIM31 | 4.37870827 | 5.648203515 | 1.61759 | 3.49174 | 0.00048 | 0.00451 |
| 454 | AQP8 | 4.25895739 | 5.63488394 | 1.87495 | 3.00534 | 0.00265 | 0.01891 |
| 680 | SPR | 16.2287461 | 5.63444912 | 1.19283 | 4.72361 | 2.3E-06 | 4.3E-05 |
| 1047 | UNC5CL | 40.5488385 | 5.631586469 | 0.86327 | 6.52355 | 6.87E-11 | 4.24E-09 |
| 1041 | MLN | 4.39160363 | 5.631523167 | 1.91761 | 2.93674 | 0.00332 | 0.02268 |
| 1267 | SERPINA7 | 4.43614937 | 5.619404631 | 1.62829 | 3.4511 | 0.00056 | 0.00515 |
| 193 | VWA2 | 4.36905265 | 5.617735658 | 1.61222 | 3.48447 | 0.00049 | 0.00463 |
| 267 | TREH | 4.28527997 | 5.611113018 | 1.83111 | 3.06433 | 0.00218 | 0.01615 |
| 1160 | SHH | 4.28720609 | 5.602312736 | 1.62786 | 3.44152 | 0.00058 | 0.00531 |
| 393 | SERPINA4 | 4.18997118 | 5.597113776 | 1.7774 | 3.14905 | 0.00164 | 0.01276 |
| 889 | LIPH | 15.7852399 | 5.596861143 | 1.16876 | 4.78871 | 1.7E-06 | 3.3E-05 |
| 1248 | ENTPD8 | 4.17552864 | 5.587751307 | 1.80988 | 3.08737 | 0.00202 | 0.01514 |
| 142 | PROX1 | 4.24953956 | 5.576527 | 1.64154 | 3.39713 | 0.00068 | 0.0061 |
| 605 | CYP4F3 | 4.28389732 | 5.57105709 | 1.65861 | 3.35888 | 0.00078 | 0.00686 |
| 1097 | CYP2W1 | 4.08874175 | 5.568558098 | 1.79385 | 3.10425 | 0.00191 | 0.01442 |
| 922 | SLC4A4 | 8.155899 | 5.564683994 | 1.42805 | 3.8967 | 9.8E-05 | 0.00115 |
| 1036 | C2 | 8.17278805 | 5.560002649 | 1.5252 | 3.64542 | 0.00027 | 0.00274 |
| 846 | ROBO2 | 4.03974028 | 5.553275024 | 1.78858 | 3.10484 | 0.0019 | 0.0144 |
| 701 | LINC01123 | 4.27523477 | 5.552443596 | 1.76396 | 3.14771 | 0.00165 | 0.01282 |
| 896 | SLC51A | 15.6028173 | 5.547523687 | 1.26248 | 4.39416 | 1.1E-05 | 0.00017 |

| | | | | | | | |
|---|---|---|---|---|---|---|---|
| 878 | ARL14 | 4.18760822 | 5.541519007 | 1.63406 | 3.39127 | 0.0007 | 0.00622 |
| 80 | ELAPOR1 | 7.98871898 | 5.538180913 | 1.44137 | 3.8423 | 0.00012 | 0.00139 |
| 338 | GJB2 | 4.0749377 | 5.533818001 | 1.62018 | 3.41556 | 0.00064 | 0.00576 |
| 893 | LINC02924 | 4.11272313 | 5.532752779 | 1.59979 | 3.45843 | 0.00054 | 0.00503 |
| 597 | STXBP2 | 14.8328199 | 5.52964448 | 1.19439 | 4.62969 | 3.7E-06 | 6.5E-05 |
| 874 | P2RY1 | 7.86660548 | 5.528106521 | 1.49327 | 3.70202 | 0.00021 | 0.00227 |
| 111 | LINC01133 | 29.3155496 | 5.504843352 | 0.8624 | 6.38316 | 1.73E-10 | 1.00E-08 |
| 83 | DENND2D | 14.52425 | 5.503972719 | 1.18756 | 4.63467 | 3.6E-06 | 6.4E-05 |
| 1020 | SLC17A1 | 4.01259291 | 5.500957883 | 1.67941 | 3.27554 | 0.00105 | 0.00877 |
| 48 | C1orf210 | 7.83525129 | 5.493281516 | 1.42397 | 3.85772 | 0.00011 | 0.00132 |
| 154 | AKR1C3 | 14.5676748 | 5.490076622 | 1.2115 | 4.53163 | 5.9E-06 | 9.8E-05 |
| 734 | CASP10 | 14.7925667 | 5.489439938 | 1.20918 | 4.53982 | 5.6E-06 | 9.5E-05 |
| 1257 | MAOA | 95.268463 | 5.488934806 | 0.57847 | 9.48879 | 2.34E-21 | 9.47E-19 |
| 533 | HOXB13 | 3.80362053 | 5.467582423 | 1.90403 | 2.87159 | 0.00408 | 0.02694 |
| 876 | ENSG00000272087 | 3.90004129 | 5.46653793 | 1.78359 | 3.0649 | 0.00218 | 0.01614 |
| 288 | ERP27 | 3.86064818 | 5.451431012 | 1.62927 | 3.34594 | 0.00082 | 0.00713 |
| 936 | ATOH1 | 3.84620563 | 5.439194357 | 1.61395 | 3.37012 | 0.00075 | 0.00661 |
| 446 | MYH11 | 21.591032 | 5.434381717 | 1.97966 | 2.7451 | 0.00605 | 0.03684 |
| 791 | SLC17A9 | 33.9502409 | 5.42892581 | 0.82498 | 6.58071 | 4.68E-11 | 2.98E-09 |
| 890 | RTP4 | 3.92729692 | 5.427001509 | 1.86502 | 2.90989 | 0.00362 | 0.02436 |
| 1200 | SPINK4 | 3.82481438 | 5.418436134 | 1.98242 | 2.73325 | 0.00627 | 0.03796 |
| 322 | RNY1P16 | 7.56303736 | 5.41444002 | 1.48208 | 3.65328 | 0.00026 | 0.00267 |
| 391 | SERPINA10 | 3.77356673 | 5.407188654 | 1.68058 | 3.21745 | 0.00129 | 0.01044 |
| 368 | SSTR1 | 3.7357813 | 5.406063022 | 1.64236 | 3.29164 | 0.001 | 0.00835 |
| 348 | LMO7 | 20.5598838 | 5.405875179 | 0.99743 | 5.41979 | 5.97E-08 | 1.7E-06 |
| 1149 | SLC13A1 | 3.67674527 | 5.403197119 | 1.84631 | 2.92648 | 0.00343 | 0.02332 |
| 395 | DEGS2 | 3.64613361 | 5.402817768 | 1.67075 | 3.23376 | 0.00122 | 0.00997 |
| 29 | CNKSR1 | 13.4594039 | 5.402117341 | 1.20992 | 4.46484 | 8E-06 | 0.00013 |
| 926 | LINC02562 | 3.8270478 | 5.397547606 | 1.71726 | 3.14311 | 0.00167 | 0.01297 |
| 404 | SPINT1-AS1 | 13.4874334 | 5.382495787 | 1.24233 | 4.33257 | 1.5E-05 | 0.00022 |
| 1084 | ADGRG6 | 7.34593479 | 5.366919492 | 1.45256 | 3.69481 | 0.00022 | 0.00233 |
| 374 | GCH1 | 7.2672612 | 5.360508056 | 1.43866 | 3.72605 | 0.00019 | 0.00209 |
| 1205 | ENSG00000235523 | 3.59683758 | 5.357376771 | 1.98028 | 2.70536 | 0.00682 | 0.04079 |
| 593 | FUT3 | 3.53904191 | 5.348496467 | 1.87179 | 2.85742 | 0.00427 | 0.02794 |
| 1100 | CARD11 | 3.48213317 | 5.346133771 | 1.86699 | 2.8635 | 0.00419 | 0.02752 |
| 748 | SP140L | 7.00597407 | 5.343694538 | 1.44611 | 3.69523 | 0.00022 | 0.00233 |
| 676 | EPCAM-DT | 3.57786816 | 5.34326474 | 1.63823 | 3.26161 | 0.00111 | 0.00918 |
| 745 | SLC19A3 | 19.9386518 | 5.339494345 | 1.03438 | 5.16205 | 2.44E-07 | 6E-06 |
| 780 | TGM2 | 46.2715838 | 5.333699244 | 0.72689 | 7.33775 | 2.17E-13 | 2.60E-11 |

| | | | | | | | |
|---|---|---|---|---|---|---|---|
| 417 | MYZAP | 6.92916738 | 5.326884872 | 1.43223 | 3.71929 | 0.0002 | 0.00214 |
| 925 | CXCL5 | 6.84274619 | 5.321218207 | 1.46614 | 3.62941 | 0.00028 | 0.00288 |
| 413 | GATM | 19.3718489 | 5.3114728 | 1.0018 | 5.30195 | 1.15E-07 | 3.1E-06 |
| 418 | GCNT3 | 19.1777802 | 5.296827486 | 1.00037 | 5.29485 | 1.19E-07 | 3.2E-06 |
| 763 | SDCBP2 | 25.3813167 | 5.295162721 | 0.92122 | 5.74799 | 9.03E-09 | 3.26E-07 |
| 577 | PLPP2 | 18.4963774 | 5.288926042 | 1.06945 | 4.94547 | 7.60E-07 | 1.6E-05 |
| 955 | TLR2 | 3.48382524 | 5.283124288 | 1.68796 | 3.12988 | 0.00175 | 0.01347 |
| 1197 | TTC39B | 38.1651273 | 5.280825326 | 0.75321 | 7.01107 | 2.37E-12 | 2.06E-10 |
| 946 | PANCR | 3.35395753 | 5.269588592 | 1.70084 | 3.09822 | 0.00195 | 0.01467 |
| 405 | SPINT1 | 37.4793756 | 5.259245751 | 0.78921 | 6.66397 | 2.67E-11 | 1.83E-09 |
| 483 | ATP2C2 | 36.9152114 | 5.25623981 | 0.77242 | 6.80492 | 1.01E-11 | 7.69E-10 |
| 399 | ENSG00000287346 | 3.26767911 | 5.254907308 | 1.95446 | 2.68867 | 0.00717 | 0.04243 |
| 980 | CCNI2 | 3.32891356 | 5.252694375 | 1.68438 | 3.11847 | 0.00182 | 0.01389 |
| 717 | LINC01806 | 3.38400233 | 5.250382364 | 1.7197 | 3.05308 | 0.00227 | 0.01662 |
| 518 | ENSG00000234477 | 3.35479719 | 5.241632272 | 1.70184 | 3.07998 | 0.00207 | 0.01543 |
| 992 | SLC26A2 | 226.481654 | 5.240289734 | 0.40178 | 13.0426 | 7.00E-39 | 3.21E-35 |
| 242 | LINC02747 | 6.68341752 | 5.237379173 | 1.5274 | 3.42894 | 0.00061 | 0.00553 |
| 984 | PITX1 | 24.4796405 | 5.233292723 | 1.00569 | 5.20366 | 1.95E-07 | 4.9E-06 |
| 762 | NEU4 | 3.21250807 | 5.225446337 | 1.73496 | 3.01185 | 0.0026 | 0.01856 |
| 803 | TRAPPC10 | 6.45394742 | 5.208675591 | 1.75234 | 2.97241 | 0.00295 | 0.02068 |
| 271 | VSIG2 | 17.9686271 | 5.20296252 | 1.12103 | 4.64121 | 3.5E-06 | 6.2E-05 |
| 256 | MMP1 | 3.29702703 | 5.197031031 | 1.84077 | 2.82329 | 0.00475 | 0.03031 |
| 958 | TMEM144 | 18.0124349 | 5.194561625 | 1.03236 | 5.03175 | 4.86E-07 | 1.1E-05 |
| 390 | IFI27 | 236.006474 | 5.192825597 | 0.43994 | 11.8035 | 3.74E-32 | 7.36E-29 |
| 57 | FOXD2-AS1 | 3.14199645 | 5.19072567 | 1.72979 | 3.00078 | 0.00269 | 0.01911 |
| 232 | BATF2 | 6.50161399 | 5.187337171 | 1.55337 | 3.33941 | 0.00084 | 0.00728 |
| 633 | KCNN4 | 3.09532343 | 5.176628079 | 1.9404 | 2.66781 | 0.00763 | 0.04449 |
| 1078 | TARID | 3.25155937 | 5.176532949 | 1.81797 | 2.84743 | 0.00441 | 0.02871 |
| 806 | ADORA2A-AS1 | 3.14413649 | 5.167608605 | 1.86075 | 2.77716 | 0.00548 | 0.03409 |
| 85 | BCL2L15 | 22.6721394 | 5.166782229 | 0.92966 | 5.55773 | 2.73E-08 | 8.61E-07 |
| 285 | BCL2L14 | 11.8031513 | 5.16461385 | 1.22312 | 4.22251 | 2.4E-05 | 0.00034 |
| 854 | SIDT1 | 3.12199896 | 5.163101188 | 1.71891 | 3.0037 | 0.00267 | 0.01898 |
| 450 | GP2 | 3.14242266 | 5.16299354 | 1.74014 | 2.96699 | 0.00301 | 0.02093 |
| 3 | C1QTNF12 | 11.4823785 | 5.152923413 | 1.27814 | 4.03158 | 5.5E-05 | 0.00071 |
| 1028 | TRIM10 | 3.19408376 | 5.150929717 | 1.75875 | 2.92875 | 0.0034 | 0.02319 |
| 273 | TMEM45B | 74.4853522 | 5.144356126 | 0.53628 | 9.59266 | 8.58E-22 | 3.94E-19 |
| 222 | CREB3L1 | 16.6762678 | 5.142465152 | 1.07871 | 4.76722 | 1.9E-06 | 3.6E-05 |
| 1057 | FAM83B | 3.19377645 | 5.14155156 | 1.73926 | 2.95617 | 0.00311 | 0.02158 |
| 468 | PLLP | 11.5322604 | 5.139483349 | 1.22003 | 4.21258 | 2.5E-05 | 0.00035 |

| | | | | | | | |
|---|---|---|---|---|---|---|---|
| 41 | GJB3 | 3.01956417 | 5.136824273 | 1.92398 | 2.6699 | 0.00759 | 0.04431 |
| 275 | ADAMTS15 | 11.3684611 | 5.130108944 | 1.22976 | 4.17164 | 3E-05 | 0.00041 |
| 741 | USP37 | 5.96795566 | 5.122032473 | 1.55978 | 3.28382 | 0.00102 | 0.00855 |
| 702 | MALL | 3.01692841 | 5.119895184 | 1.72055 | 2.97574 | 0.00292 | 0.02049 |
| 713 | KYNU | 3.05928847 | 5.1091521 | 1.80985 | 2.82296 | 0.00476 | 0.03032 |
| 725 | SLC40A1 | 51.3983542 | 5.10832953 | 0.67425 | 7.57629 | 3.56E-14 | 5.15E-12 |
| 1101 | ICA1 | 16.6730007 | 5.106639738 | 1.02644 | 4.97511 | 6.52E-07 | 1.4E-05 |
| 625 | CYP2S1 | 11.1605493 | 5.096617486 | 1.31353 | 3.88009 | 0.0001 | 0.00122 |
| 863 | ALDH1L1 | 2.95337826 | 5.088717887 | 1.93006 | 2.63656 | 0.00838 | 0.04742 |
| 809 | GAL3ST1 | 5.83654077 | 5.08631552 | 1.56344 | 3.25329 | 0.00114 | 0.00942 |
| 1279 | ERCC00154 | 2.98138917 | 5.085671211 | 1.91935 | 2.64968 | 0.00806 | 0.04622 |
| 492 | RPH3AL-AS2 | 2.9534844 | 5.084402472 | 1.72696 | 2.94414 | 0.00324 | 0.02226 |
| 257 | CASP5 | 2.98461567 | 5.078992036 | 1.75621 | 2.89203 | 0.00383 | 0.02555 |
| 964 | HPGD | 27.5071755 | 5.077122883 | 0.93057 | 5.45593 | 4.87E-08 | 1.4E-06 |
| 1105 | LINC02889 | 3.03263682 | 5.073579741 | 1.77292 | 2.86171 | 0.00421 | 0.02763 |
| 1053 | ENSG00000235122 | 6.03316914 | 5.06745226 | 1.58843 | 3.19022 | 0.00142 | 0.01135 |
| 27 | NCMAP | 2.95421792 | 5.06280141 | 1.80964 | 2.79769 | 0.00515 | 0.03236 |
| 1172 | ENSG00000253414 | 2.88266547 | 5.036117652 | 1.83637 | 2.74243 | 0.0061 | 0.03706 |
| 1122 | ENSG00000274080 | 2.86530374 | 5.035401801 | 1.83552 | 2.74332 | 0.00608 | 0.03701 |
| 534 | TMEM92 | 20.9843548 | 5.027678036 | 0.89528 | 5.61575 | 1.96E-08 | 6.43E-07 |
| 279 | PLEKHG6 | 26.3016635 | 5.021491711 | 0.88098 | 5.6999 | 1.20E-08 | 4.19E-07 |
| 347 | KLF5 | 114.900402 | 5.014144042 | 0.43403 | 11.5525 | 7.17E-31 | 1.10E-27 |
| 865 | MGLL | 47.6020926 | 5.009060589 | 0.69083 | 7.25083 | 4.14E-13 | 4.39E-11 |
| 1108 | HOXA3 | 5.52274141 | 5.006683613 | 1.63432 | 3.06346 | 0.00219 | 0.01617 |
| 58 | FOXD2 | 10.1626842 | 5.005678653 | 1.31526 | 3.80583 | 0.00014 | 0.00157 |
| 1094 | PDGFA | 20.1425809 | 4.978934502 | 0.97858 | 5.08791 | 3.62E-07 | 8.4E-06 |
| 613 | CEBPA-DT | 2.75608476 | 4.968614608 | 1.78313 | 2.78646 | 0.00533 | 0.03329 |
| 203 | ANO9 | 116.999888 | 4.968483901 | 0.52158 | 9.52581 | 1.64E-21 | 7.05E-19 |
| 258 | CASP1 | 10.4041311 | 4.965063007 | 1.27219 | 3.90277 | 9.5E-05 | 0.00113 |
| 319 | NTN4 | 10.3928863 | 4.961733901 | 1.32099 | 3.75607 | 0.00017 | 0.0019 |
| 882 | LRRC31 | 2.74558948 | 4.952167402 | 1.76949 | 2.79864 | 0.00513 | 0.03232 |
| 1023 | H2BC9 | 2.71768472 | 4.951186481 | 1.85388 | 2.67071 | 0.00757 | 0.04427 |
| 687 | TRABD2A | 30.3191548 | 4.948820349 | 0.80854 | 6.12067 | 9.32E-10 | 4.44E-08 |
| 138 | G0S2 | 20.166995 | 4.94328074 | 1.04109 | 4.74819 | 2.1E-06 | 3.9E-05 |
| 930 | PRKG2 | 15.1705712 | 4.942277093 | 1.25571 | 3.93586 | 8.3E-05 | 0.001 |
| 557 | SLC16A3 | 29.0830117 | 4.942105828 | 0.89545 | 5.51913 | 3.41E-08 | 1E-06 |
| 1204 | GDA | 49.0486869 | 4.918507468 | 0.72935 | 6.74372 | 1.54E-11 | 1.11E-09 |
| 62 | TTC22 | 14.7356168 | 4.914213956 | 1.02202 | 4.80834 | 1.5E-06 | 3E-05 |
| 509 | LGALS9 | 24.7157664 | 4.912432101 | 0.83908 | 5.85457 | 4.78E-09 | 1.90E-07 |

| | | | | | | | |
|---|---|---|---|---|---|---|---|
| 1001 | FGFR4 | 14.4083946 | 4.912188387 | 1.06363 | 4.61832 | 3.9E-06 | 6.8E-05 |
| 706 | TMEM37 | 14.4645551 | 4.906180593 | 1.08409 | 4.52561 | 6E-06 | 0.0001 |
| 785 | TNNC2 | 2.59878624 | 4.901778124 | 1.84319 | 2.65939 | 0.00783 | 0.04522 |
| 482 | WFDC1 | 5.11458688 | 4.898719314 | 1.54807 | 3.16441 | 0.00155 | 0.01225 |
| 1167 | DMTN | 14.4630943 | 4.893008141 | 1.085 | 4.50967 | 6.5E-06 | 0.00011 |
| 1070 | ENSG00000253194 | 5.1470358 | 4.855420479 | 1.52511 | 3.18365 | 0.00145 | 0.01156 |
| 1262 | IL2RG | 24.0281863 | 4.848634117 | 0.96106 | 5.04509 | 4.53E-07 | 1E-05 |
| 816 | MFNG | 2.46025598 | 4.829806767 | 1.82768 | 2.64259 | 0.00823 | 0.04675 |
| 104 | IL6R | 5.08967909 | 4.820991625 | 1.56184 | 3.08673 | 0.00202 | 0.01514 |
| 74 | GBP3 | 23.6632115 | 4.820775705 | 0.92834 | 5.19291 | 2.07E-07 | 5.2E-06 |
| 1005 | BTNL9 | 2.45147453 | 4.819384525 | 1.8201 | 2.64787 | 0.0081 | 0.04635 |
| 328 | OAS1 | 69.1468079 | 4.818117054 | 0.55971 | 8.60827 | 7.42E-18 | 1.96E-15 |
| 424 | CTXND1 | 4.90878425 | 4.813516732 | 1.79094 | 2.68771 | 0.00719 | 0.04251 |
| 611 | TM6SF2 | 9.03473229 | 4.804566101 | 1.25936 | 3.81508 | 0.00014 | 0.00152 |
| 530 | HOXB-AS4 | 4.80847673 | 4.796729039 | 1.80615 | 2.65577 | 0.00791 | 0.04557 |
| 635 | CBLC | 13.5837657 | 4.792338019 | 1.06873 | 4.48416 | 7.3E-06 | 0.00012 |
| 949 | UGT8 | 36.438522 | 4.776135266 | 0.79429 | 6.0131 | 1.82E-09 | 8.27E-08 |
| 947 | PITX2 | 30.6375029 | 4.759124769 | 0.71597 | 6.64706 | 2.99E-11 | 2.04E-09 |
| 794 | CBR3-AS1 | 4.78002898 | 4.746887964 | 1.53698 | 3.08846 | 0.00201 | 0.0151 |
| 881 | MECOM | 29.9701869 | 4.723129364 | 0.69703 | 6.77606 | 1.23E-11 | 9.20E-10 |
| 849 | NFKBIZ | 107.329189 | 4.72084174 | 0.64082 | 7.36686 | 1.75E-13 | 2.17E-11 |
| 157 | ST8SIA6-AS1 | 4.75446995 | 4.714012831 | 1.58331 | 2.97731 | 0.00291 | 0.0204 |
| 71 | MCOLN3 | 8.72811092 | 4.701036261 | 1.576 | 2.98289 | 0.00286 | 0.0201 |
| 646 | SPIB | 16.3758168 | 4.677647626 | 1.15357 | 4.05494 | 5E-05 | 0.00065 |
| 487 | FENDRR | 24.9338302 | 4.669131553 | 0.8609 | 5.42357 | 5.84E-08 | 1.7E-06 |
| 1130 | SAMD9L | 4.41034235 | 4.664940183 | 1.54382 | 3.02168 | 0.00251 | 0.01804 |
| 989 | SH3RF2 | 24.649604 | 4.662232218 | 0.80967 | 5.75817 | 8.50E-09 | 3.15E-07 |
| 550 | EVPL | 11.8671 | 4.658576916 | 1.29823 | 3.58841 | 0.00033 | 0.00331 |
| 247 | SLCO2B1 | 4.34331101 | 4.650701301 | 1.5651 | 2.9715 | 0.00296 | 0.02073 |
| 532 | LINC02086 | 16.3037955 | 4.643088499 | 1.09811 | 4.22825 | 2.4E-05 | 0.00033 |
| 318 | TMCC3 | 7.91418403 | 4.618166726 | 1.3868 | 3.33009 | 0.00087 | 0.00747 |
| 547 | LLGL2 | 125.058916 | 4.61431862 | 0.40715 | 11.3332 | 8.99E-30 | 1.24E-26 |
| 385 | TC2N | 15.6631322 | 4.610989514 | 1.0211 | 4.51572 | 6.3E-06 | 0.0001 |
| 842 | CACNA1D | 8.02225777 | 4.600259982 | 1.30937 | 3.51334 | 0.00044 | 0.00421 |
| 1177 | TPD52 | 56.6133798 | 4.584444001 | 0.61358 | 7.47167 | 7.92E-14 | 1.05E-11 |
| 199 | PRAP1 | 4.19985317 | 4.584412201 | 1.6974 | 2.70084 | 0.00692 | 0.04126 |
| 114 | F11R | 28.0489357 | 4.579457566 | 0.76171 | 6.0121 | 1.83E-09 | 8.29E-08 |
| 792 | HELZ2 | 7.68615997 | 4.575398998 | 1.31706 | 3.47395 | 0.00051 | 0.00478 |
| 1243 | LCN12 | 7.68222546 | 4.573431499 | 1.40244 | 3.26105 | 0.00111 | 0.00919 |

| | | | | | | | |
|---|---|---|---|---|---|---|---|
| 353 | F7 | 11.6390062 | 4.544408992 | 1.2414 | 3.66073 | 0.00025 | 0.0026 |
| 1258 | SLC38A5 | 15.1388496 | 4.534310187 | 0.92628 | 4.89517 | 9.82E-07 | 2E-05 |
| 20 | CAMK2N1 | 102.627709 | 4.532223761 | 0.61408 | 7.38051 | 1.58E-13 | 1.97E-11 |
| 917 | ENSG00000286093 | 3.9951786 | 4.513209727 | 1.64544 | 2.74286 | 0.00609 | 0.03704 |
| 670 | QPCT | 11.3761763 | 4.511512986 | 1.09215 | 4.13084 | 3.6E-05 | 0.00049 |
| 1025 | HLA-F | 4.03544554 | 4.504784465 | 1.65776 | 2.7174 | 0.00658 | 0.03964 |
| 5 | TNFRSF14 | 68.0566975 | 4.502460398 | 0.54572 | 8.2505 | 1.58E-16 | 3.39E-14 |
| 1276 | PNCK | 30.6907449 | 4.497249244 | 0.91097 | 4.93678 | 7.94E-07 | 1.7E-05 |
| 1110 | HOXA9 | 18.5247038 | 4.484876848 | 0.89961 | 4.98537 | 6.18E-07 | 1.3E-05 |
| 24 | GALE | 42.8465873 | 4.483935676 | 0.66984 | 6.69406 | 2.17E-11 | 1.52E-09 |
| 350 | SLITRK6 | 3.88994268 | 4.474055312 | 1.67091 | 2.67761 | 0.00741 | 0.04351 |
| 899 | MELTF | 14.6977751 | 4.473215426 | 1.04487 | 4.28111 | 1.9E-05 | 0.00027 |
| 905 | RN7SL589P | 3.86141753 | 4.463012761 | 1.70707 | 2.61443 | 0.00894 | 0.04999 |
| 299 | GPD1 | 14.5845377 | 4.462614839 | 0.98989 | 4.50819 | 6.5E-06 | 0.00011 |
| 119 | FMO4 | 3.9486714 | 4.462364329 | 1.58503 | 2.81532 | 0.00487 | 0.03095 |
| 700 | SULT1C2 | 45.5773623 | 4.45509296 | 1.55708 | 2.86119 | 0.00422 | 0.02766 |
| 535 | ABCC3 | 60.5728198 | 4.449234018 | 0.55554 | 8.0088 | 1.16E-15 | 2.18E-13 |
| 523 | ARHGAP27 | 53.5535768 | 4.445834271 | 0.53779 | 8.2669 | 1.37E-16 | 3.00E-14 |
| 721 | SPC25 | 3.79381234 | 4.4454451 | 1.60671 | 2.7668 | 0.00566 | 0.035 |
| 571 | RAB27B | 3.88179972 | 4.426045054 | 1.63239 | 2.71139 | 0.0067 | 0.04027 |
| 1232 | KIF12 | 17.8655413 | 4.408481975 | 0.966 | 4.56364 | 5E-06 | 8.7E-05 |
| 210 | SLC22A18 | 33.9958571 | 4.406375923 | 0.67067 | 6.57009 | 5.03E-11 | 3.18E-09 |
| 755 | MLPH | 7.11375827 | 4.402467992 | 1.35762 | 3.24279 | 0.00118 | 0.00971 |
| 218 | CAT | 13.6781852 | 4.40166294 | 1.0932 | 4.02641 | 5.7E-05 | 0.00072 |
| 888 | EHHADH | 10.8614398 | 4.400665314 | 1.18448 | 3.71527 | 0.0002 | 0.00217 |
| 1150 | AKR1B10 | 7.07322729 | 4.399744562 | 1.67542 | 2.62606 | 0.00864 | 0.04857 |
| 560 | PIEZO2 | 3.77137539 | 4.390538192 | 1.62839 | 2.69624 | 0.00701 | 0.04173 |
| 972 | ELOVL7 | 18.0638684 | 4.390332667 | 1.12687 | 3.89605 | 9.8E-05 | 0.00115 |
| 552 | TMC6 | 10.1552806 | 4.382619248 | 1.13933 | 3.84665 | 0.00012 | 0.00137 |
| 185 | COL17A1 | 23.4903123 | 4.381712169 | 0.81733 | 5.36101 | 8.28E-08 | 2.3E-06 |
| 870 | PLS1 | 31.0295021 | 4.381415928 | 0.65248 | 6.71503 | 1.88E-11 | 1.33E-09 |
| 311 | TAC3 | 6.94269022 | 4.376962708 | 1.36299 | 3.2113 | 0.00132 | 0.01065 |
| 90 | HMGCS2 | 134.188121 | 4.368024797 | 0.92748 | 4.70954 | 2.5E-06 | 4.6E-05 |
| 1129 | SAMD9 | 6.92030707 | 4.363034227 | 1.38457 | 3.15119 | 0.00163 | 0.01269 |
| 132 | RAB29 | 16.943513 | 4.353911192 | 0.92564 | 4.70367 | 2.6E-06 | 4.7E-05 |
| 476 | ESRP2 | 9.92822288 | 4.350742819 | 1.16599 | 3.73137 | 0.00019 | 0.00205 |
| 1024 | PRSS16 | 3.69530881 | 4.34326159 | 1.64339 | 2.64286 | 0.00822 | 0.04675 |
| 1247 | NRARP | 26.2489325 | 4.342122765 | 0.75532 | 5.74874 | 8.99E-09 | 3.26E-07 |
| 733 | SATB2-AS1 | 16.6126381 | 4.340871419 | 1.00852 | 4.30419 | 1.7E-05 | 0.00025 |

| | | | | | | | |
|---|---|---|---|---|---|---|---|
| 314 | GLI1 | 6.54195541 | 4.336539472 | 1.47025 | 2.94952 | 0.00318 | 0.02197 |
| 735 | CASP8 | 17.0106891 | 4.334081359 | 0.95755 | 4.52622 | 6E-06 | 0.0001 |
| 719 | SCN9A | 3.52419385 | 4.321049876 | 1.62957 | 2.65165 | 0.00801 | 0.04603 |
| 120 | CRYZL2P | 22.9294781 | 4.307930946 | 0.9084 | 4.74232 | 2.1E-06 | 4E-05 |
| 813 | APOL6 | 26.9234094 | 4.30292603 | 0.80421 | 5.35049 | 8.77E-08 | 2.4E-06 |
| 51 | CCDC17 | 3.5259546 | 4.300989909 | 1.61079 | 2.67011 | 0.00758 | 0.04431 |
| 184 | NEURL1 | 16.103415 | 4.297423303 | 0.91462 | 4.6986 | 2.6E-06 | 4.8E-05 |
| 822 | FAM83F | 9.87856042 | 4.294449317 | 1.12526 | 3.81641 | 0.00014 | 0.00152 |
| 726 | C2orf88 | 6.3771293 | 4.294400369 | 1.35863 | 3.16084 | 0.00157 | 0.01236 |
| 307 | HOXC6 | 9.44689626 | 4.285372056 | 1.31846 | 3.25028 | 0.00115 | 0.00949 |
| 163 | PRKG1 | 9.94583609 | 4.277618207 | 1.14152 | 3.74729 | 0.00018 | 0.00195 |
| 581 | EFNA2 | 12.8280939 | 4.268257467 | 0.95873 | 4.45197 | 8.5E-06 | 0.00014 |
| 662 | LRATD1 | 94.1304634 | 4.259205315 | 0.46327 | 9.19373 | 3.79E-20 | 1.27E-17 |
| 884 | PLD1 | 31.675188 | 4.258963536 | 0.76061 | 5.5994 | 2.15E-08 | 6.92E-07 |
| 598 | AP1M2 | 18.9232396 | 4.254538312 | 0.79218 | 5.37068 | 7.84E-08 | 2.2E-06 |
| 478 | MARVELD3 | 34.9297951 | 4.254261167 | 0.61524 | 6.91476 | 4.69E-12 | 3.71E-10 |
| 612 | SLC7A9 | 18.8476775 | 4.250750047 | 0.98931 | 4.29667 | 1.7E-05 | 0.00025 |
| 967 | OTULINL | 24.2242857 | 4.234271297 | 0.76822 | 5.51176 | 3.55E-08 | 1.1E-06 |
| 352 | SLC15A1 | 36.6745601 | 4.215444066 | 0.75238 | 5.60281 | 2.11E-08 | 6.83E-07 |
| 1166 | SH2D4A | 6.01151625 | 4.194693735 | 1.38748 | 3.02325 | 0.0025 | 0.01795 |
| 17 | NBL1 | 38.8495808 | 4.194136678 | 0.63308 | 6.62502 | 3.47E-11 | 2.30E-09 |
| 830 | VILL | 20.9469275 | 4.19314981 | 0.74545 | 5.62498 | 1.86E-08 | 6.19E-07 |
| 1261 | HEPH | 106.878726 | 4.190717486 | 0.47453 | 8.83122 | 1.04E-18 | 3.03E-16 |
| 603 | ADGRE5 | 14.7944641 | 4.183430745 | 0.88173 | 4.74458 | 2.1E-06 | 4E-05 |
| 895 | MUC4 | 8.87690236 | 4.169707849 | 1.16648 | 3.57459 | 0.00035 | 0.00346 |
| 771 | INSM1 | 5.73801258 | 4.162553195 | 1.4013 | 2.9705 | 0.00297 | 0.02076 |
| 1165 | MTUS1 | 5.73080917 | 4.158035617 | 1.38489 | 3.00244 | 0.00268 | 0.01903 |
| 866 | CLDN18 | 5.94633582 | 4.149993696 | 1.57647 | 2.63246 | 0.00848 | 0.0479 |
| 69 | IFI44L | 44.9705848 | 4.145338105 | 0.70097 | 5.91368 | 3.35E-09 | 1.40E-07 |
| 619 | KCNK6 | 17.4118925 | 4.125865582 | 0.78928 | 5.22739 | 1.72E-07 | 4.4E-06 |
| 651 | CDC42EP5 | 22.9712967 | 4.124774784 | 0.73522 | 5.61024 | 2.02E-08 | 6.59E-07 |
| 375 | LGALS3 | 844.961694 | 4.12458671 | 0.74913 | 5.50581 | 3.67E-08 | 1.1E-06 |
| 160 | MPP7 | 8.70019838 | 4.117607462 | 1.1626 | 3.54171 | 0.0004 | 0.00384 |
| 843 | CHDH | 20.2593151 | 4.111104131 | 0.87679 | 4.68882 | 2.7E-06 | 5E-05 |
| 654 | TMEM238 | 25.6869851 | 4.109663048 | 0.70303 | 5.84562 | 5.05E-09 | 1.98E-07 |
| 747 | PID1 | 5.76235479 | 4.105525635 | 1.38209 | 2.97053 | 0.00297 | 0.02076 |
| 1093 | FRMD1 | 5.67524946 | 4.102953614 | 1.54957 | 2.6478 | 0.0081 | 0.04635 |
| 496 | RAP1GAP2 | 5.68284942 | 4.089968371 | 1.44778 | 2.82499 | 0.00473 | 0.03021 |
| 77 | SLC44A3 | 22.306558 | 4.074627867 | 0.73861 | 5.51663 | 3.46E-08 | 1E-06 |

| | | | | | | | |
|---|---|---|---|---|---|---|---|
| 746 | CCL20 | 5.41743025 | 4.066625631 | 1.42831 | 2.84716 | 0.00441 | 0.02872 |
| 286 | GPRC5A | 19.0813182 | 4.044167956 | 0.76552 | 5.2829 | 1.27E-07 | 3.4E-06 |
| 853 | PLCXD2 | 5.5923115 | 4.041762331 | 1.45659 | 2.77481 | 0.00552 | 0.0343 |
| 634 | PVR | 18.9405673 | 4.034990861 | 0.78855 | 5.11699 | 3.10E-07 | 7.4E-06 |
| 245 | STARD10 | 53.9406285 | 4.03274236 | 0.49552 | 8.13833 | 4.01E-16 | 8.24E-14 |
| 207 | EPS8L2 | 72.7547273 | 4.028834396 | 0.49778 | 8.0936 | 5.79E-16 | 1.16E-13 |
| 383 | FLVCR2 | 5.46586575 | 4.011911003 | 1.4662 | 2.73626 | 0.00621 | 0.03766 |
| 555 | PCYT2 | 16.3210336 | 4.010705812 | 0.81295 | 4.93352 | 8.08E-07 | 1.7E-05 |
| 1215 | TRIM14 | 19.0775056 | 4.006860227 | 0.77548 | 5.16697 | 2.38E-07 | 5.9E-06 |
| 217 | DEPDC7 | 13.4311568 | 4.00111725 | 0.93615 | 4.27399 | 1.9E-05 | 0.00028 |
| 1180 | CA13 | 10.727116 | 3.999988953 | 1.12527 | 3.5547 | 0.00038 | 0.00368 |
| 8 | ESPN | 5.32144935 | 3.996606128 | 1.49029 | 2.68177 | 0.00732 | 0.04309 |
| 781 | SGK2 | 10.630135 | 3.984914847 | 0.97909 | 4.07003 | 4.7E-05 | 0.00061 |
| 1266 | NOX1 | 20.4051329 | 3.95739348 | 0.80954 | 4.88847 | 1E-06 | 2.1E-05 |
| 933 | HSD17B11 | 62.5742176 | 3.947040793 | 0.51609 | 7.64794 | 2.04E-14 | 3.16E-12 |
| 703 | TMEM87B | 12.5944372 | 3.942425394 | 0.90017 | 4.37965 | 1.2E-05 | 0.00018 |
| 935 | GPRIN3 | 7.48640932 | 3.941194533 | 1.21137 | 3.25351 | 0.00114 | 0.00942 |
| 1088 | ZDHHC14 | 7.61213521 | 3.916856575 | 1.15679 | 3.38598 | 0.00071 | 0.00631 |
| 1139 | MUC17 | 17.6847093 | 3.913688838 | 0.92945 | 4.21077 | 2.5E-05 | 0.00035 |
| 1138 | MUC12 | 17.6756392 | 3.911181081 | 1.18633 | 3.29686 | 0.00098 | 0.00822 |
| 304 | KRT8 | 141.666673 | 3.910417165 | 0.38165 | 10.2461 | 1.23E-24 | 9.98E-22 |
| 301 | ENSG00000257298 | 7.34789182 | 3.910017918 | 1.28024 | 3.05414 | 0.00226 | 0.01657 |
| 817 | CARD10 | 25.2375633 | 3.897432106 | 0.67872 | 5.74232 | 9.34E-09 | 3.35E-07 |
| 1193 | DGAT1 | 138.014688 | 3.883407254 | 0.46162 | 8.41258 | 4.01E-17 | 9.20E-15 |
| 1127 | STEAP1 | 12.4428373 | 3.881973415 | 0.9441 | 4.11183 | 3.9E-05 | 0.00052 |
| 588 | TJP3 | 29.4428514 | 3.87569889 | 0.62165 | 6.23456 | 4.53E-10 | 2.33E-08 |
| 976 | GLRX | 29.4983969 | 3.858353788 | 0.67166 | 5.74448 | 9.22E-09 | 3.31E-07 |
| 1066 | CD24 | 133.762098 | 3.856306596 | 1.17004 | 3.29589 | 0.00098 | 0.00825 |
| 1196 | SLC1A1 | 14.5167714 | 3.842621245 | 0.88458 | 4.34399 | 1.4E-05 | 0.00021 |
| 837 | MST1R | 16.5025784 | 3.819215975 | 0.88377 | 4.3215 | 1.5E-05 | 0.00023 |
| 235 | EHBP1L1 | 11.5820798 | 3.814927184 | 1.01021 | 3.77636 | 0.00016 | 0.00176 |
| 6 | ARHGEF16 | 29.8377542 | 3.812482534 | 0.68067 | 5.60106 | 2.13E-08 | 6.87E-07 |
| 1231 | HDHD3 | 66.7247852 | 3.810518244 | 0.50889 | 7.4879 | 7.00E-14 | 9.45E-12 |
| 583 | ADAMTSL5 | 7.28012013 | 3.794469844 | 1.26795 | 2.9926 | 0.00277 | 0.01959 |
| 150 | SIPA1L2 | 16.1211974 | 3.788179936 | 0.91225 | 4.15258 | 3.3E-05 | 0.00044 |
| 255 | PIWIL4 | 9.10852781 | 3.784231187 | 1.31397 | 2.88001 | 0.00398 | 0.02635 |
| 249 | CAPN5 | 46.9489352 | 3.783781755 | 0.60226 | 6.28267 | 3.33E-10 | 1.76E-08 |
| 303 | ACVRL1 | 13.8687408 | 3.783497108 | 0.97121 | 3.89567 | 9.8E-05 | 0.00115 |
| 788 | BCAS1 | 28.284624 | 3.782469392 | 0.67516 | 5.60232 | 2.11E-08 | 6.84E-07 |

| | | | | | | | |
|---|---|---|---|---|---|---|---|
| 66 | LINC01359 | 7.19343938 | 3.77958372 | 1.27344 | 2.96802 | 0.003 | 0.0209 |
| 682 | ACTG2 | 25.5793896 | 3.778706737 | 1.01184 | 3.7345 | 0.00019 | 0.00203 |
| 666 | KHK | 16.0188292 | 3.771763228 | 0.8824 | 4.27444 | 1.9E-05 | 0.00028 |
| 277 | B4GALNT3 | 16.0080081 | 3.766870377 | 0.90397 | 4.16705 | 3.1E-05 | 0.00042 |
| 302 | SMAGP | 16.0731766 | 3.7668091 | 1.0307 | 3.6546 | 0.00026 | 0.00266 |
| 931 | PLAC8 | 20.9395061 | 3.762118671 | 0.73826 | 5.09589 | 3.47E-07 | 8.1E-06 |
| 565 | DSC2 | 44.8857586 | 3.75230724 | 0.50616 | 7.41331 | 1.23E-13 | 1.59E-11 |
| 1201 | AQP7 | 13.613615 | 3.747453773 | 1.04615 | 3.58215 | 0.00034 | 0.00337 |
| 326 | SH2B3 | 13.0915482 | 3.742864748 | 0.97732 | 3.82971 | 0.00013 | 0.00145 |
| 164 | PBLD | 42.5303417 | 3.734413917 | 0.73139 | 5.1059 | 3.29E-07 | 7.8E-06 |
| 441 | DNASE1 | 60.3775304 | 3.728438326 | 0.49425 | 7.54369 | 4.57E-14 | 6.48E-12 |
| 295 | PKP2 | 33.6351991 | 3.721798339 | 0.59756 | 6.22833 | 4.71E-10 | 2.40E-08 |
| 574 | TNFRSF11A | 41.3612029 | 3.708004626 | 0.61833 | 5.99678 | 2.01E-09 | 8.97E-08 |
| 1102 | TSPAN13 | 39.945023 | 3.701205794 | 0.64903 | 5.70266 | 1.18E-08 | 4.13E-07 |
| 852 | DPPA4 | 8.26330803 | 3.684897916 | 1.2266 | 3.00415 | 0.00266 | 0.01896 |
| 461 | MVP | 49.4216768 | 3.670629436 | 0.54331 | 6.75603 | 1.42E-11 | 1.03E-09 |
| 585 | TLE2 | 12.5439367 | 3.659242819 | 0.92163 | 3.97039 | 7.2E-05 | 0.00089 |
| 819 | SH3BP1 | 16.6549638 | 3.656773319 | 0.75947 | 4.81493 | 1.5E-06 | 2.9E-05 |
| 331 | ACADS | 40.0996711 | 3.653592461 | 0.56435 | 6.47402 | 9.54E-11 | 5.76E-09 |
| 94 | MTMR11 | 87.2329599 | 3.651373335 | 0.42742 | 8.54277 | 1.31E-17 | 3.34E-15 |
| 1118 | COBL | 10.784997 | 3.647503076 | 1.09964 | 3.31699 | 0.00091 | 0.00776 |
| 1063 | NT5E | 14.6266569 | 3.645292381 | 0.83874 | 4.34617 | 1.4E-05 | 0.00021 |
| 848 | ABI3BP | 6.33789265 | 3.642642964 | 1.23816 | 2.94199 | 0.00326 | 0.02238 |
| 339 | CRYL1 | 46.0243433 | 3.64031544 | 0.73529 | 4.95086 | 7.39E-07 | 1.6E-05 |
| 425 | ABHD17C | 36.040613 | 3.638581979 | 0.66216 | 5.49498 | 3.91E-08 | 1.2E-06 |
| 903 | FGFR3 | 39.0170728 | 3.633819904 | 0.71234 | 5.10124 | 3.37E-07 | 7.9E-06 |
| 1187 | MAL2 | 48.802347 | 3.633382696 | 0.53657 | 6.77146 | 1.27E-11 | 9.39E-10 |
| 359 | SLC7A7 | 21.2919372 | 3.633172064 | 0.90395 | 4.01922 | 5.8E-05 | 0.00074 |
| 1011 | DSP | 59.665836 | 3.622960654 | 0.60471 | 5.99124 | 2.08E-09 | 9.25E-08 |
| 449 | TMC5 | 21.0972321 | 3.618484263 | 0.73971 | 4.89176 | 9.99E-07 | 2.1E-05 |
| 691 | VAMP8 | 47.8710172 | 3.612660844 | 0.49404 | 7.31243 | 2.62E-13 | 2.94E-11 |
| 465 | PYCARD | 8.20170636 | 3.602565724 | 1.07635 | 3.34702 | 0.00082 | 0.00711 |
| 580 | ARID3A | 86.3090588 | 3.601523266 | 0.4223 | 8.5283 | 1.49E-17 | 3.72E-15 |
| 64 | L1TD1 | 5.83180791 | 3.57969437 | 1.33699 | 2.67742 | 0.00742 | 0.04351 |
| 1186 | TNFRSF11B | 8.14454623 | 3.568214534 | 1.09506 | 3.25847 | 0.00112 | 0.00926 |
| 469 | DOK4 | 16.0438634 | 3.565787822 | 0.82015 | 4.34771 | 1.4E-05 | 0.00021 |
| 657 | UBE2CP5 | 5.95347552 | 3.536289602 | 1.30605 | 2.70763 | 0.00678 | 0.0406 |
| 479 | NPIPB15 | 9.60962732 | 3.535226534 | 0.97987 | 3.60787 | 0.00031 | 0.00311 |
| 1043 | MAPK13 | 46.5307578 | 3.53324404 | 0.49955 | 7.07289 | 1.52E-12 | 1.37E-10 |

| | | | | | | | |
|---|---|---|---|---|---|---|---|
| 229 | MYRF | 13.9000109 | 3.533225005 | 1.00881 | 3.50236 | 0.00046 | 0.00437 |
| 513 | DHRS11 | 77.5198818 | 3.51591061 | 0.40348 | 8.71398 | 2.93E-18 | 8.08E-16 |
| 233 | NAALADL1 | 21.9447922 | 3.511514334 | 0.92145 | 3.81084 | 0.00014 | 0.00155 |
| 988 | ARHGAP26 | 21.1971965 | 3.49220598 | 0.69748 | 5.0069 | 5.53E-07 | 1.2E-05 |
| 444 | CPPED1 | 9.49678612 | 3.492006334 | 0.96892 | 3.60403 | 0.00031 | 0.00314 |
| 582 | REEP6 | 48.3835223 | 3.490477265 | 0.65442 | 5.3337 | 9.62E-08 | 2.6E-06 |
| 1071 | MAN1A1 | 20.8702065 | 3.472542883 | 0.68707 | 5.05416 | 4.32E-07 | 9.8E-06 |
| 832 | CDCP1 | 24.2675773 | 3.460820285 | 0.60952 | 5.67795 | 1.36E-08 | 4.68E-07 |
| 1081 | MAP3K5 | 5.46162151 | 3.460210305 | 1.30557 | 2.65034 | 0.00804 | 0.04615 |
| 70 | IFI44 | 25.0016432 | 3.453516898 | 0.8122 | 4.25207 | 2.1E-05 | 0.0003 |
| 1062 | TENT5A | 20.2349188 | 3.445499801 | 0.69384 | 4.96581 | 6.84E-07 | 1.5E-05 |
| 438 | IL32 | 24.0517377 | 3.434404492 | 0.67955 | 5.05393 | 4.33E-07 | 9.8E-06 |
| 266 | MPZL3 | 5.43493569 | 3.43321565 | 1.19654 | 2.86928 | 0.00411 | 0.0271 |
| 995 | SLC36A1 | 27.5082335 | 3.432140417 | 0.61166 | 5.61122 | 2.01E-08 | 6.59E-07 |
| 360 | PCK2 | 69.020867 | 3.424936119 | 0.56748 | 6.03532 | 1.59E-09 | 7.31E-08 |
| 216 | PRRG4 | 11.3156296 | 3.411825294 | 1.0245 | 3.33024 | 0.00087 | 0.00747 |
| 1199 | LRRC19 | 32.6834676 | 3.410467585 | 0.58822 | 5.79795 | 6.71E-09 | 2.57E-07 |
| 928 | BMP2K | 7.41872617 | 3.410452896 | 1.09794 | 3.10623 | 0.00189 | 0.01435 |
| 508 | FAM83G | 9.12234063 | 3.404422392 | 1.09407 | 3.11172 | 0.00186 | 0.01416 |
| 124 | INAVA | 7.02493387 | 3.384610504 | 1.12048 | 3.02069 | 0.00252 | 0.01808 |
| 14 | MST1P2 | 7.38644429 | 3.376327898 | 1.21841 | 2.77109 | 0.00559 | 0.03464 |
| 297 | VDR | 10.483343 | 3.376143957 | 1.00618 | 3.35541 | 0.00079 | 0.00692 |
| 1244 | SAPCD2 | 10.3119734 | 3.375869005 | 0.94191 | 3.58408 | 0.00034 | 0.00335 |
| 885 | TNFSF10 | 34.2503961 | 3.367368786 | 0.73226 | 4.59862 | 4.3E-06 | 7.5E-05 |
| 595 | SH2D3A | 15.6179716 | 3.365969361 | 0.77934 | 4.31902 | 1.6E-05 | 0.00023 |
| 838 | HYAL1 | 6.69859332 | 3.362833099 | 1.18504 | 2.83775 | 0.00454 | 0.02937 |
| 220 | PRR5L | 11.8362641 | 3.360855012 | 0.98688 | 3.40552 | 0.00066 | 0.00594 |
| 448 | ABCC6P1 | 17.804267 | 3.357667853 | 0.8504 | 3.94834 | 7.9E-05 | 0.00096 |
| 987 | PCDH1 | 10.3345709 | 3.357535554 | 1.0072 | 3.33354 | 0.00086 | 0.00742 |
| 75 | GBP2 | 8.57847627 | 3.356114345 | 0.99518 | 3.37239 | 0.00075 | 0.00658 |
| 663 | SDC1 | 33.6115716 | 3.352658574 | 0.67132 | 4.9941 | 5.91E-07 | 1.3E-05 |
| 632 | ETHE1 | 58.3083077 | 3.34362734 | 0.47101 | 7.09881 | 1.26E-12 | 1.16E-10 |
| 787 | SALL4 | 6.93690137 | 3.338499286 | 1.13564 | 2.93976 | 0.00328 | 0.0225 |
| 971 | HMGCS1 | 6.86550769 | 3.338293192 | 1.14355 | 2.91922 | 0.00351 | 0.02379 |
| 1015 | MBOAT1 | 44.0367813 | 3.333581959 | 0.53792 | 6.19712 | 5.75E-10 | 2.85E-08 |
| 982 | SHROOM1 | 15.5271923 | 3.329004009 | 0.82829 | 4.01915 | 5.8E-05 | 0.00074 |
| 1072 | PKIB | 10.4793391 | 3.325335402 | 1.04037 | 3.19629 | 0.00139 | 0.01114 |
| 1123 | CLDN4 | 92.1073487 | 3.32438211 | 0.44112 | 7.53626 | 4.84E-14 | 6.80E-12 |
| 253 | NAALAD2 | 8.56327845 | 3.321490047 | 1.02005 | 3.25621 | 0.00113 | 0.00933 |

| | | | | | | | |
|---|---|---|---|---|---|---|---|
| 316 | KITLG | 6.834195 | 3.318033174 | 1.10956 | 2.99039 | 0.00279 | 0.01969 |
| 1229 | SNX30 | 8.48402733 | 3.317266899 | 0.97933 | 3.38729 | 0.00071 | 0.00629 |
| 900 | BDH1 | 35.994754 | 3.31148743 | 0.53112 | 6.23488 | 4.52E-10 | 2.33E-08 |
| 886 | B3GNT5 | 8.56204435 | 3.297550466 | 1.08884 | 3.02851 | 0.00246 | 0.01771 |
| 306 | PRR13 | 66.7034065 | 3.290082903 | 0.49106 | 6.70002 | 2.08E-11 | 1.47E-09 |
| 573 | NEDD4L | 44.3430943 | 3.286337672 | 0.55218 | 5.9516 | 2.66E-09 | 1.15E-07 |
| 1080 | MTFR2 | 7.00533557 | 3.285241768 | 1.25475 | 2.61825 | 0.00884 | 0.04953 |
| 1017 | TDP2 | 63.6403554 | 3.281924563 | 0.47418 | 6.92132 | 4.47E-12 | 3.58E-10 |
| 1111 | HOXA10 | 6.43409786 | 3.273014225 | 1.14323 | 2.86295 | 0.0042 | 0.02755 |
| 736 | FZD5 | 16.1985205 | 3.259936804 | 0.75092 | 4.34126 | 1.4E-05 | 0.00021 |
| 655 | ZNF524 | 7.91283865 | 3.24878378 | 1.05543 | 3.07815 | 0.00208 | 0.01551 |
| 953 | FHDC1 | 7.90113102 | 3.248583752 | 1.07087 | 3.03358 | 0.00242 | 0.01746 |
| 840 | GLYCTK | 43.0017531 | 3.236500227 | 0.57838 | 5.59585 | 2.20E-08 | 7.05E-07 |
| 778 | ACSS2 | 45.1588698 | 3.23380822 | 0.46376 | 6.97298 | 3.10E-12 | 2.61E-10 |
| 912 | SMIM14 | 38.5302815 | 3.225921298 | 0.58197 | 5.54309 | 2.97E-08 | 9.17E-07 |
| 292 | PDE3A | 16.3006288 | 3.224506124 | 0.74033 | 4.35552 | 1.3E-05 | 0.0002 |
| 1120 | ASL | 57.5043704 | 3.206505054 | 0.53834 | 5.95623 | 2.58E-09 | 1.13E-07 |
| 148 | TMEM63A | 27.0273632 | 3.202690947 | 0.69614 | 4.60065 | 4.2E-06 | 7.4E-05 |
| 738 | CPS1 | 89.1320827 | 3.197225963 | 0.60182 | 5.3126 | 1.08E-07 | 2.9E-06 |
| 1085 | STXBP5-AS1 | 6.62123229 | 3.196286708 | 1.17264 | 2.72572 | 0.00642 | 0.03871 |
| 704 | PSD4 | 19.9769764 | 3.181929445 | 0.68545 | 4.64213 | 3.4E-06 | 6.2E-05 |
| 1068 | MICAL1 | 35.586187 | 3.158627804 | 0.56245 | 5.61583 | 1.96E-08 | 6.43E-07 |
| 30 | SFN | 22.1491532 | 3.158566725 | 0.86711 | 3.64265 | 0.00027 | 0.00276 |
| 128 | FMOD | 11.173903 | 3.157141755 | 1.05649 | 2.98833 | 0.00281 | 0.01982 |
| 1091 | ENSG00000285730 | 6.16677247 | 3.153608311 | 1.17322 | 2.68799 | 0.00719 | 0.0425 |
| 126 | ELF3-AS1 | 6.17830241 | 3.153323144 | 1.14291 | 2.75904 | 0.0058 | 0.03566 |
| 329 | VSIG10 | 16.8594496 | 3.15029623 | 0.73376 | 4.29337 | 1.8E-05 | 0.00026 |
| 135 | PFKFB2 | 35.6401309 | 3.147706698 | 0.53603 | 5.87226 | 4.30E-09 | 1.76E-07 |
| 570 | MYO5B | 24.2259412 | 3.131856635 | 0.60638 | 5.16488 | 2.41E-07 | 5.9E-06 |
| 542 | \P27P1-BPTFP1-KF | 29.4363156 | 3.130519516 | 0.68642 | 4.56064 | 5.1E-06 | 8.7E-05 |
| 793 | NRIP1 | 11.9260156 | 3.121561368 | 0.79379 | 3.93247 | 8.4E-05 | 0.00101 |
| 107 | MUC1 | 67.6424556 | 3.111858135 | 0.62418 | 4.98553 | 6.18E-07 | 1.3E-05 |
| 759 | GPR35 | 16.0401862 | 3.110880648 | 0.75481 | 4.12144 | 3.8E-05 | 0.0005 |
| 715 | PLA2R1 | 19.3675886 | 3.108017287 | 0.6907 | 4.49978 | 6.8E-06 | 0.00011 |
| 474 | ELMO3 | 17.992112 | 3.10615454 | 0.72607 | 4.27803 | 1.9E-05 | 0.00027 |
| 798 | MX1 | 80.2365532 | 3.105480139 | 0.57508 | 5.40008 | 6.66E-08 | 1.9E-06 |
| 1238 | TMEM141 | 92.5163283 | 3.094446945 | 0.4328 | 7.14982 | 8.69E-13 | 8.48E-11 |
| 1021 | TRIM38 | 41.0625763 | 3.09415022 | 0.64543 | 4.79393 | 1.6E-06 | 3.2E-05 |
| 1274 | GPC3 | 107.189169 | 3.091333752 | 0.49672 | 6.22344 | 4.86E-10 | 2.45E-08 |

| | | | | | | | |
|---|---|---|---|---|---|---|---|
| 36 | SERINC2 | 47.9316956 | 3.088862401 | 0.46706 | 6.61339 | 3.76E-11 | 2.44E-09 |
| 472 | CES2 | 86.3888021 | 3.084141831 | 0.44103 | 6.9931 | 2.69E-12 | 2.29E-10 |
| 679 | MXD1 | 24.0881652 | 3.083497378 | 0.77965 | 3.95499 | 7.7E-05 | 0.00093 |
| 488 | FOXF1 | 5.76763959 | 3.08269145 | 1.13452 | 2.71717 | 0.00658 | 0.03964 |
| 796 | B3GALT5 | 33.2340077 | 3.081341083 | 0.59557 | 5.17373 | 2.29E-07 | 5.7E-06 |
| 281 | LTBR | 27.6813531 | 3.072206295 | 0.56706 | 5.41777 | 6.03E-08 | 1.7E-06 |
| 455 | IL4R | 11.4840493 | 3.069922387 | 1.00983 | 3.04004 | 0.00237 | 0.01716 |
| 1208 | GCNT1 | 52.7845782 | 3.069603854 | 0.58531 | 5.24443 | 1.57E-07 | 4E-06 |
| 300 | LIMA1 | 61.5742876 | 3.064241287 | 0.44572 | 6.87488 | 6.20E-12 | 4.85E-10 |
| 501 | SLC52A1 | 7.16643852 | 3.06305503 | 1.15754 | 2.64617 | 0.00814 | 0.04649 |
| 724 | TFPI | 14.7688218 | 3.060876532 | 0.94929 | 3.2244 | 0.00126 | 0.01025 |
| 1089 | SYNJ2 | 15.6108009 | 3.043675406 | 0.77275 | 3.93875 | 8.2E-05 | 0.00099 |
| 1270 | KLHL13 | 14.6438014 | 3.036659413 | 0.7786 | 3.90016 | 9.6E-05 | 0.00114 |
| 37 | TINAGL1 | 12.5957243 | 3.035923768 | 0.81391 | 3.73003 | 0.00019 | 0.00206 |
| 617 | SDHAF1 | 7.20455597 | 3.031895627 | 1.09933 | 2.75794 | 0.00582 | 0.03574 |
| 470 | MMP15 | 41.4723971 | 3.017783482 | 0.52065 | 5.79618 | 6.78E-09 | 2.58E-07 |
| 1222 | KLF4 | 22.89745 | 3.011443382 | 0.66561 | 4.52432 | 6.1E-06 | 0.0001 |
| 712 | HNMT | 25.8507217 | 3.005313187 | 0.62196 | 4.83199 | 1.4E-06 | 2.7E-05 |
| 246 | KCNE3 | 10.9619128 | 2.988182172 | 0.90268 | 3.31036 | 0.00093 | 0.00791 |
| 904 | ZFYVE28 | 8.22623947 | 2.98240601 | 1.1192 | 2.66476 | 0.0077 | 0.04472 |
| 875 | MME | 11.1867594 | 2.981056314 | 1.02656 | 2.90394 | 0.00368 | 0.02476 |
| 568 | LIPG | 17.459574 | 2.976306985 | 0.70955 | 4.19462 | 2.7E-05 | 0.00038 |
| 139 | IRF6 | 11.3147581 | 2.958062094 | 1.01669 | 2.9095 | 0.00362 | 0.02437 |
| 35 | EPB41 | 13.6394633 | 2.952508366 | 0.98518 | 2.99693 | 0.00273 | 0.01934 |
| 1236 | AGPAT2 | 90.619442 | 2.938176313 | 0.3778 | 7.77712 | 7.42E-15 | 1.25E-12 |
| 1158 | PRKAG2 | 10.4778512 | 2.937748315 | 0.91233 | 3.22004 | 0.00128 | 0.01037 |
| 548 | MYO15B | 52.3162411 | 2.920414559 | 0.58806 | 4.96617 | 6.83E-07 | 1.5E-05 |
| 355 | GRTP1 | 18.8303412 | 2.916738515 | 0.79463 | 3.67056 | 0.00024 | 0.00252 |
| 970 | SELENOP | 14.7000598 | 2.914287639 | 0.77978 | 3.73733 | 0.00019 | 0.00201 |
| 709 | CCDC115 | 14.7106572 | 2.904771368 | 0.79503 | 3.65365 | 0.00026 | 0.00266 |
| 945 | CFI | 10.6540077 | 2.904435979 | 0.93458 | 3.10773 | 0.00189 | 0.0143 |
| 293 | FAR2 | 11.6638429 | 2.893614097 | 0.79001 | 3.66277 | 0.00025 | 0.00259 |
| 489 | CYBA | 42.6777408 | 2.88762689 | 0.6064 | 4.76195 | 1.9E-06 | 3.7E-05 |
| 991 | CARMN | 25.1096594 | 2.882307998 | 0.81975 | 3.51608 | 0.00044 | 0.00418 |
| 957 | GASK1B | 19.2918896 | 2.872707174 | 0.6472 | 4.43868 | 9.1E-06 | 0.00014 |
| 294 | RESF1 | 27.4060367 | 2.871401931 | 0.61805 | 4.64589 | 3.4E-06 | 6.1E-05 |
| 913 | ATP8A1 | 14.3764048 | 2.87079583 | 0.80521 | 3.5653 | 0.00036 | 0.00356 |
| 47 | TMEM125 | 10.5639881 | 2.869703297 | 0.99045 | 2.89738 | 0.00376 | 0.0252 |
| 333 | P2RX4 | 27.7907515 | 2.865883992 | 0.64645 | 4.43327 | 9.3E-06 | 0.00015 |

| | | | | | | | |
|---|---|---|---|---|---|---|---|
| 1162 | PRAG1 | 7.55177108 | 2.863367823 | 0.99735 | 2.87099 | 0.00409 | 0.02697 |
| 1173 | ZNF703 | 26.6792649 | 2.857157613 | 0.59385 | 4.81124 | 1.5E-06 | 3E-05 |
| 366 | MIPOL1 | 11.8778561 | 2.854130104 | 0.97723 | 2.92064 | 0.00349 | 0.02372 |
| 211 | SPON1 | 16.9975049 | 2.853975618 | 0.72831 | 3.91862 | 8.9E-05 | 0.00107 |
| 416 | SEMA6D | 21.6649816 | 2.853389742 | 0.61294 | 4.65522 | 3.2E-06 | 5.9E-05 |
| 1073 | NCOA7 | 35.6929655 | 2.850700944 | 0.77281 | 3.68875 | 0.00023 | 0.00238 |
| 97 | CGN | 39.721754 | 2.840223015 | 0.51854 | 5.47733 | 4.32E-08 | 1.3E-06 |
| 836 | MST1 | 35.9139618 | 2.838024098 | 0.70274 | 4.0385 | 5.4E-05 | 0.00069 |
| 814 | TST | 63.4868507 | 2.837362274 | 0.40274 | 7.04523 | 1.85E-12 | 1.65E-10 |
| 342 | SLC46A3 | 21.763521 | 2.830134882 | 0.84899 | 3.33355 | 0.00086 | 0.00742 |
| 16 | ARHGEF10L | 56.0036508 | 2.823008502 | 0.44673 | 6.3192 | 2.63E-10 | 1.43E-08 |
| 511 | MYO1D | 43.0697619 | 2.812913715 | 0.49066 | 5.73296 | 9.87E-09 | 3.52E-07 |
| 1250 | REPS2 | 10.2670656 | 2.810229771 | 1.04194 | 2.69712 | 0.00699 | 0.04164 |
| 305 | KRT18 | 75.5271213 | 2.809294418 | 0.40474 | 6.94104 | 3.89E-12 | 3.16E-10 |
| 1170 | EPHX2 | 76.3903028 | 2.804521543 | 0.47091 | 5.95548 | 2.59E-09 | 1.13E-07 |
| 841 | PRKCD | 21.7740578 | 2.804461232 | 0.62215 | 4.5077 | 6.6E-06 | 0.00011 |
| 1269 | AMOT | 21.0177142 | 2.791790738 | 0.62793 | 4.44604 | 8.7E-06 | 0.00014 |
| 98 | S100A10 | 339.337455 | 2.788337707 | 0.49407 | 5.64359 | 1.67E-08 | 5.66E-07 |
| 674 | OXER1 | 10.9869547 | 2.780797349 | 0.82107 | 3.38679 | 0.00071 | 0.0063 |
| 394 | GSKIP | 16.4063524 | 2.780068226 | 0.72637 | 3.82736 | 0.00013 | 0.00146 |
| 962 | SH3RF1 | 20.6601757 | 2.77682344 | 0.60878 | 4.56133 | 5.1E-06 | 8.7E-05 |
| 254 | FUT4 | 17.7531267 | 2.770284537 | 0.86031 | 3.2201 | 0.00128 | 0.01037 |
| 1032 | HLA-C | 99.1283405 | 2.769231434 | 0.42281 | 6.54955 | 5.77E-11 | 3.60E-09 |
| 829 | CTDSPL | 19.0535566 | 2.764845561 | 0.72228 | 3.82797 | 0.00013 | 0.00146 |
| 810 | TCN2 | 15.1268198 | 2.753829051 | 0.77101 | 3.57171 | 0.00035 | 0.00348 |
| 858 | DTX3L | 22.6302934 | 2.750247647 | 0.57635 | 4.77186 | 1.8E-06 | 3.5E-05 |
| 1083 | ABRACL | 40.8896419 | 2.749428857 | 0.5143 | 5.34598 | 8.99E-08 | 2.5E-06 |
| 685 | HK2 | 28.7509997 | 2.748604111 | 0.7602 | 3.61565 | 0.0003 | 0.00302 |
| 1045 | ETV7 | 8.58322971 | 2.733488674 | 1.01558 | 2.69155 | 0.00711 | 0.04221 |
| 710 | PTPN18 | 42.0001108 | 2.716194133 | 0.45802 | 5.93025 | 3.02E-09 | 1.29E-07 |
| 466 | TOX3 | 12.5326463 | 2.71473858 | 0.88592 | 3.06432 | 0.00218 | 0.01615 |
| 53 | MAST2 | 28.8872728 | 2.712491514 | 0.55352 | 4.90046 | 9.56E-07 | 2E-05 |
| 1141 | CLDN15 | 27.3635603 | 2.706519393 | 0.60503 | 4.47334 | 7.7E-06 | 0.00012 |
| 76 | SLC44A3-AS1 | 8.16617603 | 2.70378009 | 1.02395 | 2.64053 | 0.00828 | 0.04692 |
| 601 | DOCK6 | 17.4412412 | 2.700099304 | 0.78 | 3.46165 | 0.00054 | 0.00497 |
| 230 | PLCB3 | 26.6454724 | 2.689913134 | 0.68599 | 3.92119 | 8.8E-05 | 0.00106 |
| 1144 | NAMPT | 36.5344464 | 2.688081547 | 0.52084 | 5.16106 | 2.46E-07 | 6E-06 |
| 226 | STX3 | 21.5774975 | 2.685385566 | 0.59585 | 4.50684 | 6.6E-06 | 0.00011 |
| 689 | RETSAT | 31.8203901 | 2.684272701 | 0.54498 | 4.92541 | 8.42E-07 | 1.8E-05 |

| | | | | | | | |
|---|---|---|---|---|---|---|---|
| 1113 | NT5C3A | 41.5192171 | 2.681480825 | 0.46583 | 5.75637 | 8.59E-09 | 3.17E-07 |
| 834 | SLC26A6 | 24.1886391 | 2.662327565 | 0.59622 | 4.46535 | 8E-06 | 0.00013 |
| 171 | IFIT3 | 10.6416404 | 2.659797084 | 0.98428 | 2.70227 | 0.00689 | 0.0411 |
| 7 | TPRG1L | 25.1223586 | 2.656869828 | 0.56618 | 4.69265 | 2.7E-06 | 5E-05 |
| 638 | SLC1A5 | 30.6055676 | 2.655057728 | 0.66149 | 4.01377 | 6E-05 | 0.00076 |
| 181 | AVPI1 | 12.589014 | 2.653632783 | 0.81963 | 3.23761 | 0.00121 | 0.00984 |
| 13 | FBLIM1 | 35.4852997 | 2.651485373 | 0.47946 | 5.53014 | 3.20E-08 | 9.79E-07 |
| 1253 | SAT1 | 303.187984 | 2.643929847 | 0.37746 | 7.0045 | 2.48E-12 | 2.15E-10 |
| 943 | HADH | 61.5671716 | 2.642425164 | 0.48248 | 5.4767 | 4.33E-08 | 1.3E-06 |
| 1241 | MAMDC4 | 28.4404311 | 2.628793356 | 0.65914 | 3.98824 | 6.7E-05 | 0.00084 |
| 1272 | ELF4 | 16.2813688 | 2.627484732 | 0.72172 | 3.64058 | 0.00027 | 0.00278 |
| 204 | RASSF7 | 33.1746055 | 2.616018346 | 0.57008 | 4.58883 | 4.5E-06 | 7.8E-05 |
| 240 | LINC02754 | 10.108419 | 2.614989972 | 0.89567 | 2.9196 | 0.0035 | 0.02378 |
| 668 | LCLAT1 | 15.0210059 | 2.61444308 | 0.82892 | 3.15402 | 0.00161 | 0.01258 |
| 457 | SULT1A1 | 65.4178197 | 2.611534605 | 0.49393 | 5.28729 | 1.24E-07 | 3.3E-06 |
| 1060 | MYO6 | 88.2097308 | 2.609894203 | 0.3979 | 6.5592 | 5.41E-11 | 3.40E-09 |
| 602 | HOOK2 | 25.7691899 | 2.609248216 | 0.59079 | 4.41651 | 1E-05 | 0.00016 |
| 537 | AKAP1 | 60.7515842 | 2.595652487 | 0.49242 | 5.27127 | 1.35E-07 | 3.5E-06 |
| 757 | PER2 | 10.7782992 | 2.58558169 | 0.84121 | 3.07363 | 0.00211 | 0.01571 |
| 101 | S100A16 | 39.2669104 | 2.585262176 | 0.48835 | 5.29389 | 1.20E-07 | 3.2E-06 |
| 1059 | SLC17A5 | 11.8607276 | 2.581031347 | 0.75708 | 3.40917 | 0.00065 | 0.00586 |
| 516 | GSDMB | 50.0512908 | 2.579413533 | 0.48529 | 5.31524 | 1.07E-07 | 2.9E-06 |
| 356 | TEP1 | 19.1270135 | 2.57875667 | 0.61198 | 4.21381 | 2.5E-05 | 0.00035 |
| 714 | LY75 | 13.4419828 | 2.575024461 | 0.96367 | 2.67209 | 0.00754 | 0.04413 |
| 1125 | DTX2 | 13.7117896 | 2.572375822 | 0.79266 | 3.24525 | 0.00117 | 0.00963 |
| 130 | PIK3C2B | 18.09703 | 2.570813374 | 0.7533 | 3.41272 | 0.00064 | 0.0058 |
| 695 | CNNM4 | 16.7113705 | 2.567660494 | 0.69369 | 3.70145 | 0.00021 | 0.00227 |
| 131 | CDK18 | 18.3975511 | 2.563485885 | 0.86297 | 2.97055 | 0.00297 | 0.02076 |
| 1006 | GMDS | 38.5453933 | 2.561089396 | 0.53235 | 4.81089 | 1.5E-06 | 3E-05 |
| 205 | MIR210HG | 21.1238565 | 2.560317382 | 0.68475 | 3.73903 | 0.00018 | 0.00201 |
| 1151 | ZC3HAV1 | 29.7714507 | 2.529639951 | 0.60783 | 4.16174 | 3.2E-05 | 0.00043 |
| 671 | GALM | 41.2961089 | 2.529576146 | 0.47205 | 5.35867 | 8.38E-08 | 2.3E-06 |
| 166 | COMTD1 | 16.4133688 | 2.523250871 | 0.71941 | 3.50737 | 0.00045 | 0.0043 |
| 833 | PFKFB4 | 18.1173041 | 2.522279248 | 0.71277 | 3.5387 | 0.0004 | 0.00388 |
| 180 | FRAT2 | 10.435516 | 2.519772607 | 0.81626 | 3.08696 | 0.00202 | 0.01514 |
| 1012 | GCNT2 | 8.24907531 | 2.515620508 | 0.96062 | 2.61875 | 0.00883 | 0.04948 |
| 291 | MGST1 | 92.9367186 | 2.514644548 | 0.5636 | 4.46178 | 8.1E-06 | 0.00013 |
| 122 | ASPM | 25.2967092 | 2.512603447 | 0.80974 | 3.10298 | 0.00192 | 0.01446 |
| 1038 | PSMB8 | 38.8484791 | 2.507111354 | 0.60713 | 4.12947 | 3.6E-05 | 0.00049 |

| | | | | | | | |
|---|---|---|---|---|---|---|---|
| 1039 | PSMB9 | 13.7882259 | 2.506853558 | 0.88719 | 2.82561 | 0.00472 | 0.03019 |
| 916 | FRYL | 60.3130856 | 2.502996191 | 0.41443 | 6.03964 | 1.54E-09 | 7.14E-08 |
| 538 | YPEL2 | 9.42933316 | 2.498995943 | 0.87749 | 2.84791 | 0.0044 | 0.02868 |
| 507 | SREBF1 | 22.5620773 | 2.493127937 | 0.67834 | 3.67532 | 0.00024 | 0.00248 |
| 484 | GSE1 | 32.4349961 | 2.492767518 | 0.68998 | 3.61282 | 0.0003 | 0.00305 |
| 1260 | CLCN5 | 22.4491192 | 2.489996691 | 0.67525 | 3.68752 | 0.00023 | 0.00239 |
| 648 | ZNF600 | 14.7720246 | 2.488134108 | 0.81392 | 3.05696 | 0.00224 | 0.01644 |
| 81 | SORT1 | 21.9343071 | 2.487718112 | 0.60623 | 4.10356 | 4.1E-05 | 0.00054 |
| 1147 | TES | 30.7640825 | 2.482629346 | 0.58218 | 4.26437 | 2E-05 | 0.00029 |
| 1198 | MLLT3 | 16.4177968 | 2.480003882 | 0.69137 | 3.58709 | 0.00033 | 0.00332 |
| 562 | ABHD3 | 18.0915208 | 2.479042238 | 0.67148 | 3.69192 | 0.00022 | 0.00235 |
| 948 | ALPK1 | 17.2544305 | 2.478336011 | 0.66357 | 3.73487 | 0.00019 | 0.00203 |
| 473 | CES3 | 10.9336631 | 2.476376117 | 0.87455 | 2.83161 | 0.00463 | 0.02976 |
| 643 | TRPM4 | 26.1989838 | 2.475296599 | 0.6084 | 4.06853 | 4.7E-05 | 0.00062 |
| 1033 | HLA-B | 150.085786 | 2.474911723 | 0.41944 | 5.90056 | 3.62E-09 | 1.49E-07 |
| 591 | STAP2 | 20.5418199 | 2.452791109 | 0.75435 | 3.25152 | 0.00115 | 0.00946 |
| 272 | ARHGAP32 | 11.7692861 | 2.433551567 | 0.86184 | 2.82368 | 0.00475 | 0.03029 |
| 38 | HDAC1 | 51.8098619 | 2.432724966 | 0.39925 | 6.09327 | 1.11E-09 | 5.16E-08 |
| 696 | MGAT4A | 16.5542095 | 2.430339027 | 0.7332 | 3.31471 | 0.00092 | 0.00782 |
| 1255 | XK | 14.4467852 | 2.429335948 | 0.78472 | 3.09579 | 0.00196 | 0.01478 |
| 1179 | CHMP4C | 9.53952132 | 2.428847365 | 0.84533 | 2.87326 | 0.00406 | 0.02683 |
| 494 | RILP | 10.8333614 | 2.423543005 | 0.87993 | 2.75423 | 0.00588 | 0.03605 |
| 1 | ISG15 | 35.7891334 | 2.420786988 | 0.55096 | 4.39378 | 1.1E-05 | 0.00017 |
| 749 | SP100 | 19.7304712 | 2.417525163 | 0.64739 | 3.73426 | 0.00019 | 0.00203 |
| 769 | RRBP1 | 79.5902733 | 2.417429581 | 0.3719 | 6.50024 | 8.02E-11 | 4.89E-09 |
| 732 | SATB2 | 17.1482002 | 2.411662274 | 0.72528 | 3.32513 | 0.00088 | 0.00757 |
| 23 | TCEA3 | 16.7910062 | 2.397119104 | 0.90649 | 2.64441 | 0.00818 | 0.04666 |
| 894 | TMEM44 | 11.4755969 | 2.38789789 | 0.80802 | 2.95524 | 0.00312 | 0.02164 |
| 802 | SLC37A1 | 37.4617305 | 2.383870137 | 0.60081 | 3.96775 | 7.3E-05 | 0.0009 |
| 1245 | TPRN | 28.3234521 | 2.383158656 | 0.50503 | 4.71881 | 2.4E-06 | 4.4E-05 |
| 208 | CRACR2B | 29.0590784 | 2.382206028 | 0.54737 | 4.35208 | 1.3E-05 | 0.0002 |
| 544 | GPRC5C | 31.5596377 | 2.375175031 | 0.5102 | 4.65543 | 3.2E-06 | 5.9E-05 |
| 1022 | H2AC6 | 11.4413872 | 2.375053464 | 0.75938 | 3.12762 | 0.00176 | 0.01356 |
| 147 | ENSG00000242861 | 13.2412678 | 2.374154939 | 0.79954 | 2.96939 | 0.00298 | 0.02082 |
| 175 | PDLIM1 | 125.348432 | 2.351691653 | 0.41876 | 5.61589 | 1.96E-08 | 6.43E-07 |
| 190 | TCF7L2 | 59.450418 | 2.342835744 | 0.40897 | 5.72856 | 1.01E-08 | 3.59E-07 |
| 1203 | MYORG | 23.4441332 | 2.339945461 | 0.5916 | 3.95526 | 7.6E-05 | 0.00093 |
| 600 | LDLR | 30.1939636 | 2.33687988 | 0.52878 | 4.4194 | 9.9E-06 | 0.00015 |
| 910 | LIAS | 9.43935093 | 2.331434191 | 0.84405 | 2.76221 | 0.00574 | 0.03542 |

| | | | | | | | |
|---|---|---|---|---|---|---|---|
| 739 | TMBIM1 | 31.8974227 | 2.328311903 | 0.55158 | 4.2212 | 2.4E-05 | 0.00034 |
| 34 | RCC1 | 9.81220272 | 2.325051988 | 0.8438 | 2.75545 | 0.00586 | 0.03593 |
| 1184 | NIPAL2 | 13.7489532 | 2.324722818 | 0.85078 | 2.73245 | 0.00629 | 0.03803 |
| 871 | PLSCR1 | 37.4777251 | 2.324404638 | 0.47584 | 4.8848 | 1E-06 | 2.1E-05 |
| 1169 | ADAM28 | 12.0482451 | 2.3214832 | 0.88683 | 2.61773 | 0.00885 | 0.04958 |
| 183 | SCD | 94.4095779 | 2.320803514 | 0.61815 | 3.75446 | 0.00017 | 0.0019 |
| 1016 | E2F3 | 17.5230474 | 2.318607171 | 0.71911 | 3.22429 | 0.00126 | 0.01025 |
| 596 | CAMSAP3 | 37.448532 | 2.314264986 | 0.47957 | 4.82568 | 1.4E-06 | 2.8E-05 |
| 337 | BRI3BP | 31.3443984 | 2.304345729 | 0.48872 | 4.71507 | 2.4E-06 | 4.5E-05 |
| 821 | APOBEC3C | 9.90266993 | 2.294969553 | 0.80035 | 2.86745 | 0.00414 | 0.02721 |
| 321 | CHPT1 | 14.4029318 | 2.294334908 | 0.71634 | 3.20284 | 0.00136 | 0.01091 |
| 1077 | STX7 | 42.04342 | 2.292576539 | 0.53902 | 4.25322 | 2.1E-05 | 0.0003 |
| 644 | FCGRT | 178.677751 | 2.286202216 | 0.39896 | 5.73039 | 1.00E-08 | 3.56E-07 |
| 1002 | ZFP62 | 15.9503076 | 2.277674158 | 0.66505 | 3.4248 | 0.00062 | 0.0056 |
| 344 | ELF1 | 17.3636014 | 2.275172555 | 0.64557 | 3.52429 | 0.00042 | 0.00407 |
| 435 | FAHD1 | 21.7059322 | 2.272294002 | 0.58815 | 3.86348 | 0.00011 | 0.00129 |
| 447 | ABCC6 | 22.8110305 | 2.268399164 | 0.76568 | 2.9626 | 0.00305 | 0.02118 |
| 649 | ZNF160 | 33.8515337 | 2.266109786 | 0.66259 | 3.42008 | 0.00063 | 0.00568 |
| 308 | SUOX | 18.4703557 | 2.261572181 | 0.65676 | 3.44353 | 0.00057 | 0.00528 |
| 975 | RHOBTB3 | 19.6136493 | 2.248153431 | 0.75168 | 2.99083 | 0.00278 | 0.01968 |
| 191 | RPS15AP30 | 9.89090068 | 2.242189392 | 0.85347 | 2.62715 | 0.00861 | 0.04845 |
| 961 | DDX60 | 14.6602048 | 2.241252007 | 0.76399 | 2.93362 | 0.00335 | 0.02286 |
| 1026 | HLA-A | 286.991817 | 2.238498029 | 0.35563 | 6.29441 | 3.09E-10 | 1.65E-08 |
| 672 | PKDCC | 17.276446 | 2.238340306 | 0.61439 | 3.6432 | 0.00027 | 0.00276 |
| 1031 | HLA-E | 62.4661889 | 2.238251949 | 0.42997 | 5.20566 | 1.93E-07 | 4.9E-06 |
| 522 | IFI35 | 14.5475922 | 2.234226542 | 0.65813 | 3.3948 | 0.00069 | 0.00614 |
| 764 | LZTS3 | 13.5792531 | 2.233642761 | 0.71796 | 3.11111 | 0.00186 | 0.01418 |
| 1086 | STXBP5 | 16.8607936 | 2.231099706 | 0.70577 | 3.16123 | 0.00157 | 0.01236 |
| 730 | MYO1B | 17.2138672 | 2.230645745 | 0.73198 | 3.0474 | 0.00231 | 0.01683 |
| 426 | ALPK3 | 19.9570794 | 2.226163233 | 0.76382 | 2.91453 | 0.00356 | 0.02408 |
| 346 | ATP7B | 14.1449071 | 2.215709635 | 0.78264 | 2.83107 | 0.00464 | 0.02978 |
| 462 | MAPK3 | 68.7811415 | 2.215415663 | 0.36893 | 6.00495 | 1.91E-09 | 8.58E-08 |
| 196 | OAT | 54.9638235 | 2.196011254 | 0.68261 | 3.2171 | 0.00129 | 0.01045 |
| 1079 | MYB | 16.9528664 | 2.194881911 | 0.7946 | 2.76226 | 0.00574 | 0.03542 |
| 1220 | SLC44A1 | 63.2198921 | 2.191139003 | 0.3958 | 5.53598 | 3.09E-08 | 9.53E-07 |
| 1185 | SYBU | 10.8179706 | 2.189245134 | 0.78893 | 2.77494 | 0.00552 | 0.0343 |
| 586 | GNA11 | 62.4134895 | 2.189197776 | 0.42228 | 5.18422 | 2.17E-07 | 5.4E-06 |
| 432 | DECR2 | 23.2380944 | 2.173446556 | 0.56433 | 3.85141 | 0.00012 | 0.00135 |
| 225 | DTX4 | 32.4393788 | 2.172797247 | 0.5951 | 3.65112 | 0.00026 | 0.00268 |

| | | | | | | | |
|---|---|---|---|---|---|---|---|
| 151 | NID1 | 22.2650764 | 2.172358672 | 0.73939 | 2.93804 | 0.0033 | 0.02261 |
| 351 | FARP1 | 14.1536803 | 2.172021273 | 0.7724 | 2.81205 | 0.00492 | 0.03121 |
| 795 | DOP1B | 12.1900929 | 2.166577886 | 0.73237 | 2.95831 | 0.00309 | 0.02146 |
| 1099 | LFNG | 14.4907765 | 2.166305851 | 0.7674 | 2.82292 | 0.00476 | 0.03032 |
| 343 | POSTN | 27.5048211 | 2.166170459 | 0.71637 | 3.0238 | 0.0025 | 0.01793 |
| 845 | FOXP1 | 63.4701916 | 2.165654783 | 0.38329 | 5.6501 | 1.60E-08 | 5.48E-07 |
| 610 | PGPEP1 | 39.7049481 | 2.162821054 | 0.49023 | 4.41188 | 1E-05 | 0.00016 |
| 839 | ABHD14B | 26.740576 | 2.160267578 | 0.57666 | 3.7462 | 0.00018 | 0.00196 |
| 767 | BMP2 | 12.2659353 | 2.159854464 | 0.768 | 2.8123 | 0.00492 | 0.0312 |
| 966 | CMBL | 57.3655747 | 2.157821498 | 0.56562 | 3.81496 | 0.00014 | 0.00152 |
| 1163 | FDFT1 | 65.8544366 | 2.152668043 | 0.53517 | 4.02237 | 5.8E-05 | 0.00073 |
| 520 | JUP | 69.5778726 | 2.148402632 | 0.36156 | 5.942 | 2.82E-09 | 1.20E-07 |
| 566 | DSG2 | 47.7566317 | 2.141069428 | 0.50689 | 4.22392 | 2.4E-05 | 0.00033 |
| 1214 | SLC35D2 | 33.6589207 | 2.141057415 | 0.61511 | 3.48078 | 0.0005 | 0.00468 |
| 389 | LYSET | 16.5734027 | 2.136264929 | 0.72541 | 2.94489 | 0.00323 | 0.02224 |
| 121 | C1orf21 | 14.4374408 | 2.122975165 | 0.6933 | 3.06213 | 0.0022 | 0.01623 |
| 515 | CISD3 | 14.1266158 | 2.116224027 | 0.76739 | 2.7577 | 0.00582 | 0.03574 |
| 909 | C4orf19 | 20.926959 | 2.115200175 | 0.5851 | 3.61513 | 0.0003 | 0.00303 |
| 221 | SYT13 | 11.0157011 | 2.114893959 | 0.76144 | 2.7775 | 0.00548 | 0.03407 |
| 1128 | STEAP2 | 24.1778555 | 2.112451443 | 0.51774 | 4.08017 | 4.5E-05 | 0.00059 |
| 963 | GALNT7 | 31.4697908 | 2.111668589 | 0.55187 | 3.82638 | 0.00013 | 0.00147 |
| 177 | ALDH18A1 | 67.4898054 | 2.111428116 | 0.3626 | 5.82299 | 5.78E-09 | 2.24E-07 |
| 1171 | ZNF395 | 15.0945446 | 2.108690999 | 0.73069 | 2.88591 | 0.0039 | 0.02594 |
| 661 | HPCAL1 | 24.4362223 | 2.103838586 | 0.63252 | 3.32614 | 0.00088 | 0.00755 |
| 826 | TTC38 | 29.8148831 | 2.09634304 | 0.52911 | 3.96198 | 7.4E-05 | 0.00091 |
| 92 | TXNIP | 115.192991 | 2.095910367 | 0.48297 | 4.33966 | 1.4E-05 | 0.00021 |
| 223 | NR1H3 | 19.2625697 | 2.088062643 | 0.66643 | 3.1332 | 0.00173 | 0.01333 |
| 543 | ABCA5 | 29.7675404 | 2.083931112 | 0.59334 | 3.51219 | 0.00044 | 0.00423 |
| 110 | IFI16 | 21.8100601 | 2.082579341 | 0.76926 | 2.70724 | 0.00678 | 0.04063 |
| 728 | GLS | 144.569768 | 2.070924209 | 0.60512 | 3.42231 | 0.00062 | 0.00564 |
| 731 | NABP1 | 32.9103016 | 2.070379979 | 0.6212 | 3.3329 | 0.00086 | 0.00742 |
| 1075 | ARHGAP18 | 20.9555575 | 2.068420419 | 0.67042 | 3.08526 | 0.00203 | 0.01521 |
| 290 | DERA | 20.0846386 | 2.06317872 | 0.69355 | 2.97479 | 0.00293 | 0.02054 |
| 463 | SEPHS2 | 26.2834673 | 2.061289621 | 0.65272 | 3.15801 | 0.00159 | 0.01246 |
| 1195 | C8orf82 | 15.3744878 | 2.060542692 | 0.7453 | 2.7647 | 0.0057 | 0.03521 |
| 1219 | PGAP4 | 14.2275927 | 2.058517581 | 0.73966 | 2.78307 | 0.00538 | 0.03359 |
| 558 | RAB40B | 14.6451322 | 2.055560238 | 0.74164 | 2.77164 | 0.00558 | 0.03459 |
| 377 | KIAA0586 | 13.4513456 | 2.055425019 | 0.75014 | 2.74007 | 0.00614 | 0.03726 |
| 406 | CHP1 | 69.2007489 | 2.044524668 | 0.39162 | 5.22075 | 1.78E-07 | 4.5E-06 |

| | | | | | | | |
|---|---|---|---|---|---|---|---|
| 1007 | SERPINB1 | 88.3218115 | 2.038504908 | 0.35786 | 5.69645 | 1.22E-08 | 4.25E-07 |
| 1153 | SLC37A3 | 23.4175534 | 2.037285628 | 0.56087 | 3.63238 | 0.00028 | 0.00286 |
| 412 | B2M | 504.594102 | 2.034759023 | 0.32893 | 6.18592 | 6.17E-10 | 3.03E-08 |
| 861 | KALRN | 19.7125954 | 2.033062103 | 0.68548 | 2.9659 | 0.00302 | 0.02099 |
| 1178 | PAG1 | 34.6528136 | 2.03271724 | 0.45802 | 4.4381 | 9.1E-06 | 0.00014 |
| 642 | FTL | 372.671956 | 2.031822052 | 0.34679 | 5.85899 | 4.66E-09 | 1.87E-07 |
| 241 | CPT1A | 85.0268131 | 2.031020589 | 0.43957 | 4.62043 | 3.8E-06 | 6.8E-05 |
| 270 | SIAE | 15.7343426 | 2.025607549 | 0.64836 | 3.12421 | 0.00178 | 0.01368 |
| 144 | C1orf115 | 21.508979 | 2.015341774 | 0.67593 | 2.98159 | 0.00287 | 0.02016 |
| 330 | MLEC | 98.442552 | 2.010589081 | 0.32143 | 6.25506 | 3.97E-10 | 2.07E-08 |
| 1278 | ERCC00062 | 37.0973587 | 2.006094105 | 0.55335 | 3.62537 | 0.00029 | 0.00292 |
| 198 | MKI67 | 45.977207 | 2.005241818 | 0.68077 | 2.94553 | 0.00322 | 0.02221 |
| 503 | ELP5 | 13.4649382 | 1.994083684 | 0.68414 | 2.91472 | 0.00356 | 0.02408 |
| 1010 | RREB1 | 43.5701394 | 1.991959561 | 0.47751 | 4.17157 | 3E-05 | 0.00041 |
| 750 | ITM2C | 240.193613 | 1.990567629 | 0.32731 | 6.08164 | 1.19E-09 | 5.53E-08 |
| 584 | PLEKHJ1 | 74.1093584 | 1.988506961 | 0.38636 | 5.1468 | 2.65E-07 | 6.4E-06 |
| 411 | CKMT1A | 14.8789207 | 1.979480015 | 0.66402 | 2.98105 | 0.00287 | 0.02019 |
| 243 | NADSYN1 | 41.5802376 | 1.974743798 | 0.41839 | 4.71981 | 2.4E-06 | 4.4E-05 |
| 1221 | TMEM38B | 19.4342697 | 1.971868876 | 0.57191 | 3.44788 | 0.00057 | 0.0052 |
| 296 | SLC38A1 | 43.9610664 | 1.968169151 | 0.50608 | 3.88902 | 0.0001 | 0.00118 |
| 1213 | PTCH1 | 16.0926074 | 1.963957478 | 0.64709 | 3.03506 | 0.0024 | 0.0174 |
| 369 | LRR1 | 14.735933 | 1.956144364 | 0.69997 | 2.7946 | 0.0052 | 0.0326 |
| 1126 | CROT | 16.8803889 | 1.945480551 | 0.66315 | 2.9337 | 0.00335 | 0.02286 |
| 952 | MGST2 | 38.9336282 | 1.943536939 | 0.45854 | 4.23857 | 2.2E-05 | 0.00032 |
| 100 | S100A6 | 405.139442 | 1.941964579 | 0.38268 | 5.07471 | 3.88E-07 | 8.9E-06 |
| 950 | USP53 | 28.2886741 | 1.936181137 | 0.51771 | 3.73989 | 0.00018 | 0.002 |
| 944 | CASP6 | 14.5321561 | 1.935791522 | 0.68622 | 2.82095 | 0.00479 | 0.03047 |
| 1209 | VPS13A | 55.5788467 | 1.932159475 | 0.54934 | 3.51725 | 0.00044 | 0.00417 |
| 1119 | GUSB | 48.5825611 | 1.931234566 | 0.42974 | 4.49394 | 7E-06 | 0.00011 |
| 280 | SCNN1A | 13.8134714 | 1.916214891 | 0.68358 | 2.8032 | 0.00506 | 0.03196 |
| 727 | NAB1 | 28.3049134 | 1.911439985 | 0.50075 | 3.81715 | 0.00014 | 0.00152 |
| 87 | IGSF3 | 27.1770059 | 1.910248385 | 0.52516 | 3.63744 | 0.00028 | 0.00281 |
| 1275 | MIR503HG | 19.2826071 | 1.902853391 | 0.72343 | 2.63034 | 0.00853 | 0.0481 |
| 698 | NCK2 | 27.6836554 | 1.897396828 | 0.56957 | 3.33128 | 0.00086 | 0.00745 |
| 1192 | GSDMD | 50.0088758 | 1.897058484 | 0.47882 | 3.96191 | 7.4E-05 | 0.00091 |
| 200 | PGGHG | 49.050651 | 1.874904931 | 0.54887 | 3.41593 | 0.00064 | 0.00575 |
| 660 | RRM2 | 41.6020149 | 1.874810715 | 0.66719 | 2.81001 | 0.00495 | 0.03138 |
| 388 | ITPK1 | 26.726839 | 1.866092695 | 0.53987 | 3.45653 | 0.00055 | 0.00506 |
| 645 | RCN3 | 17.3571653 | 1.860998539 | 0.68043 | 2.73505 | 0.00624 | 0.03778 |

| | | | | | | | |
|---|---|---|---|---|---|---|---|
| 458 | QPRT | 61.4343733 | 1.854746872 | 0.41522 | 4.46693 | 7.9E-06 | 0.00013 |
| 1132 | BAIAP2L1 | 28.4736401 | 1.854638578 | 0.54926 | 3.37663 | 0.00073 | 0.0065 |
| 973 | CWC27 | 14.7293477 | 1.8525205 | 0.69608 | 2.66136 | 0.00778 | 0.04502 |
| 1254 | PDK3 | 42.1453975 | 1.852112133 | 0.49201 | 3.76439 | 0.00017 | 0.00184 |
| 159 | ACBD5 | 21.2317182 | 1.850313818 | 0.5903 | 3.13455 | 0.00172 | 0.01329 |
| 554 | SLC25A10 | 16.2596163 | 1.846328373 | 0.67117 | 2.75091 | 0.00594 | 0.03629 |
| 443 | ATF7IP2 | 30.2694032 | 1.846088088 | 0.55412 | 3.33155 | 0.00086 | 0.00745 |
| 1052 | CD2AP | 40.1267663 | 1.838101474 | 0.60454 | 3.0405 | 0.00236 | 0.01714 |
| 1175 | CHCHD7 | 20.4026258 | 1.836944031 | 0.56217 | 3.26758 | 0.00108 | 0.009 |
| 575 | CNDP2 | 59.148336 | 1.83688443 | 0.52683 | 3.48668 | 0.00049 | 0.00459 |
| 313 | STAT6 | 64.180224 | 1.834987678 | 0.51851 | 3.53893 | 0.0004 | 0.00388 |
| 423 | CTSH | 31.5826028 | 1.83263577 | 0.59367 | 3.08695 | 0.00202 | 0.01514 |
| 268 | SLC37A4 | 55.04697 | 1.831992156 | 0.46012 | 3.98159 | 6.8E-05 | 0.00085 |
| 932 | CDS1 | 52.3996924 | 1.831922665 | 0.55873 | 3.27875 | 0.00104 | 0.00869 |
| 428 | PDE8A | 31.5514844 | 1.831152554 | 0.48766 | 3.75498 | 0.00017 | 0.0019 |
| 775 | CST3 | 346.949586 | 1.827291175 | 0.40045 | 4.56307 | 5E-06 | 8.7E-05 |
| 942 | CENPE | 24.2125091 | 1.820227907 | 0.68781 | 2.64642 | 0.00813 | 0.04649 |
| 1207 | PCSK5 | 27.159582 | 1.813492706 | 0.59526 | 3.04658 | 0.00231 | 0.01687 |
| 807 | GUCD1 | 35.2437828 | 1.81193186 | 0.65867 | 2.75089 | 0.00594 | 0.03629 |
| 941 | SLC39A8 | 19.2166676 | 1.809397611 | 0.62966 | 2.87361 | 0.00406 | 0.02682 |
| 549 | ITGB4 | 18.3654565 | 1.80938727 | 0.67768 | 2.66999 | 0.00759 | 0.04431 |
| 1061 | SH3BGRL2 | 39.2902921 | 1.808619486 | 0.51716 | 3.49721 | 0.00047 | 0.00444 |
| 737 | IDH1 | 52.3855813 | 1.799097783 | 0.4191 | 4.29281 | 1.8E-05 | 0.00026 |
| 790 | RAB22A | 19.4894167 | 1.798463517 | 0.6171 | 2.9144 | 0.00356 | 0.02408 |
| 1008 | SERPINB6 | 86.2876997 | 1.79844079 | 0.36024 | 4.99239 | 5.96E-07 | 1.3E-05 |
| 659 | ADI1 | 43.5775909 | 1.794877509 | 0.45039 | 3.98516 | 6.7E-05 | 0.00084 |
| 42 | PPIE | 16.4761198 | 1.79327104 | 0.67801 | 2.64491 | 0.00817 | 0.04663 |
| 238 | SSH3 | 21.104912 | 1.790777993 | 0.64924 | 2.75828 | 0.00581 | 0.03572 |
| 1190 | MTSS1 | 23.5148884 | 1.790537315 | 0.61877 | 2.89369 | 0.00381 | 0.02542 |
| 1090 | RNASET2 | 20.0813011 | 1.789887357 | 0.58637 | 3.05249 | 0.00227 | 0.01664 |
| 1223 | EPB41L4B | 29.4413373 | 1.789085501 | 0.57572 | 3.10755 | 0.00189 | 0.0143 |
| 589 | ZBTB7A | 64.0653079 | 1.787979651 | 0.44087 | 4.0556 | 5E-05 | 0.00065 |
| 379 | PLEKHG3 | 30.2276066 | 1.785151258 | 0.53938 | 3.30966 | 0.00093 | 0.00793 |
| 1049 | KLC4 | 24.5235981 | 1.784754635 | 0.5787 | 3.08408 | 0.00204 | 0.01524 |
| 491 | SPIRE2 | 46.6975943 | 1.783358339 | 0.42789 | 4.16779 | 3.1E-05 | 0.00042 |
| 614 | CEBPG | 19.3749835 | 1.782752935 | 0.56447 | 3.15829 | 0.00159 | 0.01246 |
| 213 | HTATIP2 | 22.5947122 | 1.781577864 | 0.57668 | 3.08939 | 0.00201 | 0.01506 |
| 665 | ITSN2 | 29.4504992 | 1.77919554 | 0.61744 | 2.88159 | 0.00396 | 0.02624 |
| 188 | PDCD4 | 65.8269429 | 1.778456609 | 0.39209 | 4.53589 | 5.7E-06 | 9.7E-05 |

| | | | | | | | |
|---|---|---|---|---|---|---|---|
| 1168 | SLC39A14 | 48.7411609 | 1.778151276 | 0.54778 | 3.24611 | 0.00117 | 0.00961 |
| 1064 | PREP | 39.5105179 | 1.775480974 | 0.42027 | 4.22458 | 2.4E-05 | 0.00033 |
| 1280 | ERCC00162 | 52.7352221 | 1.771398402 | 0.52113 | 3.39913 | 0.00068 | 0.00606 |
| 1224 | PTPN3 | 24.6590622 | 1.76723576 | 0.64738 | 2.72981 | 0.00634 | 0.03829 |
| 227 | TKFC | 33.6201376 | 1.764271465 | 0.49372 | 3.57344 | 0.00035 | 0.00347 |
| 1273 | IGSF1 | 48.3433028 | 1.763581868 | 0.55546 | 3.17498 | 0.0015 | 0.01187 |
| 433 | MCRIP2 | 30.2280809 | 1.76260584 | 0.59666 | 2.95413 | 0.00314 | 0.0217 |
| 422 | CSK | 22.8407508 | 1.761415204 | 0.62961 | 2.79764 | 0.00515 | 0.03236 |
| 1174 | TACC1 | 36.4259493 | 1.761247382 | 0.47178 | 3.73323 | 0.00019 | 0.00204 |
| 1087 | RMND1 | 14.6793207 | 1.759619471 | 0.65401 | 2.69051 | 0.00713 | 0.04225 |
| 278 | CCND2 | 138.703048 | 1.759416789 | 0.35492 | 4.95719 | 7.15E-07 | 1.5E-05 |
| 434 | MRPS34 | 30.2688744 | 1.756734563 | 0.4971 | 3.53396 | 0.00041 | 0.00394 |
| 182 | DNMBP | 31.4066243 | 1.754400428 | 0.52564 | 3.33766 | 0.00084 | 0.00732 |
| 490 | ACSF3 | 18.0434626 | 1.75340811 | 0.62067 | 2.82502 | 0.00473 | 0.03021 |
| 186 | GSTO1 | 54.4041193 | 1.753000492 | 0.44195 | 3.96649 | 7.3E-05 | 0.0009 |
| 1096 | COX19 | 22.8657647 | 1.752397566 | 0.5657 | 3.09773 | 0.00195 | 0.01469 |
| 1124 | TMEM120A | 34.3550927 | 1.747485536 | 0.58544 | 2.9849 | 0.00284 | 0.02001 |
| 55 | FAAH | 22.4207576 | 1.747397708 | 0.55398 | 3.15425 | 0.00161 | 0.01258 |
| 705 | DBI | 124.243907 | 1.746987672 | 0.30285 | 5.76855 | 8.00E-09 | 2.98E-07 |
| 868 | ZBTB38 | 23.8170507 | 1.7450787 | 0.54986 | 3.17369 | 0.00151 | 0.0119 |
| 495 | TLCD2 | 24.1325872 | 1.742938966 | 0.54941 | 3.17236 | 0.00151 | 0.01194 |
| 327 | ALDH2 | 61.5744952 | 1.730245841 | 0.39505 | 4.3798 | 1.2E-05 | 0.00018 |
| 252 | TMEM135 | 14.8341151 | 1.729334596 | 0.65051 | 2.65843 | 0.00785 | 0.04532 |
| 178 | TM9SF3 | 134.459664 | 1.728429997 | 0.32069 | 5.38975 | 7.06E-08 | 2E-06 |
| 934 | FAM13A | 37.6656368 | 1.724290494 | 0.47157 | 3.65645 | 0.00026 | 0.00265 |
| 658 | SH3YL1 | 33.7393966 | 1.716036508 | 0.49025 | 3.50034 | 0.00046 | 0.0044 |
| 161 | RASSF4 | 18.8532908 | 1.713440174 | 0.65329 | 2.62281 | 0.00872 | 0.04893 |
| 140 | LPGAT1 | 25.0699206 | 1.707178278 | 0.63342 | 2.69516 | 0.00704 | 0.04184 |
| 115 | TSTD1 | 28.3907285 | 1.706135392 | 0.50608 | 3.37129 | 0.00075 | 0.0066 |
| 86 | ATP1A1 | 165.151613 | 1.704996187 | 0.36431 | 4.68007 | 2.9E-06 | 5.2E-05 |
| 67 | SLC35D1 | 17.3045442 | 1.704912251 | 0.63971 | 2.66514 | 0.0077 | 0.04471 |
| 761 | D2HGDH | 44.3231819 | 1.70409278 | 0.49069 | 3.47282 | 0.00052 | 0.0048 |
| 607 | NR2F6 | 91.9664002 | 1.703647973 | 0.41731 | 4.08248 | 4.5E-05 | 0.00058 |
| 61 | ECHDC2 | 64.1504373 | 1.701698456 | 0.42499 | 4.00409 | 6.2E-05 | 0.00078 |
| 50 | AKR1A1 | 57.7416059 | 1.698221299 | 0.38802 | 4.3766 | 1.2E-05 | 0.00018 |
| 56 | MKNK1 | 29.8726615 | 1.692989739 | 0.47495 | 3.56456 | 0.00036 | 0.00357 |
| 349 | NDFIP2 | 19.9067933 | 1.690426821 | 0.58552 | 2.88705 | 0.00389 | 0.02588 |
| 545 | NHERF1 | 74.6416161 | 1.687340223 | 0.41254 | 4.09016 | 4.3E-05 | 0.00057 |
| 1152 | PARP12 | 23.3692638 | 1.683261256 | 0.50827 | 3.31177 | 0.00093 | 0.00788 |

| | | | | | | | |
|---|---|---|---|---|---|---|---|
| 902 | DGKQ | 26.5888648 | 1.67825466 | 0.60405 | 2.77836 | 0.00546 | 0.03399 |
| 1009 | SLC22A23 | 22.7190994 | 1.675454498 | 0.57164 | 2.93097 | 0.00338 | 0.02303 |
| 1240 | RABL6 | 84.6618639 | 1.673563965 | 0.35682 | 4.69017 | 2.7E-06 | 5E-05 |
| 897 | NCBP2AS2 | 23.4021213 | 1.669137049 | 0.53165 | 3.13954 | 0.00169 | 0.0131 |
| 152 | IDI1 | 59.2134748 | 1.669010327 | 0.45427 | 3.67403 | 0.00024 | 0.00249 |
| 561 | IMPA2 | 38.4679501 | 1.663500818 | 0.5052 | 3.29276 | 0.00099 | 0.00833 |
| 1277 | SLC6A8 | 29.2483627 | 1.662707612 | 0.521 | 3.19137 | 0.00142 | 0.01131 |
| 323 | APPL2 | 33.7243223 | 1.662302603 | 0.52912 | 3.14162 | 0.00168 | 0.01301 |
| 758 | RNPEPL1 | 23.7901375 | 1.661842192 | 0.57521 | 2.88908 | 0.00386 | 0.02572 |
| 1048 | FOXP4 | 24.1806508 | 1.661288347 | 0.54753 | 3.03415 | 0.00241 | 0.01744 |
| 722 | CDCA7 | 34.8527967 | 1.661005564 | 0.46894 | 3.54205 | 0.0004 | 0.00384 |
| 1159 | INSIG1 | 16.3588501 | 1.660844582 | 0.63122 | 2.63115 | 0.00851 | 0.04802 |
| 569 | ACAA2 | 72.9701726 | 1.657455388 | 0.44547 | 3.72073 | 0.0002 | 0.00213 |
| 808 | XBP1 | 71.5278981 | 1.654052125 | 0.4573 | 3.61696 | 0.0003 | 0.00301 |
| 743 | DNPEP | 24.8692294 | 1.645267601 | 0.50363 | 3.2668 | 0.00109 | 0.00902 |
| 32 | IFI6 | 134.898049 | 1.64258886 | 0.50103 | 3.2784 | 0.00104 | 0.0087 |
| 202 | SIGIRR | 17.0488047 | 1.641674005 | 0.58411 | 2.81058 | 0.00495 | 0.03134 |
| 103 | TPM3 | 28.4170376 | 1.641584555 | 0.54177 | 3.03002 | 0.00245 | 0.01765 |
| 146 | LBR | 32.4602032 | 1.638913941 | 0.55969 | 2.92824 | 0.00341 | 0.02321 |
| 334 | ORAI1 | 15.8592907 | 1.633755672 | 0.60515 | 2.69976 | 0.00694 | 0.04138 |
| 106 | SLC50A1 | 28.6865342 | 1.629639606 | 0.51999 | 3.134 | 0.00172 | 0.0133 |
| 880 | GOLIM4 | 40.9447243 | 1.629506195 | 0.4172 | 3.9058 | 9.4E-05 | 0.00112 |
| 1074 | PTPRK | 27.227105 | 1.626573962 | 0.52006 | 3.12768 | 0.00176 | 0.01356 |
| 623 | BLVRB | 27.7011287 | 1.621244084 | 0.50252 | 3.22625 | 0.00125 | 0.0102 |
| 823 | SREBF2 | 68.2121275 | 1.618684709 | 0.36368 | 4.45081 | 8.6E-06 | 0.00014 |
| 96 | SELENBP1 | 56.6443853 | 1.614035252 | 0.36715 | 4.39606 | 1.1E-05 | 0.00017 |
| 911 | UGDH | 44.6492103 | 1.611727228 | 0.42259 | 3.81392 | 0.00014 | 0.00153 |
| 576 | HSBP1L1 | 41.8543817 | 1.610652835 | 0.46636 | 3.45368 | 0.00055 | 0.0051 |
| 869 | XRN1 | 23.5949013 | 1.594218929 | 0.553 | 2.88284 | 0.00394 | 0.02615 |
| 1226 | TXN | 88.761161 | 1.592158327 | 0.37262 | 4.27289 | 1.9E-05 | 0.00028 |
| 365 | FAM177A1 | 45.9306852 | 1.590610787 | 0.41197 | 3.86102 | 0.00011 | 0.0013 |
| 616 | LSR | 44.1630117 | 1.589981693 | 0.41482 | 3.83292 | 0.00013 | 0.00144 |
| 141 | FLVCR1 | 39.467107 | 1.589113473 | 0.56816 | 2.79693 | 0.00516 | 0.03242 |
| 599 | DNM2 | 73.9440565 | 1.587976743 | 0.41399 | 3.83576 | 0.00013 | 0.00142 |
| 815 | MPST | 38.2541453 | 1.587929027 | 0.42161 | 3.76633 | 0.00017 | 0.00183 |
| 49 | PTPRF | 144.234104 | 1.579144835 | 0.32951 | 4.79241 | 1.6E-06 | 3.2E-05 |
| 460 | KIF22 | 32.7123263 | 1.562055274 | 0.58121 | 2.68757 | 0.0072 | 0.04251 |
| 1259 | EBP | 45.9541872 | 1.55865372 | 0.39938 | 3.90265 | 9.5E-05 | 0.00113 |
| 1030 | TRIM26 | 27.9719092 | 1.54968731 | 0.50152 | 3.08996 | 0.002 | 0.01504 |

| | | | | | | | |
|---|---|---|---|---|---|---|---|
| 887 | POLR2H | 39.0536422 | 1.546939201 | 0.4375 | 3.53588 | 0.00041 | 0.00392 |
| 686 | SUCLG1 | 92.8628161 | 1.546436325 | 0.32124 | 4.81399 | 1.5E-06 | 2.9E-05 |
| 224 | CTNND1 | 89.9425043 | 1.544765466 | 0.33864 | 4.56174 | 5.1E-06 | 8.7E-05 |
| 784 | SERINC3 | 31.4866552 | 1.538405917 | 0.45259 | 3.39912 | 0.00068 | 0.00606 |
| 1014 | MCUR1 | 26.6579761 | 1.537792988 | 0.48606 | 3.16378 | 0.00156 | 0.01227 |
| 386 | LGMN | 22.959447 | 1.529213789 | 0.57338 | 2.66703 | 0.00765 | 0.04453 |
| 401 | LPCAT4 | 26.2949028 | 1.527934727 | 0.58037 | 2.63267 | 0.00847 | 0.0479 |
| 289 | EPS8 | 31.4823183 | 1.527520463 | 0.56847 | 2.68708 | 0.00721 | 0.04256 |
| 1228 | PTBP3 | 43.8124508 | 1.527347622 | 0.44649 | 3.42079 | 0.00062 | 0.00567 |
| 1210 | GOLM1 | 226.299384 | 1.524363884 | 0.27467 | 5.54984 | 2.86E-08 | 8.93E-07 |
| 857 | FAM162A | 56.5954795 | 1.523626495 | 0.39357 | 3.8713 | 0.00011 | 0.00126 |
| 688 | TGOLN2 | 40.8611816 | 1.519144947 | 0.49925 | 3.04287 | 0.00234 | 0.01704 |
| 46 | PPCS | 41.0396008 | 1.515914648 | 0.50332 | 3.01181 | 0.0026 | 0.01856 |
| 979 | MAN2A1 | 29.9091926 | 1.514089509 | 0.46701 | 3.24206 | 0.00119 | 0.00972 |
| 398 | CKB | 206.798671 | 1.509736629 | 0.56067 | 2.69272 | 0.00709 | 0.04208 |
| 1133 | ZNF789 | 19.6007645 | 1.507051089 | 0.57255 | 2.63217 | 0.00848 | 0.0479 |
| 493 | MYO1C | 39.1469101 | 1.504667618 | 0.42282 | 3.55864 | 0.00037 | 0.00364 |
| 1034 | CLIC1 | 67.0701171 | 1.50061953 | 0.44534 | 3.36961 | 0.00075 | 0.00662 |
| 1058 | DST | 61.8407086 | 1.500323459 | 0.458 | 3.27579 | 0.00105 | 0.00877 |
| 22 | LUZP1 | 24.2908155 | 1.497788208 | 0.52614 | 2.84674 | 0.00442 | 0.02873 |
| 117 | MPC2 | 70.6107808 | 1.497595158 | 0.40302 | 3.71597 | 0.0002 | 0.00216 |
| 974 | F2RL1 | 22.5954203 | 1.497542009 | 0.52583 | 2.84797 | 0.0044 | 0.02868 |
| 824 | PACSIN2 | 22.6425122 | 1.497525807 | 0.55254 | 2.71026 | 0.00672 | 0.04039 |
| 927 | CCNG2 | 38.8746564 | 1.488016272 | 0.54991 | 2.70595 | 0.00681 | 0.04076 |
| 693 | CHMP3 | 40.4020221 | 1.487457058 | 0.49948 | 2.97801 | 0.0029 | 0.02038 |
| 1067 | CD164 | 239.441718 | 1.486418032 | 0.40824 | 3.64108 | 0.00027 | 0.00278 |
| 370 | RPL36AL | 88.8460593 | 1.48550607 | 0.35487 | 4.18607 | 2.8E-05 | 0.00039 |
| 960 | MSMO1 | 36.3347881 | 1.484864507 | 0.53016 | 2.8008 | 0.0051 | 0.03214 |
| 983 | SAR1B | 45.6060547 | 1.477146175 | 0.52053 | 2.83778 | 0.00454 | 0.02937 |
| 604 | ILVBL | 50.2463235 | 1.474571065 | 0.44524 | 3.31183 | 0.00093 | 0.00788 |
| 88 | TTF2 | 26.6689087 | 1.459962007 | 0.49586 | 2.94428 | 0.00324 | 0.02226 |
| 400 | INF2 | 36.2801581 | 1.45770133 | 0.53065 | 2.74703 | 0.00601 | 0.03669 |
| 999 | PANK3 | 34.8654191 | 1.457565597 | 0.44131 | 3.3028 | 0.00096 | 0.0081 |
| 770 | SNX5 | 74.5418299 | 1.450417773 | 0.42899 | 3.38097 | 0.00072 | 0.00642 |
| 179 | LCOR | 41.942035 | 1.448757098 | 0.52317 | 2.76916 | 0.00562 | 0.03481 |
| 421 | CLK3 | 34.4896024 | 1.44583402 | 0.47174 | 3.06491 | 0.00218 | 0.01614 |
| 502 | TXNDC17 | 80.993991 | 1.44347891 | 0.42752 | 3.37642 | 0.00073 | 0.0065 |
| 786 | SULF2 | 37.6216632 | 1.436935459 | 0.44626 | 3.21992 | 0.00128 | 0.01037 |
| 1281 | ERCC00165 | 73.0692771 | 1.427610226 | 0.5325 | 2.68096 | 0.00734 | 0.04316 |

| | | | | | | | |
|---|---|---|---|---|---|---|---|
| 892 | LPP | 90.7964834 | 1.426454535 | 0.41993 | 3.39688 | 0.00068 | 0.0061 |
| 1227 | PTGR1 | 61.9912021 | 1.424484762 | 0.48912 | 2.91237 | 0.00359 | 0.0242 |
| 776 | PYGB | 60.7087502 | 1.422187532 | 0.46648 | 3.04879 | 0.0023 | 0.01679 |
| 65 | RAVER2 | 28.5256956 | 1.421982052 | 0.53633 | 2.65133 | 0.00802 | 0.04606 |
| 768 | PLCB4 | 48.4523771 | 1.42142134 | 0.4045 | 3.51406 | 0.00044 | 0.0042 |
| 977 | CAST | 51.1438705 | 1.419407148 | 0.38996 | 3.63985 | 0.00027 | 0.00279 |
| 859 | PARP14 | 46.3213466 | 1.41779437 | 0.53364 | 2.65684 | 0.00789 | 0.04548 |
| 420 | DPP8 | 32.5197148 | 1.416584566 | 0.50435 | 2.80873 | 0.00497 | 0.03149 |
| 1264 | PBDC1 | 46.5494896 | 1.409145501 | 0.45743 | 3.08054 | 0.00207 | 0.01541 |
| 804 | PFKL | 93.6568585 | 1.407404252 | 0.35715 | 3.94062 | 8.1E-05 | 0.00098 |
| 937 | METAP1 | 37.1843987 | 1.398863323 | 0.4151 | 3.36993 | 0.00075 | 0.00661 |
| 410 | TMEM87A | 37.8825206 | 1.397448802 | 0.44714 | 3.12532 | 0.00178 | 0.01365 |
| 234 | CAPN1 | 62.0952604 | 1.392478557 | 0.38545 | 3.61256 | 0.0003 | 0.00305 |
| 1018 | GMNN | 41.2040936 | 1.390667993 | 0.45349 | 3.06661 | 0.00216 | 0.01608 |
| 52 | GPBP1L1 | 39.1463656 | 1.381959954 | 0.48595 | 2.8438 | 0.00446 | 0.02894 |
| 996 | CLINT1 | 48.9418792 | 1.380810204 | 0.48266 | 2.86082 | 0.00423 | 0.02768 |
| 779 | EIF6 | 59.3743418 | 1.379912041 | 0.37508 | 3.67894 | 0.00023 | 0.00245 |
| 772 | RALGAPA2 | 32.9029113 | 1.372324314 | 0.46394 | 2.95798 | 0.0031 | 0.02147 |
| 215 | LGR4 | 39.4151535 | 1.364041061 | 0.47304 | 2.88356 | 0.00393 | 0.0261 |
| 539 | VMP1 | 103.775171 | 1.35869207 | 0.44053 | 3.08424 | 0.00204 | 0.01524 |
| 805 | SLC25A1 | 47.2254579 | 1.357109885 | 0.37653 | 3.60423 | 0.00031 | 0.00314 |
| 40 | AK2 | 45.7809277 | 1.353787292 | 0.42926 | 3.15376 | 0.00161 | 0.01259 |
| 860 | HACD2 | 25.0280577 | 1.350453571 | 0.4878 | 2.76844 | 0.00563 | 0.03487 |
| 187 | ADD3 | 72.5381297 | 1.348342176 | 0.43278 | 3.11555 | 0.00184 | 0.01401 |
| 78 | SLC35A3 | 35.5145953 | 1.337840885 | 0.42815 | 3.12473 | 0.00178 | 0.01367 |
| 362 | PSME1 | 127.757091 | 1.327026702 | 0.40222 | 3.29929 | 0.00097 | 0.00817 |
| 324 | ARPC3 | 69.8614236 | 1.321056458 | 0.40396 | 3.27025 | 0.00107 | 0.00893 |
| 118 | ATP1B1 | 38.3552857 | 1.319447365 | 0.44171 | 2.98712 | 0.00282 | 0.01987 |
| 73 | LMO4 | 89.0855448 | 1.316394409 | 0.46556 | 2.82755 | 0.00469 | 0.03003 |
| 108 | RUSC1 | 34.9213625 | 1.313059903 | 0.44483 | 2.9518 | 0.00316 | 0.02185 |
| 361 | DCAF11 | 32.289339 | 1.31122549 | 0.44827 | 2.92506 | 0.00344 | 0.02342 |
| 250 | PAK1 | 28.4348343 | 1.307398858 | 0.47233 | 2.76798 | 0.00564 | 0.03489 |
| 354 | LAMP1 | 74.1996918 | 1.302520817 | 0.34846 | 3.73794 | 0.00019 | 0.00201 |
| 692 | IMMT | 37.2873217 | 1.299431479 | 0.44851 | 2.89725 | 0.00376 | 0.0252 |
| 777 | NOL4L | 57.5823205 | 1.297313837 | 0.40293 | 3.21971 | 0.00128 | 0.01037 |
| 1256 | CASK | 77.7025141 | 1.2951387 | 0.40478 | 3.19963 | 0.00138 | 0.01102 |
| 678 | GFPT1 | 70.5334567 | 1.291901972 | 0.41474 | 3.11498 | 0.00184 | 0.01403 |
| 156 | NET1 | 84.178846 | 1.290102336 | 0.47356 | 2.72424 | 0.00645 | 0.03887 |
| 872 | SERP1 | 61.3383143 | 1.287098966 | 0.40309 | 3.19306 | 0.00141 | 0.01125 |

| | | | | | | | |
|---|---|---|---|---|---|---|---|
| 1251 | PDHA1 | 66.2978688 | 1.266865243 | 0.37228 | 3.40298 | 0.00067 | 0.00599 |
| 954 | TRIM2 | 62.4698812 | 1.265485315 | 0.39282 | 3.2215 | 0.00128 | 0.01033 |
| 153 | KLF6 | 43.3774723 | 1.262433002 | 0.41269 | 3.05907 | 0.00222 | 0.01637 |
| 382 | NUMB | 36.3731927 | 1.261568058 | 0.47296 | 2.66738 | 0.00764 | 0.04451 |
| 505 | TMEM256 | 26.7049885 | 1.261545265 | 0.47765 | 2.64117 | 0.00826 | 0.04687 |
| 1082 | PERP | 32.7415545 | 1.259895165 | 0.46827 | 2.69056 | 0.00713 | 0.04225 |
| 371 | ARF6 | 63.2631855 | 1.249005286 | 0.39565 | 3.15686 | 0.00159 | 0.01249 |
| 1092 | AFDN | 100.429308 | 1.242386763 | 0.38654 | 3.21409 | 0.00131 | 0.01056 |
| 430 | IQGAP1 | 42.6551203 | 1.241421268 | 0.46005 | 2.69842 | 0.00697 | 0.04149 |
| 828 | GOLGA4 | 112.231276 | 1.240232123 | 0.43357 | 2.8605 | 0.00423 | 0.02769 |
| 901 | GAK | 63.1261463 | 1.206801488 | 0.43168 | 2.79562 | 0.00518 | 0.03251 |
| 445 | PDXDC1 | 50.3373743 | 1.2046942 | 0.40326 | 2.98737 | 0.00281 | 0.01987 |
| 84 | RHOC | 107.711467 | 1.197242677 | 0.38297 | 3.1262 | 0.00177 | 0.01361 |
| 1217 | GALNT12 | 40.565245 | 1.192505134 | 0.4398 | 2.71149 | 0.0067 | 0.04027 |
| 1116 | TBRG4 | 44.1856173 | 1.188837199 | 0.41962 | 2.8331 | 0.00461 | 0.02966 |
| 673 | EML4 | 60.7156145 | 1.16878729 | 0.4222 | 2.76833 | 0.00563 | 0.03487 |
| 729 | STAT1 | 51.949285 | 1.159931163 | 0.38053 | 3.04822 | 0.0023 | 0.0168 |
| 1135 | LAMTOR4 | 91.9464801 | 1.15894045 | 0.36872 | 3.14312 | 0.00167 | 0.01297 |
| 656 | EPN1 | 42.4149201 | 1.158067571 | 0.41037 | 2.82201 | 0.00477 | 0.03039 |
| 431 | PGAP6 | 35.0995906 | 1.152553042 | 0.43123 | 2.6727 | 0.00752 | 0.04407 |
| 33 | ATP5IF1 | 207.867263 | 1.148947307 | 0.25245 | 4.55125 | 5.3E-06 | 9.1E-05 |
| 500 | KIF1C | 51.2696303 | 1.118443057 | 0.42091 | 2.65722 | 0.00788 | 0.04547 |
| 373 | ERO1A | 45.2062394 | 1.115932421 | 0.42228 | 2.64265 | 0.00823 | 0.04675 |
| 317 | BTG1 | 75.0959963 | 1.096384693 | 0.40456 | 2.71004 | 0.00673 | 0.0404 |
| 556 | SIRT7 | 38.7242054 | 1.09636229 | 0.41313 | 2.65378 | 0.00796 | 0.04579 |
| 1154 | GSTK1 | 63.6023883 | 1.090430198 | 0.40979 | 2.66092 | 0.00779 | 0.04506 |
| 765 | MAVS | 50.7879661 | 1.087649818 | 0.3782 | 2.87587 | 0.00403 | 0.02668 |
| 835 | IP6K2 | 109.009627 | 1.079725885 | 0.32107 | 3.36295 | 0.00077 | 0.00676 |
| 363 | PSME2 | 99.9399099 | 1.073052073 | 0.33011 | 3.25058 | 0.00115 | 0.00948 |
| 335 | MLXIP | 78.0863978 | 1.068206098 | 0.36865 | 2.89759 | 0.00376 | 0.0252 |
| 236 | CFL1 | 412.043322 | 1.068163511 | 0.27969 | 3.81906 | 0.00013 | 0.00151 |
| 167 | ANXA11 | 50.8627458 | 1.063756911 | 0.36875 | 2.88476 | 0.00392 | 0.02602 |
| 325 | VPS29 | 66.2976865 | 1.052954355 | 0.34059 | 3.09159 | 0.00199 | 0.01496 |
| 1040 | TAPBP | 87.4514148 | 1.050532304 | 0.39824 | 2.63795 | 0.00834 | 0.04726 |
| 559 | ANKRD12 | 88.3268077 | 1.012427101 | 0.37626 | 2.69078 | 0.00713 | 0.04225 |
| 1252 | RPS6KA3 | 69.0003262 | 1.007309253 | 0.35914 | 2.8048 | 0.00503 | 0.03181 |
| 287 | DYNLL1 | 132.977031 | -1.004289113 | 0.30939 | -3.246 | 0.00117 | 0.00961 |
| 136 | PSAP | 166.313209 | -1.037294081 | 0.27397 | -3.7861 | 0.00015 | 0.00169 |
| 712 | ATXN10 | 59.8907527 | -1.052996555 | 0.37573 | -2.8025 | 0.00507 | 0.032 |

| | | | | | | | |
|---|---|---|---|---|---|---|---|
| 1120 | TCEAL9 | 80.1656925 | -1.064182283 | 0.37713 | -2.8218 | 0.00478 | 0.0304 |
| 647 | COPS9 | 51.532223 | -1.067183187 | 0.35364 | -3.0177 | 0.00255 | 0.01822 |
| 806 | MRFAP1 | 91.4826816 | -1.07174892 | 0.34329 | -3.122 | 0.0018 | 0.01376 |
| 1073 | ATP6V1G1 | 63.5948497 | -1.072472801 | 0.35072 | -3.0579 | 0.00223 | 0.01642 |
| 851 | DAP | 47.9839105 | -1.077769846 | 0.40441 | -2.665 | 0.0077 | 0.04471 |
| 380 | RCN2 | 65.6627317 | -1.081620212 | 0.36674 | -2.9493 | 0.00319 | 0.02197 |
| 605 | GALNT5 | 55.1766129 | -1.088975453 | 0.4025 | -2.7055 | 0.00682 | 0.04079 |
| 250 | CBX5 | 78.1129481 | -1.10040447 | 0.40326 | -2.7288 | 0.00636 | 0.03839 |
| 158 | PRDX3 | 68.3623801 | -1.105784611 | 0.42257 | -2.6168 | 0.00888 | 0.0497 |
| 549 | PTOV1 | 85.6033547 | -1.107461408 | 0.35691 | -3.103 | 0.00192 | 0.01446 |
| 317 | DAD1 | 87.4344068 | -1.108995928 | 0.33728 | -3.2881 | 0.00101 | 0.00845 |
| 8 | PARK7 | 83.4481464 | -1.116373527 | 0.32761 | -3.4076 | 0.00066 | 0.00589 |
| 940 | DYNLT1 | 49.8263721 | -1.123409886 | 0.38189 | -2.9417 | 0.00326 | 0.02239 |
| 365 | CAPN15 | 27.7666894 | -1.123740715 | 0.43337 | 2.59302 | 0.00951 | 0.04464 |
| 182 | FADS1 | 30.671105 | -1.128015695 | 0.39592 | 2.8491 | 0.00438 | 0.02425 |
| 767 | OPA1 | 42.3836277 | -1.128362415 | 0.35703 | 3.16046 | 0.00158 | 0.01063 |
| 67 | AHCYL1 | 72.3025027 | -1.130160109 | 0.39746 | -2.8435 | 0.00446 | 0.02895 |
| 884 | HLA-A | 127.765581 | -1.131767773 | 0.34851 | 3.24749 | 0.00116 | 0.00823 |
| 258 | CDK2AP1 | 91.3640954 | -1.133449841 | 0.2781 | 4.07572 | 4.6E-05 | 0.00055 |
| 452 | SPOP | 22.4373874 | -1.134145855 | 0.43949 | 2.58058 | 0.00986 | 0.04608 |
| 728 | IMPDH2 | 55.0497787 | -1.134368644 | 0.32614 | 3.47811 | 0.0005 | 0.00411 |
| 502 | LMNB2 | 38.9090241 | -1.135717955 | 0.41978 | 2.70552 | 0.00682 | 0.03457 |
| 773 | NSD2 | 50.5957465 | -1.135731927 | 0.41697 | 2.72375 | 0.00645 | 0.03312 |
| 680 | PCNT | 36.2361451 | -1.136577029 | 0.35504 | 3.20127 | 0.00137 | 0.00946 |
| 690 | PRMT2 | 52.6466994 | -1.139501093 | 0.40341 | -2.8247 | 0.00473 | 0.03023 |
| 25 | KHDRBS1 | 92.3902567 | -1.144850739 | 0.24639 | 4.64652 | 3.4E-06 | 5.6E-05 |
| 338 | DUT | 70.1088563 | -1.145765456 | 0.31147 | 3.67862 | 0.00023 | 0.0022 |
| 663 | CSE1L | 45.5075297 | -1.147324444 | 0.34787 | 3.29814 | 0.00097 | 0.00705 |
| 1018 | GPAT4 | 23.7446831 | -1.151517048 | 0.42013 | 2.74086 | 0.00613 | 0.03179 |
| 957 | ZMIZ2 | 44.5543101 | -1.155843328 | 0.36156 | 3.19679 | 0.00139 | 0.00956 |
| 742 | GPX1 | 73.9520385 | -1.157874663 | 0.40794 | -2.8383 | 0.00454 | 0.02934 |
| 525 | LSM14A | 57.3057145 | -1.158735145 | 0.43075 | -2.69 | 0.00714 | 0.04229 |
| 318 | RBM23 | 66.054873 | -1.159639533 | 0.38041 | -3.0484 | 0.0023 | 0.0168 |
| 260 | CCT2 | 95.2193465 | -1.160426697 | 0.33408 | -3.4735 | 0.00051 | 0.00479 |
| 997 | AGAP3 | 57.7308393 | -1.160956826 | 0.33638 | 3.45131 | 0.00056 | 0.00447 |
| 457 | MTMR4 | 33.3261964 | -1.161915575 | 0.38504 | 3.01761 | 0.00255 | 0.01562 |
| 790 | PAICS | 99.282139 | -1.163605831 | 0.27199 | 4.2782 | 1.9E-05 | 0.00025 |
| 424 | PELP1 | 33.5432669 | -1.168356398 | 0.3727 | 3.13486 | 0.00172 | 0.01138 |
| 944 | KLHL7 | 43.9568137 | -1.173004207 | 0.34315 | 3.4183 | 0.00063 | 0.00495 |

| | | | | | | | |
|---|---|---|---|---|---|---|---|
| 475 | ALYREF | 30.8079725 | -1.173589624 | 0.39293 | 2.9868 | 0.00282 | 0.01694 |
| 152 | SMC3 | 33.9752718 | -1.174522975 | 0.38517 | 3.04935 | 0.00229 | 0.01442 |
| 451 | TMEM98 | 91.946362 | -1.174766634 | 0.44152 | -2.6607 | 0.0078 | 0.04507 |
| 1104 | RBM10 | 37.3639956 | -1.175422277 | 0.3508 | 3.35066 | 0.00081 | 0.00605 |
| 229 | ATN1 | 49.8444739 | -1.176131481 | 0.36719 | -3.203 | 0.00136 | 0.01091 |
| 561 | PPP3R1 | 27.8023456 | -1.179498758 | 0.457 | 2.58098 | 0.00985 | 0.04606 |
| 361 | IDH2 | 108.671065 | -1.184683486 | 0.22688 | 5.22165 | 1.8E-07 | 4.4E-06 |
| 35 | UTP11 | 26.7998357 | -1.186390595 | 0.46006 | 2.57879 | 0.00991 | 0.04625 |
| 961 | GUSB | 27.0812077 | -1.187738178 | 0.41653 | 2.85149 | 0.00435 | 0.02411 |
| 732 | PBRM1 | 39.2955228 | -1.189488116 | 0.38029 | 3.12783 | 0.00176 | 0.0116 |
| 569 | IMMT | 28.9441678 | -1.191153086 | 0.40603 | 2.93366 | 0.00335 | 0.01943 |
| 1019 | SNHG6 | 45.3760266 | -1.192043128 | 0.43184 | -2.7604 | 0.00577 | 0.03555 |
| 485 | SNRPD1 | 49.9141122 | -1.192755528 | 0.38278 | 3.11606 | 0.00183 | 0.012 |
| 611 | GLS | 72.7233201 | -1.192875197 | 0.28786 | 4.14394 | 3.4E-05 | 0.00043 |
| 795 | WDFY3 | 22.7019176 | -1.193083171 | 0.44382 | 2.68823 | 0.00718 | 0.03601 |
| 621 | EPHA4 | 28.345324 | -1.19486802 | 0.43595 | 2.74087 | 0.00613 | 0.03179 |
| 151 | MXI1 | 25.3597552 | -1.19512985 | 0.4295 | 2.78264 | 0.00539 | 0.02872 |
| 1022 | BPNT2 | 36.9193586 | -1.196367611 | 0.38972 | 3.06979 | 0.00214 | 0.01361 |
| 306 | MAX | 34.9074481 | -1.196406296 | 0.39863 | 3.00131 | 0.00269 | 0.01631 |
| 539 | PAFAH1B3 | 39.8138584 | -1.19683354 | 0.37754 | 3.17005 | 0.00152 | 0.01031 |
| 524 | UBA2 | 51.0208854 | -1.197493815 | 0.33614 | 3.56254 | 0.00037 | 0.00318 |
| 145 | NOLC1 | 52.7899981 | -1.198861228 | 0.34522 | 3.47279 | 0.00052 | 0.00418 |
| 616 | LANCL1 | 41.5252992 | -1.198966159 | 0.34714 | 3.45388 | 0.00055 | 0.00444 |
| 639 | MFF | 39.8029447 | -1.199601213 | 0.43569 | -2.7534 | 0.0059 | 0.03613 |
| 111 | CDC42BPA | 44.4860443 | -1.199861131 | 0.34278 | 3.50042 | 0.00046 | 0.00385 |
| 153 | TRUB1 | 28.6275365 | -1.20083033 | 0.39457 | 3.04338 | 0.00234 | 0.01465 |
| 774 | ATP2C1 | 43.5994206 | -1.203058774 | 0.44959 | -2.6759 | 0.00745 | 0.04367 |
| 714 | PLXNB2 | 98.227825 | -1.203298922 | 0.39722 | -3.0293 | 0.00245 | 0.01767 |
| 305 | MTHFD1 | 37.5095741 | -1.204423085 | 0.40005 | 3.0107 | 0.00261 | 0.01591 |
| 661 | TOMM34 | 35.3596655 | -1.206054384 | 0.38287 | 3.15007 | 0.00163 | 0.01093 |
| 594 | RIF1 | 39.6835906 | -1.206080366 | 0.35768 | 3.37191 | 0.00075 | 0.00567 |
| 38 | CTPS1 | 30.5759646 | -1.207260319 | 0.3887 | 3.10589 | 0.0019 | 0.01234 |
| 751 | RBP1 | 214.933467 | -1.207723547 | 0.27356 | 4.41491 | 1E-05 | 0.00015 |
| 805 | SNHG8 | 104.327975 | -1.212823081 | 0.23367 | 5.19036 | 2.1E-07 | 5E-06 |
| 1035 | ZNF706 | 32.6283209 | -1.214742419 | 0.39552 | 3.07129 | 0.00213 | 0.01356 |
| 530 | POLR2I | 39.9588259 | -1.215057257 | 0.39957 | -3.0409 | 0.00236 | 0.01713 |
| 536 | EID2 | 20.7948424 | -1.2195777 | 0.46042 | 2.64881 | 0.00808 | 0.03933 |
| 162 | ZNF195 | 24.9313088 | -1.229308628 | 0.47146 | 2.60748 | 0.00912 | 0.04336 |
| 41 | NASP | 78.463619 | -1.230819367 | 0.38135 | 3.22751 | 0.00125 | 0.00874 |

| | | | | | | | |
|---|---|---|---|---|---|---|---|
| 285 | IRS2 | 74.7682435 | -1.233584955 | 0.31647 | 3.89789 | 9.7E-05 | 0.00105 |
| 313 | PPP4R3A | 25.1047801 | -1.235073804 | 0.42751 | 2.88901 | 0.00386 | 0.02176 |
| 1135 | HMGB3 | 58.3296916 | -1.237355926 | 0.36839 | 3.35878 | 0.00078 | 0.0059 |
| 664 | ZFAS1 | 111.69947 | -1.242285987 | 0.26437 | 4.699 | 2.6E-06 | 4.5E-05 |
| 709 | NUP210 | 37.0490958 | -1.244300834 | 0.39908 | 3.11792 | 0.00182 | 0.01194 |
| 464 | CBX1 | 41.1320746 | -1.250565386 | 0.44468 | -2.8123 | 0.00492 | 0.0312 |
| 765 | YEATS2 | 43.0066413 | -1.250593795 | 0.3854 | 3.24494 | 0.00117 | 0.00829 |
| 780 | QDPR | 19.9513922 | -1.251278338 | 0.47811 | 2.61712 | 0.00887 | 0.04233 |
| 374 | UBN1 | 27.1375422 | -1.253678328 | 0.40651 | 3.08397 | 0.00204 | 0.01313 |
| 885 | TUBB | 486.958182 | -1.255600288 | 0.24951 | 5.03221 | 4.8E-07 | 1E-05 |
| 408 | EXOSC6 | 37.8075313 | -1.259018093 | 0.35492 | 3.54738 | 0.00039 | 0.00334 |
| 487 | RNF138 | 22.0561876 | -1.261142023 | 0.46341 | 2.72146 | 0.0065 | 0.03332 |
| 176 | ZFP91 | 22.6179685 | -1.267959668 | 0.45883 | 2.76345 | 0.00572 | 0.03016 |
| 175 | CKAP5 | 40.2066761 | -1.269472761 | 0.35498 | 3.57614 | 0.00035 | 0.00305 |
| 155 | MCMBP | 29.0503587 | -1.269540775 | 0.47274 | 2.6855 | 0.00724 | 0.0362 |
| 21 | SNHG12 | 26.2344853 | -1.271703754 | 0.46823 | 2.71596 | 0.00661 | 0.03379 |
| 685 | POFUT2 | 33.0967737 | -1.275208861 | 0.46432 | -2.7464 | 0.00603 | 0.03672 |
| 347 | CEP128 | 25.5420238 | -1.275465107 | 0.48282 | -2.6417 | 0.00825 | 0.04684 |
| 135 | HELLS | 92.9384732 | -1.27783285 | 0.41149 | 3.1054 | 0.0019 | 0.01236 |
| 183 | CSKMT | 22.5077536 | -1.279014258 | 0.48604 | 2.63152 | 0.0085 | 0.04092 |
| 538 | ERCC1 | 28.1844119 | -1.279340741 | 0.47647 | -2.685 | 0.00725 | 0.04271 |
| 626 | PTMA | 162.12023 | -1.280712477 | 0.27448 | 4.6659 | 3.1E-06 | 5.2E-05 |
| 71 | TTF2 | 20.29797 | -1.281652427 | 0.45151 | 2.83859 | 0.00453 | 0.02496 |
| 692 | GTPBP1 | 23.0924348 | -1.281786445 | 0.44209 | 2.89939 | 0.00374 | 0.0212 |
| 586 | POLR2D | 26.0411956 | -1.285140571 | 0.4297 | 2.99077 | 0.00278 | 0.01675 |
| 517 | PRKACA | 25.2140122 | -1.285324311 | 0.48916 | -2.6276 | 0.0086 | 0.0484 |
| 1006 | PSD3 | 18.0824775 | -1.290201 | 0.50578 | 2.5509 | 0.01074 | 0.04924 |
| 406 | CDIPT | 39.3308299 | -1.290520134 | 0.44073 | -2.9282 | 0.00341 | 0.02321 |
| 724 | TMEM158 | 19.1963746 | -1.296957827 | 0.50384 | 2.57414 | 0.01005 | 0.04677 |
| 381 | XPO6 | 34.6584843 | -1.297751157 | 0.383 | 3.38843 | 0.0007 | 0.0054 |
| 825 | NIPBL-DT | 36.3252642 | -1.297961077 | 0.41639 | 3.11714 | 0.00183 | 0.01196 |
| 1063 | C9orf72 | 23.6917075 | -1.298419457 | 0.49173 | 2.64051 | 0.00828 | 0.04007 |
| 682 | YDJC | 26.1790737 | -1.301173342 | 0.42148 | 3.08715 | 0.00202 | 0.01301 |
| 360 | TJP1 | 31.4982845 | -1.30273841 | 0.49642 | -2.6243 | 0.00868 | 0.04878 |
| 581 | DDX18 | 35.3221773 | -1.304460473 | 0.36326 | 3.59094 | 0.00033 | 0.00292 |
| 686 | YWHAH | 43.9163459 | -1.304576514 | 0.40509 | 3.22049 | 0.00128 | 0.0089 |
| 960 | NIPSNAP2 | 35.4412172 | -1.306238171 | 0.49312 | -2.6489 | 0.00807 | 0.04629 |
| 1018 | SDCBP | 51.2546086 | -1.309768068 | 0.41135 | -3.1841 | 0.00145 | 0.01155 |
| 75 | SEMA6C | 24.6897324 | -1.313284739 | 0.46111 | 2.84809 | 0.0044 | 0.02429 |

| | | | | | | | |
|---|---|---|---|---|---|---|---|
| 978 | PCOLCE | 40.2558285 | -1.315191721 | 0.48521 | 2.71054 | 0.00672 | 0.03418 |
| 411 | TERF2IP | 59.5768297 | -1.3163336 | 0.32131 | 4.09676 | 4.2E-05 | 0.00051 |
| 879 | GMNN | 32.9983781 | -1.318591685 | 0.43514 | 3.03026 | 0.00244 | 0.01516 |
| 261 | PSPC1 | 23.7865336 | -1.320267186 | 0.46685 | 2.828 | 0.00468 | 0.02565 |
| 1100 | USP11 | 95.2622081 | -1.322926642 | 0.45315 | -2.9194 | 0.00351 | 0.02379 |
| 1148 | MTMR1 | 37.9650082 | -1.325141201 | 0.47234 | -2.8055 | 0.00502 | 0.03178 |
| 874 | PXDC1 | 30.6868995 | -1.328703857 | 0.45882 | 2.89591 | 0.00378 | 0.02141 |
| 70 | HIPK1 | 34.2165833 | -1.329844894 | 0.4607 | -2.8866 | 0.00389 | 0.02591 |
| 194 | TAF1D | 28.8371215 | -1.330523097 | 0.42084 | 3.16161 | 0.00157 | 0.01059 |
| 658 | TMEM230 | 32.3118398 | -1.330832367 | 0.46862 | -2.8399 | 0.00451 | 0.02922 |
| 934 | PHACTR2 | 51.8274081 | -1.335923125 | 0.42372 | -3.1528 | 0.00162 | 0.01262 |
| 7 | EXOSC10 | 25.0833467 | -1.339238921 | 0.42635 | 3.14119 | 0.00168 | 0.01119 |
| 474 | PRKAR1A | 98.4486721 | -1.341547297 | 0.35246 | -3.8062 | 0.00014 | 0.00157 |
| 466 | MSI2 | 73.5588541 | -1.341566432 | 0.33353 | -4.0224 | 5.8E-05 | 0.00073 |
| 833 | HMGCR | 22.0232215 | -1.344633562 | 0.45514 | 2.95432 | 0.00313 | 0.01842 |
| 903 | MCM3 | 65.1335207 | -1.344878072 | 0.38123 | 3.52772 | 0.00042 | 0.00354 |
| 428 | POLR2A | 22.9874533 | -1.345868199 | 0.46998 | 2.86368 | 0.00419 | 0.02333 |
| 85 | PBX1 | 50.2637938 | -1.34885473 | 0.36593 | 3.68608 | 0.00023 | 0.00215 |
| 282 | RPL6 | 69.3838308 | -1.34936413 | 0.34743 | -3.8839 | 0.0001 | 0.0012 |
| 433 | HSBP1 | 142.071383 | -1.349532378 | 0.32274 | -4.1815 | 2.9E-05 | 0.0004 |
| 716 | SLC4A7 | 26.3695583 | -1.351455932 | 0.49536 | 2.72825 | 0.00637 | 0.03275 |
| 914 | HMGN3 | 101.633122 | -1.354625198 | 0.36159 | -3.7463 | 0.00018 | 0.00196 |
| 399 | NUDT21 | 29.7192951 | -1.356179473 | 0.4251 | 3.19024 | 0.00142 | 0.00975 |
| 162 | GLRX3 | 33.9520458 | -1.358311462 | 0.48155 | -2.8207 | 0.00479 | 0.03047 |
| 98 | PLEKHA6 | 45.9761405 | -1.358670658 | 0.37313 | 3.64132 | 0.00027 | 0.00248 |
| 1079 | FBXW2 | 51.2837613 | -1.358674842 | 0.30793 | 4.41235 | 1E-05 | 0.00015 |
| 786 | COMMD2 | 27.4599465 | -1.360780354 | 0.48827 | -2.7869 | 0.00532 | 0.03326 |
| 730 | PCBP4 | 36.8526634 | -1.365304734 | 0.37969 | 3.59586 | 0.00032 | 0.00287 |
| 884 | IK | 24.8698568 | -1.366190295 | 0.49629 | -2.7528 | 0.00591 | 0.03617 |
| 883 | CXXC5 | 29.7310417 | -1.367166816 | 0.50799 | -2.6913 | 0.00712 | 0.04222 |
| 456 | AKAP1 | 24.993343 | -1.367701977 | 0.45097 | 3.03279 | 0.00242 | 0.01508 |
| 1008 | ATP6V1B2 | 40.9934991 | -1.36829699 | 0.39136 | 3.49629 | 0.00047 | 0.00389 |
| 577 | EPAS1 | 64.3128584 | -1.373144336 | 0.34811 | -3.9446 | 8E-05 | 0.00097 |
| 376 | PKM | 382.473667 | -1.392326646 | 0.36298 | -3.8358 | 0.00013 | 0.00142 |
| 211 | CENATAC | 31.7686356 | -1.39521821 | 0.49199 | -2.8359 | 0.00457 | 0.02952 |
| 356 | HERC2P2 | 26.455102 | -1.398441216 | 0.49069 | -2.8499 | 0.00437 | 0.02854 |
| 152 | SHTN1 | 49.7956715 | -1.40239018 | 0.42603 | -3.2918 | 0.001 | 0.00835 |
| 876 | C5orf15 | 31.7968313 | -1.405277942 | 0.47636 | -2.95 | 0.00318 | 0.02194 |
| 1065 | TLE4 | 31.4808294 | -1.405573594 | 0.49104 | -2.8624 | 0.0042 | 0.02758 |

| | | | | | | | |
|---|---|---|---|---|---|---|---|
| 363 | ZNF106 | 43.4865231 | -1.407885596 | 0.47849 | -2.9424 | 0.00326 | 0.02237 |
| 10 | SRM | 52.1878168 | -1.408235851 | 0.40468 | -3.4799 | 0.0005 | 0.00469 |
| 274 | RPL18AP3 | 24.3463575 | -1.413882869 | 0.53649 | -2.6355 | 0.0084 | 0.04755 |
| 400 | LITAF | 53.3858113 | -1.419478478 | 0.48719 | -2.9136 | 0.00357 | 0.02413 |
| 293 | CDK2AP1 | 41.8973287 | -1.425186656 | 0.46515 | -3.0639 | 0.00218 | 0.01616 |
| 964 | HSPB1 | 66.2044529 | -1.431378683 | 0.41854 | -3.4199 | 0.00063 | 0.00568 |
| 580 | MCFD2 | 34.9343324 | -1.438438121 | 0.44583 | -3.2264 | 0.00125 | 0.0102 |
| 559 | ZSCAN18 | 31.0884024 | -1.443492675 | 0.50917 | -2.835 | 0.00458 | 0.02954 |
| 831 | ANXA5 | 74.0844388 | -1.446421833 | 0.36294 | -3.9853 | 6.7E-05 | 0.00084 |
| 424 | PRMT7 | 32.2950019 | -1.450128959 | 0.51157 | -2.8346 | 0.00459 | 0.02954 |
| 1099 | NDUFB11 | 47.3233964 | -1.466345468 | 0.46676 | -3.1416 | 0.00168 | 0.01301 |
| 21 | SELENON | 55.1512544 | -1.475109472 | 0.4886 | -3.019 | 0.00254 | 0.01816 |
| 581 | ASB3 | 22.2097013 | -1.476575555 | 0.52969 | -2.7876 | 0.00531 | 0.0332 |
| 357 | NDN | 57.4869389 | -1.495531736 | 0.43354 | -3.4496 | 0.00056 | 0.00517 |
| 1035 | EMC2 | 33.0690387 | -1.498174344 | 0.49268 | -3.0409 | 0.00236 | 0.01713 |
| 155 | RAB11FIP2 | 25.5060662 | -1.512382321 | 0.52192 | -2.8978 | 0.00376 | 0.0252 |
| 1053 | NFIB | 48.2204698 | -1.513350123 | 0.49225 | -3.0743 | 0.00211 | 0.01569 |
| 359 | SNHG14 | 112.090774 | -1.520597381 | 0.30182 | -5.0382 | 4.7E-07 | 1E-05 |
| 825 | PDLIM5 | 80.6355673 | -1.529222631 | 0.37022 | -4.1306 | 3.6E-05 | 0.00049 |
| 248 | EIF4B | 26.1907315 | -1.532813459 | 0.51386 | -2.983 | 0.00285 | 0.0201 |
| 43 | BTF3L4 | 19.9217899 | -1.535519473 | 0.57148 | -2.6869 | 0.00721 | 0.04256 |
| 548 | MED25 | 36.9904366 | -1.538344527 | 0.44809 | -3.4331 | 0.0006 | 0.00546 |
| 743 | DAG1 | 38.5656716 | -1.541660138 | 0.54499 | -2.8288 | 0.00467 | 0.02995 |
| 903 | RGL2 | 43.2172967 | -1.546839195 | 0.5092 | -3.0378 | 0.00238 | 0.01726 |
| 1092 | ACOT9 | 29.1225406 | -1.548285929 | 0.48782 | -3.1739 | 0.0015 | 0.0119 |
| 540 | DMWD | 17.2804429 | -1.550709965 | 0.5774 | -2.6857 | 0.00724 | 0.04268 |
| 1020 | CSPP1 | 37.2464499 | -1.554522406 | 0.47964 | -3.241 | 0.00119 | 0.00975 |
| 233 | RIMKLB | 17.2473139 | -1.560013147 | 0.58089 | -2.6856 | 0.00724 | 0.04268 |
| 788 | GMPS | 21.6744543 | -1.568444158 | 0.56758 | -2.7634 | 0.00572 | 0.03533 |
| 516 | NFIX | 32.0055168 | -1.589181018 | 0.59572 | -2.6677 | 0.00764 | 0.04449 |
| 772 | RUVBL1 | 31.3564955 | -1.596467208 | 0.47285 | -3.3762 | 0.00073 | 0.0065 |
| 30 | MAP7D1 | 30.8840572 | -1.602292073 | 0.58322 | -2.7473 | 0.00601 | 0.03667 |
| 918 | PM20D2 | 43.8277113 | -1.602440436 | 0.52132 | -3.0738 | 0.00211 | 0.01571 |
| 527 | GRAMD1A | 27.2926358 | -1.603217124 | 0.54954 | -2.9174 | 0.00353 | 0.02392 |
| 135 | HK1 | 40.2707886 | -1.609566657 | 0.59554 | -2.7027 | 0.00688 | 0.04109 |
| 1055 | NOL6 | 18.585638 | -1.610120901 | 0.58715 | -2.7423 | 0.0061 | 0.03706 |
| 727 | NGLY1 | 39.011679 | -1.613409779 | 0.44453 | -3.6295 | 0.00028 | 0.00288 |
| 951 | SCRN1 | 38.6157858 | -1.615480272 | 0.53498 | -3.0197 | 0.00253 | 0.01813 |
| 758 | PDZRN3 | 25.6729761 | -1.617016591 | 0.48059 | -3.3647 | 0.00077 | 0.00673 |

| | | | | | | | |
|---|---|---|---|---|---|---|---|
| 264 | NAP1L1 | 164.532513 | -1.622325638 | 0.41202 | -3.9375 | 8.2E-05 | 0.00099 |
| 1149 | CD99L2 | 20.6609661 | -1.627802489 | 0.53583 | -3.0379 | 0.00238 | 0.01726 |
| 455 | NT5C3B | 22.207219 | -1.628758043 | 0.53513 | -3.0437 | 0.00234 | 0.01702 |
| 1031 | LAPTM4B | 73.1790383 | -1.630660944 | 0.36894 | -4.4198 | 9.9E-06 | 0.00015 |
| 704 | CBX6 | 35.5832011 | -1.630963785 | 0.50305 | -3.2422 | 0.00119 | 0.00972 |
| 198 | PRCP | 25.6415148 | -1.631591852 | 0.53783 | -3.0337 | 0.00242 | 0.01746 |
| 134 | ARID5B | 53.8585524 | -1.633165719 | 0.40679 | -4.0148 | 5.9E-05 | 0.00075 |
| 388 | NR2F2 | 66.5688879 | -1.638746854 | 0.48874 | -3.353 | 0.0008 | 0.00697 |
| 532 | PLD3 | 82.0576831 | -1.639711626 | 0.37655 | -4.3546 | 1.3E-05 | 0.0002 |
| 1096 | RPGR | 58.7820008 | -1.641793786 | 0.46961 | -3.4961 | 0.00047 | 0.00445 |
| 147 | TRIM8 | 33.3942073 | -1.646600967 | 0.51377 | -3.2049 | 0.00135 | 0.01086 |
| 1122 | TCEAL4 | 56.5480991 | -1.655650017 | 0.37826 | -4.377 | 1.2E-05 | 0.00018 |
| 345 | IRF2BPL | 37.5122123 | -1.658073938 | 0.44384 | -3.7357 | 0.00019 | 0.00202 |
| 688 | COL6A1 | 214.075609 | -1.668131001 | 0.38144 | -4.3732 | 1.2E-05 | 0.00019 |
| 167 | IFITM2 | 155.770643 | -1.671109492 | 0.29922 | -5.5849 | 2.3E-08 | 7.5E-07 |
| 607 | RBMS1 | 30.966767 | -1.673018024 | 0.57531 | -2.908 | 0.00364 | 0.02448 |
| 94 | TMEM9 | 51.1041983 | -1.673149624 | 0.42828 | -3.9067 | 9.4E-05 | 0.00111 |
| 812 | STIM2 | 55.8948794 | -1.674855522 | 0.41765 | -4.0102 | 6.1E-05 | 0.00077 |
| 869 | ADGRV1 | 31.1870355 | -1.678322417 | 0.55993 | -2.9974 | 0.00272 | 0.01932 |
| 372 | MAP2K1 | 17.2120938 | -1.680409898 | 0.61186 | -2.7464 | 0.00603 | 0.03672 |
| 1106 | MAGED2 | 169.080705 | -1.680914319 | 0.32535 | -5.1665 | 2.4E-07 | 5.9E-06 |
| 343 | NEK9 | 19.0599302 | -1.689704127 | 0.60952 | -2.7722 | 0.00557 | 0.03455 |
| 1071 | ZNF483 | 21.3889486 | -1.694074561 | 0.57215 | -2.9609 | 0.00307 | 0.02129 |
| 108 | RAB3GAP2 | 75.5310574 | -1.703486556 | 0.39033 | -4.3643 | 1.3E-05 | 0.00019 |
| 809 | CPEB2 | 23.7276332 | -1.704962193 | 0.64533 | -2.642 | 0.00824 | 0.04682 |
| 1023 | CA2 | 75.5615793 | -1.705560029 | 0.58873 | -2.897 | 0.00377 | 0.0252 |
| 843 | SLC25A4 | 34.1140216 | -1.71725002 | 0.57122 | -3.0063 | 0.00264 | 0.01886 |
| 574 | PREPL | 26.492511 | -1.724049077 | 0.54124 | -3.1854 | 0.00145 | 0.01152 |
| 994 | CALD1 | 80.3909078 | -1.728561587 | 0.58572 | -2.9512 | 0.00317 | 0.02187 |
| 902 | HSPA1A | 25.3071156 | -1.729814608 | 0.54132 | -3.1955 | 0.0014 | 0.01116 |
| 460 | HDAC5 | 14.8665272 | -1.734875997 | 0.65084 | -2.6656 | 0.00769 | 0.04467 |
| 936 | PPIL4 | 18.9931062 | -1.737897643 | 0.64387 | -2.6991 | 0.00695 | 0.04144 |
| 703 | CBY1 | 30.5202117 | -1.738063271 | 0.55909 | -3.1087 | 0.00188 | 0.01426 |
| 970 | PEG10 | 15.3120061 | -1.740068776 | 0.65226 | -2.6678 | 0.00764 | 0.04449 |
| 480 | SEC14L1 | 14.3124856 | -1.740559894 | 0.63119 | -2.7576 | 0.00582 | 0.03574 |
| 450 | COPRS | 28.4398241 | -1.743116905 | 0.62622 | -2.7835 | 0.00538 | 0.03356 |
| 336 | PCNX4 | 37.3737713 | -1.743298318 | 0.4439 | -3.9273 | 8.6E-05 | 0.00103 |
| 243 | HDAC7 | 28.6971375 | -1.745578295 | 0.63261 | -2.7593 | 0.00579 | 0.03565 |
| 554 | NAT14 | 14.9337923 | -1.746168432 | 0.65004 | -2.6862 | 0.00723 | 0.04263 |

| | | | | | | | |
|---|---|---|---|---|---|---|---|
| 407 | YPEL3 | 31.130229 | -1.754125091 | 0.50694 | -3.4602 | 0.00054 | 0.005 |
| 634 | TTLL4 | 38.6817014 | -1.756262203 | 0.65167 | -2.695 | 0.00704 | 0.04185 |
| 1088 | WWC3 | 23.3988286 | -1.759243016 | 0.6042 | -2.9117 | 0.00359 | 0.02424 |
| 618 | FRZB | 25.7867048 | -1.759987869 | 0.61342 | -2.8691 | 0.00412 | 0.0271 |
| 543 | INAFM1 | 24.1712393 | -1.76229706 | 0.53722 | -3.2804 | 0.00104 | 0.00865 |
| 222 | WNK1 | 54.6810162 | -1.764660622 | 0.45503 | -3.8781 | 0.00011 | 0.00123 |
| 963 | RHBDD2 | 41.874116 | -1.765583829 | 0.47751 | -3.6975 | 0.00022 | 0.00231 |
| 337 | SYNE2 | 51.1133273 | -1.773863596 | 0.48109 | -3.6871 | 0.00023 | 0.00239 |
| 610 | STK39 | 49.1664065 | -1.777806949 | 0.44796 | -3.9687 | 7.2E-05 | 0.00089 |
| 1027 | CIBAR1 | 21.2641798 | -1.77871356 | 0.58212 | -3.0556 | 0.00225 | 0.0165 |
| 411 | CPNE2 | 27.0948688 | -1.779821472 | 0.56784 | -3.1343 | 0.00172 | 0.01329 |
| 948 | KLHL7 | 18.3197203 | -1.788243165 | 0.59956 | -2.9826 | 0.00286 | 0.02011 |
| 168 | IFITM3 | 435.72999 | -1.791544671 | 0.35711 | -5.0168 | 5.3E-07 | 1.2E-05 |
| 878 | TXNDC15 | 19.6420498 | -1.794349915 | 0.5685 | -3.1563 | 0.0016 | 0.01251 |
| 509 | NFIC | 34.8912904 | -1.797386515 | 0.57536 | -3.124 | 0.00178 | 0.01368 |
| 38 | SLC2A1 | 83.626345 | -1.800373739 | 0.63247 | -2.8466 | 0.00442 | 0.02873 |
| 362 | GOLGA8B | 79.6735252 | -1.804835611 | 0.52654 | -3.4277 | 0.00061 | 0.00555 |
| 833 | SCOC | 39.9491211 | -1.806759814 | 0.51189 | -3.5296 | 0.00042 | 0.004 |
| 1107 | FAM104B | 14.9655616 | -1.807310918 | 0.68249 | -2.6481 | 0.00809 | 0.04635 |
| 232 | PEX5 | 17.5551677 | -1.808093037 | 0.62008 | -2.9159 | 0.00355 | 0.02401 |
| 37 | P3H1 | 19.6468416 | -1.812931652 | 0.68538 | -2.6452 | 0.00817 | 0.04661 |
| 999 | HIPK2 | 31.4237301 | -1.816248178 | 0.58367 | -3.1118 | 0.00186 | 0.01416 |
| 482 | SLC26A11 | 25.5089742 | -1.821324666 | 0.52084 | -3.4969 | 0.00047 | 0.00444 |
| 795 | SKIL | 31.7054655 | -1.823460869 | 0.46668 | -3.9073 | 9.3E-05 | 0.00111 |
| 667 | SYS1 | 14.7411742 | -1.830324649 | 0.63805 | -2.8686 | 0.00412 | 0.02714 |
| 225 | IFFO1 | 12.9010411 | -1.833168836 | 0.67109 | -2.7316 | 0.0063 | 0.03811 |
| 58 | GCLM | 14.9222305 | -1.837360046 | 0.63991 | -2.8713 | 0.00409 | 0.02696 |
| 324 | IPO4 | 13.6152885 | -1.838806948 | 0.69729 | -2.6371 | 0.00836 | 0.04736 |
| 383 | MFGE8 | 23.8483024 | -1.839988011 | 0.54881 | -3.3527 | 0.0008 | 0.00697 |
| 500 | TPGS2 | 79.8220325 | -1.841528009 | 0.3754 | -4.9055 | 9.3E-07 | 1.9E-05 |
| 1093 | APOO | 22.9885147 | -1.843752177 | 0.57953 | -3.1815 | 0.00147 | 0.01163 |
| 726 | NR1D2 | 15.9906955 | -1.849194678 | 0.5958 | -3.1037 | 0.00191 | 0.01443 |
| 361 | SCG5 | 15.7370894 | -1.849871092 | 0.69949 | -2.6446 | 0.00818 | 0.04665 |
| 981 | AASS | 31.4592437 | -1.851220627 | 0.51769 | -3.5759 | 0.00035 | 0.00344 |
| 823 | MAPK10 | 29.2462776 | -1.855559667 | 0.56714 | -3.2718 | 0.00107 | 0.00889 |
| 445 | MYH10 | 52.6222357 | -1.857700352 | 0.37962 | -4.8936 | 9.9E-07 | 2E-05 |
| 719 | SRGAP3 | 11.0449059 | -1.860496771 | 0.70252 | -2.6483 | 0.00809 | 0.04635 |
| 888 | ANXA6 | 39.341834 | -1.86186861 | 0.48858 | -3.8107 | 0.00014 | 0.00155 |
| 566 | IFT172 | 17.8652029 | -1.871178879 | 0.64759 | -2.8895 | 0.00386 | 0.02571 |

| | | | | | | | |
|---|---|---|---|---|---|---|---|
| 408 | VKORC1 | 28.0479022 | -1.87277973 | 0.48869 | -3.8323 | 0.00013 | 0.00144 |
| 677 | MRPS6 | 127.841075 | -1.874193194 | 0.33774 | -5.5492 | 2.9E-08 | 8.9E-07 |
| 606 | TANC1 | 29.6667688 | -1.881399993 | 0.49239 | -3.8209 | 0.00013 | 0.0015 |
| 1127 | COL4A5 | 89.6181886 | -1.882871842 | 0.4266 | -4.4136 | 1E-05 | 0.00016 |
| 1069 | CTSV | 18.828728 | -1.894061026 | 0.6779 | -2.794 | 0.00521 | 0.03263 |
| 468 | DHX40 | 36.6620535 | -1.894257333 | 0.46665 | -4.0592 | 4.9E-05 | 0.00064 |
| 973 | TRIP6 | 40.6355782 | -1.895787336 | 0.56754 | -3.3404 | 0.00084 | 0.00727 |
| 1077 | BBLN | 25.2557015 | -1.896325886 | 0.56666 | -3.3465 | 0.00082 | 0.00712 |
| 1115 | GPRASP2 | 15.374308 | -1.899794585 | 0.70984 | -2.6764 | 0.00744 | 0.04362 |
| 288 | UNC119B | 12.6328999 | -1.901086767 | 0.69243 | -2.7455 | 0.00604 | 0.03681 |
| 583 | FAM161A | 20.0286348 | -1.902425897 | 0.70401 | -2.7023 | 0.00689 | 0.0411 |
| 105 | PTPN14 | 12.1750727 | -1.908002681 | 0.71519 | -2.6678 | 0.00763 | 0.04449 |
| 173 | CD81 | 126.615896 | -1.909028135 | 0.34819 | -5.4827 | 4.2E-08 | 1.2E-06 |
| 499 | SLC39A6 | 39.962761 | -1.919962347 | 0.46809 | -4.1017 | 4.1E-05 | 0.00054 |
| 968 | CDK14 | 18.0867613 | -1.930507445 | 0.73569 | -2.6241 | 0.00869 | 0.04879 |
| 659 | TMX4 | 18.9999061 | -1.934438316 | 0.62407 | -3.0997 | 0.00194 | 0.01461 |
| 986 | CALU | 79.3709216 | -1.934721292 | 0.37852 | -5.1112 | 3.2E-07 | 7.6E-06 |
| 537 | APOE | 20.6178668 | -1.936306221 | 0.69151 | -2.8001 | 0.00511 | 0.03219 |
| 993 | AKR1B1 | 16.4433413 | -1.936690392 | 0.72906 | -2.6564 | 0.0079 | 0.04552 |
| 1078 | ODF2 | 31.7332203 | -1.937060006 | 0.56525 | -3.4269 | 0.00061 | 0.00556 |
| 921 | AK9 | 11.4872738 | -1.941059617 | 0.71688 | -2.7076 | 0.00678 | 0.0406 |
| 894 | DBN1 | 28.3470736 | -1.944866772 | 0.66129 | -2.941 | 0.00327 | 0.02242 |
| 528 | PSENEN | 45.1754736 | -1.947143723 | 0.51344 | -3.7923 | 0.00015 | 0.00165 |
| 325 | NFATC4 | 22.2522589 | -1.954664486 | 0.61471 | -3.1798 | 0.00147 | 0.01168 |
| 928 | HSF2 | 19.0971785 | -1.955652504 | 0.61686 | -3.1704 | 0.00152 | 0.01202 |
| 1000 | AGK | 13.6869844 | -1.961404998 | 0.71285 | -2.7515 | 0.00593 | 0.03625 |
| 672 | LAMA5 | 83.2621211 | -1.971698527 | 0.35172 | -5.6059 | 2.1E-08 | 6.7E-07 |
| 965 | CCDC146 | 16.2844273 | -1.975837206 | 0.65449 | -3.0189 | 0.00254 | 0.01816 |
| 175 | APBB1 | 21.0052397 | -1.977556651 | 0.55557 | -3.5595 | 0.00037 | 0.00363 |
| 458 | TUBG2 | 13.7066676 | -1.9855388 | 0.69748 | -2.8467 | 0.00442 | 0.02873 |
| 191 | TRMT112 | 48.1682205 | -1.989363572 | 0.51687 | -3.8489 | 0.00012 | 0.00136 |
| 729 | CRTAP | 31.2277991 | -1.990692581 | 0.46074 | -4.3206 | 1.6E-05 | 0.00023 |
| 625 | SUMO1 | 12.2208341 | -1.995166185 | 0.72492 | -2.7523 | 0.00592 | 0.0362 |
| 257 | XPOT | 31.5337369 | -1.996163215 | 0.57066 | -3.498 | 0.00047 | 0.00443 |
| 646 | TRAF3IP1 | 10.8581684 | -1.999012173 | 0.75636 | -2.6429 | 0.00822 | 0.04675 |
| 873 | GRAMD2B | 42.6883238 | -1.999023693 | 0.58342 | -3.4264 | 0.00061 | 0.00557 |
| 911 | ELOVL5 | 50.2516817 | -2.002980948 | 0.43522 | -4.6022 | 4.2E-06 | 7.4E-05 |
| 561 | MBOAT2 | 29.624176 | -2.003801129 | 0.54724 | -3.6616 | 0.00025 | 0.0026 |
| 358 | SNRPN | 41.7663857 | -2.009929903 | 0.49668 | -4.0467 | 5.2E-05 | 0.00067 |

| | | | | | | | |
|---|---|---|---|---|---|---|---|
| 838 | CPE | 67.80029 | -2.011057695 | 0.44271 | -4.5426 | 5.6E-06 | 9.4E-05 |
| 214 | GRAMD1B | 26.9013839 | -2.015848155 | 0.64128 | -3.1435 | 0.00167 | 0.01297 |
| 1051 | RCL1 | 14.047501 | -2.015912313 | 0.71639 | -2.814 | 0.00489 | 0.03106 |
| 1004 | KCNH2 | 17.347236 | -2.016052379 | 0.74713 | -2.6984 | 0.00697 | 0.04149 |
| 488 | TUBB6 | 29.3521329 | -2.025673586 | 0.66255 | -3.0574 | 0.00223 | 0.01643 |
| 110 | DEGS1 | 21.4542453 | -2.033347776 | 0.64324 | -3.1611 | 0.00157 | 0.01236 |
| 508 | SLC39A3 | 11.6556608 | -2.036219371 | 0.77032 | -2.6434 | 0.00821 | 0.04675 |
| 1038 | LRATD2 | 27.0493782 | -2.041242651 | 0.61811 | -3.3024 | 0.00096 | 0.0081 |
| 190 | PLAAT3 | 25.3530316 | -2.045673521 | 0.6753 | -3.0293 | 0.00245 | 0.01767 |
| 384 | ABHD2 | 64.8548732 | -2.047721645 | 0.38841 | -5.2721 | 1.3E-07 | 3.5E-06 |
| 441 | NLGN2 | 29.026094 | -2.05081665 | 0.58648 | -3.4968 | 0.00047 | 0.00444 |
| 751 | GLT8D1 | 23.9119993 | -2.054512018 | 0.65591 | -3.1323 | 0.00173 | 0.01336 |
| 470 | AXIN2 | 26.2173583 | -2.057846227 | 0.68482 | -3.005 | 0.00266 | 0.01892 |
| 436 | COTL1 | 54.0383748 | -2.063104059 | 0.53712 | -3.841 | 0.00012 | 0.0014 |
| 707 | TSPO | 24.3279711 | -2.064559687 | 0.66438 | -3.1075 | 0.00189 | 0.0143 |
| 462 | DCAKD | 17.1356075 | -2.071621887 | 0.66478 | -3.1163 | 0.00183 | 0.01399 |
| 777 | AMOTL2 | 23.6954723 | -2.082645974 | 0.53883 | -3.8651 | 0.00011 | 0.00129 |
| 927 | GJA1 | 31.3734974 | -2.085910439 | 0.61561 | -3.3883 | 0.0007 | 0.00627 |
| 265 | OSBPL8 | 37.375479 | -2.093458725 | 0.50956 | -4.1084 | 4E-05 | 0.00053 |
| 565 | SLC5A6 | 62.9895737 | -2.093486322 | 0.37508 | -5.5814 | 2.4E-08 | 7.6E-07 |
| 1144 | ATP11C | 68.7746556 | -2.093818558 | 0.40863 | -5.1239 | 3E-07 | 7.2E-06 |
| 739 | P4HTM | 22.0421309 | -2.096897174 | 0.59879 | -3.5019 | 0.00046 | 0.00437 |
| 454 | FKBP10 | 11.2006728 | -2.102493174 | 0.74035 | -2.8399 | 0.00451 | 0.02922 |
| 73 | NOTCH2 | 31.9275007 | -2.10270112 | 0.64672 | -3.2513 | 0.00115 | 0.00946 |
| 394 | IFT140 | 13.7721119 | -2.10354361 | 0.77118 | -2.7277 | 0.00638 | 0.0385 |
| 371 | TRIP4 | 9.45395004 | -2.106897603 | 0.78961 | -2.6683 | 0.00762 | 0.04448 |
| 502 | POLI | 18.5647616 | -2.109433329 | 0.6983 | -3.0208 | 0.00252 | 0.01808 |
| 1050 | GLIS3 | 28.01994 | -2.111061517 | 0.69236 | -3.0491 | 0.0023 | 0.01679 |
| 626 | ICA1L | 8.96836799 | -2.111589584 | 0.78644 | -2.685 | 0.00725 | 0.04271 |
| 644 | SH3BP4 | 11.02266 | -2.112706164 | 0.75966 | -2.7811 | 0.00542 | 0.03373 |
| 171 | SLC25A22 | 12.0701897 | -2.115342026 | 0.77124 | -2.7428 | 0.00609 | 0.03704 |
| 1032 | RPL30 | 21.3736466 | -2.120558074 | 0.71844 | -2.9516 | 0.00316 | 0.02185 |
| 146 | CUEDC2 | 35.347896 | -2.125355369 | 0.5219 | -4.0724 | 4.7E-05 | 0.00061 |
| 1009 | TUSC3 | 93.5764945 | -2.125858612 | 0.36372 | -5.8447 | 5.1E-09 | 2E-07 |
| 253 | GDF11 | 14.2189844 | -2.129623339 | 0.677 | -3.1457 | 0.00166 | 0.01288 |
| 1133 | PLS3 | 221.342709 | -2.144349407 | 0.51412 | -4.1709 | 3E-05 | 0.00041 |
| 272 | IKBIP | 10.3076418 | -2.14447197 | 0.79182 | -2.7083 | 0.00676 | 0.04056 |
| 621 | SLC39A10 | 21.7448145 | -2.144939398 | 0.57769 | -3.7129 | 0.0002 | 0.00219 |
| 764 | CCDC80 | 12.4540254 | -2.14816946 | 0.75111 | -2.86 | 0.00424 | 0.02772 |

| | | | | | | | |
|---|---|---|---|---|---|---|---|
| 331 | BMP4 | 64.903029 | -2.158796273 | 0.68623 | -3.1459 | 0.00166 | 0.01288 |
| 770 | SLC41A3 | 15.7610208 | -2.159353149 | 0.72878 | -2.963 | 0.00305 | 0.02117 |
| 263 | PHLDA1 | 14.630384 | -2.160726914 | 0.74859 | -2.8864 | 0.0039 | 0.02591 |
| 835 | DCLK2 | 11.5765383 | -2.16301093 | 0.72481 | -2.9842 | 0.00284 | 0.02004 |
| 204 | RDX | 31.5195028 | -2.165238912 | 0.49934 | -4.3362 | 1.4E-05 | 0.00022 |
| 787 | PFN2 | 147.4027 | -2.16597516 | 0.44424 | -4.8757 | 1.1E-06 | 2.2E-05 |
| 1123 | TCEAL3 | 21.7216269 | -2.169502592 | 0.60281 | -3.599 | 0.00032 | 0.0032 |
| 1139 | RTL8C | 101.125071 | -2.175227213 | 0.32871 | -6.6175 | 3.7E-11 | 2.4E-09 |
| 41 | BTBD19 | 12.499665 | -2.182952589 | 0.7933 | -2.7517 | 0.00593 | 0.03624 |
| 421 | RANBP10 | 9.98044322 | -2.185650114 | 0.75046 | -2.9124 | 0.00359 | 0.0242 |
| 11 | FBXO44 | 12.5260808 | -2.187976354 | 0.77321 | -2.8297 | 0.00466 | 0.02988 |
| 1016 | FGFR1 | 30.5209496 | -2.189302606 | 0.63831 | -3.4298 | 0.0006 | 0.00552 |
| 864 | ZFYVE16 | 20.877402 | -2.197238227 | 0.57236 | -3.8389 | 0.00012 | 0.00141 |
| 78 | PBXIP1 | 41.3968621 | -2.202182264 | 0.46239 | -4.7626 | 1.9E-06 | 3.7E-05 |
| 1086 | NPDC1 | 24.7806639 | -2.202297964 | 0.5996 | -3.673 | 0.00024 | 0.0025 |
| 578 | RHOQ | 22.2395959 | -2.205385182 | 0.72407 | -3.0458 | 0.00232 | 0.0169 |
| 740 | LAMB2 | 56.9581705 | -2.205607267 | 0.39537 | -5.5785 | 2.4E-08 | 7.7E-07 |
| 633 | STK36 | 54.9813352 | -2.20795342 | 0.44067 | -5.0104 | 5.4E-07 | 1.2E-05 |
| 765 | ZBTB20 | 15.037668 | -2.211396202 | 0.74995 | -2.9487 | 0.00319 | 0.022 |
| 1124 | MORC4 | 17.3731024 | -2.232212664 | 0.77051 | -2.8971 | 0.00377 | 0.0252 |
| 816 | PDGFRA | 47.7543022 | -2.232682538 | 0.48916 | -4.5643 | 5E-06 | 8.6E-05 |
| 514 | ADAMTS10 | 12.890571 | -2.238490276 | 0.84104 | -2.6616 | 0.00778 | 0.04502 |
| 1087 | NSMF | 44.7799634 | -2.240155985 | 0.44259 | -5.0614 | 4.2E-07 | 9.5E-06 |
| 1136 | SMARCA1 | 48.9795104 | -2.242545648 | 0.49554 | -4.5255 | 6E-06 | 0.0001 |
| 95 | NAV1 | 21.8305147 | -2.250321499 | 0.64259 | -3.5019 | 0.00046 | 0.00437 |
| 410 | BBS2 | 13.8927587 | -2.252362195 | 0.73809 | -3.0516 | 0.00228 | 0.01667 |
| 969 | FZD1 | 14.7705493 | -2.254323942 | 0.66321 | -3.3991 | 0.00068 | 0.00606 |
| 114 | CEP170 | 7.89709089 | -2.255962372 | 0.8462 | -2.666 | 0.00768 | 0.04463 |
| 526 | WTIP | 8.43618063 | -2.256909969 | 0.85882 | -2.6279 | 0.00859 | 0.04838 |
| 930 | LAMA2 | 9.26104571 | -2.259499785 | 0.83876 | -2.6939 | 0.00706 | 0.04195 |
| 546 | PLEKHA4 | 16.6604607 | -2.261138512 | 0.65458 | -3.4543 | 0.00055 | 0.0051 |
| 322 | EFS | 10.2661511 | -2.264578326 | 0.84499 | -2.68 | 0.00736 | 0.04326 |
| 294 | SLC15A4 | 29.975814 | -2.265051463 | 0.56744 | -3.9917 | 6.6E-05 | 0.00082 |
| 142 | MYOF | 18.1558623 | -2.27066892 | 0.71299 | -3.1847 | 0.00145 | 0.01154 |
| 650 | NRSN2 | 16.7627425 | -2.272608433 | 0.65876 | -3.4498 | 0.00056 | 0.00517 |
| 188 | TMEM109 | 36.7151509 | -2.284828438 | 0.44918 | -5.0866 | 3.6E-07 | 8.4E-06 |
| 756 | FRMD4B | 9.31156987 | -2.285160691 | 0.81537 | -2.8026 | 0.00507 | 0.032 |
| 512 | MLLT1 | 62.1406696 | -2.286130576 | 0.37924 | -6.0281 | 1.7E-09 | 7.6E-08 |
| 54 | WLS | 77.9425372 | -2.286177992 | 0.35103 | -6.5128 | 7.4E-11 | 4.5E-09 |

| | | | | | | | |
|---|---|---|---|---|---|---|---|
| 848 | LPCAT1 | 21.5694623 | -2.289673752 | 0.70972 | -3.2262 | 0.00125 | 0.0102 |
| 28 | ADPRS | 26.1848418 | -2.291377722 | 0.55563 | -4.1239 | 3.7E-05 | 0.0005 |
| 837 | PDGFC | 15.0235781 | -2.291652763 | 0.756 | -3.0313 | 0.00244 | 0.01758 |
| 856 | SPEF2 | 23.5058219 | -2.291799638 | 0.64941 | -3.5291 | 0.00042 | 0.004 |
| 953 | DPY19L1 | 25.5054481 | -2.292788206 | 0.52113 | -4.3996 | 1.1E-05 | 0.00017 |
| 12 | AGTRAP | 20.564767 | -2.295549528 | 0.68358 | -3.3581 | 0.00078 | 0.00687 |
| 978 | PRKAR2B | 18.3795658 | -2.296647324 | 0.78191 | -2.9372 | 0.00331 | 0.02266 |
| 47 | NFIA | 147.089459 | -2.302049732 | 0.34634 | -6.6467 | 3E-11 | 2E-09 |
| 562 | WDR35 | 15.2510661 | -2.304573278 | 0.77686 | -2.9665 | 0.00301 | 0.02096 |
| 1118 | BEX2 | 37.1018467 | -2.311394291 | 0.49215 | -4.6965 | 2.6E-06 | 4.9E-05 |
| 15 | CROCC | 25.6011911 | -2.314253633 | 0.5868 | -3.9439 | 8E-05 | 0.00097 |
| 696 | CCDC157 | 9.98525484 | -2.320112809 | 0.82263 | -2.8204 | 0.0048 | 0.03049 |
| 958 | IGFBP3 | 28.7158475 | -2.325926644 | 0.70346 | -3.3064 | 0.00095 | 0.00801 |
| 213 | MIR100HG | 22.2934899 | -2.32781997 | 0.71592 | -3.2515 | 0.00115 | 0.00946 |
| 1070 | CTNNAL1 | 11.4106709 | -2.328163435 | 0.84286 | -2.7622 | 0.00574 | 0.03542 |
| 207 | DIXDC1 | 9.81658372 | -2.333079245 | 0.88946 | -2.623 | 0.00872 | 0.04892 |
| 1015 | DDHD2 | 25.5878404 | -2.334222082 | 0.55116 | -4.2351 | 2.3E-05 | 0.00032 |
| 945 | FSCN1 | 35.1167428 | -2.344344692 | 0.63585 | -3.687 | 0.00023 | 0.00239 |
| 1037 | TMEM65 | 22.2044697 | -2.351350794 | 0.69546 | -3.381 | 0.00072 | 0.00642 |
| 328 | CFL2 | 9.90022127 | -2.35847453 | 0.87239 | -2.7035 | 0.00686 | 0.04101 |
| 24 | ADGRB2 | 12.1663863 | -2.358728834 | 0.76454 | -3.0852 | 0.00203 | 0.01521 |
| 7 | KLHL21 | 17.4378042 | -2.364476905 | 0.71947 | -3.2864 | 0.00101 | 0.00849 |
| 148 | ARL3 | 25.3330905 | -2.365579903 | 0.54548 | -4.3367 | 1.4E-05 | 0.00021 |
| 797 | EPHB3 | 11.0574739 | -2.366289294 | 0.79106 | -2.9913 | 0.00278 | 0.01966 |
| 1080 | LINC00963 | 9.04063792 | -2.368071213 | 0.90196 | -2.6255 | 0.00865 | 0.04863 |
| 1091 | AP1S2 | 47.1333326 | -2.369655499 | 0.4693 | -5.0494 | 4.4E-07 | 1E-05 |
| 202 | DYNC2H1 | 27.5949494 | -2.372091076 | 0.54536 | -4.3496 | 1.4E-05 | 0.0002 |
| 49 | CACHD1 | 43.7251726 | -2.373010481 | 0.4651 | -5.1022 | 3.4E-07 | 7.9E-06 |
| 349 | IFI27L1 | 25.2452527 | -2.373388976 | 0.56198 | -4.2233 | 2.4E-05 | 0.00034 |
| 763 | IFT57 | 20.5789439 | -2.375408626 | 0.67897 | -3.4985 | 0.00047 | 0.00442 |
| 675 | MIR99AHG | 24.4327767 | -2.378012881 | 0.58946 | -4.0343 | 5.5E-05 | 0.0007 |
| 815 | SPATA18 | 10.4038973 | -2.381966506 | 0.79609 | -2.9921 | 0.00277 | 0.01961 |
| 244 | COL2A1 | 24.741612 | -2.383617032 | 0.89822 | -2.6537 | 0.00796 | 0.04579 |
| 819 | ADGRL3 | 17.8080241 | -2.392776085 | 0.70037 | -3.4164 | 0.00063 | 0.00575 |
| 414 | KATNB1 | 11.519233 | -2.396320606 | 0.8637 | -2.7745 | 0.00553 | 0.03432 |
| 997 | PTN | 11.8748535 | -2.396459453 | 0.89751 | -2.6701 | 0.00758 | 0.04431 |
| 234 | BCAT1 | 34.594278 | -2.406338734 | 0.48752 | -4.9358 | 8E-07 | 1.7E-05 |
| 1002 | ATP6V0E2 | 8.78813159 | -2.408229999 | 0.8506 | -2.8312 | 0.00464 | 0.02978 |
| 599 | RABL2A | 21.4505898 | -2.408486297 | 0.69178 | -3.4816 | 0.0005 | 0.00467 |

| | | | | | | | |
|---|---|---|---|---|---|---|---|
| 416 | CDH11 | 27.1482624 | -2.411826373 | 0.65018 | -3.7095 | 0.00021 | 0.00221 |
| 1109 | SLC16A2 | 20.4517646 | -2.411872902 | 0.58998 | -4.088 | 4.4E-05 | 0.00057 |
| 874 | FBN2 | 30.329961 | -2.412141389 | 0.64869 | -3.7185 | 0.0002 | 0.00214 |
| 176 | RIC3 | 15.2249557 | -2.412325368 | 0.71945 | -3.353 | 0.0008 | 0.00697 |
| 700 | IFT27 | 17.9600173 | -2.413445261 | 0.71452 | -3.3777 | 0.00073 | 0.00648 |
| 113 | FAM89A | 25.6360569 | -2.416851947 | 0.61172 | -3.9509 | 7.8E-05 | 0.00095 |
| 26 | NCDN | 17.0251671 | -2.41844477 | 0.72876 | -3.3186 | 0.0009 | 0.00772 |
| 531 | LINC00665 | 14.0668206 | -2.420798484 | 0.67701 | -3.5757 | 0.00035 | 0.00345 |
| 1085 | CAMSAP1 | 17.2383195 | -2.424114691 | 0.68316 | -3.5484 | 0.00039 | 0.00376 |
| 180 | NAV2 | 14.7886287 | -2.432426193 | 0.87477 | -2.7807 | 0.00542 | 0.03377 |
| 588 | WDR54 | 20.7926662 | -2.441813697 | 0.85812 | -2.8455 | 0.00443 | 0.02879 |
| 824 | PKD2 | 15.2221823 | -2.446392573 | 0.72455 | -3.3764 | 0.00073 | 0.0065 |
| 1119 | TCEAL7 | 14.4065289 | -2.456895901 | 0.70931 | -3.4638 | 0.00053 | 0.00495 |
| 1049 | RFX3 | 23.3959971 | -2.46039299 | 0.58008 | -4.2415 | 2.2E-05 | 0.00031 |
| 539 | DMPK | 11.3123837 | -2.469587921 | 0.78068 | -3.1634 | 0.00156 | 0.01228 |
| 290 | TMEM120B | 13.5500944 | -2.473578376 | 0.83363 | -2.9672 | 0.003 | 0.02093 |
| 1101 | TIMP1 | 106.262568 | -2.473778621 | 0.436 | -5.6738 | 1.4E-08 | 4.8E-07 |
| 92 | XPR1 | 13.6868801 | -2.475710058 | 0.8088 | -3.061 | 0.00221 | 0.01629 |
| 145 | DPCD | 22.3335515 | -2.476165689 | 0.62741 | -3.9466 | 7.9E-05 | 0.00096 |
| 906 | CCDC167 | 13.1307687 | -2.477262858 | 0.72987 | -3.3941 | 0.00069 | 0.00616 |
| 793 | EGFEM1P | 12.3633123 | -2.479105134 | 0.87375 | -2.8373 | 0.00455 | 0.0294 |
| 830 | MCUB | 9.49008108 | -2.48025652 | 0.81369 | -3.0482 | 0.0023 | 0.0168 |
| 875 | PDLIM4 | 28.363236 | -2.480772186 | 0.5607 | -4.4244 | 9.7E-06 | 0.00015 |
| 490 | RBBP8 | 31.9052158 | -2.487014447 | 0.57972 | -4.29 | 1.8E-05 | 0.00026 |
| 1039 | DNAAF11 | 19.547731 | -2.487885064 | 0.76812 | -3.2389 | 0.0012 | 0.00981 |
| 839 | PALLD | 46.2629061 | -2.488042784 | 0.69333 | -3.5885 | 0.00033 | 0.00331 |
| 221 | JAM3 | 15.201819 | -2.489464711 | 0.74165 | -3.3567 | 0.00079 | 0.00689 |
| 344 | TGFB3 | 20.2194648 | -2.491729243 | 0.66245 | -3.7614 | 0.00017 | 0.00186 |
| 922 | WASF1 | 16.0401694 | -2.495780113 | 0.80262 | -3.1095 | 0.00187 | 0.01425 |
| 992 | PODXL | 17.3332727 | -2.495792954 | 0.81642 | -3.057 | 0.00224 | 0.01644 |
| 966 | MAGI2-AS3 | 22.2651058 | -2.501792984 | 0.67956 | -3.6815 | 0.00023 | 0.00243 |
| 71 | TRIM45 | 18.004574 | -2.501851195 | 0.70646 | -3.5414 | 0.0004 | 0.00385 |
| 210 | IFT46 | 15.8667758 | -2.504312758 | 0.77403 | -3.2354 | 0.00121 | 0.00991 |
| 678 | SLC5A3 | 79.9509535 | -2.507092047 | 0.46841 | -5.3523 | 8.7E-08 | 2.4E-06 |
| 299 | SACS | 14.0884714 | -2.513228099 | 0.85671 | -2.9336 | 0.00335 | 0.02286 |
| 390 | CHSY1 | 15.8033945 | -2.513504378 | 0.75674 | -3.3215 | 0.0009 | 0.00765 |
| 943 | QKI | 47.3260517 | -2.513601007 | 0.53937 | -4.6603 | 3.2E-06 | 5.7E-05 |
| 217 | PKNOX2 | 20.2097037 | -2.514613633 | 0.78314 | -3.2109 | 0.00132 | 0.01065 |
| 1094 | DMD | 75.6649104 | -2.5204571 | 0.45883 | -5.4932 | 3.9E-08 | 1.2E-06 |

| | | | | | | | |
|---|---|---|---|---|---|---|---|
| 192 | LTBP3 | 20.4825318 | -2.521701916 | 0.68885 | -3.6607 | 0.00025 | 0.0026 |
| 352 | EML1 | 15.0988611 | -2.524995087 | 0.76189 | -3.3141 | 0.00092 | 0.00782 |
| 377 | BBS4 | 20.3385262 | -2.525623879 | 0.61227 | -4.125 | 3.7E-05 | 0.0005 |
| 637 | OBSL1 | 58.6414997 | -2.527981192 | 0.43856 | -5.7642 | 8.2E-09 | 3E-07 |
| 209 | FXYD6 | 20.3863315 | -2.528334956 | 0.68901 | -3.6695 | 0.00024 | 0.00253 |
| 45 | PRKAA2 | 10.1303603 | -2.538981869 | 0.82315 | -3.0845 | 0.00204 | 0.01523 |
| 941 | AGPAT4 | 15.2766836 | -2.541850805 | 0.67013 | -3.793 | 0.00015 | 0.00165 |
| 9 | TMEM201 | 9.04189706 | -2.544583136 | 0.81741 | -3.113 | 0.00185 | 0.01411 |
| 438 | NXN | 18.429739 | -2.546348427 | 0.60369 | -4.218 | 2.5E-05 | 0.00034 |
| 511 | RFX2 | 17.1969339 | -2.557314251 | 0.68735 | -3.7206 | 0.0002 | 0.00213 |
| 140 | ZNF503 | 16.3428398 | -2.563032229 | 0.92112 | -2.7825 | 0.00539 | 0.03364 |
| 298 | LATS2 | 11.0174736 | -2.571297836 | 0.87286 | -2.9458 | 0.00322 | 0.0222 |
| 510 | CAPS | 22.4965913 | -2.575694302 | 0.80315 | -3.207 | 0.00134 | 0.01079 |
| 75 | SV2A | 17.358347 | -2.581491302 | 0.77446 | -3.3333 | 0.00086 | 0.00742 |
| 1132 | LRCH2 | 9.59019147 | -2.582812841 | 0.95051 | -2.7173 | 0.00658 | 0.03964 |
| 706 | POLR3H | 7.18749338 | -2.592901446 | 0.96796 | -2.6787 | 0.00739 | 0.04339 |
| 53 | GADD45A | 11.0879839 | -2.593222945 | 0.86592 | -2.9948 | 0.00275 | 0.01946 |
| 701 | KCTD17 | 15.8800082 | -2.5991999 | 0.67245 | -3.8653 | 0.00011 | 0.00129 |
| 381 | LINGO1 | 7.43283195 | -2.600465115 | 0.92698 | -2.8053 | 0.00503 | 0.03178 |
| 962 | ELN | 24.8601014 | -2.605411232 | 0.77236 | -3.3733 | 0.00074 | 0.00656 |
| 20 | STPG1 | 8.2022914 | -2.60741467 | 0.87966 | -2.9641 | 0.00304 | 0.0211 |
| 1068 | CTSL | 14.5787756 | -2.61581947 | 0.71683 | -3.6492 | 0.00026 | 0.0027 |
| 520 | JAK3 | 8.20042672 | -2.619667354 | 0.90313 | -2.9007 | 0.00372 | 0.025 |
| 130 | CSGALNACT2 | 14.8563214 | -2.621958811 | 0.7273 | -3.6051 | 0.00031 | 0.00313 |
| 369 | PYGO1 | 11.0551719 | -2.624511504 | 0.80998 | -3.2402 | 0.00119 | 0.00977 |
| 616 | CHN1 | 14.9260272 | -2.625406951 | 0.71395 | -3.6773 | 0.00024 | 0.00247 |
| 715 | DENND6B | 16.2971112 | -2.626500679 | 0.67198 | -3.9086 | 9.3E-05 | 0.00111 |
| 1005 | SMARCD3 | 16.7010525 | -2.627835535 | 0.74083 | -3.5471 | 0.00039 | 0.00378 |
| 208 | CADM1 | 26.4593363 | -2.633616861 | 0.68957 | -3.8192 | 0.00013 | 0.00151 |
| 744 | SEMA3F | 6.92408558 | -2.634012424 | 0.99618 | -2.6441 | 0.00819 | 0.04668 |
| 1114 | GPRASP1 | 10.5175053 | -2.649207253 | 0.89963 | -2.9448 | 0.00323 | 0.02224 |
| 780 | TRPC1 | 6.96409968 | -2.660420075 | 0.95618 | -2.7823 | 0.0054 | 0.03364 |
| 853 | FBXL7 | 13.4103613 | -2.668789836 | 0.85036 | -3.1384 | 0.0017 | 0.01314 |
| 85 | PEA15 | 32.1572644 | -2.669798811 | 0.61328 | -4.3533 | 1.3E-05 | 0.0002 |
| 755 | PTPRG | 19.8867096 | -2.671146459 | 0.63838 | -4.1843 | 2.9E-05 | 0.00039 |
| 972 | PILRA | 8.78479314 | -2.674746606 | 0.84307 | -3.1726 | 0.00151 | 0.01194 |
| 348 | STON2 | 19.8467836 | -2.676229896 | 0.65559 | -4.0822 | 4.5E-05 | 0.00058 |
| 735 | POMGNT2 | 14.0424135 | -2.678456681 | 0.80479 | -3.3281 | 0.00087 | 0.00751 |
| 415 | NDRG4 | 9.9344173 | -2.682673049 | 0.9462 | -2.8352 | 0.00458 | 0.02954 |

| | | | | | | | |
|---|---|---|---|---|---|---|---|
| 420 | ENKD1 | 12.178 | -2.683886822 | 0.84764 | -3.1663 | 0.00154 | 0.01218 |
| 769 | HEG1 | 16.8578549 | -2.687094144 | 0.70619 | -3.805 | 0.00014 | 0.00158 |
| 560 | PXDN | 39.6607419 | -2.694387116 | 0.49269 | -5.4687 | 4.5E-08 | 1.3E-06 |
| 736 | TMEM42 | 11.1006019 | -2.697789714 | 0.86771 | -3.1091 | 0.00188 | 0.01426 |
| 1006 | GALNT11 | 41.3458468 | -2.699329093 | 0.68011 | -3.969 | 7.2E-05 | 0.00089 |
| 258 | WIF1 | 15.7698292 | -2.70374287 | 0.99618 | -2.7141 | 0.00665 | 0.03997 |
| 841 | WDR17 | 7.97116987 | -2.7043298 | 1.03404 | -2.6153 | 0.00891 | 0.04988 |
| 386 | ZNF710-AS1 | 8.7104031 | -2.705690231 | 0.94989 | -2.8484 | 0.00439 | 0.02866 |
| 1045 | IQANK1 | 14.1150407 | -2.712605397 | 0.76432 | -3.549 | 0.00039 | 0.00376 |
| 897 | CD83 | 21.6113329 | -2.72147231 | 0.58897 | -4.6208 | 3.8E-06 | 6.8E-05 |
| 1121 | BEX3 | 184.933243 | -2.725804564 | 0.33492 | -8.1386 | 4E-16 | 8.2E-14 |
| 107 | SLC30A10 | 8.49976859 | -2.727290622 | 0.85155 | -3.2027 | 0.00136 | 0.01091 |
| 1103 | PRAF2 | 35.3568077 | -2.729283513 | 0.58522 | -4.6637 | 3.1E-06 | 5.7E-05 |
| 663 | SNTA1 | 11.2363329 | -2.730788316 | 0.93439 | -2.9225 | 0.00347 | 0.02358 |
| 151 | GFRA1 | 9.56986256 | -2.733663418 | 0.89358 | -3.0592 | 0.00222 | 0.01637 |
| 240 | BICD1 | 20.7121611 | -2.745534032 | 0.70637 | -3.8868 | 0.0001 | 0.00119 |
| 737 | LZTFL1 | 19.3051708 | -2.748946066 | 0.66533 | -4.1317 | 3.6E-05 | 0.00049 |
| 861 | GPX8 | 17.8838686 | -2.749733675 | 0.67979 | -4.045 | 5.2E-05 | 0.00067 |
| 469 | TANC2 | 17.4759503 | -2.750238093 | 0.71838 | -3.8284 | 0.00013 | 0.00146 |
| 1067 | DAPK1 | 16.1245492 | -2.756978861 | 0.94026 | -2.9321 | 0.00337 | 0.02296 |
| 713 | CELSR1 | 8.41744341 | -2.760649441 | 0.98948 | -2.79 | 0.00527 | 0.03299 |
| 266 | CSRP2 | 11.3852927 | -2.764062587 | 0.92642 | -2.9836 | 0.00285 | 0.02008 |
| 887 | PDGFRB | 33.3262801 | -2.765397636 | 0.81081 | -3.4107 | 0.00065 | 0.00584 |
| 1140 | ZNF449 | 14.1640331 | -2.768578721 | 0.71039 | -3.8973 | 9.7E-05 | 0.00115 |
| 1033 | FZD6 | 28.2858645 | -2.782246835 | 0.6091 | -4.5678 | 4.9E-06 | 8.5E-05 |
| 227 | P3H3 | 9.92344337 | -2.784895739 | 0.8368 | -3.328 | 0.00087 | 0.00751 |
| 22 | UBXN11 | 31.3414292 | -2.786628386 | 0.60034 | -4.6418 | 3.5E-06 | 6.2E-05 |
| 329 | SAV1 | 8.59084977 | -2.787337343 | 1.05971 | -2.6303 | 0.00853 | 0.0481 |
| 279 | IFT81 | 11.4929758 | -2.78902975 | 0.81084 | -3.4397 | 0.00058 | 0.00534 |
| 425 | MTSS2 | 27.0317479 | -2.789453555 | 0.68109 | -4.0956 | 4.2E-05 | 0.00055 |
| 571 | CDC42EP3 | 17.1803285 | -2.7899369 | 0.80773 | -3.454 | 0.00055 | 0.0051 |
| 496 | B4GALT6 | 14.3992747 | -2.791456488 | 0.81752 | -3.4146 | 0.00064 | 0.00578 |
| 698 | TIMP3 | 62.8140332 | -2.797072495 | 0.43862 | -6.3769 | 1.8E-10 | 1E-08 |
| 14 | EPHA2 | 7.05020594 | -2.797638043 | 0.9845 | -2.8417 | 0.00449 | 0.02909 |
| 467 | TSPOAP1 | 24.8228027 | -2.798285888 | 0.70395 | -3.9751 | 7E-05 | 0.00088 |
| 19 | ID3 | 53.3108886 | -2.798413745 | 0.44937 | -6.2274 | 4.7E-10 | 2.4E-08 |
| 863 | ADAMTS6 | 18.0505221 | -2.801003241 | 0.83092 | -3.371 | 0.00075 | 0.0066 |
| 5 | PRDM16 | 26.5398412 | -2.801379572 | 0.70626 | -3.9665 | 7.3E-05 | 0.0009 |
| 1108 | MSN | 28.7231937 | -2.804101592 | 0.65269 | -4.2962 | 1.7E-05 | 0.00025 |

| | | | | | | | |
|---|---|---|---|---|---|---|---|
| 1126 | PRPS1 | 64.1568005 | -2.804724077 | 0.38211 | -7.34 | 2.1E-13 | 2.6E-11 |
| 1 | SCNN1D | 7.13087197 | -2.819136606 | 0.93729 | -3.0078 | 0.00263 | 0.01878 |
| 308 | SMAD9 | 17.4728954 | -2.820309642 | 0.71013 | -3.9715 | 7.1E-05 | 0.00089 |
| 1112 | ARMCX2 | 39.6083034 | -2.822555167 | 0.52053 | -5.4225 | 5.9E-08 | 1.7E-06 |
| 276 | CRY1 | 19.1157989 | -2.827180257 | 0.75656 | -3.7369 | 0.00019 | 0.00202 |
| 768 | FSTL1 | 165.599951 | -2.830765375 | 0.2795 | -10.128 | 4.2E-24 | 2.9E-21 |
| 196 | UVRAG | 21.2086544 | -2.831893159 | 0.79577 | -3.5587 | 0.00037 | 0.00364 |
| 340 | PNMA1 | 46.7484668 | -2.833267432 | 0.60897 | -4.6526 | 3.3E-06 | 5.9E-05 |
| 975 | IFT22 | 19.3945477 | -2.835049457 | 0.73993 | -3.8315 | 0.00013 | 0.00144 |
| 23 | PTPRU | 7.93825464 | -2.835212181 | 0.93637 | -3.0279 | 0.00246 | 0.01773 |
| 350 | C14orf132 | 8.33524654 | -2.835363734 | 0.9363 | -3.0283 | 0.00246 | 0.01771 |
| 1046 | NRBP2 | 48.1576543 | -2.838577325 | 0.48467 | -5.8567 | 4.7E-09 | 1.9E-07 |
| 2 | MXRA8 | 46.1246581 | -2.848495305 | 0.50099 | -5.6857 | 1.3E-08 | 4.5E-07 |
| 611 | BBS5 | 12.4239922 | -2.848766137 | 0.83559 | -3.4093 | 0.00065 | 0.00586 |
| 855 | CDH6 | 21.5363139 | -2.850761743 | 0.86416 | -3.2989 | 0.00097 | 0.00818 |
| 716 | SHANK3 | 10.4062029 | -2.851881886 | 0.84333 | -3.3817 | 0.00072 | 0.00641 |
| 99 | DSTYK | 21.5811863 | -2.854745847 | 0.60456 | -4.722 | 2.3E-06 | 4.4E-05 |
| 681 | GET1 | 30.0957248 | -2.856969256 | 0.59648 | -4.7897 | 1.7E-06 | 3.3E-05 |
| 947 | HDAC9 | 13.3226508 | -2.860808619 | 0.79637 | -3.5923 | 0.00033 | 0.00327 |
| 199 | SLC36A4 | 12.5157483 | -2.861681908 | 0.82529 | -3.4675 | 0.00053 | 0.00488 |
| 419 | RIPOR1 | 6.327402 | -2.866172175 | 1.08526 | -2.641 | 0.00827 | 0.04688 |
| 339 | DNAL1 | 11.4243729 | -2.868960373 | 0.80362 | -3.57 | 0.00036 | 0.0035 |
| 301 | SLC7A1 | 27.0016492 | -2.871005566 | 0.56554 | -5.0765 | 3.8E-07 | 8.8E-06 |
| 699 | APOL4 | 6.53313235 | -2.874823147 | 1.04125 | -2.7609 | 0.00576 | 0.03551 |
| 226 | GPR162 | 12.9830009 | -2.881669026 | 0.80659 | -3.5727 | 0.00035 | 0.00348 |
| 44 | GPX7 | 22.1381463 | -2.885482019 | 0.62229 | -4.6369 | 3.5E-06 | 6.3E-05 |
| 121 | VIM | 221.837904 | -2.906195004 | 0.53027 | -5.4806 | 4.2E-08 | 1.3E-06 |
| 172 | TSPAN4 | 37.4944614 | -2.907937062 | 0.52425 | -5.5468 | 2.9E-08 | 9E-07 |
| 443 | EFNB3 | 10.7596567 | -2.908520183 | 0.96836 | -3.0035 | 0.00267 | 0.01898 |
| 912 | COL9A1 | 31.4916617 | -2.90852567 | 0.88027 | -3.3041 | 0.00095 | 0.00807 |
| 544 | C5AR1 | 16.022327 | -2.912898824 | 0.74074 | -3.9324 | 8.4E-05 | 0.00101 |
| 890 | CYFIP2 | 24.7362857 | -2.924547856 | 0.55297 | -5.2888 | 1.2E-07 | 3.3E-06 |
| 895 | NEDD9 | 24.8892669 | -2.930075737 | 0.59239 | -4.9462 | 7.6E-07 | 1.6E-05 |
| 1007 | TDRP | 18.9666432 | -2.933533339 | 0.64159 | -4.5723 | 4.8E-06 | 8.4E-05 |
| 929 | TPD52L1 | 40.1197248 | -2.934583038 | 0.76533 | -3.8344 | 0.00013 | 0.00143 |
| 542 | PNMA8A | 15.8086731 | -2.937545378 | 0.74271 | -3.9552 | 7.6E-05 | 0.00093 |
| 164 | LRRC27 | 6.07186712 | -2.937686225 | 1.09172 | -2.6909 | 0.00713 | 0.04225 |
| 128 | BAMBI | 17.0232715 | -2.944058474 | 0.64946 | -4.5331 | 5.8E-06 | 9.8E-05 |
| 178 | AMPD3 | 6.4545142 | -2.946410948 | 1.08772 | -2.7088 | 0.00675 | 0.04053 |

| | | | | | | | |
|---|---|---|---|---|---|---|---|
| 392 | METRN | 8.5371029 | -2.95159797 | 0.88736 | -3.3263 | 0.00088 | 0.00755 |
| 718 | LRRN1 | 18.0435228 | -2.95735549 | 0.89227 | -3.3144 | 0.00092 | 0.00782 |
| 13 | PDPN | 31.4586733 | -2.959084217 | 0.61191 | -4.8358 | 1.3E-06 | 2.7E-05 |
| 396 | CLUAP1 | 12.9580793 | -2.960114388 | 0.76594 | -3.8647 | 0.00011 | 0.00129 |
| 398 | EMP2 | 18.8573038 | -2.984626431 | 0.66409 | -4.4943 | 7E-06 | 0.00011 |
| 312 | SLITRK5 | 6.93443638 | -2.985735906 | 1.06551 | -2.8022 | 0.00508 | 0.03202 |
| 320 | SLC7A8 | 21.3947254 | -2.995268214 | 0.82592 | -3.6266 | 0.00029 | 0.00291 |
| 697 | SEC14L2 | 13.1760476 | -2.995876024 | 0.84377 | -3.5506 | 0.00038 | 0.00374 |
| 638 | SLC4A3 | 10.9380008 | -3.000929042 | 0.84011 | -3.5721 | 0.00035 | 0.00348 |
| 779 | RBP1 | 74.0075478 | -3.007589564 | 0.42142 | -7.1369 | 9.5E-13 | 9.1E-11 |
| 1047 | ARHGAP39 | 6.23049078 | -3.016544994 | 1.0578 | -2.8517 | 0.00435 | 0.0284 |
| 471 | CEP112 | 8.13891139 | -3.019375173 | 1.06913 | -2.8241 | 0.00474 | 0.03026 |
| 457 | CAVIN1 | 22.3566916 | -3.020592461 | 0.6899 | -4.3783 | 1.2E-05 | 0.00018 |
| 627 | NRP2 | 15.5859801 | -3.021193974 | 0.9856 | -3.0653 | 0.00217 | 0.01614 |
| 679 | CLIC6 | 110.919062 | -3.02625428 | 0.4358 | -6.9442 | 3.8E-12 | 3.1E-10 |
| 595 | POU3F3 | 8.79145024 | -3.027688085 | 1.06803 | -2.8348 | 0.00458 | 0.02954 |
| 977 | ORAI2 | 16.3720717 | -3.033932303 | 0.73807 | -4.1106 | 3.9E-05 | 0.00052 |
| 346 | DIO2 | 12.7766763 | -3.034849434 | 1.04667 | -2.8995 | 0.00374 | 0.02508 |
| 459 | ARL4D | 8.80693538 | -3.042536771 | 0.94141 | -3.2319 | 0.00123 | 0.01002 |
| 759 | CNTN3 | 19.8541596 | -3.04807284 | 0.71377 | -4.2704 | 2E-05 | 0.00028 |
| 1008 | TRMT9B | 27.9897 | -3.062764009 | 0.84254 | -3.6352 | 0.00028 | 0.00283 |
| 169 | B4GALNT4 | 15.9434604 | -3.072050778 | 0.76887 | -3.9956 | 6.5E-05 | 0.00081 |
| 319 | AJUBA | 12.770788 | -3.077387831 | 0.87495 | -3.5172 | 0.00044 | 0.00417 |
| 449 | B9D1 | 26.8880638 | -3.078285653 | 0.60156 | -5.1172 | 3.1E-07 | 7.4E-06 |
| 582 | EFEMP1 | 13.090754 | -3.084603316 | 0.94546 | -3.2625 | 0.0011 | 0.00915 |
| 939 | SYNE1 | 20.9206476 | -3.093211422 | 0.69727 | -4.4361 | 9.2E-06 | 0.00014 |
| 286 | MSI1 | 18.9024507 | -3.093726804 | 0.67488 | -4.5841 | 4.6E-06 | 7.9E-05 |
| 802 | IQCG | 19.5662075 | -3.095152097 | 0.73396 | -4.217 | 2.5E-05 | 0.00034 |
| 237 | RASSF8 | 15.0401923 | -3.0971633 | 0.72607 | -4.2657 | 2E-05 | 0.00028 |
| 847 | ZDHHC11 | 7.70078654 | -3.107017504 | 1.05625 | -2.9416 | 0.00327 | 0.02239 |
| 247 | SLC4A8 | 13.1161854 | -3.114696923 | 0.86936 | -3.5827 | 0.00034 | 0.00337 |
| 86 | RGS5 | 19.009422 | -3.117543489 | 0.86611 | -3.5995 | 0.00032 | 0.00319 |
| 61 | CNN3 | 87.4625811 | -3.119517329 | 0.49662 | -6.2815 | 3.4E-10 | 1.8E-08 |
| 230 | C1S | 31.0052987 | -3.122247977 | 0.58574 | -5.3305 | 9.8E-08 | 2.7E-06 |
| 882 | STING1 | 20.0371557 | -3.127067738 | 0.88958 | -3.5152 | 0.00044 | 0.00419 |
| 412 | CX3CL1 | 10.7657003 | -3.128881695 | 0.94832 | -3.2994 | 0.00097 | 0.00817 |
| 373 | CORO2B | 9.03741347 | -3.131961255 | 0.98652 | -3.1748 | 0.0015 | 0.01187 |
| 730 | FBXL2 | 8.98687048 | -3.132367585 | 1.02429 | -3.0581 | 0.00223 | 0.01642 |
| 519 | SLC27A1 | 11.8604036 | -3.136911911 | 0.7618 | -4.1178 | 3.8E-05 | 0.00051 |

| | | | | | | | |
|---|---|---|---|---|---|---|---|
| 379 | STRA6 | 9.92340563 | -3.141788154 | 0.85503 | -3.6745 | 0.00024 | 0.00249 |
| 836 | MAP9 | 29.9565458 | -3.142872647 | 0.62229 | -5.0505 | 4.4E-07 | 9.9E-06 |
| 456 | ODAD4 | 10.6771694 | -3.142920688 | 1.02051 | -3.0798 | 0.00207 | 0.01544 |
| 129 | FZD8 | 48.7581702 | -3.144134186 | 0.65911 | -4.7703 | 1.8E-06 | 3.6E-05 |
| 810 | CC2D2A | 16.0867796 | -3.145300936 | 0.85311 | -3.6869 | 0.00023 | 0.00239 |
| 1012 | RHOBTB2 | 6.68278602 | -3.146461844 | 1.01213 | -3.1087 | 0.00188 | 0.01426 |
| 615 | CYBRD1 | 15.0434695 | -3.149696394 | 0.90282 | -3.4887 | 0.00049 | 0.00456 |
| 461 | FAM171A2 | 9.64583872 | -3.149756767 | 0.87839 | -3.5858 | 0.00034 | 0.00333 |
| 477 | CDR2L | 7.58879244 | -3.152343484 | 1.08535 | -2.9044 | 0.00368 | 0.02473 |
| 822 | SCD5 | 11.8125326 | -3.152423867 | 0.87073 | -3.6204 | 0.00029 | 0.00298 |
| 497 | ASXL3 | 17.2415252 | -3.15424112 | 1.00959 | -3.1243 | 0.00178 | 0.01368 |
| 778 | DZIP1L | 9.74890872 | -3.155078508 | 0.93992 | -3.3568 | 0.00079 | 0.00689 |
| 218 | FEZ1 | 8.8181857 | -3.166962788 | 0.89712 | -3.5301 | 0.00042 | 0.00399 |
| 653 | EBF4 | 29.0031101 | -3.167562745 | 0.74779 | -4.2359 | 2.3E-05 | 0.00032 |
| 184 | PRDM11 | 11.2038021 | -3.170265085 | 0.89464 | -3.5436 | 0.00039 | 0.00382 |
| 447 | GAS7 | 11.7535821 | -3.178280661 | 0.952 | -3.3385 | 0.00084 | 0.0073 |
| 116 | CELF2 | 43.6567737 | -3.183695755 | 0.47997 | -6.6331 | 3.3E-11 | 2.2E-09 |
| 1025 | TMEM64 | 17.2798696 | -3.185282501 | 0.66412 | -4.7962 | 1.6E-06 | 3.2E-05 |
| 1111 | NAP1L3 | 13.9939524 | -3.190504453 | 0.88047 | -3.6237 | 0.00029 | 0.00294 |
| 50 | LEPR | 6.09666087 | -3.195495307 | 1.21361 | -2.633 | 0.00846 | 0.04787 |
| 899 | RBM24 | 6.61186112 | -3.195762354 | 1.14409 | -2.7933 | 0.00522 | 0.03268 |
| 251 | ZNF385A | 5.65173154 | -3.197510636 | 1.12503 | -2.8421 | 0.00448 | 0.02906 |
| 56 | TTLL7 | 16.1672922 | -3.199728479 | 0.74307 | -4.3061 | 1.7E-05 | 0.00024 |
| 796 | ABCC5 | 33.8644478 | -3.200494738 | 0.54121 | -5.9136 | 3.3E-09 | 1.4E-07 |
| 170 | LRRC56 | 4.65390865 | -3.209057804 | 1.20268 | -2.6683 | 0.00762 | 0.04448 |
| 573 | MORN2 | 26.5950069 | -3.211183408 | 0.68575 | -4.6827 | 2.8E-06 | 5.2E-05 |
| 368 | DNAAF4 | 5.12297551 | -3.218009409 | 1.19898 | -2.6839 | 0.00728 | 0.04283 |
| 919 | ANKRD6 | 5.51123712 | -3.715579999 | 1.30352 | -2.8504 | 0.00437 | 0.02851 |
| 585 | ANTXR1 | 29.8520741 | -3.717116263 | 0.57015 | -6.5195 | 7.1E-11 | 4.3E-09 |
| 624 | PGAP1 | 55.6965867 | -3.718177385 | 0.56702 | -6.5574 | 5.5E-11 | 3.4E-09 |
| 901 | ID4 | 18.5418601 | -3.723198231 | 0.78781 | -4.726 | 2.3E-06 | 4.3E-05 |
| 946 | SOSTDC1 | 14.0030913 | -3.72581202 | 0.91222 | -4.0843 | 4.4E-05 | 0.00058 |
| 950 | JAZF1 | 6.69451678 | -3.726617672 | 1.11879 | -3.3309 | 0.00087 | 0.00746 |
| 1138 | RTL8B | 8.87008942 | -3.726666195 | 1.03865 | -3.588 | 0.00033 | 0.00332 |
| 1042 | LY6H | 4.70375562 | -3.731816262 | 1.35585 | -2.7524 | 0.00592 | 0.0362 |
| 224 | TSPAN9 | 13.1165467 | -3.734473302 | 0.78981 | -4.7283 | 2.3E-06 | 4.3E-05 |
| 881 | REEP2 | 14.8392722 | -3.741451581 | 0.83107 | -4.502 | 6.7E-06 | 0.00011 |
| 893 | PRR7 | 7.18249752 | -3.74725116 | 1.16244 | -3.2236 | 0.00127 | 0.01027 |
| 463 | LRRC46 | 8.55764857 | -3.763261722 | 1.24311 | -3.0273 | 0.00247 | 0.01773 |

| | | | | | | | |
|---|---|---|---|---|---|---|---|
| 382 | ARNT2 | 13.3194262 | -3.767386062 | 0.85034 | -4.4304 | 9.4E-06 | 0.00015 |
| 533 | PRX | 4.78190697 | -3.768320471 | 1.33391 | -2.825 | 0.00473 | 0.03021 |
| 74 | LIX1L | 17.3644821 | -3.776654213 | 0.71372 | -5.2915 | 1.2E-07 | 3.3E-06 |
| 597 | SULT1C4 | 11.719667 | -3.777172153 | 1.00005 | -3.777 | 0.00016 | 0.00175 |
| 84 | ATP1A2 | 14.3226415 | -3.781516998 | 0.88313 | -4.2819 | 1.9E-05 | 0.00027 |
| 428 | TMEM231 | 29.5708653 | -3.784428188 | 0.68231 | -5.5465 | 2.9E-08 | 9E-07 |
| 270 | RMST | 22.7621384 | -3.787146482 | 0.70017 | -5.4089 | 6.3E-08 | 1.8E-06 |
| 711 | RIBC2 | 6.06298514 | -3.791144153 | 1.26252 | -3.0028 | 0.00267 | 0.01901 |
| 1084 | PIERCE1 | 13.7502849 | -3.806772708 | 0.83917 | -4.5364 | 5.7E-06 | 9.7E-05 |
| 494 | CDH2 | 41.1480382 | -3.808201641 | 0.72081 | -5.2832 | 1.3E-07 | 3.4E-06 |
| 1001 | CNTNAP2 | 31.9183245 | -3.820278923 | 0.67708 | -5.6423 | 1.7E-08 | 5.7E-07 |
| 671 | BMP7 | 23.2068649 | -3.840393326 | 0.65845 | -5.8324 | 5.5E-09 | 2.1E-07 |
| 670 | DOK5 | 7.79303487 | -3.841453884 | 1.16809 | -3.2886 | 0.00101 | 0.00844 |
| 138 | SPOCK2 | 7.79847182 | -3.846382093 | 1.12701 | -3.4129 | 0.00064 | 0.0058 |
| 613 | ENSG00000239467 | 8.48720798 | -3.852749483 | 1.04607 | -3.6831 | 0.00023 | 0.00242 |
| 35 | CCDC30 | 11.6163747 | -3.857812833 | 0.88761 | -4.3463 | 1.4E-05 | 0.00021 |
| 189 | TMEM132A | 13.8173523 | -3.860392242 | 0.81453 | -4.7394 | 2.1E-06 | 4E-05 |
| 1145 | LINC00632 | 45.2236063 | -3.863343487 | 0.53592 | -7.2087 | 5.6E-13 | 5.8E-11 |
| 1076 | AK1 | 7.88396739 | -3.864717914 | 1.20109 | -3.2177 | 0.00129 | 0.01044 |
| 203 | GUCY1A2 | 12.3780011 | -3.867009881 | 0.93825 | -4.1215 | 3.8E-05 | 0.0005 |
| 556 | ZNF667 | 4.87071001 | -3.884812128 | 1.36796 | -2.8399 | 0.00451 | 0.02922 |
| 680 | RIPPLY3 | 7.65170292 | -3.885980584 | 1.40848 | -2.759 | 0.0058 | 0.03566 |
| 529 | APLP1 | 11.2475623 | -3.888895641 | 0.9666 | -4.0233 | 5.7E-05 | 0.00073 |
| 115 | AKT3 | 7.92172539 | -3.894315081 | 1.17701 | -3.3086 | 0.00094 | 0.00795 |
| 402 | SYT17 | 15.6679749 | -3.896692056 | 0.91219 | -4.2718 | 1.9E-05 | 0.00028 |
| 513 | CTXN1 | 16.0991926 | -3.903512179 | 0.84214 | -4.6352 | 3.6E-06 | 6.4E-05 |
| 472 | ARSG | 6.43209157 | -3.915592412 | 1.3196 | -2.9673 | 0.003 | 0.02093 |
| 747 | CACNA2D2 | 6.54521378 | -3.915674547 | 1.19221 | -3.2844 | 0.00102 | 0.00854 |
| 81 | KIRREL1 | 8.56070134 | -3.928654929 | 1.03032 | -3.8131 | 0.00014 | 0.00153 |
| 687 | COL18A1 | 10.533006 | -3.929441852 | 0.91737 | -4.2834 | 1.8E-05 | 0.00027 |
| 212 | TLCD5 | 7.62698794 | -3.941731757 | 1.14776 | -3.4343 | 0.00059 | 0.00544 |
| 370 | CGNL1 | 27.0756971 | -3.945585573 | 0.67559 | -5.8402 | 5.2E-09 | 2E-07 |
| 160 | CPXM2 | 24.0417477 | -3.94715031 | 0.79381 | -4.9724 | 6.6E-07 | 1.4E-05 |
| 295 | GLT1D1 | 5.69231823 | -3.949708354 | 1.28893 | -3.0643 | 0.00218 | 0.01615 |
| 31 | OSCP1 | 11.5957361 | -3.951127447 | 1.01249 | -3.9024 | 9.5E-05 | 0.00113 |
| 159 | FGFR2 | 57.4127096 | -3.955701495 | 0.46488 | -8.5091 | 1.8E-17 | 4.3E-15 |
| 957 | GLI3 | 9.06823189 | -3.973194291 | 1.12442 | -3.5336 | 0.00041 | 0.00395 |
| 1014 | PURG | 7.51148397 | -3.975534872 | 1.1082 | -3.5874 | 0.00033 | 0.00332 |
| 865 | FAM151B-DT | 3.74785567 | -3.995619093 | 1.46907 | -2.7198 | 0.00653 | 0.03938 |

| | | | | | | | |
|---|---|---|---|---|---|---|---|
| 900 | CAP2 | 6.9926229 | -3.998523649 | 1.15458 | -3.4632 | 0.00053 | 0.00495 |
| 161 | FANK1 | 9.89380588 | -3.998889285 | 1.03103 | -3.8785 | 0.00011 | 0.00123 |
| 163 | DPYSL4 | 8.35582446 | -4.004436122 | 1.31296 | -3.0499 | 0.00229 | 0.01675 |
| 1072 | WHRN | 17.7342106 | -4.005867249 | 0.77012 | -5.2016 | 2E-07 | 5E-06 |
| 1146 | AFF2 | 11.1811214 | -4.021492309 | 1.03042 | -3.9028 | 9.5E-05 | 0.00113 |
| 651 | SDCBP2-AS1 | 3.80185361 | -4.022114758 | 1.50226 | -2.6774 | 0.00742 | 0.04351 |
| 395 | ZSCAN32 | 6.25035155 | -4.026270967 | 1.22499 | -3.2868 | 0.00101 | 0.00848 |
| 995 | WDR91 | 13.9177685 | -4.029283612 | 0.94861 | -4.2475 | 2.2E-05 | 0.00031 |
| 669 | BCAS4 | 9.58200246 | -4.030094547 | 1.01104 | -3.9861 | 6.7E-05 | 0.00084 |
| 1116 | BEX1 | 36.1663199 | -4.03368234 | 0.79676 | -5.0626 | 4.1E-07 | 9.4E-06 |
| 635 | CFAP65 | 12.1411436 | -4.035289381 | 1.02016 | -3.9556 | 7.6E-05 | 0.00093 |
| 666 | NNAT | 41.1650239 | -4.035289477 | 0.72953 | -5.5313 | 3.2E-08 | 9.7E-07 |
| 915 | TPBG | 63.0026715 | -4.044880691 | 0.59695 | -6.7759 | 1.2E-11 | 9.2E-10 |
| 195 | MAP6 | 23.3321948 | -4.050324835 | 0.74589 | -5.4302 | 5.6E-08 | 1.6E-06 |
| 351 | AK7 | 7.49563103 | -4.054721657 | 1.1384 | -3.5618 | 0.00037 | 0.0036 |
| 722 | RFTN1 | 17.2494562 | -4.070376626 | 0.76849 | -5.2966 | 1.2E-07 | 3.2E-06 |
| 632 | ERBB4 | 11.2598283 | -4.070486668 | 1.0036 | -4.0559 | 5E-05 | 0.00065 |
| 547 | CPT1C | 15.0786522 | -4.082488215 | 0.79281 | -5.1494 | 2.6E-07 | 6.3E-06 |
| 259 | MDM1 | 14.0790181 | -4.086495976 | 1.04424 | -3.9134 | 9.1E-05 | 0.00109 |
| 877 | TCF7 | 15.0315267 | -4.08829053 | 0.84106 | -4.8609 | 1.2E-06 | 2.4E-05 |
| 389 | ADAMTS17 | 4.03033024 | -4.107327724 | 1.42078 | -2.8909 | 0.00384 | 0.02562 |
| 17 | ALDH4A1 | 5.68354454 | -4.10772848 | 1.30213 | -3.1546 | 0.00161 | 0.01257 |
| 550 | LRRC4B | 6.27113416 | -4.109239702 | 1.23888 | -3.3169 | 0.00091 | 0.00776 |
| 852 | DNAH5 | 12.5679577 | -4.110288685 | 1.01853 | -4.0355 | 5.4E-05 | 0.0007 |
| 1026 | NECAB1 | 20.3110437 | -4.117537727 | 0.84872 | -4.8514 | 1.2E-06 | 2.5E-05 |
| 693 | LRRC74B | 4.3786119 | -4.119414529 | 1.43431 | -2.8721 | 0.00408 | 0.02692 |
| 998 | DGKI | 5.73321353 | -4.121964783 | 1.33154 | -3.0956 | 0.00196 | 0.01478 |
| 289 | MORN3 | 5.75786529 | -4.131303378 | 1.35265 | -3.0542 | 0.00226 | 0.01657 |
| 487 | GNAL | 4.37977251 | -4.132504163 | 1.41771 | -2.9149 | 0.00356 | 0.02407 |
| 281 | CUX2 | 4.0723444 | -4.132969579 | 1.48158 | -2.7896 | 0.00528 | 0.03302 |
| 1154 | PNMA3 | 6.99784415 | -4.13480362 | 1.2047 | -3.4322 | 0.0006 | 0.00548 |
| 674 | STMN3 | 15.540498 | -4.140694134 | 0.86115 | -4.8083 | 1.5E-06 | 3E-05 |
| 185 | MAPK8IP1 | 6.1118965 | -4.141188895 | 1.23008 | -3.3666 | 0.00076 | 0.00668 |
| 150 | ATRNL1 | 7.69576395 | -4.151087682 | 1.19482 | -3.4742 | 0.00051 | 0.00478 |
| 60 | ARHGAP29 | 13.8014347 | -4.153728956 | 0.8613 | -4.8226 | 1.4E-06 | 2.8E-05 |
| 273 | SLC17A8 | 19.8642801 | -4.15479651 | 0.77503 | -5.3608 | 8.3E-08 | 2.3E-06 |
| 691 | CLTCL1 | 4.2267907 | -4.177797703 | 1.57772 | -2.648 | 0.0081 | 0.04635 |
| 660 | DZANK1 | 13.1581845 | -4.186389015 | 1.01894 | -4.1086 | 4E-05 | 0.00053 |
| 302 | KATNAL1 | 6.89623521 | -4.194947714 | 1.17414 | -3.5728 | 0.00035 | 0.00348 |

| | | | | | | | |
|---|---|---|---|---|---|---|---|
| 628 | ZDBF2 | 7.98411548 | -4.195763182 | 1.13222 | -3.7058 | 0.00021 | 0.00224 |
| 567 | FNDC4 | 8.01682394 | -4.208457887 | 1.14232 | -3.6841 | 0.00023 | 0.00241 |
| 834 | CLGN | 4.29494406 | -4.212973025 | 1.4543 | -2.8969 | 0.00377 | 0.0252 |
| 892 | MSX2 | 13.3951007 | -4.217524916 | 0.9305 | -4.5325 | 5.8E-06 | 9.8E-05 |
| 187 | SERPING1 | 23.7881022 | -4.239465912 | 0.71664 | -5.9157 | 3.3E-09 | 1.4E-07 |
| 137 | CHST3 | 6.19851169 | -4.242328413 | 1.27795 | -3.3196 | 0.0009 | 0.0077 |
| 938 | CCDC170 | 16.537609 | -4.24870698 | 0.88726 | -4.7886 | 1.7E-06 | 3.3E-05 |
| 904 | SYNGAP1 | 20.9164779 | -4.253093728 | 0.74023 | -5.7456 | 9.2E-09 | 3.3E-07 |
| 731 | DLEC1 | 5.01826353 | -4.255174084 | 1.50489 | -2.8276 | 0.00469 | 0.03003 |
| 476 | BTBD17 | 8.01110303 | -4.269690472 | 1.18446 | -3.6048 | 0.00031 | 0.00314 |
| 387 | SLCO3A1 | 13.2065307 | -4.270743092 | 0.9197 | -4.6436 | 3.4E-06 | 6.2E-05 |
| 355 | CRIP2 | 43.4044911 | -4.273211715 | 0.62559 | -6.8307 | 8.5E-12 | 6.5E-10 |
| 262 | CAPS2 | 4.72425681 | -4.274191795 | 1.37268 | -3.1138 | 0.00185 | 0.01408 |
| 593 | TSGA10 | 4.50027831 | -4.283609193 | 1.4818 | -2.8908 | 0.00384 | 0.02562 |
| 931 | CCN2 | 71.6194398 | -4.286769623 | 0.64781 | -6.6173 | 3.7E-11 | 2.4E-09 |
| 859 | SLC1A3 | 21.8370045 | -4.294933328 | 0.94692 | -4.5357 | 5.7E-06 | 9.7E-05 |
| 334 | ARMH4 | 4.56292828 | -4.303971521 | 1.45002 | -2.9682 | 0.003 | 0.02089 |
| 1061 | ENTREP1 | 6.49537377 | -4.307263689 | 1.29372 | -3.3294 | 0.00087 | 0.00748 |
| 558 | ZNF772 | 4.89827347 | -4.307955419 | 1.3794 | -3.1231 | 0.00179 | 0.01371 |
| 989 | MEST | 87.1971136 | -4.310638154 | 0.43696 | -9.865 | 5.9E-23 | 3.4E-20 |
| 1043 | RHPN1 | 23.2561447 | -4.310777957 | 0.73439 | -5.8699 | 4.4E-09 | 1.8E-07 |
| 749 | DOCK3 | 15.7950495 | -4.312170257 | 0.84752 | -5.088 | 3.6E-07 | 8.4E-06 |
| 55 | RPE65 | 10.8515249 | -4.323604716 | 0.97276 | -4.4447 | 8.8E-06 | 0.00014 |
| 132 | SLC16A9 | 39.5124783 | -4.339066974 | 0.59249 | -7.3234 | 2.4E-13 | 2.8E-11 |
| 924 | FAM229B | 18.1072474 | -4.350638874 | 0.83233 | -5.227 | 1.7E-07 | 4.4E-06 |
| 3 | MRPL20-AS1 | 3.15624805 | -4.372490956 | 1.65377 | -2.644 | 0.00819 | 0.04668 |
| 622 | DNAH7 | 9.01456483 | -4.372903975 | 1.06227 | -4.1166 | 3.8E-05 | 0.00051 |
| 77 | KCNN3 | 5.1338096 | -4.37780076 | 1.38636 | -3.1578 | 0.00159 | 0.01247 |
| 156 | CASC2 | 3.03293115 | -4.379129499 | 1.64222 | -2.6666 | 0.00766 | 0.04457 |
| 728 | NEK10 | 3.04399278 | -4.389207835 | 1.62198 | -2.7061 | 0.00681 | 0.04076 |
| 109 | MARK1 | 33.1144549 | -4.40749878 | 0.72148 | -6.109 | 1E-09 | 4.7E-08 |
| 228 | ENO2 | 26.6595105 | -4.407898897 | 0.81351 | -5.4184 | 6E-08 | 1.7E-06 |
| 655 | PRNP | 86.8397522 | -4.413623187 | 1.32758 | -3.3246 | 0.00089 | 0.00757 |
| 255 | GIHCG | 14.61717 | -4.415465971 | 0.88619 | -4.9825 | 6.3E-07 | 1.4E-05 |
| 652 | ENSG00000226644 | 3.12904833 | -4.426807278 | 1.60617 | -2.7561 | 0.00585 | 0.03587 |
| 913 | LINC00472 | 36.9130638 | -4.427639221 | 0.58795 | -7.5306 | 5.1E-14 | 7E-12 |
| 705 | SYNGR1 | 20.7736209 | -4.429973474 | 0.84076 | -5.269 | 1.4E-07 | 3.6E-06 |
| 300 | AMER2 | 3.2902803 | -4.43266439 | 1.69068 | -2.6218 | 0.00875 | 0.04905 |
| 59 | ABCA4 | 22.3018626 | -4.438011656 | 0.78909 | -5.6242 | 1.9E-08 | 6.2E-07 |

| | | | | | | | |
|---|---|---|---|---|---|---|---|
| 721 | FBLN2 | 13.1209721 | -4.439784088 | 1.01389 | -4.379 | 1.2E-05 | 0.00018 |
| 684 | ADARB1 | 20.8076171 | -4.440608676 | 0.85995 | -5.1638 | 2.4E-07 | 5.9E-06 |
| 990 | ENSG00000270953 | 3.16116958 | -4.445399791 | 1.67449 | -2.6548 | 0.00794 | 0.04568 |
| 64 | WDR47 | 7.52609969 | -4.455545005 | 1.21935 | -3.654 | 0.00026 | 0.00266 |
| 709 | SHISAL1 | 7.17000829 | -4.461053292 | 1.24946 | -3.5704 | 0.00036 | 0.0035 |
| 967 | CFAP69 | 19.2650402 | -4.461448805 | 0.80255 | -5.5591 | 2.7E-08 | 8.6E-07 |
| 333 | OTX2-AS1 | 13.2988351 | -4.461846167 | 0.97169 | -4.5918 | 4.4E-06 | 7.7E-05 |
| 668 | PLTP | 32.198718 | -4.467292926 | 0.76377 | -5.849 | 4.9E-09 | 2E-07 |
| 1135 | AKAP14 | 9.99115823 | -4.470652398 | 1.24753 | -3.5836 | 0.00034 | 0.00336 |
| 139 | CFAP70 | 5.10869265 | -4.475173061 | 1.46724 | -3.0501 | 0.00229 | 0.01675 |
| 393 | BAIAP3 | 3.27055006 | -4.49222885 | 1.62911 | -2.7575 | 0.00582 | 0.03574 |
| 907 | MDGA1 | 3.26969434 | -4.496504241 | 1.60678 | -2.7985 | 0.00513 | 0.03233 |
| 367 | MYO5A | 9.93784348 | -4.521542736 | 1.07582 | -4.2029 | 2.6E-05 | 0.00036 |
| 36 | CLDN19 | 3.48218782 | -4.522765743 | 1.56234 | -2.8949 | 0.00379 | 0.02535 |
| 656 | PRND | 12.3651492 | -4.524394004 | 0.99373 | -4.5529 | 5.3E-06 | 9E-05 |
| 954 | MATCAP2 | 8.18385087 | -4.526315887 | 1.14426 | -3.9557 | 7.6E-05 | 0.00093 |
| 750 | TNNC1 | 3.35181141 | -4.530431679 | 1.59196 | -2.8458 | 0.00443 | 0.02878 |
| 201 | CFAP300 | 8.62072154 | -4.53129226 | 1.20539 | -3.7592 | 0.00017 | 0.00187 |
| 879 | CXCL14 | 193.856599 | -4.536105771 | 0.41477 | -10.937 | 7.7E-28 | 7.6E-25 |
| 1150 | CSAG1 | 12.8793905 | -4.540759919 | 0.95755 | -4.7421 | 2.1E-06 | 4E-05 |
| 323 | MYH7 | 3.53377617 | -4.543709472 | 1.58122 | -2.8735 | 0.00406 | 0.02682 |
| 753 | IL17RD | 29.8297468 | -4.553165622 | 0.71663 | -6.3536 | 2.1E-10 | 1.2E-08 |
| 636 | GLB1L | 3.45165059 | -4.55400657 | 1.56857 | -2.9033 | 0.00369 | 0.0248 |
| 423 | SLC7A6 | 79.6810156 | -4.554695359 | 0.48308 | -9.4284 | 4.2E-21 | 1.6E-18 |
| 342 | LTBP2 | 7.62514727 | -4.564973315 | 1.37308 | -3.3246 | 0.00089 | 0.00757 |
| 93 | COLGALT2 | 3.43023937 | -4.565058518 | 1.59891 | -2.8551 | 0.0043 | 0.02811 |
| 206 | CRYAB | 18.0181604 | -4.567733821 | 0.90364 | -5.0548 | 4.3E-07 | 9.8E-06 |
| 1030 | TSPYL5 | 6.30486321 | -4.588048521 | 1.37723 | -3.3314 | 0.00086 | 0.00745 |
| 536 | CADM4 | 13.6277388 | -4.588615292 | 0.97061 | -4.7275 | 2.3E-06 | 4.3E-05 |
| 97 | PRELP | 3.51143428 | -4.598312136 | 1.58143 | -2.9077 | 0.00364 | 0.02449 |
| 69 | WNT2B | 14.1165836 | -4.600677438 | 1.13849 | -4.041 | 5.3E-05 | 0.00068 |
| 238 | PTHLH | 5.57478818 | -4.603403499 | 1.41238 | -3.2593 | 0.00112 | 0.00924 |
| 216 | ROBO3 | 7.9746227 | -4.604331409 | 1.20326 | -3.8266 | 0.00013 | 0.00147 |
| 827 | TACR3 | 3.72719307 | -4.625127908 | 1.68702 | -2.7416 | 0.00611 | 0.03712 |
| 601 | ARHGEF4 | 20.1621678 | -4.628537083 | 0.82741 | -5.594 | 2.2E-08 | 7.1E-07 |
| 326 | LINC01551 | 3.74001063 | -4.636863552 | 1.63676 | -2.8329 | 0.00461 | 0.02966 |
| 609 | SLC38A11 | 48.8583986 | -4.637352667 | 0.60432 | -7.6737 | 1.7E-14 | 2.6E-12 |
| 988 | CPA4 | 3.60343043 | -4.641871876 | 1.58052 | -2.9369 | 0.00331 | 0.02267 |
| 645 | IQCA1 | 16.2534136 | -4.655501084 | 0.89986 | -5.1736 | 2.3E-07 | 5.7E-06 |

| | | | | | | | |
|---|---|---|---|---|---|---|---|
| 492 | ZNF521 | 16.5369656 | -4.676869697 | 0.87173 | -5.3651 | 8.1E-08 | 2.3E-06 |
| 177 | STK33 | 19.1961767 | -4.681901655 | 0.84373 | -5.5491 | 2.9E-08 | 8.9E-07 |
| 1010 | LZTS1 | 3.71987042 | -4.682411874 | 1.67588 | -2.794 | 0.00521 | 0.03263 |
| 316 | SOX1 | 3.73822142 | -4.691473809 | 1.75587 | -2.6719 | 0.00754 | 0.04414 |
| 694 | RAB36 | 23.4956798 | -4.691537162 | 0.73819 | -6.3555 | 2.1E-10 | 1.2E-08 |
| 98 | LRRN2 | 15.1690728 | -4.704248688 | 0.86989 | -5.4079 | 6.4E-08 | 1.8E-06 |
| 102 | ENSG00000286619 | 18.0430008 | -4.708231495 | 0.84507 | -5.5714 | 2.5E-08 | 8E-07 |
| 284 | LINC00173 | 3.85750575 | -4.723529343 | 1.6098 | -2.9342 | 0.00334 | 0.02285 |
| 617 | PDE11A | 14.9498269 | -4.72577348 | 0.93142 | -5.0737 | 3.9E-07 | 8.9E-06 |
| 101 | SLC41A1 | 17.1521732 | -4.744136913 | 0.94059 | -5.0438 | 4.6E-07 | 1E-05 |
| 16 | KLHDC7A | 6.17389683 | -4.75357792 | 1.3664 | -3.4789 | 0.0005 | 0.00471 |
| 34 | COL9A2 | 159.857754 | -4.760837986 | 0.46884 | -10.154 | 3.2E-24 | 2.3E-21 |
| 88 | ILDR2 | 15.0494613 | -4.764491032 | 0.94554 | -5.0389 | 4.7E-07 | 1E-05 |
| 535 | GRIK5 | 8.89944126 | -4.766200573 | 1.16982 | -4.0743 | 4.6E-05 | 0.0006 |
| 126 | ENKUR | 6.30140655 | -4.77085897 | 1.35611 | -3.5181 | 0.00043 | 0.00416 |
| 277 | ABTB3 | 4.00120915 | -4.771482647 | 1.62128 | -2.943 | 0.00325 | 0.02233 |
| 1074 | BRINP1 | 3.95739235 | -4.776244473 | 1.51274 | -3.1573 | 0.00159 | 0.01248 |
| 932 | EYA4 | 8.94990539 | -4.779833813 | 1.16691 | -4.0961 | 4.2E-05 | 0.00055 |
| 76 | CA14 | 14.846387 | -4.788277786 | 0.98095 | -4.8813 | 1.1E-06 | 2.2E-05 |
| 794 | LINC02082 | 4.04471886 | -4.793143264 | 1.5052 | -3.1844 | 0.00145 | 0.01155 |
| 818 | IGFBP7-AS1 | 6.38919854 | -4.796776437 | 1.43888 | -3.3337 | 0.00086 | 0.00742 |
| 600 | CCDC74B | 2.37798662 | -4.806786448 | 1.80913 | -2.657 | 0.00789 | 0.04548 |
| 335 | RTN1 | 12.0405703 | -4.809382207 | 1.04715 | -4.5928 | 4.4E-06 | 7.7E-05 |
| 826 | DDIT4L | 14.8148837 | -4.81832179 | 1.12798 | -4.2716 | 1.9E-05 | 0.00028 |
| 305 | ENSG00000287650 | 4.06071051 | -4.819354689 | 1.57906 | -3.052 | 0.00227 | 0.01666 |
| 223 | WNT5B | 4.10054493 | -4.833100554 | 1.51977 | -3.1802 | 0.00147 | 0.01167 |
| 541 | PNMA8C | 2.43577876 | -4.838930128 | 1.83092 | -2.6429 | 0.00822 | 0.04675 |
| 866 | RASGRF2 | 2.44181914 | -4.842239989 | 1.83326 | -2.6413 | 0.00826 | 0.04687 |
| 803 | PDE6B-AS1 | 4.13593149 | -4.846599742 | 1.77287 | -2.7338 | 0.00626 | 0.03791 |
| 179 | DKK3 | 33.8562882 | -4.860095142 | 0.73182 | -6.6411 | 3.1E-11 | 2.1E-09 |
| 506 | NFATC1 | 4.39205359 | -4.879617956 | 1.50137 | -3.2501 | 0.00115 | 0.00949 |
| 131 | RASGEF1A | 12.5977919 | -4.881862094 | 1.12975 | -4.3212 | 1.6E-05 | 0.00023 |
| 821 | LINC01088 | 6.81829305 | -4.890913561 | 1.27323 | -3.8413 | 0.00012 | 0.0014 |
| 375 | LRRC49 | 6.83256165 | -4.892937163 | 1.31583 | -3.7185 | 0.0002 | 0.00214 |
| 485 | CLUL1 | 2.53499784 | -4.897467765 | 1.79323 | -2.7311 | 0.00631 | 0.03816 |
| 241 | PDZRN4 | 4.29406691 | -4.898046208 | 1.49894 | -3.2677 | 0.00108 | 0.009 |
| 1129 | CHRDL1 | 2.53071339 | -4.899951435 | 1.84114 | -2.6614 | 0.00778 | 0.04502 |
| 592 | FER1L5 | 2.54103821 | -4.900619604 | 1.78951 | -2.7385 | 0.00617 | 0.03742 |
| 306 | CCNA1 | 2.54707859 | -4.903772283 | 1.78892 | -2.7412 | 0.00612 | 0.03715 |

| | | | | | | | |
|---|---|---|---|---|---|---|---|
| 905 | ZBTB9 | 71.1025172 | -4.905180067 | 0.47421 | -10.344 | 4.5E-25 | 3.8E-22 |
| 734 | ZNF662 | 4.33832725 | -4.916568423 | 1.49206 | -3.2952 | 0.00098 | 0.00826 |
| 614 | GAD1 | 2.58332087 | -4.923014127 | 1.87425 | -2.6267 | 0.00862 | 0.0485 |
| 551 | PRKCG | 4.53388215 | -4.928176653 | 1.5456 | -3.1885 | 0.00143 | 0.01141 |
| 641 | DNER | 24.2354431 | -4.934283532 | 0.83486 | -5.9103 | 3.4E-09 | 1.4E-07 |
| 321 | SLC22A17 | 84.7312888 | -4.935064249 | 0.5072 | -9.7301 | 2.2E-22 | 1.1E-19 |
| 589 | SEMA4F | 2.6007497 | -4.936688032 | 1.77293 | -2.7845 | 0.00536 | 0.03348 |
| 587 | C2orf81 | 2.60989198 | -4.941376317 | 1.77638 | -2.7817 | 0.00541 | 0.03369 |
| 40 | CFAP57 | 4.59930718 | -4.951120691 | 1.47463 | -3.3575 | 0.00079 | 0.00688 |
| 127 | ODAD2 | 2.63715539 | -4.955178353 | 1.75096 | -2.83 | 0.00466 | 0.02987 |
| 271 | PAFAH1B2P2 | 2.64303235 | -4.958098049 | 1.79004 | -2.7698 | 0.00561 | 0.03475 |
| 857 | ENSG00000279737 | 2.64923615 | -4.96122775 | 1.74992 | -2.8351 | 0.00458 | 0.02954 |
| 1041 | LYNX1 | 4.50709239 | -4.971560784 | 1.47486 | -3.3709 | 0.00075 | 0.0066 |
| 505 | LINC00305 | 2.68270336 | -4.97815649 | 1.78409 | -2.7903 | 0.00527 | 0.03297 |
| 760 | CADM2 | 2.68286677 | -4.97865041 | 1.88136 | -2.6463 | 0.00814 | 0.04649 |
| 1036 | ENPP2 | 44.7745111 | -4.990731503 | 0.73776 | -6.7647 | 1.3E-11 | 9.7E-10 |
| 523 | CERS1 | 2.71188612 | -4.997260153 | 1.74337 | -2.8664 | 0.00415 | 0.02728 |
| 524 | PBX4 | 4.59541326 | -4.998017974 | 1.47478 | -3.389 | 0.0007 | 0.00626 |
| 493 | ENSG00000279439 | 2.72413029 | -5.003321661 | 1.729 | -2.8938 | 0.00381 | 0.02542 |
| 291 | CFAP251 | 13.7570888 | -5.010468172 | 1.09442 | -4.5782 | 4.7E-06 | 8.1E-05 |
| 720 | ATP2B2 | 24.5167865 | -5.017597064 | 0.79791 | -6.2884 | 3.2E-10 | 1.7E-08 |
| 522 | COMP | 2.7606994 | -5.021156421 | 1.74716 | -2.8739 | 0.00405 | 0.02681 |
| 1081 | PTGES | 7.48133929 | -5.024362281 | 1.29418 | -3.8823 | 0.0001 | 0.00121 |
| 63 | COL11A1 | 27.8554721 | -5.029841732 | 0.76926 | -6.5385 | 6.2E-11 | 3.9E-09 |
| 307 | SERTM1 | 2.78518775 | -5.03340544 | 1.89991 | -2.6493 | 0.00807 | 0.04626 |
| 980 | CTTNBP2 | 4.77410419 | -5.03800982 | 1.50089 | -3.3567 | 0.00079 | 0.00689 |
| 4 | PRDM16-DT | 32.4339588 | -5.046168299 | 0.69982 | -7.2106 | 5.6E-13 | 5.7E-11 |
| 784 | TM4SF18 | 4.74989596 | -5.051809649 | 1.48978 | -3.391 | 0.0007 | 0.00622 |
| 501 | MAPK4 | 2.83024546 | -5.05465657 | 1.80742 | -2.7966 | 0.00516 | 0.03244 |
| 748 | C3orf18 | 4.88049948 | -5.088787881 | 1.44634 | -3.5184 | 0.00043 | 0.00416 |
| 771 | CFAP100 | 17.9253938 | -5.09968432 | 1.03287 | -4.9374 | 7.9E-07 | 1.7E-05 |
| 1089 | CLCN4 | 11.2503279 | -5.102576362 | 1.08593 | -4.6988 | 2.6E-06 | 4.8E-05 |
| 916 | SNAP91 | 8.74080539 | -5.113375565 | 1.28421 | -3.9817 | 6.8E-05 | 0.00085 |
| 515 | COL5A3 | 31.6738897 | -5.117304588 | 0.71582 | -7.1489 | 8.7E-13 | 8.5E-11 |
| 935 | ADGB | 2.97415679 | -5.129539351 | 1.69 | -3.0352 | 0.0024 | 0.0174 |
| 269 | CFAP54 | 3.02574513 | -5.152467181 | 1.69325 | -3.0429 | 0.00234 | 0.01704 |
| 444 | TMEM88 | 5.10428169 | -5.152628358 | 1.45056 | -3.5522 | 0.00038 | 0.00372 |
| 120 | RPP38-DT | 3.05023348 | -5.163836378 | 1.82581 | -2.8282 | 0.00468 | 0.02999 |
| 976 | COL26A1 | 11.623541 | -5.165770612 | 1.22073 | -4.2317 | 2.3E-05 | 0.00032 |

| | | | | | | | |
|---|---|---|---|---|---|---|---|
| 141 | DYDC2 | 3.07112968 | -5.172587149 | 1.73972 | -2.9732 | 0.00295 | 0.02064 |
| 391 | RGS11 | 3.0883951 | -5.184038153 | 1.66132 | -3.1204 | 0.00181 | 0.01381 |
| 708 | EFCAB6 | 3.0883951 | -5.184038153 | 1.66132 | -3.1204 | 0.00181 | 0.01381 |
| 166 | KNDC1 | 3.09737396 | -5.187840792 | 1.69337 | -3.0636 | 0.00219 | 0.01617 |
| 717 | IL5RA | 3.1287584 | -5.198658652 | 1.88279 | -2.7611 | 0.00576 | 0.0355 |
| 1079 | LINC01503 | 5.27572492 | -5.201909957 | 1.41442 | -3.6778 | 0.00024 | 0.00246 |
| 860 | CDC20B | 8.30063577 | -5.202078372 | 1.42368 | -3.654 | 0.00026 | 0.00266 |
| 418 | TPPP3 | 35.9058443 | -5.202716951 | 0.76484 | -6.8024 | 1E-11 | 7.8E-10 |
| 920 | ARMC2 | 5.368355 | -5.211654233 | 1.54774 | -3.3673 | 0.00076 | 0.00667 |
| 754 | FAM107A | 16.476057 | -5.22382709 | 0.95946 | -5.4445 | 5.2E-08 | 1.5E-06 |
| 25 | FNDC5 | 11.9016727 | -5.233341904 | 1.39923 | -3.7402 | 0.00018 | 0.002 |
| 39 | CFAP144 | 12.3523916 | -5.245429304 | 1.16843 | -4.4893 | 7.1E-06 | 0.00012 |
| 643 | ARL4C | 79.8471585 | -5.247633249 | 0.54084 | -9.7027 | 2.9E-22 | 1.4E-19 |
| 304 | NBEA | 16.3540142 | -5.26119127 | 1.00891 | -5.2147 | 1.8E-07 | 4.7E-06 |
| 792 | WDR49 | 3.26646593 | -5.2629012 | 1.65428 | -3.1814 | 0.00147 | 0.01163 |
| 773 | EFCAB12 | 5.5113194 | -5.26551074 | 1.49847 | -3.5139 | 0.00044 | 0.0042 |
| 557 | ZNF667-AS1 | 3.27854669 | -5.267896021 | 1.67614 | -3.1429 | 0.00167 | 0.01298 |
| 842 | ACSL1 | 30.4855353 | -5.272709794 | 0.77074 | -6.8411 | 7.9E-12 | 6.1E-10 |
| 807 | ENSG00000287117 | 3.29079086 | -5.272907252 | 1.66815 | -3.1609 | 0.00157 | 0.01236 |
| 481 | CCDC40 | 20.753631 | -5.274557709 | 0.90206 | -5.8472 | 5E-09 | 2E-07 |
| 314 | ZIC5 | 36.1181208 | -5.282683695 | 1.98298 | -2.664 | 0.00772 | 0.04478 |
| 1062 | MAMDC2 | 3.31805428 | -5.284154633 | 1.70158 | -3.1054 | 0.0019 | 0.01438 |
| 808 | LINC01085 | 3.34152349 | -5.297227858 | 1.63585 | -3.2382 | 0.0012 | 0.00983 |
| 776 | RAB6B | 9.52354035 | -5.317229652 | 1.31221 | -4.0521 | 5.1E-05 | 0.00065 |
| 926 | SLC35F1 | 5.75523332 | -5.318695269 | 1.42137 | -3.742 | 0.00018 | 0.00199 |
| 1011 | PHYHIP | 3.39605032 | -5.318793993 | 1.63806 | -3.247 | 0.00117 | 0.00959 |
| 814 | KCTD8 | 3.40796766 | -5.323765265 | 1.73119 | -3.0752 | 0.0021 | 0.01566 |
| 1075 | MORN5 | 3.41417146 | -5.325985223 | 1.6604 | -3.2076 | 0.00134 | 0.01077 |
| 117 | ECHDC3 | 3.47196359 | -5.349374076 | 1.77797 | -3.0087 | 0.00262 | 0.01873 |
| 695 | C22orf15 | 3.47714826 | -5.354127178 | 1.62264 | -3.2996 | 0.00097 | 0.00817 |
| 1110 | PCDH11X | 3.49526939 | -5.361005366 | 1.61253 | -3.3246 | 0.00089 | 0.00757 |
| 186 | LARGE2 | 5.91658734 | -5.372503348 | 1.39525 | -3.8506 | 0.00012 | 0.00135 |
| 219 | NTM | 5.97852528 | -5.373022034 | 1.38965 | -3.8665 | 0.00011 | 0.00128 |
| 261 | KCNMB4 | 5.92203239 | -5.374553437 | 1.37642 | -3.9047 | 9.4E-05 | 0.00112 |
| 733 | CCDC13-AS1 | 3.54995964 | -5.382077365 | 1.66893 | -3.2249 | 0.00126 | 0.01024 |
| 1029 | GEM | 10.607716 | -5.382834848 | 1.39685 | -3.8536 | 0.00012 | 0.00134 |
| 664 | GDF5 | 3.57016354 | -5.392997647 | 1.59545 | -3.3802 | 0.00072 | 0.00643 |
| 256 | C12orf56 | 3.58930381 | -5.39723419 | 1.71776 | -3.142 | 0.00168 | 0.01301 |
| 745 | SLC38A3 | 6.25063903 | -5.414498115 | 1.55053 | -3.492 | 0.00048 | 0.00451 |

| | | | | | | | |
|---|---|---|---|---|---|---|---|
| 275 | CHST11 | 9.7474972 | -5.41488757 | 1.22914 | -4.4054 | 1.1E-05 | 0.00016 |
| 908 | TREM1 | 3.64574998 | -5.421240159 | 1.704 | -3.1815 | 0.00147 | 0.01163 |
| 686 | ENSG00000273796 | 3.65387313 | -5.427310078 | 1.62993 | -3.3298 | 0.00087 | 0.00747 |
| 403 | ENSG00000259871 | 9.73346132 | -5.428452182 | 1.48005 | -3.6677 | 0.00024 | 0.00254 |
| 1113 | TCEAL2 | 6.15479346 | -5.429000628 | 1.3784 | -3.9386 | 8.2E-05 | 0.00099 |
| 568 | TOGARAM2 | 3.69439984 | -5.439159616 | 1.68394 | -3.23 | 0.00124 | 0.01008 |
| 33 | RSPO1 | 3.68648462 | -5.441965251 | 1.73607 | -3.1346 | 0.00172 | 0.01329 |
| 452 | CCL2 | 48.427201 | -5.443479454 | 1.49085 | -3.6513 | 0.00026 | 0.00268 |
| 898 | GMPR | 10.243246 | -5.445062247 | 1.21463 | -4.4829 | 7.4E-06 | 0.00012 |
| 801 | CEP19 | 3.71668651 | -5.453491068 | 1.61487 | -3.3771 | 0.00073 | 0.0065 |
| 623 | C2orf66 | 6.25358874 | -5.454275373 | 1.3913 | -3.9203 | 8.8E-05 | 0.00106 |
| 267 | SLC6A15 | 6.32140364 | -5.456079893 | 1.38028 | -3.9529 | 7.7E-05 | 0.00094 |
| 925 | RSPH4A | 10.7495276 | -5.465690124 | 1.22861 | -4.4487 | 8.6E-06 | 0.00014 |
| 197 | FAM181B | 10.77267 | -5.470385505 | 1.22557 | -4.4636 | 8.1E-06 | 0.00013 |
| 292 | LRRC43 | 3.78137474 | -5.473546441 | 1.66675 | -3.284 | 0.00102 | 0.00855 |
| 100 | BLACAT1 | 3.77758055 | -5.47535402 | 1.57532 | -3.4757 | 0.00051 | 0.00476 |
| 483 | BAIAP2 | 3.80500738 | -5.485021759 | 1.57977 | -3.472 | 0.00052 | 0.00481 |
| 1066 | GAS1 | 3.83831117 | -5.496591619 | 1.56368 | -3.5152 | 0.00044 | 0.00419 |
| 236 | RASSF8-AS1 | 6.59180596 | -5.497497023 | 1.35112 | -4.0689 | 4.7E-05 | 0.00062 |
| 249 | MAP3K12 | 3.86867648 | -5.507135412 | 1.57365 | -3.4996 | 0.00047 | 0.00441 |
| 799 | OSTN | 3.89283799 | -5.515495323 | 1.58992 | -3.469 | 0.00052 | 0.00486 |
| 971 | TMEM130 | 3.89818607 | -5.520594054 | 1.61061 | -3.4276 | 0.00061 | 0.00555 |
| 805 | MSX1 | 49.8102426 | -5.533285651 | 0.64665 | -8.5569 | 1.2E-17 | 3E-15 |
| 42 | DMRTA2 | 3.9363477 | -5.536573323 | 1.60901 | -3.441 | 0.00058 | 0.00532 |
| 1137 | CCDC160 | 10.6558091 | -5.546249668 | 1.26871 | -4.3716 | 1.2E-05 | 0.00019 |
| 82 | ACKR1 | 3.97467274 | -5.551411038 | 1.74274 | -3.1854 | 0.00145 | 0.01152 |
| 133 | MRLN | 4.00499354 | -5.555207853 | 1.78922 | -3.1048 | 0.0019 | 0.0144 |
| 417 | RRAD | 6.72241483 | -5.556704481 | 1.61163 | -3.4479 | 0.00057 | 0.0052 |
| 399 | TVP23A | 4.00932249 | -5.561363843 | 1.55286 | -3.5814 | 0.00034 | 0.00338 |
| 629 | MDH1B | 15.5259837 | -5.570405954 | 1.05367 | -5.2867 | 1.2E-07 | 3.3E-06 |
| 341 | BBOF1 | 11.7434081 | -5.57207949 | 1.23283 | -4.5197 | 6.2E-06 | 0.0001 |
| 303 | KL | 4.05262427 | -5.572880033 | 1.62753 | -3.4241 | 0.00062 | 0.00561 |
| 828 | DKK2 | 22.6314017 | -5.583673169 | 0.93939 | -5.9439 | 2.8E-09 | 1.2E-07 |
| 631 | UNC80 | 6.88748861 | -5.597755667 | 1.3925 | -4.0199 | 5.8E-05 | 0.00074 |
| 413 | DRC7 | 4.12457994 | -5.599184135 | 1.55664 | -3.597 | 0.00032 | 0.00322 |
| 313 | ENSG00000286757 | 6.92240686 | -5.602174978 | 1.33705 | -4.1899 | 2.8E-05 | 0.00038 |
| 498 | DTNA | 16.3543058 | -5.611940812 | 1.01883 | -5.5082 | 3.6E-08 | 1.1E-06 |
| 385 | WDR93 | 4.24192015 | -5.639756417 | 1.54865 | -3.6417 | 0.00027 | 0.00277 |
| 586 | SLC4A5 | 7.2312694 | -5.654114595 | 1.35975 | -4.1582 | 3.2E-05 | 0.00044 |

| | | | | | | | |
|---|---|---|---|---|---|---|---|
| 106 | SPATA17 | 7.34274116 | -5.657058886 | 1.36747 | -4.1369 | 3.5E-05 | 0.00048 |
| 1098 | CHST7 | 7.18122686 | -5.65923572 | 1.43928 | -3.932 | 8.4E-05 | 0.00101 |
| 563 | KLHL29 | 16.556768 | -5.661566428 | 1.04129 | -5.437 | 5.4E-08 | 1.6E-06 |
| 427 | CLEC18B | 4.31873364 | -5.669549612 | 1.546 | -3.6672 | 0.00025 | 0.00255 |
| 1153 | ZNF185 | 7.27544663 | -5.673375551 | 1.34535 | -4.217 | 2.5E-05 | 0.00034 |
| 1064 | ANXA1 | 46.5686262 | -5.673893723 | 0.73934 | -7.6743 | 1.7E-14 | 2.6E-12 |
| 68 | PIFO | 59.8158262 | -5.675948543 | 0.71232 | -7.9682 | 1.6E-15 | 3E-13 |
| 757 | MITF | 32.731414 | -5.68632366 | 0.81541 | -6.9736 | 3.1E-12 | 2.6E-10 |
| 29 | COL8A2 | 7.34446381 | -5.6880142 | 1.33674 | -4.2551 | 2.1E-05 | 0.0003 |
| 174 | ART5 | 4.39582946 | -5.690537213 | 1.55115 | -3.6686 | 0.00024 | 0.00254 |
| 933 | SLC2A12 | 4.42896984 | -5.701121699 | 1.63803 | -3.4805 | 0.0005 | 0.00469 |
| 917 | CFAP206 | 4.42223716 | -5.701307984 | 1.50549 | -3.787 | 0.00015 | 0.00169 |
| 845 | C4orf47 | 18.077812 | -5.721790702 | 1.06866 | -5.3542 | 8.6E-08 | 2.4E-06 |
| 1052 | TYRP1 | 4.50488311 | -5.724977244 | 1.7313 | -3.3068 | 0.00094 | 0.008 |
| 439 | SERPINF1 | 90.9491424 | -5.740068703 | 0.65694 | -8.7376 | 2.4E-18 | 6.7E-16 |
| 91 | ANKRD45 | 4.55165813 | -5.742643967 | 1.49175 | -3.8496 | 0.00012 | 0.00136 |
| 584 | OTX1 | 4.5568428 | -5.746835831 | 1.55175 | -3.7034 | 0.00021 | 0.00226 |
| 1024 | CALB1 | 24.6384752 | -5.756156852 | 1.9014 | -3.0273 | 0.00247 | 0.01773 |
| 1056 | C9orf24 | 32.9657058 | -5.760923903 | 1.0352 | -5.565 | 2.6E-08 | 8.3E-07 |
| 1143 | ZIC3 | 12.6643769 | -5.763954052 | 1.17543 | -4.9037 | 9.4E-07 | 2E-05 |
| 409 | CHD9NB | 12.2453831 | -5.764558544 | 1.31267 | -4.3915 | 1.1E-05 | 0.00017 |
| 602 | CCDC74A | 12.5551737 | -5.784286721 | 1.16442 | -4.9675 | 6.8E-07 | 1.5E-05 |
| 144 | ENSG00000280660 | 8.03279354 | -5.821302833 | 1.35755 | -4.2881 | 1.8E-05 | 0.00026 |
| 149 | CFAP43 | 20.0170878 | -5.823449934 | 1.05989 | -5.4944 | 3.9E-08 | 1.2E-06 |
| 182 | FJX1 | 4.84119221 | -5.83059798 | 1.49057 | -3.9117 | 9.2E-05 | 0.00109 |
| 1131 | SNORA35 | 4.85209042 | -5.836164623 | 1.47366 | -3.9603 | 7.5E-05 | 0.00092 |
| 268 | LRRIQ1 | 54.8053286 | -5.83944453 | 0.67502 | -8.6508 | 5.1E-18 | 1.4E-15 |
| 283 | CFAP73 | 4.92298245 | -5.853462094 | 1.54792 | -3.7815 | 0.00016 | 0.00172 |
| 862 | CCNO | 24.9567262 | -5.868903321 | 0.98625 | -5.9507 | 2.7E-09 | 1.2E-07 |
| 575 | LINC01833 | 13.7007336 | -5.870185953 | 1.14562 | -5.124 | 3E-07 | 7.2E-06 |
| 181 | DCDC1 | 4.96975747 | -5.870807445 | 1.47764 | -3.9731 | 7.1E-05 | 0.00088 |
| 783 | CP | 27.0695415 | -5.881336019 | 0.97661 | -6.0222 | 1.7E-09 | 7.8E-08 |
| 820 | CDKL2 | 5.02395746 | -5.88522604 | 1.51263 | -3.8907 | 1E-04 | 0.00118 |
| 90 | F5 | 8.58099492 | -5.889142729 | 1.4416 | -4.0851 | 4.4E-05 | 0.00058 |
| 285 | HSPB8 | 5.04946494 | -5.890040652 | 1.5431 | -3.817 | 0.00014 | 0.00152 |
| 662 | BPIFB4 | 5.0981148 | -5.903904692 | 1.66051 | -3.5555 | 0.00038 | 0.00368 |
| 437 | CIBAR2 | 5.10917643 | -5.90782703 | 1.48419 | -3.9805 | 6.9E-05 | 0.00086 |
| 27 | TEKT2 | 5.10933984 | -5.907925995 | 1.4866 | -3.9741 | 7.1E-05 | 0.00088 |
| 1105 | CENPVL3 | 5.13113626 | -5.918004008 | 1.53376 | -3.8585 | 0.00011 | 0.00132 |

| | | | | | | | |
|---|---|---|---|---|---|---|---|
| 89 | CCDC181 | 5.14958423 | -5.923176221 | 1.46595 | -4.0405 | 5.3E-05 | 0.00068 |
| 123 | NEBL | 5.1920303 | -5.934132414 | 1.46015 | -4.0641 | 4.8E-05 | 0.00063 |
| 157 | LINC02944 | 5.24790309 | -5.94624847 | 1.4858 | -4.0021 | 6.3E-05 | 0.00079 |
| 183 | ENSG00000246250 | 8.85725719 | -5.952749978 | 1.34184 | -4.4362 | 9.2E-06 | 0.00014 |
| 62 | PALMD | 8.91532945 | -5.972156582 | 1.28578 | -4.6448 | 3.4E-06 | 6.1E-05 |
| 246 | AQP5 | 5.37026455 | -5.98164633 | 1.43137 | -4.179 | 2.9E-05 | 0.0004 |
| 453 | GAS2L2 | 5.3861395 | -5.984057033 | 1.60054 | -3.7388 | 0.00018 | 0.00201 |
| 374 | KIF23-AS1 | 5.39528178 | -5.98626316 | 1.5738 | -3.8037 | 0.00014 | 0.00158 |
| 594 | LONRF2 | 9.0567342 | -5.995897627 | 1.28626 | -4.6615 | 3.1E-06 | 5.7E-05 |
| 789 | MLF1 | 21.1327066 | -6.016438421 | 1.02755 | -5.8551 | 4.8E-09 | 1.9E-07 |
| 591 | TEKT4 | 5.55245641 | -6.027071468 | 1.52869 | -3.9426 | 8.1E-05 | 0.00098 |
| 112 | OBSCN-AS1 | 9.37806269 | -6.047210253 | 1.26535 | -4.7791 | 1.8E-06 | 3.4E-05 |
| 431 | DYNLRB2 | 5.65755244 | -6.054239994 | 1.50262 | -4.0291 | 5.6E-05 | 0.00071 |
| 982 | FEZF1 | 5.68032936 | -6.063553335 | 1.47369 | -4.1145 | 3.9E-05 | 0.00052 |
| 846 | PLEKHG4B | 9.53507391 | -6.071014213 | 1.25979 | -4.8191 | 1.4E-06 | 2.9E-05 |
| 79 | THBS3-AS1 | 15.607692 | -6.102274021 | 1.15329 | -5.2912 | 1.2E-07 | 3.3E-06 |
| 32 | DNALI1 | 31.2255008 | -6.155172197 | 0.91184 | -6.7503 | 1.5E-11 | 1.1E-09 |
| 949 | GPNMB | 16.5947673 | -6.163254485 | 1.16694 | -5.2815 | 1.3E-07 | 3.4E-06 |
| 475 | DNAI2 | 6.22303047 | -6.192459662 | 1.4897 | -4.1569 | 3.2E-05 | 0.00044 |
| 165 | CFAP46 | 10.5047481 | -6.211485642 | 1.26876 | -4.8957 | 9.8E-07 | 2E-05 |
| 959 | C7orf57 | 10.6059333 | -6.21781714 | 1.27426 | -4.8796 | 1.1E-06 | 2.2E-05 |
| 885 | KCTD16 | 6.35085892 | -6.221126288 | 1.60973 | -3.8647 | 0.00011 | 0.00129 |
| 604 | LRP1B | 6.35727064 | -6.226438404 | 1.44056 | -4.3222 | 1.5E-05 | 0.00023 |
| 553 | DNAAF3 | 6.38816483 | -6.231027031 | 1.39757 | -4.4585 | 8.3E-06 | 0.00013 |
| 868 | LINC00461 | 17.5119229 | -6.238734799 | 1.12709 | -5.5353 | 3.1E-08 | 9.6E-07 |
| 891 | KCNIP1 | 6.45750884 | -6.247544002 | 1.40665 | -4.4414 | 8.9E-06 | 0.00014 |
| 422 | TSNAXIP1 | 6.49619935 | -6.255557998 | 1.45775 | -4.2912 | 1.8E-05 | 0.00026 |
| 1095 | CFAP47 | 6.70614487 | -6.299853389 | 1.58754 | -3.9683 | 7.2E-05 | 0.0009 |
| 791 | SERPINI2 | 6.74773522 | -6.308858468 | 1.45275 | -4.3427 | 1.4E-05 | 0.00021 |
| 143 | LGI1 | 26.813432 | -6.331859356 | 0.95706 | -6.6159 | 3.7E-11 | 2.4E-09 |
| 552 | CACNG8 | 6.86389288 | -6.334736768 | 1.36448 | -4.6426 | 3.4E-06 | 6.2E-05 |
| 804 | DOK7 | 6.90013516 | -6.341900233 | 1.39352 | -4.551 | 5.3E-06 | 9.1E-05 |
| 220 | IGSF9B | 6.91931993 | -6.349052322 | 1.37848 | -4.6058 | 4.1E-06 | 7.3E-05 |
| 429 | CNTNAP4 | 11.617694 | -6.361119091 | 1.45042 | -4.3857 | 1.2E-05 | 0.00018 |
| 752 | CACNA2D3 | 6.9836427 | -6.361547951 | 1.38254 | -4.6014 | 4.2E-06 | 7.4E-05 |
| 434 | NECAB2 | 6.97625635 | -6.361591336 | 1.40797 | -4.5183 | 6.2E-06 | 0.0001 |
| 724 | EFHB | 7.03073867 | -6.367754252 | 1.45312 | -4.3821 | 1.2E-05 | 0.00018 |
| 858 | CAPSL | 7.10563282 | -6.383348348 | 1.41458 | -4.5125 | 6.4E-06 | 0.00011 |
| 297 | ENSG00000274373 | 11.9815376 | -6.384994914 | 1.23868 | -5.1547 | 2.5E-07 | 6.2E-06 |

| | | | | | | | |
|---|---|---|---|---|---|---|---|
| 432 | PKD1L2 | 7.11273683 | -6.387782406 | 1.34838 | -4.7374 | 2.2E-06 | 4.1E-05 |
| 83 | CFAP45 | 11.887672 | -6.390965145 | 1.25182 | -5.1054 | 3.3E-07 | 7.8E-06 |
| 732 | CCK | 7.1335944 | -6.392725919 | 1.55741 | -4.1047 | 4E-05 | 0.00054 |
| 495 | TTR | 7505.41531 | -6.406068833 | 0.9211 | -6.9548 | 3.5E-12 | 2.9E-10 |
| 608 | SLC4A10 | 20.4326658 | -6.422066357 | 1.12731 | -5.6968 | 1.2E-08 | 4.3E-07 |
| 896 | RNF182 | 12.198637 | -6.431027507 | 1.2247 | -5.2511 | 1.5E-07 | 3.9E-06 |
| 850 | ROPN1L | 7.37602663 | -6.438185646 | 1.35314 | -4.758 | 2E-06 | 3.7E-05 |
| 448 | DRC3 | 12.6366659 | -6.480409889 | 1.26938 | -5.1052 | 3.3E-07 | 7.8E-06 |
| 1082 | CFAP77 | 7.76824715 | -6.511887558 | 1.44671 | -4.5012 | 6.8E-06 | 0.00011 |
| 1022 | EYA1 | 7.83881822 | -6.527868126 | 1.35358 | -4.8227 | 1.4E-06 | 2.8E-05 |
| 673 | COL9A3 | 7.87436819 | -6.53513385 | 1.43931 | -4.5405 | 5.6E-06 | 9.5E-05 |
| 193 | CTSF | 8.00741994 | -6.557851341 | 1.31394 | -4.991 | 6E-07 | 1.3E-05 |
| 572 | CYP1B1 | 13.3234309 | -6.558958688 | 1.44594 | -4.5361 | 5.7E-06 | 9.7E-05 |
| 649 | KIF1A | 22.2017452 | -6.559103328 | 1.10051 | -5.9601 | 2.5E-09 | 1.1E-07 |
| 1021 | SULF1 | 153.889325 | -6.568091698 | 0.55327 | -11.871 | 1.7E-32 | 3.8E-29 |
| 57 | DNAI3 | 8.08141386 | -6.569166205 | 1.38394 | -4.7467 | 2.1E-06 | 3.9E-05 |
| 545 | ODAD1 | 31.0343589 | -6.576541032 | 0.99909 | -6.5825 | 4.6E-11 | 3E-09 |
| 603 | MAP3K19 | 8.17945039 | -6.587844474 | 1.31605 | -5.0058 | 5.6E-07 | 1.2E-05 |
| 1060 | BANCR | 8.18206204 | -6.588390451 | 1.3683 | -4.815 | 1.5E-06 | 2.9E-05 |
| 503 | RAX | 8.20324644 | -6.593332858 | 1.36023 | -4.8472 | 1.3E-06 | 2.5E-05 |
| 1090 | EGFL6 | 8.24728503 | -6.601964298 | 1.31809 | -5.0087 | 5.5E-07 | 1.2E-05 |
| 103 | SYT14 | 8.32993098 | -6.613867051 | 1.34215 | -4.9278 | 8.3E-07 | 1.7E-05 |
| 435 | DNAAF1 | 42.2377252 | -6.623601161 | 0.92328 | -7.174 | 7.3E-13 | 7.2E-11 |
| 612 | CFAP210 | 13.9499386 | -6.62437769 | 1.23398 | -5.3683 | 7.9E-08 | 2.2E-06 |
| 798 | GMNC | 8.43016918 | -6.630219485 | 1.36583 | -4.8544 | 1.2E-06 | 2.4E-05 |
| 689 | S100B | 8.46865764 | -6.637472348 | 1.33073 | -4.9878 | 6.1E-07 | 1.3E-05 |
| 996 | SLC13A4 | 101.549525 | -6.641228456 | 1.49534 | -4.4413 | 8.9E-06 | 0.00014 |
| 886 | PPP2R2B | 8.50249619 | -6.646568964 | 1.40034 | -4.7464 | 2.1E-06 | 3.9E-05 |
| 1054 | LINC01239 | 8.63950554 | -6.66708333 | 1.33402 | -4.9977 | 5.8E-07 | 1.3E-05 |
| 974 | SERPINE1 | 14.9541492 | -6.714373047 | 1.37694 | -4.8763 | 1.1E-06 | 2.2E-05 |
| 909 | RSPH9 | 8.95940494 | -6.718707253 | 1.32282 | -5.0791 | 3.8E-07 | 8.7E-06 |
| 767 | CFAP91 | 14.8823595 | -6.718987621 | 1.19638 | -5.6161 | 2E-08 | 6.4E-07 |
| 800 | ATP13A5 | 9.06829517 | -6.735977914 | 1.47094 | -4.5794 | 4.7E-06 | 8.1E-05 |
| 489 | LDLRAD4 | 9.1774736 | -6.75285046 | 1.36257 | -4.9559 | 7.2E-07 | 1.5E-05 |
| 983 | FEZF1-AS1 | 9.18323165 | -6.757105201 | 1.29168 | -5.2312 | 1.7E-07 | 4.3E-06 |
| 80 | SYT11 | 24.518021 | -6.762068172 | 1.09695 | -6.1644 | 7.1E-10 | 3.4E-08 |
| 746 | ZMYND10 | 9.36125798 | -6.781049842 | 1.37563 | -4.9294 | 8.2E-07 | 1.7E-05 |
| 278 | FOXN4 | 9.49398877 | -6.804304289 | 1.27558 | -5.3343 | 9.6E-08 | 2.6E-06 |
| 315 | ZIC2 | 66.8850017 | -6.828162664 | 1.34555 | -5.0746 | 3.9E-07 | 8.9E-06 |

| | | | | | | | |
|---|---|---|---|---|---|---|---|
| 404 | VWA3A | 26.6803201 | -6.880617898 | 1.10714 | -6.2148 | 5.1E-10 | 2.6E-08 |
| 654 | SPEF1 | 10.0352608 | -6.881628432 | 1.32631 | -5.1886 | 2.1E-07 | 5.3E-06 |
| 1134 | ZCCHC12 | 10.0164918 | -6.882748242 | 1.39762 | -4.9246 | 8.5E-07 | 1.8E-05 |
| 790 | ZBBX | 10.2992874 | -6.919716847 | 1.28725 | -5.3756 | 7.6E-08 | 2.2E-06 |
| 18 | UBXN10 | 10.4194026 | -6.936619811 | 1.31724 | -5.266 | 1.4E-07 | 3.6E-06 |
| 1048 | DMRT3 | 10.5686621 | -6.95894077 | 1.25332 | -5.5524 | 2.8E-08 | 8.9E-07 |
| 124 | SPAG6 | 10.7703596 | -6.983696781 | 1.43093 | -4.8805 | 1.1E-06 | 2.2E-05 |
| 944 | C6orf118 | 18.1636367 | -7.007260708 | 1.24177 | -5.643 | 1.7E-08 | 5.7E-07 |
| 598 | NPHP1 | 18.6735635 | -7.03714066 | 1.17189 | -6.0049 | 1.9E-09 | 8.6E-08 |
| 961 | GALNT17 | 18.6352824 | -7.044924234 | 1.28879 | -5.4663 | 4.6E-08 | 1.4E-06 |
| 576 | SIX3 | 31.2941548 | -7.075131738 | 1.05793 | -6.6877 | 2.3E-11 | 1.6E-09 |
| 478 | FOXJ1 | 44.1257665 | -7.087725685 | 0.96612 | -7.3363 | 2.2E-13 | 2.6E-11 |
| 952 | AQP1 | 32.664845 | -7.104767983 | 1.17919 | -6.0251 | 1.7E-09 | 7.7E-08 |
| 984 | GRM8 | 11.870056 | -7.125460163 | 1.23498 | -5.7697 | 7.9E-09 | 3E-07 |
| 87 | LMX1A | 19.9292906 | -7.133870512 | 1.15845 | -6.1581 | 7.4E-10 | 3.5E-08 |
| 72 | SPAG17 | 12.1924037 | -7.163548006 | 1.25949 | -5.6877 | 1.3E-08 | 4.5E-07 |
| 473 | SLC16A6 | 20.7560887 | -7.190958 | 1.35766 | -5.2966 | 1.2E-07 | 3.2E-06 |
| 849 | CFAP90 | 12.610503 | -7.212591025 | 1.23188 | -5.855 | 4.8E-09 | 1.9E-07 |
| 153 | EMX2OS | 12.7830237 | -7.231882806 | 1.23706 | -5.846 | 5E-09 | 2E-07 |
| 817 | IGFBP7 | 408.320998 | -7.251713845 | 0.97069 | -7.4707 | 8E-14 | 1E-11 |
| 440 | TEKT1 | 13.1874884 | -7.275986124 | 1.26785 | -5.7388 | 9.5E-09 | 3.4E-07 |
| 518 | NWD1 | 13.3700457 | -7.295655068 | 1.43046 | -5.1002 | 3.4E-07 | 7.9E-06 |
| 1057 | DNAI1 | 13.4257996 | -7.301670387 | 1.3225 | -5.5211 | 3.4E-08 | 1E-06 |
| 1151 | MAGEA12 | 13.653465 | -7.328338579 | 1.23178 | -5.9494 | 2.7E-09 | 1.2E-07 |
| 426 | HYDIN | 14.0340505 | -7.365594057 | 1.30595 | -5.64 | 1.7E-08 | 5.7E-07 |
| 870 | FAM81B | 14.7291129 | -7.435510512 | 1.29333 | -5.7491 | 9E-09 | 3.3E-07 |
| 692 | CLDN5 | 56.8902919 | -7.468031346 | 1.01858 | -7.3318 | 2.3E-13 | 2.6E-11 |
| 446 | CFAP52 | 15.1908409 | -7.479869404 | 1.30851 | -5.7163 | 1.1E-08 | 3.8E-07 |
| 871 | CDO1 | 41.5427839 | -7.484682066 | 1.06244 | -7.0448 | 1.9E-12 | 1.6E-10 |
| 596 | ECRG4 | 15.3742598 | -7.49726686 | 1.28389 | -5.8395 | 5.2E-09 | 2E-07 |
| 880 | NME5 | 15.4581773 | -7.506567206 | 1.19426 | -6.2855 | 3.3E-10 | 1.7E-08 |
| 205 | HOATZ | 15.4976404 | -7.508786333 | 1.28129 | -5.8603 | 4.6E-09 | 1.9E-07 |
| 1125 | FRMPD3 | 15.5228655 | -7.511645156 | 1.22066 | -6.1538 | 7.6E-10 | 3.6E-08 |
| 640 | DAW1 | 15.8050133 | -7.538027749 | 1.21186 | -6.2202 | 5E-10 | 2.5E-08 |
| 702 | FAM227A | 15.9689651 | -7.554009146 | 1.19241 | -6.3351 | 2.4E-10 | 1.3E-08 |
| 194 | FOLR1 | 28.7551223 | -7.674132971 | 1.29145 | -5.9423 | 2.8E-09 | 1.2E-07 |
| 1034 | RSPO2 | 28.7953872 | -7.67923436 | 1.20342 | -6.3812 | 1.8E-10 | 1E-08 |
| 154 | EMX2 | 19.774834 | -7.86365682 | 1.23639 | -6.3602 | 2E-10 | 1.1E-08 |
| 122 | SLC39A12 | 20.0765619 | -7.882529162 | 1.2243 | -6.4384 | 1.2E-10 | 7.2E-09 |

| | | | | | | | |
|---|---|---|---|---|---|---|---|
| 96 | CHI3L1 | 20.9737881 | -7.948687429 | 1.30043 | -6.1123 | 9.8E-10 | 4.7E-08 |
| 65 | CFAP276 | 21.1003392 | -7.954303906 | 1.23233 | -6.4547 | 1.1E-10 | 6.5E-09 |
| 564 | DRC1 | 22.0133068 | -8.015407691 | 1.23803 | -6.4743 | 9.5E-11 | 5.8E-09 |
| 1017 | CLXN | 38.4665872 | -8.089904376 | 1.16914 | -6.9196 | 4.5E-12 | 3.6E-10 |
| 51 | DYNLT5 | 27.2246794 | -8.322319348 | 1.25625 | -6.6247 | 3.5E-11 | 2.3E-09 |
| 1044 | MAPK15 | 74.6361336 | -8.374699402 | 1.06488 | -7.8645 | 3.7E-15 | 6.6E-13 |
| 125 | ARMC3 | 28.3712698 | -8.381727337 | 1.18757 | -7.0579 | 1.7E-12 | 1.5E-10 |
| 442 | ATP1B2 | 28.6281982 | -8.395215997 | 1.14861 | -7.309 | 2.7E-13 | 3E-11 |
| 781 | ZIC4 | 30.419038 | -8.483671607 | 1.12278 | -7.5559 | 4.2E-14 | 6E-12 |
| 1058 | SPAG8 | 30.987731 | -8.509064186 | 1.19105 | -7.1442 | 9.1E-13 | 8.7E-11 |
| 1063 | TRPM3 | 191.316081 | -8.712783914 | 0.77911 | -11.183 | 4.9E-29 | 5.7E-26 |
| 430 | VAT1L | 60.9738762 | -8.76203422 | 1.1347 | -7.7219 | 1.1E-14 | 1.9E-12 |
| 642 | ECEL1 | 40.6215305 | -8.901235696 | 1.94527 | -4.5758 | 4.7E-06 | 8.2E-05 |
| 682 | PCP4 | 54.5783576 | -9.327699546 | 1.10224 | -8.4625 | 2.6E-17 | 6.2E-15 |
| 782 | ZIC1 | 76.6514263 | -9.817582137 | 1.09585 | -8.9589 | 3.3E-19 | 1E-16 |
| 1128 | ENSG00000261409 | 76.9630836 | -9.823503997 | 1.10109 | -8.9216 | 4.6E-19 | 1.4E-16 |
| 332 | OTX2 | 141.498555 | -9.977666118 | 1.06428 | -9.3751 | 6.9E-21 | 2.6E-18 |
| 840 | GPM6A | 93.7871265 | -10.10779046 | 1.08582 | -9.3089 | 1.3E-20 | 4.7E-18 |
| 465 | WFIKKN2 | 175.958135 | -11.01589124 | 2.36537 | -4.6572 | 3.2E-06 | 5.8E-05 |
| 1130 | HTR2C | 716.973897 | -13.04260085 | 1.05587 | -12.352 | 4.7E-35 | 1.3E-31 |

TableS5

| Pathways upregulated in Undiff compared to Gland_A | | | |
|---|---|---|---|
| **KEGG pathway** | **No. of genes** | **%** | **p-value** |
| DNA replication | 14 | 1 | 1.50E-07 |
| Cell cycle | 28 | 2.1 | 1.90E-06 |
| Synaptic vesicle cycle | 17 | 1.3 | 2.60E-05 |
| Motor proteins | 26 | 1.9 | 5.50E-04 |
| Dopaminergic synapse | 19 | 1.4 | 1.80E-03 |
| Cholinergic synapse | 17 | 1.3 | 2.10E-03 |
| Purine metabolism | 18 | 1.3 | 3.10E-03 |
| Morphine addiction | 14 | 1 | 4.90E-03 |
| Human immunodeficiency virus 1 infection | 24 | 1.8 | 8.70E-03 |
| Endocrine resistance | 14 | 1 | 9.20E-03 |
| MAPK signaling pathway | 31 | 2.3 | 1.00E-02 |
| Cell adhesion molecules | 19 | 1.4 | 1.10E-02 |
| Axon guidance | 21 | 1.5 | 1.20E-02 |
| Nicotine addiction | 8 | 0.6 | 1.20E-02 |
| Mismatch repair | 6 | 0.4 | 1.30E-02 |
| Glutamatergic synapse | 15 | 1.1 | 1.50E-02 |
| Ras signaling pathway | 25 | 1.8 | 1.60E-02 |
| ErbB signaling pathway | 12 | 0.9 | 1.90E-02 |
| Regulation of actin cytoskeleton | 24 | 1.8 | 2.00E-02 |
| Insulin secretion | 12 | 0.9 | 2.00E-02 |
| Carbon metabolism | 14 | 1 | 3.10E-02 |
| Epithelial cell signaling in Helicobacter pylori infection | 10 | 0.7 | 3.30E-02 |
| Focal adhesion | 21 | 1.5 | 3.50E-02 |
| Bladder cancer | 7 | 0.5 | 4.40E-02 |
| HIF-1 signaling pathway | 13 | 1 | 4.50E-02 |
| Rap1 signaling pathway | 21 | 1.5 | 4.70E-02 |
| Salmonella infection | 24 | 1.8 | 4.70E-02 |
| Biosynthesis of amino acids | 10 | 0.7 | 4.90E-02 |
| Glioma | 10 | 0.7 | 4.90E-02 |
| Endocrine and other factor-regulated calcium reabsorption | 8 | 0.6 | 5.00E-02 |

TableS6

| Pathways upregulated in Gland_A compared to Undiff | | | |
|---|---|---|---|
| **KEGG pathway** | **No. of genes** | **%** | **p-value** |
| Metabolic pathways | 281 | 13.4 | 1.50E-14 |
| ECM-receptor interaction | 30 | 1.4 | 3.10E-07 |
| Focal adhesion | 49 | 2.3 | 3.10E-06 |
| Human papillomavirus infection | 66 | 3.1 | 4.70E-05 |
| Pathogenic Escherichia coli infection | 44 | 2.1 | 7.90E-05 |
| Proteoglycans in cancer | 45 | 2.1 | 9.90E-05 |
| PPAR signaling pathway | 22 | 1 | 1.60E-04 |
| Insulin resistance | 28 | 1.3 | 1.70E-04 |
| Leukocyte transendothelial migration | 29 | 1.4 | 1.80E-04 |
| Tight junction | 38 | 1.8 | 2.30E-04 |
| Drug metabolism - cytochrome P450 | 21 | 1 | 2.60E-04 |
| Glycerolipid metabolism | 19 | 0.9 | 3.50E-04 |
| Fat digestion and absorption | 15 | 0.7 | 3.60E-04 |
| Fructose and mannose metabolism | 13 | 0.6 | 4.10E-04 |
| Sulfur metabolism | 7 | 0.3 | 4.30E-04 |
| Pathways in cancer | 91 | 4.3 | 5.20E-04 |
| Mineral absorption | 18 | 0.9 | 5.70E-04 |
| Lipid and atherosclerosis | 44 | 2.1 | 5.90E-04 |
| PI3K-Akt signaling pathway | 65 | 3.1 | 0.00061 |
| Lysosome | 30 | 1.4 | 9.90E-04 |
| Biosynthesis of nucleotide sugars | 13 | 0.6 | 9.90E-04 |
| Protein digestion and absorption | 25 | 1.2 | 1.10E-03 |
| Sphingolipid metabolism | 16 | 0.8 | 1.20E-03 |
| Regulation of actin cytoskeleton | 45 | 2.1 | 1.20E-03 |
| Adherens junction | 23 | 1.1 | 0.0014 |
| Amino sugar and nucleotide sugar metabolism | 15 | 0.7 | 1.60E-03 |
| Arrhythmogenic right ventricular cardiomyopathy | 20 | 1 | 1.70E-03 |
| TNF signaling pathway | 26 | 1.2 | 2.20E-03 |
| Glycosphingolipid biosynthesis - lacto and neolacto series | 10 | 0.5 | 3.40E-03 |
| Axon guidance | 36 | 1.7 | 3.70E-03 |
| AGE-RAGE signaling pathway in diabetic complications | 23 | 1.1 | 3.80E-03 |
| Hepatitis C | 32 | 1.5 | 4.00E-03 |
| Fatty acid degradation | 13 | 0.6 | 4.10E-03 |
| Metabolism of xenobiotics by cytochrome P450 | 19 | 0.9 | 5.00E-03 |
| NOD-like receptor signaling pathway | 36 | 1.7 | 5.20E-03 |
| Salmonella infection | 45 | 2.1 | 6.30E-03 |

| | | | |
|---|---|---|---|
| Bladder cancer | 12 | 0.6 | 0.0081 |
| Mucin type O-glycan biosynthesis | 11 | 0.5 | 8.80E-03 |
| beta-Alanine metabolism | 10 | 0.5 | 9.40E-03 |
| Various types of N-glycan biosynthesis | 12 | 0.6 | 9.80E-03 |
| Amoebiasis | 22 | 1 | 1.00E-02 |
| Galactose metabolism | 10 | 0.5 | 1.20E-02 |
| Propanoate metabolism | 10 | 0.5 | 1.20E-02 |
| Nitrogen metabolism | 7 | 0.3 | 1.20E-02 |
| Histidine metabolism | 8 | 0.4 | 1.30E-02 |
| Arachidonic acid metabolism | 15 | 0.7 | 1.30E-02 |
| Glycolysis / Gluconeogenesis | 16 | 0.8 | 1.30E-02 |
| Arginine and proline metabolism | 13 | 0.6 | 1.50E-02 |
| Necroptosis | 30 | 1.4 | 1.60E-02 |
| Chemical carcinogenesis - DNA adducts | 16 | 0.8 | 1.70E-02 |
| Glutathione metabolism | 14 | 0.7 | 1.70E-02 |
| Epstein-Barr virus infection | 36 | 1.7 | 1.80E-02 |
| TGF-beta signaling pathway | 22 | 1 | 1.90E-02 |
| Phospholipase D signaling pathway | 28 | 1.3 | 1.90E-02 |
| Apoptosis | 26 | 1.2 | 2.20E-02 |
| Ferroptosis | 11 | 0.5 | 2.20E-02 |
| Pyruvate metabolism | 12 | 0.6 | 2.30E-02 |
| Cell adhesion molecules | 29 | 1.4 | 2.30E-02 |
| C-type lectin receptor signaling pathway | 21 | 1 | 2.40E-02 |
| Tryptophan metabolism | 11 | 0.5 | 2.60E-02 |
| Valine, leucine and isoleucine degradation | 12 | 0.6 | 2.60E-02 |
| Platinum drug resistance | 16 | 0.8 | 0.027 |
| Diabetic cardiomyopathy | 35 | 1.7 | 3.10E-02 |
| Maturity onset diabetes of the young | 8 | 0.4 | 3.20E-02 |
| Peroxisome | 17 | 0.8 | 3.60E-02 |
| Butanoate metabolism | 8 | 0.4 | 3.80E-02 |
| Fatty acid metabolism | 13 | 0.6 | 3.90E-02 |
| Chemical carcinogenesis - reactive oxygen species | 37 | 1.8 | 4.50E-02 |
| Fluid shear stress and atherosclerosis | 25 | 1.2 | 4.70E-02 |
| Toxoplasmosis | 21 | 1 | 4.90E-02 |

TableS7

| Pathways upregulated in Undiff compared to Gland_B | | | |
|---|---|---|---|
| **KEGG pathway** | **No. of genes** | **%** | **p-value** |
| Cell cycle | 32 | 3 | 4.00E-11 |
| Cellular senescence | 20 | 1.9 | 3.20E-04 |
| DNA replication | 9 | 0.8 | 3.50E-04 |
| Glutamatergic synapse | 16 | 1.5 | 6.60E-04 |
| Base excision repair | 9 | 0.8 | 1.40E-03 |
| Synaptic vesicle cycle | 12 | 1.1 | 1.80E-03 |
| Amphetamine addiction | 11 | 1 | 2.30E-03 |
| Nicotine addiction | 8 | 0.8 | 3.60E-03 |
| Motor proteins | 20 | 1.9 | 4.10E-03 |
| Cholinergic synapse | 14 | 1.3 | 4.60E-03 |
| Axon guidance | 19 | 1.8 | 4.90E-03 |
| Cell adhesion molecules | 17 | 1.6 | 5.90E-03 |
| Epstein-Barr virus infection | 20 | 1.9 | 6.70E-03 |
| Circadian entrainment | 12 | 1.1 | 9.70E-03 |
| Endocrine resistance | 12 | 1.1 | 1.00E-02 |
| MAPK signaling pathway | 26 | 2.5 | 1.10E-02 |
| Purine metabolism | 14 | 1.3 | 1.30E-02 |
| Chronic myeloid leukemia | 10 | 0.9 | 1.40E-02 |
| GABAergic synapse | 11 | 1 | 1.40E-02 |
| Th1 and Th2 cell differentiation | 11 | 1 | 1.80E-02 |
| Human immunodeficiency virus 1 infection | 19 | 1.8 | 2.20E-02 |
| Spinocerebellar ataxia | 14 | 1.3 | 2.90E-02 |
| Carbon metabolism | 12 | 1.1 | 3.10E-02 |
| Human T-cell leukemia virus 1 infection | 19 | 1.8 | 3.30E-02 |
| Oocyte meiosis | 13 | 1.2 | 3.40E-02 |
| Glioma | 9 | 0.8 | 3.50E-02 |
| Human cytomegalovirus infection | 19 | 1.8 | 3.70E-02 |
| Leishmaniasis | 9 | 0.8 | 4.00E-02 |
| Small cell lung cancer | 10 | 0.9 | 4.30E-02 |
| Oxytocin signaling pathway | 14 | 1.3 | 4.90E-02 |

TableS8

| Pathways upregulated in Gland_B compared to Undiff | | | |
|---|---|---|---|
| **KEGG pathway** | **No. of genes** | **%** | **p-value** |
| ECM-receptor interaction | 33 | 2 | 6.40.E-14 |
| Human papillomavirus infection | 65 | 4 | 1.00.E-11 |
| Focal adhesion | 47 | 2.9 | 4.00.E-11 |
| Hippo signaling pathway | 37 | 2.3 | 4.00.E-09 |
| Protein digestion and absorption | 29 | 1.8 | 4.00.E-09 |
| Proteoglycans in cancer | 42 | 2.6 | 2.30.E-08 |
| PI3K-Akt signaling pathway | 60 | 3.6 | 2.60.E-08 |
| Pathways in cancer | 78 | 4.7 | 9.70.E-08 |
| Signaling pathways regulating pluripotency of stem cells | 32 | 1.9 | 1.90.E-07 |
| Wnt signaling pathway | 35 | 2.1 | 4.30.E-07 |
| Arrhythmogenic right ventricular cardiomyopathy | 21 | 1.3 | 1.50.E-06 |
| TGF-beta signaling pathway | 24 | 1.5 | 9.90.E-06 |
| Axon guidance | 32 | 1.9 | 3.70.E-05 |
| Breast cancer | 27 | 1.6 | 8.20.E-05 |
| Basal cell carcinoma | 16 | 1 | 9.10.E-05 |
| AGE-RAGE signaling pathway in diabetic complications | 21 | 1.3 | 9.50.E-05 |
| Gastric cancer | 27 | 1.6 | 1.00.E-04 |
| Adherens junction | 20 | 1.2 | 1.00.E-04 |
| Hepatocellular carcinoma | 29 | 1.8 | 1.30.E-04 |
| Prostate cancer | 20 | 1.2 | 1.90.E-04 |
| Regulation of actin cytoskeleton | 35 | 2.1 | 2.70.E-04 |
| Tight junction | 27 | 1.6 | 8.00.E-04 |
| Melanogenesis | 18 | 1.1 | 2.40.E-03 |
| Cellular senescence | 24 | 1.5 | 2.80.E-03 |
| Salmonella infection | 33 | 2 | 4.50.E-03 |
| Hypertrophic cardiomyopathy | 16 | 1 | 4.60.E-03 |
| Small cell lung cancer | 16 | 1 | 5.70.E-03 |
| Calcium signaling pathway | 33 | 2 | 5.70.E-03 |
| Amoebiasis | 17 | 1 | 6.40.E-03 |
| Aldosterone-regulated sodium reabsorption | 9 | 0.5 | 7.50.E-03 |
| Complement and coagulation cascades | 15 | 0.9 | 7.50.E-03 |
| Lysosome | 20 | 1.2 | 8.00.E-03 |
| Dilated cardiomyopathy | 16 | 1 | 8.40.E-03 |
| Other types of O-glycan biosynthesis | 10 | 0.6 | 1.10.E-02 |
| Kaposi sarcoma-associated herpesvirus infection | 26 | 1.6 | 1.10.E-02 |
| Apoptosis | 20 | 1.2 | 1.10.E-02 |

| | | | |
|---|---|---|---|
| mTOR signaling pathway | 22 | 1.3 | 1.20.E-02 |
| Endometrial cancer | 11 | 0.7 | 1.50.E-02 |
| Hepatitis B | 22 | 1.3 | 1.70.E-02 |
| Colorectal cancer | 14 | 0.9 | 1.80.E-02 |
| Leukocyte transendothelial migration | 17 | 1 | 1.80.E-02 |
| Motor proteins | 25 | 1.5 | 1.90.E-02 |
| Transcriptional misregulation in cancer | 25 | 1.5 | 1.90.E-02 |
| Mucin type O-glycan biosynthesis | 8 | 0.5 | 2.10.E-02 |
| EGFR tyrosine kinase inhibitor resistance | 13 | 0.8 | 2.10.E-02 |
| Pathogenic Escherichia coli infection | 25 | 1.5 | 2.30.E-02 |
| Hepatitis C | 21 | 1.3 | 2.40.E-02 |
| MAPK signaling pathway | 35 | 2.1 | 2.40.E-02 |
| Pathways of neurodegeneration - multiple diseases | 51 | 3.1 | 2.40.E-02 |
| Thyroid cancer | 8 | 0.5 | 2.50.E-02 |
| Melanoma | 12 | 0.7 | 2.60.E-02 |
| Human T-cell leukemia virus 1 infection | 27 | 1.6 | 2.90.E-02 |
| Th1 and Th2 cell differentiation | 14 | 0.9 | 2.90.E-02 |
| Thyroid hormone signaling pathway | 17 | 1 | 3.00.E-02 |
| Protein processing in endoplasmic reticulum | 22 | 1.3 | 3.50.E-02 |
| Pancreatic cancer | 12 | 0.7 | 3.70.E-02 |
| Acute myeloid leukemia | 11 | 0.7 | 3.80.E-02 |
| Cell adhesion molecules | 20 | 1.2 | 4.30.E-02 |
| Mineral absorption | 10 | 0.6 | 4.60.E-02 |
| Cholesterol metabolism | 9 | 0.5 | 4.60.E-02 |

TableS9

| Pathways upregulated in Gland_A compared to Gland_B | | | |
|---|---|---|---|
| **KEGG pathway** | **No. of genes** | **%** | **p-value** |
| Metabolic pathways | 185 | 15.1 | 5.60E-15 |
| PPAR signaling pathway | 19 | 1.5 | 3.10E-06 |
| Sulfur metabolism | 7 | 0.6 | 2.00E-05 |
| Glycerolipid metabolism | 16 | 1.3 | 2.30E-05 |
| Fat digestion and absorption | 13 | 1.1 | 2.70E-05 |
| Drug metabolism - cytochrome P450 | 17 | 1.4 | 3.10E-05 |
| Fructose and mannose metabolism | 11 | 0.9 | 7.90E-05 |
| Glycolysis / Gluconeogenesis | 14 | 1.1 | 6.90E-04 |
| Proximal tubule bicarbonate reclamation | 8 | 0.7 | 7.60E-04 |
| Mineral absorption | 13 | 1.1 | 8.30E-04 |
| Chemical carcinogenesis - DNA adducts | 14 | 1.1 | 9.20E-04 |
| Metabolism of xenobiotics by cytochrome P450 | 15 | 1.2 | 9.80E-04 |
| Maturity onset diabetes of the young | 8 | 0.7 | 1.70E-03 |
| Glycosphingolipid biosynthesis - lacto and neolacto series | 8 | 0.7 | 2.10E-03 |
| Protein digestion and absorption | 17 | 1.4 | 2.20E-03 |
| Arginine and proline metabolism | 11 | 0.9 | 2.20E-03 |
| Retinol metabolism | 13 | 1.1 | 2.60E-03 |
| Carbohydrate digestion and absorption | 10 | 0.8 | 4.90E-03 |
| Pathogenic Escherichia coli infection | 25 | 2 | 5.90E-03 |
| Glutathione metabolism | 11 | 0.9 | 6.00E-03 |
| Tryptophan metabolism | 9 | 0.7 | 8.10E-03 |
| Insulin resistance | 16 | 1.3 | 8.50E-03 |
| Fatty acid degradation | 9 | 0.7 | 9.40E-03 |
| NOD-like receptor signaling pathway | 23 | 1.9 | 1.10E-02 |
| Cortisol synthesis and secretion | 11 | 0.9 | 1.50E-02 |
| Tight junction | 21 | 1.7 | 1.50E-02 |
| Fatty acid metabolism | 10 | 0.8 | 1.70E-02 |
| Valine, leucine and isoleucine degradation | 9 | 0.7 | 1.80E-02 |
| Renin-angiotensin system | 6 | 0.5 | 2.00E-02 |
| Antigen processing and presentation | 12 | 1 | 2.00E-02 |
| Glycerophospholipid metabolism | 14 | 1.1 | 2.10E-02 |
| Bile secretion | 13 | 1.1 | 2.20E-02 |
| Galactose metabolism | 7 | 0.6 | 2.20E-02 |
| Vitamin digestion and absorption | 6 | 0.5 | 2.40E-02 |
| Drug metabolism - other enzymes | 12 | 1 | 2.40E-02 |
| Arachidonic acid metabolism | 10 | 0.8 | 2.60E-02 |

| | | | |
|---|---|---|---|
| Nitrogen metabolism | 5 | 0.4 | 2.80E-02 |
| Lipid and atherosclerosis | 24 | 2 | 2.90E-02 |
| Butanoate metabolism | 6 | 0.5 | 3.80E-02 |
| Pentose and glucuronate interconversions | 7 | 0.6 | 3.80E-02 |
| Aldosterone-regulated sodium reabsorption | 7 | 0.6 | 4.30E-02 |
| Pyruvate metabolism | 8 | 0.7 | 4.50E-02 |
| AMPK signaling pathway | 15 | 1.2 | 4.50E-02 |

TableS10

| Pathways upregulated in Gland_B compared to Gland_A | | | |
|---|---|---|---|
| **KEGG pathway** | **No. of genes** | **%** | **p-value** |
| Wnt signaling pathway | 23 | 2.1 | 1.40E-05 |
| Protein digestion and absorption | 17 | 1.5 | 2.10E-05 |
| Signaling pathways regulating pluripotency of stem cells | 20 | 1.8 | 3.50E-05 |
| Hippo signaling pathway | 19 | 1.7 | 3.70E-04 |
| Human papillomavirus infection | 30 | 2.7 | 8.00E-04 |
| Central carbon metabolism in cancer | 11 | 1 | 1.40E-03 |
| Basal cell carcinoma | 10 | 0.9 | 2.50E-03 |
| Gap junction | 12 | 1.1 | 2.50E-03 |
| ECM-receptor interaction | 12 | 1.1 | 2.80E-03 |
| Proximal tubule bicarbonate reclamation | 6 | 0.5 | 3.70E-03 |
| MAPK signaling pathway | 25 | 2.3 | 7.60E-03 |
| Motor proteins | 18 | 1.6 | 8.90E-03 |
| Endocrine and other factor-regulated calcium reabsorption | 8 | 0.7 | 1.10E-02 |
| Focal adhesion | 18 | 1.6 | 1.40E-02 |
| Glycosaminoglycan biosynthesis - chondroitin sulfate / dermatan sulfate | 5 | 0.5 | 1.50E-02 |
| Nucleotide metabolism | 10 | 0.9 | 1.70E-02 |
| Melanogenesis | 11 | 1 | 1.90E-02 |
| Thyroid hormone synthesis | 9 | 0.8 | 2.30E-02 |
| Axon guidance | 16 | 1.5 | 2.30E-02 |
| Hypertrophic cardiomyopathy | 10 | 0.9 | 2.40E-02 |
| Thyroid hormone signaling pathway | 12 | 1.1 | 2.60E-02 |
| Arrhythmogenic right ventricular cardiomyopathy | 9 | 0.8 | 2.70E-02 |
| Calcium signaling pathway | 20 | 1.8 | 2.80E-02 |
| Cellular senescence | 14 | 1.3 | 3.10E-02 |
| Dilated cardiomyopathy | 10 | 0.9 | 3.50E-02 |
| Adipocytokine signaling pathway | 8 | 0.7 | 4.20E-02 |
| Glutamatergic synapse | 11 | 1 | 4.30E-02 |
| Glutathione metabolism | 7 | 0.6 | 4.90E-02 |
| Melanoma | 8 | 0.7 | 5.00E-02 |

TableS11

| Pathways upregulated in Undiff compared to Gland_A | | | |
|---|---|---|---|
| **GO Pathway** | **No. of genes** | **%** | **p-value** |
| Nervous system development | 332 | 24.0 | 8.33E-52 |
| Neurogenesis | 218 | 16.0 | 3.79E-36 |
| Generation of neurons | 195 | 14.3 | 1.41E-34 |
| Neuron differentiation | 180 | 13.2 | 5.65E-31 |
| Neuron development | 153 | 11.2 | 1.73E-29 |
| Neuron projection development | 134 | 9.9 | 2.82E-29 |
| Cell projection morphogenesis | 101 | 7.4 | 6.49E-24 |
| Plasma membrane bounded cell projection organization | 167 | 12.3 | 1.75E-22 |
| Plasma membrane bounded cell projection morphogenesis | 97 | 7.1 | 3.65E-22 |
| Cell projection organization | 171 | 12.6 | 5.2E-22 |
| Neuron projection morphogenesis | 96 | 7.1 | 6.47E-22 |
| Modulation of chemical synaptic transmission | 97 | 7.1 | 1.93E-21 |
| Regulation of trans-synaptic signaling | 97 | 7.1 | 2.21E-21 |
| Cell morphogenesis | 116 | 8.5 | 3.92E-20 |
| Cell morphogenesis involved in neuron differentiation | 86 | 6.3 | 2.2E-19 |
| Axon development | 82 | 6.0 | 6.93E-19 |
| Regulation of cellular component organization | 268 | 19.7 | 8.47E-19 |
| Regulation of neuron projection development | 80 | 5.9 | 6.97E-17 |
| Central nervous system development | 136 | 10.0 | 2.75E-16 |
| Axonogenesis | 71 | 5.2 | 3.7E-16 |
| Brain development | 109 | 8.0 | 8.65E-16 |
| Trans-synaptic signaling | 77 | 5.7 | 1.02E-15 |
| Head development | 113 | 8.3 | 1.67E-15 |
| Synaptic signaling | 80 | 5.9 | 2.43E-15 |
| Regulation of plasma membrane bounded cell projection organization | 98 | 7.2 | 2.7E-15 |
| Chemical synaptic transmission | 73 | 5.4 | 9E-15 |
| Anterograde trans-synaptic signaling | 73 | 5.4 | 9E-15 |
| Behavior | 96 | 7.1 | 9.23E-15 |
| Regulation of cell projection organization | 98 | 7.2 | 1.37E-14 |
| Synapse organization | 67 | 4.9 | 1.48E-14 |
| Regulation of localization | 220 | 16.2 | 3.49E-14 |
| Mitotic cell cycle | 90 | 6.6 | 5.29E-14 |
| Mitotic cell cycle process | 81 | 6.0 | 8.61E-14 |
| Cell development | 232 | 17.1 | 3.52E-13 |
| Regulation of synapse structure or activity | 59 | 4.3 | 3.86E-13 |
| Regulation of synapse organization | 58 | 4.3 | 5.15E-13 |
| Cell junction organization | 84 | 6.2 | 5.39E-13 |
| Cell cycle process | 113 | 8.3 | 6.7E-12 |
| Regulation of cell communication | 327 | 24.0 | 1.24E-11 |
| Regulation of signaling | 326 | 24.0 | 1.54E-11 |
| Cell cycle | 120 | 8.8 | 1.62E-11 |
| Regulation of biological quality | 287 | 21.1 | 1.79E-11 |
| Regulation of postsynapse organization | 35 | 2.6 | 4.05E-11 |
| Forebrain development | 65 | 4.8 | 5.28E-11 |
| Hindbrain development | 36 | 2.7 | 1.23E-10 |
| Regulation of cellular localization | 121 | 8.9 | 1.41E-10 |
| Cell migration | 112 | 8.2 | 1.52E-10 |
| Vesicle-mediated transport in synapse | 37 | 2.7 | 1.98E-10 |
| Synaptic vesicle cycle | 35 | 2.6 | 2.56E-10 |
| Microtubule cytoskeleton organization | 80 | 5.9 | 3.25E-10 |
| Telencephalon development | 49 | 3.6 | 4.87E-10 |
| Neuron migration | 33 | 2.4 | 6.54E-10 |
| Cytoskeleton organization | 139 | 10.2 | 9.62E-10 |
| Regulation of transport | 169 | 12.4 | 1.11E-09 |
| Regulation of protein localization | 108 | 7.9 | 1.58E-09 |
| Regulation of synapse assembly | 34 | 2.5 | 4.51E-09 |
| Learning | 33 | 2.4 | 5.77E-09 |
| Cell-cell signaling | 98 | 7.2 | 7.12E-09 |
| Cellular localization | 245 | 18.0 | 8.34E-09 |
| Organelle localization | 72 | 5.3 | 1.24E-08 |
| Cerebellum development | 26 | 1.9 | 1.83E-08 |
| Negative regulation of cellular component organization | 86 | 6.3 | 2.57E-08 |
| Microtubule-based process | 101 | 7.4 | 3.13E-08 |
| Negative regulation of neuron projection development | 28 | 2.1 | 4.23E-08 |
| Dendrite development | 26 | 1.9 | 7.81E-08 |
| Learning or memory | 44 | 3.2 | 8.64E-08 |
| Negative regulation of cell projection organization | 34 | 2.5 | 8.72E-08 |
| Metencephalon development | 26 | 1.9 | 1.1E-07 |
| Cell motility | 121 | 8.9 | 1.24E-07 |
| Regulation of synaptic plasticity | 38 | 2.8 | 1.57E-07 |
| Regulation of neurogenesis | 53 | 3.9 | 1.61E-07 |
| Cognition | 48 | 3.5 | 1.65E-07 |
| Locomotory behavior | 36 | 2.7 | 1.84E-07 |
| Regulation of nervous system development | 60 | 4.4 | 2.37E-07 |

| | | | |
|---|---|---|---|
| Axon guidance | 38 | 2.8 | 2.43E-07 |
| Positive regulation of cell projection organization | 51 | 3.8 | 2.55E-07 |
| Neuron projection guidance | 38 | 2.8 | 2.71E-07 |
| Regulation of axonogenesis | 28 | 2.1 | 3.62E-07 |
| Gliogenesis | 43 | 3.2 | 3.8E-07 |
| Central nervous system neuron differentiation | 35 | 2.6 | 3.94E-07 |
| Signal release from synapse | 21 | 1.5 | 4.25E-07 |
| Neurotransmitter secretion | 21 | 1.5 | 4.25E-07 |
| Cellular macromolecule localization | 183 | 13.5 | 5.34E-07 |
| Vesicle localization | 33 | 2.4 | 7.26E-07 |
| Cell communication | 414 | 30.4 | 7.59E-07 |
| Embryo development | 108 | 7.9 | 8.06E-07 |
| Synapse assembly | 26 | 1.9 | 8.29E-07 |
| Establishment of organelle localization | 55 | 4.0 | 1.01E-06 |
| Regulation of neurotransmitter secretion | 20 | 1.5 | 1.57E-06 |
| Regulation of postsynapse assembly | 14 | 1.0 | 1.69E-06 |
| Regulation of synaptic vesicle exocytosis | 15 | 1.1 | 1.81E-06 |
| Establishment of localization in cell | 154 | 11.3 | 1.99E-06 |
| Secretory granule localization | 10 | 0.7 | 2.04E-06 |
| Telencephalon cell migration | 17 | 1.3 | 2.07E-06 |
| Organelle organization | 261 | 19.2 | 2.07E-06 |
| Protein localization | 179 | 13.2 | 2.38E-06 |
| Signaling | 407 | 29.9 | 2.61E-06 |
| Positive regulation of cellular component organization | 112 | 8.2 | 2.92E-06 |
| Pallium development | 32 | 2.4 | 0.0000033 |
| Regulation of microtubule cytoskeleton organization | 29 | 2.1 | 3.47E-06 |
| Regulation of cellular component biogenesis | 105 | 7.7 | 3.58E-06 |
| Forebrain cell migration | 17 | 1.3 | 3.88E-06 |
| Postsynapse organization | 23 | 1.7 | 4.42E-06 |
| Adult behavior | 27 | 2.0 | 5.04E-06 |
| Positive regulation of protein localization | 60 | 4.4 | 5.07E-06 |
| Macromolecule localization | 208 | 15.3 | 5.47E-06 |
| Dense core granule localization | 9 | 0.7 | 5.54E-06 |
| Regulation of microtubule polymerization or depolymerization | 21 | 1.5 | 6.52E-06 |
| Microtubule polymerization or depolymerization | 16 | 1.2 | 6.72E-06 |
| Cell junction assembly | 41 | 3.0 | 7.27E-06 |
| Dna replication | 32 | 2.4 | 7.44E-06 |
| Regulation of cell junction assembly | 38 | 2.8 | 7.57E-06 |
| Regulation of neurotransmitter transport | 21 | 1.5 | 7.58E-06 |
| Regulation of anatomical structure morphogenesis | 92 | 6.8 | 7.83E-06 |
| Regulation of secretion | 70 | 5.1 | 8.58E-06 |
| Synaptic vesicle exocytosis | 16 | 1.2 | 0.0000101 |
| Regulation of cell differentiation | 149 | 11.0 | 0.0000114 |
| Signal release | 30 | 2.2 | 0.0000114 |
| Export from cell | 58 | 4.3 | 0.0000121 |
| Protein localization to synapse | 16 | 1.2 | 0.0000122 |
| Regulation of organelle organization | 113 | 8.3 | 0.0000131 |
| Cell division | 62 | 4.6 | 0.0000132 |
| Embryo development ending in birth or egg hatching | 73 | 5.4 | 0.0000143 |
| Synaptic vesicle localization | 14 | 1.0 | 0.0000162 |
| Regulation of cytoskeleton organization | 62 | 4.6 | 0.0000166 |
| Negative regulation of microtubule polymerization or depolymerization | 13 | 1.0 | 0.0000174 |
| Protein localization to cell junction | 19 | 1.4 | 0.0000201 |
| Regulation of secretion by cell | 64 | 4.7 | 0.0000203 |
| Presynaptic modulation of chemical synaptic transmission | 12 | 0.9 | 0.0000227 |
| Motor neuron axon guidance | 10 | 0.7 | 0.0000231 |
| Regulation of dendritic spine development | 15 | 1.1 | 0.0000259 |
| Regulation of neuronal synaptic plasticity | 15 | 1.1 | 0.0000259 |
| Neurotransmitter transport | 24 | 1.8 | 0.0000284 |
| Cytoskeleton-dependent intracellular transport | 30 | 2.2 | 0.0000299 |
| Secretion by cell | 51 | 3.8 | 0.0000309 |
| Hippocampus development | 19 | 1.4 | 0.0000312 |
| Regulation of neuron migration | 14 | 1.0 | 0.0000374 |
| Secretion | 61 | 4.5 | 0.0000375 |
| Exocytic process | 15 | 1.1 | 0.0000376 |
| Synapse maturation | 8 | 0.6 | 0.0000401 |
| Cerebral cortex development | 23 | 1.7 | 0.0000434 |
| Cell adhesion | 94 | 6.9 | 0.000044 |
| Mitotic cell cycle checkpoint signaling | 21 | 1.5 | 0.00005 |
| Dna metabolic process | 78 | 5.7 | 0.0000523 |
| Mitotic cell cycle phase transition | 24 | 1.8 | 0.000055 |
| Regulation of receptor localization to synapse | 10 | 0.7 | 0.0000596 |
| Regulation of protein localization to synapse | 10 | 0.7 | 0.0000596 |
| Microtubule depolymerization | 8 | 0.6 | 0.0000621 |
| Cerebral cortex cell migration | 13 | 1.0 | 0.0000654 |
| Vesicle docking | 14 | 1.0 | 0.0000663 |
| Cellular component biogenesis | 234 | 17.2 | 0.0000682 |

| | | | |
|---|---|---|---|
| Positive regulation of nervous system development | 38 | 2.8 | 0.0000703 |
| Cell cycle phase transition | 25 | 1.8 | 0.0000743 |
| Regulation of dendritic spine morphogenesis | 12 | 0.9 | 0.0000748 |
| Establishment of vesicle localization | 27 | 2.0 | 0.000077 |
| Regulation of microtubule depolymerization | 11 | 0.8 | 0.0000807 |
| Vesicle docking involved in exocytosis | 11 | 0.8 | 0.0000807 |
| Regulation of neuron differentiation | 28 | 2.1 | 0.000088 |
| Supramolecular fiber organization | 65 | 4.8 | 0.0000885 |
| Central nervous system neuron development | 18 | 1.3 | 0.0000946 |
| Negative regulation of microtubule depolymerization | 10 | 0.7 | 0.000104 |
| Axo-dendritic transport | 16 | 1.2 | 0.000109 |
| Regulation of cell morphogenesis | 33 | 2.4 | 0.000118 |
| Regulation of membrane potential | 52 | 3.8 | 0.000124 |
| Negative regulation of mitotic cell cycle phase transition | 26 | 1.9 | 0.000127 |
| Protein depolymerization | 11 | 0.8 | 0.000129 |
| Positive regulation of neuron projection development | 25 | 1.8 | 0.000132 |
| Regulation of neurotransmitter receptor localization to postsynaptic specialization membrane | 8 | 0.6 | 0.000135 |
| Post-anal tail morphogenesis | 8 | 0.6 | 0.000135 |
| Regulation of protein localization to membrane | 28 | 2.1 | 0.000147 |
| Vesicle-mediated transport | 118 | 8.7 | 0.000156 |
| Negative regulation of organelle organization | 41 | 3.0 | 0.000157 |
| Limbic system development | 21 | 1.5 | 0.000157 |
| Regulation of regulated secretory pathway | 21 | 1.5 | 0.000174 |
| Chordate embryonic development | 67 | 4.9 | 0.00018 |
| Intracellular signal transduction | 151 | 11.1 | 0.000184 |
| Axonal transport | 14 | 1.0 | 0.000185 |
| Associative learning | 17 | 1.3 | 0.000214 |
| Regulation of microtubule-based process | 35 | 2.6 | 0.000227 |
| Positive regulation of transport | 83 | 6.1 | 0.000235 |
| Dendrite morphogenesis | 14 | 1.0 | 0.000253 |
| Glial cell differentiation | 30 | 2.2 | 0.000256 |
| Negative regulation of mitotic cell cycle | 30 | 2.2 | 0.000256 |
| Dna strand elongation involved in dna replication | 7 | 0.5 | 0.000269 |
| Transport along microtubule | 25 | 1.8 | 0.00027 |
| Organelle transport along microtubule | 16 | 1.2 | 0.000288 |
| Glial cell migration | 12 | 0.9 | 0.000298 |
| Intracellular mrna localization | 6 | 0.4 | 0.000314 |
| Double-strand break repair | 29 | 2.1 | 0.000323 |
| Negative regulation of protein modification process | 37 | 2.7 | 0.000327 |
| Adult locomotory behavior | 16 | 1.2 | 0.000328 |
| Cell cycle checkpoint signaling | 23 | 1.7 | 0.000328 |
| Exocytosis | 31 | 2.3 | 0.000335 |
| Negative regulation of cell cycle phase transition | 31 | 2.3 | 0.000335 |
| Cellular component assembly | 211 | 15.5 | 0.000352 |
| Positive regulation of developmental process | 122 | 9.0 | 0.000369 |
| Negative regulation of supramolecular fiber organization | 24 | 1.8 | 0.000371 |
| Peripheral nervous system development | 16 | 1.2 | 0.000372 |
| Neuron projection extension | 16 | 1.2 | 0.000372 |
| Central nervous system projection neuron axonogenesis | 9 | 0.7 | 0.000378 |
| Regulation of exocytosis | 26 | 1.9 | 0.000382 |
| Dna damage response | 78 | 5.7 | 0.000383 |
| Dna repair | 56 | 4.1 | 0.000419 |
| Organelle localization by membrane tethering | 15 | 1.1 | 0.000443 |
| Regulation of guanyl-nucleotide exchange factor activity | 5 | 0.4 | 0.000451 |
| Regulation of establishment of protein localization | 57 | 4.2 | 0.000472 |
| Negative regulation of dna-templated transcription | 112 | 8.2 | 0.000478 |
| Cerebral cortex radial glia-guided migration | 9 | 0.7 | 0.000482 |
| Telencephalon glial cell migration | 9 | 0.7 | 0.000482 |
| Regulation of cell growth | 45 | 3.3 | 0.000493 |
| Nerve development | 17 | 1.3 | 0.000495 |
| Regulation of extent of cell growth | 17 | 1.3 | 0.000495 |
| Positive regulation of neurogenesis | 31 | 2.3 | 0.000504 |
| Synaptic vesicle clustering | 6 | 0.4 | 0.00051 |
| Calcium ion-regulated exocytosis of neurotransmitter | 6 | 0.4 | 0.00051 |
| Double-strand break repair via break-induced replication | 6 | 0.4 | 0.00051 |
| Regulation of dendrite development | 17 | 1.3 | 0.000553 |
| Cell migration in hindbrain | 7 | 0.5 | 0.00056 |
| Synaptic vesicle priming | 7 | 0.5 | 0.00056 |
| Cell-cell adhesion | 56 | 4.1 | 0.000588 |
| Presynapse organization | 9 | 0.7 | 0.000609 |
| Anterograde axonal transport | 11 | 0.8 | 0.000632 |
| Regulation of axon extension | 15 | 1.1 | 0.000648 |
| Cerebral cortex radially oriented cell migration | 10 | 0.7 | 0.000648 |
| Negative regulation of rna biosynthetic process | 112 | 8.2 | 0.000668 |
| Regulation of vesicle-mediated transport | 56 | 4.1 | 0.000716 |
| Regulation of protein depolymerization | 16 | 1.2 | 0.000759 |
| Phosphorylation | 46 | 3.4 | 0.000788 |

| | | | |
|---|---|---|---|
| Neuromuscular process controlling balance | 12 | 0.9 | 0.000797 |
| Regulation of system process | 58 | 4.3 | 0.000826 |
| Nuclear dna replication | 8 | 0.6 | 0.000826 |
| Developmental maturation | 35 | 2.6 | 0.000827 |
| Maintenance of presynaptic active zone structure | 5 | 0.4 | 0.000857 |
| Small gtpase-mediated signal transduction | 32 | 2.4 | 0.000867 |
| Regulation of signal transduction | 246 | 18.1 | 0.000906 |
| Positive regulation of establishment of protein localization | 38 | 2.8 | 0.000911 |
| Cellular component maintenance | 13 | 1.0 | 0.000918 |
| Chromosome organization | 47 | 3.5 | 0.000923 |
| Animal organ morphogenesis | 89 | 6.5 | 0.000933 |
| Developmental growth involved in morphogenesis | 21 | 1.5 | 0.00096 |
| Import into cell | 67 | 4.9 | 0.001 |
| Cell junction maintenance | 11 | 0.8 | 0.00104 |
| Cell cycle dna replication | 8 | 0.6 | 0.00106 |
| Ear development | 29 | 2.1 | 0.00111 |
| Positive regulation of cellular component biogenesis | 52 | 3.8 | 0.00112 |
| Neuronal action potential | 9 | 0.7 | 0.00115 |
| Regulated exocytosis | 20 | 1.5 | 0.00121 |
| Regulation of filopodium assembly | 11 | 0.8 | 0.00122 |
| Positive regulation of cell differentiation | 83 | 6.1 | 0.00124 |
| Sensory organ development | 60 | 4.4 | 0.00124 |
| Regulation of cell development | 78 | 5.7 | 0.00125 |
| Regulation of g2/m transition of mitotic cell cycle | 17 | 1.3 | 0.00128 |
| Mitotic nuclear division | 23 | 1.7 | 0.00133 |
| Regulation of supramolecular fiber organization | 43 | 3.2 | 0.00134 |
| Sister chromatid segregation | 21 | 1.5 | 0.00134 |
| Notch signaling pathway | 18 | 1.3 | 0.00135 |
| Negative regulation of protein depolymerization | 13 | 1.0 | 0.00136 |
| Dna strand elongation | 7 | 0.5 | 0.00139 |
| Ribonucleoside monophosphate biosynthetic process | 9 | 0.7 | 0.0014 |
| Vesicle cytoskeletal trafficking | 12 | 0.9 | 0.00142 |
| Negative regulation of protein-containing complex disassembly | 14 | 1.0 | 0.00143 |
| Spinal cord development | 16 | 1.2 | 0.00144 |
| Negative regulation of cytoskeleton organization | 22 | 1.6 | 0.00145 |
| Dense core granule transport | 5 | 0.4 | 0.00147 |
| Dense core granule cytoskeletal transport | 5 | 0.4 | 0.00147 |
| Purine nucleobase biosynthetic process | 5 | 0.4 | 0.00147 |
| Negative regulation of phosphate metabolic process | 31 | 2.3 | 0.00151 |
| Negative regulation of phosphorus metabolic process | 31 | 2.3 | 0.00151 |
| Regulation of postsynaptic membrane potential | 17 | 1.3 | 0.00155 |
| Nucleoside monophosphate biosynthetic process | 10 | 0.7 | 0.00159 |
| Positive regulation of dendritic spine development | 10 | 0.7 | 0.00159 |
| Membrane docking | 15 | 1.1 | 0.00162 |
| Regulation of cell cycle g2/m phase transition | 18 | 1.3 | 0.00162 |
| Regulation of dna replication | 18 | 1.3 | 0.00162 |
| Negative regulation of axonogenesis | 11 | 0.8 | 0.00165 |
| Gmp biosynthetic process | 6 | 0.4 | 0.00165 |
| Negative regulation of rna metabolic process | 117 | 8.6 | 0.00168 |
| Positive regulation of cell morphogenesis | 9 | 0.7 | 0.00169 |
| Developmental cell growth | 18 | 1.3 | 0.00177 |
| Regulation of transcription by rna polymerase ii | 209 | 15.4 | 0.00179 |
| Presynapse assembly | 7 | 0.5 | 0.00182 |
| Regulation of synaptic vesicle membrane organization | 7 | 0.5 | 0.00182 |
| Regulation of synaptic vesicle fusion to presynaptic active zone membrane | 7 | 0.5 | 0.00182 |
| Purine nucleoside monophosphate biosynthetic process | 7 | 0.5 | 0.00182 |
| Inner ear development | 26 | 1.9 | 0.00185 |
| Glutamate receptor signaling pathway | 11 | 0.8 | 0.0019 |
| Cerebellar cortex development | 11 | 0.8 | 0.0019 |
| Positive regulation of nucleobase-containing compound metabolic process | 165 | 12.1 | 0.00193 |
| Neuron fate commitment | 13 | 1.0 | 0.00196 |
| Regulation of binding | 26 | 1.9 | 0.00197 |
| Nephron development | 20 | 1.5 | 0.00199 |
| Cell cycle g1/s phase transition | 14 | 1.0 | 0.00201 |
| Calcium-ion regulated exocytosis | 9 | 0.7 | 0.00203 |
| Central nervous system neuron axonogenesis | 9 | 0.7 | 0.00203 |
| Mitotic spindle checkpoint signaling | 8 | 0.6 | 0.00205 |
| Maintenance of synapse structure | 8 | 0.6 | 0.00205 |
| Mitotic spindle assembly checkpoint signaling | 8 | 0.6 | 0.00205 |
| Spindle assembly checkpoint signaling | 8 | 0.6 | 0.00205 |
| Regulation of mitotic cell cycle | 51 | 3.8 | 0.00206 |
| Negative regulation of cell cycle | 40 | 2.9 | 0.0021 |
| Cell growth | 18 | 1.3 | 0.00211 |
| Anatomical structure formation involved in morphogenesis | 87 | 6.4 | 0.00214 |
| Regulation of mitotic cell cycle phase transition | 38 | 2.8 | 0.00214 |
| Vesicle transport along microtubule | 10 | 0.7 | 0.00218 |
| Regulation of dna-templated dna replication initiation | 6 | 0.4 | 0.00228 |

| | | | |
|---|---|---|---|
| Mitotic sister chromatid segregation | 20 | 1.5 | 0.00232 |
| Presynaptic active zone organization | 5 | 0.4 | 0.00233 |
| Postsynaptic membrane assembly | 5 | 0.4 | 0.00233 |
| Hindbrain radial glia guided cell migration | 5 | 0.4 | 0.00233 |
| Spindle checkpoint signaling | 8 | 0.6 | 0.00251 |
| Motor behavior | 8 | 0.6 | 0.00251 |
| Negative regulation of neurogenesis | 19 | 1.4 | 0.00252 |
| Renal system development | 35 | 2.6 | 0.00253 |
| Negative regulation of nucleobase-containing compound metabolic process | 125 | 9.2 | 0.00254 |
| Regulation of cell cycle | 95 | 7.0 | 0.00262 |
| Sensory organ morphogenesis | 32 | 2.4 | 0.0028 |
| Regulation of growth | 58 | 4.3 | 0.00284 |
| Metal ion transport | 61 | 4.5 | 0.00289 |
| Regulation of dna metabolic process | 49 | 3.6 | 0.00291 |
| Response to light stimulus | 35 | 2.6 | 0.00291 |
| Negative regulation of neuron projection regeneration | 6 | 0.4 | 0.00307 |
| Intracellular signaling cassette | 75 | 5.5 | 0.0031 |
| Regulation of molecular function | 119 | 8.7 | 0.00313 |
| Negative regulation of cell cycle process | 32 | 2.4 | 0.00327 |
| Regulation of peptide hormone secretion | 25 | 1.8 | 0.0034 |
| Regulation of protein-containing complex disassembly | 18 | 1.3 | 0.00347 |
| Presynaptic membrane organization | 5 | 0.4 | 0.00348 |
| Regulation of gtpase activity | 22 | 1.6 | 0.00366 |
| Negative regulation of mitotic sister chromatid segregation | 8 | 0.6 | 0.00366 |
| Positive regulation of filopodium assembly | 8 | 0.6 | 0.00366 |
| Negative regulation of sister chromatid segregation | 8 | 0.6 | 0.00366 |
| Negative regulation of mitotic metaphase/anaphase transition | 8 | 0.6 | 0.00366 |
| Negative regulation of mitotic sister chromatid separation | 8 | 0.6 | 0.00366 |
| Visual behavior | 11 | 0.8 | 0.0037 |
| Social behavior | 11 | 0.8 | 0.0037 |
| Biological process involved in intraspecies interaction between organisms | 11 | 0.8 | 0.0037 |
| Establishment of cell polarity | 19 | 1.4 | 0.00371 |
| Memory | 18 | 1.3 | 0.00375 |
| Rna localization | 24 | 1.8 | 0.0038 |
| Synaptic vesicle recycling | 12 | 0.9 | 0.00383 |
| Modulation of excitatory postsynaptic potential | 10 | 0.7 | 0.0039 |
| Segmentation | 15 | 1.1 | 0.00392 |
| Dna recombination | 29 | 2.1 | 0.004 |
| Layer formation in cerebral cortex | 6 | 0.4 | 0.00403 |
| Response to l-glutamate | 6 | 0.4 | 0.00403 |
| Nucleobase biosynthetic process | 6 | 0.4 | 0.00403 |
| Regulation of peptide secretion | 25 | 1.8 | 0.00407 |
| Negative regulation of guanyl-nucleotide exchange factor activity | 4 | 0.3 | 0.00419 |
| Positive regulation of inhibitory postsynaptic potential | 4 | 0.3 | 0.00419 |
| Synaptic vesicle endocytosis | 11 | 0.8 | 0.00419 |
| Neural nucleus development | 12 | 0.9 | 0.00428 |
| Mitotic spindle organization | 15 | 1.1 | 0.00429 |
| Dna-templated dna replication | 19 | 1.4 | 0.0043 |
| Negative regulation of axon extension | 8 | 0.6 | 0.00437 |
| Establishment of mitochondrion localization | 8 | 0.6 | 0.00437 |
| Regulation of protein modification process | 79 | 5.8 | 0.00442 |
| Regulation of insulin secretion | 22 | 1.6 | 0.00445 |
| Visual learning | 10 | 0.7 | 0.00446 |
| G2/m transition of mitotic cell cycle | 10 | 0.7 | 0.00446 |
| Retina layer formation | 7 | 0.5 | 0.00453 |
| Negative regulation of protein phosphorylation | 24 | 1.8 | 0.00456 |
| Regulation of peptide transport | 25 | 1.8 | 0.00458 |
| Negative regulation of nervous system development | 19 | 1.4 | 0.00462 |
| G1/s transition of mitotic cell cycle | 13 | 1.0 | 0.0047 |
| Chromatin remodeling | 92 | 6.8 | 0.00471 |
| Membrane biogenesis | 12 | 0.9 | 0.00478 |
| Mitotic dna damage checkpoint signaling | 12 | 0.9 | 0.00478 |
| Positive regulation of dna-templated transcription | 132 | 9.7 | 0.00492 |
| Protein phosphorylation | 39 | 2.9 | 0.00505 |
| Regulation of vesicle fusion | 8 | 0.6 | 0.00517 |
| Negative regulation of chromosome separation | 8 | 0.6 | 0.00517 |
| Positive regulation of dendrite morphogenesis | 8 | 0.6 | 0.00517 |
| Regulation of protein targeting to membrane | 8 | 0.6 | 0.00517 |
| Negative regulation of metaphase/anaphase transition of cell cycle | 8 | 0.6 | 0.00517 |
| Regulation of intracellular transport | 30 | 2.2 | 0.00534 |
| Positive regulation of rna biosynthetic process | 132 | 9.7 | 0.0054 |
| Microtubule cytoskeleton organization involved in mitosis | 18 | 1.3 | 0.00547 |
| Corpus callosum development | 7 | 0.5 | 0.00552 |
| Regulation of phosphate metabolic process | 71 | 5.2 | 0.00559 |
| Protein localization to cell periphery | 27 | 2.0 | 0.00563 |
| Response to radiation | 43 | 3.2 | 0.00565 |
| Response to mechanical stimulus | 26 | 1.9 | 0.0057 |

| | | | |
|---|---|---|---|
| Regulation of phosphorus metabolic process | 71 | 5.2 | 0.00571 |
| Negative regulation of phosphorylation | 25 | 1.8 | 0.00575 |
| Aorta development | 11 | 0.8 | 0.00594 |
| Renal tubule development | 14 | 1.0 | 0.00596 |
| Positive regulation of plasma membrane bounded cell projection assembly | 16 | 1.2 | 0.00608 |
| Cell-cell junction organization | 22 | 1.6 | 0.00608 |
| Positive regulation of excitatory postsynaptic potential | 8 | 0.6 | 0.00608 |
| Negative regulation of chromosome segregation | 8 | 0.6 | 0.00608 |
| Positive regulation of transcription by rna polymerase ii | 101 | 7.4 | 0.00622 |
| Nuclear chromosome segregation | 27 | 2.0 | 0.00625 |
| Neuromuscular process | 20 | 1.5 | 0.00628 |
| Positive regulation of dna metabolic process | 31 | 2.3 | 0.00631 |
| Chromatin organization | 102 | 7.5 | 0.00641 |
| Signal transduction | 353 | 25.9 | 0.00647 |
| Phosphate-containing compound metabolic process | 119 | 8.7 | 0.00648 |
| Adherens junction organization | 10 | 0.7 | 0.00653 |
| Regulation of protein transport | 44 | 3.2 | 0.00656 |
| Positive regulation of dendritic spine morphogenesis | 6 | 0.4 | 0.00658 |
| Purine ribonucleoside monophosphate biosynthetic process | 6 | 0.4 | 0.00658 |
| Proteoglycan metabolic process | 15 | 1.1 | 0.00659 |
| Presynaptic endocytosis | 11 | 0.8 | 0.00664 |
| Regulation of dendrite morphogenesis | 11 | 0.8 | 0.00664 |
| Dentate gyrus development | 7 | 0.5 | 0.00666 |
| Positive regulation of g2/m transition of mitotic cell cycle | 7 | 0.5 | 0.00666 |
| Microtubule-based transport | 25 | 1.8 | 0.00678 |
| Synaptic vesicle docking | 5 | 0.4 | 0.00681 |
| Centriole-centriole cohesion | 5 | 0.4 | 0.00681 |
| Phosphorus metabolic process | 119 | 8.7 | 0.00685 |
| Regulation of dna-templated transcription | 259 | 19.0 | 0.00689 |
| Modulation of inhibitory postsynaptic potential | 4 | 0.3 | 0.00699 |
| Neuronal dense core vesicle exocytosis | 4 | 0.3 | 0.00699 |
| Presynaptic dense core vesicle exocytosis | 4 | 0.3 | 0.00699 |
| Positive regulation of protein localization to centrosome | 4 | 0.3 | 0.00699 |
| Rhythmic process | 30 | 2.2 | 0.00711 |
| Epithelium development | 93 | 6.8 | 0.00715 |
| Mitotic dna integrity checkpoint signaling | 12 | 0.9 | 0.00722 |
| Monoatomic ion transport | 86 | 6.3 | 0.00735 |
| Cellular senescence | 11 | 0.8 | 0.0074 |
| Neuron cellular homeostasis | 11 | 0.8 | 0.0074 |
| Negative regulation of binding | 14 | 1.0 | 0.00773 |
| Protein localization to chromosome | 15 | 1.1 | 0.00775 |
| Cellular response to stress | 132 | 9.7 | 0.00786 |
| Calcium-dependent cell-cell adhesion via plasma membrane cell adhesion molecules | 9 | 0.7 | 0.00795 |
| Positive regulation of protein targeting to membrane | 7 | 0.5 | 0.00796 |
| Innervation | 7 | 0.5 | 0.00796 |
| Postsynaptic membrane organization | 7 | 0.5 | 0.00796 |
| Regulation of rna biosynthetic process | 259 | 19.0 | 0.00809 |
| Regulation of rna splicing | 22 | 1.6 | 0.00817 |
| Embryonic organ development | 44 | 3.2 | 0.00824 |
| Positive regulation of sodium ion transport | 8 | 0.6 | 0.00825 |
| Response to cocaine | 10 | 0.7 | 0.00827 |
| Cell cycle g2/m phase transition | 10 | 0.7 | 0.00827 |
| Positive regulation of macromolecule metabolic process | 244 | 17.9 | 0.00829 |
| Positive regulation of secretion by cell | 31 | 2.3 | 0.00831 |
| Endocytosis | 48 | 3.5 | 0.00834 |
| Centrosome cycle | 14 | 1.0 | 0.00841 |
| Positive regulation of secretion | 33 | 2.4 | 0.00848 |
| Regulation of protein localization to cell periphery | 19 | 1.4 | 0.00851 |
| Kidney development | 32 | 2.4 | 0.00871 |
| Sulfur compound biosynthetic process | 19 | 1.4 | 0.00907 |
| Regulation of microtubule polymerization | 11 | 0.8 | 0.00912 |
| Membrane depolarization | 10 | 0.7 | 0.00926 |
| Cell fate commitment | 28 | 2.1 | 0.00953 |
| Regulation of nervous system process | 16 | 1.2 | 0.00955 |
| Proteoglycan biosynthetic process | 12 | 0.9 | 0.00963 |
| Monoatomic cation transport | 71 | 5.2 | 0.00965 |
| Regulation of nucleobase-containing compound metabolic process | 302 | 22.2 | 0.00972 |
| Regulation of postsynaptic specialization assembly | 6 | 0.4 | 0.0101 |
| Retrograde axonal transport | 6 | 0.4 | 0.0101 |
| Regulation of protein polymerization | 23 | 1.7 | 0.0101 |
| Regulation of hormone secretion | 28 | 2.1 | 0.0102 |
| Double-strand break repair via homologous recombination | 16 | 1.2 | 0.0103 |
| Regulation of developmental growth | 33 | 2.4 | 0.0104 |
| Negative regulation of chromosome organization | 12 | 0.9 | 0.0106 |
| Somite development | 12 | 0.9 | 0.0106 |
| Negative regulation of cell differentiation | 64 | 4.7 | 0.0106 |
| Spontaneous synaptic transmission | 4 | 0.3 | 0.0107 |

| | | | |
|---|---|---|---|
| Corticospinal tract morphogenesis | 4 | 0.3 | 0.0107 |
| Regulation of monoatomic ion transport | 44 | 3.2 | 0.0108 |
| Positive regulation of metabolic process | 264 | 19.4 | 0.011 |
| Positive regulation of cell cycle g2/m phase transition | 7 | 0.5 | 0.0111 |
| Dendritic spine development | 7 | 0.5 | 0.0111 |
| Neurofilament bundle assembly | 3 | 0.2 | 0.0112 |
| Neuroligin clustering involved in postsynaptic membrane assembly | 3 | 0.2 | 0.0112 |
| Postsynaptic intermediate filament cytoskeleton organization | 3 | 0.2 | 0.0112 |
| Negative regulation of developmental process | 85 | 6.3 | 0.0115 |
| Regulation of blood circulation | 28 | 2.1 | 0.0116 |
| Regulation of protein-containing complex assembly | 41 | 3.0 | 0.0116 |
| Developmental growth | 43 | 3.2 | 0.0116 |
| Growth | 43 | 3.2 | 0.0116 |
| Negative regulation of axon regeneration | 5 | 0.4 | 0.0118 |
| Base-excision repair, gap-filling | 5 | 0.4 | 0.0118 |
| Response to nitrogen compound | 88 | 6.5 | 0.0125 |
| Establishment or maintenance of cell polarity | 23 | 1.7 | 0.0125 |
| Negative regulation of mitotic nuclear division | 8 | 0.6 | 0.0125 |
| Regulation of glial cell differentiation | 12 | 0.9 | 0.0126 |
| Chromosome segregation | 34 | 2.5 | 0.0127 |
| Regulation of phosphorylation | 59 | 4.3 | 0.0129 |
| Attachment of spindle microtubules to kinetochore | 7 | 0.5 | 0.013 |
| Postsynaptic specialization organization | 7 | 0.5 | 0.013 |
| Protein localization to cell surface | 7 | 0.5 | 0.013 |
| Negative regulation of protein metabolic process | 63 | 4.6 | 0.013 |
| Cell population proliferation | 63 | 4.6 | 0.0131 |
| Nephron epithelium development | 15 | 1.1 | 0.0131 |
| Tissue morphogenesis | 52 | 3.8 | 0.0132 |
| Negative regulation of signaling | 122 | 9.0 | 0.0133 |
| Embryonic organ morphogenesis | 31 | 2.3 | 0.0133 |
| Negative regulation of molecular function | 51 | 3.8 | 0.0134 |
| Negative regulation of cell communication | 122 | 9.0 | 0.0136 |
| Regulation of transmembrane transport | 44 | 3.2 | 0.0139 |
| Regulation of cell cycle phase transition | 43 | 3.2 | 0.0142 |
| Actin filament-based process | 55 | 4.0 | 0.0143 |
| Potassium ion transport | 21 | 1.5 | 0.0144 |
| Regulation of small gtpase mediated signal transduction | 31 | 2.3 | 0.0144 |
| Regulation of neural precursor cell proliferation | 14 | 1.0 | 0.0145 |
| Protein localization to organelle | 67 | 4.9 | 0.0146 |
| Regulation of protein phosphorylation | 55 | 4.0 | 0.0147 |
| Postsynaptic cytoskeleton organization | 6 | 0.4 | 0.0147 |
| Cell differentiation in hindbrain | 6 | 0.4 | 0.0147 |
| Regulation of dopamine secretion | 6 | 0.4 | 0.0147 |
| Peptidyl-serine modification | 13 | 1.0 | 0.0148 |
| Heart morphogenesis | 28 | 2.1 | 0.0149 |
| Neuron cell-cell adhesion | 5 | 0.4 | 0.0149 |
| Negative regulation of cell cycle g1/s phase transition | 12 | 0.9 | 0.0149 |
| Ionotropic glutamate receptor signaling pathway | 7 | 0.5 | 0.015 |
| Regulation of circadian rhythm | 15 | 1.1 | 0.0151 |
| Dense core granule exocytosis | 4 | 0.3 | 0.0153 |
| Sensory perception of sound | 20 | 1.5 | 0.0154 |
| Recombinational repair | 16 | 1.2 | 0.0154 |
| Apoptotic process | 89 | 6.5 | 0.0155 |
| Regulation of neuron apoptotic process | 27 | 2.0 | 0.0155 |
| Somitogenesis | 10 | 0.7 | 0.0156 |
| Positive regulation of binding | 14 | 1.0 | 0.0156 |
| In utero embryonic development | 38 | 2.8 | 0.0158 |
| Regulation of dna-templated dna replication | 8 | 0.6 | 0.0161 |
| Regulation of catecholamine secretion | 8 | 0.6 | 0.0161 |
| Synaptic vesicle transport | 8 | 0.6 | 0.0161 |
| Renal tubule morphogenesis | 11 | 0.8 | 0.0161 |
| Mitochondrion localization | 9 | 0.7 | 0.0162 |
| Cell-cell adhesion via plasma-membrane adhesion molecules | 27 | 2.0 | 0.0162 |
| Cell-cell junction assembly | 16 | 1.2 | 0.0164 |
| Regulation of heart contraction | 23 | 1.7 | 0.0168 |
| Regulation of cell cycle process | 65 | 4.8 | 0.0169 |
| Regulation of g1/s transition of mitotic cell cycle | 20 | 1.5 | 0.0171 |
| Sensory perception of mechanical stimulus | 22 | 1.6 | 0.0173 |
| Positive regulation of neuron differentiation | 13 | 1.0 | 0.0173 |
| Response to auditory stimulus | 7 | 0.5 | 0.0173 |
| Wnt signaling pathway, planar cell polarity pathway | 7 | 0.5 | 0.0173 |
| Postsynapse assembly | 6 | 0.4 | 0.0174 |
| Action potential | 16 | 1.2 | 0.0175 |
| Negative regulation of g1/s transition of mitotic cell cycle | 11 | 0.8 | 0.0176 |
| Forebrain neuron differentiation | 11 | 0.8 | 0.0176 |
| Cochlea development | 9 | 0.7 | 0.018 |
| Mitotic spindle assembly | 9 | 0.7 | 0.018 |

| | | | |
|---|---|---|---|
| Spindle organization | 20 | 1.5 | 0.0181 |
| Negative regulation of neuron apoptotic process | 20 | 1.5 | 0.0181 |
| Potassium ion transmembrane transport | 19 | 1.4 | 0.0184 |
| Organelle fission | 34 | 2.5 | 0.0185 |
| Regulation of cellular component size | 34 | 2.5 | 0.0185 |
| Attachment of mitotic spindle microtubules to kinetochore | 5 | 0.4 | 0.0186 |
| Synaptic vesicle membrane organization | 5 | 0.4 | 0.0186 |
| Membrane assembly | 10 | 0.7 | 0.0189 |
| Metanephros development | 12 | 0.9 | 0.0191 |
| Signal transduction in response to dna damage | 17 | 1.3 | 0.0198 |
| Nucleobase metabolic process | 7 | 0.5 | 0.0198 |
| Adult walking behavior | 7 | 0.5 | 0.0198 |
| Regulation of neuron projection regeneration | 7 | 0.5 | 0.0198 |
| Regulation of chromosome organization | 25 | 1.8 | 0.0198 |
| Embryonic morphogenesis | 52 | 3.8 | 0.0199 |
| Negative regulation of transcription by rna polymerase ii | 77 | 5.7 | 0.0201 |
| Glycosaminoglycan biosynthetic process | 8 | 0.6 | 0.0204 |
| Homophilic cell adhesion via plasma membrane adhesion molecules | 19 | 1.4 | 0.0205 |
| Negative regulation of rna splicing | 6 | 0.4 | 0.0205 |
| Inhibitory synapse assembly | 6 | 0.4 | 0.0205 |
| Purine ribonucleotide catabolic process | 12 | 0.9 | 0.0206 |
| Dna integrity checkpoint signaling | 14 | 1.0 | 0.0206 |
| Positive regulation of synapse assembly | 10 | 0.7 | 0.0207 |
| Cranial nerve development | 10 | 0.7 | 0.0207 |
| Transmission of nerve impulse | 10 | 0.7 | 0.0207 |
| Regulation of organelle transport along microtubule | 4 | 0.3 | 0.0208 |
| Presynaptic membrane assembly | 4 | 0.3 | 0.0208 |
| Metanephric part of ureteric bud development | 3 | 0.2 | 0.0214 |
| Third ventricle development | 3 | 0.2 | 0.0214 |
| Interneuron axon guidance | 3 | 0.2 | 0.0214 |
| Radial glia guided migration of purkinje cell | 3 | 0.2 | 0.0214 |
| Leading strand elongation | 3 | 0.2 | 0.0214 |
| Protein localization to membrane | 43 | 3.2 | 0.022 |
| Peptidyl-amino acid modification | 34 | 2.5 | 0.0221 |
| Nephron tubule development | 12 | 0.9 | 0.0222 |
| Kidney epithelium development | 17 | 1.3 | 0.0222 |
| Response to amino acid | 16 | 1.2 | 0.0224 |
| Postreplication repair | 7 | 0.5 | 0.0226 |
| Regulation of locomotion | 94 | 6.9 | 0.0226 |
| Cytoplasmic microtubule organization | 10 | 0.7 | 0.0226 |
| Ganglion development | 5 | 0.4 | 0.0227 |
| Microtubule polymerization | 8 | 0.6 | 0.0228 |
| Cerebellum morphogenesis | 8 | 0.6 | 0.0228 |
| Regulation of plasma membrane bounded cell projection assembly | 22 | 1.6 | 0.023 |
| Negative regulation of cell development | 28 | 2.1 | 0.0233 |
| Establishment of mitochondrion localization, microtubule-mediated | 6 | 0.4 | 0.0239 |
| Blastocyst hatching | 6 | 0.4 | 0.0239 |
| Organism emergence from protective structure | 6 | 0.4 | 0.0239 |
| Mitochondrion transport along microtubule | 6 | 0.4 | 0.0239 |
| Regulation of long-term neuronal synaptic plasticity | 6 | 0.4 | 0.0239 |
| Postsynaptic density organization | 6 | 0.4 | 0.0239 |
| Hatching | 6 | 0.4 | 0.0239 |
| Regulation of actin filament-based process | 37 | 2.7 | 0.024 |
| Regulation of synaptic transmission, glutamatergic | 11 | 0.8 | 0.0245 |
| Positive regulation of axonogenesis | 11 | 0.8 | 0.0245 |
| Dna damage checkpoint signaling | 13 | 1.0 | 0.0247 |
| Positive regulation of hormone secretion | 17 | 1.3 | 0.0249 |
| Tube morphogenesis | 59 | 4.3 | 0.025 |
| Protein-containing complex organization | 120 | 8.8 | 0.0251 |
| Inner ear morphogenesis | 14 | 1.0 | 0.0252 |
| Microtubule organizing center organization | 14 | 1.0 | 0.0252 |
| Regulation of cell projection assembly | 22 | 1.6 | 0.0253 |
| Lamellipodium organization | 8 | 0.6 | 0.0254 |
| Neuron projection organization | 8 | 0.6 | 0.0254 |
| Base-excision repair | 8 | 0.6 | 0.0254 |
| Cardiac muscle hypertrophy | 7 | 0.5 | 0.0256 |
| Positive regulation of insulin secretion | 12 | 0.9 | 0.0258 |
| Response to abiotic stimulus | 91 | 6.7 | 0.0261 |
| Response to acid chemical | 17 | 1.3 | 0.0263 |
| Negative regulation of catalytic activity | 28 | 2.1 | 0.0264 |
| Regulation of mitotic sister chromatid segregation | 9 | 0.7 | 0.0266 |
| Regulation of cyclin-dependent protein serine/threonine kinase activity | 9 | 0.7 | 0.0266 |
| Positive regulation of protein transport | 26 | 1.9 | 0.0267 |
| Regulation of catalytic activity | 70 | 5.1 | 0.0267 |
| Nephron tubule morphogenesis | 10 | 0.7 | 0.0269 |
| Imp biosynthetic process | 4 | 0.3 | 0.0273 |
| Regulation of synaptic vesicle priming | 4 | 0.3 | 0.0273 |

| | | | |
|---|---|---|---|
| Dendrite extension | 4 | 0.3 | 0.0273 |
| Purine nucleobase metabolic process | 5 | 0.4 | 0.0274 |
| Nuclear body organization | 5 | 0.4 | 0.0274 |
| Gmp metabolic process | 6 | 0.4 | 0.0276 |
| Regulation of dendrite extension | 6 | 0.4 | 0.0276 |
| Neuromuscular synaptic transmission | 6 | 0.4 | 0.0276 |
| Dna replication initiation | 6 | 0.4 | 0.0276 |
| Positive regulation of epithelial cell proliferation | 23 | 1.7 | 0.0278 |
| Regulation of cell motility | 90 | 6.6 | 0.0281 |
| Receptor clustering | 8 | 0.6 | 0.0282 |
| Synaptic transmission, glutamatergic | 8 | 0.6 | 0.0282 |
| Response to alkaloid | 13 | 1.0 | 0.0283 |
| Regulation of multicellular organismal development | 115 | 8.5 | 0.0283 |
| Nuclear division | 31 | 2.3 | 0.0284 |
| Ras protein signal transduction | 11 | 0.8 | 0.0286 |
| Protein localization to plasma membrane | 21 | 1.5 | 0.0287 |
| Striated muscle hypertrophy | 7 | 0.5 | 0.0288 |
| Forebrain neuron development | 7 | 0.5 | 0.0288 |
| Cerebellar cortex morphogenesis | 7 | 0.5 | 0.0288 |
| Regulation of cytosolic calcium ion concentration | 9 | 0.7 | 0.0292 |
| Response to amyloid-beta | 9 | 0.7 | 0.0292 |
| Regulation of protein localization to nucleus | 17 | 1.3 | 0.0292 |
| Tissue development | 137 | 10.1 | 0.0295 |
| Regulation of mitotic metaphase/anaphase transition | 12 | 0.9 | 0.0297 |
| Regulation of protein secretion | 27 | 2.0 | 0.0297 |
| Positive regulation of cell development | 41 | 3.0 | 0.03 |
| Regulation of g protein-coupled receptor signaling pathway | 18 | 1.3 | 0.0301 |
| Cell death | 90 | 6.6 | 0.0301 |
| Artery development | 13 | 1.0 | 0.0302 |
| Positive regulation of peptide hormone secretion | 14 | 1.0 | 0.0303 |
| Positive regulation of rna metabolic process | 140 | 10.3 | 0.0307 |
| Establishment of protein localization | 103 | 7.6 | 0.0309 |
| Negative regulation of nuclear division | 8 | 0.6 | 0.0312 |
| Hindbrain morphogenesis | 8 | 0.6 | 0.0312 |
| Regulation of erythrocyte differentiation | 8 | 0.6 | 0.0312 |
| Negative regulation of kinase activity | 16 | 1.2 | 0.0315 |
| Positive regulation of synaptic transmission | 18 | 1.3 | 0.0316 |
| Nephron epithelium morphogenesis | 10 | 0.7 | 0.0317 |
| Excitatory postsynaptic potential | 10 | 0.7 | 0.0317 |
| Positive regulation of oligodendrocyte differentiation | 6 | 0.4 | 0.0317 |
| Negative regulation of nucleocytoplasmic transport | 6 | 0.4 | 0.0317 |
| Cerebellar purkinje cell layer development | 6 | 0.4 | 0.0317 |
| Mitochondrial genome maintenance | 6 | 0.4 | 0.0317 |
| Negative regulation of dna binding | 6 | 0.4 | 0.0317 |
| Regulation of dna biosynthetic process | 12 | 0.9 | 0.0318 |
| Respiratory gaseous exchange by respiratory system | 9 | 0.7 | 0.0319 |
| Regulation of cell migration | 85 | 6.3 | 0.0321 |
| Long-term memory | 7 | 0.5 | 0.0323 |
| Pattern specification process | 41 | 3.0 | 0.0326 |
| Regulation of monoatomic ion transmembrane transporter activity | 18 | 1.3 | 0.0332 |
| Peptidyl-serine phosphorylation | 11 | 0.8 | 0.0332 |
| Maintenance of cell number | 15 | 1.1 | 0.0338 |
| Ribonucleotide catabolic process | 12 | 0.9 | 0.034 |
| Protein-containing complex disassembly | 17 | 1.3 | 0.0342 |
| Nuclear cell cycle dna replication initiation | 3 | 0.2 | 0.0342 |
| Skeletal muscle satellite cell maintenance involved in skeletal muscle regeneration | 3 | 0.2 | 0.0342 |
| Retrograde neuronal dense core vesicle transport | 3 | 0.2 | 0.0342 |
| Hindbrain maturation | 3 | 0.2 | 0.0342 |
| Mitotic dna replication initiation | 3 | 0.2 | 0.0342 |
| Cell cycle dna replication initiation | 3 | 0.2 | 0.0342 |
| Gephyrin clustering involved in postsynaptic density assembly | 3 | 0.2 | 0.0342 |
| Positive regulation of peptide secretion | 14 | 1.0 | 0.0342 |
| Regulation of monoatomic ion transmembrane transport | 32 | 2.4 | 0.0344 |
| Regulation of metal ion transport | 36 | 2.7 | 0.0346 |
| Regulation of telomerase rna localization to cajal body | 4 | 0.3 | 0.0348 |
| Regulation of protein localization to centrosome | 4 | 0.3 | 0.0348 |
| Reelin-mediated signaling pathway | 4 | 0.3 | 0.0348 |
| Regulation of neurotrophin trk receptor signaling pathway | 4 | 0.3 | 0.0348 |
| Positive regulation of epithelial to mesenchymal transition | 9 | 0.7 | 0.0348 |
| Programmed cell death | 89 | 6.5 | 0.0355 |
| Muscle hypertrophy | 7 | 0.5 | 0.0361 |
| Regulation of metaphase/anaphase transition of cell cycle | 12 | 0.9 | 0.0363 |
| Regulation of sister chromatid segregation | 13 | 1.0 | 0.0365 |
| Chromosome localization | 13 | 1.0 | 0.0365 |
| Regulation of cell size | 19 | 1.4 | 0.037 |
| Negative regulation of neuron differentiation | 10 | 0.7 | 0.037 |
| Nephron morphogenesis | 10 | 0.7 | 0.037 |

| | | | |
|---|---|---|---|
| Cellular homeostasis | 58 | 4.3 | 0.0373 |
| Axon extension | 8 | 0.6 | 0.0378 |
| Response to uv | 17 | 1.3 | 0.0378 |
| Ribonucleoside monophosphate metabolic process | 9 | 0.7 | 0.0379 |
| Forebrain generation of neurons | 11 | 0.8 | 0.0383 |
| Positive regulation of gtpase activity | 14 | 1.0 | 0.0384 |
| Homeostatic process | 114 | 8.4 | 0.0387 |
| Purine nucleotide catabolic process | 12 | 0.9 | 0.0388 |
| Regulation of hormone levels | 47 | 3.5 | 0.0388 |
| Cellular response to nitrogen compound | 53 | 3.9 | 0.0396 |
| Negative regulation of protein kinase activity | 15 | 1.1 | 0.0398 |
| Regulation of receptor-mediated endocytosis | 15 | 1.1 | 0.0398 |
| Negative regulation of g2/m transition of mitotic cell cycle | 10 | 0.7 | 0.0399 |
| Intracellular transport | 94 | 6.9 | 0.04 |
| Walking behavior | 7 | 0.5 | 0.0401 |
| Ribose phosphate metabolic process | 30 | 2.2 | 0.0403 |
| Calcium ion transport | 26 | 1.9 | 0.0405 |
| Regulation of cell cycle g1/s phase transition | 21 | 1.5 | 0.0406 |
| Metaphase chromosome alignment | 11 | 0.8 | 0.041 |
| Response to endogenous stimulus | 97 | 7.1 | 0.0416 |
| Nucleotide metabolic process | 39 | 2.9 | 0.0419 |
| Ear morphogenesis | 15 | 1.1 | 0.042 |
| Axon ensheathment | 15 | 1.1 | 0.042 |
| Ensheathment of neurons | 15 | 1.1 | 0.042 |
| Localization within membrane | 48 | 3.5 | 0.0427 |
| Chaperone-mediated protein folding | 10 | 0.7 | 0.043 |
| Dephosphorylation | 16 | 1.2 | 0.043 |
| Amp biosynthetic process | 4 | 0.3 | 0.0431 |
| Microtubule anchoring at microtubule organizing center | 4 | 0.3 | 0.0431 |
| Neurotransmitter-gated ion channel clustering | 4 | 0.3 | 0.0431 |
| Protein localization to condensed chromosome | 4 | 0.3 | 0.0431 |
| Protein localization to kinetochore | 4 | 0.3 | 0.0431 |
| Neural precursor cell proliferation | 13 | 1.0 | 0.0437 |
| Monoatomic ion transmembrane transport | 68 | 5.0 | 0.0439 |
| Regulation of postsynaptic density organization | 5 | 0.4 | 0.0447 |
| Positive regulation of dna biosynthetic process | 8 | 0.6 | 0.0452 |
| Protein sumoylation | 8 | 0.6 | 0.0452 |
| Aminoglycan biosynthetic process | 8 | 0.6 | 0.0452 |
| Establishment of rna localization | 18 | 1.3 | 0.0461 |
| Negative regulation of cell cycle g2/m phase transition | 10 | 0.7 | 0.0462 |
| Response to vitamin d | 6 | 0.4 | 0.0462 |
| Regulation of axon regeneration | 6 | 0.4 | 0.0462 |
| Establishment of chromosome localization | 12 | 0.9 | 0.0468 |
| Negative regulation of signal transduction | 110 | 8.1 | 0.0469 |
| Regulation of rna metabolic process | 272 | 20.0 | 0.0474 |
| Protein targeting to mitochondrion | 9 | 0.7 | 0.0481 |
| Regulation of transmembrane transporter activity | 18 | 1.3 | 0.0482 |
| Purine nucleotide metabolic process | 34 | 2.5 | 0.0489 |
| Somatic cell dna recombination | 7 | 0.5 | 0.049 |
| Somatic diversification of immune receptors via germline recombination within a single locus | 7 | 0.5 | 0.049 |
| Spontaneous neurotransmitter secretion | 3 | 0.2 | 0.0492 |
| B cell lineage commitment | 3 | 0.2 | 0.0492 |
| Positive regulation of catecholamine metabolic process | 3 | 0.2 | 0.0492 |
| Positive regulation of dopamine metabolic process | 3 | 0.2 | 0.0492 |
| 'de novo' imp biosynthetic process | 3 | 0.2 | 0.0492 |
| Regulation of axon diameter | 3 | 0.2 | 0.0492 |
| Postsynaptic density protein 95 clustering | 3 | 0.2 | 0.0492 |
| Synaptic vesicle uncoating | 3 | 0.2 | 0.0492 |
| Dna synthesis involved in dna repair | 8 | 0.6 | 0.0493 |
| Response to nerve growth factor | 8 | 0.6 | 0.0493 |
| Somatic diversification of immune receptors | 8 | 0.6 | 0.0493 |
| Morphogenesis of an epithelium | 40 | 2.9 | 0.05 |

TableS12

| Pathways upregulated in Gland_A compared to Undiff | | | |
|---|---|---|---|
| **GO Pathway** | **No. of genes** | **%** | **p-value** |
| Lipid metabolic process | 246 | 11.74 | 2.69E-28 |
| Tissue development | 313 | 14.94 | 2.12E-23 |
| Homeostatic process | 263 | 12.55 | 1.13E-20 |
| Response to stress | 514 | 24.53 | 3.02E-20 |
| Epithelium development | 206 | 9.83 | 7.47E-19 |
| Epithelial cell differentiation | 138 | 6.59 | 5.97E-18 |
| Response to external stimulus | 318 | 15.18 | 7.21E-18 |
| Oxoacid metabolic process | 158 | 7.54 | 4.14E-17 |
| Cell adhesion | 180 | 8.59 | 4.79E-17 |
| Regulation of signaling | 504 | 24.06 | 1.67E-16 |
| Regulation of cell adhesion | 159 | 7.59 | 1.89E-16 |
| Cellular response to chemical stimulus | 313 | 14.94 | 1.99E-16 |
| Organic acid metabolic process | 158 | 7.54 | 2.24E-16 |
| Regulation of cell communication | 503 | 24.01 | 3.56E-16 |
| Actin filament organization | 77 | 3.68 | 4.41E-16 |
| Biological process involved in interspecies interaction between organisms | 268 | 12.79 | 6.1E-16 |
| Carboxylic acid metabolic process | 152 | 7.26 | 7.68E-16 |
| Chemical homeostasis | 174 | 8.31 | 1.17E-15 |
| Regulation of localization | 319 | 15.23 | 1.39E-15 |
| Response to endogenous stimulus | 215 | 10.26 | 2.43E-15 |
| Regulation of signal transduction | 447 | 21.34 | 2.63E-15 |
| Phosphate-containing compound metabolic process | 242 | 11.55 | 2.79E-15 |
| Phosphorus metabolic process | 242 | 11.55 | 3.58E-15 |
| Response to external biotic stimulus | 244 | 11.65 | 4.89E-15 |
| Response to biotic stimulus | 249 | 11.89 | 5.18E-15 |
| Positive regulation of signaling | 286 | 13.65 | 5.32E-15 |
| Response to other organism | 243 | 11.6 | 7.28E-15 |
| Response to oxygen-containing compound | 259 | 12.36 | 1.51E-14 |
| Positive regulation of cell communication | 283 | 13.51 | 2.34E-14 |
| Tube development | 167 | 7.97 | 2.47E-14 |
| Actin cytoskeleton organization | 115 | 5.49 | 3.6E-14 |
| Positive regulation of signal transduction | 258 | 12.32 | 3.64E-14 |
| Regulation of cell motility | 194 | 9.26 | 6.9E-14 |
| Regulation of protein localization | 166 | 7.92 | 8.19E-14 |
| Cell-cell junction organization | 55 | 2.63 | 9.27E-14 |
| Regulation of locomotion | 199 | 9.5 | 9.85E-14 |
| Supramolecular fiber organization | 123 | 5.87 | 1.06E-13 |
| Regulation of cell migration | 185 | 8.83 | 1.08E-13 |
| Regulation of biological quality | 429 | 20.48 | 1.52E-13 |
| Actin filament-based process | 123 | 5.87 | 2.13E-13 |
| Regulation of cellular localization | 179 | 8.54 | 2.78E-13 |
| Positive regulation of response to stimulus | 341 | 16.28 | 3.32E-13 |
| Response to hormone | 153 | 7.3 | 4.72E-13 |
| Defense response | 242 | 11.55 | 6.23E-13 |
| Monocarboxylic acid metabolic process | 106 | 5.06 | 6.9E-13 |
| Regulation of response to external stimulus | 194 | 9.26 | 7.16E-13 |
| Gland development | 94 | 4.49 | 7.54E-13 |
| Intracellular signal transduction | 268 | 12.79 | 1.23E-12 |
| Response to cytokine | 156 | 7.45 | 3.49E-12 |
| Response to wounding | 96 | 4.58 | 3.75E-12 |
| Regulation of intracellular signal transduction | 299 | 14.27 | 4.3E-12 |
| Response to peptide | 157 | 7.49 | 7.07E-12 |
| Defense response to other organism | 182 | 8.69 | 1.15E-11 |
| Catabolic process | 295 | 14.08 | 1.61E-11 |
| Regulation of hormone levels | 107 | 5.11 | 2.55E-11 |
| Organophosphate metabolic process | 165 | 7.88 | 2.83E-11 |
| Tube morphogenesis | 127 | 6.06 | 3.29E-11 |
| Actin filament bundle organization | 28 | 1.34 | 3.6E-11 |
| Cell-cell adhesion | 106 | 5.06 | 3.63E-11 |
| Blood vessel development | 105 | 5.01 | 4.21E-11 |
| Cellular catabolic process | 70 | 3.34 | 4.92E-11 |
| Extracellular matrix organization | 66 | 3.15 | 6.01E-11 |
| Extracellular structure organization | 66 | 3.15 | 7.04E-11 |
| Cell development | 321 | 15.32 | 7.29E-11 |
| Regulation of actin filament-based process | 84 | 4.01 | 8.06E-11 |

| | | | |
|---|---|---|---|
| Positive regulation of cell adhesion | 98 | 4.68 | 8.18E-11 |
| External encapsulating structure organization | 66 | 3.15 | 8.25E-11 |
| Fatty acid metabolic process | 73 | 3.48 | 8.77E-11 |
| Positive regulation of biosynthetic process | 367 | 17.52 | 9.27E-11 |
| Circulatory system development | 159 | 7.59 | 9.97E-11 |
| Wound healing | 75 | 3.58 | 1.08E-10 |
| Positive regulation of multicellular organismal process | 254 | 12.12 | 1.12E-10 |
| Small molecule catabolic process | 78 | 3.72 | 1.31E-10 |
| Regulation of cellular component organization | 344 | 16.42 | 1.32E-10 |
| Regulation of transport | 246 | 11.74 | 1.44E-10 |
| Positive regulation of cell motility | 116 | 5.54 | 1.45E-10 |
| Positive regulation of locomotion | 118 | 5.63 | 1.56E-10 |
| Cell junction organization | 106 | 5.06 | 1.61E-10 |
| Alcohol metabolic process | 71 | 3.39 | 1.68E-10 |
| Negative regulation of multicellular organismal process | 194 | 9.26 | 1.7E-10 |
| Epithelial cell development | 52 | 2.48 | 2.39E-10 |
| Actin filament bundle assembly | 26 | 1.24 | 2.83E-10 |
| Respiratory tube development | 51 | 2.43 | 2.97E-10 |
| Positive regulation of cell migration | 111 | 5.3 | 3.08E-10 |
| Multicellular organismal-level homeostasis | 117 | 5.58 | 3.2E-10 |
| Regulation of cell population proliferation | 258 | 12.32 | 3.86E-10 |
| Vasculature development | 105 | 5.01 | 4.21E-10 |
| Lipid catabolic process | 69 | 3.29 | 4.24E-10 |
| Positive regulation of intracellular signal transduction | 182 | 8.69 | 4.54E-10 |
| Cell migration | 154 | 7.35 | 5.12E-10 |
| Amide metabolic process | 83 | 3.96 | 5.24E-10 |
| Innate immune response | 139 | 6.63 | 6.31E-10 |
| Positive regulation of macromolecule biosynthetic process | 348 | 16.61 | 7.81E-10 |
| Positive regulation of protein localization | 96 | 4.58 | 8.66E-10 |
| Positive regulation of metabolic process | 465 | 22.2 | 8.84E-10 |
| Carbohydrate derivative metabolic process | 149 | 7.11 | 1.1E-09 |
| Apoptotic signaling pathway | 71 | 3.39 | 1.14E-09 |
| Anatomical structure formation involved in morphogenesis | 159 | 7.59 | 1.26E-09 |
| Digestive system development | 42 | 2 | 1.27E-09 |
| Regulation of protein localization to membrane | 50 | 2.39 | 1.35E-09 |
| Lung development | 49 | 2.34 | 1.41E-09 |
| Response to type ii interferon | 35 | 1.67 | 1.59E-09 |
| Circulatory system process | 98 | 4.68 | 1.6E-09 |
| Cell-cell junction assembly | 38 | 1.81 | 1.66E-09 |
| Steroid metabolic process | 60 | 2.86 | 1.68E-09 |
| Cellular response to endogenous stimulus | 156 | 7.45 | 1.75E-09 |
| Hemopoiesis | 121 | 5.78 | 1.8E-09 |
| Positive regulation of developmental process | 209 | 9.98 | 1.81E-09 |
| Digestive system process | 29 | 1.38 | 2E-09 |
| Positive regulation of gene expression | 189 | 9.02 | 2.15E-09 |
| Regulation of actin cytoskeleton organization | 74 | 3.53 | 2.16E-09 |
| Positive regulation of molecular function | 138 | 6.59 | 2.61E-09 |
| Intestinal absorption | 18 | 0.86 | 2.81E-09 |
| Defense response to symbiont | 150 | 7.16 | 3.54E-09 |
| Regulation of phosphate metabolic process | 134 | 6.4 | 3.83E-09 |
| Response to lipid | 143 | 6.83 | 4.03E-09 |
| Regulation of immune system process | 244 | 11.65 | 4.05E-09 |
| Regulation of phosphorus metabolic process | 134 | 6.4 | 4.12E-09 |
| Lipid biosynthetic process | 105 | 5.01 | 4.27E-09 |
| Lipid homeostasis | 48 | 2.29 | 4.33E-09 |
| Cell death | 176 | 8.4 | 4.54E-09 |
| Response to metal ion | 71 | 3.39 | 4.75E-09 |
| Regulation of actin filament organization | 61 | 2.91 | 4.84E-09 |
| Enzyme-linked receptor protein signaling pathway | 111 | 5.3 | 4.97E-09 |
| Regulation of body fluid levels | 75 | 3.58 | 5.08E-09 |
| Cellular response to cytokine stimulus | 132 | 6.3 | 5.23E-09 |
| Programmed cell death | 175 | 8.35 | 5.29E-09 |
| Digestion | 35 | 1.67 | 5.43E-09 |
| Respiratory system development | 52 | 2.48 | 5.45E-09 |
| Small molecule biosynthetic process | 89 | 4.25 | 5.54E-09 |
| Regulation of molecular function | 212 | 10.12 | 5.8E-09 |
| Macromolecule localization | 326 | 15.56 | 5.87E-09 |
| Response to growth factor | 101 | 4.82 | 5.91E-09 |

| | | | |
|---|---|---|---|
| Regulation of programmed cell death | 232 | 11.07 | 7.24E-09 |
| Regulation of small molecule metabolic process | 70 | 3.34 | 8.83E-09 |
| Negative regulation of cell adhesion | 66 | 3.15 | 8.93E-09 |
| Digestive tract development | 38 | 1.81 | 9.77E-09 |
| Response to virus | 78 | 3.72 | 1.05E-08 |
| Negative regulation of response to stimulus | 257 | 12.27 | 1.15E-08 |
| Vesicle-mediated transport | 195 | 9.31 | 1.28E-08 |
| Apoptotic process | 168 | 8.02 | 1.43E-08 |
| Immune system process | 326 | 15.56 | 1.54E-08 |
| Positive regulation of phosphatidylinositol 3-kinase/protein kinase b signal transduction | 48 | 2.29 | 1.68E-08 |
| Response to peptide hormone | 82 | 3.91 | 1.74E-08 |
| Blood vessel morphogenesis | 84 | 4.01 | 1.75E-08 |
| Monocarboxylic acid catabolic process | 33 | 1.58 | 1.92E-08 |
| Negative regulation of response to external stimulus | 84 | 4.01 | 1.94E-08 |
| Cellular response to growth factor stimulus | 95 | 4.53 | 1.94E-08 |
| Animal organ morphogenesis | 154 | 7.35 | 1.99E-08 |
| Positive regulation of transport | 139 | 6.63 | 2.07E-08 |
| Defense response to virus | 61 | 2.91 | 2.15E-08 |
| Response to xenobiotic stimulus | 84 | 4.01 | 2.38E-08 |
| Response to nitrogen compound | 164 | 7.83 | 2.58E-08 |
| Carbohydrate metabolic process | 85 | 4.06 | 2.76E-08 |
| Cell surface receptor signaling pathway | 288 | 13.75 | 2.77E-08 |
| Glycosylation | 54 | 2.58 | 2.79E-08 |
| Macromolecule glycosylation | 51 | 2.43 | 2.83E-08 |
| Protein glycosylation | 51 | 2.43 | 2.83E-08 |
| Regulation of protein transport | 84 | 4.01 | 4.3E-08 |
| Cell junction assembly | 62 | 2.96 | 4.63E-08 |
| Signal transduction | 592 | 28.26 | 4.98E-08 |
| Regulation of lipid metabolic process | 68 | 3.25 | 5.34E-08 |
| Regulation of establishment of protein localization | 97 | 4.63 | 5.99E-08 |
| Regulation of apoptotic process | 222 | 10.6 | 6.09E-08 |
| Response to bacterium | 121 | 5.78 | 6.44E-08 |
| Cellular response to oxygen-containing compound | 168 | 8.02 | 6.52E-08 |
| Regulation of defense response | 140 | 6.68 | 6.9E-08 |
| Response to toxic substance | 55 | 2.63 | 7.53E-08 |
| Positive regulation of response to external stimulus | 102 | 4.87 | 7.7E-08 |
| Cell communication | 630 | 30.07 | 7.71E-08 |
| Positive regulation of macromolecule metabolic process | 417 | 19.9 | 7.89E-08 |
| Regulation of viral process | 39 | 1.86 | 8.22E-08 |
| Regulation of actin polymerization or depolymerization | 38 | 1.81 | 1.03E-07 |
| Regulation of cytokine production | 139 | 6.63 | 1.04E-07 |
| Regulation of cell-substrate adhesion | 50 | 2.39 | 1.16E-07 |
| Nucleobase-containing small molecule metabolic process | 104 | 4.96 | 1.27E-07 |
| Response to organophosphorus | 36 | 1.72 | 1.33E-07 |
| Blood circulation | 80 | 3.82 | 1.4E-07 |
| Regulation of phosphatidylinositol 3-kinase/protein kinase b signal transduction | 59 | 2.82 | 1.43E-07 |
| Negative regulation of programmed cell death | 148 | 7.06 | 1.45E-07 |
| Regulation of actin filament length | 38 | 1.81 | 1.76E-07 |
| Hormone metabolic process | 47 | 2.24 | 1.85E-07 |
| Import into cell | 114 | 5.44 | 1.87E-07 |
| Signaling | 622 | 29.69 | 1.9E-07 |
| Cell surface receptor protein tyrosine kinase signaling pathway | 79 | 3.77 | 1.97E-07 |
| Regulation of cell differentiation | 228 | 10.88 | 2.01E-07 |
| Negative regulation of viral process | 25 | 1.19 | 2.04E-07 |
| Regulation of protein localization to plasma membrane | 32 | 1.53 | 2.08E-07 |
| Organic acid catabolic process | 50 | 2.39 | 2.66E-07 |
| Carboxylic acid catabolic process | 50 | 2.39 | 2.66E-07 |
| Regulation of multicellular organismal development | 210 | 10.02 | 2.72E-07 |
| Regulation of response to stress | 208 | 9.93 | 2.78E-07 |
| Organic anion transport | 75 | 3.58 | 2.97E-07 |
| Embryo development | 156 | 7.45 | 3.11E-07 |
| Myeloid cell differentiation | 61 | 2.91 | 3.13E-07 |
| Cellular response to tumor necrosis factor | 38 | 1.81 | 3.52E-07 |
| Positive regulation of phosphorus metabolic process | 85 | 4.06 | 3.71E-07 |
| Positive regulation of phosphate metabolic process | 85 | 4.06 | 3.71E-07 |
| Cholesterol homeostasis | 31 | 1.48 | 3.83E-07 |
| Regulation of cell-cell adhesion | 88 | 4.2 | 3.94E-07 |
| Regulation of viral life cycle | 33 | 1.58 | 4.52E-07 |

| | | | |
|---|---|---|---|
| Sterol homeostasis | 31 | 1.48 | 4.69E-07 |
| Regulation of anatomical structure morphogenesis | 137 | 6.54 | 4.73E-07 |
| Skeletal system development | 88 | 4.2 | 5.07E-07 |
| Triglyceride metabolic process | 24 | 1.15 | 5.08E-07 |
| Nucleoside phosphate metabolic process | 98 | 4.68 | 5.1E-07 |
| Positive regulation of cell differentiation | 139 | 6.63 | 5.26E-07 |
| Fatty acid catabolic process | 27 | 1.29 | 5.3E-07 |
| Mammary gland development | 33 | 1.58 | 5.46E-07 |
| Negative regulation of immune system process | 95 | 4.53 | 5.87E-07 |
| Positive regulation of supramolecular fiber organization | 41 | 1.96 | 6.1E-07 |
| Embryonic organ development | 81 | 3.87 | 6.82E-07 |
| Regulation of secretion | 103 | 4.92 | 6.94E-07 |
| Response to purine-containing compound | 37 | 1.77 | 7.35E-07 |
| Regulation of protein localization to cell periphery | 37 | 1.77 | 7.35E-07 |
| Connective tissue development | 50 | 2.39 | 7.54E-07 |
| Response to camp | 27 | 1.29 | 8.31E-07 |
| Positive regulation of protein transport | 53 | 2.53 | 8.32E-07 |
| Embryonic morphogenesis | 98 | 4.68 | 8.71E-07 |
| Lipid transport | 66 | 3.15 | 9.23E-07 |
| Export from cell | 85 | 4.06 | 9.5E-07 |
| Negative regulation of apoptotic process | 140 | 6.68 | 9.79E-07 |
| Cellular response to type ii interferon | 26 | 1.24 | 9.94E-07 |
| Lipid localization | 66 | 3.15 | 1.02E-06 |
| Sulfur compound metabolic process | 61 | 2.91 | 1.04E-06 |
| Secretion | 92 | 4.39 | 0.0000011 |
| Endocytosis | 87 | 4.15 | 1.13E-06 |
| Regulation of immune response | 154 | 7.35 | 1.14E-06 |
| Positive regulation of transcription by rna polymerase ii | 174 | 8.31 | 1.23E-06 |
| Acylglycerol metabolic process | 28 | 1.34 | 0.0000013 |
| Regulation of protein phosphorylation | 101 | 4.82 | 1.33E-06 |
| Positive regulation of defense response | 79 | 3.77 | 1.42E-06 |
| Response to ketone | 47 | 2.24 | 1.42E-06 |
| Cellular macromolecule localization | 264 | 12.6 | 1.51E-06 |
| Regulation of phosphorylation | 107 | 5.11 | 1.54E-06 |
| Regulation of blood pressure | 44 | 2.1 | 1.58E-06 |
| Organophosphate biosynthetic process | 94 | 4.49 | 1.59E-06 |
| Neutral lipid metabolic process | 28 | 1.34 | 1.59E-06 |
| Protein localization | 263 | 12.55 | 1.61E-06 |
| Regulation of epithelial cell proliferation | 73 | 3.48 | 1.75E-06 |
| Cell activation | 118 | 5.63 | 0.0000018 |
| Carbohydrate homeostasis | 47 | 2.24 | 1.83E-06 |
| Regulation of actin filament depolymerization | 19 | 0.91 | 1.84E-06 |
| Response to carbohydrate | 42 | 2 | 0.000002 |
| Actin polymerization or depolymerization | 21 | 1 | 2.08E-06 |
| Response to tumor necrosis factor | 40 | 1.91 | 2.51E-06 |
| Regulation of response to wounding | 40 | 1.91 | 2.51E-06 |
| Positive regulation of cellular component organization | 161 | 7.68 | 2.61E-06 |
| Glycerolipid metabolic process | 65 | 3.1 | 2.77E-06 |
| Renal system development | 60 | 2.86 | 2.78E-06 |
| Regulation of cytoskeleton organization | 90 | 4.3 | 2.86E-06 |
| Liver development | 34 | 1.62 | 0.0000029 |
| Positive regulation of cytoskeleton organization | 41 | 1.96 | 2.96E-06 |
| Organic acid biosynthetic process | 55 | 2.63 | 3.05E-06 |
| Epithelial tube branching involved in lung morphogenesis | 13 | 0.62 | 3.07E-06 |
| Icosanoid metabolic process | 30 | 1.43 | 3.22E-06 |
| Angiogenesis | 64 | 3.05 | 0.0000036 |
| Cell motility | 165 | 7.88 | 3.66E-06 |
| Regulation of supramolecular fiber organization | 71 | 3.39 | 3.69E-06 |
| Glucose homeostasis | 46 | 2.2 | 3.86E-06 |
| Positive regulation of programmed cell death | 95 | 4.53 | 0.000004 |
| Kidney development | 58 | 2.77 | 4.04E-06 |
| Extracellular matrix assembly | 14 | 0.67 | 4.06E-06 |
| Intracellular chemical homeostasis | 95 | 4.53 | 0.0000043 |
| Regulation of wound healing | 33 | 1.58 | 4.34E-06 |
| Response to nutrient levels | 85 | 4.06 | 4.59E-06 |
| Hepaticobiliary system development | 34 | 1.62 | 4.65E-06 |
| Positive regulation of cell population proliferation | 144 | 6.87 | 4.84E-06 |
| Response to abiotic stimulus | 162 | 7.73 | 5.16E-06 |

| | | | |
|---|---|---|---|
| Response to fatty acid | 20 | 0.95 | 5.18E-06 |
| Monoatomic ion homeostasis | 86 | 4.11 | 5.19E-06 |
| Positive regulation of immune system process | 151 | 7.21 | 5.19E-06 |
| Purine-containing compound metabolic process | 84 | 4.01 | 5.31E-06 |
| Positive regulation of canonical nf-kappab signal transduction | 46 | 2.2 | 5.51E-06 |
| Response to oxygen levels | 63 | 3.01 | 5.53E-06 |
| Monosaccharide metabolic process | 41 | 1.96 | 5.73E-06 |
| Glycoprotein biosynthetic process | 54 | 2.58 | 5.95E-06 |
| Regulation of extracellular matrix organization | 22 | 1.05 | 6.24E-06 |
| Leukocyte differentiation | 73 | 3.48 | 6.33E-06 |
| Epidermis development | 62 | 2.96 | 6.42E-06 |
| Regulation of cellular component biogenesis | 149 | 7.11 | 6.51E-06 |
| Morphogenesis of an epithelium | 78 | 3.72 | 0.0000066 |
| Regulation of microvillus organization | 9 | 0.43 | 6.62E-06 |
| Glomerulus development | 20 | 0.95 | 0.0000067 |
| Glycosphingolipid metabolic process | 20 | 0.95 | 0.0000067 |
| Positive regulation of establishment of protein localization | 61 | 2.91 | 0.0000068 |
| Glycoprotein metabolic process | 63 | 3.01 | 7.27E-06 |
| Pancreas development | 22 | 1.05 | 7.84E-06 |
| Monoatomic ion transport | 146 | 6.97 | 7.85E-06 |
| Response to transforming growth factor beta | 37 | 1.77 | 8.12E-06 |
| Negative regulation of cell-cell adhesion | 43 | 2.05 | 8.25E-06 |
| Regulation of actin filament polymerization | 31 | 1.48 | 8.29E-06 |
| Regulation of catalytic activity | 126 | 6.01 | 8.32E-06 |
| Peptide metabolic process | 20 | 0.95 | 8.61E-06 |
| Cellular response to lipid | 88 | 4.2 | 8.67E-06 |
| Phosphorylation | 73 | 3.48 | 8.75E-06 |
| Regulation of inflammatory response | 75 | 3.58 | 8.86E-06 |
| Xenobiotic metabolic process | 32 | 1.53 | 8.89E-06 |
| Intracellular signaling cassette | 124 | 5.92 | 9.35E-06 |
| Tissue morphogenesis | 92 | 4.39 | 9.78E-06 |
| Apical junction assembly | 19 | 0.91 | 9.91E-06 |
| Cellular homeostasis | 107 | 5.11 | 0.0000101 |
| Carboxylic acid biosynthetic process | 53 | 2.53 | 0.0000103 |
| Positive regulation of inflammatory response | 36 | 1.72 | 0.0000105 |
| Regulation of hydrolase activity | 58 | 2.77 | 0.0000119 |
| Lung morphogenesis | 17 | 0.81 | 0.0000123 |
| Negative regulation of signaling | 207 | 9.88 | 0.0000127 |
| Monoatomic cation homeostasis | 83 | 3.96 | 0.0000127 |
| Tight junction organization | 19 | 0.91 | 0.0000128 |
| Negative regulation of cell communication | 207 | 9.88 | 0.0000131 |
| Regulation of response to biotic stimulus | 91 | 4.34 | 0.0000135 |
| Epidermal growth factor receptor signaling pathway | 26 | 1.24 | 0.0000138 |
| Transition metal ion transport | 26 | 1.24 | 0.0000138 |
| Epithelial cell proliferation | 38 | 1.81 | 0.000014 |
| Homotypic cell-cell adhesion | 20 | 0.95 | 0.000014 |
| Regulation of ketone metabolic process | 32 | 1.53 | 0.000014 |
| Intestinal epithelial cell differentiation | 12 | 0.57 | 0.0000142 |
| Antigen processing and presentation of peptide antigen | 21 | 1 | 0.0000147 |
| Plasma membrane organization | 35 | 1.67 | 0.0000154 |
| Cell morphogenesis | 109 | 5.2 | 0.0000155 |
| Nitrogen compound transport | 205 | 9.79 | 0.0000156 |
| Inflammatory response | 93 | 4.44 | 0.000016 |
| Regulation of protein metabolic process | 236 | 11.26 | 0.0000161 |
| Lipoprotein localization | 11 | 0.53 | 0.0000163 |
| Negative regulation of viral genome replication | 17 | 0.81 | 0.0000164 |
| Primary alcohol catabolic process | 10 | 0.48 | 0.0000168 |
| Negative regulation of protein localization to plasma membrane | 13 | 0.62 | 0.0000172 |
| Cellular localization | 335 | 15.99 | 0.0000175 |
| Response to insulin | 50 | 2.39 | 0.0000178 |
| Response to monosaccharide | 36 | 1.72 | 0.0000179 |
| Lipid modification | 36 | 1.72 | 0.0000179 |
| Cartilage development | 38 | 1.81 | 0.000018 |
| Response to steroid hormone | 53 | 2.53 | 0.0000186 |
| Positive regulation of dna-templated transcription | 216 | 10.31 | 0.0000193 |
| Positive regulation of protein metabolic process | 139 | 6.63 | 0.0000195 |
| Carbohydrate derivative biosynthetic process | 88 | 4.2 | 0.0000196 |
| Protein processing | 39 | 1.86 | 0.0000202 |

| | | | |
|---|---|---|---|
| Regulation of secretion by cell | 90 | 4.3 | 0.0000203 |
| Regulation of substrate adhesion-dependent cell spreading | 19 | 0.91 | 0.0000209 |
| Positive regulation of actin filament bundle assembly | 19 | 0.91 | 0.0000209 |
| Positive regulation of rna biosynthetic process | 216 | 10.31 | 0.0000211 |
| Response to hypoxia | 55 | 2.63 | 0.0000221 |
| Myeloid leukocyte differentiation | 35 | 1.67 | 0.0000231 |
| Response to decreased oxygen levels | 57 | 2.72 | 0.0000239 |
| Positive regulation of cytokine production | 82 | 3.91 | 0.0000242 |
| Cellular response to nitrogen compound | 97 | 4.63 | 0.0000247 |
| Cholesterol metabolic process | 29 | 1.38 | 0.0000254 |
| Cellular response to hormone stimulus | 88 | 4.2 | 0.0000255 |
| Antigen processing and presentation | 27 | 1.29 | 0.0000258 |
| Negative regulation of leukocyte cell-cell adhesion | 34 | 1.62 | 0.0000261 |
| Regulation of peptide transport | 42 | 2 | 0.0000269 |
| Aldehyde metabolic process | 20 | 0.95 | 0.0000276 |
| Positive regulation by symbiont of entry into host | 9 | 0.43 | 0.0000277 |
| Positive regulation of viral entry into host cell | 9 | 0.43 | 0.0000277 |
| Regulation of monoatomic anion transport | 9 | 0.43 | 0.0000277 |
| Unsaturated fatty acid metabolic process | 26 | 1.24 | 0.0000278 |
| Negative regulation of cell activation | 44 | 2.1 | 0.0000281 |
| Regulation of lipid biosynthetic process | 40 | 1.91 | 0.0000284 |
| Cell-substrate adhesion | 38 | 1.81 | 0.0000294 |
| Response to nutrient | 35 | 1.67 | 0.00003 |
| Regulation of protein-containing complex assembly | 71 | 3.39 | 0.0000326 |
| Intrinsic apoptotic signaling pathway | 37 | 1.77 | 0.0000336 |
| Regulation of coagulation | 21 | 1 | 0.0000345 |
| Positive regulation of apoptotic process | 89 | 4.25 | 0.0000345 |
| Regulation of immune effector process | 69 | 3.29 | 0.0000347 |
| Cytokine-mediated signaling pathway | 72 | 3.44 | 0.0000347 |
| Negative regulation of protein localization to cell periphery | 13 | 0.62 | 0.0000359 |
| Regulation of vesicle-mediated transport | 86 | 4.11 | 0.0000365 |
| Anatomical structure homeostasis | 45 | 2.15 | 0.000037 |
| Tissue homeostasis | 45 | 2.15 | 0.000037 |
| Positive regulation of immune response | 104 | 4.96 | 0.0000378 |
| Heart development | 90 | 4.3 | 0.0000386 |
| Response to hexose | 34 | 1.62 | 0.0000387 |
| Liposaccharide metabolic process | 26 | 1.24 | 0.0000387 |
| Fatty acid beta-oxidation | 18 | 0.86 | 0.0000396 |
| Multicellular organismal-level chemical homeostasis | 22 | 1.05 | 0.0000412 |
| Regulation of cell development | 122 | 5.82 | 0.0000435 |
| Positive regulation of phosphorylation | 70 | 3.34 | 0.0000438 |
| Positive regulation of cell-substrate adhesion | 29 | 1.38 | 0.0000462 |
| Lung alveolus development | 17 | 0.81 | 0.0000465 |
| Regulation of protein secretion | 50 | 2.39 | 0.0000469 |
| Cellular response to peptide hormone stimulus | 53 | 2.53 | 0.000047 |
| Regulation of t cell proliferation | 38 | 1.81 | 0.0000471 |
| Positive regulation of rho protein signal transduction | 12 | 0.57 | 0.0000483 |
| Erbb signaling pathway | 27 | 1.29 | 0.0000487 |
| Regulation of anatomical structure size | 80 | 3.82 | 0.0000489 |
| Regulation of canonical nf-kappab signal transduction | 55 | 2.63 | 0.0000493 |
| Lung epithelium development | 14 | 0.67 | 0.0000494 |
| Hexose metabolic process | 35 | 1.67 | 0.0000495 |
| Cellular response to transforming growth factor beta stimulus | 34 | 1.62 | 0.0000499 |
| Intestinal epithelial cell development | 9 | 0.43 | 0.0000506 |
| Regulation of protein modification process | 128 | 6.11 | 0.0000512 |
| Negative regulation of cell population proliferation | 114 | 5.44 | 0.0000518 |
| Sterol metabolic process | 30 | 1.43 | 0.0000549 |
| Secondary alcohol metabolic process | 30 | 1.43 | 0.0000549 |
| Regulation of mapk cascade | 100 | 4.77 | 0.0000555 |
| Detoxification | 32 | 1.53 | 0.0000566 |
| Regulation of cellular component size | 60 | 2.86 | 0.0000577 |
| Regulation of leukocyte cell-cell adhesion | 65 | 3.1 | 0.000059 |
| Regulation of alpha-beta t cell activation | 28 | 1.34 | 0.0000595 |
| Embryonic placenta development | 22 | 1.05 | 0.0000598 |
| Regulation of small gtpase mediated signal transduction | 54 | 2.58 | 0.0000625 |
| Negative regulation of protein-containing complex assembly | 31 | 1.48 | 0.000064 |
| Protein localization to plasma membrane | 39 | 1.86 | 0.0000642 |
| Regulation of dna-binding transcription factor activity | 56 | 2.67 | 0.0000647 |

| | | | |
|---|---|---|---|
| Blood coagulation | 38 | 1.81 | 0.0000662 |
| Negative regulation of signal transduction | 191 | 9.12 | 0.0000665 |
| Negative regulation of protein localization to membrane | 14 | 0.67 | 0.0000667 |
| Cholesterol biosynthetic process | 14 | 0.67 | 0.0000667 |
| Secondary alcohol biosynthetic process | 14 | 0.67 | 0.0000667 |
| Cellular response to fatty acid | 14 | 0.67 | 0.0000667 |
| Regulation of adaptive immune response | 42 | 2 | 0.0000706 |
| Organic hydroxy compound transport | 35 | 1.67 | 0.0000712 |
| Positive regulation of dna-binding transcription factor activity | 38 | 1.81 | 0.000074 |
| Negative regulation of response to wounding | 24 | 1.15 | 0.0000743 |
| Alcohol biosynthetic process | 24 | 1.15 | 0.0000743 |
| Lipoprotein transport | 10 | 0.48 | 0.0000774 |
| Glycolipid catabolic process | 10 | 0.48 | 0.0000774 |
| Regulation of integrin-mediated signaling pathway | 10 | 0.48 | 0.0000774 |
| Negative regulation of wound healing | 20 | 0.95 | 0.0000776 |
| Response to oxidative stress | 62 | 2.96 | 0.0000794 |
| Inorganic ion homeostasis | 71 | 3.39 | 0.000082 |
| Coagulation | 38 | 1.81 | 0.0000827 |
| Steroid biosynthetic process | 24 | 1.15 | 0.0000874 |
| Morphogenesis of a branching epithelium | 34 | 1.62 | 0.0000919 |
| Glycolipid metabolic process | 25 | 1.19 | 0.0000936 |
| Tight junction assembly | 17 | 0.81 | 0.0000943 |
| Primary alcohol metabolic process | 23 | 1.1 | 0.0000948 |
| Chordate embryonic development | 97 | 4.63 | 0.0000948 |
| Positive regulation of secretion | 55 | 2.63 | 0.0000965 |
| Regulation of viral entry into host cell | 12 | 0.57 | 0.0000984 |
| Regulation of peptide secretion | 40 | 1.91 | 0.000105 |
| Positive regulation of cellular component biogenesis | 79 | 3.77 | 0.000107 |
| Negative regulation of cellular component organization | 104 | 4.96 | 0.000111 |
| Positive regulation of protein localization to nucleus | 23 | 1.1 | 0.000112 |
| Leukocyte activation | 94 | 4.49 | 0.000114 |
| Prostanoid metabolic process | 14 | 0.67 | 0.000118 |
| Cellular response to fibroblast growth factor stimulus | 29 | 1.38 | 0.000122 |
| Regulation of interleukin-1 production | 29 | 1.38 | 0.000122 |
| Mononuclear cell differentiation | 60 | 2.86 | 0.000125 |
| Hemostasis | 38 | 1.81 | 0.000127 |
| Negative regulation of phosphorus metabolic process | 47 | 2.24 | 0.000127 |
| Negative regulation of phosphate metabolic process | 47 | 2.24 | 0.000127 |
| Extrinsic apoptotic signaling pathway | 26 | 1.24 | 0.000132 |
| Membrane lipid catabolic process | 15 | 0.72 | 0.000133 |
| Sterol biosynthetic process | 15 | 0.72 | 0.000133 |
| Monoatomic anion transport | 33 | 1.58 | 0.000133 |
| Positive regulation of protein phosphorylation | 65 | 3.1 | 0.000141 |
| Morphogenesis of a branching structure | 35 | 1.67 | 0.000142 |
| Negative regulation of secretion | 35 | 1.67 | 0.000142 |
| Regulation of insulin secretion | 35 | 1.67 | 0.000142 |
| Regulation of biological process involved in symbiotic interaction | 16 | 0.76 | 0.000142 |
| Lipid oxidation | 22 | 1.05 | 0.000143 |
| Cellular response to xenobiotic stimulus | 39 | 1.86 | 0.000149 |
| Desmosome organization | 8 | 0.38 | 0.000153 |
| Fatty acid oxidation | 21 | 1 | 0.000154 |
| Regulation of microvillus length | 6 | 0.29 | 0.000156 |
| Renal system process | 28 | 1.34 | 0.000157 |
| Protein transport | 149 | 7.11 | 0.000161 |
| Regulation of cell activation | 95 | 4.53 | 0.000162 |
| Regulation of peptide hormone secretion | 39 | 1.86 | 0.000165 |
| Secretion by cell | 68 | 3.25 | 0.000168 |
| Regulation of extrinsic apoptotic signaling pathway | 33 | 1.58 | 0.000168 |
| Positive regulation of cell-cell adhesion | 56 | 2.67 | 0.000169 |
| Positive regulation of small gtpase mediated signal transduction | 19 | 0.91 | 0.000172 |
| Embryo development ending in birth or egg hatching | 98 | 4.68 | 0.000174 |
| Actomyosin structure organization | 27 | 1.29 | 0.000178 |
| Cytoskeleton organization | 169 | 8.07 | 0.00018 |
| Ceramide metabolic process | 23 | 1.1 | 0.00018 |
| Carbohydrate derivative catabolic process | 39 | 1.86 | 0.000182 |
| Organophosphate ester transport | 34 | 1.62 | 0.000183 |
| Response to starvation | 42 | 2 | 0.000188 |
| Phospholipid transport | 24 | 1.15 | 0.000189 |

| | | | |
|---|---|---|---|
| Columnar/cuboidal epithelial cell development | 11 | 0.53 | 0.000192 |
| Protein autoprocessing | 11 | 0.53 | 0.000192 |
| Response to molecule of bacterial origin | 59 | 2.82 | 0.000195 |
| Cell population proliferation | 104 | 4.96 | 0.000196 |
| Transmembrane transport | 168 | 8.02 | 0.000199 |
| Regulation of leukocyte proliferation | 48 | 2.29 | 0.000201 |
| Positive regulation of nf-kappab transcription factor activity | 26 | 1.24 | 0.000201 |
| Negative regulation of t cell activation | 29 | 1.38 | 0.000203 |
| Regulation of growth | 90 | 4.3 | 0.000213 |
| Protein localization to cell periphery | 43 | 2.05 | 0.000213 |
| Sphingolipid metabolic process | 32 | 1.53 | 0.000217 |
| Collagen-activated signaling pathway | 17 | 0.81 | 0.000222 |
| Amide biosynthetic process | 31 | 1.48 | 0.000222 |
| Phospholipid metabolic process | 59 | 2.82 | 0.000225 |
| Negative regulation of actin filament depolymerization | 13 | 0.62 | 0.000228 |
| Actin filament polymerization | 13 | 0.62 | 0.000228 |
| Regulation of homotypic cell-cell adhesion | 13 | 0.62 | 0.000228 |
| Cellular response to nutrient levels | 44 | 2.1 | 0.00024 |
| Regulation of response to cytokine stimulus | 37 | 1.77 | 0.000246 |
| Regulation of alpha-beta t cell proliferation | 14 | 0.67 | 0.000255 |
| Elastic fiber assembly | 7 | 0.33 | 0.000256 |
| Regulation of blood coagulation | 18 | 0.86 | 0.000261 |
| Response to thyroid hormone | 10 | 0.48 | 0.000263 |
| Positive regulation of lipid metabolic process | 28 | 1.34 | 0.000263 |
| Nicotinamide nucleotide metabolic process | 28 | 1.34 | 0.000263 |
| Pyridine nucleotide metabolic process | 28 | 1.34 | 0.000263 |
| Ethanol catabolic process | 8 | 0.38 | 0.000263 |
| Negative regulation of coagulation | 15 | 0.72 | 0.000269 |
| Bicellular tight junction assembly | 15 | 0.72 | 0.000269 |
| Regulation of protein depolymerization | 22 | 1.05 | 0.00027 |
| Cellular response to camp | 16 | 0.76 | 0.000274 |
| Iron ion transport | 16 | 0.76 | 0.000274 |
| Regulation of viral genome replication | 20 | 0.95 | 0.000275 |
| Regulation of chemotaxis | 40 | 1.91 | 0.00028 |
| Response to glucose | 31 | 1.48 | 0.00028 |
| Regulation of innate immune response | 74 | 3.53 | 0.000283 |
| Positive regulation of secretion by cell | 50 | 2.39 | 0.000284 |
| Membrane lipid metabolic process | 38 | 1.81 | 0.000284 |
| Negative regulation of protein metabolic process | 103 | 4.92 | 0.000288 |
| Pyridine-containing compound metabolic process | 29 | 1.38 | 0.000293 |
| Positive regulation of transporter activity | 19 | 0.91 | 0.000296 |
| Regulation of production of molecular mediator of immune response | 37 | 1.77 | 0.0003 |
| Monoatomic ion transmembrane transport | 117 | 5.58 | 0.0003 |
| Response to alcohol | 48 | 2.29 | 0.000303 |
| Nervous system development | 275 | 13.13 | 0.000304 |
| Regulation of mononuclear cell proliferation | 44 | 2.1 | 0.000311 |
| Positive regulation of small molecule metabolic process | 31 | 1.48 | 0.000314 |
| Labyrinthine layer development | 14 | 0.67 | 0.000324 |
| Exocrine system development | 14 | 0.67 | 0.000324 |
| Nucleotide-sugar metabolic process | 14 | 0.67 | 0.000324 |
| Positive regulation of immune effector process | 47 | 2.24 | 0.000325 |
| Response to fibroblast growth factor | 29 | 1.38 | 0.00033 |
| Regulation of cell shape | 29 | 1.38 | 0.00033 |
| Regulation of cytokine-mediated signaling pathway | 35 | 1.67 | 0.000331 |
| Regulation of erk1 and erk2 cascade | 50 | 2.39 | 0.000332 |
| Epithelial tube morphogenesis | 53 | 2.53 | 0.000333 |
| Actin filament capping | 12 | 0.57 | 0.000336 |
| Collagen-activated tyrosine kinase receptor signaling pathway | 16 | 0.76 | 0.000337 |
| Barbed-end actin filament capping | 9 | 0.43 | 0.000347 |
| Peptide catabolic process | 9 | 0.43 | 0.000347 |
| Lipoprotein metabolic process | 19 | 0.91 | 0.000351 |
| Regulation of leukocyte migration | 42 | 2 | 0.000355 |
| Intracellular receptor signaling pathway | 37 | 1.77 | 0.000364 |
| Nucleotide-sugar biosynthetic process | 11 | 0.53 | 0.000366 |
| Positive regulation of epithelial cell proliferation | 40 | 1.91 | 0.000368 |
| Regulation of hemostasis | 18 | 0.86 | 0.000376 |
| Response to lipopolysaccharide | 55 | 2.63 | 0.000382 |
| Prostaglandin metabolic process | 13 | 0.62 | 0.000382 |

| | | | |
|---|---|---|---|
| Regulation of actin filament bundle assembly | 24 | 1.15 | 0.000385 |
| Gland morphogenesis | 24 | 1.15 | 0.000385 |
| Regulation of endocytosis | 52 | 2.48 | 0.000387 |
| Regulation of lymphocyte proliferation | 43 | 2.05 | 0.000395 |
| Placenta development | 30 | 1.43 | 0.000405 |
| Negative regulation of transport | 71 | 3.39 | 0.000405 |
| Intestinal hexose absorption | 5 | 0.24 | 0.000416 |
| Positive regulation of stress fiber assembly | 15 | 0.72 | 0.000416 |
| Positive regulation of endocytosis | 32 | 1.53 | 0.00042 |
| Positive regulation of vasculature development | 36 | 1.72 | 0.000423 |
| Ethanol metabolic process | 8 | 0.38 | 0.000429 |
| Myelination | 27 | 1.29 | 0.000435 |
| Cellular response to carbohydrate stimulus | 24 | 1.15 | 0.000441 |
| Response to growth hormone | 12 | 0.57 | 0.000442 |
| Regulation of cell morphogenesis | 43 | 2.05 | 0.000467 |
| Regulation of ventricular cardiac muscle cell action potential | 7 | 0.33 | 0.000469 |
| Estrogen metabolic process | 13 | 0.62 | 0.000488 |
| Antigen processing and presentation of peptide antigen via mhc class i | 13 | 0.62 | 0.000488 |
| Endocrine system development | 27 | 1.29 | 0.000491 |
| Response to endoplasmic reticulum stress | 41 | 1.96 | 0.000491 |
| Cellular response to starvation | 35 | 1.67 | 0.000492 |
| Regulation of systemic arterial blood pressure | 24 | 1.15 | 0.000503 |
| Developmental growth involved in morphogenesis | 29 | 1.38 | 0.000523 |
| Platelet-derived growth factor receptor signaling pathway | 20 | 0.95 | 0.000523 |
| Lung cell differentiation | 10 | 0.48 | 0.000524 |
| Lung epithelial cell differentiation | 10 | 0.48 | 0.000524 |
| Negative regulation of lymphocyte activation | 33 | 1.58 | 0.000546 |
| Regulation of cell projection organization | 94 | 4.49 | 0.000546 |
| Axon ensheathment | 27 | 1.29 | 0.000552 |
| Ensheathment of neurons | 27 | 1.29 | 0.000552 |
| Negative regulation of defense response | 51 | 2.43 | 0.000559 |
| Viral process | 45 | 2.15 | 0.000561 |
| Protein phosphorylation | 60 | 2.86 | 0.000566 |
| Regulation of angiogenesis | 57 | 2.72 | 0.000567 |
| Animal organ regeneration | 17 | 0.81 | 0.000571 |
| Bicarbonate transport | 12 | 0.57 | 0.000574 |
| Epithelial structure maintenance | 12 | 0.57 | 0.000574 |
| Developmental growth | 69 | 3.29 | 0.000577 |
| Growth | 69 | 3.29 | 0.000577 |
| Muscle structure development | 77 | 3.68 | 0.000587 |
| Regulation of vasculature development | 58 | 2.77 | 0.000598 |
| Positive regulation of catalytic activity | 74 | 3.53 | 0.000603 |
| Establishment or maintenance of bipolar cell polarity | 15 | 0.72 | 0.000626 |
| Establishment or maintenance of apical/basal cell polarity | 15 | 0.72 | 0.000626 |
| Negative regulation of intracellular signal transduction | 103 | 4.92 | 0.000628 |
| Maintenance of location | 21 | 1 | 0.000632 |
| Regulation of cytokine production involved in immune response | 26 | 1.24 | 0.000635 |
| Regulation of adaptive immune response based on somatic recombination of immune receptors built from immunoglobulin superfamily domains | 37 | 1.77 | 0.000636 |
| Regulation of t cell activation | 61 | 2.91 | 0.000646 |
| Positive regulation of angiogenesis | 35 | 1.67 | 0.000655 |
| Regeneration | 31 | 1.48 | 0.000665 |
| Epithelial cell-cell adhesion | 8 | 0.38 | 0.000668 |
| Regulation of plasma membrane organization | 8 | 0.38 | 0.000668 |
| Positive regulation of response to biotic stimulus | 54 | 2.58 | 0.000675 |
| Metal ion transport | 92 | 4.39 | 0.000677 |
| Cellular component maintenance | 17 | 0.81 | 0.00068 |
| Skin development | 47 | 2.24 | 0.00071 |
| Nucleotide metabolic process | 67 | 3.2 | 0.000715 |
| Podocyte differentiation | 10 | 0.48 | 0.000719 |
| Renal filtration cell differentiation | 10 | 0.48 | 0.000719 |
| Regulation of systemic arterial blood pressure by renin-angiotensin | 10 | 0.48 | 0.000719 |
| Mammary gland epithelium development | 16 | 0.76 | 0.000725 |
| Biological process involved in interaction with host | 29 | 1.38 | 0.000726 |
| Labyrinthine layer morphogenesis | 9 | 0.43 | 0.000737 |
| Stress fiber assembly | 9 | 0.43 | 0.000737 |
| Contractile actin filament bundle assembly | 9 | 0.43 | 0.000737 |
| Modulation by symbiont of entry into host | 12 | 0.57 | 0.000737 |

| | | | |
|---|---|---|---|
| Positive regulation of alpha-beta t cell activation | 18 | 0.86 | 0.00074 |
| Regulation of lymphocyte activation | 76 | 3.63 | 0.000748 |
| Carboxylic acid transport | 49 | 2.34 | 0.000758 |
| Negative regulation of actin filament polymerization | 15 | 0.72 | 0.000761 |
| Tumor necrosis factor-mediated signaling pathway | 15 | 0.72 | 0.000761 |
| Establishment or maintenance of epithelial cell apical/basal polarity | 14 | 0.67 | 0.00078 |
| Response to fructose | 6 | 0.29 | 0.00079 |
| Galactolipid metabolic process | 6 | 0.29 | 0.00079 |
| Monoatomic cation transport | 112 | 5.35 | 0.000794 |
| Midgut development | 7 | 0.33 | 0.0008 |
| Establishment of protein localization | 169 | 8.07 | 0.000802 |
| Positive regulation of protein secretion | 29 | 1.38 | 0.000807 |
| Organic acid transport | 49 | 2.34 | 0.000814 |
| Peripheral nervous system development | 20 | 0.95 | 0.000817 |
| Regulation of rho protein signal transduction | 20 | 0.95 | 0.000817 |
| Regulation of amide metabolic process | 20 | 0.95 | 0.000817 |
| Muscle cell differentiation | 47 | 2.24 | 0.000823 |
| Cellular response to chemical stress | 47 | 2.24 | 0.000823 |
| Epidermal cell differentiation | 37 | 1.77 | 0.000831 |
| Intracellular monoatomic ion homeostasis | 66 | 3.15 | 0.000835 |
| Regulation of plasma membrane bounded cell projection organization | 91 | 4.34 | 0.000835 |
| Positive regulation of hydrolase activity | 34 | 1.62 | 0.000838 |
| Regulation of t cell mediated immunity | 23 | 1.1 | 0.000845 |
| Purine nucleotide metabolic process | 59 | 2.82 | 0.000846 |
| Regulation of system process | 83 | 3.96 | 0.000849 |
| Positive regulation of rna metabolic process | 228 | 10.88 | 0.00085 |
| Regulation of reactive oxygen species metabolic process | 30 | 1.43 | 0.000856 |
| Positive regulation of t cell mediated immunity | 18 | 0.86 | 0.000868 |
| Response to calcium ion | 27 | 1.29 | 0.000872 |
| Neuron projection development | 100 | 4.77 | 0.000886 |
| Regulation of hormone secretion | 44 | 2.1 | 0.000892 |
| Regulation of lipid localization | 34 | 1.62 | 0.000919 |
| Intracellular glucose homeostasis | 25 | 1.19 | 0.000925 |
| Nephron development | 28 | 1.34 | 0.000936 |
| Regulation of nucleotide metabolic process | 20 | 0.95 | 0.000943 |
| Myeloid leukocyte activation | 30 | 1.43 | 0.000948 |
| Regulation of tumor necrosis factor-mediated signaling pathway | 17 | 0.81 | 0.000952 |
| Terpenoid metabolic process | 22 | 1.05 | 0.000963 |
| Transforming growth factor beta receptor signaling pathway | 22 | 1.05 | 0.000963 |
| Schwann cell development | 13 | 0.62 | 0.000965 |
| Plasma membrane phospholipid scrambling | 10 | 0.48 | 0.000969 |
| Antigen processing and presentation of endogenous peptide antigen | 10 | 0.48 | 0.000969 |
| Epithelial cell maturation | 10 | 0.48 | 0.000969 |
| Glomerular epithelial cell differentiation | 10 | 0.48 | 0.000969 |
| Cell-matrix adhesion | 27 | 1.29 | 0.000974 |
| Regulation of carbohydrate metabolic process | 33 | 1.58 | 0.000976 |
| In utero embryonic development | 61 | 2.91 | 0.000982 |
| Regulation of hepatocyte proliferation | 8 | 0.38 | 0.001 |
| Glycosphingolipid catabolic process | 8 | 0.38 | 0.001 |
| Stem cell differentiation | 34 | 1.62 | 0.00101 |
| Carbohydrate catabolic process | 26 | 1.24 | 0.00101 |
| Positive regulation of reactive oxygen species metabolic process | 18 | 0.86 | 0.00101 |
| Maintenance of gastrointestinal epithelium | 9 | 0.43 | 0.00103 |
| Endomembrane system organization | 82 | 3.91 | 0.00104 |
| Negative regulation of leukocyte activation | 36 | 1.72 | 0.00106 |
| Membrane organization | 104 | 4.96 | 0.00107 |
| Nucleoside phosphate biosynthetic process | 48 | 2.29 | 0.00109 |
| Negative regulation of protein localization | 39 | 1.86 | 0.0011 |
| Positive regulation of protein modification process | 80 | 3.82 | 0.00111 |
| Regulation of chloride transport | 5 | 0.24 | 0.00115 |
| Bundle of his cell-purkinje myocyte adhesion involved in cell communication | 5 | 0.24 | 0.00115 |
| Hemidesmosome assembly | 5 | 0.24 | 0.00115 |
| Embryonic organ morphogenesis | 49 | 2.34 | 0.00115 |
| Monoatomic anion transmembrane transport | 28 | 1.34 | 0.00116 |
| Nad metabolic process | 13 | 0.62 | 0.00119 |
| Negative regulation of blood coagulation | 13 | 0.62 | 0.00119 |
| Positive regulation of apoptotic signaling pathway | 29 | 1.38 | 0.00122 |
| Antiviral innate immune response | 16 | 0.76 | 0.00122 |

| | | | |
|---|---|---|---|
| Neuron development | 119 | 5.68 | 0.00123 |
| Negative regulation of protein modification process | 49 | 2.34 | 0.00124 |
| Regulation of stress fiber assembly | 21 | 1 | 0.00125 |
| Positive regulation of intracellular protein transport | 21 | 1 | 0.00125 |
| Cellular response to stress | 205 | 9.79 | 0.00128 |
| Intracellular monoatomic cation homeostasis | 64 | 3.05 | 0.00131 |
| Response to type i interferon | 15 | 0.72 | 0.00132 |
| Positive regulation of innate immune response | 49 | 2.34 | 0.00132 |
| Alpha-beta t cell activation | 24 | 1.15 | 0.00135 |
| Positive regulation of interleukin-1 production | 18 | 0.86 | 0.00137 |
| Embryonic skeletal system development | 26 | 1.24 | 0.0014 |
| Negative regulation of chemotaxis | 14 | 0.67 | 0.0014 |
| Regulation of leukocyte activation | 83 | 3.96 | 0.00141 |
| Protein o-linked glycosylation | 21 | 1 | 0.00142 |
| Antigen processing and presentation of endogenous antigen | 11 | 0.53 | 0.00142 |
| Negative regulation of developmental process | 133 | 6.35 | 0.00144 |
| Positive regulation of interleukin-1 beta production | 16 | 0.76 | 0.00144 |
| Bronchus development | 6 | 0.29 | 0.00145 |
| Trachea development | 8 | 0.38 | 0.00145 |
| Regulation of systemic arterial blood pressure by circulatory renin-angiotensin | 8 | 0.38 | 0.00145 |
| Negative regulation of hemostasis | 13 | 0.62 | 0.00146 |
| Semaphorin-plexin signaling pathway | 13 | 0.62 | 0.00146 |
| Regulation of proteolysis | 62 | 2.96 | 0.00146 |
| Negative regulation of secretion by cell | 29 | 1.38 | 0.00149 |
| Regulation of intracellular protein transport | 29 | 1.38 | 0.00149 |
| Defense response to bacterium | 53 | 2.53 | 0.00153 |
| Regulation of steroid metabolic process | 23 | 1.1 | 0.00155 |
| Symbiont entry into host | 23 | 1.1 | 0.00155 |
| Regulation of interleukin-1 beta production | 23 | 1.1 | 0.00155 |
| Symbiont entry into host cell | 23 | 1.1 | 0.00155 |
| Protein localization to extracellular region | 26 | 1.24 | 0.00156 |
| Carbohydrate transmembrane transport | 15 | 0.72 | 0.00157 |
| Protein n-linked glycosylation | 18 | 0.86 | 0.00159 |
| Neural crest cell development | 18 | 0.86 | 0.00159 |
| Regulation of apoptotic signaling pathway | 60 | 2.86 | 0.00161 |
| Regulation of lipid transport | 29 | 1.38 | 0.00164 |
| Female pregnancy | 32 | 1.53 | 0.00164 |
| Cellular response to metal ion | 32 | 1.53 | 0.00164 |
| Regulation of platelet aggregation | 10 | 0.48 | 0.00168 |
| Brain-derived neurotrophic factor receptor signaling pathway | 14 | 0.67 | 0.00169 |
| Regulation of t cell receptor signaling pathway | 14 | 0.67 | 0.00169 |
| Phospholipid translocation | 14 | 0.67 | 0.00169 |
| Regulation of carbohydrate catabolic process | 16 | 0.76 | 0.00169 |
| Regulation of protein localization to nucleus | 28 | 1.34 | 0.00173 |
| Regulation of mononuclear cell migration | 28 | 1.34 | 0.00173 |
| Positive regulation of protein localization to membrane | 22 | 1.05 | 0.00179 |
| Regulation of fatty acid metabolic process | 22 | 1.05 | 0.00179 |
| Renal absorption | 11 | 0.53 | 0.0018 |
| Regulation of kinase activity | 58 | 2.77 | 0.00182 |
| Antigen processing and presentation of exogenous peptide antigen | 12 | 0.57 | 0.00182 |
| Sphingolipid catabolic process | 12 | 0.57 | 0.00182 |
| Cellular response to external stimulus | 18 | 0.86 | 0.00183 |
| Vascular process in circulatory system | 45 | 2.15 | 0.00183 |
| Positive regulation of cytokine production involved in immune response | 19 | 0.91 | 0.00185 |
| Olefinic compound metabolic process | 28 | 1.34 | 0.00191 |
| Phospholipid biosynthetic process | 39 | 1.86 | 0.00191 |
| Positive regulation of alpha-beta t cell proliferation | 9 | 0.43 | 0.00191 |
| Positive regulation of peptide hormone secretion | 23 | 1.1 | 0.00195 |
| Transforming growth factor beta receptor superfamily signaling pathway | 32 | 1.53 | 0.00196 |
| Negative regulation of cell development | 45 | 2.15 | 0.00196 |
| Glycosylceramide metabolic process | 7 | 0.33 | 0.00197 |
| Regulation of protein-containing complex disassembly | 25 | 1.19 | 0.00201 |
| Regulation of actomyosin structure organization | 22 | 1.05 | 0.00202 |
| Cellular response to ketone | 22 | 1.05 | 0.00202 |
| Positive regulation of production of molecular mediator of immune response | 27 | 1.29 | 0.00202 |
| Epoxygenase p450 pathway | 8 | 0.38 | 0.00205 |
| Response to mechanical stimulus | 38 | 1.81 | 0.00206 |
| Response to hydrogen peroxide | 21 | 1 | 0.00206 |

| | | | |
|---|---|---|---|
| Negative regulation of mapk cascade | 31 | 1.48 | 0.00209 |
| Maintenance of location in cell | 18 | 0.86 | 0.0021 |
| Vesicle budding from membrane | 18 | 0.86 | 0.0021 |
| Regulation of purine nucleotide metabolic process | 19 | 0.91 | 0.00211 |
| Cytokinetic process | 13 | 0.62 | 0.00215 |
| Neuron projection morphogenesis | 72 | 3.44 | 0.00215 |
| Pyridine-containing compound biosynthetic process | 10 | 0.48 | 0.00216 |
| Glomerular epithelium development | 10 | 0.48 | 0.00216 |
| Inorganic anion transport | 29 | 1.38 | 0.00217 |
| Export across plasma membrane | 15 | 0.72 | 0.00219 |
| Regulation of leukocyte chemotaxis | 25 | 1.19 | 0.00223 |
| Regulation of digestive system process | 12 | 0.57 | 0.00224 |
| Positive regulation of protein import into nucleus | 12 | 0.57 | 0.00224 |
| Immune response | 216 | 10.31 | 0.00225 |
| Establishment of skin barrier | 11 | 0.53 | 0.00226 |
| Smooth muscle cell differentiation | 11 | 0.53 | 0.00226 |
| Intrinsic apoptotic signaling pathway in response to endoplasmic reticulum stress | 11 | 0.53 | 0.00226 |
| Maintenance of protein location | 16 | 0.76 | 0.00231 |
| Homeostasis of number of cells | 47 | 2.24 | 0.00232 |
| Fibroblast growth factor receptor signaling pathway | 21 | 1 | 0.00233 |
| Positive regulation of adaptive immune response | 26 | 1.24 | 0.00235 |
| Golgi vesicle transport | 46 | 2.2 | 0.00236 |
| Regulation of erbb signaling pathway | 17 | 0.81 | 0.00238 |
| Regulation of purine nucleotide catabolic process | 14 | 0.67 | 0.00239 |
| Regulation of nucleotide catabolic process | 14 | 0.67 | 0.00239 |
| Regulation of glycolytic process | 14 | 0.67 | 0.00239 |
| Anterior/posterior pattern specification | 36 | 1.72 | 0.0024 |
| Regulation of protein kinase activity | 54 | 2.58 | 0.0024 |
| Endothelial cell migration | 18 | 0.86 | 0.00241 |
| Positive regulation of peptide secretion | 23 | 1.1 | 0.00244 |
| Golgi ribbon formation | 6 | 0.29 | 0.00245 |
| Regulation of basement membrane organization | 6 | 0.29 | 0.00245 |
| Protein folding in endoplasmic reticulum | 6 | 0.29 | 0.00245 |
| Bradykinin catabolic process | 5 | 0.24 | 0.00248 |
| Desmosome assembly | 5 | 0.24 | 0.00248 |
| Cardiac muscle cell-cardiac muscle cell adhesion | 5 | 0.24 | 0.00248 |
| Regulation of plasma membrane repair | 5 | 0.24 | 0.00248 |
| Cell morphogenesis involved in neuron differentiation | 66 | 3.15 | 0.00249 |
| Host-mediated suppression of symbiont invasion | 9 | 0.43 | 0.00253 |
| Antigen processing and presentation of endogenous peptide antigen via mhc class i | 9 | 0.43 | 0.00253 |
| Regulation of cell growth | 60 | 2.86 | 0.00255 |
| Positive regulation of cytokine-mediated signaling pathway | 15 | 0.72 | 0.00257 |
| Schwann cell differentiation | 13 | 0.62 | 0.00258 |
| Regulation of systemic arterial blood pressure mediated by a chemical signal | 13 | 0.62 | 0.00258 |
| Regulation of cholesterol metabolic process | 13 | 0.62 | 0.00258 |
| Platelet-derived growth factor receptor-alpha signaling pathway | 13 | 0.62 | 0.00258 |
| Ameboidal-type cell migration | 26 | 1.24 | 0.00259 |
| Columnar/cuboidal epithelial cell differentiation | 21 | 1 | 0.00262 |
| Cell projection morphogenesis | 73 | 3.48 | 0.00266 |
| Positive regulation of response to cytokine stimulus | 16 | 0.76 | 0.00268 |
| Dicarboxylic acid metabolic process | 20 | 0.95 | 0.00268 |
| Positive regulation of insulin secretion | 19 | 0.91 | 0.00273 |
| Plasma membrane bounded cell projection morphogenesis | 72 | 3.44 | 0.00275 |
| Regulation of cardiac muscle cell action potential | 10 | 0.48 | 0.00275 |
| Cell surface receptor protein serine/threonine kinase signaling pathway | 33 | 1.58 | 0.00278 |
| Negative regulation of protein phosphorylation | 34 | 1.62 | 0.00279 |
| Positive regulation of proteolysis | 37 | 1.77 | 0.00279 |
| Response to interferon-beta | 11 | 0.53 | 0.0028 |
| Multi-multicellular organism process | 35 | 1.67 | 0.0028 |
| Triglyceride biosynthetic process | 8 | 0.38 | 0.00282 |
| Regulation of viral transcription | 8 | 0.38 | 0.00282 |
| T cell activation | 52 | 2.48 | 0.00282 |
| Regulation of chondrocyte differentiation | 14 | 0.67 | 0.00283 |
| Monosaccharide catabolic process | 14 | 0.67 | 0.00283 |
| Actin filament severing | 7 | 0.33 | 0.00289 |
| Regulation of adipose tissue development | 7 | 0.33 | 0.00289 |
| Neurogenesis | 173 | 8.26 | 0.00291 |
| Skeletal system morphogenesis | 38 | 1.81 | 0.00299 |

| | | | |
|---|---|---|---|
| Limb development | 33 | 1.58 | 0.00301 |
| Appendage development | 33 | 1.58 | 0.00301 |
| Bone development | 34 | 1.62 | 0.00302 |
| Cellular detoxification | 23 | 1.1 | 0.00302 |
| Antigen processing and presentation of exogenous antigen | 13 | 0.62 | 0.00308 |
| Negative regulation of alpha-beta t cell activation | 13 | 0.62 | 0.00308 |
| Response to epidermal growth factor | 13 | 0.62 | 0.00308 |
| Negative regulation of protein depolymerization | 16 | 0.76 | 0.00309 |
| Inorganic ion transmembrane transport | 101 | 4.82 | 0.0031 |
| Ephrin receptor signaling pathway | 18 | 0.86 | 0.00313 |
| Regulation of atp metabolic process | 17 | 0.81 | 0.00314 |
| Regulation of muscle hypertrophy | 17 | 0.81 | 0.00314 |
| Negative regulation of t cell proliferation | 17 | 0.81 | 0.00314 |
| Long-chain fatty acid metabolic process | 22 | 1.05 | 0.00316 |
| Positive regulation of transmembrane receptor protein serine/threonine kinase signaling pathway | 22 | 1.05 | 0.00316 |
| Negative regulation of cell projection organization | 32 | 1.53 | 0.00324 |
| Positive regulation of viral life cycle | 9 | 0.43 | 0.00329 |
| Embryonic placenta morphogenesis | 9 | 0.43 | 0.00329 |
| Pyridine nucleotide biosynthetic process | 9 | 0.43 | 0.00329 |
| Nicotinamide nucleotide biosynthetic process | 9 | 0.43 | 0.00329 |
| Modulation of process of another organism | 23 | 1.1 | 0.00336 |
| Positive regulation of lipid biosynthetic process | 20 | 0.95 | 0.0034 |
| Cellular response to hexose stimulus | 20 | 0.95 | 0.0034 |
| Cellular response to fructose stimulus | 4 | 0.19 | 0.00343 |
| Intestinal d-glucose absorption | 4 | 0.19 | 0.00343 |
| Neuropeptide catabolic process | 4 | 0.19 | 0.00343 |
| Positive regulation of plasma membrane repair | 4 | 0.19 | 0.00343 |
| Endocrine pancreas development | 11 | 0.53 | 0.00344 |
| Glycosphingolipid biosynthetic process | 11 | 0.53 | 0.00344 |
| Antigen processing and presentation of peptide or polysaccharide antigen via mhc class ii | 11 | 0.53 | 0.00344 |
| Establishment of endothelial barrier | 10 | 0.48 | 0.00346 |
| Positive regulation of protein localization to plasma membrane | 15 | 0.72 | 0.00348 |
| Carbohydrate transport | 15 | 0.72 | 0.00348 |
| Neural crest cell differentiation | 19 | 0.91 | 0.00349 |
| Diterpenoid metabolic process | 19 | 0.91 | 0.00349 |
| Cellular response to vascular endothelial growth factor stimulus | 19 | 0.91 | 0.00349 |
| Heart process | 22 | 1.05 | 0.00352 |
| Isoprenoid metabolic process | 24 | 1.15 | 0.00353 |
| Stem cell development | 18 | 0.86 | 0.00356 |
| Regulation of muscle system process | 41 | 1.96 | 0.00356 |
| Activation of immune response | 59 | 2.82 | 0.00363 |
| Response to corticosteroid | 28 | 1.34 | 0.00364 |
| Regulation of t cell mediated cytotoxicity | 13 | 0.62 | 0.00365 |
| Positive regulation of actin filament polymerization | 13 | 0.62 | 0.00365 |
| Actin filament-based movement | 21 | 1 | 0.00367 |
| Negative regulation of response to biotic stimulus | 25 | 1.19 | 0.00367 |
| Viral life cycle | 31 | 1.48 | 0.00378 |
| Biological process involved in symbiotic interaction | 32 | 1.53 | 0.0038 |
| Cellular response to insulin stimulus | 33 | 1.58 | 0.00381 |
| Glomerular basement membrane development | 6 | 0.29 | 0.00387 |
| Regulation of intestinal lipid absorption | 6 | 0.29 | 0.00387 |
| Positive regulation of integrin-mediated signaling pathway | 6 | 0.29 | 0.00387 |
| Collagen biosynthetic process | 6 | 0.29 | 0.00387 |
| Positive regulation of extrinsic apoptotic signaling pathway | 14 | 0.67 | 0.00389 |
| Positive regulation of receptor-mediated endocytosis | 14 | 0.67 | 0.00389 |
| Lipid translocation | 14 | 0.67 | 0.00389 |
| Peptidyl-tyrosine modification | 14 | 0.67 | 0.00389 |
| Regulation of leukocyte differentiation | 51 | 2.43 | 0.00389 |
| Positive regulation of lipid localization | 22 | 1.05 | 0.00391 |
| Cellular response to epidermal growth factor stimulus | 12 | 0.57 | 0.00395 |
| Triglyceride homeostasis | 12 | 0.57 | 0.00395 |
| Regulation of cellular response to growth factor stimulus | 56 | 2.67 | 0.00398 |
| Regulation of nervous system development | 65 | 3.1 | 0.00403 |
| Regulation of plasma lipoprotein particle levels | 15 | 0.72 | 0.00403 |
| Regulation of membrane lipid distribution | 15 | 0.72 | 0.00403 |
| Regulation of protein import into nucleus | 15 | 0.72 | 0.00403 |
| Regulation of interleukin-10 production | 15 | 0.72 | 0.00403 |
| Cellular response to mechanical stimulus | 17 | 0.81 | 0.00408 |

| | | | |
|---|---|---|---|
| Glycerolipid biosynthetic process | 35 | 1.67 | 0.00408 |
| Regulation of peptidase activity | 16 | 0.76 | 0.00409 |
| Negative regulation of viral transcription | 7 | 0.33 | 0.0041 |
| Regulation of metallopeptidase activity | 7 | 0.33 | 0.0041 |
| Lipid digestion | 7 | 0.33 | 0.0041 |
| Purine ribonucleoside bisphosphate metabolic process | 7 | 0.33 | 0.0041 |
| 3'-phosphoadenosine 5'-phosphosulfate metabolic process | 7 | 0.33 | 0.0041 |
| Branch elongation of an epithelium | 7 | 0.33 | 0.0041 |
| Mammary gland epithelial cell differentiation | 7 | 0.33 | 0.0041 |
| Regulation of vasculogenesis | 7 | 0.33 | 0.0041 |
| Developmental cell growth | 23 | 1.1 | 0.00412 |
| Chloride transport | 23 | 1.1 | 0.00412 |
| Regulation of t cell cytokine production | 11 | 0.53 | 0.00419 |
| O-glycan processing | 11 | 0.53 | 0.00419 |
| Regulation of hormone metabolic process | 11 | 0.53 | 0.00419 |
| Ruffle organization | 9 | 0.43 | 0.00421 |
| Cellular response to monosaccharide stimulus | 20 | 0.95 | 0.00427 |
| Pyroptotic inflammatory response | 10 | 0.48 | 0.00431 |
| Hexose catabolic process | 13 | 0.62 | 0.00431 |
| Negative regulation of immune response | 35 | 1.67 | 0.00439 |
| Gastrulation | 30 | 1.43 | 0.00442 |
| Post-embryonic development | 19 | 0.91 | 0.00442 |
| Response to glucocorticoid | 25 | 1.19 | 0.00444 |
| Regulation of nucleocytoplasmic transport | 23 | 1.1 | 0.00455 |
| Retinoid metabolic process | 18 | 0.86 | 0.00455 |
| Kidney epithelium development | 26 | 1.24 | 0.00455 |
| Regulation of non-canonical nf-kappab signal transduction | 21 | 1 | 0.00456 |
| Galactosylceramide metabolic process | 5 | 0.24 | 0.00457 |
| Ire1-mediated unfolded protein response | 5 | 0.24 | 0.00457 |
| Brush border assembly | 5 | 0.24 | 0.00457 |
| Renal protein absorption | 5 | 0.24 | 0.00457 |
| Regulation of alpha-beta t cell differentiation | 17 | 0.81 | 0.00464 |
| Cellular response to corticosteroid stimulus | 15 | 0.72 | 0.00465 |
| Regulation of hemopoiesis | 62 | 2.96 | 0.00468 |
| Acylglycerol homeostasis | 12 | 0.57 | 0.00472 |
| Positive regulation of nitric oxide biosynthetic process | 12 | 0.57 | 0.00472 |
| Negative chemotaxis | 12 | 0.57 | 0.00472 |
| Cellular response to toxic substance | 24 | 1.15 | 0.00472 |
| Inorganic anion transmembrane transport | 24 | 1.15 | 0.00472 |
| Negative regulation of innate immune response | 20 | 0.95 | 0.00477 |
| Multi-organism reproductive process | 33 | 1.58 | 0.00479 |
| Branching morphogenesis of an epithelial tube | 25 | 1.19 | 0.00486 |
| Regulated exocytosis | 25 | 1.19 | 0.00486 |
| Response to ethanol | 25 | 1.19 | 0.00486 |
| Positive regulation of mapk cascade | 66 | 3.15 | 0.00494 |
| Polarized epithelial cell differentiation | 8 | 0.38 | 0.005 |
| Cell differentiation involved in embryonic placenta development | 8 | 0.38 | 0.005 |
| Striated muscle cell differentiation | 36 | 1.72 | 0.00501 |
| Negative regulation of small molecule metabolic process | 23 | 1.1 | 0.00502 |
| Cell growth | 23 | 1.1 | 0.00502 |
| Protein secretion | 23 | 1.1 | 0.00502 |
| Cell-substrate junction assembly | 11 | 0.53 | 0.00507 |
| Regulation of systemic arterial blood pressure by hormone | 11 | 0.53 | 0.00507 |
| Negative regulation of phosphorylation | 35 | 1.67 | 0.00507 |
| Glial cell development | 24 | 1.15 | 0.00519 |
| Regulation of cardiac muscle contraction | 17 | 0.81 | 0.00525 |
| Negative regulation of locomotion | 59 | 2.82 | 0.00526 |
| Negative regulation of growth | 37 | 1.77 | 0.0053 |
| Axon guidance | 37 | 1.77 | 0.0053 |
| Intracellular zinc ion homeostasis | 10 | 0.48 | 0.00531 |
| Establishment of epithelial cell polarity | 10 | 0.48 | 0.00531 |
| Regulation of mesenchymal cell proliferation | 10 | 0.48 | 0.00531 |
| Negative regulation of cell junction assembly | 10 | 0.48 | 0.00531 |
| Vesicle cargo loading | 10 | 0.48 | 0.00531 |
| Regulation of biomineral tissue development | 20 | 0.95 | 0.00531 |
| Tissue remodeling | 22 | 1.05 | 0.00531 |
| Neutral lipid biosynthetic process | 9 | 0.43 | 0.00533 |
| Multicellular organismal-level iron ion homeostasis | 9 | 0.43 | 0.00533 |

| | | | |
|---|---|---|---|
| Acylglycerol biosynthetic process | 9 | 0.43 | 0.00533 |
| Regulation of membrane protein ectodomain proteolysis | 9 | 0.43 | 0.00533 |
| Renal system process involved in regulation of systemic arterial blood pressure | 9 | 0.43 | 0.00533 |
| Positive regulation of peptidase activity | 9 | 0.43 | 0.00533 |
| Regulation of cardiac muscle hypertrophy | 16 | 0.76 | 0.00533 |
| Sprouting angiogenesis | 15 | 0.72 | 0.00534 |
| Regulation of leukocyte apoptotic process | 19 | 0.91 | 0.00555 |
| Xenobiotic transport | 12 | 0.57 | 0.00559 |
| Positive regulation of epithelial cell migration | 12 | 0.57 | 0.00559 |
| Positive regulation of nitric oxide metabolic process | 12 | 0.57 | 0.00559 |
| Substrate adhesion-dependent cell spreading | 12 | 0.57 | 0.00559 |
| Regulation of plasminogen activation | 7 | 0.33 | 0.00565 |
| Purine nucleoside bisphosphate metabolic process | 7 | 0.33 | 0.00565 |
| Nucleoside bisphosphate metabolic process | 7 | 0.33 | 0.00565 |
| Ribonucleoside bisphosphate metabolic process | 7 | 0.33 | 0.00565 |
| Neuron projection guidance | 37 | 1.77 | 0.00567 |
| Positive regulation of adaptive immune response based on somatic recombination of immune receptors built from immunoglobulin superfamily domains | 24 | 1.15 | 0.00569 |
| Positive regulation of actin filament depolymerization | 6 | 0.29 | 0.00578 |
| Branching involved in labyrinthine layer morphogenesis | 6 | 0.29 | 0.00578 |
| Regulation of binding | 34 | 1.62 | 0.00591 |
| Hematopoietic or lymphoid organ development | 20 | 0.95 | 0.00591 |
| Post-golgi vesicle-mediated transport | 20 | 0.95 | 0.00591 |
| Adherens junction organization | 13 | 0.62 | 0.00591 |
| Regulation of protein polymerization | 33 | 1.58 | 0.00596 |
| Vasculogenesis | 16 | 0.76 | 0.00606 |
| Modified amino acid metabolic process | 30 | 1.43 | 0.00606 |
| Epithelial cell differentiation involved in kidney development | 11 | 0.53 | 0.00608 |
| Regulation of heart rate by cardiac conduction | 11 | 0.53 | 0.00608 |
| Cell adhesion mediated by integrin | 11 | 0.53 | 0.00608 |
| Positive regulation of interleukin-10 production | 11 | 0.53 | 0.00608 |
| Positive regulation of t cell mediated cytotoxicity | 11 | 0.53 | 0.00608 |
| Polyol metabolic process | 19 | 0.91 | 0.00619 |
| Organic cation transport | 19 | 0.91 | 0.00619 |
| Neuron differentiation | 140 | 6.68 | 0.0062 |
| Host-mediated perturbation of symbiont process | 21 | 1 | 0.00621 |
| Chloride transmembrane transport | 21 | 1 | 0.00621 |
| Positive regulation of kinase activity | 35 | 1.67 | 0.00625 |
| Positive regulation of intracellular transport | 25 | 1.19 | 0.00636 |
| Positive regulation of leukocyte cell-cell adhesion | 43 | 2.05 | 0.00643 |
| Chondrocyte differentiation | 18 | 0.86 | 0.00645 |
| Signal transduction by p53 class mediator | 18 | 0.86 | 0.00645 |
| Ossification | 45 | 2.15 | 0.00646 |
| Sodium ion transport | 32 | 1.53 | 0.00647 |
| Neural tube development | 27 | 1.29 | 0.00651 |
| Positive regulation of nucleobase-containing compound metabolic process | 241 | 11.5 | 0.00657 |
| Vascular endothelial growth factor receptor-1 signaling pathway | 12 | 0.57 | 0.00659 |
| Regulation of transferase activity | 63 | 3.01 | 0.00661 |
| Androgen metabolic process | 9 | 0.43 | 0.00666 |
| Regulation of cell-matrix adhesion | 23 | 1.1 | 0.00666 |
| Establishment of protein localization to extracellular region | 23 | 1.1 | 0.00666 |
| Endoderm development | 17 | 0.81 | 0.00668 |
| Regulation of heart contraction | 34 | 1.62 | 0.00679 |
| Cd4-positive, alpha-beta t cell activation | 16 | 0.76 | 0.00686 |
| Platelet-derived growth factor receptor-beta signaling pathway | 13 | 0.62 | 0.00687 |
| Positive regulation of alpha-beta t cell differentiation | 13 | 0.62 | 0.00687 |
| Regulation of cell-substrate junction organization | 15 | 0.72 | 0.00697 |
| Cellular response to steroid hormone stimulus | 27 | 1.29 | 0.00706 |
| Receptor-mediated endocytosis | 29 | 1.38 | 0.00708 |
| Gliogenesis | 43 | 2.05 | 0.00723 |
| Cell-substrate junction organization | 11 | 0.53 | 0.00725 |
| Response to progesterone | 11 | 0.53 | 0.00725 |
| Regulation of transmembrane receptor protein serine/threonine kinase signaling pathway | 48 | 2.29 | 0.0074 |
| Negative regulation of molecular function | 76 | 3.63 | 0.00747 |
| Alpha-beta t cell differentiation | 17 | 0.81 | 0.0075 |
| Necroptotic process | 7 | 0.33 | 0.00758 |
| Actin filament network formation | 5 | 0.24 | 0.0076 |
| Positive regulation of mononuclear cell proliferation | 26 | 1.24 | 0.00762 |

| | | | |
|---|---|---|---|
| Glycerophospholipid metabolic process | 43 | 2.05 | 0.00766 |
| Negative regulation of response to endoplasmic reticulum stress | 12 | 0.57 | 0.00772 |
| Regulation of endoplasmic reticulum unfolded protein response | 10 | 0.48 | 0.00783 |
| Endocrine process | 13 | 0.62 | 0.00795 |
| Regulation of epithelial cell apoptotic process | 13 | 0.62 | 0.00795 |
| Morphogenesis of an epithelial sheet | 13 | 0.62 | 0.00795 |
| Propionate metabolic process | 4 | 0.19 | 0.00796 |
| Liver regeneration | 9 | 0.43 | 0.00822 |
| Antigen processing and presentation of exogenous peptide antigen via mhc class ii | 9 | 0.43 | 0.00822 |
| Regulation of sodium ion transmembrane transporter activity | 9 | 0.43 | 0.00822 |
| Negative regulation of t cell receptor signaling pathway | 9 | 0.43 | 0.00822 |
| Epiboly involved in wound healing | 9 | 0.43 | 0.00822 |
| Wound healing, spreading of cells | 9 | 0.43 | 0.00822 |
| Carbohydrate phosphorylation | 8 | 0.38 | 0.00824 |
| Axis elongation | 8 | 0.38 | 0.00824 |
| Nad biosynthetic process | 8 | 0.38 | 0.00824 |
| Fructose metabolic process | 6 | 0.29 | 0.00828 |
| Positive regulation of protein localization to endosome | 6 | 0.29 | 0.00828 |
| Negative regulation of substrate adhesion-dependent cell spreading | 6 | 0.29 | 0.00828 |
| Hepatocyte apoptotic process | 6 | 0.29 | 0.00828 |
| Bundle of his cell to purkinje myocyte communication | 6 | 0.29 | 0.00828 |
| Regulation of intestinal absorption | 6 | 0.29 | 0.00828 |
| Regulation of cell projection size | 6 | 0.29 | 0.00828 |
| Regulation of leukocyte mediated immunity | 40 | 1.91 | 0.00829 |
| Nephron epithelium development | 21 | 1 | 0.00832 |
| Regulation of cd4-positive, alpha-beta t cell activation | 17 | 0.81 | 0.00841 |
| Positive regulation of cold-induced thermogenesis | 19 | 0.91 | 0.0085 |
| Cellular response to biotic stimulus | 39 | 1.86 | 0.0085 |
| Positive regulation of substrate adhesion-dependent cell spreading | 11 | 0.53 | 0.00858 |
| Alcohol catabolic process | 11 | 0.53 | 0.00858 |
| Signaling receptor ligand precursor processing | 11 | 0.53 | 0.00858 |
| Fat cell differentiation | 23 | 1.1 | 0.00873 |
| Positive regulation of t cell proliferation | 20 | 0.95 | 0.00886 |
| Phosphatidylcholine metabolic process | 15 | 0.72 | 0.00898 |
| Negative regulation of protein polymerization | 15 | 0.72 | 0.00898 |
| Cellular response to stem cell factor stimulus | 12 | 0.57 | 0.00899 |
| Kit signaling pathway | 12 | 0.57 | 0.00899 |
| Macrophage colony-stimulating factor signaling pathway | 12 | 0.57 | 0.00899 |
| Cell junction maintenance | 12 | 0.57 | 0.00899 |
| Response to stem cell factor | 12 | 0.57 | 0.00899 |
| Oligosaccharide metabolic process | 14 | 0.67 | 0.00913 |
| Vascular endothelial growth factor signaling pathway | 14 | 0.67 | 0.00913 |
| Peptidyl-tyrosine phosphorylation | 13 | 0.62 | 0.00916 |
| Leukocyte migration | 38 | 1.81 | 0.00926 |
| Regulation of blood circulation | 40 | 1.91 | 0.00933 |
| Endothelial cell proliferation | 10 | 0.48 | 0.00939 |
| Protein-lipid complex remodeling | 10 | 0.48 | 0.00939 |
| Plasma lipoprotein particle remodeling | 10 | 0.48 | 0.00939 |
| Positive regulation of cell development | 63 | 3.01 | 0.00952 |
| Pattern recognition receptor signaling pathway | 23 | 1.1 | 0.00952 |
| Negative regulation of neuron projection development | 23 | 1.1 | 0.00952 |
| Activation of innate immune response | 29 | 1.38 | 0.00955 |
| Monocarboxylic acid transport | 26 | 1.24 | 0.0097 |
| Lymphocyte activation | 69 | 3.29 | 0.00971 |
| Cell projection organization | 151 | 7.21 | 0.00971 |
| Regulation of sodium ion transport | 16 | 0.76 | 0.0098 |
| Sterol transport | 16 | 0.76 | 0.0098 |
| Negative regulation of erk1 and erk2 cascade | 16 | 0.76 | 0.0098 |
| Antigen processing and presentation of peptide antigen via mhc class ib | 7 | 0.33 | 0.00995 |
| Cell-cell junction maintenance | 7 | 0.33 | 0.00995 |
| Epiboly | 9 | 0.43 | 0.01 |
| Response to ph | 11 | 0.53 | 0.0101 |
| Adipose tissue development | 11 | 0.53 | 0.0101 |
| Formation of primary germ layer | 22 | 1.05 | 0.0102 |
| Amyloid-beta clearance | 8 | 0.38 | 0.0103 |
| Smooth muscle tissue development | 8 | 0.38 | 0.0103 |
| Regulation of receptor recycling | 8 | 0.38 | 0.0103 |
| Positive regulation of leukocyte proliferation | 28 | 1.34 | 0.0104 |

| | | | |
|---|---|---|---|
| Interferon-mediated signaling pathway | 14 | 0.67 | 0.0104 |
| Collagen fibril organization | 14 | 0.67 | 0.0104 |
| Cell differentiation involved in kidney development | 12 | 0.57 | 0.0104 |
| Glial cell migration | 12 | 0.57 | 0.0104 |
| Negative regulation of cell differentiation | 94 | 4.49 | 0.0105 |
| Positive regulation of leukocyte migration | 26 | 1.24 | 0.0105 |
| Regulation of cell junction assembly | 40 | 1.91 | 0.0105 |
| Positive regulation of hormone secretion | 25 | 1.19 | 0.0105 |
| Hair cell differentiation | 13 | 0.62 | 0.0105 |
| Mesenchymal cell migration | 13 | 0.62 | 0.0105 |
| Negative regulation of leukocyte apoptotic process | 13 | 0.62 | 0.0105 |
| Regulation of protein binding | 20 | 0.95 | 0.0107 |
| Regulation of heart rate | 20 | 0.95 | 0.0107 |
| Erythrocyte differentiation | 20 | 0.95 | 0.0107 |
| Negative regulation of protein-containing complex disassembly | 16 | 0.76 | 0.011 |
| Maintenance of blood-brain barrier | 10 | 0.48 | 0.0112 |
| Limb morphogenesis | 26 | 1.24 | 0.0113 |
| Appendage morphogenesis | 26 | 1.24 | 0.0113 |
| Response to acid chemical | 25 | 1.19 | 0.0114 |
| Regulation of cold-induced thermogenesis | 25 | 1.19 | 0.0114 |
| Positive regulation of lymphocyte proliferation | 25 | 1.19 | 0.0114 |
| Generation of neurons | 146 | 6.97 | 0.0114 |
| Phosphatidylinositol 3-kinase/protein kinase b signal transduction | 15 | 0.72 | 0.0114 |
| Cell-cell adhesion via plasma-membrane adhesion molecules | 39 | 1.86 | 0.0114 |
| Positive regulation of t cell activation | 39 | 1.86 | 0.0114 |
| Intestinal lipid absorption | 6 | 0.29 | 0.0114 |
| Regulation of protein localization to endosome | 6 | 0.29 | 0.0114 |
| Galactose metabolic process | 6 | 0.29 | 0.0114 |
| Urea metabolic process | 6 | 0.29 | 0.0114 |
| Platelet activation | 19 | 0.91 | 0.0114 |
| Negative regulation of extrinsic apoptotic signaling pathway | 19 | 0.91 | 0.0114 |
| Zinc ion import across plasma membrane | 5 | 0.24 | 0.0117 |
| Peroxisome proliferator activated receptor signaling pathway | 5 | 0.24 | 0.0117 |
| Regulation of glucocorticoid biosynthetic process | 5 | 0.24 | 0.0117 |
| Positive regulation of protein localization to early endosome | 5 | 0.24 | 0.0117 |
| Positive regulation of adipose tissue development | 5 | 0.24 | 0.0117 |
| Regulation of protein localization to early endosome | 5 | 0.24 | 0.0117 |
| Cellular response to sterol depletion | 5 | 0.24 | 0.0117 |
| Positive regulation of non-canonical nf-kappab signal transduction | 14 | 0.67 | 0.0118 |
| Positive regulation of nucleocytoplasmic transport | 14 | 0.67 | 0.0118 |
| Integrin-mediated signaling pathway | 20 | 0.95 | 0.0118 |
| Multicellular organism growth | 20 | 0.95 | 0.0118 |
| Positive regulation of tumor necrosis factor production | 20 | 0.95 | 0.0118 |
| Receptor metabolic process | 11 | 0.53 | 0.0118 |
| Epithelial cell apoptotic process | 11 | 0.53 | 0.0118 |
| Positive regulation of lymphocyte activation | 48 | 2.29 | 0.0118 |
| Establishment of vesicle localization | 29 | 1.38 | 0.0118 |
| Mesenchymal cell differentiation | 28 | 1.34 | 0.012 |
| Maintenance of protein location in cell | 13 | 0.62 | 0.012 |
| Anterior/posterior axis specification | 13 | 0.62 | 0.012 |
| Osteoclast differentiation | 13 | 0.62 | 0.012 |
| Regulation of neurogenesis | 53 | 2.53 | 0.012 |
| Mrna transcription | 12 | 0.57 | 0.012 |
| Icosanoid biosynthetic process | 12 | 0.57 | 0.012 |
| Antigen processing and presentation of peptide antigen via mhc class ii | 9 | 0.43 | 0.0122 |
| Positive regulation of transforming growth factor beta receptor signaling pathway | 9 | 0.43 | 0.0122 |
| Salivary gland development | 9 | 0.43 | 0.0122 |
| Positive regulation of cellular response to transforming growth factor beta stimulus | 9 | 0.43 | 0.0122 |
| Embryonic digestive tract development | 9 | 0.43 | 0.0122 |
| Regulation of stem cell proliferation | 18 | 0.86 | 0.0122 |
| Positive regulation of exocytosis | 16 | 0.76 | 0.0122 |
| Response to amino acid | 23 | 1.1 | 0.0123 |
| Positive regulation of endothelial cell migration | 23 | 1.1 | 0.0123 |
| Myeloid cell homeostasis | 25 | 1.19 | 0.0123 |
| Response to topologically incorrect protein | 24 | 1.15 | 0.0123 |
| Plasma membrane bounded cell projection organization | 144 | 6.87 | 0.0124 |
| Regulation of cd8-positive, alpha-beta t cell activation | 7 | 0.33 | 0.0128 |
| Mammary gland alveolus development | 7 | 0.33 | 0.0128 |

| | | | |
|---|---|---|---|
| Calcium-independent cell-cell adhesion via plasma membrane cell-adhesion molecules | 7 | 0.33 | 0.0128 |
| Sulfation | 7 | 0.33 | 0.0128 |
| Arginine metabolic process | 7 | 0.33 | 0.0128 |
| Ceramide catabolic process | 7 | 0.33 | 0.0128 |
| Surfactant homeostasis | 7 | 0.33 | 0.0128 |
| Positive regulation of chondrocyte differentiation | 7 | 0.33 | 0.0128 |
| Mammary gland lobule development | 7 | 0.33 | 0.0128 |
| Regulation of antigen receptor-mediated signaling pathway | 15 | 0.72 | 0.0128 |
| Regulation of steroid biosynthetic process | 17 | 0.81 | 0.0129 |
| Notch signaling pathway | 21 | 1 | 0.0131 |
| Nadp metabolic process | 10 | 0.48 | 0.0132 |
| Positive regulation of fatty acid metabolic process | 10 | 0.48 | 0.0132 |
| Protein-containing complex remodeling | 10 | 0.48 | 0.0132 |
| Dendritic cell differentiation | 10 | 0.48 | 0.0132 |
| Cortical actin cytoskeleton organization | 10 | 0.48 | 0.0132 |
| Negative regulation of cell migration | 53 | 2.53 | 0.0132 |
| Regulation of monoatomic ion transport | 63 | 3.01 | 0.0132 |
| Positive regulation of chemotaxis | 25 | 1.19 | 0.0132 |
| Cellular response to hydrogen peroxide | 14 | 0.67 | 0.0133 |
| Negative regulation of cell-substrate adhesion | 14 | 0.67 | 0.0133 |
| Regulation of nitric oxide biosynthetic process | 14 | 0.67 | 0.0133 |
| Cholesterol transport | 14 | 0.67 | 0.0133 |
| Regulation of generation of precursor metabolites and energy | 24 | 1.15 | 0.0133 |
| Regulation of bone mineralization | 16 | 0.76 | 0.0136 |
| Positive regulation of cd4-positive, alpha-beta t cell activation | 11 | 0.53 | 0.0137 |
| Regulation of fatty acid biosynthetic process | 12 | 0.57 | 0.0138 |
| Neutral amino acid transport | 12 | 0.57 | 0.0138 |
| Sphingolipid biosynthetic process | 19 | 0.91 | 0.0138 |
| Regulation of notch signaling pathway | 19 | 0.91 | 0.0138 |
| Negative regulation of supramolecular fiber organization | 27 | 1.29 | 0.014 |
| Erythrocyte homeostasis | 21 | 1 | 0.0143 |
| Protein metabolic process | 379 | 18.09 | 0.0145 |
| Positive regulation of cartilage development | 9 | 0.43 | 0.0146 |
| Protein localization to golgi apparatus | 9 | 0.43 | 0.0146 |
| Regulation of transforming growth factor beta production | 9 | 0.43 | 0.0146 |
| Peptide antigen assembly with mhc class i protein complex | 4 | 0.19 | 0.0148 |
| Negative regulation of ire1-mediated unfolded protein response | 4 | 0.19 | 0.0148 |
| Mhc class i protein complex assembly | 4 | 0.19 | 0.0148 |
| Response to gram-positive bacterium | 4 | 0.19 | 0.0148 |
| Positive regulation of ion transmembrane transporter activity | 14 | 0.67 | 0.0149 |
| Regulation of epidermal growth factor receptor signaling pathway | 14 | 0.67 | 0.0149 |
| Regulation of epithelial cell migration | 14 | 0.67 | 0.0149 |
| Cardiac conduction | 14 | 0.67 | 0.0149 |
| Response to temperature stimulus | 27 | 1.29 | 0.015 |
| Positive regulation of leukocyte mediated immunity | 26 | 1.24 | 0.0152 |
| Oligosaccharide catabolic process | 6 | 0.29 | 0.0153 |
| Regulation of receptor binding | 6 | 0.29 | 0.0153 |
| Non-proteinogenic amino acid biosynthetic process | 6 | 0.29 | 0.0153 |
| Regulation of steroid hormone biosynthetic process | 6 | 0.29 | 0.0153 |
| Non-proteinogenic amino acid metabolic process | 13 | 0.62 | 0.0155 |
| Regulation of intracellular transport | 40 | 1.91 | 0.0155 |
| Negative regulation of muscle hypertrophy | 10 | 0.48 | 0.0155 |
| Spleen development | 10 | 0.48 | 0.0155 |
| Response to estradiol | 21 | 1 | 0.0155 |
| Negative regulation of immune effector process | 23 | 1.1 | 0.0156 |
| Negative regulation of hemopoiesis | 22 | 1.05 | 0.0156 |
| Cellular response to oxidative stress | 34 | 1.62 | 0.0156 |
| Protein maturation | 71 | 3.39 | 0.0157 |
| Negative regulation of natural killer cell mediated cytotoxicity | 8 | 0.38 | 0.0157 |
| Positive regulation of t cell cytokine production | 8 | 0.38 | 0.0157 |
| Positive regulation of mesenchymal cell proliferation | 8 | 0.38 | 0.0157 |
| Cellular response to macrophage colony-stimulating factor stimulus | 12 | 0.57 | 0.0158 |
| Regulation of platelet activation | 12 | 0.57 | 0.0158 |
| Negative regulation of t cell differentiation | 12 | 0.57 | 0.0158 |
| Positive regulation of lipid transport | 17 | 0.81 | 0.0158 |
| Monosaccharide transmembrane transport | 11 | 0.53 | 0.0158 |
| Response to fluid shear stress | 11 | 0.53 | 0.0158 |
| Mrna transcription by rna polymerase ii | 11 | 0.53 | 0.0158 |

| | | | |
|---|---|---|---|
| Regulation of cartilage development | 15 | 0.72 | 0.016 |
| Positive regulation of protein localization to cell periphery | 15 | 0.72 | 0.016 |
| Cell activation involved in immune response | 33 | 1.58 | 0.0161 |
| Establishment of apical/basal cell polarity | 7 | 0.33 | 0.0162 |
| Biogenic amine catabolic process | 7 | 0.33 | 0.0162 |
| Response to hyperoxia | 7 | 0.33 | 0.0162 |
| Amine catabolic process | 7 | 0.33 | 0.0162 |
| Regulation of translation in response to stress | 7 | 0.33 | 0.0162 |
| N-glycan processing | 7 | 0.33 | 0.0162 |
| Negative regulation of endoplasmic reticulum unfolded protein response | 7 | 0.33 | 0.0162 |
| Protein polymerization | 19 | 0.91 | 0.0166 |
| Regulation of nitric oxide metabolic process | 14 | 0.67 | 0.0168 |
| Positive regulation of tumor necrosis factor superfamily cytokine production | 20 | 0.95 | 0.0168 |
| Negative regulation of leukocyte differentiation | 21 | 1 | 0.0169 |
| Regulation of mesenchymal stem cell differentiation | 5 | 0.24 | 0.017 |
| Trachea morphogenesis | 5 | 0.24 | 0.017 |
| Protein modification process | 209 | 9.98 | 0.017 |
| Cartilage development involved in endochondral bone morphogenesis | 9 | 0.43 | 0.0173 |
| Regulation of protein targeting to membrane | 9 | 0.43 | 0.0173 |
| Nucleobase metabolic process | 9 | 0.43 | 0.0173 |
| Negative regulation of erbb signaling pathway | 9 | 0.43 | 0.0173 |
| Organophosphate catabolic process | 30 | 1.43 | 0.0173 |
| Negative regulation of cell motility | 54 | 2.58 | 0.0174 |
| Regulation of tissue remodeling | 17 | 0.81 | 0.0174 |
| Regulation of response to endoplasmic reticulum stress | 17 | 0.81 | 0.0174 |
| Positive regulation of transferase activity | 38 | 1.81 | 0.0174 |
| Monoatomic cation transmembrane transport | 87 | 4.15 | 0.0174 |
| Arachidonate metabolic process | 13 | 0.62 | 0.0174 |
| Amino acid biosynthetic process | 15 | 0.72 | 0.0178 |
| Mechanoreceptor differentiation | 15 | 0.72 | 0.0178 |
| Receptor internalization | 15 | 0.72 | 0.0178 |
| Sulfur compound biosynthetic process | 25 | 1.19 | 0.0178 |
| Positive regulation of leukocyte activation | 51 | 2.43 | 0.0179 |
| Hepatocyte growth factor receptor signaling pathway | 12 | 0.57 | 0.018 |
| Protein-lipid complex organization | 12 | 0.57 | 0.018 |
| Cortical cytoskeleton organization | 12 | 0.57 | 0.018 |
| Response to reactive oxygen species | 28 | 1.34 | 0.0181 |
| Positive regulation of lymphocyte mediated immunity | 23 | 1.1 | 0.0182 |
| Platelet aggregation | 11 | 0.53 | 0.0182 |
| Odontogenesis | 21 | 1 | 0.0183 |
| Vesicle localization | 30 | 1.43 | 0.0184 |
| Cellular response to molecule of bacterial origin | 34 | 1.62 | 0.0186 |
| Mesodermal cell differentiation | 8 | 0.38 | 0.019 |
| Negative regulation of natural killer cell mediated immunity | 8 | 0.38 | 0.019 |
| Cellular response to amino acid stimulus | 17 | 0.81 | 0.0191 |
| Primary neural tube formation | 17 | 0.81 | 0.0191 |
| Nucleoside diphosphate metabolic process | 17 | 0.81 | 0.0191 |
| Neural tube formation | 18 | 0.86 | 0.0195 |
| Heart contraction | 18 | 0.86 | 0.0195 |
| Endochondral bone morphogenesis | 13 | 0.62 | 0.0196 |
| Intrinsic apoptotic signaling pathway in response to dna damage | 15 | 0.72 | 0.0197 |
| Cellular component assembly involved in morphogenesis | 22 | 1.05 | 0.0198 |
| Cellular anatomical entity morphogenesis | 22 | 1.05 | 0.0198 |
| Negative regulation of inflammatory response | 32 | 1.53 | 0.0198 |
| Cellular response to lipopolysaccharide | 32 | 1.53 | 0.0198 |
| Insulin receptor signaling pathway | 20 | 0.95 | 0.0198 |
| Regulation of mitochondrion organization | 20 | 0.95 | 0.0198 |
| Fatty acid biosynthetic process | 21 | 1 | 0.0198 |
| Response to unfolded protein | 21 | 1 | 0.0198 |
| Positive regulation of phospholipid metabolic process | 6 | 0.29 | 0.02 |
| Copii-coated vesicle cargo loading | 6 | 0.29 | 0.02 |
| Establishment of epithelial cell apical/basal polarity | 6 | 0.29 | 0.02 |
| Lactone metabolic process | 6 | 0.29 | 0.02 |
| Embryonic digestive tract morphogenesis | 6 | 0.29 | 0.02 |
| Phospholipid homeostasis | 6 | 0.29 | 0.02 |
| Antigen processing and presentation of endogenous peptide antigen via mhc class i via er pathway, tap-independent | 6 | 0.29 | 0.02 |
| Cytokine production | 6 | 0.29 | 0.02 |

| | | | |
|---|---|---|---|
| T cell differentiation | 29 | 1.38 | 0.0201 |
| Response to salt | 7 | 0.33 | 0.0201 |
| Detoxification of inorganic compound | 7 | 0.33 | 0.0201 |
| Cellular response to growth hormone stimulus | 7 | 0.33 | 0.0201 |
| Proteinogenic amino acid biosynthetic process | 12 | 0.57 | 0.0204 |
| Neural crest cell migration | 12 | 0.57 | 0.0204 |
| Regulation of cardiac muscle cell contraction | 9 | 0.43 | 0.0204 |
| Regulation of fatty acid oxidation | 9 | 0.43 | 0.0204 |
| Glandular epithelial cell differentiation | 9 | 0.43 | 0.0204 |
| Dna damage response, signal transduction by p53 class mediator | 9 | 0.43 | 0.0204 |
| Glycosyl compound metabolic process | 16 | 0.76 | 0.0205 |
| Vascular transport | 16 | 0.76 | 0.0205 |
| Transport across blood-brain barrier | 16 | 0.76 | 0.0205 |
| Positive regulation of carbohydrate metabolic process | 16 | 0.76 | 0.0205 |
| Osteoblast differentiation | 24 | 1.15 | 0.0208 |
| Positive regulation of fat cell differentiation | 14 | 0.67 | 0.0209 |
| Leukocyte activation involved in immune response | 32 | 1.53 | 0.0209 |
| Negative regulation of lymphocyte proliferation | 17 | 0.81 | 0.021 |
| Cellular response to acid chemical | 18 | 0.86 | 0.0213 |
| Regulation of tumor necrosis factor production | 29 | 1.38 | 0.0214 |
| Import across plasma membrane | 26 | 1.24 | 0.0215 |
| Positive regulation of protein kinase activity | 31 | 1.48 | 0.0216 |
| Positive regulation of interleukin-8 production | 13 | 0.62 | 0.022 |
| Regulation of focal adhesion assembly | 13 | 0.62 | 0.022 |
| Negative regulation of lymphocyte differentiation | 13 | 0.62 | 0.022 |
| Regulation of cell-substrate junction assembly | 13 | 0.62 | 0.022 |
| Axonogenesis | 51 | 2.43 | 0.0222 |
| Negative regulation of epithelial cell proliferation | 27 | 1.29 | 0.0225 |
| Cellular response to environmental stimulus | 46 | 2.2 | 0.0225 |
| Cellular response to abiotic stimulus | 46 | 2.2 | 0.0225 |
| Nucleoside phosphate catabolic process | 23 | 1.1 | 0.0227 |
| Negative regulation of nucleotide metabolic process | 8 | 0.38 | 0.0227 |
| Negative regulation of mirna transcription | 8 | 0.38 | 0.0227 |
| Establishment or maintenance of cell polarity | 31 | 1.48 | 0.0229 |
| Embryonic skeletal system morphogenesis | 17 | 0.81 | 0.023 |
| Regulation of smooth muscle cell migration | 17 | 0.81 | 0.023 |
| Ceramide biosynthetic process | 12 | 0.57 | 0.023 |
| Regulation of glucose metabolic process | 18 | 0.86 | 0.0232 |
| Endothelin receptor signaling pathway | 5 | 0.24 | 0.0235 |
| Regulation of glucocorticoid metabolic process | 5 | 0.24 | 0.0235 |
| Regulation of ferroptosis | 5 | 0.24 | 0.0235 |
| Columnar/cuboidal epithelial cell maturation | 5 | 0.24 | 0.0235 |
| Cardiac neural crest cell differentiation involved in heart development | 5 | 0.24 | 0.0235 |
| Response to sterol depletion | 5 | 0.24 | 0.0235 |
| Negative regulation of receptor binding | 5 | 0.24 | 0.0235 |
| Cardiac neural crest cell development involved in heart development | 5 | 0.24 | 0.0235 |
| Skin morphogenesis | 5 | 0.24 | 0.0235 |
| Self proteolysis | 5 | 0.24 | 0.0235 |
| Cellular response to thyroid hormone stimulus | 5 | 0.24 | 0.0235 |
| Actin filament depolymerization | 5 | 0.24 | 0.0235 |
| Positive regulation of vasculogenesis | 5 | 0.24 | 0.0235 |
| Positive regulation of hepatocyte proliferation | 5 | 0.24 | 0.0235 |
| Negative regulation of protein localization to cell surface | 5 | 0.24 | 0.0235 |
| Hormone catabolic process | 5 | 0.24 | 0.0235 |
| Cellular response to type i interferon | 11 | 0.53 | 0.0237 |
| Regulation of membrane depolarization | 11 | 0.53 | 0.0237 |
| Regulation of sodium ion transmembrane transport | 11 | 0.53 | 0.0237 |
| Small gtpase-mediated signal transduction | 38 | 1.81 | 0.0238 |
| Mature b cell differentiation | 9 | 0.43 | 0.0239 |
| Response to copper ion | 9 | 0.43 | 0.0239 |
| Blood vessel endothelial cell migration | 9 | 0.43 | 0.0239 |
| Regulation of fatty acid transport | 9 | 0.43 | 0.0239 |
| Sodium ion transmembrane transport | 24 | 1.15 | 0.024 |
| Vitamin e metabolic process | 4 | 0.19 | 0.024 |
| Lung lobe morphogenesis | 4 | 0.19 | 0.024 |
| Glycosylceramide biosynthetic process | 4 | 0.19 | 0.024 |
| Citrulline metabolic process | 4 | 0.19 | 0.024 |
| Blood vessel endothelial cell proliferation involved in sprouting angiogenesis | 4 | 0.19 | 0.024 |

| | | | |
|---|---|---|---|
| Ceramide 1-phosphate transport | 4 | 0.19 | 0.024 |
| Adherens junction maintenance | 4 | 0.19 | 0.024 |
| Positive regulation of metallopeptidase activity | 4 | 0.19 | 0.024 |
| Bud elongation involved in lung branching | 4 | 0.19 | 0.024 |
| Interleukin-4-mediated signaling pathway | 4 | 0.19 | 0.024 |
| Lung lobe development | 4 | 0.19 | 0.024 |
| Response to testosterone | 10 | 0.48 | 0.0241 |
| Nucleoside monophosphate metabolic process | 15 | 0.72 | 0.0241 |
| Regulation of mirna transcription | 15 | 0.72 | 0.0241 |
| Insulin-like growth factor receptor signaling pathway | 15 | 0.72 | 0.0241 |
| L-amino acid metabolic process | 26 | 1.24 | 0.0246 |
| Collagen metabolic process | 13 | 0.62 | 0.0246 |
| Plasma lipoprotein particle clearance | 7 | 0.33 | 0.0247 |
| T-helper 1 type immune response | 7 | 0.33 | 0.0247 |
| Establishment of monopolar cell polarity | 7 | 0.33 | 0.0247 |
| Negative regulation of mononuclear cell proliferation | 17 | 0.81 | 0.0251 |
| Cellular response to glucose stimulus | 17 | 0.81 | 0.0251 |
| Negative regulation of leukocyte proliferation | 18 | 0.86 | 0.0253 |
| Regulation of striated muscle contraction | 18 | 0.86 | 0.0253 |
| Negative regulation of platelet activation | 6 | 0.29 | 0.0255 |
| Nitrogen cycle metabolic process | 6 | 0.29 | 0.0255 |
| Chondrocyte proliferation | 6 | 0.29 | 0.0255 |
| Regulation of extracellular matrix disassembly | 6 | 0.29 | 0.0255 |
| Fatty acid homeostasis | 6 | 0.29 | 0.0255 |
| Keratan sulfate proteoglycan metabolic process | 6 | 0.29 | 0.0255 |
| Regulation of epithelial cell differentiation | 27 | 1.29 | 0.0255 |
| Axon development | 57 | 2.72 | 0.0256 |
| Regulation of pattern recognition receptor signaling pathway | 29 | 1.38 | 0.0257 |
| Regulation of myeloid leukocyte mediated immunity | 12 | 0.57 | 0.0258 |
| Response to macrophage colony-stimulating factor | 12 | 0.57 | 0.0258 |
| Maternal process involved in female pregnancy | 12 | 0.57 | 0.0258 |
| Regulation of endothelial cell migration | 34 | 1.62 | 0.026 |
| Inorganic cation transmembrane transport | 84 | 4.01 | 0.0261 |
| Tube formation | 23 | 1.1 | 0.0261 |
| Lung goblet cell differentiation | 3 | 0.14 | 0.0267 |
| Substance p catabolic process | 3 | 0.14 | 0.0267 |
| Fructose catabolic process to hydroxyacetone phosphate and glyceraldehyde-3-phosphate | 3 | 0.14 | 0.0267 |
| Fructose catabolic process | 3 | 0.14 | 0.0267 |
| Sesquiterpenoid metabolic process | 3 | 0.14 | 0.0267 |
| Mesodermal cell fate determination | 3 | 0.14 | 0.0267 |
| Terminal web assembly | 3 | 0.14 | 0.0267 |
| Regulation of dense core granule biogenesis | 3 | 0.14 | 0.0267 |
| Galactolipid catabolic process | 3 | 0.14 | 0.0267 |
| Bile acid and bile salt transport | 8 | 0.38 | 0.027 |
| Neutrophil activation | 8 | 0.38 | 0.027 |
| Retinoic acid metabolic process | 8 | 0.38 | 0.027 |
| Negative regulation of catalytic activity | 40 | 1.91 | 0.027 |
| Regulation of muscle contraction | 27 | 1.29 | 0.0272 |
| Regulation of mirna metabolic process | 17 | 0.81 | 0.0274 |
| Sulfur compound transport | 13 | 0.62 | 0.0274 |
| Positive regulation of viral process | 13 | 0.62 | 0.0274 |
| Aortic valve development | 10 | 0.48 | 0.0276 |
| Hexose transmembrane transport | 10 | 0.48 | 0.0276 |
| Positive regulation of lamellipodium organization | 9 | 0.43 | 0.0277 |
| Positive regulation of atp metabolic process | 9 | 0.43 | 0.0277 |
| Lymphocyte apoptotic process | 9 | 0.43 | 0.0277 |
| Positive regulation of organelle organization | 64 | 3.05 | 0.0279 |
| Morphogenesis of embryonic epithelium | 23 | 1.1 | 0.028 |
| Cytoplasmic pattern recognition receptor signaling pathway | 14 | 0.67 | 0.0285 |
| Positive regulation of cell activation | 52 | 2.48 | 0.0285 |
| Phagocytosis | 25 | 1.19 | 0.0286 |
| Regulation of tumor necrosis factor superfamily cytokine production | 29 | 1.38 | 0.0289 |
| Embryonic epithelial tube formation | 20 | 0.95 | 0.0294 |
| Positive regulation of erk1 and erk2 cascade | 30 | 1.43 | 0.0297 |
| Hematopoietic progenitor cell differentiation | 18 | 0.86 | 0.0298 |
| Canonical wnt signaling pathway | 18 | 0.86 | 0.0298 |
| Organ growth | 18 | 0.86 | 0.0298 |
| Amine metabolic process | 17 | 0.81 | 0.0298 |

| | | | |
|---|---|---|---|
| Regulation of hormone biosynthetic process | 7 | 0.33 | 0.0299 |
| Positive regulation of protein processing | 7 | 0.33 | 0.0299 |
| Establishment or maintenance of monopolar cell polarity | 7 | 0.33 | 0.0299 |
| Endothelial cell development | 11 | 0.53 | 0.0303 |
| Plasma lipoprotein particle organization | 11 | 0.53 | 0.0303 |
| L-amino acid biosynthetic process | 11 | 0.53 | 0.0303 |
| Erk1 and erk2 cascade | 11 | 0.53 | 0.0303 |
| Rho protein signal transduction | 13 | 0.62 | 0.0304 |
| Regulation of lymphocyte mediated immunity | 30 | 1.43 | 0.0314 |
| Intracellular iron ion homeostasis | 14 | 0.67 | 0.0314 |
| Body fluid secretion | 14 | 0.67 | 0.0314 |
| Type ii interferon-mediated signaling pathway | 5 | 0.24 | 0.0314 |
| Mucus secretion | 5 | 0.24 | 0.0314 |
| Maintenance of protein localization in endoplasmic reticulum | 5 | 0.24 | 0.0314 |
| Negative regulation of homotypic cell-cell adhesion | 5 | 0.24 | 0.0314 |
| Negative regulation of peptidyl-threonine phosphorylation | 5 | 0.24 | 0.0314 |
| Very-low-density lipoprotein particle assembly | 5 | 0.24 | 0.0314 |
| Fatty acid beta-oxidation using acyl-coa oxidase | 5 | 0.24 | 0.0314 |
| Positive regulation of sodium ion transmembrane transporter activity | 5 | 0.24 | 0.0314 |
| Keratinocyte development | 5 | 0.24 | 0.0314 |
| Angiotensin maturation | 5 | 0.24 | 0.0314 |
| Response to cold | 10 | 0.48 | 0.0314 |
| Plasma membrane invagination | 10 | 0.48 | 0.0314 |
| Immune system development | 26 | 1.24 | 0.0316 |
| Negative regulation of mirna metabolic process | 8 | 0.38 | 0.0317 |
| Positive regulation of extracellular matrix organization | 8 | 0.38 | 0.0317 |
| Viral protein processing | 8 | 0.38 | 0.0317 |
| Glycoside metabolic process | 6 | 0.29 | 0.0319 |
| Antigen processing and presentation of endogenous peptide antigen via mhc class i via er pathway | 6 | 0.29 | 0.0319 |
| Cellular response to aldehyde | 6 | 0.29 | 0.0319 |
| Antigen processing and presentation of endogenous peptide antigen via mhc class ib | 6 | 0.29 | 0.0319 |
| Nucleoside monophosphate catabolic process | 6 | 0.29 | 0.0319 |
| Heart morphogenesis | 38 | 1.81 | 0.032 |
| Negative regulation of cardiac muscle hypertrophy | 9 | 0.43 | 0.032 |
| Endothelium development | 18 | 0.86 | 0.0323 |
| Glutathione metabolic process | 12 | 0.57 | 0.0323 |
| Endoplasmic reticulum unfolded protein response | 12 | 0.57 | 0.0323 |
| Acyl-coa metabolic process | 16 | 0.76 | 0.0323 |
| Membrane docking | 16 | 0.76 | 0.0323 |
| Thioester metabolic process | 16 | 0.76 | 0.0323 |
| Alpha-amino acid biosynthetic process | 13 | 0.62 | 0.0337 |
| Regulation of action potential | 13 | 0.62 | 0.0337 |
| Regulation of morphogenesis of an epithelium | 13 | 0.62 | 0.0337 |
| Face development | 11 | 0.53 | 0.0341 |
| Retinol metabolic process | 11 | 0.53 | 0.0341 |
| Negative regulation of interleukin-1 production | 11 | 0.53 | 0.0341 |
| Regulation of lymphocyte differentiation | 32 | 1.53 | 0.0344 |
| Immune response-activating signaling pathway | 45 | 2.15 | 0.0345 |
| Negative regulation of lipid metabolic process | 19 | 0.91 | 0.0345 |
| Negative regulation of rna metabolic process | 160 | 7.64 | 0.0348 |
| Response to retinoic acid | 18 | 0.86 | 0.0349 |
| Cellular response to alcohol | 18 | 0.86 | 0.0349 |
| Positive regulation of protein-containing complex assembly | 30 | 1.43 | 0.035 |
| Regulation of membrane potential | 60 | 2.86 | 0.0351 |
| Purine-containing compound biosynthetic process | 34 | 1.62 | 0.0354 |
| Regulation of t cell differentiation | 28 | 1.34 | 0.0355 |
| Gap junction assembly | 4 | 0.19 | 0.0356 |
| Ganglioside catabolic process | 4 | 0.19 | 0.0356 |
| Putrescine metabolic process | 4 | 0.19 | 0.0356 |
| Protein retention in er lumen | 4 | 0.19 | 0.0356 |
| Positive regulation of mammary gland epithelial cell proliferation | 4 | 0.19 | 0.0356 |
| Interleukin-27-mediated signaling pathway | 4 | 0.19 | 0.0356 |
| Copper ion transmembrane transport | 4 | 0.19 | 0.0356 |
| Positive regulation of plasminogen activation | 4 | 0.19 | 0.0356 |
| Negative regulation of membrane protein ectodomain proteolysis | 4 | 0.19 | 0.0356 |
| Nadph oxidation | 4 | 0.19 | 0.0356 |
| Necroptotic signaling pathway | 4 | 0.19 | 0.0356 |
| Male sex differentiation | 26 | 1.24 | 0.0356 |

| | | | |
|---|---|---|---|
| Monocarboxylic acid biosynthetic process | 26 | 1.24 | 0.0356 |
| Endodermal cell differentiation | 10 | 0.48 | 0.0356 |
| Mammary gland morphogenesis | 10 | 0.48 | 0.0356 |
| Prostate gland development | 10 | 0.48 | 0.0356 |
| Response to angiotensin | 10 | 0.48 | 0.0356 |
| Body morphogenesis | 10 | 0.48 | 0.0356 |
| Peptide biosynthetic process | 7 | 0.33 | 0.0357 |
| Filopodium assembly | 7 | 0.33 | 0.0357 |
| Regulation of receptor-mediated endocytosis | 21 | 1 | 0.0358 |
| Positive regulation of leukocyte differentiation | 29 | 1.38 | 0.0362 |
| Positive regulation of hemopoiesis | 29 | 1.38 | 0.0362 |
| Regulation of cellular response to transforming growth factor beta stimulus | 27 | 1.29 | 0.0366 |
| Negative regulation of antigen receptor-mediated signaling pathway | 9 | 0.43 | 0.0367 |
| Animal organ formation | 9 | 0.43 | 0.0367 |
| Epithelial cell morphogenesis | 9 | 0.43 | 0.0367 |
| Copii-coated vesicle budding | 9 | 0.43 | 0.0367 |
| Peptide hormone processing | 9 | 0.43 | 0.0367 |
| Generation of precursor metabolites and energy | 50 | 2.39 | 0.0369 |
| Negative regulation of epithelial cell apoptotic process | 8 | 0.38 | 0.037 |
| Positive regulation of amide metabolic process | 8 | 0.38 | 0.037 |
| Transepithelial transport | 8 | 0.38 | 0.037 |
| Lymphocyte differentiation | 39 | 1.86 | 0.0371 |
| Negative regulation of lipid biosynthetic process | 13 | 0.62 | 0.0372 |
| Positive regulation of neuron apoptotic process | 13 | 0.62 | 0.0372 |
| Exocytosis | 34 | 1.62 | 0.0372 |
| Circadian rhythm | 22 | 1.05 | 0.0374 |
| Proteinogenic amino acid metabolic process | 26 | 1.24 | 0.0378 |
| Vacuolar transport | 26 | 1.24 | 0.0378 |
| Positive regulation of wnt signaling pathway | 24 | 1.15 | 0.0378 |
| Stem cell proliferation | 14 | 0.67 | 0.0379 |
| Cellular response to unfolded protein | 14 | 0.67 | 0.0379 |
| Cellular response to calcium ion | 14 | 0.67 | 0.0379 |
| Regulation of mitochondrial membrane potential | 14 | 0.67 | 0.0379 |
| Regulation of lamellipodium organization | 11 | 0.53 | 0.0382 |
| Cellular oxidant detoxification | 16 | 0.76 | 0.0382 |
| Cellular response to topologically incorrect protein | 16 | 0.76 | 0.0382 |
| Innate immune response-activating signaling pathway | 23 | 1.1 | 0.0389 |
| Development of primary male sexual characteristics | 23 | 1.1 | 0.0389 |
| Carbohydrate biosynthetic process | 20 | 0.95 | 0.0392 |
| Regulation of defense response to bacterium | 6 | 0.29 | 0.0392 |
| Positive regulation of protein depolymerization | 6 | 0.29 | 0.0392 |
| Behavioral response to pain | 6 | 0.29 | 0.0392 |
| Fibroblast migration | 6 | 0.29 | 0.0392 |
| Production of molecular mediator involved in inflammatory response | 6 | 0.29 | 0.0392 |
| Positive regulation of homotypic cell-cell adhesion | 6 | 0.29 | 0.0392 |
| Protein localization to cell-cell junction | 6 | 0.29 | 0.0392 |
| Regulation of cd4-positive, alpha-beta t cell differentiation | 12 | 0.57 | 0.0399 |
| Cd4-positive, alpha-beta t cell differentiation | 12 | 0.57 | 0.0399 |
| Regulation of gtpase activity | 26 | 1.24 | 0.0401 |
| Regulation of interleukin-6 production | 26 | 1.24 | 0.0401 |
| Leukocyte apoptotic process | 10 | 0.48 | 0.0402 |
| Neuron recognition | 10 | 0.48 | 0.0402 |
| Response to dexamethasone | 10 | 0.48 | 0.0402 |
| Regulation of leukocyte degranulation | 10 | 0.48 | 0.0402 |
| Positive regulation of protein binding | 10 | 0.48 | 0.0402 |
| Mesenchyme development | 39 | 1.86 | 0.0405 |
| Regulation of organelle organization | 137 | 6.54 | 0.0406 |
| Regulation of dendritic cell differentiation | 5 | 0.24 | 0.0407 |
| L-ascorbic acid metabolic process | 5 | 0.24 | 0.0407 |
| Urea cycle | 5 | 0.24 | 0.0407 |
| Neutrophil activation involved in immune response | 5 | 0.24 | 0.0407 |
| Interleukin-6-mediated signaling pathway | 5 | 0.24 | 0.0407 |
| Regulation of angiotensin levels in blood | 5 | 0.24 | 0.0407 |
| Deoxyribonucleoside monophosphate catabolic process | 5 | 0.24 | 0.0407 |
| Xenobiotic export from cell | 5 | 0.24 | 0.0407 |
| Myofibril assembly | 13 | 0.62 | 0.0409 |
| Regulation of muscle adaptation | 17 | 0.81 | 0.0411 |
| Axis specification | 17 | 0.81 | 0.0411 |

| | | | |
|---|---|---|---|
| Negative regulation of developmental growth | 17 | 0.81 | 0.0411 |
| Positive regulation of autophagy | 25 | 1.19 | 0.0414 |
| Protein localization to endoplasmic reticulum | 14 | 0.67 | 0.0414 |
| Glycolipid biosynthetic process | 14 | 0.67 | 0.0414 |
| Negative regulation of mrna catabolic process | 14 | 0.67 | 0.0414 |
| Negative regulation of protein secretion | 14 | 0.67 | 0.0414 |
| Pyruvate metabolic process | 14 | 0.67 | 0.0414 |
| Ribonucleotide catabolic process | 16 | 0.76 | 0.0415 |
| Ribonucleoside diphosphate metabolic process | 15 | 0.72 | 0.0416 |
| Metanephros development | 15 | 0.72 | 0.0416 |
| Regulation of catabolic process | 133 | 6.35 | 0.0416 |
| Lactation | 9 | 0.43 | 0.0419 |
| Regulation of programmed necrotic cell death | 9 | 0.43 | 0.0419 |
| Regulation of epithelial to mesenchymal transition | 20 | 0.95 | 0.042 |
| Regulation of bicellular tight junction assembly | 7 | 0.33 | 0.0423 |
| Programmed necrotic cell death | 7 | 0.33 | 0.0423 |
| Myeloid dendritic cell activation | 7 | 0.33 | 0.0423 |
| Positive regulation of protein maturation | 7 | 0.33 | 0.0423 |
| Antigen processing and presentation via mhc class ib | 7 | 0.33 | 0.0423 |
| Negative regulation of regulated secretory pathway | 7 | 0.33 | 0.0423 |
| Vitamin transport | 11 | 0.53 | 0.0426 |
| Sensory perception of mechanical stimulus | 29 | 1.38 | 0.0426 |
| Negative regulation of leukocyte mediated cytotoxicity | 8 | 0.38 | 0.0428 |
| Er-nucleus signaling pathway | 8 | 0.38 | 0.0428 |
| Brown fat cell differentiation | 8 | 0.38 | 0.0428 |
| Intrinsic apoptotic signaling pathway in response to dna damage by p53 class mediator | 8 | 0.38 | 0.0428 |
| Negative regulation of rna biosynthetic process | 148 | 7.06 | 0.044 |
| Regulation of lymphocyte apoptotic process | 12 | 0.57 | 0.0441 |
| Glycerophospholipid biosynthetic process | 28 | 1.34 | 0.0442 |
| Response to alkaloid | 17 | 0.81 | 0.0444 |
| Negative regulation of dna-templated transcription | 147 | 7.02 | 0.0448 |
| Negative regulation of protein transport | 20 | 0.95 | 0.0449 |
| Regulation of dna binding | 13 | 0.62 | 0.0449 |
| Polyol biosynthetic process | 10 | 0.48 | 0.0452 |
| Digestive tract morphogenesis | 10 | 0.48 | 0.0452 |
| Neural tube closure | 15 | 0.72 | 0.0452 |
| Endothelial cell differentiation | 15 | 0.72 | 0.0452 |
| Purine ribonucleotide catabolic process | 15 | 0.72 | 0.0452 |
| Vascular endothelial growth factor receptor signaling pathway | 14 | 0.67 | 0.0452 |
| Regulation of neuron projection development | 57 | 2.72 | 0.0453 |
| Membrane lipid biosynthetic process | 22 | 1.05 | 0.0454 |
| Camera-type eye development | 47 | 2.24 | 0.046 |
| Regulation of myeloid leukocyte differentiation | 21 | 1 | 0.0467 |
| Negative regulation of nucleobase-containing compound metabolic process | 172 | 8.21 | 0.0467 |
| Positive regulation of purine nucleotide metabolic process | 9 | 0.43 | 0.0475 |
| Myoblast differentiation | 9 | 0.43 | 0.0475 |
| Positive regulation of nucleotide metabolic process | 9 | 0.43 | 0.0475 |
| Cell communication involved in cardiac conduction | 9 | 0.43 | 0.0475 |
| Regulation of actin filament-based movement | 9 | 0.43 | 0.0475 |
| Sodium ion homeostasis | 9 | 0.43 | 0.0475 |
| Purine ribonucleoside monophosphate metabolic process | 9 | 0.43 | 0.0475 |
| Stress response to metal ion | 6 | 0.29 | 0.0475 |
| Regulation of phospholipid biosynthetic process | 6 | 0.29 | 0.0475 |
| Response to interferon-alpha | 6 | 0.29 | 0.0475 |
| Positive regulation of type i interferon-mediated signaling pathway | 6 | 0.29 | 0.0475 |
| Regulation of ketone biosynthetic process | 6 | 0.29 | 0.0475 |
| Transcytosis | 6 | 0.29 | 0.0475 |
| Regulation of chemokine (c-x-c motif) ligand 2 production | 6 | 0.29 | 0.0475 |
| Artery development | 17 | 0.81 | 0.0478 |
| Response to interleukin-1 | 17 | 0.81 | 0.0478 |
| Regulation of interleukin-8 production | 17 | 0.81 | 0.0478 |
| Regulation of ossification | 20 | 0.95 | 0.048 |
| Ribonucleotide metabolic process | 42 | 2 | 0.0482 |
| Negative regulation of nervous system development | 22 | 1.05 | 0.0483 |
| Purine nucleotide catabolic process | 16 | 0.76 | 0.0485 |
| Membrane assembly | 12 | 0.57 | 0.0485 |
| Presynaptic endocytosis | 12 | 0.57 | 0.0485 |
| Exocytic process | 12 | 0.57 | 0.0485 |

| | | | |
|---|---|---|---|
| Tube closure | 15 | 0.72 | 0.049 |
| Cellular response to fluid shear stress | 8 | 0.38 | 0.0491 |
| Regulation of necroptotic process | 8 | 0.38 | 0.0491 |
| Leukotriene metabolic process | 8 | 0.38 | 0.0491 |
| Positive regulation of insulin secretion involved in cellular response to glucose stimulus | 8 | 0.38 | 0.0491 |
| Inner ear receptor cell differentiation | 13 | 0.62 | 0.0492 |
| Positive regulation of leukocyte mediated cytotoxicity | 13 | 0.62 | 0.0492 |
| Glucose metabolic process | 19 | 0.91 | 0.0492 |
| Nucleotide catabolic process | 19 | 0.91 | 0.0492 |
| Organelle localization by membrane tethering | 14 | 0.67 | 0.0493 |
| Positive regulation of lipid catabolic process | 7 | 0.33 | 0.0495 |
| Protein-lipid complex assembly | 7 | 0.33 | 0.0495 |
| Endochondral bone growth | 7 | 0.33 | 0.0495 |
| Endoplasmic reticulum calcium ion homeostasis | 7 | 0.33 | 0.0495 |
| Cellular response to ph | 7 | 0.33 | 0.0495 |
| Autophagy | 44 | 2.1 | 0.0496 |
| Process utilizing autophagic mechanism | 44 | 2.1 | 0.0496 |
| Regulation of cd8-positive, alpha-beta t cell proliferation | 4 | 0.19 | 0.0497 |
| Response to zinc ion starvation | 4 | 0.19 | 0.0497 |
| Cellular response to zinc ion starvation | 4 | 0.19 | 0.0497 |
| Daunorubicin metabolic process | 4 | 0.19 | 0.0497 |
| Limb bud formation | 4 | 0.19 | 0.0497 |
| Long-chain fatty acid catabolic process | 4 | 0.19 | 0.0497 |
| Allantoin metabolic process | 4 | 0.19 | 0.0497 |
| Exocrine pancreas development | 4 | 0.19 | 0.0497 |
| L-serine transport | 4 | 0.19 | 0.0497 |
| Srebp signaling pathway | 4 | 0.19 | 0.0497 |
| Galactose catabolic process | 4 | 0.19 | 0.0497 |
| Regulation of iron ion transport | 4 | 0.19 | 0.0497 |
| Lysosomal transport | 21 | 1 | 0.0497 |
| Renin secretion into blood stream | 3 | 0.14 | 0.0499 |
| Astrocyte activation involved in immune response | 3 | 0.14 | 0.0499 |
| Regulation of isoprenoid metabolic process | 3 | 0.14 | 0.0499 |
| Activation of janus kinase activity | 3 | 0.14 | 0.0499 |
| Cholesterol biosynthetic process via lathosterol | 3 | 0.14 | 0.0499 |
| Dgmp catabolic process | 3 | 0.14 | 0.0499 |
| Negative regulation of very-low-density lipoprotein particle remodeling | 3 | 0.14 | 0.0499 |
| Co-translational protein modification | 3 | 0.14 | 0.0499 |
| Cholesterol biosynthetic process via desmosterol | 3 | 0.14 | 0.0499 |
| Langerhans cell differentiation | 3 | 0.14 | 0.0499 |
| Immunoglobulin transcytosis in epithelial cells | 3 | 0.14 | 0.0499 |
| Urinary bladder development | 3 | 0.14 | 0.0499 |
| Actomyosin contractile ring assembly actin filament organization | 3 | 0.14 | 0.0499 |
| Lobar bronchus epithelium development | 3 | 0.14 | 0.0499 |
| Positive regulation of rna polymerase ii regulatory region sequence-specific dna binding | 3 | 0.14 | 0.0499 |
| Negative regulation of extracellular matrix constituent secretion | 3 | 0.14 | 0.0499 |
| Renal response to blood flow involved in circulatory renin-angiotensin regulation of systemic arterial blood pressure | 3 | 0.14 | 0.0499 |
| Negative regulation of monoatomic anion transport | 3 | 0.14 | 0.0499 |

TableS13

| Pathways upregulated in Undiff compared to Gland_B | | | |
|---|---|---|---|
| **GO Pathway** | **No. of genes** | **%** | **p-value** |
| Nervous system development | 246 | 23.08 | 3.87E-33 |
| Neuron differentiation | 144 | 13.51 | 3.34E-25 |
| Generation of neurons | 150 | 14.07 | 3.35E-25 |
| Neurogenesis | 164 | 15.38 | 1.27E-24 |
| Neuron development | 123 | 11.54 | 3.05E-24 |
| Mitotic cell cycle process | 83 | 7.79 | 6.36E-21 |
| Neuron projection development | 101 | 9.47 | 2.01E-20 |
| Mitotic cell cycle | 89 | 8.35 | 5.72E-20 |
| Neuron projection morphogenesis | 75 | 7.04 | 8.39E-17 |
| Cell projection morphogenesis | 76 | 7.13 | 9.68E-17 |
| Plasma membrane bounded cell projection morphogenesis | 75 | 7.04 | 2.53E-16 |
| Cell cycle process | 105 | 9.85 | 1.53E-15 |
| Cell cycle | 111 | 10.41 | 4.01E-15 |
| Cell morphogenesis | 89 | 8.35 | 1.52E-14 |
| Cell morphogenesis involved in neuron differentiation | 64 | 6 | 4.21E-13 |
| Regulation of cellular component organization | 206 | 19.32 | 4.37E-13 |
| Modulation of chemical synaptic transmission | 69 | 6.47 | 6.79E-13 |
| Regulation of trans-synaptic signaling | 69 | 6.47 | 7.36E-13 |
| Synaptic signaling | 63 | 5.91 | 4.79E-12 |
| Central nervous system development | 105 | 9.85 | 4.93E-12 |
| Trans-synaptic signaling | 60 | 5.63 | 5.04E-12 |
| Anterograde trans-synaptic signaling | 58 | 5.44 | 7.4E-12 |
| Chemical synaptic transmission | 58 | 5.44 | 7.4E-12 |
| Head development | 88 | 8.26 | 8.2E-12 |
| Brain development | 84 | 7.88 | 1.12E-11 |
| Axon development | 58 | 5.44 | 2.72E-11 |
| Plasma membrane bounded cell projection organization | 114 | 10.69 | 5.21E-11 |
| Axonogenesis | 52 | 4.88 | 1.17E-10 |
| Behavior | 73 | 6.85 | 1.51E-10 |
| Cell development | 182 | 17.07 | 3.03E-10 |
| Regulation of neuron projection development | 57 | 5.35 | 3.19E-10 |
| Cell projection organization | 115 | 10.79 | 3.96E-10 |
| Dna metabolic process | 80 | 7.5 | 1.17E-09 |
| Dna replication | 34 | 3.19 | 3.71E-09 |
| Mitotic sister chromatid segregation | 28 | 2.63 | 3.83E-09 |
| Sister chromatid segregation | 28 | 2.63 | 6.95E-09 |
| Mitotic nuclear division | 30 | 2.81 | 8.58E-09 |
| Organelle localization | 61 | 5.72 | 1.73E-08 |
| Cytoskeleton organization | 112 | 10.51 | 1.79E-08 |
| Dna damage response | 80 | 7.5 | 1.95E-08 |
| Cell division | 60 | 5.63 | 2.66E-08 |
| Microtubule cytoskeleton organization involved in mitosis | 25 | 2.35 | 6.33E-08 |
| Forebrain development | 49 | 4.6 | 8.58E-08 |
| Mitotic spindle organization | 21 | 1.97 | 1.03E-07 |
| Regulation of localization | 159 | 14.92 | 1.05E-07 |
| Chromosome organization | 51 | 4.78 | 1.08E-07 |
| Dna repair | 58 | 5.44 | 1.1E-07 |
| Telencephalon development | 38 | 3.56 | 1.12E-07 |
| Regulation of neuron differentiation | 30 | 2.81 | 1.13E-07 |
| Nuclear chromosome segregation | 34 | 3.19 | 1.84E-07 |
| Negative regulation of dna-templated transcription | 107 | 10.04 | 2.09E-07 |
| Regulation of plasma membrane bounded cell projection organization | 65 | 6.1 | 2.27E-07 |
| Cell-cell signaling | 78 | 7.32 | 2.38E-07 |
| Spindle organization | 28 | 2.63 | 2.59E-07 |
| Regulation of synapse structure or activity | 40 | 3.75 | 3.1E-07 |
| Negative regulation of rna biosynthetic process | 107 | 10.04 | 3.21E-07 |

| | | | |
|---|---|---|---|
| Establishment of organelle localization | 48 | 4.5 | 3.77E-07 |
| Negative regulation of nucleobase-containing compound metabolic process | 121 | 11.35 | 4.58E-07 |
| Regulation of cell differentiation | 129 | 12.1 | 4.86E-07 |
| Regulation of synapse organization | 39 | 3.66 | 4.86E-07 |
| Regulation of transport | 130 | 12.2 | 4.99E-07 |
| Vesicle-mediated transport in synapse | 27 | 2.53 | 5.2E-07 |
| Regulation of cell projection organization | 65 | 6.1 | 5.54E-07 |
| Regulation of signaling | 245 | 22.98 | 6.03E-07 |
| Organelle fission | 42 | 3.94 | 6.15E-07 |
| Regulation of neurogenesis | 44 | 4.13 | 6.34E-07 |
| Synapse organization | 43 | 4.03 | 6.51E-07 |
| Regulation of nervous system development | 50 | 4.69 | 6.6E-07 |
| Regulation of cell communication | 245 | 22.98 | 7.03E-07 |
| Chromosome segregation | 41 | 3.85 | 7.73E-07 |
| Cell junction organization | 57 | 5.35 | 9.08E-07 |
| Regulation of membrane potential | 50 | 4.69 | 1.01E-06 |
| Synaptic vesicle cycle | 25 | 2.35 | 0.0000012 |
| Neurotransmitter secretion | 18 | 1.69 | 1.35E-06 |
| Signal release from synapse | 18 | 1.69 | 1.35E-06 |
| Nuclear division | 39 | 3.66 | 1.41E-06 |
| Cognition | 39 | 3.66 | 1.63E-06 |
| Regulation of secretion | 61 | 5.72 | 1.88E-06 |
| Regulation of secretion by cell | 57 | 5.35 | 2.02E-06 |
| Regulation of biological quality | 212 | 19.89 | 2.14E-06 |
| Negative regulation of rna metabolic process | 110 | 10.32 | 2.19E-06 |
| Axon guidance | 31 | 2.91 | 2.26E-06 |
| Nuclear dna replication | 10 | 0.94 | 2.39E-06 |
| Neuron projection guidance | 31 | 2.91 | 2.46E-06 |
| Regulation of cell cycle | 92 | 8.63 | 2.53E-06 |
| Dna-templated dna replication | 23 | 2.16 | 2.75E-06 |
| Negative regulation of transcription by rna polymerase ii | 81 | 7.6 | 3.09E-06 |
| Synaptic vesicle exocytosis | 15 | 1.41 | 3.15E-06 |
| Cell cycle dna replication | 10 | 0.94 | 3.43E-06 |
| Microtubule cytoskeleton organization | 57 | 5.35 | 4.44E-06 |
| Dna replication initiation | 10 | 0.94 | 4.83E-06 |
| Learning | 24 | 2.25 | 5.15E-06 |
| Regulation of system process | 56 | 5.25 | 5.48E-06 |
| Positive regulation of developmental process | 109 | 10.23 | 5.78E-06 |
| Neurotransmitter transport | 22 | 2.06 | 7.78E-06 |
| Regulation of postsynaptic membrane potential | 19 | 1.78 | 8.74E-06 |
| Establishment of localization in cell | 124 | 11.63 | 8.84E-06 |
| Regulation of mitotic cell cycle phase transition | 39 | 3.66 | 9.55E-06 |
| Calcium ion-regulated exocytosis of neurotransmitter | 7 | 0.66 | 0.00001 |
| Regulation of synapse assembly | 24 | 2.25 | 0.0000105 |
| Signal release | 26 | 2.44 | 0.0000115 |
| Dense core granule localization | 8 | 0.75 | 0.0000135 |
| Regulation of cell cycle process | 67 | 6.29 | 0.0000147 |
| Neuron migration | 22 | 2.06 | 0.0000148 |
| Learning or memory | 33 | 3.1 | 0.0000153 |
| Positive regulation of neuron differentiation | 17 | 1.59 | 0.0000195 |
| Locomotory behavior | 27 | 2.53 | 0.0000238 |
| Regulation of mitotic cell cycle | 49 | 4.6 | 0.0000313 |
| Regulation of synaptic plasticity | 28 | 2.63 | 0.0000357 |
| Regulation of axonogenesis | 21 | 1.97 | 0.0000357 |
| Positive regulation of nervous system development | 33 | 3.1 | 0.000042 |
| Regulation of dendrite extension | 9 | 0.84 | 0.0000431 |
| Positive regulation of cell differentiation | 75 | 7.04 | 0.0000432 |
| Secretory granule localization | 8 | 0.75 | 0.0000434 |
| Dna recombination | 30 | 2.81 | 0.0000435 |

| | | | |
|---|---|---|---|
| Positive regulation of intracellular steroid hormone receptor signaling pathway | 7 | 0.66 | 0.0000478 |
| Central nervous system neuron differentiation | 26 | 2.44 | 0.0000517 |
| Regulation of nucleobase-containing compound metabolic process | 262 | 24.58 | 0.0000581 |
| Positive regulation of dna-templated transcription | 119 | 11.16 | 0.0000606 |
| Hindbrain development | 22 | 2.06 | 0.0000626 |
| Regulation of cellular localization | 82 | 7.69 | 0.0000642 |
| Pallium development | 25 | 2.35 | 0.0000646 |
| Regulation of dna-templated transcription | 225 | 21.11 | 0.0000671 |
| Positive regulation of rna biosynthetic process | 119 | 11.16 | 0.0000682 |
| Cellular response to stress | 120 | 11.26 | 0.000072 |
| Positive regulation of transcription by rna polymerase ii | 93 | 8.72 | 0.000073 |
| Vesicle docking involved in exocytosis | 10 | 0.94 | 0.0000732 |
| Regulation of dna-templated dna replication initiation | 7 | 0.66 | 0.0000732 |
| Regulation of regulated secretory pathway | 19 | 1.78 | 0.0000758 |
| Positive regulation of macromolecule metabolic process | 213 | 19.98 | 0.0000782 |
| Positive regulation of nucleobase-containing compound metabolic process | 143 | 13.41 | 0.0000797 |
| Organelle organization | 203 | 19.04 | 0.000081 |
| Regulation of rna biosynthetic process | 225 | 21.11 | 0.000085 |
| Regulation of chromosome organization | 28 | 2.63 | 0.0000858 |
| Regulation of cell cycle phase transition | 44 | 4.13 | 0.0000922 |
| Attachment of mitotic spindle microtubules to kinetochore | 7 | 0.66 | 0.000108 |
| Double-strand break repair | 26 | 2.44 | 0.000109 |
| Regulation of cell cycle g2/m phase transition | 18 | 1.69 | 0.00011 |
| Positive regulation of macromolecule biosynthetic process | 171 | 16.04 | 0.000112 |
| Positive regulation of dendrite extension | 8 | 0.75 | 0.000113 |
| Regulation of dna metabolic process | 46 | 4.32 | 0.000114 |
| Regulation of anatomical structure morphogenesis | 72 | 6.75 | 0.000117 |
| Cerebellum development | 17 | 1.59 | 0.000117 |
| Spindle assembly | 17 | 1.59 | 0.000131 |
| Positive regulation of metabolic process | 229 | 21.48 | 0.000131 |
| Regulation of chromosome segregation | 19 | 1.78 | 0.000149 |
| Regulation of postsynapse organization | 20 | 1.88 | 0.00016 |
| Potassium ion transport | 23 | 2.16 | 0.000161 |
| Modulation of excitatory postsynaptic potential | 11 | 1.03 | 0.000161 |
| Regulation of glial cell differentiation | 14 | 1.31 | 0.000174 |
| Microtubule-based process | 71 | 6.66 | 0.000194 |
| Regulation of exocytosis | 23 | 2.16 | 0.000202 |
| Regulation of transcription by rna polymerase ii | 176 | 16.51 | 0.000214 |
| Regulation of cell development | 68 | 6.38 | 0.000215 |
| Regulation of cell junction assembly | 29 | 2.72 | 0.000216 |
| Regulation of cell growth | 39 | 3.66 | 0.000222 |
| Cell migration | 73 | 6.85 | 0.000245 |
| Potassium ion transmembrane transport | 21 | 1.97 | 0.000252 |
| Action potential | 18 | 1.69 | 0.000264 |
| Synaptic vesicle docking | 6 | 0.56 | 0.000264 |
| Dendrite development | 17 | 1.59 | 0.000265 |
| Mitotic cell cycle checkpoint signaling | 17 | 1.59 | 0.000265 |
| Negative regulation of neuron projection development | 18 | 1.69 | 0.000289 |
| Adult behavior | 20 | 1.88 | 0.00029 |
| Positive regulation of excitatory postsynaptic potential | 9 | 0.84 | 0.000296 |
| Exocytic process | 12 | 1.13 | 0.000302 |
| Positive regulation of neurogenesis | 27 | 2.53 | 0.000307 |
| Regulation of g2/m transition of mitotic cell cycle | 16 | 1.5 | 0.000319 |
| Metencephalon development | 17 | 1.59 | 0.000321 |
| Positive regulation of glial cell differentiation | 10 | 0.94 | 0.000411 |
| Positive regulation of oligodendrocyte differentiation | 8 | 0.75 | 0.000412 |
| Cell cycle g2/m phase transition | 11 | 1.03 | 0.000418 |
| Mitotic cell cycle phase transition | 19 | 1.78 | 0.000425 |
| Cell cycle phase transition | 20 | 1.88 | 0.000431 |

| | | | |
|---|---|---|---|
| Cerebral cortex development | 18 | 1.69 | 0.000447 |
| Cellular localization | 175 | 16.42 | 0.000449 |
| Regulation of synaptic transmission, glutamatergic | 13 | 1.22 | 0.00046 |
| Negative regulation of cellular component organization | 58 | 5.44 | 0.000505 |
| Regulation of synaptic vesicle membrane organization | 7 | 0.66 | 0.000529 |
| Regulation of synaptic vesicle fusion to presynaptic active zone membrane | 7 | 0.66 | 0.000529 |
| Regulation of organelle organization | 86 | 8.07 | 0.000536 |
| Mitotic metaphase chromosome alignment | 11 | 1.03 | 0.000556 |
| Gmp biosynthetic process | 6 | 0.56 | 0.000568 |
| Positive regulation of biosynthetic process | 173 | 16.23 | 0.000568 |
| Regulated exocytosis | 18 | 1.69 | 0.000575 |
| Positive regulation of rna metabolic process | 126 | 11.82 | 0.000597 |
| Protein depolymerization | 9 | 0.84 | 0.000631 |
| Vesicle docking | 11 | 1.03 | 0.000638 |
| Regulation of rna metabolic process | 237 | 22.23 | 0.00065 |
| Excitatory postsynaptic potential | 12 | 1.13 | 0.000661 |
| Associative learning | 14 | 1.31 | 0.000684 |
| Establishment of cell polarity | 18 | 1.69 | 0.000732 |
| Establishment or maintenance of cell polarity | 23 | 2.16 | 0.00075 |
| Regulation of synaptic vesicle exocytosis | 10 | 0.94 | 0.00077 |
| Attachment of spindle microtubules to kinetochore | 8 | 0.75 | 0.000781 |
| Positive regulation of calcium ion-dependent exocytosis | 6 | 0.56 | 0.000792 |
| Cell cycle checkpoint signaling | 19 | 1.78 | 0.000793 |
| Positive regulation of cellular component organization | 82 | 7.69 | 0.000802 |
| Postsynapse organization | 16 | 1.5 | 0.000814 |
| Vesicle localization | 22 | 2.06 | 0.000838 |
| Ionotropic glutamate receptor signaling pathway | 8 | 0.75 | 0.00095 |
| Regulation of microtubule cytoskeleton organization | 20 | 1.88 | 0.000965 |
| Metaphase chromosome alignment | 13 | 1.22 | 0.001 |
| Hippocampus development | 14 | 1.31 | 0.00102 |
| Embryo development ending in birth or egg hatching | 54 | 5.07 | 0.00104 |
| Regulation of nervous system process | 16 | 1.5 | 0.00105 |
| Regulation of dna replication | 16 | 1.5 | 0.00105 |
| Cell communication | 311 | 29.17 | 0.00106 |
| Aorta development | 11 | 1.03 | 0.00107 |
| Mitotic dna replication initiation | 4 | 0.38 | 0.0011 |
| Cell cycle dna replication initiation | 4 | 0.38 | 0.0011 |
| Nuclear cell cycle dna replication initiation | 4 | 0.38 | 0.0011 |
| Double-strand break repair via homologous recombination | 16 | 1.5 | 0.00114 |
| Exocytosis | 25 | 2.35 | 0.00115 |
| Regulation of transmembrane transport | 41 | 3.85 | 0.00115 |
| Mitotic spindle assembly | 10 | 0.94 | 0.00118 |
| Signaling | 308 | 28.89 | 0.0012 |
| Negative regulation of cell projection organization | 21 | 1.97 | 0.00121 |
| Regulation of neurotransmitter secretion | 13 | 1.22 | 0.00123 |
| Cell junction assembly | 29 | 2.72 | 0.00126 |
| Protein complex oligomerization | 27 | 2.53 | 0.00127 |
| Regulation of protein localization | 69 | 6.47 | 0.00128 |
| Circulatory system process | 44 | 4.13 | 0.00129 |
| Developmental maturation | 29 | 2.72 | 0.00131 |
| Glutamate receptor signaling pathway | 10 | 0.94 | 0.00135 |
| Export from cell | 42 | 3.94 | 0.00135 |
| Axonal transport | 11 | 1.03 | 0.00137 |
| Regulation of vesicle fusion | 8 | 0.75 | 0.00137 |
| Synaptic vesicle priming | 6 | 0.56 | 0.00143 |
| Presynaptic membrane organization | 5 | 0.47 | 0.00146 |
| Mitotic dna replication checkpoint signaling | 5 | 0.47 | 0.00146 |
| Positive regulation of synaptic transmission | 19 | 1.78 | 0.0015 |
| Chemical synaptic transmission, postsynaptic | 12 | 1.13 | 0.00163 |

| | | | |
|---|---|---|---|
| Spindle localization | 11 | 1.03 | 0.00173 |
| Regulation of cell morphogenesis | 25 | 2.35 | 0.00176 |
| Positive regulation of secretion | 30 | 2.81 | 0.00183 |
| Recombinational repair | 16 | 1.5 | 0.00185 |
| Neuron maturation | 9 | 0.84 | 0.0019 |
| Base-excision repair | 9 | 0.84 | 0.0019 |
| Positive regulation of regulated secretory pathway | 9 | 0.84 | 0.0019 |
| Artery development | 14 | 1.31 | 0.00193 |
| Regulation of mitotic metaphase/anaphase transition | 13 | 1.22 | 0.002 |
| Organelle localization by membrane tethering | 12 | 1.13 | 0.002 |
| Embryo development | 75 | 7.04 | 0.0021 |
| Appendage morphogenesis | 18 | 1.69 | 0.00219 |
| Limb morphogenesis | 18 | 1.69 | 0.00219 |
| Anterograde axonal transport | 9 | 0.84 | 0.00219 |
| Neuron recognition | 9 | 0.84 | 0.00219 |
| Establishment of spindle orientation | 9 | 0.84 | 0.00219 |
| Positive regulation of secretion by cell | 28 | 2.63 | 0.00221 |
| Calcium-ion regulated exocytosis | 8 | 0.75 | 0.00227 |
| Phosphorus metabolic process | 100 | 9.38 | 0.00239 |
| Dna replication checkpoint signaling | 6 | 0.56 | 0.00239 |
| Purine ribonucleoside monophosphate biosynthetic process | 6 | 0.56 | 0.00239 |
| Neural nucleus development | 11 | 1.03 | 0.0024 |
| Positive regulation of transport | 64 | 6 | 0.00243 |
| Cellular component assembly | 165 | 15.48 | 0.0025 |
| Chordate embryonic development | 51 | 4.78 | 0.00259 |
| Regulation of metaphase/anaphase transition of cell cycle | 13 | 1.22 | 0.00262 |
| Synapse assembly | 16 | 1.5 | 0.00268 |
| Regulation of cellular component biogenesis | 74 | 6.94 | 0.00282 |
| Central nervous system neuron development | 13 | 1.22 | 0.00286 |
| Embryonic skeletal system development | 16 | 1.5 | 0.00288 |
| Positive regulation of intracellular estrogen receptor signaling pathway | 5 | 0.47 | 0.00292 |
| Regulation of growth | 48 | 4.5 | 0.00293 |
| Forebrain generation of neurons | 12 | 1.13 | 0.00295 |
| Regulation of amine transport | 12 | 1.13 | 0.00295 |
| Dna strand elongation | 6 | 0.56 | 0.00301 |
| Maintenance of synapse structure | 7 | 0.66 | 0.00301 |
| Cellular component biogenesis | 178 | 16.7 | 0.00301 |
| Phosphate-containing compound metabolic process | 99 | 9.29 | 0.00305 |
| Positive regulation of microtubule polymerization or depolymerization | 8 | 0.75 | 0.00307 |
| Chromatin organization | 85 | 7.97 | 0.00316 |
| Positive regulation of cell projection organization | 32 | 3 | 0.00321 |
| Negative regulation of mitotic cell cycle phase transition | 19 | 1.78 | 0.00323 |
| Regulation of spindle organization | 9 | 0.84 | 0.00324 |
| Cell motility | 81 | 7.6 | 0.00327 |
| Negative regulation of g2/m transition of mitotic cell cycle | 11 | 1.03 | 0.00328 |
| Regulation of g1/s transition of mitotic cell cycle | 19 | 1.78 | 0.00343 |
| Chromatin remodeling | 76 | 7.13 | 0.00352 |
| Regulation of dendritic spine development | 10 | 0.94 | 0.00354 |
| Establishment of spindle localization | 10 | 0.94 | 0.00354 |
| Regulation of neuronal synaptic plasticity | 10 | 0.94 | 0.00354 |
| Positive regulation of cell cycle g2/m phase transition | 7 | 0.66 | 0.00358 |
| Presynaptic dense core vesicle exocytosis | 4 | 0.38 | 0.00358 |
| Regulation of guanyl-nucleotide exchange factor activity | 4 | 0.38 | 0.00358 |
| Neuronal dense core vesicle exocytosis | 4 | 0.38 | 0.00358 |
| 'de novo' xmp biosynthetic process | 4 | 0.38 | 0.00358 |
| Xmp metabolic process | 4 | 0.38 | 0.00358 |
| Xmp biosynthetic process | 4 | 0.38 | 0.00358 |
| Forebrain neuron differentiation | 11 | 1.03 | 0.00362 |
| Regulation of insulin secretion | 19 | 1.78 | 0.00364 |

| | | | |
|---|---|---|---|
| Metal ion transport | 50 | 4.69 | 0.00367 |
| G2/m transition of mitotic cell cycle | 9 | 0.84 | 0.00367 |
| Establishment of chromosome localization | 13 | 1.22 | 0.00368 |
| Nucleoside phosphate metabolic process | 46 | 4.32 | 0.00371 |
| Purine nucleoside monophosphate biosynthetic process | 6 | 0.56 | 0.00374 |
| Regulation of postsynaptic specialization assembly | 6 | 0.56 | 0.00374 |
| Secretion by cell | 36 | 3.38 | 0.00384 |
| Regulation of hormone secretion | 25 | 2.35 | 0.00389 |
| Response to nitrogen compound | 75 | 7.04 | 0.0039 |
| Mitotic dna replication | 5 | 0.47 | 0.00393 |
| Microtubule polymerization or depolymerization | 10 | 0.94 | 0.00394 |
| Regulation of microtubule-based process | 26 | 2.44 | 0.00397 |
| Negative regulation of cell cycle g2/m phase transition | 11 | 1.03 | 0.00399 |
| Mitotic dna integrity checkpoint signaling | 11 | 1.03 | 0.00399 |
| Muscle system process | 27 | 2.53 | 0.00399 |
| Nerve development | 13 | 1.22 | 0.00399 |
| Regulation of peptide hormone secretion | 21 | 1.97 | 0.00413 |
| Synaptic vesicle localization | 9 | 0.84 | 0.00414 |
| Positive regulation of cell cycle phase transition | 15 | 1.41 | 0.00421 |
| Peripheral nervous system development | 12 | 1.13 | 0.00423 |
| Establishment of vesicle localization | 19 | 1.78 | 0.00433 |
| Embryonic appendage morphogenesis | 15 | 1.41 | 0.00452 |
| Embryonic limb morphogenesis | 15 | 1.41 | 0.00452 |
| Regulation of catecholamine secretion | 8 | 0.75 | 0.00468 |
| Regulation of dna-templated dna replication | 8 | 0.75 | 0.00468 |
| Regulation of monoatomic ion transport | 38 | 3.56 | 0.00483 |
| Regulation of peptide secretion | 21 | 1.97 | 0.00483 |
| Blood circulation | 36 | 3.38 | 0.00493 |
| Peripheral nervous system neuron development | 5 | 0.47 | 0.00515 |
| Regulation of peptide transport | 21 | 1.97 | 0.00535 |
| Protein localization to synapse | 10 | 0.94 | 0.00537 |
| Membrane docking | 12 | 1.13 | 0.00546 |
| Regulation of microtubule polymerization or depolymerization | 13 | 1.22 | 0.00547 |
| Negative regulation of cell cycle | 32 | 3 | 0.00552 |
| Spontaneous synaptic transmission | 4 | 0.38 | 0.00553 |
| Import into cell | 52 | 4.88 | 0.00558 |
| Signal transduction in response to dna damage | 16 | 1.5 | 0.00561 |
| Nucleobase-containing small molecule metabolic process | 47 | 4.41 | 0.00562 |
| Negative regulation of chromosome organization | 11 | 1.03 | 0.00577 |
| Axo-dendritic transport | 11 | 1.03 | 0.00577 |
| Establishment of mitochondrion localization | 7 | 0.66 | 0.00577 |
| Mitotic g2/m transition checkpoint | 9 | 0.84 | 0.00583 |
| Receptor localization to synapse | 9 | 0.84 | 0.00583 |
| Regulation of neurotransmitter transport | 13 | 1.22 | 0.0059 |
| Positive regulation of mitotic cell cycle phase transition | 13 | 1.22 | 0.0059 |
| Regulation of dendritic spine morphogenesis | 8 | 0.75 | 0.00605 |
| Regulation of rna splicing | 19 | 1.78 | 0.00637 |
| Positive regulation of cell cycle process | 25 | 2.35 | 0.00637 |
| Nucleotide metabolic process | 36 | 3.38 | 0.00645 |
| Cardiac muscle cell action potential | 9 | 0.84 | 0.0065 |
| Regulation of neuron migration | 9 | 0.84 | 0.0065 |
| Negative regulation of developmental process | 71 | 6.66 | 0.00656 |
| Dna strand elongation involved in dna replication | 5 | 0.47 | 0.0066 |
| Peripheral nervous system neuron differentiation | 5 | 0.47 | 0.0066 |
| Regulation of protein secretion | 25 | 2.35 | 0.00667 |
| Positive regulation of microtubule polymerization | 7 | 0.66 | 0.00668 |
| Regulation of sister chromatid segregation | 13 | 1.22 | 0.00684 |
| Chromosome localization | 13 | 1.22 | 0.00684 |
| G1/s transition of mitotic cell cycle | 11 | 1.03 | 0.00686 |

| | | | |
|---|---|---|---|
| Neurofilament bundle assembly | 3 | 0.28 | 0.00702 |
| Positive regulation of g2/mi transition of meiotic cell cycle | 3 | 0.28 | 0.00702 |
| Postsynaptic intermediate filament cytoskeleton organization | 3 | 0.28 | 0.00702 |
| Muscle contraction | 22 | 2.06 | 0.00721 |
| Regulation of cyclin-dependent protein serine/threonine kinase activity | 9 | 0.84 | 0.00722 |
| Ligand-gated ion channel signaling pathway | 8 | 0.75 | 0.0077 |
| Regulation of mitotic spindle organization | 8 | 0.75 | 0.0077 |
| Regulation of oligodendrocyte differentiation | 8 | 0.75 | 0.0077 |
| Dense core granule transport | 4 | 0.38 | 0.00799 |
| Dense core granule exocytosis | 4 | 0.38 | 0.00799 |
| Dense core granule cytoskeletal transport | 4 | 0.38 | 0.00799 |
| Protein-containing complex organization | 101 | 9.47 | 0.00808 |
| Cell cycle g1/s phase transition | 11 | 1.03 | 0.00812 |
| Positive regulation of mitotic cell cycle | 15 | 1.41 | 0.00815 |
| Regulation of developmental growth | 28 | 2.63 | 0.00816 |
| Protein heterotetramerization | 5 | 0.47 | 0.0083 |
| Negative regulation of neuron projection regeneration | 5 | 0.47 | 0.0083 |
| Regulation of cell cycle g1/s phase transition | 20 | 1.88 | 0.0083 |
| Ear development | 22 | 2.06 | 0.0083 |
| Animal organ morphogenesis | 68 | 6.38 | 0.0084 |
| Dna integrity checkpoint signaling | 13 | 1.22 | 0.00847 |
| Positive regulation of cell cycle | 29 | 2.72 | 0.00882 |
| Visual behavior | 9 | 0.84 | 0.00885 |
| Cellular response to nitrogen compound | 47 | 4.41 | 0.0089 |
| Regulation of vesicle-mediated transport | 42 | 3.94 | 0.00891 |
| Appendage development | 19 | 1.78 | 0.00913 |
| Limb development | 19 | 1.78 | 0.00913 |
| Regulation of monoatomic ion transmembrane transport | 29 | 2.72 | 0.00913 |
| Negative regulation of dna metabolic process | 15 | 1.41 | 0.0092 |
| Corpus callosum development | 6 | 0.56 | 0.00935 |
| Motor neuron axon guidance | 6 | 0.56 | 0.00935 |
| Meiotic chromosome segregation | 12 | 1.13 | 0.0094 |
| Inner ear development | 20 | 1.88 | 0.00958 |
| Neuron fate commitment | 10 | 0.94 | 0.0102 |
| Microtubule depolymerization | 5 | 0.47 | 0.0103 |
| Regulation of postsynaptic density assembly | 5 | 0.47 | 0.0103 |
| Serine family amino acid biosynthetic process | 5 | 0.47 | 0.0103 |
| Regulation of gliogenesis | 13 | 1.22 | 0.0104 |
| Negative regulation of cell cycle phase transition | 22 | 2.06 | 0.0104 |
| Negative regulation of mitotic cell cycle | 21 | 1.97 | 0.0104 |
| Gliogenesis | 25 | 2.35 | 0.0104 |
| Sulfur compound biosynthetic process | 16 | 1.5 | 0.0107 |
| Gmp metabolic process | 6 | 0.56 | 0.0109 |
| Positive regulation of g2/m transition of mitotic cell cycle | 6 | 0.56 | 0.0109 |
| Mesenchymal cell proliferation | 6 | 0.56 | 0.0109 |
| Neuromuscular synaptic transmission | 6 | 0.56 | 0.0109 |
| Regulation of fat cell proliferation | 4 | 0.38 | 0.011 |
| Presynaptic membrane assembly | 4 | 0.38 | 0.011 |
| Regulation of heart contraction | 20 | 1.88 | 0.011 |
| Adult locomotory behavior | 11 | 1.03 | 0.0111 |
| Positive regulation of gliogenesis | 10 | 0.94 | 0.0111 |
| Regulation of calcium ion-dependent exocytosis | 7 | 0.66 | 0.0114 |
| Regulation of dendrite development | 12 | 1.13 | 0.0116 |
| Protein tetramerization | 12 | 1.13 | 0.0116 |
| Monoatomic ion transport | 69 | 6.47 | 0.0117 |
| Limbic system development | 14 | 1.31 | 0.0118 |
| Positive regulation of multicellular organismal process | 109 | 10.23 | 0.0118 |
| B cell activation | 19 | 1.78 | 0.0122 |
| Meiotic spindle assembly | 5 | 0.47 | 0.0125 |

| | | | |
|---|---|---|---|
| Regulation of circadian sleep/wake cycle | 5 | 0.47 | 0.0125 |
| Tetrahydrofolate metabolic process | 5 | 0.47 | 0.0125 |
| Innervation | 6 | 0.56 | 0.0127 |
| Presynaptic modulation of chemical synaptic transmission | 7 | 0.66 | 0.0128 |
| Cell-cell adhesion via plasma-membrane adhesion molecules | 23 | 2.16 | 0.013 |
| Visual learning | 8 | 0.75 | 0.0133 |
| Regulation of signal transduction | 188 | 17.64 | 0.0134 |
| Negative regulation of transport | 36 | 3.38 | 0.0135 |
| Positive regulation of meiotic cell cycle phase transition | 3 | 0.28 | 0.0136 |
| Negative regulation of cell proliferation involved in contact inhibition | 3 | 0.28 | 0.0136 |
| Sensory organ development | 45 | 4.22 | 0.0136 |
| Negative regulation of nervous system development | 15 | 1.41 | 0.0138 |
| Secretion | 41 | 3.85 | 0.0138 |
| Regulation of behavior | 10 | 0.94 | 0.0142 |
| Regulation of chromosome separation | 10 | 0.94 | 0.0142 |
| Response to radiation | 34 | 3.19 | 0.0146 |
| Imp biosynthetic process | 4 | 0.38 | 0.0146 |
| Dna modification | 6 | 0.56 | 0.0146 |
| Mitotic spindle assembly checkpoint signaling | 6 | 0.56 | 0.0146 |
| Positive regulation of isotype switching | 6 | 0.56 | 0.0146 |
| Mitotic spindle checkpoint signaling | 6 | 0.56 | 0.0146 |
| Spindle assembly checkpoint signaling | 6 | 0.56 | 0.0146 |
| Mitochondrion localization | 8 | 0.75 | 0.0147 |
| Cell adhesion | 65 | 6.1 | 0.0153 |
| Regulation of establishment of protein localization | 41 | 3.85 | 0.0153 |
| Telencephalon cell migration | 9 | 0.84 | 0.0154 |
| Actin filament-based process | 45 | 4.22 | 0.0157 |
| Regulation of blood circulation | 23 | 2.16 | 0.0158 |
| Roof of mouth development | 11 | 1.03 | 0.016 |
| Interstrand cross-link repair | 7 | 0.66 | 0.0161 |
| Notch signaling pathway | 13 | 1.22 | 0.0161 |
| Spindle checkpoint signaling | 6 | 0.56 | 0.0167 |
| Positive regulation of chromosome segregation | 6 | 0.56 | 0.0167 |
| Regulation of multicellular organismal development | 95 | 8.91 | 0.017 |
| Purine-containing compound metabolic process | 38 | 3.56 | 0.0173 |
| Protein homooligomerization | 19 | 1.78 | 0.0175 |
| Positive regulation of cell development | 35 | 3.28 | 0.0175 |
| Monoatomic ion transmembrane transport | 58 | 5.44 | 0.0176 |
| Negative regulation of protein-containing complex disassembly | 10 | 0.94 | 0.0179 |
| Skeletal system development | 38 | 3.56 | 0.0179 |
| Embryonic morphogenesis | 43 | 4.03 | 0.0181 |
| Regulation of microtubule polymerization | 9 | 0.84 | 0.0182 |
| Renal tubule development | 11 | 1.03 | 0.0184 |
| Protein localization to chromosome | 12 | 1.13 | 0.0184 |
| Sno(s)rna 3'-end processing | 4 | 0.38 | 0.0187 |
| Synaptic vesicle clustering | 4 | 0.38 | 0.0187 |
| Double-strand break repair via break-induced replication | 4 | 0.38 | 0.0187 |
| Heparan sulfate proteoglycan biosynthetic process, enzymatic modification | 4 | 0.38 | 0.0187 |
| Presynapse organization | 6 | 0.56 | 0.0191 |
| Negative regulation of microtubule depolymerization | 6 | 0.56 | 0.0191 |
| Protein heterooligomerization | 6 | 0.56 | 0.0191 |
| Regulation of metal ion transport | 31 | 2.91 | 0.0191 |
| Membraneless organelle assembly | 29 | 2.72 | 0.0192 |
| Cell-cell adhesion | 40 | 3.75 | 0.0194 |
| Regulation of cellular response to stress | 39 | 3.66 | 0.0196 |
| Embryonic skeletal system morphogenesis | 11 | 1.03 | 0.0196 |
| Forebrain cell migration | 9 | 0.84 | 0.0197 |
| Negative regulation of neurogenesis | 14 | 1.31 | 0.0197 |
| Negative regulation of cell differentiation | 51 | 4.78 | 0.0199 |

| | | | |
|---|---|---|---|
| Regulation of postsynapse assembly | 7 | 0.66 | 0.02 |
| Tissue development | 112 | 10.51 | 0.0205 |
| Calcium ion transmembrane import into cytosol | 11 | 1.03 | 0.021 |
| Presynapse assembly | 5 | 0.47 | 0.021 |
| Regulation of postsynaptic density organization | 5 | 0.47 | 0.021 |
| Negative regulation of cell cycle process | 24 | 2.25 | 0.0211 |
| Vesicle-mediated transport | 84 | 7.88 | 0.0213 |
| Positive regulation of filopodium assembly | 6 | 0.56 | 0.0216 |
| Negative regulation of mitotic sister chromatid separation | 6 | 0.56 | 0.0216 |
| Negative regulation of sister chromatid segregation | 6 | 0.56 | 0.0216 |
| Negative regulation of mitotic sister chromatid segregation | 6 | 0.56 | 0.0216 |
| Negative regulation of mitotic metaphase/anaphase transition | 6 | 0.56 | 0.0216 |
| Intracellular signal transduction | 109 | 10.23 | 0.0217 |
| Regulation of dna repair | 20 | 1.88 | 0.0218 |
| Monoatomic cation transport | 56 | 5.25 | 0.0219 |
| Homophilic cell adhesion via plasma membrane adhesion molecules | 16 | 1.5 | 0.0219 |
| Protein-containing complex disassembly | 15 | 1.41 | 0.022 |
| Female meiotic nuclear division | 7 | 0.66 | 0.0222 |
| Nucleoside monophosphate biosynthetic process | 7 | 0.66 | 0.0222 |
| Regulation of axon extension | 10 | 0.94 | 0.0222 |
| Negative regulation of protein depolymerization | 9 | 0.84 | 0.023 |
| Neuromuscular process controlling balance | 8 | 0.75 | 0.0232 |
| Membrane depolarization | 8 | 0.75 | 0.0232 |
| Organophosphate metabolic process | 65 | 6.1 | 0.0232 |
| Amp biosynthetic process | 4 | 0.38 | 0.0235 |
| Positive regulation of synaptic vesicle fusion to presynaptic active zone membrane | 4 | 0.38 | 0.0235 |
| Calcium-dependent activation of synaptic vesicle fusion | 4 | 0.38 | 0.0235 |
| Eyelid development in camera-type eye | 4 | 0.38 | 0.0235 |
| Positive regulation of synaptic vesicle membrane organization | 4 | 0.38 | 0.0235 |
| Regulation of astrocyte differentiation | 6 | 0.56 | 0.0243 |
| Negative regulation of microtubule polymerization or depolymerization | 7 | 0.66 | 0.0244 |
| Membrane depolarization during action potential | 5 | 0.47 | 0.0245 |
| Regulation of excitatory synapse assembly | 5 | 0.47 | 0.0245 |
| Regulation of arf protein signal transduction | 5 | 0.47 | 0.0245 |
| Mitotic dna damage checkpoint signaling | 9 | 0.84 | 0.0248 |
| Response to amyloid-beta | 8 | 0.75 | 0.0252 |
| Negative regulation of secretion by cell | 15 | 1.41 | 0.0254 |
| Nucleoside phosphate catabolic process | 14 | 1.31 | 0.0255 |
| Negative regulation of neuron differentiation | 9 | 0.84 | 0.0267 |
| Calcium-dependent cell-cell adhesion via plasma membrane cell adhesion molecules | 7 | 0.66 | 0.0269 |
| Spinal cord development | 11 | 1.03 | 0.0269 |
| Rhythmic process | 23 | 2.16 | 0.0271 |
| Regulation of mesenchymal cell proliferation | 6 | 0.56 | 0.0272 |
| Negative regulation of chromosome separation | 6 | 0.56 | 0.0272 |
| Neuronal action potential | 6 | 0.56 | 0.0272 |
| Negative regulation of metaphase/anaphase transition of cell cycle | 6 | 0.56 | 0.0272 |
| Adult walking behavior | 6 | 0.56 | 0.0272 |
| Pteridine-containing compound metabolic process | 6 | 0.56 | 0.0272 |
| Tube morphogenesis | 48 | 4.5 | 0.0282 |
| Postsynaptic cytoskeleton organization | 5 | 0.47 | 0.0282 |
| Glutamine metabolic process | 5 | 0.47 | 0.0282 |
| Regulation of cellular response to oxidative stress | 5 | 0.47 | 0.0282 |
| Positive regulation of cellular component biogenesis | 37 | 3.47 | 0.0286 |
| Regulation of chromosome condensation | 4 | 0.38 | 0.0288 |
| Sno(s)rna processing | 4 | 0.38 | 0.0288 |
| Negative regulation of dendritic spine development | 4 | 0.38 | 0.0288 |
| Regulation of meiotic cell cycle phase transition | 4 | 0.38 | 0.0288 |

| | | | |
|---|---|---|---|
| Synaptic transmission, glutamatergic | 7 | 0.66 | 0.0295 |
| Sister chromatid cohesion | 7 | 0.66 | 0.0295 |
| Negative regulation of macromolecule metabolic process | 179 | 16.79 | 0.0295 |
| Positive regulation of mirna transcription | 8 | 0.75 | 0.0297 |
| Ribonucleotide metabolic process | 25 | 2.35 | 0.03 |
| Neuroepithelial cell differentiation | 6 | 0.56 | 0.0304 |
| Ribonucleoside monophosphate biosynthetic process | 6 | 0.56 | 0.0304 |
| Regulation of spindle assembly | 6 | 0.56 | 0.0304 |
| Postreplication repair | 6 | 0.56 | 0.0304 |
| Negative regulation of chromosome segregation | 6 | 0.56 | 0.0304 |
| Negative regulation of signaling | 96 | 9.01 | 0.0305 |
| Regulation of stem cell population maintenance | 9 | 0.84 | 0.0308 |
| Neuron projection extension | 10 | 0.94 | 0.0309 |
| Memory | 13 | 1.22 | 0.0312 |
| Response to oxygen-containing compound | 100 | 9.38 | 0.0315 |
| Negative regulation of guanyl-nucleotide exchange factor activity | 3 | 0.28 | 0.0318 |
| Positive regulation of inhibitory postsynaptic potential | 3 | 0.28 | 0.0318 |
| Synaptic vesicle uncoating | 3 | 0.28 | 0.0318 |
| 'de novo' imp biosynthetic process | 3 | 0.28 | 0.0318 |
| Spontaneous neurotransmitter secretion | 3 | 0.28 | 0.0318 |
| Vesicle cytoskeletal trafficking | 8 | 0.75 | 0.0321 |
| Regulation of cell cycle checkpoint | 7 | 0.66 | 0.0323 |
| Regulation of erythrocyte differentiation | 7 | 0.66 | 0.0323 |
| Insulin secretion | 7 | 0.66 | 0.0323 |
| Purine nucleotide metabolic process | 29 | 2.72 | 0.0323 |
| U2-type prespliceosome assembly | 5 | 0.47 | 0.0323 |
| Ventricular septum development | 9 | 0.84 | 0.033 |
| Positive regulation of synaptic transmission, glutamatergic | 6 | 0.56 | 0.0338 |
| Regulation of response to oxidative stress | 6 | 0.56 | 0.0338 |
| Protein localization to postsynaptic membrane | 6 | 0.56 | 0.0338 |
| Regulation of intracellular estrogen receptor signaling pathway | 6 | 0.56 | 0.0338 |
| Negative regulation of secretion | 16 | 1.5 | 0.034 |
| Positive regulation of hormone secretion | 14 | 1.31 | 0.0341 |
| Regulation of double-strand break repair | 14 | 1.31 | 0.0341 |
| Protein-containing complex assembly | 87 | 8.16 | 0.0345 |
| Regulation of circadian sleep/wake cycle, sleep | 4 | 0.38 | 0.0347 |
| Interneuron migration | 4 | 0.38 | 0.0347 |
| Actin cytoskeleton organization | 39 | 3.66 | 0.035 |
| Regulation of kinase activity | 29 | 2.72 | 0.0353 |
| Meiotic nuclear division | 16 | 1.5 | 0.0355 |
| Meiotic spindle organization | 5 | 0.47 | 0.0368 |
| Regulation of synapse maturation | 5 | 0.47 | 0.0368 |
| Glial cell differentiation | 19 | 1.78 | 0.037 |
| Regulation of protein transport | 33 | 3.1 | 0.037 |
| Feeding behavior | 10 | 0.94 | 0.0371 |
| Positive regulation of transmembrane transport | 18 | 1.69 | 0.0372 |
| Regulation of immunoglobulin mediated immune response | 8 | 0.75 | 0.0373 |
| Regulation of b cell mediated immunity | 8 | 0.75 | 0.0373 |
| Establishment of mitotic spindle orientation | 6 | 0.56 | 0.0374 |
| Mitotic g2 dna damage checkpoint signaling | 6 | 0.56 | 0.0374 |
| Protein localization to postsynapse | 6 | 0.56 | 0.0374 |
| Forebrain neuron development | 6 | 0.56 | 0.0374 |
| Regulation of microtubule depolymerization | 6 | 0.56 | 0.0374 |
| Proteoglycan metabolic process | 11 | 1.03 | 0.038 |
| Positive regulation of binding | 11 | 1.03 | 0.038 |
| Hormone transport | 11 | 1.03 | 0.038 |
| Positive regulation of cell growth | 15 | 1.41 | 0.0381 |
| Nucleotide catabolic process | 12 | 1.13 | 0.0382 |
| Striated muscle contraction | 12 | 1.13 | 0.0382 |

| | | | |
|---|---|---|---|
| Response to light stimulus | 25 | 2.35 | 0.0383 |
| Ribose phosphate metabolic process | 25 | 2.35 | 0.0383 |
| Cell junction maintenance | 7 | 0.66 | 0.0383 |
| Negative regulation of cell communication | 95 | 8.91 | 0.0392 |
| Positive regulation of neuron projection development | 15 | 1.41 | 0.0397 |
| Regulation of dendrite morphogenesis | 8 | 0.75 | 0.0401 |
| Proteoglycan biosynthetic process | 9 | 0.84 | 0.0402 |
| Positive regulation of axonogenesis | 9 | 0.84 | 0.0402 |
| Negative regulation of organelle organization | 26 | 2.44 | 0.0408 |
| Regulation of isotype switching | 6 | 0.56 | 0.0412 |
| Long-term memory | 6 | 0.56 | 0.0412 |
| Sno(s)rna metabolic process | 4 | 0.38 | 0.0412 |
| Spindle assembly involved in female meiosis | 4 | 0.38 | 0.0412 |
| Neuron cell-cell adhesion | 4 | 0.38 | 0.0412 |
| Regulation of monoatomic cation transmembrane transport | 24 | 2.25 | 0.0415 |
| Establishment of mitochondrion localization, microtubule-mediated | 5 | 0.47 | 0.0416 |
| Mitochondrion transport along microtubule | 5 | 0.47 | 0.0416 |
| Cerebral cortex neuron differentiation | 5 | 0.47 | 0.0416 |
| Granulocyte differentiation | 5 | 0.47 | 0.0416 |
| Glial cell migration | 7 | 0.66 | 0.0416 |
| Regulation of filopodium assembly | 7 | 0.66 | 0.0416 |
| Developmental cell growth | 12 | 1.13 | 0.0422 |
| Purine ribonucleotide metabolic process | 23 | 2.16 | 0.0423 |
| Regulation of phosphate metabolic process | 53 | 4.97 | 0.0426 |
| Regulation of phosphorus metabolic process | 53 | 4.97 | 0.0427 |
| Tissue morphogenesis | 40 | 3.75 | 0.0427 |
| Regulation of receptor internalization | 9 | 0.84 | 0.0428 |
| Cranial nerve development | 8 | 0.75 | 0.043 |
| Neuron cellular homeostasis | 8 | 0.75 | 0.043 |
| Regulation of g2/mi transition of meiotic cell cycle | 3 | 0.28 | 0.0431 |
| Modulation of inhibitory postsynaptic potential | 3 | 0.28 | 0.0431 |
| Cell fate commitment | 21 | 1.97 | 0.044 |
| Protein localization to cell junction | 10 | 0.94 | 0.0442 |
| Protein sumoylation | 7 | 0.66 | 0.045 |
| Serine family amino acid metabolic process | 6 | 0.56 | 0.0452 |
| Regulation of organic acid transport | 9 | 0.84 | 0.0456 |
| Jnk cascade | 9 | 0.84 | 0.0456 |
| Cytosolic transport | 13 | 1.22 | 0.0457 |
| Negative regulation of phosphate metabolic process | 21 | 1.97 | 0.0458 |
| Negative regulation of phosphorus metabolic process | 21 | 1.97 | 0.0458 |
| Response to nutrient levels | 36 | 3.38 | 0.0464 |
| Cell growth | 12 | 1.13 | 0.0465 |
| Cytoskeleton-dependent intracellular transport | 17 | 1.59 | 0.0466 |
| Calcium ion import across plasma membrane | 5 | 0.47 | 0.0467 |
| Positive regulation of mesenchymal cell proliferation | 5 | 0.47 | 0.0467 |
| Folic acid-containing compound metabolic process | 5 | 0.47 | 0.0467 |
| Hormone secretion | 10 | 0.94 | 0.0467 |
| Nucleoside phosphate biosynthetic process | 23 | 2.16 | 0.048 |
| Cellular response to oxygen-containing compound | 70 | 6.57 | 0.0482 |
| Segment specification | 4 | 0.38 | 0.0483 |
| Synapse maturation | 4 | 0.38 | 0.0483 |
| Membrane depolarization during cardiac muscle cell action potential | 4 | 0.38 | 0.0483 |
| Positive regulation of exocytosis | 9 | 0.84 | 0.0484 |
| Postsynaptic signal transduction | 7 | 0.66 | 0.0486 |
| Negative regulation of axonogenesis | 7 | 0.66 | 0.0486 |
| Regulation of phosphorylation | 45 | 4.22 | 0.0492 |
| Dendrite morphogenesis | 8 | 0.75 | 0.0493 |
| Tube development | 59 | 5.53 | 0.0494 |
| Negative regulation of cell development | 22 | 2.06 | 0.0494 |

| | | | |
|---|---|---|---|
| Calcium ion import | 6 | 0.56 | 0.0495 |
| Negative regulation of mitotic nuclear division | 6 | 0.56 | 0.0495 |
| Walking behavior | 6 | 0.56 | 0.0495 |

TableS14

| Pathways upregulated in Gland_B compared to Undiff | | | |
|---|---|---|---|
| GO Pathway | No. of genes | % | p-value |
| Cell projection organization | 247 | 15.04 | 2.68E-45 |
| Plasma membrane bounded cell projection organization | 240 | 14.62 | 3.07E-45 |
| Cilium organization | 120 | 7.31 | 9.78E-41 |
| Cilium assembly | 114 | 6.94 | 1.46E-40 |
| Cell projection assembly | 124 | 7.55 | 1.78E-35 |
| Plasma membrane bounded cell projection assembly | 122 | 7.43 | 1.96E-35 |
| Circulatory system development | 191 | 11.63 | 2.52E-35 |
| Tissue development | 289 | 17.6 | 2.04E-34 |
| Cell development | 333 | 20.28 | 1.93E-32 |
| Cell adhesion | 182 | 11.08 | 2.38E-30 |
| Cell motility | 203 | 12.36 | 6.01E-29 |
| Cilium movement | 77 | 4.69 | 1.16E-27 |
| Microtubule-based movement | 105 | 6.39 | 7.89E-25 |
| Heart development | 123 | 7.49 | 1.1E-24 |
| Axoneme assembly | 47 | 2.86 | 1.37E-23 |
| Epithelium development | 183 | 11.14 | 4.73E-23 |
| Microtubule bundle formation | 51 | 3.11 | 1.65E-21 |
| Nervous system development | 297 | 18.09 | 2.26E-21 |
| Regulation of signaling | 424 | 25.82 | 6.29E-21 |
| Microtubule-based process | 152 | 9.26 | 1.49E-20 |
| Regulation of cell communication | 423 | 25.76 | 1.69E-20 |
| Regulation of signal transduction | 381 | 23.2 | 2.31E-20 |
| Tube development | 153 | 9.32 | 9.54E-20 |
| Cilium-dependent cell motility | 59 | 3.59 | 2.82E-19 |
| Cilium or flagellum-dependent cell motility | 59 | 3.59 | 2.82E-19 |
| Animal organ morphogenesis | 159 | 9.68 | 2.84E-19 |
| Cytoskeleton organization | 191 | 11.63 | 5.07E-19 |
| Cilium movement involved in cell motility | 58 | 3.53 | 6.57E-19 |
| Generation of neurons | 182 | 11.08 | 7.05E-19 |
| Organelle assembly | 148 | 9.01 | 9.99E-19 |
| Neurogenesis | 202 | 12.3 | 1.44E-18 |
| Anatomical structure formation involved in morphogenesis | 157 | 9.56 | 1.79E-18 |
| Motile cilium assembly | 36 | 2.19 | 4.5E-18 |
| Enzyme-linked receptor protein signaling pathway | 115 | 7 | 5.77E-18 |
| Extracellular matrix organization | 69 | 4.2 | 6.26E-18 |
| Extracellular structure organization | 69 | 4.2 | 7.61E-18 |
| External encapsulating structure organization | 69 | 4.2 | 9.23E-18 |
| Regulation of cell motility | 171 | 10.41 | 1.3E-17 |
| Neuron differentiation | 170 | 10.35 | 1.62E-17 |
| Cell surface receptor signaling pathway | 269 | 16.38 | 3.39E-17 |
| Cell morphogenesis | 124 | 7.55 | 3.63E-17 |
| Regulation of locomotion | 174 | 10.6 | 4.44E-17 |
| Response to growth factor | 104 | 6.33 | 5.39E-17 |
| Cell-cell adhesion | 102 | 6.21 | 8.68E-17 |
| Tube morphogenesis | 119 | 7.25 | 3.4E-16 |
| Sperm motility | 52 | 3.17 | 3.82E-16 |
| Flagellated sperm motility | 52 | 3.17 | 3.82E-16 |
| Cell junction organization | 102 | 6.21 | 6.9E-16 |
| Embryonic morphogenesis | 106 | 6.46 | 8.34E-16 |
| Epithelial cilium movement involved in extracellular fluid movement | 26 | 1.58 | 8.4E-16 |
| Response to endogenous stimulus | 179 | 10.9 | 1.16E-15 |
| Regulation of cell migration | 159 | 9.68 | 1.44E-15 |
| Vasculature development | 101 | 6.15 | 2.49E-15 |
| Cell migration | 143 | 8.71 | 3.43E-15 |
| Sensory organ development | 107 | 6.52 | 3.63E-15 |
| Central nervous system development | 152 | 9.26 | 4.21E-15 |
| Extracellular transport | 26 | 1.58 | 4.81E-15 |
| Positive regulation of developmental process | 191 | 11.63 | 5.22E-15 |
| Cellular response to growth factor stimulus | 96 | 5.85 | 5.58E-15 |
| Pattern specification process | 88 | 5.36 | 6.64E-15 |
| Blood vessel development | 97 | 5.91 | 1.06E-14 |
| Embryo development | 153 | 9.32 | 1.47E-14 |

| | | | |
|---|---|---|---|
| Left/right pattern formation | 44 | 2.68 | 1.84E-14 |
| Tissue morphogenesis | 101 | 6.15 | 1.93E-14 |
| Regulation of multicellular organismal development | 198 | 12.06 | 3.64E-14 |
| Positive regulation of cell motility | 106 | 6.46 | 5.45E-14 |
| Cellular response to endogenous stimulus | 143 | 8.71 | 5.53E-14 |
| Visual system development | 79 | 4.81 | 7.01E-14 |
| Muscle structure development | 93 | 5.66 | 7.05E-14 |
| Determination of left/right symmetry | 42 | 2.56 | 7.68E-14 |
| Positive regulation of cell migration | 102 | 6.21 | 9.81E-14 |
| Positive regulation of locomotion | 107 | 6.52 | 1.1E-13 |
| Eye development | 78 | 4.75 | 1.22E-13 |
| Microtubule-based transport | 53 | 3.23 | 1.73E-13 |
| Sensory system development | 79 | 4.81 | 1.74E-13 |
| Determination of bilateral symmetry | 42 | 2.56 | 4.26E-13 |
| Specification of symmetry | 42 | 2.56 | 4.26E-13 |
| Organelle organization | 346 | 21.07 | 4.34E-13 |
| Positive regulation of signaling | 227 | 13.82 | 4.77E-13 |
| Regionalization | 78 | 4.75 | 4.97E-13 |
| Growth | 83 | 5.05 | 5.47E-13 |
| Developmental growth | 83 | 5.05 | 5.47E-13 |
| Response to wounding | 83 | 5.05 | 6.14E-13 |
| Camera-type eye development | 70 | 4.26 | 1.23E-12 |
| Positive regulation of cell communication | 225 | 13.7 | 1.37E-12 |
| Connective tissue development | 54 | 3.29 | 1.68E-12 |
| Neuron projection development | 112 | 6.82 | 2.59E-12 |
| Regulation of anatomical structure morphogenesis | 130 | 7.92 | 3.62E-12 |
| Neuron development | 130 | 7.92 | 3.96E-12 |
| Positive regulation of signal transduction | 204 | 12.42 | 4.42E-12 |
| Cellular component assembly | 297 | 18.09 | 5.55E-12 |
| Respiratory system development | 50 | 3.05 | 7.46E-12 |
| Neuron projection morphogenesis | 86 | 5.24 | 7.75E-12 |
| Epithelial cell differentiation | 102 | 6.21 | 8.45E-12 |
| Morphogenesis of an epithelium | 81 | 4.93 | 9.55E-12 |
| Plasma membrane bounded cell projection morphogenesis | 86 | 5.24 | 1.3E-11 |
| Skeletal system development | 86 | 5.24 | 1.3E-11 |
| Regulation of cell population proliferation | 216 | 13.15 | 1.32E-11 |
| Regulation of cell differentiation | 203 | 12.36 | 1.36E-11 |
| Regulation of cellular component organization | 282 | 17.17 | 1.95E-11 |
| Head development | 117 | 7.13 | 2.16E-11 |
| Cell projection morphogenesis | 86 | 5.24 | 2.9E-11 |
| Cell morphogenesis involved in neuron differentiation | 79 | 4.81 | 2.96E-11 |
| Positive regulation of cellular component organization | 150 | 9.14 | 4.31E-11 |
| Negative regulation of apoptotic process | 130 | 7.92 | 5.55E-11 |
| Regulation of wnt signaling pathway | 66 | 4.02 | 5.71E-11 |
| Cell communication | 520 | 31.67 | 7.04E-11 |
| Microtubule cytoskeleton organization | 93 | 5.66 | 7.71E-11 |
| Positive regulation of multicellular organismal process | 207 | 12.61 | 1.03E-10 |
| Renal system development | 61 | 3.71 | 1.16E-10 |
| Lung development | 44 | 2.68 | 1.5E-10 |
| Negative regulation of programmed cell death | 132 | 8.04 | 1.64E-10 |
| Kidney development | 59 | 3.59 | 2.34E-10 |
| Respiratory tube development | 44 | 2.68 | 2.96E-10 |
| Blood vessel morphogenesis | 75 | 4.57 | 3.06E-10 |
| Axon development | 73 | 4.45 | 4.06E-10 |
| Homeostatic process | 183 | 11.14 | 4.62E-10 |
| Response to hormone | 118 | 7.19 | 5.45E-10 |
| Wound healing | 62 | 3.78 | 6.07E-10 |
| Cell surface receptor protein serine/threonine kinase signaling pathway | 43 | 2.62 | 6.25E-10 |
| Axonogenesis | 66 | 4.02 | 6.51E-10 |
| Epithelial cell development | 44 | 2.68 | 6.7E-10 |
| Sperm flagellum assembly | 22 | 1.34 | 7.6E-10 |
| Regulation of cell adhesion | 114 | 6.94 | 7.93E-10 |
| Heart morphogenesis | 53 | 3.23 | 8.9E-10 |
| Signaling | 509 | 31 | 9.86E-10 |
| Cell junction assembly | 56 | 3.41 | 1.34E-09 |

| | | | |
|---|---|---|---|
| Signal transduction | 480 | 29.23 | 1.77E-09 |
| Regulation of programmed cell death | 191 | 11.63 | 1.96E-09 |
| Wnt signaling pathway | 54 | 3.29 | 2.12E-09 |
| Platelet-derived growth factor receptor signaling pathway | 26 | 1.58 | 2.49E-09 |
| Positive regulation of response to stimulus | 259 | 15.77 | 2.7E-09 |
| Regulation of apoptotic process | 186 | 11.33 | 2.81E-09 |
| Cell surface receptor protein tyrosine kinase signaling pathway | 71 | 4.32 | 2.97E-09 |
| Transforming growth factor beta receptor superfamily signaling pathway | 40 | 2.44 | 3.56E-09 |
| Response to abiotic stimulus | 145 | 8.83 | 3.93E-09 |
| Cellular component biogenesis | 305 | 18.57 | 4.04E-09 |
| Cell-cell junction organization | 40 | 2.44 | 4.18E-09 |
| Response to transforming growth factor beta | 38 | 2.31 | 5E-09 |
| Muscle cell differentiation | 53 | 3.23 | 5.91E-09 |
| Regulation of canonical wnt signaling pathway | 52 | 3.17 | 6.2E-09 |
| Brain development | 104 | 6.33 | 6.64E-09 |
| Gland development | 70 | 4.26 | 7.06E-09 |
| Supramolecular fiber organization | 90 | 5.48 | 7.72E-09 |
| Regulation of cell projection organization | 95 | 5.79 | 8E-09 |
| Circulatory system process | 80 | 4.87 | 8.02E-09 |
| Cellular localization | 288 | 17.54 | 8.11E-09 |
| Regulation of plasma membrane bounded cell projection organization | 93 | 5.66 | 9.58E-09 |
| Cartilage development | 39 | 2.38 | 1.19E-08 |
| Positive regulation of cell differentiation | 120 | 7.31 | 1.3E-08 |
| Cellular response to transforming growth factor beta stimulus | 36 | 2.19 | 1.89E-08 |
| Epithelial tube morphogenesis | 56 | 3.41 | 1.91E-08 |
| Response to oxygen-containing compound | 188 | 11.45 | 2.1E-08 |
| Cell-substrate adhesion | 39 | 2.38 | 2.19E-08 |
| Axon guidance | 45 | 2.74 | 2.49E-08 |
| Cellular response to fibroblast growth factor stimulus | 32 | 1.95 | 2.5E-08 |
| Positive regulation of cell-substrate adhesion | 31 | 1.89 | 2.6E-08 |
| Response to fibroblast growth factor | 33 | 2.01 | 2.82E-08 |
| Neuron projection guidance | 45 | 2.74 | 2.84E-08 |
| Regulation of localization | 229 | 13.95 | 3.04E-08 |
| Forebrain development | 66 | 4.02 | 4.32E-08 |
| Collagen fibril organization | 21 | 1.28 | 4.47E-08 |
| Regulation of intracellular signal transduction | 224 | 13.64 | 5.61E-08 |
| Sperm axoneme assembly | 15 | 0.91 | 6.08E-08 |
| Embryonic organ morphogenesis | 53 | 3.23 | 6.36E-08 |
| Canonical wnt signaling pathway | 27 | 1.64 | 7.18E-08 |
| Appendage development | 39 | 2.38 | 8E-08 |
| Limb development | 39 | 2.38 | 8E-08 |
| Angiogenesis | 58 | 3.53 | 8.24E-08 |
| Negative regulation of cell population proliferation | 104 | 6.33 | 9.08E-08 |
| Intraciliary transport | 18 | 1.1 | 9.59E-08 |
| Macromolecule localization | 256 | 15.59 | 1.04E-07 |
| Regulation of molecular function | 167 | 10.17 | 1.07E-07 |
| Sensory organ morphogenesis | 50 | 3.05 | 1.11E-07 |
| Glomerulus development | 20 | 1.22 | 1.39E-07 |
| Collagen-activated tyrosine kinase receptor signaling pathway | 19 | 1.16 | 1.4E-07 |
| Chordate embryonic development | 90 | 5.48 | 1.45E-07 |
| Protein localization | 217 | 13.22 | 1.56E-07 |
| Gland morphogenesis | 27 | 1.64 | 1.58E-07 |
| Gastrulation | 36 | 2.19 | 1.6E-07 |
| Cellular response to chemical stimulus | 218 | 13.28 | 1.67E-07 |
| Response to peptide | 115 | 7 | 1.67E-07 |
| Cell population proliferation | 97 | 5.91 | 1.89E-07 |
| Axonemal dynein complex assembly | 16 | 0.97 | 1.93E-07 |
| Cellular macromolecule localization | 217 | 13.22 | 2.15E-07 |
| Neuron recognition | 17 | 1.04 | 2.28E-07 |
| Skin development | 49 | 2.98 | 2.29E-07 |
| Limb morphogenesis | 33 | 2.01 | 2.71E-07 |
| Appendage morphogenesis | 33 | 2.01 | 2.71E-07 |
| Embryo development ending in birth or egg hatching | 91 | 5.54 | 2.82E-07 |
| Regulation of cellular response to growth factor stimulus | 60 | 3.65 | 2.93E-07 |
| Hemopoiesis | 93 | 5.66 | 2.97E-07 |

| | | | |
|---|---|---|---|
| Retina development in camera-type eye | 35 | 2.13 | 3.71E-07 |
| Response to cytokine | 112 | 6.82 | 3.92E-07 |
| Actin filament-based process | 85 | 5.18 | 4.05E-07 |
| Embryonic organ development | 68 | 4.14 | 4.26E-07 |
| Collagen-activated signaling pathway | 19 | 1.16 | 4.3E-07 |
| Regulation of cellular localization | 126 | 7.67 | 4.57E-07 |
| Nephron development | 31 | 1.89 | 4.64E-07 |
| Eye morphogenesis | 34 | 2.07 | 4.91E-07 |
| Positive regulation of intracellular signal transduction | 137 | 8.34 | 5.1E-07 |
| Telencephalon development | 48 | 2.92 | 5.17E-07 |
| Regulation of cell-substrate adhesion | 41 | 2.5 | 5.68E-07 |
| Positive regulation of phosphate metabolic process | 70 | 4.26 | 6.08E-07 |
| Positive regulation of phosphorus metabolic process | 70 | 4.26 | 6.08E-07 |
| Regulation of bone mineralization | 22 | 1.34 | 6.19E-07 |
| Actin cytoskeleton organization | 77 | 4.69 | 6.71E-07 |
| Protein phosphorylation | 59 | 3.59 | 7.12E-07 |
| Cell-cell adhesion via plasma-membrane adhesion molecules | 45 | 2.74 | 7.41E-07 |
| Synapse organization | 57 | 3.47 | 7.46E-07 |
| Negative regulation of multicellular organismal process | 144 | 8.77 | 7.57E-07 |
| Regulation of biological quality | 310 | 18.88 | 8.12E-07 |
| Embryonic heart tube morphogenesis | 21 | 1.28 | 9.07E-07 |
| Negative regulation of response to stimulus | 198 | 12.06 | 9.97E-07 |
| Negative regulation of developmental process | 123 | 7.49 | 1.11E-06 |
| Positive regulation of chemotaxis | 31 | 1.89 | 1.13E-06 |
| Digestive system development | 31 | 1.89 | 1.13E-06 |
| Chemical homeostasis | 115 | 7 | 0.0000012 |
| Ossification | 49 | 2.98 | 1.25E-06 |
| Regulation of transmembrane receptor protein serine/threonine kinase signaling pathway | 52 | 3.17 | 1.33E-06 |
| Striated muscle cell differentiation | 40 | 2.44 | 1.35E-06 |
| Brain-derived neurotrophic factor receptor signaling pathway | 17 | 1.04 | 1.38E-06 |
| Negative regulation of wnt signaling pathway | 35 | 2.13 | 1.58E-06 |
| Gliogenesis | 47 | 2.86 | 1.69E-06 |
| Extracellular matrix assembly | 13 | 0.79 | 1.79E-06 |
| Odontogenesis | 27 | 1.64 | 1.82E-06 |
| Regulation of phosphatidylinositol 3-kinase/protein kinase b signal transduction | 47 | 2.86 | 1.86E-06 |
| Vascular endothelial growth factor receptor-1 signaling pathway | 16 | 0.97 | 1.86E-06 |
| Positive regulation of cell population proliferation | 119 | 7.25 | 1.89E-06 |
| Ear development | 41 | 2.5 | 1.94E-06 |
| Epidermal growth factor receptor signaling pathway | 24 | 1.46 | 1.97E-06 |
| Tissue homeostasis | 41 | 2.5 | 2.15E-06 |
| Anatomical structure homeostasis | 41 | 2.5 | 2.15E-06 |
| Regulation of biomineral tissue development | 24 | 1.46 | 2.35E-06 |
| Platelet-derived growth factor receptor-beta signaling pathway | 17 | 1.04 | 2.36E-06 |
| Hepatocyte growth factor receptor signaling pathway | 17 | 1.04 | 2.36E-06 |
| Epidermis development | 53 | 3.23 | 2.38E-06 |
| Positive regulation of phosphorylation | 62 | 3.78 | 2.44E-06 |
| Platelet-derived growth factor receptor-alpha signaling pathway | 16 | 0.97 | 2.48E-06 |
| Regulation of protein localization | 112 | 6.82 | 2.56E-06 |
| Regulation of mapk cascade | 87 | 5.3 | 2.68E-06 |
| Regulation of phosphorylation | 87 | 5.3 | 2.85E-06 |
| Spermatid differentiation | 41 | 2.5 | 2.95E-06 |
| Negative regulation of cell differentiation | 93 | 5.66 | 0.000003 |
| Positive regulation of cell adhesion | 69 | 4.2 | 3.06E-06 |
| Regulation of epithelial cell proliferation | 60 | 3.65 | 3.13E-06 |
| Spermatid development | 40 | 2.44 | 3.21E-06 |
| Regulation of nervous system development | 65 | 3.96 | 3.21E-06 |
| Response to stem cell factor | 16 | 0.97 | 3.27E-06 |
| Kit signaling pathway | 16 | 0.97 | 3.27E-06 |
| Cellular response to stem cell factor stimulus | 16 | 0.97 | 3.27E-06 |
| Macrophage colony-stimulating factor signaling pathway | 16 | 0.97 | 3.27E-06 |
| Negative regulation of canonical wnt signaling pathway | 30 | 1.83 | 3.46E-06 |
| Embryonic heart tube development | 22 | 1.34 | 3.75E-06 |
| Regulation of cell development | 104 | 6.33 | 3.75E-06 |
| Neural retina development | 21 | 1.28 | 3.86E-06 |
| Apoptotic process | 127 | 7.73 | 4.01E-06 |

| | | | |
|---|---|---|---|
| Positive regulation of angiogenesis | 35 | 2.13 | 4.14E-06 |
| Vascular endothelial growth factor signaling pathway | 18 | 1.1 | 0.0000044 |
| Cellular response to vascular endothelial growth factor stimulus | 22 | 1.34 | 0.0000045 |
| Digestive tract development | 28 | 1.71 | 0.0000046 |
| Cellular response to hormone stimulus | 75 | 4.57 | 4.82E-06 |
| Regulation of phosphate metabolic process | 99 | 6.03 | 4.94E-06 |
| Regulation of erk1 and erk2 cascade | 47 | 2.86 | 5.12E-06 |
| Developmental cell growth | 26 | 1.58 | 5.15E-06 |
| Regulation of phosphorus metabolic process | 99 | 6.03 | 5.35E-06 |
| Erbb signaling pathway | 25 | 1.52 | 5.78E-06 |
| Vascular endothelial growth factor receptor signaling pathway | 20 | 1.22 | 5.78E-06 |
| Positive regulation of vasculature development | 35 | 2.13 | 5.84E-06 |
| Developmental growth involved in morphogenesis | 29 | 1.77 | 6.02E-06 |
| Phosphorylation | 61 | 3.71 | 6.21E-06 |
| Response to peptide hormone | 61 | 3.71 | 6.66E-06 |
| Insulin receptor signaling pathway | 25 | 1.52 | 6.74E-06 |
| Positive regulation of molecular function | 101 | 6.15 | 6.81E-06 |
| Reproductive process | 172 | 10.48 | 6.82E-06 |
| Cell growth | 26 | 1.58 | 6.93E-06 |
| Camera-type eye morphogenesis | 28 | 1.71 | 6.98E-06 |
| Insulin-like growth factor receptor signaling pathway | 20 | 1.22 | 7.03E-06 |
| Mesenchymal cell differentiation | 32 | 1.95 | 7.87E-06 |
| Cell death | 130 | 7.92 | 7.89E-06 |
| Regulation of protein localization to membrane | 35 | 2.13 | 8.15E-06 |
| Fibroblast growth factor receptor signaling pathway | 23 | 1.4 | 8.32E-06 |
| Protein localization to cilium | 18 | 1.1 | 8.52E-06 |
| Integrin-mediated signaling pathway | 24 | 1.46 | 8.81E-06 |
| Cellular response to macrophage colony-stimulating factor stimulus | 16 | 0.97 | 9.14E-06 |
| Regulation of angiogenesis | 53 | 3.23 | 9.44E-06 |
| Programmed cell death | 129 | 7.86 | 9.69E-06 |
| Regulation of protein phosphorylation | 80 | 4.87 | 9.69E-06 |
| Positive regulation of phosphatidylinositol 3-kinase/protein kinase b signal transduction | 35 | 2.13 | 0.0000101 |
| Regulation of neuron projection development | 62 | 3.78 | 0.0000102 |
| Negative regulation of signaling | 167 | 10.17 | 0.0000108 |
| Response to steroid hormone | 45 | 2.74 | 0.0000108 |
| Regulation of cellular component biogenesis | 120 | 7.31 | 0.000011 |
| Negative regulation of cell communication | 167 | 10.17 | 0.0000111 |
| Negative regulation of signal transduction | 158 | 9.62 | 0.0000111 |
| Regulation of actin filament-based process | 57 | 3.47 | 0.0000112 |
| Inner ear development | 36 | 2.19 | 0.0000115 |
| Chondrocyte differentiation | 21 | 1.28 | 0.0000116 |
| Positive regulation of transmembrane receptor protein serine/threonine kinase signaling pathway | 24 | 1.46 | 0.000012 |
| Regulation of ossification | 26 | 1.58 | 0.0000123 |
| Multicellular organismal-level homeostasis | 82 | 4.99 | 0.0000124 |
| Response to mechanical stimulus | 38 | 2.31 | 0.000013 |
| Organ growth | 23 | 1.4 | 0.0000134 |
| Cellular response to cytokine stimulus | 96 | 5.85 | 0.000014 |
| Cell-cell junction assembly | 26 | 1.58 | 0.0000141 |
| Regulation of chemotaxis | 37 | 2.25 | 0.0000143 |
| Positive regulation of protein phosphorylation | 57 | 3.47 | 0.0000149 |
| Cellular response to bmp stimulus | 22 | 1.34 | 0.0000149 |
| Response to bmp | 22 | 1.34 | 0.0000149 |
| Muscle tissue development | 52 | 3.17 | 0.0000157 |
| Positive regulation of cellular component biogenesis | 68 | 4.14 | 0.000016 |
| Bmp signaling pathway | 20 | 1.22 | 0.0000176 |
| Regeneration | 30 | 1.83 | 0.0000179 |
| Regulation of vasculature development | 53 | 3.23 | 0.0000186 |
| Heart looping | 18 | 1.1 | 0.0000192 |
| Smoothened signaling pathway | 21 | 1.28 | 0.0000193 |
| Blood circulation | 60 | 3.65 | 0.0000199 |
| Stem cell differentiation | 33 | 2.01 | 0.0000203 |
| Formation of primary germ layer | 25 | 1.52 | 0.0000215 |
| Osteoblast differentiation | 28 | 1.71 | 0.0000224 |
| Response to macrophage colony-stimulating factor | 16 | 0.97 | 0.000023 |

| | | | |
|---|---|---|---|
| Cell recognition | 27 | 1.64 | 0.0000234 |
| Regulation of cilium movement | 14 | 0.85 | 0.0000236 |
| Positive regulation of wnt signaling pathway | 29 | 1.77 | 0.000024 |
| Neural tube development | 29 | 1.77 | 0.000024 |
| Positive regulation of cell projection organization | 52 | 3.17 | 0.0000264 |
| Muscle cell development | 31 | 1.89 | 0.0000265 |
| Transforming growth factor beta receptor signaling pathway | 22 | 1.34 | 0.0000279 |
| Positive regulation of programmed cell death | 75 | 4.57 | 0.0000283 |
| Bone development | 34 | 2.07 | 0.0000313 |
| Mammary gland development | 25 | 1.52 | 0.0000322 |
| Cell-substrate junction assembly | 13 | 0.79 | 0.000033 |
| Embryonic eye morphogenesis | 12 | 0.73 | 0.0000335 |
| Response to oxygen levels | 50 | 3.05 | 0.0000366 |
| Intraciliary anterograde transport | 9 | 0.55 | 0.0000368 |
| Regulation of cell morphogenesis | 39 | 2.38 | 0.0000394 |
| Regulation of neurogenesis | 53 | 3.23 | 0.0000408 |
| Response to stress | 338 | 20.58 | 0.0000428 |
| Cell adhesion mediated by integrin | 13 | 0.79 | 0.000043 |
| Kidney epithelium development | 27 | 1.64 | 0.0000432 |
| Mammary gland epithelium development | 16 | 0.97 | 0.0000433 |
| Behavior | 79 | 4.81 | 0.0000439 |
| In utero embryonic development | 55 | 3.35 | 0.0000468 |
| Cellular response to peptide hormone stimulus | 44 | 2.68 | 0.0000537 |
| Regulation of catalytic activity | 99 | 6.03 | 0.0000556 |
| Cell-substrate junction organization | 13 | 0.79 | 0.0000556 |
| Roof of mouth development | 20 | 1.22 | 0.0000559 |
| Determination of heart left/right asymmetry | 18 | 1.1 | 0.0000578 |
| Biomineral tissue development | 22 | 1.34 | 0.0000578 |
| Response to nitrogen compound | 119 | 7.25 | 0.0000582 |
| Bone growth | 11 | 0.67 | 0.0000595 |
| Positive regulation of apoptotic process | 72 | 4.38 | 0.0000609 |
| Lung morphogenesis | 14 | 0.85 | 0.0000621 |
| Stem cell development | 19 | 1.16 | 0.0000623 |
| Monoatomic ion homeostasis | 67 | 4.08 | 0.0000632 |
| Morphogenesis of an epithelial sheet | 15 | 0.91 | 0.0000647 |
| Leukocyte differentiation | 57 | 3.47 | 0.000069 |
| Cardiocyte differentiation | 24 | 1.46 | 0.000072 |
| Regulation of osteoblast differentiation | 28 | 1.71 | 0.0000798 |
| Regulation of integrin-mediated signaling pathway | 9 | 0.55 | 0.0000865 |
| Multicellular organism growth | 22 | 1.34 | 0.0000872 |
| Protein localization to cell periphery | 37 | 2.25 | 0.0000906 |
| Embryonic appendage morphogenesis | 24 | 1.46 | 0.0000924 |
| Embryonic limb morphogenesis | 24 | 1.46 | 0.0000924 |
| Regulation of transport | 171 | 10.41 | 0.0000934 |
| Regulation of cell junction assembly | 40 | 2.44 | 0.0000952 |
| Positive regulation of mapk cascade | 61 | 3.71 | 0.0000992 |
| Positive regulation of cell junction assembly | 22 | 1.34 | 0.0000996 |
| Cellular process involved in reproduction in multicellular organism | 56 | 3.41 | 0.000101 |
| Monoatomic cation homeostasis | 65 | 3.96 | 0.000103 |
| Cellular response to oxygen-containing compound | 122 | 7.43 | 0.000105 |
| Epithelial tube branching involved in lung morphogenesis | 10 | 0.61 | 0.000105 |
| Response to ketone | 35 | 2.13 | 0.000105 |
| Regulation of protein metabolic process | 184 | 11.21 | 0.000112 |
| Cellular homeostasis | 83 | 5.05 | 0.000116 |
| Regulation of extracellular matrix organization | 17 | 1.04 | 0.000124 |
| Cell-matrix adhesion | 25 | 1.52 | 0.000124 |
| Regulation of cytoskeleton organization | 68 | 4.14 | 0.000129 |
| Developmental process involved in reproduction | 117 | 7.13 | 0.000132 |
| Vascular process in circulatory system | 41 | 2.5 | 0.000135 |
| Inner dynein arm assembly | 8 | 0.49 | 0.000137 |
| Response to decreased oxygen levels | 45 | 2.74 | 0.000143 |
| Endochondral bone growth | 10 | 0.61 | 0.000144 |
| Regulation of non-canonical wnt signaling pathway | 10 | 0.61 | 0.000144 |
| Positive regulation of fibroblast proliferation | 14 | 0.85 | 0.000146 |
| Face development | 14 | 0.85 | 0.000146 |

| | | | |
|---|---|---|---|
| Protein-containing complex localization | 28 | 1.71 | 0.00015 |
| Regulation of animal organ morphogenesis | 18 | 1.1 | 0.000153 |
| Cardiac muscle tissue development | 32 | 1.95 | 0.00016 |
| Cytoskeleton-dependent intracellular transport | 32 | 1.95 | 0.00016 |
| Glial cell development | 24 | 1.46 | 0.000167 |
| Positive regulation of leukocyte chemotaxis | 20 | 1.22 | 0.000178 |
| Ephrin receptor signaling pathway | 18 | 1.1 | 0.000178 |
| Positive regulation of erk1 and erk2 cascade | 32 | 1.95 | 0.000191 |
| Adrenal gland development | 10 | 0.61 | 0.000195 |
| Inorganic ion homeostasis | 57 | 3.47 | 0.000201 |
| Regulation of cilium beat frequency | 8 | 0.49 | 0.000209 |
| Tissue remodeling | 22 | 1.34 | 0.000213 |
| Regulation of supramolecular fiber organization | 53 | 3.23 | 0.000216 |
| Response to lipid | 98 | 5.97 | 0.000223 |
| Positive regulation of metabolic process | 334 | 20.34 | 0.000234 |
| Positive regulation of canonical wnt signaling pathway | 22 | 1.34 | 0.00024 |
| Skeletal system morphogenesis | 35 | 2.13 | 0.000242 |
| Forebrain regionalization | 9 | 0.55 | 0.000255 |
| Membrane protein ectodomain proteolysis | 9 | 0.55 | 0.000255 |
| Cellular response to insulin stimulus | 31 | 1.89 | 0.000256 |
| Intracellular chemical homeostasis | 71 | 4.32 | 0.000264 |
| Germ cell development | 51 | 3.11 | 0.000264 |
| Response to progesterone | 12 | 0.73 | 0.000264 |
| Morphogenesis of embryonic epithelium | 25 | 1.52 | 0.000268 |
| Regulation of microtubule-based movement | 15 | 0.91 | 0.00029 |
| Regulation of response to wounding | 29 | 1.77 | 0.000292 |
| Glial cell differentiation | 34 | 2.07 | 0.000296 |
| Protein localization to plasma membrane | 31 | 1.89 | 0.000304 |
| Embryonic camera-type eye development | 11 | 0.67 | 0.000308 |
| Tissue regeneration | 16 | 0.97 | 0.000308 |
| Regulation of body fluid levels | 49 | 2.98 | 0.000308 |
| Epithelial tube formation | 23 | 1.4 | 0.000316 |
| Negative regulation of dna-templated transcription | 133 | 8.1 | 0.000319 |
| Regulation of cellular response to transforming growth factor beta stimulus | 29 | 1.77 | 0.00032 |
| Neuromuscular junction development | 12 | 0.73 | 0.000328 |
| Response to metal ion | 46 | 2.8 | 0.000338 |
| Embryonic epithelial tube formation | 22 | 1.34 | 0.00034 |
| Wound healing, spreading of cells | 10 | 0.61 | 0.000342 |
| Epiboly involved in wound healing | 10 | 0.61 | 0.000342 |
| Regulation of protein maturation | 15 | 0.91 | 0.000343 |
| Response to oxidative stress | 49 | 2.98 | 0.000348 |
| Mononuclear cell differentiation | 48 | 2.92 | 0.000352 |
| Transport along microtubule | 28 | 1.71 | 0.000357 |
| Regulation of wound healing | 24 | 1.46 | 0.00036 |
| Mesenchyme development | 40 | 2.44 | 0.000362 |
| Symbiont entry into host | 21 | 1.28 | 0.000363 |
| Symbiont entry into host cell | 21 | 1.28 | 0.000363 |
| Regulation of smoothened signaling pathway | 18 | 1.1 | 0.000363 |
| Neuron migration | 25 | 1.52 | 0.000364 |
| Positive regulation of cell-matrix adhesion | 14 | 0.85 | 0.000374 |
| Olfactory lobe development | 11 | 0.67 | 0.00039 |
| Ameboidal-type cell migration | 24 | 1.46 | 0.0004 |
| Response to acid chemical | 25 | 1.52 | 0.000403 |
| Positive regulation of bone mineralization | 12 | 0.73 | 0.000405 |
| Regulation of growth | 72 | 4.38 | 0.000419 |
| Positive regulation of protein modification process | 67 | 4.08 | 0.000428 |
| Morphogenesis of a branching epithelium | 27 | 1.64 | 0.000437 |
| Regulation of protein modification process | 99 | 6.03 | 0.000442 |
| Response to corticosteroid | 26 | 1.58 | 0.000443 |
| Epiboly | 10 | 0.61 | 0.000444 |
| Neuron projection regeneration | 10 | 0.61 | 0.000444 |
| Face morphogenesis | 10 | 0.61 | 0.000444 |
| Detoxification | 25 | 1.52 | 0.000445 |
| Regulation of actin cytoskeleton organization | 47 | 2.86 | 0.000455 |
| Negative regulation of rna biosynthetic process | 133 | 8.1 | 0.000455 |

| | | | |
|---|---|---|---|
| Embryonic camera-type eye morphogenesis | 9 | 0.55 | 0.000472 |
| Cellular component maintenance | 15 | 0.91 | 0.000474 |
| Intracellular monoatomic ion homeostasis | 55 | 3.35 | 0.000483 |
| Pancreas development | 16 | 0.97 | 0.000486 |
| Head morphogenesis | 11 | 0.67 | 0.000489 |
| Regulation of wnt signaling pathway, planar cell polarity pathway | 7 | 0.43 | 0.000506 |
| Cytokine-mediated signaling pathway | 55 | 3.35 | 0.000507 |
| Locomotion | 48 | 2.92 | 0.000533 |
| Regulation of hydrolase activity | 43 | 2.62 | 0.000538 |
| Positive regulation of endothelial cell migration | 23 | 1.4 | 0.000539 |
| Neural crest cell differentiation | 18 | 1.1 | 0.000541 |
| Regulation of fibroblast proliferation | 18 | 1.1 | 0.000541 |
| Response to alcohol | 39 | 2.38 | 0.000548 |
| Regulation of transforming growth factor beta receptor signaling pathway | 28 | 1.71 | 0.000559 |
| Ventricular system development | 10 | 0.61 | 0.00057 |
| Positive regulation of protein localization | 61 | 3.71 | 0.000576 |
| Negative regulation of cell development | 39 | 2.38 | 0.000581 |
| Neural tube formation | 19 | 1.16 | 0.000585 |
| Regulation of epithelial to mesenchymal transition | 22 | 1.34 | 0.000589 |
| Positive regulation of protein metabolic process | 105 | 6.39 | 0.000589 |
| Mammary gland morphogenesis | 12 | 0.73 | 0.000604 |
| Body morphogenesis | 12 | 0.73 | 0.000604 |
| Epithelial cell proliferation | 28 | 1.71 | 0.00061 |
| Cellular response to amino acid stimulus | 18 | 1.1 | 0.000615 |
| Cellular oxidant detoxification | 18 | 1.1 | 0.000615 |
| Export across plasma membrane | 14 | 0.85 | 0.000622 |
| Actin filament organization | 39 | 2.38 | 0.000622 |
| Lens development in camera-type eye | 17 | 1.04 | 0.000637 |
| Regulation of protein localization to cell periphery | 25 | 1.52 | 0.000653 |
| Negative regulation of cell adhesion | 42 | 2.56 | 0.000654 |
| Response to toxic substance | 36 | 2.19 | 0.000656 |
| Cellular response to acid chemical | 19 | 1.16 | 0.00066 |
| Negative regulation of nervous system development | 24 | 1.46 | 0.00066 |
| Positive regulation of biomineral tissue development | 13 | 0.79 | 0.000684 |
| Regulation of dna-binding transcription factor activity | 43 | 2.62 | 0.000685 |
| Response to insulin | 37 | 2.25 | 0.000708 |
| Cellular response to nitrogen compound | 73 | 4.45 | 0.00071 |
| System process | 204 | 12.42 | 0.00071 |
| Transport across blood-brain barrier | 17 | 1.04 | 0.000727 |
| Metanephros development | 17 | 1.04 | 0.000727 |
| Vascular transport | 17 | 1.04 | 0.000727 |
| Retina homeostasis | 14 | 0.85 | 0.00073 |
| Substrate adhesion-dependent cell spreading | 12 | 0.73 | 0.000731 |
| Negative regulation of extrinsic apoptotic signaling pathway | 19 | 1.16 | 0.000742 |
| Myofibril assembly | 15 | 0.91 | 0.000748 |
| Regulation of organelle organization | 122 | 7.43 | 0.000752 |
| Dendrite self-avoidance | 7 | 0.43 | 0.000759 |
| Positive regulation of non-canonical wnt signaling pathway | 7 | 0.43 | 0.000759 |
| Positive regulation of catalytic activity | 60 | 3.65 | 0.000759 |
| Positive regulation of bicellular tight junction assembly | 6 | 0.37 | 0.000766 |
| Regulation of mesenchymal stem cell differentiation | 6 | 0.37 | 0.000766 |
| Regulation of odontoblast differentiation | 6 | 0.37 | 0.000766 |
| Elastic fiber assembly | 6 | 0.37 | 0.000766 |
| Cellular response to tumor necrosis factor | 25 | 1.52 | 0.000786 |
| Response to glucocorticoid | 23 | 1.4 | 0.000805 |
| Response to xenobiotic stimulus | 55 | 3.35 | 0.000813 |
| Negative regulation of epithelial cell proliferation | 27 | 1.64 | 0.000815 |
| Striated muscle tissue development | 44 | 2.68 | 0.000815 |
| Glomerular epithelial cell differentiation | 9 | 0.55 | 0.00082 |
| Endothelial cell differentiation | 17 | 1.04 | 0.000827 |
| Artery development | 19 | 1.16 | 0.000834 |
| Regulation of extrinsic apoptotic signaling pathway | 26 | 1.58 | 0.00084 |
| Myeloid cell differentiation | 41 | 2.5 | 0.000847 |
| Bone mineralization | 14 | 0.85 | 0.000854 |
| Neural crest cell development | 16 | 0.97 | 0.000855 |

| | | | |
|---|---|---|---|
| Positive regulation of osteoblast differentiation | 16 | 0.97 | 0.000855 |
| Branching morphogenesis of an epithelial tube | 23 | 1.4 | 0.000888 |
| Regulation of small gtpase mediated signal transduction | 41 | 2.5 | 0.0009 |
| Intracellular monoatomic cation homeostasis | 53 | 3.23 | 0.00091 |
| Positive regulation of neuron projection development | 26 | 1.58 | 0.000917 |
| Neuron projection extension | 17 | 1.04 | 0.000939 |
| Peripheral nervous system development | 17 | 1.04 | 0.000939 |
| Regulation of plasma membrane repair | 5 | 0.3 | 0.000941 |
| Regulation of synapse structure or activity | 42 | 2.56 | 0.000945 |
| Positive regulation of ossification | 13 | 0.79 | 0.000958 |
| Cellular response to toxic substance | 22 | 1.34 | 0.000983 |
| Regulation of blood pressure | 30 | 1.83 | 0.000991 |
| Positive regulation of small gtpase mediated signal transduction | 15 | 0.91 | 0.000998 |
| Muscle organ development | 42 | 2.56 | 0.001 |
| Intracellular signaling cassette | 91 | 5.54 | 0.00102 |
| Positive regulation of epithelial cell proliferation | 32 | 1.95 | 0.00103 |
| Regulation of system process | 67 | 4.08 | 0.00105 |
| Morphogenesis of a polarized epithelium | 12 | 0.73 | 0.00105 |
| Schwann cell differentiation | 12 | 0.73 | 0.00105 |
| Cell fate commitment | 36 | 2.19 | 0.00106 |
| Regulation of animal organ formation | 9 | 0.55 | 0.00106 |
| Nephron epithelium development | 20 | 1.22 | 0.00107 |
| Endocrine system development | 22 | 1.34 | 0.00108 |
| Branch elongation of an epithelium | 7 | 0.43 | 0.0011 |
| Cerebrospinal fluid circulation | 7 | 0.43 | 0.0011 |
| Epithelial cell-cell adhesion | 7 | 0.43 | 0.0011 |
| Regulation of endothelial cell migration | 33 | 2.01 | 0.00112 |
| Peptidyl-tyrosine phosphorylation | 13 | 0.79 | 0.00113 |
| Olfactory bulb development | 10 | 0.61 | 0.00113 |
| Morphogenesis of a branching structure | 27 | 1.64 | 0.00114 |
| Positive regulation of supramolecular fiber organization | 27 | 1.64 | 0.00114 |
| Negative regulation of neuron apoptotic process | 27 | 1.64 | 0.00114 |
| Collagen metabolic process | 14 | 0.85 | 0.00115 |
| Positive regulation of gene expression | 126 | 7.67 | 0.00118 |
| Negative regulation of rna metabolic process | 139 | 8.47 | 0.00118 |
| Regulation of hormone levels | 64 | 3.9 | 0.00119 |
| Actomyosin structure organization | 21 | 1.28 | 0.0012 |
| Intracellular signal transduction | 172 | 10.48 | 0.00122 |
| Skin morphogenesis | 6 | 0.37 | 0.00123 |
| Lateral ventricle development | 6 | 0.37 | 0.00123 |
| Positive regulation of integrin-mediated signaling pathway | 6 | 0.37 | 0.00123 |
| Peptidyl-threonine modification | 12 | 0.73 | 0.00125 |
| Placenta development | 24 | 1.46 | 0.00126 |
| Positive regulation of biosynthetic process | 250 | 15.23 | 0.00131 |
| Response to amino acid | 22 | 1.34 | 0.00132 |
| Response to hypoxia | 40 | 2.44 | 0.00132 |
| Mesenchymal cell migration | 13 | 0.79 | 0.00132 |
| Peptidyl-tyrosine modification | 13 | 0.79 | 0.00132 |
| Cardiac septum development | 20 | 1.22 | 0.00132 |
| Homotypic cell-cell adhesion | 14 | 0.85 | 0.00133 |
| Positive regulation of bmp signaling pathway | 11 | 0.67 | 0.00135 |
| Monoatomic cation transport | 89 | 5.42 | 0.00141 |
| Pallium development | 29 | 1.77 | 0.00142 |
| Positive regulation of nervous system development | 39 | 2.38 | 0.00142 |
| Tube formation | 23 | 1.4 | 0.00142 |
| Endoderm development | 16 | 0.97 | 0.00144 |
| Response to tumor necrosis factor | 27 | 1.64 | 0.00145 |
| Regulation of keratinocyte proliferation | 12 | 0.73 | 0.00148 |
| Outer dynein arm assembly | 8 | 0.49 | 0.00152 |
| Apical junction assembly | 13 | 0.79 | 0.00154 |
| Regulation of protein processing | 13 | 0.79 | 0.00154 |
| Regulation of plasminogen activation | 7 | 0.43 | 0.00154 |
| Positive regulation of kinase activity | 31 | 1.89 | 0.00157 |
| Phosphate-containing compound metabolic process | 146 | 8.89 | 0.00157 |
| Sensory perception of light stimulus | 32 | 1.95 | 0.00158 |

| | | | |
|---|---|---|---|
| Regulation of bmp signaling pathway | 21 | 1.28 | 0.00161 |
| Neural precursor cell proliferation | 19 | 1.16 | 0.00161 |
| Dendrite development | 20 | 1.22 | 0.00162 |
| Schwann cell development | 11 | 0.67 | 0.00162 |
| Adipose tissue development | 11 | 0.67 | 0.00162 |
| Astrocyte activation involved in immune response | 4 | 0.24 | 0.00162 |
| Negative regulation of prostatic bud formation | 4 | 0.24 | 0.00162 |
| Positive regulation of plasma membrane repair | 4 | 0.24 | 0.00162 |
| Odontogenesis of dentin-containing tooth | 16 | 0.97 | 0.00162 |
| Import into cell | 77 | 4.69 | 0.00169 |
| Synaptic membrane adhesion | 9 | 0.55 | 0.0017 |
| Glomerular epithelium development | 9 | 0.55 | 0.0017 |
| Regulation of apoptotic signaling pathway | 49 | 2.98 | 0.0017 |
| Regulation of cartilage development | 15 | 0.91 | 0.00171 |
| Regulation of coagulation | 15 | 0.91 | 0.00171 |
| Primary neural tube formation | 17 | 1.04 | 0.00171 |
| Phosphorus metabolic process | 146 | 8.89 | 0.00173 |
| Regulation of neuron differentiation | 28 | 1.71 | 0.00174 |
| Negative regulation of neurogenesis | 22 | 1.34 | 0.00174 |
| Endothelial cell development | 12 | 0.73 | 0.00174 |
| Membranous septum morphogenesis | 5 | 0.3 | 0.00177 |
| Tight junction organization | 13 | 0.79 | 0.00178 |
| Regulation of neuron apoptotic process | 35 | 2.13 | 0.00178 |
| Striated muscle cell development | 23 | 1.4 | 0.00185 |
| Negative regulation of apoptotic signaling pathway | 33 | 2.01 | 0.00193 |
| Glomerulus vasculature development | 8 | 0.49 | 0.00198 |
| Axis elongation | 8 | 0.49 | 0.00198 |
| Apoptotic signaling pathway | 42 | 2.56 | 0.00199 |
| Positive regulation of endocytosis | 25 | 1.52 | 0.00202 |
| Astrocyte differentiation | 14 | 0.85 | 0.00202 |
| Positive regulation of response to external stimulus | 67 | 4.08 | 0.00205 |
| Membrane protein proteolysis | 10 | 0.61 | 0.00208 |
| Regulation of epithelial cell differentiation | 26 | 1.58 | 0.00208 |
| Regulation of synapse organization | 40 | 2.44 | 0.00209 |
| Trachea development | 7 | 0.43 | 0.00211 |
| Respiratory system process | 9 | 0.55 | 0.00212 |
| Regulation of protein-containing complex disassembly | 21 | 1.28 | 0.00213 |
| Fat cell differentiation | 21 | 1.28 | 0.00213 |
| Renal system process | 21 | 1.28 | 0.00213 |
| Columnar/cuboidal epithelial cell differentiation | 18 | 1.1 | 0.00218 |
| Response to external stimulus | 191 | 11.63 | 0.00221 |
| Chemotaxis | 43 | 2.62 | 0.00225 |
| Semaphorin-plexin signaling pathway | 11 | 0.67 | 0.00229 |
| Endodermal cell differentiation | 11 | 0.67 | 0.00229 |
| Regulation of hemostasis | 14 | 0.85 | 0.00231 |
| Neural tube closure | 16 | 0.97 | 0.00232 |
| Regulation of leukocyte chemotaxis | 21 | 1.28 | 0.00234 |
| Regulation of kinase activity | 47 | 2.86 | 0.00234 |
| Monoatomic ion transport | 105 | 6.39 | 0.00236 |
| Visual perception | 31 | 1.89 | 0.00236 |
| Lipid metabolic process | 122 | 7.43 | 0.00239 |
| Brain morphogenesis | 10 | 0.61 | 0.0025 |
| Taxis | 43 | 2.62 | 0.00253 |
| Secondary palate development | 8 | 0.49 | 0.00253 |
| Regulation of cell growth | 49 | 2.98 | 0.00253 |
| Regulation of actin filament organization | 36 | 2.19 | 0.00254 |
| Localization within membrane | 64 | 3.9 | 0.00254 |
| Cellular anatomical entity morphogenesis | 21 | 1.28 | 0.00256 |
| Cellular component assembly involved in morphogenesis | 21 | 1.28 | 0.00256 |
| Regulation of protein localization to nucleus | 23 | 1.4 | 0.0026 |
| Tube closure | 16 | 0.97 | 0.0026 |
| Ureteric bud development | 16 | 0.97 | 0.0026 |
| Establishment of endothelial barrier | 9 | 0.55 | 0.00261 |
| Cardiac chamber development | 26 | 1.58 | 0.00262 |
| T cell differentiation | 27 | 1.64 | 0.00266 |

| | | | |
|---|---|---|---|
| Positive regulation of canonical nf-kappab signal transduction | 31 | 1.89 | 0.00269 |
| Regulation of mesenchymal cell apoptotic process | 6 | 0.37 | 0.00274 |
| Nitrogen compound transport | 150 | 9.14 | 0.00279 |
| Female pregnancy | 26 | 1.58 | 0.00282 |
| Cell-cell junction maintenance | 7 | 0.43 | 0.00283 |
| Mesonephric epithelium development | 16 | 0.97 | 0.0029 |
| Mesonephric tubule development | 16 | 0.97 | 0.0029 |
| Regulation of cell shape | 22 | 1.34 | 0.00294 |
| Positive regulation of mononuclear cell migration | 17 | 1.04 | 0.00295 |
| Endothelium development | 18 | 1.1 | 0.00296 |
| Glomerulus vasculature morphogenesis | 5 | 0.3 | 0.003 |
| Protein localization to ciliary transition zone | 5 | 0.3 | 0.003 |
| Glomerular capillary formation | 5 | 0.3 | 0.003 |
| Embryonic skeletal system development | 21 | 1.28 | 0.00305 |
| Monoatomic ion transmembrane transport | 89 | 5.42 | 0.00312 |
| Establishment of planar polarity | 8 | 0.49 | 0.00319 |
| Establishment of tissue polarity | 8 | 0.49 | 0.00319 |
| Kidney vasculature development | 8 | 0.49 | 0.00319 |
| Renal filtration cell differentiation | 8 | 0.49 | 0.00319 |
| Renal system vasculature development | 8 | 0.49 | 0.00319 |
| Podocyte differentiation | 8 | 0.49 | 0.00319 |
| Regulation of morphogenesis of a branching structure | 12 | 0.73 | 0.0032 |
| Regulation of chondrocyte differentiation | 12 | 0.73 | 0.0032 |
| Neural crest cell migration | 12 | 0.73 | 0.0032 |
| Regulation of protein localization to plasma membrane | 19 | 1.16 | 0.00322 |
| Positive regulation of macromolecule biosynthetic process | 236 | 14.37 | 0.00325 |
| Viral life cycle | 26 | 1.58 | 0.00327 |
| Mesoderm formation | 14 | 0.85 | 0.00337 |
| Myeloid leukocyte differentiation | 24 | 1.46 | 0.0034 |
| Response to carbohydrate | 27 | 1.64 | 0.00353 |
| Monoatomic cation transmembrane transport | 74 | 4.51 | 0.00355 |
| Regulation of actin filament-based movement | 10 | 0.61 | 0.00356 |
| Non-canonical wnt signaling pathway | 11 | 0.67 | 0.0037 |
| Bone remodeling | 11 | 0.67 | 0.0037 |
| Regulation of cell-matrix adhesion | 20 | 1.22 | 0.00375 |
| Negative regulation of wound healing | 14 | 0.85 | 0.0038 |
| Positive regulation of amyloid fibril formation | 4 | 0.24 | 0.00383 |
| Mammary gland epithelial cell proliferation | 6 | 0.37 | 0.00386 |
| Regulation of mesenchymal cell proliferation | 9 | 0.55 | 0.00387 |
| Cartilage development involved in endochondral bone morphogenesis | 9 | 0.55 | 0.00387 |
| Positive regulation of leukocyte migration | 23 | 1.4 | 0.00387 |
| Regulation of actin filament bundle assembly | 18 | 1.1 | 0.00396 |
| Endoplasmic reticulum calcium ion homeostasis | 8 | 0.49 | 0.00399 |
| Chondrocyte development | 8 | 0.49 | 0.00399 |
| Midbrain development | 16 | 0.97 | 0.00401 |
| Regulation of gtpase activity | 25 | 1.52 | 0.00401 |
| Vesicle-mediated transport | 129 | 7.86 | 0.00407 |
| Negative regulation of nucleobase-containing compound metabolic process | 146 | 8.89 | 0.0042 |
| Response to axon injury | 12 | 0.73 | 0.00422 |
| Positive regulation of receptor-mediated endocytosis | 12 | 0.73 | 0.00422 |
| Cellular detoxification | 19 | 1.16 | 0.00423 |
| Mesoderm morphogenesis | 14 | 0.85 | 0.00428 |
| Response to epidermal growth factor | 11 | 0.67 | 0.0043 |
| Cell junction maintenance | 11 | 0.67 | 0.0043 |
| Retrograde vesicle-mediated transport, golgi to endoplasmic reticulum | 11 | 0.67 | 0.0043 |
| Regulation of reproductive process | 28 | 1.71 | 0.00432 |
| Export from cell | 56 | 3.41 | 0.00441 |
| Regulation of fat cell differentiation | 22 | 1.34 | 0.00441 |
| Negative regulation of transcription by rna polymerase ii | 96 | 5.85 | 0.00442 |
| Mesonephros development | 16 | 0.97 | 0.00444 |
| Regulation of cardiac muscle cell contraction | 9 | 0.55 | 0.00465 |
| Renal absorption | 9 | 0.55 | 0.00465 |
| Regulation of glucocorticoid biosynthetic process | 5 | 0.3 | 0.0047 |
| Regulation of anatomical structure size | 57 | 3.47 | 0.00473 |
| Notochord development | 7 | 0.43 | 0.00477 |

| | | | |
|---|---|---|---|
| Non-motile cilium assembly | 12 | 0.73 | 0.00483 |
| Regulation of protein depolymerization | 16 | 0.97 | 0.00492 |
| Homophilic cell adhesion via plasma membrane adhesion molecules | 24 | 1.46 | 0.00493 |
| Positive regulation of hydrolase activity | 26 | 1.58 | 0.00497 |
| Negative regulation of cell projection organization | 26 | 1.58 | 0.00497 |
| Cell differentiation involved in kidney development | 11 | 0.67 | 0.00497 |
| Glial cell migration | 11 | 0.67 | 0.00497 |
| Negative regulation of cell activation | 30 | 1.83 | 0.00511 |
| Biological process involved in interaction with host | 22 | 1.34 | 0.00515 |
| Regulation of blood coagulation | 13 | 0.79 | 0.00517 |
| Regulation of morphogenesis of an epithelium | 13 | 0.79 | 0.00517 |
| Forebrain cell migration | 13 | 0.79 | 0.00517 |
| Cell activation | 80 | 4.87 | 0.00518 |
| Multicellular organismal reproductive process | 104 | 6.33 | 0.0052 |
| Glomerular epithelial cell development | 6 | 0.37 | 0.00526 |
| Branching involved in salivary gland morphogenesis | 6 | 0.37 | 0.00526 |
| Regulation of steroid hormone biosynthetic process | 6 | 0.37 | 0.00526 |
| Embryonic skeletal joint development | 6 | 0.37 | 0.00526 |
| Cardiac septum morphogenesis | 14 | 0.85 | 0.00537 |
| Mechanoreceptor differentiation | 14 | 0.85 | 0.00537 |
| Cardiac muscle cell differentiation | 16 | 0.97 | 0.00543 |
| Regulation of stem cell differentiation | 15 | 0.91 | 0.00545 |
| Epithelial to mesenchymal transition | 15 | 0.91 | 0.00545 |
| Positive regulation of transport | 89 | 5.42 | 0.00545 |
| Positive regulation of macromolecule metabolic process | 294 | 17.9 | 0.00549 |
| Positive regulation of epithelial to mesenchymal transition | 12 | 0.73 | 0.00549 |
| Modulation of chemical synaptic transmission | 58 | 3.53 | 0.00552 |
| Multi-multicellular organism process | 28 | 1.71 | 0.00559 |
| Intrinsic apoptotic signaling pathway | 25 | 1.52 | 0.00569 |
| Regulation of muscle system process | 33 | 2.01 | 0.00571 |
| Regulation of lipid metabolic process | 41 | 2.5 | 0.00572 |
| Regulation of proteolysis | 48 | 2.92 | 0.00572 |
| Regulation of trans-synaptic signaling | 58 | 3.53 | 0.00579 |
| Bone morphogenesis | 17 | 1.04 | 0.00587 |
| Positive regulation of neuron differentiation | 16 | 0.97 | 0.00599 |
| Regulation of leukocyte apoptotic process | 16 | 0.97 | 0.00599 |
| Myoblast fusion | 8 | 0.49 | 0.006 |
| Basement membrane organization | 8 | 0.49 | 0.006 |
| Multi-organism reproductive process | 27 | 1.64 | 0.00602 |
| Positive regulation of transferase activity | 33 | 2.01 | 0.00603 |
| Negative regulation of protein processing | 7 | 0.43 | 0.00605 |
| Positive regulation of cell development | 52 | 3.17 | 0.00618 |
| Cellular response to abiotic stimulus | 40 | 2.44 | 0.00623 |
| Cellular response to environmental stimulus | 40 | 2.44 | 0.00623 |
| Response to light stimulus | 39 | 2.38 | 0.00646 |
| Artery morphogenesis | 13 | 0.79 | 0.00654 |
| Positive regulation of protein-containing complex disassembly | 9 | 0.55 | 0.00658 |
| Skeletal muscle tissue regeneration | 9 | 0.55 | 0.00658 |
| Organic cation transport | 16 | 0.97 | 0.00659 |
| Cardiac chamber morphogenesis | 20 | 1.22 | 0.0067 |
| Cellular response to epidermal growth factor stimulus | 10 | 0.61 | 0.00671 |
| Exocrine system development | 10 | 0.61 | 0.00671 |
| Blood coagulation | 26 | 1.58 | 0.0069 |
| Renal system vasculature morphogenesis | 5 | 0.3 | 0.00694 |
| Nail development | 5 | 0.3 | 0.00694 |
| Regulation of basement membrane organization | 5 | 0.3 | 0.00694 |
| Prostate gland growth | 5 | 0.3 | 0.00694 |
| Kidney vasculature morphogenesis | 5 | 0.3 | 0.00694 |
| Regulation of plasma membrane organization | 6 | 0.37 | 0.007 |
| Actin filament bundle assembly | 12 | 0.73 | 0.00705 |
| Lymphocyte differentiation | 35 | 2.13 | 0.00705 |
| Regulation of prostatic bud formation | 4 | 0.24 | 0.00722 |
| Axonemal central apparatus assembly | 4 | 0.24 | 0.00722 |
| Determination of pancreatic left/right asymmetry | 4 | 0.24 | 0.00722 |
| Positive regulation of fat cell differentiation | 13 | 0.79 | 0.00732 |

| | | | |
|---|---|---|---|
| Regulation of endocytosis | 38 | 2.31 | 0.00738 |
| Regulation of protein kinase activity | 42 | 2.56 | 0.00767 |
| Peptidyl-amino acid modification | 42 | 2.56 | 0.00767 |
| Positive regulation of protein localization to membrane | 17 | 1.04 | 0.00768 |
| Maintenance of blood-brain barrier | 9 | 0.55 | 0.00774 |
| Regulation of homotypic cell-cell adhesion | 9 | 0.55 | 0.00774 |
| Astrocyte development | 9 | 0.55 | 0.00774 |
| Negative regulation of hemostasis | 10 | 0.61 | 0.00775 |
| Prostate gland development | 10 | 0.61 | 0.00775 |
| Protein localization to membrane | 53 | 3.23 | 0.00776 |
| Coagulation | 26 | 1.58 | 0.00783 |
| Negative regulation of response to wounding | 16 | 0.97 | 0.00794 |
| Urogenital system development | 12 | 0.73 | 0.00795 |
| Kidney morphogenesis | 15 | 0.91 | 0.00813 |
| Inner ear receptor cell differentiation | 13 | 0.79 | 0.00817 |
| Regulation of peptidase activity | 13 | 0.79 | 0.00817 |
| Positive regulation of response to wounding | 14 | 0.85 | 0.00822 |
| Cognition | 39 | 2.38 | 0.00825 |
| Negative regulation of locomotion | 47 | 2.86 | 0.00829 |
| Viral process | 33 | 2.01 | 0.00832 |
| Neural tube patterning | 8 | 0.49 | 0.0087 |
| Positive regulation of protein kinase activity | 27 | 1.64 | 0.00871 |
| Positive regulation of neurogenesis | 31 | 1.89 | 0.00884 |
| Photoreceptor cell maintenance | 10 | 0.61 | 0.00892 |
| Modified amino acid transport | 9 | 0.55 | 0.00905 |
| L-amino acid transport | 14 | 0.85 | 0.00909 |
| Left/right axis specification | 6 | 0.37 | 0.0091 |
| Cardiac cell development | 13 | 0.79 | 0.0091 |
| Cellular response to lipid | 59 | 3.59 | 0.00915 |
| Response to radiation | 49 | 2.98 | 0.00931 |
| Negative regulation of protein maturation | 7 | 0.43 | 0.00933 |
| Axon regeneration | 7 | 0.43 | 0.00933 |
| Regulation of hormone biosynthetic process | 7 | 0.43 | 0.00933 |
| Regulation of leukocyte migration | 30 | 1.83 | 0.0096 |
| Endoderm formation | 11 | 0.67 | 0.00964 |
| Response to ethanol | 20 | 1.22 | 0.00979 |
| Negative regulation of protein metabolic process | 75 | 4.57 | 0.00979 |
| Response to monosaccharide | 23 | 1.4 | 0.0098 |
| Regulation of glucocorticoid metabolic process | 5 | 0.3 | 0.0098 |
| Mesenchymal cell apoptotic process | 5 | 0.3 | 0.0098 |
| Cellular response to thyroid hormone stimulus | 5 | 0.3 | 0.0098 |
| Glomerular basement membrane development | 5 | 0.3 | 0.0098 |
| Collagen biosynthetic process | 5 | 0.3 | 0.0098 |
| Glomerulus morphogenesis | 5 | 0.3 | 0.0098 |
| Regulation of ventricular cardiac muscle cell action potential | 5 | 0.3 | 0.0098 |
| Metanephric glomerulus development | 5 | 0.3 | 0.0098 |
| Regulation of cellular component size | 41 | 2.5 | 0.0098 |
| Regulation of reactive oxygen species metabolic process | 22 | 1.34 | 0.00985 |
| Regulation of actin polymerization or depolymerization | 21 | 1.28 | 0.00985 |
| Regulation of microtubule-based process | 34 | 2.07 | 0.00987 |
| Inorganic cation transmembrane transport | 70 | 4.26 | 0.01 |
| Hemostasis | 26 | 1.58 | 0.01 |
| Heart valve morphogenesis | 12 | 0.73 | 0.01 |
| Negative regulation of cellular component organization | 74 | 4.51 | 0.01 |
| Mammary gland duct morphogenesis | 8 | 0.49 | 0.0103 |
| Response to hydrogen peroxide | 16 | 0.97 | 0.0104 |
| Central nervous system neuron differentiation | 27 | 1.64 | 0.0104 |
| Peptidyl-threonine phosphorylation | 9 | 0.55 | 0.0105 |
| Regulation of fibroblast growth factor receptor signaling pathway | 9 | 0.55 | 0.0105 |
| Regulation of lipid biosynthetic process | 26 | 1.58 | 0.0106 |
| Metal ion transport | 68 | 4.14 | 0.0107 |
| Tight junction assembly | 11 | 0.67 | 0.0109 |
| Inner ear receptor cell development | 11 | 0.67 | 0.0109 |
| Cellular response to glucocorticoid stimulus | 11 | 0.67 | 0.0109 |
| Regulation of transferase activity | 50 | 3.05 | 0.0109 |

| | | | |
|---|---|---|---|
| Epidermal cell differentiation | 27 | 1.64 | 0.011 |
| Response to activity | 14 | 0.85 | 0.011 |
| Alpha-beta t cell differentiation | 14 | 0.85 | 0.011 |
| Regulation of muscle cell differentiation | 23 | 1.4 | 0.0112 |
| Actin filament bundle organization | 12 | 0.73 | 0.0112 |
| Cellular response to corticosteroid stimulus | 12 | 0.73 | 0.0112 |
| Memory | 19 | 1.16 | 0.0112 |
| Myelination | 19 | 1.16 | 0.0112 |
| Response to hexose | 22 | 1.34 | 0.0113 |
| Platelet activation | 16 | 0.97 | 0.0113 |
| Regulation of mononuclear cell migration | 21 | 1.28 | 0.0113 |
| Regulation of lymphocyte chemotaxis | 7 | 0.43 | 0.0114 |
| Retina layer formation | 7 | 0.43 | 0.0114 |
| Smooth muscle tissue development | 7 | 0.43 | 0.0114 |
| Amyloid-beta clearance | 7 | 0.43 | 0.0114 |
| Spermatogenesis | 72 | 4.38 | 0.0116 |
| Pericardium development | 6 | 0.37 | 0.0116 |
| Paraxial mesoderm development | 6 | 0.37 | 0.0116 |
| Response to type ii interferon | 17 | 1.04 | 0.0117 |
| Determination of liver left/right asymmetry | 4 | 0.24 | 0.0119 |
| Chorio-allantoic fusion | 4 | 0.24 | 0.0119 |
| Bud elongation involved in lung branching | 4 | 0.24 | 0.0119 |
| Response to heparin | 4 | 0.24 | 0.0119 |
| Regulation of actin filament length | 21 | 1.28 | 0.0121 |
| Epithelial cell differentiation involved in kidney development | 9 | 0.55 | 0.0122 |
| Developmental induction | 8 | 0.49 | 0.0122 |
| Regulation of cilium-dependent cell motility | 8 | 0.49 | 0.0122 |
| Response to muscle activity | 8 | 0.49 | 0.0122 |
| Wnt signaling pathway, planar cell polarity pathway | 8 | 0.49 | 0.0122 |
| Regulation of cilium movement involved in cell motility | 8 | 0.49 | 0.0122 |
| Negative regulation of leukocyte apoptotic process | 11 | 0.67 | 0.0122 |
| Positive regulation of reactive oxygen species metabolic process | 13 | 0.79 | 0.0124 |
| Regulation of cilium assembly | 13 | 0.79 | 0.0124 |
| Dendrite morphogenesis | 12 | 0.73 | 0.0125 |
| Heart process | 17 | 1.04 | 0.0126 |
| Ear morphogenesis | 19 | 1.16 | 0.013 |
| Ensheathment of neurons | 19 | 1.16 | 0.013 |
| Axon ensheathment | 19 | 1.16 | 0.013 |
| Liver development | 20 | 1.22 | 0.013 |
| Regulation of macrophage migration | 10 | 0.61 | 0.0132 |
| Axon extension | 10 | 0.61 | 0.0132 |
| Endocardial cushion development | 10 | 0.61 | 0.0132 |
| Negative regulation of coagulation | 10 | 0.61 | 0.0132 |
| Keratinocyte development | 5 | 0.3 | 0.0133 |
| Telencephalon regionalization | 5 | 0.3 | 0.0133 |
| Positive regulation of actin filament depolymerization | 5 | 0.3 | 0.0133 |
| Aromatic amino acid transport | 5 | 0.3 | 0.0133 |
| Extracellular regulation of signal transduction | 5 | 0.3 | 0.0133 |
| Extracellular negative regulation of signal transduction | 5 | 0.3 | 0.0133 |
| Trans-synaptic signaling, modulating synaptic transmission | 5 | 0.3 | 0.0133 |
| Transmembrane transport | 122 | 7.43 | 0.0134 |
| Regulation of receptor signaling pathway via jak-stat | 13 | 0.79 | 0.0137 |
| Myotube differentiation | 13 | 0.79 | 0.0137 |
| Regulation of bicellular tight junction assembly | 7 | 0.43 | 0.0137 |
| Cell proliferation in forebrain | 7 | 0.43 | 0.0137 |
| Cardiac conduction | 12 | 0.73 | 0.0138 |
| Cellular response to retinoic acid | 12 | 0.73 | 0.0138 |
| Retina morphogenesis in camera-type eye | 12 | 0.73 | 0.0138 |
| Inner ear receptor cell stereocilium organization | 9 | 0.55 | 0.014 |
| Syncytium formation | 9 | 0.55 | 0.014 |
| Positive regulation of proteolysis | 28 | 1.71 | 0.0141 |
| Hematopoietic progenitor cell differentiation | 16 | 0.97 | 0.0145 |
| Positive regulation of lymphocyte chemotaxis | 6 | 0.37 | 0.0145 |
| Protein localization to cell-cell junction | 6 | 0.37 | 0.0145 |
| Positive regulation of synaptic transmission | 22 | 1.34 | 0.0146 |

| | | | |
|---|---|---|---|
| Extrinsic apoptotic signaling pathway | 17 | 1.04 | 0.0148 |
| Regulation of axonogenesis | 20 | 1.22 | 0.0149 |
| Protein autophosphorylation | 20 | 1.22 | 0.0149 |
| Cerebral cortex cell migration | 10 | 0.61 | 0.0149 |
| Semi-lunar valve development | 10 | 0.61 | 0.0149 |
| Negative regulation of leukocyte activation | 26 | 1.58 | 0.015 |
| Positive regulation of organelle organization | 53 | 3.23 | 0.0156 |
| Lipid transport | 40 | 2.44 | 0.0156 |
| Hepaticobiliary system development | 20 | 1.22 | 0.0159 |
| Cerebral cortex development | 19 | 1.16 | 0.016 |
| Sterol metabolic process | 19 | 1.16 | 0.016 |
| Negative regulation of lymphocyte activation | 23 | 1.4 | 0.0162 |
| Positive regulation of cardiac neural crest cell migration involved in outflow tract morphogenesis | 3 | 0.18 | 0.0162 |
| Radial spoke assembly | 3 | 0.18 | 0.0162 |
| Regulation of cardiac neural crest cell migration involved in outflow tract morphogenesis | 3 | 0.18 | 0.0162 |
| Lens induction in camera-type eye | 3 | 0.18 | 0.0162 |
| Mesenchymal cell proliferation | 7 | 0.43 | 0.0164 |
| Lipid localization | 40 | 2.44 | 0.0164 |
| Heart valve development | 13 | 0.79 | 0.0165 |
| Cellular response to mechanical stimulus | 13 | 0.79 | 0.0165 |
| Regulation of stress fiber assembly | 15 | 0.91 | 0.0165 |
| Positive regulation of animal organ morphogenesis | 8 | 0.49 | 0.0166 |
| Positive t cell selection | 8 | 0.49 | 0.0166 |
| Regulation of fatty acid oxidation | 8 | 0.49 | 0.0166 |
| Dopaminergic neuron differentiation | 8 | 0.49 | 0.0166 |
| Positive regulation of stress fiber assembly | 10 | 0.61 | 0.0168 |
| T cell activation | 39 | 2.38 | 0.017 |
| Regulation of glycoprotein metabolic process | 11 | 0.67 | 0.0171 |
| Amino acid transport | 19 | 1.16 | 0.0172 |
| Regulation of steroid biosynthetic process | 14 | 0.85 | 0.0174 |
| Negative regulation of supramolecular fiber organization | 22 | 1.34 | 0.0175 |
| Regulation of synapse assembly | 22 | 1.34 | 0.0175 |
| Intraciliary retrograde transport | 5 | 0.3 | 0.0176 |
| Negative regulation of synapse organization | 5 | 0.3 | 0.0176 |
| Epithelial cilium movement involved in determination of left/right asymmetry | 5 | 0.3 | 0.0176 |
| Negative regulation of long-term synaptic potentiation | 5 | 0.3 | 0.0176 |
| Desmosome organization | 5 | 0.3 | 0.0176 |
| Neuron projection fasciculation | 6 | 0.37 | 0.0179 |
| Calcium-independent cell-cell adhesion via plasma membrane cell-adhesion molecules | 6 | 0.37 | 0.0179 |
| Axonal fasciculation | 6 | 0.37 | 0.0179 |
| Metanephric nephron epithelium development | 6 | 0.37 | 0.0179 |
| Negative regulation of plasminogen activation | 4 | 0.24 | 0.018 |
| Positive regulation of neuroepithelial cell differentiation | 4 | 0.24 | 0.018 |
| Determination of digestive tract left/right asymmetry | 4 | 0.24 | 0.018 |
| Hard palate development | 4 | 0.24 | 0.018 |
| Chondrocyte development involved in endochondral bone morphogenesis | 4 | 0.24 | 0.018 |
| Cellular response to external stimulus | 13 | 0.79 | 0.0181 |
| Regulation of mirna transcription | 13 | 0.79 | 0.0181 |
| Regulation of actomyosin structure organization | 16 | 0.97 | 0.0183 |
| Learning or memory | 33 | 2.01 | 0.0183 |
| Limbic system development | 18 | 1.1 | 0.0185 |
| Positive regulation of cytoskeleton organization | 24 | 1.46 | 0.0187 |
| Male gamete generation | 72 | 4.38 | 0.0188 |
| Negative regulation of axonogenesis | 10 | 0.61 | 0.0189 |
| Positive regulation of actin filament bundle assembly | 11 | 0.67 | 0.019 |
| Response to mineralocorticoid | 8 | 0.49 | 0.0192 |
| Response to dietary excess | 8 | 0.49 | 0.0192 |
| Smooth muscle cell differentiation | 8 | 0.49 | 0.0192 |
| Response to reactive oxygen species | 23 | 1.4 | 0.0192 |
| Salivary gland morphogenesis | 7 | 0.43 | 0.0193 |
| Positive regulation of rho protein signal transduction | 7 | 0.43 | 0.0193 |
| Regulation of membrane protein ectodomain proteolysis | 7 | 0.43 | 0.0193 |
| Endocytosis | 54 | 3.29 | 0.0194 |
| Steroid metabolic process | 31 | 1.89 | 0.0197 |

| | | | |
|---|---|---|---|
| Regulation of protein binding | 16 | 0.97 | 0.0197 |
| Biological process involved in symbiotic interaction | 24 | 1.46 | 0.0198 |
| Outflow tract morphogenesis | 13 | 0.79 | 0.0198 |
| Notch signaling pathway | 17 | 1.04 | 0.0198 |
| Alpha-beta t cell activation | 17 | 1.04 | 0.0198 |
| Inorganic ion transmembrane transport | 76 | 4.63 | 0.0201 |
| Negative regulation of cell migration | 42 | 2.56 | 0.0204 |
| Negative regulation of neuron differentiation | 12 | 0.73 | 0.0205 |
| Sarcomere organization | 9 | 0.55 | 0.0206 |
| Negative regulation of blood coagulation | 9 | 0.55 | 0.0206 |
| Labyrinthine layer development | 9 | 0.55 | 0.0206 |
| Regulation of hemopoiesis | 47 | 2.86 | 0.0207 |
| Amino acid transmembrane transport | 15 | 0.91 | 0.021 |
| Endochondral bone morphogenesis | 11 | 0.67 | 0.021 |
| Positive chemotaxis | 10 | 0.61 | 0.0211 |
| Lung alveolus development | 10 | 0.61 | 0.0211 |
| Mesenchyme morphogenesis | 10 | 0.61 | 0.0211 |
| Cell-cell signaling | 82 | 4.99 | 0.0216 |
| Establishment of protein localization | 124 | 7.55 | 0.0217 |
| Labyrinthine layer morphogenesis | 6 | 0.37 | 0.0218 |
| Positive regulation of macrophage chemotaxis | 6 | 0.37 | 0.0218 |
| Heterotypic cell-cell adhesion | 8 | 0.49 | 0.022 |
| Positive regulation of morphogenesis of an epithelium | 8 | 0.49 | 0.022 |
| Negative regulation of growth | 28 | 1.71 | 0.0222 |
| Circadian rhythm | 19 | 1.16 | 0.0223 |
| Response to corticosterone | 5 | 0.3 | 0.0225 |
| Podocyte development | 5 | 0.3 | 0.0225 |
| Galactose metabolic process | 5 | 0.3 | 0.0225 |
| Negative regulation of dna damage response, signal transduction by p53 class mediator | 5 | 0.3 | 0.0225 |
| Cardiac ventricle development | 18 | 1.1 | 0.0226 |
| Negative regulation of neuron projection development | 18 | 1.1 | 0.0226 |
| Negative regulation of mirna transcription | 7 | 0.43 | 0.0227 |
| Lens fiber cell differentiation | 7 | 0.43 | 0.0227 |
| Membrane raft organization | 7 | 0.43 | 0.0227 |
| Positive regulation of macrophage migration | 7 | 0.43 | 0.0227 |
| Axis specification | 15 | 0.91 | 0.0227 |
| Regulation of systemic arterial blood pressure | 16 | 0.97 | 0.0228 |
| Regulation of sprouting angiogenesis | 11 | 0.67 | 0.0232 |
| Negative regulation of lymphocyte differentiation | 11 | 0.67 | 0.0232 |
| Neuron projection organization | 9 | 0.55 | 0.0232 |
| Response to angiotensin | 9 | 0.55 | 0.0232 |
| Negative chemotaxis | 9 | 0.55 | 0.0232 |
| Regulation of response to external stimulus | 110 | 6.7 | 0.0233 |
| Regulation of glycoprotein biosynthetic process | 10 | 0.61 | 0.0235 |
| Negative regulation of cell cycle g1/s phase transition | 13 | 0.79 | 0.0235 |
| Negative regulation of cell motility | 43 | 2.62 | 0.0236 |
| Regulation of cell-cell adhesion | 53 | 3.23 | 0.0238 |
| Protein transport | 105 | 6.39 | 0.0238 |
| Hindbrain development | 21 | 1.28 | 0.0241 |
| Negative regulation of cytoskeleton organization | 21 | 1.28 | 0.0241 |
| Regulation of receptor-mediated endocytosis | 18 | 1.1 | 0.0241 |
| Maintenance of location | 14 | 0.85 | 0.0242 |
| Organic anion transport | 44 | 2.68 | 0.0243 |
| Positive regulation of dna-templated transcription | 150 | 9.14 | 0.0245 |
| Lung epithelium development | 8 | 0.49 | 0.0251 |
| Cell-cell fusion | 8 | 0.49 | 0.0251 |
| Syncytium formation by plasma membrane fusion | 8 | 0.49 | 0.0251 |
| Keratinocyte differentiation | 19 | 1.16 | 0.0253 |
| Limb bud formation | 4 | 0.24 | 0.0256 |
| Regulation of hair follicle maturation | 4 | 0.24 | 0.0256 |
| Positive regulation of wnt signaling pathway, planar cell polarity pathway | 4 | 0.24 | 0.0256 |
| Bone trabecula formation | 4 | 0.24 | 0.0256 |
| Iris morphogenesis | 4 | 0.24 | 0.0256 |
| Mesenchymal-epithelial cell signaling | 4 | 0.24 | 0.0256 |
| Positive regulation of chondrocyte proliferation | 4 | 0.24 | 0.0256 |

| | | | |
|---|---|---|---|
| Negative regulation of morphogenesis of an epithelium | 4 | 0.24 | 0.0256 |
| Telencephalon cell migration | 11 | 0.67 | 0.0256 |
| Positive regulation of wound healing | 11 | 0.67 | 0.0256 |
| Regulation of peptidyl-serine phosphorylation | 11 | 0.67 | 0.0256 |
| Forebrain generation of neurons | 13 | 0.79 | 0.0256 |
| Positive regulation of rna biosynthetic process | 150 | 9.14 | 0.0256 |
| Negative regulation of molecular function | 58 | 3.53 | 0.0258 |
| Regulation of cytokine production | 86 | 5.24 | 0.0261 |
| Positive regulation of epithelial cell migration | 9 | 0.55 | 0.0261 |
| Membrane lipid catabolic process | 9 | 0.55 | 0.0261 |
| Renal filtration | 6 | 0.37 | 0.0261 |
| Smad protein signal transduction | 6 | 0.37 | 0.0261 |
| Positive regulation of stem cell differentiation | 6 | 0.37 | 0.0261 |
| Tooth mineralization | 6 | 0.37 | 0.0261 |
| Post-embryonic development | 14 | 0.85 | 0.0261 |
| Regulation of epithelial cell apoptotic process | 10 | 0.61 | 0.0261 |
| Regulation of steroid metabolic process | 16 | 0.97 | 0.0263 |
| Myelination in peripheral nervous system | 7 | 0.43 | 0.0264 |
| Positive regulation of sprouting angiogenesis | 7 | 0.43 | 0.0264 |
| Peripheral nervous system axon ensheathment | 7 | 0.43 | 0.0264 |
| Regulation of organelle assembly | 28 | 1.71 | 0.0268 |
| Organic acid transmembrane transport | 19 | 1.16 | 0.0269 |
| Negative regulation of reproductive process | 12 | 0.73 | 0.0269 |
| Ventricular septum development | 12 | 0.73 | 0.0269 |
| Positive regulation of lipid metabolic process | 18 | 1.1 | 0.0274 |
| Negative regulation of transmembrane receptor protein serine/threonine kinase signaling pathway | 25 | 1.52 | 0.0275 |
| Mesoderm development | 22 | 1.34 | 0.0277 |
| Organic acid transport | 34 | 2.07 | 0.0277 |
| Embryonic placenta development | 13 | 0.79 | 0.0278 |
| L-alpha-amino acid transmembrane transport | 11 | 0.67 | 0.0281 |
| Cardiac muscle cell development | 11 | 0.67 | 0.0281 |
| Positive regulation of protein localization to nucleus | 14 | 0.85 | 0.0282 |
| Regulation of cytokine-mediated signaling pathway | 23 | 1.4 | 0.0282 |
| Negative regulation of cell fate commitment | 5 | 0.3 | 0.0283 |
| Positive regulation of endothelial cell chemotaxis | 5 | 0.3 | 0.0283 |
| Wound healing, spreading of epidermal cells | 5 | 0.3 | 0.0283 |
| Regulation of neural precursor cell proliferation | 15 | 0.91 | 0.0283 |
| Regulation of cell activation | 65 | 3.96 | 0.0283 |
| Epithelial cell morphogenesis | 8 | 0.49 | 0.0285 |
| Animal organ formation | 8 | 0.49 | 0.0285 |
| Regulation of hormone metabolic process | 8 | 0.49 | 0.0285 |
| Regulation of canonical nf-kappab signal transduction | 35 | 2.13 | 0.0286 |
| Digestive tract morphogenesis | 9 | 0.55 | 0.0292 |
| Platelet aggregation | 9 | 0.55 | 0.0292 |
| Single fertilization | 22 | 1.34 | 0.0292 |
| Negative regulation of protein localization | 27 | 1.64 | 0.0293 |
| Cd4-positive, alpha-beta t cell activation | 12 | 0.73 | 0.0294 |
| Adult behavior | 20 | 1.22 | 0.0295 |
| Response to glucose | 20 | 1.22 | 0.0295 |
| Presynapse organization | 7 | 0.43 | 0.0305 |
| Positive regulation of extracellular matrix organization | 7 | 0.43 | 0.0305 |
| Negative regulation of mirna metabolic process | 7 | 0.43 | 0.0305 |
| Negative regulation of microtubule depolymerization | 7 | 0.43 | 0.0305 |
| Regulation of cardiac muscle cell action potential | 7 | 0.43 | 0.0305 |
| Regulation of developmental growth | 36 | 2.19 | 0.0306 |
| Apoptotic mitochondrial changes | 11 | 0.67 | 0.0308 |
| Circadian regulation of gene expression | 11 | 0.67 | 0.0308 |
| Regulation of muscle contraction | 22 | 1.34 | 0.0308 |
| Fusion of virus membrane with host plasma membrane | 3 | 0.18 | 0.0308 |
| Cardiac muscle tissue regeneration | 3 | 0.18 | 0.0308 |
| Cellular response to heparin | 3 | 0.18 | 0.0308 |
| Sequestering of tgfbeta in extracellular matrix | 3 | 0.18 | 0.0308 |
| Negative regulation of apoptotic process in bone marrow cell | 3 | 0.18 | 0.0308 |
| Embryonic nail plate morphogenesis | 3 | 0.18 | 0.0308 |

| | | | |
|---|---|---|---|
| Membrane fusion involved in viral entry into host cell | 3 | 0.18 | 0.0308 |
| Response to thyroid hormone | 6 | 0.37 | 0.031 |
| Prostate gland epithelium morphogenesis | 6 | 0.37 | 0.031 |
| Negative regulation of keratinocyte proliferation | 6 | 0.37 | 0.031 |
| Negative regulation of dna-binding transcription factor activity | 18 | 1.1 | 0.031 |
| Secretion | 56 | 3.41 | 0.0313 |
| Regulation of vesicle-mediated transport | 56 | 3.41 | 0.0318 |
| Regulation of cell projection assembly | 25 | 1.52 | 0.0319 |
| Hormone metabolic process | 25 | 1.52 | 0.0319 |
| Maternal process involved in female pregnancy | 10 | 0.61 | 0.0319 |
| Negative regulation of secretion | 22 | 1.34 | 0.0325 |
| Response to lipopolysaccharide | 37 | 2.25 | 0.0326 |
| Actin filament-based movement | 15 | 0.91 | 0.0326 |
| Regulation of mirna metabolic process | 14 | 0.85 | 0.0327 |
| Negative regulation of leukocyte cell-cell adhesion | 20 | 1.22 | 0.033 |
| Endomembrane system organization | 58 | 3.53 | 0.0336 |
| Regulation of epithelial cell migration | 11 | 0.67 | 0.0337 |
| Amide transport | 19 | 1.16 | 0.034 |
| Mucociliary clearance | 4 | 0.24 | 0.0345 |
| Regulation of cilium beat frequency involved in ciliary motility | 4 | 0.24 | 0.0345 |
| Cell proliferation involved in kidney development | 4 | 0.24 | 0.0345 |
| Regulation of glomerular filtration | 4 | 0.24 | 0.0345 |
| Negative regulation of axon extension involved in axon guidance | 4 | 0.24 | 0.0345 |
| Protein processing | 23 | 1.4 | 0.0346 |
| Regulation of intracellular transport | 31 | 1.89 | 0.0346 |
| Mammary gland epithelial cell differentiation | 5 | 0.3 | 0.0348 |
| Type b pancreatic cell development | 5 | 0.3 | 0.0348 |
| Positive regulation of fatty acid oxidation | 5 | 0.3 | 0.0348 |
| Regulation of metallopeptidase activity | 5 | 0.3 | 0.0348 |
| Cellular response to steroid hormone stimulus | 20 | 1.22 | 0.0348 |
| Pigmentation | 15 | 0.91 | 0.0349 |
| Response to retinoic acid | 15 | 0.91 | 0.0349 |
| Salivary gland development | 7 | 0.43 | 0.0349 |
| Regulation of macrophage chemotaxis | 7 | 0.43 | 0.0349 |
| Negative regulation of epithelial cell apoptotic process | 7 | 0.43 | 0.0349 |
| Metanephric nephron development | 7 | 0.43 | 0.0349 |
| Negative regulation of protein localization to cell periphery | 7 | 0.43 | 0.0349 |
| Respiratory gaseous exchange by respiratory system | 10 | 0.61 | 0.0351 |
| Cellular response to metal ion | 22 | 1.34 | 0.036 |
| Regulation of postsynapse organization | 19 | 1.16 | 0.036 |
| Bicellular tight junction assembly | 9 | 0.55 | 0.0361 |
| Regulation of sodium ion transmembrane transport | 9 | 0.55 | 0.0361 |
| Embryo implantation | 9 | 0.55 | 0.0361 |
| Regulation of endocrine process | 8 | 0.49 | 0.0363 |
| Cell communication involved in cardiac conduction | 8 | 0.49 | 0.0363 |
| Regulation of endothelial cell chemotaxis | 6 | 0.37 | 0.0364 |
| Amyloid precursor protein metabolic process | 6 | 0.37 | 0.0364 |
| Metanephric epithelium development | 6 | 0.37 | 0.0364 |
| Negative regulation of fibroblast growth factor receptor signaling pathway | 6 | 0.37 | 0.0364 |
| Decidualization | 6 | 0.37 | 0.0364 |
| Regulation of appetite | 6 | 0.37 | 0.0364 |
| Locomotory behavior | 25 | 1.52 | 0.0367 |
| Regulation of lymphocyte differentiation | 26 | 1.58 | 0.0368 |
| Negative regulation of t cell activation | 18 | 1.1 | 0.0371 |
| Membrane organization | 74 | 4.51 | 0.0373 |
| Regulation of receptor internalization | 12 | 0.73 | 0.0376 |
| Sterol transport | 12 | 0.73 | 0.0376 |
| Negative regulation of erk1 and erk2 cascade | 12 | 0.73 | 0.0376 |
| Regulation of sodium ion transport | 12 | 0.73 | 0.0376 |
| Hair cycle process | 13 | 0.79 | 0.0379 |
| Multicellular organismal response to stress | 13 | 0.79 | 0.0379 |
| Molting cycle process | 13 | 0.79 | 0.0379 |
| Positive regulation of gtpase activity | 16 | 0.97 | 0.0391 |
| Placenta blood vessel development | 7 | 0.43 | 0.0398 |
| Brown fat cell differentiation | 7 | 0.43 | 0.0398 |

| | | | |
|---|---|---|---|
| Positive regulation of cartilage development | 7 | 0.43 | 0.0398 |
| Negative regulation of protein depolymerization | 11 | 0.67 | 0.04 |
| Humoral immune response mediated by circulating immunoglobulin | 9 | 0.55 | 0.04 |
| Lipid homeostasis | 23 | 1.4 | 0.0401 |
| Positive regulation of cytokine production | 52 | 3.17 | 0.0404 |
| Regulation of calcium-mediated signaling | 14 | 0.85 | 0.0404 |
| Positive regulation of smad protein signal transduction | 8 | 0.49 | 0.0406 |
| Ventricular septum morphogenesis | 8 | 0.49 | 0.0406 |
| Carboxylic acid transport | 33 | 2.01 | 0.0412 |
| Organic hydroxy compound transport | 21 | 1.28 | 0.0414 |
| Response to nutrient levels | 52 | 3.17 | 0.0416 |
| Carboxylic acid transmembrane transport | 18 | 1.1 | 0.0416 |
| Regulation of response to food | 5 | 0.3 | 0.0422 |
| Heart formation | 5 | 0.3 | 0.0422 |
| Cd4-positive, alpha-beta t cell lineage commitment | 5 | 0.3 | 0.0422 |
| Chondrocyte proliferation | 5 | 0.3 | 0.0422 |
| Positive regulation of membrane protein ectodomain proteolysis | 5 | 0.3 | 0.0422 |
| Specification of animal organ identity | 5 | 0.3 | 0.0422 |
| Cardiac myofibril assembly | 5 | 0.3 | 0.0422 |
| Preganglionic parasympathetic fiber development | 5 | 0.3 | 0.0422 |
| Cell surface receptor signaling pathway via jak-stat | 10 | 0.61 | 0.0422 |
| Release of cytochrome c from mitochondria | 6 | 0.37 | 0.0424 |
| Prostate gland morphogenesis | 6 | 0.37 | 0.0424 |
| Positive regulation of transcription by rna polymerase ii | 112 | 6.82 | 0.0432 |
| Heart contraction | 14 | 0.85 | 0.0433 |
| Regulation of smooth muscle cell proliferation | 21 | 1.28 | 0.0436 |
| Positive regulation of reproductive process | 13 | 0.79 | 0.0438 |
| Regulation of myeloid cell differentiation | 26 | 1.58 | 0.0439 |
| Response to topologically incorrect protein | 18 | 1.1 | 0.044 |
| Floor plate development | 4 | 0.24 | 0.0448 |
| Negative regulation of mesenchymal cell apoptotic process | 4 | 0.24 | 0.0448 |
| Dendritic spine maintenance | 4 | 0.24 | 0.0448 |
| Positive regulation of membrane depolarization | 4 | 0.24 | 0.0448 |
| Trachea morphogenesis | 4 | 0.24 | 0.0448 |
| Regulation of glial cell apoptotic process | 4 | 0.24 | 0.0448 |
| Regulation of glomerular mesangial cell proliferation | 4 | 0.24 | 0.0448 |
| Sequestering of extracellular ligand from receptor | 4 | 0.24 | 0.0448 |
| Olfactory behavior | 4 | 0.24 | 0.0448 |
| Dendritic spine organization | 7 | 0.43 | 0.0451 |
| Vesicle cargo loading | 7 | 0.43 | 0.0451 |
| Camera-type eye photoreceptor cell differentiation | 7 | 0.43 | 0.0451 |
| Maternal placenta development | 7 | 0.43 | 0.0451 |
| Positive regulation of substrate adhesion-dependent cell spreading | 8 | 0.49 | 0.0453 |
| Receptor-mediated endocytosis | 21 | 1.28 | 0.0458 |
| Cd4-positive, alpha-beta t cell differentiation | 10 | 0.61 | 0.0461 |
| Cellular pigmentation | 10 | 0.61 | 0.0461 |
| Regulation of substrate adhesion-dependent cell spreading | 10 | 0.61 | 0.0461 |
| Membrane lipid metabolic process | 24 | 1.46 | 0.0461 |
| Regulation of muscle adaptation | 14 | 0.85 | 0.0463 |
| Regulation of jnk cascade | 18 | 1.1 | 0.0465 |
| Skin epidermis development | 16 | 0.97 | 0.0469 |
| Leukocyte migration | 28 | 1.71 | 0.047 |
| Nephron morphogenesis | 11 | 0.67 | 0.047 |
| Actin-mediated cell contraction | 12 | 0.73 | 0.0473 |
| Muscle organ morphogenesis | 12 | 0.73 | 0.0473 |
| Regulation of lymphocyte activation | 52 | 3.17 | 0.0479 |
| Synapse assembly | 17 | 1.04 | 0.0481 |
| Regulation of cell cycle g1/s phase transition | 24 | 1.46 | 0.0482 |
| Regulation of plasma membrane bounded cell projection assembly | 24 | 1.46 | 0.0482 |
| Inner ear morphogenesis | 15 | 0.91 | 0.0482 |
| Sodium ion transport | 23 | 1.4 | 0.0483 |
| Regulation of actin filament depolymerization | 9 | 0.55 | 0.0485 |
| Regulation of smooth muscle cell differentiation | 9 | 0.55 | 0.0485 |
| Response to molecule of bacterial origin | 38 | 2.31 | 0.0488 |
| Cell surface receptor signaling pathway involved in heart development | 3 | 0.18 | 0.0489 |

| | | | |
|---|---|---|---|
| Positive regulation of mesenchymal stem cell differentiation | 3 | 0.18 | 0.0489 |
| Chondrocyte hypertrophy | 3 | 0.18 | 0.0489 |
| Positive regulation of glutamate receptor signaling pathway | 3 | 0.18 | 0.0489 |
| Negative regulation of glucocorticoid metabolic process | 3 | 0.18 | 0.0489 |
| Regulation of apoptotic process in bone marrow cell | 3 | 0.18 | 0.0489 |
| Chondroblast differentiation | 3 | 0.18 | 0.0489 |
| Negative regulation of glucocorticoid biosynthetic process | 3 | 0.18 | 0.0489 |
| Regulation of branching involved in prostate gland morphogenesis | 3 | 0.18 | 0.0489 |
| Tendon development | 3 | 0.18 | 0.0489 |
| Negative regulation of chondrocyte differentiation | 6 | 0.37 | 0.0489 |
| Embryonic placenta morphogenesis | 6 | 0.37 | 0.0489 |
| Positive regulation of collagen biosynthetic process | 6 | 0.37 | 0.0489 |
| Outflow tract septum morphogenesis | 6 | 0.37 | 0.0489 |
| Pulmonary valve development | 6 | 0.37 | 0.0489 |
| Positive regulation of apoptotic signaling pathway | 19 | 1.16 | 0.0497 |
| Regulation of nucleocytoplasmic transport | 16 | 0.97 | 0.0498 |

TableS15

| Pathways upregulated in Gland_A compared to Gland_B | | | |
|---|---|---|---|
| **GO Pathway** | **No. of genes** | **%** | **p-value** |
| Lipid metabolic process | 162 | 13.19 | 5.86E-24 |
| Organic acid metabolic process | 112 | 9.12 | 1.03E-17 |
| Oxoacid metabolic process | 111 | 9.04 | 1.28E-17 |
| Carboxylic acid metabolic process | 106 | 8.63 | 3.68E-16 |
| Monocarboxylic acid metabolic process | 77 | 6.27 | 4.38E-14 |
| Organophosphate metabolic process | 116 | 9.45 | 4.02E-13 |
| Small molecule biosynthetic process | 70 | 5.7 | 1.07E-12 |
| Alcohol metabolic process | 54 | 4.4 | 3.31E-12 |
| Regulation of hormone levels | 76 | 6.19 | 4.45E-12 |
| Homeostatic process | 153 | 12.46 | 5.83E-12 |
| Response to biotic stimulus | 155 | 12.62 | 9.38E-12 |
| Gland development | 64 | 5.21 | 1.41E-11 |
| Response to external biotic stimulus | 151 | 12.3 | 1.76E-11 |
| Biological process involved in interspecies interaction between organisms | 163 | 13.27 | 2.05E-11 |
| Organic anion transport | 61 | 4.97 | 2.7E-11 |
| Response to other organism | 150 | 12.21 | 2.93E-11 |
| Phosphate-containing compound metabolic process | 148 | 12.05 | 4.14E-11 |
| Phosphorus metabolic process | 148 | 12.05 | 4.98E-11 |
| Small molecule catabolic process | 56 | 4.56 | 5.81E-11 |
| Glycerolipid metabolic process | 55 | 4.48 | 9.54E-11 |
| Response to external stimulus | 186 | 15.15 | 1.11E-10 |
| Epithelial cell differentiation | 81 | 6.6 | 1.24E-10 |
| Steroid metabolic process | 45 | 3.66 | 1.67E-10 |
| Epithelium development | 118 | 9.61 | 2.43E-10 |
| Digestion | 28 | 2.28 | 4.19E-10 |
| Lipid biosynthetic process | 74 | 6.03 | 5.44E-10 |
| Digestive system process | 23 | 1.87 | 7.44E-10 |
| Regulation of transport | 157 | 12.79 | 9.36E-10 |
| Sulfur compound metabolic process | 49 | 3.99 | 1.13E-09 |
| Regulation of localization | 186 | 15.15 | 1.89E-09 |
| Regulation of small molecule metabolic process | 50 | 4.07 | 3.84E-09 |
| Triglyceride metabolic process | 21 | 1.71 | 5.03E-09 |
| Regulation of protein transport | 60 | 4.89 | 5.49E-09 |
| Fatty acid metabolic process | 48 | 3.91 | 8.19E-09 |
| Transmembrane transport | 125 | 10.18 | 1.02E-08 |
| Regulation of lipid metabolic process | 49 | 3.99 | 1.2E-08 |
| Chemical homeostasis | 99 | 8.06 | 1.3E-08 |
| Cell-cell junction organization | 33 | 2.69 | 1.63E-08 |
| Tissue development | 163 | 13.27 | 1.68E-08 |
| Lipid homeostasis | 34 | 2.77 | 1.69E-08 |
| Intestinal absorption | 14 | 1.14 | 1.73E-08 |
| Cellular catabolic process | 45 | 3.66 | 1.9E-08 |
| Monosaccharide metabolic process | 34 | 2.77 | 1.92E-08 |
| Actin filament organization | 43 | 3.5 | 2.02E-08 |

| | | | |
|---|---|---|---|
| Actin filament bundle organization | 19 | 1.55 | 2.06E-08 |
| Cellular response to chemical stimulus | 176 | 14.33 | 2.09E-08 |
| Regulation of protein localization | 97 | 7.9 | 2.16E-08 |
| Cholesterol homeostasis | 25 | 2.04 | 2.24E-08 |
| Defense response | 143 | 11.64 | 2.29E-08 |
| Regulation of establishment of protein localization | 67 | 5.46 | 2.41E-08 |
| Regulation of response to external stimulus | 115 | 9.36 | 2.42E-08 |
| Sterol homeostasis | 25 | 2.04 | 2.67E-08 |
| Defense response to other organism | 110 | 8.96 | 3.4E-08 |
| Carbohydrate metabolic process | 58 | 4.72 | 4.27E-08 |
| Positive regulation of transport | 91 | 7.41 | 5.31E-08 |
| Acylglycerol metabolic process | 23 | 1.87 | 5.52E-08 |
| Xenobiotic metabolic process | 27 | 2.2 | 6.28E-08 |
| Neutral lipid metabolic process | 23 | 1.87 | 6.63E-08 |
| Response to oxygen-containing compound | 147 | 11.97 | 7.22E-08 |
| Actin filament-based process | 71 | 5.78 | 8.63E-08 |
| Actin cytoskeleton organization | 65 | 5.29 | 1.05E-07 |
| Regulation of cellular localization | 103 | 8.39 | 1.12E-07 |
| Hexose metabolic process | 30 | 2.44 | 1.19E-07 |
| Carboxylic acid biosynthetic process | 41 | 3.34 | 1.52E-07 |
| Response to stress | 277 | 22.56 | 1.57E-07 |
| Organic acid biosynthetic process | 41 | 3.34 | 2.01E-07 |
| Regulation of defense response | 91 | 7.41 | 2.01E-07 |
| Actin filament bundle assembly | 17 | 1.38 | 2.92E-07 |
| Response to xenobiotic stimulus | 55 | 4.48 | 3.01E-07 |
| Monoatomic ion transport | 100 | 8.14 | 3.04E-07 |
| Carboxylic acid transport | 42 | 3.42 | 3.11E-07 |
| Organic acid transport | 42 | 3.42 | 3.39E-07 |
| Circulatory system process | 61 | 4.97 | 0.0000004 |
| Import into cell | 75 | 6.11 | 4.18E-07 |
| Regulation of biological quality | 243 | 19.79 | 4.39E-07 |
| Response to bacterium | 78 | 6.35 | 4.75E-07 |
| Innate immune response | 84 | 6.84 | 5.95E-07 |
| Organic hydroxy compound transport | 29 | 2.36 | 5.95E-07 |
| Regulation of secretion | 69 | 5.62 | 6.08E-07 |
| Response to lipid | 88 | 7.17 | 6.4E-07 |
| Positive regulation of secretion | 43 | 3.5 | 6.93E-07 |
| Multicellular organismal-level homeostasis | 70 | 5.7 | 8.09E-07 |
| Catabolic process | 170 | 13.84 | 9.14E-07 |
| Amide metabolic process | 50 | 4.07 | 1.11E-06 |
| Defense response to symbiont | 91 | 7.41 | 1.23E-06 |
| Hormone metabolic process | 32 | 2.61 | 1.26E-06 |
| Phospholipid metabolic process | 46 | 3.75 | 1.27E-06 |
| Monocarboxylic acid catabolic process | 22 | 1.79 | 1.28E-06 |
| Cholesterol biosynthetic process | 13 | 1.06 | 0.0000013 |
| Secondary alcohol biosynthetic process | 13 | 1.06 | 0.0000013 |
| Lipid transport | 45 | 3.66 | 1.48E-06 |

| | | | |
|---|---|---|---|
| Regulation of cell adhesion | 82 | 6.68 | 1.52E-06 |
| Lipid localization | 45 | 3.66 | 1.59E-06 |
| Sterol biosynthetic process | 14 | 1.14 | 1.72E-06 |
| Positive regulation of response to stimulus | 190 | 15.47 | 1.98E-06 |
| Liver development | 25 | 2.04 | 2.02E-06 |
| Lipid catabolic process | 41 | 3.34 | 2.02E-06 |
| Cholesterol metabolic process | 23 | 1.87 | 2.14E-06 |
| Regulation of microvillus organization | 8 | 0.65 | 2.32E-06 |
| Positive regulation of cell communication | 154 | 12.54 | 2.36E-06 |
| Immune system process | 195 | 15.88 | 2.45E-06 |
| Positive regulation of signaling | 154 | 12.54 | 2.55E-06 |
| Alcohol biosynthetic process | 20 | 1.63 | 2.87E-06 |
| Secondary alcohol metabolic process | 24 | 1.95 | 0.0000029 |
| Sterol metabolic process | 24 | 1.95 | 0.0000029 |
| Hepaticobiliary system development | 25 | 2.04 | 2.93E-06 |
| Positive regulation of protein transport | 36 | 2.93 | 2.93E-06 |
| Organophosphate biosynthetic process | 62 | 5.05 | 3.28E-06 |
| Positive regulation of secretion by cell | 39 | 3.18 | 3.58E-06 |
| Regulation of secretion by cell | 62 | 5.05 | 3.96E-06 |
| Regulation of lipid biosynthetic process | 30 | 2.44 | 4.02E-06 |
| Carboxylic acid catabolic process | 33 | 2.69 | 4.06E-06 |
| Organic acid catabolic process | 33 | 2.69 | 4.06E-06 |
| Positive regulation of defense response | 52 | 4.23 | 5.04E-06 |
| Regulation of body fluid levels | 45 | 3.66 | 5.34E-06 |
| Positive regulation of molecular function | 81 | 6.6 | 6.16E-06 |
| Negative regulation of response to external stimulus | 51 | 4.15 | 0.0000067 |
| Metal ion transport | 67 | 5.46 | 7.73E-06 |
| Regulation of signaling | 269 | 21.91 | 8.49E-06 |
| Respiratory tube development | 29 | 2.36 | 8.67E-06 |
| Nucleobase-containing small molecule metabolic process | 64 | 5.21 | 9.36E-06 |
| Positive regulation of protein localization | 55 | 4.48 | 0.0000106 |
| Regulation of protein secretion | 36 | 2.93 | 0.0000109 |
| Positive regulation of signal transduction | 137 | 11.16 | 0.0000113 |
| Regulation of inflammatory response | 50 | 4.07 | 0.0000134 |
| Regulation of cell communication | 268 | 21.82 | 0.0000139 |
| Tube development | 86 | 7 | 0.0000154 |
| Regulation of plasma lipoprotein particle levels | 15 | 1.22 | 0.0000155 |
| Regulation of ketone metabolic process | 23 | 1.87 | 0.0000158 |
| Lung development | 28 | 2.28 | 0.0000168 |
| Digestive system development | 24 | 1.95 | 0.0000169 |
| Transition metal ion transport | 19 | 1.55 | 0.0000174 |
| Cellular response to xenobiotic stimulus | 29 | 2.36 | 0.0000238 |
| Positive regulation of inflammatory response | 25 | 2.04 | 0.0000241 |
| Defense response to virus | 36 | 2.93 | 0.0000248 |
| Epithelial structure maintenance | 11 | 0.9 | 0.0000277 |
| Purine-containing compound metabolic process | 54 | 4.4 | 0.0000287 |
| Positive regulation of cell adhesion | 53 | 4.32 | 0.0000297 |

| | | | |
|---|---|---|---|
| Regulation of insulin secretion | 26 | 2.12 | 0.0000326 |
| Primary alcohol catabolic process | 8 | 0.65 | 0.0000334 |
| Regulation of response to biotic stimulus | 59 | 4.8 | 0.0000335 |
| Endocytosis | 54 | 4.4 | 0.0000344 |
| Blood circulation | 48 | 3.91 | 0.0000344 |
| Regulation of lipid transport | 24 | 1.95 | 0.0000351 |
| Digestive tract development | 22 | 1.79 | 0.0000359 |
| Regulation of cell population proliferation | 144 | 11.73 | 0.0000366 |
| Regulation of peptide transport | 29 | 2.36 | 0.0000398 |
| Positive regulation of multicellular organismal process | 140 | 11.4 | 0.0000416 |
| Response to fatty acid | 14 | 1.14 | 0.0000422 |
| Icosanoid metabolic process | 20 | 1.63 | 0.0000432 |
| Glycerolipid biosynthetic process | 29 | 2.36 | 0.0000432 |
| Phosphatidylcholine metabolic process | 15 | 1.22 | 0.0000436 |
| Cell-cell junction assembly | 21 | 1.71 | 0.0000469 |
| Response to endogenous stimulus | 108 | 8.79 | 0.0000482 |
| Steroid biosynthetic process | 18 | 1.47 | 0.0000488 |
| Regulation of immune system process | 138 | 11.24 | 0.0000519 |
| Glucose metabolic process | 20 | 1.63 | 0.0000546 |
| Regulation of molecular function | 120 | 9.77 | 0.0000546 |
| Nucleoside phosphate metabolic process | 59 | 4.8 | 0.0000561 |
| Organophosphate ester transport | 25 | 2.04 | 0.0000572 |
| Regulation of hormone secretion | 33 | 2.69 | 0.0000633 |
| Desmosome organization | 7 | 0.57 | 0.0000657 |
| Fructose metabolic process | 7 | 0.57 | 0.0000657 |
| Response to hormone | 78 | 6.35 | 0.0000662 |
| Monoatomic ion transmembrane transport | 78 | 6.35 | 0.0000662 |
| Response to virus | 44 | 3.58 | 0.0000665 |
| Regulation of peptide hormone secretion | 28 | 2.28 | 0.0000681 |
| Positive regulation of establishment of protein localization | 39 | 3.18 | 0.0000687 |
| Response to purine-containing compound | 23 | 1.87 | 0.0000691 |
| Terpenoid metabolic process | 18 | 1.47 | 0.0000714 |
| Monocarboxylic acid transport | 23 | 1.87 | 0.0000762 |
| Response to toxic substance | 32 | 2.61 | 0.0000786 |
| Regulation of steroid metabolic process | 19 | 1.55 | 0.0000794 |
| Primary alcohol metabolic process | 17 | 1.38 | 0.00008 |
| Positive regulation of transcription by rna polymerase ii | 104 | 8.47 | 0.0000801 |
| Positive regulation of response to external stimulus | 59 | 4.8 | 0.000081 |
| Pancreas development | 15 | 1.22 | 0.0000811 |
| Regulation of peptide secretion | 28 | 2.28 | 0.0000868 |
| Regulation of lipid localization | 26 | 2.12 | 0.0000871 |
| Regulation of cholesterol metabolic process | 12 | 0.98 | 0.0000881 |
| Positive regulation of lipid localization | 19 | 1.55 | 0.0000892 |
| Protein processing | 26 | 2.12 | 0.0000948 |
| Phospholipid biosynthetic process | 30 | 2.44 | 0.000097 |
| Regulation of signal transduction | 233 | 18.97 | 0.0000974 |
| Regulation of cell-cell adhesion | 52 | 4.23 | 0.0001 |

| | | | |
|---|---|---|---|
| Inorganic ion transmembrane transport | 71 | 5.78 | 0.000105 |
| Modified amino acid metabolic process | 25 | 2.04 | 0.000107 |
| Negative regulation of viral process | 15 | 1.22 | 0.000109 |
| Regulation of cytokine production | 80 | 6.51 | 0.000112 |
| Organic cation transport | 17 | 1.38 | 0.000117 |
| Mammary gland development | 20 | 1.63 | 0.000118 |
| Positive regulation of immune response | 66 | 5.37 | 0.00012 |
| Carbohydrate derivative metabolic process | 81 | 6.6 | 0.000123 |
| Regulation of vesicle-mediated transport | 55 | 4.48 | 0.000125 |
| Regulation of innate immune response | 50 | 4.07 | 0.000129 |
| Inorganic anion transport | 23 | 1.87 | 0.000134 |
| Regulation of immune response | 91 | 7.41 | 0.000136 |
| Response to nutrient levels | 52 | 4.23 | 0.000139 |
| Bundle of his cell-purkinje myocyte adhesion involved in cell communication | 5 | 0.41 | 0.00014 |
| Glycerophospholipid metabolic process | 34 | 2.77 | 0.00014 |
| Sodium ion transmembrane transport | 22 | 1.79 | 0.000148 |
| Positive regulation of protein secretion | 22 | 1.79 | 0.000148 |
| Negative regulation of defense response | 36 | 2.93 | 0.00015 |
| Plasma lipoprotein particle organization | 12 | 0.98 | 0.000154 |
| Respiratory system development | 28 | 2.28 | 0.000161 |
| Bicarbonate transport | 10 | 0.81 | 0.00017 |
| Regulation of response to stress | 120 | 9.77 | 0.000171 |
| Monoatomic anion transport | 23 | 1.87 | 0.000175 |
| Cell-cell adhesion | 54 | 4.4 | 0.000183 |
| Negative regulation of transport | 48 | 3.91 | 0.000185 |
| Cell junction organization | 55 | 4.48 | 0.000185 |
| Positive regulation of lipid transport | 16 | 1.3 | 0.000192 |
| Regulation of viral process | 22 | 1.79 | 0.000195 |
| Maintenance of gastrointestinal epithelium | 8 | 0.65 | 0.000201 |
| Supramolecular fiber organization | 59 | 4.8 | 0.000206 |
| Positive regulation of biosynthetic process | 199 | 16.21 | 0.000209 |
| Protein-lipid complex remodeling | 10 | 0.81 | 0.000212 |
| Plasma lipoprotein particle remodeling | 10 | 0.81 | 0.000212 |
| Negative regulation of innate immune response | 17 | 1.38 | 0.000213 |
| Fatty acid catabolic process | 16 | 1.3 | 0.000218 |
| Secretion | 54 | 4.4 | 0.000218 |
| Lung alveolus development | 12 | 0.98 | 0.000218 |
| Positive regulation of macromolecule biosynthetic process | 191 | 15.55 | 0.000219 |
| Positive regulation of programmed cell death | 57 | 4.64 | 0.000223 |
| Apoptotic signaling pathway | 37 | 3.01 | 0.00023 |
| Positive regulation of lipid metabolic process | 20 | 1.63 | 0.000239 |
| Protein-lipid complex organization | 12 | 0.98 | 0.000257 |
| Negative regulation of small molecule metabolic process | 19 | 1.55 | 0.00026 |
| Negative regulation of response to biotic stimulus | 20 | 1.63 | 0.000289 |
| Regulation of leukocyte degranulation | 11 | 0.9 | 0.000295 |
| Cell junction assembly | 34 | 2.77 | 0.000297 |
| Iron ion transport | 12 | 0.98 | 0.000303 |

| | | | |
|---|---|---|---|
| Regulation of blood pressure | 26 | 2.12 | 0.00031 |
| Regulation of microvillus length | 5 | 0.41 | 0.000311 |
| Cardiac muscle cell-cardiac muscle cell adhesion | 5 | 0.41 | 0.000311 |
| Programmed cell death | 95 | 7.74 | 0.000319 |
| Protein-containing complex remodeling | 10 | 0.81 | 0.000321 |
| Cellular response to fatty acid | 10 | 0.81 | 0.000321 |
| Regulation of ventricular cardiac muscle cell action potential | 6 | 0.49 | 0.000326 |
| Negative regulation of protein localization | 28 | 2.28 | 0.000329 |
| Negative regulation of phosphate metabolic process | 31 | 2.52 | 0.00033 |
| Negative regulation of phosphorus metabolic process | 31 | 2.52 | 0.00033 |
| Cellular response to lipid | 53 | 4.32 | 0.000332 |
| Unsaturated fatty acid metabolic process | 17 | 1.38 | 0.000333 |
| Regulation of protein localization to plasma membrane | 18 | 1.47 | 0.000345 |
| Response to organophosphorus | 20 | 1.63 | 0.000348 |
| Phosphatidylcholine biosynthetic process | 8 | 0.65 | 0.000356 |
| Cell death | 95 | 7.74 | 0.000366 |
| Detoxification | 21 | 1.71 | 0.000372 |
| Positive regulation of immune system process | 89 | 7.25 | 0.000376 |
| Regulation of cellular component size | 38 | 3.09 | 0.000381 |
| Regulation of wound healing | 20 | 1.63 | 0.000381 |
| Monoatomic cation transport | 73 | 5.94 | 0.000385 |
| Isoprenoid metabolic process | 19 | 1.55 | 0.000385 |
| Negative regulation of protein localization to cell periphery | 9 | 0.73 | 0.0004 |
| Negative regulation of response to cytokine stimulus | 17 | 1.38 | 0.000412 |
| Animal organ morphogenesis | 84 | 6.84 | 0.000429 |
| Amine metabolic process | 16 | 1.3 | 0.000437 |
| Epidermis development | 37 | 3.01 | 0.000448 |
| Carboxylic acid transmembrane transport | 20 | 1.63 | 0.000456 |
| Cellular detoxification | 18 | 1.47 | 0.000465 |
| Regulation of phosphate metabolic process | 71 | 5.78 | 0.000494 |
| Inorganic anion transmembrane transport | 19 | 1.55 | 0.000509 |
| Carbohydrate catabolic process | 19 | 1.55 | 0.000509 |
| Regulation of phosphorus metabolic process | 71 | 5.78 | 0.000516 |
| Regulation of protein localization to cell periphery | 21 | 1.71 | 0.000523 |
| Vesicle-mediated transport | 106 | 8.63 | 0.000525 |
| Positive regulation of innate immune response | 34 | 2.77 | 0.000528 |
| Response to ketone | 27 | 2.2 | 0.000539 |
| Sulfur compound biosynthetic process | 21 | 1.71 | 0.000568 |
| Regulation of t cell mediated cytotoxicity | 11 | 0.9 | 0.000581 |
| Positive regulation of cell-cell adhesion | 36 | 2.93 | 0.000587 |
| Organic acid transmembrane transport | 20 | 1.63 | 0.000592 |
| Brush border assembly | 5 | 0.41 | 0.000594 |
| Positive regulation of apoptotic process | 54 | 4.4 | 0.000612 |
| Regulation of cellular component organization | 183 | 14.9 | 0.00063 |
| Regulation of anatomical structure size | 49 | 3.99 | 0.00063 |
| Cellular response to oxygen-containing compound | 92 | 7.49 | 0.000661 |
| Positive regulation of exocytosis | 14 | 1.14 | 0.000665 |

| | | | |
|---|---|---|---|
| Pyridine nucleotide metabolic process | 19 | 1.55 | 0.000667 |
| Nicotinamide nucleotide metabolic process | 19 | 1.55 | 0.000667 |
| Positive regulation of cell differentiation | 78 | 6.35 | 0.000668 |
| Hexose catabolic process | 11 | 0.9 | 0.00068 |
| Regulation of immune effector process | 42 | 3.42 | 0.000681 |
| Intracellular chemical homeostasis | 55 | 4.48 | 0.000716 |
| Regulation of response to wounding | 23 | 1.87 | 0.000743 |
| Negative regulation of cytokine-mediated signaling pathway | 16 | 1.3 | 0.000744 |
| Fatty acid biosynthetic process | 18 | 1.47 | 0.000746 |
| Response to wounding | 45 | 3.66 | 0.000749 |
| Bundle of his cell to purkinje myocyte communication | 6 | 0.49 | 0.000749 |
| Regulation of cell projection size | 6 | 0.49 | 0.000749 |
| Glandular epithelial cell differentiation | 9 | 0.73 | 0.00075 |
| Antigen processing and presentation of endogenous peptide antigen | 8 | 0.65 | 0.000756 |
| Protein autoprocessing | 8 | 0.65 | 0.000756 |
| Antigen processing and presentation of peptide antigen | 13 | 1.06 | 0.00076 |
| Monosaccharide biosynthetic process | 11 | 0.9 | 0.000793 |
| Phospholipid translocation | 11 | 0.9 | 0.000793 |
| Inflammatory response | 55 | 4.48 | 0.000803 |
| Cell migration | 79 | 6.43 | 0.000815 |
| Positive regulation of hormone secretion | 20 | 1.63 | 0.000828 |
| Triglyceride biosynthetic process | 7 | 0.57 | 0.000843 |
| Lipoprotein transport | 7 | 0.57 | 0.000843 |
| Defense response to bacterium | 36 | 2.93 | 0.000844 |
| Response to camp | 15 | 1.22 | 0.000889 |
| Regulation of viral life cycle | 18 | 1.47 | 0.000894 |
| Phospholipid transport | 16 | 1.3 | 0.00091 |
| Regulation of catalytic activity | 73 | 5.94 | 0.000956 |
| Carbohydrate transport | 12 | 0.98 | 0.000972 |
| Sodium ion transport | 24 | 1.95 | 0.000978 |
| Regulation of chemokine production | 16 | 1.3 | 0.001 |
| Renal system development | 34 | 2.77 | 0.00101 |
| Galactolipid metabolic process | 5 | 0.41 | 0.00102 |
| Regulation of response to cytokine stimulus | 24 | 1.95 | 0.00105 |
| Ethanol catabolic process | 6 | 0.49 | 0.00107 |
| Regulation of actin filament-based process | 40 | 3.26 | 0.00108 |
| Regulation of erbb signaling pathway | 13 | 1.06 | 0.0011 |
| Lipoprotein localization | 7 | 0.57 | 0.0011 |
| Actin filament-based movement | 16 | 1.3 | 0.00111 |
| Regulation of membrane lipid distribution | 12 | 0.98 | 0.00111 |
| Response to molecule of bacterial origin | 37 | 3.01 | 0.00111 |
| Gluconeogenesis | 10 | 0.81 | 0.00112 |
| Regulation of protein localization to membrane | 24 | 1.95 | 0.00112 |
| Olefinic compound metabolic process | 20 | 1.63 | 0.00114 |
| Export from cell | 47 | 3.83 | 0.00117 |
| Apoptotic process | 89 | 7.25 | 0.00119 |
| Regulation of fatty acid metabolic process | 16 | 1.3 | 0.00122 |

| | | | |
|---|---|---|---|
| Monosaccharide catabolic process | 11 | 0.9 | 0.00123 |
| Positive regulation of metabolic process | 251 | 20.44 | 0.00127 |
| Tissue homeostasis | 27 | 2.2 | 0.0013 |
| Anatomical structure homeostasis | 27 | 2.2 | 0.0013 |
| Pyridine-containing compound metabolic process | 19 | 1.55 | 0.00131 |
| Regulation of phospholipid metabolic process | 9 | 0.73 | 0.00131 |
| Response to nutrient | 21 | 1.71 | 0.00133 |
| Import across plasma membrane | 21 | 1.71 | 0.00133 |
| Positive regulation of response to biotic stimulus | 35 | 2.85 | 0.00134 |
| Positive regulation of immune effector process | 30 | 2.44 | 0.00134 |
| Epithelial cell development | 24 | 1.95 | 0.00136 |
| Cellular response to toxic substance | 18 | 1.47 | 0.00137 |
| Liver regeneration | 8 | 0.65 | 0.00145 |
| Negative regulation of protein localization to plasma membrane | 8 | 0.65 | 0.00145 |
| Positive regulation of lipid biosynthetic process | 15 | 1.22 | 0.00148 |
| Ethanol metabolic process | 6 | 0.49 | 0.00149 |
| Intracellular signal transduction | 133 | 10.83 | 0.00151 |
| Embryonic organ development | 44 | 3.58 | 0.00152 |
| Regulation of cell migration | 87 | 7.08 | 0.00152 |
| Positive regulation of dna-templated transcription | 125 | 10.18 | 0.00152 |
| Positive regulation of rna biosynthetic process | 125 | 10.18 | 0.00155 |
| Negative regulation of protein localization to membrane | 9 | 0.73 | 0.00156 |
| Wound healing | 35 | 2.85 | 0.00156 |
| Regulation of blood coagulation | 12 | 0.98 | 0.00161 |
| Lipid translocation | 11 | 0.9 | 0.00161 |
| Regulation of myeloid leukocyte mediated immunity | 11 | 0.9 | 0.00161 |
| Regulation of steroid biosynthetic process | 14 | 1.14 | 0.00161 |
| Cell proliferation involved in kidney development | 5 | 0.41 | 0.00163 |
| Cellular response to sterol depletion | 5 | 0.41 | 0.00163 |
| Regulation of intracellular protein transport | 20 | 1.63 | 0.00166 |
| Response to insulin | 29 | 2.36 | 0.00167 |
| Regulation of cell motility | 91 | 7.41 | 0.00172 |
| Antigen processing and presentation | 16 | 1.3 | 0.00176 |
| Cellular homeostasis | 61 | 4.97 | 0.00176 |
| Hexose biosynthetic process | 10 | 0.81 | 0.00177 |
| Positive regulation of insulin secretion | 14 | 1.14 | 0.00179 |
| Immune response | 134 | 10.91 | 0.00179 |
| Negative regulation of keratinocyte proliferation | 7 | 0.57 | 0.00181 |
| Aldehyde metabolic process | 12 | 0.98 | 0.00182 |
| Glycerophospholipid biosynthetic process | 23 | 1.87 | 0.00182 |
| Positive regulation of dna-binding transcription factor activity | 23 | 1.87 | 0.00182 |
| Carbohydrate transmembrane transport | 11 | 0.9 | 0.00184 |
| Positive regulation of intracellular signal transduction | 92 | 7.49 | 0.00188 |
| Monoatomic anion transmembrane transport | 19 | 1.55 | 0.00193 |
| Leukocyte differentiation | 41 | 3.34 | 0.00193 |
| Vascular process in circulatory system | 30 | 2.44 | 0.00195 |
| Regulation of hydrolase activity | 33 | 2.69 | 0.00196 |

| | | | |
|---|---|---|---|
| Purine ribonucleoside bisphosphate metabolic process | 6 | 0.49 | 0.00201 |
| Phospholipid homeostasis | 6 | 0.49 | 0.00201 |
| 3'-phosphoadenosine 5'-phosphosulfate metabolic process | 6 | 0.49 | 0.00201 |
| Cytokine production | 6 | 0.49 | 0.00201 |
| Regulation of hemostasis | 12 | 0.98 | 0.00204 |
| Regulation of small gtpase mediated signal transduction | 32 | 2.61 | 0.00209 |
| Positive regulation of t cell mediated cytotoxicity | 9 | 0.73 | 0.00217 |
| Regulation of heart rate by cardiac conduction | 9 | 0.73 | 0.00217 |
| Cell communication involved in cardiac conduction | 9 | 0.73 | 0.00217 |
| Antigen processing and presentation of peptide antigen via mhc class i | 9 | 0.73 | 0.00217 |
| Carbohydrate biosynthetic process | 17 | 1.38 | 0.00218 |
| Kidney development | 32 | 2.61 | 0.00219 |
| Regulation of actin polymerization or depolymerization | 19 | 1.55 | 0.00224 |
| Carbohydrate derivative catabolic process | 24 | 1.95 | 0.00226 |
| Positive regulation of leukocyte degranulation | 7 | 0.57 | 0.00227 |
| Embryo development | 84 | 6.84 | 0.00227 |
| Response to peptide hormone | 41 | 3.34 | 0.00236 |
| Mammary gland epithelium development | 11 | 0.9 | 0.00237 |
| Hemopoiesis | 60 | 4.89 | 0.00239 |
| Epidermal cell differentiation | 24 | 1.95 | 0.0024 |
| Positive regulation of cytokine production | 47 | 3.83 | 0.00242 |
| Regulation of cytokine-mediated signaling pathway | 22 | 1.79 | 0.00243 |
| Choline transport | 5 | 0.41 | 0.00244 |
| Regulation of locomotion | 93 | 7.57 | 0.00248 |
| Positive regulation of peptide hormone secretion | 16 | 1.3 | 0.00249 |
| Regulation of digestive system process | 9 | 0.73 | 0.00253 |
| Pyroptotic inflammatory response | 8 | 0.65 | 0.00255 |
| Negative regulation of protein transport | 17 | 1.38 | 0.00256 |
| Tube morphogenesis | 60 | 4.89 | 0.00257 |
| Regulation of leukocyte mediated cytotoxicity | 15 | 1.22 | 0.00259 |
| Cytosolic transport | 18 | 1.47 | 0.00259 |
| Purine nucleoside bisphosphate metabolic process | 6 | 0.49 | 0.00265 |
| Nucleoside bisphosphate metabolic process | 6 | 0.49 | 0.00265 |
| Ribonucleoside bisphosphate metabolic process | 6 | 0.49 | 0.00265 |
| Negative regulation of cell adhesion | 32 | 2.61 | 0.00267 |
| Cell adhesion | 78 | 6.35 | 0.00271 |
| Regulation of actin filament length | 19 | 1.55 | 0.00278 |
| Antigen processing and presentation of endogenous peptide antigen via mhc class i | 7 | 0.57 | 0.00281 |
| Positive regulation of cell population proliferation | 80 | 6.51 | 0.00286 |
| Regulation of exocytosis | 22 | 1.79 | 0.00294 |
| Positive regulation of peptide secretion | 16 | 1.3 | 0.00294 |
| Alcohol catabolic process | 9 | 0.73 | 0.00294 |
| Signaling receptor ligand precursor processing | 9 | 0.73 | 0.00294 |
| Response to cytokine | 73 | 5.94 | 0.00299 |
| Regulation of regulated secretory pathway | 17 | 1.38 | 0.003 |
| Negative regulation of protein phosphorylation | 23 | 1.87 | 0.003 |

| | | | |
|---|---|---|---|
| Regulation of intracellular signal transduction | 149 | 12.13 | 0.00302 |
| Negative regulation of response to stimulus | 135 | 10.99 | 0.00303 |
| Response to metal ion | 34 | 2.77 | 0.00304 |
| Antigen processing and presentation of endogenous antigen | 8 | 0.65 | 0.00304 |
| Negative regulation of erbb signaling pathway | 8 | 0.65 | 0.00304 |
| Positive regulation of insulin secretion involved in cellular response to glucose stimulus | 8 | 0.65 | 0.00304 |
| Neutral amino acid transport | 10 | 0.81 | 0.00307 |
| Positive regulation of alpha-beta t cell activation | 12 | 0.98 | 0.00318 |
| Anterior/posterior pattern specification | 24 | 1.95 | 0.00321 |
| Cellular response to cytokine stimulus | 66 | 5.37 | 0.0033 |
| Peptide metabolic process | 11 | 0.9 | 0.00338 |
| Actin polymerization or depolymerization | 11 | 0.9 | 0.00338 |
| Regulation of cholesterol storage | 6 | 0.49 | 0.00344 |
| Regulation of systemic arterial blood pressure by renin-angiotensin | 7 | 0.57 | 0.00344 |
| Negative regulation of establishment of protein localization | 18 | 1.47 | 0.00348 |
| Dicarboxylic acid metabolic process | 14 | 1.14 | 0.00349 |
| Polyol metabolic process | 14 | 1.14 | 0.00349 |
| Myelination | 17 | 1.38 | 0.00349 |
| Glyceraldehyde-3-phosphate metabolic process | 5 | 0.41 | 0.0035 |
| Response to sterol depletion | 5 | 0.41 | 0.0035 |
| Self proteolysis | 5 | 0.41 | 0.0035 |
| Glycosyl compound metabolic process | 13 | 1.06 | 0.00355 |
| Vascular transport | 13 | 1.06 | 0.00355 |
| Transport across blood-brain barrier | 13 | 1.06 | 0.00355 |
| Morphogenesis of a branching epithelium | 20 | 1.63 | 0.00357 |
| Regulation of macrophage derived foam cell differentiation | 8 | 0.65 | 0.00359 |
| Monoatomic ion homeostasis | 47 | 3.83 | 0.0036 |
| Regulation of t cell mediated immunity | 15 | 1.22 | 0.00365 |
| Response to carbohydrate | 22 | 1.79 | 0.00375 |
| Endocrine system development | 17 | 1.38 | 0.00376 |
| Myeloid cell differentiation | 31 | 2.52 | 0.00378 |
| Positive regulation of interleukin-1 beta production | 11 | 0.9 | 0.00379 |
| Lipid modification | 20 | 1.63 | 0.00381 |
| Response to monosaccharide | 20 | 1.63 | 0.00381 |
| Retinoid metabolic process | 13 | 1.06 | 0.00391 |
| Carbohydrate derivative transport | 13 | 1.06 | 0.00391 |
| Regulation of coagulation | 12 | 0.98 | 0.00392 |
| Negative regulation of blood coagulation | 9 | 0.73 | 0.00392 |
| Response to starvation | 25 | 2.04 | 0.00401 |
| Ensheathment of neurons | 17 | 1.38 | 0.00405 |
| Axon ensheathment | 17 | 1.38 | 0.00405 |
| Mononuclear cell differentiation | 35 | 2.85 | 0.0041 |
| Regulation of protein-containing complex assembly | 40 | 3.26 | 0.0041 |
| Regulation of carbohydrate metabolic process | 21 | 1.71 | 0.00416 |
| Negative regulation of natural killer cell mediated cytotoxicity | 7 | 0.57 | 0.00417 |
| Establishment of skin barrier | 8 | 0.65 | 0.00422 |

| | | | |
|---|---|---|---|
| Animal organ regeneration | 11 | 0.9 | 0.00424 |
| Homotypic cell-cell adhesion | 11 | 0.9 | 0.00424 |
| Regulation of lipid catabolic process | 11 | 0.9 | 0.00424 |
| Sulfur compound transport | 11 | 0.9 | 0.00424 |
| Amino acid biosynthetic process | 12 | 0.98 | 0.00433 |
| Monoatomic cation transmembrane transport | 58 | 4.72 | 0.00434 |
| Antigen processing and presentation of peptide antigen via mhc class ib | 6 | 0.49 | 0.00437 |
| Epoxygenase p450 pathway | 6 | 0.49 | 0.00437 |
| Phosphate ion transport | 6 | 0.49 | 0.00437 |
| Positive regulation of macromolecule metabolic process | 226 | 18.4 | 0.0045 |
| Negative regulation of hemostasis | 9 | 0.73 | 0.0045 |
| Response to peptide | 73 | 5.94 | 0.00452 |
| Response to abiotic stimulus | 89 | 7.25 | 0.00454 |
| Cellular response to endogenous stimulus | 77 | 6.27 | 0.00456 |
| Negative regulation of multicellular organismal process | 95 | 7.74 | 0.00459 |
| Response to lipopolysaccharide | 33 | 2.69 | 0.00464 |
| Regulation of systemic arterial blood pressure | 15 | 1.22 | 0.00466 |
| Regulation of interleukin-1 production | 17 | 1.38 | 0.00468 |
| Biogenic amine metabolic process | 13 | 1.06 | 0.0047 |
| Pyruvate metabolic process | 12 | 0.98 | 0.00478 |
| Epithelial fluid transport | 5 | 0.41 | 0.00483 |
| Very-low-density lipoprotein particle assembly | 5 | 0.41 | 0.00483 |
| Phosphate ion transmembrane transport | 5 | 0.41 | 0.00483 |
| Negative regulation of homotypic cell-cell adhesion | 5 | 0.41 | 0.00483 |
| Cell activation | 63 | 5.13 | 0.00491 |
| Acylglycerol biosynthetic process | 7 | 0.57 | 0.00501 |
| Xenobiotic catabolic process | 7 | 0.57 | 0.00501 |
| Negative regulation of natural killer cell mediated immunity | 7 | 0.57 | 0.00501 |
| Neutral lipid biosynthetic process | 7 | 0.57 | 0.00501 |
| Monocarboxylic acid biosynthetic process | 20 | 1.63 | 0.00521 |
| Cardiac conduction | 11 | 0.9 | 0.00525 |
| Regulation of epidermal growth factor receptor signaling pathway | 11 | 0.9 | 0.00525 |
| Positive regulation of developmental process | 105 | 8.55 | 0.00527 |
| Purine nucleotide metabolic process | 36 | 2.93 | 0.00532 |
| Regulation of phospholipid catabolic process | 4 | 0.33 | 0.00533 |
| Cell proliferation involved in metanephros development | 4 | 0.33 | 0.00533 |
| Desmosome assembly | 4 | 0.33 | 0.00533 |
| Regulation of actin filament organization | 28 | 2.28 | 0.00543 |
| Regulation of interleukin-1 beta production | 15 | 1.22 | 0.00545 |
| Regulation of epithelial cell proliferation | 38 | 3.09 | 0.0055 |
| Intracellular monoatomic ion homeostasis | 40 | 3.26 | 0.00554 |
| Embryonic morphogenesis | 51 | 4.15 | 0.00562 |
| Thioester metabolic process | 13 | 1.06 | 0.00562 |
| Acyl-coa metabolic process | 13 | 1.06 | 0.00562 |
| Regulation of production of molecular mediator of immune response | 22 | 1.79 | 0.00562 |
| Lung epithelium development | 8 | 0.65 | 0.00571 |
| Regulation of homotypic cell-cell adhesion | 8 | 0.65 | 0.00571 |

| | | | |
|---|---|---|---|
| Regulation of leukocyte mediated immunity | 27 | 2.2 | 0.00584 |
| Regulation of cell killing | 15 | 1.22 | 0.00589 |
| Positive regulation of gene expression | 94 | 7.65 | 0.00596 |
| Regulation of platelet aggregation | 7 | 0.57 | 0.00597 |
| Diterpenoid metabolic process | 13 | 1.06 | 0.00613 |
| Nitrogen compound transport | 114 | 9.28 | 0.00619 |
| Sensory perception of mechanical stimulus | 22 | 1.79 | 0.00628 |
| Regulation of cytokine production involved in immune response | 16 | 1.3 | 0.0063 |
| Post-golgi vesicle-mediated transport | 14 | 1.14 | 0.0063 |
| Regulation of viral genome replication | 12 | 0.98 | 0.00636 |
| Monoatomic cation homeostasis | 45 | 3.66 | 0.00639 |
| Regulation of growth | 52 | 4.23 | 0.00641 |
| Regulation of intestinal absorption | 5 | 0.41 | 0.00647 |
| Negative regulation of immune response | 23 | 1.87 | 0.00657 |
| Inorganic cation transmembrane transport | 56 | 4.56 | 0.00659 |
| Regulation of systemic arterial blood pressure mediated by a chemical signal | 9 | 0.73 | 0.00661 |
| Regulation of t cell proliferation | 21 | 1.71 | 0.00665 |
| Amine catabolic process | 6 | 0.49 | 0.00676 |
| Biogenic amine catabolic process | 6 | 0.49 | 0.00676 |
| Triglyceride catabolic process | 6 | 0.49 | 0.00676 |
| Negative regulation of protein modification process | 30 | 2.44 | 0.00696 |
| Negative regulation of secretion | 20 | 1.63 | 0.00702 |
| Morphogenesis of a branching structure | 20 | 1.63 | 0.00702 |
| Negative regulation of peptidyl-tyrosine phosphorylation | 7 | 0.57 | 0.00705 |
| Amide biosynthetic process | 18 | 1.47 | 0.00726 |
| Negative regulation of phosphorylation | 23 | 1.87 | 0.00729 |
| Regulation of anatomical structure morphogenesis | 71 | 5.78 | 0.00736 |
| Antigen processing and presentation of exogenous antigen | 9 | 0.73 | 0.00746 |
| Negative regulation of coagulation | 9 | 0.73 | 0.00746 |
| Negative regulation of viral genome replication | 9 | 0.73 | 0.00746 |
| Regulation of systemic arterial blood pressure by hormone | 8 | 0.65 | 0.00757 |
| Multicellular organismal-level chemical homeostasis | 12 | 0.98 | 0.00761 |
| Regulation of actin filament polymerization | 16 | 1.3 | 0.00776 |
| Cellular response to hormone stimulus | 48 | 3.91 | 0.00779 |
| Regulation of alcohol biosynthetic process | 10 | 0.81 | 0.00782 |
| Positive regulation of leukocyte mediated cytotoxicity | 11 | 0.9 | 0.00783 |
| Epithelial cell proliferation | 20 | 1.63 | 0.00787 |
| Inorganic ion import across plasma membrane | 14 | 1.14 | 0.00796 |
| Inorganic cation import across plasma membrane | 14 | 1.14 | 0.00796 |
| Galactosylceramide metabolic process | 4 | 0.33 | 0.00817 |
| Cellular response to tumor necrosis factor | 18 | 1.47 | 0.00821 |
| Negative regulation of tissue remodeling | 6 | 0.49 | 0.00824 |
| Response to tumor necrosis factor | 20 | 1.63 | 0.00832 |
| Positive regulation by symbiont of entry into host | 5 | 0.41 | 0.00843 |
| Positive regulation of viral entry into host cell | 5 | 0.41 | 0.00843 |
| Regulation of phagocytosis | 14 | 1.14 | 0.00859 |
| Positive regulation of chemokine production | 11 | 0.9 | 0.00861 |

| | | | |
|---|---|---|---|
| Antigen processing and presentation of exogenous peptide antigen | 8 | 0.65 | 0.00865 |
| Estrogen metabolic process | 8 | 0.65 | 0.00865 |
| Signal transduction | 321 | 26.14 | 0.00865 |
| Positive regulation of small molecule metabolic process | 18 | 1.47 | 0.00873 |
| Regulation of cellular component biogenesis | 80 | 6.51 | 0.00912 |
| Positive regulation of production of molecular mediator of immune response | 17 | 1.38 | 0.00912 |
| Fructose catabolic process to hydroxyacetone phosphate and glyceraldehyde-3-phosphate | 3 | 0.24 | 0.00924 |
| Lung goblet cell differentiation | 3 | 0.24 | 0.00924 |
| Sulfide oxidation, using sulfide:quinone oxidoreductase | 3 | 0.24 | 0.00924 |
| Fructose catabolic process | 3 | 0.24 | 0.00924 |
| Sulfide oxidation | 3 | 0.24 | 0.00924 |
| Negative regulation of wound healing | 11 | 0.9 | 0.00944 |
| Cell motility | 88 | 7.17 | 0.00953 |
| Regulation of programmed cell death | 117 | 9.53 | 0.00961 |
| Negative regulation of chemokine production | 7 | 0.57 | 0.00964 |
| Macromolecule localization | 168 | 13.68 | 0.00975 |
| Response to estradiol | 15 | 1.22 | 0.00976 |
| Cellular response to nutrient levels | 25 | 2.04 | 0.00983 |
| Acyl-coa biosynthetic process | 8 | 0.65 | 0.00984 |
| Thioester biosynthetic process | 8 | 0.65 | 0.00984 |
| Regulation of apoptotic process | 114 | 9.28 | 0.00988 |
| Relaxation of muscle | 6 | 0.49 | 0.00993 |
| Plasma lipoprotein particle clearance | 6 | 0.49 | 0.00993 |
| Response to thyroid hormone | 6 | 0.49 | 0.00993 |
| Regulation of actin cytoskeleton organization | 33 | 2.69 | 0.01 |
| Negative regulation of cell-cell adhesion | 22 | 1.79 | 0.0101 |
| Cell surface receptor signaling pathway | 149 | 12.13 | 0.0101 |
| Positive regulation of cell killing | 11 | 0.9 | 0.0103 |
| Positive regulation of t cell mediated immunity | 11 | 0.9 | 0.0103 |
| Positive regulation of catalytic activity | 43 | 3.5 | 0.0105 |
| Regulation of actin filament depolymerization | 9 | 0.73 | 0.0105 |
| Retinol metabolic process | 9 | 0.73 | 0.0105 |
| Positive regulation of cytokine production involved in immune response | 12 | 0.98 | 0.0107 |
| Monoacylglycerol metabolic process | 5 | 0.41 | 0.0107 |
| Plasma membrane organization | 18 | 1.47 | 0.011 |
| Response to hexose | 18 | 1.47 | 0.011 |
| Regulation of leukocyte migration | 24 | 1.95 | 0.0111 |
| Intracellular monoatomic cation homeostasis | 38 | 3.09 | 0.0111 |
| Neutral lipid catabolic process | 7 | 0.57 | 0.0112 |
| Acylglycerol catabolic process | 7 | 0.57 | 0.0112 |
| Negative regulation of leukocyte mediated cytotoxicity | 7 | 0.57 | 0.0112 |
| Regulation of nucleocytoplasmic transport | 15 | 1.22 | 0.0112 |
| Regulation of alpha-beta t cell activation | 15 | 1.22 | 0.0112 |
| Nucleotide metabolic process | 39 | 3.18 | 0.0112 |
| Lipoprotein metabolic process | 11 | 0.9 | 0.0113 |
| Positive regulation of cell motility | 52 | 4.23 | 0.0114 |

| | | | |
|---|---|---|---|
| Positive regulation of cell migration | 50 | 4.07 | 0.0114 |
| Positive regulation of locomotion | 53 | 4.32 | 0.0116 |
| Negative regulation of epithelial cell proliferation | 19 | 1.55 | 0.0116 |
| Adherens junction organization | 9 | 0.73 | 0.0117 |
| Response to fructose | 4 | 0.33 | 0.0117 |
| Srebp signaling pathway | 4 | 0.33 | 0.0117 |
| Carbohydrate phosphorylation | 6 | 0.49 | 0.0118 |
| Plasma lipoprotein particle assembly | 6 | 0.49 | 0.0118 |
| Long-chain fatty-acyl-coa metabolic process | 6 | 0.49 | 0.0118 |
| Positive regulation of alpha-beta t cell proliferation | 6 | 0.49 | 0.0118 |
| Embryonic organ morphogenesis | 29 | 2.36 | 0.0119 |
| Actomyosin structure organization | 15 | 1.22 | 0.0119 |
| Response to type ii interferon | 14 | 1.14 | 0.0123 |
| Long-chain fatty acid metabolic process | 14 | 1.14 | 0.0123 |
| Regulation of sterol transport | 11 | 0.9 | 0.0123 |
| Regulation of cholesterol transport | 11 | 0.9 | 0.0123 |
| Regulation of nucleotide metabolic process | 12 | 0.98 | 0.0126 |
| Cellular response to type ii interferon | 12 | 0.98 | 0.0126 |
| Regulation of triglyceride metabolic process | 8 | 0.65 | 0.0126 |
| Regulation of alpha-beta t cell proliferation | 8 | 0.65 | 0.0126 |
| Modified amino acid biosynthetic process | 8 | 0.65 | 0.0126 |
| Establishment of epithelial cell polarity | 7 | 0.57 | 0.0128 |
| Positive regulation of alpha-beta t cell differentiation | 9 | 0.73 | 0.013 |
| Myeloid leukocyte differentiation | 18 | 1.47 | 0.0131 |
| Response to mechanical stimulus | 23 | 1.87 | 0.0131 |
| Positive regulation of interleukin-1 production | 11 | 0.9 | 0.0134 |
| Lipid digestion | 5 | 0.41 | 0.0134 |
| Sulfate transmembrane transport | 5 | 0.41 | 0.0134 |
| Positive regulation of phospholipid metabolic process | 5 | 0.41 | 0.0134 |
| Epithelial cell-cell adhesion | 5 | 0.41 | 0.0134 |
| Mammary gland epithelial cell differentiation | 5 | 0.41 | 0.0134 |
| Antigen processing and presentation of endogenous peptide antigen via mhc class i via er pathway, tap-independent | 5 | 0.41 | 0.0134 |
| Regulation of system process | 48 | 3.91 | 0.0134 |
| Cellular response to starvation | 20 | 1.63 | 0.0135 |
| Response to hydrogen peroxide | 13 | 1.06 | 0.0135 |
| Negative regulation of immune system process | 47 | 3.83 | 0.0136 |
| Cell communication | 341 | 27.77 | 0.0137 |
| Defense response to gram-positive bacterium | 16 | 1.3 | 0.0138 |
| Regulation of dna-binding transcription factor activity | 30 | 2.44 | 0.0139 |
| Glycosyl compound catabolic process | 6 | 0.49 | 0.014 |
| Intestinal epithelial cell differentiation | 6 | 0.49 | 0.014 |
| Lung cell differentiation | 6 | 0.49 | 0.014 |
| Lung epithelial cell differentiation | 6 | 0.49 | 0.014 |
| Positive regulation of cellular response to insulin stimulus | 6 | 0.49 | 0.014 |
| Cellular component maintenance | 10 | 0.81 | 0.0141 |
| Cholesterol transport | 10 | 0.81 | 0.0141 |

| | | | |
|---|---|---|---|
| Cellular response to hydrogen peroxide | 10 | 0.81 | 0.0141 |
| Nad metabolic process | 8 | 0.65 | 0.0141 |
| Triglyceride homeostasis | 8 | 0.65 | 0.0141 |
| Cellular response to lipoprotein particle stimulus | 8 | 0.65 | 0.0141 |
| Endocrine process | 9 | 0.73 | 0.0144 |
| Activation of innate immune response | 19 | 1.55 | 0.0144 |
| Sterol transport | 11 | 0.9 | 0.0146 |
| Regulation of sodium ion transport | 11 | 0.9 | 0.0146 |
| Regulation of fatty acid oxidation | 7 | 0.57 | 0.0147 |
| Cellular response to peptide hormone stimulus | 28 | 2.28 | 0.0149 |
| Skeletal system development | 43 | 3.5 | 0.015 |
| Negative regulation of lipid metabolic process | 14 | 1.14 | 0.0151 |
| Extrinsic apoptotic signaling pathway | 14 | 1.14 | 0.0151 |
| Anatomical structure formation involved in morphogenesis | 75 | 6.11 | 0.0152 |
| Alpha-amino acid biosynthetic process | 10 | 0.81 | 0.0154 |
| Hematopoietic or lymphoid organ development | 13 | 1.06 | 0.0155 |
| Columnar/cuboidal epithelial cell differentiation | 13 | 1.06 | 0.0155 |
| Acylglycerol homeostasis | 8 | 0.65 | 0.0158 |
| Positive regulation of regulated secretory pathway | 8 | 0.65 | 0.0158 |
| Regulation of cellular response to insulin stimulus | 11 | 0.9 | 0.0158 |
| Regulation of tissue remodeling | 12 | 0.98 | 0.0158 |
| Cellular response to chemical stress | 27 | 2.2 | 0.0158 |
| Tight junction assembly | 9 | 0.73 | 0.0159 |
| Zinc ion import across plasma membrane | 4 | 0.33 | 0.0161 |
| Osmosensory signaling pathway | 4 | 0.33 | 0.0161 |
| Lung secretory cell differentiation | 4 | 0.33 | 0.0161 |
| Positive regulation of interleukin-18 production | 4 | 0.33 | 0.0161 |
| Regulation of glucocorticoid biosynthetic process | 4 | 0.33 | 0.0161 |
| Positive regulation of adipose tissue development | 4 | 0.33 | 0.0161 |
| Positive regulation of phosphate metabolic process | 41 | 3.34 | 0.0163 |
| Positive regulation of phosphorus metabolic process | 41 | 3.34 | 0.0163 |
| Regulation of cell differentiation | 117 | 9.53 | 0.0164 |
| Keratinocyte proliferation | 6 | 0.49 | 0.0164 |
| Antigen processing and presentation via mhc class ib | 6 | 0.49 | 0.0164 |
| Epithelial tube branching involved in lung morphogenesis | 6 | 0.49 | 0.0164 |
| Fatty acid homeostasis | 5 | 0.41 | 0.0165 |
| Negative regulation of platelet activation | 5 | 0.41 | 0.0165 |
| Regulation of hepatocyte proliferation | 5 | 0.41 | 0.0165 |
| Negative regulation of cell killing | 7 | 0.57 | 0.0167 |
| Regulation of tumor necrosis factor-mediated signaling pathway | 10 | 0.81 | 0.0168 |
| Regulation of endocytosis | 29 | 2.36 | 0.0168 |
| Nucleoside diphosphate metabolic process | 12 | 0.98 | 0.017 |
| Signaling | 337 | 27.44 | 0.017 |
| Regulation of cardiac muscle contraction | 11 | 0.9 | 0.0171 |
| Response to nitrogen compound | 80 | 6.51 | 0.0173 |
| Response to glucose | 17 | 1.38 | 0.0174 |
| Renal system process | 15 | 1.22 | 0.0174 |

| | | | |
|---|---|---|---|
| Hair cell differentiation | 9 | 0.73 | 0.0175 |
| Immunoglobulin transcytosis in epithelial cells | 3 | 0.24 | 0.0178 |
| Ethanolamine transport | 3 | 0.24 | 0.0178 |
| Intestinal d-glucose absorption | 3 | 0.24 | 0.0178 |
| Lobar bronchus epithelium development | 3 | 0.24 | 0.0178 |
| Neuropeptide catabolic process | 3 | 0.24 | 0.0178 |
| Negative regulation of very-low-density lipoprotein particle remodeling | 3 | 0.24 | 0.0178 |
| Secretion by cell | 37 | 3.01 | 0.0178 |
| Positive regulation of nf-kappab transcription factor activity | 14 | 1.14 | 0.0183 |
| Leukocyte activation | 51 | 4.15 | 0.0184 |
| Pattern recognition receptor signaling pathway | 15 | 1.22 | 0.0185 |
| Epithelial cell maturation | 6 | 0.49 | 0.0191 |
| Protein-lipid complex assembly | 6 | 0.49 | 0.0191 |
| Apical junction assembly | 9 | 0.73 | 0.0192 |
| Regulation of d-glucose import | 9 | 0.73 | 0.0192 |
| Notch signaling pathway | 14 | 1.14 | 0.0195 |
| Fatty acid transport | 12 | 0.98 | 0.0197 |
| Negative regulation of cell population proliferation | 61 | 4.97 | 0.0198 |
| Regulation of protein phosphorylation | 50 | 4.07 | 0.0199 |
| Regulation of animal organ morphogenesis | 11 | 0.9 | 0.02 |
| Antigen processing and presentation of endogenous peptide antigen via mhc class ib | 5 | 0.41 | 0.02 |
| Cellular response to aldehyde | 5 | 0.41 | 0.02 |
| Nucleoside catabolic process | 5 | 0.41 | 0.02 |
| Regulation of systemic arterial blood pressure by circulatory renin-angiotensin | 5 | 0.41 | 0.02 |
| Antigen processing and presentation of endogenous peptide antigen via mhc class i via er pathway | 5 | 0.41 | 0.02 |
| Tissue morphogenesis | 47 | 3.83 | 0.0202 |
| Glucose homeostasis | 22 | 1.79 | 0.0203 |
| Positive regulation of t cell proliferation | 13 | 1.06 | 0.0203 |
| Extracellular matrix organization | 26 | 2.12 | 0.0209 |
| Positive regulation of mirna transcription | 9 | 0.73 | 0.0211 |
| Tight junction organization | 9 | 0.73 | 0.0211 |
| Glutathione metabolic process | 9 | 0.73 | 0.0211 |
| Regulation of blood circulation | 25 | 2.04 | 0.0211 |
| Carbohydrate homeostasis | 22 | 1.79 | 0.0211 |
| Negative regulation of platelet aggregation | 4 | 0.33 | 0.0212 |
| Fluid transport | 7 | 0.57 | 0.0213 |
| Actin filament polymerization | 7 | 0.57 | 0.0213 |
| Negative regulation of actin filament depolymerization | 7 | 0.57 | 0.0213 |
| Endocrine pancreas development | 7 | 0.57 | 0.0213 |
| Phosphatidic acid metabolic process | 7 | 0.57 | 0.0213 |
| Antigen processing and presentation of peptide or polysaccharide antigen via mhc class ii | 7 | 0.57 | 0.0213 |
| Myeloid leukocyte activation | 17 | 1.38 | 0.0215 |
| Fatty acid oxidation | 11 | 0.9 | 0.0215 |
| Schwann cell differentiation | 8 | 0.65 | 0.0217 |
| Extracellular structure organization | 26 | 2.12 | 0.0218 |

| | | | |
|---|---|---|---|
| Regulation of heart contraction | 21 | 1.71 | 0.022 |
| Acetyl-coa metabolic process | 6 | 0.49 | 0.022 |
| Embryonic skeletal system development | 15 | 1.22 | 0.022 |
| Cell development | 156 | 12.7 | 0.0221 |
| Regulation of smooth muscle cell proliferation | 18 | 1.47 | 0.0221 |
| Regulation of phosphorylation | 53 | 4.32 | 0.0225 |
| External encapsulating structure organization | 26 | 2.12 | 0.0226 |
| Response to alcohol | 26 | 2.12 | 0.0226 |
| Positive regulation of phosphatidylinositol 3-kinase/protein kinase b signal transduction | 20 | 1.63 | 0.0228 |
| Positive regulation of canonical nf-kappab signal transduction | 22 | 1.79 | 0.023 |
| Export across plasma membrane | 9 | 0.73 | 0.023 |
| Nucleoside metabolic process | 9 | 0.73 | 0.023 |
| Regulation of heart rate | 13 | 1.06 | 0.0231 |
| Gland morphogenesis | 13 | 1.06 | 0.0231 |
| Protein localization to extracellular region | 15 | 1.22 | 0.0233 |
| Chloride transport | 14 | 1.14 | 0.0233 |
| Positive regulation of hydrolase activity | 19 | 1.55 | 0.0236 |
| Blood vessel development | 44 | 3.58 | 0.0237 |
| Cellular localization | 182 | 14.82 | 0.0239 |
| Cell-cell junction maintenance | 5 | 0.41 | 0.0239 |
| Protein localization to cell-cell junction | 5 | 0.41 | 0.0239 |
| Glycerol metabolic process | 5 | 0.41 | 0.0239 |
| Barbed-end actin filament capping | 5 | 0.41 | 0.0239 |
| Peptide catabolic process | 5 | 0.41 | 0.0239 |
| Regulation of hormone metabolic process | 7 | 0.57 | 0.0239 |
| O-glycan processing | 7 | 0.57 | 0.0239 |
| Peptide hormone processing | 7 | 0.57 | 0.0239 |
| Regulation of t cell cytokine production | 7 | 0.57 | 0.0239 |
| Bicellular tight junction assembly | 8 | 0.65 | 0.024 |
| Positive regulation of intracellular protein transport | 12 | 0.98 | 0.0241 |
| Ribonucleoside diphosphate metabolic process | 11 | 0.9 | 0.0249 |
| Enzyme-linked receptor protein signaling pathway | 50 | 4.07 | 0.025 |
| Regulation of insulin secretion involved in cellular response to glucose stimulus | 9 | 0.73 | 0.0251 |
| Arachidonate metabolic process | 9 | 0.73 | 0.0251 |
| Positive regulation of actin filament bundle assembly | 9 | 0.73 | 0.0251 |
| Negative regulation of nucleotide metabolic process | 6 | 0.49 | 0.0252 |
| Regulation of mast cell degranulation | 6 | 0.49 | 0.0252 |
| Negative regulation of epidermal growth factor receptor signaling pathway | 6 | 0.49 | 0.0252 |
| Positive regulation of leukocyte cell-cell adhesion | 26 | 2.12 | 0.0253 |
| Regulation of extracellular matrix organization | 10 | 0.81 | 0.0253 |
| Positive regulation of rna metabolic process | 129 | 10.5 | 0.0254 |
| Innate immune response-activating signaling pathway | 16 | 1.3 | 0.0256 |
| Negative regulation of response to wounding | 12 | 0.98 | 0.0258 |
| Chloride transmembrane transport | 13 | 1.06 | 0.0262 |
| Regulation of keratinocyte proliferation | 8 | 0.65 | 0.0264 |
| Prostaglandin metabolic process | 7 | 0.57 | 0.0267 |

| | | | |
|---|---|---|---|
| Regulation of mononuclear cell migration | 16 | 1.3 | 0.0269 |
| Regulation of intestinal lipid absorption | 4 | 0.33 | 0.0271 |
| Hormone catabolic process | 4 | 0.33 | 0.0271 |
| Regulation of glucocorticoid metabolic process | 4 | 0.33 | 0.0271 |
| Negative regulation of receptor binding | 4 | 0.33 | 0.0271 |
| Inorganic ion homeostasis | 37 | 3.01 | 0.0271 |
| Vasculature development | 45 | 3.66 | 0.0273 |
| Purine ribonucleoside diphosphate metabolic process | 10 | 0.81 | 0.0273 |
| Body fluid secretion | 10 | 0.81 | 0.0273 |
| Purine nucleoside diphosphate metabolic process | 10 | 0.81 | 0.0273 |
| Glycosphingolipid metabolic process | 9 | 0.73 | 0.0273 |
| Positive regulation of cold-induced thermogenesis | 12 | 0.98 | 0.0275 |
| Negative regulation of signaling | 109 | 8.88 | 0.0276 |
| Homeostasis of number of cells | 27 | 2.2 | 0.0276 |
| Sulfation | 5 | 0.41 | 0.0282 |
| Surfactant homeostasis | 5 | 0.41 | 0.0282 |
| Regulation of phospholipid biosynthetic process | 5 | 0.41 | 0.0282 |
| Regulation of chemokine (c-x-c motif) ligand 2 production | 5 | 0.41 | 0.0282 |
| Positive regulation of superoxide anion generation | 5 | 0.41 | 0.0282 |
| High-density lipoprotein particle remodeling | 5 | 0.41 | 0.0282 |
| Transcytosis | 5 | 0.41 | 0.0282 |
| Response to oxidative stress | 32 | 2.61 | 0.0283 |
| Glycoside transport | 3 | 0.24 | 0.0285 |
| Lobar bronchus development | 3 | 0.24 | 0.0285 |
| Propionate metabolic process | 3 | 0.24 | 0.0285 |
| Quinolinate metabolic process | 3 | 0.24 | 0.0285 |
| Cytokine precursor processing | 3 | 0.24 | 0.0285 |
| Udp-n-acetylglucosamine transmembrane transport | 3 | 0.24 | 0.0285 |
| Ductus arteriosus closure | 3 | 0.24 | 0.0285 |
| Antigen processing and presentation of exogenous peptide antigen via mhc class ib | 3 | 0.24 | 0.0285 |
| Intestinal hexose absorption | 3 | 0.24 | 0.0285 |
| Transepithelial water transport | 3 | 0.24 | 0.0285 |
| Regulation of purine nucleotide metabolic process | 11 | 0.9 | 0.0287 |
| Bile acid and bile salt transport | 6 | 0.49 | 0.0287 |
| Cell-cell adhesion mediated by cadherin | 6 | 0.49 | 0.0287 |
| Neutrophil activation | 6 | 0.49 | 0.0287 |
| Retinoic acid metabolic process | 6 | 0.49 | 0.0287 |
| Regulation of extrinsic apoptotic signaling pathway | 17 | 1.38 | 0.0289 |
| Icosanoid biosynthetic process | 8 | 0.65 | 0.029 |
| Response to steroid hormone | 26 | 2.12 | 0.0292 |
| Positive regulation of protein localization to cell periphery | 10 | 0.81 | 0.0295 |
| Positive regulation of reactive oxygen species metabolic process | 10 | 0.81 | 0.0295 |
| Heart process | 13 | 1.06 | 0.0295 |
| Blood coagulation | 19 | 1.55 | 0.0296 |
| Cytoskeleton organization | 92 | 7.49 | 0.0296 |
| Positive regulation of t cell activation | 24 | 1.95 | 0.0297 |

| | | | |
|---|---|---|---|
| Oligosaccharide metabolic process | 9 | 0.73 | 0.0297 |
| Positive regulation of protein localization to plasma membrane | 9 | 0.73 | 0.0297 |
| Regulation of intracellular transport | 25 | 2.04 | 0.0297 |
| Positive regulation of interleukin-10 production | 7 | 0.57 | 0.0297 |
| Prostanoid metabolic process | 7 | 0.57 | 0.0297 |
| Intrinsic apoptotic signaling pathway | 18 | 1.47 | 0.0307 |
| Female pregnancy | 18 | 1.47 | 0.0307 |
| Chordate embryonic development | 51 | 4.15 | 0.0312 |
| Positive regulation of type ii interferon production | 10 | 0.81 | 0.0317 |
| Regulation of atp metabolic process | 10 | 0.81 | 0.0317 |
| Regulation of adaptive immune response | 21 | 1.71 | 0.0321 |
| Regulation of protein import into nucleus | 9 | 0.73 | 0.0322 |
| Coagulation | 19 | 1.55 | 0.0323 |
| Organophosphate catabolic process | 19 | 1.55 | 0.0323 |
| Regulation of cardiac muscle cell action potential | 6 | 0.49 | 0.0325 |
| Lipid oxidation | 11 | 0.9 | 0.0328 |
| Positive regulation of apoptotic signaling pathway | 16 | 1.3 | 0.0329 |
| Response to progesterone | 7 | 0.57 | 0.033 |
| Positive regulation of protein import into nucleus | 7 | 0.57 | 0.033 |
| Regulation of low-density lipoprotein particle clearance | 5 | 0.41 | 0.033 |
| Stress fiber assembly | 5 | 0.41 | 0.033 |
| Positive regulation of triglyceride metabolic process | 5 | 0.41 | 0.033 |
| Positive regulation of macrophage derived foam cell differentiation | 5 | 0.41 | 0.033 |
| Negative regulation of extracellular matrix organization | 5 | 0.41 | 0.033 |
| Contractile actin filament bundle assembly | 5 | 0.41 | 0.033 |
| Nucleobase-containing small molecule catabolic process | 5 | 0.41 | 0.033 |
| Morphogenesis of an epithelium | 38 | 3.09 | 0.0332 |
| Activation of immune response | 34 | 2.77 | 0.0333 |
| Protein glycosylation | 21 | 1.71 | 0.0334 |
| Macromolecule glycosylation | 21 | 1.71 | 0.0334 |
| Mucus secretion | 4 | 0.33 | 0.0338 |
| Negative regulation of leukocyte degranulation | 4 | 0.33 | 0.0338 |
| Amine transport | 4 | 0.33 | 0.0338 |
| Lysosomal protein catabolic process | 4 | 0.33 | 0.0338 |
| Angiotensin maturation | 4 | 0.33 | 0.0338 |
| Epithelial tube morphogenesis | 28 | 2.28 | 0.0338 |
| Response to osmotic stress | 10 | 0.81 | 0.0341 |
| Embryonic pattern specification | 10 | 0.81 | 0.0341 |
| Regulation of platelet activation | 8 | 0.65 | 0.0346 |
| Regulation of carbohydrate catabolic process | 9 | 0.73 | 0.0348 |
| Embryo development ending in birth or egg hatching | 52 | 4.23 | 0.0348 |
| Regulation of striated muscle contraction | 12 | 0.98 | 0.0353 |
| Positive regulation of cellular component biogenesis | 41 | 3.34 | 0.0355 |
| Negative regulation of secretion by cell | 16 | 1.3 | 0.0362 |
| Positive regulation of pattern recognition receptor signaling pathway | 10 | 0.81 | 0.0365 |
| Embryonic digestive tract development | 6 | 0.49 | 0.0365 |
| Regulation of superoxide metabolic process | 6 | 0.49 | 0.0365 |

| | | | |
|---|---|---|---|
| Transepithelial transport | 6 | 0.49 | 0.0365 |
| Negative regulation of cell communication | 108 | 8.79 | 0.0367 |
| Glycoprotein metabolic process | 30 | 2.44 | 0.0371 |
| Regulation of notch signaling pathway | 12 | 0.98 | 0.0374 |
| Cellular response to nitrogen compound | 49 | 3.99 | 0.0376 |
| Regulation of purine nucleotide catabolic process | 8 | 0.65 | 0.0377 |
| Regulation of lipid storage | 8 | 0.65 | 0.0377 |
| Regulation of nucleotide catabolic process | 8 | 0.65 | 0.0377 |
| Regulation of glycolytic process | 8 | 0.65 | 0.0377 |
| Regulation of cholesterol efflux | 8 | 0.65 | 0.0377 |
| Vitamin transport | 8 | 0.65 | 0.0377 |
| Regulation of natural killer cell mediated cytotoxicity | 8 | 0.65 | 0.0377 |
| Tumor necrosis factor-mediated signaling pathway | 8 | 0.65 | 0.0377 |
| Negative regulation of actin filament polymerization | 8 | 0.65 | 0.0377 |
| Hemostasis | 19 | 1.55 | 0.0382 |
| Cell differentiation involved in embryonic placenta development | 5 | 0.41 | 0.0382 |
| Regulation of superoxide anion generation | 5 | 0.41 | 0.0382 |
| Alditol metabolic process | 5 | 0.41 | 0.0382 |
| Regulation of leukocyte chemotaxis | 14 | 1.14 | 0.0384 |
| Protein localization to cell periphery | 22 | 1.79 | 0.0385 |
| Regulation of alpha-beta t cell differentiation | 10 | 0.81 | 0.0391 |
| Regulation of mirna transcription | 10 | 0.81 | 0.0391 |
| Cellular response to external stimulus | 10 | 0.81 | 0.0391 |
| Negative regulation of inflammatory response | 20 | 1.63 | 0.0395 |
| Cellular oxidant detoxification | 11 | 0.9 | 0.0397 |
| Maintenance of location | 11 | 0.9 | 0.0397 |
| Protein localization to plasma membrane | 19 | 1.55 | 0.0398 |
| Response to ph | 7 | 0.57 | 0.0401 |
| Schwann cell development | 7 | 0.57 | 0.0401 |
| Negative regulation of protein kinase activity | 14 | 1.14 | 0.0404 |
| Maintenance of protein location | 9 | 0.73 | 0.0405 |
| Negative regulation of tumor necrosis factor-mediated signaling pathway | 6 | 0.49 | 0.0409 |
| Regulation of transforming growth factor beta production | 6 | 0.49 | 0.0409 |
| Phosphatidic acid biosynthetic process | 6 | 0.49 | 0.0409 |
| Cellular response to camp | 8 | 0.65 | 0.041 |
| Long-chain fatty acid transport | 8 | 0.65 | 0.041 |
| Proteinogenic amino acid biosynthetic process | 8 | 0.65 | 0.041 |
| Cardiac muscle cell action potential | 8 | 0.65 | 0.041 |
| Leukocyte activation involved in immune response | 20 | 1.63 | 0.041 |
| Kidney epithelium development | 15 | 1.22 | 0.0412 |
| L-ascorbic acid metabolic process | 4 | 0.33 | 0.0412 |
| Hepatocyte differentiation | 4 | 0.33 | 0.0412 |
| Regulation of angiotensin levels in blood | 4 | 0.33 | 0.0412 |
| Hepatocyte apoptotic process | 4 | 0.33 | 0.0412 |
| Acetyl-coa biosynthetic process | 4 | 0.33 | 0.0412 |
| Positive regulation of triglyceride biosynthetic process | 4 | 0.33 | 0.0412 |
| Regulation of dendritic cell differentiation | 4 | 0.33 | 0.0412 |

| | | | |
|---|---|---|---|
| Regulation of microvillus assembly | 3 | 0.24 | 0.0412 |
| Regulation of chloride transport | 3 | 0.24 | 0.0412 |
| Galactosylceramide biosynthetic process | 3 | 0.24 | 0.0412 |
| Deoxyribonucleoside catabolic process | 3 | 0.24 | 0.0412 |
| Mhc class i protein complex assembly | 3 | 0.24 | 0.0412 |
| Protection from natural killer cell mediated cytotoxicity | 3 | 0.24 | 0.0412 |
| Regulation of very-low-density lipoprotein particle remodeling | 3 | 0.24 | 0.0412 |
| Negative regulation of golgi to plasma membrane protein transport | 3 | 0.24 | 0.0412 |
| Peptide antigen assembly with mhc class i protein complex | 3 | 0.24 | 0.0412 |
| Regulation of somitogenesis | 3 | 0.24 | 0.0412 |
| Hemidesmosome assembly | 3 | 0.24 | 0.0412 |
| Negative regulation of type b pancreatic cell apoptotic process | 3 | 0.24 | 0.0412 |
| Farnesyl diphosphate metabolic process | 3 | 0.24 | 0.0412 |
| Galactolipid biosynthetic process | 3 | 0.24 | 0.0412 |
| Alpha-beta t cell activation | 13 | 1.06 | 0.0414 |
| B cell differentiation | 13 | 1.06 | 0.0414 |
| Ribonucleotide catabolic process | 11 | 0.9 | 0.0422 |
| Cellular response to biotic stimulus | 23 | 1.87 | 0.0423 |
| Regulation of canonical nf-kappab signal transduction | 27 | 2.2 | 0.0427 |
| Regulation of kinase activity | 32 | 2.61 | 0.043 |
| Intracellular receptor signaling pathway | 19 | 1.55 | 0.0431 |
| Synaptic vesicle recycling | 9 | 0.73 | 0.0435 |
| Positive regulation of leukocyte mediated immunity | 16 | 1.3 | 0.0435 |
| Modulation of process of another organism | 13 | 1.06 | 0.0436 |
| Positive regulation of insulin receptor signaling pathway | 5 | 0.41 | 0.0439 |
| Regulation of heterotypic cell-cell adhesion | 5 | 0.41 | 0.0439 |
| T-helper 1 type immune response | 5 | 0.41 | 0.0439 |
| Exocrine system development | 7 | 0.57 | 0.044 |
| Epithelial cell apoptotic process | 7 | 0.57 | 0.044 |
| Liposaccharide metabolic process | 12 | 0.98 | 0.0445 |
| Positive regulation of protein localization to membrane | 12 | 0.98 | 0.0445 |
| Positive regulation of protein localization to nucleus | 11 | 0.9 | 0.0449 |
| Amide transport | 15 | 1.22 | 0.0453 |
| Positive regulation of cytoskeleton organization | 18 | 1.47 | 0.0453 |
| Sensory perception of sound | 17 | 1.38 | 0.0455 |
| Actin filament capping | 6 | 0.49 | 0.0456 |
| Intracellular zinc ion homeostasis | 6 | 0.49 | 0.0456 |
| Regulation of phosphatidylinositol 3-kinase/protein kinase b signal transduction | 25 | 2.04 | 0.0459 |
| Neutrophil migration | 9 | 0.73 | 0.0467 |
| Cellular response to ketone | 12 | 0.98 | 0.047 |
| Regulation of cold-induced thermogenesis | 15 | 1.22 | 0.0474 |
| Cell activation involved in immune response | 20 | 1.63 | 0.0478 |
| Glycerolipid catabolic process | 8 | 0.65 | 0.0481 |
| Leukocyte cell-cell adhesion | 8 | 0.65 | 0.0481 |
| Positive regulation of glial cell differentiation | 7 | 0.57 | 0.0481 |
| Prostate gland development | 7 | 0.57 | 0.0481 |
| Mammary gland morphogenesis | 7 | 0.57 | 0.0481 |

| | | | |
|---|---|---|---|
| Positive regulation of cd4-positive, alpha-beta t cell activation | 7 | 0.57 | 0.0481 |
| Skin epidermis development | 13 | 1.06 | 0.0484 |
| Glycine transport | 4 | 0.33 | 0.0494 |
| Relaxation of cardiac muscle | 4 | 0.33 | 0.0494 |
| Regulation of response to type ii interferon | 4 | 0.33 | 0.0494 |
| Regulation of phospholipid transport | 4 | 0.33 | 0.0494 |
| Ribonucleoside monophosphate catabolic process | 4 | 0.33 | 0.0494 |
| Fructose 6-phosphate metabolic process | 4 | 0.33 | 0.0494 |
| Protein catabolic process in the vacuole | 4 | 0.33 | 0.0494 |
| Regulation of type ii interferon-mediated signaling pathway | 4 | 0.33 | 0.0494 |
| Azole transmembrane transport | 4 | 0.33 | 0.0494 |
| Tertiary alcohol metabolic process | 4 | 0.33 | 0.0494 |
| Positive regulation of ruffle assembly | 4 | 0.33 | 0.0494 |
| Intestinal lipid absorption | 4 | 0.33 | 0.0494 |
| Negative regulation of low-density lipoprotein particle clearance | 4 | 0.33 | 0.0494 |
| Glycosylceramide metabolic process | 4 | 0.33 | 0.0494 |
| Regulation of monoatomic anion transport | 4 | 0.33 | 0.0494 |
| Urea metabolic process | 4 | 0.33 | 0.0494 |
| Positive regulation of phospholipid transport | 4 | 0.33 | 0.0494 |
| Myeloid cell homeostasis | 15 | 1.22 | 0.0496 |
| Biological process involved in interaction with host | 15 | 1.22 | 0.0496 |
| Regulation of lymphocyte proliferation | 22 | 1.79 | 0.0497 |
| Cellular response to abiotic stimulus | 28 | 2.28 | 0.0497 |
| Cellular response to environmental stimulus | 28 | 2.28 | 0.0497 |
| Nad biosynthetic process | 5 | 0.41 | 0.05 |
| Regulation of icosanoid secretion | 5 | 0.41 | 0.05 |
| Myelin assembly | 5 | 0.41 | 0.05 |

TableS16

| Pathways upregulated in Gland_B compared to Gland_A | | | |
|---|---|---|---|
| **GO Pathway** | **No. of genes** | **%** | **p-value** |
| Cilium organization | 90 | 8.17 | 7.11E-34 |
| Cilium assembly | 85 | 7.71 | 3.83E-33 |
| Cell projection organization | 170 | 15.43 | 9.94E-33 |
| Plasma membrane bounded cell projection organization | 164 | 14.88 | 5.2E-32 |
| Cell projection assembly | 94 | 8.53 | 7.85E-31 |
| Plasma membrane bounded cell projection assembly | 91 | 8.26 | 1.09E-29 |
| Microtubule-based process | 131 | 11.89 | 6E-28 |
| Microtubule-based movement | 87 | 7.89 | 6.06E-27 |
| Cilium movement | 62 | 5.63 | 2.72E-26 |
| Axoneme assembly | 40 | 3.63 | 1.56E-23 |
| Motile cilium assembly | 35 | 3.18 | 9.96E-23 |
| Microtubule bundle formation | 43 | 3.9 | 1.02E-21 |
| Nervous system development | 216 | 19.6 | 1.64E-20 |
| Cell motility | 135 | 12.25 | 2.46E-19 |
| Cilium-dependent cell motility | 48 | 4.36 | 5.64E-19 |
| Cilium or flagellum-dependent cell motility | 48 | 4.36 | 5.64E-19 |
| Cilium movement involved in cell motility | 47 | 4.26 | 1.78E-18 |
| Microtubule cytoskeleton organization | 85 | 7.71 | 8.86E-18 |
| Circulatory system development | 114 | 10.34 | 2.31E-17 |
| Cell development | 206 | 18.69 | 2.24E-16 |
| Generation of neurons | 131 | 11.89 | 2.81E-16 |
| Pattern specification process | 71 | 6.44 | 3.29E-16 |
| Cytoskeleton organization | 136 | 12.34 | 5.91E-16 |
| Sperm motility | 42 | 3.81 | 7.49E-16 |
| Flagellated sperm motility | 42 | 3.81 | 7.49E-16 |
| Neurogenesis | 143 | 12.98 | 2.22E-15 |
| Microtubule-based transport | 45 | 4.08 | 3.93E-15 |
| Organelle assembly | 105 | 9.53 | 4.22E-15 |
| Heart development | 77 | 6.99 | 6.4E-14 |
| Organelle organization | 250 | 22.69 | 7.41E-14 |
| Regionalization | 62 | 5.63 | 9.83E-14 |
| Neuron differentiation | 118 | 10.71 | 1.81E-13 |
| Sperm flagellum assembly | 22 | 2 | 3.2E-13 |
| Sensory organ development | 78 | 7.08 | 3.4E-13 |
| Epithelial cilium movement involved in extracellular fluid movement | 20 | 1.81 | 7.13E-13 |
| Extracellular transport | 20 | 1.81 | 2.5E-12 |
| Eye development | 58 | 5.26 | 3.72E-12 |
| Left/right pattern formation | 33 | 2.99 | 3.87E-12 |
| Visual system development | 58 | 5.26 | 5.51E-12 |
| Determination of left/right symmetry | 32 | 2.9 | 5.53E-12 |
| Camera-type eye development | 53 | 4.81 | 1.03E-11 |
| Sensory system development | 58 | 5.26 | 1.08E-11 |
| Central nervous system development | 105 | 9.53 | 1.19E-11 |
| Animal organ morphogenesis | 102 | 9.26 | 1.74E-11 |
| Determination of bilateral symmetry | 32 | 2.9 | 2E-11 |
| Specification of symmetry | 32 | 2.9 | 2E-11 |
| Tube development | 96 | 8.71 | 5.18E-11 |
| Tissue development | 154 | 13.97 | 3.97E-10 |
| Axonemal dynein complex assembly | 16 | 1.45 | 7.48E-10 |
| Head development | 82 | 7.44 | 2.75E-09 |
| Cellular response to endogenous stimulus | 94 | 8.53 | 3.16E-09 |
| Cellular component assembly | 201 | 18.24 | 3.41E-09 |
| Tube morphogenesis | 74 | 6.72 | 5.44E-09 |

| | | | |
|---|---|---|---|
| Regulation of signaling | 259 | 23.5 | 5.81E-09 |
| Spermatid differentiation | 37 | 3.36 | 5.93E-09 |
| Regulation of cell communication | 259 | 23.5 | 7.02E-09 |
| Epithelium development | 103 | 9.35 | 7.07E-09 |
| Spermatid development | 36 | 3.27 | 8.52E-09 |
| Intraciliary transport | 16 | 1.45 | 1.58E-08 |
| Cellular component biogenesis | 213 | 19.33 | 1.6E-08 |
| Embryo development | 97 | 8.8 | 1.72E-08 |
| Embryonic organ morphogenesis | 42 | 3.81 | 1.78E-08 |
| Brain development | 76 | 6.9 | 1.99E-08 |
| Skeletal system development | 58 | 5.26 | 2.83E-08 |
| Regulation of plasma membrane bounded cell projection organization | 68 | 6.17 | 3.55E-08 |
| Embryonic morphogenesis | 64 | 5.81 | 3.7E-08 |
| Regulation of cell projection organization | 69 | 6.26 | 4.27E-08 |
| Sperm axoneme assembly | 13 | 1.18 | 4.45E-08 |
| Homeostatic process | 125 | 11.34 | 6.75E-08 |
| Anatomical structure formation involved in morphogenesis | 91 | 8.26 | 7.04E-08 |
| Neuron development | 84 | 7.62 | 1.27E-07 |
| Response to endogenous stimulus | 108 | 9.8 | 1.62E-07 |
| Tissue morphogenesis | 61 | 5.54 | 1.8E-07 |
| Cell migration | 86 | 7.8 | 1.91E-07 |
| Embryonic organ development | 52 | 4.72 | 2.01E-07 |
| Cell morphogenesis | 71 | 6.44 | 2.54E-07 |
| Regulation of cellular component organization | 183 | 16.61 | 3.94E-07 |
| Cytoskeleton-dependent intracellular transport | 30 | 2.72 | 4.04E-07 |
| Heart morphogenesis | 36 | 3.27 | 4.76E-07 |
| Regulation of anatomical structure morphogenesis | 82 | 7.44 | 4.81E-07 |
| Enzyme-linked receptor protein signaling pathway | 63 | 5.72 | 7.5E-07 |
| Respiratory system development | 31 | 2.81 | 7.66E-07 |
| Modulation of chemical synaptic transmission | 55 | 4.99 | 7.8E-07 |
| Forebrain development | 47 | 4.26 | 8.11E-07 |
| Regulation of trans-synaptic signaling | 55 | 4.99 | 8.24E-07 |
| Neuron projection development | 69 | 6.26 | 0.0000011 |
| Epithelial tube morphogenesis | 39 | 3.54 | 1.37E-06 |
| Cellular homeostasis | 67 | 6.08 | 1.41E-06 |
| Cell junction organization | 57 | 5.17 | 1.44E-06 |
| Regulation of signal transduction | 220 | 19.96 | 1.57E-06 |
| Sensory organ morphogenesis | 36 | 3.27 | 1.65E-06 |
| Lung development | 28 | 2.54 | 0.0000017 |
| Intracellular chemical homeostasis | 59 | 5.35 | 1.79E-06 |
| Respiratory tube development | 28 | 2.54 | 2.52E-06 |
| Transport along microtubule | 26 | 2.36 | 2.58E-06 |
| Retina development in camera-type eye | 26 | 2.36 | 2.58E-06 |
| Cellular process involved in reproduction in multicellular organism | 46 | 4.17 | 2.82E-06 |
| Cellular localization | 191 | 17.33 | 3.05E-06 |
| Regulation of synapse structure or activity | 38 | 3.45 | 3.11E-06 |
| Neuron projection morphogenesis | 52 | 4.72 | 3.24E-06 |
| Morphogenesis of an epithelium | 49 | 4.45 | 3.26E-06 |
| Renal system development | 38 | 3.45 | 0.0000033 |
| Germ cell development | 43 | 3.9 | 4.18E-06 |
| Plasma membrane bounded cell projection morphogenesis | 52 | 4.72 | 4.27E-06 |
| Wnt signaling pathway | 35 | 3.18 | 4.41E-06 |
| Cell morphogenesis involved in neuron differentiation | 48 | 4.36 | 0.0000049 |
| Regulation of neuron projection development | 47 | 4.26 | 0.0000055 |
| Synapse organization | 41 | 3.72 | 5.67E-06 |
| Cartilage development | 26 | 2.36 | 5.78E-06 |

| | | | |
|---|---|---|---|
| Regulation of wnt signaling pathway | 40 | 3.63 | 6.35E-06 |
| Appendage development | 27 | 2.45 | 6.43E-06 |
| Limb development | 27 | 2.45 | 6.43E-06 |
| Cell projection morphogenesis | 52 | 4.72 | 0.0000065 |
| Vasculature development | 55 | 4.99 | 7.07E-06 |
| Cellular response to hormone stimulus | 55 | 4.99 | 7.07E-06 |
| Cell adhesion | 82 | 7.44 | 7.38E-06 |
| Intracellular monoatomic ion homeostasis | 46 | 4.17 | 7.72E-06 |
| Reproductive process | 122 | 11.07 | 8.21E-06 |
| Monoatomic ion homeostasis | 52 | 4.72 | 8.89E-06 |
| Inner dynein arm assembly | 8 | 0.73 | 9.49E-06 |
| Kidney development | 36 | 3.27 | 9.65E-06 |
| Axonogenesis | 41 | 3.72 | 0.00001 |
| Monoatomic cation homeostasis | 51 | 4.63 | 0.0000101 |
| Chondrocyte differentiation | 17 | 1.54 | 0.0000101 |
| Growth | 47 | 4.26 | 0.0000102 |
| Developmental growth | 47 | 4.26 | 0.0000102 |
| Cell surface receptor signaling pathway | 153 | 13.88 | 0.0000112 |
| Chemical homeostasis | 80 | 7.26 | 0.0000115 |
| Regulation of synapse organization | 36 | 3.27 | 0.0000119 |
| Epithelial cell development | 27 | 2.45 | 0.0000121 |
| Regulation of cilium movement | 12 | 1.09 | 0.0000121 |
| Developmental process involved in reproduction | 88 | 7.99 | 0.0000127 |
| Connective tissue development | 30 | 2.72 | 0.0000138 |
| Limb morphogenesis | 23 | 2.09 | 0.0000142 |
| Appendage morphogenesis | 23 | 2.09 | 0.0000142 |
| Regulation of cilium beat frequency | 8 | 0.73 | 0.0000149 |
| Regulation of cell motility | 92 | 8.35 | 0.000015 |
| Muscle structure development | 51 | 4.63 | 0.0000177 |
| Spermatogenesis | 63 | 5.72 | 0.0000191 |
| Intraciliary anterograde transport | 8 | 0.73 | 0.0000225 |
| Intracellular monoatomic cation homeostasis | 44 | 3.99 | 0.0000227 |
| Regulation of locomotion | 94 | 8.53 | 0.0000232 |
| Telencephalon development | 33 | 2.99 | 0.0000233 |
| Central nervous system neuron differentiation | 27 | 2.45 | 0.0000239 |
| Transport across blood-brain barrier | 16 | 1.45 | 0.0000243 |
| Vascular transport | 16 | 1.45 | 0.0000243 |
| Positive regulation of developmental process | 107 | 9.71 | 0.000029 |
| Protein-containing complex localization | 23 | 2.09 | 0.0000338 |
| Neural precursor cell proliferation | 18 | 1.63 | 0.0000353 |
| Regulation of biological quality | 207 | 18.78 | 0.0000357 |
| Male gamete generation | 63 | 5.72 | 0.0000385 |
| Muscle tissue development | 38 | 3.45 | 0.0000387 |
| Chordate embryonic development | 59 | 5.35 | 0.000039 |
| Blood vessel development | 51 | 4.63 | 0.0000417 |
| Cellular response to growth factor stimulus | 50 | 4.54 | 0.0000435 |
| Extracellular matrix organization | 32 | 2.9 | 0.0000438 |
| Eye morphogenesis | 23 | 2.09 | 0.0000444 |
| Extracellular structure organization | 32 | 2.9 | 0.0000466 |
| Embryo development ending in birth or egg hatching | 60 | 5.44 | 0.0000472 |
| External encapsulating structure organization | 32 | 2.9 | 0.00005 |
| Axon development | 43 | 3.9 | 0.0000506 |
| Heart looping | 14 | 1.27 | 0.0000507 |
| Transforming growth factor beta receptor superfamily signaling pathway | 24 | 2.18 | 0.0000519 |
| Blood vessel morphogenesis | 44 | 3.99 | 0.0000531 |
| Cellular macromolecule localization | 142 | 12.89 | 0.0000582 |

| | | | |
|---|---|---|---|
| Neuron migration | 21 | 1.91 | 0.0000586 |
| Circulatory system process | 49 | 4.45 | 0.000075 |
| Regulation of microtubule-based movement | 13 | 1.18 | 0.0000782 |
| Positive regulation of cellular component organization | 88 | 7.99 | 0.00008 |
| Regulation of localization | 146 | 13.25 | 0.0000811 |
| Camera-type eye morphogenesis | 20 | 1.81 | 0.0000847 |
| Response to growth factor | 51 | 4.63 | 0.0000979 |
| Metanephros development | 15 | 1.36 | 0.0000982 |
| Inorganic ion homeostasis | 43 | 3.9 | 0.0000988 |
| Response to hormone | 70 | 6.35 | 0.0000997 |
| Regulation of microtubule-based process | 31 | 2.81 | 0.000105 |
| Protein localization | 140 | 12.7 | 0.000107 |
| Embryonic heart tube morphogenesis | 14 | 1.27 | 0.00012 |
| Determination of heart left/right asymmetry | 14 | 1.27 | 0.00012 |
| Outer dynein arm assembly | 8 | 0.73 | 0.000124 |
| Forebrain regionalization | 8 | 0.73 | 0.000124 |
| Positive regulation of cell projection organization | 37 | 3.36 | 0.000125 |
| Striated muscle tissue development | 35 | 3.18 | 0.000129 |
| Respiratory system process | 9 | 0.82 | 0.000135 |
| Cell surface receptor protein serine/threonine kinase signaling pathway | 24 | 2.18 | 0.000137 |
| Macromolecule localization | 164 | 14.88 | 0.000141 |
| Neural tube development | 21 | 1.91 | 0.000145 |
| System process | 146 | 13.25 | 0.000158 |
| Epithelial cell differentiation | 56 | 5.08 | 0.000173 |
| Embryonic heart tube development | 15 | 1.36 | 0.000181 |
| Negative regulation of wnt signaling pathway | 23 | 2.09 | 0.000183 |
| Regulation of postsynapse organization | 20 | 1.81 | 0.000193 |
| Cell population proliferation | 61 | 5.54 | 0.000204 |
| Regulation of canonical wnt signaling pathway | 30 | 2.72 | 0.00023 |
| Behavior | 55 | 4.99 | 0.000231 |
| Embryonic limb morphogenesis | 18 | 1.63 | 0.000237 |
| Embryonic appendage morphogenesis | 18 | 1.63 | 0.000237 |
| Gland development | 41 | 3.72 | 0.000237 |
| Cardiac muscle tissue development | 24 | 2.18 | 0.000247 |
| Regulation of cell differentiation | 117 | 10.62 | 0.000251 |
| Regulation of transport | 118 | 10.71 | 0.000262 |
| Multicellular organismal reproductive process | 80 | 7.26 | 0.00027 |
| Cell proliferation in forebrain | 8 | 0.73 | 0.000279 |
| Carboxylic acid transmembrane transport | 19 | 1.72 | 0.00028 |
| Epithelial tube formation | 18 | 1.63 | 0.000285 |
| Regulation of cell migration | 82 | 7.44 | 0.000306 |
| Organic acid transmembrane transport | 19 | 1.72 | 0.000363 |
| Tube formation | 19 | 1.72 | 0.000363 |
| Regulation of cytoskeleton organization | 48 | 4.36 | 0.000391 |
| Collagen fibril organization | 12 | 1.09 | 0.000393 |
| Odontogenesis | 17 | 1.54 | 0.000497 |
| Gamete generation | 69 | 6.26 | 0.000506 |
| Regulation of organelle organization | 87 | 7.89 | 0.000539 |
| Neural tube formation | 15 | 1.36 | 0.00054 |
| Multicellular organismal-level homeostasis | 54 | 4.9 | 0.000547 |
| Organic anion transport | 38 | 3.45 | 0.000551 |
| Metal ion transport | 54 | 4.9 | 0.000615 |
| Negative regulation of canonical wnt signaling pathway | 19 | 1.72 | 0.00064 |
| Bone development | 23 | 2.09 | 0.000658 |
| Actin filament-based process | 52 | 4.72 | 0.000674 |
| Regulation of synaptic plasticity | 25 | 2.27 | 0.000687 |

| | | | |
|---|---|---|---|
| Brain morphogenesis | 9 | 0.82 | 0.000694 |
| Regulation of cellular response to growth factor stimulus | 36 | 3.27 | 0.000698 |
| Cellular response to peptide hormone stimulus | 30 | 2.72 | 0.000702 |
| Respiratory gaseous exchange by respiratory system | 11 | 1 | 0.000714 |
| Sexual reproduction | 84 | 7.62 | 0.00072 |
| Cognition | 32 | 2.9 | 0.000773 |
| Regulation of nervous system development | 41 | 3.72 | 0.000778 |
| Primary neural tube formation | 14 | 1.27 | 0.000783 |
| Smoothened signaling pathway | 14 | 1.27 | 0.000783 |
| Regulation of cellular component biogenesis | 78 | 7.08 | 0.000785 |
| Gliogenesis | 29 | 2.63 | 0.000796 |
| Muscle cell differentiation | 29 | 2.63 | 0.000796 |
| Skeletal system morphogenesis | 25 | 2.27 | 0.000872 |
| Canonical wnt signaling pathway | 15 | 1.36 | 0.000877 |
| Regulation of multicellular organismal development | 105 | 9.53 | 0.000894 |
| Ear development | 25 | 2.27 | 0.00104 |
| Morphogenesis of embryonic epithelium | 18 | 1.63 | 0.00108 |
| Nephron development | 18 | 1.63 | 0.00108 |
| Inner ear development | 23 | 2.09 | 0.00109 |
| Tissue homeostasis | 25 | 2.27 | 0.0011 |
| Anatomical structure homeostasis | 25 | 2.27 | 0.0011 |
| Lens development in camera-type eye | 13 | 1.18 | 0.00114 |
| Regulation of neurogenesis | 35 | 3.18 | 0.00114 |
| Negative regulation of protein modification process | 30 | 2.72 | 0.00117 |
| Kidney epithelium development | 18 | 1.63 | 0.00117 |
| Embryonic epithelial tube formation | 16 | 1.45 | 0.00122 |
| Gland morphogenesis | 15 | 1.36 | 0.00126 |
| Learning or memory | 28 | 2.54 | 0.0013 |
| Positive regulation of nervous system development | 29 | 2.63 | 0.00133 |
| Regulation of dendrite morphogenesis | 11 | 1 | 0.00135 |
| Positive regulation of cell differentiation | 69 | 6.26 | 0.00138 |
| Neural tube closure | 13 | 1.18 | 0.00139 |
| Embryonic eye morphogenesis | 8 | 0.73 | 0.00149 |
| Response to peptide hormone | 38 | 3.45 | 0.00152 |
| Locomotory exploration behavior | 6 | 0.54 | 0.00153 |
| Renal system vasculature morphogenesis | 5 | 0.45 | 0.00154 |
| Kidney vasculature morphogenesis | 5 | 0.45 | 0.00154 |
| Tube closure | 13 | 1.18 | 0.00154 |
| Sensory perception of light stimulus | 24 | 2.18 | 0.00156 |
| Embryonic cranial skeleton morphogenesis | 9 | 0.82 | 0.00156 |
| Regulation of transmembrane receptor protein serine/threonine kinase signaling pathway | 31 | 2.81 | 0.00156 |
| Positive regulation of cell communication | 123 | 11.16 | 0.00157 |
| Regulation of dendrite development | 14 | 1.27 | 0.00169 |
| Positive regulation of signaling | 123 | 11.16 | 0.00169 |
| Carboxylic acid transport | 29 | 2.63 | 0.00169 |
| Regulation of smoothened signaling pathway | 13 | 1.18 | 0.0017 |
| Monoatomic cation transport | 63 | 5.72 | 0.00171 |
| L-alpha-amino acid transmembrane transport | 11 | 1 | 0.00172 |
| Organic acid transport | 29 | 2.63 | 0.00176 |
| Axon guidance | 24 | 2.18 | 0.00194 |
| Monoatomic cation transmembrane transport | 54 | 4.9 | 0.00199 |
| Positive regulation of neuron projection development | 19 | 1.72 | 0.00202 |
| Neuron projection guidance | 24 | 2.18 | 0.00205 |
| Bicarbonate transport | 8 | 0.73 | 0.0021 |
| Limbic system development | 16 | 1.45 | 0.00212 |

| | | | |
|---|---|---|---|
| Columnar/cuboidal epithelial cell differentiation | 14 | 1.27 | 0.00219 |
| Forebrain ventricular zone progenitor cell division | 4 | 0.36 | 0.00222 |
| Axonemal central apparatus assembly | 4 | 0.36 | 0.00222 |
| Epithelial cell maturation | 7 | 0.64 | 0.00223 |
| Regulation of system process | 47 | 4.26 | 0.00224 |
| Insulin-like growth factor receptor signaling pathway | 12 | 1.09 | 0.00225 |
| Roof of mouth development | 13 | 1.18 | 0.00225 |
| Skeletal muscle tissue regeneration | 8 | 0.73 | 0.00246 |
| Sodium ion transport | 21 | 1.91 | 0.00249 |
| Organ growth | 14 | 1.27 | 0.0026 |
| Actin cytoskeleton organization | 45 | 4.08 | 0.0026 |
| Positive regulation of locomotion | 51 | 4.63 | 0.00265 |
| Adrenal gland development | 7 | 0.64 | 0.0027 |
| Salivary gland morphogenesis | 7 | 0.64 | 0.0027 |
| Supramolecular fiber organization | 49 | 4.45 | 0.0027 |
| Positive regulation of cell motility | 50 | 4.54 | 0.00271 |
| Visual perception | 23 | 2.09 | 0.00273 |
| L-amino acid transport | 12 | 1.09 | 0.00273 |
| Lung epithelium development | 8 | 0.73 | 0.00287 |
| Embryonic skeletal system morphogenesis | 13 | 1.18 | 0.00295 |
| Cell surface receptor protein tyrosine kinase signaling pathway | 37 | 3.36 | 0.00301 |
| Transmembrane transport | 89 | 8.08 | 0.00304 |
| Aromatic amino acid transport | 5 | 0.45 | 0.00308 |
| Monoatomic ion transmembrane transport | 63 | 5.72 | 0.00313 |
| Central nervous system neuron development | 13 | 1.18 | 0.00321 |
| Positive regulation of cellular component biogenesis | 42 | 3.81 | 0.00321 |
| Neural retina development | 12 | 1.09 | 0.0033 |
| Regulation of supramolecular fiber organization | 35 | 3.18 | 0.0033 |
| Animal organ formation | 8 | 0.73 | 0.00333 |
| Amide transport | 17 | 1.54 | 0.00343 |
| Negative regulation of post-translational protein modification | 13 | 1.18 | 0.0035 |
| Muscle cell development | 19 | 1.72 | 0.00355 |
| Monoatomic ion transport | 73 | 6.62 | 0.00364 |
| Positive regulation of erk1 and erk2 cascade | 21 | 1.91 | 0.00371 |
| Determination of liver left/right asymmetry | 4 | 0.36 | 0.00374 |
| Stem cell differentiation | 20 | 1.81 | 0.00375 |
| Embryonic digit morphogenesis | 10 | 0.91 | 0.0039 |
| Retina homeostasis | 10 | 0.91 | 0.0039 |
| Positive regulation of signal transduction | 109 | 9.89 | 0.00395 |
| Cranial skeletal system development | 11 | 1 | 0.00401 |
| Regulation of mesenchymal cell apoptotic process | 5 | 0.45 | 0.00414 |
| Negative regulation of long-term synaptic potentiation | 5 | 0.45 | 0.00414 |
| Sodium ion transmembrane transport | 17 | 1.54 | 0.00417 |
| Negative regulation of cell migration | 33 | 2.99 | 0.00433 |
| Import across plasma membrane | 18 | 1.63 | 0.00449 |
| Angiogenesis | 31 | 2.81 | 0.00452 |
| Presynapse organization | 7 | 0.64 | 0.00454 |
| Ureteric bud development | 12 | 1.09 | 0.00472 |
| Regulation of epithelial to mesenchymal transition | 15 | 1.36 | 0.00479 |
| Amino acid transmembrane transport | 13 | 1.18 | 0.00486 |
| Regulation of actin filament-based process | 34 | 3.09 | 0.0049 |
| Face development | 9 | 0.82 | 0.00508 |
| Brain-derived neurotrophic factor receptor signaling pathway | 9 | 0.82 | 0.00508 |
| Regulation of bmp signaling pathway | 15 | 1.36 | 0.00513 |
| Mesonephric epithelium development | 12 | 1.09 | 0.00514 |
| Mesonephric tubule development | 12 | 1.09 | 0.00514 |

| | | | |
|---|---|---|---|
| Ventricular system development | 7 | 0.64 | 0.00531 |
| Salivary gland development | 7 | 0.64 | 0.00531 |
| Cell communication | 308 | 27.95 | 0.00535 |
| Inorganic cation transmembrane transport | 51 | 4.63 | 0.00538 |
| Regulation of cellular localization | 73 | 6.62 | 0.00551 |
| Regulation of phosphate metabolic process | 59 | 5.35 | 0.00561 |
| Regulation of protein modification process | 65 | 5.9 | 0.00567 |
| Transforming growth factor beta receptor signaling pathway | 13 | 1.18 | 0.00568 |
| Regulation of glutamate receptor signaling pathway | 4 | 0.36 | 0.00576 |
| Membranous septum morphogenesis | 4 | 0.36 | 0.00576 |
| Cellular response to insulin stimulus | 20 | 1.81 | 0.00581 |
| Regulation of phosphorus metabolic process | 59 | 5.35 | 0.00599 |
| Insulin receptor signaling pathway | 14 | 1.27 | 0.00605 |
| Regulation of cilium movement involved in cell motility | 7 | 0.64 | 0.00619 |
| Regulation of cilium-dependent cell motility | 7 | 0.64 | 0.00619 |
| Response to auditory stimulus | 7 | 0.64 | 0.00619 |
| Negative regulation of cartilage development | 7 | 0.64 | 0.00619 |
| Regulation of heart contraction | 21 | 1.91 | 0.00626 |
| Negative regulation of signal transduction | 97 | 8.8 | 0.00633 |
| Regulation of microtubule cytoskeleton organization | 18 | 1.63 | 0.00637 |
| Pallium development | 20 | 1.81 | 0.00645 |
| Protein localization to cilium | 10 | 0.91 | 0.0065 |
| Regulation of dendritic spine morphogenesis | 8 | 0.73 | 0.00654 |
| Cellular response to vascular endothelial growth factor stimulus | 12 | 1.09 | 0.00658 |
| Branching involved in salivary gland morphogenesis | 5 | 0.45 | 0.00695 |
| Cellular hyperosmotic response | 5 | 0.45 | 0.00695 |
| Inorganic ion transmembrane transport | 56 | 5.08 | 0.00696 |
| Regulation of chondrocyte differentiation | 9 | 0.82 | 0.00708 |
| Mesenchymal cell differentiation | 18 | 1.63 | 0.00713 |
| Mesonephros development | 12 | 1.09 | 0.00713 |
| Neuron fate specification | 7 | 0.64 | 0.00716 |
| Lens induction in camera-type eye | 3 | 0.27 | 0.00723 |
| Exocrine system development | 8 | 0.73 | 0.00739 |
| Negative regulation of cell motility | 33 | 2.99 | 0.00745 |
| Regulation of intracellular transport | 25 | 2.27 | 0.00752 |
| Cellular response to leukemia inhibitory factor | 13 | 1.18 | 0.00765 |
| Positive regulation of cell migration | 46 | 4.17 | 0.00767 |
| Collagen-activated tyrosine kinase receptor signaling pathway | 9 | 0.82 | 0.00786 |
| Nucleotide metabolic process | 36 | 3.27 | 0.00787 |
| Positive regulation of neurogenesis | 23 | 2.09 | 0.00799 |
| Negative regulation of locomotion | 34 | 3.09 | 0.008 |
| Cardiac chamber morphogenesis | 15 | 1.36 | 0.00812 |
| Embryonic skeletal system development | 15 | 1.36 | 0.00812 |
| Cell fate commitment | 24 | 2.18 | 0.00828 |
| Glomerular capillary formation | 4 | 0.36 | 0.00832 |
| Cranial suture morphogenesis | 4 | 0.36 | 0.00832 |
| Glomerulus vasculature morphogenesis | 4 | 0.36 | 0.00832 |
| Embryonic camera-type eye morphogenesis | 6 | 0.54 | 0.00842 |
| Regulation of phosphorylation | 50 | 4.54 | 0.00862 |
| Mesenchymal cell migration | 9 | 0.82 | 0.0087 |
| Regulation of cell morphogenesis | 23 | 2.09 | 0.00873 |
| Cerebrospinal fluid circulation | 5 | 0.45 | 0.00873 |
| Response to leukemia inhibitory factor | 13 | 1.18 | 0.00881 |
| Photoreceptor cell maintenance | 8 | 0.73 | 0.00933 |
| Neuron recognition | 8 | 0.73 | 0.00933 |
| Response to abiotic stimulus | 78 | 7.08 | 0.00938 |

| | | | |
|---|---|---|---|
| Embryonic camera-type eye development | 7 | 0.64 | 0.00942 |
| Non-motile cilium assembly | 9 | 0.82 | 0.00961 |
| Forebrain generation of neurons | 11 | 1 | 0.0097 |
| Regulation of cell population proliferation | 114 | 10.34 | 0.00978 |
| Regulation of actin cytoskeleton organization | 30 | 2.72 | 0.00982 |
| Regulation of transforming growth factor beta receptor signaling pathway | 18 | 1.63 | 0.00982 |
| Cardiac chamber development | 18 | 1.63 | 0.00982 |
| Negative regulation of response to stimulus | 117 | 10.62 | 0.00988 |
| Negative regulation of chondrocyte differentiation | 6 | 0.54 | 0.00992 |
| Kidney vasculature development | 6 | 0.54 | 0.00992 |
| Renal system vasculature development | 6 | 0.54 | 0.00992 |
| Vascular process in circulatory system | 25 | 2.27 | 0.01 |
| Tissue regeneration | 10 | 0.91 | 0.0103 |
| Vascular endothelial growth factor receptor-1 signaling pathway | 8 | 0.73 | 0.0104 |
| Bmp signaling pathway | 11 | 1 | 0.0105 |
| Central nervous system neuron axonogenesis | 7 | 0.64 | 0.0107 |
| Heart formation | 5 | 0.45 | 0.0108 |
| Spinal cord patterning | 5 | 0.45 | 0.0108 |
| Specification of animal organ identity | 5 | 0.45 | 0.0108 |
| Branching morphogenesis of an epithelial tube | 15 | 1.36 | 0.011 |
| Striated muscle cell differentiation | 21 | 1.91 | 0.011 |
| Cellular response to bmp stimulus | 12 | 1.09 | 0.0112 |
| Response to bmp | 12 | 1.09 | 0.0112 |
| Neuron fate commitment | 10 | 0.91 | 0.0112 |
| Regulation of cilium beat frequency involved in ciliary motility | 4 | 0.36 | 0.0114 |
| Regulation of cellular response to transforming growth factor beta stimulus | 18 | 1.63 | 0.0114 |
| Positive regulation of mapk cascade | 37 | 3.36 | 0.0115 |
| Positive regulation of protein localization | 39 | 3.54 | 0.0115 |
| Chondrocyte development | 6 | 0.54 | 0.0116 |
| Bone remodeling | 8 | 0.73 | 0.0116 |
| Lung morphogenesis | 8 | 0.73 | 0.0116 |
| Platelet-derived growth factor receptor-alpha signaling pathway | 8 | 0.73 | 0.0116 |
| Collagen-activated signaling pathway | 9 | 0.82 | 0.0116 |
| Export across plasma membrane | 9 | 0.82 | 0.0116 |
| Locomotory behavior | 20 | 1.81 | 0.0121 |
| Endothelial cell differentiation | 11 | 1 | 0.0122 |
| Negative regulation of cell population proliferation | 56 | 5.08 | 0.0123 |
| Response to wounding | 36 | 3.27 | 0.0123 |
| Signaling | 301 | 27.31 | 0.0128 |
| Regulation of neuronal synaptic plasticity | 9 | 0.82 | 0.0128 |
| Regulation of dendritic spine development | 9 | 0.82 | 0.0128 |
| Cellular response to stem cell factor stimulus | 8 | 0.73 | 0.0129 |
| Macrophage colony-stimulating factor signaling pathway | 8 | 0.73 | 0.0129 |
| Response to stem cell factor | 8 | 0.73 | 0.0129 |
| Kit signaling pathway | 8 | 0.73 | 0.0129 |
| Cellular response to osmotic stress | 8 | 0.73 | 0.0129 |
| Purine nucleobase metabolic process | 5 | 0.45 | 0.0131 |
| Nephron epithelium development | 13 | 1.18 | 0.0132 |
| Purine-containing compound metabolic process | 39 | 3.54 | 0.0132 |
| Response to mechanical stimulus | 21 | 1.91 | 0.0132 |
| Subpallium development | 6 | 0.54 | 0.0134 |
| Phosphorylation | 34 | 3.09 | 0.0138 |
| Muscle organ development | 27 | 2.45 | 0.0139 |
| Embryonic nail plate morphogenesis | 3 | 0.27 | 0.014 |
| Frontal suture morphogenesis | 3 | 0.27 | 0.014 |
| Metanephric part of ureteric bud development | 3 | 0.27 | 0.014 |

| | | | |
|---|---|---|---|
| Negative regulation of prostatic bud formation | 3 | 0.27 | 0.014 |
| Glomerulus development | 9 | 0.82 | 0.014 |
| Positive regulation of wnt signaling pathway | 16 | 1.45 | 0.0143 |
| Regulation of cartilage development | 10 | 0.91 | 0.0143 |
| Negative regulation of signaling | 100 | 9.07 | 0.0144 |
| Negative regulation of cell communication | 100 | 9.07 | 0.0144 |
| Negative regulation of protein metabolic process | 52 | 4.72 | 0.0147 |
| Protein phosphorylation | 31 | 2.81 | 0.0147 |
| Manchette assembly | 4 | 0.36 | 0.0152 |
| Ventral spinal cord interneuron fate commitment | 4 | 0.36 | 0.0152 |
| Cell fate commitment involved in pattern specification | 4 | 0.36 | 0.0152 |
| Vascular endothelial growth factor signaling pathway | 9 | 0.82 | 0.0153 |
| Ribose phosphate metabolic process | 27 | 2.45 | 0.0154 |
| Exploration behavior | 7 | 0.64 | 0.0154 |
| Central nervous system projection neuron axonogenesis | 6 | 0.54 | 0.0155 |
| Lens fiber cell differentiation | 6 | 0.54 | 0.0155 |
| Cell communication by electrical coupling | 5 | 0.45 | 0.0158 |
| Chondroitin sulfate biosynthetic process | 5 | 0.45 | 0.0158 |
| Endocrine system development | 14 | 1.27 | 0.0167 |
| Regulation of cell development | 59 | 5.35 | 0.0167 |
| Embryonic pattern specification | 10 | 0.91 | 0.0168 |
| Regulation of neurotransmitter transport | 12 | 1.09 | 0.0168 |
| Negative regulation of phosphate metabolic process | 23 | 2.09 | 0.0168 |
| Negative regulation of phosphorus metabolic process | 23 | 2.09 | 0.0168 |
| Positive regulation of multicellular organismal process | 109 | 9.89 | 0.0173 |
| Regulation of molecular function | 93 | 8.44 | 0.0174 |
| Midbrain development | 11 | 1 | 0.0176 |
| Regulation of protein metabolic process | 114 | 10.34 | 0.0176 |
| Single fertilization | 17 | 1.54 | 0.0176 |
| Negative regulation of cellular component organization | 51 | 4.63 | 0.0177 |
| Zinc ion transmembrane transport | 6 | 0.54 | 0.0177 |
| Peptidyl-amino acid modification | 29 | 2.63 | 0.0177 |
| Protein localization to organelle | 55 | 4.99 | 0.018 |
| Negative regulation of dna-templated transcription | 83 | 7.53 | 0.0181 |
| Vascular endothelial growth factor receptor signaling pathway | 10 | 0.91 | 0.0181 |
| Response to insulin | 23 | 2.09 | 0.0182 |
| Nucleoside phosphate metabolic process | 43 | 3.9 | 0.0187 |
| Metanephric nephron epithelium development | 5 | 0.45 | 0.0187 |
| Vitamin transmembrane transport | 5 | 0.45 | 0.0187 |
| Negative regulation of protein modification by small protein conjugation or removal | 11 | 1 | 0.0188 |
| Regulation of erk1 and erk2 cascade | 25 | 2.27 | 0.019 |
| Cellular response to macrophage colony-stimulating factor stimulus | 8 | 0.73 | 0.0191 |
| Negative regulation of cell development | 24 | 2.18 | 0.0191 |
| Actin filament-based movement | 12 | 1.09 | 0.0191 |
| Regulation of neuron differentiation | 18 | 1.63 | 0.0193 |
| Negative regulation of protein ubiquitination | 10 | 0.91 | 0.0195 |
| Lateral ventricle development | 4 | 0.36 | 0.0195 |
| Mesenchymal cell apoptotic process | 4 | 0.36 | 0.0195 |
| Glomerulus morphogenesis | 4 | 0.36 | 0.0195 |
| Metanephric glomerulus development | 4 | 0.36 | 0.0195 |
| Negative regulation of cell adhesion mediated by integrin | 4 | 0.36 | 0.0195 |
| Regulation of cell junction assembly | 23 | 2.09 | 0.0196 |
| Cell junction assembly | 25 | 2.27 | 0.0197 |
| Chondroitin sulfate proteoglycan metabolic process | 6 | 0.54 | 0.0202 |
| Zinc ion transport | 6 | 0.54 | 0.0202 |

| | | | |
|---|---|---|---|
| Face morphogenesis | 6 | 0.54 | 0.0202 |
| Ribonucleotide metabolic process | 26 | 2.36 | 0.0203 |
| Signal transduction | 281 | 25.5 | 0.0205 |
| Cerebral cortex development | 14 | 1.27 | 0.0208 |
| Hepatocyte growth factor receptor signaling pathway | 8 | 0.73 | 0.0209 |
| Platelet-derived growth factor receptor-beta signaling pathway | 8 | 0.73 | 0.0209 |
| Positive regulation of bmp signaling pathway | 7 | 0.64 | 0.0213 |
| Cellular response to chemical stimulus | 124 | 11.25 | 0.0214 |
| Negative regulation of apoptotic process | 63 | 5.72 | 0.0217 |
| Amino acid transport | 14 | 1.27 | 0.0219 |
| Negative regulation of neurogenesis | 14 | 1.27 | 0.0219 |
| Lens morphogenesis in camera-type eye | 5 | 0.45 | 0.022 |
| Tendon development | 3 | 0.27 | 0.0225 |
| Ampa glutamate receptor clustering | 3 | 0.27 | 0.0225 |
| Glutamate receptor clustering | 3 | 0.27 | 0.0225 |
| Cell surface receptor signaling pathway involved in heart development | 3 | 0.27 | 0.0225 |
| Positive regulation of glutamate receptor signaling pathway | 3 | 0.27 | 0.0225 |
| Positive regulation of transmembrane transport | 19 | 1.72 | 0.0226 |
| Negative regulation of rna biosynthetic process | 83 | 7.53 | 0.0226 |
| Purine nucleotide metabolic process | 30 | 2.72 | 0.0227 |
| Glial cell differentiation | 20 | 1.81 | 0.023 |
| Negative regulation of supramolecular fiber organization | 16 | 1.45 | 0.0234 |
| Morphogenesis of a branching epithelium | 16 | 1.45 | 0.0234 |
| Positive regulation of dendritic spine development | 7 | 0.64 | 0.0236 |
| Glycosaminoglycan biosynthetic process | 7 | 0.64 | 0.0236 |
| Formation of primary germ layer | 13 | 1.18 | 0.0238 |
| Actomyosin structure organization | 13 | 1.18 | 0.0238 |
| Cell-cell adhesion | 40 | 3.63 | 0.024 |
| Cell-cell junction organization | 17 | 1.54 | 0.0243 |
| Auditory behavior | 4 | 0.36 | 0.0244 |
| Telencephalon regionalization | 4 | 0.36 | 0.0244 |
| Short-term memory | 4 | 0.36 | 0.0244 |
| Neurotransmitter-gated ion channel clustering | 4 | 0.36 | 0.0244 |
| Ventral spinal cord interneuron differentiation | 4 | 0.36 | 0.0244 |
| Artery morphogenesis | 9 | 0.82 | 0.0248 |
| Peptidyl-tyrosine phosphorylation | 8 | 0.73 | 0.0249 |
| Nervous system process | 93 | 8.44 | 0.0252 |
| In utero embryonic development | 31 | 2.81 | 0.0252 |
| Ameboidal-type cell migration | 14 | 1.27 | 0.0256 |
| Nucleobase-containing small molecule metabolic process | 44 | 3.99 | 0.0256 |
| Chondroitin sulfate metabolic process | 5 | 0.45 | 0.0256 |
| Striatum development | 5 | 0.45 | 0.0256 |
| Response to transforming growth factor beta | 16 | 1.45 | 0.0257 |
| Response to muscle activity | 6 | 0.54 | 0.0257 |
| Regulation of presynapse organization | 6 | 0.54 | 0.0257 |
| Positive regulation of cell junction assembly | 12 | 1.09 | 0.0257 |
| Ephrin receptor signaling pathway | 10 | 0.91 | 0.0258 |
| Response to activity | 10 | 0.91 | 0.0258 |
| Anterior/posterior pattern specification | 19 | 1.72 | 0.0265 |
| Positive regulation of fat cell differentiation | 9 | 0.82 | 0.0268 |
| Calcium ion homeostasis | 20 | 1.81 | 0.0269 |
| Cell recognition | 14 | 1.27 | 0.0269 |
| Response to macrophage colony-stimulating factor | 8 | 0.73 | 0.027 |
| Peptidyl-tyrosine modification | 8 | 0.73 | 0.027 |
| Blood circulation | 33 | 2.99 | 0.0272 |

| | | | |
|---|---|---|---|
| Positive regulation of transmembrane receptor protein serine/threonine kinase signaling pathway | 12 | 1.09 | 0.0273 |
| Platelet-derived growth factor receptor signaling pathway | 10 | 0.91 | 0.0276 |
| Stem cell development | 10 | 0.91 | 0.0276 |
| Mammary gland development | 13 | 1.18 | 0.028 |
| Negative regulation of phosphorylation | 19 | 1.72 | 0.0287 |
| Purine ribonucleotide metabolic process | 24 | 2.18 | 0.0287 |
| Nucleobase metabolic process | 6 | 0.54 | 0.0288 |
| Regulation of mesenchymal cell proliferation | 6 | 0.54 | 0.0288 |
| Spinal cord development | 11 | 1 | 0.0294 |
| Hyperosmotic response | 5 | 0.45 | 0.0296 |
| Regulation of cell shape | 14 | 1.27 | 0.0297 |
| Intraciliary retrograde transport | 4 | 0.36 | 0.0299 |
| Lipid import into cell | 4 | 0.36 | 0.0299 |
| Protein localization to nuclear envelope | 4 | 0.36 | 0.0299 |
| Epithelial cilium movement involved in determination of left/right asymmetry | 4 | 0.36 | 0.0299 |
| L-ascorbic acid metabolic process | 4 | 0.36 | 0.0299 |
| Negative regulation of programmed cell death | 64 | 5.81 | 0.0303 |
| Positive regulation of transport | 58 | 5.26 | 0.0306 |
| Establishment of localization in cell | 103 | 9.35 | 0.0309 |
| Mesoderm formation | 9 | 0.82 | 0.0309 |
| Receptor clustering | 7 | 0.64 | 0.0314 |
| Regulation of long-term synaptic potentiation | 7 | 0.64 | 0.0314 |
| Regulation of neurotransmitter secretion | 10 | 0.91 | 0.0314 |
| Positive regulation of epithelial to mesenchymal transition | 8 | 0.73 | 0.0318 |
| Regulation of transmembrane transport | 35 | 3.18 | 0.0321 |
| Regulation of blood circulation | 22 | 2 | 0.0322 |
| Renal system process | 13 | 1.18 | 0.0327 |
| Regulation of prostatic bud formation | 3 | 0.27 | 0.0327 |
| Determination of pancreatic left/right asymmetry | 3 | 0.27 | 0.0327 |
| Trunk segmentation | 3 | 0.27 | 0.0327 |
| L-histidine transmembrane transport | 3 | 0.27 | 0.0327 |
| Metanephric loop of henle development | 3 | 0.27 | 0.0327 |
| Trunk neural crest cell migration | 3 | 0.27 | 0.0327 |
| Negative regulation of nervous system development | 14 | 1.27 | 0.0327 |
| Regulation of axonogenesis | 14 | 1.27 | 0.0327 |
| Positive regulation of cell development | 34 | 3.09 | 0.033 |
| Transition metal ion transport | 11 | 1 | 0.033 |
| Forebrain neuron differentiation | 9 | 0.82 | 0.0332 |
| Neuron projection extension | 10 | 0.91 | 0.0335 |
| Nephron tubule development | 10 | 0.91 | 0.0335 |
| Glomerulus vasculature development | 5 | 0.45 | 0.0339 |
| Metanephric epithelium development | 5 | 0.45 | 0.0339 |
| Chondroitin sulfate proteoglycan biosynthetic process | 5 | 0.45 | 0.0339 |
| Cardiac septum development | 12 | 1.09 | 0.034 |
| Cardiac ventricle development | 13 | 1.18 | 0.0343 |
| Cellular response to toxic substance | 13 | 1.18 | 0.0343 |
| Negative regulation of transmembrane receptor protein serine/threonine kinase signaling pathway | 18 | 1.63 | 0.0345 |
| Regulation of organelle assembly | 20 | 1.81 | 0.0348 |
| Negative regulation of extrinsic apoptotic signaling pathway | 11 | 1 | 0.035 |
| Mesoderm morphogenesis | 9 | 0.82 | 0.0355 |
| Pancreas development | 9 | 0.82 | 0.0355 |
| Pyrimidine nucleotide biosynthetic process | 6 | 0.54 | 0.0356 |
| Striated muscle cell development | 14 | 1.27 | 0.0359 |

| | | | |
|---|---|---|---|
| Ossification | 24 | 2.18 | 0.036 |
| Cell communication by electrical coupling involved in cardiac conduction | 4 | 0.36 | 0.036 |
| Mechanosensory behavior | 4 | 0.36 | 0.036 |
| Negative regulation of transforming growth factor beta receptor signaling pathway | 13 | 1.18 | 0.0361 |
| Negative regulation of rna metabolic process | 87 | 7.89 | 0.0363 |
| Artery development | 11 | 1 | 0.037 |
| Negative regulation of cytoskeleton organization | 15 | 1.36 | 0.0371 |
| Cellular response to transforming growth factor beta stimulus | 15 | 1.36 | 0.0371 |
| Phosphate-containing compound metabolic process | 92 | 8.35 | 0.0371 |
| Regulation of protein localization | 61 | 5.54 | 0.0374 |
| Schwann cell differentiation | 7 | 0.64 | 0.0374 |
| Morphogenesis of a polarized epithelium | 7 | 0.64 | 0.0374 |
| Cell differentiation in spinal cord | 7 | 0.64 | 0.0374 |
| Cellular response to nitrogen compound | 44 | 3.99 | 0.0376 |
| Kidney morphogenesis | 10 | 0.91 | 0.0378 |
| Ear morphogenesis | 13 | 1.18 | 0.0379 |
| Morphogenesis of a branching structure | 16 | 1.45 | 0.0379 |
| Negative regulation of cell differentiation | 50 | 4.54 | 0.0381 |
| Regulation of lymphocyte chemotaxis | 5 | 0.45 | 0.0385 |
| Retina layer formation | 5 | 0.45 | 0.0385 |
| Regeneration | 15 | 1.36 | 0.0388 |
| Regulation of neural precursor cell proliferation | 11 | 1 | 0.0391 |
| Bone morphogenesis | 11 | 1 | 0.0391 |
| Osteoblast differentiation | 14 | 1.27 | 0.0394 |
| Cardiac muscle tissue growth | 6 | 0.54 | 0.0394 |
| Amino acid import across plasma membrane | 6 | 0.54 | 0.0394 |
| Phosphorus metabolic process | 92 | 8.35 | 0.0397 |
| Bone mineralization | 8 | 0.73 | 0.0398 |
| Urogenital system development | 8 | 0.73 | 0.0398 |
| Positive regulation of synaptic transmission | 15 | 1.36 | 0.0404 |
| Cardiac septum morphogenesis | 9 | 0.82 | 0.0405 |
| Regulation of myoblast differentiation | 9 | 0.82 | 0.0405 |
| Regulation of sodium ion transmembrane transport | 7 | 0.64 | 0.0407 |
| Cellular response to fibroblast growth factor stimulus | 13 | 1.18 | 0.0417 |
| Regulation of body fluid levels | 28 | 2.54 | 0.0422 |
| Neural crest cell differentiation | 10 | 0.91 | 0.0426 |
| Glomerular epithelial cell development | 4 | 0.36 | 0.0428 |
| Spinal cord dorsal/ventral patterning | 4 | 0.36 | 0.0428 |
| Head morphogenesis | 6 | 0.54 | 0.0434 |
| Maintenance of blood-brain barrier | 6 | 0.54 | 0.0434 |
| Epithelial tube branching involved in lung morphogenesis | 5 | 0.45 | 0.0435 |
| Regulation of bicellular tight junction assembly | 5 | 0.45 | 0.0435 |
| Gtp metabolic process | 5 | 0.45 | 0.0435 |
| Response to tumor cell | 5 | 0.45 | 0.0435 |
| Endothelium development | 11 | 1 | 0.0436 |
| Regulation of protein phosphorylation | 43 | 3.9 | 0.0437 |
| Semi-lunar valve development | 7 | 0.64 | 0.0441 |
| Cerebral cortex cell migration | 7 | 0.64 | 0.0441 |
| Glial cell migration | 7 | 0.64 | 0.0441 |
| Ventral spinal cord interneuron specification | 3 | 0.27 | 0.0443 |
| Mammary gland formation | 3 | 0.27 | 0.0443 |
| Establishment of left/right asymmetry | 3 | 0.27 | 0.0443 |
| Imitative learning | 3 | 0.27 | 0.0443 |
| Cell fate specification involved in pattern specification | 3 | 0.27 | 0.0443 |
| Metanephric glomerulus vasculature development | 3 | 0.27 | 0.0443 |

| | | | |
|---|---|---|---|
| Sympathetic neuron projection extension | 3 | 0.27 | 0.0443 |
| Metanephric distal tubule development | 3 | 0.27 | 0.0443 |
| Vocal learning | 3 | 0.27 | 0.0443 |
| Regulation of autophagy | 28 | 2.54 | 0.0444 |
| Regulation of dna-binding transcription factor activity | 25 | 2.27 | 0.045 |
| Cellular oxidant detoxification | 10 | 0.91 | 0.0451 |
| Regulation of synapse assembly | 15 | 1.36 | 0.0458 |
| Transmission of nerve impulse | 8 | 0.73 | 0.0459 |
| Biomineral tissue development | 11 | 1 | 0.0459 |
| Regulation of sodium ion transport | 9 | 0.82 | 0.046 |
| Neural crest cell development | 9 | 0.82 | 0.046 |
| Developmental cell growth | 12 | 1.09 | 0.0461 |
| Negative regulation of organelle organization | 26 | 2.36 | 0.0464 |
| Regulation of protein localization to membrane | 17 | 1.54 | 0.0465 |
| Cerebral cortex radially oriented cell migration | 6 | 0.54 | 0.0477 |
| Heart growth | 6 | 0.54 | 0.0477 |
| Modified amino acid transport | 6 | 0.54 | 0.0477 |
| Renal tubule development | 10 | 0.91 | 0.0477 |
| Post-embryonic development | 10 | 0.91 | 0.0477 |
| Endothelial cell development | 7 | 0.64 | 0.0477 |
| Aminoglycan biosynthetic process | 7 | 0.64 | 0.0477 |
| Regulation of neurotransmitter receptor activity | 5 | 0.45 | 0.0488 |
| Protein localization to membrane | 34 | 3.09 | 0.0488 |
| Nucleoside monophosphate metabolic process | 9 | 0.82 | 0.0489 |
| Actin filament bundle organization | 8 | 0.73 | 0.0491 |
| Type b pancreatic cell development | 4 | 0.36 | 0.05 |
| Lactone metabolic process | 4 | 0.36 | 0.05 |

TableS17

| Marker genes for each cluster | | | | | | |
|---|---|---|---|---|---|---|
| gene | cluster | average log2FoldChange | pct.1 | pct.2 | p-value | adjusted p-value |
| ITGBL1 | 0 | 3.145816803 | 0.602 | 0.053 | 0 | 0 |
| PDE1C | 0 | 2.630128709 | 0.752 | 0.128 | 0 | 0 |
| AL008633.1 | 0 | 2.355177536 | 0.967 | 0.608 | 0 | 0 |
| VGLL3 | 0 | 2.188805199 | 0.686 | 0.159 | 0 | 0 |
| KCNN2 | 0 | 2.148102457 | 0.663 | 0.151 | 0 | 0 |
| FGF14 | 0 | 1.928091378 | 0.934 | 0.543 | 0 | 0 |
| LINC02398 | 0 | 1.839969681 | 0.425 | 0.059 | 0 | 0 |
| FAP | 0 | 1.766580249 | 0.525 | 0.093 | 0 | 0 |
| LAMA4 | 0 | 1.714651271 | 0.747 | 0.337 | 0 | 0 |
| SORCS3 | 0 | 1.677674881 | 0.768 | 0.283 | 0 | 0 |
| AFF3 | 0 | 1.641481071 | 0.912 | 0.483 | 0 | 0 |
| AK5 | 0 | 1.626616166 | 0.48 | 0.121 | 0 | 0 |
| LRRC4C | 0 | 1.624639231 | 0.782 | 0.427 | 0 | 0 |
| LIMCH1 | 0 | 1.542120808 | 0.9 | 0.596 | 0 | 0 |
| NALCN | 0 | 1.470226493 | 0.544 | 0.094 | 0 | 0 |
| GPC6 | 0 | 1.464951226 | 0.988 | 0.747 | 0 | 0 |
| CFAP58 | 0 | 1.454506155 | 0.597 | 0.175 | 0 | 0 |
| CA10 | 0 | 1.377415715 | 0.617 | 0.196 | 0 | 0 |
| PPFIA2 | 0 | 1.340608153 | 0.935 | 0.641 | 0 | 0 |
| SLC22A3 | 0 | 1.257800592 | 0.687 | 0.294 | 0 | 0 |
| ARHGEF28 | 0 | 1.248324166 | 0.643 | 0.239 | 0 | 0 |
| OTUD7A | 0 | 1.20480296 | 0.814 | 0.468 | 0 | 0 |
| TBX15 | 0 | 1.190122298 | 0.816 | 0.419 | 0 | 0 |
| FAM13A | 0 | 1.174852178 | 0.82 | 0.454 | 0 | 0 |
| TCF4 | 0 | 1.169342914 | 0.989 | 0.849 | 0 | 0 |
| GLDC | 0 | 1.161175091 | 0.582 | 0.153 | 0 | 0 |
| KCNB2 | 0 | 1.05229019 | 0.899 | 0.412 | 0 | 0 |
| IKZF3 | 0 | 1.014477229 | 0.552 | 0.153 | 0 | 0 |
| IGF2BP1 | 0 | -0.930591211 | 0.041 | 0.424 | 0 | 0 |
| CDH2 | 0 | -1.411194083 | 0.173 | 0.595 | 0 | 0 |
| COBLL1 | 0 | -1.462410964 | 0.095 | 0.498 | 0 | 0 |
| GABRB3 | 0 | -1.528303208 | 0.096 | 0.49 | 0 | 0 |
| NBEA | 0 | -1.70623001 | 0.47 | 0.798 | 0 | 0 |
| UBE2E2 | 0 | -1.733700645 | 0.277 | 0.731 | 0 | 0 |
| IL1RAPL1 | 0 | -2.474796211 | 0.36 | 0.777 | 0 | 0 |
| HS6ST3 | 0 | -2.958893525 | 0.22 | 0.62 | 0 | 0 |
| ADAMTSL1 | 0 | -3.037051124 | 0.185 | 0.66 | 0 | 0 |
| NRG3 | 0 | -3.704579441 | 0.098 | 0.614 | 0 | 0 |
| MME | 0 | 1.184915654 | 0.441 | 0.087 | 3.197e-321 | 1.17030896e-316 |
| MIR99AHG | 0 | 1.118301984 | 0.906 | 0.597 | 4.1e-321 | 1.5013237e-316 |
| PAK3 | 0 | 0.948991046 | 0.964 | 0.83 | 3.122653e-318 | 1.1432344835e-313 |
| RIMS2 | 0 | -2.304202884 | 0.172 | 0.531 | 1.0948479822e-313 | 0 |
| SYT14 | 0 | -1.369691186 | 0.185 | 0.592 | 2.82323945099e-312 | 1.03E-307 |
| AC024901.1 | 0 | 1.504290773 | 0.417 | 0.084 | 5.012971302713e-312 | 1.84E-307 |
| PDE3A | 0 | 1.226387461 | 0.826 | 0.464 | 7.792055573637e-312 | 2.85E-307 |
| IGF2BP2 | 0 | -1.55418463 | 0.148 | 0.535 | 4.67E-308 | 1.71E-303 |
| LAMA2 | 0 | 1.464588216 | 0.669 | 0.279 | 7.14E-308 | 2.61E-303 |
| PLPP4 | 0 | 1.266875068 | 0.613 | 0.272 | 3.29E-305 | 1.2E-300 |
| CDH12 | 0 | 1.689222679 | 0.663 | 0.292 | 1.33E-303 | 4.86E-299 |
| IRAK3 | 0 | 0.951814789 | 0.759 | 0.383 | 3.07E-303 | 1.12E-298 |
| TFAP2B | 0 | 1.086252161 | 0.504 | 0.129 | 6.7E-301 | 2.45E-296 |
| PIEZO2 | 0 | 1.095200213 | 0.662 | 0.26 | 1.58E-297 | 5.77E-293 |
| WDR11 | 0 | 1.094875994 | 0.829 | 0.613 | 7.2E-294 | 2.64E-289 |
| LUZP2 | 0 | 1.293390657 | 0.817 | 0.45 | 1.93E-293 | 7.05E-289 |
| CADPS | 0 | -1.689454446 | 0.197 | 0.575 | 3.92E-291 | 1.43E-286 |

| | | | | | | |
|---|---|---|---|---|---|---|
| COL3A1 | 0 | 0.69316956 | 0.55 | 0.165 | 1.83E-289 | 6.7E-285 |
| C4orf19 | 0 | -1.222689079 | 0.059 | 0.405 | 7.49E-289 | 2.74E-284 |
| HS3ST4 | 0 | -1.976508036 | 0.087 | 0.441 | 1.19E-288 | 4.37E-284 |
| LDB2 | 0 | -1.865379239 | 0.089 | 0.437 | 7.68E-287 | 2.81E-282 |
| NKAIN3 | 0 | 1.159887217 | 0.889 | 0.601 | 3.95E-284 | 1.44E-279 |
| DPP4 | 0 | 1.099954292 | 0.401 | 0.082 | 1.53E-282 | 5.58E-278 |
| WDR11-AS1 | 0 | 1.155381129 | 0.556 | 0.231 | 1.64E-281 | 6.01E-277 |
| GHR | 0 | 1.14395548 | 0.456 | 0.11 | 2.54E-280 | 9.29E-276 |
| PPP2R2B | 0 | 1.254463939 | 0.846 | 0.569 | 6.07E-280 | 2.22E-275 |
| COL24A1 | 0 | 1.234496036 | 0.429 | 0.098 | 1.95E-272 | 7.15E-268 |
| ESRRG | 0 | 1.340170655 | 0.667 | 0.317 | 4.49E-271 | 1.64E-266 |
| KCNH1 | 0 | 1.558153611 | 0.503 | 0.173 | 3.45E-269 | 1.26E-264 |
| DPP10 | 0 | -1.447226511 | 0.356 | 0.707 | 2.01E-268 | 7.35E-264 |
| ZSWIM5 | 0 | -1.140628683 | 0.128 | 0.488 | 2.1E-262 | 7.68E-258 |
| NCALD | 0 | 1.144749963 | 0.784 | 0.479 | 1.04E-260 | 3.79E-256 |
| SORBS1 | 0 | 1.001043088 | 0.763 | 0.437 | 1.98E-260 | 7.24E-256 |
| CELF2 | 0 | -1.133794262 | 0.366 | 0.723 | 1.72E-255 | 6.31E-251 |
| SIPA1L1 | 0 | 0.923762552 | 0.909 | 0.715 | 3.44E-255 | 1.26E-250 |
| EFNA5 | 0 | -1.569730803 | 0.47 | 0.742 | 4.73E-255 | 1.73E-250 |
| KCNJ6 | 0 | 1.13715512 | 0.433 | 0.108 | 4.09E-254 | 1.5E-249 |
| SYN3 | 0 | -1.712888583 | 0.181 | 0.521 | 8.16E-253 | 2.99E-248 |
| HMGA2 | 0 | 0.998991854 | 0.689 | 0.322 | 1.32E-250 | 4.82E-246 |
| SH3GL2 | 0 | 1.153701208 | 0.635 | 0.298 | 3.17E-249 | 1.16E-244 |
| C1orf61 | 0 | 1.143915854 | 0.528 | 0.211 | 3.2E-249 | 1.17E-244 |
| PLCL1 | 0 | 1.183243172 | 0.681 | 0.318 | 3.26E-249 | 1.2E-244 |
| SNTG1 | 0 | -2.334354653 | 0.15 | 0.481 | 6.71E-248 | 2.46E-243 |
| LINC00461 | 0 | 0.994912891 | 0.74 | 0.406 | 8.15E-248 | 2.98E-243 |
| CFH | 0 | 1.166673742 | 0.35 | 0.064 | 1.03E-243 | 3.77E-239 |
| ADGRL3 | 0 | -1.371609437 | 0.651 | 0.88 | 1.42E-243 | 5.19E-239 |
| CLCN5 | 0 | 1.003721387 | 0.661 | 0.347 | 6.09E-243 | 2.23E-238 |
| AC073050.1 | 0 | -1.539707441 | 0.076 | 0.397 | 1.01E-242 | 3.69E-238 |
| FAT1 | 0 | 1.009787111 | 0.802 | 0.537 | 1.38E-242 | 5.04E-238 |
| SEMA5A | 0 | 1.768349854 | 0.592 | 0.333 | 6.69E-242 | 2.45E-237 |
| XACT | 0 | -2.894240763 | 0.051 | 0.33 | 2.42E-241 | 8.85E-237 |
| SDK1 | 0 | 1.312434986 | 0.957 | 0.883 | 2.35E-240 | 8.59E-236 |
| DOCK3 | 0 | -1.420690895 | 0.075 | 0.37 | 5.32E-240 | 1.95E-235 |
| PCDH11X | 0 | 1.606272462 | 0.504 | 0.166 | 1.15E-239 | 4.2E-235 |
| NDST3 | 0 | 1.185913154 | 0.708 | 0.384 | 7.25E-237 | 2.65E-232 |
| FRMD4A | 0 | 0.918897035 | 0.944 | 0.766 | 1.17E-234 | 4.27E-230 |
| DNAJC1 | 0 | 1.072463062 | 0.748 | 0.499 | 1.98E-233 | 7.25E-229 |
| TNMD | 0 | 0.6990695 | 0.22 | 0.014 | 1.58E-232 | 5.77E-228 |
| NAALAD2 | 0 | 0.859008723 | 0.463 | 0.148 | 5.17E-229 | 1.89E-224 |
| RAB3C | 0 | -1.315829866 | 0.186 | 0.525 | 1.13E-228 | 4.13E-224 |
| DLC1 | 0 | 0.953624596 | 0.509 | 0.168 | 1.86E-228 | 6.81E-224 |
| NELL2 | 0 | -1.378619168 | 0.055 | 0.339 | 1.73E-227 | 6.34E-223 |
| FIRRE | 0 | -1.394619652 | 0.087 | 0.394 | 4.86E-225 | 1.78E-220 |
| ZBTB20 | 0 | 0.944406135 | 0.922 | 0.733 | 6.86E-224 | 2.51E-219 |
| EPS8 | 0 | 1.051732821 | 0.773 | 0.511 | 1.16E-223 | 4.24E-219 |
| UNC5D | 0 | 0.844272539 | 0.824 | 0.478 | 1.17E-222 | 4.28E-218 |
| NFIB | 0 | 1.016916915 | 0.935 | 0.791 | 2.54E-221 | 9.32E-217 |
| MIR137HG | 0 | -1.307744843 | 0.05 | 0.331 | 9.17E-220 | 3.36E-215 |
| RASGRF2 | 0 | 1.240157484 | 0.373 | 0.103 | 1.38E-216 | 5.04E-212 |
| SEMA3E | 0 | -1.406795838 | 0.089 | 0.386 | 3.16E-216 | 1.16E-211 |
| MTUS2 | 0 | -0.978630404 | 0.027 | 0.284 | 6.43E-214 | 2.36E-209 |
| AL110292.1 | 0 | 1.383212756 | 0.198 | 0.019 | 2.05E-212 | 7.52E-208 |
| PDE10A | 0 | -1.117568453 | 0.056 | 0.333 | 9.05E-211 | 3.31E-206 |
| SPATS2L | 0 | 1.023036546 | 0.684 | 0.383 | 9.55E-210 | 3.49E-205 |
| COL25A1 | 0 | -1.555375282 | 0.165 | 0.483 | 3.68E-209 | 1.35E-204 |

| | | | | | | |
|---|---|---|---|---|---|---|
| CTTNBP2 | 0 | -1.081164784 | 0.084 | 0.381 | 2.8E-208 | 1.02E-203 |
| ADAMTS20 | 0 | -1.188694366 | 0.047 | 0.31 | 2.76E-205 | 1.01E-200 |
| LINC02263 | 0 | 1.150521027 | 0.265 | 0.042 | 1.08E-204 | 3.96E-200 |
| FREM1 | 0 | 0.835538248 | 0.487 | 0.169 | 8.37E-204 | 3.06E-199 |
| HS3ST3B1 | 0 | 0.872694848 | 0.529 | 0.205 | 2.91E-202 | 1.07E-197 |
| PTGER3 | 0 | 0.963014005 | 0.617 | 0.299 | 1.65E-200 | 6.06E-196 |
| CNTN5 | 0 | 2.044374852 | 0.308 | 0.072 | 5.55E-200 | 2.03E-195 |
| RMST | 0 | -1.50536101 | 0.261 | 0.582 | 1.96E-199 | 7.19E-195 |
| DNAH14 | 0 | -1.089630652 | 0.303 | 0.609 | 6.15E-199 | 2.25E-194 |
| SLC35F1 | 0 | -1.093335789 | 0.095 | 0.387 | 3.49E-197 | 1.28E-192 |
| CACNA1B | 0 | -1.102686375 | 0.071 | 0.342 | 1.03E-196 | 3.76E-192 |
| PTPN14 | 0 | 0.850740848 | 0.511 | 0.197 | 3.94E-196 | 1.44E-191 |
| ASTN1 | 0 | -0.813169156 | 0.056 | 0.325 | 4.51E-195 | 1.65E-190 |
| DOK5 | 0 | 0.970844822 | 0.728 | 0.472 | 5.68E-195 | 2.08E-190 |
| UST | 0 | -1.135161137 | 0.082 | 0.356 | 9.36E-195 | 3.43E-190 |
| AC023282.1 | 0 | 0.934624634 | 0.51 | 0.209 | 5.98E-194 | 2.19E-189 |
| GRIP1 | 0 | -1.177923038 | 0.142 | 0.447 | 5.21E-192 | 1.91E-187 |
| NFIX | 0 | 0.750968895 | 0.697 | 0.374 | 1.29E-191 | 4.73E-187 |
| IL1RAPL2 | 0 | -2.220855123 | 0.024 | 0.252 | 1.09E-190 | 3.99E-186 |
| LRMP | 0 | 1.139230537 | 0.205 | 0.021 | 1.43E-190 | 5.24E-186 |
| PSME4 | 0 | 0.894640518 | 0.881 | 0.748 | 1.02E-189 | 3.75E-185 |
| MAPK10 | 0 | -1.265081458 | 0.053 | 0.296 | 2.63E-186 | 9.64E-182 |
| PTCHD4 | 0 | -1.19013339 | 0.068 | 0.326 | 3.3E-185 | 1.21E-180 |
| CHN1 | 0 | -0.966673784 | 0.425 | 0.683 | 5.01E-185 | 1.83E-180 |
| NHS | 0 | -1.187898489 | 0.25 | 0.564 | 3.01E-182 | 1.1E-177 |
| ABCB1 | 0 | -0.903041695 | 0.054 | 0.309 | 8.01E-182 | 2.93E-177 |
| PRKG1 | 0 | 1.071273442 | 0.847 | 0.687 | 6.48E-181 | 2.37E-176 |
| PPP1R14C | 0 | 1.001828396 | 0.659 | 0.361 | 2.44E-179 | 8.92E-175 |
| NRP1 | 0 | -1.107012309 | 0.121 | 0.389 | 3.68E-179 | 1.35E-174 |
| DNM3OS | 0 | 0.900797447 | 0.559 | 0.251 | 1.43E-178 | 5.24E-174 |
| RYR2 | 0 | -1.603088399 | 0.041 | 0.275 | 6.44E-178 | 2.36E-173 |
| KANK1 | 0 | 0.947871616 | 0.71 | 0.427 | 1.98E-177 | 7.25E-173 |
| THRB | 0 | -1.34548155 | 0.117 | 0.389 | 8.79E-177 | 3.22E-172 |
| MEIS2 | 0 | 0.371526339 | 0.767 | 0.447 | 1.23E-175 | 4.51E-171 |
| DANT2 | 0 | -1.118137841 | 0.129 | 0.408 | 1.54E-175 | 5.63E-171 |
| NKAIN2 | 0 | -2.025015572 | 0.188 | 0.452 | 1.31E-174 | 4.81E-170 |
| ASTN2 | 0 | 1.013468925 | 0.682 | 0.445 | 3.09E-173 | 1.13E-168 |
| NRXN3 | 0 | -1.931988761 | 0.153 | 0.436 | 5.35E-172 | 1.96E-167 |
| PTPRN2 | 0 | -1.716590586 | 0.09 | 0.347 | 4.3E-171 | 1.57E-166 |
| FN1 | 0 | 1.006530499 | 0.58 | 0.281 | 4.97E-171 | 1.82E-166 |
| MSC-AS1 | 0 | -0.831509118 | 0.012 | 0.207 | 1.73E-170 | 6.32E-166 |
| SEMA3A | 0 | -1.565394745 | 0.097 | 0.367 | 3.75E-170 | 1.37E-165 |
| TOX3 | 0 | -0.921263804 | 0.105 | 0.384 | 4.83E-170 | 1.77E-165 |
| TMEM161B-AS1 | 0 | 0.844954002 | 0.794 | 0.608 | 1.91E-169 | 6.98E-165 |
| KCND2 | 0 | 1.269591671 | 0.72 | 0.467 | 2.37E-169 | 8.67E-165 |
| CDCP1 | 0 | 0.798322047 | 0.366 | 0.116 | 2.57E-169 | 9.41E-165 |
| SCN9A | 0 | -0.851727019 | 0.014 | 0.21 | 2.02E-168 | 7.41E-164 |
| MOCOS | 0 | 0.592457471 | 0.271 | 0.058 | 2.1E-168 | 7.68E-164 |
| LIN28B | 0 | -0.954112348 | 0.298 | 0.579 | 5.98E-168 | 2.19E-163 |
| EYA1 | 0 | -1.127460187 | 0.102 | 0.37 | 1.18E-167 | 4.32E-163 |
| PAM | 0 | -1.109118416 | 0.631 | 0.806 | 1.34E-167 | 4.9E-163 |
| LINC02197 | 0 | 0.861396038 | 0.35 | 0.097 | 1.73E-167 | 6.35E-163 |
| PPARG | 0 | -0.862132389 | 0.092 | 0.358 | 1.34E-166 | 4.89E-162 |
| CADPS2 | 0 | 0.377170564 | 0.417 | 0.161 | 6.75E-166 | 2.47E-161 |
| FAR2 | 0 | 1.109034881 | 0.557 | 0.315 | 3.87E-165 | 1.42E-160 |
| RGL1 | 0 | 0.593319247 | 0.893 | 0.658 | 5.75E-165 | 2.11E-160 |
| KLF12 | 0 | 0.804702082 | 0.783 | 0.585 | 1.77E-162 | 6.47E-158 |
| DGKI | 0 | -0.957010515 | 0.066 | 0.304 | 2.83E-162 | 1.04E-157 |

| | | | | | | |
|---|---|---|---|---|---|---|
| GLI3 | 0 | 0.671941698 | 0.56 | 0.257 | 2.84E-162 | 1.04E-157 |
| AC011586.2 | 0 | 1.153318172 | 0.231 | 0.041 | 3.39E-162 | 1.24E-157 |
| ADAMTS12 | 0 | -1.703246047 | 0.237 | 0.464 | 2.06E-160 | 7.55E-156 |
| CAMK4 | 0 | 0.792195485 | 0.504 | 0.214 | 2.06E-160 | 7.56E-156 |
| AEBP2 | 0 | 0.820738843 | 0.823 | 0.683 | 8.9E-160 | 3.26E-155 |
| LRRTM4 | 0 | -1.3426455 | 0.378 | 0.657 | 3E-159 | 1.1E-154 |
| AL691420.1 | 0 | -1.27252137 | 0.008 | 0.186 | 3.59E-159 | 1.31E-154 |
| SLC4A7 | 0 | 0.641685176 | 0.883 | 0.704 | 8.71E-159 | 3.19E-154 |
| TRIM71 | 0 | -0.738392458 | 0.079 | 0.322 | 2.1E-158 | 7.7E-154 |
| SIL1 | 0 | 0.76936144 | 0.789 | 0.612 | 2.75E-158 | 1.01E-153 |
| NMNAT2 | 0 | -0.855149899 | 0.128 | 0.399 | 1.09E-157 | 3.99E-153 |
| PPP1R9A | 0 | -0.980304798 | 0.029 | 0.226 | 1.68E-156 | 6.14E-152 |
| ANO4 | 0 | -1.4560418 | 0.445 | 0.639 | 2.86E-156 | 1.05E-151 |
| GLIS3 | 0 | -1.40458356 | 0.085 | 0.312 | 6.5E-155 | 2.38E-150 |
| CDK14 | 0 | 0.577838997 | 0.91 | 0.724 | 1.18E-154 | 4.31E-150 |
| DCDC2 | 0 | 0.892925512 | 0.209 | 0.034 | 1.05E-152 | 3.83E-148 |
| LRRC7 | 0 | -1.790085644 | 0.174 | 0.372 | 1.28E-152 | 4.7E-148 |
| HCN1 | 0 | -0.903827804 | 0.02 | 0.211 | 3.12E-152 | 1.14E-147 |
| LHFPL6 | 0 | 0.933446175 | 0.585 | 0.304 | 3.18E-152 | 1.16E-147 |
| CACNB4 | 0 | 0.348585644 | 0.756 | 0.462 | 5.37E-152 | 1.97E-147 |
| AL162718.1 | 0 | -0.724330192 | 0.011 | 0.187 | 4.13E-151 | 1.51E-146 |
| TDRD3 | 0 | 0.743614659 | 0.767 | 0.584 | 2.26E-150 | 8.27E-146 |
| ARPP21 | 0 | -1.337786712 | 0.142 | 0.385 | 2.95E-150 | 1.08E-145 |
| NSUN7 | 0 | 0.67918862 | 0.433 | 0.173 | 5.4E-150 | 1.98E-145 |
| ANKS1B | 0 | 0.747756941 | 0.912 | 0.739 | 7.72E-150 | 2.83E-145 |
| AC009803.1 | 0 | -0.649786333 | 0.008 | 0.177 | 2.11E-149 | 7.71E-145 |
| MCTP2 | 0 | -1.396741994 | 0.097 | 0.332 | 1.31E-148 | 4.78E-144 |
| GAS7 | 0 | -0.961301573 | 0.074 | 0.307 | 1.32E-148 | 4.82E-144 |
| EBF1 | 0 | -1.316183126 | 0.182 | 0.445 | 1.16E-147 | 4.23E-143 |
| PRSS12 | 0 | 0.600807213 | 0.939 | 0.741 | 3.82E-146 | 1.4E-141 |
| ADCY2 | 0 | 0.296562413 | 0.464 | 0.203 | 2.69E-145 | 9.85E-141 |
| C13orf42 | 0 | -0.646732879 | 0.01 | 0.18 | 8.35E-145 | 3.06E-140 |
| TFDP2 | 0 | 0.662350141 | 0.698 | 0.433 | 3.58E-144 | 1.31E-139 |
| DPYSL3 | 0 | -0.844124922 | 0.187 | 0.45 | 5.14E-143 | 1.88E-138 |
| SHISA6 | 0 | -0.863459777 | 0.029 | 0.221 | 6.12E-143 | 2.24E-138 |
| PLPPR1 | 0 | -0.973549458 | 0.05 | 0.253 | 9.98E-143 | 3.65E-138 |
| PCLO | 0 | -1.08293514 | 0.447 | 0.673 | 1.08E-142 | 3.95E-138 |
| EFHD1 | 0 | 0.594188519 | 0.283 | 0.074 | 1.36E-142 | 4.96E-138 |
| FMN2 | 0 | -0.90493113 | 0.164 | 0.429 | 3.35E-142 | 1.23E-137 |
| XKR4 | 0 | -0.981689047 | 0.042 | 0.245 | 4.15E-142 | 1.52E-137 |
| PAG1 | 0 | 0.804897346 | 0.48 | 0.235 | 5.79E-142 | 2.12E-137 |
| MPP6 | 0 | 0.898670586 | 0.751 | 0.604 | 7.19E-142 | 2.63E-137 |
| UNC13C | 0 | 1.456079535 | 0.192 | 0.038 | 9.86E-142 | 3.61E-137 |
| ZNF804B | 0 | -1.876187886 | 0.044 | 0.241 | 3.49E-141 | 1.28E-136 |
| CUX2 | 0 | 0.700314632 | 0.493 | 0.218 | 6.6E-141 | 2.42E-136 |
| SYCP2 | 0 | -1.264016682 | 0.074 | 0.288 | 6.62E-141 | 2.42E-136 |
| ITGA8 | 0 | 0.479293626 | 0.444 | 0.184 | 2.91E-139 | 1.07E-134 |
| VAV3 | 0 | -0.724883399 | 0.028 | 0.215 | 1.25E-138 | 4.57E-134 |
| GRID2 | 0 | 0.987456475 | 0.762 | 0.504 | 1.35E-138 | 4.93E-134 |
| DNAH10 | 0 | 0.81987975 | 0.653 | 0.394 | 3.54E-138 | 1.3E-133 |
| CXADR | 0 | -0.678900162 | 0.156 | 0.411 | 1.76E-137 | 6.45E-133 |
| RTN1 | 0 | -0.610517208 | 0.026 | 0.21 | 1.95E-137 | 7.12E-133 |
| SOX2-OT | 0 | -2.083620515 | 0.064 | 0.261 | 2.6E-137 | 9.53E-133 |
| CRISPLD1 | 0 | 1.435100599 | 0.646 | 0.436 | 7.86E-137 | 2.88E-132 |
| SPOCK1 | 0 | 1.030015437 | 0.54 | 0.29 | 2.47E-136 | 9.04E-132 |
| STXBP5L | 0 | -1.143112801 | 0.147 | 0.372 | 2.52E-136 | 9.21E-132 |
| LSAMP | 0 | -0.933646486 | 0.578 | 0.802 | 9.01E-136 | 3.3E-131 |
| MIR100HG | 0 | -1.03599382 | 0.668 | 0.775 | 3.05E-135 | 1.12E-130 |

| | | | | | | |
|---|---|---|---|---|---|---|
| MYO16 | 0 | 1.160818399 | 0.47 | 0.256 | 3.16E-135 | 1.16E-130 |
| ROBO1 | 0 | 0.558217002 | 0.979 | 0.869 | 5.05E-135 | 1.85E-130 |
| MPDZ | 0 | 0.621521744 | 0.812 | 0.645 | 5.2E-135 | 1.9E-130 |
| ERBB4 | 0 | -1.801436546 | 0.124 | 0.352 | 3.18E-133 | 1.16E-128 |
| CYP4X1 | 0 | -0.667679539 | 0.023 | 0.2 | 1.1E-132 | 4.01E-128 |
| ANK3 | 0 | -0.928265766 | 0.147 | 0.402 | 2.08E-132 | 7.63E-128 |
| BRINP3 | 0 | -1.519675441 | 0.339 | 0.578 | 3.18E-132 | 1.16E-127 |
| CCND2 | 0 | 0.641796218 | 0.495 | 0.246 | 3.32E-132 | 1.22E-127 |
| ZC3H12C | 0 | 0.784602797 | 0.728 | 0.534 | 9.81E-132 | 3.59E-127 |
| PLAGL1 | 0 | -0.906714531 | 0.185 | 0.427 | 4.39E-131 | 1.61E-126 |
| KCNIP4 | 0 | 0.299740906 | 0.692 | 0.421 | 4.41E-131 | 1.61E-126 |
| LINC01768 | 0 | -0.737656745 | 0.043 | 0.235 | 6.97E-131 | 2.55E-126 |
| PRAME | 0 | -0.43273552 | 0.014 | 0.177 | 1.58E-130 | 5.77E-126 |
| NTF3 | 0 | 0.807841606 | 0.316 | 0.108 | 2.04E-130 | 7.47E-126 |
| GULP1 | 0 | 0.366261081 | 0.705 | 0.427 | 2.77E-130 | 1.02E-125 |
| GUCY1A1 | 0 | 0.818741212 | 0.424 | 0.193 | 1.26E-129 | 4.6E-125 |
| MCC | 0 | 0.600363558 | 0.906 | 0.753 | 1.81E-129 | 6.62E-125 |
| MACROD2 | 0 | -0.986529528 | 0.397 | 0.639 | 1.83E-129 | 6.7E-125 |
| FBN2 | 0 | 1.31005522 | 0.502 | 0.328 | 1.86E-129 | 6.82E-125 |
| NCOA1 | 0 | 0.634474354 | 0.815 | 0.638 | 2.49E-129 | 9.12E-125 |
| MYO1D | 0 | -0.641805745 | 0.069 | 0.281 | 2.68E-129 | 9.82E-125 |
| LDLRAD4 | 0 | 0.780000079 | 0.684 | 0.418 | 3.73E-127 | 1.37E-122 |
| TSHZ3 | 0 | 0.641181386 | 0.68 | 0.416 | 3.5E-125 | 1.28E-120 |
| APBA1 | 0 | -1.04028494 | 0.282 | 0.504 | 5.09E-125 | 1.86E-120 |
| SORBS2 | 0 | -1.123542595 | 0.019 | 0.17 | 5.33E-124 | 1.95E-119 |
| GNG2 | 0 | -0.64739666 | 0.034 | 0.212 | 1.16E-123 | 4.26E-119 |
| TMEM163 | 0 | -0.777797205 | 0.031 | 0.203 | 2.57E-123 | 9.41E-119 |
| EFR3B | 0 | 0.644920703 | 0.632 | 0.383 | 1.48E-122 | 5.41E-118 |
| FIGN | 0 | -0.951018631 | 0.167 | 0.407 | 2.05E-122 | 7.51E-118 |
| GABRB2 | 0 | -0.964202941 | 0.138 | 0.374 | 2.75E-122 | 1.01E-117 |
| IGSF21 | 0 | 0.657019776 | 0.429 | 0.195 | 1.54E-121 | 5.63E-117 |
| SNAP91 | 0 | -0.659930412 | 0.051 | 0.238 | 2.37E-121 | 8.68E-117 |
| RGS7 | 0 | -0.98517107 | 0.067 | 0.257 | 3.12E-121 | 1.14E-116 |
| ANO5 | 0 | -0.719699143 | 0.095 | 0.313 | 3.31E-121 | 1.21E-116 |
| SYT1 | 0 | 0.513872504 | 0.895 | 0.681 | 9.05E-121 | 3.31E-116 |
| IL12RB2 | 0 | -1.223877652 | 0.148 | 0.341 | 1.07E-120 | 3.9E-116 |
| GLCCI1 | 0 | 0.786614334 | 0.696 | 0.53 | 1.96E-120 | 7.16E-116 |
| DNAH11 | 0 | -1.290720769 | 0.118 | 0.314 | 1.96E-120 | 7.17E-116 |
| GLT1D1 | 0 | -0.539191816 | 0.038 | 0.214 | 2.29E-120 | 8.4E-116 |
| GRB14 | 0 | -0.440175033 | 0.01 | 0.156 | 2.48E-120 | 9.07E-116 |
| FAM160A1 | 0 | -0.635506187 | 0.027 | 0.197 | 3.77E-120 | 1.38E-115 |
| FBXL7 | 0 | 0.504364743 | 0.946 | 0.792 | 4.75E-120 | 1.74E-115 |
| PPM1E | 0 | 0.656541245 | 0.845 | 0.692 | 1.11E-119 | 4.06E-115 |
| STEAP2 | 0 | -0.383290789 | 0.014 | 0.164 | 3.21E-119 | 1.17E-114 |
| CLECL1 | 0 | 0.601322779 | 0.135 | 0.013 | 6.7E-119 | 2.45E-114 |
| IGF1R | 0 | -0.646865957 | 0.681 | 0.843 | 8.67E-119 | 3.17E-114 |
| DACH2 | 0 | -1.589880003 | 0.33 | 0.5 | 1.31E-118 | 4.78E-114 |
| AC083906.3 | 0 | 0.378359554 | 0.167 | 0.025 | 1.32E-118 | 4.84E-114 |
| NTN1 | 0 | 0.580877308 | 0.287 | 0.088 | 1.59E-118 | 5.83E-114 |
| INTS6L | 0 | 0.652076708 | 0.546 | 0.314 | 1.87E-118 | 6.84E-114 |
| FAM184A | 0 | -0.744988491 | 0.26 | 0.5 | 3.54E-118 | 1.3E-113 |
| AHCYL2 | 0 | -0.91079089 | 0.562 | 0.708 | 3.59E-118 | 1.32E-113 |
| KIF21A | 0 | 0.680019966 | 0.636 | 0.399 | 4.94E-118 | 1.81E-113 |
| SNCAIP | 0 | 0.926676847 | 0.349 | 0.145 | 8.82E-118 | 3.23E-113 |
| HOXD3 | 0 | 0.449189407 | 0.305 | 0.101 | 4.36E-117 | 1.59E-112 |
| ZEB2 | 0 | 0.420623476 | 0.764 | 0.506 | 6.52E-117 | 2.39E-112 |
| KAZN | 0 | 0.282018646 | 0.985 | 0.864 | 7.87E-117 | 2.88E-112 |
| ADAMTS3 | 0 | 0.904161322 | 0.631 | 0.415 | 3.39E-116 | 1.24E-111 |

| | | | | | | |
|---|---|---|---|---|---|---|
| CLEC2D | 0 | 0.634320855 | 0.606 | 0.402 | 4.83E-116 | 1.77E-111 |
| CDH8 | 0 | -0.684644405 | 0.021 | 0.178 | 9.84E-116 | 3.6E-111 |
| ST6GAL1 | 0 | -0.877845951 | 0.212 | 0.435 | 1.08E-115 | 3.95E-111 |
| AC012150.1 | 0 | 0.411365524 | 0.128 | 0.012 | 1.11E-115 | 4.06E-111 |
| ZNF711 | 0 | 0.525711886 | 0.75 | 0.536 | 1.34E-115 | 4.9E-111 |
| IGFBP2 | 0 | 0.68886201 | 0.673 | 0.467 | 1.47E-115 | 5.4E-111 |
| SSX2IP | 0 | 0.638617949 | 0.662 | 0.448 | 1.8E-115 | 6.59E-111 |
| STK33 | 0 | -0.896581002 | 0.042 | 0.208 | 3.63E-115 | 1.33E-110 |
| RIMBP2 | 0 | -1.367100549 | 0.024 | 0.172 | 3.75E-115 | 1.37E-110 |
| SHANK2 | 0 | -0.797220244 | 0.078 | 0.276 | 8.6E-115 | 3.15E-110 |
| AKAP6 | 0 | -0.694371993 | 0.086 | 0.296 | 1.21E-114 | 4.44E-110 |
| COL4A2 | 0 | 0.459976801 | 0.516 | 0.261 | 1.64E-114 | 6E-110 |
| NID1 | 0 | 0.571227433 | 0.452 | 0.215 | 3.5E-114 | 1.28E-109 |
| NTM | 0 | 0.381642442 | 0.599 | 0.341 | 4.21E-114 | 1.54E-109 |
| SULF2 | 0 | 0.605938909 | 0.342 | 0.133 | 6.33E-114 | 2.32E-109 |
| BICC1 | 0 | 0.650738803 | 0.471 | 0.23 | 1.13E-113 | 4.14E-109 |
| ZNF827 | 0 | 0.621165739 | 0.697 | 0.484 | 1.69E-113 | 6.17E-109 |
| PALLD | 0 | -0.969268455 | 0.262 | 0.477 | 4.1E-113 | 1.5E-108 |
| NEO1 | 0 | 0.628004712 | 0.701 | 0.489 | 8.18E-113 | 3E-108 |
| CD200 | 0 | -0.416106415 | 0.036 | 0.207 | 1.65E-112 | 6.06E-108 |
| FTO | 0 | 0.61000048 | 0.895 | 0.777 | 2.29E-112 | 8.38E-108 |
| GPR176 | 0 | 0.62323529 | 0.566 | 0.334 | 2.64E-112 | 9.67E-108 |
| PHYHIPL | 0 | -0.570640665 | 0.038 | 0.208 | 2.91E-112 | 1.07E-107 |
| THSD7B | 0 | -1.09751298 | 0.083 | 0.264 | 3.46E-112 | 1.27E-107 |
| AC063938.1 | 0 | -0.630414877 | 0.01 | 0.146 | 3.75E-112 | 1.37E-107 |
| STAT1 | 0 | 0.745132514 | 0.417 | 0.225 | 4.43E-112 | 1.62E-107 |
| PDGFD | 0 | 0.805106722 | 0.327 | 0.124 | 1.33E-111 | 4.89E-107 |
| ZFAND6 | 0 | 0.565570747 | 0.78 | 0.605 | 1.6E-111 | 5.86E-107 |
| ELL2 | 0 | -1.075016266 | 0.066 | 0.237 | 2.15E-111 | 7.86E-107 |
| EML5 | 0 | -0.618582528 | 0.043 | 0.208 | 3.28E-111 | 1.2E-106 |
| TENM4 | 0 | -0.84096335 | 0.201 | 0.434 | 4.59E-111 | 1.68E-106 |
| MAN1C1 | 0 | 0.563020287 | 0.381 | 0.168 | 8.24E-111 | 3.02E-106 |
| RBMX | 0 | 0.479678089 | 0.892 | 0.798 | 1.02E-110 | 3.73E-106 |
| APCDD1 | 0 | 0.699457185 | 0.473 | 0.241 | 1.16E-110 | 4.24E-106 |
| PDGFRA | 0 | 0.644236853 | 0.341 | 0.13 | 1.53E-110 | 5.62E-106 |
| AC069228.1 | 0 | 0.534820256 | 0.318 | 0.121 | 2.89E-110 | 1.06E-105 |
| FNDC1 | 0 | 0.36308364 | 0.124 | 0.011 | 3.16E-110 | 1.16E-105 |
| RUNDC3B | 0 | -0.551017142 | 0.037 | 0.204 | 7.4E-110 | 2.71E-105 |
| AP002954.2 | 0 | -0.786626023 | 0.007 | 0.135 | 1.12E-109 | 4.12E-105 |
| AC040168.1 | 0 | -0.664262633 | 0.1 | 0.308 | 3.16E-109 | 1.16E-104 |
| RAPGEF4 | 0 | -0.858451696 | 0.234 | 0.471 | 4.96E-109 | 1.82E-104 |
| TENM1 | 0 | -0.797995777 | 0.022 | 0.175 | 5.34E-109 | 1.96E-104 |
| RBM47 | 0 | -1.032320928 | 0.028 | 0.17 | 1.94E-108 | 7.12E-104 |
| RAP1GAP2 | 0 | -1.30453118 | 0.115 | 0.314 | 3.32E-107 | 1.21E-102 |
| PDLIM5 | 0 | -0.741579594 | 0.228 | 0.46 | 3.67E-107 | 1.34E-102 |
| UNC79 | 0 | -0.877241098 | 0.198 | 0.402 | 5.38E-107 | 1.97E-102 |
| ARHGAP6 | 0 | 0.493509163 | 0.536 | 0.287 | 1.27E-106 | 4.64E-102 |
| AGBL4 | 0 | -1.204235012 | 0.086 | 0.257 | 1.53E-106 | 5.58E-102 |
| ZNF385B | 0 | -0.979983874 | 0.172 | 0.396 | 5.94E-106 | 2.17E-101 |
| BBX | 0 | 0.50683483 | 0.823 | 0.646 | 7.53E-106 | 2.76E-101 |
| FNDC3A | 0 | -0.669819923 | 0.733 | 0.832 | 1.05E-105 | 3.86E-101 |
| NEDD4L | 0 | 0.515193538 | 0.934 | 0.824 | 1.64E-105 | 6.02E-101 |
| MAP7D2 | 0 | -0.565756777 | 0.132 | 0.346 | 2.07E-105 | 7.59E-101 |
| PKIA | 0 | 0.646376265 | 0.61 | 0.38 | 3.31E-105 | 1.21E-100 |
| HTR7 | 0 | -0.59837914 | 0.022 | 0.166 | 3.47E-105 | 1.27E-100 |
| MSR1 | 0 | 0.659434418 | 0.149 | 0.023 | 3.86E-105 | 1.41E-100 |
| PRR16 | 0 | 0.654555288 | 0.186 | 0.039 | 4.87E-105 | 1.78E-100 |
| SLC22A15 | 0 | -0.780132873 | 0.166 | 0.368 | 3.32E-104 | 1.22E-99 |

| | | | | | | |
|---|---|---|---|---|---|---|
| DNER | 0 | -0.861785445 | 0.233 | 0.454 | 6.79E-104 | 2.49E-99 |
| MPPED2 | 0 | -0.668901335 | 0.058 | 0.238 | 9.46E-104 | 3.47E-99 |
| LINC01102 | 0 | -0.418341756 | 0.007 | 0.128 | 1.13E-103 | 4.15E-99 |
| RBFOX1 | 0 | 0.568998009 | 0.864 | 0.683 | 1.24E-103 | 4.54E-99 |
| NR2F1 | 0 | -0.683263499 | 0.084 | 0.269 | 1.28E-103 | 4.7E-99 |
| NRG1 | 0 | -1.708979778 | 0.165 | 0.342 | 2.19E-103 | 8.03E-99 |
| ANKRD6 | 0 | -0.630247192 | 0.148 | 0.365 | 3.41E-103 | 1.25E-98 |
| PDE7B | 0 | -1.159497734 | 0.089 | 0.268 | 1.35E-102 | 4.95E-98 |
| LARGE1 | 0 | 0.484940927 | 0.438 | 0.209 | 1.37E-102 | 5.03E-98 |
| SLC12A2 | 0 | 0.73665598 | 0.602 | 0.407 | 3.58E-102 | 1.31E-97 |
| SMOC1 | 0 | 0.453595335 | 0.319 | 0.12 | 4.58E-102 | 1.68E-97 |
| NPSR1-AS1 | 0 | -0.914185035 | 0.054 | 0.224 | 6.05E-102 | 2.21E-97 |
| ZMAT4 | 0 | 0.89225897 | 0.649 | 0.476 | 8.92E-102 | 3.27E-97 |
| KCNH7 | 0 | -0.936674127 | 0.031 | 0.181 | 9.96E-102 | 3.65E-97 |
| LRIG3 | 0 | 0.621250003 | 0.394 | 0.188 | 1.02E-101 | 3.73E-97 |
| DENND2A | 0 | -0.714034135 | 0.185 | 0.39 | 1.3E-101 | 4.75E-97 |
| LRFN5 | 0 | -1.190968074 | 0.313 | 0.519 | 2.75E-101 | 1.01E-96 |
| TPD52 | 0 | -0.74922909 | 0.204 | 0.423 | 3.5E-101 | 1.28E-96 |
| CRYBG3 | 0 | 0.681842803 | 0.468 | 0.253 | 1.17E-100 | 4.29E-96 |
| MEG8 | 0 | 0.667738367 | 0.71 | 0.518 | 1.43E-100 | 5.23E-96 |
| DACH1 | 0 | 0.484248872 | 0.963 | 0.856 | 2.92E-100 | 1.07E-95 |
| GRIA4 | 0 | -0.858209142 | 0.162 | 0.378 | 7.72E-100 | 2.82E-95 |
| RIC3 | 0 | -0.398887406 | 0.044 | 0.208 | 1.34E-99 | 4.91E-95 |
| DYNC1I1 | 0 | -0.752470857 | 0.188 | 0.41 | 2.19E-99 | 8.03E-95 |
| ARHGAP10 | 0 | 0.498048508 | 0.678 | 0.444 | 3.92E-99 | 1.43E-94 |
| FERMT2 | 0 | 0.714963028 | 0.563 | 0.38 | 4.6E-99 | 1.68E-94 |
| EEF1AKNMT | 0 | 0.410956678 | 0.21 | 0.06 | 8.26E-99 | 3.02E-94 |
| VAMP4 | 0 | 0.503632766 | 0.357 | 0.168 | 8.62E-99 | 3.16E-94 |
| NLRC5 | 0 | 0.483208748 | 0.293 | 0.107 | 1.02E-98 | 3.73E-94 |
| CACNA1A | 0 | -0.545165188 | 0.391 | 0.219 | 1.68E-98 | 6.13E-94 |
| LINC02315 | 0 | 0.348688491 | 0.917 | 0.731 | 1.96E-98 | 7.19E-94 |
| ROR1 | 0 | -1.077720105 | 0.125 | 0.304 | 3.31E-98 | 1.21E-93 |
| KCNH8 | 0 | 0.788126506 | 0.727 | 0.595 | 1.74E-97 | 6.37E-93 |
| B4GALT6 | 0 | -0.671451376 | 0.144 | 0.35 | 2.59E-97 | 9.5E-93 |
| PROM1 | 0 | -0.456221664 | 0.012 | 0.138 | 1.08E-96 | 3.96E-92 |
| EPB41L3 | 0 | -0.605619466 | 0.145 | 0.353 | 1.16E-96 | 4.23E-92 |
| MCTP1 | 0 | 0.888857456 | 0.434 | 0.24 | 2.02E-96 | 7.39E-92 |
| BICDL1 | 0 | -0.566119882 | 0.132 | 0.337 | 2.57E-96 | 9.4E-92 |
| EPHA5 | 0 | -1.101941856 | 0.126 | 0.312 | 2.63E-96 | 9.63E-92 |
| MYO10 | 0 | -0.684027984 | 0.182 | 0.393 | 2.87E-96 | 1.05E-91 |
| GALNT7 | 0 | 0.639318299 | 0.706 | 0.55 | 1.25E-95 | 4.57E-91 |
| C9orf135 | 0 | -0.416947809 | 0.008 | 0.124 | 2.44E-95 | 8.94E-91 |
| CSMD3 | 0 | 0.872994568 | 0.484 | 0.26 | 2.85E-95 | 1.04E-90 |
| VCL | 0 | 0.730973023 | 0.857 | 0.746 | 8.39E-95 | 3.07E-90 |
| AC006972.1 | 0 | -0.497751672 | 0.013 | 0.137 | 1.17E-94 | 4.28E-90 |
| NLGN4X | 0 | 0.437102568 | 0.869 | 0.679 | 1.59E-94 | 5.83E-90 |
| CAMK1D | 0 | -0.714092253 | 0.524 | 0.692 | 1.93E-94 | 7.07E-90 |
| STMN2 | 0 | -0.567550318 | 0.053 | 0.217 | 8.23E-94 | 3.01E-89 |
| GABRA5 | 0 | -0.321294969 | 0.008 | 0.123 | 1.06E-93 | 3.9E-89 |
| FAM189A1 | 0 | -0.714386269 | 0.681 | 0.699 | 1.95E-93 | 7.16E-89 |
| AC068587.4 | 0 | 0.634921709 | 0.499 | 0.281 | 3.1E-93 | 1.14E-88 |
| ELOVL2 | 0 | -0.449460876 | 0.039 | 0.185 | 3.34E-93 | 1.22E-88 |
| ID1 | 0 | 0.477736971 | 0.242 | 0.078 | 4.34E-93 | 1.59E-88 |
| DNM3 | 0 | 0.491906347 | 0.637 | 0.4 | 5.02E-93 | 1.84E-88 |
| GRK3 | 0 | -0.613492365 | 0.299 | 0.52 | 1.03E-92 | 3.77E-88 |
| COL4A5 | 0 | -1.295434809 | 0.025 | 0.145 | 1.4E-92 | 5.13E-88 |
| AC093765.3 | 0 | 0.433097535 | 0.144 | 0.025 | 1.79E-92 | 6.55E-88 |
| ATP8A1 | 0 | 0.550267246 | 0.573 | 0.344 | 1.89E-92 | 6.9E-88 |

| | | | | | | |
|---|---|---|---|---|---|---|
| GAP43 | 0 | -0.659123914 | 0.096 | 0.274 | 2.39E-92 | 8.74E-88 |
| RNF128 | 0 | -0.557207935 | 0.037 | 0.182 | 1.12E-91 | 4.12E-87 |
| HS3ST5 | 0 | -0.881807171 | 0.618 | 0.618 | 1.82E-91 | 6.65E-87 |
| BAIAP2L1 | 0 | -0.389147751 | 0.022 | 0.154 | 2.03E-91 | 7.45E-87 |
| RAD51B | 0 | 0.545542448 | 0.8 | 0.623 | 2.2E-91 | 8.07E-87 |
| C2CD2 | 0 | 0.484940635 | 0.271 | 0.104 | 2.22E-91 | 8.12E-87 |
| UNC80 | 0 | -0.633384009 | 0.086 | 0.258 | 3.67E-91 | 1.35E-86 |
| SLC2A13 | 0 | 0.83174126 | 0.566 | 0.4 | 4.12E-91 | 1.51E-86 |
| PTPRD | 0 | -0.456425647 | 0.923 | 0.807 | 4.82E-91 | 1.77E-86 |
| CACNG8 | 0 | -0.302890568 | 0.007 | 0.117 | 5.83E-91 | 2.13E-86 |
| RAP2A | 0 | 0.580414665 | 0.428 | 0.258 | 1.31E-90 | 4.78E-86 |
| DSP | 0 | -0.475337344 | 0.008 | 0.119 | 1.38E-90 | 5.06E-86 |
| CDH11 | 0 | 0.352235601 | 0.466 | 0.251 | 2.01E-90 | 7.38E-86 |
| ERBIN | 0 | 0.475274207 | 0.84 | 0.723 | 4.43E-90 | 1.62E-85 |
| EDNRA | 0 | 0.475276048 | 0.246 | 0.081 | 5.25E-90 | 1.92E-85 |
| ISM1 | 0 | 0.432982885 | 0.146 | 0.027 | 6.03E-90 | 2.21E-85 |
| GABRG3 | 0 | -0.551541859 | 0.01 | 0.124 | 7.4E-90 | 2.71E-85 |
| BRINP2 | 0 | -0.498478366 | 0.018 | 0.143 | 9.01E-90 | 3.3E-85 |
| ARHGAP8 | 0 | -0.582693248 | 0.087 | 0.263 | 1.48E-89 | 5.42E-85 |
| SSBP2 | 0 | 0.440536808 | 0.91 | 0.771 | 1.75E-89 | 6.39E-85 |
| DAB1 | 0 | 0.543731231 | 0.815 | 0.631 | 4.87E-89 | 1.78E-84 |
| SCN7A | 0 | -0.318472157 | 0.247 | 0.097 | 2.26E-88 | 8.29E-84 |
| ZEB1 | 0 | 0.61206529 | 0.687 | 0.509 | 3.69E-88 | 1.35E-83 |
| POU2F3 | 0 | 0.511564123 | 0.468 | 0.259 | 1.53E-87 | 5.6E-83 |
| EPB41L4A | 0 | -0.520877203 | 0.02 | 0.144 | 1.76E-87 | 6.46E-83 |
| MIR4713HG | 0 | -0.509569453 | 0.022 | 0.149 | 1.96E-87 | 7.16E-83 |
| KIAA1217 | 0 | -0.98862203 | 0.182 | 0.363 | 2.05E-87 | 7.5E-83 |
| RBMS3 | 0 | 0.451292532 | 0.913 | 0.758 | 4.68E-87 | 1.71E-82 |
| SYNE1 | 0 | -0.901643121 | 0.369 | 0.518 | 2.22E-86 | 8.12E-82 |
| ANOS1 | 0 | 0.761672593 | 0.597 | 0.378 | 2.23E-86 | 8.16E-82 |
| DPY19L3 | 0 | 0.506681131 | 0.619 | 0.445 | 4.09E-86 | 1.5E-81 |
| RASGEF1B | 0 | -0.869076125 | 0.187 | 0.379 | 4.39E-86 | 1.61E-81 |
| LINC01163 | 0 | 0.644185857 | 0.174 | 0.047 | 7.21E-86 | 2.64E-81 |
| EPHA3 | 0 | -1.172502035 | 0.105 | 0.261 | 1.03E-85 | 3.77E-81 |
| NRXN1 | 0 | -0.622204668 | 0.456 | 0.655 | 1.05E-85 | 3.83E-81 |
| SYNDIG1 | 0 | 0.579999659 | 0.707 | 0.517 | 1.16E-85 | 4.23E-81 |
| ADAM23 | 0 | -0.718104939 | 0.216 | 0.416 | 1.27E-85 | 4.64E-81 |
| RORA | 0 | -0.708121071 | 0.716 | 0.831 | 1.63E-85 | 5.97E-81 |
| MAP7D3 | 0 | 0.52663202 | 0.378 | 0.201 | 1.79E-85 | 6.56E-81 |
| TNS3 | 0 | -0.65783552 | 0.115 | 0.291 | 2.01E-85 | 7.37E-81 |
| CBFA2T3 | 0 | 0.437673375 | 0.291 | 0.116 | 2.07E-85 | 7.57E-81 |
| FGD6 | 0 | -0.684725869 | 0.2 | 0.382 | 2.89E-85 | 1.06E-80 |
| ZNF804A | 0 | 0.915239028 | 0.378 | 0.194 | 3.48E-85 | 1.27E-80 |
| KCNMB2 | 0 | -1.195347749 | 0.149 | 0.32 | 6.84E-85 | 2.5E-80 |
| SH3RF3 | 0 | 0.459404112 | 0.417 | 0.21 | 8.29E-85 | 3.04E-80 |
| RBPMS | 0 | -0.546900821 | 0.016 | 0.133 | 8.78E-85 | 3.22E-80 |
| RALYL | 0 | 0.437400004 | 0.635 | 0.41 | 2.27E-84 | 8.32E-80 |
| LHFPL3 | 0 | 0.470081937 | 0.723 | 0.502 | 2.35E-84 | 8.6E-80 |
| CTNNA1 | 0 | 0.524165882 | 0.766 | 0.62 | 2.4E-84 | 8.78E-80 |
| ADGRD1 | 0 | 0.555989742 | 0.393 | 0.199 | 2.62E-84 | 9.59E-80 |
| AOPEP | 0 | 0.454174932 | 0.846 | 0.718 | 2.63E-84 | 9.62E-80 |
| CHD9 | 0 | 0.414170036 | 0.952 | 0.908 | 2.76E-84 | 1.01E-79 |
| MCF2L | 0 | 0.520116606 | 0.398 | 0.209 | 2.85E-84 | 1.04E-79 |
| STK3 | 0 | 0.450494383 | 0.652 | 0.431 | 4.12E-84 | 1.51E-79 |
| CLVS2 | 0 | -0.561473608 | 0.01 | 0.118 | 5.67E-84 | 2.08E-79 |
| AC106799.2 | 0 | -0.393810605 | 0.008 | 0.115 | 5.83E-84 | 2.13E-79 |
| KIF5C | 0 | -0.626000676 | 0.158 | 0.345 | 8.16E-84 | 2.99E-79 |
| XRN1 | 0 | 0.503859877 | 0.765 | 0.632 | 1.28E-83 | 4.7E-79 |

| | | | | | | |
|---|---|---|---|---|---|---|
| PURPL | 0 | -0.714218174 | 0.033 | 0.164 | 2.06E-83 | 7.56E-79 |
| CELF4 | 0 | -0.626361781 | 0.156 | 0.342 | 2.14E-83 | 7.83E-79 |
| AC019197.1 | 0 | -0.488343427 | 0.028 | 0.157 | 3.34E-83 | 1.22E-78 |
| GNAO1 | 0 | -0.565093386 | 0.303 | 0.52 | 5.95E-83 | 2.18E-78 |
| SRGAP1 | 0 | 0.542338241 | 0.576 | 0.367 | 8.01E-83 | 2.93E-78 |
| GALNT17 | 0 | -0.779700425 | 0.072 | 0.235 | 1.12E-82 | 4.1E-78 |
| CPEB1 | 0 | -0.371252851 | 0.021 | 0.143 | 1.12E-82 | 4.1E-78 |
| PELI2 | 0 | -0.589487977 | 0.274 | 0.491 | 1.48E-82 | 5.44E-78 |
| PTPRM | 0 | -0.931331672 | 0.267 | 0.43 | 1.61E-82 | 5.9E-78 |
| TMEFF2 | 0 | -0.66160458 | 0.045 | 0.187 | 1.71E-82 | 6.25E-78 |
| WDR35 | 0 | 0.542256719 | 0.598 | 0.415 | 1.93E-82 | 7.06E-78 |
| ZHX2 | 0 | 0.565375938 | 0.418 | 0.234 | 2.45E-82 | 8.97E-78 |
| MIRLET7BHG | 0 | 0.429003303 | 0.807 | 0.616 | 2.55E-82 | 9.33E-78 |
| WASF3 | 0 | -0.444907214 | 0.066 | 0.219 | 3.04E-82 | 1.11E-77 |
| IDI2-AS1 | 0 | -0.510516107 | 0.04 | 0.175 | 4.52E-82 | 1.66E-77 |
| CLSTN2 | 0 | -0.819275963 | 0.198 | 0.401 | 6.1E-82 | 2.23E-77 |
| FAM184B | 0 | -0.508350013 | 0.08 | 0.239 | 1.02E-81 | 3.73E-77 |
| DOCK8 | 0 | -0.58630835 | 0.008 | 0.107 | 2.24E-81 | 8.21E-77 |
| TSPAN9 | 0 | 0.524831938 | 0.535 | 0.333 | 4.17E-81 | 1.53E-76 |
| JMJD1C | 0 | 0.347318769 | 0.98 | 0.931 | 4.9E-81 | 1.79E-76 |
| SLC17A8 | 0 | -0.854628544 | 0.156 | 0.342 | 7.32E-81 | 2.68E-76 |
| KCNK2 | 0 | 0.761136365 | 0.416 | 0.237 | 7.92E-81 | 2.9E-76 |
| SLC7A14 | 0 | -0.58360468 | 0.023 | 0.144 | 8.86E-81 | 3.24E-76 |
| TOX2 | 0 | -0.372719156 | 0.016 | 0.131 | 1.02E-80 | 3.75E-76 |
| STARD13 | 0 | 0.256984707 | 0.748 | 0.534 | 2E-80 | 7.31E-76 |
| NFASC | 0 | -0.543662182 | 0.07 | 0.222 | 2.38E-80 | 8.7E-76 |
| DSCAM | 0 | -0.948646429 | 0.03 | 0.152 | 2.96E-80 | 1.08E-75 |
| METTL24 | 0 | 0.394918765 | 0.204 | 0.063 | 5.63E-80 | 2.06E-75 |
| ARFGEF3 | 0 | -0.654373963 | 0.318 | 0.518 | 7E-80 | 2.56E-75 |
| L1TD1 | 0 | -0.607294037 | 0.108 | 0.272 | 9.16E-80 | 3.35E-75 |
| ADAMTS6 | 0 | -1.1104924 | 0.068 | 0.184 | 1.14E-79 | 4.19E-75 |
| ANKRD18A | 0 | -0.395882229 | 0.052 | 0.199 | 1.17E-79 | 4.27E-75 |
| ADAMTS19 | 0 | -0.807934451 | 0.364 | 0.535 | 1.37E-79 | 5E-75 |
| GAB2 | 0 | -0.541308427 | 0.171 | 0.368 | 2.08E-79 | 7.62E-75 |
| HIF3A | 0 | -0.426778998 | 0.034 | 0.164 | 2.11E-79 | 7.72E-75 |
| CTCFL | 0 | -0.2821424 | 0.01 | 0.114 | 3.9E-79 | 1.43E-74 |
| SYT17 | 0 | 0.378983442 | 0.224 | 0.076 | 6.99E-79 | 2.56E-74 |
| ADGRL2 | 0 | -0.924735571 | 0.538 | 0.669 | 1.49E-78 | 5.44E-74 |
| ADGRB3 | 0 | -0.614269066 | 0.384 | 0.6 | 2.12E-78 | 7.77E-74 |
| EGFR | 0 | 0.367750754 | 0.396 | 0.203 | 2.23E-78 | 8.18E-74 |
| PHF14 | 0 | 0.389129113 | 0.95 | 0.881 | 5.1E-78 | 1.87E-73 |
| TEAD1 | 0 | 0.395367596 | 0.945 | 0.883 | 5.53E-78 | 2.02E-73 |
| LINC00937 | 0 | -0.531454473 | 0.081 | 0.231 | 7.61E-78 | 2.79E-73 |
| AL050309.1 | 0 | -0.504762495 | 0.025 | 0.145 | 1.02E-77 | 3.73E-73 |
| PCDH7 | 0 | -1.192300924 | 0.336 | 0.465 | 1.32E-77 | 4.83E-73 |
| SH3GL3 | 0 | 0.571466298 | 0.666 | 0.478 | 1.35E-77 | 4.96E-73 |
| TET1 | 0 | 0.446243078 | 0.563 | 0.348 | 2.11E-77 | 7.72E-73 |
| EGR1 | 0 | 0.508294561 | 0.222 | 0.076 | 2.6E-77 | 9.51E-73 |
| EZH2 | 0 | 0.45974744 | 0.822 | 0.665 | 2.81E-77 | 1.03E-72 |
| KLHL1 | 0 | -0.36376181 | 0.158 | 0.08 | 3.87E-77 | 1.42E-72 |
| VWDE | 0 | -0.507443119 | 0.251 | 0.459 | 1.96E-76 | 7.18E-72 |
| LINC02428 | 0 | -0.382508461 | 0.01 | 0.109 | 2.58E-76 | 9.44E-72 |
| DAAM2 | 0 | -0.791907171 | 0.108 | 0.26 | 2.88E-76 | 1.06E-71 |
| CHM | 0 | 0.493392348 | 0.771 | 0.647 | 5.64E-76 | 2.06E-71 |
| AL136418.1 | 0 | -0.494614619 | 0.017 | 0.123 | 7.43E-76 | 2.72E-71 |
| SOX5 | 0 | -0.692871324 | 0.717 | 0.623 | 9.53E-76 | 3.49E-71 |
| ERC2 | 0 | -0.690814042 | 0.316 | 0.506 | 1.46E-75 | 5.34E-71 |
| PSD3 | 0 | 0.47048031 | 0.901 | 0.801 | 2.09E-75 | 7.64E-71 |

| | | | | | | |
|---|---|---|---|---|---|---|
| SFMBT2 | 0 | -0.709872262 | 0.367 | 0.543 | 2.98E-75 | 1.09E-70 |
| PRKCB | 0 | -0.525965098 | 0.023 | 0.138 | 3.38E-75 | 1.24E-70 |
| MYO5B | 0 | -0.312795146 | 0.011 | 0.111 | 2.22E-74 | 8.14E-70 |
| NR6A1 | 0 | -0.591802672 | 0.245 | 0.422 | 3.14E-74 | 1.15E-69 |
| SEMA3C | 0 | -0.763688789 | 0.036 | 0.157 | 3.71E-74 | 1.36E-69 |
| CHRM2 | 0 | -0.791111817 | 0.018 | 0.127 | 3.81E-74 | 1.39E-69 |
| ZYG11A | 0 | -0.509606342 | 0.17 | 0.356 | 4.26E-74 | 1.56E-69 |
| KLHL32 | 0 | -0.477238952 | 0.068 | 0.216 | 5.43E-74 | 1.99E-69 |
| NPAS2 | 0 | -0.448347633 | 0.066 | 0.213 | 1.29E-73 | 4.74E-69 |
| TRMT9B | 0 | 0.739064705 | 0.339 | 0.176 | 1.54E-73 | 5.65E-69 |
| TENM3 | 0 | -0.501684695 | 0.83 | 0.763 | 1.8E-73 | 6.58E-69 |
| MARCH1 | 0 | -1.075862216 | 0.181 | 0.346 | 2.58E-73 | 9.46E-69 |
| HLA-A | 0 | 0.447045818 | 0.353 | 0.179 | 2.78E-73 | 1.02E-68 |
| HTR2A | 0 | 0.320533073 | 0.147 | 0.037 | 3.13E-73 | 1.15E-68 |
| HHIP | 0 | -0.503811487 | 0.033 | 0.154 | 4.19E-73 | 1.53E-68 |
| TMEM65 | 0 | 0.461021932 | 0.373 | 0.191 | 5.03E-73 | 1.84E-68 |
| LIFR | 0 | 0.300852411 | 0.774 | 0.578 | 6.72E-73 | 2.46E-68 |
| NPAS3 | 0 | -1.360917893 | 0.147 | 0.284 | 8.25E-73 | 3.02E-68 |
| ZMYM2 | 0 | 0.355082369 | 0.94 | 0.882 | 9.2E-73 | 3.37E-68 |
| PHIP | 0 | 0.439370165 | 0.921 | 0.854 | 9.9E-73 | 3.62E-68 |
| APOE | 0 | 0.298040279 | 0.132 | 0.028 | 1.4E-72 | 5.14E-68 |
| FBXL2 | 0 | -0.49206787 | 0.108 | 0.273 | 1.78E-72 | 6.5E-68 |
| ZBTB7C | 0 | 0.563726607 | 0.218 | 0.089 | 3.21E-72 | 1.18E-67 |
| ADGRG2 | 0 | -0.679458773 | 0.084 | 0.219 | 4.56E-72 | 1.67E-67 |
| CDH20 | 0 | -0.535261282 | 0.021 | 0.131 | 6.09E-72 | 2.23E-67 |
| HLA-B | 0 | 0.345249031 | 0.241 | 0.092 | 6.18E-72 | 2.26E-67 |
| SKAP2 | 0 | -0.616487157 | 0.077 | 0.224 | 7.71E-72 | 2.82E-67 |
| THSD4 | 0 | -0.582903626 | 0.061 | 0.203 | 1.39E-71 | 5.07E-67 |
| CNTN4 | 0 | 1.016346125 | 0.23 | 0.117 | 2.24E-71 | 8.21E-67 |
| PHACTR1 | 0 | -0.675773888 | 0.19 | 0.375 | 2.34E-71 | 8.58E-67 |
| LINC01776 | 0 | -0.533316174 | 0.082 | 0.232 | 2.46E-71 | 9.02E-67 |
| FMNL3 | 0 | 0.420882428 | 0.334 | 0.165 | 2.67E-71 | 9.76E-67 |
| EEPD1 | 0 | -0.610952165 | 0.182 | 0.353 | 2.95E-71 | 1.08E-66 |
| ZDBF2 | 0 | -0.420336972 | 0.083 | 0.233 | 3.35E-71 | 1.23E-66 |
| NPNT | 0 | -0.549946364 | 0.009 | 0.101 | 3.46E-71 | 1.27E-66 |
| CGNL1 | 0 | -0.361017447 | 0.01 | 0.106 | 3.5E-71 | 1.28E-66 |
| FOXP1 | 0 | 0.436253413 | 0.89 | 0.783 | 4.02E-71 | 1.47E-66 |
| ASXL3 | 0 | -0.645812916 | 0.357 | 0.528 | 1.21E-70 | 4.43E-66 |
| TRPS1 | 0 | 0.663317394 | 0.704 | 0.542 | 1.3E-70 | 4.74E-66 |
| CABLES1 | 0 | -0.404066903 | 0.07 | 0.216 | 1.65E-70 | 6.02E-66 |
| AL390957.1 | 0 | -1.237643331 | 0.052 | 0.172 | 1.74E-70 | 6.38E-66 |
| AC093817.2 | 0 | 0.614324269 | 0.199 | 0.066 | 1.93E-70 | 7.05E-66 |
| PARD3 | 0 | 0.309050786 | 0.99 | 0.956 | 3.25E-70 | 1.19E-65 |
| EFNB2 | 0 | -0.395544566 | 0.059 | 0.198 | 4.59E-70 | 1.68E-65 |
| TAFA1 | 0 | -0.815636897 | 0.11 | 0.273 | 4.68E-70 | 1.71E-65 |
| RUNX1T1 | 0 | 0.45525559 | 0.827 | 0.668 | 5.34E-70 | 1.96E-65 |
| CUL4B | 0 | 0.493055094 | 0.518 | 0.387 | 6.08E-70 | 2.23E-65 |
| EPHA7 | 0 | -0.841668731 | 0.188 | 0.366 | 6.25E-70 | 2.29E-65 |
| PLAG1 | 0 | -0.339706953 | 0.026 | 0.138 | 1.07E-69 | 3.93E-65 |
| CTNND2 | 0 | -0.655566576 | 0.671 | 0.762 | 1.22E-69 | 4.45E-65 |
| FAM162B | 0 | 0.273516177 | 0.162 | 0.045 | 1.27E-69 | 4.63E-65 |
| KIAA0408 | 0 | -0.441717567 | 0.1 | 0.255 | 1.29E-69 | 4.72E-65 |
| FRMD5 | 0 | -0.9239792 | 0.295 | 0.454 | 1.34E-69 | 4.92E-65 |
| ACYP2 | 0 | 0.511446903 | 0.64 | 0.483 | 3.19E-69 | 1.17E-64 |
| YAP1 | 0 | 0.410351568 | 0.474 | 0.278 | 4.34E-69 | 1.59E-64 |
| KIAA1324 | 0 | -0.584938234 | 0.021 | 0.116 | 6.18E-69 | 2.26E-64 |
| ZNF521 | 0 | -0.839804827 | 0.065 | 0.176 | 6.6E-69 | 2.42E-64 |
| GABPA | 0 | 0.484620021 | 0.411 | 0.259 | 6.7E-69 | 2.45E-64 |

| | | | | | | |
|---|---|---|---|---|---|---|
| FLI1 | 0 | -0.326344841 | 0.013 | 0.111 | 7.68E-69 | 2.81E-64 |
| AUTS2 | 0 | -0.916747966 | 0.482 | 0.472 | 9.91E-69 | 3.63E-64 |
| ANTXR2 | 0 | 0.507123927 | 0.36 | 0.203 | 1.97E-68 | 7.2E-64 |
| NPSR1 | 0 | -1.021358633 | 0.076 | 0.213 | 2.81E-68 | 1.03E-63 |
| ANKH | 0 | -0.513605944 | 0.098 | 0.248 | 3.16E-68 | 1.16E-63 |
| MAPT | 0 | -0.427689378 | 0.047 | 0.17 | 5.49E-68 | 2.01E-63 |
| FUT9 | 0 | -0.434827337 | 0.02 | 0.123 | 9.63E-68 | 3.53E-63 |
| AP001981.2 | 0 | 0.294797535 | 0.139 | 0.035 | 9.79E-68 | 3.58E-63 |
| DACT1 | 0 | 0.40696153 | 0.203 | 0.077 | 9.95E-68 | 3.64E-63 |
| MBNL3 | 0 | 0.598343082 | 0.599 | 0.445 | 1.23E-67 | 4.51E-63 |
| LURAP1L | 0 | 0.537988408 | 0.329 | 0.173 | 1.64E-67 | 6.01E-63 |
| EHBP1 | 0 | 0.406999785 | 0.885 | 0.803 | 1.91E-67 | 6.98E-63 |
| LRRTM3 | 0 | -0.615009951 | 0.068 | 0.207 | 2E-67 | 7.33E-63 |
| TANC2 | 0 | 0.37579227 | 0.823 | 0.671 | 2.09E-67 | 7.65E-63 |
| MIR222HG | 0 | -0.424424754 | 0.095 | 0.249 | 3.08E-67 | 1.13E-62 |
| R3HDM1 | 0 | 0.426539297 | 0.91 | 0.851 | 5.28E-67 | 1.93E-62 |
| FSTL4 | 0 | -0.434332051 | 0.033 | 0.15 | 6.08E-67 | 2.23E-62 |
| ARID2 | 0 | 0.382115807 | 0.881 | 0.787 | 7.91E-67 | 2.9E-62 |
| MYO7A | 0 | 0.326436476 | 0.17 | 0.058 | 8.4E-67 | 3.07E-62 |
| SLC38A4 | 0 | 0.669341103 | 0.279 | 0.133 | 1.03E-66 | 3.77E-62 |
| EYA2 | 0 | -0.513146416 | 0.046 | 0.171 | 1.07E-66 | 3.91E-62 |
| GLIS1 | 0 | 0.278556788 | 0.102 | 0.017 | 1.1E-66 | 4.01E-62 |
| SRPX | 0 | 0.432730193 | 0.352 | 0.181 | 1.27E-66 | 4.64E-62 |
| AL355499.1 | 0 | -0.934515478 | 0.026 | 0.132 | 1.43E-66 | 5.23E-62 |
| ATG10 | 0 | 0.466908167 | 0.697 | 0.534 | 1.44E-66 | 5.29E-62 |
| SH3BGRL | 0 | 0.480573429 | 0.513 | 0.348 | 2.87E-66 | 1.05E-61 |
| RGS17 | 0 | -0.445852383 | 0.074 | 0.216 | 3.96E-66 | 1.45E-61 |
| LIN7A | 0 | 0.527031522 | 0.486 | 0.313 | 5.4E-66 | 1.98E-61 |
| PDGFRB | 0 | 0.420111701 | 0.206 | 0.075 | 6.25E-66 | 2.29E-61 |
| CNTN6 | 0 | 0.88830635 | 0.28 | 0.138 | 6.93E-66 | 2.54E-61 |
| FOXN3 | 0 | 0.486719256 | 0.961 | 0.92 | 7.77E-66 | 2.84E-61 |
| SLC16A9 | 0 | -0.347935575 | 0.071 | 0.211 | 1.52E-65 | 5.57E-61 |
| AC005162.3 | 0 | 0.294440113 | 0.131 | 0.033 | 1.89E-65 | 6.91E-61 |
| NTN4 | 0 | 0.439279956 | 0.298 | 0.141 | 2.5E-65 | 9.14E-61 |
| CTDSPL | 0 | 0.464604939 | 0.628 | 0.475 | 2.88E-65 | 1.06E-60 |
| CCDC93 | 0 | 0.459133158 | 0.435 | 0.292 | 2.89E-65 | 1.06E-60 |
| CBFB | 0 | 0.430031981 | 0.622 | 0.471 | 3.83E-65 | 1.4E-60 |
| AC009133.1 | 0 | 0.515941634 | 0.192 | 0.08 | 4.92E-65 | 1.8E-60 |
| SLCO5A1 | 0 | -0.479168833 | 0.059 | 0.187 | 5.47E-65 | 2E-60 |
| AL162411.1 | 0 | 0.335273896 | 0.205 | 0.075 | 6.57E-65 | 2.41E-60 |
| NSD3 | 0 | 0.378345668 | 0.828 | 0.734 | 1E-64 | 3.67E-60 |
| AC004158.1 | 0 | 0.460470483 | 0.294 | 0.144 | 1.23E-64 | 4.5E-60 |
| PTCHD1-AS | 0 | -0.716912272 | 0.038 | 0.152 | 1.32E-64 | 4.81E-60 |
| BCL11A | 0 | -0.56402547 | 0.164 | 0.319 | 1.42E-64 | 5.21E-60 |
| COL27A1 | 0 | 0.504521382 | 0.51 | 0.335 | 1.46E-64 | 5.33E-60 |
| ANGPT1 | 0 | -0.710903666 | 0.457 | 0.625 | 2.12E-64 | 7.76E-60 |
| SAMD12 | 0 | -0.37747056 | 0.013 | 0.102 | 2.2E-64 | 8.05E-60 |
| AL592156.2 | 0 | -0.450939531 | 0.018 | 0.116 | 2.94E-64 | 1.08E-59 |
| SYT10 | 0 | -0.53984831 | 0.048 | 0.156 | 3.1E-64 | 1.14E-59 |
| SLC16A7 | 0 | 0.571662745 | 0.453 | 0.312 | 3.16E-64 | 1.16E-59 |
| AC025809.1 | 0 | 0.322570738 | 0.212 | 0.079 | 4.1E-64 | 1.5E-59 |
| AL157944.1 | 0 | -0.846510506 | 0.027 | 0.127 | 4.38E-64 | 1.6E-59 |
| RASAL2 | 0 | -0.510718464 | 0.408 | 0.593 | 4.47E-64 | 1.64E-59 |
| AC105285.1 | 0 | 0.402394545 | 0.216 | 0.093 | 5.92E-64 | 2.17E-59 |
| SDC2 | 0 | 0.487868924 | 0.497 | 0.315 | 1.2E-63 | 4.41E-59 |
| GCNT2 | 0 | -0.33609731 | 0.021 | 0.12 | 1.54E-63 | 5.65E-59 |
| CTNNA2 | 0 | 0.606657175 | 0.627 | 0.435 | 1.82E-63 | 6.68E-59 |
| TMEM63A | 0 | 0.367609462 | 0.227 | 0.094 | 1.88E-63 | 6.87E-59 |

| | | | | | | |
|---|---|---|---|---|---|---|
| ERBB3 | 0 | -0.37945942 | 0.027 | 0.134 | 2.03E-63 | 7.44E-59 |
| SYNPO2 | 0 | 0.612421702 | 0.389 | 0.219 | 2.45E-63 | 8.97E-59 |
| PRTG | 0 | -0.751178932 | 0.026 | 0.121 | 3.23E-63 | 1.18E-58 |
| TBX20 | 0 | 0.493725808 | 0.377 | 0.217 | 4.27E-63 | 1.56E-58 |
| PTPRK | 0 | 0.502106931 | 0.88 | 0.779 | 5.55E-63 | 2.03E-58 |
| ITGAV | 0 | -0.640407868 | 0.423 | 0.584 | 6.06E-63 | 2.22E-58 |
| SLC35D1 | 0 | 0.442040596 | 0.365 | 0.221 | 7.61E-63 | 2.79E-58 |
| LINGO2 | 0 | -0.565240235 | 0.013 | 0.102 | 7.94E-63 | 2.91E-58 |
| CPQ | 0 | 0.438185795 | 0.415 | 0.237 | 8.11E-63 | 2.97E-58 |
| SLC9A7 | 0 | -0.390574901 | 0.059 | 0.185 | 3E-62 | 1.1E-57 |
| UNC13A | 0 | -0.364206968 | 0.049 | 0.166 | 3.28E-62 | 1.2E-57 |
| AOX1 | 0 | 0.303806592 | 0.112 | 0.025 | 4.21E-62 | 1.54E-57 |
| MAP7 | 0 | -0.639509527 | 0.487 | 0.631 | 4.72E-62 | 1.73E-57 |
| GPAM | 0 | 0.425029218 | 0.284 | 0.141 | 4.79E-62 | 1.76E-57 |
| MEG3 | 0 | 0.488553344 | 0.568 | 0.398 | 5.42E-62 | 1.98E-57 |
| PCDH15 | 0 | -1.148181652 | 0.055 | 0.168 | 6.75E-62 | 2.47E-57 |
| RCAN2 | 0 | -0.457977702 | 0.027 | 0.129 | 1.01E-61 | 3.68E-57 |
| NCKAP5 | 0 | 0.797342557 | 0.512 | 0.369 | 1.47E-61 | 5.4E-57 |
| DMGDH | 0 | 0.466519021 | 0.434 | 0.261 | 2.42E-61 | 8.88E-57 |
| SOGA3 | 0 | -0.411421356 | 0.11 | 0.261 | 2.47E-61 | 9.05E-57 |
| TLE1 | 0 | -0.487965013 | 0.143 | 0.303 | 2.76E-61 | 1.01E-56 |
| RERG | 0 | 0.627937141 | 0.288 | 0.157 | 4.43E-61 | 1.62E-56 |
| PLCH1 | 0 | 0.736699079 | 0.408 | 0.288 | 4.88E-61 | 1.78E-56 |
| COL21A1 | 0 | -0.537087426 | 0.015 | 0.105 | 5.53E-61 | 2.03E-56 |
| GAB1 | 0 | -0.582220824 | 0.625 | 0.685 | 7.8E-61 | 2.85E-56 |
| ITPR3 | 0 | -0.271916185 | 0.032 | 0.14 | 1.04E-60 | 3.81E-56 |
| ADAM11 | 0 | 0.256361835 | 0.134 | 0.036 | 1.26E-60 | 4.61E-56 |
| F3 | 0 | -0.637879212 | 0.032 | 0.128 | 1.47E-60 | 5.39E-56 |
| LINC01515 | 0 | -0.621550851 | 0.093 | 0.227 | 3.24E-60 | 1.18E-55 |
| LINC01170 | 0 | -0.683384424 | 0.489 | 0.621 | 3.89E-60 | 1.43E-55 |
| CDH6 | 0 | -0.639791967 | 0.223 | 0.381 | 4.44E-60 | 1.63E-55 |
| GRM7 | 0 | -0.925763587 | 0.055 | 0.17 | 5.02E-60 | 1.84E-55 |
| ENOX1 | 0 | -0.649316313 | 0.422 | 0.577 | 9.61E-60 | 3.52E-55 |
| HOOK1 | 0 | -0.391356303 | 0.103 | 0.248 | 1.07E-59 | 3.93E-55 |
| DOCK9 | 0 | -0.420147289 | 0.08 | 0.213 | 1.67E-59 | 6.12E-55 |
| DOCK4 | 0 | 0.600738342 | 0.715 | 0.585 | 2E-59 | 7.32E-55 |
| FBN1 | 0 | 0.434862842 | 0.285 | 0.143 | 5.51E-59 | 2.02E-54 |
| GREB1L | 0 | -0.776547893 | 0.029 | 0.125 | 7.52E-59 | 2.75E-54 |
| UTRN | 0 | -0.682372033 | 0.613 | 0.625 | 7.84E-59 | 2.87E-54 |
| VASH2 | 0 | 0.392554262 | 0.233 | 0.121 | 8.42E-59 | 3.08E-54 |
| MIR503HG | 0 | -0.530199322 | 0.072 | 0.198 | 1.13E-58 | 4.13E-54 |
| VEZT | 0 | -0.471276868 | 0.581 | 0.704 | 1.18E-58 | 4.32E-54 |
| SPRED1 | 0 | 0.413354556 | 0.538 | 0.364 | 1.22E-58 | 4.47E-54 |
| PTPN13 | 0 | -0.563311301 | 0.058 | 0.173 | 1.26E-58 | 4.61E-54 |
| SLITRK5 | 0 | 0.375994902 | 0.181 | 0.067 | 1.34E-58 | 4.9E-54 |
| FAM168A | 0 | 0.372177118 | 0.776 | 0.643 | 2.84E-58 | 1.04E-53 |
| PPFIA2-AS1 | 0 | 0.2698353 | 0.164 | 0.056 | 2.87E-58 | 1.05E-53 |
| AC004540.1 | 0 | 0.390242638 | 0.299 | 0.151 | 3.36E-58 | 1.23E-53 |
| FOXP2 | 0 | -1.306668992 | 0.179 | 0.298 | 3.59E-58 | 1.32E-53 |
| KLF3 | 0 | -0.328820591 | 0.065 | 0.189 | 4.24E-58 | 1.55E-53 |
| AC079781.5 | 0 | 0.446117536 | 0.525 | 0.396 | 5.73E-58 | 2.1E-53 |
| FRMD4B | 0 | -0.53989936 | 0.305 | 0.485 | 5.86E-58 | 2.14E-53 |
| VPS13C | 0 | 0.356907166 | 0.83 | 0.702 | 6.19E-58 | 2.27E-53 |
| RYR3 | 0 | -0.576590977 | 0.035 | 0.139 | 7.36E-58 | 2.69E-53 |
| LDHA | 0 | -0.332965501 | 0.034 | 0.139 | 8.16E-58 | 2.99E-53 |
| LBR | 0 | -0.526150692 | 0.235 | 0.383 | 1.02E-57 | 3.72E-53 |
| NWD2 | 0 | -0.487108811 | 0.057 | 0.175 | 1.19E-57 | 4.36E-53 |
| PTPRG | 0 | -0.50666581 | 0.856 | 0.835 | 1.28E-57 | 4.68E-53 |

| | | | | | | |
|---|---|---|---|---|---|---|
| RAB3B | 0 | -0.384695702 | 0.021 | 0.114 | 1.3E-57 | 4.75E-53 |
| EYS | 0 | -0.408652688 | 0.346 | 0.206 | 1.34E-57 | 4.9E-53 |
| NCOR2 | 0 | 0.384870631 | 0.703 | 0.554 | 1.73E-57 | 6.32E-53 |
| TRPM3 | 0 | -1.88853536 | 0.123 | 0.232 | 2.81E-57 | 1.03E-52 |
| KRAS | 0 | -0.624524866 | 0.289 | 0.405 | 3.07E-57 | 1.12E-52 |
| TBC1D5 | 0 | 0.395317315 | 0.916 | 0.843 | 3.92E-57 | 1.44E-52 |
| SAMD11 | 0 | 0.400158974 | 0.228 | 0.101 | 4.55E-57 | 1.66E-52 |
| PHACTR2 | 0 | -0.588534834 | 0.465 | 0.611 | 6.88E-57 | 2.52E-52 |
| CFAP47 | 0 | -1.100498899 | 0.025 | 0.111 | 9.66E-57 | 3.54E-52 |
| TRAF5 | 0 | 0.37822793 | 0.373 | 0.218 | 1.34E-56 | 4.89E-52 |
| RPGR | 0 | 0.390557125 | 0.57 | 0.398 | 1.58E-56 | 5.78E-52 |
| SOBP | 0 | 0.508511347 | 0.864 | 0.741 | 1.73E-56 | 6.34E-52 |
| FNBP1 | 0 | 0.352525443 | 0.766 | 0.618 | 1.77E-56 | 6.49E-52 |
| MIR181A2HG | 0 | 0.348078914 | 0.395 | 0.227 | 2.09E-56 | 7.64E-52 |
| GLS | 0 | 0.412338216 | 0.851 | 0.756 | 2.21E-56 | 8.09E-52 |
| MBP | 0 | -0.336778258 | 0.046 | 0.158 | 2.26E-56 | 8.29E-52 |
| SLC22A23 | 0 | -0.548517324 | 0.237 | 0.404 | 2.52E-56 | 9.22E-52 |
| RPS6KA6 | 0 | -0.413717982 | 0.105 | 0.242 | 2.99E-56 | 1.09E-51 |
| SHB | 0 | -0.422622614 | 0.136 | 0.283 | 4.98E-56 | 1.82E-51 |
| GRM8 | 0 | -1.350910907 | 0.083 | 0.181 | 6.9E-56 | 2.53E-51 |
| RABGAP1L | 0 | 0.476876635 | 0.828 | 0.716 | 6.93E-56 | 2.54E-51 |
| SLFN11 | 0 | 0.278844157 | 0.198 | 0.077 | 7.46E-56 | 2.73E-51 |
| GLRB | 0 | -0.315559197 | 0.038 | 0.142 | 1.4E-55 | 5.13E-51 |
| APLP2 | 0 | 0.400540287 | 0.587 | 0.446 | 1.72E-55 | 6.29E-51 |
| MTURN | 0 | 0.348293384 | 0.332 | 0.182 | 1.99E-55 | 7.27E-51 |
| LINC01184 | 0 | 0.446421226 | 0.53 | 0.386 | 2.07E-55 | 7.57E-51 |
| NBAS | 0 | 0.375555565 | 0.877 | 0.767 | 2.25E-55 | 8.24E-51 |
| SPATA13 | 0 | -0.514836815 | 0.138 | 0.27 | 2.97E-55 | 1.09E-50 |
| PDGFC | 0 | -0.670666984 | 0.159 | 0.305 | 3.41E-55 | 1.25E-50 |
| RERE | 0 | 0.265654831 | 0.986 | 0.964 | 5.3E-55 | 1.94E-50 |
| AP001831.1 | 0 | -0.320764805 | 0.052 | 0.166 | 7.59E-55 | 2.78E-50 |
| ZNF462 | 0 | -0.436140979 | 0.325 | 0.49 | 8.32E-55 | 3.05E-50 |
| EPCAM | 0 | -0.322781422 | 0.022 | 0.112 | 1.35E-54 | 4.94E-50 |
| EPHA5-AS1 | 0 | -0.309137955 | 0.038 | 0.144 | 2.05E-54 | 7.51E-50 |
| APP | 0 | 0.353075619 | 0.904 | 0.854 | 3.87E-54 | 1.42E-49 |
| AL354771.1 | 0 | -0.40889828 | 0.059 | 0.17 | 5.35E-54 | 1.96E-49 |
| KIF26B | 0 | 0.536717268 | 0.629 | 0.464 | 5.47E-54 | 2E-49 |
| SH3BP4 | 0 | -0.422116063 | 0.216 | 0.381 | 1.36E-53 | 4.97E-49 |
| AL139351.3 | 0 | 0.258956939 | 0.115 | 0.029 | 2.14E-53 | 7.82E-49 |
| SCN1A | 0 | -0.523634542 | 0.031 | 0.128 | 2.62E-53 | 9.6E-49 |
| COL5A2 | 0 | -0.361549853 | 0.54 | 0.444 | 2.68E-53 | 9.82E-49 |
| MGAT4C | 0 | -0.719019491 | 0.426 | 0.576 | 2.81E-53 | 1.03E-48 |
| MEGF6 | 0 | 0.287397474 | 0.197 | 0.078 | 3.21E-53 | 1.18E-48 |
| AC026704.1 | 0 | -0.365748151 | 0.044 | 0.149 | 3.31E-53 | 1.21E-48 |
| AC092821.3 | 0 | 0.383226635 | 0.395 | 0.249 | 4.83E-53 | 1.77E-48 |
| DCHS2 | 0 | -0.359975398 | 0.031 | 0.13 | 6.73E-53 | 2.47E-48 |
| DLX6-AS1 | 0 | -0.872429369 | 0.147 | 0.253 | 1.42E-52 | 5.19E-48 |
| RAB3IP | 0 | -0.32900005 | 0.06 | 0.172 | 1.71E-52 | 6.25E-48 |
| MORC3 | 0 | 0.413253395 | 0.64 | 0.507 | 2.56E-52 | 9.36E-48 |
| MKX | 0 | -0.815121342 | 0.278 | 0.4 | 2.64E-52 | 9.68E-48 |
| HIBADH | 0 | 0.41277261 | 0.608 | 0.456 | 2.91E-52 | 1.06E-47 |
| AGPAT4 | 0 | -0.411841868 | 0.24 | 0.399 | 3.04E-52 | 1.11E-47 |
| AP1S2 | 0 | 0.450357607 | 0.632 | 0.491 | 4.09E-52 | 1.5E-47 |
| HPCAL1 | 0 | 0.40097403 | 0.44 | 0.277 | 4.69E-52 | 1.72E-47 |
| CADM2 | 0 | -0.534843299 | 0.327 | 0.497 | 4.9E-52 | 1.8E-47 |
| PLD5 | 0 | 0.870926967 | 0.301 | 0.233 | 5.17E-52 | 1.89E-47 |
| MAF | 0 | -0.311308789 | 0.059 | 0.175 | 9.92E-52 | 3.63E-47 |
| DCLK1 | 0 | -0.561371643 | 0.511 | 0.632 | 1.23E-51 | 4.5E-47 |

| | | | | | | |
|---|---|---|---|---|---|---|
| AL022068.1 | 0 | -0.705866984 | 0.046 | 0.138 | 1.53E-51 | 5.59E-47 |
| PDE8A | 0 | 0.428530464 | 0.735 | 0.616 | 1.57E-51 | 5.75E-47 |
| BRINP1 | 0 | 0.681261204 | 0.515 | 0.382 | 1.91E-51 | 6.97E-47 |
| HECW2 | 0 | 0.307261564 | 0.49 | 0.319 | 1.97E-51 | 7.22E-47 |
| NCOA3 | 0 | -0.461697774 | 0.508 | 0.624 | 2.35E-51 | 8.61E-47 |
| LSM6 | 0 | 0.315310585 | 0.236 | 0.118 | 2.4E-51 | 8.78E-47 |
| PCDH10 | 0 | -0.425082747 | 0.055 | 0.164 | 3.8E-51 | 1.39E-46 |
| DCX | 0 | -0.427576493 | 0.088 | 0.216 | 3.84E-51 | 1.4E-46 |
| LINC00632 | 0 | 0.289311944 | 0.507 | 0.335 | 4.97E-51 | 1.82E-46 |
| NELL1 | 0 | -0.725103263 | 0.205 | 0.355 | 5.91E-51 | 2.16E-46 |
| OPTN | 0 | -0.333479816 | 0.079 | 0.201 | 6.38E-51 | 2.34E-46 |
| LINC01091 | 0 | -0.413899355 | 0.019 | 0.101 | 7.06E-51 | 2.58E-46 |
| CFAP70 | 0 | -0.36734645 | 0.025 | 0.112 | 7.09E-51 | 2.6E-46 |
| SHROOM4 | 0 | -0.551660847 | 0.552 | 0.626 | 7.97E-51 | 2.92E-46 |
| TCF3 | 0 | 0.352501454 | 0.482 | 0.323 | 8.15E-51 | 2.98E-46 |
| GRIA1 | 0 | 0.660609665 | 0.446 | 0.338 | 8.83E-51 | 3.23E-46 |
| TMTC1 | 0 | -0.46710296 | 0.057 | 0.169 | 1.42E-50 | 5.21E-46 |
| TES | 0 | -0.291881484 | 0.034 | 0.131 | 1.59E-50 | 5.83E-46 |
| MBNL2 | 0 | -0.48636546 | 0.312 | 0.449 | 1.95E-50 | 7.15E-46 |
| ZNF438 | 0 | 0.437317544 | 0.39 | 0.247 | 3.72E-50 | 1.36E-45 |
| ASCC3 | 0 | 0.380265229 | 0.834 | 0.752 | 4.68E-50 | 1.71E-45 |
| LHFPL4 | 0 | -0.300377001 | 0.041 | 0.139 | 5.6E-50 | 2.05E-45 |
| ZDHHC21 | 0 | 0.341890141 | 0.777 | 0.661 | 7.15E-50 | 2.62E-45 |
| ROBO2 | 0 | 0.288511853 | 0.82 | 0.669 | 7.76E-50 | 2.84E-45 |
| AL390800.1 | 0 | -0.560572601 | 0.514 | 0.52 | 7.83E-50 | 2.87E-45 |
| CNTLN | 0 | 0.393949855 | 0.786 | 0.677 | 1.02E-49 | 3.72E-45 |
| FSTL5 | 0 | -0.686886691 | 0.06 | 0.171 | 1.2E-49 | 4.38E-45 |
| USP32 | 0 | 0.369149176 | 0.756 | 0.658 | 1.55E-49 | 5.69E-45 |
| ADARB2 | 0 | -0.960109472 | 0.02 | 0.101 | 1.64E-49 | 6.01E-45 |
| ARHGAP32 | 0 | 0.375224706 | 0.798 | 0.67 | 3.88E-49 | 1.42E-44 |
| AL589740.1 | 0 | 0.657167944 | 0.318 | 0.177 | 4.24E-49 | 1.55E-44 |
| CDYL | 0 | 0.411815658 | 0.791 | 0.69 | 4.76E-49 | 1.74E-44 |
| RHBDL3 | 0 | 0.353500688 | 0.413 | 0.253 | 7.57E-49 | 2.77E-44 |
| AC092902.4 | 0 | 0.324249987 | 0.25 | 0.125 | 7.84E-49 | 2.87E-44 |
| GCNT1 | 0 | -0.367948557 | 0.031 | 0.118 | 9.76E-49 | 3.58E-44 |
| DTNA | 0 | 0.651108381 | 0.769 | 0.667 | 1.09E-48 | 3.99E-44 |
| DGKB | 0 | -1.031788366 | 0.223 | 0.311 | 1.09E-48 | 4.01E-44 |
| SPAG9 | 0 | 0.291224771 | 0.89 | 0.818 | 1.65E-48 | 6.06E-44 |
| NEAT1 | 0 | -0.775025166 | 0.921 | 0.843 | 2.4E-48 | 8.78E-44 |
| FGD5 | 0 | -0.34648283 | 0.099 | 0.224 | 2.7E-48 | 9.9E-44 |
| LINC01934 | 0 | -1.250663266 | 0.046 | 0.126 | 2.78E-48 | 1.02E-43 |
| KIAA1549L | 0 | -0.475820715 | 0.129 | 0.259 | 3.19E-48 | 1.17E-43 |
| CCDC81 | 0 | -0.267015515 | 0.024 | 0.106 | 3.57E-48 | 1.31E-43 |
| STXBP6 | 0 | 0.370133051 | 0.108 | 0.029 | 4.28E-48 | 1.57E-43 |
| IGSF9B | 0 | -0.287274545 | 0.057 | 0.165 | 4.29E-48 | 1.57E-43 |
| IQGAP2 | 0 | -0.569872788 | 0.331 | 0.429 | 4.45E-48 | 1.63E-43 |
| SREBF2 | 0 | -0.502105165 | 0.404 | 0.536 | 5.02E-48 | 1.84E-43 |
| CTNNB1 | 0 | 0.403994584 | 0.732 | 0.63 | 6.08E-48 | 2.23E-43 |
| NGF | 0 | 0.376856882 | 0.203 | 0.092 | 7.88E-48 | 2.89E-43 |
| PREX1 | 0 | 0.340041138 | 0.321 | 0.18 | 8.69E-48 | 3.18E-43 |
| SHROOM3 | 0 | -0.386025777 | 0.031 | 0.119 | 9.85E-48 | 3.6E-43 |
| AC114971.1 | 0 | 0.43447598 | 0.123 | 0.041 | 9.93E-48 | 3.63E-43 |
| GRK5 | 0 | -0.454809364 | 0.226 | 0.372 | 1.13E-47 | 4.15E-43 |
| LRMDA | 0 | -0.733385501 | 0.054 | 0.14 | 1.86E-47 | 6.82E-43 |
| MAGI1 | 0 | -0.442376947 | 0.863 | 0.866 | 2.13E-47 | 7.81E-43 |
| MYO1B | 0 | 0.38457199 | 0.686 | 0.566 | 2.25E-47 | 8.23E-43 |
| HMCN1 | 0 | 0.563003336 | 0.314 | 0.174 | 2.34E-47 | 8.55E-43 |
| SYNPR | 0 | 0.571340108 | 0.449 | 0.323 | 2.49E-47 | 9.12E-43 |

| | | | | | | |
|---|---|---|---|---|---|---|
| GRAMD2B | 0 | -0.364264351 | 0.079 | 0.189 | 2.53E-47 | 9.26E-43 |
| SVEP1 | 0 | -0.520428239 | 0.284 | 0.297 | 2.61E-47 | 9.56E-43 |
| VGLL4 | 0 | 0.318746497 | 0.699 | 0.541 | 2.66E-47 | 9.74E-43 |
| CMTM7 | 0 | 0.3306039 | 0.273 | 0.141 | 3.25E-47 | 1.19E-42 |
| NAAA | 0 | 0.431992246 | 0.376 | 0.261 | 3.56E-47 | 1.3E-42 |
| IMMP2L | 0 | 0.302054764 | 0.894 | 0.79 | 3.67E-47 | 1.34E-42 |
| ACSL4 | 0 | -0.473227496 | 0.471 | 0.598 | 4.73E-47 | 1.73E-42 |
| AC018731.1 | 0 | -0.590346564 | 0.601 | 0.588 | 4.99E-47 | 1.83E-42 |
| FOCAD | 0 | 0.360934495 | 0.792 | 0.68 | 5.3E-47 | 1.94E-42 |
| PAX3 | 0 | -0.311802467 | 0.168 | 0.127 | 5.57E-47 | 2.04E-42 |
| RHOBTB3 | 0 | 0.3157147 | 0.579 | 0.418 | 7.01E-47 | 2.56E-42 |
| GSR | 0 | -0.419734437 | 0.199 | 0.335 | 7.75E-47 | 2.84E-42 |
| BARD1 | 0 | 0.390755164 | 0.491 | 0.337 | 8.61E-47 | 3.15E-42 |
| LMO4 | 0 | 0.470593151 | 0.347 | 0.232 | 9.05E-47 | 3.31E-42 |
| STK32B | 0 | -0.496849053 | 0.425 | 0.566 | 9.9E-47 | 3.62E-42 |
| SUPT3H | 0 | -0.440207098 | 0.645 | 0.764 | 1.06E-46 | 3.89E-42 |
| MTSS1 | 0 | -0.492923249 | 0.105 | 0.227 | 1.24E-46 | 4.55E-42 |
| CSRNP3 | 0 | -0.566186506 | 0.265 | 0.4 | 1.24E-46 | 4.55E-42 |
| CDH7 | 0 | -0.738910179 | 0.207 | 0.318 | 1.53E-46 | 5.6E-42 |
| PDE3B | 0 | 0.404906932 | 0.735 | 0.596 | 1.66E-46 | 6.09E-42 |
| FAM126A | 0 | 0.367118138 | 0.702 | 0.563 | 1.76E-46 | 6.45E-42 |
| SCN3A | 0 | -0.32921324 | 0.164 | 0.308 | 1.83E-46 | 6.7E-42 |
| ADGRA3 | 0 | 0.340628066 | 0.767 | 0.646 | 2.42E-46 | 8.87E-42 |
| TBC1D1 | 0 | 0.388526262 | 0.569 | 0.445 | 2.46E-46 | 9.01E-42 |
| HS3ST3A1 | 0 | 0.416536345 | 0.359 | 0.212 | 3.4E-46 | 1.24E-41 |
| TOP2B | 0 | 0.329044549 | 0.825 | 0.741 | 3.5E-46 | 1.28E-41 |
| PHACTR3 | 0 | -0.378737963 | 0.052 | 0.153 | 3.84E-46 | 1.41E-41 |
| MAML2 | 0 | 0.429378273 | 0.634 | 0.473 | 3.94E-46 | 1.44E-41 |
| MEIS1 | 0 | -0.625527472 | 0.268 | 0.387 | 4.09E-46 | 1.5E-41 |
| NIPAL2 | 0 | 0.30359306 | 0.173 | 0.072 | 4.34E-46 | 1.59E-41 |
| CPNE5 | 0 | 0.418661437 | 0.202 | 0.096 | 4.34E-46 | 1.59E-41 |
| LINC00511 | 0 | -0.572224498 | 0.186 | 0.331 | 4.39E-46 | 1.61E-41 |
| MEGF10 | 0 | 0.382721755 | 0.517 | 0.353 | 6.53E-46 | 2.39E-41 |
| AFDN | 0 | 0.363313446 | 0.857 | 0.768 | 6.54E-46 | 2.4E-41 |
| PTK7 | 0 | 0.351458557 | 0.493 | 0.342 | 6.69E-46 | 2.45E-41 |
| LMO7 | 0 | -0.646937314 | 0.456 | 0.553 | 6.91E-46 | 2.53E-41 |
| AFF2 | 0 | -0.655910876 | 0.612 | 0.659 | 7.02E-46 | 2.57E-41 |
| DUXAP8 | 0 | -0.252252903 | 0.038 | 0.13 | 9.83E-46 | 3.6E-41 |
| CRIM1 | 0 | -0.435983364 | 0.23 | 0.379 | 9.91E-46 | 3.63E-41 |
| E2F3 | 0 | 0.392144846 | 0.674 | 0.568 | 1.19E-45 | 4.35E-41 |
| AF117829.1 | 0 | 0.383398828 | 0.466 | 0.318 | 1.22E-45 | 4.45E-41 |
| CASC15 | 0 | -0.533428376 | 0.571 | 0.574 | 1.5E-45 | 5.48E-41 |
| KCNN3 | 0 | 0.388864306 | 0.215 | 0.109 | 1.92E-45 | 7.04E-41 |
| SPATA17 | 0 | -0.429716615 | 0.036 | 0.117 | 2.01E-45 | 7.35E-41 |
| SH3KBP1 | 0 | 0.392838753 | 0.769 | 0.642 | 2.33E-45 | 8.54E-41 |
| LANCL2 | 0 | 0.379218187 | 0.372 | 0.255 | 3.82E-45 | 1.4E-40 |
| CSNK1A1 | 0 | 0.312774441 | 0.894 | 0.824 | 4.7E-45 | 1.72E-40 |
| ITPR1 | 0 | -0.537432284 | 0.344 | 0.409 | 1.3E-44 | 4.75E-40 |
| PLEKHA7 | 0 | -0.42836376 | 0.109 | 0.227 | 1.33E-44 | 4.88E-40 |
| CPE | 0 | 0.363628163 | 0.725 | 0.591 | 1.62E-44 | 5.92E-40 |
| KLHDC8A | 0 | 0.310742398 | 0.257 | 0.133 | 2.05E-44 | 7.52E-40 |
| NYAP2 | 0 | -0.366405523 | 0.025 | 0.102 | 2.59E-44 | 9.49E-40 |
| IDH2 | 0 | 0.35528001 | 0.345 | 0.227 | 2.68E-44 | 9.81E-40 |
| ABCA5 | 0 | -0.430701478 | 0.277 | 0.432 | 2.76E-44 | 1.01E-39 |
| LINC01956 | 0 | 0.311917371 | 0.194 | 0.085 | 2.94E-44 | 1.08E-39 |
| TLN2 | 0 | 0.413121041 | 0.56 | 0.409 | 3.58E-44 | 1.31E-39 |
| ENC1 | 0 | 0.426096161 | 0.308 | 0.179 | 5.21E-44 | 1.91E-39 |
| FARP1 | 0 | 0.331702725 | 0.811 | 0.694 | 7.49E-44 | 2.74E-39 |

| | | | | | | |
|---|---|---|---|---|---|---|
| LRRC8B | 0 | -0.49776071 | 0.337 | 0.463 | 9.54E-44 | 3.49E-39 |
| LGMN | 0 | 0.295705293 | 0.257 | 0.132 | 1.18E-43 | 4.3E-39 |
| RNF24 | 0 | 0.331598065 | 0.658 | 0.511 | 1.28E-43 | 4.67E-39 |
| TRAF3 | 0 | 0.394440543 | 0.551 | 0.419 | 1.28E-43 | 4.69E-39 |
| ANKRD36 | 0 | 0.362212536 | 0.863 | 0.788 | 1.6E-43 | 5.86E-39 |
| GNG4 | 0 | -0.44659855 | 0.225 | 0.363 | 1.93E-43 | 7.08E-39 |
| NREP | 0 | 0.363088618 | 0.509 | 0.378 | 2.16E-43 | 7.92E-39 |
| OTX2 | 0 | -0.438492737 | 0.038 | 0.119 | 2.66E-43 | 9.73E-39 |
| EPHA6 | 0 | -0.550434551 | 0.047 | 0.142 | 3.09E-43 | 1.13E-38 |
| TBC1D23 | 0 | -0.428509649 | 0.403 | 0.524 | 4.3E-43 | 1.58E-38 |
| FAM155A | 0 | 0.350200233 | 0.871 | 0.752 | 6.7E-43 | 2.45E-38 |
| PPP2R3A | 0 | -0.37718123 | 0.247 | 0.394 | 7.04E-43 | 2.58E-38 |
| DLGAP1 | 0 | 0.29431384 | 0.906 | 0.805 | 7.57E-43 | 2.77E-38 |
| PCOLCE2 | 0 | 0.400373238 | 0.442 | 0.312 | 8.68E-43 | 3.18E-38 |
| TDRP | 0 | -0.256032364 | 0.054 | 0.151 | 9.02E-43 | 3.3E-38 |
| NOVA1 | 0 | 0.513838746 | 0.692 | 0.6 | 1.1E-42 | 4.04E-38 |
| AL353747.4 | 0 | -0.333251518 | 0.099 | 0.216 | 1.38E-42 | 5.07E-38 |
| RRAS2 | 0 | -0.378880198 | 0.183 | 0.319 | 1.47E-42 | 5.37E-38 |
| NBPF1 | 0 | 0.322076264 | 0.439 | 0.306 | 1.53E-42 | 5.59E-38 |
| AGK | 0 | 0.369066197 | 0.452 | 0.347 | 1.66E-42 | 6.08E-38 |
| VPS13A | 0 | -0.5479472 | 0.585 | 0.678 | 2.3E-42 | 8.42E-38 |
| EML6 | 0 | -0.432189216 | 0.152 | 0.281 | 2.74E-42 | 1E-37 |
| DCDC1 | 0 | -0.48327174 | 0.032 | 0.103 | 3.09E-42 | 1.13E-37 |
| ITGA7 | 0 | 0.276748026 | 0.198 | 0.094 | 3.21E-42 | 1.17E-37 |
| NETO1 | 0 | 0.499260512 | 0.25 | 0.139 | 3.28E-42 | 1.2E-37 |
| FAT3 | 0 | -0.589157585 | 0.293 | 0.431 | 4.21E-42 | 1.54E-37 |
| MAGI2-AS3 | 0 | 0.311484649 | 0.307 | 0.182 | 5.64E-42 | 2.07E-37 |
| TNIK | 0 | -0.581791845 | 0.264 | 0.399 | 5.67E-42 | 2.07E-37 |
| AGMO | 0 | -0.751114325 | 0.053 | 0.14 | 6.97E-42 | 2.55E-37 |
| PATJ | 0 | -0.462984333 | 0.344 | 0.466 | 7.08E-42 | 2.59E-37 |
| AC061958.1 | 0 | 0.394702165 | 0.413 | 0.269 | 8.16E-42 | 2.99E-37 |
| BCO2 | 0 | -0.279833597 | 0.049 | 0.142 | 9.29E-42 | 3.4E-37 |
| LINC01876 | 0 | -0.574003155 | 0.406 | 0.549 | 1.04E-41 | 3.82E-37 |
| MSRA | 0 | 0.421058139 | 0.509 | 0.386 | 1.18E-41 | 4.31E-37 |
| KNTC1 | 0 | 0.371955137 | 0.581 | 0.46 | 1.32E-41 | 4.83E-37 |
| POLA1 | 0 | 0.368117672 | 0.894 | 0.825 | 1.34E-41 | 4.92E-37 |
| DPYSL5 | 0 | -0.407632181 | 0.14 | 0.258 | 1.5E-41 | 5.49E-37 |
| MIER1 | 0 | 0.353659982 | 0.576 | 0.45 | 1.52E-41 | 5.56E-37 |
| ST8SIA4 | 0 | -0.302460108 | 0.05 | 0.145 | 1.66E-41 | 6.06E-37 |
| C5orf17 | 0 | 0.553895526 | 0.165 | 0.074 | 1.72E-41 | 6.29E-37 |
| STK39 | 0 | -0.454651263 | 0.581 | 0.666 | 3.18E-41 | 1.16E-36 |
| MEF2C-AS2 | 0 | 0.33321383 | 0.188 | 0.089 | 3.21E-41 | 1.17E-36 |
| SEPTIN6 | 0 | 0.276371366 | 0.28 | 0.152 | 3.38E-41 | 1.24E-36 |
| EYA4 | 0 | -0.337185361 | 0.246 | 0.187 | 3.75E-41 | 1.37E-36 |
| WSB1 | 0 | -0.553959576 | 0.747 | 0.805 | 4.48E-41 | 1.64E-36 |
| ATP8A2 | 0 | 0.26885366 | 0.417 | 0.271 | 5.7E-41 | 2.09E-36 |
| ZEB1-AS1 | 0 | 0.306854892 | 0.145 | 0.057 | 6.92E-41 | 2.54E-36 |
| PXYLP1 | 0 | -0.291302656 | 0.09 | 0.2 | 8.97E-41 | 3.28E-36 |
| OLFM2 | 0 | -0.283628172 | 0.054 | 0.15 | 9.54E-41 | 3.49E-36 |
| CNKSR2 | 0 | -0.549649821 | 0.687 | 0.651 | 1.26E-40 | 4.6E-36 |
| PDZRN3 | 0 | -0.569144661 | 0.898 | 0.906 | 1.35E-40 | 4.93E-36 |
| CRMP1 | 0 | -0.40377596 | 0.234 | 0.361 | 1.44E-40 | 5.28E-36 |
| NOTCH2 | 0 | 0.452198023 | 0.407 | 0.27 | 1.46E-40 | 5.35E-36 |
| LINC01414 | 0 | -0.640912765 | 0.032 | 0.104 | 1.66E-40 | 6.09E-36 |
| WDR66 | 0 | -0.307936185 | 0.057 | 0.152 | 1.67E-40 | 6.11E-36 |
| PLEKHA8 | 0 | 0.336111892 | 0.459 | 0.327 | 1.8E-40 | 6.58E-36 |
| SOX6 | 0 | -0.708413788 | 0.154 | 0.275 | 1.82E-40 | 6.68E-36 |
| DCP2 | 0 | 0.349505657 | 0.663 | 0.542 | 1.92E-40 | 7.04E-36 |

| | | | | | | |
|---|---|---|---|---|---|---|
| PKNOX2 | 0 | -0.424274325 | 0.144 | 0.259 | 3.19E-40 | 1.17E-35 |
| HLA-C | 0 | 0.258342028 | 0.197 | 0.092 | 3.3E-40 | 1.21E-35 |
| C12orf56 | 0 | 0.317404538 | 0.228 | 0.121 | 3.33E-40 | 1.22E-35 |
| KCTD16 | 0 | -0.471458883 | 0.087 | 0.196 | 3.68E-40 | 1.35E-35 |
| AKAP9 | 0 | -0.390970895 | 0.681 | 0.758 | 5.98E-40 | 2.19E-35 |
| FAM171B | 0 | -0.300647508 | 0.067 | 0.167 | 6.96E-40 | 2.55E-35 |
| CPLANE1 | 0 | 0.329290749 | 0.684 | 0.554 | 7.8E-40 | 2.86E-35 |
| PKD2 | 0 | 0.31528326 | 0.634 | 0.501 | 8.8E-40 | 3.22E-35 |
| CYBRD1 | 0 | 0.257311556 | 0.187 | 0.086 | 9.6E-40 | 3.51E-35 |
| SNX25 | 0 | 0.317206745 | 0.722 | 0.594 | 9.75E-40 | 3.57E-35 |
| PDS5B | 0 | 0.287104648 | 0.888 | 0.825 | 9.79E-40 | 3.59E-35 |
| FBXL17 | 0 | 0.316172272 | 0.832 | 0.739 | 1.18E-39 | 4.32E-35 |
| RALGAPA2 | 0 | -0.412218165 | 0.49 | 0.615 | 1.19E-39 | 4.34E-35 |
| TACR3 | 0 | 0.334866266 | 0.139 | 0.052 | 1.28E-39 | 4.68E-35 |
| MIAT | 0 | -0.290183132 | 0.044 | 0.132 | 1.34E-39 | 4.92E-35 |
| FZD6 | 0 | -0.291144635 | 0.142 | 0.268 | 1.47E-39 | 5.39E-35 |
| SYTL4 | 0 | 0.284924565 | 0.203 | 0.099 | 1.53E-39 | 5.6E-35 |
| SLC8A1 | 0 | -0.60726727 | 0.239 | 0.367 | 1.65E-39 | 6.06E-35 |
| STXBP5 | 0 | -0.396009707 | 0.563 | 0.65 | 1.74E-39 | 6.39E-35 |
| PALM2-AKAP2 | 0 | -0.435313302 | 0.409 | 0.52 | 2.88E-39 | 1.05E-34 |
| SEPTIN10 | 0 | 0.306657461 | 0.355 | 0.225 | 4.71E-39 | 1.72E-34 |
| CDC14B | 0 | 0.316443408 | 0.336 | 0.203 | 6.4E-39 | 2.34E-34 |
| ZMYND8 | 0 | -0.42624401 | 0.499 | 0.61 | 7.7E-39 | 2.82E-34 |
| LINC02595 | 0 | -0.272770023 | 0.046 | 0.13 | 1.16E-38 | 4.25E-34 |
| EBF3 | 0 | 0.489399714 | 0.345 | 0.213 | 1.19E-38 | 4.37E-34 |
| KIAA1671 | 0 | -0.370415013 | 0.098 | 0.196 | 1.44E-38 | 5.25E-34 |
| NCAN | 0 | 0.357896915 | 0.301 | 0.182 | 1.45E-38 | 5.31E-34 |
| TMEM132C | 0 | -0.324935691 | 0.15 | 0.11 | 2E-38 | 7.34E-34 |
| CNTNAP3B | 0 | 0.370247327 | 0.405 | 0.266 | 2.26E-38 | 8.29E-34 |
| TACC2 | 0 | -0.316134891 | 0.121 | 0.239 | 2.39E-38 | 8.74E-34 |
| MINAR1 | 0 | 0.284728173 | 0.246 | 0.131 | 3.7E-38 | 1.35E-33 |
| ARNTL | 0 | 0.300174509 | 0.224 | 0.117 | 3.93E-38 | 1.44E-33 |
| CNTNAP3 | 0 | -0.362494791 | 0.082 | 0.178 | 4.19E-38 | 1.53E-33 |
| PLCB1 | 0 | -0.403826091 | 0.418 | 0.568 | 4.3E-38 | 1.58E-33 |
| PMEPA1 | 0 | 0.272964332 | 0.321 | 0.19 | 5.01E-38 | 1.84E-33 |
| ZC4H2 | 0 | -0.334350048 | 0.056 | 0.147 | 7.53E-38 | 2.76E-33 |
| ITGB8 | 0 | -0.672432458 | 0.272 | 0.31 | 7.69E-38 | 2.81E-33 |
| ANKRD10 | 0 | 0.257635071 | 0.761 | 0.639 | 7.82E-38 | 2.86E-33 |
| SDK2 | 0 | -0.400796183 | 0.061 | 0.148 | 8.85E-38 | 3.24E-33 |
| PDZD2 | 0 | 0.44749207 | 0.479 | 0.351 | 9.36E-38 | 3.43E-33 |
| CENPP | 0 | -0.596419762 | 0.65 | 0.709 | 9.77E-38 | 3.58E-33 |
| SYBU | 0 | -0.469234923 | 0.316 | 0.45 | 1E-37 | 3.68E-33 |
| RNF157 | 0 | -0.318368928 | 0.165 | 0.291 | 1.15E-37 | 4.19E-33 |
| SAMD3 | 0 | -0.445657083 | 0.042 | 0.112 | 1.48E-37 | 5.42E-33 |
| SLIT1 | 0 | -0.316556582 | 0.049 | 0.136 | 1.52E-37 | 5.55E-33 |
| HTR2C | 0 | -3.799368909 | 0.072 | 0.102 | 1.58E-37 | 5.78E-33 |
| GPR156 | 0 | -0.359393128 | 0.144 | 0.26 | 2E-37 | 7.31E-33 |
| EPB41L2 | 0 | 0.384821772 | 0.7 | 0.623 | 2.28E-37 | 8.36E-33 |
| AC074286.1 | 0 | -0.315569933 | 0.049 | 0.138 | 2.43E-37 | 8.9E-33 |
| CD47 | 0 | 0.325702312 | 0.497 | 0.368 | 2.85E-37 | 1.04E-32 |
| MYRIP | 0 | 0.287443688 | 0.755 | 0.619 | 2.92E-37 | 1.07E-32 |
| C5 | 0 | 0.285283316 | 0.19 | 0.097 | 4.25E-37 | 1.55E-32 |
| DPPA4 | 0 | -0.398160341 | 0.36 | 0.465 | 5.08E-37 | 1.86E-32 |
| PACRG | 0 | -0.582624643 | 0.044 | 0.118 | 5.39E-37 | 1.97E-32 |
| PIP5K1B | 0 | -0.61923452 | 0.228 | 0.3 | 5.64E-37 | 2.06E-32 |
| GGH | 0 | 0.299223845 | 0.188 | 0.095 | 6.48E-37 | 2.37E-32 |
| MAGI2 | 0 | 0.269234631 | 0.784 | 0.666 | 7.75E-37 | 2.84E-32 |
| RPS6KA3 | 0 | -0.599594366 | 0.529 | 0.604 | 8.6E-37 | 3.15E-32 |

| | | | | | | |
|---|---|---|---|---|---|---|
| REEP1 | 0 | -0.280984589 | 0.065 | 0.158 | 8.88E-37 | 3.25E-32 |
| WWC2 | 0 | 0.333722059 | 0.503 | 0.36 | 9.23E-37 | 3.38E-32 |
| PRKN | 0 | 0.399850727 | 0.621 | 0.503 | 1.02E-36 | 3.74E-32 |
| AL591519.1 | 0 | -0.361612945 | 0.076 | 0.177 | 1.04E-36 | 3.82E-32 |
| LRP1B | 0 | -1.022637576 | 0.124 | 0.229 | 1.05E-36 | 3.84E-32 |
| BMPR2 | 0 | 0.295548087 | 0.871 | 0.806 | 1.11E-36 | 4.07E-32 |
| CNTN1 | 0 | -0.629190283 | 0.19 | 0.28 | 1.69E-36 | 6.2E-32 |
| SUB1 | 0 | 0.300010772 | 0.437 | 0.314 | 1.75E-36 | 6.41E-32 |
| AC138305.1 | 0 | -0.583840704 | 0.092 | 0.193 | 1.76E-36 | 6.45E-32 |
| SEMA6D | 0 | -0.270258327 | 0.727 | 0.65 | 1.97E-36 | 7.22E-32 |
| DENND1B | 0 | 0.375296173 | 0.413 | 0.288 | 2.04E-36 | 7.46E-32 |
| MEF2C | 0 | 0.259502753 | 0.454 | 0.311 | 2.1E-36 | 7.68E-32 |
| CSPP1 | 0 | -0.398160962 | 0.412 | 0.53 | 2.14E-36 | 7.82E-32 |
| ME3 | 0 | -0.398863859 | 0.221 | 0.347 | 2.22E-36 | 8.14E-32 |
| TLE4 | 0 | 0.349824582 | 0.581 | 0.445 | 2.53E-36 | 9.26E-32 |
| GKAP1 | 0 | 0.35711593 | 0.307 | 0.205 | 3.66E-36 | 1.34E-31 |
| AXIN2 | 0 | -0.289322784 | 0.104 | 0.205 | 3.94E-36 | 1.44E-31 |
| SIMC1 | 0 | 0.333822655 | 0.545 | 0.421 | 4.29E-36 | 1.57E-31 |
| MET | 0 | -0.346193964 | 0.123 | 0.237 | 4.78E-36 | 1.75E-31 |
| LINC01619 | 0 | 0.383016013 | 0.424 | 0.296 | 5.31E-36 | 1.94E-31 |
| UNC5C | 0 | -0.306271406 | 0.424 | 0.571 | 5.74E-36 | 2.1E-31 |
| ARNTL2 | 0 | -0.271110943 | 0.059 | 0.149 | 6.25E-36 | 2.29E-31 |
| PLPPR4 | 0 | 0.482683841 | 0.363 | 0.251 | 7.18E-36 | 2.63E-31 |
| SPRY4-AS1 | 0 | 0.353444654 | 0.249 | 0.14 | 7.24E-36 | 2.65E-31 |
| PIP4K2A | 0 | 0.31434911 | 0.677 | 0.555 | 8.2E-36 | 3E-31 |
| DAPK2 | 0 | 0.270503341 | 0.305 | 0.182 | 8.71E-36 | 3.19E-31 |
| RNF212 | 0 | -0.280597744 | 0.126 | 0.241 | 1.14E-35 | 4.18E-31 |
| PRKD2 | 0 | 0.316762797 | 0.246 | 0.139 | 1.15E-35 | 4.22E-31 |
| DOP1B | 0 | 0.31788245 | 0.588 | 0.454 | 1.31E-35 | 4.8E-31 |
| VEGFC | 0 | 0.471677503 | 0.298 | 0.181 | 1.32E-35 | 4.82E-31 |
| RAPGEF1 | 0 | -0.326382557 | 0.301 | 0.437 | 1.43E-35 | 5.24E-31 |
| UACA | 0 | 0.302567801 | 0.478 | 0.336 | 1.44E-35 | 5.29E-31 |
| NIPA1 | 0 | -0.304949396 | 0.149 | 0.265 | 1.5E-35 | 5.47E-31 |
| FIP1L1 | 0 | 0.318326409 | 0.635 | 0.545 | 1.5E-35 | 5.51E-31 |
| UNC13B | 0 | -0.487342674 | 0.501 | 0.555 | 1.69E-35 | 6.18E-31 |
| USP25 | 0 | -0.374230368 | 0.447 | 0.574 | 1.89E-35 | 6.91E-31 |
| PRKCA | 0 | -0.487264814 | 0.222 | 0.335 | 2.7E-35 | 9.88E-31 |
| CFAP69 | 0 | -0.33577927 | 0.043 | 0.117 | 3.03E-35 | 1.11E-30 |
| SATL1 | 0 | 0.25120522 | 0.159 | 0.075 | 3.12E-35 | 1.14E-30 |
| SRGAP3 | 0 | 0.31767667 | 0.646 | 0.512 | 3.25E-35 | 1.19E-30 |
| SEMA5B | 0 | -0.2664609 | 0.061 | 0.15 | 4.95E-35 | 1.81E-30 |
| DIP2B | 0 | 0.29454372 | 0.762 | 0.683 | 4.97E-35 | 1.82E-30 |
| OSBPL10 | 0 | -0.473630978 | 0.29 | 0.423 | 5.31E-35 | 1.94E-30 |
| GPD2 | 0 | 0.351540493 | 0.501 | 0.389 | 7.51E-35 | 2.75E-30 |
| KHDRBS3 | 0 | -0.562035301 | 0.166 | 0.274 | 7.91E-35 | 2.89E-30 |
| MTA3 | 0 | -0.374823912 | 0.64 | 0.705 | 8.84E-35 | 3.24E-30 |
| LRCH1 | 0 | 0.308109665 | 0.696 | 0.594 | 1.19E-34 | 4.36E-30 |
| SATB1-AS1 | 0 | -0.281250013 | 0.102 | 0.208 | 1.41E-34 | 5.17E-30 |
| PLEKHA5 | 0 | 0.326820386 | 0.892 | 0.839 | 1.43E-34 | 5.24E-30 |
| OSBPL11 | 0 | 0.373934008 | 0.4 | 0.308 | 1.47E-34 | 5.4E-30 |
| ANO6 | 0 | 0.302986821 | 0.67 | 0.57 | 2.46E-34 | 9.01E-30 |
| CAPN7 | 0 | 0.306472978 | 0.56 | 0.451 | 2.51E-34 | 9.2E-30 |
| HACE1 | 0 | -0.422487985 | 0.563 | 0.64 | 2.92E-34 | 1.07E-29 |
| TBC1D8 | 0 | -0.313275394 | 0.13 | 0.237 | 3.03E-34 | 1.11E-29 |
| SCN8A | 0 | 0.34352591 | 0.724 | 0.641 | 3.58E-34 | 1.31E-29 |
| GRIK2 | 0 | 0.333094731 | 0.386 | 0.253 | 3.93E-34 | 1.44E-29 |
| SPTBN4 | 0 | -0.284736687 | 0.046 | 0.123 | 3.93E-34 | 1.44E-29 |
| RFX2 | 0 | -0.455494063 | 0.17 | 0.244 | 4.14E-34 | 1.52E-29 |

| | | | | | | |
|---|---|---|---|---|---|---|
| ETV5 | 0 | 0.262073964 | 0.208 | 0.107 | 4.23E-34 | 1.55E-29 |
| GAS2L3 | 0 | 0.309314128 | 0.362 | 0.241 | 4.51E-34 | 1.65E-29 |
| CYP7B1 | 0 | -0.523636034 | 0.279 | 0.405 | 5.18E-34 | 1.9E-29 |
| STRBP | 0 | -0.376980728 | 0.445 | 0.568 | 7.34E-34 | 2.69E-29 |
| KCTD8 | 0 | -0.424590821 | 0.08 | 0.174 | 7.55E-34 | 2.76E-29 |
| PCDH19 | 0 | -0.485439315 | 0.123 | 0.227 | 7.79E-34 | 2.85E-29 |
| KCNQ3 | 0 | -0.550857438 | 0.309 | 0.419 | 8.63E-34 | 3.16E-29 |
| PRDM5 | 0 | 0.33174557 | 0.405 | 0.287 | 8.94E-34 | 3.27E-29 |
| FAM171A1 | 0 | 0.313891211 | 0.628 | 0.536 | 9.83E-34 | 3.6E-29 |
| PLEKHH2 | 0 | 0.263801857 | 0.249 | 0.138 | 1.14E-33 | 4.16E-29 |
| FAM219A | 0 | -0.262279667 | 0.097 | 0.194 | 1.17E-33 | 4.29E-29 |
| MOK | 0 | -0.323829753 | 0.076 | 0.16 | 1.23E-33 | 4.51E-29 |
| BCL2 | 0 | 0.365171862 | 0.275 | 0.163 | 1.36E-33 | 4.98E-29 |
| YLPM1 | 0 | 0.283322724 | 0.676 | 0.589 | 1.4E-33 | 5.13E-29 |
| DARS | 0 | 0.299645436 | 0.645 | 0.53 | 1.82E-33 | 6.65E-29 |
| PDE4DIP | 0 | -0.344468618 | 0.25 | 0.365 | 1.91E-33 | 7E-29 |
| ABCC1 | 0 | -0.424409693 | 0.424 | 0.518 | 2.3E-33 | 8.42E-29 |
| GFOD1 | 0 | 0.331218266 | 0.422 | 0.3 | 2.56E-33 | 9.37E-29 |
| SLAIN2 | 0 | 0.314037204 | 0.596 | 0.487 | 2.64E-33 | 9.68E-29 |
| DCBLD1 | 0 | -0.289489061 | 0.113 | 0.217 | 3.22E-33 | 1.18E-28 |
| PIK3AP1 | 0 | -0.310933648 | 0.09 | 0.186 | 3.8E-33 | 1.39E-28 |
| PAWR | 0 | -0.337109482 | 0.261 | 0.392 | 4.07E-33 | 1.49E-28 |
| SCFD2 | 0 | 0.361029328 | 0.662 | 0.569 | 5.55E-33 | 2.03E-28 |
| CCDC30 | 0 | -0.353390865 | 0.076 | 0.153 | 7.19E-33 | 2.63E-28 |
| BCR | 0 | -0.301613014 | 0.165 | 0.279 | 8.09E-33 | 2.96E-28 |
| POU5F1 | 0 | 0.269943915 | 0.578 | 0.444 | 8.78E-33 | 3.22E-28 |
| RCOR1 | 0 | 0.276973057 | 0.754 | 0.655 | 9.48E-33 | 3.47E-28 |
| KITLG | 0 | -0.293608534 | 0.057 | 0.141 | 1.14E-32 | 4.16E-28 |
| HS6ST2 | 0 | -0.403042852 | 0.61 | 0.69 | 1.3E-32 | 4.75E-28 |
| STK26 | 0 | -0.26858811 | 0.086 | 0.182 | 1.45E-32 | 5.31E-28 |
| FAM178B | 0 | -0.366568174 | 0.039 | 0.108 | 1.5E-32 | 5.48E-28 |
| CLYBL | 0 | -0.379730155 | 0.194 | 0.309 | 1.85E-32 | 6.76E-28 |
| PIK3R3 | 0 | 0.316188418 | 0.589 | 0.458 | 1.95E-32 | 7.15E-28 |
| TRAK2 | 0 | 0.28736014 | 0.354 | 0.234 | 2.14E-32 | 7.84E-28 |
| NTNG1 | 0 | 0.434488129 | 0.561 | 0.442 | 2.2E-32 | 8.06E-28 |
| OSBPL6 | 0 | 0.363468141 | 0.538 | 0.429 | 2.22E-32 | 8.12E-28 |
| KIRREL3 | 0 | -0.655960314 | 0.115 | 0.206 | 2.52E-32 | 9.21E-28 |
| ZNF91 | 0 | -0.272750686 | 0.165 | 0.282 | 3.28E-32 | 1.2E-27 |
| SPEF2 | 0 | -0.425727536 | 0.089 | 0.172 | 3.4E-32 | 1.24E-27 |
| MRC2 | 0 | 0.275122666 | 0.323 | 0.202 | 4.08E-32 | 1.49E-27 |
| TIAM1 | 0 | 0.47621136 | 0.687 | 0.576 | 4.21E-32 | 1.54E-27 |
| KIAA1958 | 0 | 0.293589469 | 0.824 | 0.74 | 5.03E-32 | 1.84E-27 |
| KIAA1324L | 0 | -0.300635066 | 0.141 | 0.253 | 6.24E-32 | 2.29E-27 |
| NEDD4 | 0 | 0.308759256 | 0.403 | 0.283 | 6.72E-32 | 2.46E-27 |
| VCAN | 0 | 0.419346071 | 0.371 | 0.435 | 7.3E-32 | 2.67E-27 |
| AQP4-AS1 | 0 | -0.263357622 | 0.053 | 0.134 | 7.65E-32 | 2.8E-27 |
| CCNH | 0 | 0.333639024 | 0.651 | 0.555 | 8.48E-32 | 3.1E-27 |
| IGSF11 | 0 | 0.36167531 | 0.215 | 0.121 | 1.03E-31 | 3.76E-27 |
| SNHG14 | 0 | -0.334535921 | 0.804 | 0.843 | 1.06E-31 | 3.89E-27 |
| TAB2 | 0 | -0.307776432 | 0.414 | 0.544 | 1.12E-31 | 4.09E-27 |
| KATNBL1 | 0 | 0.28588901 | 0.576 | 0.476 | 1.17E-31 | 4.3E-27 |
| TBL1X | 0 | -0.38163344 | 0.736 | 0.791 | 1.4E-31 | 5.13E-27 |
| OPCML | 0 | -0.48340594 | 0.253 | 0.381 | 1.4E-31 | 5.13E-27 |
| HES1 | 0 | 0.315892242 | 0.121 | 0.049 | 1.49E-31 | 5.45E-27 |
| UFL1-AS1 | 0 | -0.456905693 | 0.059 | 0.125 | 1.54E-31 | 5.62E-27 |
| TTC3 | 0 | 0.267182309 | 0.909 | 0.864 | 1.61E-31 | 5.89E-27 |
| MEF2C-AS1 | 0 | 0.301020269 | 0.332 | 0.212 | 1.65E-31 | 6.06E-27 |
| RNF152 | 0 | 0.263487976 | 0.148 | 0.068 | 1.81E-31 | 6.64E-27 |

| | | | | | | |
|---|---|---|---|---|---|---|
| ZNF814 | 0 | -0.265218456 | 0.128 | 0.232 | 2.07E-31 | 7.57E-27 |
| HMGCS1 | 0 | -0.415525918 | 0.113 | 0.21 | 2.2E-31 | 8.06E-27 |
| NEB | 0 | -0.524043631 | 0.073 | 0.151 | 2.27E-31 | 8.32E-27 |
| TDRD7 | 0 | 0.286156327 | 0.263 | 0.17 | 2.31E-31 | 8.47E-27 |
| ARNT2 | 0 | 0.311005884 | 0.483 | 0.367 | 2.75E-31 | 1.01E-26 |
| LNX1 | 0 | 0.277894655 | 0.191 | 0.102 | 2.88E-31 | 1.05E-26 |
| LRCH3 | 0 | 0.275021128 | 0.699 | 0.591 | 3.94E-31 | 1.44E-26 |
| DNM1 | 0 | 0.293706365 | 0.399 | 0.278 | 4.13E-31 | 1.51E-26 |
| NAB1 | 0 | 0.280575022 | 0.396 | 0.28 | 4.95E-31 | 1.81E-26 |
| AC107223.1 | 0 | 0.269660063 | 0.22 | 0.12 | 6.03E-31 | 2.21E-26 |
| PXDN | 0 | 0.349681676 | 0.311 | 0.202 | 7.25E-31 | 2.66E-26 |
| THOC1 | 0 | 0.289036461 | 0.594 | 0.498 | 9.48E-31 | 3.47E-26 |
| MAMLD1 | 0 | -0.388539314 | 0.227 | 0.347 | 1.28E-30 | 4.69E-26 |
| HSPA12A | 0 | -0.328162806 | 0.201 | 0.31 | 1.34E-30 | 4.92E-26 |
| GARNL3 | 0 | 0.285858099 | 0.311 | 0.199 | 1.5E-30 | 5.51E-26 |
| CACNB2 | 0 | -0.561988852 | 0.73 | 0.733 | 1.65E-30 | 6.03E-26 |
| ADGRV1 | 0 | 0.583175949 | 0.648 | 0.699 | 1.69E-30 | 6.18E-26 |
| LDLR | 0 | -0.276133422 | 0.056 | 0.135 | 1.73E-30 | 6.33E-26 |
| AC135782.1 | 0 | 0.26575698 | 0.214 | 0.117 | 1.77E-30 | 6.48E-26 |
| PRIM2 | 0 | 0.293970409 | 0.699 | 0.615 | 2.25E-30 | 8.25E-26 |
| SUN1 | 0 | 0.269886922 | 0.653 | 0.55 | 2.28E-30 | 8.34E-26 |
| NECTIN3 | 0 | 0.317895681 | 0.681 | 0.568 | 2.31E-30 | 8.45E-26 |
| LPIN1 | 0 | -0.262142485 | 0.14 | 0.244 | 3.2E-30 | 1.17E-25 |
| PDIA6 | 0 | 0.279312358 | 0.363 | 0.248 | 3.51E-30 | 1.29E-25 |
| SERPINB6 | 0 | -0.289895996 | 0.149 | 0.257 | 4.2E-30 | 1.54E-25 |
| ADAMTS9 | 0 | 0.419385927 | 0.419 | 0.36 | 4.39E-30 | 1.61E-25 |
| ADGRL1 | 0 | -0.273946043 | 0.162 | 0.271 | 5.32E-30 | 1.95E-25 |
| PIAS1 | 0 | -0.323668382 | 0.67 | 0.755 | 5.76E-30 | 2.11E-25 |
| VMP1 | 0 | -0.439707342 | 0.533 | 0.62 | 6.06E-30 | 2.22E-25 |
| FAM124A | 0 | -0.327420551 | 0.172 | 0.276 | 6.09E-30 | 2.23E-25 |
| MYT1L | 0 | -0.500245386 | 0.069 | 0.154 | 6.13E-30 | 2.24E-25 |
| AC108690.1 | 0 | 0.368635236 | 0.157 | 0.11 | 6.38E-30 | 2.33E-25 |
| SNCA | 0 | -0.420855182 | 0.2 | 0.304 | 6.58E-30 | 2.41E-25 |
| LINC00599 | 0 | -0.288073496 | 0.094 | 0.183 | 7.59E-30 | 2.78E-25 |
| MED12L | 0 | 0.380995764 | 0.581 | 0.47 | 7.75E-30 | 2.84E-25 |
| RFC3 | 0 | -0.413041447 | 0.394 | 0.504 | 8E-30 | 2.93E-25 |
| GAREM1 | 0 | 0.307668522 | 0.368 | 0.247 | 8.29E-30 | 3.03E-25 |
| BACH1 | 0 | 0.310032939 | 0.547 | 0.435 | 8.47E-30 | 3.1E-25 |
| ZNF516 | 0 | 0.318214822 | 0.627 | 0.521 | 8.64E-30 | 3.16E-25 |
| ETV1 | 0 | -0.546887483 | 0.18 | 0.207 | 9.92E-30 | 3.63E-25 |
| PWWP3A | 0 | 0.283734539 | 0.438 | 0.324 | 1.04E-29 | 3.81E-25 |
| GCLM | 0 | -0.376076827 | 0.096 | 0.182 | 1.25E-29 | 4.57E-25 |
| TMOD3 | 0 | -0.297408411 | 0.255 | 0.373 | 1.66E-29 | 6.07E-25 |
| TTC8 | 0 | -0.301250551 | 0.172 | 0.28 | 1.79E-29 | 6.54E-25 |
| PARM1 | 0 | 0.319908633 | 0.278 | 0.172 | 1.9E-29 | 6.96E-25 |
| PPP1R13B | 0 | -0.322030716 | 0.18 | 0.288 | 2.59E-29 | 9.47E-25 |
| PDCD4 | 0 | 0.314366891 | 0.469 | 0.377 | 3.05E-29 | 1.12E-24 |
| AC006148.1 | 0 | -0.382792465 | 0.092 | 0.184 | 3.26E-29 | 1.19E-24 |
| WIPF1 | 0 | 0.271677499 | 0.256 | 0.154 | 3.45E-29 | 1.26E-24 |
| DMD | 0 | -0.927699092 | 0.658 | 0.678 | 3.53E-29 | 1.29E-24 |
| GCLC | 0 | -0.657491445 | 0.176 | 0.227 | 3.83E-29 | 1.4E-24 |
| WEE1 | 0 | -0.364744542 | 0.377 | 0.453 | 4.32E-29 | 1.58E-24 |
| AL157778.1 | 0 | -0.694598997 | 0.167 | 0.268 | 5.4E-29 | 1.98E-24 |
| PLEKHG4B | 0 | 0.261186673 | 0.304 | 0.193 | 5.58E-29 | 2.04E-24 |
| SRRM3 | 0 | -0.253243075 | 0.095 | 0.188 | 6.71E-29 | 2.46E-24 |
| TMEM131L | 0 | 0.286014015 | 0.648 | 0.533 | 6.95E-29 | 2.55E-24 |
| GMPS | 0 | 0.276062509 | 0.622 | 0.515 | 1.01E-28 | 3.68E-24 |
| SCP2 | 0 | -0.357928642 | 0.293 | 0.4 | 1.05E-28 | 3.86E-24 |

| | | | | | | |
|---|---|---|---|---|---|---|
| SFXN5 | 0 | 0.287436242 | 0.391 | 0.278 | 1.37E-28 | 5.01E-24 |
| ARRB1 | 0 | 0.256421158 | 0.22 | 0.128 | 1.68E-28 | 6.15E-24 |
| HULC | 0 | -0.780311701 | 0.363 | 0.435 | 1.91E-28 | 7E-24 |
| MTHFD1L | 0 | -0.367901554 | 0.484 | 0.582 | 2.13E-28 | 7.81E-24 |
| PCDH9 | 0 | -0.820349056 | 0.287 | 0.37 | 2.43E-28 | 8.9E-24 |
| VPS26A | 0 | 0.284655895 | 0.372 | 0.273 | 2.86E-28 | 1.05E-23 |
| GPI | 0 | -0.420835901 | 0.189 | 0.292 | 3.64E-28 | 1.33E-23 |
| MAML3 | 0 | -0.258578847 | 0.637 | 0.546 | 4.08E-28 | 1.49E-23 |
| CTNNA3 | 0 | -0.542307756 | 0.114 | 0.209 | 4.11E-28 | 1.5E-23 |
| AC008014.1 | 0 | -0.430369492 | 0.284 | 0.382 | 4.47E-28 | 1.63E-23 |
| TASOR2 | 0 | -0.28391317 | 0.577 | 0.685 | 4.71E-28 | 1.73E-23 |
| DGKH | 0 | -0.352458102 | 0.684 | 0.709 | 4.83E-28 | 1.77E-23 |
| ITPR2 | 0 | -0.44249601 | 0.261 | 0.379 | 5.46E-28 | 2E-23 |
| CARMIL1 | 0 | -0.403019845 | 0.259 | 0.348 | 5.85E-28 | 2.14E-23 |
| PER3 | 0 | 0.267633211 | 0.272 | 0.168 | 6.01E-28 | 2.2E-23 |
| GOLIM4 | 0 | 0.280479458 | 0.589 | 0.485 | 7.77E-28 | 2.84E-23 |
| DPY19L2 | 0 | 0.280805481 | 0.47 | 0.362 | 8.31E-28 | 3.04E-23 |
| NEBL | 0 | -0.306619858 | 0.13 | 0.23 | 8.47E-28 | 3.1E-23 |
| FGD4 | 0 | -0.427692741 | 0.357 | 0.445 | 9.5E-28 | 3.48E-23 |
| GPM6A | 0 | -0.727578847 | 0.15 | 0.219 | 1.06E-27 | 3.88E-23 |
| MEGF11 | 0 | 0.269747936 | 0.4 | 0.282 | 1.14E-27 | 4.18E-23 |
| PCNX1 | 0 | 0.25419092 | 0.805 | 0.733 | 1.14E-27 | 4.18E-23 |
| ZNF280D | 0 | -0.278611057 | 0.229 | 0.345 | 1.26E-27 | 4.63E-23 |
| RIN2 | 0 | 0.262929825 | 0.445 | 0.324 | 1.33E-27 | 4.86E-23 |
| ADARB1 | 0 | 0.361207036 | 0.363 | 0.254 | 1.77E-27 | 6.46E-23 |
| TCF20 | 0 | -0.289580271 | 0.34 | 0.459 | 2.4E-27 | 8.79E-23 |
| AC119673.2 | 0 | 0.267309014 | 0.462 | 0.355 | 2.44E-27 | 8.95E-23 |
| IPO9 | 0 | 0.253813267 | 0.678 | 0.587 | 2.58E-27 | 9.43E-23 |
| AKAP7 | 0 | 0.29885345 | 0.314 | 0.212 | 2.85E-27 | 1.04E-22 |
| ARHGAP44 | 0 | -0.411538934 | 0.34 | 0.418 | 3.27E-27 | 1.2E-22 |
| MYO3A | 0 | -0.785515462 | 0.138 | 0.197 | 3.48E-27 | 1.27E-22 |
| ARHGAP21 | 0 | 0.265336993 | 0.628 | 0.522 | 3.5E-27 | 1.28E-22 |
| LINC01122 | 0 | -0.547239461 | 0.205 | 0.296 | 4.88E-27 | 1.79E-22 |
| SPINK5 | 0 | -0.280011487 | 0.092 | 0.178 | 5.56E-27 | 2.04E-22 |
| SH3BGRL2 | 0 | 0.274926042 | 0.236 | 0.154 | 6.17E-27 | 2.26E-22 |
| PRKAG2 | 0 | 0.312229393 | 0.517 | 0.405 | 6.22E-27 | 2.28E-22 |
| RFX3 | 0 | -0.408965991 | 0.812 | 0.823 | 7.37E-27 | 2.7E-22 |
| DSE | 0 | 0.313200688 | 0.409 | 0.317 | 8.64E-27 | 3.16E-22 |
| TMPO | 0 | 0.298838919 | 0.579 | 0.489 | 9.3E-27 | 3.41E-22 |
| VDAC3 | 0 | -0.310047301 | 0.242 | 0.346 | 1.36E-26 | 4.98E-22 |
| SOX11 | 0 | 0.293386307 | 0.401 | 0.295 | 1.47E-26 | 5.37E-22 |
| DZIP1 | 0 | 0.263852709 | 0.55 | 0.443 | 1.72E-26 | 6.3E-22 |
| RDX | 0 | 0.275339236 | 0.636 | 0.552 | 1.74E-26 | 6.36E-22 |
| AC023469.1 | 0 | -0.511484917 | 0.304 | 0.372 | 1.8E-26 | 6.58E-22 |
| PRRT2 | 0 | 0.319988598 | 0.474 | 0.373 | 1.8E-26 | 6.61E-22 |
| MIR646HG | 0 | -0.305745095 | 0.135 | 0.109 | 1.93E-26 | 7.06E-22 |
| PTBP3 | 0 | -0.271647107 | 0.293 | 0.413 | 1.97E-26 | 7.2E-22 |
| FERMT1 | 0 | -0.334396484 | 0.135 | 0.224 | 2.05E-26 | 7.49E-22 |
| AP3B1 | 0 | 0.259991373 | 0.714 | 0.616 | 2.17E-26 | 7.96E-22 |
| AC090578.1 | 0 | 0.348282805 | 0.344 | 0.249 | 2.2E-26 | 8.05E-22 |
| GALNTL6 | 0 | -0.656307781 | 0.178 | 0.278 | 2.28E-26 | 8.36E-22 |
| FAM49B | 0 | -0.281897308 | 0.573 | 0.674 | 2.4E-26 | 8.79E-22 |
| TENT5A | 0 | 0.264036939 | 0.196 | 0.109 | 2.46E-26 | 9.02E-22 |
| WNK2 | 0 | 0.319728019 | 0.659 | 0.556 | 2.71E-26 | 9.91E-22 |
| LPGAT1 | 0 | 0.285479724 | 0.562 | 0.455 | 3.74E-26 | 1.37E-21 |
| EXPH5 | 0 | -0.291377836 | 0.096 | 0.183 | 4.55E-26 | 1.67E-21 |
| FOXO1 | 0 | -0.296073449 | 0.141 | 0.239 | 5.39E-26 | 1.97E-21 |
| PRH1 | 0 | 0.310583382 | 0.54 | 0.444 | 7.77E-26 | 2.85E-21 |

| | | | | | | |
|---|---|---|---|---|---|---|
| NBEAL1 | 0 | 0.285472131 | 0.528 | 0.432 | 9.63E-26 | 3.52E-21 |
| SRRM4 | 0 | -0.293168786 | 0.116 | 0.209 | 9.83E-26 | 3.6E-21 |
| SMC2 | 0 | 0.305363954 | 0.407 | 0.306 | 1.05E-25 | 3.85E-21 |
| CSMD1 | 0 | -0.666084263 | 0.408 | 0.46 | 1.2E-25 | 4.39E-21 |
| SORCS1 | 0 | -0.389064016 | 0.16 | 0.259 | 1.23E-25 | 4.49E-21 |
| WLS | 0 | -0.355424974 | 0.358 | 0.445 | 1.39E-25 | 5.11E-21 |
| SKIL | 0 | 0.274780017 | 0.692 | 0.589 | 1.5E-25 | 5.51E-21 |
| PPP2R5C | 0 | 0.255189118 | 0.625 | 0.542 | 1.77E-25 | 6.46E-21 |
| NLGN1 | 0 | -0.269055464 | 0.854 | 0.785 | 1.95E-25 | 7.12E-21 |
| PLXDC2 | 0 | 0.251773535 | 0.747 | 0.637 | 2.35E-25 | 8.59E-21 |
| WDR17 | 0 | -0.329485324 | 0.271 | 0.382 | 2.5E-25 | 9.15E-21 |
| LPAR1 | 0 | -0.518346089 | 0.161 | 0.224 | 2.75E-25 | 1.01E-20 |
| BMP6 | 0 | 0.316022827 | 0.156 | 0.086 | 2.86E-25 | 1.05E-20 |
| AL591501.1 | 0 | -0.593796322 | 0.106 | 0.117 | 2.87E-25 | 1.05E-20 |
| JADE3 | 0 | -0.271495538 | 0.274 | 0.389 | 2.91E-25 | 1.07E-20 |
| GTDC1 | 0 | 0.298849561 | 0.564 | 0.46 | 3.04E-25 | 1.11E-20 |
| NR3C2 | 0 | -0.550335736 | 0.221 | 0.296 | 3.11E-25 | 1.14E-20 |
| RBFOX3 | 0 | 0.36496865 | 0.267 | 0.174 | 3.61E-25 | 1.32E-20 |
| TMEM164 | 0 | -0.354246748 | 0.419 | 0.519 | 4.34E-25 | 1.59E-20 |
| NETO2 | 0 | 0.298599297 | 0.602 | 0.503 | 4.38E-25 | 1.6E-20 |
| ELAVL3 | 0 | -0.314462515 | 0.105 | 0.189 | 4.44E-25 | 1.62E-20 |
| TOM1L1 | 0 | -0.26529019 | 0.215 | 0.319 | 4.45E-25 | 1.63E-20 |
| C1GALT1 | 0 | -0.25221604 | 0.156 | 0.254 | 4.47E-25 | 1.64E-20 |
| TARID | 0 | 0.31430445 | 0.118 | 0.065 | 6.13E-25 | 2.25E-20 |
| SIPA1L3 | 0 | -0.583111328 | 0.496 | 0.537 | 6.51E-25 | 2.38E-20 |
| RPGRIP1L | 0 | 0.311933782 | 0.499 | 0.398 | 7E-25 | 2.56E-20 |
| CPEB3 | 0 | -0.33736809 | 0.319 | 0.419 | 8.15E-25 | 2.99E-20 |
| ANKRD36B | 0 | 0.290329574 | 0.542 | 0.462 | 8.5E-25 | 3.11E-20 |
| BCAT1 | 0 | -0.276274539 | 0.121 | 0.211 | 1.11E-24 | 4.07E-20 |
| PPARGC1A | 0 | -0.360784493 | 0.108 | 0.189 | 1.21E-24 | 4.43E-20 |
| PTPRJ | 0 | -0.423676036 | 0.461 | 0.538 | 1.62E-24 | 5.92E-20 |
| AC004917.1 | 0 | 0.348218038 | 0.14 | 0.081 | 1.79E-24 | 6.54E-20 |
| GPR135 | 0 | 0.254338819 | 0.324 | 0.242 | 1.8E-24 | 6.58E-20 |
| LEPR | 0 | -0.408799958 | 0.082 | 0.152 | 2.18E-24 | 7.99E-20 |
| ARL6IP6 | 0 | 0.269661168 | 0.554 | 0.464 | 3.13E-24 | 1.15E-19 |
| BCLAF3 | 0 | -0.324484806 | 0.352 | 0.437 | 3.6E-24 | 1.32E-19 |
| PKP4 | 0 | 0.264320368 | 0.768 | 0.682 | 4.25E-24 | 1.56E-19 |
| FEZ1 | 0 | -0.27440053 | 0.153 | 0.247 | 4.35E-24 | 1.59E-19 |
| LIMD1 | 0 | 0.265636168 | 0.442 | 0.334 | 4.69E-24 | 1.72E-19 |
| SMYD3 | 0 | 0.273839266 | 0.952 | 0.915 | 6.39E-24 | 2.34E-19 |
| HS3ST1 | 0 | 0.291083948 | 0.25 | 0.177 | 6.51E-24 | 2.38E-19 |
| AL033504.1 | 0 | -0.321768595 | 0.494 | 0.612 | 6.73E-24 | 2.46E-19 |
| CHPT1 | 0 | -0.354902442 | 0.547 | 0.573 | 7.66E-24 | 2.8E-19 |
| GDAP1 | 0 | -0.354861462 | 0.241 | 0.34 | 9.21E-24 | 3.37E-19 |
| PTAR1 | 0 | -0.297755886 | 0.397 | 0.492 | 9.26E-24 | 3.39E-19 |
| FZD3 | 0 | -0.320904931 | 0.201 | 0.288 | 9.34E-24 | 3.42E-19 |
| CDK17 | 0 | 0.266059943 | 0.65 | 0.561 | 1E-23 | 3.67E-19 |
| KIAA0586 | 0 | 0.251863515 | 0.464 | 0.356 | 1.47E-23 | 5.36E-19 |
| CUL1 | 0 | 0.254893994 | 0.638 | 0.553 | 1.52E-23 | 5.58E-19 |
| NDRG1 | 0 | -0.5033409 | 0.068 | 0.129 | 1.55E-23 | 5.66E-19 |
| RPS6KA5 | 0 | 0.279746342 | 0.477 | 0.379 | 1.62E-23 | 5.93E-19 |
| EDEM3 | 0 | 0.262680393 | 0.43 | 0.335 | 1.75E-23 | 6.41E-19 |
| GPM6B | 0 | 0.304254673 | 0.702 | 0.605 | 1.88E-23 | 6.87E-19 |
| WASF1 | 0 | 0.266394867 | 0.509 | 0.43 | 2.03E-23 | 7.44E-19 |
| MRE11 | 0 | 0.258771951 | 0.542 | 0.458 | 2.27E-23 | 8.33E-19 |
| TMEM87B | 0 | 0.268630412 | 0.345 | 0.25 | 2.67E-23 | 9.77E-19 |
| GK | 0 | -0.324482267 | 0.214 | 0.302 | 2.93E-23 | 1.07E-18 |
| DIRC3 | 0 | -0.263536037 | 0.137 | 0.23 | 3.11E-23 | 1.14E-18 |

| | | | | | | |
|---|---|---|---|---|---|---|
| ATP11C | 0 | -0.294894684 | 0.714 | 0.718 | 3.16E-23 | 1.16E-18 |
| KCTD13 | 0 | 0.250986002 | 0.395 | 0.297 | 4.02E-23 | 1.47E-18 |
| CDC42BPA | 0 | 0.294473392 | 0.769 | 0.773 | 4.53E-23 | 1.66E-18 |
| MARK1 | 0 | -0.412806379 | 0.456 | 0.524 | 4.59E-23 | 1.68E-18 |
| ADCY3 | 0 | 0.258267298 | 0.369 | 0.272 | 5.95E-23 | 2.18E-18 |
| LANCL3 | 0 | -0.296570856 | 0.159 | 0.253 | 6.34E-23 | 2.32E-18 |
| PRRG1 | 0 | -0.25125539 | 0.281 | 0.392 | 7.56E-23 | 2.77E-18 |
| NAALADL2 | 0 | 0.27243996 | 0.491 | 0.384 | 8.02E-23 | 2.94E-18 |
| RAPH1 | 0 | 0.282473817 | 0.487 | 0.408 | 9.45E-23 | 3.46E-18 |
| SERGEF | 0 | -0.303445526 | 0.342 | 0.447 | 9.63E-23 | 3.53E-18 |
| RNASEH2B | 0 | 0.260791598 | 0.405 | 0.316 | 1.08E-22 | 3.96E-18 |
| PAPSS1 | 0 | 0.254164852 | 0.475 | 0.379 | 1.22E-22 | 4.45E-18 |
| PAK5 | 0 | -0.390646026 | 0.433 | 0.457 | 1.52E-22 | 5.56E-18 |
| ITFG1 | 0 | 0.342562898 | 0.822 | 0.778 | 1.52E-22 | 5.58E-18 |
| KCTD1 | 0 | -0.332964484 | 0.261 | 0.336 | 1.53E-22 | 5.6E-18 |
| DLEU2 | 0 | 0.284029103 | 0.786 | 0.728 | 1.55E-22 | 5.67E-18 |
| SLC4A5 | 0 | -0.258353467 | 0.057 | 0.114 | 1.63E-22 | 5.99E-18 |
| PLXNA2 | 0 | -0.309243218 | 0.306 | 0.415 | 1.96E-22 | 7.17E-18 |
| NOL4 | 0 | -0.449296947 | 0.283 | 0.345 | 2.66E-22 | 9.74E-18 |
| STAC | 0 | -0.354465806 | 0.147 | 0.235 | 2.94E-22 | 1.08E-17 |
| CHST11 | 0 | -0.329045489 | 0.553 | 0.634 | 2.97E-22 | 1.09E-17 |
| ATP6V0A1 | 0 | -0.255604013 | 0.258 | 0.361 | 3.19E-22 | 1.17E-17 |
| FRAS1 | 0 | -0.48392714 | 0.339 | 0.392 | 3.25E-22 | 1.19E-17 |
| EPB41L5 | 0 | 0.266617609 | 0.558 | 0.476 | 3.56E-22 | 1.31E-17 |
| DENND4A | 0 | -0.341038308 | 0.532 | 0.608 | 3.6E-22 | 1.32E-17 |
| FMNL2 | 0 | -0.277514477 | 0.861 | 0.88 | 4.7E-22 | 1.72E-17 |
| FHOD3 | 0 | -0.31865969 | 0.24 | 0.334 | 5.48E-22 | 2.01E-17 |
| CCNE2 | 0 | 0.259946979 | 0.219 | 0.138 | 5.58E-22 | 2.04E-17 |
| TBC1D32 | 0 | 0.252221565 | 0.622 | 0.552 | 8.54E-22 | 3.13E-17 |
| SLC35F3 | 0 | -0.263669485 | 0.062 | 0.13 | 9.26E-22 | 3.39E-17 |
| NFIA | 0 | -0.291412835 | 0.902 | 0.89 | 9.35E-22 | 3.42E-17 |
| EPM2A | 0 | 0.287341961 | 0.334 | 0.262 | 1.15E-21 | 4.21E-17 |
| CENPK | 0 | 0.275211284 | 0.554 | 0.469 | 1.5E-21 | 5.51E-17 |
| RFLNA | 0 | 0.253949916 | 0.323 | 0.231 | 1.67E-21 | 6.12E-17 |
| GRIN3A | 0 | -0.484965263 | 0.2 | 0.28 | 1.69E-21 | 6.19E-17 |
| PRKACB | 0 | 0.251818614 | 0.658 | 0.57 | 1.71E-21 | 6.28E-17 |
| HOMER1 | 0 | 0.270096884 | 0.561 | 0.467 | 2.49E-21 | 9.13E-17 |
| SLC16A10 | 0 | -0.482126437 | 0.176 | 0.248 | 2.68E-21 | 9.82E-17 |
| CROCC | 0 | -0.321211672 | 0.385 | 0.468 | 2.82E-21 | 1.03E-16 |
| ITSN1 | 0 | 0.255647681 | 0.661 | 0.581 | 3.35E-21 | 1.23E-16 |
| EPHB1 | 0 | -0.312648987 | 0.111 | 0.182 | 3.58E-21 | 1.31E-16 |
| PLCB4 | 0 | -0.338511558 | 0.624 | 0.691 | 3.79E-21 | 1.39E-16 |
| ERICH1 | 0 | -0.25676406 | 0.202 | 0.29 | 4.23E-21 | 1.55E-16 |
| ACAP2 | 0 | -0.316010468 | 0.618 | 0.683 | 4.24E-21 | 1.55E-16 |
| POLQ | 0 | -0.295436455 | 0.247 | 0.349 | 4.59E-21 | 1.68E-16 |
| ZFPM2 | 0 | 0.304822793 | 0.381 | 0.277 | 6.04E-21 | 2.21E-16 |
| LGR4 | 0 | -0.317531824 | 0.371 | 0.475 | 7.03E-21 | 2.58E-16 |
| ZFAND5 | 0 | 0.285538511 | 0.496 | 0.429 | 8.55E-21 | 3.13E-16 |
| SIK3 | 0 | -0.314934246 | 0.614 | 0.692 | 8.75E-21 | 3.2E-16 |
| BORCS5 | 0 | 0.252982904 | 0.414 | 0.332 | 1.05E-20 | 3.83E-16 |
| MAP1B | 0 | -0.304879645 | 0.681 | 0.74 | 1.06E-20 | 3.89E-16 |
| MYLIP | 0 | -0.259016216 | 0.121 | 0.199 | 1.06E-20 | 3.89E-16 |
| STK17B | 0 | -0.277830972 | 0.098 | 0.167 | 1.09E-20 | 4E-16 |
| MDGA2 | 0 | -0.52079963 | 0.048 | 0.106 | 1.32E-20 | 4.82E-16 |
| ACVR2A | 0 | -0.377470185 | 0.394 | 0.482 | 1.43E-20 | 5.22E-16 |
| TMSB4X | 0 | -0.251074537 | 0.185 | 0.278 | 1.57E-20 | 5.76E-16 |
| MEST | 0 | -0.30300099 | 0.09 | 0.154 | 1.81E-20 | 6.63E-16 |
| MAP3K4 | 0 | -0.302985557 | 0.447 | 0.521 | 1.87E-20 | 6.83E-16 |

| | | | | | | |
|---|---|---|---|---|---|---|
| TRPC6 | 0 | -0.464637403 | 0.055 | 0.107 | 1.91E-20 | 6.98E-16 |
| UBE2D1 | 0 | 0.251386811 | 0.325 | 0.24 | 1.96E-20 | 7.16E-16 |
| TUNAR | 0 | -0.291189816 | 0.052 | 0.108 | 2.27E-20 | 8.31E-16 |
| ZFP64 | 0 | -0.252852975 | 0.219 | 0.313 | 2.93E-20 | 1.07E-15 |
| NR2F2-AS1 | 0 | -0.386638622 | 0.059 | 0.103 | 3.31E-20 | 1.21E-15 |
| LINC02307 | 0 | -0.254037148 | 0.083 | 0.155 | 3.55E-20 | 1.3E-15 |
| SESN3 | 0 | 0.286493275 | 0.477 | 0.379 | 3.8E-20 | 1.39E-15 |
| LIPE-AS1 | 0 | -0.291601905 | 0.257 | 0.352 | 3.91E-20 | 1.43E-15 |
| AL356108.1 | 0 | -0.592460808 | 0.073 | 0.127 | 4.07E-20 | 1.49E-15 |
| TARS | 0 | -0.268152122 | 0.262 | 0.348 | 4.58E-20 | 1.68E-15 |
| JAZF1 | 0 | -0.363970191 | 0.71 | 0.741 | 5.33E-20 | 1.95E-15 |
| GALNT18 | 0 | -0.344964735 | 0.334 | 0.424 | 5.61E-20 | 2.05E-15 |
| ACACA | 0 | -0.266437555 | 0.705 | 0.748 | 5.72E-20 | 2.09E-15 |
| PDE1A | 0 | 0.518239394 | 0.264 | 0.177 | 6.92E-20 | 2.53E-15 |
| NEK7 | 0 | 0.263869316 | 0.678 | 0.595 | 7.86E-20 | 2.88E-15 |
| KIAA1211 | 0 | -0.270273667 | 0.646 | 0.716 | 8.83E-20 | 3.23E-15 |
| ZNF608 | 0 | -0.324609482 | 0.598 | 0.652 | 9.12E-20 | 3.34E-15 |
| CDON | 0 | -0.321048339 | 0.265 | 0.36 | 1.12E-19 | 4.11E-15 |
| MYBL1 | 0 | -0.284693031 | 0.182 | 0.27 | 1.23E-19 | 4.51E-15 |
| FAT4 | 0 | -0.387802656 | 0.241 | 0.32 | 1.36E-19 | 4.99E-15 |
| ARHGAP26 | 0 | -0.267933202 | 0.123 | 0.199 | 1.4E-19 | 5.12E-15 |
| LEF1 | 0 | -0.261081956 | 0.071 | 0.137 | 1.41E-19 | 5.15E-15 |
| CMIP | 0 | -0.270433481 | 0.522 | 0.612 | 1.63E-19 | 5.95E-15 |
| CEP135 | 0 | -0.272218988 | 0.294 | 0.382 | 1.73E-19 | 6.34E-15 |
| CDHR3 | 0 | -0.262124867 | 0.322 | 0.424 | 1.97E-19 | 7.22E-15 |
| SPOPL | 0 | 0.275222508 | 0.432 | 0.355 | 3.22E-19 | 1.18E-14 |
| COL18A1 | 0 | 0.287822466 | 0.185 | 0.111 | 3.65E-19 | 1.34E-14 |
| UBE3D | 0 | 0.264964163 | 0.529 | 0.454 | 4.14E-19 | 1.51E-14 |
| SLC24A2 | 0 | 0.340457647 | 0.463 | 0.419 | 4.21E-19 | 1.54E-14 |
| PPM1L | 0 | 0.259880626 | 0.591 | 0.492 | 4.33E-19 | 1.59E-14 |
| TBC1D19 | 0 | -0.254453219 | 0.267 | 0.361 | 4.51E-19 | 1.65E-14 |
| LINC01473 | 0 | 0.253134432 | 0.633 | 0.544 | 4.72E-19 | 1.73E-14 |
| MAGI3 | 0 | -0.317731636 | 0.4 | 0.471 | 5.14E-19 | 1.88E-14 |
| CTIF | 0 | -0.267908089 | 0.358 | 0.447 | 5.2E-19 | 1.9E-14 |
| MICAL3 | 0 | -0.287594108 | 0.407 | 0.487 | 6E-19 | 2.2E-14 |
| HFM1 | 0 | 0.254346856 | 0.41 | 0.337 | 6.42E-19 | 2.35E-14 |
| SPAG17 | 0 | 0.414040049 | 0.345 | 0.269 | 6.71E-19 | 2.46E-14 |
| ADCY9 | 0 | -0.276962196 | 0.307 | 0.405 | 7.62E-19 | 2.79E-14 |
| POLR3G | 0 | -0.273855401 | 0.219 | 0.283 | 8.73E-19 | 3.2E-14 |
| PHLDB2 | 0 | -0.344626553 | 0.177 | 0.253 | 1.21E-18 | 4.42E-14 |
| AHR | 0 | -0.374669309 | 0.124 | 0.193 | 1.22E-18 | 4.48E-14 |
| YWHAE | 0 | -0.28140145 | 0.545 | 0.626 | 1.33E-18 | 4.88E-14 |
| LAMA3 | 0 | -0.337046284 | 0.155 | 0.233 | 1.47E-18 | 5.37E-14 |
| SPATA5 | 0 | 0.25293287 | 0.671 | 0.605 | 2.47E-18 | 9.04E-14 |
| GMDS | 0 | -0.324789812 | 0.459 | 0.435 | 2.51E-18 | 9.19E-14 |
| NDFIP1 | 0 | -0.266243869 | 0.347 | 0.431 | 2.75E-18 | 1.01E-13 |
| ANO2 | 0 | -0.383741192 | 0.155 | 0.208 | 2.79E-18 | 1.02E-13 |
| WDR45B | 0 | -0.265614576 | 0.243 | 0.33 | 4.01E-18 | 1.47E-13 |
| SLC30A8 | 0 | 0.251566219 | 0.102 | 0.05 | 4.06E-18 | 1.49E-13 |
| CACNA1C | 0 | -0.54479961 | 0.434 | 0.499 | 4.19E-18 | 1.53E-13 |
| PTPRO | 0 | -0.44373808 | 0.431 | 0.446 | 4.21E-18 | 1.54E-13 |
| GAS2 | 0 | -0.252495232 | 0.061 | 0.111 | 5.09E-18 | 1.86E-13 |
| AC009315.1 | 0 | -0.324478556 | 0.347 | 0.435 | 6.85E-18 | 2.51E-13 |
| APBB2 | 0 | -0.306744095 | 0.406 | 0.484 | 6.99E-18 | 2.56E-13 |
| ENO1 | 0 | -0.355644014 | 0.413 | 0.493 | 8.26E-18 | 3.02E-13 |
| NEK11 | 0 | -0.26898378 | 0.093 | 0.149 | 8.41E-18 | 3.08E-13 |
| ENAH | 0 | -0.294177279 | 0.686 | 0.732 | 9.64E-18 | 3.53E-13 |
| NEUROD1 | 0 | -0.288008611 | 0.173 | 0.184 | 1.08E-17 | 3.95E-13 |

| | | | | | | |
|---|---|---|---|---|---|---|
| MVB12B | 0 | -0.280002398 | 0.394 | 0.431 | 1.2E-17 | 4.39E-13 |
| TSPAN7 | 0 | 0.253443059 | 0.61 | 0.512 | 1.22E-17 | 4.48E-13 |
| INSR | 0 | -0.363290927 | 0.228 | 0.267 | 1.24E-17 | 4.53E-13 |
| GALNT11 | 0 | -0.256774539 | 0.363 | 0.448 | 1.35E-17 | 4.95E-13 |
| PLS3 | 0 | -0.355389282 | 0.35 | 0.425 | 1.45E-17 | 5.3E-13 |
| RAPGEF2 | 0 | -0.296958557 | 0.642 | 0.66 | 1.67E-17 | 6.11E-13 |
| PCAT1 | 0 | -0.313939365 | 0.105 | 0.098 | 1.68E-17 | 6.16E-13 |
| DNAH7 | 0 | -0.499661598 | 0.177 | 0.178 | 1.7E-17 | 6.23E-13 |
| CADM1 | 0 | -0.288759459 | 0.369 | 0.469 | 2.5E-17 | 9.16E-13 |
| PCBP3 | 0 | -0.322555252 | 0.201 | 0.268 | 4.83E-17 | 1.77E-12 |
| GLRA2 | 0 | -0.315502175 | 0.052 | 0.104 | 5.04E-17 | 1.84E-12 |
| CHST9 | 0 | -0.436815873 | 0.13 | 0.159 | 5.42E-17 | 1.98E-12 |
| PLCL2 | 0 | -0.299686619 | 0.542 | 0.609 | 5.81E-17 | 2.13E-12 |
| PDE11A | 0 | -0.288439658 | 0.155 | 0.202 | 5.91E-17 | 2.16E-12 |
| LINC00907 | 0 | 0.552685878 | 0.309 | 0.231 | 8.72E-17 | 3.19E-12 |
| ANKRD50 | 0 | -0.286349463 | 0.255 | 0.338 | 1.13E-16 | 4.15E-12 |
| RUNX2 | 0 | -0.286887953 | 0.091 | 0.156 | 1.28E-16 | 4.68E-12 |
| SAT1 | 0 | 0.575351937 | 0.169 | 0.139 | 1.53E-16 | 5.62E-12 |
| EDIL3 | 0 | -0.449020845 | 0.604 | 0.626 | 1.61E-16 | 5.9E-12 |
| ACTG1 | 0 | -0.273623002 | 0.377 | 0.465 | 2.69E-16 | 9.84E-12 |
| TBC1D4 | 0 | -0.301510167 | 0.348 | 0.42 | 3.2E-16 | 1.17E-11 |
| MALAT1 | 0 | -0.45851453 | 1 | 0.998 | 4.35E-16 | 1.59E-11 |
| CREB5 | 0 | -0.315180756 | 0.152 | 0.225 | 4.58E-16 | 1.68E-11 |
| LINC01376 | 0 | -0.269248363 | 0.109 | 0.168 | 4.93E-16 | 1.81E-11 |
| AKAP12 | 0 | -0.339056323 | 0.154 | 0.218 | 4.94E-16 | 1.81E-11 |
| GRIA3 | 0 | 0.275202639 | 0.236 | 0.164 | 5.39E-16 | 1.97E-11 |
| PEX5L | 0 | -0.273921227 | 0.766 | 0.763 | 5.68E-16 | 2.08E-11 |
| MALRD1 | 0 | -0.483204957 | 0.229 | 0.27 | 9.53E-16 | 3.49E-11 |
| SLC7A6 | 0 | -0.313175081 | 0.2 | 0.263 | 1.49E-15 | 5.46E-11 |
| TSPAN18 | 0 | -0.275959597 | 0.219 | 0.3 | 1.84E-15 | 6.75E-11 |
| ZFHX4 | 0 | -0.353167818 | 0.37 | 0.454 | 2.09E-15 | 7.65E-11 |
| NFAT5 | 0 | -0.285489138 | 0.786 | 0.8 | 2.15E-15 | 7.88E-11 |
| RPS6KA2 | 0 | -0.302994932 | 0.269 | 0.348 | 2.21E-15 | 8.11E-11 |
| FCHSD2 | 0 | -0.251106056 | 0.716 | 0.756 | 2.25E-15 | 8.25E-11 |
| SUGCT | 0 | -0.288397323 | 0.342 | 0.424 | 2.33E-15 | 8.53E-11 |
| BMERB1 | 0 | -0.292085868 | 0.436 | 0.486 | 2.35E-15 | 8.61E-11 |
| PLEKHG1 | 0 | -0.278333492 | 0.096 | 0.149 | 2.58E-15 | 9.44E-11 |
| CCDC102B | 0 | 0.637943014 | 0.179 | 0.123 | 3.02E-15 | 1.11E-10 |
| SOX4 | 0 | -0.284604532 | 0.218 | 0.296 | 3.19E-15 | 1.17E-10 |
| MAN1A1 | 0 | -0.400362207 | 0.113 | 0.141 | 3.25E-15 | 1.19E-10 |
| SMARCA2 | 0 | 0.2548395 | 0.491 | 0.405 | 4.32E-15 | 1.58E-10 |
| COLEC12 | 0 | -0.336949288 | 0.102 | 0.146 | 6.12E-15 | 2.24E-10 |
| ARRDC3-AS1 | 0 | -0.626666201 | 0.26 | 0.293 | 7.82E-15 | 2.86E-10 |
| TMEM232 | 0 | -0.338463307 | 0.104 | 0.147 | 9.17E-15 | 3.36E-10 |
| AGAP1 | 0 | -0.254193561 | 0.851 | 0.841 | 9.49E-15 | 3.47E-10 |
| FMN1 | 0 | -0.334223657 | 0.071 | 0.123 | 1.44E-14 | 5.27E-10 |
| SHC3 | 0 | -0.310905072 | 0.308 | 0.375 | 1.68E-14 | 6.16E-10 |
| HSP90AA1 | 0 | -0.658733974 | 0.405 | 0.476 | 2.42E-14 | 8.86E-10 |
| CSGALNACT1 | 0 | -0.266278221 | 0.083 | 0.14 | 2.77E-14 | 1.01E-09 |
| THSD7A | 0 | -0.321477763 | 0.373 | 0.394 | 3.28E-14 | 1.2E-09 |
| GPR158 | 0 | 0.342920287 | 0.178 | 0.126 | 6.33E-14 | 2.32E-09 |
| COL1A2 | 0 | -0.313891131 | 0.134 | 0.123 | 6.33E-14 | 2.32E-09 |
| SLC1A3 | 0 | -0.259320321 | 0.084 | 0.117 | 8.84E-14 | 3.24E-09 |
| HSPH1 | 0 | -0.506654158 | 0.219 | 0.288 | 1.04E-13 | 3.79E-09 |
| BMPER | 0 | -0.352439372 | 0.161 | 0.232 | 1.23E-13 | 4.52E-09 |
| RPL13A | 0 | -0.266878898 | 0.502 | 0.574 | 1.75E-13 | 6.39E-09 |
| PTCH1 | 0 | -0.265697952 | 0.142 | 0.199 | 1.91E-13 | 7.01E-09 |
| ARHGAP28 | 0 | -0.341609832 | 0.208 | 0.274 | 2.18E-13 | 0.000000008 |

| | | | | | | |
|---|---|---|---|---|---|---|
| PTPRZ1 | 0 | -0.362833078 | 0.384 | 0.417 | 2.5E-13 | 9.14E-09 |
| IVNS1ABP | 0 | -0.290589752 | 0.478 | 0.518 | 2.5E-13 | 9.14E-09 |
| AL589693.1 | 0 | -0.433455992 | 0.063 | 0.108 | 3.07E-13 | 1.12E-08 |
| GALNT13 | 0 | 0.309931032 | 0.741 | 0.687 | 4.25E-13 | 1.56E-08 |
| PHF21B | 0 | -0.252186803 | 0.331 | 0.393 | 1.16E-12 | 4.25E-08 |
| AC109466.1 | 0 | -0.595808338 | 0.078 | 0.126 | 1.39E-12 | 5.09E-08 |
| ERICH3 | 0 | -0.253986415 | 0.112 | 0.151 | 1.94E-12 | 7.09E-08 |
| MIR4435-2HG | 0 | 0.254874954 | 0.272 | 0.22 | 2.46E-12 | 8.99E-08 |
| SEZ6L | 0 | -0.313412578 | 0.066 | 0.107 | 2.54E-12 | 9.29E-08 |
| PDK3 | 0 | -0.25064001 | 0.377 | 0.445 | 2.6E-12 | 9.53E-08 |
| PFKFB4 | 0 | -0.255796822 | 0.11 | 0.158 | 2.63E-12 | 9.62E-08 |
| UBE2D2 | 0 | -0.250928455 | 0.548 | 0.608 | 3.45E-12 | 0.000000126 |
| SPECC1 | 0 | -0.260845481 | 0.497 | 0.563 | 3.74E-12 | 0.000000137 |
| STOX2 | 0 | -0.262245705 | 0.27 | 0.32 | 6.41E-12 | 0.000000235 |
| KCNJ3 | 0 | -0.374904832 | 0.111 | 0.154 | 8.85E-12 | 0.000000324 |
| SLC39A11 | 0 | -0.279219621 | 0.476 | 0.523 | 1.08E-11 | 0.000000395 |
| DLG2 | 0 | -0.614823825 | 0.61 | 0.645 | 1.55E-11 | 0.000000567 |
| PGK1 | 0 | -0.396670956 | 0.3 | 0.355 | 1.58E-11 | 0.000000577 |
| ITGA1 | 0 | -0.301561898 | 0.116 | 0.131 | 2.57E-11 | 0.000000942 |
| AC093298.2 | 0 | -0.597941081 | 0.17 | 0.196 | 3.17E-11 | 0.00000116 |
| AC068286.2 | 0 | -0.758690142 | 0.124 | 0.141 | 3.96E-11 | 0.00000145 |
| SSBP3 | 0 | -0.252659138 | 0.523 | 0.567 | 5.8E-11 | 0.00000212 |
| AL138720.1 | 0 | -0.279522969 | 0.131 | 0.174 | 1.22E-10 | 0.00000445 |
| PLOD2 | 0 | -0.375540526 | 0.245 | 0.302 | 1.97E-10 | 0.00000721 |
| CDK6 | 0 | -0.270150687 | 0.11 | 0.156 | 2.93E-10 | 0.0000107 |
| SLC4A4 | 0 | 0.296301562 | 0.206 | 0.16 | 3.4E-10 | 0.0000124 |
| HDAC9 | 0 | -0.354649009 | 0.555 | 0.583 | 1.39E-09 | 0.0000509 |
| CNTNAP5 | 0 | -0.382408554 | 0.242 | 0.289 | 1.79E-09 | 0.0000656 |
| LAMB1 | 0 | 0.294532173 | 0.429 | 0.371 | 3.02E-09 | 0.000111 |
| HIF1A-AS3 | 0 | -0.281782943 | 0.111 | 0.137 | 8.74E-09 | 0.00032 |
| KCNMA1 | 0 | 0.307841074 | 0.308 | 0.28 | 1.02E-08 | 0.000373 |
| SGO1-AS1 | 0 | -0.343167391 | 0.148 | 0.2 | 2.25E-08 | 0.000822 |
| PDK1 | 0 | -0.274370417 | 0.349 | 0.393 | 2.29E-08 | 0.000839 |
| SLC2A3 | 0 | -0.284676881 | 0.112 | 0.148 | 6.22E-08 | 0.002278822 |
| IQCJ-SCHIP1 | 0 | -0.555828771 | 0.158 | 0.179 | 0.000000116 | 0.004234323 |
| PRICKLE1 | 0 | -0.342010187 | 0.378 | 0.424 | 0.000000642 | 0.023517216 |
| BNC2 | 0 | -0.33161611 | 0.176 | 0.207 | 0.00000261 | 0.095559971 |
| PBX3 | 0 | -0.294864133 | 0.49 | 0.512 | 0.000104 | 1 |
| TTN | 0 | -0.271891324 | 0.15 | 0.152 | 0.002579397 | 1 |
| XACT | 1 | 2.716671433 | 0.666 | 0.113 | 0 | 0 |
| ADAMTS12 | 1 | 2.571841836 | 0.778 | 0.275 | 0 | 0 |
| SYCP2 | 1 | 2.279503103 | 0.645 | 0.09 | 0 | 0 |
| CSMD1 | 1 | 2.24259324 | 0.782 | 0.335 | 0 | 0 |
| IL12RB2 | 1 | 2.165284395 | 0.657 | 0.164 | 0 | 0 |
| MCTP2 | 1 | 2.087869701 | 0.63 | 0.145 | 0 | 0 |
| AL691420.1 | 1 | 2.016910212 | 0.457 | 0.029 | 0 | 0 |
| THRB | 1 | 2.016364127 | 0.665 | 0.195 | 0 | 0 |
| MKX | 1 | 1.916325725 | 0.719 | 0.249 | 0 | 0 |
| LDB2 | 1 | 1.892896272 | 0.658 | 0.232 | 0 | 0 |
| HS3ST5 | 1 | 1.817905948 | 0.867 | 0.537 | 0 | 0 |
| COL25A1 | 1 | 1.812429923 | 0.708 | 0.288 | 0 | 0 |
| SLC17A8 | 1 | 1.794862409 | 0.617 | 0.181 | 0 | 0 |
| NPSR1 | 1 | 1.786201192 | 0.461 | 0.08 | 0 | 0 |
| AP002954.2 | 1 | 1.602016058 | 0.363 | 0.011 | 0 | 0 |
| AHCYL2 | 1 | 1.567355681 | 0.839 | 0.61 | 0 | 0 |
| CACNB4 | 1 | 1.512958451 | 0.833 | 0.454 | 0 | 0 |
| RMST | 1 | 1.511300441 | 0.831 | 0.378 | 0 | 0 |
| AL162718.1 | 1 | 1.49388913 | 0.507 | 0.016 | 0 | 0 |

| | | | | | | |
|---|---|---|---|---|---|---|
| DAAM2 | 1 | 1.45709453 | 0.529 | 0.114 | 0 | 0 |
| ADGRG2 | 1 | 1.42608322 | 0.489 | 0.08 | 0 | 0 |
| SLC22A15 | 1 | 1.390376603 | 0.621 | 0.209 | 0 | 0 |
| KRAS | 1 | 1.383264792 | 0.65 | 0.281 | 0 | 0 |
| ADAM23 | 1 | 1.362497375 | 0.648 | 0.265 | 0 | 0 |
| LINC01768 | 1 | 1.356151713 | 0.504 | 0.074 | 0 | 0 |
| UST | 1 | 1.323647266 | 0.6 | 0.172 | 0 | 0 |
| EEPD1 | 1 | 1.303838742 | 0.616 | 0.202 | 0 | 0 |
| SEMA3E | 1 | 1.28811179 | 0.636 | 0.191 | 0 | 0 |
| C13orf42 | 1 | 1.261933838 | 0.45 | 0.027 | 0 | 0 |
| AC009803.1 | 1 | 1.261910211 | 0.447 | 0.025 | 0 | 0 |
| AC063938.1 | 1 | 1.252385896 | 0.368 | 0.022 | 0 | 0 |
| CHPT1 | 1 | 1.207123014 | 0.783 | 0.495 | 0 | 0 |
| HTR7 | 1 | 1.138951476 | 0.394 | 0.037 | 0 | 0 |
| IDI2-AS1 | 1 | 1.132830141 | 0.41 | 0.047 | 0 | 0 |
| PDE4D | 1 | -1.357348573 | 0.647 | 0.916 | 0 | 0 |
| MAP2 | 1 | -1.742427727 | 0.197 | 0.659 | 0 | 0 |
| MIR99AHG | 1 | -1.928754302 | 0.395 | 0.781 | 0 | 0 |
| GPC6 | 1 | -2.204340298 | 0.484 | 0.925 | 0 | 0 |
| KCNB2 | 1 | -2.536258081 | 0.143 | 0.685 | 0 | 0 |
| HCN1 | 1 | 1.326109128 | 0.429 | 0.067 | 5.395e-321 | 1.9752355e-316 |
| GABRB2 | 1 | 1.467236145 | 0.619 | 0.204 | 1.15788e-318 | 4.239123131e-314 |
| UBE2E2 | 1 | 1.230794398 | 0.829 | 0.526 | 4.6212237e-317 | 1.691876227045e-312 |
| APBA1 | 1 | 1.454948568 | 0.697 | 0.356 | 8.2863004e-316 | 3.0336974348473e-311 |
| ARHGAP24 | 1 | -1.988953385 | 0.106 | 0.51 | 4.398171549723e-312 | 1.61E-307 |
| GLT1D1 | 1 | 0.92706276 | 0.431 | 0.076 | 6.96E-307 | 2.55E-302 |
| FGF14 | 1 | -1.80983639 | 0.399 | 0.74 | 1.37E-302 | 5.02E-298 |
| NRXN3 | 1 | 2.256964396 | 0.595 | 0.275 | 3.23E-302 | 1.18E-297 |
| TSHZ2 | 1 | 1.257401793 | 0.906 | 0.761 | 1.08E-301 | 3.96E-297 |
| FGD6 | 1 | 1.168140252 | 0.618 | 0.235 | 2.75E-299 | 1.01E-294 |
| SFMBT2 | 1 | 1.279209284 | 0.739 | 0.412 | 8.26E-299 | 3.03E-294 |
| NCALD | 1 | -1.48859236 | 0.257 | 0.668 | 3.63E-297 | 1.33E-292 |
| KANK1 | 1 | -1.342635297 | 0.193 | 0.612 | 2.25E-292 | 8.25E-288 |
| ADAMTSL1 | 1 | 1.307295763 | 0.828 | 0.423 | 3.1E-287 | 1.13E-282 |
| LUZP2 | 1 | -2.186251518 | 0.282 | 0.646 | 1.84E-286 | 6.74E-282 |
| L1TD1 | 1 | 1.061940813 | 0.508 | 0.132 | 3.73E-283 | 1.37E-278 |
| AC006972.1 | 1 | 0.914850614 | 0.318 | 0.031 | 1.9E-281 | 6.96E-277 |
| IQGAP2 | 1 | 1.224688604 | 0.669 | 0.313 | 5.74E-281 | 2.1E-276 |
| FAM13A | 1 | -1.006738496 | 0.238 | 0.664 | 1.13E-277 | 4.12E-273 |
| LINC01102 | 1 | 0.825891837 | 0.298 | 0.026 | 2.46E-276 | 9E-272 |
| NXN | 1 | -1.147288355 | 0.108 | 0.507 | 3.89E-275 | 1.42E-270 |
| UNC5D | 1 | -1.936944677 | 0.316 | 0.663 | 2.58E-274 | 9.43E-270 |
| MIR100HG | 1 | 1.199607559 | 0.884 | 0.698 | 4.96E-273 | 1.82E-268 |
| DPPA4 | 1 | 0.993694344 | 0.695 | 0.35 | 5.33E-273 | 1.95E-268 |
| RYR2 | 1 | 1.507298395 | 0.484 | 0.118 | 2.06E-270 | 7.55E-266 |
| GAB1 | 1 | 1.044639461 | 0.826 | 0.616 | 2.23E-270 | 8.15E-266 |
| CDH12 | 1 | -2.046326684 | 0.118 | 0.491 | 4.61E-270 | 1.69E-265 |
| TMEM108 | 1 | 1.288900494 | 0.814 | 0.603 | 6.31E-268 | 2.31E-263 |
| AFF3 | 1 | -1.485290185 | 0.332 | 0.696 | 8E-268 | 2.93E-263 |
| ZSWIM5 | 1 | 1.111081297 | 0.665 | 0.292 | 4.3E-267 | 1.57E-262 |
| RALYL | 1 | -2.3853993 | 0.215 | 0.559 | 1.18E-266 | 4.31E-262 |
| PLCL1 | 1 | -1.435067743 | 0.126 | 0.52 | 1.2E-264 | 4.4E-260 |
| AL355499.1 | 1 | 1.828319889 | 0.303 | 0.036 | 4.75E-264 | 1.74E-259 |
| DPYSL3 | 1 | 1.076423235 | 0.65 | 0.284 | 1.62E-259 | 5.95E-255 |
| F3 | 1 | 1.330134427 | 0.294 | 0.037 | 2.06E-257 | 7.55E-253 |
| MGAT4C | 1 | 1.228323898 | 0.803 | 0.445 | 2.63E-257 | 9.64E-253 |
| NFAT5 | 1 | 0.952041943 | 0.89 | 0.765 | 1E-256 | 3.66E-252 |
| PCCA | 1 | -0.976126498 | 0.349 | 0.73 | 2.05E-255 | 7.49E-251 |

| | | | | | | |
|---|---|---|---|---|---|---|
| MTMR7 | 1 | -1.084100075 | 0.077 | 0.437 | 3.47E-254 | 1.27E-249 |
| DNM3OS | 1 | -1.166431033 | 0.07 | 0.428 | 3.75E-253 | 1.37E-248 |
| C9orf135 | 1 | 0.813254488 | 0.284 | 0.028 | 9.89E-253 | 3.62E-248 |
| DNAH11 | 1 | 1.57553361 | 0.516 | 0.173 | 5.64E-248 | 2.07E-243 |
| AL390800.1 | 1 | 1.261655681 | 0.758 | 0.441 | 1.32E-247 | 4.84E-243 |
| ROBO1 | 1 | -1.371913019 | 0.816 | 0.929 | 3.78E-247 | 1.38E-242 |
| LDLRAD4 | 1 | -1.556563295 | 0.207 | 0.588 | 2.9E-244 | 1.06E-239 |
| ZNF385B | 1 | 1.295624792 | 0.608 | 0.241 | 9.88E-244 | 3.62E-239 |
| NR2F1 | 1 | 1.092342014 | 0.455 | 0.137 | 7.13E-243 | 2.61E-238 |
| AL136418.1 | 1 | 0.951108735 | 0.285 | 0.03 | 1.71E-241 | 6.27E-237 |
| NEAT1 | 1 | 1.442291716 | 0.919 | 0.848 | 1.57E-240 | 5.74E-236 |
| EPHA7 | 1 | 1.200684489 | 0.599 | 0.222 | 3.48E-240 | 1.27E-235 |
| LRRC8B | 1 | 1.040266233 | 0.67 | 0.347 | 5.67E-239 | 2.07E-234 |
| DNM3 | 1 | -1.182130014 | 0.182 | 0.562 | 8.84E-238 | 3.24E-233 |
| FAM189A1 | 1 | 1.013484363 | 0.879 | 0.633 | 6.31E-235 | 2.31E-230 |
| IL1RAPL2 | 1 | 1.980261215 | 0.43 | 0.107 | 4.48E-233 | 1.64E-228 |
| PCDH7 | 1 | 1.24397824 | 0.689 | 0.342 | 4.4E-230 | 1.61E-225 |
| RAPGEF4 | 1 | 1.124408584 | 0.669 | 0.315 | 1.49E-229 | 5.44E-225 |
| IDO1 | 1 | 0.721566068 | 0.209 | 0.009 | 6.85E-228 | 2.51E-223 |
| ANO2 | 1 | 1.162517145 | 0.427 | 0.116 | 7.64E-228 | 2.8E-223 |
| DGKB | 1 | -1.879210388 | 0.052 | 0.361 | 4.19E-226 | 1.53E-221 |
| HECW2 | 1 | -1.175044933 | 0.112 | 0.452 | 5.87E-226 | 2.15E-221 |
| EXOC6B | 1 | -0.94529206 | 0.457 | 0.769 | 9.93E-224 | 3.64E-219 |
| AC018731.1 | 1 | 1.401081695 | 0.783 | 0.529 | 1.61E-217 | 5.9E-213 |
| MYBPC1 | 1 | 0.633602812 | 0.213 | 0.012 | 1.02E-214 | 3.74E-210 |
| ST6GALNAC3 | 1 | -1.319288129 | 0.043 | 0.339 | 1.63E-212 | 5.96E-208 |
| LSAMP | 1 | 1.177033251 | 0.885 | 0.689 | 1.78E-210 | 6.51E-206 |
| GLI3 | 1 | -1.092152692 | 0.099 | 0.424 | 1.99E-210 | 7.29E-206 |
| ANO4 | 1 | 1.129539631 | 0.835 | 0.501 | 3.37E-210 | 1.24E-205 |
| SLIT2 | 1 | -1.716162122 | 0.548 | 0.777 | 4.02E-209 | 1.47E-204 |
| STK17B | 1 | 0.854663172 | 0.355 | 0.08 | 1.49E-207 | 5.46E-203 |
| MAML3 | 1 | -1.473118884 | 0.332 | 0.65 | 3.26E-207 | 1.19E-202 |
| TIAM2 | 1 | 0.918202073 | 0.864 | 0.754 | 6.49E-206 | 2.38E-201 |
| AC026704.1 | 1 | 0.815094068 | 0.305 | 0.058 | 9.26E-206 | 3.39E-201 |
| ZEB2 | 1 | -1.372547962 | 0.335 | 0.66 | 1.12E-202 | 4.1E-198 |
| CD70 | 1 | 0.465127166 | 0.205 | 0.012 | 1.35E-202 | 4.94E-198 |
| PRAME | 1 | 0.619750412 | 0.333 | 0.064 | 6.18E-202 | 2.26E-197 |
| ZNF385D | 1 | -1.336060967 | 0.32 | 0.655 | 9.03E-202 | 3.31E-197 |
| SPINK5 | 1 | 0.729643032 | 0.368 | 0.083 | 2.38E-201 | 8.7E-197 |
| JMJD1C | 1 | -0.656257241 | 0.886 | 0.965 | 4.34E-201 | 1.59E-196 |
| LAMA2 | 1 | -1.561461195 | 0.153 | 0.469 | 9.23E-201 | 3.38E-196 |
| SIPA1L1 | 1 | -0.990172592 | 0.596 | 0.828 | 2.76E-200 | 1.01E-195 |
| SYN3 | 1 | 0.85817457 | 0.697 | 0.334 | 3.48E-199 | 1.27E-194 |
| KCNIP4 | 1 | -2.039832452 | 0.279 | 0.571 | 1.35E-198 | 4.95E-194 |
| GLIS3 | 1 | -1.584978957 | 0.042 | 0.313 | 1.64E-198 | 6E-194 |
| DLGAP1 | 1 | -1.123606209 | 0.677 | 0.886 | 4.41E-198 | 1.62E-193 |
| ZYG11A | 1 | 0.809608458 | 0.555 | 0.22 | 5.14E-198 | 1.88E-193 |
| EGFR | 1 | -1.002827609 | 0.046 | 0.328 | 6.69E-197 | 2.45E-192 |
| BTNL9 | 1 | 0.598519084 | 0.229 | 0.022 | 2.19E-196 | 8E-192 |
| CCNO | 1 | 0.617168547 | 0.216 | 0.017 | 1.48E-195 | 5.41E-191 |
| EFNA5 | 1 | 0.951481208 | 0.84 | 0.606 | 2.46E-193 | 8.99E-189 |
| EGLN3 | 1 | 0.890266899 | 0.477 | 0.173 | 5.02E-193 | 1.84E-188 |
| BCLAF3 | 1 | 0.833178216 | 0.622 | 0.344 | 2.84E-191 | 1.04E-186 |
| SATB2 | 1 | -1.117606626 | 0.224 | 0.536 | 9.5E-191 | 3.48E-186 |
| TFAP2C | 1 | 0.504360153 | 0.228 | 0.023 | 2.14E-190 | 7.84E-186 |
| MCTP1 | 1 | -1.365500398 | 0.077 | 0.367 | 4.43E-190 | 1.62E-185 |
| NCOA1 | 1 | -0.901217185 | 0.475 | 0.758 | 4.93E-190 | 1.8E-185 |
| EXOC2 | 1 | 0.975014151 | 0.813 | 0.626 | 2.44E-189 | 8.92E-185 |

| | | | | | | |
|---|---|---|---|---|---|---|
| RGL1 | 1 | -1.08852023 | 0.548 | 0.784 | 3.75E-189 | 1.37E-184 |
| PIK3AP1 | 1 | 0.750316917 | 0.365 | 0.091 | 1.07E-188 | 3.9E-184 |
| CACNG8 | 1 | 0.529506427 | 0.25 | 0.031 | 2.48E-188 | 9.09E-184 |
| OSBPL3 | 1 | 0.971043419 | 0.665 | 0.39 | 2.06E-187 | 7.56E-183 |
| MAP7 | 1 | 1.011709677 | 0.778 | 0.528 | 2.09E-186 | 7.63E-182 |
| NPSR1-AS1 | 1 | 0.86039931 | 0.392 | 0.104 | 3.19E-186 | 1.17E-181 |
| GK | 1 | 0.866957188 | 0.482 | 0.209 | 1.51E-183 | 5.52E-179 |
| ARHGAP32 | 1 | -0.859334953 | 0.503 | 0.773 | 2.05E-183 | 7.5E-179 |
| NEGR1 | 1 | -2.429490694 | 0.442 | 0.621 | 2.61E-183 | 9.56E-179 |
| SYBU | 1 | 0.977057258 | 0.652 | 0.333 | 9.51E-183 | 3.48E-178 |
| GAP43 | 1 | 0.960063812 | 0.438 | 0.152 | 1.13E-182 | 4.14E-178 |
| FAM160A1 | 1 | 0.855849077 | 0.35 | 0.082 | 3.28E-182 | 1.2E-177 |
| TRMT9B | 1 | -0.915600606 | 0.031 | 0.286 | 8.09E-182 | 2.96E-177 |
| CNTN1 | 1 | -1.202335732 | 0.05 | 0.321 | 2.21E-181 | 8.09E-177 |
| HS6ST3 | 1 | -0.958493308 | 0.701 | 0.441 | 9.97E-181 | 3.65E-176 |
| AL117329.1 | 1 | 1.10277618 | 0.202 | 0.018 | 3.67E-180 | 1.34E-175 |
| AL008633.1 | 1 | -2.198281801 | 0.579 | 0.754 | 6.03E-180 | 2.21E-175 |
| PDE1C | 1 | -1.625220611 | 0.09 | 0.379 | 4.01E-179 | 1.47E-174 |
| SNCA | 1 | 0.951364284 | 0.498 | 0.201 | 6.51E-179 | 2.38E-174 |
| SKAP2 | 1 | 0.953906125 | 0.393 | 0.113 | 6.68E-179 | 2.45E-174 |
| CNTNAP5 | 1 | -1.609053388 | 0.071 | 0.342 | 2.32E-178 | 8.49E-174 |
| NRG3 | 1 | 0.882294181 | 0.725 | 0.381 | 3.33E-178 | 1.22E-173 |
| SPOCK1 | 1 | -1.132826686 | 0.126 | 0.44 | 7.19E-178 | 2.63E-173 |
| B4GALT6 | 1 | 1.004363693 | 0.493 | 0.225 | 2.16E-176 | 7.9E-172 |
| SHC3 | 1 | 0.969033581 | 0.57 | 0.286 | 1.23E-175 | 4.51E-171 |
| TBC1D23 | 1 | 0.820396296 | 0.672 | 0.43 | 2.84E-175 | 1.04E-170 |
| ANOS1 | 1 | -1.396350377 | 0.222 | 0.513 | 1.01E-174 | 3.69E-170 |
| SDC2 | 1 | -0.82557508 | 0.133 | 0.445 | 2.1E-172 | 7.69E-168 |
| AC006058.1 | 1 | 0.597778178 | 0.273 | 0.049 | 2.66E-172 | 9.72E-168 |
| MARCH1 | 1 | 1.128432441 | 0.53 | 0.223 | 2.7E-170 | 9.89E-166 |
| HDAC2-AS2 | 1 | 0.907426992 | 0.714 | 0.44 | 3.82E-170 | 1.4E-165 |
| TMEM178B | 1 | -1.035045601 | 0.262 | 0.582 | 4.33E-170 | 1.59E-165 |
| MTA3 | 1 | 0.755742532 | 0.805 | 0.648 | 6.54E-170 | 2.39E-165 |
| FERMT1 | 1 | 0.845329863 | 0.401 | 0.132 | 6.61E-170 | 2.42E-165 |
| CENPP | 1 | 1.138778453 | 0.794 | 0.658 | 7.28E-170 | 2.67E-165 |
| LANCL3 | 1 | 0.844747299 | 0.44 | 0.157 | 1.41E-169 | 5.16E-165 |
| FLI1 | 1 | 0.553815778 | 0.237 | 0.032 | 1.18E-168 | 4.33E-164 |
| KIRREL3 | 1 | 1.104699475 | 0.381 | 0.114 | 2.87E-168 | 1.05E-163 |
| DANT2 | 1 | 0.795423616 | 0.573 | 0.247 | 4.1E-168 | 1.5E-163 |
| PPP1R14C | 1 | -0.996811766 | 0.206 | 0.526 | 1.23E-166 | 4.5E-162 |
| COBLL1 | 1 | 1.007612392 | 0.611 | 0.307 | 5.18E-166 | 1.9E-161 |
| FAM124A | 1 | 0.769578776 | 0.452 | 0.179 | 1.11E-165 | 4.06E-161 |
| PHACTR1 | 1 | -1.135130903 | 0.108 | 0.392 | 1.38E-165 | 5.04E-161 |
| NAALAD2 | 1 | -0.725699339 | 0.046 | 0.301 | 1.79E-165 | 6.54E-161 |
| ABCC1 | 1 | 0.851763352 | 0.689 | 0.426 | 1.9E-165 | 6.95E-161 |
| ABCD3 | 1 | 0.834816022 | 0.778 | 0.596 | 2E-165 | 7.31E-161 |
| DACH2 | 1 | -2.266502436 | 0.282 | 0.506 | 8.22E-165 | 3.01E-160 |
| AC105460.1 | 1 | 0.476776127 | 0.166 | 0.01 | 1.31E-164 | 4.8E-160 |
| STXBP5L | 1 | -1.313039221 | 0.105 | 0.373 | 1.47E-164 | 5.37E-160 |
| DOCK10 | 1 | -1.185781664 | 0.152 | 0.451 | 9.62E-164 | 3.52E-159 |
| AL390957.1 | 1 | 1.889650661 | 0.297 | 0.085 | 1.13E-163 | 4.14E-159 |
| FRMD5 | 1 | -1.525338269 | 0.204 | 0.474 | 2.63E-163 | 9.62E-159 |
| CTNNA2 | 1 | -1.397803015 | 0.287 | 0.557 | 2.91E-163 | 1.07E-158 |
| BCL11A | 1 | 0.771374139 | 0.502 | 0.2 | 5.48E-163 | 2.01E-158 |
| MEIS2 | 1 | -0.90453259 | 0.292 | 0.62 | 1.85E-162 | 6.77E-158 |
| KIF21A | 1 | -0.915921742 | 0.239 | 0.542 | 4.56E-162 | 1.67E-157 |
| CPQ | 1 | -0.792827928 | 0.082 | 0.355 | 6.44E-162 | 2.36E-157 |
| ITPR3 | 1 | 0.510140447 | 0.278 | 0.054 | 7.4E-162 | 2.71E-157 |

| | | | | | | |
|---|---|---|---|---|---|---|
| GLYATL2 | 1 | -0.881673341 | 0.116 | 0.402 | 1.99E-161 | 7.3E-157 |
| STOX2 | 1 | -0.829151746 | 0.093 | 0.375 | 9.53E-161 | 3.49E-156 |
| FRMD3 | 1 | -0.956493856 | 0.143 | 0.437 | 3.48E-160 | 1.27E-155 |
| CADPS2 | 1 | -1.230758303 | 0.053 | 0.294 | 4.02E-160 | 1.47E-155 |
| IL1RAPL1 | 1 | 0.66839589 | 0.853 | 0.593 | 3.26E-159 | 1.19E-154 |
| TRIM71 | 1 | 0.792328271 | 0.465 | 0.183 | 2E-158 | 7.33E-154 |
| CDK14 | 1 | -0.695168309 | 0.605 | 0.835 | 5.38E-158 | 1.97E-153 |
| ADGRB3 | 1 | 1.031409953 | 0.736 | 0.472 | 1.8E-157 | 6.58E-153 |
| DAPK1 | 1 | 0.860140578 | 0.796 | 0.651 | 4.61E-157 | 1.69E-152 |
| TRPC1 | 1 | 0.849864446 | 0.696 | 0.495 | 1.01E-156 | 3.7E-152 |
| UTRN | 1 | 1.154136586 | 0.753 | 0.579 | 4.8E-156 | 1.76E-151 |
| CXADR | 1 | 0.736203538 | 0.559 | 0.265 | 5.67E-156 | 2.08E-151 |
| STK32B | 1 | 0.8245048 | 0.737 | 0.456 | 8.16E-156 | 2.99E-151 |
| FIRRE | 1 | 0.911452233 | 0.533 | 0.231 | 6.05E-155 | 2.22E-150 |
| MAST4 | 1 | -1.02906582 | 0.155 | 0.431 | 1.19E-154 | 4.36E-150 |
| MIR4713HG | 1 | 0.657642765 | 0.28 | 0.058 | 1.51E-154 | 5.55E-150 |
| KCNMB2 | 1 | -1.750701497 | 0.094 | 0.328 | 2.06E-154 | 7.53E-150 |
| AL138720.1 | 1 | 0.909051094 | 0.347 | 0.101 | 2.19E-154 | 8E-150 |
| NCOA3 | 1 | 0.752290895 | 0.749 | 0.538 | 3.69E-154 | 1.35E-149 |
| CDH11 | 1 | -1.368660951 | 0.108 | 0.38 | 6.85E-154 | 2.51E-149 |
| DCHS2 | 1 | 0.621037814 | 0.261 | 0.049 | 9.83E-154 | 3.6E-149 |
| AGAP1 | 1 | -0.73974109 | 0.709 | 0.888 | 4.96E-153 | 1.81E-148 |
| SLC2A13 | 1 | -0.967443881 | 0.221 | 0.522 | 1.72E-152 | 6.29E-148 |
| HS3ST4 | 1 | 0.927638579 | 0.573 | 0.263 | 2.56E-152 | 9.38E-148 |
| SGCZ | 1 | 0.880703423 | 0.781 | 0.479 | 3.66E-152 | 1.34E-147 |
| CACNA1A | 1 | -1.703304091 | 0.076 | 0.331 | 7.73E-151 | 2.83E-146 |
| TCF4 | 1 | -0.822419824 | 0.85 | 0.903 | 2.22E-150 | 8.13E-146 |
| FBN2 | 1 | -1.3419616 | 0.163 | 0.448 | 2.57E-150 | 9.42E-146 |
| CLVS2 | 1 | 0.942317246 | 0.232 | 0.04 | 2.68E-150 | 9.82E-146 |
| AL157944.1 | 1 | 1.271105432 | 0.25 | 0.048 | 3.64E-150 | 1.33E-145 |
| KLRG2 | 1 | 0.378946911 | 0.175 | 0.015 | 7.35E-150 | 2.69E-145 |
| FIGN | 1 | 1.083176534 | 0.541 | 0.271 | 1.06E-149 | 3.87E-145 |
| AL353747.4 | 1 | 0.690391006 | 0.371 | 0.121 | 1.08E-149 | 3.97E-145 |
| CAMK4 | 1 | -0.793336308 | 0.094 | 0.365 | 2.38E-149 | 8.7E-145 |
| MYO16 | 1 | -1.34040415 | 0.131 | 0.379 | 2.9E-148 | 1.06E-143 |
| FAP | 1 | -1.248983329 | 0.045 | 0.274 | 3.61E-148 | 1.32E-143 |
| FOXP2 | 1 | -1.636621417 | 0.074 | 0.326 | 7.04E-148 | 2.58E-143 |
| COL4A2 | 1 | -0.855118146 | 0.126 | 0.402 | 1.44E-147 | 5.26E-143 |
| CHN1 | 1 | 0.782563423 | 0.765 | 0.558 | 1.58E-147 | 5.8E-143 |
| DOCK3 | 1 | -1.178205476 | 0.096 | 0.347 | 1.86E-147 | 6.79E-143 |
| MIR503HG | 1 | 0.723586103 | 0.348 | 0.101 | 1.94E-147 | 7.11E-143 |
| PARM1 | 1 | -0.723471185 | 0.034 | 0.257 | 5.94E-147 | 2.17E-142 |
| CDH13 | 1 | 0.861717211 | 0.204 | 0.03 | 6.74E-147 | 2.47E-142 |
| POLR3G | 1 | 0.724503563 | 0.464 | 0.2 | 1.75E-146 | 6.43E-142 |
| AC023469.1 | 1 | 1.110001787 | 0.538 | 0.291 | 2.2E-146 | 8.04E-142 |
| NPAS3 | 1 | -1.869183565 | 0.079 | 0.298 | 5.57E-146 | 2.04E-141 |
| BICC1 | 1 | -0.830747759 | 0.099 | 0.365 | 8.38E-146 | 3.07E-141 |
| AL591519.1 | 1 | 0.803728486 | 0.327 | 0.089 | 9.59E-146 | 3.51E-141 |
| NEO1 | 1 | -0.834548718 | 0.351 | 0.615 | 1.3E-145 | 4.75E-141 |
| PIP5K1B | 1 | -1.278748559 | 0.097 | 0.339 | 3.2E-145 | 1.17E-140 |
| AL024495.1 | 1 | 0.709939214 | 0.176 | 0.019 | 8.37E-145 | 3.06E-140 |
| ELAVL2 | 1 | -0.98046782 | 0.169 | 0.452 | 1.72E-144 | 6.28E-140 |
| GPC4 | 1 | 0.946764199 | 0.397 | 0.144 | 4.67E-144 | 1.71E-139 |
| ARHGAP8 | 1 | 0.770935716 | 0.403 | 0.15 | 6E-144 | 2.2E-139 |
| AL033504.1 | 1 | 0.9236592 | 0.763 | 0.518 | 1.75E-143 | 6.4E-139 |
| MYO16-AS1 | 1 | 0.594016734 | 0.172 | 0.018 | 1.03E-142 | 3.76E-138 |
| PEX1 | 1 | 0.735066046 | 0.566 | 0.348 | 1.72E-142 | 6.31E-138 |
| DLC1 | 1 | -1.134982325 | 0.079 | 0.327 | 4.54E-142 | 1.66E-137 |

| | | | | | | |
|---|---|---|---|---|---|---|
| LIMCH1 | 1 | -1.120996098 | 0.508 | 0.741 | 6.25E-142 | 2.29E-137 |
| EBF3 | 1 | -0.978387943 | 0.066 | 0.311 | 1.1E-141 | 4.04E-137 |
| NELL1 | 1 | 1.111012586 | 0.515 | 0.246 | 3.04E-141 | 1.11E-136 |
| ATP8A2 | 1 | -0.868065483 | 0.115 | 0.378 | 5.14E-140 | 1.88E-135 |
| STEAP2 | 1 | 0.520293827 | 0.28 | 0.068 | 9.71E-140 | 3.56E-135 |
| ENPP2 | 1 | -0.666385635 | 0.043 | 0.267 | 1.09E-139 | 4E-135 |
| PIEZO2 | 1 | -1.053491218 | 0.176 | 0.441 | 1.69E-139 | 6.18E-135 |
| CATSPERE | 1 | 0.871918984 | 0.422 | 0.181 | 1.7E-139 | 6.23E-135 |
| PXDN | 1 | -0.641031284 | 0.057 | 0.291 | 1.72E-138 | 6.3E-134 |
| PPARG | 1 | 0.69315584 | 0.496 | 0.211 | 2.09E-137 | 7.67E-133 |
| CASK | 1 | -0.643431123 | 0.757 | 0.892 | 6.75E-137 | 2.47E-132 |
| NKAIN3 | 1 | -1.109147776 | 0.511 | 0.74 | 1.5E-136 | 5.5E-132 |
| RNASET2 | 1 | 0.517582882 | 0.314 | 0.089 | 2.13E-136 | 7.81E-132 |
| TNFRSF19 | 1 | -0.761229315 | 0.063 | 0.299 | 2.17E-136 | 7.95E-132 |
| RAB3C | 1 | 1.007852618 | 0.619 | 0.365 | 4.01E-136 | 1.47E-131 |
| CMTM6 | 1 | 0.566472767 | 0.328 | 0.099 | 6.07E-136 | 2.22E-131 |
| MALAT1 | 1 | 0.9157421 | 0.994 | 1 | 7.4E-136 | 2.71E-131 |
| SLC7A11 | 1 | 0.720422474 | 0.188 | 0.026 | 3.41E-135 | 1.25E-130 |
| ADAMTSL3 | 1 | 0.874778198 | 0.482 | 0.224 | 2.33E-134 | 8.52E-130 |
| ESRG | 1 | 0.38731687 | 0.134 | 0.007 | 3.57E-134 | 1.31E-129 |
| GHR | 1 | -0.811218547 | 0.045 | 0.263 | 6E-133 | 2.2E-128 |
| BRINP2 | 1 | 0.747344615 | 0.255 | 0.058 | 9.44E-133 | 3.45E-128 |
| NR1H4 | 1 | 0.715561471 | 0.191 | 0.028 | 4.59E-132 | 1.68E-127 |
| KCNJ6 | 1 | -0.861953232 | 0.041 | 0.254 | 4.78E-132 | 1.75E-127 |
| ST6GAL1 | 1 | 0.964554967 | 0.511 | 0.325 | 3.19E-131 | 1.17E-126 |
| TMEM14A | 1 | 0.498914001 | 0.294 | 0.082 | 3.46E-131 | 1.27E-126 |
| FAM184A | 1 | 0.792149699 | 0.622 | 0.368 | 3.81E-131 | 1.4E-126 |
| MYBL1 | 1 | 0.764153668 | 0.429 | 0.185 | 6.24E-131 | 2.28E-126 |
| PDE7B | 1 | 0.901784277 | 0.41 | 0.153 | 1.42E-130 | 5.19E-126 |
| AL589740.1 | 1 | -1.128974652 | 0.054 | 0.271 | 2E-130 | 7.34E-126 |
| MAGI2 | 1 | -0.842010311 | 0.53 | 0.755 | 2.03E-130 | 7.42E-126 |
| SH3RF3 | 1 | -0.701076847 | 0.086 | 0.329 | 4.01E-130 | 1.47E-125 |
| SORCS3 | 1 | -0.900959918 | 0.216 | 0.49 | 8.25E-130 | 3.02E-125 |
| AC092422.1 | 1 | 1.047476771 | 0.325 | 0.108 | 9.21E-130 | 3.37E-125 |
| GMDS | 1 | -1.009636606 | 0.244 | 0.506 | 1.33E-129 | 4.87E-125 |
| TMEM220 | 1 | 0.592720534 | 0.385 | 0.144 | 1.41E-129 | 5.16E-125 |
| ITPR1 | 1 | 0.866406267 | 0.586 | 0.327 | 1.49E-129 | 5.47E-125 |
| DENND1B | 1 | -0.729983109 | 0.135 | 0.386 | 7.47E-129 | 2.73E-124 |
| SCP2 | 1 | 0.74852994 | 0.541 | 0.313 | 1.33E-128 | 4.87E-124 |
| LRRTM3 | 1 | 0.871313314 | 0.34 | 0.11 | 1.52E-128 | 5.57E-124 |
| EMP1 | 1 | 0.474176332 | 0.224 | 0.042 | 1.91E-128 | 7E-124 |
| NOL4 | 1 | -1.174618248 | 0.156 | 0.383 | 2.02E-128 | 7.41E-124 |
| ENPP1 | 1 | 0.865397785 | 0.549 | 0.29 | 2.49E-128 | 9.12E-124 |
| GRIA4 | 1 | -1.178508523 | 0.135 | 0.375 | 3.59E-128 | 1.31E-123 |
| PCDH11X | 1 | -1.691287284 | 0.089 | 0.321 | 4.24E-128 | 1.55E-123 |
| AC008014.1 | 1 | 0.882083276 | 0.529 | 0.296 | 2.54E-127 | 9.3E-123 |
| WEE1 | 1 | 0.745819665 | 0.605 | 0.374 | 3.37E-127 | 1.23E-122 |
| RNF150 | 1 | -0.772808711 | 0.188 | 0.458 | 4.49E-126 | 1.64E-121 |
| AC040168.1 | 1 | 0.75199902 | 0.437 | 0.186 | 5.57E-126 | 2.04E-121 |
| SEMA6D | 1 | -1.009088774 | 0.511 | 0.725 | 6.97E-126 | 2.55E-121 |
| KALRN | 1 | -0.849206568 | 0.205 | 0.473 | 7.28E-126 | 2.66E-121 |
| CDH23 | 1 | 0.770187743 | 0.464 | 0.211 | 1.09E-125 | 4.01E-121 |
| VEZT | 1 | 0.645188494 | 0.785 | 0.631 | 1.28E-125 | 4.7E-121 |
| RASGRP1 | 1 | 0.661389753 | 0.412 | 0.158 | 1.77E-125 | 6.49E-121 |
| RALGPS1 | 1 | -0.612945464 | 0.099 | 0.338 | 1.09E-124 | 3.99E-120 |
| CASZ1 | 1 | 0.685150936 | 0.444 | 0.19 | 1.17E-124 | 4.27E-120 |
| CTNND2 | 1 | 0.816913789 | 0.825 | 0.707 | 2.92E-124 | 1.07E-119 |
| GSR | 1 | 0.697120237 | 0.471 | 0.239 | 4.91E-124 | 1.8E-119 |

| | | | | | | |
|---|---|---|---|---|---|---|
| TESK2 | 1 | 0.665222218 | 0.352 | 0.129 | 1.23E-123 | 4.52E-119 |
| CHD7 | 1 | -0.623961918 | 0.669 | 0.848 | 2.26E-123 | 8.28E-119 |
| GPC5 | 1 | 0.873938357 | 0.195 | 0.032 | 5.98E-123 | 2.19E-118 |
| FNDC3A | 1 | 0.682859586 | 0.853 | 0.787 | 9.33E-123 | 3.42E-118 |
| PLEKHA6 | 1 | 0.767144288 | 0.628 | 0.414 | 1.77E-122 | 6.49E-118 |
| RABGAP1L | 1 | -0.790057944 | 0.614 | 0.792 | 2.86E-122 | 1.05E-117 |
| EYS | 1 | -2.319657755 | 0.094 | 0.296 | 4.85E-122 | 1.77E-117 |
| ADCY2 | 1 | -1.199173294 | 0.102 | 0.336 | 2.82E-121 | 1.03E-116 |
| SYNPO2 | 1 | 0.662547416 | 0.466 | 0.203 | 4.16E-121 | 1.52E-116 |
| DYNC1I1 | 1 | 0.869348198 | 0.539 | 0.283 | 6.64E-121 | 2.43E-116 |
| SLC44A1 | 1 | 0.663479255 | 0.812 | 0.651 | 7.77E-121 | 2.85E-116 |
| MBNL2 | 1 | 0.67520575 | 0.608 | 0.345 | 1.72E-120 | 6.31E-116 |
| LINC02470 | 1 | -1.106747115 | 0.105 | 0.322 | 6.07E-120 | 2.22E-115 |
| CASC15 | 1 | 0.64624803 | 0.771 | 0.509 | 6.57E-120 | 2.41E-115 |
| SHB | 1 | 0.630343993 | 0.425 | 0.18 | 1.26E-119 | 4.62E-115 |
| LINC02428 | 1 | 0.516732538 | 0.207 | 0.039 | 1.39E-119 | 5.1E-115 |
| TRIM22 | 1 | 0.691376139 | 0.468 | 0.226 | 3.33E-119 | 1.22E-114 |
| AC007326.5 | 1 | 0.435606277 | 0.117 | 0.006 | 3.35E-119 | 1.23E-114 |
| ZBTB18 | 1 | 0.606567404 | 0.393 | 0.166 | 3.49E-119 | 1.28E-114 |
| ANKRD18A | 1 | 0.552397783 | 0.321 | 0.103 | 3.95E-119 | 1.45E-114 |
| CACHD1 | 1 | 0.576926331 | 0.767 | 0.521 | 6.65E-119 | 2.43E-114 |
| FHOD3 | 1 | -0.833949788 | 0.133 | 0.364 | 6.77E-119 | 2.48E-114 |
| UNC5C | 1 | 0.836011067 | 0.719 | 0.467 | 6.82E-119 | 2.5E-114 |
| IVNS1ABP | 1 | 0.789706347 | 0.617 | 0.47 | 9.16E-119 | 3.35E-114 |
| ZNF804A | 1 | -1.458594599 | 0.081 | 0.301 | 1.19E-118 | 4.36E-114 |
| NRP1 | 1 | 0.796003631 | 0.507 | 0.248 | 1.87E-118 | 6.86E-114 |
| SORCS1 | 1 | -0.931360788 | 0.07 | 0.283 | 2.69E-118 | 9.84E-114 |
| STK32A | 1 | 0.489316854 | 0.185 | 0.03 | 3.68E-118 | 1.35E-113 |
| PTPRO | 1 | 0.914628871 | 0.605 | 0.389 | 5.79E-118 | 2.12E-113 |
| TLL2 | 1 | 0.645115253 | 0.324 | 0.104 | 7.32E-118 | 2.68E-113 |
| SCMH1 | 1 | -0.648781335 | 0.454 | 0.693 | 1.33E-117 | 4.88E-113 |
| GRIA2 | 1 | -1.162225135 | 0.569 | 0.688 | 2.5E-117 | 9.15E-113 |
| ZNF532 | 1 | 0.679415699 | 0.776 | 0.643 | 2.79E-117 | 1.02E-112 |
| GMDS-DT | 1 | -0.720301994 | 0.093 | 0.32 | 3.89E-117 | 1.42E-112 |
| SVIL | 1 | 0.809651732 | 0.706 | 0.47 | 1.15E-116 | 4.21E-112 |
| SLC26A7 | 1 | 0.858515995 | 0.188 | 0.032 | 1.94E-116 | 7.1E-112 |
| LINC02398 | 1 | -1.21897356 | 0.031 | 0.208 | 2.1E-116 | 7.69E-112 |
| SAMD5 | 1 | 0.687609846 | 0.786 | 0.598 | 2.8E-116 | 1.02E-111 |
| TRIM9 | 1 | 0.789006428 | 0.599 | 0.367 | 3.43E-116 | 1.25E-111 |
| MPPED2 | 1 | -0.731044077 | 0.04 | 0.233 | 6.71E-116 | 2.46E-111 |
| SMOC1 | 1 | -0.538007753 | 0.035 | 0.223 | 7.11E-116 | 2.6E-111 |
| ARHGEF28 | 1 | -0.925507257 | 0.178 | 0.413 | 1.02E-115 | 3.72E-111 |
| SNCAIP | 1 | -0.90052162 | 0.053 | 0.253 | 1.18E-115 | 4.32E-111 |
| AC068286.2 | 1 | 1.788208823 | 0.263 | 0.095 | 2.11E-115 | 7.73E-111 |
| AL356108.1 | 1 | 1.130288353 | 0.249 | 0.067 | 3.69E-115 | 1.35E-110 |
| KIF16B | 1 | -0.570975358 | 0.123 | 0.363 | 4.17E-115 | 1.53E-110 |
| DDIT4 | 1 | 0.544831339 | 0.238 | 0.059 | 4.79E-115 | 1.75E-110 |
| AC068587.4 | 1 | -0.792155844 | 0.157 | 0.405 | 1.97E-114 | 7.2E-110 |
| SH3GL2 | 1 | -0.859166848 | 0.206 | 0.457 | 2.41E-114 | 8.82E-110 |
| ACAP2 | 1 | 0.679769262 | 0.753 | 0.635 | 4.2E-114 | 1.54E-109 |
| GRM3 | 1 | 0.573180905 | 0.159 | 0.021 | 4.58E-114 | 1.68E-109 |
| OSTF1 | 1 | 0.503490459 | 0.237 | 0.06 | 3.29E-113 | 1.2E-108 |
| ANKRD29 | 1 | 0.53768169 | 0.205 | 0.044 | 8.84E-113 | 3.24E-108 |
| ALOX15 | 1 | 0.412409072 | 0.126 | 0.01 | 1.18E-112 | 4.33E-108 |
| MAGI1 | 1 | 0.605371853 | 0.853 | 0.869 | 1.47E-112 | 5.38E-108 |
| AC120193.1 | 1 | 0.672772938 | 0.276 | 0.079 | 1.98E-112 | 7.26E-108 |
| ERICH3 | 1 | -0.61174844 | 0.017 | 0.179 | 2.64E-112 | 9.67E-108 |
| YAP1 | 1 | -0.75668706 | 0.15 | 0.394 | 2.82E-112 | 1.03E-107 |

| | | | | | | |
|---|---|---|---|---|---|---|
| ITGBL1 | 1 | -1.824515172 | 0.065 | 0.259 | 4.53E-112 | 1.66E-107 |
| ALCAM | 1 | -1.174015563 | 0.155 | 0.389 | 5.82E-112 | 2.13E-107 |
| AUTS2 | 1 | -1.339651025 | 0.353 | 0.514 | 1.04E-111 | 3.83E-107 |
| AC006148.1 | 1 | 0.813188216 | 0.312 | 0.107 | 1.38E-111 | 5.05E-107 |
| LBR | 1 | 0.757783076 | 0.484 | 0.293 | 1.53E-111 | 5.6E-107 |
| PDE1B | 1 | 0.337914604 | 0.16 | 0.022 | 2.08E-111 | 7.63E-107 |
| PRKCA | 1 | -0.820039207 | 0.129 | 0.359 | 3.46E-111 | 1.27E-106 |
| ADD3-AS1 | 1 | 0.536660281 | 0.228 | 0.054 | 3.56E-111 | 1.3E-106 |
| SPATS2L | 1 | -0.923791783 | 0.286 | 0.53 | 1.29E-110 | 4.72E-106 |
| GNG2 | 1 | -0.624440173 | 0.029 | 0.203 | 1.82E-110 | 6.65E-106 |
| CHST9 | 1 | -1.06095231 | 0.027 | 0.191 | 2.31E-110 | 8.47E-106 |
| ITGB8 | 1 | -1.098462422 | 0.139 | 0.351 | 6.7E-110 | 2.45E-105 |
| EZH2 | 1 | -0.669398247 | 0.581 | 0.753 | 8.9E-110 | 3.26E-105 |
| TENM3 | 1 | 0.671724615 | 0.862 | 0.757 | 1.2E-109 | 4.38E-105 |
| BMERB1 | 1 | 0.707840447 | 0.637 | 0.418 | 1.57E-109 | 5.73E-105 |
| PRELID2 | 1 | -0.712019663 | 0.14 | 0.382 | 1.58E-109 | 5.78E-105 |
| SORBS1 | 1 | -0.764060017 | 0.348 | 0.591 | 2.76E-109 | 1.01E-104 |
| TBX3 | 1 | 0.644768636 | 0.348 | 0.141 | 3.44E-109 | 1.26E-104 |
| BBX | 1 | -0.616176288 | 0.532 | 0.751 | 6.31E-109 | 2.31E-104 |
| COL3A1 | 1 | -1.050216847 | 0.106 | 0.331 | 9.04E-109 | 3.31E-104 |
| EYA4 | 1 | -1.235624356 | 0.066 | 0.249 | 2.57E-108 | 9.41E-104 |
| PAX3 | 1 | -0.915889666 | 0.018 | 0.178 | 7.74E-108 | 2.83E-103 |
| TRPM3 | 1 | -2.589153638 | 0.067 | 0.244 | 1.47E-107 | 5.39E-103 |
| MAF | 1 | 0.488560036 | 0.291 | 0.092 | 8.41E-107 | 3.08E-102 |
| HIF3A | 1 | 0.526990767 | 0.271 | 0.079 | 8.55E-107 | 3.13E-102 |
| CADPS | 1 | 0.945725281 | 0.639 | 0.409 | 1.31E-106 | 4.8E-102 |
| FUT8 | 1 | 0.630545642 | 0.866 | 0.8 | 2.11E-106 | 7.72E-102 |
| IDH3A | 1 | 0.544325446 | 0.357 | 0.154 | 2.94E-106 | 1.08E-101 |
| SVEP1 | 1 | -0.907550759 | 0.126 | 0.348 | 3.28E-106 | 1.2E-101 |
| LIN28B | 1 | 0.621861148 | 0.691 | 0.435 | 3.34E-106 | 1.22E-101 |
| SPECC1 | 1 | 0.743117656 | 0.69 | 0.496 | 5.79E-106 | 2.12E-101 |
| ZNF814 | 1 | 0.513078957 | 0.368 | 0.148 | 9.81E-106 | 3.59E-101 |
| CYP4X1 | 1 | 0.684951907 | 0.301 | 0.1 | 1.26E-105 | 4.6E-101 |
| DMD | 1 | 0.537816533 | 0.82 | 0.624 | 1.53E-105 | 5.62E-101 |
| SORBS2 | 1 | -1.063779203 | 0.014 | 0.163 | 1.92E-105 | 7.02E-101 |
| EXPH5 | 1 | 0.684551885 | 0.3 | 0.111 | 3.38E-105 | 1.24E-100 |
| CD109 | 1 | 0.519952414 | 0.288 | 0.091 | 3.92E-105 | 1.44E-100 |
| ST18 | 1 | -1.096151817 | 0.365 | 0.548 | 4.63E-105 | 1.7E-100 |
| SLC10A4 | 1 | 0.485825561 | 0.223 | 0.057 | 5.09E-105 | 1.86E-100 |
| PRDM5 | 1 | -0.625308327 | 0.149 | 0.378 | 7.91E-105 | 2.9E-100 |
| SCN7A | 1 | -1.38601737 | 0.021 | 0.179 | 1.02E-104 | 3.75E-100 |
| AL592156.2 | 1 | 0.742041353 | 0.207 | 0.049 | 1.96E-104 | 7.19E-100 |
| AL354809.1 | 1 | 0.427598728 | 0.26 | 0.074 | 3.79E-104 | 1.39E-99 |
| SLC22A23 | 1 | 0.739954272 | 0.535 | 0.298 | 9.35E-104 | 3.42E-99 |
| ANKRD44 | 1 | -0.632020767 | 0.152 | 0.383 | 1.1E-102 | 4.04E-98 |
| ATP11A | 1 | -0.612727911 | 0.293 | 0.541 | 1.48E-102 | 5.43E-98 |
| HS3ST3B1 | 1 | -0.760715654 | 0.13 | 0.353 | 2E-102 | 7.34E-98 |
| NLGN4X | 1 | -0.760986827 | 0.591 | 0.78 | 2.32E-102 | 8.51E-98 |
| LAMA4 | 1 | -1.216683122 | 0.413 | 0.469 | 2.36E-102 | 8.66E-98 |
| BCL2 | 1 | -0.64095722 | 0.053 | 0.241 | 2.65E-102 | 9.7E-98 |
| ATXN1 | 1 | 0.570025906 | 0.832 | 0.648 | 4.21E-102 | 1.54E-97 |
| FARP1 | 1 | -0.617761345 | 0.576 | 0.778 | 4.6E-102 | 1.68E-97 |
| MAP4K3 | 1 | 0.574064596 | 0.912 | 0.874 | 4.87E-102 | 1.78E-97 |
| MAGI3 | 1 | -0.732849489 | 0.275 | 0.508 | 4.92E-102 | 1.8E-97 |
| ZNF91 | 1 | 0.503770113 | 0.425 | 0.191 | 6.25E-102 | 2.29E-97 |
| FREM1 | 1 | -1.040207726 | 0.109 | 0.31 | 1.05E-101 | 3.85E-97 |
| BACH2 | 1 | -0.747199382 | 0.202 | 0.446 | 2.55E-101 | 9.32E-97 |
| PDGFRA | 1 | -0.695235478 | 0.052 | 0.236 | 7.66E-101 | 2.8E-96 |

| | | | | | | |
|---|---|---|---|---|---|---|
| TTC3 | 1 | -0.519699596 | 0.799 | 0.902 | 9.23E-101 | 3.38E-96 |
| MBOAT2 | 1 | -0.598015545 | 0.273 | 0.52 | 1.44E-100 | 5.27E-96 |
| NMNAT2 | 1 | 0.775968592 | 0.484 | 0.268 | 2.1E-100 | 7.7E-96 |
| TAOK3 | 1 | 0.689589649 | 0.591 | 0.371 | 2.25E-100 | 8.22E-96 |
| NKAIN2 | 1 | -1.692955807 | 0.285 | 0.406 | 2.31E-100 | 8.47E-96 |
| NEUROD1 | 1 | -0.621036632 | 0.045 | 0.225 | 3.76E-100 | 1.38E-95 |
| LINC00907 | 1 | -1.390704902 | 0.124 | 0.296 | 5.43E-100 | 1.99E-95 |
| AC007255.1 | 1 | 0.347173089 | 0.159 | 0.026 | 5.81E-100 | 2.13E-95 |
| SESTD1 | 1 | -0.619542087 | 0.314 | 0.554 | 1.9E-99 | 6.97E-95 |
| KCNQ3 | 1 | 0.832803967 | 0.571 | 0.328 | 2.16E-99 | 7.92E-95 |
| ANK1 | 1 | -0.638207998 | 0.056 | 0.239 | 2.38E-99 | 8.71E-95 |
| GAS7 | 1 | 0.329961578 | 0.405 | 0.185 | 2.52E-99 | 9.21E-95 |
| TFDP2 | 1 | -0.631951846 | 0.328 | 0.569 | 2.98E-99 | 1.09E-94 |
| TANC2 | 1 | -0.662326343 | 0.564 | 0.764 | 3.09E-99 | 1.13E-94 |
| RBFOX1 | 1 | -0.722509422 | 0.695 | 0.748 | 6.48E-98 | 2.37E-93 |
| HPSE2 | 1 | 0.537518479 | 0.43 | 0.201 | 7.93E-98 | 2.9E-93 |
| MYLIP | 1 | 0.544048642 | 0.33 | 0.126 | 9.79E-98 | 3.59E-93 |
| CHRNA7 | 1 | -0.604728755 | 0.085 | 0.285 | 1.18E-97 | 4.33E-93 |
| GPRC5D-AS1 | 1 | 0.321588875 | 0.178 | 0.035 | 1.54E-97 | 5.66E-93 |
| GPR1 | 1 | 0.342183221 | 0.134 | 0.017 | 2.08E-97 | 7.6E-93 |
| TMEM178A | 1 | 0.698972523 | 0.624 | 0.42 | 2.58E-97 | 9.44E-93 |
| MME | 1 | -0.620153647 | 0.052 | 0.234 | 3E-97 | 1.1E-92 |
| GRID2 | 1 | -1.340260639 | 0.463 | 0.616 | 8.06E-97 | 2.95E-92 |
| AC108690.1 | 1 | -0.565369826 | 0.015 | 0.159 | 1.36E-96 | 4.96E-92 |
| DIP2C | 1 | -0.540850631 | 0.604 | 0.801 | 1.37E-96 | 5E-92 |
| NECTIN3 | 1 | 0.7201455 | 0.739 | 0.555 | 2.29E-96 | 8.4E-92 |
| TMEM163 | 1 | -0.714768643 | 0.033 | 0.193 | 3.64E-96 | 1.33E-91 |
| UNC79 | 1 | -0.864622176 | 0.183 | 0.395 | 6.24E-96 | 2.28E-91 |
| TMTC2 | 1 | -0.928837536 | 0.347 | 0.563 | 8.65E-96 | 3.17E-91 |
| SLC7A14 | 1 | 0.929382347 | 0.201 | 0.079 | 1.92E-95 | 7.02E-91 |
| CFH | 1 | -0.774252909 | 0.028 | 0.185 | 3.55E-95 | 1.3E-90 |
| MET | 1 | 0.61103177 | 0.361 | 0.153 | 3.67E-95 | 1.35E-90 |
| MIR124-2HG | 1 | -0.442572709 | 0.019 | 0.167 | 4.13E-95 | 1.51E-90 |
| NDST3 | 1 | -0.954020607 | 0.338 | 0.523 | 4.89E-95 | 1.79E-90 |
| CDKAL1 | 1 | -0.776621312 | 0.725 | 0.858 | 7.34E-95 | 2.69E-90 |
| TMOD3 | 1 | 0.55194253 | 0.506 | 0.285 | 7.88E-95 | 2.89E-90 |
| NHS | 1 | 0.996495661 | 0.643 | 0.419 | 3.8E-94 | 1.39E-89 |
| ADAMTS6 | 1 | -1.195576175 | 0.036 | 0.188 | 6.09E-94 | 2.23E-89 |
| GRIP1 | 1 | 0.728622723 | 0.532 | 0.303 | 1.15E-93 | 4.21E-89 |
| KLF12 | 1 | -0.700336661 | 0.492 | 0.691 | 1.35E-93 | 4.93E-89 |
| NCOA2 | 1 | -0.614220337 | 0.506 | 0.686 | 1.39E-93 | 5.09E-89 |
| ITGA4 | 1 | -1.182825983 | 0.343 | 0.514 | 1.44E-93 | 5.27E-89 |
| RPS6KA2 | 1 | 0.630878012 | 0.503 | 0.267 | 1.85E-93 | 6.78E-89 |
| CPLANE1 | 1 | -0.583959827 | 0.431 | 0.644 | 4.09E-93 | 1.5E-88 |
| AL109930.1 | 1 | 0.531145391 | 0.43 | 0.208 | 5.98E-93 | 2.19E-88 |
| MAPK10 | 1 | -0.995762741 | 0.09 | 0.27 | 9.58E-93 | 3.51E-88 |
| NRCAM | 1 | -0.68382883 | 0.219 | 0.446 | 1E-92 | 3.68E-88 |
| DACH1 | 1 | -0.582303528 | 0.795 | 0.917 | 1.26E-92 | 4.6E-88 |
| ANKRD33B | 1 | 0.608817366 | 0.311 | 0.119 | 1.91E-92 | 7E-88 |
| SNTG1 | 1 | -1.520277302 | 0.242 | 0.432 | 2.38E-92 | 8.71E-88 |
| MSI2 | 1 | -0.587273882 | 0.78 | 0.882 | 3.44E-92 | 1.26E-87 |
| ARRDC3-AS1 | 1 | 1.178037175 | 0.432 | 0.235 | 3.82E-92 | 1.4E-87 |
| UBE2G1 | 1 | 0.571400445 | 0.731 | 0.575 | 8.73E-92 | 3.19E-87 |
| NEB | 1 | 0.838624196 | 0.256 | 0.087 | 1.02E-91 | 3.74E-87 |
| MACF1 | 1 | -0.451847518 | 0.758 | 0.89 | 1.67E-91 | 6.12E-87 |
| SLITRK2 | 1 | 0.300160866 | 0.132 | 0.018 | 2.47E-91 | 9.06E-87 |
| EIF1AX | 1 | 0.577068276 | 0.597 | 0.391 | 4.69E-91 | 1.72E-86 |
| HOTAIRM1 | 1 | 0.432349353 | 0.174 | 0.036 | 9.6E-91 | 3.51E-86 |

| | | | | | | |
|---|---|---|---|---|---|---|
| GAREM1 | 1 | -0.722429777 | 0.137 | 0.329 | 1.36E-90 | 4.97E-86 |
| NAV3 | 1 | -1.109302052 | 0.232 | 0.434 | 2.78E-90 | 1.02E-85 |
| SBF2 | 1 | -0.54425465 | 0.628 | 0.793 | 2.85E-90 | 1.04E-85 |
| TNFSF11 | 1 | 0.280543659 | 0.11 | 0.01 | 4.15E-90 | 1.52E-85 |
| HMGA2 | 1 | -0.886768844 | 0.26 | 0.483 | 4.36E-90 | 1.6E-85 |
| KIAA1549L | 1 | -0.656589206 | 0.081 | 0.267 | 9.34E-90 | 3.42E-85 |
| ZFP64 | 1 | 0.545503329 | 0.442 | 0.236 | 1.25E-89 | 4.56E-85 |
| GARNL3 | 1 | -0.490288936 | 0.088 | 0.278 | 2.28E-89 | 8.33E-85 |
| CLEC2A | 1 | -0.543856644 | 0.022 | 0.164 | 2.89E-89 | 1.06E-84 |
| VCAN | 1 | -1.096459899 | 0.248 | 0.471 | 3.29E-89 | 1.2E-84 |
| DNAJC1 | 1 | -0.727244622 | 0.414 | 0.622 | 4.84E-89 | 1.77E-84 |
| KIAA0319L | 1 | 0.515988109 | 0.681 | 0.522 | 5.2E-89 | 1.9E-84 |
| AL445623.2 | 1 | -0.556886569 | 0.063 | 0.24 | 5.45E-89 | 1.99E-84 |
| TRIM36 | 1 | 0.378627705 | 0.168 | 0.034 | 6.16E-89 | 2.25E-84 |
| NBAS | 1 | 0.651005717 | 0.836 | 0.787 | 8.46E-89 | 3.1E-84 |
| ZNF680 | 1 | 0.481312595 | 0.406 | 0.193 | 1.13E-88 | 4.12E-84 |
| TSPAN5 | 1 | -0.697954641 | 0.329 | 0.547 | 1.47E-88 | 5.36E-84 |
| NR3C2 | 1 | 1.006179891 | 0.415 | 0.229 | 2.55E-88 | 9.34E-84 |
| CELF2 | 1 | 0.598871135 | 0.767 | 0.572 | 6.57E-88 | 2.41E-83 |
| RAP1GDS1 | 1 | 0.553419989 | 0.765 | 0.636 | 1.12E-87 | 4.12E-83 |
| SH3BGRL | 1 | -0.588632115 | 0.233 | 0.449 | 1.84E-87 | 6.73E-83 |
| PKIA | 1 | -0.658697058 | 0.282 | 0.5 | 2.03E-87 | 7.43E-83 |
| PBX3 | 1 | -0.832391361 | 0.332 | 0.563 | 3E-87 | 1.1E-82 |
| TARS | 1 | 0.547377867 | 0.473 | 0.275 | 3.01E-87 | 1.1E-82 |
| C1orf21 | 1 | -0.607949549 | 0.151 | 0.363 | 3.38E-87 | 1.24E-82 |
| PTAR1 | 1 | 0.560200806 | 0.598 | 0.421 | 4.66E-87 | 1.7E-82 |
| FAM155A | 1 | 0.666423513 | 0.876 | 0.757 | 4.92E-87 | 1.8E-82 |
| ZNF506 | 1 | 0.399456624 | 0.279 | 0.1 | 5.81E-87 | 2.13E-82 |
| ACSS3 | 1 | -0.576095256 | 0.065 | 0.24 | 7.17E-87 | 2.63E-82 |
| FREM2 | 1 | 0.499640188 | 0.169 | 0.036 | 1.87E-86 | 6.83E-82 |
| BEND6 | 1 | 0.31622072 | 0.12 | 0.015 | 2.92E-86 | 1.07E-81 |
| ITGAV | 1 | 0.660522816 | 0.686 | 0.49 | 3.11E-86 | 1.14E-81 |
| INPP4B | 1 | -0.786228427 | 0.046 | 0.206 | 3.9E-86 | 1.43E-81 |
| AKNAD1 | 1 | 0.510591581 | 0.398 | 0.191 | 4.08E-86 | 1.49E-81 |
| RBMS3 | 1 | -0.682040578 | 0.732 | 0.825 | 4.18E-86 | 1.53E-81 |
| CCDC81 | 1 | 0.408919559 | 0.19 | 0.048 | 5.12E-86 | 1.87E-81 |
| HSBP1L1 | 1 | 0.303618377 | 0.172 | 0.038 | 5.59E-86 | 2.05E-81 |
| HECTD1 | 1 | 0.487897108 | 0.852 | 0.784 | 6.37E-86 | 2.33E-81 |
| COX20 | 1 | 0.446314171 | 0.35 | 0.154 | 7.46E-86 | 2.73E-81 |
| AL357054.2 | 1 | 0.524228492 | 0.357 | 0.157 | 1.27E-85 | 4.65E-81 |
| IFI27 | 1 | 0.340532709 | 0.164 | 0.034 | 1.38E-85 | 5.04E-81 |
| ARAP2 | 1 | 0.620039968 | 0.581 | 0.353 | 1.79E-85 | 6.57E-81 |
| AMACR | 1 | 0.38363915 | 0.202 | 0.056 | 2.84E-85 | 1.04E-80 |
| MPDZ | 1 | -0.56526846 | 0.577 | 0.731 | 3.84E-85 | 1.41E-80 |
| HIPK2 | 1 | -0.602412332 | 0.292 | 0.508 | 4.33E-85 | 1.59E-80 |
| BRINP3 | 1 | -1.187944282 | 0.494 | 0.514 | 5.69E-85 | 2.08E-80 |
| NETO1 | 1 | -0.584216492 | 0.048 | 0.211 | 7.99E-85 | 2.93E-80 |
| RRAGD | 1 | 0.42330833 | 0.313 | 0.125 | 9.07E-85 | 3.32E-80 |
| TNIK | 1 | -0.805773451 | 0.197 | 0.413 | 9.64E-85 | 3.53E-80 |
| AMOTL1 | 1 | -0.507324435 | 0.189 | 0.403 | 1.23E-84 | 4.5E-80 |
| CDK6 | 1 | -0.682809426 | 0.033 | 0.179 | 1.28E-84 | 4.67E-80 |
| ELMOD1 | 1 | 0.639094852 | 0.566 | 0.345 | 1.62E-84 | 5.91E-80 |
| SULF1 | 1 | -1.071068763 | 0.164 | 0.356 | 1.77E-84 | 6.46E-80 |
| APAF1 | 1 | 0.567544481 | 0.482 | 0.286 | 1.8E-84 | 6.6E-80 |
| C4orf19 | 1 | 0.530389634 | 0.475 | 0.249 | 1.91E-84 | 6.98E-80 |
| VEGFC | 1 | -0.741416629 | 0.08 | 0.259 | 2.29E-84 | 8.38E-80 |
| ACTN2 | 1 | 0.49612944 | 0.228 | 0.072 | 3.04E-84 | 1.11E-79 |
| AMPH | 1 | -0.479601022 | 0.05 | 0.213 | 6.73E-84 | 2.46E-79 |

| | | | | | | |
|---|---|---|---|---|---|---|
| GRK5 | 1 | 0.543698709 | 0.498 | 0.276 | 8.29E-84 | 3.04E-79 |
| VDAC3 | 1 | 0.550236179 | 0.462 | 0.268 | 9.88E-84 | 3.62E-79 |
| CNTN5 | 1 | -1.629129907 | 0.033 | 0.175 | 1.58E-83 | 5.78E-79 |
| MYO1D | 1 | -0.596395107 | 0.089 | 0.263 | 3.15E-83 | 1.15E-78 |
| SMG1 | 1 | 0.506119864 | 0.848 | 0.765 | 3.21E-83 | 1.17E-78 |
| TBC1D5 | 1 | -0.539062996 | 0.781 | 0.891 | 7.54E-83 | 2.76E-78 |
| KIAA1328 | 1 | -0.57605332 | 0.46 | 0.656 | 9.87E-83 | 3.61E-78 |
| COL18A1 | 1 | -0.515682372 | 0.027 | 0.167 | 1.05E-82 | 3.83E-78 |
| LAMC2 | 1 | 0.252630407 | 0.103 | 0.01 | 1.38E-82 | 5.05E-78 |
| NLRP2 | 1 | 0.315940448 | 0.191 | 0.049 | 1.39E-82 | 5.08E-78 |
| PCDH9 | 1 | -1.30226834 | 0.206 | 0.391 | 1.92E-82 | 7.02E-78 |
| YWHAZ | 1 | 0.547068553 | 0.713 | 0.561 | 1.99E-82 | 7.27E-78 |
| STK33 | 1 | -0.797184863 | 0.047 | 0.197 | 2.89E-82 | 1.06E-77 |
| AC024901.1 | 1 | -0.919570432 | 0.063 | 0.218 | 4.3E-82 | 1.57E-77 |
| PTPRZ1 | 1 | -0.943070417 | 0.265 | 0.454 | 6.34E-82 | 2.32E-77 |
| SUGCT | 1 | 0.636461311 | 0.556 | 0.35 | 6.38E-82 | 2.34E-77 |
| DLG2 | 1 | 1.174033883 | 0.68 | 0.621 | 8.77E-82 | 3.21E-77 |
| MYO6 | 1 | -0.511566423 | 0.131 | 0.328 | 1.89E-81 | 6.93E-77 |
| PTPRK | 1 | -0.664057634 | 0.699 | 0.844 | 1.93E-81 | 7.07E-77 |
| ROBO2 | 1 | -0.878307322 | 0.61 | 0.746 | 2.97E-81 | 1.09E-76 |
| C2CD2 | 1 | -0.405735257 | 0.039 | 0.189 | 7.63E-81 | 2.79E-76 |
| KCNN2 | 1 | -1.100686972 | 0.167 | 0.342 | 9.46E-81 | 3.46E-76 |
| SYT10 | 1 | -0.569334552 | 0.024 | 0.158 | 9.81E-81 | 3.59E-76 |
| PHYHIPL | 1 | -0.501588869 | 0.044 | 0.197 | 1.48E-80 | 5.43E-76 |
| C6orf141 | 1 | 0.274947476 | 0.132 | 0.022 | 1.6E-80 | 5.85E-76 |
| AC026765.3 | 1 | 0.290000402 | 0.128 | 0.02 | 2.03E-80 | 7.43E-76 |
| VPS54 | 1 | 0.585736065 | 0.682 | 0.553 | 2.28E-80 | 8.35E-76 |
| TTLL7 | 1 | -0.630185385 | 0.32 | 0.52 | 4.8E-80 | 1.76E-75 |
| TLE1 | 1 | 0.499026648 | 0.413 | 0.205 | 5.9E-80 | 2.16E-75 |
| CA12 | 1 | 0.469241567 | 0.401 | 0.194 | 6.57E-80 | 2.4E-75 |
| GLCE | 1 | 0.585800589 | 0.534 | 0.334 | 1.1E-79 | 4.02E-75 |
| STK3 | 1 | -0.611491309 | 0.341 | 0.545 | 1.5E-79 | 5.5E-75 |
| NTM | 1 | 0.912137907 | 0.53 | 0.378 | 1.53E-79 | 5.61E-75 |
| DENND4C | 1 | 0.520682563 | 0.777 | 0.675 | 1.94E-79 | 7.11E-75 |
| ABLIM1 | 1 | -0.588980841 | 0.104 | 0.287 | 2.4E-79 | 8.78E-75 |
| LINC02263 | 1 | -0.694221578 | 0.014 | 0.136 | 2.55E-79 | 9.34E-75 |
| SHROOM4 | 1 | 0.655748851 | 0.728 | 0.564 | 2.77E-79 | 1.01E-74 |
| MIR181A1HG | 1 | -0.657574415 | 0.316 | 0.527 | 4.84E-79 | 1.77E-74 |
| RARB | 1 | 0.890055599 | 0.382 | 0.222 | 6.59E-79 | 2.41E-74 |
| GSAP | 1 | 0.420132484 | 0.252 | 0.09 | 1.35E-78 | 4.94E-74 |
| PDE1A | 1 | -1.393425128 | 0.079 | 0.243 | 1.37E-78 | 5.03E-74 |
| ZMYM4 | 1 | 0.434070804 | 0.863 | 0.843 | 3.59E-78 | 1.31E-73 |
| CLMP | 1 | -0.435912505 | 0.034 | 0.176 | 3.87E-78 | 1.42E-73 |
| PKD1L1 | 1 | 0.296764216 | 0.101 | 0.011 | 4.86E-78 | 1.78E-73 |
| PDGFD | 1 | -0.768789126 | 0.065 | 0.222 | 5.57E-78 | 2.04E-73 |
| PXYLP1 | 1 | 0.44683807 | 0.299 | 0.125 | 6.14E-78 | 2.25E-73 |
| IRS2 | 1 | 0.618228739 | 0.539 | 0.35 | 7.25E-78 | 2.65E-73 |
| MINAR1 | 1 | -0.420749977 | 0.049 | 0.201 | 1.57E-77 | 5.75E-73 |
| MAN1A1 | 1 | -0.801740701 | 0.031 | 0.166 | 1.71E-77 | 6.27E-73 |
| HUNK | 1 | -0.447499296 | 0.042 | 0.19 | 2.01E-77 | 7.37E-73 |
| GPI | 1 | 0.73140472 | 0.389 | 0.221 | 4.23E-77 | 1.55E-72 |
| KCNG3 | 1 | 0.462907109 | 0.296 | 0.121 | 6.58E-77 | 2.41E-72 |
| NEBL | 1 | 0.29610815 | 0.333 | 0.158 | 2.84E-76 | 1.04E-71 |
| TSPAN18 | 1 | 0.46212588 | 0.434 | 0.226 | 8.92E-76 | 3.27E-71 |
| ZNF493 | 1 | 0.379748293 | 0.284 | 0.114 | 1.15E-75 | 4.2E-71 |
| AC093298.2 | 1 | 1.174239374 | 0.314 | 0.147 | 1.24E-75 | 4.53E-71 |
| ACVR2A | 1 | 0.688162765 | 0.591 | 0.413 | 1.31E-75 | 4.81E-71 |
| AGMO | 1 | -0.917820295 | 0.022 | 0.145 | 1.83E-75 | 6.69E-71 |

| | | | | | | |
|---|---|---|---|---|---|---|
| KIAA1217 | 1 | 0.630594993 | 0.472 | 0.258 | 1.88E-75 | 6.87E-71 |
| KIAA1324L | 1 | -0.496962187 | 0.09 | 0.263 | 1.98E-75 | 7.27E-71 |
| KCNT2 | 1 | -0.588430764 | 0.579 | 0.747 | 2.24E-75 | 8.19E-71 |
| LURAP1L | 1 | -0.562570908 | 0.09 | 0.26 | 2.3E-75 | 8.42E-71 |
| SLC38A4 | 1 | -0.70139534 | 0.06 | 0.212 | 2.72E-75 | 9.97E-71 |
| LINC02197 | 1 | -0.726793743 | 0.058 | 0.207 | 2.88E-75 | 1.05E-70 |
| CNTNAP3 | 1 | 0.475767393 | 0.278 | 0.109 | 4.72E-75 | 1.73E-70 |
| MAOB | 1 | 0.444261918 | 0.221 | 0.075 | 5.31E-75 | 1.94E-70 |
| PARD3B | 1 | -0.841665953 | 0.334 | 0.512 | 6.06E-75 | 2.22E-70 |
| ZC4H2 | 1 | -0.442605189 | 0.024 | 0.152 | 6.2E-75 | 2.27E-70 |
| SMARCA2 | 1 | -0.594899994 | 0.272 | 0.481 | 6.71E-75 | 2.45E-70 |
| CCDC144NL-AS1 | 1 | -0.561525032 | 0.736 | 0.865 | 9.26E-75 | 3.39E-70 |
| SEZ6L | 1 | -0.797908501 | 0.013 | 0.122 | 9.83E-75 | 3.6E-70 |
| SLC8A1 | 1 | -0.863744007 | 0.187 | 0.377 | 1.09E-74 | 3.99E-70 |
| DGKI | 1 | -0.648483707 | 0.102 | 0.279 | 1.14E-74 | 4.16E-70 |
| RAD51B | 1 | -0.601014378 | 0.535 | 0.72 | 1.27E-74 | 4.65E-70 |
| STK26 | 1 | 0.430500937 | 0.282 | 0.113 | 1.27E-74 | 4.67E-70 |
| SLFN11 | 1 | -0.273141244 | 0.021 | 0.142 | 1.92E-74 | 7.03E-70 |
| ST6GALNAC5 | 1 | -0.894380497 | 0.231 | 0.423 | 2.91E-74 | 1.06E-69 |
| LHFPL2 | 1 | -0.501425978 | 0.103 | 0.277 | 3.43E-74 | 1.26E-69 |
| NAALADL2 | 1 | -0.796892994 | 0.291 | 0.455 | 3.92E-74 | 1.43E-69 |
| EBF1 | 1 | -1.031172641 | 0.242 | 0.411 | 4.2E-74 | 1.54E-69 |
| CPEB3 | 1 | 0.575004457 | 0.526 | 0.345 | 5.53E-74 | 2.03E-69 |
| DDAH1 | 1 | -0.557718323 | 0.202 | 0.405 | 5.54E-74 | 2.03E-69 |
| KLHL32 | 1 | 0.554902047 | 0.302 | 0.132 | 5.9E-74 | 2.16E-69 |
| TSPAN7 | 1 | -0.650561057 | 0.41 | 0.583 | 6.67E-74 | 2.44E-69 |
| LAMB1 | 1 | -0.724996068 | 0.244 | 0.435 | 1E-73 | 3.68E-69 |
| BCO2 | 1 | 0.442128423 | 0.223 | 0.08 | 1.41E-73 | 5.17E-69 |
| MVB12B | 1 | -0.544614691 | 0.266 | 0.471 | 2.27E-73 | 8.31E-69 |
| AC096747.1 | 1 | 0.372552704 | 0.205 | 0.067 | 2.52E-73 | 9.23E-69 |
| LAMC1 | 1 | 0.514251798 | 0.673 | 0.475 | 3.11E-73 | 1.14E-68 |
| MYO10 | 1 | 0.345668324 | 0.491 | 0.281 | 3.28E-73 | 1.2E-68 |
| PLEKHB2 | 1 | 0.465933873 | 0.42 | 0.229 | 3.52E-73 | 1.29E-68 |
| PCDH17 | 1 | 0.485783465 | 0.171 | 0.046 | 3.6E-73 | 1.32E-68 |
| AK5 | 1 | -0.880933869 | 0.106 | 0.264 | 3.9E-73 | 1.43E-68 |
| PTPN14 | 1 | -0.585490222 | 0.146 | 0.334 | 4.07E-73 | 1.49E-68 |
| PDE11A | 1 | -0.549552503 | 0.069 | 0.227 | 4.69E-73 | 1.72E-68 |
| COL1A2 | 1 | -0.734789033 | 0.028 | 0.158 | 6.1E-73 | 2.23E-68 |
| RGS7 | 1 | -0.782428372 | 0.082 | 0.241 | 6.3E-73 | 2.31E-68 |
| GRIP2 | 1 | 0.428404801 | 0.291 | 0.122 | 6.56E-73 | 2.4E-68 |
| DPP6 | 1 | 0.821536835 | 0.163 | 0.053 | 7.62E-73 | 2.79E-68 |
| HSPA12A | 1 | 0.455329348 | 0.43 | 0.229 | 8.09E-73 | 2.96E-68 |
| SLC4A7 | 1 | -0.506454372 | 0.664 | 0.785 | 1.79E-72 | 6.55E-68 |
| LARP4B | 1 | 0.44858329 | 0.786 | 0.693 | 1.85E-72 | 6.79E-68 |
| ELF2 | 1 | 0.487987647 | 0.848 | 0.774 | 1.9E-72 | 6.94E-68 |
| AC016831.7 | 1 | 0.353619762 | 0.265 | 0.102 | 2.79E-72 | 1.02E-67 |
| LINC00937 | 1 | 0.599574217 | 0.304 | 0.15 | 3.13E-72 | 1.15E-67 |
| TOX2 | 1 | 0.3877792 | 0.205 | 0.063 | 3.52E-72 | 1.29E-67 |
| NALCN | 1 | -0.68101637 | 0.097 | 0.265 | 3.86E-72 | 1.41E-67 |
| AC073050.1 | 1 | 0.661177868 | 0.461 | 0.253 | 4.1E-72 | 1.5E-67 |
| PDE3B | 1 | -0.684752858 | 0.559 | 0.661 | 4.64E-72 | 1.7E-67 |
| PLCE1 | 1 | -0.459192235 | 0.058 | 0.209 | 5.65E-72 | 2.07E-67 |
| MITF | 1 | -0.569994006 | 0.071 | 0.227 | 5.93E-72 | 2.17E-67 |
| CERKL | 1 | -0.428254155 | 0.033 | 0.167 | 7.41E-72 | 2.71E-67 |
| WDR44 | 1 | 0.510678647 | 0.653 | 0.484 | 8.07E-72 | 2.96E-67 |
| PRKACB | 1 | -0.551681372 | 0.456 | 0.64 | 1.14E-71 | 4.19E-67 |
| NBEA | 1 | 0.77723755 | 0.808 | 0.669 | 1.98E-71 | 7.25E-67 |
| XKR4 | 1 | -0.706096899 | 0.068 | 0.225 | 2.24E-71 | 8.21E-67 |

| | | | | | | |
|---|---|---|---|---|---|---|
| LINC02232 | 1 | 0.470378675 | 0.124 | 0.022 | 2.97E-71 | 1.09E-66 |
| DNAH2 | 1 | 0.386326095 | 0.263 | 0.102 | 3.74E-71 | 1.37E-66 |
| TIAM1 | 1 | -0.666444613 | 0.502 | 0.643 | 4.82E-71 | 1.76E-66 |
| GABRA2 | 1 | -0.765189962 | 0.018 | 0.133 | 4.94E-71 | 1.81E-66 |
| PTCHD4 | 1 | 0.725851794 | 0.395 | 0.205 | 5.63E-71 | 2.06E-66 |
| ZNF589 | 1 | 0.481471772 | 0.52 | 0.326 | 6.2E-71 | 2.27E-66 |
| NSMCE3 | 1 | 0.358315422 | 0.217 | 0.075 | 6.57E-71 | 2.41E-66 |
| SKIL | 1 | 0.55092058 | 0.729 | 0.583 | 7.36E-71 | 2.69E-66 |
| EYA2 | 1 | 0.316332383 | 0.251 | 0.097 | 7.73E-71 | 2.83E-66 |
| ATP11C | 1 | 0.538282914 | 0.775 | 0.698 | 8.22E-71 | 3.01E-66 |
| LINC01122 | 1 | 0.888498772 | 0.395 | 0.229 | 9.17E-71 | 3.36E-66 |
| PTPRM | 1 | -0.892092358 | 0.263 | 0.422 | 9.24E-71 | 3.38E-66 |
| DPP4 | 1 | -0.591858274 | 0.061 | 0.211 | 1.05E-70 | 3.86E-66 |
| SCFD2 | 1 | -0.59843475 | 0.471 | 0.636 | 1.35E-70 | 4.95E-66 |
| SEPTIN6 | 1 | -0.356700085 | 0.07 | 0.228 | 1.47E-70 | 5.39E-66 |
| FSTL1 | 1 | 0.467377344 | 0.453 | 0.258 | 1.9E-70 | 6.96E-66 |
| RBFOX3 | 1 | -0.538840483 | 0.078 | 0.241 | 2E-70 | 7.32E-66 |
| RAP1GAP2 | 1 | -1.215833843 | 0.148 | 0.292 | 2.83E-70 | 1.04E-65 |
| MAML2 | 1 | -0.942353649 | 0.436 | 0.547 | 2.93E-70 | 1.07E-65 |
| AC093817.2 | 1 | -0.807896936 | 0.018 | 0.132 | 3.57E-70 | 1.31E-65 |
| ZFHX4 | 1 | 0.281276999 | 0.589 | 0.378 | 4.07E-70 | 1.49E-65 |
| FTX | 1 | 0.318768008 | 0.96 | 0.994 | 4.36E-70 | 1.6E-65 |
| TMEM65 | 1 | -0.457128419 | 0.111 | 0.287 | 5.47E-70 | 2E-65 |
| SCN9A | 1 | 0.424810558 | 0.277 | 0.113 | 5.65E-70 | 2.07E-65 |
| TMF1 | 1 | 0.504790061 | 0.504 | 0.323 | 5.71E-70 | 2.09E-65 |
| CNTN4 | 1 | -0.820027238 | 0.044 | 0.184 | 5.71E-70 | 2.09E-65 |
| NUP50-DT | 1 | 0.443562515 | 0.324 | 0.155 | 7.25E-70 | 2.65E-65 |
| HACE1 | 1 | 0.601111007 | 0.714 | 0.587 | 8.12E-70 | 2.97E-65 |
| AC004540.1 | 1 | -0.428323582 | 0.076 | 0.232 | 1.23E-69 | 4.49E-65 |
| PDE4B | 1 | -1.050199418 | 0.039 | 0.162 | 1.34E-69 | 4.9E-65 |
| GREB1 | 1 | -0.426135878 | 0.022 | 0.139 | 1.35E-69 | 4.94E-65 |
| KCND2 | 1 | -0.931098341 | 0.476 | 0.561 | 1.95E-69 | 7.15E-65 |
| TFRC | 1 | 0.515016713 | 0.43 | 0.251 | 2.02E-69 | 7.38E-65 |
| ARL15 | 1 | -0.581908639 | 0.561 | 0.705 | 2.13E-69 | 7.8E-65 |
| BAZ2B | 1 | -0.512687514 | 0.881 | 0.927 | 3.14E-69 | 1.15E-64 |
| TSPAN9 | 1 | -0.519417576 | 0.242 | 0.44 | 3.37E-69 | 1.24E-64 |
| AL445123.2 | 1 | 0.557959518 | 0.372 | 0.191 | 3.96E-69 | 1.45E-64 |
| RELN | 1 | 0.543909688 | 0.193 | 0.059 | 7.02E-69 | 2.57E-64 |
| FMNL3 | 1 | -0.393706682 | 0.09 | 0.254 | 7.5E-69 | 2.75E-64 |
| ACTR3C | 1 | 0.307642078 | 0.205 | 0.066 | 1.02E-68 | 3.73E-64 |
| AP001636.3 | 1 | -0.393408026 | 0.026 | 0.146 | 1.06E-68 | 3.88E-64 |
| LRRC7 | 1 | -1.233254486 | 0.233 | 0.341 | 1.11E-68 | 4.05E-64 |
| BCHE | 1 | -0.492436407 | 0.028 | 0.152 | 1.37E-68 | 5.02E-64 |
| GRAMD1B | 1 | -0.474172201 | 0.062 | 0.211 | 2.04E-68 | 7.48E-64 |
| SHANK2 | 1 | 0.567072668 | 0.357 | 0.174 | 2.29E-68 | 8.38E-64 |
| ASXL3 | 1 | 0.569124569 | 0.623 | 0.431 | 2.44E-68 | 8.92E-64 |
| ITGA1 | 1 | -0.631462359 | 0.033 | 0.158 | 3.33E-68 | 1.22E-63 |
| ESYT2 | 1 | 0.692115268 | 0.661 | 0.519 | 3.78E-68 | 1.38E-63 |
| LHFPL3 | 1 | -0.788730788 | 0.46 | 0.601 | 4.72E-68 | 1.73E-63 |
| COL27A1 | 1 | -0.636496154 | 0.251 | 0.429 | 5.56E-68 | 2.04E-63 |
| MAMLD1 | 1 | -0.636165922 | 0.181 | 0.355 | 6.11E-68 | 2.24E-63 |
| XYLT1 | 1 | -0.5407597 | 0.04 | 0.17 | 1.21E-67 | 4.44E-63 |
| EVL | 1 | -0.492131696 | 0.426 | 0.606 | 2.56E-67 | 9.37E-63 |
| NIPA1 | 1 | 0.449387749 | 0.365 | 0.189 | 2.65E-67 | 9.7E-63 |
| MTUS2 | 1 | 0.579433724 | 0.344 | 0.166 | 2.77E-67 | 1.01E-62 |
| GLB1L3 | 1 | -0.30588114 | 0.013 | 0.117 | 3.46E-67 | 1.27E-62 |
| POLA1 | 1 | -0.608864654 | 0.803 | 0.858 | 4.43E-67 | 1.62E-62 |
| VSIG10 | 1 | 0.475158994 | 0.375 | 0.197 | 5.84E-67 | 2.14E-62 |

| | | | | | | |
|---|---|---|---|---|---|---|
| EDA | 1 | -0.828321329 | 0.158 | 0.322 | 9.4E-67 | 3.44E-62 |
| MBP | 1 | 0.354657133 | 0.238 | 0.089 | 1.03E-66 | 3.77E-62 |
| NFASC | 1 | -0.493798775 | 0.066 | 0.215 | 1.62E-66 | 5.95E-62 |
| RAB3IP | 1 | -0.354376176 | 0.04 | 0.172 | 1.7E-66 | 6.22E-62 |
| TSC22D1 | 1 | -0.582894193 | 0.641 | 0.732 | 1.97E-66 | 7.2E-62 |
| CECR2 | 1 | -0.592745575 | 0.175 | 0.357 | 2.11E-66 | 7.72E-62 |
| KLF3 | 1 | 0.391229102 | 0.269 | 0.115 | 2.77E-66 | 1.01E-61 |
| CDC16 | 1 | 0.455254998 | 0.583 | 0.41 | 2.89E-66 | 1.06E-61 |
| ETV1 | 1 | -0.815112169 | 0.088 | 0.235 | 3.15E-66 | 1.15E-61 |
| AC011476.3 | 1 | 0.29653654 | 0.167 | 0.047 | 4.82E-66 | 1.76E-61 |
| PEX5L | 1 | 0.389499725 | 0.883 | 0.725 | 7.58E-66 | 2.77E-61 |
| AC117453.1 | 1 | -0.313997129 | 0.008 | 0.102 | 7.99E-66 | 2.93E-61 |
| TPD52 | 1 | 0.513081181 | 0.512 | 0.31 | 8.55E-66 | 3.13E-61 |
| LINC01170 | 1 | 0.4994201 | 0.735 | 0.534 | 9.11E-66 | 3.33E-61 |
| ARSJ | 1 | -0.474302437 | 0.025 | 0.141 | 1.22E-65 | 4.48E-61 |
| PPHLN1 | 1 | 0.501686613 | 0.737 | 0.632 | 1.32E-65 | 4.82E-61 |
| ADGRA3 | 1 | -0.476767128 | 0.562 | 0.72 | 1.4E-65 | 5.12E-61 |
| UNC13B | 1 | -0.660499698 | 0.422 | 0.578 | 2.16E-65 | 7.89E-61 |
| TAFA1 | 1 | 0.580760964 | 0.363 | 0.181 | 2.2E-65 | 8.05E-61 |
| CSGALNACT1 | 1 | 0.533095354 | 0.233 | 0.087 | 3.67E-65 | 1.34E-60 |
| LRP1B | 1 | -1.232405036 | 0.094 | 0.233 | 4.36E-65 | 1.6E-60 |
| GALNT2 | 1 | -0.521148148 | 0.269 | 0.445 | 4.78E-65 | 1.75E-60 |
| NNT | 1 | 0.513877596 | 0.558 | 0.454 | 5.07E-65 | 1.86E-60 |
| ZFAS1 | 1 | 0.556415694 | 0.465 | 0.303 | 9.11E-65 | 3.33E-60 |
| PIAS2 | 1 | 0.522065824 | 0.694 | 0.588 | 9.51E-65 | 3.48E-60 |
| AL157778.1 | 1 | -0.490389393 | 0.285 | 0.223 | 9.74E-65 | 3.57E-60 |
| LINC02307 | 1 | 0.455366342 | 0.248 | 0.097 | 1.05E-64 | 3.86E-60 |
| EPB41L4A | 1 | -0.482376915 | 0.024 | 0.136 | 1.08E-64 | 3.95E-60 |
| PAK1 | 1 | 0.508698554 | 0.693 | 0.534 | 1.11E-64 | 4.06E-60 |
| HULC | 1 | 1.096225729 | 0.512 | 0.382 | 1.47E-64 | 5.38E-60 |
| ELOVL4 | 1 | 0.388124365 | 0.229 | 0.088 | 1.49E-64 | 5.44E-60 |
| LINC-PINT | 1 | 0.271590552 | 0.195 | 0.063 | 1.64E-64 | 6E-60 |
| UNC80 | 1 | -0.528574718 | 0.09 | 0.247 | 2E-64 | 7.33E-60 |
| RERE | 1 | -0.362304208 | 0.942 | 0.979 | 2.39E-64 | 8.76E-60 |
| TRERF1 | 1 | 0.455819218 | 0.717 | 0.599 | 3.01E-64 | 1.1E-59 |
| LGR4 | 1 | 0.526345172 | 0.589 | 0.398 | 3.32E-64 | 1.21E-59 |
| LINC00276 | 1 | 1.310818112 | 0.168 | 0.055 | 3.82E-64 | 1.4E-59 |
| CEP85L | 1 | 0.504808687 | 0.66 | 0.514 | 3.93E-64 | 1.44E-59 |
| KCNH7 | 1 | -0.792946889 | 0.042 | 0.169 | 4.54E-64 | 1.66E-59 |
| SCARB2 | 1 | -0.436774343 | 0.26 | 0.456 | 5E-64 | 1.83E-59 |
| RNF212 | 1 | 0.412003765 | 0.338 | 0.166 | 5.59E-64 | 2.05E-59 |
| CREB3L2 | 1 | -0.445580026 | 0.127 | 0.292 | 6.08E-64 | 2.22E-59 |
| RAPGEF2 | 1 | 0.580718068 | 0.747 | 0.625 | 8.53E-64 | 3.12E-59 |
| JAZF1 | 1 | 0.42763531 | 0.814 | 0.705 | 1.09E-63 | 3.99E-59 |
| CBFA2T3 | 1 | -0.350068476 | 0.059 | 0.201 | 1.19E-63 | 4.37E-59 |
| GLP1R | 1 | 0.317532201 | 0.161 | 0.046 | 1.38E-63 | 5.04E-59 |
| SNAP25 | 1 | -0.479625674 | 0.161 | 0.34 | 1.43E-63 | 5.23E-59 |
| AC061958.1 | 1 | -0.56616978 | 0.184 | 0.352 | 1.44E-63 | 5.26E-59 |
| CLIP4 | 1 | -0.407121281 | 0.173 | 0.357 | 2.01E-63 | 7.36E-59 |
| NEFL | 1 | 0.29540038 | 0.164 | 0.046 | 2.32E-63 | 8.51E-59 |
| CTCFL | 1 | 0.316558187 | 0.177 | 0.054 | 3.2E-63 | 1.17E-58 |
| PPP1R9A | 1 | -0.702463857 | 0.069 | 0.202 | 3.2E-63 | 1.17E-58 |
| MAP4K3-DT | 1 | 0.469082524 | 0.533 | 0.347 | 3.21E-63 | 1.18E-58 |
| GABRG3 | 1 | 0.481896122 | 0.188 | 0.06 | 3.25E-63 | 1.19E-58 |
| GLCCI1 | 1 | -0.714560857 | 0.46 | 0.616 | 3.29E-63 | 1.2E-58 |
| ADARB1 | 1 | -0.558959484 | 0.159 | 0.327 | 3.47E-63 | 1.27E-58 |
| MBNL3 | 1 | -0.601271751 | 0.36 | 0.532 | 4.94E-63 | 1.81E-58 |
| NDFIP1 | 1 | 0.500566287 | 0.529 | 0.367 | 5.99E-63 | 2.19E-58 |

| | | | | | | |
|---|---|---|---|---|---|---|
| FBXL7 | 1 | -0.560655632 | 0.796 | 0.849 | 7.48E-63 | 2.74E-58 |
| ERICH1 | 1 | 0.440911619 | 0.397 | 0.222 | 8.19E-63 | 3E-58 |
| PLEKHH2 | 1 | -0.426239243 | 0.063 | 0.205 | 1.17E-62 | 4.29E-58 |
| RYR3 | 1 | -0.624862419 | 0.028 | 0.135 | 1.2E-62 | 4.4E-58 |
| LINC01776 | 1 | 0.502259231 | 0.314 | 0.148 | 1.84E-62 | 6.72E-58 |
| SRGAP1 | 1 | -0.488273292 | 0.278 | 0.476 | 1.94E-62 | 7.11E-58 |
| AGBL4 | 1 | -0.93931319 | 0.162 | 0.222 | 2.55E-62 | 9.33E-58 |
| NUP93 | 1 | 0.475908929 | 0.499 | 0.327 | 2.64E-62 | 9.65E-58 |
| ITPR2 | 1 | -0.742628646 | 0.224 | 0.384 | 2.89E-62 | 1.06E-57 |
| DENND2A | 1 | 0.570442515 | 0.457 | 0.29 | 3.17E-62 | 1.16E-57 |
| PAPSS2 | 1 | 0.456067404 | 0.281 | 0.13 | 3.43E-62 | 1.25E-57 |
| KLHL29 | 1 | -0.720377259 | 0.529 | 0.645 | 6.28E-62 | 2.3E-57 |
| SSH2 | 1 | -0.463683526 | 0.707 | 0.832 | 6.84E-62 | 2.5E-57 |
| PCBP3 | 1 | -0.548573738 | 0.125 | 0.289 | 9.37E-62 | 3.43E-57 |
| APBA2 | 1 | 0.503310157 | 0.683 | 0.536 | 1.05E-61 | 3.84E-57 |
| TSHZ3 | 1 | -0.623850242 | 0.36 | 0.536 | 1.31E-61 | 4.78E-57 |
| RIMKLA | 1 | 0.299580189 | 0.194 | 0.066 | 1.79E-61 | 6.55E-57 |
| NAPEPLD | 1 | 0.483413765 | 0.475 | 0.317 | 2.44E-61 | 8.95E-57 |
| FAM204A | 1 | 0.489922724 | 0.494 | 0.329 | 2.7E-61 | 9.9E-57 |
| KCNJ2 | 1 | 0.268235726 | 0.135 | 0.033 | 3.06E-61 | 1.12E-56 |
| PRTG | 1 | -0.731699473 | 0.018 | 0.118 | 4.04E-61 | 1.48E-56 |
| RETREG1 | 1 | 0.488487305 | 0.283 | 0.135 | 5.73E-61 | 2.1E-56 |
| DOCK8 | 1 | -0.543798622 | 0.01 | 0.101 | 6.67E-61 | 2.44E-56 |
| SPATA6 | 1 | -0.495329352 | 0.076 | 0.205 | 6.95E-61 | 2.55E-56 |
| LINC01414 | 1 | -0.707971279 | 0.013 | 0.106 | 7.35E-61 | 2.69E-56 |
| DCDC2 | 1 | -0.519844155 | 0.013 | 0.109 | 8.72E-61 | 3.19E-56 |
| ZNF43 | 1 | 0.407570343 | 0.41 | 0.228 | 9.51E-61 | 3.48E-56 |
| PPM1L | 1 | -0.578345817 | 0.396 | 0.56 | 9.63E-61 | 3.52E-56 |
| CDH20 | 1 | -0.507997383 | 0.02 | 0.125 | 1.49E-60 | 5.45E-56 |
| PARD6G-AS1 | 1 | 0.303972337 | 0.191 | 0.064 | 1.7E-60 | 6.23E-56 |
| LIFR | 1 | 0.61034218 | 0.757 | 0.595 | 1.99E-60 | 7.29E-56 |
| KLHL1 | 1 | -0.888243485 | 0.022 | 0.129 | 2.05E-60 | 7.51E-56 |
| ATP8A1 | 1 | -0.750337166 | 0.311 | 0.442 | 2.25E-60 | 8.25E-56 |
| ANXA6 | 1 | -0.304938355 | 0.072 | 0.217 | 2.53E-60 | 9.26E-56 |
| FBN1 | 1 | -0.428976391 | 0.076 | 0.219 | 2.61E-60 | 9.56E-56 |
| RBM47 | 1 | -0.770156511 | 0.089 | 0.142 | 2.77E-60 | 1.01E-55 |
| LDHA | 1 | 0.411839391 | 0.207 | 0.077 | 3.04E-60 | 1.11E-55 |
| GPR143 | 1 | 0.312163722 | 0.165 | 0.05 | 3.05E-60 | 1.12E-55 |
| LINC01550 | 1 | 0.27593876 | 0.152 | 0.042 | 3.64E-60 | 1.33E-55 |
| GPR158 | 1 | -0.593555544 | 0.048 | 0.172 | 3.79E-60 | 1.39E-55 |
| DNAH6 | 1 | -0.503355683 | 0.01 | 0.1 | 3.85E-60 | 1.41E-55 |
| MYO5A | 1 | 0.469967823 | 0.709 | 0.553 | 3.86E-60 | 1.41E-55 |
| OPTN | 1 | 0.408783111 | 0.281 | 0.128 | 5.3E-60 | 1.94E-55 |
| SEPTIN9 | 1 | -0.523327936 | 0.329 | 0.506 | 8.03E-60 | 2.94E-55 |
| PTGFRN | 1 | 0.390614114 | 0.471 | 0.281 | 8.92E-60 | 3.27E-55 |
| SASH1 | 1 | 0.533868929 | 0.697 | 0.518 | 1.04E-59 | 3.8E-55 |
| RIMS1 | 1 | -0.752356141 | 0.326 | 0.473 | 1.53E-59 | 5.6E-55 |
| SOBP | 1 | 0.421649194 | 0.849 | 0.753 | 2.47E-59 | 9.03E-55 |
| METTL24 | 1 | -0.35171136 | 0.024 | 0.13 | 2.78E-59 | 1.02E-54 |
| ANXA1 | 1 | 0.299997871 | 0.146 | 0.04 | 4.72E-59 | 1.73E-54 |
| HIBADH | 1 | -0.489401469 | 0.365 | 0.544 | 6.05E-59 | 2.21E-54 |
| COL5A2 | 1 | 0.703879099 | 0.604 | 0.429 | 6.99E-59 | 2.56E-54 |
| DCBLD1 | 1 | 0.438769197 | 0.298 | 0.151 | 8.62E-59 | 3.16E-54 |
| CYTOR | 1 | 0.273919748 | 0.263 | 0.116 | 1.18E-58 | 4.32E-54 |
| SLC22A3 | 1 | -0.644571656 | 0.284 | 0.448 | 1.69E-58 | 6.19E-54 |
| SLC7A2 | 1 | -0.42122579 | 0.042 | 0.164 | 1.7E-58 | 6.22E-54 |
| VGLL3 | 1 | -0.952118483 | 0.216 | 0.342 | 2.03E-58 | 7.44E-54 |
| SNHG3 | 1 | 0.408619399 | 0.315 | 0.156 | 2.04E-58 | 7.46E-54 |

| | | | | | | |
|---|---|---|---|---|---|---|
| ANKH | 1 | -0.498041447 | 0.099 | 0.239 | 2.15E-58 | 7.87E-54 |
| ROR1 | 1 | -0.910481277 | 0.161 | 0.283 | 2.56E-58 | 9.37E-54 |
| C5orf17 | 1 | -0.59518705 | 0.022 | 0.126 | 3.11E-58 | 1.14E-53 |
| DPF3 | 1 | 0.401134621 | 0.406 | 0.226 | 3.12E-58 | 1.14E-53 |
| DOK5 | 1 | -0.674380813 | 0.469 | 0.571 | 3.31E-58 | 1.21E-53 |
| LRATD1 | 1 | 0.330960176 | 0.238 | 0.097 | 3.86E-58 | 1.41E-53 |
| PTPRD | 1 | 0.296249465 | 0.918 | 0.815 | 5.03E-58 | 1.84E-53 |
| LIN7A | 1 | -0.498060672 | 0.228 | 0.407 | 5.28E-58 | 1.93E-53 |
| SOX6 | 1 | -0.925257757 | 0.176 | 0.261 | 6.45E-58 | 2.36E-53 |
| FAM189A2 | 1 | 0.261825283 | 0.123 | 0.028 | 7.55E-58 | 2.77E-53 |
| COL4A5 | 1 | -1.151428804 | 0.036 | 0.134 | 9.53E-58 | 3.49E-53 |
| PRPF18 | 1 | 0.414916951 | 0.343 | 0.191 | 1.19E-57 | 4.35E-53 |
| LRMDA | 1 | -0.79617727 | 0.038 | 0.141 | 1.92E-57 | 7.03E-53 |
| SPAG9 | 1 | -0.375131128 | 0.773 | 0.86 | 2.53E-57 | 9.25E-53 |
| EHBP1 | 1 | -0.415537788 | 0.742 | 0.854 | 2.98E-57 | 1.09E-52 |
| SYNE1 | 1 | 0.758583235 | 0.581 | 0.441 | 3.69E-57 | 1.35E-52 |
| MSC-AS1 | 1 | -0.597818038 | 0.064 | 0.179 | 3.78E-57 | 1.38E-52 |
| RNF19A | 1 | 0.494457097 | 0.624 | 0.468 | 4.77E-57 | 1.74E-52 |
| SHROOM3 | 1 | -0.412265708 | 0.019 | 0.118 | 4.92E-57 | 1.8E-52 |
| FUT9 | 1 | -0.404609713 | 0.019 | 0.117 | 6.09E-57 | 2.23E-52 |
| PLEKHA1 | 1 | -0.418790993 | 0.221 | 0.395 | 7.06E-57 | 2.58E-52 |
| PTGER3 | 1 | -0.636490434 | 0.29 | 0.424 | 1.01E-56 | 3.72E-52 |
| APCDD1L-DT | 1 | 0.373205823 | 0.157 | 0.05 | 1.27E-56 | 4.63E-52 |
| LINC02133 | 1 | 0.252590532 | 0.111 | 0.023 | 1.34E-56 | 4.89E-52 |
| GNAQ | 1 | -0.448630596 | 0.629 | 0.757 | 1.34E-56 | 4.92E-52 |
| ITGA8 | 1 | -0.99496045 | 0.167 | 0.288 | 1.54E-56 | 5.64E-52 |
| ZC2HC1A | 1 | -0.449119381 | 0.261 | 0.436 | 1.9E-56 | 6.96E-52 |
| STK17A | 1 | 0.348084031 | 0.242 | 0.105 | 2.34E-56 | 8.56E-52 |
| PRSS12 | 1 | 0.41053542 | 0.897 | 0.766 | 2.63E-56 | 9.64E-52 |
| RASGRF2 | 1 | -0.716846409 | 0.085 | 0.212 | 3.05E-56 | 1.12E-51 |
| FAR2 | 1 | -0.741228456 | 0.274 | 0.421 | 3.11E-56 | 1.14E-51 |
| BRIP1 | 1 | -0.638283925 | 0.484 | 0.587 | 3.2E-56 | 1.17E-51 |
| SLC5A4-AS1 | 1 | -0.308257514 | 0.04 | 0.158 | 3.36E-56 | 1.23E-51 |
| SLC9A6 | 1 | 0.383457985 | 0.372 | 0.205 | 4.13E-56 | 1.51E-51 |
| BACH1 | 1 | -0.53646327 | 0.348 | 0.506 | 4.33E-56 | 1.59E-51 |
| NELL2 | 1 | -0.799309636 | 0.162 | 0.288 | 6.54E-56 | 2.4E-51 |
| ABCC5 | 1 | 0.467378647 | 0.479 | 0.342 | 9.11E-56 | 3.34E-51 |
| GPCPD1 | 1 | 0.499731007 | 0.675 | 0.537 | 1.27E-55 | 4.66E-51 |
| ZNF521 | 1 | -0.767361512 | 0.055 | 0.173 | 1.32E-55 | 4.85E-51 |
| REPS2 | 1 | 0.565543476 | 0.446 | 0.291 | 1.6E-55 | 5.86E-51 |
| NMU | 1 | 0.485696532 | 0.377 | 0.211 | 1.66E-55 | 6.07E-51 |
| CROCC | 1 | 0.495375943 | 0.561 | 0.406 | 2.84E-55 | 1.04E-50 |
| DCLK1 | 1 | 0.401519331 | 0.737 | 0.551 | 3.06E-55 | 1.12E-50 |
| EPHA3 | 1 | -1.004229105 | 0.127 | 0.245 | 3.12E-55 | 1.14E-50 |
| SIMC1 | 1 | -0.421633954 | 0.32 | 0.501 | 3.28E-55 | 1.2E-50 |
| MYT1 | 1 | 0.29870102 | 0.181 | 0.062 | 3.46E-55 | 1.27E-50 |
| ARHGAP31 | 1 | 0.394141103 | 0.355 | 0.188 | 3.6E-55 | 1.32E-50 |
| LIN28B-AS1 | 1 | 0.261851598 | 0.169 | 0.055 | 4.04E-55 | 1.48E-50 |
| SATB1-AS1 | 1 | 0.390153623 | 0.293 | 0.14 | 4.14E-55 | 1.52E-50 |
| GLRB | 1 | -0.311039292 | 0.031 | 0.139 | 5.82E-55 | 2.13E-50 |
| ICA1L | 1 | -0.368924192 | 0.081 | 0.219 | 6.62E-55 | 2.42E-50 |
| HDAC2 | 1 | 0.509036129 | 0.673 | 0.567 | 8.35E-55 | 3.06E-50 |
| SLF2 | 1 | -0.438720344 | 0.325 | 0.492 | 9.85E-55 | 3.61E-50 |
| LINC01515 | 1 | -0.635832844 | 0.099 | 0.217 | 1.28E-54 | 4.7E-50 |
| PTPN13 | 1 | -0.553305326 | 0.05 | 0.169 | 1.29E-54 | 4.71E-50 |
| ASCC3 | 1 | -0.448361964 | 0.696 | 0.801 | 1.31E-54 | 4.79E-50 |
| AC105916.1 | 1 | -0.309875799 | 0.027 | 0.131 | 1.49E-54 | 5.44E-50 |
| COX7A2L | 1 | 0.374565746 | 0.277 | 0.133 | 1.58E-54 | 5.8E-50 |

| | | | | | | |
|---|---|---|---|---|---|---|
| MEG3 | 1 | 0.542544264 | 0.581 | 0.404 | 1.62E-54 | 5.92E-50 |
| CHD2 | 1 | 0.432524871 | 0.846 | 0.773 | 1.92E-54 | 7.02E-50 |
| LRIG3 | 1 | -0.472243192 | 0.134 | 0.284 | 2.09E-54 | 7.65E-50 |
| STARD9 | 1 | -0.410242548 | 0.246 | 0.42 | 2.37E-54 | 8.67E-50 |
| SIPA1L3 | 1 | -0.757593848 | 0.423 | 0.558 | 2.54E-54 | 9.29E-50 |
| AC002460.2 | 1 | 0.451181953 | 0.128 | 0.034 | 3.06E-54 | 1.12E-49 |
| NAV1 | 1 | -0.469558463 | 0.28 | 0.456 | 3.29E-54 | 1.2E-49 |
| SH3BP4 | 1 | 0.405745261 | 0.471 | 0.289 | 3.31E-54 | 1.21E-49 |
| CRYM | 1 | -0.388192819 | 0.024 | 0.124 | 4.36E-54 | 1.6E-49 |
| RNF217 | 1 | -0.432497705 | 0.265 | 0.442 | 6.78E-54 | 2.48E-49 |
| SPIDR | 1 | -0.433341814 | 0.648 | 0.78 | 6.92E-54 | 2.54E-49 |
| STAT6 | 1 | 0.326136841 | 0.283 | 0.135 | 7.48E-54 | 2.74E-49 |
| AC109466.1 | 1 | -1.13757314 | 0.036 | 0.137 | 1.48E-53 | 5.42E-49 |
| AC117461.1 | 1 | -0.316954303 | 0.018 | 0.113 | 1.86E-53 | 6.82E-49 |
| ADAMTS19 | 1 | 0.597590564 | 0.62 | 0.442 | 2.02E-53 | 7.38E-49 |
| PDGFRB | 1 | -0.356026703 | 0.032 | 0.14 | 2.87E-53 | 1.05E-48 |
| SGCD | 1 | 0.661437354 | 0.608 | 0.447 | 3.35E-53 | 1.23E-48 |
| ADGRL3 | 1 | -0.800993659 | 0.797 | 0.82 | 3.77E-53 | 1.38E-48 |
| PTPRN2 | 1 | -1.253617006 | 0.172 | 0.306 | 4.1E-53 | 1.5E-48 |
| NADK | 1 | 0.368019478 | 0.346 | 0.188 | 4.14E-53 | 1.51E-48 |
| ALMS1 | 1 | -0.432298462 | 0.375 | 0.532 | 5.3E-53 | 1.94E-48 |
| PRKCI | 1 | 0.434308783 | 0.593 | 0.44 | 6E-53 | 2.2E-48 |
| CEMIP2 | 1 | -0.507097192 | 0.219 | 0.39 | 8.15E-53 | 2.99E-48 |
| TBX15 | 1 | -0.562285107 | 0.405 | 0.575 | 8.77E-53 | 3.21E-48 |
| ARHGEF10 | 1 | -0.381383921 | 0.15 | 0.311 | 8.89E-53 | 3.25E-48 |
| TUNAR | 1 | -0.400359023 | 0.021 | 0.114 | 1.35E-52 | 4.94E-48 |
| ARL5A | 1 | 0.502723513 | 0.525 | 0.408 | 1.61E-52 | 5.89E-48 |
| CPED1 | 1 | -0.460274296 | 0.013 | 0.1 | 3.01E-52 | 1.1E-47 |
| MEGF6 | 1 | -0.26377586 | 0.032 | 0.139 | 3.08E-52 | 1.13E-47 |
| NDRG1 | 1 | 0.801696262 | 0.181 | 0.089 | 3.37E-52 | 1.23E-47 |
| L3MBTL4 | 1 | 0.588155102 | 0.65 | 0.492 | 8.53E-52 | 3.12E-47 |
| SLC25A3 | 1 | 0.396135848 | 0.401 | 0.25 | 1.3E-51 | 4.77E-47 |
| PKIB | 1 | 0.501141091 | 0.518 | 0.351 | 1.49E-51 | 5.45E-47 |
| CAMK2D | 1 | -0.487178905 | 0.45 | 0.612 | 2.08E-51 | 7.61E-47 |
| PHACTR3 | 1 | -0.39297262 | 0.041 | 0.151 | 2.22E-51 | 8.12E-47 |
| UBE3A | 1 | 0.370907861 | 0.784 | 0.698 | 2.29E-51 | 8.39E-47 |
| SYNE2 | 1 | -0.469391241 | 0.621 | 0.731 | 2.69E-51 | 9.85E-47 |
| LINC01163 | 1 | -0.428864094 | 0.016 | 0.106 | 3.06E-51 | 1.12E-46 |
| CNTLN | 1 | -0.464265955 | 0.615 | 0.739 | 3.41E-51 | 1.25E-46 |
| CBLB | 1 | -0.579147504 | 0.508 | 0.617 | 3.51E-51 | 1.28E-46 |
| OTUD7A | 1 | -0.579450277 | 0.472 | 0.599 | 4.19E-51 | 1.53E-46 |
| AC104365.1 | 1 | 0.503106055 | 0.151 | 0.058 | 5.87E-51 | 2.15E-46 |
| CEP135 | 1 | 0.419142107 | 0.478 | 0.317 | 8.4E-51 | 3.08E-46 |
| ZNF438 | 1 | -0.462924155 | 0.172 | 0.326 | 8.66E-51 | 3.17E-46 |
| GAPDH | 1 | 0.494888 | 0.365 | 0.217 | 1.01E-50 | 3.71E-46 |
| AR | 1 | 0.289537748 | 0.105 | 0.024 | 1.02E-50 | 3.74E-46 |
| ELOVL2 | 1 | -0.345283411 | 0.054 | 0.172 | 1.21E-50 | 4.43E-46 |
| DLEU2 | 1 | -0.511542951 | 0.701 | 0.759 | 1.27E-50 | 4.67E-46 |
| BBS9 | 1 | -0.528477846 | 0.356 | 0.502 | 1.62E-50 | 5.91E-46 |
| PRKG2 | 1 | 0.284705595 | 0.168 | 0.058 | 1.72E-50 | 6.31E-46 |
| KCNK12 | 1 | 0.254256351 | 0.159 | 0.052 | 2.07E-50 | 7.57E-46 |
| MEG8 | 1 | 0.651398023 | 0.699 | 0.532 | 2.09E-50 | 7.64E-46 |
| RSPRY1 | 1 | 0.388628121 | 0.49 | 0.323 | 2.36E-50 | 8.64E-46 |
| ALDH1A2 | 1 | 0.325278528 | 0.145 | 0.045 | 2.94E-50 | 1.08E-45 |
| CRYBG3 | 1 | -0.561227889 | 0.215 | 0.348 | 3.09E-50 | 1.13E-45 |
| STIM2 | 1 | -0.473818588 | 0.368 | 0.501 | 3.49E-50 | 1.28E-45 |
| CACNB2 | 1 | 0.33988144 | 0.827 | 0.702 | 3.6E-50 | 1.32E-45 |
| IRS1 | 1 | -0.335007351 | 0.068 | 0.194 | 3.7E-50 | 1.36E-45 |

| | | | | | | |
|---|---|---|---|---|---|---|
| MOCOS | 1 | -0.312690573 | 0.039 | 0.145 | 4.66E-50 | 1.71E-45 |
| NID1 | 1 | -0.436322824 | 0.167 | 0.321 | 4.94E-50 | 1.81E-45 |
| SON | 1 | 0.38386202 | 0.761 | 0.657 | 5.07E-50 | 1.86E-45 |
| VAV2 | 1 | -0.444639059 | 0.198 | 0.361 | 5.39E-50 | 1.97E-45 |
| GALNT13 | 1 | -0.569491353 | 0.652 | 0.719 | 5.42E-50 | 1.98E-45 |
| LBX1-AS1 | 1 | 0.284406164 | 0.246 | 0.111 | 7.19E-50 | 2.63E-45 |
| LRFN5 | 1 | -0.899113257 | 0.371 | 0.489 | 9.45E-50 | 3.46E-45 |
| MIR646HG | 1 | -0.65776668 | 0.04 | 0.141 | 9.93E-50 | 3.64E-45 |
| ESRRG | 1 | -0.750834328 | 0.317 | 0.451 | 1.04E-49 | 3.8E-45 |
| B4GALT5 | 1 | 0.422084632 | 0.528 | 0.383 | 1.86E-49 | 6.81E-45 |
| RBM25 | 1 | 0.372822999 | 0.798 | 0.683 | 2.54E-49 | 9.31E-45 |
| MANBA | 1 | -0.325647785 | 0.098 | 0.236 | 2.7E-49 | 9.9E-45 |
| HBEGF | 1 | -0.318245518 | 0.04 | 0.148 | 3.03E-49 | 1.11E-44 |
| LINC02018 | 1 | -0.323213176 | 0.042 | 0.151 | 3.13E-49 | 1.15E-44 |
| PCNX4 | 1 | 0.432799348 | 0.637 | 0.517 | 3.23E-49 | 1.18E-44 |
| TMX1 | 1 | 0.373395336 | 0.437 | 0.279 | 3.56E-49 | 1.31E-44 |
| PHLPP1 | 1 | -0.414911337 | 0.346 | 0.52 | 3.63E-49 | 1.33E-44 |
| CFAP58 | 1 | -0.632295096 | 0.198 | 0.329 | 4.18E-49 | 1.53E-44 |
| IRAK3 | 1 | -0.433008124 | 0.372 | 0.531 | 5.12E-49 | 1.88E-44 |
| SEPTIN10 | 1 | -0.360654025 | 0.151 | 0.299 | 5.96E-49 | 2.18E-44 |
| ENO1 | 1 | 0.579324939 | 0.575 | 0.436 | 8.32E-49 | 3.05E-44 |
| MAP7D2 | 1 | 0.339182232 | 0.41 | 0.243 | 9.52E-49 | 3.49E-44 |
| SLC2A1 | 1 | 0.471649041 | 0.2 | 0.101 | 9.71E-49 | 3.55E-44 |
| EFHD1 | 1 | -0.30116837 | 0.049 | 0.162 | 1.53E-48 | 5.6E-44 |
| CDKN2B-AS1 | 1 | -0.526808712 | 0.221 | 0.358 | 1.75E-48 | 6.42E-44 |
| NPAS2 | 1 | 0.390202937 | 0.276 | 0.136 | 1.82E-48 | 6.65E-44 |
| NRXN1 | 1 | -0.545695524 | 0.583 | 0.603 | 2.31E-48 | 8.47E-44 |
| CACNA1D | 1 | -0.586539134 | 0.333 | 0.45 | 2.39E-48 | 8.76E-44 |
| ANKDD1A | 1 | 0.385232481 | 0.26 | 0.129 | 3.26E-48 | 1.19E-43 |
| SLC4A10 | 1 | -0.522482819 | 0.021 | 0.107 | 4.21E-48 | 1.54E-43 |
| MARCH8 | 1 | -0.363503247 | 0.082 | 0.209 | 4.41E-48 | 1.61E-43 |
| AL591501.1 | 1 | -0.897197628 | 0.041 | 0.137 | 4.76E-48 | 1.74E-43 |
| DLX6-AS1 | 1 | -0.8235167 | 0.139 | 0.249 | 4.85E-48 | 1.78E-43 |
| TPST1 | 1 | -0.471867218 | 0.293 | 0.46 | 7.34E-48 | 2.69E-43 |
| AAK1 | 1 | 0.427135224 | 0.628 | 0.499 | 7.81E-48 | 2.86E-43 |
| SNHG1 | 1 | 0.406515986 | 0.418 | 0.271 | 9.18E-48 | 3.36E-43 |
| BMPR1B | 1 | -0.639716715 | 0.21 | 0.355 | 9.73E-48 | 3.56E-43 |
| CCDC144A | 1 | 0.365001362 | 0.29 | 0.149 | 1.12E-47 | 4.09E-43 |
| SLC39A8 | 1 | -0.289534588 | 0.038 | 0.142 | 1.15E-47 | 4.23E-43 |
| COL24A1 | 1 | -0.648205891 | 0.098 | 0.225 | 1.2E-47 | 4.4E-43 |
| SLC25A48 | 1 | 0.29188279 | 0.143 | 0.045 | 1.71E-47 | 6.27E-43 |
| ASAP2 | 1 | -0.48902203 | 0.277 | 0.425 | 2.28E-47 | 8.36E-43 |
| NIPBL | 1 | -0.306526024 | 0.829 | 0.902 | 3.16E-47 | 1.16E-42 |
| TFAP2B | 1 | -0.562732433 | 0.14 | 0.268 | 3.41E-47 | 1.25E-42 |
| ARFGEF3 | 1 | 0.318622501 | 0.595 | 0.417 | 3.59E-47 | 1.32E-42 |
| ARHGAP22 | 1 | 0.318681675 | 0.206 | 0.087 | 3.7E-47 | 1.35E-42 |
| NTNG1 | 1 | 0.478255058 | 0.601 | 0.436 | 3.73E-47 | 1.37E-42 |
| LNPK | 1 | -0.375989777 | 0.308 | 0.479 | 3.89E-47 | 1.42E-42 |
| ARID1B | 1 | -0.349067653 | 0.924 | 0.968 | 4.11E-47 | 1.5E-42 |
| KCTD1 | 1 | 0.495495991 | 0.417 | 0.281 | 4.17E-47 | 1.53E-42 |
| DHX29 | 1 | 0.426707883 | 0.428 | 0.297 | 5.15E-47 | 1.88E-42 |
| VPS8 | 1 | 0.434815414 | 0.673 | 0.579 | 6.47E-47 | 2.37E-42 |
| LINC00536 | 1 | -0.774231522 | 0.026 | 0.118 | 6.68E-47 | 2.45E-42 |
| ZMPSTE24 | 1 | 0.328392851 | 0.298 | 0.159 | 6.87E-47 | 2.51E-42 |
| SPTLC2 | 1 | 0.417223712 | 0.472 | 0.332 | 8.03E-47 | 2.94E-42 |
| AL049828.1 | 1 | -0.486385478 | 0.13 | 0.258 | 8.35E-47 | 3.06E-42 |
| SPRY4-AS1 | 1 | -0.438076487 | 0.082 | 0.201 | 8.58E-47 | 3.14E-42 |
| MMS22L | 1 | -0.460802544 | 0.525 | 0.629 | 9.98E-47 | 3.65E-42 |

| | | | | | | |
|---|---|---|---|---|---|---|
| ANKRD6 | 1 | 0.446233044 | 0.422 | 0.263 | 1.03E-46 | 3.78E-42 |
| NACC2 | 1 | 0.382495516 | 0.356 | 0.204 | 1.09E-46 | 4.01E-42 |
| ROCK2 | 1 | -0.377480196 | 0.581 | 0.718 | 1.13E-46 | 4.15E-42 |
| GNAL | 1 | -0.336974281 | 0.043 | 0.148 | 1.18E-46 | 4.32E-42 |
| LRRC8C | 1 | 0.356199345 | 0.265 | 0.133 | 1.6E-46 | 5.87E-42 |
| DOK6 | 1 | -0.580716545 | 0.545 | 0.658 | 1.64E-46 | 6.01E-42 |
| PCNX2 | 1 | -0.408798125 | 0.387 | 0.54 | 1.92E-46 | 7.04E-42 |
| KANSL1 | 1 | -0.30181222 | 0.849 | 0.917 | 2.02E-46 | 7.4E-42 |
| GAB2 | 1 | 0.560014194 | 0.42 | 0.275 | 2.49E-46 | 9.12E-42 |
| COL6A1 | 1 | -0.385549763 | 0.104 | 0.235 | 2.93E-46 | 1.07E-41 |
| ITPRID2 | 1 | -0.394683819 | 0.211 | 0.359 | 3.06E-46 | 1.12E-41 |
| SGO1-AS1 | 1 | -0.862791091 | 0.097 | 0.214 | 3.46E-46 | 1.27E-41 |
| CSRNP3 | 1 | -0.524370107 | 0.247 | 0.398 | 5.18E-46 | 1.9E-41 |
| PIR | 1 | 0.333789937 | 0.252 | 0.121 | 9.63E-46 | 3.53E-41 |
| ARHGAP6 | 1 | -0.671100478 | 0.252 | 0.394 | 1.01E-45 | 3.71E-41 |
| ZNF160 | 1 | 0.339634013 | 0.372 | 0.221 | 1.03E-45 | 3.78E-41 |
| SPATA5 | 1 | -0.451527801 | 0.534 | 0.654 | 1.1E-45 | 4.04E-41 |
| CADM2 | 1 | -0.751644394 | 0.453 | 0.446 | 1.11E-45 | 4.07E-41 |
| PDLIM5 | 1 | 0.457196292 | 0.522 | 0.351 | 1.12E-45 | 4.11E-41 |
| PGAP1 | 1 | -0.4216466 | 0.615 | 0.732 | 1.18E-45 | 4.33E-41 |
| LINC01456 | 1 | 0.314222846 | 0.221 | 0.099 | 1.22E-45 | 4.45E-41 |
| TBC1D8 | 1 | 0.369526958 | 0.312 | 0.172 | 1.24E-45 | 4.56E-41 |
| LRRFIP1 | 1 | -0.44206465 | 0.401 | 0.538 | 1.3E-45 | 4.77E-41 |
| MTF2 | 1 | 0.386825724 | 0.731 | 0.63 | 1.41E-45 | 5.18E-41 |
| PPFIA2 | 1 | -0.602799858 | 0.678 | 0.741 | 1.51E-45 | 5.53E-41 |
| SYT17 | 1 | -0.259809673 | 0.041 | 0.144 | 1.56E-45 | 5.73E-41 |
| GUCY1A2 | 1 | -0.605645482 | 0.285 | 0.434 | 1.59E-45 | 5.83E-41 |
| PLEKHG4B | 1 | -0.326934968 | 0.12 | 0.259 | 1.66E-45 | 6.08E-41 |
| PRPF40A | 1 | 0.411479361 | 0.713 | 0.617 | 1.93E-45 | 7.06E-41 |
| NIPAL2 | 1 | -0.262859653 | 0.03 | 0.124 | 1.94E-45 | 7.09E-41 |
| AKAP13 | 1 | -0.432745533 | 0.49 | 0.573 | 2.03E-45 | 7.42E-41 |
| AC135782.1 | 1 | -0.293572534 | 0.059 | 0.173 | 2.64E-45 | 9.67E-41 |
| RIMBP2 | 1 | -1.067989269 | 0.087 | 0.143 | 3.02E-45 | 1.1E-40 |
| IL6ST | 1 | -0.38410284 | 0.085 | 0.209 | 3.44E-45 | 1.26E-40 |
| DPYD | 1 | -0.627635603 | 0.325 | 0.437 | 3.48E-45 | 1.27E-40 |
| RIMS2 | 1 | -0.522885867 | 0.422 | 0.429 | 3.88E-45 | 1.42E-40 |
| TEX9 | 1 | -0.399779365 | 0.11 | 0.233 | 3.97E-45 | 1.45E-40 |
| GKAP1 | 1 | -0.386631492 | 0.133 | 0.267 | 4.13E-45 | 1.51E-40 |
| CEP112 | 1 | -0.540249251 | 0.3 | 0.429 | 4.22E-45 | 1.54E-40 |
| EVI5 | 1 | -0.445358935 | 0.473 | 0.585 | 5.02E-45 | 1.84E-40 |
| ZSWIM6 | 1 | -0.384716626 | 0.806 | 0.836 | 5.54E-45 | 2.03E-40 |
| KDM4A | 1 | 0.34752146 | 0.403 | 0.253 | 5.67E-45 | 2.08E-40 |
| SNX25 | 1 | 0.413127734 | 0.731 | 0.599 | 7.6E-45 | 2.78E-40 |
| GALNT18 | 1 | 0.434290786 | 0.525 | 0.357 | 9.31E-45 | 3.41E-40 |
| UNC13C | 1 | -0.875327557 | 0.025 | 0.101 | 9.6E-45 | 3.52E-40 |
| ZNF536 | 1 | -0.296148508 | 0.862 | 0.809 | 1.3E-44 | 4.77E-40 |
| COL5A1 | 1 | -0.463041134 | 0.06 | 0.167 | 1.31E-44 | 4.79E-40 |
| CFAP47 | 1 | -1.028466023 | 0.024 | 0.107 | 1.39E-44 | 5.09E-40 |
| DOP1A | 1 | -0.384438324 | 0.399 | 0.547 | 1.59E-44 | 5.84E-40 |
| RIPOR2 | 1 | -0.44372871 | 0.242 | 0.406 | 1.7E-44 | 6.23E-40 |
| AATK | 1 | -0.294052624 | 0.077 | 0.199 | 1.79E-44 | 6.57E-40 |
| MIR181A2HG | 1 | -0.415727765 | 0.168 | 0.31 | 2.24E-44 | 8.21E-40 |
| REV1 | 1 | -0.370601759 | 0.46 | 0.608 | 2.34E-44 | 8.56E-40 |
| ZEB1 | 1 | -0.512707723 | 0.544 | 0.566 | 2.44E-44 | 8.93E-40 |
| DMXL2 | 1 | -0.393035652 | 0.473 | 0.607 | 2.5E-44 | 9.14E-40 |
| XK | 1 | 0.267687408 | 0.194 | 0.081 | 2.52E-44 | 9.22E-40 |
| ZNF730 | 1 | 0.322322963 | 0.319 | 0.174 | 2.55E-44 | 9.33E-40 |
| CHRM2 | 1 | -0.671495619 | 0.03 | 0.117 | 2.78E-44 | 1.02E-39 |

| | | | | | | |
|---|---|---|---|---|---|---|
| SDCBP | 1 | 0.33186454 | 0.268 | 0.136 | 2.87E-44 | 1.05E-39 |
| TRAF3 | 1 | -0.427674942 | 0.347 | 0.493 | 3.34E-44 | 1.22E-39 |
| SYNJ2 | 1 | -0.491378869 | 0.104 | 0.219 | 3.38E-44 | 1.24E-39 |
| SLC10A7 | 1 | -0.419971971 | 0.209 | 0.349 | 3.46E-44 | 1.27E-39 |
| LRRC4C | 1 | -0.798581005 | 0.479 | 0.546 | 3.5E-44 | 1.28E-39 |
| TRPC6 | 1 | 0.432217148 | 0.172 | 0.066 | 3.6E-44 | 1.32E-39 |
| VPS13B | 1 | -0.355629373 | 0.762 | 0.847 | 4.12E-44 | 1.51E-39 |
| LINC01934 | 1 | -1.274506546 | 0.04 | 0.123 | 4.49E-44 | 1.65E-39 |
| OPCML | 1 | 0.715388764 | 0.453 | 0.309 | 5.55E-44 | 2.03E-39 |
| FEM1C | 1 | 0.366400981 | 0.247 | 0.128 | 6.44E-44 | 2.36E-39 |
| MAGED1 | 1 | 0.372267363 | 0.704 | 0.574 | 7.06E-44 | 2.58E-39 |
| TMEM132C | 1 | 0.578956214 | 0.197 | 0.097 | 7.12E-44 | 2.61E-39 |
| ABCA1 | 1 | 0.319600794 | 0.38 | 0.225 | 7.32E-44 | 2.68E-39 |
| CDC14A | 1 | -0.397289352 | 0.202 | 0.343 | 7.7E-44 | 2.82E-39 |
| CHCHD6 | 1 | -0.346284217 | 0.15 | 0.292 | 1.03E-43 | 3.79E-39 |
| EPHB1 | 1 | 0.426498037 | 0.257 | 0.13 | 1.2E-43 | 4.39E-39 |
| GABRB3 | 1 | 0.406467485 | 0.503 | 0.335 | 1.61E-43 | 5.88E-39 |
| AGTPBP1 | 1 | -0.394862512 | 0.506 | 0.629 | 1.63E-43 | 5.95E-39 |
| GAS5 | 1 | 0.40218472 | 0.473 | 0.326 | 2.25E-43 | 8.24E-39 |
| SEPTIN7 | 1 | 0.387072569 | 0.749 | 0.66 | 2.78E-43 | 1.02E-38 |
| DNER | 1 | 0.418524721 | 0.517 | 0.349 | 2.85E-43 | 1.04E-38 |
| COL4A1 | 1 | -0.405533821 | 0.056 | 0.161 | 3.41E-43 | 1.25E-38 |
| KDM6A | 1 | 0.32605956 | 0.902 | 0.892 | 3.43E-43 | 1.26E-38 |
| SIK2 | 1 | -0.399538566 | 0.437 | 0.583 | 3.6E-43 | 1.32E-38 |
| E2F7 | 1 | -0.412213258 | 0.099 | 0.217 | 4.22E-43 | 1.55E-38 |
| TM2D1 | 1 | 0.38206006 | 0.515 | 0.373 | 4.57E-43 | 1.67E-38 |
| ZFYVE16 | 1 | -0.447405304 | 0.488 | 0.6 | 4.67E-43 | 1.71E-38 |
| LPAR1 | 1 | -0.616685508 | 0.112 | 0.236 | 5.26E-43 | 1.93E-38 |
| LRCH3 | 1 | -0.369306788 | 0.513 | 0.658 | 5.61E-43 | 2.05E-38 |
| EPB41 | 1 | -0.458409092 | 0.395 | 0.511 | 5.75E-43 | 2.1E-38 |
| TKT | 1 | 0.331541268 | 0.259 | 0.131 | 5.82E-43 | 2.13E-38 |
| CCDC88A | 1 | -0.435807764 | 0.547 | 0.671 | 5.9E-43 | 2.16E-38 |
| NTN4 | 1 | -0.385665777 | 0.095 | 0.216 | 7.07E-43 | 2.59E-38 |
| OSBP2 | 1 | -0.310122251 | 0.107 | 0.236 | 8.08E-43 | 2.96E-38 |
| YTHDC1 | 1 | 0.371484711 | 0.675 | 0.564 | 8.11E-43 | 2.97E-38 |
| EOGT | 1 | 0.294985555 | 0.265 | 0.135 | 9.18E-43 | 3.36E-38 |
| PKD2 | 1 | -0.366556809 | 0.428 | 0.575 | 9.42E-43 | 3.45E-38 |
| PDK3 | 1 | 0.466088403 | 0.525 | 0.393 | 1.06E-42 | 3.87E-38 |
| EFR3B | 1 | 0.347193544 | 0.579 | 0.414 | 1.26E-42 | 4.61E-38 |
| FAF1 | 1 | -0.327345505 | 0.77 | 0.848 | 1.27E-42 | 4.66E-38 |
| ICA1 | 1 | -0.389646408 | 0.12 | 0.248 | 1.33E-42 | 4.88E-38 |
| NREP | 1 | -0.371016296 | 0.304 | 0.452 | 1.61E-42 | 5.89E-38 |
| NRG1 | 1 | -1.146334959 | 0.258 | 0.301 | 1.7E-42 | 6.22E-38 |
| CHRM3 | 1 | 0.492212945 | 0.545 | 0.377 | 2.05E-42 | 7.51E-38 |
| KCNH8 | 1 | -0.551552487 | 0.571 | 0.654 | 2.3E-42 | 8.44E-38 |
| BRINP1 | 1 | -0.627684252 | 0.31 | 0.457 | 2.33E-42 | 8.53E-38 |
| PDK1 | 1 | 0.583793423 | 0.479 | 0.348 | 2.39E-42 | 8.75E-38 |
| GTF2A2 | 1 | 0.266895182 | 0.193 | 0.084 | 2.42E-42 | 8.86E-38 |
| ME3 | 1 | -0.445907629 | 0.211 | 0.343 | 2.53E-42 | 9.27E-38 |
| DMXL1 | 1 | -0.38119131 | 0.424 | 0.564 | 2.55E-42 | 9.35E-38 |
| OSBPL1A | 1 | -0.415187361 | 0.326 | 0.477 | 2.56E-42 | 9.36E-38 |
| PLS1 | 1 | 0.357413942 | 0.312 | 0.172 | 2.64E-42 | 9.67E-38 |
| LARP4 | 1 | 0.38954777 | 0.589 | 0.461 | 2.73E-42 | 1E-37 |
| OSBPL8 | 1 | -0.375828695 | 0.523 | 0.661 | 3.87E-42 | 1.42E-37 |
| MYT1L | 1 | -0.613775492 | 0.052 | 0.155 | 5.43E-42 | 1.99E-37 |
| KIF6 | 1 | 0.38873826 | 0.149 | 0.054 | 6.42E-42 | 2.35E-37 |
| CLVS1 | 1 | -0.492022678 | 0.115 | 0.237 | 6.59E-42 | 2.41E-37 |
| ZC3H12C | 1 | -0.492998399 | 0.505 | 0.618 | 6.84E-42 | 2.51E-37 |

| | | | | | | |
|---|---|---|---|---|---|---|
| PLXDC2 | 1 | -0.458701981 | 0.658 | 0.672 | 6.93E-42 | 2.54E-37 |
| RASSF8 | 1 | 0.455498866 | 0.606 | 0.469 | 7.33E-42 | 2.68E-37 |
| TENM2 | 1 | -0.625006086 | 0.528 | 0.641 | 7.7E-42 | 2.82E-37 |
| AFDN | 1 | -0.372441216 | 0.742 | 0.811 | 1.01E-41 | 3.68E-37 |
| CAMSAP2 | 1 | -0.362865333 | 0.51 | 0.64 | 1.01E-41 | 3.71E-37 |
| GAS2 | 1 | -0.355413621 | 0.035 | 0.117 | 1.01E-41 | 3.71E-37 |
| SEMA5A | 1 | -0.927639186 | 0.341 | 0.429 | 1.02E-41 | 3.73E-37 |
| ZDBF2 | 1 | 0.272543785 | 0.292 | 0.156 | 1.03E-41 | 3.77E-37 |
| ACTR3B | 1 | 0.388550615 | 0.53 | 0.387 | 1.08E-41 | 3.94E-37 |
| CSMD2 | 1 | 0.45017298 | 0.424 | 0.269 | 1.08E-41 | 3.97E-37 |
| KCTD3 | 1 | -0.380360734 | 0.356 | 0.506 | 1.1E-41 | 4.01E-37 |
| FBXW2 | 1 | 0.350309364 | 0.488 | 0.353 | 1.19E-41 | 4.36E-37 |
| SRRM4 | 1 | -0.401504829 | 0.091 | 0.212 | 1.28E-41 | 4.7E-37 |
| REEP1 | 1 | -0.272714233 | 0.053 | 0.157 | 1.59E-41 | 5.82E-37 |
| ENAH | 1 | 0.372285688 | 0.788 | 0.696 | 1.75E-41 | 6.39E-37 |
| LNX1 | 1 | -0.291527909 | 0.051 | 0.153 | 1.92E-41 | 7.02E-37 |
| MAPRE2 | 1 | 0.437534186 | 0.443 | 0.303 | 2.02E-41 | 7.38E-37 |
| BEX2 | 1 | 0.268190246 | 0.156 | 0.061 | 2.12E-41 | 7.77E-37 |
| DGKH | 1 | 0.472826162 | 0.776 | 0.678 | 2.42E-41 | 8.85E-37 |
| SEC61A2 | 1 | 0.292023183 | 0.309 | 0.172 | 2.79E-41 | 1.02E-36 |
| AHDC1 | 1 | -0.285800212 | 0.093 | 0.214 | 2.99E-41 | 1.09E-36 |
| TC2N | 1 | 0.2769958 | 0.169 | 0.067 | 3.48E-41 | 1.27E-36 |
| CACNA2D1 | 1 | -0.861253768 | 0.817 | 0.866 | 3.5E-41 | 1.28E-36 |
| C12orf56 | 1 | -0.290848305 | 0.068 | 0.179 | 3.51E-41 | 1.28E-36 |
| MORC4 | 1 | -0.256040034 | 0.083 | 0.201 | 3.74E-41 | 1.37E-36 |
| RPS18 | 1 | 0.494549724 | 0.474 | 0.327 | 4.07E-41 | 1.49E-36 |
| CD200 | 1 | 0.293113396 | 0.253 | 0.127 | 4.77E-41 | 1.75E-36 |
| ZNF704 | 1 | 0.395623363 | 0.704 | 0.589 | 4.82E-41 | 1.76E-36 |
| PCMTD2 | 1 | 0.368849731 | 0.43 | 0.301 | 5.4E-41 | 1.98E-36 |
| KAT2B | 1 | -0.338166531 | 0.179 | 0.323 | 5.63E-41 | 2.06E-36 |
| SPATA13 | 1 | -0.472768914 | 0.152 | 0.258 | 5.75E-41 | 2.11E-36 |
| SLC25A16 | 1 | 0.369999157 | 0.417 | 0.281 | 6.73E-41 | 2.46E-36 |
| AGPAT3 | 1 | -0.342441415 | 0.265 | 0.421 | 7.29E-41 | 2.67E-36 |
| PLEKHA5 | 1 | -0.407897381 | 0.808 | 0.869 | 7.36E-41 | 2.7E-36 |
| PARVA | 1 | 0.377470218 | 0.504 | 0.357 | 7.68E-41 | 2.81E-36 |
| ZNF804B | 1 | -1.256503161 | 0.118 | 0.205 | 8.03E-41 | 2.94E-36 |
| SLC6A15 | 1 | -0.43334658 | 0.121 | 0.241 | 1.07E-40 | 3.93E-36 |
| FMNL2 | 1 | 0.396491505 | 0.899 | 0.867 | 1.13E-40 | 4.13E-36 |
| TPM3 | 1 | 0.35504263 | 0.328 | 0.198 | 1.13E-40 | 4.16E-36 |
| RNF144A | 1 | -0.417351772 | 0.404 | 0.536 | 1.32E-40 | 4.83E-36 |
| OXCT1 | 1 | -0.322325489 | 0.115 | 0.241 | 1.33E-40 | 4.87E-36 |
| NXPH2 | 1 | -0.299720827 | 0.043 | 0.14 | 1.7E-40 | 6.21E-36 |
| WWTR1 | 1 | -0.578108137 | 0.332 | 0.458 | 1.92E-40 | 7.04E-36 |
| FAT4 | 1 | -0.50371622 | 0.197 | 0.33 | 2.21E-40 | 8.09E-36 |
| HIBCH | 1 | 0.384682985 | 0.656 | 0.56 | 2.61E-40 | 9.54E-36 |
| RHBDL3 | 1 | -0.413205737 | 0.192 | 0.334 | 3.16E-40 | 1.16E-35 |
| UFL1-AS1 | 1 | -0.497400143 | 0.044 | 0.126 | 3.16E-40 | 1.16E-35 |
| PRKD3 | 1 | -0.377855101 | 0.3 | 0.445 | 3.29E-40 | 1.2E-35 |
| NGLY1 | 1 | 0.447274984 | 0.56 | 0.446 | 4.4E-40 | 1.61E-35 |
| EBF4 | 1 | -0.331407433 | 0.108 | 0.229 | 4.53E-40 | 1.66E-35 |
| TNFAIP8 | 1 | -0.276312201 | 0.027 | 0.109 | 5.39E-40 | 1.97E-35 |
| MYO3A | 1 | -0.886083986 | 0.138 | 0.194 | 5.54E-40 | 2.03E-35 |
| UBE2E1 | 1 | 0.382673612 | 0.686 | 0.587 | 5.56E-40 | 2.04E-35 |
| NFIX | 1 | -0.471115088 | 0.368 | 0.5 | 5.88E-40 | 2.15E-35 |
| NSD3 | 1 | -0.32802159 | 0.696 | 0.782 | 6.09E-40 | 2.23E-35 |
| EPCAM | 1 | 0.356307452 | 0.156 | 0.063 | 7.81E-40 | 2.86E-35 |
| AC090945.1 | 1 | 0.30287066 | 0.276 | 0.147 | 8.1E-40 | 2.97E-35 |
| CPM | 1 | 0.305294531 | 0.147 | 0.054 | 9.64E-40 | 3.53E-35 |

| | | | | | | |
|---|---|---|---|---|---|---|
| CCDC91 | 1 | -0.376667678 | 0.652 | 0.75 | 1.03E-39 | 3.78E-35 |
| GOLM1 | 1 | -0.347867246 | 0.233 | 0.377 | 1.11E-39 | 4.05E-35 |
| ARHGAP10 | 1 | -0.461400651 | 0.43 | 0.538 | 1.22E-39 | 4.47E-35 |
| HTR2C | 1 | -3.855774318 | 0.035 | 0.112 | 1.25E-39 | 4.59E-35 |
| HOMER2 | 1 | 0.399542736 | 0.413 | 0.279 | 1.33E-39 | 4.87E-35 |
| TUT4 | 1 | 0.338440668 | 0.889 | 0.857 | 1.69E-39 | 6.17E-35 |
| MAX | 1 | 0.352722697 | 0.407 | 0.259 | 1.7E-39 | 6.21E-35 |
| PPM1B | 1 | 0.372613131 | 0.617 | 0.477 | 1.95E-39 | 7.13E-35 |
| CLCN3 | 1 | 0.377174299 | 0.613 | 0.491 | 1.98E-39 | 7.26E-35 |
| SRPX | 1 | -0.340317162 | 0.133 | 0.262 | 2.74E-39 | 1E-34 |
| TAF1D | 1 | 0.374827213 | 0.739 | 0.656 | 2.88E-39 | 1.05E-34 |
| HECTD4 | 1 | 0.353997999 | 0.8 | 0.74 | 3.35E-39 | 1.23E-34 |
| IPO7 | 1 | 0.381586657 | 0.647 | 0.534 | 3.41E-39 | 1.25E-34 |
| HDAC4 | 1 | -0.371123491 | 0.579 | 0.707 | 4.57E-39 | 1.67E-34 |
| AC025809.1 | 1 | -0.255852476 | 0.046 | 0.141 | 5.49E-39 | 2.01E-34 |
| IGDCC3 | 1 | -0.383058096 | 0.166 | 0.288 | 5.83E-39 | 2.14E-34 |
| TXLNG | 1 | 0.361818871 | 0.501 | 0.362 | 6.84E-39 | 2.5E-34 |
| HSD17B4 | 1 | -0.339561989 | 0.207 | 0.342 | 7.02E-39 | 2.57E-34 |
| ADAM9 | 1 | 0.419597388 | 0.45 | 0.327 | 9.06E-39 | 3.32E-34 |
| SECISBP2L | 1 | 0.402954334 | 0.549 | 0.419 | 1.07E-38 | 3.91E-34 |
| KAZN | 1 | -0.360336152 | 0.904 | 0.897 | 1.31E-38 | 4.8E-34 |
| ZNF714 | 1 | 0.332586815 | 0.437 | 0.29 | 1.35E-38 | 4.94E-34 |
| PPP4R3A | 1 | 0.35071574 | 0.715 | 0.61 | 1.36E-38 | 4.96E-34 |
| FBXL17 | 1 | -0.353843988 | 0.695 | 0.789 | 1.48E-38 | 5.4E-34 |
| PARP14 | 1 | -0.290705623 | 0.094 | 0.211 | 1.51E-38 | 5.54E-34 |
| GYG2 | 1 | -0.300425382 | 0.067 | 0.17 | 1.59E-38 | 5.83E-34 |
| ADAMTS3 | 1 | -0.655718473 | 0.408 | 0.5 | 1.61E-38 | 5.88E-34 |
| SLC16A9 | 1 | 0.275508132 | 0.266 | 0.14 | 1.81E-38 | 6.61E-34 |
| ECHDC2 | 1 | 0.35992589 | 0.521 | 0.379 | 1.81E-38 | 6.62E-34 |
| SWAP70 | 1 | 0.304447886 | 0.265 | 0.142 | 1.85E-38 | 6.76E-34 |
| NSMCE2 | 1 | -0.344030969 | 0.613 | 0.729 | 1.98E-38 | 7.24E-34 |
| HERC4 | 1 | 0.364477933 | 0.797 | 0.697 | 2.05E-38 | 7.5E-34 |
| SYDE2 | 1 | -0.309981712 | 0.109 | 0.232 | 2.36E-38 | 8.63E-34 |
| RPL13A | 1 | 0.467807506 | 0.648 | 0.522 | 2.51E-38 | 9.19E-34 |
| SLC9A7 | 1 | -0.31565541 | 0.069 | 0.174 | 3.38E-38 | 1.24E-33 |
| AP001831.1 | 1 | 0.312932156 | 0.215 | 0.106 | 3.48E-38 | 1.27E-33 |
| HOXD3 | 1 | -0.306915901 | 0.078 | 0.186 | 3.49E-38 | 1.28E-33 |
| SLC38A2 | 1 | 0.396882119 | 0.561 | 0.416 | 3.56E-38 | 1.3E-33 |
| KHDRBS3 | 1 | -0.583721654 | 0.159 | 0.27 | 3.89E-38 | 1.42E-33 |
| RERG | 1 | -0.545195696 | 0.126 | 0.217 | 4.36E-38 | 1.6E-33 |
| RSF1 | 1 | 0.339844439 | 0.742 | 0.666 | 4.54E-38 | 1.66E-33 |
| SMYD3 | 1 | -0.29608126 | 0.874 | 0.942 | 5.58E-38 | 2.04E-33 |
| OLFM2 | 1 | 0.284180518 | 0.206 | 0.096 | 6.52E-38 | 2.39E-33 |
| EPS8 | 1 | -0.510946521 | 0.524 | 0.608 | 7.08E-38 | 2.59E-33 |
| WWOX | 1 | -0.472381322 | 0.712 | 0.792 | 7.96E-38 | 2.91E-33 |
| RBFOX2 | 1 | -0.336235253 | 0.806 | 0.859 | 8.74E-38 | 3.2E-33 |
| LPP | 1 | -0.36834114 | 0.642 | 0.761 | 9.64E-38 | 3.53E-33 |
| EDNRA | 1 | -0.44819672 | 0.056 | 0.153 | 1.18E-37 | 4.33E-33 |
| TRIB2 | 1 | -0.297932054 | 0.105 | 0.217 | 1.21E-37 | 4.42E-33 |
| CCZ1 | 1 | 0.303346485 | 0.321 | 0.192 | 1.63E-37 | 5.98E-33 |
| SOX4 | 1 | 0.479034138 | 0.37 | 0.242 | 1.95E-37 | 7.15E-33 |
| DFFA | 1 | 0.292100517 | 0.267 | 0.144 | 2.14E-37 | 7.83E-33 |
| PKNOX2 | 1 | -0.424242154 | 0.138 | 0.254 | 2.37E-37 | 8.66E-33 |
| KATNAL2 | 1 | 0.298381314 | 0.284 | 0.158 | 2.48E-37 | 9.08E-33 |
| GSE1 | 1 | -0.423385696 | 0.667 | 0.749 | 2.62E-37 | 9.58E-33 |
| SNHG19 | 1 | 0.297641741 | 0.239 | 0.122 | 2.78E-37 | 1.02E-32 |
| LEPR | 1 | -0.493043533 | 0.065 | 0.154 | 2.96E-37 | 1.09E-32 |
| SLC7A14-AS1 | 1 | 0.279885799 | 0.112 | 0.036 | 3E-37 | 1.1E-32 |

| | | | | | | |
|---|---|---|---|---|---|---|
| GTDC1 | 1 | -0.445134571 | 0.402 | 0.519 | 3.18E-37 | 1.17E-32 |
| PKP4 | 1 | 0.402634831 | 0.77 | 0.686 | 3.42E-37 | 1.25E-32 |
| ZFAND4 | 1 | -0.280651077 | 0.09 | 0.2 | 3.62E-37 | 1.32E-32 |
| MID1 | 1 | 0.260579454 | 0.72 | 0.57 | 3.84E-37 | 1.4E-32 |
| MNAT1 | 1 | -0.299205854 | 0.669 | 0.774 | 4.04E-37 | 1.48E-32 |
| PCAT1 | 1 | -0.597410631 | 0.042 | 0.119 | 4.66E-37 | 1.71E-32 |
| LRRTM4 | 1 | -0.630031609 | 0.6 | 0.569 | 4.71E-37 | 1.72E-32 |
| SNX29 | 1 | -0.413457194 | 0.306 | 0.433 | 4.74E-37 | 1.74E-32 |
| RNF165 | 1 | -0.308148997 | 0.101 | 0.214 | 5.58E-37 | 2.04E-32 |
| DYNC1I2 | 1 | 0.425361348 | 0.713 | 0.628 | 6.12E-37 | 2.24E-32 |
| VPS13A | 1 | -0.437701337 | 0.641 | 0.655 | 6.75E-37 | 2.47E-32 |
| GRIN3A | 1 | -0.602443361 | 0.208 | 0.273 | 6.76E-37 | 2.48E-32 |
| IFNAR2 | 1 | 0.32103753 | 0.341 | 0.21 | 7.18E-37 | 2.63E-32 |
| BMP6 | 1 | -0.35736179 | 0.04 | 0.128 | 8.08E-37 | 2.96E-32 |
| HMCN1 | 1 | -0.646127183 | 0.129 | 0.243 | 9.6E-37 | 3.52E-32 |
| UNC5B | 1 | 0.2881271 | 0.193 | 0.089 | 1.21E-36 | 4.44E-32 |
| UGP2 | 1 | 0.373946644 | 0.363 | 0.238 | 1.54E-36 | 5.63E-32 |
| PRKG1 | 1 | -0.775724808 | 0.673 | 0.753 | 1.54E-36 | 5.63E-32 |
| MALRD1 | 1 | -0.753726817 | 0.163 | 0.29 | 1.63E-36 | 5.95E-32 |
| GPR156 | 1 | 0.360989137 | 0.325 | 0.194 | 1.68E-36 | 6.16E-32 |
| SLC4A4 | 1 | -0.405877549 | 0.09 | 0.2 | 1.73E-36 | 6.35E-32 |
| FAM162A | 1 | 0.607381337 | 0.287 | 0.186 | 1.76E-36 | 6.44E-32 |
| ATL3 | 1 | 0.346411229 | 0.413 | 0.275 | 2.11E-36 | 7.72E-32 |
| STK39 | 1 | 0.384027949 | 0.699 | 0.623 | 2.44E-36 | 8.92E-32 |
| MCF2L | 1 | -0.36190646 | 0.168 | 0.294 | 2.56E-36 | 9.39E-32 |
| KIAA1549 | 1 | -0.358166643 | 0.325 | 0.459 | 2.69E-36 | 9.85E-32 |
| ARHGEF10L | 1 | -0.335741549 | 0.131 | 0.246 | 2.72E-36 | 9.94E-32 |
| DHRSX | 1 | 0.374523218 | 0.72 | 0.632 | 2.78E-36 | 1.02E-31 |
| AFTPH | 1 | 0.392164071 | 0.509 | 0.391 | 2.8E-36 | 1.03E-31 |
| INVS | 1 | -0.363522797 | 0.459 | 0.584 | 3.12E-36 | 1.14E-31 |
| LRRC1 | 1 | -0.275855078 | 0.08 | 0.185 | 3.14E-36 | 1.15E-31 |
| AC011586.2 | 1 | -0.527031467 | 0.034 | 0.116 | 3.35E-36 | 1.23E-31 |
| GREB1L | 1 | -0.68335396 | 0.038 | 0.117 | 3.55E-36 | 1.3E-31 |
| CCSER1 | 1 | 0.528434642 | 0.686 | 0.578 | 3.87E-36 | 1.42E-31 |
| KIAA0895 | 1 | 0.28151492 | 0.197 | 0.093 | 4.04E-36 | 1.48E-31 |
| ENOX1 | 1 | 0.497573966 | 0.617 | 0.504 | 4.22E-36 | 1.54E-31 |
| MBNL1 | 1 | 0.326998367 | 0.929 | 0.909 | 4.26E-36 | 1.56E-31 |
| AC023469.2 | 1 | 0.255882088 | 0.172 | 0.075 | 4.36E-36 | 1.6E-31 |
| LRPPRC | 1 | 0.338873357 | 0.829 | 0.771 | 5.24E-36 | 1.92E-31 |
| WDR7 | 1 | -0.349172385 | 0.423 | 0.548 | 5.63E-36 | 2.06E-31 |
| SNHG8 | 1 | 0.332164489 | 0.302 | 0.188 | 6.1E-36 | 2.23E-31 |
| SYNPR | 1 | -0.675623847 | 0.285 | 0.384 | 6.19E-36 | 2.27E-31 |
| APBB2 | 1 | 0.404154663 | 0.549 | 0.433 | 6.62E-36 | 2.42E-31 |
| LPIN1 | 1 | 0.316837225 | 0.307 | 0.184 | 6.71E-36 | 2.46E-31 |
| FNBP4 | 1 | 0.321671112 | 0.782 | 0.687 | 7.98E-36 | 2.92E-31 |
| RFX2 | 1 | -0.460828693 | 0.149 | 0.247 | 8.01E-36 | 2.93E-31 |
| AOPEP | 1 | -0.361982657 | 0.698 | 0.774 | 8.4E-36 | 3.07E-31 |
| ACTR3 | 1 | 0.376685965 | 0.544 | 0.417 | 8.62E-36 | 3.15E-31 |
| ZNF423 | 1 | -0.520972248 | 0.33 | 0.436 | 9.99E-36 | 3.66E-31 |
| TTC7B | 1 | -0.330652799 | 0.29 | 0.428 | 1.31E-35 | 4.81E-31 |
| MDFIC | 1 | -0.297902035 | 0.048 | 0.139 | 1.47E-35 | 5.37E-31 |
| PID1 | 1 | -0.506092072 | 0.579 | 0.584 | 1.73E-35 | 6.35E-31 |
| LAMP2 | 1 | 0.362572753 | 0.385 | 0.259 | 1.78E-35 | 6.53E-31 |
| CDC14B | 1 | -0.396574595 | 0.149 | 0.271 | 1.86E-35 | 6.81E-31 |
| TENM1 | 1 | 0.345062489 | 0.208 | 0.105 | 1.93E-35 | 7.05E-31 |
| FRMD4A | 1 | -0.482792667 | 0.786 | 0.828 | 2.04E-35 | 7.48E-31 |
| AP001347.1 | 1 | 0.27512094 | 0.467 | 0.318 | 2.12E-35 | 7.75E-31 |
| USP9X | 1 | 0.350466137 | 0.877 | 0.848 | 2.16E-35 | 7.91E-31 |

| | | | | | | |
|---|---|---|---|---|---|---|
| EPB41L1 | 1 | 0.298378269 | 0.228 | 0.119 | 2.31E-35 | 8.44E-31 |
| AL596087.2 | 1 | 0.588380221 | 0.292 | 0.176 | 2.34E-35 | 8.56E-31 |
| ATRNL1 | 1 | -0.681107298 | 0.788 | 0.813 | 3.09E-35 | 1.13E-30 |
| TMSB4X | 1 | 0.409868033 | 0.343 | 0.221 | 3.3E-35 | 1.21E-30 |
| CDHR3 | 1 | 0.345409964 | 0.503 | 0.359 | 3.74E-35 | 1.37E-30 |
| MTREX | 1 | 0.357410766 | 0.646 | 0.567 | 4.2E-35 | 1.54E-30 |
| PMEPA1 | 1 | -0.380810614 | 0.137 | 0.258 | 4.36E-35 | 1.6E-30 |
| AQP4-AS1 | 1 | 0.313480143 | 0.187 | 0.086 | 4.61E-35 | 1.69E-30 |
| RTN1 | 1 | -0.367273565 | 0.084 | 0.181 | 5.16E-35 | 1.89E-30 |
| AC092821.3 | 1 | -0.337854168 | 0.204 | 0.319 | 5.64E-35 | 2.06E-30 |
| NR2F2-AS1 | 1 | -0.460712425 | 0.032 | 0.109 | 7.03E-35 | 2.57E-30 |
| FGF13 | 1 | -0.65973343 | 0.371 | 0.469 | 7.11E-35 | 2.6E-30 |
| HSPA9 | 1 | 0.376439427 | 0.448 | 0.314 | 8.2E-35 | 3E-30 |
| ATP2B2 | 1 | -0.32402575 | 0.045 | 0.128 | 8.47E-35 | 3.1E-30 |
| PTPN12 | 1 | 0.333696824 | 0.749 | 0.654 | 8.56E-35 | 3.13E-30 |
| AFAP1 | 1 | 0.295329407 | 0.588 | 0.438 | 9.48E-35 | 3.47E-30 |
| DAAM1 | 1 | -0.420737154 | 0.413 | 0.532 | 1.11E-34 | 4.05E-30 |
| FAM107B | 1 | -0.28325364 | 0.098 | 0.209 | 1.14E-34 | 4.18E-30 |
| SOX5 | 1 | 0.344888425 | 0.721 | 0.626 | 1.21E-34 | 4.43E-30 |
| JAK1 | 1 | 0.348645326 | 0.673 | 0.557 | 1.65E-34 | 6.04E-30 |
| APC | 1 | -0.375936116 | 0.428 | 0.557 | 1.73E-34 | 6.34E-30 |
| FCHO2 | 1 | -0.33989511 | 0.274 | 0.408 | 1.96E-34 | 7.18E-30 |
| RTKN2 | 1 | -0.327835277 | 0.094 | 0.201 | 2.18E-34 | 7.98E-30 |
| EIF4A2 | 1 | 0.342559583 | 0.647 | 0.549 | 2.47E-34 | 9.05E-30 |
| MYEF2 | 1 | -0.349923507 | 0.414 | 0.538 | 2.56E-34 | 9.39E-30 |
| GRK3 | 1 | 0.370492873 | 0.563 | 0.421 | 2.87E-34 | 1.05E-29 |
| SHOX2 | 1 | -0.288040486 | 0.096 | 0.207 | 3.21E-34 | 1.18E-29 |
| N4BP2L2 | 1 | 0.326331124 | 0.804 | 0.744 | 3.74E-34 | 1.37E-29 |
| SYT14 | 1 | 0.36926083 | 0.588 | 0.438 | 3.87E-34 | 1.42E-29 |
| COCH | 1 | -0.341870681 | 0.146 | 0.263 | 3.9E-34 | 1.43E-29 |
| DDX3X | 1 | 0.348614802 | 0.609 | 0.497 | 4.4E-34 | 1.61E-29 |
| UBA3 | 1 | 0.331397251 | 0.491 | 0.355 | 4.5E-34 | 1.65E-29 |
| RASAL2 | 1 | -0.413619736 | 0.47 | 0.562 | 5.22E-34 | 1.91E-29 |
| CDH4 | 1 | 0.531744927 | 0.69 | 0.597 | 5.34E-34 | 1.96E-29 |
| C3orf70 | 1 | 0.306007906 | 0.367 | 0.231 | 5.43E-34 | 1.99E-29 |
| GNB1 | 1 | 0.326734753 | 0.816 | 0.735 | 5.76E-34 | 2.11E-29 |
| CCND3 | 1 | 0.349242087 | 0.31 | 0.187 | 6.24E-34 | 2.28E-29 |
| PLPP4 | 1 | -0.505961285 | 0.311 | 0.39 | 6.8E-34 | 2.49E-29 |
| SLC13A3 | 1 | 0.290141355 | 0.24 | 0.128 | 7.13E-34 | 2.61E-29 |
| KIRREL1 | 1 | 0.324510316 | 0.548 | 0.408 | 7.72E-34 | 2.83E-29 |
| OXR1 | 1 | 0.337263304 | 0.743 | 0.63 | 9.21E-34 | 3.37E-29 |
| CRTC3 | 1 | -0.312879604 | 0.279 | 0.417 | 1.04E-33 | 3.81E-29 |
| ZNF431 | 1 | 0.305909849 | 0.461 | 0.319 | 1.08E-33 | 3.97E-29 |
| MTSS1 | 1 | -0.466747045 | 0.118 | 0.215 | 1.21E-33 | 4.44E-29 |
| NEK10 | 1 | -0.303641697 | 0.053 | 0.142 | 1.27E-33 | 4.66E-29 |
| BICD1 | 1 | -0.305894276 | 0.742 | 0.824 | 1.31E-33 | 4.8E-29 |
| DDX24 | 1 | 0.284279922 | 0.332 | 0.205 | 1.73E-33 | 6.35E-29 |
| AC004158.1 | 1 | -0.334976103 | 0.106 | 0.214 | 1.94E-33 | 7.1E-29 |
| EPHA4 | 1 | -0.408631058 | 0.501 | 0.616 | 2.48E-33 | 9.08E-29 |
| AC004917.1 | 1 | 0.259726285 | 0.167 | 0.075 | 4.7E-33 | 1.72E-28 |
| SCAPER | 1 | -0.325154307 | 0.59 | 0.692 | 6.29E-33 | 2.3E-28 |
| RCAN2 | 1 | 0.400726212 | 0.17 | 0.077 | 7.69E-33 | 2.82E-28 |
| URGCP | 1 | 0.314869643 | 0.483 | 0.351 | 8.55E-33 | 3.13E-28 |
| NF1 | 1 | -0.276355471 | 0.866 | 0.92 | 8.84E-33 | 3.24E-28 |
| SUMF1 | 1 | -0.387399272 | 0.36 | 0.484 | 9.05E-33 | 3.31E-28 |
| CAMTA1 | 1 | -0.393308893 | 0.664 | 0.725 | 9.07E-33 | 3.32E-28 |
| PAN3 | 1 | 0.298624026 | 0.841 | 0.783 | 9.1E-33 | 3.33E-28 |
| EFCAB2 | 1 | -0.36394363 | 0.226 | 0.349 | 1.14E-32 | 4.18E-28 |

| | | | | | | |
|---|---|---|---|---|---|---|
| FAM135A | 1 | -0.339763232 | 0.475 | 0.582 | 1.16E-32 | 4.26E-28 |
| DIXDC1 | 1 | 0.278055378 | 0.351 | 0.22 | 1.2E-32 | 4.38E-28 |
| PRPF38B | 1 | 0.297558879 | 0.42 | 0.289 | 1.21E-32 | 4.42E-28 |
| VPS50 | 1 | -0.334445941 | 0.284 | 0.411 | 1.34E-32 | 4.91E-28 |
| CTNNAL1 | 1 | -0.27074238 | 0.169 | 0.295 | 1.36E-32 | 4.96E-28 |
| TMEM135 | 1 | -0.366584894 | 0.434 | 0.546 | 1.45E-32 | 5.32E-28 |
| FBXL2 | 1 | 0.291318324 | 0.321 | 0.194 | 1.64E-32 | 6.02E-28 |
| SPRED1 | 1 | -0.390732269 | 0.332 | 0.441 | 1.67E-32 | 6.12E-28 |
| AL157392.3 | 1 | 0.266858102 | 0.26 | 0.147 | 1.92E-32 | 7.04E-28 |
| SCAMP1 | 1 | 0.35327704 | 0.448 | 0.324 | 1.93E-32 | 7.07E-28 |
| NDUFAF2 | 1 | -0.367148274 | 0.231 | 0.354 | 2.11E-32 | 7.74E-28 |
| NR2F1-AS1 | 1 | 0.406421208 | 0.56 | 0.432 | 2.74E-32 | 1E-27 |
| GPHN | 1 | -0.270278818 | 0.876 | 0.938 | 2.82E-32 | 1.03E-27 |
| ERBB4 | 1 | 0.260052975 | 0.388 | 0.253 | 3.04E-32 | 1.11E-27 |
| POU2F3 | 1 | 0.29984416 | 0.424 | 0.285 | 3.49E-32 | 1.28E-27 |
| AL354771.1 | 1 | -0.343741963 | 0.077 | 0.158 | 3.51E-32 | 1.29E-27 |
| ABTB2 | 1 | -0.385356327 | 0.091 | 0.186 | 3.62E-32 | 1.33E-27 |
| LINC01876 | 1 | -0.403594354 | 0.512 | 0.506 | 3.85E-32 | 1.41E-27 |
| PBX1 | 1 | 0.285808698 | 0.864 | 0.846 | 4.17E-32 | 1.53E-27 |
| UHMK1 | 1 | 0.303090106 | 0.323 | 0.202 | 4.5E-32 | 1.65E-27 |
| MKLN1 | 1 | 0.354966671 | 0.86 | 0.818 | 4.7E-32 | 1.72E-27 |
| FAM228B | 1 | -0.298296497 | 0.142 | 0.252 | 4.85E-32 | 1.77E-27 |
| UBE3D | 1 | -0.377655906 | 0.382 | 0.506 | 5.28E-32 | 1.93E-27 |
| TMEFF1 | 1 | -0.377332581 | 0.271 | 0.394 | 6.21E-32 | 2.27E-27 |
| ZNF277 | 1 | 0.338912882 | 0.484 | 0.352 | 7.63E-32 | 2.79E-27 |
| IGSF11 | 1 | -0.364454232 | 0.077 | 0.171 | 8.53E-32 | 3.12E-27 |
| CYYR1 | 1 | -0.265675489 | 0.03 | 0.103 | 8.58E-32 | 3.14E-27 |
| BOC | 1 | -0.395114015 | 0.124 | 0.228 | 9.26E-32 | 3.39E-27 |
| MT-CO1 | 1 | -0.349452507 | 0.361 | 0.49 | 1.01E-31 | 3.69E-27 |
| MRPS21 | 1 | 0.280167472 | 0.334 | 0.211 | 1.13E-31 | 4.15E-27 |
| SCAI | 1 | -0.322679861 | 0.416 | 0.546 | 1.17E-31 | 4.29E-27 |
| STRN | 1 | 0.321112917 | 0.599 | 0.479 | 1.24E-31 | 4.54E-27 |
| ULK2 | 1 | -0.335304235 | 0.315 | 0.44 | 1.25E-31 | 4.57E-27 |
| MAP3K1 | 1 | -0.432216586 | 0.128 | 0.232 | 1.37E-31 | 5.03E-27 |
| DOCK11 | 1 | -0.441167142 | 0.497 | 0.54 | 1.55E-31 | 5.66E-27 |
| ZBTB20 | 1 | -0.499563494 | 0.78 | 0.79 | 2.06E-31 | 7.54E-27 |
| AF117829.1 | 1 | -0.373536767 | 0.277 | 0.389 | 2.07E-31 | 7.58E-27 |
| SEPTIN11 | 1 | -0.347536206 | 0.382 | 0.511 | 2.27E-31 | 8.32E-27 |
| MED12L | 1 | -0.421393087 | 0.406 | 0.534 | 3.1E-31 | 1.13E-26 |
| GCLC | 1 | -0.691098594 | 0.152 | 0.232 | 3.12E-31 | 1.14E-26 |
| SOX2-OT | 1 | 0.452534327 | 0.291 | 0.176 | 4.45E-31 | 1.63E-26 |
| FNBP1 | 1 | 0.32767596 | 0.743 | 0.634 | 4.55E-31 | 1.67E-26 |
| ACTR6 | 1 | 0.269979678 | 0.248 | 0.141 | 4.62E-31 | 1.69E-26 |
| SUSD4 | 1 | -0.425354062 | 0.188 | 0.29 | 5.04E-31 | 1.84E-26 |
| PRRG1 | 1 | 0.315821492 | 0.463 | 0.326 | 5.28E-31 | 1.93E-26 |
| SLTM | 1 | 0.333511364 | 0.672 | 0.573 | 5.59E-31 | 2.05E-26 |
| ALDH1L2 | 1 | 0.25744948 | 0.262 | 0.149 | 5.88E-31 | 2.15E-26 |
| GULP1 | 1 | -0.523950141 | 0.462 | 0.522 | 6.55E-31 | 2.4E-26 |
| GSTO2 | 1 | 0.272161734 | 0.325 | 0.203 | 6.59E-31 | 2.41E-26 |
| ZEB1-AS1 | 1 | -0.258539879 | 0.029 | 0.1 | 7.65E-31 | 2.8E-26 |
| LINC00882 | 1 | -0.344974331 | 0.043 | 0.12 | 7.89E-31 | 2.89E-26 |
| CLEC2D | 1 | -0.375812493 | 0.375 | 0.489 | 1.1E-30 | 4.02E-26 |
| DSCAM | 1 | 0.54359714 | 0.189 | 0.094 | 1.13E-30 | 4.13E-26 |
| PM20D2 | 1 | 0.268504734 | 0.29 | 0.174 | 1.13E-30 | 4.14E-26 |
| PGK1 | 1 | 0.627099203 | 0.41 | 0.316 | 1.23E-30 | 4.5E-26 |
| SLC3A2 | 1 | 0.302054602 | 0.324 | 0.207 | 1.28E-30 | 4.68E-26 |
| NBEAL1 | 1 | -0.348469275 | 0.376 | 0.486 | 1.54E-30 | 5.64E-26 |
| SPHKAP | 1 | -0.447020992 | 0.052 | 0.135 | 1.91E-30 | 6.98E-26 |

| | | | | | | |
|---|---|---|---|---|---|---|
| TRAF5 | 1 | -0.288695836 | 0.175 | 0.291 | 1.96E-30 | 7.16E-26 |
| IPO5 | 1 | 0.380726422 | 0.51 | 0.411 | 2.07E-30 | 7.57E-26 |
| CCNY | 1 | -0.281595723 | 0.658 | 0.747 | 2.19E-30 | 8.01E-26 |
| AL022068.1 | 1 | -0.617294406 | 0.064 | 0.127 | 2.25E-30 | 8.23E-26 |
| PRKCB | 1 | -0.35176226 | 0.045 | 0.124 | 2.32E-30 | 8.48E-26 |
| BARD1 | 1 | -0.34700973 | 0.291 | 0.411 | 2.37E-30 | 8.69E-26 |
| KIF5C | 1 | -0.352455463 | 0.202 | 0.32 | 2.49E-30 | 9.1E-26 |
| PHLDB2 | 1 | -0.450675261 | 0.175 | 0.249 | 2.76E-30 | 1.01E-25 |
| DOCK5 | 1 | -0.3819887 | 0.277 | 0.399 | 2.91E-30 | 1.06E-25 |
| MSRB3 | 1 | -0.285651443 | 0.043 | 0.116 | 2.96E-30 | 1.08E-25 |
| SNX4 | 1 | 0.318614868 | 0.483 | 0.362 | 2.97E-30 | 1.09E-25 |
| RPS12 | 1 | 0.362748813 | 0.387 | 0.261 | 3.15E-30 | 1.15E-25 |
| DDHD1 | 1 | 0.328890451 | 0.637 | 0.54 | 3.66E-30 | 1.34E-25 |
| SNHG4 | 1 | 0.27153808 | 0.239 | 0.134 | 3.98E-30 | 1.46E-25 |
| ADAMTS16 | 1 | -0.465709466 | 0.192 | 0.288 | 4.37E-30 | 1.6E-25 |
| CCDC102B | 1 | -0.981302153 | 0.082 | 0.158 | 5.21E-30 | 1.91E-25 |
| PDE9A | 1 | -0.311512109 | 0.178 | 0.297 | 5.22E-30 | 1.91E-25 |
| TBL1XR1 | 1 | -0.263623247 | 0.809 | 0.873 | 5.49E-30 | 2.01E-25 |
| RIF1 | 1 | 0.327364565 | 0.813 | 0.766 | 5.76E-30 | 2.11E-25 |
| CDK17 | 1 | -0.287474643 | 0.493 | 0.617 | 5.82E-30 | 2.13E-25 |
| TPCN1 | 1 | -0.30599116 | 0.102 | 0.198 | 6.41E-30 | 2.35E-25 |
| COL6A3 | 1 | -0.53416643 | 0.061 | 0.137 | 6.56E-30 | 2.4E-25 |
| AL138828.1 | 1 | 0.320274537 | 0.146 | 0.063 | 6.77E-30 | 2.48E-25 |
| TMEM117 | 1 | -0.472438608 | 0.52 | 0.598 | 8.28E-30 | 3.03E-25 |
| CCNE2 | 1 | -0.268200981 | 0.087 | 0.185 | 8.53E-30 | 3.12E-25 |
| IMMP1L | 1 | -0.353314583 | 0.385 | 0.487 | 8.54E-30 | 3.13E-25 |
| TMEM232 | 1 | -0.440989582 | 0.081 | 0.152 | 8.91E-30 | 3.26E-25 |
| IMMP2L | 1 | -0.358757803 | 0.76 | 0.839 | 9.59E-30 | 3.51E-25 |
| GSTCD | 1 | 0.367912727 | 0.563 | 0.447 | 1.01E-29 | 3.71E-25 |
| ACSL4 | 1 | 0.364825304 | 0.637 | 0.537 | 1.12E-29 | 4.11E-25 |
| GALNT7 | 1 | -0.412383958 | 0.533 | 0.615 | 1.27E-29 | 4.65E-25 |
| CSNK2A1 | 1 | 0.303702929 | 0.557 | 0.431 | 1.3E-29 | 4.77E-25 |
| FRMD4B | 1 | -0.473458114 | 0.363 | 0.456 | 1.32E-29 | 4.83E-25 |
| P4HA1 | 1 | 0.708223776 | 0.524 | 0.469 | 1.4E-29 | 5.12E-25 |
| BCAS3 | 1 | -0.390990919 | 0.593 | 0.678 | 1.45E-29 | 5.32E-25 |
| BCL7A | 1 | -0.292649405 | 0.15 | 0.261 | 1.48E-29 | 5.43E-25 |
| FRAS1 | 1 | -0.552617242 | 0.293 | 0.404 | 1.53E-29 | 5.6E-25 |
| RSRC1 | 1 | -0.297455282 | 0.749 | 0.827 | 1.69E-29 | 6.19E-25 |
| TUT7 | 1 | 0.303018705 | 0.408 | 0.288 | 1.81E-29 | 6.62E-25 |
| RSRC2 | 1 | 0.328955449 | 0.583 | 0.474 | 1.86E-29 | 6.79E-25 |
| EYA1 | 1 | 0.371894516 | 0.39 | 0.261 | 1.99E-29 | 7.28E-25 |
| PNPLA8 | 1 | 0.309666642 | 0.373 | 0.251 | 2.06E-29 | 7.53E-25 |
| SERPINE2 | 1 | -0.417360696 | 0.081 | 0.17 | 2.18E-29 | 8E-25 |
| CHCHD3 | 1 | 0.302009969 | 0.761 | 0.701 | 2.65E-29 | 9.71E-25 |
| EFNB2 | 1 | 0.263541383 | 0.238 | 0.132 | 3.04E-29 | 1.11E-24 |
| DST | 1 | 0.292095571 | 0.879 | 0.875 | 3.06E-29 | 1.12E-24 |
| TBC1D16 | 1 | -0.332927447 | 0.187 | 0.303 | 3.91E-29 | 1.43E-24 |
| RPL12 | 1 | 0.32949346 | 0.276 | 0.172 | 4E-29 | 1.46E-24 |
| PCNX1 | 1 | -0.278221752 | 0.701 | 0.771 | 4.02E-29 | 1.47E-24 |
| OTUD3 | 1 | 0.259728345 | 0.276 | 0.173 | 4.02E-29 | 1.47E-24 |
| ZNF544 | 1 | 0.298672721 | 0.48 | 0.357 | 4.05E-29 | 1.48E-24 |
| CEP126 | 1 | -0.349372265 | 0.22 | 0.326 | 4.25E-29 | 1.56E-24 |
| TFG | 1 | 0.314254124 | 0.542 | 0.429 | 4.83E-29 | 1.77E-24 |
| CDC42BPA | 1 | -0.360877786 | 0.728 | 0.786 | 5.11E-29 | 1.87E-24 |
| METAP1D | 1 | -0.279272291 | 0.15 | 0.261 | 5.24E-29 | 1.92E-24 |
| HFM1 | 1 | -0.350785932 | 0.286 | 0.382 | 5.44E-29 | 1.99E-24 |
| SHROOM2 | 1 | 0.341030573 | 0.455 | 0.337 | 5.67E-29 | 2.08E-24 |
| SLC44A5 | 1 | -0.402432312 | 0.045 | 0.12 | 6.32E-29 | 2.31E-24 |

| | | | | | | |
|---|---|---|---|---|---|---|
| ANO10 | 1 | -0.3851453 | 0.395 | 0.498 | 6.45E-29 | 2.36E-24 |
| RPS24 | 1 | 0.395834623 | 0.544 | 0.414 | 6.74E-29 | 2.47E-24 |
| RAB6A | 1 | 0.321290567 | 0.662 | 0.569 | 7.79E-29 | 2.85E-24 |
| THUMPD3-AS1 | 1 | 0.29972006 | 0.496 | 0.371 | 7.8E-29 | 2.86E-24 |
| C2CD5 | 1 | -0.316783189 | 0.605 | 0.698 | 8.18E-29 | 2.99E-24 |
| RANBP17 | 1 | -0.303204579 | 0.665 | 0.759 | 8.43E-29 | 3.09E-24 |
| ACTR2 | 1 | 0.313189251 | 0.425 | 0.312 | 8.48E-29 | 3.11E-24 |
| CCSER2 | 1 | -0.314861964 | 0.437 | 0.552 | 8.5E-29 | 3.11E-24 |
| ALG8 | 1 | 0.307989139 | 0.402 | 0.294 | 8.76E-29 | 3.21E-24 |
| TXNDC16 | 1 | -0.345986987 | 0.239 | 0.357 | 8.98E-29 | 3.29E-24 |
| FTO | 1 | -0.425958585 | 0.749 | 0.831 | 9.38E-29 | 3.43E-24 |
| TXNIP | 1 | 0.317770448 | 0.148 | 0.071 | 9.88E-29 | 3.62E-24 |
| INTS6L | 1 | -0.370216416 | 0.304 | 0.407 | 1.08E-28 | 3.96E-24 |
| HK2 | 1 | 0.377397089 | 0.2 | 0.113 | 1.09E-28 | 3.99E-24 |
| MT-CO2 | 1 | -0.307210383 | 0.271 | 0.394 | 1.38E-28 | 5.05E-24 |
| REEP3 | 1 | -0.322952905 | 0.357 | 0.465 | 1.43E-28 | 5.22E-24 |
| ZNRF2 | 1 | -0.312145132 | 0.251 | 0.366 | 1.44E-28 | 5.27E-24 |
| FANCL | 1 | -0.332333823 | 0.551 | 0.647 | 1.68E-28 | 6.16E-24 |
| RPL41 | 1 | 0.431312923 | 0.632 | 0.506 | 1.71E-28 | 6.27E-24 |
| KCTD13 | 1 | 0.289511483 | 0.421 | 0.294 | 1.78E-28 | 6.53E-24 |
| KSR1 | 1 | -0.330563536 | 0.302 | 0.426 | 1.79E-28 | 6.56E-24 |
| ZNRF1 | 1 | -0.312148963 | 0.332 | 0.445 | 1.84E-28 | 6.73E-24 |
| STXBP4 | 1 | -0.346395839 | 0.392 | 0.498 | 2.24E-28 | 8.21E-24 |
| PGAM1 | 1 | 0.282007121 | 0.251 | 0.149 | 2.45E-28 | 8.98E-24 |
| SLCO3A1 | 1 | -0.378158292 | 0.191 | 0.3 | 2.68E-28 | 9.8E-24 |
| INSR | 1 | -0.452671703 | 0.184 | 0.279 | 2.75E-28 | 1.01E-23 |
| PDGFC | 1 | -0.500506105 | 0.184 | 0.288 | 2.88E-28 | 1.06E-23 |
| HOMER1 | 1 | 0.414465306 | 0.567 | 0.47 | 3.08E-28 | 1.13E-23 |
| ROR2 | 1 | -0.45694633 | 0.348 | 0.446 | 3.11E-28 | 1.14E-23 |
| RBBP7 | 1 | 0.327343308 | 0.589 | 0.473 | 3.34E-28 | 1.22E-23 |
| ZNF10 | 1 | 0.288777661 | 0.369 | 0.249 | 3.44E-28 | 1.26E-23 |
| ELP4 | 1 | -0.401870028 | 0.363 | 0.421 | 3.45E-28 | 1.26E-23 |
| DIRC3 | 1 | 0.290595493 | 0.289 | 0.175 | 3.47E-28 | 1.27E-23 |
| EDIL3 | 1 | -0.477644955 | 0.539 | 0.646 | 3.63E-28 | 1.33E-23 |
| SLC20A1 | 1 | -0.251390948 | 0.207 | 0.329 | 3.72E-28 | 1.36E-23 |
| WDR11-AS1 | 1 | -0.342820814 | 0.315 | 0.328 | 4.18E-28 | 1.53E-23 |
| SNX8 | 1 | 0.314769415 | 0.379 | 0.257 | 4.25E-28 | 1.55E-23 |
| CDK19 | 1 | 0.343984032 | 0.742 | 0.661 | 4.32E-28 | 1.58E-23 |
| EEF1A1 | 1 | 0.415016276 | 0.651 | 0.553 | 5.76E-28 | 2.11E-23 |
| MEIS1 | 1 | 0.324032652 | 0.446 | 0.322 | 6.53E-28 | 2.39E-23 |
| FAM126A | 1 | -0.299881414 | 0.522 | 0.63 | 6.58E-28 | 2.41E-23 |
| MAN1C1 | 1 | -0.308596914 | 0.152 | 0.255 | 7.24E-28 | 2.65E-23 |
| CEP290 | 1 | -0.308066181 | 0.273 | 0.396 | 7.39E-28 | 2.7E-23 |
| KLHL15 | 1 | 0.297498139 | 0.26 | 0.16 | 7.44E-28 | 2.72E-23 |
| GPC3 | 1 | 0.361735913 | 0.32 | 0.204 | 8.09E-28 | 2.96E-23 |
| CRADD | 1 | -0.372430663 | 0.333 | 0.439 | 8.73E-28 | 3.2E-23 |
| CHD1 | 1 | 0.336217212 | 0.737 | 0.694 | 1.2E-27 | 4.39E-23 |
| WLS | 1 | 0.335932106 | 0.516 | 0.389 | 1.28E-27 | 4.7E-23 |
| IKBIP | 1 | 0.25676168 | 0.249 | 0.145 | 1.32E-27 | 4.85E-23 |
| CPEB2 | 1 | -0.419193203 | 0.295 | 0.377 | 1.33E-27 | 4.86E-23 |
| TBC1D19 | 1 | -0.324473511 | 0.261 | 0.358 | 1.58E-27 | 5.77E-23 |
| MGAT4A | 1 | -0.274192344 | 0.113 | 0.213 | 1.86E-27 | 6.79E-23 |
| AC107223.1 | 1 | -0.287303024 | 0.082 | 0.171 | 2.15E-27 | 7.86E-23 |
| MEST | 1 | -0.335093558 | 0.075 | 0.155 | 2.19E-27 | 8.01E-23 |
| GRM8 | 1 | -1.116169844 | 0.099 | 0.17 | 2.22E-27 | 8.12E-23 |
| BTBD9 | 1 | -0.345766442 | 0.504 | 0.591 | 2.28E-27 | 8.33E-23 |
| NSMAF | 1 | 0.32390132 | 0.547 | 0.436 | 2.34E-27 | 8.57E-23 |
| KDM7A | 1 | 0.378390866 | 0.45 | 0.326 | 2.36E-27 | 8.64E-23 |

| | | | | | | |
|---|---|---|---|---|---|---|
| TOX | 1 | -0.453101803 | 0.64 | 0.685 | 2.56E-27 | 9.38E-23 |
| EFCAB6 | 1 | -0.298975635 | 0.095 | 0.164 | 2.89E-27 | 1.06E-22 |
| CANX | 1 | 0.330485577 | 0.463 | 0.347 | 2.93E-27 | 1.07E-22 |
| CCDC14 | 1 | -0.283690999 | 0.655 | 0.731 | 3.11E-27 | 1.14E-22 |
| DENR | 1 | 0.296648495 | 0.398 | 0.283 | 3.18E-27 | 1.16E-22 |
| RIMKLB | 1 | 0.286989218 | 0.522 | 0.404 | 3.44E-27 | 1.26E-22 |
| ZNF280C | 1 | 0.33228251 | 0.486 | 0.374 | 3.93E-27 | 1.44E-22 |
| KIF5B | 1 | 0.304707431 | 0.6 | 0.481 | 4.09E-27 | 1.5E-22 |
| SAMD3 | 1 | -0.398945985 | 0.045 | 0.107 | 4.45E-27 | 1.63E-22 |
| FAM149A | 1 | -0.288469167 | 0.226 | 0.349 | 4.83E-27 | 1.77E-22 |
| PSMA6 | 1 | 0.31125917 | 0.477 | 0.363 | 5.15E-27 | 1.89E-22 |
| CHSY3 | 1 | 0.493454229 | 0.78 | 0.688 | 5.24E-27 | 1.92E-22 |
| ITM2B | 1 | 0.285032585 | 0.325 | 0.216 | 5.54E-27 | 2.03E-22 |
| MAP1B | 1 | 0.329384677 | 0.785 | 0.703 | 6.01E-27 | 2.2E-22 |
| GRM7 | 1 | -0.749314024 | 0.09 | 0.152 | 7.56E-27 | 2.77E-22 |
| LMF1 | 1 | -0.25456599 | 0.104 | 0.194 | 8.4E-27 | 3.08E-22 |
| EVA1C | 1 | 0.260892308 | 0.187 | 0.098 | 8.74E-27 | 3.2E-22 |
| L3MBTL3 | 1 | -0.259236138 | 0.092 | 0.184 | 1.03E-26 | 3.78E-22 |
| JAK2 | 1 | 0.389491707 | 0.377 | 0.276 | 1.08E-26 | 3.94E-22 |
| VAMP4 | 1 | -0.278804416 | 0.149 | 0.246 | 1.08E-26 | 3.94E-22 |
| PAIP1 | 1 | 0.285117006 | 0.37 | 0.255 | 1.21E-26 | 4.43E-22 |
| WDR60 | 1 | 0.30018557 | 0.504 | 0.376 | 1.33E-26 | 4.87E-22 |
| TTBK2 | 1 | -0.294748928 | 0.352 | 0.472 | 1.34E-26 | 4.92E-22 |
| PTCHD1-AS | 1 | 0.460905047 | 0.185 | 0.097 | 1.38E-26 | 5.07E-22 |
| ERC2 | 1 | -0.298000915 | 0.453 | 0.45 | 1.39E-26 | 5.07E-22 |
| VPS13C | 1 | -0.270710884 | 0.666 | 0.763 | 1.4E-26 | 5.13E-22 |
| CALD1 | 1 | 0.351556913 | 0.512 | 0.4 | 1.47E-26 | 5.37E-22 |
| BCKDHB | 1 | 0.340328146 | 0.57 | 0.467 | 1.55E-26 | 5.66E-22 |
| FAM78B | 1 | 0.372548526 | 0.359 | 0.26 | 1.56E-26 | 5.7E-22 |
| KHDC4 | 1 | 0.283103478 | 0.486 | 0.369 | 1.58E-26 | 5.8E-22 |
| AC002377.1 | 1 | -0.319814272 | 0.067 | 0.146 | 1.67E-26 | 6.11E-22 |
| DYNC2H1 | 1 | -0.361661908 | 0.328 | 0.373 | 1.68E-26 | 6.13E-22 |
| SLC2A3 | 1 | 0.50165396 | 0.204 | 0.117 | 1.74E-26 | 6.36E-22 |
| VWC2 | 1 | 0.336184003 | 0.211 | 0.116 | 1.82E-26 | 6.65E-22 |
| ZNF827 | 1 | -0.308133664 | 0.523 | 0.553 | 1.91E-26 | 6.98E-22 |
| EXT1 | 1 | -0.385053561 | 0.653 | 0.703 | 2.17E-26 | 7.95E-22 |
| ENTPD1-AS1 | 1 | -0.327324704 | 0.283 | 0.377 | 2.31E-26 | 8.44E-22 |
| LPGAT1 | 1 | -0.308197737 | 0.399 | 0.514 | 2.4E-26 | 8.8E-22 |
| AC016042.1 | 1 | -0.323242584 | 0.172 | 0.255 | 2.55E-26 | 9.33E-22 |
| DDX21 | 1 | 0.331523953 | 0.372 | 0.262 | 2.58E-26 | 9.44E-22 |
| GOLIM4 | 1 | -0.258755013 | 0.422 | 0.545 | 2.66E-26 | 9.73E-22 |
| AKAP6 | 1 | -0.446719943 | 0.155 | 0.261 | 2.76E-26 | 1.01E-21 |
| AK4 | 1 | 0.315623419 | 0.347 | 0.236 | 2.98E-26 | 1.09E-21 |
| IKZF3 | 1 | -0.335858465 | 0.192 | 0.293 | 4.02E-26 | 1.47E-21 |
| FSIP2-AS1 | 1 | -0.271298619 | 0.074 | 0.154 | 4.25E-26 | 1.56E-21 |
| ZFAND5 | 1 | 0.355129634 | 0.526 | 0.423 | 4.3E-26 | 1.57E-21 |
| CCDC171 | 1 | -0.351105241 | 0.36 | 0.468 | 4.35E-26 | 1.59E-21 |
| SH3D19 | 1 | -0.348169761 | 0.515 | 0.599 | 4.92E-26 | 1.8E-21 |
| PARP8 | 1 | -0.273836077 | 0.241 | 0.362 | 4.93E-26 | 1.8E-21 |
| CACNA1B | 1 | -0.389956444 | 0.186 | 0.289 | 5.24E-26 | 1.92E-21 |
| KIF20B | 1 | 0.326921559 | 0.405 | 0.287 | 5.25E-26 | 1.92E-21 |
| THSD4 | 1 | 0.305641474 | 0.238 | 0.137 | 5.52E-26 | 2.02E-21 |
| LINC01619 | 1 | -0.368315392 | 0.255 | 0.359 | 6.25E-26 | 2.29E-21 |
| UHRF1BP1L | 1 | 0.312565781 | 0.552 | 0.453 | 6.32E-26 | 2.31E-21 |
| DOCK7 | 1 | -0.288080738 | 0.53 | 0.637 | 6.32E-26 | 2.31E-21 |
| PROM1 | 1 | 0.28606395 | 0.164 | 0.081 | 6.34E-26 | 2.32E-21 |
| FBXW7 | 1 | -0.428779233 | 0.709 | 0.748 | 6.35E-26 | 2.33E-21 |
| BAZ1B | 1 | 0.268746897 | 0.709 | 0.596 | 6.39E-26 | 2.34E-21 |

| | | | | | | |
|---|---|---|---|---|---|---|
| TRPM7 | 1 | 0.29933118 | 0.801 | 0.741 | 6.64E-26 | 2.43E-21 |
| TTC6 | 1 | -0.455475782 | 0.1 | 0.184 | 7.42E-26 | 2.72E-21 |
| AC008591.1 | 1 | 0.351108564 | 0.531 | 0.401 | 7.9E-26 | 2.89E-21 |
| NKD1 | 1 | -0.419578499 | 0.095 | 0.187 | 8.08E-26 | 2.96E-21 |
| SCN1A | 1 | 0.49893873 | 0.159 | 0.081 | 8.78E-26 | 3.21E-21 |
| RABGGTB | 1 | 0.253020097 | 0.369 | 0.251 | 8.79E-26 | 3.22E-21 |
| SERPINB6 | 1 | 0.304612774 | 0.308 | 0.199 | 8.89E-26 | 3.25E-21 |
| NR3C1 | 1 | -0.369593589 | 0.288 | 0.388 | 9.85E-26 | 3.61E-21 |
| REL | 1 | 0.267918424 | 0.274 | 0.173 | 1.01E-25 | 3.7E-21 |
| SPAG17 | 1 | -0.309459895 | 0.323 | 0.281 | 1.29E-25 | 4.72E-21 |
| KBTBD12 | 1 | 0.285282231 | 0.297 | 0.191 | 1.29E-25 | 4.74E-21 |
| NTN1 | 1 | -0.367920162 | 0.081 | 0.166 | 1.33E-25 | 4.87E-21 |
| SRGAP3 | 1 | -0.346634137 | 0.467 | 0.578 | 1.67E-25 | 6.11E-21 |
| MEGF11 | 1 | 0.345201856 | 0.404 | 0.288 | 1.71E-25 | 6.25E-21 |
| TIPRL | 1 | 0.256325985 | 0.249 | 0.153 | 1.89E-25 | 6.9E-21 |
| YWHAQ | 1 | 0.316411101 | 0.603 | 0.502 | 2.09E-25 | 7.66E-21 |
| SEC63 | 1 | 0.309457333 | 0.709 | 0.646 | 2.19E-25 | 8E-21 |
| PTBP3 | 1 | 0.314260167 | 0.464 | 0.351 | 2.47E-25 | 9.03E-21 |
| RHOQ | 1 | 0.264070731 | 0.374 | 0.256 | 2.6E-25 | 9.53E-21 |
| MIS18BP1 | 1 | 0.26184537 | 0.621 | 0.493 | 2.65E-25 | 9.72E-21 |
| FNBP1L | 1 | -0.257335872 | 0.615 | 0.714 | 3.05E-25 | 1.12E-20 |
| CADM1 | 1 | -0.476773358 | 0.411 | 0.45 | 3.95E-25 | 1.45E-20 |
| MGST1 | 1 | 0.311972688 | 0.212 | 0.123 | 3.99E-25 | 1.46E-20 |
| SESN3 | 1 | -0.439932801 | 0.387 | 0.414 | 4.9E-25 | 1.8E-20 |
| RPS2 | 1 | 0.317614468 | 0.46 | 0.352 | 5.16E-25 | 1.89E-20 |
| RPLP0 | 1 | 0.304118647 | 0.511 | 0.396 | 6.12E-25 | 2.24E-20 |
| PRH1 | 1 | -0.323853306 | 0.392 | 0.498 | 6.9E-25 | 2.53E-20 |
| VPS37B | 1 | 0.289133696 | 0.452 | 0.334 | 8.67E-25 | 3.18E-20 |
| TTC37 | 1 | -0.255983887 | 0.561 | 0.672 | 8.94E-25 | 3.27E-20 |
| PELI2 | 1 | -0.328902519 | 0.381 | 0.444 | 9.1E-25 | 3.33E-20 |
| RMND1 | 1 | 0.270166049 | 0.401 | 0.285 | 9.82E-25 | 3.6E-20 |
| EIF3B | 1 | 0.307732855 | 0.46 | 0.36 | 1.03E-24 | 3.77E-20 |
| AC023282.1 | 1 | -0.382778203 | 0.247 | 0.312 | 1.06E-24 | 3.89E-20 |
| CBFB | 1 | -0.275712324 | 0.436 | 0.54 | 1.16E-24 | 4.26E-20 |
| PKM | 1 | 0.319205465 | 0.557 | 0.465 | 1.3E-24 | 4.74E-20 |
| PIGN | 1 | -0.300747181 | 0.267 | 0.371 | 1.38E-24 | 5.07E-20 |
| LDLRAD3 | 1 | -0.335821005 | 0.483 | 0.575 | 1.4E-24 | 5.11E-20 |
| GPD2 | 1 | -0.32127586 | 0.345 | 0.446 | 1.4E-24 | 5.12E-20 |
| EPN2 | 1 | -0.274145938 | 0.361 | 0.478 | 1.78E-24 | 6.5E-20 |
| OSBPL10 | 1 | -0.457092844 | 0.316 | 0.407 | 1.94E-24 | 7.09E-20 |
| CPNE5 | 1 | -0.278350557 | 0.067 | 0.146 | 2.25E-24 | 8.23E-20 |
| RPS6KA6 | 1 | -0.305249787 | 0.137 | 0.223 | 2.27E-24 | 8.3E-20 |
| C1orf61 | 1 | -0.439635723 | 0.247 | 0.32 | 2.39E-24 | 8.77E-20 |
| LRCH2 | 1 | 0.3465725 | 0.753 | 0.679 | 2.49E-24 | 9.13E-20 |
| ARHGAP21 | 1 | -0.292500133 | 0.473 | 0.579 | 2.72E-24 | 9.95E-20 |
| SLCO5A1 | 1 | -0.319170843 | 0.09 | 0.169 | 2.76E-24 | 1.01E-19 |
| TMLHE | 1 | -0.250403422 | 0.172 | 0.272 | 3.27E-24 | 1.2E-19 |
| FAM135B | 1 | 0.331415211 | 0.461 | 0.34 | 3.62E-24 | 1.32E-19 |
| WASF3 | 1 | -0.27115234 | 0.11 | 0.196 | 4.05E-24 | 1.48E-19 |
| DYNC1H1 | 1 | 0.337231274 | 0.678 | 0.592 | 4.09E-24 | 1.5E-19 |
| EIF2AK3 | 1 | 0.289950342 | 0.344 | 0.236 | 4.18E-24 | 1.53E-19 |
| RAB28 | 1 | -0.287767793 | 0.388 | 0.491 | 4.27E-24 | 1.56E-19 |
| TTN | 1 | -0.599304443 | 0.096 | 0.169 | 4.34E-24 | 1.59E-19 |
| EPHA5 | 1 | -0.459432255 | 0.27 | 0.254 | 4.46E-24 | 1.63E-19 |
| ANTXR2 | 1 | -0.306749118 | 0.173 | 0.273 | 4.76E-24 | 1.74E-19 |
| PPP3CA | 1 | -0.317510445 | 0.795 | 0.854 | 4.96E-24 | 1.82E-19 |
| SNHG14 | 1 | -0.281223507 | 0.792 | 0.845 | 5.58E-24 | 2.04E-19 |
| GOLGB1 | 1 | 0.268371632 | 0.539 | 0.422 | 5.85E-24 | 2.14E-19 |

| | | | | | | |
|---|---|---|---|---|---|---|
| CLINT1 | 1 | 0.267109082 | 0.539 | 0.428 | 6.08E-24 | 2.23E-19 |
| RASGEF1B | 1 | 0.298522259 | 0.414 | 0.294 | 6.35E-24 | 2.33E-19 |
| RDX | 1 | -0.276494826 | 0.507 | 0.598 | 6.38E-24 | 2.34E-19 |
| RAB3B | 1 | -0.280813983 | 0.05 | 0.1 | 6.41E-24 | 2.35E-19 |
| ATP2B1 | 1 | -0.348634804 | 0.348 | 0.455 | 6.58E-24 | 2.41E-19 |
| RAB5A | 1 | 0.307294417 | 0.45 | 0.347 | 7.26E-24 | 2.66E-19 |
| RPL6 | 1 | 0.349474743 | 0.488 | 0.376 | 8.44E-24 | 3.09E-19 |
| PAK3 | 1 | -0.325220174 | 0.837 | 0.879 | 8.89E-24 | 3.26E-19 |
| PARN | 1 | 0.299491317 | 0.525 | 0.425 | 9.22E-24 | 3.37E-19 |
| TMEM41B | 1 | 0.313007895 | 0.549 | 0.437 | 9.46E-24 | 3.46E-19 |
| FMN1 | 1 | -0.404507589 | 0.053 | 0.125 | 9.81E-24 | 3.59E-19 |
| EDEM3 | 1 | -0.280253536 | 0.288 | 0.387 | 1.01E-23 | 3.7E-19 |
| QSER1 | 1 | 0.287287743 | 0.762 | 0.688 | 1.07E-23 | 3.91E-19 |
| REV3L | 1 | -0.318537887 | 0.838 | 0.874 | 1.16E-23 | 4.25E-19 |
| DPY19L3 | 1 | -0.297271717 | 0.425 | 0.518 | 1.19E-23 | 4.35E-19 |
| NFATC2 | 1 | -0.289527473 | 0.098 | 0.178 | 1.39E-23 | 5.1E-19 |
| TLK1 | 1 | -0.303057929 | 0.65 | 0.727 | 1.42E-23 | 5.21E-19 |
| FARS2 | 1 | -0.282942201 | 0.555 | 0.642 | 1.5E-23 | 5.47E-19 |
| PTN | 1 | -0.305284422 | 0.041 | 0.105 | 1.54E-23 | 5.66E-19 |
| PCDH19 | 1 | -0.479515799 | 0.141 | 0.215 | 1.56E-23 | 5.72E-19 |
| SNED1 | 1 | -0.278162839 | 0.055 | 0.124 | 1.66E-23 | 6.08E-19 |
| AC119673.2 | 1 | -0.284223586 | 0.32 | 0.407 | 2.01E-23 | 7.36E-19 |
| YLPM1 | 1 | -0.255022012 | 0.562 | 0.631 | 2.37E-23 | 8.67E-19 |
| DAB1 | 1 | -0.324121795 | 0.622 | 0.704 | 2.38E-23 | 8.7E-19 |
| STMN2 | 1 | -0.29295106 | 0.103 | 0.192 | 2.51E-23 | 9.21E-19 |
| UBE2O | 1 | 0.262868289 | 0.479 | 0.362 | 2.74E-23 | 1E-18 |
| NAPB | 1 | 0.267706411 | 0.379 | 0.28 | 2.97E-23 | 1.09E-18 |
| FYN | 1 | -0.333730477 | 0.616 | 0.674 | 3.56E-23 | 1.3E-18 |
| DHX15 | 1 | 0.256396778 | 0.791 | 0.74 | 4.15E-23 | 1.52E-18 |
| TET1 | 1 | -0.345531047 | 0.332 | 0.435 | 4.72E-23 | 1.73E-18 |
| TTC14 | 1 | 0.261418502 | 0.637 | 0.544 | 4.82E-23 | 1.77E-18 |
| MDGA2 | 1 | -0.59645488 | 0.042 | 0.105 | 4.83E-23 | 1.77E-18 |
| SCYL2 | 1 | 0.30227728 | 0.501 | 0.409 | 5.15E-23 | 1.88E-18 |
| PEX14 | 1 | 0.282858125 | 0.49 | 0.382 | 5.74E-23 | 2.1E-18 |
| RPL5 | 1 | 0.301238856 | 0.488 | 0.378 | 6.37E-23 | 2.33E-18 |
| SNX9 | 1 | 0.331293514 | 0.417 | 0.333 | 6.65E-23 | 2.43E-18 |
| KCTD8 | 1 | -0.676367359 | 0.11 | 0.159 | 6.76E-23 | 2.48E-18 |
| RBM39 | 1 | 0.251663879 | 0.884 | 0.852 | 6.87E-23 | 2.52E-18 |
| PRKCE | 1 | -0.35315364 | 0.441 | 0.546 | 7.03E-23 | 2.57E-18 |
| PLEKHA7 | 1 | -0.346566573 | 0.133 | 0.213 | 7.57E-23 | 2.77E-18 |
| SLC12A2 | 1 | -0.374341225 | 0.401 | 0.484 | 8.09E-23 | 2.96E-18 |
| CCDC6 | 1 | 0.292373608 | 0.561 | 0.45 | 8.43E-23 | 3.09E-18 |
| GAPVD1 | 1 | 0.270894726 | 0.535 | 0.439 | 9.61E-23 | 3.52E-18 |
| PDE5A | 1 | -0.342638929 | 0.205 | 0.279 | 1.14E-22 | 4.16E-18 |
| CPNE8 | 1 | -0.34821518 | 0.127 | 0.174 | 1.68E-22 | 6.15E-18 |
| SETBP1 | 1 | -0.371147113 | 0.525 | 0.598 | 1.72E-22 | 6.31E-18 |
| HIVEP1 | 1 | -0.285606689 | 0.419 | 0.521 | 2.09E-22 | 7.65E-18 |
| LRCH1 | 1 | -0.281886207 | 0.569 | 0.641 | 2.16E-22 | 7.91E-18 |
| NOTCH2 | 1 | -0.35931601 | 0.241 | 0.332 | 2.21E-22 | 8.1E-18 |
| PARP1 | 1 | 0.261359178 | 0.365 | 0.26 | 2.23E-22 | 8.18E-18 |
| ANP32B | 1 | 0.256311168 | 0.443 | 0.331 | 2.26E-22 | 8.28E-18 |
| TUBGCP3 | 1 | -0.262014491 | 0.393 | 0.494 | 2.4E-22 | 8.79E-18 |
| PFKFB4 | 1 | 0.385452592 | 0.187 | 0.13 | 2.52E-22 | 9.22E-18 |
| ST3GAL6 | 1 | -0.293798187 | 0.221 | 0.308 | 2.87E-22 | 1.05E-17 |
| HSPA4 | 1 | 0.269489141 | 0.396 | 0.287 | 2.94E-22 | 1.08E-17 |
| SIL1 | 1 | -0.309672839 | 0.661 | 0.664 | 3.03E-22 | 1.11E-17 |
| DNAH10 | 1 | -0.390801926 | 0.412 | 0.487 | 3.35E-22 | 1.23E-17 |
| STXBP5 | 1 | 0.299051702 | 0.66 | 0.613 | 3.36E-22 | 1.23E-17 |

| | | | | | | |
|---|---|---|---|---|---|---|
| ZRANB2-AS2 | 1 | -0.26133208 | 0.122 | 0.212 | 3.67E-22 | 1.34E-17 |
| MINDY3 | 1 | 0.271584587 | 0.493 | 0.389 | 3.83E-22 | 1.4E-17 |
| RESF1 | 1 | 0.271994234 | 0.278 | 0.184 | 4.03E-22 | 1.48E-17 |
| ITGB1 | 1 | 0.335389862 | 0.631 | 0.541 | 4.12E-22 | 1.51E-17 |
| PLPPR1 | 1 | -0.434762208 | 0.132 | 0.215 | 4.29E-22 | 1.57E-17 |
| ABL1 | 1 | -0.284572688 | 0.515 | 0.599 | 4.36E-22 | 1.59E-17 |
| CLSTN2 | 1 | 0.606244404 | 0.406 | 0.321 | 4.66E-22 | 1.71E-17 |
| NUTM2A-AS1 | 1 | -0.25581611 | 0.481 | 0.581 | 4.92E-22 | 1.8E-17 |
| GUCY1A1 | 1 | -0.414402778 | 0.213 | 0.275 | 5.25E-22 | 1.92E-17 |
| MAP4K4 | 1 | -0.2867912 | 0.831 | 0.861 | 5.37E-22 | 1.97E-17 |
| EEA1 | 1 | -0.284828967 | 0.417 | 0.516 | 5.73E-22 | 2.1E-17 |
| TJP1 | 1 | 0.286672887 | 0.754 | 0.69 | 8.92E-22 | 3.27E-17 |
| ATE1 | 1 | -0.260182275 | 0.581 | 0.656 | 9.9E-22 | 3.62E-17 |
| SYNJ1 | 1 | 0.253607861 | 0.446 | 0.336 | 1.04E-21 | 3.82E-17 |
| RPL35A | 1 | 0.342978632 | 0.334 | 0.234 | 1.25E-21 | 4.58E-17 |
| USP3 | 1 | 0.290399166 | 0.751 | 0.677 | 1.3E-21 | 4.75E-17 |
| PIK3C3 | 1 | 0.282466077 | 0.654 | 0.579 | 1.31E-21 | 4.81E-17 |
| HHIP | 1 | -0.299938562 | 0.067 | 0.136 | 1.34E-21 | 4.9E-17 |
| GPR137C | 1 | -0.331544395 | 0.378 | 0.446 | 1.35E-21 | 4.93E-17 |
| PPM1E | 1 | -0.323149851 | 0.697 | 0.748 | 1.36E-21 | 4.99E-17 |
| KCNJ3 | 1 | -0.481698615 | 0.097 | 0.156 | 1.66E-21 | 6.07E-17 |
| RPL24 | 1 | 0.320661527 | 0.278 | 0.186 | 2.11E-21 | 7.73E-17 |
| CHM | 1 | -0.27865229 | 0.648 | 0.694 | 2.15E-21 | 7.87E-17 |
| GNB4 | 1 | -0.26759022 | 0.12 | 0.203 | 2.22E-21 | 8.14E-17 |
| NLGN1 | 1 | -0.381130439 | 0.845 | 0.792 | 2.28E-21 | 8.36E-17 |
| ADGRA2 | 1 | -0.260436109 | 0.09 | 0.162 | 2.34E-21 | 8.57E-17 |
| AC108169.1 | 1 | 0.399313603 | 0.195 | 0.116 | 2.52E-21 | 9.23E-17 |
| MAP2K4 | 1 | -0.253083026 | 0.478 | 0.574 | 2.59E-21 | 9.48E-17 |
| LARGE1 | 1 | -0.323219808 | 0.202 | 0.299 | 2.78E-21 | 1.02E-16 |
| HHAT | 1 | 0.279459121 | 0.351 | 0.25 | 2.99E-21 | 1.09E-16 |
| AMD1 | 1 | 0.319570126 | 0.444 | 0.363 | 3E-21 | 1.1E-16 |
| NWD2 | 1 | -0.328875961 | 0.096 | 0.156 | 3.1E-21 | 1.13E-16 |
| WDPCP | 1 | -0.274881316 | 0.512 | 0.599 | 3.28E-21 | 1.2E-16 |
| PJA2 | 1 | 0.313676778 | 0.587 | 0.487 | 3.67E-21 | 1.34E-16 |
| MGAT5 | 1 | -0.303876065 | 0.398 | 0.497 | 3.82E-21 | 1.4E-16 |
| LRIG1 | 1 | -0.313347875 | 0.124 | 0.205 | 4.05E-21 | 1.48E-16 |
| HACD3 | 1 | 0.279152469 | 0.58 | 0.478 | 4.26E-21 | 1.56E-16 |
| RPS14 | 1 | 0.347296566 | 0.384 | 0.282 | 4.35E-21 | 1.59E-16 |
| RPGR | 1 | -0.306592365 | 0.392 | 0.466 | 4.69E-21 | 1.72E-16 |
| NCOR2 | 1 | -0.273967756 | 0.529 | 0.619 | 4.95E-21 | 1.81E-16 |
| EPC1 | 1 | 0.262187022 | 0.794 | 0.732 | 6.06E-21 | 2.22E-16 |
| RPS27A | 1 | 0.379903981 | 0.417 | 0.312 | 6.27E-21 | 2.29E-16 |
| MED14 | 1 | 0.254240774 | 0.701 | 0.623 | 7.12E-21 | 2.61E-16 |
| ATG4C | 1 | -0.252926588 | 0.226 | 0.321 | 7.72E-21 | 2.83E-16 |
| CDON | 1 | 0.280165387 | 0.416 | 0.305 | 8.18E-21 | 3E-16 |
| TRA2A | 1 | 0.271531542 | 0.762 | 0.709 | 9.05E-21 | 3.31E-16 |
| KIAA1211 | 1 | 0.270462968 | 0.724 | 0.687 | 1E-20 | 3.66E-16 |
| NPM1 | 1 | 0.302408797 | 0.464 | 0.369 | 1.09E-20 | 4.01E-16 |
| SNHG5 | 1 | 0.344451361 | 0.346 | 0.262 | 1.24E-20 | 4.54E-16 |
| RHOBTB1 | 1 | 0.270367744 | 0.327 | 0.226 | 1.25E-20 | 4.58E-16 |
| KDM3A | 1 | 0.47698311 | 0.477 | 0.397 | 1.26E-20 | 4.62E-16 |
| HS6ST2 | 1 | 0.30261523 | 0.735 | 0.644 | 1.31E-20 | 4.81E-16 |
| CLHC1 | 1 | -0.265841113 | 0.238 | 0.327 | 1.33E-20 | 4.86E-16 |
| STAT3 | 1 | 0.254931124 | 0.528 | 0.416 | 1.36E-20 | 4.99E-16 |
| PSEN1 | 1 | 0.262346341 | 0.432 | 0.333 | 1.46E-20 | 5.35E-16 |
| TIA1 | 1 | -0.257298078 | 0.545 | 0.632 | 1.48E-20 | 5.42E-16 |
| IFT80 | 1 | -0.261846142 | 0.385 | 0.481 | 1.67E-20 | 6.12E-16 |
| KLHL22 | 1 | -0.252695128 | 0.254 | 0.354 | 1.73E-20 | 6.35E-16 |

| | | | | | | |
|---|---|---|---|---|---|---|
| METTL15 | 1 | -0.260148204 | 0.6 | 0.653 | 1.94E-20 | 7.12E-16 |
| POT1-AS1 | 1 | -0.257291614 | 0.075 | 0.143 | 1.97E-20 | 7.2E-16 |
| TPM1 | 1 | -0.367466941 | 0.266 | 0.347 | 2.4E-20 | 8.77E-16 |
| ACYP2 | 1 | -0.288081473 | 0.49 | 0.541 | 2.56E-20 | 9.37E-16 |
| MORF4L2 | 1 | 0.264751362 | 0.625 | 0.524 | 2.77E-20 | 1.01E-15 |
| PPP2R2B | 1 | -0.250703235 | 0.566 | 0.676 | 2.88E-20 | 1.05E-15 |
| RACK1 | 1 | 0.261382172 | 0.39 | 0.287 | 3.4E-20 | 1.25E-15 |
| GDAP1 | 1 | -0.367940044 | 0.247 | 0.332 | 3.57E-20 | 1.31E-15 |
| ADAM22 | 1 | -0.297249844 | 0.341 | 0.428 | 3.76E-20 | 1.38E-15 |
| MGMT | 1 | -0.293294996 | 0.348 | 0.436 | 3.8E-20 | 1.39E-15 |
| TNRC6A | 1 | 0.259363627 | 0.839 | 0.812 | 3.96E-20 | 1.45E-15 |
| VEGFA | 1 | 0.398368242 | 0.353 | 0.355 | 4.2E-20 | 1.54E-15 |
| KIAA1671 | 1 | -0.279303237 | 0.121 | 0.183 | 5.3E-20 | 1.94E-15 |
| LCLAT1 | 1 | 0.265763157 | 0.653 | 0.558 | 6.03E-20 | 2.21E-15 |
| RADX | 1 | 0.29128329 | 0.579 | 0.465 | 6.14E-20 | 2.25E-15 |
| GART | 1 | 0.250476335 | 0.516 | 0.414 | 6.39E-20 | 2.34E-15 |
| VOPP1 | 1 | -0.261590332 | 0.265 | 0.356 | 8.15E-20 | 2.98E-15 |
| PPP4R1 | 1 | 0.270339417 | 0.614 | 0.542 | 8.34E-20 | 3.05E-15 |
| XRN1 | 1 | -0.252952196 | 0.615 | 0.688 | 1.21E-19 | 4.43E-15 |
| CNKSR2 | 1 | -0.431472714 | 0.65 | 0.665 | 1.32E-19 | 4.82E-15 |
| FAT1 | 1 | -0.335026298 | 0.574 | 0.627 | 1.52E-19 | 5.56E-15 |
| GLMN | 1 | 0.274980855 | 0.56 | 0.48 | 1.59E-19 | 5.81E-15 |
| LINC01578 | 1 | 0.302853237 | 0.616 | 0.535 | 1.63E-19 | 5.95E-15 |
| PRIM2 | 1 | -0.250005975 | 0.59 | 0.656 | 1.9E-19 | 6.95E-15 |
| MALT1 | 1 | 0.267189231 | 0.368 | 0.272 | 1.94E-19 | 7.09E-15 |
| CPXM2 | 1 | -0.267201528 | 0.044 | 0.1 | 2.16E-19 | 7.9E-15 |
| ANKRD26 | 1 | 0.264071708 | 0.661 | 0.597 | 3E-19 | 1.1E-14 |
| CUL3 | 1 | 0.285640404 | 0.733 | 0.672 | 3.04E-19 | 1.11E-14 |
| ACVR1 | 1 | 0.314006864 | 0.457 | 0.368 | 3.15E-19 | 1.15E-14 |
| ATP6V1A | 1 | 0.264474503 | 0.37 | 0.276 | 3.55E-19 | 1.3E-14 |
| BCLAF1 | 1 | 0.293499229 | 0.732 | 0.667 | 3.94E-19 | 1.44E-14 |
| DCX | 1 | 0.250636333 | 0.244 | 0.157 | 4.12E-19 | 1.51E-14 |
| FZD3 | 1 | -0.294611883 | 0.211 | 0.28 | 4.15E-19 | 1.52E-14 |
| B2M | 1 | 0.302955192 | 0.304 | 0.213 | 4.55E-19 | 1.67E-14 |
| WASHC4 | 1 | 0.266742517 | 0.669 | 0.584 | 4.66E-19 | 1.71E-14 |
| FGFR1 | 1 | 0.260345627 | 0.439 | 0.333 | 4.69E-19 | 1.72E-14 |
| RUNX1 | 1 | -0.445418803 | 0.076 | 0.131 | 4.76E-19 | 1.74E-14 |
| ANKRD37 | 1 | 0.283138535 | 0.116 | 0.061 | 6.34E-19 | 2.32E-14 |
| INSIG2 | 1 | 0.266535448 | 0.246 | 0.266 | 6.86E-19 | 2.51E-14 |
| DDX1 | 1 | 0.270486639 | 0.515 | 0.435 | 7.25E-19 | 2.65E-14 |
| SAT1 | 1 | -0.484232617 | 0.101 | 0.163 | 7.53E-19 | 2.76E-14 |
| WDR43 | 1 | 0.295856528 | 0.55 | 0.473 | 7.73E-19 | 2.83E-14 |
| TRIM24 | 1 | -0.288555107 | 0.433 | 0.516 | 8.1E-19 | 2.97E-14 |
| RIN2 | 1 | -0.269450772 | 0.293 | 0.38 | 8.24E-19 | 3.02E-14 |
| YME1L1 | 1 | 0.25732326 | 0.589 | 0.492 | 9.05E-19 | 3.31E-14 |
| RPL37 | 1 | 0.345867556 | 0.449 | 0.356 | 9.54E-19 | 3.49E-14 |
| CMTM8 | 1 | -0.277222562 | 0.09 | 0.154 | 9.59E-19 | 3.51E-14 |
| TMEM131L | 1 | -0.256605361 | 0.501 | 0.588 | 1E-18 | 3.67E-14 |
| NAAA | 1 | -0.266091684 | 0.234 | 0.314 | 1E-18 | 3.68E-14 |
| RPRD2 | 1 | 0.261326424 | 0.668 | 0.597 | 1.04E-18 | 3.79E-14 |
| PLD5 | 1 | -0.320373061 | 0.281 | 0.244 | 1.12E-18 | 4.1E-14 |
| RFX3 | 1 | -0.345602252 | 0.781 | 0.833 | 1.26E-18 | 4.61E-14 |
| ANK3 | 1 | 0.412042723 | 0.405 | 0.304 | 1.48E-18 | 5.43E-14 |
| SPAG16 | 1 | -0.319222086 | 0.231 | 0.296 | 1.8E-18 | 6.57E-14 |
| DGKD | 1 | 0.27901539 | 0.45 | 0.36 | 1.95E-18 | 7.14E-14 |
| ANLN | 1 | 0.35319163 | 0.337 | 0.249 | 2.09E-18 | 7.67E-14 |
| RPL11 | 1 | 0.322583662 | 0.384 | 0.295 | 2.18E-18 | 7.99E-14 |
| WDR26 | 1 | 0.255318815 | 0.552 | 0.46 | 2.29E-18 | 8.39E-14 |

| | | | | | | |
|---|---|---|---|---|---|---|
| SLC16A1 | 1 | 0.298227682 | 0.362 | 0.282 | 2.49E-18 | 9.1E-14 |
| SPPL2A | 1 | 0.259488711 | 0.41 | 0.325 | 2.51E-18 | 9.2E-14 |
| CD2AP | 1 | 0.257168877 | 0.646 | 0.56 | 2.71E-18 | 9.91E-14 |
| APCDD1 | 1 | -0.350195016 | 0.277 | 0.318 | 4.35E-18 | 1.59E-13 |
| PDZRN3 | 1 | -0.290588434 | 0.881 | 0.911 | 4.8E-18 | 1.76E-13 |
| PSME4 | 1 | -0.263048189 | 0.752 | 0.797 | 5.29E-18 | 1.94E-13 |
| RPL10A | 1 | 0.273502063 | 0.351 | 0.259 | 5.54E-18 | 2.03E-13 |
| TXNRD1 | 1 | 0.267006893 | 0.415 | 0.322 | 6.5E-18 | 2.38E-13 |
| KDM3B | 1 | 0.257647437 | 0.542 | 0.449 | 6.57E-18 | 2.41E-13 |
| RAI14 | 1 | -0.270247112 | 0.396 | 0.497 | 7.1E-18 | 2.6E-13 |
| CHKA | 1 | -0.261716602 | 0.293 | 0.354 | 9.03E-18 | 3.31E-13 |
| ARHGAP28 | 1 | 0.40440717 | 0.322 | 0.233 | 9.55E-18 | 3.5E-13 |
| TPT1 | 1 | 0.272336185 | 0.328 | 0.237 | 1.01E-17 | 3.71E-13 |
| MRPS28 | 1 | -0.250975783 | 0.373 | 0.438 | 1.23E-17 | 4.51E-13 |
| SLC35F1 | 1 | -0.323250141 | 0.287 | 0.308 | 1.28E-17 | 4.7E-13 |
| DCP2 | 1 | 0.267296943 | 0.643 | 0.556 | 1.49E-17 | 5.46E-13 |
| RPL26 | 1 | 0.310428107 | 0.389 | 0.299 | 1.64E-17 | 6.01E-13 |
| PABPC1 | 1 | 0.270878122 | 0.713 | 0.64 | 1.96E-17 | 7.16E-13 |
| RHOBTB3 | 1 | -0.276950622 | 0.396 | 0.486 | 2.13E-17 | 7.79E-13 |
| NFIA | 1 | -0.489705012 | 0.908 | 0.889 | 2.29E-17 | 8.38E-13 |
| RPL27A | 1 | 0.280021145 | 0.337 | 0.25 | 2.42E-17 | 8.86E-13 |
| MACROD2 | 1 | 0.490286735 | 0.61 | 0.556 | 2.79E-17 | 1.02E-12 |
| RPS9 | 1 | 0.254277357 | 0.39 | 0.294 | 2.92E-17 | 1.07E-12 |
| ZBTB7C | 1 | -0.276759176 | 0.08 | 0.141 | 2.98E-17 | 1.09E-12 |
| LYPD6 | 1 | 0.270373124 | 0.425 | 0.324 | 3.08E-17 | 1.13E-12 |
| CCNL1 | 1 | 0.254345465 | 0.577 | 0.49 | 3.31E-17 | 1.21E-12 |
| RPL7 | 1 | 0.292473139 | 0.273 | 0.196 | 3.39E-17 | 1.24E-12 |
| SNAP91 | 1 | -0.264187296 | 0.126 | 0.203 | 3.79E-17 | 1.39E-12 |
| SLC35F3 | 1 | -0.330378251 | 0.072 | 0.122 | 4.47E-17 | 1.63E-12 |
| CRMP1 | 1 | 0.26312243 | 0.396 | 0.301 | 4.65E-17 | 1.7E-12 |
| SERGEF | 1 | 0.253387396 | 0.492 | 0.393 | 5.5E-17 | 2.01E-12 |
| RPL4 | 1 | 0.259828369 | 0.339 | 0.253 | 5.61E-17 | 2.05E-12 |
| RPS16 | 1 | 0.269848629 | 0.389 | 0.296 | 6.2E-17 | 2.27E-12 |
| FGF12 | 1 | 0.378328416 | 0.753 | 0.731 | 6.77E-17 | 2.48E-12 |
| ARHGAP26 | 1 | -0.254603525 | 0.121 | 0.195 | 8.35E-17 | 3.06E-12 |
| FANCI | 1 | -0.268144524 | 0.345 | 0.418 | 9.13E-17 | 3.34E-12 |
| ILRUN | 1 | 0.27320289 | 0.421 | 0.348 | 9.38E-17 | 3.43E-12 |
| VCL | 1 | -0.348960901 | 0.758 | 0.785 | 1.12E-16 | 4.08E-12 |
| ADGRD1 | 1 | -0.289726326 | 0.211 | 0.269 | 1.26E-16 | 4.6E-12 |
| DAPK2 | 1 | 0.27228083 | 0.278 | 0.198 | 1.49E-16 | 5.45E-12 |
| RALGAPA1 | 1 | 0.257905424 | 0.768 | 0.726 | 1.75E-16 | 6.41E-12 |
| PACRG | 1 | -0.290482083 | 0.102 | 0.095 | 1.86E-16 | 6.79E-12 |
| LHFPL6 | 1 | -0.287722719 | 0.39 | 0.383 | 2.06E-16 | 7.52E-12 |
| NHSL1 | 1 | 0.261731513 | 0.649 | 0.555 | 2.12E-16 | 7.76E-12 |
| CENPF | 1 | 0.275001508 | 0.266 | 0.185 | 2.35E-16 | 8.6E-12 |
| FKBP5 | 1 | -0.257717571 | 0.258 | 0.338 | 2.59E-16 | 9.48E-12 |
| RPL28 | 1 | 0.363342014 | 0.381 | 0.298 | 3.04E-16 | 1.11E-11 |
| RPS8 | 1 | 0.336682835 | 0.529 | 0.441 | 3.45E-16 | 1.26E-11 |
| PTEN | 1 | -0.293081528 | 0.662 | 0.714 | 4.34E-16 | 1.59E-11 |
| SERBP1 | 1 | 0.256266418 | 0.555 | 0.484 | 4.49E-16 | 1.64E-11 |
| FRY | 1 | 0.272388155 | 0.634 | 0.553 | 4.56E-16 | 1.67E-11 |
| ZMYND8 | 1 | 0.250248809 | 0.636 | 0.559 | 5.24E-16 | 1.92E-11 |
| CMIP | 1 | -0.260351363 | 0.529 | 0.604 | 5.8E-16 | 2.12E-11 |
| BNIP3 | 1 | 0.325316516 | 0.23 | 0.197 | 8.17E-16 | 2.99E-11 |
| NCKAP5 | 1 | -0.510474564 | 0.37 | 0.424 | 9.1E-16 | 3.33E-11 |
| MARK1 | 1 | -0.356764506 | 0.473 | 0.515 | 1.01E-15 | 3.71E-11 |
| PPP2R2A | 1 | 0.261238187 | 0.712 | 0.649 | 1.38E-15 | 5.05E-11 |
| NTF3 | 1 | -0.32670459 | 0.121 | 0.184 | 1.39E-15 | 5.09E-11 |

| | | | | | | |
|---|---|---|---|---|---|---|
| RPL7A | 1 | 0.282605397 | 0.453 | 0.362 | 1.63E-15 | 5.98E-11 |
| CAMK1D | 1 | -0.271032599 | 0.614 | 0.652 | 1.68E-15 | 6.16E-11 |
| FAM214A | 1 | -0.264744182 | 0.345 | 0.406 | 1.87E-15 | 6.83E-11 |
| CRISPLD1 | 1 | -0.30903501 | 0.558 | 0.477 | 2.26E-15 | 8.27E-11 |
| HPCAL1 | 1 | 0.260035998 | 0.394 | 0.301 | 2.64E-15 | 9.68E-11 |
| DNAH7 | 1 | -0.374252348 | 0.19 | 0.173 | 2.69E-15 | 9.86E-11 |
| RPS25 | 1 | 0.271248034 | 0.383 | 0.293 | 2.8E-15 | 1.03E-10 |
| PAG1 | 1 | -0.301855729 | 0.264 | 0.319 | 3.04E-15 | 1.11E-10 |
| RPL13 | 1 | 0.311048849 | 0.489 | 0.397 | 4.07E-15 | 1.49E-10 |
| HMGCR | 1 | 0.264784869 | 0.268 | 0.196 | 4.95E-15 | 1.81E-10 |
| GARS | 1 | 0.257164719 | 0.325 | 0.262 | 5.16E-15 | 1.89E-10 |
| GLDC | 1 | -0.268289246 | 0.226 | 0.294 | 5.25E-15 | 1.92E-10 |
| DTNA | 1 | -0.315326469 | 0.64 | 0.715 | 5.31E-15 | 1.94E-10 |
| CNTN6 | 1 | -0.534411124 | 0.149 | 0.188 | 5.5E-15 | 2.02E-10 |
| MEF2A | 1 | 0.281047896 | 0.811 | 0.807 | 7.23E-15 | 2.65E-10 |
| GPM6A | 1 | -0.543009774 | 0.15 | 0.215 | 7.69E-15 | 2.81E-10 |
| MOB1B | 1 | -0.264432058 | 0.35 | 0.426 | 8.68E-15 | 3.18E-10 |
| TMEM165 | 1 | 0.2748706 | 0.605 | 0.535 | 9.3E-15 | 3.41E-10 |
| RPL3 | 1 | 0.262449031 | 0.409 | 0.321 | 9.49E-15 | 3.48E-10 |
| RPL37A | 1 | 0.347802719 | 0.555 | 0.475 | 9.6E-15 | 3.51E-10 |
| TXNL1 | 1 | 0.266772843 | 0.395 | 0.323 | 9.99E-15 | 3.66E-10 |
| PTPRR | 1 | 0.282182238 | 0.102 | 0.058 | 1.09E-14 | 3.99E-10 |
| RPS3A | 1 | 0.352473357 | 0.403 | 0.328 | 1.19E-14 | 4.37E-10 |
| RALGPS2 | 1 | -0.258084686 | 0.344 | 0.375 | 1.99E-14 | 7.3E-10 |
| MIR137HG | 1 | -0.352692136 | 0.204 | 0.265 | 2.48E-14 | 9.1E-10 |
| ARIH1 | 1 | 0.252364318 | 0.811 | 0.759 | 2.83E-14 | 1.04E-09 |
| FGFR2 | 1 | -0.318003073 | 0.346 | 0.404 | 2.91E-14 | 1.06E-09 |
| RPL18A | 1 | 0.319346813 | 0.282 | 0.208 | 3.2E-14 | 1.17E-09 |
| PALLD | 1 | 0.313491997 | 0.47 | 0.397 | 3.61E-14 | 1.32E-09 |
| LRP8 | 1 | 0.27794709 | 0.346 | 0.275 | 5.05E-14 | 1.85E-09 |
| CDH10 | 1 | -0.378727529 | 0.305 | 0.336 | 8.75E-14 | 3.2E-09 |
| RPS6 | 1 | 0.321093413 | 0.424 | 0.35 | 9.82E-14 | 3.6E-09 |
| ARHGEF3 | 1 | 0.274295216 | 0.519 | 0.439 | 1.27E-13 | 4.66E-09 |
| RND3 | 1 | 0.348784109 | 0.37 | 0.321 | 1.36E-13 | 4.99E-09 |
| AL589693.1 | 1 | -0.466627276 | 0.059 | 0.107 | 1.56E-13 | 5.7E-09 |
| RAI2 | 1 | 0.263062636 | 0.204 | 0.139 | 1.61E-13 | 5.9E-09 |
| SNTG2 | 1 | 0.291680508 | 0.298 | 0.219 | 3.47E-13 | 1.27E-08 |
| HIF1A-AS3 | 1 | 0.34096317 | 0.172 | 0.115 | 3.66E-13 | 1.34E-08 |
| RPS28 | 1 | 0.292122936 | 0.34 | 0.275 | 6.38E-13 | 2.34E-08 |
| RPS19 | 1 | 0.307488253 | 0.461 | 0.384 | 6.68E-13 | 2.45E-08 |
| FMN2 | 1 | -0.29986638 | 0.305 | 0.368 | 6.7E-13 | 2.45E-08 |
| RPS7 | 1 | 0.268073814 | 0.349 | 0.28 | 7.11E-13 | 0.000000026 |
| RPS29 | 1 | 0.298751292 | 0.357 | 0.293 | 7.33E-13 | 2.68E-08 |
| RPS23 | 1 | 0.285655686 | 0.373 | 0.296 | 9.58E-13 | 3.51E-08 |
| RPL35 | 1 | 0.26946521 | 0.299 | 0.226 | 9.72E-13 | 3.56E-08 |
| PLPP1 | 1 | -0.307604155 | 0.337 | 0.349 | 1.23E-12 | 4.49E-08 |
| MARCH3 | 1 | -0.253862301 | 0.161 | 0.208 | 1.74E-12 | 6.39E-08 |
| FNDC3B | 1 | -0.251233512 | 0.66 | 0.706 | 1.75E-12 | 0.000000064 |
| TDRD3 | 1 | -0.251660689 | 0.609 | 0.645 | 1.77E-12 | 6.48E-08 |
| RYBP | 1 | 0.255718849 | 0.567 | 0.499 | 2.05E-12 | 0.000000075 |
| PRICKLE1 | 1 | 0.444591756 | 0.462 | 0.394 | 3.47E-12 | 0.000000127 |
| RPS11 | 1 | 0.306137026 | 0.352 | 0.272 | 4.16E-12 | 0.000000152 |
| TAF3 | 1 | 0.269346922 | 0.532 | 0.479 | 4.29E-12 | 0.000000157 |
| RPS4X | 1 | 0.288464619 | 0.35 | 0.276 | 5.13E-12 | 0.000000188 |
| PCDH15 | 1 | -0.551055378 | 0.096 | 0.148 | 5.44E-12 | 0.000000199 |
| CALN1 | 1 | -0.265991281 | 0.063 | 0.102 | 5.87E-12 | 0.000000215 |
| PDE3A | 1 | -0.280639498 | 0.57 | 0.567 | 8.39E-12 | 0.000000307 |
| MAN2A1 | 1 | 0.308913549 | 0.503 | 0.441 | 1.31E-11 | 0.00000048 |

| | | | | | | |
|---|---|---|---|---|---|---|
| KDM5B | 1 | 0.310181521 | 0.467 | 0.394 | 1.4E-11 | 0.000000513 |
| AL513166.1 | 1 | -0.291053417 | 0.217 | 0.262 | 1.74E-11 | 0.000000638 |
| WSB1 | 1 | 0.333044817 | 0.809 | 0.782 | 3.89E-11 | 0.00000142 |
| ARHGAP42 | 1 | -0.259652876 | 0.34 | 0.383 | 4.74E-11 | 0.00000173 |
| RPLP1 | 1 | 0.25548408 | 0.504 | 0.429 | 5.78E-11 | 0.00000212 |
| ASIC2 | 1 | -0.251979852 | 0.222 | 0.283 | 7.29E-11 | 0.00000267 |
| TMEFF2 | 1 | 0.28065215 | 0.19 | 0.131 | 1.25E-10 | 0.00000459 |
| THSD7B | 1 | -0.408488471 | 0.188 | 0.219 | 1.35E-10 | 0.00000495 |
| RLF | 1 | 0.282377784 | 0.515 | 0.517 | 2.01E-10 | 0.00000736 |
| RPS20 | 1 | 0.261457032 | 0.379 | 0.313 | 2.4E-10 | 0.00000879 |
| CDH7 | 1 | -0.264518969 | 0.252 | 0.297 | 4.33E-10 | 0.0000158 |
| LAMA3 | 1 | 0.301399242 | 0.232 | 0.204 | 5.45E-10 | 0.0000199 |
| CACNA1C | 1 | -0.338908322 | 0.49 | 0.477 | 4.38E-09 | 0.00016 |
| COL19A1 | 1 | -0.280264741 | 0.091 | 0.107 | 1.28E-08 | 0.00047 |
| ZFHX3 | 1 | -0.253889656 | 0.568 | 0.586 | 1.52E-08 | 0.000557 |
| SEMA3C | 1 | -0.274656958 | 0.118 | 0.124 | 2.34E-08 | 0.000856 |
| RPL32 | 1 | 0.274412149 | 0.277 | 0.222 | 0.00000006 | 0.002198269 |
| JAKMIP2 | 1 | 0.269380966 | 0.389 | 0.341 | 8.98E-08 | 0.003289236 |
| AC019197.1 | 1 | 0.257725996 | 0.148 | 0.11 | 0.000000946 | 0.034640907 |
| THSD7A | 1 | -0.31146662 | 0.348 | 0.4 | 0.00000128 | 0.046927598 |
| CSMD3 | 1 | -0.336410914 | 0.283 | 0.338 | 0.00000409 | 0.149559508 |
| KCTD16 | 1 | -0.267153206 | 0.14 | 0.173 | 0.00000667 | 0.244270053 |
| HSPH1 | 1 | 0.250795904 | 0.299 | 0.258 | 0.000142 | 1 |
| GBE1 | 1 | 0.300646933 | 0.339 | 0.312 | 0.000237 | 1 |
| PDE10A | 1 | 0.260764625 | 0.278 | 0.245 | 0.0004 | 1 |
| ASPH | 1 | -0.250737293 | 0.69 | 0.704 | 0.000711 | 1 |
| HS6ST3 | 2 | 3.691021919 | 0.995 | 0.392 | 0 | 0 |
| DACH2 | 2 | 3.316109707 | 0.998 | 0.325 | 0 | 0 |
| LRRC7 | 2 | 3.228809011 | 0.934 | 0.173 | 0 | 0 |
| ZNF804B | 2 | 3.00296702 | 0.673 | 0.072 | 0 | 0 |
| NKAIN2 | 2 | 2.961129507 | 0.908 | 0.254 | 0 | 0 |
| NEGR1 | 2 | 2.887123598 | 0.99 | 0.482 | 0 | 0 |
| LINC01934 | 2 | 2.772827958 | 0.382 | 0.039 | 0 | 0 |
| SNTG1 | 2 | 2.529902275 | 0.888 | 0.27 | 0 | 0 |
| FRMD5 | 2 | 2.47564322 | 0.966 | 0.28 | 0 | 0 |
| KCNMB2 | 2 | 2.340768299 | 0.712 | 0.17 | 0 | 0 |
| DLX6-AS1 | 2 | 2.31935705 | 0.724 | 0.107 | 0 | 0 |
| KCNIP4 | 2 | 2.254721523 | 0.963 | 0.393 | 0 | 0 |
| NELL2 | 2 | 2.251512058 | 0.84 | 0.123 | 0 | 0 |
| LRRTM4 | 2 | 2.235804945 | 0.973 | 0.486 | 0 | 0 |
| NRG3 | 2 | 2.198496178 | 0.989 | 0.345 | 0 | 0 |
| ARPP21 | 2 | 2.192868711 | 0.82 | 0.199 | 0 | 0 |
| CADM2 | 2 | 2.086974981 | 0.898 | 0.344 | 0 | 0 |
| LINC02470 | 2 | 2.078899219 | 0.792 | 0.149 | 0 | 0 |
| SLIT2 | 2 | 2.07794459 | 0.996 | 0.658 | 0 | 0 |
| EPHA5 | 2 | 2.0743855 | 0.663 | 0.165 | 0 | 0 |
| CNTNAP5 | 2 | 2.070953089 | 0.82 | 0.15 | 0 | 0 |
| RALYL | 2 | 2.068304075 | 0.959 | 0.364 | 0 | 0 |
| THSD7B | 2 | 2.043597967 | 0.687 | 0.102 | 0 | 0 |
| ARHGAP24 | 2 | 2.008121472 | 0.84 | 0.312 | 0 | 0 |
| ITGA4 | 2 | 2.006964258 | 0.834 | 0.389 | 0 | 0 |
| RIMS2 | 2 | 1.952434646 | 0.975 | 0.302 | 0 | 0 |
| GRIN3A | 2 | 1.95114739 | 0.738 | 0.147 | 0 | 0 |
| LRFN5 | 2 | 1.948620277 | 0.836 | 0.373 | 0 | 0 |
| BRINP3 | 2 | 1.939622635 | 0.86 | 0.428 | 0 | 0 |
| CHRM2 | 2 | 1.902505497 | 0.417 | 0.022 | 0 | 0 |
| SVEP1 | 2 | 1.892849397 | 0.753 | 0.188 | 0 | 0 |
| STXBP5L | 2 | 1.889989135 | 0.867 | 0.178 | 0 | 0 |

| | | | | | | |
|---|---|---|---|---|---|---|
| PHACTR1 | 2 | 1.818314835 | 0.846 | 0.202 | 0 | 0 |
| CNKSR2 | 2 | 1.816528475 | 0.947 | 0.596 | 0 | 0 |
| UNC79 | 2 | 1.786501755 | 0.883 | 0.219 | 0 | 0 |
| ST18 | 2 | 1.780867725 | 0.941 | 0.403 | 0 | 0 |
| PIP5K1B | 2 | 1.776462412 | 0.769 | 0.167 | 0 | 0 |
| NLGN1 | 2 | 1.766023217 | 0.993 | 0.762 | 0 | 0 |
| ADAMTSL1 | 2 | 1.765653284 | 0.892 | 0.438 | 0 | 0 |
| CDH7 | 2 | 1.746861349 | 0.702 | 0.191 | 0 | 0 |
| GLIS3 | 2 | 1.729222796 | 0.664 | 0.151 | 0 | 0 |
| NRXN1 | 2 | 1.712032857 | 0.991 | 0.507 | 0 | 0 |
| HS3ST4 | 2 | 1.706501261 | 0.769 | 0.24 | 0 | 0 |
| UNC5D | 2 | 1.689302461 | 0.98 | 0.486 | 0 | 0 |
| ATRNL1 | 2 | 1.652960976 | 0.988 | 0.765 | 0 | 0 |
| RIMS1 | 2 | 1.650504788 | 0.879 | 0.336 | 0 | 0 |
| GRIA4 | 2 | 1.634533816 | 0.791 | 0.207 | 0 | 0 |
| DGKI | 2 | 1.631566452 | 0.77 | 0.113 | 0 | 0 |
| GRIA2 | 2 | 1.623669176 | 0.996 | 0.581 | 0 | 0 |
| ADAMTS20 | 2 | 1.617118268 | 0.699 | 0.127 | 0 | 0 |
| EBF1 | 2 | 1.606386441 | 0.851 | 0.259 | 0 | 0 |
| NFIA | 2 | 1.575560971 | 0.996 | 0.87 | 0 | 0 |
| SEMA6D | 2 | 1.574072084 | 0.959 | 0.607 | 0 | 0 |
| PCDH19 | 2 | 1.566286539 | 0.551 | 0.115 | 0 | 0 |
| AFF2 | 2 | 1.561896603 | 0.909 | 0.585 | 0 | 0 |
| PCLO | 2 | 1.555270408 | 0.949 | 0.529 | 0 | 0 |
| MIR137HG | 2 | 1.551100141 | 0.641 | 0.16 | 0 | 0 |
| ELAVL2 | 2 | 1.543594677 | 0.845 | 0.276 | 0 | 0 |
| PTPRZ1 | 2 | 1.543356332 | 0.826 | 0.312 | 0 | 0 |
| CNTN1 | 2 | 1.541176561 | 0.734 | 0.144 | 0 | 0 |
| PID1 | 2 | 1.531587784 | 0.921 | 0.505 | 0 | 0 |
| CACNA1B | 2 | 1.520522737 | 0.755 | 0.151 | 0 | 0 |
| FMN2 | 2 | 1.50635707 | 0.807 | 0.248 | 0 | 0 |
| UNC13B | 2 | 1.505873476 | 0.847 | 0.469 | 0 | 0 |
| SYT10 | 2 | 1.500902711 | 0.509 | 0.037 | 0 | 0 |
| ANO4 | 2 | 1.495839705 | 0.98 | 0.492 | 0 | 0 |
| VWDE | 2 | 1.483303032 | 0.78 | 0.311 | 0 | 0 |
| ZNF385D | 2 | 1.468297528 | 0.939 | 0.488 | 0 | 0 |
| DOCK3 | 2 | 1.467878906 | 0.791 | 0.169 | 0 | 0 |
| ERC2 | 2 | 1.463419812 | 0.854 | 0.358 | 0 | 0 |
| PAK5 | 2 | 1.453871517 | 0.82 | 0.365 | 0 | 0 |
| CTTNBP2 | 2 | 1.453066597 | 0.725 | 0.196 | 0 | 0 |
| TMEM163 | 2 | 1.448409201 | 0.596 | 0.052 | 0 | 0 |
| LHFPL3 | 2 | 1.423586061 | 0.933 | 0.482 | 0 | 0 |
| FGF13 | 2 | 1.415344375 | 0.855 | 0.351 | 0 | 0 |
| DOCK11 | 2 | 1.403100981 | 0.898 | 0.445 | 0 | 0 |
| CTNNA2 | 2 | 1.40274755 | 0.943 | 0.387 | 0 | 0 |
| XKR4 | 2 | 1.394539997 | 0.57 | 0.099 | 0 | 0 |
| PTPRD | 2 | 1.391923013 | 0.988 | 0.807 | 0 | 0 |
| ZNF536 | 2 | 1.372937014 | 0.991 | 0.783 | 0 | 0 |
| CACNA1D | 2 | 1.333416916 | 0.818 | 0.33 | 0 | 0 |
| PRKCB | 2 | 1.308588038 | 0.434 | 0.029 | 0 | 0 |
| MAP2 | 2 | 1.305750315 | 0.946 | 0.453 | 0 | 0 |
| KIAA1549L | 2 | 1.305295774 | 0.644 | 0.125 | 0 | 0 |
| PDE11A | 2 | 1.299966249 | 0.544 | 0.107 | 0 | 0 |
| FHOD3 | 2 | 1.291169725 | 0.727 | 0.21 | 0 | 0 |
| RBFOX1 | 2 | 1.251902267 | 0.977 | 0.679 | 0 | 0 |
| DPP10 | 2 | 1.250722009 | 0.925 | 0.532 | 0 | 0 |
| SNAP91 | 2 | 1.244905187 | 0.593 | 0.09 | 0 | 0 |
| HHIP | 2 | 1.242610296 | 0.449 | 0.043 | 0 | 0 |

| | | | | | | |
|---|---|---|---|---|---|---|
| RGL1 | 2 | 1.241882636 | 0.947 | 0.676 | 0 | 0 |
| ADGRL3 | 2 | 1.225353486 | 0.993 | 0.773 | 0 | 0 |
| ME3 | 2 | 1.220693061 | 0.686 | 0.225 | 0 | 0 |
| DCLK1 | 2 | 1.218627035 | 0.949 | 0.516 | 0 | 0 |
| CDH20 | 2 | 1.217254747 | 0.407 | 0.028 | 0 | 0 |
| SORCS1 | 2 | 1.213317084 | 0.621 | 0.141 | 0 | 0 |
| PEX5L | 2 | 1.204840292 | 0.992 | 0.711 | 0 | 0 |
| CELF4 | 2 | 1.20483075 | 0.678 | 0.199 | 0 | 0 |
| NOL4 | 2 | 1.195309653 | 0.747 | 0.231 | 0 | 0 |
| RUNDC3B | 2 | 1.189850993 | 0.518 | 0.072 | 0 | 0 |
| SYT14 | 2 | 1.187529109 | 0.869 | 0.384 | 0 | 0 |
| FUT9 | 2 | 1.168488002 | 0.395 | 0.024 | 0 | 0 |
| NBEA | 2 | 1.152218523 | 0.969 | 0.642 | 0 | 0 |
| STMN2 | 2 | 1.149199173 | 0.581 | 0.075 | 0 | 0 |
| CAMTA1 | 2 | 1.133658829 | 0.892 | 0.668 | 0 | 0 |
| TMEM178B | 2 | 1.080242635 | 0.891 | 0.414 | 0 | 0 |
| MAPT | 2 | 1.031659408 | 0.472 | 0.057 | 0 | 0 |
| AKAP13 | 2 | -1.485294528 | 0.136 | 0.648 | 0 | 0 |
| TENM3 | 2 | -1.569111342 | 0.452 | 0.858 | 0 | 0 |
| STARD13 | 2 | -1.845967716 | 0.202 | 0.686 | 0 | 0 |
| SVIL | 2 | -1.860679174 | 0.127 | 0.62 | 0 | 0 |
| GULP1 | 2 | -1.923427115 | 0.083 | 0.604 | 0 | 0 |
| UTRN | 2 | -1.981570007 | 0.223 | 0.713 | 0 | 0 |
| ZBTB20 | 2 | -1.984016623 | 0.467 | 0.861 | 0 | 0 |
| COL5A2 | 2 | -2.085609791 | 0.059 | 0.567 | 0 | 0 |
| TCF7L2 | 2 | -2.094457362 | 0.084 | 0.702 | 0 | 0 |
| TRPS1 | 2 | -2.112227653 | 0.117 | 0.697 | 0 | 0 |
| CASC15 | 2 | -2.119304695 | 0.194 | 0.66 | 0 | 0 |
| SOX5 | 2 | -2.461105953 | 0.231 | 0.746 | 0 | 0 |
| LAMA4 | 2 | -2.487716979 | 0.059 | 0.546 | 0 | 0 |
| MAML2 | 2 | -2.516204185 | 0.075 | 0.622 | 0 | 0 |
| NEAT1 | 2 | -2.621220463 | 0.653 | 0.914 | 0 | 0 |
| DMD | 2 | -3.576218924 | 0.333 | 0.75 | 0 | 0 |
| MYT1L | 2 | 1.443768665 | 0.451 | 0.056 | 5e-324 | 1.8088e-319 |
| TENM2 | 2 | 1.249255397 | 0.934 | 0.539 | 9e-323 | 3.255883e-318 |
| DPYSL5 | 2 | 1.019814484 | 0.601 | 0.137 | 1.527e-321 | 5.5892653e-317 |
| SLC35F1 | 2 | 1.222454593 | 0.695 | 0.213 | 9.39e-321 | 3.4385739e-316 |
| ANOS1 | 2 | 1.234177312 | 0.823 | 0.354 | 3.451e-318 | 1.26344529135e-313 |
| CACNB4 | 2 | -2.104460752 | 0.198 | 0.627 | 2.50191187e-316 | 9.159749546094e-312 |
| MAMLD1 | 2 | 1.255026819 | 0.683 | 0.227 | 1.020251265e-315 | 3.735241906732e-311 |
| NTM | 2 | -2.211242551 | 0.057 | 0.498 | 4.494696705e-315 | 1.64555341062243e-310 |
| DCLK2 | 2 | 1.038056202 | 0.957 | 0.675 | 2.4847693526e-313 | 0 |
| DGKB | 2 | 0.620504033 | 0.669 | 0.197 | 6.9483513937e-313 | 2.54E-308 |
| ASTN1 | 2 | 1.105436778 | 0.617 | 0.162 | 0 | 4.69E-304 |
| SYN3 | 2 | 1.534737182 | 0.791 | 0.339 | 9.56E-308 | 3.5E-303 |
| SCN3A | 2 | 1.015794019 | 0.647 | 0.179 | 6.6E-302 | 2.42E-297 |
| ANK1 | 2 | 1.052015796 | 0.546 | 0.113 | 3.34E-301 | 1.22E-296 |
| RAB3C | 2 | 1.005431635 | 0.819 | 0.337 | 1.68E-299 | 6.16E-295 |
| MID1 | 2 | -1.760364724 | 0.256 | 0.687 | 7.82E-298 | 2.86E-293 |
| DNER | 2 | 1.206822045 | 0.77 | 0.303 | 1.23E-297 | 4.51E-293 |
| MEGF10 | 2 | -1.743656842 | 0.054 | 0.48 | 4.77E-296 | 1.75E-291 |
| AC073050.1 | 2 | 1.453589348 | 0.677 | 0.218 | 5.8E-296 | 2.12E-291 |
| WWTR1 | 2 | -1.593118307 | 0.072 | 0.509 | 2.81E-295 | 1.03E-290 |
| KHDRBS3 | 2 | 1.530763003 | 0.569 | 0.167 | 4.92E-295 | 1.8E-290 |
| ABCB1 | 2 | 1.268448505 | 0.583 | 0.156 | 3.74E-292 | 1.37E-287 |
| LHFPL6 | 2 | -1.644481799 | 0.044 | 0.463 | 1.34E-286 | 4.91E-282 |
| ZNF827 | 2 | -1.252424244 | 0.178 | 0.63 | 2.39E-280 | 8.75E-276 |
| CYP7B1 | 2 | 1.404300667 | 0.69 | 0.295 | 8.25E-279 | 3.02E-274 |

| | | | | | | |
|---|---|---|---|---|---|---|
| AL157778.1 | 2 | 1.959800094 | 0.51 | 0.176 | 5.09E-278 | 1.86E-273 |
| SLCO5A1 | 2 | 1.040085123 | 0.457 | 0.079 | 3.63E-277 | 1.33E-272 |
| PHACTR3 | 2 | 0.884562453 | 0.415 | 0.057 | 3.95E-275 | 1.44E-270 |
| SRRM4 | 2 | 0.9040725 | 0.512 | 0.106 | 8.65E-275 | 3.17E-270 |
| KIF5C | 2 | 0.962463355 | 0.659 | 0.206 | 1.67E-274 | 6.13E-270 |
| RREB1 | 2 | -1.037182952 | 0.027 | 0.416 | 4.08E-274 | 1.49E-269 |
| PCBP3 | 2 | 1.177793233 | 0.59 | 0.171 | 5.75E-274 | 2.1E-269 |
| EPHA3 | 2 | 1.587704514 | 0.546 | 0.14 | 1.27E-273 | 4.66E-269 |
| KLHL29 | 2 | 1.116048462 | 0.9 | 0.551 | 7.16E-273 | 2.62E-268 |
| PCDH9 | 2 | 1.969658045 | 0.648 | 0.277 | 4.44E-272 | 1.62E-267 |
| MPPED2 | 2 | 0.858351078 | 0.514 | 0.111 | 1.63E-270 | 5.97E-266 |
| ADAMTS19 | 2 | 1.084302007 | 0.838 | 0.405 | 2.35E-267 | 8.62E-263 |
| ATP8A2 | 2 | 1.114530357 | 0.668 | 0.232 | 3.7E-264 | 1.36E-259 |
| AC074286.1 | 2 | 0.977833652 | 0.382 | 0.05 | 1.67E-262 | 6.11E-258 |
| YAP1 | 2 | -1.286625975 | 0.029 | 0.404 | 1.38E-261 | 5.04E-257 |
| FAT3 | 2 | 1.208779435 | 0.75 | 0.309 | 9.31E-261 | 3.41E-256 |
| NLGN4X | 2 | 1.029502408 | 0.949 | 0.684 | 2.8E-260 | 1.02E-255 |
| PHYHIPL | 2 | 0.879835054 | 0.466 | 0.089 | 2.82E-259 | 1.03E-254 |
| NYAP2 | 2 | 1.025022098 | 0.314 | 0.026 | 5.87E-258 | 2.15E-253 |
| CALD1 | 2 | -1.302434521 | 0.098 | 0.503 | 9.05E-255 | 3.31E-250 |
| STK3 | 2 | -1.269093575 | 0.16 | 0.572 | 2.19E-254 | 8.01E-250 |
| SSBP2 | 2 | -1.349136627 | 0.574 | 0.865 | 3.52E-254 | 1.29E-249 |
| DYNC2H1 | 2 | -1.267720767 | 0.051 | 0.434 | 2.39E-251 | 8.75E-247 |
| ARHGAP44 | 2 | 1.108517003 | 0.718 | 0.322 | 3.35E-251 | 1.23E-246 |
| ENPP2 | 2 | 0.966317784 | 0.54 | 0.137 | 7.62E-251 | 2.79E-246 |
| GPHN | 2 | 0.802766535 | 0.983 | 0.909 | 1.82E-250 | 6.67E-246 |
| MALRD1 | 2 | 1.717708988 | 0.574 | 0.186 | 3.69E-249 | 1.35E-244 |
| TUNAR | 2 | 1.057878821 | 0.33 | 0.037 | 5.38E-249 | 1.97E-244 |
| SYT1 | 2 | 0.933060974 | 0.975 | 0.689 | 4.39E-248 | 1.61E-243 |
| AL008633.1 | 2 | -3.46958269 | 0.522 | 0.755 | 1.63E-246 | 5.96E-242 |
| PDZRN3 | 2 | 0.937688411 | 0.988 | 0.884 | 2.91E-246 | 1.06E-241 |
| GNAO1 | 2 | 0.979435405 | 0.793 | 0.38 | 1.99E-245 | 7.28E-241 |
| FLT1 | 2 | 0.717451308 | 0.292 | 0.021 | 5.98E-245 | 2.19E-240 |
| GABRB3 | 2 | 0.965676535 | 0.734 | 0.294 | 3.78E-243 | 1.39E-238 |
| TET1 | 2 | -1.264553383 | 0.093 | 0.483 | 1E-241 | 3.67E-237 |
| FAM189A1 | 2 | 1.006389748 | 0.943 | 0.636 | 2.35E-241 | 8.6E-237 |
| PTCHD4 | 2 | 1.091648618 | 0.585 | 0.175 | 4.21E-239 | 1.54E-234 |
| MBNL2 | 2 | -1.148833518 | 0.09 | 0.483 | 7.89E-239 | 2.89E-234 |
| PDE10A | 2 | 0.880123611 | 0.586 | 0.177 | 2.16E-236 | 7.91E-232 |
| CNTN3 | 2 | 1.074537864 | 0.818 | 0.42 | 6.05E-235 | 2.21E-230 |
| FAM184B | 2 | 0.910315517 | 0.497 | 0.123 | 8.71E-235 | 3.19E-230 |
| WWC2 | 2 | -1.165123874 | 0.091 | 0.473 | 2.61E-231 | 9.56E-227 |
| FRMD3 | 2 | 1.012430369 | 0.705 | 0.287 | 2.9E-231 | 1.06E-226 |
| VAV3 | 2 | 0.976116843 | 0.45 | 0.095 | 4E-231 | 1.46E-226 |
| CDKN2B-AS1 | 2 | 1.086785135 | 0.641 | 0.252 | 5.3E-230 | 1.94E-225 |
| SUSD4 | 2 | 0.931760123 | 0.595 | 0.19 | 2.35E-229 | 8.62E-225 |
| CDH2 | 2 | 0.87923408 | 0.813 | 0.395 | 1.27E-228 | 4.65E-224 |
| AL445623.2 | 2 | 0.934157843 | 0.5 | 0.127 | 2.51E-228 | 9.19E-224 |
| RGS7 | 2 | 0.915549688 | 0.508 | 0.131 | 8.05E-228 | 2.95E-223 |
| ELAVL3 | 2 | 0.749530546 | 0.453 | 0.099 | 4.92E-227 | 1.8E-222 |
| MTMR7 | 2 | 0.904165557 | 0.689 | 0.27 | 9.29E-227 | 3.4E-222 |
| SOGA3 | 2 | 0.84926058 | 0.528 | 0.146 | 1.25E-226 | 4.59E-222 |
| ARHGAP42 | 2 | -1.166622971 | 0.07 | 0.442 | 8.45E-226 | 3.09E-221 |
| LINC00632 | 2 | -1.284628449 | 0.08 | 0.454 | 1.1E-225 | 4.04E-221 |
| SYTL5 | 2 | 1.071734612 | 0.645 | 0.24 | 3.91E-225 | 1.43E-220 |
| PARD3B | 2 | -1.610815986 | 0.154 | 0.541 | 2.46E-224 | 9E-220 |
| RAPGEF5 | 2 | 0.882009483 | 0.831 | 0.443 | 4.64E-224 | 1.7E-219 |
| SGCD | 2 | -1.597008131 | 0.161 | 0.561 | 1.07E-223 | 3.92E-219 |

| | | | | | | |
|---|---|---|---|---|---|---|
| TSPAN7 | 2 | 0.943772888 | 0.842 | 0.471 | 1.28E-223 | 4.68E-219 |
| KIAA1217 | 2 | -1.60737169 | 0.035 | 0.374 | 2.79E-223 | 1.02E-218 |
| PTPN14 | 2 | -1.101728573 | 0.022 | 0.349 | 5.83E-223 | 2.13E-218 |
| CDH11 | 2 | -1.792543201 | 0.038 | 0.377 | 1.35E-221 | 4.95E-217 |
| AL592463.1 | 2 | 0.476224459 | 0.205 | 0.004 | 5.31E-221 | 1.94E-216 |
| EDIL3 | 2 | 1.132061855 | 0.888 | 0.558 | 5.51E-221 | 2.02E-216 |
| GRM7 | 2 | 1.890109829 | 0.369 | 0.083 | 3.43E-220 | 1.26E-215 |
| BRIP1 | 2 | 1.035355378 | 0.845 | 0.496 | 6.76E-220 | 2.48E-215 |
| CADPS | 2 | 0.297333996 | 0.789 | 0.392 | 1.73E-219 | 6.34E-215 |
| GLI3 | 2 | -1.235034193 | 0.058 | 0.41 | 1.8E-219 | 6.57E-215 |
| EGFR | 2 | 1.107701269 | 0.567 | 0.188 | 5.38E-219 | 1.97E-214 |
| CLVS1 | 2 | 1.060389729 | 0.505 | 0.139 | 3.01E-218 | 1.1E-213 |
| DANT2 | 2 | -1.42837599 | 0.046 | 0.392 | 5.72E-218 | 2.09E-213 |
| GRID2 | 2 | 1.236250159 | 0.86 | 0.514 | 5.97E-217 | 2.19E-212 |
| TCF7L1 | 2 | -1.116182645 | 0.037 | 0.371 | 1.13E-216 | 4.15E-212 |
| SLC1A2 | 2 | 0.882965664 | 0.602 | 0.209 | 1.54E-216 | 5.63E-212 |
| TTLL7 | 2 | 0.96900882 | 0.767 | 0.403 | 4.52E-215 | 1.65E-210 |
| SLC8A1 | 2 | 1.245022195 | 0.638 | 0.26 | 1.08E-213 | 3.96E-209 |
| AC117453.1 | 2 | 0.710183354 | 0.292 | 0.03 | 5.7E-213 | 2.09E-208 |
| TMEFF1 | 2 | 0.910071447 | 0.68 | 0.291 | 6.38E-213 | 2.34E-208 |
| SESN3 | 2 | -1.36124151 | 0.109 | 0.476 | 3.39E-212 | 1.24E-207 |
| LINC00599 | 2 | 0.769761566 | 0.431 | 0.095 | 2.32E-211 | 8.48E-207 |
| RIPOR2 | 2 | 0.781063013 | 0.699 | 0.289 | 2.32E-211 | 8.5E-207 |
| IL1RAPL1 | 2 | 0.399242147 | 0.944 | 0.591 | 6.71E-210 | 2.46E-205 |
| MALAT1 | 2 | -1.194203039 | 1 | 0.998 | 2.31E-209 | 8.44E-205 |
| SPHKAP | 2 | 1.036958558 | 0.358 | 0.058 | 7.2E-209 | 2.64E-204 |
| GLYATL2 | 2 | 0.941260044 | 0.646 | 0.26 | 2.01E-208 | 7.34E-204 |
| PPARG | 2 | 0.976437324 | 0.595 | 0.209 | 3.76E-208 | 1.38E-203 |
| ARHGAP6 | 2 | -1.415378773 | 0.07 | 0.425 | 3.85E-207 | 1.41E-202 |
| CNTNAP3B | 2 | -1.165790866 | 0.04 | 0.367 | 9.15E-207 | 3.35E-202 |
| KCNIP4-IT1 | 2 | 0.649512686 | 0.289 | 0.031 | 1.22E-206 | 4.46E-202 |
| FIRRE | 2 | -1.524525193 | 0.038 | 0.367 | 1.67E-206 | 6.12E-202 |
| MIR124-2HG | 2 | 0.751288214 | 0.385 | 0.072 | 1.81E-205 | 6.62E-201 |
| ADAM22 | 2 | 0.871726427 | 0.718 | 0.336 | 1.88E-205 | 6.88E-201 |
| CLSTN2 | 2 | 0.914641943 | 0.668 | 0.267 | 2.27E-205 | 8.32E-201 |
| MAP4K4 | 2 | 0.791913935 | 0.957 | 0.83 | 1.85E-204 | 6.76E-200 |
| GDAP1 | 2 | 1.086882732 | 0.605 | 0.244 | 3.48E-204 | 1.27E-199 |
| PCDH7 | 2 | -2.14938749 | 0.137 | 0.494 | 1.76E-203 | 6.46E-199 |
| ADCY2 | 2 | -1.610002584 | 0.025 | 0.336 | 7.24E-203 | 2.65E-198 |
| ITPR1 | 2 | -1.216844106 | 0.098 | 0.458 | 1.83E-202 | 6.7E-198 |
| AKAP6 | 2 | 0.91579637 | 0.536 | 0.166 | 4.12E-202 | 1.51E-197 |
| SEMA3E | 2 | -1.487757362 | 0.039 | 0.36 | 2.47E-201 | 9.03E-197 |
| PCDH10 | 2 | 0.873803175 | 0.382 | 0.075 | 6.42E-200 | 2.35E-195 |
| LINC01414 | 2 | 1.043611876 | 0.293 | 0.035 | 2.73E-199 | 9.98E-195 |
| UACA | 2 | -1.054629066 | 0.092 | 0.443 | 2.08E-198 | 7.62E-194 |
| CPLX2 | 2 | 0.750175572 | 0.443 | 0.109 | 2.61E-198 | 9.56E-194 |
| GNG4 | 2 | 0.8581644 | 0.634 | 0.252 | 2.73E-198 | 9.98E-194 |
| ACSL1 | 2 | 0.940726057 | 0.676 | 0.317 | 4.15E-197 | 1.52E-192 |
| NFASC | 2 | 0.59825562 | 0.444 | 0.117 | 5.74E-195 | 2.1E-190 |
| CASK | 2 | 0.737778053 | 0.957 | 0.836 | 1.45E-194 | 5.31E-190 |
| ZNF519 | 2 | 0.855081504 | 0.687 | 0.347 | 5.41E-194 | 1.98E-189 |
| TSPAN5 | 2 | 0.879717963 | 0.794 | 0.424 | 9.52E-194 | 3.49E-189 |
| FGD4 | 2 | -1.206212423 | 0.141 | 0.483 | 2.5E-193 | 9.15E-189 |
| ABCD3 | 2 | -1.142350337 | 0.384 | 0.7 | 3.69E-193 | 1.35E-188 |
| MSC-AS1 | 2 | 0.789842208 | 0.407 | 0.092 | 7.73E-193 | 2.83E-188 |
| DNM3 | 2 | 0.812237815 | 0.773 | 0.399 | 2.29E-192 | 8.39E-188 |
| MYO16 | 2 | 0.724569098 | 0.63 | 0.246 | 5.51E-192 | 2.02E-187 |
| SHISA6 | 2 | 1.157783314 | 0.418 | 0.108 | 1.76E-191 | 6.43E-187 |

| | | | | | | |
|---|---|---|---|---|---|---|
| KAZN | 2 | 0.862560104 | 0.992 | 0.878 | 7.17E-191 | 2.62E-186 |
| NECTIN3 | 2 | -1.42699137 | 0.351 | 0.658 | 4.76E-190 | 1.74E-185 |
| THSD7A | 2 | 1.157130346 | 0.683 | 0.32 | 3.27E-189 | 1.2E-184 |
| GALNT14 | 2 | 0.911087326 | 0.535 | 0.183 | 5.85E-189 | 2.14E-184 |
| SRRM3 | 2 | 0.702416471 | 0.422 | 0.102 | 8.37E-188 | 3.07E-183 |
| ERICH3 | 2 | 0.984811783 | 0.381 | 0.084 | 3.01E-187 | 1.1E-182 |
| PLCH1 | 2 | -1.290816585 | 0.063 | 0.382 | 5.74E-187 | 2.1E-182 |
| LINC01170 | 2 | 0.855230376 | 0.867 | 0.518 | 5.06E-185 | 1.85E-180 |
| VGLL3 | 2 | -1.785695645 | 0.059 | 0.369 | 1.39E-184 | 5.08E-180 |
| GALNT13 | 2 | 0.793092293 | 0.928 | 0.651 | 2.78E-184 | 1.02E-179 |
| RBFOX3 | 2 | 0.762872443 | 0.474 | 0.138 | 8.14E-184 | 2.98E-179 |
| BAZ2B | 2 | 0.736926558 | 0.967 | 0.904 | 1.42E-182 | 5.21E-178 |
| NRIP1 | 2 | -1.10588672 | 0.346 | 0.67 | 2.2E-182 | 8.06E-178 |
| CA10 | 2 | -1.54025239 | 0.066 | 0.375 | 3.09E-182 | 1.13E-177 |
| MIR99AHG | 2 | -1.797628051 | 0.548 | 0.718 | 5.35E-182 | 1.96E-177 |
| SYNPO2 | 2 | -1.192842283 | 0.032 | 0.322 | 5.77E-181 | 2.11E-176 |
| ZFHX4 | 2 | -1.290703113 | 0.153 | 0.493 | 3.1E-180 | 1.13E-175 |
| SYNE2 | 2 | 0.791479483 | 0.91 | 0.656 | 9.83E-180 | 3.6E-175 |
| AC018731.1 | 2 | 0.474167516 | 0.88 | 0.525 | 6.35E-179 | 2.32E-174 |
| ATXN1 | 2 | -1.195842881 | 0.485 | 0.741 | 7.37E-178 | 2.7E-173 |
| DNAJC6 | 2 | 0.895151628 | 0.704 | 0.345 | 1.26E-177 | 4.61E-173 |
| SYT2 | 2 | 0.693077352 | 0.253 | 0.029 | 2.13E-177 | 7.81E-173 |
| AC009264.1 | 2 | 0.493806806 | 0.196 | 0.01 | 2.34E-177 | 8.57E-173 |
| BICDL1 | 2 | 0.788339446 | 0.566 | 0.211 | 8.11E-177 | 2.97E-172 |
| MAP3K5 | 2 | 0.869372339 | 0.608 | 0.254 | 9.46E-177 | 3.46E-172 |
| PDE1C | 2 | -1.906248001 | 0.064 | 0.364 | 2.2E-176 | 8.05E-172 |
| FN1 | 2 | -1.662063366 | 0.102 | 0.429 | 4.37E-176 | 1.6E-171 |
| MIR924HG | 2 | 0.900730906 | 0.899 | 0.676 | 1.39E-174 | 5.09E-170 |
| PDE3A | 2 | -1.553907418 | 0.299 | 0.63 | 2.11E-174 | 7.71E-170 |
| KIAA0408 | 2 | 0.714364176 | 0.48 | 0.148 | 7.16E-174 | 2.62E-169 |
| PLPPR1 | 2 | 0.911009378 | 0.46 | 0.134 | 1.15E-173 | 4.22E-169 |
| NHSL1 | 2 | -1.144205959 | 0.302 | 0.641 | 5.03E-173 | 1.84E-168 |
| KCND2 | 2 | 0.686151408 | 0.828 | 0.474 | 7.34E-173 | 2.69E-168 |
| AC093523.1 | 2 | 0.755811777 | 0.225 | 0.019 | 8.23E-173 | 3.01E-168 |
| MAP7D2 | 2 | 0.750913065 | 0.569 | 0.219 | 8.4E-173 | 3.07E-168 |
| PELI2 | 2 | 0.865105971 | 0.715 | 0.362 | 1.41E-172 | 5.15E-168 |
| BCL11B | 2 | 0.892768953 | 0.709 | 0.359 | 2.79E-172 | 1.02E-167 |
| UST | 2 | -1.188675759 | 0.04 | 0.331 | 9.8E-172 | 3.59E-167 |
| LIN28B | 2 | 0.846517933 | 0.779 | 0.433 | 2.5E-171 | 9.14E-167 |
| SNAP25 | 2 | 0.743962623 | 0.586 | 0.23 | 3.13E-170 | 1.15E-165 |
| RYR3 | 2 | 0.80746747 | 0.323 | 0.06 | 7.49E-169 | 2.74E-164 |
| ZFHX3 | 2 | -1.160868688 | 0.317 | 0.643 | 1.71E-168 | 6.26E-164 |
| GNG12 | 2 | -0.739325266 | 0.03 | 0.308 | 1.88E-168 | 6.9E-164 |
| NETO1 | 2 | 0.678377381 | 0.418 | 0.114 | 1.24E-166 | 4.53E-162 |
| AC008591.1 | 2 | 0.994834733 | 0.721 | 0.367 | 1.74E-166 | 6.36E-162 |
| LUZP2 | 2 | 0.993469682 | 0.829 | 0.493 | 3.96E-166 | 1.45E-161 |
| AC016042.1 | 2 | 0.834924235 | 0.491 | 0.176 | 1.04E-164 | 3.82E-160 |
| ZEB1 | 2 | -1.128460509 | 0.276 | 0.626 | 1.51E-164 | 5.52E-160 |
| MYO10 | 2 | -1.010151833 | 0.082 | 0.39 | 1.71E-164 | 6.26E-160 |
| TLN2 | 2 | -1.116695404 | 0.191 | 0.513 | 1.76E-164 | 6.46E-160 |
| EZH2 | 2 | 0.797391493 | 0.889 | 0.67 | 5.58E-164 | 2.04E-159 |
| ABCA1 | 2 | -0.952603246 | 0.036 | 0.315 | 1.06E-163 | 3.9E-159 |
| ITCH | 2 | -0.849217687 | 0.417 | 0.714 | 1.96E-163 | 7.17E-159 |
| ZHX2 | 2 | -0.833736994 | 0.049 | 0.341 | 2.01E-163 | 7.37E-159 |
| IQGAP2 | 2 | -1.185492343 | 0.142 | 0.46 | 2.9E-163 | 1.06E-158 |
| KCNQ3 | 2 | 0.765776336 | 0.68 | 0.32 | 8.54E-163 | 3.13E-158 |
| NAV3 | 2 | 1.113324752 | 0.648 | 0.324 | 1.25E-162 | 4.59E-158 |
| DIAPH3 | 2 | 0.971599031 | 0.679 | 0.349 | 1.89E-162 | 6.92E-158 |

| | | | | | | |
|---|---|---|---|---|---|---|
| LGR5 | 2 | 0.71889531 | 0.251 | 0.032 | 2.53E-162 | 9.28E-158 |
| TOX | 2 | 0.710594759 | 0.907 | 0.62 | 1.55E-161 | 5.67E-157 |
| ATAD2 | 2 | 0.892191119 | 0.754 | 0.447 | 2.47E-161 | 9.04E-157 |
| LHFPL4 | 2 | 0.574169686 | 0.321 | 0.063 | 1.23E-160 | 4.5E-156 |
| ARID5B | 2 | -1.08911651 | 0.144 | 0.473 | 1.48E-160 | 5.43E-156 |
| COCH | 2 | 0.756095557 | 0.494 | 0.174 | 6.76E-160 | 2.47E-155 |
| AFF1 | 2 | -0.983122578 | 0.157 | 0.47 | 1.15E-159 | 4.2E-155 |
| FRAS1 | 2 | 1.200936501 | 0.63 | 0.319 | 2.97E-159 | 1.09E-154 |
| PDE4DIP | 2 | 0.774594588 | 0.584 | 0.274 | 5.03E-159 | 1.84E-154 |
| REEP1 | 2 | 0.632714056 | 0.353 | 0.08 | 5.27E-159 | 1.93E-154 |
| NRCAM | 2 | 0.668922889 | 0.681 | 0.323 | 7.38E-159 | 2.7E-154 |
| RBM39 | 2 | -0.729015374 | 0.74 | 0.888 | 1.53E-158 | 5.61E-154 |
| GUCY1A2 | 2 | 0.82323927 | 0.681 | 0.333 | 2.99E-158 | 1.09E-153 |
| ZFPM2 | 2 | -1.544330512 | 0.079 | 0.359 | 4.31E-158 | 1.58E-153 |
| CDH6 | 2 | 0.90790452 | 0.604 | 0.274 | 9.36E-158 | 3.43E-153 |
| RBMS1 | 2 | -1.062595332 | 0.292 | 0.605 | 9.76E-158 | 3.57E-153 |
| AC009315.1 | 2 | 0.863286609 | 0.679 | 0.348 | 2.12E-156 | 7.76E-152 |
| KCNH7 | 2 | 1.150989264 | 0.36 | 0.086 | 3.44E-156 | 1.26E-151 |
| AL354771.1 | 2 | 0.758603507 | 0.357 | 0.088 | 4.94E-156 | 1.81E-151 |
| LMO3 | 2 | 0.809113342 | 0.242 | 0.032 | 5.69E-156 | 2.08E-151 |
| CADPS2 | 2 | -1.278856499 | 0.027 | 0.282 | 6.66E-156 | 2.44E-151 |
| AMPH | 2 | 0.693348692 | 0.414 | 0.117 | 9.91E-156 | 3.63E-151 |
| SASH1 | 2 | -1.169396008 | 0.303 | 0.621 | 4.57E-155 | 1.67E-150 |
| CHRNA7 | 2 | 0.814229952 | 0.489 | 0.178 | 1.26E-153 | 4.61E-149 |
| MCTP2 | 2 | -1.630935768 | 0.052 | 0.312 | 2.24E-153 | 8.19E-149 |
| FREM1 | 2 | -1.287573172 | 0.045 | 0.31 | 2.91E-153 | 1.06E-148 |
| SOX6 | 2 | -1.578787774 | 0.034 | 0.288 | 3.28E-153 | 1.2E-148 |
| TXNDC16 | 2 | 0.850088151 | 0.584 | 0.269 | 2.58E-152 | 9.46E-148 |
| NOTCH2 | 2 | -0.96038048 | 0.077 | 0.363 | 9.79E-152 | 3.59E-147 |
| DSCAML1 | 2 | 0.748534436 | 0.702 | 0.353 | 3.89E-151 | 1.42E-146 |
| TFAP2B | 2 | -1.007553967 | 0.029 | 0.285 | 5.14E-151 | 1.88E-146 |
| SLC6A15 | 2 | 0.698048162 | 0.462 | 0.154 | 1.36E-150 | 4.97E-146 |
| NMNAT2 | 2 | 0.599285238 | 0.596 | 0.257 | 3.65E-150 | 1.34E-145 |
| ALCAM | 2 | -1.562529433 | 0.096 | 0.385 | 8.24E-150 | 3.02E-145 |
| PGBD5 | 2 | 0.708549233 | 0.569 | 0.241 | 8.41E-150 | 3.08E-145 |
| CLEC2A | 2 | 0.75504921 | 0.343 | 0.08 | 8.77E-150 | 3.21E-145 |
| GPR137C | 2 | 0.849998168 | 0.678 | 0.372 | 4.88E-149 | 1.79E-144 |
| FOXO1 | 2 | -0.858321523 | 0.017 | 0.255 | 6.59E-149 | 2.41E-144 |
| DLGAP1 | 2 | 0.576355581 | 0.975 | 0.802 | 8.87E-149 | 3.25E-144 |
| FGD5 | 2 | 0.669875998 | 0.426 | 0.133 | 1.29E-148 | 4.71E-144 |
| CRYBG3 | 2 | -0.948929274 | 0.082 | 0.369 | 5.27E-148 | 1.93E-143 |
| MAST1 | 2 | 0.547073156 | 0.317 | 0.068 | 2.12E-147 | 7.77E-143 |
| MYO1E | 2 | -0.751951699 | 0.056 | 0.33 | 4.43E-147 | 1.62E-142 |
| DDX17 | 2 | -0.683730507 | 0.761 | 0.881 | 5.3E-147 | 1.94E-142 |
| FAP | 2 | -1.360003159 | 0.024 | 0.262 | 1.86E-146 | 6.82E-142 |
| KCTD8 | 2 | 1.493016045 | 0.356 | 0.099 | 1.34E-145 | 4.9E-141 |
| PLAGL1 | 2 | 1.076308139 | 0.589 | 0.304 | 3.88E-145 | 1.42E-140 |
| CHD9 | 2 | -0.708212975 | 0.859 | 0.935 | 1.11E-144 | 4.07E-140 |
| PDE7B | 2 | -1.390158427 | 0.022 | 0.261 | 1.98E-144 | 7.26E-140 |
| EPB41 | 2 | 0.74028547 | 0.727 | 0.426 | 2.44E-144 | 8.93E-140 |
| REST | 2 | -0.663910629 | 0.025 | 0.269 | 3.09E-144 | 1.13E-139 |
| TAFA1 | 2 | 1.390402434 | 0.467 | 0.171 | 6.02E-144 | 2.2E-139 |
| DOCK1 | 2 | -0.893312234 | 0.453 | 0.721 | 6.76E-144 | 2.47E-139 |
| DTL | 2 | 0.798036711 | 0.651 | 0.326 | 6.91E-144 | 2.53E-139 |
| PLXDC2 | 2 | 0.800215993 | 0.872 | 0.622 | 1.04E-143 | 3.81E-139 |
| KCNS3 | 2 | 0.584992997 | 0.293 | 0.058 | 5.11E-143 | 1.87E-138 |
| CDH10 | 2 | 1.146630285 | 0.556 | 0.276 | 5.93E-142 | 2.17E-137 |
| AC104073.4 | 2 | 0.532763429 | 0.283 | 0.054 | 6.15E-142 | 2.25E-137 |

| | | | | | | |
|---|---|---|---|---|---|---|
| SKAP1 | 2 | 0.647459242 | 0.248 | 0.04 | 1.09E-141 | 3.98E-137 |
| SLC38A1 | 2 | 0.703225473 | 0.797 | 0.517 | 1.86E-141 | 6.81E-137 |
| COL3A1 | 2 | -1.513649497 | 0.062 | 0.325 | 2.79E-141 | 1.02E-136 |
| PON2 | 2 | -0.611345828 | 0.024 | 0.263 | 3.95E-141 | 1.45E-136 |
| HS3ST3A1 | 2 | -0.99516482 | 0.046 | 0.302 | 5.93E-141 | 2.17E-136 |
| NFAT5 | 2 | -0.944030717 | 0.664 | 0.826 | 6.43E-141 | 2.35E-136 |
| CRIM1 | 2 | 0.881284698 | 0.566 | 0.283 | 7.97E-141 | 2.92E-136 |
| ARAP2 | 2 | -1.089395523 | 0.176 | 0.463 | 1.97E-140 | 7.21E-136 |
| EYA4 | 2 | -1.405773199 | 0.02 | 0.247 | 2.19E-140 | 8.03E-136 |
| MDGA2 | 2 | 1.413672659 | 0.263 | 0.049 | 3.46E-140 | 1.27E-135 |
| LAMC1 | 2 | -0.969318662 | 0.275 | 0.581 | 4.45E-140 | 1.63E-135 |
| LTBP1 | 2 | -1.05068941 | 0.091 | 0.367 | 5.34E-140 | 1.95E-135 |
| CAMK4 | 2 | 0.687000766 | 0.562 | 0.237 | 1.34E-139 | 4.91E-135 |
| POLA1 | 2 | 0.698133846 | 0.938 | 0.823 | 5.61E-139 | 2.05E-134 |
| RFC3 | 2 | 0.84144186 | 0.701 | 0.42 | 1.37E-138 | 5.02E-134 |
| ADAMTS16 | 2 | 0.769470004 | 0.519 | 0.205 | 1.69E-138 | 6.18E-134 |
| BCL11A | 2 | -0.849419609 | 0.059 | 0.324 | 2.47E-138 | 9.06E-134 |
| AL121578.3 | 2 | 0.72789629 | 0.456 | 0.163 | 2.99E-138 | 1.09E-133 |
| EPHA4 | 2 | 0.888240329 | 0.772 | 0.545 | 6.63E-138 | 2.43E-133 |
| RAD18 | 2 | 0.748508146 | 0.727 | 0.481 | 7.52E-138 | 2.75E-133 |
| ITGA8 | 2 | -1.442570512 | 0.055 | 0.305 | 8.8E-138 | 3.22E-133 |
| ITGBL1 | 2 | -2.178618852 | 0.029 | 0.254 | 3.17E-137 | 1.16E-132 |
| ALK | 2 | 0.583858355 | 0.243 | 0.038 | 3.88E-137 | 1.42E-132 |
| SIL1 | 2 | -1.011319427 | 0.475 | 0.706 | 8.71E-137 | 3.19E-132 |
| FANCI | 2 | 0.705203077 | 0.659 | 0.34 | 1.87E-135 | 6.83E-131 |
| L3MBTL4 | 2 | 0.620876496 | 0.792 | 0.471 | 3.17E-135 | 1.16E-130 |
| LIFR | 2 | -1.291572723 | 0.431 | 0.681 | 8.66E-135 | 3.17E-130 |
| LRIG3 | 2 | -0.765656612 | 0.046 | 0.294 | 1.61E-134 | 5.9E-130 |
| AUTS2 | 2 | -1.69324701 | 0.278 | 0.52 | 2.71E-134 | 9.92E-130 |
| CTNNA1 | 2 | -0.854994689 | 0.44 | 0.713 | 5.06E-134 | 1.85E-129 |
| ABLIM1 | 2 | -0.837718565 | 0.044 | 0.288 | 6.34E-134 | 2.32E-129 |
| CACHD1 | 2 | -1.146356629 | 0.4 | 0.623 | 1.54E-133 | 5.63E-129 |
| KCNH1 | 2 | -1.420697692 | 0.065 | 0.315 | 3.05E-133 | 1.12E-128 |
| WDR35 | 2 | -0.852955709 | 0.222 | 0.524 | 1.49E-132 | 5.44E-128 |
| ANKH | 2 | 0.544880373 | 0.436 | 0.152 | 1.06E-131 | 3.88E-127 |
| LINC01278 | 2 | -0.841927577 | 0.106 | 0.384 | 1.18E-131 | 4.32E-127 |
| HELLS | 2 | 0.678051311 | 0.834 | 0.584 | 1.71E-131 | 6.27E-127 |
| ZNF91 | 2 | -0.647737223 | 0.046 | 0.294 | 1.81E-131 | 6.63E-127 |
| ETV1 | 2 | -1.064467679 | 0.021 | 0.24 | 3.53E-131 | 1.29E-126 |
| NRP1 | 2 | -1.100386171 | 0.097 | 0.361 | 4.62E-131 | 1.69E-126 |
| BLM | 2 | 0.737189881 | 0.533 | 0.241 | 8.56E-131 | 3.13E-126 |
| ROBO2 | 2 | 0.461740006 | 0.919 | 0.665 | 2.59E-130 | 9.47E-126 |
| RTKN2 | 2 | 0.657878876 | 0.393 | 0.124 | 3.48E-130 | 1.28E-125 |
| HBEGF | 2 | 0.57823827 | 0.315 | 0.077 | 3.79E-130 | 1.39E-125 |
| TNS3 | 2 | -0.858639812 | 0.047 | 0.285 | 1.48E-128 | 5.43E-124 |
| BCL7A | 2 | 0.691422869 | 0.46 | 0.181 | 5.6E-128 | 2.05E-123 |
| AC093898.1 | 2 | 0.67964151 | 0.227 | 0.036 | 1.19E-127 | 4.35E-123 |
| CDH8 | 2 | 0.881592631 | 0.325 | 0.088 | 1.22E-127 | 4.46E-123 |
| ACSS3 | 2 | -0.746232014 | 0.022 | 0.238 | 8.56E-127 | 3.13E-122 |
| LINC01473 | 2 | -0.901923616 | 0.33 | 0.625 | 9.83E-127 | 3.6E-122 |
| ELOVL2 | 2 | 0.529475439 | 0.345 | 0.097 | 2.16E-126 | 7.91E-122 |
| ELP4 | 2 | -0.899914331 | 0.172 | 0.46 | 3.15E-126 | 1.15E-121 |
| C1orf21 | 2 | 0.67466659 | 0.56 | 0.253 | 5.76E-125 | 2.11E-120 |
| SLIT1 | 2 | 0.515729433 | 0.294 | 0.069 | 6.9E-125 | 2.53E-120 |
| OLFM3 | 2 | 0.523480898 | 0.16 | 0.012 | 9.54E-125 | 3.49E-120 |
| GLRA1 | 2 | 0.372334328 | 0.162 | 0.013 | 8.92E-124 | 3.27E-119 |
| RAPGEF2 | 2 | -0.94711624 | 0.451 | 0.702 | 9.07E-124 | 3.32E-119 |
| C2CD5 | 2 | 0.644559838 | 0.822 | 0.641 | 2.12E-123 | 7.77E-119 |

| | | | | | | |
|---|---|---|---|---|---|---|
| GRIN1 | 2 | 0.300008621 | 0.13 | 0.005 | 5.97E-123 | 2.18E-118 |
| AASS | 2 | -0.714745495 | 0.063 | 0.312 | 8.79E-123 | 3.22E-118 |
| MARK1 | 2 | 0.544373763 | 0.754 | 0.447 | 1.04E-122 | 3.82E-118 |
| GLRB | 2 | 0.599739919 | 0.29 | 0.071 | 1.13E-122 | 4.15E-118 |
| CNKSR3 | 2 | -0.776217115 | 0.081 | 0.337 | 3.21E-122 | 1.18E-117 |
| MYRIP | 2 | 0.622845551 | 0.878 | 0.608 | 5.3E-122 | 1.94E-117 |
| INTS6L | 2 | -0.830697924 | 0.16 | 0.432 | 8.39E-122 | 3.07E-117 |
| POLQ | 2 | 0.716485235 | 0.565 | 0.264 | 9.55E-122 | 3.5E-117 |
| APBA1 | 2 | 0.331744213 | 0.69 | 0.382 | 1.33E-121 | 4.86E-117 |
| AL589740.1 | 2 | 0.901019154 | 0.443 | 0.166 | 1.84E-121 | 6.75E-117 |
| ANO5 | 2 | 0.614233694 | 0.485 | 0.196 | 3.92E-121 | 1.43E-116 |
| HMCN1 | 2 | -1.388373758 | 0.037 | 0.256 | 4.14E-121 | 1.52E-116 |
| PLS3 | 2 | -0.927644434 | 0.175 | 0.456 | 5.14E-121 | 1.88E-116 |
| MINAR1 | 2 | 0.577680733 | 0.371 | 0.116 | 8.22E-121 | 3.01E-116 |
| SEZ6 | 2 | 0.424706707 | 0.221 | 0.036 | 1.35E-120 | 4.93E-116 |
| KDM5B | 2 | -0.857133109 | 0.181 | 0.465 | 2.31E-120 | 8.44E-116 |
| AF233439.1 | 2 | 0.433277317 | 0.237 | 0.043 | 1.24E-119 | 4.55E-115 |
| RPS6KA6 | 2 | 0.649561329 | 0.419 | 0.152 | 1.31E-119 | 4.8E-115 |
| GAB1 | 2 | -0.966544045 | 0.494 | 0.707 | 1.63E-119 | 5.95E-115 |
| PDGFC | 2 | 1.073046351 | 0.467 | 0.216 | 1.82E-119 | 6.68E-115 |
| MEG8 | 2 | -0.321377893 | 0.756 | 0.531 | 2.49E-119 | 9.13E-115 |
| AL513166.1 | 2 | 0.945928603 | 0.457 | 0.203 | 3.03E-119 | 1.11E-114 |
| TTC8 | 2 | -0.643446997 | 0.056 | 0.293 | 7.18E-119 | 2.63E-114 |
| PHACTR2 | 2 | 0.840976486 | 0.77 | 0.522 | 8.97E-119 | 3.28E-114 |
| KIF15 | 2 | 0.615980331 | 0.506 | 0.215 | 1.18E-118 | 4.31E-114 |
| GLB1L3 | 2 | 0.464362335 | 0.257 | 0.054 | 1.72E-118 | 6.31E-114 |
| LMO7 | 2 | 0.901715656 | 0.716 | 0.481 | 2.1E-118 | 7.69E-114 |
| SPATA6 | 2 | -0.650503769 | 0.015 | 0.209 | 2.36E-118 | 8.65E-114 |
| ARHGAP10 | 2 | -0.945022343 | 0.299 | 0.56 | 4.85E-118 | 1.78E-113 |
| XACT | 2 | -2.424196859 | 0.068 | 0.291 | 4.98E-118 | 1.82E-113 |
| KLHL1 | 2 | 1.561334117 | 0.226 | 0.074 | 9.48E-118 | 3.47E-113 |
| LINC01933 | 2 | 0.570797536 | 0.202 | 0.029 | 1.31E-117 | 4.78E-113 |
| LINC02197 | 2 | -0.910131608 | 0.013 | 0.206 | 1.44E-117 | 5.25E-113 |
| GAS7 | 2 | -1.008532293 | 0.056 | 0.281 | 1.61E-117 | 5.9E-113 |
| NSUN7 | 2 | -0.709704442 | 0.057 | 0.292 | 1.75E-117 | 6.41E-113 |
| L1TD1 | 2 | -0.796840353 | 0.045 | 0.266 | 2.25E-117 | 8.25E-113 |
| CCND2 | 2 | -0.74185061 | 0.114 | 0.365 | 3.99E-117 | 1.46E-112 |
| ZSWIM6 | 2 | 0.599547571 | 0.931 | 0.805 | 5.85E-117 | 2.14E-112 |
| PRKG1 | 2 | -1.499596081 | 0.613 | 0.761 | 8.91E-117 | 3.26E-112 |
| PCDH11X | 2 | -1.949169264 | 0.075 | 0.307 | 1.38E-116 | 5.05E-112 |
| CACNG5 | 2 | 0.478499865 | 0.238 | 0.046 | 1.52E-116 | 5.56E-112 |
| HDAC9 | 2 | 0.841394753 | 0.792 | 0.525 | 2.83E-116 | 1.04E-111 |
| AC096576.2 | 2 | 0.359667583 | 0.164 | 0.016 | 6.01E-116 | 2.2E-111 |
| PARP14 | 2 | -0.571804408 | 0.019 | 0.22 | 8.43E-116 | 3.08E-111 |
| NEBL | 2 | -1.047280973 | 0.033 | 0.24 | 1.07E-115 | 3.93E-111 |
| MITF | 2 | -0.797780346 | 0.025 | 0.226 | 1.27E-115 | 4.66E-111 |
| DDR2 | 2 | -0.655500549 | 0.034 | 0.246 | 1.96E-115 | 7.16E-111 |
| LINC00907 | 2 | -1.63864099 | 0.074 | 0.295 | 3.78E-114 | 1.38E-109 |
| BCAS3 | 2 | -0.862205512 | 0.467 | 0.701 | 8.31E-114 | 3.04E-109 |
| CDKL5 | 2 | 0.649251311 | 0.646 | 0.352 | 8.34E-114 | 3.05E-109 |
| TDRD10 | 2 | 0.471598396 | 0.257 | 0.056 | 1.17E-113 | 4.28E-109 |
| EXO1 | 2 | 0.611983126 | 0.444 | 0.174 | 1.53E-113 | 5.6E-109 |
| VGLL4 | 2 | -0.770365404 | 0.37 | 0.637 | 2.13E-113 | 7.79E-109 |
| IGF1R | 2 | 0.586280805 | 0.934 | 0.765 | 2.54E-113 | 9.28E-109 |
| DAAM2 | 2 | -0.984287362 | 0.043 | 0.256 | 3.06E-113 | 1.12E-108 |
| CDK1 | 2 | 0.577042781 | 0.415 | 0.153 | 3.15E-113 | 1.15E-108 |
| LMNB1 | 2 | 0.650581871 | 0.704 | 0.436 | 4.41E-113 | 1.61E-108 |
| BASP1 | 2 | 0.653908661 | 0.698 | 0.413 | 5.16E-113 | 1.89E-108 |

| | | | | | | |
|---|---|---|---|---|---|---|
| SH3GL3 | 2 | 0.629499277 | 0.768 | 0.478 | 5.8E-113 | 2.12E-108 |
| AC096576.3 | 2 | 0.293165539 | 0.138 | 0.009 | 9.47E-113 | 3.47E-108 |
| RAMP1 | 2 | 0.40413113 | 0.193 | 0.028 | 1.39E-112 | 5.1E-108 |
| MIR4435-2HG | 2 | -0.797117795 | 0.054 | 0.277 | 2.32E-112 | 8.48E-108 |
| GALNT16 | 2 | 0.489147875 | 0.28 | 0.07 | 5.96E-112 | 2.18E-107 |
| MIR503HG | 2 | -0.659375957 | 0.012 | 0.196 | 2.94E-111 | 1.08E-106 |
| CHSY3 | 2 | 0.685974941 | 0.892 | 0.669 | 3.08E-111 | 1.13E-106 |
| KCNH8 | 2 | 0.461065869 | 0.853 | 0.583 | 3.81E-111 | 1.39E-106 |
| ANO2 | 2 | -0.892254483 | 0.028 | 0.23 | 4.54E-111 | 1.66E-106 |
| FGFR1 | 2 | -0.777707786 | 0.153 | 0.406 | 6.29E-111 | 2.3E-106 |
| TMEFF2 | 2 | 0.634998265 | 0.336 | 0.102 | 8.48E-111 | 3.11E-106 |
| DNAH7 | 2 | -1.020718991 | 0.021 | 0.213 | 1.21E-110 | 4.43E-106 |
| FRMD4A | 2 | 0.457793408 | 0.959 | 0.785 | 1.43E-110 | 5.25E-106 |
| NR3C1 | 2 | -0.851789454 | 0.15 | 0.412 | 1.78E-110 | 6.52E-106 |
| TMEM132E | 2 | 0.31055326 | 0.154 | 0.015 | 2.11E-110 | 7.73E-106 |
| LAMA3 | 2 | 0.654428218 | 0.424 | 0.162 | 2.31E-110 | 8.46E-106 |
| GMDS-DT | 2 | -0.792112615 | 0.075 | 0.308 | 2.84E-110 | 1.04E-105 |
| NETO2 | 2 | 0.662607726 | 0.738 | 0.484 | 3.11E-110 | 1.14E-105 |
| C4orf19 | 2 | 0.897565839 | 0.504 | 0.259 | 4.99E-110 | 1.83E-105 |
| ARHGAP31 | 2 | -0.638128899 | 0.049 | 0.271 | 2.65E-109 | 9.69E-105 |
| AC105383.1 | 2 | 0.513678682 | 0.231 | 0.046 | 3.14E-109 | 1.15E-104 |
| NCAPG | 2 | 0.649090067 | 0.496 | 0.221 | 7.43E-109 | 2.72E-104 |
| CPNE8 | 2 | -0.718931007 | 0.015 | 0.196 | 1.05E-108 | 3.85E-104 |
| TIAM1 | 2 | 0.512595784 | 0.822 | 0.559 | 1.33E-108 | 4.86E-104 |
| ADGRV1 | 2 | -1.574061823 | 0.625 | 0.698 | 1.49E-108 | 5.45E-104 |
| GRIA3 | 2 | -0.718414234 | 0.025 | 0.221 | 2.38E-108 | 8.71E-104 |
| IMMP2L | 2 | -0.865222526 | 0.688 | 0.85 | 2.49E-108 | 9.13E-104 |
| AC009899.1 | 2 | 0.49628168 | 0.23 | 0.046 | 3.32E-108 | 1.21E-103 |
| NPHP3 | 2 | -0.629067798 | 0.12 | 0.373 | 7.83E-108 | 2.87E-103 |
| EXT1 | 2 | -0.845373094 | 0.501 | 0.734 | 1.81E-107 | 6.64E-103 |
| KIF4A | 2 | 0.613227001 | 0.395 | 0.144 | 3.49E-107 | 1.28E-102 |
| AC079148.3 | 2 | 0.443824314 | 0.17 | 0.021 | 8.56E-107 | 3.13E-102 |
| ERBIN | 2 | -0.730230657 | 0.597 | 0.793 | 1.43E-106 | 5.25E-102 |
| INPP5F | 2 | 0.610419558 | 0.601 | 0.338 | 1.71E-106 | 6.28E-102 |
| FGF9 | 2 | 0.33017143 | 0.166 | 0.02 | 1.74E-106 | 6.37E-102 |
| PTPRM | 2 | -1.242338206 | 0.193 | 0.426 | 2.55E-106 | 9.33E-102 |
| SNX9 | 2 | -0.730455151 | 0.151 | 0.4 | 3.29E-106 | 1.2E-101 |
| TAOK3 | 2 | -0.842480829 | 0.214 | 0.473 | 5.12E-106 | 1.87E-101 |
| DOK6 | 2 | 0.616410102 | 0.833 | 0.584 | 5.13E-106 | 1.88E-101 |
| EPB41L4A | 2 | 0.716054147 | 0.273 | 0.071 | 1.26E-105 | 4.61E-101 |
| FAM110B | 2 | -0.870996136 | 0.191 | 0.448 | 4.01E-105 | 1.47E-100 |
| PGAP1 | 2 | 0.662955227 | 0.839 | 0.672 | 4.32E-105 | 1.58E-100 |
| SCG3 | 2 | 0.369345529 | 0.236 | 0.052 | 8.76E-105 | 3.21E-100 |
| AC135782.1 | 2 | 0.486849482 | 0.33 | 0.103 | 9.65E-105 | 3.53E-100 |
| STXBP1 | 2 | 0.597863208 | 0.55 | 0.277 | 1.03E-104 | 3.76E-100 |
| NCKAP5 | 2 | -1.425737567 | 0.231 | 0.452 | 1.56E-104 | 5.73E-100 |
| LINC01201 | 2 | 0.473612692 | 0.243 | 0.054 | 1.61E-104 | 5.9E-100 |
| RARB | 2 | -1.016554807 | 0.075 | 0.304 | 1.75E-104 | 6.42E-100 |
| PLSCR4 | 2 | -0.501651811 | 0.013 | 0.187 | 4.12E-104 | 1.51E-99 |
| DLC1 | 2 | -1.174540395 | 0.088 | 0.307 | 9.31E-104 | 3.41E-99 |
| NFATC2 | 2 | -0.546229742 | 0.014 | 0.192 | 9.34E-104 | 3.42E-99 |
| RNF144A | 2 | 0.597406707 | 0.718 | 0.454 | 1.41E-103 | 5.15E-99 |
| TGFBR2 | 2 | -0.498742925 | 0.008 | 0.173 | 2.07E-103 | 7.57E-99 |
| BCKDHB | 2 | -0.758088212 | 0.282 | 0.54 | 1.14E-102 | 4.17E-98 |
| UNC80 | 2 | 0.427758495 | 0.407 | 0.162 | 1.41E-102 | 5.18E-98 |
| SYNJ2 | 2 | -0.717715562 | 0.034 | 0.227 | 1.49E-102 | 5.46E-98 |
| ATAD5 | 2 | 0.618335779 | 0.729 | 0.479 | 1.79E-102 | 6.57E-98 |
| ACTL6B | 2 | 0.327450785 | 0.177 | 0.026 | 2.7E-102 | 9.88E-98 |

| | | | | | | |
|---|---|---|---|---|---|---|
| ANKRD34C-AS1 | 2 | 0.365838122 | 0.134 | 0.011 | 4.18E-102 | 1.53E-97 |
| RTN3 | 2 | 0.563200594 | 0.84 | 0.643 | 4.37E-102 | 1.6E-97 |
| COL24A1 | 2 | -0.957934415 | 0.036 | 0.23 | 1.27E-101 | 4.66E-97 |
| NAV2 | 2 | -0.973281869 | 0.271 | 0.514 | 5.14E-101 | 1.88E-96 |
| PPFIBP1 | 2 | -0.850665711 | 0.188 | 0.438 | 5.7E-101 | 2.09E-96 |
| PCNX2 | 2 | 0.636599602 | 0.692 | 0.458 | 5.81E-101 | 2.13E-96 |
| CNTNAP3 | 2 | -0.53364332 | 0.012 | 0.182 | 2.7E-100 | 9.87E-96 |
| ANKS1B | 2 | -1.229343175 | 0.752 | 0.798 | 3.87E-100 | 1.42E-95 |
| STOX2 | 2 | 0.415813015 | 0.522 | 0.256 | 5.31E-100 | 1.94E-95 |
| KIAA1671 | 2 | -0.576411921 | 0.022 | 0.201 | 8.19E-100 | 3E-95 |
| AC106799.2 | 2 | 0.608032801 | 0.227 | 0.051 | 8.28E-100 | 3.03E-95 |
| CMTM7 | 2 | -0.518216924 | 0.026 | 0.214 | 1.22E-99 | 4.46E-95 |
| ESRRG | 2 | -1.175842761 | 0.222 | 0.463 | 1.33E-99 | 4.88E-95 |
| C13orf42 | 2 | -0.607948426 | 0.005 | 0.16 | 2.12E-99 | 7.76E-95 |
| FLT4 | 2 | 0.429030403 | 0.253 | 0.062 | 2.74E-99 | 1E-94 |
| FRRS1L | 2 | 0.391170883 | 0.2 | 0.037 | 3.86E-99 | 1.41E-94 |
| PLCL2 | 2 | 0.64511765 | 0.779 | 0.546 | 4.18E-99 | 1.53E-94 |
| SUCLG2 | 2 | -0.708050495 | 0.257 | 0.518 | 7.64E-99 | 2.8E-94 |
| STRBP | 2 | 0.64501489 | 0.715 | 0.49 | 1.02E-98 | 3.74E-94 |
| MMS22L | 2 | 0.596500146 | 0.792 | 0.561 | 1.04E-98 | 3.8E-94 |
| SNX25 | 2 | -0.869703831 | 0.491 | 0.664 | 1.83E-98 | 6.69E-94 |
| CYTOR | 2 | -0.520713823 | 0.014 | 0.184 | 2.73E-98 | 9.99E-94 |
| CCDC150 | 2 | 0.654131572 | 0.502 | 0.25 | 3.06E-98 | 1.12E-93 |
| NPAS3 | 2 | -1.729816865 | 0.079 | 0.282 | 6.68E-98 | 2.44E-93 |
| RHBDL3 | 2 | 0.479961414 | 0.518 | 0.249 | 1.53E-97 | 5.61E-93 |
| SEMA5A | 2 | -1.583893332 | 0.23 | 0.449 | 3.85E-97 | 1.41E-92 |
| IQSEC3 | 2 | 0.283506564 | 0.16 | 0.021 | 4.41E-97 | 1.61E-92 |
| LRIG1 | 2 | -0.614187007 | 0.033 | 0.22 | 5.27E-97 | 1.93E-92 |
| TOP2A | 2 | 0.654618339 | 0.443 | 0.19 | 6.55E-97 | 2.4E-92 |
| HS6ST2 | 2 | 0.526743724 | 0.858 | 0.622 | 1.43E-96 | 5.24E-92 |
| IGF2BP2 | 2 | -1.087638613 | 0.287 | 0.454 | 1.54E-96 | 5.65E-92 |
| AP3B2 | 2 | 0.42828811 | 0.285 | 0.082 | 3.89E-96 | 1.42E-91 |
| ZGRF1 | 2 | 0.607593703 | 0.587 | 0.326 | 5.01E-96 | 1.83E-91 |
| PAX3 | 2 | -0.963339658 | 0.01 | 0.168 | 5.91E-96 | 2.16E-91 |
| GLRA2 | 2 | 0.700529159 | 0.235 | 0.056 | 6.88E-96 | 2.52E-91 |
| EFR3B | 2 | -0.791087042 | 0.264 | 0.498 | 8.08E-96 | 2.96E-91 |
| LDB2 | 2 | -1.367982179 | 0.157 | 0.378 | 9.39E-96 | 3.44E-91 |
| CHN1 | 2 | 0.487503636 | 0.819 | 0.56 | 1.59E-95 | 5.82E-91 |
| AC093298.2 | 2 | -1.671948214 | 0.037 | 0.223 | 1.88E-95 | 6.9E-91 |
| RTN1 | 2 | 0.387758944 | 0.329 | 0.117 | 2.07E-95 | 7.56E-91 |
| HACE1 | 2 | 0.564194436 | 0.797 | 0.577 | 2.57E-95 | 9.4E-91 |
| EPHA5-AS1 | 2 | 0.49866646 | 0.273 | 0.077 | 3.07E-95 | 1.12E-90 |
| BNC2 | 2 | -0.990694834 | 0.04 | 0.234 | 4.05E-95 | 1.48E-90 |
| HECW1 | 2 | 0.428754866 | 0.185 | 0.032 | 4.14E-95 | 1.51E-90 |
| SPAG17 | 2 | -1.066060453 | 0.12 | 0.33 | 6.08E-95 | 2.23E-90 |
| NPSR1 | 2 | -1.244315956 | 0.029 | 0.207 | 7.82E-95 | 2.86E-90 |
| MTMR1 | 2 | 0.591229732 | 0.66 | 0.435 | 8.63E-95 | 3.16E-90 |
| NLK | 2 | -0.875116733 | 0.513 | 0.699 | 1.04E-94 | 3.8E-90 |
| VPS13A | 2 | 0.449561947 | 0.838 | 0.608 | 1.07E-94 | 3.92E-90 |
| DUXAP8 | 2 | 0.431731403 | 0.257 | 0.068 | 2.2E-94 | 8.05E-90 |
| CASZ1 | 2 | -0.676781158 | 0.078 | 0.293 | 4.26E-94 | 1.56E-89 |
| BICD1 | 2 | 0.508398804 | 0.912 | 0.779 | 4.34E-94 | 1.59E-89 |
| RAB11FIP4 | 2 | 0.45026883 | 0.295 | 0.09 | 5.15E-94 | 1.89E-89 |
| BBS9 | 2 | -0.812464126 | 0.277 | 0.51 | 1.34E-93 | 4.91E-89 |
| FNDC3B | 2 | -0.779416228 | 0.56 | 0.726 | 1.57E-93 | 5.73E-89 |
| SEC24D | 2 | -0.65316969 | 0.083 | 0.292 | 1.93E-93 | 7.08E-89 |
| TFDP2 | 2 | -0.723870768 | 0.301 | 0.558 | 1.94E-93 | 7.09E-89 |
| LGR4 | 2 | -0.774131897 | 0.236 | 0.493 | 2.69E-93 | 9.83E-89 |

| | | | | | | |
|---|---|---|---|---|---|---|
| SEMA6A | 2 | -0.831771929 | 0.296 | 0.515 | 4.28E-93 | 1.57E-88 |
| ARNTL2 | 2 | 0.482238264 | 0.286 | 0.086 | 4.32E-93 | 1.58E-88 |
| NWD2 | 2 | 0.637812958 | 0.312 | 0.102 | 6.06E-93 | 2.22E-88 |
| IL6ST | 2 | -0.560186025 | 0.031 | 0.212 | 6.83E-93 | 2.5E-88 |
| REV3L | 2 | 0.561790398 | 0.933 | 0.85 | 1.16E-92 | 4.26E-88 |
| NRSN1 | 2 | 0.265314449 | 0.13 | 0.012 | 1.33E-92 | 4.87E-88 |
| ARRDC3-AS1 | 2 | -1.525202079 | 0.11 | 0.323 | 1.41E-92 | 5.17E-88 |
| CATSPERE | 2 | -0.753634969 | 0.077 | 0.278 | 1.85E-92 | 6.76E-88 |
| HS3ST5 | 2 | -1.106013889 | 0.585 | 0.626 | 3.04E-92 | 1.11E-87 |
| CLCN5 | 2 | -0.858577996 | 0.25 | 0.48 | 3.23E-92 | 1.18E-87 |
| ST8SIA4 | 2 | 0.546360712 | 0.277 | 0.081 | 3.4E-92 | 1.24E-87 |
| LRRC4C | 2 | -1.347067826 | 0.37 | 0.566 | 3.47E-92 | 1.27E-87 |
| RYR2 | 2 | -1.361637248 | 0.053 | 0.243 | 4.35E-92 | 1.59E-87 |
| SKA2 | 2 | 0.607069824 | 0.62 | 0.375 | 5.09E-92 | 1.86E-87 |
| SMPD3 | 2 | 0.374019948 | 0.211 | 0.046 | 5.23E-92 | 1.91E-87 |
| RASGRF2 | 2 | -0.946506239 | 0.036 | 0.214 | 1.21E-91 | 4.44E-87 |
| DPH6 | 2 | -0.698674625 | 0.186 | 0.439 | 1.34E-91 | 4.91E-87 |
| WSB1 | 2 | -0.90387271 | 0.68 | 0.813 | 1.57E-91 | 5.74E-87 |
| SCN8A | 2 | 0.447854692 | 0.854 | 0.622 | 1.68E-91 | 6.15E-87 |
| CUX2 | 2 | -1.014089966 | 0.118 | 0.338 | 2.28E-91 | 8.35E-87 |
| HNRNPC | 2 | -0.50909212 | 0.822 | 0.897 | 3.36E-91 | 1.23E-86 |
| PALM2-AKAP2 | 2 | 0.538138835 | 0.698 | 0.439 | 3.82E-91 | 1.4E-86 |
| LINC01515 | 2 | -0.820142152 | 0.044 | 0.221 | 1.76E-90 | 6.45E-86 |
| VIM | 2 | -0.645232206 | 0.092 | 0.31 | 1.77E-90 | 6.49E-86 |
| ROR1 | 2 | -1.15152138 | 0.09 | 0.29 | 2.76E-90 | 1.01E-85 |
| MRC2 | 2 | -0.603642052 | 0.072 | 0.275 | 3.34E-90 | 1.22E-85 |
| PTPN13 | 2 | -0.703434281 | 0.014 | 0.168 | 4.81E-90 | 1.76E-85 |
| AKAP7 | 2 | -0.58663351 | 0.073 | 0.28 | 1.05E-89 | 3.85E-85 |
| HERC4 | 2 | -0.6305086 | 0.573 | 0.756 | 1.73E-89 | 6.32E-85 |
| SMAD3 | 2 | -0.619985721 | 0.143 | 0.384 | 2.42E-89 | 8.88E-85 |
| LINC00937 | 2 | 0.445563019 | 0.372 | 0.145 | 2.53E-89 | 9.25E-85 |
| DIAPH2 | 2 | -0.730349175 | 0.888 | 0.909 | 3.67E-89 | 1.34E-84 |
| ANXA2 | 2 | -0.65463427 | 0.096 | 0.308 | 4.47E-89 | 1.64E-84 |
| SYCP2 | 2 | -1.229209417 | 0.083 | 0.259 | 5.43E-89 | 1.99E-84 |
| FRMD4B | 2 | 0.463929311 | 0.652 | 0.382 | 8.44E-89 | 3.09E-84 |
| SLC1A1 | 2 | 0.347331414 | 0.156 | 0.023 | 9.82E-89 | 3.59E-84 |
| SPRED1 | 2 | -0.722602792 | 0.223 | 0.458 | 1.73E-88 | 6.34E-84 |
| AL390800.1 | 2 | 0.506565092 | 0.734 | 0.469 | 1.83E-88 | 6.7E-84 |
| FOXN3 | 2 | 0.441913186 | 0.984 | 0.92 | 2.36E-88 | 8.63E-84 |
| TNRC6B | 2 | -0.533997033 | 0.773 | 0.874 | 2.51E-88 | 9.19E-84 |
| NGLY1 | 2 | -0.733551637 | 0.29 | 0.516 | 2.51E-88 | 9.19E-84 |
| AL691420.1 | 2 | -1.159812004 | 0.013 | 0.162 | 3.95E-88 | 1.45E-83 |
| NAAA | 2 | 0.465306487 | 0.502 | 0.247 | 5.68E-88 | 2.08E-83 |
| BRINP1 | 2 | 0.550762456 | 0.637 | 0.371 | 6.09E-88 | 2.23E-83 |
| ARHGEF26 | 2 | -0.614986168 | 0.102 | 0.317 | 7.34E-88 | 2.69E-83 |
| RBBP8 | 2 | 0.572264175 | 0.715 | 0.48 | 9.26E-88 | 3.39E-83 |
| FAM126A | 2 | 0.59582036 | 0.772 | 0.565 | 1.16E-87 | 4.24E-83 |
| DCAF6 | 2 | -0.679088578 | 0.362 | 0.592 | 1.33E-87 | 4.88E-83 |
| KDM6A | 2 | -0.544035996 | 0.837 | 0.908 | 3.03E-87 | 1.11E-82 |
| HMGA2 | 2 | -1.060543399 | 0.263 | 0.466 | 4.1E-87 | 1.5E-82 |
| ITGB8 | 2 | -1.068182041 | 0.128 | 0.338 | 4.55E-87 | 1.67E-82 |
| ST8SIA1 | 2 | 0.379254424 | 0.205 | 0.047 | 4.77E-87 | 1.75E-82 |
| ADAMTSL3 | 2 | -0.875459563 | 0.124 | 0.325 | 5.56E-87 | 2.04E-82 |
| ADAMTS6 | 2 | -1.216091646 | 0.022 | 0.18 | 6.2E-87 | 2.27E-82 |
| GSN | 2 | -0.558243114 | 0.069 | 0.263 | 7.68E-87 | 2.81E-82 |
| CYP4X1 | 2 | 0.557783878 | 0.317 | 0.111 | 1.96E-86 | 7.16E-82 |
| CFH | 2 | -0.855001752 | 0.021 | 0.176 | 2.24E-86 | 8.19E-82 |
| TOX3 | 2 | 0.452675399 | 0.508 | 0.256 | 3.21E-86 | 1.18E-81 |

| | | | | | | |
|---|---|---|---|---|---|---|
| BOC | 2 | -0.636619067 | 0.051 | 0.237 | 3.8E-86 | 1.39E-81 |
| AEBP2 | 2 | -0.803613116 | 0.595 | 0.753 | 4.15E-86 | 1.52E-81 |
| FAT4 | 2 | 0.706501532 | 0.492 | 0.253 | 4.32E-86 | 1.58E-81 |
| DACH1 | 2 | 0.468118926 | 0.976 | 0.867 | 4.34E-86 | 1.59E-81 |
| FAM13A | 2 | -0.870278627 | 0.405 | 0.595 | 4.36E-86 | 1.6E-81 |
| FGD6 | 2 | -0.774615302 | 0.154 | 0.37 | 6.33E-86 | 2.32E-81 |
| PTGER3 | 2 | 0.524217346 | 0.605 | 0.342 | 7.38E-86 | 2.7E-81 |
| PPP1R14C | 2 | 0.488153918 | 0.663 | 0.398 | 9.29E-86 | 3.4E-81 |
| TPT1-AS1 | 2 | -0.629023064 | 0.214 | 0.446 | 1.43E-85 | 5.24E-81 |
| CDK6 | 2 | -0.725327099 | 0.018 | 0.172 | 2.01E-85 | 7.37E-81 |
| FBXL2 | 2 | -0.616206386 | 0.069 | 0.261 | 2.12E-85 | 7.76E-81 |
| AC009803.1 | 2 | -0.57161622 | 0.01 | 0.156 | 2.34E-85 | 8.56E-81 |
| ELF1 | 2 | -0.627450977 | 0.111 | 0.333 | 3.85E-85 | 1.41E-80 |
| CHD7 | 2 | 0.501891107 | 0.905 | 0.78 | 4.05E-85 | 1.48E-80 |
| ELL2 | 2 | -1.022522354 | 0.045 | 0.221 | 4.19E-85 | 1.53E-80 |
| ZNF367 | 2 | 0.480997912 | 0.425 | 0.19 | 6.36E-85 | 2.33E-80 |
| IL1RAPL2 | 2 | -2.044633697 | 0.05 | 0.218 | 9.37E-85 | 3.43E-80 |
| PPP2R2B | 2 | -0.360444297 | 0.8 | 0.615 | 1.16E-84 | 4.25E-80 |
| RRM2 | 2 | 0.440275117 | 0.406 | 0.176 | 2.4E-84 | 8.77E-80 |
| RALGPS1 | 2 | 0.521864028 | 0.475 | 0.234 | 4.57E-84 | 1.67E-79 |
| CDC14B | 2 | -0.673444508 | 0.082 | 0.278 | 5.18E-84 | 1.9E-79 |
| SYBU | 2 | 0.455254393 | 0.624 | 0.363 | 7.28E-84 | 2.67E-79 |
| PPARGC1A | 2 | -0.636962645 | 0.031 | 0.197 | 7.56E-84 | 2.77E-79 |
| MDFIC | 2 | -0.437874882 | 0.007 | 0.142 | 8.26E-84 | 3.02E-79 |
| CAMK1D | 2 | 0.442834647 | 0.83 | 0.6 | 1.31E-83 | 4.78E-79 |
| DNA2 | 2 | 0.546048259 | 0.569 | 0.32 | 1.74E-83 | 6.38E-79 |
| DGKH | 2 | 0.514547624 | 0.855 | 0.667 | 1.87E-83 | 6.86E-79 |
| TLE4 | 2 | -0.813310323 | 0.307 | 0.525 | 2.39E-83 | 8.76E-79 |
| EML4 | 2 | -0.661882563 | 0.525 | 0.717 | 2.54E-83 | 9.3E-79 |
| SLC4A4 | 2 | 0.504579977 | 0.346 | 0.133 | 2.87E-83 | 1.05E-78 |
| GPC4 | 2 | -0.741089092 | 0.057 | 0.24 | 3.08E-83 | 1.13E-78 |
| GRAMD2B | 2 | -0.491159744 | 0.026 | 0.187 | 3.67E-83 | 1.34E-78 |
| NBAS | 2 | -0.75908609 | 0.691 | 0.824 | 4.65E-83 | 1.7E-78 |
| KIAA1211 | 2 | 0.503071278 | 0.857 | 0.659 | 5.98E-83 | 2.19E-78 |
| SULF2 | 2 | -0.602066624 | 0.049 | 0.227 | 6.03E-83 | 2.21E-78 |
| MAN2A1 | 2 | -0.809871549 | 0.281 | 0.496 | 6.15E-83 | 2.25E-78 |
| PKP4 | 2 | -0.833988458 | 0.575 | 0.737 | 6.34E-83 | 2.32E-78 |
| AL162718.1 | 2 | -0.631949537 | 0.015 | 0.164 | 7.41E-83 | 2.71E-78 |
| SPC25 | 2 | 0.529274211 | 0.338 | 0.13 | 7.9E-83 | 2.89E-78 |
| TMEM179 | 2 | 0.307734156 | 0.147 | 0.022 | 1.06E-82 | 3.87E-78 |
| CDH23 | 2 | -0.727079486 | 0.107 | 0.311 | 1.1E-82 | 4.01E-78 |
| RAB3IP | 2 | 0.465920769 | 0.299 | 0.103 | 1.36E-82 | 4.99E-78 |
| FAM149A | 2 | 0.529794119 | 0.522 | 0.272 | 1.69E-82 | 6.17E-78 |
| FHIT | 2 | -0.760792153 | 0.589 | 0.771 | 2.08E-82 | 7.62E-78 |
| NCAM1 | 2 | 0.352786328 | 0.955 | 0.805 | 2.11E-82 | 7.73E-78 |
| FAM160A1 | 2 | -0.649461483 | 0.022 | 0.177 | 3.91E-82 | 1.43E-77 |
| CRMP1 | 2 | 0.526284694 | 0.525 | 0.279 | 4.06E-82 | 1.49E-77 |
| PDE4B | 2 | -1.149018233 | 0.015 | 0.159 | 6.78E-82 | 2.48E-77 |
| SH3GL2 | 2 | 0.342877399 | 0.599 | 0.348 | 7.65E-82 | 2.8E-77 |
| SCN1A | 2 | 0.619546394 | 0.24 | 0.068 | 8.53E-82 | 3.12E-77 |
| CBLB | 2 | -0.823774055 | 0.416 | 0.631 | 9.21E-82 | 3.37E-77 |
| PDE4D | 2 | 0.510673086 | 0.939 | 0.83 | 1.16E-81 | 4.24E-77 |
| ARSJ | 2 | -0.529167232 | 0.005 | 0.137 | 1.26E-81 | 4.6E-77 |
| ECE2 | 2 | 0.317835879 | 0.161 | 0.028 | 1.56E-81 | 5.71E-77 |
| YES1 | 2 | -0.607660702 | 0.183 | 0.413 | 1.64E-81 | 6.01E-77 |
| GLDC | 2 | -0.701819196 | 0.111 | 0.315 | 1.76E-81 | 6.46E-77 |
| ARID1B | 2 | 0.483944515 | 0.983 | 0.951 | 2.18E-81 | 7.97E-77 |
| AC108690.1 | 2 | -0.577009192 | 0.011 | 0.149 | 2.19E-81 | 8.01E-77 |

| | | | | | | |
|---|---|---|---|---|---|---|
| CUX1 | 2 | -0.648893884 | 0.678 | 0.811 | 2.51E-81 | 9.17E-77 |
| MAP3K1 | 2 | -0.676400171 | 0.058 | 0.24 | 3.9E-81 | 1.43E-76 |
| FAM135B | 2 | 0.58715355 | 0.575 | 0.322 | 1.01E-80 | 3.7E-76 |
| AP3B1 | 2 | -0.595879748 | 0.473 | 0.684 | 1.18E-80 | 4.32E-76 |
| TENT5A | 2 | -0.701786714 | 0.015 | 0.161 | 1.41E-80 | 5.17E-76 |
| EPS15 | 2 | -0.611692479 | 0.433 | 0.657 | 1.74E-80 | 6.38E-76 |
| ATF7IP2 | 2 | -0.617298675 | 0.089 | 0.289 | 2.14E-80 | 7.83E-76 |
| MYO1D | 2 | 0.469863403 | 0.403 | 0.178 | 3.19E-80 | 1.17E-75 |
| KIF1A | 2 | 0.43111039 | 0.334 | 0.128 | 3.24E-80 | 1.18E-75 |
| DAAM1 | 2 | 0.73670685 | 0.653 | 0.468 | 4.41E-80 | 1.62E-75 |
| TENM1 | 2 | -0.808420091 | 0.014 | 0.157 | 5.27E-80 | 1.93E-75 |
| SMARCA1 | 2 | -0.691060889 | 0.517 | 0.706 | 5.46E-80 | 2E-75 |
| FBN2 | 2 | -0.363791781 | 0.521 | 0.346 | 9.85E-80 | 3.61E-75 |
| HECW2 | 2 | 0.30439805 | 0.566 | 0.323 | 9.87E-80 | 3.61E-75 |
| NXPH2 | 2 | 0.41045098 | 0.261 | 0.083 | 1E-79 | 3.68E-75 |
| EDNRA | 2 | -0.616298364 | 0.014 | 0.155 | 2.51E-79 | 9.19E-75 |
| CHCHD6 | 2 | 0.533912568 | 0.436 | 0.216 | 2.85E-79 | 1.04E-74 |
| PLCB1 | 2 | 0.616650081 | 0.721 | 0.48 | 3.24E-79 | 1.19E-74 |
| CHD2 | 2 | -0.601160944 | 0.694 | 0.813 | 3.76E-79 | 1.38E-74 |
| RALGPS2 | 2 | -0.716941448 | 0.188 | 0.409 | 3.97E-79 | 1.45E-74 |
| ACYP2 | 2 | -0.699196905 | 0.343 | 0.571 | 4.19E-79 | 1.53E-74 |
| ZNF521 | 2 | -0.940783456 | 0.025 | 0.171 | 4.59E-79 | 1.68E-74 |
| MYLIP | 2 | -0.492060691 | 0.039 | 0.208 | 6.12E-79 | 2.24E-74 |
| RAD51AP1 | 2 | 0.431966487 | 0.341 | 0.135 | 1.15E-78 | 4.2E-74 |
| EFHC2 | 2 | -0.513305902 | 0.018 | 0.166 | 1.25E-78 | 4.57E-74 |
| UBE2E2 | 2 | 0.272135672 | 0.793 | 0.556 | 2.18E-78 | 7.97E-74 |
| CDKN2A | 2 | 0.498211033 | 0.522 | 0.282 | 2.44E-78 | 8.92E-74 |
| OSBPL3 | 2 | -0.788752931 | 0.293 | 0.496 | 3.05E-78 | 1.12E-73 |
| KCNN2 | 2 | -1.241277394 | 0.16 | 0.331 | 4.01E-78 | 1.47E-73 |
| SLC4A7 | 2 | 0.437942015 | 0.893 | 0.724 | 9.23E-78 | 3.38E-73 |
| COLEC12 | 2 | -0.654164067 | 0.019 | 0.16 | 1.37E-77 | 5.03E-73 |
| NFATC4 | 2 | -0.371611874 | 0.021 | 0.166 | 1.98E-77 | 7.26E-73 |
| JPH1 | 2 | 0.507165438 | 0.316 | 0.122 | 2.61E-77 | 9.54E-73 |
| NEIL3 | 2 | 0.496110174 | 0.327 | 0.126 | 3.62E-77 | 1.33E-72 |
| KIF2A | 2 | 0.525863504 | 0.756 | 0.572 | 3.81E-77 | 1.39E-72 |
| MIR181A2HG | 2 | -0.648989508 | 0.118 | 0.311 | 4.37E-77 | 1.6E-72 |
| PLEKHG1 | 2 | -0.558227993 | 0.018 | 0.16 | 4.67E-77 | 1.71E-72 |
| WDR76 | 2 | 0.489072616 | 0.546 | 0.302 | 7.09E-77 | 2.59E-72 |
| PRKN | 2 | -0.747604716 | 0.359 | 0.578 | 8.15E-77 | 2.98E-72 |
| MIER1 | 2 | -0.610848111 | 0.308 | 0.528 | 1.63E-76 | 5.96E-72 |
| ZFAND6 | 2 | -0.645152373 | 0.511 | 0.689 | 1.96E-76 | 7.17E-72 |
| PCSK5 | 2 | 0.67592493 | 0.479 | 0.258 | 3.29E-76 | 1.2E-71 |
| RBM47 | 2 | -0.968172417 | 0.018 | 0.154 | 6.31E-76 | 2.31E-71 |
| MIR9-3HG | 2 | 0.351466915 | 0.244 | 0.074 | 6.45E-76 | 2.36E-71 |
| ADGRG2 | 2 | -0.740177147 | 0.05 | 0.21 | 8.9E-76 | 3.26E-71 |
| BCHE | 2 | -0.5587776 | 0.012 | 0.146 | 1.34E-75 | 4.92E-71 |
| TAPT1-AS1 | 2 | -0.490966058 | 0.066 | 0.247 | 1.63E-75 | 5.99E-71 |
| WDR60 | 2 | -0.595669956 | 0.229 | 0.449 | 1.72E-75 | 6.29E-71 |
| TTC28 | 2 | 0.37974887 | 0.989 | 0.958 | 3.05E-75 | 1.12E-70 |
| GABRA5 | 2 | 0.441049574 | 0.214 | 0.062 | 3.93E-75 | 1.44E-70 |
| LINC00511 | 2 | -1.052485773 | 0.134 | 0.325 | 5.68E-75 | 2.08E-70 |
| MTUS2 | 2 | 0.461279953 | 0.383 | 0.17 | 6.48E-75 | 2.37E-70 |
| AC117461.1 | 2 | 0.465129292 | 0.218 | 0.06 | 6.86E-75 | 2.51E-70 |
| AGMO | 2 | -0.942026381 | 0.009 | 0.139 | 7.02E-75 | 2.57E-70 |
| NCAPG2 | 2 | 0.488542648 | 0.562 | 0.322 | 1.24E-74 | 4.55E-70 |
| CCDC18-AS1 | 2 | -0.577249294 | 0.179 | 0.397 | 2.01E-74 | 7.34E-70 |
| MAGI1 | 2 | -0.640597713 | 0.842 | 0.87 | 2.02E-74 | 7.38E-70 |
| YBX3 | 2 | -0.473250471 | 0.086 | 0.279 | 2.15E-74 | 7.87E-70 |

| | | | | | | |
|---|---|---|---|---|---|---|
| APBB2 | 2 | -0.698105598 | 0.301 | 0.498 | 2.43E-74 | 8.89E-70 |
| COL18A1 | 2 | -0.532165875 | 0.018 | 0.159 | 3.02E-74 | 1.11E-69 |
| MAP3K15 | 2 | -0.619931193 | 0.041 | 0.2 | 3.2E-74 | 1.17E-69 |
| MAML3 | 2 | -1.053288881 | 0.439 | 0.603 | 3.42E-74 | 1.25E-69 |
| PAN3 | 2 | -0.536211814 | 0.704 | 0.819 | 6.21E-74 | 2.27E-69 |
| GRK3 | 2 | 0.533710844 | 0.641 | 0.413 | 7.33E-74 | 2.68E-69 |
| ANXA4 | 2 | -0.479643228 | 0.084 | 0.275 | 1.04E-73 | 3.81E-69 |
| CHST11 | 2 | 0.521623235 | 0.779 | 0.572 | 1.6E-73 | 5.84E-69 |
| LINC01956 | 2 | -0.445493899 | 0.012 | 0.141 | 1.66E-73 | 6.08E-69 |
| PARP8 | 2 | -0.753216904 | 0.175 | 0.369 | 2.06E-73 | 7.54E-69 |
| PBK | 2 | 0.412319358 | 0.262 | 0.087 | 2.12E-73 | 7.75E-69 |
| SON | 2 | -0.550998015 | 0.542 | 0.715 | 2.21E-73 | 8.1E-69 |
| PRSS23 | 2 | 0.438081665 | 0.236 | 0.071 | 2.64E-73 | 9.68E-69 |
| IGSF9B | 2 | 0.407648443 | 0.282 | 0.1 | 2.86E-73 | 1.05E-68 |
| INSR | 2 | -0.736255349 | 0.109 | 0.289 | 3.12E-73 | 1.14E-68 |
| RAB39A | 2 | 0.344961791 | 0.22 | 0.063 | 4.17E-73 | 1.53E-68 |
| DPP4 | 2 | -0.676257514 | 0.049 | 0.203 | 5.15E-73 | 1.89E-68 |
| APC | 2 | 0.616070213 | 0.676 | 0.491 | 7.72E-73 | 2.83E-68 |
| ANKRD10 | 2 | -0.570525987 | 0.535 | 0.706 | 8.02E-73 | 2.94E-68 |
| DLEU2 | 2 | 0.462770021 | 0.867 | 0.717 | 9.1E-73 | 3.33E-68 |
| SESTD1 | 2 | 0.54343711 | 0.659 | 0.457 | 9.27E-73 | 3.39E-68 |
| VCAN | 2 | -0.793095933 | 0.488 | 0.4 | 1.28E-72 | 4.68E-68 |
| PAM | 2 | 0.359150164 | 0.898 | 0.723 | 1.29E-72 | 4.71E-68 |
| LINC02263 | 2 | -0.719228168 | 0.007 | 0.129 | 1.56E-72 | 5.7E-68 |
| HS3ST1 | 2 | 0.384558447 | 0.364 | 0.16 | 2.12E-72 | 7.76E-68 |
| AOPEP | 2 | -0.584148676 | 0.648 | 0.78 | 2.48E-72 | 9.09E-68 |
| HPCAL1 | 2 | -0.640553534 | 0.156 | 0.363 | 2.62E-72 | 9.61E-68 |
| HLF | 2 | 0.298666296 | 0.134 | 0.021 | 4.92E-72 | 1.8E-67 |
| SIDT1 | 2 | 0.354446341 | 0.148 | 0.027 | 7.97E-72 | 2.92E-67 |
| ARMC9 | 2 | -0.578695855 | 0.147 | 0.35 | 8.2E-72 | 3E-67 |
| RIN2 | 2 | -0.558094052 | 0.181 | 0.399 | 2.64E-71 | 9.66E-67 |
| NT5DC1 | 2 | -0.662975495 | 0.22 | 0.433 | 2.65E-71 | 9.69E-67 |
| FTX | 2 | -0.436160785 | 0.988 | 0.985 | 2.7E-71 | 9.9E-67 |
| EFCAB6 | 2 | -0.478412968 | 0.028 | 0.174 | 3.51E-71 | 1.29E-66 |
| KCNT2 | 2 | 0.534866419 | 0.836 | 0.676 | 5.2E-71 | 1.9E-66 |
| KCNQ2 | 2 | 0.424630417 | 0.369 | 0.163 | 7.48E-71 | 2.74E-66 |
| ETV5 | 2 | -0.393405604 | 0.022 | 0.162 | 7.87E-71 | 2.88E-66 |
| NEK7 | 2 | -0.640120811 | 0.454 | 0.657 | 8.25E-71 | 3.02E-66 |
| PDE7A | 2 | 0.476892682 | 0.755 | 0.579 | 1.05E-70 | 3.84E-66 |
| ZNF385B | 2 | -1.122151142 | 0.192 | 0.363 | 1.11E-70 | 4.07E-66 |
| SERPINE2 | 2 | -0.595004965 | 0.03 | 0.175 | 1.14E-70 | 4.18E-66 |
| GALNT18 | 2 | 0.577087887 | 0.582 | 0.356 | 1.83E-70 | 6.68E-66 |
| PARM1 | 2 | 0.404593823 | 0.367 | 0.165 | 3.28E-70 | 1.2E-65 |
| FAM227A | 2 | -0.388852636 | 0.022 | 0.161 | 4.2E-70 | 1.54E-65 |
| CDC42BPA | 2 | 0.475923579 | 0.877 | 0.748 | 4.73E-70 | 1.73E-65 |
| DLG1 | 2 | -0.549552251 | 0.635 | 0.785 | 5.12E-70 | 1.87E-65 |
| ZNF254 | 2 | -0.555694236 | 0.205 | 0.421 | 5.49E-70 | 2.01E-65 |
| FAM214A | 2 | -0.619177977 | 0.228 | 0.428 | 5.62E-70 | 2.06E-65 |
| CAMKV | 2 | 0.316897044 | 0.199 | 0.053 | 6.09E-70 | 2.23E-65 |
| B3GAT1 | 2 | 0.306787024 | 0.188 | 0.047 | 7.2E-70 | 2.64E-65 |
| MAFB | 2 | 0.288193729 | 0.146 | 0.027 | 9.44E-70 | 3.45E-65 |
| TMEM145 | 2 | 0.28079885 | 0.154 | 0.031 | 1.31E-69 | 4.78E-65 |
| DEPDC1B | 2 | 0.459638412 | 0.362 | 0.161 | 1.36E-69 | 4.98E-65 |
| KCNJ6 | 2 | -0.798883424 | 0.082 | 0.229 | 1.59E-69 | 5.81E-65 |
| AF117829.1 | 2 | -0.613204955 | 0.204 | 0.397 | 1.79E-69 | 6.54E-65 |
| UNC13A | 2 | 0.400330926 | 0.276 | 0.1 | 1.83E-69 | 6.69E-65 |
| SEMA3C | 2 | -0.795612946 | 0.017 | 0.146 | 1.83E-69 | 6.72E-65 |
| ATP8A1 | 2 | -0.872630452 | 0.271 | 0.442 | 1.9E-69 | 6.96E-65 |

| | | | | | | |
|---|---|---|---|---|---|---|
| DNM3OS | 2 | 0.288184028 | 0.52 | 0.299 | 2.14E-69 | 7.83E-65 |
| SUGCT | 2 | -0.64645892 | 0.221 | 0.442 | 2.24E-69 | 8.21E-65 |
| EBF4 | 2 | 0.452438817 | 0.363 | 0.162 | 2.8E-69 | 1.02E-64 |
| EFNA5 | 2 | -0.968572646 | 0.629 | 0.671 | 2.91E-69 | 1.06E-64 |
| RAPGEF4 | 2 | -0.735670159 | 0.374 | 0.409 | 3.11E-69 | 1.14E-64 |
| COL4A5 | 2 | -1.24768972 | 0.015 | 0.132 | 3.23E-69 | 1.18E-64 |
| EFHC1 | 2 | -0.567578556 | 0.115 | 0.299 | 3.31E-69 | 1.21E-64 |
| BCAR3 | 2 | -0.539464091 | 0.092 | 0.276 | 4.46E-69 | 1.63E-64 |
| SERINC5 | 2 | -0.661178865 | 0.133 | 0.316 | 5.22E-69 | 1.91E-64 |
| RAP2A | 2 | -0.595527563 | 0.153 | 0.342 | 5.98E-69 | 2.19E-64 |
| MBNL1 | 2 | 0.665440458 | 0.957 | 0.904 | 7.96E-69 | 2.91E-64 |
| LRMDA | 2 | -0.863668848 | 0.016 | 0.138 | 8.03E-69 | 2.94E-64 |
| AC006972.1 | 2 | -0.470994508 | 0.007 | 0.123 | 1.01E-68 | 3.7E-64 |
| MMP24 | 2 | 0.340203707 | 0.185 | 0.048 | 1.06E-68 | 3.88E-64 |
| STK32B | 2 | 0.313739842 | 0.714 | 0.482 | 1.92E-68 | 7.05E-64 |
| GPBP1L1 | 2 | -0.533913325 | 0.299 | 0.516 | 2.14E-68 | 7.84E-64 |
| GABRA2 | 2 | -0.794794509 | 0.009 | 0.127 | 2.23E-68 | 8.17E-64 |
| ADAMTS12 | 2 | -1.403013734 | 0.272 | 0.428 | 2.86E-68 | 1.05E-63 |
| ARMC4 | 2 | -0.403176869 | 0.02 | 0.154 | 3.2E-68 | 1.17E-63 |
| SLC1A3 | 2 | -0.483192527 | 0.01 | 0.13 | 4.05E-68 | 1.48E-63 |
| EML6 | 2 | 0.500045859 | 0.415 | 0.205 | 7.75E-68 | 2.84E-63 |
| SORBS2 | 2 | -1.000668173 | 0.026 | 0.149 | 8.2E-68 | 3E-63 |
| TMEM63A | 2 | -0.371763439 | 0.022 | 0.158 | 1.24E-67 | 4.52E-63 |
| COL6A1 | 2 | -0.518714213 | 0.069 | 0.234 | 1.37E-67 | 5.03E-63 |
| ZNF516 | 2 | -0.614472167 | 0.399 | 0.587 | 1.61E-67 | 5.9E-63 |
| USP31 | 2 | 0.499017443 | 0.492 | 0.285 | 1.8E-67 | 6.59E-63 |
| KDM2A | 2 | -0.518211323 | 0.647 | 0.799 | 2.23E-67 | 8.17E-63 |
| NRXN3 | 2 | -1.717322212 | 0.267 | 0.374 | 3.74E-67 | 1.37E-62 |
| PTPRU | 2 | 0.322371372 | 0.199 | 0.056 | 4.17E-67 | 1.53E-62 |
| ABL1 | 2 | -0.595793653 | 0.422 | 0.615 | 4.35E-67 | 1.59E-62 |
| ACSL4 | 2 | 0.530571353 | 0.722 | 0.524 | 4.87E-67 | 1.78E-62 |
| TMEM87B | 2 | -0.53943164 | 0.124 | 0.313 | 5.04E-67 | 1.85E-62 |
| CHRM3 | 2 | -1.184437904 | 0.262 | 0.454 | 5.51E-67 | 2.02E-62 |
| ASRGL1 | 2 | 0.382807432 | 0.288 | 0.111 | 6.64E-67 | 2.43E-62 |
| KIAA1328 | 2 | 0.523406561 | 0.74 | 0.578 | 7.23E-67 | 2.65E-62 |
| POLR3G | 2 | -0.570974494 | 0.117 | 0.299 | 7.54E-67 | 2.76E-62 |
| N4BP2L2 | 2 | -0.511692781 | 0.645 | 0.785 | 1.03E-66 | 3.78E-62 |
| PHLDB2 | 2 | -0.711762428 | 0.092 | 0.262 | 1.06E-66 | 3.88E-62 |
| AGTPBP1 | 2 | 0.471788628 | 0.75 | 0.564 | 1.41E-66 | 5.16E-62 |
| E2F7 | 2 | 0.487208851 | 0.344 | 0.152 | 1.98E-66 | 7.25E-62 |
| LINC02315 | 2 | 0.419186755 | 0.918 | 0.754 | 2.3E-66 | 8.41E-62 |
| BACH2 | 2 | 0.435980972 | 0.574 | 0.343 | 2.43E-66 | 8.91E-62 |
| KANK1 | 2 | 0.291571533 | 0.695 | 0.466 | 2.59E-66 | 9.49E-62 |
| NPFFR2 | 2 | 0.347276898 | 0.104 | 0.013 | 4.19E-66 | 1.53E-61 |
| CAP2 | 2 | -0.528818129 | 0.115 | 0.304 | 4.57E-66 | 1.67E-61 |
| AC063938.1 | 2 | -0.563633545 | 0.011 | 0.129 | 5.66E-66 | 2.07E-61 |
| LINC01876 | 2 | 0.38619965 | 0.695 | 0.465 | 8.25E-66 | 3.02E-61 |
| KCNK10 | 2 | 0.355136602 | 0.138 | 0.026 | 9.54E-66 | 3.49E-61 |
| ST5 | 2 | -0.606778761 | 0.147 | 0.331 | 1.01E-65 | 3.7E-61 |
| ITPKB | 2 | -0.372856484 | 0.022 | 0.151 | 1.02E-65 | 3.74E-61 |
| AP002954.2 | 2 | -0.71762105 | 0.007 | 0.119 | 1.03E-65 | 3.79E-61 |
| TBC1D16 | 2 | -0.523859078 | 0.121 | 0.31 | 1.6E-65 | 5.87E-61 |
| CENPP | 2 | 0.256294497 | 0.852 | 0.655 | 1.65E-65 | 6.04E-61 |
| NPSR1-AS1 | 2 | -0.857074527 | 0.052 | 0.203 | 1.89E-65 | 6.94E-61 |
| ZNF184 | 2 | 0.401675428 | 0.208 | 0.065 | 1.91E-65 | 7E-61 |
| GALNTL6 | 2 | 1.261553474 | 0.37 | 0.222 | 2.85E-65 | 1.04E-60 |
| TMEM220 | 2 | -0.444210684 | 0.069 | 0.234 | 2.95E-65 | 1.08E-60 |
| GLT1D1 | 2 | -0.470837664 | 0.044 | 0.191 | 3.67E-65 | 1.34E-60 |

| | | | | | | |
|---|---|---|---|---|---|---|
| MAP7 | 2 | 0.355432593 | 0.77 | 0.548 | 3.74E-65 | 1.37E-60 |
| TTC23 | 2 | -0.377467813 | 0.06 | 0.226 | 3.74E-65 | 1.37E-60 |
| CSMD3 | 2 | -1.193124882 | 0.192 | 0.355 | 4.57E-65 | 1.67E-60 |
| AC040168.1 | 2 | 0.453978486 | 0.417 | 0.209 | 5.41E-65 | 1.98E-60 |
| TANC1 | 2 | -0.653672121 | 0.386 | 0.562 | 6.1E-65 | 2.23E-60 |
| ADD3 | 2 | -0.583415507 | 0.275 | 0.482 | 7.41E-65 | 2.71E-60 |
| RCOR1 | 2 | -0.574105241 | 0.559 | 0.713 | 8.79E-65 | 3.22E-60 |
| KIF1B | 2 | 0.429196427 | 0.824 | 0.722 | 1.05E-64 | 3.83E-60 |
| COL5A1 | 2 | -0.552808748 | 0.031 | 0.166 | 1.63E-64 | 5.98E-60 |
| JADE3 | 2 | 0.486838662 | 0.531 | 0.315 | 1.86E-64 | 6.8E-60 |
| CNRIP1 | 2 | 0.363369465 | 0.267 | 0.099 | 1.88E-64 | 6.9E-60 |
| FNBP1 | 2 | -0.577722053 | 0.537 | 0.689 | 2.65E-64 | 9.71E-60 |
| BMP6 | 2 | 0.411665499 | 0.23 | 0.078 | 2.82E-64 | 1.03E-59 |
| AL591501.1 | 2 | -1.002626406 | 0.017 | 0.136 | 2.93E-64 | 1.07E-59 |
| NEDD9 | 2 | -0.363282083 | 0.018 | 0.146 | 3.11E-64 | 1.14E-59 |
| MBP | 2 | -0.398149466 | 0.021 | 0.15 | 3.9E-64 | 1.43E-59 |
| MELK | 2 | 0.393055629 | 0.446 | 0.234 | 4.3E-64 | 1.58E-59 |
| AL596087.2 | 2 | 0.364624283 | 0.359 | 0.169 | 5.12E-64 | 1.88E-59 |
| RIN3 | 2 | -0.326315508 | 0.022 | 0.153 | 5.98E-64 | 2.19E-59 |
| FGFR2 | 2 | -0.763626534 | 0.24 | 0.425 | 7.67E-64 | 2.81E-59 |
| AC016831.7 | 2 | -0.342802577 | 0.029 | 0.168 | 8.06E-64 | 2.95E-59 |
| PPFIBP2 | 2 | -0.285385666 | 0.004 | 0.106 | 9.87E-64 | 3.61E-59 |
| ZKSCAN1 | 2 | -0.567902184 | 0.27 | 0.474 | 1.02E-63 | 3.74E-59 |
| NCALD | 2 | 0.383421367 | 0.747 | 0.526 | 1.07E-63 | 3.92E-59 |
| RSRC1 | 2 | 0.463338021 | 0.889 | 0.79 | 1.12E-63 | 4.1E-59 |
| LGMN | 2 | -0.429497432 | 0.046 | 0.197 | 1.27E-63 | 4.63E-59 |
| GPATCH2 | 2 | -0.575864308 | 0.282 | 0.474 | 1.41E-63 | 5.18E-59 |
| HSPA12A | 2 | -0.528507363 | 0.13 | 0.312 | 1.68E-63 | 6.17E-59 |
| EBF3 | 2 | 0.44525144 | 0.418 | 0.212 | 1.83E-63 | 6.71E-59 |
| RNF157 | 2 | 0.443950785 | 0.421 | 0.216 | 1.98E-63 | 7.24E-59 |
| TBC1D30 | 2 | 0.28030114 | 0.199 | 0.06 | 2.04E-63 | 7.47E-59 |
| TM9SF3 | 2 | 0.502033333 | 0.733 | 0.587 | 2.38E-63 | 8.71E-59 |
| TIAM2 | 2 | -0.699098141 | 0.736 | 0.791 | 2.62E-63 | 9.61E-59 |
| CENPU | 2 | 0.452427193 | 0.482 | 0.267 | 3.64E-63 | 1.33E-58 |
| PDCD4 | 2 | -0.537185306 | 0.241 | 0.441 | 4.12E-63 | 1.51E-58 |
| CTNND1 | 2 | -0.554554201 | 0.483 | 0.654 | 6.91E-63 | 2.53E-58 |
| NALCN | 2 | -0.736296399 | 0.108 | 0.251 | 9.84E-63 | 3.6E-58 |
| CIP2A | 2 | 0.414246844 | 0.473 | 0.259 | 1.3E-62 | 4.75E-58 |
| MAP1B | 2 | 0.386991573 | 0.866 | 0.69 | 1.37E-62 | 5.01E-58 |
| KIRREL3 | 2 | -0.979130657 | 0.066 | 0.205 | 1.83E-62 | 6.71E-58 |
| VPS54 | 2 | -0.603003034 | 0.446 | 0.616 | 1.89E-62 | 6.91E-58 |
| CRTAC1 | 2 | 0.269845686 | 0.145 | 0.031 | 3.11E-62 | 1.14E-57 |
| HHAT | 2 | -0.527102478 | 0.13 | 0.308 | 3.32E-62 | 1.22E-57 |
| CSMD1 | 2 | -1.247630668 | 0.43 | 0.448 | 4.47E-62 | 1.64E-57 |
| CPNE4 | 2 | 0.348294436 | 0.164 | 0.041 | 4.52E-62 | 1.65E-57 |
| SPINK5 | 2 | -0.461337486 | 0.037 | 0.179 | 4.88E-62 | 1.79E-57 |
| F3 | 2 | -0.63627389 | 0.01 | 0.121 | 5.72E-62 | 2.09E-57 |
| FAM111B | 2 | 0.396058727 | 0.301 | 0.124 | 6.21E-62 | 2.27E-57 |
| TSHZ3 | 2 | 0.51575759 | 0.672 | 0.451 | 6.44E-62 | 2.36E-57 |
| AC023469.1 | 2 | 0.324452632 | 0.528 | 0.312 | 6.55E-62 | 2.4E-57 |
| MEG3 | 2 | -0.297243719 | 0.522 | 0.43 | 8.7E-62 | 3.18E-57 |
| PCMTD2 | 2 | -0.466523423 | 0.175 | 0.369 | 1.06E-61 | 3.87E-57 |
| FANCA | 2 | 0.415578181 | 0.633 | 0.413 | 1.55E-61 | 5.67E-57 |
| PLXNA2 | 2 | 0.507024481 | 0.557 | 0.343 | 1.6E-61 | 5.87E-57 |
| LRP4 | 2 | -0.356599802 | 0.014 | 0.128 | 1.67E-61 | 6.12E-57 |
| BMPER | 2 | 0.641831612 | 0.369 | 0.175 | 1.83E-61 | 6.69E-57 |
| FTO | 2 | -0.817001551 | 0.792 | 0.816 | 1.9E-61 | 6.96E-57 |
| GAREM1 | 2 | -0.675632648 | 0.147 | 0.313 | 2.33E-61 | 8.52E-57 |

| | | | | | | |
|---|---|---|---|---|---|---|
| TCF12 | 2 | -0.468427316 | 0.893 | 0.927 | 4.04E-61 | 1.48E-56 |
| CELF5 | 2 | 0.308320399 | 0.178 | 0.048 | 4.9E-61 | 1.8E-56 |
| KIAA1324L | 2 | 0.498520289 | 0.376 | 0.185 | 6.95E-61 | 2.54E-56 |
| MCM10 | 2 | 0.337274172 | 0.351 | 0.164 | 7.47E-61 | 2.73E-56 |
| NTN1 | 2 | -0.581462103 | 0.036 | 0.171 | 7.69E-61 | 2.82E-56 |
| PYGL | 2 | -0.32556861 | 0.01 | 0.121 | 8.29E-61 | 3.03E-56 |
| SGO1-AS1 | 2 | -1.055535956 | 0.069 | 0.212 | 9.48E-61 | 3.47E-56 |
| AL138720.1 | 2 | -0.630684307 | 0.047 | 0.188 | 1.17E-60 | 4.3E-56 |
| RANBP17 | 2 | 0.473950851 | 0.838 | 0.712 | 1.24E-60 | 4.53E-56 |
| POU2F1 | 2 | 0.470286048 | 0.89 | 0.819 | 1.25E-60 | 4.58E-56 |
| FHL1 | 2 | 0.424922314 | 0.401 | 0.203 | 1.34E-60 | 4.9E-56 |
| KSR1 | 2 | 0.464772426 | 0.569 | 0.356 | 1.62E-60 | 5.95E-56 |
| CUL4A | 2 | -0.513678399 | 0.307 | 0.496 | 3.04E-60 | 1.11E-55 |
| NOTCH2NLC | 2 | -0.330812885 | 0.034 | 0.171 | 3.26E-60 | 1.19E-55 |
| GPR156 | 2 | 0.477882215 | 0.381 | 0.191 | 3.43E-60 | 1.26E-55 |
| COL19A1 | 2 | -0.633821076 | 0.012 | 0.124 | 3.51E-60 | 1.28E-55 |
| C4orf47 | 2 | -0.426772684 | 0.017 | 0.137 | 3.72E-60 | 1.36E-55 |
| RAD51B | 2 | -0.675864002 | 0.549 | 0.703 | 4.29E-60 | 1.57E-55 |
| LEF1 | 2 | -0.489953185 | 0.02 | 0.14 | 5.25E-60 | 1.92E-55 |
| FLI1 | 2 | -0.333387674 | 0.004 | 0.101 | 6.81E-60 | 2.49E-55 |
| SS18 | 2 | -0.501308456 | 0.334 | 0.527 | 6.91E-60 | 2.53E-55 |
| FEZ1 | 2 | 0.454968661 | 0.373 | 0.185 | 7.73E-60 | 2.83E-55 |
| HULC | 2 | 0.591749324 | 0.578 | 0.377 | 8.32E-60 | 3.05E-55 |
| MME | 2 | -0.702128086 | 0.086 | 0.213 | 8.86E-60 | 3.24E-55 |
| MGAT5B | 2 | 0.254898946 | 0.165 | 0.043 | 1.01E-59 | 3.72E-55 |
| HSPA4L | 2 | 0.461081456 | 0.364 | 0.179 | 1.06E-59 | 3.87E-55 |
| ST3GAL6 | 2 | 0.463496592 | 0.453 | 0.248 | 1.28E-59 | 4.7E-55 |
| TRABD2B | 2 | -0.406734952 | 0.017 | 0.135 | 1.6E-59 | 5.86E-55 |
| TMEM168 | 2 | -0.280833767 | 0.007 | 0.111 | 1.65E-59 | 6.03E-55 |
| RNF150 | 2 | 0.363178981 | 0.569 | 0.351 | 1.86E-59 | 6.83E-55 |
| DLX5 | 2 | 0.30953745 | 0.19 | 0.056 | 2.7E-59 | 9.9E-55 |
| CELSR3 | 2 | 0.299354927 | 0.18 | 0.051 | 2.97E-59 | 1.09E-54 |
| SMC4 | 2 | 0.444964049 | 0.736 | 0.546 | 3.7E-59 | 1.36E-54 |
| KCNC2 | 2 | 0.325770069 | 0.125 | 0.024 | 5.62E-59 | 2.06E-54 |
| ADAMTS3 | 2 | 0.443930268 | 0.655 | 0.437 | 5.94E-59 | 2.18E-54 |
| SYNE1 | 2 | -0.795365298 | 0.32 | 0.511 | 6.32E-59 | 2.31E-54 |
| FANCB | 2 | 0.426976447 | 0.401 | 0.207 | 7.6E-59 | 2.78E-54 |
| EVI5 | 2 | -0.590241762 | 0.407 | 0.592 | 7.88E-59 | 2.89E-54 |
| ADD3-AS1 | 2 | -0.356544995 | 0.011 | 0.117 | 9.33E-59 | 3.42E-54 |
| MYBL2 | 2 | 0.339365999 | 0.28 | 0.114 | 1.17E-58 | 4.29E-54 |
| RFX2 | 2 | -0.629067838 | 0.101 | 0.251 | 1.19E-58 | 4.35E-54 |
| AFAP1 | 2 | -0.582118114 | 0.312 | 0.512 | 1.24E-58 | 4.54E-54 |
| AC092939.1 | 2 | 0.427940145 | 0.328 | 0.154 | 1.3E-58 | 4.77E-54 |
| ITGB5 | 2 | -0.335638117 | 0.014 | 0.126 | 1.31E-58 | 4.81E-54 |
| MCTP1 | 2 | 0.656023419 | 0.463 | 0.258 | 1.35E-58 | 4.96E-54 |
| CDH4 | 2 | 0.51532383 | 0.773 | 0.584 | 1.5E-58 | 5.47E-54 |
| USP34 | 2 | -0.397899879 | 0.882 | 0.927 | 1.7E-58 | 6.21E-54 |
| KIF18B | 2 | 0.40535562 | 0.263 | 0.102 | 2.25E-58 | 8.23E-54 |
| PRRX1 | 2 | -0.515676007 | 0.013 | 0.124 | 2.67E-58 | 9.78E-54 |
| SLFN13 | 2 | 0.404609874 | 0.263 | 0.105 | 3.4E-58 | 1.25E-53 |
| ABAT | 2 | -0.388881101 | 0.031 | 0.162 | 5.08E-58 | 1.86E-53 |
| DLX6 | 2 | 0.332474923 | 0.183 | 0.054 | 5.43E-58 | 1.99E-53 |
| RADX | 2 | 0.328803049 | 0.669 | 0.452 | 5.49E-58 | 2.01E-53 |
| DSP | 2 | -0.430169413 | 0.006 | 0.105 | 6.27E-58 | 2.3E-53 |
| PRKCH | 2 | -0.346361536 | 0.018 | 0.134 | 6.66E-58 | 2.44E-53 |
| LINC00271 | 2 | -0.379755739 | 0.028 | 0.157 | 6.7E-58 | 2.45E-53 |
| C5orf17 | 2 | -0.642008448 | 0.014 | 0.121 | 6.8E-58 | 2.49E-53 |
| CPT1A | 2 | -0.327322214 | 0.022 | 0.145 | 9.31E-58 | 3.41E-53 |

| | | | | | | |
|---|---|---|---|---|---|---|
| CARMIL1 | 2 | -0.646371875 | 0.183 | 0.354 | 9.47E-58 | 3.47E-53 |
| PTBP2 | 2 | 0.370914725 | 0.947 | 0.888 | 9.51E-58 | 3.48E-53 |
| ONECUT2 | 2 | 0.281728623 | 0.158 | 0.04 | 9.57E-58 | 3.5E-53 |
| MAP4K3 | 2 | -0.489790715 | 0.862 | 0.889 | 1E-57 | 3.67E-53 |
| PLEKHA5 | 2 | 0.388918076 | 0.928 | 0.837 | 1E-57 | 3.67E-53 |
| AC011586.2 | 2 | -0.763075381 | 0.012 | 0.115 | 1.06E-57 | 3.88E-53 |
| KIF21A | 2 | 0.315328225 | 0.644 | 0.427 | 1.16E-57 | 4.25E-53 |
| LDLRAD4 | 2 | -1.018997859 | 0.455 | 0.504 | 1.39E-57 | 5.08E-53 |
| IFT74 | 2 | -0.397027486 | 0.064 | 0.216 | 1.5E-57 | 5.5E-53 |
| LMO4 | 2 | -0.588306502 | 0.131 | 0.295 | 1.64E-57 | 6.01E-53 |
| KCNK1 | 2 | -0.2951655 | 0.007 | 0.107 | 1.64E-57 | 6.02E-53 |
| TDRD3 | 2 | -0.585008068 | 0.514 | 0.665 | 1.66E-57 | 6.06E-53 |
| DNAH11 | 2 | -1.116850664 | 0.163 | 0.279 | 1.87E-57 | 6.85E-53 |
| TJP1 | 2 | -0.534639025 | 0.579 | 0.735 | 1.89E-57 | 6.92E-53 |
| CCAR1 | 2 | -0.474110303 | 0.618 | 0.756 | 2.01E-57 | 7.35E-53 |
| WDR11 | 2 | -0.716451361 | 0.582 | 0.697 | 2.13E-57 | 7.82E-53 |
| NRG2 | 2 | 0.319541773 | 0.163 | 0.043 | 2.17E-57 | 7.96E-53 |
| ASPH | 2 | -0.782065746 | 0.615 | 0.721 | 2.44E-57 | 8.95E-53 |
| BSN | 2 | 0.296412055 | 0.234 | 0.085 | 2.61E-57 | 9.57E-53 |
| SDK1 | 2 | -0.829531375 | 0.844 | 0.919 | 2.86E-57 | 1.05E-52 |
| RIC1 | 2 | -0.516794641 | 0.417 | 0.599 | 3.02E-57 | 1.11E-52 |
| ANKRD36 | 2 | 0.477670046 | 0.887 | 0.792 | 3.74E-57 | 1.37E-52 |
| RRBP1 | 2 | -0.360403471 | 0.036 | 0.169 | 4.01E-57 | 1.47E-52 |
| RUNX1 | 2 | -0.645638736 | 0.022 | 0.14 | 4.57E-57 | 1.67E-52 |
| ZNF609 | 2 | -0.523200348 | 0.669 | 0.778 | 5.75E-57 | 2.11E-52 |
| CHEK1 | 2 | 0.443146033 | 0.503 | 0.3 | 6.26E-57 | 2.29E-52 |
| FOXP2 | 2 | -1.468643245 | 0.142 | 0.292 | 7.41E-57 | 2.71E-52 |
| CCND3 | 2 | -0.486284512 | 0.089 | 0.247 | 1.01E-56 | 3.71E-52 |
| FRMD6 | 2 | -0.581203364 | 0.105 | 0.267 | 1.2E-56 | 4.38E-52 |
| PUM1 | 2 | -0.417503892 | 0.819 | 0.885 | 1.26E-56 | 4.6E-52 |
| FAM155A | 2 | -0.308997211 | 0.897 | 0.761 | 1.49E-56 | 5.44E-52 |
| AK5 | 2 | -0.880597312 | 0.117 | 0.25 | 1.62E-56 | 5.93E-52 |
| SCMH1 | 2 | 0.463162761 | 0.751 | 0.608 | 1.83E-56 | 6.7E-52 |
| FMNL2 | 2 | 0.369588815 | 0.95 | 0.857 | 1.95E-56 | 7.15E-52 |
| HOXC6 | 2 | -0.302462725 | 0.019 | 0.134 | 2.02E-56 | 7.4E-52 |
| RBM25 | 2 | -0.483153637 | 0.59 | 0.739 | 2.19E-56 | 8.03E-52 |
| HOXC4 | 2 | -0.304953212 | 0.025 | 0.146 | 2.21E-56 | 8.09E-52 |
| FMN1 | 2 | -0.696492652 | 0.018 | 0.128 | 2.23E-56 | 8.17E-52 |
| MOK | 2 | -0.429161819 | 0.035 | 0.159 | 3.12E-56 | 1.14E-51 |
| DDX60L | 2 | -0.294144398 | 0.006 | 0.103 | 3.28E-56 | 1.2E-51 |
| CSMD2 | 2 | 0.327901639 | 0.469 | 0.27 | 3.47E-56 | 1.27E-51 |
| FLNB | 2 | -0.58701396 | 0.337 | 0.507 | 3.75E-56 | 1.37E-51 |
| CCDC191 | 2 | -0.312394632 | 0.021 | 0.138 | 3.87E-56 | 1.42E-51 |
| RBMS2 | 2 | -0.358490277 | 0.042 | 0.179 | 3.96E-56 | 1.45E-51 |
| NAALADL2 | 2 | -0.766376372 | 0.264 | 0.45 | 4.3E-56 | 1.57E-51 |
| AIM2 | 2 | 0.291818669 | 0.18 | 0.054 | 5.17E-56 | 1.89E-51 |
| WRN | 2 | -0.510297435 | 0.493 | 0.664 | 5.8E-56 | 2.12E-51 |
| TC2N | 2 | -0.294924894 | 0.01 | 0.111 | 6.06E-56 | 2.22E-51 |
| AC012501.2 | 2 | -0.309625125 | 0.014 | 0.122 | 8.66E-56 | 3.17E-51 |
| OGT | 2 | -0.467888876 | 0.535 | 0.688 | 8.94E-56 | 3.27E-51 |
| SPRY4-AS1 | 2 | -0.520011091 | 0.059 | 0.198 | 9.04E-56 | 3.31E-51 |
| CACNA2D3 | 2 | 0.782460342 | 0.637 | 0.49 | 9.18E-56 | 3.36E-51 |
| CCDC148 | 2 | -0.395280292 | 0.027 | 0.149 | 9.3E-56 | 3.41E-51 |
| SH3D19 | 2 | -0.60111594 | 0.456 | 0.607 | 1.31E-55 | 4.81E-51 |
| AL022068.1 | 2 | -0.730642604 | 0.021 | 0.132 | 1.41E-55 | 5.16E-51 |
| COL1A2 | 2 | -0.772081602 | 0.029 | 0.148 | 1.63E-55 | 5.98E-51 |
| MPP5 | 2 | -0.40157926 | 0.098 | 0.264 | 1.64E-55 | 6.02E-51 |
| MAST4 | 2 | 0.583696376 | 0.517 | 0.328 | 1.74E-55 | 6.37E-51 |

| | | | | | | |
|---|---|---|---|---|---|---|
| PLPP3 | 2 | -0.411044172 | 0.034 | 0.161 | 1.75E-55 | 6.41E-51 |
| RSRP1 | 2 | -0.47481343 | 0.355 | 0.54 | 2.25E-55 | 8.24E-51 |
| GPR63 | 2 | 0.34576852 | 0.231 | 0.086 | 2.41E-55 | 8.84E-51 |
| PLCE1 | 2 | 0.317685305 | 0.308 | 0.14 | 2.59E-55 | 9.49E-51 |
| UBE2H | 2 | -0.543167402 | 0.412 | 0.567 | 2.99E-55 | 1.1E-50 |
| EMP1 | 2 | -0.284344109 | 0.007 | 0.105 | 3.17E-55 | 1.16E-50 |
| NKAIN3 | 2 | -1.118347816 | 0.611 | 0.7 | 3.3E-55 | 1.21E-50 |
| NEXMIF | 2 | 0.47951108 | 0.318 | 0.146 | 3.89E-55 | 1.42E-50 |
| AC138627.1 | 2 | -0.736346694 | 0.053 | 0.189 | 4.25E-55 | 1.56E-50 |
| TBX15 | 2 | -0.609565554 | 0.51 | 0.539 | 4.57E-55 | 1.67E-50 |
| ESCO2 | 2 | 0.317458078 | 0.211 | 0.072 | 4.97E-55 | 1.82E-50 |
| LYN | 2 | -0.364698507 | 0.027 | 0.148 | 5.01E-55 | 1.83E-50 |
| PDGFD | 2 | -0.670067332 | 0.063 | 0.211 | 5.48E-55 | 2.01E-50 |
| DISC1 | 2 | -0.506348568 | 0.137 | 0.311 | 6.29E-55 | 2.3E-50 |
| TRIM66 | 2 | -0.355426973 | 0.064 | 0.214 | 7.06E-55 | 2.58E-50 |
| TEX15 | 2 | 0.324219 | 0.176 | 0.052 | 7.51E-55 | 2.75E-50 |
| PRTG | 2 | -0.735995504 | 0.012 | 0.112 | 7.76E-55 | 2.84E-50 |
| CNNM1 | 2 | 0.303329228 | 0.251 | 0.099 | 8.22E-55 | 3.01E-50 |
| CADM1 | 2 | -0.874469487 | 0.311 | 0.47 | 8.6E-55 | 3.15E-50 |
| ATP2B4 | 2 | -0.53675989 | 0.337 | 0.518 | 9.1E-55 | 3.33E-50 |
| TMEM26 | 2 | 0.289077146 | 0.131 | 0.028 | 9.83E-55 | 3.6E-50 |
| FAM169A | 2 | 0.37993159 | 0.254 | 0.101 | 1.18E-54 | 4.32E-50 |
| FRMPD3 | 2 | 0.312283759 | 0.232 | 0.086 | 1.29E-54 | 4.71E-50 |
| PRC1 | 2 | 0.348538745 | 0.285 | 0.122 | 2.3E-54 | 8.43E-50 |
| STIL | 2 | 0.425121951 | 0.424 | 0.232 | 2.41E-54 | 8.82E-50 |
| KLF12 | 2 | -0.645136899 | 0.546 | 0.664 | 2.57E-54 | 9.42E-50 |
| STS | 2 | 0.558426022 | 0.488 | 0.305 | 2.93E-54 | 1.07E-49 |
| PROM1 | 2 | -0.403862362 | 0.015 | 0.121 | 3.11E-54 | 1.14E-49 |
| XYLT1 | 2 | -0.554843254 | 0.044 | 0.159 | 3.38E-54 | 1.24E-49 |
| ERBB3 | 2 | -0.396922825 | 0.016 | 0.123 | 3.66E-54 | 1.34E-49 |
| PAK1 | 2 | 0.388742247 | 0.732 | 0.536 | 3.72E-54 | 1.36E-49 |
| DLG3 | 2 | 0.401069332 | 0.448 | 0.254 | 3.88E-54 | 1.42E-49 |
| WDR27 | 2 | -0.520845711 | 0.329 | 0.502 | 3.89E-54 | 1.42E-49 |
| WDR78 | 2 | -0.446452045 | 0.03 | 0.149 | 4.51E-54 | 1.65E-49 |
| MKX | 2 | -0.452249204 | 0.431 | 0.349 | 4.62E-54 | 1.69E-49 |
| COBLL1 | 2 | 0.377764201 | 0.55 | 0.343 | 4.68E-54 | 1.71E-49 |
| OSMR | 2 | -0.28803061 | 0.007 | 0.1 | 5.09E-54 | 1.86E-49 |
| DECR1 | 2 | -0.374564365 | 0.072 | 0.221 | 5.93E-54 | 2.17E-49 |
| PLPP1 | 2 | -0.792140609 | 0.211 | 0.377 | 6E-54 | 2.2E-49 |
| WDR17 | 2 | 0.504197967 | 0.511 | 0.313 | 6.68E-54 | 2.45E-49 |
| BRCA1 | 2 | 0.4342242 | 0.496 | 0.296 | 7.12E-54 | 2.61E-49 |
| PKNOX2 | 2 | 0.283145477 | 0.37 | 0.192 | 7.39E-54 | 2.7E-49 |
| CDR2 | 2 | -0.467603785 | 0.13 | 0.296 | 8.64E-54 | 3.16E-49 |
| LINC02398 | 2 | -1.033786956 | 0.077 | 0.184 | 9.85E-54 | 3.6E-49 |
| OSMR-AS1 | 2 | -0.274878467 | 0.008 | 0.103 | 1.14E-53 | 4.19E-49 |
| TSPAN18 | 2 | -0.554524273 | 0.141 | 0.308 | 1.46E-53 | 5.34E-49 |
| ASIC2 | 2 | 0.327541644 | 0.424 | 0.233 | 1.79E-53 | 6.55E-49 |
| CLSPN | 2 | 0.387259173 | 0.42 | 0.228 | 1.8E-53 | 6.59E-49 |
| CEP112 | 2 | -0.640763979 | 0.253 | 0.43 | 2.05E-53 | 7.52E-49 |
| AP001636.3 | 2 | 0.404430696 | 0.234 | 0.089 | 2.22E-53 | 8.14E-49 |
| CENPK | 2 | 0.308802721 | 0.663 | 0.454 | 2.48E-53 | 9.08E-49 |
| SATB1-AS1 | 2 | 0.411222713 | 0.315 | 0.146 | 2.72E-53 | 9.97E-49 |
| SUPT3H | 2 | 0.487198932 | 0.836 | 0.705 | 2.79E-53 | 1.02E-48 |
| NOTCH1 | 2 | -0.346482631 | 0.018 | 0.127 | 2.98E-53 | 1.09E-48 |
| HEATR5A | 2 | -0.45302723 | 0.144 | 0.321 | 3.09E-53 | 1.13E-48 |
| DIP2B | 2 | -0.526475153 | 0.617 | 0.727 | 3.98E-53 | 1.46E-48 |
| CYP27C1 | 2 | -0.283467827 | 0.012 | 0.113 | 4.44E-53 | 1.63E-48 |
| AC053513.1 | 2 | 0.414365351 | 0.403 | 0.223 | 5.02E-53 | 1.84E-48 |

| | | | | | | |
|---|---|---|---|---|---|---|
| ATCAY | 2 | 0.253349811 | 0.157 | 0.044 | 5.6E-53 | 2.05E-48 |
| DCDC2 | 2 | -0.520313502 | 0.008 | 0.103 | 5.95E-53 | 2.18E-48 |
| IGF2BP1 | 2 | 0.389460702 | 0.475 | 0.276 | 6.27E-53 | 2.29E-48 |
| MKI67 | 2 | 0.327421905 | 0.281 | 0.123 | 7.19E-53 | 2.63E-48 |
| SRSF5 | 2 | -0.451793996 | 0.357 | 0.536 | 8.34E-53 | 3.05E-48 |
| TRIM2 | 2 | -0.576103119 | 0.549 | 0.661 | 8.98E-53 | 3.29E-48 |
| NR3C2 | 2 | -0.740998368 | 0.133 | 0.307 | 9.75E-53 | 3.57E-48 |
| FAM49B | 2 | 0.424423311 | 0.767 | 0.616 | 1.02E-52 | 3.75E-48 |
| NSD2 | 2 | 0.397710035 | 0.8 | 0.657 | 1.04E-52 | 3.8E-48 |
| NBEAL1 | 2 | -0.513528911 | 0.317 | 0.492 | 1.05E-52 | 3.83E-48 |
| VDR | 2 | 0.286369045 | 0.133 | 0.031 | 1.41E-52 | 5.16E-48 |
| EVA1C | 2 | 0.307459558 | 0.238 | 0.093 | 1.53E-52 | 5.61E-48 |
| EGLN3 | 2 | -0.553476815 | 0.126 | 0.276 | 1.63E-52 | 5.97E-48 |
| AC008543.1 | 2 | 0.274143636 | 0.205 | 0.071 | 1.73E-52 | 6.34E-48 |
| ENOX1 | 2 | 0.394253253 | 0.698 | 0.494 | 1.74E-52 | 6.37E-48 |
| IQCK | 2 | -0.372220444 | 0.061 | 0.201 | 2.06E-52 | 7.53E-48 |
| TRIM71 | 2 | 0.275124529 | 0.401 | 0.218 | 2.34E-52 | 8.55E-48 |
| PTPRS | 2 | 0.393811019 | 0.722 | 0.546 | 2.85E-52 | 1.04E-47 |
| STRN3 | 2 | 0.536362245 | 0.853 | 0.793 | 3.08E-52 | 1.13E-47 |
| LINC01376 | 2 | -0.458981467 | 0.046 | 0.175 | 3.58E-52 | 1.31E-47 |
| PDGFRA | 2 | -0.623567509 | 0.089 | 0.214 | 3.62E-52 | 1.33E-47 |
| RGS3 | 2 | 0.372317001 | 0.26 | 0.108 | 3.65E-52 | 1.33E-47 |
| PDE5A | 2 | 0.37669557 | 0.411 | 0.226 | 3.97E-52 | 1.45E-47 |
| RIOX2 | 2 | -0.315386769 | 0.038 | 0.166 | 4.18E-52 | 1.53E-47 |
| ATG10 | 2 | -0.517004331 | 0.433 | 0.615 | 4.53E-52 | 1.66E-47 |
| ZFP36L1 | 2 | -0.304967249 | 0.013 | 0.113 | 6.77E-52 | 2.48E-47 |
| CPXM2 | 2 | -0.395937754 | 0.009 | 0.104 | 6.87E-52 | 2.52E-47 |
| PLAG1 | 2 | -0.305203961 | 0.018 | 0.126 | 7.51E-52 | 2.75E-47 |
| TMEM67 | 2 | -0.485219793 | 0.133 | 0.295 | 1.05E-51 | 3.84E-47 |
| PRKCE | 2 | -0.58210231 | 0.386 | 0.551 | 1.05E-51 | 3.85E-47 |
| AHCYL2 | 2 | -0.694847096 | 0.622 | 0.676 | 1.43E-51 | 5.25E-47 |
| SYDE2 | 2 | 0.389480639 | 0.343 | 0.169 | 1.45E-51 | 5.3E-47 |
| STAT6 | 2 | -0.345723746 | 0.062 | 0.196 | 1.9E-51 | 6.96E-47 |
| CSGALNACT1 | 2 | -0.51442374 | 0.028 | 0.145 | 1.93E-51 | 7.05E-47 |
| SAT1 | 2 | -0.80476107 | 0.046 | 0.171 | 2.18E-51 | 7.98E-47 |
| ANKRA2 | 2 | -0.413875061 | 0.103 | 0.26 | 2.68E-51 | 9.8E-47 |
| FAXC | 2 | 0.325328414 | 0.257 | 0.107 | 3.37E-51 | 1.23E-46 |
| NUF2 | 2 | 0.386145492 | 0.293 | 0.133 | 3.4E-51 | 1.24E-46 |
| IRS1 | 2 | 0.359802087 | 0.294 | 0.133 | 3.98E-51 | 1.46E-46 |
| AHR | 2 | -0.603831709 | 0.067 | 0.197 | 4.39E-51 | 1.61E-46 |
| ZC4H2 | 2 | 0.371105599 | 0.237 | 0.094 | 5.16E-51 | 1.89E-46 |
| CNOT6L | 2 | 0.449245823 | 0.65 | 0.476 | 6.31E-51 | 2.31E-46 |
| ARHGAP32 | 2 | 0.384912813 | 0.827 | 0.679 | 7.51E-51 | 2.75E-46 |
| KNL1 | 2 | 0.363446445 | 0.358 | 0.182 | 7.56E-51 | 2.77E-46 |
| FNBP4 | 2 | -0.440204237 | 0.607 | 0.734 | 8.48E-51 | 3.11E-46 |
| TUSC3 | 2 | -0.500551187 | 0.6 | 0.71 | 9.77E-51 | 3.58E-46 |
| MERTK | 2 | -0.315913219 | 0.025 | 0.136 | 1.34E-50 | 4.92E-46 |
| C6orf62 | 2 | -0.399140922 | 0.145 | 0.312 | 1.35E-50 | 4.94E-46 |
| UBE2E3 | 2 | 0.464363561 | 0.801 | 0.68 | 1.4E-50 | 5.11E-46 |
| XPNPEP3 | 2 | -0.397884236 | 0.109 | 0.27 | 1.85E-50 | 6.78E-46 |
| OSTF1 | 2 | -0.298587518 | 0.018 | 0.123 | 1.92E-50 | 7.03E-46 |
| AC092422.1 | 2 | -0.755902678 | 0.062 | 0.184 | 1.96E-50 | 7.16E-46 |
| ROR2 | 2 | 0.299878804 | 0.586 | 0.384 | 1.97E-50 | 7.22E-46 |
| MBNL3 | 2 | 0.423975598 | 0.646 | 0.454 | 2.08E-50 | 7.63E-46 |
| FNDC5 | 2 | 0.288200558 | 0.179 | 0.057 | 2.31E-50 | 8.45E-46 |
| SRSF6 | 2 | -0.431849821 | 0.233 | 0.408 | 2.68E-50 | 9.81E-46 |
| SIK3 | 2 | -0.530929265 | 0.559 | 0.695 | 2.8E-50 | 1.03E-45 |
| PLXNA4 | 2 | 0.312441197 | 0.194 | 0.067 | 2.83E-50 | 1.03E-45 |

| | | | | | | |
|---|---|---|---|---|---|---|
| CFAP47 | 2 | -1.045198492 | 0.011 | 0.104 | 3.35E-50 | 1.23E-45 |
| C12orf56 | 2 | 0.282899786 | 0.276 | 0.123 | 3.46E-50 | 1.27E-45 |
| NCAPH | 2 | 0.274781694 | 0.212 | 0.078 | 3.81E-50 | 1.39E-45 |
| ST6GALNAC3 | 2 | -0.844972701 | 0.169 | 0.288 | 4.51E-50 | 1.65E-45 |
| DMXL2 | 2 | 0.411595913 | 0.708 | 0.544 | 5.01E-50 | 1.83E-45 |
| SLC17A8 | 2 | -0.414904503 | 0.341 | 0.276 | 5.14E-50 | 1.88E-45 |
| PIKFYVE | 2 | -0.452002641 | 0.376 | 0.552 | 5.73E-50 | 2.1E-45 |
| MKLN1 | 2 | -0.444520215 | 0.778 | 0.84 | 6.28E-50 | 2.3E-45 |
| AC107980.1 | 2 | 0.34202486 | 0.225 | 0.091 | 6.38E-50 | 2.33E-45 |
| CDC73 | 2 | -0.46846459 | 0.431 | 0.608 | 6.38E-50 | 2.34E-45 |
| CIT | 2 | 0.417752082 | 0.483 | 0.29 | 6.66E-50 | 2.44E-45 |
| CPNE7 | 2 | 0.304175737 | 0.177 | 0.057 | 6.78E-50 | 2.48E-45 |
| MAOB | 2 | -0.331146077 | 0.022 | 0.131 | 7.49E-50 | 2.74E-45 |
| PDGFRB | 2 | -0.404542239 | 0.024 | 0.133 | 9.31E-50 | 3.41E-45 |
| GOLGA4 | 2 | -0.464582459 | 0.589 | 0.71 | 9.46E-50 | 3.46E-45 |
| RBMS3 | 2 | 0.354869809 | 0.908 | 0.778 | 9.87E-50 | 3.61E-45 |
| KCNC1 | 2 | 0.250731471 | 0.172 | 0.054 | 1.19E-49 | 4.37E-45 |
| CEP83 | 2 | -0.542402479 | 0.335 | 0.507 | 1.4E-49 | 5.13E-45 |
| B3GLCT | 2 | 0.442371212 | 0.438 | 0.267 | 1.51E-49 | 5.51E-45 |
| SPATA17 | 2 | -0.45007148 | 0.015 | 0.112 | 1.75E-49 | 6.39E-45 |
| EYS | 2 | -1.951504745 | 0.162 | 0.266 | 2.03E-49 | 7.42E-45 |
| CSAD | 2 | -0.40123546 | 0.138 | 0.299 | 2.14E-49 | 7.85E-45 |
| PVT1 | 2 | 0.377827094 | 0.902 | 0.787 | 2.27E-49 | 8.3E-45 |
| ZNF721 | 2 | -0.467584386 | 0.459 | 0.62 | 2.46E-49 | 8.99E-45 |
| ANTXR1 | 2 | -0.539056885 | 0.337 | 0.499 | 2.55E-49 | 9.35E-45 |
| PPP1R13B | 2 | 0.402435629 | 0.402 | 0.223 | 2.85E-49 | 1.04E-44 |
| CBFA2T2 | 2 | -0.519194182 | 0.502 | 0.67 | 2.85E-49 | 1.04E-44 |
| GHR | 2 | -0.658934982 | 0.095 | 0.236 | 2.9E-49 | 1.06E-44 |
| NTF3 | 2 | -0.575757489 | 0.062 | 0.193 | 3.1E-49 | 1.13E-44 |
| RBPMS | 2 | -0.489130054 | 0.018 | 0.118 | 4.22E-49 | 1.55E-44 |
| CACNB2 | 2 | -0.801771954 | 0.698 | 0.74 | 5.6E-49 | 2.05E-44 |
| RACGAP1 | 2 | 0.343042703 | 0.305 | 0.144 | 6.12E-49 | 2.24E-44 |
| CD44 | 2 | -0.350002659 | 0.014 | 0.112 | 6.21E-49 | 2.27E-44 |
| FAM120A | 2 | -0.440422556 | 0.418 | 0.587 | 7.15E-49 | 2.62E-44 |
| RILPL1 | 2 | -0.394341286 | 0.099 | 0.249 | 1.08E-48 | 3.94E-44 |
| GJC1 | 2 | 0.373127683 | 0.458 | 0.27 | 1.27E-48 | 4.65E-44 |
| CTBP2 | 2 | -0.473538722 | 0.714 | 0.82 | 1.47E-48 | 5.39E-44 |
| ABCC1 | 2 | -0.557025183 | 0.351 | 0.523 | 1.58E-48 | 5.8E-44 |
| KIF11 | 2 | 0.384319388 | 0.378 | 0.202 | 1.81E-48 | 6.63E-44 |
| FBXL13 | 2 | -0.371923657 | 0.047 | 0.17 | 1.93E-48 | 7.07E-44 |
| ADAM33 | 2 | -0.306328265 | 0.011 | 0.103 | 1.96E-48 | 7.18E-44 |
| SORT1 | 2 | 0.419008465 | 0.443 | 0.264 | 2.05E-48 | 7.5E-44 |
| CFAP69 | 2 | -0.383059944 | 0.016 | 0.114 | 2.06E-48 | 7.56E-44 |
| ITGA1 | 2 | -0.609506811 | 0.038 | 0.147 | 2.83E-48 | 1.04E-43 |
| SAMD4A | 2 | -0.703237026 | 0.245 | 0.419 | 3.06E-48 | 1.12E-43 |
| PIBF1 | 2 | -0.450907917 | 0.588 | 0.713 | 3.28E-48 | 1.2E-43 |
| COL27A1 | 2 | -0.622125352 | 0.256 | 0.415 | 3.56E-48 | 1.3E-43 |
| TMBIM4 | 2 | -0.389164006 | 0.13 | 0.29 | 3.61E-48 | 1.32E-43 |
| HSPG2 | 2 | -0.336788117 | 0.034 | 0.147 | 3.62E-48 | 1.32E-43 |
| HYMAI | 2 | 0.362139055 | 0.16 | 0.058 | 4.05E-48 | 1.48E-43 |
| SLAIN2 | 2 | -0.487541892 | 0.391 | 0.548 | 4.62E-48 | 1.69E-43 |
| CACNA1A | 2 | -1.380999123 | 0.207 | 0.283 | 4.84E-48 | 1.77E-43 |
| SERPINI1 | 2 | 0.325544437 | 0.14 | 0.041 | 5.07E-48 | 1.86E-43 |
| YPEL2 | 2 | -0.373516423 | 0.059 | 0.19 | 5.72E-48 | 2.09E-43 |
| IFT88 | 2 | -0.477345636 | 0.245 | 0.417 | 6.04E-48 | 2.21E-43 |
| STXBP3 | 2 | -0.398678702 | 0.146 | 0.311 | 6.18E-48 | 2.26E-43 |
| HNRNPH1 | 2 | -0.38992108 | 0.772 | 0.851 | 6.26E-48 | 2.29E-43 |
| SLC18B1 | 2 | -0.290803013 | 0.038 | 0.159 | 6.49E-48 | 2.38E-43 |

| | | | | | | |
|---|---|---|---|---|---|---|
| AL162411.1 | 2 | -0.295904152 | 0.024 | 0.133 | 7.16E-48 | 2.62E-43 |
| SHROOM3 | 2 | -0.408157157 | 0.016 | 0.111 | 9.93E-48 | 3.64E-43 |
| C1orf112 | 2 | 0.39849059 | 0.43 | 0.253 | 1.06E-47 | 3.87E-43 |
| EPSTI1 | 2 | -0.320916052 | 0.012 | 0.106 | 1.31E-47 | 4.78E-43 |
| PACRGL | 2 | -0.41856857 | 0.158 | 0.318 | 1.53E-47 | 5.6E-43 |
| PKIA | 2 | 0.409573828 | 0.602 | 0.411 | 1.74E-47 | 6.38E-43 |
| FANK1 | 2 | -0.309082151 | 0.011 | 0.1 | 1.83E-47 | 6.71E-43 |
| VPS13C | 2 | -0.455381952 | 0.65 | 0.76 | 1.91E-47 | 7E-43 |
| ITSN2 | 2 | -0.447305963 | 0.21 | 0.385 | 1.92E-47 | 7.01E-43 |
| PHLDB1 | 2 | -0.317264379 | 0.043 | 0.162 | 1.95E-47 | 7.15E-43 |
| PBX3 | 2 | 0.767508733 | 0.639 | 0.475 | 2.38E-47 | 8.7E-43 |
| SPEF2 | 2 | -0.521385833 | 0.051 | 0.17 | 3.02E-47 | 1.11E-42 |
| SLC35G2 | 2 | 0.358372786 | 0.36 | 0.19 | 3.63E-47 | 1.33E-42 |
| LINC00536 | 2 | -0.846024352 | 0.017 | 0.114 | 4.09E-47 | 1.5E-42 |
| HIF3A | 2 | -0.383893534 | 0.036 | 0.147 | 4.33E-47 | 1.59E-42 |
| TMOD2 | 2 | 0.288751831 | 0.214 | 0.082 | 5.11E-47 | 1.87E-42 |
| UNC5C | 2 | -0.420500531 | 0.631 | 0.505 | 5.54E-47 | 2.03E-42 |
| RELN | 2 | -0.517800023 | 0.015 | 0.11 | 5.8E-47 | 2.12E-42 |
| DPY19L3 | 2 | -0.470427874 | 0.369 | 0.524 | 8.05E-47 | 2.95E-42 |
| ARHGEF28 | 2 | -0.66165946 | 0.237 | 0.383 | 8.28E-47 | 3.03E-42 |
| ASB3 | 2 | -0.455665064 | 0.531 | 0.667 | 8.84E-47 | 3.24E-42 |
| FAM114A1 | 2 | -0.260346583 | 0.017 | 0.11 | 1.23E-46 | 4.49E-42 |
| PCOLCE2 | 2 | 0.36848446 | 0.504 | 0.314 | 1.27E-46 | 4.65E-42 |
| NCOR2 | 2 | -0.494284618 | 0.483 | 0.623 | 1.48E-46 | 5.43E-42 |
| GBE1 | 2 | -0.621116775 | 0.186 | 0.35 | 1.6E-46 | 5.85E-42 |
| SREK1 | 2 | -0.413665154 | 0.557 | 0.701 | 1.67E-46 | 6.13E-42 |
| PRKD2 | 2 | -0.390479612 | 0.063 | 0.195 | 1.76E-46 | 6.45E-42 |
| KLHL7 | 2 | 0.430904147 | 0.64 | 0.469 | 1.77E-46 | 6.48E-42 |
| SMYD3 | 2 | 0.328011806 | 0.965 | 0.916 | 1.79E-46 | 6.55E-42 |
| DOP1B | 2 | -0.512909316 | 0.374 | 0.52 | 1.83E-46 | 6.71E-42 |
| PLEKHB2 | 2 | -0.411667743 | 0.149 | 0.305 | 2.04E-46 | 7.48E-42 |
| CCNL1 | 2 | -0.45283506 | 0.378 | 0.542 | 2.17E-46 | 7.93E-42 |
| AL136418.1 | 2 | -0.44152741 | 0.015 | 0.11 | 2.18E-46 | 7.97E-42 |
| AHI1 | 2 | -0.473161559 | 0.588 | 0.733 | 2.2E-46 | 8.06E-42 |
| HBP1 | 2 | -0.414819014 | 0.114 | 0.264 | 2.46E-46 | 9.01E-42 |
| RNF213 | 2 | -0.519360426 | 0.312 | 0.472 | 2.92E-46 | 1.07E-41 |
| C1orf61 | 2 | -0.63131953 | 0.23 | 0.319 | 3.21E-46 | 1.18E-41 |
| RASAL2 | 2 | 0.336052972 | 0.693 | 0.504 | 3.37E-46 | 1.23E-41 |
| KRAS | 2 | -0.65377201 | 0.264 | 0.397 | 3.48E-46 | 1.27E-41 |
| OGA | 2 | -0.435571151 | 0.573 | 0.721 | 3.54E-46 | 1.3E-41 |
| AL138828.1 | 2 | -0.469108033 | 0.011 | 0.1 | 3.66E-46 | 1.34E-41 |
| KYAT3 | 2 | -0.390226089 | 0.122 | 0.277 | 3.75E-46 | 1.37E-41 |
| RCAN2 | 2 | -0.443485802 | 0.02 | 0.118 | 3.76E-46 | 1.38E-41 |
| SKAP2 | 2 | -0.616409922 | 0.082 | 0.205 | 4.14E-46 | 1.52E-41 |
| TRAF3IP2-AS1 | 2 | 0.418081076 | 0.728 | 0.576 | 4.38E-46 | 1.61E-41 |
| P4HA1 | 2 | -0.704652202 | 0.338 | 0.515 | 4.85E-46 | 1.78E-41 |
| IL12RB2 | 2 | -0.877715553 | 0.232 | 0.298 | 4.87E-46 | 1.78E-41 |
| WNK2 | 2 | 0.292670414 | 0.737 | 0.551 | 5.14E-46 | 1.88E-41 |
| PDE8A | 2 | -0.540866895 | 0.539 | 0.676 | 5.57E-46 | 2.04E-41 |
| CCDC30 | 2 | -0.429751429 | 0.039 | 0.152 | 5.75E-46 | 2.1E-41 |
| ULK4 | 2 | -0.542403694 | 0.303 | 0.463 | 6.84E-46 | 2.5E-41 |
| PPP1R12A | 2 | -0.441474705 | 0.668 | 0.768 | 7.62E-46 | 2.79E-41 |
| PARD3 | 2 | -0.388649628 | 0.969 | 0.965 | 8.25E-46 | 3.02E-41 |
| PDE6B | 2 | 0.255104882 | 0.12 | 0.034 | 9.19E-46 | 3.36E-41 |
| COG5 | 2 | -0.445414417 | 0.541 | 0.681 | 9.33E-46 | 3.42E-41 |
| ANLN | 2 | 0.326864737 | 0.415 | 0.237 | 9.55E-46 | 3.5E-41 |
| ALG13 | 2 | -0.449640464 | 0.573 | 0.699 | 1.04E-45 | 3.8E-41 |
| SLC4A10 | 2 | -0.544938657 | 0.014 | 0.103 | 1.12E-45 | 4.12E-41 |

| | | | | | | |
|---|---|---|---|---|---|---|
| XRN1 | 2 | -0.464106893 | 0.566 | 0.694 | 1.2E-45 | 4.38E-41 |
| FUT10 | 2 | -0.402095298 | 0.139 | 0.297 | 1.4E-45 | 5.12E-41 |
| EPHA6 | 2 | -0.651743546 | 0.031 | 0.134 | 1.52E-45 | 5.55E-41 |
| ADAMTS19-AS1 | 2 | 0.26662139 | 0.15 | 0.045 | 1.7E-45 | 6.24E-41 |
| SLC35D1 | 2 | -0.43661445 | 0.146 | 0.29 | 1.76E-45 | 6.45E-41 |
| FAXDC2 | 2 | -0.265230859 | 0.02 | 0.117 | 1.79E-45 | 6.54E-41 |
| E2F1 | 2 | 0.275681101 | 0.254 | 0.112 | 2.34E-45 | 8.57E-41 |
| JAZF1 | 2 | -0.564889983 | 0.705 | 0.738 | 2.51E-45 | 9.18E-41 |
| GABPB1-AS1 | 2 | -0.446459467 | 0.396 | 0.565 | 2.51E-45 | 9.19E-41 |
| CFAP70 | 2 | -0.363941481 | 0.012 | 0.103 | 2.59E-45 | 9.47E-41 |
| IQCA1 | 2 | -0.289806994 | 0.025 | 0.129 | 2.6E-45 | 9.51E-41 |
| SLC35B3 | 2 | 0.451493554 | 0.52 | 0.361 | 2.81E-45 | 1.03E-40 |
| RAD9A | 2 | -0.429755553 | 0.103 | 0.242 | 2.85E-45 | 1.04E-40 |
| CFAP44 | 2 | -0.542960009 | 0.221 | 0.382 | 3.1E-45 | 1.13E-40 |
| KALRN | 2 | 0.340605027 | 0.562 | 0.371 | 3.3E-45 | 1.21E-40 |
| MTMR10 | 2 | -0.435536392 | 0.216 | 0.382 | 4.33E-45 | 1.59E-40 |
| GRIK4 | 2 | 0.406209314 | 0.389 | 0.219 | 4.83E-45 | 1.77E-40 |
| ARHGAP12 | 2 | -0.52435247 | 0.436 | 0.581 | 5.09E-45 | 1.86E-40 |
| FCHSD2 | 2 | 0.380201174 | 0.847 | 0.721 | 5.56E-45 | 2.04E-40 |
| TEAD1 | 2 | -0.434448336 | 0.852 | 0.912 | 5.71E-45 | 2.09E-40 |
| CYTH3 | 2 | -0.395480249 | 0.101 | 0.249 | 6.16E-45 | 2.26E-40 |
| ZEB2 | 2 | -0.42148988 | 0.66 | 0.562 | 6.72E-45 | 2.46E-40 |
| NEK11 | 2 | -0.429870556 | 0.044 | 0.154 | 7.65E-45 | 2.8E-40 |
| XKR6 | 2 | 0.408634461 | 0.67 | 0.494 | 7.88E-45 | 2.88E-40 |
| SIPA1L2 | 2 | 0.393969883 | 0.586 | 0.402 | 1.01E-44 | 3.71E-40 |
| SENP6 | 2 | -0.384567828 | 0.758 | 0.84 | 1.13E-44 | 4.13E-40 |
| RASSF3 | 2 | -0.448963687 | 0.149 | 0.299 | 1.14E-44 | 4.16E-40 |
| RCAN3 | 2 | -0.282978966 | 0.027 | 0.129 | 1.15E-44 | 4.22E-40 |
| CLYBL | 2 | -0.491248111 | 0.149 | 0.305 | 1.33E-44 | 4.86E-40 |
| AP000787.1 | 2 | -0.285716254 | 0.026 | 0.127 | 1.39E-44 | 5.08E-40 |
| GRIK1 | 2 | 0.42550477 | 0.16 | 0.052 | 1.48E-44 | 5.43E-40 |
| FRYL | 2 | 0.362232731 | 0.9 | 0.841 | 1.63E-44 | 5.96E-40 |
| ROCK2 | 2 | 0.382089042 | 0.791 | 0.659 | 1.7E-44 | 6.22E-40 |
| TRIP12 | 2 | -0.401785205 | 0.656 | 0.757 | 1.79E-44 | 6.56E-40 |
| CRYM | 2 | -0.385270498 | 0.022 | 0.117 | 2.35E-44 | 8.59E-40 |
| TACC2 | 2 | 0.288507351 | 0.336 | 0.175 | 2.66E-44 | 9.73E-40 |
| TMEM91 | 2 | -0.362722544 | 0.108 | 0.256 | 2.75E-44 | 1.01E-39 |
| ENOSF1 | 2 | -0.45704325 | 0.489 | 0.625 | 3.08E-44 | 1.13E-39 |
| LHFPL2 | 2 | -0.460073298 | 0.124 | 0.26 | 3.46E-44 | 1.27E-39 |
| TFAP2E | 2 | -0.275797424 | 0.024 | 0.125 | 3.9E-44 | 1.43E-39 |
| ETV6 | 2 | -0.475617575 | 0.353 | 0.524 | 3.96E-44 | 1.45E-39 |
| CREB3L2 | 2 | -0.44106631 | 0.137 | 0.278 | 3.99E-44 | 1.46E-39 |
| SNCAIP | 2 | -0.720808161 | 0.119 | 0.224 | 4.67E-44 | 1.71E-39 |
| SLC5A4-AS1 | 2 | 0.366776925 | 0.237 | 0.104 | 5.07E-44 | 1.86E-39 |
| SGK3 | 2 | -0.429004619 | 0.211 | 0.368 | 5.09E-44 | 1.86E-39 |
| DYNC2LI1 | 2 | -0.331345361 | 0.082 | 0.221 | 5.31E-44 | 1.94E-39 |
| MIR181A1HG | 2 | 0.318975246 | 0.63 | 0.44 | 6.04E-44 | 2.21E-39 |
| IDI2-AS1 | 2 | -0.444478246 | 0.047 | 0.156 | 6.33E-44 | 2.32E-39 |
| NEDD4 | 2 | -0.427329258 | 0.185 | 0.349 | 6.85E-44 | 2.51E-39 |
| ATP1B3 | 2 | 0.401783751 | 0.545 | 0.387 | 8.66E-44 | 3.17E-39 |
| DCX | 2 | 0.281388538 | 0.299 | 0.151 | 9.44E-44 | 3.46E-39 |
| CAMK2D | 2 | -0.606435214 | 0.467 | 0.596 | 9.95E-44 | 3.64E-39 |
| ZMYND8 | 2 | -0.512629043 | 0.466 | 0.604 | 9.99E-44 | 3.66E-39 |
| MAP7D3 | 2 | -0.411466392 | 0.13 | 0.28 | 1.1E-43 | 4.04E-39 |
| UHRF1 | 2 | 0.303621868 | 0.398 | 0.227 | 1.19E-43 | 4.37E-39 |
| EGR1 | 2 | -0.433670863 | 0.034 | 0.138 | 1.37E-43 | 5.02E-39 |
| SLC9A7 | 2 | 0.273964125 | 0.264 | 0.122 | 1.39E-43 | 5.09E-39 |
| FAT1 | 2 | -0.588618206 | 0.515 | 0.636 | 1.51E-43 | 5.54E-39 |

| | | | | | | |
|---|---|---|---|---|---|---|
| PRELID2 | 2 | 0.56549467 | 0.458 | 0.291 | 1.57E-43 | 5.74E-39 |
| CCNE2 | 2 | 0.283610054 | 0.279 | 0.134 | 1.73E-43 | 6.32E-39 |
| LINC00461 | 2 | -0.635388986 | 0.414 | 0.523 | 2E-43 | 7.32E-39 |
| MAP4K3-DT | 2 | -0.467301753 | 0.259 | 0.423 | 2.23E-43 | 8.17E-39 |
| EVL | 2 | 0.381790996 | 0.688 | 0.533 | 2.35E-43 | 8.59E-39 |
| C11orf49 | 2 | -0.451279631 | 0.42 | 0.575 | 2.38E-43 | 8.7E-39 |
| AC024901.1 | 2 | -0.810438477 | 0.096 | 0.199 | 2.53E-43 | 9.27E-39 |
| EPDR1 | 2 | 0.275884615 | 0.196 | 0.075 | 2.78E-43 | 1.02E-38 |
| ADGRA2 | 2 | -0.384121166 | 0.05 | 0.166 | 3.18E-43 | 1.16E-38 |
| NR2F1 | 2 | -0.540171005 | 0.115 | 0.238 | 3.49E-43 | 1.28E-38 |
| ZDHHC21 | 2 | -0.44761069 | 0.614 | 0.713 | 3.51E-43 | 1.28E-38 |
| STUM | 2 | -0.281602262 | 0.02 | 0.115 | 3.69E-43 | 1.35E-38 |
| AGPS | 2 | 0.40897746 | 0.688 | 0.546 | 4E-43 | 1.46E-38 |
| SEMA3D | 2 | -0.423518237 | 0.021 | 0.118 | 4.06E-43 | 1.49E-38 |
| PARN | 2 | -0.423782151 | 0.314 | 0.481 | 5.8E-43 | 2.12E-38 |
| ITPR2 | 2 | 0.296014956 | 0.493 | 0.311 | 5.95E-43 | 2.18E-38 |
| TNFRSF19 | 2 | -0.572276176 | 0.142 | 0.263 | 6.59E-43 | 2.41E-38 |
| PDE3B | 2 | 0.403465977 | 0.772 | 0.605 | 8.4E-43 | 3.07E-38 |
| CAST | 2 | -0.375114539 | 0.115 | 0.255 | 8.5E-43 | 3.11E-38 |
| RRM1 | 2 | 0.34693768 | 0.427 | 0.253 | 9.32E-43 | 3.41E-38 |
| TCTN1 | 2 | -0.296014709 | 0.05 | 0.167 | 9.4E-43 | 3.44E-38 |
| ZMAT4 | 2 | -0.280498379 | 0.616 | 0.506 | 9.96E-43 | 3.64E-38 |
| PCNX4 | 2 | -0.45297877 | 0.429 | 0.573 | 1.06E-42 | 3.89E-38 |
| SLC22A15 | 2 | -0.599082579 | 0.221 | 0.331 | 1.1E-42 | 4.02E-38 |
| CYFIP2 | 2 | 0.305567668 | 0.254 | 0.115 | 1.1E-42 | 4.02E-38 |
| CACNG8 | 2 | -0.255438945 | 0.015 | 0.101 | 1.32E-42 | 4.84E-38 |
| PTPRR | 2 | 0.327622055 | 0.153 | 0.049 | 1.59E-42 | 5.82E-38 |
| CEP128 | 2 | 0.294039428 | 0.958 | 0.917 | 1.71E-42 | 6.27E-38 |
| CAVIN4 | 2 | 0.255110701 | 0.172 | 0.061 | 1.75E-42 | 6.4E-38 |
| DDX31 | 2 | -0.369065588 | 0.161 | 0.318 | 2.73E-42 | 1E-37 |
| ARID2 | 2 | -0.406738529 | 0.746 | 0.83 | 3.06E-42 | 1.12E-37 |
| EVC | 2 | -0.296919326 | 0.064 | 0.19 | 3.3E-42 | 1.21E-37 |
| PHKA1 | 2 | 0.341570512 | 0.431 | 0.258 | 3.53E-42 | 1.29E-37 |
| KDM7A | 2 | -0.481237255 | 0.22 | 0.388 | 3.82E-42 | 1.4E-37 |
| CDC16 | 2 | -0.410106126 | 0.317 | 0.484 | 4.09E-42 | 1.5E-37 |
| SSH2 | 2 | 0.327151385 | 0.891 | 0.781 | 4.89E-42 | 1.79E-37 |
| SEZ6L2 | 2 | 0.272720872 | 0.227 | 0.097 | 6.53E-42 | 2.39E-37 |
| PPFIA1 | 2 | -0.469391299 | 0.436 | 0.582 | 7.72E-42 | 2.83E-37 |
| TBC1D5 | 2 | 0.324901458 | 0.925 | 0.85 | 7.86E-42 | 2.88E-37 |
| ZNF106 | 2 | -0.403765404 | 0.175 | 0.324 | 9.33E-42 | 3.42E-37 |
| AC138150.1 | 2 | -0.314177905 | 0.071 | 0.2 | 9.49E-42 | 3.47E-37 |
| SPIRE1 | 2 | 0.358106567 | 0.635 | 0.465 | 9.75E-42 | 3.57E-37 |
| MTUS1 | 2 | -0.323258817 | 0.025 | 0.122 | 1.01E-41 | 3.7E-37 |
| STAT2 | 2 | -0.399499759 | 0.141 | 0.283 | 1.1E-41 | 4.04E-37 |
| ZNF586 | 2 | -0.390817424 | 0.125 | 0.262 | 1.59E-41 | 5.83E-37 |
| PCAT1 | 2 | -0.629461887 | 0.025 | 0.117 | 1.64E-41 | 6.02E-37 |
| DGKE | 2 | 0.337572236 | 0.338 | 0.181 | 1.76E-41 | 6.43E-37 |
| PTPRO | 2 | -0.664321855 | 0.405 | 0.451 | 1.96E-41 | 7.16E-37 |
| ATF7 | 2 | -0.419127591 | 0.304 | 0.465 | 2.15E-41 | 7.87E-37 |
| APCDD1 | 2 | -0.654478568 | 0.208 | 0.331 | 2.29E-41 | 8.39E-37 |
| DTWD1 | 2 | -0.402301079 | 0.229 | 0.392 | 2.71E-41 | 9.92E-37 |
| RIC3 | 2 | 0.283639786 | 0.277 | 0.134 | 2.97E-41 | 1.09E-36 |
| TLL2 | 2 | -0.528511049 | 0.07 | 0.179 | 3.78E-41 | 1.38E-36 |
| GTSE1 | 2 | 0.265459794 | 0.195 | 0.077 | 4.29E-41 | 1.57E-36 |
| CDC6 | 2 | 0.306588347 | 0.246 | 0.112 | 4.34E-41 | 1.59E-36 |
| ECT2 | 2 | 0.414684981 | 0.422 | 0.261 | 4.41E-41 | 1.61E-36 |
| DNAAF4 | 2 | -0.26083667 | 0.02 | 0.11 | 4.74E-41 | 1.73E-36 |
| THRB | 2 | -0.783249891 | 0.257 | 0.322 | 5.3E-41 | 1.94E-36 |

| | | | | | | |
|---|---|---|---|---|---|---|
| AC026704.1 | 2 | -0.360657485 | 0.036 | 0.138 | 5.42E-41 | 1.98E-36 |
| NAV1 | 2 | 0.320721918 | 0.56 | 0.379 | 5.94E-41 | 2.18E-36 |
| LINC01006 | 2 | -0.35437098 | 0.076 | 0.201 | 6.29E-41 | 2.3E-36 |
| CACNA1C | 2 | -0.578969671 | 0.529 | 0.469 | 6.32E-41 | 2.31E-36 |
| BTBD3 | 2 | -0.411938471 | 0.254 | 0.414 | 6.82E-41 | 2.5E-36 |
| AUH | 2 | -0.444676506 | 0.375 | 0.533 | 6.84E-41 | 2.5E-36 |
| ZNF292 | 2 | 0.312195939 | 0.924 | 0.877 | 7.44E-41 | 2.72E-36 |
| SNRNP70 | 2 | -0.421518666 | 0.356 | 0.515 | 7.73E-41 | 2.83E-36 |
| MAP4 | 2 | -0.429038211 | 0.644 | 0.74 | 9.78E-41 | 3.58E-36 |
| ACVR1 | 2 | -0.505067071 | 0.269 | 0.417 | 9.9E-41 | 3.63E-36 |
| GRIA1 | 2 | -0.735507549 | 0.296 | 0.386 | 1.13E-40 | 4.13E-36 |
| RBM5 | 2 | -0.399620934 | 0.468 | 0.621 | 1.15E-40 | 4.22E-36 |
| LURAP1L | 2 | 0.308174056 | 0.344 | 0.189 | 1.22E-40 | 4.45E-36 |
| RPGRIP1L | 2 | -0.59523028 | 0.317 | 0.452 | 1.31E-40 | 4.79E-36 |
| SF3B1 | 2 | -0.414204118 | 0.492 | 0.633 | 1.47E-40 | 5.39E-36 |
| BDNF-AS | 2 | -0.272477165 | 0.031 | 0.131 | 1.56E-40 | 5.72E-36 |
| GCA | 2 | -0.320658396 | 0.067 | 0.186 | 1.63E-40 | 5.98E-36 |
| TICRR | 2 | 0.291706969 | 0.305 | 0.157 | 1.64E-40 | 6.01E-36 |
| PPP4R3B | 2 | -0.417906637 | 0.459 | 0.598 | 2.09E-40 | 7.65E-36 |
| EXTL2 | 2 | -0.332900791 | 0.122 | 0.267 | 2.32E-40 | 8.5E-36 |
| AFDN | 2 | 0.342261762 | 0.868 | 0.777 | 2.37E-40 | 8.68E-36 |
| SLK | 2 | -0.38615735 | 0.179 | 0.335 | 2.37E-40 | 8.69E-36 |
| PHF21A | 2 | -0.415277825 | 0.579 | 0.703 | 2.43E-40 | 8.9E-36 |
| ZNF217 | 2 | -0.28713572 | 0.049 | 0.162 | 2.71E-40 | 9.94E-36 |
| SCML2 | 2 | 0.38097445 | 0.577 | 0.406 | 2.84E-40 | 1.04E-35 |
| PTPRJ | 2 | -0.610189339 | 0.44 | 0.533 | 3.06E-40 | 1.12E-35 |
| LIMCH1 | 2 | -0.557065008 | 0.741 | 0.67 | 3.34E-40 | 1.22E-35 |
| DCAF8 | 2 | -0.372885812 | 0.249 | 0.413 | 4.49E-40 | 1.64E-35 |
| UBL3 | 2 | -0.495116158 | 0.256 | 0.408 | 4.61E-40 | 1.69E-35 |
| GNAI1 | 2 | 0.440356507 | 0.705 | 0.557 | 4.67E-40 | 1.71E-35 |
| PLK4 | 2 | 0.30011588 | 0.301 | 0.154 | 4.79E-40 | 1.75E-35 |
| PLEKHA4 | 2 | -0.277800926 | 0.036 | 0.138 | 4.83E-40 | 1.77E-35 |
| FBXO9 | 2 | -0.357841804 | 0.184 | 0.341 | 5.24E-40 | 1.92E-35 |
| FBN1 | 2 | -0.435346887 | 0.093 | 0.205 | 5.4E-40 | 1.98E-35 |
| PRIM2 | 2 | 0.368805294 | 0.751 | 0.614 | 6.22E-40 | 2.28E-35 |
| TMEM108 | 2 | -0.642226286 | 0.651 | 0.656 | 7.18E-40 | 2.63E-35 |
| LYPD6 | 2 | 0.413902737 | 0.488 | 0.317 | 7.28E-40 | 2.67E-35 |
| SNCA | 2 | -0.547710143 | 0.157 | 0.301 | 7.47E-40 | 2.73E-35 |
| TES | 2 | -0.297682062 | 0.027 | 0.12 | 8.09E-40 | 2.96E-35 |
| ST3GAL1 | 2 | -0.33983637 | 0.064 | 0.185 | 9.26E-40 | 3.39E-35 |
| NOTCH3 | 2 | -0.301990255 | 0.022 | 0.11 | 9.53E-40 | 3.49E-35 |
| LINC00174 | 2 | -0.375229046 | 0.159 | 0.304 | 9.61E-40 | 3.52E-35 |
| MYO19 | 2 | 0.376982095 | 0.597 | 0.432 | 9.82E-40 | 3.59E-35 |
| DNAJC1 | 2 | -0.638404119 | 0.488 | 0.59 | 1.01E-39 | 3.71E-35 |
| AC022424.1 | 2 | 0.281649368 | 0.185 | 0.072 | 1.04E-39 | 3.81E-35 |
| AC138305.1 | 2 | -0.706964668 | 0.072 | 0.185 | 1.07E-39 | 3.91E-35 |
| RPS6KA2 | 2 | -0.558232361 | 0.205 | 0.353 | 1.29E-39 | 4.72E-35 |
| TLE1 | 2 | -0.491541477 | 0.153 | 0.28 | 1.32E-39 | 4.85E-35 |
| NUMB | 2 | -0.400033923 | 0.466 | 0.606 | 1.43E-39 | 5.24E-35 |
| CTCFL | 2 | 0.259261631 | 0.173 | 0.064 | 1.49E-39 | 5.45E-35 |
| FANCD2 | 2 | 0.304241307 | 0.368 | 0.209 | 1.8E-39 | 6.58E-35 |
| CD109 | 2 | -0.349454498 | 0.049 | 0.161 | 1.82E-39 | 6.65E-35 |
| ITGA6 | 2 | -0.306760297 | 0.03 | 0.128 | 1.85E-39 | 6.78E-35 |
| RORA | 2 | 0.473189034 | 0.832 | 0.79 | 2.16E-39 | 7.89E-35 |
| EPHB2 | 2 | 0.307955857 | 0.32 | 0.17 | 2.35E-39 | 8.59E-35 |
| WDR26 | 2 | -0.406233248 | 0.356 | 0.511 | 2.53E-39 | 9.26E-35 |
| CERS6 | 2 | 0.345381746 | 0.885 | 0.799 | 2.62E-39 | 9.61E-35 |
| EVC2 | 2 | -0.318960174 | 0.107 | 0.244 | 3.07E-39 | 1.12E-34 |

| | | | | | | |
|---|---|---|---|---|---|---|
| DTNA | 2 | 0.383639553 | 0.809 | 0.67 | 3.27E-39 | 1.2E-34 |
| PTGFRN | 2 | -0.423740826 | 0.213 | 0.354 | 3.73E-39 | 1.36E-34 |
| DNMT1 | 2 | 0.353543754 | 0.611 | 0.443 | 4E-39 | 1.46E-34 |
| USP32 | 2 | -0.437866055 | 0.608 | 0.704 | 5.02E-39 | 1.84E-34 |
| BCL6 | 2 | 0.258792779 | 0.163 | 0.059 | 5.12E-39 | 1.87E-34 |
| EP300 | 2 | -0.40904598 | 0.496 | 0.635 | 5.55E-39 | 2.03E-34 |
| FOXP1 | 2 | -0.396037236 | 0.744 | 0.83 | 6.85E-39 | 2.51E-34 |
| LDHA | 2 | -0.318206437 | 0.031 | 0.126 | 6.9E-39 | 2.53E-34 |
| TTC21B | 2 | 0.525741238 | 0.66 | 0.569 | 6.98E-39 | 2.56E-34 |
| HIST1H4C | 2 | 0.337144246 | 0.281 | 0.142 | 7.06E-39 | 2.58E-34 |
| SMCHD1 | 2 | 0.312256167 | 0.885 | 0.821 | 7.2E-39 | 2.64E-34 |
| DDX58 | 2 | -0.334324077 | 0.025 | 0.118 | 8.04E-39 | 2.94E-34 |
| XYLB | 2 | 0.337720265 | 0.434 | 0.27 | 1.16E-38 | 4.24E-34 |
| SDC2 | 2 | 0.350684555 | 0.509 | 0.335 | 1.3E-38 | 4.75E-34 |
| FOXK1 | 2 | -0.434967013 | 0.322 | 0.465 | 1.36E-38 | 4.99E-34 |
| NKD1 | 2 | -0.660688583 | 0.076 | 0.185 | 1.37E-38 | 5.01E-34 |
| ATF7IP | 2 | -0.456478803 | 0.42 | 0.561 | 2.04E-38 | 7.48E-34 |
| TRPM3 | 2 | -2.028369747 | 0.153 | 0.212 | 2.18E-38 | 7.98E-34 |
| AC061958.1 | 2 | 0.378218495 | 0.446 | 0.279 | 2.3E-38 | 8.42E-34 |
| MTCL1 | 2 | 0.303365443 | 0.356 | 0.201 | 2.49E-38 | 9.11E-34 |
| SPTBN2 | 2 | 0.307015875 | 0.311 | 0.166 | 2.82E-38 | 1.03E-33 |
| ADAM23 | 2 | -0.35960171 | 0.383 | 0.353 | 3.16E-38 | 1.16E-33 |
| RIMBP2 | 2 | -1.096750979 | 0.063 | 0.145 | 3.29E-38 | 1.2E-33 |
| AGPAT5 | 2 | 0.383957881 | 0.615 | 0.46 | 3.48E-38 | 1.27E-33 |
| PPP6R2 | 2 | -0.403574236 | 0.531 | 0.659 | 4.09E-38 | 1.5E-33 |
| NFYC | 2 | -0.378971046 | 0.182 | 0.322 | 4.1E-38 | 1.5E-33 |
| LYPLAL1 | 2 | -0.410139229 | 0.315 | 0.465 | 4.27E-38 | 1.56E-33 |
| CYBRD1 | 2 | -0.273030524 | 0.039 | 0.133 | 4.67E-38 | 1.71E-33 |
| GNAS | 2 | -0.425665603 | 0.496 | 0.622 | 4.71E-38 | 1.72E-33 |
| AC079142.1 | 2 | -0.451935569 | 0.486 | 0.642 | 6.15E-38 | 2.25E-33 |
| TMF1 | 2 | -0.417215843 | 0.248 | 0.395 | 6.33E-38 | 2.32E-33 |
| SMIM14 | 2 | -0.349022552 | 0.104 | 0.235 | 6.84E-38 | 2.51E-33 |
| RHOQ | 2 | -0.36348779 | 0.166 | 0.313 | 7.1E-38 | 2.6E-33 |
| PDSS2 | 2 | -0.409190462 | 0.491 | 0.644 | 7.62E-38 | 2.79E-33 |
| PELI1 | 2 | -0.497321117 | 0.186 | 0.312 | 8.34E-38 | 3.05E-33 |
| TNPO1 | 2 | -0.389298995 | 0.584 | 0.687 | 9.71E-38 | 3.56E-33 |
| ENOX2 | 2 | -0.459971475 | 0.208 | 0.361 | 9.75E-38 | 3.57E-33 |
| SNAPC1 | 2 | -0.310652367 | 0.078 | 0.2 | 1.03E-37 | 3.78E-33 |
| LINC02615 | 2 | -0.310035881 | 0.054 | 0.166 | 1.1E-37 | 4.04E-33 |
| CYYR1 | 2 | -0.307579597 | 0.017 | 0.1 | 1.17E-37 | 4.28E-33 |
| TPD52L1 | 2 | -0.339413223 | 0.047 | 0.153 | 1.18E-37 | 4.32E-33 |
| EIF2AK4 | 2 | -0.355075036 | 0.17 | 0.321 | 1.18E-37 | 4.33E-33 |
| PLSCR1 | 2 | -0.41814655 | 0.169 | 0.298 | 1.25E-37 | 4.59E-33 |
| METTL15 | 2 | -0.426674723 | 0.537 | 0.664 | 1.26E-37 | 4.61E-33 |
| NBPF26 | 2 | -0.308922865 | 0.09 | 0.216 | 1.28E-37 | 4.67E-33 |
| MIR100HG | 2 | 0.358027421 | 0.855 | 0.719 | 1.3E-37 | 4.77E-33 |
| DLEU1 | 2 | -0.566915616 | 0.224 | 0.344 | 1.5E-37 | 5.48E-33 |
| FBXL4 | 2 | -0.4415246 | 0.411 | 0.552 | 1.62E-37 | 5.92E-33 |
| TRIM37 | 2 | 0.364603807 | 0.562 | 0.406 | 1.64E-37 | 6E-33 |
| EEPD1 | 2 | -0.560299633 | 0.217 | 0.324 | 1.7E-37 | 6.24E-33 |
| GRIK2 | 2 | -0.852528381 | 0.176 | 0.318 | 1.85E-37 | 6.76E-33 |
| CEMIP2 | 2 | 0.377564349 | 0.484 | 0.317 | 1.89E-37 | 6.93E-33 |
| AKAP1 | 2 | 0.314599947 | 0.298 | 0.159 | 1.92E-37 | 7.04E-33 |
| NID1 | 2 | -0.452286543 | 0.209 | 0.301 | 1.94E-37 | 7.11E-33 |
| ASPM | 2 | 0.280038401 | 0.365 | 0.213 | 1.97E-37 | 7.22E-33 |
| LINC01578 | 2 | -0.407610474 | 0.434 | 0.583 | 2.07E-37 | 7.58E-33 |
| CEP164 | 2 | -0.353855967 | 0.132 | 0.268 | 2.29E-37 | 8.38E-33 |
| RHOBTB1 | 2 | -0.423538546 | 0.141 | 0.276 | 2.51E-37 | 9.19E-33 |

| | | | | | | |
|---|---|---|---|---|---|---|
| CNOT2 | 2 | -0.374284663 | 0.721 | 0.804 | 2.59E-37 | 9.48E-33 |
| AL354809.1 | 2 | -0.365452864 | 0.041 | 0.138 | 2.67E-37 | 9.78E-33 |
| RPS6KC1 | 2 | -0.440878802 | 0.399 | 0.545 | 2.75E-37 | 1.01E-32 |
| PRAG1 | 2 | -0.354237792 | 0.047 | 0.15 | 2.83E-37 | 1.04E-32 |
| ESYT2 | 2 | -0.532866543 | 0.46 | 0.576 | 3.11E-37 | 1.14E-32 |
| MON2 | 2 | -0.375801201 | 0.662 | 0.772 | 3.47E-37 | 1.27E-32 |
| IBTK | 2 | -0.416495079 | 0.334 | 0.486 | 3.72E-37 | 1.36E-32 |
| GEN1 | 2 | 0.349055357 | 0.548 | 0.383 | 3.77E-37 | 1.38E-32 |
| MARCH3 | 2 | -0.4742939 | 0.111 | 0.216 | 4.32E-37 | 1.58E-32 |
| CLMN | 2 | -0.39199086 | 0.093 | 0.215 | 4.63E-37 | 1.7E-32 |
| PIK3C3 | 2 | -0.41639948 | 0.505 | 0.619 | 4.65E-37 | 1.7E-32 |
| LINC00472 | 2 | -0.428721945 | 0.117 | 0.239 | 5.16E-37 | 1.89E-32 |
| ATP11A | 2 | 0.336295973 | 0.617 | 0.448 | 6.14E-37 | 2.25E-32 |
| BTBD9 | 2 | -0.453009441 | 0.453 | 0.597 | 6.92E-37 | 2.53E-32 |
| VPS37B | 2 | -0.383950019 | 0.237 | 0.392 | 7.59E-37 | 2.78E-32 |
| SRSF11 | 2 | -0.298909834 | 0.842 | 0.901 | 8.85E-37 | 3.24E-32 |
| UBE2G2 | 2 | -0.379286444 | 0.296 | 0.45 | 9.71E-37 | 3.55E-32 |
| PACRG | 2 | -0.61180677 | 0.029 | 0.112 | 1.04E-36 | 3.8E-32 |
| SATB2 | 2 | 0.556831293 | 0.569 | 0.435 | 1.04E-36 | 3.81E-32 |
| DIO2 | 2 | 0.465010549 | 0.421 | 0.282 | 1.08E-36 | 3.94E-32 |
| SLC7A1 | 2 | 0.327627149 | 0.428 | 0.266 | 1.14E-36 | 4.17E-32 |
| EIF4A2 | 2 | -0.385770497 | 0.467 | 0.597 | 1.19E-36 | 4.35E-32 |
| SEMA3A | 2 | -0.49335417 | 0.382 | 0.267 | 1.23E-36 | 4.51E-32 |
| PRDM2 | 2 | 0.459649502 | 0.667 | 0.545 | 1.23E-36 | 4.52E-32 |
| HMGN3 | 2 | -0.401102551 | 0.234 | 0.374 | 1.51E-36 | 5.54E-32 |
| KIF13A | 2 | -0.449497857 | 0.466 | 0.615 | 1.58E-36 | 5.8E-32 |
| CASC2 | 2 | -0.277150151 | 0.027 | 0.115 | 1.61E-36 | 5.88E-32 |
| ACVR2A | 2 | -0.536359425 | 0.345 | 0.482 | 1.78E-36 | 6.52E-32 |
| PCBP2 | 2 | -0.375792846 | 0.512 | 0.642 | 1.8E-36 | 6.58E-32 |
| PDLIM5 | 2 | -0.579816362 | 0.295 | 0.416 | 2.06E-36 | 7.54E-32 |
| DAB1 | 2 | 0.321234911 | 0.801 | 0.657 | 2.57E-36 | 9.42E-32 |
| EFR3A | 2 | -0.405508472 | 0.317 | 0.465 | 2.79E-36 | 1.02E-31 |
| MAN1C1 | 2 | -0.401113709 | 0.13 | 0.252 | 3.13E-36 | 1.15E-31 |
| AC068286.2 | 2 | -1.104618984 | 0.055 | 0.155 | 3.33E-36 | 1.22E-31 |
| STAT3 | 2 | -0.443471448 | 0.326 | 0.47 | 3.34E-36 | 1.22E-31 |
| AKT3 | 2 | 0.320806071 | 0.907 | 0.821 | 3.52E-36 | 1.29E-31 |
| WNT5B | 2 | -0.375828691 | 0.043 | 0.139 | 3.55E-36 | 1.3E-31 |
| FANCL | 2 | 0.372254477 | 0.726 | 0.6 | 3.74E-36 | 1.37E-31 |
| TMEM132C | 2 | -0.6371108 | 0.045 | 0.139 | 3.92E-36 | 1.44E-31 |
| AP002026.1 | 2 | -0.260605338 | 0.02 | 0.101 | 4.32E-36 | 1.58E-31 |
| ZC3H12C | 2 | 0.273824566 | 0.721 | 0.56 | 4.39E-36 | 1.61E-31 |
| CNTN5 | 2 | -1.398725666 | 0.062 | 0.158 | 4.4E-36 | 1.61E-31 |
| SF1 | 2 | -0.36664772 | 0.442 | 0.582 | 5.36E-36 | 1.96E-31 |
| LINC02614 | 2 | -0.278302434 | 0.055 | 0.161 | 5.57E-36 | 2.04E-31 |
| NUP210 | 2 | 0.31588879 | 0.446 | 0.284 | 6.15E-36 | 2.25E-31 |
| WASF3 | 2 | 0.316803475 | 0.286 | 0.149 | 6.71E-36 | 2.46E-31 |
| C3orf70 | 2 | 0.345054719 | 0.391 | 0.236 | 7.15E-36 | 2.62E-31 |
| FAM204A | 2 | -0.412708345 | 0.26 | 0.395 | 8.43E-36 | 3.09E-31 |
| LRBA | 2 | -0.357225399 | 0.876 | 0.892 | 9.1E-36 | 3.33E-31 |
| IP6K2 | 2 | -0.394086518 | 0.378 | 0.522 | 9.46E-36 | 3.46E-31 |
| PDE1A | 2 | 0.946404456 | 0.302 | 0.179 | 1.01E-35 | 3.69E-31 |
| CHPT1 | 2 | -0.488515995 | 0.524 | 0.575 | 1.15E-35 | 4.2E-31 |
| HTR7 | 2 | -0.439225874 | 0.045 | 0.143 | 1.19E-35 | 4.37E-31 |
| CFAP36 | 2 | -0.28242505 | 0.084 | 0.206 | 1.25E-35 | 4.59E-31 |
| FKBP5 | 2 | 0.319864065 | 0.451 | 0.288 | 1.32E-35 | 4.83E-31 |
| THADA | 2 | -0.405170705 | 0.479 | 0.613 | 1.45E-35 | 5.31E-31 |
| ASTN2 | 2 | -0.674361519 | 0.444 | 0.529 | 1.79E-35 | 6.57E-31 |
| SIPA1L1 | 2 | -0.522170656 | 0.731 | 0.781 | 1.94E-35 | 7.11E-31 |

| | | | | | | |
|---|---|---|---|---|---|---|
| WDR62 | 2 | 0.270469665 | 0.214 | 0.097 | 2.32E-35 | 8.48E-31 |
| PIEZO2 | 2 | -0.580356975 | 0.328 | 0.387 | 2.39E-35 | 8.76E-31 |
| CERK | 2 | 0.307206668 | 0.388 | 0.237 | 3.35E-35 | 1.23E-30 |
| CCDC136 | 2 | 0.258553872 | 0.272 | 0.139 | 3.46E-35 | 1.27E-30 |
| ZNF160 | 2 | -0.33980781 | 0.15 | 0.282 | 3.48E-35 | 1.27E-30 |
| SOX2-OT | 2 | -0.689208752 | 0.263 | 0.191 | 3.69E-35 | 1.35E-30 |
| ZSCAN5A | 2 | -0.351350669 | 0.135 | 0.266 | 3.71E-35 | 1.36E-30 |
| PALLD | 2 | -0.640322827 | 0.379 | 0.423 | 4.1E-35 | 1.5E-30 |
| AC008014.1 | 2 | -0.554371012 | 0.247 | 0.378 | 4.35E-35 | 1.59E-30 |
| ATP10A | 2 | 0.497665458 | 0.388 | 0.259 | 4.4E-35 | 1.61E-30 |
| CENPI | 2 | 0.264221909 | 0.3 | 0.162 | 4.77E-35 | 1.75E-30 |
| SLC22A3 | 2 | -0.516247857 | 0.396 | 0.41 | 5.05E-35 | 1.85E-30 |
| CNTLN | 2 | 0.35145231 | 0.804 | 0.687 | 5.25E-35 | 1.92E-30 |
| SPG11 | 2 | -0.379999695 | 0.373 | 0.524 | 5.39E-35 | 1.97E-30 |
| KDM4C | 2 | -0.433318379 | 0.518 | 0.635 | 5.69E-35 | 2.08E-30 |
| ZNF75D | 2 | -0.2714159 | 0.057 | 0.165 | 6.02E-35 | 2.2E-30 |
| CYTL1 | 2 | -0.251975923 | 0.028 | 0.113 | 6.38E-35 | 2.33E-30 |
| GNAL | 2 | 0.286467233 | 0.218 | 0.101 | 7E-35 | 2.56E-30 |
| CC2D2A | 2 | -0.354343481 | 0.158 | 0.288 | 7.02E-35 | 2.57E-30 |
| CIRBP | 2 | -0.304329926 | 0.116 | 0.244 | 7.36E-35 | 2.69E-30 |
| AC113383.1 | 2 | 0.441512246 | 0.304 | 0.172 | 7.68E-35 | 2.81E-30 |
| TMEM170B | 2 | 0.277516244 | 0.221 | 0.103 | 9.48E-35 | 3.47E-30 |
| CABLES1 | 2 | 0.32683457 | 0.282 | 0.149 | 9.48E-35 | 3.47E-30 |
| SERGEF | 2 | 0.410530082 | 0.529 | 0.391 | 1.02E-34 | 3.72E-30 |
| CBR4 | 2 | -0.394913129 | 0.454 | 0.583 | 1.08E-34 | 3.94E-30 |
| ANKRD36C | 2 | 0.332998603 | 0.762 | 0.633 | 1.11E-34 | 4.08E-30 |
| BCLAF3 | 2 | -0.430803257 | 0.324 | 0.433 | 1.13E-34 | 4.12E-30 |
| DDX39B | 2 | -0.367390684 | 0.379 | 0.517 | 1.24E-34 | 4.53E-30 |
| VSIG10 | 2 | -0.381455101 | 0.137 | 0.265 | 1.4E-34 | 5.13E-30 |
| CAPN2 | 2 | -0.322232354 | 0.114 | 0.235 | 1.53E-34 | 5.6E-30 |
| PIK3R3 | 2 | 0.367213045 | 0.621 | 0.467 | 1.79E-34 | 6.54E-30 |
| EIF4G3 | 2 | -0.307813169 | 0.92 | 0.938 | 1.97E-34 | 7.2E-30 |
| TUBA1A | 2 | 0.301916694 | 0.365 | 0.216 | 2E-34 | 7.34E-30 |
| NMU | 2 | 0.272289646 | 0.373 | 0.224 | 2.01E-34 | 7.36E-30 |
| SLMAP | 2 | -0.396548995 | 0.491 | 0.617 | 2.03E-34 | 7.42E-30 |
| TOP2B | 2 | -0.397626459 | 0.702 | 0.78 | 2.1E-34 | 7.68E-30 |
| CEP152 | 2 | 0.329613188 | 0.44 | 0.284 | 2.27E-34 | 8.29E-30 |
| TMEM181 | 2 | 0.346957722 | 0.596 | 0.466 | 2.47E-34 | 9.06E-30 |
| ARHGEF3 | 2 | -0.474573041 | 0.35 | 0.483 | 3.24E-34 | 1.19E-29 |
| CD99 | 2 | -0.366870293 | 0.182 | 0.319 | 3.41E-34 | 1.25E-29 |
| ADD2 | 2 | 0.316672284 | 0.419 | 0.264 | 3.48E-34 | 1.27E-29 |
| ALDH1L2 | 2 | -0.344158261 | 0.085 | 0.198 | 3.94E-34 | 1.44E-29 |
| MCF2L | 2 | -0.416336448 | 0.169 | 0.285 | 4.42E-34 | 1.62E-29 |
| STX17 | 2 | -0.30903065 | 0.114 | 0.24 | 4.47E-34 | 1.64E-29 |
| ZNF44 | 2 | -0.259157265 | 0.057 | 0.163 | 4.69E-34 | 1.72E-29 |
| ADCY5 | 2 | 0.256293582 | 0.13 | 0.043 | 4.89E-34 | 1.79E-29 |
| OLA1 | 2 | 0.317295972 | 0.846 | 0.755 | 4.94E-34 | 1.81E-29 |
| HMBOX1 | 2 | -0.367502559 | 0.488 | 0.619 | 5.1E-34 | 1.87E-29 |
| KIAA0825 | 2 | -0.429438269 | 0.125 | 0.249 | 5.48E-34 | 2.01E-29 |
| DENND2A | 2 | -0.498889299 | 0.238 | 0.352 | 5.48E-34 | 2.01E-29 |
| C9orf135 | 2 | -0.313710595 | 0.024 | 0.106 | 6.05E-34 | 2.22E-29 |
| IGF2R | 2 | -0.379127619 | 0.298 | 0.432 | 6.15E-34 | 2.25E-29 |
| EXOC6 | 2 | 0.362449644 | 0.684 | 0.544 | 6.19E-34 | 2.27E-29 |
| TPCN1 | 2 | -0.361318635 | 0.086 | 0.195 | 7.11E-34 | 2.6E-29 |
| FKBP10 | 2 | -0.310456428 | 0.135 | 0.269 | 7.49E-34 | 2.74E-29 |
| RAP1B | 2 | -0.397030637 | 0.413 | 0.548 | 8.58E-34 | 3.14E-29 |
| SLC24A2 | 2 | 0.257097573 | 0.563 | 0.402 | 8.85E-34 | 3.24E-29 |
| MYCBP2 | 2 | 0.339306357 | 0.856 | 0.791 | 9.75E-34 | 3.57E-29 |

| | | | | | | |
|---|---|---|---|---|---|---|
| GGA1 | 2 | -0.308776613 | 0.116 | 0.24 | 1.06E-33 | 3.88E-29 |
| MMP17 | 2 | -0.270869529 | 0.044 | 0.139 | 1.09E-33 | 3.99E-29 |
| KLHL23 | 2 | 0.34721608 | 0.507 | 0.357 | 1.22E-33 | 4.45E-29 |
| NIN | 2 | 0.337324553 | 0.662 | 0.514 | 1.22E-33 | 4.45E-29 |
| PRPF4B | 2 | -0.350592906 | 0.585 | 0.706 | 1.64E-33 | 6E-29 |
| ACAD11 | 2 | -0.34295258 | 0.153 | 0.289 | 1.65E-33 | 6.03E-29 |
| PJA2 | 2 | -0.430756245 | 0.401 | 0.537 | 1.9E-33 | 6.96E-29 |
| TENT2 | 2 | -0.344256903 | 0.662 | 0.758 | 2.14E-33 | 7.83E-29 |
| ABTB2 | 2 | -0.418867848 | 0.075 | 0.183 | 2.48E-33 | 9.08E-29 |
| AL157944.1 | 2 | -0.732617369 | 0.034 | 0.113 | 2.51E-33 | 9.18E-29 |
| CCDC102B | 2 | -1.020606491 | 0.062 | 0.157 | 2.54E-33 | 9.32E-29 |
| MGLL | 2 | -0.373478127 | 0.096 | 0.215 | 2.89E-33 | 1.06E-28 |
| SULF1 | 2 | -0.695601662 | 0.196 | 0.335 | 2.95E-33 | 1.08E-28 |
| EYA1 | 2 | -0.748940511 | 0.201 | 0.313 | 3.01E-33 | 1.1E-28 |
| OFD1 | 2 | -0.391500082 | 0.341 | 0.469 | 3.08E-33 | 1.13E-28 |
| AXIN2 | 2 | 0.30836438 | 0.28 | 0.152 | 3.09E-33 | 1.13E-28 |
| CDC7 | 2 | 0.316835375 | 0.476 | 0.318 | 3.26E-33 | 1.19E-28 |
| PPP1R21 | 2 | -0.369570081 | 0.238 | 0.377 | 3.59E-33 | 1.31E-28 |
| CRISPLD1 | 2 | -1.115821996 | 0.412 | 0.516 | 3.75E-33 | 1.37E-28 |
| SH3BP4 | 2 | 0.361582483 | 0.46 | 0.304 | 3.78E-33 | 1.38E-28 |
| RBL1 | 2 | 0.312791417 | 0.55 | 0.387 | 3.82E-33 | 1.4E-28 |
| RNF168 | 2 | 0.295381209 | 0.567 | 0.405 | 4.27E-33 | 1.56E-28 |
| TVP23C | 2 | -0.361253263 | 0.199 | 0.331 | 4.41E-33 | 1.62E-28 |
| ZNF337-AS1 | 2 | -0.35875977 | 0.185 | 0.323 | 4.45E-33 | 1.63E-28 |
| EXOC6B | 2 | 0.284052898 | 0.8 | 0.668 | 4.52E-33 | 1.66E-28 |
| PRAME | 2 | -0.289839745 | 0.051 | 0.148 | 4.75E-33 | 1.74E-28 |
| ZFAND3 | 2 | -0.310897887 | 0.872 | 0.912 | 4.95E-33 | 1.81E-28 |
| HIBCH | 2 | -0.396133817 | 0.477 | 0.608 | 5.19E-33 | 1.9E-28 |
| ARMCX4 | 2 | -0.354508165 | 0.179 | 0.317 | 5.25E-33 | 1.92E-28 |
| TRIM9 | 2 | -0.396331862 | 0.417 | 0.426 | 5.42E-33 | 1.99E-28 |
| VRK1 | 2 | 0.29751325 | 0.638 | 0.48 | 5.74E-33 | 2.1E-28 |
| APTX | 2 | -0.333059059 | 0.213 | 0.359 | 5.95E-33 | 2.18E-28 |
| MAP9 | 2 | -0.275478355 | 0.069 | 0.179 | 6.01E-33 | 2.2E-28 |
| STK17B | 2 | -0.380919815 | 0.067 | 0.166 | 7.29E-33 | 2.67E-28 |
| TRIB2 | 2 | 0.312279465 | 0.295 | 0.165 | 7.8E-33 | 2.85E-28 |
| TPM1 | 2 | -0.550636349 | 0.24 | 0.347 | 7.93E-33 | 2.9E-28 |
| ACTR3B | 2 | -0.394284409 | 0.312 | 0.447 | 8.06E-33 | 2.95E-28 |
| HECTD4 | 2 | -0.369597053 | 0.69 | 0.77 | 9.44E-33 | 3.46E-28 |
| GCNT2 | 2 | -0.288384577 | 0.027 | 0.106 | 9.54E-33 | 3.49E-28 |
| FAM178B | 2 | -0.383335425 | 0.025 | 0.102 | 1.17E-32 | 4.28E-28 |
| NR6A1 | 2 | 0.383221969 | 0.494 | 0.343 | 1.26E-32 | 4.62E-28 |
| CAMSAP2 | 2 | 0.329164305 | 0.717 | 0.583 | 1.29E-32 | 4.7E-28 |
| SNX24 | 2 | -0.336073197 | 0.109 | 0.227 | 1.36E-32 | 4.96E-28 |
| DIXDC1 | 2 | -0.362644988 | 0.154 | 0.275 | 1.44E-32 | 5.28E-28 |
| TUT7 | 2 | -0.343210758 | 0.204 | 0.343 | 1.54E-32 | 5.63E-28 |
| LRCH3 | 2 | -0.380798447 | 0.547 | 0.639 | 1.58E-32 | 5.8E-28 |
| FYN | 2 | 0.254835424 | 0.771 | 0.634 | 1.6E-32 | 5.86E-28 |
| SEC31A | 2 | -0.388899197 | 0.52 | 0.634 | 1.6E-32 | 5.87E-28 |
| OTX2 | 2 | -0.421938002 | 0.032 | 0.11 | 1.71E-32 | 6.27E-28 |
| PPP2R5A | 2 | -0.313814055 | 0.106 | 0.226 | 2.04E-32 | 7.45E-28 |
| PSEN1 | 2 | -0.365532042 | 0.25 | 0.382 | 2.29E-32 | 8.37E-28 |
| AGL | 2 | -0.387991624 | 0.238 | 0.382 | 2.39E-32 | 8.73E-28 |
| VPS13B | 2 | -0.332766558 | 0.784 | 0.836 | 2.41E-32 | 8.82E-28 |
| AL390957.1 | 2 | -1.068367756 | 0.063 | 0.154 | 2.57E-32 | 9.41E-28 |
| AC118549.1 | 2 | -0.374961386 | 0.589 | 0.704 | 2.89E-32 | 1.06E-27 |
| LSAMP | 2 | -0.606866901 | 0.782 | 0.727 | 3.03E-32 | 1.11E-27 |
| PABPN1 | 2 | -0.349778767 | 0.337 | 0.484 | 3.09E-32 | 1.13E-27 |
| NME7 | 2 | -0.349050874 | 0.271 | 0.422 | 3.31E-32 | 1.21E-27 |

| | | | | | | |
|---|---|---|---|---|---|---|
| WHRN | 2 | -0.305821349 | 0.092 | 0.201 | 3.52E-32 | 1.29E-27 |
| VEGFA | 2 | -0.547522975 | 0.233 | 0.382 | 3.79E-32 | 1.39E-27 |
| ACIN1 | 2 | -0.336942071 | 0.324 | 0.475 | 4.51E-32 | 1.65E-27 |
| MSRA | 2 | -0.474862638 | 0.314 | 0.447 | 4.73E-32 | 1.73E-27 |
| ADGRL1 | 2 | 0.254226836 | 0.356 | 0.213 | 5.34E-32 | 1.95E-27 |
| PHC3 | 2 | -0.359861049 | 0.502 | 0.628 | 5.86E-32 | 2.15E-27 |
| TNRC6C | 2 | 0.340325884 | 0.716 | 0.583 | 6.17E-32 | 2.26E-27 |
| MIA2 | 2 | -0.385364463 | 0.241 | 0.375 | 6.41E-32 | 2.35E-27 |
| LARP4B | 2 | -0.340077229 | 0.657 | 0.729 | 7.39E-32 | 2.7E-27 |
| OTUD7A | 2 | -0.414325742 | 0.58 | 0.565 | 7.72E-32 | 2.83E-27 |
| AGAP1 | 2 | 0.266917341 | 0.909 | 0.829 | 7.72E-32 | 2.83E-27 |
| NSD1 | 2 | -0.360300784 | 0.713 | 0.78 | 7.97E-32 | 2.92E-27 |
| ASAP1 | 2 | -0.365577014 | 0.819 | 0.86 | 8.03E-32 | 2.94E-27 |
| TM2D1 | 2 | -0.362965197 | 0.297 | 0.433 | 8.14E-32 | 2.98E-27 |
| CHM | 2 | -0.413073281 | 0.603 | 0.701 | 9.43E-32 | 3.45E-27 |
| AP1S2 | 2 | -0.435648248 | 0.43 | 0.555 | 9.46E-32 | 3.46E-27 |
| SMC2 | 2 | 0.302289246 | 0.46 | 0.306 | 9.98E-32 | 3.65E-27 |
| RALGAPA2 | 2 | 0.307127675 | 0.689 | 0.554 | 1.4E-31 | 5.12E-27 |
| CDRT4 | 2 | -0.30408951 | 0.107 | 0.224 | 1.49E-31 | 5.46E-27 |
| R3HDM1 | 2 | 0.268368828 | 0.926 | 0.854 | 1.67E-31 | 6.11E-27 |
| LPAR1 | 2 | -0.61125525 | 0.138 | 0.221 | 1.7E-31 | 6.23E-27 |
| GLCE | 2 | -0.462848453 | 0.277 | 0.408 | 1.82E-31 | 6.65E-27 |
| HMGA1P4 | 2 | -0.253156349 | 0.029 | 0.11 | 2.25E-31 | 8.22E-27 |
| USP47 | 2 | -0.344440688 | 0.623 | 0.723 | 2.48E-31 | 9.07E-27 |
| ATP6V1B2 | 2 | 0.298581438 | 0.325 | 0.191 | 2.67E-31 | 9.78E-27 |
| CHKA | 2 | -0.408821859 | 0.24 | 0.362 | 3.08E-31 | 1.13E-26 |
| HNRNPA2B1 | 2 | -0.293216366 | 0.808 | 0.866 | 3.6E-31 | 1.32E-26 |
| IGFBP2 | 2 | -0.530511887 | 0.454 | 0.543 | 3.84E-31 | 1.41E-26 |
| SCFD1 | 2 | -0.363897525 | 0.513 | 0.636 | 3.94E-31 | 1.44E-26 |
| KCNN3 | 2 | -0.331946905 | 0.064 | 0.157 | 4.11E-31 | 1.5E-26 |
| MMP16 | 2 | 0.324448273 | 0.82 | 0.704 | 4.19E-31 | 1.53E-26 |
| NCAM2 | 2 | 0.337973177 | 0.147 | 0.058 | 4.42E-31 | 1.62E-26 |
| MBD5 | 2 | -0.316944475 | 0.889 | 0.9 | 4.75E-31 | 1.74E-26 |
| MSH2 | 2 | 0.309463731 | 0.644 | 0.502 | 5.05E-31 | 1.85E-26 |
| CD2AP | 2 | -0.386223588 | 0.495 | 0.601 | 5.43E-31 | 1.99E-26 |
| ANKRD13B | 2 | 0.262717902 | 0.207 | 0.099 | 5.62E-31 | 2.06E-26 |
| TMPO | 2 | 0.303127547 | 0.638 | 0.487 | 5.87E-31 | 2.15E-26 |
| MAGI3 | 2 | 0.280089709 | 0.579 | 0.421 | 7.03E-31 | 2.58E-26 |
| DPPA4 | 2 | -0.331705798 | 0.414 | 0.439 | 9.53E-31 | 3.49E-26 |
| ZYG11A | 2 | 0.31379263 | 0.424 | 0.275 | 1E-30 | 3.66E-26 |
| GMNN | 2 | 0.269369823 | 0.321 | 0.187 | 1.03E-30 | 3.76E-26 |
| SLC25A36 | 2 | -0.362032025 | 0.505 | 0.627 | 1.03E-30 | 3.76E-26 |
| IRAK1BP1 | 2 | -0.313309337 | 0.148 | 0.277 | 1.08E-30 | 3.94E-26 |
| PRMT3 | 2 | -0.387876376 | 0.286 | 0.424 | 1.1E-30 | 4.04E-26 |
| COL6A2 | 2 | -0.321265537 | 0.064 | 0.155 | 1.13E-30 | 4.14E-26 |
| ZNF331 | 2 | 0.259903788 | 0.325 | 0.19 | 1.2E-30 | 4.4E-26 |
| AGBL4 | 2 | -0.746035996 | 0.186 | 0.213 | 1.33E-30 | 4.88E-26 |
| PPP1R9A | 2 | -0.579746297 | 0.101 | 0.185 | 1.35E-30 | 4.93E-26 |
| IKZF3 | 2 | -0.40432805 | 0.177 | 0.289 | 1.38E-30 | 5.05E-26 |
| CCNL2 | 2 | -0.366360776 | 0.383 | 0.512 | 1.53E-30 | 5.59E-26 |
| ARGLU1 | 2 | -0.316067275 | 0.769 | 0.84 | 1.55E-30 | 5.68E-26 |
| CEP192 | 2 | 0.327113046 | 0.695 | 0.597 | 1.56E-30 | 5.7E-26 |
| INTU | 2 | -0.360317727 | 0.201 | 0.334 | 1.74E-30 | 6.38E-26 |
| GANC | 2 | -0.321578241 | 0.135 | 0.26 | 2.07E-30 | 7.57E-26 |
| CDC42SE2 | 2 | -0.381242086 | 0.334 | 0.468 | 2.19E-30 | 8.02E-26 |
| MORF4L2 | 2 | -0.372046683 | 0.447 | 0.572 | 2.3E-30 | 8.43E-26 |
| VRK2 | 2 | -0.372133188 | 0.217 | 0.346 | 2.36E-30 | 8.64E-26 |
| DOCK5 | 2 | 0.371205952 | 0.481 | 0.343 | 2.37E-30 | 8.66E-26 |

| | | | | | | |
|---|---|---|---|---|---|---|
| CREBRF | 2 | -0.324895479 | 0.101 | 0.212 | 2.89E-30 | 1.06E-25 |
| JAM3 | 2 | -0.375286062 | 0.296 | 0.435 | 4.03E-30 | 1.47E-25 |
| RNF128 | 2 | -0.410672392 | 0.062 | 0.158 | 4.05E-30 | 1.48E-25 |
| GTF2I | 2 | -0.289750806 | 0.87 | 0.906 | 4.14E-30 | 1.51E-25 |
| TCF25 | 2 | -0.362668536 | 0.365 | 0.499 | 4.69E-30 | 1.72E-25 |
| SMAD1 | 2 | -0.363234103 | 0.254 | 0.393 | 5.21E-30 | 1.91E-25 |
| CCDC146 | 2 | -0.404695803 | 0.143 | 0.26 | 5.34E-30 | 1.96E-25 |
| LIX1-AS1 | 2 | -0.256345438 | 0.035 | 0.12 | 5.6E-30 | 2.05E-25 |
| NUBPL | 2 | -0.349057217 | 0.375 | 0.525 | 5.62E-30 | 2.06E-25 |
| UBR2 | 2 | -0.365209952 | 0.462 | 0.581 | 5.64E-30 | 2.06E-25 |
| SUPT20H | 2 | -0.355405793 | 0.411 | 0.545 | 5.68E-30 | 2.08E-25 |
| EMC2 | 2 | -0.328852838 | 0.199 | 0.322 | 5.81E-30 | 2.13E-25 |
| IQCJ-SCHIP1 | 2 | -0.699740009 | 0.09 | 0.192 | 6.2E-30 | 2.27E-25 |
| RAB30 | 2 | 0.266697144 | 0.215 | 0.107 | 6.68E-30 | 2.45E-25 |
| TMEM232 | 2 | -0.450557921 | 0.059 | 0.152 | 6.82E-30 | 2.5E-25 |
| NCAPD3 | 2 | 0.321336943 | 0.483 | 0.336 | 7.28E-30 | 2.66E-25 |
| SRPK2 | 2 | -0.326064403 | 0.779 | 0.851 | 7.4E-30 | 2.71E-25 |
| ZMYM2 | 2 | -0.30987578 | 0.872 | 0.905 | 7.53E-30 | 2.76E-25 |
| CHIC1 | 2 | 0.364158203 | 0.505 | 0.389 | 7.61E-30 | 2.79E-25 |
| HIVEP2 | 2 | 0.299446149 | 0.555 | 0.401 | 7.88E-30 | 2.89E-25 |
| KCTD13 | 2 | -0.361546609 | 0.219 | 0.349 | 8.44E-30 | 3.09E-25 |
| ARID3A | 2 | -0.331717471 | 0.076 | 0.169 | 8.89E-30 | 3.26E-25 |
| CSNK1G1 | 2 | -0.357191113 | 0.425 | 0.563 | 8.99E-30 | 3.29E-25 |
| ZMAT1 | 2 | -0.274932705 | 0.07 | 0.168 | 9.16E-30 | 3.35E-25 |
| SHROOM2 | 2 | -0.38486662 | 0.263 | 0.39 | 1E-29 | 3.68E-25 |
| JAG1 | 2 | -0.396069909 | 0.069 | 0.163 | 1.1E-29 | 4.04E-25 |
| SNHG25 | 2 | -0.356611281 | 0.208 | 0.343 | 1.11E-29 | 4.07E-25 |
| REPS2 | 2 | -0.459448895 | 0.238 | 0.35 | 1.16E-29 | 4.24E-25 |
| ENPP1 | 2 | -0.518657899 | 0.286 | 0.37 | 1.16E-29 | 4.25E-25 |
| SLC44A1 | 2 | -0.337846267 | 0.691 | 0.691 | 1.22E-29 | 4.45E-25 |
| UFL1 | 2 | -0.298271095 | 0.161 | 0.293 | 1.23E-29 | 4.49E-25 |
| DENND1B | 2 | -0.424412998 | 0.229 | 0.346 | 1.27E-29 | 4.65E-25 |
| PARPBP | 2 | 0.281408962 | 0.385 | 0.243 | 1.32E-29 | 4.84E-25 |
| STAT1 | 2 | -0.469040855 | 0.192 | 0.301 | 1.33E-29 | 4.87E-25 |
| SNHG4 | 2 | -0.284771907 | 0.076 | 0.179 | 1.34E-29 | 4.91E-25 |
| CDCP1 | 2 | -0.425328382 | 0.108 | 0.206 | 1.34E-29 | 4.91E-25 |
| CABIN1 | 2 | -0.349238316 | 0.279 | 0.407 | 1.36E-29 | 4.96E-25 |
| NAPEPLD | 2 | -0.370050113 | 0.253 | 0.379 | 1.37E-29 | 5.02E-25 |
| IRAK3 | 2 | -0.269638541 | 0.505 | 0.489 | 1.41E-29 | 5.16E-25 |
| OGFRL1 | 2 | 0.253200569 | 0.249 | 0.131 | 1.42E-29 | 5.19E-25 |
| BRCA2 | 2 | 0.267487665 | 0.384 | 0.243 | 1.45E-29 | 5.31E-25 |
| CFAP58 | 2 | -0.364505311 | 0.303 | 0.295 | 1.57E-29 | 5.73E-25 |
| STRN | 2 | -0.375960811 | 0.409 | 0.531 | 1.85E-29 | 6.77E-25 |
| CENPJ | 2 | 0.271266998 | 0.397 | 0.255 | 1.9E-29 | 6.96E-25 |
| AKAP9 | 2 | -0.382145028 | 0.665 | 0.751 | 1.97E-29 | 7.22E-25 |
| NFIX | 2 | -0.426702864 | 0.354 | 0.493 | 1.97E-29 | 7.22E-25 |
| KLF3 | 2 | -0.27453785 | 0.075 | 0.171 | 2.01E-29 | 7.35E-25 |
| SLC8A2 | 2 | -0.326920922 | 0.076 | 0.172 | 2.03E-29 | 7.45E-25 |
| STX18 | 2 | -0.347166099 | 0.242 | 0.374 | 2.13E-29 | 7.79E-25 |
| TRIP11 | 2 | -0.343789967 | 0.287 | 0.424 | 2.44E-29 | 8.92E-25 |
| CDK12 | 2 | -0.341392345 | 0.338 | 0.472 | 2.61E-29 | 9.57E-25 |
| GLS | 2 | -0.390308301 | 0.753 | 0.791 | 2.95E-29 | 1.08E-24 |
| ENTPD5 | 2 | -0.273216311 | 0.075 | 0.176 | 3.06E-29 | 1.12E-24 |
| PRKAA2 | 2 | 0.253286735 | 0.283 | 0.158 | 3.11E-29 | 1.14E-24 |
| RUNX1T1 | 2 | 0.35039439 | 0.813 | 0.691 | 3.29E-29 | 1.2E-24 |
| VMP1 | 2 | -0.501951355 | 0.524 | 0.611 | 3.31E-29 | 1.21E-24 |
| ADD1 | 2 | -0.335804079 | 0.383 | 0.523 | 3.31E-29 | 1.21E-24 |
| GLMN | 2 | 0.334314091 | 0.611 | 0.474 | 3.47E-29 | 1.27E-24 |

| | | | | | | |
|---|---|---|---|---|---|---|
| RBBP6 | 2 | -0.356643932 | 0.375 | 0.501 | 3.94E-29 | 1.44E-24 |
| RTTN | 2 | 0.305483999 | 0.638 | 0.492 | 4.01E-29 | 1.47E-24 |
| PAG1 | 2 | -0.507165114 | 0.221 | 0.325 | 4.34E-29 | 1.59E-24 |
| GTF2F2 | 2 | -0.388106168 | 0.34 | 0.465 | 5.54E-29 | 2.03E-24 |
| ATAD2B | 2 | -0.333328423 | 0.525 | 0.641 | 5.87E-29 | 2.15E-24 |
| COL6A3 | 2 | -0.574391661 | 0.062 | 0.131 | 6.31E-29 | 2.31E-24 |
| ULK1 | 2 | -0.332644492 | 0.186 | 0.302 | 6.56E-29 | 2.4E-24 |
| GPATCH8 | 2 | 0.294906514 | 0.871 | 0.805 | 6.56E-29 | 2.4E-24 |
| SPG7 | 2 | -0.358031138 | 0.357 | 0.49 | 6.71E-29 | 2.46E-24 |
| ICK | 2 | -0.263912062 | 0.092 | 0.202 | 7.37E-29 | 2.7E-24 |
| NBPF11 | 2 | -0.301031358 | 0.182 | 0.308 | 8.55E-29 | 3.13E-24 |
| BBS4 | 2 | -0.271016771 | 0.107 | 0.22 | 8.71E-29 | 3.19E-24 |
| GLI2 | 2 | -0.444823967 | 0.21 | 0.339 | 8.79E-29 | 3.22E-24 |
| USP40 | 2 | -0.333911114 | 0.219 | 0.35 | 9.14E-29 | 3.35E-24 |
| MIPEP | 2 | -0.326638968 | 0.162 | 0.286 | 9.85E-29 | 3.61E-24 |
| DOK5 | 2 | -0.409813027 | 0.564 | 0.541 | 1.06E-28 | 3.88E-24 |
| LETMD1 | 2 | -0.309387924 | 0.169 | 0.292 | 1.07E-28 | 3.92E-24 |
| SLC20A2 | 2 | -0.380136698 | 0.354 | 0.477 | 1.09E-28 | 3.98E-24 |
| GCC2 | 2 | -0.35954962 | 0.309 | 0.439 | 1.11E-28 | 4.07E-24 |
| PNISR | 2 | -0.296927572 | 0.757 | 0.839 | 1.11E-28 | 4.08E-24 |
| YTHDC1 | 2 | -0.340229811 | 0.493 | 0.614 | 1.4E-28 | 5.13E-24 |
| FERMT2 | 2 | -0.482188991 | 0.338 | 0.454 | 1.4E-28 | 5.13E-24 |
| RPGR | 2 | -0.4089992 | 0.368 | 0.466 | 1.4E-28 | 5.14E-24 |
| AL162253.2 | 2 | -0.255836337 | 0.054 | 0.143 | 1.46E-28 | 5.35E-24 |
| PITRM1 | 2 | 0.299547147 | 0.296 | 0.179 | 1.5E-28 | 5.5E-24 |
| GAS2 | 2 | -0.314630648 | 0.034 | 0.111 | 1.5E-28 | 5.5E-24 |
| ANKRD17 | 2 | -0.277900818 | 0.909 | 0.924 | 1.6E-28 | 5.85E-24 |
| KCNIP1 | 2 | -0.316360103 | 0.038 | 0.12 | 1.66E-28 | 6.09E-24 |
| SPATS2L | 2 | -0.318813634 | 0.35 | 0.498 | 1.8E-28 | 6.61E-24 |
| MTMR2 | 2 | -0.391192827 | 0.266 | 0.39 | 1.83E-28 | 6.69E-24 |
| ATP2A2 | 2 | 0.314876838 | 0.768 | 0.655 | 1.85E-28 | 6.78E-24 |
| ATR | 2 | -0.368426223 | 0.496 | 0.61 | 1.89E-28 | 6.94E-24 |
| TRIM22 | 2 | -0.390659256 | 0.192 | 0.307 | 2.08E-28 | 7.63E-24 |
| NLRC5 | 2 | -0.306830094 | 0.079 | 0.179 | 2.11E-28 | 7.74E-24 |
| SLC38A6 | 2 | -0.332054142 | 0.208 | 0.333 | 2.25E-28 | 8.24E-24 |
| TGIF2 | 2 | -0.287457835 | 0.109 | 0.223 | 2.86E-28 | 1.05E-23 |
| TIA1 | 2 | -0.334270204 | 0.511 | 0.633 | 2.88E-28 | 1.06E-23 |
| SEZ6L | 2 | -0.633505426 | 0.051 | 0.105 | 2.99E-28 | 1.09E-23 |
| WAC | 2 | -0.320984566 | 0.761 | 0.836 | 3.34E-28 | 1.22E-23 |
| SDK2 | 2 | -0.380972215 | 0.053 | 0.139 | 3.44E-28 | 1.26E-23 |
| ACTN1 | 2 | -0.36046347 | 0.176 | 0.295 | 4.16E-28 | 1.52E-23 |
| PIP5K1A | 2 | -0.348952984 | 0.436 | 0.559 | 4.66E-28 | 1.7E-23 |
| MARCH8 | 2 | -0.322937322 | 0.093 | 0.197 | 4.83E-28 | 1.77E-23 |
| TWIST1 | 2 | -0.40440027 | 0.096 | 0.192 | 4.89E-28 | 1.79E-23 |
| MED15 | 2 | -0.336389368 | 0.271 | 0.389 | 5.36E-28 | 1.96E-23 |
| SDCCAG8 | 2 | -0.359476918 | 0.451 | 0.579 | 5.41E-28 | 1.98E-23 |
| CPEB2 | 2 | 0.402658103 | 0.456 | 0.334 | 5.51E-28 | 2.02E-23 |
| TRAPPC12 | 2 | -0.335907118 | 0.314 | 0.441 | 5.7E-28 | 2.09E-23 |
| MOB3B | 2 | -0.343588619 | 0.114 | 0.226 | 5.82E-28 | 2.13E-23 |
| PARVA | 2 | -0.368572707 | 0.292 | 0.416 | 5.97E-28 | 2.18E-23 |
| ULK2 | 2 | 0.330079338 | 0.519 | 0.384 | 6.02E-28 | 2.2E-23 |
| RAD50 | 2 | -0.335124928 | 0.38 | 0.518 | 6.52E-28 | 2.39E-23 |
| SALL4 | 2 | -0.322228803 | 0.181 | 0.288 | 7.32E-28 | 2.68E-23 |
| BEND7 | 2 | -0.37863811 | 0.156 | 0.278 | 7.61E-28 | 2.79E-23 |
| HOOK1 | 2 | 0.265757042 | 0.309 | 0.183 | 8.22E-28 | 3.01E-23 |
| ANXA5 | 2 | -0.260275495 | 0.076 | 0.174 | 8.34E-28 | 3.05E-23 |
| SLC25A21 | 2 | -0.299463609 | 0.058 | 0.153 | 8.69E-28 | 3.18E-23 |
| TJP2 | 2 | -0.446066793 | 0.305 | 0.413 | 9.01E-28 | 3.3E-23 |

| | | | | | | |
|---|---|---|---|---|---|---|
| SCAPER | 2 | 0.320345613 | 0.747 | 0.648 | 9.27E-28 | 3.39E-23 |
| SNED1 | 2 | -0.32412055 | 0.046 | 0.121 | 1.1E-27 | 4.03E-23 |
| SLC25A37 | 2 | -0.353255023 | 0.199 | 0.314 | 1.46E-27 | 5.35E-23 |
| STK39 | 2 | -0.408925443 | 0.612 | 0.648 | 1.61E-27 | 5.91E-23 |
| TXNIP | 2 | -0.265407279 | 0.03 | 0.104 | 1.62E-27 | 5.95E-23 |
| GNG12-AS1 | 2 | -0.322824193 | 0.091 | 0.191 | 1.68E-27 | 6.16E-23 |
| BCL2L13 | 2 | -0.311234867 | 0.194 | 0.325 | 1.73E-27 | 6.34E-23 |
| ORMDL1 | 2 | -0.312249338 | 0.24 | 0.37 | 1.74E-27 | 6.36E-23 |
| BBX | 2 | -0.374872261 | 0.66 | 0.705 | 1.83E-27 | 6.69E-23 |
| NCK2 | 2 | -0.364374288 | 0.37 | 0.502 | 1.92E-27 | 7.03E-23 |
| FOXN2 | 2 | -0.382597135 | 0.257 | 0.37 | 2.01E-27 | 7.37E-23 |
| PIR | 2 | -0.264182062 | 0.072 | 0.172 | 2.06E-27 | 7.55E-23 |
| PARVB | 2 | -0.311637413 | 0.137 | 0.252 | 2.3E-27 | 8.41E-23 |
| ITFG1 | 2 | 0.318587569 | 0.85 | 0.777 | 2.41E-27 | 8.81E-23 |
| ARRB1 | 2 | -0.28616467 | 0.076 | 0.172 | 2.55E-27 | 9.33E-23 |
| ZFP82 | 2 | 0.250717487 | 0.213 | 0.111 | 2.75E-27 | 1.01E-22 |
| RAD54B | 2 | 0.259859133 | 0.345 | 0.214 | 3.72E-27 | 1.36E-22 |
| DDX3X | 2 | -0.324303086 | 0.43 | 0.546 | 4.5E-27 | 1.65E-22 |
| MAF | 2 | -0.277594224 | 0.066 | 0.159 | 4.75E-27 | 1.74E-22 |
| BCO2 | 2 | -0.269195193 | 0.048 | 0.13 | 4.8E-27 | 1.76E-22 |
| MCUB | 2 | -0.333911818 | 0.11 | 0.219 | 4.93E-27 | 1.8E-22 |
| PICALM | 2 | -0.35783491 | 0.635 | 0.726 | 5.15E-27 | 1.88E-22 |
| ABCA3 | 2 | 0.290115074 | 0.362 | 0.229 | 5.62E-27 | 2.06E-22 |
| ID3 | 2 | -0.259585367 | 0.033 | 0.11 | 5.63E-27 | 2.06E-22 |
| CEP85L | 2 | -0.404010995 | 0.491 | 0.564 | 5.7E-27 | 2.09E-22 |
| GALNT11 | 2 | 0.304634771 | 0.536 | 0.398 | 5.82E-27 | 2.13E-22 |
| GOLGB1 | 2 | -0.329992655 | 0.347 | 0.474 | 6.64E-27 | 2.43E-22 |
| ATXN3 | 2 | -0.339894205 | 0.25 | 0.372 | 6.87E-27 | 2.51E-22 |
| AC079781.5 | 2 | -0.371537981 | 0.34 | 0.454 | 8.08E-27 | 2.96E-22 |
| DENND5B | 2 | 0.307121423 | 0.616 | 0.48 | 8.9E-27 | 3.26E-22 |
| VPS8 | 2 | -0.38254864 | 0.543 | 0.616 | 9.34E-27 | 3.42E-22 |
| SNHG3 | 2 | -0.301208059 | 0.11 | 0.215 | 1.1E-26 | 4.03E-22 |
| NRDC | 2 | -0.330120885 | 0.434 | 0.561 | 1.22E-26 | 4.48E-22 |
| UNC5B | 2 | -0.251563137 | 0.049 | 0.13 | 1.23E-26 | 4.51E-22 |
| DNMT3A | 2 | -0.336228721 | 0.247 | 0.37 | 1.26E-26 | 4.62E-22 |
| EDA | 2 | -0.58918974 | 0.224 | 0.295 | 1.36E-26 | 4.99E-22 |
| TTC6 | 2 | -0.499150283 | 0.088 | 0.181 | 1.37E-26 | 5.02E-22 |
| DPYSL2 | 2 | 0.266859336 | 0.626 | 0.483 | 1.38E-26 | 5.07E-22 |
| BMPR1B | 2 | -0.589229287 | 0.228 | 0.34 | 1.42E-26 | 5.21E-22 |
| LEPR | 2 | -0.455191221 | 0.061 | 0.148 | 1.42E-26 | 5.22E-22 |
| RALGAPA1 | 2 | -0.334333703 | 0.658 | 0.754 | 1.46E-26 | 5.33E-22 |
| ADAM17 | 2 | -0.337438239 | 0.333 | 0.459 | 1.47E-26 | 5.37E-22 |
| PDZD2 | 2 | -0.507669444 | 0.334 | 0.4 | 1.7E-26 | 6.22E-22 |
| FUBP1 | 2 | -0.294191788 | 0.629 | 0.73 | 1.83E-26 | 6.68E-22 |
| CA5B | 2 | -0.325126147 | 0.202 | 0.322 | 1.87E-26 | 6.83E-22 |
| KIAA2026 | 2 | -0.341665837 | 0.38 | 0.504 | 1.89E-26 | 6.92E-22 |
| MCM9 | 2 | -0.314083615 | 0.191 | 0.31 | 1.98E-26 | 7.26E-22 |
| SETX | 2 | -0.322679649 | 0.453 | 0.582 | 2.42E-26 | 8.86E-22 |
| VPS35L | 2 | -0.300325358 | 0.251 | 0.383 | 2.55E-26 | 9.34E-22 |
| ZNF438 | 2 | -0.367844129 | 0.191 | 0.31 | 2.6E-26 | 9.54E-22 |
| COQ4 | 2 | -0.252528578 | 0.089 | 0.186 | 2.66E-26 | 9.73E-22 |
| HSDL2 | 2 | -0.296158347 | 0.153 | 0.271 | 2.72E-26 | 9.97E-22 |
| TULP4 | 2 | 0.310420255 | 0.653 | 0.54 | 3.26E-26 | 1.19E-21 |
| C2orf92 | 2 | -0.254767171 | 0.073 | 0.169 | 3.45E-26 | 1.26E-21 |
| COP1 | 2 | 0.26736191 | 0.869 | 0.813 | 3.55E-26 | 1.3E-21 |
| GCLC | 2 | -0.691721594 | 0.147 | 0.227 | 3.61E-26 | 1.32E-21 |
| USP8 | 2 | -0.300633183 | 0.313 | 0.45 | 3.67E-26 | 1.34E-21 |
| WDR19 | 2 | -0.311148153 | 0.162 | 0.276 | 3.76E-26 | 1.38E-21 |

| | | | | | | |
|---|---|---|---|---|---|---|
| AC112722.1 | 2 | -0.280445052 | 0.088 | 0.184 | 3.78E-26 | 1.39E-21 |
| LRPPRC | 2 | -0.332293777 | 0.75 | 0.794 | 3.98E-26 | 1.46E-21 |
| MCM6 | 2 | 0.268096933 | 0.563 | 0.418 | 4.11E-26 | 1.51E-21 |
| ZMYM1 | 2 | -0.291840564 | 0.15 | 0.265 | 4.75E-26 | 1.74E-21 |
| ZNF644 | 2 | -0.322947796 | 0.671 | 0.755 | 4.92E-26 | 1.8E-21 |
| GOPC | 2 | -0.253178887 | 0.062 | 0.152 | 5.14E-26 | 1.88E-21 |
| CTNNAL1 | 2 | 0.273842516 | 0.37 | 0.239 | 5.18E-26 | 1.9E-21 |
| SRRM2 | 2 | -0.355893496 | 0.556 | 0.653 | 5.28E-26 | 1.93E-21 |
| GXYLT1 | 2 | 0.313238235 | 0.395 | 0.269 | 6.35E-26 | 2.32E-21 |
| PER3 | 2 | 0.254042654 | 0.296 | 0.176 | 7E-26 | 2.56E-21 |
| KIRREL1 | 2 | 0.289087942 | 0.554 | 0.416 | 7.34E-26 | 2.69E-21 |
| PRRG1 | 2 | 0.332070914 | 0.463 | 0.336 | 8.18E-26 | 3E-21 |
| CSNK1D | 2 | -0.296109278 | 0.193 | 0.314 | 8.51E-26 | 3.12E-21 |
| TNIK | 2 | -0.488041502 | 0.337 | 0.365 | 1.04E-25 | 3.82E-21 |
| PLPPR5 | 2 | 0.282625963 | 0.238 | 0.13 | 1.17E-25 | 4.29E-21 |
| KAT6B | 2 | -0.329697765 | 0.659 | 0.748 | 1.42E-25 | 5.2E-21 |
| FBXW8 | 2 | -0.337469349 | 0.288 | 0.397 | 1.5E-25 | 5.48E-21 |
| UPF2 | 2 | -0.324387684 | 0.429 | 0.553 | 1.51E-25 | 5.54E-21 |
| TCAF1 | 2 | 0.310120469 | 0.352 | 0.241 | 1.56E-25 | 5.73E-21 |
| LARP4 | 2 | -0.345359939 | 0.405 | 0.513 | 1.77E-25 | 6.5E-21 |
| WASHC4 | 2 | -0.327444743 | 0.512 | 0.626 | 1.79E-25 | 6.55E-21 |
| DAPK1 | 2 | -0.373299979 | 0.691 | 0.685 | 2.04E-25 | 7.45E-21 |
| ZNF518A | 2 | -0.32274485 | 0.624 | 0.706 | 2.12E-25 | 7.77E-21 |
| USP9X | 2 | -0.327306356 | 0.815 | 0.864 | 2.68E-25 | 9.82E-21 |
| PPP2R2A | 2 | -0.36841754 | 0.586 | 0.683 | 2.73E-25 | 9.99E-21 |
| MYO3A | 2 | -0.674292885 | 0.172 | 0.182 | 2.8E-25 | 1.03E-20 |
| LARGE1 | 2 | -0.435375448 | 0.207 | 0.291 | 2.88E-25 | 1.05E-20 |
| AKAP8L | 2 | -0.321363692 | 0.235 | 0.352 | 3.05E-25 | 1.12E-20 |
| IDH1 | 2 | -0.34680862 | 0.133 | 0.239 | 3.11E-25 | 1.14E-20 |
| ZFX | 2 | -0.337498944 | 0.378 | 0.501 | 3.15E-25 | 1.15E-20 |
| AC104596.1 | 2 | -0.272599491 | 0.05 | 0.135 | 3.33E-25 | 1.22E-20 |
| HDX | 2 | 0.282561846 | 0.367 | 0.239 | 3.59E-25 | 1.31E-20 |
| NIPBL | 2 | 0.280145505 | 0.917 | 0.877 | 3.84E-25 | 1.41E-20 |
| MTM1 | 2 | 0.337023075 | 0.368 | 0.256 | 4.08E-25 | 1.49E-20 |
| DPYSL3 | 2 | -0.323367615 | 0.373 | 0.375 | 4.28E-25 | 1.57E-20 |
| EPC1 | 2 | -0.313582271 | 0.674 | 0.764 | 4.32E-25 | 1.58E-20 |
| CTPS2 | 2 | -0.318738966 | 0.469 | 0.568 | 4.96E-25 | 1.81E-20 |
| PAFAH1B1 | 2 | 0.27456835 | 0.847 | 0.773 | 5.13E-25 | 1.88E-20 |
| ATP10D | 2 | -0.250543347 | 0.069 | 0.159 | 5.65E-25 | 2.07E-20 |
| GPAM | 2 | -0.306287457 | 0.105 | 0.2 | 5.71E-25 | 2.09E-20 |
| DENND1A | 2 | -0.318570426 | 0.669 | 0.742 | 5.74E-25 | 2.1E-20 |
| ZBTB18 | 2 | -0.299221294 | 0.136 | 0.242 | 6.03E-25 | 2.21E-20 |
| THUMPD3-AS1 | 2 | -0.310102192 | 0.298 | 0.426 | 6.06E-25 | 2.22E-20 |
| GABBR1 | 2 | -0.299146619 | 0.182 | 0.294 | 6.6E-25 | 2.41E-20 |
| WIPF1 | 2 | -0.310868057 | 0.106 | 0.201 | 7.01E-25 | 2.57E-20 |
| AFF4 | 2 | -0.340570764 | 0.635 | 0.724 | 7.32E-25 | 2.68E-20 |
| FMR1 | 2 | 0.325285183 | 0.589 | 0.478 | 7.84E-25 | 2.87E-20 |
| WDR73 | 2 | -0.256528154 | 0.111 | 0.211 | 7.85E-25 | 2.87E-20 |
| ST3GAL4 | 2 | -0.306289603 | 0.128 | 0.229 | 8.01E-25 | 2.93E-20 |
| TNKS2 | 2 | -0.31627653 | 0.383 | 0.499 | 8.39E-25 | 3.07E-20 |
| PTCD3 | 2 | -0.29104656 | 0.312 | 0.444 | 1.14E-24 | 4.17E-20 |
| TMEM161B-AS1 | 2 | -0.422804817 | 0.586 | 0.679 | 1.14E-24 | 4.18E-20 |
| CARNMT1 | 2 | -0.294324402 | 0.178 | 0.29 | 1.18E-24 | 4.34E-20 |
| DHX32 | 2 | -0.294714273 | 0.173 | 0.289 | 1.46E-24 | 5.35E-20 |
| CD47 | 2 | -0.33913064 | 0.313 | 0.426 | 1.5E-24 | 5.48E-20 |
| NARS2 | 2 | -0.342044567 | 0.382 | 0.501 | 1.53E-24 | 5.6E-20 |
| XKR9 | 2 | 0.25921009 | 0.204 | 0.109 | 1.53E-24 | 5.6E-20 |
| FAF1 | 2 | 0.258537027 | 0.866 | 0.82 | 1.54E-24 | 5.64E-20 |

| | | | | | | |
|---|---|---|---|---|---|---|
| USP49 | 2 | 0.299434965 | 0.393 | 0.272 | 1.56E-24 | 5.71E-20 |
| SLF1 | 2 | 0.280444428 | 0.576 | 0.441 | 1.69E-24 | 6.18E-20 |
| ETNK1 | 2 | 0.254458942 | 0.876 | 0.796 | 2.04E-24 | 7.48E-20 |
| PAK3 | 2 | -0.360661765 | 0.901 | 0.861 | 2.05E-24 | 7.52E-20 |
| OSBPL10 | 2 | -0.453232767 | 0.36 | 0.39 | 2.2E-24 | 8.04E-20 |
| VAMP4 | 2 | -0.285800556 | 0.141 | 0.241 | 2.2E-24 | 8.07E-20 |
| LUC7L | 2 | -0.321120769 | 0.515 | 0.624 | 2.22E-24 | 8.14E-20 |
| DOCK6 | 2 | -0.290910606 | 0.101 | 0.201 | 2.26E-24 | 8.26E-20 |
| MYEF2 | 2 | 0.262888109 | 0.618 | 0.482 | 2.32E-24 | 8.48E-20 |
| EIF3E | 2 | -0.334178599 | 0.458 | 0.56 | 2.35E-24 | 8.62E-20 |
| SLC4A5 | 2 | -0.284402563 | 0.04 | 0.111 | 2.48E-24 | 9.07E-20 |
| LCOR | 2 | 0.253370818 | 0.84 | 0.786 | 2.69E-24 | 9.84E-20 |
| SFPQ | 2 | -0.296621009 | 0.679 | 0.749 | 2.75E-24 | 1.01E-19 |
| DSCAM | 2 | -0.6749191 | 0.054 | 0.132 | 2.83E-24 | 1.04E-19 |
| IFT81 | 2 | -0.320995031 | 0.262 | 0.385 | 2.95E-24 | 1.08E-19 |
| TRMT9B | 2 | -0.471344522 | 0.187 | 0.232 | 2.96E-24 | 1.08E-19 |
| INTS3 | 2 | -0.270859303 | 0.165 | 0.275 | 3.29E-24 | 1.21E-19 |
| FUS | 2 | -0.309401294 | 0.709 | 0.789 | 3.3E-24 | 1.21E-19 |
| DPY19L4 | 2 | -0.298572117 | 0.202 | 0.32 | 3.43E-24 | 1.25E-19 |
| PLEKHH2 | 2 | -0.33778626 | 0.098 | 0.187 | 3.54E-24 | 1.3E-19 |
| LIMA1 | 2 | -0.263874671 | 0.078 | 0.169 | 4.21E-24 | 1.54E-19 |
| UQCRC2 | 2 | -0.279878411 | 0.253 | 0.379 | 4.24E-24 | 1.55E-19 |
| TFCP2 | 2 | -0.324730753 | 0.491 | 0.592 | 4.45E-24 | 1.63E-19 |
| PIP4K2A | 2 | 0.298671499 | 0.689 | 0.567 | 4.45E-24 | 1.63E-19 |
| TPR | 2 | -0.336320797 | 0.609 | 0.703 | 4.46E-24 | 1.63E-19 |
| CHST15 | 2 | 0.254992538 | 0.2 | 0.104 | 4.57E-24 | 1.67E-19 |
| ST6GAL1 | 2 | -0.293209793 | 0.404 | 0.363 | 4.85E-24 | 1.78E-19 |
| TMOD3 | 2 | -0.32689522 | 0.251 | 0.359 | 5.26E-24 | 1.93E-19 |
| EYA2 | 2 | -0.327563048 | 0.063 | 0.151 | 5.37E-24 | 1.97E-19 |
| E2F5 | 2 | -0.265765999 | 0.111 | 0.214 | 5.45E-24 | 2E-19 |
| SCAF11 | 2 | -0.301909985 | 0.572 | 0.667 | 5.72E-24 | 2.1E-19 |
| SKIL | 2 | -0.414831627 | 0.583 | 0.627 | 6.52E-24 | 2.39E-19 |
| DTD1 | 2 | 0.298975192 | 0.37 | 0.25 | 6.62E-24 | 2.42E-19 |
| STX18-AS1 | 2 | -0.344961062 | 0.282 | 0.404 | 6.9E-24 | 2.53E-19 |
| ZDHHC13 | 2 | 0.278823412 | 0.349 | 0.23 | 7.6E-24 | 2.78E-19 |
| SIN3B | 2 | -0.271063694 | 0.143 | 0.25 | 7.78E-24 | 2.85E-19 |
| KIAA1109 | 2 | -0.306543418 | 0.638 | 0.721 | 8.21E-24 | 3E-19 |
| LNPK | 2 | 0.318027967 | 0.541 | 0.413 | 8.73E-24 | 3.2E-19 |
| DNMBP | 2 | -0.254548826 | 0.112 | 0.213 | 9.51E-24 | 3.48E-19 |
| ATP2B1 | 2 | -0.438393763 | 0.353 | 0.446 | 9.8E-24 | 3.59E-19 |
| FIGN | 2 | -0.570576862 | 0.263 | 0.354 | 1.07E-23 | 3.92E-19 |
| AGO3 | 2 | -0.322199963 | 0.626 | 0.715 | 1.08E-23 | 3.96E-19 |
| TRIT1 | 2 | -0.279123338 | 0.183 | 0.297 | 1.09E-23 | 3.99E-19 |
| LMNA | 2 | -0.300245317 | 0.176 | 0.281 | 1.17E-23 | 4.3E-19 |
| FER | 2 | -0.374457158 | 0.809 | 0.816 | 1.19E-23 | 4.37E-19 |
| HOMER2 | 2 | -0.332137746 | 0.234 | 0.329 | 1.25E-23 | 4.57E-19 |
| TMEM87A | 2 | -0.316207261 | 0.391 | 0.511 | 1.29E-23 | 4.71E-19 |
| MSI2 | 2 | -0.34285254 | 0.847 | 0.859 | 1.32E-23 | 4.85E-19 |
| PLEKHA7 | 2 | -0.386976554 | 0.129 | 0.208 | 1.56E-23 | 5.7E-19 |
| VEGFC | 2 | 0.343813538 | 0.31 | 0.193 | 1.71E-23 | 6.27E-19 |
| NEB | 2 | -0.508200676 | 0.066 | 0.143 | 1.77E-23 | 6.46E-19 |
| ANKRD50 | 2 | -0.368326055 | 0.231 | 0.334 | 1.77E-23 | 6.49E-19 |
| TSEN2 | 2 | -0.288411209 | 0.217 | 0.336 | 1.91E-23 | 7.01E-19 |
| RYK | 2 | -0.299412544 | 0.388 | 0.513 | 1.95E-23 | 7.16E-19 |
| TBC1D4 | 2 | 0.271292887 | 0.509 | 0.374 | 2E-23 | 7.33E-19 |
| TUBGCP3 | 2 | 0.289336728 | 0.567 | 0.447 | 2.02E-23 | 7.38E-19 |
| SMC3 | 2 | 0.287941879 | 0.507 | 0.379 | 2.02E-23 | 7.4E-19 |
| CYTH1 | 2 | 0.302619714 | 0.56 | 0.453 | 2.71E-23 | 9.91E-19 |

| | | | | | | |
|---|---|---|---|---|---|---|
| PALD1 | 2 | 0.278268557 | 0.283 | 0.174 | 2.77E-23 | 1.01E-18 |
| AC093817.2 | 2 | -0.617171439 | 0.064 | 0.114 | 2.97E-23 | 1.09E-18 |
| KIF26B | 2 | -0.515717606 | 0.525 | 0.509 | 3.17E-23 | 1.16E-18 |
| METTL24 | 2 | -0.265093742 | 0.045 | 0.117 | 3.45E-23 | 1.26E-18 |
| ARNTL | 2 | -0.274082362 | 0.083 | 0.163 | 3.52E-23 | 1.29E-18 |
| OTUD4 | 2 | -0.311288826 | 0.302 | 0.413 | 3.56E-23 | 1.3E-18 |
| MED13 | 2 | -0.271857431 | 0.738 | 0.806 | 4.21E-23 | 1.54E-18 |
| PRKACB | 2 | 0.330541911 | 0.68 | 0.576 | 4.37E-23 | 1.6E-18 |
| RHOBTB3 | 2 | -0.358791378 | 0.404 | 0.478 | 4.5E-23 | 1.65E-18 |
| PMEPA1 | 2 | -0.40303386 | 0.162 | 0.243 | 4.54E-23 | 1.66E-18 |
| ZXDC | 2 | -0.287545168 | 0.292 | 0.41 | 4.59E-23 | 1.68E-18 |
| CBFB | 2 | -0.339682633 | 0.429 | 0.534 | 4.69E-23 | 1.72E-18 |
| DENND4A | 2 | 0.270101563 | 0.688 | 0.563 | 5.15E-23 | 1.88E-18 |
| CTTN | 2 | -0.290303003 | 0.296 | 0.404 | 5.82E-23 | 2.13E-18 |
| BPTF | 2 | -0.256866675 | 0.822 | 0.854 | 6.27E-23 | 2.29E-18 |
| HSP90AA1 | 2 | 0.519380163 | 0.558 | 0.432 | 6.52E-23 | 2.39E-18 |
| ZNF532 | 2 | -0.323058696 | 0.656 | 0.68 | 6.79E-23 | 2.49E-18 |
| COL25A1 | 2 | -0.578657483 | 0.4 | 0.389 | 6.98E-23 | 2.56E-18 |
| FGFR1OP2 | 2 | -0.278877764 | 0.191 | 0.3 | 7.31E-23 | 2.68E-18 |
| PRPF38B | 2 | -0.278831717 | 0.225 | 0.343 | 7.63E-23 | 2.79E-18 |
| ATP5F1C | 2 | -0.268788237 | 0.22 | 0.337 | 8.18E-23 | 3E-18 |
| SP4 | 2 | 0.303484899 | 0.378 | 0.265 | 1.07E-22 | 3.92E-18 |
| SSBP3 | 2 | -0.392028405 | 0.495 | 0.568 | 1.16E-22 | 4.25E-18 |
| DNAJC24 | 2 | -0.253653984 | 0.124 | 0.226 | 1.22E-22 | 4.48E-18 |
| UBAP2L | 2 | -0.300476255 | 0.346 | 0.454 | 1.25E-22 | 4.58E-18 |
| RERG | 2 | -0.472180202 | 0.131 | 0.209 | 1.34E-22 | 4.9E-18 |
| PRKRIP1 | 2 | -0.300056219 | 0.325 | 0.442 | 1.39E-22 | 5.08E-18 |
| DHTKD1 | 2 | -0.293876032 | 0.161 | 0.264 | 1.6E-22 | 5.85E-18 |
| ANAPC16 | 2 | -0.25523176 | 0.185 | 0.3 | 1.62E-22 | 5.94E-18 |
| ZMYM4 | 2 | -0.297397744 | 0.807 | 0.857 | 1.66E-22 | 6.06E-18 |
| SCN9A | 2 | -0.427463573 | 0.085 | 0.169 | 1.79E-22 | 6.56E-18 |
| STX16 | 2 | -0.284417892 | 0.371 | 0.49 | 1.94E-22 | 7.12E-18 |
| ELMOD1 | 2 | -0.38860049 | 0.379 | 0.403 | 2.2E-22 | 8.05E-18 |
| C11orf58 | 2 | -0.290623647 | 0.304 | 0.416 | 2.21E-22 | 8.08E-18 |
| SMAD9 | 2 | 0.251178101 | 0.501 | 0.369 | 2.22E-22 | 8.11E-18 |
| KIFAP3 | 2 | 0.295558295 | 0.505 | 0.386 | 2.33E-22 | 8.55E-18 |
| SIK2 | 2 | 0.263591851 | 0.649 | 0.524 | 2.52E-22 | 9.23E-18 |
| SOAT1 | 2 | -0.255183193 | 0.088 | 0.173 | 2.64E-22 | 9.67E-18 |
| DGLUCY | 2 | -0.296801342 | 0.172 | 0.279 | 2.86E-22 | 1.05E-17 |
| CLK4 | 2 | -0.26535236 | 0.188 | 0.297 | 3.29E-22 | 1.2E-17 |
| PDS5A | 2 | -0.291572791 | 0.764 | 0.827 | 3.64E-22 | 1.33E-17 |
| TTN | 2 | -0.638444152 | 0.098 | 0.164 | 4.3E-22 | 1.57E-17 |
| TEX9 | 2 | -0.301094775 | 0.124 | 0.221 | 4.42E-22 | 1.62E-17 |
| LRRC40 | 2 | 0.283660335 | 0.521 | 0.416 | 4.6E-22 | 1.68E-17 |
| GLCCI1 | 2 | -0.435411648 | 0.578 | 0.578 | 4.63E-22 | 1.69E-17 |
| LINC02018 | 2 | -0.270770489 | 0.064 | 0.138 | 5.09E-22 | 1.86E-17 |
| CLHC1 | 2 | -0.304440906 | 0.221 | 0.324 | 5.12E-22 | 1.87E-17 |
| FARS2 | 2 | -0.314837961 | 0.54 | 0.639 | 5.15E-22 | 1.88E-17 |
| TOP3A | 2 | 0.262835621 | 0.449 | 0.325 | 5.5E-22 | 2.01E-17 |
| TANC2 | 2 | -0.331496184 | 0.732 | 0.711 | 7.2E-22 | 2.64E-17 |
| MGST1 | 2 | -0.323758092 | 0.082 | 0.159 | 7.21E-22 | 2.64E-17 |
| ANO10 | 2 | 0.329232287 | 0.566 | 0.451 | 7.56E-22 | 2.77E-17 |
| PTPRG | 2 | 0.316696529 | 0.888 | 0.83 | 7.76E-22 | 2.84E-17 |
| JAK1 | 2 | -0.341787265 | 0.508 | 0.603 | 7.88E-22 | 2.89E-17 |
| SOS1 | 2 | -0.362257801 | 0.75 | 0.801 | 8.16E-22 | 2.99E-17 |
| SH3PXD2A | 2 | -0.312478629 | 0.295 | 0.382 | 9.16E-22 | 3.35E-17 |
| DSE | 2 | -0.319452741 | 0.247 | 0.366 | 9.18E-22 | 3.36E-17 |
| GPD2 | 2 | 0.281278935 | 0.524 | 0.397 | 9.73E-22 | 3.56E-17 |

| | | | | | | |
|---|---|---|---|---|---|---|
| STAT5B | 2 | 0.281446277 | 0.536 | 0.419 | 9.79E-22 | 3.58E-17 |
| MAU2 | 2 | -0.277022445 | 0.218 | 0.331 | 9.83E-22 | 3.6E-17 |
| MAPK8 | 2 | 0.269414015 | 0.811 | 0.737 | 1.01E-21 | 3.71E-17 |
| SMG1 | 2 | -0.29778063 | 0.731 | 0.798 | 1.03E-21 | 3.78E-17 |
| COMMD1 | 2 | -0.309149864 | 0.391 | 0.503 | 1.06E-21 | 3.87E-17 |
| LMCD1-AS1 | 2 | -0.39385435 | 0.218 | 0.326 | 1.19E-21 | 4.35E-17 |
| DENR | 2 | -0.283023574 | 0.222 | 0.332 | 1.23E-21 | 4.49E-17 |
| ACACA | 2 | 0.261939463 | 0.818 | 0.717 | 1.24E-21 | 4.53E-17 |
| PEX1 | 2 | -0.331848532 | 0.33 | 0.418 | 1.26E-21 | 4.63E-17 |
| ATG7 | 2 | -0.308867795 | 0.406 | 0.521 | 1.27E-21 | 4.64E-17 |
| TDRD7 | 2 | -0.273083833 | 0.128 | 0.212 | 1.45E-21 | 5.31E-17 |
| TTC7B | 2 | 0.279327595 | 0.492 | 0.372 | 1.65E-21 | 6.03E-17 |
| CKMT2-AS1 | 2 | -0.297095159 | 0.302 | 0.419 | 1.72E-21 | 6.28E-17 |
| KIAA0319L | 2 | -0.296548559 | 0.483 | 0.579 | 1.74E-21 | 6.36E-17 |
| PABPC1 | 2 | -0.302114957 | 0.594 | 0.672 | 1.76E-21 | 6.43E-17 |
| MARCH1 | 2 | -0.76650546 | 0.227 | 0.315 | 1.8E-21 | 6.59E-17 |
| TTC14 | 2 | -0.276149554 | 0.483 | 0.586 | 1.86E-21 | 6.82E-17 |
| IWS1 | 2 | -0.29095319 | 0.259 | 0.369 | 1.94E-21 | 7.08E-17 |
| IGSF11 | 2 | -0.344105552 | 0.083 | 0.163 | 1.96E-21 | 7.16E-17 |
| GARS-DT | 2 | -0.277877461 | 0.239 | 0.345 | 2.12E-21 | 7.78E-17 |
| SLC38A4 | 2 | -0.257259317 | 0.2 | 0.169 | 2.28E-21 | 8.33E-17 |
| ANKIB1 | 2 | -0.388799541 | 0.775 | 0.813 | 2.42E-21 | 8.85E-17 |
| SYMPK | 2 | -0.267080325 | 0.202 | 0.308 | 2.51E-21 | 9.19E-17 |
| EPS15L1 | 2 | -0.266458386 | 0.323 | 0.446 | 2.61E-21 | 9.57E-17 |
| TTC28-AS1 | 2 | -0.288697227 | 0.163 | 0.26 | 2.65E-21 | 9.7E-17 |
| APOLD1 | 2 | 0.311837282 | 0.217 | 0.124 | 2.7E-21 | 9.89E-17 |
| ZNF26 | 2 | -0.258593826 | 0.211 | 0.324 | 2.9E-21 | 1.06E-16 |
| SLC25A26 | 2 | -0.261242305 | 0.237 | 0.358 | 2.92E-21 | 1.07E-16 |
| ATXN7L1 | 2 | -0.307405521 | 0.487 | 0.593 | 3.08E-21 | 1.13E-16 |
| KCNJ3 | 2 | -0.542662495 | 0.096 | 0.152 | 3.12E-21 | 1.14E-16 |
| RRAS2 | 2 | 0.288661062 | 0.372 | 0.258 | 3.85E-21 | 1.41E-16 |
| TTC17 | 2 | -0.264614298 | 0.74 | 0.796 | 4.16E-21 | 1.52E-16 |
| CTNNA3 | 2 | -0.588210737 | 0.112 | 0.198 | 4.21E-21 | 1.54E-16 |
| ZFC3H1 | 2 | -0.298807203 | 0.503 | 0.603 | 4.68E-21 | 1.71E-16 |
| TCAIM | 2 | -0.284392171 | 0.204 | 0.307 | 5.17E-21 | 1.89E-16 |
| AP3S1 | 2 | 0.267168193 | 0.417 | 0.299 | 5.44E-21 | 1.99E-16 |
| RSPRY1 | 2 | -0.288927469 | 0.278 | 0.383 | 5.49E-21 | 2.01E-16 |
| AK9 | 2 | -0.262315651 | 0.083 | 0.162 | 6.62E-21 | 2.42E-16 |
| SUCLG2-AS1 | 2 | -0.260697354 | 0.086 | 0.174 | 7.25E-21 | 2.65E-16 |
| ASXL3 | 2 | 0.301590353 | 0.582 | 0.455 | 7.43E-21 | 2.72E-16 |
| FSTL1 | 2 | -0.29393134 | 0.222 | 0.325 | 8.42E-21 | 3.08E-16 |
| LAMB1 | 2 | -0.450924781 | 0.356 | 0.395 | 8.44E-21 | 3.09E-16 |
| AL365295.1 | 2 | -0.276725449 | 0.04 | 0.101 | 8.57E-21 | 3.14E-16 |
| FOCAD | 2 | -0.299377604 | 0.661 | 0.724 | 8.64E-21 | 3.16E-16 |
| ARHGEF9 | 2 | -0.281856496 | 0.186 | 0.279 | 9.11E-21 | 3.33E-16 |
| TUG1 | 2 | -0.26823179 | 0.215 | 0.321 | 9.68E-21 | 3.54E-16 |
| BMPR2 | 2 | -0.32833368 | 0.798 | 0.831 | 9.76E-21 | 3.57E-16 |
| SLC16A10 | 2 | -0.316631307 | 0.247 | 0.222 | 1.1E-20 | 4.03E-16 |
| GOLM1 | 2 | 0.267062784 | 0.439 | 0.319 | 1.13E-20 | 4.15E-16 |
| RAP1GDS1 | 2 | -0.307992816 | 0.585 | 0.687 | 1.29E-20 | 4.73E-16 |
| TMTC1 | 2 | -0.409939101 | 0.08 | 0.15 | 1.37E-20 | 5.02E-16 |
| SQSTM1 | 2 | -0.330507432 | 0.157 | 0.251 | 1.41E-20 | 5.17E-16 |
| MBTPS1 | 2 | -0.263368292 | 0.258 | 0.362 | 1.43E-20 | 5.23E-16 |
| CLCN3 | 2 | -0.312032888 | 0.442 | 0.539 | 1.55E-20 | 5.66E-16 |
| LENG8 | 2 | -0.258025838 | 0.209 | 0.318 | 1.77E-20 | 6.48E-16 |
| KATNAL2 | 2 | -0.252557139 | 0.115 | 0.206 | 1.8E-20 | 6.59E-16 |
| DYNC1LI1 | 2 | 0.298675629 | 0.436 | 0.324 | 1.85E-20 | 6.76E-16 |
| USP53 | 2 | -0.276955193 | 0.178 | 0.287 | 1.86E-20 | 6.82E-16 |

| | | | | | | |
|---|---|---|---|---|---|---|
| ARHGAP5 | 2 | -0.320026591 | 0.437 | 0.541 | 2.1E-20 | 7.7E-16 |
| RAB5A | 2 | -0.293753578 | 0.282 | 0.393 | 2.13E-20 | 7.79E-16 |
| FNDC3A | 2 | 0.259822246 | 0.853 | 0.792 | 2.31E-20 | 8.46E-16 |
| TFB1M | 2 | -0.289964504 | 0.311 | 0.423 | 2.35E-20 | 8.6E-16 |
| FBXO17 | 2 | -0.2940155 | 0.158 | 0.254 | 2.4E-20 | 8.8E-16 |
| ADARB1 | 2 | -0.411790423 | 0.215 | 0.302 | 2.41E-20 | 8.83E-16 |
| TYW1 | 2 | -0.279749854 | 0.36 | 0.474 | 2.45E-20 | 8.96E-16 |
| AC009120.2 | 2 | -0.299703309 | 0.228 | 0.327 | 2.64E-20 | 9.68E-16 |
| ANKRD26 | 2 | -0.302342216 | 0.535 | 0.631 | 2.79E-20 | 1.02E-15 |
| NCOA1 | 2 | -0.336740219 | 0.673 | 0.692 | 2.86E-20 | 1.05E-15 |
| LINC01004 | 2 | -0.281580927 | 0.4 | 0.515 | 2.89E-20 | 1.06E-15 |
| C21orf58 | 2 | 0.252769923 | 0.335 | 0.223 | 2.92E-20 | 1.07E-15 |
| XPO6 | 2 | 0.250371218 | 0.613 | 0.5 | 2.93E-20 | 1.07E-15 |
| CAPN7 | 2 | -0.285859081 | 0.407 | 0.5 | 2.97E-20 | 1.09E-15 |
| MIA3 | 2 | -0.266759927 | 0.217 | 0.322 | 3.04E-20 | 1.11E-15 |
| PRPF40A | 2 | -0.295598158 | 0.59 | 0.652 | 3.18E-20 | 1.16E-15 |
| PUM3 | 2 | -0.323909808 | 0.493 | 0.585 | 3.22E-20 | 1.18E-15 |
| AMMECR1 | 2 | -0.310888311 | 0.251 | 0.356 | 3.32E-20 | 1.22E-15 |
| TBCK | 2 | -0.276683465 | 0.661 | 0.743 | 3.43E-20 | 1.26E-15 |
| BABAM2 | 2 | -0.277088614 | 0.483 | 0.6 | 3.64E-20 | 1.33E-15 |
| AP3S2 | 2 | -0.255021809 | 0.228 | 0.341 | 4.48E-20 | 1.64E-15 |
| ZFAND5 | 2 | -0.362331809 | 0.391 | 0.461 | 4.78E-20 | 1.75E-15 |
| SFSWAP | 2 | -0.276615738 | 0.404 | 0.512 | 6.64E-20 | 2.43E-15 |
| SMOC1 | 2 | -0.251922139 | 0.159 | 0.181 | 6.8E-20 | 2.49E-15 |
| SUB1 | 2 | -0.283564572 | 0.266 | 0.369 | 7.35E-20 | 2.69E-15 |
| RICTOR | 2 | -0.290072621 | 0.669 | 0.725 | 7.58E-20 | 2.77E-15 |
| JMY | 2 | -0.270634034 | 0.254 | 0.369 | 7.76E-20 | 2.84E-15 |
| HOMER1 | 2 | -0.374039244 | 0.437 | 0.507 | 7.94E-20 | 2.91E-15 |
| HSPA4 | 2 | -0.276742407 | 0.225 | 0.334 | 8.53E-20 | 3.12E-15 |
| COX10-AS1 | 2 | -0.297930639 | 0.228 | 0.333 | 8.53E-20 | 3.12E-15 |
| PNPLA8 | 2 | -0.279251351 | 0.199 | 0.3 | 8.6E-20 | 3.15E-15 |
| TMEM14B | 2 | -0.318798706 | 0.382 | 0.476 | 9.15E-20 | 3.35E-15 |
| TRPC1 | 2 | -0.353586809 | 0.511 | 0.552 | 9.59E-20 | 3.51E-15 |
| AL445123.2 | 2 | -0.357433927 | 0.172 | 0.25 | 9.63E-20 | 3.53E-15 |
| HOOK3 | 2 | -0.310431199 | 0.467 | 0.555 | 9.71E-20 | 3.56E-15 |
| AC021231.1 | 2 | -0.25822628 | 0.069 | 0.148 | 1.07E-19 | 3.93E-15 |
| SNX7 | 2 | -0.307479206 | 0.118 | 0.195 | 1.14E-19 | 4.16E-15 |
| GPR107 | 2 | -0.263526803 | 0.251 | 0.348 | 1.19E-19 | 4.36E-15 |
| ARL5A | 2 | -0.33574984 | 0.362 | 0.454 | 1.21E-19 | 4.44E-15 |
| ADGRD1 | 2 | -0.382667788 | 0.186 | 0.27 | 1.27E-19 | 4.65E-15 |
| TFG | 2 | -0.282316909 | 0.373 | 0.476 | 1.34E-19 | 4.89E-15 |
| ST13 | 2 | -0.278606589 | 0.233 | 0.332 | 1.43E-19 | 5.25E-15 |
| ZFYVE16 | 2 | -0.358667592 | 0.521 | 0.584 | 1.51E-19 | 5.53E-15 |
| CSNK2A1 | 2 | -0.28427368 | 0.389 | 0.479 | 1.59E-19 | 5.83E-15 |
| FARP2 | 2 | -0.272698296 | 0.238 | 0.347 | 1.62E-19 | 5.93E-15 |
| MAPKAP1 | 2 | -0.280585986 | 0.443 | 0.553 | 1.67E-19 | 6.12E-15 |
| KDM4B | 2 | -0.349066961 | 0.411 | 0.51 | 1.68E-19 | 6.16E-15 |
| SECISBP2L | 2 | -0.312776439 | 0.37 | 0.47 | 1.87E-19 | 6.86E-15 |
| NDRG1 | 2 | -0.489058677 | 0.06 | 0.123 | 2.12E-19 | 7.75E-15 |
| NISCH | 2 | -0.270092722 | 0.242 | 0.35 | 2.12E-19 | 7.76E-15 |
| AC107223.1 | 2 | -0.283456971 | 0.085 | 0.164 | 2.18E-19 | 7.99E-15 |
| MEF2C | 2 | -0.386291627 | 0.268 | 0.372 | 2.28E-19 | 8.35E-15 |
| TAX1BP1 | 2 | -0.283896368 | 0.372 | 0.479 | 2.36E-19 | 8.63E-15 |
| LINC00882 | 2 | -0.314836755 | 0.054 | 0.112 | 2.41E-19 | 8.84E-15 |
| ALS2 | 2 | -0.265004116 | 0.285 | 0.394 | 2.43E-19 | 8.9E-15 |
| HIF1A | 2 | -0.320054683 | 0.436 | 0.534 | 2.48E-19 | 9.08E-15 |
| UBE4B | 2 | -0.312921876 | 0.491 | 0.564 | 2.63E-19 | 9.63E-15 |
| STAM2 | 2 | -0.261673189 | 0.177 | 0.277 | 2.7E-19 | 9.87E-15 |

| | | | | | | |
|---|---|---|---|---|---|---|
| PWWP3A | 2 | -0.273982834 | 0.269 | 0.377 | 2.78E-19 | 1.02E-14 |
| PHYKPL | 2 | -0.258522312 | 0.207 | 0.313 | 3.04E-19 | 1.11E-14 |
| ANKDD1A | 2 | -0.254896837 | 0.095 | 0.176 | 3.06E-19 | 1.12E-14 |
| GNB4 | 2 | -0.294736372 | 0.119 | 0.198 | 3.14E-19 | 1.15E-14 |
| SDHAF3 | 2 | -0.263858129 | 0.173 | 0.275 | 3.25E-19 | 1.19E-14 |
| TMEM164 | 2 | 0.287823373 | 0.576 | 0.471 | 3.44E-19 | 1.26E-14 |
| CUL4B | 2 | -0.323651963 | 0.359 | 0.44 | 3.81E-19 | 1.4E-14 |
| BRWD1 | 2 | -0.262560583 | 0.793 | 0.832 | 4.38E-19 | 1.6E-14 |
| LRP12 | 2 | 0.262973052 | 0.505 | 0.394 | 4.44E-19 | 1.63E-14 |
| GMDS | 2 | -0.509195081 | 0.393 | 0.453 | 5.02E-19 | 1.84E-14 |
| WASF2 | 2 | -0.283494014 | 0.251 | 0.352 | 6.04E-19 | 2.21E-14 |
| FANCC | 2 | 0.295903141 | 0.586 | 0.505 | 6.68E-19 | 2.45E-14 |
| AL589693.1 | 2 | 0.451761914 | 0.155 | 0.081 | 6.98E-19 | 2.55E-14 |
| AQP4-AS1 | 2 | -0.250476446 | 0.055 | 0.123 | 7.07E-19 | 2.59E-14 |
| GUCY1A1 | 2 | -0.448191788 | 0.208 | 0.271 | 8.45E-19 | 3.09E-14 |
| RCOR3 | 2 | -0.266147253 | 0.509 | 0.607 | 9.24E-19 | 3.38E-14 |
| TASP1 | 2 | -0.288609381 | 0.444 | 0.534 | 9.44E-19 | 3.46E-14 |
| UBE2E1 | 2 | -0.293459836 | 0.539 | 0.628 | 1.02E-18 | 3.74E-14 |
| GRM8 | 2 | -0.63341627 | 0.192 | 0.143 | 1.07E-18 | 3.94E-14 |
| ZNF326 | 2 | -0.274420836 | 0.375 | 0.47 | 1.09E-18 | 3.97E-14 |
| CUL3 | 2 | -0.267063954 | 0.621 | 0.702 | 1.13E-18 | 4.13E-14 |
| MEIS2 | 2 | -0.432650253 | 0.53 | 0.541 | 1.18E-18 | 4.3E-14 |
| MAP3K2 | 2 | -0.300738605 | 0.509 | 0.608 | 1.24E-18 | 4.52E-14 |
| ZNRF3 | 2 | -0.338640797 | 0.322 | 0.421 | 1.27E-18 | 4.66E-14 |
| KDM5A | 2 | -0.276592122 | 0.442 | 0.55 | 1.29E-18 | 4.71E-14 |
| MTREX | 2 | -0.295444635 | 0.515 | 0.603 | 1.47E-18 | 5.39E-14 |
| EIF2S3 | 2 | -0.275678711 | 0.428 | 0.518 | 1.5E-18 | 5.5E-14 |
| ARHGAP8 | 2 | -0.319149196 | 0.179 | 0.22 | 1.51E-18 | 5.51E-14 |
| KHDC4 | 2 | -0.255851055 | 0.309 | 0.419 | 1.54E-18 | 5.63E-14 |
| ZBTB44 | 2 | -0.265305344 | 0.329 | 0.442 | 1.62E-18 | 5.94E-14 |
| GPM6B | 2 | -0.381054172 | 0.618 | 0.637 | 1.65E-18 | 6.03E-14 |
| SAMD11 | 2 | -0.262305083 | 0.079 | 0.151 | 1.66E-18 | 6.09E-14 |
| ID1 | 2 | -0.269485172 | 0.075 | 0.137 | 1.67E-18 | 6.11E-14 |
| SRRM1 | 2 | -0.292368968 | 0.519 | 0.607 | 1.8E-18 | 6.58E-14 |
| CHD1 | 2 | -0.318203354 | 0.657 | 0.716 | 1.95E-18 | 7.14E-14 |
| ANTXR2 | 2 | -0.303055874 | 0.173 | 0.265 | 2.03E-18 | 7.43E-14 |
| CYFIP1 | 2 | -0.275508699 | 0.275 | 0.378 | 2.07E-18 | 7.58E-14 |
| ORC5 | 2 | -0.268472353 | 0.216 | 0.319 | 2.18E-18 | 7.99E-14 |
| L3MBTL3 | 2 | -0.261282376 | 0.098 | 0.176 | 2.42E-18 | 8.86E-14 |
| RC3H2 | 2 | -0.280620723 | 0.425 | 0.526 | 2.73E-18 | 9.98E-14 |
| MYSM1 | 2 | -0.268366509 | 0.409 | 0.518 | 2.73E-18 | 9.98E-14 |
| CALN1 | 2 | -0.326577481 | 0.044 | 0.103 | 2.82E-18 | 1.03E-13 |
| MINDY3 | 2 | -0.263326519 | 0.338 | 0.432 | 2.92E-18 | 1.07E-13 |
| SESN1 | 2 | 0.280397204 | 0.58 | 0.467 | 3.13E-18 | 1.15E-13 |
| DHX36 | 2 | -0.271758516 | 0.623 | 0.708 | 3.16E-18 | 1.16E-13 |
| PTK7 | 2 | -0.305471136 | 0.322 | 0.4 | 3.5E-18 | 1.28E-13 |
| IREB2 | 2 | -0.259158057 | 0.441 | 0.546 | 3.82E-18 | 1.4E-13 |
| HIF1A-AS3 | 2 | -0.384453205 | 0.076 | 0.142 | 3.85E-18 | 1.41E-13 |
| PHLPP2 | 2 | -0.251381485 | 0.193 | 0.282 | 3.98E-18 | 1.46E-13 |
| JARID2 | 2 | -0.299197037 | 0.609 | 0.69 | 4.12E-18 | 1.51E-13 |
| GAS5 | 2 | -0.271561054 | 0.275 | 0.382 | 4.21E-18 | 1.54E-13 |
| PLPPR4 | 2 | -0.38799935 | 0.243 | 0.292 | 5.06E-18 | 1.85E-13 |
| ZFAS1 | 2 | -0.305463971 | 0.268 | 0.359 | 5.56E-18 | 2.04E-13 |
| LNPEP | 2 | -0.271594395 | 0.262 | 0.359 | 5.65E-18 | 2.07E-13 |
| XIAP | 2 | -0.272022943 | 0.368 | 0.476 | 5.74E-18 | 2.1E-13 |
| ME1 | 2 | -0.272426719 | 0.088 | 0.158 | 6.02E-18 | 2.2E-13 |
| PPP4R3A | 2 | -0.266479649 | 0.562 | 0.653 | 6.03E-18 | 2.21E-13 |
| CACNA2D1 | 2 | -0.442738466 | 0.918 | 0.84 | 6.99E-18 | 2.56E-13 |

| | | | | | | |
|---|---|---|---|---|---|---|
| USO1 | 2 | -0.281328998 | 0.377 | 0.46 | 7.1E-18 | 2.6E-13 |
| LINC01184 | 2 | -0.313576575 | 0.343 | 0.448 | 7.13E-18 | 2.61E-13 |
| NTNG1 | 2 | -0.404142146 | 0.45 | 0.482 | 7.23E-18 | 2.65E-13 |
| PIGN | 2 | -0.276928649 | 0.26 | 0.365 | 7.34E-18 | 2.69E-13 |
| ANKRD11 | 2 | -0.272754995 | 0.789 | 0.839 | 7.78E-18 | 2.85E-13 |
| ZNF804A | 2 | -0.619423644 | 0.246 | 0.247 | 7.81E-18 | 2.86E-13 |
| MIR4713HG | 2 | -0.287091041 | 0.057 | 0.125 | 8.1E-18 | 2.97E-13 |
| AL033504.1 | 2 | -0.409050255 | 0.585 | 0.577 | 8.28E-18 | 3.03E-13 |
| GCNT1 | 2 | -0.269742452 | 0.048 | 0.103 | 9.01E-18 | 3.3E-13 |
| RASGEF1B | 2 | -0.530172719 | 0.257 | 0.339 | 9.7E-18 | 3.55E-13 |
| AZIN1-AS1 | 2 | -0.272619752 | 0.173 | 0.261 | 9.7E-18 | 3.55E-13 |
| DPYD | 2 | -0.450803753 | 0.354 | 0.422 | 9.81E-18 | 3.59E-13 |
| POU2F3 | 2 | -0.278609244 | 0.276 | 0.329 | 1.04E-17 | 3.82E-13 |
| MRPS27 | 2 | -0.251643347 | 0.266 | 0.372 | 1.08E-17 | 3.97E-13 |
| MACF1 | 2 | -0.2659096 | 0.836 | 0.862 | 1.34E-17 | 4.89E-13 |
| DHX29 | 2 | -0.284997723 | 0.264 | 0.344 | 1.58E-17 | 5.79E-13 |
| CMC1 | 2 | -0.263433242 | 0.263 | 0.368 | 1.61E-17 | 5.9E-13 |
| KLHL2 | 2 | 0.250340141 | 0.394 | 0.286 | 1.61E-17 | 5.9E-13 |
| PTPN12 | 2 | -0.278405755 | 0.638 | 0.687 | 1.61E-17 | 5.9E-13 |
| MALT1 | 2 | -0.279349427 | 0.224 | 0.311 | 1.67E-17 | 6.13E-13 |
| EXPH5 | 2 | -0.320200622 | 0.098 | 0.171 | 1.72E-17 | 6.29E-13 |
| PMP22 | 2 | -0.278212742 | 0.131 | 0.209 | 1.86E-17 | 6.81E-13 |
| DHRSX | 2 | -0.320425164 | 0.602 | 0.665 | 1.87E-17 | 6.86E-13 |
| SAMD3 | 2 | -0.355170854 | 0.046 | 0.102 | 1.92E-17 | 7.04E-13 |
| TSHZ1 | 2 | 0.261741124 | 0.492 | 0.382 | 2.08E-17 | 7.62E-13 |
| PRICKLE2 | 2 | -0.329091558 | 0.289 | 0.351 | 2.54E-17 | 9.31E-13 |
| CA12 | 2 | -0.345335738 | 0.18 | 0.26 | 2.74E-17 | 1E-12 |
| LINC01768 | 2 | -0.280882339 | 0.168 | 0.182 | 2.79E-17 | 1.02E-12 |
| DDX6 | 2 | -0.263696688 | 0.383 | 0.485 | 2.85E-17 | 1.04E-12 |
| SPATA5 | 2 | 0.25695821 | 0.696 | 0.608 | 2.98E-17 | 1.09E-12 |
| SPTBN1 | 2 | -0.282262603 | 0.679 | 0.69 | 3.51E-17 | 1.28E-12 |
| CHST9 | 2 | -0.478511639 | 0.14 | 0.153 | 3.73E-17 | 1.36E-12 |
| PPP1R10 | 2 | -0.255275554 | 0.236 | 0.336 | 3.76E-17 | 1.37E-12 |
| FERMT1 | 2 | -0.329815537 | 0.137 | 0.212 | 3.84E-17 | 1.41E-12 |
| MDM4 | 2 | -0.255119139 | 0.486 | 0.587 | 3.85E-17 | 1.41E-12 |
| PDCD11 | 2 | -0.2551029 | 0.189 | 0.277 | 4.01E-17 | 1.47E-12 |
| SCN7A | 2 | -0.947259346 | 0.115 | 0.146 | 4.19E-17 | 1.53E-12 |
| STK24 | 2 | -0.279403125 | 0.395 | 0.49 | 4.49E-17 | 1.64E-12 |
| RAB31 | 2 | -0.287286152 | 0.183 | 0.276 | 4.64E-17 | 1.7E-12 |
| ADGRA3 | 2 | -0.288391274 | 0.642 | 0.69 | 4.94E-17 | 1.81E-12 |
| GREB1L | 2 | -0.369196427 | 0.116 | 0.093 | 5.09E-17 | 1.86E-12 |
| GK | 2 | -0.28851142 | 0.232 | 0.287 | 5.2E-17 | 1.9E-12 |
| PAPOLA | 2 | -0.260331246 | 0.573 | 0.65 | 5.7E-17 | 2.09E-12 |
| TRAM2 | 2 | -0.250993477 | 0.169 | 0.248 | 6.33E-17 | 2.32E-12 |
| KCNMB4 | 2 | -0.306207874 | 0.174 | 0.256 | 6.88E-17 | 2.52E-12 |
| GPCPD1 | 2 | -0.298668519 | 0.499 | 0.587 | 7.26E-17 | 2.66E-12 |
| CRNDE | 2 | -0.282681155 | 0.236 | 0.324 | 8.14E-17 | 2.98E-12 |
| WDR12 | 2 | -0.250856461 | 0.214 | 0.309 | 8.21E-17 | 3E-12 |
| SEM1 | 2 | 0.272898835 | 0.378 | 0.291 | 8.68E-17 | 3.18E-12 |
| SORCS2 | 2 | -0.358687342 | 0.197 | 0.267 | 8.7E-17 | 3.19E-12 |
| DDX21 | 2 | -0.272719383 | 0.215 | 0.306 | 9.17E-17 | 3.36E-12 |
| PLEKHA6 | 2 | -0.320480413 | 0.424 | 0.477 | 9.23E-17 | 3.38E-12 |
| AC025159.1 | 2 | -0.291591227 | 0.11 | 0.185 | 9.3E-17 | 3.4E-12 |
| ZNF608 | 2 | -0.339202864 | 0.582 | 0.649 | 9.83E-17 | 3.6E-12 |
| ANKRD36B | 2 | 0.251742421 | 0.57 | 0.465 | 9.83E-17 | 3.6E-12 |
| FAM171A1 | 2 | -0.281499253 | 0.522 | 0.572 | 1.06E-16 | 3.88E-12 |
| LINC01619 | 2 | 0.318951072 | 0.409 | 0.316 | 1.16E-16 | 4.24E-12 |
| INTS6 | 2 | -0.310786812 | 0.481 | 0.569 | 1.19E-16 | 4.37E-12 |

| | | | | | | |
|---|---|---|---|---|---|---|
| TESK2 | 2 | -0.282540631 | 0.126 | 0.197 | 1.22E-16 | 4.48E-12 |
| LINC01572 | 2 | -0.311187132 | 0.467 | 0.553 | 1.37E-16 | 5.01E-12 |
| SIPA1L3 | 2 | -0.467056287 | 0.521 | 0.526 | 1.41E-16 | 5.15E-12 |
| ARFIP1 | 2 | -0.261795579 | 0.415 | 0.513 | 1.42E-16 | 5.2E-12 |
| KLHL24 | 2 | -0.331654319 | 0.412 | 0.491 | 1.45E-16 | 5.3E-12 |
| ZNF423 | 2 | -0.408793922 | 0.36 | 0.421 | 1.52E-16 | 5.57E-12 |
| MEF2A | 2 | -0.303003664 | 0.776 | 0.815 | 1.61E-16 | 5.88E-12 |
| KCNK2 | 2 | -0.330099767 | 0.289 | 0.289 | 1.81E-16 | 6.63E-12 |
| MICU1 | 2 | -0.283442744 | 0.434 | 0.518 | 1.81E-16 | 6.64E-12 |
| WDR11-AS1 | 2 | -0.340974239 | 0.298 | 0.331 | 2.25E-16 | 8.24E-12 |
| FAM124A | 2 | -0.275272937 | 0.199 | 0.257 | 2.46E-16 | 8.99E-12 |
| SPAG16 | 2 | -0.323250006 | 0.218 | 0.295 | 2.47E-16 | 9.06E-12 |
| MAN1A1 | 2 | -0.45873579 | 0.108 | 0.138 | 2.7E-16 | 9.87E-12 |
| OSBPL11 | 2 | -0.287389438 | 0.292 | 0.344 | 3.03E-16 | 1.11E-11 |
| AZIN1 | 2 | -0.27114405 | 0.24 | 0.331 | 3.31E-16 | 1.21E-11 |
| KATNBL1 | 2 | -0.25358316 | 0.429 | 0.523 | 3.4E-16 | 1.25E-11 |
| PCF11 | 2 | -0.255473127 | 0.247 | 0.344 | 3.46E-16 | 1.27E-11 |
| AL163541.1 | 2 | 0.278714663 | 0.141 | 0.076 | 3.63E-16 | 1.33E-11 |
| FAR2 | 2 | -0.398272912 | 0.386 | 0.385 | 3.77E-16 | 1.38E-11 |
| TSHZ2 | 2 | -0.290394712 | 0.827 | 0.79 | 4.58E-16 | 1.68E-11 |
| LYST | 2 | -0.314318392 | 0.309 | 0.407 | 5.04E-16 | 1.85E-11 |
| ANKUB1 | 2 | -0.262951087 | 0.186 | 0.272 | 5.13E-16 | 1.88E-11 |
| AC068587.4 | 2 | -0.437107201 | 0.29 | 0.356 | 5.25E-16 | 1.92E-11 |
| SMAD6 | 2 | 0.256325316 | 0.381 | 0.278 | 5.5E-16 | 2.02E-11 |
| TMEM65 | 2 | -0.285291518 | 0.18 | 0.258 | 6.02E-16 | 2.2E-11 |
| MORC3 | 2 | -0.292721076 | 0.507 | 0.554 | 6.03E-16 | 2.21E-11 |
| MEF2C-AS1 | 2 | -0.276189435 | 0.173 | 0.264 | 6.34E-16 | 2.32E-11 |
| SGCZ | 2 | 0.48419634 | 0.601 | 0.542 | 6.38E-16 | 2.34E-11 |
| UFL1-AS1 | 2 | -0.370571323 | 0.058 | 0.117 | 7.5E-16 | 2.75E-11 |
| GRB10 | 2 | -0.356347786 | 0.324 | 0.4 | 7.99E-16 | 2.93E-11 |
| DZIP3 | 2 | -0.266808576 | 0.295 | 0.385 | 8.08E-16 | 2.96E-11 |
| TRAPPC8 | 2 | -0.263151564 | 0.52 | 0.584 | 8.94E-16 | 3.27E-11 |
| LPP | 2 | -0.311801314 | 0.707 | 0.737 | 1.1E-15 | 4.02E-11 |
| AC004917.1 | 2 | -0.290456905 | 0.051 | 0.109 | 1.14E-15 | 4.17E-11 |
| FAM53B | 2 | -0.250954368 | 0.234 | 0.325 | 1.16E-15 | 4.25E-11 |
| GRK5 | 2 | -0.303305266 | 0.263 | 0.346 | 1.26E-15 | 4.62E-11 |
| ANP32B | 2 | -0.264640122 | 0.293 | 0.373 | 1.3E-15 | 4.76E-11 |
| ICE2 | 2 | -0.25070797 | 0.327 | 0.424 | 1.41E-15 | 5.17E-11 |
| TMEM178A | 2 | -0.250527817 | 0.467 | 0.471 | 1.43E-15 | 5.25E-11 |
| SRGAP1 | 2 | -0.32946323 | 0.367 | 0.441 | 1.63E-15 | 5.95E-11 |
| AKNAD1 | 2 | -0.25989757 | 0.183 | 0.255 | 1.74E-15 | 6.37E-11 |
| FBXL7 | 2 | 0.269330945 | 0.892 | 0.823 | 1.85E-15 | 6.76E-11 |
| MEGF11 | 2 | -0.310470259 | 0.252 | 0.331 | 1.89E-15 | 6.93E-11 |
| PKN2-AS1 | 2 | -0.354916887 | 0.07 | 0.132 | 1.91E-15 | 6.98E-11 |
| ABCC5 | 2 | -0.26923022 | 0.323 | 0.388 | 2.56E-15 | 9.38E-11 |
| TRAPPC9 | 2 | -0.295129252 | 0.657 | 0.708 | 2.96E-15 | 1.08E-10 |
| SHLD2 | 2 | -0.253771623 | 0.392 | 0.492 | 4E-15 | 1.46E-10 |
| CDH12 | 2 | -0.539258681 | 0.353 | 0.409 | 4.19E-15 | 1.54E-10 |
| CAPRIN2 | 2 | -0.261356683 | 0.222 | 0.305 | 4.21E-15 | 1.54E-10 |
| MPDZ | 2 | -0.269653488 | 0.662 | 0.7 | 4.24E-15 | 1.55E-10 |
| URI1 | 2 | 0.258303186 | 0.799 | 0.74 | 4.49E-15 | 1.64E-10 |
| HIPK2 | 2 | -0.290966028 | 0.389 | 0.47 | 5.51E-15 | 2.02E-10 |
| LIN7A | 2 | -0.259916774 | 0.355 | 0.365 | 5.61E-15 | 2.05E-10 |
| NHS | 2 | -0.29189146 | 0.515 | 0.464 | 6.45E-15 | 2.36E-10 |
| PTEN | 2 | -0.340480093 | 0.657 | 0.712 | 6.46E-15 | 2.37E-10 |
| RSRC2 | 2 | -0.251039594 | 0.43 | 0.518 | 7E-15 | 2.56E-10 |
| MCU | 2 | -0.252075177 | 0.454 | 0.549 | 9.33E-15 | 3.42E-10 |
| CCDC7 | 2 | -0.253226641 | 0.266 | 0.364 | 1.05E-14 | 3.83E-10 |

| | | | | | | |
|---|---|---|---|---|---|---|
| ELOVL5 | 2 | 0.250600663 | 0.378 | 0.291 | 1.17E-14 | 4.3E-10 |
| FHDC1 | 2 | -0.259256615 | 0.118 | 0.178 | 1.2E-14 | 4.41E-10 |
| ENC1 | 2 | -0.283845287 | 0.197 | 0.22 | 1.38E-14 | 5.05E-10 |
| AC009133.1 | 2 | -0.271942534 | 0.07 | 0.122 | 1.47E-14 | 5.4E-10 |
| HDAC2-AS2 | 2 | -0.274260818 | 0.498 | 0.509 | 1.51E-14 | 5.52E-10 |
| ZNF280C | 2 | -0.256292156 | 0.322 | 0.419 | 1.55E-14 | 5.68E-10 |
| DISP1 | 2 | -0.294091578 | 0.299 | 0.389 | 1.96E-14 | 7.19E-10 |
| DPF3 | 2 | -0.264967004 | 0.227 | 0.28 | 2.16E-14 | 7.91E-10 |
| WARS2-AS1 | 2 | -0.2538543 | 0.323 | 0.422 | 2.25E-14 | 8.23E-10 |
| SND1 | 2 | -0.258150345 | 0.673 | 0.718 | 3.13E-14 | 1.15E-09 |
| CSRNP3 | 2 | -0.332993124 | 0.337 | 0.366 | 3.26E-14 | 1.19E-09 |
| AL591519.1 | 2 | -0.356479846 | 0.113 | 0.156 | 3.9E-14 | 1.43E-09 |
| SCUBE2 | 2 | -0.344644975 | 0.133 | 0.179 | 3.95E-14 | 1.44E-09 |
| PKM | 2 | -0.253981242 | 0.427 | 0.502 | 4.03E-14 | 1.47E-09 |
| VAV2 | 2 | -0.256515098 | 0.31 | 0.324 | 4.12E-14 | 1.51E-09 |
| MYO5A | 2 | -0.275044474 | 0.544 | 0.602 | 5.15E-14 | 1.88E-09 |
| ZNF280D | 2 | -0.252541276 | 0.246 | 0.326 | 6.16E-14 | 2.25E-09 |
| UBE2D2 | 2 | -0.275451616 | 0.537 | 0.603 | 6.75E-14 | 2.47E-09 |
| GSE1 | 2 | -0.352868237 | 0.709 | 0.734 | 6.95E-14 | 2.54E-09 |
| MAP2K5 | 2 | -0.25266823 | 0.435 | 0.516 | 7.07E-14 | 2.59E-09 |
| SLC39A11 | 2 | -0.327124483 | 0.445 | 0.524 | 8.44E-14 | 3.09E-09 |
| ENTPD1-AS1 | 2 | -0.26341123 | 0.286 | 0.369 | 8.59E-14 | 3.15E-09 |
| MAP3K4 | 2 | -0.29257246 | 0.454 | 0.51 | 9.32E-14 | 3.41E-09 |
| SNX8 | 2 | -0.263203733 | 0.22 | 0.302 | 1.05E-13 | 3.83E-09 |
| SLC2A3 | 2 | -0.414983096 | 0.091 | 0.149 | 1.23E-13 | 4.52E-09 |
| PIAS2 | 2 | -0.265833291 | 0.565 | 0.625 | 1.28E-13 | 4.68E-09 |
| TAFA2 | 2 | -0.272634916 | 0.062 | 0.117 | 1.34E-13 | 4.91E-09 |
| RGS12 | 2 | -0.253988241 | 0.402 | 0.488 | 1.4E-13 | 5.13E-09 |
| TBL1X | 2 | 0.279315583 | 0.776 | 0.774 | 1.49E-13 | 5.44E-09 |
| COPA | 2 | -0.251052483 | 0.418 | 0.497 | 1.96E-13 | 7.19E-09 |
| RPL41 | 2 | -0.269608512 | 0.467 | 0.553 | 2.15E-13 | 7.86E-09 |
| GCLM | 2 | -0.298355324 | 0.116 | 0.167 | 2.35E-13 | 8.61E-09 |
| STT3B | 2 | -0.280900724 | 0.647 | 0.703 | 2.39E-13 | 8.74E-09 |
| FSTL5 | 2 | -0.531013351 | 0.088 | 0.151 | 3.05E-13 | 1.12E-08 |
| NEK1 | 2 | -0.25186709 | 0.435 | 0.518 | 3.17E-13 | 1.16E-08 |
| ADAM9 | 2 | -0.262994928 | 0.292 | 0.372 | 3.35E-13 | 1.23E-08 |
| ANKRD12 | 2 | -0.260320394 | 0.609 | 0.669 | 3.5E-13 | 1.28E-08 |
| ID2 | 2 | -0.335802496 | 0.105 | 0.174 | 3.51E-13 | 1.29E-08 |
| FNIP1 | 2 | -0.285066061 | 0.58 | 0.632 | 3.61E-13 | 1.32E-08 |
| GSR | 2 | -0.26413285 | 0.247 | 0.307 | 3.62E-13 | 1.33E-08 |
| ACSL3 | 2 | -0.268354476 | 0.323 | 0.391 | 3.89E-13 | 1.42E-08 |
| HSPA9 | 2 | -0.250958702 | 0.285 | 0.361 | 4.38E-13 | 0.000000016 |
| ITGB1 | 2 | -0.270269441 | 0.502 | 0.577 | 4.44E-13 | 1.62E-08 |
| AC104041.1 | 2 | -0.350895852 | 0.462 | 0.493 | 4.47E-13 | 1.64E-08 |
| AGK | 2 | -0.253331717 | 0.317 | 0.391 | 4.77E-13 | 1.74E-08 |
| DCP2 | 2 | -0.262806683 | 0.527 | 0.589 | 5.95E-13 | 2.18E-08 |
| AL355499.1 | 2 | -0.608037512 | 0.076 | 0.107 | 6.37E-13 | 2.33E-08 |
| DOCK4 | 2 | -0.330259389 | 0.631 | 0.62 | 8.05E-13 | 2.95E-08 |
| RABGEF1 | 2 | -0.257538523 | 0.625 | 0.689 | 8.15E-13 | 2.98E-08 |
| ANKRD33B | 2 | -0.266828872 | 0.114 | 0.178 | 8.75E-13 | 0.000000032 |
| LRRC8B | 2 | -0.256963044 | 0.416 | 0.429 | 9.36E-13 | 3.43E-08 |
| KDM3A | 2 | -0.322681341 | 0.344 | 0.433 | 1.34E-12 | 4.91E-08 |
| SLC16A1 | 2 | -0.256303736 | 0.243 | 0.315 | 1.42E-12 | 5.19E-08 |
| TMEM165 | 2 | -0.265426251 | 0.495 | 0.565 | 1.42E-12 | 5.19E-08 |
| HDAC4 | 2 | -0.271379178 | 0.63 | 0.687 | 1.75E-12 | 6.39E-08 |
| MLLT3 | 2 | -0.333536703 | 0.818 | 0.818 | 2.61E-12 | 9.54E-08 |
| PHF6 | 2 | -0.251942824 | 0.428 | 0.496 | 2.7E-12 | 9.87E-08 |
| CSPP1 | 2 | -0.268220978 | 0.44 | 0.509 | 2.76E-12 | 0.000000101 |

| | | | | | | |
|---|---|---|---|---|---|---|
| RNF19A | 2 | -0.270924449 | 0.479 | 0.512 | 3.13E-12 | 0.000000114 |
| THSD4 | 2 | -0.370385359 | 0.123 | 0.171 | 3.13E-12 | 0.000000115 |
| COL4A1 | 2 | -0.279934931 | 0.092 | 0.145 | 3.32E-12 | 0.000000121 |
| USP25 | 2 | -0.251876534 | 0.465 | 0.554 | 3.89E-12 | 0.000000142 |
| PSME4 | 2 | -0.361453702 | 0.769 | 0.79 | 5.18E-12 | 0.00000019 |
| PFKFB4 | 2 | -0.285080878 | 0.11 | 0.151 | 5.47E-12 | 0.0000002 |
| GSTCD | 2 | -0.274213032 | 0.42 | 0.489 | 5.67E-12 | 0.000000208 |
| LRRFIP1 | 2 | -0.266986601 | 0.46 | 0.515 | 8.77E-12 | 0.000000321 |
| SFMBT2 | 2 | -0.292279091 | 0.48 | 0.495 | 1.59E-11 | 0.000000581 |
| GPC3 | 2 | -0.530061394 | 0.195 | 0.241 | 1.81E-11 | 0.000000664 |
| MGMT | 2 | -0.255354842 | 0.351 | 0.429 | 1.88E-11 | 0.000000687 |
| BCL2 | 2 | -0.284191138 | 0.166 | 0.202 | 1.99E-11 | 0.000000727 |
| WWOX | 2 | -0.351556121 | 0.731 | 0.781 | 2.42E-11 | 0.000000885 |
| CPEB3 | 2 | -0.25381952 | 0.344 | 0.4 | 3.68E-11 | 0.00000135 |
| RPS6KA3 | 2 | -0.379116406 | 0.573 | 0.585 | 4.6E-11 | 0.00000169 |
| RASSF8 | 2 | -0.276278803 | 0.468 | 0.511 | 5.12E-11 | 0.00000188 |
| CCDC85A | 2 | -0.254623845 | 0.066 | 0.112 | 5.28E-11 | 0.00000193 |
| ST6GALNAC5 | 2 | -0.420706186 | 0.369 | 0.377 | 6.04E-11 | 0.00000221 |
| GAB2 | 2 | -0.286731562 | 0.285 | 0.317 | 1.79E-10 | 0.00000655 |
| TSC22D1 | 2 | -0.287544457 | 0.685 | 0.716 | 2.05E-10 | 0.00000751 |
| GPI | 2 | -0.295783946 | 0.221 | 0.272 | 2.23E-10 | 0.00000817 |
| ATP9A | 2 | -0.253694981 | 0.363 | 0.42 | 2.27E-10 | 0.00000831 |
| STK33 | 2 | -0.337041224 | 0.146 | 0.164 | 2.35E-10 | 0.00000862 |
| PTPRK | 2 | 0.270456996 | 0.832 | 0.803 | 2.51E-10 | 0.00000919 |
| TSPAN9 | 2 | -0.267090966 | 0.339 | 0.404 | 2.79E-10 | 0.0000102 |
| CDON | 2 | -0.251377464 | 0.302 | 0.339 | 4E-10 | 0.0000146 |
| SETBP1 | 2 | -0.28028793 | 0.562 | 0.584 | 4.71E-10 | 0.0000173 |
| MEIS1 | 2 | -0.275015182 | 0.362 | 0.35 | 5.14E-10 | 0.0000188 |
| KCNQ1OT1 | 2 | -0.280775809 | 0.598 | 0.596 | 5.34E-10 | 0.0000196 |
| PLOD2 | 2 | -0.359993146 | 0.236 | 0.297 | 1.18E-09 | 0.0000431 |
| SEPTIN9 | 2 | -0.251761491 | 0.425 | 0.471 | 1.63E-09 | 0.0000595 |
| BZW2 | 2 | -0.263790309 | 0.376 | 0.41 | 1.57E-08 | 0.000573 |
| SLC12A2 | 2 | -0.255836664 | 0.45 | 0.466 | 1.61E-08 | 0.000588 |
| SHC3 | 2 | -0.255349079 | 0.321 | 0.364 | 2.06E-08 | 0.000753 |
| AC109466.1 | 2 | -0.678124708 | 0.082 | 0.12 | 2.65E-08 | 0.000968 |
| PRICKLE1 | 2 | -0.345384459 | 0.405 | 0.412 | 2.87E-08 | 0.001051964 |
| ENO1 | 2 | -0.272790249 | 0.43 | 0.479 | 4.12E-08 | 0.001509922 |
| FAM162A | 2 | -0.279322843 | 0.166 | 0.221 | 4.69E-08 | 0.001716273 |
| QKI | 2 | -0.270164179 | 0.771 | 0.789 | 0.00000032 | 0.011729542 |
| CDKAL1 | 2 | -0.278589126 | 0.861 | 0.817 | 0.000000444 | 0.016244009 |
| ADGRL2 | 2 | -0.36841382 | 0.617 | 0.635 | 0.00000161 | 0.058968391 |
| CNTN6 | 2 | 0.279439981 | 0.222 | 0.169 | 0.00000215 | 0.078768707 |
| FBXW7 | 2 | -0.296479968 | 0.723 | 0.742 | 0.00000633 | 0.231911476 |
| HSPH1 | 2 | 0.359457808 | 0.278 | 0.265 | 0.0000114 | 0.41695944 |
| LAMA2 | 2 | -0.28091458 | 0.353 | 0.4 | 0.0000934 | 1 |
| HPSE2 | 2 | 0.317413609 | 0.27 | 0.254 | 0.000185 | 1 |
| NRG1 | 2 | -0.397290133 | 0.292 | 0.29 | 0.003711586 | 1 |
| ADARB2 | 3 | 2.506209895 | 0.469 | 0.027 | 0 | 0 |
| DMD | 3 | 2.499682855 | 0.948 | 0.636 | 0 | 0 |
| CDH19 | 3 | 2.235763568 | 0.476 | 0.028 | 0 | 0 |
| PAX3 | 3 | 2.033530157 | 0.601 | 0.079 | 0 | 0 |
| LPAR1 | 3 | 1.883601682 | 0.681 | 0.144 | 0 | 0 |
| FAM178B | 3 | 1.454097322 | 0.486 | 0.036 | 0 | 0 |
| GABRA2 | 3 | 1.877162847 | 0.516 | 0.051 | 0 | 2.25E-304 |
| WWTR1 | 3 | 1.702898221 | 0.831 | 0.375 | 2.29E-308 | 8.38E-304 |
| AGMO | 3 | 2.185282088 | 0.515 | 0.063 | 2.11E-299 | 7.71E-295 |
| DOCK10 | 3 | 1.842088046 | 0.748 | 0.329 | 1.05E-283 | 3.83E-279 |
| MAML2 | 3 | 1.732958069 | 0.9 | 0.47 | 1.7E-281 | 6.22E-277 |

| | | | | | | |
|---|---|---|---|---|---|---|
| MEOX2 | 3 | 1.564458719 | 0.425 | 0.029 | 6.61E-279 | 2.42E-274 |
| ADCY2 | 3 | 2.132669611 | 0.692 | 0.224 | 6.24E-273 | 2.29E-268 |
| CDH11 | 3 | 1.690255542 | 0.781 | 0.253 | 1.68E-271 | 6.16E-267 |
| PDE4B | 3 | 2.179974427 | 0.521 | 0.081 | 6.65E-271 | 2.43E-266 |
| ANGPT1 | 3 | 1.77982154 | 0.88 | 0.537 | 2.09E-267 | 7.64E-263 |
| PPFIBP1 | 3 | 1.418247673 | 0.744 | 0.345 | 1.42E-255 | 5.2E-251 |
| ZEB2 | 3 | 1.629508811 | 0.876 | 0.542 | 1.44E-230 | 5.28E-226 |
| NRG3 | 3 | -3.58851595 | 0.064 | 0.517 | 5.98E-223 | 2.19E-218 |
| PDE7B | 3 | 1.751188459 | 0.615 | 0.164 | 5.4E-222 | 1.98E-217 |
| PTPRM | 3 | 1.462605487 | 0.787 | 0.33 | 2.57E-218 | 9.4E-214 |
| ITGA8 | 3 | 1.984232718 | 0.583 | 0.216 | 1.91E-209 | 7.01E-205 |
| SEMA3C | 3 | 1.457745906 | 0.456 | 0.079 | 6.63E-199 | 2.43E-194 |
| WSB1 | 3 | 1.149770643 | 0.943 | 0.768 | 3.84E-194 | 1.4E-189 |
| FGF14 | 3 | -1.95662201 | 0.256 | 0.708 | 5.44E-194 | 1.99E-189 |
| RBFOX1 | 3 | -1.892865127 | 0.381 | 0.781 | 1.35E-192 | 4.93E-188 |
| GAS7 | 3 | 1.531540327 | 0.577 | 0.195 | 1.98E-192 | 7.25E-188 |
| KCNB2 | 3 | -2.302716819 | 0.171 | 0.602 | 6.06E-190 | 2.22E-185 |
| NEGR1 | 3 | -3.34778971 | 0.199 | 0.626 | 1.32E-189 | 4.82E-185 |
| VWDE | 3 | -1.701837109 | 0.035 | 0.446 | 1.09E-188 | 3.99E-184 |
| L3MBTL4 | 3 | -1.605916142 | 0.131 | 0.583 | 1.29E-188 | 4.74E-184 |
| ATP10B | 3 | 1.025526057 | 0.244 | 0.009 | 6.48E-183 | 2.37E-178 |
| EDNRB | 3 | 1.177506571 | 0.324 | 0.034 | 1.62E-181 | 5.92E-177 |
| GULP1 | 3 | 1.41511987 | 0.799 | 0.469 | 2.73E-179 | 1E-174 |
| BCHE | 3 | 1.132261461 | 0.436 | 0.08 | 4.35E-179 | 1.59E-174 |
| COL5A2 | 3 | 1.306199542 | 0.826 | 0.426 | 2.53E-178 | 9.27E-174 |
| AL162493.1 | 3 | 1.031652467 | 0.332 | 0.034 | 9.04E-178 | 3.31E-173 |
| RIMS2 | 3 | -2.718898817 | 0.092 | 0.471 | 1.84E-175 | 6.75E-171 |
| ATRNL1 | 3 | -1.878428162 | 0.537 | 0.842 | 2.04E-173 | 7.46E-169 |
| LAMA4 | 3 | 0.851462378 | 0.846 | 0.404 | 2.3E-171 | 8.43E-167 |
| IL1RAPL1 | 3 | 1.777281299 | 0.888 | 0.627 | 7.73E-171 | 2.83E-166 |
| AL442636.1 | 3 | 0.831076876 | 0.331 | 0.036 | 7.94E-171 | 2.91E-166 |
| ACSS3 | 3 | 1.160014315 | 0.526 | 0.155 | 1.88E-168 | 6.88E-164 |
| AUTS2 | 3 | 1.034927829 | 0.817 | 0.43 | 1.41E-165 | 5.15E-161 |
| ST18 | 3 | -1.951027174 | 0.137 | 0.551 | 1.53E-165 | 5.6E-161 |
| PCDH15 | 3 | 2.189770208 | 0.41 | 0.1 | 2.77E-165 | 1.01E-160 |
| FAM189A1 | 3 | -1.676967707 | 0.352 | 0.738 | 9.13E-165 | 3.34E-160 |
| TCF7L2 | 3 | 1.133572717 | 0.884 | 0.548 | 7.76E-163 | 2.84E-158 |
| ANO4 | 3 | -2.141586628 | 0.242 | 0.628 | 1.24E-158 | 4.55E-154 |
| AL008633.1 | 3 | 1.165866645 | 0.951 | 0.68 | 1.22E-155 | 4.45E-151 |
| STARD13 | 3 | 1.344232045 | 0.797 | 0.569 | 2.46E-154 | 8.99E-150 |
| AC109466.1 | 3 | 2.24882847 | 0.374 | 0.079 | 3.56E-154 | 1.3E-149 |
| UNC5D | 3 | -2.118182909 | 0.238 | 0.622 | 4.71E-153 | 1.72E-148 |
| PTPRD | 3 | -1.480023234 | 0.628 | 0.868 | 8.07E-152 | 2.95E-147 |
| AC138305.1 | 3 | 1.563354948 | 0.443 | 0.127 | 3.34E-151 | 1.22E-146 |
| SNX7 | 3 | 1.053286948 | 0.415 | 0.15 | 4.98E-151 | 1.82E-146 |
| ARHGAP42 | 3 | 1.141418311 | 0.684 | 0.332 | 6.93E-150 | 2.54E-145 |
| NKAIN3 | 3 | 1.313667857 | 0.92 | 0.653 | 1.64E-148 | 6.01E-144 |
| CPED1 | 3 | 1.155030377 | 0.327 | 0.046 | 1.76E-147 | 6.43E-143 |
| AL390800.1 | 3 | -1.53397946 | 0.168 | 0.564 | 3.07E-144 | 1.12E-139 |
| FGF13 | 3 | -1.920103236 | 0.117 | 0.488 | 3.51E-144 | 1.29E-139 |
| NBEA | 3 | -1.664079982 | 0.466 | 0.734 | 9.21E-144 | 3.37E-139 |
| DSCAML1 | 3 | -1.214757101 | 0.084 | 0.462 | 1.17E-142 | 4.28E-138 |
| ERC2 | 3 | -1.346890031 | 0.122 | 0.494 | 4.39E-142 | 1.61E-137 |
| LINC00511 | 3 | 1.51204914 | 0.526 | 0.259 | 1.36E-141 | 4.99E-137 |
| YAP1 | 3 | 0.788571144 | 0.695 | 0.287 | 2.5E-140 | 9.16E-136 |
| MEGF10 | 3 | 1.239373861 | 0.704 | 0.361 | 6.66E-140 | 2.44E-135 |
| ZFHX4 | 3 | 1.114607347 | 0.711 | 0.393 | 2.03E-138 | 7.43E-134 |
| MLLT3 | 3 | 1.133437737 | 0.911 | 0.806 | 3.46E-138 | 1.27E-133 |

| | | | | | | |
|---|---|---|---|---|---|---|
| RAPGEF5 | 3 | -1.266181056 | 0.178 | 0.56 | 3.69E-138 | 1.35E-133 |
| CASC15 | 3 | 1.0098222 | 0.873 | 0.534 | 2.38E-136 | 8.7E-132 |
| MIR99AHG | 3 | 1.20846202 | 0.912 | 0.657 | 2.9E-136 | 1.06E-131 |
| AC108690.1 | 3 | 0.786746021 | 0.402 | 0.087 | 1.4E-135 | 5.14E-131 |
| ERBB3 | 3 | 0.972625422 | 0.345 | 0.072 | 1.45E-135 | 5.29E-131 |
| FTO | 3 | 1.129081328 | 0.912 | 0.798 | 1.75E-135 | 6.4E-131 |
| CSMD2 | 3 | -1.158807416 | 0.025 | 0.344 | 5.78E-135 | 2.12E-130 |
| SYT1 | 3 | -1.209336596 | 0.432 | 0.783 | 1.73E-134 | 6.33E-130 |
| MARK1 | 3 | -1.223286685 | 0.166 | 0.548 | 1.16E-133 | 4.24E-129 |
| SOX5 | 3 | 1.707318474 | 0.818 | 0.628 | 4.93E-133 | 1.81E-128 |
| TNIK | 3 | 1.352904161 | 0.63 | 0.325 | 1.15E-132 | 4.21E-128 |
| AL136018.1 | 3 | 0.525722621 | 0.182 | 0.007 | 6.21E-132 | 2.27E-127 |
| GPNMB | 3 | 0.792257206 | 0.282 | 0.038 | 1.37E-131 | 5E-127 |
| TTLL7 | 3 | -1.103599495 | 0.143 | 0.514 | 2.25E-131 | 8.23E-127 |
| PCDH7 | 3 | 1.596094139 | 0.733 | 0.388 | 2.32E-131 | 8.5E-127 |
| AC104041.1 | 3 | -0.997355293 | 0.147 | 0.532 | 2.59E-131 | 9.47E-127 |
| DLGAP1 | 3 | 1.086652332 | 0.964 | 0.817 | 3.67E-131 | 1.34E-126 |
| FRAS1 | 3 | -1.437189595 | 0.069 | 0.417 | 6.07E-131 | 2.22E-126 |
| DSCAM | 3 | 1.434264613 | 0.38 | 0.083 | 5.38E-130 | 1.97E-125 |
| HELLS | 3 | -1.068213856 | 0.312 | 0.672 | 1.09E-128 | 3.99E-124 |
| RAB3C | 3 | -1.502833857 | 0.123 | 0.467 | 5.34E-128 | 1.96E-123 |
| AL139383.1 | 3 | 1.334023967 | 0.267 | 0.053 | 6.48E-128 | 2.37E-123 |
| KIF21A | 3 | -1.116780608 | 0.147 | 0.509 | 1.29E-127 | 4.72E-123 |
| HMCN1 | 3 | 1.425463388 | 0.517 | 0.175 | 3.68E-127 | 1.35E-122 |
| PLP1 | 3 | 0.578905698 | 0.204 | 0.014 | 8.35E-127 | 3.06E-122 |
| SOX10 | 3 | 0.509028629 | 0.172 | 0.007 | 1.69E-126 | 6.18E-122 |
| LINC00536 | 3 | 1.863542361 | 0.313 | 0.067 | 1.95E-126 | 7.16E-122 |
| KCNQ3 | 3 | -1.340990323 | 0.085 | 0.427 | 1.53E-125 | 5.58E-121 |
| JAZF1 | 3 | 1.07310744 | 0.84 | 0.718 | 2.31E-125 | 8.47E-121 |
| FAM184A | 3 | -1.077608198 | 0.121 | 0.471 | 3.24E-125 | 1.18E-120 |
| VCAN | 3 | 1.116908602 | 0.722 | 0.376 | 2.1E-124 | 7.68E-120 |
| EBF1 | 3 | -1.886764734 | 0.078 | 0.407 | 3.08E-124 | 1.13E-119 |
| SERPINE2 | 3 | 1.239532502 | 0.378 | 0.118 | 2.99E-123 | 1.09E-118 |
| GASK1B | 3 | 0.662625127 | 0.244 | 0.026 | 6.28E-123 | 2.3E-118 |
| CTNNA2 | 3 | -1.663382084 | 0.194 | 0.529 | 4.37E-122 | 1.6E-117 |
| CTNND2 | 3 | -1.296316868 | 0.496 | 0.767 | 5.84E-121 | 2.14E-116 |
| GPR149 | 3 | 1.377529491 | 0.287 | 0.051 | 1.17E-120 | 4.3E-116 |
| MEG8 | 3 | -1.136303061 | 0.239 | 0.617 | 1.39E-120 | 5.08E-116 |
| STXBP5L | 3 | -1.511935002 | 0.042 | 0.341 | 2.71E-120 | 9.94E-116 |
| NOL4 | 3 | -1.48557229 | 0.057 | 0.363 | 2.93E-120 | 1.07E-115 |
| GABRB3 | 3 | -1.361697636 | 0.088 | 0.414 | 4.62E-120 | 1.69E-115 |
| FNDC3B | 3 | 0.873671141 | 0.861 | 0.673 | 3.37E-118 | 1.24E-113 |
| LINC02315 | 3 | -1.051629827 | 0.552 | 0.815 | 4.34E-118 | 1.59E-113 |
| APBA1 | 3 | -1.341230055 | 0.139 | 0.479 | 1.55E-117 | 5.67E-113 |
| SYNE2 | 3 | -1.099896763 | 0.449 | 0.737 | 3.89E-116 | 1.42E-111 |
| KIF26B | 3 | -1.283323268 | 0.19 | 0.554 | 6.49E-116 | 2.38E-111 |
| KIF5C | 3 | -0.940681959 | 0.033 | 0.324 | 9.1E-116 | 3.33E-111 |
| ADGRL3 | 3 | 1.337888447 | 0.927 | 0.799 | 1.96E-114 | 7.16E-110 |
| NMNAT2 | 3 | -1.042906807 | 0.052 | 0.356 | 1.46E-113 | 5.34E-109 |
| SGO1-AS1 | 3 | 1.657165484 | 0.437 | 0.152 | 2.27E-113 | 8.32E-109 |
| PTPRJ | 3 | 1.111699323 | 0.676 | 0.494 | 3.87E-113 | 1.42E-108 |
| LINC01876 | 3 | -1.321575455 | 0.203 | 0.547 | 5.49E-113 | 2.01E-108 |
| HS3ST4 | 3 | -1.795246164 | 0.064 | 0.375 | 1.43E-112 | 5.25E-108 |
| CYP7B1 | 3 | -1.272910079 | 0.087 | 0.405 | 4.95E-112 | 1.81E-107 |
| OSBPL10 | 3 | 1.039967263 | 0.639 | 0.352 | 6.4E-112 | 2.34E-107 |
| RIMS1 | 3 | -1.299394204 | 0.138 | 0.476 | 6.62E-112 | 2.42E-107 |
| KCNIP4 | 3 | -1.567251572 | 0.194 | 0.539 | 7.97E-112 | 2.92E-107 |
| RBMS1 | 3 | 0.876131143 | 0.791 | 0.515 | 9.24E-112 | 3.38E-107 |

| | | | | | | |
|---|---|---|---|---|---|---|
| UNC79 | 3 | -1.317082033 | 0.085 | 0.377 | 2.29E-110 | 8.4E-106 |
| CDKN2B-AS1 | 3 | -1.022373501 | 0.061 | 0.359 | 3.18E-110 | 1.16E-105 |
| PLPP1 | 3 | 1.184754957 | 0.589 | 0.315 | 1.2E-109 | 4.41E-105 |
| RHOJ | 3 | 0.687572173 | 0.263 | 0.041 | 2.52E-109 | 9.22E-105 |
| PLD1 | 3 | 0.707793866 | 0.259 | 0.039 | 1.36E-108 | 4.98E-104 |
| TIAM1 | 3 | -1.164883266 | 0.327 | 0.645 | 2.52E-108 | 9.24E-104 |
| LDLRAD4 | 3 | 1.272270116 | 0.715 | 0.466 | 5.72E-108 | 2.09E-103 |
| VPS13A | 3 | -1.119510815 | 0.379 | 0.687 | 1.07E-107 | 3.91E-103 |
| PPARG | 3 | -1.024126347 | 0.035 | 0.313 | 1.32E-107 | 4.83E-103 |
| BZW2 | 3 | 0.937968084 | 0.62 | 0.376 | 1.52E-107 | 5.56E-103 |
| NR3C1 | 3 | 0.949236101 | 0.636 | 0.328 | 1.8E-107 | 6.6E-103 |
| PLPP4 | 3 | -1.064852328 | 0.089 | 0.407 | 2.5E-107 | 9.16E-103 |
| GSN | 3 | 0.782436258 | 0.49 | 0.192 | 7E-107 | 2.56E-102 |
| ANKRD36 | 3 | -0.93035751 | 0.637 | 0.832 | 7.19E-107 | 2.63E-102 |
| CDH2 | 3 | -1.023815115 | 0.169 | 0.513 | 1.99E-106 | 7.3E-102 |
| SCN9A | 3 | 1.166095056 | 0.383 | 0.124 | 4.44E-106 | 1.63E-101 |
| PLA2G4A | 3 | 0.739047243 | 0.292 | 0.057 | 1.02E-105 | 3.75E-101 |
| SORCS3 | 3 | -1.367581949 | 0.132 | 0.46 | 1.52E-105 | 5.55E-101 |
| PIP5K1B | 3 | -1.395322703 | 0.038 | 0.31 | 1.72E-105 | 6.29E-101 |
| GRIN3A | 3 | -1.260134252 | 0.026 | 0.287 | 2.81E-105 | 1.03E-100 |
| DOCK3 | 3 | -1.335619216 | 0.046 | 0.316 | 6.82E-105 | 2.5E-100 |
| RGS21 | 3 | 0.848167837 | 0.216 | 0.029 | 1.28E-104 | 4.7E-100 |
| PID1 | 3 | -1.511478057 | 0.306 | 0.619 | 3.87E-104 | 1.42E-99 |
| OSMR | 3 | 0.666078713 | 0.284 | 0.057 | 4.09E-104 | 1.5E-99 |
| AHR | 3 | 0.952889685 | 0.433 | 0.139 | 5.93E-104 | 2.17E-99 |
| MAP3K1 | 3 | 1.04850833 | 0.437 | 0.176 | 2.05E-103 | 7.5E-99 |
| C2orf80 | 3 | 0.571875972 | 0.2 | 0.021 | 2.55E-103 | 9.34E-99 |
| CHN1 | 3 | -0.940478634 | 0.308 | 0.648 | 3.1E-103 | 1.13E-98 |
| BICDL1 | 3 | -0.810455235 | 0.037 | 0.309 | 3.47E-103 | 1.27E-98 |
| CACNA1B | 3 | -1.090502576 | 0.032 | 0.294 | 5.15E-102 | 1.89E-97 |
| ROBO1 | 3 | 0.998351005 | 0.978 | 0.891 | 2.49E-101 | 9.1E-97 |
| ADAMTSL1 | 3 | -2.390033786 | 0.353 | 0.545 | 2.61E-101 | 9.55E-97 |
| ALCAM | 3 | 1.140896855 | 0.625 | 0.293 | 2.86E-101 | 1.05E-96 |
| SASH1 | 3 | 0.892997233 | 0.793 | 0.532 | 1.03E-100 | 3.76E-96 |
| CLSTN2 | 3 | -1.391639589 | 0.081 | 0.376 | 1.32E-100 | 4.84E-96 |
| SH3GL2 | 3 | -1.210065752 | 0.126 | 0.43 | 2.64E-100 | 9.68E-96 |
| ARID5B | 3 | 0.888077741 | 0.695 | 0.375 | 2.79E-100 | 1.02E-95 |
| AC073050.1 | 3 | -1.440034709 | 0.057 | 0.336 | 3.3E-100 | 1.21E-95 |
| LIFR | 3 | 0.931094121 | 0.864 | 0.605 | 2.67E-99 | 9.77E-95 |
| CTTNBP2 | 3 | -1.059184564 | 0.051 | 0.327 | 3.4E-99 | 1.24E-94 |
| DPYSL5 | 3 | -0.766099048 | 0.014 | 0.251 | 4.45E-99 | 1.63E-94 |
| MAP7D2 | 3 | -0.736341645 | 0.043 | 0.315 | 5.09E-99 | 1.86E-94 |
| CDK6 | 3 | 0.976835379 | 0.38 | 0.112 | 5.29E-99 | 1.94E-94 |
| GPR137C | 3 | -0.997146036 | 0.152 | 0.465 | 1.37E-98 | 5.01E-94 |
| IDH1 | 3 | 0.867159617 | 0.439 | 0.191 | 3.25E-98 | 1.19E-93 |
| UBE2E2 | 3 | -1.220187853 | 0.346 | 0.633 | 6.66E-98 | 2.44E-93 |
| CADPS | 3 | -1.468131849 | 0.193 | 0.501 | 1.02E-97 | 3.73E-93 |
| GNAO1 | 3 | -0.984520029 | 0.181 | 0.493 | 2.13E-97 | 7.8E-93 |
| DOCK11 | 3 | -1.264242718 | 0.274 | 0.563 | 2.96E-97 | 1.08E-92 |
| POSTN | 3 | 1.115229798 | 0.171 | 0.015 | 3.6E-97 | 1.32E-92 |
| DNER | 3 | -1.103785768 | 0.115 | 0.426 | 1.48E-96 | 5.41E-92 |
| LINC02470 | 3 | -1.279315624 | 0.041 | 0.298 | 4.49E-96 | 1.64E-91 |
| POLA1 | 3 | -1.003277644 | 0.74 | 0.858 | 2.03E-95 | 7.45E-91 |
| PLCL2 | 3 | -0.982544228 | 0.313 | 0.626 | 5.21E-95 | 1.91E-90 |
| ABCB1 | 3 | -0.904689631 | 0.022 | 0.263 | 1.53E-94 | 5.6E-90 |
| CAMK4 | 3 | -0.992712444 | 0.059 | 0.329 | 2.65E-94 | 9.71E-90 |
| ADAMTS20 | 3 | -1.111830474 | 0.023 | 0.262 | 7.46E-94 | 2.73E-89 |
| CAMK2D | 3 | 0.838864095 | 0.793 | 0.543 | 1.25E-93 | 4.58E-89 |

| | | | | | | |
|---|---|---|---|---|---|---|
| LPP | 3 | 0.779672748 | 0.875 | 0.713 | 1.75E-93 | 6.4E-89 |
| CBLB | 3 | 0.96699236 | 0.746 | 0.57 | 2.04E-93 | 7.47E-89 |
| RHBDL3 | 3 | -0.886249033 | 0.058 | 0.33 | 2.04E-93 | 7.47E-89 |
| CRYM | 3 | 0.837716676 | 0.293 | 0.074 | 4.03E-93 | 1.48E-88 |
| COL11A1 | 3 | 1.179721793 | 0.868 | 0.692 | 6.3E-93 | 2.31E-88 |
| KCND2 | 3 | -1.480481435 | 0.268 | 0.576 | 3.36E-92 | 1.23E-87 |
| ARHGAP44 | 3 | -0.9421318 | 0.134 | 0.43 | 5.92E-92 | 2.17E-87 |
| MIR924HG | 3 | -0.928713465 | 0.476 | 0.749 | 9.88E-92 | 3.62E-87 |
| RADX | 3 | -0.980839675 | 0.208 | 0.53 | 1.16E-91 | 4.24E-87 |
| STK3 | 3 | 0.714505407 | 0.757 | 0.461 | 4.41E-91 | 1.61E-86 |
| TOX | 3 | -0.783510018 | 0.389 | 0.711 | 9.46E-91 | 3.47E-86 |
| HS6ST3 | 3 | -2.124125716 | 0.24 | 0.539 | 2.74E-90 | 1E-85 |
| TSPAN7 | 3 | -1.007400765 | 0.277 | 0.575 | 3.38E-90 | 1.24E-85 |
| CCSER1 | 3 | -0.871289686 | 0.316 | 0.643 | 4.09E-90 | 1.5E-85 |
| ABTB2 | 3 | 0.887903671 | 0.385 | 0.134 | 1.84E-89 | 6.73E-85 |
| NHS | 3 | -1.310428267 | 0.209 | 0.508 | 2.33E-89 | 8.54E-85 |
| RMST | 3 | -1.667328752 | 0.239 | 0.522 | 6E-89 | 2.2E-84 |
| MITF | 3 | 0.968432739 | 0.405 | 0.161 | 3.55E-88 | 1.3E-83 |
| EYA1 | 3 | 1.075551879 | 0.541 | 0.26 | 8.44E-88 | 3.09E-83 |
| PDE4D | 3 | 1.179729657 | 0.944 | 0.838 | 9.12E-88 | 3.34E-83 |
| SYTL5 | 3 | -0.972881003 | 0.074 | 0.347 | 1.44E-87 | 5.29E-83 |
| DDR2 | 3 | 0.668337463 | 0.459 | 0.173 | 3.94E-87 | 1.44E-82 |
| SYBU | 3 | -1.055332333 | 0.152 | 0.445 | 4.11E-87 | 1.51E-82 |
| NLK | 3 | 0.810458645 | 0.818 | 0.644 | 4.81E-87 | 1.76E-82 |
| WDR17 | 3 | -0.854657168 | 0.106 | 0.382 | 5.45E-87 | 1.99E-82 |
| IFI16 | 3 | 0.545599839 | 0.215 | 0.033 | 6.64E-87 | 2.43E-82 |
| BASP1 | 3 | -0.837980457 | 0.194 | 0.502 | 1.2E-86 | 4.38E-82 |
| FAM135B | 3 | -0.87164711 | 0.112 | 0.403 | 7.55E-86 | 2.77E-81 |
| P3H2 | 3 | 0.902485665 | 0.238 | 0.047 | 8.53E-86 | 3.12E-81 |
| GLYATL2 | 3 | -0.885114294 | 0.091 | 0.363 | 2.31E-85 | 8.45E-81 |
| CACNA2D1 | 3 | -1.351820489 | 0.762 | 0.866 | 2.66E-85 | 9.75E-81 |
| LINC02821 | 3 | 0.535834811 | 0.188 | 0.026 | 1.23E-84 | 4.51E-80 |
| MMP26 | 3 | 0.866803173 | 0.258 | 0.056 | 2.23E-84 | 8.15E-80 |
| OSMR-AS1 | 3 | 0.551399636 | 0.27 | 0.062 | 1.14E-83 | 4.16E-79 |
| FRY | 3 | -0.96950785 | 0.314 | 0.606 | 1.53E-83 | 5.62E-79 |
| TMEM168 | 3 | 0.498838655 | 0.283 | 0.066 | 1.83E-83 | 6.7E-79 |
| CACNA1D | 3 | -1.061285068 | 0.174 | 0.453 | 2.01E-83 | 7.37E-79 |
| BCAS3 | 3 | 0.740808829 | 0.823 | 0.636 | 2.14E-83 | 7.85E-79 |
| CNTNAP3B | 3 | 0.979092754 | 0.529 | 0.277 | 3.25E-83 | 1.19E-78 |
| HS3ST5 | 3 | -0.755667666 | 0.338 | 0.655 | 4.03E-83 | 1.48E-78 |
| ADAM23 | 3 | -0.999521785 | 0.118 | 0.39 | 7.43E-83 | 2.72E-78 |
| PRRX1 | 3 | 0.736755426 | 0.303 | 0.077 | 8.85E-83 | 3.24E-78 |
| COL19A1 | 3 | 1.143719626 | 0.288 | 0.079 | 1.25E-82 | 4.59E-78 |
| SLC38A1 | 3 | -0.825780204 | 0.321 | 0.602 | 3.88E-82 | 1.42E-77 |
| LIN28B | 3 | -0.962960597 | 0.237 | 0.531 | 6.42E-82 | 2.35E-77 |
| PGBD5 | 3 | -0.707189175 | 0.074 | 0.332 | 8.3E-82 | 3.04E-77 |
| BRIP1 | 3 | -0.941236906 | 0.296 | 0.596 | 8.5E-82 | 3.11E-77 |
| FAT3 | 3 | -1.134619764 | 0.14 | 0.424 | 8.53E-82 | 3.12E-77 |
| TSPAN5 | 3 | -0.988912348 | 0.237 | 0.527 | 1.38E-81 | 5.05E-77 |
| IGSF21 | 3 | -0.719329728 | 0.048 | 0.29 | 1.39E-81 | 5.08E-77 |
| PLPPR1 | 3 | -0.952430436 | 0.014 | 0.218 | 1.45E-81 | 5.3E-77 |
| SEMA3E | 3 | 1.087104885 | 0.541 | 0.269 | 2.37E-81 | 8.68E-77 |
| ERBB4 | 3 | -2.0499348 | 0.077 | 0.313 | 2.49E-81 | 9.1E-77 |
| VMP1 | 3 | 0.956768642 | 0.712 | 0.58 | 3.06E-81 | 1.12E-76 |
| TET1 | 3 | 0.728274354 | 0.672 | 0.376 | 4.43E-81 | 1.62E-76 |
| SIL1 | 3 | 0.719563469 | 0.835 | 0.641 | 6.69E-81 | 2.45E-76 |
| SLC16A10 | 3 | -0.888939125 | 0.03 | 0.252 | 2.02E-80 | 7.4E-76 |
| CA12 | 3 | -0.731754869 | 0.04 | 0.272 | 2.72E-80 | 9.97E-76 |

| | | | | | | |
|---|---|---|---|---|---|---|
| LSAMP | 3 | 0.733304487 | 0.925 | 0.713 | 2.88E-80 | 1.06E-75 |
| FER | 3 | 0.779578522 | 0.89 | 0.805 | 3.21E-80 | 1.18E-75 |
| CECR2 | 3 | -0.886069225 | 0.082 | 0.342 | 3.51E-80 | 1.29E-75 |
| SH3GL3 | 3 | -1.032103123 | 0.298 | 0.562 | 6.9E-80 | 2.53E-75 |
| SNAP25 | 3 | -0.73487777 | 0.073 | 0.326 | 9.25E-80 | 3.39E-75 |
| RAPGEF4 | 3 | -1.172330127 | 0.17 | 0.432 | 1.01E-79 | 3.68E-75 |
| MAP7 | 3 | -0.836013853 | 0.318 | 0.625 | 1.24E-79 | 4.55E-75 |
| DLX6-AS1 | 3 | -1.27764738 | 0.031 | 0.247 | 1.42E-79 | 5.21E-75 |
| SBF2 | 3 | 0.632985738 | 0.884 | 0.735 | 1.51E-79 | 5.52E-75 |
| FSTL5 | 3 | 1.516244749 | 0.313 | 0.117 | 1.55E-79 | 5.67E-75 |
| TGFBI | 3 | 0.588123957 | 0.193 | 0.029 | 1.59E-79 | 5.82E-75 |
| MAP3K5 | 3 | -0.781456164 | 0.088 | 0.35 | 1.95E-79 | 7.15E-75 |
| PLS3 | 3 | 0.902376581 | 0.611 | 0.377 | 2.11E-79 | 7.71E-75 |
| MAPK10 | 3 | -1.151603528 | 0.033 | 0.251 | 3.22E-79 | 1.18E-74 |
| LRRTM4 | 3 | -1.206291283 | 0.311 | 0.611 | 6.38E-79 | 2.33E-74 |
| ROR2 | 3 | -0.686649825 | 0.164 | 0.456 | 7.58E-79 | 2.77E-74 |
| CFAP58 | 3 | -1.004643163 | 0.077 | 0.325 | 8.57E-79 | 3.14E-74 |
| AC023282.1 | 3 | -0.840022911 | 0.072 | 0.325 | 8.79E-79 | 3.22E-74 |
| GALNT14 | 3 | -0.72784893 | 0.043 | 0.275 | 1.68E-78 | 6.17E-74 |
| COL6A3 | 3 | 0.935929896 | 0.313 | 0.093 | 1.95E-78 | 7.13E-74 |
| SOBP | 3 | -0.943990839 | 0.637 | 0.795 | 2.92E-78 | 1.07E-73 |
| ENPP2 | 3 | -0.738169169 | 0.025 | 0.236 | 3.09E-78 | 1.13E-73 |
| GNG4 | 3 | -0.774165241 | 0.093 | 0.353 | 3.99E-78 | 1.46E-73 |
| MEF2C | 3 | 0.872658689 | 0.591 | 0.321 | 4.03E-78 | 1.47E-73 |
| ZBTB20 | 3 | 0.672659857 | 0.925 | 0.77 | 6.29E-78 | 2.3E-73 |
| CDH7 | 3 | -1.088911634 | 0.067 | 0.315 | 7.59E-78 | 2.78E-73 |
| ST5 | 3 | 0.730163118 | 0.535 | 0.266 | 1.04E-77 | 3.82E-73 |
| CDKN2A | 3 | -0.689676061 | 0.1 | 0.356 | 2.17E-77 | 7.93E-73 |
| FRMD4A | 3 | -0.897480809 | 0.74 | 0.828 | 2.89E-77 | 1.06E-72 |
| GNG2 | 3 | 0.825053998 | 0.361 | 0.134 | 2.99E-77 | 1.09E-72 |
| CENPP | 3 | -1.136458922 | 0.517 | 0.714 | 3.28E-77 | 1.2E-72 |
| LRRC7 | 3 | -1.858410451 | 0.102 | 0.343 | 3.75E-77 | 1.37E-72 |
| ART3 | 3 | 0.66060612 | 0.224 | 0.046 | 3.81E-77 | 1.4E-72 |
| ZFHX3 | 3 | 0.792797758 | 0.775 | 0.557 | 9.33E-77 | 3.42E-72 |
| BRINP3 | 3 | 1.157169404 | 0.731 | 0.48 | 1.23E-76 | 4.51E-72 |
| CCDC144NL-AS1 | 3 | 0.646104266 | 0.916 | 0.822 | 1.86E-76 | 6.8E-72 |
| ATAD2 | 3 | -0.896846767 | 0.25 | 0.537 | 2.04E-76 | 7.47E-72 |
| DGKI | 3 | -0.934893006 | 0.042 | 0.261 | 2.44E-76 | 8.94E-72 |
| MIR181A2HG | 3 | 0.735868544 | 0.509 | 0.245 | 2.5E-76 | 9.15E-72 |
| AFF3 | 3 | -1.174500092 | 0.396 | 0.634 | 4.35E-76 | 1.59E-71 |
| ARHGAP6 | 3 | 0.826610179 | 0.613 | 0.326 | 1.03E-75 | 3.78E-71 |
| HPSE2 | 3 | -1.831307108 | 0.059 | 0.283 | 1.15E-75 | 4.22E-71 |
| SLC17A8 | 3 | -1.192244615 | 0.075 | 0.316 | 1.31E-75 | 4.78E-71 |
| GNB4 | 3 | 0.744904348 | 0.371 | 0.158 | 6.65E-75 | 2.44E-70 |
| RGS7 | 3 | -1.004336189 | 0.024 | 0.225 | 9.68E-75 | 3.54E-70 |
| AFDN | 3 | -0.753311636 | 0.617 | 0.817 | 1.04E-74 | 3.8E-70 |
| PDE3B | 3 | -0.843486875 | 0.381 | 0.669 | 1.04E-74 | 3.82E-70 |
| AC138627.1 | 3 | 1.349316376 | 0.336 | 0.142 | 1.13E-74 | 4.15E-70 |
| DNAJC6 | 3 | -0.842709761 | 0.165 | 0.444 | 1.35E-74 | 4.95E-70 |
| MIR181A1HG | 3 | 0.719799551 | 0.72 | 0.444 | 2.6E-74 | 9.52E-70 |
| COCH | 3 | -0.620034891 | 0.04 | 0.259 | 3.62E-74 | 1.32E-69 |
| SOGA3 | 3 | -0.622651192 | 0.033 | 0.241 | 4.25E-74 | 1.56E-69 |
| MAML3 | 3 | 0.732981749 | 0.791 | 0.544 | 4.28E-74 | 1.57E-69 |
| FRMD3 | 3 | -0.975217032 | 0.141 | 0.394 | 5.39E-74 | 1.97E-69 |
| LDLRAD3 | 3 | 0.720785596 | 0.73 | 0.529 | 1.17E-73 | 4.28E-69 |
| BBS9 | 3 | 0.659943244 | 0.711 | 0.434 | 1.18E-73 | 4.33E-69 |
| TMEM108 | 3 | -1.086084901 | 0.461 | 0.68 | 1.4E-73 | 5.11E-69 |
| HDAC2-AS2 | 3 | -0.802149331 | 0.251 | 0.54 | 3.2E-73 | 1.17E-68 |

| | | | | | | |
|---|---|---|---|---|---|---|
| CSMD3 | 3 | 1.079393846 | 0.556 | 0.295 | 4.67E-73 | 1.71E-68 |
| AC009315.1 | 3 | -0.791754521 | 0.165 | 0.442 | 2.29E-72 | 8.39E-68 |
| ANK1 | 3 | -0.724296876 | 0.024 | 0.216 | 3.29E-72 | 1.2E-67 |
| TRIM38 | 3 | 0.507787445 | 0.225 | 0.048 | 4.28E-72 | 1.57E-67 |
| PPFIA1 | 3 | 0.76169213 | 0.684 | 0.538 | 4.59E-72 | 1.68E-67 |
| NRCAM | 3 | -0.802529324 | 0.152 | 0.421 | 5.49E-72 | 2.01E-67 |
| DTL | 3 | -0.725728573 | 0.145 | 0.418 | 6.16E-72 | 2.26E-67 |
| ABCA1 | 3 | 0.875352881 | 0.475 | 0.236 | 6.78E-72 | 2.48E-67 |
| SRRM4 | 3 | -0.640113989 | 0.015 | 0.204 | 1.08E-71 | 3.95E-67 |
| SESN3 | 3 | 1.04046575 | 0.59 | 0.384 | 1.2E-71 | 4.38E-67 |
| SCML2 | 3 | -0.691531405 | 0.202 | 0.468 | 2.37E-71 | 8.66E-67 |
| PTCHD4 | 3 | -1.027301385 | 0.056 | 0.277 | 3.88E-71 | 1.42E-66 |
| SLIT2 | 3 | -1.448032301 | 0.614 | 0.735 | 3.91E-71 | 1.43E-66 |
| NEUROD1 | 3 | -0.726256699 | 0.016 | 0.202 | 4.41E-71 | 1.62E-66 |
| GAP43 | 3 | -0.73057427 | 0.037 | 0.246 | 6.53E-71 | 2.39E-66 |
| SLC35F1 | 3 | -0.71328103 | 0.085 | 0.331 | 7.44E-71 | 2.73E-66 |
| DIAPH3 | 3 | -1.059695648 | 0.173 | 0.442 | 1.08E-70 | 3.96E-66 |
| RBFOX3 | 3 | -0.836018524 | 0.025 | 0.223 | 1.45E-70 | 5.29E-66 |
| KCNMB2 | 3 | -1.705088391 | 0.092 | 0.294 | 1.54E-70 | 5.65E-66 |
| PTPRN2 | 3 | -1.577308415 | 0.076 | 0.299 | 2.36E-70 | 8.63E-66 |
| SPRED1 | 3 | 0.717785788 | 0.639 | 0.385 | 2.49E-70 | 9.13E-66 |
| EBF3 | 3 | -1.066965943 | 0.057 | 0.276 | 1.36E-69 | 4.98E-65 |
| EGFR | 3 | -0.789521339 | 0.06 | 0.285 | 3.17E-69 | 1.16E-64 |
| PTGER3 | 3 | -0.84043376 | 0.157 | 0.421 | 3.99E-69 | 1.46E-64 |
| TCF7L1 | 3 | 0.617324207 | 0.556 | 0.277 | 5.15E-69 | 1.88E-64 |
| SLC1A2 | 3 | -0.715996453 | 0.079 | 0.309 | 6.44E-69 | 2.36E-64 |
| PKIB | 3 | -0.771730251 | 0.159 | 0.422 | 6.79E-69 | 2.48E-64 |
| MYO3A | 3 | -1.230216203 | 0.019 | 0.201 | 9.16E-69 | 3.35E-64 |
| MEG3 | 3 | -0.704048908 | 0.201 | 0.48 | 1.23E-68 | 4.5E-64 |
| THSD7B | 3 | -1.159066676 | 0.039 | 0.234 | 1.57E-68 | 5.75E-64 |
| MMS22L | 3 | -0.693199087 | 0.361 | 0.635 | 2.33E-68 | 8.52E-64 |
| THSD7A | 3 | -1.397740258 | 0.16 | 0.417 | 2.59E-68 | 9.48E-64 |
| PRKAG2 | 3 | -0.69890454 | 0.196 | 0.468 | 7.58E-68 | 2.78E-63 |
| FHOD3 | 3 | -0.774784448 | 0.094 | 0.335 | 1.14E-67 | 4.16E-63 |
| XKR4 | 3 | -0.948846673 | 0.022 | 0.208 | 1.23E-67 | 4.49E-63 |
| NELL2 | 3 | -1.133967312 | 0.07 | 0.281 | 1.45E-67 | 5.29E-63 |
| DLEU1 | 3 | 0.92472999 | 0.478 | 0.301 | 1.61E-67 | 5.9E-63 |
| ANKIB1 | 3 | 0.749289514 | 0.88 | 0.796 | 3.11E-67 | 1.14E-62 |
| FANCI | 3 | -0.733747093 | 0.167 | 0.43 | 4E-67 | 1.46E-62 |
| AFAP1L2 | 3 | 0.38312199 | 0.164 | 0.024 | 5.53E-67 | 2.02E-62 |
| PRRT2 | 3 | -0.676228676 | 0.171 | 0.432 | 6.19E-67 | 2.27E-62 |
| BACH1 | 3 | 0.747028317 | 0.657 | 0.442 | 6.3E-67 | 2.31E-62 |
| DISC1 | 3 | 0.692432224 | 0.476 | 0.253 | 6.78E-67 | 2.48E-62 |
| IGF2BP2 | 3 | 0.722151154 | 0.666 | 0.392 | 2.43E-66 | 8.89E-62 |
| ELAVL2 | 3 | -0.915080383 | 0.157 | 0.412 | 2.51E-66 | 9.21E-62 |
| PTEN | 3 | 0.886503161 | 0.788 | 0.69 | 4.01E-66 | 1.47E-61 |
| AC018731.1 | 3 | -1.246361147 | 0.384 | 0.618 | 4.21E-66 | 1.54E-61 |
| EZH2 | 3 | -0.859662519 | 0.551 | 0.731 | 4.78E-66 | 1.75E-61 |
| CENPK | 3 | -0.649025029 | 0.249 | 0.525 | 5.34E-66 | 1.95E-61 |
| ARHGAP10 | 3 | 0.807103678 | 0.687 | 0.489 | 7.63E-66 | 2.8E-61 |
| SLCO3A1 | 3 | -0.710752211 | 0.073 | 0.299 | 9.68E-66 | 3.54E-61 |
| AP001347.1 | 3 | -0.648211498 | 0.131 | 0.384 | 1.8E-65 | 6.59E-61 |
| PHYHIPL | 3 | -0.561061404 | 0.011 | 0.179 | 2.07E-65 | 7.58E-61 |
| SAMD4A | 3 | 0.974909059 | 0.571 | 0.363 | 2.13E-65 | 7.81E-61 |
| GNG7 | 3 | 0.476077539 | 0.192 | 0.039 | 2.6E-65 | 9.52E-61 |
| TMEM135 | 3 | 0.666057682 | 0.68 | 0.498 | 3.24E-65 | 1.19E-60 |
| DCLK1 | 3 | -0.898058634 | 0.368 | 0.627 | 7.43E-65 | 2.72E-60 |
| RBBP8 | 3 | -0.7441945 | 0.3 | 0.553 | 7.64E-65 | 2.8E-60 |

| | | | | | | |
|---|---|---|---|---|---|---|
| ARPP21 | 3 | -1.269301954 | 0.125 | 0.34 | 1.06E-64 | 3.89E-60 |
| PARM1 | 3 | -0.65592708 | 0.032 | 0.225 | 1.33E-64 | 4.89E-60 |
| FOXO1 | 3 | 0.723352892 | 0.425 | 0.183 | 1.34E-64 | 4.89E-60 |
| IGSF11 | 3 | 0.677702776 | 0.34 | 0.123 | 1.57E-64 | 5.73E-60 |
| KCNJ6 | 3 | -0.8643827 | 0.034 | 0.223 | 1.87E-64 | 6.86E-60 |
| PHLDB1 | 3 | 0.495787994 | 0.327 | 0.115 | 2.59E-64 | 9.49E-60 |
| ANKRD50 | 3 | 0.756321998 | 0.481 | 0.293 | 7.56E-64 | 2.77E-59 |
| GPR176 | 3 | -0.716476555 | 0.177 | 0.43 | 8.87E-64 | 3.25E-59 |
| PDZRN3 | 3 | 0.823736962 | 0.955 | 0.897 | 9.19E-64 | 3.36E-59 |
| PHACTR2 | 3 | -0.915688954 | 0.387 | 0.592 | 1.13E-63 | 4.14E-59 |
| CDH4 | 3 | -0.903798082 | 0.402 | 0.648 | 1.88E-63 | 6.89E-59 |
| AF279873.3 | 3 | 0.71104871 | 0.119 | 0.021 | 2.73E-63 | 1E-58 |
| IMMP2L | 3 | 0.678504455 | 0.93 | 0.805 | 4.23E-63 | 1.55E-58 |
| AC097528.1 | 3 | 1.107148452 | 0.125 | 0.014 | 4.57E-63 | 1.67E-58 |
| ABCA3 | 3 | -0.587875335 | 0.066 | 0.278 | 4.61E-63 | 1.69E-58 |
| DPY19L2 | 3 | -0.620358981 | 0.171 | 0.422 | 7.22E-63 | 2.64E-58 |
| AL035446.2 | 3 | 0.503554696 | 0.128 | 0.014 | 1.17E-62 | 4.28E-58 |
| MCM6 | 3 | -0.646351724 | 0.216 | 0.475 | 1.43E-62 | 5.24E-58 |
| SPATA6 | 3 | 0.732591011 | 0.354 | 0.15 | 4.54E-62 | 1.66E-57 |
| NMU | 3 | -0.639677639 | 0.064 | 0.276 | 5.52E-62 | 2.02E-57 |
| LINC00599 | 3 | -0.480737599 | 0.013 | 0.176 | 7.11E-62 | 2.6E-57 |
| ADD2 | 3 | -0.531002295 | 0.091 | 0.319 | 9.35E-62 | 3.42E-57 |
| PRKCE | 3 | 0.642021037 | 0.722 | 0.494 | 9.75E-62 | 3.57E-57 |
| NETO2 | 3 | -0.734728185 | 0.313 | 0.56 | 1.1E-61 | 4.04E-57 |
| AL121578.3 | 3 | -0.599171277 | 0.044 | 0.24 | 1.33E-61 | 4.86E-57 |
| GPM6B | 3 | 0.724628551 | 0.761 | 0.617 | 1.44E-61 | 5.27E-57 |
| ASTN1 | 3 | -0.657811615 | 0.062 | 0.271 | 2.08E-61 | 7.6E-57 |
| STMN2 | 3 | -0.610659445 | 0.019 | 0.189 | 2.49E-61 | 9.1E-57 |
| PLEKHA6 | 3 | -0.690493423 | 0.237 | 0.497 | 3.58E-61 | 1.31E-56 |
| CDCA7 | 3 | -0.593112718 | 0.187 | 0.444 | 3.58E-61 | 1.31E-56 |
| NFASC | 3 | -0.601205638 | 0.024 | 0.198 | 4.18E-61 | 1.53E-56 |
| RPS6KA6 | 3 | -0.544456627 | 0.036 | 0.224 | 6.14E-61 | 2.25E-56 |
| PTPN14 | 3 | 0.560621288 | 0.518 | 0.258 | 6.67E-61 | 2.44E-56 |
| ELAVL3 | 3 | -0.570238764 | 0.017 | 0.184 | 7.3E-61 | 2.67E-56 |
| NFIB | 3 | -0.968485886 | 0.726 | 0.846 | 7.82E-61 | 2.86E-56 |
| MYO16 | 3 | -1.177284329 | 0.12 | 0.344 | 1.03E-60 | 3.77E-56 |
| CACNA2D3 | 3 | -0.513021142 | 0.279 | 0.549 | 1.06E-60 | 3.89E-56 |
| AL353784.1 | 3 | 0.306286849 | 0.114 | 0.01 | 1.06E-60 | 3.89E-56 |
| KCNH7 | 3 | -0.957442414 | 0.007 | 0.154 | 1.88E-60 | 6.88E-56 |
| MRC2 | 3 | 0.556080068 | 0.451 | 0.209 | 2.4E-60 | 8.78E-56 |
| ROR1 | 3 | 0.391528013 | 0.469 | 0.225 | 2.44E-60 | 8.94E-56 |
| MKX | 3 | -1.080234188 | 0.153 | 0.392 | 3.08E-60 | 1.13E-55 |
| NEO1 | 3 | -0.746828438 | 0.346 | 0.577 | 4.4E-60 | 1.61E-55 |
| PCDH11X | 3 | 1.015735766 | 0.485 | 0.235 | 4.65E-60 | 1.7E-55 |
| DGKB | 3 | -1.414276963 | 0.1 | 0.31 | 5.27E-60 | 1.93E-55 |
| ARHGAP12 | 3 | 0.668440207 | 0.7 | 0.535 | 5.6E-60 | 2.05E-55 |
| RTN1 | 3 | -0.540546457 | 0.014 | 0.176 | 1.01E-59 | 3.71E-55 |
| RFTN2 | 3 | 0.438318276 | 0.165 | 0.029 | 1.07E-59 | 3.91E-55 |
| MED12L | 3 | -0.716822683 | 0.271 | 0.532 | 1.11E-59 | 4.05E-55 |
| C4orf19 | 3 | -0.878202922 | 0.118 | 0.329 | 1.11E-59 | 4.07E-55 |
| TCF12 | 3 | 0.475764453 | 0.96 | 0.916 | 1.12E-59 | 4.08E-55 |
| GRIP1 | 3 | -0.960809413 | 0.153 | 0.386 | 1.22E-59 | 4.48E-55 |
| PARD3 | 3 | 0.490123718 | 0.99 | 0.963 | 1.24E-59 | 4.53E-55 |
| WWC2 | 3 | 0.580320197 | 0.629 | 0.372 | 1.41E-59 | 5.18E-55 |
| STAT3 | 3 | 0.603509331 | 0.647 | 0.417 | 1.93E-59 | 7.06E-55 |
| AC040168.1 | 3 | -0.709285399 | 0.069 | 0.271 | 2.2E-59 | 8.06E-55 |
| MGLL | 3 | 0.703411754 | 0.362 | 0.171 | 2.22E-59 | 8.13E-55 |
| KCNG3 | 3 | -0.467718165 | 0.018 | 0.183 | 7.06E-59 | 2.59E-54 |

| | | | | | | |
|---|---|---|---|---|---|---|
| NEXMIF | 3 | -0.604005663 | 0.025 | 0.198 | 1.05E-58 | 3.86E-54 |
| NOVA1 | 3 | -0.822341233 | 0.414 | 0.654 | 1.11E-58 | 4.08E-54 |
| NEBL | 3 | -0.88797619 | 0.038 | 0.223 | 1.81E-58 | 6.62E-54 |
| SYN3 | 3 | -1.230465069 | 0.236 | 0.447 | 2.32E-58 | 8.51E-54 |
| KIAA0408 | 3 | -0.545269631 | 0.045 | 0.232 | 3.36E-58 | 1.23E-53 |
| GLI3 | 3 | 0.516866825 | 0.575 | 0.314 | 4.68E-58 | 1.71E-53 |
| ADAMTS19 | 3 | -0.9437309 | 0.267 | 0.514 | 5.11E-58 | 1.87E-53 |
| PCOLCE2 | 3 | -0.646173267 | 0.142 | 0.376 | 6.79E-58 | 2.48E-53 |
| AK5 | 3 | -0.999435943 | 0.06 | 0.246 | 7.83E-58 | 2.87E-53 |
| DCX | 3 | -0.609030955 | 0.026 | 0.199 | 7.9E-58 | 2.89E-53 |
| PHLDB2 | 3 | 0.83614657 | 0.401 | 0.208 | 9E-58 | 3.3E-53 |
| LMNB1 | 3 | -0.601152064 | 0.261 | 0.515 | 1.04E-57 | 3.82E-53 |
| SMC4 | 3 | -0.711391105 | 0.374 | 0.609 | 1.85E-57 | 6.77E-53 |
| SHANK2 | 3 | -0.748716746 | 0.05 | 0.241 | 5.67E-57 | 2.08E-52 |
| TBX20 | 3 | -0.527255703 | 0.078 | 0.288 | 6.01E-57 | 2.2E-52 |
| ZNF717 | 3 | 0.444900094 | 0.222 | 0.059 | 8.55E-57 | 3.13E-52 |
| GPM6A | 3 | -1.134372157 | 0.042 | 0.219 | 8.71E-57 | 3.19E-52 |
| CADPS2 | 3 | -1.073615095 | 0.077 | 0.255 | 1.1E-56 | 4.04E-52 |
| ZFPM2 | 3 | 0.639144222 | 0.531 | 0.278 | 1.11E-56 | 4.06E-52 |
| CAMTA1 | 3 | -0.789537662 | 0.577 | 0.727 | 1.27E-56 | 4.67E-52 |
| NOTCH2 | 3 | 0.585895608 | 0.531 | 0.281 | 1.31E-56 | 4.8E-52 |
| AL162718.1 | 3 | -0.619805206 | 0.01 | 0.153 | 1.39E-56 | 5.08E-52 |
| FAM149A | 3 | -0.593775924 | 0.117 | 0.345 | 1.86E-56 | 6.8E-52 |
| GUCY1A1 | 3 | -0.272573653 | 0.09 | 0.282 | 2.16E-56 | 7.89E-52 |
| KCNQ2 | 3 | -0.46397122 | 0.04 | 0.223 | 2.39E-56 | 8.76E-52 |
| NEAT1 | 3 | 0.73899843 | 0.957 | 0.853 | 2.55E-56 | 9.34E-52 |
| AC016042.1 | 3 | -0.595138268 | 0.063 | 0.257 | 3.47E-56 | 1.27E-51 |
| AMPH | 3 | -0.525739014 | 0.025 | 0.192 | 3.95E-56 | 1.45E-51 |
| IL6ST | 3 | 0.632945576 | 0.362 | 0.155 | 4.34E-56 | 1.59E-51 |
| PMEPA1 | 3 | 0.761145935 | 0.4 | 0.206 | 4.34E-56 | 1.59E-51 |
| FANCA | 3 | -0.575760975 | 0.232 | 0.483 | 4.45E-56 | 1.63E-51 |
| PDE11A | 3 | -0.658562711 | 0.036 | 0.208 | 4.92E-56 | 1.8E-51 |
| ANKS1B | 3 | 0.757761266 | 0.632 | 0.809 | 5.45E-56 | 1.99E-51 |
| MIR124-2HG | 3 | -0.441216677 | 0.007 | 0.147 | 1.2E-55 | 4.38E-51 |
| CADM2 | 3 | -1.352788286 | 0.252 | 0.473 | 2.23E-55 | 8.18E-51 |
| RIC3 | 3 | -0.404383727 | 0.022 | 0.179 | 3.57E-55 | 1.31E-50 |
| SHROOM4 | 3 | 0.701014265 | 0.765 | 0.584 | 3.85E-55 | 1.41E-50 |
| GOPC | 3 | 0.501594818 | 0.302 | 0.114 | 4.04E-55 | 1.48E-50 |
| PLPP3 | 3 | 0.650393329 | 0.296 | 0.117 | 4.48E-55 | 1.64E-50 |
| ACSL1 | 3 | -0.716222503 | 0.188 | 0.41 | 7.2E-55 | 2.64E-50 |
| CHRM3 | 3 | 1.106333561 | 0.586 | 0.397 | 7.26E-55 | 2.66E-50 |
| ZNF521 | 3 | 0.328724341 | 0.305 | 0.123 | 8.2E-55 | 3E-50 |
| VAV3 | 3 | -0.638171043 | 0.023 | 0.179 | 8.74E-55 | 3.2E-50 |
| PPM1E | 3 | -0.614858423 | 0.548 | 0.76 | 9.17E-55 | 3.36E-50 |
| FRMD4B | 3 | -0.864187057 | 0.227 | 0.46 | 1.23E-54 | 4.51E-50 |
| AC040174.1 | 3 | 0.448534402 | 0.183 | 0.042 | 1.33E-54 | 4.88E-50 |
| FHL1 | 3 | -0.480768239 | 0.065 | 0.262 | 1.62E-54 | 5.94E-50 |
| SFMBT1 | 3 | -0.605747759 | 0.213 | 0.454 | 1.89E-54 | 6.91E-50 |
| PAK1 | 3 | -0.647865661 | 0.36 | 0.601 | 4.34E-54 | 1.59E-49 |
| SNAP91 | 3 | -0.577430519 | 0.033 | 0.203 | 4.84E-54 | 1.77E-49 |
| DOK5 | 3 | 0.725343175 | 0.719 | 0.523 | 6.33E-54 | 2.32E-49 |
| SYT10 | 3 | -0.583087437 | 0.007 | 0.14 | 8.4E-54 | 3.07E-49 |
| RTN3 | 3 | -0.627540014 | 0.502 | 0.703 | 1.95E-53 | 7.13E-49 |
| AL513166.1 | 3 | -0.722782332 | 0.077 | 0.273 | 1.98E-53 | 7.25E-49 |
| CPQ | 3 | 0.608454007 | 0.489 | 0.262 | 2.17E-53 | 7.94E-49 |
| MTUS2 | 3 | -0.759880271 | 0.051 | 0.23 | 2.39E-53 | 8.76E-49 |
| NAAA | 3 | -0.625878375 | 0.112 | 0.318 | 2.55E-53 | 9.34E-49 |
| SRRM3 | 3 | -0.466324406 | 0.023 | 0.18 | 3.16E-53 | 1.16E-48 |

| | | | | | | |
|---|---|---|---|---|---|---|
| KANK1 | 3 | -0.68560788 | 0.294 | 0.537 | 3.28E-53 | 1.2E-48 |
| ANKRD36C | 3 | -0.618154725 | 0.476 | 0.68 | 3.49E-53 | 1.28E-48 |
| CREB5 | 3 | 0.730638998 | 0.396 | 0.179 | 4.1E-53 | 1.5E-48 |
| KCTD8 | 3 | -1.137857464 | 0.017 | 0.163 | 4.76E-53 | 1.74E-48 |
| CACNA1A | 3 | -1.510358539 | 0.095 | 0.291 | 6.13E-53 | 2.24E-48 |
| MTSS1 | 3 | -0.652854049 | 0.039 | 0.211 | 6.51E-53 | 2.38E-48 |
| LHFPL3 | 3 | -0.944126743 | 0.371 | 0.591 | 7.17E-53 | 2.63E-48 |
| ERICH3 | 3 | -0.560627627 | 0.012 | 0.156 | 7.31E-53 | 2.67E-48 |
| COL1A2 | 3 | 0.5989762 | 0.293 | 0.104 | 9.56E-53 | 3.5E-48 |
| FMN2 | 3 | -0.805093245 | 0.158 | 0.378 | 1.01E-52 | 3.68E-48 |
| STS | 3 | -0.736538803 | 0.156 | 0.363 | 1.07E-52 | 3.91E-48 |
| SCN8A | 3 | -0.64225252 | 0.48 | 0.689 | 1.25E-52 | 4.57E-48 |
| SDK1 | 3 | -0.82420387 | 0.86 | 0.911 | 1.61E-52 | 5.89E-48 |
| ATXN1 | 3 | 0.537831077 | 0.858 | 0.672 | 1.88E-52 | 6.89E-48 |
| ZNF804B | 3 | -1.690659066 | 0.037 | 0.203 | 2.57E-52 | 9.39E-48 |
| GPHN | 3 | -0.553746985 | 0.877 | 0.928 | 3.34E-52 | 1.22E-47 |
| TBX15 | 3 | 0.478748834 | 0.751 | 0.505 | 3.95E-52 | 1.45E-47 |
| PPP2R2A | 3 | 0.562829013 | 0.778 | 0.65 | 8.58E-52 | 3.14E-47 |
| CPLX2 | 3 | -0.462914971 | 0.03 | 0.19 | 9.62E-52 | 3.52E-47 |
| SHC4 | 3 | 0.350151128 | 0.122 | 0.016 | 1.07E-51 | 3.92E-47 |
| CALD1 | 3 | 0.54189007 | 0.641 | 0.4 | 1.18E-51 | 4.33E-47 |
| TTN | 3 | 1.34291324 | 0.256 | 0.138 | 1.3E-51 | 4.77E-47 |
| ZNRF2 | 3 | -0.570764439 | 0.146 | 0.363 | 1.34E-51 | 4.92E-47 |
| PRKG1 | 3 | 0.667362407 | 0.879 | 0.714 | 1.36E-51 | 4.98E-47 |
| TENT5A | 3 | 0.625410419 | 0.303 | 0.112 | 1.63E-51 | 5.98E-47 |
| ZNF536 | 3 | -0.907305212 | 0.749 | 0.832 | 1.78E-51 | 6.5E-47 |
| ROBO2 | 3 | 1.068647913 | 0.836 | 0.696 | 2.54E-51 | 9.29E-47 |
| TENM4 | 3 | -0.736526053 | 0.167 | 0.392 | 2.55E-51 | 9.35E-47 |
| RUNDC3B | 3 | -0.520332203 | 0.023 | 0.173 | 2.58E-51 | 9.45E-47 |
| C12orf56 | 3 | -0.40036601 | 0.019 | 0.169 | 3.69E-51 | 1.35E-46 |
| ITGB5 | 3 | 0.501222325 | 0.25 | 0.087 | 3.78E-51 | 1.38E-46 |
| AC093817.2 | 3 | 1.174851551 | 0.235 | 0.087 | 6.5E-51 | 2.38E-46 |
| GDAP1 | 3 | -0.741829714 | 0.133 | 0.334 | 7.01E-51 | 2.57E-46 |
| EXT1 | 3 | 0.629416155 | 0.819 | 0.674 | 8.25E-51 | 3.02E-46 |
| WDR76 | 3 | -0.543898603 | 0.15 | 0.373 | 9.05E-51 | 3.31E-46 |
| ARAP2 | 3 | 0.680951935 | 0.603 | 0.384 | 1.04E-50 | 3.82E-46 |
| NFIA | 3 | -0.882716292 | 0.877 | 0.896 | 1.23E-50 | 4.51E-46 |
| NPAS3 | 3 | 0.502792919 | 0.445 | 0.218 | 1.49E-50 | 5.46E-46 |
| LINC00937 | 3 | -0.544836724 | 0.041 | 0.207 | 1.86E-50 | 6.8E-46 |
| SLC1A3 | 3 | 0.511793893 | 0.262 | 0.088 | 2.22E-50 | 8.13E-46 |
| NSD2 | 3 | -0.563658221 | 0.499 | 0.707 | 2.23E-50 | 8.17E-46 |
| DOK6 | 3 | -0.332787324 | 0.416 | 0.658 | 2.29E-50 | 8.38E-46 |
| ITGA4 | 3 | -1.228414349 | 0.325 | 0.491 | 3.16E-50 | 1.16E-45 |
| STAC | 3 | 0.582840078 | 0.181 | 0.213 | 3.67E-50 | 1.34E-45 |
| RASGRP1 | 3 | -0.632297243 | 0.061 | 0.241 | 3.78E-50 | 1.38E-45 |
| AGBL4 | 3 | -1.099053785 | 0.058 | 0.227 | 4.8E-50 | 1.76E-45 |
| CADM1 | 3 | 0.79520858 | 0.547 | 0.426 | 6.02E-50 | 2.2E-45 |
| PLCB4 | 3 | 0.627859843 | 0.742 | 0.663 | 6.82E-50 | 2.5E-45 |
| IL33 | 3 | 0.698366922 | 0.112 | 0.016 | 8.63E-50 | 3.16E-45 |
| SP100 | 3 | 0.35265346 | 0.165 | 0.036 | 9.25E-50 | 3.39E-45 |
| NID1 | 3 | 0.604414075 | 0.474 | 0.259 | 9.62E-50 | 3.52E-45 |
| PTGIS | 3 | 0.309168028 | 0.123 | 0.019 | 9.88E-50 | 3.62E-45 |
| MPP6 | 3 | -0.853413696 | 0.484 | 0.668 | 1E-49 | 3.66E-45 |
| KCNMB4 | 3 | 0.65965851 | 0.402 | 0.219 | 1.04E-49 | 3.79E-45 |
| CEP170 | 3 | 0.52536284 | 0.874 | 0.757 | 1.26E-49 | 4.61E-45 |
| NETO1 | 3 | -0.626782859 | 0.033 | 0.189 | 1.29E-49 | 4.71E-45 |
| MEF2C-AS1 | 3 | 0.613408063 | 0.433 | 0.222 | 1.32E-49 | 4.83E-45 |
| DMXL2 | 3 | -0.613638969 | 0.38 | 0.6 | 1.36E-49 | 4.96E-45 |

| | | | | | | |
|---|---|---|---|---|---|---|
| AC068756.1 | 3 | 0.296664733 | 0.148 | 0.028 | 1.36E-49 | 4.99E-45 |
| DTNA | 3 | -0.837755845 | 0.522 | 0.719 | 1.4E-49 | 5.13E-45 |
| VGLL4 | 3 | 0.536454752 | 0.744 | 0.567 | 1.67E-49 | 6.11E-45 |
| TFAP2A | 3 | 0.277721456 | 0.118 | 0.016 | 2.82E-49 | 1.03E-44 |
| TES | 3 | 0.463504814 | 0.25 | 0.084 | 2.9E-49 | 1.06E-44 |
| GLDC | 3 | -0.675909239 | 0.111 | 0.298 | 3.31E-49 | 1.21E-44 |
| COL4A2 | 3 | 0.712228065 | 0.499 | 0.313 | 3.41E-49 | 1.25E-44 |
| LOXL4 | 3 | 0.405243507 | 0.137 | 0.028 | 3.6E-49 | 1.32E-44 |
| MYT1L | 3 | -0.895927643 | 0.014 | 0.145 | 4.54E-49 | 1.66E-44 |
| NCAPG | 3 | -0.635712379 | 0.099 | 0.295 | 4.77E-49 | 1.75E-44 |
| ZMAT4 | 3 | -0.698107566 | 0.313 | 0.554 | 4.87E-49 | 1.78E-44 |
| MTMR1 | 3 | -0.570893282 | 0.285 | 0.502 | 8.93E-49 | 3.27E-44 |
| HECW2 | 3 | -0.781948004 | 0.181 | 0.393 | 1.04E-48 | 3.82E-44 |
| LRRC4C | 3 | 0.344498337 | 0.741 | 0.502 | 1.05E-48 | 3.86E-44 |
| NXPH2 | 3 | -0.421559645 | 0.007 | 0.13 | 1.37E-48 | 5.01E-44 |
| RYR2 | 3 | -1.334278756 | 0.063 | 0.227 | 1.48E-48 | 5.41E-44 |
| RPGRIP1L | 3 | 0.685411901 | 0.566 | 0.409 | 1.5E-48 | 5.48E-44 |
| AL133346.1 | 3 | 0.632760568 | 0.312 | 0.125 | 1.62E-48 | 5.93E-44 |
| STARD13-AS | 3 | 0.382965835 | 0.214 | 0.072 | 2.06E-48 | 7.56E-44 |
| GPC3 | 3 | -0.978702694 | 0.073 | 0.253 | 2.07E-48 | 7.56E-44 |
| CRIM1 | 3 | -0.678453168 | 0.154 | 0.359 | 3.31E-48 | 1.21E-43 |
| FRMD5 | 3 | -0.919955521 | 0.216 | 0.433 | 3.63E-48 | 1.33E-43 |
| NWD2 | 3 | -0.556665565 | 0.017 | 0.157 | 3.99E-48 | 1.46E-43 |
| POLQ | 3 | -0.546588981 | 0.133 | 0.344 | 4.62E-48 | 1.69E-43 |
| ZYG11A | 3 | -0.605791542 | 0.13 | 0.325 | 8.7E-48 | 3.19E-43 |
| MAP2 | 3 | -0.871268892 | 0.355 | 0.57 | 9.11E-48 | 3.34E-43 |
| C1orf21 | 3 | -0.610290586 | 0.13 | 0.334 | 9.4E-48 | 3.44E-43 |
| ADGRG2 | 3 | -0.696422286 | 0.041 | 0.198 | 1.36E-47 | 5E-43 |
| ANKRD36B | 3 | -0.598731629 | 0.293 | 0.51 | 1.61E-47 | 5.9E-43 |
| NUP210 | 3 | -0.485824022 | 0.13 | 0.339 | 1.7E-47 | 6.21E-43 |
| SLC35D2 | 3 | 0.434995545 | 0.254 | 0.091 | 1.79E-47 | 6.55E-43 |
| KIF15 | 3 | -0.566878693 | 0.098 | 0.291 | 1.93E-47 | 7.08E-43 |
| WDHD1 | 3 | -0.528967188 | 0.225 | 0.454 | 1.96E-47 | 7.16E-43 |
| UNC80 | 3 | -0.627475126 | 0.06 | 0.227 | 2.04E-47 | 7.48E-43 |
| AP3B2 | 3 | -0.323036815 | 0.009 | 0.135 | 2.1E-47 | 7.69E-43 |
| MAST1 | 3 | -0.323768441 | 0.007 | 0.129 | 2.84E-47 | 1.04E-42 |
| SATB1-AS1 | 3 | -0.480853469 | 0.038 | 0.195 | 3.12E-47 | 1.14E-42 |
| SYT14 | 3 | -0.620800769 | 0.269 | 0.501 | 3.28E-47 | 1.2E-42 |
| ANKRD18A | 3 | -0.439378868 | 0.029 | 0.173 | 3.43E-47 | 1.25E-42 |
| FAM160A1 | 3 | -0.552217791 | 0.021 | 0.164 | 3.52E-47 | 1.29E-42 |
| KDM5B | 3 | 0.53013112 | 0.606 | 0.386 | 4.41E-47 | 1.61E-42 |
| WDFY3 | 3 | -0.528744895 | 0.851 | 0.903 | 4.92E-47 | 1.8E-42 |
| PCDH19 | 3 | -0.786008416 | 0.055 | 0.215 | 5.07E-47 | 1.86E-42 |
| AC093298.2 | 3 | 1.190996992 | 0.354 | 0.167 | 5.71E-47 | 2.09E-42 |
| SLC4A7 | 3 | -0.600518559 | 0.62 | 0.773 | 6.27E-47 | 2.29E-42 |
| AL356124.1 | 3 | 0.674349084 | 0.375 | 0.195 | 6.56E-47 | 2.4E-42 |
| TNC | 3 | 0.640890392 | 0.108 | 0.014 | 8.25E-47 | 3.02E-42 |
| TMPRSS5 | 3 | 0.29622422 | 0.134 | 0.026 | 8.64E-47 | 3.16E-42 |
| BPTF | 3 | 0.44864759 | 0.92 | 0.839 | 9.41E-47 | 3.44E-42 |
| KNTC1 | 3 | -0.502888705 | 0.287 | 0.522 | 9.85E-47 | 3.61E-42 |
| COBLL1 | 3 | -0.867402238 | 0.21 | 0.404 | 1.14E-46 | 4.17E-42 |
| PCBP3 | 3 | -0.691875406 | 0.091 | 0.27 | 1.18E-46 | 4.33E-42 |
| KCNH1 | 3 | -1.126984401 | 0.113 | 0.288 | 1.38E-46 | 5.07E-42 |
| DGCR9 | 3 | -0.404289268 | 0.019 | 0.159 | 1.44E-46 | 5.26E-42 |
| ANKRD10 | 3 | 0.52174687 | 0.777 | 0.661 | 1.94E-46 | 7.12E-42 |
| WEE1 | 3 | -0.598464864 | 0.237 | 0.456 | 1.96E-46 | 7.19E-42 |
| RABGAP1L | 3 | -0.744303568 | 0.655 | 0.761 | 1.98E-46 | 7.25E-42 |
| SAT1 | 3 | 0.571184368 | 0.31 | 0.126 | 2.22E-46 | 8.13E-42 |

| | | | | | | |
|---|---|---|---|---|---|---|
| PALM2-AKAP2 | 3 | 0.650839107 | 0.623 | 0.47 | 2.24E-46 | 8.18E-42 |
| CPNE8 | 3 | -0.584047467 | 0.03 | 0.179 | 2.25E-46 | 8.23E-42 |
| RAB31 | 3 | 0.601530159 | 0.43 | 0.236 | 2.46E-46 | 9.01E-42 |
| AL138828.1 | 3 | 0.380689725 | 0.215 | 0.067 | 2.59E-46 | 9.49E-42 |
| AC063944.1 | 3 | 0.491598163 | 0.118 | 0.021 | 2.75E-46 | 1.01E-41 |
| ASXL3 | 3 | -0.658785375 | 0.277 | 0.505 | 2.76E-46 | 1.01E-41 |
| AC002377.1 | 3 | 0.690940659 | 0.255 | 0.11 | 2.76E-46 | 1.01E-41 |
| GPC6 | 3 | -0.662991097 | 0.794 | 0.819 | 2.78E-46 | 1.02E-41 |
| NLGN1 | 3 | -0.91477706 | 0.722 | 0.816 | 3.63E-46 | 1.33E-41 |
| UACA | 3 | 0.50343211 | 0.578 | 0.351 | 4.76E-46 | 1.74E-41 |
| VIM | 3 | 0.608738392 | 0.439 | 0.247 | 5.29E-46 | 1.94E-41 |
| SCN3A | 3 | -0.67489417 | 0.098 | 0.289 | 5.79E-46 | 2.12E-41 |
| PELI2 | 3 | -0.574091782 | 0.228 | 0.454 | 6.06E-46 | 2.22E-41 |
| ALPK1 | 3 | 0.353651764 | 0.194 | 0.054 | 7.11E-46 | 2.6E-41 |
| GLCCI1 | 3 | 0.735782343 | 0.704 | 0.562 | 9.68E-46 | 3.55E-41 |
| JAKMIP2 | 3 | -0.530965018 | 0.163 | 0.378 | 1.01E-45 | 3.71E-41 |
| XKR6 | 3 | -0.618773202 | 0.332 | 0.552 | 1.06E-45 | 3.88E-41 |
| AC009435.1 | 3 | 0.765585988 | 0.105 | 0.016 | 1.27E-45 | 4.65E-41 |
| CENPU | 3 | -0.525904049 | 0.13 | 0.33 | 1.68E-45 | 6.15E-41 |
| SCN7A | 3 | -1.267997253 | 0.021 | 0.156 | 1.74E-45 | 6.38E-41 |
| TPD52L1 | 3 | 0.546539585 | 0.274 | 0.115 | 1.9E-45 | 6.97E-41 |
| PPP1R12A | 3 | 0.517007723 | 0.82 | 0.74 | 2.35E-45 | 8.61E-41 |
| TRPS1 | 3 | 0.463606388 | 0.781 | 0.564 | 3.41E-45 | 1.25E-40 |
| RFC3 | 3 | -0.669042682 | 0.286 | 0.497 | 5.66E-45 | 2.07E-40 |
| TCERG1L | 3 | 0.482684201 | 0.131 | 0.024 | 7.59E-45 | 2.78E-40 |
| KIF1A | 3 | -0.405343793 | 0.034 | 0.183 | 8.79E-45 | 3.22E-40 |
| GRB10 | 3 | 0.592152377 | 0.567 | 0.362 | 9.29E-45 | 3.4E-40 |
| KCNQ5 | 3 | 1.020481573 | 0.603 | 0.502 | 1.06E-44 | 3.87E-40 |
| ANKH | 3 | -0.55750946 | 0.062 | 0.223 | 1.24E-44 | 4.54E-40 |
| HBEGF | 3 | -0.401030068 | 0.012 | 0.136 | 1.55E-44 | 5.67E-40 |
| LINC02018 | 3 | 0.357713029 | 0.275 | 0.105 | 2.06E-44 | 7.53E-40 |
| KIAA1755 | 3 | 0.367621857 | 0.16 | 0.038 | 2.13E-44 | 7.81E-40 |
| MCTP1 | 3 | -0.961054079 | 0.133 | 0.317 | 2.35E-44 | 8.62E-40 |
| MMP16 | 3 | -0.449780416 | 0.547 | 0.749 | 2.5E-44 | 9.15E-40 |
| SP110 | 3 | 0.329492974 | 0.191 | 0.054 | 2.57E-44 | 9.43E-40 |
| CERKL | 3 | -0.455759922 | 0.018 | 0.149 | 2.64E-44 | 9.66E-40 |
| STEAP2 | 3 | -0.318655782 | 0.012 | 0.135 | 3.71E-44 | 1.36E-39 |
| GALNT18 | 3 | -0.605109564 | 0.207 | 0.423 | 4.22E-44 | 1.54E-39 |
| SLC25A37 | 3 | 0.584401385 | 0.441 | 0.274 | 4.31E-44 | 1.58E-39 |
| ITGA1 | 3 | 0.89739932 | 0.253 | 0.111 | 4.81E-44 | 1.76E-39 |
| CCDC68 | 3 | 0.335063061 | 0.131 | 0.024 | 4.86E-44 | 1.78E-39 |
| RHOBTB3 | 3 | 0.626969174 | 0.598 | 0.447 | 5.55E-44 | 2.03E-39 |
| SUSD4 | 3 | -0.558249344 | 0.099 | 0.287 | 6.01E-44 | 2.2E-39 |
| RNF213 | 3 | 0.579285209 | 0.609 | 0.42 | 6.61E-44 | 2.42E-39 |
| IL12RB2 | 3 | -0.999716587 | 0.129 | 0.306 | 6.65E-44 | 2.43E-39 |
| LINC01456 | 3 | -0.457205353 | 0.015 | 0.144 | 8.29E-44 | 3.03E-39 |
| KAZN | 3 | -0.405100296 | 0.881 | 0.901 | 1.08E-43 | 3.97E-39 |
| SF3B1 | 3 | 0.455327264 | 0.747 | 0.589 | 1.13E-43 | 4.13E-39 |
| AC022075.1 | 3 | 0.285369075 | 0.113 | 0.018 | 1.55E-43 | 5.66E-39 |
| ICA1 | 3 | 0.56887571 | 0.37 | 0.196 | 2.67E-43 | 9.78E-39 |
| STXBP1 | 3 | -0.495395497 | 0.154 | 0.351 | 2.77E-43 | 1.01E-38 |
| ZNF207 | 3 | 0.415792289 | 0.82 | 0.728 | 2.99E-43 | 1.1E-38 |
| LHFPL4 | 3 | -0.340588397 | 0.009 | 0.124 | 3.12E-43 | 1.14E-38 |
| EPB41 | 3 | -0.651329694 | 0.311 | 0.504 | 4.69E-43 | 1.72E-38 |
| ARHGAP18 | 3 | 0.695004489 | 0.59 | 0.438 | 5.68E-43 | 2.08E-38 |
| MEGF11 | 3 | -0.453091782 | 0.139 | 0.339 | 6.92E-43 | 2.53E-38 |
| ELMO1 | 3 | 0.684749074 | 0.708 | 0.53 | 8.29E-43 | 3.03E-38 |
| SPTBN1 | 3 | 0.592515648 | 0.774 | 0.676 | 9.05E-43 | 3.31E-38 |

| | | | | | | |
|---|---|---|---|---|---|---|
| TMEM65 | 3 | -0.535227434 | 0.09 | 0.264 | 9.74E-43 | 3.57E-38 |
| DNA2 | 3 | -0.51846963 | 0.182 | 0.39 | 1.26E-42 | 4.63E-38 |
| KANSL1 | 3 | 0.394139359 | 0.943 | 0.895 | 1.3E-42 | 4.77E-38 |
| PARP8 | 3 | 0.536929456 | 0.524 | 0.308 | 1.35E-42 | 4.95E-38 |
| STAT2 | 3 | 0.486472113 | 0.427 | 0.235 | 1.41E-42 | 5.14E-38 |
| TENM2 | 3 | -0.729176539 | 0.431 | 0.636 | 1.44E-42 | 5.26E-38 |
| ANXA2 | 3 | 0.574074676 | 0.428 | 0.248 | 1.5E-42 | 5.48E-38 |
| AKAP13 | 3 | 0.401512667 | 0.745 | 0.528 | 1.78E-42 | 6.51E-38 |
| ZNF730 | 3 | -0.483851179 | 0.064 | 0.228 | 1.81E-42 | 6.61E-38 |
| TANC2 | 3 | 0.69038758 | 0.789 | 0.705 | 2.19E-42 | 8.01E-38 |
| TOPBP1 | 3 | -0.539711452 | 0.372 | 0.57 | 2.4E-42 | 8.79E-38 |
| LINC01473 | 3 | 0.548883322 | 0.734 | 0.549 | 3.09E-42 | 1.13E-37 |
| PURPL | 3 | -0.6753358 | 0.017 | 0.141 | 6.29E-42 | 2.3E-37 |
| IGDCC3 | 3 | -0.482590377 | 0.098 | 0.279 | 6.33E-42 | 2.32E-37 |
| NAALADL2 | 3 | 0.689032992 | 0.586 | 0.393 | 7.04E-42 | 2.58E-37 |
| FRMD6 | 3 | 0.588816945 | 0.405 | 0.215 | 7.54E-42 | 2.76E-37 |
| AC008591.1 | 3 | -0.787509519 | 0.246 | 0.458 | 7.75E-42 | 2.84E-37 |
| GRM1 | 3 | 1.16959367 | 0.148 | 0.08 | 7.89E-42 | 2.89E-37 |
| SPRY4-AS1 | 3 | 0.579689379 | 0.329 | 0.151 | 8.92E-42 | 3.26E-37 |
| MIS18BP1 | 3 | -0.572743026 | 0.351 | 0.547 | 9.71E-42 | 3.55E-37 |
| GALNT16 | 3 | -0.312726419 | 0.011 | 0.122 | 1.15E-41 | 4.19E-37 |
| EML5 | 3 | -0.512260961 | 0.036 | 0.176 | 1.17E-41 | 4.27E-37 |
| TMPO | 3 | -0.515402882 | 0.329 | 0.54 | 1.18E-41 | 4.32E-37 |
| ATP11C | 3 | -0.583866531 | 0.575 | 0.735 | 1.32E-41 | 4.82E-37 |
| HS6ST2 | 3 | -0.340149468 | 0.473 | 0.692 | 1.41E-41 | 5.15E-37 |
| ZSWIM7 | 3 | 0.455845345 | 0.408 | 0.221 | 1.42E-41 | 5.19E-37 |
| MBTD1 | 3 | 0.442454489 | 0.747 | 0.573 | 1.8E-41 | 6.58E-37 |
| CLSPN | 3 | -0.469710468 | 0.102 | 0.284 | 2.07E-41 | 7.57E-37 |
| ZNF711 | 3 | -0.523755955 | 0.423 | 0.621 | 2.2E-41 | 8.04E-37 |
| PMP22 | 3 | 0.554961953 | 0.34 | 0.175 | 2.31E-41 | 8.47E-37 |
| SHTN1 | 3 | 0.612413305 | 0.749 | 0.678 | 2.88E-41 | 1.05E-36 |
| TOP2A | 3 | -0.615354518 | 0.085 | 0.257 | 3.35E-41 | 1.23E-36 |
| PHKA1 | 3 | -0.547557545 | 0.131 | 0.311 | 3.71E-41 | 1.36E-36 |
| ARL15 | 3 | 0.541086244 | 0.784 | 0.655 | 3.86E-41 | 1.41E-36 |
| SLC22A23 | 3 | -0.609123571 | 0.179 | 0.379 | 4.97E-41 | 1.82E-36 |
| ARNTL | 3 | 0.47765994 | 0.291 | 0.129 | 5.79E-41 | 2.12E-36 |
| WDR11-AS1 | 3 | -0.622232315 | 0.156 | 0.347 | 6.98E-41 | 2.56E-36 |
| CHEK1 | 3 | -0.485938771 | 0.163 | 0.361 | 7.07E-41 | 2.59E-36 |
| RGL1 | 3 | -0.753221348 | 0.671 | 0.733 | 7.85E-41 | 2.88E-36 |
| HULC | 3 | -1.117921703 | 0.262 | 0.434 | 8.88E-41 | 3.25E-36 |
| BMP6 | 3 | -0.443499918 | 0.009 | 0.119 | 1.02E-40 | 3.74E-36 |
| RBM47 | 3 | -0.885465675 | 0.023 | 0.142 | 1.28E-40 | 4.68E-36 |
| RAPGEF2 | 3 | 0.541720196 | 0.786 | 0.638 | 1.49E-40 | 5.44E-36 |
| AC008415.1 | 3 | 0.589364341 | 0.12 | 0.023 | 1.81E-40 | 6.63E-36 |
| ZFHX4-AS1 | 3 | 0.341443694 | 0.163 | 0.043 | 1.83E-40 | 6.7E-36 |
| GREB1 | 3 | 0.27560972 | 0.242 | 0.093 | 2.13E-40 | 7.81E-36 |
| CLVS1 | 3 | -0.593255326 | 0.066 | 0.225 | 2.28E-40 | 8.36E-36 |
| RIC1 | 3 | 0.474008097 | 0.702 | 0.548 | 2.69E-40 | 9.83E-36 |
| SEMA3A | 3 | 1.28337139 | 0.353 | 0.28 | 3.29E-40 | 1.21E-35 |
| MPPED2 | 3 | 0.563228726 | 0.349 | 0.165 | 3.57E-40 | 1.31E-35 |
| SEMA3D | 3 | 0.621466007 | 0.229 | 0.083 | 4.81E-40 | 1.76E-35 |
| GTF2F2 | 3 | 0.522121099 | 0.583 | 0.423 | 5.58E-40 | 2.04E-35 |
| FGD5 | 3 | -0.403307596 | 0.053 | 0.205 | 6.29E-40 | 2.3E-35 |
| EIF1AX | 3 | -0.526971605 | 0.27 | 0.464 | 6.42E-40 | 2.35E-35 |
| LPCAT2 | 3 | 0.274355622 | 0.158 | 0.04 | 6.44E-40 | 2.36E-35 |
| TAFA1 | 3 | -1.232089433 | 0.085 | 0.244 | 6.44E-40 | 2.36E-35 |
| COL27A1 | 3 | 0.577639051 | 0.565 | 0.362 | 6.44E-40 | 2.36E-35 |
| NF1 | 3 | 0.389509934 | 0.948 | 0.901 | 6.86E-40 | 2.51E-35 |

| | | | | | | |
|---|---|---|---|---|---|---|
| IQGAP2 | 3 | -0.706017322 | 0.235 | 0.422 | 7.38E-40 | 2.7E-35 |
| CHRM2 | 3 | -0.764190834 | 0.006 | 0.107 | 9.85E-40 | 3.6E-35 |
| DCLK2 | 3 | -0.637868741 | 0.671 | 0.735 | 1.11E-39 | 4.06E-35 |
| AC003659.1 | 3 | 0.268904991 | 0.154 | 0.039 | 1.16E-39 | 4.23E-35 |
| GAB1 | 3 | 0.517428445 | 0.807 | 0.649 | 1.38E-39 | 5.07E-35 |
| BICD1 | 3 | -0.544040667 | 0.714 | 0.815 | 1.44E-39 | 5.28E-35 |
| PARP14 | 3 | 0.442680178 | 0.343 | 0.162 | 1.45E-39 | 5.31E-35 |
| POLR2F | 3 | 0.577951426 | 0.235 | 0.112 | 1.47E-39 | 5.39E-35 |
| KIF4A | 3 | -0.50475605 | 0.057 | 0.209 | 1.96E-39 | 7.18E-35 |
| TLCD4 | 3 | -0.338949168 | 0.02 | 0.143 | 2.09E-39 | 7.63E-35 |
| IFI44 | 3 | 0.343068517 | 0.133 | 0.029 | 2.54E-39 | 9.31E-35 |
| RGS17 | 3 | -0.466490742 | 0.047 | 0.191 | 2.78E-39 | 1.02E-34 |
| RAP1GAP2 | 3 | -1.107349401 | 0.108 | 0.276 | 3.01E-39 | 1.1E-34 |
| FOXN3 | 3 | -0.527546841 | 0.9 | 0.936 | 3.25E-39 | 1.19E-34 |
| AC074286.1 | 3 | -0.426591842 | 0.013 | 0.125 | 3.25E-39 | 1.19E-34 |
| RREB1 | 3 | 0.464887079 | 0.524 | 0.319 | 3.37E-39 | 1.23E-34 |
| FAXDC2 | 3 | 0.34608973 | 0.228 | 0.082 | 4.14E-39 | 1.52E-34 |
| ITGBL1 | 3 | -1.583260756 | 0.097 | 0.226 | 4.59E-39 | 1.68E-34 |
| SREBF2 | 3 | -0.615664915 | 0.326 | 0.52 | 4.81E-39 | 1.76E-34 |
| APPBP2 | 3 | 0.482718668 | 0.611 | 0.435 | 4.94E-39 | 1.81E-34 |
| CACNA1C | 3 | -0.781905601 | 0.295 | 0.504 | 5.76E-39 | 2.11E-34 |
| FCHSD2 | 3 | -0.50430915 | 0.609 | 0.762 | 6.01E-39 | 2.2E-34 |
| CA3 | 3 | 0.369677022 | 0.133 | 0.031 | 6.6E-39 | 2.42E-34 |
| PLCB1 | 3 | -0.782864051 | 0.372 | 0.545 | 8.72E-39 | 3.19E-34 |
| CDK1 | 3 | -0.427184749 | 0.064 | 0.22 | 9.11E-39 | 3.34E-34 |
| LINC01768 | 3 | -0.579036058 | 0.053 | 0.196 | 9.48E-39 | 3.47E-34 |
| RIMBP2 | 3 | -1.137772938 | 0.024 | 0.143 | 1.13E-38 | 4.14E-34 |
| SGCZ | 3 | -0.909814942 | 0.397 | 0.573 | 1.24E-38 | 4.55E-34 |
| LBX1-AS1 | 3 | -0.33406744 | 0.029 | 0.159 | 1.29E-38 | 4.73E-34 |
| HACE1 | 3 | -0.630186037 | 0.486 | 0.635 | 1.31E-38 | 4.78E-34 |
| ARNTL2 | 3 | -0.356338759 | 0.018 | 0.137 | 1.38E-38 | 5.04E-34 |
| GALNT2 | 3 | 0.566783973 | 0.533 | 0.385 | 1.38E-38 | 5.05E-34 |
| CDC14B | 3 | 0.580740631 | 0.403 | 0.22 | 1.73E-38 | 6.32E-34 |
| CUX2 | 3 | -0.534129601 | 0.134 | 0.318 | 1.83E-38 | 6.71E-34 |
| LINC02516 | 3 | 0.304212279 | 0.116 | 0.024 | 1.84E-38 | 6.73E-34 |
| EFCAB13 | 3 | 0.493276154 | 0.387 | 0.226 | 1.95E-38 | 7.15E-34 |
| NRXN1 | 3 | -0.758577118 | 0.428 | 0.62 | 2.34E-38 | 8.56E-34 |
| ARHGAP8 | 3 | -0.412227436 | 0.071 | 0.231 | 2.43E-38 | 8.89E-34 |
| DLG3 | 3 | -0.450590452 | 0.131 | 0.311 | 2.47E-38 | 9.06E-34 |
| FUT9 | 3 | -0.414361107 | 0.006 | 0.105 | 2.61E-38 | 9.56E-34 |
| USP32 | 3 | 0.496254913 | 0.801 | 0.671 | 2.87E-38 | 1.05E-33 |
| AL157778.1 | 3 | -1.118231303 | 0.099 | 0.257 | 2.9E-38 | 1.06E-33 |
| ANOS1 | 3 | -0.865967063 | 0.296 | 0.46 | 2.98E-38 | 1.09E-33 |
| FAM184B | 3 | -0.466910019 | 0.061 | 0.21 | 3.56E-38 | 1.3E-33 |
| ITGA2 | 3 | 0.30567808 | 0.125 | 0.026 | 3.76E-38 | 1.38E-33 |
| COL4A1 | 3 | 0.508556814 | 0.269 | 0.118 | 5.14E-38 | 1.88E-33 |
| ZDBF2 | 3 | -0.417398139 | 0.058 | 0.207 | 6.51E-38 | 2.38E-33 |
| TBC1D1 | 3 | -0.521930246 | 0.309 | 0.503 | 6.69E-38 | 2.45E-33 |
| KANK4 | 3 | 0.481820201 | 0.156 | 0.042 | 7.48E-38 | 2.74E-33 |
| CDC7 | 3 | -0.471701929 | 0.183 | 0.369 | 7.9E-38 | 2.89E-33 |
| PAM | 3 | -0.811804727 | 0.673 | 0.767 | 8.98E-38 | 3.29E-33 |
| PRICKLE1 | 3 | 0.737699327 | 0.564 | 0.391 | 9.17E-38 | 3.36E-33 |
| PIK3R3 | 3 | -0.575478326 | 0.319 | 0.519 | 1.13E-37 | 4.14E-33 |
| SUCLG2 | 3 | 0.452465921 | 0.641 | 0.447 | 1.3E-37 | 4.76E-33 |
| CIP2A | 3 | -0.507150621 | 0.142 | 0.319 | 1.3E-37 | 4.77E-33 |
| HHIP | 3 | -0.482908757 | 0.02 | 0.132 | 1.38E-37 | 5.06E-33 |
| TBC1D4 | 3 | -0.623272417 | 0.233 | 0.421 | 1.49E-37 | 5.44E-33 |
| NOS1AP | 3 | -0.492694978 | 0.156 | 0.342 | 1.87E-37 | 6.83E-33 |

| | | | | | | |
|---|---|---|---|---|---|---|
| ZNF516 | 3 | 0.456366822 | 0.71 | 0.531 | 1.91E-37 | 6.99E-33 |
| LINC00882 | 3 | 0.624020489 | 0.217 | 0.086 | 2.65E-37 | 9.72E-33 |
| TVP23C | 3 | 0.44229219 | 0.475 | 0.285 | 3.07E-37 | 1.13E-32 |
| WWC1 | 3 | -0.447436646 | 0.171 | 0.363 | 3.55E-37 | 1.3E-32 |
| AC120193.1 | 3 | -0.523295686 | 0.022 | 0.141 | 3.58E-37 | 1.31E-32 |
| FSTL4 | 3 | -0.454323425 | 0.016 | 0.129 | 4.19E-37 | 1.53E-32 |
| FHIT | 3 | 0.476299121 | 0.862 | 0.721 | 4.2E-37 | 1.54E-32 |
| PCLO | 3 | -0.444552392 | 0.427 | 0.631 | 4.24E-37 | 1.55E-32 |
| PDZD2 | 3 | -0.531184841 | 0.211 | 0.411 | 4.77E-37 | 1.75E-32 |
| WDR44 | 3 | -0.526496278 | 0.353 | 0.548 | 5.13E-37 | 1.88E-32 |
| HCN1 | 3 | -0.671270511 | 0.038 | 0.171 | 5.99E-37 | 2.19E-32 |
| NCAPG2 | 3 | -0.505866809 | 0.199 | 0.389 | 6.89E-37 | 2.52E-32 |
| FMR1 | 3 | -0.542256284 | 0.349 | 0.518 | 7.09E-37 | 2.6E-32 |
| REEP1 | 3 | -0.34822569 | 0.024 | 0.145 | 7.68E-37 | 2.81E-32 |
| BBX | 3 | 0.496723309 | 0.814 | 0.682 | 8.69E-37 | 3.18E-32 |
| TMEM178B | 3 | -0.662517493 | 0.324 | 0.527 | 9.11E-37 | 3.34E-32 |
| PTPN4 | 3 | -0.51927653 | 0.492 | 0.651 | 9.36E-37 | 3.43E-32 |
| ATF7IP2 | 3 | 0.530712694 | 0.402 | 0.232 | 1.11E-36 | 4.06E-32 |
| ITPR2 | 3 | 0.702317051 | 0.486 | 0.327 | 1.24E-36 | 4.54E-32 |
| CELF4 | 3 | -0.567457111 | 0.134 | 0.308 | 1.27E-36 | 4.66E-32 |
| CCDC18-AS1 | 3 | 0.46503216 | 0.529 | 0.334 | 1.27E-36 | 4.66E-32 |
| ATP2B2 | 3 | -0.37120117 | 0.012 | 0.12 | 1.35E-36 | 4.93E-32 |
| EYA2 | 3 | -0.441697049 | 0.026 | 0.149 | 1.46E-36 | 5.34E-32 |
| EVA1C | 3 | -0.310736797 | 0.018 | 0.133 | 1.48E-36 | 5.43E-32 |
| ELK3 | 3 | 0.308897516 | 0.203 | 0.071 | 1.54E-36 | 5.62E-32 |
| TRPC6 | 3 | 0.822881699 | 0.131 | 0.087 | 1.61E-36 | 5.91E-32 |
| COL3A1 | 3 | 0.713390105 | 0.444 | 0.254 | 1.64E-36 | 5.99E-32 |
| SYCP2 | 3 | -1.00092588 | 0.101 | 0.243 | 1.77E-36 | 6.48E-32 |
| AL589740.1 | 3 | -0.875607781 | 0.089 | 0.234 | 1.86E-36 | 6.8E-32 |
| ARHGAP26 | 3 | 0.548491741 | 0.317 | 0.159 | 1.87E-36 | 6.85E-32 |
| MAPT | 3 | -0.412230244 | 0.029 | 0.148 | 2.42E-36 | 8.86E-32 |
| ADGRG6 | 3 | 0.515831149 | 0.179 | 0.057 | 2.94E-36 | 1.08E-31 |
| PDE1C | 3 | -0.893389124 | 0.306 | 0.308 | 2.97E-36 | 1.09E-31 |
| E2F7 | 3 | -0.437967356 | 0.059 | 0.205 | 3.47E-36 | 1.27E-31 |
| SSH2 | 3 | 0.501789163 | 0.827 | 0.798 | 3.65E-36 | 1.34E-31 |
| SETBP1 | 3 | -0.546964843 | 0.405 | 0.602 | 4.28E-36 | 1.57E-31 |
| GABRB1 | 3 | 0.273420999 | 0.142 | 0.038 | 4.52E-36 | 1.65E-31 |
| CPS1 | 3 | 0.257336805 | 0.133 | 0.031 | 5.06E-36 | 1.85E-31 |
| ACSS1 | 3 | 0.350958113 | 0.223 | 0.085 | 5.22E-36 | 1.91E-31 |
| SMC2 | 3 | -0.468242373 | 0.171 | 0.356 | 5.84E-36 | 2.14E-31 |
| SLC22A15 | 3 | -0.66406663 | 0.166 | 0.329 | 6.05E-36 | 2.22E-31 |
| PDE4DIP | 3 | -0.482512933 | 0.173 | 0.353 | 7.47E-36 | 2.73E-31 |
| MSH2 | 3 | -0.487615334 | 0.363 | 0.55 | 8.3E-36 | 3.04E-31 |
| MED14 | 3 | -0.481691908 | 0.5 | 0.661 | 9.47E-36 | 3.47E-31 |
| DOCK1 | 3 | 0.41875765 | 0.817 | 0.652 | 9.77E-36 | 3.58E-31 |
| CNNM1 | 3 | -0.296529976 | 0.023 | 0.141 | 9.8E-36 | 3.59E-31 |
| THRB | 3 | -0.815424848 | 0.152 | 0.331 | 1.01E-35 | 3.7E-31 |
| SYNJ2 | 3 | 0.585273611 | 0.334 | 0.172 | 1.51E-35 | 5.54E-31 |
| C1orf198 | 3 | 0.272502946 | 0.155 | 0.045 | 2E-35 | 7.31E-31 |
| MVB12B | 3 | 0.513210423 | 0.561 | 0.402 | 2.46E-35 | 9E-31 |
| BICC1 | 3 | -0.565972232 | 0.143 | 0.32 | 2.68E-35 | 9.81E-31 |
| LINC00907 | 3 | 0.376069342 | 0.419 | 0.232 | 2.69E-35 | 9.84E-31 |
| FZD6 | 3 | -0.39611172 | 0.094 | 0.249 | 2.92E-35 | 1.07E-30 |
| PEX5L | 3 | -0.679585191 | 0.664 | 0.777 | 3.04E-35 | 1.11E-30 |
| MTMR7 | 3 | -0.589911297 | 0.191 | 0.369 | 3.13E-35 | 1.14E-30 |
| XACT | 3 | -1.29196694 | 0.115 | 0.267 | 3.24E-35 | 1.19E-30 |
| MAGED1 | 3 | -0.504118001 | 0.451 | 0.626 | 3.39E-35 | 1.24E-30 |
| DLC1 | 3 | 0.742301892 | 0.419 | 0.246 | 3.54E-35 | 1.3E-30 |

| | | | | | | |
|---|---|---|---|---|---|---|
| CACNG4 | 3 | -0.318912319 | 0.022 | 0.137 | 3.71E-35 | 1.36E-30 |
| MAP3K21 | 3 | -0.294324676 | 0.014 | 0.121 | 3.81E-35 | 1.39E-30 |
| ENPP1 | 3 | -0.592350276 | 0.19 | 0.375 | 3.87E-35 | 1.42E-30 |
| IKZF3 | 3 | -0.478566345 | 0.118 | 0.288 | 4.12E-35 | 1.51E-30 |
| GK | 3 | -0.489688044 | 0.129 | 0.296 | 4.36E-35 | 1.6E-30 |
| TCF4 | 3 | -0.622295514 | 0.889 | 0.89 | 4.72E-35 | 1.73E-30 |
| FANCL | 3 | -0.530391652 | 0.481 | 0.642 | 4.91E-35 | 1.8E-30 |
| AC011246.1 | 3 | 0.792998597 | 0.505 | 0.352 | 5.88E-35 | 2.15E-30 |
| CHRNA7 | 3 | -0.510451395 | 0.098 | 0.254 | 7.02E-35 | 2.57E-30 |
| TWIST1 | 3 | 0.488592186 | 0.321 | 0.156 | 7.61E-35 | 2.79E-30 |
| HIST1H2AC | 3 | 0.486165475 | 0.187 | 0.065 | 8.36E-35 | 3.06E-30 |
| EPB41L4B | 3 | -0.30799777 | 0.025 | 0.141 | 9.47E-35 | 3.47E-30 |
| MAP1B | 3 | -0.444760525 | 0.565 | 0.744 | 1.04E-34 | 3.81E-30 |
| ME1 | 3 | 0.481425687 | 0.279 | 0.128 | 1.06E-34 | 3.88E-30 |
| LYPD6 | 3 | -0.585801738 | 0.193 | 0.369 | 1.32E-34 | 4.84E-30 |
| BLM | 3 | -0.464870461 | 0.141 | 0.316 | 1.78E-34 | 6.53E-30 |
| RBMX | 3 | -0.416556644 | 0.728 | 0.838 | 1.89E-34 | 6.92E-30 |
| ANKRD44 | 3 | 0.568389471 | 0.471 | 0.307 | 1.96E-34 | 7.19E-30 |
| UNC13A | 3 | -0.347901121 | 0.028 | 0.146 | 2.56E-34 | 9.37E-30 |
| MIR4435-2HG | 3 | 0.581636741 | 0.382 | 0.216 | 2.61E-34 | 9.56E-30 |
| C9orf135 | 3 | -0.334118437 | 0.008 | 0.102 | 2.71E-34 | 9.93E-30 |
| ANKRD28 | 3 | 0.50355354 | 0.734 | 0.629 | 2.85E-34 | 1.04E-29 |
| AC019197.1 | 3 | -0.423848191 | 0.021 | 0.132 | 3.04E-34 | 1.11E-29 |
| HUNK | 3 | -0.388009003 | 0.039 | 0.168 | 3.28E-34 | 1.2E-29 |
| ATAD5 | 3 | -0.446098141 | 0.352 | 0.549 | 3.45E-34 | 1.26E-29 |
| CDR2 | 3 | 0.461536486 | 0.408 | 0.246 | 3.65E-34 | 1.34E-29 |
| FAXC | 3 | -0.30371256 | 0.029 | 0.149 | 3.77E-34 | 1.38E-29 |
| TUNAR | 3 | -0.381736505 | 0.008 | 0.102 | 3.8E-34 | 1.39E-29 |
| AC011416.4 | 3 | 0.299783057 | 0.146 | 0.041 | 4.07E-34 | 1.49E-29 |
| ADGRA2 | 3 | 0.42785668 | 0.279 | 0.127 | 4.67E-34 | 1.71E-29 |
| KDR | 3 | 0.369892274 | 0.102 | 0.02 | 5.34E-34 | 1.95E-29 |
| XK | 3 | -0.257369943 | 0.015 | 0.121 | 5.89E-34 | 2.16E-29 |
| ANLN | 3 | -0.51703149 | 0.126 | 0.289 | 6.57E-34 | 2.41E-29 |
| FAM111B | 3 | -0.353785015 | 0.042 | 0.172 | 7.48E-34 | 2.74E-29 |
| CDH10 | 3 | -0.698724417 | 0.173 | 0.349 | 7.53E-34 | 2.76E-29 |
| OTX2 | 3 | -0.444396096 | 0.01 | 0.107 | 8.64E-34 | 3.16E-29 |
| DUXAP8 | 3 | -0.280142905 | 0.014 | 0.115 | 8.68E-34 | 3.18E-29 |
| KLHL1 | 3 | -0.902926209 | 0.014 | 0.114 | 8.78E-34 | 3.21E-29 |
| OPCML | 3 | -0.946504458 | 0.193 | 0.364 | 1.16E-33 | 4.25E-29 |
| ABI3BP | 3 | 0.661444344 | 0.11 | 0.037 | 1.29E-33 | 4.73E-29 |
| DIP2B | 3 | 0.479253277 | 0.792 | 0.695 | 1.58E-33 | 5.78E-29 |
| AC069277.1 | 3 | 0.453747305 | 0.177 | 0.058 | 1.63E-33 | 5.96E-29 |
| CACNB2 | 3 | -0.72796677 | 0.632 | 0.746 | 1.64E-33 | 6E-29 |
| POLE | 3 | -0.39239713 | 0.197 | 0.383 | 1.65E-33 | 6.06E-29 |
| KCNS3 | 3 | -0.328722001 | 0.013 | 0.114 | 1.94E-33 | 7.1E-29 |
| ARRDC3-AS1 | 3 | 1.023460138 | 0.403 | 0.268 | 2.08E-33 | 7.63E-29 |
| STX8 | 3 | 0.453029566 | 0.634 | 0.46 | 2.65E-33 | 9.7E-29 |
| MCM10 | 3 | -0.330496785 | 0.071 | 0.215 | 3.49E-33 | 1.28E-28 |
| SLC38A4 | 3 | 0.689105955 | 0.294 | 0.159 | 3.51E-33 | 1.29E-28 |
| ARHGEF4 | 3 | -0.387085664 | 0.094 | 0.25 | 3.78E-33 | 1.39E-28 |
| TOM1L2 | 3 | 0.45217214 | 0.462 | 0.298 | 4.21E-33 | 1.54E-28 |
| FKBP5 | 3 | -0.470151568 | 0.164 | 0.339 | 4.57E-33 | 1.67E-28 |
| EPB41L5 | 3 | -0.498215421 | 0.347 | 0.519 | 4.97E-33 | 1.82E-28 |
| JARID2 | 3 | 0.427213014 | 0.778 | 0.662 | 5.16E-33 | 1.89E-28 |
| TXNDC16 | 3 | -0.526750366 | 0.178 | 0.348 | 7.26E-33 | 2.66E-28 |
| TMEFF1 | 3 | -0.540331084 | 0.214 | 0.383 | 7.52E-33 | 2.75E-28 |
| SMAD9 | 3 | -0.51086968 | 0.234 | 0.414 | 7.71E-33 | 2.82E-28 |
| SMC1A | 3 | -0.425686528 | 0.238 | 0.419 | 9.61E-33 | 3.52E-28 |

| | | | | | | |
|---|---|---|---|---|---|---|
| MIR137HG | 3 | -0.81902881 | 0.123 | 0.266 | 1.33E-32 | 4.87E-28 |
| RBL1 | 3 | -0.414138377 | 0.251 | 0.44 | 1.37E-32 | 5.01E-28 |
| KNL1 | 3 | -0.398635518 | 0.083 | 0.232 | 1.52E-32 | 5.56E-28 |
| PRIM2 | 3 | -0.478896744 | 0.501 | 0.658 | 1.83E-32 | 6.7E-28 |
| ZNF280D | 3 | 0.430998383 | 0.463 | 0.291 | 1.94E-32 | 7.11E-28 |
| CNTN3 | 3 | -0.708935384 | 0.343 | 0.514 | 2.12E-32 | 7.75E-28 |
| KCNN2 | 3 | -0.92442918 | 0.167 | 0.316 | 2.22E-32 | 8.11E-28 |
| ARMC4 | 3 | -0.341039375 | 0.029 | 0.142 | 2.29E-32 | 8.37E-28 |
| LIFR-AS1 | 3 | 0.391749825 | 0.216 | 0.093 | 2.5E-32 | 9.15E-28 |
| PRKCB | 3 | -0.471264343 | 0.016 | 0.116 | 2.73E-32 | 9.98E-28 |
| ANXA1 | 3 | 0.580618054 | 0.146 | 0.056 | 2.75E-32 | 1.01E-27 |
| RNF157 | 3 | -0.366805283 | 0.112 | 0.273 | 2.85E-32 | 1.04E-27 |
| CDCP1 | 3 | -0.458377038 | 0.065 | 0.204 | 3E-32 | 1.1E-27 |
| SHOX2 | 3 | 0.664056753 | 0.295 | 0.164 | 3.01E-32 | 1.1E-27 |
| IGSF9B | 3 | -0.306858823 | 0.032 | 0.148 | 3.64E-32 | 1.33E-27 |
| NRP1 | 3 | 0.362565036 | 0.478 | 0.29 | 3.7E-32 | 1.35E-27 |
| CHST8 | 3 | -0.419825191 | 0.053 | 0.185 | 3.76E-32 | 1.38E-27 |
| ANTXR1 | 3 | 0.433052515 | 0.624 | 0.448 | 3.79E-32 | 1.39E-27 |
| STON2 | 3 | 0.282209165 | 0.307 | 0.356 | 4.21E-32 | 1.54E-27 |
| TNFRSF19 | 3 | 0.628170389 | 0.369 | 0.224 | 4.42E-32 | 1.62E-27 |
| RANBP17 | 3 | -0.487400596 | 0.627 | 0.75 | 4.46E-32 | 1.63E-27 |
| TMEM163 | 3 | -0.54282368 | 0.043 | 0.168 | 4.47E-32 | 1.64E-27 |
| BMPR1B | 3 | 0.684278698 | 0.451 | 0.302 | 4.73E-32 | 1.73E-27 |
| AP001172.1 | 3 | 0.291978347 | 0.15 | 0.045 | 6.43E-32 | 2.35E-27 |
| GPC4 | 3 | -0.585287971 | 0.082 | 0.222 | 6.47E-32 | 2.37E-27 |
| DYNC2H1 | 3 | 0.353545198 | 0.533 | 0.34 | 7.84E-32 | 2.87E-27 |
| AC023469.1 | 3 | -0.734922503 | 0.204 | 0.371 | 8.29E-32 | 3.03E-27 |
| ZNF367 | 3 | -0.378073589 | 0.098 | 0.251 | 8.51E-32 | 3.12E-27 |
| AXL | 3 | 0.261951324 | 0.147 | 0.043 | 8.54E-32 | 3.12E-27 |
| RAB3IP | 3 | -0.343706816 | 0.041 | 0.152 | 8.6E-32 | 3.15E-27 |
| RALYL | 3 | -0.868354239 | 0.336 | 0.493 | 8.65E-32 | 3.17E-27 |
| EPHA4 | 3 | -0.615538279 | 0.451 | 0.605 | 9.02E-32 | 3.3E-27 |
| PRKDC | 3 | -0.396071639 | 0.725 | 0.842 | 1.19E-31 | 4.34E-27 |
| BAIAP2L1 | 3 | -0.314641645 | 0.022 | 0.128 | 1.3E-31 | 4.75E-27 |
| TRIM71 | 3 | -0.464261233 | 0.115 | 0.27 | 1.3E-31 | 4.77E-27 |
| SOX2-OT | 3 | -1.607890928 | 0.092 | 0.219 | 1.42E-31 | 5.19E-27 |
| SLFN13 | 3 | -0.314862742 | 0.033 | 0.148 | 1.48E-31 | 5.42E-27 |
| RGS3 | 3 | -0.340055122 | 0.036 | 0.15 | 2.07E-31 | 7.59E-27 |
| YPEL2 | 3 | 0.416688383 | 0.294 | 0.149 | 2.31E-31 | 8.47E-27 |
| MTCL1 | 3 | -0.407412056 | 0.099 | 0.246 | 2.51E-31 | 9.18E-27 |
| CYYR1 | 3 | 0.308760122 | 0.192 | 0.071 | 2.51E-31 | 9.19E-27 |
| MYT1 | 3 | -0.263215935 | 0.011 | 0.102 | 3E-31 | 1.1E-26 |
| PLXNA2 | 3 | -0.511111534 | 0.227 | 0.404 | 3.39E-31 | 1.24E-26 |
| IQSEC1 | 3 | -0.32482236 | 0.226 | 0.409 | 3.39E-31 | 1.24E-26 |
| PSIP1 | 3 | -0.452306519 | 0.402 | 0.572 | 3.46E-31 | 1.27E-26 |
| VPS13B | 3 | 0.394142119 | 0.892 | 0.817 | 3.95E-31 | 1.45E-26 |
| HDAC9 | 3 | -0.527490958 | 0.414 | 0.596 | 4.01E-31 | 1.47E-26 |
| SLC39A8 | 3 | -0.305386704 | 0.023 | 0.128 | 4.82E-31 | 1.77E-26 |
| B4GALNT3 | 3 | -0.300374934 | 0.025 | 0.133 | 5.54E-31 | 2.03E-26 |
| GEN1 | 3 | -0.443128664 | 0.258 | 0.434 | 5.76E-31 | 2.11E-26 |
| STEAP1B | 3 | -0.420891675 | 0.076 | 0.213 | 5.78E-31 | 2.12E-26 |
| CHD9 | 3 | 0.34398448 | 0.951 | 0.917 | 6.44E-31 | 2.36E-26 |
| GLB1L3 | 3 | -0.264927859 | 0.011 | 0.102 | 6.76E-31 | 2.47E-26 |
| ASAP2 | 3 | 0.569836582 | 0.483 | 0.376 | 8E-31 | 2.93E-26 |
| KIAA1324L | 3 | 0.315326067 | 0.37 | 0.201 | 9.03E-31 | 3.3E-26 |
| CAPN5 | 3 | 0.350805788 | 0.181 | 0.068 | 9.6E-31 | 3.52E-26 |
| KBTBD12 | 3 | -0.377616554 | 0.088 | 0.233 | 1.15E-30 | 4.2E-26 |
| AL596087.2 | 3 | -0.692213292 | 0.088 | 0.22 | 1.21E-30 | 4.43E-26 |

| | | | | | | |
|---|---|---|---|---|---|---|
| CSMD1 | 3 | -0.711338106 | 0.279 | 0.466 | 1.23E-30 | 4.5E-26 |
| RTTN | 3 | -0.493890645 | 0.383 | 0.537 | 1.5E-30 | 5.5E-26 |
| QKI | 3 | 0.45968427 | 0.856 | 0.776 | 1.51E-30 | 5.51E-26 |
| ASPM | 3 | -0.518540399 | 0.107 | 0.259 | 1.56E-30 | 5.7E-26 |
| AP000829.1 | 3 | -0.481762995 | 0.135 | 0.295 | 1.57E-30 | 5.76E-26 |
| MSC-AS1 | 3 | -0.553992129 | 0.046 | 0.165 | 1.68E-30 | 6.16E-26 |
| AL355499.1 | 3 | -0.907876459 | 0.019 | 0.112 | 1.68E-30 | 6.16E-26 |
| SLC6A15 | 3 | -0.47737004 | 0.086 | 0.228 | 1.72E-30 | 6.29E-26 |
| E2F1 | 3 | -0.285372429 | 0.037 | 0.152 | 1.76E-30 | 6.46E-26 |
| EVI5 | 3 | 0.41180953 | 0.702 | 0.539 | 1.87E-30 | 6.85E-26 |
| NUSAP1 | 3 | -0.395063621 | 0.102 | 0.252 | 2.12E-30 | 7.77E-26 |
| BCAR3 | 3 | 0.459311273 | 0.371 | 0.225 | 2.28E-30 | 8.36E-26 |
| B4GALT6 | 3 | -0.560687321 | 0.16 | 0.308 | 2.89E-30 | 1.06E-25 |
| LIN7A | 3 | 0.49875092 | 0.501 | 0.345 | 3.14E-30 | 1.15E-25 |
| PRIMA1 | 3 | 0.279344752 | 0.112 | 0.029 | 3.29E-30 | 1.21E-25 |
| GABRB2 | 3 | -0.638534661 | 0.16 | 0.325 | 3.61E-30 | 1.32E-25 |
| SLC7A14 | 3 | -0.470961706 | 0.02 | 0.12 | 3.77E-30 | 1.38E-25 |
| ADAMTS9 | 3 | -0.594734064 | 0.226 | 0.397 | 4.04E-30 | 1.48E-25 |
| AP002954.2 | 3 | -0.624723034 | 0.015 | 0.108 | 4.07E-30 | 1.49E-25 |
| JADE3 | 3 | -0.413110238 | 0.201 | 0.376 | 4.36E-30 | 1.6E-25 |
| EXO1 | 3 | -0.424209444 | 0.095 | 0.241 | 4.51E-30 | 1.65E-25 |
| RUNX1T1 | 3 | -0.58074151 | 0.614 | 0.727 | 4.58E-30 | 1.68E-25 |
| MCMBP | 3 | -0.433484115 | 0.284 | 0.459 | 4.63E-30 | 1.7E-25 |
| SORL1 | 3 | -0.327602908 | 0.043 | 0.163 | 6.18E-30 | 2.26E-25 |
| DEK | 3 | -0.4494817 | 0.379 | 0.549 | 6.36E-30 | 2.33E-25 |
| AC135782.1 | 3 | -0.340903134 | 0.042 | 0.159 | 6.71E-30 | 2.46E-25 |
| LBR | 3 | -0.515766935 | 0.206 | 0.358 | 7.75E-30 | 2.84E-25 |
| KLHL32 | 3 | -0.42674459 | 0.061 | 0.188 | 8.59E-30 | 3.14E-25 |
| CNTN4 | 3 | -0.879652502 | 0.047 | 0.163 | 8.61E-30 | 3.15E-25 |
| GALNT13 | 3 | -0.608212638 | 0.586 | 0.718 | 1.26E-29 | 4.61E-25 |
| AL355612.1 | 3 | 0.351491649 | 0.178 | 0.068 | 1.47E-29 | 5.38E-25 |
| AL445623.2 | 3 | -0.447262302 | 0.078 | 0.212 | 1.93E-29 | 7.05E-25 |
| NFATC1 | 3 | 0.291642651 | 0.151 | 0.049 | 2.23E-29 | 8.15E-25 |
| YAF2 | 3 | -0.455252114 | 0.412 | 0.57 | 2.26E-29 | 8.28E-25 |
| REV3L | 3 | -0.455455461 | 0.815 | 0.872 | 2.72E-29 | 9.96E-25 |
| PLCH1 | 3 | -0.627863012 | 0.188 | 0.34 | 2.89E-29 | 1.06E-24 |
| SATB2 | 3 | 0.505714305 | 0.616 | 0.439 | 3.07E-29 | 1.12E-24 |
| CYP4X1 | 3 | -0.467765382 | 0.045 | 0.163 | 3.08E-29 | 1.13E-24 |
| AC104389.5 | 3 | 0.291988517 | 0.171 | 0.061 | 3.13E-29 | 1.15E-24 |
| SLC8A2 | 3 | 0.473888567 | 0.265 | 0.14 | 3.25E-29 | 1.19E-24 |
| ZGRF1 | 3 | -0.48408875 | 0.23 | 0.394 | 3.45E-29 | 1.26E-24 |
| AP001636.3 | 3 | -0.334600017 | 0.025 | 0.128 | 3.5E-29 | 1.28E-24 |
| ITPRID2 | 3 | 0.436892967 | 0.424 | 0.309 | 3.75E-29 | 1.37E-24 |
| MINAR1 | 3 | -0.339726862 | 0.054 | 0.178 | 3.91E-29 | 1.43E-24 |
| CNTNAP5 | 3 | -0.50265452 | 0.136 | 0.294 | 3.95E-29 | 1.45E-24 |
| REST | 3 | 0.407491884 | 0.366 | 0.205 | 4.39E-29 | 1.61E-24 |
| WNK2 | 3 | -0.450267649 | 0.432 | 0.605 | 5.7E-29 | 2.09E-24 |
| DECR1 | 3 | 0.346681082 | 0.329 | 0.175 | 5.79E-29 | 2.12E-24 |
| CHAF1A | 3 | -0.308852359 | 0.131 | 0.285 | 5.86E-29 | 2.14E-24 |
| PRKD1 | 3 | 0.665540611 | 0.704 | 0.655 | 6.48E-29 | 2.37E-24 |
| NCAPD3 | 3 | -0.394548377 | 0.212 | 0.384 | 6.77E-29 | 2.48E-24 |
| MT-CO3 | 3 | -0.538484738 | 0.254 | 0.432 | 7.59E-29 | 2.78E-24 |
| NR2C1 | 3 | 0.435538602 | 0.574 | 0.453 | 7.88E-29 | 2.89E-24 |
| SYDE2 | 3 | -0.428190089 | 0.088 | 0.216 | 8.7E-29 | 3.18E-24 |
| DDX58 | 3 | 0.376446656 | 0.211 | 0.087 | 8.73E-29 | 3.2E-24 |
| CPT1A | 3 | 0.336479038 | 0.238 | 0.107 | 9.02E-29 | 3.3E-24 |
| PIK3R1 | 3 | 0.477129533 | 0.472 | 0.323 | 1.02E-28 | 3.73E-24 |
| ASTN2 | 3 | -0.565478674 | 0.362 | 0.533 | 1.11E-28 | 4.07E-24 |

| | | | | | | |
|---|---|---|---|---|---|---|
| MCM3 | 3 | -0.348336048 | 0.178 | 0.345 | 1.13E-28 | 4.13E-24 |
| SYNPO | 3 | 0.255651742 | 0.142 | 0.045 | 1.2E-28 | 4.38E-24 |
| UHRF1 | 3 | -0.318317048 | 0.125 | 0.277 | 1.26E-28 | 4.62E-24 |
| CNOT2 | 3 | 0.366435794 | 0.862 | 0.779 | 1.4E-28 | 5.13E-24 |
| GABRA5 | 3 | -0.252043096 | 0.012 | 0.1 | 1.42E-28 | 5.21E-24 |
| CDC42BPA | 3 | -0.507796175 | 0.709 | 0.78 | 1.53E-28 | 5.61E-24 |
| GRID2 | 3 | -1.028843343 | 0.476 | 0.592 | 1.76E-28 | 6.44E-24 |
| GRK3 | 3 | -0.480272649 | 0.311 | 0.475 | 1.79E-28 | 6.56E-24 |
| ZSWIM5 | 3 | -0.556951073 | 0.242 | 0.402 | 1.81E-28 | 6.62E-24 |
| AC107980.1 | 3 | -0.250671659 | 0.026 | 0.128 | 1.83E-28 | 6.68E-24 |
| RBM39 | 3 | 0.309894819 | 0.915 | 0.853 | 1.87E-28 | 6.85E-24 |
| POLE2 | 3 | -0.352530141 | 0.175 | 0.34 | 2.04E-28 | 7.47E-24 |
| LAMB1 | 3 | 0.44238503 | 0.541 | 0.368 | 2.14E-28 | 7.83E-24 |
| SRPK1 | 3 | -0.435099638 | 0.468 | 0.627 | 2.15E-28 | 7.88E-24 |
| GRAMD2B | 3 | 0.370142875 | 0.284 | 0.14 | 2.24E-28 | 8.18E-24 |
| AL356108.1 | 3 | -0.73963905 | 0.026 | 0.122 | 2.3E-28 | 8.43E-24 |
| WDR11 | 3 | -0.579222339 | 0.615 | 0.683 | 2.35E-28 | 8.59E-24 |
| ASRGL1 | 3 | -0.268802191 | 0.042 | 0.157 | 2.37E-28 | 8.68E-24 |
| PLS1 | 3 | -0.413692826 | 0.088 | 0.222 | 2.84E-28 | 1.04E-23 |
| LRFN5 | 3 | 0.406581809 | 0.622 | 0.439 | 3.58E-28 | 1.31E-23 |
| SEZ6L | 3 | -0.636641262 | 0.015 | 0.106 | 3.67E-28 | 1.34E-23 |
| PRPSAP1 | 3 | 0.404575677 | 0.36 | 0.227 | 3.96E-28 | 1.45E-23 |
| ATP8A2 | 3 | -0.430822846 | 0.169 | 0.332 | 4.19E-28 | 1.53E-23 |
| PAK5 | 3 | -0.609983399 | 0.314 | 0.468 | 4.44E-28 | 1.63E-23 |
| MCM4 | 3 | -0.369439091 | 0.138 | 0.291 | 4.67E-28 | 1.71E-23 |
| CUL4B | 3 | -0.4624121 | 0.284 | 0.443 | 4.78E-28 | 1.75E-23 |
| CPT1C | 3 | -0.278241124 | 0.046 | 0.162 | 4.96E-28 | 1.82E-23 |
| KHDRBS3 | 3 | -0.674447433 | 0.128 | 0.257 | 5E-28 | 1.83E-23 |
| GRM7 | 3 | -0.906349571 | 0.045 | 0.149 | 5.9E-28 | 2.16E-23 |
| CCND2 | 3 | -0.418429353 | 0.175 | 0.337 | 6.42E-28 | 2.35E-23 |
| JMJD1C | 3 | 0.318483736 | 0.976 | 0.942 | 7.1E-28 | 2.6E-23 |
| FNIP1 | 3 | 0.446964907 | 0.709 | 0.611 | 7.11E-28 | 2.6E-23 |
| BCL11B | 3 | -0.531532183 | 0.277 | 0.443 | 8.58E-28 | 3.14E-23 |
| SYNPR | 3 | 0.872021544 | 0.448 | 0.348 | 8.67E-28 | 3.18E-23 |
| IPO9 | 3 | -0.418097719 | 0.48 | 0.63 | 8.97E-28 | 3.28E-23 |
| SPART | 3 | 0.344699797 | 0.284 | 0.151 | 9.25E-28 | 3.39E-23 |
| CERS6 | 3 | -0.409984626 | 0.749 | 0.824 | 1.02E-27 | 3.73E-23 |
| MELK | 3 | -0.383824434 | 0.137 | 0.291 | 1.03E-27 | 3.78E-23 |
| PDE10A | 3 | -0.55119349 | 0.122 | 0.27 | 1.06E-27 | 3.87E-23 |
| ZNF827 | 3 | 0.435430976 | 0.674 | 0.529 | 1.07E-27 | 3.9E-23 |
| ETV5 | 3 | 0.333420175 | 0.256 | 0.121 | 1.13E-27 | 4.15E-23 |
| MCM8 | 3 | -0.381653554 | 0.199 | 0.365 | 1.2E-27 | 4.39E-23 |
| PHIP | 3 | -0.353371177 | 0.807 | 0.882 | 1.26E-27 | 4.61E-23 |
| C20orf194 | 3 | 0.406348107 | 0.606 | 0.451 | 1.31E-27 | 4.79E-23 |
| ST3GAL1 | 3 | 0.406174697 | 0.281 | 0.147 | 1.31E-27 | 4.8E-23 |
| RNF168 | 3 | -0.425391575 | 0.288 | 0.455 | 1.43E-27 | 5.24E-23 |
| LINC02050 | 3 | 0.301560125 | 0.112 | 0.03 | 1.53E-27 | 5.58E-23 |
| PDE7A | 3 | -0.444253527 | 0.483 | 0.629 | 1.87E-27 | 6.86E-23 |
| RRM1 | 3 | -0.387773935 | 0.151 | 0.303 | 1.99E-27 | 7.27E-23 |
| SPTBN2 | 3 | -0.302192742 | 0.078 | 0.208 | 2.09E-27 | 7.65E-23 |
| KCNH8 | 3 | -0.55778636 | 0.516 | 0.649 | 2.41E-27 | 8.84E-23 |
| VCAN-AS1 | 3 | 0.254002912 | 0.154 | 0.052 | 2.47E-27 | 9.04E-23 |
| EEPD1 | 3 | -0.586409335 | 0.186 | 0.319 | 2.76E-27 | 1.01E-22 |
| IVNS1ABP | 3 | -0.520891862 | 0.376 | 0.524 | 2.88E-27 | 1.06E-22 |
| ALOX12-AS1 | 3 | 0.404406237 | 0.302 | 0.157 | 3.35E-27 | 1.22E-22 |
| AC068587.4 | 3 | -0.415615284 | 0.197 | 0.363 | 3.36E-27 | 1.23E-22 |
| EPHB2 | 3 | -0.359003336 | 0.082 | 0.213 | 3.6E-27 | 1.32E-22 |
| TFAP2B | 3 | 0.488769807 | 0.378 | 0.218 | 3.88E-27 | 1.42E-22 |

| | | | | | | |
|---|---|---|---|---|---|---|
| GMDS-DT | 3 | 0.33221829 | 0.411 | 0.245 | 4.07E-27 | 1.49E-22 |
| MBNL2 | 3 | 0.410929028 | 0.564 | 0.39 | 4.32E-27 | 1.58E-22 |
| TAOK3 | 3 | 0.394813321 | 0.575 | 0.405 | 4.43E-27 | 1.62E-22 |
| SKA2 | 3 | -0.436783558 | 0.276 | 0.44 | 4.61E-27 | 1.69E-22 |
| NUBPL | 3 | 0.387101885 | 0.645 | 0.478 | 4.76E-27 | 1.74E-22 |
| CRMP1 | 3 | -0.423449561 | 0.184 | 0.343 | 4.83E-27 | 1.77E-22 |
| AC119674.1 | 3 | 0.348278176 | 0.161 | 0.059 | 5.42E-27 | 1.98E-22 |
| BUB1B | 3 | -0.344335283 | 0.061 | 0.183 | 5.53E-27 | 2.02E-22 |
| KIF2A | 3 | -0.448543131 | 0.476 | 0.623 | 7.24E-27 | 2.65E-22 |
| DNMT1 | 3 | -0.402764857 | 0.327 | 0.494 | 7.62E-27 | 2.79E-22 |
| PER3 | 3 | -0.372343143 | 0.083 | 0.213 | 7.81E-27 | 2.86E-22 |
| PRAG1 | 3 | 0.424942148 | 0.244 | 0.116 | 9.95E-27 | 3.64E-22 |
| RPS6KA2 | 3 | -0.530682015 | 0.188 | 0.343 | 1.06E-26 | 3.89E-22 |
| SVEP1 | 3 | -0.700026977 | 0.185 | 0.308 | 1.21E-26 | 4.42E-22 |
| BRCA2 | 3 | -0.373613363 | 0.136 | 0.286 | 1.3E-26 | 4.77E-22 |
| ZC4H2 | 3 | -0.378348825 | 0.032 | 0.132 | 1.3E-26 | 4.77E-22 |
| MERTK | 3 | 0.321843804 | 0.225 | 0.101 | 1.33E-26 | 4.85E-22 |
| PTPRO | 3 | 0.683245624 | 0.518 | 0.432 | 1.42E-26 | 5.2E-22 |
| MON2 | 3 | 0.36849151 | 0.841 | 0.74 | 1.45E-26 | 5.3E-22 |
| MGMT | 3 | 0.327924338 | 0.572 | 0.394 | 1.63E-26 | 5.96E-22 |
| LRP8 | 3 | -0.384353634 | 0.158 | 0.31 | 1.65E-26 | 6.03E-22 |
| PICALM | 3 | 0.394294563 | 0.776 | 0.701 | 1.74E-26 | 6.39E-22 |
| CCNE2 | 3 | -0.335614718 | 0.057 | 0.175 | 2.05E-26 | 7.5E-22 |
| C3orf70 | 3 | -0.393207535 | 0.134 | 0.282 | 2.28E-26 | 8.35E-22 |
| AC138150.1 | 3 | 0.319988931 | 0.302 | 0.159 | 2.54E-26 | 9.28E-22 |
| POU5F1 | 3 | 0.549380953 | 0.586 | 0.469 | 2.71E-26 | 9.9E-22 |
| LMO4 | 3 | 0.411521188 | 0.406 | 0.246 | 2.72E-26 | 9.98E-22 |
| ZKSCAN1 | 3 | 0.396360321 | 0.576 | 0.417 | 2.94E-26 | 1.08E-21 |
| SULF1 | 3 | 0.437869739 | 0.459 | 0.29 | 3.23E-26 | 1.18E-21 |
| ECT2 | 3 | -0.421021308 | 0.162 | 0.307 | 3.67E-26 | 1.34E-21 |
| PDS5B | 3 | -0.37851082 | 0.757 | 0.854 | 3.95E-26 | 1.45E-21 |
| ZFR2 | 3 | -0.27014078 | 0.04 | 0.148 | 4.01E-26 | 1.47E-21 |
| BRINP2 | 3 | -0.387811855 | 0.024 | 0.117 | 4.15E-26 | 1.52E-21 |
| TTC27 | 3 | 0.353395601 | 0.452 | 0.288 | 4.76E-26 | 1.74E-21 |
| RAD18 | 3 | -0.461948862 | 0.411 | 0.542 | 4.89E-26 | 1.79E-21 |
| TSHZ3 | 3 | -0.501466589 | 0.348 | 0.511 | 4.97E-26 | 1.82E-21 |
| CTNNA3 | 3 | 0.998647443 | 0.244 | 0.173 | 5.19E-26 | 1.9E-21 |
| OSBPL9 | 3 | 0.397398064 | 0.677 | 0.558 | 5.59E-26 | 2.05E-21 |
| ZNF608 | 3 | 0.439792412 | 0.736 | 0.623 | 6.39E-26 | 2.34E-21 |
| LUZP2 | 3 | -0.293063348 | 0.653 | 0.544 | 6.65E-26 | 2.43E-21 |
| SEPTIN9 | 3 | 0.425579166 | 0.598 | 0.445 | 6.67E-26 | 2.44E-21 |
| RNF128 | 3 | -0.444727169 | 0.049 | 0.152 | 7.64E-26 | 2.8E-21 |
| C8orf37-AS1 | 3 | 0.49488029 | 0.168 | 0.072 | 8.46E-26 | 3.1E-21 |
| NCOA1 | 3 | 0.37449857 | 0.806 | 0.674 | 8.59E-26 | 3.14E-21 |
| C21orf58 | 3 | -0.361695699 | 0.119 | 0.26 | 9.63E-26 | 3.53E-21 |
| TFDP1 | 3 | -0.316630407 | 0.288 | 0.46 | 9.79E-26 | 3.58E-21 |
| SLC35G2 | 3 | -0.327352739 | 0.102 | 0.237 | 1.04E-25 | 3.82E-21 |
| MT-CO1 | 3 | -0.540466422 | 0.312 | 0.477 | 1.11E-25 | 4.05E-21 |
| AP002336.2 | 3 | 0.335741445 | 0.231 | 0.133 | 1.11E-25 | 4.07E-21 |
| ST6GALNAC3 | 3 | 0.579188652 | 0.392 | 0.25 | 1.24E-25 | 4.55E-21 |
| LRCH3 | 3 | 0.381619532 | 0.733 | 0.608 | 1.29E-25 | 4.71E-21 |
| SCN1A | 3 | -0.527506033 | 0.021 | 0.11 | 1.34E-25 | 4.9E-21 |
| MYB | 3 | -0.290489909 | 0.075 | 0.199 | 1.36E-25 | 4.98E-21 |
| ZFP36L1 | 3 | 0.312051624 | 0.194 | 0.081 | 1.44E-25 | 5.26E-21 |
| AC092683.1 | 3 | -0.45560212 | 0.229 | 0.376 | 1.53E-25 | 5.6E-21 |
| KIF18A | 3 | -0.320773651 | 0.058 | 0.173 | 1.64E-25 | 6.01E-21 |
| EML4 | 3 | 0.403787426 | 0.755 | 0.672 | 1.7E-25 | 6.22E-21 |
| RACGAP1 | 3 | -0.303444287 | 0.068 | 0.188 | 1.7E-25 | 6.23E-21 |

| | | | | | | |
|---|---|---|---|---|---|---|
| SMAD3 | 3 | 0.403629785 | 0.482 | 0.321 | 1.8E-25 | 6.59E-21 |
| PIEZO2 | 3 | 0.584497027 | 0.495 | 0.36 | 1.97E-25 | 7.22E-21 |
| SMCHD1 | 3 | -0.348621204 | 0.768 | 0.841 | 2.02E-25 | 7.41E-21 |
| PLEKHH2 | 3 | 0.422297175 | 0.284 | 0.156 | 2.12E-25 | 7.75E-21 |
| AL354771.1 | 3 | 0.445527718 | 0.242 | 0.125 | 2.23E-25 | 8.17E-21 |
| DPF3 | 3 | -0.343644486 | 0.14 | 0.287 | 2.44E-25 | 8.92E-21 |
| IBTK | 3 | 0.377022334 | 0.573 | 0.442 | 2.59E-25 | 9.47E-21 |
| SLF1 | 3 | -0.396038026 | 0.321 | 0.485 | 2.91E-25 | 1.07E-20 |
| IFIH1 | 3 | 0.388508018 | 0.142 | 0.053 | 3.1E-25 | 1.13E-20 |
| EFNA5 | 3 | 0.617145629 | 0.769 | 0.649 | 3.55E-25 | 1.3E-20 |
| ZCCHC24 | 3 | 0.259265107 | 0.172 | 0.067 | 4.1E-25 | 1.5E-20 |
| FBXL2 | 3 | 0.377467564 | 0.359 | 0.208 | 4.27E-25 | 1.56E-20 |
| GALK2 | 3 | 0.400590431 | 0.537 | 0.417 | 4.27E-25 | 1.56E-20 |
| FGF12 | 3 | 0.7866539 | 0.791 | 0.729 | 4.35E-25 | 1.59E-20 |
| ARFGEF3 | 3 | -0.511921669 | 0.319 | 0.479 | 4.41E-25 | 1.61E-20 |
| FMN1 | 3 | 0.567445658 | 0.212 | 0.094 | 4.68E-25 | 1.71E-20 |
| RBM10 | 3 | -0.33053922 | 0.241 | 0.405 | 4.88E-25 | 1.79E-20 |
| CPEB1 | 3 | -0.281889285 | 0.026 | 0.119 | 5.25E-25 | 1.92E-20 |
| CHM | 3 | 0.362559863 | 0.773 | 0.671 | 5.45E-25 | 1.99E-20 |
| LINC01122 | 3 | -0.626499901 | 0.146 | 0.286 | 5.57E-25 | 2.04E-20 |
| FLCN | 3 | 0.300762369 | 0.239 | 0.119 | 5.95E-25 | 2.18E-20 |
| KIF11 | 3 | -0.391751652 | 0.116 | 0.25 | 6.23E-25 | 2.28E-20 |
| RRM2 | 3 | -0.31643535 | 0.102 | 0.234 | 6.3E-25 | 2.31E-20 |
| ZNF519 | 3 | -0.413212862 | 0.273 | 0.428 | 6.6E-25 | 2.42E-20 |
| HMBOX1 | 3 | 0.354041477 | 0.704 | 0.58 | 7E-25 | 2.56E-20 |
| ZNF83 | 3 | -0.376988894 | 0.251 | 0.409 | 7.02E-25 | 2.57E-20 |
| DENND4C | 3 | -0.426880917 | 0.604 | 0.712 | 7.08E-25 | 2.59E-20 |
| LINC00632 | 3 | 0.353151614 | 0.536 | 0.365 | 7.08E-25 | 2.59E-20 |
| LITAF | 3 | 0.333171529 | 0.22 | 0.108 | 9.11E-25 | 3.34E-20 |
| INSYN2B | 3 | 0.319721964 | 0.165 | 0.065 | 9.34E-25 | 3.42E-20 |
| ZNF100 | 3 | -0.282339577 | 0.087 | 0.215 | 9.74E-25 | 3.57E-20 |
| MAN1A1 | 3 | -0.685921675 | 0.052 | 0.143 | 9.79E-25 | 3.58E-20 |
| CDON | 3 | -0.473685535 | 0.201 | 0.349 | 1.09E-24 | 4.01E-20 |
| KLHDC8A | 3 | -0.296890444 | 0.065 | 0.182 | 1.22E-24 | 4.48E-20 |
| GREB1L | 3 | -0.660623747 | 0.022 | 0.107 | 1.23E-24 | 4.51E-20 |
| LINC01934 | 3 | -1.242059902 | 0.031 | 0.112 | 1.27E-24 | 4.64E-20 |
| SMARCA1 | 3 | 0.415922512 | 0.759 | 0.66 | 1.31E-24 | 4.78E-20 |
| DLGAP1-AS5 | 3 | 0.270508239 | 0.166 | 0.069 | 1.31E-24 | 4.8E-20 |
| MBD5 | 3 | 0.337894532 | 0.931 | 0.893 | 1.39E-24 | 5.1E-20 |
| RAD54B | 3 | -0.331339158 | 0.122 | 0.254 | 1.41E-24 | 5.17E-20 |
| RALGPS1 | 3 | -0.388157501 | 0.157 | 0.295 | 1.53E-24 | 5.61E-20 |
| LINC02197 | 3 | -0.601752996 | 0.077 | 0.182 | 1.77E-24 | 6.49E-20 |
| RPTOR | 3 | 0.357436667 | 0.694 | 0.585 | 1.89E-24 | 6.92E-20 |
| LYPLAL1 | 3 | 0.361238197 | 0.573 | 0.419 | 2.07E-24 | 7.56E-20 |
| STK33 | 3 | -0.630832138 | 0.082 | 0.171 | 2.16E-24 | 7.9E-20 |
| ZMAT3 | 3 | 0.363510789 | 0.292 | 0.162 | 2.2E-24 | 8.06E-20 |
| ZNF718 | 3 | -0.380105602 | 0.337 | 0.493 | 2.6E-24 | 9.51E-20 |
| STIL | 3 | -0.386326563 | 0.143 | 0.284 | 2.65E-24 | 9.72E-20 |
| GRM8 | 3 | -1.119179587 | 0.06 | 0.165 | 2.79E-24 | 1.02E-19 |
| LAMA3 | 3 | -0.474743264 | 0.098 | 0.225 | 3.04E-24 | 1.11E-19 |
| ITGB8 | 3 | 0.375561482 | 0.441 | 0.28 | 3.27E-24 | 1.2E-19 |
| MYBL2 | 3 | -0.255947499 | 0.05 | 0.158 | 3.42E-24 | 1.25E-19 |
| PLAT | 3 | 0.328466861 | 0.115 | 0.035 | 3.6E-24 | 1.32E-19 |
| LIMA1 | 3 | 0.340349514 | 0.268 | 0.137 | 3.83E-24 | 1.4E-19 |
| SCN1A-AS1 | 3 | 0.290233979 | 0.144 | 0.058 | 3.92E-24 | 1.43E-19 |
| SSX2IP | 3 | -0.463505803 | 0.378 | 0.527 | 4.06E-24 | 1.49E-19 |
| BSN | 3 | -0.255447238 | 0.031 | 0.124 | 4.38E-24 | 1.6E-19 |
| COL4A5 | 3 | -0.769409452 | 0.109 | 0.11 | 4.87E-24 | 1.78E-19 |

| | | | | | | |
|---|---|---|---|---|---|---|
| GRB14 | 3 | -0.279387472 | 0.031 | 0.125 | 5.24E-24 | 1.92E-19 |
| PCNX2 | 3 | -0.42505548 | 0.375 | 0.519 | 5.33E-24 | 1.95E-19 |
| HELZ | 3 | 0.363224273 | 0.749 | 0.657 | 5.79E-24 | 2.12E-19 |
| TBC1D9 | 3 | -0.362463802 | 0.229 | 0.387 | 6.08E-24 | 2.23E-19 |
| LHFPL2 | 3 | 0.410395643 | 0.352 | 0.219 | 6.53E-24 | 2.39E-19 |
| CABLES1 | 3 | -0.333129772 | 0.071 | 0.187 | 7.79E-24 | 2.85E-19 |
| WHRN | 3 | -0.302026947 | 0.077 | 0.194 | 8.22E-24 | 3.01E-19 |
| DTWD1 | 3 | 0.357359592 | 0.485 | 0.346 | 8.38E-24 | 3.07E-19 |
| BTBD9 | 3 | 0.372683748 | 0.699 | 0.553 | 8.44E-24 | 3.09E-19 |
| ARHGAP31 | 3 | 0.383218813 | 0.354 | 0.213 | 8.9E-24 | 3.26E-19 |
| CCDC150 | 3 | -0.390181446 | 0.168 | 0.314 | 9.2E-24 | 3.37E-19 |
| CAPRIN2 | 3 | 0.394466642 | 0.393 | 0.276 | 9.68E-24 | 3.54E-19 |
| GPR156 | 3 | -0.374744196 | 0.112 | 0.241 | 9.76E-24 | 3.57E-19 |
| ABLIM2 | 3 | -0.286419874 | 0.053 | 0.161 | 1.01E-23 | 3.71E-19 |
| CDH20 | 3 | -0.420285144 | 0.023 | 0.109 | 1.07E-23 | 3.91E-19 |
| SRSF10 | 3 | -0.38058129 | 0.504 | 0.628 | 1.1E-23 | 4.04E-19 |
| MSI2 | 3 | 0.311320877 | 0.921 | 0.849 | 1.11E-23 | 4.07E-19 |
| SLC9A6 | 3 | -0.319937843 | 0.127 | 0.262 | 1.26E-23 | 4.61E-19 |
| SLC9A9 | 3 | 0.286388654 | 0.195 | 0.085 | 1.32E-23 | 4.84E-19 |
| TNS3 | 3 | 0.397876849 | 0.373 | 0.223 | 1.35E-23 | 4.93E-19 |
| SLC44A3 | 3 | -0.251991162 | 0.023 | 0.109 | 1.37E-23 | 5E-19 |
| NSRP1 | 3 | 0.334553323 | 0.78 | 0.66 | 1.42E-23 | 5.19E-19 |
| FAM126A | 3 | -0.472649521 | 0.507 | 0.616 | 1.65E-23 | 6.02E-19 |
| SLC35B3 | 3 | -0.425718163 | 0.273 | 0.406 | 1.65E-23 | 6.06E-19 |
| OSBPL6 | 3 | -0.298278747 | 0.314 | 0.48 | 1.75E-23 | 6.41E-19 |
| CDC73 | 3 | 0.357260237 | 0.691 | 0.56 | 1.77E-23 | 6.48E-19 |
| IDS | 3 | -0.331939794 | 0.113 | 0.241 | 1.79E-23 | 6.55E-19 |
| SLC16A9 | 3 | -0.310113274 | 0.072 | 0.184 | 1.88E-23 | 6.87E-19 |
| TOX3 | 3 | -0.454907997 | 0.173 | 0.32 | 2.25E-23 | 8.24E-19 |
| POLA2 | 3 | -0.264038138 | 0.115 | 0.247 | 2.3E-23 | 8.43E-19 |
| KIAA1211 | 3 | -0.437685439 | 0.59 | 0.71 | 3.08E-23 | 1.13E-18 |
| PSMA1 | 3 | -0.375763429 | 0.342 | 0.499 | 3.23E-23 | 1.18E-18 |
| PLAGL1 | 3 | -0.602310553 | 0.232 | 0.373 | 3.9E-23 | 1.43E-18 |
| CREB3L2 | 3 | 0.364715043 | 0.379 | 0.235 | 4.04E-23 | 1.48E-18 |
| RAI2 | 3 | -0.329426481 | 0.06 | 0.167 | 4.04E-23 | 1.48E-18 |
| HIF1A-AS3 | 3 | 0.530248959 | 0.212 | 0.119 | 4.31E-23 | 1.58E-18 |
| PYGL | 3 | 0.286233825 | 0.197 | 0.087 | 4.34E-23 | 1.59E-18 |
| KAT2A | 3 | 0.306161497 | 0.207 | 0.097 | 4.41E-23 | 1.61E-18 |
| POTEF | 3 | 0.343777186 | 0.355 | 0.217 | 4.63E-23 | 1.7E-18 |
| MYLIP | 3 | 0.340852336 | 0.292 | 0.161 | 4.74E-23 | 1.74E-18 |
| FRYL | 3 | -0.307623932 | 0.831 | 0.855 | 4.97E-23 | 1.82E-18 |
| KLHL23 | 3 | -0.379621503 | 0.26 | 0.401 | 5.31E-23 | 1.94E-18 |
| AC092691.1 | 3 | -0.54461032 | 0.023 | 0.105 | 5.48E-23 | 2.01E-18 |
| FANCD2 | 3 | -0.28863424 | 0.121 | 0.254 | 5.67E-23 | 2.08E-18 |
| PRKAR2B | 3 | -0.367892026 | 0.144 | 0.282 | 5.93E-23 | 2.17E-18 |
| CNTLN | 3 | -0.403976433 | 0.613 | 0.721 | 6.21E-23 | 2.27E-18 |
| ST3GAL4 | 3 | 0.379796537 | 0.311 | 0.197 | 6.38E-23 | 2.34E-18 |
| AFF2 | 3 | -0.652255772 | 0.561 | 0.657 | 6.49E-23 | 2.38E-18 |
| CNR1 | 3 | 0.35653602 | 0.134 | 0.054 | 6.56E-23 | 2.4E-18 |
| DLG2 | 3 | -0.908244453 | 0.573 | 0.643 | 6.6E-23 | 2.42E-18 |
| AL137009.1 | 3 | -0.291690136 | 0.1 | 0.224 | 6.83E-23 | 2.5E-18 |
| SCRN1 | 3 | -0.304029809 | 0.113 | 0.24 | 7.3E-23 | 2.67E-18 |
| SLAIN1 | 3 | -0.43057422 | 0.193 | 0.325 | 7.35E-23 | 2.69E-18 |
| CSE1L | 3 | -0.354404149 | 0.35 | 0.508 | 7.37E-23 | 2.7E-18 |
| ID1 | 3 | -0.300214137 | 0.039 | 0.137 | 7.85E-23 | 2.87E-18 |
| KCNT2 | 3 | -0.478988433 | 0.636 | 0.715 | 8.06E-23 | 2.95E-18 |
| SLC27A1 | 3 | 0.270448711 | 0.183 | 0.078 | 8.27E-23 | 3.03E-18 |
| MTURN | 3 | -0.313766381 | 0.114 | 0.239 | 8.75E-23 | 3.2E-18 |

| | | | | | | |
|---|---|---|---|---|---|---|
| STRBP | 3 | -0.440630955 | 0.413 | 0.548 | 9.81E-23 | 3.59E-18 |
| MTMR2 | 3 | 0.427889945 | 0.472 | 0.353 | 1.06E-22 | 3.9E-18 |
| GKAP1 | 3 | -0.348091204 | 0.121 | 0.249 | 1.14E-22 | 4.17E-18 |
| AFF4 | 3 | 0.339150334 | 0.787 | 0.697 | 1.34E-22 | 4.9E-18 |
| WDR41 | 3 | 0.332194814 | 0.369 | 0.228 | 1.42E-22 | 5.2E-18 |
| PTMA | 3 | -0.346988532 | 0.268 | 0.425 | 1.58E-22 | 5.77E-18 |
| TMEM63C | 3 | -0.260217411 | 0.051 | 0.149 | 1.6E-22 | 5.87E-18 |
| NFATC2 | 3 | 0.400676491 | 0.26 | 0.145 | 1.62E-22 | 5.94E-18 |
| EDIL3 | 3 | 0.614663377 | 0.717 | 0.607 | 1.63E-22 | 5.95E-18 |
| AC061958.1 | 3 | -0.38968451 | 0.183 | 0.327 | 1.64E-22 | 6E-18 |
| LYN | 3 | 0.341905789 | 0.218 | 0.113 | 1.68E-22 | 6.15E-18 |
| AC004158.1 | 3 | 0.315697677 | 0.307 | 0.172 | 1.82E-22 | 6.67E-18 |
| GRAMD1B | 3 | -0.410922458 | 0.08 | 0.187 | 1.82E-22 | 6.67E-18 |
| PLEKHG1 | 3 | 0.426666103 | 0.235 | 0.12 | 1.95E-22 | 7.14E-18 |
| AC069437.1 | 3 | 0.281787603 | 0.132 | 0.056 | 2.04E-22 | 7.46E-18 |
| ACSL4 | 3 | -0.440795651 | 0.435 | 0.578 | 2.06E-22 | 7.53E-18 |
| GMPS | 3 | -0.372183664 | 0.411 | 0.563 | 2.13E-22 | 7.8E-18 |
| PDZD8 | 3 | -0.384282335 | 0.3 | 0.443 | 2.21E-22 | 8.1E-18 |
| SGIP1 | 3 | -0.335413078 | 0.023 | 0.105 | 2.23E-22 | 8.18E-18 |
| BOC | 3 | 0.372388528 | 0.321 | 0.187 | 2.79E-22 | 1.02E-17 |
| CCDC6 | 3 | -0.441944397 | 0.359 | 0.492 | 2.83E-22 | 1.04E-17 |
| BEND7 | 3 | 0.449750284 | 0.362 | 0.242 | 2.99E-22 | 1.1E-17 |
| LINC02307 | 3 | -0.34771892 | 0.047 | 0.146 | 3.07E-22 | 1.12E-17 |
| TMEM87B | 3 | 0.390206508 | 0.39 | 0.263 | 3.17E-22 | 1.16E-17 |
| CDKL5 | 3 | -0.447755452 | 0.28 | 0.424 | 3.29E-22 | 1.2E-17 |
| TNFRSF10B | 3 | 0.357880811 | 0.309 | 0.178 | 3.65E-22 | 1.34E-17 |
| CACNB4 | 3 | -0.546986591 | 0.538 | 0.548 | 3.66E-22 | 1.34E-17 |
| EFEMP1 | 3 | 0.280696795 | 0.121 | 0.042 | 3.9E-22 | 1.43E-17 |
| AL022068.1 | 3 | -0.627682234 | 0.037 | 0.121 | 3.9E-22 | 1.43E-17 |
| EPHA5 | 3 | -0.732577299 | 0.141 | 0.273 | 3.93E-22 | 1.44E-17 |
| EPHB1 | 3 | -0.411547872 | 0.067 | 0.174 | 4.19E-22 | 1.53E-17 |
| AP000787.1 | 3 | 0.342816576 | 0.194 | 0.097 | 4.45E-22 | 1.63E-17 |
| NRXN3 | 3 | -1.241827367 | 0.263 | 0.366 | 4.79E-22 | 1.75E-17 |
| DYNC1I2 | 3 | -0.435855009 | 0.567 | 0.66 | 4.97E-22 | 1.82E-17 |
| LANCL3 | 3 | -0.422058465 | 0.119 | 0.24 | 5.01E-22 | 1.83E-17 |
| VEGFA | 3 | 0.562046984 | 0.476 | 0.338 | 5.03E-22 | 1.84E-17 |
| EML6 | 3 | -0.353424699 | 0.129 | 0.259 | 5.04E-22 | 1.85E-17 |
| MAP3K9 | 3 | -0.254333068 | 0.066 | 0.175 | 5.41E-22 | 1.98E-17 |
| RICTOR | 3 | 0.36554775 | 0.778 | 0.706 | 5.48E-22 | 2.01E-17 |
| PLEKHA4 | 3 | 0.325684235 | 0.209 | 0.107 | 5.56E-22 | 2.03E-17 |
| GINS1 | 3 | -0.291938569 | 0.137 | 0.271 | 5.83E-22 | 2.13E-17 |
| TRPM3 | 3 | -0.925789661 | 0.097 | 0.214 | 7.19E-22 | 2.63E-17 |
| GUCY1A2 | 3 | -0.348678625 | 0.26 | 0.415 | 7.42E-22 | 2.72E-17 |
| GRID1 | 3 | -0.374515478 | 0.04 | 0.135 | 7.62E-22 | 2.79E-17 |
| CLCN5 | 3 | -0.49533844 | 0.314 | 0.453 | 9.16E-22 | 3.35E-17 |
| MT-CO2 | 3 | -0.504612873 | 0.234 | 0.381 | 9.88E-22 | 3.62E-17 |
| HIVEP3 | 3 | -0.430776111 | 0.165 | 0.287 | 1.1E-21 | 4.03E-17 |
| PLEKHG4B | 3 | -0.316121987 | 0.113 | 0.239 | 1.39E-21 | 5.09E-17 |
| CSRNP3 | 3 | 0.396841899 | 0.494 | 0.344 | 1.4E-21 | 5.12E-17 |
| ABCC4 | 3 | -0.336435671 | 0.167 | 0.306 | 1.45E-21 | 5.3E-17 |
| HBP1 | 3 | 0.352710773 | 0.353 | 0.221 | 1.86E-21 | 6.82E-17 |
| AASS | 3 | 0.299492394 | 0.396 | 0.249 | 1.98E-21 | 7.24E-17 |
| STK26 | 3 | -0.303544569 | 0.067 | 0.166 | 2.17E-21 | 7.95E-17 |
| PRC1 | 3 | -0.259206613 | 0.061 | 0.165 | 2.43E-21 | 8.91E-17 |
| SASS6 | 3 | -0.334188133 | 0.191 | 0.332 | 2.56E-21 | 9.36E-17 |
| RIPOR2 | 3 | 0.440255892 | 0.319 | 0.372 | 2.79E-21 | 1.02E-16 |
| H2AFY | 3 | -0.337990253 | 0.349 | 0.489 | 3.16E-21 | 1.16E-16 |
| CDRT4 | 3 | 0.321817154 | 0.312 | 0.188 | 3.2E-21 | 1.17E-16 |

| | | | | | | |
|---|---|---|---|---|---|---|
| PBK | 3 | -0.255784624 | 0.039 | 0.131 | 3.23E-21 | 1.18E-16 |
| TRIM2 | 3 | 0.483705407 | 0.723 | 0.629 | 3.37E-21 | 1.23E-16 |
| PATJ | 3 | -0.416188608 | 0.298 | 0.448 | 3.45E-21 | 1.26E-16 |
| EYS | 3 | -1.62323931 | 0.19 | 0.254 | 3.63E-21 | 1.33E-16 |
| PLEKHA5 | 3 | -0.337366846 | 0.782 | 0.863 | 3.63E-21 | 1.33E-16 |
| PSME4 | 3 | -0.573679229 | 0.72 | 0.795 | 4.02E-21 | 1.47E-16 |
| CASP8AP2 | 3 | -0.330084382 | 0.342 | 0.492 | 4.05E-21 | 1.48E-16 |
| TAOK1 | 3 | 0.305533831 | 0.787 | 0.724 | 4.27E-21 | 1.56E-16 |
| ZNF43 | 3 | -0.318722767 | 0.154 | 0.288 | 4.52E-21 | 1.65E-16 |
| POU2F3 | 3 | -0.293782564 | 0.193 | 0.335 | 5.02E-21 | 1.84E-16 |
| MARCH1 | 3 | 0.940864705 | 0.389 | 0.287 | 5.08E-21 | 1.86E-16 |
| TSHZ2 | 3 | -0.560814367 | 0.833 | 0.792 | 5.26E-21 | 1.93E-16 |
| C2CD5 | 3 | -0.362084284 | 0.599 | 0.685 | 5.76E-21 | 2.11E-16 |
| YES1 | 3 | 0.315734706 | 0.505 | 0.352 | 6.28E-21 | 2.3E-16 |
| OSBPL11 | 3 | -0.4154935 | 0.229 | 0.348 | 6.3E-21 | 2.31E-16 |
| AUH | 3 | 0.361114859 | 0.613 | 0.489 | 6.53E-21 | 2.39E-16 |
| AC025159.1 | 3 | 0.335858927 | 0.282 | 0.157 | 7.01E-21 | 2.57E-16 |
| DNM3 | 3 | -0.500821905 | 0.358 | 0.483 | 7.22E-21 | 2.64E-16 |
| FANCC | 3 | -0.399678859 | 0.404 | 0.535 | 7.39E-21 | 2.7E-16 |
| TLE2 | 3 | 0.27302707 | 0.17 | 0.074 | 7.53E-21 | 2.76E-16 |
| NBAS | 3 | -0.496634126 | 0.778 | 0.802 | 7.79E-21 | 2.85E-16 |
| AC004917.1 | 3 | -0.370433663 | 0.028 | 0.107 | 7.98E-21 | 2.92E-16 |
| CNTN1 | 3 | 0.764985265 | 0.349 | 0.242 | 8.11E-21 | 2.97E-16 |
| TPT1-AS1 | 3 | 0.350465542 | 0.525 | 0.387 | 9.04E-21 | 3.31E-16 |
| KIF1B | 3 | -0.386572381 | 0.662 | 0.751 | 9.12E-21 | 3.34E-16 |
| MT-CYB | 3 | -0.389445195 | 0.124 | 0.249 | 9.19E-21 | 3.37E-16 |
| RSRP1 | 3 | 0.326125577 | 0.624 | 0.49 | 9.52E-21 | 3.48E-16 |
| TRPC1 | 3 | -0.457404635 | 0.444 | 0.558 | 9.64E-21 | 3.53E-16 |
| RRBP1 | 3 | 0.274479115 | 0.249 | 0.13 | 9.71E-21 | 3.55E-16 |
| SLC4A8 | 3 | -0.267716399 | 0.057 | 0.156 | 9.99E-21 | 3.66E-16 |
| EYA4 | 3 | -0.823600086 | 0.113 | 0.216 | 1.13E-20 | 4.15E-16 |
| AC079142.1 | 3 | 0.363967098 | 0.723 | 0.598 | 1.14E-20 | 4.18E-16 |
| GOLGA4 | 3 | 0.305060221 | 0.784 | 0.675 | 1.16E-20 | 4.26E-16 |
| R3HDM1 | 3 | -0.336234719 | 0.805 | 0.876 | 1.23E-20 | 4.48E-16 |
| SNX25 | 3 | 0.494007297 | 0.692 | 0.623 | 1.26E-20 | 4.62E-16 |
| USP31 | 3 | -0.35888261 | 0.211 | 0.339 | 1.35E-20 | 4.93E-16 |
| RUNX1 | 3 | 0.518617667 | 0.214 | 0.105 | 1.38E-20 | 5.05E-16 |
| DOCK7 | 3 | 0.334673711 | 0.684 | 0.601 | 1.41E-20 | 5.16E-16 |
| POLR3G | 3 | 0.384237784 | 0.384 | 0.249 | 1.49E-20 | 5.46E-16 |
| BCL7A | 3 | -0.360642498 | 0.134 | 0.246 | 1.5E-20 | 5.49E-16 |
| ZFAND6 | 3 | 0.365014118 | 0.745 | 0.644 | 1.59E-20 | 5.84E-16 |
| ITPR1 | 3 | -0.453185651 | 0.318 | 0.4 | 1.59E-20 | 5.84E-16 |
| ADGRL2 | 3 | 0.474161178 | 0.744 | 0.617 | 1.71E-20 | 6.27E-16 |
| CCDC50 | 3 | 0.361267417 | 0.521 | 0.411 | 1.77E-20 | 6.49E-16 |
| PHACTR1 | 3 | -0.51893545 | 0.27 | 0.329 | 1.79E-20 | 6.56E-16 |
| BRCA1 | 3 | -0.285817394 | 0.207 | 0.35 | 1.93E-20 | 7.05E-16 |
| RASA3 | 3 | 0.265012239 | 0.179 | 0.081 | 2.11E-20 | 7.72E-16 |
| ZNF438 | 3 | 0.319714651 | 0.417 | 0.271 | 2.51E-20 | 9.17E-16 |
| HDGFL3 | 3 | -0.349116675 | 0.323 | 0.463 | 2.51E-20 | 9.2E-16 |
| ELF1 | 3 | 0.381456632 | 0.417 | 0.275 | 2.64E-20 | 9.65E-16 |
| KCNK2 | 3 | -0.40264337 | 0.169 | 0.304 | 2.64E-20 | 9.66E-16 |
| TM9SF3 | 3 | -0.389216758 | 0.512 | 0.628 | 2.69E-20 | 9.84E-16 |
| SCML1 | 3 | -0.322069984 | 0.149 | 0.27 | 2.76E-20 | 1.01E-15 |
| SOS1 | 3 | 0.416899758 | 0.86 | 0.782 | 2.88E-20 | 1.05E-15 |
| ATL3 | 3 | -0.350494955 | 0.195 | 0.324 | 3.43E-20 | 1.26E-15 |
| NBR1 | 3 | 0.309552166 | 0.301 | 0.178 | 3.53E-20 | 1.29E-15 |
| ST6GALNAC5 | 3 | -0.669001979 | 0.254 | 0.391 | 3.76E-20 | 1.38E-15 |
| ATP2A2 | 3 | -0.386578495 | 0.572 | 0.689 | 3.77E-20 | 1.38E-15 |

| | | | | | | |
|---|---|---|---|---|---|---|
| RNLS | 3 | -0.25347842 | 0.031 | 0.114 | 3.84E-20 | 1.4E-15 |
| TMEM132C | 3 | -0.405105241 | 0.041 | 0.132 | 3.96E-20 | 1.45E-15 |
| SLIT1 | 3 | -0.319639351 | 0.037 | 0.12 | 4.21E-20 | 1.54E-15 |
| MMP17 | 3 | 0.318967854 | 0.213 | 0.11 | 4.47E-20 | 1.64E-15 |
| CTNNA1 | 3 | 0.309002476 | 0.77 | 0.648 | 4.5E-20 | 1.65E-15 |
| ARHGEF28 | 3 | -0.531876588 | 0.242 | 0.37 | 6.09E-20 | 2.23E-15 |
| PRMT2 | 3 | 0.324561238 | 0.443 | 0.317 | 6.55E-20 | 2.4E-15 |
| HS3ST1 | 3 | -0.320051688 | 0.097 | 0.211 | 6.61E-20 | 2.42E-15 |
| PPP1R9A | 3 | -0.542016925 | 0.091 | 0.18 | 6.7E-20 | 2.45E-15 |
| DDX60L | 3 | 0.288560956 | 0.167 | 0.074 | 7.05E-20 | 2.58E-15 |
| ITCH | 3 | 0.289532324 | 0.758 | 0.646 | 7.34E-20 | 2.69E-15 |
| RFWD3 | 3 | -0.297385267 | 0.205 | 0.344 | 7.55E-20 | 2.76E-15 |
| HOXD3 | 3 | -0.292820659 | 0.07 | 0.171 | 7.79E-20 | 2.85E-15 |
| PDE3A | 3 | 0.351936548 | 0.701 | 0.551 | 8.08E-20 | 2.96E-15 |
| TRIP12 | 3 | 0.309609914 | 0.814 | 0.728 | 8.92E-20 | 3.27E-15 |
| MAP4 | 3 | 0.341224091 | 0.792 | 0.713 | 8.99E-20 | 3.29E-15 |
| DCHS2 | 3 | -0.314179434 | 0.032 | 0.11 | 1.71E-19 | 6.25E-15 |
| SPAG1 | 3 | -0.297633425 | 0.053 | 0.142 | 1.72E-19 | 6.29E-15 |
| MYO9A | 3 | 0.314225216 | 0.777 | 0.666 | 1.73E-19 | 6.34E-15 |
| GARNL3 | 3 | 0.342820273 | 0.334 | 0.218 | 1.76E-19 | 6.44E-15 |
| SPTBN4 | 3 | -0.26691158 | 0.03 | 0.11 | 1.77E-19 | 6.49E-15 |
| ASXL1 | 3 | 0.299194932 | 0.723 | 0.622 | 1.78E-19 | 6.51E-15 |
| APCDD1 | 3 | 0.562009672 | 0.394 | 0.297 | 1.78E-19 | 6.52E-15 |
| SPC25 | 3 | -0.312835229 | 0.077 | 0.181 | 1.83E-19 | 6.68E-15 |
| PBX1 | 3 | -0.414424403 | 0.821 | 0.854 | 1.84E-19 | 6.74E-15 |
| FANCB | 3 | -0.313129056 | 0.137 | 0.257 | 1.88E-19 | 6.88E-15 |
| PEAK1 | 3 | 0.405212522 | 0.828 | 0.772 | 1.93E-19 | 7.07E-15 |
| GAREM1 | 3 | 0.388706402 | 0.401 | 0.267 | 2E-19 | 7.32E-15 |
| RAD51B | 3 | 0.343392761 | 0.787 | 0.66 | 2.03E-19 | 7.43E-15 |
| SPINK5 | 3 | -0.309813558 | 0.064 | 0.164 | 2.15E-19 | 7.87E-15 |
| ATP6V1C1 | 3 | 0.339198027 | 0.391 | 0.281 | 2.32E-19 | 8.51E-15 |
| L1TD1 | 3 | 0.255576776 | 0.339 | 0.21 | 2.5E-19 | 9.14E-15 |
| DOCK4 | 3 | -0.424210514 | 0.516 | 0.636 | 2.51E-19 | 9.19E-15 |
| EML1 | 3 | -0.340354719 | 0.277 | 0.419 | 2.52E-19 | 9.24E-15 |
| PITPNC1 | 3 | 0.426392193 | 0.652 | 0.537 | 2.64E-19 | 9.67E-15 |
| COL6A2 | 3 | 0.297643979 | 0.233 | 0.126 | 2.99E-19 | 1.09E-14 |
| AC092422.1 | 3 | -0.576538139 | 0.074 | 0.173 | 3.22E-19 | 1.18E-14 |
| SFMBT2 | 3 | -0.517358575 | 0.428 | 0.5 | 3.28E-19 | 1.2E-14 |
| CKMT2-AS1 | 3 | 0.326752731 | 0.507 | 0.383 | 3.51E-19 | 1.29E-14 |
| TOGARAM1 | 3 | -0.338773655 | 0.313 | 0.453 | 4.18E-19 | 1.53E-14 |
| RTKN2 | 3 | -0.335228824 | 0.085 | 0.186 | 4.4E-19 | 1.61E-14 |
| TRAF3 | 3 | -0.329851515 | 0.329 | 0.474 | 4.97E-19 | 1.82E-14 |
| TUBGCP3 | 3 | -0.358793932 | 0.356 | 0.484 | 5.14E-19 | 1.88E-14 |
| NEIL3 | 3 | -0.330459841 | 0.075 | 0.175 | 5.25E-19 | 1.92E-14 |
| TPM4 | 3 | 0.354140361 | 0.468 | 0.335 | 5.39E-19 | 1.97E-14 |
| DPH6-DT | 3 | 0.306043835 | 0.222 | 0.115 | 6.15E-19 | 2.25E-14 |
| SIK3 | 3 | 0.368461086 | 0.757 | 0.658 | 6.39E-19 | 2.34E-14 |
| DACH2 | 3 | -0.912351481 | 0.348 | 0.464 | 6.75E-19 | 2.47E-14 |
| CPE | 3 | -0.299548005 | 0.5 | 0.646 | 7.13E-19 | 2.61E-14 |
| SCCPDH | 3 | 0.271415899 | 0.259 | 0.147 | 7.38E-19 | 2.7E-14 |
| NALCN | 3 | -0.496760823 | 0.135 | 0.235 | 7.56E-19 | 2.77E-14 |
| NDFIP2 | 3 | 0.393133049 | 0.332 | 0.217 | 7.78E-19 | 2.85E-14 |
| CSGALNACT1 | 3 | 0.426602799 | 0.215 | 0.111 | 7.99E-19 | 2.93E-14 |
| AL589693.1 | 3 | 0.880882646 | 0.146 | 0.088 | 8.27E-19 | 3.03E-14 |
| COMMD10 | 3 | 0.341880668 | 0.691 | 0.577 | 8.48E-19 | 3.1E-14 |
| NLGN4X | 3 | -0.416032313 | 0.705 | 0.737 | 8.71E-19 | 3.19E-14 |
| PHF21B | 3 | -0.352053117 | 0.251 | 0.392 | 9.52E-19 | 3.49E-14 |
| LGR4 | 3 | 0.338342245 | 0.574 | 0.428 | 1.01E-18 | 3.7E-14 |

| | | | | | | |
|---|---|---|---|---|---|---|
| SLC30A8 | 3 | 0.325614276 | 0.138 | 0.056 | 1.02E-18 | 3.74E-14 |
| PAGR1 | 3 | -0.268500568 | 0.12 | 0.236 | 1.04E-18 | 3.8E-14 |
| TLK2 | 3 | 0.307898568 | 0.694 | 0.6 | 1.1E-18 | 4.02E-14 |
| GXYLT1 | 3 | -0.319889757 | 0.187 | 0.306 | 1.11E-18 | 4.06E-14 |
| AGPAT5 | 3 | -0.368742268 | 0.377 | 0.504 | 1.17E-18 | 4.27E-14 |
| MT-ND4 | 3 | -0.310257625 | 0.144 | 0.268 | 1.19E-18 | 4.36E-14 |
| XYLB | 3 | -0.298318309 | 0.186 | 0.316 | 1.27E-18 | 4.67E-14 |
| TRIM9 | 3 | -0.361085053 | 0.297 | 0.441 | 1.3E-18 | 4.76E-14 |
| BABAM2 | 3 | 0.315049115 | 0.683 | 0.564 | 1.55E-18 | 5.69E-14 |
| CNKSR2 | 3 | -0.579347604 | 0.557 | 0.675 | 1.56E-18 | 5.73E-14 |
| NKD1 | 3 | 0.604761309 | 0.244 | 0.154 | 1.63E-18 | 5.95E-14 |
| HLA-A | 3 | -0.326095915 | 0.131 | 0.242 | 1.65E-18 | 6.03E-14 |
| SNTG2 | 3 | -0.466565468 | 0.134 | 0.252 | 1.72E-18 | 6.28E-14 |
| SLC4A4 | 3 | -0.345211207 | 0.082 | 0.185 | 1.81E-18 | 6.62E-14 |
| ANKHD1 | 3 | -0.281806864 | 0.708 | 0.788 | 1.82E-18 | 6.66E-14 |
| NAV3 | 3 | -0.676219879 | 0.277 | 0.399 | 1.94E-18 | 7.11E-14 |
| PROM1 | 3 | -0.284403746 | 0.032 | 0.111 | 2.08E-18 | 7.61E-14 |
| GCA | 3 | 0.283686175 | 0.262 | 0.151 | 2.25E-18 | 8.22E-14 |
| WDR70 | 3 | 0.297577961 | 0.764 | 0.662 | 2.46E-18 | 9.02E-14 |
| DANT2 | 3 | -0.447587149 | 0.209 | 0.343 | 2.74E-18 | 1E-13 |
| TRIM16 | 3 | 0.266362179 | 0.288 | 0.171 | 2.79E-18 | 1.02E-13 |
| NSUN7 | 3 | -0.373538388 | 0.152 | 0.26 | 2.8E-18 | 1.02E-13 |
| HIPK3 | 3 | 0.349318464 | 0.531 | 0.445 | 2.87E-18 | 1.05E-13 |
| GINS3 | 3 | 0.272252984 | 0.117 | 0.061 | 3.09E-18 | 1.13E-13 |
| MT-ATP6 | 3 | -0.42349928 | 0.16 | 0.282 | 3.12E-18 | 1.14E-13 |
| GMNN | 3 | -0.289155753 | 0.112 | 0.225 | 3.19E-18 | 1.17E-13 |
| HS3ST3A1 | 3 | 0.444697943 | 0.365 | 0.24 | 3.28E-18 | 1.2E-13 |
| LRRN1 | 3 | -0.305714657 | 0.099 | 0.208 | 3.32E-18 | 1.21E-13 |
| HACD3 | 3 | -0.324982577 | 0.378 | 0.519 | 3.51E-18 | 1.29E-13 |
| HERC4 | 3 | 0.285353533 | 0.812 | 0.71 | 3.75E-18 | 1.37E-13 |
| TLK1 | 3 | -0.364923404 | 0.637 | 0.717 | 4.14E-18 | 1.51E-13 |
| RFC4 | 3 | -0.260594125 | 0.135 | 0.254 | 4.68E-18 | 1.71E-13 |
| ZNF814 | 3 | -0.274795762 | 0.108 | 0.214 | 4.99E-18 | 1.83E-13 |
| TBC1D16 | 3 | 0.328249229 | 0.388 | 0.26 | 5.36E-18 | 1.96E-13 |
| PACRG | 3 | -0.533722292 | 0.039 | 0.104 | 5.46E-18 | 2E-13 |
| TOX2 | 3 | 0.323945512 | 0.178 | 0.088 | 5.47E-18 | 2E-13 |
| CPXM2 | 3 | 0.383407721 | 0.16 | 0.077 | 5.55E-18 | 2.03E-13 |
| RNF217 | 3 | 0.354470612 | 0.51 | 0.384 | 5.55E-18 | 2.03E-13 |
| PRPS2 | 3 | -0.275490126 | 0.133 | 0.245 | 5.61E-18 | 2.06E-13 |
| RB1CC1 | 3 | 0.300908895 | 0.667 | 0.554 | 5.8E-18 | 2.12E-13 |
| DAPK2 | 3 | -0.29304675 | 0.117 | 0.231 | 6.05E-18 | 2.22E-13 |
| OSBP2 | 3 | -0.28781621 | 0.107 | 0.217 | 6.25E-18 | 2.29E-13 |
| GLCE | 3 | -0.427949971 | 0.305 | 0.394 | 6.48E-18 | 2.37E-13 |
| SPAG17 | 3 | 0.349798237 | 0.412 | 0.275 | 6.83E-18 | 2.5E-13 |
| TUT4 | 3 | -0.290469937 | 0.82 | 0.871 | 7.28E-18 | 2.67E-13 |
| USP3 | 3 | -0.371855959 | 0.62 | 0.705 | 7.6E-18 | 2.78E-13 |
| CCDC138 | 3 | -0.32183427 | 0.397 | 0.535 | 8.7E-18 | 3.19E-13 |
| PHF20L1 | 3 | 0.298162298 | 0.605 | 0.522 | 9.81E-18 | 3.59E-13 |
| TJP1 | 3 | 0.305494355 | 0.789 | 0.695 | 9.88E-18 | 3.62E-13 |
| UHRF1BP1L | 3 | -0.322135382 | 0.36 | 0.492 | 9.99E-18 | 3.66E-13 |
| AIG1 | 3 | -0.353221477 | 0.278 | 0.399 | 1.01E-17 | 3.69E-13 |
| DACH1 | 3 | -0.274877112 | 0.836 | 0.894 | 1.03E-17 | 3.75E-13 |
| SKP2 | 3 | -0.272254965 | 0.133 | 0.247 | 1.08E-17 | 3.96E-13 |
| PIGN | 3 | 0.308089276 | 0.466 | 0.33 | 1.09E-17 | 4E-13 |
| ANK3 | 3 | 0.543089264 | 0.419 | 0.317 | 1.12E-17 | 4.11E-13 |
| VPS37B | 3 | 0.29657259 | 0.483 | 0.348 | 1.15E-17 | 4.2E-13 |
| CBX5 | 3 | -0.292832106 | 0.27 | 0.405 | 1.22E-17 | 4.47E-13 |
| C5orf17 | 3 | 0.324450905 | 0.183 | 0.09 | 1.33E-17 | 4.86E-13 |

| | | | | | | |
|---|---|---|---|---|---|---|
| VGLL3 | 3 | -0.385854388 | 0.353 | 0.306 | 1.37E-17 | 5.02E-13 |
| PLAG1 | 3 | 0.282281939 | 0.188 | 0.095 | 1.49E-17 | 5.45E-13 |
| LINC00461 | 3 | -0.468937962 | 0.392 | 0.517 | 1.55E-17 | 5.66E-13 |
| RPS27L | 3 | 0.289542398 | 0.286 | 0.173 | 1.59E-17 | 5.81E-13 |
| DGKH | 3 | -0.346578801 | 0.598 | 0.715 | 1.65E-17 | 6.03E-13 |
| SLC12A2 | 3 | -0.47358644 | 0.362 | 0.476 | 1.73E-17 | 6.34E-13 |
| TPD52 | 3 | -0.460001236 | 0.248 | 0.375 | 1.74E-17 | 6.38E-13 |
| PALLD | 3 | -0.543282551 | 0.328 | 0.426 | 1.76E-17 | 6.44E-13 |
| AKAP12 | 3 | 0.540067557 | 0.251 | 0.193 | 1.82E-17 | 6.67E-13 |
| KDM4C | 3 | 0.304593192 | 0.698 | 0.602 | 1.83E-17 | 6.72E-13 |
| NGF | 3 | 0.319479166 | 0.207 | 0.113 | 2.05E-17 | 7.52E-13 |
| CNPY2 | 3 | 0.322519935 | 0.328 | 0.224 | 2.57E-17 | 9.42E-13 |
| PTPRG-AS1 | 3 | -0.275075461 | 0.115 | 0.224 | 2.64E-17 | 9.65E-13 |
| FBLN1 | 3 | -0.250181698 | 0.123 | 0.235 | 2.73E-17 | 1E-12 |
| CENPF | 3 | -0.338837088 | 0.11 | 0.217 | 2.76E-17 | 1.01E-12 |
| ETV6 | 3 | 0.359100418 | 0.602 | 0.478 | 2.82E-17 | 1.03E-12 |
| RASGEF1B | 3 | 0.569595633 | 0.39 | 0.315 | 2.92E-17 | 1.07E-12 |
| LDB2 | 3 | -0.743460323 | 0.271 | 0.345 | 3.07E-17 | 1.12E-12 |
| LIN9 | 3 | -0.312068069 | 0.213 | 0.337 | 3.1E-17 | 1.13E-12 |
| IDI2-AS1 | 3 | -0.354124093 | 0.066 | 0.145 | 3.29E-17 | 1.2E-12 |
| NSMCE2 | 3 | 0.290676978 | 0.793 | 0.688 | 3.46E-17 | 1.27E-12 |
| LINC00342 | 3 | -0.340218352 | 0.345 | 0.476 | 3.55E-17 | 1.3E-12 |
| TPX2 | 3 | -0.270328186 | 0.084 | 0.182 | 3.67E-17 | 1.34E-12 |
| CEP192 | 3 | -0.324008445 | 0.527 | 0.627 | 3.76E-17 | 1.38E-12 |
| PLK4 | 3 | -0.275543752 | 0.093 | 0.193 | 3.87E-17 | 1.42E-12 |
| TACC1 | 3 | -0.325886787 | 0.19 | 0.307 | 3.88E-17 | 1.42E-12 |
| GNG12-AS1 | 3 | 0.331206069 | 0.256 | 0.162 | 4.17E-17 | 1.53E-12 |
| MTF2 | 3 | -0.320996315 | 0.577 | 0.665 | 4.45E-17 | 1.63E-12 |
| CDH23 | 3 | -0.311023912 | 0.165 | 0.287 | 4.6E-17 | 1.68E-12 |
| SLC36A4 | 3 | -0.328231997 | 0.335 | 0.453 | 4.61E-17 | 1.69E-12 |
| STX16 | 3 | 0.286402146 | 0.571 | 0.454 | 4.62E-17 | 1.69E-12 |
| SERGEF | 3 | -0.370827283 | 0.305 | 0.431 | 4.71E-17 | 1.72E-12 |
| MIR646HG | 3 | -0.555483719 | 0.051 | 0.125 | 4.84E-17 | 1.77E-12 |
| AKAP10 | 3 | 0.291882217 | 0.656 | 0.557 | 4.9E-17 | 1.79E-12 |
| CREBRF | 3 | 0.297794632 | 0.285 | 0.179 | 5.03E-17 | 1.84E-12 |
| FBXO7 | 3 | 0.272285638 | 0.313 | 0.2 | 5.1E-17 | 1.87E-12 |
| TTC23 | 3 | 0.250216127 | 0.299 | 0.181 | 5.77E-17 | 2.11E-12 |
| ZC3H12C | 3 | -0.455110122 | 0.512 | 0.6 | 6.02E-17 | 2.2E-12 |
| NDC80 | 3 | -0.261110137 | 0.07 | 0.163 | 6.03E-17 | 2.21E-12 |
| LAMC1 | 3 | 0.297954455 | 0.646 | 0.508 | 6.97E-17 | 2.55E-12 |
| SSBP2 | 3 | 0.349777579 | 0.881 | 0.802 | 7.32E-17 | 2.68E-12 |
| GPR137B | 3 | 0.315162548 | 0.346 | 0.233 | 7.57E-17 | 2.77E-12 |
| DIP2C | 3 | 0.338214027 | 0.82 | 0.744 | 7.76E-17 | 2.84E-12 |
| MEST | 3 | -0.312798836 | 0.06 | 0.146 | 7.86E-17 | 2.88E-12 |
| SLC25A36 | 3 | 0.299320768 | 0.704 | 0.592 | 8.08E-17 | 2.96E-12 |
| PRTG | 3 | -0.536456423 | 0.032 | 0.101 | 8.42E-17 | 3.08E-12 |
| TUBB | 3 | -0.279086225 | 0.151 | 0.266 | 8.54E-17 | 3.13E-12 |
| MFHAS1 | 3 | -0.330315638 | 0.185 | 0.303 | 8.72E-17 | 3.19E-12 |
| SPHKAP | 3 | -0.297789001 | 0.044 | 0.123 | 1.17E-16 | 4.29E-12 |
| AC093879.1 | 3 | 0.324832353 | 0.252 | 0.176 | 1.18E-16 | 4.31E-12 |
| ADAMTS6 | 3 | -0.638107257 | 0.139 | 0.152 | 1.2E-16 | 4.41E-12 |
| CNTNAP3 | 3 | 0.350631888 | 0.237 | 0.139 | 1.21E-16 | 4.41E-12 |
| RTN4 | 3 | 0.33362373 | 0.757 | 0.709 | 1.3E-16 | 4.75E-12 |
| MAMLD1 | 3 | -0.445311424 | 0.236 | 0.322 | 1.31E-16 | 4.81E-12 |
| DENND1B | 3 | 0.322847351 | 0.438 | 0.309 | 1.34E-16 | 4.91E-12 |
| CYTOR | 3 | 0.32378069 | 0.244 | 0.14 | 1.36E-16 | 4.98E-12 |
| ABI2 | 3 | -0.312811741 | 0.569 | 0.677 | 1.36E-16 | 4.99E-12 |
| MB21D2 | 3 | -0.37191134 | 0.191 | 0.297 | 1.44E-16 | 5.28E-12 |

| | | | | | | |
|---|---|---|---|---|---|---|
| AHCYL2 | 3 | -0.482854799 | 0.66 | 0.667 | 1.46E-16 | 5.33E-12 |
| SYNJ1 | 3 | -0.311340584 | 0.254 | 0.377 | 1.5E-16 | 5.47E-12 |
| MYO6 | 3 | -0.339480915 | 0.183 | 0.292 | 1.52E-16 | 5.55E-12 |
| MYO3B | 3 | 0.495534078 | 0.11 | 0.06 | 1.69E-16 | 6.2E-12 |
| FLVCR1 | 3 | -0.295504928 | 0.252 | 0.378 | 1.78E-16 | 6.51E-12 |
| KAT2B | 3 | 0.356451616 | 0.365 | 0.278 | 1.8E-16 | 6.6E-12 |
| DPH6 | 3 | 0.302789342 | 0.512 | 0.376 | 1.83E-16 | 6.71E-12 |
| EXPH5 | 3 | -0.343829769 | 0.078 | 0.168 | 1.89E-16 | 6.93E-12 |
| NEDD9 | 3 | 0.26091191 | 0.208 | 0.111 | 2E-16 | 7.32E-12 |
| MCTS1 | 3 | -0.26872613 | 0.155 | 0.262 | 2.23E-16 | 8.17E-12 |
| COL5A1 | 3 | 0.341633498 | 0.229 | 0.129 | 2.3E-16 | 8.43E-12 |
| CEP152 | 3 | -0.32293113 | 0.211 | 0.326 | 2.38E-16 | 8.71E-12 |
| FUT8 | 3 | -0.397042849 | 0.785 | 0.82 | 2.58E-16 | 9.45E-12 |
| AC005562.1 | 3 | 0.267122765 | 0.34 | 0.22 | 2.72E-16 | 9.97E-12 |
| ZNF804A | 3 | 0.997531853 | 0.325 | 0.237 | 2.76E-16 | 1.01E-11 |
| COLEC12 | 3 | -0.387714674 | 0.059 | 0.143 | 2.79E-16 | 1.02E-11 |
| CA5B | 3 | 0.309845252 | 0.396 | 0.287 | 2.88E-16 | 1.05E-11 |
| FARP1 | 3 | 0.32543648 | 0.802 | 0.718 | 3.39E-16 | 1.24E-11 |
| FOCAD | 3 | -0.321174171 | 0.618 | 0.725 | 3.51E-16 | 1.28E-11 |
| LTBP1 | 3 | 0.308696 | 0.43 | 0.3 | 3.52E-16 | 1.29E-11 |
| F3 | 3 | -0.482070807 | 0.04 | 0.108 | 3.55E-16 | 1.3E-11 |
| RAB30-DT | 3 | 0.287888841 | 0.391 | 0.278 | 3.82E-16 | 1.4E-11 |
| TJP2 | 3 | -0.352175643 | 0.28 | 0.407 | 4.03E-16 | 1.48E-11 |
| HSPA4L | 3 | -0.278335397 | 0.122 | 0.225 | 4.07E-16 | 1.49E-11 |
| SLBP | 3 | -0.272104695 | 0.128 | 0.236 | 4.28E-16 | 1.57E-11 |
| MSH6 | 3 | -0.281943626 | 0.207 | 0.329 | 4.39E-16 | 1.61E-11 |
| PPL | 3 | -0.29782818 | 0.057 | 0.136 | 4.8E-16 | 1.76E-11 |
| EXOC4 | 3 | 0.275533474 | 0.905 | 0.875 | 5.32E-16 | 1.95E-11 |
| CDH12 | 3 | -0.538906042 | 0.291 | 0.413 | 5.7E-16 | 2.09E-11 |
| CTDSPL2 | 3 | -0.316174364 | 0.531 | 0.642 | 5.98E-16 | 2.19E-11 |
| CKAP2 | 3 | -0.281649884 | 0.142 | 0.249 | 6.25E-16 | 2.29E-11 |
| PHYKPL | 3 | 0.285287363 | 0.394 | 0.28 | 6.54E-16 | 2.4E-11 |
| SH3RF3 | 3 | -0.302081384 | 0.166 | 0.283 | 6.61E-16 | 2.42E-11 |
| MIRLET7BHG | 3 | 0.364693248 | 0.767 | 0.659 | 6.66E-16 | 2.44E-11 |
| ABAT | 3 | 0.299112323 | 0.224 | 0.126 | 6.81E-16 | 2.49E-11 |
| MCF2L | 3 | -0.321918215 | 0.163 | 0.276 | 6.93E-16 | 2.54E-11 |
| UCK2 | 3 | -0.330622158 | 0.331 | 0.449 | 7.49E-16 | 2.74E-11 |
| CXADR | 3 | -0.304466658 | 0.228 | 0.352 | 8.07E-16 | 2.95E-11 |
| CARMIL1 | 3 | 0.254340494 | 0.438 | 0.307 | 8.94E-16 | 3.27E-11 |
| FSIP2 | 3 | -0.250379293 | 0.105 | 0.206 | 9.13E-16 | 3.34E-11 |
| LINC00630 | 3 | 0.297568697 | 0.433 | 0.315 | 1.07E-15 | 3.9E-11 |
| C1orf112 | 3 | -0.27431534 | 0.183 | 0.299 | 1.23E-15 | 4.51E-11 |
| TTLL5 | 3 | -0.32558105 | 0.53 | 0.614 | 1.37E-15 | 5.02E-11 |
| CRPPA | 3 | 0.295351108 | 0.334 | 0.22 | 1.56E-15 | 5.69E-11 |
| XPO6 | 3 | -0.311037279 | 0.419 | 0.535 | 1.59E-15 | 5.84E-11 |
| TRABD2B | 3 | 0.352755873 | 0.178 | 0.105 | 1.77E-15 | 6.49E-11 |
| USP37 | 3 | -0.312343482 | 0.393 | 0.514 | 1.85E-15 | 6.76E-11 |
| WASF1 | 3 | -0.340187946 | 0.347 | 0.467 | 1.95E-15 | 7.14E-11 |
| FKBP10 | 3 | 0.254017326 | 0.348 | 0.23 | 1.97E-15 | 7.22E-11 |
| METAP1D | 3 | 0.274819283 | 0.337 | 0.221 | 1.98E-15 | 7.26E-11 |
| CTDSPL | 3 | -0.318057863 | 0.411 | 0.534 | 2.45E-15 | 8.96E-11 |
| NUFIP2 | 3 | 0.287049711 | 0.537 | 0.446 | 2.72E-15 | 9.97E-11 |
| EPHA3 | 3 | -0.762749087 | 0.143 | 0.225 | 2.8E-15 | 1.02E-10 |
| NSD3 | 3 | -0.285047207 | 0.698 | 0.769 | 2.82E-15 | 1.03E-10 |
| SLMAP | 3 | 0.292749711 | 0.668 | 0.584 | 3.01E-15 | 1.1E-10 |
| ZDHHC20 | 3 | -0.320841546 | 0.365 | 0.47 | 3.08E-15 | 1.13E-10 |
| EDA | 3 | -0.338133688 | 0.179 | 0.295 | 3.17E-15 | 1.16E-10 |
| RFX3 | 3 | -0.427550566 | 0.782 | 0.825 | 3.29E-15 | 1.21E-10 |

| | | | | | | |
|---|---|---|---|---|---|---|
| SFI1 | 3 | -0.269173269 | 0.177 | 0.289 | 3.44E-15 | 1.26E-10 |
| HMGN2 | 3 | -0.310838073 | 0.28 | 0.402 | 3.54E-15 | 1.3E-10 |
| ADAMTSL3 | 3 | 0.448651172 | 0.385 | 0.274 | 3.61E-15 | 1.32E-10 |
| GPD2 | 3 | -0.335699683 | 0.32 | 0.434 | 3.7E-15 | 1.35E-10 |
| BRD8 | 3 | 0.282716938 | 0.357 | 0.251 | 3.85E-15 | 1.41E-10 |
| COMMD1 | 3 | 0.27870045 | 0.593 | 0.468 | 4.02E-15 | 1.47E-10 |
| HDAC8 | 3 | -0.311820948 | 0.63 | 0.717 | 4.15E-15 | 1.52E-10 |
| AKAP6 | 3 | -0.51021749 | 0.148 | 0.246 | 4.2E-15 | 1.54E-10 |
| HMGCR | 3 | -0.312488134 | 0.125 | 0.225 | 4.47E-15 | 1.63E-10 |
| DNAJC7 | 3 | 0.270570771 | 0.375 | 0.266 | 4.52E-15 | 1.65E-10 |
| TTC28 | 3 | -0.265543161 | 0.967 | 0.964 | 4.85E-15 | 1.78E-10 |
| WWOX | 3 | -0.448535202 | 0.737 | 0.777 | 4.95E-15 | 1.81E-10 |
| KCNMA1 | 3 | -0.265761337 | 0.186 | 0.301 | 5E-15 | 1.83E-10 |
| PKN2-AS1 | 3 | 0.265627799 | 0.203 | 0.11 | 5.06E-15 | 1.85E-10 |
| SNHG4 | 3 | 0.263921144 | 0.246 | 0.149 | 5.14E-15 | 1.88E-10 |
| DYNC1LI1 | 3 | -0.316076631 | 0.246 | 0.358 | 5.7E-15 | 2.09E-10 |
| KIF20B | 3 | -0.290647138 | 0.21 | 0.33 | 6.32E-15 | 2.32E-10 |
| ELL2 | 3 | -0.628872019 | 0.115 | 0.197 | 6.58E-15 | 2.41E-10 |
| DAAM1 | 3 | -0.377698647 | 0.398 | 0.516 | 6.95E-15 | 2.54E-10 |
| EFR3B | 3 | -0.316259582 | 0.344 | 0.469 | 7.02E-15 | 2.57E-10 |
| FST | 3 | 0.251775659 | 0.15 | 0.072 | 7.75E-15 | 2.84E-10 |
| PCAT1 | 3 | -0.51688682 | 0.043 | 0.107 | 7.85E-15 | 2.87E-10 |
| ID2 | 3 | -0.390221419 | 0.081 | 0.171 | 7.86E-15 | 2.88E-10 |
| DBF4 | 3 | -0.293028164 | 0.187 | 0.3 | 8.46E-15 | 3.1E-10 |
| MARCH3 | 3 | 0.382683792 | 0.279 | 0.186 | 8.9E-15 | 3.26E-10 |
| KYAT3 | 3 | 0.250240833 | 0.352 | 0.235 | 9.24E-15 | 3.38E-10 |
| GLMN | 3 | -0.326713055 | 0.416 | 0.51 | 9.89E-15 | 3.62E-10 |
| CEP126 | 3 | -0.339550543 | 0.21 | 0.312 | 9.93E-15 | 3.63E-10 |
| SYNRG | 3 | 0.282213958 | 0.495 | 0.388 | 9.97E-15 | 3.65E-10 |
| NUP58 | 3 | -0.301310334 | 0.517 | 0.623 | 1.08E-14 | 3.97E-10 |
| APOLD1 | 3 | -0.322804774 | 0.068 | 0.151 | 1.09E-14 | 4.01E-10 |
| MTHFD1 | 3 | -0.273255454 | 0.232 | 0.349 | 1.11E-14 | 4.06E-10 |
| CDK12 | 3 | 0.285447819 | 0.543 | 0.435 | 1.11E-14 | 4.07E-10 |
| AOPEP | 3 | 0.28115682 | 0.813 | 0.748 | 1.13E-14 | 4.14E-10 |
| KCNJ3 | 3 | -0.521736069 | 0.075 | 0.15 | 1.18E-14 | 4.31E-10 |
| HPRT1 | 3 | -0.293318441 | 0.317 | 0.437 | 1.18E-14 | 4.33E-10 |
| RHOBTB1 | 3 | 0.33070139 | 0.336 | 0.24 | 1.21E-14 | 4.41E-10 |
| CNOT6L | 3 | -0.324244515 | 0.414 | 0.521 | 1.25E-14 | 4.58E-10 |
| ABI1 | 3 | 0.297319597 | 0.591 | 0.506 | 1.3E-14 | 4.76E-10 |
| SCP2 | 3 | -0.349579076 | 0.289 | 0.379 | 1.36E-14 | 4.98E-10 |
| BCKDHB | 3 | 0.301606296 | 0.596 | 0.479 | 1.36E-14 | 5E-10 |
| ARHGAP11B | 3 | -0.310082601 | 0.174 | 0.282 | 1.38E-14 | 5.07E-10 |
| RASSF3 | 3 | 0.327731298 | 0.363 | 0.259 | 1.4E-14 | 5.11E-10 |
| ELOVL6 | 3 | -0.340901043 | 0.335 | 0.445 | 1.5E-14 | 5.51E-10 |
| ACACA | 3 | -0.323900015 | 0.665 | 0.745 | 1.51E-14 | 5.51E-10 |
| SMC5 | 3 | -0.282663234 | 0.578 | 0.665 | 1.52E-14 | 5.55E-10 |
| PRKAA2 | 3 | -0.250614059 | 0.099 | 0.192 | 1.52E-14 | 5.57E-10 |
| ZC2HC1A | 3 | 0.262530825 | 0.505 | 0.379 | 1.55E-14 | 5.68E-10 |
| PARPBP | 3 | -0.275953549 | 0.172 | 0.282 | 1.58E-14 | 5.79E-10 |
| WASF3 | 3 | -0.273900866 | 0.096 | 0.185 | 1.65E-14 | 6.03E-10 |
| PHF6 | 3 | 0.33358497 | 0.563 | 0.473 | 1.69E-14 | 6.18E-10 |
| TGFBR2 | 3 | 0.297739668 | 0.227 | 0.132 | 1.73E-14 | 6.35E-10 |
| LINC01170 | 3 | 0.431434288 | 0.681 | 0.57 | 1.85E-14 | 6.76E-10 |
| SLC9A7 | 3 | 0.354526279 | 0.205 | 0.141 | 1.9E-14 | 6.97E-10 |
| CASK | 3 | -0.288879288 | 0.816 | 0.864 | 1.94E-14 | 7.12E-10 |
| SDK2 | 3 | 0.286331569 | 0.203 | 0.112 | 2.18E-14 | 7.99E-10 |
| OBI1 | 3 | -0.279922808 | 0.21 | 0.313 | 2.43E-14 | 8.9E-10 |
| PIKFYVE | 3 | 0.280828278 | 0.611 | 0.508 | 2.43E-14 | 8.9E-10 |

| | | | | | | |
|---|---|---|---|---|---|---|
| CCDC85A | 3 | 0.330665656 | 0.11 | 0.102 | 2.48E-14 | 9.09E-10 |
| FZD3 | 3 | -0.326637471 | 0.172 | 0.275 | 2.49E-14 | 9.1E-10 |
| GPRIN3 | 3 | -0.27857532 | 0.265 | 0.388 | 2.57E-14 | 9.42E-10 |
| ZDHHC21 | 3 | 0.341137265 | 0.738 | 0.689 | 2.61E-14 | 9.54E-10 |
| LRRC37B | 3 | 0.27362864 | 0.341 | 0.244 | 2.71E-14 | 9.91E-10 |
| SERPINB9 | 3 | 0.257630706 | 0.166 | 0.085 | 2.71E-14 | 9.94E-10 |
| ZNF106 | 3 | 0.29171435 | 0.392 | 0.284 | 2.75E-14 | 1.01E-09 |
| KIF13A | 3 | 0.319850968 | 0.671 | 0.576 | 3.08E-14 | 1.13E-09 |
| AC092821.3 | 3 | -0.294563362 | 0.206 | 0.302 | 3.18E-14 | 1.16E-09 |
| NUP205 | 3 | -0.290480631 | 0.391 | 0.512 | 3.19E-14 | 1.17E-09 |
| METTL15 | 3 | 0.275517516 | 0.71 | 0.631 | 3.25E-14 | 1.19E-09 |
| ZMYND8 | 3 | 0.325228327 | 0.66 | 0.567 | 3.36E-14 | 1.23E-09 |
| AC012501.2 | 3 | 0.255857328 | 0.159 | 0.094 | 3.46E-14 | 1.27E-09 |
| MID1 | 3 | 0.507057668 | 0.695 | 0.595 | 3.53E-14 | 1.29E-09 |
| MME | 3 | -0.36802891 | 0.108 | 0.2 | 3.62E-14 | 1.32E-09 |
| LRCH2 | 3 | 0.362585139 | 0.736 | 0.692 | 3.69E-14 | 1.35E-09 |
| SMC3 | 3 | -0.297571601 | 0.297 | 0.416 | 3.92E-14 | 1.44E-09 |
| DPP4 | 3 | -0.348322478 | 0.093 | 0.184 | 3.97E-14 | 1.45E-09 |
| RIF1 | 3 | -0.276642023 | 0.739 | 0.783 | 4.13E-14 | 1.51E-09 |
| HSPA12A | 3 | 0.312753076 | 0.365 | 0.267 | 4.14E-14 | 1.52E-09 |
| EXOC2 | 3 | -0.420053669 | 0.613 | 0.679 | 4.33E-14 | 1.58E-09 |
| KIAA1549L | 3 | -0.317185301 | 0.132 | 0.233 | 4.55E-14 | 1.67E-09 |
| RFX2 | 3 | -0.392957666 | 0.153 | 0.232 | 5.51E-14 | 2.02E-09 |
| ASAP1 | 3 | 0.272096479 | 0.898 | 0.846 | 5.56E-14 | 2.04E-09 |
| ETS1 | 3 | 0.289747543 | 0.256 | 0.158 | 5.58E-14 | 2.04E-09 |
| MYRIP | 3 | 0.516251591 | 0.703 | 0.652 | 5.91E-14 | 2.16E-09 |
| ARFIP1 | 3 | 0.278377434 | 0.594 | 0.481 | 5.93E-14 | 2.17E-09 |
| GCNT1 | 3 | -0.269798814 | 0.037 | 0.1 | 6.34E-14 | 2.32E-09 |
| GOSR1 | 3 | 0.268722901 | 0.507 | 0.4 | 6.95E-14 | 2.54E-09 |
| ATP8A1 | 3 | -0.512147784 | 0.326 | 0.421 | 7.74E-14 | 2.83E-09 |
| SESN1 | 3 | -0.387396298 | 0.404 | 0.499 | 7.94E-14 | 2.91E-09 |
| CCDC88A | 3 | -0.362122382 | 0.575 | 0.649 | 8.33E-14 | 3.05E-09 |
| NR6A1 | 3 | -0.387207437 | 0.288 | 0.382 | 8.93E-14 | 3.27E-09 |
| HIBADH | 3 | 0.320615837 | 0.587 | 0.489 | 9.04E-14 | 3.31E-09 |
| AC063938.1 | 3 | -0.37409394 | 0.052 | 0.114 | 9.78E-14 | 3.58E-09 |
| QSER1 | 3 | -0.264086854 | 0.629 | 0.716 | 9.78E-14 | 3.58E-09 |
| FGD4 | 3 | 0.362675878 | 0.504 | 0.408 | 9.83E-14 | 3.6E-09 |
| RUNX2 | 3 | 0.596391117 | 0.203 | 0.129 | 9.91E-14 | 3.63E-09 |
| SGK3 | 3 | 0.263315456 | 0.439 | 0.326 | 1.14E-13 | 4.16E-09 |
| OLA1 | 3 | -0.289804398 | 0.711 | 0.78 | 1.16E-13 | 4.23E-09 |
| UBR2 | 3 | 0.253399316 | 0.656 | 0.546 | 1.16E-13 | 4.26E-09 |
| MBNL3 | 3 | -0.397245411 | 0.396 | 0.502 | 1.19E-13 | 4.34E-09 |
| BMP2K | 3 | -0.357261768 | 0.473 | 0.568 | 1.25E-13 | 4.59E-09 |
| SLCO5A1 | 3 | -0.32825967 | 0.082 | 0.159 | 1.29E-13 | 4.72E-09 |
| GRIA4 | 3 | -0.516470264 | 0.26 | 0.323 | 1.35E-13 | 4.94E-09 |
| LRBA | 3 | -0.286946583 | 0.866 | 0.892 | 1.39E-13 | 5.08E-09 |
| PHTF1 | 3 | -0.303166498 | 0.222 | 0.325 | 1.44E-13 | 5.28E-09 |
| ACAP2 | 3 | -0.322650988 | 0.597 | 0.673 | 1.47E-13 | 5.39E-09 |
| SUPT16H | 3 | -0.288003614 | 0.339 | 0.447 | 1.53E-13 | 5.59E-09 |
| SLC25A26 | 3 | 0.25291583 | 0.44 | 0.322 | 1.54E-13 | 5.62E-09 |
| AC090825.1 | 3 | 0.316885523 | 0.354 | 0.257 | 1.57E-13 | 5.75E-09 |
| MTHFD1L | 3 | 0.3417325 | 0.611 | 0.546 | 1.65E-13 | 6.05E-09 |
| FEZ1 | 3 | -0.294826162 | 0.148 | 0.229 | 1.73E-13 | 6.34E-09 |
| TMEM164 | 3 | -0.331612335 | 0.403 | 0.502 | 1.81E-13 | 6.63E-09 |
| SORCS2 | 3 | -0.38067727 | 0.163 | 0.266 | 1.82E-13 | 6.65E-09 |
| XPR1 | 3 | -0.319408704 | 0.416 | 0.519 | 1.83E-13 | 6.68E-09 |
| ABHD3 | 3 | -0.285543924 | 0.136 | 0.232 | 2.15E-13 | 7.86E-09 |
| HDAC4 | 3 | 0.300918324 | 0.73 | 0.669 | 2.32E-13 | 8.48E-09 |

| | | | | | | |
|---|---|---|---|---|---|---|
| KIF6 | 3 | 0.401140841 | 0.127 | 0.071 | 2.33E-13 | 8.54E-09 |
| GCLM | 3 | -0.339666192 | 0.092 | 0.166 | 2.42E-13 | 8.87E-09 |
| GPR158 | 3 | -0.278547953 | 0.07 | 0.151 | 2.46E-13 | 9.02E-09 |
| ATP1B3 | 3 | -0.274963226 | 0.318 | 0.43 | 2.54E-13 | 9.3E-09 |
| MAPKAP1 | 3 | 0.267046722 | 0.608 | 0.523 | 2.64E-13 | 9.65E-09 |
| EPB41L4A | 3 | -0.324802894 | 0.049 | 0.116 | 2.72E-13 | 9.94E-09 |
| USP40 | 3 | 0.276793114 | 0.418 | 0.314 | 2.81E-13 | 1.03E-08 |
| PVT1 | 3 | 0.287871324 | 0.838 | 0.805 | 3.06E-13 | 1.12E-08 |
| CLHC1 | 3 | 0.269253765 | 0.403 | 0.292 | 3.35E-13 | 1.23E-08 |
| CEP95 | 3 | 0.27275904 | 0.589 | 0.523 | 3.55E-13 | 0.000000013 |
| MAGI3 | 3 | -0.372481067 | 0.368 | 0.461 | 3.56E-13 | 0.000000013 |
| FAF1 | 3 | -0.27138064 | 0.809 | 0.831 | 3.58E-13 | 1.31E-08 |
| GRIK2 | 3 | 0.568766131 | 0.386 | 0.279 | 3.67E-13 | 1.34E-08 |
| ZNF385B | 3 | 0.518242124 | 0.426 | 0.319 | 3.74E-13 | 1.37E-08 |
| KIAA1217 | 3 | 0.333565898 | 0.415 | 0.297 | 3.74E-13 | 1.37E-08 |
| EPN2 | 3 | 0.283259356 | 0.545 | 0.437 | 3.78E-13 | 1.38E-08 |
| ERBIN | 3 | 0.304992746 | 0.819 | 0.748 | 4.26E-13 | 1.56E-08 |
| MYBL1 | 3 | -0.259030863 | 0.155 | 0.257 | 4.64E-13 | 0.000000017 |
| SCUBE2 | 3 | 0.362817688 | 0.252 | 0.16 | 4.74E-13 | 1.73E-08 |
| NUF2 | 3 | -0.250832685 | 0.09 | 0.172 | 4.78E-13 | 1.75E-08 |
| RPS21 | 3 | -0.29475795 | 0.183 | 0.285 | 5.05E-13 | 1.85E-08 |
| NMT1 | 3 | 0.253178362 | 0.38 | 0.276 | 5.28E-13 | 1.93E-08 |
| SPATS2L | 3 | -0.458882653 | 0.409 | 0.478 | 5.63E-13 | 2.06E-08 |
| N4BP2 | 3 | -0.296463564 | 0.451 | 0.536 | 5.88E-13 | 2.15E-08 |
| NBEAL1 | 3 | 0.275629154 | 0.557 | 0.447 | 6.03E-13 | 2.21E-08 |
| ZNF91 | 3 | -0.252129264 | 0.159 | 0.26 | 6.29E-13 | 0.000000023 |
| NFIX | 3 | 0.371064465 | 0.565 | 0.455 | 6.46E-13 | 2.36E-08 |
| MORC3 | 3 | 0.276647084 | 0.633 | 0.534 | 6.48E-13 | 2.37E-08 |
| NAP1L4 | 3 | -0.269563013 | 0.585 | 0.679 | 6.67E-13 | 2.44E-08 |
| SORBS2 | 3 | -0.580035216 | 0.072 | 0.133 | 7.21E-13 | 2.64E-08 |
| SLC7A1 | 3 | -0.283977602 | 0.207 | 0.308 | 7.26E-13 | 2.66E-08 |
| OSBPL1A | 3 | 0.279516929 | 0.541 | 0.427 | 7.71E-13 | 2.82E-08 |
| MT-ND3 | 3 | -0.251457008 | 0.087 | 0.172 | 7.73E-13 | 2.83E-08 |
| AL353747.4 | 3 | -0.276889518 | 0.108 | 0.192 | 8.09E-13 | 2.96E-08 |
| GALNTL6 | 3 | -0.556139246 | 0.16 | 0.261 | 8.52E-13 | 3.12E-08 |
| TMEM178A | 3 | -0.346568528 | 0.371 | 0.483 | 8.64E-13 | 3.16E-08 |
| HSF2 | 3 | -0.271956695 | 0.231 | 0.325 | 8.65E-13 | 3.17E-08 |
| TSC22D1 | 3 | -0.329251508 | 0.641 | 0.719 | 9.21E-13 | 3.37E-08 |
| KLHL24 | 3 | 0.337225505 | 0.549 | 0.467 | 9.91E-13 | 3.63E-08 |
| TMEM181 | 3 | -0.278728001 | 0.41 | 0.501 | 1.01E-12 | 3.69E-08 |
| PEX1 | 3 | -0.294564873 | 0.337 | 0.41 | 1.03E-12 | 3.78E-08 |
| HMGN3 | 3 | 0.273561067 | 0.438 | 0.336 | 1.06E-12 | 3.88E-08 |
| NNT | 3 | -0.325279685 | 0.402 | 0.489 | 1.18E-12 | 4.33E-08 |
| RFFL | 3 | 0.25591764 | 0.321 | 0.223 | 1.42E-12 | 5.19E-08 |
| NCAM1 | 3 | -0.256801796 | 0.83 | 0.834 | 1.44E-12 | 5.27E-08 |
| SEMA6D | 3 | -0.397175968 | 0.603 | 0.681 | 1.52E-12 | 5.55E-08 |
| MTA3 | 3 | -0.29910741 | 0.642 | 0.692 | 1.69E-12 | 0.000000062 |
| SLC7A6 | 3 | -0.352345807 | 0.173 | 0.254 | 1.76E-12 | 6.44E-08 |
| DDHD1 | 3 | -0.288410674 | 0.48 | 0.575 | 1.78E-12 | 6.53E-08 |
| DERA | 3 | -0.287115399 | 0.288 | 0.398 | 2.04E-12 | 7.47E-08 |
| RAB1A | 3 | 0.262946946 | 0.552 | 0.448 | 2.08E-12 | 7.62E-08 |
| MYEF2 | 3 | -0.258645831 | 0.407 | 0.52 | 2.15E-12 | 7.86E-08 |
| NCBP3 | 3 | 0.259837611 | 0.412 | 0.312 | 2.31E-12 | 8.47E-08 |
| KRAS | 3 | -0.390837091 | 0.331 | 0.377 | 2.33E-12 | 8.55E-08 |
| NR2F1 | 3 | -0.366295333 | 0.159 | 0.223 | 2.39E-12 | 8.76E-08 |
| SOX6 | 3 | 0.383396273 | 0.334 | 0.228 | 2.44E-12 | 8.94E-08 |
| NCKAP5 | 3 | 0.365606009 | 0.506 | 0.398 | 2.5E-12 | 9.16E-08 |
| WIPF2 | 3 | 0.266719436 | 0.404 | 0.309 | 2.52E-12 | 9.22E-08 |

| | | | | | | |
|---|---|---|---|---|---|---|
| CEP295 | 3 | -0.267445472 | 0.335 | 0.443 | 2.59E-12 | 9.48E-08 |
| NAA16 | 3 | -0.276501628 | 0.524 | 0.606 | 2.84E-12 | 0.000000104 |
| GCH1 | 3 | -0.263699796 | 0.098 | 0.179 | 2.85E-12 | 0.000000104 |
| BACH2 | 3 | 0.348532677 | 0.488 | 0.373 | 2.93E-12 | 0.000000107 |
| GBE1 | 3 | 0.348059882 | 0.404 | 0.308 | 3.09E-12 | 0.000000113 |
| ACIN1 | 3 | 0.258858295 | 0.535 | 0.436 | 3.24E-12 | 0.000000119 |
| SAE1 | 3 | -0.250716296 | 0.289 | 0.398 | 3.27E-12 | 0.00000012 |
| ACTN1 | 3 | 0.286997755 | 0.352 | 0.263 | 3.31E-12 | 0.000000121 |
| SH3BP4 | 3 | -0.310697162 | 0.239 | 0.346 | 3.51E-12 | 0.000000129 |
| DHX36 | 3 | 0.251385811 | 0.763 | 0.683 | 3.53E-12 | 0.000000129 |
| CENPE | 3 | -0.316577871 | 0.116 | 0.205 | 3.58E-12 | 0.000000131 |
| TBC1D12 | 3 | 0.296305211 | 0.369 | 0.276 | 3.62E-12 | 0.000000133 |
| PARVB | 3 | 0.276125953 | 0.309 | 0.22 | 3.87E-12 | 0.000000142 |
| TBC1D5 | 3 | -0.272104414 | 0.86 | 0.864 | 3.87E-12 | 0.000000142 |
| SORT1 | 3 | -0.275977058 | 0.217 | 0.308 | 4.06E-12 | 0.000000149 |
| NCAM2 | 3 | 0.504724466 | 0.125 | 0.068 | 4.19E-12 | 0.000000153 |
| ARHGEF3 | 3 | 0.295493389 | 0.549 | 0.447 | 4.25E-12 | 0.000000156 |
| RNF138 | 3 | -0.265211218 | 0.276 | 0.381 | 4.27E-12 | 0.000000156 |
| ICA1L | 3 | -0.252024471 | 0.114 | 0.194 | 4.3E-12 | 0.000000157 |
| BRINP1 | 3 | -0.319286177 | 0.318 | 0.434 | 4.37E-12 | 0.00000016 |
| RBBP7 | 3 | -0.281772445 | 0.418 | 0.512 | 4.5E-12 | 0.000000165 |
| TRIM24 | 3 | -0.316382047 | 0.409 | 0.507 | 4.59E-12 | 0.000000168 |
| VEGFC | 3 | 0.304522858 | 0.305 | 0.203 | 4.63E-12 | 0.000000169 |
| INPP4B | 3 | 0.689806648 | 0.22 | 0.159 | 5.14E-12 | 0.000000188 |
| KIF13B | 3 | 0.259670149 | 0.337 | 0.244 | 5.34E-12 | 0.000000196 |
| VCL | 3 | 0.266207203 | 0.846 | 0.769 | 5.38E-12 | 0.000000197 |
| SIPA1L2 | 3 | -0.297406942 | 0.339 | 0.449 | 5.48E-12 | 0.000000201 |
| CTNND1 | 3 | 0.257426637 | 0.706 | 0.611 | 5.57E-12 | 0.000000204 |
| RPS6KC1 | 3 | 0.255529912 | 0.598 | 0.507 | 5.9E-12 | 0.000000216 |
| PPP1R14C | 3 | -0.467412679 | 0.38 | 0.456 | 6.1E-12 | 0.000000223 |
| VPS13C | 3 | 0.251494087 | 0.785 | 0.733 | 7.46E-12 | 0.000000273 |
| GLT1D1 | 3 | -0.266247785 | 0.096 | 0.172 | 7.54E-12 | 0.000000276 |
| RALGAPA2 | 3 | -0.315821021 | 0.509 | 0.588 | 7.62E-12 | 0.000000279 |
| TRMT9B | 3 | -0.325733726 | 0.143 | 0.234 | 7.89E-12 | 0.000000289 |
| LRP6 | 3 | 0.279127751 | 0.636 | 0.572 | 7.96E-12 | 0.000000291 |
| NUP153 | 3 | -0.270865379 | 0.502 | 0.591 | 8.18E-12 | 0.000000299 |
| PTPN13 | 3 | -0.381949164 | 0.08 | 0.147 | 9.19E-12 | 0.000000336 |
| ASIC2 | 3 | 0.442655305 | 0.302 | 0.264 | 1.01E-11 | 0.000000372 |
| STK39 | 3 | -0.345435351 | 0.576 | 0.65 | 1.06E-11 | 0.000000388 |
| FSD1L | 3 | -0.267017501 | 0.361 | 0.453 | 1.1E-11 | 0.000000401 |
| SPPL3 | 3 | 0.266971882 | 0.663 | 0.582 | 1.15E-11 | 0.00000042 |
| EPM2A | 3 | -0.313490549 | 0.205 | 0.293 | 1.25E-11 | 0.000000457 |
| CHSY3 | 3 | -0.303672649 | 0.725 | 0.709 | 1.28E-11 | 0.00000047 |
| HFM1 | 3 | -0.28946721 | 0.268 | 0.37 | 1.31E-11 | 0.000000479 |
| TMEM161B-AS1 | 3 | -0.351588153 | 0.616 | 0.668 | 1.45E-11 | 0.000000529 |
| RPS6KA3 | 3 | -0.32888836 | 0.49 | 0.595 | 1.51E-11 | 0.000000552 |
| PLOD2 | 3 | 0.386259743 | 0.377 | 0.274 | 1.58E-11 | 0.000000578 |
| LIPE-AS1 | 3 | -0.314792535 | 0.246 | 0.335 | 1.59E-11 | 0.000000581 |
| NPAT | 3 | -0.278453062 | 0.326 | 0.425 | 1.67E-11 | 0.000000611 |
| EPHA7 | 3 | -0.376874286 | 0.222 | 0.326 | 1.91E-11 | 0.000000698 |
| TRMT11 | 3 | 0.31664665 | 0.697 | 0.641 | 1.95E-11 | 0.000000716 |
| CLEC2A | 3 | -0.278113914 | 0.066 | 0.138 | 2.09E-11 | 0.000000766 |
| EEA1 | 3 | 0.263979364 | 0.569 | 0.482 | 2.19E-11 | 0.000000801 |
| NDUFAF2 | 3 | 0.2868867 | 0.409 | 0.313 | 2.19E-11 | 0.000000802 |
| UNC5C | 3 | 0.587974874 | 0.609 | 0.518 | 2.47E-11 | 0.000000905 |
| TNRC6A | 3 | -0.260729999 | 0.768 | 0.825 | 2.6E-11 | 0.000000952 |
| EXOC6 | 3 | -0.292639577 | 0.499 | 0.58 | 2.68E-11 | 0.00000098 |
| KDM4B | 3 | 0.307076796 | 0.575 | 0.481 | 2.78E-11 | 0.00000102 |

| | | | | | | |
|---|---|---|---|---|---|---|
| MGA | 3 | -0.25184713 | 0.612 | 0.687 | 2.99E-11 | 0.0000011 |
| KLHL7 | 3 | -0.298943258 | 0.442 | 0.509 | 3.11E-11 | 0.00000114 |
| APAF1 | 3 | -0.280290637 | 0.257 | 0.344 | 3.19E-11 | 0.00000117 |
| TSHZ1 | 3 | -0.305555451 | 0.314 | 0.414 | 3.23E-11 | 0.00000118 |
| AC108169.1 | 3 | -0.313029487 | 0.075 | 0.143 | 3.39E-11 | 0.00000124 |
| DLG5 | 3 | -0.256459817 | 0.296 | 0.395 | 3.39E-11 | 0.00000124 |
| SLC2A3 | 3 | 0.366195592 | 0.206 | 0.129 | 3.71E-11 | 0.00000136 |
| VPS8 | 3 | 0.285457061 | 0.67 | 0.594 | 4.04E-11 | 0.00000148 |
| IPO5 | 3 | -0.267087823 | 0.342 | 0.448 | 4.32E-11 | 0.00000158 |
| KIRREL1 | 3 | -0.257587579 | 0.35 | 0.454 | 4.33E-11 | 0.00000158 |
| RPS3 | 3 | -0.286642005 | 0.395 | 0.48 | 4.73E-11 | 0.00000173 |
| CDHR3 | 3 | -0.29640229 | 0.306 | 0.406 | 5.1E-11 | 0.00000187 |
| RASGRF2 | 3 | -0.295065997 | 0.109 | 0.19 | 5.53E-11 | 0.00000202 |
| TEX41 | 3 | 0.347305394 | 0.127 | 0.065 | 5.87E-11 | 0.00000215 |
| FAM171A1 | 3 | -0.288821363 | 0.483 | 0.573 | 5.98E-11 | 0.00000219 |
| SMARCA2 | 3 | -0.328403097 | 0.35 | 0.44 | 6.39E-11 | 0.00000234 |
| FBXL20 | 3 | 0.250108411 | 0.644 | 0.587 | 6.98E-11 | 0.00000255 |
| AGTPBP1 | 3 | -0.280768902 | 0.528 | 0.608 | 7.34E-11 | 0.00000269 |
| ANK2 | 3 | 0.432016281 | 0.839 | 0.84 | 7.79E-11 | 0.00000285 |
| CRADD | 3 | 0.300342785 | 0.502 | 0.401 | 7.89E-11 | 0.00000289 |
| DNAJC1 | 3 | -0.393210954 | 0.543 | 0.575 | 8.05E-11 | 0.00000295 |
| PPAT | 3 | -0.262283148 | 0.203 | 0.29 | 8.36E-11 | 0.00000306 |
| SLC35F3 | 3 | 0.322015148 | 0.171 | 0.102 | 9.47E-11 | 0.00000347 |
| AKT3 | 3 | -0.271056142 | 0.794 | 0.842 | 9.53E-11 | 0.00000349 |
| MYCBP2 | 3 | -0.255288133 | 0.753 | 0.81 | 1.05E-10 | 0.00000383 |
| ACTR3B | 3 | -0.263864954 | 0.355 | 0.431 | 1.33E-10 | 0.00000485 |
| TMTC4 | 3 | 0.263530935 | 0.332 | 0.257 | 1.43E-10 | 0.00000524 |
| OSBPL3 | 3 | 0.256953989 | 0.549 | 0.446 | 1.57E-10 | 0.00000574 |
| NR3C2 | 3 | -0.389010866 | 0.193 | 0.285 | 1.6E-10 | 0.00000586 |
| UNC13B | 3 | -0.390745902 | 0.501 | 0.545 | 1.98E-10 | 0.00000724 |
| NTM | 3 | -0.398425335 | 0.324 | 0.428 | 2E-10 | 0.00000731 |
| ARHGAP24 | 3 | -0.398471288 | 0.378 | 0.415 | 2.01E-10 | 0.00000735 |
| AFAP1 | 3 | 0.26767066 | 0.563 | 0.463 | 2.06E-10 | 0.00000753 |
| SEMA5A | 3 | -0.372786381 | 0.47 | 0.4 | 2.2E-10 | 0.00000804 |
| TENT4A | 3 | -0.264914475 | 0.367 | 0.463 | 2.29E-10 | 0.00000838 |
| CALM2 | 3 | -0.261829599 | 0.438 | 0.526 | 2.42E-10 | 0.00000885 |
| CNTN6 | 3 | -0.558440096 | 0.112 | 0.187 | 2.52E-10 | 0.00000922 |
| RBMS3 | 3 | 0.264086527 | 0.869 | 0.794 | 2.78E-10 | 0.0000102 |
| DNAH7 | 3 | -0.576411906 | 0.134 | 0.183 | 2.89E-10 | 0.0000106 |
| GCLC | 3 | -0.566320818 | 0.162 | 0.219 | 2.95E-10 | 0.0000108 |
| SPOCK1 | 3 | 0.465234595 | 0.449 | 0.351 | 3.2E-10 | 0.0000117 |
| CHCHD3 | 3 | -0.25112272 | 0.647 | 0.725 | 3.39E-10 | 0.0000124 |
| DHX9 | 3 | -0.251995303 | 0.604 | 0.672 | 3.44E-10 | 0.0000126 |
| CCNL2 | 3 | 0.260465152 | 0.556 | 0.479 | 4.09E-10 | 0.000015 |
| MAP7D3 | 3 | -0.256994182 | 0.183 | 0.261 | 4.43E-10 | 0.0000162 |
| LIMS1 | 3 | 0.282283391 | 0.56 | 0.481 | 4.83E-10 | 0.0000177 |
| LRP5 | 3 | 0.265317222 | 0.363 | 0.283 | 5.55E-10 | 0.0000203 |
| PRRG1 | 3 | -0.267590906 | 0.288 | 0.369 | 6.69E-10 | 0.0000245 |
| TMTC2 | 3 | 0.361155657 | 0.583 | 0.501 | 6.91E-10 | 0.0000253 |
| GTDC1 | 3 | 0.292527577 | 0.574 | 0.479 | 7E-10 | 0.0000256 |
| RNF125 | 3 | 0.301940449 | 0.196 | 0.147 | 7.16E-10 | 0.0000262 |
| LEPR | 3 | -0.287296144 | 0.124 | 0.133 | 7.99E-10 | 0.0000293 |
| EEF1A1 | 3 | -0.278884166 | 0.492 | 0.588 | 9.17E-10 | 0.0000336 |
| APBA2 | 3 | -0.267590631 | 0.514 | 0.58 | 9.87E-10 | 0.0000361 |
| HDAC2 | 3 | -0.279420699 | 0.543 | 0.599 | 1.27E-09 | 0.0000465 |
| FBN2 | 3 | -0.442685207 | 0.296 | 0.389 | 1.36E-09 | 0.0000499 |
| HSP90AA1 | 3 | -0.608978523 | 0.369 | 0.467 | 1.51E-09 | 0.0000552 |
| ITFG1 | 3 | -0.301483713 | 0.772 | 0.793 | 1.55E-09 | 0.0000566 |

| | | | | | | |
|---|---|---|---|---|---|---|
| CPEB3 | 3 | -0.279438981 | 0.31 | 0.4 | 1.65E-09 | 0.0000605 |
| TLE4 | 3 | -0.313397701 | 0.417 | 0.493 | 1.66E-09 | 0.0000606 |
| CDH8 | 3 | -0.262265789 | 0.074 | 0.14 | 1.66E-09 | 0.0000606 |
| RAP1A | 3 | 0.251916308 | 0.527 | 0.453 | 2.07E-09 | 0.0000758 |
| DENND2A | 3 | -0.28372436 | 0.249 | 0.341 | 2.23E-09 | 0.0000815 |
| SYNPO2 | 3 | -0.357467927 | 0.201 | 0.277 | 3.16E-09 | 0.000116 |
| KIAA1328 | 3 | -0.275080519 | 0.549 | 0.616 | 3.23E-09 | 0.000118 |
| PRKCA | 3 | 0.375654082 | 0.374 | 0.293 | 3.6E-09 | 0.000132 |
| DENND4A | 3 | -0.260602467 | 0.544 | 0.591 | 3.64E-09 | 0.000133 |
| KLHL29 | 3 | -0.358147761 | 0.564 | 0.623 | 3.64E-09 | 0.000133 |
| FGFR2 | 3 | -0.31341553 | 0.306 | 0.401 | 0.000000004 | 0.000146 |
| FOXO3 | 3 | 0.251678485 | 0.589 | 0.499 | 4.39E-09 | 0.000161 |
| LINC01776 | 3 | 0.269922577 | 0.26 | 0.18 | 5.26E-09 | 0.000193 |
| SLC16A7 | 3 | -0.299096487 | 0.282 | 0.362 | 5.38E-09 | 0.000197 |
| HMGCS1 | 3 | -0.327781053 | 0.131 | 0.188 | 6.03E-09 | 0.000221 |
| ADAMTS12 | 3 | -0.523913642 | 0.403 | 0.398 | 6.07E-09 | 0.000222 |
| EHBP1 | 3 | 0.294567982 | 0.85 | 0.823 | 6.08E-09 | 0.000223 |
| ARHGEF26 | 3 | 0.259292321 | 0.334 | 0.269 | 6.83E-09 | 0.00025 |
| RPGR | 3 | -0.278852541 | 0.391 | 0.455 | 7.53E-09 | 0.000276 |
| CLIP2 | 3 | 0.305495866 | 0.374 | 0.304 | 1.05E-08 | 0.000386 |
| RPS16 | 3 | -0.250036143 | 0.273 | 0.325 | 1.23E-08 | 0.000452 |
| C13orf42 | 3 | -0.26586635 | 0.079 | 0.137 | 1.47E-08 | 0.000537 |
| UFL1-AS1 | 3 | -0.339100917 | 0.076 | 0.11 | 1.61E-08 | 0.00059 |
| PCSK5 | 3 | -0.306658406 | 0.231 | 0.308 | 1.71E-08 | 0.000627 |
| TANC1 | 3 | 0.27870204 | 0.59 | 0.521 | 1.72E-08 | 0.000629 |
| AC009803.1 | 3 | -0.267705238 | 0.078 | 0.135 | 1.99E-08 | 0.000727 |
| DNAH11 | 3 | -0.437168214 | 0.188 | 0.267 | 2.11E-08 | 0.000772 |
| MAGI1 | 3 | -0.278701987 | 0.836 | 0.869 | 2.13E-08 | 0.000779 |
| SNCA | 3 | -0.257763293 | 0.203 | 0.283 | 2.86E-08 | 0.001045799 |
| MIR222HG | 3 | 0.276474327 | 0.254 | 0.198 | 2.87E-08 | 0.001049147 |
| RPS4X | 3 | -0.264254419 | 0.228 | 0.303 | 2.87E-08 | 0.001052168 |
| NCALD | 3 | -0.327208254 | 0.514 | 0.574 | 0.000000033 | 0.001209361 |
| DDAH1 | 3 | 0.282322366 | 0.422 | 0.347 | 4.05E-08 | 0.001484171 |
| JAG1 | 3 | -0.253622611 | 0.091 | 0.152 | 4.73E-08 | 0.001731583 |
| TTC6 | 3 | -0.299560922 | 0.107 | 0.171 | 7.11E-08 | 0.002602225 |
| MAGI2 | 3 | 0.306363211 | 0.724 | 0.697 | 8.83E-08 | 0.003234267 |
| ZNF385D | 3 | 0.354596637 | 0.622 | 0.566 | 9.26E-08 | 0.003388929 |
| CDKAL1 | 3 | -0.446768155 | 0.813 | 0.827 | 0.000000104 | 0.003795357 |
| NTN1 | 3 | 0.267001485 | 0.205 | 0.138 | 0.000000104 | 0.003804285 |
| GHR | 3 | -0.27241163 | 0.146 | 0.218 | 0.000000129 | 0.004736861 |
| GRIA1 | 3 | 0.329309916 | 0.436 | 0.36 | 0.000000147 | 0.005390413 |
| SIPA1L3 | 3 | -0.443596236 | 0.479 | 0.531 | 0.000000183 | 0.006710048 |
| TTC21B | 3 | -0.331659951 | 0.549 | 0.59 | 0.000000208 | 0.007614446 |
| ITGAV | 3 | 0.359011352 | 0.553 | 0.536 | 0.000000238 | 0.008697623 |
| APC | 3 | -0.256915105 | 0.476 | 0.532 | 0.000000278 | 0.010162104 |
| PPP2R2B | 3 | -0.271129148 | 0.593 | 0.657 | 0.000000286 | 0.010488811 |
| LRMDA | 3 | -0.411280592 | 0.08 | 0.12 | 0.000000298 | 0.01092325 |
| SORCS1 | 3 | 0.395761899 | 0.284 | 0.223 | 0.000000326 | 0.011936581 |
| PLPPR4 | 3 | 0.26006692 | 0.326 | 0.278 | 0.00000037 | 0.013554163 |
| MAST4 | 3 | -0.293432208 | 0.313 | 0.37 | 0.000000404 | 0.014789011 |
| SLC44A5 | 3 | -0.290627169 | 0.06 | 0.107 | 0.000000465 | 0.017010315 |
| DPYD | 3 | 0.316724087 | 0.477 | 0.4 | 0.000000486 | 0.017788015 |
| CFAP44 | 3 | -0.281220741 | 0.306 | 0.358 | 0.000000635 | 0.0232386 |
| AL591501.1 | 3 | -0.530597192 | 0.085 | 0.117 | 0.0000008 | 0.029273265 |
| RPL13A | 3 | -0.260236836 | 0.492 | 0.561 | 0.00000102 | 0.037240394 |
| CEP85L | 3 | 0.258459483 | 0.584 | 0.546 | 0.00000143 | 0.052267243 |
| GLIS3 | 3 | 0.437295957 | 0.311 | 0.238 | 0.00000177 | 0.064620789 |
| DYNC1I1 | 3 | -0.281619685 | 0.303 | 0.352 | 0.00000311 | 0.113713565 |

| | | | | | | |
|---|---|---|---|---|---|---|
| ASPH | 3 | -0.377644965 | 0.672 | 0.705 | 0.00000402 | 0.147111588 |
| BMPER | 3 | 0.382619068 | 0.257 | 0.205 | 0.00000412 | 0.150844753 |
| ADGRV1 | 3 | 0.485173264 | 0.66 | 0.687 | 0.00000419 | 0.153456129 |
| EPS8 | 3 | -0.293217873 | 0.557 | 0.591 | 0.00000443 | 0.162195021 |
| MEIS1 | 3 | -0.32477057 | 0.329 | 0.355 | 0.00000492 | 0.180182137 |
| PRELID2 | 3 | -0.267542713 | 0.302 | 0.325 | 0.0000188 | 0.688821475 |
| CHPT1 | 3 | -0.253317048 | 0.522 | 0.571 | 0.0000391 | 1 |
| PTCHD1-AS | 3 | -0.320382233 | 0.087 | 0.123 | 0.000217 | 1 |
| CNTN5 | 3 | -0.753179651 | 0.109 | 0.144 | 0.000544 | 1 |
| MCTP2 | 3 | -0.339335298 | 0.23 | 0.268 | 0.000664 | 1 |
| KIRREL3 | 3 | 0.252018977 | 0.169 | 0.181 | 0.000799 | 1 |
| NELL1 | 3 | -0.273101884 | 0.298 | 0.314 | 0.001868929 | 1 |
| AC068286.2 | 3 | -0.449632266 | 0.12 | 0.138 | 0.003027099 | 1 |
| KCTD16 | 3 | 0.29873424 | 0.173 | 0.163 | 0.00986479 | 1 |
| HTR2C | 4 | 6.548429114 | 0.391 | 0.059 | 0 | 0 |
| AC092957.1 | 4 | 4.084034661 | 0.419 | 0.025 | 0 | 0 |
| COL4A5 | 4 | 3.363302642 | 0.756 | 0.036 | 0 | 0 |
| AUTS2 | 4 | 2.62737504 | 0.927 | 0.422 | 0 | 0 |
| AC233296.1 | 4 | 2.502918595 | 0.504 | 0.016 | 0 | 0 |
| AL137139.2 | 4 | 2.465700629 | 0.444 | 0.018 | 0 | 0 |
| PRTG | 4 | 2.438645852 | 0.636 | 0.031 | 0 | 0 |
| ZNF521 | 4 | 2.371404819 | 0.688 | 0.081 | 0 | 0 |
| MAPK10 | 4 | 2.10089283 | 0.719 | 0.169 | 0 | 0 |
| COL4A6 | 4 | 2.067860553 | 0.488 | 0.006 | 0 | 0 |
| STK33 | 4 | 2.058247433 | 0.629 | 0.106 | 0 | 0 |
| VAT1L | 4 | 1.938022955 | 0.407 | 0.011 | 0 | 0 |
| PPP1R9A | 4 | 1.863284039 | 0.69 | 0.109 | 0 | 0 |
| PEX5L | 4 | -3.104780625 | 0.216 | 0.827 | 0 | 0 |
| IGF2BP2 | 4 | 1.641534256 | 0.912 | 0.367 | 3.883e-321 | 1.42173546e-316 |
| HYDIN | 4 | 2.024525532 | 0.42 | 0.035 | 1.495240663e-314 | [illegible].47422559080433e-310 |
| PTPN13 | 4 | 1.662047364 | 0.615 | 0.085 | 1.44915055246e-313 | 0 |
| GRM8 | 4 | 3.049893494 | 0.379 | 0.126 | 0 | 8.4E-305 |
| CFAP54 | 4 | 2.400351264 | 0.379 | 0.024 | 4.21E-308 | 1.54E-303 |
| LRRIQ1 | 4 | 1.9432771 | 0.401 | 0.022 | 1.72E-304 | 6.29E-300 |
| MYRIP | 4 | -2.486822765 | 0.144 | 0.718 | 2E-292 | 7.31E-288 |
| NR2F2-AS1 | 4 | 1.761185451 | 0.479 | 0.045 | 2.55E-288 | 9.35E-284 |
| TRPM3 | 4 | 3.620469357 | 0.569 | 0.158 | 4.21E-283 | 1.54E-278 |
| PRSS12 | 4 | -2.340997525 | 0.38 | 0.846 | 1.15E-274 | 4.21E-270 |
| ARMC3 | 4 | 1.912090287 | 0.332 | 0.022 | 3.07E-273 | 1.12E-268 |
| DNAH7 | 4 | 2.162295211 | 0.415 | 0.15 | 1.16E-265 | 4.25E-261 |
| WDR49 | 4 | 2.258886645 | 0.329 | 0.017 | 9.13E-256 | 3.34E-251 |
| PACRG | 4 | 1.981603846 | 0.415 | 0.06 | 2.18E-255 | 7.98E-251 |
| PRDM16 | 4 | 2.134237816 | 0.378 | 0.036 | 3.22E-255 | 1.18E-250 |
| FGF12 | 4 | -2.124390822 | 0.272 | 0.79 | 5.36E-254 | 1.96E-249 |
| GPM6A | 4 | 2.274472729 | 0.518 | 0.162 | 1.03E-252 | 3.78E-248 |
| CFAP47 | 4 | 2.878762525 | 0.38 | 0.052 | 3.18E-251 | 1.17E-246 |
| ADAMTS6 | 4 | 2.132168759 | 0.557 | 0.104 | 2.23E-249 | 8.16E-245 |
| CFAP43 | 4 | 1.751365684 | 0.347 | 0.033 | 9.76E-247 | 3.57E-242 |
| CLIC6 | 4 | 1.440320896 | 0.301 | 0.009 | 7.19E-246 | 2.63E-241 |
| IGFBP7 | 4 | 1.378974751 | 0.33 | 0.015 | 1.47E-241 | 5.39E-237 |
| BMP7 | 4 | 1.196797967 | 0.374 | 0.024 | 2.86E-241 | 1.05E-236 |
| LGI1 | 4 | 0.93526158 | 0.308 | 0.008 | 3.34E-241 | 1.22E-236 |
| TENM4 | 4 | 1.591816431 | 0.74 | 0.323 | 1.09E-240 | 4E-236 |
| MBNL1 | 4 | -1.78117827 | 0.718 | 0.937 | 1.21E-240 | 4.44E-236 |
| XYLT1 | 4 | 1.479849515 | 0.524 | 0.093 | 2.56E-238 | 9.39E-234 |
| NHSL2 | 4 | 1.21339155 | 0.396 | 0.028 | 3.41E-236 | 1.25E-231 |
| PCAT1 | 4 | 1.960330118 | 0.397 | 0.065 | 5.65E-236 | 2.07E-231 |
| AC018731.1 | 4 | -2.419894436 | 0.12 | 0.646 | 3.77E-235 | 1.38E-230 |

| | | | | | | |
|---|---|---|---|---|---|---|
| LMX1A | 4 | 1.418014693 | 0.347 | 0.02 | 1.62E-234 | 5.94E-230 |
| MAATS1 | 4 | 1.003348874 | 0.341 | 0.018 | 9.61E-234 | 3.52E-229 |
| ST6GALNAC3 | 4 | 1.791643949 | 0.683 | 0.218 | 3.87E-232 | 1.42E-227 |
| SHROOM3 | 4 | 1.326981262 | 0.441 | 0.053 | 6.24E-227 | 2.29E-222 |
| RBPMS | 4 | 1.409972633 | 0.456 | 0.058 | 2.4E-226 | 8.8E-222 |
| CFAP221 | 4 | 0.940009918 | 0.307 | 0.011 | 1.38E-224 | 5.06E-220 |
| CFAP61 | 4 | 1.144872589 | 0.309 | 0.014 | 2.75E-223 | 1.01E-218 |
| DCDC1 | 4 | 1.646052656 | 0.373 | 0.049 | 5.65E-221 | 2.07E-216 |
| ERBB4 | 4 | 2.43753927 | 0.631 | 0.246 | 1.44E-220 | 5.28E-216 |
| SLC7A6 | 4 | 1.263797087 | 0.418 | 0.225 | 3.9E-219 | 1.43E-214 |
| SPATA17 | 4 | 1.4615064 | 0.363 | 0.063 | 2.7E-215 | 9.87E-211 |
| LRRC9 | 4 | 1.483239376 | 0.295 | 0.02 | 1.33E-214 | 4.88E-210 |
| SLC16A10 | 4 | 1.636500768 | 0.4 | 0.207 | 3.93E-213 | 1.44E-208 |
| SERPINI2 | 4 | 1.269536645 | 0.278 | 0.008 | 4.69E-213 | 1.72E-208 |
| AC021504.1 | 4 | 1.25440631 | 0.393 | 0.037 | 9.01E-212 | 3.3E-207 |
| CHSY3 | 4 | -1.689584563 | 0.266 | 0.762 | 1.8E-211 | 6.6E-207 |
| TTR | 4 | 1.759544198 | 0.33 | 0.044 | 8.84E-211 | 3.24E-206 |
| SPEF2 | 4 | 1.492908258 | 0.42 | 0.117 | 9.04E-211 | 3.31E-206 |
| PPFIA2 | 4 | -1.831577484 | 0.306 | 0.774 | 2.15E-209 | 7.86E-205 |
| TMEM232 | 4 | 1.565749566 | 0.436 | 0.1 | 2.27E-208 | 8.33E-204 |
| ADGB | 4 | 1.257236498 | 0.246 | 0.004 | 1.52E-207 | 5.57E-203 |
| KCNB2 | 4 | -2.528239993 | 0.126 | 0.601 | 8.18E-205 | 2.99E-200 |
| DNAAF1 | 4 | 0.856117241 | 0.255 | 0.005 | 8.22E-204 | 3.01E-199 |
| CFAP69 | 4 | 1.26274324 | 0.394 | 0.061 | 1.47E-203 | 5.36E-199 |
| CFAP46 | 4 | 1.140832396 | 0.286 | 0.014 | 1.47E-202 | 5.37E-198 |
| MIR646HG | 4 | 1.844394145 | 0.332 | 0.092 | 7.22E-202 | 2.64E-197 |
| RBFOX1 | 4 | -2.01345261 | 0.344 | 0.78 | 9.32E-202 | 3.41E-197 |
| ZC3H12B | 4 | 0.921688283 | 0.287 | 0.012 | 6.08E-201 | 2.23E-196 |
| GREB1L | 4 | 2.160179349 | 0.365 | 0.066 | 1E-200 | 3.68E-196 |
| ZIC1 | 4 | 1.0299652 | 0.319 | 0.02 | 5.75E-200 | 2.11E-195 |
| VWA5B1 | 4 | 1.065266088 | 0.288 | 0.014 | 9.84E-200 | 3.6E-195 |
| CFAP70 | 4 | 1.252013969 | 0.355 | 0.055 | 2.65E-195 | 9.7E-191 |
| LINC01170 | 4 | -1.984945594 | 0.146 | 0.634 | 9.6E-195 | 3.51E-190 |
| AL050309.1 | 4 | 1.295395529 | 0.447 | 0.072 | 1.46E-193 | 5.34E-189 |
| VWA3A | 4 | 1.05315499 | 0.257 | 0.009 | 1.15E-192 | 4.21E-188 |
| AKAP13 | 4 | 1.090567965 | 0.851 | 0.518 | 5.97E-191 | 2.19E-186 |
| GALNT13 | 4 | -1.767081112 | 0.316 | 0.747 | 1.42E-190 | 5.2E-186 |
| DANT2 | 4 | 1.293067122 | 0.745 | 0.279 | 1.93E-190 | 7.06E-186 |
| DNAH6 | 4 | 1.569595448 | 0.342 | 0.047 | 3.42E-190 | 1.25E-185 |
| SLC39A12 | 4 | 0.728808521 | 0.214 | 0.002 | 9.82E-188 | 3.59E-183 |
| GRIA2 | 4 | -2.442487178 | 0.278 | 0.702 | 2.92E-186 | 1.07E-181 |
| EDIL3 | 4 | -1.757117318 | 0.19 | 0.669 | 4.51E-186 | 1.65E-181 |
| DYNC2H1 | 4 | 1.286961827 | 0.696 | 0.324 | 6.33E-186 | 2.32E-181 |
| DRC1 | 4 | 0.674926345 | 0.244 | 0.006 | 1.18E-185 | 4.34E-181 |
| CCDC40 | 4 | 1.058340724 | 0.359 | 0.047 | 1.62E-185 | 5.93E-181 |
| ZBBX | 4 | 0.991452524 | 0.261 | 0.012 | 7.65E-185 | 2.8E-180 |
| SLC38A11 | 4 | 1.788991509 | 0.28 | 0.02 | 9.76E-184 | 3.57E-179 |
| MIRLET7BHG | 4 | -1.315766707 | 0.253 | 0.719 | 2.32E-182 | 8.5E-178 |
| CCDC173 | 4 | 0.971102067 | 0.282 | 0.019 | 1.34E-181 | 4.92E-177 |
| AC011029.1 | 4 | 0.682172275 | 0.222 | 0.003 | 5.06E-181 | 1.85E-176 |
| LUZP2 | 4 | -2.644306886 | 0.158 | 0.602 | 7.23E-180 | 2.65E-175 |
| OTX2 | 4 | 1.346892552 | 0.378 | 0.063 | 8.22E-180 | 3.01E-175 |
| NEDD4L | 4 | -1.200889365 | 0.556 | 0.89 | 2.03E-178 | 7.45E-174 |
| OTX2-AS1 | 4 | 1.864214592 | 0.345 | 0.045 | 2.29E-178 | 8.39E-174 |
| PKNOX2 | 4 | 1.234392578 | 0.548 | 0.189 | 5.35E-177 | 1.96E-172 |
| LRP1B | 4 | 2.324768268 | 0.47 | 0.168 | 1.35E-176 | 4.95E-172 |
| CCDC30 | 4 | 1.214545882 | 0.421 | 0.097 | 4.5E-176 | 1.65E-171 |
| SPAG8 | 4 | 0.811614666 | 0.252 | 0.011 | 4.94E-176 | 1.81E-171 |

| | | | | | | |
|---|---|---|---|---|---|---|
| DNAI1 | 4 | 0.962777607 | 0.286 | 0.021 | 5.07E-175 | 1.85E-170 |
| SORBS2 | 4 | 1.021119537 | 0.457 | 0.088 | 6.87E-175 | 2.51E-170 |
| ZNF423 | 4 | 1.403180986 | 0.712 | 0.375 | 7.18E-175 | 2.63E-170 |
| WDR78 | 4 | 1.192087602 | 0.381 | 0.097 | 9.82E-175 | 3.6E-170 |
| NEK11 | 4 | 1.204759431 | 0.391 | 0.103 | 1.54E-174 | 5.63E-170 |
| C6orf118 | 4 | 0.755169317 | 0.267 | 0.014 | 3.01E-174 | 1.1E-169 |
| AGBL4 | 4 | 1.90130709 | 0.491 | 0.175 | 3.39E-174 | 1.24E-169 |
| CFAP52 | 4 | 0.799919206 | 0.24 | 0.008 | 7.82E-173 | 2.86E-168 |
| NXN | 4 | 1.499596856 | 0.738 | 0.371 | 4.76E-172 | 1.74E-167 |
| SOX2-OT | 4 | 2.647468918 | 0.467 | 0.174 | 1.31E-171 | 4.79E-167 |
| FGF14 | 4 | -2.162650173 | 0.309 | 0.696 | 4.14E-171 | 1.52E-166 |
| MOK | 4 | 1.098902637 | 0.433 | 0.101 | 4.24E-171 | 1.55E-166 |
| TBX15 | 4 | -1.537953456 | 0.124 | 0.581 | 2.09E-170 | 7.65E-166 |
| ZIC4 | 4 | 0.481185758 | 0.198 | 0.002 | 7.71E-169 | 2.82E-164 |
| SDK2 | 4 | 1.224897637 | 0.437 | 0.086 | 1.13E-168 | 4.14E-164 |
| AC069277.1 | 4 | 1.580899358 | 0.343 | 0.04 | 1.29E-168 | 4.73E-164 |
| TCTEX1D1 | 4 | 0.754540423 | 0.232 | 0.009 | 1.31E-168 | 4.79E-164 |
| FZD3 | 4 | 1.131630902 | 0.55 | 0.23 | 6.8E-168 | 2.49E-163 |
| AC016152.1 | 4 | 1.827426101 | 0.297 | 0.035 | 1.82E-167 | 6.66E-163 |
| SPATA13 | 4 | 1.188355737 | 0.549 | 0.195 | 3.18E-167 | 1.16E-162 |
| NCAM1 | 4 | -0.732496568 | 0.474 | 0.875 | 2.25E-166 | 8.23E-162 |
| CFAP157 | 4 | 0.682457997 | 0.225 | 0.007 | 1.05E-165 | 3.85E-161 |
| RGL1 | 4 | -1.655478224 | 0.446 | 0.759 | 1.5E-165 | 5.49E-161 |
| SGCZ | 4 | -2.890996736 | 0.201 | 0.594 | 3.87E-165 | 1.42E-160 |
| MAP3K19 | 4 | 0.681804719 | 0.221 | 0.006 | 5.39E-165 | 1.97E-160 |
| CECR2 | 4 | 1.23466775 | 0.679 | 0.269 | 1.34E-164 | 4.91E-160 |
| PLEKHG1 | 4 | 1.073883867 | 0.47 | 0.095 | 1.66E-164 | 6.08E-160 |
| ZIC5 | 4 | 0.512756776 | 0.197 | 0.002 | 4.2E-164 | 1.54E-159 |
| CEP112 | 4 | 1.218906953 | 0.705 | 0.361 | 5.75E-164 | 2.11E-159 |
| AL035425.3 | 4 | 0.499181466 | 0.188 | 0.001 | 1.02E-163 | 3.73E-159 |
| RADX | 4 | -1.485768727 | 0.11 | 0.537 | 1.17E-162 | 4.3E-158 |
| ZC4H2 | 4 | 1.034124935 | 0.441 | 0.083 | 3.26E-162 | 1.19E-157 |
| NEK5 | 4 | 0.846004986 | 0.256 | 0.016 | 5.06E-162 | 1.85E-157 |
| AC110023.1 | 4 | 1.335202133 | 0.215 | 0.006 | 2.57E-161 | 9.41E-157 |
| BTBD11 | 4 | 0.827500338 | 0.245 | 0.012 | 5.73E-161 | 2.1E-156 |
| ATP13A4 | 4 | 0.843160987 | 0.251 | 0.016 | 1.09E-160 | 3.97E-156 |
| KIAA1217 | 4 | 1.470326351 | 0.654 | 0.271 | 1.39E-160 | 5.08E-156 |
| DCC | 4 | 3.208784706 | 0.261 | 0.021 | 4.19E-160 | 1.53E-155 |
| FLRT2 | 4 | 1.519152856 | 0.35 | 0.053 | 2.39E-159 | 8.76E-155 |
| LHFPL3 | 4 | -2.088803727 | 0.205 | 0.608 | 9.24E-159 | 3.38E-154 |
| LIFR | 4 | -1.761278147 | 0.285 | 0.675 | 5.27E-158 | 1.93E-153 |
| PREX2 | 4 | 1.28074965 | 0.345 | 0.046 | 1.44E-157 | 5.27E-153 |
| CASC1 | 4 | 0.866223329 | 0.28 | 0.023 | 3.28E-157 | 1.2E-152 |
| GRAMD1B | 4 | 1.168431867 | 0.487 | 0.139 | 1.15E-156 | 4.22E-152 |
| SLC4A5 | 4 | 1.063208567 | 0.339 | 0.07 | 1.88E-156 | 6.87E-152 |
| SLC13A4 | 4 | 0.881144371 | 0.246 | 0.019 | 8.27E-156 | 3.03E-151 |
| SLIT3 | 4 | 1.599039842 | 0.349 | 0.046 | 1.28E-155 | 4.69E-151 |
| FANK1 | 4 | 0.970312541 | 0.335 | 0.055 | 3.89E-155 | 1.42E-150 |
| HS3ST5 | 4 | -1.70038959 | 0.222 | 0.664 | 1.48E-154 | 5.41E-150 |
| FAM189A1 | 4 | -1.804795628 | 0.393 | 0.728 | 2.52E-154 | 9.23E-150 |
| CGNL1 | 4 | 1.023980025 | 0.334 | 0.048 | 3.78E-154 | 1.39E-149 |
| CFAP44 | 4 | 1.171510715 | 0.531 | 0.331 | 4.63E-154 | 1.7E-149 |
| MSI2 | 4 | 0.946193164 | 0.927 | 0.849 | 5.86E-154 | 2.15E-149 |
| SPATA6 | 4 | 0.93625434 | 0.521 | 0.133 | 1.02E-153 | 3.72E-149 |
| GMNC | 4 | 1.114371604 | 0.22 | 0.012 | 3.22E-153 | 1.18E-148 |
| COLEC12 | 4 | 1.407469852 | 0.424 | 0.1 | 7.48E-153 | 2.74E-148 |
| DAW1 | 4 | 0.435111395 | 0.176 | 0.001 | 6.33E-152 | 2.32E-147 |
| EFHC1 | 4 | 1.032052321 | 0.48 | 0.24 | 6.54E-151 | 2.39E-146 |

| | | | | | | |
|---|---|---|---|---|---|---|
| LINC01278 | 4 | 1.12948386 | 0.626 | 0.298 | 8.17E-151 | 2.99E-146 |
| ANO4 | 4 | -2.316211637 | 0.3 | 0.616 | 1.43E-150 | 5.22E-146 |
| NPHP1 | 4 | 0.776907367 | 0.328 | 0.043 | 1.57E-150 | 5.77E-146 |
| IRAK3 | 4 | -1.186323484 | 0.115 | 0.535 | 3.73E-150 | 1.37E-145 |
| SULF1 | 4 | 1.755832604 | 0.499 | 0.287 | 2.87E-149 | 1.05E-144 |
| PCCA | 4 | 0.933085605 | 0.857 | 0.611 | 8.28E-149 | 3.03E-144 |
| ARMC2 | 4 | 0.967519734 | 0.439 | 0.14 | 2.3E-148 | 8.41E-144 |
| TEKT1 | 4 | 0.558165471 | 0.183 | 0.003 | 2.36E-147 | 8.63E-143 |
| PAK3 | 4 | -1.161009009 | 0.659 | 0.893 | 4.61E-147 | 1.69E-142 |
| GALNT17 | 4 | 1.679328492 | 0.455 | 0.157 | 1.15E-146 | 4.2E-142 |
| LINC01515 | 4 | 1.119971728 | 0.535 | 0.148 | 1.6E-146 | 5.87E-142 |
| SYT1 | 4 | -1.413723327 | 0.439 | 0.778 | 1.74E-146 | 6.39E-142 |
| FHAD1 | 4 | 0.640345807 | 0.267 | 0.022 | 2.73E-146 | 9.99E-142 |
| SPAG16 | 4 | 1.163023696 | 0.52 | 0.253 | 3.87E-146 | 1.42E-141 |
| CACNB4 | 4 | -1.852966126 | 0.196 | 0.588 | 5.24E-146 | 1.92E-141 |
| PTGER3 | 4 | -1.372353753 | 0.05 | 0.43 | 9.86E-146 | 3.61E-141 |
| FYB2 | 4 | 0.785015146 | 0.237 | 0.014 | 2.97E-145 | 1.09E-140 |
| COL5A3 | 4 | 0.576180972 | 0.207 | 0.007 | 4.56E-144 | 1.67E-139 |
| DNAH12 | 4 | 1.317433854 | 0.247 | 0.022 | 3.87E-143 | 1.42E-138 |
| ABLIM1 | 4 | 1.115059644 | 0.569 | 0.204 | 6.17E-143 | 2.26E-138 |
| FAM81B | 4 | 0.466096704 | 0.169 | 0.002 | 8.49E-143 | 3.11E-138 |
| CCDC114 | 4 | 0.524230686 | 0.204 | 0.007 | 4.66E-142 | 1.71E-137 |
| ZFYVE16 | 4 | 1.02812049 | 0.688 | 0.559 | 1.88E-141 | 6.88E-137 |
| SLC44A1 | 4 | -1.15088519 | 0.355 | 0.73 | 2.1E-141 | 7.69E-137 |
| STOX2 | 4 | 1.14815016 | 0.605 | 0.271 | 2.13E-141 | 7.81E-137 |
| ZIC2 | 4 | 0.440243647 | 0.178 | 0.003 | 3.3E-141 | 1.21E-136 |
| ATP2B2 | 4 | 1.037499295 | 0.299 | 0.086 | 5.63E-140 | 2.06E-135 |
| LEF1 | 4 | 1.039822734 | 0.409 | 0.084 | 6.37E-140 | 2.33E-135 |
| PTPRM | 4 | 1.258294917 | 0.714 | 0.345 | 1.43E-139 | 5.24E-135 |
| AC073114.1 | 4 | 1.598887477 | 0.279 | 0.038 | 2.58E-139 | 9.45E-135 |
| LRRC6 | 4 | 0.91320566 | 0.334 | 0.072 | 2.83E-139 | 1.04E-134 |
| FGFR2 | 4 | 1.25737606 | 0.603 | 0.366 | 2.88E-138 | 1.06E-133 |
| MAPK15 | 4 | 0.503630324 | 0.209 | 0.009 | 4.11E-138 | 1.5E-133 |
| PARD3B | 4 | 1.251951996 | 0.752 | 0.436 | 1.15E-137 | 4.23E-133 |
| KCNIP4 | 4 | -2.529883521 | 0.178 | 0.536 | 8.68E-137 | 3.18E-132 |
| NME5 | 4 | 0.614164793 | 0.27 | 0.025 | 1.34E-136 | 4.89E-132 |
| GRID2 | 4 | -2.282771077 | 0.232 | 0.619 | 2.13E-136 | 7.79E-132 |
| KCNH8 | 4 | -1.261088311 | 0.262 | 0.676 | 2.69E-136 | 9.87E-132 |
| NLGN1 | 4 | -1.695305261 | 0.568 | 0.832 | 1.66E-135 | 6.08E-131 |
| CNKSR2 | 4 | -1.621411072 | 0.34 | 0.698 | 2.83E-135 | 1.04E-130 |
| COL1A2 | 4 | 1.687669602 | 0.395 | 0.095 | 3.79E-135 | 1.39E-130 |
| EFHC2 | 4 | 1.090323311 | 0.332 | 0.116 | 4.98E-135 | 1.82E-130 |
| SCN8A | 4 | -1.132997418 | 0.339 | 0.703 | 5.59E-135 | 2.05E-130 |
| ST18 | 4 | -1.477191102 | 0.139 | 0.545 | 1.07E-134 | 3.93E-130 |
| AL008633.1 | 4 | -3.535975698 | 0.627 | 0.721 | 1.48E-134 | 5.41E-130 |
| DLGAP1 | 4 | -1.360016115 | 0.607 | 0.861 | 1.67E-134 | 6.11E-130 |
| PVT1 | 4 | -1.037538935 | 0.546 | 0.839 | 7.43E-133 | 2.72E-128 |
| KIAA2012 | 4 | 0.595797395 | 0.193 | 0.007 | 1.03E-132 | 3.79E-128 |
| YAP1 | 4 | 1.005741606 | 0.688 | 0.293 | 2.51E-132 | 9.2E-128 |
| BRINP3 | 4 | -1.796445333 | 0.155 | 0.55 | 5.91E-132 | 2.16E-127 |
| SH3GL3 | 4 | -1.354876301 | 0.183 | 0.572 | 6.39E-132 | 2.34E-127 |
| CFAP299 | 4 | 1.349392419 | 0.229 | 0.02 | 1.04E-131 | 3.83E-127 |
| FMN1 | 4 | 1.29133983 | 0.379 | 0.076 | 6.17E-131 | 2.26E-126 |
| WDR63 | 4 | 0.644042723 | 0.231 | 0.018 | 3.29E-130 | 1.2E-125 |
| AC091078.1 | 4 | 0.931390437 | 0.232 | 0.023 | 5.84E-129 | 2.14E-124 |
| LRGUK | 4 | 0.8477442 | 0.339 | 0.065 | 6.91E-129 | 2.53E-124 |
| TEX9 | 4 | 0.947844149 | 0.494 | 0.169 | 1.11E-128 | 4.06E-124 |
| SORCS3 | 4 | -1.633880782 | 0.089 | 0.461 | 6.49E-128 | 2.38E-123 |

| | | | | | | |
|---|---|---|---|---|---|---|
| ZNF536 | 4 | -1.583200509 | 0.641 | 0.843 | 1.11E-127 | 4.06E-123 |
| PRKCQ | 4 | 0.537425919 | 0.213 | 0.012 | 1.11E-127 | 4.07E-123 |
| PPP1R14C | 4 | -1.439943463 | 0.113 | 0.486 | 1.16E-127 | 4.26E-123 |
| COL18A1 | 4 | 0.88503823 | 0.425 | 0.099 | 1.67E-127 | 6.12E-123 |
| SYNDIG1 | 4 | -1.226751101 | 0.228 | 0.612 | 1.72E-127 | 6.29E-123 |
| ANKS1B | 4 | -1.41891646 | 0.528 | 0.819 | 2.84E-127 | 1.04E-122 |
| MARK1 | 4 | 1.218382709 | 0.619 | 0.491 | 5.48E-127 | 2.01E-122 |
| ANGPT1 | 4 | -1.521734002 | 0.219 | 0.618 | 7.25E-127 | 2.65E-122 |
| PCP4 | 4 | 0.411889069 | 0.157 | 0.002 | 1.88E-126 | 6.89E-122 |
| FAM135B | 4 | -1.327035299 | 0.058 | 0.405 | 5.64E-126 | 2.06E-121 |
| CFAP74 | 4 | 0.459435314 | 0.167 | 0.004 | 7.28E-126 | 2.67E-121 |
| MROH7 | 4 | 0.829299577 | 0.294 | 0.054 | 8.81E-126 | 3.23E-121 |
| LINC00305 | 4 | 0.381785921 | 0.138 | 0 | 1.14E-125 | 4.17E-121 |
| RBM47 | 4 | 1.565113339 | 0.387 | 0.099 | 1.87E-125 | 6.84E-121 |
| AC105916.1 | 4 | 0.908620556 | 0.331 | 0.079 | 3.68E-125 | 1.35E-120 |
| SOX6 | 4 | 1.605313742 | 0.536 | 0.206 | 4.57E-125 | 1.67E-120 |
| TMEM67 | 4 | 0.985465599 | 0.527 | 0.234 | 4.58E-125 | 1.68E-120 |
| NLGN4X | 4 | -1.262829072 | 0.437 | 0.768 | 5.02E-125 | 1.84E-120 |
| PPP1R42 | 4 | 0.470704745 | 0.175 | 0.005 | 1.17E-124 | 4.28E-120 |
| OCA2 | 4 | 0.838682838 | 0.292 | 0.039 | 2.3E-124 | 8.41E-120 |
| RALYL | 4 | -2.431251222 | 0.154 | 0.512 | 3.4E-124 | 1.24E-119 |
| SLC4A10 | 4 | 1.354691097 | 0.295 | 0.062 | 1.57E-123 | 5.75E-119 |
| LINC01376 | 4 | 1.051191358 | 0.389 | 0.123 | 2.33E-123 | 8.53E-119 |
| EPB41L4A | 4 | 0.902668103 | 0.379 | 0.077 | 6.46E-123 | 2.36E-118 |
| NEGR1 | 4 | -3.050361602 | 0.317 | 0.607 | 7.9E-123 | 2.89E-118 |
| MKX | 4 | -1.541111569 | 0.057 | 0.4 | 6.16E-122 | 2.26E-117 |
| CFAP100 | 4 | 0.398122116 | 0.187 | 0.008 | 7.35E-122 | 2.69E-117 |
| CA2 | 4 | 0.686296608 | 0.216 | 0.016 | 1.8E-121 | 6.58E-117 |
| DKK2 | 4 | 1.633852263 | 0.28 | 0.04 | 4.82E-121 | 1.76E-116 |
| CRB2 | 4 | 0.383193778 | 0.163 | 0.004 | 5.77E-121 | 2.11E-116 |
| AL161757.4 | 4 | 0.571801273 | 0.197 | 0.012 | 8.57E-121 | 3.14E-116 |
| COL6A1 | 4 | 0.939534015 | 0.499 | 0.169 | 1.12E-120 | 4.1E-116 |
| AL157396.1 | 4 | 0.377208658 | 0.162 | 0.004 | 1.49E-120 | 5.44E-116 |
| COL26A1 | 4 | 0.805962866 | 0.269 | 0.034 | 1.65E-120 | 6.03E-116 |
| MEG8 | 4 | -1.238442312 | 0.225 | 0.613 | 2.01E-120 | 7.37E-116 |
| RSPO2 | 4 | 1.01610805 | 0.185 | 0.009 | 2.8E-120 | 1.03E-115 |
| LINC01088 | 4 | 1.079040456 | 0.274 | 0.048 | 5.91E-120 | 2.16E-115 |
| EFCAB1 | 4 | 0.575092996 | 0.203 | 0.013 | 7.16E-120 | 2.62E-115 |
| ITGA4 | 4 | -1.761076422 | 0.146 | 0.509 | 5.07E-119 | 1.86E-114 |
| PRKCA | 4 | 1.135641827 | 0.585 | 0.27 | 1.03E-118 | 3.77E-114 |
| CCDC180 | 4 | 0.770515738 | 0.328 | 0.073 | 1.1E-118 | 4.03E-114 |
| COL9A1 | 4 | 0.724295201 | 0.218 | 0.019 | 1.33E-118 | 4.87E-114 |
| MKRN3 | 4 | 0.377474128 | 0.153 | 0.003 | 4.81E-118 | 1.76E-113 |
| RFX2 | 4 | 1.030230188 | 0.512 | 0.19 | 1.32E-117 | 4.81E-113 |
| ABCA4 | 4 | 0.999449008 | 0.272 | 0.038 | 1.59E-117 | 5.84E-113 |
| EMX2OS | 4 | 0.420631633 | 0.17 | 0.006 | 1.78E-117 | 6.53E-113 |
| MDFIC | 4 | 0.856235803 | 0.378 | 0.086 | 1.86E-117 | 6.8E-113 |
| KCNQ5 | 4 | -1.673659302 | 0.191 | 0.551 | 8.12E-117 | 2.97E-112 |
| SERINC5 | 4 | 1.056876649 | 0.553 | 0.251 | 6.39E-116 | 2.34E-111 |
| EXT1 | 4 | 0.957666558 | 0.869 | 0.67 | 8.11E-116 | 2.97E-111 |
| CUX1 | 4 | 0.871463966 | 0.92 | 0.771 | 9.92E-116 | 3.63E-111 |
| SLC22A3 | 4 | -1.259125449 | 0.091 | 0.444 | 4.47E-115 | 1.64E-110 |
| SPAG6 | 4 | 0.558432523 | 0.212 | 0.016 | 6.08E-115 | 2.23E-110 |
| POU2F3 | 4 | -0.897549684 | 0.038 | 0.351 | 1.54E-114 | 5.65E-110 |
| SIX3 | 4 | 0.305633456 | 0.13 | 0.001 | 1.84E-114 | 6.75E-110 |
| KLHL13 | 4 | 1.20958646 | 0.321 | 0.061 | 3.53E-114 | 1.29E-109 |
| ST6GAL2 | 4 | 0.849849366 | 0.228 | 0.025 | 3.66E-114 | 1.34E-109 |
| ROBO1 | 4 | -0.932570424 | 0.682 | 0.926 | 3.86E-114 | 1.41E-109 |

| | | | | | | |
|---|---|---|---|---|---|---|
| NPAS3 | 4 | 1.193200903 | 0.573 | 0.207 | 3.94E-114 | 1.44E-109 |
| CCDC60 | 4 | 0.604340981 | 0.198 | 0.015 | 4.06E-114 | 1.49E-109 |
| C5AR1 | 4 | 0.707676636 | 0.25 | 0.031 | 4.6E-114 | 1.68E-109 |
| ULK4 | 4 | 0.992405056 | 0.615 | 0.412 | 5.29E-114 | 1.94E-109 |
| LRIG1 | 4 | 1.073687788 | 0.438 | 0.156 | 1.09E-113 | 3.98E-109 |
| COL11A1 | 4 | -1.709028603 | 0.48 | 0.739 | 1.56E-113 | 5.7E-109 |
| C4orf47 | 4 | 0.941266156 | 0.355 | 0.087 | 2.11E-113 | 7.72E-109 |
| AKAP9 | 4 | 0.813730274 | 0.842 | 0.723 | 2.31E-113 | 8.46E-109 |
| MEIS2 | 4 | 1.264935347 | 0.562 | 0.537 | 3.08E-113 | 1.13E-108 |
| LRP2 | 4 | 0.643232821 | 0.145 | 0.003 | 3.23E-113 | 1.18E-108 |
| FAM155A | 4 | -1.1152436 | 0.504 | 0.819 | 1.47E-112 | 5.36E-108 |
| ILDR2 | 4 | 0.447042845 | 0.181 | 0.01 | 2.66E-112 | 9.74E-108 |
| AC234582.1 | 4 | 0.662610658 | 0.317 | 0.062 | 5.64E-112 | 2.07E-107 |
| PPP2R2B | 4 | -1.708857887 | 0.406 | 0.677 | 9.53E-112 | 3.49E-107 |
| RFX4 | 4 | 1.339652399 | 0.184 | 0.025 | 1.06E-111 | 3.87E-107 |
| AL583808.1 | 4 | 0.649992083 | 0.154 | 0.005 | 1.09E-111 | 3.98E-107 |
| HSPG2 | 4 | 0.770269614 | 0.382 | 0.096 | 3.08E-111 | 1.13E-106 |
| CSMD1 | 4 | -2.496182917 | 0.15 | 0.479 | 5.95E-111 | 2.18E-106 |
| TCF7L1 | 4 | 0.961041253 | 0.618 | 0.273 | 1.33E-110 | 4.87E-106 |
| CDK6 | 4 | 1.022152278 | 0.422 | 0.111 | 2.42E-110 | 8.84E-106 |
| BBOX1-AS1 | 4 | 0.878811589 | 0.266 | 0.039 | 3.25E-110 | 1.19E-105 |
| TSC22D1 | 4 | 0.938835075 | 0.788 | 0.701 | 4.16E-110 | 1.52E-105 |
| SLC9C2 | 4 | 0.529771048 | 0.218 | 0.02 | 1.17E-109 | 4.27E-105 |
| AL390800.1 | 4 | -1.468610921 | 0.212 | 0.554 | 1.72E-109 | 6.28E-105 |
| CEP83 | 4 | 0.906474129 | 0.672 | 0.452 | 5.93E-109 | 2.17E-104 |
| ARAP2 | 4 | -1.071083482 | 0.098 | 0.445 | 6.92E-109 | 2.53E-104 |
| GLI3 | 4 | 1.022874639 | 0.655 | 0.308 | 7.94E-109 | 2.91E-104 |
| TNFAIP8 | 4 | 0.808304056 | 0.308 | 0.064 | 3.51E-108 | 1.29E-103 |
| SH3GL2 | 4 | -1.206936269 | 0.091 | 0.43 | 3.81E-108 | 1.4E-103 |
| SNRPN | 4 | 0.669874008 | 0.277 | 0.046 | 1.01E-107 | 3.71E-103 |
| GALNT10 | 4 | 0.836126869 | 0.366 | 0.104 | 1.51E-107 | 5.52E-103 |
| DGKH | 4 | -1.010969837 | 0.395 | 0.737 | 3.81E-107 | 1.4E-102 |
| IQUB | 4 | 0.537248991 | 0.216 | 0.021 | 4.55E-107 | 1.66E-102 |
| TSBP1-AS1 | 4 | 0.661642196 | 0.198 | 0.016 | 4.88E-107 | 1.79E-102 |
| SLC44A5 | 4 | 1.183682969 | 0.339 | 0.074 | 5.64E-107 | 2.06E-102 |
| LIMCH1 | 4 | -1.389945043 | 0.385 | 0.718 | 5.95E-107 | 2.18E-102 |
| ZFHX3 | 4 | 1.057874953 | 0.816 | 0.555 | 6.51E-107 | 2.38E-102 |
| CADM1 | 4 | 1.111261202 | 0.737 | 0.406 | 1.72E-106 | 6.3E-102 |
| LINC02470 | 4 | -1.359971225 | 0.022 | 0.297 | 1.89E-106 | 6.9E-102 |
| IL17RD | 4 | 0.716284785 | 0.322 | 0.066 | 3.82E-106 | 1.4E-101 |
| STIM2 | 4 | 0.865248604 | 0.587 | 0.454 | 4.67E-106 | 1.71E-101 |
| MEIS1 | 4 | 1.278261358 | 0.541 | 0.331 | 2.87E-105 | 1.05E-100 |
| BBS9 | 4 | 0.941553601 | 0.717 | 0.437 | 5.68E-105 | 2.08E-100 |
| RSPH1 | 4 | 0.429812864 | 0.158 | 0.006 | 1.97E-104 | 7.22E-100 |
| HOXB3 | 4 | 0.77290945 | 0.24 | 0.037 | 3.27E-104 | 1.2E-99 |
| NWD1 | 4 | 0.666087574 | 0.19 | 0.015 | 3.31E-104 | 1.21E-99 |
| KCND2 | 4 | -1.672558213 | 0.244 | 0.575 | 3.75E-104 | 1.37E-99 |
| TDRP | 4 | 0.654028114 | 0.375 | 0.094 | 6.28E-104 | 2.3E-99 |
| PID1 | 4 | -1.464690843 | 0.29 | 0.617 | 6.62E-104 | 2.43E-99 |
| CDKN2A | 4 | -0.809252126 | 0.052 | 0.358 | 9.03E-104 | 3.31E-99 |
| PROM1 | 4 | 0.735789996 | 0.343 | 0.074 | 9.24E-104 | 3.38E-99 |
| CNTNAP2 | 4 | 0.840588881 | 0.239 | 0.031 | 1.24E-103 | 4.53E-99 |
| DNAH5 | 4 | 0.579992931 | 0.205 | 0.018 | 1.5E-103 | 5.49E-99 |
| RUNX1T1 | 4 | -0.74687865 | 0.393 | 0.751 | 2.58E-103 | 9.46E-99 |
| EFCAB6 | 4 | 0.865193694 | 0.377 | 0.12 | 3.11E-103 | 1.14E-98 |
| TMEM178A | 4 | -1.053348289 | 0.162 | 0.506 | 3.91E-103 | 1.43E-98 |
| TRIM9 | 4 | -0.976045053 | 0.118 | 0.46 | 4.75E-103 | 1.74E-98 |
| ARFGEF3 | 4 | -1.115127046 | 0.159 | 0.495 | 1.14E-102 | 4.18E-98 |

| | | | | | | |
|---|---|---|---|---|---|---|
| SYBU | 4 | -1.167809339 | 0.113 | 0.446 | 1.32E-102 | 4.84E-98 |
| ZBTB9 | 4 | 0.523513612 | 0.21 | 0.022 | 1.46E-102 | 5.34E-98 |
| SPECC1 | 4 | -1.170452517 | 0.26 | 0.577 | 2.34E-102 | 8.56E-98 |
| AC023469.1 | 4 | -1.26491578 | 0.071 | 0.384 | 4.97E-102 | 1.82E-97 |
| AC009271.1 | 4 | 0.772057128 | 0.204 | 0.022 | 1.4E-101 | 5.13E-97 |
| ZNF385D | 4 | -1.17182851 | 0.252 | 0.609 | 1.47E-101 | 5.38E-97 |
| FUT8 | 4 | -0.978321684 | 0.62 | 0.839 | 3.3E-101 | 1.21E-96 |
| TRPV4 | 4 | 0.497585084 | 0.183 | 0.014 | 3.81E-101 | 1.4E-96 |
| NKD1 | 4 | 1.161381364 | 0.436 | 0.133 | 5.77E-101 | 2.11E-96 |
| CFAP206 | 4 | 0.515149625 | 0.2 | 0.018 | 9.32E-101 | 3.41E-96 |
| SPTLC3 | 4 | 0.498117432 | 0.16 | 0.008 | 1.02E-100 | 3.72E-96 |
| PRDM16-DT | 4 | 0.339812821 | 0.14 | 0.004 | 1.17E-100 | 4.28E-96 |
| ANKFN1 | 4 | 0.556727378 | 0.173 | 0.012 | 1.6E-100 | 5.85E-96 |
| ELMOD1 | 4 | -1.084315173 | 0.105 | 0.433 | 2.33E-100 | 8.55E-96 |
| TMEM117 | 4 | 0.991941228 | 0.571 | 0.58 | 3.34E-100 | 1.22E-95 |
| ADAMTSL1 | 4 | -2.107759646 | 0.229 | 0.557 | 1.37E-99 | 5.03E-95 |
| NOL4 | 4 | -1.433518622 | 0.065 | 0.358 | 2.02E-99 | 7.39E-95 |
| LINC01833 | 4 | 0.251673722 | 0.119 | 0.001 | 3.04E-99 | 1.11E-94 |
| DOCK3 | 4 | 1.302795156 | 0.49 | 0.261 | 8.98E-99 | 3.29E-94 |
| L3MBTL4 | 4 | -0.944911589 | 0.21 | 0.568 | 9.26E-99 | 3.39E-94 |
| FAM227A | 4 | 0.762052878 | 0.36 | 0.109 | 3.46E-98 | 1.27E-93 |
| NPY1R | 4 | 0.707419715 | 0.205 | 0.024 | 9.9E-98 | 3.62E-93 |
| AC114316.1 | 4 | 0.54131463 | 0.143 | 0.005 | 1.63E-97 | 5.97E-93 |
| CCDC175 | 4 | 0.454871651 | 0.196 | 0.018 | 3.69E-97 | 1.35E-92 |
| LINC00648 | 4 | 0.457364538 | 0.2 | 0.019 | 5E-97 | 1.83E-92 |
| DNAI2 | 4 | 0.436683543 | 0.159 | 0.008 | 5.24E-97 | 1.92E-92 |
| GRIN2B | 4 | 0.677736464 | 0.171 | 0.012 | 6.08E-97 | 2.23E-92 |
| ATRNL1 | 4 | -1.459801659 | 0.712 | 0.818 | 1.93E-96 | 7.05E-92 |
| CDKN2B-AS1 | 4 | -0.993901328 | 0.061 | 0.355 | 3.26E-96 | 1.19E-91 |
| GREB1 | 4 | 0.95436925 | 0.284 | 0.09 | 3.71E-96 | 1.36E-91 |
| FIRRE | 4 | 0.780774297 | 0.616 | 0.269 | 3.89E-96 | 1.43E-91 |
| TTC29 | 4 | 0.454138635 | 0.15 | 0.007 | 6.71E-96 | 2.46E-91 |
| NRXN1 | 4 | -1.487064632 | 0.302 | 0.632 | 1.27E-95 | 4.65E-91 |
| CDH2 | 4 | 0.999234638 | 0.732 | 0.443 | 4.61E-95 | 1.69E-90 |
| SYTL5 | 4 | -1.101330241 | 0.057 | 0.345 | 7.78E-95 | 2.85E-90 |
| GPHN | 4 | -0.79096981 | 0.876 | 0.928 | 1.36E-94 | 4.97E-90 |
| CCDC146 | 4 | 0.944055087 | 0.465 | 0.212 | 1.63E-94 | 5.97E-90 |
| TAFA2 | 4 | 0.994374001 | 0.301 | 0.084 | 1.85E-94 | 6.77E-90 |
| SOBP | 4 | -1.104420233 | 0.632 | 0.793 | 3.64E-94 | 1.33E-89 |
| LDB2 | 4 | 0.678577007 | 0.649 | 0.301 | 1.04E-93 | 3.79E-89 |
| TPBG | 4 | 0.563856284 | 0.259 | 0.043 | 2.47E-93 | 9.03E-89 |
| DOK5 | 4 | -1.243351198 | 0.282 | 0.576 | 4.54E-93 | 1.66E-88 |
| NFIB | 4 | -1.122798198 | 0.689 | 0.849 | 6.51E-93 | 2.38E-88 |
| CFAP45 | 4 | 0.29481459 | 0.141 | 0.006 | 2.12E-92 | 7.76E-88 |
| LMO7 | 4 | -1.21945942 | 0.239 | 0.558 | 2.93E-92 | 1.07E-87 |
| IQCK | 4 | 0.714454194 | 0.424 | 0.146 | 2.98E-92 | 1.09E-87 |
| IGSF1 | 4 | 0.569459195 | 0.194 | 0.021 | 3.63E-92 | 1.33E-87 |
| SDK1 | 4 | -1.443343863 | 0.806 | 0.916 | 4.01E-92 | 1.47E-87 |
| DNAH14 | 4 | 0.880429446 | 0.775 | 0.491 | 1.15E-91 | 4.2E-87 |
| ARHGAP29 | 4 | 0.786538562 | 0.264 | 0.049 | 9.44E-91 | 3.46E-86 |
| GPRIN3 | 4 | -0.855046681 | 0.097 | 0.405 | 1.58E-90 | 5.77E-86 |
| RBM20 | 4 | 0.779107053 | 0.168 | 0.014 | 2.38E-90 | 8.72E-86 |
| ACOT11 | 4 | 0.74983543 | 0.327 | 0.117 | 4.44E-90 | 1.62E-85 |
| CFAP77 | 4 | 0.508046582 | 0.183 | 0.018 | 7.48E-90 | 2.74E-85 |
| ATP13A5 | 4 | 0.372677066 | 0.121 | 0.003 | 7.97E-90 | 2.92E-85 |
| PCLO | 4 | -1.338432021 | 0.39 | 0.633 | 9.34E-90 | 3.42E-85 |
| TBC1D19 | 4 | 0.770315083 | 0.573 | 0.306 | 9.61E-90 | 3.52E-85 |
| JHY | 4 | 0.407989905 | 0.184 | 0.017 | 9.7E-90 | 3.55E-85 |

| | | | | | | |
|---|---|---|---|---|---|---|
| AC231759.2 | 4 | 0.367201832 | 0.122 | 0.003 | 1.02E-89 | 3.73E-85 |
| MAP7 | 4 | -1.166937546 | 0.314 | 0.621 | 1.69E-89 | 6.19E-85 |
| VGLL3 | 4 | -1.633886506 | 0.065 | 0.34 | 2.61E-89 | 9.54E-85 |
| RASAL2 | 4 | 0.840104103 | 0.765 | 0.513 | 2.79E-89 | 1.02E-84 |
| GPR176 | 4 | -0.862276512 | 0.123 | 0.433 | 5.45E-89 | 1.99E-84 |
| SNHG14 | 4 | 0.744939489 | 0.893 | 0.825 | 1.23E-88 | 4.5E-84 |
| AL158055.1 | 4 | 0.442070395 | 0.188 | 0.02 | 1.3E-88 | 4.75E-84 |
| POU5F1 | 4 | -0.948776162 | 0.209 | 0.514 | 1.88E-88 | 6.88E-84 |
| PCOLCE2 | 4 | -0.850494376 | 0.09 | 0.379 | 1.91E-88 | 7E-84 |
| AC105411.1 | 4 | 0.840041219 | 0.159 | 0.012 | 1.93E-88 | 7.08E-84 |
| LINC01239 | 4 | 0.761085592 | 0.184 | 0.02 | 1.96E-88 | 7.19E-84 |
| MAP6 | 4 | 0.674002239 | 0.27 | 0.056 | 2.26E-88 | 8.28E-84 |
| DLEU2 | 4 | -1.011250311 | 0.516 | 0.771 | 2.68E-88 | 9.81E-84 |
| RIMS2 | 4 | -1.95694324 | 0.207 | 0.453 | 3.57E-88 | 1.31E-83 |
| VWDE | 4 | -1.258872092 | 0.127 | 0.43 | 3.78E-88 | 1.38E-83 |
| SLC7A8 | 4 | 0.422874264 | 0.189 | 0.019 | 5.08E-88 | 1.86E-83 |
| RABL2A | 4 | 0.561655868 | 0.268 | 0.057 | 6.3E-88 | 2.31E-83 |
| NID2 | 4 | 0.448370413 | 0.156 | 0.01 | 6.44E-88 | 2.36E-83 |
| SPAG1 | 4 | 0.785354153 | 0.312 | 0.111 | 7.65E-88 | 2.8E-83 |
| PLEKHA7 | 4 | 0.910436382 | 0.428 | 0.166 | 9.76E-88 | 3.57E-83 |
| AC104041.1 | 4 | 1.046763756 | 0.672 | 0.466 | 1.79E-87 | 6.54E-83 |
| MIR924HG | 4 | -0.655047468 | 0.425 | 0.752 | 2.86E-87 | 1.05E-82 |
| AC008591.1 | 4 | -1.293465517 | 0.158 | 0.465 | 3.7E-87 | 1.35E-82 |
| CSPP1 | 4 | 0.807229795 | 0.646 | 0.479 | 5.64E-87 | 2.07E-82 |
| SYNPR | 4 | -1.134538876 | 0.092 | 0.39 | 7.5E-87 | 2.74E-82 |
| FSIP1 | 4 | 0.548612481 | 0.265 | 0.051 | 8.97E-87 | 3.28E-82 |
| PDZRN4 | 4 | 1.170793178 | 0.281 | 0.057 | 1.25E-86 | 4.56E-82 |
| MECOM | 4 | 1.530391459 | 0.225 | 0.042 | 2.61E-86 | 9.56E-82 |
| SLC1A3 | 4 | 0.924003073 | 0.313 | 0.084 | 4.33E-86 | 1.59E-81 |
| DNAH10 | 4 | -1.167864787 | 0.197 | 0.5 | 5.33E-86 | 1.95E-81 |
| RAB3GAP2 | 4 | 0.765632091 | 0.696 | 0.583 | 6.5E-86 | 2.38E-81 |
| CHPT1 | 4 | -1.017980825 | 0.308 | 0.595 | 1.26E-85 | 4.61E-81 |
| SNTB1 | 4 | 0.736619552 | 0.265 | 0.053 | 1.3E-85 | 4.75E-81 |
| ADGRB3 | 4 | -1.110915554 | 0.247 | 0.571 | 2.18E-85 | 7.99E-81 |
| NTRK2 | 4 | 1.036792444 | 0.257 | 0.049 | 3.58E-85 | 1.31E-80 |
| GABRA3 | 4 | 0.456009653 | 0.181 | 0.019 | 8.28E-85 | 3.03E-80 |
| AKAP14 | 4 | 0.390857564 | 0.152 | 0.01 | 9.29E-85 | 3.4E-80 |
| GRIP1 | 4 | 1.127408442 | 0.62 | 0.329 | 1.8E-84 | 6.59E-80 |
| PPFIBP2 | 4 | 0.573265446 | 0.288 | 0.064 | 1.83E-84 | 6.7E-80 |
| LINC02550 | 4 | 0.382962867 | 0.149 | 0.009 | 2.19E-84 | 8.01E-80 |
| ELF2 | 4 | -0.811206137 | 0.619 | 0.812 | 2.93E-84 | 1.07E-79 |
| KIF9 | 4 | 0.571915551 | 0.248 | 0.048 | 5.25E-84 | 1.92E-79 |
| KIAA1671 | 4 | 0.673340398 | 0.42 | 0.139 | 6.92E-84 | 2.53E-79 |
| FBLN1 | 4 | 0.814903223 | 0.426 | 0.199 | 8.7E-84 | 3.18E-79 |
| UGT8 | 4 | -0.784325867 | 0.071 | 0.35 | 2.35E-83 | 8.62E-79 |
| CPE | 4 | -0.91089368 | 0.377 | 0.659 | 5.25E-83 | 1.92E-78 |
| NAV2 | 4 | 1.092423036 | 0.657 | 0.447 | 6.72E-83 | 2.46E-78 |
| AC060834.2 | 4 | 0.392362863 | 0.126 | 0.005 | 7.2E-83 | 2.64E-78 |
| MAK | 4 | 0.625248887 | 0.316 | 0.09 | 1.19E-82 | 4.34E-78 |
| ROR2 | 4 | 1.021986515 | 0.609 | 0.4 | 2.36E-82 | 8.62E-78 |
| EPAS1 | 4 | 0.347045156 | 0.135 | 0.007 | 2.41E-82 | 8.82E-78 |
| DNAAF4 | 4 | 0.586248071 | 0.295 | 0.07 | 3.78E-82 | 1.38E-77 |
| IKZF3 | 4 | -0.78258379 | 0.048 | 0.294 | 4.63E-82 | 1.69E-77 |
| CACNA2D1 | 4 | -0.924450668 | 0.657 | 0.877 | 5.55E-82 | 2.03E-77 |
| PRICKLE2 | 4 | 0.867288075 | 0.537 | 0.316 | 1.06E-81 | 3.87E-77 |
| SHISA9 | 4 | 0.780134758 | 0.169 | 0.016 | 1.19E-81 | 4.34E-77 |
| FMNL2 | 4 | -0.784603424 | 0.735 | 0.891 | 1.57E-81 | 5.76E-77 |
| LINC01091 | 4 | 1.044652562 | 0.251 | 0.057 | 1.67E-81 | 6.11E-77 |

| | | | | | | |
|---|---|---|---|---|---|---|
| ADAMTS3 | 4 | -1.383197234 | 0.229 | 0.506 | 2.26E-81 | 8.28E-77 |
| SORBS1 | 4 | -0.871681928 | 0.249 | 0.564 | 3.16E-81 | 1.16E-76 |
| DAB1 | 4 | -0.85322527 | 0.42 | 0.715 | 3.48E-81 | 1.27E-76 |
| CYYR1 | 4 | 0.687484418 | 0.28 | 0.062 | 3.66E-81 | 1.34E-76 |
| PPM1H | 4 | 0.580062384 | 0.202 | 0.028 | 4.68E-81 | 1.71E-76 |
| KCNIP1 | 4 | 0.852015102 | 0.295 | 0.083 | 4.97E-81 | 1.82E-76 |
| GRIN3A | 4 | -1.210902094 | 0.043 | 0.282 | 5.62E-81 | 2.06E-76 |
| SLC2A12 | 4 | 0.482767756 | 0.174 | 0.022 | 6.51E-81 | 2.38E-76 |
| GTF2IRD1 | 4 | 0.788395788 | 0.656 | 0.418 | 6.89E-81 | 2.52E-76 |
| SLC25A27 | 4 | 0.534663738 | 0.261 | 0.054 | 8.25E-81 | 3.02E-76 |
| OTUD7A | 4 | -1.088309625 | 0.308 | 0.598 | 1.04E-80 | 3.81E-76 |
| LINC01876 | 4 | 0.991518913 | 0.591 | 0.498 | 1.07E-80 | 3.92E-76 |
| PAK5 | 4 | -1.149808669 | 0.207 | 0.478 | 1.09E-80 | 3.97E-76 |
| TSPAN7 | 4 | -1.035601406 | 0.277 | 0.571 | 1.31E-80 | 4.81E-76 |
| RAPGEF5 | 4 | -0.999390423 | 0.255 | 0.546 | 2.56E-80 | 9.36E-76 |
| DNAJC6 | 4 | -0.99044374 | 0.16 | 0.441 | 2.75E-80 | 1.01E-75 |
| ZMAT4 | 4 | -0.873729615 | 0.236 | 0.56 | 3.72E-80 | 1.36E-75 |
| PXDN | 4 | 0.733321175 | 0.496 | 0.203 | 3.87E-80 | 1.42E-75 |
| GFRA1 | 4 | 0.888292035 | 0.232 | 0.05 | 4.09E-80 | 1.5E-75 |
| ZNRF3 | 4 | 0.837055157 | 0.615 | 0.378 | 6.49E-80 | 2.38E-75 |
| SHANK2 | 4 | 1.074631585 | 0.428 | 0.195 | 7.78E-80 | 2.85E-75 |
| STK39 | 4 | 0.798456316 | 0.752 | 0.629 | 8.25E-80 | 3.02E-75 |
| FAR2 | 4 | -1.086054782 | 0.125 | 0.415 | 1.04E-79 | 3.8E-75 |
| RAD9A | 4 | 0.757149084 | 0.435 | 0.191 | 1.39E-79 | 5.1E-75 |
| MCC | 4 | -0.919541808 | 0.607 | 0.819 | 2.4E-79 | 8.78E-75 |
| UNC5D | 4 | -1.548508586 | 0.437 | 0.594 | 3.45E-79 | 1.26E-74 |
| CSF2RA | 4 | 0.260976587 | 0.116 | 0.004 | 1.11E-78 | 4.08E-74 |
| CERS6 | 4 | -0.756042158 | 0.645 | 0.835 | 1.83E-78 | 6.69E-74 |
| AGAP1 | 4 | 0.740638318 | 0.934 | 0.833 | 2.03E-78 | 7.44E-74 |
| AC023282.1 | 4 | -0.930205291 | 0.072 | 0.322 | 2.1E-78 | 7.68E-74 |
| FBXL13 | 4 | 0.700690639 | 0.358 | 0.122 | 4.52E-78 | 1.65E-73 |
| AC104461.1 | 4 | 0.358828387 | 0.137 | 0.009 | 4.59E-78 | 1.68E-73 |
| FOXP2 | 4 | 0.583933214 | 0.529 | 0.233 | 7.61E-78 | 2.79E-73 |
| MEG3 | 4 | -0.804782063 | 0.172 | 0.479 | 7.76E-78 | 2.84E-73 |
| NAALADL2 | 4 | 0.977896282 | 0.655 | 0.387 | 1.01E-77 | 3.69E-73 |
| CFAP99 | 4 | 0.254111067 | 0.114 | 0.004 | 2.05E-77 | 7.49E-73 |
| NUP62CL | 4 | 0.475024014 | 0.201 | 0.03 | 3.08E-77 | 1.13E-72 |
| SLC16A7 | 4 | -0.878987424 | 0.108 | 0.381 | 4.2E-77 | 1.54E-72 |
| CROCC2 | 4 | 0.638484169 | 0.129 | 0.008 | 6.73E-77 | 2.46E-72 |
| DOCK11 | 4 | -1.154478769 | 0.271 | 0.559 | 6.79E-77 | 2.49E-72 |
| GLI2 | 4 | 0.928386465 | 0.518 | 0.291 | 1.05E-76 | 3.85E-72 |
| RSRC1 | 4 | -0.753763317 | 0.651 | 0.827 | 2.15E-76 | 7.86E-72 |
| MGAT4C | 4 | -1.334087374 | 0.264 | 0.564 | 3.59E-76 | 1.32E-71 |
| SHROOM4 | 4 | -1.072803955 | 0.359 | 0.633 | 5.35E-76 | 1.96E-71 |
| DNALI1 | 4 | 0.31983313 | 0.171 | 0.019 | 5.5E-76 | 2.02E-71 |
| FAM13C | 4 | 0.402651408 | 0.193 | 0.027 | 9.7E-76 | 3.55E-71 |
| PTPRD | 4 | -0.8977549 | 0.798 | 0.846 | 1.05E-75 | 3.85E-71 |
| PWRN1 | 4 | 0.573125885 | 0.165 | 0.017 | 1.45E-75 | 5.29E-71 |
| CUEDC1 | 4 | 0.673933556 | 0.405 | 0.153 | 4.61E-75 | 1.69E-70 |
| WNT5B | 4 | 0.806390623 | 0.315 | 0.099 | 8.9E-75 | 3.26E-70 |
| OXR1 | 4 | -0.973655417 | 0.416 | 0.686 | 2.5E-74 | 9.14E-70 |
| SORCS2 | 4 | 0.975612192 | 0.467 | 0.23 | 3.73E-74 | 1.37E-69 |
| MRPS6 | 4 | 0.692772745 | 0.427 | 0.211 | 4.38E-74 | 1.6E-69 |
| AL022068.1 | 4 | 0.488869512 | 0.314 | 0.088 | 4.63E-74 | 1.7E-69 |
| PDE4D | 4 | -0.748507782 | 0.702 | 0.867 | 5.24E-74 | 1.92E-69 |
| NDST3 | 4 | -0.995730438 | 0.206 | 0.509 | 7.13E-74 | 2.61E-69 |
| PROS1 | 4 | 0.456127099 | 0.201 | 0.031 | 1.33E-73 | 4.85E-69 |
| MYOF | 4 | 0.459606408 | 0.172 | 0.02 | 2.48E-73 | 9.07E-69 |

| | | | | | | |
|---|---|---|---|---|---|---|
| PTPRC | 4 | 0.312438261 | 0.105 | 0.003 | 2.97E-73 | 1.09E-68 |
| HDAC2-AS2 | 4 | -0.920775144 | 0.239 | 0.538 | 5.72E-73 | 2.1E-68 |
| CDH4 | 4 | -0.95643514 | 0.353 | 0.65 | 5.84E-73 | 2.14E-68 |
| IL12RB2 | 4 | -1.191812255 | 0.063 | 0.311 | 1.24E-72 | 4.55E-68 |
| TMEM231 | 4 | 0.558637927 | 0.243 | 0.057 | 1.37E-72 | 5.01E-68 |
| CRISPLD1 | 4 | -1.918339148 | 0.274 | 0.523 | 1.38E-72 | 5.05E-68 |
| EXOC2 | 4 | -1.052934622 | 0.527 | 0.689 | 1.43E-72 | 5.22E-68 |
| PLPP4 | 4 | -1.019296553 | 0.145 | 0.397 | 1.66E-72 | 6.06E-68 |
| ELMO1 | 4 | -1.006112367 | 0.297 | 0.58 | 2.57E-72 | 9.43E-68 |
| SLC5A3 | 4 | 0.669509799 | 0.397 | 0.185 | 2.66E-72 | 9.76E-68 |
| AGBL1 | 4 | 0.617888439 | 0.152 | 0.015 | 4.32E-72 | 1.58E-67 |
| CADPS2 | 4 | 1.099992216 | 0.384 | 0.217 | 4.78E-72 | 1.75E-67 |
| FSIP2 | 4 | 0.753041646 | 0.42 | 0.169 | 7.14E-72 | 2.61E-67 |
| ABCB1 | 4 | -0.87777703 | 0.041 | 0.258 | 7.64E-72 | 2.8E-67 |
| FGFR1 | 4 | 0.776236556 | 0.593 | 0.332 | 8.06E-72 | 2.95E-67 |
| GALNT5 | 4 | 0.342983549 | 0.15 | 0.014 | 8.38E-72 | 3.07E-67 |
| TPM1 | 4 | 0.958518058 | 0.544 | 0.302 | 8.92E-72 | 3.26E-67 |
| DOP1B | 4 | -0.724887472 | 0.229 | 0.523 | 1.38E-71 | 5.04E-67 |
| HOXA3 | 4 | 0.5067915 | 0.144 | 0.014 | 2.9E-71 | 1.06E-66 |
| EFCAB2 | 4 | 0.737633773 | 0.55 | 0.292 | 2.94E-71 | 1.08E-66 |
| MYO10 | 4 | 0.764223201 | 0.596 | 0.302 | 3.04E-71 | 1.11E-66 |
| PDE3B | 4 | -1.025755933 | 0.404 | 0.663 | 3.49E-71 | 1.28E-66 |
| HS6ST3 | 4 | -2.365131366 | 0.444 | 0.512 | 4.74E-71 | 1.73E-66 |
| USH2A | 4 | 0.592425751 | 0.183 | 0.026 | 5.19E-71 | 1.9E-66 |
| LINC02018 | 4 | 0.696164098 | 0.311 | 0.103 | 7.78E-71 | 2.85E-66 |
| ZNF609 | 4 | 0.624270271 | 0.88 | 0.743 | 8.4E-71 | 3.08E-66 |
| FRMPD4 | 4 | 0.821596617 | 0.159 | 0.02 | 9.5E-71 | 3.48E-66 |
| ZYG11A | 4 | -0.737674565 | 0.079 | 0.328 | 1.19E-70 | 4.35E-66 |
| LRRC4C | 4 | -1.125601861 | 0.271 | 0.559 | 1.47E-70 | 5.37E-66 |
| CELSR1 | 4 | 0.622203952 | 0.229 | 0.048 | 2.14E-70 | 7.83E-66 |
| TGFBR2 | 4 | 0.674440064 | 0.349 | 0.119 | 2.21E-70 | 8.09E-66 |
| ARHGEF28 | 4 | -0.90459099 | 0.113 | 0.383 | 2.32E-70 | 8.51E-66 |
| PDZRN3 | 4 | -0.977516204 | 0.784 | 0.917 | 2.35E-70 | 8.62E-66 |
| CASC9 | 4 | 0.619515118 | 0.226 | 0.044 | 2.66E-70 | 9.74E-66 |
| CACNA1D | 4 | -0.770419181 | 0.165 | 0.451 | 2.7E-70 | 9.88E-66 |
| ENPP1 | 4 | -0.85685598 | 0.113 | 0.382 | 4.02E-70 | 1.47E-65 |
| BBS5 | 4 | 0.494240641 | 0.292 | 0.079 | 5.19E-70 | 1.9E-65 |
| TRIM24 | 4 | 0.838797183 | 0.662 | 0.477 | 5.85E-70 | 2.14E-65 |
| RPE65 | 4 | 0.304020062 | 0.145 | 0.014 | 7.91E-70 | 2.89E-65 |
| ZNF487 | 4 | 0.414638485 | 0.193 | 0.031 | 1.36E-69 | 4.99E-65 |
| BBOF1 | 4 | 0.4536624 | 0.231 | 0.048 | 1.58E-69 | 5.78E-65 |
| IL1RAPL1 | 4 | -1.329699835 | 0.546 | 0.67 | 1.8E-69 | 6.58E-65 |
| GLS | 4 | -0.789106788 | 0.645 | 0.8 | 2.79E-69 | 1.02E-64 |
| SLC16A6 | 4 | 0.356429427 | 0.149 | 0.016 | 2.81E-69 | 1.03E-64 |
| DLEC1 | 4 | 0.324296612 | 0.134 | 0.011 | 4.42E-69 | 1.62E-64 |
| ZFP36L1 | 4 | 0.581055562 | 0.276 | 0.073 | 9.92E-69 | 3.63E-64 |
| TACC2 | 4 | 0.686442401 | 0.422 | 0.18 | 1.35E-68 | 4.95E-64 |
| CFAP58 | 4 | -0.895464777 | 0.077 | 0.322 | 1.4E-68 | 5.12E-64 |
| C11orf49 | 4 | 0.651702757 | 0.726 | 0.525 | 1.45E-68 | 5.32E-64 |
| SHC3 | 4 | -0.898176833 | 0.124 | 0.382 | 1.53E-68 | 5.59E-64 |
| ARL4C | 4 | 0.345712567 | 0.17 | 0.022 | 1.78E-68 | 6.52E-64 |
| PRKAG2 | 4 | -0.808061551 | 0.185 | 0.466 | 2.07E-68 | 7.6E-64 |
| ZBTB46 | 4 | 0.640800805 | 0.253 | 0.063 | 2.11E-68 | 7.71E-64 |
| RPGR | 4 | 0.704025188 | 0.551 | 0.436 | 2.65E-68 | 9.72E-64 |
| LINC01980 | 4 | 0.3408718 | 0.143 | 0.014 | 3.27E-68 | 1.2E-63 |
| INTU | 4 | 0.685330052 | 0.528 | 0.284 | 4.26E-68 | 1.56E-63 |
| AGPAT4 | 4 | 0.672465249 | 0.59 | 0.326 | 5.3E-68 | 1.94E-63 |
| AL121578.3 | 4 | -0.640940483 | 0.029 | 0.239 | 5.44E-68 | 1.99E-63 |

| | | | | | | |
|---|---|---|---|---|---|---|
| ARMC9 | 4 | 0.67722577 | 0.537 | 0.286 | 6.7E-68 | 2.45E-63 |
| EFHB | 4 | 0.278801749 | 0.119 | 0.007 | 1.52E-67 | 5.57E-63 |
| ANOS1 | 4 | -1.21671622 | 0.219 | 0.467 | 1.78E-67 | 6.52E-63 |
| LINC02296 | 4 | 0.386064298 | 0.11 | 0.006 | 1.98E-67 | 7.26E-63 |
| WDR11-AS1 | 4 | -0.877070401 | 0.109 | 0.35 | 2.91E-67 | 1.07E-62 |
| UBASH3B | 4 | 0.349882977 | 0.158 | 0.019 | 3.57E-67 | 1.31E-62 |
| WDR66 | 4 | 0.693186237 | 0.31 | 0.103 | 4.66E-67 | 1.71E-62 |
| GABRB2 | 4 | -1.119721994 | 0.088 | 0.331 | 6.58E-67 | 2.41E-62 |
| LINC00472 | 4 | 0.77397917 | 0.41 | 0.194 | 1.02E-66 | 3.75E-62 |
| NHS | 4 | 0.643908305 | 0.732 | 0.444 | 1.52E-66 | 5.55E-62 |
| SATB2 | 4 | -0.925673569 | 0.226 | 0.487 | 1.62E-66 | 5.94E-62 |
| UST | 4 | 0.688373414 | 0.531 | 0.248 | 1.89E-66 | 6.92E-62 |
| SYNGAP1 | 4 | 0.603969207 | 0.428 | 0.196 | 2.51E-66 | 9.18E-62 |
| AC233976.1 | 4 | 0.431715689 | 0.202 | 0.037 | 4.24E-66 | 1.55E-61 |
| CCDC170 | 4 | 0.424823412 | 0.177 | 0.027 | 4.97E-66 | 1.82E-61 |
| MIR99AHG | 4 | 0.574440563 | 0.889 | 0.663 | 5.42E-66 | 1.98E-61 |
| KCND3 | 4 | 0.376014636 | 0.111 | 0.006 | 5.42E-66 | 1.98E-61 |
| DOCK10 | 4 | -1.145706433 | 0.149 | 0.404 | 5.59E-66 | 2.04E-61 |
| ZNF19 | 4 | 0.285079849 | 0.156 | 0.019 | 5.92E-66 | 2.17E-61 |
| ANKDD1B | 4 | 0.400428909 | 0.177 | 0.028 | 9.25E-66 | 3.39E-61 |
| FRMD4A | 4 | -0.786718885 | 0.737 | 0.827 | 1.08E-65 | 3.95E-61 |
| DCLK2 | 4 | -0.856435766 | 0.565 | 0.746 | 1.39E-65 | 5.08E-61 |
| ZNF462 | 4 | 0.664635487 | 0.67 | 0.416 | 1.57E-65 | 5.75E-61 |
| AC092939.1 | 4 | -0.483982455 | 0.017 | 0.206 | 2E-65 | 7.34E-61 |
| DACH1 | 4 | -0.61979179 | 0.754 | 0.902 | 2.25E-65 | 8.24E-61 |
| MERTK | 4 | 0.554998793 | 0.304 | 0.094 | 3.44E-65 | 1.26E-60 |
| ARHGAP20 | 4 | 0.30517834 | 0.131 | 0.011 | 3.51E-65 | 1.28E-60 |
| DENND2A | 4 | 0.70613084 | 0.578 | 0.302 | 3.63E-65 | 1.33E-60 |
| EVI5 | 4 | 0.678025152 | 0.71 | 0.54 | 5.68E-65 | 2.08E-60 |
| KALRN | 4 | 1.00208775 | 0.593 | 0.385 | 7.73E-65 | 2.83E-60 |
| SNCAIP | 4 | 0.460369586 | 0.432 | 0.178 | 1.19E-64 | 4.34E-60 |
| TTLL9 | 4 | 0.570464065 | 0.259 | 0.073 | 1.2E-64 | 4.38E-60 |
| NT5DC1 | 4 | 0.764869616 | 0.463 | 0.385 | 1.42E-64 | 5.21E-60 |
| ST8SIA1 | 4 | 0.619445536 | 0.233 | 0.058 | 1.5E-64 | 5.48E-60 |
| ROR1 | 4 | 1.03029824 | 0.49 | 0.225 | 1.84E-64 | 6.74E-60 |
| MUSK | 4 | 0.356733376 | 0.126 | 0.011 | 2.48E-64 | 9.07E-60 |
| PPARG | 4 | -0.875862956 | 0.076 | 0.305 | 2.99E-64 | 1.09E-59 |
| TACR3 | 4 | 0.953705238 | 0.236 | 0.059 | 3.11E-64 | 1.14E-59 |
| WDR11 | 4 | -0.949704886 | 0.527 | 0.692 | 6.39E-64 | 2.34E-59 |
| RIMBP2 | 4 | -0.303998025 | 0.262 | 0.114 | 7.31E-64 | 2.68E-59 |
| TBX20 | 4 | -0.64377541 | 0.059 | 0.287 | 8.12E-64 | 2.97E-59 |
| ABCD3 | 4 | -0.849100233 | 0.493 | 0.658 | 9.77E-64 | 3.58E-59 |
| PRLR | 4 | 0.57816657 | 0.199 | 0.041 | 1.09E-63 | 4.01E-59 |
| AL442636.1 | 4 | 0.554762234 | 0.231 | 0.052 | 1.85E-63 | 6.77E-59 |
| TTC23L | 4 | 0.336907963 | 0.171 | 0.025 | 2.09E-63 | 7.67E-59 |
| AC114689.3 | 4 | 0.914204649 | 0.22 | 0.057 | 3.9E-63 | 1.43E-58 |
| CCDC13 | 4 | 0.257675788 | 0.121 | 0.009 | 4.07E-63 | 1.49E-58 |
| TSGA10 | 4 | 0.680457866 | 0.333 | 0.132 | 4.45E-63 | 1.63E-58 |
| CSMD3 | 4 | -1.366867315 | 0.11 | 0.349 | 5.1E-63 | 1.87E-58 |
| KIF27 | 4 | 0.658655828 | 0.4 | 0.2 | 8.14E-63 | 2.98E-58 |
| SMAD3 | 4 | 0.679718307 | 0.566 | 0.313 | 9.09E-63 | 3.33E-58 |
| TBC1D5 | 4 | -0.673433713 | 0.763 | 0.875 | 9.49E-63 | 3.47E-58 |
| CFAP300 | 4 | 0.262668631 | 0.145 | 0.018 | 1.07E-62 | 3.9E-58 |
| COL5A1 | 4 | 0.975300092 | 0.284 | 0.124 | 1.72E-62 | 6.31E-58 |
| CCDC148 | 4 | 0.621700554 | 0.317 | 0.105 | 2.13E-62 | 7.8E-58 |
| DNAH9 | 4 | 0.405719409 | 0.125 | 0.012 | 2.34E-62 | 8.58E-58 |
| MACF1 | 4 | 0.553746324 | 0.943 | 0.847 | 3.19E-62 | 1.17E-57 |
| TLK1 | 4 | 0.643220621 | 0.747 | 0.704 | 3.58E-62 | 1.31E-57 |

| | | | | | | |
|---|---|---|---|---|---|---|
| POLA1 | 4 | -0.807759564 | 0.71 | 0.86 | 7.69E-62 | 2.82E-57 |
| AL365295.1 | 4 | 0.778089995 | 0.233 | 0.073 | 9.02E-62 | 3.3E-57 |
| ROBO2 | 4 | -0.911121277 | 0.494 | 0.738 | 1.13E-61 | 4.14E-57 |
| SAMD12 | 4 | 0.480218685 | 0.236 | 0.058 | 1.23E-61 | 4.52E-57 |
| MSRB3 | 4 | 0.626965511 | 0.274 | 0.078 | 1.26E-61 | 4.62E-57 |
| CCDC178 | 4 | 0.613942096 | 0.201 | 0.041 | 1.49E-61 | 5.47E-57 |
| RGS22 | 4 | 0.563429295 | 0.205 | 0.045 | 1.75E-61 | 6.41E-57 |
| MNS1 | 4 | 0.396676875 | 0.216 | 0.047 | 1.76E-61 | 6.44E-57 |
| RMST | 4 | 0.932965599 | 0.73 | 0.461 | 1.78E-61 | 6.52E-57 |
| FBXL2 | 4 | 0.65877702 | 0.443 | 0.2 | 2.41E-61 | 8.83E-57 |
| CENPP | 4 | -0.982513571 | 0.503 | 0.713 | 2.63E-61 | 9.64E-57 |
| AL049828.2 | 4 | 0.254495012 | 0.107 | 0.006 | 3.13E-61 | 1.14E-56 |
| ZDBF2 | 4 | 0.642517012 | 0.368 | 0.169 | 1.06E-60 | 3.88E-56 |
| CYP46A1 | 4 | 0.317825391 | 0.137 | 0.015 | 1.19E-60 | 4.34E-56 |
| BCAT1 | 4 | 0.650785809 | 0.397 | 0.16 | 1.27E-60 | 4.66E-56 |
| SYN3 | 4 | -1.148642057 | 0.193 | 0.45 | 1.39E-60 | 5.1E-56 |
| AC105052.5 | 4 | 0.269342478 | 0.136 | 0.015 | 2.06E-60 | 7.55E-56 |
| TNS3 | 4 | 0.881403105 | 0.443 | 0.217 | 2.16E-60 | 7.91E-56 |
| MIR100HG | 4 | -1.042022896 | 0.651 | 0.755 | 3.07E-60 | 1.12E-55 |
| CACHD1 | 4 | 0.632032028 | 0.781 | 0.559 | 4.28E-60 | 1.57E-55 |
| ITGA6 | 4 | 0.509850752 | 0.293 | 0.089 | 4.9E-60 | 1.79E-55 |
| MTMR7 | 4 | -0.80643212 | 0.137 | 0.373 | 5.35E-60 | 1.96E-55 |
| MAP3K5 | 4 | -0.717726558 | 0.105 | 0.345 | 6.12E-60 | 2.24E-55 |
| MSI1 | 4 | 0.500706371 | 0.299 | 0.098 | 7.36E-60 | 2.69E-55 |
| TMEM108 | 4 | -1.042649102 | 0.468 | 0.676 | 7.66E-60 | 2.8E-55 |
| FOXN3 | 4 | -0.755805229 | 0.874 | 0.939 | 1.29E-59 | 4.71E-55 |
| MPP6 | 4 | -0.924381875 | 0.465 | 0.667 | 1.48E-59 | 5.42E-55 |
| AC125613.1 | 4 | 0.376452885 | 0.155 | 0.023 | 1.71E-59 | 6.24E-55 |
| ZBTB16 | 4 | 0.500815537 | 0.114 | 0.014 | 2.99E-59 | 1.09E-54 |
| BMP5 | 4 | 0.749083429 | 0.107 | 0.009 | 3.46E-59 | 1.27E-54 |
| PIP4K2A | 4 | -0.671393099 | 0.347 | 0.618 | 3.53E-59 | 1.29E-54 |
| PSD3 | 4 | -0.647911229 | 0.683 | 0.847 | 3.91E-59 | 1.43E-54 |
| ITGBL1 | 4 | -1.942365246 | 0.042 | 0.231 | 4.02E-59 | 1.47E-54 |
| GMDS-DT | 4 | 0.626500304 | 0.497 | 0.237 | 4.28E-59 | 1.57E-54 |
| NEK10 | 4 | 0.741280509 | 0.281 | 0.101 | 5.5E-59 | 2.01E-54 |
| SLC38A4 | 4 | -0.778485123 | 0.018 | 0.193 | 6.36E-59 | 2.33E-54 |
| MVB12B | 4 | 0.658197625 | 0.631 | 0.396 | 1.26E-58 | 4.62E-54 |
| TRPC4 | 4 | 0.469791693 | 0.135 | 0.015 | 1.57E-58 | 5.75E-54 |
| NAAA | 4 | -0.686086289 | 0.096 | 0.317 | 1.75E-58 | 6.42E-54 |
| TET1 | 4 | 0.592091491 | 0.656 | 0.382 | 1.92E-58 | 7.01E-54 |
| CDON | 4 | 0.758856682 | 0.534 | 0.309 | 1.98E-58 | 7.25E-54 |
| NTNG1 | 4 | -0.911660236 | 0.247 | 0.503 | 2.17E-58 | 7.93E-54 |
| MLLT3 | 4 | -0.617564501 | 0.628 | 0.84 | 3.22E-58 | 1.18E-53 |
| ITGB8 | 4 | 0.355779146 | 0.537 | 0.271 | 3.51E-58 | 1.28E-53 |
| MCM6 | 4 | -0.61560036 | 0.206 | 0.473 | 5.83E-58 | 2.13E-53 |
| DRC3 | 4 | 0.291629813 | 0.159 | 0.024 | 8.68E-58 | 3.18E-53 |
| STOX1 | 4 | 0.522983613 | 0.288 | 0.094 | 1.07E-57 | 3.92E-53 |
| PKIB | 4 | -0.667987075 | 0.161 | 0.418 | 1.12E-57 | 4.1E-53 |
| TMEM178B | 4 | -1.011824897 | 0.291 | 0.528 | 1.46E-57 | 5.34E-53 |
| PLXNC1 | 4 | 0.532339177 | 0.23 | 0.059 | 1.65E-57 | 6.03E-53 |
| ITPKB | 4 | 0.532635629 | 0.311 | 0.106 | 1.93E-57 | 7.07E-53 |
| IFT57 | 4 | 0.403741578 | 0.272 | 0.08 | 2.24E-57 | 8.21E-53 |
| AL139815.1 | 4 | 0.353975328 | 0.146 | 0.021 | 3.07E-57 | 1.13E-52 |
| CADPS | 4 | -1.030901419 | 0.229 | 0.493 | 4.77E-57 | 1.75E-52 |
| COL12A1 | 4 | 0.620209752 | 0.101 | 0.007 | 5.17E-57 | 1.89E-52 |
| CUX2 | 4 | 0.843826323 | 0.52 | 0.271 | 5.62E-57 | 2.06E-52 |
| RPS6KA2 | 4 | 0.744406634 | 0.548 | 0.3 | 5.93E-57 | 2.17E-52 |
| ARPP21 | 4 | -1.039502403 | 0.106 | 0.339 | 6.21E-57 | 2.27E-52 |

| | | | | | | |
|---|---|---|---|---|---|---|
| SEMA6D | 4 | -0.799100967 | 0.463 | 0.697 | 1.12E-56 | 4.1E-52 |
| RTN3 | 4 | -0.666544506 | 0.499 | 0.7 | 1.51E-56 | 5.52E-52 |
| AF117829.1 | 4 | 0.645633343 | 0.566 | 0.337 | 1.82E-56 | 6.67E-52 |
| BOC | 4 | 0.873396369 | 0.373 | 0.183 | 2.15E-56 | 7.89E-52 |
| ATP1A2 | 4 | 0.294742993 | 0.139 | 0.018 | 3.02E-56 | 1.11E-51 |
| UNC13B | 4 | -0.886436661 | 0.356 | 0.561 | 3.1E-56 | 1.14E-51 |
| SLC30A10 | 4 | 0.368511785 | 0.153 | 0.024 | 3.16E-56 | 1.16E-51 |
| RAB3C | 4 | -0.915182433 | 0.202 | 0.453 | 3.5E-56 | 1.28E-51 |
| STPG2-AS1 | 4 | 0.359488781 | 0.108 | 0.009 | 3.91E-56 | 1.43E-51 |
| KBTBD12 | 4 | -0.533563473 | 0.043 | 0.237 | 4.22E-56 | 1.55E-51 |
| AK8 | 4 | 0.355646133 | 0.174 | 0.032 | 5.97E-56 | 2.19E-51 |
| C8orf34 | 4 | 0.484873193 | 0.144 | 0.021 | 9.34E-56 | 3.42E-51 |
| TOX | 4 | -0.738633709 | 0.473 | 0.697 | 9.9E-56 | 3.62E-51 |
| SGK3 | 4 | 0.619022719 | 0.526 | 0.317 | 1.11E-55 | 4.06E-51 |
| STK32B | 4 | -0.835250303 | 0.298 | 0.551 | 1.14E-55 | 4.18E-51 |
| SLC16A12 | 4 | 0.537845091 | 0.188 | 0.038 | 1.23E-55 | 4.5E-51 |
| NELL1 | 4 | -0.804751782 | 0.104 | 0.336 | 1.4E-55 | 5.13E-51 |
| ANK3 | 4 | 0.803299181 | 0.563 | 0.302 | 1.64E-55 | 5.99E-51 |
| TTC25 | 4 | 0.3589914 | 0.167 | 0.029 | 2.15E-55 | 7.86E-51 |
| MCTP2 | 4 | -1.328715234 | 0.082 | 0.285 | 2.19E-55 | 8.03E-51 |
| TAFA1 | 4 | -1.578490556 | 0.056 | 0.246 | 2.35E-55 | 8.6E-51 |
| TCTN1 | 4 | 0.513379668 | 0.318 | 0.125 | 2.57E-55 | 9.39E-51 |
| MAP4K3 | 4 | -0.711342573 | 0.811 | 0.892 | 2.63E-55 | 9.62E-51 |
| LINGO2 | 4 | 0.818022535 | 0.229 | 0.059 | 2.88E-55 | 1.05E-50 |
| TMTC2 | 4 | 0.914086559 | 0.682 | 0.49 | 2.92E-55 | 1.07E-50 |
| IQCE | 4 | 0.445579311 | 0.256 | 0.076 | 3.43E-55 | 1.26E-50 |
| ZNF117 | 4 | 0.439482481 | 0.263 | 0.081 | 4.99E-55 | 1.83E-50 |
| KCNK2 | 4 | -0.83981804 | 0.091 | 0.312 | 5.65E-55 | 2.07E-50 |
| PLXNB1 | 4 | 0.522980876 | 0.333 | 0.127 | 7.93E-55 | 2.9E-50 |
| USP2-AS1 | 4 | 0.426185796 | 0.234 | 0.062 | 1.73E-54 | 6.32E-50 |
| NRG3 | 4 | -0.808474634 | 0.556 | 0.455 | 1.94E-54 | 7.12E-50 |
| MOB3B | 4 | 0.60557444 | 0.407 | 0.182 | 2.03E-54 | 7.43E-50 |
| NMU | 4 | -0.613727788 | 0.065 | 0.273 | 2.04E-54 | 7.46E-50 |
| B3GALT5 | 4 | 0.262095364 | 0.122 | 0.013 | 2.06E-54 | 7.53E-50 |
| ANKRD6 | 4 | 0.642175997 | 0.526 | 0.276 | 2.08E-54 | 7.63E-50 |
| DAB2 | 4 | 0.300576697 | 0.168 | 0.03 | 2.25E-54 | 8.23E-50 |
| SLC38A6 | 4 | 0.580150822 | 0.511 | 0.286 | 2.28E-54 | 8.34E-50 |
| RFX3 | 4 | 0.642595574 | 0.854 | 0.816 | 2.55E-54 | 9.34E-50 |
| PITX2 | 4 | 0.635519455 | 0.169 | 0.048 | 3.77E-54 | 1.38E-49 |
| FARS2 | 4 | 0.562041077 | 0.751 | 0.606 | 4.27E-54 | 1.56E-49 |
| ICA1L | 4 | 0.579640746 | 0.376 | 0.163 | 4.69E-54 | 1.72E-49 |
| DCLK1 | 4 | -0.964516707 | 0.409 | 0.618 | 6.27E-54 | 2.3E-49 |
| P3H2 | 4 | 0.53259225 | 0.215 | 0.052 | 6.51E-54 | 2.38E-49 |
| FTO | 4 | -0.979373646 | 0.728 | 0.821 | 7.45E-54 | 2.73E-49 |
| CACNA2D4 | 4 | 0.254935638 | 0.114 | 0.011 | 1.01E-53 | 3.69E-49 |
| PTPRG | 4 | 0.684546683 | 0.905 | 0.834 | 1.02E-53 | 3.73E-49 |
| EYA4 | 4 | 0.385081571 | 0.403 | 0.181 | 1.22E-53 | 4.47E-49 |
| DTNA | 4 | -1.168130737 | 0.544 | 0.714 | 1.23E-53 | 4.49E-49 |
| CCDC191 | 4 | 0.455728277 | 0.29 | 0.096 | 1.27E-53 | 4.64E-49 |
| Z93403.1 | 4 | 0.651622237 | 0.108 | 0.014 | 1.37E-53 | 5E-49 |
| VCL | 4 | -0.833139081 | 0.634 | 0.795 | 1.39E-53 | 5.1E-49 |
| PCCA-DT | 4 | 0.303929565 | 0.151 | 0.024 | 1.45E-53 | 5.3E-49 |
| MYEF2 | 4 | 0.567829849 | 0.673 | 0.488 | 1.61E-53 | 5.89E-49 |
| SKIL | 4 | -0.710114788 | 0.428 | 0.641 | 3E-53 | 1.1E-48 |
| PALMD | 4 | 0.349902131 | 0.175 | 0.034 | 3.09E-53 | 1.13E-48 |
| ATAD2 | 4 | -0.789032584 | 0.277 | 0.53 | 3.59E-53 | 1.31E-48 |
| AC017074.1 | 4 | 0.265259462 | 0.101 | 0.008 | 4.86E-53 | 1.78E-48 |
| PALLD | 4 | 0.802011308 | 0.629 | 0.39 | 4.87E-53 | 1.78E-48 |

| | | | | | | |
|---|---|---|---|---|---|---|
| TMEM168 | 4 | 0.380509206 | 0.251 | 0.073 | 4.89E-53 | 1.79E-48 |
| OSBPL8 | 4 | 0.630247111 | 0.737 | 0.614 | 6.11E-53 | 2.24E-48 |
| DLX6-AS1 | 4 | -1.074351776 | 0.05 | 0.242 | 6.6E-53 | 2.42E-48 |
| ADGRD1 | 4 | -0.581005266 | 0.068 | 0.276 | 6.8E-53 | 2.49E-48 |
| CAPS2 | 4 | 0.547028883 | 0.309 | 0.112 | 7.54E-53 | 2.76E-48 |
| ANKRD45 | 4 | 0.26404394 | 0.137 | 0.019 | 1E-52 | 3.67E-48 |
| COL1A1 | 4 | 0.971921085 | 0.201 | 0.065 | 1.19E-52 | 4.37E-48 |
| ELAVL2 | 4 | -0.851297107 | 0.173 | 0.407 | 1.5E-52 | 5.49E-48 |
| GRIA4 | 4 | -1.052355249 | 0.118 | 0.339 | 1.81E-52 | 6.62E-48 |
| DACH2 | 4 | -1.159703158 | 0.5 | 0.445 | 2.16E-52 | 7.92E-48 |
| BRIP1 | 4 | -0.705343419 | 0.328 | 0.588 | 2.52E-52 | 9.23E-48 |
| DPPA4 | 4 | -0.663086293 | 0.214 | 0.46 | 2.53E-52 | 9.27E-48 |
| ARHGAP39 | 4 | 0.430122825 | 0.312 | 0.111 | 2.55E-52 | 9.34E-48 |
| CDYL2 | 4 | 0.306202434 | 0.142 | 0.021 | 2.83E-52 | 1.04E-47 |
| SLC20A2 | 4 | 0.602435433 | 0.606 | 0.437 | 2.92E-52 | 1.07E-47 |
| AC090015.1 | 4 | 0.37128989 | 0.114 | 0.013 | 2.97E-52 | 1.09E-47 |
| PDLIM7 | 4 | 0.478205762 | 0.276 | 0.091 | 3.05E-52 | 1.12E-47 |
| SSBP3 | 4 | 0.722902903 | 0.694 | 0.538 | 3.38E-52 | 1.24E-47 |
| ASAP1 | 4 | -0.540809163 | 0.765 | 0.862 | 5.82E-52 | 2.13E-47 |
| CNTN1 | 4 | -0.888612841 | 0.072 | 0.275 | 5.89E-52 | 2.16E-47 |
| IRS2 | 4 | -0.724279953 | 0.192 | 0.42 | 9.2E-52 | 3.37E-47 |
| DNAL1 | 4 | 0.453482995 | 0.281 | 0.095 | 1.02E-51 | 3.73E-47 |
| KCNQ3 | 4 | -0.764408857 | 0.171 | 0.413 | 1.3E-51 | 4.75E-47 |
| ACSL1 | 4 | -0.726157693 | 0.199 | 0.406 | 1.34E-51 | 4.91E-47 |
| ODF2 | 4 | 0.567693854 | 0.363 | 0.162 | 1.78E-51 | 6.53E-47 |
| GALNTL6 | 4 | -1.153746072 | 0.068 | 0.27 | 1.83E-51 | 6.7E-47 |
| IFT74 | 4 | 0.546478203 | 0.37 | 0.166 | 2.69E-51 | 9.84E-47 |
| ACSS3 | 4 | 0.459641566 | 0.4 | 0.174 | 3.49E-51 | 1.28E-46 |
| ST8SIA5 | 4 | 0.28875922 | 0.131 | 0.018 | 3.66E-51 | 1.34E-46 |
| AC106798.1 | 4 | 0.440810887 | 0.141 | 0.022 | 3.97E-51 | 1.45E-46 |
| SYN2 | 4 | 0.350854796 | 0.129 | 0.017 | 4.83E-51 | 1.77E-46 |
| PGM5 | 4 | 0.578059535 | 0.166 | 0.032 | 6.04E-51 | 2.21E-46 |
| DCP2 | 4 | -0.666242206 | 0.388 | 0.599 | 6.18E-51 | 2.26E-46 |
| TOGARAM2 | 4 | 0.300519674 | 0.151 | 0.026 | 7.54E-51 | 2.76E-46 |
| MAP3K8 | 4 | 0.321740392 | 0.159 | 0.029 | 9.39E-51 | 3.44E-46 |
| NOTCH3 | 4 | 0.605031143 | 0.232 | 0.077 | 1.05E-50 | 3.84E-46 |
| TLN2 | 4 | 0.697999175 | 0.644 | 0.431 | 1.25E-50 | 4.58E-46 |
| LTBP1 | 4 | 0.874042719 | 0.51 | 0.293 | 1.51E-50 | 5.52E-46 |
| DDR2 | 4 | 0.570036768 | 0.405 | 0.183 | 1.77E-50 | 6.49E-46 |
| CLHC1 | 4 | 0.552026974 | 0.5 | 0.282 | 2.22E-50 | 8.12E-46 |
| IGSF21 | 4 | -0.61836158 | 0.084 | 0.283 | 2.27E-50 | 8.33E-46 |
| MALRD1 | 4 | -1.814443299 | 0.119 | 0.275 | 2.47E-50 | 9.03E-46 |
| CCDC141 | 4 | 0.308635675 | 0.109 | 0.011 | 2.56E-50 | 9.36E-46 |
| MID1 | 4 | 0.601051004 | 0.804 | 0.584 | 2.63E-50 | 9.64E-46 |
| KCNMB2 | 4 | -1.393771973 | 0.106 | 0.29 | 3.06E-50 | 1.12E-45 |
| SPAG17 | 4 | 0.848878646 | 0.379 | 0.281 | 3.08E-50 | 1.13E-45 |
| POU2F1 | 4 | -0.619960307 | 0.735 | 0.844 | 4.91E-50 | 1.8E-45 |
| LINC02197 | 4 | -0.774607049 | 0.025 | 0.187 | 5.55E-50 | 2.03E-45 |
| CASC15 | 4 | 0.864635545 | 0.738 | 0.554 | 5.8E-50 | 2.13E-45 |
| LRP4 | 4 | 0.586095635 | 0.246 | 0.091 | 6.32E-50 | 2.31E-45 |
| NR2F2 | 4 | 0.435959783 | 0.226 | 0.064 | 7.12E-50 | 2.61E-45 |
| AL162493.1 | 4 | 0.777594255 | 0.202 | 0.053 | 9.26E-50 | 3.39E-45 |
| F5 | 4 | 0.340738662 | 0.131 | 0.019 | 1.03E-49 | 3.77E-45 |
| SLC38A1 | 4 | -0.593730582 | 0.342 | 0.596 | 1.27E-49 | 4.64E-45 |
| TFAP2B | 4 | -0.679457919 | 0.062 | 0.257 | 1.4E-49 | 5.13E-45 |
| KCNJ6 | 4 | -0.802438794 | 0.044 | 0.22 | 1.49E-49 | 5.45E-45 |
| NETO2 | 4 | -0.658934752 | 0.314 | 0.557 | 1.49E-49 | 5.45E-45 |
| FREM2 | 4 | 0.533061879 | 0.207 | 0.053 | 1.76E-49 | 6.43E-45 |

| | | | | | | |
|---|---|---|---|---|---|---|
| HACE1 | 4 | -0.733298123 | 0.447 | 0.638 | 1.77E-49 | 6.48E-45 |
| MAP7D2 | 4 | -0.582261161 | 0.098 | 0.305 | 1.91E-49 | 7E-45 |
| TRIM22 | 4 | -0.628270299 | 0.105 | 0.306 | 1.97E-49 | 7.22E-45 |
| BCL2 | 4 | 0.685404679 | 0.386 | 0.173 | 2E-49 | 7.31E-45 |
| IQCG | 4 | 0.463068812 | 0.281 | 0.103 | 2.16E-49 | 7.91E-45 |
| AFF1 | 4 | 0.63059889 | 0.62 | 0.388 | 2.48E-49 | 9.07E-45 |
| GLYATL2 | 4 | -0.694366136 | 0.134 | 0.355 | 2.68E-49 | 9.82E-45 |
| FAM184B | 4 | -0.549483285 | 0.041 | 0.211 | 2.75E-49 | 1.01E-44 |
| LRCH2 | 4 | -0.693886424 | 0.558 | 0.713 | 2.83E-49 | 1.04E-44 |
| STRIP2 | 4 | 0.540730713 | 0.214 | 0.084 | 3.2E-49 | 1.17E-44 |
| CACNB2 | 4 | -0.771990839 | 0.534 | 0.755 | 3.25E-49 | 1.19E-44 |
| HS3ST4 | 4 | -1.398714447 | 0.187 | 0.357 | 3.28E-49 | 1.2E-44 |
| RBM41 | 4 | 0.546559776 | 0.612 | 0.417 | 5.99E-49 | 2.19E-44 |
| CDCP1 | 4 | -0.592393817 | 0.036 | 0.206 | 9.15E-49 | 3.35E-44 |
| STRA6 | 4 | 0.303824871 | 0.141 | 0.023 | 1.03E-48 | 3.76E-44 |
| RBPMS2 | 4 | 0.256842252 | 0.126 | 0.017 | 1.03E-48 | 3.79E-44 |
| LRRN2 | 4 | 0.399271914 | 0.244 | 0.074 | 1.2E-48 | 4.39E-44 |
| AK5 | 4 | -0.934596897 | 0.059 | 0.244 | 1.34E-48 | 4.91E-44 |
| MAML2 | 4 | 0.302238879 | 0.744 | 0.494 | 1.46E-48 | 5.33E-44 |
| PKDCC | 4 | 0.329284041 | 0.17 | 0.036 | 1.54E-48 | 5.65E-44 |
| TMEM132D | 4 | 0.859749917 | 0.138 | 0.023 | 1.76E-48 | 6.44E-44 |
| PTN | 4 | 0.794430404 | 0.212 | 0.075 | 1.85E-48 | 6.78E-44 |
| WDR19 | 4 | 0.492004688 | 0.46 | 0.23 | 1.89E-48 | 6.91E-44 |
| LHFPL2 | 4 | 0.600598507 | 0.422 | 0.213 | 1.92E-48 | 7.04E-44 |
| CLEC2A | 4 | -0.532275659 | 0.007 | 0.143 | 1.97E-48 | 7.23E-44 |
| LAMB1 | 4 | 0.741295687 | 0.534 | 0.371 | 1.99E-48 | 7.28E-44 |
| COLQ | 4 | 0.47720353 | 0.275 | 0.102 | 2.24E-48 | 8.21E-44 |
| FSIP2-AS1 | 4 | 0.539071721 | 0.302 | 0.115 | 2.39E-48 | 8.75E-44 |
| SSH2 | 4 | -0.621225313 | 0.697 | 0.814 | 2.43E-48 | 8.9E-44 |
| OSBPL3 | 4 | -0.783081838 | 0.298 | 0.476 | 2.48E-48 | 9.08E-44 |
| CLMN | 4 | 0.626310087 | 0.345 | 0.174 | 2.53E-48 | 9.28E-44 |
| GNAO1 | 4 | -0.714426778 | 0.251 | 0.481 | 2.88E-48 | 1.06E-43 |
| IFT88 | 4 | 0.590075827 | 0.565 | 0.364 | 3.45E-48 | 1.26E-43 |
| LINC01798 | 4 | 0.386421562 | 0.106 | 0.012 | 4.78E-48 | 1.75E-43 |
| MEF2C | 4 | -0.645553537 | 0.152 | 0.375 | 5.23E-48 | 1.92E-43 |
| COL2A1 | 4 | 0.463498114 | 0.186 | 0.044 | 5.96E-48 | 2.18E-43 |
| AC113383.1 | 4 | -0.564224725 | 0.041 | 0.215 | 6.04E-48 | 2.21E-43 |
| LRRC4B | 4 | 0.380067267 | 0.149 | 0.027 | 6.76E-48 | 2.48E-43 |
| PBX4 | 4 | 0.455789962 | 0.243 | 0.075 | 8.27E-48 | 3.03E-43 |
| PDLIM5 | 4 | 0.671771152 | 0.564 | 0.373 | 8.55E-48 | 3.13E-43 |
| TSHZ2 | 4 | -0.913038725 | 0.731 | 0.804 | 8.61E-48 | 3.15E-43 |
| CPNE8 | 4 | 0.50294079 | 0.346 | 0.141 | 1.14E-47 | 4.16E-43 |
| LAMA5 | 4 | 0.451227936 | 0.241 | 0.077 | 1.14E-47 | 4.18E-43 |
| ZDHHC15 | 4 | 0.344327719 | 0.197 | 0.049 | 1.33E-47 | 4.88E-43 |
| IQCJ-SCHIP1 | 4 | 1.691034838 | 0.248 | 0.164 | 1.37E-47 | 5.01E-43 |
| MAPK4 | 4 | 0.404040762 | 0.163 | 0.036 | 1.41E-47 | 5.17E-43 |
| NALCN | 4 | -0.772231713 | 0.062 | 0.243 | 1.54E-47 | 5.63E-43 |
| COL9A2 | 4 | 0.449327945 | 0.214 | 0.06 | 1.65E-47 | 6.06E-43 |
| PDGFRB | 4 | 0.493806015 | 0.275 | 0.094 | 1.77E-47 | 6.48E-43 |
| ASAP2 | 4 | 0.587678558 | 0.591 | 0.365 | 1.78E-47 | 6.52E-43 |
| PTCH1 | 4 | 0.724779486 | 0.326 | 0.166 | 2.01E-47 | 7.35E-43 |
| MYO16 | 4 | -0.940029048 | 0.127 | 0.34 | 2.38E-47 | 8.71E-43 |
| NEUROD1 | 4 | -0.638986231 | 0.033 | 0.198 | 5.15E-47 | 1.88E-42 |
| ANKIB1 | 4 | -0.658165181 | 0.717 | 0.816 | 7.56E-47 | 2.77E-42 |
| BASP1-AS1 | 4 | 0.426987909 | 0.193 | 0.049 | 8.29E-47 | 3.04E-42 |
| LATS2 | 4 | 0.412801729 | 0.243 | 0.076 | 8.53E-47 | 3.12E-42 |
| CPEB4 | 4 | 0.550900923 | 0.34 | 0.158 | 9.86E-47 | 3.61E-42 |
| LINC00632 | 4 | 0.661938335 | 0.56 | 0.364 | 1.03E-46 | 3.77E-42 |

| | | | | | | |
|---|---|---|---|---|---|---|
| RHOJ | 4 | 0.41493914 | 0.199 | 0.051 | 1.28E-46 | 4.68E-42 |
| KCNH1 | 4 | -1.038167066 | 0.096 | 0.288 | 1.71E-46 | 6.26E-42 |
| POU6F2 | 4 | 0.554698768 | 0.181 | 0.043 | 1.75E-46 | 6.4E-42 |
| AC135782.1 | 4 | -0.419841368 | 0.018 | 0.16 | 1.75E-46 | 6.41E-42 |
| RAD18 | 4 | -0.60623405 | 0.323 | 0.551 | 1.96E-46 | 7.18E-42 |
| CA10 | 4 | -0.702032377 | 0.124 | 0.34 | 2.3E-46 | 8.4E-42 |
| AC096570.1 | 4 | 0.60740318 | 0.103 | 0.013 | 2.44E-46 | 8.93E-42 |
| APP | 4 | -0.512230587 | 0.768 | 0.88 | 2.62E-46 | 9.59E-42 |
| USP54 | 4 | 0.579814681 | 0.486 | 0.285 | 3.47E-46 | 1.27E-41 |
| DYNC1I2 | 4 | -0.67598666 | 0.489 | 0.667 | 3.98E-46 | 1.46E-41 |
| LRRC7 | 4 | -1.471293139 | 0.163 | 0.332 | 4.24E-46 | 1.55E-41 |
| CACNA1E | 4 | 0.424927393 | 0.116 | 0.016 | 6.19E-46 | 2.27E-41 |
| MAP2K5 | 4 | 0.578127563 | 0.625 | 0.487 | 1.17E-45 | 4.28E-41 |
| ADAMTSL3 | 4 | -0.73021676 | 0.109 | 0.308 | 1.28E-45 | 4.7E-41 |
| CACNA2D3 | 4 | -0.985404109 | 0.327 | 0.54 | 1.37E-45 | 5.02E-41 |
| AC069228.1 | 4 | -0.439382275 | 0.033 | 0.195 | 1.48E-45 | 5.41E-41 |
| AL049828.1 | 4 | 0.715858526 | 0.41 | 0.205 | 1.54E-45 | 5.65E-41 |
| KIAA0556 | 4 | 0.48045006 | 0.39 | 0.191 | 1.58E-45 | 5.77E-41 |
| MYO1B | 4 | -0.528489734 | 0.381 | 0.626 | 1.62E-45 | 5.93E-41 |
| MINDY4 | 4 | 0.443929382 | 0.277 | 0.105 | 2.22E-45 | 8.12E-41 |
| TP53BP1 | 4 | 0.521813533 | 0.621 | 0.402 | 2.46E-45 | 8.99E-41 |
| CAMK4 | 4 | -0.736670689 | 0.121 | 0.318 | 2.63E-45 | 9.64E-41 |
| AL357054.2 | 4 | -0.541808559 | 0.053 | 0.223 | 3.78E-45 | 1.39E-40 |
| NOTCH2 | 4 | 0.479830124 | 0.519 | 0.285 | 3.85E-45 | 1.41E-40 |
| MAGI1 | 4 | 0.490962849 | 0.927 | 0.858 | 4.47E-45 | 1.64E-40 |
| FAM149A | 4 | -0.642452033 | 0.144 | 0.339 | 4.5E-45 | 1.65E-40 |
| DIP2C | 4 | 0.555232066 | 0.86 | 0.741 | 4.56E-45 | 1.67E-40 |
| STXBP5L | 4 | -0.99380859 | 0.171 | 0.323 | 5.21E-45 | 1.91E-40 |
| BNC2 | 4 | 1.183356086 | 0.301 | 0.186 | 8.84E-45 | 3.24E-40 |
| LRMDA | 4 | 0.66177221 | 0.274 | 0.097 | 9.71E-45 | 3.55E-40 |
| PARD3 | 4 | -0.496490585 | 0.953 | 0.967 | 1.03E-44 | 3.78E-40 |
| RNF220 | 4 | 0.810823105 | 0.172 | 0.07 | 1.12E-44 | 4.1E-40 |
| PTPRK | 4 | -0.839310987 | 0.735 | 0.817 | 1.29E-44 | 4.74E-40 |
| ARPC5 | 4 | -0.463406384 | 0.108 | 0.305 | 1.93E-44 | 7.08E-40 |
| EPCAM-DT | 4 | 0.531113901 | 0.219 | 0.08 | 2.59E-44 | 9.47E-40 |
| FREM1 | 4 | 1.246164371 | 0.42 | 0.242 | 2.68E-44 | 9.8E-40 |
| BRINP1 | 4 | -0.959436576 | 0.246 | 0.441 | 3E-44 | 1.1E-39 |
| IDS | 4 | 0.531405117 | 0.391 | 0.207 | 3.01E-44 | 1.1E-39 |
| OSBPL1A | 4 | 0.574750313 | 0.619 | 0.419 | 3.14E-44 | 1.15E-39 |
| CCDC39 | 4 | 0.400317575 | 0.187 | 0.052 | 3.78E-44 | 1.38E-39 |
| FOXO1 | 4 | 0.600157721 | 0.397 | 0.189 | 3.95E-44 | 1.45E-39 |
| FAM78B | 4 | -0.603675187 | 0.114 | 0.304 | 4.54E-44 | 1.66E-39 |
| DMD | 4 | -1.558335512 | 0.67 | 0.673 | 4.55E-44 | 1.67E-39 |
| SSX2IP | 4 | -0.647915376 | 0.337 | 0.53 | 4.84E-44 | 1.77E-39 |
| WEE1 | 4 | -0.592869329 | 0.228 | 0.454 | 5.06E-44 | 1.85E-39 |
| VCAN | 4 | 0.702196232 | 0.625 | 0.392 | 6.97E-44 | 2.55E-39 |
| PDE3A | 4 | -0.831170736 | 0.371 | 0.591 | 7.08E-44 | 2.59E-39 |
| SYNPO2 | 4 | -0.777384185 | 0.095 | 0.288 | 8.29E-44 | 3.03E-39 |
| DENND5B | 4 | 0.539655677 | 0.673 | 0.486 | 9.13E-44 | 3.34E-39 |
| RIN2 | 4 | 0.536833492 | 0.559 | 0.335 | 1.05E-43 | 3.86E-39 |
| MIAT | 4 | 0.628484899 | 0.216 | 0.094 | 1.1E-43 | 4.04E-39 |
| HIP1 | 4 | 0.670711627 | 0.544 | 0.356 | 1.28E-43 | 4.69E-39 |
| SRPX | 4 | -0.493912021 | 0.067 | 0.249 | 1.44E-43 | 5.29E-39 |
| INPP4B | 4 | 0.665040836 | 0.343 | 0.146 | 1.59E-43 | 5.81E-39 |
| NR6A1 | 4 | 0.683597304 | 0.522 | 0.354 | 1.64E-43 | 6.02E-39 |
| LINC01768 | 4 | -0.611484399 | 0.037 | 0.196 | 1.78E-43 | 6.53E-39 |
| CDCA7 | 4 | -0.552688097 | 0.217 | 0.437 | 1.82E-43 | 6.67E-39 |
| MED25 | 4 | 0.446377451 | 0.307 | 0.125 | 1.92E-43 | 7.04E-39 |

| | | | | | | |
|---|---|---|---|---|---|---|
| NPSR1-AS1 | 4 | -0.895004306 | 0.037 | 0.191 | 1.95E-43 | 7.14E-39 |
| HIPK2 | 4 | 0.556520931 | 0.645 | 0.433 | 2.13E-43 | 7.81E-39 |
| PIBF1 | 4 | 0.496127226 | 0.79 | 0.678 | 2.82E-43 | 1.03E-38 |
| C1orf61 | 4 | -0.55500494 | 0.119 | 0.324 | 3.05E-43 | 1.12E-38 |
| IQCH | 4 | 0.46201104 | 0.272 | 0.1 | 3.67E-43 | 1.34E-38 |
| SNX25 | 4 | -0.749875027 | 0.48 | 0.649 | 3.95E-43 | 1.44E-38 |
| SMC4 | 4 | -0.576376754 | 0.373 | 0.606 | 4.23E-43 | 1.55E-38 |
| AL512662.2 | 4 | 0.286001647 | 0.119 | 0.02 | 6.42E-43 | 2.35E-38 |
| PPP3CA | 4 | -0.549788521 | 0.747 | 0.85 | 6.61E-43 | 2.42E-38 |
| ZNF280D | 4 | 0.51447057 | 0.504 | 0.289 | 6.75E-43 | 2.47E-38 |
| ZNF586 | 4 | 0.522028634 | 0.413 | 0.216 | 1.01E-42 | 3.69E-38 |
| SAXO2 | 4 | 0.347803297 | 0.169 | 0.04 | 1.05E-42 | 3.83E-38 |
| CDH11 | 4 | 0.821834816 | 0.519 | 0.29 | 1.24E-42 | 4.53E-38 |
| ZSCAN5A | 4 | 0.489225558 | 0.423 | 0.22 | 1.34E-42 | 4.91E-38 |
| PTCHD1-AS | 4 | 0.845657873 | 0.271 | 0.101 | 1.42E-42 | 5.22E-38 |
| SHANK3 | 4 | 0.353986263 | 0.159 | 0.039 | 1.58E-42 | 5.77E-38 |
| SCUBE3 | 4 | 0.258915432 | 0.103 | 0.014 | 1.6E-42 | 5.87E-38 |
| FRMD5 | 4 | -1.174843603 | 0.274 | 0.423 | 1.68E-42 | 6.15E-38 |
| PRMT7 | 4 | 0.480948744 | 0.405 | 0.214 | 2.04E-42 | 7.46E-38 |
| SNX24 | 4 | 0.470035402 | 0.386 | 0.184 | 2.2E-42 | 8.05E-38 |
| SPATA6L | 4 | 0.451567958 | 0.237 | 0.088 | 2.29E-42 | 8.4E-38 |
| MYO3B | 4 | 0.480919928 | 0.187 | 0.052 | 2.4E-42 | 8.8E-38 |
| ETNK1 | 4 | -0.521674108 | 0.708 | 0.823 | 2.71E-42 | 9.93E-38 |
| IRAK1BP1 | 4 | 0.50536099 | 0.438 | 0.232 | 2.91E-42 | 1.06E-37 |
| RAPGEF4 | 4 | -0.926992427 | 0.245 | 0.42 | 3E-42 | 1.1E-37 |
| FBRSL1 | 4 | 0.476599658 | 0.426 | 0.221 | 3.04E-42 | 1.11E-37 |
| WDR76 | 4 | -0.5546249 | 0.156 | 0.369 | 3.07E-42 | 1.12E-37 |
| COL6A3 | 4 | 0.987592961 | 0.232 | 0.105 | 4.5E-42 | 1.65E-37 |
| HELLS | 4 | -0.584285727 | 0.44 | 0.653 | 4.65E-42 | 1.7E-37 |
| MEST | 4 | 0.383529259 | 0.297 | 0.117 | 4.69E-42 | 1.72E-37 |
| MAP9 | 4 | 0.430044356 | 0.326 | 0.139 | 4.72E-42 | 1.73E-37 |
| NOX4 | 4 | 0.506023883 | 0.158 | 0.036 | 5.16E-42 | 1.89E-37 |
| WDR60 | 4 | 0.512072884 | 0.578 | 0.388 | 5.33E-42 | 1.95E-37 |
| RABL2B | 4 | 0.334848181 | 0.232 | 0.076 | 5.96E-42 | 2.18E-37 |
| PDGFRA | 4 | 0.689329584 | 0.354 | 0.172 | 6.12E-42 | 2.24E-37 |
| GPC3 | 4 | 1.292974677 | 0.353 | 0.218 | 6.7E-42 | 2.45E-37 |
| DNM3OS | 4 | -0.741834067 | 0.154 | 0.362 | 7.2E-42 | 2.64E-37 |
| EHMT1 | 4 | 0.492465065 | 0.838 | 0.703 | 8E-42 | 2.93E-37 |
| PCYT1B | 4 | 0.316959357 | 0.152 | 0.034 | 8.52E-42 | 3.12E-37 |
| ROBO3 | 4 | 0.712584663 | 0.138 | 0.032 | 1E-41 | 3.66E-37 |
| CSMD2 | 4 | -0.717680067 | 0.135 | 0.327 | 1.06E-41 | 3.87E-37 |
| PLCH1 | 4 | 0.598432344 | 0.506 | 0.301 | 1.26E-41 | 4.6E-37 |
| SLIT2 | 4 | -1.39850441 | 0.692 | 0.724 | 1.27E-41 | 4.65E-37 |
| CYP4X1 | 4 | -0.569265976 | 0.024 | 0.164 | 1.78E-41 | 6.52E-37 |
| SLCO3A1 | 4 | 0.690287853 | 0.399 | 0.259 | 1.82E-41 | 6.68E-37 |
| ZNF91 | 4 | 0.385588359 | 0.44 | 0.226 | 2.02E-41 | 7.39E-37 |
| ARHGAP4 | 4 | -0.397429398 | 0.051 | 0.216 | 2.41E-41 | 8.81E-37 |
| PAK1 | 4 | -0.612805682 | 0.408 | 0.592 | 2.62E-41 | 9.61E-37 |
| SPSB4 | 4 | 0.440225995 | 0.272 | 0.104 | 2.85E-41 | 1.04E-36 |
| CNTNAP5 | 4 | -0.804214683 | 0.104 | 0.295 | 3.8E-41 | 1.39E-36 |
| MYO1D | 4 | 0.488374636 | 0.406 | 0.199 | 3.93E-41 | 1.44E-36 |
| MIR137HG | 4 | -0.818192119 | 0.085 | 0.269 | 4.23E-41 | 1.55E-36 |
| PPIL6 | 4 | 0.323020874 | 0.162 | 0.039 | 4.36E-41 | 1.6E-36 |
| PHC1 | 4 | 0.440093235 | 0.374 | 0.183 | 4.69E-41 | 1.72E-36 |
| IGFBP2 | 4 | -0.71602114 | 0.349 | 0.547 | 5.31E-41 | 1.94E-36 |
| TMTC1 | 4 | 0.726017841 | 0.291 | 0.119 | 6.59E-41 | 2.41E-36 |
| GRIN2A | 4 | 0.559745323 | 0.104 | 0.017 | 7.67E-41 | 2.81E-36 |
| TTN | 4 | 0.654244913 | 0.31 | 0.133 | 8.03E-41 | 2.94E-36 |

| | | | | | | |
|---|---|---|---|---|---|---|
| STK36 | 4 | 0.345745115 | 0.272 | 0.103 | 8.34E-41 | 3.05E-36 |
| SGO1-AS1 | 4 | 0.691012744 | 0.359 | 0.165 | 8.4E-41 | 3.08E-36 |
| NETO1 | 4 | -0.600567011 | 0.037 | 0.187 | 9.17E-41 | 3.36E-36 |
| RNF168 | 4 | -0.530795154 | 0.248 | 0.457 | 1.06E-40 | 3.9E-36 |
| CNKSR3 | 4 | 0.591981693 | 0.464 | 0.269 | 1.12E-40 | 4.11E-36 |
| RASGRP1 | 4 | -0.599132009 | 0.067 | 0.238 | 1.17E-40 | 4.3E-36 |
| LINC02263 | 4 | -0.680522507 | 0.005 | 0.118 | 1.24E-40 | 4.54E-36 |
| AFF2 | 4 | -0.933805321 | 0.52 | 0.66 | 1.3E-40 | 4.76E-36 |
| ARHGAP18 | 4 | -0.723170338 | 0.28 | 0.476 | 1.34E-40 | 4.92E-36 |
| THSD7B | 4 | -0.567976192 | 0.061 | 0.229 | 1.68E-40 | 6.14E-36 |
| PLAG1 | 4 | 0.399372865 | 0.251 | 0.089 | 1.81E-40 | 6.64E-36 |
| TEX41 | 4 | 0.574622308 | 0.197 | 0.058 | 2.46E-40 | 9.01E-36 |
| GFOD1 | 4 | -0.461448763 | 0.15 | 0.356 | 2.57E-40 | 9.4E-36 |
| ICA1 | 4 | -0.49079106 | 0.064 | 0.234 | 2.95E-40 | 1.08E-35 |
| RIPOR2 | 4 | -0.7280356 | 0.182 | 0.387 | 2.98E-40 | 1.09E-35 |
| CLYBL | 4 | 0.64387169 | 0.421 | 0.259 | 3.47E-40 | 1.27E-35 |
| LIN28B | 4 | -0.61044507 | 0.295 | 0.521 | 4.26E-40 | 1.56E-35 |
| PSME4 | 4 | -0.731584023 | 0.706 | 0.795 | 4.52E-40 | 1.65E-35 |
| URI1 | 4 | -0.556002702 | 0.631 | 0.765 | 5.96E-40 | 2.18E-35 |
| PPP3CC | 4 | 0.534362529 | 0.387 | 0.251 | 7.85E-40 | 2.87E-35 |
| MINAR1 | 4 | -0.423005713 | 0.035 | 0.179 | 1.01E-39 | 3.68E-35 |
| DOCK5 | 4 | -0.667005083 | 0.199 | 0.388 | 1.01E-39 | 3.71E-35 |
| AC013652.1 | 4 | 0.42098729 | 0.13 | 0.027 | 1.48E-39 | 5.42E-35 |
| MTMR8 | 4 | 0.320037566 | 0.187 | 0.053 | 1.78E-39 | 6.52E-35 |
| ID4 | 4 | 0.390082348 | 0.153 | 0.036 | 1.81E-39 | 6.62E-35 |
| ZNF717 | 4 | 0.267180458 | 0.199 | 0.064 | 1.83E-39 | 6.71E-35 |
| ZC3H12C | 4 | -0.629846151 | 0.408 | 0.611 | 1.94E-39 | 7.1E-35 |
| MEF2C-AS1 | 4 | -0.557986198 | 0.088 | 0.265 | 1.95E-39 | 7.12E-35 |
| OLA1 | 4 | -0.50527579 | 0.678 | 0.783 | 1.96E-39 | 7.19E-35 |
| COTL1 | 4 | 0.412975491 | 0.217 | 0.073 | 1.97E-39 | 7.2E-35 |
| FBN2 | 4 | 0.352376776 | 0.582 | 0.355 | 1.98E-39 | 7.24E-35 |
| COL23A1 | 4 | 0.665521271 | 0.156 | 0.045 | 2.06E-39 | 7.53E-35 |
| SIL1 | 4 | -0.688965815 | 0.535 | 0.678 | 2.41E-39 | 8.84E-35 |
| SLC12A7 | 4 | 0.26451608 | 0.142 | 0.03 | 2.54E-39 | 9.32E-35 |
| NCAN | 4 | -0.445794614 | 0.065 | 0.234 | 3.04E-39 | 1.11E-34 |
| PHF14 | 4 | -0.451533363 | 0.85 | 0.907 | 3.13E-39 | 1.15E-34 |
| COL21A1 | 4 | 0.53344145 | 0.204 | 0.065 | 3.24E-39 | 1.19E-34 |
| PIK3CB | 4 | 0.537114248 | 0.503 | 0.348 | 3.43E-39 | 1.26E-34 |
| NKAIN3 | 4 | -1.310165316 | 0.596 | 0.694 | 3.55E-39 | 1.3E-34 |
| MARCH2 | 4 | 0.305749021 | 0.188 | 0.054 | 3.75E-39 | 1.37E-34 |
| WHRN | 4 | 0.467609224 | 0.319 | 0.164 | 3.88E-39 | 1.42E-34 |
| PDE1C | 4 | -1.243795412 | 0.17 | 0.324 | 4E-39 | 1.46E-34 |
| DENND6B | 4 | 0.259444477 | 0.15 | 0.034 | 5.4E-39 | 1.98E-34 |
| EZH2 | 4 | -0.666872233 | 0.563 | 0.728 | 5.5E-39 | 2.01E-34 |
| MYBL1 | 4 | -0.528273391 | 0.088 | 0.263 | 5.83E-39 | 2.13E-34 |
| SH3D19 | 4 | 0.513831689 | 0.63 | 0.573 | 6.53E-39 | 2.39E-34 |
| PLCB1 | 4 | 0.681362981 | 0.669 | 0.508 | 7.61E-39 | 2.79E-34 |
| HS3ST3B1 | 4 | -0.703438928 | 0.135 | 0.317 | 8.02E-39 | 2.94E-34 |
| DDX17 | 4 | 0.37070244 | 0.929 | 0.851 | 8.04E-39 | 2.94E-34 |
| LINC00461 | 4 | -0.302524414 | 0.301 | 0.526 | 1.09E-38 | 4E-34 |
| RFTN2 | 4 | 0.414957901 | 0.143 | 0.033 | 1.15E-38 | 4.2E-34 |
| ITGB5 | 4 | 0.397514402 | 0.247 | 0.089 | 1.22E-38 | 4.48E-34 |
| ENOX1 | 4 | -0.670101862 | 0.342 | 0.554 | 1.29E-38 | 4.73E-34 |
| TTC21A | 4 | 0.300237968 | 0.166 | 0.043 | 1.37E-38 | 5.01E-34 |
| AASS | 4 | 0.478336658 | 0.452 | 0.244 | 1.73E-38 | 6.34E-34 |
| ATP10A | 4 | -0.56674537 | 0.114 | 0.303 | 1.74E-38 | 6.37E-34 |
| ARMCX4 | 4 | 0.489718077 | 0.457 | 0.272 | 2E-38 | 7.31E-34 |
| HUNK | 4 | 0.590333313 | 0.268 | 0.14 | 2.85E-38 | 1.04E-33 |

| | | | | | | |
|---|---|---|---|---|---|---|
| STPG2 | 4 | 0.545283467 | 0.223 | 0.093 | 3.05E-38 | 1.12E-33 |
| PGBD5 | 4 | -0.482766292 | 0.129 | 0.323 | 3.15E-38 | 1.15E-33 |
| SEMA3E | 4 | -0.847240222 | 0.141 | 0.318 | 4.76E-38 | 1.74E-33 |
| CAB39L | 4 | 0.427693198 | 0.261 | 0.104 | 4.77E-38 | 1.75E-33 |
| PLPPR4 | 4 | -0.581824028 | 0.114 | 0.303 | 5.71E-38 | 2.09E-33 |
| KCNN2 | 4 | -1.074361248 | 0.143 | 0.317 | 6.03E-38 | 2.21E-33 |
| ASTN2 | 4 | -0.646314712 | 0.339 | 0.534 | 6.07E-38 | 2.22E-33 |
| REST | 4 | 0.4656716 | 0.397 | 0.204 | 6.5E-38 | 2.38E-33 |
| TNS1 | 4 | 0.412350407 | 0.192 | 0.06 | 6.69E-38 | 2.45E-33 |
| ZFAND6 | 4 | -0.575612652 | 0.522 | 0.671 | 6.71E-38 | 2.46E-33 |
| NHSL1 | 4 | 0.556416329 | 0.745 | 0.559 | 7.23E-38 | 2.65E-33 |
| NKAIN2 | 4 | -1.425789993 | 0.219 | 0.394 | 8.33E-38 | 3.05E-33 |
| LINC01967 | 4 | 0.348175859 | 0.1 | 0.016 | 8.35E-38 | 3.06E-33 |
| RNF180 | 4 | 0.360470352 | 0.205 | 0.065 | 8.99E-38 | 3.29E-33 |
| GAB1 | 4 | -0.662854629 | 0.559 | 0.68 | 9.5E-38 | 3.48E-33 |
| AL596087.2 | 4 | -0.715950444 | 0.062 | 0.221 | 9.56E-38 | 3.5E-33 |
| HS3ST1 | 4 | -0.447275075 | 0.057 | 0.215 | 9.75E-38 | 3.57E-33 |
| ALPK1 | 4 | 0.322568082 | 0.19 | 0.056 | 9.98E-38 | 3.65E-33 |
| IGF2BP1 | 4 | 0.504796494 | 0.494 | 0.292 | 1E-37 | 3.67E-33 |
| AC040168.1 | 4 | -0.575670763 | 0.092 | 0.265 | 1.02E-37 | 3.74E-33 |
| AL353747.4 | 4 | -0.419695218 | 0.046 | 0.198 | 1.08E-37 | 3.96E-33 |
| FAM107B | 4 | 0.526927098 | 0.33 | 0.165 | 1.19E-37 | 4.37E-33 |
| GOPC | 4 | 0.376080103 | 0.287 | 0.118 | 1.25E-37 | 4.59E-33 |
| CTNNA3 | 4 | 0.876611889 | 0.331 | 0.164 | 1.43E-37 | 5.22E-33 |
| ARHGEF10 | 4 | 0.504036697 | 0.446 | 0.251 | 1.86E-37 | 6.8E-33 |
| LYPD6 | 4 | -0.629089394 | 0.177 | 0.369 | 1.92E-37 | 7.04E-33 |
| ITGA9 | 4 | 0.441976062 | 0.183 | 0.057 | 2.23E-37 | 8.15E-33 |
| MAMDC2 | 4 | 0.309290939 | 0.175 | 0.049 | 2.25E-37 | 8.24E-33 |
| SPARC | 4 | 0.303707579 | 0.169 | 0.045 | 2.36E-37 | 8.63E-33 |
| TIAM2 | 4 | -0.567930215 | 0.672 | 0.793 | 2.44E-37 | 8.93E-33 |
| ZNF385B | 4 | -0.533753705 | 0.154 | 0.352 | 2.61E-37 | 9.55E-33 |
| ACACB | 4 | 0.425184177 | 0.317 | 0.144 | 2.99E-37 | 1.09E-32 |
| HOTAIRM1 | 4 | 0.47708579 | 0.184 | 0.057 | 3.14E-37 | 1.15E-32 |
| CASC2 | 4 | 0.434857987 | 0.23 | 0.083 | 3.58E-37 | 1.31E-32 |
| MAGI2 | 4 | 0.639598699 | 0.732 | 0.696 | 3.59E-37 | 1.31E-32 |
| CABCOCO1 | 4 | 0.252729882 | 0.1 | 0.015 | 4.31E-37 | 1.58E-32 |
| LAMA4 | 4 | -0.775323308 | 0.261 | 0.477 | 4.33E-37 | 1.59E-32 |
| LMNB1 | 4 | -0.535880899 | 0.297 | 0.508 | 4.57E-37 | 1.67E-32 |
| SYTL3 | 4 | 0.321148377 | 0.138 | 0.031 | 4.68E-37 | 1.71E-32 |
| EPS8 | 4 | -0.672471774 | 0.417 | 0.607 | 4.9E-37 | 1.79E-32 |
| ASIC2 | 4 | -0.597072508 | 0.108 | 0.287 | 5.99E-37 | 2.19E-32 |
| KRAS | 4 | -0.709371524 | 0.212 | 0.39 | 7.08E-37 | 2.59E-32 |
| COCH | 4 | -0.489654222 | 0.085 | 0.251 | 7.54E-37 | 2.76E-32 |
| BCL11B | 4 | -0.600305109 | 0.244 | 0.445 | 9.97E-37 | 3.65E-32 |
| PDGFC | 4 | 0.480872976 | 0.447 | 0.242 | 1.15E-36 | 4.22E-32 |
| UNC5C | 4 | -0.75373363 | 0.334 | 0.551 | 1.35E-36 | 4.93E-32 |
| GRAMD2B | 4 | 0.47817073 | 0.308 | 0.14 | 1.36E-36 | 4.97E-32 |
| RGMA | 4 | 0.423474709 | 0.143 | 0.044 | 1.59E-36 | 5.81E-32 |
| GAS2L3 | 4 | -0.524421534 | 0.118 | 0.294 | 2.13E-36 | 7.78E-32 |
| AP000317.1 | 4 | 0.259438653 | 0.152 | 0.038 | 2.32E-36 | 8.48E-32 |
| ARHGAP24 | 4 | -0.896188511 | 0.34 | 0.419 | 2.5E-36 | 9.15E-32 |
| ABAT | 4 | 0.424271543 | 0.286 | 0.12 | 3.01E-36 | 1.1E-31 |
| CHSY1 | 4 | 0.514483548 | 0.487 | 0.313 | 3.07E-36 | 1.12E-31 |
| SP3 | 4 | -0.500906093 | 0.52 | 0.674 | 3.07E-36 | 1.12E-31 |
| GXYLT2 | 4 | 0.357471985 | 0.193 | 0.061 | 3.55E-36 | 1.3E-31 |
| DNAH3 | 4 | 0.412542686 | 0.104 | 0.021 | 3.82E-36 | 1.4E-31 |
| DTL | 4 | -0.613118994 | 0.206 | 0.407 | 4.31E-36 | 1.58E-31 |
| JAZF1 | 4 | -0.645086597 | 0.598 | 0.747 | 4.39E-36 | 1.61E-31 |

| | | | | | | |
|---|---|---|---|---|---|---|
| IGF2BP3 | 4 | -0.413331562 | 0.802 | 0.895 | 4.55E-36 | 1.67E-31 |
| GRIA1 | 4 | -0.651969065 | 0.188 | 0.39 | 4.91E-36 | 1.8E-31 |
| IL1RAPL2 | 4 | 0.786904681 | 0.347 | 0.168 | 5.07E-36 | 1.86E-31 |
| PON2 | 4 | 0.477203222 | 0.38 | 0.2 | 5.09E-36 | 1.86E-31 |
| EIF4G3 | 4 | 0.359831347 | 0.973 | 0.931 | 5.44E-36 | 1.99E-31 |
| LINC02398 | 4 | -1.001503398 | 0.043 | 0.178 | 5.76E-36 | 2.11E-31 |
| ZDHHC14 | 4 | -0.549739462 | 0.26 | 0.466 | 6.23E-36 | 2.28E-31 |
| RNF182 | 4 | 0.261907723 | 0.155 | 0.04 | 6.55E-36 | 2.4E-31 |
| DPP4 | 4 | -0.608074967 | 0.05 | 0.188 | 6.84E-36 | 2.5E-31 |
| SLC45A4 | 4 | 0.361810047 | 0.25 | 0.096 | 6.93E-36 | 2.54E-31 |
| NPSR1 | 4 | -0.988980669 | 0.045 | 0.188 | 7.24E-36 | 2.65E-31 |
| ZFP36L2 | 4 | 0.36516382 | 0.183 | 0.055 | 7.65E-36 | 2.8E-31 |
| CNTN3 | 4 | -0.927644725 | 0.343 | 0.512 | 9.38E-36 | 3.43E-31 |
| HS6ST2 | 4 | -0.581201279 | 0.503 | 0.685 | 9.67E-36 | 3.54E-31 |
| ATF7IP | 4 | 0.481744133 | 0.692 | 0.517 | 9.81E-36 | 3.59E-31 |
| LPCAT2 | 4 | 0.303483228 | 0.158 | 0.042 | 1.03E-35 | 3.77E-31 |
| AIM2 | 4 | 0.314978456 | 0.198 | 0.064 | 1.12E-35 | 4.1E-31 |
| EBF2 | 4 | 0.917383597 | 0.113 | 0.04 | 1.17E-35 | 4.28E-31 |
| PHF21B | 4 | 0.574362094 | 0.546 | 0.356 | 1.2E-35 | 4.38E-31 |
| SPPL2B | 4 | 0.37398233 | 0.338 | 0.161 | 1.23E-35 | 4.49E-31 |
| ADARB1 | 4 | 0.591571242 | 0.451 | 0.267 | 1.33E-35 | 4.87E-31 |
| ARHGAP6 | 4 | 0.882055201 | 0.525 | 0.34 | 1.46E-35 | 5.36E-31 |
| RORA | 4 | 0.545581198 | 0.881 | 0.788 | 1.51E-35 | 5.53E-31 |
| LURAP1L | 4 | -0.492450086 | 0.073 | 0.235 | 1.52E-35 | 5.56E-31 |
| MOCOS | 4 | -0.346091418 | 0.017 | 0.131 | 1.61E-35 | 5.91E-31 |
| C19orf18 | 4 | 0.252057817 | 0.125 | 0.027 | 1.69E-35 | 6.17E-31 |
| SLC23A2 | 4 | 0.496354048 | 0.436 | 0.319 | 1.71E-35 | 6.25E-31 |
| IFT43 | 4 | 0.403032482 | 0.374 | 0.188 | 1.81E-35 | 6.62E-31 |
| DTX4 | 4 | 0.30451113 | 0.149 | 0.039 | 1.84E-35 | 6.72E-31 |
| DYNC2LI1 | 4 | 0.397651202 | 0.359 | 0.176 | 1.88E-35 | 6.88E-31 |
| ITGA8 | 4 | -0.958322968 | 0.105 | 0.276 | 1.89E-35 | 6.92E-31 |
| COL6A2 | 4 | 0.556665605 | 0.253 | 0.125 | 2.15E-35 | 7.85E-31 |
| MDM1 | 4 | 0.392259244 | 0.286 | 0.124 | 2.31E-35 | 8.47E-31 |
| ANK1 | 4 | -0.559387432 | 0.06 | 0.21 | 2.47E-35 | 9.05E-31 |
| NCALD | 4 | -0.786592496 | 0.442 | 0.581 | 2.64E-35 | 9.66E-31 |
| MLF1 | 4 | 0.332241944 | 0.231 | 0.084 | 2.95E-35 | 1.08E-30 |
| MIS18BP1 | 4 | -0.38822263 | 0.331 | 0.547 | 3.34E-35 | 1.22E-30 |
| GUCY1A1 | 4 | -0.395934228 | 0.105 | 0.277 | 3.38E-35 | 1.24E-30 |
| DCN | 4 | 0.702352913 | 0.157 | 0.062 | 3.4E-35 | 1.24E-30 |
| EYS | 4 | -1.854431253 | 0.111 | 0.262 | 3.43E-35 | 1.26E-30 |
| LINC00310 | 4 | 0.273925525 | 0.166 | 0.046 | 4.61E-35 | 1.69E-30 |
| AC024901.1 | 4 | -0.83211988 | 0.054 | 0.194 | 4.77E-35 | 1.74E-30 |
| UNC79 | 4 | -0.753557202 | 0.183 | 0.362 | 5.02E-35 | 1.84E-30 |
| PPM1E | 4 | -0.419491531 | 0.565 | 0.755 | 5.68E-35 | 2.08E-30 |
| TIAM1 | 4 | -0.549867626 | 0.421 | 0.63 | 5.82E-35 | 2.13E-30 |
| GPATCH2 | 4 | 0.492730376 | 0.59 | 0.421 | 6.28E-35 | 2.3E-30 |
| EGLN3 | 4 | -0.519327127 | 0.1 | 0.265 | 7.81E-35 | 2.86E-30 |
| SHTN1 | 4 | -0.422677826 | 0.501 | 0.707 | 8.21E-35 | 3.01E-30 |
| SLC4A7 | 4 | -0.519847301 | 0.634 | 0.77 | 8.44E-35 | 3.09E-30 |
| TTC12 | 4 | 0.331494747 | 0.22 | 0.081 | 9.85E-35 | 3.61E-30 |
| PRKD1 | 4 | -0.676904269 | 0.521 | 0.677 | 1.03E-34 | 3.78E-30 |
| FAAH2 | 4 | 0.426385867 | 0.207 | 0.072 | 1.04E-34 | 3.8E-30 |
| CYFIP2 | 4 | 0.366984734 | 0.287 | 0.124 | 1.14E-34 | 4.16E-30 |
| SRSF5 | 4 | 0.433000988 | 0.628 | 0.488 | 1.34E-34 | 4.91E-30 |
| IFT140 | 4 | 0.427659209 | 0.33 | 0.165 | 1.44E-34 | 5.27E-30 |
| LANCL3 | 4 | -0.542523422 | 0.087 | 0.242 | 1.59E-34 | 5.84E-30 |
| SLC1A2 | 4 | -0.509613861 | 0.122 | 0.301 | 1.99E-34 | 7.27E-30 |
| ZNRF1 | 4 | 0.478373982 | 0.584 | 0.398 | 2.02E-34 | 7.4E-30 |

| | | | | | | |
|---|---|---|---|---|---|---|
| RAB11FIP1 | 4 | 0.287035675 | 0.166 | 0.047 | 2.42E-34 | 8.86E-30 |
| SETBP1 | 4 | 0.635718069 | 0.698 | 0.566 | 2.51E-34 | 9.19E-30 |
| POFUT2 | 4 | 0.257455171 | 0.208 | 0.072 | 2.7E-34 | 9.89E-30 |
| MME | 4 | -0.531172992 | 0.056 | 0.204 | 2.83E-34 | 1.04E-29 |
| COL25A1 | 4 | -0.806262599 | 0.216 | 0.411 | 2.9E-34 | 1.06E-29 |
| AC010148.1 | 4 | 0.254644884 | 0.104 | 0.019 | 3.09E-34 | 1.13E-29 |
| WSCD1 | 4 | 0.302809375 | 0.125 | 0.028 | 3.21E-34 | 1.17E-29 |
| ARHGEF9 | 4 | 0.429978758 | 0.431 | 0.242 | 3.3E-34 | 1.21E-29 |
| AC119673.2 | 4 | 0.486812637 | 0.524 | 0.37 | 3.49E-34 | 1.28E-29 |
| CNTN6 | 4 | -0.913094358 | 0.052 | 0.193 | 3.69E-34 | 1.35E-29 |
| SGCD | 4 | -0.775601243 | 0.319 | 0.506 | 5.41E-34 | 1.98E-29 |
| LCORL | 4 | -0.485448565 | 0.645 | 0.774 | 5.71E-34 | 2.09E-29 |
| AC016042.1 | 4 | -0.508950065 | 0.096 | 0.251 | 6.01E-34 | 2.2E-29 |
| AC092691.1 | 4 | 0.985241217 | 0.193 | 0.084 | 6.42E-34 | 2.35E-29 |
| CDRT4 | 4 | 0.425257123 | 0.361 | 0.184 | 6.46E-34 | 2.36E-29 |
| MAP4K4 | 4 | -0.434823467 | 0.76 | 0.865 | 6.87E-34 | 2.51E-29 |
| GALNT14 | 4 | -0.466220173 | 0.097 | 0.266 | 7.15E-34 | 2.62E-29 |
| PTPRZ1 | 4 | -0.697882939 | 0.229 | 0.428 | 7.3E-34 | 2.67E-29 |
| CHKA | 4 | 0.46804728 | 0.511 | 0.319 | 7.31E-34 | 2.68E-29 |
| BAZ2B | 4 | -0.464512083 | 0.889 | 0.919 | 7.33E-34 | 2.68E-29 |
| LCA5 | 4 | 0.418542355 | 0.262 | 0.114 | 7.83E-34 | 2.87E-29 |
| GCH1 | 4 | -0.429860679 | 0.046 | 0.184 | 7.93E-34 | 2.9E-29 |
| DGKB | 4 | -0.924591105 | 0.135 | 0.303 | 8.32E-34 | 3.05E-29 |
| GPR173 | 4 | 0.484270207 | 0.657 | 0.482 | 8.41E-34 | 3.08E-29 |
| ADAMTS12 | 4 | -1.086763982 | 0.373 | 0.402 | 8.8E-34 | 3.22E-29 |
| ARHGAP10 | 4 | -0.605164843 | 0.335 | 0.532 | 9.49E-34 | 3.48E-29 |
| CCDC57 | 4 | 0.493110363 | 0.578 | 0.408 | 9.91E-34 | 3.63E-29 |
| MAP3K15 | 4 | -0.546356193 | 0.051 | 0.184 | 1.09E-33 | 3.97E-29 |
| EPHA3 | 4 | 1.127719584 | 0.333 | 0.202 | 1.19E-33 | 4.36E-29 |
| LINC00630 | 4 | 0.448165953 | 0.497 | 0.309 | 1.19E-33 | 4.37E-29 |
| E2F3 | 4 | -0.478065303 | 0.423 | 0.619 | 1.77E-33 | 6.47E-29 |
| ENPEP | 4 | 0.295614532 | 0.151 | 0.04 | 1.77E-33 | 6.5E-29 |
| NFIA | 4 | -0.549920163 | 0.779 | 0.907 | 1.8E-33 | 6.59E-29 |
| POLR3G | 4 | -0.507671383 | 0.123 | 0.281 | 1.86E-33 | 6.81E-29 |
| AHNAK | 4 | 0.375972677 | 0.183 | 0.06 | 1.98E-33 | 7.26E-29 |
| SLIT1 | 4 | 0.501593338 | 0.13 | 0.109 | 2.01E-33 | 7.38E-29 |
| PRR5 | 4 | 0.257643257 | 0.143 | 0.036 | 2.05E-33 | 7.51E-29 |
| ADAM22 | 4 | -0.534320494 | 0.233 | 0.427 | 2.16E-33 | 7.91E-29 |
| MBNL3 | 4 | -0.627242672 | 0.323 | 0.509 | 2.19E-33 | 8.01E-29 |
| WDR27 | 4 | 0.481121614 | 0.611 | 0.454 | 2.75E-33 | 1.01E-28 |
| FXYD6 | 4 | 0.325464151 | 0.177 | 0.059 | 2.89E-33 | 1.06E-28 |
| DAPK2 | 4 | -0.423923567 | 0.077 | 0.234 | 2.98E-33 | 1.09E-28 |
| NWD2 | 4 | -0.487028455 | 0.029 | 0.154 | 3.01E-33 | 1.1E-28 |
| CDH7 | 4 | -0.79748416 | 0.131 | 0.304 | 3.23E-33 | 1.18E-28 |
| ITSN1 | 4 | 0.492112456 | 0.737 | 0.589 | 3.28E-33 | 1.2E-28 |
| NBAS | 4 | -0.67974516 | 0.725 | 0.807 | 3.5E-33 | 1.28E-28 |
| GABRA2 | 4 | -0.717637296 | 0.012 | 0.116 | 5.11E-33 | 1.87E-28 |
| NLRC5 | 4 | -0.388566068 | 0.041 | 0.174 | 5.17E-33 | 1.89E-28 |
| SNAP25 | 4 | -0.491988205 | 0.138 | 0.315 | 5.34E-33 | 1.96E-28 |
| SNED1 | 4 | 0.455326991 | 0.236 | 0.092 | 5.73E-33 | 2.1E-28 |
| FRMPD3 | 4 | 0.398748943 | 0.241 | 0.099 | 6.68E-33 | 2.45E-28 |
| PIEZO2 | 4 | -0.400323918 | 0.202 | 0.396 | 6.86E-33 | 2.51E-28 |
| AKAP12 | 4 | 0.619369184 | 0.343 | 0.183 | 7.17E-33 | 2.63E-28 |
| ASPH | 4 | -0.765483634 | 0.59 | 0.714 | 9.01E-33 | 3.3E-28 |
| EXOC6B | 4 | 0.431121998 | 0.828 | 0.677 | 9.09E-33 | 3.33E-28 |
| AK9 | 4 | 0.418618188 | 0.288 | 0.131 | 9.75E-33 | 3.57E-28 |
| ZMYND8 | 4 | 0.50824909 | 0.721 | 0.561 | 9.91E-33 | 3.63E-28 |
| ATL3 | 4 | -0.454464965 | 0.16 | 0.326 | 1.05E-32 | 3.85E-28 |

| | | | | | | |
|---|---|---|---|---|---|---|
| KCTD2 | 4 | 0.319147161 | 0.176 | 0.057 | 1.2E-32 | 4.4E-28 |
| POTEF | 4 | 0.406701418 | 0.399 | 0.214 | 1.24E-32 | 4.55E-28 |
| CNNM1 | 4 | -0.285192924 | 0.022 | 0.14 | 1.34E-32 | 4.91E-28 |
| ADPRHL2 | 4 | 0.34658623 | 0.205 | 0.074 | 1.42E-32 | 5.18E-28 |
| PLS1 | 4 | -0.450571837 | 0.073 | 0.222 | 1.79E-32 | 6.54E-28 |
| ZEB1 | 4 | 0.604315254 | 0.726 | 0.541 | 1.83E-32 | 6.71E-28 |
| CBY1 | 4 | 0.341055597 | 0.253 | 0.104 | 1.9E-32 | 6.94E-28 |
| RGS6 | 4 | 0.69880582 | 0.103 | 0.027 | 2.05E-32 | 7.5E-28 |
| ADAMTS19 | 4 | -0.604226058 | 0.304 | 0.507 | 2.05E-32 | 7.52E-28 |
| PKIA | 4 | -0.60102532 | 0.286 | 0.465 | 2.24E-32 | 8.2E-28 |
| AC023480.1 | 4 | 0.336763079 | 0.155 | 0.044 | 2.28E-32 | 8.34E-28 |
| RND3 | 4 | -0.618219052 | 0.17 | 0.352 | 2.43E-32 | 8.91E-28 |
| SNX10 | 4 | -0.347412363 | 0.043 | 0.177 | 2.78E-32 | 1.02E-27 |
| PTPN21 | 4 | 0.336341776 | 0.28 | 0.122 | 2.82E-32 | 1.03E-27 |
| RBMS2 | 4 | 0.385098083 | 0.298 | 0.137 | 2.95E-32 | 1.08E-27 |
| GLIS3 | 4 | 0.671250044 | 0.405 | 0.228 | 3.87E-32 | 1.42E-27 |
| FAM126A | 4 | -0.59088016 | 0.46 | 0.62 | 4.33E-32 | 1.59E-27 |
| GALNT7 | 4 | -0.551920328 | 0.444 | 0.612 | 4.78E-32 | 1.75E-27 |
| SYNE1 | 4 | 0.634828627 | 0.55 | 0.466 | 4.88E-32 | 1.79E-27 |
| SLC24A3 | 4 | 0.480089635 | 0.12 | 0.035 | 5.15E-32 | 1.88E-27 |
| AKAP8L | 4 | 0.443579699 | 0.497 | 0.31 | 5.52E-32 | 2.02E-27 |
| FRAS1 | 4 | 0.616237009 | 0.545 | 0.357 | 5.8E-32 | 2.12E-27 |
| KCTD16 | 4 | 0.577355815 | 0.312 | 0.147 | 6.03E-32 | 2.21E-27 |
| ACAD11 | 4 | 0.442785293 | 0.422 | 0.245 | 6.3E-32 | 2.31E-27 |
| TTN-AS1 | 4 | 0.460804814 | 0.61 | 0.432 | 6.66E-32 | 2.44E-27 |
| CASTOR3 | 4 | 0.502109851 | 0.454 | 0.275 | 7.84E-32 | 2.87E-27 |
| INSR | 4 | 0.42572774 | 0.425 | 0.236 | 7.93E-32 | 2.9E-27 |
| HCN1 | 4 | -0.650933111 | 0.041 | 0.169 | 8.97E-32 | 3.28E-27 |
| KCNK1 | 4 | 0.320012207 | 0.207 | 0.075 | 1E-31 | 3.66E-27 |
| GAD1 | 4 | 0.348195328 | 0.109 | 0.023 | 1.67E-31 | 6.13E-27 |
| EDA | 4 | 0.555132398 | 0.45 | 0.262 | 1.7E-31 | 6.24E-27 |
| UVRAG | 4 | 0.420238447 | 0.662 | 0.594 | 1.92E-31 | 7.05E-27 |
| ANTXR1 | 4 | 0.410713664 | 0.637 | 0.449 | 2.06E-31 | 7.56E-27 |
| NBPF14 | 4 | 0.292154562 | 0.204 | 0.073 | 2.12E-31 | 7.76E-27 |
| CTNNA2 | 4 | -0.455756688 | 0.315 | 0.511 | 2.13E-31 | 7.81E-27 |
| MAGI3 | 4 | 0.487101904 | 0.623 | 0.431 | 2.32E-31 | 8.5E-27 |
| TMEM47 | 4 | 0.268940883 | 0.142 | 0.038 | 2.47E-31 | 9.06E-27 |
| LRRC49 | 4 | 0.462068435 | 0.371 | 0.209 | 2.57E-31 | 9.4E-27 |
| GLDC | 4 | -0.555061385 | 0.13 | 0.294 | 2.58E-31 | 9.45E-27 |
| ANKRD36 | 4 | -0.484978576 | 0.768 | 0.814 | 2.96E-31 | 1.08E-26 |
| TRPC1 | 4 | -0.58516656 | 0.417 | 0.559 | 3.14E-31 | 1.15E-26 |
| KCNG3 | 4 | -0.363425492 | 0.045 | 0.178 | 3.2E-31 | 1.17E-26 |
| HIF3A | 4 | 0.42303704 | 0.259 | 0.111 | 3.47E-31 | 1.27E-26 |
| ARHGEF4 | 4 | 0.532620003 | 0.363 | 0.217 | 3.68E-31 | 1.35E-26 |
| CBFA2T3 | 4 | -0.376284076 | 0.047 | 0.18 | 3.76E-31 | 1.38E-26 |
| RNF213-AS1 | 4 | 0.407181001 | 0.342 | 0.184 | 3.77E-31 | 1.38E-26 |
| FERMT1 | 4 | -0.508861104 | 0.075 | 0.213 | 3.97E-31 | 1.45E-26 |
| WDFY3 | 4 | -0.406023772 | 0.847 | 0.903 | 4.4E-31 | 1.61E-26 |
| GNG2 | 4 | 0.299360783 | 0.299 | 0.145 | 5.01E-31 | 1.83E-26 |
| ATP8A1 | 4 | -0.594001513 | 0.246 | 0.429 | 5.06E-31 | 1.85E-26 |
| BAZ1A | 4 | -0.451977323 | 0.429 | 0.616 | 5.97E-31 | 2.19E-26 |
| B3GNTL1 | 4 | 0.296904287 | 0.264 | 0.114 | 6.09E-31 | 2.23E-26 |
| RAB3IP | 4 | 0.363759203 | 0.277 | 0.124 | 6.16E-31 | 2.26E-26 |
| CDC14B | 4 | 0.451337118 | 0.405 | 0.222 | 6.45E-31 | 2.36E-26 |
| RHPN2 | 4 | 0.331802392 | 0.194 | 0.069 | 6.68E-31 | 2.44E-26 |
| FER | 4 | -0.540705915 | 0.753 | 0.822 | 6.7E-31 | 2.45E-26 |
| MAP3K1 | 4 | 0.329327086 | 0.361 | 0.188 | 6.76E-31 | 2.48E-26 |
| RALGPS2 | 4 | 0.432619117 | 0.545 | 0.347 | 7.17E-31 | 2.63E-26 |

| | | | | | | |
|---|---|---|---|---|---|---|
| SNTG1 | 4 | -1.415617965 | 0.285 | 0.397 | 8.74E-31 | 3.2E-26 |
| CIT | 4 | -0.442677627 | 0.165 | 0.345 | 9.27E-31 | 3.39E-26 |
| METTL24 | 4 | 0.440033994 | 0.226 | 0.09 | 1.08E-30 | 3.95E-26 |
| RFPL1S | 4 | -0.390103139 | 0.09 | 0.241 | 1.13E-30 | 4.15E-26 |
| MIPEP | 4 | 0.460190078 | 0.41 | 0.246 | 1.19E-30 | 4.37E-26 |
| AC087500.1 | 4 | 0.306679186 | 0.194 | 0.069 | 1.33E-30 | 4.88E-26 |
| SYT17 | 4 | 0.31096515 | 0.248 | 0.104 | 1.42E-30 | 5.2E-26 |
| ASCC3 | 4 | -0.462938471 | 0.685 | 0.786 | 1.74E-30 | 6.38E-26 |
| DPP10 | 4 | -0.3872459 | 0.471 | 0.621 | 1.8E-30 | 6.6E-26 |
| THSD4 | 4 | 0.71036909 | 0.284 | 0.148 | 1.96E-30 | 7.17E-26 |
| MPP7 | 4 | 0.262740196 | 0.124 | 0.03 | 2.04E-30 | 7.46E-26 |
| PATJ | 4 | 0.572569818 | 0.547 | 0.417 | 2.08E-30 | 7.61E-26 |
| AL162718.1 | 4 | -0.5060985 | 0.031 | 0.149 | 2.18E-30 | 7.97E-26 |
| TCF7 | 4 | 0.27586934 | 0.176 | 0.058 | 2.72E-30 | 9.97E-26 |
| TTC26 | 4 | 0.352251192 | 0.278 | 0.126 | 2.83E-30 | 1.04E-25 |
| LRRC8B | 4 | -0.554217523 | 0.263 | 0.445 | 2.85E-30 | 1.05E-25 |
| TVP23A | 4 | 0.312786667 | 0.15 | 0.045 | 2.91E-30 | 1.07E-25 |
| C5orf17 | 4 | 0.482888345 | 0.22 | 0.087 | 3.84E-30 | 1.41E-25 |
| COL3A1 | 4 | 1.077426578 | 0.264 | 0.277 | 4.35E-30 | 1.59E-25 |
| UPRT | 4 | 0.311191027 | 0.212 | 0.081 | 4.42E-30 | 1.62E-25 |
| CSNK1A1 | 4 | -0.437226194 | 0.763 | 0.854 | 4.56E-30 | 1.67E-25 |
| AC011287.1 | 4 | 0.796848338 | 0.14 | 0.039 | 4.72E-30 | 1.73E-25 |
| GPC6 | 4 | -0.562725072 | 0.802 | 0.818 | 5.75E-30 | 2.11E-25 |
| RNF128 | 4 | -0.467697871 | 0.034 | 0.152 | 6E-30 | 2.2E-25 |
| ADAMTS16 | 4 | -0.336851241 | 0.122 | 0.28 | 6.45E-30 | 2.36E-25 |
| ECHDC2 | 4 | -0.43229457 | 0.251 | 0.433 | 6.76E-30 | 2.48E-25 |
| SHISA5 | 4 | 0.341651513 | 0.177 | 0.062 | 6.86E-30 | 2.51E-25 |
| FBN1 | 4 | 0.401479348 | 0.332 | 0.167 | 1.02E-29 | 3.73E-25 |
| BTBD9 | 4 | 0.424688375 | 0.72 | 0.553 | 1.12E-29 | 4.09E-25 |
| KLHL4 | 4 | 0.467024945 | 0.132 | 0.036 | 1.14E-29 | 4.16E-25 |
| VRK1 | 4 | -0.488812187 | 0.345 | 0.529 | 1.15E-29 | 4.21E-25 |
| GPR158 | 4 | -0.581108622 | 0.038 | 0.153 | 1.27E-29 | 4.64E-25 |
| SEMA3C | 4 | 0.862549654 | 0.246 | 0.108 | 1.39E-29 | 5.09E-25 |
| TLE4 | 4 | 0.576330664 | 0.623 | 0.468 | 1.5E-29 | 5.48E-25 |
| TCF4 | 4 | -0.438893221 | 0.897 | 0.889 | 1.62E-29 | 5.91E-25 |
| LAMA1 | 4 | 0.486774078 | 0.156 | 0.062 | 1.67E-29 | 6.12E-25 |
| LINC01619 | 4 | -0.537171867 | 0.178 | 0.351 | 1.7E-29 | 6.22E-25 |
| RAP2A | 4 | -0.497982646 | 0.172 | 0.322 | 1.74E-29 | 6.38E-25 |
| TTC28 | 4 | -0.36553791 | 0.944 | 0.966 | 1.75E-29 | 6.41E-25 |
| APBA1 | 4 | -0.748532788 | 0.318 | 0.454 | 1.82E-29 | 6.67E-25 |
| SLC17A8 | 4 | -0.850716371 | 0.161 | 0.303 | 1.83E-29 | 6.7E-25 |
| DLG5 | 4 | 0.445611267 | 0.52 | 0.368 | 1.91E-29 | 6.98E-25 |
| FAM160A1 | 4 | 0.387515171 | 0.285 | 0.132 | 1.91E-29 | 6.99E-25 |
| PTGFRN | 4 | -0.458427108 | 0.182 | 0.344 | 2.2E-29 | 8.05E-25 |
| APBA2 | 4 | -0.425183445 | 0.399 | 0.592 | 2.33E-29 | 8.55E-25 |
| ATP8A2 | 4 | -0.602761939 | 0.175 | 0.329 | 2.45E-29 | 8.98E-25 |
| ALMS1 | 4 | 0.450238539 | 0.632 | 0.477 | 2.45E-29 | 8.99E-25 |
| PCGF5 | 4 | 0.325291773 | 0.197 | 0.073 | 2.69E-29 | 9.84E-25 |
| CERS4 | 4 | 0.403641121 | 0.249 | 0.111 | 2.79E-29 | 1.02E-24 |
| EPHA5 | 4 | -0.860019087 | 0.122 | 0.274 | 3.1E-29 | 1.13E-24 |
| FNDC5 | 4 | 0.322893083 | 0.189 | 0.068 | 3.11E-29 | 1.14E-24 |
| PRKDC | 4 | -0.411873037 | 0.775 | 0.835 | 3.35E-29 | 1.22E-24 |
| LRRN1 | 4 | 0.54441743 | 0.325 | 0.181 | 3.66E-29 | 1.34E-24 |
| POTEI | 4 | 0.334020604 | 0.238 | 0.102 | 3.67E-29 | 1.34E-24 |
| TLE1 | 4 | 0.563645931 | 0.401 | 0.24 | 4.02E-29 | 1.47E-24 |
| SNRNP70 | 4 | 0.431380995 | 0.626 | 0.469 | 4.18E-29 | 1.53E-24 |
| MEGF11 | 4 | -0.469865642 | 0.163 | 0.334 | 4.49E-29 | 1.64E-24 |
| SVEP1 | 4 | -0.655347348 | 0.147 | 0.31 | 5.03E-29 | 1.84E-24 |

| | | | | | | |
|---|---|---|---|---|---|---|
| CHRM3 | 4 | -1.019703663 | 0.292 | 0.433 | 5.59E-29 | 2.05E-24 |
| CDH12 | 4 | -0.977490748 | 0.261 | 0.415 | 5.61E-29 | 2.06E-24 |
| AL354771.1 | 4 | -0.360031572 | 0.033 | 0.15 | 5.94E-29 | 2.18E-24 |
| TJP2 | 4 | -0.504180881 | 0.241 | 0.41 | 6.66E-29 | 2.44E-24 |
| GRIP2 | 4 | -0.324489324 | 0.048 | 0.176 | 6.77E-29 | 2.48E-24 |
| DGKI | 4 | 0.518185353 | 0.393 | 0.218 | 6.82E-29 | 2.5E-24 |
| CARMIL3 | 4 | -0.302605255 | 0.038 | 0.149 | 6.93E-29 | 2.54E-24 |
| MAMDC2-AS1 | 4 | 0.25440877 | 0.145 | 0.043 | 8.4E-29 | 3.08E-24 |
| FAT4 | 4 | 0.560174345 | 0.454 | 0.279 | 8.43E-29 | 3.09E-24 |
| NCAPG2 | 4 | -0.456674769 | 0.205 | 0.386 | 8.47E-29 | 3.1E-24 |
| ATP2A2 | 4 | -0.522565581 | 0.581 | 0.687 | 9E-29 | 3.29E-24 |
| AC108169.1 | 4 | -0.452375383 | 0.032 | 0.147 | 9.35E-29 | 3.42E-24 |
| CDC14A | 4 | 0.419716347 | 0.466 | 0.29 | 1.04E-28 | 3.81E-24 |
| DPYD | 4 | -0.471736387 | 0.243 | 0.428 | 1.11E-28 | 4.05E-24 |
| TSPOAP1 | 4 | 0.259470206 | 0.218 | 0.087 | 1.25E-28 | 4.59E-24 |
| C3orf67 | 4 | 0.631143306 | 0.325 | 0.244 | 1.31E-28 | 4.8E-24 |
| HLA-A | 4 | -0.411853565 | 0.099 | 0.244 | 1.39E-28 | 5.1E-24 |
| TRAF3IP1 | 4 | 0.426799501 | 0.35 | 0.2 | 1.56E-28 | 5.71E-24 |
| LPIN2 | 4 | 0.42695841 | 0.442 | 0.274 | 1.68E-28 | 6.13E-24 |
| PCSK6 | 4 | 0.367941169 | 0.132 | 0.04 | 1.75E-28 | 6.41E-24 |
| CASC4 | 4 | 0.425583755 | 0.623 | 0.473 | 1.81E-28 | 6.63E-24 |
| BBS4 | 4 | 0.326094792 | 0.347 | 0.182 | 1.84E-28 | 6.74E-24 |
| DNMT3A | 4 | 0.444825529 | 0.488 | 0.331 | 2.32E-28 | 8.5E-24 |
| LINC00271 | 4 | 0.407956601 | 0.259 | 0.119 | 2.39E-28 | 8.76E-24 |
| PLPP3 | 4 | 0.308948923 | 0.268 | 0.122 | 2.41E-28 | 8.81E-24 |
| NME7 | 4 | 0.41581302 | 0.54 | 0.377 | 2.56E-28 | 9.38E-24 |
| AC107223.1 | 4 | 0.50472876 | 0.266 | 0.136 | 2.81E-28 | 1.03E-23 |
| GNG4 | 4 | -0.499082246 | 0.176 | 0.34 | 2.95E-28 | 1.08E-23 |
| PDE7B | 4 | -0.936409874 | 0.126 | 0.227 | 3.06E-28 | 1.12E-23 |
| AL691420.1 | 4 | -0.915240591 | 0.036 | 0.146 | 3.48E-28 | 1.27E-23 |
| TTYH1 | 4 | 0.324417676 | 0.157 | 0.057 | 3.78E-28 | 1.38E-23 |
| SGMS2 | 4 | 0.265314418 | 0.182 | 0.065 | 3.8E-28 | 1.39E-23 |
| NFATC1 | 4 | 0.272522901 | 0.156 | 0.05 | 4.1E-28 | 1.5E-23 |
| SOX13 | 4 | 0.409772685 | 0.269 | 0.135 | 5.34E-28 | 1.96E-23 |
| VWC2 | 4 | -0.459218631 | 0.035 | 0.151 | 5.51E-28 | 2.02E-23 |
| FBXO36 | 4 | 0.362588452 | 0.249 | 0.114 | 5.71E-28 | 2.09E-23 |
| KDM2B | 4 | 0.477471047 | 0.463 | 0.319 | 6.3E-28 | 2.31E-23 |
| TVP23C | 4 | 0.381986011 | 0.463 | 0.288 | 6.43E-28 | 2.35E-23 |
| CARMN | 4 | 0.425912705 | 0.108 | 0.027 | 6.54E-28 | 2.4E-23 |
| SLC25A48 | 4 | 0.393590185 | 0.168 | 0.058 | 6.86E-28 | 2.51E-23 |
| AC011246.1 | 4 | -0.567734176 | 0.209 | 0.389 | 6.98E-28 | 2.55E-23 |
| CCNL2 | 4 | 0.410454215 | 0.627 | 0.472 | 9.92E-28 | 3.63E-23 |
| ZNF254 | 4 | 0.431320586 | 0.538 | 0.363 | 1.06E-27 | 3.87E-23 |
| TPT1-AS1 | 4 | 0.425236017 | 0.561 | 0.384 | 1.17E-27 | 4.29E-23 |
| FAM111B | 4 | -0.315983661 | 0.047 | 0.17 | 1.23E-27 | 4.52E-23 |
| ATG10 | 4 | -0.507951864 | 0.444 | 0.597 | 1.25E-27 | 4.58E-23 |
| EXO1 | 4 | -0.367754213 | 0.093 | 0.24 | 1.27E-27 | 4.66E-23 |
| CELF2 | 4 | 0.554464875 | 0.681 | 0.613 | 1.39E-27 | 5.1E-23 |
| DYNC1I1 | 4 | -0.57216236 | 0.194 | 0.363 | 1.44E-27 | 5.28E-23 |
| WDPCP | 4 | 0.442669692 | 0.703 | 0.563 | 1.54E-27 | 5.62E-23 |
| CYTH2 | 4 | 0.291776349 | 0.218 | 0.091 | 1.87E-27 | 6.83E-23 |
| LINC02614 | 4 | 0.330650966 | 0.27 | 0.126 | 2.07E-27 | 7.59E-23 |
| FGD5 | 4 | -0.313638274 | 0.067 | 0.202 | 2.16E-27 | 7.9E-23 |
| NCOA2 | 4 | 0.428380411 | 0.764 | 0.628 | 2.24E-27 | 8.2E-23 |
| IFT122 | 4 | 0.332295907 | 0.303 | 0.151 | 2.3E-27 | 8.4E-23 |
| VGLL4 | 4 | 0.405304243 | 0.729 | 0.571 | 2.39E-27 | 8.76E-23 |
| SLC35B3 | 4 | -0.476240253 | 0.251 | 0.406 | 2.54E-27 | 9.32E-23 |
| ANKRD33B | 4 | -0.414678128 | 0.054 | 0.179 | 2.64E-27 | 9.67E-23 |

| | | | | | | |
|---|---|---|---|---|---|---|
| PDE8A | 4 | -0.490130485 | 0.532 | 0.664 | 3.09E-27 | 1.13E-22 |
| KIAA1522 | 4 | 0.259167041 | 0.169 | 0.058 | 3.17E-27 | 1.16E-22 |
| PRRX1 | 4 | 0.666481177 | 0.157 | 0.097 | 3.35E-27 | 1.23E-22 |
| CTXND1 | 4 | -0.34322796 | 0.026 | 0.129 | 3.38E-27 | 1.24E-22 |
| C14orf39 | 4 | 0.302160744 | 0.168 | 0.06 | 3.45E-27 | 1.26E-22 |
| SMC2 | 4 | -0.435856573 | 0.185 | 0.352 | 3.77E-27 | 1.38E-22 |
| SESN1 | 4 | -0.532111009 | 0.349 | 0.504 | 3.84E-27 | 1.41E-22 |
| HECTD1 | 4 | -0.466483698 | 0.756 | 0.806 | 3.85E-27 | 1.41E-22 |
| AC061958.1 | 4 | -0.370392989 | 0.163 | 0.328 | 3.86E-27 | 1.41E-22 |
| CMIP | 4 | 0.483223073 | 0.702 | 0.572 | 3.98E-27 | 1.46E-22 |
| ADAMTS10 | 4 | 0.322253626 | 0.203 | 0.085 | 4.34E-27 | 1.59E-22 |
| TBC1D23 | 4 | -0.502523562 | 0.361 | 0.504 | 4.45E-27 | 1.63E-22 |
| AC006230.1 | 4 | -0.299823066 | 0.038 | 0.154 | 4.47E-27 | 1.63E-22 |
| TEAD2 | 4 | 0.317938896 | 0.17 | 0.065 | 4.61E-27 | 1.69E-22 |
| TRABD2B | 4 | 0.481255188 | 0.22 | 0.101 | 4.66E-27 | 1.71E-22 |
| AL355499.1 | 4 | -0.902267481 | 0.018 | 0.111 | 4.93E-27 | 1.8E-22 |
| NEIL3 | 4 | -0.412944635 | 0.056 | 0.176 | 5.07E-27 | 1.86E-22 |
| KIAA0825 | 4 | 0.530769503 | 0.354 | 0.211 | 5.29E-27 | 1.94E-22 |
| LMNA | 4 | 0.403537728 | 0.403 | 0.245 | 6.07E-27 | 2.22E-22 |
| LMLN | 4 | 0.25465591 | 0.22 | 0.092 | 6.29E-27 | 2.3E-22 |
| ALDH1L2 | 4 | -0.362499183 | 0.062 | 0.19 | 6.5E-27 | 2.38E-22 |
| CDH23 | 4 | -0.530706263 | 0.15 | 0.287 | 6.64E-27 | 2.43E-22 |
| RASA3 | 4 | 0.35411486 | 0.189 | 0.081 | 6.74E-27 | 2.47E-22 |
| FAP | 4 | -0.872427021 | 0.111 | 0.23 | 7.36E-27 | 2.69E-22 |
| C4orf19 | 4 | -0.572186474 | 0.238 | 0.313 | 7.52E-27 | 2.75E-22 |
| SHOX2 | 4 | -0.334107393 | 0.064 | 0.193 | 8.04E-27 | 2.94E-22 |
| LINC00571 | 4 | 0.308000834 | 0.189 | 0.071 | 8.12E-27 | 2.97E-22 |
| MYO5C | 4 | 0.310509203 | 0.234 | 0.101 | 8.31E-27 | 3.04E-22 |
| CRPPA | 4 | 0.507990708 | 0.36 | 0.219 | 8.34E-27 | 3.05E-22 |
| LHFPL6 | 4 | 0.546726881 | 0.547 | 0.366 | 8.71E-27 | 3.19E-22 |
| RANBP10 | 4 | 0.406979574 | 0.404 | 0.245 | 8.97E-27 | 3.28E-22 |
| UBE2G1 | 4 | -0.464324727 | 0.49 | 0.628 | 9.99E-27 | 3.66E-22 |
| GLB1L3 | 4 | -0.266454337 | 0.013 | 0.101 | 1.07E-26 | 3.91E-22 |
| RSPH3 | 4 | 0.300523842 | 0.228 | 0.098 | 1.24E-26 | 4.55E-22 |
| ARID3A | 4 | 0.478469602 | 0.254 | 0.14 | 1.25E-26 | 4.57E-22 |
| RAB11FIP3 | 4 | 0.472551455 | 0.527 | 0.387 | 1.38E-26 | 5.07E-22 |
| ZFAND5 | 4 | -0.532767128 | 0.342 | 0.46 | 1.49E-26 | 5.44E-22 |
| CEP89 | 4 | 0.371200432 | 0.297 | 0.153 | 1.57E-26 | 5.75E-22 |
| NCKAP5 | 4 | 0.644340359 | 0.56 | 0.393 | 1.58E-26 | 5.79E-22 |
| KCNQ1OT1 | 4 | 0.603382459 | 0.677 | 0.587 | 1.64E-26 | 6.02E-22 |
| BACH2 | 4 | 0.595149198 | 0.46 | 0.377 | 1.76E-26 | 6.45E-22 |
| ATXN2 | 4 | 0.387581483 | 0.832 | 0.731 | 1.79E-26 | 6.57E-22 |
| CSRNP3 | 4 | 0.429882608 | 0.525 | 0.342 | 1.86E-26 | 6.81E-22 |
| MBD6 | 4 | 0.268393345 | 0.215 | 0.089 | 1.88E-26 | 6.88E-22 |
| DAB2IP | 4 | 0.403404936 | 0.325 | 0.194 | 2.15E-26 | 7.87E-22 |
| MAML3 | 4 | 0.532523505 | 0.722 | 0.555 | 2.32E-26 | 8.48E-22 |
| SYT14 | 4 | -0.62323702 | 0.373 | 0.486 | 2.4E-26 | 8.78E-22 |
| PTBP2 | 4 | -0.285242334 | 0.864 | 0.903 | 2.82E-26 | 1.03E-21 |
| LBX1-AS1 | 4 | -0.279719803 | 0.041 | 0.156 | 3.06E-26 | 1.12E-21 |
| SPATA7 | 4 | 0.355607801 | 0.349 | 0.191 | 3.07E-26 | 1.12E-21 |
| LRRTM4 | 4 | -0.74486346 | 0.581 | 0.576 | 3.15E-26 | 1.15E-21 |
| MEGF10 | 4 | -0.611482585 | 0.257 | 0.417 | 3.68E-26 | 1.35E-21 |
| L3MBTL3 | 4 | 0.348700081 | 0.293 | 0.146 | 4.25E-26 | 1.56E-21 |
| B2M | 4 | -0.405426925 | 0.108 | 0.25 | 4.29E-26 | 1.57E-21 |
| ACADVL | 4 | 0.288505736 | 0.237 | 0.105 | 4.3E-26 | 1.58E-21 |
| SCD5 | 4 | 0.461401774 | 0.373 | 0.231 | 4.45E-26 | 1.63E-21 |
| H2AFY2 | 4 | 0.358082902 | 0.233 | 0.12 | 5.7E-26 | 2.09E-21 |
| TTLL3 | 4 | 0.338284767 | 0.347 | 0.19 | 6.47E-26 | 2.37E-21 |

| | | | | | | |
|---|---|---|---|---|---|---|
| PRMT2 | 4 | 0.379970213 | 0.485 | 0.314 | 6.59E-26 | 2.41E-21 |
| PRICKLE1 | 4 | -0.394601148 | 0.252 | 0.429 | 6.74E-26 | 2.47E-21 |
| ARID5B | 4 | 0.506744891 | 0.561 | 0.395 | 7.8E-26 | 2.86E-21 |
| CC2D2A | 4 | 0.387039793 | 0.404 | 0.248 | 7.82E-26 | 2.86E-21 |
| SLC8A2 | 4 | -0.331732439 | 0.048 | 0.167 | 8.69E-26 | 3.18E-21 |
| XACT | 4 | 0.752496891 | 0.4 | 0.232 | 8.91E-26 | 3.26E-21 |
| IDI2-AS1 | 4 | -0.43043351 | 0.045 | 0.146 | 9.32E-26 | 3.41E-21 |
| R3HDM1 | 4 | -0.440682499 | 0.821 | 0.873 | 9.48E-26 | 3.47E-21 |
| AC068051.1 | 4 | 0.388660992 | 0.162 | 0.057 | 9.79E-26 | 3.58E-21 |
| MYO6 | 4 | 0.451041681 | 0.422 | 0.263 | 1.02E-25 | 3.74E-21 |
| TCF12 | 4 | 0.395967274 | 0.941 | 0.919 | 1.13E-25 | 4.13E-21 |
| CNNM2 | 4 | 0.47328188 | 0.432 | 0.286 | 1.2E-25 | 4.4E-21 |
| ERICH3 | 4 | -0.444671082 | 0.043 | 0.151 | 1.23E-25 | 4.49E-21 |
| TNFRSF19 | 4 | 0.624674809 | 0.355 | 0.227 | 1.23E-25 | 4.52E-21 |
| LUC7L2 | 4 | -0.372945185 | 0.676 | 0.775 | 1.25E-25 | 4.57E-21 |
| TMEM132C | 4 | 0.832257523 | 0.153 | 0.118 | 1.47E-25 | 5.38E-21 |
| CDC7 | 4 | -0.344362797 | 0.197 | 0.365 | 1.52E-25 | 5.58E-21 |
| AC093817.2 | 4 | -0.715617993 | 0.021 | 0.114 | 1.67E-25 | 6.1E-21 |
| PIP5K1B | 4 | 0.722254304 | 0.368 | 0.269 | 1.68E-25 | 6.14E-21 |
| LIN7A | 4 | -0.495991331 | 0.222 | 0.379 | 1.84E-25 | 6.72E-21 |
| CCDC102B | 4 | 1.222098118 | 0.24 | 0.127 | 1.86E-25 | 6.81E-21 |
| FAM114A1 | 4 | 0.318563534 | 0.197 | 0.08 | 1.94E-25 | 7.09E-21 |
| LRATD1 | 4 | -0.27902502 | 0.035 | 0.143 | 1.98E-25 | 7.24E-21 |
| PTPRO | 4 | -0.63489841 | 0.295 | 0.459 | 2.25E-25 | 8.23E-21 |
| ADD3 | 4 | 0.444594392 | 0.592 | 0.426 | 2.38E-25 | 8.7E-21 |
| OSBPL6 | 4 | -0.464296674 | 0.307 | 0.479 | 2.39E-25 | 8.74E-21 |
| GOLIM4 | 4 | -0.449125033 | 0.399 | 0.528 | 2.62E-25 | 9.59E-21 |
| LARGE1 | 4 | 0.607276937 | 0.345 | 0.267 | 2.8E-25 | 1.02E-20 |
| APCDD1 | 4 | -0.406235561 | 0.166 | 0.325 | 2.8E-25 | 1.03E-20 |
| IQCA1 | 4 | 0.251885013 | 0.221 | 0.097 | 3.3E-25 | 1.21E-20 |
| AC007091.1 | 4 | 0.266490526 | 0.132 | 0.041 | 3.92E-25 | 1.43E-20 |
| IGF1R | 4 | 0.340777139 | 0.902 | 0.784 | 4.18E-25 | 1.53E-20 |
| CCDC81 | 4 | 0.311187515 | 0.184 | 0.071 | 4.55E-25 | 1.67E-20 |
| CEP290 | 4 | 0.423826198 | 0.515 | 0.348 | 4.65E-25 | 1.7E-20 |
| DNAH11 | 4 | 0.653783401 | 0.372 | 0.244 | 4.69E-25 | 1.72E-20 |
| SCP2 | 4 | -0.494565909 | 0.262 | 0.381 | 4.79E-25 | 1.76E-20 |
| TIA1 | 4 | 0.379766052 | 0.738 | 0.596 | 4.98E-25 | 1.82E-20 |
| SUSD4 | 4 | -0.475916562 | 0.132 | 0.281 | 5.39E-25 | 1.97E-20 |
| GSE1 | 4 | 0.532033851 | 0.801 | 0.721 | 5.68E-25 | 2.08E-20 |
| INPP5A | 4 | 0.370505845 | 0.335 | 0.188 | 5.85E-25 | 2.14E-20 |
| LINC01473 | 4 | 0.412714976 | 0.716 | 0.553 | 5.9E-25 | 2.16E-20 |
| PPP2R3A | 4 | 0.36710953 | 0.509 | 0.334 | 6.03E-25 | 2.21E-20 |
| MFSD14C | 4 | 0.375930378 | 0.614 | 0.462 | 6.05E-25 | 2.22E-20 |
| MCMBP | 4 | -0.400263219 | 0.288 | 0.456 | 6.16E-25 | 2.25E-20 |
| STIM1 | 4 | 0.381393616 | 0.405 | 0.243 | 6.29E-25 | 2.3E-20 |
| KIAA1958 | 4 | -0.453336865 | 0.673 | 0.774 | 6.36E-25 | 2.33E-20 |
| KDM5B | 4 | 0.291682272 | 0.574 | 0.393 | 6.6E-25 | 2.42E-20 |
| CLSPN | 4 | -0.391593721 | 0.129 | 0.279 | 6.88E-25 | 2.52E-20 |
| ARGLU1 | 4 | 0.336125247 | 0.902 | 0.818 | 8.33E-25 | 3.05E-20 |
| RBM6 | 4 | 0.328332401 | 0.92 | 0.84 | 8.58E-25 | 3.14E-20 |
| GOLGA8B | 4 | 0.409108197 | 0.604 | 0.46 | 9.29E-25 | 3.4E-20 |
| PHIP | 4 | -0.351765347 | 0.795 | 0.882 | 9.77E-25 | 3.58E-20 |
| TBC1D16 | 4 | 0.445889081 | 0.408 | 0.259 | 9.96E-25 | 3.65E-20 |
| ATAD5 | 4 | -0.344724164 | 0.364 | 0.545 | 1.15E-24 | 4.21E-20 |
| CCNE2 | 4 | -0.325445203 | 0.054 | 0.173 | 1.16E-24 | 4.25E-20 |
| RILPL1 | 4 | 0.405288778 | 0.348 | 0.206 | 1.18E-24 | 4.31E-20 |
| MYO5A | 4 | -0.4627069 | 0.471 | 0.605 | 1.21E-24 | 4.41E-20 |
| ST3GAL6 | 4 | -0.408320891 | 0.149 | 0.302 | 1.29E-24 | 4.73E-20 |

| | | | | | | |
|---|---|---|---|---|---|---|
| SLC10A4 | 4 | -0.265167859 | 0.018 | 0.107 | 1.3E-24 | 4.75E-20 |
| COQ4 | 4 | 0.356875103 | 0.293 | 0.153 | 1.39E-24 | 5.08E-20 |
| STK3 | 4 | 0.409928912 | 0.652 | 0.477 | 1.39E-24 | 5.1E-20 |
| ABCA5 | 4 | -0.55357641 | 0.271 | 0.401 | 1.42E-24 | 5.21E-20 |
| RNF175 | 4 | 0.333730351 | 0.173 | 0.07 | 1.48E-24 | 5.41E-20 |
| STX18-AS1 | 4 | 0.407307392 | 0.53 | 0.364 | 1.49E-24 | 5.44E-20 |
| GAS7 | 4 | 0.36727443 | 0.384 | 0.223 | 1.49E-24 | 5.45E-20 |
| PXDNL | 4 | 0.475949924 | 0.16 | 0.071 | 1.53E-24 | 5.6E-20 |
| ADAMTS17 | 4 | 0.507897888 | 0.222 | 0.111 | 2.11E-24 | 7.71E-20 |
| ST7 | 4 | -0.402610713 | 0.312 | 0.482 | 2.17E-24 | 7.93E-20 |
| TOPBP1 | 4 | -0.372681668 | 0.397 | 0.564 | 2.46E-24 | 9E-20 |
| KLHL2 | 4 | -0.374720988 | 0.167 | 0.323 | 2.46E-24 | 9.01E-20 |
| ARHGEF17 | 4 | 0.341087353 | 0.218 | 0.1 | 2.84E-24 | 1.04E-19 |
| FCHO2 | 4 | 0.388596953 | 0.522 | 0.358 | 3.1E-24 | 1.13E-19 |
| CNOT6L | 4 | -0.450782494 | 0.405 | 0.521 | 3.12E-24 | 1.14E-19 |
| FAM110B | 4 | 0.576069812 | 0.528 | 0.385 | 3.19E-24 | 1.17E-19 |
| NT5C2 | 4 | 0.405437637 | 0.652 | 0.532 | 3.49E-24 | 1.28E-19 |
| NCAPG | 4 | -0.415640416 | 0.138 | 0.288 | 3.68E-24 | 1.35E-19 |
| MTHFD1 | 4 | -0.380869118 | 0.196 | 0.351 | 4.01E-24 | 1.47E-19 |
| WASF3 | 4 | 0.295419432 | 0.304 | 0.16 | 4.33E-24 | 1.58E-19 |
| LINC01414 | 4 | 0.989222172 | 0.129 | 0.078 | 4.35E-24 | 1.59E-19 |
| CACNA1B | 4 | -0.499408489 | 0.131 | 0.279 | 4.67E-24 | 1.71E-19 |
| ZFHX4-AS1 | 4 | 0.536808432 | 0.113 | 0.05 | 5.09E-24 | 1.86E-19 |
| PMP22 | 4 | -0.385826599 | 0.08 | 0.207 | 6.45E-24 | 2.36E-19 |
| TECPR2 | 4 | 0.383195976 | 0.418 | 0.268 | 6.54E-24 | 2.39E-19 |
| USP3 | 4 | -0.413773204 | 0.6 | 0.706 | 6.7E-24 | 2.45E-19 |
| AL138720.1 | 4 | -0.464061766 | 0.057 | 0.174 | 6.74E-24 | 2.47E-19 |
| C7orf50 | 4 | 0.352376808 | 0.379 | 0.225 | 8.44E-24 | 3.09E-19 |
| ARL6IP6 | 4 | -0.424191124 | 0.356 | 0.505 | 8.57E-24 | 3.14E-19 |
| FHDC1 | 4 | -0.33215748 | 0.06 | 0.179 | 9.01E-24 | 3.3E-19 |
| THADA | 4 | 0.366821685 | 0.671 | 0.578 | 9.25E-24 | 3.39E-19 |
| JAG1 | 4 | 0.46266976 | 0.256 | 0.132 | 9.83E-24 | 3.6E-19 |
| COMMD1 | 4 | 0.381107456 | 0.614 | 0.467 | 1.11E-23 | 4.06E-19 |
| CACNG4 | 4 | 0.375629748 | 0.231 | 0.112 | 1.15E-23 | 4.2E-19 |
| ZFHX4 | 4 | 0.549969215 | 0.584 | 0.412 | 1.2E-23 | 4.39E-19 |
| SLC35F1 | 4 | 0.670640597 | 0.437 | 0.287 | 1.21E-23 | 4.43E-19 |
| FOXP4 | 4 | 0.254826087 | 0.153 | 0.058 | 1.25E-23 | 4.59E-19 |
| IGDCC4 | 4 | 0.298858788 | 0.21 | 0.093 | 1.36E-23 | 4.97E-19 |
| ATP2B1 | 4 | 0.493269244 | 0.569 | 0.412 | 1.46E-23 | 5.35E-19 |
| AC034195.1 | 4 | 0.416632965 | 0.167 | 0.063 | 1.51E-23 | 5.52E-19 |
| RAPGEF1 | 4 | 0.390718394 | 0.541 | 0.382 | 1.55E-23 | 5.68E-19 |
| HBEGF | 4 | -0.337794317 | 0.033 | 0.132 | 1.68E-23 | 6.15E-19 |
| CENPK | 4 | -0.295298981 | 0.337 | 0.511 | 1.72E-23 | 6.28E-19 |
| ABHD2 | 4 | 0.367607941 | 0.309 | 0.176 | 1.86E-23 | 6.82E-19 |
| BMPER | 4 | -0.55649945 | 0.094 | 0.225 | 1.89E-23 | 6.91E-19 |
| SOX2 | 4 | 0.283991617 | 0.181 | 0.075 | 1.92E-23 | 7.05E-19 |
| HDAC6 | 4 | 0.264964619 | 0.207 | 0.09 | 1.99E-23 | 7.28E-19 |
| MPP5 | 4 | 0.331993122 | 0.372 | 0.217 | 2.01E-23 | 7.36E-19 |
| NLK | 4 | -0.516406462 | 0.597 | 0.672 | 2.13E-23 | 7.78E-19 |
| PLPP1 | 4 | 0.40947997 | 0.499 | 0.329 | 2.21E-23 | 8.1E-19 |
| PRIM2 | 4 | -0.42379826 | 0.557 | 0.649 | 2.52E-23 | 9.23E-19 |
| CCDC186 | 4 | 0.268901991 | 0.284 | 0.146 | 2.67E-23 | 9.79E-19 |
| PLCE1 | 4 | 0.455812408 | 0.292 | 0.158 | 2.85E-23 | 1.04E-18 |
| DZANK1 | 4 | 0.255854593 | 0.265 | 0.131 | 3.1E-23 | 1.13E-18 |
| TM9SF3 | 4 | -0.417695548 | 0.519 | 0.626 | 3.16E-23 | 1.16E-18 |
| MORC3 | 4 | -0.475282612 | 0.427 | 0.559 | 3.19E-23 | 1.17E-18 |
| SPON1 | 4 | 0.452741836 | 0.13 | 0.043 | 3.23E-23 | 1.18E-18 |
| CELF4 | 4 | -0.346243918 | 0.154 | 0.304 | 3.46E-23 | 1.27E-18 |

| | | | | | | |
|---|---|---|---|---|---|---|
| ZNF367 | 4 | -0.289887521 | 0.111 | 0.248 | 4.27E-23 | 1.56E-18 |
| MOSPD1 | 4 | 0.289935829 | 0.235 | 0.112 | 4.51E-23 | 1.65E-18 |
| XPO6 | 4 | -0.38563337 | 0.384 | 0.537 | 5E-23 | 1.83E-18 |
| HSF2 | 4 | 0.378054388 | 0.426 | 0.301 | 5.09E-23 | 1.86E-18 |
| CLMP | 4 | 0.456255235 | 0.214 | 0.133 | 5.11E-23 | 1.87E-18 |
| RNF24 | 4 | -0.348335511 | 0.397 | 0.571 | 5.45E-23 | 2E-18 |
| FIGN | 4 | 0.470361916 | 0.485 | 0.32 | 5.67E-23 | 2.08E-18 |
| EHBP1 | 4 | -0.365967192 | 0.761 | 0.834 | 6.36E-23 | 2.33E-18 |
| CENPU | 4 | -0.35216354 | 0.17 | 0.323 | 6.54E-23 | 2.4E-18 |
| NGF | 4 | -0.308852131 | 0.034 | 0.134 | 7.1E-23 | 2.6E-18 |
| ARMC4 | 4 | 0.322216389 | 0.243 | 0.116 | 9.53E-23 | 3.49E-18 |
| SIN3B | 4 | 0.353595845 | 0.358 | 0.215 | 1.24E-22 | 4.53E-18 |
| IFT172 | 4 | 0.337874701 | 0.395 | 0.243 | 1.32E-22 | 4.84E-18 |
| SIK3 | 4 | 0.337369273 | 0.783 | 0.657 | 1.4E-22 | 5.12E-18 |
| PAN3 | 4 | 0.321316633 | 0.865 | 0.789 | 1.43E-22 | 5.23E-18 |
| SLC9A7 | 4 | 0.359221147 | 0.266 | 0.135 | 1.53E-22 | 5.61E-18 |
| HLA-B | 4 | -0.288118653 | 0.045 | 0.145 | 1.69E-22 | 6.2E-18 |
| AP002954.2 | 4 | -0.605050801 | 0.025 | 0.106 | 2.01E-22 | 7.35E-18 |
| NPEPL1 | 4 | 0.288318768 | 0.279 | 0.146 | 2.35E-22 | 8.61E-18 |
| ELL2 | 4 | -0.764325077 | 0.103 | 0.198 | 2.37E-22 | 8.69E-18 |
| COL4A1 | 4 | 0.572673564 | 0.16 | 0.132 | 2.48E-22 | 9.09E-18 |
| NECTIN3 | 4 | -0.583756543 | 0.52 | 0.61 | 2.5E-22 | 9.14E-18 |
| PCM1 | 4 | 0.33590096 | 0.855 | 0.791 | 2.51E-22 | 9.18E-18 |
| NRG1 | 4 | 0.304644062 | 0.435 | 0.274 | 2.52E-22 | 9.23E-18 |
| WWOX | 4 | 0.429425191 | 0.824 | 0.766 | 2.89E-22 | 1.06E-17 |
| LRCH1 | 4 | -0.336801998 | 0.485 | 0.639 | 2.95E-22 | 1.08E-17 |
| EXTL2 | 4 | 0.337322888 | 0.37 | 0.225 | 2.96E-22 | 1.08E-17 |
| KLHL5 | 4 | 0.411933391 | 0.304 | 0.198 | 2.97E-22 | 1.09E-17 |
| HS6ST1 | 4 | 0.413423069 | 0.239 | 0.137 | 3.25E-22 | 1.19E-17 |
| FGFR1OP | 4 | 0.373034739 | 0.534 | 0.383 | 3.26E-22 | 1.2E-17 |
| GNA12 | 4 | 0.380748232 | 0.446 | 0.299 | 3.38E-22 | 1.24E-17 |
| NPHP4 | 4 | 0.348676475 | 0.356 | 0.212 | 3.46E-22 | 1.26E-17 |
| KCNT2 | 4 | -0.435069611 | 0.64 | 0.713 | 3.47E-22 | 1.27E-17 |
| EBF3 | 4 | 0.251616009 | 0.157 | 0.262 | 3.49E-22 | 1.28E-17 |
| SPATA18 | 4 | 0.287334965 | 0.118 | 0.037 | 3.51E-22 | 1.29E-17 |
| NOL4L | 4 | 0.373587651 | 0.284 | 0.161 | 3.78E-22 | 1.38E-17 |
| SPG7 | 4 | 0.353797207 | 0.607 | 0.449 | 3.86E-22 | 1.41E-17 |
| HIVEP2 | 4 | -0.484515916 | 0.315 | 0.443 | 3.93E-22 | 1.44E-17 |
| BORCS5 | 4 | -0.390256876 | 0.231 | 0.37 | 4.19E-22 | 1.53E-17 |
| MKLN1 | 4 | -0.401128309 | 0.762 | 0.836 | 4.68E-22 | 1.71E-17 |
| BCAS3 | 4 | 0.366538753 | 0.774 | 0.644 | 5.06E-22 | 1.85E-17 |
| SENP7 | 4 | 0.384205193 | 0.725 | 0.61 | 5.15E-22 | 1.89E-17 |
| MECP2 | 4 | 0.369926878 | 0.497 | 0.357 | 6.07E-22 | 2.22E-17 |
| KLF3-AS1 | 4 | 0.311351081 | 0.179 | 0.077 | 6.42E-22 | 2.35E-17 |
| PABPN1 | 4 | 0.357863334 | 0.589 | 0.442 | 6.47E-22 | 2.37E-17 |
| AC093298.2 | 4 | -1.178256941 | 0.122 | 0.196 | 7.04E-22 | 2.58E-17 |
| GABRB3 | 4 | -0.530179172 | 0.244 | 0.392 | 7.23E-22 | 2.65E-17 |
| GPD2 | 4 | -0.422335447 | 0.294 | 0.436 | 7.52E-22 | 2.75E-17 |
| TBC1D4 | 4 | -0.41880393 | 0.255 | 0.416 | 7.67E-22 | 2.81E-17 |
| AL157778.1 | 4 | -0.832324681 | 0.12 | 0.252 | 7.79E-22 | 2.85E-17 |
| OPHN1 | 4 | 0.580091357 | 0.506 | 0.387 | 8.21E-22 | 3.01E-17 |
| DGKD | 4 | 0.398705184 | 0.486 | 0.37 | 8.36E-22 | 3.06E-17 |
| AC092279.1 | 4 | 0.258547798 | 0.237 | 0.116 | 8.67E-22 | 3.18E-17 |
| ADGRA3 | 4 | 0.358173819 | 0.757 | 0.672 | 9.66E-22 | 3.54E-17 |
| KDM4B | 4 | 0.414146424 | 0.613 | 0.477 | 1.2E-21 | 4.41E-17 |
| SMO | 4 | 0.356981647 | 0.274 | 0.152 | 1.22E-21 | 4.46E-17 |
| EFR3B | 4 | -0.45260807 | 0.33 | 0.469 | 1.33E-21 | 4.86E-17 |
| CBLB | 4 | 0.473135833 | 0.719 | 0.576 | 1.37E-21 | 5.03E-17 |

| | | | | | | |
|---|---|---|---|---|---|---|
| LRRC1 | 4 | 0.30882931 | 0.277 | 0.145 | 1.44E-21 | 5.28E-17 |
| SPAG9 | 4 | -0.343230785 | 0.778 | 0.846 | 1.45E-21 | 5.31E-17 |
| ID2 | 4 | 0.344320581 | 0.28 | 0.147 | 1.49E-21 | 5.46E-17 |
| EXD3 | 4 | 0.318788014 | 0.26 | 0.138 | 1.58E-21 | 5.78E-17 |
| APAF1 | 4 | -0.39994737 | 0.21 | 0.348 | 1.64E-21 | 5.99E-17 |
| KIF9-AS1 | 4 | 0.397745317 | 0.368 | 0.227 | 1.66E-21 | 6.09E-17 |
| CAST | 4 | 0.345326836 | 0.355 | 0.215 | 1.84E-21 | 6.72E-17 |
| ADCY2 | 4 | -0.797493794 | 0.163 | 0.291 | 1.93E-21 | 7.05E-17 |
| DCX | 4 | 0.56149506 | 0.284 | 0.167 | 2.05E-21 | 7.49E-17 |
| NTF3 | 4 | -0.501223142 | 0.099 | 0.176 | 2.25E-21 | 8.24E-17 |
| OBSL1 | 4 | 0.287150116 | 0.272 | 0.142 | 2.37E-21 | 8.67E-17 |
| DNAAF5 | 4 | 0.291958325 | 0.309 | 0.171 | 2.43E-21 | 8.91E-17 |
| EFCAB7 | 4 | 0.322416443 | 0.292 | 0.161 | 2.46E-21 | 9E-17 |
| AC109466.1 | 4 | 0.291781258 | 0.212 | 0.101 | 2.48E-21 | 9.07E-17 |
| CATSPERE | 4 | -0.498137164 | 0.137 | 0.252 | 2.56E-21 | 9.39E-17 |
| LNX1 | 4 | -0.288038431 | 0.04 | 0.138 | 2.81E-21 | 1.03E-16 |
| RC3H1 | 4 | 0.344412886 | 0.709 | 0.619 | 2.83E-21 | 1.04E-16 |
| FAM135A | 4 | 0.352297985 | 0.649 | 0.545 | 2.86E-21 | 1.05E-16 |
| CPLANE1 | 4 | 0.334587118 | 0.716 | 0.577 | 2.95E-21 | 1.08E-16 |
| CEMIP2 | 4 | 0.480970268 | 0.483 | 0.332 | 3.11E-21 | 1.14E-16 |
| CSAD | 4 | 0.35996669 | 0.397 | 0.254 | 3.17E-21 | 1.16E-16 |
| STK17B | 4 | -0.367643935 | 0.053 | 0.158 | 3.2E-21 | 1.17E-16 |
| LENG8 | 4 | 0.345315664 | 0.433 | 0.282 | 3.36E-21 | 1.23E-16 |
| LAMP2 | 4 | -0.362067347 | 0.168 | 0.304 | 3.45E-21 | 1.26E-16 |
| FNBP1 | 4 | -0.378256378 | 0.546 | 0.674 | 3.6E-21 | 1.32E-16 |
| GLCCI1 | 4 | -0.55776887 | 0.549 | 0.581 | 3.7E-21 | 1.36E-16 |
| RUFY2 | 4 | 0.383163149 | 0.55 | 0.406 | 3.9E-21 | 1.43E-16 |
| PACS2 | 4 | 0.365271613 | 0.394 | 0.251 | 3.93E-21 | 1.44E-16 |
| NBPF26 | 4 | 0.290178894 | 0.317 | 0.178 | 3.97E-21 | 1.45E-16 |
| AC046134.2 | 4 | 0.389125084 | 0.431 | 0.29 | 4.01E-21 | 1.47E-16 |
| MYC | 4 | -0.38264825 | 0.173 | 0.303 | 4.23E-21 | 1.55E-16 |
| PRKCB | 4 | -0.337619039 | 0.026 | 0.114 | 4.33E-21 | 1.59E-16 |
| PHACTR1 | 4 | -0.594878636 | 0.21 | 0.335 | 4.74E-21 | 1.74E-16 |
| AXIN2 | 4 | 0.330022149 | 0.291 | 0.163 | 4.93E-21 | 1.81E-16 |
| VEZT | 4 | -0.402166831 | 0.574 | 0.68 | 4.97E-21 | 1.82E-16 |
| MCM3 | 4 | -0.342234467 | 0.192 | 0.341 | 5.39E-21 | 1.97E-16 |
| ATN1 | 4 | 0.32164705 | 0.324 | 0.189 | 5.71E-21 | 2.09E-16 |
| WDR73 | 4 | 0.309732119 | 0.315 | 0.178 | 6.02E-21 | 2.2E-16 |
| LAMA3 | 4 | -0.510167482 | 0.116 | 0.221 | 6.09E-21 | 2.23E-16 |
| IFT81 | 4 | 0.359259351 | 0.5 | 0.346 | 7E-21 | 2.56E-16 |
| PNISR | 4 | 0.296144431 | 0.88 | 0.817 | 7.08E-21 | 2.59E-16 |
| HECW2 | 4 | -0.305827014 | 0.231 | 0.384 | 7.18E-21 | 2.63E-16 |
| ST5 | 4 | 0.438103311 | 0.428 | 0.282 | 7.84E-21 | 2.87E-16 |
| PLD1 | 4 | 0.354610413 | 0.147 | 0.055 | 8.33E-21 | 3.05E-16 |
| FBXO16 | 4 | 0.275743335 | 0.208 | 0.096 | 8.51E-21 | 3.11E-16 |
| CFLAR | 4 | 0.255592799 | 0.234 | 0.114 | 8.91E-21 | 3.26E-16 |
| LINC01934 | 4 | -1.222583099 | 0.043 | 0.11 | 9.59E-21 | 3.51E-16 |
| AC009803.1 | 4 | -0.39855157 | 0.043 | 0.138 | 1.01E-20 | 3.69E-16 |
| SLC44A2 | 4 | 0.349302072 | 0.22 | 0.116 | 1.08E-20 | 3.94E-16 |
| LINC01456 | 4 | 0.393420293 | 0.236 | 0.117 | 1.1E-20 | 4.02E-16 |
| STAC | 4 | -0.5088691 | 0.109 | 0.221 | 1.14E-20 | 4.18E-16 |
| SLC25A21 | 4 | 0.423516564 | 0.237 | 0.123 | 1.16E-20 | 4.26E-16 |
| EPHA5-AS1 | 4 | -0.273800827 | 0.032 | 0.123 | 1.44E-20 | 5.28E-16 |
| REPS2 | 4 | -0.472133167 | 0.221 | 0.341 | 1.44E-20 | 5.28E-16 |
| FN1 | 4 | -0.399335189 | 0.23 | 0.384 | 1.5E-20 | 5.49E-16 |
| NIPSNAP2 | 4 | 0.329904459 | 0.468 | 0.321 | 1.52E-20 | 5.56E-16 |
| WDR90 | 4 | 0.270113124 | 0.24 | 0.12 | 1.56E-20 | 5.71E-16 |
| GALNT11 | 4 | 0.382984691 | 0.487 | 0.416 | 1.58E-20 | 5.79E-16 |

| | | | | | | |
|---|---|---|---|---|---|---|
| TNIK | 4 | 0.295597097 | 0.504 | 0.343 | 1.58E-20 | 5.79E-16 |
| SENP6 | 4 | 0.289264002 | 0.9 | 0.816 | 1.66E-20 | 6.09E-16 |
| KANK4 | 4 | 0.40790163 | 0.126 | 0.047 | 1.73E-20 | 6.33E-16 |
| RGMB | 4 | 0.322560345 | 0.122 | 0.072 | 1.83E-20 | 6.7E-16 |
| THRB | 4 | -0.691446609 | 0.268 | 0.315 | 1.9E-20 | 6.94E-16 |
| BZW2 | 4 | -0.475959568 | 0.307 | 0.415 | 1.94E-20 | 7.1E-16 |
| PRRG1 | 4 | -0.409506325 | 0.244 | 0.373 | 2E-20 | 7.33E-16 |
| RAD54B | 4 | -0.307390859 | 0.124 | 0.252 | 2.03E-20 | 7.42E-16 |
| NEDD9 | 4 | 0.289952262 | 0.226 | 0.11 | 2.18E-20 | 7.99E-16 |
| PHLPP1 | 4 | 0.40254193 | 0.595 | 0.463 | 2.2E-20 | 8.04E-16 |
| CNTNAP3 | 4 | 0.280428127 | 0.263 | 0.138 | 2.2E-20 | 8.06E-16 |
| KMT5B | 4 | 0.356465105 | 0.609 | 0.466 | 2.21E-20 | 8.1E-16 |
| GNAI1 | 4 | -0.443256373 | 0.467 | 0.598 | 2.3E-20 | 8.41E-16 |
| BTG1 | 4 | -0.280599204 | 0.067 | 0.178 | 2.31E-20 | 8.44E-16 |
| AKAP6 | 4 | 0.793604352 | 0.342 | 0.223 | 2.53E-20 | 9.26E-16 |
| TTC8 | 4 | 0.295697226 | 0.381 | 0.234 | 2.58E-20 | 9.46E-16 |
| AHDC1 | 4 | 0.308990952 | 0.303 | 0.17 | 2.59E-20 | 9.48E-16 |
| RAI1 | 4 | 0.389634046 | 0.473 | 0.335 | 2.75E-20 | 1.01E-15 |
| CHRM2 | 4 | -0.66157259 | 0.03 | 0.103 | 2.78E-20 | 1.02E-15 |
| CHD7 | 4 | -0.330486725 | 0.746 | 0.81 | 2.85E-20 | 1.04E-15 |
| EYA1 | 4 | 0.659687091 | 0.418 | 0.278 | 2.94E-20 | 1.08E-15 |
| NFIX | 4 | -0.389061643 | 0.327 | 0.484 | 3.08E-20 | 1.13E-15 |
| YEATS2 | 4 | -0.321756481 | 0.657 | 0.764 | 3.16E-20 | 1.16E-15 |
| MAP4K3-DT | 4 | -0.40716304 | 0.274 | 0.406 | 3.31E-20 | 1.21E-15 |
| SLC25A13 | 4 | 0.369350124 | 0.59 | 0.46 | 3.31E-20 | 1.21E-15 |
| RIMS1 | 4 | -0.592947765 | 0.354 | 0.447 | 3.35E-20 | 1.23E-15 |
| AL445623.2 | 4 | -0.285040201 | 0.091 | 0.209 | 3.69E-20 | 1.35E-15 |
| CHEK1 | 4 | -0.307033345 | 0.205 | 0.353 | 3.71E-20 | 1.36E-15 |
| KAZN | 4 | -0.491864229 | 0.87 | 0.902 | 3.72E-20 | 1.36E-15 |
| GULP1 | 4 | 0.343072336 | 0.64 | 0.492 | 3.83E-20 | 1.4E-15 |
| DRAXIN | 4 | 0.269930809 | 0.121 | 0.052 | 4.28E-20 | 1.57E-15 |
| PIK3AP1 | 4 | -0.315070179 | 0.063 | 0.17 | 4.32E-20 | 1.58E-15 |
| KIAA1324L | 4 | 0.363777681 | 0.346 | 0.206 | 4.4E-20 | 1.61E-15 |
| ANKRD18A | 4 | -0.301110024 | 0.062 | 0.167 | 4.52E-20 | 1.65E-15 |
| MLLT1 | 4 | 0.302407807 | 0.312 | 0.18 | 4.56E-20 | 1.67E-15 |
| PAG1 | 4 | -0.460308937 | 0.186 | 0.319 | 4.58E-20 | 1.68E-15 |
| MTA1 | 4 | 0.311043418 | 0.306 | 0.178 | 4.69E-20 | 1.72E-15 |
| FBLN7 | 4 | 0.260163624 | 0.113 | 0.043 | 4.94E-20 | 1.81E-15 |
| FANCI | 4 | -0.366301022 | 0.262 | 0.416 | 4.98E-20 | 1.82E-15 |
| LINC01138 | 4 | 0.328228533 | 0.206 | 0.104 | 5.21E-20 | 1.91E-15 |
| APOO | 4 | 0.325065134 | 0.34 | 0.202 | 5.46E-20 | 2E-15 |
| PRDM2 | 4 | -0.442164044 | 0.457 | 0.581 | 5.59E-20 | 2.05E-15 |
| ATP7B | 4 | 0.264480239 | 0.141 | 0.054 | 5.75E-20 | 2.11E-15 |
| SLC7A14 | 4 | -0.470528515 | 0.033 | 0.118 | 5.8E-20 | 2.13E-15 |
| PCDH7 | 4 | -0.587205838 | 0.465 | 0.423 | 5.97E-20 | 2.19E-15 |
| NID1 | 4 | -0.302455754 | 0.159 | 0.298 | 6.28E-20 | 2.3E-15 |
| ITFG1 | 4 | -0.409584728 | 0.756 | 0.795 | 6.41E-20 | 2.35E-15 |
| CAMK2B | 4 | -0.310616174 | 0.066 | 0.162 | 6.48E-20 | 2.37E-15 |
| SON | 4 | 0.321381036 | 0.769 | 0.672 | 6.96E-20 | 2.55E-15 |
| PPP1R12C | 4 | 0.300859858 | 0.247 | 0.13 | 7.59E-20 | 2.78E-15 |
| DLG1 | 4 | 0.350064002 | 0.832 | 0.749 | 7.7E-20 | 2.82E-15 |
| TTLL4 | 4 | 0.257734673 | 0.272 | 0.146 | 1.02E-19 | 3.75E-15 |
| PSMA6 | 4 | -0.37037246 | 0.283 | 0.404 | 1.03E-19 | 3.77E-15 |
| RIF1 | 4 | -0.331675506 | 0.705 | 0.786 | 1.04E-19 | 3.8E-15 |
| NAPEPLD | 4 | -0.385660861 | 0.234 | 0.37 | 1.16E-19 | 4.25E-15 |
| KIAA0753 | 4 | 0.298818724 | 0.405 | 0.26 | 1.26E-19 | 4.62E-15 |
| TMEM91 | 4 | 0.302999212 | 0.353 | 0.214 | 1.33E-19 | 4.86E-15 |
| MITF | 4 | 0.368606396 | 0.308 | 0.175 | 1.33E-19 | 4.88E-15 |

| | | | | | | |
|---|---|---|---|---|---|---|
| NKAIN1 | 4 | 0.383771014 | 0.141 | 0.104 | 1.35E-19 | 4.93E-15 |
| CD47 | 4 | -0.333690795 | 0.272 | 0.42 | 1.39E-19 | 5.1E-15 |
| TAPT1-AS1 | 4 | 0.372848191 | 0.323 | 0.2 | 1.4E-19 | 5.12E-15 |
| EFNA5 | 4 | -0.552814204 | 0.551 | 0.676 | 1.43E-19 | 5.22E-15 |
| AC004158.1 | 4 | -0.341469287 | 0.087 | 0.199 | 1.44E-19 | 5.28E-15 |
| GLMN | 4 | -0.397030482 | 0.39 | 0.512 | 1.59E-19 | 5.84E-15 |
| ANXA6 | 4 | 0.28368103 | 0.297 | 0.168 | 1.77E-19 | 6.49E-15 |
| RAB28 | 4 | 0.366647045 | 0.585 | 0.451 | 1.78E-19 | 6.52E-15 |
| ARHGAP11B | 4 | -0.309536117 | 0.149 | 0.283 | 1.79E-19 | 6.54E-15 |
| RNF130 | 4 | 0.389582964 | 0.757 | 0.672 | 1.89E-19 | 6.92E-15 |
| RUNX1 | 4 | 0.74463354 | 0.187 | 0.11 | 1.95E-19 | 7.14E-15 |
| LRIG3 | 4 | 0.279714658 | 0.374 | 0.233 | 2.2E-19 | 8.06E-15 |
| TRAK1 | 4 | 0.381211276 | 0.557 | 0.421 | 2.3E-19 | 8.44E-15 |
| PLCL1 | 4 | -0.54647089 | 0.301 | 0.437 | 2.5E-19 | 9.14E-15 |
| CDK10 | 4 | 0.302064083 | 0.298 | 0.175 | 2.62E-19 | 9.58E-15 |
| SFMBT1 | 4 | -0.399514564 | 0.302 | 0.441 | 2.74E-19 | 1E-14 |
| DECR1 | 4 | 0.283997853 | 0.311 | 0.179 | 2.75E-19 | 1.01E-14 |
| ARMH4 | 4 | 0.329694087 | 0.319 | 0.196 | 2.76E-19 | 1.01E-14 |
| NQO2 | 4 | 0.319205894 | 0.201 | 0.103 | 2.78E-19 | 1.02E-14 |
| CCPG1 | 4 | 0.26459019 | 0.255 | 0.134 | 2.87E-19 | 1.05E-14 |
| TPD52L2 | 4 | 0.270898797 | 0.253 | 0.135 | 2.93E-19 | 1.07E-14 |
| PDS5B | 4 | -0.338513347 | 0.786 | 0.85 | 3.21E-19 | 1.18E-14 |
| MOB3A | 4 | 0.257403648 | 0.175 | 0.077 | 3.37E-19 | 1.23E-14 |
| CPLX2 | 4 | -0.296666078 | 0.074 | 0.183 | 3.9E-19 | 1.43E-14 |
| NSMAF | 4 | -0.384892835 | 0.349 | 0.476 | 3.92E-19 | 1.43E-14 |
| AP3D1 | 4 | 0.338688945 | 0.397 | 0.258 | 4.29E-19 | 1.57E-14 |
| ATXN3 | 4 | 0.363371206 | 0.471 | 0.335 | 4.43E-19 | 1.62E-14 |
| ZER1 | 4 | 0.300993822 | 0.327 | 0.193 | 4.99E-19 | 1.83E-14 |
| NAP1L4 | 4 | -0.347003363 | 0.569 | 0.68 | 5.17E-19 | 1.89E-14 |
| RNF19A | 4 | -0.380191951 | 0.384 | 0.52 | 5.52E-19 | 2.02E-14 |
| AGMO | 4 | -0.732008305 | 0.048 | 0.122 | 5.62E-19 | 2.06E-14 |
| CYP7B1 | 4 | -0.441533904 | 0.375 | 0.368 | 5.74E-19 | 2.1E-14 |
| ENO4 | 4 | 0.285901777 | 0.239 | 0.144 | 6.7E-19 | 2.45E-14 |
| AP001636.3 | 4 | -0.273192743 | 0.036 | 0.125 | 6.77E-19 | 2.48E-14 |
| ZNF587 | 4 | 0.283008973 | 0.264 | 0.145 | 6.8E-19 | 2.49E-14 |
| CEP126 | 4 | 0.35619059 | 0.349 | 0.295 | 6.88E-19 | 2.52E-14 |
| CACNA2D2 | 4 | -0.352230796 | 0.183 | 0.298 | 6.9E-19 | 2.53E-14 |
| ANLN | 4 | -0.314929283 | 0.152 | 0.284 | 7.47E-19 | 2.74E-14 |
| AC104596.1 | 4 | 0.376361982 | 0.214 | 0.108 | 8.01E-19 | 2.93E-14 |
| ME3 | 4 | -0.416975789 | 0.216 | 0.322 | 8.05E-19 | 2.95E-14 |
| AC004917.1 | 4 | -0.34864058 | 0.028 | 0.106 | 8.21E-19 | 3.01E-14 |
| AL356108.1 | 4 | -0.656193948 | 0.036 | 0.12 | 8.5E-19 | 3.11E-14 |
| NCOA3 | 4 | -0.407888741 | 0.465 | 0.605 | 8.73E-19 | 3.2E-14 |
| CFAP36 | 4 | 0.28213798 | 0.297 | 0.17 | 9.24E-19 | 3.38E-14 |
| UBE2E3 | 4 | -0.395819666 | 0.641 | 0.71 | 9.55E-19 | 3.49E-14 |
| SYT10 | 4 | -0.415957687 | 0.047 | 0.134 | 1.01E-18 | 3.68E-14 |
| TEAD1 | 4 | -0.304177104 | 0.855 | 0.907 | 1.03E-18 | 3.78E-14 |
| RUFY1 | 4 | 0.322405941 | 0.449 | 0.309 | 1.23E-18 | 4.51E-14 |
| HTR7 | 4 | -0.39115795 | 0.044 | 0.134 | 1.3E-18 | 4.77E-14 |
| LINC00536 | 4 | -0.722883796 | 0.029 | 0.103 | 1.36E-18 | 4.97E-14 |
| RRAS2 | 4 | 0.251355021 | 0.409 | 0.265 | 1.37E-18 | 5.01E-14 |
| SCN7A | 4 | -1.009016855 | 0.059 | 0.15 | 1.39E-18 | 5.11E-14 |
| TSPAN18 | 4 | 0.484600136 | 0.369 | 0.266 | 1.4E-18 | 5.14E-14 |
| MMS22L | 4 | -0.381736992 | 0.483 | 0.618 | 1.46E-18 | 5.34E-14 |
| BBS2 | 4 | 0.283862283 | 0.263 | 0.149 | 1.56E-18 | 5.7E-14 |
| LYRM4 | 4 | 0.359530823 | 0.483 | 0.349 | 1.69E-18 | 6.17E-14 |
| ZFPM2 | 4 | 0.779337452 | 0.361 | 0.301 | 1.83E-18 | 6.68E-14 |
| ARL13B | 4 | 0.326244827 | 0.451 | 0.309 | 1.84E-18 | 6.74E-14 |

| | | | | | | |
|---|---|---|---|---|---|---|
| KLHL29 | 4 | -0.617117537 | 0.509 | 0.629 | 1.85E-18 | 6.78E-14 |
| GALNT2 | 4 | 0.352014105 | 0.529 | 0.387 | 1.91E-18 | 7.01E-14 |
| MBOAT2 | 4 | 0.357067795 | 0.58 | 0.445 | 2.02E-18 | 7.38E-14 |
| DIS3L2 | 4 | 0.320311114 | 0.603 | 0.478 | 2.07E-18 | 7.58E-14 |
| HOMER2 | 4 | -0.371392392 | 0.197 | 0.325 | 2.09E-18 | 7.65E-14 |
| PIK3R3 | 4 | -0.362405748 | 0.363 | 0.511 | 2.29E-18 | 8.37E-14 |
| PHLDB1 | 4 | 0.253674909 | 0.244 | 0.128 | 2.3E-18 | 8.43E-14 |
| ZNF721 | 4 | 0.330414588 | 0.701 | 0.577 | 2.31E-18 | 8.44E-14 |
| PLXNA4 | 4 | 0.451498446 | 0.114 | 0.088 | 2.34E-18 | 8.58E-14 |
| CDH10 | 4 | -0.586805623 | 0.214 | 0.342 | 2.35E-18 | 8.61E-14 |
| RHOBTB3 | 4 | -0.283425877 | 0.327 | 0.48 | 2.39E-18 | 8.76E-14 |
| RELL1 | 4 | -0.31000194 | 0.138 | 0.262 | 2.42E-18 | 8.87E-14 |
| RBPJ | 4 | 0.306596542 | 0.751 | 0.652 | 2.5E-18 | 9.15E-14 |
| DHX15 | 4 | -0.31177717 | 0.679 | 0.761 | 2.52E-18 | 9.22E-14 |
| PRKCZ | 4 | 0.280657078 | 0.201 | 0.102 | 2.69E-18 | 9.84E-14 |
| PDK3 | 4 | -0.427502684 | 0.302 | 0.439 | 2.76E-18 | 1.01E-13 |
| CLSTN1 | 4 | 0.354826074 | 0.483 | 0.351 | 3.06E-18 | 1.12E-13 |
| GRIK5 | 4 | 0.291024251 | 0.217 | 0.111 | 3.25E-18 | 1.19E-13 |
| UPF3B | 4 | 0.338477766 | 0.425 | 0.29 | 3.34E-18 | 1.22E-13 |
| NARS2 | 4 | 0.34576311 | 0.591 | 0.466 | 3.4E-18 | 1.25E-13 |
| XIAP | 4 | 0.35594875 | 0.573 | 0.442 | 3.41E-18 | 1.25E-13 |
| GCA | 4 | -0.26864062 | 0.073 | 0.174 | 3.49E-18 | 1.28E-13 |
| ZFP90 | 4 | 0.301496693 | 0.299 | 0.182 | 4.48E-18 | 1.64E-13 |
| BCR | 4 | 0.342731732 | 0.363 | 0.233 | 4.6E-18 | 1.68E-13 |
| LINC00907 | 4 | -0.734514741 | 0.159 | 0.264 | 4.6E-18 | 1.68E-13 |
| LINC00511 | 4 | -0.651695263 | 0.206 | 0.299 | 4.71E-18 | 1.72E-13 |
| CASD1 | 4 | 0.351578901 | 0.543 | 0.424 | 5.24E-18 | 1.92E-13 |
| EPHB2 | 4 | 0.372580601 | 0.274 | 0.189 | 5.29E-18 | 1.94E-13 |
| RNF152 | 4 | 0.274860074 | 0.178 | 0.081 | 5.37E-18 | 1.96E-13 |
| TMEM65 | 4 | 0.325444468 | 0.366 | 0.229 | 5.42E-18 | 1.98E-13 |
| MFSD6 | 4 | -0.256511689 | 0.082 | 0.19 | 5.59E-18 | 2.05E-13 |
| LCLAT1 | 4 | -0.357695691 | 0.464 | 0.595 | 5.85E-18 | 2.14E-13 |
| C1GALT1 | 4 | 0.32299067 | 0.338 | 0.213 | 6.14E-18 | 2.25E-13 |
| DNMBP | 4 | 0.269153234 | 0.309 | 0.181 | 6.3E-18 | 2.31E-13 |
| DSTYK | 4 | 0.284482505 | 0.316 | 0.19 | 6.67E-18 | 2.44E-13 |
| RBM25 | 4 | 0.315319204 | 0.803 | 0.701 | 6.73E-18 | 2.46E-13 |
| ADGRL3 | 4 | -0.426737986 | 0.869 | 0.808 | 6.79E-18 | 2.49E-13 |
| WLS | 4 | 0.387263804 | 0.537 | 0.407 | 7.21E-18 | 2.64E-13 |
| RAPGEF2 | 4 | 0.255355119 | 0.771 | 0.641 | 7.48E-18 | 2.74E-13 |
| OBI1 | 4 | -0.32474509 | 0.192 | 0.314 | 7.55E-18 | 2.76E-13 |
| SEMA3A | 4 | 1.006052644 | 0.348 | 0.282 | 8.21E-18 | 3E-13 |
| CTBP2 | 4 | 0.361274508 | 0.875 | 0.792 | 8.32E-18 | 3.05E-13 |
| HULC | 4 | -0.834353443 | 0.337 | 0.423 | 8.53E-18 | 3.12E-13 |
| LINC00882 | 4 | 0.371237965 | 0.188 | 0.091 | 8.54E-18 | 3.13E-13 |
| LRRC23 | 4 | 0.272844369 | 0.209 | 0.104 | 8.61E-18 | 3.15E-13 |
| TRPS1 | 4 | 0.32523518 | 0.719 | 0.574 | 8.67E-18 | 3.17E-13 |
| ZNF75D | 4 | 0.268692524 | 0.246 | 0.133 | 9.15E-18 | 3.35E-13 |
| RYR3 | 4 | 0.760185712 | 0.176 | 0.101 | 9.5E-18 | 3.48E-13 |
| SCARB2 | 4 | -0.331118478 | 0.286 | 0.422 | 9.9E-18 | 3.62E-13 |
| SCN9A | 4 | -0.458347409 | 0.066 | 0.164 | 9.99E-18 | 3.66E-13 |
| KIAA0355 | 4 | 0.306409025 | 0.46 | 0.318 | 1.01E-17 | 3.69E-13 |
| FRYL | 4 | -0.29408223 | 0.801 | 0.858 | 1.01E-17 | 3.7E-13 |
| ANKRD36B | 4 | -0.365305422 | 0.369 | 0.498 | 1.02E-17 | 3.74E-13 |
| NEK7 | 4 | -0.438235206 | 0.54 | 0.628 | 1.12E-17 | 4.1E-13 |
| SSBP2 | 4 | 0.333130934 | 0.889 | 0.802 | 1.12E-17 | 4.12E-13 |
| DNAJC1 | 4 | -0.479133763 | 0.498 | 0.58 | 1.21E-17 | 4.44E-13 |
| HNRNPDL | 4 | 0.302293804 | 0.722 | 0.622 | 1.22E-17 | 4.48E-13 |
| GOLGA4 | 4 | 0.299218949 | 0.77 | 0.678 | 1.24E-17 | 4.54E-13 |

| | | | | | | |
|---|---|---|---|---|---|---|
| SRRM2 | 4 | 0.342310811 | 0.72 | 0.625 | 1.27E-17 | 4.64E-13 |
| ERI3 | 4 | 0.306557875 | 0.412 | 0.275 | 1.28E-17 | 4.7E-13 |
| AC245297.3 | 4 | 0.254808643 | 0.271 | 0.152 | 1.37E-17 | 5.02E-13 |
| SF3B1 | 4 | 0.325241924 | 0.704 | 0.596 | 1.42E-17 | 5.21E-13 |
| PPARGC1A | 4 | 0.370138497 | 0.264 | 0.155 | 1.43E-17 | 5.22E-13 |
| AL033504.1 | 4 | -0.487437696 | 0.495 | 0.588 | 1.43E-17 | 5.24E-13 |
| LSAMP | 4 | -0.372867955 | 0.785 | 0.732 | 1.5E-17 | 5.5E-13 |
| DISP1 | 4 | 0.302819353 | 0.505 | 0.357 | 1.6E-17 | 5.87E-13 |
| CHD4 | 4 | 0.319791339 | 0.498 | 0.359 | 1.72E-17 | 6.28E-13 |
| SEPTIN7 | 4 | -0.364106769 | 0.593 | 0.692 | 1.82E-17 | 6.66E-13 |
| MT-CO1 | 4 | 0.750138626 | 0.533 | 0.449 | 1.88E-17 | 6.89E-13 |
| FLNA | 4 | 0.322440436 | 0.313 | 0.189 | 1.96E-17 | 7.17E-13 |
| CARMIL1 | 4 | 0.44281906 | 0.44 | 0.309 | 2.07E-17 | 7.59E-13 |
| PRKAA2 | 4 | 0.290182535 | 0.292 | 0.169 | 2.09E-17 | 7.64E-13 |
| CLEC2D | 4 | -0.384484967 | 0.369 | 0.472 | 2.1E-17 | 7.7E-13 |
| LASP1 | 4 | 0.271191109 | 0.218 | 0.113 | 2.22E-17 | 8.14E-13 |
| AL109930.1 | 4 | -0.340487127 | 0.156 | 0.275 | 2.4E-17 | 8.77E-13 |
| CEP192 | 4 | -0.340342532 | 0.518 | 0.627 | 2.49E-17 | 9.12E-13 |
| ATG16L1 | 4 | 0.275910192 | 0.279 | 0.162 | 2.5E-17 | 9.15E-13 |
| PALM | 4 | 0.2836534 | 0.183 | 0.094 | 2.51E-17 | 9.2E-13 |
| VAV2 | 4 | 0.348778547 | 0.447 | 0.307 | 2.53E-17 | 9.28E-13 |
| SLBP | 4 | -0.286791211 | 0.119 | 0.236 | 2.66E-17 | 9.75E-13 |
| IL6ST | 4 | 0.396755667 | 0.285 | 0.166 | 2.78E-17 | 1.02E-12 |
| PIK3C2B | 4 | 0.310850658 | 0.204 | 0.119 | 2.78E-17 | 1.02E-12 |
| USP34 | 4 | 0.26006498 | 0.942 | 0.916 | 2.79E-17 | 1.02E-12 |
| CDK1 | 4 | -0.293331966 | 0.101 | 0.214 | 3.07E-17 | 1.13E-12 |
| RAP1GDS1 | 4 | -0.368201442 | 0.6 | 0.676 | 3.2E-17 | 1.17E-12 |
| TOM1L2 | 4 | 0.322764757 | 0.442 | 0.302 | 3.27E-17 | 1.2E-12 |
| AC074286.1 | 4 | -0.250505833 | 0.037 | 0.121 | 3.28E-17 | 1.2E-12 |
| ECT2 | 4 | -0.335367171 | 0.178 | 0.304 | 3.5E-17 | 1.28E-12 |
| FBN3 | 4 | 0.288720207 | 0.137 | 0.065 | 3.5E-17 | 1.28E-12 |
| RAB40C | 4 | 0.290464771 | 0.214 | 0.113 | 4.3E-17 | 1.57E-12 |
| BICDL1 | 4 | -0.363534455 | 0.192 | 0.287 | 4.44E-17 | 1.63E-12 |
| WDHD1 | 4 | -0.263387206 | 0.298 | 0.442 | 4.53E-17 | 1.66E-12 |
| CASK | 4 | 0.266764792 | 0.929 | 0.85 | 4.57E-17 | 1.67E-12 |
| LINC01572 | 4 | 0.401979021 | 0.607 | 0.529 | 4.6E-17 | 1.68E-12 |
| TTC21B | 4 | -0.465479086 | 0.533 | 0.592 | 4.61E-17 | 1.69E-12 |
| CPXM2 | 4 | 0.28515768 | 0.169 | 0.077 | 4.85E-17 | 1.77E-12 |
| TBX3 | 4 | -0.303436127 | 0.094 | 0.203 | 5.44E-17 | 1.99E-12 |
| ARID1B | 4 | -0.372199716 | 0.941 | 0.959 | 5.47E-17 | 2E-12 |
| AL358232.2 | 4 | 0.344830486 | 0.201 | 0.109 | 5.81E-17 | 2.13E-12 |
| AL807742.1 | 4 | -0.288127535 | 0.052 | 0.144 | 6.49E-17 | 2.37E-12 |
| EIF1AX | 4 | -0.362234456 | 0.332 | 0.454 | 6.9E-17 | 2.52E-12 |
| MTMR1 | 4 | -0.323125252 | 0.403 | 0.486 | 7.27E-17 | 2.66E-12 |
| ODF2L | 4 | 0.351515393 | 0.606 | 0.482 | 7.67E-17 | 2.81E-12 |
| ZNF713 | 4 | 0.300523841 | 0.261 | 0.15 | 7.69E-17 | 2.81E-12 |
| NECTIN2 | 4 | 0.261943346 | 0.176 | 0.084 | 8.13E-17 | 2.98E-12 |
| FNDC3A | 4 | -0.404909911 | 0.776 | 0.807 | 8.23E-17 | 3.01E-12 |
| LDLRAD4 | 4 | 0.358575389 | 0.615 | 0.481 | 9E-17 | 3.3E-12 |
| NPHP3 | 4 | 0.2873225 | 0.452 | 0.311 | 9.43E-17 | 3.45E-12 |
| TMEM123 | 4 | 0.271890467 | 0.311 | 0.187 | 9.66E-17 | 3.54E-12 |
| LRFN5 | 4 | -0.663991442 | 0.448 | 0.461 | 1E-16 | 3.67E-12 |
| FBXW7 | 4 | -0.466128013 | 0.678 | 0.746 | 1.02E-16 | 3.75E-12 |
| AC011586.2 | 4 | -0.576575077 | 0.032 | 0.103 | 1.08E-16 | 3.95E-12 |
| AC092821.3 | 4 | -0.290841394 | 0.176 | 0.304 | 1.13E-16 | 4.14E-12 |
| FAM92A | 4 | 0.271173505 | 0.303 | 0.184 | 1.15E-16 | 4.22E-12 |
| DIAPH1 | 4 | -0.351973941 | 0.334 | 0.438 | 1.22E-16 | 4.46E-12 |
| MCM8 | 4 | -0.256261286 | 0.224 | 0.36 | 1.26E-16 | 4.61E-12 |

| | | | | | | |
|---|---|---|---|---|---|---|
| SMYD3 | 4 | -0.358238533 | 0.9 | 0.928 | 1.37E-16 | 5.02E-12 |
| TPST2 | 4 | 0.267916704 | 0.184 | 0.093 | 1.38E-16 | 5.05E-12 |
| CDC42BPB | 4 | 0.314978624 | 0.567 | 0.428 | 1.4E-16 | 5.11E-12 |
| KIAA1109 | 4 | 0.27059103 | 0.794 | 0.695 | 1.4E-16 | 5.13E-12 |
| SLC6A15 | 4 | 0.537410531 | 0.292 | 0.202 | 1.57E-16 | 5.73E-12 |
| PHTF1 | 4 | 0.332357327 | 0.425 | 0.3 | 1.74E-16 | 6.38E-12 |
| TNKS2 | 4 | 0.297172803 | 0.585 | 0.465 | 1.75E-16 | 6.42E-12 |
| SYCP2 | 4 | -0.731387615 | 0.165 | 0.233 | 1.76E-16 | 6.46E-12 |
| PACRGL | 4 | 0.277089769 | 0.409 | 0.274 | 1.78E-16 | 6.51E-12 |
| YPEL1 | 4 | 0.316757468 | 0.23 | 0.136 | 1.86E-16 | 6.82E-12 |
| AP2A2 | 4 | 0.342605577 | 0.512 | 0.396 | 2.07E-16 | 7.6E-12 |
| AMMECR1 | 4 | 0.357902804 | 0.454 | 0.323 | 2.11E-16 | 7.73E-12 |
| HEG1 | 4 | 0.334539463 | 0.27 | 0.162 | 2.27E-16 | 8.32E-12 |
| TTC39B | 4 | 0.320434294 | 0.23 | 0.129 | 2.56E-16 | 9.38E-12 |
| PEAK1 | 4 | -0.357997523 | 0.697 | 0.788 | 2.61E-16 | 9.57E-12 |
| ZNF334 | 4 | 0.255722504 | 0.315 | 0.193 | 2.63E-16 | 9.64E-12 |
| ANP32B | 4 | -0.323343044 | 0.249 | 0.371 | 2.65E-16 | 9.69E-12 |
| GPCPD1 | 4 | -0.344843106 | 0.467 | 0.583 | 2.8E-16 | 1.03E-11 |
| TNPO1 | 4 | 0.306923559 | 0.744 | 0.659 | 3.13E-16 | 1.15E-11 |
| NFYC | 4 | 0.258531425 | 0.419 | 0.282 | 3.24E-16 | 1.19E-11 |
| ARHGAP8 | 4 | 0.354666198 | 0.323 | 0.2 | 3.27E-16 | 1.2E-11 |
| AFF3 | 4 | -0.439712856 | 0.485 | 0.621 | 3.32E-16 | 1.22E-11 |
| ARNTL | 4 | -0.276475341 | 0.068 | 0.157 | 3.35E-16 | 1.23E-11 |
| EDNRA | 4 | 0.519205609 | 0.159 | 0.125 | 3.45E-16 | 1.26E-11 |
| RBBP8 | 4 | -0.373903067 | 0.428 | 0.535 | 3.69E-16 | 1.35E-11 |
| MTURN | 4 | -0.255452452 | 0.125 | 0.236 | 3.74E-16 | 1.37E-11 |
| TPD52 | 4 | -0.417181798 | 0.243 | 0.373 | 3.83E-16 | 1.4E-11 |
| ARSB | 4 | 0.28256318 | 0.262 | 0.152 | 5.13E-16 | 1.88E-11 |
| NBPF1 | 4 | -0.289987404 | 0.234 | 0.357 | 5.49E-16 | 2.01E-11 |
| DBN1 | 4 | 0.282927895 | 0.24 | 0.139 | 5.59E-16 | 2.05E-11 |
| PPP6R2 | 4 | 0.309677607 | 0.73 | 0.624 | 5.66E-16 | 2.07E-11 |
| ASXL3 | 4 | 0.371713178 | 0.579 | 0.467 | 5.8E-16 | 2.12E-11 |
| FAM168A | 4 | -0.325119851 | 0.592 | 0.691 | 6.06E-16 | 2.22E-11 |
| TRMT9B | 4 | 0.279522928 | 0.334 | 0.21 | 6.39E-16 | 2.34E-11 |
| UCHL5 | 4 | -0.31313148 | 0.279 | 0.387 | 6.56E-16 | 2.4E-11 |
| SACS | 4 | 0.363643788 | 0.448 | 0.323 | 6.86E-16 | 2.51E-11 |
| ILF3 | 4 | 0.338007803 | 0.692 | 0.582 | 7.07E-16 | 2.59E-11 |
| IBTK | 4 | 0.320028011 | 0.569 | 0.445 | 7.15E-16 | 2.62E-11 |
| ADCY3 | 4 | -0.296139172 | 0.188 | 0.313 | 8.04E-16 | 2.94E-11 |
| PRPF40B | 4 | 0.310996919 | 0.317 | 0.206 | 8.14E-16 | 2.98E-11 |
| CTDSPL | 4 | -0.339705866 | 0.411 | 0.532 | 8.29E-16 | 3.03E-11 |
| DGLUCY | 4 | 0.318737979 | 0.37 | 0.247 | 8.58E-16 | 3.14E-11 |
| HPCAL1 | 4 | -0.344984804 | 0.212 | 0.337 | 8.64E-16 | 3.16E-11 |
| WWC2 | 4 | 0.314462351 | 0.528 | 0.387 | 8.81E-16 | 3.23E-11 |
| ITPR2 | 4 | 0.360605822 | 0.276 | 0.353 | 9.63E-16 | 3.52E-11 |
| GPM6B | 4 | -0.264166715 | 0.51 | 0.648 | 1.04E-15 | 3.79E-11 |
| CD109 | 4 | -0.262024156 | 0.059 | 0.149 | 1.06E-15 | 3.88E-11 |
| TNRC6A | 4 | 0.300927908 | 0.856 | 0.814 | 1.12E-15 | 4.11E-11 |
| CORO2B | 4 | 0.349025616 | 0.342 | 0.227 | 1.19E-15 | 4.36E-11 |
| KPNB1 | 4 | -0.316954022 | 0.475 | 0.594 | 1.2E-15 | 4.4E-11 |
| UNC5B | 4 | 0.27156318 | 0.201 | 0.105 | 1.27E-15 | 4.65E-11 |
| KIF15 | 4 | -0.263269618 | 0.162 | 0.282 | 1.29E-15 | 4.73E-11 |
| MPPED2 | 4 | 0.495663555 | 0.207 | 0.183 | 1.32E-15 | 4.83E-11 |
| MEF2A | 4 | -0.348104087 | 0.759 | 0.814 | 1.55E-15 | 5.66E-11 |
| SH3RF1 | 4 | -0.346721864 | 0.438 | 0.554 | 1.69E-15 | 6.17E-11 |
| PSMA1 | 4 | -0.352525845 | 0.39 | 0.492 | 1.75E-15 | 6.42E-11 |
| APBB2 | 4 | 0.402490146 | 0.563 | 0.45 | 1.76E-15 | 6.44E-11 |
| KNL1 | 4 | -0.281408055 | 0.116 | 0.226 | 1.99E-15 | 7.28E-11 |

| | | | | | | |
|---|---|---|---|---|---|---|
| TMEM161B | 4 | -0.314912681 | 0.409 | 0.525 | 2.04E-15 | 7.48E-11 |
| CUL4B | 4 | -0.357577331 | 0.348 | 0.434 | 2.28E-15 | 8.35E-11 |
| MTMR10 | 4 | -0.321310731 | 0.246 | 0.363 | 2.46E-15 | 8.99E-11 |
| TSHZ3 | 4 | -0.326569055 | 0.366 | 0.507 | 2.49E-15 | 9.12E-11 |
| USP25 | 4 | 0.306598828 | 0.655 | 0.523 | 2.61E-15 | 9.57E-11 |
| CDKAL1 | 4 | -0.512330346 | 0.795 | 0.829 | 2.69E-15 | 9.85E-11 |
| AEBP2 | 4 | -0.457792904 | 0.639 | 0.734 | 2.76E-15 | 1.01E-10 |
| SAMD5 | 4 | -0.344300179 | 0.562 | 0.654 | 2.89E-15 | 1.06E-10 |
| PRKN | 4 | 0.353156028 | 0.64 | 0.526 | 2.9E-15 | 1.06E-10 |
| TNRC18 | 4 | 0.376890704 | 0.635 | 0.527 | 3.27E-15 | 1.2E-10 |
| PCBP3 | 4 | -0.345026187 | 0.149 | 0.261 | 3.29E-15 | 1.21E-10 |
| LANCL2 | 4 | -0.305467833 | 0.194 | 0.3 | 3.42E-15 | 1.25E-10 |
| CIP2A | 4 | -0.324957537 | 0.192 | 0.311 | 3.47E-15 | 1.27E-10 |
| LRP6 | 4 | 0.306948784 | 0.674 | 0.568 | 3.58E-15 | 1.31E-10 |
| TFB1M | 4 | -0.330109385 | 0.299 | 0.414 | 3.74E-15 | 1.37E-10 |
| BTBD2 | 4 | 0.287745607 | 0.25 | 0.15 | 3.76E-15 | 1.38E-10 |
| AL390957.1 | 4 | -0.898291612 | 0.066 | 0.145 | 3.92E-15 | 1.43E-10 |
| NRP2 | 4 | 0.343726948 | 0.166 | 0.099 | 3.99E-15 | 1.46E-10 |
| MGAT5 | 4 | 0.36868808 | 0.49 | 0.471 | 4.5E-15 | 1.65E-10 |
| CSNK1G2 | 4 | 0.309057539 | 0.262 | 0.157 | 4.76E-15 | 1.74E-10 |
| MT-ATP8 | 4 | 0.270572638 | 0.12 | 0.058 | 4.89E-15 | 1.79E-10 |
| CTNND2 | 4 | 0.319887748 | 0.763 | 0.733 | 4.92E-15 | 1.8E-10 |
| C11orf58 | 4 | 0.312890102 | 0.502 | 0.383 | 5.23E-15 | 1.91E-10 |
| REV3L | 4 | -0.340656968 | 0.798 | 0.873 | 5.48E-15 | 2.01E-10 |
| MT-CYB | 4 | 0.494614559 | 0.314 | 0.225 | 5.64E-15 | 2.06E-10 |
| PHACTR2 | 4 | 0.355773715 | 0.621 | 0.563 | 5.79E-15 | 2.12E-10 |
| ANKS3 | 4 | 0.256882307 | 0.322 | 0.204 | 5.95E-15 | 2.18E-10 |
| TMEM220 | 4 | -0.28270248 | 0.112 | 0.214 | 6.39E-15 | 2.34E-10 |
| RRBP1 | 4 | 0.336006818 | 0.224 | 0.135 | 6.46E-15 | 2.37E-10 |
| ZNF277 | 4 | -0.321479067 | 0.283 | 0.396 | 6.97E-15 | 2.55E-10 |
| PRDM15 | 4 | -0.267693483 | 0.151 | 0.253 | 7.36E-15 | 2.69E-10 |
| ADGRA2 | 4 | 0.353724379 | 0.2 | 0.138 | 7.37E-15 | 2.7E-10 |
| HOMER1 | 4 | -0.414779216 | 0.423 | 0.502 | 7.41E-15 | 2.71E-10 |
| BAZ1B | 4 | -0.292625904 | 0.52 | 0.636 | 7.63E-15 | 2.79E-10 |
| TENM2 | 4 | -0.517090171 | 0.553 | 0.62 | 8.74E-15 | 3.2E-10 |
| AC068587.4 | 4 | 0.46039732 | 0.464 | 0.33 | 8.79E-15 | 3.22E-10 |
| ZNF804B | 4 | -0.5177698 | 0.095 | 0.194 | 8.9E-15 | 3.26E-10 |
| SH3PXD2B | 4 | 0.315668647 | 0.349 | 0.252 | 9.01E-15 | 3.3E-10 |
| AC009315.1 | 4 | 0.410819199 | 0.52 | 0.397 | 9.09E-15 | 3.33E-10 |
| C13orf42 | 4 | -0.350779153 | 0.057 | 0.139 | 9.33E-15 | 3.42E-10 |
| CREB3L2 | 4 | 0.284073172 | 0.363 | 0.239 | 9.46E-15 | 3.46E-10 |
| EXTL3 | 4 | 0.276995818 | 0.223 | 0.125 | 9.68E-15 | 3.54E-10 |
| MGAT4A | 4 | -0.273703288 | 0.099 | 0.199 | 1.03E-14 | 3.76E-10 |
| SLC30A8 | 4 | 0.28454384 | 0.134 | 0.057 | 1.03E-14 | 3.77E-10 |
| REEP5 | 4 | -0.286982483 | 0.182 | 0.293 | 1.07E-14 | 3.92E-10 |
| HHIP | 4 | -0.3013852 | 0.049 | 0.127 | 1.14E-14 | 4.16E-10 |
| JPH1 | 4 | -0.255462395 | 0.076 | 0.168 | 1.19E-14 | 4.34E-10 |
| CRTC1 | 4 | 0.321820765 | 0.413 | 0.292 | 1.21E-14 | 4.42E-10 |
| PLXDC2 | 4 | -0.533566199 | 0.665 | 0.669 | 1.25E-14 | 4.59E-10 |
| CTTNBP2 | 4 | -0.327336017 | 0.307 | 0.294 | 1.31E-14 | 4.78E-10 |
| COPG2 | 4 | 0.297410334 | 0.37 | 0.249 | 1.36E-14 | 4.96E-10 |
| CCDC144A | 4 | -0.252958092 | 0.096 | 0.194 | 1.37E-14 | 5.01E-10 |
| FST | 4 | 0.365616097 | 0.121 | 0.076 | 1.39E-14 | 5.11E-10 |
| RGS20 | 4 | 0.271497656 | 0.116 | 0.081 | 1.41E-14 | 5.15E-10 |
| NDUFAF6 | 4 | -0.306263734 | 0.351 | 0.466 | 1.44E-14 | 5.27E-10 |
| SCAI | 4 | 0.320930709 | 0.621 | 0.502 | 1.45E-14 | 5.31E-10 |
| TOGARAM1 | 4 | 0.30628926 | 0.528 | 0.427 | 1.5E-14 | 5.48E-10 |
| ZNF292 | 4 | -0.27628431 | 0.877 | 0.886 | 1.5E-14 | 5.5E-10 |

| | | | | | | |
|---|---|---|---|---|---|---|
| STT3B | 4 | -0.312117423 | 0.597 | 0.704 | 1.52E-14 | 5.58E-10 |
| PRRC2B | 4 | 0.267835957 | 0.856 | 0.788 | 1.55E-14 | 5.69E-10 |
| TOX3 | 4 | 0.459456763 | 0.316 | 0.302 | 1.74E-14 | 6.39E-10 |
| ATM | 4 | -0.299389913 | 0.668 | 0.752 | 1.75E-14 | 6.39E-10 |
| SHISA6 | 4 | -0.452805632 | 0.105 | 0.173 | 1.83E-14 | 6.69E-10 |
| SLAIN2 | 4 | -0.338264527 | 0.423 | 0.529 | 1.88E-14 | 6.9E-10 |
| HPRT1 | 4 | -0.306685092 | 0.311 | 0.436 | 1.9E-14 | 6.95E-10 |
| CTNND1 | 4 | 0.270881601 | 0.72 | 0.611 | 1.91E-14 | 6.97E-10 |
| FHOD3 | 4 | 0.292527543 | 0.423 | 0.294 | 1.91E-14 | 7E-10 |
| IL1RAP | 4 | -0.306207492 | 0.085 | 0.176 | 1.96E-14 | 7.16E-10 |
| LMBR1 | 4 | -0.317886748 | 0.567 | 0.659 | 1.97E-14 | 7.19E-10 |
| NAMPT | 4 | -0.384126897 | 0.206 | 0.312 | 1.99E-14 | 7.3E-10 |
| GOLGA8A | 4 | 0.295406376 | 0.732 | 0.621 | 2.08E-14 | 7.61E-10 |
| TRDMT1 | 4 | 0.259011565 | 0.257 | 0.152 | 2.18E-14 | 7.98E-10 |
| AC006148.1 | 4 | -0.334662856 | 0.075 | 0.166 | 2.35E-14 | 8.6E-10 |
| RESF1 | 4 | 0.286292583 | 0.307 | 0.196 | 2.4E-14 | 8.79E-10 |
| ALCAM | 4 | -0.624536137 | 0.239 | 0.342 | 2.45E-14 | 8.98E-10 |
| TXNDC16 | 4 | -0.354632264 | 0.226 | 0.34 | 2.57E-14 | 9.42E-10 |
| RPS6KA6 | 4 | 0.311538782 | 0.303 | 0.191 | 2.72E-14 | 9.98E-10 |
| SLC24A2 | 4 | -0.471170934 | 0.325 | 0.444 | 2.82E-14 | 1.03E-09 |
| RBBP7 | 4 | -0.304621044 | 0.396 | 0.514 | 3.06E-14 | 1.12E-09 |
| SLC35F5 | 4 | 0.274126169 | 0.532 | 0.402 | 3.2E-14 | 1.17E-09 |
| IGDCC3 | 4 | 0.345409712 | 0.302 | 0.253 | 3.26E-14 | 1.19E-09 |
| NCAPD3 | 4 | -0.303347809 | 0.266 | 0.375 | 3.31E-14 | 1.21E-09 |
| SRSF6 | 4 | 0.282157754 | 0.481 | 0.363 | 3.34E-14 | 1.22E-09 |
| AOPEP | 4 | 0.299223214 | 0.822 | 0.747 | 3.54E-14 | 1.3E-09 |
| LGMN | 4 | -0.289161781 | 0.09 | 0.177 | 3.77E-14 | 1.38E-09 |
| ARHGAP44 | 4 | -0.415319011 | 0.314 | 0.405 | 3.81E-14 | 1.39E-09 |
| TMEM132B | 4 | 0.45067882 | 0.135 | 0.071 | 3.84E-14 | 1.41E-09 |
| NT5C3A | 4 | -0.317246052 | 0.328 | 0.444 | 3.96E-14 | 1.45E-09 |
| BCL11A | 4 | 0.426748247 | 0.366 | 0.264 | 3.99E-14 | 1.46E-09 |
| POLQ | 4 | -0.286735956 | 0.21 | 0.333 | 4E-14 | 1.47E-09 |
| TENT5A | 4 | -0.441039065 | 0.06 | 0.143 | 4.02E-14 | 1.47E-09 |
| ASPM | 4 | -0.36532809 | 0.143 | 0.253 | 4.17E-14 | 1.53E-09 |
| ENOX2 | 4 | 0.462335541 | 0.435 | 0.321 | 4.36E-14 | 1.6E-09 |
| SMARCA2 | 4 | -0.391638468 | 0.339 | 0.44 | 4.39E-14 | 1.61E-09 |
| ARHGEF6 | 4 | -0.263246634 | 0.077 | 0.162 | 4.73E-14 | 1.73E-09 |
| SH3KBP1 | 4 | -0.296481787 | 0.566 | 0.691 | 5.17E-14 | 1.89E-09 |
| GLCE | 4 | -0.373588531 | 0.309 | 0.392 | 5.21E-14 | 1.91E-09 |
| MMP16 | 4 | -0.319841985 | 0.658 | 0.733 | 5.33E-14 | 1.95E-09 |
| GRK4 | 4 | -0.285851415 | 0.193 | 0.305 | 5.58E-14 | 2.04E-09 |
| GOLGB1 | 4 | 0.279160558 | 0.558 | 0.438 | 5.6E-14 | 2.05E-09 |
| NBEAL1 | 4 | 0.270944171 | 0.577 | 0.446 | 5.65E-14 | 2.07E-09 |
| NUBPL | 4 | 0.302957977 | 0.605 | 0.485 | 5.87E-14 | 2.15E-09 |
| IGFBP5 | 4 | 0.257957219 | 0.108 | 0.045 | 6.07E-14 | 2.22E-09 |
| ARRDC3-AS1 | 4 | -0.773277917 | 0.298 | 0.282 | 6.18E-14 | 2.26E-09 |
| PDK1 | 4 | -0.424834328 | 0.275 | 0.393 | 6.34E-14 | 2.32E-09 |
| PDE10A | 4 | 0.456913341 | 0.355 | 0.241 | 6.39E-14 | 2.34E-09 |
| USP46 | 4 | 0.250850462 | 0.291 | 0.184 | 6.76E-14 | 2.48E-09 |
| CYTH3 | 4 | 0.34072425 | 0.319 | 0.21 | 6.85E-14 | 2.51E-09 |
| CAMSAP1 | 4 | 0.301735938 | 0.478 | 0.361 | 7.05E-14 | 2.58E-09 |
| ACTN2 | 4 | -0.254101181 | 0.046 | 0.117 | 7.17E-14 | 2.63E-09 |
| SNX9 | 4 | 0.299011243 | 0.467 | 0.341 | 7.35E-14 | 2.69E-09 |
| GART | 4 | -0.283430676 | 0.347 | 0.45 | 7.7E-14 | 2.82E-09 |
| DMXL1 | 4 | 0.290131313 | 0.629 | 0.518 | 7.76E-14 | 2.84E-09 |
| ADGRL1 | 4 | 0.301425291 | 0.341 | 0.228 | 8.15E-14 | 2.98E-09 |
| MAP7D3 | 4 | -0.315534435 | 0.168 | 0.262 | 8.36E-14 | 3.06E-09 |
| PRTFDC1 | 4 | -0.280374193 | 0.189 | 0.302 | 8.53E-14 | 3.12E-09 |

| | | | | | | |
|---|---|---|---|---|---|---|
| KIF2A | 4 | -0.322706823 | 0.525 | 0.616 | 8.84E-14 | 3.24E-09 |
| ZNF235 | 4 | 0.250953412 | 0.33 | 0.214 | 9.4E-14 | 3.44E-09 |
| TTC28-AS1 | 4 | 0.25539236 | 0.347 | 0.229 | 9.63E-14 | 3.53E-09 |
| SLC22A15 | 4 | -0.430988603 | 0.229 | 0.319 | 1.01E-13 | 3.71E-09 |
| TDRD7 | 4 | -0.257085619 | 0.121 | 0.206 | 1.07E-13 | 3.93E-09 |
| PIP5K1A | 4 | 0.307195809 | 0.635 | 0.525 | 1.08E-13 | 3.97E-09 |
| STXBP5 | 4 | -0.332037549 | 0.552 | 0.633 | 1.1E-13 | 4.04E-09 |
| UBE2D1 | 4 | -0.3108738 | 0.181 | 0.274 | 1.14E-13 | 4.16E-09 |
| ANKRD27 | 4 | 0.25049466 | 0.285 | 0.178 | 1.24E-13 | 4.52E-09 |
| PBX3 | 4 | 0.553202119 | 0.56 | 0.5 | 1.3E-13 | 4.76E-09 |
| IGSF11 | 4 | -0.344255273 | 0.081 | 0.156 | 1.36E-13 | 4.99E-09 |
| JARID2 | 4 | 0.31697323 | 0.75 | 0.666 | 1.39E-13 | 5.08E-09 |
| PPCDC | 4 | 0.328282851 | 0.288 | 0.196 | 1.39E-13 | 5.09E-09 |
| PARPBP | 4 | -0.261513729 | 0.169 | 0.281 | 1.4E-13 | 5.12E-09 |
| USP32 | 4 | -0.33058754 | 0.637 | 0.692 | 1.4E-13 | 5.13E-09 |
| PAPSS2 | 4 | -0.279242681 | 0.088 | 0.176 | 1.4E-13 | 5.13E-09 |
| PJA2 | 4 | -0.340557908 | 0.44 | 0.52 | 1.43E-13 | 5.23E-09 |
| MT-CO3 | 4 | 0.590936418 | 0.451 | 0.407 | 1.44E-13 | 5.28E-09 |
| PLD5 | 4 | -0.669866363 | 0.172 | 0.262 | 1.48E-13 | 5.41E-09 |
| SOS1 | 4 | -0.347870974 | 0.767 | 0.794 | 1.49E-13 | 5.46E-09 |
| CEP162 | 4 | 0.272069507 | 0.435 | 0.314 | 1.57E-13 | 5.73E-09 |
| NAV3 | 4 | 0.834648984 | 0.464 | 0.375 | 1.63E-13 | 5.95E-09 |
| CLCN3 | 4 | -0.312596384 | 0.441 | 0.53 | 1.64E-13 | 0.000000006 |
| MSL1 | 4 | 0.250021185 | 0.376 | 0.256 | 1.65E-13 | 6.05E-09 |
| ITGB1 | 4 | -0.343869909 | 0.475 | 0.573 | 1.69E-13 | 6.2E-09 |
| EPB41L2 | 4 | -0.263697588 | 0.537 | 0.658 | 1.71E-13 | 6.25E-09 |
| GRK3 | 4 | -0.336885199 | 0.386 | 0.464 | 1.74E-13 | 6.39E-09 |
| KIAA2026 | 4 | 0.297507355 | 0.567 | 0.471 | 1.77E-13 | 6.49E-09 |
| YTHDC2 | 4 | -0.28569868 | 0.636 | 0.654 | 1.79E-13 | 6.56E-09 |
| SFRP1 | 4 | 0.268966866 | 0.161 | 0.082 | 1.8E-13 | 6.57E-09 |
| AC139887.2 | 4 | 0.310841996 | 0.354 | 0.26 | 1.85E-13 | 6.76E-09 |
| MTA3 | 4 | -0.310962623 | 0.637 | 0.692 | 1.89E-13 | 6.91E-09 |
| USP37 | 4 | -0.273116322 | 0.392 | 0.512 | 2.04E-13 | 7.46E-09 |
| FSTL5 | 4 | -0.534094077 | 0.066 | 0.148 | 2.16E-13 | 7.9E-09 |
| GOLM1 | 4 | 0.270111435 | 0.454 | 0.328 | 2.16E-13 | 7.91E-09 |
| LPP | 4 | 0.350475822 | 0.798 | 0.724 | 2.24E-13 | 8.2E-09 |
| CNN3 | 4 | 0.294509471 | 0.38 | 0.273 | 2.34E-13 | 8.56E-09 |
| CHST9 | 4 | -0.573391401 | 0.08 | 0.159 | 2.39E-13 | 8.75E-09 |
| DIAPH3 | 4 | -0.356317335 | 0.302 | 0.423 | 2.44E-13 | 8.92E-09 |
| TOP2A | 4 | -0.347720651 | 0.142 | 0.248 | 2.55E-13 | 9.33E-09 |
| AMPH | 4 | 0.272036357 | 0.266 | 0.162 | 2.6E-13 | 9.5E-09 |
| PFKP | 4 | -0.308876175 | 0.279 | 0.386 | 2.82E-13 | 1.03E-08 |
| ITGAV | 4 | -0.415969188 | 0.45 | 0.548 | 2.93E-13 | 1.07E-08 |
| SCN1A | 4 | -0.419022329 | 0.037 | 0.107 | 2.97E-13 | 1.09E-08 |
| CMSS1 | 4 | -0.316250806 | 0.649 | 0.715 | 3.1E-13 | 1.14E-08 |
| FANCB | 4 | -0.250310822 | 0.147 | 0.254 | 3.28E-13 | 0.000000012 |
| ZMIZ1 | 4 | 0.340858253 | 0.544 | 0.463 | 3.31E-13 | 1.21E-08 |
| ADAM12 | 4 | 0.421659151 | 0.106 | 0.064 | 3.43E-13 | 1.25E-08 |
| SHROOM2 | 4 | 0.303983781 | 0.472 | 0.354 | 3.7E-13 | 1.35E-08 |
| MAPKBP1 | 4 | 0.251785838 | 0.308 | 0.198 | 3.87E-13 | 1.42E-08 |
| PCDH19 | 4 | -0.477092084 | 0.151 | 0.202 | 3.87E-13 | 1.42E-08 |
| PSMD14 | 4 | -0.291446024 | 0.45 | 0.526 | 3.89E-13 | 1.43E-08 |
| KCNJ3 | 4 | -0.418769492 | 0.072 | 0.15 | 4.12E-13 | 1.51E-08 |
| PPL | 4 | -0.276885693 | 0.06 | 0.135 | 4.13E-13 | 1.51E-08 |
| ADCY9 | 4 | 0.356257456 | 0.463 | 0.366 | 4.25E-13 | 1.56E-08 |
| TULP3 | 4 | 0.273333213 | 0.329 | 0.221 | 4.34E-13 | 1.59E-08 |
| SMCHD1 | 4 | -0.274866086 | 0.798 | 0.837 | 4.81E-13 | 1.76E-08 |
| AC025159.1 | 4 | 0.31694366 | 0.261 | 0.161 | 4.85E-13 | 1.78E-08 |

| | | | | | | |
|---|---|---|---|---|---|---|
| RBM5 | 4 | 0.268449301 | 0.686 | 0.582 | 4.99E-13 | 1.83E-08 |
| STAT1 | 4 | -0.379164806 | 0.196 | 0.29 | 5.01E-13 | 1.83E-08 |
| LAMC1 | 4 | -0.336847106 | 0.474 | 0.529 | 5.05E-13 | 1.85E-08 |
| TESK2 | 4 | -0.267262966 | 0.1 | 0.193 | 5.47E-13 | 0.00000002 |
| MYO19 | 4 | -0.291979175 | 0.363 | 0.475 | 5.69E-13 | 2.08E-08 |
| CABLES1 | 4 | 0.281321565 | 0.265 | 0.163 | 5.95E-13 | 2.18E-08 |
| MAU2 | 4 | 0.284709138 | 0.409 | 0.299 | 6.43E-13 | 2.35E-08 |
| SYNE2 | 4 | 0.269286524 | 0.797 | 0.693 | 6.64E-13 | 2.43E-08 |
| ADSS | 4 | -0.287708203 | 0.29 | 0.383 | 6.66E-13 | 2.44E-08 |
| WNK3 | 4 | 0.267245612 | 0.357 | 0.243 | 6.81E-13 | 2.49E-08 |
| VDAC3 | 4 | -0.298015886 | 0.238 | 0.325 | 7.08E-13 | 2.59E-08 |
| SNTG2 | 4 | 0.478998054 | 0.298 | 0.232 | 7.18E-13 | 2.63E-08 |
| AL136418.1 | 4 | -0.287795659 | 0.033 | 0.1 | 7.35E-13 | 2.69E-08 |
| TTC7A | 4 | 0.25750401 | 0.234 | 0.146 | 7.83E-13 | 2.87E-08 |
| PTCHD4 | 4 | -0.397492692 | 0.158 | 0.263 | 8.01E-13 | 2.93E-08 |
| PPHLN1 | 4 | -0.309775124 | 0.589 | 0.666 | 8.05E-13 | 2.95E-08 |
| FAM162A | 4 | -0.434072302 | 0.156 | 0.217 | 8.1E-13 | 2.96E-08 |
| GPAM | 4 | -0.278518791 | 0.112 | 0.19 | 9.35E-13 | 3.42E-08 |
| FAM120B | 4 | 0.258145576 | 0.591 | 0.477 | 9.67E-13 | 3.54E-08 |
| YWHAZ | 4 | -0.294528954 | 0.535 | 0.606 | 1.03E-12 | 3.76E-08 |
| DZIP1L | 4 | 0.256937547 | 0.312 | 0.21 | 1.08E-12 | 3.97E-08 |
| RNASEH2B | 4 | -0.276527675 | 0.239 | 0.354 | 1.09E-12 | 3.99E-08 |
| TBL1XR1 | 4 | 0.275559346 | 0.89 | 0.853 | 1.11E-12 | 4.05E-08 |
| PPP2R2A | 4 | -0.294506654 | 0.611 | 0.671 | 1.14E-12 | 4.17E-08 |
| RHBDL3 | 4 | 0.261442971 | 0.263 | 0.303 | 1.14E-12 | 4.18E-08 |
| STRN3 | 4 | -0.346662984 | 0.782 | 0.807 | 1.16E-12 | 4.25E-08 |
| AGPS | 4 | -0.295970664 | 0.504 | 0.581 | 1.18E-12 | 4.32E-08 |
| PDE5A | 4 | 0.439211522 | 0.26 | 0.261 | 1.19E-12 | 4.34E-08 |
| OBSCN | 4 | 0.279566432 | 0.435 | 0.324 | 1.19E-12 | 4.37E-08 |
| NNT | 4 | -0.309371032 | 0.403 | 0.488 | 1.21E-12 | 4.42E-08 |
| SERGEF | 4 | -0.328690927 | 0.339 | 0.426 | 1.22E-12 | 4.46E-08 |
| MT-CO2 | 4 | 0.575014312 | 0.404 | 0.359 | 1.24E-12 | 4.55E-08 |
| ST3GAL2 | 4 | 0.316492342 | 0.351 | 0.255 | 1.3E-12 | 4.77E-08 |
| ERO1A | 4 | -0.29599928 | 0.178 | 0.287 | 1.37E-12 | 5.01E-08 |
| RNF216 | 4 | 0.287105488 | 0.629 | 0.525 | 1.37E-12 | 5.01E-08 |
| MAN2A1 | 4 | 0.320342093 | 0.567 | 0.443 | 1.52E-12 | 5.56E-08 |
| ZDHHC21 | 4 | -0.287022772 | 0.647 | 0.7 | 1.66E-12 | 6.07E-08 |
| CSNK1D | 4 | 0.251797969 | 0.394 | 0.279 | 1.79E-12 | 6.55E-08 |
| RBMS1 | 4 | 0.301518211 | 0.646 | 0.535 | 1.82E-12 | 6.68E-08 |
| GATAD2B | 4 | 0.307281198 | 0.771 | 0.673 | 1.9E-12 | 6.95E-08 |
| SCML2 | 4 | -0.270151301 | 0.33 | 0.45 | 1.94E-12 | 7.09E-08 |
| RNF150 | 4 | 0.433639491 | 0.487 | 0.381 | 1.97E-12 | 7.22E-08 |
| WWP2 | 4 | 0.261209647 | 0.368 | 0.257 | 2E-12 | 7.32E-08 |
| NONO | 4 | 0.283519379 | 0.496 | 0.397 | 2.07E-12 | 7.58E-08 |
| UCHL3 | 4 | -0.27431739 | 0.214 | 0.301 | 2.09E-12 | 7.67E-08 |
| ABR | 4 | 0.329397975 | 0.45 | 0.34 | 2.15E-12 | 7.86E-08 |
| STK38 | 4 | 0.267899152 | 0.481 | 0.365 | 2.16E-12 | 7.91E-08 |
| GRB10 | 4 | 0.307438737 | 0.494 | 0.373 | 2.16E-12 | 7.91E-08 |
| FBXW2 | 4 | -0.270621365 | 0.303 | 0.396 | 2.16E-12 | 7.92E-08 |
| XPNPEP3 | 4 | 0.255302773 | 0.339 | 0.229 | 2.17E-12 | 7.95E-08 |
| FOXK1 | 4 | 0.31020595 | 0.534 | 0.427 | 2.2E-12 | 8.04E-08 |
| AC073050.1 | 4 | -0.435576525 | 0.285 | 0.306 | 2.34E-12 | 8.56E-08 |
| PER3 | 4 | -0.254737776 | 0.115 | 0.208 | 2.47E-12 | 9.03E-08 |
| BRINP2 | 4 | -0.303375442 | 0.046 | 0.114 | 2.47E-12 | 9.05E-08 |
| SPOCK1 | 4 | -0.463545718 | 0.345 | 0.364 | 2.48E-12 | 9.07E-08 |
| ZNF730 | 4 | 0.318397703 | 0.249 | 0.205 | 2.59E-12 | 9.47E-08 |
| GABPA | 4 | -0.304854238 | 0.226 | 0.312 | 2.68E-12 | 0.000000098 |
| HECTD2 | 4 | 0.279705406 | 0.415 | 0.306 | 2.72E-12 | 9.96E-08 |

| | | | | | | |
|---|---|---|---|---|---|---|
| FAM117B | 4 | 0.292304371 | 0.487 | 0.369 | 2.74E-12 | 0.0000001 |
| PDE4DIP | 4 | -0.271449219 | 0.282 | 0.338 | 2.75E-12 | 0.000000101 |
| PCDH9 | 4 | -0.394180138 | 0.241 | 0.358 | 2.78E-12 | 0.000000102 |
| EGFR | 4 | -0.361199364 | 0.176 | 0.269 | 2.88E-12 | 0.000000105 |
| MAMLD1 | 4 | -0.406153431 | 0.228 | 0.322 | 2.94E-12 | 0.000000108 |
| ENO1 | 4 | -0.386692944 | 0.392 | 0.479 | 3.09E-12 | 0.000000113 |
| MGLL | 4 | -0.261076079 | 0.109 | 0.202 | 3.12E-12 | 0.000000114 |
| ANK2 | 4 | -0.340369095 | 0.772 | 0.848 | 3.17E-12 | 0.000000116 |
| AP001011.1 | 4 | -0.254684976 | 0.179 | 0.272 | 3.27E-12 | 0.00000012 |
| MT-ND5 | 4 | 0.332357122 | 0.182 | 0.116 | 3.52E-12 | 0.000000129 |
| PLCB4 | 4 | 0.310403183 | 0.744 | 0.664 | 3.82E-12 | 0.00000014 |
| UMAD1 | 4 | -0.296967931 | 0.446 | 0.529 | 3.9E-12 | 0.000000143 |
| TAOK3 | 4 | -0.330746188 | 0.355 | 0.433 | 3.98E-12 | 0.000000146 |
| GCLC | 4 | -0.384561216 | 0.244 | 0.209 | 4.04E-12 | 0.000000148 |
| SKAP2 | 4 | 0.400584777 | 0.233 | 0.176 | 4.08E-12 | 0.000000149 |
| ACTR3B | 4 | -0.298154207 | 0.348 | 0.431 | 4.11E-12 | 0.000000151 |
| KCTD13 | 4 | -0.283552439 | 0.238 | 0.335 | 4.14E-12 | 0.000000151 |
| CALD1 | 4 | 0.465910918 | 0.535 | 0.415 | 4.15E-12 | 0.000000152 |
| ADD1 | 4 | 0.273809332 | 0.594 | 0.485 | 4.24E-12 | 0.000000155 |
| CBL | 4 | 0.298517092 | 0.44 | 0.343 | 4.3E-12 | 0.000000157 |
| UBE2H | 4 | 0.288355163 | 0.63 | 0.527 | 4.35E-12 | 0.000000159 |
| SEPTIN9 | 4 | 0.315247729 | 0.561 | 0.451 | 4.62E-12 | 0.000000169 |
| RGS12 | 4 | 0.25182281 | 0.578 | 0.459 | 4.7E-12 | 0.000000172 |
| AC063938.1 | 4 | -0.288294311 | 0.046 | 0.114 | 4.78E-12 | 0.000000175 |
| IPMK | 4 | -0.273719407 | 0.188 | 0.284 | 5.06E-12 | 0.000000185 |
| KIAA1841 | 4 | 0.264590774 | 0.451 | 0.334 | 5.07E-12 | 0.000000186 |
| USP6NL | 4 | 0.295912421 | 0.515 | 0.422 | 5.08E-12 | 0.000000186 |
| EFHD1 | 4 | -0.251111947 | 0.066 | 0.142 | 5.11E-12 | 0.000000187 |
| ATP2C1 | 4 | 0.274229237 | 0.645 | 0.554 | 5.15E-12 | 0.000000189 |
| NCOA7 | 4 | -0.253368432 | 0.151 | 0.25 | 5.24E-12 | 0.000000192 |
| FGD4 | 4 | 0.281894337 | 0.53 | 0.407 | 5.41E-12 | 0.000000198 |
| ULK1 | 4 | 0.261657512 | 0.378 | 0.269 | 5.61E-12 | 0.000000205 |
| MT-ATP6 | 4 | 0.43214851 | 0.304 | 0.264 | 5.69E-12 | 0.000000208 |
| TMPO | 4 | -0.301026851 | 0.426 | 0.526 | 5.84E-12 | 0.000000214 |
| ATF7IP2 | 4 | 0.333095712 | 0.347 | 0.24 | 5.86E-12 | 0.000000215 |
| PDE9A | 4 | 0.351842212 | 0.353 | 0.258 | 6.27E-12 | 0.00000023 |
| HMGN3 | 4 | 0.290180001 | 0.436 | 0.338 | 6.32E-12 | 0.000000231 |
| DOT1L | 4 | 0.263045389 | 0.385 | 0.273 | 6.46E-12 | 0.000000237 |
| NUP58 | 4 | -0.277093148 | 0.536 | 0.619 | 6.59E-12 | 0.000000241 |
| ZBED3-AS1 | 4 | 0.258628403 | 0.311 | 0.213 | 7.11E-12 | 0.00000026 |
| SINHCAF | 4 | 0.264509707 | 0.483 | 0.367 | 7.11E-12 | 0.00000026 |
| HPSE2 | 4 | -0.328723577 | 0.166 | 0.267 | 7.48E-12 | 0.000000274 |
| MTX2 | 4 | -0.279552195 | 0.553 | 0.634 | 7.65E-12 | 0.00000028 |
| PITPNA | 4 | 0.252040217 | 0.314 | 0.22 | 8.39E-12 | 0.000000307 |
| ZNF710 | 4 | 0.266450618 | 0.26 | 0.175 | 8.48E-12 | 0.00000031 |
| UBE2R2 | 4 | 0.29090273 | 0.759 | 0.667 | 8.51E-12 | 0.000000312 |
| PLPPR1 | 4 | -0.425787961 | 0.131 | 0.202 | 8.51E-12 | 0.000000312 |
| FBXL20 | 4 | 0.297956443 | 0.688 | 0.583 | 8.58E-12 | 0.000000314 |
| CCAR1 | 4 | 0.267220721 | 0.802 | 0.722 | 8.6E-12 | 0.000000315 |
| GDAP1 | 4 | -0.354497848 | 0.216 | 0.322 | 8.68E-12 | 0.000000318 |
| TUSC3 | 4 | 0.251265366 | 0.725 | 0.685 | 9.02E-12 | 0.00000033 |
| VAMP4 | 4 | -0.250646553 | 0.151 | 0.231 | 9.47E-12 | 0.000000347 |
| ZBTB18 | 4 | -0.258822449 | 0.142 | 0.231 | 9.52E-12 | 0.000000348 |
| PGK1 | 4 | -0.51177408 | 0.277 | 0.346 | 9.82E-12 | 0.000000359 |
| AATF | 4 | -0.299171986 | 0.34 | 0.427 | 1.02E-11 | 0.000000375 |
| GALNT18 | 4 | -0.322307783 | 0.301 | 0.409 | 1.03E-11 | 0.000000377 |
| IQSEC1 | 4 | 0.345223509 | 0.47 | 0.379 | 1.15E-11 | 0.00000042 |
| DNMT1 | 4 | -0.252557164 | 0.369 | 0.487 | 1.19E-11 | 0.000000435 |

| | | | | | | |
|---|---|---|---|---|---|---|
| PTEN | 4 | -0.336100846 | 0.649 | 0.707 | 1.49E-11 | 0.000000545 |
| EFCAB13 | 4 | -0.263599197 | 0.159 | 0.255 | 1.72E-11 | 0.000000629 |
| LRRFIP1 | 4 | 0.311147369 | 0.582 | 0.495 | 1.77E-11 | 0.000000647 |
| FEZ2 | 4 | 0.25566396 | 0.37 | 0.272 | 1.97E-11 | 0.000000723 |
| PALM2-AKAP2 | 4 | -0.266019467 | 0.385 | 0.5 | 2.07E-11 | 0.000000758 |
| STARD13 | 4 | -0.493723892 | 0.531 | 0.603 | 2.12E-11 | 0.000000777 |
| CSNK1E | 4 | 0.253982062 | 0.452 | 0.34 | 2.17E-11 | 0.000000796 |
| THUMPD2 | 4 | -0.290012953 | 0.402 | 0.489 | 2.25E-11 | 0.000000823 |
| ATP1B1 | 4 | -0.297820993 | 0.194 | 0.277 | 2.29E-11 | 0.000000837 |
| MDM4 | 4 | 0.274340121 | 0.639 | 0.56 | 2.61E-11 | 0.000000956 |
| NCOA6 | 4 | 0.251142561 | 0.564 | 0.456 | 2.71E-11 | 0.000000993 |
| AC090517.5 | 4 | 0.26191617 | 0.184 | 0.113 | 3.11E-11 | 0.00000114 |
| FGD6 | 4 | -0.361313186 | 0.277 | 0.336 | 3.11E-11 | 0.00000114 |
| EGR1 | 4 | -0.286884158 | 0.057 | 0.125 | 3.12E-11 | 0.00000114 |
| CADM2 | 4 | -0.782264469 | 0.392 | 0.454 | 3.3E-11 | 0.00000121 |
| AC079142.1 | 4 | 0.29295841 | 0.697 | 0.603 | 3.3E-11 | 0.00000121 |
| ANKRD11 | 4 | 0.25462711 | 0.87 | 0.825 | 3.33E-11 | 0.00000122 |
| TRIP4 | 4 | 0.27410364 | 0.309 | 0.22 | 3.39E-11 | 0.00000124 |
| BCAR3 | 4 | 0.352111998 | 0.324 | 0.232 | 3.47E-11 | 0.00000127 |
| INTS7 | 4 | -0.263300945 | 0.365 | 0.466 | 3.58E-11 | 0.00000131 |
| LINC02315 | 4 | -0.30625235 | 0.724 | 0.791 | 3.66E-11 | 0.00000134 |
| RYR2 | 4 | -0.488073335 | 0.202 | 0.209 | 3.9E-11 | 0.00000143 |
| SLC35F3 | 4 | 0.439736941 | 0.168 | 0.103 | 3.93E-11 | 0.00000144 |
| CCNH | 4 | -0.295497711 | 0.514 | 0.591 | 3.99E-11 | 0.00000146 |
| PWWP2A | 4 | 0.257838841 | 0.451 | 0.348 | 4.33E-11 | 0.00000159 |
| MIR4713HG | 4 | 0.329232706 | 0.157 | 0.108 | 4.53E-11 | 0.00000166 |
| DAAM2 | 4 | -0.46808731 | 0.149 | 0.224 | 4.61E-11 | 0.00000169 |
| TTYH3 | 4 | 0.263011024 | 0.283 | 0.19 | 4.62E-11 | 0.00000169 |
| TMEM163 | 4 | 0.36355332 | 0.137 | 0.155 | 4.64E-11 | 0.0000017 |
| ZNF251 | 4 | 0.263089312 | 0.376 | 0.278 | 4.98E-11 | 0.00000182 |
| AC092683.1 | 4 | -0.283602231 | 0.276 | 0.368 | 5.27E-11 | 0.00000193 |
| FAM228B | 4 | 0.25966188 | 0.31 | 0.215 | 5.34E-11 | 0.00000195 |
| ZNF804A | 4 | -0.544380446 | 0.159 | 0.257 | 5.5E-11 | 0.00000201 |
| FGGY | 4 | 0.251521896 | 0.322 | 0.22 | 5.53E-11 | 0.00000203 |
| SYDE2 | 4 | -0.27468053 | 0.145 | 0.208 | 5.57E-11 | 0.00000204 |
| LUC7L | 4 | 0.257424309 | 0.69 | 0.593 | 5.68E-11 | 0.00000208 |
| EYA2 | 4 | 0.402331415 | 0.154 | 0.133 | 6.02E-11 | 0.0000022 |
| TARS | 4 | -0.265588552 | 0.247 | 0.332 | 6.16E-11 | 0.00000226 |
| ADAM19 | 4 | 0.267722282 | 0.142 | 0.09 | 6.34E-11 | 0.00000232 |
| PDE8B | 4 | 0.259208228 | 0.112 | 0.059 | 6.7E-11 | 0.00000245 |
| RAB31 | 4 | -0.253005453 | 0.17 | 0.269 | 6.9E-11 | 0.00000253 |
| NPAT | 4 | -0.263973629 | 0.324 | 0.424 | 7.14E-11 | 0.00000262 |
| ACTN4 | 4 | 0.282594235 | 0.458 | 0.353 | 7.2E-11 | 0.00000263 |
| MCTP1 | 4 | -0.488450224 | 0.232 | 0.303 | 7.28E-11 | 0.00000266 |
| ARHGEF26 | 4 | -0.270603403 | 0.207 | 0.285 | 7.34E-11 | 0.00000269 |
| DEK | 4 | -0.276753761 | 0.441 | 0.54 | 7.75E-11 | 0.00000284 |
| PUS7 | 4 | -0.281500747 | 0.347 | 0.452 | 8.47E-11 | 0.0000031 |
| VEGFA | 4 | -0.295122386 | 0.392 | 0.35 | 8.63E-11 | 0.00000316 |
| TMEM14B | 4 | 0.304828771 | 0.545 | 0.448 | 8.95E-11 | 0.00000328 |
| VEGFC | 4 | -0.420745523 | 0.145 | 0.223 | 9.34E-11 | 0.00000342 |
| ATP11C | 4 | 0.263301432 | 0.718 | 0.717 | 9.94E-11 | 0.00000364 |
| PRKRIP1 | 4 | 0.271511973 | 0.51 | 0.41 | 1.01E-10 | 0.00000369 |
| MT-ND3 | 4 | 0.336580955 | 0.209 | 0.157 | 1.02E-10 | 0.00000373 |
| RNF115 | 4 | 0.266375471 | 0.497 | 0.399 | 1.03E-10 | 0.00000378 |
| SATB1-AS1 | 4 | -0.257096267 | 0.107 | 0.185 | 1.04E-10 | 0.00000382 |
| MT-ND4 | 4 | 0.470251674 | 0.314 | 0.246 | 1.1E-10 | 0.00000404 |
| SPHKAP | 4 | -0.40130516 | 0.061 | 0.12 | 1.14E-10 | 0.00000417 |
| WNK2 | 4 | -0.306310396 | 0.526 | 0.592 | 1.14E-10 | 0.00000417 |

| | | | | | | |
|---|---|---|---|---|---|---|
| STAU2 | 4 | 0.258236363 | 0.698 | 0.61 | 1.14E-10 | 0.00000417 |
| AL589740.1 | 4 | -0.380667027 | 0.14 | 0.227 | 1.16E-10 | 0.00000424 |
| SPTAN1 | 4 | 0.260536156 | 0.525 | 0.425 | 1.19E-10 | 0.00000437 |
| PCGF3 | 4 | 0.255487177 | 0.432 | 0.329 | 1.27E-10 | 0.00000464 |
| MAST4 | 4 | 0.32057565 | 0.452 | 0.353 | 1.28E-10 | 0.0000047 |
| HDAC2 | 4 | -0.268702597 | 0.497 | 0.604 | 1.29E-10 | 0.00000473 |
| STRBP | 4 | 0.317587279 | 0.612 | 0.523 | 1.3E-10 | 0.00000475 |
| SFMBT2 | 4 | -0.434843084 | 0.448 | 0.497 | 1.36E-10 | 0.00000499 |
| ELAVL1 | 4 | 0.25013707 | 0.372 | 0.275 | 1.37E-10 | 0.00000503 |
| MT-ND1 | 4 | 0.321731162 | 0.239 | 0.159 | 1.5E-10 | 0.00000548 |
| ARMCX5-GPRASP2 | 4 | 0.274855079 | 0.392 | 0.298 | 1.58E-10 | 0.00000577 |
| OLFM2 | 4 | 0.2773855 | 0.181 | 0.116 | 1.79E-10 | 0.00000655 |
| MAD1L1 | 4 | 0.257777283 | 0.573 | 0.475 | 1.8E-10 | 0.00000658 |
| DCUN1D5 | 4 | -0.258626893 | 0.302 | 0.393 | 1.85E-10 | 0.00000677 |
| UCK2 | 4 | -0.289762643 | 0.349 | 0.445 | 1.86E-10 | 0.0000068 |
| LINC01776 | 4 | -0.258324518 | 0.112 | 0.198 | 1.88E-10 | 0.00000689 |
| AL356124.1 | 4 | -0.333405162 | 0.143 | 0.224 | 1.92E-10 | 0.00000702 |
| INSIG2 | 4 | -0.333878761 | 0.204 | 0.268 | 1.92E-10 | 0.00000703 |
| CHCHD3 | 4 | -0.252458918 | 0.666 | 0.722 | 2.21E-10 | 0.0000081 |
| TTLL5 | 4 | 0.263439724 | 0.668 | 0.597 | 2.3E-10 | 0.00000844 |
| MTF2 | 4 | -0.250290465 | 0.583 | 0.663 | 2.57E-10 | 0.00000941 |
| CLIP1 | 4 | 0.345568625 | 0.512 | 0.424 | 2.6E-10 | 0.00000952 |
| ZFAS1 | 4 | -0.292618494 | 0.262 | 0.352 | 2.8E-10 | 0.0000102 |
| PARP14 | 4 | 0.265210413 | 0.264 | 0.173 | 2.88E-10 | 0.0000106 |
| DENND4C | 4 | -0.26831611 | 0.657 | 0.705 | 3.42E-10 | 0.0000125 |
| NOTCH1 | 4 | 0.275821857 | 0.168 | 0.1 | 3.62E-10 | 0.0000133 |
| DOCK7 | 4 | 0.257368501 | 0.696 | 0.601 | 3.75E-10 | 0.0000137 |
| USO1 | 4 | 0.259313595 | 0.509 | 0.437 | 3.89E-10 | 0.0000142 |
| VPS13A | 4 | -0.373712022 | 0.635 | 0.653 | 3.89E-10 | 0.0000143 |
| RPL26 | 4 | -0.278055423 | 0.252 | 0.329 | 3.91E-10 | 0.0000143 |
| AL133346.1 | 4 | 0.331172222 | 0.22 | 0.138 | 4.22E-10 | 0.0000154 |
| TANC1 | 4 | 0.27765306 | 0.603 | 0.52 | 4.42E-10 | 0.0000162 |
| PIGL | 4 | 0.25238607 | 0.488 | 0.383 | 4.45E-10 | 0.0000163 |
| PLCL2 | 4 | -0.251478617 | 0.503 | 0.6 | 4.72E-10 | 0.0000173 |
| ARID1A | 4 | 0.255330266 | 0.674 | 0.582 | 5.07E-10 | 0.0000186 |
| ABCA3 | 4 | -0.268625965 | 0.182 | 0.262 | 5.08E-10 | 0.0000186 |
| AL139260.1 | 4 | 0.252339245 | 0.226 | 0.15 | 5.13E-10 | 0.0000188 |
| GNAQ | 4 | 0.277559531 | 0.79 | 0.718 | 5.3E-10 | 0.0000194 |
| ENTPD1-AS1 | 4 | 0.280202251 | 0.421 | 0.346 | 5.35E-10 | 0.0000196 |
| RAP1GAP2 | 4 | -0.753932724 | 0.199 | 0.263 | 5.38E-10 | 0.0000197 |
| FAM184A | 4 | -0.315870685 | 0.371 | 0.437 | 5.4E-10 | 0.0000198 |
| RAI2 | 4 | -0.256602755 | 0.1 | 0.161 | 5.77E-10 | 0.0000211 |
| SPIRE1 | 4 | 0.253470528 | 0.593 | 0.486 | 5.91E-10 | 0.0000216 |
| PPP3R1 | 4 | -0.252734592 | 0.52 | 0.6 | 6.32E-10 | 0.0000232 |
| RUNDC3B | 4 | -0.270459982 | 0.101 | 0.162 | 6.34E-10 | 0.0000232 |
| SMAD9 | 4 | -0.2510559 | 0.3 | 0.405 | 6.81E-10 | 0.0000249 |
| PLEKHH2 | 4 | 0.291701799 | 0.249 | 0.161 | 7.75E-10 | 0.0000284 |
| CAPZA2 | 4 | -0.264822056 | 0.4 | 0.461 | 8.18E-10 | 0.00003 |
| MCF2L2 | 4 | 0.256842954 | 0.13 | 0.079 | 8.42E-10 | 0.0000308 |
| GK | 4 | -0.297387534 | 0.209 | 0.284 | 8.79E-10 | 0.0000322 |
| BARD1 | 4 | -0.258641459 | 0.29 | 0.392 | 8.81E-10 | 0.0000322 |
| USP12 | 4 | -0.25679091 | 0.406 | 0.499 | 8.83E-10 | 0.0000323 |
| PDE11A | 4 | -0.299995403 | 0.132 | 0.195 | 8.88E-10 | 0.0000325 |
| LINC00937 | 4 | -0.275686354 | 0.122 | 0.195 | 9.48E-10 | 0.0000347 |
| KIF1B | 4 | 0.263661107 | 0.809 | 0.733 | 1.05E-09 | 0.0000386 |
| FHIT | 4 | -0.301291624 | 0.707 | 0.741 | 1.06E-09 | 0.0000389 |
| ELAVL4 | 4 | 0.307893982 | 0.1 | 0.058 | 1.16E-09 | 0.0000425 |
| SLC4A8 | 4 | 0.261746748 | 0.212 | 0.136 | 1.18E-09 | 0.000043 |

| | | | | | | |
|---|---|---|---|---|---|---|
| HBS1L | 4 | -0.260284634 | 0.372 | 0.45 | 1.27E-09 | 0.0000466 |
| JAM3 | 4 | 0.250237494 | 0.501 | 0.398 | 1.27E-09 | 0.0000466 |
| GPI | 4 | -0.363520265 | 0.2 | 0.269 | 1.31E-09 | 0.000048 |
| PEX1 | 4 | -0.283137518 | 0.34 | 0.408 | 1.52E-09 | 0.0000556 |
| FAT3 | 4 | 0.287664667 | 0.386 | 0.392 | 1.6E-09 | 0.0000585 |
| ZNF43 | 4 | 0.275123043 | 0.357 | 0.263 | 1.68E-09 | 0.0000616 |
| ARHGEF7 | 4 | 0.254942011 | 0.629 | 0.564 | 1.73E-09 | 0.0000633 |
| LRRC8D | 4 | 0.298099085 | 0.417 | 0.355 | 2.03E-09 | 0.0000743 |
| DNAJC5 | 4 | 0.268964322 | 0.381 | 0.291 | 2.12E-09 | 0.0000776 |
| IPO5 | 4 | -0.259218036 | 0.354 | 0.445 | 2.23E-09 | 0.0000816 |
| ZNF407 | 4 | -0.261993579 | 0.663 | 0.733 | 2.34E-09 | 0.0000857 |
| F3 | 4 | -0.303550851 | 0.101 | 0.1 | 2.39E-09 | 0.0000874 |
| FRMD3 | 4 | -0.328413157 | 0.319 | 0.37 | 2.42E-09 | 0.0000887 |
| MAPK8IP3 | 4 | 0.250537397 | 0.433 | 0.34 | 2.62E-09 | 0.0000961 |
| SIK2 | 4 | -0.278185754 | 0.495 | 0.553 | 2.68E-09 | 0.0000981 |
| SUFU | 4 | 0.251214508 | 0.351 | 0.266 | 2.79E-09 | 0.000102 |
| MAP4 | 4 | 0.254147297 | 0.767 | 0.717 | 2.81E-09 | 0.000103 |
| CACNA1A | 4 | -0.715063826 | 0.271 | 0.268 | 2.83E-09 | 0.000104 |
| AC138305.1 | 4 | -0.423173329 | 0.098 | 0.171 | 3.02E-09 | 0.000111 |
| EPHB1 | 4 | 0.344411975 | 0.201 | 0.157 | 3.48E-09 | 0.000128 |
| CDK14 | 4 | 0.272298259 | 0.827 | 0.772 | 3.77E-09 | 0.000138 |
| AC098829.1 | 4 | 0.25289339 | 0.163 | 0.102 | 3.93E-09 | 0.000144 |
| LINC02428 | 4 | 0.292399811 | 0.129 | 0.074 | 4.23E-09 | 0.000155 |
| IVNS1ABP | 4 | -0.307382304 | 0.458 | 0.512 | 4.32E-09 | 0.000158 |
| NDRG1 | 4 | -0.438687181 | 0.065 | 0.117 | 4.32E-09 | 0.000158 |
| RPS6KA3 | 4 | -0.423830291 | 0.554 | 0.586 | 4.83E-09 | 0.000177 |
| AC009120.2 | 4 | 0.26633277 | 0.391 | 0.299 | 5.26E-09 | 0.000193 |
| LAMA2 | 4 | -0.343429177 | 0.349 | 0.396 | 5.48E-09 | 0.000201 |
| FAT1 | 4 | -0.364602988 | 0.58 | 0.618 | 5.72E-09 | 0.000209295 |
| UFL1-AS1 | 4 | 0.260401479 | 0.167 | 0.099 | 5.93E-09 | 0.000217 |
| SIPA1L1 | 4 | -0.36932172 | 0.739 | 0.775 | 6.35E-09 | 0.000233 |
| PLEKHA6 | 4 | -0.279741058 | 0.396 | 0.475 | 7.25E-09 | 0.000265 |
| NCS1 | 4 | 0.282186085 | 0.281 | 0.208 | 7.78E-09 | 0.000285 |
| RHBDD1 | 4 | -0.259878439 | 0.286 | 0.362 | 8.62E-09 | 0.000316 |
| HMGA2 | 4 | 0.548306682 | 0.456 | 0.425 | 8.78E-09 | 0.000321 |
| BNIP3 | 4 | -0.268711261 | 0.135 | 0.213 | 8.96E-09 | 0.000328 |
| DOCK9 | 4 | 0.280548986 | 0.192 | 0.173 | 1.11E-08 | 0.000405 |
| SMARCA4 | 4 | 0.290983861 | 0.619 | 0.54 | 1.14E-08 | 0.000418 |
| CPEB3 | 4 | -0.260251233 | 0.327 | 0.397 | 1.27E-08 | 0.000465 |
| KCNMA1 | 4 | -0.261202237 | 0.207 | 0.298 | 1.29E-08 | 0.000474 |
| AC068286.2 | 4 | -0.787167225 | 0.09 | 0.142 | 1.31E-08 | 0.00048 |
| LINC01320 | 4 | 0.407932835 | 0.1 | 0.06 | 1.39E-08 | 0.000509 |
| GAPDH | 4 | -0.261894446 | 0.182 | 0.262 | 1.53E-08 | 0.00056 |
| RPS18 | 4 | -0.280247334 | 0.299 | 0.37 | 1.62E-08 | 0.000591 |
| SEPTIN11 | 4 | 0.286144213 | 0.541 | 0.472 | 1.62E-08 | 0.000594 |
| AL445123.2 | 4 | -0.283148815 | 0.17 | 0.243 | 1.66E-08 | 0.000609 |
| CFH | 4 | -0.322707888 | 0.085 | 0.154 | 1.69E-08 | 0.00062 |
| SRGAP3 | 4 | 0.329616752 | 0.601 | 0.545 | 1.77E-08 | 0.000648 |
| HDAC9 | 4 | 0.295017992 | 0.56 | 0.576 | 1.82E-08 | 0.000665 |
| PRDM5 | 4 | 0.252163912 | 0.406 | 0.312 | 1.82E-08 | 0.000665 |
| CDK19 | 4 | -0.25808628 | 0.62 | 0.688 | 1.93E-08 | 0.000705 |
| UBE2D2 | 4 | 0.286588897 | 0.65 | 0.584 | 2.25E-08 | 0.000822 |
| PTPRN2 | 4 | -0.251425923 | 0.339 | 0.265 | 2.62E-08 | 0.000959 |
| MT-ND2 | 4 | 0.268979369 | 0.191 | 0.132 | 0.000000028 | 0.001025848 |
| EEPD1 | 4 | -0.327796431 | 0.229 | 0.312 | 2.83E-08 | 0.001036645 |
| FRMD6 | 4 | -0.254996366 | 0.163 | 0.246 | 2.97E-08 | 0.001085758 |
| EPHA6 | 4 | 0.440870639 | 0.173 | 0.108 | 3.43E-08 | 0.001256965 |
| SNX7 | 4 | -0.253705122 | 0.126 | 0.186 | 3.45E-08 | 0.001261989 |

| | | | | | | |
|---|---|---|---|---|---|---|
| KHDRBS3 | 4 | -0.384796209 | 0.176 | 0.25 | 3.64E-08 | 0.00133426 |
| MIR181A2HG | 4 | 0.255904458 | 0.358 | 0.266 | 3.83E-08 | 0.001403598 |
| ESYT2 | 4 | -0.296657159 | 0.493 | 0.561 | 4.25E-08 | 0.001556304 |
| FNDC3B | 4 | 0.276289545 | 0.741 | 0.69 | 4.25E-08 | 0.001557394 |
| LINC00894 | 4 | 0.267111519 | 0.426 | 0.349 | 4.45E-08 | 0.001629786 |
| NPAS2 | 4 | 0.278408417 | 0.226 | 0.164 | 5.37E-08 | 0.001965208 |
| CPED1 | 4 | 0.39040249 | 0.113 | 0.075 | 6.89E-08 | 0.00252408 |
| SKI | 4 | 0.250032233 | 0.453 | 0.369 | 7.66E-08 | 0.00280478 |
| CRYBG3 | 4 | -0.282367446 | 0.259 | 0.322 | 7.97E-08 | 0.002917203 |
| CNIH3 | 4 | 0.260777896 | 0.231 | 0.178 | 8.55E-08 | 0.003130436 |
| MTSS1 | 4 | 0.281678338 | 0.262 | 0.183 | 0.000000109 | 0.004004971 |
| MAPRE2 | 4 | -0.255882488 | 0.27 | 0.345 | 0.000000111 | 0.004073699 |
| DSCAM | 4 | -0.320921257 | 0.065 | 0.123 | 0.000000115 | 0.004222333 |
| ANO10 | 4 | 0.316372006 | 0.505 | 0.469 | 0.000000119 | 0.004374508 |
| KDM4C | 4 | -0.26286909 | 0.559 | 0.62 | 0.000000126 | 0.004628287 |
| MYO3A | 4 | -0.596465894 | 0.139 | 0.185 | 0.000000128 | 0.00470269 |
| OSBPL11 | 4 | -0.260056024 | 0.271 | 0.341 | 0.000000142 | 0.005213512 |
| RASSF3 | 4 | 0.265708796 | 0.344 | 0.262 | 0.000000152 | 0.005552228 |
| RGS7 | 4 | -0.341328296 | 0.152 | 0.208 | 0.000000163 | 0.005955411 |
| OPCML | 4 | -0.412282454 | 0.337 | 0.345 | 0.000000168 | 0.006147026 |
| FSTL4 | 4 | 0.346165625 | 0.171 | 0.11 | 0.000000185 | 0.006757089 |
| PLEKHG4B | 4 | 0.253668835 | 0.288 | 0.217 | 0.000000219 | 0.008030845 |
| BICC1 | 4 | 0.322510134 | 0.376 | 0.291 | 0.000000237 | 0.008682062 |
| EPB41 | 4 | 0.276385098 | 0.482 | 0.482 | 0.000000312 | 0.011435502 |
| MARCH1 | 4 | -0.369641251 | 0.223 | 0.307 | 0.000000338 | 0.012362307 |
| GPR137C | 4 | -0.260417449 | 0.358 | 0.437 | 0.000000359 | 0.013130151 |
| CACNA1C | 4 | -0.382027614 | 0.412 | 0.488 | 0.000000399 | 0.014603794 |
| TMEM161B-AS1 | 4 | -0.294880304 | 0.606 | 0.668 | 0.000000415 | 0.015206493 |
| SCFD2 | 4 | -0.269280703 | 0.556 | 0.6 | 0.000000417 | 0.015272341 |
| PHLDB2 | 4 | 0.292543949 | 0.293 | 0.223 | 0.000000469 | 0.017160807 |
| RPL23A | 4 | -0.257433481 | 0.287 | 0.345 | 0.000000487 | 0.0178146 |
| AC138627.1 | 4 | -0.438995692 | 0.112 | 0.17 | 0.000000496 | 0.018173732 |
| PLAGL1 | 4 | 0.297202862 | 0.431 | 0.348 | 0.000000557 | 0.020381303 |
| FBXL4 | 4 | -0.261799501 | 0.479 | 0.531 | 0.000000601 | 0.022012462 |
| ADGRG2 | 4 | -0.318194122 | 0.134 | 0.186 | 0.00000106 | 0.03879974 |
| P4HA1 | 4 | -0.35683372 | 0.475 | 0.483 | 0.00000108 | 0.039431218 |
| SAMD4A | 4 | -0.313049573 | 0.343 | 0.392 | 0.00000111 | 0.040458358 |
| PAM | 4 | -0.328219177 | 0.744 | 0.757 | 0.00000169 | 0.061871082 |
| ARHGAP42 | 4 | 0.256892325 | 0.447 | 0.364 | 0.00000292 | 0.106748594 |
| PKN2-AS1 | 4 | -0.291744907 | 0.078 | 0.125 | 0.00000301 | 0.110284547 |
| HSP90AA1 | 4 | -0.643884401 | 0.438 | 0.458 | 0.00000316 | 0.11570314 |
| CLSTN2 | 4 | -0.271992054 | 0.29 | 0.348 | 0.00000405 | 0.148220365 |
| PCDH11X | 4 | -0.436809949 | 0.246 | 0.266 | 0.00000419 | 0.153340825 |
| PDE1A | 4 | -0.72156387 | 0.153 | 0.208 | 0.00000441 | 0.161336127 |
| AHCYL2 | 4 | -0.256142686 | 0.7 | 0.662 | 0.0000084 | 0.307457988 |
| ADAMTS9-AS2 | 4 | 0.280605962 | 0.375 | 0.302 | 0.00000867 | 0.317366369 |
| FYN | 4 | 0.297841458 | 0.682 | 0.657 | 0.0000109 | 0.39842273 |
| ENAH | 4 | 0.268536059 | 0.737 | 0.716 | 0.0000126 | 0.462308893 |
| AL591519.1 | 4 | -0.251406943 | 0.098 | 0.153 | 0.0000143 | 0.524188037 |
| TENM3 | 4 | 0.253588817 | 0.84 | 0.776 | 0.0000189 | 0.693478104 |
| SLC2A13 | 4 | -0.297570193 | 0.438 | 0.449 | 0.00002 | 0.730929445 |
| GABRG3 | 4 | 0.291345562 | 0.113 | 0.089 | 0.0000233 | 0.851626005 |
| MAN1A1 | 4 | -0.253087197 | 0.132 | 0.133 | 0.0000316 | 1 |
| PTPRS | 4 | 0.261180989 | 0.61 | 0.575 | 0.0000353 | 1 |
| NEB | 4 | 0.326770822 | 0.131 | 0.128 | 0.0000433 | 1 |
| PDGFD | 4 | 0.276469897 | 0.237 | 0.177 | 0.000065 | 1 |
| DAPK1 | 4 | -0.252934742 | 0.636 | 0.692 | 0.0000728 | 1 |
| CNTN4 | 4 | -0.352575031 | 0.124 | 0.153 | 0.0000737 | 1 |

| | | | | | | |
|---|---|---|---|---|---|---|
| PLS3 | 4 | 0.29028173 | 0.453 | 0.398 | 0.0000739 | 1 |
| MTUS2 | 4 | 0.393060137 | 0.259 | 0.204 | 0.0000752 | 1 |
| NRP1 | 4 | 0.290563832 | 0.345 | 0.308 | 0.0000898 | 1 |
| PARM1 | 4 | 0.332470525 | 0.24 | 0.198 | 0.000135 | 1 |
| TENM1 | 4 | 0.372695385 | 0.16 | 0.127 | 0.000248 | 1 |
| THSD7A | 4 | 0.522352645 | 0.396 | 0.387 | 0.000256 | 1 |
| GUCY1A2 | 4 | 0.327432772 | 0.41 | 0.396 | 0.000278 | 1 |
| ADK | 4 | 0.417878383 | 0.794 | 0.783 | 0.000344 | 1 |
| KANK1 | 4 | -0.268254046 | 0.476 | 0.512 | 0.000459 | 1 |
| KIF26B | 4 | 0.30665552 | 0.479 | 0.516 | 0.000598 | 1 |
| MAP2 | 4 | 0.445965999 | 0.563 | 0.543 | 0.000602 | 1 |
| MALAT1 | 4 | -0.343363069 | 0.999 | 0.998 | 0.003109974 | 1 |
| AL591501.1 | 4 | -0.279601839 | 0.116 | 0.113 | 0.004350021 | 1 |
| EYS | 5 | 4.411969967 | 0.991 | 0.199 | 0 | 0 |
| CACNA1A | 5 | 3.969518507 | 0.985 | 0.223 | 0 | 0 |
| RIMBP2 | 5 | 3.912997714 | 0.991 | 0.074 | 0 | 0 |
| PTPRN2 | 5 | 3.79994539 | 0.978 | 0.228 | 0 | 0 |
| RAP1GAP2 | 5 | 3.679769066 | 0.994 | 0.209 | 0 | 0 |
| FOXP2 | 5 | 3.447977519 | 0.985 | 0.218 | 0 | 0 |
| MYO3A | 5 | 3.440018239 | 0.958 | 0.13 | 0 | 0 |
| MACC1 | 5 | 3.393064721 | 0.928 | 0.01 | 0 | 0 |
| DGKB | 5 | 3.306990057 | 0.963 | 0.242 | 0 | 0 |
| AL591501.1 | 5 | 3.176528581 | 0.794 | 0.07 | 0 | 0 |
| RFX6 | 5 | 3.153989879 | 0.91 | 0.006 | 0 | 0 |
| NRG1 | 5 | 3.127359323 | 0.963 | 0.248 | 0 | 0 |
| SLC14A2 | 5 | 3.113892657 | 0.626 | 0.016 | 0 | 0 |
| EYA4 | 5 | 3.108967888 | 0.974 | 0.155 | 0 | 0 |
| NPAS3 | 5 | 3.08619697 | 0.954 | 0.199 | 0 | 0 |
| RASSF6 | 5 | 3.014061744 | 0.883 | 0.005 | 0 | 0 |
| GCLC | 5 | 3.012759236 | 0.873 | 0.17 | 0 | 0 |
| SORBS2 | 5 | 2.976473702 | 0.945 | 0.074 | 0 | 0 |
| LINC01811 | 5 | 2.960452553 | 0.806 | 0.016 | 0 | 0 |
| CACNA2D1 | 5 | 2.95579504 | 0.994 | 0.845 | 0 | 0 |
| ELL2 | 5 | 2.874928468 | 0.943 | 0.139 | 0 | 0 |
| ETV1 | 5 | 2.847846245 | 0.921 | 0.153 | 0 | 0 |
| CHST9 | 5 | 2.810720584 | 0.89 | 0.104 | 0 | 0 |
| MAN1A1 | 5 | 2.748351415 | 0.873 | 0.085 | 0 | 0 |
| ITGB8 | 5 | 2.694025025 | 0.927 | 0.258 | 0 | 0 |
| CACNA1C | 5 | 2.689596609 | 0.978 | 0.448 | 0 | 0 |
| LRMDA | 5 | 2.673390043 | 0.831 | 0.07 | 0 | 0 |
| AL022068.1 | 5 | 2.650297557 | 0.694 | 0.074 | 0 | 0 |
| SIPA1L3 | 5 | 2.644081891 | 0.982 | 0.496 | 0 | 0 |
| SEZ6L | 5 | 2.641090289 | 0.837 | 0.048 | 0 | 0 |
| KIAA1324 | 5 | 2.595652733 | 0.884 | 0.038 | 0 | 0 |
| KCNJ3 | 5 | 2.555532999 | 0.861 | 0.096 | 0 | 0 |
| A1CF | 5 | 2.522410588 | 0.828 | 0.005 | 0 | 0 |
| DOCK8 | 5 | 2.503199298 | 0.839 | 0.03 | 0 | 0 |
| HEPACAM2 | 5 | 2.467872698 | 0.756 | 0.004 | 0 | 0 |
| ROR1 | 5 | 2.460430173 | 0.923 | 0.21 | 0 | 0 |
| CDKAL1 | 5 | 2.444866737 | 0.974 | 0.816 | 0 | 0 |
| ST6GALNAC5 | 5 | 2.387230416 | 0.921 | 0.341 | 0 | 0 |
| RBM47 | 5 | 2.367811251 | 0.877 | 0.081 | 0 | 0 |
| KCNB2 | 5 | 2.349661275 | 0.991 | 0.524 | 0 | 0 |
| ASPH | 5 | 2.349257969 | 0.993 | 0.682 | 0 | 0 |
| GMDS | 5 | 2.336523118 | 0.91 | 0.412 | 0 | 0 |
| CACNB2 | 5 | 2.29949259 | 0.998 | 0.715 | 0 | 0 |
| NPNT | 5 | 2.286520004 | 0.752 | 0.031 | 0 | 0 |
| RPS6KA3 | 5 | 2.253956041 | 0.95 | 0.559 | 0 | 0 |

| | | | | | | |
|---|---|---|---|---|---|---|
| SAMD3 | 5 | 2.229056358 | 0.756 | 0.049 | 0 | 0 |
| ALDH1A1 | 5 | 2.188793744 | 0.716 | 0.01 | 0 | 0 |
| NEUROD1 | 5 | 2.186879685 | 0.835 | 0.139 | 0 | 0 |
| MAML3 | 5 | 2.147752758 | 0.976 | 0.546 | 0 | 0 |
| SCGN | 5 | 2.142690031 | 0.728 | 0.008 | 0 | 0 |
| AL445584.2 | 5 | 2.127269293 | 0.69 | 0.007 | 0 | 0 |
| RNF128 | 5 | 2.09027415 | 0.815 | 0.097 | 0 | 0 |
| VPS13A | 5 | 2.067238273 | 0.976 | 0.631 | 0 | 0 |
| INSR | 5 | 2.051988867 | 0.862 | 0.217 | 0 | 0 |
| FBXW7 | 5 | 1.968515003 | 0.96 | 0.725 | 0 | 0 |
| MYOM1 | 5 | 1.95074712 | 0.655 | 0.03 | 0 | 0 |
| MLXIPL | 5 | 1.943448837 | 0.712 | 0.009 | 0 | 0 |
| RAB3B | 5 | 1.931109965 | 0.741 | 0.046 | 0 | 0 |
| HNF4G | 5 | 1.90852711 | 0.661 | 0.013 | 0 | 0 |
| GCNT1 | 5 | 1.775866621 | 0.71 | 0.054 | 0 | 0 |
| GIPR | 5 | 1.712195277 | 0.593 | 0.009 | 0 | 0 |
| KIAA1211L | 5 | 1.550889914 | 0.609 | 0.022 | 0 | 0 |
| AC107220.1 | 5 | 1.482046052 | 0.479 | 0.002 | 0 | 0 |
| S100Z | 5 | 1.441411189 | 0.503 | 0.004 | 0 | 0 |
| LINC02717 | 5 | 1.344107985 | 0.492 | 0.002 | 0 | 0 |
| GATA4 | 5 | 1.343183488 | 0.517 | 0.005 | 0 | 0 |
| FBXL7 | 5 | -3.085068187 | 0.117 | 0.882 | 0 | 0 |
| KAZN | 5 | -3.125964801 | 0.261 | 0.94 | 0 | 0 |
| SIM1 | 5 | 1.560398671 | 0.56 | 0.012 | 3.8e-322 | 1.3927943e-317 |
| PTPRD | 5 | -3.181585651 | 0.156 | 0.885 | 3.264e-320 | 1.19490896e-315 |
| COL21A1 | 5 | 2.073183569 | 0.679 | 0.041 | 4.0657e-320 | 1.48848105e-315 |
| ABCC8 | 5 | 2.025095919 | 0.587 | 0.027 | 4.489692e-317 | 1.6437211789e-312 |
| PURPL | 5 | 2.135616029 | 0.765 | 0.085 | 1.25401222e-316 | 4.59106415668e-312 |
| ZNF536 | 5 | -3.170114762 | 0.121 | 0.867 | 1.7838258e-316 | 6.530764610555e-312 |
| HNF4A | 5 | 1.264522282 | 0.499 | 0.005 | 4.165759675e-315 | 1.5251262746925e-310 |
| LINC00907 | 5 | 2.287593433 | 0.917 | 0.211 | 1.01340480783e-312 | 3.71E-308 |
| UFL1-AS1 | 5 | 2.126864985 | 0.684 | 0.069 | 2.27160870669e-311 | 8.32E-307 |
| LEPR | 5 | 2.139651738 | 0.705 | 0.095 | 8.06E-306 | 2.95E-301 |
| TCF4 | 5 | -2.682103645 | 0.336 | 0.925 | 1.23E-305 | 4.52E-301 |
| SAMD12 | 5 | 1.645170956 | 0.644 | 0.04 | 9.47E-304 | 3.47E-299 |
| GCLM | 5 | 1.921078864 | 0.723 | 0.121 | 6.54E-303 | 2.39E-298 |
| ANXA13 | 5 | 1.230932266 | 0.451 | 0.002 | 3.14E-300 | 1.15E-295 |
| DSP | 5 | 1.851690864 | 0.659 | 0.05 | 2.45E-299 | 8.97E-295 |
| CXXC4 | 5 | 1.544599128 | 0.567 | 0.021 | 3.8E-299 | 1.39E-294 |
| PAM | 5 | 1.942766432 | 0.965 | 0.742 | 9.57E-299 | 3.5E-294 |
| LRRC53 | 5 | 1.723035171 | 0.455 | 0.004 | 7.04E-298 | 2.58E-293 |
| AC112487.1 | 5 | 1.414785358 | 0.512 | 0.011 | 3.23E-297 | 1.18E-292 |
| CPNE8 | 5 | 1.884226329 | 0.782 | 0.122 | 3.32E-297 | 1.21E-292 |
| MCOLN3 | 5 | 1.435030242 | 0.593 | 0.029 | 3.39E-297 | 1.24E-292 |
| ATP8A1 | 5 | 1.953833418 | 0.905 | 0.378 | 4.2E-295 | 1.54E-290 |
| CA8 | 5 | 1.658494499 | 0.503 | 0.009 | 6.42E-293 | 2.35E-288 |
| RIMS2 | 5 | 2.140144307 | 0.993 | 0.391 | 4.32E-292 | 1.58E-287 |
| AGBL4 | 5 | 2.187956934 | 0.84 | 0.167 | 7.1E-291 | 2.6E-286 |
| RGS7 | 5 | 2.170151303 | 0.811 | 0.163 | 2.96E-288 | 1.09E-283 |
| DPYD | 5 | 1.900445752 | 0.914 | 0.377 | 5.14E-287 | 1.88E-282 |
| ALCAM | 5 | 2.013449314 | 0.952 | 0.291 | 9.23E-287 | 3.38E-282 |
| PROX1 | 5 | 1.316484722 | 0.536 | 0.017 | 9.5E-284 | 3.48E-279 |
| LINC02006 | 5 | 1.583941032 | 0.539 | 0.019 | 1.6E-277 | 5.86E-273 |
| NOL4 | 5 | 2.092272342 | 0.905 | 0.29 | 2.36E-275 | 8.64E-271 |
| RBMS3 | 5 | -2.775642955 | 0.16 | 0.844 | 1.65E-274 | 6.06E-270 |
| TMOD1 | 5 | 1.719855359 | 0.545 | 0.029 | 3.14E-274 | 1.15E-269 |
| FRMD4A | 5 | -2.818898868 | 0.191 | 0.858 | 2.59E-273 | 9.49E-269 |
| SLC39A11 | 5 | 1.696796611 | 0.868 | 0.486 | 1.9E-272 | 6.96E-268 |

| | | | | | | |
|---|---|---|---|---|---|---|
| LINC02315 | 5 | -2.70766733 | 0.125 | 0.827 | 1.77E-270 | 6.49E-266 |
| MTSS1 | 5 | 1.880763762 | 0.756 | 0.155 | 2.49E-267 | 9.11E-263 |
| MIR217HG | 5 | 1.184120168 | 0.439 | 0.005 | 6.56E-265 | 2.4E-260 |
| PAX5 | 5 | 1.196342397 | 0.475 | 0.011 | 1.27E-264 | 4.66E-260 |
| CDH1 | 5 | 1.467491199 | 0.554 | 0.027 | 1.72E-264 | 6.3E-260 |
| PRSS12 | 5 | -3.131108256 | 0.178 | 0.838 | 3.38E-262 | 1.24E-257 |
| KIF12 | 5 | 1.128954049 | 0.429 | 0.005 | 1.23E-261 | 4.49E-257 |
| DSC3 | 5 | 1.210102227 | 0.435 | 0.006 | 7.71E-260 | 2.82E-255 |
| RGL3 | 5 | 1.198375002 | 0.44 | 0.007 | 6.96E-257 | 2.55E-252 |
| AL391117.1 | 5 | 1.834036354 | 0.479 | 0.018 | 1.7E-256 | 6.22E-252 |
| HNF1A-AS1 | 5 | 1.371336278 | 0.453 | 0.009 | 3.04E-256 | 1.11E-251 |
| UNC80 | 5 | 1.757555798 | 0.787 | 0.171 | 1.13E-255 | 4.14E-251 |
| FRMD4B | 5 | 1.789083297 | 0.916 | 0.402 | 1.79E-253 | 6.54E-249 |
| KCNH6 | 5 | 1.218426425 | 0.455 | 0.011 | 6.19E-253 | 2.27E-248 |
| NEBL | 5 | 1.978765546 | 0.804 | 0.163 | 1.1E-252 | 4.04E-248 |
| TSHZ2 | 5 | -2.87006361 | 0.163 | 0.837 | 1.89E-250 | 6.93E-246 |
| SCN7A | 5 | 3.566827957 | 0.56 | 0.113 | 2.23E-247 | 8.15E-243 |
| EML5 | 5 | 1.627044559 | 0.727 | 0.124 | 3.13E-247 | 1.15E-242 |
| ELP4 | 5 | 1.712461047 | 0.848 | 0.378 | 1.58E-246 | 5.78E-242 |
| TTC6 | 5 | 1.840975538 | 0.743 | 0.126 | 4.07E-243 | 1.49E-238 |
| DCLK2 | 5 | -2.482463496 | 0.095 | 0.768 | 1.14E-242 | 4.18E-238 |
| SOX6 | 5 | 1.758966627 | 0.87 | 0.2 | 2.96E-241 | 1.08E-236 |
| CADPS | 5 | 1.815863837 | 0.972 | 0.433 | 3.59E-240 | 1.31E-235 |
| MAP3K4 | 5 | 1.449423035 | 0.873 | 0.476 | 1.72E-239 | 6.29E-235 |
| SI | 5 | 1.656014261 | 0.369 | 0.002 | 1.77E-238 | 6.46E-234 |
| PLPPR1 | 5 | 1.878544031 | 0.776 | 0.158 | 2.72E-238 | 9.95E-234 |
| TRPM3 | 5 | 1.412784757 | 0.829 | 0.16 | 3.24E-238 | 1.19E-233 |
| MIR100HG | 5 | -2.706655316 | 0.11 | 0.785 | 2.87E-237 | 1.05E-232 |
| RFX2 | 5 | 1.676267148 | 0.793 | 0.187 | 3.11E-236 | 1.14E-231 |
| ADAMTS6 | 5 | 2.088089697 | 0.738 | 0.113 | 2.63E-235 | 9.62E-231 |
| NKX2-2 | 5 | 0.993572302 | 0.356 | 0.002 | 1.26E-233 | 4.62E-229 |
| CMTM8 | 5 | 1.584073772 | 0.672 | 0.104 | 1.45E-233 | 5.3E-229 |
| OTOGL | 5 | 1.33706185 | 0.411 | 0.01 | 9.49E-232 | 3.47E-227 |
| ANKH | 5 | 1.569465206 | 0.756 | 0.17 | 1.31E-231 | 4.8E-227 |
| GAS2 | 5 | 1.548212509 | 0.585 | 0.066 | 6.21E-229 | 2.27E-224 |
| SPTSSB | 5 | 1.017754941 | 0.367 | 0.003 | 8.68E-229 | 3.18E-224 |
| MARCH4 | 5 | 1.360111405 | 0.527 | 0.036 | 1.89E-228 | 6.92E-224 |
| RAPGEF4 | 5 | 1.663440283 | 0.923 | 0.369 | 3.54E-228 | 1.3E-223 |
| NEURL1 | 5 | 1.241120086 | 0.517 | 0.038 | 6.3E-227 | 2.31E-222 |
| TENM1 | 5 | 1.619266044 | 0.695 | 0.094 | 8E-227 | 2.93E-222 |
| SLC16A12 | 5 | 1.418338061 | 0.497 | 0.025 | 3.98E-226 | 1.46E-221 |
| KSR2 | 5 | 1.332202838 | 0.541 | 0.043 | 8.58E-226 | 3.14E-221 |
| FAM155A | 5 | -2.945286614 | 0.213 | 0.823 | 8.1E-224 | 2.96E-219 |
| RTN1 | 5 | 1.554616011 | 0.675 | 0.124 | 4.6E-223 | 1.68E-218 |
| PLAC8 | 5 | 1.061299648 | 0.385 | 0.007 | 5.58E-220 | 2.04E-215 |
| SYT1 | 5 | -2.853109066 | 0.147 | 0.781 | 1.12E-219 | 4.1E-215 |
| STXBP5L | 5 | 1.681648588 | 0.897 | 0.269 | 1.84E-217 | 6.73E-213 |
| PROX1-AS1 | 5 | 1.04571758 | 0.36 | 0.004 | 4.25E-217 | 1.56E-212 |
| AC090679.2 | 5 | 1.229506128 | 0.33 | 0.001 | 4.46E-217 | 1.63E-212 |
| LDB3 | 5 | 1.139897241 | 0.426 | 0.016 | 2.62E-216 | 9.58E-212 |
| ARSJ | 5 | 1.565355235 | 0.622 | 0.08 | 5.52E-216 | 2.02E-211 |
| ROBO2 | 5 | -3.510186364 | 0.116 | 0.751 | 5.98E-214 | 2.19E-209 |
| LLGL2 | 5 | 1.196128282 | 0.483 | 0.029 | 1.71E-213 | 6.26E-209 |
| MIRLET7BHG | 5 | -1.911215972 | 0.066 | 0.71 | 1.31E-212 | 4.79E-208 |
| COL11A1 | 5 | -2.973387019 | 0.112 | 0.751 | 1.31E-212 | 4.81E-208 |
| MATN2 | 5 | 1.267611666 | 0.543 | 0.053 | 7.92E-209 | 2.9E-204 |
| GCNT2 | 5 | 1.357751322 | 0.554 | 0.062 | 1.21E-208 | 4.45E-204 |
| KCNH7 | 5 | 1.683436292 | 0.681 | 0.103 | 1.31E-204 | 4.8E-200 |

| | | | | | | |
|---|---|---|---|---|---|---|
| RFX3 | 5 | 1.491841888 | 0.945 | 0.812 | 1.95E-204 | 7.13E-200 |
| CHSY3 | 5 | -2.781807854 | 0.127 | 0.748 | 5.16E-204 | 1.89E-199 |
| CADPS2 | 5 | 1.871107163 | 0.793 | 0.199 | 2.62E-203 | 9.58E-199 |
| AC093866.1 | 5 | 1.244391815 | 0.383 | 0.011 | 1.64E-202 | 6.02E-198 |
| SOBP | 5 | -2.083784583 | 0.228 | 0.812 | 1.19E-200 | 4.37E-196 |
| CSRNP3 | 5 | 1.626245558 | 0.826 | 0.331 | 1.23E-200 | 4.51E-196 |
| EBF1 | 5 | 1.592128829 | 0.932 | 0.333 | 3.43E-200 | 1.26E-195 |
| KHDRBS2 | 5 | 1.203796219 | 0.44 | 0.02 | 4.85E-200 | 1.78E-195 |
| COBL | 5 | 1.123578308 | 0.457 | 0.026 | 2.37E-199 | 8.67E-195 |
| PTPRG | 5 | -2.318486446 | 0.365 | 0.871 | 2.64E-199 | 9.67E-195 |
| GAD2 | 5 | 1.278277982 | 0.484 | 0.034 | 8.78E-199 | 3.21E-194 |
| ABCA5 | 5 | 1.490096492 | 0.839 | 0.359 | 8.99E-198 | 3.29E-193 |
| MAP3K15 | 5 | 1.549729362 | 0.686 | 0.137 | 1.92E-197 | 7.03E-193 |
| MSC-AS1 | 5 | 1.743957715 | 0.639 | 0.12 | 5.12E-197 | 1.88E-192 |
| AC005062.1 | 5 | 0.948265698 | 0.347 | 0.006 | 1.49E-196 | 5.44E-192 |
| CERKL | 5 | 1.494114162 | 0.611 | 0.104 | 1.53E-196 | 5.59E-192 |
| TPST1 | 5 | 1.554558388 | 0.828 | 0.393 | 8.58E-195 | 3.14E-190 |
| CCDC88A | 5 | 1.362081231 | 0.895 | 0.624 | 3.57E-194 | 1.31E-189 |
| PPP1R9A | 5 | 1.479287089 | 0.717 | 0.134 | 1.1E-192 | 4.02E-188 |
| LINC01515 | 5 | 1.669884715 | 0.706 | 0.155 | 2.83E-192 | 1.04E-187 |
| NFASC | 5 | 1.464756615 | 0.684 | 0.146 | 4.87E-192 | 1.78E-187 |
| TPD52 | 5 | 1.553762784 | 0.855 | 0.328 | 9.87E-191 | 3.61E-186 |
| GLRA3 | 5 | 0.964050183 | 0.374 | 0.011 | 6.18E-189 | 2.26E-184 |
| ARFGEF3 | 5 | 1.598208121 | 0.835 | 0.436 | 2.61E-188 | 9.56E-184 |
| HECW2 | 5 | 1.602323337 | 0.813 | 0.34 | 1.12E-187 | 4.11E-183 |
| SLIT2 | 5 | -3.334194739 | 0.172 | 0.756 | 5.66E-186 | 2.07E-181 |
| RIPPLY3 | 5 | 1.087858613 | 0.461 | 0.04 | 4.66E-185 | 1.71E-180 |
| ANKRD30B | 5 | 1.251752388 | 0.389 | 0.018 | 1.18E-184 | 4.33E-180 |
| CHKA | 5 | 1.323632227 | 0.769 | 0.311 | 2.92E-184 | 1.07E-179 |
| GRB14 | 5 | 1.247141944 | 0.574 | 0.085 | 3.17E-183 | 1.16E-178 |
| VIL1 | 5 | 0.908385089 | 0.345 | 0.009 | 6.65E-183 | 2.43E-178 |
| AL591378.1 | 5 | 0.883234273 | 0.317 | 0.005 | 1.08E-181 | 3.94E-177 |
| MEST | 5 | 1.562415205 | 0.565 | 0.108 | 2.65E-181 | 9.69E-177 |
| KIAA1671 | 5 | 1.354040442 | 0.655 | 0.137 | 8.03E-180 | 2.94E-175 |
| PALLD | 5 | 1.594779322 | 0.835 | 0.388 | 1.18E-179 | 4.32E-175 |
| CDH17 | 5 | 0.843788563 | 0.297 | 0.003 | 1.86E-177 | 6.83E-173 |
| KCNMB2 | 5 | 1.379205983 | 0.833 | 0.235 | 5.95E-177 | 2.18E-172 |
| MAGI3 | 5 | 1.366433019 | 0.873 | 0.424 | 1.2E-176 | 4.41E-172 |
| LINC02197 | 5 | 1.572029383 | 0.683 | 0.137 | 7.82E-176 | 2.86E-171 |
| RGS17 | 5 | 1.413004932 | 0.629 | 0.146 | 1.55E-175 | 5.66E-171 |
| NWD2 | 5 | 1.44815025 | 0.626 | 0.11 | 3.19E-175 | 1.17E-170 |
| ADGRL2 | 5 | 1.744322593 | 0.93 | 0.612 | 2.17E-174 | 7.95E-170 |
| FAM214A | 5 | 1.310464192 | 0.793 | 0.365 | 3.38E-174 | 1.24E-169 |
| PPP2R2B | 5 | -2.669105454 | 0.101 | 0.684 | 4.35E-173 | 1.59E-168 |
| PLXDC2 | 5 | -2.334066987 | 0.125 | 0.703 | 1.52E-172 | 5.57E-168 |
| ISL1 | 5 | 0.951225636 | 0.361 | 0.014 | 1.9E-172 | 6.96E-168 |
| NPC1L1 | 5 | 0.762846669 | 0.283 | 0.002 | 2.79E-172 | 1.02E-167 |
| DGKG | 5 | 1.388982408 | 0.409 | 0.032 | 3.82E-172 | 1.4E-167 |
| MIR7-3HG | 5 | 0.925660628 | 0.314 | 0.007 | 4.25E-172 | 1.56E-167 |
| MAPK10 | 5 | 1.416723658 | 0.761 | 0.192 | 2.43E-171 | 8.88E-167 |
| FRK | 5 | 1.124182519 | 0.473 | 0.044 | 4.92E-171 | 1.8E-166 |
| ARX | 5 | 0.833643289 | 0.305 | 0.005 | 8.11E-170 | 2.97E-165 |
| ZEB2 | 5 | -2.374320732 | 0.05 | 0.614 | 2.5E-169 | 9.14E-165 |
| AL133325.3 | 5 | 0.872997872 | 0.356 | 0.016 | 4.73E-169 | 1.73E-164 |
| TPCN1 | 5 | 1.309379133 | 0.609 | 0.146 | 8.59E-169 | 3.14E-164 |
| TBC1D4 | 5 | 1.421753532 | 0.826 | 0.372 | 4.16E-168 | 1.52E-163 |
| PRUNE2 | 5 | 1.065575075 | 0.413 | 0.029 | 1.51E-167 | 5.52E-163 |
| MYO1D | 5 | 1.296669157 | 0.721 | 0.188 | 9.23E-167 | 3.38E-162 |

| | | | | | | |
|---|---|---|---|---|---|---|
| TMPRSS6 | 5 | 0.748714538 | 0.283 | 0.004 | 1.91E-166 | 6.98E-162 |
| IL1RAPL1 | 5 | -3.394928751 | 0.134 | 0.69 | 5.74E-166 | 2.1E-161 |
| SEC24D | 5 | 1.397053495 | 0.67 | 0.226 | 7.56E-166 | 2.77E-161 |
| FAM241A | 5 | 0.931398044 | 0.33 | 0.01 | 6.8E-165 | 2.49E-160 |
| ZSWIM6 | 5 | -1.645722335 | 0.398 | 0.856 | 8.13E-165 | 2.98E-160 |
| VWA5B2 | 5 | 1.002418243 | 0.42 | 0.035 | 1.67E-164 | 6.13E-160 |
| ESRP1 | 5 | 0.81129374 | 0.314 | 0.008 | 1.84E-163 | 6.75E-159 |
| NLGN1 | 5 | -2.607185591 | 0.385 | 0.832 | 8.44E-163 | 3.09E-158 |
| ABCC2 | 5 | 0.857966829 | 0.338 | 0.012 | 9.1E-163 | 3.33E-158 |
| WNK2 | 5 | -1.608915516 | 0.061 | 0.619 | 1.09E-162 | 4E-158 |
| GABBR2 | 5 | 1.114187962 | 0.389 | 0.027 | 6.75E-162 | 2.47E-157 |
| SHROOM4 | 5 | -1.931900929 | 0.077 | 0.638 | 1.04E-161 | 3.81E-157 |
| GABRB3 | 5 | 1.382274309 | 0.873 | 0.345 | 3.58E-161 | 1.31E-156 |
| EFNA5 | 5 | 1.448565859 | 0.961 | 0.644 | 4.53E-161 | 1.66E-156 |
| PARD3B | 5 | 1.534026358 | 0.894 | 0.441 | 2.43E-160 | 8.9E-156 |
| SNTG1 | 5 | 0.597484475 | 0.919 | 0.351 | 2.78E-160 | 1.02E-155 |
| MYO10 | 5 | 1.308288245 | 0.818 | 0.301 | 7.36E-160 | 2.69E-155 |
| CAMK1D | 5 | 1.304268082 | 0.912 | 0.626 | 1.95E-159 | 7.12E-155 |
| AC018731.1 | 5 | -2.636060035 | 0.075 | 0.625 | 1.87E-157 | 6.85E-153 |
| SYNJ2 | 5 | 1.276244596 | 0.679 | 0.159 | 5.81E-157 | 2.13E-152 |
| KCNJ6 | 5 | 1.233764054 | 0.71 | 0.169 | 2E-156 | 7.31E-152 |
| RGS9 | 5 | 0.967123306 | 0.382 | 0.027 | 2.12E-156 | 7.75E-152 |
| NT5DC3 | 5 | 1.167585118 | 0.561 | 0.108 | 2.79E-156 | 1.02E-151 |
| VAV2 | 5 | 1.40385835 | 0.705 | 0.297 | 5.1E-156 | 1.87E-151 |
| EDA | 5 | 1.713617949 | 0.736 | 0.252 | 9.99E-155 | 3.66E-150 |
| SVOP | 5 | 0.999543284 | 0.437 | 0.044 | 2.35E-154 | 8.6E-150 |
| ELAVL4 | 5 | 1.027554167 | 0.437 | 0.039 | 2.86E-154 | 1.05E-149 |
| ADORA2A-AS1 | 5 | 0.766021839 | 0.283 | 0.006 | 5E-152 | 1.83E-147 |
| PID1 | 5 | -2.492457041 | 0.073 | 0.616 | 2.12E-151 | 7.78E-147 |
| RPH3AL | 5 | 1.115384951 | 0.457 | 0.057 | 5.04E-151 | 1.85E-146 |
| AC004704.1 | 5 | 1.349912523 | 0.306 | 0.01 | 5.44E-151 | 1.99E-146 |
| ANGPT1 | 5 | -2.486073333 | 0.072 | 0.609 | 8.8E-151 | 3.22E-146 |
| RADX | 5 | 1.246861016 | 0.908 | 0.466 | 1.37E-150 | 5.03E-146 |
| FBXL4 | 5 | 1.177118438 | 0.813 | 0.508 | 2.5E-150 | 9.16E-146 |
| CHGA | 5 | 1.117973191 | 0.338 | 0.02 | 7.7E-150 | 2.82E-145 |
| RABGAP1L | 5 | 1.136614163 | 0.967 | 0.735 | 8.77E-150 | 3.21E-145 |
| DOK5 | 5 | -1.924756466 | 0.051 | 0.577 | 4.15E-149 | 1.52E-144 |
| EPB41L3 | 5 | 1.396078087 | 0.686 | 0.268 | 4.2E-149 | 1.54E-144 |
| ITGA1 | 5 | 1.235238538 | 0.572 | 0.098 | 8.75E-149 | 3.2E-144 |
| TFPI | 5 | 1.011251267 | 0.38 | 0.026 | 4.51E-148 | 1.65E-143 |
| MEG8 | 5 | -2.43079748 | 0.075 | 0.605 | 6.32E-148 | 2.31E-143 |
| SLC9C1 | 5 | 0.929569311 | 0.319 | 0.014 | 8.87E-148 | 3.25E-143 |
| RALGPS2 | 5 | 1.386758173 | 0.747 | 0.343 | 1.29E-147 | 4.71E-143 |
| C5orf66 | 5 | 1.160188073 | 0.45 | 0.056 | 6.07E-146 | 2.22E-141 |
| MARCH8 | 5 | 1.217534563 | 0.571 | 0.152 | 6.72E-146 | 2.46E-141 |
| GRIA2 | 5 | -3.108437313 | 0.22 | 0.687 | 7.82E-146 | 2.86E-141 |
| SPTBN4 | 5 | 1.193640386 | 0.486 | 0.076 | 8.52E-146 | 3.12E-141 |
| TIAM2 | 5 | -1.812125209 | 0.36 | 0.808 | 2.99E-145 | 1.09E-140 |
| ANO3 | 5 | 1.101949974 | 0.358 | 0.021 | 4.69E-145 | 1.72E-140 |
| CARMIL1 | 5 | 1.263689945 | 0.774 | 0.293 | 5.34E-145 | 1.95E-140 |
| USH1C | 5 | 0.612838282 | 0.259 | 0.004 | 1.77E-144 | 6.48E-140 |
| PDK4 | 5 | 1.214152709 | 0.334 | 0.022 | 3.21E-144 | 1.17E-139 |
| LINC00616 | 5 | 0.817928216 | 0.255 | 0.004 | 3.3E-144 | 1.21E-139 |
| DENND4A | 5 | 1.182668615 | 0.877 | 0.567 | 1.26E-143 | 4.61E-139 |
| LUZP2 | 5 | -3.261653917 | 0.079 | 0.587 | 3.97E-143 | 1.45E-138 |
| GRID2 | 5 | -3.430260407 | 0.092 | 0.61 | 6.1E-143 | 2.23E-138 |
| RBFOX2 | 5 | 0.989134968 | 0.954 | 0.839 | 6.23E-143 | 2.28E-138 |
| PEX5L | 5 | -2.06912871 | 0.694 | 0.768 | 1.46E-142 | 5.34E-138 |

| | | | | | | |
|---|---|---|---|---|---|---|
| RAB26 | 5 | 1.002821845 | 0.451 | 0.056 | 4.93E-142 | 1.8E-137 |
| PHLDB2 | 5 | 1.221455627 | 0.712 | 0.2 | 5.85E-142 | 2.14E-137 |
| FGD4 | 5 | 1.260000236 | 0.846 | 0.392 | 1.1E-141 | 4.03E-137 |
| DCLK1 | 5 | -2.175221521 | 0.108 | 0.628 | 1.81E-141 | 6.63E-137 |
| AL160408.3 | 5 | 0.875798831 | 0.229 | 0.002 | 4.8E-140 | 1.76E-135 |
| EPHA6 | 5 | 1.430613937 | 0.528 | 0.088 | 2.2E-139 | 8.05E-135 |
| LINC01170 | 5 | -2.421128303 | 0.097 | 0.614 | 2.33E-139 | 8.52E-135 |
| TBX15 | 5 | -1.98030973 | 0.062 | 0.564 | 2.84E-139 | 1.04E-134 |
| PLEKHA7 | 5 | 1.179067632 | 0.655 | 0.164 | 3.37E-139 | 1.23E-134 |
| CASC15 | 5 | -2.40888089 | 0.092 | 0.604 | 3.56E-139 | 1.3E-134 |
| AL008633.1 | 5 | -4.532141212 | 0.299 | 0.738 | 3.6E-139 | 1.32E-134 |
| PTPN13 | 5 | 1.101997987 | 0.582 | 0.111 | 6.5E-139 | 2.38E-134 |
| PPFIA2 | 5 | -2.241205157 | 0.332 | 0.751 | 8.25E-139 | 3.02E-134 |
| AC084816.1 | 5 | 1.123877764 | 0.35 | 0.024 | 8.4E-139 | 3.08E-134 |
| SGCZ | 5 | -3.476537886 | 0.075 | 0.584 | 1.48E-137 | 5.41E-133 |
| UNC5D | 5 | -3.085751749 | 0.112 | 0.608 | 1.51E-137 | 5.53E-133 |
| IMMP1L | 5 | 1.173401842 | 0.802 | 0.44 | 9.95E-137 | 3.64E-132 |
| SNX29 | 5 | 1.243394271 | 0.743 | 0.38 | 1.15E-136 | 4.23E-132 |
| LSAMP | 5 | -2.676781008 | 0.319 | 0.764 | 8.42E-136 | 3.08E-131 |
| LHFPL3 | 5 | -2.595961825 | 0.099 | 0.596 | 1.1E-135 | 4.03E-131 |
| ZNF532 | 5 | -1.466655915 | 0.193 | 0.707 | 1.89E-135 | 6.93E-131 |
| LRRC4C | 5 | -2.441782993 | 0.059 | 0.56 | 2.37E-135 | 8.66E-131 |
| GPR39 | 5 | 0.926165654 | 0.323 | 0.018 | 8.61E-135 | 3.15E-130 |
| RCAN2 | 5 | 1.186374879 | 0.492 | 0.075 | 9.73E-135 | 3.56E-130 |
| RYR2 | 5 | 1.91033861 | 0.646 | 0.18 | 1.95E-134 | 7.15E-130 |
| SYNDIG1 | 5 | -1.642075965 | 0.09 | 0.603 | 2.02E-134 | 7.41E-130 |
| MIA2 | 5 | 1.135722277 | 0.734 | 0.325 | 1.88E-133 | 6.87E-129 |
| PDE3B | 5 | 1.191487626 | 0.943 | 0.617 | 2.84E-133 | 1.04E-128 |
| NFIA | 5 | -1.866272602 | 0.708 | 0.906 | 3.62E-133 | 1.32E-128 |
| PLCXD3 | 5 | 0.98602113 | 0.356 | 0.026 | 4.47E-133 | 1.64E-128 |
| ZMAT4 | 5 | -2.144333239 | 0.064 | 0.556 | 1.02E-132 | 3.72E-128 |
| SLC29A4 | 5 | 0.939329419 | 0.402 | 0.053 | 3.8E-132 | 1.39E-127 |
| TSPAN2 | 5 | 0.779681446 | 0.332 | 0.024 | 6.44E-132 | 2.36E-127 |
| CYP4F3 | 5 | 0.606672389 | 0.235 | 0.003 | 1.3E-131 | 4.77E-127 |
| ATXN1 | 5 | -1.762990234 | 0.237 | 0.723 | 4.82E-131 | 1.77E-126 |
| GCK | 5 | 0.713487654 | 0.299 | 0.015 | 8.77E-131 | 3.21E-126 |
| LINC00461 | 5 | -1.590856661 | 0.05 | 0.532 | 1.05E-129 | 3.84E-125 |
| TENM2 | 5 | -2.484568733 | 0.152 | 0.642 | 2.14E-129 | 7.84E-125 |
| CDH9 | 5 | 0.943830697 | 0.299 | 0.014 | 2.22E-129 | 8.12E-125 |
| GPR158 | 5 | 1.197715645 | 0.565 | 0.114 | 2.39E-129 | 8.76E-125 |
| GNG2 | 5 | 1.214901261 | 0.583 | 0.133 | 4.76E-129 | 1.74E-124 |
| MEIS2 | 5 | -2.046408088 | 0.084 | 0.569 | 1.61E-128 | 5.89E-124 |
| KCND2 | 5 | -2.636282641 | 0.077 | 0.57 | 2.38E-127 | 8.71E-123 |
| CACNB4 | 5 | -2.162471145 | 0.086 | 0.577 | 2.66E-127 | 9.72E-123 |
| CD36 | 5 | 1.282923009 | 0.42 | 0.054 | 3.77E-127 | 1.38E-122 |
| SLC7A2 | 5 | 1.252021784 | 0.534 | 0.109 | 6.25E-127 | 2.29E-122 |
| ANO4 | 5 | -2.591041255 | 0.125 | 0.613 | 1.98E-126 | 7.24E-122 |
| CFAP47 | 5 | 0.374220228 | 0.424 | 0.065 | 2.91E-126 | 1.07E-121 |
| SLC44A3 | 5 | 0.953731989 | 0.464 | 0.075 | 6.17E-126 | 2.26E-121 |
| LRBA | 5 | 0.931263105 | 0.978 | 0.883 | 2E-125 | 7.32E-121 |
| XKR4 | 5 | 1.217315254 | 0.633 | 0.158 | 4.41E-125 | 1.61E-120 |
| SETBP1 | 5 | 1.123203545 | 0.901 | 0.559 | 6.35E-125 | 2.33E-120 |
| NDNF | 5 | 0.651295062 | 0.262 | 0.009 | 7.79E-125 | 2.85E-120 |
| MYRIP | 5 | -2.16954587 | 0.235 | 0.685 | 3.14E-124 | 1.15E-119 |
| ADGRB3 | 5 | -2.379935596 | 0.086 | 0.566 | 5.21E-124 | 1.91E-119 |
| SYT7 | 5 | 0.921974406 | 0.394 | 0.052 | 9.53E-124 | 3.49E-119 |
| PPP3CA | 5 | 1.055686201 | 0.947 | 0.832 | 7.15E-123 | 2.62E-118 |
| PPL | 5 | 1.22644895 | 0.483 | 0.104 | 8.21E-123 | 3.01E-118 |

| | | | | | | |
|---|---|---|---|---|---|---|
| TMEM108 | 5 | -1.958402373 | 0.22 | 0.683 | 2.36E-122 | 8.63E-118 |
| NFIB | 5 | -1.814879753 | 0.501 | 0.854 | 9.23E-122 | 3.38E-117 |
| WWOX | 5 | 1.30749603 | 0.947 | 0.761 | 9.7E-122 | 3.55E-117 |
| NPSR1-AS1 | 5 | 1.656385184 | 0.58 | 0.149 | 3.18E-121 | 1.16E-116 |
| HIPK2 | 5 | 1.096526484 | 0.804 | 0.432 | 5.01E-121 | 1.84E-116 |
| DOCK11 | 5 | -1.785223333 | 0.083 | 0.558 | 6.83E-121 | 2.5E-116 |
| STK33 | 5 | 0.904702024 | 0.578 | 0.134 | 9.7E-121 | 3.55E-116 |
| CDH12 | 5 | 0.554264078 | 0.866 | 0.369 | 4.27E-120 | 1.56E-115 |
| AC104123.1 | 5 | 0.987066962 | 0.371 | 0.044 | 4.46E-120 | 1.63E-115 |
| MOB1B | 5 | 1.180121456 | 0.686 | 0.39 | 4.94E-120 | 1.81E-115 |
| GMDS-DT | 5 | 1.322387605 | 0.659 | 0.239 | 2.57E-119 | 9.39E-115 |
| ANTXR1 | 5 | -1.221825585 | 0.04 | 0.496 | 6.17E-119 | 2.26E-114 |
| MIR137HG | 5 | 1.135405919 | 0.705 | 0.22 | 7.98E-119 | 2.92E-114 |
| MIR181A1HG | 5 | -1.522404853 | 0.046 | 0.503 | 2.42E-118 | 8.88E-114 |
| AC087280.2 | 5 | 0.831929257 | 0.325 | 0.026 | 2.67E-118 | 9.79E-114 |
| ATP2B1 | 5 | 1.166771638 | 0.782 | 0.406 | 4.97E-118 | 1.82E-113 |
| TENM3 | 5 | 1.102547991 | 0.978 | 0.77 | 7.63E-118 | 2.79E-113 |
| ERN1 | 5 | 0.918989999 | 0.396 | 0.06 | 8.5E-118 | 3.11E-113 |
| GRIA4 | 5 | 1.330994338 | 0.765 | 0.287 | 5.03E-117 | 1.84E-112 |
| CACHD1 | 5 | -1.704709865 | 0.138 | 0.61 | 5.11E-117 | 1.87E-112 |
| EPSTI1 | 5 | 0.902825189 | 0.44 | 0.066 | 1.25E-116 | 4.56E-112 |
| TESC | 5 | 0.749825973 | 0.294 | 0.018 | 6.36E-116 | 2.33E-111 |
| NDST3 | 5 | -1.85374226 | 0.055 | 0.504 | 1.18E-115 | 4.34E-111 |
| LINC00511 | 5 | 1.185085237 | 0.739 | 0.261 | 1.38E-115 | 5.06E-111 |
| SLC27A2 | 5 | 0.692488155 | 0.292 | 0.018 | 2.07E-115 | 7.59E-111 |
| CALN1 | 5 | 1.366873516 | 0.409 | 0.072 | 3.62E-115 | 1.33E-110 |
| KCTD3 | 5 | 1.038370389 | 0.785 | 0.449 | 8.64E-115 | 3.16E-110 |
| DNAH10 | 5 | -1.754512975 | 0.048 | 0.496 | 9.38E-115 | 3.43E-110 |
| PTPRS | 5 | -1.29802957 | 0.134 | 0.608 | 9.89E-115 | 3.62E-110 |
| AC061961.1 | 5 | 0.622685755 | 0.266 | 0.014 | 1.09E-114 | 3.98E-110 |
| GALNT3 | 5 | 0.77912129 | 0.345 | 0.035 | 1.22E-114 | 4.45E-110 |
| NTNG1 | 5 | -1.765739057 | 0.055 | 0.503 | 7.45E-114 | 2.73E-109 |
| TNS3 | 5 | 1.067303803 | 0.672 | 0.213 | 7.61E-114 | 2.79E-109 |
| ATP9A | 5 | 1.060346686 | 0.783 | 0.385 | 3.8E-113 | 1.39E-108 |
| FHL5 | 5 | 0.480905455 | 0.206 | 0.004 | 9.59E-113 | 3.51E-108 |
| MEG3 | 5 | -1.511702072 | 0.039 | 0.474 | 3.04E-112 | 1.11E-107 |
| TMEM164 | 5 | 1.117176302 | 0.749 | 0.474 | 1.79E-111 | 6.57E-107 |
| MAP3K21 | 5 | 0.906840949 | 0.457 | 0.086 | 1.93E-111 | 7.08E-107 |
| AL390800.1 | 5 | -1.926833701 | 0.092 | 0.546 | 2.89E-111 | 1.06E-106 |
| TMEM178A | 5 | -1.335797099 | 0.051 | 0.497 | 3.24E-111 | 1.19E-106 |
| LAMA4 | 5 | -2.244866807 | 0.044 | 0.481 | 4.44E-111 | 1.63E-106 |
| DDC | 5 | 0.468896591 | 0.18 | 0.001 | 5.78E-111 | 2.12E-106 |
| BAIAP2L1 | 5 | 0.968338044 | 0.466 | 0.093 | 9.2E-111 | 3.37E-106 |
| NRXN3 | 5 | 0.403379417 | 0.807 | 0.325 | 1.26E-110 | 4.61E-106 |
| ADAMTS3 | 5 | -2.031403018 | 0.068 | 0.504 | 1.46E-110 | 5.34E-106 |
| SH3GL3 | 5 | -1.634978205 | 0.11 | 0.559 | 1.48E-110 | 5.41E-106 |
| MYO5A | 5 | -1.325561016 | 0.161 | 0.618 | 1.54E-110 | 5.64E-106 |
| STK32B | 5 | -1.646031647 | 0.099 | 0.552 | 7.44E-110 | 2.72E-105 |
| BTG2 | 5 | 0.93784604 | 0.29 | 0.03 | 1.13E-109 | 4.12E-105 |
| TSHZ3 | 5 | -1.494683831 | 0.07 | 0.52 | 1.14E-109 | 4.16E-105 |
| IRAK3 | 5 | -1.32333263 | 0.073 | 0.519 | 2.43E-109 | 8.89E-105 |
| LINC01060 | 5 | 1.081017896 | 0.352 | 0.04 | 2.96E-109 | 1.08E-104 |
| CHRM3 | 5 | -2.16404965 | 0.026 | 0.443 | 3.57E-109 | 1.31E-104 |
| NKAIN3 | 5 | -2.849346464 | 0.328 | 0.707 | 7.84E-109 | 2.87E-104 |
| ASCL1 | 5 | 0.733588483 | 0.187 | 0.002 | 9.78E-109 | 3.58E-104 |
| SH3KBP1 | 5 | -1.430360124 | 0.272 | 0.705 | 1.13E-108 | 4.15E-104 |
| MYO5B | 5 | 0.903681549 | 0.404 | 0.062 | 1.14E-108 | 4.17E-104 |
| GPM6B | 5 | -1.530069642 | 0.22 | 0.66 | 4.35E-108 | 1.59E-103 |

| | | | | | | |
|---|---|---|---|---|---|---|
| MARCH3 | 5 | 1.159246397 | 0.572 | 0.172 | 4.56E-108 | 1.67E-103 |
| TNIK | 5 | 1.025624851 | 0.793 | 0.332 | 7.13E-108 | 2.61E-103 |
| ABCA4 | 5 | 0.954151786 | 0.363 | 0.043 | 8.58E-108 | 3.14E-103 |
| ABLIM1 | 5 | 1.087128678 | 0.65 | 0.216 | 1.26E-107 | 4.63E-103 |
| LINC02595 | 5 | 0.932799951 | 0.444 | 0.084 | 2.44E-107 | 8.92E-103 |
| UNC5C | 5 | -2.511428426 | 0.119 | 0.555 | 2.57E-107 | 9.41E-103 |
| WASF3 | 5 | 1.006602008 | 0.543 | 0.151 | 2.59E-107 | 9.49E-103 |
| BASP1 | 5 | -1.322716852 | 0.061 | 0.492 | 6.18E-107 | 2.26E-102 |
| ELOVL2 | 5 | 0.979826008 | 0.505 | 0.12 | 7.73E-107 | 2.83E-102 |
| POU5F1 | 5 | -1.242197463 | 0.068 | 0.509 | 1.58E-106 | 5.77E-102 |
| UNC79 | 5 | 0.968742098 | 0.787 | 0.315 | 1.86E-106 | 6.81E-102 |
| FIRRE | 5 | 1.137995027 | 0.743 | 0.277 | 1.95E-106 | 7.14E-102 |
| CYP4F12 | 5 | 0.549428453 | 0.209 | 0.005 | 2.05E-106 | 7.49E-102 |
| BMERB1 | 5 | -1.217117652 | 0.062 | 0.498 | 2.82E-106 | 1.03E-101 |
| IRF2BP2 | 5 | 0.899740862 | 0.457 | 0.104 | 5.08E-106 | 1.86E-101 |
| SPATA13 | 5 | 1.071255507 | 0.635 | 0.206 | 7.92E-106 | 2.9E-101 |
| TTC8 | 5 | 1.01612882 | 0.631 | 0.224 | 9.5E-106 | 3.48E-101 |
| DPP10 | 5 | 1.013244266 | 0.941 | 0.584 | 1.14E-105 | 4.18E-101 |
| CRHBP | 5 | 0.506970969 | 0.189 | 0.003 | 1.4E-105 | 5.12E-101 |
| TRAPPC9 | 5 | 0.990133386 | 0.895 | 0.686 | 2.87E-104 | 1.05E-99 |
| CNTN3 | 5 | -1.854446401 | 0.081 | 0.521 | 2.88E-104 | 1.05E-99 |
| RNF8 | 5 | 0.954754951 | 0.477 | 0.133 | 4.66E-104 | 1.71E-99 |
| DNAH6 | 5 | 0.685459379 | 0.383 | 0.058 | 4.7E-104 | 1.72E-99 |
| PPARGC1A | 5 | 1.295068349 | 0.532 | 0.142 | 7.81E-104 | 2.86E-99 |
| MYO5C | 5 | 0.920997439 | 0.444 | 0.094 | 1.45E-103 | 5.3E-99 |
| CBFA2T2 | 5 | 0.960119903 | 0.884 | 0.623 | 1.91E-103 | 7E-99 |
| ELMO1 | 5 | -1.665849746 | 0.145 | 0.577 | 3.82E-103 | 1.4E-98 |
| MAL2 | 5 | 0.469173622 | 0.198 | 0.004 | 4.03E-103 | 1.48E-98 |
| EXOC6B | 5 | 0.982009337 | 0.901 | 0.679 | 9.35E-103 | 3.42E-98 |
| LRRFIP1 | 5 | 1.023791845 | 0.787 | 0.486 | 9.86E-103 | 3.61E-98 |
| AFF3 | 5 | -2.063099772 | 0.213 | 0.632 | 1.29E-102 | 4.72E-98 |
| ARHGEF10L | 5 | 1.000984844 | 0.563 | 0.196 | 2.1E-102 | 7.7E-98 |
| AGMO | 5 | 1.02264093 | 0.464 | 0.092 | 2.31E-102 | 8.46E-98 |
| SYDE2 | 5 | 1.014129621 | 0.576 | 0.178 | 2.58E-102 | 9.43E-98 |
| AFF1 | 5 | 1.016891942 | 0.78 | 0.388 | 6.42E-102 | 2.35E-97 |
| PIAS1 | 5 | 0.949229891 | 0.916 | 0.719 | 1.52E-101 | 5.56E-97 |
| GRAMD2B | 5 | 0.966596833 | 0.517 | 0.134 | 3.9E-101 | 1.43E-96 |
| RMST | 5 | -2.659416799 | 0.103 | 0.514 | 4.56E-101 | 1.67E-96 |
| GPR173 | 5 | -1.134688607 | 0.097 | 0.526 | 2.75E-100 | 1.01E-95 |
| FOXN3 | 5 | -1.303948228 | 0.815 | 0.939 | 3.14E-100 | 1.15E-95 |
| DPPA4 | 5 | -1.162790409 | 0.05 | 0.459 | 4.6E-100 | 1.68E-95 |
| TTLL6 | 5 | 0.565786136 | 0.213 | 0.008 | 6.91E-100 | 2.53E-95 |
| EPS8 | 5 | -1.670309338 | 0.178 | 0.613 | 7.22E-100 | 2.64E-95 |
| ZNF215 | 5 | 0.579831186 | 0.242 | 0.014 | 9.48E-100 | 3.47E-95 |
| FAM83F | 5 | 0.504726273 | 0.189 | 0.004 | 1.1E-99 | 4.02E-95 |
| NEDD4L | 5 | -1.330736188 | 0.635 | 0.87 | 1.27E-99 | 4.65E-95 |
| FAT1 | 5 | -1.490158346 | 0.213 | 0.639 | 1.33E-99 | 4.88E-95 |
| RALYL | 5 | -2.89727447 | 0.084 | 0.5 | 2.37E-99 | 8.68E-95 |
| XG | 5 | 0.673779417 | 0.257 | 0.018 | 2.63E-99 | 9.62E-95 |
| SMARCA2 | 5 | 1.060602517 | 0.794 | 0.406 | 3.59E-99 | 1.31E-94 |
| PTPRO | 5 | -1.56568515 | 0.053 | 0.467 | 3.81E-99 | 1.4E-94 |
| TMEM45B | 5 | 0.622120669 | 0.196 | 0.006 | 4.23E-99 | 1.55E-94 |
| WLS | 5 | -1.1442447 | 0.042 | 0.444 | 2.24E-98 | 8.19E-94 |
| BCL11B | 5 | -1.319797649 | 0.044 | 0.449 | 2.87E-98 | 1.05E-93 |
| ABCC9 | 5 | 0.707282001 | 0.301 | 0.028 | 3.18E-98 | 1.16E-93 |
| LIFR | 5 | -1.876814317 | 0.261 | 0.659 | 4.91E-98 | 1.8E-93 |
| XBP1 | 5 | 0.90754109 | 0.286 | 0.032 | 6.88E-98 | 2.52E-93 |
| F5 | 5 | 0.601838411 | 0.253 | 0.016 | 2.51E-97 | 9.17E-93 |

| | | | | | | |
|---|---|---|---|---|---|---|
| KIF19 | 5 | 0.593021093 | 0.211 | 0.009 | 3.54E-97 | 1.3E-92 |
| LRFN5 | 5 | -2.031681989 | 0.066 | 0.485 | 3.6E-97 | 1.32E-92 |
| CPEB1 | 5 | 0.943397273 | 0.428 | 0.088 | 5.84E-97 | 2.14E-92 |
| SHROOM3 | 5 | 0.727295867 | 0.413 | 0.073 | 6.14E-97 | 2.25E-92 |
| HPN | 5 | 0.647034497 | 0.233 | 0.013 | 7.62E-97 | 2.79E-92 |
| PCDH7 | 5 | -2.544681933 | 0.053 | 0.451 | 1.95E-96 | 7.13E-92 |
| NRCAM | 5 | 1.068260331 | 0.763 | 0.366 | 2.14E-96 | 7.85E-92 |
| MAP4K3 | 5 | -1.188743288 | 0.705 | 0.895 | 2.19E-96 | 8.01E-92 |
| MTMR7 | 5 | 1.086939715 | 0.723 | 0.325 | 3.55E-96 | 1.3E-91 |
| LINC01091 | 5 | 0.990149444 | 0.376 | 0.058 | 4.34E-96 | 1.59E-91 |
| DLG2 | 5 | -2.477195257 | 0.262 | 0.659 | 9.05E-96 | 3.31E-91 |
| AGFG2 | 5 | 0.76358134 | 0.323 | 0.045 | 1.23E-95 | 4.5E-91 |
| SLC22A3 | 5 | -1.56725524 | 0.04 | 0.431 | 1.25E-95 | 4.56E-91 |
| SNED1 | 5 | 1.039344817 | 0.422 | 0.087 | 1.54E-95 | 5.64E-91 |
| UNC13A | 5 | 0.930627246 | 0.472 | 0.111 | 3.38E-95 | 1.24E-90 |
| DNAH14 | 5 | 1.118004456 | 0.822 | 0.501 | 3.7E-95 | 1.36E-90 |
| AC006369.1 | 5 | 0.566068776 | 0.16 | 0.001 | 4.08E-95 | 1.49E-90 |
| CARD11 | 5 | 0.834760146 | 0.336 | 0.042 | 4.38E-95 | 1.6E-90 |
| TMEM200A | 5 | 0.843093584 | 0.36 | 0.056 | 6.86E-95 | 2.51E-90 |
| PDXDC1 | 5 | 0.945330294 | 0.719 | 0.42 | 7.72E-95 | 2.83E-90 |
| MEGF10 | 5 | -1.637109098 | 0.039 | 0.423 | 1.66E-94 | 6.09E-90 |
| RGS4 | 5 | 0.828268183 | 0.183 | 0.004 | 1.79E-94 | 6.56E-90 |
| CAMKK1 | 5 | 0.904253427 | 0.429 | 0.096 | 2.03E-94 | 7.44E-90 |
| CRISPLD1 | 5 | -2.13177925 | 0.101 | 0.522 | 2.8E-94 | 1.02E-89 |
| SSTR2 | 5 | 0.746748794 | 0.312 | 0.04 | 3.33E-94 | 1.22E-89 |
| ANOS1 | 5 | -1.843780655 | 0.064 | 0.466 | 3.42E-94 | 1.25E-89 |
| PALM2-AKAP2 | 5 | -1.215589617 | 0.097 | 0.513 | 5.79E-94 | 2.12E-89 |
| EYA2 | 5 | 1.027063067 | 0.483 | 0.112 | 6.53E-94 | 2.39E-89 |
| VCL | 5 | -1.546495232 | 0.44 | 0.8 | 8.6E-94 | 3.15E-89 |
| AC112770.1 | 5 | 0.886229359 | 0.158 | 0.002 | 1.2E-93 | 4.39E-89 |
| KIF5C | 5 | 0.993320916 | 0.684 | 0.266 | 1.67E-93 | 6.12E-89 |
| EFCAB6 | 5 | 0.930035587 | 0.494 | 0.125 | 6.36E-93 | 2.33E-88 |
| TOX3 | 5 | 1.02785325 | 0.705 | 0.277 | 8.38E-93 | 3.07E-88 |
| NRP1 | 5 | 1.039709374 | 0.716 | 0.286 | 8.46E-93 | 3.1E-88 |
| SEPTIN9 | 5 | 1.066961754 | 0.76 | 0.443 | 8.83E-93 | 3.23E-88 |
| GYG2 | 5 | 0.926137707 | 0.475 | 0.123 | 1.01E-92 | 3.71E-88 |
| LINGO2 | 5 | 1.101708321 | 0.371 | 0.057 | 1.2E-92 | 4.38E-88 |
| MIR222HG | 5 | 0.926335644 | 0.572 | 0.181 | 1.82E-92 | 6.66E-88 |
| MBP | 5 | 0.870687622 | 0.461 | 0.104 | 2.44E-92 | 8.93E-88 |
| SLC24A2 | 5 | -1.669751944 | 0.064 | 0.456 | 3.98E-92 | 1.46E-87 |
| BAIAP3 | 5 | 0.573942341 | 0.231 | 0.014 | 6.65E-92 | 2.44E-87 |
| CXCL6 | 5 | 0.435897756 | 0.143 | 0 | 7.74E-92 | 2.83E-87 |
| CHST8 | 5 | 0.987182278 | 0.528 | 0.147 | 1.09E-91 | 3.99E-87 |
| IL4R | 5 | 0.671427679 | 0.294 | 0.032 | 2.45E-91 | 8.97E-87 |
| PHYHIPL | 5 | 0.94154201 | 0.512 | 0.137 | 3.34E-91 | 1.22E-86 |
| CHST11 | 5 | -1.258575919 | 0.213 | 0.636 | 3.96E-91 | 1.45E-86 |
| AC009315.1 | 5 | -1.26355335 | 0.046 | 0.433 | 6.5E-91 | 2.38E-86 |
| MTUS1 | 5 | 0.897357552 | 0.411 | 0.084 | 1.11E-90 | 4.07E-86 |
| AC078864.1 | 5 | 0.3576671 | 0.152 | 0.001 | 2.27E-90 | 8.32E-86 |
| REV3L | 5 | 0.857249683 | 0.967 | 0.859 | 2.64E-90 | 9.66E-86 |
| ENPEP | 5 | 0.726707268 | 0.301 | 0.036 | 3.49E-90 | 1.28E-85 |
| ZKSCAN1 | 5 | 0.975624779 | 0.712 | 0.418 | 4.23E-90 | 1.55E-85 |
| ITPR1 | 5 | 1.051176804 | 0.743 | 0.368 | 8.03E-90 | 2.94E-85 |
| MGAT4C | 5 | -2.34681551 | 0.152 | 0.557 | 1.01E-89 | 3.7E-85 |
| CTNNA2 | 5 | -2.270654356 | 0.132 | 0.514 | 1.32E-89 | 4.83E-85 |
| FBN2 | 5 | -1.859409698 | 0.031 | 0.401 | 3.26E-89 | 1.2E-84 |
| AL033504.1 | 5 | -1.782630626 | 0.202 | 0.602 | 4.51E-89 | 1.65E-84 |
| SGSM1 | 5 | 0.678943777 | 0.257 | 0.022 | 4.78E-89 | 1.75E-84 |

| | | | | | | |
|---|---|---|---|---|---|---|
| ATRNL1 | 5 | -1.970231937 | 0.585 | 0.821 | 1.54E-88 | 5.65E-84 |
| KLHL29 | 5 | -1.686664715 | 0.237 | 0.641 | 1.87E-88 | 6.84E-84 |
| FAAH2 | 5 | 0.824038224 | 0.374 | 0.068 | 2.32E-88 | 8.51E-84 |
| ADAMTS5 | 5 | 0.664301199 | 0.273 | 0.026 | 2.43E-88 | 8.88E-84 |
| SEC31A | 5 | 0.921522315 | 0.85 | 0.597 | 5.68E-88 | 2.08E-83 |
| LINC01876 | 5 | 1.104662689 | 0.833 | 0.487 | 8.37E-88 | 3.06E-83 |
| ELMOD1 | 5 | -1.262602803 | 0.046 | 0.422 | 9.24E-88 | 3.38E-83 |
| KLF5 | 5 | 0.611614302 | 0.25 | 0.02 | 6.1E-87 | 2.23E-82 |
| AC008591.1 | 5 | -1.637641011 | 0.075 | 0.456 | 7.89E-87 | 2.89E-82 |
| PGM5 | 5 | 0.711617586 | 0.286 | 0.031 | 8.91E-87 | 3.26E-82 |
| MIR99AHG | 5 | -2.26938733 | 0.466 | 0.7 | 1.09E-86 | 3.99E-82 |
| TLCD4 | 5 | 0.849038806 | 0.44 | 0.109 | 1.28E-86 | 4.7E-82 |
| ADAMTS9 | 5 | -1.2471482 | 0.033 | 0.399 | 1.53E-86 | 5.6E-82 |
| CACNA1B | 5 | 0.971167879 | 0.655 | 0.239 | 2.17E-86 | 7.96E-82 |
| CADM2 | 5 | -2.196517992 | 0.079 | 0.471 | 2.21E-86 | 8.09E-82 |
| SEC11C | 5 | 0.640568977 | 0.316 | 0.045 | 2.68E-86 | 9.82E-82 |
| BAIAP2L2 | 5 | 0.626361883 | 0.251 | 0.025 | 2.88E-86 | 1.05E-81 |
| GAREM1 | 5 | 1.085432827 | 0.637 | 0.259 | 3.37E-86 | 1.23E-81 |
| LINC00632 | 5 | -1.287503192 | 0.04 | 0.406 | 5.54E-86 | 2.03E-81 |
| PIEZO2 | 5 | -1.432686643 | 0.037 | 0.397 | 5.69E-86 | 2.08E-81 |
| HOOK3 | 5 | 0.851294665 | 0.791 | 0.523 | 7.25E-86 | 2.65E-81 |
| FAM189A1 | 5 | -1.796018117 | 0.396 | 0.713 | 1.36E-85 | 4.99E-81 |
| TET1 | 5 | -1.198606958 | 0.055 | 0.433 | 1.61E-85 | 5.88E-81 |
| PNPLA7 | 5 | 0.807768618 | 0.33 | 0.058 | 1.89E-85 | 6.93E-81 |
| DNER | 5 | -1.509805452 | 0.046 | 0.412 | 1.97E-85 | 7.21E-81 |
| UTRN | 5 | 0.868093659 | 0.916 | 0.603 | 2.26E-85 | 8.28E-81 |
| ATP8B4 | 5 | 0.603418029 | 0.244 | 0.02 | 5.37E-85 | 1.97E-80 |
| LAMA2 | 5 | -1.767216578 | 0.044 | 0.414 | 6.59E-85 | 2.41E-80 |
| ZNF423 | 5 | -1.253254849 | 0.059 | 0.432 | 6.65E-85 | 2.44E-80 |
| MCC | 5 | -1.317607168 | 0.525 | 0.815 | 1.14E-84 | 4.18E-80 |
| TPK1 | 5 | 0.812558803 | 0.354 | 0.065 | 2.88E-84 | 1.06E-79 |
| AUTS2 | 5 | -1.975643763 | 0.114 | 0.498 | 2.73E-83 | 9.98E-79 |
| SND1 | 5 | 0.876842167 | 0.848 | 0.701 | 2.78E-83 | 1.02E-78 |
| TBC1D30 | 5 | 0.771883428 | 0.358 | 0.069 | 4.49E-83 | 1.64E-78 |
| DPEP1 | 5 | 0.402823029 | 0.161 | 0.004 | 5.27E-83 | 1.93E-78 |
| GPR176 | 5 | -1.074668249 | 0.055 | 0.423 | 6.01E-83 | 2.2E-78 |
| LRCH2 | 5 | -1.192528072 | 0.361 | 0.719 | 6.24E-83 | 2.29E-78 |
| TLE4 | 5 | -1.294119192 | 0.119 | 0.508 | 8.44E-83 | 3.09E-78 |
| DIAPH2 | 5 | 0.799220942 | 0.98 | 0.9 | 8.44E-83 | 3.09E-78 |
| PAK5 | 5 | -1.416856526 | 0.09 | 0.473 | 8.61E-83 | 3.15E-78 |
| DNM3OS | 5 | -1.275016372 | 0.02 | 0.361 | 9.49E-83 | 3.48E-78 |
| GCH1 | 5 | 0.999487598 | 0.481 | 0.149 | 1.99E-82 | 7.28E-78 |
| ARNTL2 | 5 | 0.850607152 | 0.428 | 0.104 | 3.67E-82 | 1.34E-77 |
| ADAMTS12 | 5 | -2.048947457 | 0.057 | 0.421 | 4.56E-82 | 1.67E-77 |
| KIRREL1 | 5 | -0.962804888 | 0.086 | 0.465 | 6.7E-82 | 2.45E-77 |
| GNAO1 | 5 | 0.879373956 | 0.811 | 0.434 | 8.43E-82 | 3.09E-77 |
| NIPSNAP1 | 5 | 0.850789012 | 0.417 | 0.118 | 9.92E-82 | 3.63E-77 |
| TTC3 | 5 | 0.72862486 | 0.982 | 0.87 | 1.1E-81 | 4.03E-77 |
| FGF5 | 5 | 0.307878279 | 0.125 | 0 | 1.57E-81 | 5.76E-77 |
| HS6ST3 | 5 | -2.990143984 | 0.158 | 0.527 | 5.83E-81 | 2.14E-76 |
| CRYBG3 | 5 | 0.963228441 | 0.67 | 0.292 | 1.36E-80 | 4.99E-76 |
| NRG3 | 5 | -2.851216054 | 0.152 | 0.485 | 1.45E-80 | 5.31E-76 |
| KCNIP4 | 5 | -2.841668643 | 0.171 | 0.52 | 1.61E-80 | 5.89E-76 |
| LINC02842 | 5 | 0.730227829 | 0.156 | 0.004 | 1.68E-80 | 6.16E-76 |
| PLPP4 | 5 | -1.266949736 | 0.042 | 0.392 | 1.72E-80 | 6.29E-76 |
| GRIA1 | 5 | -1.479326427 | 0.042 | 0.39 | 2.17E-80 | 7.95E-76 |
| SNAP25 | 5 | 0.912553724 | 0.651 | 0.274 | 2.34E-80 | 8.58E-76 |
| SAMD5 | 5 | -1.510762569 | 0.294 | 0.667 | 2.89E-80 | 1.06E-75 |

| | | | | | | |
|---|---|---|---|---|---|---|
| SYN3 | 5 | -1.897055351 | 0.079 | 0.445 | 3.23E-80 | 1.18E-75 |
| EXOC2 | 5 | -1.370728472 | 0.332 | 0.694 | 4.43E-80 | 1.62E-75 |
| LINC01725 | 5 | 0.690311273 | 0.257 | 0.026 | 4.54E-80 | 1.66E-75 |
| RASGEF1B | 5 | 0.957053013 | 0.686 | 0.3 | 5.01E-80 | 1.84E-75 |
| DZIP3 | 5 | 0.907474921 | 0.661 | 0.349 | 6.02E-80 | 2.2E-75 |
| CNGA3 | 5 | 0.416193596 | 0.165 | 0.005 | 9.31E-80 | 3.41E-75 |
| DOCK4 | 5 | -1.500067398 | 0.268 | 0.645 | 1.16E-79 | 4.25E-75 |
| CRYBG1 | 5 | 0.796101105 | 0.319 | 0.053 | 1.26E-79 | 4.63E-75 |
| TMC4 | 5 | 0.494540338 | 0.18 | 0.008 | 1.6E-79 | 5.85E-75 |
| RNF24 | 5 | -1.144360456 | 0.185 | 0.577 | 5.36E-79 | 1.96E-74 |
| LHFPL6 | 5 | -1.437760326 | 0.05 | 0.407 | 6.54E-79 | 2.4E-74 |
| ITGA4 | 5 | -1.869693379 | 0.127 | 0.494 | 7.92E-79 | 2.9E-74 |
| DOCK10 | 5 | -1.518359255 | 0.05 | 0.398 | 1.16E-78 | 4.23E-74 |
| PTPRZ1 | 5 | -1.679572159 | 0.066 | 0.43 | 1.54E-78 | 5.64E-74 |
| KCNQ3 | 5 | -1.475275364 | 0.059 | 0.409 | 2.33E-78 | 8.52E-74 |
| PARD3 | 5 | -0.901724845 | 0.897 | 0.97 | 3.14E-78 | 1.15E-73 |
| AL078600.1 | 5 | 0.463594817 | 0.174 | 0.007 | 2.4E-77 | 8.78E-73 |
| PLS1 | 5 | 0.88015793 | 0.536 | 0.185 | 2.49E-77 | 9.12E-73 |
| SREBF2 | 5 | 0.983424772 | 0.785 | 0.479 | 2.59E-77 | 9.49E-73 |
| COL4A2 | 5 | -1.091743696 | 0.028 | 0.354 | 5.08E-77 | 1.86E-72 |
| FAF1 | 5 | 0.734737969 | 0.947 | 0.821 | 5.3E-77 | 1.94E-72 |
| POU2F1 | 5 | -1.012974724 | 0.565 | 0.849 | 6.3E-77 | 2.31E-72 |
| USP25 | 5 | 0.882887585 | 0.809 | 0.52 | 9.06E-77 | 3.32E-72 |
| FHIT | 5 | 0.970885867 | 0.917 | 0.726 | 1.12E-76 | 4.1E-72 |
| CSMD1 | 5 | -3.004469483 | 0.116 | 0.466 | 1.56E-76 | 5.72E-72 |
| HNF1A | 5 | 0.340327005 | 0.139 | 0.002 | 1.96E-76 | 7.16E-72 |
| FYN | 5 | -1.165197481 | 0.316 | 0.682 | 2.89E-76 | 1.06E-71 |
| ICA1 | 5 | 0.847913213 | 0.556 | 0.195 | 9.49E-76 | 3.47E-71 |
| CAMK2D | 5 | 0.909424996 | 0.829 | 0.556 | 1.35E-75 | 4.93E-71 |
| MCTP2 | 5 | 0.431027381 | 0.606 | 0.242 | 1.41E-75 | 5.15E-71 |
| TANC1 | 5 | -1.099741334 | 0.172 | 0.552 | 1.73E-75 | 6.32E-71 |
| CEP126 | 5 | 0.935137143 | 0.611 | 0.28 | 1.92E-75 | 7.03E-71 |
| CALCOCO2 | 5 | 0.733379615 | 0.374 | 0.097 | 2.74E-75 | 1E-70 |
| TRAK1 | 5 | 0.857234821 | 0.712 | 0.417 | 3.88E-75 | 1.42E-70 |
| KCNQ1OT1 | 5 | -1.220754448 | 0.24 | 0.62 | 7.25E-75 | 2.66E-70 |
| LINC00261 | 5 | 0.370792711 | 0.138 | 0.002 | 1.1E-74 | 4.04E-70 |
| ST18 | 5 | 0.479404219 | 0.866 | 0.48 | 2E-74 | 7.34E-70 |
| CDC42SE2 | 5 | 0.870898999 | 0.71 | 0.426 | 2.65E-74 | 9.7E-70 |
| APBA2 | 5 | -1.067543591 | 0.222 | 0.595 | 3.8E-74 | 1.39E-69 |
| TBC1D8B | 5 | 0.692550433 | 0.349 | 0.074 | 3.81E-74 | 1.39E-69 |
| SRGAP3 | 5 | -1.151617689 | 0.198 | 0.573 | 4.44E-74 | 1.63E-69 |
| NR0B1 | 5 | 0.36988865 | 0.127 | 0.001 | 4.95E-74 | 1.81E-69 |
| GRIA3 | 5 | 0.857523483 | 0.517 | 0.163 | 6.53E-74 | 2.39E-69 |
| EFR3B | 5 | -1.088853517 | 0.112 | 0.476 | 7.82E-74 | 2.86E-69 |
| DDX17 | 5 | 0.66974417 | 0.96 | 0.852 | 1.35E-73 | 4.96E-69 |
| RIPOR2 | 5 | -1.284973491 | 0.051 | 0.386 | 1.82E-73 | 6.68E-69 |
| TTC39A | 5 | 0.349562578 | 0.149 | 0.004 | 2.06E-73 | 7.54E-69 |
| VGLL3 | 5 | -1.81350147 | 0.026 | 0.329 | 3.55E-73 | 1.3E-68 |
| DOK6 | 5 | 0.887930908 | 0.881 | 0.615 | 3.84E-73 | 1.4E-68 |
| DACH2 | 5 | -2.960671485 | 0.134 | 0.471 | 6.81E-73 | 2.49E-68 |
| PRKG1 | 5 | -2.091461177 | 0.501 | 0.748 | 7.88E-73 | 2.89E-68 |
| TUBA4A | 5 | 0.489803335 | 0.191 | 0.012 | 8E-73 | 2.93E-68 |
| AC023469.1 | 5 | -1.392658463 | 0.046 | 0.372 | 1.23E-72 | 4.52E-68 |
| SULF1 | 5 | -1.569366024 | 0.022 | 0.327 | 1.82E-72 | 6.66E-68 |
| CDH11 | 5 | -1.688684425 | 0.024 | 0.332 | 2.8E-72 | 1.03E-67 |
| SIK3 | 5 | 0.852076835 | 0.853 | 0.658 | 8.85E-72 | 3.24E-67 |
| AC011246.1 | 5 | -1.61645447 | 0.059 | 0.39 | 1.21E-71 | 4.43E-67 |
| MYC | 5 | 0.915258411 | 0.527 | 0.274 | 2.81E-71 | 1.03E-66 |

| | | | | | | |
|---|---|---|---|---|---|---|
| SMOC2 | 5 | 0.682262229 | 0.228 | 0.022 | 3.05E-71 | 1.12E-66 |
| OCA2 | 5 | 0.793912788 | 0.305 | 0.05 | 3.55E-71 | 1.3E-66 |
| PLXNC1 | 5 | 0.697374772 | 0.323 | 0.061 | 3.74E-71 | 1.37E-66 |
| FNIP2 | 5 | 0.823243449 | 0.615 | 0.311 | 8.8E-71 | 3.22E-66 |
| ITSN1 | 5 | -0.958281739 | 0.248 | 0.627 | 1.4E-70 | 5.13E-66 |
| FNBP1 | 5 | -0.986788528 | 0.321 | 0.683 | 1.54E-70 | 5.64E-66 |
| FOXA1 | 5 | 0.371988631 | 0.158 | 0.006 | 1.72E-70 | 6.28E-66 |
| DUSP16 | 5 | 0.850409594 | 0.615 | 0.289 | 2.03E-70 | 7.43E-66 |
| FTX | 5 | -0.685746509 | 0.987 | 0.985 | 2.44E-70 | 8.92E-66 |
| MEF2C | 5 | -1.079280016 | 0.048 | 0.372 | 5.62E-70 | 2.06E-65 |
| HIP1R | 5 | 0.718725271 | 0.365 | 0.093 | 6.39E-70 | 2.34E-65 |
| WDR11-AS1 | 5 | -1.036069841 | 0.033 | 0.343 | 8.17E-70 | 2.99E-65 |
| ADD2 | 5 | -0.748097527 | 0.017 | 0.311 | 1.17E-69 | 4.27E-65 |
| AL109613.1 | 5 | 0.378886173 | 0.139 | 0.004 | 1.51E-69 | 5.54E-65 |
| UBE2E3 | 5 | -1.070725191 | 0.382 | 0.723 | 2.53E-69 | 9.25E-65 |
| TJP2 | 5 | 0.86220405 | 0.708 | 0.372 | 4.26E-69 | 1.56E-64 |
| AMPD3 | 5 | 0.472392661 | 0.213 | 0.019 | 4.29E-69 | 1.57E-64 |
| CDC42EP3 | 5 | 0.630632416 | 0.27 | 0.041 | 4.93E-69 | 1.81E-64 |
| AC023424.3 | 5 | 0.78850474 | 0.404 | 0.146 | 5.13E-69 | 1.88E-64 |
| ADAMTS19 | 5 | -1.638740265 | 0.16 | 0.507 | 6.16E-69 | 2.25E-64 |
| PDLIM5 | 5 | 0.89732388 | 0.723 | 0.372 | 7.21E-69 | 2.64E-64 |
| MKX | 5 | -1.564889085 | 0.061 | 0.384 | 8.43E-69 | 3.09E-64 |
| SERPINF2 | 5 | 0.402045376 | 0.15 | 0.006 | 1.88E-68 | 6.9E-64 |
| TOX | 5 | 0.872572678 | 0.914 | 0.659 | 2.54E-68 | 9.28E-64 |
| TOGARAM2 | 5 | 0.572463553 | 0.229 | 0.026 | 3.59E-68 | 1.31E-63 |
| KLB | 5 | 0.359053226 | 0.138 | 0.004 | 5.07E-68 | 1.86E-63 |
| CUX2 | 5 | -1.325551821 | 0.022 | 0.315 | 5.77E-68 | 2.11E-63 |
| FGF13 | 5 | -1.910713732 | 0.134 | 0.465 | 7.39E-68 | 2.71E-63 |
| TENT5C | 5 | 0.419634598 | 0.139 | 0.005 | 8.16E-68 | 2.99E-63 |
| SNTB1 | 5 | 0.765553242 | 0.316 | 0.059 | 9.52E-68 | 3.49E-63 |
| AC023282.1 | 5 | -1.025651577 | 0.022 | 0.314 | 9.92E-68 | 3.63E-63 |
| ABL1 | 5 | 0.813065769 | 0.818 | 0.563 | 1.03E-67 | 3.77E-63 |
| POU2F3 | 5 | -0.874419392 | 0.033 | 0.337 | 1.25E-67 | 4.59E-63 |
| BDKRB2 | 5 | 0.590613385 | 0.202 | 0.02 | 3.37E-67 | 1.23E-62 |
| CDH6 | 5 | -1.146077913 | 0.042 | 0.354 | 4.83E-67 | 1.77E-62 |
| GUCY2C | 5 | 0.717175437 | 0.158 | 0.01 | 5.43E-67 | 1.99E-62 |
| ERC1 | 5 | 0.674606754 | 0.952 | 0.856 | 6.85E-67 | 2.51E-62 |
| CCDC198 | 5 | 0.340731008 | 0.13 | 0.003 | 1.38E-66 | 5.07E-62 |
| CLMP | 5 | 0.816800211 | 0.424 | 0.123 | 1.87E-66 | 6.86E-62 |
| HS3ST5 | 5 | 0.491663441 | 0.923 | 0.599 | 2.86E-66 | 1.05E-61 |
| TMCO3 | 5 | 0.786079986 | 0.549 | 0.262 | 2.87E-66 | 1.05E-61 |
| LTBP1 | 5 | -1.107522967 | 0.033 | 0.333 | 3E-66 | 1.1E-61 |
| TCF20 | 5 | 0.791309206 | 0.677 | 0.408 | 5.85E-66 | 2.14E-61 |
| ZFHX4 | 5 | -1.311789001 | 0.117 | 0.45 | 5.87E-66 | 2.15E-61 |
| ERICH3 | 5 | 0.771245675 | 0.431 | 0.121 | 6.06E-66 | 2.22E-61 |
| LINC01814 | 5 | 0.455947639 | 0.193 | 0.016 | 8.95E-66 | 3.28E-61 |
| BMPR1B | 5 | 1.077607632 | 0.644 | 0.298 | 9.26E-66 | 3.39E-61 |
| HS3ST4 | 5 | -1.860058571 | 0.048 | 0.358 | 1.23E-65 | 4.52E-61 |
| COL5A2 | 5 | -1.569996658 | 0.165 | 0.491 | 1.24E-65 | 4.53E-61 |
| DCAF6 | 5 | 0.804011779 | 0.743 | 0.536 | 1.3E-65 | 4.75E-61 |
| ABCA10 | 5 | 0.740899351 | 0.341 | 0.074 | 1.39E-65 | 5.1E-61 |
| EML6 | 5 | 0.886266678 | 0.556 | 0.224 | 1.43E-65 | 5.25E-61 |
| FAM167A | 5 | 0.379671996 | 0.149 | 0.006 | 1.77E-65 | 6.47E-61 |
| TCF7L1 | 5 | -1.009618325 | 0.033 | 0.326 | 2.09E-65 | 7.65E-61 |
| RAP1GAP | 5 | 0.532946783 | 0.229 | 0.036 | 2.13E-65 | 7.78E-61 |
| USH2A | 5 | 0.758701189 | 0.235 | 0.03 | 2.2E-65 | 8.05E-61 |
| PTGER3 | 5 | -1.24406939 | 0.09 | 0.41 | 2.28E-65 | 8.34E-61 |
| CCDC144NL-AS1 | 5 | -1.085297217 | 0.635 | 0.846 | 3.46E-65 | 1.27E-60 |

| | | | | | | |
|---|---|---|---|---|---|---|
| TBL1X | 5 | 0.776274486 | 0.925 | 0.765 | 3.98E-65 | 1.46E-60 |
| ACSL1 | 5 | -0.984153537 | 0.083 | 0.404 | 4.75E-65 | 1.74E-60 |
| SLCO3A1 | 5 | 0.922595384 | 0.576 | 0.254 | 1.44E-64 | 5.28E-60 |
| MAP4K4 | 5 | -0.980078676 | 0.688 | 0.865 | 1.68E-64 | 6.13E-60 |
| ARHGEF28 | 5 | 0.717334465 | 0.703 | 0.333 | 1.7E-64 | 6.24E-60 |
| SATB2 | 5 | 0.828942836 | 0.761 | 0.44 | 2.11E-64 | 7.72E-60 |
| FOXA2 | 5 | 0.282130784 | 0.105 | 0.001 | 2.34E-64 | 8.57E-60 |
| GRB10 | 5 | -0.951406291 | 0.079 | 0.405 | 3.12E-64 | 1.14E-59 |
| E6 | 5 | 0.650507057 | 0.286 | 0.055 | 4.15E-64 | 1.52E-59 |
| CBLC | 5 | 0.30918523 | 0.117 | 0.002 | 4.29E-64 | 1.57E-59 |
| MBNL1 | 5 | -1.26185112 | 0.817 | 0.92 | 6.13E-64 | 2.24E-59 |
| PRKCE | 5 | 0.905641102 | 0.789 | 0.503 | 1.26E-63 | 4.6E-59 |
| FSTL1 | 5 | -0.737878089 | 0.031 | 0.323 | 1.3E-63 | 4.75E-59 |
| PCSK1 | 5 | 0.494503135 | 0.185 | 0.016 | 1.41E-63 | 5.17E-59 |
| ATP8A2 | 5 | -1.083185221 | 0.04 | 0.331 | 1.5E-63 | 5.49E-59 |
| EBF3 | 5 | 0.393562095 | 0.569 | 0.231 | 1.59E-63 | 5.84E-59 |
| KCNK2 | 5 | -1.088945614 | 0.024 | 0.306 | 1.72E-63 | 6.3E-59 |
| ADAMTSL1 | 5 | -1.835553308 | 0.582 | 0.519 | 1.97E-63 | 7.2E-59 |
| CALY | 5 | 0.464076233 | 0.187 | 0.016 | 2.02E-63 | 7.39E-59 |
| CDK19 | 5 | -0.970210666 | 0.371 | 0.701 | 2.43E-63 | 8.89E-59 |
| KLHL7 | 5 | -0.930434887 | 0.174 | 0.522 | 2.64E-63 | 9.67E-59 |
| NTM | 5 | -1.734378182 | 0.112 | 0.435 | 3.41E-63 | 1.25E-58 |
| SPOCK1 | 5 | -1.711580782 | 0.073 | 0.381 | 3.85E-63 | 1.41E-58 |
| CCND2 | 5 | -0.838498524 | 0.042 | 0.336 | 5.02E-63 | 1.84E-58 |
| FAM135A | 5 | 0.750400297 | 0.76 | 0.543 | 5.64E-63 | 2.06E-58 |
| PROC | 5 | 0.331622178 | 0.15 | 0.007 | 7.73E-63 | 2.83E-58 |
| NEGR1 | 5 | -0.984908162 | 0.853 | 0.56 | 1.15E-62 | 4.2E-58 |
| ATP2A3 | 5 | 0.462894307 | 0.193 | 0.018 | 1.2E-62 | 4.41E-58 |
| FGFR1 | 5 | -0.887407296 | 0.064 | 0.378 | 1.22E-62 | 4.48E-58 |
| TMEFF1 | 5 | -0.916618523 | 0.068 | 0.383 | 1.27E-62 | 4.66E-58 |
| UST | 5 | -1.16222362 | 0.022 | 0.293 | 1.53E-62 | 5.61E-58 |
| AQP4-AS1 | 5 | 0.655886306 | 0.374 | 0.094 | 2.09E-62 | 7.64E-58 |
| DLGAP1 | 5 | -1.44103705 | 0.842 | 0.834 | 2.44E-62 | 8.95E-58 |
| SORCS3 | 5 | -1.562231453 | 0.116 | 0.442 | 2.93E-62 | 1.07E-57 |
| EPB41L2 | 5 | -1.073280355 | 0.35 | 0.664 | 3.17E-62 | 1.16E-57 |
| TRIM22 | 5 | -0.81991667 | 0.024 | 0.302 | 3.3E-62 | 1.21E-57 |
| PKP2 | 5 | 0.433258426 | 0.18 | 0.014 | 3.64E-62 | 1.33E-57 |
| TNRC6C | 5 | -0.921480581 | 0.286 | 0.628 | 3.74E-62 | 1.37E-57 |
| OSBPL3 | 5 | -1.100212186 | 0.143 | 0.478 | 4.31E-62 | 1.58E-57 |
| PPM1H | 5 | 0.533274051 | 0.242 | 0.033 | 5.31E-62 | 1.95E-57 |
| DNM3 | 5 | -1.285260027 | 0.163 | 0.488 | 7.57E-62 | 2.77E-57 |
| AP1S2 | 5 | -1.003541052 | 0.217 | 0.552 | 7.8E-62 | 2.85E-57 |
| LDLRAD1 | 5 | 0.293782063 | 0.106 | 0.001 | 9.05E-62 | 3.31E-57 |
| CLN8 | 5 | 0.587008452 | 0.305 | 0.067 | 9.46E-62 | 3.46E-57 |
| DAB1 | 5 | -1.251571319 | 0.391 | 0.703 | 1.03E-61 | 3.78E-57 |
| KLHL32 | 5 | 0.783597323 | 0.464 | 0.155 | 1.04E-61 | 3.82E-57 |
| GAS8 | 5 | 0.526552265 | 0.272 | 0.047 | 1.25E-61 | 4.57E-57 |
| SYTL5 | 5 | -1.111004434 | 0.04 | 0.333 | 1.28E-61 | 4.67E-57 |
| C4orf19 | 5 | 0.695749385 | 0.64 | 0.283 | 1.76E-61 | 6.44E-57 |
| EPHA7 | 5 | -1.485188295 | 0.04 | 0.332 | 2.07E-61 | 7.59E-57 |
| COL3A1 | 5 | -1.521962739 | 0.02 | 0.292 | 2.2E-61 | 8.07E-57 |
| LHFPL4 | 5 | 0.709540221 | 0.363 | 0.095 | 3.49E-61 | 1.28E-56 |
| RHBDL3 | 5 | -0.951792534 | 0.033 | 0.316 | 3.62E-61 | 1.33E-56 |
| APCDD1 | 5 | -1.017111519 | 0.037 | 0.325 | 4.62E-61 | 1.69E-56 |
| CABIN1 | 5 | 0.737702177 | 0.642 | 0.367 | 5.46E-61 | 2E-56 |
| TNNI3K | 5 | 0.549354724 | 0.235 | 0.033 | 5.86E-61 | 2.15E-56 |
| MGST2 | 5 | 0.613865681 | 0.286 | 0.058 | 6.82E-61 | 2.5E-56 |
| GLT8D2 | 5 | 0.539127327 | 0.224 | 0.029 | 7.56E-61 | 2.77E-56 |

| | | | | | | |
|---|---|---|---|---|---|---|
| PTPN14 | 5 | -0.957284197 | 0.028 | 0.305 | 7.62E-61 | 2.79E-56 |
| SOX11 | 5 | -0.818686179 | 0.048 | 0.343 | 7.9E-61 | 2.89E-56 |
| KRT19 | 5 | 0.495956275 | 0.141 | 0.006 | 1.41E-60 | 5.16E-56 |
| OPRD1 | 5 | 0.531896159 | 0.213 | 0.027 | 1.85E-60 | 6.78E-56 |
| DSG2 | 5 | 0.508975311 | 0.226 | 0.031 | 2.57E-60 | 9.39E-56 |
| GLI2 | 5 | -0.91748278 | 0.044 | 0.332 | 3.2E-60 | 1.17E-55 |
| FBXW8 | 5 | 0.748628815 | 0.6 | 0.362 | 4.17E-60 | 1.53E-55 |
| TMTC1 | 5 | 0.608254454 | 0.411 | 0.119 | 6.04E-60 | 2.21E-55 |
| DZIP1 | 5 | -0.787130008 | 0.152 | 0.495 | 7.01E-60 | 2.56E-55 |
| ARHGEF9 | 5 | 0.746733881 | 0.525 | 0.245 | 7.53E-60 | 2.76E-55 |
| LINC01278 | 5 | -0.904182179 | 0.055 | 0.35 | 7.8E-60 | 2.85E-55 |
| PDE3A | 5 | -1.283488243 | 0.244 | 0.589 | 1.65E-59 | 6.04E-55 |
| NR5A2 | 5 | 0.46975123 | 0.149 | 0.008 | 1.68E-59 | 6.14E-55 |
| CEP170 | 5 | -0.85777645 | 0.528 | 0.786 | 1.72E-59 | 6.31E-55 |
| NHS | 5 | 0.698808281 | 0.787 | 0.454 | 1.97E-59 | 7.23E-55 |
| ARHGAP6 | 5 | -1.353995212 | 0.075 | 0.377 | 2.03E-59 | 7.43E-55 |
| C1orf61 | 5 | -1.005260519 | 0.039 | 0.319 | 2.48E-59 | 9.06E-55 |
| ECE1 | 5 | 0.685739742 | 0.453 | 0.156 | 2.88E-59 | 1.05E-54 |
| TRIM55 | 5 | 0.333836234 | 0.117 | 0.003 | 4.91E-59 | 1.8E-54 |
| CA10 | 5 | -1.458592247 | 0.05 | 0.335 | 6.09E-59 | 2.23E-54 |
| USP54 | 5 | 0.789015833 | 0.578 | 0.288 | 6.75E-59 | 2.47E-54 |
| TLN2 | 5 | -1.083595907 | 0.149 | 0.472 | 7.41E-59 | 2.71E-54 |
| TENM4 | 5 | -1.172939663 | 0.086 | 0.384 | 8.03E-59 | 2.94E-54 |
| FRMD5 | 5 | -0.34001815 | 0.65 | 0.392 | 8.8E-59 | 3.22E-54 |
| HACD2 | 5 | 0.732849958 | 0.461 | 0.191 | 1.09E-58 | 3.98E-54 |
| ARHGAP18 | 5 | -1.165132759 | 0.167 | 0.474 | 1.11E-58 | 4.05E-54 |
| CCDC125 | 5 | 0.751770216 | 0.457 | 0.185 | 1.3E-58 | 4.75E-54 |
| PLCL1 | 5 | -1.423310576 | 0.125 | 0.442 | 1.36E-58 | 4.98E-54 |
| SLC39A14 | 5 | 0.722298844 | 0.382 | 0.124 | 1.52E-58 | 5.56E-54 |
| ANPEP | 5 | 0.511438388 | 0.174 | 0.016 | 1.99E-58 | 7.3E-54 |
| DANT2 | 5 | 0.713559963 | 0.657 | 0.306 | 2.22E-58 | 8.14E-54 |
| NKAIN2 | 5 | -2.205566308 | 0.09 | 0.395 | 2.52E-58 | 9.24E-54 |
| DOCK3 | 5 | 0.503875196 | 0.598 | 0.265 | 2.65E-58 | 9.71E-54 |
| MEIS1 | 5 | -1.065500484 | 0.07 | 0.37 | 2.98E-58 | 1.09E-53 |
| SCG2 | 5 | 0.338123367 | 0.123 | 0.004 | 3.09E-58 | 1.13E-53 |
| ZFPM2 | 5 | -1.49453975 | 0.044 | 0.324 | 4.1E-58 | 1.5E-53 |
| OSBPL10 | 5 | 0.866957948 | 0.692 | 0.365 | 4.68E-58 | 1.71E-53 |
| SEMA5A | 5 | -1.751365977 | 0.116 | 0.427 | 4.81E-58 | 1.76E-53 |
| SMAD6 | 5 | -0.80989308 | 0.037 | 0.314 | 6.65E-58 | 2.43E-53 |
| COX10 | 5 | 0.734501271 | 0.582 | 0.295 | 8.78E-58 | 3.21E-53 |
| ATP10A | 5 | -0.86694318 | 0.029 | 0.299 | 9.15E-58 | 3.35E-53 |
| PCDH11X | 5 | -2.306176288 | 0.024 | 0.279 | 1.06E-57 | 3.9E-53 |
| CFAP58 | 5 | -1.097914361 | 0.037 | 0.313 | 1.12E-57 | 4.09E-53 |
| SPECC1L | 5 | 0.681815396 | 0.778 | 0.578 | 1.13E-57 | 4.13E-53 |
| ARHGAP24 | 5 | -1.680854521 | 0.127 | 0.429 | 1.23E-57 | 4.49E-53 |
| KCNN2 | 5 | -1.594064441 | 0.042 | 0.315 | 1.56E-57 | 5.72E-53 |
| KCNJ16 | 5 | 0.253597762 | 0.105 | 0.002 | 2.87E-57 | 1.05E-52 |
| NID1 | 5 | -0.804606042 | 0.031 | 0.3 | 3.14E-57 | 1.15E-52 |
| SYNE1 | 5 | 0.688856176 | 0.778 | 0.456 | 3.25E-57 | 1.19E-52 |
| AASS | 5 | -0.756930674 | 0.022 | 0.281 | 3.87E-57 | 1.42E-52 |
| LARGE1 | 5 | -0.854442775 | 0.029 | 0.291 | 4.11E-57 | 1.5E-52 |
| ZC3H12C | 5 | -1.198174779 | 0.328 | 0.607 | 4.16E-57 | 1.52E-52 |
| CSMD3 | 5 | -1.737132111 | 0.066 | 0.341 | 4.83E-57 | 1.77E-52 |
| ADAMTS9-AS2 | 5 | -0.976842585 | 0.046 | 0.327 | 7.59E-57 | 2.78E-52 |
| CACNA1H | 5 | 0.436832429 | 0.187 | 0.02 | 9.03E-57 | 3.31E-52 |
| IL1RAP | 5 | 0.786427146 | 0.422 | 0.15 | 1.03E-56 | 3.78E-52 |
| PARP8 | 5 | 0.812116443 | 0.644 | 0.313 | 1.15E-56 | 4.2E-52 |
| TUSC3 | 5 | 0.711264892 | 0.844 | 0.679 | 1.23E-56 | 4.49E-52 |

| | | | | | | |
|---|---|---|---|---|---|---|
| PRKCH | 5 | 0.63501725 | 0.36 | 0.096 | 1.34E-56 | 4.92E-52 |
| PSD3 | 5 | -0.956225947 | 0.631 | 0.842 | 1.5E-56 | 5.48E-52 |
| SERPINB1 | 5 | 0.405455454 | 0.189 | 0.021 | 2.15E-56 | 7.86E-52 |
| PHTF1 | 5 | 0.762721265 | 0.58 | 0.296 | 2.38E-56 | 8.71E-52 |
| HS3ST3A1 | 5 | -0.986926335 | 0.018 | 0.269 | 2.4E-56 | 8.78E-52 |
| RIMS1 | 5 | -1.398679017 | 0.152 | 0.455 | 2.97E-56 | 1.09E-51 |
| CDKL5 | 5 | -0.959156051 | 0.116 | 0.426 | 3.82E-56 | 1.4E-51 |
| YAP1 | 5 | -1.03350493 | 0.066 | 0.351 | 3.97E-56 | 1.45E-51 |
| EPOR | 5 | 0.318818167 | 0.127 | 0.005 | 4.05E-56 | 1.48E-51 |
| CDYL | 5 | -0.954560932 | 0.47 | 0.735 | 4.31E-56 | 1.58E-51 |
| NAV3 | 5 | -1.555556567 | 0.105 | 0.402 | 4.71E-56 | 1.72E-51 |
| FXYD3 | 5 | 0.288755217 | 0.114 | 0.003 | 4.81E-56 | 1.76E-51 |
| SPECC1 | 5 | -1.107439862 | 0.248 | 0.563 | 6.16E-56 | 2.25E-51 |
| ADAMTSL3 | 5 | -1.048878816 | 0.037 | 0.303 | 6.25E-56 | 2.29E-51 |
| DOCK9 | 5 | 0.737145384 | 0.446 | 0.157 | 7.88E-56 | 2.89E-51 |
| AC019197.1 | 5 | 0.870177336 | 0.349 | 0.105 | 7.89E-56 | 2.89E-51 |
| TRMT9B | 5 | 0.623701818 | 0.525 | 0.204 | 1.02E-55 | 3.75E-51 |
| KLHL3 | 5 | 0.634817676 | 0.35 | 0.1 | 1.26E-55 | 4.61E-51 |
| ABCG2 | 5 | 0.29612356 | 0.121 | 0.004 | 1.72E-55 | 6.29E-51 |
| FER | 5 | -0.979373441 | 0.62 | 0.827 | 1.81E-55 | 6.64E-51 |
| CYB5R1 | 5 | 0.534831607 | 0.242 | 0.045 | 2.07E-55 | 7.59E-51 |
| DEPTOR | 5 | 0.596076237 | 0.259 | 0.047 | 2.35E-55 | 8.61E-51 |
| ITGA8 | 5 | -1.353183627 | 0.022 | 0.274 | 2.47E-55 | 9.03E-51 |
| GLCCI1 | 5 | -1.226444872 | 0.305 | 0.595 | 2.78E-55 | 1.02E-50 |
| SLC12A2 | 5 | -1.032584696 | 0.161 | 0.483 | 2.88E-55 | 1.06E-50 |
| BACE1 | 5 | 0.638691396 | 0.378 | 0.13 | 3.49E-55 | 1.28E-50 |
| CD46 | 5 | 0.684059796 | 0.738 | 0.515 | 3.66E-55 | 1.34E-50 |
| EFCC1 | 5 | 0.360669322 | 0.13 | 0.006 | 4.78E-55 | 1.75E-50 |
| FUT8 | 5 | -1.036911557 | 0.631 | 0.828 | 5.24E-55 | 1.92E-50 |
| SPINK1 | 5 | 0.318666663 | 0.125 | 0.005 | 5.7E-55 | 2.09E-50 |
| TBX20 | 5 | -0.790212231 | 0.026 | 0.279 | 5.87E-55 | 2.15E-50 |
| ADAM23 | 5 | -1.051587728 | 0.084 | 0.376 | 7.51E-55 | 2.75E-50 |
| RCAN3 | 5 | 0.654429227 | 0.323 | 0.096 | 9.5E-55 | 3.48E-50 |
| RAB11FIP1 | 5 | 0.650397029 | 0.246 | 0.047 | 1.21E-54 | 4.43E-50 |
| SLC1A2 | 5 | -0.824007039 | 0.037 | 0.298 | 1.64E-54 | 5.99E-50 |
| HOOK1 | 5 | 0.687194088 | 0.484 | 0.189 | 1.8E-54 | 6.6E-50 |
| RARB | 5 | -1.123933142 | 0.024 | 0.276 | 2.05E-54 | 7.51E-50 |
| APBA1 | 5 | -1.26184841 | 0.154 | 0.458 | 2.36E-54 | 8.65E-50 |
| DCDC1 | 5 | 0.496265661 | 0.294 | 0.069 | 2.85E-54 | 1.04E-49 |
| DSC2 | 5 | 0.420862706 | 0.187 | 0.021 | 6.07E-54 | 2.22E-49 |
| ARPP21 | 5 | -1.429604704 | 0.057 | 0.332 | 8.03E-54 | 2.94E-49 |
| SLC17A8 | 5 | -1.314811277 | 0.04 | 0.304 | 1.02E-53 | 3.72E-49 |
| SCD5 | 5 | -0.666980109 | 0.02 | 0.261 | 1.66E-53 | 6.08E-49 |
| LARP1B | 5 | 0.74643101 | 0.741 | 0.518 | 1.68E-53 | 6.16E-49 |
| LINC00299 | 5 | 0.475833784 | 0.18 | 0.02 | 2.18E-53 | 7.99E-49 |
| OXR1 | 5 | -1.243878011 | 0.415 | 0.673 | 2.28E-53 | 8.36E-49 |
| TRERF1 | 5 | -0.842063924 | 0.339 | 0.647 | 2.45E-53 | 8.95E-49 |
| CDH23 | 5 | -0.85325251 | 0.031 | 0.288 | 3.39E-53 | 1.24E-48 |
| SLC19A3 | 5 | 0.372803455 | 0.145 | 0.01 | 3.4E-53 | 1.25E-48 |
| RND3 | 5 | -0.987390365 | 0.072 | 0.35 | 3.74E-53 | 1.37E-48 |
| LINC02470 | 5 | -1.207752766 | 0.029 | 0.284 | 5.45E-53 | 2E-48 |
| PTPRM | 5 | 0.894883336 | 0.675 | 0.364 | 5.47E-53 | 2E-48 |
| SUSD4 | 5 | -0.831587921 | 0.028 | 0.281 | 5.72E-53 | 2.09E-48 |
| BACH2 | 5 | -1.09092602 | 0.108 | 0.404 | 5.94E-53 | 2.17E-48 |
| SPAG17 | 5 | -1.166583787 | 0.048 | 0.307 | 6.52E-53 | 2.39E-48 |
| SH3BP4 | 5 | -0.8195652 | 0.068 | 0.35 | 7.26E-53 | 2.66E-48 |
| PLCB1 | 5 | -1.318187363 | 0.255 | 0.542 | 7.97E-53 | 2.92E-48 |
| AL353576.1 | 5 | 0.339464004 | 0.139 | 0.009 | 1.03E-52 | 3.78E-48 |

| | | | | | | |
|---|---|---|---|---|---|---|
| MRC2 | 5 | -0.680210032 | 0.017 | 0.251 | 1.05E-52 | 3.84E-48 |
| CD99 | 5 | 0.732128947 | 0.541 | 0.277 | 2.05E-52 | 7.51E-48 |
| CELF3 | 5 | 0.460733428 | 0.202 | 0.029 | 2.65E-52 | 9.69E-48 |
| EPHX2 | 5 | 0.517780152 | 0.242 | 0.048 | 2.69E-52 | 9.86E-48 |
| TRIM71 | 5 | -0.729274128 | 0.022 | 0.267 | 3.17E-52 | 1.16E-47 |
| CAMK2N1 | 5 | 0.494100838 | 0.244 | 0.045 | 3.19E-52 | 1.17E-47 |
| PXDN | 5 | -0.729558574 | 0.015 | 0.248 | 3.29E-52 | 1.2E-47 |
| FAM135B | 5 | -1.143942307 | 0.101 | 0.387 | 5.93E-52 | 2.17E-47 |
| ZFAND4 | 5 | 0.664382533 | 0.413 | 0.157 | 6.36E-52 | 2.33E-47 |
| IGF1R | 5 | -0.904991282 | 0.576 | 0.81 | 6.9E-52 | 2.53E-47 |
| MALRD1 | 5 | -2.068958986 | 0.028 | 0.273 | 6.91E-52 | 2.53E-47 |
| TIAM1 | 5 | 0.697016936 | 0.859 | 0.592 | 7.39E-52 | 2.7E-47 |
| SKIL | 5 | -0.975281393 | 0.338 | 0.637 | 7.73E-52 | 2.83E-47 |
| NOTCH2 | 5 | -0.912150502 | 0.061 | 0.325 | 8.51E-52 | 3.12E-47 |
| XPR1 | 5 | 0.704337395 | 0.734 | 0.492 | 9.14E-52 | 3.35E-47 |
| ADGRG5 | 5 | 0.309469138 | 0.121 | 0.006 | 1.06E-51 | 3.88E-47 |
| ATP11C | 5 | -0.946431056 | 0.486 | 0.732 | 1.65E-51 | 6.05E-47 |
| MALAT1 | 5 | 0.592657616 | 0.998 | 0.998 | 1.89E-51 | 6.91E-47 |
| PDZRN3 | 5 | -1.216719275 | 0.866 | 0.906 | 2.73E-51 | 9.98E-47 |
| SLC38A11 | 5 | 0.302441884 | 0.204 | 0.037 | 2.92E-51 | 1.07E-46 |
| ESRP2 | 5 | 0.264536301 | 0.103 | 0.003 | 3E-51 | 1.1E-46 |
| LDLRAD4 | 5 | -1.659067091 | 0.257 | 0.51 | 3.62E-51 | 1.32E-46 |
| MIR181A2HG | 5 | -0.771379257 | 0.037 | 0.291 | 3.89E-51 | 1.42E-46 |
| PLPPR4 | 5 | -0.927042433 | 0.042 | 0.299 | 5.05E-51 | 1.85E-46 |
| TUB | 5 | 0.561165715 | 0.272 | 0.065 | 5.46E-51 | 2E-46 |
| FAM81A | 5 | 0.501990911 | 0.255 | 0.051 | 5.46E-51 | 2E-46 |
| RIPOR3 | 5 | 0.457177734 | 0.187 | 0.023 | 5.68E-51 | 2.08E-46 |
| JAKMIP2 | 5 | 0.763422701 | 0.635 | 0.335 | 5.76E-51 | 2.11E-46 |
| DENND1B | 5 | 0.709279427 | 0.615 | 0.305 | 9.82E-51 | 3.6E-46 |
| SYNPO2 | 5 | -1.034004629 | 0.035 | 0.283 | 1.08E-50 | 3.96E-46 |
| PLEKHF2 | 5 | 0.440055438 | 0.202 | 0.029 | 1.13E-50 | 4.13E-46 |
| TMEM38B | 5 | 0.5196108 | 0.301 | 0.073 | 1.41E-50 | 5.15E-46 |
| NPIPB15 | 5 | 0.530746354 | 0.277 | 0.068 | 1.53E-50 | 5.62E-46 |
| IL6ST | 5 | 0.663177654 | 0.442 | 0.162 | 1.67E-50 | 6.13E-46 |
| AFAP1 | 5 | -0.89205968 | 0.187 | 0.493 | 2.12E-50 | 7.75E-46 |
| ZNF704 | 5 | -0.985937736 | 0.36 | 0.634 | 2.34E-50 | 8.58E-46 |
| KCNH1 | 5 | -1.407190223 | 0.037 | 0.283 | 2.68E-50 | 9.83E-46 |
| HMGA1P4 | 5 | 0.696622128 | 0.294 | 0.082 | 2.84E-50 | 1.04E-45 |
| AL109930.1 | 5 | -0.688636328 | 0.033 | 0.277 | 3.21E-50 | 1.18E-45 |
| RERE | 5 | 0.591077482 | 0.985 | 0.969 | 3.83E-50 | 1.4E-45 |
| LRRC7 | 5 | -2.000566255 | 0.07 | 0.331 | 4.28E-50 | 1.57E-45 |
| THSD7A | 5 | -1.732531725 | 0.123 | 0.405 | 6.17E-50 | 2.26E-45 |
| DNAH5 | 5 | 0.550985531 | 0.196 | 0.027 | 7.08E-50 | 2.59E-45 |
| SVEP1 | 5 | -1.115821384 | 0.053 | 0.309 | 8.4E-50 | 3.07E-45 |
| CSMD2 | 5 | -1.012658506 | 0.062 | 0.323 | 1.05E-49 | 3.84E-45 |
| VCAN | 5 | -1.588149308 | 0.154 | 0.433 | 1.19E-49 | 4.35E-45 |
| CGN | 5 | 0.352428367 | 0.154 | 0.015 | 1.56E-49 | 5.73E-45 |
| SPOCK3 | 5 | 0.571063104 | 0.176 | 0.02 | 1.69E-49 | 6.19E-45 |
| RAB31 | 5 | -0.731005345 | 0.033 | 0.273 | 1.82E-49 | 6.67E-45 |
| PTPN12 | 5 | -0.805486774 | 0.415 | 0.695 | 1.83E-49 | 6.69E-45 |
| PLD5 | 5 | -1.338338914 | 0.029 | 0.267 | 1.84E-49 | 6.75E-45 |
| RAB3IP | 5 | 0.618647706 | 0.383 | 0.124 | 2.14E-49 | 7.83E-45 |
| ANKRD18A | 5 | 0.62560671 | 0.409 | 0.14 | 2.25E-49 | 8.26E-45 |
| AL157778.1 | 5 | -1.513926447 | 0.022 | 0.252 | 2.72E-49 | 9.97E-45 |
| PPIL2 | 5 | 0.655131098 | 0.492 | 0.234 | 2.99E-49 | 1.1E-44 |
| NALCN | 5 | -0.909938013 | 0.015 | 0.237 | 3.26E-49 | 1.2E-44 |
| ROCK1 | 5 | 0.645441586 | 0.793 | 0.586 | 3.38E-49 | 1.24E-44 |
| QPCT | 5 | 0.28598102 | 0.121 | 0.006 | 3.74E-49 | 1.37E-44 |

| | | | | | | |
|---|---|---|---|---|---|---|
| HS3ST3B1 | 5 | -0.969167689 | 0.061 | 0.314 | 5.73E-49 | 2.1E-44 |
| SNX25 | 5 | -1.00577098 | 0.371 | 0.648 | 8.35E-49 | 3.06E-44 |
| C15orf41 | 5 | -0.935627575 | 0.279 | 0.563 | 8.7E-49 | 3.18E-44 |
| SNHG14 | 5 | 0.630520114 | 0.923 | 0.826 | 8.99E-49 | 3.29E-44 |
| NRN1 | 5 | 0.260915322 | 0.121 | 0.006 | 1.03E-48 | 3.77E-44 |
| SRPX | 5 | -0.66744369 | 0.018 | 0.244 | 1.13E-48 | 4.12E-44 |
| CRKL | 5 | 0.610062006 | 0.36 | 0.122 | 1.28E-48 | 4.68E-44 |
| SNTG2 | 5 | -1.043483977 | 0.022 | 0.253 | 1.32E-48 | 4.85E-44 |
| MAOA | 5 | 0.345237678 | 0.13 | 0.009 | 1.46E-48 | 5.36E-44 |
| SPRED1 | 5 | -0.896283313 | 0.156 | 0.431 | 1.5E-48 | 5.49E-44 |
| OXCT1 | 5 | 0.68974856 | 0.462 | 0.194 | 1.95E-48 | 7.14E-44 |
| GPR160 | 5 | 0.323202277 | 0.141 | 0.011 | 1.99E-48 | 7.28E-44 |
| ADK | 5 | 0.629186906 | 0.919 | 0.775 | 2.59E-48 | 9.49E-44 |
| ITGBL1 | 5 | -2.121087448 | 0.013 | 0.224 | 2.65E-48 | 9.7E-44 |
| HMGA2 | 5 | -1.168561021 | 0.15 | 0.446 | 2.99E-48 | 1.09E-43 |
| MTA3 | 5 | -0.833630139 | 0.428 | 0.703 | 4.05E-48 | 1.48E-43 |
| PTPN18 | 5 | 0.533462518 | 0.275 | 0.071 | 4.13E-48 | 1.51E-43 |
| NFIX | 5 | -0.910423277 | 0.183 | 0.486 | 5.18E-48 | 1.9E-43 |
| NMU | 5 | -0.740817959 | 0.031 | 0.266 | 6.19E-48 | 2.27E-43 |
| TFAP2B | 5 | -0.937485762 | 0.026 | 0.25 | 7.26E-48 | 2.66E-43 |
| ADAMTS16 | 5 | -0.964948197 | 0.039 | 0.278 | 8.44E-48 | 3.09E-43 |
| TMEM132D | 5 | 1.107885541 | 0.176 | 0.025 | 1.08E-47 | 3.95E-43 |
| HMCN1 | 5 | -1.346256486 | 0.013 | 0.228 | 1.52E-47 | 5.57E-43 |
| PLA2G12A | 5 | 0.612024576 | 0.382 | 0.154 | 2.02E-47 | 7.4E-43 |
| PITX1 | 5 | 0.327717978 | 0.128 | 0.008 | 2.04E-47 | 7.45E-43 |
| ERBB4 | 5 | -2.159267365 | 0.059 | 0.301 | 2.2E-47 | 8.07E-43 |
| ENAH | 5 | -0.837267071 | 0.483 | 0.733 | 2.39E-47 | 8.75E-43 |
| APPL2 | 5 | 0.690538716 | 0.607 | 0.356 | 2.51E-47 | 9.17E-43 |
| EGFR | 5 | -0.912079353 | 0.04 | 0.273 | 2.65E-47 | 9.69E-43 |
| PDE1C | 5 | -1.683531044 | 0.07 | 0.323 | 2.75E-47 | 1.01E-42 |
| PIP5K1B | 5 | -1.311641381 | 0.057 | 0.293 | 3.05E-47 | 1.12E-42 |
| FAT3 | 5 | -1.270265854 | 0.138 | 0.408 | 3.17E-47 | 1.16E-42 |
| FSTL4 | 5 | 0.664619621 | 0.345 | 0.102 | 3.4E-47 | 1.24E-42 |
| METTL15 | 5 | 0.673325962 | 0.806 | 0.629 | 3.94E-47 | 1.44E-42 |
| PKNOX2 | 5 | -0.7096066 | 0.02 | 0.239 | 3.98E-47 | 1.46E-42 |
| AHI1 | 5 | 0.644819417 | 0.87 | 0.696 | 4.42E-47 | 1.62E-42 |
| ESRRG | 5 | -1.236486856 | 0.152 | 0.435 | 5.36E-47 | 1.96E-42 |
| FAT4 | 5 | -0.899214521 | 0.059 | 0.313 | 5.94E-47 | 2.17E-42 |
| ZNF385D | 5 | -1.502466898 | 0.334 | 0.588 | 6.59E-47 | 2.41E-42 |
| LIMK2 | 5 | 0.629414588 | 0.417 | 0.173 | 6.91E-47 | 2.53E-42 |
| SRSF11 | 5 | 0.503022925 | 0.967 | 0.885 | 6.94E-47 | 2.54E-42 |
| BHMT | 5 | 0.305678384 | 0.112 | 0.006 | 7.33E-47 | 2.68E-42 |
| PLIN5 | 5 | 0.311780549 | 0.117 | 0.007 | 9.46E-47 | 3.46E-42 |
| AC127502.2 | 5 | 0.523678077 | 0.292 | 0.083 | 1.1E-46 | 4.01E-42 |
| ASIC2 | 5 | -0.913565592 | 0.044 | 0.283 | 1.16E-46 | 4.26E-42 |
| SUMF1 | 5 | 0.814164855 | 0.644 | 0.441 | 1.43E-46 | 5.22E-42 |
| PATJ | 5 | 0.688582354 | 0.692 | 0.414 | 1.55E-46 | 5.66E-42 |
| SEMA3A | 5 | 0.679480319 | 0.572 | 0.271 | 1.57E-46 | 5.75E-42 |
| EML1 | 5 | -0.78120152 | 0.138 | 0.419 | 1.85E-46 | 6.76E-42 |
| GALNT13 | 5 | 0.610830325 | 0.914 | 0.689 | 1.89E-46 | 6.9E-42 |
| SFI1 | 5 | 0.657445451 | 0.508 | 0.261 | 1.94E-46 | 7.09E-42 |
| ZNRF2 | 5 | 0.711565442 | 0.582 | 0.322 | 1.99E-46 | 7.3E-42 |
| EPCAM | 5 | 0.426477855 | 0.29 | 0.073 | 2.16E-46 | 7.91E-42 |
| INSM1 | 5 | 0.404066033 | 0.176 | 0.023 | 2.29E-46 | 8.37E-42 |
| TMEM178B | 5 | 0.470466486 | 0.794 | 0.485 | 2.8E-46 | 1.02E-41 |
| OTUD7A | 5 | -1.130329885 | 0.325 | 0.583 | 3.08E-46 | 1.13E-41 |
| SLC5A4-AS1 | 5 | 0.621201271 | 0.352 | 0.115 | 3.45E-46 | 1.26E-41 |
| PLEKHG3 | 5 | 0.51571492 | 0.295 | 0.082 | 3.5E-46 | 1.28E-41 |

| | | | | | | |
|---|---|---|---|---|---|---|
| TESMIN | 5 | 0.429223003 | 0.228 | 0.043 | 3.72E-46 | 1.36E-41 |
| RBM26 | 5 | 0.538973848 | 0.912 | 0.832 | 4.58E-46 | 1.68E-41 |
| PLEKHA1 | 5 | 0.667210068 | 0.598 | 0.336 | 4.81E-46 | 1.76E-41 |
| SOGA1 | 5 | -0.755499869 | 0.127 | 0.408 | 4.83E-46 | 1.77E-41 |
| PDGFC | 5 | -1.001966846 | 0.04 | 0.277 | 5.36E-46 | 1.96E-41 |
| IDS | 5 | 0.628032274 | 0.475 | 0.21 | 5.6E-46 | 2.05E-41 |
| STARD10 | 5 | 0.260797581 | 0.121 | 0.007 | 6.08E-46 | 2.22E-41 |
| TRNP1 | 5 | 0.39547082 | 0.16 | 0.02 | 6.09E-46 | 2.23E-41 |
| STAT2 | 5 | -0.575308395 | 0.035 | 0.271 | 7.11E-46 | 2.6E-41 |
| SLC38A1 | 5 | -0.809106376 | 0.288 | 0.587 | 7.52E-46 | 2.75E-41 |
| NIN | 5 | -0.824918932 | 0.281 | 0.558 | 7.64E-46 | 2.8E-41 |
| PCED1B | 5 | 0.667351062 | 0.242 | 0.067 | 1.1E-45 | 4.04E-41 |
| RGS3 | 5 | 0.628671292 | 0.363 | 0.122 | 1.19E-45 | 4.35E-41 |
| SLC25A27 | 5 | 0.483393205 | 0.266 | 0.063 | 1.24E-45 | 4.55E-41 |
| TP53I13 | 5 | 0.457481383 | 0.198 | 0.035 | 1.27E-45 | 4.65E-41 |
| FAM124A | 5 | -0.640298345 | 0.031 | 0.26 | 1.53E-45 | 5.61E-41 |
| DMKN | 5 | 0.341714807 | 0.145 | 0.014 | 1.6E-45 | 5.87E-41 |
| LINC00472 | 5 | -0.634114906 | 0.017 | 0.229 | 2.17E-45 | 7.95E-41 |
| CNKSR3 | 5 | 0.747982735 | 0.541 | 0.273 | 2.6E-45 | 9.5E-41 |
| PAWR | 5 | 0.713289719 | 0.596 | 0.338 | 3.11E-45 | 1.14E-40 |
| FBLN1 | 5 | -0.598053516 | 0.02 | 0.235 | 3.33E-45 | 1.22E-40 |
| CNTN1 | 5 | -1.165945905 | 0.039 | 0.268 | 4E-45 | 1.47E-40 |
| PDGFRA | 5 | -0.755440059 | 0.007 | 0.203 | 4.53E-45 | 1.66E-40 |
| CNKSR2 | 5 | -1.329231259 | 0.448 | 0.675 | 5.35E-45 | 1.96E-40 |
| NLGN4X | 5 | -1.040425926 | 0.561 | 0.745 | 7.12E-45 | 2.61E-40 |
| RIC3 | 5 | 0.586473699 | 0.393 | 0.146 | 7.43E-45 | 2.72E-40 |
| DYNC1I2 | 5 | -0.854503564 | 0.393 | 0.665 | 7.56E-45 | 2.77E-40 |
| GPM6A | 5 | -1.214962842 | 0.011 | 0.211 | 8.03E-45 | 2.94E-40 |
| TNRC18 | 5 | -0.75438494 | 0.251 | 0.556 | 8.79E-45 | 3.22E-40 |
| CA12 | 5 | 0.921595828 | 0.455 | 0.232 | 8.92E-45 | 3.27E-40 |
| NCAN | 5 | -0.616524455 | 0.02 | 0.229 | 9E-45 | 3.29E-40 |
| SRBD1 | 5 | 0.743591853 | 0.721 | 0.524 | 1.08E-44 | 3.96E-40 |
| CDYL2 | 5 | 0.392710374 | 0.176 | 0.024 | 1.39E-44 | 5.1E-40 |
| SPIRE2 | 5 | 0.595393797 | 0.369 | 0.136 | 1.45E-44 | 5.31E-40 |
| RGL1 | 5 | -1.133105738 | 0.554 | 0.737 | 1.47E-44 | 5.38E-40 |
| MYO16 | 5 | -1.379670379 | 0.088 | 0.333 | 1.47E-44 | 5.4E-40 |
| SEMA3E | 5 | 0.26900222 | 0.563 | 0.283 | 1.76E-44 | 6.44E-40 |
| LIMCH1 | 5 | -1.320424457 | 0.576 | 0.69 | 1.79E-44 | 6.55E-40 |
| GREB1L | 5 | 0.285971644 | 0.295 | 0.085 | 1.89E-44 | 6.93E-40 |
| SMOC1 | 5 | 0.71942 | 0.42 | 0.162 | 2.12E-44 | 7.77E-40 |
| HPSE2 | 5 | -1.859871116 | 0.046 | 0.27 | 2.53E-44 | 9.27E-40 |
| TRIM2 | 5 | -0.883532161 | 0.387 | 0.657 | 2.66E-44 | 9.75E-40 |
| NOX4 | 5 | 0.491392546 | 0.213 | 0.038 | 3.19E-44 | 1.17E-39 |
| GAS7 | 5 | -0.969651852 | 0.033 | 0.253 | 3.32E-44 | 1.22E-39 |
| ABCA3 | 5 | 0.683718273 | 0.497 | 0.238 | 3.37E-44 | 1.23E-39 |
| SLC4A7 | 5 | -0.823992052 | 0.56 | 0.768 | 4.67E-44 | 1.71E-39 |
| ASTN1 | 5 | -0.737998917 | 0.037 | 0.261 | 5.48E-44 | 2.01E-39 |
| VEGFC | 5 | -0.894757628 | 0.02 | 0.227 | 5.81E-44 | 2.13E-39 |
| ONECUT2 | 5 | 0.522539851 | 0.235 | 0.051 | 6.65E-44 | 2.44E-39 |
| BCR | 5 | 0.677989872 | 0.472 | 0.232 | 6.94E-44 | 2.54E-39 |
| PAIP2B | 5 | 0.351419338 | 0.154 | 0.018 | 8.18E-44 | 3E-39 |
| TIMP2 | 5 | -0.589993085 | 0.039 | 0.268 | 8.57E-44 | 3.14E-39 |
| NR2F1-AS1 | 5 | -0.972368534 | 0.206 | 0.48 | 8.92E-44 | 3.27E-39 |
| ZBTB16 | 5 | 0.265639009 | 0.143 | 0.017 | 9.89E-44 | 3.62E-39 |
| ADAMTS20 | 5 | -1.049857061 | 0.031 | 0.247 | 1.29E-43 | 4.72E-39 |
| ARL15 | 5 | 0.652638889 | 0.85 | 0.658 | 1.31E-43 | 4.79E-39 |
| ITPR2 | 5 | 0.504694045 | 0.631 | 0.327 | 1.41E-43 | 5.18E-39 |
| PCLO | 5 | 0.486227315 | 0.864 | 0.591 | 1.46E-43 | 5.34E-39 |

| | | | | | | |
|---|---|---|---|---|---|---|
| MPC2 | 5 | 0.434493879 | 0.217 | 0.044 | 1.53E-43 | 5.59E-39 |
| LINC00884 | 5 | 0.309132843 | 0.125 | 0.01 | 1.72E-43 | 6.29E-39 |
| SNCAIP | 5 | -0.908171488 | 0.015 | 0.216 | 1.8E-43 | 6.59E-39 |
| HSD17B11 | 5 | 0.522516378 | 0.29 | 0.086 | 1.82E-43 | 6.67E-39 |
| SSBP3 | 5 | 0.71671472 | 0.754 | 0.541 | 2.07E-43 | 7.57E-39 |
| STX1A | 5 | 0.468789425 | 0.235 | 0.053 | 2.19E-43 | 8.03E-39 |
| ELF1 | 5 | 0.690338489 | 0.547 | 0.275 | 2.41E-43 | 8.83E-39 |
| PREX1 | 5 | -0.550038468 | 0.022 | 0.234 | 2.47E-43 | 9.04E-39 |
| STAG2 | 5 | -0.666288768 | 0.655 | 0.84 | 2.72E-43 | 9.96E-39 |
| KIAA0319 | 5 | 0.438487538 | 0.161 | 0.021 | 4.92E-43 | 1.8E-38 |
| FAM126A | 5 | -0.928087776 | 0.369 | 0.618 | 5.08E-43 | 1.86E-38 |
| OPRM1 | 5 | 0.384116138 | 0.13 | 0.016 | 6.82E-43 | 2.5E-38 |
| TRIB2 | 5 | -0.471567079 | 0.009 | 0.201 | 6.83E-43 | 2.5E-38 |
| SMC4 | 5 | 0.755953996 | 0.776 | 0.569 | 7.83E-43 | 2.87E-38 |
| MSI2 | 5 | -0.814937395 | 0.745 | 0.864 | 8.21E-43 | 3E-38 |
| PSD | 5 | 0.48571632 | 0.231 | 0.055 | 8.51E-43 | 3.11E-38 |
| SCARB2 | 5 | 0.630557914 | 0.655 | 0.392 | 1.04E-42 | 3.79E-38 |
| SYNPR | 5 | -1.300281525 | 0.125 | 0.374 | 1.22E-42 | 4.47E-38 |
| SLC35G2 | 5 | 0.616926942 | 0.455 | 0.207 | 1.44E-42 | 5.27E-38 |
| DLX6-AS1 | 5 | -1.260228776 | 0.028 | 0.235 | 1.52E-42 | 5.55E-38 |
| EPHA5 | 5 | -1.303180578 | 0.048 | 0.272 | 2.14E-42 | 7.82E-38 |
| ZNF710 | 5 | 0.652950092 | 0.4 | 0.17 | 2.36E-42 | 8.65E-38 |
| NT5C3A | 5 | 0.629344735 | 0.659 | 0.418 | 2.7E-42 | 9.88E-38 |
| PVR | 5 | 0.539723046 | 0.306 | 0.1 | 3.06E-42 | 1.12E-37 |
| FYB2 | 5 | 0.393771041 | 0.178 | 0.028 | 3.73E-42 | 1.37E-37 |
| PLEKHA8 | 5 | -0.651622719 | 0.114 | 0.381 | 3.81E-42 | 1.39E-37 |
| PRKAR2B | 5 | -0.671277098 | 0.051 | 0.28 | 5.3E-42 | 1.94E-37 |
| KCTD12 | 5 | 0.482460044 | 0.215 | 0.048 | 7.72E-42 | 2.83E-37 |
| CASTOR2 | 5 | 0.652843784 | 0.385 | 0.166 | 7.97E-42 | 2.92E-37 |
| REST | 5 | -0.583613899 | 0.028 | 0.236 | 8.7E-42 | 3.18E-37 |
| SULT4A1 | 5 | 0.540361009 | 0.314 | 0.11 | 1.26E-41 | 4.6E-37 |
| RNF43 | 5 | 0.335346349 | 0.154 | 0.019 | 1.38E-41 | 5.04E-37 |
| LYPD6 | 5 | -0.860788563 | 0.112 | 0.364 | 1.39E-41 | 5.09E-37 |
| SIRPA | 5 | 0.321136951 | 0.145 | 0.016 | 1.46E-41 | 5.34E-37 |
| PSMF1 | 5 | 0.574807458 | 0.413 | 0.193 | 1.48E-41 | 5.43E-37 |
| MED25 | 5 | 0.550286531 | 0.358 | 0.13 | 1.69E-41 | 6.19E-37 |
| ST3GAL5 | 5 | 0.609076632 | 0.334 | 0.122 | 1.82E-41 | 6.68E-37 |
| SORD | 5 | 0.482651169 | 0.262 | 0.073 | 1.88E-41 | 6.9E-37 |
| TCP11L2 | 5 | 0.510871636 | 0.246 | 0.067 | 1.9E-41 | 6.96E-37 |
| PTCHD4 | 5 | -1.069998445 | 0.046 | 0.265 | 2.18E-41 | 7.99E-37 |
| WDR11 | 5 | -0.973910197 | 0.451 | 0.69 | 2.52E-41 | 9.22E-37 |
| DGKI | 5 | -0.879947266 | 0.033 | 0.249 | 2.54E-41 | 9.3E-37 |
| HIST1H2BJ | 5 | 0.671957814 | 0.121 | 0.016 | 2.79E-41 | 1.02E-36 |
| DYNC2H1 | 5 | 0.672953102 | 0.631 | 0.345 | 2.82E-41 | 1.03E-36 |
| AFF2 | 5 | -1.235320178 | 0.44 | 0.659 | 3.38E-41 | 1.24E-36 |
| TEX2 | 5 | 0.557729388 | 0.396 | 0.157 | 3.39E-41 | 1.24E-36 |
| YTHDC2 | 5 | -0.750384091 | 0.413 | 0.668 | 4.85E-41 | 1.77E-36 |
| LMO4 | 5 | -0.721899303 | 0.051 | 0.278 | 5.02E-41 | 1.84E-36 |
| ANK1 | 5 | 0.569627239 | 0.44 | 0.178 | 5.29E-41 | 1.94E-36 |
| DNAJC12 | 5 | 0.441903502 | 0.187 | 0.035 | 7.48E-41 | 2.74E-36 |
| PCSK5 | 5 | -0.829680162 | 0.075 | 0.313 | 7.7E-41 | 2.82E-36 |
| ABHD3 | 5 | 0.624241966 | 0.453 | 0.206 | 8.44E-41 | 3.09E-36 |
| AL645568.1 | 5 | 0.608821802 | 0.382 | 0.162 | 9.02E-41 | 3.3E-36 |
| AL121578.3 | 5 | -0.651351819 | 0.029 | 0.23 | 1.04E-40 | 3.79E-36 |
| OCLN | 5 | 0.501961967 | 0.273 | 0.088 | 1.04E-40 | 3.81E-36 |
| WWC3 | 5 | -0.654323261 | 0.136 | 0.397 | 1.05E-40 | 3.86E-36 |
| CMIP | 5 | 0.64437195 | 0.761 | 0.575 | 1.08E-40 | 3.96E-36 |
| PLOD2 | 5 | 1.045794349 | 0.514 | 0.271 | 1.19E-40 | 4.34E-36 |

| | | | | | | |
|---|---|---|---|---|---|---|
| MSH5 | 5 | 0.63550871 | 0.486 | 0.258 | 1.29E-40 | 4.71E-36 |
| PPM1L | 5 | 0.715887649 | 0.721 | 0.507 | 1.4E-40 | 5.14E-36 |
| EPHA3 | 5 | -1.227208487 | 0.024 | 0.228 | 1.46E-40 | 5.34E-36 |
| GRIN3A | 5 | -1.095181598 | 0.05 | 0.271 | 1.86E-40 | 6.82E-36 |
| HSD17B7 | 5 | 0.528217385 | 0.281 | 0.086 | 1.88E-40 | 6.87E-36 |
| ARHGAP21 | 5 | -0.756716864 | 0.295 | 0.569 | 2.01E-40 | 7.35E-36 |
| PMM2 | 5 | 0.605443744 | 0.508 | 0.257 | 2.06E-40 | 7.53E-36 |
| CHM | 5 | -0.736866044 | 0.466 | 0.697 | 2.35E-40 | 8.61E-36 |
| AL513188.1 | 5 | 0.333389393 | 0.183 | 0.031 | 2.58E-40 | 9.43E-36 |
| NTF3 | 5 | -0.707491313 | 0.006 | 0.179 | 3.04E-40 | 1.11E-35 |
| NFIC | 5 | -0.684731462 | 0.103 | 0.358 | 3.42E-40 | 1.25E-35 |
| AKAP12 | 5 | -0.738299735 | 0.018 | 0.211 | 3.53E-40 | 1.29E-35 |
| IL20RA | 5 | 0.307464086 | 0.132 | 0.013 | 3.59E-40 | 1.31E-35 |
| CAMTA1 | 5 | -0.914783231 | 0.528 | 0.722 | 3.68E-40 | 1.35E-35 |
| FAP | 5 | -1.176058849 | 0.028 | 0.23 | 3.87E-40 | 1.42E-35 |
| SIPA1L1 | 5 | 0.565645115 | 0.923 | 0.762 | 4.78E-40 | 1.75E-35 |
| SORCS1 | 5 | -0.979636256 | 0.037 | 0.243 | 5.5E-40 | 2.01E-35 |
| GALNT18 | 5 | -0.87875098 | 0.152 | 0.414 | 5.76E-40 | 2.11E-35 |
| PDE8B | 5 | 0.533550273 | 0.231 | 0.054 | 5.82E-40 | 2.13E-35 |
| FSIP2 | 5 | -0.584343818 | 0.017 | 0.206 | 6.36E-40 | 2.33E-35 |
| COL24A1 | 5 | -0.930284423 | 0.017 | 0.205 | 7.66E-40 | 2.8E-35 |
| TBX3 | 5 | -0.56299324 | 0.017 | 0.203 | 7.84E-40 | 2.87E-35 |
| SMC5-AS1 | 5 | 0.535819904 | 0.35 | 0.128 | 8.56E-40 | 3.13E-35 |
| DISP1 | 5 | -0.744377824 | 0.128 | 0.388 | 1.01E-39 | 3.7E-35 |
| MARCH1 | 5 | -1.521468396 | 0.092 | 0.312 | 1.02E-39 | 3.75E-35 |
| MAPK1 | 5 | 0.597313179 | 0.769 | 0.587 | 1.03E-39 | 3.79E-35 |
| L1TD1 | 5 | -0.740635148 | 0.033 | 0.237 | 1.12E-39 | 4.12E-35 |
| LRP11 | 5 | 0.491211561 | 0.275 | 0.083 | 1.28E-39 | 4.69E-35 |
| SPTBN1 | 5 | -0.778299299 | 0.466 | 0.702 | 1.33E-39 | 4.87E-35 |
| UBE2G1 | 5 | -0.756620296 | 0.38 | 0.629 | 1.44E-39 | 5.26E-35 |
| AL357052.1 | 5 | 0.262060299 | 0.103 | 0.006 | 1.44E-39 | 5.28E-35 |
| HULC | 5 | -1.394022789 | 0.178 | 0.429 | 1.59E-39 | 5.81E-35 |
| ANO2 | 5 | -0.838022622 | 0.017 | 0.204 | 1.83E-39 | 6.69E-35 |
| HID1 | 5 | 0.327593747 | 0.154 | 0.021 | 2.33E-39 | 8.51E-35 |
| HES6 | 5 | 0.600121088 | 0.266 | 0.084 | 2.62E-39 | 9.58E-35 |
| CAMK4 | 5 | -0.911737958 | 0.088 | 0.311 | 2.78E-39 | 1.02E-34 |
| SYT14 | 5 | 0.377916959 | 0.745 | 0.457 | 2.81E-39 | 1.03E-34 |
| MORC3 | 5 | -0.7803607 | 0.29 | 0.561 | 2.89E-39 | 1.06E-34 |
| SLC25A33 | 5 | 0.552927746 | 0.347 | 0.127 | 3E-39 | 1.1E-34 |
| AC090204.1 | 5 | 0.528776266 | 0.248 | 0.066 | 3.37E-39 | 1.23E-34 |
| DLG5 | 5 | 0.639961386 | 0.596 | 0.37 | 3.41E-39 | 1.25E-34 |
| PKP4 | 5 | -0.913097192 | 0.501 | 0.72 | 3.82E-39 | 1.4E-34 |
| AL390783.1 | 5 | 0.434407505 | 0.262 | 0.072 | 4.25E-39 | 1.55E-34 |
| FTO | 5 | -1.038350021 | 0.675 | 0.82 | 4.65E-39 | 1.7E-34 |
| SLC44A1 | 5 | -0.825032257 | 0.49 | 0.704 | 5.51E-39 | 2.02E-34 |
| AL356124.1 | 5 | -0.692809527 | 0.028 | 0.228 | 5.76E-39 | 2.11E-34 |
| FERMT2 | 5 | -0.85248191 | 0.187 | 0.448 | 6E-39 | 2.2E-34 |
| PAG1 | 5 | -0.824564767 | 0.088 | 0.319 | 7.52E-39 | 2.75E-34 |
| PDE7B | 5 | -1.267406636 | 0.031 | 0.228 | 7.93E-39 | 2.9E-34 |
| RAB11A | 5 | 0.613539569 | 0.47 | 0.253 | 8.68E-39 | 3.18E-34 |
| MAN2A1 | 5 | 0.637489268 | 0.699 | 0.44 | 1.07E-38 | 3.9E-34 |
| ENC1 | 5 | -0.684250672 | 0.029 | 0.228 | 1.09E-38 | 3.98E-34 |
| ELOVL6 | 5 | 0.667303468 | 0.622 | 0.421 | 1.14E-38 | 4.18E-34 |
| PER2 | 5 | 0.507892818 | 0.286 | 0.093 | 1.65E-38 | 6.05E-34 |
| TAFA1 | 5 | -1.690979284 | 0.039 | 0.238 | 1.73E-38 | 6.32E-34 |
| PCDH9 | 5 | -1.710890431 | 0.132 | 0.36 | 1.73E-38 | 6.32E-34 |
| C11orf49 | 5 | 0.59022394 | 0.736 | 0.534 | 1.76E-38 | 6.43E-34 |
| KCNK9 | 5 | 0.297376639 | 0.114 | 0.009 | 2.12E-38 | 7.77E-34 |

| | | | | | | |
|---|---|---|---|---|---|---|
| FAM184A | 5 | 0.593138834 | 0.684 | 0.414 | 2.33E-38 | 8.54E-34 |
| LINC01184 | 5 | -0.733561761 | 0.182 | 0.444 | 2.65E-38 | 9.69E-34 |
| ARRDC3-AS1 | 5 | -1.560230727 | 0.077 | 0.297 | 2.71E-38 | 9.93E-34 |
| MMUT | 5 | 0.600272935 | 0.383 | 0.171 | 2.74E-38 | 1E-33 |
| CMTM4 | 5 | 0.649518742 | 0.459 | 0.232 | 2.96E-38 | 1.08E-33 |
| DPYSL5 | 5 | -0.670049907 | 0.037 | 0.236 | 3.54E-38 | 1.3E-33 |
| HS3ST1 | 5 | -0.554174293 | 0.02 | 0.21 | 3.62E-38 | 1.33E-33 |
| USP32 | 5 | -0.736823596 | 0.457 | 0.701 | 3.77E-38 | 1.38E-33 |
| GRIK4 | 5 | -0.605118582 | 0.048 | 0.264 | 3.93E-38 | 1.44E-33 |
| PKIA | 5 | -0.907288061 | 0.213 | 0.461 | 4.29E-38 | 1.57E-33 |
| PERP | 5 | 0.303375717 | 0.114 | 0.009 | 4.39E-38 | 1.61E-33 |
| BEND7 | 5 | -0.644620204 | 0.051 | 0.269 | 4.85E-38 | 1.78E-33 |
| ARHGAP31 | 5 | -0.604523894 | 0.037 | 0.242 | 5.4E-38 | 1.98E-33 |
| MEF2C-AS1 | 5 | -0.699848709 | 0.05 | 0.259 | 5.5E-38 | 2.01E-33 |
| ELOVL7 | 5 | 0.344780173 | 0.143 | 0.018 | 5.72E-38 | 2.09E-33 |
| LINC01122 | 5 | -0.968332919 | 0.064 | 0.283 | 6.47E-38 | 2.37E-33 |
| SYCP2 | 5 | -1.179455094 | 0.037 | 0.238 | 6.9E-38 | 2.53E-33 |
| SCN8A | 5 | -0.796390939 | 0.494 | 0.676 | 6.96E-38 | 2.55E-33 |
| E7 | 5 | 0.301109704 | 0.13 | 0.014 | 7.01E-38 | 2.57E-33 |
| GULP1 | 5 | -1.07407051 | 0.272 | 0.522 | 1.03E-37 | 3.78E-33 |
| PAK1 | 5 | -0.816676538 | 0.365 | 0.586 | 1.06E-37 | 3.88E-33 |
| IGF2BP3 | 5 | -0.641719533 | 0.767 | 0.893 | 1.18E-37 | 4.3E-33 |
| AP001347.1 | 5 | -0.728993303 | 0.121 | 0.37 | 1.45E-37 | 5.31E-33 |
| LANCL3 | 5 | -0.673348741 | 0.037 | 0.238 | 1.72E-37 | 6.29E-33 |
| C1orf21 | 5 | 0.629776923 | 0.565 | 0.294 | 2.27E-37 | 8.3E-33 |
| IRS2 | 5 | -0.795740854 | 0.16 | 0.412 | 2.29E-37 | 8.37E-33 |
| C6orf118 | 5 | 0.272513819 | 0.169 | 0.032 | 2.31E-37 | 8.46E-33 |
| ECPAS | 5 | 0.581505962 | 0.723 | 0.504 | 2.42E-37 | 8.88E-33 |
| PLEKHG4B | 5 | -0.568910729 | 0.04 | 0.236 | 2.62E-37 | 9.59E-33 |
| GAP43 | 5 | -0.72291297 | 0.037 | 0.234 | 3.19E-37 | 1.17E-32 |
| XACT | 5 | -2.366321713 | 0.061 | 0.261 | 3.76E-37 | 1.38E-32 |
| MYO6 | 5 | 0.575275638 | 0.527 | 0.263 | 4.49E-37 | 1.64E-32 |
| ST8SIA3 | 5 | 0.285797989 | 0.134 | 0.016 | 4.65E-37 | 1.7E-32 |
| FLNA | 5 | -0.480832798 | 0.024 | 0.213 | 4.97E-37 | 1.82E-32 |
| MYO18B | 5 | 0.258262946 | 0.123 | 0.012 | 5.28E-37 | 1.93E-32 |
| DCP2 | 5 | -0.738274671 | 0.341 | 0.592 | 5.66E-37 | 2.07E-32 |
| EIF4G3 | 5 | -0.545187731 | 0.892 | 0.938 | 5.91E-37 | 2.16E-32 |
| CPE | 5 | 0.607946428 | 0.835 | 0.616 | 6.39E-37 | 2.34E-32 |
| HK1 | 5 | -0.512405578 | 0.061 | 0.283 | 6.53E-37 | 2.39E-32 |
| MCF2L | 5 | 0.574281532 | 0.508 | 0.248 | 9.84E-37 | 3.6E-32 |
| TUFT1 | 5 | 0.322675669 | 0.158 | 0.024 | 1.02E-36 | 3.72E-32 |
| MYH9 | 5 | 0.675551201 | 0.464 | 0.257 | 1.25E-36 | 4.59E-32 |
| STARD13 | 5 | -1.020474335 | 0.483 | 0.603 | 1.6E-36 | 5.86E-32 |
| BMPER | 5 | -0.991583304 | 0.031 | 0.223 | 1.73E-36 | 6.33E-32 |
| AC024901.1 | 5 | -0.995097736 | 0.017 | 0.19 | 1.98E-36 | 7.25E-32 |
| KCNT2 | 5 | 0.553895219 | 0.883 | 0.694 | 2.1E-36 | 7.68E-32 |
| AC068587.4 | 5 | 0.519616958 | 0.604 | 0.327 | 2.14E-36 | 7.83E-32 |
| DYNC1I1 | 5 | 0.755431761 | 0.561 | 0.332 | 2.21E-36 | 8.1E-32 |
| PELI2 | 5 | 0.59359385 | 0.683 | 0.412 | 2.26E-36 | 8.27E-32 |
| DAPK1 | 5 | -0.857882839 | 0.486 | 0.699 | 2.29E-36 | 8.39E-32 |
| DMTN | 5 | 0.281009967 | 0.106 | 0.009 | 2.42E-36 | 8.86E-32 |
| GPR156 | 5 | -0.585619697 | 0.039 | 0.239 | 2.57E-36 | 9.41E-32 |
| LINC00571 | 5 | 0.530866849 | 0.248 | 0.073 | 3.04E-36 | 1.11E-31 |
| VPS37B | 5 | -0.635306907 | 0.134 | 0.378 | 3.21E-36 | 1.18E-31 |
| RPGRIP1L | 5 | -0.852587074 | 0.2 | 0.441 | 3.54E-36 | 1.29E-31 |
| SPIDR | 5 | 0.571939688 | 0.879 | 0.739 | 3.95E-36 | 1.44E-31 |
| DAPK2 | 5 | -0.537602042 | 0.033 | 0.23 | 4.08E-36 | 1.49E-31 |
| ABHD16A | 5 | 0.449592809 | 0.246 | 0.07 | 4.14E-36 | 1.52E-31 |

| | | | | | | |
|---|---|---|---|---|---|---|
| CDK8 | 5 | 0.584982673 | 0.734 | 0.543 | 4.74E-36 | 1.73E-31 |
| NCAM1 | 5 | -0.741355939 | 0.75 | 0.839 | 5.47E-36 | 2E-31 |
| TASP1 | 5 | 0.594132404 | 0.71 | 0.505 | 5.57E-36 | 2.04E-31 |
| RASSF8 | 5 | -0.844620354 | 0.275 | 0.517 | 5.6E-36 | 2.05E-31 |
| CEMIP2 | 5 | -0.828071612 | 0.127 | 0.362 | 7.42E-36 | 2.72E-31 |
| LNPEP | 5 | 0.58603303 | 0.547 | 0.327 | 7.43E-36 | 2.72E-31 |
| RAB27B | 5 | 0.393914943 | 0.149 | 0.022 | 8.32E-36 | 3.04E-31 |
| COBLL1 | 5 | 0.445662508 | 0.642 | 0.365 | 1.04E-35 | 3.8E-31 |
| TOM1L1 | 5 | 0.577410974 | 0.486 | 0.276 | 1.17E-35 | 4.28E-31 |
| FGD5 | 5 | -0.492416957 | 0.02 | 0.199 | 1.33E-35 | 4.87E-31 |
| PCDH19 | 5 | -0.852030103 | 0.026 | 0.208 | 1.38E-35 | 5.05E-31 |
| DPF3 | 5 | -0.646377214 | 0.068 | 0.283 | 1.44E-35 | 5.27E-31 |
| IQCH | 5 | 0.531311101 | 0.308 | 0.106 | 1.45E-35 | 5.31E-31 |
| PLEKHH1 | 5 | 0.425192725 | 0.218 | 0.055 | 1.93E-35 | 7.05E-31 |
| SAMD4A | 5 | 0.599951944 | 0.637 | 0.371 | 2E-35 | 7.31E-31 |
| ITGAV | 5 | -0.931706761 | 0.325 | 0.551 | 2.03E-35 | 7.45E-31 |
| MMP16 | 5 | -0.819098855 | 0.53 | 0.738 | 2.16E-35 | 7.9E-31 |
| SLC6A6 | 5 | 0.539244849 | 0.323 | 0.133 | 2.3E-35 | 8.41E-31 |
| HDAC9 | 5 | -1.16227067 | 0.429 | 0.584 | 2.48E-35 | 9.09E-31 |
| LAMC1 | 5 | -0.854893535 | 0.31 | 0.537 | 2.48E-35 | 9.1E-31 |
| EIF2S3 | 5 | 0.555216372 | 0.666 | 0.491 | 2.5E-35 | 9.16E-31 |
| ZNF711 | 5 | -0.679712576 | 0.361 | 0.613 | 2.6E-35 | 9.52E-31 |
| MRAS | 5 | -0.419640432 | 0.015 | 0.186 | 2.77E-35 | 1.01E-30 |
| ARHGEF3 | 5 | -0.792937074 | 0.226 | 0.473 | 2.84E-35 | 1.04E-30 |
| WFS1 | 5 | 0.336712862 | 0.16 | 0.029 | 2.88E-35 | 1.05E-30 |
| BACH1 | 5 | -0.801031396 | 0.244 | 0.481 | 3.06E-35 | 1.12E-30 |
| TRPC1 | 5 | -0.783028863 | 0.31 | 0.56 | 3.23E-35 | 1.18E-30 |
| VWA5B1 | 5 | 0.258940456 | 0.161 | 0.035 | 3.62E-35 | 1.32E-30 |
| DLG1 | 5 | 0.534605262 | 0.877 | 0.75 | 3.95E-35 | 1.44E-30 |
| ANXA7 | 5 | 0.509054248 | 0.36 | 0.147 | 5.09E-35 | 1.86E-30 |
| ITGB1 | 5 | 0.589928089 | 0.723 | 0.553 | 5.99E-35 | 2.19E-30 |
| TSGA10 | 5 | 0.558075136 | 0.35 | 0.14 | 6.04E-35 | 2.21E-30 |
| KCTD16 | 5 | 0.564106252 | 0.38 | 0.151 | 6.55E-35 | 2.4E-30 |
| ABHD2 | 5 | 0.554530742 | 0.387 | 0.177 | 7.2E-35 | 2.64E-30 |
| CASZ1 | 5 | -0.660920375 | 0.059 | 0.265 | 7.4E-35 | 2.71E-30 |
| PPP1R21 | 5 | 0.612155649 | 0.558 | 0.338 | 7.72E-35 | 2.83E-30 |
| LINC02398 | 5 | -1.151902984 | 0.013 | 0.174 | 7.75E-35 | 2.84E-30 |
| CACNA2D2 | 5 | -0.554657196 | 0.077 | 0.3 | 7.76E-35 | 2.84E-30 |
| RAB3C | 5 | -0.383816762 | 0.552 | 0.419 | 8.63E-35 | 3.16E-30 |
| SLC39A8 | 5 | 0.486629428 | 0.305 | 0.104 | 9.43E-35 | 3.45E-30 |
| PMP22 | 5 | -0.553281567 | 0.026 | 0.205 | 1.04E-34 | 3.81E-30 |
| HUNK | 5 | 0.499881439 | 0.363 | 0.14 | 1.09E-34 | 3.98E-30 |
| GPC4 | 5 | -0.729899544 | 0.031 | 0.217 | 1.11E-34 | 4.05E-30 |
| MARVELD2 | 5 | 0.324140254 | 0.141 | 0.022 | 1.22E-34 | 4.47E-30 |
| NDFIP2 | 5 | 0.586590251 | 0.444 | 0.217 | 1.41E-34 | 5.17E-30 |
| KCNMB2-AS1 | 5 | 0.418184438 | 0.226 | 0.057 | 1.97E-34 | 7.22E-30 |
| GRB7 | 5 | 0.326800782 | 0.116 | 0.014 | 2.1E-34 | 7.7E-30 |
| PIP4K2A | 5 | -0.73439943 | 0.363 | 0.605 | 2.16E-34 | 7.9E-30 |
| DESI1 | 5 | 0.461619268 | 0.319 | 0.117 | 2.28E-34 | 8.35E-30 |
| IGSF21 | 5 | -0.655557611 | 0.064 | 0.275 | 2.38E-34 | 8.7E-30 |
| SUCLG1 | 5 | 0.520403854 | 0.42 | 0.204 | 2.46E-34 | 9.02E-30 |
| CHGB | 5 | 0.315974384 | 0.136 | 0.019 | 2.56E-34 | 9.37E-30 |
| C12orf75 | 5 | 0.512212779 | 0.338 | 0.132 | 2.99E-34 | 1.09E-29 |
| PURG | 5 | -0.441462513 | 0.018 | 0.186 | 3E-34 | 1.1E-29 |
| FASTKD1 | 5 | 0.59785628 | 0.585 | 0.386 | 3.64E-34 | 1.33E-29 |
| CTTNBP2 | 5 | -0.906809609 | 0.097 | 0.308 | 3.76E-34 | 1.38E-29 |
| SCAPER | 5 | 0.548064445 | 0.822 | 0.657 | 3.78E-34 | 1.38E-29 |
| TNR | 5 | 0.274946648 | 0.112 | 0.011 | 3.84E-34 | 1.41E-29 |

| | | | | | | |
|---|---|---|---|---|---|---|
| BMP2K | 5 | -0.718874441 | 0.323 | 0.572 | 3.91E-34 | 1.43E-29 |
| PDE8A | 5 | 0.523809791 | 0.828 | 0.639 | 4.03E-34 | 1.48E-29 |
| KIRREL3 | 5 | -0.995495672 | 0.018 | 0.19 | 4.11E-34 | 1.5E-29 |
| AL357054.2 | 5 | -0.579665397 | 0.033 | 0.217 | 4.63E-34 | 1.7E-29 |
| HOXD3 | 5 | -0.443648804 | 0.011 | 0.169 | 5.37E-34 | 1.97E-29 |
| AC093298.2 | 5 | -1.639188151 | 0.026 | 0.199 | 5.72E-34 | 2.1E-29 |
| PTPN3 | 5 | 0.547994719 | 0.341 | 0.154 | 5.84E-34 | 2.14E-29 |
| AL591519.1 | 5 | -0.644477477 | 0.006 | 0.157 | 6.09E-34 | 2.23E-29 |
| ADGRV1 | 5 | -0.854716804 | 0.848 | 0.674 | 6.11E-34 | 2.24E-29 |
| TWIST1 | 5 | -0.599280602 | 0.018 | 0.185 | 8.72E-34 | 3.19E-29 |
| THSD7B | 5 | -1.060404584 | 0.037 | 0.223 | 1.05E-33 | 3.85E-29 |
| MPPED2 | 5 | -0.72172225 | 0.026 | 0.196 | 1.08E-33 | 3.96E-29 |
| PSMA1 | 5 | 0.574842783 | 0.677 | 0.469 | 1.19E-33 | 4.34E-29 |
| BCL2L13 | 5 | 0.576639869 | 0.508 | 0.287 | 1.2E-33 | 4.41E-29 |
| OFD1 | 5 | 0.591312753 | 0.626 | 0.433 | 1.38E-33 | 5.05E-29 |
| ABCB1 | 5 | 0.313339343 | 0.455 | 0.222 | 1.43E-33 | 5.24E-29 |
| SLAIN1 | 5 | 0.60982661 | 0.521 | 0.296 | 1.45E-33 | 5.31E-29 |
| RAPGEF5 | 5 | 0.525955747 | 0.75 | 0.501 | 1.71E-33 | 6.27E-29 |
| AL596087.2 | 5 | -0.804158424 | 0.031 | 0.216 | 1.71E-33 | 6.27E-29 |
| CFAP70 | 5 | 0.296958227 | 0.244 | 0.076 | 1.75E-33 | 6.42E-29 |
| MOK | 5 | 0.483537672 | 0.334 | 0.123 | 1.94E-33 | 7.09E-29 |
| STXBP4 | 5 | 0.613161289 | 0.642 | 0.461 | 2.16E-33 | 7.92E-29 |
| ANKIB1 | 5 | -0.7522069 | 0.666 | 0.815 | 2.47E-33 | 9.03E-29 |
| NCKAP5 | 5 | -1.310055168 | 0.207 | 0.424 | 2.5E-33 | 9.16E-29 |
| YEATS2 | 5 | -0.596397698 | 0.561 | 0.765 | 2.59E-33 | 9.48E-29 |
| DNAJC5 | 5 | 0.577638407 | 0.508 | 0.287 | 2.73E-33 | 1E-28 |
| KIF13A | 5 | -0.727132523 | 0.363 | 0.601 | 3.57E-33 | 1.31E-28 |
| CCDC92 | 5 | -0.478447641 | 0.066 | 0.276 | 3.59E-33 | 1.31E-28 |
| PARVA | 5 | -0.637698076 | 0.169 | 0.407 | 3.8E-33 | 1.39E-28 |
| TENT5A | 5 | 0.447999787 | 0.327 | 0.122 | 3.94E-33 | 1.44E-28 |
| FMNL3 | 5 | -0.503983913 | 0.04 | 0.225 | 3.95E-33 | 1.45E-28 |
| RBFOX3 | 5 | -0.833585004 | 0.039 | 0.211 | 5.54E-33 | 2.03E-28 |
| AL353747.4 | 5 | -0.491512132 | 0.022 | 0.192 | 6.24E-33 | 2.28E-28 |
| ID4 | 5 | 0.435669785 | 0.183 | 0.039 | 8.97E-33 | 3.28E-28 |
| HOMER1 | 5 | -0.76801018 | 0.266 | 0.509 | 9.24E-33 | 3.38E-28 |
| TRAF5 | 5 | -0.510565587 | 0.068 | 0.275 | 1.05E-32 | 3.85E-28 |
| AACS | 5 | 0.511436469 | 0.345 | 0.142 | 1.06E-32 | 3.87E-28 |
| AC073050.1 | 5 | -1.212636528 | 0.105 | 0.317 | 1.2E-32 | 4.41E-28 |
| ARHGAP26 | 5 | 0.535894682 | 0.385 | 0.164 | 1.22E-32 | 4.47E-28 |
| ANO9 | 5 | 0.305225411 | 0.117 | 0.014 | 1.23E-32 | 4.49E-28 |
| KIAA1257 | 5 | 0.371491333 | 0.18 | 0.038 | 1.23E-32 | 4.51E-28 |
| KCNQ2 | 5 | -0.480377459 | 0.033 | 0.212 | 1.31E-32 | 4.8E-28 |
| TBC1D8 | 5 | 0.540347258 | 0.415 | 0.193 | 1.59E-32 | 5.82E-28 |
| CBR3-AS1 | 5 | 0.346320109 | 0.156 | 0.029 | 1.86E-32 | 6.8E-28 |
| APLP2 | 5 | -0.637740798 | 0.253 | 0.501 | 1.9E-32 | 6.97E-28 |
| SLC12A8 | 5 | 0.447079266 | 0.24 | 0.073 | 1.95E-32 | 7.14E-28 |
| QKI | 5 | -0.770460345 | 0.644 | 0.794 | 2.09E-32 | 7.66E-28 |
| HDAC7 | 5 | -0.413842451 | 0.031 | 0.209 | 2.26E-32 | 8.27E-28 |
| BSN | 5 | 0.472754924 | 0.288 | 0.102 | 2.89E-32 | 1.06E-27 |
| IFIH1 | 5 | 0.353078484 | 0.211 | 0.054 | 2.93E-32 | 1.07E-27 |
| DMXL2 | 5 | 0.547351358 | 0.754 | 0.563 | 3.56E-32 | 1.3E-27 |
| TAOK3 | 5 | -0.748998708 | 0.202 | 0.439 | 3.97E-32 | 1.45E-27 |
| SLC43A3 | 5 | 0.486656745 | 0.312 | 0.124 | 4.28E-32 | 1.57E-27 |
| LMX1A | 5 | 0.337439752 | 0.182 | 0.046 | 4.45E-32 | 1.63E-27 |
| ZNF521 | 5 | -0.894030125 | 0.007 | 0.153 | 4.48E-32 | 1.64E-27 |
| GHR | 5 | -0.799302006 | 0.04 | 0.221 | 4.51E-32 | 1.65E-27 |
| PGAP1 | 5 | -0.776348455 | 0.565 | 0.712 | 4.61E-32 | 1.69E-27 |
| DNAJC1 | 5 | 0.438921663 | 0.8 | 0.556 | 5E-32 | 1.83E-27 |

| | | | | | | |
|---|---|---|---|---|---|---|
| RHOH | 5 | 0.340565027 | 0.143 | 0.025 | 5.78E-32 | 2.12E-27 |
| IKZF2 | 5 | 0.488936366 | 0.277 | 0.095 | 6.14E-32 | 2.25E-27 |
| RASGRP1 | 5 | -0.659382091 | 0.046 | 0.232 | 6.24E-32 | 2.29E-27 |
| PROS1 | 5 | 0.392262784 | 0.182 | 0.04 | 6.28E-32 | 2.3E-27 |
| NLK | 5 | -0.87385481 | 0.495 | 0.675 | 6.91E-32 | 2.53E-27 |
| MIR646HG | 5 | -0.698556054 | 0 | 0.124 | 7.05E-32 | 2.58E-27 |
| PCBD2 | 5 | 0.556906836 | 0.448 | 0.232 | 7.19E-32 | 2.63E-27 |
| DMGDH | 5 | -0.716834496 | 0.114 | 0.324 | 9.47E-32 | 3.47E-27 |
| MAGI2 | 5 | -0.928054476 | 0.538 | 0.711 | 9.71E-32 | 3.55E-27 |
| PTGFRN | 5 | -0.584491345 | 0.119 | 0.341 | 9.71E-32 | 3.55E-27 |
| CNTN6 | 5 | -1.026482752 | 0.022 | 0.189 | 9.82E-32 | 3.6E-27 |
| LINC00599 | 5 | -0.460973351 | 0.015 | 0.167 | 9.86E-32 | 3.61E-27 |
| IGFBP2 | 5 | -0.80501212 | 0.299 | 0.541 | 1.11E-31 | 4.08E-27 |
| MGAT4A | 5 | 0.487579124 | 0.4 | 0.175 | 1.12E-31 | 4.08E-27 |
| GSE1 | 5 | 0.550416756 | 0.875 | 0.72 | 1.14E-31 | 4.19E-27 |
| NOS1AP | 5 | -0.638979889 | 0.119 | 0.334 | 1.15E-31 | 4.23E-27 |
| SPINK5 | 5 | -0.489469548 | 0.011 | 0.162 | 1.2E-31 | 4.4E-27 |
| NKD1 | 5 | -0.771366406 | 0.017 | 0.174 | 1.22E-31 | 4.48E-27 |
| MAPK8 | 5 | -0.613749543 | 0.589 | 0.762 | 1.23E-31 | 4.49E-27 |
| DAAM2 | 5 | -0.829927066 | 0.042 | 0.227 | 1.3E-31 | 4.77E-27 |
| CYBRD1 | 5 | -0.317401937 | 0 | 0.123 | 1.43E-31 | 5.24E-27 |
| UBE2E2 | 5 | 0.360538399 | 0.829 | 0.586 | 1.59E-31 | 5.81E-27 |
| PI4KA | 5 | 0.569388578 | 0.712 | 0.541 | 1.67E-31 | 6.1E-27 |
| PPM1E | 5 | -0.778673381 | 0.607 | 0.744 | 1.68E-31 | 6.15E-27 |
| RBMX | 5 | -0.57179762 | 0.701 | 0.833 | 1.98E-31 | 7.25E-27 |
| NDST4 | 5 | 0.3686453 | 0.183 | 0.044 | 2.18E-31 | 7.97E-27 |
| FGFR4 | 5 | 0.255812655 | 0.105 | 0.011 | 2.29E-31 | 8.38E-27 |
| LDLRAD3 | 5 | -0.714531583 | 0.341 | 0.566 | 2.33E-31 | 8.51E-27 |
| IQGAP2 | 5 | 0.505098414 | 0.642 | 0.385 | 2.78E-31 | 1.02E-26 |
| AL445123.2 | 5 | -0.628291587 | 0.059 | 0.247 | 3.72E-31 | 1.36E-26 |
| EBF4 | 5 | -0.473344517 | 0.033 | 0.21 | 3.9E-31 | 1.43E-26 |
| PRKAA2 | 5 | 0.469722366 | 0.389 | 0.168 | 4.87E-31 | 1.78E-26 |
| SLC4A4 | 5 | -0.707072723 | 0.022 | 0.183 | 5.12E-31 | 1.87E-26 |
| P2RX4 | 5 | 0.319179856 | 0.18 | 0.04 | 5.27E-31 | 1.93E-26 |
| NUP210 | 5 | 0.56552774 | 0.53 | 0.301 | 5.6E-31 | 2.05E-26 |
| SHROOM2 | 5 | -0.605921422 | 0.15 | 0.38 | 5.71E-31 | 2.09E-26 |
| NECTIN3 | 5 | -0.997179781 | 0.457 | 0.61 | 6.78E-31 | 2.48E-26 |
| BRINP1 | 5 | -1.078443143 | 0.244 | 0.432 | 7.13E-31 | 2.61E-26 |
| BTRC | 5 | -0.657635583 | 0.508 | 0.706 | 7.39E-31 | 2.7E-26 |
| DAB2IP | 5 | -0.453549507 | 0.04 | 0.218 | 8.34E-31 | 3.05E-26 |
| LRRN1 | 5 | -0.531458686 | 0.033 | 0.206 | 8.76E-31 | 3.21E-26 |
| RPS6KA6 | 5 | 0.549698986 | 0.407 | 0.189 | 9.6E-31 | 3.51E-26 |
| CEACAM19 | 5 | 0.423847471 | 0.211 | 0.062 | 9.69E-31 | 3.55E-26 |
| RREB1 | 5 | 0.592578816 | 0.558 | 0.329 | 1.04E-30 | 3.81E-26 |
| NELL2 | 5 | -1.062747716 | 0.084 | 0.268 | 1.07E-30 | 3.93E-26 |
| CAPZB | 5 | 0.544511634 | 0.664 | 0.505 | 1.11E-30 | 4.05E-26 |
| CAMKMT | 5 | 0.560219223 | 0.818 | 0.685 | 1.14E-30 | 4.16E-26 |
| CATSPER2 | 5 | 0.474835336 | 0.341 | 0.149 | 1.24E-30 | 4.56E-26 |
| ADD3 | 5 | 0.611087811 | 0.617 | 0.432 | 1.33E-30 | 4.85E-26 |
| SATB1-AS1 | 5 | -0.507889889 | 0.026 | 0.187 | 1.48E-30 | 5.4E-26 |
| CCDC136 | 5 | -0.388921798 | 0.018 | 0.173 | 1.69E-30 | 6.2E-26 |
| GNAZ | 5 | 0.334825443 | 0.185 | 0.044 | 1.8E-30 | 6.58E-26 |
| RASA4 | 5 | 0.356476565 | 0.18 | 0.043 | 1.81E-30 | 6.61E-26 |
| POU6F1 | 5 | -0.455308448 | 0.055 | 0.244 | 1.86E-30 | 6.8E-26 |
| FBXO17 | 5 | 0.577105807 | 0.429 | 0.224 | 1.87E-30 | 6.83E-26 |
| TASOR2 | 5 | 0.543136592 | 0.765 | 0.647 | 1.88E-30 | 6.87E-26 |
| GTDC1 | 5 | -0.76222961 | 0.288 | 0.503 | 1.97E-30 | 7.21E-26 |
| REEP1 | 5 | 0.502696604 | 0.31 | 0.12 | 2.04E-30 | 7.48E-26 |

| | | | | | | |
|---|---|---|---|---|---|---|
| CACUL1 | 5 | 0.555031123 | 0.582 | 0.375 | 2.1E-30 | 7.69E-26 |
| TDRD3 | 5 | -0.712877495 | 0.44 | 0.649 | 2.11E-30 | 7.71E-26 |
| EPS8L2 | 5 | 0.263949419 | 0.116 | 0.015 | 2.3E-30 | 8.41E-26 |
| CD2AP | 5 | 0.531409834 | 0.728 | 0.571 | 2.33E-30 | 8.53E-26 |
| ARID3A | 5 | 0.567870572 | 0.332 | 0.14 | 2.54E-30 | 9.29E-26 |
| SCN3A | 5 | -0.792518797 | 0.088 | 0.278 | 2.67E-30 | 9.78E-26 |
| GABRB2 | 5 | -1.054672633 | 0.116 | 0.318 | 2.88E-30 | 1.05E-25 |
| SLC18A2 | 5 | 0.482154135 | 0.101 | 0.013 | 3.08E-30 | 1.13E-25 |
| TMEM184B | 5 | 0.526521142 | 0.378 | 0.187 | 3.5E-30 | 1.28E-25 |
| VASH2 | 5 | -0.403309661 | 0.015 | 0.162 | 3.55E-30 | 1.3E-25 |
| NEO1 | 5 | 0.611199124 | 0.708 | 0.54 | 3.61E-30 | 1.32E-25 |
| RHOBTB1 | 5 | -0.550495766 | 0.066 | 0.263 | 3.77E-30 | 1.38E-25 |
| CASTOR3 | 5 | -0.544205453 | 0.095 | 0.307 | 3.9E-30 | 1.43E-25 |
| GNG4 | 5 | 0.553567518 | 0.55 | 0.309 | 4.36E-30 | 1.6E-25 |
| PIK3R3 | 5 | -0.754272482 | 0.286 | 0.509 | 4.6E-30 | 1.69E-25 |
| PRAME | 5 | -0.367436928 | 0.006 | 0.138 | 5.15E-30 | 1.89E-25 |
| SLC37A1 | 5 | 0.414804663 | 0.242 | 0.081 | 5.51E-30 | 2.02E-25 |
| BMPR2 | 5 | -0.628564311 | 0.694 | 0.833 | 5.81E-30 | 2.13E-25 |
| GLT1D1 | 5 | -0.479355902 | 0.018 | 0.173 | 6.02E-30 | 2.2E-25 |
| ATP6V0A1 | 5 | 0.546939477 | 0.525 | 0.319 | 6.09E-30 | 2.23E-25 |
| OPCML | 5 | 1.009759995 | 0.501 | 0.334 | 6.19E-30 | 2.27E-25 |
| PTBP3 | 5 | 0.555094541 | 0.589 | 0.365 | 6.41E-30 | 2.35E-25 |
| ZDHHC21 | 5 | -0.662769453 | 0.521 | 0.706 | 6.71E-30 | 2.46E-25 |
| PHLDB3 | 5 | 0.391357621 | 0.193 | 0.052 | 7.42E-30 | 2.72E-25 |
| TSPAN7 | 5 | 0.470516173 | 0.761 | 0.526 | 7.48E-30 | 2.74E-25 |
| GALNT17 | 5 | -0.973569757 | 0.033 | 0.198 | 7.8E-30 | 2.86E-25 |
| TTC28 | 5 | -0.489121637 | 0.945 | 0.965 | 9.13E-30 | 3.34E-25 |
| ANKRD33B | 5 | -0.516465373 | 0.02 | 0.176 | 9.74E-30 | 3.56E-25 |
| GPCPD1 | 5 | -0.665479354 | 0.371 | 0.584 | 1.15E-29 | 4.23E-25 |
| LRRTM3 | 5 | -0.722427271 | 0.02 | 0.176 | 1.2E-29 | 4.39E-25 |
| LRRC28 | 5 | 0.581885839 | 0.514 | 0.331 | 1.25E-29 | 4.57E-25 |
| AC135782.1 | 5 | -0.40956817 | 0.011 | 0.154 | 1.58E-29 | 5.79E-25 |
| PAX3 | 5 | -0.87080393 | 0.009 | 0.147 | 1.7E-29 | 6.22E-25 |
| AC069228.1 | 5 | -0.452723717 | 0.028 | 0.188 | 1.85E-29 | 6.76E-25 |
| PISD | 5 | 0.499799867 | 0.404 | 0.2 | 1.9E-29 | 6.95E-25 |
| LYST | 5 | 0.56515341 | 0.589 | 0.376 | 2.1E-29 | 7.71E-25 |
| LINC01768 | 5 | -0.614954069 | 0.028 | 0.189 | 2.2E-29 | 8.07E-25 |
| CCDC3 | 5 | -0.550027192 | 0.05 | 0.223 | 2.48E-29 | 9.09E-25 |
| ST8SIA2 | 5 | -0.356429984 | 0.009 | 0.149 | 2.58E-29 | 9.46E-25 |
| MAP1B | 5 | -0.769482708 | 0.569 | 0.733 | 2.81E-29 | 1.03E-24 |
| ARHGAP5 | 5 | 0.532826218 | 0.659 | 0.513 | 2.84E-29 | 1.04E-24 |
| XRCC6 | 5 | 0.517733435 | 0.477 | 0.265 | 2.92E-29 | 1.07E-24 |
| UVRAG | 5 | 0.503321731 | 0.75 | 0.591 | 3.12E-29 | 1.14E-24 |
| HLA-A | 5 | -0.50462492 | 0.057 | 0.24 | 4.07E-29 | 1.49E-24 |
| AAK1 | 5 | -0.624747615 | 0.316 | 0.545 | 4.09E-29 | 1.5E-24 |
| RPS6KA2 | 5 | -0.71466021 | 0.125 | 0.338 | 4.17E-29 | 1.53E-24 |
| COL25A1 | 5 | -1.36790573 | 0.224 | 0.402 | 4.33E-29 | 1.58E-24 |
| WNK1 | 5 | 0.511957076 | 0.848 | 0.741 | 4.35E-29 | 1.59E-24 |
| L3MBTL4 | 5 | -0.931539183 | 0.378 | 0.54 | 4.82E-29 | 1.77E-24 |
| RNF130 | 5 | -0.623845166 | 0.506 | 0.692 | 4.84E-29 | 1.77E-24 |
| MVK | 5 | 0.300412577 | 0.147 | 0.03 | 5.04E-29 | 1.84E-24 |
| BCHE | 5 | -0.525130381 | 0.004 | 0.129 | 5.34E-29 | 1.96E-24 |
| TMEM161B-AS1 | 5 | -0.761596963 | 0.472 | 0.674 | 5.4E-29 | 1.98E-24 |
| FEZ1 | 5 | -0.491711861 | 0.05 | 0.231 | 5.43E-29 | 1.99E-24 |
| CDKN2B-AS1 | 5 | 0.487617324 | 0.554 | 0.31 | 6.6E-29 | 2.42E-24 |
| TOP2B | 5 | -0.604931091 | 0.626 | 0.774 | 8.07E-29 | 2.96E-24 |
| NFATC2IP | 5 | 0.553885532 | 0.534 | 0.347 | 8.26E-29 | 3.02E-24 |
| NETO1 | 5 | -0.630790606 | 0.026 | 0.18 | 8.26E-29 | 3.03E-24 |

| | | | | | | |
|---|---|---|---|---|---|---|
| COL1A2 | 5 | -0.778559215 | 0.006 | 0.134 | 8.39E-29 | 3.07E-24 |
| LINC00667 | 5 | 0.408234901 | 0.237 | 0.079 | 8.56E-29 | 3.14E-24 |
| FLNB | 5 | 0.569870373 | 0.67 | 0.463 | 1.07E-28 | 3.93E-24 |
| IL1RAPL2 | 5 | -1.714003458 | 0.035 | 0.196 | 1.23E-28 | 4.5E-24 |
| THSD4 | 5 | 0.48515653 | 0.354 | 0.15 | 1.24E-28 | 4.53E-24 |
| TBC1D5 | 5 | 0.453599715 | 0.943 | 0.859 | 1.35E-28 | 4.94E-24 |
| NMNAT3 | 5 | 0.462138131 | 0.33 | 0.14 | 1.36E-28 | 4.97E-24 |
| CFH | 5 | -0.760493659 | 0.013 | 0.155 | 1.37E-28 | 5.02E-24 |
| NELL1 | 5 | -1.160656942 | 0.134 | 0.323 | 1.44E-28 | 5.26E-24 |
| JMJD1C | 5 | 0.393564884 | 0.978 | 0.943 | 1.48E-28 | 5.41E-24 |
| PTPRN | 5 | 0.275676236 | 0.138 | 0.025 | 1.71E-28 | 6.24E-24 |
| E2F3 | 5 | -0.634484265 | 0.387 | 0.612 | 1.72E-28 | 6.3E-24 |
| SLC39A10 | 5 | -0.584495633 | 0.235 | 0.468 | 1.76E-28 | 6.46E-24 |
| GSN | 5 | -0.506307189 | 0.057 | 0.237 | 1.86E-28 | 6.83E-24 |
| SH3D19 | 5 | 0.531531326 | 0.763 | 0.567 | 2.16E-28 | 7.91E-24 |
| MSRA | 5 | -0.740390731 | 0.228 | 0.434 | 2.27E-28 | 8.32E-24 |
| COL6A2 | 5 | -0.406609766 | 0.011 | 0.146 | 2.55E-28 | 9.33E-24 |
| LAMP2 | 5 | -0.508569202 | 0.099 | 0.302 | 2.97E-28 | 1.09E-23 |
| LINC01138 | 5 | 0.427736544 | 0.277 | 0.104 | 3.03E-28 | 1.11E-23 |
| SSX2IP | 5 | -0.691830908 | 0.306 | 0.523 | 3.07E-28 | 1.12E-23 |
| CDC14B | 5 | -0.588230698 | 0.066 | 0.253 | 3.42E-28 | 1.25E-23 |
| LBX1-AS1 | 5 | -0.36151819 | 0.013 | 0.152 | 3.43E-28 | 1.26E-23 |
| ANXA2 | 5 | -0.550665383 | 0.084 | 0.28 | 4.33E-28 | 1.58E-23 |
| RGS22 | 5 | 0.455066936 | 0.193 | 0.053 | 4.59E-28 | 1.68E-23 |
| ACLY | 5 | 0.537772422 | 0.47 | 0.281 | 5.04E-28 | 1.85E-23 |
| CFAP299 | 5 | 0.256828727 | 0.15 | 0.035 | 5.2E-28 | 1.9E-23 |
| WWC2 | 5 | -0.710034022 | 0.194 | 0.415 | 5.66E-28 | 2.07E-23 |
| QDPR | 5 | 0.440385785 | 0.264 | 0.101 | 5.72E-28 | 2.09E-23 |
| CAMSAP3 | 5 | 0.270449318 | 0.11 | 0.016 | 5.82E-28 | 2.13E-23 |
| PLK2 | 5 | 0.485251947 | 0.172 | 0.05 | 6.1E-28 | 2.23E-23 |
| MTCL1 | 5 | -0.506552181 | 0.062 | 0.24 | 6.52E-28 | 2.39E-23 |
| OSBPL5 | 5 | -0.325876519 | 0.011 | 0.146 | 6.96E-28 | 2.55E-23 |
| RAD51B | 5 | 0.538068164 | 0.833 | 0.664 | 7.16E-28 | 2.62E-23 |
| VGLL4 | 5 | -0.669381829 | 0.404 | 0.599 | 7.43E-28 | 2.72E-23 |
| NWD1 | 5 | 0.332165817 | 0.139 | 0.026 | 8.16E-28 | 2.99E-23 |
| ZFX | 5 | 0.538050559 | 0.635 | 0.468 | 8.39E-28 | 3.07E-23 |
| NTN1 | 5 | -0.585630736 | 0.015 | 0.154 | 8.69E-28 | 3.18E-23 |
| TMEM59 | 5 | 0.50919067 | 0.394 | 0.213 | 9.26E-28 | 3.39E-23 |
| TBCD | 5 | -0.598995442 | 0.372 | 0.599 | 9.91E-28 | 3.63E-23 |
| SDK1 | 5 | -0.910941899 | 0.961 | 0.901 | 1.02E-27 | 3.72E-23 |
| NCALD | 5 | -0.785200031 | 0.55 | 0.568 | 1.1E-27 | 4.04E-23 |
| AL138720.1 | 5 | -0.646062374 | 0.024 | 0.17 | 1.11E-27 | 4.07E-23 |
| RNF150 | 5 | 0.654235234 | 0.571 | 0.38 | 1.15E-27 | 4.2E-23 |
| NFATC2 | 5 | 0.404376919 | 0.345 | 0.147 | 1.23E-27 | 4.49E-23 |
| NFKB1 | 5 | 0.544158106 | 0.475 | 0.291 | 1.44E-27 | 5.28E-23 |
| HOXC4 | 5 | -0.315187317 | 0.006 | 0.131 | 1.48E-27 | 5.42E-23 |
| PPP1CB | 5 | 0.503531975 | 0.585 | 0.41 | 1.59E-27 | 5.81E-23 |
| AC006148.1 | 5 | -0.621731622 | 0.02 | 0.166 | 1.6E-27 | 5.87E-23 |
| MICAL3 | 5 | 0.558477432 | 0.62 | 0.454 | 1.75E-27 | 6.41E-23 |
| RASAL2 | 5 | 0.56740503 | 0.716 | 0.528 | 2E-27 | 7.33E-23 |
| KCNMA1 | 5 | -0.877310905 | 0.103 | 0.3 | 2.04E-27 | 7.47E-23 |
| AC022893.2 | 5 | 0.252212452 | 0.106 | 0.015 | 2.16E-27 | 7.92E-23 |
| MYT1L | 5 | -0.882068986 | 0.013 | 0.137 | 2.3E-27 | 8.42E-23 |
| GRK5 | 5 | 0.621833165 | 0.517 | 0.318 | 2.46E-27 | 9E-23 |
| MET | 5 | -0.566275763 | 0.048 | 0.214 | 2.48E-27 | 9.09E-23 |
| AC108690.1 | 5 | -0.510828857 | 0.006 | 0.131 | 2.54E-27 | 9.28E-23 |
| PDSS2 | 5 | 0.495992086 | 0.787 | 0.604 | 2.54E-27 | 9.3E-23 |
| XYLT1 | 5 | -0.567091679 | 0.013 | 0.146 | 2.55E-27 | 9.33E-23 |

| | | | | | | |
|---|---|---|---|---|---|---|
| SCUBE2 | 5 | -0.602727739 | 0.028 | 0.18 | 2.8E-27 | 1.02E-22 |
| C20orf194 | 5 | -0.493568037 | 0.248 | 0.483 | 2.9E-27 | 1.06E-22 |
| TCEA3 | 5 | 0.372433125 | 0.207 | 0.064 | 2.92E-27 | 1.07E-22 |
| ZNF644 | 5 | 0.499444654 | 0.831 | 0.733 | 2.94E-27 | 1.08E-22 |
| SASS6 | 5 | 0.553733741 | 0.497 | 0.304 | 2.99E-27 | 1.09E-22 |
| SYT4 | 5 | 0.357712386 | 0.15 | 0.032 | 3.21E-27 | 1.17E-22 |
| MAP2 | 5 | -0.893100637 | 0.514 | 0.547 | 3.35E-27 | 1.23E-22 |
| AHCYL2 | 5 | -0.878062761 | 0.549 | 0.674 | 3.4E-27 | 1.24E-22 |
| AC006230.1 | 5 | -0.356292894 | 0.013 | 0.15 | 3.58E-27 | 1.31E-22 |
| DPP10-AS1 | 5 | 0.317388437 | 0.204 | 0.058 | 3.6E-27 | 1.32E-22 |
| SMO | 5 | -0.34529155 | 0.024 | 0.173 | 3.65E-27 | 1.34E-22 |
| PELI1 | 5 | -0.634379512 | 0.106 | 0.3 | 3.83E-27 | 1.4E-22 |
| COLEC12 | 5 | -0.598305191 | 0.011 | 0.141 | 4.16E-27 | 1.52E-22 |
| AP001636.3 | 5 | 0.578069188 | 0.262 | 0.107 | 4.54E-27 | 1.66E-22 |
| DIXDC1 | 5 | -0.457798345 | 0.075 | 0.264 | 4.88E-27 | 1.79E-22 |
| CNTNAP3B | 5 | -0.791064938 | 0.132 | 0.317 | 5.25E-27 | 1.92E-22 |
| ST13 | 5 | 0.52704109 | 0.481 | 0.303 | 5.43E-27 | 1.99E-22 |
| POLA1 | 5 | 0.552906488 | 0.919 | 0.84 | 5.54E-27 | 2.03E-22 |
| MBD5 | 5 | -0.529573428 | 0.811 | 0.903 | 5.59E-27 | 2.05E-22 |
| ARSE | 5 | 0.28534904 | 0.13 | 0.024 | 6.58E-27 | 2.41E-22 |
| MAST4 | 5 | 0.401751752 | 0.587 | 0.349 | 6.61E-27 | 2.42E-22 |
| AL596214.1 | 5 | 0.375083121 | 0.198 | 0.058 | 7.71E-27 | 2.82E-22 |
| MROH1 | 5 | 0.501607561 | 0.426 | 0.227 | 7.86E-27 | 2.88E-22 |
| C12orf56 | 5 | -0.398905653 | 0.02 | 0.16 | 8.31E-27 | 3.04E-22 |
| SLC1A3 | 5 | -0.454175906 | 0.002 | 0.115 | 9.01E-27 | 3.3E-22 |
| LINC01776 | 5 | -0.592368455 | 0.04 | 0.198 | 9.75E-27 | 3.57E-22 |
| MED14 | 5 | -0.577202708 | 0.473 | 0.653 | 9.76E-27 | 3.57E-22 |
| NFXL1 | 5 | 0.526626777 | 0.521 | 0.344 | 1.03E-26 | 3.78E-22 |
| SLCO5A1 | 5 | -0.523703096 | 0.02 | 0.158 | 1.19E-26 | 4.35E-22 |
| OLFM2 | 5 | -0.388065054 | 0.007 | 0.13 | 1.21E-26 | 4.44E-22 |
| PTPRK | 5 | 0.408181376 | 0.925 | 0.801 | 1.41E-26 | 5.17E-22 |
| NRIP1 | 5 | 0.500604838 | 0.772 | 0.599 | 1.56E-26 | 5.72E-22 |
| GPRC5B | 5 | -0.397818502 | 0.044 | 0.206 | 1.63E-26 | 5.98E-22 |
| TLL2 | 5 | -0.603167822 | 0.024 | 0.167 | 1.65E-26 | 6.04E-22 |
| ZXDC | 5 | 0.503898482 | 0.541 | 0.378 | 1.89E-26 | 6.91E-22 |
| PROM1 | 5 | 0.474580394 | 0.259 | 0.091 | 2.06E-26 | 7.53E-22 |
| ZNF804B | 5 | -1.723570922 | 0.039 | 0.193 | 2.22E-26 | 8.13E-22 |
| ZNF589 | 5 | -0.505212971 | 0.171 | 0.387 | 2.24E-26 | 8.2E-22 |
| NUP93 | 5 | -0.555368081 | 0.172 | 0.382 | 2.34E-26 | 8.58E-22 |
| CNTN5 | 5 | -1.630897819 | 0.017 | 0.148 | 2.99E-26 | 1.09E-21 |
| SLFN11 | 5 | -0.283364153 | 0.004 | 0.119 | 3.18E-26 | 1.16E-21 |
| CRYZ | 5 | 0.402950897 | 0.261 | 0.1 | 3.3E-26 | 1.21E-21 |
| MIR124-2HG | 5 | -0.415365626 | 0.011 | 0.138 | 3.34E-26 | 1.22E-21 |
| ZNF326 | 5 | 0.482508058 | 0.574 | 0.445 | 3.61E-26 | 1.32E-21 |
| AC016042.1 | 5 | -0.557928487 | 0.077 | 0.245 | 3.82E-26 | 1.4E-21 |
| AC092422.1 | 5 | -0.787502985 | 0.026 | 0.17 | 3.91E-26 | 1.43E-21 |
| GRIP2 | 5 | -0.398426261 | 0.026 | 0.172 | 4.2E-26 | 1.54E-21 |
| TXNDC11 | 5 | 0.512885131 | 0.374 | 0.198 | 4.49E-26 | 1.65E-21 |
| EPHA4 | 5 | -0.783819261 | 0.407 | 0.599 | 4.6E-26 | 1.69E-21 |
| KDM7A | 5 | 0.575808969 | 0.55 | 0.344 | 5.04E-26 | 1.84E-21 |
| PHACTR2 | 5 | 0.516364126 | 0.752 | 0.557 | 5.31E-26 | 1.95E-21 |
| FOXP1 | 5 | 0.415098424 | 0.923 | 0.807 | 5.81E-26 | 2.13E-21 |
| NLRC5 | 5 | -0.434738888 | 0.026 | 0.169 | 6.91E-26 | 2.53E-21 |
| FANCC | 5 | 0.505412929 | 0.697 | 0.509 | 7.37E-26 | 2.7E-21 |
| MIR4435-2HG | 5 | -0.644243766 | 0.072 | 0.246 | 7.44E-26 | 2.72E-21 |
| FRMD3 | 5 | 0.494126586 | 0.583 | 0.351 | 7.67E-26 | 2.81E-21 |
| LRP1 | 5 | -0.426574777 | 0.068 | 0.246 | 7.97E-26 | 2.92E-21 |
| ST3GAL4 | 5 | -0.439256558 | 0.053 | 0.22 | 8.73E-26 | 3.2E-21 |

| | | | | | | |
|---|---|---|---|---|---|---|
| CASK | 5 | -0.581448193 | 0.776 | 0.864 | 8.9E-26 | 3.26E-21 |
| KBTBD12 | 5 | -0.472460857 | 0.057 | 0.227 | 8.92E-26 | 3.27E-21 |
| ARHGEF6 | 5 | -0.404792905 | 0.022 | 0.161 | 9.34E-26 | 3.42E-21 |
| AC245297.3 | 5 | 0.448684051 | 0.327 | 0.154 | 9.69E-26 | 3.55E-21 |
| FGF12 | 5 | -0.703922869 | 0.806 | 0.732 | 1.13E-25 | 4.15E-21 |
| HECTD1 | 5 | -0.612092308 | 0.706 | 0.807 | 1.16E-25 | 4.24E-21 |
| ANKRD50 | 5 | -0.575158067 | 0.132 | 0.326 | 1.19E-25 | 4.36E-21 |
| C3orf70 | 5 | -0.546843205 | 0.092 | 0.276 | 1.22E-25 | 4.47E-21 |
| ARAP3 | 5 | -0.309866095 | 0.009 | 0.132 | 1.24E-25 | 4.53E-21 |
| AKNA | 5 | -0.378674447 | 0.051 | 0.219 | 1.31E-25 | 4.81E-21 |
| ERO1B | 5 | 0.566682843 | 0.261 | 0.12 | 1.42E-25 | 5.22E-21 |
| GLDC | 5 | -0.641361914 | 0.108 | 0.288 | 1.47E-25 | 5.39E-21 |
| SLC7A5 | 5 | 0.403071042 | 0.226 | 0.078 | 1.5E-25 | 5.48E-21 |
| COL4A1 | 5 | -0.479831085 | 0.017 | 0.143 | 1.5E-25 | 5.48E-21 |
| PHF3 | 5 | 0.467123159 | 0.738 | 0.622 | 1.53E-25 | 5.58E-21 |
| NMT2 | 5 | -0.519783105 | 0.136 | 0.34 | 1.58E-25 | 5.77E-21 |
| TMEM263 | 5 | 0.409333041 | 0.325 | 0.142 | 1.59E-25 | 5.83E-21 |
| MITF | 5 | 0.339545733 | 0.371 | 0.177 | 1.63E-25 | 5.95E-21 |
| LPAR1 | 5 | -0.80193673 | 0.053 | 0.215 | 1.67E-25 | 6.11E-21 |
| SLC41A2 | 5 | 0.409237889 | 0.345 | 0.154 | 1.72E-25 | 6.31E-21 |
| CHST15 | 5 | -0.356936674 | 0.009 | 0.129 | 1.72E-25 | 6.31E-21 |
| FBN1 | 5 | -0.498993524 | 0.039 | 0.193 | 1.87E-25 | 6.85E-21 |
| MPDZ | 5 | -0.609508401 | 0.532 | 0.703 | 2.04E-25 | 7.47E-21 |
| NLGN2 | 5 | -0.286375295 | 0.009 | 0.133 | 2.05E-25 | 7.51E-21 |
| HS2ST1 | 5 | 0.510235424 | 0.738 | 0.634 | 2.13E-25 | 7.8E-21 |
| ANK2 | 5 | -0.755835298 | 0.877 | 0.838 | 2.16E-25 | 7.91E-21 |
| PKIB | 5 | 0.558785519 | 0.585 | 0.379 | 2.51E-25 | 9.18E-21 |
| LINC02307 | 5 | -0.469219972 | 0.013 | 0.142 | 2.56E-25 | 9.39E-21 |
| PCOLCE2 | 5 | -0.619964365 | 0.172 | 0.361 | 2.62E-25 | 9.59E-21 |
| PLEKHA5 | 5 | -0.64313818 | 0.793 | 0.858 | 2.64E-25 | 9.66E-21 |
| TMEM65 | 5 | 0.488292337 | 0.442 | 0.231 | 2.68E-25 | 9.83E-21 |
| PRRX1 | 5 | -0.496546641 | 0.002 | 0.11 | 2.79E-25 | 1.02E-20 |
| KCNMB4 | 5 | -0.53218907 | 0.073 | 0.251 | 2.81E-25 | 1.03E-20 |
| TSPAN5 | 5 | 0.394095024 | 0.71 | 0.479 | 2.82E-25 | 1.03E-20 |
| GLYATL1 | 5 | 0.322515047 | 0.128 | 0.03 | 2.86E-25 | 1.05E-20 |
| FKBP5 | 5 | 0.524737393 | 0.514 | 0.306 | 3.22E-25 | 1.18E-20 |
| AC131571.1 | 5 | 0.30078424 | 0.165 | 0.043 | 3.27E-25 | 1.2E-20 |
| APP | 5 | -0.541516718 | 0.796 | 0.873 | 3.4E-25 | 1.25E-20 |
| CCDC144A | 5 | -0.393256311 | 0.039 | 0.193 | 3.5E-25 | 1.28E-20 |
| NUAK1 | 5 | 0.457679947 | 0.242 | 0.096 | 3.55E-25 | 1.3E-20 |
| CATSPERE | 5 | -0.630270573 | 0.077 | 0.251 | 3.82E-25 | 1.4E-20 |
| IQCJ-SCHIP1 | 5 | -0.860126629 | 0.033 | 0.182 | 3.86E-25 | 1.41E-20 |
| PLS3 | 5 | 0.389688895 | 0.62 | 0.39 | 3.88E-25 | 1.42E-20 |
| MINAR1 | 5 | -0.421028407 | 0.029 | 0.173 | 3.9E-25 | 1.43E-20 |
| BRINP3 | 5 | -1.293707056 | 0.479 | 0.511 | 4.2E-25 | 1.54E-20 |
| ARIH2 | 5 | 0.483381266 | 0.639 | 0.462 | 4.39E-25 | 1.61E-20 |
| CHPT1 | 5 | -0.744537237 | 0.426 | 0.575 | 4.69E-25 | 1.72E-20 |
| MGMT | 5 | 0.540760927 | 0.611 | 0.402 | 4.76E-25 | 1.74E-20 |
| RBP1 | 5 | -0.3726507 | 0.037 | 0.19 | 4.92E-25 | 1.8E-20 |
| TWSG1 | 5 | -0.437691329 | 0.072 | 0.248 | 5.4E-25 | 1.98E-20 |
| MBNL2 | 5 | 0.476788306 | 0.622 | 0.396 | 5.52E-25 | 2.02E-20 |
| KLHDC8A | 5 | -0.413726023 | 0.033 | 0.178 | 5.68E-25 | 2.08E-20 |
| DNAH11 | 5 | -1.154598884 | 0.099 | 0.268 | 5.79E-25 | 2.12E-20 |
| CPNE5 | 5 | -0.434331056 | 0.011 | 0.134 | 5.85E-25 | 2.14E-20 |
| SPHKAP | 5 | -0.619630552 | 0.007 | 0.121 | 6.31E-25 | 2.31E-20 |
| DACT1 | 5 | -0.343396714 | 0.006 | 0.12 | 7.29E-25 | 2.67E-20 |
| YPEL2 | 5 | 0.522327287 | 0.325 | 0.156 | 7.54E-25 | 2.76E-20 |
| FOXK1 | 5 | 0.515921638 | 0.611 | 0.427 | 7.54E-25 | 2.76E-20 |

| | | | | | | |
|---|---|---|---|---|---|---|
| COL18A1 | 5 | -0.474961126 | 0.013 | 0.14 | 8.72E-25 | 3.19E-20 |
| CEP112 | 5 | 0.506152545 | 0.6 | 0.384 | 8.79E-25 | 3.22E-20 |
| PMEPA1 | 5 | -0.552007262 | 0.066 | 0.238 | 9.06E-25 | 3.32E-20 |
| CD109 | 5 | -0.423442048 | 0.018 | 0.147 | 9.71E-25 | 3.56E-20 |
| ST3GAL1 | 5 | -0.391232753 | 0.028 | 0.172 | 1.04E-24 | 3.8E-20 |
| VOPP1 | 5 | 0.501520824 | 0.517 | 0.322 | 1.05E-24 | 3.85E-20 |
| CPEB3 | 5 | 0.51526646 | 0.576 | 0.378 | 1.07E-24 | 3.9E-20 |
| FOCAD | 5 | -0.592204441 | 0.582 | 0.721 | 1.14E-24 | 4.17E-20 |
| RETREG1 | 5 | -0.45772059 | 0.033 | 0.181 | 1.15E-24 | 4.22E-20 |
| NGF | 5 | -0.397696407 | 0.011 | 0.131 | 1.22E-24 | 4.48E-20 |
| RALGPS1 | 5 | 0.540052267 | 0.453 | 0.268 | 1.3E-24 | 4.76E-20 |
| AP000829.1 | 5 | 0.570158909 | 0.47 | 0.264 | 1.34E-24 | 4.9E-20 |
| BICD1 | 5 | -0.630321317 | 0.695 | 0.811 | 1.35E-24 | 4.93E-20 |
| FREM1 | 5 | -0.929036781 | 0.095 | 0.271 | 1.38E-24 | 5.06E-20 |
| SLC25A53 | 5 | 0.434994505 | 0.294 | 0.134 | 1.43E-24 | 5.24E-20 |
| SPSB4 | 5 | -0.315129518 | 0.009 | 0.129 | 2.11E-24 | 7.72E-20 |
| ZSWIM5 | 5 | 0.397640342 | 0.598 | 0.37 | 2.18E-24 | 7.98E-20 |
| PCDH10 | 5 | -0.48059105 | 0.015 | 0.14 | 2.27E-24 | 8.32E-20 |
| RGS12 | 5 | -0.534429218 | 0.268 | 0.485 | 2.3E-24 | 8.41E-20 |
| SYBU | 5 | -0.808477034 | 0.235 | 0.423 | 2.7E-24 | 9.89E-20 |
| RAP2A | 5 | -0.557399865 | 0.13 | 0.318 | 2.78E-24 | 1.02E-19 |
| BCAT2 | 5 | 0.401986147 | 0.218 | 0.085 | 2.82E-24 | 1.03E-19 |
| STXBP1 | 5 | 0.478308319 | 0.51 | 0.317 | 2.92E-24 | 1.07E-19 |
| SMAD3 | 5 | -0.579745979 | 0.16 | 0.351 | 2.98E-24 | 1.09E-19 |
| COL4A5 | 5 | -1.141956524 | 0.006 | 0.117 | 3.02E-24 | 1.11E-19 |
| PRKD3 | 5 | 0.480472962 | 0.591 | 0.397 | 3.51E-24 | 1.28E-19 |
| RFTN1 | 5 | -0.381246646 | 0.024 | 0.161 | 3.63E-24 | 1.33E-19 |
| EPHA5-AS1 | 5 | -0.346887753 | 0.007 | 0.12 | 3.71E-24 | 1.36E-19 |
| DDR2 | 5 | -0.49694117 | 0.055 | 0.216 | 3.94E-24 | 1.44E-19 |
| AC023469.2 | 5 | -0.278261273 | 0.002 | 0.105 | 3.97E-24 | 1.45E-19 |
| TANC2 | 5 | -0.722275558 | 0.587 | 0.723 | 3.99E-24 | 1.46E-19 |
| RAPGEF1 | 5 | 0.507089099 | 0.567 | 0.387 | 4.01E-24 | 1.47E-19 |
| NHSL1 | 5 | 0.353258181 | 0.778 | 0.565 | 4.88E-24 | 1.79E-19 |
| LINC01456 | 5 | -0.466297816 | 0.013 | 0.136 | 5.31E-24 | 1.95E-19 |
| ARHGAP28 | 5 | -0.693795258 | 0.09 | 0.265 | 5.45E-24 | 1.99E-19 |
| DRAIC | 5 | 0.34482013 | 0.136 | 0.036 | 5.6E-24 | 2.05E-19 |
| RBMS1 | 5 | -0.687813268 | 0.358 | 0.559 | 5.63E-24 | 2.06E-19 |
| IL13RA1 | 5 | 0.388703037 | 0.183 | 0.059 | 5.79E-24 | 2.12E-19 |
| NBPF1 | 5 | -0.462866277 | 0.156 | 0.356 | 6.01E-24 | 2.2E-19 |
| MYH10 | 5 | -0.490698258 | 0.554 | 0.748 | 6.01E-24 | 2.2E-19 |
| RNF13 | 5 | 0.490672252 | 0.617 | 0.447 | 6.23E-24 | 2.28E-19 |
| LINC02802 | 5 | 0.298265869 | 0.165 | 0.046 | 7.2E-24 | 2.64E-19 |
| CFAP300 | 5 | 0.256799757 | 0.125 | 0.025 | 7.71E-24 | 2.82E-19 |
| LINC00937 | 5 | -0.50947942 | 0.046 | 0.197 | 7.82E-24 | 2.86E-19 |
| LINC00894 | 5 | -0.528337379 | 0.165 | 0.369 | 8.18E-24 | 2.99E-19 |
| CRNDE | 5 | -0.49909721 | 0.127 | 0.319 | 8.71E-24 | 3.19E-19 |
| NUMB | 5 | 0.439026664 | 0.734 | 0.57 | 8.75E-24 | 3.2E-19 |
| GRAMD4 | 5 | 0.439625137 | 0.321 | 0.152 | 8.8E-24 | 3.22E-19 |
| MAML2 | 5 | -0.751137103 | 0.58 | 0.516 | 9.05E-24 | 3.31E-19 |
| EMB | 5 | 0.424030373 | 0.305 | 0.139 | 9.13E-24 | 3.34E-19 |
| NREP | 5 | -0.545221756 | 0.226 | 0.428 | 1.03E-23 | 3.78E-19 |
| EXTL3 | 5 | 0.400551566 | 0.292 | 0.126 | 1.06E-23 | 3.86E-19 |
| GJC1 | 5 | -0.546416651 | 0.13 | 0.316 | 1.14E-23 | 4.17E-19 |
| ADGRL3 | 5 | -1.003625223 | 0.8 | 0.815 | 1.19E-23 | 4.37E-19 |
| ADCY1 | 5 | 0.444817857 | 0.29 | 0.123 | 1.24E-23 | 4.54E-19 |
| RBFOX1 | 5 | -0.475391075 | 0.873 | 0.726 | 1.27E-23 | 4.65E-19 |
| EDIL3 | 5 | -0.369246027 | 0.741 | 0.612 | 1.4E-23 | 5.13E-19 |
| DENND2A | 5 | 0.455796255 | 0.534 | 0.318 | 1.42E-23 | 5.19E-19 |

| | | | | | | |
|---|---|---|---|---|---|---|
| DYNC1H1 | 5 | -0.528466889 | 0.418 | 0.626 | 1.44E-23 | 5.28E-19 |
| LRRC37B | 5 | -0.423788392 | 0.092 | 0.266 | 1.46E-23 | 5.33E-19 |
| SLC10A4 | 5 | -0.284076882 | 0.002 | 0.104 | 1.48E-23 | 5.42E-19 |
| AC092939.1 | 5 | -0.391834567 | 0.044 | 0.196 | 1.57E-23 | 5.75E-19 |
| UBE2K | 5 | 0.409519759 | 0.851 | 0.776 | 1.6E-23 | 5.84E-19 |
| ARHGAP1 | 5 | 0.414620636 | 0.229 | 0.093 | 1.69E-23 | 6.2E-19 |
| RAPGEF2 | 5 | -0.723701883 | 0.523 | 0.663 | 1.92E-23 | 7.05E-19 |
| OTX2 | 5 | -0.466963933 | 0.002 | 0.101 | 1.97E-23 | 7.22E-19 |
| EFHC2 | 5 | -0.45834418 | 0.018 | 0.146 | 2.01E-23 | 7.36E-19 |
| SLC16A2 | 5 | -0.466427354 | 0.167 | 0.372 | 2.09E-23 | 7.64E-19 |
| ZNF100 | 5 | 0.452669321 | 0.352 | 0.191 | 2.12E-23 | 7.76E-19 |
| NCOA3 | 5 | -0.606650582 | 0.42 | 0.601 | 2.15E-23 | 7.87E-19 |
| KCTD8 | 5 | -1.033499174 | 0.026 | 0.154 | 2.18E-23 | 7.98E-19 |
| CTNNB1 | 5 | -0.591814322 | 0.472 | 0.672 | 2.2E-23 | 8.07E-19 |
| TES | 5 | 0.428921541 | 0.248 | 0.094 | 2.48E-23 | 9.08E-19 |
| AXIN2 | 5 | -0.376996112 | 0.039 | 0.185 | 2.9E-23 | 1.06E-18 |
| STEAP1B | 5 | -0.469683645 | 0.051 | 0.207 | 2.91E-23 | 1.06E-18 |
| MACROD2 | 5 | 0.429446803 | 0.763 | 0.557 | 2.95E-23 | 1.08E-18 |
| ASIC4 | 5 | 0.264780439 | 0.121 | 0.024 | 3E-23 | 1.1E-18 |
| FOXO1 | 5 | 0.425201503 | 0.396 | 0.199 | 3.23E-23 | 1.18E-18 |
| PPARA | 5 | 0.461441825 | 0.273 | 0.117 | 3.41E-23 | 1.25E-18 |
| SIL1 | 5 | -0.718247001 | 0.547 | 0.671 | 3.48E-23 | 1.27E-18 |
| AP004608.1 | 5 | 0.25500679 | 0.15 | 0.037 | 3.68E-23 | 1.35E-18 |
| EFHD1 | 5 | -0.406853001 | 0.018 | 0.142 | 3.71E-23 | 1.36E-18 |
| KIF13B | 5 | 0.435931102 | 0.442 | 0.243 | 3.71E-23 | 1.36E-18 |
| TECPR1 | 5 | 0.41798736 | 0.29 | 0.135 | 3.73E-23 | 1.36E-18 |
| ALPK3 | 5 | 0.287770443 | 0.127 | 0.03 | 3.91E-23 | 1.43E-18 |
| PGRMC2 | 5 | 0.393100443 | 0.29 | 0.137 | 4.05E-23 | 1.48E-18 |
| CYTOR | 5 | -0.418965377 | 0.026 | 0.16 | 4.2E-23 | 1.54E-18 |
| KLHL24 | 5 | -0.62954382 | 0.292 | 0.488 | 4.34E-23 | 1.59E-18 |
| TMSB15B | 5 | 0.386331808 | 0.292 | 0.132 | 4.4E-23 | 1.61E-18 |
| DCAF10 | 5 | 0.487936359 | 0.594 | 0.442 | 4.49E-23 | 1.65E-18 |
| HDAC8 | 5 | -0.533194391 | 0.576 | 0.715 | 4.83E-23 | 1.77E-18 |
| ETFDH | 5 | 0.502438685 | 0.389 | 0.218 | 5.1E-23 | 1.87E-18 |
| TMEM132C | 5 | -0.645477539 | 0.011 | 0.128 | 5.1E-23 | 1.87E-18 |
| PLCH1 | 5 | -0.863604643 | 0.165 | 0.332 | 5.22E-23 | 1.91E-18 |
| AGAP1 | 5 | 0.386960414 | 0.938 | 0.838 | 5.39E-23 | 1.97E-18 |
| LZTS1 | 5 | -0.373545781 | 0.035 | 0.178 | 5.39E-23 | 1.97E-18 |
| MAMDC2-AS1 | 5 | 0.317983814 | 0.163 | 0.046 | 5.42E-23 | 1.99E-18 |
| PPP2R1B | 5 | 0.440478563 | 0.378 | 0.197 | 5.53E-23 | 2.02E-18 |
| SRSF10 | 5 | 0.424520118 | 0.725 | 0.607 | 5.8E-23 | 2.12E-18 |
| SCG3 | 5 | 0.35872149 | 0.218 | 0.078 | 6.02E-23 | 2.2E-18 |
| CYB5R4 | 5 | 0.467469695 | 0.433 | 0.265 | 6.72E-23 | 2.46E-18 |
| TBC1D23 | 5 | -0.613577984 | 0.316 | 0.5 | 6.83E-23 | 2.5E-18 |
| TTC39B | 5 | 0.443857224 | 0.29 | 0.13 | 6.96E-23 | 2.55E-18 |
| AC004158.1 | 5 | -0.480641384 | 0.05 | 0.196 | 7.13E-23 | 2.61E-18 |
| RNF144A | 5 | -0.509389469 | 0.305 | 0.516 | 7.14E-23 | 2.61E-18 |
| SEMA5B | 5 | -0.34143703 | 0.013 | 0.132 | 7.84E-23 | 2.87E-18 |
| AC025809.1 | 5 | -0.315852275 | 0.011 | 0.124 | 8.67E-23 | 3.17E-18 |
| LINC02302 | 5 | -0.252885844 | 0.007 | 0.117 | 8.96E-23 | 3.28E-18 |
| YTHDC1 | 5 | 0.460221943 | 0.714 | 0.583 | 9.2E-23 | 3.37E-18 |
| AKAP7 | 5 | -0.506147121 | 0.092 | 0.251 | 9.4E-23 | 3.44E-18 |
| LCOR | 5 | -0.509022599 | 0.672 | 0.804 | 9.64E-23 | 3.53E-18 |
| CYP4X1 | 5 | -0.549642653 | 0.026 | 0.157 | 9.79E-23 | 3.58E-18 |
| CLIC4 | 5 | -0.54092706 | 0.176 | 0.371 | 1.04E-22 | 3.82E-18 |
| CAMSAP2 | 5 | -0.536954516 | 0.426 | 0.619 | 1.06E-22 | 3.89E-18 |
| AC009803.1 | 5 | -0.477529312 | 0.015 | 0.136 | 1.11E-22 | 4.06E-18 |
| PARD6G | 5 | -0.397161521 | 0.062 | 0.22 | 1.18E-22 | 4.32E-18 |

| | | | | | | |
|---|---|---|---|---|---|---|
| KCNK1 | 5 | 0.334272897 | 0.226 | 0.08 | 1.21E-22 | 4.43E-18 |
| SASH1 | 5 | -0.826163261 | 0.413 | 0.571 | 1.21E-22 | 4.45E-18 |
| MXRA5 | 5 | -0.390481113 | 0.013 | 0.128 | 1.28E-22 | 4.68E-18 |
| CYB5R3 | 5 | 0.3688426 | 0.198 | 0.071 | 1.38E-22 | 5.03E-18 |
| ANK3 | 5 | -0.99588177 | 0.176 | 0.339 | 1.4E-22 | 5.13E-18 |
| ACER3 | 5 | -0.496256706 | 0.163 | 0.352 | 1.41E-22 | 5.17E-18 |
| SORBS1 | 5 | -0.723925032 | 0.38 | 0.541 | 1.42E-22 | 5.2E-18 |
| JUN | 5 | 0.34241358 | 0.16 | 0.043 | 1.54E-22 | 5.65E-18 |
| SQSTM1 | 5 | 0.582423514 | 0.356 | 0.226 | 1.69E-22 | 6.18E-18 |
| ABHD18 | 5 | 0.480032649 | 0.604 | 0.437 | 1.74E-22 | 6.37E-18 |
| AK9 | 5 | 0.478970957 | 0.292 | 0.138 | 1.76E-22 | 6.44E-18 |
| ADH5 | 5 | 0.440456806 | 0.369 | 0.208 | 1.77E-22 | 6.47E-18 |
| RPS3 | 5 | 0.489238379 | 0.613 | 0.461 | 1.8E-22 | 6.59E-18 |
| OXNAD1 | 5 | 0.429540237 | 0.429 | 0.243 | 1.8E-22 | 6.59E-18 |
| ERBB3 | 5 | 0.339335121 | 0.246 | 0.094 | 1.82E-22 | 6.67E-18 |
| AL162718.1 | 5 | -0.547810225 | 0.022 | 0.144 | 1.93E-22 | 7.08E-18 |
| AC090578.1 | 5 | -0.627191538 | 0.114 | 0.287 | 2.09E-22 | 7.65E-18 |
| ZCCHC7 | 5 | 0.463469717 | 0.872 | 0.826 | 2.15E-22 | 7.88E-18 |
| EWSR1 | 5 | 0.423663099 | 0.789 | 0.669 | 2.36E-22 | 8.63E-18 |
| NOTCH2NLC | 5 | -0.318989239 | 0.024 | 0.153 | 2.4E-22 | 8.77E-18 |
| TMEM63C | 5 | 0.381601647 | 0.29 | 0.128 | 2.42E-22 | 8.87E-18 |
| FAM171A1 | 5 | -0.549861446 | 0.389 | 0.574 | 2.45E-22 | 8.97E-18 |
| RIMKLB | 5 | -0.473447265 | 0.24 | 0.445 | 2.46E-22 | 9.01E-18 |
| PDGFRB | 5 | -0.405040568 | 0.011 | 0.12 | 2.46E-22 | 9.02E-18 |
| MRTFA | 5 | 0.451525128 | 0.75 | 0.645 | 2.48E-22 | 9.09E-18 |
| AKAP13 | 5 | 0.393634521 | 0.747 | 0.54 | 2.52E-22 | 9.21E-18 |
| AC118549.1 | 5 | 0.455245134 | 0.798 | 0.675 | 2.53E-22 | 9.26E-18 |
| C1GALT1 | 5 | 0.411195969 | 0.406 | 0.215 | 2.56E-22 | 9.38E-18 |
| IRX2 | 5 | 0.428560727 | 0.279 | 0.126 | 2.62E-22 | 9.6E-18 |
| OSBPL8 | 5 | -0.60177313 | 0.475 | 0.637 | 2.7E-22 | 9.88E-18 |
| ITGA2 | 5 | 0.266801597 | 0.136 | 0.031 | 2.78E-22 | 1.02E-17 |
| CORO1C | 5 | -0.542218261 | 0.218 | 0.409 | 2.93E-22 | 1.07E-17 |
| SNAP91 | 5 | -0.527804818 | 0.048 | 0.193 | 2.94E-22 | 1.08E-17 |
| GFPT1 | 5 | 0.47405448 | 0.479 | 0.303 | 2.98E-22 | 1.09E-17 |
| ZFAND5 | 5 | -0.619127173 | 0.268 | 0.46 | 2.98E-22 | 1.09E-17 |
| GFOD1 | 5 | -0.531294977 | 0.16 | 0.346 | 3E-22 | 1.1E-17 |
| RAB27A | 5 | 0.464140253 | 0.391 | 0.228 | 3.28E-22 | 1.2E-17 |
| AC105460.2 | 5 | 0.325142531 | 0.112 | 0.03 | 3.63E-22 | 1.33E-17 |
| RBM39 | 5 | 0.340561071 | 0.956 | 0.854 | 3.73E-22 | 1.36E-17 |
| TMEFF2 | 5 | -0.618816703 | 0.026 | 0.154 | 3.87E-22 | 1.42E-17 |
| AC019205.1 | 5 | -0.408747153 | 0.103 | 0.281 | 3.89E-22 | 1.43E-17 |
| ATP13A3 | 5 | 0.50388788 | 0.708 | 0.531 | 3.97E-22 | 1.45E-17 |
| AMOTL1 | 5 | 0.462305234 | 0.536 | 0.338 | 4.01E-22 | 1.47E-17 |
| SHISA6 | 5 | -0.664987354 | 0.039 | 0.174 | 4.35E-22 | 1.59E-17 |
| JAZF1 | 5 | -0.714106495 | 0.609 | 0.74 | 4.43E-22 | 1.62E-17 |
| AC096747.1 | 5 | -0.271172325 | 0.006 | 0.107 | 4.81E-22 | 1.76E-17 |
| TTC39C | 5 | -0.492774038 | 0.248 | 0.454 | 4.93E-22 | 1.81E-17 |
| MMP17 | 5 | -0.302137967 | 0.013 | 0.128 | 5.27E-22 | 1.93E-17 |
| SORCS2 | 5 | -0.61503934 | 0.097 | 0.264 | 5.39E-22 | 1.97E-17 |
| ARID1B | 5 | -0.50137513 | 0.93 | 0.959 | 5.59E-22 | 2.05E-17 |
| FGF2 | 5 | -0.338044167 | 0.033 | 0.17 | 5.71E-22 | 2.09E-17 |
| LRP12 | 5 | -0.492155897 | 0.226 | 0.427 | 5.96E-22 | 2.18E-17 |
| MMP26 | 5 | 0.554999504 | 0.207 | 0.071 | 6.3E-22 | 2.31E-17 |
| VWC2 | 5 | -0.54765628 | 0.026 | 0.146 | 6.53E-22 | 2.39E-17 |
| DNAH2 | 5 | -0.349779444 | 0.024 | 0.149 | 6.57E-22 | 2.41E-17 |
| COL19A1 | 5 | -0.585173458 | 0.006 | 0.109 | 6.69E-22 | 2.45E-17 |
| STK24 | 5 | 0.478930337 | 0.618 | 0.463 | 7.19E-22 | 2.63E-17 |
| PHACTR3 | 5 | -0.424159967 | 0.015 | 0.131 | 7.25E-22 | 2.65E-17 |

| | | | | | | |
|---|---|---|---|---|---|---|
| HOXC6 | 5 | -0.292301938 | 0.011 | 0.119 | 8.35E-22 | 3.06E-17 |
| BTBD9 | 5 | 0.515464605 | 0.71 | 0.561 | 8.76E-22 | 3.21E-17 |
| AL589740.1 | 5 | -0.940942665 | 0.09 | 0.226 | 9.15E-22 | 3.35E-17 |
| NSG1 | 5 | -0.276409114 | 0.011 | 0.122 | 9.42E-22 | 3.45E-17 |
| GDAP1 | 5 | 0.331692941 | 0.506 | 0.299 | 1.03E-21 | 3.79E-17 |
| KLHDC10 | 5 | 0.444058398 | 0.554 | 0.385 | 1.09E-21 | 3.97E-17 |
| CNTNAP3 | 5 | -0.406056804 | 0.028 | 0.158 | 1.1E-21 | 4.04E-17 |
| LINC01934 | 5 | -1.294648697 | 0.007 | 0.109 | 1.1E-21 | 4.04E-17 |
| ANKRD18B | 5 | 0.329119129 | 0.189 | 0.064 | 1.13E-21 | 4.14E-17 |
| GRAMD1B | 5 | -0.50425434 | 0.046 | 0.183 | 1.13E-21 | 4.14E-17 |
| DPH6 | 5 | 0.527981456 | 0.554 | 0.382 | 1.17E-21 | 4.3E-17 |
| TSNARE1 | 5 | 0.394548201 | 0.277 | 0.125 | 1.18E-21 | 4.31E-17 |
| PRRC2C | 5 | 0.399971017 | 0.872 | 0.774 | 1.24E-21 | 4.53E-17 |
| ARHGAP10 | 5 | 0.475958525 | 0.677 | 0.501 | 1.25E-21 | 4.59E-17 |
| HMGN2 | 5 | 0.487361305 | 0.565 | 0.377 | 1.28E-21 | 4.7E-17 |
| CLEC2A | 5 | -0.485612738 | 0.02 | 0.136 | 1.28E-21 | 4.7E-17 |
| FN1 | 5 | -1.031719407 | 0.209 | 0.378 | 1.31E-21 | 4.78E-17 |
| ZFPM1 | 5 | 0.310211923 | 0.156 | 0.044 | 1.44E-21 | 5.29E-17 |
| XXYLT1 | 5 | -0.470096062 | 0.165 | 0.354 | 1.49E-21 | 5.47E-17 |
| MN1 | 5 | -0.271489055 | 0.006 | 0.107 | 1.53E-21 | 5.61E-17 |
| CERS6 | 5 | -0.521838523 | 0.734 | 0.82 | 1.67E-21 | 6.1E-17 |
| NR3C1 | 5 | 0.473592964 | 0.556 | 0.351 | 1.69E-21 | 6.2E-17 |
| SAMD11 | 5 | -0.385207706 | 0.022 | 0.145 | 1.73E-21 | 6.35E-17 |
| EFCAB2 | 5 | 0.465780312 | 0.495 | 0.307 | 1.76E-21 | 6.43E-17 |
| FBXL5 | 5 | -0.501756069 | 0.305 | 0.497 | 1.91E-21 | 7E-17 |
| STOX2 | 5 | -0.636863913 | 0.143 | 0.316 | 1.97E-21 | 7.21E-17 |
| INPP5F | 5 | -0.494211045 | 0.213 | 0.399 | 2.03E-21 | 7.44E-17 |
| NAALADL2 | 5 | -0.793910805 | 0.284 | 0.423 | 2.08E-21 | 7.61E-17 |
| KCTD1 | 5 | -0.581898671 | 0.158 | 0.325 | 2.11E-21 | 7.74E-17 |
| COX5A | 5 | 0.370697221 | 0.277 | 0.124 | 2.17E-21 | 7.96E-17 |
| WDR17 | 5 | 0.448165037 | 0.541 | 0.338 | 2.38E-21 | 8.72E-17 |
| TMEM63A | 5 | -0.334545608 | 0.02 | 0.14 | 2.44E-21 | 8.95E-17 |
| CCND1 | 5 | -0.396554067 | 0.048 | 0.192 | 2.45E-21 | 8.97E-17 |
| ZNF433-AS1 | 5 | -0.274553723 | 0.013 | 0.125 | 2.66E-21 | 9.72E-17 |
| MEF2C-AS2 | 5 | -0.348251386 | 0.013 | 0.124 | 2.68E-21 | 9.8E-17 |
| SOX13 | 5 | -0.349333301 | 0.028 | 0.156 | 2.72E-21 | 9.97E-17 |
| SRRM3 | 5 | 0.32935699 | 0.323 | 0.151 | 2.88E-21 | 1.05E-16 |
| CAMK2B | 5 | 0.400004731 | 0.308 | 0.142 | 3.05E-21 | 1.12E-16 |
| PCDH15 | 5 | -1.158096388 | 0.026 | 0.143 | 3.06E-21 | 1.12E-16 |
| ATP1B1 | 5 | 0.478762619 | 0.422 | 0.259 | 3.1E-21 | 1.14E-16 |
| AC079781.5 | 5 | -0.486055633 | 0.244 | 0.445 | 3.45E-21 | 1.26E-16 |
| PRICKLE2 | 5 | -0.584635819 | 0.171 | 0.35 | 3.48E-21 | 1.27E-16 |
| TAPT1 | 5 | 0.432865295 | 0.371 | 0.209 | 3.58E-21 | 1.31E-16 |
| ST6GALNAC3 | 5 | -0.90198913 | 0.123 | 0.275 | 3.75E-21 | 1.37E-16 |
| DUXAP8 | 5 | -0.278856643 | 0.007 | 0.11 | 3.81E-21 | 1.4E-16 |
| KCNG3 | 5 | -0.398548121 | 0.039 | 0.172 | 3.82E-21 | 1.4E-16 |
| CNTNAP5 | 5 | -1.108172229 | 0.145 | 0.284 | 4.16E-21 | 1.52E-16 |
| LINC01128 | 5 | 0.29120297 | 0.189 | 0.065 | 4.16E-21 | 1.52E-16 |
| STUM | 5 | 0.332264845 | 0.233 | 0.088 | 4.22E-21 | 1.55E-16 |
| SHOX2 | 5 | -0.512394301 | 0.048 | 0.188 | 4.34E-21 | 1.59E-16 |
| HBEGF | 5 | -0.380355838 | 0.017 | 0.128 | 4.49E-21 | 1.64E-16 |
| CPLX1 | 5 | 0.288829871 | 0.136 | 0.035 | 4.59E-21 | 1.68E-16 |
| AL691420.1 | 5 | -0.991032446 | 0.026 | 0.141 | 4.76E-21 | 1.74E-16 |
| MEGF6 | 5 | -0.289564105 | 0.011 | 0.119 | 4.95E-21 | 1.81E-16 |
| PLXNB1 | 5 | 0.412593511 | 0.292 | 0.139 | 5.42E-21 | 1.99E-16 |
| ANAPC5 | 5 | 0.418606748 | 0.672 | 0.508 | 5.56E-21 | 2.04E-16 |
| IL6R | 5 | 0.340675667 | 0.16 | 0.051 | 5.77E-21 | 2.11E-16 |
| DDHD1 | 5 | -0.515466475 | 0.382 | 0.575 | 5.84E-21 | 2.14E-16 |

| | | | | | | |
|---|---|---|---|---|---|---|
| MSRB3 | 5 | -0.336588612 | 0.006 | 0.104 | 6.14E-21 | 2.25E-16 |
| EGLN3 | 5 | -0.558630547 | 0.106 | 0.257 | 6.2E-21 | 2.27E-16 |
| MSN | 5 | -0.290949179 | 0.022 | 0.144 | 6.43E-21 | 2.35E-16 |
| CCDC14 | 5 | 0.413531165 | 0.772 | 0.708 | 6.6E-21 | 2.42E-16 |
| DLG3 | 5 | -0.444446642 | 0.125 | 0.301 | 6.71E-21 | 2.46E-16 |
| CRTC1 | 5 | 0.495756531 | 0.455 | 0.295 | 7.42E-21 | 2.72E-16 |
| MYBL1 | 5 | -0.54550127 | 0.095 | 0.254 | 7.59E-21 | 2.78E-16 |
| SLC8A2 | 5 | -0.40039123 | 0.033 | 0.162 | 7.63E-21 | 2.79E-16 |
| CSNK1A1 | 5 | -0.448954412 | 0.747 | 0.851 | 7.77E-21 | 2.84E-16 |
| VWDE | 5 | -0.690599186 | 0.389 | 0.4 | 8.12E-21 | 2.97E-16 |
| LINC00536 | 5 | -0.824209408 | 0.006 | 0.101 | 8.48E-21 | 3.1E-16 |
| MOB3B | 5 | -0.45922 | 0.064 | 0.214 | 8.5E-21 | 3.11E-16 |
| CPT1C | 5 | -0.314032359 | 0.029 | 0.156 | 8.75E-21 | 3.2E-16 |
| LINC01956 | 5 | -0.367671387 | 0.013 | 0.123 | 9.02E-21 | 3.3E-16 |
| ZNF519 | 5 | -0.519863011 | 0.237 | 0.421 | 9.32E-21 | 3.41E-16 |
| RPGR | 5 | -0.558966016 | 0.27 | 0.459 | 9.62E-21 | 3.52E-16 |
| KIF7 | 5 | -0.299854698 | 0.029 | 0.156 | 9.99E-21 | 3.66E-16 |
| NEK8 | 5 | 0.267455942 | 0.127 | 0.034 | 1.02E-20 | 3.75E-16 |
| EPDR1 | 5 | -0.250575246 | 0.006 | 0.104 | 1.07E-20 | 3.9E-16 |
| FAM168A | 5 | -0.519108557 | 0.532 | 0.691 | 1.09E-20 | 4E-16 |
| BAZ2B | 5 | -0.551476646 | 0.872 | 0.919 | 1.12E-20 | 4.1E-16 |
| LMCD1-AS1 | 5 | 0.523419016 | 0.49 | 0.294 | 1.14E-20 | 4.16E-16 |
| UBTD1 | 5 | -0.267596578 | 0.007 | 0.109 | 1.32E-20 | 4.83E-16 |
| IMMP2L | 5 | 0.492079694 | 0.899 | 0.815 | 1.39E-20 | 5.09E-16 |
| KLHL1 | 5 | -0.851528311 | 0.007 | 0.108 | 1.44E-20 | 5.28E-16 |
| NBPF26 | 5 | -0.346130356 | 0.055 | 0.202 | 1.49E-20 | 5.44E-16 |
| MXI1 | 5 | -0.487188135 | 0.182 | 0.37 | 1.53E-20 | 5.6E-16 |
| KCNH8 | 5 | -0.669139425 | 0.62 | 0.634 | 1.66E-20 | 6.07E-16 |
| ACOT9 | 5 | -0.345254265 | 0.053 | 0.198 | 1.7E-20 | 6.24E-16 |
| LINC02615 | 5 | 0.460685772 | 0.268 | 0.137 | 1.75E-20 | 6.4E-16 |
| WRN | 5 | 0.463866013 | 0.739 | 0.625 | 1.83E-20 | 6.69E-16 |
| LIMA1 | 5 | 0.406478179 | 0.305 | 0.143 | 1.83E-20 | 6.7E-16 |
| SKAP2 | 5 | -0.631169767 | 0.055 | 0.19 | 2.08E-20 | 7.61E-16 |
| CPXM1 | 5 | -0.303666978 | 0.026 | 0.149 | 2.2E-20 | 8.04E-16 |
| TCF3 | 5 | -0.47337551 | 0.196 | 0.38 | 3.02E-20 | 1.11E-15 |
| LINC00290 | 5 | -0.262754359 | 0.009 | 0.112 | 3.13E-20 | 1.15E-15 |
| ZFYVE9 | 5 | 0.450424184 | 0.635 | 0.533 | 3.48E-20 | 1.28E-15 |
| DNAJC24 | 5 | 0.413156706 | 0.363 | 0.197 | 3.64E-20 | 1.33E-15 |
| LRP8 | 5 | -0.436026385 | 0.128 | 0.303 | 3.68E-20 | 1.35E-15 |
| ATF7IP | 5 | -0.560979916 | 0.383 | 0.544 | 3.7E-20 | 1.36E-15 |
| RNF34 | 5 | 0.396231278 | 0.295 | 0.156 | 3.71E-20 | 1.36E-15 |
| TFAP2E | 5 | -0.281620879 | 0.011 | 0.112 | 3.79E-20 | 1.39E-15 |
| LMCD1 | 5 | 0.325019913 | 0.204 | 0.075 | 3.95E-20 | 1.45E-15 |
| HOMER2 | 5 | -0.495879293 | 0.154 | 0.322 | 3.97E-20 | 1.45E-15 |
| ATE1 | 5 | 0.417860281 | 0.771 | 0.629 | 4.07E-20 | 1.49E-15 |
| ADAM19 | 5 | -0.275878591 | 0.006 | 0.101 | 4.25E-20 | 1.56E-15 |
| SSBP4 | 5 | 0.36828476 | 0.303 | 0.15 | 4.29E-20 | 1.57E-15 |
| NOP58 | 5 | 0.449396309 | 0.675 | 0.533 | 4.35E-20 | 1.59E-15 |
| CLIP2 | 5 | -0.486132198 | 0.143 | 0.323 | 4.49E-20 | 1.64E-15 |
| NBEA | 5 | -0.280759043 | 0.842 | 0.694 | 4.64E-20 | 1.7E-15 |
| ATP2B2 | 5 | -0.368600062 | 0.011 | 0.114 | 4.92E-20 | 1.8E-15 |
| FARP1 | 5 | 0.36023516 | 0.855 | 0.72 | 5.11E-20 | 1.87E-15 |
| HLA-B | 5 | -0.328077443 | 0.024 | 0.142 | 5.33E-20 | 1.95E-15 |
| EPB41L4A | 5 | -0.447100601 | 0.013 | 0.115 | 5.46E-20 | 2E-15 |
| AKT3 | 5 | 0.395063792 | 0.908 | 0.832 | 5.55E-20 | 2.03E-15 |
| CYP27C1 | 5 | -0.26453541 | 0.006 | 0.1 | 6.55E-20 | 2.4E-15 |
| SPPL2B | 5 | 0.398507472 | 0.317 | 0.17 | 6.69E-20 | 2.45E-15 |
| PITPNC1 | 5 | -0.628348291 | 0.389 | 0.561 | 7.06E-20 | 2.58E-15 |

| | | | | | | |
|---|---|---|---|---|---|---|
| HLA-C | 5 | -0.304383495 | 0.018 | 0.129 | 7.6E-20 | 2.78E-15 |
| SPOPL | 5 | 0.433522233 | 0.55 | 0.366 | 7.61E-20 | 2.79E-15 |
| EFNB2 | 5 | -0.384330667 | 0.037 | 0.166 | 7.77E-20 | 2.85E-15 |
| ELF2 | 5 | -0.534519006 | 0.697 | 0.798 | 7.95E-20 | 2.91E-15 |
| MEIS3 | 5 | -0.265243661 | 0.017 | 0.127 | 7.96E-20 | 2.91E-15 |
| ADPRHL2 | 5 | 0.292690916 | 0.213 | 0.08 | 8.47E-20 | 3.1E-15 |
| LINC02263 | 5 | -0.631545681 | 0.011 | 0.112 | 8.98E-20 | 3.29E-15 |
| SEMA4D | 5 | -0.442661575 | 0.167 | 0.344 | 1.04E-19 | 3.8E-15 |
| NTN4 | 5 | -0.443316069 | 0.055 | 0.195 | 1.08E-19 | 3.95E-15 |
| ARAP1 | 5 | 0.369243275 | 0.206 | 0.108 | 1.08E-19 | 3.95E-15 |
| SGO2 | 5 | 0.408893841 | 0.229 | 0.106 | 1.08E-19 | 3.96E-15 |
| CREB5 | 5 | -0.652859953 | 0.07 | 0.212 | 1.21E-19 | 4.44E-15 |
| FAM160A1 | 5 | 0.470463238 | 0.299 | 0.138 | 1.38E-19 | 5.04E-15 |
| AC063938.1 | 5 | -0.503875017 | 0.013 | 0.113 | 1.41E-19 | 5.15E-15 |
| PDE4B | 5 | -0.98137045 | 0.026 | 0.139 | 1.44E-19 | 5.27E-15 |
| KCNIP1 | 5 | -0.369376946 | 0.011 | 0.111 | 1.56E-19 | 5.72E-15 |
| C13orf42 | 5 | -0.47938041 | 0.024 | 0.138 | 1.57E-19 | 5.75E-15 |
| SLC25A24 | 5 | -0.379526066 | 0.099 | 0.26 | 1.61E-19 | 5.91E-15 |
| RICTOR | 5 | 0.392430836 | 0.835 | 0.707 | 1.64E-19 | 6.01E-15 |
| HSPB1 | 5 | 0.674419536 | 0.182 | 0.071 | 1.67E-19 | 6.1E-15 |
| BCAS4 | 5 | 0.276648242 | 0.147 | 0.044 | 1.71E-19 | 6.25E-15 |
| ZNF395 | 5 | 0.313091076 | 0.154 | 0.048 | 1.77E-19 | 6.47E-15 |
| RHOQ | 5 | -0.428995115 | 0.132 | 0.295 | 1.79E-19 | 6.54E-15 |
| CSGALNACT1 | 5 | -0.463666636 | 0.018 | 0.13 | 1.79E-19 | 6.55E-15 |
| LNX2 | 5 | 0.408565148 | 0.27 | 0.131 | 1.83E-19 | 6.69E-15 |
| SBF2 | 5 | 0.416513825 | 0.833 | 0.747 | 1.94E-19 | 7.12E-15 |
| ROBO1 | 5 | -0.58825053 | 0.987 | 0.896 | 1.98E-19 | 7.24E-15 |
| TMEM51 | 5 | 0.3544363 | 0.283 | 0.127 | 2.14E-19 | 7.85E-15 |
| EVI5 | 5 | 0.396106429 | 0.723 | 0.547 | 2.14E-19 | 7.85E-15 |
| USP3 | 5 | 0.464608478 | 0.765 | 0.69 | 2.16E-19 | 7.93E-15 |
| DMC1 | 5 | 0.309009409 | 0.156 | 0.049 | 2.19E-19 | 8.01E-15 |
| AC074286.1 | 5 | -0.403256751 | 0.017 | 0.118 | 2.24E-19 | 8.19E-15 |
| ITFG1 | 5 | -0.563931004 | 0.701 | 0.797 | 2.27E-19 | 8.3E-15 |
| KIAA1211 | 5 | -0.606437963 | 0.606 | 0.702 | 2.45E-19 | 8.96E-15 |
| PLCL2 | 5 | 0.410196285 | 0.754 | 0.579 | 2.48E-19 | 9.07E-15 |
| APH1B | 5 | 0.377530627 | 0.262 | 0.121 | 2.51E-19 | 9.17E-15 |
| ANXA4 | 5 | 0.404892345 | 0.406 | 0.229 | 2.53E-19 | 9.26E-15 |
| NAV1 | 5 | 0.474520927 | 0.583 | 0.402 | 2.63E-19 | 9.64E-15 |
| PIP4P2 | 5 | 0.364789636 | 0.261 | 0.118 | 2.81E-19 | 1.03E-14 |
| HACE1 | 5 | -0.589209131 | 0.479 | 0.627 | 2.81E-19 | 1.03E-14 |
| RNF182 | 5 | 0.28212542 | 0.154 | 0.046 | 2.96E-19 | 1.08E-14 |
| SULF2 | 5 | -0.459721764 | 0.062 | 0.202 | 2.99E-19 | 1.09E-14 |
| COCH | 5 | 0.394208835 | 0.406 | 0.223 | 3.02E-19 | 1.11E-14 |
| PLXNA2 | 5 | -0.620364924 | 0.228 | 0.393 | 3.12E-19 | 1.14E-14 |
| RANBP2 | 5 | 0.419090516 | 0.626 | 0.498 | 3.15E-19 | 1.15E-14 |
| TTYH3 | 5 | -0.355455254 | 0.064 | 0.209 | 3.4E-19 | 1.25E-14 |
| CYTL1 | 5 | -0.257077053 | 0.007 | 0.103 | 3.41E-19 | 1.25E-14 |
| CD164 | 5 | 0.392385178 | 0.314 | 0.176 | 3.66E-19 | 1.34E-14 |
| GLI3 | 5 | -0.65841968 | 0.224 | 0.352 | 3.8E-19 | 1.39E-14 |
| C7orf50 | 5 | 0.422398689 | 0.383 | 0.232 | 4.11E-19 | 1.5E-14 |
| JAM2 | 5 | -0.277081883 | 0.009 | 0.105 | 4.26E-19 | 1.56E-14 |
| PTER | 5 | 0.325232881 | 0.2 | 0.077 | 4.31E-19 | 1.58E-14 |
| KLF13 | 5 | -0.366215139 | 0.062 | 0.204 | 4.38E-19 | 1.6E-14 |
| SNAP25-AS1 | 5 | 0.279294664 | 0.211 | 0.08 | 4.62E-19 | 1.69E-14 |
| HSP90AA1 | 5 | 1.044674858 | 0.58 | 0.448 | 4.62E-19 | 1.69E-14 |
| AC109466.1 | 5 | -1.15291994 | 0.017 | 0.119 | 4.67E-19 | 1.71E-14 |
| PDE7A | 5 | -0.499966679 | 0.47 | 0.621 | 4.81E-19 | 1.76E-14 |
| HSPA4L | 5 | 0.423543432 | 0.374 | 0.203 | 4.82E-19 | 1.76E-14 |

| | | | | | | |
|---|---|---|---|---|---|---|
| ADAMTS2 | 5 | 0.319296564 | 0.213 | 0.081 | 5.11E-19 | 1.87E-14 |
| MT-CO1 | 5 | -0.583142754 | 0.273 | 0.47 | 5.45E-19 | 2E-14 |
| NPAS2 | 5 | -0.436245721 | 0.05 | 0.178 | 6.37E-19 | 2.33E-14 |
| CDH4 | 5 | -0.794196432 | 0.591 | 0.621 | 6.79E-19 | 2.48E-14 |
| SPAG9 | 5 | -0.463488999 | 0.736 | 0.846 | 8.09E-19 | 2.96E-14 |
| NUDT4 | 5 | 0.392362645 | 0.29 | 0.153 | 8.46E-19 | 3.1E-14 |
| HSDL2 | 5 | 0.411390251 | 0.396 | 0.239 | 8.85E-19 | 3.24E-14 |
| HABP4 | 5 | -0.332098195 | 0.048 | 0.179 | 8.93E-19 | 3.27E-14 |
| SLC29A1 | 5 | 0.271949403 | 0.152 | 0.049 | 9.12E-19 | 3.34E-14 |
| AC006972.1 | 5 | -0.407461243 | 0.011 | 0.107 | 9.78E-19 | 3.58E-14 |
| CNTN4 | 5 | -0.955969821 | 0.04 | 0.157 | 1E-18 | 3.68E-14 |
| AHR | 5 | -0.576056382 | 0.053 | 0.181 | 1.04E-18 | 3.79E-14 |
| FAM149A | 5 | 0.420997786 | 0.497 | 0.307 | 1.04E-18 | 3.81E-14 |
| RANBP17 | 5 | 0.400104386 | 0.848 | 0.729 | 1.06E-18 | 3.87E-14 |
| ZNF217 | 5 | -0.300799171 | 0.029 | 0.148 | 1.06E-18 | 3.87E-14 |
| ABCB10 | 5 | 0.398943103 | 0.343 | 0.198 | 1.07E-18 | 3.91E-14 |
| KIAA0825 | 5 | -0.480351222 | 0.084 | 0.235 | 1.07E-18 | 3.93E-14 |
| SGPL1 | 5 | -0.347875373 | 0.07 | 0.209 | 1.08E-18 | 3.94E-14 |
| CECR2 | 5 | 0.275567915 | 0.488 | 0.301 | 1.08E-18 | 3.96E-14 |
| AC008050.1 | 5 | 0.276568279 | 0.105 | 0.025 | 1.24E-18 | 4.53E-14 |
| RAB8B | 5 | -0.521921052 | 0.163 | 0.33 | 1.25E-18 | 4.56E-14 |
| KDM5B | 5 | -0.523636448 | 0.235 | 0.423 | 1.25E-18 | 4.58E-14 |
| THRB | 5 | -0.606130612 | 0.328 | 0.309 | 1.29E-18 | 4.72E-14 |
| BAZ1A | 5 | -0.480731494 | 0.428 | 0.607 | 1.29E-18 | 4.73E-14 |
| CREB3L2 | 5 | 0.441026599 | 0.406 | 0.242 | 1.34E-18 | 4.9E-14 |
| ID2 | 5 | 0.670996914 | 0.288 | 0.153 | 1.35E-18 | 4.94E-14 |
| MAGI1 | 5 | 0.314307106 | 0.945 | 0.86 | 1.35E-18 | 4.96E-14 |
| VPS13C | 5 | 0.385110152 | 0.853 | 0.732 | 1.43E-18 | 5.24E-14 |
| LHFPL2 | 5 | 0.431770433 | 0.396 | 0.224 | 1.44E-18 | 5.26E-14 |
| STAT3 | 5 | -0.482762993 | 0.27 | 0.455 | 1.47E-18 | 5.39E-14 |
| FBLN7 | 5 | 0.262822183 | 0.149 | 0.044 | 1.48E-18 | 5.42E-14 |
| NOTCH1 | 5 | -0.315654728 | 0.013 | 0.113 | 1.5E-18 | 5.49E-14 |
| TSPAN18 | 5 | -0.521326508 | 0.127 | 0.286 | 1.53E-18 | 5.61E-14 |
| PLCE1 | 5 | 0.328972104 | 0.325 | 0.162 | 1.62E-18 | 5.93E-14 |
| P4HB | 5 | 0.441394502 | 0.369 | 0.222 | 1.63E-18 | 5.98E-14 |
| WASF2 | 5 | 0.396005353 | 0.499 | 0.322 | 1.67E-18 | 6.11E-14 |
| AF117829.1 | 5 | -0.495108132 | 0.198 | 0.372 | 1.71E-18 | 6.25E-14 |
| HTR7 | 5 | -0.465641627 | 0.022 | 0.131 | 1.9E-18 | 6.94E-14 |
| ZFHX3 | 5 | -0.665578379 | 0.437 | 0.591 | 1.92E-18 | 7.03E-14 |
| RFLNA | 5 | -0.432180934 | 0.108 | 0.267 | 1.97E-18 | 7.21E-14 |
| ZDHHC9 | 5 | 0.341908202 | 0.279 | 0.131 | 2.07E-18 | 7.59E-14 |
| SACS | 5 | -0.451863235 | 0.172 | 0.346 | 2.08E-18 | 7.61E-14 |
| WDFY3 | 5 | 0.334739182 | 0.936 | 0.895 | 2.26E-18 | 8.27E-14 |
| UBE2D2 | 5 | 0.406877146 | 0.697 | 0.584 | 2.29E-18 | 8.4E-14 |
| PLEKHG1 | 5 | -0.453156029 | 0.028 | 0.14 | 2.39E-18 | 8.75E-14 |
| FAM174B | 5 | 0.380885706 | 0.284 | 0.136 | 2.45E-18 | 8.97E-14 |
| PDZD2 | 5 | -0.567744604 | 0.215 | 0.399 | 2.73E-18 | 1E-13 |
| ACAP3 | 5 | 0.400647848 | 0.339 | 0.191 | 2.77E-18 | 1.01E-13 |
| ADORA1 | 5 | -0.257066345 | 0.013 | 0.112 | 2.87E-18 | 1.05E-13 |
| AL133346.1 | 5 | -0.512472746 | 0.037 | 0.154 | 3.07E-18 | 1.12E-13 |
| ZHX3 | 5 | 0.4252515 | 0.464 | 0.307 | 3.14E-18 | 1.15E-13 |
| EDEM3 | 5 | 0.425805124 | 0.492 | 0.354 | 3.21E-18 | 1.17E-13 |
| PRR14L | 5 | 0.464163745 | 0.494 | 0.348 | 3.36E-18 | 1.23E-13 |
| MIR4713HG | 5 | -0.418470756 | 0.017 | 0.119 | 3.64E-18 | 1.33E-13 |
| GLIS3 | 5 | -0.965278237 | 0.169 | 0.252 | 3.64E-18 | 1.33E-13 |
| CDH8 | 5 | -0.521839227 | 0.028 | 0.139 | 3.68E-18 | 1.35E-13 |
| ARPC5 | 5 | -0.375433248 | 0.13 | 0.295 | 3.93E-18 | 1.44E-13 |
| ANKDD1A | 5 | -0.357336098 | 0.046 | 0.169 | 3.93E-18 | 1.44E-13 |

| | | | | | | |
|---|---|---|---|---|---|---|
| GNAL | 5 | -0.366610677 | 0.024 | 0.129 | 3.99E-18 | 1.46E-13 |
| SEC61A2 | 5 | -0.327097449 | 0.072 | 0.214 | 4.03E-18 | 1.48E-13 |
| XRN2 | 5 | -0.424723037 | 0.466 | 0.638 | 4.04E-18 | 1.48E-13 |
| AC093817.2 | 5 | -0.736731917 | 0.017 | 0.11 | 4.28E-18 | 1.57E-13 |
| XKR6 | 5 | 0.434568967 | 0.69 | 0.517 | 4.4E-18 | 1.61E-13 |
| SLC25A20 | 5 | 0.257254152 | 0.145 | 0.044 | 4.72E-18 | 1.73E-13 |
| SLC13A3 | 5 | -0.337005979 | 0.04 | 0.163 | 4.74E-18 | 1.74E-13 |
| MED27 | 5 | 0.400238576 | 0.541 | 0.427 | 5.16E-18 | 1.89E-13 |
| GNPAT | 5 | 0.401767277 | 0.404 | 0.257 | 5.19E-18 | 1.9E-13 |
| RABL6 | 5 | 0.447774125 | 0.506 | 0.366 | 5.75E-18 | 2.1E-13 |
| ASB13 | 5 | 0.291355099 | 0.178 | 0.066 | 5.77E-18 | 2.11E-13 |
| KCNAB3 | 5 | -0.270253605 | 0.026 | 0.137 | 5.97E-18 | 2.19E-13 |
| SOX2-OT | 5 | -1.728985879 | 0.105 | 0.211 | 5.98E-18 | 2.19E-13 |
| MAP2K6 | 5 | -0.457794959 | 0.117 | 0.274 | 6.49E-18 | 2.38E-13 |
| SLC9B2 | 5 | 0.375974723 | 0.305 | 0.162 | 6.57E-18 | 2.4E-13 |
| RGS2 | 5 | 0.465213938 | 0.18 | 0.078 | 6.7E-18 | 2.45E-13 |
| R3HDM1 | 5 | -0.504726178 | 0.796 | 0.872 | 6.94E-18 | 2.54E-13 |
| CFLAR | 5 | 0.364039204 | 0.242 | 0.119 | 7.05E-18 | 2.58E-13 |
| AL356108.1 | 5 | -0.734559907 | 0.017 | 0.117 | 7.12E-18 | 2.61E-13 |
| CBFB | 5 | 0.401446389 | 0.675 | 0.504 | 7.29E-18 | 2.67E-13 |
| EML4 | 5 | 0.410111776 | 0.774 | 0.676 | 7.3E-18 | 2.67E-13 |
| CUL4B | 5 | -0.484528556 | 0.264 | 0.435 | 7.36E-18 | 2.69E-13 |
| NAV2 | 5 | 0.345497177 | 0.65 | 0.457 | 7.51E-18 | 2.75E-13 |
| SOX4 | 5 | -0.444764088 | 0.121 | 0.283 | 7.65E-18 | 2.8E-13 |
| GPR137C | 5 | -0.604716937 | 0.273 | 0.439 | 7.71E-18 | 2.82E-13 |
| ENOSF1 | 5 | 0.401796769 | 0.732 | 0.591 | 7.74E-18 | 2.83E-13 |
| PPFIBP2 | 5 | 0.32410884 | 0.204 | 0.079 | 7.96E-18 | 2.91E-13 |
| WIPI1 | 5 | 0.303333851 | 0.202 | 0.08 | 8.67E-18 | 3.18E-13 |
| SFMBT2 | 5 | -0.631559409 | 0.319 | 0.503 | 8.68E-18 | 3.18E-13 |
| CCDC12 | 5 | 0.370198213 | 0.299 | 0.156 | 8.77E-18 | 3.21E-13 |
| KDM4A | 5 | -0.378923803 | 0.136 | 0.3 | 9.39E-18 | 3.44E-13 |
| SCG5 | 5 | 0.33865717 | 0.255 | 0.127 | 9.52E-18 | 3.48E-13 |
| CDK14 | 5 | -0.525223166 | 0.675 | 0.785 | 9.72E-18 | 3.56E-13 |
| CLMN | 5 | -0.419255004 | 0.066 | 0.2 | 9.77E-18 | 3.58E-13 |
| ELAVL3 | 5 | -0.440190239 | 0.048 | 0.172 | 1.01E-17 | 3.7E-13 |
| C11orf54 | 5 | 0.402601411 | 0.365 | 0.223 | 1.04E-17 | 3.81E-13 |
| KDM4C | 5 | 0.389871031 | 0.763 | 0.604 | 1.05E-17 | 3.85E-13 |
| CARMIL3 | 5 | -0.296900721 | 0.035 | 0.144 | 1.08E-17 | 3.95E-13 |
| SCN1A | 5 | -0.565950649 | 0.013 | 0.105 | 1.1E-17 | 4.03E-13 |
| AC011586.2 | 5 | -0.68939655 | 0.013 | 0.101 | 1.12E-17 | 4.09E-13 |
| PFKFB3 | 5 | 0.515842763 | 0.464 | 0.306 | 1.2E-17 | 4.38E-13 |
| INVS | 5 | 0.408476537 | 0.684 | 0.545 | 1.21E-17 | 4.42E-13 |
| DSCAM | 5 | -0.870579067 | 0.024 | 0.123 | 1.21E-17 | 4.43E-13 |
| PHLDB1 | 5 | -0.280682909 | 0.031 | 0.147 | 1.23E-17 | 4.51E-13 |
| KIAA1328 | 5 | 0.442825221 | 0.732 | 0.6 | 1.28E-17 | 4.68E-13 |
| AL157944.1 | 5 | -0.749206115 | 0.011 | 0.104 | 1.39E-17 | 5.07E-13 |
| AC006058.1 | 5 | -0.281354009 | 0.015 | 0.11 | 1.44E-17 | 5.26E-13 |
| LRRC8B | 5 | -0.59718021 | 0.29 | 0.435 | 1.49E-17 | 5.44E-13 |
| MAOB | 5 | -0.318965648 | 0.018 | 0.117 | 1.59E-17 | 5.83E-13 |
| SOS1 | 5 | -0.515019196 | 0.705 | 0.797 | 1.67E-17 | 6.1E-13 |
| PTGES3 | 5 | 0.415803174 | 0.624 | 0.48 | 1.71E-17 | 6.26E-13 |
| AMMECR1 | 5 | 0.441594406 | 0.486 | 0.327 | 1.76E-17 | 6.45E-13 |
| CPSF6 | 5 | -0.40464465 | 0.765 | 0.852 | 1.78E-17 | 6.51E-13 |
| AEBP2 | 5 | -0.615423389 | 0.651 | 0.728 | 1.84E-17 | 6.72E-13 |
| USP53 | 5 | 0.40189858 | 0.428 | 0.256 | 1.86E-17 | 6.81E-13 |
| DOCK1 | 5 | 0.390307659 | 0.802 | 0.663 | 1.96E-17 | 7.17E-13 |
| SLC17A9 | 5 | 0.287076912 | 0.132 | 0.046 | 1.98E-17 | 7.26E-13 |
| HIRA | 5 | 0.320664954 | 0.257 | 0.12 | 1.99E-17 | 7.28E-13 |

| | | | | | | |
|---|---|---|---|---|---|---|
| PRKCB | 5 | -0.456249802 | 0.017 | 0.11 | 2.08E-17 | 7.61E-13 |
| LMAN1 | 5 | 0.407652876 | 0.42 | 0.276 | 2.25E-17 | 8.24E-13 |
| CTC1 | 5 | 0.300675521 | 0.185 | 0.071 | 2.34E-17 | 8.55E-13 |
| BOC | 5 | -0.490502906 | 0.075 | 0.21 | 2.41E-17 | 8.84E-13 |
| BRINP2 | 5 | -0.408036108 | 0.017 | 0.112 | 2.47E-17 | 9.03E-13 |
| SYCP2L | 5 | 0.334405203 | 0.25 | 0.111 | 2.65E-17 | 9.72E-13 |
| WIPF1 | 5 | -0.351748041 | 0.059 | 0.191 | 2.76E-17 | 1.01E-12 |
| CPEB4 | 5 | 0.361624587 | 0.323 | 0.167 | 2.94E-17 | 1.08E-12 |
| ADGRA2 | 5 | -0.352899273 | 0.035 | 0.151 | 2.98E-17 | 1.09E-12 |
| MARK4 | 5 | -0.35055568 | 0.117 | 0.275 | 3.01E-17 | 1.1E-12 |
| CCDC150 | 5 | 0.456861417 | 0.457 | 0.287 | 3.02E-17 | 1.11E-12 |
| PRDM5 | 5 | 0.407259386 | 0.49 | 0.311 | 3.04E-17 | 1.11E-12 |
| SFMBT1 | 5 | 0.404685314 | 0.594 | 0.416 | 3.14E-17 | 1.15E-12 |
| PTPN21 | 5 | 0.324402688 | 0.273 | 0.13 | 3.33E-17 | 1.22E-12 |
| FRMPD3 | 5 | -0.282707601 | 0.02 | 0.119 | 3.33E-17 | 1.22E-12 |
| MIPOL1 | 5 | 0.395181561 | 0.683 | 0.528 | 3.43E-17 | 1.26E-12 |
| MTMR10 | 5 | -0.442874199 | 0.196 | 0.361 | 3.44E-17 | 1.26E-12 |
| GALNT7 | 5 | -0.638447443 | 0.484 | 0.602 | 3.78E-17 | 1.39E-12 |
| AC004917.1 | 5 | -0.406231521 | 0.013 | 0.104 | 3.81E-17 | 1.4E-12 |
| MAPK14 | 5 | 0.386272789 | 0.563 | 0.421 | 4.31E-17 | 1.58E-12 |
| BMP6 | 5 | -0.403793828 | 0.017 | 0.112 | 4.48E-17 | 1.64E-12 |
| ESYT1 | 5 | 0.329831449 | 0.207 | 0.091 | 4.72E-17 | 1.73E-12 |
| IFT74 | 5 | 0.354439139 | 0.334 | 0.178 | 4.77E-17 | 1.75E-12 |
| TIPARP | 5 | 0.329606329 | 0.237 | 0.11 | 4.89E-17 | 1.79E-12 |
| FANCL | 5 | 0.387300124 | 0.745 | 0.615 | 4.91E-17 | 1.8E-12 |
| ARHGEF17 | 5 | -0.267621473 | 0.018 | 0.118 | 5.04E-17 | 1.85E-12 |
| CNNM1 | 5 | 0.357305168 | 0.253 | 0.12 | 5.23E-17 | 1.91E-12 |
| NAALAD2 | 5 | -0.48946629 | 0.121 | 0.246 | 5.33E-17 | 1.95E-12 |
| RALGAPA1 | 5 | 0.372025159 | 0.811 | 0.732 | 5.74E-17 | 2.1E-12 |
| TBC1D22A | 5 | 0.394724359 | 0.771 | 0.661 | 5.76E-17 | 2.11E-12 |
| SH3RF1 | 5 | 0.411720318 | 0.684 | 0.533 | 5.85E-17 | 2.14E-12 |
| AGO3 | 5 | 0.357706752 | 0.794 | 0.692 | 5.85E-17 | 2.14E-12 |
| SCAMP5 | 5 | 0.330042804 | 0.246 | 0.116 | 5.87E-17 | 2.15E-12 |
| PCM1 | 5 | 0.352326878 | 0.873 | 0.792 | 5.93E-17 | 2.17E-12 |
| EIF5 | 5 | 0.383273171 | 0.409 | 0.248 | 5.94E-17 | 2.17E-12 |
| POLE2 | 5 | 0.359213766 | 0.488 | 0.31 | 7.01E-17 | 2.57E-12 |
| BNC2 | 5 | 0.476713004 | 0.35 | 0.188 | 7.36E-17 | 2.69E-12 |
| PALD1 | 5 | -0.358409691 | 0.07 | 0.202 | 7.38E-17 | 2.7E-12 |
| EVA1C | 5 | -0.29884769 | 0.024 | 0.126 | 7.41E-17 | 2.71E-12 |
| MYO1E | 5 | 0.389217319 | 0.44 | 0.269 | 7.78E-17 | 2.85E-12 |
| PPP1R14C | 5 | -0.589022873 | 0.45 | 0.447 | 7.82E-17 | 2.86E-12 |
| MVB12B | 5 | 0.483828951 | 0.558 | 0.412 | 7.94E-17 | 2.91E-12 |
| AC092691.1 | 5 | -0.563948066 | 0.011 | 0.101 | 8.08E-17 | 2.96E-12 |
| CNN3 | 5 | -0.418658484 | 0.141 | 0.294 | 8.1E-17 | 2.97E-12 |
| NACC2 | 5 | -0.402076985 | 0.103 | 0.25 | 8.34E-17 | 3.05E-12 |
| GAB2 | 5 | 0.270749146 | 0.479 | 0.3 | 8.48E-17 | 3.1E-12 |
| MIR924HG | 5 | -0.704351381 | 0.681 | 0.72 | 8.71E-17 | 3.19E-12 |
| FMNL2 | 5 | -0.520587454 | 0.818 | 0.878 | 8.87E-17 | 3.25E-12 |
| CUL1 | 5 | -0.443282638 | 0.422 | 0.587 | 9.28E-17 | 3.4E-12 |
| DUSP10 | 5 | 0.292963569 | 0.149 | 0.054 | 1.02E-16 | 3.75E-12 |
| MAPT | 5 | -0.363961439 | 0.031 | 0.141 | 1.19E-16 | 4.35E-12 |
| METTL24 | 5 | -0.315736898 | 0.017 | 0.109 | 1.32E-16 | 4.84E-12 |
| PHTF2 | 5 | 0.326307788 | 0.8 | 0.751 | 1.35E-16 | 4.94E-12 |
| NETO2 | 5 | -0.530855619 | 0.376 | 0.541 | 1.36E-16 | 4.97E-12 |
| RUNX1 | 5 | -0.572483663 | 0.024 | 0.124 | 1.43E-16 | 5.23E-12 |
| PTCH1 | 5 | 0.330940972 | 0.328 | 0.173 | 1.44E-16 | 5.29E-12 |
| AC107223.1 | 5 | -0.399198628 | 0.046 | 0.156 | 1.45E-16 | 5.29E-12 |
| AC012501.2 | 5 | -0.271729176 | 0.015 | 0.107 | 1.49E-16 | 5.47E-12 |

| | | | | | | |
|---|---|---|---|---|---|---|
| HPS4 | 5 | 0.365545971 | 0.356 | 0.208 | 1.54E-16 | 5.63E-12 |
| ARHGEF38 | 5 | 0.253951847 | 0.13 | 0.038 | 1.54E-16 | 5.64E-12 |
| NIPAL3 | 5 | 0.358978242 | 0.235 | 0.109 | 1.57E-16 | 5.74E-12 |
| RFX7 | 5 | -0.449582377 | 0.633 | 0.747 | 1.61E-16 | 5.89E-12 |
| MIAT | 5 | -0.295692743 | 0.017 | 0.112 | 1.65E-16 | 6.02E-12 |
| TPGS2 | 5 | 0.331386084 | 0.262 | 0.155 | 1.72E-16 | 6.29E-12 |
| STX3 | 5 | 0.315657287 | 0.231 | 0.116 | 1.74E-16 | 6.37E-12 |
| HHIP | 5 | -0.434066246 | 0.028 | 0.125 | 1.78E-16 | 6.51E-12 |
| XYLB | 5 | 0.370536383 | 0.453 | 0.291 | 1.98E-16 | 7.24E-12 |
| GPR89B | 5 | 0.363705068 | 0.288 | 0.162 | 2E-16 | 7.31E-12 |
| FLT4 | 5 | -0.257509086 | 0.013 | 0.103 | 2E-16 | 7.34E-12 |
| GUCY1A2 | 5 | -0.711335572 | 0.279 | 0.405 | 2.03E-16 | 7.43E-12 |
| URI1 | 5 | -0.461390085 | 0.618 | 0.759 | 2.1E-16 | 7.69E-12 |
| CHKB | 5 | 0.313365365 | 0.261 | 0.127 | 2.2E-16 | 8.07E-12 |
| CAPRIN2 | 5 | -0.399777915 | 0.141 | 0.299 | 2.28E-16 | 8.36E-12 |
| MECOM | 5 | -0.325420273 | 0.121 | 0.057 | 2.32E-16 | 8.51E-12 |
| ZC2HC1A | 5 | -0.461447934 | 0.237 | 0.403 | 2.36E-16 | 8.64E-12 |
| SPATA6 | 5 | 0.252694195 | 0.312 | 0.164 | 2.4E-16 | 8.8E-12 |
| SDK2 | 5 | -0.42170562 | 0.028 | 0.129 | 2.42E-16 | 8.84E-12 |
| FRMPD4 | 5 | 0.326822905 | 0.114 | 0.03 | 2.43E-16 | 8.9E-12 |
| SCYL2 | 5 | -0.408924472 | 0.264 | 0.442 | 2.44E-16 | 8.94E-12 |
| RSPH3 | 5 | 0.329045194 | 0.228 | 0.104 | 2.5E-16 | 9.15E-12 |
| JADE3 | 5 | 0.383551907 | 0.521 | 0.345 | 2.65E-16 | 9.71E-12 |
| SPTAN1 | 5 | -0.418588867 | 0.27 | 0.446 | 2.78E-16 | 1.02E-11 |
| AC005224.2 | 5 | -0.262851069 | 0.018 | 0.111 | 2.8E-16 | 1.03E-11 |
| CCNL1 | 5 | 0.388010862 | 0.617 | 0.505 | 3E-16 | 1.1E-11 |
| ADAT2 | 5 | 0.343032366 | 0.261 | 0.132 | 3.03E-16 | 1.11E-11 |
| MBOAT2 | 5 | 0.441234471 | 0.611 | 0.45 | 3.17E-16 | 1.16E-11 |
| BCLAF3 | 5 | -0.465637961 | 0.257 | 0.423 | 3.3E-16 | 1.21E-11 |
| AC112722.1 | 5 | -0.315843378 | 0.051 | 0.174 | 3.41E-16 | 1.25E-11 |
| AP002026.1 | 5 | 0.334711992 | 0.196 | 0.079 | 3.58E-16 | 1.31E-11 |
| WHRN | 5 | -0.333295756 | 0.062 | 0.188 | 3.65E-16 | 1.33E-11 |
| ARMC4 | 5 | 0.356640893 | 0.257 | 0.121 | 3.85E-16 | 1.41E-11 |
| STX17 | 5 | 0.37735291 | 0.349 | 0.208 | 3.88E-16 | 1.42E-11 |
| TPM2 | 5 | -0.251681823 | 0.022 | 0.122 | 3.92E-16 | 1.43E-11 |
| PLEKHB2 | 5 | -0.369409804 | 0.132 | 0.285 | 3.95E-16 | 1.44E-11 |
| ANKRD10 | 5 | -0.432565711 | 0.536 | 0.683 | 3.95E-16 | 1.45E-11 |
| SMAD1 | 5 | -0.439407845 | 0.213 | 0.377 | 4.02E-16 | 1.47E-11 |
| KIAA1217 | 5 | -0.558207064 | 0.341 | 0.309 | 4.11E-16 | 1.51E-11 |
| AC009879.4 | 5 | 0.278914016 | 0.141 | 0.048 | 4.12E-16 | 1.51E-11 |
| SLC9A9 | 5 | -0.272321109 | 0.015 | 0.103 | 4.27E-16 | 1.56E-11 |
| INTS10 | 5 | 0.376606534 | 0.514 | 0.362 | 4.31E-16 | 1.58E-11 |
| WDFY3-AS2 | 5 | 0.339431906 | 0.217 | 0.098 | 4.33E-16 | 1.58E-11 |
| VPS8 | 5 | -0.480837465 | 0.457 | 0.612 | 4.59E-16 | 1.68E-11 |
| ERGIC1 | 5 | -0.347135866 | 0.174 | 0.339 | 4.85E-16 | 1.78E-11 |
| AMPD2 | 5 | 0.292851631 | 0.154 | 0.067 | 4.86E-16 | 1.78E-11 |
| CHRNA7 | 5 | -0.375306094 | 0.101 | 0.245 | 4.86E-16 | 1.78E-11 |
| PLPP1 | 5 | -0.71275222 | 0.231 | 0.354 | 4.98E-16 | 1.82E-11 |
| FAM53B | 5 | -0.389478318 | 0.16 | 0.318 | 5.13E-16 | 1.88E-11 |
| SLITRK5 | 5 | -0.298263011 | 0.015 | 0.105 | 5.16E-16 | 1.89E-11 |
| LRRC1 | 5 | 0.308515953 | 0.297 | 0.15 | 5.28E-16 | 1.93E-11 |
| PIK3CA | 5 | -0.461514283 | 0.363 | 0.523 | 5.34E-16 | 1.96E-11 |
| OTUD4 | 5 | 0.39498905 | 0.532 | 0.383 | 5.45E-16 | 2E-11 |
| MYCBP2 | 5 | -0.404634353 | 0.703 | 0.81 | 5.46E-16 | 2E-11 |
| NEDD9 | 5 | -0.296626825 | 0.026 | 0.128 | 5.51E-16 | 2.02E-11 |
| PTBP2 | 5 | -0.382366516 | 0.828 | 0.904 | 6.03E-16 | 2.21E-11 |
| ARHGEF12 | 5 | -0.40857639 | 0.624 | 0.751 | 6.22E-16 | 2.28E-11 |
| ACBD6 | 5 | 0.363070731 | 0.745 | 0.659 | 6.38E-16 | 2.34E-11 |

| | | | | | | |
|---|---|---|---|---|---|---|
| KHDC4 | 5 | 0.382755133 | 0.539 | 0.389 | 6.46E-16 | 2.36E-11 |
| KDM2B | 5 | -0.398988544 | 0.18 | 0.344 | 6.67E-16 | 2.44E-11 |
| AP001831.1 | 5 | -0.319193694 | 0.037 | 0.139 | 7.19E-16 | 2.63E-11 |
| PHF20L1 | 5 | -0.417390382 | 0.372 | 0.541 | 7.21E-16 | 2.64E-11 |
| GFOD2 | 5 | 0.352247999 | 0.27 | 0.137 | 7.35E-16 | 2.69E-11 |
| BDH1 | 5 | 0.287786182 | 0.211 | 0.092 | 7.37E-16 | 2.7E-11 |
| CARM1 | 5 | -0.321585096 | 0.139 | 0.295 | 7.41E-16 | 2.71E-11 |
| LITAF | 5 | -0.271383527 | 0.026 | 0.127 | 7.61E-16 | 2.79E-11 |
| DMD | 5 | -1.465621218 | 0.664 | 0.673 | 7.92E-16 | 2.9E-11 |
| DBN1 | 5 | -0.273015809 | 0.042 | 0.157 | 8.06E-16 | 2.95E-11 |
| FSIP2-AS1 | 5 | -0.322636904 | 0.033 | 0.141 | 8.31E-16 | 3.04E-11 |
| CACNG4 | 5 | 0.341509973 | 0.246 | 0.116 | 8.44E-16 | 3.09E-11 |
| ARFGEF2 | 5 | 0.386296872 | 0.583 | 0.441 | 8.6E-16 | 3.15E-11 |
| FGFR2 | 5 | -0.638471625 | 0.257 | 0.399 | 8.95E-16 | 3.28E-11 |
| AL807742.1 | 5 | -0.431111851 | 0.042 | 0.14 | 9.1E-16 | 3.33E-11 |
| DCHS2 | 5 | -0.359706199 | 0.018 | 0.107 | 9.18E-16 | 3.36E-11 |
| CDH7 | 5 | -0.787560557 | 0.198 | 0.292 | 9.59E-16 | 3.51E-11 |
| WNT5B | 5 | -0.364793156 | 0.028 | 0.127 | 1.07E-15 | 3.91E-11 |
| KCTD13 | 5 | -0.404148219 | 0.178 | 0.334 | 1.08E-15 | 3.94E-11 |
| PDE1A | 5 | -0.33779933 | 0.319 | 0.195 | 1.09E-15 | 4E-11 |
| ENO2 | 5 | 0.358962139 | 0.286 | 0.151 | 1.1E-15 | 4.03E-11 |
| IMMT | 5 | 0.385912733 | 0.534 | 0.398 | 1.12E-15 | 4.09E-11 |
| GOPC | 5 | 0.266785668 | 0.264 | 0.127 | 1.13E-15 | 4.14E-11 |
| MAN1C1 | 5 | -0.416065007 | 0.108 | 0.237 | 1.13E-15 | 4.15E-11 |
| SESN2 | 5 | 0.257645286 | 0.139 | 0.047 | 1.18E-15 | 4.3E-11 |
| CISD1 | 5 | 0.323261602 | 0.284 | 0.148 | 1.21E-15 | 4.44E-11 |
| C5orf17 | 5 | -0.547134606 | 0.017 | 0.106 | 1.23E-15 | 4.51E-11 |
| CEPT1 | 5 | 0.364645019 | 0.501 | 0.373 | 1.24E-15 | 4.54E-11 |
| VSTM2L | 5 | 0.282843779 | 0.108 | 0.032 | 1.26E-15 | 4.62E-11 |
| AKNAD1 | 5 | -0.403833852 | 0.114 | 0.25 | 1.35E-15 | 4.96E-11 |
| SUB1 | 5 | -0.401416672 | 0.202 | 0.359 | 1.37E-15 | 5.01E-11 |
| CCSER1 | 5 | -0.690171971 | 0.604 | 0.605 | 1.45E-15 | 5.3E-11 |
| AC108169.1 | 5 | -0.433947179 | 0.035 | 0.142 | 1.45E-15 | 5.3E-11 |
| ST3GAL6 | 5 | 0.412225581 | 0.439 | 0.277 | 1.45E-15 | 5.32E-11 |
| EPRS | 5 | 0.410209652 | 0.439 | 0.313 | 1.46E-15 | 5.34E-11 |
| RAD18 | 5 | -0.453619361 | 0.372 | 0.537 | 1.51E-15 | 5.54E-11 |
| CDK10 | 5 | 0.372035525 | 0.319 | 0.18 | 1.53E-15 | 5.61E-11 |
| LMTK2 | 5 | 0.38904302 | 0.475 | 0.342 | 1.55E-15 | 5.67E-11 |
| GLYATL2 | 5 | 0.566902329 | 0.457 | 0.324 | 1.57E-15 | 5.74E-11 |
| ABCA1 | 5 | -0.566955079 | 0.145 | 0.271 | 1.65E-15 | 6.05E-11 |
| SS18L2 | 5 | 0.312184017 | 0.218 | 0.107 | 1.77E-15 | 6.48E-11 |
| NXPH2 | 5 | -0.354267939 | 0.028 | 0.122 | 1.77E-15 | 6.48E-11 |
| UBA1 | 5 | -0.357038416 | 0.244 | 0.416 | 1.91E-15 | 7E-11 |
| BCL2L1 | 5 | 0.319870436 | 0.244 | 0.122 | 2.02E-15 | 7.38E-11 |
| CRAMP1 | 5 | -0.329904757 | 0.123 | 0.27 | 2.16E-15 | 7.91E-11 |
| TAF4B | 5 | 0.417984212 | 0.385 | 0.249 | 2.26E-15 | 8.27E-11 |
| KIF18A | 5 | 0.391839245 | 0.277 | 0.152 | 2.31E-15 | 8.48E-11 |
| POLR3G | 5 | -0.455874625 | 0.145 | 0.272 | 2.57E-15 | 9.42E-11 |
| GGT7 | 5 | -0.29095865 | 0.094 | 0.231 | 2.61E-15 | 9.56E-11 |
| PPP1R3E | 5 | 0.27627237 | 0.196 | 0.084 | 2.68E-15 | 9.8E-11 |
| GPC3 | 5 | -0.866379052 | 0.119 | 0.24 | 2.77E-15 | 1.01E-10 |
| DPY19L3 | 5 | -0.436458221 | 0.35 | 0.505 | 2.79E-15 | 1.02E-10 |
| SLC2A3 | 5 | -0.425553289 | 0.037 | 0.144 | 2.83E-15 | 1.03E-10 |
| C14orf39 | 5 | 0.259167703 | 0.169 | 0.064 | 2.93E-15 | 1.07E-10 |
| ROR2 | 5 | -0.557270605 | 0.262 | 0.432 | 3.01E-15 | 1.1E-10 |
| STON2 | 5 | 0.407103373 | 0.505 | 0.341 | 3.19E-15 | 1.17E-10 |
| SKAP1 | 5 | 0.279684102 | 0.183 | 0.072 | 3.2E-15 | 1.17E-10 |
| SRGAP2 | 5 | -0.341842559 | 0.187 | 0.35 | 3.22E-15 | 1.18E-10 |

| | | | | | | |
|---|---|---|---|---|---|---|
| ARHGEF26 | 5 | 0.361922666 | 0.431 | 0.267 | 3.42E-15 | 1.25E-10 |
| RFC5 | 5 | 0.309753621 | 0.257 | 0.127 | 3.44E-15 | 1.26E-10 |
| CACNG5 | 5 | 0.253203074 | 0.187 | 0.075 | 3.48E-15 | 1.27E-10 |
| SNU13 | 5 | 0.297457583 | 0.222 | 0.102 | 3.61E-15 | 1.32E-10 |
| DRG1 | 5 | 0.319293279 | 0.325 | 0.179 | 3.74E-15 | 1.37E-10 |
| Z83843.1 | 5 | -0.405295287 | 0.339 | 0.496 | 3.86E-15 | 1.41E-10 |
| ERC2 | 5 | -0.590330963 | 0.374 | 0.456 | 4.4E-15 | 1.61E-10 |
| SLC8A1-AS1 | 5 | -0.407753445 | 0.031 | 0.126 | 4.67E-15 | 1.71E-10 |
| MCEE | 5 | 0.334541675 | 0.261 | 0.133 | 4.73E-15 | 1.73E-10 |
| RASGRF2 | 5 | -0.70352081 | 0.079 | 0.187 | 4.82E-15 | 1.77E-10 |
| PPP4R4 | 5 | 0.293062627 | 0.239 | 0.11 | 4.84E-15 | 1.77E-10 |
| STAM | 5 | 0.392505165 | 0.468 | 0.348 | 4.88E-15 | 1.79E-10 |
| NOVA1 | 5 | -0.60894108 | 0.545 | 0.632 | 4.91E-15 | 1.8E-10 |
| STK17B | 5 | -0.389155037 | 0.046 | 0.154 | 4.97E-15 | 1.82E-10 |
| SSH2 | 5 | 0.272191397 | 0.899 | 0.795 | 5.34E-15 | 1.95E-10 |
| PCBP3 | 5 | 0.297815146 | 0.4 | 0.239 | 5.48E-15 | 2.01E-10 |
| CDK17 | 5 | -0.459837377 | 0.437 | 0.596 | 5.55E-15 | 2.03E-10 |
| ZNF292 | 5 | -0.353803977 | 0.815 | 0.89 | 5.89E-15 | 2.16E-10 |
| ASTN2 | 5 | -0.695688802 | 0.407 | 0.52 | 5.99E-15 | 2.19E-10 |
| DIP2C | 5 | -0.460822687 | 0.653 | 0.759 | 6.02E-15 | 2.2E-10 |
| UBAP2 | 5 | 0.370349375 | 0.747 | 0.636 | 6.23E-15 | 2.28E-10 |
| CCDC30 | 5 | 0.311072019 | 0.255 | 0.123 | 6.23E-15 | 2.28E-10 |
| SRCIN1 | 5 | -0.327161893 | 0.077 | 0.202 | 6.74E-15 | 2.47E-10 |
| IDI2-AS1 | 5 | -0.413673419 | 0.042 | 0.142 | 6.97E-15 | 2.55E-10 |
| COL6A1 | 5 | -0.409975073 | 0.086 | 0.211 | 7.17E-15 | 2.63E-10 |
| ASXL3 | 5 | -0.566107391 | 0.361 | 0.486 | 7.2E-15 | 2.63E-10 |
| RAB6A | 5 | -0.42571486 | 0.446 | 0.601 | 7.36E-15 | 2.7E-10 |
| GPR137B | 5 | -0.346944594 | 0.114 | 0.255 | 7.38E-15 | 2.7E-10 |
| PRSS23 | 5 | -0.301122578 | 0.018 | 0.107 | 7.87E-15 | 2.88E-10 |
| PDK3 | 5 | 0.405332146 | 0.571 | 0.416 | 7.87E-15 | 2.88E-10 |
| TUG1 | 5 | 0.371332349 | 0.402 | 0.295 | 7.89E-15 | 2.89E-10 |
| LIFR-AS1 | 5 | -0.25156689 | 0.02 | 0.112 | 8.14E-15 | 2.98E-10 |
| REPS2 | 5 | -0.517948941 | 0.191 | 0.338 | 8.29E-15 | 3.04E-10 |
| PCBP1-AS1 | 5 | -0.426469096 | 0.35 | 0.507 | 8.93E-15 | 3.27E-10 |
| GPSM2 | 5 | -0.428946147 | 0.16 | 0.299 | 9.13E-15 | 3.34E-10 |
| IFT172 | 5 | -0.32925121 | 0.123 | 0.267 | 9.45E-15 | 3.46E-10 |
| RAB10 | 5 | 0.374699046 | 0.699 | 0.582 | 1.02E-14 | 3.72E-10 |
| WDR34 | 5 | 0.262972475 | 0.185 | 0.078 | 1.05E-14 | 3.85E-10 |
| ZNRF1 | 5 | -0.384024181 | 0.259 | 0.428 | 1.12E-14 | 4.1E-10 |
| TBC1D32 | 5 | -0.45050311 | 0.437 | 0.581 | 1.12E-14 | 4.11E-10 |
| FAM118A | 5 | 0.30930376 | 0.262 | 0.142 | 1.13E-14 | 4.12E-10 |
| CALM2 | 5 | -0.425781339 | 0.374 | 0.525 | 1.17E-14 | 4.29E-10 |
| SLC16A9 | 5 | -0.325326013 | 0.064 | 0.178 | 1.2E-14 | 4.39E-10 |
| KITLG | 5 | 0.31568291 | 0.235 | 0.109 | 1.2E-14 | 4.41E-10 |
| LPCAT3 | 5 | 0.331046685 | 0.297 | 0.174 | 1.31E-14 | 4.79E-10 |
| HNRNPLL | 5 | -0.389255737 | 0.281 | 0.443 | 1.47E-14 | 5.39E-10 |
| TUBB3 | 5 | -0.257614473 | 0.024 | 0.117 | 1.51E-14 | 5.52E-10 |
| EBLN3P | 5 | 0.317860478 | 0.251 | 0.134 | 1.54E-14 | 5.63E-10 |
| SFPQ | 5 | -0.369822775 | 0.617 | 0.744 | 1.58E-14 | 5.77E-10 |
| TSPAN3 | 5 | -0.342464841 | 0.172 | 0.325 | 1.62E-14 | 5.93E-10 |
| SUGCT | 5 | -0.470342218 | 0.251 | 0.41 | 1.62E-14 | 5.95E-10 |
| LDHA | 5 | 0.380211136 | 0.211 | 0.102 | 1.66E-14 | 6.07E-10 |
| SGCD | 5 | 0.477491729 | 0.64 | 0.477 | 1.73E-14 | 6.33E-10 |
| AP002954.2 | 5 | -0.568407309 | 0.017 | 0.103 | 1.83E-14 | 6.7E-10 |
| AL355499.1 | 5 | -0.842549559 | 0.02 | 0.107 | 1.85E-14 | 6.79E-10 |
| KIF26B | 5 | -0.646755741 | 0.424 | 0.517 | 1.96E-14 | 7.19E-10 |
| LRRFIP2 | 5 | 0.371220705 | 0.556 | 0.446 | 1.98E-14 | 7.23E-10 |
| INPPL1 | 5 | -0.255994388 | 0.057 | 0.172 | 2.05E-14 | 7.51E-10 |

| | | | | | | |
|---|---|---|---|---|---|---|
| ANKRD44 | 5 | 0.342451426 | 0.483 | 0.316 | 2.23E-14 | 8.17E-10 |
| XPNPEP3 | 5 | 0.395156432 | 0.347 | 0.233 | 2.29E-14 | 8.38E-10 |
| TMEM14B | 5 | -0.426257586 | 0.306 | 0.468 | 2.31E-14 | 8.45E-10 |
| UBE3D | 5 | 0.338049787 | 0.629 | 0.466 | 2.46E-14 | 9.02E-10 |
| GGA1 | 5 | 0.338149304 | 0.314 | 0.211 | 2.5E-14 | 9.15E-10 |
| PTPRG-AS1 | 5 | 0.360348687 | 0.332 | 0.204 | 2.51E-14 | 9.19E-10 |
| FMR1 | 5 | -0.415236985 | 0.341 | 0.509 | 2.54E-14 | 9.32E-10 |
| AC097376.3 | 5 | 0.402919906 | 0.339 | 0.233 | 2.56E-14 | 9.36E-10 |
| ZNF91 | 5 | 0.343832116 | 0.394 | 0.239 | 2.63E-14 | 9.62E-10 |
| AFMID | 5 | 0.366752217 | 0.262 | 0.138 | 2.67E-14 | 9.76E-10 |
| SLFN13 | 5 | 0.32724218 | 0.255 | 0.127 | 2.72E-14 | 9.95E-10 |
| TIMP3 | 5 | -0.279746333 | 0.017 | 0.102 | 2.77E-14 | 1.01E-09 |
| SHB | 5 | -0.396424563 | 0.116 | 0.248 | 2.94E-14 | 1.08E-09 |
| ATG7 | 5 | 0.414412809 | 0.615 | 0.492 | 2.97E-14 | 1.09E-09 |
| VPS37A | 5 | 0.365812039 | 0.437 | 0.302 | 3E-14 | 1.1E-09 |
| KIF1B | 5 | -0.407902673 | 0.607 | 0.75 | 3.16E-14 | 1.16E-09 |
| PXYLP1 | 5 | -0.285107063 | 0.059 | 0.175 | 3.24E-14 | 1.19E-09 |
| DDX5 | 5 | -0.394554731 | 0.606 | 0.742 | 3.61E-14 | 1.32E-09 |
| BICDL1 | 5 | 0.302120785 | 0.429 | 0.268 | 3.64E-14 | 1.33E-09 |
| SOAT1 | 5 | 0.311091319 | 0.281 | 0.149 | 3.81E-14 | 1.4E-09 |
| PPM1K | 5 | 0.291850059 | 0.209 | 0.097 | 3.93E-14 | 1.44E-09 |
| AC068286.2 | 5 | -1.005677053 | 0.042 | 0.142 | 3.93E-14 | 1.44E-09 |
| REC8 | 5 | 0.258362666 | 0.18 | 0.08 | 4.32E-14 | 1.58E-09 |
| HCN1 | 5 | -0.605019531 | 0.055 | 0.162 | 4.38E-14 | 1.6E-09 |
| VPS53 | 5 | 0.331727055 | 0.536 | 0.485 | 4.68E-14 | 1.71E-09 |
| MAGI2-AS3 | 5 | -0.30715292 | 0.095 | 0.226 | 4.7E-14 | 1.72E-09 |
| AP002495.1 | 5 | 0.378915807 | 0.413 | 0.285 | 4.73E-14 | 1.73E-09 |
| KLHL22 | 5 | 0.385574915 | 0.435 | 0.322 | 4.87E-14 | 1.78E-09 |
| CYSTM1 | 5 | 0.309205904 | 0.262 | 0.141 | 4.94E-14 | 1.81E-09 |
| ATXN10 | 5 | 0.336201572 | 0.771 | 0.709 | 4.97E-14 | 1.82E-09 |
| ARNTL | 5 | -0.325334635 | 0.051 | 0.154 | 5.18E-14 | 1.89E-09 |
| HSPA8 | 5 | 0.420503644 | 0.281 | 0.162 | 5.73E-14 | 2.1E-09 |
| RCC2 | 5 | -0.365122642 | 0.31 | 0.474 | 5.76E-14 | 2.11E-09 |
| SVIL | 5 | -0.650805474 | 0.494 | 0.53 | 5.84E-14 | 2.14E-09 |
| MCM3AP | 5 | -0.346689732 | 0.255 | 0.417 | 6.11E-14 | 2.24E-09 |
| SEMA3C | 5 | -0.569729282 | 0.031 | 0.128 | 6.38E-14 | 2.34E-09 |
| LAPTM4A | 5 | -0.285584307 | 0.079 | 0.203 | 6.39E-14 | 2.34E-09 |
| RAI2 | 5 | -0.323904113 | 0.051 | 0.162 | 6.53E-14 | 2.39E-09 |
| CHRM2 | 5 | -0.657554954 | 0.022 | 0.1 | 6.59E-14 | 2.41E-09 |
| EME2 | 5 | 0.261077704 | 0.171 | 0.075 | 6.85E-14 | 2.51E-09 |
| RILPL1 | 5 | -0.272723727 | 0.099 | 0.229 | 7.35E-14 | 2.69E-09 |
| EFHC1 | 5 | 0.344728108 | 0.409 | 0.255 | 7.57E-14 | 2.77E-09 |
| BICC1 | 5 | 0.387260042 | 0.451 | 0.29 | 7.68E-14 | 2.81E-09 |
| AGO4 | 5 | -0.373119475 | 0.237 | 0.394 | 7.76E-14 | 2.84E-09 |
| IL12RB2 | 5 | -0.699851571 | 0.277 | 0.286 | 7.83E-14 | 2.87E-09 |
| TRABD2B | 5 | -0.329442987 | 0.028 | 0.119 | 7.96E-14 | 2.91E-09 |
| ZNF33B | 5 | 0.355370338 | 0.422 | 0.271 | 8.02E-14 | 2.94E-09 |
| CTNNA1 | 5 | -0.457490844 | 0.541 | 0.67 | 8.09E-14 | 2.96E-09 |
| PPP2R5C | 5 | -0.392552761 | 0.42 | 0.576 | 8.38E-14 | 3.07E-09 |
| GPT2 | 5 | -0.271089391 | 0.048 | 0.153 | 8.81E-14 | 3.23E-09 |
| SDC2 | 5 | -0.511498858 | 0.27 | 0.374 | 8.96E-14 | 3.28E-09 |
| SH3BGRL2 | 5 | -0.321969014 | 0.07 | 0.184 | 8.98E-14 | 3.29E-09 |
| CELF2 | 5 | -0.316624998 | 0.686 | 0.615 | 9.04E-14 | 3.31E-09 |
| DZIP1L | 5 | -0.321421125 | 0.099 | 0.229 | 9.09E-14 | 3.33E-09 |
| CNNM2 | 5 | 0.373420978 | 0.448 | 0.292 | 9.18E-14 | 3.36E-09 |
| INSIG1 | 5 | 0.258484931 | 0.136 | 0.05 | 9.23E-14 | 3.38E-09 |
| RYR3 | 5 | -0.512912785 | 0.028 | 0.114 | 9.33E-14 | 3.42E-09 |
| ZCCHC2 | 5 | 0.295473927 | 0.244 | 0.12 | 9.39E-14 | 3.44E-09 |

| | | | | | | |
|---|---|---|---|---|---|---|
| SEC23B | 5 | 0.313308547 | 0.286 | 0.164 | 1E-13 | 3.67E-09 |
| BRSK2 | 5 | 0.305402773 | 0.207 | 0.096 | 1.02E-13 | 3.75E-09 |
| SSBP2 | 5 | -0.500701798 | 0.833 | 0.809 | 1.04E-13 | 3.83E-09 |
| MDK | 5 | -0.265706842 | 0.046 | 0.15 | 1.05E-13 | 3.86E-09 |
| SOCS7 | 5 | -0.29780424 | 0.095 | 0.224 | 1.08E-13 | 3.97E-09 |
| HPCAL1 | 5 | -0.469599834 | 0.191 | 0.333 | 1.1E-13 | 4.04E-09 |
| LDLR | 5 | 0.391440527 | 0.222 | 0.105 | 1.16E-13 | 4.26E-09 |
| SLC25A26 | 5 | 0.391717887 | 0.47 | 0.327 | 1.17E-13 | 4.27E-09 |
| UBE2D1 | 5 | -0.381750432 | 0.145 | 0.272 | 1.19E-13 | 4.34E-09 |
| PPARG | 5 | -0.551718436 | 0.165 | 0.289 | 1.19E-13 | 4.37E-09 |
| SHTN1 | 5 | -0.440880792 | 0.62 | 0.69 | 1.2E-13 | 4.38E-09 |
| AC015871.1 | 5 | -0.289661714 | 0.083 | 0.204 | 1.23E-13 | 4.5E-09 |
| COL6A3 | 5 | -0.567250369 | 0.035 | 0.124 | 1.27E-13 | 4.66E-09 |
| ZNF618 | 5 | 0.378152574 | 0.657 | 0.552 | 1.36E-13 | 4.97E-09 |
| SUFU | 5 | -0.341569531 | 0.143 | 0.283 | 1.37E-13 | 5.02E-09 |
| PSPC1 | 5 | -0.37381408 | 0.651 | 0.766 | 1.41E-13 | 5.15E-09 |
| ASAP2 | 5 | 0.408106322 | 0.516 | 0.38 | 1.41E-13 | 5.17E-09 |
| CDH20 | 5 | -0.44007124 | 0.022 | 0.104 | 1.47E-13 | 5.38E-09 |
| TRIT1 | 5 | 0.330372544 | 0.406 | 0.268 | 1.48E-13 | 5.44E-09 |
| BDP1 | 5 | 0.353761421 | 0.73 | 0.638 | 1.5E-13 | 5.49E-09 |
| ROCK2 | 5 | -0.419146434 | 0.594 | 0.69 | 1.55E-13 | 5.68E-09 |
| SOCS6 | 5 | 0.277341573 | 0.213 | 0.104 | 1.78E-13 | 6.52E-09 |
| TRMT11 | 5 | 0.349942481 | 0.747 | 0.641 | 1.81E-13 | 6.61E-09 |
| EPHB1 | 5 | -0.409087136 | 0.059 | 0.168 | 1.92E-13 | 7.02E-09 |
| CENPQ | 5 | 0.283961866 | 0.246 | 0.124 | 1.95E-13 | 7.13E-09 |
| EP300 | 5 | 0.352301673 | 0.71 | 0.603 | 1.97E-13 | 7.2E-09 |
| SHC3 | 5 | -0.543650951 | 0.262 | 0.362 | 2.06E-13 | 7.54E-09 |
| LMO7 | 5 | -0.601153105 | 0.499 | 0.527 | 2.09E-13 | 7.65E-09 |
| RBPJ | 5 | 0.37757802 | 0.767 | 0.655 | 2.1E-13 | 7.68E-09 |
| AKT2 | 5 | 0.317749277 | 0.385 | 0.252 | 2.12E-13 | 7.77E-09 |
| RAB39A | 5 | 0.2806049 | 0.193 | 0.086 | 2.22E-13 | 8.13E-09 |
| FBXO7 | 5 | 0.320207621 | 0.334 | 0.205 | 2.28E-13 | 8.35E-09 |
| AGO1 | 5 | -0.368985573 | 0.233 | 0.384 | 2.37E-13 | 8.68E-09 |
| CTNND1 | 5 | 0.336738492 | 0.728 | 0.615 | 2.44E-13 | 8.95E-09 |
| MPP6 | 5 | -0.615562749 | 0.549 | 0.653 | 2.47E-13 | 9.03E-09 |
| ATXN7 | 5 | 0.344036371 | 0.464 | 0.332 | 2.48E-13 | 9.08E-09 |
| NEB | 5 | -0.441325817 | 0.037 | 0.134 | 2.63E-13 | 9.64E-09 |
| SDCBP | 5 | -0.260741929 | 0.062 | 0.175 | 2.66E-13 | 9.74E-09 |
| ME3 | 5 | -0.472340275 | 0.211 | 0.317 | 2.76E-13 | 1.01E-08 |
| CCDC146 | 5 | -0.431607248 | 0.121 | 0.246 | 2.78E-13 | 1.02E-08 |
| RCOR1 | 5 | -0.441701451 | 0.576 | 0.691 | 2.78E-13 | 1.02E-08 |
| COLGALT2 | 5 | -0.348237703 | 0.059 | 0.163 | 2.96E-13 | 1.08E-08 |
| PHF21B | 5 | -0.449326952 | 0.239 | 0.384 | 3.08E-13 | 1.13E-08 |
| TOX2 | 5 | -0.282845308 | 0.02 | 0.103 | 3.1E-13 | 1.14E-08 |
| TTC14 | 5 | -0.370910435 | 0.429 | 0.576 | 3.17E-13 | 1.16E-08 |
| LRRC6 | 5 | -0.263633745 | 0.02 | 0.104 | 3.18E-13 | 1.16E-08 |
| CBY1 | 5 | 0.300087366 | 0.217 | 0.113 | 3.26E-13 | 1.19E-08 |
| PI4K2B | 5 | 0.324997163 | 0.347 | 0.216 | 3.29E-13 | 0.000000012 |
| TBL1XR1 | 5 | 0.32289235 | 0.89 | 0.855 | 3.39E-13 | 1.24E-08 |
| EIF4E3 | 5 | -0.274433968 | 0.051 | 0.156 | 3.58E-13 | 1.31E-08 |
| ATP8B1 | 5 | 0.383002819 | 0.27 | 0.15 | 3.79E-13 | 1.39E-08 |
| GTF2I | 5 | -0.304300527 | 0.861 | 0.902 | 3.81E-13 | 1.39E-08 |
| TMSB4X | 5 | -0.380281063 | 0.127 | 0.259 | 3.81E-13 | 0.000000014 |
| HSPD1 | 5 | 0.461323965 | 0.545 | 0.436 | 3.85E-13 | 1.41E-08 |
| SH3GL2 | 5 | -0.383941895 | 0.424 | 0.393 | 3.92E-13 | 1.44E-08 |
| POU6F2 | 5 | 0.324222492 | 0.141 | 0.052 | 4.05E-13 | 1.48E-08 |
| FDPS | 5 | 0.324616278 | 0.294 | 0.178 | 4.1E-13 | 0.000000015 |
| FIG4 | 5 | 0.331154351 | 0.31 | 0.197 | 4.16E-13 | 1.52E-08 |

| | | | | | | |
|---|---|---|---|---|---|---|
| RFPL1S | 5 | 0.337211393 | 0.343 | 0.218 | 4.17E-13 | 1.52E-08 |
| ABAT | 5 | 0.335799826 | 0.248 | 0.13 | 4.28E-13 | 1.57E-08 |
| CTNNA3 | 5 | -0.61673596 | 0.073 | 0.188 | 4.36E-13 | 1.59E-08 |
| PRKACB | 5 | -0.416376909 | 0.45 | 0.605 | 4.47E-13 | 1.64E-08 |
| CENPC | 5 | 0.351020839 | 0.528 | 0.399 | 4.68E-13 | 1.71E-08 |
| IDH2 | 5 | -0.34366385 | 0.136 | 0.269 | 4.74E-13 | 1.74E-08 |
| VPS13B | 5 | 0.292285028 | 0.866 | 0.823 | 4.8E-13 | 1.76E-08 |
| APAF1 | 5 | -0.35248206 | 0.194 | 0.343 | 4.95E-13 | 1.81E-08 |
| COQ8A | 5 | 0.272642389 | 0.217 | 0.108 | 5.02E-13 | 1.84E-08 |
| TNFRSF21 | 5 | 0.262388734 | 0.207 | 0.096 | 5.31E-13 | 1.94E-08 |
| ATRX | 5 | -0.284817933 | 0.892 | 0.908 | 6E-13 | 2.19E-08 |
| ADGRB2 | 5 | -0.279469776 | 0.07 | 0.174 | 6.1E-13 | 2.23E-08 |
| KIF3C | 5 | -0.263422336 | 0.05 | 0.154 | 6.16E-13 | 2.25E-08 |
| FRMD6 | 5 | 0.467798272 | 0.31 | 0.232 | 6.29E-13 | 0.000000023 |
| ANKRD17 | 5 | 0.264381803 | 0.945 | 0.92 | 6.33E-13 | 2.32E-08 |
| GABBR1 | 5 | -0.315596012 | 0.143 | 0.281 | 6.45E-13 | 2.36E-08 |
| GCC2 | 5 | 0.355320454 | 0.538 | 0.407 | 6.63E-13 | 2.43E-08 |
| RPS9 | 5 | 0.339952553 | 0.446 | 0.309 | 6.67E-13 | 2.44E-08 |
| ACTN2 | 5 | -0.284533409 | 0.028 | 0.115 | 6.68E-13 | 2.44E-08 |
| CLEC2D | 5 | -0.43913713 | 0.336 | 0.469 | 6.72E-13 | 2.46E-08 |
| NT5DC1 | 5 | 0.320195479 | 0.541 | 0.383 | 7E-13 | 2.56E-08 |
| L3MBTL3 | 5 | -0.333092654 | 0.064 | 0.167 | 7.13E-13 | 2.61E-08 |
| CTIF | 5 | -0.299308575 | 0.272 | 0.431 | 7.46E-13 | 2.73E-08 |
| PON2 | 5 | -0.325489455 | 0.103 | 0.226 | 7.5E-13 | 2.75E-08 |
| BRI3BP | 5 | 0.296801733 | 0.229 | 0.126 | 7.62E-13 | 2.79E-08 |
| GCA | 5 | 0.291777538 | 0.284 | 0.156 | 7.92E-13 | 0.000000029 |
| RUNX1T1 | 5 | -0.511989442 | 0.618 | 0.72 | 8.01E-13 | 2.93E-08 |
| CNOT2 | 5 | -0.384568673 | 0.73 | 0.792 | 8.19E-13 | 0.00000003 |
| MT-CO3 | 5 | -0.513841945 | 0.264 | 0.421 | 8.53E-13 | 3.12E-08 |
| PDLIM1 | 5 | 0.258942825 | 0.215 | 0.102 | 8.65E-13 | 3.17E-08 |
| RPL3 | 5 | 0.355363077 | 0.455 | 0.336 | 8.8E-13 | 3.22E-08 |
| TSPAN14 | 5 | 0.288533552 | 0.295 | 0.166 | 8.86E-13 | 3.24E-08 |
| IGSF3 | 5 | -0.298006258 | 0.092 | 0.208 | 8.88E-13 | 3.25E-08 |
| QSOX2 | 5 | 0.333371318 | 0.314 | 0.189 | 9.15E-13 | 3.35E-08 |
| GBE1 | 5 | 0.36850707 | 0.457 | 0.31 | 9.28E-13 | 0.000000034 |
| SMCO4 | 5 | 0.25133169 | 0.178 | 0.081 | 9.37E-13 | 3.43E-08 |
| DGCR8 | 5 | 0.339653173 | 0.492 | 0.375 | 9.38E-13 | 3.43E-08 |
| TEX9 | 5 | 0.299860879 | 0.334 | 0.194 | 9.53E-13 | 3.49E-08 |
| WDPCP | 5 | -0.432849038 | 0.455 | 0.586 | 9.54E-13 | 3.49E-08 |
| ETFA | 5 | 0.369753261 | 0.492 | 0.37 | 9.55E-13 | 0.000000035 |
| CYP7B1 | 5 | -0.620688648 | 0.303 | 0.373 | 9.69E-13 | 3.55E-08 |
| DIO2 | 5 | -0.475134644 | 0.182 | 0.316 | 9.94E-13 | 3.64E-08 |
| PDCD4 | 5 | 0.357921589 | 0.532 | 0.395 | 1.05E-12 | 3.86E-08 |
| DSTN | 5 | 0.294585725 | 0.261 | 0.144 | 1.24E-12 | 4.55E-08 |
| ZNF124 | 5 | -0.390932496 | 0.312 | 0.46 | 1.27E-12 | 4.67E-08 |
| CALD1 | 5 | -0.553714611 | 0.306 | 0.435 | 1.3E-12 | 4.77E-08 |
| SDCCAG8 | 5 | 0.348851053 | 0.686 | 0.547 | 1.36E-12 | 4.97E-08 |
| SUSD6 | 5 | 0.293378071 | 0.294 | 0.168 | 1.38E-12 | 5.06E-08 |
| RNFT2 | 5 | 0.314078752 | 0.248 | 0.137 | 1.39E-12 | 0.000000051 |
| PLA2G6 | 5 | 0.272678578 | 0.139 | 0.067 | 1.4E-12 | 5.12E-08 |
| AGBL3 | 5 | 0.27760608 | 0.169 | 0.094 | 1.57E-12 | 5.73E-08 |
| GRSF1 | 5 | 0.352231829 | 0.396 | 0.273 | 1.58E-12 | 5.77E-08 |
| ELAVL2 | 5 | -0.41943214 | 0.389 | 0.382 | 1.59E-12 | 5.82E-08 |
| ACADM | 5 | 0.364257628 | 0.477 | 0.357 | 1.65E-12 | 6.04E-08 |
| ANAPC4 | 5 | 0.3239913 | 0.433 | 0.303 | 1.65E-12 | 6.05E-08 |
| GPR89A | 5 | 0.253741676 | 0.248 | 0.129 | 1.67E-12 | 6.13E-08 |
| NFAT5 | 5 | -0.459785534 | 0.783 | 0.797 | 1.75E-12 | 6.42E-08 |
| LASP1 | 5 | 0.317064591 | 0.213 | 0.118 | 1.77E-12 | 6.49E-08 |

| | | | | | | |
|---|---|---|---|---|---|---|
| CYTH1 | 5 | -0.388057033 | 0.338 | 0.481 | 1.82E-12 | 6.66E-08 |
| FARSB | 5 | 0.346933784 | 0.561 | 0.42 | 1.85E-12 | 6.76E-08 |
| JAG1 | 5 | -0.388202917 | 0.053 | 0.151 | 1.85E-12 | 6.77E-08 |
| CD200 | 5 | 0.273184214 | 0.277 | 0.15 | 1.9E-12 | 6.95E-08 |
| ABCD3 | 5 | -0.489629194 | 0.587 | 0.645 | 1.91E-12 | 6.98E-08 |
| TTLL5 | 5 | 0.338286591 | 0.672 | 0.6 | 1.94E-12 | 0.000000071 |
| WDR37 | 5 | 0.341972497 | 0.442 | 0.342 | 1.95E-12 | 7.14E-08 |
| PTCHD1-AS | 5 | -0.593788726 | 0.037 | 0.124 | 2.08E-12 | 0.000000076 |
| ERMP1 | 5 | 0.270799882 | 0.226 | 0.117 | 2.11E-12 | 7.74E-08 |
| GPR135 | 5 | -0.331769097 | 0.145 | 0.273 | 2.12E-12 | 7.76E-08 |
| RTN3 | 5 | -0.3932037 | 0.606 | 0.684 | 2.18E-12 | 7.96E-08 |
| LARP4B | 5 | 0.301153871 | 0.8 | 0.71 | 2.18E-12 | 7.98E-08 |
| CAB39 | 5 | -0.364322134 | 0.231 | 0.372 | 2.21E-12 | 8.08E-08 |
| LIPE-AS1 | 5 | 0.407459094 | 0.413 | 0.319 | 2.31E-12 | 8.47E-08 |
| STK10 | 5 | 0.289956657 | 0.277 | 0.152 | 2.38E-12 | 8.72E-08 |
| COX11 | 5 | 0.299398738 | 0.308 | 0.189 | 2.46E-12 | 9.01E-08 |
| UAP1 | 5 | 0.27852787 | 0.226 | 0.119 | 2.52E-12 | 9.21E-08 |
| EXOG | 5 | 0.313611958 | 0.354 | 0.231 | 2.55E-12 | 9.35E-08 |
| CHN1 | 5 | -0.450015817 | 0.565 | 0.611 | 2.56E-12 | 9.38E-08 |
| CAND1 | 5 | 0.334326567 | 0.598 | 0.497 | 2.61E-12 | 9.56E-08 |
| PRXL2A | 5 | 0.302877881 | 0.261 | 0.156 | 2.62E-12 | 0.000000096 |
| NPSR1 | 5 | -0.902006184 | 0.123 | 0.177 | 2.77E-12 | 0.000000101 |
| ID1 | 5 | -0.306493225 | 0.039 | 0.131 | 2.78E-12 | 0.000000102 |
| GRK3 | 5 | 0.347835174 | 0.596 | 0.447 | 2.78E-12 | 0.000000102 |
| CHDH | 5 | 0.258809242 | 0.152 | 0.068 | 2.91E-12 | 0.000000106 |
| MYO18A | 5 | 0.274652817 | 0.211 | 0.111 | 2.95E-12 | 0.000000108 |
| BCL11A | 5 | 0.311295133 | 0.415 | 0.265 | 2.95E-12 | 0.000000108 |
| C4orf47 | 5 | -0.336762348 | 0.033 | 0.12 | 3.05E-12 | 0.000000111 |
| RIN2 | 5 | -0.358702474 | 0.22 | 0.367 | 3.4E-12 | 0.000000125 |
| CABLES1 | 5 | -0.342719043 | 0.077 | 0.18 | 3.48E-12 | 0.000000127 |
| ZNF141 | 5 | 0.329019302 | 0.506 | 0.405 | 3.8E-12 | 0.000000139 |
| DOP1B | 5 | 0.335817719 | 0.631 | 0.484 | 4.05E-12 | 0.000000148 |
| EIF3L | 5 | 0.2964401 | 0.406 | 0.263 | 4.45E-12 | 0.000000163 |
| LRCH1 | 5 | 0.35198061 | 0.71 | 0.618 | 4.57E-12 | 0.000000167 |
| AP1S3 | 5 | 0.28437944 | 0.215 | 0.108 | 4.68E-12 | 0.000000171 |
| INTU | 5 | -0.378586082 | 0.189 | 0.317 | 4.92E-12 | 0.00000018 |
| CBX6 | 5 | 0.268251846 | 0.207 | 0.114 | 4.95E-12 | 0.000000181 |
| CTTNBP2NL | 5 | -0.308070871 | 0.138 | 0.268 | 5.57E-12 | 0.000000204 |
| SMARCA4 | 5 | -0.353118029 | 0.406 | 0.558 | 5.68E-12 | 0.000000208 |
| TFDP2 | 5 | 0.333187645 | 0.648 | 0.501 | 5.74E-12 | 0.00000021 |
| FAM110B | 5 | 0.293455831 | 0.545 | 0.39 | 5.99E-12 | 0.000000219 |
| PKM | 5 | 0.327195236 | 0.617 | 0.48 | 6.13E-12 | 0.000000224 |
| BTG3 | 5 | 0.315400599 | 0.389 | 0.25 | 6.18E-12 | 0.000000226 |
| DEPDC5 | 5 | 0.338893626 | 0.431 | 0.305 | 6.22E-12 | 0.000000228 |
| LANCL1 | 5 | -0.344855491 | 0.248 | 0.393 | 6.35E-12 | 0.000000233 |
| CXADR | 5 | -0.411117101 | 0.218 | 0.345 | 6.38E-12 | 0.000000233 |
| ATXN3 | 5 | -0.34591317 | 0.215 | 0.358 | 6.69E-12 | 0.000000245 |
| AUH | 5 | 0.340509034 | 0.631 | 0.495 | 7.14E-12 | 0.000000261 |
| SLC44A2 | 5 | -0.250689827 | 0.04 | 0.133 | 7.22E-12 | 0.000000264 |
| MT-CO2 | 5 | -0.338618813 | 0.226 | 0.373 | 7.31E-12 | 0.000000268 |
| QSER1 | 5 | -0.379293388 | 0.613 | 0.712 | 7.35E-12 | 0.000000269 |
| LRRC23 | 5 | 0.26391236 | 0.215 | 0.109 | 7.58E-12 | 0.000000277 |
| PAQR3 | 5 | -0.264167669 | 0.077 | 0.185 | 7.85E-12 | 0.000000287 |
| RCBTB2 | 5 | -0.252343 | 0.064 | 0.169 | 8.1E-12 | 0.000000296 |
| PCNX1 | 5 | -0.345043026 | 0.655 | 0.76 | 8.32E-12 | 0.000000305 |
| PDZD8 | 5 | 0.326164679 | 0.536 | 0.419 | 8.56E-12 | 0.000000313 |
| REEP3 | 5 | 0.345787552 | 0.527 | 0.432 | 8.8E-12 | 0.000000322 |
| FOXJ3 | 5 | 0.296023572 | 0.765 | 0.682 | 8.87E-12 | 0.000000325 |

| | | | | | | |
|---|---|---|---|---|---|---|
| LINC01505 | 5 | 0.288616904 | 0.143 | 0.056 | 9.06E-12 | 0.000000332 |
| CEP44 | 5 | 0.321803524 | 0.385 | 0.271 | 9.73E-12 | 0.000000356 |
| FCHO2 | 5 | 0.293952138 | 0.519 | 0.366 | 9.83E-12 | 0.00000036 |
| KCNS3 | 5 | -0.257370345 | 0.026 | 0.107 | 1.02E-11 | 0.000000374 |
| ZNF286A | 5 | -0.30325795 | 0.147 | 0.275 | 1.04E-11 | 0.00000038 |
| AC105916.1 | 5 | -0.268440625 | 0.029 | 0.11 | 1.09E-11 | 0.0000004 |
| HMGCS1 | 5 | 0.343064977 | 0.303 | 0.174 | 1.13E-11 | 0.000000413 |
| FAM169A | 5 | -0.269079426 | 0.044 | 0.135 | 1.14E-11 | 0.000000418 |
| LUZP1 | 5 | -0.361745849 | 0.165 | 0.28 | 1.16E-11 | 0.000000426 |
| ID3 | 5 | -0.257398609 | 0.022 | 0.1 | 1.19E-11 | 0.000000436 |
| UGDH | 5 | 0.294767459 | 0.217 | 0.121 | 1.22E-11 | 0.000000445 |
| PSMG4 | 5 | -0.267744781 | 0.097 | 0.214 | 1.22E-11 | 0.000000448 |
| MTM1 | 5 | -0.327183441 | 0.154 | 0.285 | 1.28E-11 | 0.00000047 |
| LAMA3 | 5 | -0.468409685 | 0.106 | 0.217 | 1.33E-11 | 0.000000488 |
| FAM185A | 5 | 0.286521684 | 0.295 | 0.175 | 1.39E-11 | 0.000000508 |
| CDKN2A | 5 | 0.259009025 | 0.468 | 0.317 | 1.4E-11 | 0.000000513 |
| LRRTM4 | 5 | -0.264872333 | 0.706 | 0.568 | 1.44E-11 | 0.000000526 |
| ZDHHC2 | 5 | 0.283284186 | 0.268 | 0.15 | 1.44E-11 | 0.000000528 |
| SLC6A15 | 5 | -0.417537875 | 0.108 | 0.218 | 1.51E-11 | 0.000000553 |
| ELOVL4 | 5 | -0.252295619 | 0.04 | 0.128 | 1.51E-11 | 0.000000553 |
| IVNS1ABP | 5 | -0.465360471 | 0.387 | 0.514 | 1.51E-11 | 0.000000554 |
| CDH2 | 5 | -0.385460829 | 0.475 | 0.473 | 1.52E-11 | 0.000000557 |
| SLAIN2 | 5 | -0.378671009 | 0.391 | 0.526 | 1.6E-11 | 0.000000584 |
| SLC19A2 | 5 | 0.269565965 | 0.206 | 0.104 | 1.61E-11 | 0.000000589 |
| MAP4 | 5 | -0.349583946 | 0.615 | 0.729 | 1.61E-11 | 0.00000059 |
| AOPEP | 5 | -0.385400987 | 0.662 | 0.761 | 1.73E-11 | 0.000000633 |
| AL050309.1 | 5 | -0.362464863 | 0.031 | 0.116 | 1.73E-11 | 0.000000634 |
| RNF215 | 5 | 0.256485166 | 0.169 | 0.084 | 1.83E-11 | 0.000000668 |
| ZNF608 | 5 | 0.343984806 | 0.736 | 0.63 | 1.83E-11 | 0.000000671 |
| PARP11 | 5 | -0.292091319 | 0.116 | 0.236 | 1.85E-11 | 0.000000679 |
| GNS | 5 | 0.276136503 | 0.233 | 0.131 | 1.86E-11 | 0.000000683 |
| NMD3 | 5 | 0.319502822 | 0.374 | 0.251 | 2.01E-11 | 0.000000734 |
| DHRSX | 5 | -0.383376055 | 0.534 | 0.661 | 2.05E-11 | 0.000000752 |
| AC092683.1 | 5 | -0.360728647 | 0.228 | 0.367 | 2.21E-11 | 0.000000811 |
| RYBP | 5 | 0.353834465 | 0.596 | 0.51 | 2.31E-11 | 0.000000845 |
| PTMA | 5 | 0.374231609 | 0.51 | 0.401 | 2.42E-11 | 0.000000887 |
| F3 | 5 | -0.498619917 | 0.028 | 0.105 | 2.43E-11 | 0.000000891 |
| FLVCR1 | 5 | 0.329849052 | 0.488 | 0.355 | 2.44E-11 | 0.000000895 |
| WASL | 5 | 0.320027799 | 0.558 | 0.457 | 2.45E-11 | 0.000000897 |
| SLK | 5 | 0.315966403 | 0.431 | 0.298 | 2.56E-11 | 0.000000937 |
| B4GALNT3 | 5 | 0.330378585 | 0.204 | 0.115 | 2.62E-11 | 0.000000961 |
| CAP2 | 5 | 0.313526276 | 0.398 | 0.26 | 2.65E-11 | 0.000000969 |
| ANKRD28 | 5 | 0.286125971 | 0.738 | 0.635 | 2.79E-11 | 0.00000102 |
| AC120193.1 | 5 | -0.414873197 | 0.044 | 0.132 | 2.79E-11 | 0.00000102 |
| MAP4K5 | 5 | -0.356182491 | 0.587 | 0.69 | 2.85E-11 | 0.00000104 |
| CCNB1IP1 | 5 | 0.297657767 | 0.301 | 0.196 | 2.88E-11 | 0.00000106 |
| FAM78B | 5 | -0.415233498 | 0.172 | 0.291 | 2.94E-11 | 0.00000108 |
| WWTR1 | 5 | -0.608212441 | 0.332 | 0.433 | 2.97E-11 | 0.00000109 |
| SPG7 | 5 | 0.339449591 | 0.567 | 0.459 | 3.06E-11 | 0.00000112 |
| FCHSD2 | 5 | -0.406612177 | 0.668 | 0.749 | 3.34E-11 | 0.00000122 |
| PPP2R5E | 5 | -0.351257848 | 0.585 | 0.697 | 3.37E-11 | 0.00000123 |
| AC015813.1 | 5 | 0.276926988 | 0.295 | 0.186 | 3.38E-11 | 0.00000124 |
| PSMA6 | 5 | -0.375449164 | 0.273 | 0.399 | 3.57E-11 | 0.00000131 |
| ENPP2 | 5 | 0.374911442 | 0.336 | 0.204 | 3.64E-11 | 0.00000133 |
| SLC6A16 | 5 | 0.280687404 | 0.255 | 0.142 | 3.72E-11 | 0.00000136 |
| RBM17 | 5 | 0.311958237 | 0.468 | 0.362 | 3.76E-11 | 0.00000138 |
| CRPPA | 5 | -0.380246605 | 0.128 | 0.24 | 3.8E-11 | 0.00000139 |
| AL390957.1 | 5 | -1.036396848 | 0.061 | 0.142 | 3.85E-11 | 0.00000141 |

| | | | | | | |
|---|---|---|---|---|---|---|
| SLC36A4 | 5 | 0.328504912 | 0.534 | 0.433 | 3.87E-11 | 0.00000142 |
| LRP6 | 5 | -0.38148176 | 0.462 | 0.586 | 3.91E-11 | 0.00000143 |
| TRAF3IP2-AS1 | 5 | -0.382562572 | 0.479 | 0.613 | 3.96E-11 | 0.00000145 |
| SEL1L3 | 5 | 0.261866981 | 0.301 | 0.176 | 4.32E-11 | 0.00000158 |
| HCFC2 | 5 | 0.330795018 | 0.332 | 0.21 | 4.32E-11 | 0.00000158 |
| CRCP | 5 | 0.281629826 | 0.27 | 0.201 | 4.44E-11 | 0.00000163 |
| CORO2B | 5 | -0.319961085 | 0.127 | 0.246 | 4.62E-11 | 0.00000169 |
| HSPA1B | 5 | 0.339537817 | 0.108 | 0.037 | 4.67E-11 | 0.00000171 |
| RECK | 5 | -0.300913897 | 0.165 | 0.293 | 4.71E-11 | 0.00000172 |
| EML2 | 5 | 0.259987613 | 0.161 | 0.079 | 4.81E-11 | 0.00000176 |
| ZNF627 | 5 | 0.257250832 | 0.178 | 0.104 | 4.84E-11 | 0.00000177 |
| AC002377.1 | 5 | -0.351407339 | 0.048 | 0.132 | 4.93E-11 | 0.0000018 |
| IGSF11 | 5 | -0.392763087 | 0.066 | 0.153 | 4.99E-11 | 0.00000183 |
| GOLGA8B | 5 | -0.367916732 | 0.343 | 0.483 | 5.08E-11 | 0.00000186 |
| THOC2 | 5 | 0.284556721 | 0.778 | 0.691 | 5.23E-11 | 0.00000192 |
| RABGAP1 | 5 | -0.343996525 | 0.567 | 0.693 | 5.47E-11 | 0.000002 |
| TDP2 | 5 | 0.251932862 | 0.211 | 0.12 | 5.5E-11 | 0.00000201 |
| ABCC4 | 5 | -0.319522386 | 0.169 | 0.298 | 5.57E-11 | 0.00000204 |
| PER3 | 5 | -0.336900282 | 0.097 | 0.205 | 5.66E-11 | 0.00000207 |
| EMP2 | 5 | -0.262842874 | 0.075 | 0.17 | 5.99E-11 | 0.00000219 |
| KIAA0319L | 5 | -0.323035779 | 0.426 | 0.57 | 6.01E-11 | 0.0000022 |
| ACTR3B | 5 | -0.369452434 | 0.301 | 0.43 | 6.09E-11 | 0.00000223 |
| PDE5A | 5 | -0.420858467 | 0.156 | 0.267 | 6.16E-11 | 0.00000226 |
| NKAIN1 | 5 | -0.279105971 | 0.035 | 0.113 | 6.31E-11 | 0.00000231 |
| MKLN1 | 5 | -0.325773037 | 0.752 | 0.833 | 6.31E-11 | 0.00000231 |
| CCDC82 | 5 | -0.300359407 | 0.178 | 0.305 | 6.69E-11 | 0.00000245 |
| ITPK1 | 5 | 0.322134258 | 0.339 | 0.223 | 6.76E-11 | 0.00000247 |
| ZC3H14 | 5 | 0.30570994 | 0.576 | 0.476 | 7.21E-11 | 0.00000264 |
| ZMYM4 | 5 | -0.356192146 | 0.782 | 0.852 | 7.35E-11 | 0.00000269 |
| RNF165 | 5 | -0.308508713 | 0.092 | 0.192 | 7.5E-11 | 0.00000274 |
| DAP | 5 | 0.303828364 | 0.336 | 0.222 | 7.96E-11 | 0.00000291 |
| DDX46 | 5 | 0.268653927 | 0.637 | 0.579 | 8.12E-11 | 0.00000297 |
| PDE11A | 5 | -0.429929905 | 0.117 | 0.193 | 8.35E-11 | 0.00000306 |
| MT-ND4 | 5 | -0.380255342 | 0.138 | 0.261 | 8.35E-11 | 0.00000306 |
| C16orf70 | 5 | 0.27343706 | 0.237 | 0.134 | 8.97E-11 | 0.00000328 |
| APOO | 5 | 0.305837885 | 0.321 | 0.21 | 8.97E-11 | 0.00000329 |
| MAT2A | 5 | 0.317327835 | 0.446 | 0.386 | 9.02E-11 | 0.0000033 |
| CC2D1A | 5 | 0.302784575 | 0.266 | 0.168 | 1.01E-10 | 0.00000369 |
| NAA20 | 5 | 0.283559784 | 0.229 | 0.129 | 1.03E-10 | 0.00000376 |
| MARK1 | 5 | -0.341508569 | 0.539 | 0.502 | 1.08E-10 | 0.00000396 |
| STAT1 | 5 | -0.456609177 | 0.172 | 0.287 | 1.09E-10 | 0.00000398 |
| CAPS2 | 5 | -0.270365998 | 0.048 | 0.138 | 1.1E-10 | 0.00000403 |
| ANKHD1 | 5 | 0.266173847 | 0.839 | 0.775 | 1.13E-10 | 0.00000415 |
| MYO1B | 5 | -0.445609925 | 0.508 | 0.607 | 1.13E-10 | 0.00000415 |
| PREPL | 5 | 0.315703832 | 0.437 | 0.313 | 1.2E-10 | 0.00000438 |
| RELCH | 5 | 0.316308834 | 0.607 | 0.507 | 1.21E-10 | 0.00000443 |
| ATP2C1 | 5 | 0.319189576 | 0.664 | 0.557 | 1.22E-10 | 0.00000448 |
| STK3 | 5 | 0.268691518 | 0.633 | 0.486 | 1.26E-10 | 0.00000462 |
| MRPS6 | 5 | -0.326338352 | 0.127 | 0.24 | 1.4E-10 | 0.00000511 |
| MED12L | 5 | 0.328546666 | 0.633 | 0.494 | 1.49E-10 | 0.00000545 |
| ABCC1 | 5 | -0.443949862 | 0.378 | 0.498 | 1.56E-10 | 0.00000571 |
| MRPL1 | 5 | 0.251484308 | 0.475 | 0.333 | 1.56E-10 | 0.00000571 |
| WDR90 | 5 | 0.265588699 | 0.213 | 0.128 | 1.58E-10 | 0.00000579 |
| SFXN1 | 5 | 0.312167354 | 0.611 | 0.485 | 1.58E-10 | 0.0000058 |
| CNIH3 | 5 | -0.309664161 | 0.088 | 0.189 | 1.6E-10 | 0.00000584 |
| PDE10A | 5 | -0.402854338 | 0.139 | 0.26 | 1.62E-10 | 0.00000593 |
| ZMYM2 | 5 | -0.304235751 | 0.844 | 0.903 | 1.65E-10 | 0.00000604 |
| MYEF2 | 5 | 0.323031093 | 0.628 | 0.5 | 1.71E-10 | 0.00000624 |

| | | | | | | |
|---|---|---|---|---|---|---|
| BUB3 | 5 | 0.26804155 | 0.286 | 0.172 | 1.71E-10 | 0.00000627 |
| DCX | 5 | -0.394050871 | 0.084 | 0.185 | 1.81E-10 | 0.00000661 |
| YES1 | 5 | -0.365383653 | 0.253 | 0.377 | 1.83E-10 | 0.0000067 |
| FAM98B | 5 | 0.268224824 | 0.231 | 0.136 | 1.85E-10 | 0.00000679 |
| RALY | 5 | -0.32402972 | 0.248 | 0.379 | 1.88E-10 | 0.00000689 |
| TTC13 | 5 | 0.311972487 | 0.42 | 0.314 | 1.89E-10 | 0.00000693 |
| RAP1A | 5 | 0.326126946 | 0.561 | 0.456 | 2.01E-10 | 0.00000737 |
| RBM25 | 5 | -0.332290549 | 0.633 | 0.716 | 2.03E-10 | 0.00000743 |
| SETDB2 | 5 | 0.288654659 | 0.295 | 0.197 | 2.21E-10 | 0.00000809 |
| CASC2 | 5 | 0.275211617 | 0.185 | 0.093 | 2.21E-10 | 0.0000081 |
| AP2A2 | 5 | -0.329179228 | 0.283 | 0.416 | 2.32E-10 | 0.00000851 |
| CCDC148 | 5 | -0.293202428 | 0.048 | 0.132 | 2.37E-10 | 0.00000869 |
| ATP11B | 5 | -0.369713001 | 0.367 | 0.497 | 2.4E-10 | 0.00000878 |
| ZNF562 | 5 | -0.271036087 | 0.134 | 0.249 | 2.5E-10 | 0.00000916 |
| SPATS2L | 5 | -0.442236757 | 0.501 | 0.468 | 2.6E-10 | 0.00000954 |
| SLC22A23 | 5 | -0.477036193 | 0.244 | 0.363 | 2.76E-10 | 0.0000101 |
| TTC7B | 5 | 0.298896908 | 0.516 | 0.386 | 2.78E-10 | 0.0000102 |
| ULK1 | 5 | -0.283441808 | 0.163 | 0.288 | 2.79E-10 | 0.0000102 |
| PKNOX1 | 5 | -0.288354603 | 0.209 | 0.342 | 2.87E-10 | 0.0000105 |
| ATF4 | 5 | 0.256734708 | 0.228 | 0.129 | 2.9E-10 | 0.0000106 |
| CYB5B | 5 | 0.314004061 | 0.389 | 0.286 | 2.9E-10 | 0.0000106 |
| KIAA0408 | 5 | 0.255057843 | 0.328 | 0.203 | 2.91E-10 | 0.0000107 |
| KLHDC4 | 5 | 0.298232318 | 0.36 | 0.26 | 3.01E-10 | 0.000011 |
| PRKX | 5 | -0.329991521 | 0.231 | 0.363 | 3.05E-10 | 0.0000112 |
| CDR2 | 5 | 0.33359947 | 0.376 | 0.257 | 3.13E-10 | 0.0000114 |
| CEP63 | 5 | 0.330424186 | 0.422 | 0.304 | 3.25E-10 | 0.0000119 |
| SPATA7 | 5 | 0.321541712 | 0.297 | 0.201 | 3.26E-10 | 0.0000119 |
| UBL3 | 5 | 0.310402677 | 0.506 | 0.372 | 3.38E-10 | 0.0000124 |
| CMTM7 | 5 | 0.352343246 | 0.264 | 0.174 | 3.43E-10 | 0.0000126 |
| SP3 | 5 | -0.346468046 | 0.55 | 0.665 | 3.49E-10 | 0.0000128 |
| DDB1 | 5 | 0.294003791 | 0.338 | 0.238 | 3.58E-10 | 0.0000131 |
| LGMN | 5 | -0.309948326 | 0.084 | 0.174 | 3.62E-10 | 0.0000133 |
| THRAP3 | 5 | 0.267722201 | 0.681 | 0.58 | 3.87E-10 | 0.0000142 |
| AC119673.2 | 5 | -0.320050496 | 0.266 | 0.393 | 3.96E-10 | 0.0000145 |
| VAV3 | 5 | -0.410060017 | 0.073 | 0.167 | 4.05E-10 | 0.0000148 |
| ANO10 | 5 | -0.397672293 | 0.356 | 0.48 | 4.07E-10 | 0.0000149 |
| NUCKS1 | 5 | -0.324956117 | 0.431 | 0.569 | 4.47E-10 | 0.0000164 |
| ZRANB2-AS2 | 5 | 0.302022131 | 0.301 | 0.183 | 4.71E-10 | 0.0000173 |
| GLCE | 5 | -0.42101183 | 0.279 | 0.39 | 4.75E-10 | 0.0000174 |
| PIK3AP1 | 5 | -0.317825695 | 0.077 | 0.164 | 4.84E-10 | 0.0000177 |
| ZC3HAV1 | 5 | 0.308040803 | 0.334 | 0.228 | 5.18E-10 | 0.000019 |
| DPY30 | 5 | 0.251417774 | 0.246 | 0.143 | 5.29E-10 | 0.0000194 |
| GOLPH3 | 5 | 0.295581586 | 0.572 | 0.468 | 5.54E-10 | 0.0000203 |
| ATF6 | 5 | 0.301372031 | 0.503 | 0.396 | 5.61E-10 | 0.0000205 |
| ZC3H8 | 5 | 0.31628683 | 0.431 | 0.332 | 5.61E-10 | 0.0000205 |
| TPST2 | 5 | 0.289915741 | 0.178 | 0.097 | 5.73E-10 | 0.000021 |
| PRICKLE1 | 5 | -0.688277961 | 0.358 | 0.414 | 5.79E-10 | 0.0000212 |
| STK38L | 5 | 0.290926282 | 0.372 | 0.249 | 5.88E-10 | 0.0000215 |
| RHOBTB3 | 5 | -0.386910445 | 0.358 | 0.471 | 5.99E-10 | 0.0000219 |
| GREB1 | 5 | -0.306123502 | 0.039 | 0.114 | 6.56E-10 | 0.000024 |
| ITPRID2 | 5 | 0.299702802 | 0.435 | 0.315 | 6.68E-10 | 0.0000244 |
| PRKAG2 | 5 | 0.328758201 | 0.561 | 0.429 | 6.74E-10 | 0.0000247 |
| KSR1 | 5 | -0.373681405 | 0.273 | 0.403 | 6.77E-10 | 0.0000248 |
| TRAP1 | 5 | 0.347844562 | 0.431 | 0.341 | 7.29E-10 | 0.0000267 |
| SLC23A2 | 5 | -0.321524398 | 0.215 | 0.339 | 8.05E-10 | 0.0000295 |
| S100PBP | 5 | 0.295170072 | 0.484 | 0.377 | 8.19E-10 | 0.00003 |
| SNX13 | 5 | 0.314754067 | 0.714 | 0.612 | 8.54E-10 | 0.0000312 |
| OBSL1 | 5 | 0.271387843 | 0.244 | 0.15 | 9.1E-10 | 0.0000333 |

| | | | | | | |
|---|---|---|---|---|---|---|
| GPAM | 5 | -0.295998112 | 0.094 | 0.188 | 9.23E-10 | 0.0000338 |
| TM7SF3 | 5 | 0.273496916 | 0.292 | 0.206 | 9.32E-10 | 0.0000341 |
| ARHGEF2 | 5 | -0.253026741 | 0.106 | 0.209 | 9.7E-10 | 0.0000355 |
| EYA3 | 5 | 0.298045358 | 0.582 | 0.482 | 9.8E-10 | 0.0000359 |
| GSPT1 | 5 | 0.288160767 | 0.472 | 0.399 | 1.04E-09 | 0.000038 |
| NPDC1 | 5 | 0.264676996 | 0.237 | 0.137 | 1.04E-09 | 0.0000383 |
| EIF1B-AS1 | 5 | -0.263702892 | 0.125 | 0.237 | 1.06E-09 | 0.0000389 |
| C19orf48 | 5 | 0.250307069 | 0.206 | 0.116 | 1.07E-09 | 0.0000393 |
| OSBPL11 | 5 | -0.379591772 | 0.222 | 0.341 | 1.08E-09 | 0.0000395 |
| PPIP5K1 | 5 | 0.257239134 | 0.211 | 0.116 | 1.08E-09 | 0.0000395 |
| PTPDC1 | 5 | 0.307557284 | 0.321 | 0.228 | 1.08E-09 | 0.0000396 |
| TOR1AIP2 | 5 | 0.281018486 | 0.312 | 0.226 | 1.13E-09 | 0.0000413 |
| CCDC112 | 5 | -0.256276677 | 0.081 | 0.178 | 1.17E-09 | 0.0000428 |
| CWC27 | 5 | 0.317297807 | 0.563 | 0.475 | 1.18E-09 | 0.0000431 |
| ARHGEF10 | 5 | -0.333103953 | 0.169 | 0.278 | 1.19E-09 | 0.0000435 |
| B4GALT5 | 5 | -0.333269459 | 0.297 | 0.426 | 1.19E-09 | 0.0000435 |
| C21orf58 | 5 | 0.28408571 | 0.356 | 0.236 | 1.2E-09 | 0.0000438 |
| DYNC1LI1 | 5 | -0.336744788 | 0.229 | 0.353 | 1.26E-09 | 0.000046 |
| MRTFB | 5 | 0.295829935 | 0.673 | 0.583 | 1.3E-09 | 0.0000477 |
| SUGP2 | 5 | -0.328554299 | 0.446 | 0.563 | 1.36E-09 | 0.0000498 |
| MALT1 | 5 | -0.314804691 | 0.182 | 0.302 | 1.39E-09 | 0.0000509 |
| KATNAL2 | 5 | -0.256403102 | 0.094 | 0.195 | 1.44E-09 | 0.0000528 |
| MOCOS | 5 | -0.254961518 | 0.05 | 0.124 | 1.5E-09 | 0.0000549 |
| RAPH1 | 5 | -0.332162863 | 0.306 | 0.439 | 1.51E-09 | 0.0000555 |
| FASTKD2 | 5 | 0.274980385 | 0.363 | 0.244 | 1.54E-09 | 0.0000562 |
| TNRC6B | 5 | 0.273645709 | 0.897 | 0.852 | 1.54E-09 | 0.0000565 |
| TPM1 | 5 | -0.484730482 | 0.248 | 0.332 | 1.64E-09 | 0.0000601 |
| CLCN5 | 5 | -0.412911691 | 0.402 | 0.44 | 1.65E-09 | 0.0000603 |
| SPIN1 | 5 | -0.311284041 | 0.536 | 0.659 | 1.66E-09 | 0.0000608 |
| MT-ATP6 | 5 | -0.386941219 | 0.158 | 0.275 | 1.79E-09 | 0.0000657 |
| ATP6V1A | 5 | 0.303891331 | 0.409 | 0.292 | 1.8E-09 | 0.0000658 |
| GABPA | 5 | -0.328866723 | 0.196 | 0.31 | 1.84E-09 | 0.0000674 |
| CPQ | 5 | -0.37473274 | 0.209 | 0.293 | 1.85E-09 | 0.0000678 |
| STT3B | 5 | 0.293544158 | 0.771 | 0.688 | 1.85E-09 | 0.0000678 |
| PPM1A | 5 | 0.291616667 | 0.481 | 0.372 | 1.98E-09 | 0.0000723 |
| CMC1 | 5 | 0.308407238 | 0.464 | 0.341 | 2.02E-09 | 0.0000738 |
| AC098829.1 | 5 | -0.257299882 | 0.039 | 0.113 | 2.04E-09 | 0.0000747 |
| RANBP3 | 5 | 0.255569401 | 0.288 | 0.195 | 2.22E-09 | 0.0000812 |
| CHEK2 | 5 | 0.253149279 | 0.22 | 0.146 | 2.26E-09 | 0.0000828 |
| DNM2 | 5 | 0.30647906 | 0.38 | 0.313 | 2.31E-09 | 0.0000845 |
| WSB1 | 5 | -0.487539759 | 0.727 | 0.792 | 2.31E-09 | 0.0000846 |
| MTURN | 5 | -0.252771568 | 0.123 | 0.232 | 2.46E-09 | 0.00009 |
| MTMR2 | 5 | -0.363049731 | 0.264 | 0.373 | 2.47E-09 | 0.0000905 |
| EFL1 | 5 | 0.277443449 | 0.372 | 0.262 | 2.54E-09 | 0.000093 |
| POLI | 5 | 0.280169201 | 0.35 | 0.243 | 2.55E-09 | 0.0000933 |
| CDCA7 | 5 | -0.348029061 | 0.317 | 0.421 | 2.55E-09 | 0.0000935 |
| ZFAND6 | 5 | -0.369720528 | 0.565 | 0.662 | 2.63E-09 | 0.0000964 |
| HDX | 5 | -0.276481564 | 0.154 | 0.27 | 2.72E-09 | 0.0000994 |
| ACVR2A | 5 | -0.46297981 | 0.369 | 0.462 | 2.73E-09 | 0.0001 |
| ARL17A | 5 | 0.27102415 | 0.382 | 0.279 | 2.74E-09 | 0.0001 |
| USP46 | 5 | -0.256344942 | 0.105 | 0.201 | 2.8E-09 | 0.000103 |
| C2CD3 | 5 | 0.307904406 | 0.457 | 0.351 | 2.83E-09 | 0.000104 |
| UBE2L3 | 5 | 0.274191673 | 0.327 | 0.22 | 2.85E-09 | 0.000104 |
| LIMD1 | 5 | -0.332960843 | 0.25 | 0.372 | 2.94E-09 | 0.000108 |
| LDB2 | 5 | -0.528487388 | 0.36 | 0.335 | 3.04E-09 | 0.000111 |
| PPM1B | 5 | -0.332034508 | 0.393 | 0.519 | 3.08E-09 | 0.000113 |
| MT-CYB | 5 | -0.384621124 | 0.132 | 0.241 | 3.14E-09 | 0.000115 |
| MINK1 | 5 | 0.279720771 | 0.358 | 0.255 | 3.18E-09 | 0.000117 |

| | | | | | | |
|---|---|---|---|---|---|---|
| FAM13B | 5 | -0.322891175 | 0.512 | 0.629 | 3.19E-09 | 0.000117 |
| RBL1 | 5 | 0.276642846 | 0.541 | 0.41 | 3.2E-09 | 0.000117 |
| ANKS1B | 5 | -0.484506542 | 0.87 | 0.784 | 3.2E-09 | 0.000117 |
| MEGF9 | 5 | 0.332479907 | 0.361 | 0.281 | 3.26E-09 | 0.000119 |
| FAM172A | 5 | -0.317644034 | 0.692 | 0.779 | 3.3E-09 | 0.000121 |
| AL354771.1 | 5 | -0.322023994 | 0.07 | 0.143 | 3.37E-09 | 0.000123 |
| APOLD1 | 5 | -0.356777049 | 0.066 | 0.146 | 3.37E-09 | 0.000123 |
| NUP214 | 5 | 0.282597117 | 0.554 | 0.487 | 3.38E-09 | 0.000123726 |
| IDH1 | 5 | -0.343367066 | 0.125 | 0.225 | 3.39E-09 | 0.000124 |
| EIF2AK4 | 5 | -0.286662318 | 0.182 | 0.3 | 3.39E-09 | 0.000124 |
| MTHFD2L | 5 | 0.28690262 | 0.343 | 0.234 | 3.47E-09 | 0.000127 |
| AL139260.1 | 5 | -0.271987524 | 0.073 | 0.163 | 3.58E-09 | 0.000131 |
| ACO2 | 5 | 0.266875692 | 0.308 | 0.209 | 3.64E-09 | 0.000133 |
| CCDC191 | 5 | 0.250708261 | 0.198 | 0.11 | 3.64E-09 | 0.000133 |
| SERPINB6 | 5 | -0.328208391 | 0.136 | 0.231 | 3.65E-09 | 0.000134 |
| MMS22L | 5 | 0.292481183 | 0.716 | 0.597 | 4.02E-09 | 0.000147 |
| PCGF3 | 5 | -0.257501041 | 0.222 | 0.347 | 4.23E-09 | 0.000155 |
| MORF4L2 | 5 | 0.310702184 | 0.611 | 0.545 | 4.39E-09 | 0.000161 |
| ARMCX5-GPRASP2 | 5 | -0.284710878 | 0.196 | 0.315 | 4.47E-09 | 0.000164 |
| RPL14 | 5 | 0.259442639 | 0.332 | 0.217 | 4.5E-09 | 0.000165 |
| C2CD5 | 5 | 0.268579723 | 0.738 | 0.671 | 4.53E-09 | 0.000166 |
| VSIG10 | 5 | 0.311127649 | 0.343 | 0.235 | 4.7E-09 | 0.000172 |
| GPBP1 | 5 | 0.281289086 | 0.752 | 0.69 | 4.79E-09 | 0.000175 |
| AARS | 5 | 0.264063806 | 0.277 | 0.189 | 4.84E-09 | 0.000177 |
| PITPNB | 5 | 0.291021091 | 0.648 | 0.548 | 4.96E-09 | 0.000182 |
| KMT2C | 5 | -0.265493516 | 0.917 | 0.927 | 5.14E-09 | 0.000188 |
| SRSF4 | 5 | 0.257754179 | 0.71 | 0.64 | 5.17E-09 | 0.000189 |
| OSBPL1A | 5 | -0.373544324 | 0.33 | 0.447 | 5.33E-09 | 0.000195 |
| KNTC1 | 5 | 0.271154366 | 0.618 | 0.487 | 5.41E-09 | 0.000198 |
| LINC00882 | 5 | -0.319124689 | 0.037 | 0.105 | 5.5E-09 | 0.000201 |
| AL354809.1 | 5 | -0.29747284 | 0.053 | 0.124 | 5.57E-09 | 0.000204 |
| CYP51A1 | 5 | 0.275702131 | 0.222 | 0.126 | 5.98E-09 | 0.000219 |
| CPEB2 | 5 | -0.393043462 | 0.251 | 0.363 | 6.11E-09 | 0.000224 |
| HDAC2 | 5 | -0.326986997 | 0.475 | 0.6 | 6.11E-09 | 0.000224 |
| FHOD3 | 5 | -0.423830788 | 0.244 | 0.311 | 6.12E-09 | 0.000224 |
| RNF19A | 5 | -0.381357956 | 0.4 | 0.513 | 6.31E-09 | 0.000231 |
| MFSD14B | 5 | 0.2815324 | 0.404 | 0.308 | 6.49E-09 | 0.000238 |
| SCFD2 | 5 | 0.328045597 | 0.673 | 0.591 | 6.55E-09 | 0.00024 |
| PPP6R2 | 5 | 0.287992465 | 0.705 | 0.631 | 6.6E-09 | 0.000242 |
| GRIP1 | 5 | -0.298870415 | 0.4 | 0.356 | 6.84E-09 | 0.000251 |
| FANCA | 5 | 0.294323588 | 0.576 | 0.447 | 6.89E-09 | 0.000252 |
| TAF1D | 5 | 0.26760743 | 0.763 | 0.671 | 6.99E-09 | 0.000256 |
| SECISBP2L | 5 | -0.323304147 | 0.334 | 0.459 | 7.01E-09 | 0.000257 |
| DPY19L2 | 5 | 0.29583334 | 0.488 | 0.387 | 7.12E-09 | 0.000261 |
| MAGED1 | 5 | -0.334471772 | 0.508 | 0.612 | 7.19E-09 | 0.000263 |
| SH3PXD2B | 5 | -0.291424294 | 0.16 | 0.269 | 7.2E-09 | 0.000264 |
| PRKAR2A | 5 | 0.298941861 | 0.51 | 0.405 | 7.4E-09 | 0.000271 |
| SLIT1 | 5 | -0.288034068 | 0.042 | 0.115 | 7.47E-09 | 0.000273 |
| IPO7 | 5 | -0.332981786 | 0.459 | 0.568 | 7.79E-09 | 0.000285 |
| KNL1 | 5 | -0.293844682 | 0.119 | 0.221 | 8.29E-09 | 0.000304 |
| HIP1 | 5 | -0.364896157 | 0.262 | 0.383 | 8.44E-09 | 0.000309 |
| ARFGAP3 | 5 | 0.278580418 | 0.303 | 0.215 | 8.44E-09 | 0.000309 |
| SEC23A | 5 | 0.298603928 | 0.409 | 0.305 | 8.44E-09 | 0.000309 |
| AGFG1 | 5 | 0.284299482 | 0.609 | 0.534 | 8.79E-09 | 0.000322 |
| NMNAT2 | 5 | -0.435189687 | 0.251 | 0.325 | 8.88E-09 | 0.000325 |
| SKP2 | 5 | 0.258153925 | 0.327 | 0.228 | 9.04E-09 | 0.000331 |
| ALDH1L2 | 5 | -0.255424093 | 0.088 | 0.182 | 9.32E-09 | 0.000341 |
| NDUFAF2 | 5 | 0.350841932 | 0.426 | 0.317 | 9.86E-09 | 0.000361 |

| | | | | | | |
|---|---|---|---|---|---|---|
| RCN2 | 5 | -0.285071623 | 0.281 | 0.403 | 1.06E-08 | 0.000386 |
| ZNF721 | 5 | 0.268898197 | 0.673 | 0.585 | 1.06E-08 | 0.000389 |
| MGRN1 | 5 | 0.262227496 | 0.332 | 0.251 | 1.07E-08 | 0.00039 |
| BRD1 | 5 | 0.267094584 | 0.47 | 0.364 | 1.07E-08 | 0.000391 |
| B4GALT6 | 5 | -0.429170476 | 0.222 | 0.295 | 1.08E-08 | 0.000394 |
| PACS1 | 5 | -0.301408874 | 0.404 | 0.534 | 1.09E-08 | 0.000399 |
| C2CD2 | 5 | -0.26869532 | 0.075 | 0.157 | 0.000000011 | 0.000404 |
| NFX1 | 5 | 0.275891534 | 0.532 | 0.427 | 1.15E-08 | 0.000421 |
| TMEM220 | 5 | -0.25832743 | 0.11 | 0.209 | 1.17E-08 | 0.000429 |
| STRADA | 5 | -0.27889078 | 0.272 | 0.392 | 1.25E-08 | 0.000459 |
| CCDC18-AS1 | 5 | 0.298580147 | 0.464 | 0.349 | 1.31E-08 | 0.000479 |
| FARP2 | 5 | 0.288099953 | 0.413 | 0.321 | 1.32E-08 | 0.000482 |
| TNRC6A | 5 | -0.284358014 | 0.776 | 0.821 | 1.33E-08 | 0.000486 |
| SIK2 | 5 | 0.264978613 | 0.646 | 0.541 | 1.34E-08 | 0.00049 |
| IKZF3 | 5 | -0.339784859 | 0.171 | 0.275 | 1.36E-08 | 0.000498 |
| ICA1L | 5 | 0.273577693 | 0.283 | 0.179 | 1.36E-08 | 0.000498 |
| BZW2 | 5 | -0.416135823 | 0.303 | 0.41 | 1.37E-08 | 0.000503 |
| STMN2 | 5 | -0.370208065 | 0.099 | 0.174 | 1.41E-08 | 0.000515 |
| AC093879.1 | 5 | -0.271192635 | 0.101 | 0.19 | 1.47E-08 | 0.000537 |
| SCFD1 | 5 | 0.269875344 | 0.675 | 0.609 | 0.000000015 | 0.00055 |
| PRH1 | 5 | -0.364844875 | 0.374 | 0.478 | 1.51E-08 | 0.000553 |
| EZR | 5 | 0.259226312 | 0.361 | 0.256 | 1.52E-08 | 0.000557 |
| ST8SIA4 | 5 | -0.301044399 | 0.059 | 0.121 | 1.53E-08 | 0.000559 |
| GSTA4 | 5 | -0.284964677 | 0.226 | 0.344 | 1.62E-08 | 0.000593 |
| HSPE1 | 5 | 0.369449454 | 0.196 | 0.124 | 1.71E-08 | 0.000626 |
| SLC38A4 | 5 | -0.503025178 | 0.108 | 0.179 | 1.73E-08 | 0.000634 |
| GANC | 5 | -0.256896296 | 0.138 | 0.243 | 1.74E-08 | 0.000636 |
| SRSF1 | 5 | 0.254425735 | 0.585 | 0.494 | 0.000000018 | 0.000659 |
| SLC35A3 | 5 | -0.259854944 | 0.191 | 0.303 | 1.84E-08 | 0.000672 |
| MRPS9 | 5 | 0.266832222 | 0.319 | 0.24 | 1.87E-08 | 0.000686 |
| NAAA | 5 | -0.358917386 | 0.196 | 0.301 | 2.08E-08 | 0.000761 |
| USP40 | 5 | 0.282053002 | 0.433 | 0.319 | 0.000000021 | 0.000769 |
| PCMTD1 | 5 | -0.291243099 | 0.552 | 0.659 | 2.14E-08 | 0.000784 |
| OPTN | 5 | -0.253244419 | 0.083 | 0.171 | 2.25E-08 | 0.000822 |
| SLC10A7 | 5 | 0.274558168 | 0.431 | 0.307 | 2.25E-08 | 0.000822 |
| TMEM117 | 5 | -0.461195 | 0.514 | 0.583 | 2.27E-08 | 0.000831 |
| MT-ND1 | 5 | -0.270662136 | 0.083 | 0.173 | 2.29E-08 | 0.000837 |
| CLYBL | 5 | 0.25259233 | 0.389 | 0.268 | 2.34E-08 | 0.000858 |
| NPRL3 | 5 | 0.282527081 | 0.382 | 0.283 | 2.44E-08 | 0.000893 |
| DSE | 5 | -0.340429885 | 0.244 | 0.35 | 2.46E-08 | 0.000901 |
| HLCS | 5 | 0.272789261 | 0.64 | 0.528 | 0.000000025 | 0.000917 |
| AHDC1 | 5 | 0.252323495 | 0.281 | 0.178 | 2.68E-08 | 0.00098 |
| MRPS25 | 5 | 0.257905055 | 0.336 | 0.261 | 0.000000027 | 0.000987 |
| PIK3CB | 5 | 0.258260305 | 0.473 | 0.357 | 2.75E-08 | 0.001005758 |
| PHKA2 | 5 | 0.269733261 | 0.488 | 0.388 | 2.77E-08 | 0.001013226 |
| CHD9 | 5 | -0.301433164 | 0.899 | 0.922 | 2.87E-08 | 0.001049772 |
| BTAF1 | 5 | 0.267466287 | 0.719 | 0.638 | 2.91E-08 | 0.001063644 |
| TTN | 5 | -0.580652456 | 0.077 | 0.156 | 3.02E-08 | 0.001104788 |
| NBAS | 5 | -0.425392173 | 0.778 | 0.8 | 3.06E-08 | 0.001122058 |
| ACVR1 | 5 | -0.365131545 | 0.297 | 0.395 | 3.08E-08 | 0.001126893 |
| ZBTB7C | 5 | -0.296063771 | 0.053 | 0.131 | 3.08E-08 | 0.001128964 |
| PPP4R1 | 5 | -0.294569503 | 0.446 | 0.567 | 3.15E-08 | 0.001152332 |
| CRIM1 | 5 | -0.406491327 | 0.242 | 0.342 | 3.16E-08 | 0.001155467 |
| CREBBP | 5 | -0.271403709 | 0.591 | 0.702 | 3.19E-08 | 0.001166126 |
| EXPH5 | 5 | -0.332898227 | 0.086 | 0.162 | 3.34E-08 | 0.001221622 |
| CACNA2D3 | 5 | -0.569287654 | 0.501 | 0.519 | 0.000000034 | 0.001243601 |
| KATNBL1 | 5 | -0.300465441 | 0.393 | 0.512 | 3.45E-08 | 0.001263603 |
| CRYM | 5 | -0.270809694 | 0.037 | 0.103 | 3.68E-08 | 0.001345514 |

| | | | | | | |
|---|---|---|---|---|---|---|
| KLHL5 | 5 | -0.251568786 | 0.117 | 0.215 | 3.86E-08 | 0.00141173 |
| MACO1 | 5 | -0.261570489 | 0.242 | 0.362 | 3.94E-08 | 0.001442203 |
| HHAT | 5 | -0.307598946 | 0.189 | 0.281 | 4.01E-08 | 0.001468638 |
| SH3RF3 | 5 | 0.325540778 | 0.38 | 0.263 | 4.04E-08 | 0.001480297 |
| SCML2 | 5 | 0.273655307 | 0.539 | 0.431 | 4.09E-08 | 0.001499107 |
| GORASP2 | 5 | 0.26668146 | 0.314 | 0.219 | 0.000000041 | 0.001500135 |
| NEXMIF | 5 | -0.3293505 | 0.103 | 0.183 | 4.13E-08 | 0.001510826 |
| SLC4A5 | 5 | -0.253520433 | 0.037 | 0.101 | 4.18E-08 | 0.001529846 |
| G2E3 | 5 | -0.29011194 | 0.367 | 0.487 | 4.21E-08 | 0.001539646 |
| EEPD1 | 5 | -0.424692058 | 0.235 | 0.308 | 4.23E-08 | 0.001550327 |
| ECHDC1 | 5 | -0.260547622 | 0.325 | 0.45 | 4.27E-08 | 0.001564374 |
| CBL | 5 | -0.29258744 | 0.248 | 0.359 | 4.39E-08 | 0.001608821 |
| WDR45B | 5 | 0.363110072 | 0.356 | 0.301 | 4.51E-08 | 0.001651962 |
| RSPRY1 | 5 | 0.260314915 | 0.453 | 0.358 | 4.56E-08 | 0.001670598 |
| DGCR9 | 5 | -0.253851073 | 0.072 | 0.147 | 4.72E-08 | 0.001727252 |
| CCDC50 | 5 | -0.305384429 | 0.323 | 0.43 | 4.74E-08 | 0.001734346 |
| OGA | 5 | -0.315332095 | 0.604 | 0.699 | 4.74E-08 | 0.001736587 |
| CDKL3 | 5 | -0.251702306 | 0.123 | 0.221 | 4.91E-08 | 0.001797622 |
| PHLPP2 | 5 | -0.255161592 | 0.167 | 0.271 | 5.08E-08 | 0.001859829 |
| SEPTIN7 | 5 | -0.317150579 | 0.618 | 0.686 | 5.23E-08 | 0.001912966 |
| CTNNAL1 | 5 | 0.267099975 | 0.369 | 0.257 | 5.43E-08 | 0.00198633 |
| PRDM2 | 5 | -0.386375943 | 0.495 | 0.572 | 5.65E-08 | 0.00206716 |
| POLR2J3 | 5 | 0.281646249 | 0.572 | 0.491 | 5.67E-08 | 0.002074992 |
| CD99L2 | 5 | 0.293932254 | 0.514 | 0.426 | 5.91E-08 | 0.002164576 |
| HIBCH | 5 | -0.29347626 | 0.486 | 0.59 | 5.98E-08 | 0.002188621 |
| RAI1 | 5 | -0.305140285 | 0.248 | 0.356 | 0.00000006 | 0.002196187 |
| AC104596.1 | 5 | -0.261806986 | 0.053 | 0.123 | 6.61E-08 | 0.002420086 |
| ARID2 | 5 | -0.29614026 | 0.754 | 0.818 | 6.75E-08 | 0.002471857 |
| DCBLD2 | 5 | -0.284090231 | 0.284 | 0.404 | 6.81E-08 | 0.002492857 |
| GRM7 | 5 | -0.715853419 | 0.072 | 0.141 | 7.65E-08 | 0.002802474 |
| WBP11 | 5 | 0.27276396 | 0.292 | 0.201 | 7.67E-08 | 0.002807971 |
| GRID1 | 5 | -0.331456087 | 0.055 | 0.129 | 8.17E-08 | 0.002991894 |
| HIST1H2AC | 5 | 0.25908595 | 0.147 | 0.075 | 8.19E-08 | 0.002996836 |
| RTTN | 5 | 0.294503904 | 0.591 | 0.515 | 8.25E-08 | 0.003019344 |
| SELENOI | 5 | 0.265707505 | 0.475 | 0.373 | 8.48E-08 | 0.003102837 |
| MYO9A | 5 | -0.279952128 | 0.602 | 0.684 | 0.000000086 | 0.003147908 |
| MCU | 5 | 0.262922591 | 0.617 | 0.526 | 0.000000086 | 0.003148032 |
| SLC20A2 | 5 | 0.277695194 | 0.543 | 0.449 | 9.24E-08 | 0.003384133 |
| TET3 | 5 | -0.250320019 | 0.301 | 0.422 | 9.34E-08 | 0.003418744 |
| CAST | 5 | 0.258359861 | 0.327 | 0.223 | 9.74E-08 | 0.003567147 |
| ZNF652 | 5 | 0.267200659 | 0.539 | 0.435 | 9.81E-08 | 0.003590555 |
| MLLT3 | 5 | -0.424282832 | 0.782 | 0.82 | 9.88E-08 | 0.003615579 |
| CRLF3 | 5 | -0.252501569 | 0.222 | 0.333 | 0.000000101 | 0.003715062 |
| SMAD2 | 5 | -0.292833072 | 0.639 | 0.703 | 0.000000103 | 0.003766682 |
| GRK4 | 5 | -0.264339952 | 0.193 | 0.3 | 0.000000104 | 0.003824647 |
| ATP11A | 5 | 0.254487513 | 0.583 | 0.473 | 0.000000109 | 0.003974054 |
| DGLUCY | 5 | 0.270477359 | 0.352 | 0.253 | 0.000000114 | 0.004158704 |
| KHDRBS1 | 5 | -0.277629655 | 0.561 | 0.642 | 0.000000114 | 0.004187684 |
| ARAP2 | 5 | -0.381903028 | 0.376 | 0.411 | 0.000000115 | 0.004206889 |
| TMED8 | 5 | 0.255601929 | 0.317 | 0.22 | 0.000000116 | 0.004260991 |
| MGLL | 5 | -0.303717724 | 0.128 | 0.197 | 0.000000118 | 0.004320051 |
| PPFIA1 | 5 | -0.341813178 | 0.462 | 0.561 | 0.00000012 | 0.004406701 |
| MED13 | 5 | -0.279218925 | 0.717 | 0.798 | 0.000000121 | 0.004443852 |
| SLC35B3 | 5 | -0.272758809 | 0.281 | 0.397 | 0.000000123 | 0.004493322 |
| FAM184B | 5 | -0.300684522 | 0.114 | 0.198 | 0.000000125 | 0.004585989 |
| CA5B | 5 | -0.274786098 | 0.207 | 0.305 | 0.000000135 | 0.004951587 |
| SLC2A1 | 5 | -0.262940656 | 0.057 | 0.13 | 0.000000135 | 0.004957289 |
| POC1B | 5 | 0.257675367 | 0.327 | 0.228 | 0.000000137 | 0.004999097 |

| | | | | | | |
|---|---|---|---|---|---|---|
| MED15 | 5 | 0.250493571 | 0.444 | 0.362 | 0.000000141 | 0.005143973 |
| KIAA1958 | 5 | -0.281934593 | 0.743 | 0.765 | 0.000000143 | 0.005225275 |
| MBNL3 | 5 | -0.357084611 | 0.475 | 0.49 | 0.000000144 | 0.005273135 |
| CTSC | 5 | -0.255427596 | 0.121 | 0.21 | 0.000000147 | 0.005368394 |
| ZFP36L2 | 5 | 0.262362199 | 0.116 | 0.065 | 0.000000148 | 0.005418946 |
| NGLY1 | 5 | -0.355339954 | 0.422 | 0.477 | 0.000000157 | 0.005757036 |
| DENND1A | 5 | -0.274912692 | 0.651 | 0.733 | 0.00000016 | 0.005859912 |
| PSIP1 | 5 | -0.318427807 | 0.455 | 0.558 | 0.000000165 | 0.006054741 |
| SFXN5 | 5 | -0.284531737 | 0.217 | 0.317 | 0.000000166 | 0.006069195 |
| AC004943.2 | 5 | 0.251234964 | 0.332 | 0.255 | 0.000000169 | 0.006198627 |
| TMTC2 | 5 | -0.446636052 | 0.514 | 0.51 | 0.00000017 | 0.006223545 |
| DOP1A | 5 | 0.266818511 | 0.594 | 0.505 | 0.00000018 | 0.006605067 |
| LMNB1 | 5 | -0.331311784 | 0.4 | 0.492 | 0.000000198 | 0.007259541 |
| OBI1 | 5 | -0.262881695 | 0.206 | 0.307 | 0.00000021 | 0.007702752 |
| WDR44 | 5 | -0.329545503 | 0.426 | 0.531 | 0.000000215 | 0.007870831 |
| FERMT1 | 5 | -0.32785464 | 0.117 | 0.204 | 0.000000225 | 0.008235448 |
| CCSER2 | 5 | -0.279633981 | 0.413 | 0.531 | 0.000000227 | 0.008308548 |
| SPG11 | 5 | 0.265088116 | 0.587 | 0.49 | 0.000000231 | 0.00847241 |
| OGT | 5 | -0.27010741 | 0.572 | 0.665 | 0.000000234 | 0.008558085 |
| G3BP2 | 5 | 0.271023289 | 0.481 | 0.388 | 0.000000239 | 0.00874246 |
| SRP54 | 5 | 0.273616449 | 0.444 | 0.354 | 0.000000259 | 0.009468536 |
| WDR26 | 5 | -0.2662827 | 0.372 | 0.489 | 0.00000026 | 0.00950264 |
| AMPH | 5 | -0.27947227 | 0.095 | 0.178 | 0.00000026 | 0.009513076 |
| DPY19L1 | 5 | -0.300406537 | 0.323 | 0.432 | 0.000000261 | 0.009547024 |
| SLC38A2 | 5 | -0.303917341 | 0.35 | 0.458 | 0.000000279 | 0.010226217 |
| DMXL1 | 5 | 0.250913026 | 0.624 | 0.524 | 0.000000284 | 0.01040306 |
| UBE2R2 | 5 | -0.283716818 | 0.582 | 0.683 | 0.000000287 | 0.01051976 |
| CCND3 | 5 | -0.25624469 | 0.128 | 0.223 | 0.000000288 | 0.010533411 |
| RAB28 | 5 | 0.264002332 | 0.556 | 0.459 | 0.000000305 | 0.011151831 |
| ZYG11A | 5 | -0.311279853 | 0.246 | 0.306 | 0.000000311 | 0.011387426 |
| RSF1 | 5 | -0.287893013 | 0.609 | 0.69 | 0.000000323 | 0.011829808 |
| ARID5B | 5 | -0.304066498 | 0.409 | 0.412 | 0.000000328 | 0.011999578 |
| ITGB3BP | 5 | 0.261479328 | 0.477 | 0.393 | 0.000000332 | 0.012150763 |
| SPEF2 | 5 | -0.364967144 | 0.084 | 0.152 | 0.000000344 | 0.01258944 |
| CHSY1 | 5 | -0.279438021 | 0.233 | 0.338 | 0.000000345 | 0.012613259 |
| SRGAP1 | 5 | -0.369073398 | 0.365 | 0.431 | 0.000000352 | 0.012876614 |
| TBC1D9 | 5 | -0.285943776 | 0.266 | 0.375 | 0.000000372 | 0.013604808 |
| NUBPL | 5 | 0.261316115 | 0.591 | 0.491 | 0.00000043 | 0.01573504 |
| LANCL2 | 5 | -0.280354064 | 0.202 | 0.295 | 0.000000444 | 0.016245371 |
| NAPEPLD | 5 | -0.298458126 | 0.281 | 0.36 | 0.000000475 | 0.017392837 |
| CLHC1 | 5 | -0.260508539 | 0.22 | 0.31 | 0.000000485 | 0.017773575 |
| DPYSL2 | 5 | -0.310601946 | 0.411 | 0.516 | 0.000000491 | 0.017970606 |
| LINC01572 | 5 | 0.303181057 | 0.622 | 0.532 | 0.000000514 | 0.018805174 |
| SPRY4-AS1 | 5 | -0.311457341 | 0.095 | 0.176 | 0.000000515 | 0.018858395 |
| AATF | 5 | -0.251848001 | 0.31 | 0.425 | 0.000000516 | 0.018903038 |
| MGAT5 | 5 | -0.34847827 | 0.389 | 0.478 | 0.000000516 | 0.018907117 |
| USP49 | 5 | -0.263436313 | 0.198 | 0.301 | 0.000000535 | 0.019603098 |
| FAM13A | 5 | -0.312056383 | 0.591 | 0.557 | 0.000000539 | 0.019720858 |
| CLSTN2 | 5 | -0.45002638 | 0.35 | 0.341 | 0.000000544 | 0.019906693 |
| ADAM9 | 5 | -0.256256639 | 0.257 | 0.364 | 0.000000571 | 0.020893864 |
| ZFP64 | 5 | -0.264589339 | 0.193 | 0.292 | 0.000000577 | 0.021108504 |
| VPS50 | 5 | 0.252917089 | 0.472 | 0.374 | 0.000000578 | 0.021176316 |
| SPIRE1 | 5 | -0.28785118 | 0.394 | 0.504 | 0.000000589 | 0.021561781 |
| BPTF | 5 | -0.262054548 | 0.782 | 0.852 | 0.000000607 | 0.0222334 |
| AC092535.4 | 5 | 0.2694965 | 0.372 | 0.273 | 0.00000064 | 0.023436411 |
| SYT10 | 5 | -0.338644451 | 0.073 | 0.128 | 0.000000652 | 0.02388744 |
| DDB2 | 5 | 0.254645162 | 0.297 | 0.238 | 0.000000675 | 0.024694317 |
| POTEF | 5 | -0.251475668 | 0.145 | 0.238 | 0.000000683 | 0.025005742 |

| | | | | | | |
|---|---|---|---|---|---|---|
| CASP8AP2 | 5 | 0.268906937 | 0.571 | 0.469 | 0.000000716 | 0.026226737 |
| SAT1 | 5 | -0.517062888 | 0.079 | 0.152 | 0.000000759 | 0.027792928 |
| MB21D2 | 5 | -0.291946638 | 0.193 | 0.291 | 0.000000761 | 0.027850661 |
| SORT1 | 5 | -0.268604145 | 0.222 | 0.302 | 0.00000081 | 0.029660473 |
| RDX | 5 | -0.302166226 | 0.51 | 0.58 | 0.00000082 | 0.030031739 |
| LRP1B | 5 | -0.810223606 | 0.174 | 0.2 | 0.000000905 | 0.033119723 |
| KPNA5 | 5 | 0.261199776 | 0.45 | 0.381 | 0.000000948 | 0.034704909 |
| CLVS1 | 5 | -0.386392047 | 0.139 | 0.212 | 0.00000096 | 0.035137559 |
| TMEM163 | 5 | -0.39827229 | 0.101 | 0.157 | 0.000000982 | 0.035942675 |
| NCOA1 | 5 | -0.35660729 | 0.642 | 0.692 | 0.000000997 | 0.03651791 |
| ARL5A | 5 | -0.317475755 | 0.365 | 0.441 | 0.000000999 | 0.036586077 |
| WDR35 | 5 | -0.320894171 | 0.4 | 0.472 | 0.000001 | 0.036655482 |
| TAX1BP1 | 5 | -0.251633578 | 0.356 | 0.466 | 0.00000103 | 0.037553617 |
| FXR1 | 5 | -0.254786751 | 0.516 | 0.61 | 0.00000103 | 0.037615534 |
| ZNF280D | 5 | -0.269954773 | 0.217 | 0.317 | 0.00000112 | 0.04099698 |
| MT-ND2 | 5 | -0.258751315 | 0.073 | 0.142 | 0.00000114 | 0.04179608 |
| CEP290 | 5 | 0.268902077 | 0.453 | 0.36 | 0.00000114 | 0.041798523 |
| BMT2 | 5 | -0.286553047 | 0.266 | 0.368 | 0.00000116 | 0.042457634 |
| KIF2A | 5 | -0.286076923 | 0.53 | 0.611 | 0.00000117 | 0.042923982 |
| DACH1 | 5 | -0.250633213 | 0.947 | 0.883 | 0.00000119 | 0.043551927 |
| ADGRA3 | 5 | -0.271935341 | 0.624 | 0.685 | 0.00000127 | 0.046432796 |
| UBA6-AS1 | 5 | -0.326344072 | 0.224 | 0.302 | 0.00000129 | 0.047307308 |
| MYO19 | 5 | 0.251427939 | 0.534 | 0.459 | 0.00000133 | 0.048871017 |
| RUFY3 | 5 | -0.255333084 | 0.538 | 0.63 | 0.00000135 | 0.049429276 |
| PAK3 | 5 | -0.326219561 | 0.886 | 0.867 | 0.00000145 | 0.052951141 |
| RNF217 | 5 | -0.287386341 | 0.299 | 0.405 | 0.00000148 | 0.054211726 |
| ZMIZ1 | 5 | -0.272596489 | 0.372 | 0.477 | 0.00000153 | 0.05594813 |
| PEX1 | 5 | -0.317406989 | 0.323 | 0.406 | 0.00000154 | 0.056534427 |
| KHDRBS3 | 5 | -0.275150984 | 0.272 | 0.241 | 0.00000156 | 0.0570117 |
| E2F7 | 5 | 0.279493728 | 0.272 | 0.183 | 0.00000156 | 0.057187972 |
| LRRC40 | 5 | -0.26285369 | 0.341 | 0.441 | 0.00000163 | 0.059696697 |
| CREM | 5 | -0.252693821 | 0.174 | 0.264 | 0.00000185 | 0.067887267 |
| ELP1 | 5 | 0.250504423 | 0.398 | 0.309 | 0.0000019 | 0.069554932 |
| FIGN | 5 | -0.443762825 | 0.299 | 0.34 | 0.00000194 | 0.070963304 |
| ST5 | 5 | -0.292766809 | 0.261 | 0.299 | 0.00000197 | 0.071958791 |
| TEX10 | 5 | -0.258838869 | 0.294 | 0.395 | 0.00000198 | 0.072484215 |
| SLC8A1 | 5 | -0.450656692 | 0.312 | 0.331 | 0.00000207 | 0.075894971 |
| SH3BGRL | 5 | -0.30446318 | 0.299 | 0.402 | 0.00000232 | 0.084787877 |
| NXN | 5 | -0.272239027 | 0.442 | 0.407 | 0.00000239 | 0.087435118 |
| GABRA2 | 5 | -0.480718775 | 0.062 | 0.108 | 0.00000275 | 0.100803341 |
| SLC16A1 | 5 | -0.255908799 | 0.209 | 0.307 | 0.00000276 | 0.101150446 |
| ADAM22 | 5 | -0.335502901 | 0.352 | 0.41 | 0.0000029 | 0.106237017 |
| MBTD1 | 5 | -0.250685553 | 0.492 | 0.599 | 0.00000293 | 0.107315264 |
| PBX1 | 5 | -0.289791353 | 0.831 | 0.852 | 0.00000345 | 0.126173994 |
| TOPBP1 | 5 | 0.251915919 | 0.618 | 0.542 | 0.00000367 | 0.134330515 |
| PPIP5K2 | 5 | -0.275562356 | 0.462 | 0.549 | 0.00000388 | 0.142087315 |
| NEAT1 | 5 | -0.701047188 | 0.866 | 0.865 | 0.00000406 | 0.148494915 |
| DAAM1 | 5 | -0.369933552 | 0.47 | 0.505 | 0.00000495 | 0.181387577 |
| SON | 5 | -0.270666987 | 0.606 | 0.687 | 0.00000514 | 0.188031694 |
| DLC1 | 5 | -0.551741347 | 0.224 | 0.269 | 0.00000546 | 0.199783246 |
| ATG2B | 5 | 0.253828801 | 0.519 | 0.439 | 0.0000056 | 0.205110339 |
| ATP1B3 | 5 | -0.255461455 | 0.328 | 0.422 | 0.00000601 | 0.219850382 |
| EPM2A | 5 | -0.30647425 | 0.218 | 0.287 | 0.00000654 | 0.239293413 |
| PPA2 | 5 | -0.262002974 | 0.576 | 0.648 | 0.00000663 | 0.24275299 |
| KIFAP3 | 5 | -0.268509294 | 0.316 | 0.414 | 0.00000666 | 0.24393803 |
| CUX1 | 5 | -0.295777822 | 0.765 | 0.788 | 0.00000704 | 0.257723957 |
| UACA | 5 | -0.31031365 | 0.312 | 0.381 | 0.00000729 | 0.267060445 |
| AC004160.1 | 5 | 0.260412076 | 0.178 | 0.122 | 0.00000747 | 0.273527853 |

| | | | | | | |
|---|---|---|---|---|---|---|
| VRK1 | 5 | -0.277921799 | 0.468 | 0.512 | 0.00000748 | 0.273969431 |
| ZNF804A | 5 | -0.255211658 | 0.297 | 0.244 | 0.00000751 | 0.275105403 |
| YWHAZ | 5 | -0.271067108 | 0.516 | 0.604 | 0.00000815 | 0.298240555 |
| RUNX2 | 5 | -0.3462538 | 0.086 | 0.14 | 0.000011 | 0.402206483 |
| FRYL | 5 | -0.25284025 | 0.818 | 0.854 | 0.000012 | 0.437989677 |
| RFC3 | 5 | -0.33428323 | 0.409 | 0.477 | 0.0000125 | 0.456773862 |
| PTPRJ | 5 | -0.411262301 | 0.444 | 0.52 | 0.0000135 | 0.495931667 |
| KIAA1324L | 5 | -0.254117644 | 0.147 | 0.225 | 0.0000141 | 0.516502419 |
| AC138627.1 | 5 | -0.490802904 | 0.114 | 0.167 | 0.0000152 | 0.557135719 |
| EXT1 | 5 | -0.317943013 | 0.653 | 0.693 | 0.0000164 | 0.600827172 |
| CENPP | 5 | -0.46681291 | 0.637 | 0.695 | 0.0000166 | 0.609337782 |
| CENPE | 5 | -0.285675687 | 0.123 | 0.2 | 0.0000171 | 0.626393168 |
| DENND4C | 5 | -0.268136468 | 0.648 | 0.703 | 0.000019 | 0.694684983 |
| GPI | 5 | 0.262506701 | 0.341 | 0.257 | 0.0000213 | 0.779900497 |
| MCMBP | 5 | -0.253012793 | 0.371 | 0.443 | 0.0000215 | 0.78799136 |
| MT-ND3 | 5 | -0.262132233 | 0.097 | 0.166 | 0.0000223 | 0.815486574 |
| JAK2 | 5 | -0.272157606 | 0.22 | 0.306 | 0.0000321 | 1 |
| LBR | 5 | -0.289102865 | 0.261 | 0.345 | 0.000039 | 1 |
| ANKRD36B | 5 | -0.266264857 | 0.415 | 0.489 | 0.000041 | 1 |
| CLIP1 | 5 | -0.259239341 | 0.354 | 0.438 | 0.0000413 | 1 |
| ADCY2 | 5 | -0.355960865 | 0.301 | 0.277 | 0.0000578 | 1 |
| LIMS1 | 5 | -0.272045908 | 0.417 | 0.495 | 0.0000635 | 1 |
| PLCB4 | 5 | -0.275300154 | 0.677 | 0.671 | 0.0000643 | 1 |
| GALNT14 | 5 | -0.286572942 | 0.189 | 0.252 | 0.0000713 | 1 |
| AC078881.1 | 5 | -0.298689137 | 0.147 | 0.205 | 0.000073 | 1 |
| MEF2A | 5 | -0.260848378 | 0.772 | 0.81 | 0.0000789 | 1 |
| PRKN | 5 | -0.307035528 | 0.47 | 0.542 | 0.0000813 | 1 |
| ANKRD37 | 5 | 0.296764465 | 0.103 | 0.073 | 0.0000874 | 1 |
| RAP1GDS1 | 5 | -0.282039188 | 0.631 | 0.67 | 0.0000988 | 1 |
| DIAPH3 | 5 | -0.318766803 | 0.327 | 0.416 | 0.000114 | 1 |
| PHKA1 | 5 | -0.26039013 | 0.237 | 0.294 | 0.000114 | 1 |
| HIF1A-AS3 | 5 | -0.31426833 | 0.086 | 0.132 | 0.000124 | 1 |
| GRIK2 | 5 | -0.560274735 | 0.253 | 0.294 | 0.000154 | 1 |
| BCL2 | 5 | -0.27737616 | 0.152 | 0.198 | 0.000175 | 1 |
| FAR2 | 5 | -0.283875326 | 0.407 | 0.383 | 0.000201 | 1 |
| PLAGL1 | 5 | -0.335973229 | 0.347 | 0.358 | 0.000203 | 1 |
| TSHZ1 | 5 | -0.290278412 | 0.338 | 0.407 | 0.00021 | 1 |
| CDC42BPA | 5 | -0.29134665 | 0.741 | 0.774 | 0.000211 | 1 |
| NAMPT | 5 | -0.301358975 | 0.239 | 0.305 | 0.000212 | 1 |
| ESYT2 | 5 | -0.287386129 | 0.497 | 0.558 | 0.000232 | 1 |
| PDK1 | 5 | -0.287561064 | 0.308 | 0.385 | 0.000239 | 1 |
| GSTCD | 5 | -0.286081883 | 0.407 | 0.48 | 0.000239 | 1 |
| ANKRD6 | 5 | -0.252588429 | 0.237 | 0.306 | 0.000271 | 1 |
| SMARCA1 | 5 | -0.27414859 | 0.644 | 0.673 | 0.000284 | 1 |
| SERPINE2 | 5 | -0.29792931 | 0.112 | 0.15 | 0.000286 | 1 |
| KRAS | 5 | -0.326653941 | 0.317 | 0.375 | 0.000295 | 1 |
| CCDC102B | 5 | -0.755781115 | 0.103 | 0.141 | 0.000321 | 1 |
| INPP4B | 5 | -0.345681104 | 0.16 | 0.167 | 0.000373 | 1 |
| LAMB1 | 5 | -0.370004032 | 0.334 | 0.391 | 0.000429 | 1 |
| BBS9 | 5 | -0.260746893 | 0.424 | 0.469 | 0.000551 | 1 |
| LPP | 5 | -0.255128589 | 0.677 | 0.735 | 0.00065 | 1 |
| HIVEP2 | 5 | -0.267438929 | 0.369 | 0.433 | 0.00101029 | 1 |
| MID1 | 5 | -0.282397317 | 0.549 | 0.611 | 0.001094368 | 1 |
| OPHN1 | 5 | -0.286936196 | 0.347 | 0.402 | 0.001525334 | 1 |
| ADARB1 | 5 | -0.257866562 | 0.239 | 0.289 | 0.003368101 | 1 |
| ADGRG2 | 5 | -0.250761697 | 0.132 | 0.183 | 0.003515809 | 1 |
| SHANK2 | 5 | -0.292806402 | 0.18 | 0.222 | 0.005389923 | 1 |
| FAM162A | 5 | -0.273329875 | 0.185 | 0.212 | 0.006651061 | 1 |

TableS18

| Motif Analysis in HPV-Positive Gland_A cells | | | | | | | |
|---|---|---|---|---|---|---|---|
| motif | gene | observed | background | percent.observed | percent.background | fold.enrichment | p.adjust |
| MA0847.2 | FOXD2 | 117 | 8810 | 49.15966387 | 22.025 | 2.231993819 | 1.90E-17 |
| MA0845.1 | FOXB1 | 108 | 9166 | 45.37815126 | 22.915 | 1.98028153 | 5.88E-12 |
| MA1487.1 | FOXE1 | 97 | 7755 | 40.75630252 | 19.3875 | 2.102194843 | 5.88E-12 |
| MA0047.3 | FOXA2 | 82 | 6374 | 34.45378151 | 15.935 | 2.162145059 | 3.27E-10 |
| MA0509.2 | RFX1 | 67 | 4689 | 28.1512605 | 11.7225 | 2.401472425 | 6.73E-10 |
| MA0032.2 | FOXC1 | 105 | 9613 | 44.11764706 | 24.0325 | 1.835749383 | 9.88E-10 |
| MA0846.1 | FOXC2 | 118 | 11550 | 49.57983193 | 28.875 | 1.717050457 | 1.34E-09 |
| MA0798.2 | RFX3 | 63 | 4460 | 26.47058824 | 11.15 | 2.374043788 | 3.74E-09 |
| MA0852.2 | FOXK1 | 67 | 4988 | 28.1512605 | 12.47 | 2.257518886 | 5.84E-09 |
| MA1683.1 | FOXA3 | 71 | 5552 | 29.83193277 | 13.88 | 2.149274695 | 1.06E-08 |
| MA0148.4 | FOXA1 | 78 | 6588 | 32.77310924 | 16.47 | 1.989866985 | 3.24E-08 |
| MA0153.2 | HNF1B | 58 | 4175 | 24.3697479 | 10.4375 | 2.334826146 | 3.60E-08 |
| MA0481.3 | FOXP1 | 72 | 5998 | 30.25210084 | 14.995 | 2.017479216 | 9.70E-08 |
| MA1606.1 | Foxf1 | 76 | 6621 | 31.93277311 | 16.5525 | 1.929181278 | 1.96E-07 |
| MA1103.2 | FOXK2 | 60 | 4675 | 25.21008403 | 11.6875 | 2.157012538 | 2.76E-07 |
| MA0042.2 | FOXI1 | 57 | 4459 | 23.94957983 | 11.1475 | 2.148426089 | 6.51E-07 |
| MA0033.2 | FOXL1 | 57 | 4459 | 23.94957983 | 11.1475 | 2.148426089 | 6.51E-07 |
| MA0848.1 | FOXO4 | 57 | 4459 | 23.94957983 | 11.1475 | 2.148426089 | 6.51E-07 |
| MA0849.1 | FOXO6 | 57 | 4459 | 23.94957983 | 11.1475 | 2.148426089 | 6.51E-07 |
| MA0850.1 | FOXP3 | 57 | 4459 | 23.94957983 | 11.1475 | 2.148426089 | 6.51E-07 |
| MA0510.2 | RFX5 | 40 | 2600 | 16.80672269 | 6.5 | 2.585649644 | 1.11E-06 |
| MA1607.1 | Foxl2 | 85 | 8180 | 35.71428571 | 20.45 | 1.746419839 | 1.11E-06 |
| MA0046.2 | HNF1A | 68 | 5918 | 28.57142857 | 14.795 | 1.931154348 | 1.11E-06 |
| MA0031.1 | FOXD1 | 48 | 3543 | 20.16806723 | 8.8575 | 2.276948036 | 1.65E-06 |
| MA0157.2 | FOXO3 | 48 | 3543 | 20.16806723 | 8.8575 | 2.276948036 | 1.65E-06 |
| MA0600.2 | RFX2 | 35 | 2417 | 14.70588235 | 6.0425 | 2.433741391 | 3.02E-05 |
| MA0818.1 | BHLHE22 | 31 | 2074 | 13.02521008 | 5.185 | 2.51209452 | 6.78E-05 |
| MA0856.1 | RXRG | 48 | 4053 | 20.16806723 | 10.1325 | 1.990433479 | 7.33E-05 |
| MA0593.1 | FOXP2 | 75 | 7647 | 31.51260504 | 19.1175 | 1.648364328 | 8.33E-05 |
| MA0817.1 | BHLHE23 | 34 | 2447 | 14.28571429 | 6.1175 | 2.33522097 | 9.21E-05 |
| MA0678.1 | OLIG2 | 33 | 2338 | 13.86554622 | 5.845 | 2.372206368 | 9.21E-05 |
| MA0512.2 | Rxra | 46 | 3953 | 19.32773109 | 9.8825 | 1.955753209 | 0.000172 |
| MA1468.1 | ATOH7 | 46 | 4028 | 19.32773109 | 10.07 | 1.919337745 | 0.000271 |
| MA1621.1 | Rbpjl | 63 | 6400 | 26.47058824 | 16 | 1.654411765 | 0.000537 |
| MA1145.1 | FOSL2::JUND(var.2) | 28 | 1998 | 11.76470588 | 4.995 | 2.355296473 | 0.000537 |
| MA1568.1 | TCF21(var.2) | 33 | 2592 | 13.86554622 | 6.48 | 2.139744787 | 0.000649 |
| MA0851.1 | Foxj3 | 78 | 8598 | 32.77310924 | 21.495 | 1.52468524 | 0.000708 |
| MA0855.1 | RXRB | 45 | 4104 | 18.90756303 | 10.26 | 1.8428424 | 0.000809 |
| MA0826.1 | OLIG1 | 30 | 2286 | 12.60504202 | 5.715 | 2.205606652 | 0.000821 |
| MA0091.1 | TAL1::TCF3 | 61 | 6298 | 25.6302521 | 15.745 | 1.627834367 | 0.00106 |
| MA0827.1 | OLIG3 | 33 | 2695 | 13.86554622 | 6.7375 | 2.057966044 | 0.001214 |
| MA1620.1 | Ptf1a(var.3) | 62 | 6481 | 26.05042017 | 16.2025 | 1.60780251 | 0.001238 |
| MA0480.1 | Foxo1 | 69 | 7485 | 28.99159664 | 18.7125 | 1.549317122 | 0.001307 |
| MA0820.1 | FIGLA | 56 | 5693 | 23.52941176 | 14.2325 | 1.653217057 | 0.001415 |
| MA0668.1 | NEUROD2 | 25 | 1877 | 10.50420168 | 4.6925 | 2.238508616 | 0.002567 |
| MA0669.1 | NEUROG2 | 37 | 3323 | 15.54621849 | 8.3075 | 1.871347395 | 0.002654 |
| MA0030.1 | FOXF2 | 49 | 4913 | 20.58823529 | 12.2825 | 1.676225141 | 0.002979 |
| MA0799.1 | RFX4 | 33 | 2866 | 13.86554622 | 7.165 | 1.935177421 | 0.003266 |
| MA0623.2 | NEUROG1 | 28 | 2274 | 11.76470588 | 5.685 | 2.069429355 | 0.003456 |

| | | | | | | | |
|---|---|---|---|---|---|---|---|
| MA1109.1 | NEUROD1 | 76 | 8882 | 31.93277311 | 22.205 | 1.438089309 | 0.004888 |
| MA0613.1 | FOXG1 | 50 | 5226 | 21.00840336 | 13.065 | 1.607991072 | 0.006246 |
| MA0835.2 | BATF3 | 55 | 5941 | 23.1092437 | 14.8525 | 1.555916088 | 0.006697 |
| MA0719.1 | RHOXF1 | 30 | 2633 | 12.60504202 | 6.5825 | 1.914932323 | 0.006972 |
| MA0745.2 | SNAI2 | 58 | 6396 | 24.3697479 | 15.99 | 1.524061782 | 0.007332 |
| MA0665.1 | MSC | 55 | 6030 | 23.1092437 | 15.075 | 1.532951489 | 0.009098 |
| MA1525.1 | NFATC4 | 57 | 6319 | 23.94957983 | 15.7975 | 1.516036071 | 0.009136 |
| MA0609.2 | CREM | 19 | 1404 | 7.983193277 | 3.51 | 2.274414039 | 0.010551 |
| MA0461.2 | Atoh1 | 34 | 3231 | 14.28571429 | 8.0775 | 1.768581156 | 0.010849 |
| MA1608.1 | Isl1 | 28 | 2490 | 11.76470588 | 6.225 | 1.889912592 | 0.011877 |
| MA1642.1 | NEUROG2(var.2) | 73 | 8750 | 30.67226891 | 21.875 | 1.402160864 | 0.011991 |
| MA0115.1 | NR1H2::RXRA | 38 | 3811 | 15.96638655 | 9.5275 | 1.675821208 | 0.013636 |
| MA0065.2 | Pparg::Rxra | 73 | 8823 | 30.67226891 | 22.0575 | 1.390559624 | 0.014719 |
| MA0712.2 | OTX2 | 40 | 4121 | 16.80672269 | 10.3025 | 1.631324697 | 0.016022 |
| MA1104.2 | GATA6 | 45 | 4831 | 18.90756303 | 12.0775 | 1.565519605 | 0.017609 |
| MA0037.3 | GATA3 | 39 | 4033 | 16.38655462 | 10.0825 | 1.625247173 | 0.019203 |
| MA1550.1 | PPARD | 41 | 4362 | 17.22689076 | 10.905 | 1.579724049 | 0.024131 |
| MA0714.1 | PITX3 | 31 | 3025 | 13.02521008 | 7.5625 | 1.722341829 | 0.024303 |
| MA0041.1 | Foxd3 | 97 | 12752 | 40.75630252 | 31.88 | 1.278428561 | 0.025714 |
| MA1131.1 | FOSL2::JUN(var.2) | 25 | 2273 | 10.50420168 | 5.6825 | 1.848517674 | 0.025714 |
| MA0891.1 | GSC2 | 24 | 2168 | 10.08403361 | 5.42 | 1.860522807 | 0.028475 |
| MA1576.1 | THRB(var.3) | 39 | 4157 | 16.38655462 | 10.3925 | 1.576767344 | 0.030048 |
| MA0484.2 | HNF4G | 46 | 5145 | 19.32773109 | 12.8625 | 1.502641873 | 0.031064 |
| MA0648.1 | GSC | 24 | 2197 | 10.08403361 | 5.4925 | 1.835964245 | 0.032255 |
| MA0894.1 | HESX1 | 28 | 2710 | 11.76470588 | 6.775 | 1.736487953 | 0.032255 |
| MA0602.1 | Arid5a | 43 | 4753 | 18.06722689 | 11.8825 | 1.520490376 | 0.033478 |
| MA0775.1 | MEIS3 | 44 | 4910 | 18.48739496 | 12.275 | 1.506101422 | 0.035159 |
| MA0743.2 | SCRT1 | 37 | 3947 | 15.54621849 | 9.8675 | 1.575497186 | 0.036133 |
| MA1524.1 | MSGN1 | 29 | 2875 | 12.18487395 | 7.1875 | 1.69528681 | 0.036133 |
| MA1585.1 | ZKSCAN1 | 25 | 2358 | 10.50420168 | 5.895 | 1.781883237 | 0.036133 |
| MA1148.1 | PPARA::RXRA | 43 | 4809 | 18.06722689 | 12.0225 | 1.50278452 | 0.038342 |
| MA1647.1 | PRDM4 | 65 | 8019 | 27.31092437 | 20.0475 | 1.362310731 | 0.038342 |
| MA0707.1 | MNX1 | 49 | 5705 | 20.58823529 | 14.2625 | 1.443522194 | 0.043394 |
| MA1127.1 | FOSB::JUN | 21 | 1895 | 8.823529412 | 4.7375 | 1.862486419 | 0.043394 |
| MA0757.1 | ONECUT3 | 49 | 5744 | 20.58823529 | 14.36 | 1.433721121 | 0.048352 |
| MA1570.1 | TFAP4(var.2) | 28 | 2823 | 11.76470588 | 7.0575 | 1.666979225 | 0.048352 |
| MA0125.1 | Nobox | 26 | 2560 | 10.92436975 | 6.4 | 1.706932773 | 0.048352 |
| MA1133.1 | JUN::JUNB(var.2) | 19 | 1674 | 7.983193277 | 4.185 | 1.907573065 | 0.048444 |

TableS19

| Patients Information for TCGA (n=299) | | |
|---|---|---|
| Age-yr | Median(range) | 46(20-88) |
| Race-no.(%) | White | 204(68.2) |
| | Asian | 20(6.7) |
| | Black or African America | 30(10.0) |
| | American indian or Alaska native | 7(2.3) |
| | Native Hawaiian or Other Pacific Islander | 2(0.7) |
| | no data | 36(12.0) |
| Histological type-no.(%) | Squamous Cell Carcinoma | 247(82.6) |
| | Adenocarcinoma | 52(17.4) |
| HPV type-no.(%) | HPV18 | 38(12.7) |
| | others-HPV | 239(79.9) |
| | HPV-negative | 19(6.4) |
| | Normal | 3(1.0) |

TableS20

| **Patients Information for ALDH1A1 immunostaining** (n=78) | | |
|---|---|---|
| Age-yr | Median(range) | 46(20-88) |
| Race-no.(%) | Asian | 78(100) |
| Histological type-no.(%) | Squamous Cell Carcinoma | 50(64.1) |
| | Adenocarcinoma | 28(35.9) |
| Clinical stage-no.(%) | I-II | 74(95) |
| | III-IV | 4(5) |

TableS21

| Information on antibodies for immunostaining | | | |
|---|---|---|---|
| antibody | dilution | pH | product information |
| CDH1 | 1:1000 | 6.0 | Abcam #40772, EP700Y, Cambridge, UK |
| KRT8 | 1:100 | 6.0 | Thermo Fisher Scientific, #MA5-16420, SP102, Waltham, MA, USA |
| KRT18 | 1:200 | 6.0 | Abcam #93741, SP69, Cambridge, UK |
| CLDN7 | 1:1000 | 9.0 | Abcam #207300, EPR18073, Cambridge, UK |
| ERBB3 | 1:100 | 9.0 | Cell signaling technology, #12708, D22C5, Danvers, MA, USA |
| FOXA1 | 1:1000 | 9.0 | Abcam #170933, EPR10881, Cambridge, UK |
| FOXA2 | 1:50 | 9.0 | Sigma-aldrich, ZRB1872, clone 1H12, St. Louis, MO, USA |
| p16 INK4a | 1:400 | 9.0 | Abcam #108349, EPR1473, Cambridge, UK |
| ALDH1A1 | 1:100 | 6.0 | Abcam #134188, EP1932Y, Cambridge, UK |
| p63 | 1:100 | 9.0 | DAKO #M7317, DAK-p63, Santa Clara, CA, USA |
| CD56 | 1:100 | 9.0 | Leica #NCL-CD56-1B6, Germany |
| SYP | 1:1500 | 6.0 | Proteintech #17785-1-AP, Rosemont, IL, USA |
| CHGA | 1:1000 | 9.0 | DAKO #A0430, Santa Clara, CA, USA |

TableS22

| Information on antibodies for immunoblotting | |
|---|---|
| antibody | product information |
| CDH1 | Beckton Dickinson, #610181,clone36, Franklin Lakes, NJ, USA |
| p16INK4A | Santa Cruz, #sc-1661, F-12, Santa Cruz, CA, USA |
| HPV18 E6 | Santa Cruz, #sc-460, C1P5, Santa Cruz, CA, USA |
| GAPDH | Thermo Fisher Scientific, #AM4300, 6C5, Waltham, MA, USA |
